# An integrated protocol for multiplexed DNA FISH and protein detection in large tissue sections

**DOI:** 10.64898/2026.05.20.726465

**Authors:** Eleanor M. O’Roberts, Pranauti Panshikar, Xiaoze Li-Wang, Christophe Avenel, Quentin Verron, Eléna Coulier, Magda Bienko, Charlotte Stadler

**Affiliations:** Department of Protein Science, Royal Institute of Technology, Stockholm, Sweden; Department of Microbiology, Tumor and Cell Biology, Karolinska Institutet, Stockholm, SE-17165, Sweden; Department of Information Technology and SciLifeLab, Uppsala University, Uppsala Sweden; Science for Life Laboratory, Solna, 17165, Sweden; Human Technopole, Viale Rita Levi-Montalcini 1, 20157, Milan, Italy; Department of Pelvic Cancer, Genitourinary Oncology and Urology Unit, Theme Cancer, Karolinska, Stockholm, Sweden

**Author notes:** Authors contributed equally (Eleanor M. O’Roberts, Pranauti Panshikar).

**Keywords:** DNA FISH, multiplexed immunofluorescence, PhenoCycler-Fusion, spatial proteomics

## Abstract

Different omics types such as genomics and proteomics all contribute to deciphering biology. Applying these omics approaches in a spatial context helps reveal biology *in situ* at a single cell level. Here we present a protocol for the combined multiplexed detection of targeted genes using DNA FISH, and proteins using multiplexed immunofluorescence. The protocol is integrated on the commercial PhenoCycler platform and generates one single dataset with gene and protein readout at a single cell level in large tissue sections, allowing for a throughput of thousands to millions of cells. The workflow can be used for characterising malignant cells in large tumor areas based on genetic aberrations, while deciphering the cellular landscape and microenvironment from multiplexed protein detection using immunofluorescence.

## 1. Introduction

Over the past decade, spatial methods such as spatial transcriptomics, *in situ* sequencing and multiplexed imaging solutions have been developed, allowing for thousands of transcripts or tens to hundreds of proteins to be explored within a single tissue section (1–4). These methods provide in depth insights into tissue microenvironments and have been used to generate extensive cell atlases (5–7), proving also to be valuable for histopathology, diagnostics and response prediction, especially cancer treatment (8,9). In a systematic review and meta analysis of ten solid tumour types from over 8000 patients, it was suggested that spatial profiling of intact tumours using multiplexed IF provides better prognostic value than gene expression profiling of dissociated material (10). For such multiplexed protein detection there are several commercial platforms and methods available, e.g. PhenoCycler (11,12), commercialized by Akoya Biosciences – the CODEX platform, Bruker - CellScape (13), Miltenyi – MACSima (3) and Lunaphore – COMET (14,15) to mention a few. Some of these platforms now also offer combined detection of proteins and transcripts in one single experimental run (16,17). Using the combination of RNA and protein detection allows for validation of single cell RNA sequencing in a spatial context where protein markers can be used to phenotype individual cells and other structural components of a tissue. Furthermore, probe-based methods used for RNA detection can be used for targets with no good antibodies available.

While RNA and protein integration has gained a lot of attention over the past years, this has not been the case for the detection of proteins together with relevant DNA loci/genes, for which no commercial platform exists to date. At the same time, the latter is likely to be more informative in diagnostics and in clinical settings in general, as it can provide information on copy number variations (CNV), gene fusions or deletions, which are clinically highly relevant. In contrast to a transcript level of a given gene that typically varies across individual cells within a tissue, gene alterations are more stable and thereby have a better potential to be used as informative biomarkers for diagnosis and response prediction.

In oncology, DNA FISH is already used as a complementary technique to immunohistochemistry for targets such as HER2, where gene amplification provides more robust information than protein expression alone in borderline cases (18,19). Integrating such gene-level readouts with multiplexed immunofluorescence in a single assay could support the classification of malignant subpopulations and their immediate microenvironment. In the study by Lu et al, it was also demonstrated that integrating multiple modalities combining biomarker approaches provides superior prognostic and predictive information than relying on one single assay (10). Similarly, adding direct DNA-level information alongside protein expression to the same cells could, in principle, further increase the robustness of biomarker panels, particularly for CNV-driven oncogenes and tumour suppressors. However, previous studies on the combinations of DNA FISH with immunostaining such as FICTION, immunoFISH and others allow only for a limited number of DNA loci and/or protein targets to be analyzed within the same sample of intact cells or tissue (20–22). Furthermore, published methods are not suitable for large tissue sections, but have rather applied high-resolution imaging to study chromosomal locations within nuclei. In oncology, and in the clinics in general, such methods are limited due to insufficient throughput.

With this in mind, we developed a protocol that can be used to simultaneously detect proteins using immunofluorescence, and specific DNA targets using DNA FISH in cells and large tissue sections. We present a proof-of-principle protocol for automation using the commercial PhenoCycler platform, which allows for large tissue samples to be scanned and DNA CNV to be quantitatively explored in downstream analysis.

## 2. Results

### 2.1 Development of an integrated protocol for DNA FISH and multiplexed immunofluorescence in cells

Here, we used the PhenoCycler-Fusion (previously CODEX) from Akoya Biosciences to carry out multiplexed omics. This system involves the use of antibodies conjugated to oligonucleotide barcodes, each of which are unique within a panel. In each cycle, the microfluidics instrument adds up to three reporters labelled with distinct fluorophores (AF488 or AF750, Atto550, AF647), which are complementary to individual barcodes. The fluorophores are imaged and the reporters are eluted, and the sample is then ready for the next round of new reporters, imaging and elution. Since this approach for protein detection is very similar to the multiple rounds of fluorescently labelled probe hybridization/stripping seen in multiplexed FISH (23–25), we decided to employ the PhenoCycler-Fusion instrument to combine multiplexed DNA and protein detection (**Fig. 1)**.

**Figure 1.**
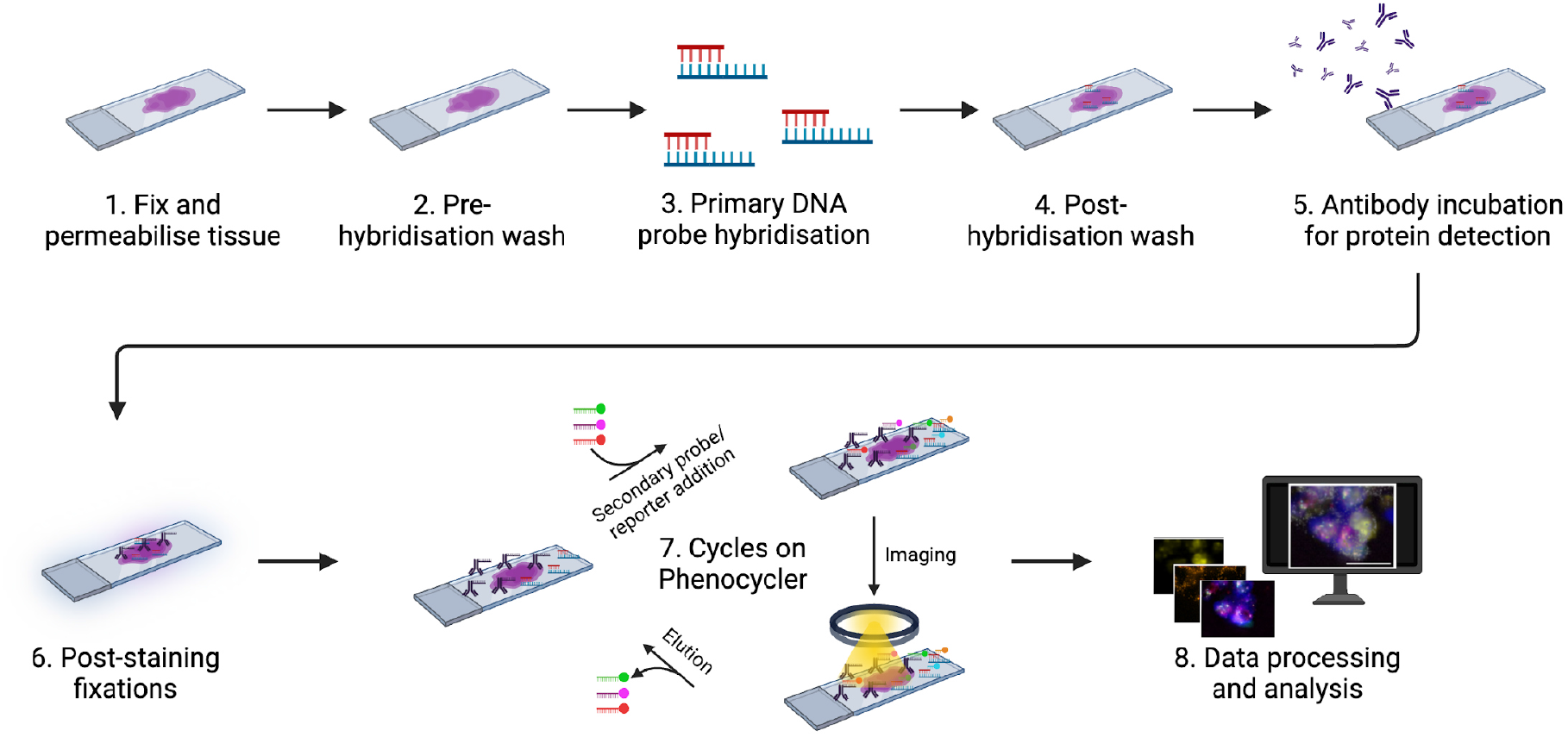
Schematic of the integrated workflow used to detect DNA and protein using the PhenoCycler-Fusion. Full details can be found in the methods section, but in brief, cells are grown on coated slides or tissue is sectioned onto slides, then fixed and permeabilised at room temperature (1). After washing the slide in hybridisation buffer (2), primary DNA FISH probes are hybridised overnight at 37°C (3). Post-hybridisation washes are carried out using SSC/0.2% tween at 56°C followed by wash buffer at room temperature (4). The panel of antibodies are incubated on the slide overnight at 4°C (5) and three post-staining fixations are carried out the next day (6). The Phenocycler-Fusion run is then set up consisting of multiple cycles. Each cycle includes hybridisation of secondary FISH probes to detect DNA, or reporters complementary to conjugated antibodies to detect proteins; imaging of fluorophores attached to secondary probes or reporters; and elution of secondary probes or reporters (7). Finally images are processed to subtract background and align cycles, and data is analysed using open source software (8).

We first tested our integrated workflow for detecting DNA and protein on cells. For DNA detection, 3 different regions of chromosome 13, as well as a probe targeting the locus of the MYC gene, were used throughout the study (**Table 1**). These probes were successfully used in previous studies and were designed taking into account probe size and oligo distribution using the iFISH pipeline (26,27). Antibodies and corresponding proteins included in this work are summarized in **Table 2**.

**Table 1.** Details of DNA FISH probes used in cell experiments.

| Target | Probe coordinates | Oligonucleotide count | Fluorophore |
| --- | --- | --- | --- |
| Ch13 region 3 | chr13:29000112-29495469 | 6000 | AF555 |
| Ch13 region 2 | chr13:53695573-54162909 | 6000 | AF647 |
| Ch13 region 1 | chr13:77193678-77674804 | 6000 | AF790 |
| Myc | chr8:127827010-128978489 | 10759 | AF647 or AF488 |

**Table 2.** Details of protein markers and antibodies used in this study.

| Marker | Clone | Supplier | Catalogue no. | Barcode | Reporter and fluorophore | Dilution | Exposure time (ms) |
| --- | --- | --- | --- | --- | --- | --- | --- |
| β-Catenin1 | 12F7 | Akoya | 4450036 | BX020 | RX020-Atto550 | 1:100 | 400 (cells) |
| CD3 | UCHT1 | Akoya | 4550103 | BX015 | RX015-AF647 | 1:100 | 200 (tissue) |
| CD4 | SK3 | Akoya | 4550105 | BX021 | RX021-AF647 | 1:100 | 200 (tissue) |
| CD11c | S-HCL-3 | Akoya | 4550107 | BX027 | RX027-AF647 | 1:150 | 200 (tissue) |
| CD31 | WM59 | Akoya | 425000<br>9 | BX032 | RX032-Atto550 | 1:150 | 200<br>(tissue) |
| CD45 | HI30 | Akoya | 415000<br>3 | BX001 | RX001-AF488 | 1:100 | 200<br>(tissue) |
| CD45RO | UCHL<br>1 | Akoya | 425002<br>3 | BX017 | RX017-Atto550 | 1:100 | 200<br>(tissue) |
| HLA-DR | L243 | Akoya | 425000<br>6 | BX026 | RX026-Atto550 | 1:150 | 200<br>(tissue) |
| HSP90B1 | CL264<br>7 | Atlas<br>Antibodi<br>es | AMAb<br>91019 | BX030 | RX030-AF647 | 1:50 | 250 (cells) |
| Keratin<br>8/18 | C51 | Akoya | 455008<br>2 | BX081 | RX081-AF647 | 1:400 | 50 (cells) |
| Ki67 | B56 | Akoya | 425001<br>9 | BX047 | RX047-Atto550 | 1:200<br>(cells)<br>1:100<br>(tissue) | 250 (cells),<br>200<br>(tissue) |
| LMNB1 | CL392<br>9 | Atlas | AMAb<br>91251 | BX026 | RX026-Atto550 | 1:50 | 500 (cells) |
| Pan-<br>cytokerati<br>n | AE-1/AE-3 | Akoya | 445002<br>0 | BX019 | RX019-AF750 (cells)<br>RX019-AF488<br>(tissue) | 1:200<br>(cells)<br>1:600<br>(tissue) | 400 (cells)<br>150<br>(tissue) |

Our workflow includes hybridisation of primary DNA FISH probes followed by antibody staining on the bench before the sample is loaded onto the PhenoCycler-Fusion instrument for cyclic addition of secondary fluorescent DNA FISH probes and reporters. Once all the rounds of imaging are complete, the data is processed in order for all the images from each cycle to be collated into a single image and analysed simultaneously in a single spatial context.

We conducted multiple optimizations of the protocol. First, we tested various stripping buffers containing increasing DMSO concentrations, starting from the lowest of 40% all the way to 70% of DMSO PhenoCycler-Fusion **(Supp. Fig. 1A-D**). We found that at least 50% of DMSO were required to elute secondary fluorescent DNA FISH probes. As shown in the **Supplementary Figure 1E**, after stripping with 40% DMSO, 89% of MYC probe signal remains compared to 23% of the signal after stripping with 50% DMSO. Although the stripping images appear to have high background in the Myc channel **(Supp. Fig. 1A-D)**, these were imaged at a higher exposure time to capture any residual signal after stripping and are not indicative of the true increase in the Myc background. This is further supported by the mean pixel intensity measurements in **(Supp. Fig. 1E)**. Re-hybridisation tests show that 50% DMSO or less maintains primary DNA FISH probe hybridisation, whereas higher concentrations of DMSO cause loss of the primary probe **(Supp. Fig. 1E**). Re-hybridisation with secondary probes after stripping with 50% DMSO yielded 94% of the original signal, whereas after stripping with 60% DMSO, re-hybridisation with secondary probe resulted in only 31% of the original signal from the MYC probe, suggesting that the primary DNA probe had been stripped **(Supp. Fig. 1E)**. We then tested whether lowering the concentration of DMSO to various levels would negatively affect the efficient elution of PhenoCycler-Fusion reporters. Complete elution of the PhenoCycler-Fusion antibody reporters is important as inefficient elution could cause false positive signals for the proteins detected in the same channel of the following cycle. **Supplementary Figure 2** shows that LMNB1 was still detected when using a stripping buffer containing 45% DMSO, as the signal to noise (SNR) level for the LMNB1 staining was similar as for the unstripped control, suggesting insufficient stripping. However, with increasing DMSO concentrations of 50 % or more, the SNR dropped compared to the control **(Supp. Fig. 2B)**. Taking these results together, we chose to use a stripping buffer containing 50% DMSO since this preserves primary DNA FISH probe hybridisation, whilst efficiently removing both secondary DNA FISH probes as well as reporters.

**Figure 2.**
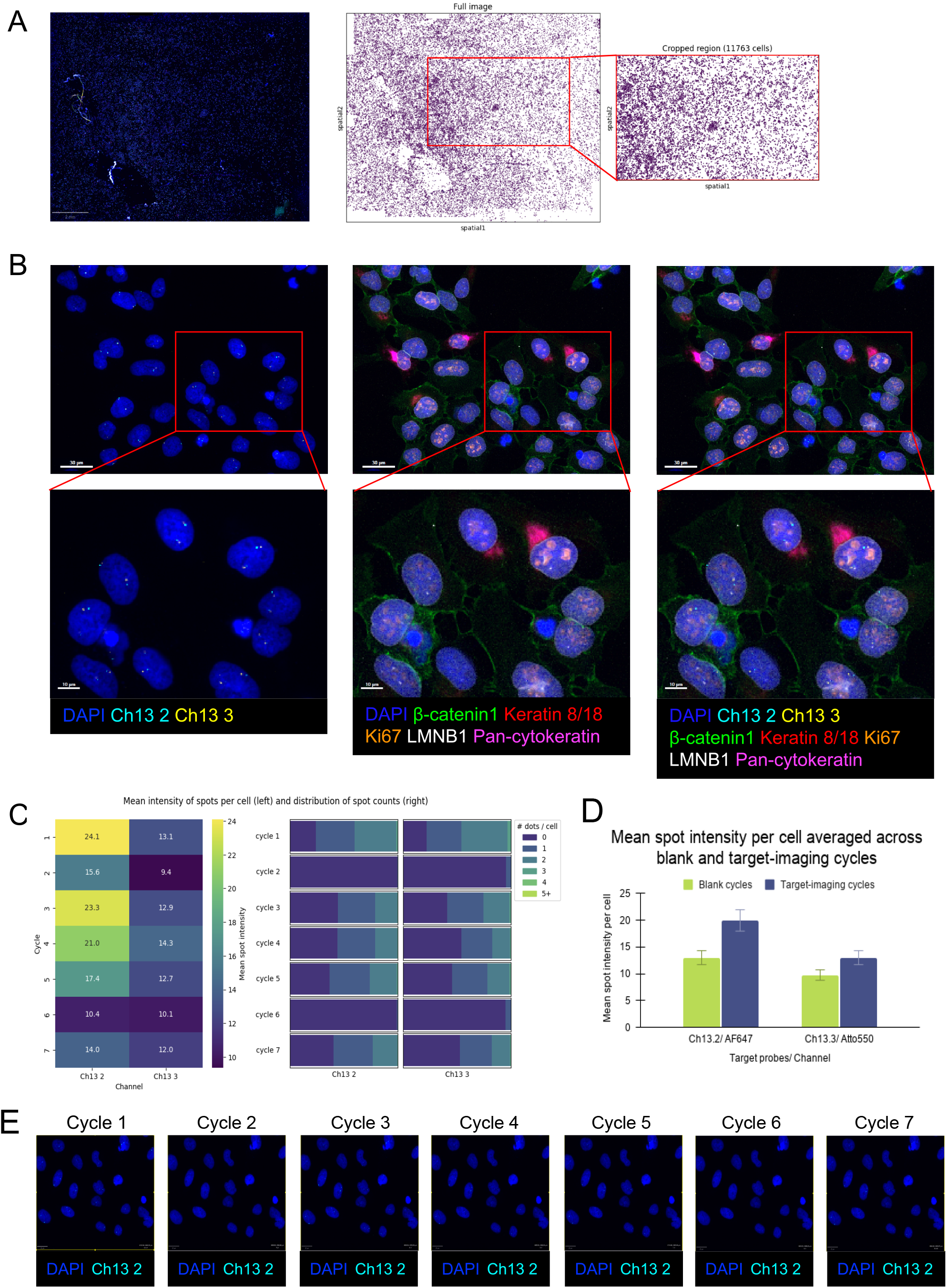
Successful integration of DNA and protein imaging in cells. (A) Left panel shows an overview of the whole imaging area for U2OS cells (scale bar represents 2 mm). DAPI shown in blue and all other markers follow the same legend as in panel (B). Right panel shows the area and segmentation mask used for analysis in panel (C). (B) DNA FISH and protein signal in U2OS cells imaged on the Phenocycler-Fusion. The red squares in the upper panel indicate the enlarged area shown for each image in the lower panel. Scale bar represents 30 µm and 10 µm. Legend indicates the colour each marker is shown in. (C) Graph of mean intensity of DNA FISH spots across imaging cycles in the left panel and distribution of the number of DNA FISH spots per cell across blank and imaging cycles in the right panel. The legend displays the colours used for the heatmap with lighter colour illustrating higher mean spot intensity or higher number of dots per cell. (D) Graph comparing the cycle-averaged mean spot intensity per cell in blank and target-imaging cycles for Ch13.2 and Ch13.3. Values for target-imaging cycles represent the average across 5 target-imaging cycles, and values for blank cycles represent the average across 2 blank cycles. (E) Representative images of DNA FISH signal across seven Phenocycler-Fusion cycles in U2OS cells. Scale bar represents 20 µm. Legend indicates the colour each marker is shown in.

The results of the integrated workflow in U2OS cells is depicted in **Figure 2**. U2OS cells were chosen for this experiment due to our prior experience working with this standard cell line in the frame of the Human Protein Atlas (www.proteinatlas.org), and five protein targets were chosen based on their expression in U2OS cells in different cellular sub-compartments (28,29). DNA FISH probes were designed using our iFISH pipeline (26), and we chose 6000 oligonucleotides per probe PhenoCycler-Fusion to secure a sensitive detection based on prior experience.

Using the setup as shown in **(Table 3**), we show a successful detection of two DNA targets together with five protein targets in an area of 16 mm x 13 mm using a 20x air objective **(Fig. 2A left panel**). While DNA FISH signal was detected in the ATTO550 and the AF647 channel (probe 2 and 3 on Chr13), no signal was detected in the AF750 channel used to detect probe 1 on Chr13. This is probably due to lower quantum yield in this wavelength and sub-optimal matching of the detection channel (AF750) with the secondary DNA probe for this target (AF790). The signal from two DNA FISH probes (Chr 13 region 2 and 3) was clearly detected in the respective channels, while protein detection was successful in all PhenoCycler-Fusion channels **(Fig. 2B)**.

**Table 3.** Cycle set up for cell experiments.

| **Cycle** | **Atto550** | **AF647** | **AF750** |
| --- | --- | --- | --- |
| Blank<br>(autofluorescence) | - | - | - |
| 1 | Ch13 region 3 | Ch13 region 2 | Ch13 region 1 |
| 2 (Blank 1) | - | - | - |
| 3 | Ch13 region 3 | Ch13 region 2 | Ch13 region 1 |
| 4 | Ch13 region 3 | Ch13 region 2 | Ch13 region 1 |
| 5 | Ch13 region 3 | Ch13 region 2 | Ch13 region 1 |
| 6 (Blank 2) | - | - | - |
| 7 | Ch13 region 3 | Ch13 region 2 | Ch13 region 1 |
| 8 | Ki67 - RX047 | HSP90B1 - RX030 | Pan-cytokeratin - RX019 |
| 9 | β-Catenin1 - RX020 | Keratin 8/18 - RX081 | - |
| 10 | LMNB1 - RX026 | - | - |
| Blank<br>(autofluorescence) | - | - | - |

We then tested the stability of the DNA FISH signal through multiple rounds of hybridization and stripping. We tested this using the PhenoCycler-Fusion with sequential re-hybridisations of secondary fluorescent DNA FISH probes across seven cycles. We found that the DNA FISH signal was still detected after seven rounds of secondary probe addition, imaging and elution in both the ATTO550 and the AF647 channels **(Fig. 2C and E**). The mean intensity of detected DNA FISH signal was maintained across cycles in the ATTO550 channel (13.1 in cycle 1, 12.0 in cycle 7) and decreased slightly across cycles in the AF647 channel (24.1 in cycle 1, 14.0 in cycle 7) (**Fig. 2C left panel**). This analysis was carried out on ∼12,000 cells, corresponding to the cells shown in **Fig. 2A right panel**. The distribution of DNA FISH spots fluctuated only very mildly across cycles, with more cells containing 2 dots (as expected since no chromosomal aberrations should be present on chromosome 13) in cycle 1 compared to other cycles (**Fig. 2C right panel)**. We also included two blank cycles in the experimental setup **(Table 3)**; these included the same three steps of hybridisation, imaging and elution, with no probe present in the solution. Consistent with our previous tests during optimization, the blank cycles confirm that elution of secondary DNA FISH probe is efficient with 50% DMSO, whilst primary DNA FISH probe hybridisation is maintained on the PhenoCycler-Fusion. **Fig. 2D** showing the comparison of mean spot intensity averaged across the blank cycles and target-imaging cycles for both the target probes.

Overall, these results show the possibility of using a commercially available system for detecting multiplexed DNA and proteins simultaneously across a large imaging area at high speed. A single cycle on the PhenoCycler-Fusion took approximately one hour. The setup requires only minimal changes to the standard protocol and while the same probe targets were used across all cycles in this work, this set-up demonstrates the potential of detecting at least 14 different DNA targets if detected in the ATTO550 and AF647 channel along with proteins.

### 2.2 Implementing the combination of DNA FISH and multiplexed immunofluorescence in tissue

Having obtained satisfactory results for the integrated protocol of DNA and protein detection from cells, we then carried out experiments on tissue samples. For this, we followed the same protocol as shown in **Figure 1** and used fresh frozen human tonsil sections (**Fig. 3A left panel)**. The same DNA targets used to optimize the protocol in cells were used for the tissue. For the protein targets, we included additional markers (CD3, CD4, CD11c, CD45, CD31, HLA-DR, CD45RO, PanCK and ki67) to explore whether we could possibly assign DNA FISH signals to individual cell types and outline different structures as tissue samples are more complex (**Table 4**). The entire sample measuring 11 mm x 11 mm was imaged. As in the case of the cells, we were able to successfully detect the same two DNA probes (Chr13 probe 2 and probe 3) detected in the ATTO550 and AF647 channels (**Fig. 3B**), while the DNA FISH probe in the AF750 channel was not detected. In the case of the proteins, the nuclear protein - Ki67 - and membrane/cytoplasmic protein - pan-cytokeratin - were well detected in tissues, illustrating that this integrated protocol can be used for detecting proteins in different cellular sub-compartments and different structures in the tissue to aid in cell phenotyping. However, the additional protein targets included in this experiment were either not detected or detected with very weak signals.

**Table 4.** Cycle set up for FF tissue experiments.

| **Cycle** | **Atto550** | **AF647** | **AF488** |
| --- | --- | --- | --- |
| 1 Blank 0<br>(autofluorescence) | - | - | - |
| 2 | Ch13 region 3 | Ch13 region 2 |  |
| 3 (Blank 1) | - | - |  |
| 4 | RX032-CD31 | RX021-CD4 |  |
| 5 | Ch13 region 3 | Ch13 region 2 |  |
| 6 | Ch13 region 3 | Ch13 region 2 |  |
| 7 | Ch13 region 3 | Ch13 region 2 |  |
| 8 (Blank 2) | - | - |  |
| 9 | Ch13 region 3 | Ch13 region 2 |  |
| 10 | Ch13 region 3 | Ch13 region 2 |  |
| 11 | Ch13 region 3 | Ch13 region 2 |  |
| 12 (Blank 3) | - | - |  |
| 13 | RX026-HLA-DR | RX027-CD11c |  |
| 14 | Ch13 region 3 | Ch13 region 2 |  |
| 15 | Ch13 region 3 | Ch13 region 2 |  |
| 16 | Ch13 region 3 | Ch13 region 2 |  |
| 17 (Blank 4) | - | - | - |
| 18 | RX017-CD45RO | - | - |
| 19 | RX047-Ki67 | RX015-CD3 | RX019-PanCK |
| 20 Blank 5<br>(autofluorescence) | - | - | - |

**Figure 3.**
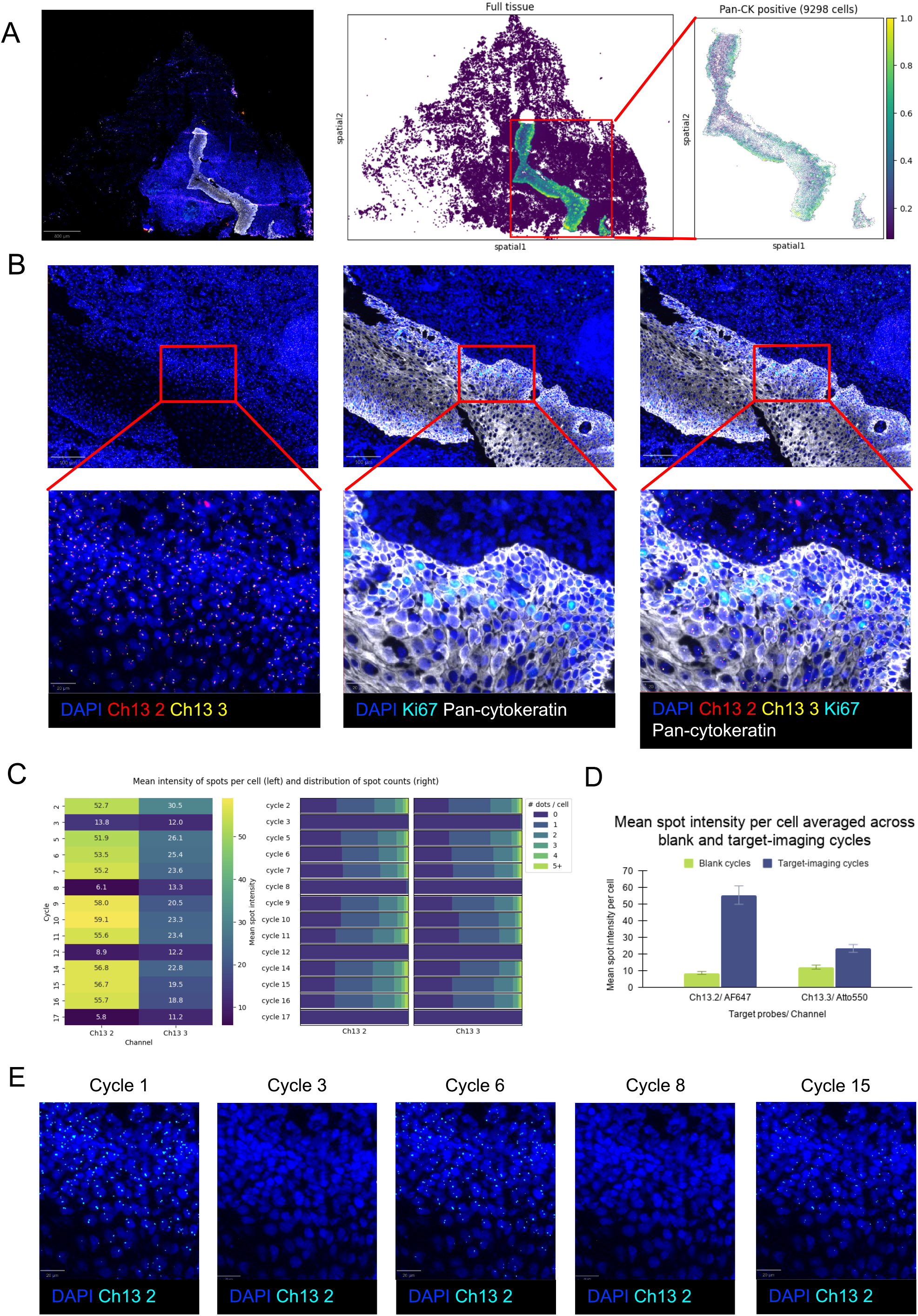
Successful integration of DNA and protein imaging in tissue. (A) Left panel shows an overview of the whole imaging area for tonsil tissue (scale bar represents 800 µm). DAPI shown in blue and all other markers follow the same legend as in panel (B). Right panel shows the area and segmentation mask used for analysis in panel (C). Legend illustrates the Pan-cytokeratin staining intensity with lighter colour indicating stronger staining. (B) DNA FISH and protein signal in fresh frozen tonsil imaged on the Phenocycler-Fusion. The red squares in the upper panel indicate the enlarged area shown for each image in the lower panel. Scale bar represents 100 µm and 20 µm. Legend indicates the colour each marker is shown in. (C) Graph of mean intensity of DNA FISH spots across imaging cycles in the left panel and distribution of the number of DNA FISH spots per cell across blank and imaging cycles in the right panel. The legend displays the colours used for the heatmap with lighter colour illustrating higher mean spot intensity or higher number of dots per cell. (D) Graph comparing the cycle-averaged mean spot intensity per cell in blank and target-imaging cycles for Ch13.2 and Ch13.3. Values for target-imaging cycles represent the average across 10 target-imaging cycles, and values for blank cycles represent the average across 4 blank cycles. (E) Representative images of DNA FISH signal across 15 Phenocycler-Fusion cycles in tonsil tissue. Scale bar represents 20 µm. Legend indicates the colour each marker is shown in.

The ability to detect DNA FISH signals across multiple cycles of imaging on the PhenoCycler-Fusion was also tested in tissue. The setup for this included sequential re-hybridisations of the secondary FISH probes (Chr 13 probe 2 and 3) on the PhenoCycler-Fusion across 15 imaging cycles. We also included three blank cycles to ensure that the signal was efficiently stripped between imaging cycles **(Table 4)**, to assure that the signal we detected was not due to a build-up of residual signals from previous cycles. Our results show that DNA FISH signals could be detected across 15 imaging cycles with no decrease in mean intensity in the AF647 channel (54.6 in cycle 2, 57.2 in cycle 16) and a slight decrease in mean intensity in the Atto550 channel (30.6 in cycle 2, 18.9 in cycle 16) (**Fig. 3C left panel and E**). The number of DNA FISH spots detected per cell across cycles was stable (**Fig. 3D**) demonstrating that our integrated protocol produces reliable DNA detection data. Overall, we see higher signal to background association in channel AF647 compared to Atto550. This quantitative analysis was carried out on a representative region of tissue, in this case pan-cytokeratin positive cells, and was based on 9298 individual cells (**Fig. 3A right panel**). Overall, our results from a tissue suggest that DNA probes plus multiple proteins could be imaged on a single tissue sample using our integrated protocol on the PhenoCycler-Fusion instrument. The successful detection of protein targets is however target-dependent and individual antibodies need to be tested/adapted.

## 3. Discussion

This is a proof-of-concept work that aims at demonstrating the possibility for simultaneous detection of targeted gene regions with DNA FISH and proteins in a large image area and at high throughput compared to traditional confocal microscopy methods. We have established a protocol for the detection of specific chromosomal regions using DNA FISH together with multiplexed immunofluorescence to add information of specific gene CNV to individual cells in a cell or tissue context. More importantly, the protocol was automated on the commercial PhenoCycler-Fusion platform, allowing for large tissue areas to be captured within one single integrated experiment. This is possible thanks to the fact that the captured images of the DNA FISH signals and protein staining are co-registered at the single cell level already during the data acquisition step. This facilitates downstream quantitative analysis from thousands to millions of cells in one experiment. By generating a single, spatially registered dataset with both DNA and protein information, the protocol provides a framework for linking genomic alterations to phenotypic states and microenvironmental context at scale, that also allows for exploring intra-tumor genetic heterogeneity.

From a practical perspective, automation is attractive because it reduces hands-on time and technical variability. In our workflow, once the FISH and antibody staining steps are completed, the subsequent fluidic exchange of reporters and imaging cycles are fully automated on the PhenoCycler-Fusion, in line with previous high-plex protein-only runs. Extending this level of automation to the DNA FISH steps themselves (for example, hybridization, post-hybridization washes and stringency adjustments performed under instrument control) would further improve standardization and throughput. While such full automation is not yet implemented here, our data demonstrate that, at minimum, the image acquisition of both modalities can be integrated into a combined DNA–protein workflow.

Technically, other instrumentation could be used for combined detection of DNA FISH probes and proteins. However, the sample preparation part and the DNA FISH detection would have to be adapted depending on the set-up of the instrument used. For example, we here used fluorescently labelled secondary probes for DNA FISH detection to mimic the cyclic process of antibody detection on the PhenoCycler-Fusion. Following the same concept as for the antibodies, all primary DNA FISH probes could be incubated in one step before loading the slide on the PhenoCycler instrument and be detected across cycles by adding only the secondary labelled reporters which allow for multiplexing. Depending on the instrumentation available, the sample preparation and the overall workflow has to be developed to be compatible with the instrumentation. For lower plex (up to ≈ 8 markers) instrumentation or slide scanners where all signals are detected simultaneously and relying on spectral unmixing, both primary or secondary labelled DNA probes could be used and combined with primary labelled or secondary antibodies for protein detection. In other instrument systems that combine staining and imaging, samples could likely be incubated with DNA FISH probes before or after the workflow on the instrument. However, achieving a single experimental run and combined image dataset might be challenging. In summary, the exact protocol for the sample prep would have to be optimized depending on the sample type and protein markers, and instrumentation capabilities.

### 3.1 Future potential of the combined protocol

Combining DNA FISH with multiplexed protein imaging has several conceptual advantages over either modality alone. Earlier work based on FICTION and related methods already demonstrated that integrating FISH with immunophenotyping enables detection of cytogenetic abnormalities in neoplastic cells, thereby increasing diagnostic specificity compared with FISH on unselected cell populations (30). Our workflow generalizes this principle to protein markers and millions of cells per section and thus has the potential to capture intratumor heterogeneity in both genotype and phenotype over large anatomical regions. An important future application is the systematic mapping of copy number variants (CNVs), amplifications and deletions onto well-defined cell states and tissue niche in tumours. In the long run, such data could be integrated with other spatial omics layers, including RNA-based assays and imaging mass spectrometry, as already explored in methods like PANINI (31) and its extensions. Due to lack of relevant samples, we were not able to demonstrate our protocol on clinically relevant cases in this work, for example detection of HER 2 gene amplification in breast cancer tumour sections. Instead, we used cell culture and available tissue sections from tonsil to demonstrate the technical possibilities of a combined sample preparation protocol for detection of genes and proteins, and to explore the sensitivity of the instrument to detect DNA FISH signals. Applying the workflow in a more relevant context would be an interesting follow-up of this study.

### 3.2 Imaging large areas efficiently – benefit and trade-off

A practical advantage of implementing the workflow on PhenoCycler-Fusion is the ability to image large tissue areas in a relatively short time (https://www.akoyabio.com/phenocycler/). This contrasts with traditional DNA FISH and immunofluorescence protocols on confocal microscopes, which are limited by small fields of view and lower acquisition speeds. For questions related to clinically relevant CNVs, intra-tumour heterogeneity or rare subclones, the ability to interrogate hundreds of thousands to millions of cells *in situ* is likely to be a major benefit. The principal trade-off is that, in the present implementation, only a single focal plane per field is acquired during the PhenoCycler-Fusion run. This can affect the sensitivity of DNA FISH detection, particularly for thick sections where individual FISH signals may fall above or below the chosen focal plane. In future work, acquiring a limited z-stack (for example, a few focal planes per field) could mitigate under-detection of FISH signals, at the cost of increased imaging time and data volume.

### 3.3 Current limitations of the present implementation

First, the number of DNA FISH probes and protein targets tested to date is limited. Our study enables high protein plex numbers by leveraging PhenoCycler-Fusion chemistry. However, we tested only a limited set of DNA targets. Additionally, we were unable to detect DNA FISH signals in the AF750 channel on the PhenoCycler-Fusion. This may be primarily because the secondary fluorescent probe used was AF790 and the PhenoCycler-Fusion does not have the optimal filter for detection in this range. As the probe tested in the AF750 channel was against target region Ch13.1, the failure in signal detection could be attributed to improper probe-target binding. However, additional experiments will be required to clearly determine the cause of the failure in signal detection. This means that, at present, the method is best suited for DNA detection in two channels only. Thankfully, thanks to our successful cycling strategies of hybridization/stripping, multiple different DNA targets could in theory be detected in a single experiment, despite relying on two channels only.

Second, the chemistry has so far been optimized and validated for a specific sample type and fixation regime (e.g. FF sections of thickness 5 µm and fixed cells). Other cell and tissue types and fixation conditions may require further optimization of antigen retrieval and hybridization. Previous work combining protein and nucleic acid detection, including PANINI and combined CODEX–RNAscope protocols, has emphasized that antigen retrieval and protease digestion need to be carefully tuned to preserve both protein epitopes and nucleic acid targets (32). The same constraints apply here and may limit direct transfer of the protocol across different laboratories and sample types without additional optimization.

Third, we have not tested the detection limit of our DNA FISH modality in the current workflow. We immediately chose a relatively large target span, selecting at least 6,000 oligos per probe (covering a few hundred kilobases of the target region) to secure strong FISH signals. Currently we do not know whether finer scale genomic alterations could be detected using our current protocol. Therefore, the potential clinical utility of DNA–protein integration should therefore be tempered by the recognition that not all classes of genomic alterations are equally accessible to this type of imaging. Standard practices for clinical FISH are often locus and assay specific but the probes used here meet the general requirement of >200 oligos per probe to obtain sufficient FISH signal (33). However the sensitivity of these probes are below the high standards required for clinical FISH probes where a sensitivity of at least 95% is required (34).

### 3.4 Concluding remarks

In summary, the integrated workflow presented here demonstrates that DNA FISH and high-plex protein imaging can be combined on an automated spatial imaging platform to produce a single dataset with gene and protein readouts at single-cell resolution over large tissue areas. This extends previous combined FISH–immunostaining approaches, which have mostly been limited to small regions and low multiplexing, and complements recent developments in protein–RNA spatial multi-omics. While the present implementation has clear limitations in terms of the number of DNA targets and the 2D imaging strategy, it provides a practical starting point for incorporating stable genomic information into high-plex spatial analyses of human samples. Future refinements to probe design, imaging and automation are likely to further increase the utility of this combined approach in both research and, potentially, clinical settings.

## 4. Methods

### 4.1 Cell culture

U2OS cells (ATCC-LGC Promochem, Borås, Sweden) were cultivated at 37 °C in a 5.0% CO_2_ humidified environment in McCoy’s 5A (modified) medium (Sigma-aldrich, MA, USA) supplemented with 2 mM L-glutamine (Sigma-aldrich, MA, USA) and 10% fetal bovine serum (VWR, PA, USA) without antibiotics.

For subsequent FISH experiments, the cells were seeded onto Poly-L-Lysine (Sigma-aldrich, MA, USA) coated Superfrost plus microscope slides (Epredia, MI, USA) at a density of 250 000 cells/ml. The cells were fixed using 4% paraformaldehyde (PFA) (Thermo Scientific, MA, USA) for 10 min then washed twice for 5 min with 125 mM glycine (Sigma-Aldrich, MA, USA) at room temperature. Cells fixed on slides were maintained in 1xPBS/0.05% NaN_3_ (Sigma-Aldrich, MA, USA) at 4°C for a maximum of one week before the FISH protocol was started.

Testing different stripping buffers was carried out using U2OS cells in fibronectin-coated glass-bottomed 96 well plates. Cells were seeded at a density of 8000 cells/well. For FISH experiments, the same procedure as described above was used. For immunostaining, cells were fixed using 4% PFA for 15 min and permeabilised using 0.5% Triton-X-100 (Sigma-Aldrich, MA, USA) for 5 min.

### 4.2 Biological Materials

Human tissue samples were collected and handled in accordance with Swedish laws and regulation. Fresh frozen anonymized residual tonsil tissue was collected after tonsillectomy performed at Sophiahemmet, Stockholm, Sweden (PhD Mattias Jangard at Sophiahemmet provided the material and Dr Lena Berglin at Karolinska Institute sectioned the material used in this study). Samples were collected under Institutional Review Board (IRB) approved protocols and de-identified in accordance with the Health Insurance Portability and Accountability Act (HIPAA). All human subjects were fully informed and explicitly asked for their consent to future research use of their samples. Fresh frozen samples were stored at -80 °C.

### 4.3 Primary FISH probe design

The FISH probes used in this study were previously described. The MYC probe was designed using the iFISH probe design pipeline, using a curated database of non-overlapping 40-mers from the hg19 human reference genome (26). The MYC probe consists of 96 oligos and spans around 14 kb of the gene locus. A sensitivity of 82.3% was calculated for the probe. The FISH probes against chromosome 13 were previously designed for another study (32), using an improved version of the iFISH probe design pipeline (available at at www.ifish4u.fht.org. Briefly, the sequences from the regions of interest were isolated in the GRCh37 reference genome. All possible 40-mers were extracted from each region, then attributed an individual weight based on their homology to the entire human genome tested using our in-house anti-alignment tool nHUSH (allowing a maximum of 32 consecutive base-pairs homology with an off-target region), their melting temperature and the existence of secondary structures. For each region, 6000 40-mers were then selected using escafish, which also integrates a pair-wise weight promoting short distances between consecutive 40-mers, while forbidding overlap. The 40-mers thus selected were then complemented with flanking 20-mers used as forward and reverse primers during probe amplification. The complete probe and primer sequences are available in **Supplementary Table 1**. An overview of the probes used in the experiments are summarized in **Table 1**.

### 4.4 Primary FISH probe production

The probe sequences were ordered as an oligonucleotide pool from Twist Biosciences and then individually selected by real-time PCR using probe-specific primers purchased from Integrated DNA Technologies. Following amplification, the products underwent DNA purification, in vitro transcription, RNA purification, and then reverse transcription to generate single-strand DNA probes, which were further purified before use. Quality control checks were performed at each stage of the protocol for eg-qPCR and NanoDrop measurements for dsDNA at intermediate steps, and a Qubit measurement for ssDNA at the final step.

### 4.5 Cells and Tissue sample preparation

Frozen tonsil samples were transferred from -80°C to -20°C and after equilibration fixed with 4% PFA for 10min. Once fixed, both cells and tissue samples were permeabilized and prepared for hybridization following a modified version of the iFISH method (PMID: 30967549). Briefly, they were washed twice 5 min with 1x PBS / 0.05% Triton X-100 (Sigma-Aldrich, MA, USA) and incubated in 1x PBS / 0.5% Triton X-100 for 20 min for permeabilization. Samples were then washed three times 5 min with 1x PBS and three times 5 min with 1x PBS / 0.05% Triton X-100. Subsequently, we rinsed them in 0.1 N HCl followed by a 5 min incubation in 0.1 N HCl at room temperature. Then, three washes with 1x PBS / 0.05% Triton X-100 followed by two 5 min washes with 1x PBS / 0.05% Triton X-100 while gently shaking at room temperature were performed. Tissue samples (but not the cells) were treated with 70ul RNaseA solution (0.1 mg/ml, in 1x PBS) by incubating at 37°C in a humidity chamber for 1 h followed by washing twice 5 min in PBS. Next, to prepare the samples for hybridization, both cells and tissue samples were rinsed once then incubated overnight in a buffer containing 50% formamide (ThermoFisher) / 2x SSC / 50 mM sodium phosphate buffer, at room temperature in the dark.

### 4.6 Single-molecule DNA FISH in cultured cells

After fixation and permeabilization, U2OS cells were incubated at 37 °C with a pre-hybridization buffer containing 50% formamide / 2x SSC / 5x Denhardt’s solution (ThermoFisher) / 50 mM sodium phosphate buffer / 1 mM EDTA (ThermoFisher) / 100 μg/mL salmon sperm DNA (Invitrogen) (pH 7.75 ± 0.25) inside a humidity chamber for 1 h. Then, the pre-hybridization buffer was replaced by the hybridization mix obtained by mixing the primary FISH probes diluted at 0.6 nM per oligonucleotide, at a ratio of 1:9 vol./vol. in a buffer consisting of 55% formamide / 2.2x SSC / 5.5x Denhardt’s solution / 55 mM sodium phosphate buffer / 1.1 mM EDTA / 111 ng/μL salmon sperm DNA / 11% w/w dextran sulfate (Thermo Fisher) (pH 7.75 ± 0.25). The hybridization mix was covered with a coverslip and sealed with Fixogum rubber cement (Marabu). Once Fixogum solidified, we denatured DNA by incubating the samples at 75 °C for 2 min 30 sec on a heating block. The denaturation parameters were determined through sample-specific optimization. The samples were then incubated for 24 h at 37 °C in a humidity chamber to allow probe hybridization. The following day, three washes of 1 min each were performed with 2x SSC / 0.2% Tween at room temperature, followed by two washes for 7 min each using 0.2x SSC / 0.2% Tween at 65 °C, and 5 min in 2x SSC at room temperature. All samples were incubated for two times 5 min in a buffer containing 25% formamide / 2x SSC before either proceeding with the hybridization of fluorescently labelled secondary FISH probes, or preparing the slides for antibody labelling and combined detection of proteins and nucleic acids.

### 4.7 Single molecule DNA FISH in Tissue

After fixation and permeabilization, tissue samples were incubated at 37°C inside a humidity chamber for 1 h in 70ul pre-hybridization buffer containing 50% formamide / 2x SSC / 5x Denhardt’s solution (ThermoFisher) / 50 mM sodium phosphate buffer / 1 mM EDTA (ThermoFisher) / 100 μg/mL salmon sperm DNA (Invitrogen) (pH 7.75 ± 0.25). Then the prehybridization buffer was replaced with by the hybridization mix obtained by mixing the primary FISH probes diluted at 0.6 nM per oligonucleotide, at a ratio of 1:9 vol./vol. in a buffer consisting of 55% formamide / 2.2x SSC / 5.5x Denhardt’s solution / 55 mM sodium phosphate buffer / 1.1 mM EDTA / 111 ng/μL salmon sperm DNA / 11% w/w dextran sulfate (Thermo Fisher) (pH 7.75 ± 0.25). The hybridization mix was covered with a coverslip and sealed with Fixogum rubber cement (Marabu). Once Fixogum solidified, we denatured DNA by incubating the samples at 75 °C for 3 min on a heating block. The denaturation parameters were determined through sample-specific optimization. The samples were then incubated for 24 h at 37 °C in a sealed humidity chamber to allow probe hybridization. The following day, samples were placed in 2xSSC/0.2% Tween to remove Fixogum and attached coverslip. Then, they were washed twice with 2XSCC/0.2% Tween at room temperature followed by two washes 5 min in prewarmed 0.2xSSC/0.2% Tween at 60°C. Next, samples were rinsed with 2X SSC at room temperature and thereafter with an RNA wash buffer consisting of 25% formamide / 2x SSC before preparing the slides for antibody labelling and combined detection of proteins and nucleic acids.

### 4.8 DNA FISH Imaging on Nikon microscope

DNA FISH fluorescence imaging was performed on a custom-built Nikon Ti-E Eclipse wide-field microscope equipped with a 100× oil-immersion objective (NA 1.45) and controlled using NIS Elements software (Nikon). Images were captured with an iXon Ultra 888 EMCCD camera (Andor Technology). For each field of view, we acquired z-stacks spanning the full cellular volume using a NIDAQ Piezo Z-drive with an interplane spacing of 200 nm. Raw image files (Nikon *.nd2 format) were converted to *.tif using nd2tool (https://github.com/elgw/nd2tool). All images were subsequently deconvolved using Deconwolf v0.4.6 (https://deconwolf.fht.org/) (35).

### 4.9 PhenoCycler Antibody Staining

The integrated protocol starts with antibody staining directly after primary DNA FISH probe hybridisation. Slides are equilibrated in staining buffer (Akoya Biosciences, MA, USA) for 20 min at room temperature prior to incubation with antibodies diluted in blocking buffer (Akoya Biosciences, MA, USA) overnight at 4 °C. The primary antibodies and dilutions used in this study are shown in **Table 2**. In-house conjugated antibodies were generated using a conjugation kit and following manufacturers instructions (Akoya Biosciences, MA, USA). Slides were then washed twice for 2 min with a staining buffer and fixed with 1.6% PFA for 10 min at room temperature. After washing with 1xPBS, a second fixation was carried out with ice-cold methanol (Sigma-aldrich, MA, USA) for 5 min. Slides were again washed in 1xPBS before a third fixation with CODEX fixative solution (Akoya Biosciences, MA, USA) for 20 min in a humidity chamber at room temperature. Slides were maintained in a storage buffer (Akoya Biosciences, MA, USA) at 4 °C prior to running on the PhenoCycler.

### 4.10 PhenoCycler Manual Addition of Reporters

For experiments testing different stripping buffers, manual addition of reporters was carried out on cells in 96 well plates. Antibody staining and post-staining fixations were carried out as detailed above prior to cells being washed three times with screening buffer (20% DMSO diluted in 1x CODEX buffer). The reporter for LMNB1 was diluted to 1:40 in reporter stock solution containing screening buffer, assay reagent and DAPI (Akoya Biosciences, MA, USA) according to manufacturer’s instructions, and added to cells for 5 min. Cells were then washed three times in screening buffer and a final wash in 1x CODEX buffer before being imaged on the Leica DMi8 microscope using the 20x objective. An exposure time of 100 ms was used for LMNB1. Stripping buffer containing different percentages of DMSO diluted in 1x CODEX buffer were then added to cells for 12 min. Cells were then washed in 1x CODEX buffer and imaged again on the Leica DMi8 microscope using the same settings to evaluate the efficiency of reporter elution.

### 4.11 PhenoCycler Multicycle Imaging

All the experiments using the PhenoCycler-Fusion were performed under the manufacturer’s instrument specifications. The operating temperature and humidity for PhenoCycler-Fusion was maintained at 20°C to 26°C and 30%-60% with no condensation, respectively, as per the manufacturer’s instructions.

Before starting the PhenoCycler run, the microfluidics were prepared by running a ‘clean instrument’ wash and a 96-well plate (product number: 7000006, Akoya Biosciences, MA, USA) containing fluorescent oligos was created. Each well of the plate contains fluorescent oligos diluted in 1x CODEX buffer, 1x buffer additive, nuclear stain and assay reagent (product number: 7000008, 7000002, Akoya Biosciences, MA, USA) according to manufacturer’s instructions. For the sequential workflow the fluorescent oligos consisted of reporters that are complementary to barcodes conjugated to antibodies, these were diluted 1:50. For the integrated workflow the fluorescent oligos included secondary DNA FISH probes diluted to 100 nM or reporters diluted 1:50. Buffers for the run were also prepared on the PhenoCycler instrument; these included 1x CODEX with 1x buffer additive (Akoya Biosciences, MA, USA), 55.5% DMSO diluted in 1x CODEX/additive (‘high DMSO buffer’), 20% DMSO diluted in 1x CODEX/additive (‘low DMSO buffer’) and distilled water.

Prior to the run the sample was left to equilibrate in 1x CODEX/additive for 10 min at room temperature before attaching the flow cell. The sample was then loaded into the sample carrier and filled with 1xCODEX/additive using a syringe. The PhenoCycler instrument is set up following the instrument wizard which includes checking microfluidic lines and priming. The sample then undergoes automated cycles of reporter/secondary probe addition, imaging, and reporter/secondary probe elution with each cycle taking approximately 1 h - 1 h 30 depending on exposure times used. Notably, the exact incubation times for each step of the multicycle imaging on the instrument are variable and entirely dependent on the number of cycles, fluorescent channels and the size of the specimen. The cycle set up for cell experiments is shown in **Table 3** and for FF tissue experiments in **Table 4**. The exposure times used for imaging antibodies are indicated in Table 2. The exposure time used for DNA FISH probes was 1000 ms and 5 ms for DAPI. The 20x objective was used for imaging.

Image processing is carried out automatically after the run following Akoya’s pipeline. This includes background subtraction using signals from blank cycles, stitching of individual tiles, and creating a qptiff file by compiling images from all the cycles into one image.

### 4.12 Data Analysis

Imaging data were exported as 8-bit compressed .qptiff files and processed using the PIPEX software pipeline (36). Cell segmentation was performed with StarDist (default parameters; nuclei diameter = 20 pixels, cytoplasmic expansion = 20 pixels), followed by extraction of mean fluorescence intensities for each marker per segmented cell.

DNA FISH signal detection was performed using Big-FISH (https://big-fish.readthedocs.io) (37). Images were denoised and filtered with a Laplacian-of-Gaussian (LoG) filter to enhance spots, local maxima were detected, and an adaptive intensity threshold was applied to remove background noise. Dense and bright regions were further corrected using the Big-FISH dense-region decomposition module with the following parameters: spot_radius = 1.5 pixels, selected based on the expected diffraction-limited spot size given the microscope point spread function and pixel size; alpha = 0.7, chosen to exclude both dim spots near the detection threshold and unusually bright spots when defining reference models for decomposition; beta = 1, retained at the default value because empirical testing showed that it appropriately distinguished clustered signals from isolated spots; and gamma = 5, selected to capture local intensity variations while remaining smaller than inter-spot distances for accurate background estimation. Finally, spots with low intensity (<100) were discarded using a threshold determined empirically through manual visual inspection of detected spots overlaid on raw images, thereby balancing sensitivity against false-positive detection of background noise.

For each cell, the total number of DNA FISH spots was quantified. The resulting per-cell data, including marker intensities and spot counts, were stored in AnnData objects for downstream analysis (38).

For each sample, we calculated mean counts, mean intensities, and total spot numbers per marker and cycle. AnnData objects were converted into TissUUmaps projects (39) for interactive visualization, preserving image layers, segmentation masks, and manually defined regions of interest. The data for this study can be explored at this url: (https://is-pla.serve.scilifelab.se/TissUUmaps_project_1.tmap?path=dna_fish). Code is available on https://github.com/BIIFSweden/6870_DNA_isPLA

## 5. Acknowledgements

This work was supported by the Spatial proteomics Unit at SciLifeLab and the specific Technology development grant from SciLifeLab, VC-2021-0042, The Swedish Research Council 2020-06182 and from the DISCERN project, HORIZON-MISS-2021-CANCER-02, grant agreement number **101096888**. Support was also provided by BioImage Informatics Unit (BIIF) at the National Bioinformatics Infrastructure Sweden (NBIS) and SciLifeLab. Tonsil samples were kindly provided by Mattias Jangard PhD, Sophiahemmet Stockholm. This work was also supported by funding from the Swedish Cancer Research Foundation (Cancerfonden, grant no. 22 2240 Pj 01 H) and the Swedish Research Council (grant. no. 2020-02657_3) to M.B. Chat GPT was used to help improve the language in the discussion part of the manuscript after author writing with relevant content.

## 8. Figure legends

**Supplementary Figure 1.** *Optimisation of stripping buffer DMSO concentration.* Representative images of DNA FISH signal before stripping, after stripping with different concentrations of DMSO and after secondary probe re-hybridisation using (A) 70% DMSO, (B) 60% DMSO, (C) 50% DMSO, and (D) 40% DMSO in the stripping buffer. (E) Graphs showing pixel intensity of original staining of hoechst, as a control, and Myc after staining, stripping and re-hybridisation with secondary probe using different concentrations of DMSO in the stripping buffer, as indicated in the graph legend.

**Supplementary Figure 2.** *Effect of stripping buffer DMSO concentration on protein detection.* (A) Representative images of LMNB1 protein signal before stripping (control) and after stripping with different concentrations of DMSO. Scale bar represents 100 µm. (B) Graph showing the signal to noise ratio of LMNB1 staining after stripping with different concentrations of DMSO and control staining.

