## Supplementary figures and images for "An integrated protocol for multiplexed DNA FISH and protein detection in large tissue sections"

### Supplementary Figure 1

#### Supplementary Figure 1

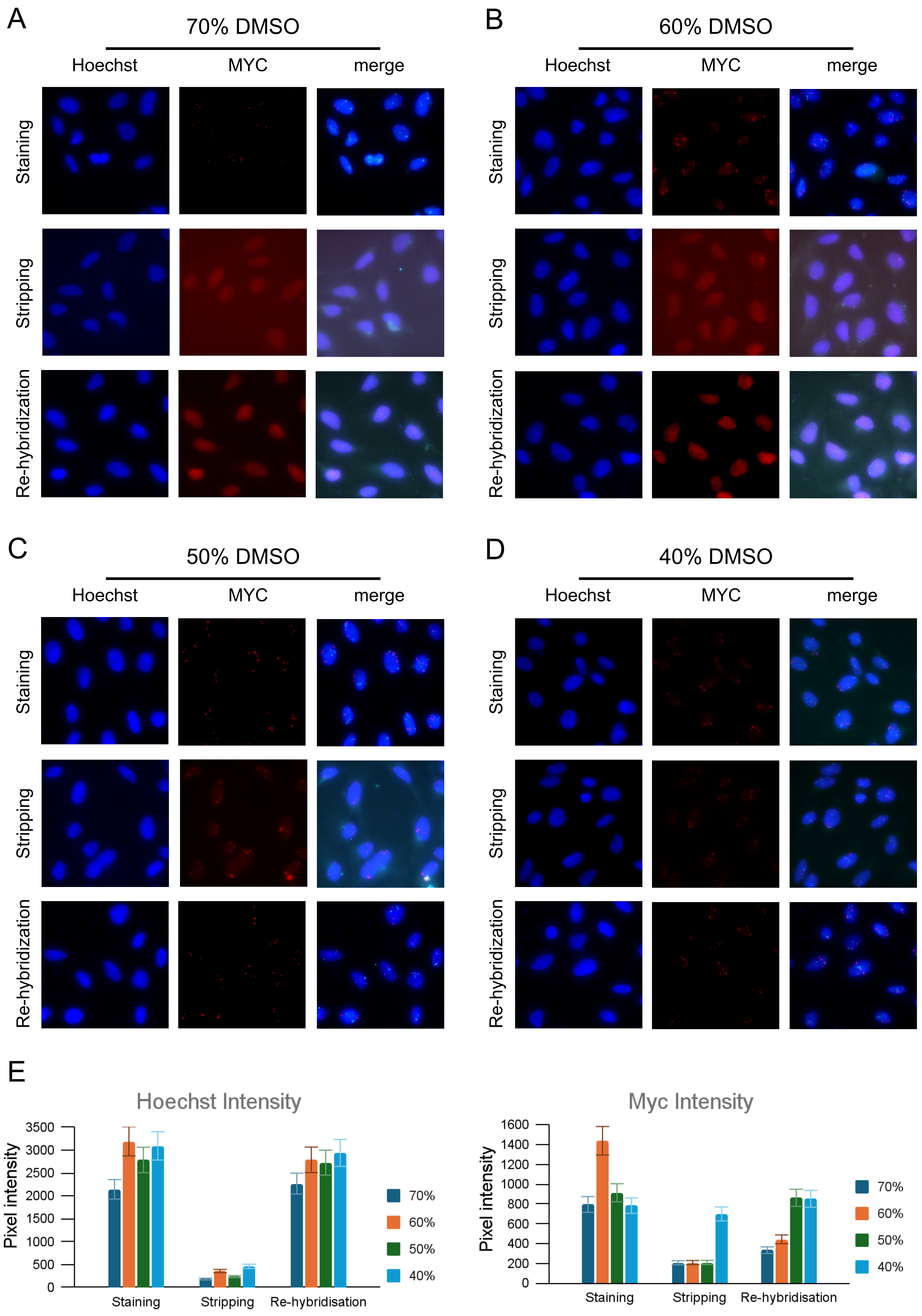

### Supplementary Figure 2

##### Supplementary Figure 2

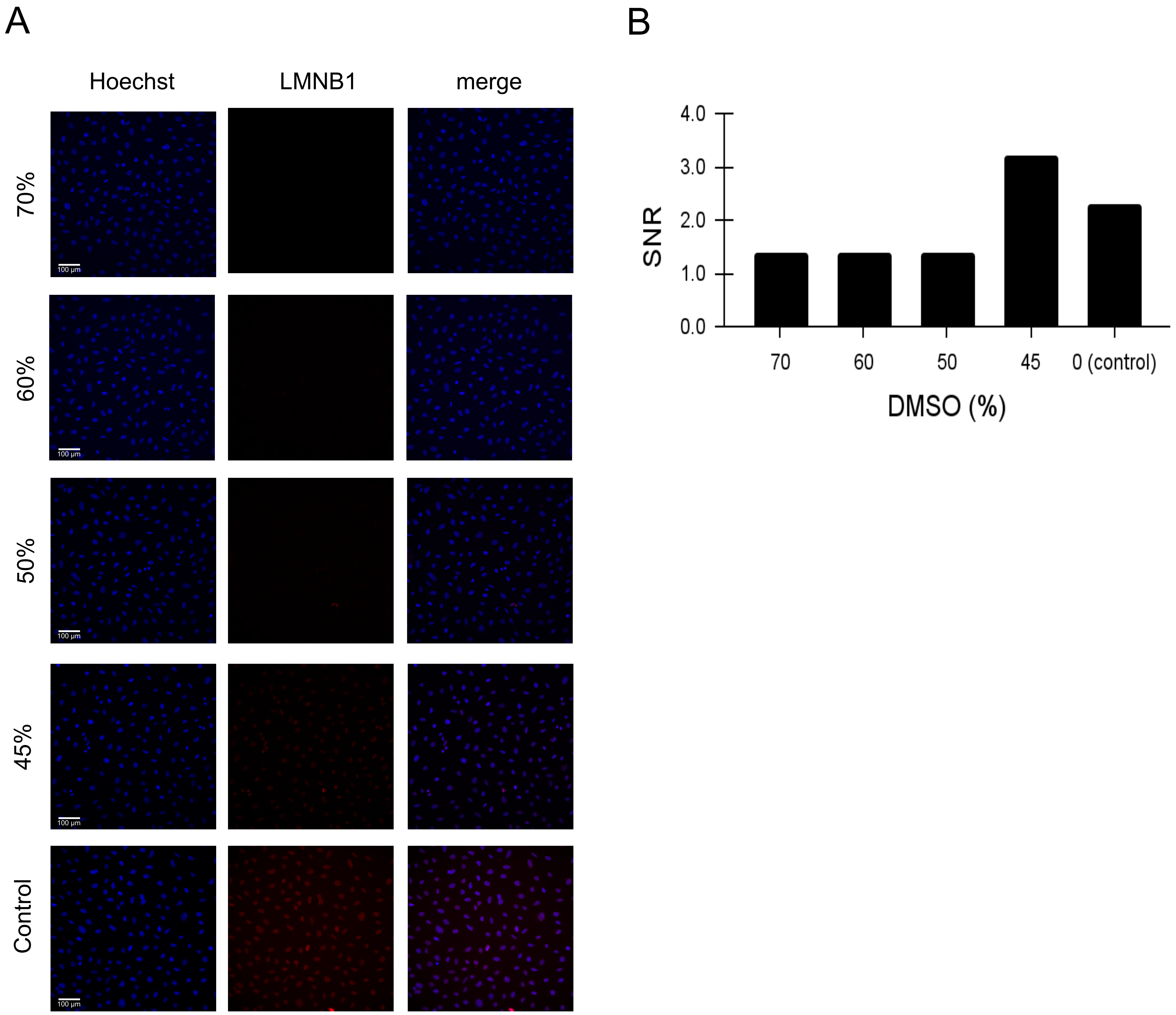
