## Supplementary Table 1 for "An integrated protocol for multiplexed DNA FISH and protein detection in large tissue sections"

Supplementary Table 1. Sequence of the oligos compos

*Upper case: target sequence. Lower case: F and R adapt

| Start-end coordinates (GRCh37) | Probe |
|---|---|
| chr13:29000065-29000105 | Ch13 region 1 |
| chr13:29000112-29000152 | Ch13 region 1 |
| chr13:29000170-29000210 | Ch13 region 1 |
| chr13:29000310-29000350 | Ch13 region 1 |
| chr13:29000380-29000420 | Ch13 region 1 |
| chr13:29000436-29000476 | Ch13 region 1 |
| chr13:29000523-29000563 | Ch13 region 1 |
| chr13:29000573-29000613 | Ch13 region 1 |
| chr13:29000632-29000672 | Ch13 region 1 |
| chr13:29000687-29000727 | Ch13 region 1 |
| chr13:29000751-29000791 | Ch13 region 1 |
| chr13:29000830-29000870 | Ch13 region 1 |
| chr13:29000875-29000915 | Ch13 region 1 |
| chr13:29000951-29000991 | Ch13 region 1 |
| chr13:29001023-29001063 | Ch13 region 1 |
| chr13:29001105-29001145 | Ch13 region 1 |
| chr13:29001175-29001215 | Ch13 region 1 |
| chr13:29001220-29001260 | Ch13 region 1 |
| chr13:29001272-29001312 | Ch13 region 1 |
| chr13:29001334-29001374 | Ch13 region 1 |
| chr13:29001410-29001450 | Ch13 region 1 |
| chr13:29001490-29001530 | Ch13 region 1 |
| chr13:29001540-29001580 | Ch13 region 1 |
| chr13:29001676-29001716 | Ch13 region 1 |
| chr13:29001729-29001769 | Ch13 region 1 |
| chr13:29001835-29001875 | Ch13 region 1 |
| chr13:29001891-29001931 | Ch13 region 1 |
| chr13:29001947-29001987 | Ch13 region 1 |
| chr13:29002020-29002060 | Ch13 region 1 |
| chr13:29002074-29002114 | Ch13 region 1 |
| chr13:29002125-29002165 | Ch13 region 1 |
| chr13:29002225-29002265 | Ch13 region 1 |
| chr13:29002324-29002364 | Ch13 region 1 |
| chr13:29002376-29002416 | Ch13 region 1 |
| chr13:29002478-29002518 | Ch13 region 1 |
| chr13:29002532-29002572 | Ch13 region 1 |
| chr13:29002586-29002626 | Ch13 region 1 |
| chr13:29002661-29002701 | Ch13 region 1 |
| chr13:29002716-29002756 | Ch13 region 1 |
| chr13:29002761-29002801 | Ch13 region 1 |
| chr13:29002862-29002902 | Ch13 region 1 |
| chr13:29002971-29003011 | Ch13 region 1 |
| chr13:29003020-29003060 | Ch13 region 1 |
| chr13:29003096-29003136 | Ch13 region 1 |
| chr13:29003145-29003185 | Ch13 region 1 |
| chr13:29003190-29003230 | Ch13 region 1 |
| chr13:29003284-29003324 | Ch13 region 1 |
| chr13:29003367-29003407 | Ch13 region 1 |

| | |
|---|---|
| chr13:29003413-29003453 | Ch13 region 1 |
| chr13:29003572-29003612 | Ch13 region 1 |
| chr13:29003619-29003659 | Ch13 region 1 |
| chr13:29003668-29003708 | Ch13 region 1 |
| chr13:29003752-29003792 | Ch13 region 1 |
| chr13:29003818-29003858 | Ch13 region 1 |
| chr13:29003905-29003945 | Ch13 region 1 |
| chr13:29003964-29004004 | Ch13 region 1 |
| chr13:29004026-29004066 | Ch13 region 1 |
| chr13:29004081-29004121 | Ch13 region 1 |
| chr13:29004129-29004169 | Ch13 region 1 |
| chr13:29004180-29004220 | Ch13 region 1 |
| chr13:29004248-29004288 | Ch13 region 1 |
| chr13:29004338-29004378 | Ch13 region 1 |
| chr13:29004420-29004460 | Ch13 region 1 |
| chr13:29004487-29004527 | Ch13 region 1 |
| chr13:29004536-29004576 | Ch13 region 1 |
| chr13:29004594-29004634 | Ch13 region 1 |
| chr13:29004683-29004723 | Ch13 region 1 |
| chr13:29004728-29004768 | Ch13 region 1 |
| chr13:29004816-29004856 | Ch13 region 1 |
| chr13:29005036-29005076 | Ch13 region 1 |
| chr13:29005103-29005143 | Ch13 region 1 |
| chr13:29005170-29005210 | Ch13 region 1 |
| chr13:29005220-29005260 | Ch13 region 1 |
| chr13:29005288-29005328 | Ch13 region 1 |
| chr13:29005357-29005397 | Ch13 region 1 |
| chr13:29005407-29005447 | Ch13 region 1 |
| chr13:29005455-29005495 | Ch13 region 1 |
| chr13:29005548-29005588 | Ch13 region 1 |
| chr13:29005595-29005635 | Ch13 region 1 |
| chr13:29005676-29005716 | Ch13 region 1 |
| chr13:29005747-29005787 | Ch13 region 1 |
| chr13:29005797-29005837 | Ch13 region 1 |
| chr13:29005856-29005896 | Ch13 region 1 |
| chr13:29005923-29005963 | Ch13 region 1 |
| chr13:29005981-29006021 | Ch13 region 1 |
| chr13:29006058-29006098 | Ch13 region 1 |
| chr13:29006131-29006171 | Ch13 region 1 |
| chr13:29006176-29006216 | Ch13 region 1 |
| chr13:29006226-29006266 | Ch13 region 1 |
| chr13:29006279-29006319 | Ch13 region 1 |
| chr13:29006324-29006364 | Ch13 region 1 |
| chr13:29006372-29006412 | Ch13 region 1 |
| chr13:29006455-29006495 | Ch13 region 1 |
| chr13:29006500-29006540 | Ch13 region 1 |
| chr13:29006572-29006612 | Ch13 region 1 |
| chr13:29006653-29006693 | Ch13 region 1 |
| chr13:29006760-29006800 | Ch13 region 1 |
| chr13:29006805-29006845 | Ch13 region 1 |
| chr13:29006882-29006922 | Ch13 region 1 |
| chr13:29006928-29006968 | Ch13 region 1 |

| | |
|---|---|
| chr13:29007038-29007078 | Ch13 region 1 |
| chr13:29007083-29007123 | Ch13 region 1 |
| chr13:29007129-29007169 | Ch13 region 1 |
| chr13:29007207-29007247 | Ch13 region 1 |
| chr13:29007267-29007307 | Ch13 region 1 |
| chr13:29007366-29007406 | Ch13 region 1 |
| chr13:29007443-29007483 | Ch13 region 1 |
| chr13:29007488-29007528 | Ch13 region 1 |
| chr13:29007559-29007599 | Ch13 region 1 |
| chr13:29007623-29007663 | Ch13 region 1 |
| chr13:29007677-29007717 | Ch13 region 1 |
| chr13:29007724-29007764 | Ch13 region 1 |
| chr13:29007788-29007828 | Ch13 region 1 |
| chr13:29007855-29007895 | Ch13 region 1 |
| chr13:29007930-29007970 | Ch13 region 1 |
| chr13:29007985-29008025 | Ch13 region 1 |
| chr13:29008056-29008096 | Ch13 region 1 |
| chr13:29008125-29008165 | Ch13 region 1 |
| chr13:29008187-29008227 | Ch13 region 1 |
| chr13:29008247-29008287 | Ch13 region 1 |
| chr13:29008297-29008337 | Ch13 region 1 |
| chr13:29008364-29008404 | Ch13 region 1 |
| chr13:29008411-29008451 | Ch13 region 1 |
| chr13:29008586-29008626 | Ch13 region 1 |
| chr13:29008635-29008675 | Ch13 region 1 |
| chr13:29008682-29008722 | Ch13 region 1 |
| chr13:29008790-29008830 | Ch13 region 1 |
| chr13:29009022-29009062 | Ch13 region 1 |
| chr13:29009101-29009141 | Ch13 region 1 |
| chr13:29009150-29009190 | Ch13 region 1 |
| chr13:29009219-29009259 | Ch13 region 1 |
| chr13:29009325-29009365 | Ch13 region 1 |
| chr13:29009375-29009415 | Ch13 region 1 |
| chr13:29009420-29009460 | Ch13 region 1 |
| chr13:29009465-29009505 | Ch13 region 1 |
| chr13:29009522-29009562 | Ch13 region 1 |
| chr13:29009568-29009608 | Ch13 region 1 |
| chr13:29009655-29009695 | Ch13 region 1 |
| chr13:29009741-29009781 | Ch13 region 1 |
| chr13:29009802-29009842 | Ch13 region 1 |
| chr13:29009887-29009927 | Ch13 region 1 |
| chr13:29009948-29009988 | Ch13 region 1 |
| chr13:29009996-29010036 | Ch13 region 1 |
| chr13:29010041-29010081 | Ch13 region 1 |
| chr13:29010120-29010160 | Ch13 region 1 |
| chr13:29010182-29010222 | Ch13 region 1 |
| chr13:29010238-29010278 | Ch13 region 1 |
| chr13:29010329-29010369 | Ch13 region 1 |
| chr13:29010389-29010429 | Ch13 region 1 |
| chr13:29010442-29010482 | Ch13 region 1 |
| chr13:29010501-29010541 | Ch13 region 1 |
| chr13:29010546-29010586 | Ch13 region 1 |

| | |
|---|---|
| chr13:29010594-29010634 | Ch13 region 1 |
| chr13:29010639-29010679 | Ch13 region 1 |
| chr13:29010701-29010741 | Ch13 region 1 |
| chr13:29010746-29010786 | Ch13 region 1 |
| chr13:29010796-29010836 | Ch13 region 1 |
| chr13:29010865-29010905 | Ch13 region 1 |
| chr13:29010938-29010978 | Ch13 region 1 |
| chr13:29011014-29011054 | Ch13 region 1 |
| chr13:29011063-29011103 | Ch13 region 1 |
| chr13:29011110-29011150 | Ch13 region 1 |
| chr13:29011203-29011243 | Ch13 region 1 |
| chr13:29011248-29011288 | Ch13 region 1 |
| chr13:29011302-29011342 | Ch13 region 1 |
| chr13:29011397-29011437 | Ch13 region 1 |
| chr13:29011444-29011484 | Ch13 region 1 |
| chr13:29011494-29011534 | Ch13 region 1 |
| chr13:29011570-29011610 | Ch13 region 1 |
| chr13:29011634-29011674 | Ch13 region 1 |
| chr13:29011679-29011719 | Ch13 region 1 |
| chr13:29011809-29011849 | Ch13 region 1 |
| chr13:29011855-29011895 | Ch13 region 1 |
| chr13:29011901-29011941 | Ch13 region 1 |
| chr13:29011984-29012024 | Ch13 region 1 |
| chr13:29012053-29012093 | Ch13 region 1 |
| chr13:29012154-29012194 | Ch13 region 1 |
| chr13:29012235-29012275 | Ch13 region 1 |
| chr13:29012319-29012359 | Ch13 region 1 |
| chr13:29012406-29012446 | Ch13 region 1 |
| chr13:29012451-29012491 | Ch13 region 1 |
| chr13:29012568-29012608 | Ch13 region 1 |
| chr13:29012615-29012655 | Ch13 region 1 |
| chr13:29012665-29012705 | Ch13 region 1 |
| chr13:29012742-29012782 | Ch13 region 1 |
| chr13:29012808-29012848 | Ch13 region 1 |
| chr13:29012889-29012929 | Ch13 region 1 |
| chr13:29012941-29012981 | Ch13 region 1 |
| chr13:29013009-29013049 | Ch13 region 1 |
| chr13:29013063-29013103 | Ch13 region 1 |
| chr13:29013112-29013152 | Ch13 region 1 |
| chr13:29013182-29013222 | Ch13 region 1 |
| chr13:29013232-29013272 | Ch13 region 1 |
| chr13:29013298-29013338 | Ch13 region 1 |
| chr13:29013348-29013388 | Ch13 region 1 |
| chr13:29013393-29013433 | Ch13 region 1 |
| chr13:29013442-29013482 | Ch13 region 1 |
| chr13:29013502-29013542 | Ch13 region 1 |
| chr13:29013549-29013589 | Ch13 region 1 |
| chr13:29013605-29013645 | Ch13 region 1 |
| chr13:29013675-29013715 | Ch13 region 1 |
| chr13:29013743-29013783 | Ch13 region 1 |
| chr13:29013789-29013829 | Ch13 region 1 |
| chr13:29013844-29013884 | Ch13 region 1 |

| | |
|---|---|
| chr13:29013903-29013943 | Ch13 region 1 |
| chr13:29013964-29014004 | Ch13 region 1 |
| chr13:29014057-29014097 | Ch13 region 1 |
| chr13:29014120-29014160 | Ch13 region 1 |
| chr13:29014167-29014207 | Ch13 region 1 |
| chr13:29014214-29014254 | Ch13 region 1 |
| chr13:29014267-29014307 | Ch13 region 1 |
| chr13:29014319-29014359 | Ch13 region 1 |
| chr13:29014384-29014424 | Ch13 region 1 |
| chr13:29014443-29014483 | Ch13 region 1 |
| chr13:29014509-29014549 | Ch13 region 1 |
| chr13:29014562-29014602 | Ch13 region 1 |
| chr13:29014613-29014653 | Ch13 region 1 |
| chr13:29014695-29014735 | Ch13 region 1 |
| chr13:29014751-29014791 | Ch13 region 1 |
| chr13:29014804-29014844 | Ch13 region 1 |
| chr13:29014879-29014919 | Ch13 region 1 |
| chr13:29014989-29015029 | Ch13 region 1 |
| chr13:29015036-29015076 | Ch13 region 1 |
| chr13:29015137-29015177 | Ch13 region 1 |
| chr13:29015236-29015276 | Ch13 region 1 |
| chr13:29015281-29015321 | Ch13 region 1 |
| chr13:29015349-29015389 | Ch13 region 1 |
| chr13:29015411-29015451 | Ch13 region 1 |
| chr13:29015484-29015524 | Ch13 region 1 |
| chr13:29015574-29015614 | Ch13 region 1 |
| chr13:29015658-29015698 | Ch13 region 1 |
| chr13:29015744-29015784 | Ch13 region 1 |
| chr13:29015897-29015937 | Ch13 region 1 |
| chr13:29015942-29015982 | Ch13 region 1 |
| chr13:29016175-29016215 | Ch13 region 1 |
| chr13:29016220-29016260 | Ch13 region 1 |
| chr13:29016293-29016333 | Ch13 region 1 |
| chr13:29016341-29016381 | Ch13 region 1 |
| chr13:29016463-29016503 | Ch13 region 1 |
| chr13:29016593-29016633 | Ch13 region 1 |
| chr13:29016663-29016703 | Ch13 region 1 |
| chr13:29016715-29016755 | Ch13 region 1 |
| chr13:29016791-29016831 | Ch13 region 1 |
| chr13:29016870-29016910 | Ch13 region 1 |
| chr13:29016915-29016955 | Ch13 region 1 |
| chr13:29017001-29017041 | Ch13 region 1 |
| chr13:29017077-29017117 | Ch13 region 1 |
| chr13:29017135-29017175 | Ch13 region 1 |
| chr13:29017205-29017245 | Ch13 region 1 |
| chr13:29017252-29017292 | Ch13 region 1 |
| chr13:29017332-29017372 | Ch13 region 1 |
| chr13:29017391-29017431 | Ch13 region 1 |
| chr13:29017445-29017485 | Ch13 region 1 |
| chr13:29017490-29017530 | Ch13 region 1 |
| chr13:29017583-29017623 | Ch13 region 1 |
| chr13:29017682-29017722 | Ch13 region 1 |

| | |
|---|---|
| chr13:29017767-29017807 | Ch13 region 1 |
| chr13:29017816-29017856 | Ch13 region 1 |
| chr13:29017869-29017909 | Ch13 region 1 |
| chr13:29017954-29017994 | Ch13 region 1 |
| chr13:29018074-29018114 | Ch13 region 1 |
| chr13:29018173-29018213 | Ch13 region 1 |
| chr13:29018237-29018277 | Ch13 region 1 |
| chr13:29018282-29018322 | Ch13 region 1 |
| chr13:29018424-29018464 | Ch13 region 1 |
| chr13:29018586-29018626 | Ch13 region 1 |
| chr13:29018631-29018671 | Ch13 region 1 |
| chr13:29018684-29018724 | Ch13 region 1 |
| chr13:29018873-29018913 | Ch13 region 1 |
| chr13:29019079-29019119 | Ch13 region 1 |
| chr13:29019244-29019284 | Ch13 region 1 |
| chr13:29019299-29019339 | Ch13 region 1 |
| chr13:29019382-29019422 | Ch13 region 1 |
| chr13:29019587-29019627 | Ch13 region 1 |
| chr13:29019696-29019736 | Ch13 region 1 |
| chr13:29019747-29019787 | Ch13 region 1 |
| chr13:29019809-29019849 | Ch13 region 1 |
| chr13:29019943-29019983 | Ch13 region 1 |
| chr13:29020085-29020125 | Ch13 region 1 |
| chr13:29020180-29020220 | Ch13 region 1 |
| chr13:29020228-29020268 | Ch13 region 1 |
| chr13:29020295-29020335 | Ch13 region 1 |
| chr13:29020387-29020427 | Ch13 region 1 |
| chr13:29020450-29020490 | Ch13 region 1 |
| chr13:29020519-29020559 | Ch13 region 1 |
| chr13:29020584-29020624 | Ch13 region 1 |
| chr13:29020783-29020823 | Ch13 region 1 |
| chr13:29020874-29020914 | Ch13 region 1 |
| chr13:29020978-29021018 | Ch13 region 1 |
| chr13:29021065-29021105 | Ch13 region 1 |
| chr13:29021115-29021155 | Ch13 region 1 |
| chr13:29021203-29021243 | Ch13 region 1 |
| chr13:29021328-29021368 | Ch13 region 1 |
| chr13:29021393-29021433 | Ch13 region 1 |
| chr13:29021612-29021652 | Ch13 region 1 |
| chr13:29021749-29021789 | Ch13 region 1 |
| chr13:29022075-29022115 | Ch13 region 1 |
| chr13:29022149-29022189 | Ch13 region 1 |
| chr13:29022228-29022268 | Ch13 region 1 |
| chr13:29022294-29022334 | Ch13 region 1 |
| chr13:29022501-29022541 | Ch13 region 1 |
| chr13:29022555-29022595 | Ch13 region 1 |
| chr13:29022682-29022722 | Ch13 region 1 |
| chr13:29022727-29022767 | Ch13 region 1 |
| chr13:29022785-29022825 | Ch13 region 1 |
| chr13:29022839-29022879 | Ch13 region 1 |
| chr13:29022923-29022963 | Ch13 region 1 |
| chr13:29023015-29023055 | Ch13 region 1 |

| | |
|---|---|
| chr13:29023094-29023134 | Ch13 region 1 |
| chr13:29023174-29023214 | Ch13 region 1 |
| chr13:29023319-29023359 | Ch13 region 1 |
| chr13:29023402-29023442 | Ch13 region 1 |
| chr13:29023474-29023514 | Ch13 region 1 |
| chr13:29023539-29023579 | Ch13 region 1 |
| chr13:29023611-29023651 | Ch13 region 1 |
| chr13:29023663-29023703 | Ch13 region 1 |
| chr13:29023731-29023771 | Ch13 region 1 |
| chr13:29023776-29023816 | Ch13 region 1 |
| chr13:29023885-29023925 | Ch13 region 1 |
| chr13:29023961-29024001 | Ch13 region 1 |
| chr13:29024010-29024050 | Ch13 region 1 |
| chr13:29024091-29024131 | Ch13 region 1 |
| chr13:29024195-29024235 | Ch13 region 1 |
| chr13:29024241-29024281 | Ch13 region 1 |
| chr13:29024325-29024365 | Ch13 region 1 |
| chr13:29024390-29024430 | Ch13 region 1 |
| chr13:29024453-29024493 | Ch13 region 1 |
| chr13:29024523-29024563 | Ch13 region 1 |
| chr13:29024587-29024627 | Ch13 region 1 |
| chr13:29024761-29024801 | Ch13 region 1 |
| chr13:29024818-29024858 | Ch13 region 1 |
| chr13:29024865-29024905 | Ch13 region 1 |
| chr13:29024914-29024954 | Ch13 region 1 |
| chr13:29024963-29025003 | Ch13 region 1 |
| chr13:29025032-29025072 | Ch13 region 1 |
| chr13:29025116-29025156 | Ch13 region 1 |
| chr13:29025218-29025258 | Ch13 region 1 |
| chr13:29025280-29025320 | Ch13 region 1 |
| chr13:29025401-29025441 | Ch13 region 1 |
| chr13:29025574-29025614 | Ch13 region 1 |
| chr13:29025653-29025693 | Ch13 region 1 |
| chr13:29025750-29025790 | Ch13 region 1 |
| chr13:29025804-29025844 | Ch13 region 1 |
| chr13:29025849-29025889 | Ch13 region 1 |
| chr13:29025908-29025948 | Ch13 region 1 |
| chr13:29025970-29026010 | Ch13 region 1 |
| chr13:29026023-29026063 | Ch13 region 1 |
| chr13:29026091-29026131 | Ch13 region 1 |
| chr13:29026208-29026248 | Ch13 region 1 |
| chr13:29026278-29026318 | Ch13 region 1 |
| chr13:29026355-29026395 | Ch13 region 1 |
| chr13:29026412-29026452 | Ch13 region 1 |
| chr13:29026474-29026514 | Ch13 region 1 |
| chr13:29026521-29026561 | Ch13 region 1 |
| chr13:29026578-29026618 | Ch13 region 1 |
| chr13:29026627-29026667 | Ch13 region 1 |
| chr13:29026676-29026716 | Ch13 region 1 |
| chr13:29026721-29026761 | Ch13 region 1 |
| chr13:29026766-29026806 | Ch13 region 1 |
| chr13:29026840-29026880 | Ch13 region 1 |

| | |
|---|---|
| chr13:29026937-29026977 | Ch13 region 1 |
| chr13:29027027-29027067 | Ch13 region 1 |
| chr13:29027108-29027148 | Ch13 region 1 |
| chr13:29027254-29027294 | Ch13 region 1 |
| chr13:29027303-29027343 | Ch13 region 1 |
| chr13:29027541-29027581 | Ch13 region 1 |
| chr13:29027589-29027629 | Ch13 region 1 |
| chr13:29027666-29027706 | Ch13 region 1 |
| chr13:29027728-29027768 | Ch13 region 1 |
| chr13:29027815-29027855 | Ch13 region 1 |
| chr13:29027878-29027918 | Ch13 region 1 |
| chr13:29027951-29027991 | Ch13 region 1 |
| chr13:29028010-29028050 | Ch13 region 1 |
| chr13:29028103-29028143 | Ch13 region 1 |
| chr13:29028192-29028232 | Ch13 region 1 |
| chr13:29028253-29028293 | Ch13 region 1 |
| chr13:29028300-29028340 | Ch13 region 1 |
| chr13:29028388-29028428 | Ch13 region 1 |
| chr13:29028516-29028556 | Ch13 region 1 |
| chr13:29028570-29028610 | Ch13 region 1 |
| chr13:29028615-29028655 | Ch13 region 1 |
| chr13:29028660-29028700 | Ch13 region 1 |
| chr13:29028718-29028758 | Ch13 region 1 |
| chr13:29028767-29028807 | Ch13 region 1 |
| chr13:29028899-29028939 | Ch13 region 1 |
| chr13:29029021-29029061 | Ch13 region 1 |
| chr13:29029245-29029285 | Ch13 region 1 |
| chr13:29029332-29029372 | Ch13 region 1 |
| chr13:29029504-29029544 | Ch13 region 1 |
| chr13:29029570-29029610 | Ch13 region 1 |
| chr13:29029686-29029726 | Ch13 region 1 |
| chr13:29029737-29029777 | Ch13 region 1 |
| chr13:29029939-29029979 | Ch13 region 1 |
| chr13:29029999-29030039 | Ch13 region 1 |
| chr13:29030051-29030091 | Ch13 region 1 |
| chr13:29030110-29030150 | Ch13 region 1 |
| chr13:29030202-29030242 | Ch13 region 1 |
| chr13:29030263-29030303 | Ch13 region 1 |
| chr13:29030357-29030397 | Ch13 region 1 |
| chr13:29030618-29030658 | Ch13 region 1 |
| chr13:29030942-29030982 | Ch13 region 1 |
| chr13:29030989-29031029 | Ch13 region 1 |
| chr13:29031064-29031104 | Ch13 region 1 |
| chr13:29031116-29031156 | Ch13 region 1 |
| chr13:29031197-29031237 | Ch13 region 1 |
| chr13:29031334-29031374 | Ch13 region 1 |
| chr13:29031392-29031432 | Ch13 region 1 |
| chr13:29031440-29031480 | Ch13 region 1 |
| chr13:29031485-29031525 | Ch13 region 1 |
| chr13:29031542-29031582 | Ch13 region 1 |
| chr13:29031632-29031672 | Ch13 region 1 |
| chr13:29031717-29031757 | Ch13 region 1 |

| | |
|---|---|
| chr13:29031763-29031803 | Ch13 region 1 |
| chr13:29031827-29031867 | Ch13 region 1 |
| chr13:29031884-29031924 | Ch13 region 1 |
| chr13:29031968-29032008 | Ch13 region 1 |
| chr13:29032017-29032057 | Ch13 region 1 |
| chr13:29032195-29032235 | Ch13 region 1 |
| chr13:29032349-29032389 | Ch13 region 1 |
| chr13:29032421-29032461 | Ch13 region 1 |
| chr13:29032476-29032516 | Ch13 region 1 |
| chr13:29032552-29032592 | Ch13 region 1 |
| chr13:29032609-29032649 | Ch13 region 1 |
| chr13:29032699-29032739 | Ch13 region 1 |
| chr13:29032756-29032796 | Ch13 region 1 |
| chr13:29032822-29032862 | Ch13 region 1 |
| chr13:29032892-29032932 | Ch13 region 1 |
| chr13:29032937-29032977 | Ch13 region 1 |
| chr13:29033016-29033056 | Ch13 region 1 |
| chr13:29033072-29033112 | Ch13 region 1 |
| chr13:29033182-29033222 | Ch13 region 1 |
| chr13:29033244-29033284 | Ch13 region 1 |
| chr13:29033335-29033375 | Ch13 region 1 |
| chr13:29033420-29033460 | Ch13 region 1 |
| chr13:29033491-29033531 | Ch13 region 1 |
| chr13:29033579-29033619 | Ch13 region 1 |
| chr13:29033626-29033666 | Ch13 region 1 |
| chr13:29033695-29033735 | Ch13 region 1 |
| chr13:29033774-29033814 | Ch13 region 1 |
| chr13:29033832-29033872 | Ch13 region 1 |
| chr13:29033926-29033966 | Ch13 region 1 |
| chr13:29033980-29034020 | Ch13 region 1 |
| chr13:29034049-29034089 | Ch13 region 1 |
| chr13:29034098-29034138 | Ch13 region 1 |
| chr13:29034148-29034188 | Ch13 region 1 |
| chr13:29034197-29034237 | Ch13 region 1 |
| chr13:29034267-29034307 | Ch13 region 1 |
| chr13:29034329-29034369 | Ch13 region 1 |
| chr13:29034390-29034430 | Ch13 region 1 |
| chr13:29034472-29034512 | Ch13 region 1 |
| chr13:29034532-29034572 | Ch13 region 1 |
| chr13:29034579-29034619 | Ch13 region 1 |
| chr13:29034638-29034678 | Ch13 region 1 |
| chr13:29034700-29034740 | Ch13 region 1 |
| chr13:29034760-29034800 | Ch13 region 1 |
| chr13:29034824-29034864 | Ch13 region 1 |
| chr13:29034901-29034941 | Ch13 region 1 |
| chr13:29034957-29034997 | Ch13 region 1 |
| chr13:29035040-29035080 | Ch13 region 1 |
| chr13:29035242-29035282 | Ch13 region 1 |
| chr13:29035292-29035332 | Ch13 region 1 |
| chr13:29035337-29035377 | Ch13 region 1 |
| chr13:29035445-29035485 | Ch13 region 1 |
| chr13:29035501-29035541 | Ch13 region 1 |

| | |
|---|---|
| chr13:29035565-29035605 | Ch13 region 1 |
| chr13:29035614-29035654 | Ch13 region 1 |
| chr13:29035682-29035722 | Ch13 region 1 |
| chr13:29035769-29035809 | Ch13 region 1 |
| chr13:29035838-29035878 | Ch13 region 1 |
| chr13:29035929-29035969 | Ch13 region 1 |
| chr13:29035977-29036017 | Ch13 region 1 |
| chr13:29036022-29036062 | Ch13 region 1 |
| chr13:29036074-29036114 | Ch13 region 1 |
| chr13:29036163-29036203 | Ch13 region 1 |
| chr13:29036233-29036273 | Ch13 region 1 |
| chr13:29036279-29036319 | Ch13 region 1 |
| chr13:29036332-29036372 | Ch13 region 1 |
| chr13:29036417-29036457 | Ch13 region 1 |
| chr13:29036478-29036518 | Ch13 region 1 |
| chr13:29036534-29036574 | Ch13 region 1 |
| chr13:29036609-29036649 | Ch13 region 1 |
| chr13:29036705-29036745 | Ch13 region 1 |
| chr13:29036774-29036814 | Ch13 region 1 |
| chr13:29036852-29036892 | Ch13 region 1 |
| chr13:29036913-29036953 | Ch13 region 1 |
| chr13:29037013-29037053 | Ch13 region 1 |
| chr13:29037095-29037135 | Ch13 region 1 |
| chr13:29037143-29037183 | Ch13 region 1 |
| chr13:29037353-29037393 | Ch13 region 1 |
| chr13:29037431-29037471 | Ch13 region 1 |
| chr13:29037500-29037540 | Ch13 region 1 |
| chr13:29037545-29037585 | Ch13 region 1 |
| chr13:29037606-29037646 | Ch13 region 1 |
| chr13:29037694-29037734 | Ch13 region 1 |
| chr13:29037781-29037821 | Ch13 region 1 |
| chr13:29037851-29037891 | Ch13 region 1 |
| chr13:29037896-29037936 | Ch13 region 1 |
| chr13:29037948-29037988 | Ch13 region 1 |
| chr13:29038020-29038060 | Ch13 region 1 |
| chr13:29038111-29038151 | Ch13 region 1 |
| chr13:29038160-29038200 | Ch13 region 1 |
| chr13:29038346-29038386 | Ch13 region 1 |
| chr13:29038433-29038473 | Ch13 region 1 |
| chr13:29038515-29038555 | Ch13 region 1 |
| chr13:29038592-29038632 | Ch13 region 1 |
| chr13:29038693-29038733 | Ch13 region 1 |
| chr13:29038748-29038788 | Ch13 region 1 |
| chr13:29038833-29038873 | Ch13 region 1 |
| chr13:29038913-29038953 | Ch13 region 1 |
| chr13:29039009-29039049 | Ch13 region 1 |
| chr13:29039070-29039110 | Ch13 region 1 |
| chr13:29039141-29039181 | Ch13 region 1 |
| chr13:29039199-29039239 | Ch13 region 1 |
| chr13:29039255-29039295 | Ch13 region 1 |
| chr13:29039307-29039347 | Ch13 region 1 |
| chr13:29039364-29039404 | Ch13 region 1 |

| | |
|---|---|
| chr13:29039440-29039480 | Ch13 region 1 |
| chr13:29039492-29039532 | Ch13 region 1 |
| chr13:29039550-29039590 | Ch13 region 1 |
| chr13:29039617-29039657 | Ch13 region 1 |
| chr13:29039672-29039712 | Ch13 region 1 |
| chr13:29039880-29039920 | Ch13 region 1 |
| chr13:29039984-29040024 | Ch13 region 1 |
| chr13:29040033-29040073 | Ch13 region 1 |
| chr13:29040079-29040119 | Ch13 region 1 |
| chr13:29040135-29040175 | Ch13 region 1 |
| chr13:29040200-29040240 | Ch13 region 1 |
| chr13:29040245-29040285 | Ch13 region 1 |
| chr13:29040298-29040338 | Ch13 region 1 |
| chr13:29040343-29040383 | Ch13 region 1 |
| chr13:29040496-29040536 | Ch13 region 1 |
| chr13:29040561-29040601 | Ch13 region 1 |
| chr13:29040637-29040677 | Ch13 region 1 |
| chr13:29040747-29040787 | Ch13 region 1 |
| chr13:29040805-29040845 | Ch13 region 1 |
| chr13:29040855-29040895 | Ch13 region 1 |
| chr13:29040915-29040955 | Ch13 region 1 |
| chr13:29040962-29041002 | Ch13 region 1 |
| chr13:29041052-29041092 | Ch13 region 1 |
| chr13:29041138-29041178 | Ch13 region 1 |
| chr13:29041208-29041248 | Ch13 region 1 |
| chr13:29041254-29041294 | Ch13 region 1 |
| chr13:29041309-29041349 | Ch13 region 1 |
| chr13:29041357-29041397 | Ch13 region 1 |
| chr13:29041419-29041459 | Ch13 region 1 |
| chr13:29041466-29041506 | Ch13 region 1 |
| chr13:29041512-29041552 | Ch13 region 1 |
| chr13:29041621-29041661 | Ch13 region 1 |
| chr13:29041700-29041740 | Ch13 region 1 |
| chr13:29041914-29041954 | Ch13 region 1 |
| chr13:29042141-29042181 | Ch13 region 1 |
| chr13:29042225-29042265 | Ch13 region 1 |
| chr13:29042270-29042310 | Ch13 region 1 |
| chr13:29042355-29042395 | Ch13 region 1 |
| chr13:29042425-29042465 | Ch13 region 1 |
| chr13:29042470-29042510 | Ch13 region 1 |
| chr13:29042548-29042588 | Ch13 region 1 |
| chr13:29042614-29042654 | Ch13 region 1 |
| chr13:29042666-29042706 | Ch13 region 1 |
| chr13:29042784-29042824 | Ch13 region 1 |
| chr13:29042855-29042895 | Ch13 region 1 |
| chr13:29042967-29043007 | Ch13 region 1 |
| chr13:29043026-29043066 | Ch13 region 1 |
| chr13:29043106-29043146 | Ch13 region 1 |
| chr13:29043175-29043215 | Ch13 region 1 |
| chr13:29043277-29043317 | Ch13 region 1 |
| chr13:29043322-29043362 | Ch13 region 1 |
| chr13:29043376-29043416 | Ch13 region 1 |

| | |
|---|---|
| chr13:29043455-29043495 | Ch13 region 1 |
| chr13:29043546-29043586 | Ch13 region 1 |
| chr13:29043591-29043631 | Ch13 region 1 |
| chr13:29043672-29043712 | Ch13 region 1 |
| chr13:29043766-29043806 | Ch13 region 1 |
| chr13:29043939-29043979 | Ch13 region 1 |
| chr13:29044158-29044198 | Ch13 region 1 |
| chr13:29044210-29044250 | Ch13 region 1 |
| chr13:29044300-29044340 | Ch13 region 1 |
| chr13:29044363-29044403 | Ch13 region 1 |
| chr13:29044457-29044497 | Ch13 region 1 |
| chr13:29044504-29044544 | Ch13 region 1 |
| chr13:29044587-29044627 | Ch13 region 1 |
| chr13:29044662-29044702 | Ch13 region 1 |
| chr13:29044707-29044747 | Ch13 region 1 |
| chr13:29044766-29044806 | Ch13 region 1 |
| chr13:29045107-29045147 | Ch13 region 1 |
| chr13:29045180-29045220 | Ch13 region 1 |
| chr13:29045238-29045278 | Ch13 region 1 |
| chr13:29045318-29045358 | Ch13 region 1 |
| chr13:29045379-29045419 | Ch13 region 1 |
| chr13:29045435-29045475 | Ch13 region 1 |
| chr13:29045558-29045598 | Ch13 region 1 |
| chr13:29045633-29045673 | Ch13 region 1 |
| chr13:29045690-29045730 | Ch13 region 1 |
| chr13:29045742-29045782 | Ch13 region 1 |
| chr13:29045791-29045831 | Ch13 region 1 |
| chr13:29045870-29045910 | Ch13 region 1 |
| chr13:29045915-29045955 | Ch13 region 1 |
| chr13:29045975-29046015 | Ch13 region 1 |
| chr13:29046038-29046078 | Ch13 region 1 |
| chr13:29046099-29046139 | Ch13 region 1 |
| chr13:29046164-29046204 | Ch13 region 1 |
| chr13:29046237-29046277 | Ch13 region 1 |
| chr13:29046321-29046361 | Ch13 region 1 |
| chr13:29046393-29046433 | Ch13 region 1 |
| chr13:29046458-29046498 | Ch13 region 1 |
| chr13:29046538-29046578 | Ch13 region 1 |
| chr13:29046584-29046624 | Ch13 region 1 |
| chr13:29046676-29046716 | Ch13 region 1 |
| chr13:29046740-29046780 | Ch13 region 1 |
| chr13:29046816-29046856 | Ch13 region 1 |
| chr13:29046914-29046954 | Ch13 region 1 |
| chr13:29046959-29046999 | Ch13 region 1 |
| chr13:29047040-29047080 | Ch13 region 1 |
| chr13:29047158-29047198 | Ch13 region 1 |
| chr13:29047262-29047302 | Ch13 region 1 |
| chr13:29047322-29047362 | Ch13 region 1 |
| chr13:29047405-29047445 | Ch13 region 1 |
| chr13:29047460-29047500 | Ch13 region 1 |
| chr13:29047759-29047799 | Ch13 region 1 |
| chr13:29047982-29048022 | Ch13 region 1 |

| | |
|---|---|
| chr13:29048071-29048111 | Ch13 region 1 |
| chr13:29048135-29048175 | Ch13 region 1 |
| chr13:29048214-29048254 | Ch13 region 1 |
| chr13:29048278-29048318 | Ch13 region 1 |
| chr13:29048339-29048379 | Ch13 region 1 |
| chr13:29048385-29048425 | Ch13 region 1 |
| chr13:29048502-29048542 | Ch13 region 1 |
| chr13:29048598-29048638 | Ch13 region 1 |
| chr13:29048738-29048778 | Ch13 region 1 |
| chr13:29048788-29048828 | Ch13 region 1 |
| chr13:29048882-29048922 | Ch13 region 1 |
| chr13:29048973-29049013 | Ch13 region 1 |
| chr13:29049040-29049080 | Ch13 region 1 |
| chr13:29049088-29049128 | Ch13 region 1 |
| chr13:29049168-29049208 | Ch13 region 1 |
| chr13:29049253-29049293 | Ch13 region 1 |
| chr13:29049373-29049413 | Ch13 region 1 |
| chr13:29049483-29049523 | Ch13 region 1 |
| chr13:29049540-29049580 | Ch13 region 1 |
| chr13:29049591-29049631 | Ch13 region 1 |
| chr13:29049670-29049710 | Ch13 region 1 |
| chr13:29049725-29049765 | Ch13 region 1 |
| chr13:29049785-29049825 | Ch13 region 1 |
| chr13:29049851-29049891 | Ch13 region 1 |
| chr13:29049923-29049963 | Ch13 region 1 |
| chr13:29049999-29050039 | Ch13 region 1 |
| chr13:29050044-29050084 | Ch13 region 1 |
| chr13:29050092-29050132 | Ch13 region 1 |
| chr13:29050174-29050214 | Ch13 region 1 |
| chr13:29050269-29050309 | Ch13 region 1 |
| chr13:29050327-29050367 | Ch13 region 1 |
| chr13:29050378-29050418 | Ch13 region 1 |
| chr13:29050431-29050471 | Ch13 region 1 |
| chr13:29050491-29050531 | Ch13 region 1 |
| chr13:29050572-29050612 | Ch13 region 1 |
| chr13:29050652-29050692 | Ch13 region 1 |
| chr13:29050713-29050753 | Ch13 region 1 |
| chr13:29050773-29050813 | Ch13 region 1 |
| chr13:29050835-29050875 | Ch13 region 1 |
| chr13:29050926-29050966 | Ch13 region 1 |
| chr13:29051015-29051055 | Ch13 region 1 |
| chr13:29051078-29051118 | Ch13 region 1 |
| chr13:29051139-29051179 | Ch13 region 1 |
| chr13:29051217-29051257 | Ch13 region 1 |
| chr13:29051283-29051323 | Ch13 region 1 |
| chr13:29051328-29051368 | Ch13 region 1 |
| chr13:29051394-29051434 | Ch13 region 1 |
| chr13:29051439-29051479 | Ch13 region 1 |
| chr13:29051494-29051534 | Ch13 region 1 |
| chr13:29051552-29051592 | Ch13 region 1 |
| chr13:29051597-29051637 | Ch13 region 1 |
| chr13:29051670-29051710 | Ch13 region 1 |

| | |
|---|---|
| chr13:29051745-29051785 | Ch13 region 1 |
| chr13:29051791-29051831 | Ch13 region 1 |
| chr13:29051865-29051905 | Ch13 region 1 |
| chr13:29051929-29051969 | Ch13 region 1 |
| chr13:29052006-29052046 | Ch13 region 1 |
| chr13:29052082-29052122 | Ch13 region 1 |
| chr13:29052158-29052198 | Ch13 region 1 |
| chr13:29052249-29052289 | Ch13 region 1 |
| chr13:29052342-29052382 | Ch13 region 1 |
| chr13:29052429-29052469 | Ch13 region 1 |
| chr13:29052492-29052532 | Ch13 region 1 |
| chr13:29052550-29052590 | Ch13 region 1 |
| chr13:29052651-29052691 | Ch13 region 1 |
| chr13:29052737-29052777 | Ch13 region 1 |
| chr13:29052789-29052829 | Ch13 region 1 |
| chr13:29052841-29052881 | Ch13 region 1 |
| chr13:29052898-29052938 | Ch13 region 1 |
| chr13:29052963-29053003 | Ch13 region 1 |
| chr13:29053009-29053049 | Ch13 region 1 |
| chr13:29053081-29053121 | Ch13 region 1 |
| chr13:29053187-29053227 | Ch13 region 1 |
| chr13:29053268-29053308 | Ch13 region 1 |
| chr13:29053314-29053354 | Ch13 region 1 |
| chr13:29053481-29053521 | Ch13 region 1 |
| chr13:29053526-29053566 | Ch13 region 1 |
| chr13:29053590-29053630 | Ch13 region 1 |
| chr13:29053678-29053718 | Ch13 region 1 |
| chr13:29053790-29053830 | Ch13 region 1 |
| chr13:29053857-29053897 | Ch13 region 1 |
| chr13:29053904-29053944 | Ch13 region 1 |
| chr13:29054030-29054070 | Ch13 region 1 |
| chr13:29054094-29054134 | Ch13 region 1 |
| chr13:29054160-29054200 | Ch13 region 1 |
| chr13:29054205-29054245 | Ch13 region 1 |
| chr13:29054277-29054317 | Ch13 region 1 |
| chr13:29054333-29054373 | Ch13 region 1 |
| chr13:29054427-29054467 | Ch13 region 1 |
| chr13:29054472-29054512 | Ch13 region 1 |
| chr13:29054580-29054620 | Ch13 region 1 |
| chr13:29054633-29054673 | Ch13 region 1 |
| chr13:29054710-29054750 | Ch13 region 1 |
| chr13:29054835-29054875 | Ch13 region 1 |
| chr13:29054884-29054924 | Ch13 region 1 |
| chr13:29054961-29055001 | Ch13 region 1 |
| chr13:29055044-29055084 | Ch13 region 1 |
| chr13:29055128-29055168 | Ch13 region 1 |
| chr13:29055186-29055226 | Ch13 region 1 |
| chr13:29055260-29055300 | Ch13 region 1 |
| chr13:29055313-29055353 | Ch13 region 1 |
| chr13:29055374-29055414 | Ch13 region 1 |
| chr13:29055419-29055459 | Ch13 region 1 |
| chr13:29055476-29055516 | Ch13 region 1 |

| | |
|---|---|
| chr13:29055544-29055584 | Ch13 region 1 |
| chr13:29055623-29055663 | Ch13 region 1 |
| chr13:29055690-29055730 | Ch13 region 1 |
| chr13:29055751-29055791 | Ch13 region 1 |
| chr13:29055828-29055868 | Ch13 region 1 |
| chr13:29055887-29055927 | Ch13 region 1 |
| chr13:29055941-29055981 | Ch13 region 1 |
| chr13:29056163-29056203 | Ch13 region 1 |
| chr13:29056208-29056248 | Ch13 region 1 |
| chr13:29056259-29056299 | Ch13 region 1 |
| chr13:29056343-29056383 | Ch13 region 1 |
| chr13:29056414-29056454 | Ch13 region 1 |
| chr13:29056650-29056690 | Ch13 region 1 |
| chr13:29056701-29056741 | Ch13 region 1 |
| chr13:29056783-29056823 | Ch13 region 1 |
| chr13:29056841-29056881 | Ch13 region 1 |
| chr13:29056903-29056943 | Ch13 region 1 |
| chr13:29056948-29056988 | Ch13 region 1 |
| chr13:29057016-29057056 | Ch13 region 1 |
| chr13:29057098-29057138 | Ch13 region 1 |
| chr13:29057157-29057197 | Ch13 region 1 |
| chr13:29057221-29057261 | Ch13 region 1 |
| chr13:29057274-29057314 | Ch13 region 1 |
| chr13:29057335-29057375 | Ch13 region 1 |
| chr13:29057417-29057457 | Ch13 region 1 |
| chr13:29057499-29057539 | Ch13 region 1 |
| chr13:29057551-29057591 | Ch13 region 1 |
| chr13:29057621-29057661 | Ch13 region 1 |
| chr13:29057691-29057731 | Ch13 region 1 |
| chr13:29057744-29057784 | Ch13 region 1 |
| chr13:29057789-29057829 | Ch13 region 1 |
| chr13:29057859-29057899 | Ch13 region 1 |
| chr13:29057922-29057962 | Ch13 region 1 |
| chr13:29058024-29058064 | Ch13 region 1 |
| chr13:29058088-29058128 | Ch13 region 1 |
| chr13:29058157-29058197 | Ch13 region 1 |
| chr13:29058202-29058242 | Ch13 region 1 |
| chr13:29058286-29058326 | Ch13 region 1 |
| chr13:29058348-29058388 | Ch13 region 1 |
| chr13:29058404-29058444 | Ch13 region 1 |
| chr13:29058469-29058509 | Ch13 region 1 |
| chr13:29058514-29058554 | Ch13 region 1 |
| chr13:29058560-29058600 | Ch13 region 1 |
| chr13:29058618-29058658 | Ch13 region 1 |
| chr13:29058680-29058720 | Ch13 region 1 |
| chr13:29058730-29058770 | Ch13 region 1 |
| chr13:29058828-29058868 | Ch13 region 1 |
| chr13:29058880-29058920 | Ch13 region 1 |
| chr13:29058938-29058978 | Ch13 region 1 |
| chr13:29059031-29059071 | Ch13 region 1 |
| chr13:29059293-29059333 | Ch13 region 1 |
| chr13:29059344-29059384 | Ch13 region 1 |

| | |
|---|---|
| chr13:29059408-29059448 | Ch13 region 1 |
| chr13:29059466-29059506 | Ch13 region 1 |
| chr13:29059541-29059581 | Ch13 region 1 |
| chr13:29059617-29059657 | Ch13 region 1 |
| chr13:29059680-29059720 | Ch13 region 1 |
| chr13:29059755-29059795 | Ch13 region 1 |
| chr13:29059841-29059881 | Ch13 region 1 |
| chr13:29059887-29059927 | Ch13 region 1 |
| chr13:29059975-29060015 | Ch13 region 1 |
| chr13:29060025-29060065 | Ch13 region 1 |
| chr13:29060093-29060133 | Ch13 region 1 |
| chr13:29060138-29060178 | Ch13 region 1 |
| chr13:29060225-29060265 | Ch13 region 1 |
| chr13:29060286-29060326 | Ch13 region 1 |
| chr13:29060362-29060402 | Ch13 region 1 |
| chr13:29060413-29060453 | Ch13 region 1 |
| chr13:29060458-29060498 | Ch13 region 1 |
| chr13:29060503-29060543 | Ch13 region 1 |
| chr13:29060576-29060616 | Ch13 region 1 |
| chr13:29060663-29060703 | Ch13 region 1 |
| chr13:29060721-29060761 | Ch13 region 1 |
| chr13:29060785-29060825 | Ch13 region 1 |
| chr13:29060855-29060895 | Ch13 region 1 |
| chr13:29060918-29060958 | Ch13 region 1 |
| chr13:29060963-29061003 | Ch13 region 1 |
| chr13:29061052-29061092 | Ch13 region 1 |
| chr13:29061184-29061224 | Ch13 region 1 |
| chr13:29061281-29061321 | Ch13 region 1 |
| chr13:29061387-29061427 | Ch13 region 1 |
| chr13:29061451-29061491 | Ch13 region 1 |
| chr13:29061505-29061545 | Ch13 region 1 |
| chr13:29061550-29061590 | Ch13 region 1 |
| chr13:29061626-29061666 | Ch13 region 1 |
| chr13:29061685-29061725 | Ch13 region 1 |
| chr13:29061748-29061788 | Ch13 region 1 |
| chr13:29061822-29061862 | Ch13 region 1 |
| chr13:29061871-29061911 | Ch13 region 1 |
| chr13:29061916-29061956 | Ch13 region 1 |
| chr13:29061980-29062020 | Ch13 region 1 |
| chr13:29062033-29062073 | Ch13 region 1 |
| chr13:29062117-29062157 | Ch13 region 1 |
| chr13:29062182-29062222 | Ch13 region 1 |
| chr13:29062228-29062268 | Ch13 region 1 |
| chr13:29062534-29062574 | Ch13 region 1 |
| chr13:29062634-29062674 | Ch13 region 1 |
| chr13:29062707-29062747 | Ch13 region 1 |
| chr13:29062752-29062792 | Ch13 region 1 |
| chr13:29062812-29062852 | Ch13 region 1 |
| chr13:29062895-29062935 | Ch13 region 1 |
| chr13:29062943-29062983 | Ch13 region 1 |
| chr13:29062990-29063030 | Ch13 region 1 |
| chr13:29063072-29063112 | Ch13 region 1 |

| | |
|---|---|
| chr13:29063139-29063179 | Ch13 region 1 |
| chr13:29063206-29063246 | Ch13 region 1 |
| chr13:29063265-29063305 | Ch13 region 1 |
| chr13:29063333-29063373 | Ch13 region 1 |
| chr13:29063401-29063441 | Ch13 region 1 |
| chr13:29063471-29063511 | Ch13 region 1 |
| chr13:29063559-29063599 | Ch13 region 1 |
| chr13:29063626-29063666 | Ch13 region 1 |
| chr13:29063728-29063768 | Ch13 region 1 |
| chr13:29063775-29063815 | Ch13 region 1 |
| chr13:29063821-29063861 | Ch13 region 1 |
| chr13:29063875-29063915 | Ch13 region 1 |
| chr13:29063934-29063974 | Ch13 region 1 |
| chr13:29064011-29064051 | Ch13 region 1 |
| chr13:29064127-29064167 | Ch13 region 1 |
| chr13:29064172-29064212 | Ch13 region 1 |
| chr13:29064218-29064258 | Ch13 region 1 |
| chr13:29064299-29064339 | Ch13 region 1 |
| chr13:29064347-29064387 | Ch13 region 1 |
| chr13:29064392-29064432 | Ch13 region 1 |
| chr13:29064464-29064504 | Ch13 region 1 |
| chr13:29064546-29064586 | Ch13 region 1 |
| chr13:29064599-29064639 | Ch13 region 1 |
| chr13:29064644-29064684 | Ch13 region 1 |
| chr13:29064689-29064729 | Ch13 region 1 |
| chr13:29064738-29064778 | Ch13 region 1 |
| chr13:29064809-29064849 | Ch13 region 1 |
| chr13:29064896-29064936 | Ch13 region 1 |
| chr13:29064964-29065004 | Ch13 region 1 |
| chr13:29065010-29065050 | Ch13 region 1 |
| chr13:29065082-29065122 | Ch13 region 1 |
| chr13:29065151-29065191 | Ch13 region 1 |
| chr13:29065215-29065255 | Ch13 region 1 |
| chr13:29065281-29065321 | Ch13 region 1 |
| chr13:29065555-29065595 | Ch13 region 1 |
| chr13:29065614-29065654 | Ch13 region 1 |
| chr13:29065670-29065710 | Ch13 region 1 |
| chr13:29065715-29065755 | Ch13 region 1 |
| chr13:29065780-29065820 | Ch13 region 1 |
| chr13:29065855-29065895 | Ch13 region 1 |
| chr13:29065907-29065947 | Ch13 region 1 |
| chr13:29066004-29066044 | Ch13 region 1 |
| chr13:29066080-29066120 | Ch13 region 1 |
| chr13:29066152-29066192 | Ch13 region 1 |
| chr13:29066223-29066263 | Ch13 region 1 |
| chr13:29066354-29066394 | Ch13 region 1 |
| chr13:29066409-29066449 | Ch13 region 1 |
| chr13:29066457-29066497 | Ch13 region 1 |
| chr13:29066526-29066566 | Ch13 region 1 |
| chr13:29066604-29066644 | Ch13 region 1 |
| chr13:29066652-29066692 | Ch13 region 1 |
| chr13:29066701-29066741 | Ch13 region 1 |

| | |
|---|---|
| chr13:29066746-29066786 | Ch13 region 1 |
| chr13:29066826-29066866 | Ch13 region 1 |
| chr13:29066873-29066913 | Ch13 region 1 |
| chr13:29066923-29066963 | Ch13 region 1 |
| chr13:29066970-29067010 | Ch13 region 1 |
| chr13:29067033-29067073 | Ch13 region 1 |
| chr13:29067139-29067179 | Ch13 region 1 |
| chr13:29067186-29067226 | Ch13 region 1 |
| chr13:29067271-29067311 | Ch13 region 1 |
| chr13:29067459-29067499 | Ch13 region 1 |
| chr13:29067517-29067557 | Ch13 region 1 |
| chr13:29067586-29067626 | Ch13 region 1 |
| chr13:29067659-29067699 | Ch13 region 1 |
| chr13:29067762-29067802 | Ch13 region 1 |
| chr13:29067810-29067850 | Ch13 region 1 |
| chr13:29067877-29067917 | Ch13 region 1 |
| chr13:29067940-29067980 | Ch13 region 1 |
| chr13:29068025-29068065 | Ch13 region 1 |
| chr13:29068080-29068120 | Ch13 region 1 |
| chr13:29068150-29068190 | Ch13 region 1 |
| chr13:29068201-29068241 | Ch13 region 1 |
| chr13:29068267-29068307 | Ch13 region 1 |
| chr13:29068358-29068398 | Ch13 region 1 |
| chr13:29068409-29068449 | Ch13 region 1 |
| chr13:29068455-29068495 | Ch13 region 1 |
| chr13:29068527-29068567 | Ch13 region 1 |
| chr13:29068597-29068637 | Ch13 region 1 |
| chr13:29068646-29068686 | Ch13 region 1 |
| chr13:29068691-29068731 | Ch13 region 1 |
| chr13:29068766-29068806 | Ch13 region 1 |
| chr13:29068811-29068851 | Ch13 region 1 |
| chr13:29068867-29068907 | Ch13 region 1 |
| chr13:29068954-29068994 | Ch13 region 1 |
| chr13:29069027-29069067 | Ch13 region 1 |
| chr13:29069108-29069148 | Ch13 region 1 |
| chr13:29069165-29069205 | Ch13 region 1 |
| chr13:29069286-29069326 | Ch13 region 1 |
| chr13:29069351-29069391 | Ch13 region 1 |
| chr13:29069414-29069454 | Ch13 region 1 |
| chr13:29069521-29069561 | Ch13 region 1 |
| chr13:29069571-29069611 | Ch13 region 1 |
| chr13:29069626-29069666 | Ch13 region 1 |
| chr13:29069711-29069751 | Ch13 region 1 |
| chr13:29069758-29069798 | Ch13 region 1 |
| chr13:29069806-29069846 | Ch13 region 1 |
| chr13:29069874-29069914 | Ch13 region 1 |
| chr13:29069920-29069960 | Ch13 region 1 |
| chr13:29069971-29070011 | Ch13 region 1 |
| chr13:29070021-29070061 | Ch13 region 1 |
| chr13:29070087-29070127 | Ch13 region 1 |
| chr13:29070150-29070190 | Ch13 region 1 |
| chr13:29070205-29070245 | Ch13 region 1 |

| | |
|---|---|
| chr13:29070283-29070323 | Ch13 region 1 |
| chr13:29070345-29070385 | Ch13 region 1 |
| chr13:29070395-29070435 | Ch13 region 1 |
| chr13:29070446-29070486 | Ch13 region 1 |
| chr13:29070533-29070573 | Ch13 region 1 |
| chr13:29070618-29070658 | Ch13 region 1 |
| chr13:29070686-29070726 | Ch13 region 1 |
| chr13:29070732-29070772 | Ch13 region 1 |
| chr13:29070823-29070863 | Ch13 region 1 |
| chr13:29070876-29070916 | Ch13 region 1 |
| chr13:29070958-29070998 | Ch13 region 1 |
| chr13:29071031-29071071 | Ch13 region 1 |
| chr13:29071111-29071151 | Ch13 region 1 |
| chr13:29071217-29071257 | Ch13 region 1 |
| chr13:29071268-29071308 | Ch13 region 1 |
| chr13:29071345-29071385 | Ch13 region 1 |
| chr13:29071390-29071430 | Ch13 region 1 |
| chr13:29071482-29071522 | Ch13 region 1 |
| chr13:29071564-29071604 | Ch13 region 1 |
| chr13:29071627-29071667 | Ch13 region 1 |
| chr13:29071672-29071712 | Ch13 region 1 |
| chr13:29071737-29071777 | Ch13 region 1 |
| chr13:29071797-29071837 | Ch13 region 1 |
| chr13:29071867-29071907 | Ch13 region 1 |
| chr13:29071913-29071953 | Ch13 region 1 |
| chr13:29072020-29072060 | Ch13 region 1 |
| chr13:29072108-29072148 | Ch13 region 1 |
| chr13:29072174-29072214 | Ch13 region 1 |
| chr13:29072245-29072285 | Ch13 region 1 |
| chr13:29072301-29072341 | Ch13 region 1 |
| chr13:29072348-29072388 | Ch13 region 1 |
| chr13:29072394-29072434 | Ch13 region 1 |
| chr13:29072457-29072497 | Ch13 region 1 |
| chr13:29072502-29072542 | Ch13 region 1 |
| chr13:29072553-29072593 | Ch13 region 1 |
| chr13:29072616-29072656 | Ch13 region 1 |
| chr13:29072712-29072752 | Ch13 region 1 |
| chr13:29072762-29072802 | Ch13 region 1 |
| chr13:29072841-29072881 | Ch13 region 1 |
| chr13:29072888-29072928 | Ch13 region 1 |
| chr13:29072941-29072981 | Ch13 region 1 |
| chr13:29072997-29073037 | Ch13 region 1 |
| chr13:29073042-29073082 | Ch13 region 1 |
| chr13:29073092-29073132 | Ch13 region 1 |
| chr13:29073167-29073207 | Ch13 region 1 |
| chr13:29073215-29073255 | Ch13 region 1 |
| chr13:29073297-29073337 | Ch13 region 1 |
| chr13:29073370-29073410 | Ch13 region 1 |
| chr13:29073431-29073471 | Ch13 region 1 |
| chr13:29073519-29073559 | Ch13 region 1 |
| chr13:29073585-29073625 | Ch13 region 1 |
| chr13:29073634-29073674 | Ch13 region 1 |

| | |
|---|---|
| chr13:29073710-29073750 | Ch13 region 1 |
| chr13:29073766-29073806 | Ch13 region 1 |
| chr13:29073853-29073893 | Ch13 region 1 |
| chr13:29073927-29073967 | Ch13 region 1 |
| chr13:29073974-29074014 | Ch13 region 1 |
| chr13:29074024-29074064 | Ch13 region 1 |
| chr13:29074088-29074128 | Ch13 region 1 |
| chr13:29074133-29074173 | Ch13 region 1 |
| chr13:29074203-29074243 | Ch13 region 1 |
| chr13:29074288-29074328 | Ch13 region 1 |
| chr13:29074366-29074406 | Ch13 region 1 |
| chr13:29074435-29074475 | Ch13 region 1 |
| chr13:29074492-29074532 | Ch13 region 1 |
| chr13:29074542-29074582 | Ch13 region 1 |
| chr13:29074604-29074644 | Ch13 region 1 |
| chr13:29074677-29074717 | Ch13 region 1 |
| chr13:29074758-29074798 | Ch13 region 1 |
| chr13:29074818-29074858 | Ch13 region 1 |
| chr13:29074933-29074973 | Ch13 region 1 |
| chr13:29074996-29075036 | Ch13 region 1 |
| chr13:29075050-29075090 | Ch13 region 1 |
| chr13:29075210-29075250 | Ch13 region 1 |
| chr13:29075385-29075425 | Ch13 region 1 |
| chr13:29075543-29075583 | Ch13 region 1 |
| chr13:29075592-29075632 | Ch13 region 1 |
| chr13:29075682-29075722 | Ch13 region 1 |
| chr13:29075770-29075810 | Ch13 region 1 |
| chr13:29075822-29075862 | Ch13 region 1 |
| chr13:29075867-29075907 | Ch13 region 1 |
| chr13:29075930-29075970 | Ch13 region 1 |
| chr13:29076027-29076067 | Ch13 region 1 |
| chr13:29076074-29076114 | Ch13 region 1 |
| chr13:29076144-29076184 | Ch13 region 1 |
| chr13:29076289-29076329 | Ch13 region 1 |
| chr13:29076359-29076399 | Ch13 region 1 |
| chr13:29076417-29076457 | Ch13 region 1 |
| chr13:29076524-29076564 | Ch13 region 1 |
| chr13:29076591-29076631 | Ch13 region 1 |
| chr13:29076659-29076699 | Ch13 region 1 |
| chr13:29076706-29076746 | Ch13 region 1 |
| chr13:29076776-29076816 | Ch13 region 1 |
| chr13:29076827-29076867 | Ch13 region 1 |
| chr13:29076996-29077036 | Ch13 region 1 |
| chr13:29077046-29077086 | Ch13 region 1 |
| chr13:29077133-29077173 | Ch13 region 1 |
| chr13:29077179-29077219 | Ch13 region 1 |
| chr13:29077230-29077270 | Ch13 region 1 |
| chr13:29077275-29077315 | Ch13 region 1 |
| chr13:29077323-29077363 | Ch13 region 1 |
| chr13:29077412-29077452 | Ch13 region 1 |
| chr13:29077478-29077518 | Ch13 region 1 |
| chr13:29077550-29077590 | Ch13 region 1 |

| | |
|---|---|
| chr13:29077618-29077658 | Ch13 region 1 |
| chr13:29077675-29077715 | Ch13 region 1 |
| chr13:29077754-29077794 | Ch13 region 1 |
| chr13:29077846-29077886 | Ch13 region 1 |
| chr13:29077910-29077950 | Ch13 region 1 |
| chr13:29077955-29077995 | Ch13 region 1 |
| chr13:29078044-29078084 | Ch13 region 1 |
| chr13:29078115-29078155 | Ch13 region 1 |
| chr13:29078178-29078218 | Ch13 region 1 |
| chr13:29078224-29078264 | Ch13 region 1 |
| chr13:29078270-29078310 | Ch13 region 1 |
| chr13:29078315-29078355 | Ch13 region 1 |
| chr13:29078364-29078404 | Ch13 region 1 |
| chr13:29078435-29078475 | Ch13 region 1 |
| chr13:29078500-29078540 | Ch13 region 1 |
| chr13:29078575-29078615 | Ch13 region 1 |
| chr13:29078620-29078660 | Ch13 region 1 |
| chr13:29078675-29078715 | Ch13 region 1 |
| chr13:29078756-29078796 | Ch13 region 1 |
| chr13:29078824-29078864 | Ch13 region 1 |
| chr13:29078940-29078980 | Ch13 region 1 |
| chr13:29079004-29079044 | Ch13 region 1 |
| chr13:29079094-29079134 | Ch13 region 1 |
| chr13:29079163-29079203 | Ch13 region 1 |
| chr13:29079286-29079326 | Ch13 region 1 |
| chr13:29079504-29079544 | Ch13 region 1 |
| chr13:29079566-29079606 | Ch13 region 1 |
| chr13:29079639-29079679 | Ch13 region 1 |
| chr13:29079700-29079740 | Ch13 region 1 |
| chr13:29079764-29079804 | Ch13 region 1 |
| chr13:29079830-29079870 | Ch13 region 1 |
| chr13:29079880-29079920 | Ch13 region 1 |
| chr13:29079949-29079989 | Ch13 region 1 |
| chr13:29080032-29080072 | Ch13 region 1 |
| chr13:29080108-29080148 | Ch13 region 1 |
| chr13:29080177-29080217 | Ch13 region 1 |
| chr13:29080243-29080283 | Ch13 region 1 |
| chr13:29080330-29080370 | Ch13 region 1 |
| chr13:29080478-29080518 | Ch13 region 1 |
| chr13:29080554-29080594 | Ch13 region 1 |
| chr13:29080632-29080672 | Ch13 region 1 |
| chr13:29080677-29080717 | Ch13 region 1 |
| chr13:29080742-29080782 | Ch13 region 1 |
| chr13:29080788-29080828 | Ch13 region 1 |
| chr13:29080879-29080919 | Ch13 region 1 |
| chr13:29080958-29080998 | Ch13 region 1 |
| chr13:29081056-29081096 | Ch13 region 1 |
| chr13:29081124-29081164 | Ch13 region 1 |
| chr13:29081247-29081287 | Ch13 region 1 |
| chr13:29081292-29081332 | Ch13 region 1 |
| chr13:29081355-29081395 | Ch13 region 1 |
| chr13:29081459-29081499 | Ch13 region 1 |

| | |
|---|---|
| chr13:29081504-29081544 | Ch13 region 1 |
| chr13:29081553-29081593 | Ch13 region 1 |
| chr13:29081627-29081667 | Ch13 region 1 |
| chr13:29081687-29081727 | Ch13 region 1 |
| chr13:29081748-29081788 | Ch13 region 1 |
| chr13:29081855-29081895 | Ch13 region 1 |
| chr13:29081903-29081943 | Ch13 region 1 |
| chr13:29082163-29082203 | Ch13 region 1 |
| chr13:29082255-29082295 | Ch13 region 1 |
| chr13:29082374-29082414 | Ch13 region 1 |
| chr13:29082442-29082482 | Ch13 region 1 |
| chr13:29082559-29082599 | Ch13 region 1 |
| chr13:29082818-29082858 | Ch13 region 1 |
| chr13:29082950-29082990 | Ch13 region 1 |
| chr13:29083069-29083109 | Ch13 region 1 |
| chr13:29083132-29083172 | Ch13 region 1 |
| chr13:29083200-29083240 | Ch13 region 1 |
| chr13:29083306-29083346 | Ch13 region 1 |
| chr13:29083426-29083466 | Ch13 region 1 |
| chr13:29083478-29083518 | Ch13 region 1 |
| chr13:29083582-29083622 | Ch13 region 1 |
| chr13:29083664-29083704 | Ch13 region 1 |
| chr13:29083722-29083762 | Ch13 region 1 |
| chr13:29083794-29083834 | Ch13 region 1 |
| chr13:29083842-29083882 | Ch13 region 1 |
| chr13:29083887-29083927 | Ch13 region 1 |
| chr13:29083985-29084025 | Ch13 region 1 |
| chr13:29084048-29084088 | Ch13 region 1 |
| chr13:29084093-29084133 | Ch13 region 1 |
| chr13:29084156-29084196 | Ch13 region 1 |
| chr13:29084230-29084270 | Ch13 region 1 |
| chr13:29084281-29084321 | Ch13 region 1 |
| chr13:29084453-29084493 | Ch13 region 1 |
| chr13:29084597-29084637 | Ch13 region 1 |
| chr13:29084651-29084691 | Ch13 region 1 |
| chr13:29084704-29084744 | Ch13 region 1 |
| chr13:29084757-29084797 | Ch13 region 1 |
| chr13:29084804-29084844 | Ch13 region 1 |
| chr13:29084859-29084899 | Ch13 region 1 |
| chr13:29084904-29084944 | Ch13 region 1 |
| chr13:29084965-29085005 | Ch13 region 1 |
| chr13:29085047-29085087 | Ch13 region 1 |
| chr13:29085125-29085165 | Ch13 region 1 |
| chr13:29085234-29085274 | Ch13 region 1 |
| chr13:29085289-29085329 | Ch13 region 1 |
| chr13:29085337-29085377 | Ch13 region 1 |
| chr13:29085424-29085464 | Ch13 region 1 |
| chr13:29085538-29085578 | Ch13 region 1 |
| chr13:29085756-29085796 | Ch13 region 1 |
| chr13:29085896-29085936 | Ch13 region 1 |
| chr13:29085959-29085999 | Ch13 region 1 |
| chr13:29086013-29086053 | Ch13 region 1 |

| | |
|---|---|
| chr13:29086140-29086180 | Ch13 region 1 |
| chr13:29086199-29086239 | Ch13 region 1 |
| chr13:29086335-29086375 | Ch13 region 1 |
| chr13:29086581-29086621 | Ch13 region 1 |
| chr13:29086639-29086679 | Ch13 region 1 |
| chr13:29086691-29086731 | Ch13 region 1 |
| chr13:29086750-29086790 | Ch13 region 1 |
| chr13:29086986-29087026 | Ch13 region 1 |
| chr13:29087051-29087091 | Ch13 region 1 |
| chr13:29087299-29087339 | Ch13 region 1 |
| chr13:29087402-29087442 | Ch13 region 1 |
| chr13:29087503-29087543 | Ch13 region 1 |
| chr13:29087685-29087725 | Ch13 region 1 |
| chr13:29087773-29087813 | Ch13 region 1 |
| chr13:29087841-29087881 | Ch13 region 1 |
| chr13:29087891-29087931 | Ch13 region 1 |
| chr13:29088032-29088072 | Ch13 region 1 |
| chr13:29088116-29088156 | Ch13 region 1 |
| chr13:29088198-29088238 | Ch13 region 1 |
| chr13:29088247-29088287 | Ch13 region 1 |
| chr13:29088310-29088350 | Ch13 region 1 |
| chr13:29088423-29088463 | Ch13 region 1 |
| chr13:29088501-29088541 | Ch13 region 1 |
| chr13:29088684-29088724 | Ch13 region 1 |
| chr13:29088731-29088771 | Ch13 region 1 |
| chr13:29088815-29088855 | Ch13 region 1 |
| chr13:29088975-29089015 | Ch13 region 1 |
| chr13:29089033-29089073 | Ch13 region 1 |
| chr13:29089102-29089142 | Ch13 region 1 |
| chr13:29089173-29089213 | Ch13 region 1 |
| chr13:29089236-29089276 | Ch13 region 1 |
| chr13:29089501-29089541 | Ch13 region 1 |
| chr13:29089553-29089593 | Ch13 region 1 |
| chr13:29089628-29089668 | Ch13 region 1 |
| chr13:29089688-29089728 | Ch13 region 1 |
| chr13:29089742-29089782 | Ch13 region 1 |
| chr13:29089791-29089831 | Ch13 region 1 |
| chr13:29089854-29089894 | Ch13 region 1 |
| chr13:29089901-29089941 | Ch13 region 1 |
| chr13:29089970-29090010 | Ch13 region 1 |
| chr13:29090020-29090060 | Ch13 region 1 |
| chr13:29090122-29090162 | Ch13 region 1 |
| chr13:29090185-29090225 | Ch13 region 1 |
| chr13:29090232-29090272 | Ch13 region 1 |
| chr13:29090307-29090347 | Ch13 region 1 |
| chr13:29090361-29090401 | Ch13 region 1 |
| chr13:29090465-29090505 | Ch13 region 1 |
| chr13:29090547-29090587 | Ch13 region 1 |
| chr13:29090680-29090720 | Ch13 region 1 |
| chr13:29090770-29090810 | Ch13 region 1 |
| chr13:29090988-29091028 | Ch13 region 1 |
| chr13:29091143-29091183 | Ch13 region 1 |

| | |
|---|---|
| chr13:29091203-29091243 | Ch13 region 1 |
| chr13:29091430-29091470 | Ch13 region 1 |
| chr13:29091542-29091582 | Ch13 region 1 |
| chr13:29091631-29091671 | Ch13 region 1 |
| chr13:29091703-29091743 | Ch13 region 1 |
| chr13:29091783-29091823 | Ch13 region 1 |
| chr13:29091844-29091884 | Ch13 region 1 |
| chr13:29091931-29091971 | Ch13 region 1 |
| chr13:29091976-29092016 | Ch13 region 1 |
| chr13:29092046-29092086 | Ch13 region 1 |
| chr13:29092119-29092159 | Ch13 region 1 |
| chr13:29092460-29092500 | Ch13 region 1 |
| chr13:29092577-29092617 | Ch13 region 1 |
| chr13:29092736-29092776 | Ch13 region 1 |
| chr13:29092801-29092841 | Ch13 region 1 |
| chr13:29093035-29093075 | Ch13 region 1 |
| chr13:29093163-29093203 | Ch13 region 1 |
| chr13:29093390-29093430 | Ch13 region 1 |
| chr13:29093435-29093475 | Ch13 region 1 |
| chr13:29093522-29093562 | Ch13 region 1 |
| chr13:29093586-29093626 | Ch13 region 1 |
| chr13:29093669-29093709 | Ch13 region 1 |
| chr13:29093714-29093754 | Ch13 region 1 |
| chr13:29093805-29093845 | Ch13 region 1 |
| chr13:29093857-29093897 | Ch13 region 1 |
| chr13:29094142-29094182 | Ch13 region 1 |
| chr13:29094325-29094365 | Ch13 region 1 |
| chr13:29094493-29094533 | Ch13 region 1 |
| chr13:29094565-29094605 | Ch13 region 1 |
| chr13:29094616-29094656 | Ch13 region 1 |
| chr13:29094666-29094706 | Ch13 region 1 |
| chr13:29094711-29094751 | Ch13 region 1 |
| chr13:29094798-29094838 | Ch13 region 1 |
| chr13:29094959-29094999 | Ch13 region 1 |
| chr13:29095010-29095050 | Ch13 region 1 |
| chr13:29095074-29095114 | Ch13 region 1 |
| chr13:29095209-29095249 | Ch13 region 1 |
| chr13:29095269-29095309 | Ch13 region 1 |
| chr13:29095444-29095484 | Ch13 region 1 |
| chr13:29095529-29095569 | Ch13 region 1 |
| chr13:29095700-29095740 | Ch13 region 1 |
| chr13:29095750-29095790 | Ch13 region 1 |
| chr13:29095995-29096035 | Ch13 region 1 |
| chr13:29096114-29096154 | Ch13 region 1 |
| chr13:29096161-29096201 | Ch13 region 1 |
| chr13:29096219-29096259 | Ch13 region 1 |
| chr13:29096310-29096350 | Ch13 region 1 |
| chr13:29096360-29096400 | Ch13 region 1 |
| chr13:29096405-29096445 | Ch13 region 1 |
| chr13:29096451-29096491 | Ch13 region 1 |
| chr13:29096506-29096546 | Ch13 region 1 |
| chr13:29096554-29096594 | Ch13 region 1 |

| | |
|---|---|
| chr13:29096614-29096654 | Ch13 region 1 |
| chr13:29096686-29096726 | Ch13 region 1 |
| chr13:29096768-29096808 | Ch13 region 1 |
| chr13:29096813-29096853 | Ch13 region 1 |
| chr13:29096892-29096932 | Ch13 region 1 |
| chr13:29096937-29096977 | Ch13 region 1 |
| chr13:29097001-29097041 | Ch13 region 1 |
| chr13:29097046-29097086 | Ch13 region 1 |
| chr13:29097091-29097131 | Ch13 region 1 |
| chr13:29097152-29097192 | Ch13 region 1 |
| chr13:29097252-29097292 | Ch13 region 1 |
| chr13:29097324-29097364 | Ch13 region 1 |
| chr13:29097434-29097474 | Ch13 region 1 |
| chr13:29097479-29097519 | Ch13 region 1 |
| chr13:29097581-29097621 | Ch13 region 1 |
| chr13:29097641-29097681 | Ch13 region 1 |
| chr13:29097738-29097778 | Ch13 region 1 |
| chr13:29097796-29097836 | Ch13 region 1 |
| chr13:29097880-29097920 | Ch13 region 1 |
| chr13:29097993-29098033 | Ch13 region 1 |
| chr13:29098186-29098226 | Ch13 region 1 |
| chr13:29098268-29098308 | Ch13 region 1 |
| chr13:29098315-29098355 | Ch13 region 1 |
| chr13:29098393-29098433 | Ch13 region 1 |
| chr13:29098473-29098513 | Ch13 region 1 |
| chr13:29098563-29098603 | Ch13 region 1 |
| chr13:29098633-29098673 | Ch13 region 1 |
| chr13:29098690-29098730 | Ch13 region 1 |
| chr13:29098749-29098789 | Ch13 region 1 |
| chr13:29098869-29098909 | Ch13 region 1 |
| chr13:29098931-29098971 | Ch13 region 1 |
| chr13:29098977-29099017 | Ch13 region 1 |
| chr13:29099022-29099062 | Ch13 region 1 |
| chr13:29099088-29099128 | Ch13 region 1 |
| chr13:29099165-29099205 | Ch13 region 1 |
| chr13:29099216-29099256 | Ch13 region 1 |
| chr13:29099267-29099307 | Ch13 region 1 |
| chr13:29099318-29099358 | Ch13 region 1 |
| chr13:29099376-29099416 | Ch13 region 1 |
| chr13:29099448-29099488 | Ch13 region 1 |
| chr13:29099536-29099576 | Ch13 region 1 |
| chr13:29099581-29099621 | Ch13 region 1 |
| chr13:29099673-29099713 | Ch13 region 1 |
| chr13:29099745-29099785 | Ch13 region 1 |
| chr13:29099797-29099837 | Ch13 region 1 |
| chr13:29099875-29099915 | Ch13 region 1 |
| chr13:29099939-29099979 | Ch13 region 1 |
| chr13:29099997-29100037 | Ch13 region 1 |
| chr13:29100065-29100105 | Ch13 region 1 |
| chr13:29100110-29100150 | Ch13 region 1 |
| chr13:29100183-29100223 | Ch13 region 1 |
| chr13:29100232-29100272 | Ch13 region 1 |

| | |
|---|---|
| chr13:29100376-29100416 | Ch13 region 1 |
| chr13:29100452-29100492 | Ch13 region 1 |
| chr13:29100568-29100608 | Ch13 region 1 |
| chr13:29100678-29100718 | Ch13 region 1 |
| chr13:29100781-29100821 | Ch13 region 1 |
| chr13:29100960-29101000 | Ch13 region 1 |
| chr13:29101228-29101268 | Ch13 region 1 |
| chr13:29101332-29101372 | Ch13 region 1 |
| chr13:29101584-29101624 | Ch13 region 1 |
| chr13:29101629-29101669 | Ch13 region 1 |
| chr13:29101687-29101727 | Ch13 region 1 |
| chr13:29101758-29101798 | Ch13 region 1 |
| chr13:29101804-29101844 | Ch13 region 1 |
| chr13:29101899-29101939 | Ch13 region 1 |
| chr13:29101987-29102027 | Ch13 region 1 |
| chr13:29102073-29102113 | Ch13 region 1 |
| chr13:29102120-29102160 | Ch13 region 1 |
| chr13:29102165-29102205 | Ch13 region 1 |
| chr13:29102248-29102288 | Ch13 region 1 |
| chr13:29102355-29102395 | Ch13 region 1 |
| chr13:29102417-29102457 | Ch13 region 1 |
| chr13:29102481-29102521 | Ch13 region 1 |
| chr13:29102552-29102592 | Ch13 region 1 |
| chr13:29102623-29102663 | Ch13 region 1 |
| chr13:29102674-29102714 | Ch13 region 1 |
| chr13:29102736-29102776 | Ch13 region 1 |
| chr13:29102814-29102854 | Ch13 region 1 |
| chr13:29102862-29102902 | Ch13 region 1 |
| chr13:29102941-29102981 | Ch13 region 1 |
| chr13:29102988-29103028 | Ch13 region 1 |
| chr13:29103033-29103073 | Ch13 region 1 |
| chr13:29103113-29103153 | Ch13 region 1 |
| chr13:29103193-29103233 | Ch13 region 1 |
| chr13:29103246-29103286 | Ch13 region 1 |
| chr13:29103292-29103332 | Ch13 region 1 |
| chr13:29103361-29103401 | Ch13 region 1 |
| chr13:29103414-29103454 | Ch13 region 1 |
| chr13:29103515-29103555 | Ch13 region 1 |
| chr13:29103611-29103651 | Ch13 region 1 |
| chr13:29103656-29103696 | Ch13 region 1 |
| chr13:29103701-29103741 | Ch13 region 1 |
| chr13:29103928-29103968 | Ch13 region 1 |
| chr13:29104068-29104108 | Ch13 region 1 |
| chr13:29104141-29104181 | Ch13 region 1 |
| chr13:29104202-29104242 | Ch13 region 1 |
| chr13:29104254-29104294 | Ch13 region 1 |
| chr13:29104306-29104346 | Ch13 region 1 |
| chr13:29104372-29104412 | Ch13 region 1 |
| chr13:29104422-29104462 | Ch13 region 1 |
| chr13:29104541-29104581 | Ch13 region 1 |
| chr13:29104598-29104638 | Ch13 region 1 |
| chr13:29104643-29104683 | Ch13 region 1 |

| | |
|---|---|
| chr13:29104721-29104761 | Ch13 region 1 |
| chr13:29104774-29104814 | Ch13 region 1 |
| chr13:29104826-29104866 | Ch13 region 1 |
| chr13:29104923-29104963 | Ch13 region 1 |
| chr13:29104992-29105032 | Ch13 region 1 |
| chr13:29105090-29105130 | Ch13 region 1 |
| chr13:29105141-29105181 | Ch13 region 1 |
| chr13:29105229-29105269 | Ch13 region 1 |
| chr13:29105309-29105349 | Ch13 region 1 |
| chr13:29105354-29105394 | Ch13 region 1 |
| chr13:29105427-29105467 | Ch13 region 1 |
| chr13:29105491-29105531 | Ch13 region 1 |
| chr13:29105569-29105609 | Ch13 region 1 |
| chr13:29105634-29105674 | Ch13 region 1 |
| chr13:29105714-29105754 | Ch13 region 1 |
| chr13:29105809-29105849 | Ch13 region 1 |
| chr13:29105955-29105995 | Ch13 region 1 |
| chr13:29106027-29106067 | Ch13 region 1 |
| chr13:29106126-29106166 | Ch13 region 1 |
| chr13:29106202-29106242 | Ch13 region 1 |
| chr13:29106283-29106323 | Ch13 region 1 |
| chr13:29106375-29106415 | Ch13 region 1 |
| chr13:29106422-29106462 | Ch13 region 1 |
| chr13:29106470-29106510 | Ch13 region 1 |
| chr13:29106524-29106564 | Ch13 region 1 |
| chr13:29106617-29106657 | Ch13 region 1 |
| chr13:29106673-29106713 | Ch13 region 1 |
| chr13:29106728-29106768 | Ch13 region 1 |
| chr13:29106816-29106856 | Ch13 region 1 |
| chr13:29106873-29106913 | Ch13 region 1 |
| chr13:29106918-29106958 | Ch13 region 1 |
| chr13:29106971-29107011 | Ch13 region 1 |
| chr13:29107085-29107125 | Ch13 region 1 |
| chr13:29107166-29107206 | Ch13 region 1 |
| chr13:29107213-29107253 | Ch13 region 1 |
| chr13:29107272-29107312 | Ch13 region 1 |
| chr13:29107330-29107370 | Ch13 region 1 |
| chr13:29107428-29107468 | Ch13 region 1 |
| chr13:29107513-29107553 | Ch13 region 1 |
| chr13:29107593-29107633 | Ch13 region 1 |
| chr13:29107642-29107682 | Ch13 region 1 |
| chr13:29107702-29107742 | Ch13 region 1 |
| chr13:29107800-29107840 | Ch13 region 1 |
| chr13:29107957-29107997 | Ch13 region 1 |
| chr13:29108002-29108042 | Ch13 region 1 |
| chr13:29108084-29108124 | Ch13 region 1 |
| chr13:29108137-29108177 | Ch13 region 1 |
| chr13:29108215-29108255 | Ch13 region 1 |
| chr13:29108261-29108301 | Ch13 region 1 |
| chr13:29108348-29108388 | Ch13 region 1 |
| chr13:29108432-29108472 | Ch13 region 1 |
| chr13:29108482-29108522 | Ch13 region 1 |

| | |
|---|---|
| chr13:29108566-29108606 | Ch13 region 1 |
| chr13:29108612-29108652 | Ch13 region 1 |
| chr13:29108727-29108767 | Ch13 region 1 |
| chr13:29108796-29108836 | Ch13 region 1 |
| chr13:29108871-29108911 | Ch13 region 1 |
| chr13:29108931-29108971 | Ch13 region 1 |
| chr13:29108984-29109024 | Ch13 region 1 |
| chr13:29109047-29109087 | Ch13 region 1 |
| chr13:29109197-29109237 | Ch13 region 1 |
| chr13:29109300-29109340 | Ch13 region 1 |
| chr13:29109390-29109430 | Ch13 region 1 |
| chr13:29109455-29109495 | Ch13 region 1 |
| chr13:29109500-29109540 | Ch13 region 1 |
| chr13:29109582-29109622 | Ch13 region 1 |
| chr13:29109703-29109743 | Ch13 region 1 |
| chr13:29109763-29109803 | Ch13 region 1 |
| chr13:29109834-29109874 | Ch13 region 1 |
| chr13:29109920-29109960 | Ch13 region 1 |
| chr13:29109968-29110008 | Ch13 region 1 |
| chr13:29110082-29110122 | Ch13 region 1 |
| chr13:29110143-29110183 | Ch13 region 1 |
| chr13:29110199-29110239 | Ch13 region 1 |
| chr13:29110275-29110315 | Ch13 region 1 |
| chr13:29110351-29110391 | Ch13 region 1 |
| chr13:29110515-29110555 | Ch13 region 1 |
| chr13:29110593-29110633 | Ch13 region 1 |
| chr13:29110655-29110695 | Ch13 region 1 |
| chr13:29110741-29110781 | Ch13 region 1 |
| chr13:29110807-29110847 | Ch13 region 1 |
| chr13:29110888-29110928 | Ch13 region 1 |
| chr13:29110934-29110974 | Ch13 region 1 |
| chr13:29111006-29111046 | Ch13 region 1 |
| chr13:29111090-29111130 | Ch13 region 1 |
| chr13:29111174-29111214 | Ch13 region 1 |
| chr13:29111281-29111321 | Ch13 region 1 |
| chr13:29111348-29111388 | Ch13 region 1 |
| chr13:29111442-29111482 | Ch13 region 1 |
| chr13:29111504-29111544 | Ch13 region 1 |
| chr13:29111577-29111617 | Ch13 region 1 |
| chr13:29111624-29111664 | Ch13 region 1 |
| chr13:29111837-29111877 | Ch13 region 1 |
| chr13:29111882-29111922 | Ch13 region 1 |
| chr13:29111999-29112039 | Ch13 region 1 |
| chr13:29112109-29112149 | Ch13 region 1 |
| chr13:29112248-29112288 | Ch13 region 1 |
| chr13:29112477-29112517 | Ch13 region 1 |
| chr13:29112547-29112587 | Ch13 region 1 |
| chr13:29112613-29112653 | Ch13 region 1 |
| chr13:29112662-29112702 | Ch13 region 1 |
| chr13:29112709-29112749 | Ch13 region 1 |
| chr13:29112781-29112821 | Ch13 region 1 |
| chr13:29112864-29112904 | Ch13 region 1 |

| | |
|---|---|
| chr13:29112913-29112953 | Ch13 region 1 |
| chr13:29112994-29113034 | Ch13 region 1 |
| chr13:29113056-29113096 | Ch13 region 1 |
| chr13:29113114-29113154 | Ch13 region 1 |
| chr13:29113225-29113265 | Ch13 region 1 |
| chr13:29113273-29113313 | Ch13 region 1 |
| chr13:29113326-29113366 | Ch13 region 1 |
| chr13:29113395-29113435 | Ch13 region 1 |
| chr13:29113528-29113568 | Ch13 region 1 |
| chr13:29113732-29113772 | Ch13 region 1 |
| chr13:29113778-29113818 | Ch13 region 1 |
| chr13:29113823-29113863 | Ch13 region 1 |
| chr13:29113873-29113913 | Ch13 region 1 |
| chr13:29113970-29114010 | Ch13 region 1 |
| chr13:29114056-29114096 | Ch13 region 1 |
| chr13:29114101-29114141 | Ch13 region 1 |
| chr13:29114178-29114218 | Ch13 region 1 |
| chr13:29114256-29114296 | Ch13 region 1 |
| chr13:29114315-29114355 | Ch13 region 1 |
| chr13:29114392-29114432 | Ch13 region 1 |
| chr13:29114444-29114484 | Ch13 region 1 |
| chr13:29114534-29114574 | Ch13 region 1 |
| chr13:29114625-29114665 | Ch13 region 1 |
| chr13:29114691-29114731 | Ch13 region 1 |
| chr13:29114741-29114781 | Ch13 region 1 |
| chr13:29114786-29114826 | Ch13 region 1 |
| chr13:29114835-29114875 | Ch13 region 1 |
| chr13:29114886-29114926 | Ch13 region 1 |
| chr13:29114942-29114982 | Ch13 region 1 |
| chr13:29115011-29115051 | Ch13 region 1 |
| chr13:29115112-29115152 | Ch13 region 1 |
| chr13:29115206-29115246 | Ch13 region 1 |
| chr13:29115257-29115297 | Ch13 region 1 |
| chr13:29115419-29115459 | Ch13 region 1 |
| chr13:29115471-29115511 | Ch13 region 1 |
| chr13:29115528-29115568 | Ch13 region 1 |
| chr13:29115597-29115637 | Ch13 region 1 |
| chr13:29115700-29115740 | Ch13 region 1 |
| chr13:29115768-29115808 | Ch13 region 1 |
| chr13:29115852-29115892 | Ch13 region 1 |
| chr13:29116001-29116041 | Ch13 region 1 |
| chr13:29116057-29116097 | Ch13 region 1 |
| chr13:29116122-29116162 | Ch13 region 1 |
| chr13:29116181-29116221 | Ch13 region 1 |
| chr13:29116344-29116384 | Ch13 region 1 |
| chr13:29116393-29116433 | Ch13 region 1 |
| chr13:29116445-29116485 | Ch13 region 1 |
| chr13:29116521-29116561 | Ch13 region 1 |
| chr13:29116576-29116616 | Ch13 region 1 |
| chr13:29116633-29116673 | Ch13 region 1 |
| chr13:29116678-29116718 | Ch13 region 1 |
| chr13:29116725-29116765 | Ch13 region 1 |

| | |
|---|---|
| chr13:29116803-29116843 | Ch13 region 1 |
| chr13:29116892-29116932 | Ch13 region 1 |
| chr13:29116946-29116986 | Ch13 region 1 |
| chr13:29116991-29117031 | Ch13 region 1 |
| chr13:29117037-29117077 | Ch13 region 1 |
| chr13:29117106-29117146 | Ch13 region 1 |
| chr13:29117172-29117212 | Ch13 region 1 |
| chr13:29117229-29117269 | Ch13 region 1 |
| chr13:29117274-29117314 | Ch13 region 1 |
| chr13:29117385-29117425 | Ch13 region 1 |
| chr13:29117456-29117496 | Ch13 region 1 |
| chr13:29117501-29117541 | Ch13 region 1 |
| chr13:29117574-29117614 | Ch13 region 1 |
| chr13:29117642-29117682 | Ch13 region 1 |
| chr13:29117699-29117739 | Ch13 region 1 |
| chr13:29117768-29117808 | Ch13 region 1 |
| chr13:29117822-29117862 | Ch13 region 1 |
| chr13:29117900-29117940 | Ch13 region 1 |
| chr13:29118049-29118089 | Ch13 region 1 |
| chr13:29118132-29118172 | Ch13 region 1 |
| chr13:29118207-29118247 | Ch13 region 1 |
| chr13:29118271-29118311 | Ch13 region 1 |
| chr13:29118345-29118385 | Ch13 region 1 |
| chr13:29118421-29118461 | Ch13 region 1 |
| chr13:29118479-29118519 | Ch13 region 1 |
| chr13:29118538-29118578 | Ch13 region 1 |
| chr13:29118628-29118668 | Ch13 region 1 |
| chr13:29118771-29118811 | Ch13 region 1 |
| chr13:29118902-29118942 | Ch13 region 1 |
| chr13:29118947-29118987 | Ch13 region 1 |
| chr13:29118992-29119032 | Ch13 region 1 |
| chr13:29119043-29119083 | Ch13 region 1 |
| chr13:29119101-29119141 | Ch13 region 1 |
| chr13:29119165-29119205 | Ch13 region 1 |
| chr13:29119210-29119250 | Ch13 region 1 |
| chr13:29119275-29119315 | Ch13 region 1 |
| chr13:29119600-29119640 | Ch13 region 1 |
| chr13:29119668-29119708 | Ch13 region 1 |
| chr13:29119721-29119761 | Ch13 region 1 |
| chr13:29119780-29119820 | Ch13 region 1 |
| chr13:29119852-29119892 | Ch13 region 1 |
| chr13:29119903-29119943 | Ch13 region 1 |
| chr13:29119973-29120013 | Ch13 region 1 |
| chr13:29120050-29120090 | Ch13 region 1 |
| chr13:29120097-29120137 | Ch13 region 1 |
| chr13:29120144-29120184 | Ch13 region 1 |
| chr13:29120189-29120229 | Ch13 region 1 |
| chr13:29120247-29120287 | Ch13 region 1 |
| chr13:29120349-29120389 | Ch13 region 1 |
| chr13:29120394-29120434 | Ch13 region 1 |
| chr13:29120439-29120479 | Ch13 region 1 |
| chr13:29120488-29120528 | Ch13 region 1 |

| | |
|---|---|
| chr13:29120553-29120593 | Ch13 region 1 |
| chr13:29120623-29120663 | Ch13 region 1 |
| chr13:29120686-29120726 | Ch13 region 1 |
| chr13:29120805-29120845 | Ch13 region 1 |
| chr13:29120981-29121021 | Ch13 region 1 |
| chr13:29121177-29121217 | Ch13 region 1 |
| chr13:29121232-29121272 | Ch13 region 1 |
| chr13:29121286-29121326 | Ch13 region 1 |
| chr13:29121376-29121416 | Ch13 region 1 |
| chr13:29121448-29121488 | Ch13 region 1 |
| chr13:29121689-29121729 | Ch13 region 1 |
| chr13:29121734-29121774 | Ch13 region 1 |
| chr13:29121786-29121826 | Ch13 region 1 |
| chr13:29121842-29121882 | Ch13 region 1 |
| chr13:29121912-29121952 | Ch13 region 1 |
| chr13:29121970-29122010 | Ch13 region 1 |
| chr13:29122032-29122072 | Ch13 region 1 |
| chr13:29122078-29122118 | Ch13 region 1 |
| chr13:29122146-29122186 | Ch13 region 1 |
| chr13:29122203-29122243 | Ch13 region 1 |
| chr13:29122279-29122319 | Ch13 region 1 |
| chr13:29122345-29122385 | Ch13 region 1 |
| chr13:29122460-29122500 | Ch13 region 1 |
| chr13:29122542-29122582 | Ch13 region 1 |
| chr13:29122587-29122627 | Ch13 region 1 |
| chr13:29122639-29122679 | Ch13 region 1 |
| chr13:29122702-29122742 | Ch13 region 1 |
| chr13:29122755-29122795 | Ch13 region 1 |
| chr13:29122819-29122859 | Ch13 region 1 |
| chr13:29122886-29122926 | Ch13 region 1 |
| chr13:29122965-29123005 | Ch13 region 1 |
| chr13:29123012-29123052 | Ch13 region 1 |
| chr13:29123086-29123126 | Ch13 region 1 |
| chr13:29123150-29123190 | Ch13 region 1 |
| chr13:29123195-29123235 | Ch13 region 1 |
| chr13:29123252-29123292 | Ch13 region 1 |
| chr13:29123329-29123369 | Ch13 region 1 |
| chr13:29123374-29123414 | Ch13 region 1 |
| chr13:29123437-29123477 | Ch13 region 1 |
| chr13:29123482-29123522 | Ch13 region 1 |
| chr13:29123548-29123588 | Ch13 region 1 |
| chr13:29123622-29123662 | Ch13 region 1 |
| chr13:29123748-29123788 | Ch13 region 1 |
| chr13:29123819-29123859 | Ch13 region 1 |
| chr13:29123869-29123909 | Ch13 region 1 |
| chr13:29124132-29124172 | Ch13 region 1 |
| chr13:29124312-29124352 | Ch13 region 1 |
| chr13:29124378-29124418 | Ch13 region 1 |
| chr13:29124468-29124508 | Ch13 region 1 |
| chr13:29124554-29124594 | Ch13 region 1 |
| chr13:29124629-29124669 | Ch13 region 1 |
| chr13:29124694-29124734 | Ch13 region 1 |

| | |
|---|---|
| chr13:29124752-29124792 | Ch13 region 1 |
| chr13:29124809-29124849 | Ch13 region 1 |
| chr13:29124858-29124898 | Ch13 region 1 |
| chr13:29124950-29124990 | Ch13 region 1 |
| chr13:29125013-29125053 | Ch13 region 1 |
| chr13:29125064-29125104 | Ch13 region 1 |
| chr13:29125117-29125157 | Ch13 region 1 |
| chr13:29125179-29125219 | Ch13 region 1 |
| chr13:29125224-29125264 | Ch13 region 1 |
| chr13:29125271-29125311 | Ch13 region 1 |
| chr13:29125336-29125376 | Ch13 region 1 |
| chr13:29125383-29125423 | Ch13 region 1 |
| chr13:29125436-29125476 | Ch13 region 1 |
| chr13:29125516-29125556 | Ch13 region 1 |
| chr13:29125599-29125639 | Ch13 region 1 |
| chr13:29125665-29125705 | Ch13 region 1 |
| chr13:29125745-29125785 | Ch13 region 1 |
| chr13:29125801-29125841 | Ch13 region 1 |
| chr13:29125875-29125915 | Ch13 region 1 |
| chr13:29125963-29126003 | Ch13 region 1 |
| chr13:29126106-29126146 | Ch13 region 1 |
| chr13:29126294-29126334 | Ch13 region 1 |
| chr13:29126380-29126420 | Ch13 region 1 |
| chr13:29126460-29126500 | Ch13 region 1 |
| chr13:29126510-29126550 | Ch13 region 1 |
| chr13:29126591-29126631 | Ch13 region 1 |
| chr13:29126673-29126713 | Ch13 region 1 |
| chr13:29126736-29126776 | Ch13 region 1 |
| chr13:29126807-29126847 | Ch13 region 1 |
| chr13:29126867-29126907 | Ch13 region 1 |
| chr13:29126960-29127000 | Ch13 region 1 |
| chr13:29127017-29127057 | Ch13 region 1 |
| chr13:29127093-29127133 | Ch13 region 1 |
| chr13:29127166-29127206 | Ch13 region 1 |
| chr13:29127221-29127261 | Ch13 region 1 |
| chr13:29127278-29127318 | Ch13 region 1 |
| chr13:29127377-29127417 | Ch13 region 1 |
| chr13:29127594-29127634 | Ch13 region 1 |
| chr13:29127669-29127709 | Ch13 region 1 |
| chr13:29127737-29127777 | Ch13 region 1 |
| chr13:29127827-29127867 | Ch13 region 1 |
| chr13:29127880-29127920 | Ch13 region 1 |
| chr13:29127936-29127976 | Ch13 region 1 |
| chr13:29127981-29128021 | Ch13 region 1 |
| chr13:29128038-29128078 | Ch13 region 1 |
| chr13:29128087-29128127 | Ch13 region 1 |
| chr13:29128132-29128172 | Ch13 region 1 |
| chr13:29128214-29128254 | Ch13 region 1 |
| chr13:29128268-29128308 | Ch13 region 1 |
| chr13:29128330-29128370 | Ch13 region 1 |
| chr13:29128396-29128436 | Ch13 region 1 |
| chr13:29128445-29128485 | Ch13 region 1 |

| | |
|---|---|
| chr13:29128564-29128604 | Ch13 region 1 |
| chr13:29128716-29128756 | Ch13 region 1 |
| chr13:29128801-29128841 | Ch13 region 1 |
| chr13:29128900-29128940 | Ch13 region 1 |
| chr13:29129005-29129045 | Ch13 region 1 |
| chr13:29129077-29129117 | Ch13 region 1 |
| chr13:29129145-29129185 | Ch13 region 1 |
| chr13:29129197-29129237 | Ch13 region 1 |
| chr13:29129261-29129301 | Ch13 region 1 |
| chr13:29129310-29129350 | Ch13 region 1 |
| chr13:29129387-29129427 | Ch13 region 1 |
| chr13:29129432-29129472 | Ch13 region 1 |
| chr13:29129484-29129524 | Ch13 region 1 |
| chr13:29129549-29129589 | Ch13 region 1 |
| chr13:29129608-29129648 | Ch13 region 1 |
| chr13:29129705-29129745 | Ch13 region 1 |
| chr13:29129812-29129852 | Ch13 region 1 |
| chr13:29129898-29129938 | Ch13 region 1 |
| chr13:29130084-29130124 | Ch13 region 1 |
| chr13:29130144-29130184 | Ch13 region 1 |
| chr13:29130216-29130256 | Ch13 region 1 |
| chr13:29130292-29130332 | Ch13 region 1 |
| chr13:29130382-29130422 | Ch13 region 1 |
| chr13:29130460-29130500 | Ch13 region 1 |
| chr13:29130531-29130571 | Ch13 region 1 |
| chr13:29130617-29130657 | Ch13 region 1 |
| chr13:29130830-29130870 | Ch13 region 1 |
| chr13:29130886-29130926 | Ch13 region 1 |
| chr13:29130937-29130977 | Ch13 region 1 |
| chr13:29131012-29131052 | Ch13 region 1 |
| chr13:29131067-29131107 | Ch13 region 1 |
| chr13:29131112-29131152 | Ch13 region 1 |
| chr13:29131192-29131232 | Ch13 region 1 |
| chr13:29131310-29131350 | Ch13 region 1 |
| chr13:29131428-29131468 | Ch13 region 1 |
| chr13:29131474-29131514 | Ch13 region 1 |
| chr13:29131529-29131569 | Ch13 region 1 |
| chr13:29131589-29131629 | Ch13 region 1 |
| chr13:29131719-29131759 | Ch13 region 1 |
| chr13:29131768-29131808 | Ch13 region 1 |
| chr13:29131837-29131877 | Ch13 region 1 |
| chr13:29131946-29131986 | Ch13 region 1 |
| chr13:29132009-29132049 | Ch13 region 1 |
| chr13:29132054-29132094 | Ch13 region 1 |
| chr13:29132111-29132151 | Ch13 region 1 |
| chr13:29132189-29132229 | Ch13 region 1 |
| chr13:29132264-29132304 | Ch13 region 1 |
| chr13:29132309-29132349 | Ch13 region 1 |
| chr13:29132417-29132457 | Ch13 region 1 |
| chr13:29132478-29132518 | Ch13 region 1 |
| chr13:29132535-29132575 | Ch13 region 1 |
| chr13:29132606-29132646 | Ch13 region 1 |

| | |
|---|---|
| chr13:29132651-29132691 | Ch13 region 1 |
| chr13:29132717-29132757 | Ch13 region 1 |
| chr13:29132762-29132802 | Ch13 region 1 |
| chr13:29132825-29132865 | Ch13 region 1 |
| chr13:29132881-29132921 | Ch13 region 1 |
| chr13:29132939-29132979 | Ch13 region 1 |
| chr13:29133023-29133063 | Ch13 region 1 |
| chr13:29133084-29133124 | Ch13 region 1 |
| chr13:29133129-29133169 | Ch13 region 1 |
| chr13:29133328-29133368 | Ch13 region 1 |
| chr13:29133489-29133529 | Ch13 region 1 |
| chr13:29133773-29133813 | Ch13 region 1 |
| chr13:29133820-29133860 | Ch13 region 1 |
| chr13:29133909-29133949 | Ch13 region 1 |
| chr13:29133985-29134025 | Ch13 region 1 |
| chr13:29134064-29134104 | Ch13 region 1 |
| chr13:29134148-29134188 | Ch13 region 1 |
| chr13:29134263-29134303 | Ch13 region 1 |
| chr13:29134335-29134375 | Ch13 region 1 |
| chr13:29134416-29134456 | Ch13 region 1 |
| chr13:29134511-29134551 | Ch13 region 1 |
| chr13:29134573-29134613 | Ch13 region 1 |
| chr13:29134619-29134659 | Ch13 region 1 |
| chr13:29134712-29134752 | Ch13 region 1 |
| chr13:29134759-29134799 | Ch13 region 1 |
| chr13:29134852-29134892 | Ch13 region 1 |
| chr13:29134916-29134956 | Ch13 region 1 |
| chr13:29134980-29135020 | Ch13 region 1 |
| chr13:29135054-29135094 | Ch13 region 1 |
| chr13:29135124-29135164 | Ch13 region 1 |
| chr13:29135178-29135218 | Ch13 region 1 |
| chr13:29135228-29135268 | Ch13 region 1 |
| chr13:29135276-29135316 | Ch13 region 1 |
| chr13:29135329-29135369 | Ch13 region 1 |
| chr13:29135472-29135512 | Ch13 region 1 |
| chr13:29135551-29135591 | Ch13 region 1 |
| chr13:29135658-29135698 | Ch13 region 1 |
| chr13:29135703-29135743 | Ch13 region 1 |
| chr13:29135851-29135891 | Ch13 region 1 |
| chr13:29135911-29135951 | Ch13 region 1 |
| chr13:29135990-29136030 | Ch13 region 1 |
| chr13:29136039-29136079 | Ch13 region 1 |
| chr13:29136090-29136130 | Ch13 region 1 |
| chr13:29136166-29136206 | Ch13 region 1 |
| chr13:29136222-29136262 | Ch13 region 1 |
| chr13:29136267-29136307 | Ch13 region 1 |
| chr13:29136351-29136391 | Ch13 region 1 |
| chr13:29136411-29136451 | Ch13 region 1 |
| chr13:29136485-29136525 | Ch13 region 1 |
| chr13:29136531-29136571 | Ch13 region 1 |
| chr13:29136585-29136625 | Ch13 region 1 |
| chr13:29136640-29136680 | Ch13 region 1 |

| | |
|---|---|
| chr13:29136735-29136775 | Ch13 region 1 |
| chr13:29136811-29136851 | Ch13 region 1 |
| chr13:29136890-29136930 | Ch13 region 1 |
| chr13:29136936-29136976 | Ch13 region 1 |
| chr13:29136983-29137023 | Ch13 region 1 |
| chr13:29137062-29137102 | Ch13 region 1 |
| chr13:29137112-29137152 | Ch13 region 1 |
| chr13:29137163-29137203 | Ch13 region 1 |
| chr13:29137237-29137277 | Ch13 region 1 |
| chr13:29137288-29137328 | Ch13 region 1 |
| chr13:29137333-29137373 | Ch13 region 1 |
| chr13:29137383-29137423 | Ch13 region 1 |
| chr13:29137455-29137495 | Ch13 region 1 |
| chr13:29137521-29137561 | Ch13 region 1 |
| chr13:29137575-29137615 | Ch13 region 1 |
| chr13:29137682-29137722 | Ch13 region 1 |
| chr13:29137772-29137812 | Ch13 region 1 |
| chr13:29137821-29137861 | Ch13 region 1 |
| chr13:29137917-29137957 | Ch13 region 1 |
| chr13:29138016-29138056 | Ch13 region 1 |
| chr13:29138065-29138105 | Ch13 region 1 |
| chr13:29138146-29138186 | Ch13 region 1 |
| chr13:29138251-29138291 | Ch13 region 1 |
| chr13:29138309-29138349 | Ch13 region 1 |
| chr13:29138393-29138433 | Ch13 region 1 |
| chr13:29138459-29138499 | Ch13 region 1 |
| chr13:29138532-29138572 | Ch13 region 1 |
| chr13:29138593-29138633 | Ch13 region 1 |
| chr13:29138638-29138678 | Ch13 region 1 |
| chr13:29138697-29138737 | Ch13 region 1 |
| chr13:29138745-29138785 | Ch13 region 1 |
| chr13:29138795-29138835 | Ch13 region 1 |
| chr13:29138889-29138929 | Ch13 region 1 |
| chr13:29138939-29138979 | Ch13 region 1 |
| chr13:29138994-29139034 | Ch13 region 1 |
| chr13:29139040-29139080 | Ch13 region 1 |
| chr13:29139118-29139158 | Ch13 region 1 |
| chr13:29139166-29139206 | Ch13 region 1 |
| chr13:29139237-29139277 | Ch13 region 1 |
| chr13:29139288-29139328 | Ch13 region 1 |
| chr13:29139347-29139387 | Ch13 region 1 |
| chr13:29139393-29139433 | Ch13 region 1 |
| chr13:29139455-29139495 | Ch13 region 1 |
| chr13:29139502-29139542 | Ch13 region 1 |
| chr13:29139568-29139608 | Ch13 region 1 |
| chr13:29139747-29139787 | Ch13 region 1 |
| chr13:29139891-29139931 | Ch13 region 1 |
| chr13:29140005-29140045 | Ch13 region 1 |
| chr13:29140151-29140191 | Ch13 region 1 |
| chr13:29140219-29140259 | Ch13 region 1 |
| chr13:29140321-29140361 | Ch13 region 1 |
| chr13:29140371-29140411 | Ch13 region 1 |

| | |
|---|---|
| chr13:29140416-29140456 | Ch13 region 1 |
| chr13:29140462-29140502 | Ch13 region 1 |
| chr13:29140527-29140567 | Ch13 region 1 |
| chr13:29140578-29140618 | Ch13 region 1 |
| chr13:29140647-29140687 | Ch13 region 1 |
| chr13:29140707-29140747 | Ch13 region 1 |
| chr13:29140807-29140847 | Ch13 region 1 |
| chr13:29140870-29140910 | Ch13 region 1 |
| chr13:29140950-29140990 | Ch13 region 1 |
| chr13:29140995-29141035 | Ch13 region 1 |
| chr13:29141065-29141105 | Ch13 region 1 |
| chr13:29141150-29141190 | Ch13 region 1 |
| chr13:29141195-29141235 | Ch13 region 1 |
| chr13:29141250-29141290 | Ch13 region 1 |
| chr13:29141305-29141345 | Ch13 region 1 |
| chr13:29141350-29141390 | Ch13 region 1 |
| chr13:29141419-29141459 | Ch13 region 1 |
| chr13:29141470-29141510 | Ch13 region 1 |
| chr13:29141533-29141573 | Ch13 region 1 |
| chr13:29141624-29141664 | Ch13 region 1 |
| chr13:29141672-29141712 | Ch13 region 1 |
| chr13:29141719-29141759 | Ch13 region 1 |
| chr13:29141771-29141811 | Ch13 region 1 |
| chr13:29141830-29141870 | Ch13 region 1 |
| chr13:29141921-29141961 | Ch13 region 1 |
| chr13:29142027-29142067 | Ch13 region 1 |
| chr13:29142081-29142121 | Ch13 region 1 |
| chr13:29142136-29142176 | Ch13 region 1 |
| chr13:29142218-29142258 | Ch13 region 1 |
| chr13:29142278-29142318 | Ch13 region 1 |
| chr13:29142334-29142374 | Ch13 region 1 |
| chr13:29142387-29142427 | Ch13 region 1 |
| chr13:29142439-29142479 | Ch13 region 1 |
| chr13:29142488-29142528 | Ch13 region 1 |
| chr13:29142580-29142620 | Ch13 region 1 |
| chr13:29142625-29142665 | Ch13 region 1 |
| chr13:29142689-29142729 | Ch13 region 1 |
| chr13:29142744-29142784 | Ch13 region 1 |
| chr13:29142812-29142852 | Ch13 region 1 |
| chr13:29142857-29142897 | Ch13 region 1 |
| chr13:29142936-29142976 | Ch13 region 1 |
| chr13:29142982-29143022 | Ch13 region 1 |
| chr13:29143027-29143067 | Ch13 region 1 |
| chr13:29143090-29143130 | Ch13 region 1 |
| chr13:29143181-29143221 | Ch13 region 1 |
| chr13:29143227-29143267 | Ch13 region 1 |
| chr13:29143356-29143396 | Ch13 region 1 |
| chr13:29143505-29143545 | Ch13 region 1 |
| chr13:29143568-29143608 | Ch13 region 1 |
| chr13:29143619-29143659 | Ch13 region 1 |
| chr13:29143702-29143742 | Ch13 region 1 |
| chr13:29143753-29143793 | Ch13 region 1 |

| | |
|---|---|
| chr13:29143831-29143871 | Ch13 region 1 |
| chr13:29143889-29143929 | Ch13 region 1 |
| chr13:29143958-29143998 | Ch13 region 1 |
| chr13:29144024-29144064 | Ch13 region 1 |
| chr13:29144099-29144139 | Ch13 region 1 |
| chr13:29144144-29144184 | Ch13 region 1 |
| chr13:29144193-29144233 | Ch13 region 1 |
| chr13:29144261-29144301 | Ch13 region 1 |
| chr13:29144332-29144372 | Ch13 region 1 |
| chr13:29144469-29144509 | Ch13 region 1 |
| chr13:29144656-29144696 | Ch13 region 1 |
| chr13:29144806-29144846 | Ch13 region 1 |
| chr13:29144964-29145004 | Ch13 region 1 |
| chr13:29145072-29145112 | Ch13 region 1 |
| chr13:29145129-29145169 | Ch13 region 1 |
| chr13:29145239-29145279 | Ch13 region 1 |
| chr13:29145284-29145324 | Ch13 region 1 |
| chr13:29145331-29145371 | Ch13 region 1 |
| chr13:29145432-29145472 | Ch13 region 1 |
| chr13:29145492-29145532 | Ch13 region 1 |
| chr13:29145577-29145617 | Ch13 region 1 |
| chr13:29145760-29145800 | Ch13 region 1 |
| chr13:29145805-29145845 | Ch13 region 1 |
| chr13:29145870-29145910 | Ch13 region 1 |
| chr13:29145936-29145976 | Ch13 region 1 |
| chr13:29145989-29146029 | Ch13 region 1 |
| chr13:29146044-29146084 | Ch13 region 1 |
| chr13:29146091-29146131 | Ch13 region 1 |
| chr13:29146159-29146199 | Ch13 region 1 |
| chr13:29146204-29146244 | Ch13 region 1 |
| chr13:29146265-29146305 | Ch13 region 1 |
| chr13:29146333-29146373 | Ch13 region 1 |
| chr13:29146384-29146424 | Ch13 region 1 |
| chr13:29146574-29146614 | Ch13 region 1 |
| chr13:29146654-29146694 | Ch13 region 1 |
| chr13:29146700-29146740 | Ch13 region 1 |
| chr13:29146795-29146835 | Ch13 region 1 |
| chr13:29146878-29146918 | Ch13 region 1 |
| chr13:29146930-29146970 | Ch13 region 1 |
| chr13:29147003-29147043 | Ch13 region 1 |
| chr13:29147073-29147113 | Ch13 region 1 |
| chr13:29147138-29147178 | Ch13 region 1 |
| chr13:29147249-29147289 | Ch13 region 1 |
| chr13:29147326-29147366 | Ch13 region 1 |
| chr13:29147405-29147445 | Ch13 region 1 |
| chr13:29147491-29147531 | Ch13 region 1 |
| chr13:29147568-29147608 | Ch13 region 1 |
| chr13:29147702-29147742 | Ch13 region 1 |
| chr13:29147755-29147795 | Ch13 region 1 |
| chr13:29147814-29147854 | Ch13 region 1 |
| chr13:29147865-29147905 | Ch13 region 1 |
| chr13:29147911-29147951 | Ch13 region 1 |

| | |
|---|---|
| chr13:29147970-29148010 | Ch13 region 1 |
| chr13:29148063-29148103 | Ch13 region 1 |
| chr13:29148109-29148149 | Ch13 region 1 |
| chr13:29148174-29148214 | Ch13 region 1 |
| chr13:29148219-29148259 | Ch13 region 1 |
| chr13:29148285-29148325 | Ch13 region 1 |
| chr13:29148330-29148370 | Ch13 region 1 |
| chr13:29148402-29148442 | Ch13 region 1 |
| chr13:29148472-29148512 | Ch13 region 1 |
| chr13:29148517-29148557 | Ch13 region 1 |
| chr13:29148563-29148603 | Ch13 region 1 |
| chr13:29148637-29148677 | Ch13 region 1 |
| chr13:29148694-29148734 | Ch13 region 1 |
| chr13:29148750-29148790 | Ch13 region 1 |
| chr13:29148852-29148892 | Ch13 region 1 |
| chr13:29148921-29148961 | Ch13 region 1 |
| chr13:29148979-29149019 | Ch13 region 1 |
| chr13:29149047-29149087 | Ch13 region 1 |
| chr13:29149101-29149141 | Ch13 region 1 |
| chr13:29149150-29149190 | Ch13 region 1 |
| chr13:29149212-29149252 | Ch13 region 1 |
| chr13:29149266-29149306 | Ch13 region 1 |
| chr13:29149321-29149361 | Ch13 region 1 |
| chr13:29149400-29149440 | Ch13 region 1 |
| chr13:29149452-29149492 | Ch13 region 1 |
| chr13:29149519-29149559 | Ch13 region 1 |
| chr13:29149574-29149614 | Ch13 region 1 |
| chr13:29149657-29149697 | Ch13 region 1 |
| chr13:29149731-29149771 | Ch13 region 1 |
| chr13:29149821-29149861 | Ch13 region 1 |
| chr13:29149974-29150014 | Ch13 region 1 |
| chr13:29150172-29150212 | Ch13 region 1 |
| chr13:29150240-29150280 | Ch13 region 1 |
| chr13:29150294-29150334 | Ch13 region 1 |
| chr13:29150339-29150379 | Ch13 region 1 |
| chr13:29150418-29150458 | Ch13 region 1 |
| chr13:29150497-29150537 | Ch13 region 1 |
| chr13:29150549-29150589 | Ch13 region 1 |
| chr13:29150620-29150660 | Ch13 region 1 |
| chr13:29150684-29150724 | Ch13 region 1 |
| chr13:29150746-29150786 | Ch13 region 1 |
| chr13:29150813-29150853 | Ch13 region 1 |
| chr13:29151017-29151057 | Ch13 region 1 |
| chr13:29151062-29151102 | Ch13 region 1 |
| chr13:29151130-29151170 | Ch13 region 1 |
| chr13:29151180-29151220 | Ch13 region 1 |
| chr13:29151255-29151295 | Ch13 region 1 |
| chr13:29151339-29151379 | Ch13 region 1 |
| chr13:29151392-29151432 | Ch13 region 1 |
| chr13:29151450-29151490 | Ch13 region 1 |
| chr13:29151514-29151554 | Ch13 region 1 |
| chr13:29151579-29151619 | Ch13 region 1 |

| | |
|---|---|
| chr13:29151660-29151700 | Ch13 region 1 |
| chr13:29151735-29151775 | Ch13 region 1 |
| chr13:29151781-29151821 | Ch13 region 1 |
| chr13:29151863-29151903 | Ch13 region 1 |
| chr13:29151933-29151973 | Ch13 region 1 |
| chr13:29152038-29152078 | Ch13 region 1 |
| chr13:29152106-29152146 | Ch13 region 1 |
| chr13:29152211-29152251 | Ch13 region 1 |
| chr13:29152294-29152334 | Ch13 region 1 |
| chr13:29152368-29152408 | Ch13 region 1 |
| chr13:29152415-29152455 | Ch13 region 1 |
| chr13:29152483-29152523 | Ch13 region 1 |
| chr13:29152545-29152585 | Ch13 region 1 |
| chr13:29152590-29152630 | Ch13 region 1 |
| chr13:29152677-29152717 | Ch13 region 1 |
| chr13:29152736-29152776 | Ch13 region 1 |
| chr13:29152790-29152830 | Ch13 region 1 |
| chr13:29152835-29152875 | Ch13 region 1 |
| chr13:29152880-29152920 | Ch13 region 1 |
| chr13:29152938-29152978 | Ch13 region 1 |
| chr13:29152999-29153039 | Ch13 region 1 |
| chr13:29153112-29153152 | Ch13 region 1 |
| chr13:29153204-29153244 | Ch13 region 1 |
| chr13:29153257-29153297 | Ch13 region 1 |
| chr13:29153302-29153342 | Ch13 region 1 |
| chr13:29153383-29153423 | Ch13 region 1 |
| chr13:29153445-29153485 | Ch13 region 1 |
| chr13:29153507-29153547 | Ch13 region 1 |
| chr13:29153556-29153596 | Ch13 region 1 |
| chr13:29153635-29153675 | Ch13 region 1 |
| chr13:29153697-29153737 | Ch13 region 1 |
| chr13:29153795-29153835 | Ch13 region 1 |
| chr13:29153840-29153880 | Ch13 region 1 |
| chr13:29153908-29153948 | Ch13 region 1 |
| chr13:29153967-29154007 | Ch13 region 1 |
| chr13:29154012-29154052 | Ch13 region 1 |
| chr13:29154067-29154107 | Ch13 region 1 |
| chr13:29154136-29154176 | Ch13 region 1 |
| chr13:29154222-29154262 | Ch13 region 1 |
| chr13:29154268-29154308 | Ch13 region 1 |
| chr13:29154315-29154355 | Ch13 region 1 |
| chr13:29154360-29154400 | Ch13 region 1 |
| chr13:29154405-29154445 | Ch13 region 1 |
| chr13:29154457-29154497 | Ch13 region 1 |
| chr13:29154541-29154581 | Ch13 region 1 |
| chr13:29154586-29154626 | Ch13 region 1 |
| chr13:29154651-29154691 | Ch13 region 1 |
| chr13:29154744-29154784 | Ch13 region 1 |
| chr13:29155010-29155050 | Ch13 region 1 |
| chr13:29155084-29155124 | Ch13 region 1 |
| chr13:29155214-29155254 | Ch13 region 1 |
| chr13:29155270-29155310 | Ch13 region 1 |

| | |
|---|---|
| chr13:29155476-29155516 | Ch13 region 1 |
| chr13:29155535-29155575 | Ch13 region 1 |
| chr13:29155627-29155667 | Ch13 region 1 |
| chr13:29155848-29155888 | Ch13 region 1 |
| chr13:29155904-29155944 | Ch13 region 1 |
| chr13:29156024-29156064 | Ch13 region 1 |
| chr13:29156094-29156134 | Ch13 region 1 |
| chr13:29156228-29156268 | Ch13 region 1 |
| chr13:29156295-29156335 | Ch13 region 1 |
| chr13:29156354-29156394 | Ch13 region 1 |
| chr13:29156430-29156470 | Ch13 region 1 |
| chr13:29156479-29156519 | Ch13 region 1 |
| chr13:29156525-29156565 | Ch13 region 1 |
| chr13:29156588-29156628 | Ch13 region 1 |
| chr13:29156634-29156674 | Ch13 region 1 |
| chr13:29156729-29156769 | Ch13 region 1 |
| chr13:29156796-29156836 | Ch13 region 1 |
| chr13:29156868-29156908 | Ch13 region 1 |
| chr13:29156932-29156972 | Ch13 region 1 |
| chr13:29157010-29157050 | Ch13 region 1 |
| chr13:29157060-29157100 | Ch13 region 1 |
| chr13:29157113-29157153 | Ch13 region 1 |
| chr13:29157176-29157216 | Ch13 region 1 |
| chr13:29157221-29157261 | Ch13 region 1 |
| chr13:29157300-29157340 | Ch13 region 1 |
| chr13:29157356-29157396 | Ch13 region 1 |
| chr13:29157455-29157495 | Ch13 region 1 |
| chr13:29157602-29157642 | Ch13 region 1 |
| chr13:29157659-29157699 | Ch13 region 1 |
| chr13:29157726-29157766 | Ch13 region 1 |
| chr13:29157785-29157825 | Ch13 region 1 |
| chr13:29157861-29157901 | Ch13 region 1 |
| chr13:29157913-29157953 | Ch13 region 1 |
| chr13:29157999-29158039 | Ch13 region 1 |
| chr13:29158053-29158093 | Ch13 region 1 |
| chr13:29158100-29158140 | Ch13 region 1 |
| chr13:29158197-29158237 | Ch13 region 1 |
| chr13:29158260-29158300 | Ch13 region 1 |
| chr13:29158307-29158347 | Ch13 region 1 |
| chr13:29158372-29158412 | Ch13 region 1 |
| chr13:29158449-29158489 | Ch13 region 1 |
| chr13:29158494-29158534 | Ch13 region 1 |
| chr13:29158704-29158744 | Ch13 region 1 |
| chr13:29158850-29158890 | Ch13 region 1 |
| chr13:29158919-29158959 | Ch13 region 1 |
| chr13:29158976-29159016 | Ch13 region 1 |
| chr13:29159022-29159062 | Ch13 region 1 |
| chr13:29159103-29159143 | Ch13 region 1 |
| chr13:29159199-29159239 | Ch13 region 1 |
| chr13:29159264-29159304 | Ch13 region 1 |
| chr13:29159309-29159349 | Ch13 region 1 |
| chr13:29159377-29159417 | Ch13 region 1 |

| | |
|---|---|
| chr13:29159454-29159494 | Ch13 region 1 |
| chr13:29159511-29159551 | Ch13 region 1 |
| chr13:29159556-29159596 | Ch13 region 1 |
| chr13:29159636-29159676 | Ch13 region 1 |
| chr13:29159693-29159733 | Ch13 region 1 |
| chr13:29159789-29159829 | Ch13 region 1 |
| chr13:29159862-29159902 | Ch13 region 1 |
| chr13:29159912-29159952 | Ch13 region 1 |
| chr13:29159959-29159999 | Ch13 region 1 |
| chr13:29160037-29160077 | Ch13 region 1 |
| chr13:29160113-29160153 | Ch13 region 1 |
| chr13:29160211-29160251 | Ch13 region 1 |
| chr13:29160256-29160296 | Ch13 region 1 |
| chr13:29160325-29160365 | Ch13 region 1 |
| chr13:29160404-29160444 | Ch13 region 1 |
| chr13:29160505-29160545 | Ch13 region 1 |
| chr13:29160552-29160592 | Ch13 region 1 |
| chr13:29160630-29160670 | Ch13 region 1 |
| chr13:29160739-29160779 | Ch13 region 1 |
| chr13:29160817-29160857 | Ch13 region 1 |
| chr13:29160920-29160960 | Ch13 region 1 |
| chr13:29160995-29161035 | Ch13 region 1 |
| chr13:29161085-29161125 | Ch13 region 1 |
| chr13:29161149-29161189 | Ch13 region 1 |
| chr13:29161219-29161259 | Ch13 region 1 |
| chr13:29161295-29161335 | Ch13 region 1 |
| chr13:29161402-29161442 | Ch13 region 1 |
| chr13:29161469-29161509 | Ch13 region 1 |
| chr13:29161581-29161621 | Ch13 region 1 |
| chr13:29161747-29161787 | Ch13 region 1 |
| chr13:29161916-29161956 | Ch13 region 1 |
| chr13:29162076-29162116 | Ch13 region 1 |
| chr13:29162135-29162175 | Ch13 region 1 |
| chr13:29162218-29162258 | Ch13 region 1 |
| chr13:29162269-29162309 | Ch13 region 1 |
| chr13:29162354-29162394 | Ch13 region 1 |
| chr13:29162513-29162553 | Ch13 region 1 |
| chr13:29162626-29162666 | Ch13 region 1 |
| chr13:29162704-29162744 | Ch13 region 1 |
| chr13:29162754-29162794 | Ch13 region 1 |
| chr13:29162802-29162842 | Ch13 region 1 |
| chr13:29162923-29162963 | Ch13 region 1 |
| chr13:29163118-29163158 | Ch13 region 1 |
| chr13:29163310-29163350 | Ch13 region 1 |
| chr13:29163433-29163473 | Ch13 region 1 |
| chr13:29163548-29163588 | Ch13 region 1 |
| chr13:29163739-29163779 | Ch13 region 1 |
| chr13:29163872-29163912 | Ch13 region 1 |
| chr13:29163944-29163984 | Ch13 region 1 |
| chr13:29164006-29164046 | Ch13 region 1 |
| chr13:29164071-29164111 | Ch13 region 1 |
| chr13:29164134-29164174 | Ch13 region 1 |

| | |
|---|---|
| chr13:29164204-29164244 | Ch13 region 1 |
| chr13:29164278-29164318 | Ch13 region 1 |
| chr13:29164324-29164364 | Ch13 region 1 |
| chr13:29164472-29164512 | Ch13 region 1 |
| chr13:29164545-29164585 | Ch13 region 1 |
| chr13:29164608-29164648 | Ch13 region 1 |
| chr13:29164685-29164725 | Ch13 region 1 |
| chr13:29164744-29164784 | Ch13 region 1 |
| chr13:29164805-29164845 | Ch13 region 1 |
| chr13:29164862-29164902 | Ch13 region 1 |
| chr13:29164951-29164991 | Ch13 region 1 |
| chr13:29165046-29165086 | Ch13 region 1 |
| chr13:29165131-29165171 | Ch13 region 1 |
| chr13:29165179-29165219 | Ch13 region 1 |
| chr13:29165266-29165306 | Ch13 region 1 |
| chr13:29165315-29165355 | Ch13 region 1 |
| chr13:29165405-29165445 | Ch13 region 1 |
| chr13:29165455-29165495 | Ch13 region 1 |
| chr13:29165517-29165557 | Ch13 region 1 |
| chr13:29165747-29165787 | Ch13 region 1 |
| chr13:29165799-29165839 | Ch13 region 1 |
| chr13:29165986-29166026 | Ch13 region 1 |
| chr13:29166237-29166277 | Ch13 region 1 |
| chr13:29166298-29166338 | Ch13 region 1 |
| chr13:29166392-29166432 | Ch13 region 1 |
| chr13:29166477-29166517 | Ch13 region 1 |
| chr13:29166583-29166623 | Ch13 region 1 |
| chr13:29166665-29166705 | Ch13 region 1 |
| chr13:29166725-29166765 | Ch13 region 1 |
| chr13:29166770-29166810 | Ch13 region 1 |
| chr13:29166838-29166878 | Ch13 region 1 |
| chr13:29166896-29166936 | Ch13 region 1 |
| chr13:29166964-29167004 | Ch13 region 1 |
| chr13:29167014-29167054 | Ch13 region 1 |
| chr13:29167152-29167192 | Ch13 region 1 |
| chr13:29167379-29167419 | Ch13 region 1 |
| chr13:29167525-29167565 | Ch13 region 1 |
| chr13:29167580-29167620 | Ch13 region 1 |
| chr13:29167683-29167723 | Ch13 region 1 |
| chr13:29167728-29167768 | Ch13 region 1 |
| chr13:29167781-29167821 | Ch13 region 1 |
| chr13:29167879-29167919 | Ch13 region 1 |
| chr13:29167953-29167993 | Ch13 region 1 |
| chr13:29168015-29168055 | Ch13 region 1 |
| chr13:29168116-29168156 | Ch13 region 1 |
| chr13:29168174-29168214 | Ch13 region 1 |
| chr13:29168341-29168381 | Ch13 region 1 |
| chr13:29168422-29168462 | Ch13 region 1 |
| chr13:29168475-29168515 | Ch13 region 1 |
| chr13:29168589-29168629 | Ch13 region 1 |
| chr13:29168680-29168720 | Ch13 region 1 |
| chr13:29168749-29168789 | Ch13 region 1 |

| | |
|---|---|
| chr13:29168807-29168847 | Ch13 region 1 |
| chr13:29168950-29168990 | Ch13 region 1 |
| chr13:29169003-29169043 | Ch13 region 1 |
| chr13:29169050-29169090 | Ch13 region 1 |
| chr13:29169217-29169257 | Ch13 region 1 |
| chr13:29169262-29169302 | Ch13 region 1 |
| chr13:29169318-29169358 | Ch13 region 1 |
| chr13:29169442-29169482 | Ch13 region 1 |
| chr13:29169491-29169531 | Ch13 region 1 |
| chr13:29169717-29169757 | Ch13 region 1 |
| chr13:29169801-29169841 | Ch13 region 1 |
| chr13:29169891-29169931 | Ch13 region 1 |
| chr13:29169936-29169976 | Ch13 region 1 |
| chr13:29170015-29170055 | Ch13 region 1 |
| chr13:29170062-29170102 | Ch13 region 1 |
| chr13:29170187-29170227 | Ch13 region 1 |
| chr13:29170297-29170337 | Ch13 region 1 |
| chr13:29170357-29170397 | Ch13 region 1 |
| chr13:29170423-29170463 | Ch13 region 1 |
| chr13:29170489-29170529 | Ch13 region 1 |
| chr13:29170562-29170602 | Ch13 region 1 |
| chr13:29170639-29170679 | Ch13 region 1 |
| chr13:29170689-29170729 | Ch13 region 1 |
| chr13:29170752-29170792 | Ch13 region 1 |
| chr13:29170809-29170849 | Ch13 region 1 |
| chr13:29170854-29170894 | Ch13 region 1 |
| chr13:29170980-29171020 | Ch13 region 1 |
| chr13:29171051-29171091 | Ch13 region 1 |
| chr13:29171110-29171150 | Ch13 region 1 |
| chr13:29171159-29171199 | Ch13 region 1 |
| chr13:29171211-29171251 | Ch13 region 1 |
| chr13:29171446-29171486 | Ch13 region 1 |
| chr13:29171491-29171531 | Ch13 region 1 |
| chr13:29171651-29171691 | Ch13 region 1 |
| chr13:29171717-29171757 | Ch13 region 1 |
| chr13:29171762-29171802 | Ch13 region 1 |
| chr13:29171829-29171869 | Ch13 region 1 |
| chr13:29171884-29171924 | Ch13 region 1 |
| chr13:29171932-29171972 | Ch13 region 1 |
| chr13:29171977-29172017 | Ch13 region 1 |
| chr13:29172038-29172078 | Ch13 region 1 |
| chr13:29172111-29172151 | Ch13 region 1 |
| chr13:29172184-29172224 | Ch13 region 1 |
| chr13:29172311-29172351 | Ch13 region 1 |
| chr13:29172399-29172439 | Ch13 region 1 |
| chr13:29172448-29172488 | Ch13 region 1 |
| chr13:29172542-29172582 | Ch13 region 1 |
| chr13:29172625-29172665 | Ch13 region 1 |
| chr13:29172689-29172729 | Ch13 region 1 |
| chr13:29172734-29172774 | Ch13 region 1 |
| chr13:29172804-29172844 | Ch13 region 1 |
| chr13:29172898-29172938 | Ch13 region 1 |

| | |
|---|---|
| chr13:29172945-29172985 | Ch13 region 1 |
| chr13:29173036-29173076 | Ch13 region 1 |
| chr13:29173101-29173141 | Ch13 region 1 |
| chr13:29173172-29173212 | Ch13 region 1 |
| chr13:29173235-29173275 | Ch13 region 1 |
| chr13:29173280-29173320 | Ch13 region 1 |
| chr13:29173338-29173378 | Ch13 region 1 |
| chr13:29173410-29173450 | Ch13 region 1 |
| chr13:29173504-29173544 | Ch13 region 1 |
| chr13:29173583-29173623 | Ch13 region 1 |
| chr13:29173647-29173687 | Ch13 region 1 |
| chr13:29173701-29173741 | Ch13 region 1 |
| chr13:29173764-29173804 | Ch13 region 1 |
| chr13:29173830-29173870 | Ch13 region 1 |
| chr13:29173878-29173918 | Ch13 region 1 |
| chr13:29173932-29173972 | Ch13 region 1 |
| chr13:29174040-29174080 | Ch13 region 1 |
| chr13:29174116-29174156 | Ch13 region 1 |
| chr13:29174193-29174233 | Ch13 region 1 |
| chr13:29174265-29174305 | Ch13 region 1 |
| chr13:29174363-29174403 | Ch13 region 1 |
| chr13:29174434-29174474 | Ch13 region 1 |
| chr13:29174496-29174536 | Ch13 region 1 |
| chr13:29174568-29174608 | Ch13 region 1 |
| chr13:29174613-29174653 | Ch13 region 1 |
| chr13:29174674-29174714 | Ch13 region 1 |
| chr13:29174721-29174761 | Ch13 region 1 |
| chr13:29174772-29174812 | Ch13 region 1 |
| chr13:29174832-29174872 | Ch13 region 1 |
| chr13:29174923-29174963 | Ch13 region 1 |
| chr13:29174970-29175010 | Ch13 region 1 |
| chr13:29175052-29175092 | Ch13 region 1 |
| chr13:29175097-29175137 | Ch13 region 1 |
| chr13:29175179-29175219 | Ch13 region 1 |
| chr13:29175224-29175264 | Ch13 region 1 |
| chr13:29175297-29175337 | Ch13 region 1 |
| chr13:29175353-29175393 | Ch13 region 1 |
| chr13:29175446-29175486 | Ch13 region 1 |
| chr13:29175523-29175563 | Ch13 region 1 |
| chr13:29175584-29175624 | Ch13 region 1 |
| chr13:29175677-29175717 | Ch13 region 1 |
| chr13:29175732-29175772 | Ch13 region 1 |
| chr13:29175798-29175838 | Ch13 region 1 |
| chr13:29175883-29175923 | Ch13 region 1 |
| chr13:29175949-29175989 | Ch13 region 1 |
| chr13:29176019-29176059 | Ch13 region 1 |
| chr13:29176065-29176105 | Ch13 region 1 |
| chr13:29176118-29176158 | Ch13 region 1 |
| chr13:29176213-29176253 | Ch13 region 1 |
| chr13:29176299-29176339 | Ch13 region 1 |
| chr13:29176391-29176431 | Ch13 region 1 |
| chr13:29176467-29176507 | Ch13 region 1 |

| | |
|---|---|
| chr13:29176553-29176593 | Ch13 region 1 |
| chr13:29176629-29176669 | Ch13 region 1 |
| chr13:29176712-29176752 | Ch13 region 1 |
| chr13:29176764-29176804 | Ch13 region 1 |
| chr13:29176829-29176869 | Ch13 region 1 |
| chr13:29176915-29176955 | Ch13 region 1 |
| chr13:29176963-29177003 | Ch13 region 1 |
| chr13:29177021-29177061 | Ch13 region 1 |
| chr13:29177073-29177113 | Ch13 region 1 |
| chr13:29177143-29177183 | Ch13 region 1 |
| chr13:29177223-29177263 | Ch13 region 1 |
| chr13:29177313-29177353 | Ch13 region 1 |
| chr13:29177381-29177421 | Ch13 region 1 |
| chr13:29177434-29177474 | Ch13 region 1 |
| chr13:29177508-29177548 | Ch13 region 1 |
| chr13:29177569-29177609 | Ch13 region 1 |
| chr13:29177624-29177664 | Ch13 region 1 |
| chr13:29177682-29177722 | Ch13 region 1 |
| chr13:29177752-29177792 | Ch13 region 1 |
| chr13:29177815-29177855 | Ch13 region 1 |
| chr13:29177878-29177918 | Ch13 region 1 |
| chr13:29177925-29177965 | Ch13 region 1 |
| chr13:29178059-29178099 | Ch13 region 1 |
| chr13:29178113-29178153 | Ch13 region 1 |
| chr13:29178196-29178236 | Ch13 region 1 |
| chr13:29178346-29178386 | Ch13 region 1 |
| chr13:29178541-29178581 | Ch13 region 1 |
| chr13:29178643-29178683 | Ch13 region 1 |
| chr13:29178688-29178728 | Ch13 region 1 |
| chr13:29178760-29178800 | Ch13 region 1 |
| chr13:29178862-29178902 | Ch13 region 1 |
| chr13:29178959-29178999 | Ch13 region 1 |
| chr13:29179029-29179069 | Ch13 region 1 |
| chr13:29179083-29179123 | Ch13 region 1 |
| chr13:29179273-29179313 | Ch13 region 1 |
| chr13:29179501-29179541 | Ch13 region 1 |
| chr13:29179831-29179871 | Ch13 region 1 |
| chr13:29179976-29180016 | Ch13 region 1 |
| chr13:29180025-29180065 | Ch13 region 1 |
| chr13:29180178-29180218 | Ch13 region 1 |
| chr13:29180261-29180301 | Ch13 region 1 |
| chr13:29180318-29180358 | Ch13 region 1 |
| chr13:29180378-29180418 | Ch13 region 1 |
| chr13:29180423-29180463 | Ch13 region 1 |
| chr13:29180493-29180533 | Ch13 region 1 |
| chr13:29180619-29180659 | Ch13 region 1 |
| chr13:29180704-29180744 | Ch13 region 1 |
| chr13:29180761-29180801 | Ch13 region 1 |
| chr13:29180809-29180849 | Ch13 region 1 |
| chr13:29180876-29180916 | Ch13 region 1 |
| chr13:29180939-29180979 | Ch13 region 1 |
| chr13:29180998-29181038 | Ch13 region 1 |

| | |
|---|---|
| chr13:29181077-29181117 | Ch13 region 1 |
| chr13:29181129-29181169 | Ch13 region 1 |
| chr13:29181215-29181255 | Ch13 region 1 |
| chr13:29181293-29181333 | Ch13 region 1 |
| chr13:29181367-29181407 | Ch13 region 1 |
| chr13:29181429-29181469 | Ch13 region 1 |
| chr13:29181516-29181556 | Ch13 region 1 |
| chr13:29181585-29181625 | Ch13 region 1 |
| chr13:29181647-29181687 | Ch13 region 1 |
| chr13:29181712-29181752 | Ch13 region 1 |
| chr13:29181804-29181844 | Ch13 region 1 |
| chr13:29181849-29181889 | Ch13 region 1 |
| chr13:29181907-29181947 | Ch13 region 1 |
| chr13:29181965-29182005 | Ch13 region 1 |
| chr13:29182055-29182095 | Ch13 region 1 |
| chr13:29182152-29182192 | Ch13 region 1 |
| chr13:29182211-29182251 | Ch13 region 1 |
| chr13:29182266-29182306 | Ch13 region 1 |
| chr13:29182350-29182390 | Ch13 region 1 |
| chr13:29182404-29182444 | Ch13 region 1 |
| chr13:29182511-29182551 | Ch13 region 1 |
| chr13:29182566-29182606 | Ch13 region 1 |
| chr13:29182648-29182688 | Ch13 region 1 |
| chr13:29182717-29182757 | Ch13 region 1 |
| chr13:29182811-29182851 | Ch13 region 1 |
| chr13:29182866-29182906 | Ch13 region 1 |
| chr13:29182920-29182960 | Ch13 region 1 |
| chr13:29182996-29183036 | Ch13 region 1 |
| chr13:29183041-29183081 | Ch13 region 1 |
| chr13:29183148-29183188 | Ch13 region 1 |
| chr13:29183193-29183233 | Ch13 region 1 |
| chr13:29183313-29183353 | Ch13 region 1 |
| chr13:29183379-29183419 | Ch13 region 1 |
| chr13:29183427-29183467 | Ch13 region 1 |
| chr13:29183518-29183558 | Ch13 region 1 |
| chr13:29183564-29183604 | Ch13 region 1 |
| chr13:29183646-29183686 | Ch13 region 1 |
| chr13:29183691-29183731 | Ch13 region 1 |
| chr13:29183743-29183783 | Ch13 region 1 |
| chr13:29183826-29183866 | Ch13 region 1 |
| chr13:29183876-29183916 | Ch13 region 1 |
| chr13:29183922-29183962 | Ch13 region 1 |
| chr13:29183973-29184013 | Ch13 region 1 |
| chr13:29184128-29184168 | Ch13 region 1 |
| chr13:29184264-29184304 | Ch13 region 1 |
| chr13:29184334-29184374 | Ch13 region 1 |
| chr13:29184470-29184510 | Ch13 region 1 |
| chr13:29184519-29184559 | Ch13 region 1 |
| chr13:29184603-29184643 | Ch13 region 1 |
| chr13:29184767-29184807 | Ch13 region 1 |
| chr13:29184812-29184852 | Ch13 region 1 |
| chr13:29184868-29184908 | Ch13 region 1 |

| | |
|---|---|
| chr13:29184913-29184953 | Ch13 region 1 |
| chr13:29184965-29185005 | Ch13 region 1 |
| chr13:29185027-29185067 | Ch13 region 1 |
| chr13:29185072-29185112 | Ch13 region 1 |
| chr13:29185153-29185193 | Ch13 region 1 |
| chr13:29185234-29185274 | Ch13 region 1 |
| chr13:29185304-29185344 | Ch13 region 1 |
| chr13:29185354-29185394 | Ch13 region 1 |
| chr13:29185457-29185497 | Ch13 region 1 |
| chr13:29185502-29185542 | Ch13 region 1 |
| chr13:29185582-29185622 | Ch13 region 1 |
| chr13:29185675-29185715 | Ch13 region 1 |
| chr13:29185740-29185780 | Ch13 region 1 |
| chr13:29185826-29185866 | Ch13 region 1 |
| chr13:29185887-29185927 | Ch13 region 1 |
| chr13:29185948-29185988 | Ch13 region 1 |
| chr13:29186042-29186082 | Ch13 region 1 |
| chr13:29186099-29186139 | Ch13 region 1 |
| chr13:29186163-29186203 | Ch13 region 1 |
| chr13:29186218-29186258 | Ch13 region 1 |
| chr13:29186292-29186332 | Ch13 region 1 |
| chr13:29186338-29186378 | Ch13 region 1 |
| chr13:29186413-29186453 | Ch13 region 1 |
| chr13:29186494-29186534 | Ch13 region 1 |
| chr13:29186540-29186580 | Ch13 region 1 |
| chr13:29186620-29186660 | Ch13 region 1 |
| chr13:29186693-29186733 | Ch13 region 1 |
| chr13:29186785-29186825 | Ch13 region 1 |
| chr13:29186858-29186898 | Ch13 region 1 |
| chr13:29186947-29186987 | Ch13 region 1 |
| chr13:29186993-29187033 | Ch13 region 1 |
| chr13:29187044-29187084 | Ch13 region 1 |
| chr13:29187094-29187134 | Ch13 region 1 |
| chr13:29187156-29187196 | Ch13 region 1 |
| chr13:29187205-29187245 | Ch13 region 1 |
| chr13:29187275-29187315 | Ch13 region 1 |
| chr13:29187345-29187385 | Ch13 region 1 |
| chr13:29187401-29187441 | Ch13 region 1 |
| chr13:29187452-29187492 | Ch13 region 1 |
| chr13:29187518-29187558 | Ch13 region 1 |
| chr13:29187563-29187603 | Ch13 region 1 |
| chr13:29187622-29187662 | Ch13 region 1 |
| chr13:29187687-29187727 | Ch13 region 1 |
| chr13:29187732-29187772 | Ch13 region 1 |
| chr13:29187810-29187850 | Ch13 region 1 |
| chr13:29187883-29187923 | Ch13 region 1 |
| chr13:29187933-29187973 | Ch13 region 1 |
| chr13:29187990-29188030 | Ch13 region 1 |
| chr13:29188051-29188091 | Ch13 region 1 |
| chr13:29188153-29188193 | Ch13 region 1 |
| chr13:29188244-29188284 | Ch13 region 1 |
| chr13:29188313-29188353 | Ch13 region 1 |

| | |
|---|---|
| chr13:29188358-29188398 | Ch13 region 1 |
| chr13:29188418-29188458 | Ch13 region 1 |
| chr13:29188488-29188528 | Ch13 region 1 |
| chr13:29188587-29188627 | Ch13 region 1 |
| chr13:29188668-29188708 | Ch13 region 1 |
| chr13:29188864-29188904 | Ch13 region 1 |
| chr13:29189013-29189053 | Ch13 region 1 |
| chr13:29189206-29189246 | Ch13 region 1 |
| chr13:29189402-29189442 | Ch13 region 1 |
| chr13:29189516-29189556 | Ch13 region 1 |
| chr13:29189570-29189610 | Ch13 region 1 |
| chr13:29189635-29189675 | Ch13 region 1 |
| chr13:29189706-29189746 | Ch13 region 1 |
| chr13:29189772-29189812 | Ch13 region 1 |
| chr13:29189829-29189869 | Ch13 region 1 |
| chr13:29189891-29189931 | Ch13 region 1 |
| chr13:29189942-29189982 | Ch13 region 1 |
| chr13:29190140-29190180 | Ch13 region 1 |
| chr13:29190212-29190252 | Ch13 region 1 |
| chr13:29190336-29190376 | Ch13 region 1 |
| chr13:29190390-29190430 | Ch13 region 1 |
| chr13:29190437-29190477 | Ch13 region 1 |
| chr13:29190500-29190540 | Ch13 region 1 |
| chr13:29190751-29190791 | Ch13 region 1 |
| chr13:29190822-29190862 | Ch13 region 1 |
| chr13:29191016-29191056 | Ch13 region 1 |
| chr13:29191070-29191110 | Ch13 region 1 |
| chr13:29191116-29191156 | Ch13 region 1 |
| chr13:29191187-29191227 | Ch13 region 1 |
| chr13:29191294-29191334 | Ch13 region 1 |
| chr13:29191360-29191400 | Ch13 region 1 |
| chr13:29191558-29191598 | Ch13 region 1 |
| chr13:29191632-29191672 | Ch13 region 1 |
| chr13:29191692-29191732 | Ch13 region 1 |
| chr13:29191740-29191780 | Ch13 region 1 |
| chr13:29191884-29191924 | Ch13 region 1 |
| chr13:29191933-29191973 | Ch13 region 1 |
| chr13:29191979-29192019 | Ch13 region 1 |
| chr13:29192036-29192076 | Ch13 region 1 |
| chr13:29192223-29192263 | Ch13 region 1 |
| chr13:29192349-29192389 | Ch13 region 1 |
| chr13:29192412-29192452 | Ch13 region 1 |
| chr13:29192488-29192528 | Ch13 region 1 |
| chr13:29192535-29192575 | Ch13 region 1 |
| chr13:29192583-29192623 | Ch13 region 1 |
| chr13:29192647-29192687 | Ch13 region 1 |
| chr13:29192693-29192733 | Ch13 region 1 |
| chr13:29192738-29192778 | Ch13 region 1 |
| chr13:29192790-29192830 | Ch13 region 1 |
| chr13:29192851-29192891 | Ch13 region 1 |
| chr13:29192928-29192968 | Ch13 region 1 |
| chr13:29192994-29193034 | Ch13 region 1 |

| | |
|---|---|
| chr13:29193065-29193105 | Ch13 region 1 |
| chr13:29193145-29193185 | Ch13 region 1 |
| chr13:29193202-29193242 | Ch13 region 1 |
| chr13:29193255-29193295 | Ch13 region 1 |
| chr13:29193302-29193342 | Ch13 region 1 |
| chr13:29193353-29193393 | Ch13 region 1 |
| chr13:29193407-29193447 | Ch13 region 1 |
| chr13:29193453-29193493 | Ch13 region 1 |
| chr13:29193529-29193569 | Ch13 region 1 |
| chr13:29193622-29193662 | Ch13 region 1 |
| chr13:29193675-29193715 | Ch13 region 1 |
| chr13:29193736-29193776 | Ch13 region 1 |
| chr13:29193797-29193837 | Ch13 region 1 |
| chr13:29193858-29193898 | Ch13 region 1 |
| chr13:29194014-29194054 | Ch13 region 1 |
| chr13:29194073-29194113 | Ch13 region 1 |
| chr13:29194137-29194177 | Ch13 region 1 |
| chr13:29194189-29194229 | Ch13 region 1 |
| chr13:29194271-29194311 | Ch13 region 1 |
| chr13:29194325-29194365 | Ch13 region 1 |
| chr13:29194395-29194435 | Ch13 region 1 |
| chr13:29194473-29194513 | Ch13 region 1 |
| chr13:29194525-29194565 | Ch13 region 1 |
| chr13:29194593-29194633 | Ch13 region 1 |
| chr13:29194653-29194693 | Ch13 region 1 |
| chr13:29194731-29194771 | Ch13 region 1 |
| chr13:29194776-29194816 | Ch13 region 1 |
| chr13:29194866-29194906 | Ch13 region 1 |
| chr13:29194940-29194980 | Ch13 region 1 |
| chr13:29194985-29195025 | Ch13 region 1 |
| chr13:29195044-29195084 | Ch13 region 1 |
| chr13:29195120-29195160 | Ch13 region 1 |
| chr13:29195189-29195229 | Ch13 region 1 |
| chr13:29195246-29195286 | Ch13 region 1 |
| chr13:29195295-29195335 | Ch13 region 1 |
| chr13:29195369-29195409 | Ch13 region 1 |
| chr13:29195456-29195496 | Ch13 region 1 |
| chr13:29195507-29195547 | Ch13 region 1 |
| chr13:29195581-29195621 | Ch13 region 1 |
| chr13:29195698-29195738 | Ch13 region 1 |
| chr13:29195768-29195808 | Ch13 region 1 |
| chr13:29195841-29195881 | Ch13 region 1 |
| chr13:29195931-29195971 | Ch13 region 1 |
| chr13:29195979-29196019 | Ch13 region 1 |
| chr13:29196025-29196065 | Ch13 region 1 |
| chr13:29196070-29196110 | Ch13 region 1 |
| chr13:29196211-29196251 | Ch13 region 1 |
| chr13:29196258-29196298 | Ch13 region 1 |
| chr13:29196330-29196370 | Ch13 region 1 |
| chr13:29196398-29196438 | Ch13 region 1 |
| chr13:29196443-29196483 | Ch13 region 1 |
| chr13:29196512-29196552 | Ch13 region 1 |

| | |
|---|---|
| chr13:29196579-29196619 | Ch13 region 1 |
| chr13:29196624-29196664 | Ch13 region 1 |
| chr13:29196697-29196737 | Ch13 region 1 |
| chr13:29196773-29196813 | Ch13 region 1 |
| chr13:29196838-29196878 | Ch13 region 1 |
| chr13:29196909-29196949 | Ch13 region 1 |
| chr13:29196954-29196994 | Ch13 region 1 |
| chr13:29197104-29197144 | Ch13 region 1 |
| chr13:29197221-29197261 | Ch13 region 1 |
| chr13:29197322-29197362 | Ch13 region 1 |
| chr13:29197385-29197425 | Ch13 region 1 |
| chr13:29197445-29197485 | Ch13 region 1 |
| chr13:29197530-29197570 | Ch13 region 1 |
| chr13:29197617-29197657 | Ch13 region 1 |
| chr13:29197669-29197709 | Ch13 region 1 |
| chr13:29197714-29197754 | Ch13 region 1 |
| chr13:29197884-29197924 | Ch13 region 1 |
| chr13:29197949-29197989 | Ch13 region 1 |
| chr13:29198027-29198067 | Ch13 region 1 |
| chr13:29198120-29198160 | Ch13 region 1 |
| chr13:29198297-29198337 | Ch13 region 1 |
| chr13:29198371-29198411 | Ch13 region 1 |
| chr13:29198544-29198584 | Ch13 region 1 |
| chr13:29198677-29198717 | Ch13 region 1 |
| chr13:29198763-29198803 | Ch13 region 1 |
| chr13:29198838-29198878 | Ch13 region 1 |
| chr13:29198926-29198966 | Ch13 region 1 |
| chr13:29198985-29199025 | Ch13 region 1 |
| chr13:29199031-29199071 | Ch13 region 1 |
| chr13:29199131-29199171 | Ch13 region 1 |
| chr13:29199203-29199243 | Ch13 region 1 |
| chr13:29199255-29199295 | Ch13 region 1 |
| chr13:29199348-29199388 | Ch13 region 1 |
| chr13:29199395-29199435 | Ch13 region 1 |
| chr13:29199458-29199498 | Ch13 region 1 |
| chr13:29199551-29199591 | Ch13 region 1 |
| chr13:29199621-29199661 | Ch13 region 1 |
| chr13:29199688-29199728 | Ch13 region 1 |
| chr13:29199738-29199778 | Ch13 region 1 |
| chr13:29199810-29199850 | Ch13 region 1 |
| chr13:29199882-29199922 | Ch13 region 1 |
| chr13:29200009-29200049 | Ch13 region 1 |
| chr13:29200253-29200293 | Ch13 region 1 |
| chr13:29200314-29200354 | Ch13 region 1 |
| chr13:29200366-29200406 | Ch13 region 1 |
| chr13:29200458-29200498 | Ch13 region 1 |
| chr13:29200504-29200544 | Ch13 region 1 |
| chr13:29200608-29200648 | Ch13 region 1 |
| chr13:29200653-29200693 | Ch13 region 1 |
| chr13:29200706-29200746 | Ch13 region 1 |
| chr13:29200766-29200806 | Ch13 region 1 |
| chr13:29200843-29200883 | Ch13 region 1 |

| | |
|---|---|
| chr13:29200890-29200930 | Ch13 region 1 |
| chr13:29200945-29200985 | Ch13 region 1 |
| chr13:29201029-29201069 | Ch13 region 1 |
| chr13:29201097-29201137 | Ch13 region 1 |
| chr13:29201163-29201203 | Ch13 region 1 |
| chr13:29201227-29201267 | Ch13 region 1 |
| chr13:29201272-29201312 | Ch13 region 1 |
| chr13:29201340-29201380 | Ch13 region 1 |
| chr13:29201387-29201427 | Ch13 region 1 |
| chr13:29201481-29201521 | Ch13 region 1 |
| chr13:29201548-29201588 | Ch13 region 1 |
| chr13:29201624-29201664 | Ch13 region 1 |
| chr13:29201684-29201724 | Ch13 region 1 |
| chr13:29201754-29201794 | Ch13 region 1 |
| chr13:29201837-29201877 | Ch13 region 1 |
| chr13:29201897-29201937 | Ch13 region 1 |
| chr13:29202083-29202123 | Ch13 region 1 |
| chr13:29202179-29202219 | Ch13 region 1 |
| chr13:29202258-29202298 | Ch13 region 1 |
| chr13:29202345-29202385 | Ch13 region 1 |
| chr13:29202418-29202458 | Ch13 region 1 |
| chr13:29202489-29202529 | Ch13 region 1 |
| chr13:29202559-29202599 | Ch13 region 1 |
| chr13:29202662-29202702 | Ch13 region 1 |
| chr13:29202707-29202747 | Ch13 region 1 |
| chr13:29202752-29202792 | Ch13 region 1 |
| chr13:29202802-29202842 | Ch13 region 1 |
| chr13:29202871-29202911 | Ch13 region 1 |
| chr13:29202930-29202970 | Ch13 region 1 |
| chr13:29202989-29203029 | Ch13 region 1 |
| chr13:29203048-29203088 | Ch13 region 1 |
| chr13:29203142-29203182 | Ch13 region 1 |
| chr13:29203187-29203227 | Ch13 region 1 |
| chr13:29203258-29203298 | Ch13 region 1 |
| chr13:29203356-29203396 | Ch13 region 1 |
| chr13:29203410-29203450 | Ch13 region 1 |
| chr13:29203515-29203555 | Ch13 region 1 |
| chr13:29203628-29203668 | Ch13 region 1 |
| chr13:29203710-29203750 | Ch13 region 1 |
| chr13:29203755-29203795 | Ch13 region 1 |
| chr13:29203829-29203869 | Ch13 region 1 |
| chr13:29203909-29203949 | Ch13 region 1 |
| chr13:29203983-29204023 | Ch13 region 1 |
| chr13:29204207-29204247 | Ch13 region 1 |
| chr13:29204381-29204421 | Ch13 region 1 |
| chr13:29204426-29204466 | Ch13 region 1 |
| chr13:29204507-29204547 | Ch13 region 1 |
| chr13:29204586-29204626 | Ch13 region 1 |
| chr13:29204647-29204687 | Ch13 region 1 |
| chr13:29204995-29205035 | Ch13 region 1 |
| chr13:29205088-29205128 | Ch13 region 1 |
| chr13:29205133-29205173 | Ch13 region 1 |

| | |
|---|---|
| chr13:29205209-29205249 | Ch13 region 1 |
| chr13:29205268-29205308 | Ch13 region 1 |
| chr13:29205327-29205367 | Ch13 region 1 |
| chr13:29205405-29205445 | Ch13 region 1 |
| chr13:29205452-29205492 | Ch13 region 1 |
| chr13:29205539-29205579 | Ch13 region 1 |
| chr13:29205632-29205672 | Ch13 region 1 |
| chr13:29205724-29205764 | Ch13 region 1 |
| chr13:29205772-29205812 | Ch13 region 1 |
| chr13:29205819-29205859 | Ch13 region 1 |
| chr13:29205868-29205908 | Ch13 region 1 |
| chr13:29205919-29205959 | Ch13 region 1 |
| chr13:29205978-29206018 | Ch13 region 1 |
| chr13:29206045-29206085 | Ch13 region 1 |
| chr13:29206141-29206181 | Ch13 region 1 |
| chr13:29206199-29206239 | Ch13 region 1 |
| chr13:29206246-29206286 | Ch13 region 1 |
| chr13:29206293-29206333 | Ch13 region 1 |
| chr13:29206347-29206387 | Ch13 region 1 |
| chr13:29206396-29206436 | Ch13 region 1 |
| chr13:29206466-29206506 | Ch13 region 1 |
| chr13:29206541-29206581 | Ch13 region 1 |
| chr13:29206728-29206768 | Ch13 region 1 |
| chr13:29206932-29206972 | Ch13 region 1 |
| chr13:29206979-29207019 | Ch13 region 1 |
| chr13:29207024-29207064 | Ch13 region 1 |
| chr13:29207093-29207133 | Ch13 region 1 |
| chr13:29207138-29207178 | Ch13 region 1 |
| chr13:29207191-29207231 | Ch13 region 1 |
| chr13:29207241-29207281 | Ch13 region 1 |
| chr13:29207323-29207363 | Ch13 region 1 |
| chr13:29207396-29207436 | Ch13 region 1 |
| chr13:29207441-29207481 | Ch13 region 1 |
| chr13:29207544-29207584 | Ch13 region 1 |
| chr13:29207602-29207642 | Ch13 region 1 |
| chr13:29208181-29208221 | Ch13 region 1 |
| chr13:29208269-29208309 | Ch13 region 1 |
| chr13:29208314-29208354 | Ch13 region 1 |
| chr13:29208369-29208409 | Ch13 region 1 |
| chr13:29208450-29208490 | Ch13 region 1 |
| chr13:29208496-29208536 | Ch13 region 1 |
| chr13:29208545-29208585 | Ch13 region 1 |
| chr13:29208591-29208631 | Ch13 region 1 |
| chr13:29208655-29208695 | Ch13 region 1 |
| chr13:29208715-29208755 | Ch13 region 1 |
| chr13:29208764-29208804 | Ch13 region 1 |
| chr13:29208827-29208867 | Ch13 region 1 |
| chr13:29208892-29208932 | Ch13 region 1 |
| chr13:29209075-29209115 | Ch13 region 1 |
| chr13:29209155-29209195 | Ch13 region 1 |
| chr13:29209263-29209303 | Ch13 region 1 |
| chr13:29209332-29209372 | Ch13 region 1 |

| | |
|---|---|
| chr13:29209400-29209440 | Ch13 region 1 |
| chr13:29209468-29209508 | Ch13 region 1 |
| chr13:29209541-29209581 | Ch13 region 1 |
| chr13:29209649-29209689 | Ch13 region 1 |
| chr13:29209731-29209771 | Ch13 region 1 |
| chr13:29209782-29209822 | Ch13 region 1 |
| chr13:29209923-29209963 | Ch13 region 1 |
| chr13:29209986-29210026 | Ch13 region 1 |
| chr13:29210072-29210112 | Ch13 region 1 |
| chr13:29210117-29210157 | Ch13 region 1 |
| chr13:29210216-29210256 | Ch13 region 1 |
| chr13:29210279-29210319 | Ch13 region 1 |
| chr13:29210327-29210367 | Ch13 region 1 |
| chr13:29210390-29210430 | Ch13 region 1 |
| chr13:29210435-29210475 | Ch13 region 1 |
| chr13:29210480-29210520 | Ch13 region 1 |
| chr13:29210590-29210630 | Ch13 region 1 |
| chr13:29210677-29210717 | Ch13 region 1 |
| chr13:29210722-29210762 | Ch13 region 1 |
| chr13:29210769-29210809 | Ch13 region 1 |
| chr13:29210836-29210876 | Ch13 region 1 |
| chr13:29210885-29210925 | Ch13 region 1 |
| chr13:29210942-29210982 | Ch13 region 1 |
| chr13:29211024-29211064 | Ch13 region 1 |
| chr13:29211087-29211127 | Ch13 region 1 |
| chr13:29211133-29211173 | Ch13 region 1 |
| chr13:29211232-29211272 | Ch13 region 1 |
| chr13:29211337-29211377 | Ch13 region 1 |
| chr13:29211393-29211433 | Ch13 region 1 |
| chr13:29211535-29211575 | Ch13 region 1 |
| chr13:29211655-29211695 | Ch13 region 1 |
| chr13:29211828-29211868 | Ch13 region 1 |
| chr13:29211913-29211953 | Ch13 region 1 |
| chr13:29211965-29212005 | Ch13 region 1 |
| chr13:29212035-29212075 | Ch13 region 1 |
| chr13:29212090-29212130 | Ch13 region 1 |
| chr13:29212178-29212218 | Ch13 region 1 |
| chr13:29212223-29212263 | Ch13 region 1 |
| chr13:29212286-29212326 | Ch13 region 1 |
| chr13:29212343-29212383 | Ch13 region 1 |
| chr13:29212389-29212429 | Ch13 region 1 |
| chr13:29212450-29212490 | Ch13 region 1 |
| chr13:29212496-29212536 | Ch13 region 1 |
| chr13:29212543-29212583 | Ch13 region 1 |
| chr13:29212615-29212655 | Ch13 region 1 |
| chr13:29212786-29212826 | Ch13 region 1 |
| chr13:29212863-29212903 | Ch13 region 1 |
| chr13:29212964-29213004 | Ch13 region 1 |
| chr13:29213015-29213055 | Ch13 region 1 |
| chr13:29213084-29213124 | Ch13 region 1 |
| chr13:29213145-29213185 | Ch13 region 1 |
| chr13:29213231-29213271 | Ch13 region 1 |

| | |
|---|---|
| chr13:29213292-29213332 | Ch13 region 1 |
| chr13:29213355-29213395 | Ch13 region 1 |
| chr13:29213447-29213487 | Ch13 region 1 |
| chr13:29213559-29213599 | Ch13 region 1 |
| chr13:29213607-29213647 | Ch13 region 1 |
| chr13:29213673-29213713 | Ch13 region 1 |
| chr13:29213755-29213795 | Ch13 region 1 |
| chr13:29213808-29213848 | Ch13 region 1 |
| chr13:29213875-29213915 | Ch13 region 1 |
| chr13:29213921-29213961 | Ch13 region 1 |
| chr13:29213999-29214039 | Ch13 region 1 |
| chr13:29214107-29214147 | Ch13 region 1 |
| chr13:29214152-29214192 | Ch13 region 1 |
| chr13:29214197-29214237 | Ch13 region 1 |
| chr13:29214399-29214439 | Ch13 region 1 |
| chr13:29214533-29214573 | Ch13 region 1 |
| chr13:29214616-29214656 | Ch13 region 1 |
| chr13:29214708-29214748 | Ch13 region 1 |
| chr13:29214760-29214800 | Ch13 region 1 |
| chr13:29214814-29214854 | Ch13 region 1 |
| chr13:29214859-29214899 | Ch13 region 1 |
| chr13:29214922-29214962 | Ch13 region 1 |
| chr13:29214967-29215007 | Ch13 region 1 |
| chr13:29215049-29215089 | Ch13 region 1 |
| chr13:29215132-29215172 | Ch13 region 1 |
| chr13:29215201-29215241 | Ch13 region 1 |
| chr13:29215270-29215310 | Ch13 region 1 |
| chr13:29215325-29215365 | Ch13 region 1 |
| chr13:29215373-29215413 | Ch13 region 1 |
| chr13:29215446-29215486 | Ch13 region 1 |
| chr13:29215506-29215546 | Ch13 region 1 |
| chr13:29215551-29215591 | Ch13 region 1 |
| chr13:29215596-29215636 | Ch13 region 1 |
| chr13:29215673-29215713 | Ch13 region 1 |
| chr13:29215719-29215759 | Ch13 region 1 |
| chr13:29215773-29215813 | Ch13 region 1 |
| chr13:29215826-29215866 | Ch13 region 1 |
| chr13:29215894-29215934 | Ch13 region 1 |
| chr13:29216023-29216063 | Ch13 region 1 |
| chr13:29216214-29216254 | Ch13 region 1 |
| chr13:29216262-29216302 | Ch13 region 1 |
| chr13:29216321-29216361 | Ch13 region 1 |
| chr13:29216377-29216417 | Ch13 region 1 |
| chr13:29216430-29216470 | Ch13 region 1 |
| chr13:29216521-29216561 | Ch13 region 1 |
| chr13:29216605-29216645 | Ch13 region 1 |
| chr13:29216650-29216690 | Ch13 region 1 |
| chr13:29216709-29216749 | Ch13 region 1 |
| chr13:29216770-29216810 | Ch13 region 1 |
| chr13:29216840-29216880 | Ch13 region 1 |
| chr13:29216915-29216955 | Ch13 region 1 |
| chr13:29216976-29217016 | Ch13 region 1 |

| | |
|---|---|
| chr13:29217084-29217124 | Ch13 region 1 |
| chr13:29217211-29217251 | Ch13 region 1 |
| chr13:29217256-29217296 | Ch13 region 1 |
| chr13:29217490-29217530 | Ch13 region 1 |
| chr13:29217564-29217604 | Ch13 region 1 |
| chr13:29217609-29217649 | Ch13 region 1 |
| chr13:29217707-29217747 | Ch13 region 1 |
| chr13:29217788-29217828 | Ch13 region 1 |
| chr13:29217848-29217888 | Ch13 region 1 |
| chr13:29217904-29217944 | Ch13 region 1 |
| chr13:29217978-29218018 | Ch13 region 1 |
| chr13:29218159-29218199 | Ch13 region 1 |
| chr13:29218216-29218256 | Ch13 region 1 |
| chr13:29218289-29218329 | Ch13 region 1 |
| chr13:29218381-29218421 | Ch13 region 1 |
| chr13:29218554-29218594 | Ch13 region 1 |
| chr13:29218626-29218666 | Ch13 region 1 |
| chr13:29218696-29218736 | Ch13 region 1 |
| chr13:29218744-29218784 | Ch13 region 1 |
| chr13:29218838-29218878 | Ch13 region 1 |
| chr13:29218885-29218925 | Ch13 region 1 |
| chr13:29218935-29218975 | Ch13 region 1 |
| chr13:29219042-29219082 | Ch13 region 1 |
| chr13:29219109-29219149 | Ch13 region 1 |
| chr13:29219175-29219215 | Ch13 region 1 |
| chr13:29219247-29219287 | Ch13 region 1 |
| chr13:29219376-29219416 | Ch13 region 1 |
| chr13:29219490-29219530 | Ch13 region 1 |
| chr13:29219628-29219668 | Ch13 region 1 |
| chr13:29219752-29219792 | Ch13 region 1 |
| chr13:29219809-29219849 | Ch13 region 1 |
| chr13:29219869-29219909 | Ch13 region 1 |
| chr13:29219950-29219990 | Ch13 region 1 |
| chr13:29220040-29220080 | Ch13 region 1 |
| chr13:29220104-29220144 | Ch13 region 1 |
| chr13:29220153-29220193 | Ch13 region 1 |
| chr13:29220228-29220268 | Ch13 region 1 |
| chr13:29220429-29220469 | Ch13 region 1 |
| chr13:29220486-29220526 | Ch13 region 1 |
| chr13:29220532-29220572 | Ch13 region 1 |
| chr13:29220647-29220687 | Ch13 region 1 |
| chr13:29220731-29220771 | Ch13 region 1 |
| chr13:29220895-29220935 | Ch13 region 1 |
| chr13:29220956-29220996 | Ch13 region 1 |
| chr13:29221086-29221126 | Ch13 region 1 |
| chr13:29221192-29221232 | Ch13 region 1 |
| chr13:29221260-29221300 | Ch13 region 1 |
| chr13:29221429-29221469 | Ch13 region 1 |
| chr13:29221490-29221530 | Ch13 region 1 |
| chr13:29221543-29221583 | Ch13 region 1 |
| chr13:29221646-29221686 | Ch13 region 1 |
| chr13:29221720-29221760 | Ch13 region 1 |

| | |
|---|---|
| chr13:29221771-29221811 | Ch13 region 1 |
| chr13:29221849-29221889 | Ch13 region 1 |
| chr13:29221921-29221961 | Ch13 region 1 |
| chr13:29221968-29222008 | Ch13 region 1 |
| chr13:29222044-29222084 | Ch13 region 1 |
| chr13:29222114-29222154 | Ch13 region 1 |
| chr13:29222169-29222209 | Ch13 region 1 |
| chr13:29222236-29222276 | Ch13 region 1 |
| chr13:29222318-29222358 | Ch13 region 1 |
| chr13:29222414-29222454 | Ch13 region 1 |
| chr13:29222463-29222503 | Ch13 region 1 |
| chr13:29222551-29222591 | Ch13 region 1 |
| chr13:29222672-29222712 | Ch13 region 1 |
| chr13:29222724-29222764 | Ch13 region 1 |
| chr13:29222771-29222811 | Ch13 region 1 |
| chr13:29222838-29222878 | Ch13 region 1 |
| chr13:29222925-29222965 | Ch13 region 1 |
| chr13:29222987-29223027 | Ch13 region 1 |
| chr13:29223054-29223094 | Ch13 region 1 |
| chr13:29223147-29223187 | Ch13 region 1 |
| chr13:29223222-29223262 | Ch13 region 1 |
| chr13:29223365-29223405 | Ch13 region 1 |
| chr13:29223437-29223477 | Ch13 region 1 |
| chr13:29223491-29223531 | Ch13 region 1 |
| chr13:29223536-29223576 | Ch13 region 1 |
| chr13:29223584-29223624 | Ch13 region 1 |
| chr13:29223707-29223747 | Ch13 region 1 |
| chr13:29223752-29223792 | Ch13 region 1 |
| chr13:29223980-29224020 | Ch13 region 1 |
| chr13:29224025-29224065 | Ch13 region 1 |
| chr13:29224092-29224132 | Ch13 region 1 |
| chr13:29224173-29224213 | Ch13 region 1 |
| chr13:29224228-29224268 | Ch13 region 1 |
| chr13:29224293-29224333 | Ch13 region 1 |
| chr13:29224378-29224418 | Ch13 region 1 |
| chr13:29224426-29224466 | Ch13 region 1 |
| chr13:29224502-29224542 | Ch13 region 1 |
| chr13:29224563-29224603 | Ch13 region 1 |
| chr13:29224657-29224697 | Ch13 region 1 |
| chr13:29224822-29224862 | Ch13 region 1 |
| chr13:29224926-29224966 | Ch13 region 1 |
| chr13:29225052-29225092 | Ch13 region 1 |
| chr13:29225238-29225278 | Ch13 region 1 |
| chr13:29225286-29225326 | Ch13 region 1 |
| chr13:29225381-29225421 | Ch13 region 1 |
| chr13:29225548-29225588 | Ch13 region 1 |
| chr13:29225674-29225714 | Ch13 region 1 |
| chr13:29225769-29225809 | Ch13 region 1 |
| chr13:29225860-29225900 | Ch13 region 1 |
| chr13:29225906-29225946 | Ch13 region 1 |
| chr13:29225967-29226007 | Ch13 region 1 |
| chr13:29226013-29226053 | Ch13 region 1 |

| | |
|---|---|
| chr13:29226100-29226140 | Ch13 region 1 |
| chr13:29226149-29226189 | Ch13 region 1 |
| chr13:29226197-29226237 | Ch13 region 1 |
| chr13:29226252-29226292 | Ch13 region 1 |
| chr13:29226317-29226357 | Ch13 region 1 |
| chr13:29226372-29226412 | Ch13 region 1 |
| chr13:29226451-29226491 | Ch13 region 1 |
| chr13:29226513-29226553 | Ch13 region 1 |
| chr13:29226583-29226623 | Ch13 region 1 |
| chr13:29226649-29226689 | Ch13 region 1 |
| chr13:29226694-29226734 | Ch13 region 1 |
| chr13:29226795-29226835 | Ch13 region 1 |
| chr13:29226840-29226880 | Ch13 region 1 |
| chr13:29226915-29226955 | Ch13 region 1 |
| chr13:29226996-29227036 | Ch13 region 1 |
| chr13:29227055-29227095 | Ch13 region 1 |
| chr13:29227135-29227175 | Ch13 region 1 |
| chr13:29227195-29227235 | Ch13 region 1 |
| chr13:29227243-29227283 | Ch13 region 1 |
| chr13:29227394-29227434 | Ch13 region 1 |
| chr13:29227445-29227485 | Ch13 region 1 |
| chr13:29227506-29227546 | Ch13 region 1 |
| chr13:29227552-29227592 | Ch13 region 1 |
| chr13:29227629-29227669 | Ch13 region 1 |
| chr13:29227720-29227760 | Ch13 region 1 |
| chr13:29227799-29227839 | Ch13 region 1 |
| chr13:29227852-29227892 | Ch13 region 1 |
| chr13:29227932-29227972 | Ch13 region 1 |
| chr13:29227978-29228018 | Ch13 region 1 |
| chr13:29228058-29228098 | Ch13 region 1 |
| chr13:29228155-29228195 | Ch13 region 1 |
| chr13:29228243-29228283 | Ch13 region 1 |
| chr13:29228323-29228363 | Ch13 region 1 |
| chr13:29228368-29228408 | Ch13 region 1 |
| chr13:29228438-29228478 | Ch13 region 1 |
| chr13:29228527-29228567 | Ch13 region 1 |
| chr13:29228592-29228632 | Ch13 region 1 |
| chr13:29228643-29228683 | Ch13 region 1 |
| chr13:29228721-29228761 | Ch13 region 1 |
| chr13:29228779-29228819 | Ch13 region 1 |
| chr13:29228824-29228864 | Ch13 region 1 |
| chr13:29228898-29228938 | Ch13 region 1 |
| chr13:29228957-29228997 | Ch13 region 1 |
| chr13:29229303-29229343 | Ch13 region 1 |
| chr13:29229348-29229388 | Ch13 region 1 |
| chr13:29229419-29229459 | Ch13 region 1 |
| chr13:29229466-29229506 | Ch13 region 1 |
| chr13:29229587-29229627 | Ch13 region 1 |
| chr13:29229699-29229739 | Ch13 region 1 |
| chr13:29229757-29229797 | Ch13 region 1 |
| chr13:29229802-29229842 | Ch13 region 1 |
| chr13:29229873-29229913 | Ch13 region 1 |

| | |
|---|---|
| chr13:29229922-29229962 | Ch13 region 1 |
| chr13:29229993-29230033 | Ch13 region 1 |
| chr13:29230085-29230125 | Ch13 region 1 |
| chr13:29230130-29230170 | Ch13 region 1 |
| chr13:29230193-29230233 | Ch13 region 1 |
| chr13:29230266-29230306 | Ch13 region 1 |
| chr13:29230359-29230399 | Ch13 region 1 |
| chr13:29230519-29230559 | Ch13 region 1 |
| chr13:29230602-29230642 | Ch13 region 1 |
| chr13:29230668-29230708 | Ch13 region 1 |
| chr13:29230765-29230805 | Ch13 region 1 |
| chr13:29230821-29230861 | Ch13 region 1 |
| chr13:29230874-29230914 | Ch13 region 1 |
| chr13:29230956-29230996 | Ch13 region 1 |
| chr13:29231002-29231042 | Ch13 region 1 |
| chr13:29231052-29231092 | Ch13 region 1 |
| chr13:29231127-29231167 | Ch13 region 1 |
| chr13:29231179-29231219 | Ch13 region 1 |
| chr13:29231232-29231272 | Ch13 region 1 |
| chr13:29231351-29231391 | Ch13 region 1 |
| chr13:29231570-29231610 | Ch13 region 1 |
| chr13:29231618-29231658 | Ch13 region 1 |
| chr13:29231664-29231704 | Ch13 region 1 |
| chr13:29231782-29231822 | Ch13 region 1 |
| chr13:29231839-29231879 | Ch13 region 1 |
| chr13:29231884-29231924 | Ch13 region 1 |
| chr13:29231955-29231995 | Ch13 region 1 |
| chr13:29232000-29232040 | Ch13 region 1 |
| chr13:29232064-29232104 | Ch13 region 1 |
| chr13:29232151-29232191 | Ch13 region 1 |
| chr13:29232197-29232237 | Ch13 region 1 |
| chr13:29232277-29232317 | Ch13 region 1 |
| chr13:29232331-29232371 | Ch13 region 1 |
| chr13:29232377-29232417 | Ch13 region 1 |
| chr13:29232422-29232462 | Ch13 region 1 |
| chr13:29232517-29232557 | Ch13 region 1 |
| chr13:29232594-29232634 | Ch13 region 1 |
| chr13:29232665-29232705 | Ch13 region 1 |
| chr13:29232751-29232791 | Ch13 region 1 |
| chr13:29232865-29232905 | Ch13 region 1 |
| chr13:29232910-29232950 | Ch13 region 1 |
| chr13:29232956-29232996 | Ch13 region 1 |
| chr13:29233028-29233068 | Ch13 region 1 |
| chr13:29233076-29233116 | Ch13 region 1 |
| chr13:29233129-29233169 | Ch13 region 1 |
| chr13:29233191-29233231 | Ch13 region 1 |
| chr13:29233243-29233283 | Ch13 region 1 |
| chr13:29233313-29233353 | Ch13 region 1 |
| chr13:29233374-29233414 | Ch13 region 1 |
| chr13:29233450-29233490 | Ch13 region 1 |
| chr13:29233550-29233590 | Ch13 region 1 |
| chr13:29233597-29233637 | Ch13 region 1 |

| | |
|---|---|
| chr13:29233709-29233749 | Ch13 region 1 |
| chr13:29233756-29233796 | Ch13 region 1 |
| chr13:29233829-29233869 | Ch13 region 1 |
| chr13:29233883-29233923 | Ch13 region 1 |
| chr13:29233951-29233991 | Ch13 region 1 |
| chr13:29233998-29234038 | Ch13 region 1 |
| chr13:29234104-29234144 | Ch13 region 1 |
| chr13:29234190-29234230 | Ch13 region 1 |
| chr13:29234235-29234275 | Ch13 region 1 |
| chr13:29234290-29234330 | Ch13 region 1 |
| chr13:29234373-29234413 | Ch13 region 1 |
| chr13:29234418-29234458 | Ch13 region 1 |
| chr13:29234537-29234577 | Ch13 region 1 |
| chr13:29234596-29234636 | Ch13 region 1 |
| chr13:29234707-29234747 | Ch13 region 1 |
| chr13:29234752-29234792 | Ch13 region 1 |
| chr13:29234813-29234853 | Ch13 region 1 |
| chr13:29234858-29234898 | Ch13 region 1 |
| chr13:29234928-29234968 | Ch13 region 1 |
| chr13:29234985-29235025 | Ch13 region 1 |
| chr13:29235054-29235094 | Ch13 region 1 |
| chr13:29235100-29235140 | Ch13 region 1 |
| chr13:29235240-29235280 | Ch13 region 1 |
| chr13:29235293-29235333 | Ch13 region 1 |
| chr13:29235403-29235443 | Ch13 region 1 |
| chr13:29235543-29235583 | Ch13 region 1 |
| chr13:29235626-29235666 | Ch13 region 1 |
| chr13:29235698-29235738 | Ch13 region 1 |
| chr13:29235745-29235785 | Ch13 region 1 |
| chr13:29235796-29235836 | Ch13 region 1 |
| chr13:29235846-29235886 | Ch13 region 1 |
| chr13:29235917-29235957 | Ch13 region 1 |
| chr13:29235962-29236002 | Ch13 region 1 |
| chr13:29236010-29236050 | Ch13 region 1 |
| chr13:29236079-29236119 | Ch13 region 1 |
| chr13:29236164-29236204 | Ch13 region 1 |
| chr13:29236211-29236251 | Ch13 region 1 |
| chr13:29236295-29236335 | Ch13 region 1 |
| chr13:29236367-29236407 | Ch13 region 1 |
| chr13:29236423-29236463 | Ch13 region 1 |
| chr13:29236524-29236564 | Ch13 region 1 |
| chr13:29236601-29236641 | Ch13 region 1 |
| chr13:29236711-29236751 | Ch13 region 1 |
| chr13:29236762-29236802 | Ch13 region 1 |
| chr13:29236814-29236854 | Ch13 region 1 |
| chr13:29236862-29236902 | Ch13 region 1 |
| chr13:29236928-29236968 | Ch13 region 1 |
| chr13:29236973-29237013 | Ch13 region 1 |
| chr13:29237026-29237066 | Ch13 region 1 |
| chr13:29237082-29237122 | Ch13 region 1 |
| chr13:29237182-29237222 | Ch13 region 1 |
| chr13:29237250-29237290 | Ch13 region 1 |

| | |
|---|---|
| chr13:29237303-29237343 | Ch13 region 1 |
| chr13:29237353-29237393 | Ch13 region 1 |
| chr13:29237498-29237538 | Ch13 region 1 |
| chr13:29237732-29237772 | Ch13 region 1 |
| chr13:29237777-29237817 | Ch13 region 1 |
| chr13:29237832-29237872 | Ch13 region 1 |
| chr13:29237911-29237951 | Ch13 region 1 |
| chr13:29237966-29238006 | Ch13 region 1 |
| chr13:29238100-29238140 | Ch13 region 1 |
| chr13:29238146-29238186 | Ch13 region 1 |
| chr13:29238260-29238300 | Ch13 region 1 |
| chr13:29238327-29238367 | Ch13 region 1 |
| chr13:29238375-29238415 | Ch13 region 1 |
| chr13:29238451-29238491 | Ch13 region 1 |
| chr13:29238533-29238573 | Ch13 region 1 |
| chr13:29238578-29238618 | Ch13 region 1 |
| chr13:29238663-29238703 | Ch13 region 1 |
| chr13:29238783-29238823 | Ch13 region 1 |
| chr13:29238842-29238882 | Ch13 region 1 |
| chr13:29238887-29238927 | Ch13 region 1 |
| chr13:29238950-29238990 | Ch13 region 1 |
| chr13:29239020-29239060 | Ch13 region 1 |
| chr13:29239088-29239128 | Ch13 region 1 |
| chr13:29239140-29239180 | Ch13 region 1 |
| chr13:29239188-29239228 | Ch13 region 1 |
| chr13:29239265-29239305 | Ch13 region 1 |
| chr13:29239380-29239420 | Ch13 region 1 |
| chr13:29239453-29239493 | Ch13 region 1 |
| chr13:29239567-29239607 | Ch13 region 1 |
| chr13:29239612-29239652 | Ch13 region 1 |
| chr13:29239657-29239697 | Ch13 region 1 |
| chr13:29239749-29239789 | Ch13 region 1 |
| chr13:29239816-29239856 | Ch13 region 1 |
| chr13:29239893-29239933 | Ch13 region 1 |
| chr13:29239938-29239978 | Ch13 region 1 |
| chr13:29240021-29240061 | Ch13 region 1 |
| chr13:29240104-29240144 | Ch13 region 1 |
| chr13:29240174-29240214 | Ch13 region 1 |
| chr13:29240268-29240308 | Ch13 region 1 |
| chr13:29240317-29240357 | Ch13 region 1 |
| chr13:29240373-29240413 | Ch13 region 1 |
| chr13:29240418-29240458 | Ch13 region 1 |
| chr13:29240481-29240521 | Ch13 region 1 |
| chr13:29240551-29240591 | Ch13 region 1 |
| chr13:29240616-29240656 | Ch13 region 1 |
| chr13:29240704-29240744 | Ch13 region 1 |
| chr13:29240770-29240810 | Ch13 region 1 |
| chr13:29240816-29240856 | Ch13 region 1 |
| chr13:29240881-29240921 | Ch13 region 1 |
| chr13:29240981-29241021 | Ch13 region 1 |
| chr13:29241026-29241066 | Ch13 region 1 |
| chr13:29241135-29241175 | Ch13 region 1 |

| | |
|---|---|
| chr13:29241202-29241242 | Ch13 region 1 |
| chr13:29241263-29241303 | Ch13 region 1 |
| chr13:29241358-29241398 | Ch13 region 1 |
| chr13:29241449-29241489 | Ch13 region 1 |
| chr13:29241518-29241558 | Ch13 region 1 |
| chr13:29241563-29241603 | Ch13 region 1 |
| chr13:29241608-29241648 | Ch13 region 1 |
| chr13:29241693-29241733 | Ch13 region 1 |
| chr13:29241779-29241819 | Ch13 region 1 |
| chr13:29241839-29241879 | Ch13 region 1 |
| chr13:29241885-29241925 | Ch13 region 1 |
| chr13:29241956-29241996 | Ch13 region 1 |
| chr13:29242026-29242066 | Ch13 region 1 |
| chr13:29242083-29242123 | Ch13 region 1 |
| chr13:29242227-29242267 | Ch13 region 1 |
| chr13:29242272-29242312 | Ch13 region 1 |
| chr13:29242398-29242438 | Ch13 region 1 |
| chr13:29242529-29242569 | Ch13 region 1 |
| chr13:29242595-29242635 | Ch13 region 1 |
| chr13:29242650-29242690 | Ch13 region 1 |
| chr13:29242714-29242754 | Ch13 region 1 |
| chr13:29242784-29242824 | Ch13 region 1 |
| chr13:29242874-29242914 | Ch13 region 1 |
| chr13:29242967-29243007 | Ch13 region 1 |
| chr13:29243069-29243109 | Ch13 region 1 |
| chr13:29243143-29243183 | Ch13 region 1 |
| chr13:29243202-29243242 | Ch13 region 1 |
| chr13:29243268-29243308 | Ch13 region 1 |
| chr13:29243322-29243362 | Ch13 region 1 |
| chr13:29243384-29243424 | Ch13 region 1 |
| chr13:29243443-29243483 | Ch13 region 1 |
| chr13:29243643-29243683 | Ch13 region 1 |
| chr13:29243752-29243792 | Ch13 region 1 |
| chr13:29243810-29243850 | Ch13 region 1 |
| chr13:29243868-29243908 | Ch13 region 1 |
| chr13:29243933-29243973 | Ch13 region 1 |
| chr13:29244078-29244118 | Ch13 region 1 |
| chr13:29244253-29244293 | Ch13 region 1 |
| chr13:29244352-29244392 | Ch13 region 1 |
| chr13:29244402-29244442 | Ch13 region 1 |
| chr13:29244477-29244517 | Ch13 region 1 |
| chr13:29244540-29244580 | Ch13 region 1 |
| chr13:29244592-29244632 | Ch13 region 1 |
| chr13:29244661-29244701 | Ch13 region 1 |
| chr13:29244744-29244784 | Ch13 region 1 |
| chr13:29244790-29244830 | Ch13 region 1 |
| chr13:29244850-29244890 | Ch13 region 1 |
| chr13:29244895-29244935 | Ch13 region 1 |
| chr13:29245218-29245258 | Ch13 region 1 |
| chr13:29245270-29245310 | Ch13 region 1 |
| chr13:29245321-29245361 | Ch13 region 1 |
| chr13:29245394-29245434 | Ch13 region 1 |

| | |
|---|---|
| chr13:29245475-29245515 | Ch13 region 1 |
| chr13:29245600-29245640 | Ch13 region 1 |
| chr13:29245664-29245704 | Ch13 region 1 |
| chr13:29245726-29245766 | Ch13 region 1 |
| chr13:29245858-29245898 | Ch13 region 1 |
| chr13:29245908-29245948 | Ch13 region 1 |
| chr13:29245985-29246025 | Ch13 region 1 |
| chr13:29246038-29246078 | Ch13 region 1 |
| chr13:29246103-29246143 | Ch13 region 1 |
| chr13:29246153-29246193 | Ch13 region 1 |
| chr13:29246201-29246241 | Ch13 region 1 |
| chr13:29246416-29246456 | Ch13 region 1 |
| chr13:29246469-29246509 | Ch13 region 1 |
| chr13:29246514-29246554 | Ch13 region 1 |
| chr13:29246566-29246606 | Ch13 region 1 |
| chr13:29246662-29246702 | Ch13 region 1 |
| chr13:29246722-29246762 | Ch13 region 1 |
| chr13:29246911-29246951 | Ch13 region 1 |
| chr13:29246957-29246997 | Ch13 region 1 |
| chr13:29247041-29247081 | Ch13 region 1 |
| chr13:29247166-29247206 | Ch13 region 1 |
| chr13:29247227-29247267 | Ch13 region 1 |
| chr13:29247491-29247531 | Ch13 region 1 |
| chr13:29247559-29247599 | Ch13 region 1 |
| chr13:29247615-29247655 | Ch13 region 1 |
| chr13:29247677-29247717 | Ch13 region 1 |
| chr13:29247740-29247780 | Ch13 region 1 |
| chr13:29247809-29247849 | Ch13 region 1 |
| chr13:29247858-29247898 | Ch13 region 1 |
| chr13:29247921-29247961 | Ch13 region 1 |
| chr13:29248006-29248046 | Ch13 region 1 |
| chr13:29248081-29248121 | Ch13 region 1 |
| chr13:29248181-29248221 | Ch13 region 1 |
| chr13:29248233-29248273 | Ch13 region 1 |
| chr13:29248340-29248380 | Ch13 region 1 |
| chr13:29248386-29248426 | Ch13 region 1 |
| chr13:29248451-29248491 | Ch13 region 1 |
| chr13:29248529-29248569 | Ch13 region 1 |
| chr13:29248621-29248661 | Ch13 region 1 |
| chr13:29248705-29248745 | Ch13 region 1 |
| chr13:29248750-29248790 | Ch13 region 1 |
| chr13:29248829-29248869 | Ch13 region 1 |
| chr13:29248874-29248914 | Ch13 region 1 |
| chr13:29248938-29248978 | Ch13 region 1 |
| chr13:29249019-29249059 | Ch13 region 1 |
| chr13:29249194-29249234 | Ch13 region 1 |
| chr13:29249242-29249282 | Ch13 region 1 |
| chr13:29249287-29249327 | Ch13 region 1 |
| chr13:29249415-29249455 | Ch13 region 1 |
| chr13:29249578-29249618 | Ch13 region 1 |
| chr13:29249624-29249664 | Ch13 region 1 |
| chr13:29249696-29249736 | Ch13 region 1 |

| | |
|---|---|
| chr13:29249746-29249786 | Ch13 region 1 |
| chr13:29249942-29249982 | Ch13 region 1 |
| chr13:29250041-29250081 | Ch13 region 1 |
| chr13:29250137-29250177 | Ch13 region 1 |
| chr13:29250429-29250469 | Ch13 region 1 |
| chr13:29250480-29250520 | Ch13 region 1 |
| chr13:29250581-29250621 | Ch13 region 1 |
| chr13:29250691-29250731 | Ch13 region 1 |
| chr13:29250749-29250789 | Ch13 region 1 |
| chr13:29250794-29250834 | Ch13 region 1 |
| chr13:29250859-29250899 | Ch13 region 1 |
| chr13:29250905-29250945 | Ch13 region 1 |
| chr13:29250996-29251036 | Ch13 region 1 |
| chr13:29251041-29251081 | Ch13 region 1 |
| chr13:29251131-29251171 | Ch13 region 1 |
| chr13:29251214-29251254 | Ch13 region 1 |
| chr13:29251297-29251337 | Ch13 region 1 |
| chr13:29251365-29251405 | Ch13 region 1 |
| chr13:29251423-29251463 | Ch13 region 1 |
| chr13:29251521-29251561 | Ch13 region 1 |
| chr13:29251636-29251676 | Ch13 region 1 |
| chr13:29251708-29251748 | Ch13 region 1 |
| chr13:29251787-29251827 | Ch13 region 1 |
| chr13:29251925-29251965 | Ch13 region 1 |
| chr13:29251996-29252036 | Ch13 region 1 |
| chr13:29252068-29252108 | Ch13 region 1 |
| chr13:29252166-29252206 | Ch13 region 1 |
| chr13:29252257-29252297 | Ch13 region 1 |
| chr13:29252308-29252348 | Ch13 region 1 |
| chr13:29252353-29252393 | Ch13 region 1 |
| chr13:29252402-29252442 | Ch13 region 1 |
| chr13:29252756-29252796 | Ch13 region 1 |
| chr13:29252820-29252860 | Ch13 region 1 |
| chr13:29253101-29253141 | Ch13 region 1 |
| chr13:29253174-29253214 | Ch13 region 1 |
| chr13:29253222-29253262 | Ch13 region 1 |
| chr13:29253321-29253361 | Ch13 region 1 |
| chr13:29253386-29253426 | Ch13 region 1 |
| chr13:29253471-29253511 | Ch13 region 1 |
| chr13:29253701-29253741 | Ch13 region 1 |
| chr13:29253759-29253799 | Ch13 region 1 |
| chr13:29253879-29253919 | Ch13 region 1 |
| chr13:29253944-29253984 | Ch13 region 1 |
| chr13:29254012-29254052 | Ch13 region 1 |
| chr13:29254098-29254138 | Ch13 region 1 |
| chr13:29254170-29254210 | Ch13 region 1 |
| chr13:29254220-29254260 | Ch13 region 1 |
| chr13:29254310-29254350 | Ch13 region 1 |
| chr13:29254355-29254395 | Ch13 region 1 |
| chr13:29254467-29254507 | Ch13 region 1 |
| chr13:29254533-29254573 | Ch13 region 1 |
| chr13:29254618-29254658 | Ch13 region 1 |

| | |
|---|---|
| chr13:29254665-29254705 | Ch13 region 1 |
| chr13:29254787-29254827 | Ch13 region 1 |
| chr13:29254867-29254907 | Ch13 region 1 |
| chr13:29254922-29254962 | Ch13 region 1 |
| chr13:29254989-29255029 | Ch13 region 1 |
| chr13:29255182-29255222 | Ch13 region 1 |
| chr13:29255238-29255278 | Ch13 region 1 |
| chr13:29255358-29255398 | Ch13 region 1 |
| chr13:29255441-29255481 | Ch13 region 1 |
| chr13:29255716-29255756 | Ch13 region 1 |
| chr13:29255998-29256038 | Ch13 region 1 |
| chr13:29256069-29256109 | Ch13 region 1 |
| chr13:29256157-29256197 | Ch13 region 1 |
| chr13:29256218-29256258 | Ch13 region 1 |
| chr13:29256263-29256303 | Ch13 region 1 |
| chr13:29256323-29256363 | Ch13 region 1 |
| chr13:29256424-29256464 | Ch13 region 1 |
| chr13:29256475-29256515 | Ch13 region 1 |
| chr13:29256533-29256573 | Ch13 region 1 |
| chr13:29256607-29256647 | Ch13 region 1 |
| chr13:29256670-29256710 | Ch13 region 1 |
| chr13:29256732-29256772 | Ch13 region 1 |
| chr13:29256784-29256824 | Ch13 region 1 |
| chr13:29256849-29256889 | Ch13 region 1 |
| chr13:29256918-29256958 | Ch13 region 1 |
| chr13:29256964-29257004 | Ch13 region 1 |
| chr13:29257097-29257137 | Ch13 region 1 |
| chr13:29257149-29257189 | Ch13 region 1 |
| chr13:29257235-29257275 | Ch13 region 1 |
| chr13:29257464-29257504 | Ch13 region 1 |
| chr13:29257634-29257674 | Ch13 region 1 |
| chr13:29257727-29257767 | Ch13 region 1 |
| chr13:29257787-29257827 | Ch13 region 1 |
| chr13:29257910-29257950 | Ch13 region 1 |
| chr13:29257998-29258038 | Ch13 region 1 |
| chr13:29258080-29258120 | Ch13 region 1 |
| chr13:29258128-29258168 | Ch13 region 1 |
| chr13:29258204-29258244 | Ch13 region 1 |
| chr13:29258305-29258345 | Ch13 region 1 |
| chr13:29258377-29258417 | Ch13 region 1 |
| chr13:29258448-29258488 | Ch13 region 1 |
| chr13:29258549-29258589 | Ch13 region 1 |
| chr13:29258602-29258642 | Ch13 region 1 |
| chr13:29258668-29258708 | Ch13 region 1 |
| chr13:29258780-29258820 | Ch13 region 1 |
| chr13:29258831-29258871 | Ch13 region 1 |
| chr13:29258937-29258977 | Ch13 region 1 |
| chr13:29259092-29259132 | Ch13 region 1 |
| chr13:29259153-29259193 | Ch13 region 1 |
| chr13:29259292-29259332 | Ch13 region 1 |
| chr13:29259446-29259486 | Ch13 region 1 |
| chr13:29259506-29259546 | Ch13 region 1 |

| | |
|---|---|
| chr13:29259585-29259625 | Ch13 region 1 |
| chr13:29259669-29259709 | Ch13 region 1 |
| chr13:29259878-29259918 | Ch13 region 1 |
| chr13:29259971-29260011 | Ch13 region 1 |
| chr13:29260131-29260171 | Ch13 region 1 |
| chr13:29260288-29260328 | Ch13 region 1 |
| chr13:29260364-29260404 | Ch13 region 1 |
| chr13:29260588-29260628 | Ch13 region 1 |
| chr13:29260731-29260771 | Ch13 region 1 |
| chr13:29260776-29260816 | Ch13 region 1 |
| chr13:29260955-29260995 | Ch13 region 1 |
| chr13:29261121-29261161 | Ch13 region 1 |
| chr13:29261325-29261365 | Ch13 region 1 |
| chr13:29261372-29261412 | Ch13 region 1 |
| chr13:29261566-29261606 | Ch13 region 1 |
| chr13:29261666-29261706 | Ch13 region 1 |
| chr13:29261797-29261837 | Ch13 region 1 |
| chr13:29261904-29261944 | Ch13 region 1 |
| chr13:29262047-29262087 | Ch13 region 1 |
| chr13:29262092-29262132 | Ch13 region 1 |
| chr13:29262281-29262321 | Ch13 region 1 |
| chr13:29262394-29262434 | Ch13 region 1 |
| chr13:29262491-29262531 | Ch13 region 1 |
| chr13:29262843-29262883 | Ch13 region 1 |
| chr13:29262947-29262987 | Ch13 region 1 |
| chr13:29262994-29263034 | Ch13 region 1 |
| chr13:29263055-29263095 | Ch13 region 1 |
| chr13:29263547-29263587 | Ch13 region 1 |
| chr13:29263713-29263753 | Ch13 region 1 |
| chr13:29264259-29264299 | Ch13 region 1 |
| chr13:29264418-29264458 | Ch13 region 1 |
| chr13:29264480-29264520 | Ch13 region 1 |
| chr13:29264526-29264566 | Ch13 region 1 |
| chr13:29264798-29264838 | Ch13 region 1 |
| chr13:29264939-29264979 | Ch13 region 1 |
| chr13:29265226-29265266 | Ch13 region 1 |
| chr13:29265365-29265405 | Ch13 region 1 |
| chr13:29265410-29265450 | Ch13 region 1 |
| chr13:29265506-29265546 | Ch13 region 1 |
| chr13:29265693-29265733 | Ch13 region 1 |
| chr13:29265773-29265813 | Ch13 region 1 |
| chr13:29265922-29265962 | Ch13 region 1 |
| chr13:29266088-29266128 | Ch13 region 1 |
| chr13:29266172-29266212 | Ch13 region 1 |
| chr13:29266355-29266395 | Ch13 region 1 |
| chr13:29266534-29266574 | Ch13 region 1 |
| chr13:29266583-29266623 | Ch13 region 1 |
| chr13:29266658-29266698 | Ch13 region 1 |
| chr13:29266826-29266866 | Ch13 region 1 |
| chr13:29266873-29266913 | Ch13 region 1 |
| chr13:29266976-29267016 | Ch13 region 1 |
| chr13:29267054-29267094 | Ch13 region 1 |

| | |
|---|---|
| chr13:29267225-29267265 | Ch13 region 1 |
| chr13:29267406-29267446 | Ch13 region 1 |
| chr13:29267490-29267530 | Ch13 region 1 |
| chr13:29267613-29267653 | Ch13 region 1 |
| chr13:29267683-29267723 | Ch13 region 1 |
| chr13:29267837-29267877 | Ch13 region 1 |
| chr13:29267978-29268018 | Ch13 region 1 |
| chr13:29268165-29268205 | Ch13 region 1 |
| chr13:29268328-29268368 | Ch13 region 1 |
| chr13:29268455-29268495 | Ch13 region 1 |
| chr13:29268587-29268627 | Ch13 region 1 |
| chr13:29268635-29268675 | Ch13 region 1 |
| chr13:29268851-29268891 | Ch13 region 1 |
| chr13:29268922-29268962 | Ch13 region 1 |
| chr13:29268967-29269007 | Ch13 region 1 |
| chr13:29269045-29269085 | Ch13 region 1 |
| chr13:29269118-29269158 | Ch13 region 1 |
| chr13:29269192-29269232 | Ch13 region 1 |
| chr13:29269274-29269314 | Ch13 region 1 |
| chr13:29269433-29269473 | Ch13 region 1 |
| chr13:29269514-29269554 | Ch13 region 1 |
| chr13:29269769-29269809 | Ch13 region 1 |
| chr13:29269846-29269886 | Ch13 region 1 |
| chr13:29269893-29269933 | Ch13 region 1 |
| chr13:29269947-29269987 | Ch13 region 1 |
| chr13:29270006-29270046 | Ch13 region 1 |
| chr13:29270072-29270112 | Ch13 region 1 |
| chr13:29270168-29270208 | Ch13 region 1 |
| chr13:29270257-29270297 | Ch13 region 1 |
| chr13:29270452-29270492 | Ch13 region 1 |
| chr13:29270597-29270637 | Ch13 region 1 |
| chr13:29271051-29271091 | Ch13 region 1 |
| chr13:29271097-29271137 | Ch13 region 1 |
| chr13:29271172-29271212 | Ch13 region 1 |
| chr13:29271236-29271276 | Ch13 region 1 |
| chr13:29271389-29271429 | Ch13 region 1 |
| chr13:29271531-29271571 | Ch13 region 1 |
| chr13:29271590-29271630 | Ch13 region 1 |
| chr13:29271668-29271708 | Ch13 region 1 |
| chr13:29271715-29271755 | Ch13 region 1 |
| chr13:29271763-29271803 | Ch13 region 1 |
| chr13:29271823-29271863 | Ch13 region 1 |
| chr13:29271893-29271933 | Ch13 region 1 |
| chr13:29271942-29271982 | Ch13 region 1 |
| chr13:29271994-29272034 | Ch13 region 1 |
| chr13:29272048-29272088 | Ch13 region 1 |
| chr13:29272093-29272133 | Ch13 region 1 |
| chr13:29272166-29272206 | Ch13 region 1 |
| chr13:29272280-29272320 | Ch13 region 1 |
| chr13:29272357-29272397 | Ch13 region 1 |
| chr13:29272481-29272521 | Ch13 region 1 |
| chr13:29272609-29272649 | Ch13 region 1 |

| | |
|---|---|
| chr13:29272803-29272843 | Ch13 region 1 |
| chr13:29272872-29272912 | Ch13 region 1 |
| chr13:29272943-29272983 | Ch13 region 1 |
| chr13:29273026-29273066 | Ch13 region 1 |
| chr13:29273097-29273137 | Ch13 region 1 |
| chr13:29273147-29273187 | Ch13 region 1 |
| chr13:29273195-29273235 | Ch13 region 1 |
| chr13:29273240-29273280 | Ch13 region 1 |
| chr13:29273290-29273330 | Ch13 region 1 |
| chr13:29273343-29273383 | Ch13 region 1 |
| chr13:29273388-29273428 | Ch13 region 1 |
| chr13:29273498-29273538 | Ch13 region 1 |
| chr13:29273573-29273613 | Ch13 region 1 |
| chr13:29273625-29273665 | Ch13 region 1 |
| chr13:29273705-29273745 | Ch13 region 1 |
| chr13:29273756-29273796 | Ch13 region 1 |
| chr13:29273801-29273841 | Ch13 region 1 |
| chr13:29273872-29273912 | Ch13 region 1 |
| chr13:29273952-29273992 | Ch13 region 1 |
| chr13:29274005-29274045 | Ch13 region 1 |
| chr13:29274076-29274116 | Ch13 region 1 |
| chr13:29274121-29274161 | Ch13 region 1 |
| chr13:29274254-29274294 | Ch13 region 1 |
| chr13:29274316-29274356 | Ch13 region 1 |
| chr13:29274368-29274408 | Ch13 region 1 |
| chr13:29274438-29274478 | Ch13 region 1 |
| chr13:29274549-29274589 | Ch13 region 1 |
| chr13:29274594-29274634 | Ch13 region 1 |
| chr13:29274683-29274723 | Ch13 region 1 |
| chr13:29274752-29274792 | Ch13 region 1 |
| chr13:29274799-29274839 | Ch13 region 1 |
| chr13:29274899-29274939 | Ch13 region 1 |
| chr13:29275028-29275068 | Ch13 region 1 |
| chr13:29275078-29275118 | Ch13 region 1 |
| chr13:29275184-29275224 | Ch13 region 1 |
| chr13:29275265-29275305 | Ch13 region 1 |
| chr13:29275326-29275366 | Ch13 region 1 |
| chr13:29275373-29275413 | Ch13 region 1 |
| chr13:29275438-29275478 | Ch13 region 1 |
| chr13:29275485-29275525 | Ch13 region 1 |
| chr13:29275545-29275585 | Ch13 region 1 |
| chr13:29275612-29275652 | Ch13 region 1 |
| chr13:29275685-29275725 | Ch13 region 1 |
| chr13:29275818-29275858 | Ch13 region 1 |
| chr13:29275897-29275937 | Ch13 region 1 |
| chr13:29275942-29275982 | Ch13 region 1 |
| chr13:29275998-29276038 | Ch13 region 1 |
| chr13:29276093-29276133 | Ch13 region 1 |
| chr13:29276285-29276325 | Ch13 region 1 |
| chr13:29276401-29276441 | Ch13 region 1 |
| chr13:29276456-29276496 | Ch13 region 1 |
| chr13:29276579-29276619 | Ch13 region 1 |

| | |
|---|---|
| chr13:29276679-29276719 | Ch13 region 1 |
| chr13:29276806-29276846 | Ch13 region 1 |
| chr13:29276856-29276896 | Ch13 region 1 |
| chr13:29276901-29276941 | Ch13 region 1 |
| chr13:29276969-29277009 | Ch13 region 1 |
| chr13:29277041-29277081 | Ch13 region 1 |
| chr13:29277120-29277160 | Ch13 region 1 |
| chr13:29277351-29277391 | Ch13 region 1 |
| chr13:29277411-29277451 | Ch13 region 1 |
| chr13:29277458-29277498 | Ch13 region 1 |
| chr13:29277554-29277594 | Ch13 region 1 |
| chr13:29277599-29277639 | Ch13 region 1 |
| chr13:29277723-29277763 | Ch13 region 1 |
| chr13:29277774-29277814 | Ch13 region 1 |
| chr13:29277882-29277922 | Ch13 region 1 |
| chr13:29277938-29277978 | Ch13 region 1 |
| chr13:29277994-29278034 | Ch13 region 1 |
| chr13:29278060-29278100 | Ch13 region 1 |
| chr13:29278138-29278178 | Ch13 region 1 |
| chr13:29278183-29278223 | Ch13 region 1 |
| chr13:29278231-29278271 | Ch13 region 1 |
| chr13:29278285-29278325 | Ch13 region 1 |
| chr13:29278349-29278389 | Ch13 region 1 |
| chr13:29278436-29278476 | Ch13 region 1 |
| chr13:29278523-29278563 | Ch13 region 1 |
| chr13:29278602-29278642 | Ch13 region 1 |
| chr13:29278663-29278703 | Ch13 region 1 |
| chr13:29278709-29278749 | Ch13 region 1 |
| chr13:29278789-29278829 | Ch13 region 1 |
| chr13:29278863-29278903 | Ch13 region 1 |
| chr13:29278922-29278962 | Ch13 region 1 |
| chr13:29278969-29279009 | Ch13 region 1 |
| chr13:29279020-29279060 | Ch13 region 1 |
| chr13:29279101-29279141 | Ch13 region 1 |
| chr13:29279158-29279198 | Ch13 region 1 |
| chr13:29279267-29279307 | Ch13 region 1 |
| chr13:29279329-29279369 | Ch13 region 1 |
| chr13:29279413-29279453 | Ch13 region 1 |
| chr13:29279489-29279529 | Ch13 region 1 |
| chr13:29279620-29279660 | Ch13 region 1 |
| chr13:29279688-29279728 | Ch13 region 1 |
| chr13:29279745-29279785 | Ch13 region 1 |
| chr13:29279812-29279852 | Ch13 region 1 |
| chr13:29279939-29279979 | Ch13 region 1 |
| chr13:29279984-29280024 | Ch13 region 1 |
| chr13:29280032-29280072 | Ch13 region 1 |
| chr13:29280113-29280153 | Ch13 region 1 |
| chr13:29280158-29280198 | Ch13 region 1 |
| chr13:29280233-29280273 | Ch13 region 1 |
| chr13:29280299-29280339 | Ch13 region 1 |
| chr13:29280344-29280384 | Ch13 region 1 |
| chr13:29280434-29280474 | Ch13 region 1 |

| | |
|---|---|
| chr13:29280595-29280635 | Ch13 region 1 |
| chr13:29280744-29280784 | Ch13 region 1 |
| chr13:29280838-29280878 | Ch13 region 1 |
| chr13:29281412-29281452 | Ch13 region 1 |
| chr13:29281530-29281570 | Ch13 region 1 |
| chr13:29281745-29281785 | Ch13 region 1 |
| chr13:29281822-29281862 | Ch13 region 1 |
| chr13:29281928-29281968 | Ch13 region 1 |
| chr13:29281975-29282015 | Ch13 region 1 |
| chr13:29282155-29282195 | Ch13 region 1 |
| chr13:29282208-29282248 | Ch13 region 1 |
| chr13:29282307-29282347 | Ch13 region 1 |
| chr13:29282367-29282407 | Ch13 region 1 |
| chr13:29282471-29282511 | Ch13 region 1 |
| chr13:29282522-29282562 | Ch13 region 1 |
| chr13:29282637-29282677 | Ch13 region 1 |
| chr13:29282703-29282743 | Ch13 region 1 |
| chr13:29282814-29282854 | Ch13 region 1 |
| chr13:29282881-29282921 | Ch13 region 1 |
| chr13:29283102-29283142 | Ch13 region 1 |
| chr13:29283148-29283188 | Ch13 region 1 |
| chr13:29283509-29283549 | Ch13 region 1 |
| chr13:29283574-29283614 | Ch13 region 1 |
| chr13:29283637-29283677 | Ch13 region 1 |
| chr13:29283728-29283768 | Ch13 region 1 |
| chr13:29283779-29283819 | Ch13 region 1 |
| chr13:29283859-29283899 | Ch13 region 1 |
| chr13:29283909-29283949 | Ch13 region 1 |
| chr13:29283954-29283994 | Ch13 region 1 |
| chr13:29284053-29284093 | Ch13 region 1 |
| chr13:29284143-29284183 | Ch13 region 1 |
| chr13:29284199-29284239 | Ch13 region 1 |
| chr13:29284244-29284284 | Ch13 region 1 |
| chr13:29284321-29284361 | Ch13 region 1 |
| chr13:29284389-29284429 | Ch13 region 1 |
| chr13:29284442-29284482 | Ch13 region 1 |
| chr13:29284509-29284549 | Ch13 region 1 |
| chr13:29284575-29284615 | Ch13 region 1 |
| chr13:29284628-29284668 | Ch13 region 1 |
| chr13:29284724-29284764 | Ch13 region 1 |
| chr13:29284771-29284811 | Ch13 region 1 |
| chr13:29284873-29284913 | Ch13 region 1 |
| chr13:29284918-29284958 | Ch13 region 1 |
| chr13:29284973-29285013 | Ch13 region 1 |
| chr13:29285046-29285086 | Ch13 region 1 |
| chr13:29285125-29285165 | Ch13 region 1 |
| chr13:29285180-29285220 | Ch13 region 1 |
| chr13:29285247-29285287 | Ch13 region 1 |
| chr13:29285322-29285362 | Ch13 region 1 |
| chr13:29285650-29285690 | Ch13 region 1 |
| chr13:29285720-29285760 | Ch13 region 1 |
| chr13:29285765-29285805 | Ch13 region 1 |

| | |
|---|---|
| chr13:29285894-29285934 | Ch13 region 1 |
| chr13:29285951-29285991 | Ch13 region 1 |
| chr13:29286008-29286048 | Ch13 region 1 |
| chr13:29286055-29286095 | Ch13 region 1 |
| chr13:29286130-29286170 | Ch13 region 1 |
| chr13:29286206-29286246 | Ch13 region 1 |
| chr13:29286272-29286312 | Ch13 region 1 |
| chr13:29286318-29286358 | Ch13 region 1 |
| chr13:29286488-29286528 | Ch13 region 1 |
| chr13:29286587-29286627 | Ch13 region 1 |
| chr13:29286654-29286694 | Ch13 region 1 |
| chr13:29286708-29286748 | Ch13 region 1 |
| chr13:29286792-29286832 | Ch13 region 1 |
| chr13:29286843-29286883 | Ch13 region 1 |
| chr13:29286940-29286980 | Ch13 region 1 |
| chr13:29286992-29287032 | Ch13 region 1 |
| chr13:29287080-29287120 | Ch13 region 1 |
| chr13:29287156-29287196 | Ch13 region 1 |
| chr13:29287246-29287286 | Ch13 region 1 |
| chr13:29287330-29287370 | Ch13 region 1 |
| chr13:29287392-29287432 | Ch13 region 1 |
| chr13:29287464-29287504 | Ch13 region 1 |
| chr13:29287540-29287580 | Ch13 region 1 |
| chr13:29287588-29287628 | Ch13 region 1 |
| chr13:29287635-29287675 | Ch13 region 1 |
| chr13:29287736-29287776 | Ch13 region 1 |
| chr13:29287802-29287842 | Ch13 region 1 |
| chr13:29287940-29287980 | Ch13 region 1 |
| chr13:29288057-29288097 | Ch13 region 1 |
| chr13:29288105-29288145 | Ch13 region 1 |
| chr13:29288297-29288337 | Ch13 region 1 |
| chr13:29288343-29288383 | Ch13 region 1 |
| chr13:29288414-29288454 | Ch13 region 1 |
| chr13:29288459-29288499 | Ch13 region 1 |
| chr13:29288579-29288619 | Ch13 region 1 |
| chr13:29288745-29288785 | Ch13 region 1 |
| chr13:29288899-29288939 | Ch13 region 1 |
| chr13:29288944-29288984 | Ch13 region 1 |
| chr13:29289001-29289041 | Ch13 region 1 |
| chr13:29289081-29289121 | Ch13 region 1 |
| chr13:29289162-29289202 | Ch13 region 1 |
| chr13:29289234-29289274 | Ch13 region 1 |
| chr13:29289319-29289359 | Ch13 region 1 |
| chr13:29289412-29289452 | Ch13 region 1 |
| chr13:29289460-29289500 | Ch13 region 1 |
| chr13:29289530-29289570 | Ch13 region 1 |
| chr13:29289600-29289640 | Ch13 region 1 |
| chr13:29289664-29289704 | Ch13 region 1 |
| chr13:29289854-29289894 | Ch13 region 1 |
| chr13:29289912-29289952 | Ch13 region 1 |
| chr13:29290024-29290064 | Ch13 region 1 |
| chr13:29290218-29290258 | Ch13 region 1 |

| | |
|---|---|
| chr13:29290342-29290382 | Ch13 region 1 |
| chr13:29290397-29290437 | Ch13 region 1 |
| chr13:29290460-29290500 | Ch13 region 1 |
| chr13:29290507-29290547 | Ch13 region 1 |
| chr13:29290554-29290594 | Ch13 region 1 |
| chr13:29290618-29290658 | Ch13 region 1 |
| chr13:29290680-29290720 | Ch13 region 1 |
| chr13:29290725-29290765 | Ch13 region 1 |
| chr13:29290770-29290810 | Ch13 region 1 |
| chr13:29290823-29290863 | Ch13 region 1 |
| chr13:29290879-29290919 | Ch13 region 1 |
| chr13:29290938-29290978 | Ch13 region 1 |
| chr13:29291041-29291081 | Ch13 region 1 |
| chr13:29291086-29291126 | Ch13 region 1 |
| chr13:29291146-29291186 | Ch13 region 1 |
| chr13:29291215-29291255 | Ch13 region 1 |
| chr13:29291290-29291330 | Ch13 region 1 |
| chr13:29291393-29291433 | Ch13 region 1 |
| chr13:29291454-29291494 | Ch13 region 1 |
| chr13:29291499-29291539 | Ch13 region 1 |
| chr13:29291554-29291594 | Ch13 region 1 |
| chr13:29291643-29291683 | Ch13 region 1 |
| chr13:29291726-29291766 | Ch13 region 1 |
| chr13:29291782-29291822 | Ch13 region 1 |
| chr13:29291829-29291869 | Ch13 region 1 |
| chr13:29291890-29291930 | Ch13 region 1 |
| chr13:29291987-29292027 | Ch13 region 1 |
| chr13:29292032-29292072 | Ch13 region 1 |
| chr13:29292094-29292134 | Ch13 region 1 |
| chr13:29292141-29292181 | Ch13 region 1 |
| chr13:29292216-29292256 | Ch13 region 1 |
| chr13:29292285-29292325 | Ch13 region 1 |
| chr13:29292332-29292372 | Ch13 region 1 |
| chr13:29292393-29292433 | Ch13 region 1 |
| chr13:29292459-29292499 | Ch13 region 1 |
| chr13:29292539-29292579 | Ch13 region 1 |
| chr13:29292613-29292653 | Ch13 region 1 |
| chr13:29292665-29292705 | Ch13 region 1 |
| chr13:29292710-29292750 | Ch13 region 1 |
| chr13:29292792-29292832 | Ch13 region 1 |
| chr13:29292848-29292888 | Ch13 region 1 |
| chr13:29292906-29292946 | Ch13 region 1 |
| chr13:29292991-29293031 | Ch13 region 1 |
| chr13:29293044-29293084 | Ch13 region 1 |
| chr13:29293131-29293171 | Ch13 region 1 |
| chr13:29293187-29293227 | Ch13 region 1 |
| chr13:29293260-29293300 | Ch13 region 1 |
| chr13:29293342-29293382 | Ch13 region 1 |
| chr13:29293392-29293432 | Ch13 region 1 |
| chr13:29293460-29293500 | Ch13 region 1 |
| chr13:29293532-29293572 | Ch13 region 1 |
| chr13:29293584-29293624 | Ch13 region 1 |

| | |
|---|---|
| chr13:29293647-29293687 | Ch13 region 1 |
| chr13:29293710-29293750 | Ch13 region 1 |
| chr13:29293766-29293806 | Ch13 region 1 |
| chr13:29293816-29293856 | Ch13 region 1 |
| chr13:29293879-29293919 | Ch13 region 1 |
| chr13:29293978-29294018 | Ch13 region 1 |
| chr13:29294029-29294069 | Ch13 region 1 |
| chr13:29294083-29294123 | Ch13 region 1 |
| chr13:29294143-29294183 | Ch13 region 1 |
| chr13:29294235-29294275 | Ch13 region 1 |
| chr13:29294309-29294349 | Ch13 region 1 |
| chr13:29294383-29294423 | Ch13 region 1 |
| chr13:29294440-29294480 | Ch13 region 1 |
| chr13:29294587-29294627 | Ch13 region 1 |
| chr13:29294834-29294874 | Ch13 region 1 |
| chr13:29294880-29294920 | Ch13 region 1 |
| chr13:29294935-29294975 | Ch13 region 1 |
| chr13:29295025-29295065 | Ch13 region 1 |
| chr13:29295130-29295170 | Ch13 region 1 |
| chr13:29295176-29295216 | Ch13 region 1 |
| chr13:29295262-29295302 | Ch13 region 1 |
| chr13:29295377-29295417 | Ch13 region 1 |
| chr13:29295472-29295512 | Ch13 region 1 |
| chr13:29295567-29295607 | Ch13 region 1 |
| chr13:29295694-29295734 | Ch13 region 1 |
| chr13:29295766-29295806 | Ch13 region 1 |
| chr13:29295896-29295936 | Ch13 region 1 |
| chr13:29296015-29296055 | Ch13 region 1 |
| chr13:29296101-29296141 | Ch13 region 1 |
| chr13:29296243-29296283 | Ch13 region 1 |
| chr13:29296481-29296521 | Ch13 region 1 |
| chr13:29296528-29296568 | Ch13 region 1 |
| chr13:29296664-29296704 | Ch13 region 1 |
| chr13:29296739-29296779 | Ch13 region 1 |
| chr13:29296839-29296879 | Ch13 region 1 |
| chr13:29296944-29296984 | Ch13 region 1 |
| chr13:29297040-29297080 | Ch13 region 1 |
| chr13:29297239-29297279 | Ch13 region 1 |
| chr13:29297387-29297427 | Ch13 region 1 |
| chr13:29297465-29297505 | Ch13 region 1 |
| chr13:29297645-29297685 | Ch13 region 1 |
| chr13:29297734-29297774 | Ch13 region 1 |
| chr13:29297795-29297835 | Ch13 region 1 |
| chr13:29297875-29297915 | Ch13 region 1 |
| chr13:29297944-29297984 | Ch13 region 1 |
| chr13:29298004-29298044 | Ch13 region 1 |
| chr13:29298070-29298110 | Ch13 region 1 |
| chr13:29298148-29298188 | Ch13 region 1 |
| chr13:29298217-29298257 | Ch13 region 1 |
| chr13:29298282-29298322 | Ch13 region 1 |
| chr13:29298346-29298386 | Ch13 region 1 |
| chr13:29298393-29298433 | Ch13 region 1 |

| | |
|---|---|
| chr13:29298438-29298478 | Ch13 region 1 |
| chr13:29298483-29298523 | Ch13 region 1 |
| chr13:29298586-29298626 | Ch13 region 1 |
| chr13:29298631-29298671 | Ch13 region 1 |
| chr13:29298751-29298791 | Ch13 region 1 |
| chr13:29298817-29298857 | Ch13 region 1 |
| chr13:29298875-29298915 | Ch13 region 1 |
| chr13:29298941-29298981 | Ch13 region 1 |
| chr13:29299032-29299072 | Ch13 region 1 |
| chr13:29299129-29299169 | Ch13 region 1 |
| chr13:29299187-29299227 | Ch13 region 1 |
| chr13:29299238-29299278 | Ch13 region 1 |
| chr13:29299314-29299354 | Ch13 region 1 |
| chr13:29299366-29299406 | Ch13 region 1 |
| chr13:29299436-29299476 | Ch13 region 1 |
| chr13:29299481-29299521 | Ch13 region 1 |
| chr13:29299553-29299593 | Ch13 region 1 |
| chr13:29299654-29299694 | Ch13 region 1 |
| chr13:29299699-29299739 | Ch13 region 1 |
| chr13:29299748-29299788 | Ch13 region 1 |
| chr13:29299863-29299903 | Ch13 region 1 |
| chr13:29299909-29299949 | Ch13 region 1 |
| chr13:29299956-29299996 | Ch13 region 1 |
| chr13:29300001-29300041 | Ch13 region 1 |
| chr13:29300070-29300110 | Ch13 region 1 |
| chr13:29300118-29300158 | Ch13 region 1 |
| chr13:29300188-29300228 | Ch13 region 1 |
| chr13:29300294-29300334 | Ch13 region 1 |
| chr13:29300386-29300426 | Ch13 region 1 |
| chr13:29300471-29300511 | Ch13 region 1 |
| chr13:29300525-29300565 | Ch13 region 1 |
| chr13:29300609-29300649 | Ch13 region 1 |
| chr13:29300662-29300702 | Ch13 region 1 |
| chr13:29300741-29300781 | Ch13 region 1 |
| chr13:29300786-29300826 | Ch13 region 1 |
| chr13:29300852-29300892 | Ch13 region 1 |
| chr13:29300951-29300991 | Ch13 region 1 |
| chr13:29300996-29301036 | Ch13 region 1 |
| chr13:29301097-29301137 | Ch13 region 1 |
| chr13:29301145-29301185 | Ch13 region 1 |
| chr13:29301202-29301242 | Ch13 region 1 |
| chr13:29301247-29301287 | Ch13 region 1 |
| chr13:29301351-29301391 | Ch13 region 1 |
| chr13:29301407-29301447 | Ch13 region 1 |
| chr13:29301472-29301512 | Ch13 region 1 |
| chr13:29301533-29301573 | Ch13 region 1 |
| chr13:29301624-29301664 | Ch13 region 1 |
| chr13:29301744-29301784 | Ch13 region 1 |
| chr13:29301799-29301839 | Ch13 region 1 |
| chr13:29301928-29301968 | Ch13 region 1 |
| chr13:29302111-29302151 | Ch13 region 1 |
| chr13:29302205-29302245 | Ch13 region 1 |

| | |
|---|---|
| chr13:29302432-29302472 | Ch13 region 1 |
| chr13:29302532-29302572 | Ch13 region 1 |
| chr13:29302640-29302680 | Ch13 region 1 |
| chr13:29302794-29302834 | Ch13 region 1 |
| chr13:29302880-29302920 | Ch13 region 1 |
| chr13:29302958-29302998 | Ch13 region 1 |
| chr13:29303028-29303068 | Ch13 region 1 |
| chr13:29303110-29303150 | Ch13 region 1 |
| chr13:29303165-29303205 | Ch13 region 1 |
| chr13:29303223-29303263 | Ch13 region 1 |
| chr13:29303286-29303326 | Ch13 region 1 |
| chr13:29303566-29303606 | Ch13 region 1 |
| chr13:29303660-29303700 | Ch13 region 1 |
| chr13:29303755-29303795 | Ch13 region 1 |
| chr13:29303826-29303866 | Ch13 region 1 |
| chr13:29303911-29303951 | Ch13 region 1 |
| chr13:29303957-29303997 | Ch13 region 1 |
| chr13:29304024-29304064 | Ch13 region 1 |
| chr13:29304104-29304144 | Ch13 region 1 |
| chr13:29304164-29304204 | Ch13 region 1 |
| chr13:29304316-29304356 | Ch13 region 1 |
| chr13:29304389-29304429 | Ch13 region 1 |
| chr13:29304645-29304685 | Ch13 region 1 |
| chr13:29304723-29304763 | Ch13 region 1 |
| chr13:29304819-29304859 | Ch13 region 1 |
| chr13:29304908-29304948 | Ch13 region 1 |
| chr13:29304991-29305031 | Ch13 region 1 |
| chr13:29305079-29305119 | Ch13 region 1 |
| chr13:29305180-29305220 | Ch13 region 1 |
| chr13:29305229-29305269 | Ch13 region 1 |
| chr13:29305294-29305334 | Ch13 region 1 |
| chr13:29305357-29305397 | Ch13 region 1 |
| chr13:29305478-29305518 | Ch13 region 1 |
| chr13:29305644-29305684 | Ch13 region 1 |
| chr13:29305696-29305736 | Ch13 region 1 |
| chr13:29305773-29305813 | Ch13 region 1 |
| chr13:29305849-29305889 | Ch13 region 1 |
| chr13:29305915-29305955 | Ch13 region 1 |
| chr13:29305994-29306034 | Ch13 region 1 |
| chr13:29306162-29306202 | Ch13 region 1 |
| chr13:29306209-29306249 | Ch13 region 1 |
| chr13:29306268-29306308 | Ch13 region 1 |
| chr13:29306325-29306365 | Ch13 region 1 |
| chr13:29306396-29306436 | Ch13 region 1 |
| chr13:29306483-29306523 | Ch13 region 1 |
| chr13:29306546-29306586 | Ch13 region 1 |
| chr13:29306611-29306651 | Ch13 region 1 |
| chr13:29306679-29306719 | Ch13 region 1 |
| chr13:29306780-29306820 | Ch13 region 1 |
| chr13:29306874-29306914 | Ch13 region 1 |
| chr13:29307014-29307054 | Ch13 region 1 |
| chr13:29307069-29307109 | Ch13 region 1 |

| | |
|---|---|
| chr13:29307138-29307178 | Ch13 region 1 |
| chr13:29307213-29307253 | Ch13 region 1 |
| chr13:29307262-29307302 | Ch13 region 1 |
| chr13:29307589-29307629 | Ch13 region 1 |
| chr13:29307692-29307732 | Ch13 region 1 |
| chr13:29307876-29307916 | Ch13 region 1 |
| chr13:29307927-29307967 | Ch13 region 1 |
| chr13:29307981-29308021 | Ch13 region 1 |
| chr13:29308054-29308094 | Ch13 region 1 |
| chr13:29308107-29308147 | Ch13 region 1 |
| chr13:29308176-29308216 | Ch13 region 1 |
| chr13:29308221-29308261 | Ch13 region 1 |
| chr13:29308278-29308318 | Ch13 region 1 |
| chr13:29308326-29308366 | Ch13 region 1 |
| chr13:29308419-29308459 | Ch13 region 1 |
| chr13:29308471-29308511 | Ch13 region 1 |
| chr13:29308664-29308704 | Ch13 region 1 |
| chr13:29308711-29308751 | Ch13 region 1 |
| chr13:29308756-29308796 | Ch13 region 1 |
| chr13:29308811-29308851 | Ch13 region 1 |
| chr13:29308881-29308921 | Ch13 region 1 |
| chr13:29309001-29309041 | Ch13 region 1 |
| chr13:29309046-29309086 | Ch13 region 1 |
| chr13:29309133-29309173 | Ch13 region 1 |
| chr13:29309185-29309225 | Ch13 region 1 |
| chr13:29309231-29309271 | Ch13 region 1 |
| chr13:29309298-29309338 | Ch13 region 1 |
| chr13:29309348-29309388 | Ch13 region 1 |
| chr13:29309399-29309439 | Ch13 region 1 |
| chr13:29309580-29309620 | Ch13 region 1 |
| chr13:29309634-29309674 | Ch13 region 1 |
| chr13:29309683-29309723 | Ch13 region 1 |
| chr13:29309736-29309776 | Ch13 region 1 |
| chr13:29309786-29309826 | Ch13 region 1 |
| chr13:29309874-29309914 | Ch13 region 1 |
| chr13:29309920-29309960 | Ch13 region 1 |
| chr13:29310011-29310051 | Ch13 region 1 |
| chr13:29310058-29310098 | Ch13 region 1 |
| chr13:29310140-29310180 | Ch13 region 1 |
| chr13:29310196-29310236 | Ch13 region 1 |
| chr13:29310260-29310300 | Ch13 region 1 |
| chr13:29310327-29310367 | Ch13 region 1 |
| chr13:29310377-29310417 | Ch13 region 1 |
| chr13:29310441-29310481 | Ch13 region 1 |
| chr13:29310524-29310564 | Ch13 region 1 |
| chr13:29310597-29310637 | Ch13 region 1 |
| chr13:29310644-29310684 | Ch13 region 1 |
| chr13:29310697-29310737 | Ch13 region 1 |
| chr13:29310749-29310789 | Ch13 region 1 |
| chr13:29310812-29310852 | Ch13 region 1 |
| chr13:29310858-29310898 | Ch13 region 1 |
| chr13:29310918-29310958 | Ch13 region 1 |

| | |
|---|---|
| chr13:29310978-29311018 | Ch13 region 1 |
| chr13:29311098-29311138 | Ch13 region 1 |
| chr13:29311256-29311296 | Ch13 region 1 |
| chr13:29311517-29311557 | Ch13 region 1 |
| chr13:29311621-29311661 | Ch13 region 1 |
| chr13:29311773-29311813 | Ch13 region 1 |
| chr13:29311854-29311894 | Ch13 region 1 |
| chr13:29311911-29311951 | Ch13 region 1 |
| chr13:29311962-29312002 | Ch13 region 1 |
| chr13:29312058-29312098 | Ch13 region 1 |
| chr13:29312230-29312270 | Ch13 region 1 |
| chr13:29312317-29312357 | Ch13 region 1 |
| chr13:29312545-29312585 | Ch13 region 1 |
| chr13:29312703-29312743 | Ch13 region 1 |
| chr13:29312945-29312985 | Ch13 region 1 |
| chr13:29313032-29313072 | Ch13 region 1 |
| chr13:29313085-29313125 | Ch13 region 1 |
| chr13:29313148-29313188 | Ch13 region 1 |
| chr13:29313217-29313257 | Ch13 region 1 |
| chr13:29313267-29313307 | Ch13 region 1 |
| chr13:29313361-29313401 | Ch13 region 1 |
| chr13:29313412-29313452 | Ch13 region 1 |
| chr13:29313545-29313585 | Ch13 region 1 |
| chr13:29313596-29313636 | Ch13 region 1 |
| chr13:29313699-29313739 | Ch13 region 1 |
| chr13:29313759-29313799 | Ch13 region 1 |
| chr13:29313818-29313858 | Ch13 region 1 |
| chr13:29314040-29314080 | Ch13 region 1 |
| chr13:29314172-29314212 | Ch13 region 1 |
| chr13:29314407-29314447 | Ch13 region 1 |
| chr13:29314452-29314492 | Ch13 region 1 |
| chr13:29314543-29314583 | Ch13 region 1 |
| chr13:29314747-29314787 | Ch13 region 1 |
| chr13:29314812-29314852 | Ch13 region 1 |
| chr13:29314898-29314938 | Ch13 region 1 |
| chr13:29314974-29315014 | Ch13 region 1 |
| chr13:29315036-29315076 | Ch13 region 1 |
| chr13:29315086-29315126 | Ch13 region 1 |
| chr13:29315175-29315215 | Ch13 region 1 |
| chr13:29315244-29315284 | Ch13 region 1 |
| chr13:29315295-29315335 | Ch13 region 1 |
| chr13:29315366-29315406 | Ch13 region 1 |
| chr13:29315441-29315481 | Ch13 region 1 |
| chr13:29315520-29315560 | Ch13 region 1 |
| chr13:29315565-29315605 | Ch13 region 1 |
| chr13:29315620-29315660 | Ch13 region 1 |
| chr13:29315717-29315757 | Ch13 region 1 |
| chr13:29315762-29315802 | Ch13 region 1 |
| chr13:29315825-29315865 | Ch13 region 1 |
| chr13:29315900-29315940 | Ch13 region 1 |
| chr13:29315947-29315987 | Ch13 region 1 |
| chr13:29316011-29316051 | Ch13 region 1 |

| | |
|---|---|
| chr13:29316189-29316229 | Ch13 region 1 |
| chr13:29316332-29316372 | Ch13 region 1 |
| chr13:29316390-29316430 | Ch13 region 1 |
| chr13:29316530-29316570 | Ch13 region 1 |
| chr13:29316594-29316634 | Ch13 region 1 |
| chr13:29316644-29316684 | Ch13 region 1 |
| chr13:29316720-29316760 | Ch13 region 1 |
| chr13:29316986-29317026 | Ch13 region 1 |
| chr13:29317148-29317188 | Ch13 region 1 |
| chr13:29317223-29317263 | Ch13 region 1 |
| chr13:29317405-29317445 | Ch13 region 1 |
| chr13:29317509-29317549 | Ch13 region 1 |
| chr13:29317558-29317598 | Ch13 region 1 |
| chr13:29317682-29317722 | Ch13 region 1 |
| chr13:29317910-29317950 | Ch13 region 1 |
| chr13:29317990-29318030 | Ch13 region 1 |
| chr13:29318066-29318106 | Ch13 region 1 |
| chr13:29318148-29318188 | Ch13 region 1 |
| chr13:29318208-29318248 | Ch13 region 1 |
| chr13:29318267-29318307 | Ch13 region 1 |
| chr13:29318314-29318354 | Ch13 region 1 |
| chr13:29318385-29318425 | Ch13 region 1 |
| chr13:29318519-29318559 | Ch13 region 1 |
| chr13:29318567-29318607 | Ch13 region 1 |
| chr13:29318643-29318683 | Ch13 region 1 |
| chr13:29318714-29318754 | Ch13 region 1 |
| chr13:29318780-29318820 | Ch13 region 1 |
| chr13:29318879-29318919 | Ch13 region 1 |
| chr13:29318944-29318984 | Ch13 region 1 |
| chr13:29318991-29319031 | Ch13 region 1 |
| chr13:29319067-29319107 | Ch13 region 1 |
| chr13:29319171-29319211 | Ch13 region 1 |
| chr13:29319224-29319264 | Ch13 region 1 |
| chr13:29319342-29319382 | Ch13 region 1 |
| chr13:29319413-29319453 | Ch13 region 1 |
| chr13:29319596-29319636 | Ch13 region 1 |
| chr13:29319747-29319787 | Ch13 region 1 |
| chr13:29319813-29319853 | Ch13 region 1 |
| chr13:29319864-29319904 | Ch13 region 1 |
| chr13:29319937-29319977 | Ch13 region 1 |
| chr13:29319994-29320034 | Ch13 region 1 |
| chr13:29320117-29320157 | Ch13 region 1 |
| chr13:29320319-29320359 | Ch13 region 1 |
| chr13:29320454-29320494 | Ch13 region 1 |
| chr13:29320599-29320639 | Ch13 region 1 |
| chr13:29320644-29320684 | Ch13 region 1 |
| chr13:29320709-29320749 | Ch13 region 1 |
| chr13:29320785-29320825 | Ch13 region 1 |
| chr13:29320851-29320891 | Ch13 region 1 |
| chr13:29320955-29320995 | Ch13 region 1 |
| chr13:29321049-29321089 | Ch13 region 1 |
| chr13:29321128-29321168 | Ch13 region 1 |

| | |
|---|---|
| chr13:29321205-29321245 | Ch13 region 1 |
| chr13:29321261-29321301 | Ch13 region 1 |
| chr13:29321326-29321366 | Ch13 region 1 |
| chr13:29321376-29321416 | Ch13 region 1 |
| chr13:29321438-29321478 | Ch13 region 1 |
| chr13:29321505-29321545 | Ch13 region 1 |
| chr13:29321551-29321591 | Ch13 region 1 |
| chr13:29321596-29321636 | Ch13 region 1 |
| chr13:29321642-29321682 | Ch13 region 1 |
| chr13:29321770-29321810 | Ch13 region 1 |
| chr13:29321960-29322000 | Ch13 region 1 |
| chr13:29322017-29322057 | Ch13 region 1 |
| chr13:29322074-29322114 | Ch13 region 1 |
| chr13:29322157-29322197 | Ch13 region 1 |
| chr13:29322215-29322255 | Ch13 region 1 |
| chr13:29322334-29322374 | Ch13 region 1 |
| chr13:29322379-29322419 | Ch13 region 1 |
| chr13:29322474-29322514 | Ch13 region 1 |
| chr13:29322524-29322564 | Ch13 region 1 |
| chr13:29322603-29322643 | Ch13 region 1 |
| chr13:29322677-29322717 | Ch13 region 1 |
| chr13:29322764-29322804 | Ch13 region 1 |
| chr13:29322856-29322896 | Ch13 region 1 |
| chr13:29322902-29322942 | Ch13 region 1 |
| chr13:29323105-29323145 | Ch13 region 1 |
| chr13:29323255-29323295 | Ch13 region 1 |
| chr13:29323300-29323340 | Ch13 region 1 |
| chr13:29323349-29323389 | Ch13 region 1 |
| chr13:29323400-29323440 | Ch13 region 1 |
| chr13:29323452-29323492 | Ch13 region 1 |
| chr13:29323538-29323578 | Ch13 region 1 |
| chr13:29323586-29323626 | Ch13 region 1 |
| chr13:29323771-29323811 | Ch13 region 1 |
| chr13:29323855-29323895 | Ch13 region 1 |
| chr13:29323992-29324032 | Ch13 region 1 |
| chr13:29324089-29324129 | Ch13 region 1 |
| chr13:29324143-29324183 | Ch13 region 1 |
| chr13:29324208-29324248 | Ch13 region 1 |
| chr13:29324273-29324313 | Ch13 region 1 |
| chr13:29324346-29324386 | Ch13 region 1 |
| chr13:29324405-29324445 | Ch13 region 1 |
| chr13:29324452-29324492 | Ch13 region 1 |
| chr13:29324546-29324586 | Ch13 region 1 |
| chr13:29324591-29324631 | Ch13 region 1 |
| chr13:29324650-29324690 | Ch13 region 1 |
| chr13:29324731-29324771 | Ch13 region 1 |
| chr13:29324841-29324881 | Ch13 region 1 |
| chr13:29324886-29324926 | Ch13 region 1 |
| chr13:29324978-29325018 | Ch13 region 1 |
| chr13:29325029-29325069 | Ch13 region 1 |
| chr13:29325074-29325114 | Ch13 region 1 |
| chr13:29325184-29325224 | Ch13 region 1 |

| | |
|---|---|
| chr13:29325230-29325270 | Ch13 region 1 |
| chr13:29325283-29325323 | Ch13 region 1 |
| chr13:29325493-29325533 | Ch13 region 1 |
| chr13:29325538-29325578 | Ch13 region 1 |
| chr13:29325588-29325628 | Ch13 region 1 |
| chr13:29325634-29325674 | Ch13 region 1 |
| chr13:29325679-29325719 | Ch13 region 1 |
| chr13:29325773-29325813 | Ch13 region 1 |
| chr13:29325828-29325868 | Ch13 region 1 |
| chr13:29325942-29325982 | Ch13 region 1 |
| chr13:29326002-29326042 | Ch13 region 1 |
| chr13:29326074-29326114 | Ch13 region 1 |
| chr13:29326154-29326194 | Ch13 region 1 |
| chr13:29326241-29326281 | Ch13 region 1 |
| chr13:29326286-29326326 | Ch13 region 1 |
| chr13:29326332-29326372 | Ch13 region 1 |
| chr13:29326377-29326417 | Ch13 region 1 |
| chr13:29326426-29326466 | Ch13 region 1 |
| chr13:29326513-29326553 | Ch13 region 1 |
| chr13:29326597-29326637 | Ch13 region 1 |
| chr13:29326680-29326720 | Ch13 region 1 |
| chr13:29326818-29326858 | Ch13 region 1 |
| chr13:29326877-29326917 | Ch13 region 1 |
| chr13:29326963-29327003 | Ch13 region 1 |
| chr13:29327039-29327079 | Ch13 region 1 |
| chr13:29327330-29327370 | Ch13 region 1 |
| chr13:29327387-29327427 | Ch13 region 1 |
| chr13:29327548-29327588 | Ch13 region 1 |
| chr13:29327725-29327765 | Ch13 region 1 |
| chr13:29327793-29327833 | Ch13 region 1 |
| chr13:29327871-29327911 | Ch13 region 1 |
| chr13:29327925-29327965 | Ch13 region 1 |
| chr13:29327975-29328015 | Ch13 region 1 |
| chr13:29328023-29328063 | Ch13 region 1 |
| chr13:29328088-29328128 | Ch13 region 1 |
| chr13:29328157-29328197 | Ch13 region 1 |
| chr13:29328204-29328244 | Ch13 region 1 |
| chr13:29328275-29328315 | Ch13 region 1 |
| chr13:29328322-29328362 | Ch13 region 1 |
| chr13:29328403-29328443 | Ch13 region 1 |
| chr13:29328497-29328537 | Ch13 region 1 |
| chr13:29328574-29328614 | Ch13 region 1 |
| chr13:29328625-29328665 | Ch13 region 1 |
| chr13:29328693-29328733 | Ch13 region 1 |
| chr13:29328762-29328802 | Ch13 region 1 |
| chr13:29328835-29328875 | Ch13 region 1 |
| chr13:29328904-29328944 | Ch13 region 1 |
| chr13:29328966-29329006 | Ch13 region 1 |
| chr13:29329017-29329057 | Ch13 region 1 |
| chr13:29329087-29329127 | Ch13 region 1 |
| chr13:29329132-29329172 | Ch13 region 1 |
| chr13:29329194-29329234 | Ch13 region 1 |

| | |
|---|---|
| chr13:29329289-29329329 | Ch13 region 1 |
| chr13:29329345-29329385 | Ch13 region 1 |
| chr13:29329405-29329445 | Ch13 region 1 |
| chr13:29329462-29329502 | Ch13 region 1 |
| chr13:29329517-29329557 | Ch13 region 1 |
| chr13:29329566-29329606 | Ch13 region 1 |
| chr13:29329617-29329657 | Ch13 region 1 |
| chr13:29329675-29329715 | Ch13 region 1 |
| chr13:29329746-29329786 | Ch13 region 1 |
| chr13:29329796-29329836 | Ch13 region 1 |
| chr13:29329860-29329900 | Ch13 region 1 |
| chr13:29329970-29330010 | Ch13 region 1 |
| chr13:29330017-29330057 | Ch13 region 1 |
| chr13:29330082-29330122 | Ch13 region 1 |
| chr13:29330255-29330295 | Ch13 region 1 |
| chr13:29330327-29330367 | Ch13 region 1 |
| chr13:29330393-29330433 | Ch13 region 1 |
| chr13:29330484-29330524 | Ch13 region 1 |
| chr13:29330530-29330570 | Ch13 region 1 |
| chr13:29330601-29330641 | Ch13 region 1 |
| chr13:29330663-29330703 | Ch13 region 1 |
| chr13:29330890-29330930 | Ch13 region 1 |
| chr13:29331075-29331115 | Ch13 region 1 |
| chr13:29331133-29331173 | Ch13 region 1 |
| chr13:29331245-29331285 | Ch13 region 1 |
| chr13:29331329-29331369 | Ch13 region 1 |
| chr13:29331388-29331428 | Ch13 region 1 |
| chr13:29331452-29331492 | Ch13 region 1 |
| chr13:29331512-29331552 | Ch13 region 1 |
| chr13:29331565-29331605 | Ch13 region 1 |
| chr13:29331715-29331755 | Ch13 region 1 |
| chr13:29331787-29331827 | Ch13 region 1 |
| chr13:29331876-29331916 | Ch13 region 1 |
| chr13:29332063-29332103 | Ch13 region 1 |
| chr13:29332162-29332202 | Ch13 region 1 |
| chr13:29332313-29332353 | Ch13 region 1 |
| chr13:29332369-29332409 | Ch13 region 1 |
| chr13:29332429-29332469 | Ch13 region 1 |
| chr13:29332512-29332552 | Ch13 region 1 |
| chr13:29332585-29332625 | Ch13 region 1 |
| chr13:29332633-29332673 | Ch13 region 1 |
| chr13:29332678-29332718 | Ch13 region 1 |
| chr13:29332914-29332954 | Ch13 region 1 |
| chr13:29333103-29333143 | Ch13 region 1 |
| chr13:29333168-29333208 | Ch13 region 1 |
| chr13:29333270-29333310 | Ch13 region 1 |
| chr13:29333402-29333442 | Ch13 region 1 |
| chr13:29333585-29333625 | Ch13 region 1 |
| chr13:29333713-29333753 | Ch13 region 1 |
| chr13:29333913-29333953 | Ch13 region 1 |
| chr13:29334044-29334084 | Ch13 region 1 |
| chr13:29334111-29334151 | Ch13 region 1 |

| | |
|---|---|
| chr13:29334166-29334206 | Ch13 region 1 |
| chr13:29334366-29334406 | Ch13 region 1 |
| chr13:29334415-29334455 | Ch13 region 1 |
| chr13:29334467-29334507 | Ch13 region 1 |
| chr13:29334552-29334592 | Ch13 region 1 |
| chr13:29334650-29334690 | Ch13 region 1 |
| chr13:29334829-29334869 | Ch13 region 1 |
| chr13:29334903-29334943 | Ch13 region 1 |
| chr13:29335043-29335083 | Ch13 region 1 |
| chr13:29335316-29335356 | Ch13 region 1 |
| chr13:29335366-29335406 | Ch13 region 1 |
| chr13:29335450-29335490 | Ch13 region 1 |
| chr13:29335500-29335540 | Ch13 region 1 |
| chr13:29335546-29335586 | Ch13 region 1 |
| chr13:29335626-29335666 | Ch13 region 1 |
| chr13:29335673-29335713 | Ch13 region 1 |
| chr13:29335734-29335774 | Ch13 region 1 |
| chr13:29335836-29335876 | Ch13 region 1 |
| chr13:29335885-29335925 | Ch13 region 1 |
| chr13:29335976-29336016 | Ch13 region 1 |
| chr13:29336022-29336062 | Ch13 region 1 |
| chr13:29336069-29336109 | Ch13 region 1 |
| chr13:29336182-29336222 | Ch13 region 1 |
| chr13:29336227-29336267 | Ch13 region 1 |
| chr13:29336299-29336339 | Ch13 region 1 |
| chr13:29336344-29336384 | Ch13 region 1 |
| chr13:29336423-29336463 | Ch13 region 1 |
| chr13:29336475-29336515 | Ch13 region 1 |
| chr13:29336557-29336597 | Ch13 region 1 |
| chr13:29336618-29336658 | Ch13 region 1 |
| chr13:29336675-29336715 | Ch13 region 1 |
| chr13:29336731-29336771 | Ch13 region 1 |
| chr13:29336789-29336829 | Ch13 region 1 |
| chr13:29336850-29336890 | Ch13 region 1 |
| chr13:29336914-29336954 | Ch13 region 1 |
| chr13:29336961-29337001 | Ch13 region 1 |
| chr13:29337012-29337052 | Ch13 region 1 |
| chr13:29337081-29337121 | Ch13 region 1 |
| chr13:29337199-29337239 | Ch13 region 1 |
| chr13:29337252-29337292 | Ch13 region 1 |
| chr13:29337331-29337371 | Ch13 region 1 |
| chr13:29337393-29337433 | Ch13 region 1 |
| chr13:29337458-29337498 | Ch13 region 1 |
| chr13:29337511-29337551 | Ch13 region 1 |
| chr13:29337586-29337626 | Ch13 region 1 |
| chr13:29337631-29337671 | Ch13 region 1 |
| chr13:29337721-29337761 | Ch13 region 1 |
| chr13:29337789-29337829 | Ch13 region 1 |
| chr13:29337836-29337876 | Ch13 region 1 |
| chr13:29337893-29337933 | Ch13 region 1 |
| chr13:29337945-29337985 | Ch13 region 1 |
| chr13:29338022-29338062 | Ch13 region 1 |

| | |
|---|---|
| chr13:29338104-29338144 | Ch13 region 1 |
| chr13:29338153-29338193 | Ch13 region 1 |
| chr13:29338198-29338238 | Ch13 region 1 |
| chr13:29338251-29338291 | Ch13 region 1 |
| chr13:29338346-29338386 | Ch13 region 1 |
| chr13:29338413-29338453 | Ch13 region 1 |
| chr13:29338541-29338581 | Ch13 region 1 |
| chr13:29338634-29338674 | Ch13 region 1 |
| chr13:29338732-29338772 | Ch13 region 1 |
| chr13:29338796-29338836 | Ch13 region 1 |
| chr13:29338991-29339031 | Ch13 region 1 |
| chr13:29339142-29339182 | Ch13 region 1 |
| chr13:29339187-29339227 | Ch13 region 1 |
| chr13:29339241-29339281 | Ch13 region 1 |
| chr13:29339316-29339356 | Ch13 region 1 |
| chr13:29339411-29339451 | Ch13 region 1 |
| chr13:29339474-29339514 | Ch13 region 1 |
| chr13:29339553-29339593 | Ch13 region 1 |
| chr13:29339602-29339642 | Ch13 region 1 |
| chr13:29339647-29339687 | Ch13 region 1 |
| chr13:29339696-29339736 | Ch13 region 1 |
| chr13:29339787-29339827 | Ch13 region 1 |
| chr13:29339842-29339882 | Ch13 region 1 |
| chr13:29339931-29339971 | Ch13 region 1 |
| chr13:29339999-29340039 | Ch13 region 1 |
| chr13:29340067-29340107 | Ch13 region 1 |
| chr13:29340132-29340172 | Ch13 region 1 |
| chr13:29340186-29340226 | Ch13 region 1 |
| chr13:29340240-29340280 | Ch13 region 1 |
| chr13:29340315-29340355 | Ch13 region 1 |
| chr13:29340361-29340401 | Ch13 region 1 |
| chr13:29340426-29340466 | Ch13 region 1 |
| chr13:29340492-29340532 | Ch13 region 1 |
| chr13:29340542-29340582 | Ch13 region 1 |
| chr13:29340587-29340627 | Ch13 region 1 |
| chr13:29340637-29340677 | Ch13 region 1 |
| chr13:29340704-29340744 | Ch13 region 1 |
| chr13:29340756-29340796 | Ch13 region 1 |
| chr13:29340833-29340873 | Ch13 region 1 |
| chr13:29340881-29340921 | Ch13 region 1 |
| chr13:29340965-29341005 | Ch13 region 1 |
| chr13:29341095-29341135 | Ch13 region 1 |
| chr13:29341156-29341196 | Ch13 region 1 |
| chr13:29341210-29341250 | Ch13 region 1 |
| chr13:29341259-29341299 | Ch13 region 1 |
| chr13:29341307-29341347 | Ch13 region 1 |
| chr13:29341391-29341431 | Ch13 region 1 |
| chr13:29341466-29341506 | Ch13 region 1 |
| chr13:29341529-29341569 | Ch13 region 1 |
| chr13:29341576-29341616 | Ch13 region 1 |
| chr13:29341691-29341731 | Ch13 region 1 |
| chr13:29341743-29341783 | Ch13 region 1 |

| | |
|---|---|
| chr13:29341817-29341857 | Ch13 region 1 |
| chr13:29341922-29341962 | Ch13 region 1 |
| chr13:29342013-29342053 | Ch13 region 1 |
| chr13:29342085-29342125 | Ch13 region 1 |
| chr13:29342181-29342221 | Ch13 region 1 |
| chr13:29342264-29342304 | Ch13 region 1 |
| chr13:29342367-29342407 | Ch13 region 1 |
| chr13:29342484-29342524 | Ch13 region 1 |
| chr13:29342568-29342608 | Ch13 region 1 |
| chr13:29342752-29342792 | Ch13 region 1 |
| chr13:29342851-29342891 | Ch13 region 1 |
| chr13:29343062-29343102 | Ch13 region 1 |
| chr13:29343187-29343227 | Ch13 region 1 |
| chr13:29343261-29343301 | Ch13 region 1 |
| chr13:29343473-29343513 | Ch13 region 1 |
| chr13:29343558-29343598 | Ch13 region 1 |
| chr13:29343665-29343705 | Ch13 region 1 |
| chr13:29343711-29343751 | Ch13 region 1 |
| chr13:29343851-29343891 | Ch13 region 1 |
| chr13:29344012-29344052 | Ch13 region 1 |
| chr13:29344096-29344136 | Ch13 region 1 |
| chr13:29344192-29344232 | Ch13 region 1 |
| chr13:29344349-29344389 | Ch13 region 1 |
| chr13:29344470-29344510 | Ch13 region 1 |
| chr13:29344662-29344702 | Ch13 region 1 |
| chr13:29344707-29344747 | Ch13 region 1 |
| chr13:29344974-29345014 | Ch13 region 1 |
| chr13:29345154-29345194 | Ch13 region 1 |
| chr13:29345382-29345422 | Ch13 region 1 |
| chr13:29345521-29345561 | Ch13 region 1 |
| chr13:29345702-29345742 | Ch13 region 1 |
| chr13:29345757-29345797 | Ch13 region 1 |
| chr13:29345831-29345871 | Ch13 region 1 |
| chr13:29345914-29345954 | Ch13 region 1 |
| chr13:29346054-29346094 | Ch13 region 1 |
| chr13:29346272-29346312 | Ch13 region 1 |
| chr13:29346423-29346463 | Ch13 region 1 |
| chr13:29346540-29346580 | Ch13 region 1 |
| chr13:29346694-29346734 | Ch13 region 1 |
| chr13:29346851-29346891 | Ch13 region 1 |
| chr13:29347013-29347053 | Ch13 region 1 |
| chr13:29347161-29347201 | Ch13 region 1 |
| chr13:29347281-29347321 | Ch13 region 1 |
| chr13:29347326-29347366 | Ch13 region 1 |
| chr13:29347410-29347450 | Ch13 region 1 |
| chr13:29347493-29347533 | Ch13 region 1 |
| chr13:29347570-29347610 | Ch13 region 1 |
| chr13:29347671-29347711 | Ch13 region 1 |
| chr13:29347828-29347868 | Ch13 region 1 |
| chr13:29347946-29347986 | Ch13 region 1 |
| chr13:29348033-29348073 | Ch13 region 1 |
| chr13:29348093-29348133 | Ch13 region 1 |

| | |
|---|---|
| chr13:29348219-29348259 | Ch13 region 1 |
| chr13:29348370-29348410 | Ch13 region 1 |
| chr13:29348438-29348478 | Ch13 region 1 |
| chr13:29348520-29348560 | Ch13 region 1 |
| chr13:29348591-29348631 | Ch13 region 1 |
| chr13:29348647-29348687 | Ch13 region 1 |
| chr13:29348710-29348750 | Ch13 region 1 |
| chr13:29348920-29348960 | Ch13 region 1 |
| chr13:29349032-29349072 | Ch13 region 1 |
| chr13:29349091-29349131 | Ch13 region 1 |
| chr13:29349209-29349249 | Ch13 region 1 |
| chr13:29349348-29349388 | Ch13 region 1 |
| chr13:29349433-29349473 | Ch13 region 1 |
| chr13:29349527-29349567 | Ch13 region 1 |
| chr13:29349637-29349677 | Ch13 region 1 |
| chr13:29349691-29349731 | Ch13 region 1 |
| chr13:29349835-29349875 | Ch13 region 1 |
| chr13:29349958-29349998 | Ch13 region 1 |
| chr13:29350024-29350064 | Ch13 region 1 |
| chr13:29350109-29350149 | Ch13 region 1 |
| chr13:29350189-29350229 | Ch13 region 1 |
| chr13:29350409-29350449 | Ch13 region 1 |
| chr13:29350592-29350632 | Ch13 region 1 |
| chr13:29350644-29350684 | Ch13 region 1 |
| chr13:29350702-29350742 | Ch13 region 1 |
| chr13:29350772-29350812 | Ch13 region 1 |
| chr13:29350838-29350878 | Ch13 region 1 |
| chr13:29351028-29351068 | Ch13 region 1 |
| chr13:29351197-29351237 | Ch13 region 1 |
| chr13:29351299-29351339 | Ch13 region 1 |
| chr13:29351456-29351496 | Ch13 region 1 |
| chr13:29351509-29351549 | Ch13 region 1 |
| chr13:29351726-29351766 | Ch13 region 1 |
| chr13:29351911-29351951 | Ch13 region 1 |
| chr13:29351969-29352009 | Ch13 region 1 |
| chr13:29352041-29352081 | Ch13 region 1 |
| chr13:29352157-29352197 | Ch13 region 1 |
| chr13:29352217-29352257 | Ch13 region 1 |
| chr13:29352264-29352304 | Ch13 region 1 |
| chr13:29352335-29352375 | Ch13 region 1 |
| chr13:29352479-29352519 | Ch13 region 1 |
| chr13:29352565-29352605 | Ch13 region 1 |
| chr13:29352615-29352655 | Ch13 region 1 |
| chr13:29352705-29352745 | Ch13 region 1 |
| chr13:29352774-29352814 | Ch13 region 1 |
| chr13:29352836-29352876 | Ch13 region 1 |
| chr13:29352882-29352922 | Ch13 region 1 |
| chr13:29352959-29352999 | Ch13 region 1 |
| chr13:29353039-29353079 | Ch13 region 1 |
| chr13:29353089-29353129 | Ch13 region 1 |
| chr13:29353147-29353187 | Ch13 region 1 |
| chr13:29353195-29353235 | Ch13 region 1 |

| | |
|---|---|
| chr13:29353284-29353324 | Ch13 region 1 |
| chr13:29353330-29353370 | Ch13 region 1 |
| chr13:29353435-29353475 | Ch13 region 1 |
| chr13:29353606-29353646 | Ch13 region 1 |
| chr13:29353728-29353768 | Ch13 region 1 |
| chr13:29353830-29353870 | Ch13 region 1 |
| chr13:29354049-29354089 | Ch13 region 1 |
| chr13:29354120-29354160 | Ch13 region 1 |
| chr13:29354199-29354239 | Ch13 region 1 |
| chr13:29354311-29354351 | Ch13 region 1 |
| chr13:29354391-29354431 | Ch13 region 1 |
| chr13:29354571-29354611 | Ch13 region 1 |
| chr13:29354640-29354680 | Ch13 region 1 |
| chr13:29354739-29354779 | Ch13 region 1 |
| chr13:29354960-29355000 | Ch13 region 1 |
| chr13:29355115-29355155 | Ch13 region 1 |
| chr13:29355265-29355305 | Ch13 region 1 |
| chr13:29355372-29355412 | Ch13 region 1 |
| chr13:29355552-29355592 | Ch13 region 1 |
| chr13:29355636-29355676 | Ch13 region 1 |
| chr13:29355810-29355850 | Ch13 region 1 |
| chr13:29355893-29355933 | Ch13 region 1 |
| chr13:29356118-29356158 | Ch13 region 1 |
| chr13:29356216-29356256 | Ch13 region 1 |
| chr13:29356306-29356346 | Ch13 region 1 |
| chr13:29356379-29356419 | Ch13 region 1 |
| chr13:29356457-29356497 | Ch13 region 1 |
| chr13:29356625-29356665 | Ch13 region 1 |
| chr13:29356840-29356880 | Ch13 region 1 |
| chr13:29357081-29357121 | Ch13 region 1 |
| chr13:29357201-29357241 | Ch13 region 1 |
| chr13:29357436-29357476 | Ch13 region 1 |
| chr13:29357481-29357521 | Ch13 region 1 |
| chr13:29357764-29357804 | Ch13 region 1 |
| chr13:29357844-29357884 | Ch13 region 1 |
| chr13:29358008-29358048 | Ch13 region 1 |
| chr13:29358185-29358225 | Ch13 region 1 |
| chr13:29358258-29358298 | Ch13 region 1 |
| chr13:29358360-29358400 | Ch13 region 1 |
| chr13:29358465-29358505 | Ch13 region 1 |
| chr13:29358515-29358555 | Ch13 region 1 |
| chr13:29358566-29358606 | Ch13 region 1 |
| chr13:29358757-29358797 | Ch13 region 1 |
| chr13:29358813-29358853 | Ch13 region 1 |
| chr13:29358937-29358977 | Ch13 region 1 |
| chr13:29359034-29359074 | Ch13 region 1 |
| chr13:29359268-29359308 | Ch13 region 1 |
| chr13:29359438-29359478 | Ch13 region 1 |
| chr13:29359687-29359727 | Ch13 region 1 |
| chr13:29359758-29359798 | Ch13 region 1 |
| chr13:29359985-29360025 | Ch13 region 1 |
| chr13:29360117-29360157 | Ch13 region 1 |

| | |
|---|---|
| chr13:29360225-29360265 | Ch13 region 1 |
| chr13:29360356-29360396 | Ch13 region 1 |
| chr13:29360450-29360490 | Ch13 region 1 |
| chr13:29360713-29360753 | Ch13 region 1 |
| chr13:29360829-29360869 | Ch13 region 1 |
| chr13:29360895-29360935 | Ch13 region 1 |
| chr13:29360941-29360981 | Ch13 region 1 |
| chr13:29361149-29361189 | Ch13 region 1 |
| chr13:29361219-29361259 | Ch13 region 1 |
| chr13:29361402-29361442 | Ch13 region 1 |
| chr13:29361501-29361541 | Ch13 region 1 |
| chr13:29361670-29361710 | Ch13 region 1 |
| chr13:29361715-29361755 | Ch13 region 1 |
| chr13:29361927-29361967 | Ch13 region 1 |
| chr13:29362066-29362106 | Ch13 region 1 |
| chr13:29362141-29362181 | Ch13 region 1 |
| chr13:29362218-29362258 | Ch13 region 1 |
| chr13:29362324-29362364 | Ch13 region 1 |
| chr13:29362489-29362529 | Ch13 region 1 |
| chr13:29362668-29362708 | Ch13 region 1 |
| chr13:29362905-29362945 | Ch13 region 1 |
| chr13:29363159-29363199 | Ch13 region 1 |
| chr13:29363264-29363304 | Ch13 region 1 |
| chr13:29363470-29363510 | Ch13 region 1 |
| chr13:29363548-29363588 | Ch13 region 1 |
| chr13:29363750-29363790 | Ch13 region 1 |
| chr13:29363926-29363966 | Ch13 region 1 |
| chr13:29364045-29364085 | Ch13 region 1 |
| chr13:29364164-29364204 | Ch13 region 1 |
| chr13:29364279-29364319 | Ch13 region 1 |
| chr13:29364337-29364377 | Ch13 region 1 |
| chr13:29364462-29364502 | Ch13 region 1 |
| chr13:29364615-29364655 | Ch13 region 1 |
| chr13:29364662-29364702 | Ch13 region 1 |
| chr13:29364765-29364805 | Ch13 region 1 |
| chr13:29364831-29364871 | Ch13 region 1 |
| chr13:29364877-29364917 | Ch13 region 1 |
| chr13:29364988-29365028 | Ch13 region 1 |
| chr13:29365107-29365147 | Ch13 region 1 |
| chr13:29365173-29365213 | Ch13 region 1 |
| chr13:29365218-29365258 | Ch13 region 1 |
| chr13:29365300-29365340 | Ch13 region 1 |
| chr13:29365370-29365410 | Ch13 region 1 |
| chr13:29365473-29365513 | Ch13 region 1 |
| chr13:29365587-29365627 | Ch13 region 1 |
| chr13:29365683-29365723 | Ch13 region 1 |
| chr13:29365751-29365791 | Ch13 region 1 |
| chr13:29365826-29365866 | Ch13 region 1 |
| chr13:29365930-29365970 | Ch13 region 1 |
| chr13:29365985-29366025 | Ch13 region 1 |
| chr13:29366042-29366082 | Ch13 region 1 |
| chr13:29366091-29366131 | Ch13 region 1 |

| | |
|---|---|
| chr13:29366139-29366179 | Ch13 region 1 |
| chr13:29366207-29366247 | Ch13 region 1 |
| chr13:29366277-29366317 | Ch13 region 1 |
| chr13:29366359-29366399 | Ch13 region 1 |
| chr13:29366431-29366471 | Ch13 region 1 |
| chr13:29366491-29366531 | Ch13 region 1 |
| chr13:29366540-29366580 | Ch13 region 1 |
| chr13:29366595-29366635 | Ch13 region 1 |
| chr13:29366676-29366716 | Ch13 region 1 |
| chr13:29366743-29366783 | Ch13 region 1 |
| chr13:29366795-29366835 | Ch13 region 1 |
| chr13:29366858-29366898 | Ch13 region 1 |
| chr13:29366907-29366947 | Ch13 region 1 |
| chr13:29366997-29367037 | Ch13 region 1 |
| chr13:29367055-29367095 | Ch13 region 1 |
| chr13:29367108-29367148 | Ch13 region 1 |
| chr13:29367154-29367194 | Ch13 region 1 |
| chr13:29367228-29367268 | Ch13 region 1 |
| chr13:29367302-29367342 | Ch13 region 1 |
| chr13:29367367-29367407 | Ch13 region 1 |
| chr13:29367420-29367460 | Ch13 region 1 |
| chr13:29367465-29367505 | Ch13 region 1 |
| chr13:29367516-29367556 | Ch13 region 1 |
| chr13:29367587-29367627 | Ch13 region 1 |
| chr13:29367642-29367682 | Ch13 region 1 |
| chr13:29367718-29367758 | Ch13 region 1 |
| chr13:29367806-29367846 | Ch13 region 1 |
| chr13:29367986-29368026 | Ch13 region 1 |
| chr13:29368032-29368072 | Ch13 region 1 |
| chr13:29368245-29368285 | Ch13 region 1 |
| chr13:29368306-29368346 | Ch13 region 1 |
| chr13:29368367-29368407 | Ch13 region 1 |
| chr13:29368454-29368494 | Ch13 region 1 |
| chr13:29368531-29368571 | Ch13 region 1 |
| chr13:29368593-29368633 | Ch13 region 1 |
| chr13:29368684-29368724 | Ch13 region 1 |
| chr13:29368742-29368782 | Ch13 region 1 |
| chr13:29368787-29368827 | Ch13 region 1 |
| chr13:29368858-29368898 | Ch13 region 1 |
| chr13:29368903-29368943 | Ch13 region 1 |
| chr13:29368978-29369018 | Ch13 region 1 |
| chr13:29369043-29369083 | Ch13 region 1 |
| chr13:29369141-29369181 | Ch13 region 1 |
| chr13:29369228-29369268 | Ch13 region 1 |
| chr13:29369304-29369344 | Ch13 region 1 |
| chr13:29369363-29369403 | Ch13 region 1 |
| chr13:29369411-29369451 | Ch13 region 1 |
| chr13:29369484-29369524 | Ch13 region 1 |
| chr13:29369660-29369700 | Ch13 region 1 |
| chr13:29369752-29369792 | Ch13 region 1 |
| chr13:29369817-29369857 | Ch13 region 1 |
| chr13:29369879-29369919 | Ch13 region 1 |

| | |
|---|---|
| chr13:29369936-29369976 | Ch13 region 1 |
| chr13:29369990-29370030 | Ch13 region 1 |
| chr13:29370038-29370078 | Ch13 region 1 |
| chr13:29370083-29370123 | Ch13 region 1 |
| chr13:29370136-29370176 | Ch13 region 1 |
| chr13:29370224-29370264 | Ch13 region 1 |
| chr13:29370296-29370336 | Ch13 region 1 |
| chr13:29370348-29370388 | Ch13 region 1 |
| chr13:29370400-29370440 | Ch13 region 1 |
| chr13:29370482-29370522 | Ch13 region 1 |
| chr13:29370558-29370598 | Ch13 region 1 |
| chr13:29370642-29370682 | Ch13 region 1 |
| chr13:29370719-29370759 | Ch13 region 1 |
| chr13:29370906-29370946 | Ch13 region 1 |
| chr13:29370982-29371022 | Ch13 region 1 |
| chr13:29371061-29371101 | Ch13 region 1 |
| chr13:29371112-29371152 | Ch13 region 1 |
| chr13:29371164-29371204 | Ch13 region 1 |
| chr13:29371240-29371280 | Ch13 region 1 |
| chr13:29371299-29371339 | Ch13 region 1 |
| chr13:29371349-29371389 | Ch13 region 1 |
| chr13:29371678-29371718 | Ch13 region 1 |
| chr13:29371732-29371772 | Ch13 region 1 |
| chr13:29371810-29371850 | Ch13 region 1 |
| chr13:29371882-29371922 | Ch13 region 1 |
| chr13:29371984-29372024 | Ch13 region 1 |
| chr13:29372063-29372103 | Ch13 region 1 |
| chr13:29372287-29372327 | Ch13 region 1 |
| chr13:29372348-29372388 | Ch13 region 1 |
| chr13:29372398-29372438 | Ch13 region 1 |
| chr13:29372475-29372515 | Ch13 region 1 |
| chr13:29372520-29372560 | Ch13 region 1 |
| chr13:29372565-29372605 | Ch13 region 1 |
| chr13:29372641-29372681 | Ch13 region 1 |
| chr13:29372742-29372782 | Ch13 region 1 |
| chr13:29372805-29372845 | Ch13 region 1 |
| chr13:29372959-29372999 | Ch13 region 1 |
| chr13:29373165-29373205 | Ch13 region 1 |
| chr13:29373211-29373251 | Ch13 region 1 |
| chr13:29373315-29373355 | Ch13 region 1 |
| chr13:29373400-29373440 | Ch13 region 1 |
| chr13:29373485-29373525 | Ch13 region 1 |
| chr13:29373561-29373601 | Ch13 region 1 |
| chr13:29373653-29373693 | Ch13 region 1 |
| chr13:29373704-29373744 | Ch13 region 1 |
| chr13:29373765-29373805 | Ch13 region 1 |
| chr13:29373951-29373991 | Ch13 region 1 |
| chr13:29374099-29374139 | Ch13 region 1 |
| chr13:29374263-29374303 | Ch13 region 1 |
| chr13:29374337-29374377 | Ch13 region 1 |
| chr13:29374553-29374593 | Ch13 region 1 |
| chr13:29374621-29374661 | Ch13 region 1 |

| | |
|---|---|
| chr13:29374670-29374710 | Ch13 region 1 |
| chr13:29374879-29374919 | Ch13 region 1 |
| chr13:29374925-29374965 | Ch13 region 1 |
| chr13:29374989-29375029 | Ch13 region 1 |
| chr13:29375047-29375087 | Ch13 region 1 |
| chr13:29375105-29375145 | Ch13 region 1 |
| chr13:29375178-29375218 | Ch13 region 1 |
| chr13:29375223-29375263 | Ch13 region 1 |
| chr13:29375315-29375355 | Ch13 region 1 |
| chr13:29375368-29375408 | Ch13 region 1 |
| chr13:29375439-29375479 | Ch13 region 1 |
| chr13:29375556-29375596 | Ch13 region 1 |
| chr13:29375601-29375641 | Ch13 region 1 |
| chr13:29375671-29375711 | Ch13 region 1 |
| chr13:29375840-29375880 | Ch13 region 1 |
| chr13:29375886-29375926 | Ch13 region 1 |
| chr13:29376025-29376065 | Ch13 region 1 |
| chr13:29376088-29376128 | Ch13 region 1 |
| chr13:29376160-29376200 | Ch13 region 1 |
| chr13:29376212-29376252 | Ch13 region 1 |
| chr13:29376261-29376301 | Ch13 region 1 |
| chr13:29376349-29376389 | Ch13 region 1 |
| chr13:29376413-29376453 | Ch13 region 1 |
| chr13:29376474-29376514 | Ch13 region 1 |
| chr13:29376522-29376562 | Ch13 region 1 |
| chr13:29376594-29376634 | Ch13 region 1 |
| chr13:29376676-29376716 | Ch13 region 1 |
| chr13:29376722-29376762 | Ch13 region 1 |
| chr13:29376831-29376871 | Ch13 region 1 |
| chr13:29376899-29376939 | Ch13 region 1 |
| chr13:29376963-29377003 | Ch13 region 1 |
| chr13:29377045-29377085 | Ch13 region 1 |
| chr13:29377090-29377130 | Ch13 region 1 |
| chr13:29377136-29377176 | Ch13 region 1 |
| chr13:29377233-29377273 | Ch13 region 1 |
| chr13:29377298-29377338 | Ch13 region 1 |
| chr13:29377374-29377414 | Ch13 region 1 |
| chr13:29377420-29377460 | Ch13 region 1 |
| chr13:29377486-29377526 | Ch13 region 1 |
| chr13:29377577-29377617 | Ch13 region 1 |
| chr13:29377640-29377680 | Ch13 region 1 |
| chr13:29377700-29377740 | Ch13 region 1 |
| chr13:29377876-29377916 | Ch13 region 1 |
| chr13:29377921-29377961 | Ch13 region 1 |
| chr13:29377980-29378020 | Ch13 region 1 |
| chr13:29378144-29378184 | Ch13 region 1 |
| chr13:29378211-29378251 | Ch13 region 1 |
| chr13:29378289-29378329 | Ch13 region 1 |
| chr13:29378334-29378374 | Ch13 region 1 |
| chr13:29378380-29378420 | Ch13 region 1 |
| chr13:29378451-29378491 | Ch13 region 1 |
| chr13:29378499-29378539 | Ch13 region 1 |

| | |
|---|---|
| chr13:29378557-29378597 | Ch13 region 1 |
| chr13:29378675-29378715 | Ch13 region 1 |
| chr13:29378732-29378772 | Ch13 region 1 |
| chr13:29378792-29378832 | Ch13 region 1 |
| chr13:29378837-29378877 | Ch13 region 1 |
| chr13:29378909-29378949 | Ch13 region 1 |
| chr13:29379018-29379058 | Ch13 region 1 |
| chr13:29379139-29379179 | Ch13 region 1 |
| chr13:29379215-29379255 | Ch13 region 1 |
| chr13:29379270-29379310 | Ch13 region 1 |
| chr13:29379349-29379389 | Ch13 region 1 |
| chr13:29379512-29379552 | Ch13 region 1 |
| chr13:29379582-29379622 | Ch13 region 1 |
| chr13:29379657-29379697 | Ch13 region 1 |
| chr13:29379705-29379745 | Ch13 region 1 |
| chr13:29379753-29379793 | Ch13 region 1 |
| chr13:29379821-29379861 | Ch13 region 1 |
| chr13:29379874-29379914 | Ch13 region 1 |
| chr13:29379947-29379987 | Ch13 region 1 |
| chr13:29380105-29380145 | Ch13 region 1 |
| chr13:29380286-29380326 | Ch13 region 1 |
| chr13:29380369-29380409 | Ch13 region 1 |
| chr13:29380424-29380464 | Ch13 region 1 |
| chr13:29380477-29380517 | Ch13 region 1 |
| chr13:29380529-29380569 | Ch13 region 1 |
| chr13:29380595-29380635 | Ch13 region 1 |
| chr13:29380657-29380697 | Ch13 region 1 |
| chr13:29380741-29380781 | Ch13 region 1 |
| chr13:29380786-29380826 | Ch13 region 1 |
| chr13:29380850-29380890 | Ch13 region 1 |
| chr13:29380907-29380947 | Ch13 region 1 |
| chr13:29380972-29381012 | Ch13 region 1 |
| chr13:29381056-29381096 | Ch13 region 1 |
| chr13:29381109-29381149 | Ch13 region 1 |
| chr13:29381190-29381230 | Ch13 region 1 |
| chr13:29381237-29381277 | Ch13 region 1 |
| chr13:29381294-29381334 | Ch13 region 1 |
| chr13:29381339-29381379 | Ch13 region 1 |
| chr13:29381389-29381429 | Ch13 region 1 |
| chr13:29381445-29381485 | Ch13 region 1 |
| chr13:29381504-29381544 | Ch13 region 1 |
| chr13:29381560-29381600 | Ch13 region 1 |
| chr13:29381637-29381677 | Ch13 region 1 |
| chr13:29381714-29381754 | Ch13 region 1 |
| chr13:29381769-29381809 | Ch13 region 1 |
| chr13:29381816-29381856 | Ch13 region 1 |
| chr13:29381871-29381911 | Ch13 region 1 |
| chr13:29381941-29381981 | Ch13 region 1 |
| chr13:29382003-29382043 | Ch13 region 1 |
| chr13:29382096-29382136 | Ch13 region 1 |
| chr13:29382145-29382185 | Ch13 region 1 |
| chr13:29382302-29382342 | Ch13 region 1 |

| | |
|---|---|
| chr13:29382347-29382387 | Ch13 region 1 |
| chr13:29382551-29382591 | Ch13 region 1 |
| chr13:29382670-29382710 | Ch13 region 1 |
| chr13:29382724-29382764 | Ch13 region 1 |
| chr13:29382788-29382828 | Ch13 region 1 |
| chr13:29382839-29382879 | Ch13 region 1 |
| chr13:29382905-29382945 | Ch13 region 1 |
| chr13:29383030-29383070 | Ch13 region 1 |
| chr13:29383105-29383145 | Ch13 region 1 |
| chr13:29383173-29383213 | Ch13 region 1 |
| chr13:29383252-29383292 | Ch13 region 1 |
| chr13:29383325-29383365 | Ch13 region 1 |
| chr13:29383465-29383505 | Ch13 region 1 |
| chr13:29383578-29383618 | Ch13 region 1 |
| chr13:29383669-29383709 | Ch13 region 1 |
| chr13:29383773-29383813 | Ch13 region 1 |
| chr13:29383825-29383865 | Ch13 region 1 |
| chr13:29383935-29383975 | Ch13 region 1 |
| chr13:29383980-29384020 | Ch13 region 1 |
| chr13:29384169-29384209 | Ch13 region 1 |
| chr13:29384280-29384320 | Ch13 region 1 |
| chr13:29384383-29384423 | Ch13 region 1 |
| chr13:29384479-29384519 | Ch13 region 1 |
| chr13:29384536-29384576 | Ch13 region 1 |
| chr13:29384625-29384665 | Ch13 region 1 |
| chr13:29384731-29384771 | Ch13 region 1 |
| chr13:29384776-29384816 | Ch13 region 1 |
| chr13:29384847-29384887 | Ch13 region 1 |
| chr13:29384946-29384986 | Ch13 region 1 |
| chr13:29385086-29385126 | Ch13 region 1 |
| chr13:29385297-29385337 | Ch13 region 1 |
| chr13:29385500-29385540 | Ch13 region 1 |
| chr13:29385545-29385585 | Ch13 region 1 |
| chr13:29385656-29385696 | Ch13 region 1 |
| chr13:29385877-29385917 | Ch13 region 1 |
| chr13:29385922-29385962 | Ch13 region 1 |
| chr13:29385990-29386030 | Ch13 region 1 |
| chr13:29386113-29386153 | Ch13 region 1 |
| chr13:29386211-29386251 | Ch13 region 1 |
| chr13:29386266-29386306 | Ch13 region 1 |
| chr13:29386467-29386507 | Ch13 region 1 |
| chr13:29386523-29386563 | Ch13 region 1 |
| chr13:29386584-29386624 | Ch13 region 1 |
| chr13:29386641-29386681 | Ch13 region 1 |
| chr13:29386700-29386740 | Ch13 region 1 |
| chr13:29386965-29387005 | Ch13 region 1 |
| chr13:29387120-29387160 | Ch13 region 1 |
| chr13:29387166-29387206 | Ch13 region 1 |
| chr13:29387233-29387273 | Ch13 region 1 |
| chr13:29387291-29387331 | Ch13 region 1 |
| chr13:29387376-29387416 | Ch13 region 1 |
| chr13:29387510-29387550 | Ch13 region 1 |

| | |
|---|---|
| chr13:29387571-29387611 | Ch13 region 1 |
| chr13:29387629-29387669 | Ch13 region 1 |
| chr13:29387696-29387736 | Ch13 region 1 |
| chr13:29387762-29387802 | Ch13 region 1 |
| chr13:29387813-29387853 | Ch13 region 1 |
| chr13:29387931-29387971 | Ch13 region 1 |
| chr13:29388098-29388138 | Ch13 region 1 |
| chr13:29388180-29388220 | Ch13 region 1 |
| chr13:29388320-29388360 | Ch13 region 1 |
| chr13:29388401-29388441 | Ch13 region 1 |
| chr13:29388480-29388520 | Ch13 region 1 |
| chr13:29388525-29388565 | Ch13 region 1 |
| chr13:29388591-29388631 | Ch13 region 1 |
| chr13:29388657-29388697 | Ch13 region 1 |
| chr13:29388719-29388759 | Ch13 region 1 |
| chr13:29388789-29388829 | Ch13 region 1 |
| chr13:29388837-29388877 | Ch13 region 1 |
| chr13:29388943-29388983 | Ch13 region 1 |
| chr13:29389042-29389082 | Ch13 region 1 |
| chr13:29389148-29389188 | Ch13 region 1 |
| chr13:29389223-29389263 | Ch13 region 1 |
| chr13:29389271-29389311 | Ch13 region 1 |
| chr13:29389322-29389362 | Ch13 region 1 |
| chr13:29389377-29389417 | Ch13 region 1 |
| chr13:29389430-29389470 | Ch13 region 1 |
| chr13:29389475-29389515 | Ch13 region 1 |
| chr13:29389541-29389581 | Ch13 region 1 |
| chr13:29389613-29389653 | Ch13 region 1 |
| chr13:29389658-29389698 | Ch13 region 1 |
| chr13:29389737-29389777 | Ch13 region 1 |
| chr13:29389782-29389822 | Ch13 region 1 |
| chr13:29389855-29389895 | Ch13 region 1 |
| chr13:29389905-29389945 | Ch13 region 1 |
| chr13:29389957-29389997 | Ch13 region 1 |
| chr13:29390063-29390103 | Ch13 region 1 |
| chr13:29390248-29390288 | Ch13 region 1 |
| chr13:29390298-29390338 | Ch13 region 1 |
| chr13:29390420-29390460 | Ch13 region 1 |
| chr13:29390565-29390605 | Ch13 region 1 |
| chr13:29390768-29390808 | Ch13 region 1 |
| chr13:29390843-29390883 | Ch13 region 1 |
| chr13:29391081-29391121 | Ch13 region 1 |
| chr13:29391160-29391200 | Ch13 region 1 |
| chr13:29391214-29391254 | Ch13 region 1 |
| chr13:29391298-29391338 | Ch13 region 1 |
| chr13:29391343-29391383 | Ch13 region 1 |
| chr13:29391434-29391474 | Ch13 region 1 |
| chr13:29391491-29391531 | Ch13 region 1 |
| chr13:29391576-29391616 | Ch13 region 1 |
| chr13:29391642-29391682 | Ch13 region 1 |
| chr13:29391699-29391739 | Ch13 region 1 |
| chr13:29391755-29391795 | Ch13 region 1 |

| | |
|---|---|
| chr13:29391818-29391858 | Ch13 region 1 |
| chr13:29391900-29391940 | Ch13 region 1 |
| chr13:29391974-29392014 | Ch13 region 1 |
| chr13:29392019-29392059 | Ch13 region 1 |
| chr13:29392066-29392106 | Ch13 region 1 |
| chr13:29392111-29392151 | Ch13 region 1 |
| chr13:29392209-29392249 | Ch13 region 1 |
| chr13:29392293-29392333 | Ch13 region 1 |
| chr13:29392354-29392394 | Ch13 region 1 |
| chr13:29392457-29392497 | Ch13 region 1 |
| chr13:29392626-29392666 | Ch13 region 1 |
| chr13:29392801-29392841 | Ch13 region 1 |
| chr13:29392897-29392937 | Ch13 region 1 |
| chr13:29392950-29392990 | Ch13 region 1 |
| chr13:29393031-29393071 | Ch13 region 1 |
| chr13:29393129-29393169 | Ch13 region 1 |
| chr13:29393192-29393232 | Ch13 region 1 |
| chr13:29393237-29393277 | Ch13 region 1 |
| chr13:29393300-29393340 | Ch13 region 1 |
| chr13:29393347-29393387 | Ch13 region 1 |
| chr13:29393398-29393438 | Ch13 region 1 |
| chr13:29393447-29393487 | Ch13 region 1 |
| chr13:29393517-29393557 | Ch13 region 1 |
| chr13:29393562-29393602 | Ch13 region 1 |
| chr13:29393610-29393650 | Ch13 region 1 |
| chr13:29393672-29393712 | Ch13 region 1 |
| chr13:29393740-29393780 | Ch13 region 1 |
| chr13:29393830-29393870 | Ch13 region 1 |
| chr13:29393891-29393931 | Ch13 region 1 |
| chr13:29393985-29394025 | Ch13 region 1 |
| chr13:29394064-29394104 | Ch13 region 1 |
| chr13:29394131-29394171 | Ch13 region 1 |
| chr13:29394220-29394260 | Ch13 region 1 |
| chr13:29394267-29394307 | Ch13 region 1 |
| chr13:29394336-29394376 | Ch13 region 1 |
| chr13:29394395-29394435 | Ch13 region 1 |
| chr13:29394469-29394509 | Ch13 region 1 |
| chr13:29394543-29394583 | Ch13 region 1 |
| chr13:29394602-29394642 | Ch13 region 1 |
| chr13:29394648-29394688 | Ch13 region 1 |
| chr13:29394709-29394749 | Ch13 region 1 |
| chr13:29394759-29394799 | Ch13 region 1 |
| chr13:29394816-29394856 | Ch13 region 1 |
| chr13:29394879-29394919 | Ch13 region 1 |
| chr13:29394926-29394966 | Ch13 region 1 |
| chr13:29394987-29395027 | Ch13 region 1 |
| chr13:29395039-29395079 | Ch13 region 1 |
| chr13:29395118-29395158 | Ch13 region 1 |
| chr13:29395166-29395206 | Ch13 region 1 |
| chr13:29395250-29395290 | Ch13 region 1 |
| chr13:29395335-29395375 | Ch13 region 1 |
| chr13:29395381-29395421 | Ch13 region 1 |

| | |
|---|---|
| chr13:29395595-29395635 | Ch13 region 1 |
| chr13:29395640-29395680 | Ch13 region 1 |
| chr13:29395749-29395789 | Ch13 region 1 |
| chr13:29395794-29395834 | Ch13 region 1 |
| chr13:29395867-29395907 | Ch13 region 1 |
| chr13:29395943-29395983 | Ch13 region 1 |
| chr13:29396013-29396053 | Ch13 region 1 |
| chr13:29396064-29396104 | Ch13 region 1 |
| chr13:29396145-29396185 | Ch13 region 1 |
| chr13:29396265-29396305 | Ch13 region 1 |
| chr13:29396310-29396350 | Ch13 region 1 |
| chr13:29396355-29396395 | Ch13 region 1 |
| chr13:29396430-29396470 | Ch13 region 1 |
| chr13:29396494-29396534 | Ch13 region 1 |
| chr13:29396558-29396598 | Ch13 region 1 |
| chr13:29396616-29396656 | Ch13 region 1 |
| chr13:29396668-29396708 | Ch13 region 1 |
| chr13:29396713-29396753 | Ch13 region 1 |
| chr13:29396779-29396819 | Ch13 region 1 |
| chr13:29396850-29396890 | Ch13 region 1 |
| chr13:29396902-29396942 | Ch13 region 1 |
| chr13:29397010-29397050 | Ch13 region 1 |
| chr13:29397084-29397124 | Ch13 region 1 |
| chr13:29397157-29397197 | Ch13 region 1 |
| chr13:29397230-29397270 | Ch13 region 1 |
| chr13:29397327-29397367 | Ch13 region 1 |
| chr13:29397379-29397419 | Ch13 region 1 |
| chr13:29397476-29397516 | Ch13 region 1 |
| chr13:29397534-29397574 | Ch13 region 1 |
| chr13:29397585-29397625 | Ch13 region 1 |
| chr13:29397670-29397710 | Ch13 region 1 |
| chr13:29397749-29397789 | Ch13 region 1 |
| chr13:29397802-29397842 | Ch13 region 1 |
| chr13:29398019-29398059 | Ch13 region 1 |
| chr13:29398065-29398105 | Ch13 region 1 |
| chr13:29398196-29398236 | Ch13 region 1 |
| chr13:29398251-29398291 | Ch13 region 1 |
| chr13:29398330-29398370 | Ch13 region 1 |
| chr13:29398411-29398451 | Ch13 region 1 |
| chr13:29398462-29398502 | Ch13 region 1 |
| chr13:29398549-29398589 | Ch13 region 1 |
| chr13:29398595-29398635 | Ch13 region 1 |
| chr13:29398640-29398680 | Ch13 region 1 |
| chr13:29398687-29398727 | Ch13 region 1 |
| chr13:29398733-29398773 | Ch13 region 1 |
| chr13:29398848-29398888 | Ch13 region 1 |
| chr13:29398926-29398966 | Ch13 region 1 |
| chr13:29398972-29399012 | Ch13 region 1 |
| chr13:29399020-29399060 | Ch13 region 1 |
| chr13:29399065-29399105 | Ch13 region 1 |
| chr13:29399132-29399172 | Ch13 region 1 |
| chr13:29399212-29399252 | Ch13 region 1 |

| | |
|---|---|
| chr13:29399270-29399310 | Ch13 region 1 |
| chr13:29399356-29399396 | Ch13 region 1 |
| chr13:29399402-29399442 | Ch13 region 1 |
| chr13:29399471-29399511 | Ch13 region 1 |
| chr13:29399525-29399565 | Ch13 region 1 |
| chr13:29399572-29399612 | Ch13 region 1 |
| chr13:29399617-29399657 | Ch13 region 1 |
| chr13:29399681-29399721 | Ch13 region 1 |
| chr13:29399734-29399774 | Ch13 region 1 |
| chr13:29399789-29399829 | Ch13 region 1 |
| chr13:29399839-29399879 | Ch13 region 1 |
| chr13:29399887-29399927 | Ch13 region 1 |
| chr13:29399938-29399978 | Ch13 region 1 |
| chr13:29400005-29400045 | Ch13 region 1 |
| chr13:29400074-29400114 | Ch13 region 1 |
| chr13:29400146-29400186 | Ch13 region 1 |
| chr13:29400335-29400375 | Ch13 region 1 |
| chr13:29400470-29400510 | Ch13 region 1 |
| chr13:29400524-29400564 | Ch13 region 1 |
| chr13:29400592-29400632 | Ch13 region 1 |
| chr13:29400641-29400681 | Ch13 region 1 |
| chr13:29400686-29400726 | Ch13 region 1 |
| chr13:29400758-29400798 | Ch13 region 1 |
| chr13:29400836-29400876 | Ch13 region 1 |
| chr13:29400888-29400928 | Ch13 region 1 |
| chr13:29400984-29401024 | Ch13 region 1 |
| chr13:29401041-29401081 | Ch13 region 1 |
| chr13:29401115-29401155 | Ch13 region 1 |
| chr13:29401173-29401213 | Ch13 region 1 |
| chr13:29401321-29401361 | Ch13 region 1 |
| chr13:29401376-29401416 | Ch13 region 1 |
| chr13:29401484-29401524 | Ch13 region 1 |
| chr13:29401549-29401589 | Ch13 region 1 |
| chr13:29401667-29401707 | Ch13 region 1 |
| chr13:29401734-29401774 | Ch13 region 1 |
| chr13:29401820-29401860 | Ch13 region 1 |
| chr13:29401976-29402016 | Ch13 region 1 |
| chr13:29402186-29402226 | Ch13 region 1 |
| chr13:29402353-29402393 | Ch13 region 1 |
| chr13:29402533-29402573 | Ch13 region 1 |
| chr13:29402780-29402820 | Ch13 region 1 |
| chr13:29402970-29403010 | Ch13 region 1 |
| chr13:29403115-29403155 | Ch13 region 1 |
| chr13:29403217-29403257 | Ch13 region 1 |
| chr13:29403307-29403347 | Ch13 region 1 |
| chr13:29403359-29403399 | Ch13 region 1 |
| chr13:29403405-29403445 | Ch13 region 1 |
| chr13:29403474-29403514 | Ch13 region 1 |
| chr13:29403675-29403715 | Ch13 region 1 |
| chr13:29403727-29403767 | Ch13 region 1 |
| chr13:29403784-29403824 | Ch13 region 1 |
| chr13:29403847-29403887 | Ch13 region 1 |

| | |
|---|---|
| chr13:29403956-29403996 | Ch13 region 1 |
| chr13:29404015-29404055 | Ch13 region 1 |
| chr13:29404076-29404116 | Ch13 region 1 |
| chr13:29404136-29404176 | Ch13 region 1 |
| chr13:29404215-29404255 | Ch13 region 1 |
| chr13:29404296-29404336 | Ch13 region 1 |
| chr13:29404363-29404403 | Ch13 region 1 |
| chr13:29404410-29404450 | Ch13 region 1 |
| chr13:29404502-29404542 | Ch13 region 1 |
| chr13:29404649-29404689 | Ch13 region 1 |
| chr13:29404737-29404777 | Ch13 region 1 |
| chr13:29404835-29404875 | Ch13 region 1 |
| chr13:29404880-29404920 | Ch13 region 1 |
| chr13:29404978-29405018 | Ch13 region 1 |
| chr13:29405116-29405156 | Ch13 region 1 |
| chr13:29405183-29405223 | Ch13 region 1 |
| chr13:29405331-29405371 | Ch13 region 1 |
| chr13:29405454-29405494 | Ch13 region 1 |
| chr13:29405508-29405548 | Ch13 region 1 |
| chr13:29405591-29405631 | Ch13 region 1 |
| chr13:29405700-29405740 | Ch13 region 1 |
| chr13:29405756-29405796 | Ch13 region 1 |
| chr13:29405815-29405855 | Ch13 region 1 |
| chr13:29406102-29406142 | Ch13 region 1 |
| chr13:29406187-29406227 | Ch13 region 1 |
| chr13:29406257-29406297 | Ch13 region 1 |
| chr13:29406308-29406348 | Ch13 region 1 |
| chr13:29406533-29406573 | Ch13 region 1 |
| chr13:29406658-29406698 | Ch13 region 1 |
| chr13:29407094-29407134 | Ch13 region 1 |
| chr13:29407162-29407202 | Ch13 region 1 |
| chr13:29407237-29407277 | Ch13 region 1 |
| chr13:29407296-29407336 | Ch13 region 1 |
| chr13:29407347-29407387 | Ch13 region 1 |
| chr13:29407471-29407511 | Ch13 region 1 |
| chr13:29407651-29407691 | Ch13 region 1 |
| chr13:29407729-29407769 | Ch13 region 1 |
| chr13:29407788-29407828 | Ch13 region 1 |
| chr13:29407921-29407961 | Ch13 region 1 |
| chr13:29407967-29408007 | Ch13 region 1 |
| chr13:29408209-29408249 | Ch13 region 1 |
| chr13:29408257-29408297 | Ch13 region 1 |
| chr13:29408341-29408381 | Ch13 region 1 |
| chr13:29408407-29408447 | Ch13 region 1 |
| chr13:29408471-29408511 | Ch13 region 1 |
| chr13:29408599-29408639 | Ch13 region 1 |
| chr13:29408758-29408798 | Ch13 region 1 |
| chr13:29408832-29408872 | Ch13 region 1 |
| chr13:29408945-29408985 | Ch13 region 1 |
| chr13:29409052-29409092 | Ch13 region 1 |
| chr13:29409158-29409198 | Ch13 region 1 |
| chr13:29409229-29409269 | Ch13 region 1 |

| | |
|---|---|
| chr13:29409339-29409379 | Ch13 region 1 |
| chr13:29409454-29409494 | Ch13 region 1 |
| chr13:29409530-29409570 | Ch13 region 1 |
| chr13:29409584-29409624 | Ch13 region 1 |
| chr13:29409646-29409686 | Ch13 region 1 |
| chr13:29409717-29409757 | Ch13 region 1 |
| chr13:29409771-29409811 | Ch13 region 1 |
| chr13:29409823-29409863 | Ch13 region 1 |
| chr13:29409869-29409909 | Ch13 region 1 |
| chr13:29409918-29409958 | Ch13 region 1 |
| chr13:29410002-29410042 | Ch13 region 1 |
| chr13:29410091-29410131 | Ch13 region 1 |
| chr13:29410136-29410176 | Ch13 region 1 |
| chr13:29410211-29410251 | Ch13 region 1 |
| chr13:29410357-29410397 | Ch13 region 1 |
| chr13:29410457-29410497 | Ch13 region 1 |
| chr13:29410522-29410562 | Ch13 region 1 |
| chr13:29410601-29410641 | Ch13 region 1 |
| chr13:29410666-29410706 | Ch13 region 1 |
| chr13:29410726-29410766 | Ch13 region 1 |
| chr13:29410787-29410827 | Ch13 region 1 |
| chr13:29410855-29410895 | Ch13 region 1 |
| chr13:29410903-29410943 | Ch13 region 1 |
| chr13:29410968-29411008 | Ch13 region 1 |
| chr13:29411031-29411071 | Ch13 region 1 |
| chr13:29411081-29411121 | Ch13 region 1 |
| chr13:29411126-29411166 | Ch13 region 1 |
| chr13:29411183-29411223 | Ch13 region 1 |
| chr13:29411244-29411284 | Ch13 region 1 |
| chr13:29411292-29411332 | Ch13 region 1 |
| chr13:29411351-29411391 | Ch13 region 1 |
| chr13:29411429-29411469 | Ch13 region 1 |
| chr13:29411474-29411514 | Ch13 region 1 |
| chr13:29411519-29411559 | Ch13 region 1 |
| chr13:29411613-29411653 | Ch13 region 1 |
| chr13:29411658-29411698 | Ch13 region 1 |
| chr13:29411703-29411743 | Ch13 region 1 |
| chr13:29411760-29411800 | Ch13 region 1 |
| chr13:29411839-29411879 | Ch13 region 1 |
| chr13:29411906-29411946 | Ch13 region 1 |
| chr13:29411978-29412018 | Ch13 region 1 |
| chr13:29412037-29412077 | Ch13 region 1 |
| chr13:29412089-29412129 | Ch13 region 1 |
| chr13:29412168-29412208 | Ch13 region 1 |
| chr13:29412221-29412261 | Ch13 region 1 |
| chr13:29412289-29412329 | Ch13 region 1 |
| chr13:29412369-29412409 | Ch13 region 1 |
| chr13:29412433-29412473 | Ch13 region 1 |
| chr13:29412498-29412538 | Ch13 region 1 |
| chr13:29412543-29412583 | Ch13 region 1 |
| chr13:29412588-29412628 | Ch13 region 1 |
| chr13:29412644-29412684 | Ch13 region 1 |

| | |
|---|---|
| chr13:29412692-29412732 | Ch13 region 1 |
| chr13:29412755-29412795 | Ch13 region 1 |
| chr13:29412829-29412869 | Ch13 region 1 |
| chr13:29412908-29412948 | Ch13 region 1 |
| chr13:29412963-29413003 | Ch13 region 1 |
| chr13:29413008-29413048 | Ch13 region 1 |
| chr13:29413058-29413098 | Ch13 region 1 |
| chr13:29413169-29413209 | Ch13 region 1 |
| chr13:29413220-29413260 | Ch13 region 1 |
| chr13:29413277-29413317 | Ch13 region 1 |
| chr13:29413401-29413441 | Ch13 region 1 |
| chr13:29413506-29413546 | Ch13 region 1 |
| chr13:29413644-29413684 | Ch13 region 1 |
| chr13:29413699-29413739 | Ch13 region 1 |
| chr13:29413762-29413802 | Ch13 region 1 |
| chr13:29413903-29413943 | Ch13 region 1 |
| chr13:29413969-29414009 | Ch13 region 1 |
| chr13:29414014-29414054 | Ch13 region 1 |
| chr13:29414059-29414099 | Ch13 region 1 |
| chr13:29414179-29414219 | Ch13 region 1 |
| chr13:29414231-29414271 | Ch13 region 1 |
| chr13:29414322-29414362 | Ch13 region 1 |
| chr13:29414375-29414415 | Ch13 region 1 |
| chr13:29414420-29414460 | Ch13 region 1 |
| chr13:29414473-29414513 | Ch13 region 1 |
| chr13:29414534-29414574 | Ch13 region 1 |
| chr13:29414585-29414625 | Ch13 region 1 |
| chr13:29414728-29414768 | Ch13 region 1 |
| chr13:29414843-29414883 | Ch13 region 1 |
| chr13:29414926-29414966 | Ch13 region 1 |
| chr13:29414971-29415011 | Ch13 region 1 |
| chr13:29415030-29415070 | Ch13 region 1 |
| chr13:29415082-29415122 | Ch13 region 1 |
| chr13:29415148-29415188 | Ch13 region 1 |
| chr13:29415231-29415271 | Ch13 region 1 |
| chr13:29415350-29415390 | Ch13 region 1 |
| chr13:29415431-29415471 | Ch13 region 1 |
| chr13:29415476-29415516 | Ch13 region 1 |
| chr13:29415532-29415572 | Ch13 region 1 |
| chr13:29415622-29415662 | Ch13 region 1 |
| chr13:29415705-29415745 | Ch13 region 1 |
| chr13:29415750-29415790 | Ch13 region 1 |
| chr13:29415970-29416010 | Ch13 region 1 |
| chr13:29416113-29416153 | Ch13 region 1 |
| chr13:29416201-29416241 | Ch13 region 1 |
| chr13:29416275-29416315 | Ch13 region 1 |
| chr13:29416345-29416385 | Ch13 region 1 |
| chr13:29416401-29416441 | Ch13 region 1 |
| chr13:29416499-29416539 | Ch13 region 1 |
| chr13:29416585-29416625 | Ch13 region 1 |
| chr13:29416668-29416708 | Ch13 region 1 |
| chr13:29416727-29416767 | Ch13 region 1 |

| | |
|---|---|
| chr13:29416791-29416831 | Ch13 region 1 |
| chr13:29416855-29416895 | Ch13 region 1 |
| chr13:29416916-29416956 | Ch13 region 1 |
| chr13:29416991-29417031 | Ch13 region 1 |
| chr13:29417053-29417093 | Ch13 region 1 |
| chr13:29417098-29417138 | Ch13 region 1 |
| chr13:29417155-29417195 | Ch13 region 1 |
| chr13:29417210-29417250 | Ch13 region 1 |
| chr13:29417255-29417295 | Ch13 region 1 |
| chr13:29417306-29417346 | Ch13 region 1 |
| chr13:29417357-29417397 | Ch13 region 1 |
| chr13:29417458-29417498 | Ch13 region 1 |
| chr13:29417512-29417552 | Ch13 region 1 |
| chr13:29417606-29417646 | Ch13 region 1 |
| chr13:29417661-29417701 | Ch13 region 1 |
| chr13:29417719-29417759 | Ch13 region 1 |
| chr13:29417815-29417855 | Ch13 region 1 |
| chr13:29417889-29417929 | Ch13 region 1 |
| chr13:29417940-29417980 | Ch13 region 1 |
| chr13:29418047-29418087 | Ch13 region 1 |
| chr13:29418099-29418139 | Ch13 region 1 |
| chr13:29418235-29418275 | Ch13 region 1 |
| chr13:29418288-29418328 | Ch13 region 1 |
| chr13:29418333-29418373 | Ch13 region 1 |
| chr13:29418429-29418469 | Ch13 region 1 |
| chr13:29418505-29418545 | Ch13 region 1 |
| chr13:29418562-29418602 | Ch13 region 1 |
| chr13:29418631-29418671 | Ch13 region 1 |
| chr13:29418681-29418721 | Ch13 region 1 |
| chr13:29418764-29418804 | Ch13 region 1 |
| chr13:29418963-29419003 | Ch13 region 1 |
| chr13:29419192-29419232 | Ch13 region 1 |
| chr13:29419304-29419344 | Ch13 region 1 |
| chr13:29419372-29419412 | Ch13 region 1 |
| chr13:29419451-29419491 | Ch13 region 1 |
| chr13:29419511-29419551 | Ch13 region 1 |
| chr13:29419556-29419596 | Ch13 region 1 |
| chr13:29419613-29419653 | Ch13 region 1 |
| chr13:29419665-29419705 | Ch13 region 1 |
| chr13:29419716-29419756 | Ch13 region 1 |
| chr13:29419763-29419803 | Ch13 region 1 |
| chr13:29419820-29419860 | Ch13 region 1 |
| chr13:29419875-29419915 | Ch13 region 1 |
| chr13:29419922-29419962 | Ch13 region 1 |
| chr13:29419967-29420007 | Ch13 region 1 |
| chr13:29420034-29420074 | Ch13 region 1 |
| chr13:29420090-29420130 | Ch13 region 1 |
| chr13:29420207-29420247 | Ch13 region 1 |
| chr13:29420262-29420302 | Ch13 region 1 |
| chr13:29420308-29420348 | Ch13 region 1 |
| chr13:29420386-29420426 | Ch13 region 1 |
| chr13:29420437-29420477 | Ch13 region 1 |

| | |
|---|---|
| chr13:29420509-29420549 | Ch13 region 1 |
| chr13:29420580-29420620 | Ch13 region 1 |
| chr13:29420660-29420700 | Ch13 region 1 |
| chr13:29420709-29420749 | Ch13 region 1 |
| chr13:29420782-29420822 | Ch13 region 1 |
| chr13:29420853-29420893 | Ch13 region 1 |
| chr13:29420919-29420959 | Ch13 region 1 |
| chr13:29420966-29421006 | Ch13 region 1 |
| chr13:29421045-29421085 | Ch13 region 1 |
| chr13:29421125-29421165 | Ch13 region 1 |
| chr13:29421170-29421210 | Ch13 region 1 |
| chr13:29421240-29421280 | Ch13 region 1 |
| chr13:29421286-29421326 | Ch13 region 1 |
| chr13:29421380-29421420 | Ch13 region 1 |
| chr13:29421457-29421497 | Ch13 region 1 |
| chr13:29421516-29421556 | Ch13 region 1 |
| chr13:29421561-29421601 | Ch13 region 1 |
| chr13:29421632-29421672 | Ch13 region 1 |
| chr13:29421680-29421720 | Ch13 region 1 |
| chr13:29421753-29421793 | Ch13 region 1 |
| chr13:29421798-29421838 | Ch13 region 1 |
| chr13:29421863-29421903 | Ch13 region 1 |
| chr13:29421925-29421965 | Ch13 region 1 |
| chr13:29422007-29422047 | Ch13 region 1 |
| chr13:29422100-29422140 | Ch13 region 1 |
| chr13:29422168-29422208 | Ch13 region 1 |
| chr13:29422248-29422288 | Ch13 region 1 |
| chr13:29422294-29422334 | Ch13 region 1 |
| chr13:29422402-29422442 | Ch13 region 1 |
| chr13:29422447-29422487 | Ch13 region 1 |
| chr13:29422497-29422537 | Ch13 region 1 |
| chr13:29422641-29422681 | Ch13 region 1 |
| chr13:29422700-29422740 | Ch13 region 1 |
| chr13:29422766-29422806 | Ch13 region 1 |
| chr13:29422815-29422855 | Ch13 region 1 |
| chr13:29422869-29422909 | Ch13 region 1 |
| chr13:29422965-29423005 | Ch13 region 1 |
| chr13:29423047-29423087 | Ch13 region 1 |
| chr13:29423109-29423149 | Ch13 region 1 |
| chr13:29423178-29423218 | Ch13 region 1 |
| chr13:29423248-29423288 | Ch13 region 1 |
| chr13:29423313-29423353 | Ch13 region 1 |
| chr13:29423378-29423418 | Ch13 region 1 |
| chr13:29423424-29423464 | Ch13 region 1 |
| chr13:29423520-29423560 | Ch13 region 1 |
| chr13:29423600-29423640 | Ch13 region 1 |
| chr13:29423646-29423686 | Ch13 region 1 |
| chr13:29423714-29423754 | Ch13 region 1 |
| chr13:29423803-29423843 | Ch13 region 1 |
| chr13:29423853-29423893 | Ch13 region 1 |
| chr13:29423927-29423967 | Ch13 region 1 |
| chr13:29423972-29424012 | Ch13 region 1 |

| | |
|---|---|
| chr13:29424032-29424072 | Ch13 region 1 |
| chr13:29424091-29424131 | Ch13 region 1 |
| chr13:29424137-29424177 | Ch13 region 1 |
| chr13:29424202-29424242 | Ch13 region 1 |
| chr13:29424264-29424304 | Ch13 region 1 |
| chr13:29424310-29424350 | Ch13 region 1 |
| chr13:29424396-29424436 | Ch13 region 1 |
| chr13:29424480-29424520 | Ch13 region 1 |
| chr13:29424580-29424620 | Ch13 region 1 |
| chr13:29424625-29424665 | Ch13 region 1 |
| chr13:29424706-29424746 | Ch13 region 1 |
| chr13:29424751-29424791 | Ch13 region 1 |
| chr13:29424810-29424850 | Ch13 region 1 |
| chr13:29424864-29424904 | Ch13 region 1 |
| chr13:29425013-29425053 | Ch13 region 1 |
| chr13:29425068-29425108 | Ch13 region 1 |
| chr13:29425115-29425155 | Ch13 region 1 |
| chr13:29425173-29425213 | Ch13 region 1 |
| chr13:29425218-29425258 | Ch13 region 1 |
| chr13:29425265-29425305 | Ch13 region 1 |
| chr13:29425332-29425372 | Ch13 region 1 |
| chr13:29425378-29425418 | Ch13 region 1 |
| chr13:29425450-29425490 | Ch13 region 1 |
| chr13:29425516-29425556 | Ch13 region 1 |
| chr13:29425596-29425636 | Ch13 region 1 |
| chr13:29425646-29425686 | Ch13 region 1 |
| chr13:29425695-29425735 | Ch13 region 1 |
| chr13:29425760-29425800 | Ch13 region 1 |
| chr13:29425828-29425868 | Ch13 region 1 |
| chr13:29425876-29425916 | Ch13 region 1 |
| chr13:29425925-29425965 | Ch13 region 1 |
| chr13:29425996-29426036 | Ch13 region 1 |
| chr13:29426052-29426092 | Ch13 region 1 |
| chr13:29426097-29426137 | Ch13 region 1 |
| chr13:29426144-29426184 | Ch13 region 1 |
| chr13:29426225-29426265 | Ch13 region 1 |
| chr13:29426275-29426315 | Ch13 region 1 |
| chr13:29426331-29426371 | Ch13 region 1 |
| chr13:29426388-29426428 | Ch13 region 1 |
| chr13:29426454-29426494 | Ch13 region 1 |
| chr13:29426541-29426581 | Ch13 region 1 |
| chr13:29426598-29426638 | Ch13 region 1 |
| chr13:29426688-29426728 | Ch13 region 1 |
| chr13:29426959-29426999 | Ch13 region 1 |
| chr13:29427123-29427163 | Ch13 region 1 |
| chr13:29427168-29427208 | Ch13 region 1 |
| chr13:29427220-29427260 | Ch13 region 1 |
| chr13:29427272-29427312 | Ch13 region 1 |
| chr13:29427317-29427357 | Ch13 region 1 |
| chr13:29427399-29427439 | Ch13 region 1 |
| chr13:29427445-29427485 | Ch13 region 1 |
| chr13:29427534-29427574 | Ch13 region 1 |

| | |
|---|---|
| chr13:29427591-29427631 | Ch13 region 1 |
| chr13:29427654-29427694 | Ch13 region 1 |
| chr13:29427705-29427745 | Ch13 region 1 |
| chr13:29427798-29427838 | Ch13 region 1 |
| chr13:29427845-29427885 | Ch13 region 1 |
| chr13:29427905-29427945 | Ch13 region 1 |
| chr13:29427973-29428013 | Ch13 region 1 |
| chr13:29428024-29428064 | Ch13 region 1 |
| chr13:29428076-29428116 | Ch13 region 1 |
| chr13:29428127-29428167 | Ch13 region 1 |
| chr13:29428212-29428252 | Ch13 region 1 |
| chr13:29428272-29428312 | Ch13 region 1 |
| chr13:29428344-29428384 | Ch13 region 1 |
| chr13:29428398-29428438 | Ch13 region 1 |
| chr13:29428461-29428501 | Ch13 region 1 |
| chr13:29428508-29428548 | Ch13 region 1 |
| chr13:29428561-29428601 | Ch13 region 1 |
| chr13:29428615-29428655 | Ch13 region 1 |
| chr13:29428708-29428748 | Ch13 region 1 |
| chr13:29428856-29428896 | Ch13 region 1 |
| chr13:29428948-29428988 | Ch13 region 1 |
| chr13:29428993-29429033 | Ch13 region 1 |
| chr13:29429064-29429104 | Ch13 region 1 |
| chr13:29429109-29429149 | Ch13 region 1 |
| chr13:29429171-29429211 | Ch13 region 1 |
| chr13:29429274-29429314 | Ch13 region 1 |
| chr13:29429348-29429388 | Ch13 region 1 |
| chr13:29429539-29429579 | Ch13 region 1 |
| chr13:29429589-29429629 | Ch13 region 1 |
| chr13:29429842-29429882 | Ch13 region 1 |
| chr13:29429936-29429976 | Ch13 region 1 |
| chr13:29430138-29430178 | Ch13 region 1 |
| chr13:29430197-29430237 | Ch13 region 1 |
| chr13:29430242-29430282 | Ch13 region 1 |
| chr13:29430325-29430365 | Ch13 region 1 |
| chr13:29430398-29430438 | Ch13 region 1 |
| chr13:29430465-29430505 | Ch13 region 1 |
| chr13:29430517-29430557 | Ch13 region 1 |
| chr13:29430569-29430609 | Ch13 region 1 |
| chr13:29430620-29430660 | Ch13 region 1 |
| chr13:29430681-29430721 | Ch13 region 1 |
| chr13:29430764-29430804 | Ch13 region 1 |
| chr13:29430845-29430885 | Ch13 region 1 |
| chr13:29430904-29430944 | Ch13 region 1 |
| chr13:29430963-29431003 | Ch13 region 1 |
| chr13:29431008-29431048 | Ch13 region 1 |
| chr13:29431093-29431133 | Ch13 region 1 |
| chr13:29431138-29431178 | Ch13 region 1 |
| chr13:29431225-29431265 | Ch13 region 1 |
| chr13:29431279-29431319 | Ch13 region 1 |
| chr13:29431324-29431364 | Ch13 region 1 |
| chr13:29431394-29431434 | Ch13 region 1 |

| | |
|---|---|
| chr13:29431439-29431479 | Ch13 region 1 |
| chr13:29431506-29431546 | Ch13 region 1 |
| chr13:29431561-29431601 | Ch13 region 1 |
| chr13:29431610-29431650 | Ch13 region 1 |
| chr13:29431710-29431750 | Ch13 region 1 |
| chr13:29431766-29431806 | Ch13 region 1 |
| chr13:29431812-29431852 | Ch13 region 1 |
| chr13:29431864-29431904 | Ch13 region 1 |
| chr13:29431940-29431980 | Ch13 region 1 |
| chr13:29431985-29432025 | Ch13 region 1 |
| chr13:29432043-29432083 | Ch13 region 1 |
| chr13:29432102-29432142 | Ch13 region 1 |
| chr13:29432163-29432203 | Ch13 region 1 |
| chr13:29432251-29432291 | Ch13 region 1 |
| chr13:29432321-29432361 | Ch13 region 1 |
| chr13:29432388-29432428 | Ch13 region 1 |
| chr13:29432452-29432492 | Ch13 region 1 |
| chr13:29432550-29432590 | Ch13 region 1 |
| chr13:29432616-29432656 | Ch13 region 1 |
| chr13:29432692-29432732 | Ch13 region 1 |
| chr13:29432743-29432783 | Ch13 region 1 |
| chr13:29432788-29432828 | Ch13 region 1 |
| chr13:29432860-29432900 | Ch13 region 1 |
| chr13:29432941-29432981 | Ch13 region 1 |
| chr13:29433018-29433058 | Ch13 region 1 |
| chr13:29433079-29433119 | Ch13 region 1 |
| chr13:29433132-29433172 | Ch13 region 1 |
| chr13:29433238-29433278 | Ch13 region 1 |
| chr13:29433284-29433324 | Ch13 region 1 |
| chr13:29433330-29433370 | Ch13 region 1 |
| chr13:29433403-29433443 | Ch13 region 1 |
| chr13:29433455-29433495 | Ch13 region 1 |
| chr13:29433528-29433568 | Ch13 region 1 |
| chr13:29433577-29433617 | Ch13 region 1 |
| chr13:29433624-29433664 | Ch13 region 1 |
| chr13:29433694-29433734 | Ch13 region 1 |
| chr13:29433739-29433779 | Ch13 region 1 |
| chr13:29433789-29433829 | Ch13 region 1 |
| chr13:29433855-29433895 | Ch13 region 1 |
| chr13:29433902-29433942 | Ch13 region 1 |
| chr13:29433981-29434021 | Ch13 region 1 |
| chr13:29434050-29434090 | Ch13 region 1 |
| chr13:29434109-29434149 | Ch13 region 1 |
| chr13:29434177-29434217 | Ch13 region 1 |
| chr13:29434249-29434289 | Ch13 region 1 |
| chr13:29434309-29434349 | Ch13 region 1 |
| chr13:29434414-29434454 | Ch13 region 1 |
| chr13:29434459-29434499 | Ch13 region 1 |
| chr13:29434532-29434572 | Ch13 region 1 |
| chr13:29434623-29434663 | Ch13 region 1 |
| chr13:29434683-29434723 | Ch13 region 1 |
| chr13:29434777-29434817 | Ch13 region 1 |

| | |
|---|---|
| chr13:29434824-29434864 | Ch13 region 1 |
| chr13:29434888-29434928 | Ch13 region 1 |
| chr13:29434963-29435003 | Ch13 region 1 |
| chr13:29435052-29435092 | Ch13 region 1 |
| chr13:29435101-29435141 | Ch13 region 1 |
| chr13:29435179-29435219 | Ch13 region 1 |
| chr13:29435251-29435291 | Ch13 region 1 |
| chr13:29435296-29435336 | Ch13 region 1 |
| chr13:29435389-29435429 | Ch13 region 1 |
| chr13:29435456-29435496 | Ch13 region 1 |
| chr13:29435507-29435547 | Ch13 region 1 |
| chr13:29435595-29435635 | Ch13 region 1 |
| chr13:29435707-29435747 | Ch13 region 1 |
| chr13:29435761-29435801 | Ch13 region 1 |
| chr13:29435820-29435860 | Ch13 region 1 |
| chr13:29435883-29435923 | Ch13 region 1 |
| chr13:29435935-29435975 | Ch13 region 1 |
| chr13:29435982-29436022 | Ch13 region 1 |
| chr13:29436034-29436074 | Ch13 region 1 |
| chr13:29436120-29436160 | Ch13 region 1 |
| chr13:29436200-29436240 | Ch13 region 1 |
| chr13:29436246-29436286 | Ch13 region 1 |
| chr13:29436297-29436337 | Ch13 region 1 |
| chr13:29436367-29436407 | Ch13 region 1 |
| chr13:29436415-29436455 | Ch13 region 1 |
| chr13:29436758-29436798 | Ch13 region 1 |
| chr13:29436814-29436854 | Ch13 region 1 |
| chr13:29436860-29436900 | Ch13 region 1 |
| chr13:29437008-29437048 | Ch13 region 1 |
| chr13:29437109-29437149 | Ch13 region 1 |
| chr13:29437188-29437228 | Ch13 region 1 |
| chr13:29437259-29437299 | Ch13 region 1 |
| chr13:29437318-29437358 | Ch13 region 1 |
| chr13:29437380-29437420 | Ch13 region 1 |
| chr13:29437430-29437470 | Ch13 region 1 |
| chr13:29437475-29437515 | Ch13 region 1 |
| chr13:29437558-29437598 | Ch13 region 1 |
| chr13:29437616-29437656 | Ch13 region 1 |
| chr13:29437661-29437701 | Ch13 region 1 |
| chr13:29437706-29437746 | Ch13 region 1 |
| chr13:29437758-29437798 | Ch13 region 1 |
| chr13:29437894-29437934 | Ch13 region 1 |
| chr13:29438124-29438164 | Ch13 region 1 |
| chr13:29438204-29438244 | Ch13 region 1 |
| chr13:29438249-29438289 | Ch13 region 1 |
| chr13:29438298-29438338 | Ch13 region 1 |
| chr13:29438347-29438387 | Ch13 region 1 |
| chr13:29438421-29438461 | Ch13 region 1 |
| chr13:29438502-29438542 | Ch13 region 1 |
| chr13:29438551-29438591 | Ch13 region 1 |
| chr13:29438599-29438639 | Ch13 region 1 |
| chr13:29438676-29438716 | Ch13 region 1 |

| | |
|---|---|
| chr13:29438721-29438761 | Ch13 region 1 |
| chr13:29438800-29438840 | Ch13 region 1 |
| chr13:29438847-29438887 | Ch13 region 1 |
| chr13:29438924-29438964 | Ch13 region 1 |
| chr13:29438969-29439009 | Ch13 region 1 |
| chr13:29439063-29439103 | Ch13 region 1 |
| chr13:29439155-29439195 | Ch13 region 1 |
| chr13:29439264-29439304 | Ch13 region 1 |
| chr13:29439319-29439359 | Ch13 region 1 |
| chr13:29439369-29439409 | Ch13 region 1 |
| chr13:29439414-29439454 | Ch13 region 1 |
| chr13:29439463-29439503 | Ch13 region 1 |
| chr13:29439515-29439555 | Ch13 region 1 |
| chr13:29439607-29439647 | Ch13 region 1 |
| chr13:29439672-29439712 | Ch13 region 1 |
| chr13:29439736-29439776 | Ch13 region 1 |
| chr13:29439816-29439856 | Ch13 region 1 |
| chr13:29439861-29439901 | Ch13 region 1 |
| chr13:29439915-29439955 | Ch13 region 1 |
| chr13:29439976-29440016 | Ch13 region 1 |
| chr13:29440027-29440067 | Ch13 region 1 |
| chr13:29440134-29440174 | Ch13 region 1 |
| chr13:29440179-29440219 | Ch13 region 1 |
| chr13:29440232-29440272 | Ch13 region 1 |
| chr13:29440277-29440317 | Ch13 region 1 |
| chr13:29440359-29440399 | Ch13 region 1 |
| chr13:29440420-29440460 | Ch13 region 1 |
| chr13:29440516-29440556 | Ch13 region 1 |
| chr13:29440574-29440614 | Ch13 region 1 |
| chr13:29440621-29440661 | Ch13 region 1 |
| chr13:29440672-29440712 | Ch13 region 1 |
| chr13:29440769-29440809 | Ch13 region 1 |
| chr13:29440865-29440905 | Ch13 region 1 |
| chr13:29440948-29440988 | Ch13 region 1 |
| chr13:29441049-29441089 | Ch13 region 1 |
| chr13:29441134-29441174 | Ch13 region 1 |
| chr13:29441190-29441230 | Ch13 region 1 |
| chr13:29441235-29441275 | Ch13 region 1 |
| chr13:29441327-29441367 | Ch13 region 1 |
| chr13:29441405-29441445 | Ch13 region 1 |
| chr13:29441537-29441577 | Ch13 region 1 |
| chr13:29441582-29441622 | Ch13 region 1 |
| chr13:29441637-29441677 | Ch13 region 1 |
| chr13:29441724-29441764 | Ch13 region 1 |
| chr13:29441817-29441857 | Ch13 region 1 |
| chr13:29441892-29441932 | Ch13 region 1 |
| chr13:29441945-29441985 | Ch13 region 1 |
| chr13:29442033-29442073 | Ch13 region 1 |
| chr13:29442101-29442141 | Ch13 region 1 |
| chr13:29442199-29442239 | Ch13 region 1 |
| chr13:29442262-29442302 | Ch13 region 1 |
| chr13:29442384-29442424 | Ch13 region 1 |

| | |
|---|---|
| chr13:29442431-29442471 | Ch13 region 1 |
| chr13:29442496-29442536 | Ch13 region 1 |
| chr13:29442548-29442588 | Ch13 region 1 |
| chr13:29442595-29442635 | Ch13 region 1 |
| chr13:29442640-29442680 | Ch13 region 1 |
| chr13:29442697-29442737 | Ch13 region 1 |
| chr13:29442763-29442803 | Ch13 region 1 |
| chr13:29442808-29442848 | Ch13 region 1 |
| chr13:29442873-29442913 | Ch13 region 1 |
| chr13:29442938-29442978 | Ch13 region 1 |
| chr13:29442985-29443025 | Ch13 region 1 |
| chr13:29443032-29443072 | Ch13 region 1 |
| chr13:29443092-29443132 | Ch13 region 1 |
| chr13:29443142-29443182 | Ch13 region 1 |
| chr13:29443233-29443273 | Ch13 region 1 |
| chr13:29443286-29443326 | Ch13 region 1 |
| chr13:29443347-29443387 | Ch13 region 1 |
| chr13:29443392-29443432 | Ch13 region 1 |
| chr13:29443438-29443478 | Ch13 region 1 |
| chr13:29443594-29443634 | Ch13 region 1 |
| chr13:29443682-29443722 | Ch13 region 1 |
| chr13:29443922-29443962 | Ch13 region 1 |
| chr13:29444139-29444179 | Ch13 region 1 |
| chr13:29444300-29444340 | Ch13 region 1 |
| chr13:29444418-29444458 | Ch13 region 1 |
| chr13:29444474-29444514 | Ch13 region 1 |
| chr13:29444520-29444560 | Ch13 region 1 |
| chr13:29444577-29444617 | Ch13 region 1 |
| chr13:29444623-29444663 | Ch13 region 1 |
| chr13:29444718-29444758 | Ch13 region 1 |
| chr13:29444776-29444816 | Ch13 region 1 |
| chr13:29444839-29444879 | Ch13 region 1 |
| chr13:29444927-29444967 | Ch13 region 1 |
| chr13:29444972-29445012 | Ch13 region 1 |
| chr13:29445081-29445121 | Ch13 region 1 |
| chr13:29445126-29445166 | Ch13 region 1 |
| chr13:29445220-29445260 | Ch13 region 1 |
| chr13:29445332-29445372 | Ch13 region 1 |
| chr13:29445377-29445417 | Ch13 region 1 |
| chr13:29445451-29445491 | Ch13 region 1 |
| chr13:29445499-29445539 | Ch13 region 1 |
| chr13:29445597-29445637 | Ch13 region 1 |
| chr13:29445689-29445729 | Ch13 region 1 |
| chr13:29445764-29445804 | Ch13 region 1 |
| chr13:29445861-29445901 | Ch13 region 1 |
| chr13:29445950-29445990 | Ch13 region 1 |
| chr13:29446020-29446060 | Ch13 region 1 |
| chr13:29446110-29446150 | Ch13 region 1 |
| chr13:29446163-29446203 | Ch13 region 1 |
| chr13:29446209-29446249 | Ch13 region 1 |
| chr13:29446296-29446336 | Ch13 region 1 |
| chr13:29446341-29446381 | Ch13 region 1 |

| | |
|---|---|
| chr13:29446390-29446430 | Ch13 region 1 |
| chr13:29446437-29446477 | Ch13 region 1 |
| chr13:29446482-29446522 | Ch13 region 1 |
| chr13:29446548-29446588 | Ch13 region 1 |
| chr13:29446629-29446669 | Ch13 region 1 |
| chr13:29446743-29446783 | Ch13 region 1 |
| chr13:29446790-29446830 | Ch13 region 1 |
| chr13:29446886-29446926 | Ch13 region 1 |
| chr13:29446953-29446993 | Ch13 region 1 |
| chr13:29447071-29447111 | Ch13 region 1 |
| chr13:29447116-29447156 | Ch13 region 1 |
| chr13:29447261-29447301 | Ch13 region 1 |
| chr13:29447306-29447346 | Ch13 region 1 |
| chr13:29447404-29447444 | Ch13 region 1 |
| chr13:29447456-29447496 | Ch13 region 1 |
| chr13:29447503-29447543 | Ch13 region 1 |
| chr13:29447578-29447618 | Ch13 region 1 |
| chr13:29447658-29447698 | Ch13 region 1 |
| chr13:29447771-29447811 | Ch13 region 1 |
| chr13:29447835-29447875 | Ch13 region 1 |
| chr13:29447885-29447925 | Ch13 region 1 |
| chr13:29448051-29448091 | Ch13 region 1 |
| chr13:29448318-29448358 | Ch13 region 1 |
| chr13:29448418-29448458 | Ch13 region 1 |
| chr13:29448518-29448558 | Ch13 region 1 |
| chr13:29448591-29448631 | Ch13 region 1 |
| chr13:29448644-29448684 | Ch13 region 1 |
| chr13:29448691-29448731 | Ch13 region 1 |
| chr13:29448766-29448806 | Ch13 region 1 |
| chr13:29448812-29448852 | Ch13 region 1 |
| chr13:29448881-29448921 | Ch13 region 1 |
| chr13:29448927-29448967 | Ch13 region 1 |
| chr13:29448998-29449038 | Ch13 region 1 |
| chr13:29449067-29449107 | Ch13 region 1 |
| chr13:29449130-29449170 | Ch13 region 1 |
| chr13:29449299-29449339 | Ch13 region 1 |
| chr13:29449523-29449563 | Ch13 region 1 |
| chr13:29449570-29449610 | Ch13 region 1 |
| chr13:29449616-29449656 | Ch13 region 1 |
| chr13:29449665-29449705 | Ch13 region 1 |
| chr13:29449711-29449751 | Ch13 region 1 |
| chr13:29449779-29449819 | Ch13 region 1 |
| chr13:29449830-29449870 | Ch13 region 1 |
| chr13:29450050-29450090 | Ch13 region 1 |
| chr13:29450117-29450157 | Ch13 region 1 |
| chr13:29450198-29450238 | Ch13 region 1 |
| chr13:29450270-29450310 | Ch13 region 1 |
| chr13:29450335-29450375 | Ch13 region 1 |
| chr13:29450391-29450431 | Ch13 region 1 |
| chr13:29450436-29450476 | Ch13 region 1 |
| chr13:29450501-29450541 | Ch13 region 1 |
| chr13:29450568-29450608 | Ch13 region 1 |

| | |
|---|---|
| chr13:29450629-29450669 | Ch13 region 1 |
| chr13:29450677-29450717 | Ch13 region 1 |
| chr13:29450722-29450762 | Ch13 region 1 |
| chr13:29450800-29450840 | Ch13 region 1 |
| chr13:29450885-29450925 | Ch13 region 1 |
| chr13:29450942-29450982 | Ch13 region 1 |
| chr13:29451040-29451080 | Ch13 region 1 |
| chr13:29451124-29451164 | Ch13 region 1 |
| chr13:29451221-29451261 | Ch13 region 1 |
| chr13:29451470-29451510 | Ch13 region 1 |
| chr13:29451560-29451600 | Ch13 region 1 |
| chr13:29451657-29451697 | Ch13 region 1 |
| chr13:29451705-29451745 | Ch13 region 1 |
| chr13:29451752-29451792 | Ch13 region 1 |
| chr13:29451820-29451860 | Ch13 region 1 |
| chr13:29451895-29451935 | Ch13 region 1 |
| chr13:29451973-29452013 | Ch13 region 1 |
| chr13:29452047-29452087 | Ch13 region 1 |
| chr13:29452124-29452164 | Ch13 region 1 |
| chr13:29452176-29452216 | Ch13 region 1 |
| chr13:29452253-29452293 | Ch13 region 1 |
| chr13:29452301-29452341 | Ch13 region 1 |
| chr13:29452457-29452497 | Ch13 region 1 |
| chr13:29452578-29452618 | Ch13 region 1 |
| chr13:29452628-29452668 | Ch13 region 1 |
| chr13:29452673-29452713 | Ch13 region 1 |
| chr13:29452890-29452930 | Ch13 region 1 |
| chr13:29452946-29452986 | Ch13 region 1 |
| chr13:29453048-29453088 | Ch13 region 1 |
| chr13:29453093-29453133 | Ch13 region 1 |
| chr13:29453158-29453198 | Ch13 region 1 |
| chr13:29453444-29453484 | Ch13 region 1 |
| chr13:29453489-29453529 | Ch13 region 1 |
| chr13:29453551-29453591 | Ch13 region 1 |
| chr13:29453599-29453639 | Ch13 region 1 |
| chr13:29453646-29453686 | Ch13 region 1 |
| chr13:29453691-29453731 | Ch13 region 1 |
| chr13:29453739-29453779 | Ch13 region 1 |
| chr13:29453829-29453869 | Ch13 region 1 |
| chr13:29453874-29453914 | Ch13 region 1 |
| chr13:29453928-29453968 | Ch13 region 1 |
| chr13:29454009-29454049 | Ch13 region 1 |
| chr13:29454058-29454098 | Ch13 region 1 |
| chr13:29454104-29454144 | Ch13 region 1 |
| chr13:29454160-29454200 | Ch13 region 1 |
| chr13:29454213-29454253 | Ch13 region 1 |
| chr13:29454284-29454324 | Ch13 region 1 |
| chr13:29454372-29454412 | Ch13 region 1 |
| chr13:29454447-29454487 | Ch13 region 1 |
| chr13:29454535-29454575 | Ch13 region 1 |
| chr13:29454580-29454620 | Ch13 region 1 |
| chr13:29454740-29454780 | Ch13 region 1 |

| | |
|---|---|
| chr13:29454793-29454833 | Ch13 region 1 |
| chr13:29454870-29454910 | Ch13 region 1 |
| chr13:29454915-29454955 | Ch13 region 1 |
| chr13:29454960-29455000 | Ch13 region 1 |
| chr13:29455079-29455119 | Ch13 region 1 |
| chr13:29455134-29455174 | Ch13 region 1 |
| chr13:29455232-29455272 | Ch13 region 1 |
| chr13:29455286-29455326 | Ch13 region 1 |
| chr13:29455421-29455461 | Ch13 region 1 |
| chr13:29455519-29455559 | Ch13 region 1 |
| chr13:29455747-29455787 | Ch13 region 1 |
| chr13:29455918-29455958 | Ch13 region 1 |
| chr13:29456017-29456057 | Ch13 region 1 |
| chr13:29456063-29456103 | Ch13 region 1 |
| chr13:29456115-29456155 | Ch13 region 1 |
| chr13:29456163-29456203 | Ch13 region 1 |
| chr13:29456332-29456372 | Ch13 region 1 |
| chr13:29456533-29456573 | Ch13 region 1 |
| chr13:29456578-29456618 | Ch13 region 1 |
| chr13:29456704-29456744 | Ch13 region 1 |
| chr13:29456939-29456979 | Ch13 region 1 |
| chr13:29457119-29457159 | Ch13 region 1 |
| chr13:29457167-29457207 | Ch13 region 1 |
| chr13:29457256-29457296 | Ch13 region 1 |
| chr13:29457313-29457353 | Ch13 region 1 |
| chr13:29457387-29457427 | Ch13 region 1 |
| chr13:29457463-29457503 | Ch13 region 1 |
| chr13:29457526-29457566 | Ch13 region 1 |
| chr13:29457582-29457622 | Ch13 region 1 |
| chr13:29457643-29457683 | Ch13 region 1 |
| chr13:29457768-29457808 | Ch13 region 1 |
| chr13:29457813-29457853 | Ch13 region 1 |
| chr13:29457878-29457918 | Ch13 region 1 |
| chr13:29457937-29457977 | Ch13 region 1 |
| chr13:29457999-29458039 | Ch13 region 1 |
| chr13:29458067-29458107 | Ch13 region 1 |
| chr13:29458184-29458224 | Ch13 region 1 |
| chr13:29458251-29458291 | Ch13 region 1 |
| chr13:29458369-29458409 | Ch13 region 1 |
| chr13:29458534-29458574 | Ch13 region 1 |
| chr13:29458588-29458628 | Ch13 region 1 |
| chr13:29458692-29458732 | Ch13 region 1 |
| chr13:29458764-29458804 | Ch13 region 1 |
| chr13:29458824-29458864 | Ch13 region 1 |
| chr13:29458888-29458928 | Ch13 region 1 |
| chr13:29459009-29459049 | Ch13 region 1 |
| chr13:29459059-29459099 | Ch13 region 1 |
| chr13:29459171-29459211 | Ch13 region 1 |
| chr13:29459230-29459270 | Ch13 region 1 |
| chr13:29459333-29459373 | Ch13 region 1 |
| chr13:29459385-29459425 | Ch13 region 1 |
| chr13:29459445-29459485 | Ch13 region 1 |

| | |
|---|---|
| chr13:29459499-29459539 | Ch13 region 1 |
| chr13:29459580-29459620 | Ch13 region 1 |
| chr13:29459627-29459667 | Ch13 region 1 |
| chr13:29459725-29459765 | Ch13 region 1 |
| chr13:29459804-29459844 | Ch13 region 1 |
| chr13:29459896-29459936 | Ch13 region 1 |
| chr13:29460005-29460045 | Ch13 region 1 |
| chr13:29460056-29460096 | Ch13 region 1 |
| chr13:29460109-29460149 | Ch13 region 1 |
| chr13:29460166-29460206 | Ch13 region 1 |
| chr13:29460249-29460289 | Ch13 region 1 |
| chr13:29460316-29460356 | Ch13 region 1 |
| chr13:29460388-29460428 | Ch13 region 1 |
| chr13:29460434-29460474 | Ch13 region 1 |
| chr13:29460512-29460552 | Ch13 region 1 |
| chr13:29460596-29460636 | Ch13 region 1 |
| chr13:29460648-29460688 | Ch13 region 1 |
| chr13:29460694-29460734 | Ch13 region 1 |
| chr13:29460743-29460783 | Ch13 region 1 |
| chr13:29460824-29460864 | Ch13 region 1 |
| chr13:29460875-29460915 | Ch13 region 1 |
| chr13:29460928-29460968 | Ch13 region 1 |
| chr13:29460986-29461026 | Ch13 region 1 |
| chr13:29461071-29461111 | Ch13 region 1 |
| chr13:29461177-29461217 | Ch13 region 1 |
| chr13:29461269-29461309 | Ch13 region 1 |
| chr13:29461343-29461383 | Ch13 region 1 |
| chr13:29461389-29461429 | Ch13 region 1 |
| chr13:29461442-29461482 | Ch13 region 1 |
| chr13:29461507-29461547 | Ch13 region 1 |
| chr13:29461585-29461625 | Ch13 region 1 |
| chr13:29461672-29461712 | Ch13 region 1 |
| chr13:29461742-29461782 | Ch13 region 1 |
| chr13:29461846-29461886 | Ch13 region 1 |
| chr13:29461925-29461965 | Ch13 region 1 |
| chr13:29461970-29462010 | Ch13 region 1 |
| chr13:29462029-29462069 | Ch13 region 1 |
| chr13:29462074-29462114 | Ch13 region 1 |
| chr13:29462120-29462160 | Ch13 region 1 |
| chr13:29462173-29462213 | Ch13 region 1 |
| chr13:29462231-29462271 | Ch13 region 1 |
| chr13:29462306-29462346 | Ch13 region 1 |
| chr13:29462355-29462395 | Ch13 region 1 |
| chr13:29462446-29462486 | Ch13 region 1 |
| chr13:29462525-29462565 | Ch13 region 1 |
| chr13:29462611-29462651 | Ch13 region 1 |
| chr13:29462838-29462878 | Ch13 region 1 |
| chr13:29462883-29462923 | Ch13 region 1 |
| chr13:29462935-29462975 | Ch13 region 1 |
| chr13:29463001-29463041 | Ch13 region 1 |
| chr13:29463082-29463122 | Ch13 region 1 |
| chr13:29463130-29463170 | Ch13 region 1 |

| | |
|---|---|
| chr13:29463202-29463242 | Ch13 region 1 |
| chr13:29463269-29463309 | Ch13 region 1 |
| chr13:29463335-29463375 | Ch13 region 1 |
| chr13:29463390-29463430 | Ch13 region 1 |
| chr13:29463435-29463475 | Ch13 region 1 |
| chr13:29463508-29463548 | Ch13 region 1 |
| chr13:29463584-29463624 | Ch13 region 1 |
| chr13:29463658-29463698 | Ch13 region 1 |
| chr13:29463724-29463764 | Ch13 region 1 |
| chr13:29463808-29463848 | Ch13 region 1 |
| chr13:29463869-29463909 | Ch13 region 1 |
| chr13:29463917-29463957 | Ch13 region 1 |
| chr13:29463981-29464021 | Ch13 region 1 |
| chr13:29464074-29464114 | Ch13 region 1 |
| chr13:29464120-29464160 | Ch13 region 1 |
| chr13:29464189-29464229 | Ch13 region 1 |
| chr13:29464249-29464289 | Ch13 region 1 |
| chr13:29464306-29464346 | Ch13 region 1 |
| chr13:29464371-29464411 | Ch13 region 1 |
| chr13:29464417-29464457 | Ch13 region 1 |
| chr13:29464481-29464521 | Ch13 region 1 |
| chr13:29464563-29464603 | Ch13 region 1 |
| chr13:29464629-29464669 | Ch13 region 1 |
| chr13:29464708-29464748 | Ch13 region 1 |
| chr13:29464871-29464911 | Ch13 region 1 |
| chr13:29464926-29464966 | Ch13 region 1 |
| chr13:29464976-29465016 | Ch13 region 1 |
| chr13:29465039-29465079 | Ch13 region 1 |
| chr13:29465084-29465124 | Ch13 region 1 |
| chr13:29465150-29465190 | Ch13 region 1 |
| chr13:29465197-29465237 | Ch13 region 1 |
| chr13:29465272-29465312 | Ch13 region 1 |
| chr13:29465327-29465367 | Ch13 region 1 |
| chr13:29465415-29465455 | Ch13 region 1 |
| chr13:29465465-29465505 | Ch13 region 1 |
| chr13:29465604-29465644 | Ch13 region 1 |
| chr13:29465665-29465705 | Ch13 region 1 |
| chr13:29465793-29465833 | Ch13 region 1 |
| chr13:29466020-29466060 | Ch13 region 1 |
| chr13:29466067-29466107 | Ch13 region 1 |
| chr13:29466131-29466171 | Ch13 region 1 |
| chr13:29466176-29466216 | Ch13 region 1 |
| chr13:29466238-29466278 | Ch13 region 1 |
| chr13:29466317-29466357 | Ch13 region 1 |
| chr13:29466363-29466403 | Ch13 region 1 |
| chr13:29466452-29466492 | Ch13 region 1 |
| chr13:29466499-29466539 | Ch13 region 1 |
| chr13:29466551-29466591 | Ch13 region 1 |
| chr13:29466608-29466648 | Ch13 region 1 |
| chr13:29466705-29466745 | Ch13 region 1 |
| chr13:29466768-29466808 | Ch13 region 1 |
| chr13:29466813-29466853 | Ch13 region 1 |

| | |
|---|---|
| chr13:29466899-29466939 | Ch13 region 1 |
| chr13:29466950-29466990 | Ch13 region 1 |
| chr13:29467046-29467086 | Ch13 region 1 |
| chr13:29467091-29467131 | Ch13 region 1 |
| chr13:29467173-29467213 | Ch13 region 1 |
| chr13:29467218-29467258 | Ch13 region 1 |
| chr13:29467265-29467305 | Ch13 region 1 |
| chr13:29467316-29467356 | Ch13 region 1 |
| chr13:29467384-29467424 | Ch13 region 1 |
| chr13:29467464-29467504 | Ch13 region 1 |
| chr13:29467533-29467573 | Ch13 region 1 |
| chr13:29467600-29467640 | Ch13 region 1 |
| chr13:29467683-29467723 | Ch13 region 1 |
| chr13:29467775-29467815 | Ch13 region 1 |
| chr13:29467828-29467868 | Ch13 region 1 |
| chr13:29467881-29467921 | Ch13 region 1 |
| chr13:29467938-29467978 | Ch13 region 1 |
| chr13:29467997-29468037 | Ch13 region 1 |
| chr13:29468050-29468090 | Ch13 region 1 |
| chr13:29468128-29468168 | Ch13 region 1 |
| chr13:29468205-29468245 | Ch13 region 1 |
| chr13:29468277-29468317 | Ch13 region 1 |
| chr13:29468394-29468434 | Ch13 region 1 |
| chr13:29468502-29468542 | Ch13 region 1 |
| chr13:29468577-29468617 | Ch13 region 1 |
| chr13:29468659-29468699 | Ch13 region 1 |
| chr13:29468711-29468751 | Ch13 region 1 |
| chr13:29468841-29468881 | Ch13 region 1 |
| chr13:29468922-29468962 | Ch13 region 1 |
| chr13:29468979-29469019 | Ch13 region 1 |
| chr13:29469082-29469122 | Ch13 region 1 |
| chr13:29469158-29469198 | Ch13 region 1 |
| chr13:29469229-29469269 | Ch13 region 1 |
| chr13:29469294-29469334 | Ch13 region 1 |
| chr13:29469342-29469382 | Ch13 region 1 |
| chr13:29469399-29469439 | Ch13 region 1 |
| chr13:29469476-29469516 | Ch13 region 1 |
| chr13:29469583-29469623 | Ch13 region 1 |
| chr13:29469683-29469723 | Ch13 region 1 |
| chr13:29469728-29469768 | Ch13 region 1 |
| chr13:29469802-29469842 | Ch13 region 1 |
| chr13:29469854-29469894 | Ch13 region 1 |
| chr13:29469947-29469987 | Ch13 region 1 |
| chr13:29469998-29470038 | Ch13 region 1 |
| chr13:29470046-29470086 | Ch13 region 1 |
| chr13:29470127-29470167 | Ch13 region 1 |
| chr13:29470190-29470230 | Ch13 region 1 |
| chr13:29470248-29470288 | Ch13 region 1 |
| chr13:29470293-29470333 | Ch13 region 1 |
| chr13:29470338-29470378 | Ch13 region 1 |
| chr13:29470583-29470623 | Ch13 region 1 |
| chr13:29470743-29470783 | Ch13 region 1 |

| | |
|---|---|
| chr13:29470958-29470998 | Ch13 region 1 |
| chr13:29471363-29471403 | Ch13 region 1 |
| chr13:29471717-29471757 | Ch13 region 1 |
| chr13:29471904-29471944 | Ch13 region 1 |
| chr13:29471979-29472019 | Ch13 region 1 |
| chr13:29472181-29472221 | Ch13 region 1 |
| chr13:29472226-29472266 | Ch13 region 1 |
| chr13:29472302-29472342 | Ch13 region 1 |
| chr13:29472383-29472423 | Ch13 region 1 |
| chr13:29472463-29472503 | Ch13 region 1 |
| chr13:29472531-29472571 | Ch13 region 1 |
| chr13:29472588-29472628 | Ch13 region 1 |
| chr13:29472660-29472700 | Ch13 region 1 |
| chr13:29472713-29472753 | Ch13 region 1 |
| chr13:29472766-29472806 | Ch13 region 1 |
| chr13:29472812-29472852 | Ch13 region 1 |
| chr13:29472901-29472941 | Ch13 region 1 |
| chr13:29472999-29473039 | Ch13 region 1 |
| chr13:29473052-29473092 | Ch13 region 1 |
| chr13:29473098-29473138 | Ch13 region 1 |
| chr13:29473179-29473219 | Ch13 region 1 |
| chr13:29473268-29473308 | Ch13 region 1 |
| chr13:29473425-29473465 | Ch13 region 1 |
| chr13:29473505-29473545 | Ch13 region 1 |
| chr13:29473831-29473871 | Ch13 region 1 |
| chr13:29473941-29473981 | Ch13 region 1 |
| chr13:29474034-29474074 | Ch13 region 1 |
| chr13:29474105-29474145 | Ch13 region 1 |
| chr13:29474304-29474344 | Ch13 region 1 |
| chr13:29474387-29474427 | Ch13 region 1 |
| chr13:29474446-29474486 | Ch13 region 1 |
| chr13:29474493-29474533 | Ch13 region 1 |
| chr13:29474569-29474609 | Ch13 region 1 |
| chr13:29474674-29474714 | Ch13 region 1 |
| chr13:29474768-29474808 | Ch13 region 1 |
| chr13:29474816-29474856 | Ch13 region 1 |
| chr13:29474864-29474904 | Ch13 region 1 |
| chr13:29474910-29474950 | Ch13 region 1 |
| chr13:29474958-29474998 | Ch13 region 1 |
| chr13:29475047-29475087 | Ch13 region 1 |
| chr13:29475121-29475161 | Ch13 region 1 |
| chr13:29475174-29475214 | Ch13 region 1 |
| chr13:29475275-29475315 | Ch13 region 1 |
| chr13:29475422-29475462 | Ch13 region 1 |
| chr13:29475547-29475587 | Ch13 region 1 |
| chr13:29475659-29475699 | Ch13 region 1 |
| chr13:29475721-29475761 | Ch13 region 1 |
| chr13:29475794-29475834 | Ch13 region 1 |
| chr13:29475848-29475888 | Ch13 region 1 |
| chr13:29475930-29475970 | Ch13 region 1 |
| chr13:29476161-29476201 | Ch13 region 1 |
| chr13:29476309-29476349 | Ch13 region 1 |

| | |
|---|---|
| chr13:29476371-29476411 | Ch13 region 1 |
| chr13:29476505-29476545 | Ch13 region 1 |
| chr13:29476573-29476613 | Ch13 region 1 |
| chr13:29476647-29476687 | Ch13 region 1 |
| chr13:29476786-29476826 | Ch13 region 1 |
| chr13:29476924-29476964 | Ch13 region 1 |
| chr13:29477029-29477069 | Ch13 region 1 |
| chr13:29477126-29477166 | Ch13 region 1 |
| chr13:29477172-29477212 | Ch13 region 1 |
| chr13:29477241-29477281 | Ch13 region 1 |
| chr13:29477289-29477329 | Ch13 region 1 |
| chr13:29477334-29477374 | Ch13 region 1 |
| chr13:29477643-29477683 | Ch13 region 1 |
| chr13:29478038-29478078 | Ch13 region 1 |
| chr13:29478194-29478234 | Ch13 region 1 |
| chr13:29478402-29478442 | Ch13 region 1 |
| chr13:29479179-29479219 | Ch13 region 1 |
| chr13:29479451-29479491 | Ch13 region 1 |
| chr13:29479951-29479991 | Ch13 region 1 |
| chr13:29480905-29480945 | Ch13 region 1 |
| chr13:29480950-29480990 | Ch13 region 1 |
| chr13:29481420-29481460 | Ch13 region 1 |
| chr13:29481498-29481538 | Ch13 region 1 |
| chr13:29481869-29481909 | Ch13 region 1 |
| chr13:29482006-29482046 | Ch13 region 1 |
| chr13:29482065-29482105 | Ch13 region 1 |
| chr13:29482138-29482178 | Ch13 region 1 |
| chr13:29482206-29482246 | Ch13 region 1 |
| chr13:29482393-29482433 | Ch13 region 1 |
| chr13:29482858-29482898 | Ch13 region 1 |
| chr13:29482995-29483035 | Ch13 region 1 |
| chr13:29483138-29483178 | Ch13 region 1 |
| chr13:29483343-29483383 | Ch13 region 1 |
| chr13:29483602-29483642 | Ch13 region 1 |
| chr13:29483699-29483739 | Ch13 region 1 |
| chr13:29483984-29484024 | Ch13 region 1 |
| chr13:29484116-29484156 | Ch13 region 1 |
| chr13:29484209-29484249 | Ch13 region 1 |
| chr13:29484272-29484312 | Ch13 region 1 |
| chr13:29484504-29484544 | Ch13 region 1 |
| chr13:29484584-29484624 | Ch13 region 1 |
| chr13:29484638-29484678 | Ch13 region 1 |
| chr13:29484712-29484752 | Ch13 region 1 |
| chr13:29484954-29484994 | Ch13 region 1 |
| chr13:29485037-29485077 | Ch13 region 1 |
| chr13:29485390-29485430 | Ch13 region 1 |
| chr13:29485591-29485631 | Ch13 region 1 |
| chr13:29485737-29485777 | Ch13 region 1 |
| chr13:29485861-29485901 | Ch13 region 1 |
| chr13:29486002-29486042 | Ch13 region 1 |
| chr13:29486058-29486098 | Ch13 region 1 |
| chr13:29486168-29486208 | Ch13 region 1 |

| | |
|---|---|
| chr13:29486213-29486253 | Ch13 region 1 |
| chr13:29486333-29486373 | Ch13 region 1 |
| chr13:29486407-29486447 | Ch13 region 1 |
| chr13:29486506-29486546 | Ch13 region 1 |
| chr13:29486681-29486721 | Ch13 region 1 |
| chr13:29486830-29486870 | Ch13 region 1 |
| chr13:29486951-29486991 | Ch13 region 1 |
| chr13:29487035-29487075 | Ch13 region 1 |
| chr13:29487198-29487238 | Ch13 region 1 |
| chr13:29487351-29487391 | Ch13 region 1 |
| chr13:29487558-29487598 | Ch13 region 1 |
| chr13:29487717-29487757 | Ch13 region 1 |
| chr13:29487783-29487823 | Ch13 region 1 |
| chr13:29487837-29487877 | Ch13 region 1 |
| chr13:29487959-29487999 | Ch13 region 1 |
| chr13:29488275-29488315 | Ch13 region 1 |
| chr13:29488325-29488365 | Ch13 region 1 |
| chr13:29488487-29488527 | Ch13 region 1 |
| chr13:29488663-29488703 | Ch13 region 1 |
| chr13:29488792-29488832 | Ch13 region 1 |
| chr13:29488900-29488940 | Ch13 region 1 |
| chr13:29489008-29489048 | Ch13 region 1 |
| chr13:29489146-29489186 | Ch13 region 1 |
| chr13:29489251-29489291 | Ch13 region 1 |
| chr13:29489427-29489467 | Ch13 region 1 |
| chr13:29489553-29489593 | Ch13 region 1 |
| chr13:29489611-29489651 | Ch13 region 1 |
| chr13:29489698-29489738 | Ch13 region 1 |
| chr13:29489859-29489899 | Ch13 region 1 |
| chr13:29489948-29489988 | Ch13 region 1 |
| chr13:29490125-29490165 | Ch13 region 1 |
| chr13:29490231-29490271 | Ch13 region 1 |
| chr13:29490343-29490383 | Ch13 region 1 |
| chr13:29490425-29490465 | Ch13 region 1 |
| chr13:29490779-29490819 | Ch13 region 1 |
| chr13:29490925-29490965 | Ch13 region 1 |
| chr13:29491024-29491064 | Ch13 region 1 |
| chr13:29491200-29491240 | Ch13 region 1 |
| chr13:29491413-29491453 | Ch13 region 1 |
| chr13:29491543-29491583 | Ch13 region 1 |
| chr13:29491720-29491760 | Ch13 region 1 |
| chr13:29491968-29492008 | Ch13 region 1 |
| chr13:29492037-29492077 | Ch13 region 1 |
| chr13:29492088-29492128 | Ch13 region 1 |
| chr13:29492196-29492236 | Ch13 region 1 |
| chr13:29492251-29492291 | Ch13 region 1 |
| chr13:29492479-29492519 | Ch13 region 1 |
| chr13:29492620-29492660 | Ch13 region 1 |
| chr13:29492717-29492757 | Ch13 region 1 |
| chr13:29492828-29492868 | Ch13 region 1 |
| chr13:29492913-29492953 | Ch13 region 1 |
| chr13:29493023-29493063 | Ch13 region 1 |

| | |
|---|---|
| chr13:29493284-29493324 | Ch13 region 1 |
| chr13:29493371-29493411 | Ch13 region 1 |
| chr13:29493521-29493561 | Ch13 region 1 |
| chr13:29493568-29493608 | Ch13 region 1 |
| chr13:29493671-29493711 | Ch13 region 1 |
| chr13:29493794-29493834 | Ch13 region 1 |
| chr13:29493911-29493951 | Ch13 region 1 |
| chr13:29493956-29493996 | Ch13 region 1 |
| chr13:29494011-29494051 | Ch13 region 1 |
| chr13:29494103-29494143 | Ch13 region 1 |
| chr13:29494259-29494299 | Ch13 region 1 |
| chr13:29494361-29494401 | Ch13 region 1 |
| chr13:29494466-29494506 | Ch13 region 1 |
| chr13:29494533-29494573 | Ch13 region 1 |
| chr13:29494612-29494652 | Ch13 region 1 |
| chr13:29494702-29494742 | Ch13 region 1 |
| chr13:29494749-29494789 | Ch13 region 1 |
| chr13:29494837-29494877 | Ch13 region 1 |
| chr13:29494962-29495002 | Ch13 region 1 |
| chr13:29495116-29495156 | Ch13 region 1 |
| chr13:29495210-29495250 | Ch13 region 1 |
| chr13:29495267-29495307 | Ch13 region 1 |
| chr13:29495330-29495370 | Ch13 region 1 |
| chr13:29495429-29495469 | Ch13 region 1 |
| chr13:53695501-53695541 | Ch13 region 2 |
| chr13:53695573-53695613 | Ch13 region 2 |
| chr13:53695626-53695666 | Ch13 region 2 |
| chr13:53695700-53695740 | Ch13 region 2 |
| chr13:53695745-53695785 | Ch13 region 2 |
| chr13:53695899-53695939 | Ch13 region 2 |
| chr13:53696013-53696053 | Ch13 region 2 |
| chr13:53696088-53696128 | Ch13 region 2 |
| chr13:53696151-53696191 | Ch13 region 2 |
| chr13:53696231-53696271 | Ch13 region 2 |
| chr13:53696280-53696320 | Ch13 region 2 |
| chr13:53696328-53696368 | Ch13 region 2 |
| chr13:53696411-53696451 | Ch13 region 2 |
| chr13:53696456-53696496 | Ch13 region 2 |
| chr13:53696604-53696644 | Ch13 region 2 |
| chr13:53696696-53696736 | Ch13 region 2 |
| chr13:53696751-53696791 | Ch13 region 2 |
| chr13:53696824-53696864 | Ch13 region 2 |
| chr13:53696872-53696912 | Ch13 region 2 |
| chr13:53696978-53697018 | Ch13 region 2 |
| chr13:53697165-53697205 | Ch13 region 2 |
| chr13:53697250-53697290 | Ch13 region 2 |
| chr13:53697347-53697387 | Ch13 region 2 |
| chr13:53697398-53697438 | Ch13 region 2 |
| chr13:53697452-53697492 | Ch13 region 2 |
| chr13:53697537-53697577 | Ch13 region 2 |
| chr13:53697601-53697641 | Ch13 region 2 |
| chr13:53697664-53697704 | Ch13 region 2 |

| | |
|---|---|
| chr13:53697751-53697791 | Ch13 region 2 |
| chr13:53697824-53697864 | Ch13 region 2 |
| chr13:53697872-53697912 | Ch13 region 2 |
| chr13:53698071-53698111 | Ch13 region 2 |
| chr13:53698286-53698326 | Ch13 region 2 |
| chr13:53698357-53698397 | Ch13 region 2 |
| chr13:53698431-53698471 | Ch13 region 2 |
| chr13:53698488-53698528 | Ch13 region 2 |
| chr13:53698544-53698584 | Ch13 region 2 |
| chr13:53698597-53698637 | Ch13 region 2 |
| chr13:53698661-53698701 | Ch13 region 2 |
| chr13:53698751-53698791 | Ch13 region 2 |
| chr13:53698797-53698837 | Ch13 region 2 |
| chr13:53698874-53698914 | Ch13 region 2 |
| chr13:53698919-53698959 | Ch13 region 2 |
| chr13:53698992-53699032 | Ch13 region 2 |
| chr13:53699047-53699087 | Ch13 region 2 |
| chr13:53699096-53699136 | Ch13 region 2 |
| chr13:53699157-53699197 | Ch13 region 2 |
| chr13:53699223-53699263 | Ch13 region 2 |
| chr13:53699275-53699315 | Ch13 region 2 |
| chr13:53699320-53699360 | Ch13 region 2 |
| chr13:53699391-53699431 | Ch13 region 2 |
| chr13:53699452-53699492 | Ch13 region 2 |
| chr13:53699523-53699563 | Ch13 region 2 |
| chr13:53699592-53699632 | Ch13 region 2 |
| chr13:53699673-53699713 | Ch13 region 2 |
| chr13:53699758-53699798 | Ch13 region 2 |
| chr13:53699825-53699865 | Ch13 region 2 |
| chr13:53699905-53699945 | Ch13 region 2 |
| chr13:53700022-53700062 | Ch13 region 2 |
| chr13:53700130-53700170 | Ch13 region 2 |
| chr13:53700299-53700339 | Ch13 region 2 |
| chr13:53700376-53700416 | Ch13 region 2 |
| chr13:53700424-53700464 | Ch13 region 2 |
| chr13:53700478-53700518 | Ch13 region 2 |
| chr13:53700527-53700567 | Ch13 region 2 |
| chr13:53700584-53700624 | Ch13 region 2 |
| chr13:53700642-53700682 | Ch13 region 2 |
| chr13:53700700-53700740 | Ch13 region 2 |
| chr13:53700758-53700798 | Ch13 region 2 |
| chr13:53700818-53700858 | Ch13 region 2 |
| chr13:53700923-53700963 | Ch13 region 2 |
| chr13:53700976-53701016 | Ch13 region 2 |
| chr13:53701057-53701097 | Ch13 region 2 |
| chr13:53701147-53701187 | Ch13 region 2 |
| chr13:53701234-53701274 | Ch13 region 2 |
| chr13:53701312-53701352 | Ch13 region 2 |
| chr13:53701405-53701445 | Ch13 region 2 |
| chr13:53701493-53701533 | Ch13 region 2 |
| chr13:53701543-53701583 | Ch13 region 2 |
| chr13:53701609-53701649 | Ch13 region 2 |

| | |
|---|---|
| chr13:53701689-53701729 | Ch13 region 2 |
| chr13:53701735-53701775 | Ch13 region 2 |
| chr13:53701821-53701861 | Ch13 region 2 |
| chr13:53701900-53701940 | Ch13 region 2 |
| chr13:53701970-53702010 | Ch13 region 2 |
| chr13:53702041-53702081 | Ch13 region 2 |
| chr13:53702100-53702140 | Ch13 region 2 |
| chr13:53702158-53702198 | Ch13 region 2 |
| chr13:53702215-53702255 | Ch13 region 2 |
| chr13:53702267-53702307 | Ch13 region 2 |
| chr13:53702327-53702367 | Ch13 region 2 |
| chr13:53702372-53702412 | Ch13 region 2 |
| chr13:53702417-53702457 | Ch13 region 2 |
| chr13:53702473-53702513 | Ch13 region 2 |
| chr13:53702545-53702585 | Ch13 region 2 |
| chr13:53702613-53702653 | Ch13 region 2 |
| chr13:53702659-53702699 | Ch13 region 2 |
| chr13:53702738-53702778 | Ch13 region 2 |
| chr13:53702783-53702823 | Ch13 region 2 |
| chr13:53702898-53702938 | Ch13 region 2 |
| chr13:53702943-53702983 | Ch13 region 2 |
| chr13:53703094-53703134 | Ch13 region 2 |
| chr13:53703155-53703195 | Ch13 region 2 |
| chr13:53703204-53703244 | Ch13 region 2 |
| chr13:53703256-53703296 | Ch13 region 2 |
| chr13:53703312-53703352 | Ch13 region 2 |
| chr13:53703381-53703421 | Ch13 region 2 |
| chr13:53703430-53703470 | Ch13 region 2 |
| chr13:53703492-53703532 | Ch13 region 2 |
| chr13:53703537-53703577 | Ch13 region 2 |
| chr13:53703606-53703646 | Ch13 region 2 |
| chr13:53703670-53703710 | Ch13 region 2 |
| chr13:53703715-53703755 | Ch13 region 2 |
| chr13:53703773-53703813 | Ch13 region 2 |
| chr13:53703869-53703909 | Ch13 region 2 |
| chr13:53703929-53703969 | Ch13 region 2 |
| chr13:53703989-53704029 | Ch13 region 2 |
| chr13:53704053-53704093 | Ch13 region 2 |
| chr13:53704098-53704138 | Ch13 region 2 |
| chr13:53704163-53704203 | Ch13 region 2 |
| chr13:53704209-53704249 | Ch13 region 2 |
| chr13:53704411-53704451 | Ch13 region 2 |
| chr13:53704553-53704593 | Ch13 region 2 |
| chr13:53704718-53704758 | Ch13 region 2 |
| chr13:53704764-53704804 | Ch13 region 2 |
| chr13:53704813-53704853 | Ch13 region 2 |
| chr13:53705057-53705097 | Ch13 region 2 |
| chr13:53705122-53705162 | Ch13 region 2 |
| chr13:53705232-53705272 | Ch13 region 2 |
| chr13:53705353-53705393 | Ch13 region 2 |
| chr13:53705428-53705468 | Ch13 region 2 |
| chr13:53705518-53705558 | Ch13 region 2 |

| | |
|---|---|
| chr13:53705573-53705613 | Ch13 region 2 |
| chr13:53705618-53705658 | Ch13 region 2 |
| chr13:53705678-53705718 | Ch13 region 2 |
| chr13:53705723-53705763 | Ch13 region 2 |
| chr13:53705802-53705842 | Ch13 region 2 |
| chr13:53705889-53705929 | Ch13 region 2 |
| chr13:53705938-53705978 | Ch13 region 2 |
| chr13:53706006-53706046 | Ch13 region 2 |
| chr13:53706061-53706101 | Ch13 region 2 |
| chr13:53706110-53706150 | Ch13 region 2 |
| chr13:53706181-53706221 | Ch13 region 2 |
| chr13:53706237-53706277 | Ch13 region 2 |
| chr13:53706284-53706324 | Ch13 region 2 |
| chr13:53706335-53706375 | Ch13 region 2 |
| chr13:53706417-53706457 | Ch13 region 2 |
| chr13:53706517-53706557 | Ch13 region 2 |
| chr13:53706582-53706622 | Ch13 region 2 |
| chr13:53706673-53706713 | Ch13 region 2 |
| chr13:53706735-53706775 | Ch13 region 2 |
| chr13:53706795-53706835 | Ch13 region 2 |
| chr13:53706841-53706881 | Ch13 region 2 |
| chr13:53706896-53706936 | Ch13 region 2 |
| chr13:53706984-53707024 | Ch13 region 2 |
| chr13:53707073-53707113 | Ch13 region 2 |
| chr13:53707186-53707226 | Ch13 region 2 |
| chr13:53707231-53707271 | Ch13 region 2 |
| chr13:53707276-53707316 | Ch13 region 2 |
| chr13:53707347-53707387 | Ch13 region 2 |
| chr13:53707468-53707508 | Ch13 region 2 |
| chr13:53707578-53707618 | Ch13 region 2 |
| chr13:53707664-53707704 | Ch13 region 2 |
| chr13:53707723-53707763 | Ch13 region 2 |
| chr13:53707796-53707836 | Ch13 region 2 |
| chr13:53707842-53707882 | Ch13 region 2 |
| chr13:53707967-53708007 | Ch13 region 2 |
| chr13:53708014-53708054 | Ch13 region 2 |
| chr13:53708071-53708111 | Ch13 region 2 |
| chr13:53708116-53708156 | Ch13 region 2 |
| chr13:53708197-53708237 | Ch13 region 2 |
| chr13:53708262-53708302 | Ch13 region 2 |
| chr13:53708383-53708423 | Ch13 region 2 |
| chr13:53708465-53708505 | Ch13 region 2 |
| chr13:53708513-53708553 | Ch13 region 2 |
| chr13:53708558-53708598 | Ch13 region 2 |
| chr13:53708639-53708679 | Ch13 region 2 |
| chr13:53708695-53708735 | Ch13 region 2 |
| chr13:53708740-53708780 | Ch13 region 2 |
| chr13:53708839-53708879 | Ch13 region 2 |
| chr13:53708905-53708945 | Ch13 region 2 |
| chr13:53708952-53708992 | Ch13 region 2 |
| chr13:53709019-53709059 | Ch13 region 2 |
| chr13:53709144-53709184 | Ch13 region 2 |

| | |
|---|---|
| chr13:53709209-53709249 | Ch13 region 2 |
| chr13:53709259-53709299 | Ch13 region 2 |
| chr13:53709312-53709352 | Ch13 region 2 |
| chr13:53709390-53709430 | Ch13 region 2 |
| chr13:53709484-53709524 | Ch13 region 2 |
| chr13:53709550-53709590 | Ch13 region 2 |
| chr13:53709607-53709647 | Ch13 region 2 |
| chr13:53709652-53709692 | Ch13 region 2 |
| chr13:53709727-53709767 | Ch13 region 2 |
| chr13:53709776-53709816 | Ch13 region 2 |
| chr13:53709827-53709867 | Ch13 region 2 |
| chr13:53709875-53709915 | Ch13 region 2 |
| chr13:53709920-53709960 | Ch13 region 2 |
| chr13:53709967-53710007 | Ch13 region 2 |
| chr13:53710016-53710056 | Ch13 region 2 |
| chr13:53710061-53710101 | Ch13 region 2 |
| chr13:53710133-53710173 | Ch13 region 2 |
| chr13:53710216-53710256 | Ch13 region 2 |
| chr13:53710291-53710331 | Ch13 region 2 |
| chr13:53710376-53710416 | Ch13 region 2 |
| chr13:53710421-53710461 | Ch13 region 2 |
| chr13:53710468-53710508 | Ch13 region 2 |
| chr13:53710526-53710566 | Ch13 region 2 |
| chr13:53710572-53710612 | Ch13 region 2 |
| chr13:53710617-53710657 | Ch13 region 2 |
| chr13:53710697-53710737 | Ch13 region 2 |
| chr13:53710778-53710818 | Ch13 region 2 |
| chr13:53710855-53710895 | Ch13 region 2 |
| chr13:53710927-53710967 | Ch13 region 2 |
| chr13:53710990-53711030 | Ch13 region 2 |
| chr13:53711067-53711107 | Ch13 region 2 |
| chr13:53711126-53711166 | Ch13 region 2 |
| chr13:53711208-53711248 | Ch13 region 2 |
| chr13:53711295-53711335 | Ch13 region 2 |
| chr13:53711344-53711384 | Ch13 region 2 |
| chr13:53711397-53711437 | Ch13 region 2 |
| chr13:53711461-53711501 | Ch13 region 2 |
| chr13:53711572-53711612 | Ch13 region 2 |
| chr13:53711654-53711694 | Ch13 region 2 |
| chr13:53711737-53711777 | Ch13 region 2 |
| chr13:53711793-53711833 | Ch13 region 2 |
| chr13:53711892-53711932 | Ch13 region 2 |
| chr13:53711947-53711987 | Ch13 region 2 |
| chr13:53712016-53712056 | Ch13 region 2 |
| chr13:53712088-53712128 | Ch13 region 2 |
| chr13:53712163-53712203 | Ch13 region 2 |
| chr13:53712210-53712250 | Ch13 region 2 |
| chr13:53712265-53712305 | Ch13 region 2 |
| chr13:53712355-53712395 | Ch13 region 2 |
| chr13:53712402-53712442 | Ch13 region 2 |
| chr13:53712469-53712509 | Ch13 region 2 |
| chr13:53712514-53712554 | Ch13 region 2 |

| | |
|---|---|
| chr13:53712596-53712636 | Ch13 region 2 |
| chr13:53712643-53712683 | Ch13 region 2 |
| chr13:53712742-53712782 | Ch13 region 2 |
| chr13:53712797-53712837 | Ch13 region 2 |
| chr13:53712844-53712884 | Ch13 region 2 |
| chr13:53712893-53712933 | Ch13 region 2 |
| chr13:53712943-53712983 | Ch13 region 2 |
| chr13:53713001-53713041 | Ch13 region 2 |
| chr13:53713054-53713094 | Ch13 region 2 |
| chr13:53713138-53713178 | Ch13 region 2 |
| chr13:53713206-53713246 | Ch13 region 2 |
| chr13:53713278-53713318 | Ch13 region 2 |
| chr13:53713344-53713384 | Ch13 region 2 |
| chr13:53713392-53713432 | Ch13 region 2 |
| chr13:53713481-53713521 | Ch13 region 2 |
| chr13:53713552-53713592 | Ch13 region 2 |
| chr13:53713653-53713693 | Ch13 region 2 |
| chr13:53713738-53713778 | Ch13 region 2 |
| chr13:53713846-53713886 | Ch13 region 2 |
| chr13:53713942-53713982 | Ch13 region 2 |
| chr13:53714014-53714054 | Ch13 region 2 |
| chr13:53714089-53714129 | Ch13 region 2 |
| chr13:53714415-53714455 | Ch13 region 2 |
| chr13:53714483-53714523 | Ch13 region 2 |
| chr13:53714555-53714595 | Ch13 region 2 |
| chr13:53714656-53714696 | Ch13 region 2 |
| chr13:53714753-53714793 | Ch13 region 2 |
| chr13:53714853-53714893 | Ch13 region 2 |
| chr13:53714898-53714938 | Ch13 region 2 |
| chr13:53714976-53715016 | Ch13 region 2 |
| chr13:53715065-53715105 | Ch13 region 2 |
| chr13:53715125-53715165 | Ch13 region 2 |
| chr13:53715185-53715225 | Ch13 region 2 |
| chr13:53715242-53715282 | Ch13 region 2 |
| chr13:53715360-53715400 | Ch13 region 2 |
| chr13:53715437-53715477 | Ch13 region 2 |
| chr13:53715482-53715522 | Ch13 region 2 |
| chr13:53715527-53715567 | Ch13 region 2 |
| chr13:53715617-53715657 | Ch13 region 2 |
| chr13:53715662-53715702 | Ch13 region 2 |
| chr13:53715707-53715747 | Ch13 region 2 |
| chr13:53715755-53715795 | Ch13 region 2 |
| chr13:53715802-53715842 | Ch13 region 2 |
| chr13:53715847-53715887 | Ch13 region 2 |
| chr13:53715926-53715966 | Ch13 region 2 |
| chr13:53716016-53716056 | Ch13 region 2 |
| chr13:53716102-53716142 | Ch13 region 2 |
| chr13:53716164-53716204 | Ch13 region 2 |
| chr13:53716211-53716251 | Ch13 region 2 |
| chr13:53716265-53716305 | Ch13 region 2 |
| chr13:53716368-53716408 | Ch13 region 2 |
| chr13:53716447-53716487 | Ch13 region 2 |

| | |
|---|---|
| chr13:53716527-53716567 | Ch13 region 2 |
| chr13:53716574-53716614 | Ch13 region 2 |
| chr13:53716632-53716672 | Ch13 region 2 |
| chr13:53716709-53716749 | Ch13 region 2 |
| chr13:53716775-53716815 | Ch13 region 2 |
| chr13:53716822-53716862 | Ch13 region 2 |
| chr13:53716880-53716920 | Ch13 region 2 |
| chr13:53716943-53716983 | Ch13 region 2 |
| chr13:53717019-53717059 | Ch13 region 2 |
| chr13:53717086-53717126 | Ch13 region 2 |
| chr13:53717251-53717291 | Ch13 region 2 |
| chr13:53717313-53717353 | Ch13 region 2 |
| chr13:53717400-53717440 | Ch13 region 2 |
| chr13:53717497-53717537 | Ch13 region 2 |
| chr13:53717542-53717582 | Ch13 region 2 |
| chr13:53717604-53717644 | Ch13 region 2 |
| chr13:53717670-53717710 | Ch13 region 2 |
| chr13:53717721-53717761 | Ch13 region 2 |
| chr13:53717780-53717820 | Ch13 region 2 |
| chr13:53717879-53717919 | Ch13 region 2 |
| chr13:53717925-53717965 | Ch13 region 2 |
| chr13:53718011-53718051 | Ch13 region 2 |
| chr13:53718062-53718102 | Ch13 region 2 |
| chr13:53718110-53718150 | Ch13 region 2 |
| chr13:53718168-53718208 | Ch13 region 2 |
| chr13:53718269-53718309 | Ch13 region 2 |
| chr13:53718319-53718359 | Ch13 region 2 |
| chr13:53718371-53718411 | Ch13 region 2 |
| chr13:53718466-53718506 | Ch13 region 2 |
| chr13:53718542-53718582 | Ch13 region 2 |
| chr13:53718616-53718656 | Ch13 region 2 |
| chr13:53718697-53718737 | Ch13 region 2 |
| chr13:53718822-53718862 | Ch13 region 2 |
| chr13:53718869-53718909 | Ch13 region 2 |
| chr13:53718946-53718986 | Ch13 region 2 |
| chr13:53718992-53719032 | Ch13 region 2 |
| chr13:53719051-53719091 | Ch13 region 2 |
| chr13:53719109-53719149 | Ch13 region 2 |
| chr13:53719180-53719220 | Ch13 region 2 |
| chr13:53719242-53719282 | Ch13 region 2 |
| chr13:53719312-53719352 | Ch13 region 2 |
| chr13:53719357-53719397 | Ch13 region 2 |
| chr13:53719417-53719457 | Ch13 region 2 |
| chr13:53719487-53719527 | Ch13 region 2 |
| chr13:53719532-53719572 | Ch13 region 2 |
| chr13:53719587-53719627 | Ch13 region 2 |
| chr13:53719732-53719772 | Ch13 region 2 |
| chr13:53719860-53719900 | Ch13 region 2 |
| chr13:53719914-53719954 | Ch13 region 2 |
| chr13:53719981-53720021 | Ch13 region 2 |
| chr13:53720026-53720066 | Ch13 region 2 |
| chr13:53720089-53720129 | Ch13 region 2 |

| | |
|---|---|
| chr13:53720137-53720177 | Ch13 region 2 |
| chr13:53720205-53720245 | Ch13 region 2 |
| chr13:53720435-53720475 | Ch13 region 2 |
| chr13:53720486-53720526 | Ch13 region 2 |
| chr13:53720537-53720577 | Ch13 region 2 |
| chr13:53720601-53720641 | Ch13 region 2 |
| chr13:53720683-53720723 | Ch13 region 2 |
| chr13:53720728-53720768 | Ch13 region 2 |
| chr13:53720776-53720816 | Ch13 region 2 |
| chr13:53720821-53720861 | Ch13 region 2 |
| chr13:53720903-53720943 | Ch13 region 2 |
| chr13:53720994-53721034 | Ch13 region 2 |
| chr13:53721058-53721098 | Ch13 region 2 |
| chr13:53721118-53721158 | Ch13 region 2 |
| chr13:53721195-53721235 | Ch13 region 2 |
| chr13:53721252-53721292 | Ch13 region 2 |
| chr13:53721322-53721362 | Ch13 region 2 |
| chr13:53721378-53721418 | Ch13 region 2 |
| chr13:53721440-53721480 | Ch13 region 2 |
| chr13:53721534-53721574 | Ch13 region 2 |
| chr13:53721582-53721622 | Ch13 region 2 |
| chr13:53721630-53721670 | Ch13 region 2 |
| chr13:53721721-53721761 | Ch13 region 2 |
| chr13:53721783-53721823 | Ch13 region 2 |
| chr13:53721837-53721877 | Ch13 region 2 |
| chr13:53721899-53721939 | Ch13 region 2 |
| chr13:53721944-53721984 | Ch13 region 2 |
| chr13:53722004-53722044 | Ch13 region 2 |
| chr13:53722068-53722108 | Ch13 region 2 |
| chr13:53722117-53722157 | Ch13 region 2 |
| chr13:53722201-53722241 | Ch13 region 2 |
| chr13:53722246-53722286 | Ch13 region 2 |
| chr13:53722321-53722361 | Ch13 region 2 |
| chr13:53722371-53722411 | Ch13 region 2 |
| chr13:53722419-53722459 | Ch13 region 2 |
| chr13:53722481-53722521 | Ch13 region 2 |
| chr13:53722530-53722570 | Ch13 region 2 |
| chr13:53722579-53722619 | Ch13 region 2 |
| chr13:53722636-53722676 | Ch13 region 2 |
| chr13:53722697-53722737 | Ch13 region 2 |
| chr13:53722919-53722959 | Ch13 region 2 |
| chr13:53722991-53723031 | Ch13 region 2 |
| chr13:53723058-53723098 | Ch13 region 2 |
| chr13:53723146-53723186 | Ch13 region 2 |
| chr13:53723191-53723231 | Ch13 region 2 |
| chr13:53723278-53723318 | Ch13 region 2 |
| chr13:53723330-53723370 | Ch13 region 2 |
| chr13:53723395-53723435 | Ch13 region 2 |
| chr13:53723480-53723520 | Ch13 region 2 |
| chr13:53723547-53723587 | Ch13 region 2 |
| chr13:53723616-53723656 | Ch13 region 2 |
| chr13:53723797-53723837 | Ch13 region 2 |

| | |
|---|---|
| chr13:53723873-53723913 | Ch13 region 2 |
| chr13:53723918-53723958 | Ch13 region 2 |
| chr13:53724018-53724058 | Ch13 region 2 |
| chr13:53724074-53724114 | Ch13 region 2 |
| chr13:53724166-53724206 | Ch13 region 2 |
| chr13:53724259-53724299 | Ch13 region 2 |
| chr13:53724341-53724381 | Ch13 region 2 |
| chr13:53724425-53724465 | Ch13 region 2 |
| chr13:53724470-53724510 | Ch13 region 2 |
| chr13:53724529-53724569 | Ch13 region 2 |
| chr13:53724592-53724632 | Ch13 region 2 |
| chr13:53724648-53724688 | Ch13 region 2 |
| chr13:53724701-53724741 | Ch13 region 2 |
| chr13:53724747-53724787 | Ch13 region 2 |
| chr13:53724792-53724832 | Ch13 region 2 |
| chr13:53724858-53724898 | Ch13 region 2 |
| chr13:53724913-53724953 | Ch13 region 2 |
| chr13:53724973-53725013 | Ch13 region 2 |
| chr13:53725031-53725071 | Ch13 region 2 |
| chr13:53725261-53725301 | Ch13 region 2 |
| chr13:53725310-53725350 | Ch13 region 2 |
| chr13:53725385-53725425 | Ch13 region 2 |
| chr13:53725522-53725562 | Ch13 region 2 |
| chr13:53725654-53725694 | Ch13 region 2 |
| chr13:53725740-53725780 | Ch13 region 2 |
| chr13:53725800-53725840 | Ch13 region 2 |
| chr13:53725862-53725902 | Ch13 region 2 |
| chr13:53725975-53726015 | Ch13 region 2 |
| chr13:53726056-53726096 | Ch13 region 2 |
| chr13:53726151-53726191 | Ch13 region 2 |
| chr13:53726196-53726236 | Ch13 region 2 |
| chr13:53726259-53726299 | Ch13 region 2 |
| chr13:53726304-53726344 | Ch13 region 2 |
| chr13:53726349-53726389 | Ch13 region 2 |
| chr13:53726394-53726434 | Ch13 region 2 |
| chr13:53726466-53726506 | Ch13 region 2 |
| chr13:53726540-53726580 | Ch13 region 2 |
| chr13:53726656-53726696 | Ch13 region 2 |
| chr13:53726797-53726837 | Ch13 region 2 |
| chr13:53726947-53726987 | Ch13 region 2 |
| chr13:53727124-53727164 | Ch13 region 2 |
| chr13:53727199-53727239 | Ch13 region 2 |
| chr13:53727244-53727284 | Ch13 region 2 |
| chr13:53727320-53727360 | Ch13 region 2 |
| chr13:53727388-53727428 | Ch13 region 2 |
| chr13:53727474-53727514 | Ch13 region 2 |
| chr13:53727545-53727585 | Ch13 region 2 |
| chr13:53727643-53727683 | Ch13 region 2 |
| chr13:53727692-53727732 | Ch13 region 2 |
| chr13:53727741-53727781 | Ch13 region 2 |
| chr13:53727821-53727861 | Ch13 region 2 |
| chr13:53727899-53727939 | Ch13 region 2 |

| | |
|---|---|
| chr13:53727944-53727984 | Ch13 region 2 |
| chr13:53727996-53728036 | Ch13 region 2 |
| chr13:53728120-53728160 | Ch13 region 2 |
| chr13:53728188-53728228 | Ch13 region 2 |
| chr13:53728301-53728341 | Ch13 region 2 |
| chr13:53728413-53728453 | Ch13 region 2 |
| chr13:53728498-53728538 | Ch13 region 2 |
| chr13:53728581-53728621 | Ch13 region 2 |
| chr13:53728629-53728669 | Ch13 region 2 |
| chr13:53728674-53728714 | Ch13 region 2 |
| chr13:53728719-53728759 | Ch13 region 2 |
| chr13:53728778-53728818 | Ch13 region 2 |
| chr13:53728865-53728905 | Ch13 region 2 |
| chr13:53728917-53728957 | Ch13 region 2 |
| chr13:53729012-53729052 | Ch13 region 2 |
| chr13:53729097-53729137 | Ch13 region 2 |
| chr13:53729165-53729205 | Ch13 region 2 |
| chr13:53729223-53729263 | Ch13 region 2 |
| chr13:53729377-53729417 | Ch13 region 2 |
| chr13:53729487-53729527 | Ch13 region 2 |
| chr13:53729532-53729572 | Ch13 region 2 |
| chr13:53729677-53729717 | Ch13 region 2 |
| chr13:53729739-53729779 | Ch13 region 2 |
| chr13:53729791-53729831 | Ch13 region 2 |
| chr13:53729866-53729906 | Ch13 region 2 |
| chr13:53729917-53729957 | Ch13 region 2 |
| chr13:53730009-53730049 | Ch13 region 2 |
| chr13:53730085-53730125 | Ch13 region 2 |
| chr13:53730150-53730190 | Ch13 region 2 |
| chr13:53730247-53730287 | Ch13 region 2 |
| chr13:53730307-53730347 | Ch13 region 2 |
| chr13:53730354-53730394 | Ch13 region 2 |
| chr13:53730432-53730472 | Ch13 region 2 |
| chr13:53730511-53730551 | Ch13 region 2 |
| chr13:53730628-53730668 | Ch13 region 2 |
| chr13:53730821-53730861 | Ch13 region 2 |
| chr13:53730868-53730908 | Ch13 region 2 |
| chr13:53730925-53730965 | Ch13 region 2 |
| chr13:53730997-53731037 | Ch13 region 2 |
| chr13:53731071-53731111 | Ch13 region 2 |
| chr13:53731146-53731186 | Ch13 region 2 |
| chr13:53731196-53731236 | Ch13 region 2 |
| chr13:53731241-53731281 | Ch13 region 2 |
| chr13:53731309-53731349 | Ch13 region 2 |
| chr13:53731365-53731405 | Ch13 region 2 |
| chr13:53731444-53731484 | Ch13 region 2 |
| chr13:53731504-53731544 | Ch13 region 2 |
| chr13:53731573-53731613 | Ch13 region 2 |
| chr13:53731629-53731669 | Ch13 region 2 |
| chr13:53731724-53731764 | Ch13 region 2 |
| chr13:53731787-53731827 | Ch13 region 2 |
| chr13:53731833-53731873 | Ch13 region 2 |

| | |
|---|---|
| chr13:53731887-53731927 | Ch13 region 2 |
| chr13:53731961-53732001 | Ch13 region 2 |
| chr13:53732038-53732078 | Ch13 region 2 |
| chr13:53732099-53732139 | Ch13 region 2 |
| chr13:53732148-53732188 | Ch13 region 2 |
| chr13:53732194-53732234 | Ch13 region 2 |
| chr13:53732252-53732292 | Ch13 region 2 |
| chr13:53732332-53732372 | Ch13 region 2 |
| chr13:53732389-53732429 | Ch13 region 2 |
| chr13:53732461-53732501 | Ch13 region 2 |
| chr13:53732518-53732558 | Ch13 region 2 |
| chr13:53732580-53732620 | Ch13 region 2 |
| chr13:53732625-53732665 | Ch13 region 2 |
| chr13:53732679-53732719 | Ch13 region 2 |
| chr13:53732761-53732801 | Ch13 region 2 |
| chr13:53732863-53732903 | Ch13 region 2 |
| chr13:53732908-53732948 | Ch13 region 2 |
| chr13:53733112-53733152 | Ch13 region 2 |
| chr13:53733177-53733217 | Ch13 region 2 |
| chr13:53733295-53733335 | Ch13 region 2 |
| chr13:53733360-53733400 | Ch13 region 2 |
| chr13:53733405-53733445 | Ch13 region 2 |
| chr13:53733476-53733516 | Ch13 region 2 |
| chr13:53733570-53733610 | Ch13 region 2 |
| chr13:53733619-53733659 | Ch13 region 2 |
| chr13:53733677-53733717 | Ch13 region 2 |
| chr13:53733751-53733791 | Ch13 region 2 |
| chr13:53733805-53733845 | Ch13 region 2 |
| chr13:53733880-53733920 | Ch13 region 2 |
| chr13:53733951-53733991 | Ch13 region 2 |
| chr13:53734004-53734044 | Ch13 region 2 |
| chr13:53734082-53734122 | Ch13 region 2 |
| chr13:53734143-53734183 | Ch13 region 2 |
| chr13:53734229-53734269 | Ch13 region 2 |
| chr13:53734303-53734343 | Ch13 region 2 |
| chr13:53734495-53734535 | Ch13 region 2 |
| chr13:53734615-53734655 | Ch13 region 2 |
| chr13:53734674-53734714 | Ch13 region 2 |
| chr13:53734819-53734859 | Ch13 region 2 |
| chr13:53734987-53735027 | Ch13 region 2 |
| chr13:53735042-53735082 | Ch13 region 2 |
| chr13:53735108-53735148 | Ch13 region 2 |
| chr13:53735155-53735195 | Ch13 region 2 |
| chr13:53735220-53735260 | Ch13 region 2 |
| chr13:53735289-53735329 | Ch13 region 2 |
| chr13:53735334-53735374 | Ch13 region 2 |
| chr13:53735405-53735445 | Ch13 region 2 |
| chr13:53735473-53735513 | Ch13 region 2 |
| chr13:53735562-53735602 | Ch13 region 2 |
| chr13:53735659-53735699 | Ch13 region 2 |
| chr13:53735709-53735749 | Ch13 region 2 |
| chr13:53735790-53735830 | Ch13 region 2 |

| | |
|---|---|
| chr13:53735835-53735875 | Ch13 region 2 |
| chr13:53735950-53735990 | Ch13 region 2 |
| chr13:53735999-53736039 | Ch13 region 2 |
| chr13:53736060-53736100 | Ch13 region 2 |
| chr13:53736150-53736190 | Ch13 region 2 |
| chr13:53736350-53736390 | Ch13 region 2 |
| chr13:53736449-53736489 | Ch13 region 2 |
| chr13:53736504-53736544 | Ch13 region 2 |
| chr13:53736557-53736597 | Ch13 region 2 |
| chr13:53736620-53736660 | Ch13 region 2 |
| chr13:53736682-53736722 | Ch13 region 2 |
| chr13:53736743-53736783 | Ch13 region 2 |
| chr13:53736838-53736878 | Ch13 region 2 |
| chr13:53736949-53736989 | Ch13 region 2 |
| chr13:53737013-53737053 | Ch13 region 2 |
| chr13:53737072-53737112 | Ch13 region 2 |
| chr13:53737140-53737180 | Ch13 region 2 |
| chr13:53737394-53737434 | Ch13 region 2 |
| chr13:53737454-53737494 | Ch13 region 2 |
| chr13:53737564-53737604 | Ch13 region 2 |
| chr13:53737646-53737686 | Ch13 region 2 |
| chr13:53737886-53737926 | Ch13 region 2 |
| chr13:53737967-53738007 | Ch13 region 2 |
| chr13:53738024-53738064 | Ch13 region 2 |
| chr13:53738076-53738116 | Ch13 region 2 |
| chr13:53738153-53738193 | Ch13 region 2 |
| chr13:53738315-53738355 | Ch13 region 2 |
| chr13:53738576-53738616 | Ch13 region 2 |
| chr13:53738705-53738745 | Ch13 region 2 |
| chr13:53738954-53738994 | Ch13 region 2 |
| chr13:53739057-53739097 | Ch13 region 2 |
| chr13:53739130-53739170 | Ch13 region 2 |
| chr13:53739188-53739228 | Ch13 region 2 |
| chr13:53739250-53739290 | Ch13 region 2 |
| chr13:53739303-53739343 | Ch13 region 2 |
| chr13:53739376-53739416 | Ch13 region 2 |
| chr13:53739472-53739512 | Ch13 region 2 |
| chr13:53739519-53739559 | Ch13 region 2 |
| chr13:53739594-53739634 | Ch13 region 2 |
| chr13:53739659-53739699 | Ch13 region 2 |
| chr13:53739720-53739760 | Ch13 region 2 |
| chr13:53739808-53739848 | Ch13 region 2 |
| chr13:53739864-53739904 | Ch13 region 2 |
| chr13:53739979-53740019 | Ch13 region 2 |
| chr13:53740042-53740082 | Ch13 region 2 |
| chr13:53740127-53740167 | Ch13 region 2 |
| chr13:53740176-53740216 | Ch13 region 2 |
| chr13:53740267-53740307 | Ch13 region 2 |
| chr13:53740341-53740381 | Ch13 region 2 |
| chr13:53740392-53740432 | Ch13 region 2 |
| chr13:53740439-53740479 | Ch13 region 2 |
| chr13:53740518-53740558 | Ch13 region 2 |

| | |
|---|---|
| chr13:53740565-53740605 | Ch13 region 2 |
| chr13:53740656-53740696 | Ch13 region 2 |
| chr13:53740711-53740751 | Ch13 region 2 |
| chr13:53740786-53740826 | Ch13 region 2 |
| chr13:53740833-53740873 | Ch13 region 2 |
| chr13:53740918-53740958 | Ch13 region 2 |
| chr13:53741002-53741042 | Ch13 region 2 |
| chr13:53741052-53741092 | Ch13 region 2 |
| chr13:53741100-53741140 | Ch13 region 2 |
| chr13:53741157-53741197 | Ch13 region 2 |
| chr13:53741231-53741271 | Ch13 region 2 |
| chr13:53741279-53741319 | Ch13 region 2 |
| chr13:53741325-53741365 | Ch13 region 2 |
| chr13:53741409-53741449 | Ch13 region 2 |
| chr13:53741462-53741502 | Ch13 region 2 |
| chr13:53741516-53741556 | Ch13 region 2 |
| chr13:53741583-53741623 | Ch13 region 2 |
| chr13:53741650-53741690 | Ch13 region 2 |
| chr13:53741711-53741751 | Ch13 region 2 |
| chr13:53741757-53741797 | Ch13 region 2 |
| chr13:53741802-53741842 | Ch13 region 2 |
| chr13:53741851-53741891 | Ch13 region 2 |
| chr13:53741902-53741942 | Ch13 region 2 |
| chr13:53741966-53742006 | Ch13 region 2 |
| chr13:53742040-53742080 | Ch13 region 2 |
| chr13:53742088-53742128 | Ch13 region 2 |
| chr13:53742143-53742183 | Ch13 region 2 |
| chr13:53742188-53742228 | Ch13 region 2 |
| chr13:53742276-53742316 | Ch13 region 2 |
| chr13:53742322-53742362 | Ch13 region 2 |
| chr13:53742478-53742518 | Ch13 region 2 |
| chr13:53742523-53742563 | Ch13 region 2 |
| chr13:53742568-53742608 | Ch13 region 2 |
| chr13:53742724-53742764 | Ch13 region 2 |
| chr13:53742802-53742842 | Ch13 region 2 |
| chr13:53742848-53742888 | Ch13 region 2 |
| chr13:53742896-53742936 | Ch13 region 2 |
| chr13:53742941-53742981 | Ch13 region 2 |
| chr13:53743023-53743063 | Ch13 region 2 |
| chr13:53743081-53743121 | Ch13 region 2 |
| chr13:53743148-53743188 | Ch13 region 2 |
| chr13:53743193-53743233 | Ch13 region 2 |
| chr13:53743243-53743283 | Ch13 region 2 |
| chr13:53743298-53743338 | Ch13 region 2 |
| chr13:53743346-53743386 | Ch13 region 2 |
| chr13:53743396-53743436 | Ch13 region 2 |
| chr13:53743501-53743541 | Ch13 region 2 |
| chr13:53743563-53743603 | Ch13 region 2 |
| chr13:53743668-53743708 | Ch13 region 2 |
| chr13:53743715-53743755 | Ch13 region 2 |
| chr13:53743785-53743825 | Ch13 region 2 |
| chr13:53743875-53743915 | Ch13 region 2 |

| | |
|---|---|
| chr13:53743956-53743996 | Ch13 region 2 |
| chr13:53744030-53744070 | Ch13 region 2 |
| chr13:53744119-53744159 | Ch13 region 2 |
| chr13:53744168-53744208 | Ch13 region 2 |
| chr13:53744313-53744353 | Ch13 region 2 |
| chr13:53744373-53744413 | Ch13 region 2 |
| chr13:53744418-53744458 | Ch13 region 2 |
| chr13:53744560-53744600 | Ch13 region 2 |
| chr13:53744611-53744651 | Ch13 region 2 |
| chr13:53744682-53744722 | Ch13 region 2 |
| chr13:53744774-53744814 | Ch13 region 2 |
| chr13:53744843-53744883 | Ch13 region 2 |
| chr13:53744918-53744958 | Ch13 region 2 |
| chr13:53744983-53745023 | Ch13 region 2 |
| chr13:53745037-53745077 | Ch13 region 2 |
| chr13:53745083-53745123 | Ch13 region 2 |
| chr13:53745167-53745207 | Ch13 region 2 |
| chr13:53745262-53745302 | Ch13 region 2 |
| chr13:53745323-53745363 | Ch13 region 2 |
| chr13:53745400-53745440 | Ch13 region 2 |
| chr13:53745445-53745485 | Ch13 region 2 |
| chr13:53745509-53745549 | Ch13 region 2 |
| chr13:53745595-53745635 | Ch13 region 2 |
| chr13:53745642-53745682 | Ch13 region 2 |
| chr13:53745702-53745742 | Ch13 region 2 |
| chr13:53745747-53745787 | Ch13 region 2 |
| chr13:53745807-53745847 | Ch13 region 2 |
| chr13:53745857-53745897 | Ch13 region 2 |
| chr13:53745907-53745947 | Ch13 region 2 |
| chr13:53745980-53746020 | Ch13 region 2 |
| chr13:53746043-53746083 | Ch13 region 2 |
| chr13:53746128-53746168 | Ch13 region 2 |
| chr13:53746173-53746213 | Ch13 region 2 |
| chr13:53746225-53746265 | Ch13 region 2 |
| chr13:53746282-53746322 | Ch13 region 2 |
| chr13:53746341-53746381 | Ch13 region 2 |
| chr13:53746387-53746427 | Ch13 region 2 |
| chr13:53746486-53746526 | Ch13 region 2 |
| chr13:53746541-53746581 | Ch13 region 2 |
| chr13:53746622-53746662 | Ch13 region 2 |
| chr13:53746698-53746738 | Ch13 region 2 |
| chr13:53746748-53746788 | Ch13 region 2 |
| chr13:53746809-53746849 | Ch13 region 2 |
| chr13:53746894-53746934 | Ch13 region 2 |
| chr13:53747024-53747064 | Ch13 region 2 |
| chr13:53747125-53747165 | Ch13 region 2 |
| chr13:53747189-53747229 | Ch13 region 2 |
| chr13:53747388-53747428 | Ch13 region 2 |
| chr13:53747495-53747535 | Ch13 region 2 |
| chr13:53747561-53747601 | Ch13 region 2 |
| chr13:53747642-53747682 | Ch13 region 2 |
| chr13:53747703-53747743 | Ch13 region 2 |

| | |
|---|---|
| chr13:53747772-53747812 | Ch13 region 2 |
| chr13:53747821-53747861 | Ch13 region 2 |
| chr13:53747889-53747929 | Ch13 region 2 |
| chr13:53747961-53748001 | Ch13 region 2 |
| chr13:53748022-53748062 | Ch13 region 2 |
| chr13:53748069-53748109 | Ch13 region 2 |
| chr13:53748139-53748179 | Ch13 region 2 |
| chr13:53748190-53748230 | Ch13 region 2 |
| chr13:53748260-53748300 | Ch13 region 2 |
| chr13:53748340-53748380 | Ch13 region 2 |
| chr13:53748496-53748536 | Ch13 region 2 |
| chr13:53748624-53748664 | Ch13 region 2 |
| chr13:53748701-53748741 | Ch13 region 2 |
| chr13:53748757-53748797 | Ch13 region 2 |
| chr13:53748833-53748873 | Ch13 region 2 |
| chr13:53748917-53748957 | Ch13 region 2 |
| chr13:53748972-53749012 | Ch13 region 2 |
| chr13:53749029-53749069 | Ch13 region 2 |
| chr13:53749082-53749122 | Ch13 region 2 |
| chr13:53749133-53749173 | Ch13 region 2 |
| chr13:53749214-53749254 | Ch13 region 2 |
| chr13:53749309-53749349 | Ch13 region 2 |
| chr13:53749385-53749425 | Ch13 region 2 |
| chr13:53749430-53749470 | Ch13 region 2 |
| chr13:53749673-53749713 | Ch13 region 2 |
| chr13:53749752-53749792 | Ch13 region 2 |
| chr13:53749828-53749868 | Ch13 region 2 |
| chr13:53749959-53749999 | Ch13 region 2 |
| chr13:53750004-53750044 | Ch13 region 2 |
| chr13:53750195-53750235 | Ch13 region 2 |
| chr13:53750331-53750371 | Ch13 region 2 |
| chr13:53750461-53750501 | Ch13 region 2 |
| chr13:53750538-53750578 | Ch13 region 2 |
| chr13:53750605-53750645 | Ch13 region 2 |
| chr13:53750701-53750741 | Ch13 region 2 |
| chr13:53750781-53750821 | Ch13 region 2 |
| chr13:53750858-53750898 | Ch13 region 2 |
| chr13:53750903-53750943 | Ch13 region 2 |
| chr13:53750983-53751023 | Ch13 region 2 |
| chr13:53751084-53751124 | Ch13 region 2 |
| chr13:53751129-53751169 | Ch13 region 2 |
| chr13:53751209-53751249 | Ch13 region 2 |
| chr13:53751269-53751309 | Ch13 region 2 |
| chr13:53751341-53751381 | Ch13 region 2 |
| chr13:53751437-53751477 | Ch13 region 2 |
| chr13:53751491-53751531 | Ch13 region 2 |
| chr13:53751564-53751604 | Ch13 region 2 |
| chr13:53751645-53751685 | Ch13 region 2 |
| chr13:53751711-53751751 | Ch13 region 2 |
| chr13:53751850-53751890 | Ch13 region 2 |
| chr13:53751943-53751983 | Ch13 region 2 |
| chr13:53752009-53752049 | Ch13 region 2 |

| | |
|---|---|
| chr13:53752092-53752132 | Ch13 region 2 |
| chr13:53752147-53752187 | Ch13 region 2 |
| chr13:53752192-53752232 | Ch13 region 2 |
| chr13:53752244-53752284 | Ch13 region 2 |
| chr13:53752354-53752394 | Ch13 region 2 |
| chr13:53752463-53752503 | Ch13 region 2 |
| chr13:53752520-53752560 | Ch13 region 2 |
| chr13:53752620-53752660 | Ch13 region 2 |
| chr13:53752693-53752733 | Ch13 region 2 |
| chr13:53752843-53752883 | Ch13 region 2 |
| chr13:53752888-53752928 | Ch13 region 2 |
| chr13:53753038-53753078 | Ch13 region 2 |
| chr13:53753222-53753262 | Ch13 region 2 |
| chr13:53753269-53753309 | Ch13 region 2 |
| chr13:53753379-53753419 | Ch13 region 2 |
| chr13:53753492-53753532 | Ch13 region 2 |
| chr13:53753541-53753581 | Ch13 region 2 |
| chr13:53753612-53753652 | Ch13 region 2 |
| chr13:53753720-53753760 | Ch13 region 2 |
| chr13:53753765-53753805 | Ch13 region 2 |
| chr13:53753848-53753888 | Ch13 region 2 |
| chr13:53753900-53753940 | Ch13 region 2 |
| chr13:53753991-53754031 | Ch13 region 2 |
| chr13:53754170-53754210 | Ch13 region 2 |
| chr13:53754259-53754299 | Ch13 region 2 |
| chr13:53754345-53754385 | Ch13 region 2 |
| chr13:53754423-53754463 | Ch13 region 2 |
| chr13:53754506-53754546 | Ch13 region 2 |
| chr13:53754624-53754664 | Ch13 region 2 |
| chr13:53754685-53754725 | Ch13 region 2 |
| chr13:53754748-53754788 | Ch13 region 2 |
| chr13:53754800-53754840 | Ch13 region 2 |
| chr13:53754849-53754889 | Ch13 region 2 |
| chr13:53754894-53754934 | Ch13 region 2 |
| chr13:53754959-53754999 | Ch13 region 2 |
| chr13:53755006-53755046 | Ch13 region 2 |
| chr13:53755122-53755162 | Ch13 region 2 |
| chr13:53755181-53755221 | Ch13 region 2 |
| chr13:53755293-53755333 | Ch13 region 2 |
| chr13:53755353-53755393 | Ch13 region 2 |
| chr13:53755402-53755442 | Ch13 region 2 |
| chr13:53755461-53755501 | Ch13 region 2 |
| chr13:53755506-53755546 | Ch13 region 2 |
| chr13:53755566-53755606 | Ch13 region 2 |
| chr13:53755623-53755663 | Ch13 region 2 |
| chr13:53755685-53755725 | Ch13 region 2 |
| chr13:53755741-53755781 | Ch13 region 2 |
| chr13:53755786-53755826 | Ch13 region 2 |
| chr13:53755878-53755918 | Ch13 region 2 |
| chr13:53755965-53756005 | Ch13 region 2 |
| chr13:53756012-53756052 | Ch13 region 2 |
| chr13:53756281-53756321 | Ch13 region 2 |

| | |
|---|---|
| chr13:53756332-53756372 | Ch13 region 2 |
| chr13:53756380-53756420 | Ch13 region 2 |
| chr13:53756433-53756473 | Ch13 region 2 |
| chr13:53756539-53756579 | Ch13 region 2 |
| chr13:53756607-53756647 | Ch13 region 2 |
| chr13:53756696-53756736 | Ch13 region 2 |
| chr13:53756745-53756785 | Ch13 region 2 |
| chr13:53756843-53756883 | Ch13 region 2 |
| chr13:53756888-53756928 | Ch13 region 2 |
| chr13:53756992-53757032 | Ch13 region 2 |
| chr13:53757072-53757112 | Ch13 region 2 |
| chr13:53757162-53757202 | Ch13 region 2 |
| chr13:53757226-53757266 | Ch13 region 2 |
| chr13:53757301-53757341 | Ch13 region 2 |
| chr13:53757346-53757386 | Ch13 region 2 |
| chr13:53757395-53757435 | Ch13 region 2 |
| chr13:53757472-53757512 | Ch13 region 2 |
| chr13:53757549-53757589 | Ch13 region 2 |
| chr13:53757599-53757639 | Ch13 region 2 |
| chr13:53757647-53757687 | Ch13 region 2 |
| chr13:53757746-53757786 | Ch13 region 2 |
| chr13:53757820-53757860 | Ch13 region 2 |
| chr13:53757900-53757940 | Ch13 region 2 |
| chr13:53757963-53758003 | Ch13 region 2 |
| chr13:53758009-53758049 | Ch13 region 2 |
| chr13:53758055-53758095 | Ch13 region 2 |
| chr13:53758150-53758190 | Ch13 region 2 |
| chr13:53758364-53758404 | Ch13 region 2 |
| chr13:53758412-53758452 | Ch13 region 2 |
| chr13:53758466-53758506 | Ch13 region 2 |
| chr13:53758525-53758565 | Ch13 region 2 |
| chr13:53758700-53758740 | Ch13 region 2 |
| chr13:53758811-53758851 | Ch13 region 2 |
| chr13:53758931-53758971 | Ch13 region 2 |
| chr13:53758985-53759025 | Ch13 region 2 |
| chr13:53759062-53759102 | Ch13 region 2 |
| chr13:53759128-53759168 | Ch13 region 2 |
| chr13:53759277-53759317 | Ch13 region 2 |
| chr13:53759553-53759593 | Ch13 region 2 |
| chr13:53759598-53759638 | Ch13 region 2 |
| chr13:53759675-53759715 | Ch13 region 2 |
| chr13:53759792-53759832 | Ch13 region 2 |
| chr13:53760077-53760117 | Ch13 region 2 |
| chr13:53760129-53760169 | Ch13 region 2 |
| chr13:53760274-53760314 | Ch13 region 2 |
| chr13:53760331-53760371 | Ch13 region 2 |
| chr13:53760376-53760416 | Ch13 region 2 |
| chr13:53760805-53760845 | Ch13 region 2 |
| chr13:53760904-53760944 | Ch13 region 2 |
| chr13:53760990-53761030 | Ch13 region 2 |
| chr13:53761043-53761083 | Ch13 region 2 |
| chr13:53761112-53761152 | Ch13 region 2 |

| | |
|---|---|
| chr13:53761198-53761238 | Ch13 region 2 |
| chr13:53761251-53761291 | Ch13 region 2 |
| chr13:53761384-53761424 | Ch13 region 2 |
| chr13:53761512-53761552 | Ch13 region 2 |
| chr13:53761604-53761644 | Ch13 region 2 |
| chr13:53761755-53761795 | Ch13 region 2 |
| chr13:53761938-53761978 | Ch13 region 2 |
| chr13:53761983-53762023 | Ch13 region 2 |
| chr13:53762028-53762068 | Ch13 region 2 |
| chr13:53762094-53762134 | Ch13 region 2 |
| chr13:53762182-53762222 | Ch13 region 2 |
| chr13:53762254-53762294 | Ch13 region 2 |
| chr13:53762307-53762347 | Ch13 region 2 |
| chr13:53762359-53762399 | Ch13 region 2 |
| chr13:53762456-53762496 | Ch13 region 2 |
| chr13:53762515-53762555 | Ch13 region 2 |
| chr13:53762618-53762658 | Ch13 region 2 |
| chr13:53762700-53762740 | Ch13 region 2 |
| chr13:53762748-53762788 | Ch13 region 2 |
| chr13:53762813-53762853 | Ch13 region 2 |
| chr13:53762859-53762899 | Ch13 region 2 |
| chr13:53762910-53762950 | Ch13 region 2 |
| chr13:53763018-53763058 | Ch13 region 2 |
| chr13:53763064-53763104 | Ch13 region 2 |
| chr13:53763111-53763151 | Ch13 region 2 |
| chr13:53763382-53763422 | Ch13 region 2 |
| chr13:53763434-53763474 | Ch13 region 2 |
| chr13:53763481-53763521 | Ch13 region 2 |
| chr13:53763615-53763655 | Ch13 region 2 |
| chr13:53763685-53763725 | Ch13 region 2 |
| chr13:53763732-53763772 | Ch13 region 2 |
| chr13:53763789-53763829 | Ch13 region 2 |
| chr13:53763907-53763947 | Ch13 region 2 |
| chr13:53763984-53764024 | Ch13 region 2 |
| chr13:53764058-53764098 | Ch13 region 2 |
| chr13:53764116-53764156 | Ch13 region 2 |
| chr13:53764178-53764218 | Ch13 region 2 |
| chr13:53764253-53764293 | Ch13 region 2 |
| chr13:53764309-53764349 | Ch13 region 2 |
| chr13:53764356-53764396 | Ch13 region 2 |
| chr13:53764451-53764491 | Ch13 region 2 |
| chr13:53764509-53764549 | Ch13 region 2 |
| chr13:53764576-53764616 | Ch13 region 2 |
| chr13:53764646-53764686 | Ch13 region 2 |
| chr13:53764706-53764746 | Ch13 region 2 |
| chr13:53764772-53764812 | Ch13 region 2 |
| chr13:53764868-53764908 | Ch13 region 2 |
| chr13:53764915-53764955 | Ch13 region 2 |
| chr13:53764994-53765034 | Ch13 region 2 |
| chr13:53765051-53765091 | Ch13 region 2 |
| chr13:53765119-53765159 | Ch13 region 2 |
| chr13:53765202-53765242 | Ch13 region 2 |

| | |
|---|---|
| chr13:53765293-53765333 | Ch13 region 2 |
| chr13:53765367-53765407 | Ch13 region 2 |
| chr13:53765435-53765475 | Ch13 region 2 |
| chr13:53765496-53765536 | Ch13 region 2 |
| chr13:53765569-53765609 | Ch13 region 2 |
| chr13:53765614-53765654 | Ch13 region 2 |
| chr13:53765699-53765739 | Ch13 region 2 |
| chr13:53765768-53765808 | Ch13 region 2 |
| chr13:53765847-53765887 | Ch13 region 2 |
| chr13:53765901-53765941 | Ch13 region 2 |
| chr13:53765953-53765993 | Ch13 region 2 |
| chr13:53766015-53766055 | Ch13 region 2 |
| chr13:53766089-53766129 | Ch13 region 2 |
| chr13:53766201-53766241 | Ch13 region 2 |
| chr13:53766259-53766299 | Ch13 region 2 |
| chr13:53766334-53766374 | Ch13 region 2 |
| chr13:53766406-53766446 | Ch13 region 2 |
| chr13:53766480-53766520 | Ch13 region 2 |
| chr13:53766542-53766582 | Ch13 region 2 |
| chr13:53766597-53766637 | Ch13 region 2 |
| chr13:53766650-53766690 | Ch13 region 2 |
| chr13:53766717-53766757 | Ch13 region 2 |
| chr13:53766828-53766868 | Ch13 region 2 |
| chr13:53766873-53766913 | Ch13 region 2 |
| chr13:53766940-53766980 | Ch13 region 2 |
| chr13:53766989-53767029 | Ch13 region 2 |
| chr13:53767037-53767077 | Ch13 region 2 |
| chr13:53767083-53767123 | Ch13 region 2 |
| chr13:53767138-53767178 | Ch13 region 2 |
| chr13:53767185-53767225 | Ch13 region 2 |
| chr13:53767272-53767312 | Ch13 region 2 |
| chr13:53767375-53767415 | Ch13 region 2 |
| chr13:53767466-53767506 | Ch13 region 2 |
| chr13:53767556-53767596 | Ch13 region 2 |
| chr13:53767632-53767672 | Ch13 region 2 |
| chr13:53767678-53767718 | Ch13 region 2 |
| chr13:53767768-53767808 | Ch13 region 2 |
| chr13:53767857-53767897 | Ch13 region 2 |
| chr13:53767940-53767980 | Ch13 region 2 |
| chr13:53767990-53768030 | Ch13 region 2 |
| chr13:53768053-53768093 | Ch13 region 2 |
| chr13:53768169-53768209 | Ch13 region 2 |
| chr13:53768253-53768293 | Ch13 region 2 |
| chr13:53768298-53768338 | Ch13 region 2 |
| chr13:53768367-53768407 | Ch13 region 2 |
| chr13:53768412-53768452 | Ch13 region 2 |
| chr13:53768474-53768514 | Ch13 region 2 |
| chr13:53768585-53768625 | Ch13 region 2 |
| chr13:53768656-53768696 | Ch13 region 2 |
| chr13:53768701-53768741 | Ch13 region 2 |
| chr13:53768784-53768824 | Ch13 region 2 |
| chr13:53768829-53768869 | Ch13 region 2 |

| | |
|---|---|
| chr13:53768874-53768914 | Ch13 region 2 |
| chr13:53769132-53769172 | Ch13 region 2 |
| chr13:53769181-53769221 | Ch13 region 2 |
| chr13:53769226-53769266 | Ch13 region 2 |
| chr13:53769273-53769313 | Ch13 region 2 |
| chr13:53769333-53769373 | Ch13 region 2 |
| chr13:53769412-53769452 | Ch13 region 2 |
| chr13:53769573-53769613 | Ch13 region 2 |
| chr13:53769818-53769858 | Ch13 region 2 |
| chr13:53769864-53769904 | Ch13 region 2 |
| chr13:53769918-53769958 | Ch13 region 2 |
| chr13:53770251-53770291 | Ch13 region 2 |
| chr13:53770437-53770477 | Ch13 region 2 |
| chr13:53770504-53770544 | Ch13 region 2 |
| chr13:53770577-53770617 | Ch13 region 2 |
| chr13:53770657-53770697 | Ch13 region 2 |
| chr13:53770762-53770802 | Ch13 region 2 |
| chr13:53770903-53770943 | Ch13 region 2 |
| chr13:53770964-53771004 | Ch13 region 2 |
| chr13:53771044-53771084 | Ch13 region 2 |
| chr13:53771092-53771132 | Ch13 region 2 |
| chr13:53771181-53771221 | Ch13 region 2 |
| chr13:53771259-53771299 | Ch13 region 2 |
| chr13:53771392-53771432 | Ch13 region 2 |
| chr13:53771469-53771509 | Ch13 region 2 |
| chr13:53771545-53771585 | Ch13 region 2 |
| chr13:53771641-53771681 | Ch13 region 2 |
| chr13:53771719-53771759 | Ch13 region 2 |
| chr13:53771801-53771841 | Ch13 region 2 |
| chr13:53771889-53771929 | Ch13 region 2 |
| chr13:53771957-53771997 | Ch13 region 2 |
| chr13:53772027-53772067 | Ch13 region 2 |
| chr13:53772106-53772146 | Ch13 region 2 |
| chr13:53772171-53772211 | Ch13 region 2 |
| chr13:53772239-53772279 | Ch13 region 2 |
| chr13:53772296-53772336 | Ch13 region 2 |
| chr13:53772341-53772381 | Ch13 region 2 |
| chr13:53772407-53772447 | Ch13 region 2 |
| chr13:53772488-53772528 | Ch13 region 2 |
| chr13:53772533-53772573 | Ch13 region 2 |
| chr13:53772592-53772632 | Ch13 region 2 |
| chr13:53772669-53772709 | Ch13 region 2 |
| chr13:53772716-53772756 | Ch13 region 2 |
| chr13:53772776-53772816 | Ch13 region 2 |
| chr13:53772826-53772866 | Ch13 region 2 |
| chr13:53772879-53772919 | Ch13 region 2 |
| chr13:53772934-53772974 | Ch13 region 2 |
| chr13:53772979-53773019 | Ch13 region 2 |
| chr13:53773028-53773068 | Ch13 region 2 |
| chr13:53773076-53773116 | Ch13 region 2 |
| chr13:53773127-53773167 | Ch13 region 2 |
| chr13:53773172-53773212 | Ch13 region 2 |

| | |
|---|---|
| chr13:53773243-53773283 | Ch13 region 2 |
| chr13:53773294-53773334 | Ch13 region 2 |
| chr13:53773342-53773382 | Ch13 region 2 |
| chr13:53773400-53773440 | Ch13 region 2 |
| chr13:53773472-53773512 | Ch13 region 2 |
| chr13:53773529-53773569 | Ch13 region 2 |
| chr13:53773608-53773648 | Ch13 region 2 |
| chr13:53773681-53773721 | Ch13 region 2 |
| chr13:53773747-53773787 | Ch13 region 2 |
| chr13:53773819-53773859 | Ch13 region 2 |
| chr13:53773884-53773924 | Ch13 region 2 |
| chr13:53773954-53773994 | Ch13 region 2 |
| chr13:53773999-53774039 | Ch13 region 2 |
| chr13:53774089-53774129 | Ch13 region 2 |
| chr13:53774157-53774197 | Ch13 region 2 |
| chr13:53774262-53774302 | Ch13 region 2 |
| chr13:53774334-53774374 | Ch13 region 2 |
| chr13:53774379-53774419 | Ch13 region 2 |
| chr13:53774446-53774486 | Ch13 region 2 |
| chr13:53774498-53774538 | Ch13 region 2 |
| chr13:53774548-53774588 | Ch13 region 2 |
| chr13:53774608-53774648 | Ch13 region 2 |
| chr13:53774657-53774697 | Ch13 region 2 |
| chr13:53774727-53774767 | Ch13 region 2 |
| chr13:53774815-53774855 | Ch13 region 2 |
| chr13:53774869-53774909 | Ch13 region 2 |
| chr13:53774915-53774955 | Ch13 region 2 |
| chr13:53774967-53775007 | Ch13 region 2 |
| chr13:53775058-53775098 | Ch13 region 2 |
| chr13:53775151-53775191 | Ch13 region 2 |
| chr13:53775198-53775238 | Ch13 region 2 |
| chr13:53775267-53775307 | Ch13 region 2 |
| chr13:53775371-53775411 | Ch13 region 2 |
| chr13:53775421-53775461 | Ch13 region 2 |
| chr13:53775511-53775551 | Ch13 region 2 |
| chr13:53775589-53775629 | Ch13 region 2 |
| chr13:53775634-53775674 | Ch13 region 2 |
| chr13:53775720-53775760 | Ch13 region 2 |
| chr13:53775799-53775839 | Ch13 region 2 |
| chr13:53775849-53775889 | Ch13 region 2 |
| chr13:53775923-53775963 | Ch13 region 2 |
| chr13:53775968-53776008 | Ch13 region 2 |
| chr13:53776040-53776080 | Ch13 region 2 |
| chr13:53776085-53776125 | Ch13 region 2 |
| chr13:53776150-53776190 | Ch13 region 2 |
| chr13:53776213-53776253 | Ch13 region 2 |
| chr13:53776288-53776328 | Ch13 region 2 |
| chr13:53776378-53776418 | Ch13 region 2 |
| chr13:53776444-53776484 | Ch13 region 2 |
| chr13:53776558-53776598 | Ch13 region 2 |
| chr13:53776616-53776656 | Ch13 region 2 |
| chr13:53776718-53776758 | Ch13 region 2 |

| | |
|---|---|
| chr13:53776798-53776838 | Ch13 region 2 |
| chr13:53776847-53776887 | Ch13 region 2 |
| chr13:53776914-53776954 | Ch13 region 2 |
| chr13:53776959-53776999 | Ch13 region 2 |
| chr13:53777037-53777077 | Ch13 region 2 |
| chr13:53777101-53777141 | Ch13 region 2 |
| chr13:53777272-53777312 | Ch13 region 2 |
| chr13:53777329-53777369 | Ch13 region 2 |
| chr13:53777374-53777414 | Ch13 region 2 |
| chr13:53777428-53777468 | Ch13 region 2 |
| chr13:53777496-53777536 | Ch13 region 2 |
| chr13:53777624-53777664 | Ch13 region 2 |
| chr13:53777679-53777719 | Ch13 region 2 |
| chr13:53777739-53777779 | Ch13 region 2 |
| chr13:53777785-53777825 | Ch13 region 2 |
| chr13:53777870-53777910 | Ch13 region 2 |
| chr13:53777931-53777971 | Ch13 region 2 |
| chr13:53778017-53778057 | Ch13 region 2 |
| chr13:53778077-53778117 | Ch13 region 2 |
| chr13:53778160-53778200 | Ch13 region 2 |
| chr13:53778222-53778262 | Ch13 region 2 |
| chr13:53778270-53778310 | Ch13 region 2 |
| chr13:53778315-53778355 | Ch13 region 2 |
| chr13:53778383-53778423 | Ch13 region 2 |
| chr13:53778469-53778509 | Ch13 region 2 |
| chr13:53778534-53778574 | Ch13 region 2 |
| chr13:53778628-53778668 | Ch13 region 2 |
| chr13:53778680-53778720 | Ch13 region 2 |
| chr13:53778765-53778805 | Ch13 region 2 |
| chr13:53778854-53778894 | Ch13 region 2 |
| chr13:53778904-53778944 | Ch13 region 2 |
| chr13:53778964-53779004 | Ch13 region 2 |
| chr13:53779009-53779049 | Ch13 region 2 |
| chr13:53779054-53779094 | Ch13 region 2 |
| chr13:53779108-53779148 | Ch13 region 2 |
| chr13:53779207-53779247 | Ch13 region 2 |
| chr13:53779283-53779323 | Ch13 region 2 |
| chr13:53779348-53779388 | Ch13 region 2 |
| chr13:53779393-53779433 | Ch13 region 2 |
| chr13:53779438-53779478 | Ch13 region 2 |
| chr13:53779488-53779528 | Ch13 region 2 |
| chr13:53779569-53779609 | Ch13 region 2 |
| chr13:53779619-53779659 | Ch13 region 2 |
| chr13:53779730-53779770 | Ch13 region 2 |
| chr13:53779819-53779859 | Ch13 region 2 |
| chr13:53779907-53779947 | Ch13 region 2 |
| chr13:53779976-53780016 | Ch13 region 2 |
| chr13:53780030-53780070 | Ch13 region 2 |
| chr13:53780098-53780138 | Ch13 region 2 |
| chr13:53780182-53780222 | Ch13 region 2 |
| chr13:53780230-53780270 | Ch13 region 2 |
| chr13:53780307-53780347 | Ch13 region 2 |

| | |
|---|---|
| chr13:53780381-53780421 | Ch13 region 2 |
| chr13:53780428-53780468 | Ch13 region 2 |
| chr13:53780509-53780549 | Ch13 region 2 |
| chr13:53780664-53780704 | Ch13 region 2 |
| chr13:53780756-53780796 | Ch13 region 2 |
| chr13:53780813-53780853 | Ch13 region 2 |
| chr13:53780864-53780904 | Ch13 region 2 |
| chr13:53781001-53781041 | Ch13 region 2 |
| chr13:53781067-53781107 | Ch13 region 2 |
| chr13:53781112-53781152 | Ch13 region 2 |
| chr13:53781170-53781210 | Ch13 region 2 |
| chr13:53781224-53781264 | Ch13 region 2 |
| chr13:53781280-53781320 | Ch13 region 2 |
| chr13:53781336-53781376 | Ch13 region 2 |
| chr13:53781404-53781444 | Ch13 region 2 |
| chr13:53781573-53781613 | Ch13 region 2 |
| chr13:53781649-53781689 | Ch13 region 2 |
| chr13:53781724-53781764 | Ch13 region 2 |
| chr13:53781771-53781811 | Ch13 region 2 |
| chr13:53781819-53781859 | Ch13 region 2 |
| chr13:53781881-53781921 | Ch13 region 2 |
| chr13:53781948-53781988 | Ch13 region 2 |
| chr13:53782041-53782081 | Ch13 region 2 |
| chr13:53782102-53782142 | Ch13 region 2 |
| chr13:53782173-53782213 | Ch13 region 2 |
| chr13:53782241-53782281 | Ch13 region 2 |
| chr13:53782305-53782345 | Ch13 region 2 |
| chr13:53782383-53782423 | Ch13 region 2 |
| chr13:53782440-53782480 | Ch13 region 2 |
| chr13:53782553-53782593 | Ch13 region 2 |
| chr13:53782599-53782639 | Ch13 region 2 |
| chr13:53782680-53782720 | Ch13 region 2 |
| chr13:53782748-53782788 | Ch13 region 2 |
| chr13:53782800-53782840 | Ch13 region 2 |
| chr13:53782870-53782910 | Ch13 region 2 |
| chr13:53782919-53782959 | Ch13 region 2 |
| chr13:53782972-53783012 | Ch13 region 2 |
| chr13:53783033-53783073 | Ch13 region 2 |
| chr13:53783078-53783118 | Ch13 region 2 |
| chr13:53783193-53783233 | Ch13 region 2 |
| chr13:53783292-53783332 | Ch13 region 2 |
| chr13:53783352-53783392 | Ch13 region 2 |
| chr13:53783400-53783440 | Ch13 region 2 |
| chr13:53783522-53783562 | Ch13 region 2 |
| chr13:53783573-53783613 | Ch13 region 2 |
| chr13:53783634-53783674 | Ch13 region 2 |
| chr13:53783729-53783769 | Ch13 region 2 |
| chr13:53783910-53783950 | Ch13 region 2 |
| chr13:53783995-53784035 | Ch13 region 2 |
| chr13:53784151-53784191 | Ch13 region 2 |
| chr13:53784275-53784315 | Ch13 region 2 |
| chr13:53784409-53784449 | Ch13 region 2 |

| | |
|---|---|
| chr13:53784517-53784557 | Ch13 region 2 |
| chr13:53784563-53784603 | Ch13 region 2 |
| chr13:53784608-53784648 | Ch13 region 2 |
| chr13:53784654-53784694 | Ch13 region 2 |
| chr13:53784722-53784762 | Ch13 region 2 |
| chr13:53784931-53784971 | Ch13 region 2 |
| chr13:53784981-53785021 | Ch13 region 2 |
| chr13:53785055-53785095 | Ch13 region 2 |
| chr13:53785139-53785179 | Ch13 region 2 |
| chr13:53785184-53785224 | Ch13 region 2 |
| chr13:53785336-53785376 | Ch13 region 2 |
| chr13:53785397-53785437 | Ch13 region 2 |
| chr13:53785450-53785490 | Ch13 region 2 |
| chr13:53785506-53785546 | Ch13 region 2 |
| chr13:53785599-53785639 | Ch13 region 2 |
| chr13:53785749-53785789 | Ch13 region 2 |
| chr13:53785822-53785862 | Ch13 region 2 |
| chr13:53785899-53785939 | Ch13 region 2 |
| chr13:53785946-53785986 | Ch13 region 2 |
| chr13:53786017-53786057 | Ch13 region 2 |
| chr13:53786080-53786120 | Ch13 region 2 |
| chr13:53786135-53786175 | Ch13 region 2 |
| chr13:53786188-53786228 | Ch13 region 2 |
| chr13:53786253-53786293 | Ch13 region 2 |
| chr13:53786340-53786380 | Ch13 region 2 |
| chr13:53786458-53786498 | Ch13 region 2 |
| chr13:53786511-53786551 | Ch13 region 2 |
| chr13:53786560-53786600 | Ch13 region 2 |
| chr13:53786691-53786731 | Ch13 region 2 |
| chr13:53786910-53786950 | Ch13 region 2 |
| chr13:53787013-53787053 | Ch13 region 2 |
| chr13:53787090-53787130 | Ch13 region 2 |
| chr13:53787219-53787259 | Ch13 region 2 |
| chr13:53787269-53787309 | Ch13 region 2 |
| chr13:53787369-53787409 | Ch13 region 2 |
| chr13:53787414-53787454 | Ch13 region 2 |
| chr13:53787485-53787525 | Ch13 region 2 |
| chr13:53787537-53787577 | Ch13 region 2 |
| chr13:53787594-53787634 | Ch13 region 2 |
| chr13:53787640-53787680 | Ch13 region 2 |
| chr13:53787696-53787736 | Ch13 region 2 |
| chr13:53787747-53787787 | Ch13 region 2 |
| chr13:53787794-53787834 | Ch13 region 2 |
| chr13:53787886-53787926 | Ch13 region 2 |
| chr13:53787931-53787971 | Ch13 region 2 |
| chr13:53787979-53788019 | Ch13 region 2 |
| chr13:53788084-53788124 | Ch13 region 2 |
| chr13:53788155-53788195 | Ch13 region 2 |
| chr13:53788223-53788263 | Ch13 region 2 |
| chr13:53788274-53788314 | Ch13 region 2 |
| chr13:53788362-53788402 | Ch13 region 2 |
| chr13:53788430-53788470 | Ch13 region 2 |

| | |
|---|---|
| chr13:53788532-53788572 | Ch13 region 2 |
| chr13:53788611-53788651 | Ch13 region 2 |
| chr13:53788669-53788709 | Ch13 region 2 |
| chr13:53788715-53788755 | Ch13 region 2 |
| chr13:53788802-53788842 | Ch13 region 2 |
| chr13:53788885-53788925 | Ch13 region 2 |
| chr13:53788941-53788981 | Ch13 region 2 |
| chr13:53789033-53789073 | Ch13 region 2 |
| chr13:53789093-53789133 | Ch13 region 2 |
| chr13:53789220-53789260 | Ch13 region 2 |
| chr13:53789298-53789338 | Ch13 region 2 |
| chr13:53789379-53789419 | Ch13 region 2 |
| chr13:53789438-53789478 | Ch13 region 2 |
| chr13:53789500-53789540 | Ch13 region 2 |
| chr13:53789545-53789585 | Ch13 region 2 |
| chr13:53789612-53789652 | Ch13 region 2 |
| chr13:53789659-53789699 | Ch13 region 2 |
| chr13:53789717-53789757 | Ch13 region 2 |
| chr13:53789762-53789802 | Ch13 region 2 |
| chr13:53789844-53789884 | Ch13 region 2 |
| chr13:53789895-53789935 | Ch13 region 2 |
| chr13:53789996-53790036 | Ch13 region 2 |
| chr13:53790081-53790121 | Ch13 region 2 |
| chr13:53790195-53790235 | Ch13 region 2 |
| chr13:53790254-53790294 | Ch13 region 2 |
| chr13:53790310-53790350 | Ch13 region 2 |
| chr13:53790355-53790395 | Ch13 region 2 |
| chr13:53790443-53790483 | Ch13 region 2 |
| chr13:53790488-53790528 | Ch13 region 2 |
| chr13:53790533-53790573 | Ch13 region 2 |
| chr13:53790594-53790634 | Ch13 region 2 |
| chr13:53790639-53790679 | Ch13 region 2 |
| chr13:53790691-53790731 | Ch13 region 2 |
| chr13:53790790-53790830 | Ch13 region 2 |
| chr13:53790876-53790916 | Ch13 region 2 |
| chr13:53790921-53790961 | Ch13 region 2 |
| chr13:53791034-53791074 | Ch13 region 2 |
| chr13:53791087-53791127 | Ch13 region 2 |
| chr13:53791137-53791177 | Ch13 region 2 |
| chr13:53791266-53791306 | Ch13 region 2 |
| chr13:53791329-53791369 | Ch13 region 2 |
| chr13:53791400-53791440 | Ch13 region 2 |
| chr13:53791445-53791485 | Ch13 region 2 |
| chr13:53791493-53791533 | Ch13 region 2 |
| chr13:53791551-53791591 | Ch13 region 2 |
| chr13:53791597-53791637 | Ch13 region 2 |
| chr13:53791688-53791728 | Ch13 region 2 |
| chr13:53791774-53791814 | Ch13 region 2 |
| chr13:53791839-53791879 | Ch13 region 2 |
| chr13:53791897-53791937 | Ch13 region 2 |
| chr13:53791957-53791997 | Ch13 region 2 |
| chr13:53792015-53792055 | Ch13 region 2 |

| | |
|---|---|
| chr13:53792061-53792101 | Ch13 region 2 |
| chr13:53792113-53792153 | Ch13 region 2 |
| chr13:53792161-53792201 | Ch13 region 2 |
| chr13:53792207-53792247 | Ch13 region 2 |
| chr13:53792301-53792341 | Ch13 region 2 |
| chr13:53792363-53792403 | Ch13 region 2 |
| chr13:53792468-53792508 | Ch13 region 2 |
| chr13:53792515-53792555 | Ch13 region 2 |
| chr13:53792766-53792806 | Ch13 region 2 |
| chr13:53792950-53792990 | Ch13 region 2 |
| chr13:53793071-53793111 | Ch13 region 2 |
| chr13:53793217-53793257 | Ch13 region 2 |
| chr13:53793274-53793314 | Ch13 region 2 |
| chr13:53793375-53793415 | Ch13 region 2 |
| chr13:53793454-53793494 | Ch13 region 2 |
| chr13:53793548-53793588 | Ch13 region 2 |
| chr13:53793682-53793722 | Ch13 region 2 |
| chr13:53793766-53793806 | Ch13 region 2 |
| chr13:53793868-53793908 | Ch13 region 2 |
| chr13:53793979-53794019 | Ch13 region 2 |
| chr13:53794064-53794104 | Ch13 region 2 |
| chr13:53794132-53794172 | Ch13 region 2 |
| chr13:53794204-53794244 | Ch13 region 2 |
| chr13:53794283-53794323 | Ch13 region 2 |
| chr13:53794340-53794380 | Ch13 region 2 |
| chr13:53794385-53794425 | Ch13 region 2 |
| chr13:53794453-53794493 | Ch13 region 2 |
| chr13:53794504-53794544 | Ch13 region 2 |
| chr13:53794591-53794631 | Ch13 region 2 |
| chr13:53794649-53794689 | Ch13 region 2 |
| chr13:53794770-53794810 | Ch13 region 2 |
| chr13:53794868-53794908 | Ch13 region 2 |
| chr13:53794967-53795007 | Ch13 region 2 |
| chr13:53795012-53795052 | Ch13 region 2 |
| chr13:53795092-53795132 | Ch13 region 2 |
| chr13:53795146-53795186 | Ch13 region 2 |
| chr13:53795218-53795258 | Ch13 region 2 |
| chr13:53795294-53795334 | Ch13 region 2 |
| chr13:53795367-53795407 | Ch13 region 2 |
| chr13:53795423-53795463 | Ch13 region 2 |
| chr13:53795491-53795531 | Ch13 region 2 |
| chr13:53795538-53795578 | Ch13 region 2 |
| chr13:53795622-53795662 | Ch13 region 2 |
| chr13:53795671-53795711 | Ch13 region 2 |
| chr13:53795858-53795898 | Ch13 region 2 |
| chr13:53795948-53795988 | Ch13 region 2 |
| chr13:53796041-53796081 | Ch13 region 2 |
| chr13:53796099-53796139 | Ch13 region 2 |
| chr13:53796157-53796197 | Ch13 region 2 |
| chr13:53796207-53796247 | Ch13 region 2 |
| chr13:53796280-53796320 | Ch13 region 2 |
| chr13:53796376-53796416 | Ch13 region 2 |

| | |
|---|---|
| chr13:53796445-53796485 | Ch13 region 2 |
| chr13:53796512-53796552 | Ch13 region 2 |
| chr13:53796605-53796645 | Ch13 region 2 |
| chr13:53796668-53796708 | Ch13 region 2 |
| chr13:53796719-53796759 | Ch13 region 2 |
| chr13:53796843-53796883 | Ch13 region 2 |
| chr13:53796899-53796939 | Ch13 region 2 |
| chr13:53796973-53797013 | Ch13 region 2 |
| chr13:53797098-53797138 | Ch13 region 2 |
| chr13:53797164-53797204 | Ch13 region 2 |
| chr13:53797217-53797257 | Ch13 region 2 |
| chr13:53797341-53797381 | Ch13 region 2 |
| chr13:53797386-53797426 | Ch13 region 2 |
| chr13:53797441-53797481 | Ch13 region 2 |
| chr13:53797497-53797537 | Ch13 region 2 |
| chr13:53797566-53797606 | Ch13 region 2 |
| chr13:53797619-53797659 | Ch13 region 2 |
| chr13:53797680-53797720 | Ch13 region 2 |
| chr13:53797725-53797765 | Ch13 region 2 |
| chr13:53797770-53797810 | Ch13 region 2 |
| chr13:53797831-53797871 | Ch13 region 2 |
| chr13:53797886-53797926 | Ch13 region 2 |
| chr13:53797962-53798002 | Ch13 region 2 |
| chr13:53798046-53798086 | Ch13 region 2 |
| chr13:53798106-53798146 | Ch13 region 2 |
| chr13:53798180-53798220 | Ch13 region 2 |
| chr13:53798246-53798286 | Ch13 region 2 |
| chr13:53798301-53798341 | Ch13 region 2 |
| chr13:53798389-53798429 | Ch13 region 2 |
| chr13:53798457-53798497 | Ch13 region 2 |
| chr13:53798868-53798908 | Ch13 region 2 |
| chr13:53798987-53799027 | Ch13 region 2 |
| chr13:53799123-53799163 | Ch13 region 2 |
| chr13:53799191-53799231 | Ch13 region 2 |
| chr13:53799340-53799380 | Ch13 region 2 |
| chr13:53799424-53799464 | Ch13 region 2 |
| chr13:53799546-53799586 | Ch13 region 2 |
| chr13:53799638-53799678 | Ch13 region 2 |
| chr13:53799777-53799817 | Ch13 region 2 |
| chr13:53799827-53799867 | Ch13 region 2 |
| chr13:53799880-53799920 | Ch13 region 2 |
| chr13:53799951-53799991 | Ch13 region 2 |
| chr13:53800032-53800072 | Ch13 region 2 |
| chr13:53800108-53800148 | Ch13 region 2 |
| chr13:53800180-53800220 | Ch13 region 2 |
| chr13:53800257-53800297 | Ch13 region 2 |
| chr13:53800302-53800342 | Ch13 region 2 |
| chr13:53800353-53800393 | Ch13 region 2 |
| chr13:53800400-53800440 | Ch13 region 2 |
| chr13:53800467-53800507 | Ch13 region 2 |
| chr13:53800711-53800751 | Ch13 region 2 |
| chr13:53800785-53800825 | Ch13 region 2 |

| | |
|---|---|
| chr13:53800918-53800958 | Ch13 region 2 |
| chr13:53800999-53801039 | Ch13 region 2 |
| chr13:53801051-53801091 | Ch13 region 2 |
| chr13:53801098-53801138 | Ch13 region 2 |
| chr13:53801163-53801203 | Ch13 region 2 |
| chr13:53801284-53801324 | Ch13 region 2 |
| chr13:53801329-53801369 | Ch13 region 2 |
| chr13:53801397-53801437 | Ch13 region 2 |
| chr13:53801530-53801570 | Ch13 region 2 |
| chr13:53801598-53801638 | Ch13 region 2 |
| chr13:53801643-53801683 | Ch13 region 2 |
| chr13:53801702-53801742 | Ch13 region 2 |
| chr13:53801849-53801889 | Ch13 region 2 |
| chr13:53801929-53801969 | Ch13 region 2 |
| chr13:53802030-53802070 | Ch13 region 2 |
| chr13:53802109-53802149 | Ch13 region 2 |
| chr13:53802156-53802196 | Ch13 region 2 |
| chr13:53802215-53802255 | Ch13 region 2 |
| chr13:53802377-53802417 | Ch13 region 2 |
| chr13:53802430-53802470 | Ch13 region 2 |
| chr13:53802512-53802552 | Ch13 region 2 |
| chr13:53802627-53802667 | Ch13 region 2 |
| chr13:53802708-53802748 | Ch13 region 2 |
| chr13:53802782-53802822 | Ch13 region 2 |
| chr13:53802827-53802867 | Ch13 region 2 |
| chr13:53802917-53802957 | Ch13 region 2 |
| chr13:53803001-53803041 | Ch13 region 2 |
| chr13:53803090-53803130 | Ch13 region 2 |
| chr13:53803202-53803242 | Ch13 region 2 |
| chr13:53803271-53803311 | Ch13 region 2 |
| chr13:53803330-53803370 | Ch13 region 2 |
| chr13:53803442-53803482 | Ch13 region 2 |
| chr13:53803504-53803544 | Ch13 region 2 |
| chr13:53803558-53803598 | Ch13 region 2 |
| chr13:53803605-53803645 | Ch13 region 2 |
| chr13:53803653-53803693 | Ch13 region 2 |
| chr13:53803699-53803739 | Ch13 region 2 |
| chr13:53803806-53803846 | Ch13 region 2 |
| chr13:53803863-53803903 | Ch13 region 2 |
| chr13:53803920-53803960 | Ch13 region 2 |
| chr13:53803978-53804018 | Ch13 region 2 |
| chr13:53804052-53804092 | Ch13 region 2 |
| chr13:53804097-53804137 | Ch13 region 2 |
| chr13:53804203-53804243 | Ch13 region 2 |
| chr13:53804254-53804294 | Ch13 region 2 |
| chr13:53804306-53804346 | Ch13 region 2 |
| chr13:53804369-53804409 | Ch13 region 2 |
| chr13:53804467-53804507 | Ch13 region 2 |
| chr13:53804523-53804563 | Ch13 region 2 |
| chr13:53804589-53804629 | Ch13 region 2 |
| chr13:53804637-53804677 | Ch13 region 2 |
| chr13:53804711-53804751 | Ch13 region 2 |

| | |
|---|---|
| chr13:53804761-53804801 | Ch13 region 2 |
| chr13:53804814-53804854 | Ch13 region 2 |
| chr13:53804897-53804937 | Ch13 region 2 |
| chr13:53804953-53804993 | Ch13 region 2 |
| chr13:53805027-53805067 | Ch13 region 2 |
| chr13:53805103-53805143 | Ch13 region 2 |
| chr13:53805155-53805195 | Ch13 region 2 |
| chr13:53805225-53805265 | Ch13 region 2 |
| chr13:53805289-53805329 | Ch13 region 2 |
| chr13:53805461-53805501 | Ch13 region 2 |
| chr13:53805517-53805557 | Ch13 region 2 |
| chr13:53805589-53805629 | Ch13 region 2 |
| chr13:53805662-53805702 | Ch13 region 2 |
| chr13:53805732-53805772 | Ch13 region 2 |
| chr13:53805806-53805846 | Ch13 region 2 |
| chr13:53805852-53805892 | Ch13 region 2 |
| chr13:53805926-53805966 | Ch13 region 2 |
| chr13:53805998-53806038 | Ch13 region 2 |
| chr13:53806067-53806107 | Ch13 region 2 |
| chr13:53806138-53806178 | Ch13 region 2 |
| chr13:53806192-53806232 | Ch13 region 2 |
| chr13:53806250-53806290 | Ch13 region 2 |
| chr13:53806295-53806335 | Ch13 region 2 |
| chr13:53806355-53806395 | Ch13 region 2 |
| chr13:53806414-53806454 | Ch13 region 2 |
| chr13:53806504-53806544 | Ch13 region 2 |
| chr13:53806575-53806615 | Ch13 region 2 |
| chr13:53806635-53806675 | Ch13 region 2 |
| chr13:53806680-53806720 | Ch13 region 2 |
| chr13:53806769-53806809 | Ch13 region 2 |
| chr13:53806843-53806883 | Ch13 region 2 |
| chr13:53806897-53806937 | Ch13 region 2 |
| chr13:53806943-53806983 | Ch13 region 2 |
| chr13:53807049-53807089 | Ch13 region 2 |
| chr13:53807098-53807138 | Ch13 region 2 |
| chr13:53807146-53807186 | Ch13 region 2 |
| chr13:53807191-53807231 | Ch13 region 2 |
| chr13:53807274-53807314 | Ch13 region 2 |
| chr13:53807378-53807418 | Ch13 region 2 |
| chr13:53807469-53807509 | Ch13 region 2 |
| chr13:53807661-53807701 | Ch13 region 2 |
| chr13:53807761-53807801 | Ch13 region 2 |
| chr13:53807849-53807889 | Ch13 region 2 |
| chr13:53807913-53807953 | Ch13 region 2 |
| chr13:53807958-53807998 | Ch13 region 2 |
| chr13:53808009-53808049 | Ch13 region 2 |
| chr13:53808063-53808103 | Ch13 region 2 |
| chr13:53808119-53808159 | Ch13 region 2 |
| chr13:53808260-53808300 | Ch13 region 2 |
| chr13:53808442-53808482 | Ch13 region 2 |
| chr13:53808488-53808528 | Ch13 region 2 |
| chr13:53808557-53808597 | Ch13 region 2 |

| | |
|---|---|
| chr13:53808618-53808658 | Ch13 region 2 |
| chr13:53808663-53808703 | Ch13 region 2 |
| chr13:53808781-53808821 | Ch13 region 2 |
| chr13:53808828-53808868 | Ch13 region 2 |
| chr13:53808894-53808934 | Ch13 region 2 |
| chr13:53808945-53808985 | Ch13 region 2 |
| chr13:53809002-53809042 | Ch13 region 2 |
| chr13:53809087-53809127 | Ch13 region 2 |
| chr13:53809153-53809193 | Ch13 region 2 |
| chr13:53809234-53809274 | Ch13 region 2 |
| chr13:53809319-53809359 | Ch13 region 2 |
| chr13:53809365-53809405 | Ch13 region 2 |
| chr13:53809445-53809485 | Ch13 region 2 |
| chr13:53809490-53809530 | Ch13 region 2 |
| chr13:53809562-53809602 | Ch13 region 2 |
| chr13:53809631-53809671 | Ch13 region 2 |
| chr13:53809679-53809719 | Ch13 region 2 |
| chr13:53809737-53809777 | Ch13 region 2 |
| chr13:53809782-53809822 | Ch13 region 2 |
| chr13:53809864-53809904 | Ch13 region 2 |
| chr13:53809918-53809958 | Ch13 region 2 |
| chr13:53810002-53810042 | Ch13 region 2 |
| chr13:53810096-53810136 | Ch13 region 2 |
| chr13:53810143-53810183 | Ch13 region 2 |
| chr13:53810228-53810268 | Ch13 region 2 |
| chr13:53810276-53810316 | Ch13 region 2 |
| chr13:53810490-53810530 | Ch13 region 2 |
| chr13:53810537-53810577 | Ch13 region 2 |
| chr13:53810661-53810701 | Ch13 region 2 |
| chr13:53810740-53810780 | Ch13 region 2 |
| chr13:53810820-53810860 | Ch13 region 2 |
| chr13:53810878-53810918 | Ch13 region 2 |
| chr13:53810924-53810964 | Ch13 region 2 |
| chr13:53811017-53811057 | Ch13 region 2 |
| chr13:53811066-53811106 | Ch13 region 2 |
| chr13:53811119-53811159 | Ch13 region 2 |
| chr13:53811173-53811213 | Ch13 region 2 |
| chr13:53811218-53811258 | Ch13 region 2 |
| chr13:53811279-53811319 | Ch13 region 2 |
| chr13:53811339-53811379 | Ch13 region 2 |
| chr13:53811398-53811438 | Ch13 region 2 |
| chr13:53811495-53811535 | Ch13 region 2 |
| chr13:53811551-53811591 | Ch13 region 2 |
| chr13:53811597-53811637 | Ch13 region 2 |
| chr13:53811666-53811706 | Ch13 region 2 |
| chr13:53811738-53811778 | Ch13 region 2 |
| chr13:53811785-53811825 | Ch13 region 2 |
| chr13:53811855-53811895 | Ch13 region 2 |
| chr13:53811913-53811953 | Ch13 region 2 |
| chr13:53812005-53812045 | Ch13 region 2 |
| chr13:53812065-53812105 | Ch13 region 2 |
| chr13:53812136-53812176 | Ch13 region 2 |

| | |
|---|---|
| chr13:53812197-53812237 | Ch13 region 2 |
| chr13:53812248-53812288 | Ch13 region 2 |
| chr13:53812296-53812336 | Ch13 region 2 |
| chr13:53812354-53812394 | Ch13 region 2 |
| chr13:53812401-53812441 | Ch13 region 2 |
| chr13:53812447-53812487 | Ch13 region 2 |
| chr13:53812549-53812589 | Ch13 region 2 |
| chr13:53812608-53812648 | Ch13 region 2 |
| chr13:53812679-53812719 | Ch13 region 2 |
| chr13:53812724-53812764 | Ch13 region 2 |
| chr13:53812807-53812847 | Ch13 region 2 |
| chr13:53812852-53812892 | Ch13 region 2 |
| chr13:53812901-53812941 | Ch13 region 2 |
| chr13:53812973-53813013 | Ch13 region 2 |
| chr13:53813020-53813060 | Ch13 region 2 |
| chr13:53813078-53813118 | Ch13 region 2 |
| chr13:53813156-53813196 | Ch13 region 2 |
| chr13:53813202-53813242 | Ch13 region 2 |
| chr13:53813286-53813326 | Ch13 region 2 |
| chr13:53813355-53813395 | Ch13 region 2 |
| chr13:53813420-53813460 | Ch13 region 2 |
| chr13:53813472-53813512 | Ch13 region 2 |
| chr13:53813559-53813599 | Ch13 region 2 |
| chr13:53813623-53813663 | Ch13 region 2 |
| chr13:53813668-53813708 | Ch13 region 2 |
| chr13:53813743-53813783 | Ch13 region 2 |
| chr13:53813794-53813834 | Ch13 region 2 |
| chr13:53813860-53813900 | Ch13 region 2 |
| chr13:53813925-53813965 | Ch13 region 2 |
| chr13:53813971-53814011 | Ch13 region 2 |
| chr13:53814021-53814061 | Ch13 region 2 |
| chr13:53814066-53814106 | Ch13 region 2 |
| chr13:53814112-53814152 | Ch13 region 2 |
| chr13:53814164-53814204 | Ch13 region 2 |
| chr13:53814215-53814255 | Ch13 region 2 |
| chr13:53814290-53814330 | Ch13 region 2 |
| chr13:53814367-53814407 | Ch13 region 2 |
| chr13:53814415-53814455 | Ch13 region 2 |
| chr13:53814469-53814509 | Ch13 region 2 |
| chr13:53814524-53814564 | Ch13 region 2 |
| chr13:53814619-53814659 | Ch13 region 2 |
| chr13:53814697-53814737 | Ch13 region 2 |
| chr13:53814771-53814811 | Ch13 region 2 |
| chr13:53814829-53814869 | Ch13 region 2 |
| chr13:53814874-53814914 | Ch13 region 2 |
| chr13:53814944-53814984 | Ch13 region 2 |
| chr13:53814999-53815039 | Ch13 region 2 |
| chr13:53815049-53815089 | Ch13 region 2 |
| chr13:53815098-53815138 | Ch13 region 2 |
| chr13:53815145-53815185 | Ch13 region 2 |
| chr13:53815209-53815249 | Ch13 region 2 |
| chr13:53815265-53815305 | Ch13 region 2 |

| | |
|---|---|
| chr13:53815340-53815380 | Ch13 region 2 |
| chr13:53815399-53815439 | Ch13 region 2 |
| chr13:53815495-53815535 | Ch13 region 2 |
| chr13:53815664-53815704 | Ch13 region 2 |
| chr13:53815713-53815753 | Ch13 region 2 |
| chr13:53815759-53815799 | Ch13 region 2 |
| chr13:53815820-53815860 | Ch13 region 2 |
| chr13:53815921-53815961 | Ch13 region 2 |
| chr13:53815967-53816007 | Ch13 region 2 |
| chr13:53816028-53816068 | Ch13 region 2 |
| chr13:53816103-53816143 | Ch13 region 2 |
| chr13:53816159-53816199 | Ch13 region 2 |
| chr13:53816218-53816258 | Ch13 region 2 |
| chr13:53816285-53816325 | Ch13 region 2 |
| chr13:53816371-53816411 | Ch13 region 2 |
| chr13:53816435-53816475 | Ch13 region 2 |
| chr13:53816483-53816523 | Ch13 region 2 |
| chr13:53816531-53816571 | Ch13 region 2 |
| chr13:53816603-53816643 | Ch13 region 2 |
| chr13:53816670-53816710 | Ch13 region 2 |
| chr13:53816722-53816762 | Ch13 region 2 |
| chr13:53816790-53816830 | Ch13 region 2 |
| chr13:53816835-53816875 | Ch13 region 2 |
| chr13:53816897-53816937 | Ch13 region 2 |
| chr13:53816950-53816990 | Ch13 region 2 |
| chr13:53816999-53817039 | Ch13 region 2 |
| chr13:53817356-53817396 | Ch13 region 2 |
| chr13:53817447-53817487 | Ch13 region 2 |
| chr13:53817500-53817540 | Ch13 region 2 |
| chr13:53817547-53817587 | Ch13 region 2 |
| chr13:53817628-53817668 | Ch13 region 2 |
| chr13:53817690-53817730 | Ch13 region 2 |
| chr13:53817762-53817802 | Ch13 region 2 |
| chr13:53817807-53817847 | Ch13 region 2 |
| chr13:53817909-53817949 | Ch13 region 2 |
| chr13:53817962-53818002 | Ch13 region 2 |
| chr13:53818019-53818059 | Ch13 region 2 |
| chr13:53818093-53818133 | Ch13 region 2 |
| chr13:53818173-53818213 | Ch13 region 2 |
| chr13:53818275-53818315 | Ch13 region 2 |
| chr13:53818320-53818360 | Ch13 region 2 |
| chr13:53818367-53818407 | Ch13 region 2 |
| chr13:53818446-53818486 | Ch13 region 2 |
| chr13:53818504-53818544 | Ch13 region 2 |
| chr13:53818583-53818623 | Ch13 region 2 |
| chr13:53818650-53818690 | Ch13 region 2 |
| chr13:53818723-53818763 | Ch13 region 2 |
| chr13:53818790-53818830 | Ch13 region 2 |
| chr13:53818861-53818901 | Ch13 region 2 |
| chr13:53818906-53818946 | Ch13 region 2 |
| chr13:53818994-53819034 | Ch13 region 2 |
| chr13:53819057-53819097 | Ch13 region 2 |

| | |
|---|---|
| chr13:53819107-53819147 | Ch13 region 2 |
| chr13:53819164-53819204 | Ch13 region 2 |
| chr13:53819229-53819269 | Ch13 region 2 |
| chr13:53819301-53819341 | Ch13 region 2 |
| chr13:53819346-53819386 | Ch13 region 2 |
| chr13:53819394-53819434 | Ch13 region 2 |
| chr13:53819488-53819528 | Ch13 region 2 |
| chr13:53819556-53819596 | Ch13 region 2 |
| chr13:53819617-53819657 | Ch13 region 2 |
| chr13:53819682-53819722 | Ch13 region 2 |
| chr13:53819745-53819785 | Ch13 region 2 |
| chr13:53819792-53819832 | Ch13 region 2 |
| chr13:53819868-53819908 | Ch13 region 2 |
| chr13:53819913-53819953 | Ch13 region 2 |
| chr13:53819997-53820037 | Ch13 region 2 |
| chr13:53820056-53820096 | Ch13 region 2 |
| chr13:53820105-53820145 | Ch13 region 2 |
| chr13:53820177-53820217 | Ch13 region 2 |
| chr13:53820248-53820288 | Ch13 region 2 |
| chr13:53820337-53820377 | Ch13 region 2 |
| chr13:53820425-53820465 | Ch13 region 2 |
| chr13:53820511-53820551 | Ch13 region 2 |
| chr13:53820576-53820616 | Ch13 region 2 |
| chr13:53820694-53820734 | Ch13 region 2 |
| chr13:53820748-53820788 | Ch13 region 2 |
| chr13:53820833-53820873 | Ch13 region 2 |
| chr13:53820885-53820925 | Ch13 region 2 |
| chr13:53820930-53820970 | Ch13 region 2 |
| chr13:53821045-53821085 | Ch13 region 2 |
| chr13:53821116-53821156 | Ch13 region 2 |
| chr13:53821171-53821211 | Ch13 region 2 |
| chr13:53821254-53821294 | Ch13 region 2 |
| chr13:53821300-53821340 | Ch13 region 2 |
| chr13:53821361-53821401 | Ch13 region 2 |
| chr13:53821406-53821446 | Ch13 region 2 |
| chr13:53821461-53821501 | Ch13 region 2 |
| chr13:53821520-53821560 | Ch13 region 2 |
| chr13:53821590-53821630 | Ch13 region 2 |
| chr13:53821640-53821680 | Ch13 region 2 |
| chr13:53821727-53821767 | Ch13 region 2 |
| chr13:53821836-53821876 | Ch13 region 2 |
| chr13:53822034-53822074 | Ch13 region 2 |
| chr13:53822105-53822145 | Ch13 region 2 |
| chr13:53822170-53822210 | Ch13 region 2 |
| chr13:53822262-53822302 | Ch13 region 2 |
| chr13:53822307-53822347 | Ch13 region 2 |
| chr13:53822377-53822417 | Ch13 region 2 |
| chr13:53822442-53822482 | Ch13 region 2 |
| chr13:53822524-53822564 | Ch13 region 2 |
| chr13:53822608-53822648 | Ch13 region 2 |
| chr13:53822665-53822705 | Ch13 region 2 |
| chr13:53822720-53822760 | Ch13 region 2 |

| | |
|---|---|
| chr13:53822785-53822825 | Ch13 region 2 |
| chr13:53822830-53822870 | Ch13 region 2 |
| chr13:53822902-53822942 | Ch13 region 2 |
| chr13:53822952-53822992 | Ch13 region 2 |
| chr13:53823000-53823040 | Ch13 region 2 |
| chr13:53823105-53823145 | Ch13 region 2 |
| chr13:53823161-53823201 | Ch13 region 2 |
| chr13:53823278-53823318 | Ch13 region 2 |
| chr13:53823344-53823384 | Ch13 region 2 |
| chr13:53823396-53823436 | Ch13 region 2 |
| chr13:53823456-53823496 | Ch13 region 2 |
| chr13:53823537-53823577 | Ch13 region 2 |
| chr13:53823623-53823663 | Ch13 region 2 |
| chr13:53823753-53823793 | Ch13 region 2 |
| chr13:53823806-53823846 | Ch13 region 2 |
| chr13:53823911-53823951 | Ch13 region 2 |
| chr13:53823982-53824022 | Ch13 region 2 |
| chr13:53824058-53824098 | Ch13 region 2 |
| chr13:53824121-53824161 | Ch13 region 2 |
| chr13:53824249-53824289 | Ch13 region 2 |
| chr13:53824294-53824334 | Ch13 region 2 |
| chr13:53824514-53824554 | Ch13 region 2 |
| chr13:53824603-53824643 | Ch13 region 2 |
| chr13:53824684-53824724 | Ch13 region 2 |
| chr13:53824820-53824860 | Ch13 region 2 |
| chr13:53824886-53824926 | Ch13 region 2 |
| chr13:53824951-53824991 | Ch13 region 2 |
| chr13:53825009-53825049 | Ch13 region 2 |
| chr13:53825055-53825095 | Ch13 region 2 |
| chr13:53825113-53825153 | Ch13 region 2 |
| chr13:53825185-53825225 | Ch13 region 2 |
| chr13:53825245-53825285 | Ch13 region 2 |
| chr13:53825332-53825372 | Ch13 region 2 |
| chr13:53825380-53825420 | Ch13 region 2 |
| chr13:53825452-53825492 | Ch13 region 2 |
| chr13:53825527-53825567 | Ch13 region 2 |
| chr13:53825595-53825635 | Ch13 region 2 |
| chr13:53825681-53825721 | Ch13 region 2 |
| chr13:53825726-53825766 | Ch13 region 2 |
| chr13:53825773-53825813 | Ch13 region 2 |
| chr13:53825834-53825874 | Ch13 region 2 |
| chr13:53825898-53825938 | Ch13 region 2 |
| chr13:53825946-53825986 | Ch13 region 2 |
| chr13:53826039-53826079 | Ch13 region 2 |
| chr13:53826085-53826125 | Ch13 region 2 |
| chr13:53826148-53826188 | Ch13 region 2 |
| chr13:53826198-53826238 | Ch13 region 2 |
| chr13:53826245-53826285 | Ch13 region 2 |
| chr13:53826346-53826386 | Ch13 region 2 |
| chr13:53826398-53826438 | Ch13 region 2 |
| chr13:53826476-53826516 | Ch13 region 2 |
| chr13:53826522-53826562 | Ch13 region 2 |

| | |
|---|---|
| chr13:53826572-53826612 | Ch13 region 2 |
| chr13:53826622-53826662 | Ch13 region 2 |
| chr13:53826676-53826716 | Ch13 region 2 |
| chr13:53826744-53826784 | Ch13 region 2 |
| chr13:53826791-53826831 | Ch13 region 2 |
| chr13:53826849-53826889 | Ch13 region 2 |
| chr13:53826915-53826955 | Ch13 region 2 |
| chr13:53826994-53827034 | Ch13 region 2 |
| chr13:53827070-53827110 | Ch13 region 2 |
| chr13:53827119-53827159 | Ch13 region 2 |
| chr13:53827166-53827206 | Ch13 region 2 |
| chr13:53827213-53827253 | Ch13 region 2 |
| chr13:53827272-53827312 | Ch13 region 2 |
| chr13:53827386-53827426 | Ch13 region 2 |
| chr13:53827443-53827483 | Ch13 region 2 |
| chr13:53827515-53827555 | Ch13 region 2 |
| chr13:53827574-53827614 | Ch13 region 2 |
| chr13:53827622-53827662 | Ch13 region 2 |
| chr13:53827685-53827725 | Ch13 region 2 |
| chr13:53827736-53827776 | Ch13 region 2 |
| chr13:53827792-53827832 | Ch13 region 2 |
| chr13:53827838-53827878 | Ch13 region 2 |
| chr13:53827903-53827943 | Ch13 region 2 |
| chr13:53828000-53828040 | Ch13 region 2 |
| chr13:53828075-53828115 | Ch13 region 2 |
| chr13:53828122-53828162 | Ch13 region 2 |
| chr13:53828191-53828231 | Ch13 region 2 |
| chr13:53828313-53828353 | Ch13 region 2 |
| chr13:53828383-53828423 | Ch13 region 2 |
| chr13:53828428-53828468 | Ch13 region 2 |
| chr13:53828478-53828518 | Ch13 region 2 |
| chr13:53828565-53828605 | Ch13 region 2 |
| chr13:53828630-53828670 | Ch13 region 2 |
| chr13:53828675-53828715 | Ch13 region 2 |
| chr13:53828720-53828760 | Ch13 region 2 |
| chr13:53828770-53828810 | Ch13 region 2 |
| chr13:53828815-53828855 | Ch13 region 2 |
| chr13:53828903-53828943 | Ch13 region 2 |
| chr13:53828976-53829016 | Ch13 region 2 |
| chr13:53829111-53829151 | Ch13 region 2 |
| chr13:53829159-53829199 | Ch13 region 2 |
| chr13:53829223-53829263 | Ch13 region 2 |
| chr13:53829321-53829361 | Ch13 region 2 |
| chr13:53829415-53829455 | Ch13 region 2 |
| chr13:53829505-53829545 | Ch13 region 2 |
| chr13:53829607-53829647 | Ch13 region 2 |
| chr13:53829671-53829711 | Ch13 region 2 |
| chr13:53829716-53829756 | Ch13 region 2 |
| chr13:53829778-53829818 | Ch13 region 2 |
| chr13:53829849-53829889 | Ch13 region 2 |
| chr13:53829917-53829957 | Ch13 region 2 |
| chr13:53829984-53830024 | Ch13 region 2 |

| | |
|---|---|
| chr13:53830043-53830083 | Ch13 region 2 |
| chr13:53830123-53830163 | Ch13 region 2 |
| chr13:53830198-53830238 | Ch13 region 2 |
| chr13:53830293-53830333 | Ch13 region 2 |
| chr13:53830344-53830384 | Ch13 region 2 |
| chr13:53830429-53830469 | Ch13 region 2 |
| chr13:53830486-53830526 | Ch13 region 2 |
| chr13:53830578-53830618 | Ch13 region 2 |
| chr13:53830630-53830670 | Ch13 region 2 |
| chr13:53830733-53830773 | Ch13 region 2 |
| chr13:53830784-53830824 | Ch13 region 2 |
| chr13:53830836-53830876 | Ch13 region 2 |
| chr13:53830896-53830936 | Ch13 region 2 |
| chr13:53830980-53831020 | Ch13 region 2 |
| chr13:53831064-53831104 | Ch13 region 2 |
| chr13:53831143-53831183 | Ch13 region 2 |
| chr13:53831189-53831229 | Ch13 region 2 |
| chr13:53831279-53831319 | Ch13 region 2 |
| chr13:53831370-53831410 | Ch13 region 2 |
| chr13:53831417-53831457 | Ch13 region 2 |
| chr13:53831462-53831502 | Ch13 region 2 |
| chr13:53831559-53831599 | Ch13 region 2 |
| chr13:53831646-53831686 | Ch13 region 2 |
| chr13:53831709-53831749 | Ch13 region 2 |
| chr13:53831764-53831804 | Ch13 region 2 |
| chr13:53831857-53831897 | Ch13 region 2 |
| chr13:53831908-53831948 | Ch13 region 2 |
| chr13:53831962-53832002 | Ch13 region 2 |
| chr13:53832031-53832071 | Ch13 region 2 |
| chr13:53832087-53832127 | Ch13 region 2 |
| chr13:53832144-53832184 | Ch13 region 2 |
| chr13:53832190-53832230 | Ch13 region 2 |
| chr13:53832278-53832318 | Ch13 region 2 |
| chr13:53832323-53832363 | Ch13 region 2 |
| chr13:53832433-53832473 | Ch13 region 2 |
| chr13:53832562-53832602 | Ch13 region 2 |
| chr13:53832641-53832681 | Ch13 region 2 |
| chr13:53832686-53832726 | Ch13 region 2 |
| chr13:53832741-53832781 | Ch13 region 2 |
| chr13:53832788-53832828 | Ch13 region 2 |
| chr13:53832850-53832890 | Ch13 region 2 |
| chr13:53832906-53832946 | Ch13 region 2 |
| chr13:53832977-53833017 | Ch13 region 2 |
| chr13:53833060-53833100 | Ch13 region 2 |
| chr13:53833107-53833147 | Ch13 region 2 |
| chr13:53833157-53833197 | Ch13 region 2 |
| chr13:53833203-53833243 | Ch13 region 2 |
| chr13:53833254-53833294 | Ch13 region 2 |
| chr13:53833331-53833371 | Ch13 region 2 |
| chr13:53833382-53833422 | Ch13 region 2 |
| chr13:53833455-53833495 | Ch13 region 2 |
| chr13:53833522-53833562 | Ch13 region 2 |

| | |
|---|---|
| chr13:53833641-53833681 | Ch13 region 2 |
| chr13:53833809-53833849 | Ch13 region 2 |
| chr13:53833864-53833904 | Ch13 region 2 |
| chr13:53833963-53834003 | Ch13 region 2 |
| chr13:53834053-53834093 | Ch13 region 2 |
| chr13:53834124-53834164 | Ch13 region 2 |
| chr13:53834173-53834213 | Ch13 region 2 |
| chr13:53834220-53834260 | Ch13 region 2 |
| chr13:53834290-53834330 | Ch13 region 2 |
| chr13:53834353-53834393 | Ch13 region 2 |
| chr13:53834399-53834439 | Ch13 region 2 |
| chr13:53834482-53834522 | Ch13 region 2 |
| chr13:53834527-53834567 | Ch13 region 2 |
| chr13:53834615-53834655 | Ch13 region 2 |
| chr13:53834674-53834714 | Ch13 region 2 |
| chr13:53834738-53834778 | Ch13 region 2 |
| chr13:53834818-53834858 | Ch13 region 2 |
| chr13:53834868-53834908 | Ch13 region 2 |
| chr13:53834949-53834989 | Ch13 region 2 |
| chr13:53835038-53835078 | Ch13 region 2 |
| chr13:53835103-53835143 | Ch13 region 2 |
| chr13:53835166-53835206 | Ch13 region 2 |
| chr13:53835240-53835280 | Ch13 region 2 |
| chr13:53835304-53835344 | Ch13 region 2 |
| chr13:53835375-53835415 | Ch13 region 2 |
| chr13:53835445-53835485 | Ch13 region 2 |
| chr13:53835490-53835530 | Ch13 region 2 |
| chr13:53835542-53835582 | Ch13 region 2 |
| chr13:53835618-53835658 | Ch13 region 2 |
| chr13:53835663-53835703 | Ch13 region 2 |
| chr13:53835743-53835783 | Ch13 region 2 |
| chr13:53835821-53835861 | Ch13 region 2 |
| chr13:53835874-53835914 | Ch13 region 2 |
| chr13:53835951-53835991 | Ch13 region 2 |
| chr13:53836032-53836072 | Ch13 region 2 |
| chr13:53836131-53836171 | Ch13 region 2 |
| chr13:53836190-53836230 | Ch13 region 2 |
| chr13:53836251-53836291 | Ch13 region 2 |
| chr13:53836325-53836365 | Ch13 region 2 |
| chr13:53836422-53836462 | Ch13 region 2 |
| chr13:53836504-53836544 | Ch13 region 2 |
| chr13:53836630-53836670 | Ch13 region 2 |
| chr13:53836698-53836738 | Ch13 region 2 |
| chr13:53836748-53836788 | Ch13 region 2 |
| chr13:53836799-53836839 | Ch13 region 2 |
| chr13:53836898-53836938 | Ch13 region 2 |
| chr13:53836953-53836993 | Ch13 region 2 |
| chr13:53837030-53837070 | Ch13 region 2 |
| chr13:53837094-53837134 | Ch13 region 2 |
| chr13:53837140-53837180 | Ch13 region 2 |
| chr13:53837202-53837242 | Ch13 region 2 |
| chr13:53837282-53837322 | Ch13 region 2 |

| | |
|---|---|
| chr13:53837327-53837367 | Ch13 region 2 |
| chr13:53837373-53837413 | Ch13 region 2 |
| chr13:53837460-53837500 | Ch13 region 2 |
| chr13:53837530-53837570 | Ch13 region 2 |
| chr13:53837623-53837663 | Ch13 region 2 |
| chr13:53837720-53837760 | Ch13 region 2 |
| chr13:53837784-53837824 | Ch13 region 2 |
| chr13:53837832-53837872 | Ch13 region 2 |
| chr13:53837897-53837937 | Ch13 region 2 |
| chr13:53837988-53838028 | Ch13 region 2 |
| chr13:53838050-53838090 | Ch13 region 2 |
| chr13:53838095-53838135 | Ch13 region 2 |
| chr13:53838196-53838236 | Ch13 region 2 |
| chr13:53838289-53838329 | Ch13 region 2 |
| chr13:53838334-53838374 | Ch13 region 2 |
| chr13:53838440-53838480 | Ch13 region 2 |
| chr13:53838507-53838547 | Ch13 region 2 |
| chr13:53838557-53838597 | Ch13 region 2 |
| chr13:53838603-53838643 | Ch13 region 2 |
| chr13:53838648-53838688 | Ch13 region 2 |
| chr13:53838718-53838758 | Ch13 region 2 |
| chr13:53838787-53838827 | Ch13 region 2 |
| chr13:53838880-53838920 | Ch13 region 2 |
| chr13:53838992-53839032 | Ch13 region 2 |
| chr13:53839042-53839082 | Ch13 region 2 |
| chr13:53839087-53839127 | Ch13 region 2 |
| chr13:53839138-53839178 | Ch13 region 2 |
| chr13:53839230-53839270 | Ch13 region 2 |
| chr13:53839279-53839319 | Ch13 region 2 |
| chr13:53839335-53839375 | Ch13 region 2 |
| chr13:53839425-53839465 | Ch13 region 2 |
| chr13:53839490-53839530 | Ch13 region 2 |
| chr13:53839599-53839639 | Ch13 region 2 |
| chr13:53839653-53839693 | Ch13 region 2 |
| chr13:53839704-53839744 | Ch13 region 2 |
| chr13:53839762-53839802 | Ch13 region 2 |
| chr13:53839823-53839863 | Ch13 region 2 |
| chr13:53839868-53839908 | Ch13 region 2 |
| chr13:53839916-53839956 | Ch13 region 2 |
| chr13:53839995-53840035 | Ch13 region 2 |
| chr13:53840056-53840096 | Ch13 region 2 |
| chr13:53840130-53840170 | Ch13 region 2 |
| chr13:53840236-53840276 | Ch13 region 2 |
| chr13:53840287-53840327 | Ch13 region 2 |
| chr13:53840348-53840388 | Ch13 region 2 |
| chr13:53840411-53840451 | Ch13 region 2 |
| chr13:53840456-53840496 | Ch13 region 2 |
| chr13:53840523-53840563 | Ch13 region 2 |
| chr13:53840589-53840629 | Ch13 region 2 |
| chr13:53840649-53840689 | Ch13 region 2 |
| chr13:53840719-53840759 | Ch13 region 2 |
| chr13:53840788-53840828 | Ch13 region 2 |

| | |
|---|---|
| chr13:53840847-53840887 | Ch13 region 2 |
| chr13:53840915-53840955 | Ch13 region 2 |
| chr13:53840987-53841027 | Ch13 region 2 |
| chr13:53841324-53841364 | Ch13 region 2 |
| chr13:53841406-53841446 | Ch13 region 2 |
| chr13:53841499-53841539 | Ch13 region 2 |
| chr13:53841725-53841765 | Ch13 region 2 |
| chr13:53841770-53841810 | Ch13 region 2 |
| chr13:53841961-53842001 | Ch13 region 2 |
| chr13:53842025-53842065 | Ch13 region 2 |
| chr13:53842130-53842170 | Ch13 region 2 |
| chr13:53842326-53842366 | Ch13 region 2 |
| chr13:53842374-53842414 | Ch13 region 2 |
| chr13:53842421-53842461 | Ch13 region 2 |
| chr13:53842470-53842510 | Ch13 region 2 |
| chr13:53842557-53842597 | Ch13 region 2 |
| chr13:53842771-53842811 | Ch13 region 2 |
| chr13:53842842-53842882 | Ch13 region 2 |
| chr13:53842901-53842941 | Ch13 region 2 |
| chr13:53842998-53843038 | Ch13 region 2 |
| chr13:53843058-53843098 | Ch13 region 2 |
| chr13:53843137-53843177 | Ch13 region 2 |
| chr13:53843188-53843228 | Ch13 region 2 |
| chr13:53843255-53843295 | Ch13 region 2 |
| chr13:53843302-53843342 | Ch13 region 2 |
| chr13:53843347-53843387 | Ch13 region 2 |
| chr13:53843394-53843434 | Ch13 region 2 |
| chr13:53843477-53843517 | Ch13 region 2 |
| chr13:53843522-53843562 | Ch13 region 2 |
| chr13:53843576-53843616 | Ch13 region 2 |
| chr13:53843621-53843661 | Ch13 region 2 |
| chr13:53843707-53843747 | Ch13 region 2 |
| chr13:53843754-53843794 | Ch13 region 2 |
| chr13:53843814-53843854 | Ch13 region 2 |
| chr13:53843911-53843951 | Ch13 region 2 |
| chr13:53844013-53844053 | Ch13 region 2 |
| chr13:53844067-53844107 | Ch13 region 2 |
| chr13:53844128-53844168 | Ch13 region 2 |
| chr13:53844174-53844214 | Ch13 region 2 |
| chr13:53844303-53844343 | Ch13 region 2 |
| chr13:53844359-53844399 | Ch13 region 2 |
| chr13:53844422-53844462 | Ch13 region 2 |
| chr13:53844467-53844507 | Ch13 region 2 |
| chr13:53844589-53844629 | Ch13 region 2 |
| chr13:53844685-53844725 | Ch13 region 2 |
| chr13:53844745-53844785 | Ch13 region 2 |
| chr13:53844790-53844830 | Ch13 region 2 |
| chr13:53844854-53844894 | Ch13 region 2 |
| chr13:53844947-53844987 | Ch13 region 2 |
| chr13:53844997-53845037 | Ch13 region 2 |
| chr13:53845065-53845105 | Ch13 region 2 |
| chr13:53845167-53845207 | Ch13 region 2 |

| | |
|---|---|
| chr13:53845212-53845252 | Ch13 region 2 |
| chr13:53845287-53845327 | Ch13 region 2 |
| chr13:53845347-53845387 | Ch13 region 2 |
| chr13:53845432-53845472 | Ch13 region 2 |
| chr13:53845480-53845520 | Ch13 region 2 |
| chr13:53845559-53845599 | Ch13 region 2 |
| chr13:53845651-53845691 | Ch13 region 2 |
| chr13:53845742-53845782 | Ch13 region 2 |
| chr13:53845848-53845888 | Ch13 region 2 |
| chr13:53845910-53845950 | Ch13 region 2 |
| chr13:53845958-53845998 | Ch13 region 2 |
| chr13:53846008-53846048 | Ch13 region 2 |
| chr13:53846058-53846098 | Ch13 region 2 |
| chr13:53846104-53846144 | Ch13 region 2 |
| chr13:53846166-53846206 | Ch13 region 2 |
| chr13:53846214-53846254 | Ch13 region 2 |
| chr13:53846317-53846357 | Ch13 region 2 |
| chr13:53846369-53846409 | Ch13 region 2 |
| chr13:53846435-53846475 | Ch13 region 2 |
| chr13:53846480-53846520 | Ch13 region 2 |
| chr13:53846535-53846575 | Ch13 region 2 |
| chr13:53846610-53846650 | Ch13 region 2 |
| chr13:53846691-53846731 | Ch13 region 2 |
| chr13:53846736-53846776 | Ch13 region 2 |
| chr13:53846783-53846823 | Ch13 region 2 |
| chr13:53846867-53846907 | Ch13 region 2 |
| chr13:53846944-53846984 | Ch13 region 2 |
| chr13:53847085-53847125 | Ch13 region 2 |
| chr13:53847162-53847202 | Ch13 region 2 |
| chr13:53847238-53847278 | Ch13 region 2 |
| chr13:53847296-53847336 | Ch13 region 2 |
| chr13:53847388-53847428 | Ch13 region 2 |
| chr13:53847469-53847509 | Ch13 region 2 |
| chr13:53847515-53847555 | Ch13 region 2 |
| chr13:53847576-53847616 | Ch13 region 2 |
| chr13:53847621-53847661 | Ch13 region 2 |
| chr13:53847678-53847718 | Ch13 region 2 |
| chr13:53847762-53847802 | Ch13 region 2 |
| chr13:53847852-53847892 | Ch13 region 2 |
| chr13:53847897-53847937 | Ch13 region 2 |
| chr13:53847962-53848002 | Ch13 region 2 |
| chr13:53848059-53848099 | Ch13 region 2 |
| chr13:53848167-53848207 | Ch13 region 2 |
| chr13:53848238-53848278 | Ch13 region 2 |
| chr13:53848285-53848325 | Ch13 region 2 |
| chr13:53848338-53848378 | Ch13 region 2 |
| chr13:53848464-53848504 | Ch13 region 2 |
| chr13:53848529-53848569 | Ch13 region 2 |
| chr13:53848574-53848614 | Ch13 region 2 |
| chr13:53848627-53848667 | Ch13 region 2 |
| chr13:53848711-53848751 | Ch13 region 2 |
| chr13:53848757-53848797 | Ch13 region 2 |

| | |
|---|---|
| chr13:53848820-53848860 | Ch13 region 2 |
| chr13:53848876-53848916 | Ch13 region 2 |
| chr13:53848943-53848983 | Ch13 region 2 |
| chr13:53849334-53849374 | Ch13 region 2 |
| chr13:53849408-53849448 | Ch13 region 2 |
| chr13:53849557-53849597 | Ch13 region 2 |
| chr13:53849638-53849678 | Ch13 region 2 |
| chr13:53849737-53849777 | Ch13 region 2 |
| chr13:53849815-53849855 | Ch13 region 2 |
| chr13:53849905-53849945 | Ch13 region 2 |
| chr13:53849988-53850028 | Ch13 region 2 |
| chr13:53850035-53850075 | Ch13 region 2 |
| chr13:53850155-53850195 | Ch13 region 2 |
| chr13:53850200-53850240 | Ch13 region 2 |
| chr13:53850253-53850293 | Ch13 region 2 |
| chr13:53850301-53850341 | Ch13 region 2 |
| chr13:53850346-53850386 | Ch13 region 2 |
| chr13:53850409-53850449 | Ch13 region 2 |
| chr13:53850458-53850498 | Ch13 region 2 |
| chr13:53850539-53850579 | Ch13 region 2 |
| chr13:53850595-53850635 | Ch13 region 2 |
| chr13:53850646-53850686 | Ch13 region 2 |
| chr13:53850719-53850759 | Ch13 region 2 |
| chr13:53850764-53850804 | Ch13 region 2 |
| chr13:53850817-53850857 | Ch13 region 2 |
| chr13:53850869-53850909 | Ch13 region 2 |
| chr13:53850922-53850962 | Ch13 region 2 |
| chr13:53851010-53851050 | Ch13 region 2 |
| chr13:53851061-53851101 | Ch13 region 2 |
| chr13:53851119-53851159 | Ch13 region 2 |
| chr13:53851173-53851213 | Ch13 region 2 |
| chr13:53851219-53851259 | Ch13 region 2 |
| chr13:53851328-53851368 | Ch13 region 2 |
| chr13:53851387-53851427 | Ch13 region 2 |
| chr13:53851440-53851480 | Ch13 region 2 |
| chr13:53851561-53851601 | Ch13 region 2 |
| chr13:53851616-53851656 | Ch13 region 2 |
| chr13:53851661-53851701 | Ch13 region 2 |
| chr13:53851752-53851792 | Ch13 region 2 |
| chr13:53851835-53851875 | Ch13 region 2 |
| chr13:53851897-53851937 | Ch13 region 2 |
| chr13:53851969-53852009 | Ch13 region 2 |
| chr13:53852027-53852067 | Ch13 region 2 |
| chr13:53852223-53852263 | Ch13 region 2 |
| chr13:53852350-53852390 | Ch13 region 2 |
| chr13:53852416-53852456 | Ch13 region 2 |
| chr13:53852468-53852508 | Ch13 region 2 |
| chr13:53852528-53852568 | Ch13 region 2 |
| chr13:53852624-53852664 | Ch13 region 2 |
| chr13:53852798-53852838 | Ch13 region 2 |
| chr13:53852926-53852966 | Ch13 region 2 |
| chr13:53853009-53853049 | Ch13 region 2 |

| | |
|---|---|
| chr13:53853054-53853094 | Ch13 region 2 |
| chr13:53853149-53853189 | Ch13 region 2 |
| chr13:53853297-53853337 | Ch13 region 2 |
| chr13:53853523-53853563 | Ch13 region 2 |
| chr13:53853626-53853666 | Ch13 region 2 |
| chr13:53853671-53853711 | Ch13 region 2 |
| chr13:53853737-53853777 | Ch13 region 2 |
| chr13:53853815-53853855 | Ch13 region 2 |
| chr13:53853860-53853900 | Ch13 region 2 |
| chr13:53853921-53853961 | Ch13 region 2 |
| chr13:53853971-53854011 | Ch13 region 2 |
| chr13:53854047-53854087 | Ch13 region 2 |
| chr13:53854113-53854153 | Ch13 region 2 |
| chr13:53854194-53854234 | Ch13 region 2 |
| chr13:53854266-53854306 | Ch13 region 2 |
| chr13:53854321-53854361 | Ch13 region 2 |
| chr13:53854369-53854409 | Ch13 region 2 |
| chr13:53854442-53854482 | Ch13 region 2 |
| chr13:53854505-53854545 | Ch13 region 2 |
| chr13:53854562-53854602 | Ch13 region 2 |
| chr13:53854612-53854652 | Ch13 region 2 |
| chr13:53854660-53854700 | Ch13 region 2 |
| chr13:53854744-53854784 | Ch13 region 2 |
| chr13:53854821-53854861 | Ch13 region 2 |
| chr13:53854873-53854913 | Ch13 region 2 |
| chr13:53854947-53854987 | Ch13 region 2 |
| chr13:53854992-53855032 | Ch13 region 2 |
| chr13:53855061-53855101 | Ch13 region 2 |
| chr13:53855113-53855153 | Ch13 region 2 |
| chr13:53855158-53855198 | Ch13 region 2 |
| chr13:53855260-53855300 | Ch13 region 2 |
| chr13:53855306-53855346 | Ch13 region 2 |
| chr13:53855395-53855435 | Ch13 region 2 |
| chr13:53855505-53855545 | Ch13 region 2 |
| chr13:53855600-53855640 | Ch13 region 2 |
| chr13:53855669-53855709 | Ch13 region 2 |
| chr13:53855747-53855787 | Ch13 region 2 |
| chr13:53855814-53855854 | Ch13 region 2 |
| chr13:53855872-53855912 | Ch13 region 2 |
| chr13:53855959-53855999 | Ch13 region 2 |
| chr13:53856175-53856215 | Ch13 region 2 |
| chr13:53856271-53856311 | Ch13 region 2 |
| chr13:53856339-53856379 | Ch13 region 2 |
| chr13:53856427-53856467 | Ch13 region 2 |
| chr13:53856494-53856534 | Ch13 region 2 |
| chr13:53856561-53856601 | Ch13 region 2 |
| chr13:53856645-53856685 | Ch13 region 2 |
| chr13:53856695-53856735 | Ch13 region 2 |
| chr13:53856812-53856852 | Ch13 region 2 |
| chr13:53856863-53856903 | Ch13 region 2 |
| chr13:53856944-53856984 | Ch13 region 2 |
| chr13:53856994-53857034 | Ch13 region 2 |

| | |
|---|---|
| chr13:53857045-53857085 | Ch13 region 2 |
| chr13:53857138-53857178 | Ch13 region 2 |
| chr13:53857207-53857247 | Ch13 region 2 |
| chr13:53857259-53857299 | Ch13 region 2 |
| chr13:53857324-53857364 | Ch13 region 2 |
| chr13:53857381-53857421 | Ch13 region 2 |
| chr13:53857456-53857496 | Ch13 region 2 |
| chr13:53857509-53857549 | Ch13 region 2 |
| chr13:53857554-53857594 | Ch13 region 2 |
| chr13:53857604-53857644 | Ch13 region 2 |
| chr13:53857650-53857690 | Ch13 region 2 |
| chr13:53857733-53857773 | Ch13 region 2 |
| chr13:53857812-53857852 | Ch13 region 2 |
| chr13:53857885-53857925 | Ch13 region 2 |
| chr13:53857971-53858011 | Ch13 region 2 |
| chr13:53858024-53858064 | Ch13 region 2 |
| chr13:53858070-53858110 | Ch13 region 2 |
| chr13:53858139-53858179 | Ch13 region 2 |
| chr13:53858205-53858245 | Ch13 region 2 |
| chr13:53858259-53858299 | Ch13 region 2 |
| chr13:53858315-53858355 | Ch13 region 2 |
| chr13:53858368-53858408 | Ch13 region 2 |
| chr13:53858730-53858770 | Ch13 region 2 |
| chr13:53858799-53858839 | Ch13 region 2 |
| chr13:53858844-53858884 | Ch13 region 2 |
| chr13:53858927-53858967 | Ch13 region 2 |
| chr13:53858991-53859031 | Ch13 region 2 |
| chr13:53859053-53859093 | Ch13 region 2 |
| chr13:53859098-53859138 | Ch13 region 2 |
| chr13:53859146-53859186 | Ch13 region 2 |
| chr13:53859224-53859264 | Ch13 region 2 |
| chr13:53859306-53859346 | Ch13 region 2 |
| chr13:53859402-53859442 | Ch13 region 2 |
| chr13:53859477-53859517 | Ch13 region 2 |
| chr13:53859611-53859651 | Ch13 region 2 |
| chr13:53859769-53859809 | Ch13 region 2 |
| chr13:53859828-53859868 | Ch13 region 2 |
| chr13:53859882-53859922 | Ch13 region 2 |
| chr13:53859927-53859967 | Ch13 region 2 |
| chr13:53860001-53860041 | Ch13 region 2 |
| chr13:53860053-53860093 | Ch13 region 2 |
| chr13:53860146-53860186 | Ch13 region 2 |
| chr13:53860238-53860278 | Ch13 region 2 |
| chr13:53860338-53860378 | Ch13 region 2 |
| chr13:53860398-53860438 | Ch13 region 2 |
| chr13:53860451-53860491 | Ch13 region 2 |
| chr13:53860513-53860553 | Ch13 region 2 |
| chr13:53860603-53860643 | Ch13 region 2 |
| chr13:53860654-53860694 | Ch13 region 2 |
| chr13:53860699-53860739 | Ch13 region 2 |
| chr13:53860761-53860801 | Ch13 region 2 |
| chr13:53860823-53860863 | Ch13 region 2 |

| | |
|---|---|
| chr13:53860886-53860926 | Ch13 region 2 |
| chr13:53860977-53861017 | Ch13 region 2 |
| chr13:53861025-53861065 | Ch13 region 2 |
| chr13:53861156-53861196 | Ch13 region 2 |
| chr13:53861230-53861270 | Ch13 region 2 |
| chr13:53861283-53861323 | Ch13 region 2 |
| chr13:53861329-53861369 | Ch13 region 2 |
| chr13:53861625-53861665 | Ch13 region 2 |
| chr13:53861670-53861710 | Ch13 region 2 |
| chr13:53861717-53861757 | Ch13 region 2 |
| chr13:53861775-53861815 | Ch13 region 2 |
| chr13:53861823-53861863 | Ch13 region 2 |
| chr13:53861883-53861923 | Ch13 region 2 |
| chr13:53861965-53862005 | Ch13 region 2 |
| chr13:53862050-53862090 | Ch13 region 2 |
| chr13:53862142-53862182 | Ch13 region 2 |
| chr13:53862187-53862227 | Ch13 region 2 |
| chr13:53862268-53862308 | Ch13 region 2 |
| chr13:53862336-53862376 | Ch13 region 2 |
| chr13:53862382-53862422 | Ch13 region 2 |
| chr13:53862477-53862517 | Ch13 region 2 |
| chr13:53862523-53862563 | Ch13 region 2 |
| chr13:53862593-53862633 | Ch13 region 2 |
| chr13:53862773-53862813 | Ch13 region 2 |
| chr13:53862851-53862891 | Ch13 region 2 |
| chr13:53862950-53862990 | Ch13 region 2 |
| chr13:53863020-53863060 | Ch13 region 2 |
| chr13:53863081-53863121 | Ch13 region 2 |
| chr13:53863154-53863194 | Ch13 region 2 |
| chr13:53863213-53863253 | Ch13 region 2 |
| chr13:53863274-53863314 | Ch13 region 2 |
| chr13:53863332-53863372 | Ch13 region 2 |
| chr13:53863396-53863436 | Ch13 region 2 |
| chr13:53863465-53863505 | Ch13 region 2 |
| chr13:53863513-53863553 | Ch13 region 2 |
| chr13:53863566-53863606 | Ch13 region 2 |
| chr13:53863634-53863674 | Ch13 region 2 |
| chr13:53863679-53863719 | Ch13 region 2 |
| chr13:53863759-53863799 | Ch13 region 2 |
| chr13:53863806-53863846 | Ch13 region 2 |
| chr13:53863892-53863932 | Ch13 region 2 |
| chr13:53863959-53863999 | Ch13 region 2 |
| chr13:53864042-53864082 | Ch13 region 2 |
| chr13:53864091-53864131 | Ch13 region 2 |
| chr13:53864166-53864206 | Ch13 region 2 |
| chr13:53864216-53864256 | Ch13 region 2 |
| chr13:53864294-53864334 | Ch13 region 2 |
| chr13:53864425-53864465 | Ch13 region 2 |
| chr13:53864596-53864636 | Ch13 region 2 |
| chr13:53864642-53864682 | Ch13 region 2 |
| chr13:53864687-53864727 | Ch13 region 2 |
| chr13:53864737-53864777 | Ch13 region 2 |

| | |
|---|---|
| chr13:53864822-53864862 | Ch13 region 2 |
| chr13:53864915-53864955 | Ch13 region 2 |
| chr13:53864960-53865000 | Ch13 region 2 |
| chr13:53865011-53865051 | Ch13 region 2 |
| chr13:53865133-53865173 | Ch13 region 2 |
| chr13:53865220-53865260 | Ch13 region 2 |
| chr13:53865265-53865305 | Ch13 region 2 |
| chr13:53865310-53865350 | Ch13 region 2 |
| chr13:53865368-53865408 | Ch13 region 2 |
| chr13:53865415-53865455 | Ch13 region 2 |
| chr13:53865461-53865501 | Ch13 region 2 |
| chr13:53865534-53865574 | Ch13 region 2 |
| chr13:53865614-53865654 | Ch13 region 2 |
| chr13:53865675-53865715 | Ch13 region 2 |
| chr13:53865773-53865813 | Ch13 region 2 |
| chr13:53865836-53865876 | Ch13 region 2 |
| chr13:53865883-53865923 | Ch13 region 2 |
| chr13:53865932-53865972 | Ch13 region 2 |
| chr13:53866018-53866058 | Ch13 region 2 |
| chr13:53866076-53866116 | Ch13 region 2 |
| chr13:53866130-53866170 | Ch13 region 2 |
| chr13:53866187-53866227 | Ch13 region 2 |
| chr13:53866257-53866297 | Ch13 region 2 |
| chr13:53866308-53866348 | Ch13 region 2 |
| chr13:53866353-53866393 | Ch13 region 2 |
| chr13:53866408-53866448 | Ch13 region 2 |
| chr13:53866465-53866505 | Ch13 region 2 |
| chr13:53866528-53866568 | Ch13 region 2 |
| chr13:53866588-53866628 | Ch13 region 2 |
| chr13:53866634-53866674 | Ch13 region 2 |
| chr13:53866689-53866729 | Ch13 region 2 |
| chr13:53866746-53866786 | Ch13 region 2 |
| chr13:53866795-53866835 | Ch13 region 2 |
| chr13:53866872-53866912 | Ch13 region 2 |
| chr13:53866950-53866990 | Ch13 region 2 |
| chr13:53867026-53867066 | Ch13 region 2 |
| chr13:53867078-53867118 | Ch13 region 2 |
| chr13:53867172-53867212 | Ch13 region 2 |
| chr13:53867237-53867277 | Ch13 region 2 |
| chr13:53867300-53867340 | Ch13 region 2 |
| chr13:53867345-53867385 | Ch13 region 2 |
| chr13:53867420-53867460 | Ch13 region 2 |
| chr13:53867518-53867558 | Ch13 region 2 |
| chr13:53867563-53867603 | Ch13 region 2 |
| chr13:53867642-53867682 | Ch13 region 2 |
| chr13:53867692-53867732 | Ch13 region 2 |
| chr13:53867742-53867782 | Ch13 region 2 |
| chr13:53867794-53867834 | Ch13 region 2 |
| chr13:53867871-53867911 | Ch13 region 2 |
| chr13:53867957-53867997 | Ch13 region 2 |
| chr13:53868019-53868059 | Ch13 region 2 |
| chr13:53868072-53868112 | Ch13 region 2 |

| | |
|---|---|
| chr13:53868153-53868193 | Ch13 region 2 |
| chr13:53868220-53868260 | Ch13 region 2 |
| chr13:53868270-53868310 | Ch13 region 2 |
| chr13:53868381-53868421 | Ch13 region 2 |
| chr13:53868439-53868479 | Ch13 region 2 |
| chr13:53868505-53868545 | Ch13 region 2 |
| chr13:53868568-53868608 | Ch13 region 2 |
| chr13:53868650-53868690 | Ch13 region 2 |
| chr13:53868696-53868736 | Ch13 region 2 |
| chr13:53868772-53868812 | Ch13 region 2 |
| chr13:53868882-53868922 | Ch13 region 2 |
| chr13:53868932-53868972 | Ch13 region 2 |
| chr13:53869003-53869043 | Ch13 region 2 |
| chr13:53869102-53869142 | Ch13 region 2 |
| chr13:53869160-53869200 | Ch13 region 2 |
| chr13:53869305-53869345 | Ch13 region 2 |
| chr13:53869350-53869390 | Ch13 region 2 |
| chr13:53869415-53869455 | Ch13 region 2 |
| chr13:53869465-53869505 | Ch13 region 2 |
| chr13:53869525-53869565 | Ch13 region 2 |
| chr13:53869570-53869610 | Ch13 region 2 |
| chr13:53869622-53869662 | Ch13 region 2 |
| chr13:53869686-53869726 | Ch13 region 2 |
| chr13:53869751-53869791 | Ch13 region 2 |
| chr13:53869796-53869836 | Ch13 region 2 |
| chr13:53869865-53869905 | Ch13 region 2 |
| chr13:53869924-53869964 | Ch13 region 2 |
| chr13:53869993-53870033 | Ch13 region 2 |
| chr13:53870112-53870152 | Ch13 region 2 |
| chr13:53870184-53870224 | Ch13 region 2 |
| chr13:53870290-53870330 | Ch13 region 2 |
| chr13:53870351-53870391 | Ch13 region 2 |
| chr13:53870396-53870436 | Ch13 region 2 |
| chr13:53870504-53870544 | Ch13 region 2 |
| chr13:53870600-53870640 | Ch13 region 2 |
| chr13:53870721-53870761 | Ch13 region 2 |
| chr13:53870926-53870966 | Ch13 region 2 |
| chr13:53870993-53871033 | Ch13 region 2 |
| chr13:53871286-53871326 | Ch13 region 2 |
| chr13:53871744-53871784 | Ch13 region 2 |
| chr13:53872163-53872203 | Ch13 region 2 |
| chr13:53872529-53872569 | Ch13 region 2 |
| chr13:53872608-53872648 | Ch13 region 2 |
| chr13:53872681-53872721 | Ch13 region 2 |
| chr13:53872741-53872781 | Ch13 region 2 |
| chr13:53872934-53872974 | Ch13 region 2 |
| chr13:53872993-53873033 | Ch13 region 2 |
| chr13:53873115-53873155 | Ch13 region 2 |
| chr13:53873192-53873232 | Ch13 region 2 |
| chr13:53873299-53873339 | Ch13 region 2 |
| chr13:53873344-53873384 | Ch13 region 2 |
| chr13:53873423-53873463 | Ch13 region 2 |

| | |
|---|---|
| chr13:53873477-53873517 | Ch13 region 2 |
| chr13:53873534-53873574 | Ch13 region 2 |
| chr13:53873591-53873631 | Ch13 region 2 |
| chr13:53873805-53873845 | Ch13 region 2 |
| chr13:53873858-53873898 | Ch13 region 2 |
| chr13:53873903-53873943 | Ch13 region 2 |
| chr13:53873951-53873991 | Ch13 region 2 |
| chr13:53873997-53874037 | Ch13 region 2 |
| chr13:53874176-53874216 | Ch13 region 2 |
| chr13:53874262-53874302 | Ch13 region 2 |
| chr13:53874431-53874471 | Ch13 region 2 |
| chr13:53874535-53874575 | Ch13 region 2 |
| chr13:53874703-53874743 | Ch13 region 2 |
| chr13:53874789-53874829 | Ch13 region 2 |
| chr13:53874858-53874898 | Ch13 region 2 |
| chr13:53874985-53875025 | Ch13 region 2 |
| chr13:53875045-53875085 | Ch13 region 2 |
| chr13:53875109-53875149 | Ch13 region 2 |
| chr13:53875182-53875222 | Ch13 region 2 |
| chr13:53875242-53875282 | Ch13 region 2 |
| chr13:53875300-53875340 | Ch13 region 2 |
| chr13:53875398-53875438 | Ch13 region 2 |
| chr13:53875483-53875523 | Ch13 region 2 |
| chr13:53875574-53875614 | Ch13 region 2 |
| chr13:53875680-53875720 | Ch13 region 2 |
| chr13:53875754-53875794 | Ch13 region 2 |
| chr13:53875840-53875880 | Ch13 region 2 |
| chr13:53875901-53875941 | Ch13 region 2 |
| chr13:53875988-53876028 | Ch13 region 2 |
| chr13:53876092-53876132 | Ch13 region 2 |
| chr13:53876137-53876177 | Ch13 region 2 |
| chr13:53876217-53876257 | Ch13 region 2 |
| chr13:53876310-53876350 | Ch13 region 2 |
| chr13:53876387-53876427 | Ch13 region 2 |
| chr13:53876460-53876500 | Ch13 region 2 |
| chr13:53876552-53876592 | Ch13 region 2 |
| chr13:53876601-53876641 | Ch13 region 2 |
| chr13:53876682-53876722 | Ch13 region 2 |
| chr13:53876825-53876865 | Ch13 region 2 |
| chr13:53876875-53876915 | Ch13 region 2 |
| chr13:53876964-53877004 | Ch13 region 2 |
| chr13:53877057-53877097 | Ch13 region 2 |
| chr13:53877110-53877150 | Ch13 region 2 |
| chr13:53877209-53877249 | Ch13 region 2 |
| chr13:53877263-53877303 | Ch13 region 2 |
| chr13:53877322-53877362 | Ch13 region 2 |
| chr13:53877385-53877425 | Ch13 region 2 |
| chr13:53877475-53877515 | Ch13 region 2 |
| chr13:53877553-53877593 | Ch13 region 2 |
| chr13:53877637-53877677 | Ch13 region 2 |
| chr13:53877682-53877722 | Ch13 region 2 |
| chr13:53877748-53877788 | Ch13 region 2 |

| | |
|---|---|
| chr13:53877822-53877862 | Ch13 region 2 |
| chr13:53877875-53877915 | Ch13 region 2 |
| chr13:53877920-53877960 | Ch13 region 2 |
| chr13:53878038-53878078 | Ch13 region 2 |
| chr13:53878087-53878127 | Ch13 region 2 |
| chr13:53878146-53878186 | Ch13 region 2 |
| chr13:53878221-53878261 | Ch13 region 2 |
| chr13:53878302-53878342 | Ch13 region 2 |
| chr13:53878390-53878430 | Ch13 region 2 |
| chr13:53878435-53878475 | Ch13 region 2 |
| chr13:53878520-53878560 | Ch13 region 2 |
| chr13:53878589-53878629 | Ch13 region 2 |
| chr13:53878634-53878674 | Ch13 region 2 |
| chr13:53878689-53878729 | Ch13 region 2 |
| chr13:53878757-53878797 | Ch13 region 2 |
| chr13:53878857-53878897 | Ch13 region 2 |
| chr13:53878974-53879014 | Ch13 region 2 |
| chr13:53879043-53879083 | Ch13 region 2 |
| chr13:53879133-53879173 | Ch13 region 2 |
| chr13:53879240-53879280 | Ch13 region 2 |
| chr13:53879389-53879429 | Ch13 region 2 |
| chr13:53879460-53879500 | Ch13 region 2 |
| chr13:53879548-53879588 | Ch13 region 2 |
| chr13:53879741-53879781 | Ch13 region 2 |
| chr13:53879816-53879856 | Ch13 region 2 |
| chr13:53879996-53880036 | Ch13 region 2 |
| chr13:53880088-53880128 | Ch13 region 2 |
| chr13:53880141-53880181 | Ch13 region 2 |
| chr13:53880226-53880266 | Ch13 region 2 |
| chr13:53880504-53880544 | Ch13 region 2 |
| chr13:53880579-53880619 | Ch13 region 2 |
| chr13:53880638-53880678 | Ch13 region 2 |
| chr13:53880713-53880753 | Ch13 region 2 |
| chr13:53880774-53880814 | Ch13 region 2 |
| chr13:53880864-53880904 | Ch13 region 2 |
| chr13:53880920-53880960 | Ch13 region 2 |
| chr13:53880980-53881020 | Ch13 region 2 |
| chr13:53881050-53881090 | Ch13 region 2 |
| chr13:53881126-53881166 | Ch13 region 2 |
| chr13:53881255-53881295 | Ch13 region 2 |
| chr13:53881332-53881372 | Ch13 region 2 |
| chr13:53881379-53881419 | Ch13 region 2 |
| chr13:53881445-53881485 | Ch13 region 2 |
| chr13:53881490-53881530 | Ch13 region 2 |
| chr13:53881572-53881612 | Ch13 region 2 |
| chr13:53881630-53881670 | Ch13 region 2 |
| chr13:53881692-53881732 | Ch13 region 2 |
| chr13:53881744-53881784 | Ch13 region 2 |
| chr13:53881795-53881835 | Ch13 region 2 |
| chr13:53881856-53881896 | Ch13 region 2 |
| chr13:53881906-53881946 | Ch13 region 2 |
| chr13:53881951-53881991 | Ch13 region 2 |

| | |
|---|---|
| chr13:53882044-53882084 | Ch13 region 2 |
| chr13:53882100-53882140 | Ch13 region 2 |
| chr13:53882254-53882294 | Ch13 region 2 |
| chr13:53882454-53882494 | Ch13 region 2 |
| chr13:53882499-53882539 | Ch13 region 2 |
| chr13:53882572-53882612 | Ch13 region 2 |
| chr13:53882633-53882673 | Ch13 region 2 |
| chr13:53882728-53882768 | Ch13 region 2 |
| chr13:53882774-53882814 | Ch13 region 2 |
| chr13:53882827-53882867 | Ch13 region 2 |
| chr13:53882877-53882917 | Ch13 region 2 |
| chr13:53882939-53882979 | Ch13 region 2 |
| chr13:53882999-53883039 | Ch13 region 2 |
| chr13:53883102-53883142 | Ch13 region 2 |
| chr13:53883149-53883189 | Ch13 region 2 |
| chr13:53883212-53883252 | Ch13 region 2 |
| chr13:53883268-53883308 | Ch13 region 2 |
| chr13:53883318-53883358 | Ch13 region 2 |
| chr13:53883394-53883434 | Ch13 region 2 |
| chr13:53883444-53883484 | Ch13 region 2 |
| chr13:53883490-53883530 | Ch13 region 2 |
| chr13:53883535-53883575 | Ch13 region 2 |
| chr13:53883587-53883627 | Ch13 region 2 |
| chr13:53883675-53883715 | Ch13 region 2 |
| chr13:53883720-53883760 | Ch13 region 2 |
| chr13:53883791-53883831 | Ch13 region 2 |
| chr13:53883869-53883909 | Ch13 region 2 |
| chr13:53883914-53883954 | Ch13 region 2 |
| chr13:53884021-53884061 | Ch13 region 2 |
| chr13:53884078-53884118 | Ch13 region 2 |
| chr13:53884130-53884170 | Ch13 region 2 |
| chr13:53884206-53884246 | Ch13 region 2 |
| chr13:53884282-53884322 | Ch13 region 2 |
| chr13:53884331-53884371 | Ch13 region 2 |
| chr13:53884428-53884468 | Ch13 region 2 |
| chr13:53884491-53884531 | Ch13 region 2 |
| chr13:53884536-53884576 | Ch13 region 2 |
| chr13:53884621-53884661 | Ch13 region 2 |
| chr13:53884677-53884717 | Ch13 region 2 |
| chr13:53884733-53884773 | Ch13 region 2 |
| chr13:53884826-53884866 | Ch13 region 2 |
| chr13:53884874-53884914 | Ch13 region 2 |
| chr13:53884973-53885013 | Ch13 region 2 |
| chr13:53885018-53885058 | Ch13 region 2 |
| chr13:53885063-53885103 | Ch13 region 2 |
| chr13:53885119-53885159 | Ch13 region 2 |
| chr13:53885195-53885235 | Ch13 region 2 |
| chr13:53885266-53885306 | Ch13 region 2 |
| chr13:53885318-53885358 | Ch13 region 2 |
| chr13:53885374-53885414 | Ch13 region 2 |
| chr13:53885456-53885496 | Ch13 region 2 |
| chr13:53885516-53885556 | Ch13 region 2 |

| | |
|---|---|
| chr13:53885582-53885622 | Ch13 region 2 |
| chr13:53885658-53885698 | Ch13 region 2 |
| chr13:53885732-53885772 | Ch13 region 2 |
| chr13:53885831-53885871 | Ch13 region 2 |
| chr13:53885876-53885916 | Ch13 region 2 |
| chr13:53885921-53885961 | Ch13 region 2 |
| chr13:53885981-53886021 | Ch13 region 2 |
| chr13:53886026-53886066 | Ch13 region 2 |
| chr13:53886101-53886141 | Ch13 region 2 |
| chr13:53886150-53886190 | Ch13 region 2 |
| chr13:53886248-53886288 | Ch13 region 2 |
| chr13:53886325-53886365 | Ch13 region 2 |
| chr13:53886383-53886423 | Ch13 region 2 |
| chr13:53886443-53886483 | Ch13 region 2 |
| chr13:53886488-53886528 | Ch13 region 2 |
| chr13:53886547-53886587 | Ch13 region 2 |
| chr13:53886643-53886683 | Ch13 region 2 |
| chr13:53886758-53886798 | Ch13 region 2 |
| chr13:53886834-53886874 | Ch13 region 2 |
| chr13:53886884-53886924 | Ch13 region 2 |
| chr13:53886938-53886978 | Ch13 region 2 |
| chr13:53887019-53887059 | Ch13 region 2 |
| chr13:53887064-53887104 | Ch13 region 2 |
| chr13:53887117-53887157 | Ch13 region 2 |
| chr13:53887167-53887207 | Ch13 region 2 |
| chr13:53887236-53887276 | Ch13 region 2 |
| chr13:53887332-53887372 | Ch13 region 2 |
| chr13:53887582-53887622 | Ch13 region 2 |
| chr13:53887659-53887699 | Ch13 region 2 |
| chr13:53887778-53887818 | Ch13 region 2 |
| chr13:53887879-53887919 | Ch13 region 2 |
| chr13:53887929-53887969 | Ch13 region 2 |
| chr13:53887979-53888019 | Ch13 region 2 |
| chr13:53888038-53888078 | Ch13 region 2 |
| chr13:53888123-53888163 | Ch13 region 2 |
| chr13:53888212-53888252 | Ch13 region 2 |
| chr13:53888262-53888302 | Ch13 region 2 |
| chr13:53888315-53888355 | Ch13 region 2 |
| chr13:53888383-53888423 | Ch13 region 2 |
| chr13:53888429-53888469 | Ch13 region 2 |
| chr13:53888489-53888529 | Ch13 region 2 |
| chr13:53888536-53888576 | Ch13 region 2 |
| chr13:53888615-53888655 | Ch13 region 2 |
| chr13:53888663-53888703 | Ch13 region 2 |
| chr13:53889069-53889109 | Ch13 region 2 |
| chr13:53889129-53889169 | Ch13 region 2 |
| chr13:53889213-53889253 | Ch13 region 2 |
| chr13:53889258-53889298 | Ch13 region 2 |
| chr13:53889303-53889343 | Ch13 region 2 |
| chr13:53889501-53889541 | Ch13 region 2 |
| chr13:53889550-53889590 | Ch13 region 2 |
| chr13:53889600-53889640 | Ch13 region 2 |

| | |
|---|---|
| chr13:53889722-53889762 | Ch13 region 2 |
| chr13:53889841-53889881 | Ch13 region 2 |
| chr13:53889973-53890013 | Ch13 region 2 |
| chr13:53890046-53890086 | Ch13 region 2 |
| chr13:53890166-53890206 | Ch13 region 2 |
| chr13:53890236-53890276 | Ch13 region 2 |
| chr13:53890296-53890336 | Ch13 region 2 |
| chr13:53890341-53890381 | Ch13 region 2 |
| chr13:53890433-53890473 | Ch13 region 2 |
| chr13:53890524-53890564 | Ch13 region 2 |
| chr13:53890612-53890652 | Ch13 region 2 |
| chr13:53890658-53890698 | Ch13 region 2 |
| chr13:53890722-53890762 | Ch13 region 2 |
| chr13:53890785-53890825 | Ch13 region 2 |
| chr13:53890862-53890902 | Ch13 region 2 |
| chr13:53890924-53890964 | Ch13 region 2 |
| chr13:53891007-53891047 | Ch13 region 2 |
| chr13:53891053-53891093 | Ch13 region 2 |
| chr13:53891135-53891175 | Ch13 region 2 |
| chr13:53891180-53891220 | Ch13 region 2 |
| chr13:53891269-53891309 | Ch13 region 2 |
| chr13:53891316-53891356 | Ch13 region 2 |
| chr13:53891436-53891476 | Ch13 region 2 |
| chr13:53891507-53891547 | Ch13 region 2 |
| chr13:53891571-53891611 | Ch13 region 2 |
| chr13:53891616-53891656 | Ch13 region 2 |
| chr13:53891671-53891711 | Ch13 region 2 |
| chr13:53891736-53891776 | Ch13 region 2 |
| chr13:53891846-53891886 | Ch13 region 2 |
| chr13:53891922-53891962 | Ch13 region 2 |
| chr13:53892005-53892045 | Ch13 region 2 |
| chr13:53892077-53892117 | Ch13 region 2 |
| chr13:53892139-53892179 | Ch13 region 2 |
| chr13:53892229-53892269 | Ch13 region 2 |
| chr13:53892279-53892319 | Ch13 region 2 |
| chr13:53892330-53892370 | Ch13 region 2 |
| chr13:53892421-53892461 | Ch13 region 2 |
| chr13:53892473-53892513 | Ch13 region 2 |
| chr13:53892543-53892583 | Ch13 region 2 |
| chr13:53892608-53892648 | Ch13 region 2 |
| chr13:53892664-53892704 | Ch13 region 2 |
| chr13:53892741-53892781 | Ch13 region 2 |
| chr13:53892786-53892826 | Ch13 region 2 |
| chr13:53892869-53892909 | Ch13 region 2 |
| chr13:53892955-53892995 | Ch13 region 2 |
| chr13:53893048-53893088 | Ch13 region 2 |
| chr13:53893096-53893136 | Ch13 region 2 |
| chr13:53893174-53893214 | Ch13 region 2 |
| chr13:53893280-53893320 | Ch13 region 2 |
| chr13:53893356-53893396 | Ch13 region 2 |
| chr13:53893421-53893461 | Ch13 region 2 |
| chr13:53893496-53893536 | Ch13 region 2 |

| | |
|---|---|
| chr13:53893563-53893603 | Ch13 region 2 |
| chr13:53893660-53893700 | Ch13 region 2 |
| chr13:53893749-53893789 | Ch13 region 2 |
| chr13:53893801-53893841 | Ch13 region 2 |
| chr13:53893890-53893930 | Ch13 region 2 |
| chr13:53893935-53893975 | Ch13 region 2 |
| chr13:53893993-53894033 | Ch13 region 2 |
| chr13:53894039-53894079 | Ch13 region 2 |
| chr13:53894098-53894138 | Ch13 region 2 |
| chr13:53894184-53894224 | Ch13 region 2 |
| chr13:53894277-53894317 | Ch13 region 2 |
| chr13:53894325-53894365 | Ch13 region 2 |
| chr13:53894373-53894413 | Ch13 region 2 |
| chr13:53894418-53894458 | Ch13 region 2 |
| chr13:53894502-53894542 | Ch13 region 2 |
| chr13:53894547-53894587 | Ch13 region 2 |
| chr13:53894641-53894681 | Ch13 region 2 |
| chr13:53894689-53894729 | Ch13 region 2 |
| chr13:53894798-53894838 | Ch13 region 2 |
| chr13:53894844-53894884 | Ch13 region 2 |
| chr13:53894916-53894956 | Ch13 region 2 |
| chr13:53894961-53895001 | Ch13 region 2 |
| chr13:53895021-53895061 | Ch13 region 2 |
| chr13:53895075-53895115 | Ch13 region 2 |
| chr13:53895120-53895160 | Ch13 region 2 |
| chr13:53895203-53895243 | Ch13 region 2 |
| chr13:53895250-53895290 | Ch13 region 2 |
| chr13:53895303-53895343 | Ch13 region 2 |
| chr13:53895372-53895412 | Ch13 region 2 |
| chr13:53895443-53895483 | Ch13 region 2 |
| chr13:53895527-53895567 | Ch13 region 2 |
| chr13:53895592-53895632 | Ch13 region 2 |
| chr13:53895652-53895692 | Ch13 region 2 |
| chr13:53895759-53895799 | Ch13 region 2 |
| chr13:53895835-53895875 | Ch13 region 2 |
| chr13:53895925-53895965 | Ch13 region 2 |
| chr13:53895987-53896027 | Ch13 region 2 |
| chr13:53896053-53896093 | Ch13 region 2 |
| chr13:53896099-53896139 | Ch13 region 2 |
| chr13:53896154-53896194 | Ch13 region 2 |
| chr13:53896233-53896273 | Ch13 region 2 |
| chr13:53896350-53896390 | Ch13 region 2 |
| chr13:53896503-53896543 | Ch13 region 2 |
| chr13:53896559-53896599 | Ch13 region 2 |
| chr13:53896643-53896683 | Ch13 region 2 |
| chr13:53896688-53896728 | Ch13 region 2 |
| chr13:53896770-53896810 | Ch13 region 2 |
| chr13:53896822-53896862 | Ch13 region 2 |
| chr13:53896898-53896938 | Ch13 region 2 |
| chr13:53896943-53896983 | Ch13 region 2 |
| chr13:53897005-53897045 | Ch13 region 2 |
| chr13:53897060-53897100 | Ch13 region 2 |

| | |
|---|---|
| chr13:53897114-53897154 | Ch13 region 2 |
| chr13:53897169-53897209 | Ch13 region 2 |
| chr13:53897256-53897296 | Ch13 region 2 |
| chr13:53897323-53897363 | Ch13 region 2 |
| chr13:53897372-53897412 | Ch13 region 2 |
| chr13:53897470-53897510 | Ch13 region 2 |
| chr13:53897519-53897559 | Ch13 region 2 |
| chr13:53897609-53897649 | Ch13 region 2 |
| chr13:53897654-53897694 | Ch13 region 2 |
| chr13:53897699-53897739 | Ch13 region 2 |
| chr13:53897744-53897784 | Ch13 region 2 |
| chr13:53897845-53897885 | Ch13 region 2 |
| chr13:53897927-53897967 | Ch13 region 2 |
| chr13:53898140-53898180 | Ch13 region 2 |
| chr13:53898222-53898262 | Ch13 region 2 |
| chr13:53898286-53898326 | Ch13 region 2 |
| chr13:53898361-53898401 | Ch13 region 2 |
| chr13:53898429-53898469 | Ch13 region 2 |
| chr13:53898477-53898517 | Ch13 region 2 |
| chr13:53898565-53898605 | Ch13 region 2 |
| chr13:53898668-53898708 | Ch13 region 2 |
| chr13:53898756-53898796 | Ch13 region 2 |
| chr13:53898819-53898859 | Ch13 region 2 |
| chr13:53898870-53898910 | Ch13 region 2 |
| chr13:53898960-53899000 | Ch13 region 2 |
| chr13:53899005-53899045 | Ch13 region 2 |
| chr13:53899071-53899111 | Ch13 region 2 |
| chr13:53899121-53899161 | Ch13 region 2 |
| chr13:53899166-53899206 | Ch13 region 2 |
| chr13:53899212-53899252 | Ch13 region 2 |
| chr13:53899338-53899378 | Ch13 region 2 |
| chr13:53899388-53899428 | Ch13 region 2 |
| chr13:53899491-53899531 | Ch13 region 2 |
| chr13:53899581-53899621 | Ch13 region 2 |
| chr13:53899715-53899755 | Ch13 region 2 |
| chr13:53899771-53899811 | Ch13 region 2 |
| chr13:53899856-53899896 | Ch13 region 2 |
| chr13:53899909-53899949 | Ch13 region 2 |
| chr13:53899954-53899994 | Ch13 region 2 |
| chr13:53900023-53900063 | Ch13 region 2 |
| chr13:53900068-53900108 | Ch13 region 2 |
| chr13:53900116-53900156 | Ch13 region 2 |
| chr13:53900519-53900559 | Ch13 region 2 |
| chr13:53900629-53900669 | Ch13 region 2 |
| chr13:53900695-53900735 | Ch13 region 2 |
| chr13:53900755-53900795 | Ch13 region 2 |
| chr13:53900800-53900840 | Ch13 region 2 |
| chr13:53900876-53900916 | Ch13 region 2 |
| chr13:53900921-53900961 | Ch13 region 2 |
| chr13:53900998-53901038 | Ch13 region 2 |
| chr13:53901117-53901157 | Ch13 region 2 |
| chr13:53901291-53901331 | Ch13 region 2 |

| | |
|---|---|
| chr13:53901362-53901402 | Ch13 region 2 |
| chr13:53901467-53901507 | Ch13 region 2 |
| chr13:53901516-53901556 | Ch13 region 2 |
| chr13:53901594-53901634 | Ch13 region 2 |
| chr13:53901639-53901679 | Ch13 region 2 |
| chr13:53901703-53901743 | Ch13 region 2 |
| chr13:53901775-53901815 | Ch13 region 2 |
| chr13:53901894-53901934 | Ch13 region 2 |
| chr13:53901988-53902028 | Ch13 region 2 |
| chr13:53902040-53902080 | Ch13 region 2 |
| chr13:53902105-53902145 | Ch13 region 2 |
| chr13:53902153-53902193 | Ch13 region 2 |
| chr13:53902238-53902278 | Ch13 region 2 |
| chr13:53902294-53902334 | Ch13 region 2 |
| chr13:53902354-53902394 | Ch13 region 2 |
| chr13:53902411-53902451 | Ch13 region 2 |
| chr13:53902508-53902548 | Ch13 region 2 |
| chr13:53903003-53903043 | Ch13 region 2 |
| chr13:53903065-53903105 | Ch13 region 2 |
| chr13:53903119-53903159 | Ch13 region 2 |
| chr13:53903175-53903215 | Ch13 region 2 |
| chr13:53903246-53903286 | Ch13 region 2 |
| chr13:53903339-53903379 | Ch13 region 2 |
| chr13:53903387-53903427 | Ch13 region 2 |
| chr13:53903445-53903485 | Ch13 region 2 |
| chr13:53903492-53903532 | Ch13 region 2 |
| chr13:53903538-53903578 | Ch13 region 2 |
| chr13:53903620-53903660 | Ch13 region 2 |
| chr13:53903668-53903708 | Ch13 region 2 |
| chr13:53903759-53903799 | Ch13 region 2 |
| chr13:53903804-53903844 | Ch13 region 2 |
| chr13:53903849-53903889 | Ch13 region 2 |
| chr13:53903911-53903951 | Ch13 region 2 |
| chr13:53904030-53904070 | Ch13 region 2 |
| chr13:53904075-53904115 | Ch13 region 2 |
| chr13:53904126-53904166 | Ch13 region 2 |
| chr13:53904205-53904245 | Ch13 region 2 |
| chr13:53904268-53904308 | Ch13 region 2 |
| chr13:53904337-53904377 | Ch13 region 2 |
| chr13:53904393-53904433 | Ch13 region 2 |
| chr13:53904455-53904495 | Ch13 region 2 |
| chr13:53904518-53904558 | Ch13 region 2 |
| chr13:53904581-53904621 | Ch13 region 2 |
| chr13:53904647-53904687 | Ch13 region 2 |
| chr13:53904694-53904734 | Ch13 region 2 |
| chr13:53904745-53904785 | Ch13 region 2 |
| chr13:53904796-53904836 | Ch13 region 2 |
| chr13:53904894-53904934 | Ch13 region 2 |
| chr13:53904964-53905004 | Ch13 region 2 |
| chr13:53905020-53905060 | Ch13 region 2 |
| chr13:53905065-53905105 | Ch13 region 2 |
| chr13:53905140-53905180 | Ch13 region 2 |

| | |
|---|---|
| chr13:53905203-53905243 | Ch13 region 2 |
| chr13:53905273-53905313 | Ch13 region 2 |
| chr13:53905318-53905358 | Ch13 region 2 |
| chr13:53905391-53905431 | Ch13 region 2 |
| chr13:53905467-53905507 | Ch13 region 2 |
| chr13:53905531-53905571 | Ch13 region 2 |
| chr13:53905629-53905669 | Ch13 region 2 |
| chr13:53905682-53905722 | Ch13 region 2 |
| chr13:53905763-53905803 | Ch13 region 2 |
| chr13:53905811-53905851 | Ch13 region 2 |
| chr13:53905873-53905913 | Ch13 region 2 |
| chr13:53905932-53905972 | Ch13 region 2 |
| chr13:53905980-53906020 | Ch13 region 2 |
| chr13:53906041-53906081 | Ch13 region 2 |
| chr13:53906099-53906139 | Ch13 region 2 |
| chr13:53906167-53906207 | Ch13 region 2 |
| chr13:53906264-53906304 | Ch13 region 2 |
| chr13:53906429-53906469 | Ch13 region 2 |
| chr13:53906495-53906535 | Ch13 region 2 |
| chr13:53906612-53906652 | Ch13 region 2 |
| chr13:53906673-53906713 | Ch13 region 2 |
| chr13:53906743-53906783 | Ch13 region 2 |
| chr13:53906828-53906868 | Ch13 region 2 |
| chr13:53906911-53906951 | Ch13 region 2 |
| chr13:53906962-53907002 | Ch13 region 2 |
| chr13:53907043-53907083 | Ch13 region 2 |
| chr13:53907100-53907140 | Ch13 region 2 |
| chr13:53907167-53907207 | Ch13 region 2 |
| chr13:53907217-53907257 | Ch13 region 2 |
| chr13:53907288-53907328 | Ch13 region 2 |
| chr13:53907339-53907379 | Ch13 region 2 |
| chr13:53907418-53907458 | Ch13 region 2 |
| chr13:53907492-53907532 | Ch13 region 2 |
| chr13:53907674-53907714 | Ch13 region 2 |
| chr13:53907780-53907820 | Ch13 region 2 |
| chr13:53907876-53907916 | Ch13 region 2 |
| chr13:53907946-53907986 | Ch13 region 2 |
| chr13:53907991-53908031 | Ch13 region 2 |
| chr13:53908079-53908119 | Ch13 region 2 |
| chr13:53908172-53908212 | Ch13 region 2 |
| chr13:53908285-53908325 | Ch13 region 2 |
| chr13:53908533-53908573 | Ch13 region 2 |
| chr13:53908589-53908629 | Ch13 region 2 |
| chr13:53908687-53908727 | Ch13 region 2 |
| chr13:53908806-53908846 | Ch13 region 2 |
| chr13:53908890-53908930 | Ch13 region 2 |
| chr13:53908935-53908975 | Ch13 region 2 |
| chr13:53909005-53909045 | Ch13 region 2 |
| chr13:53909097-53909137 | Ch13 region 2 |
| chr13:53909220-53909260 | Ch13 region 2 |
| chr13:53909269-53909309 | Ch13 region 2 |
| chr13:53909328-53909368 | Ch13 region 2 |

| | |
|---|---|
| chr13:53909412-53909452 | Ch13 region 2 |
| chr13:53909462-53909502 | Ch13 region 2 |
| chr13:53909637-53909677 | Ch13 region 2 |
| chr13:53909758-53909798 | Ch13 region 2 |
| chr13:53909912-53909952 | Ch13 region 2 |
| chr13:53910046-53910086 | Ch13 region 2 |
| chr13:53910107-53910147 | Ch13 region 2 |
| chr13:53910168-53910208 | Ch13 region 2 |
| chr13:53910213-53910253 | Ch13 region 2 |
| chr13:53910288-53910328 | Ch13 region 2 |
| chr13:53910344-53910384 | Ch13 region 2 |
| chr13:53910500-53910540 | Ch13 region 2 |
| chr13:53910576-53910616 | Ch13 region 2 |
| chr13:53910637-53910677 | Ch13 region 2 |
| chr13:53910745-53910785 | Ch13 region 2 |
| chr13:53910797-53910837 | Ch13 region 2 |
| chr13:53910842-53910882 | Ch13 region 2 |
| chr13:53910922-53910962 | Ch13 region 2 |
| chr13:53910967-53911007 | Ch13 region 2 |
| chr13:53911035-53911075 | Ch13 region 2 |
| chr13:53911081-53911121 | Ch13 region 2 |
| chr13:53911171-53911211 | Ch13 region 2 |
| chr13:53911217-53911257 | Ch13 region 2 |
| chr13:53911285-53911325 | Ch13 region 2 |
| chr13:53911331-53911371 | Ch13 region 2 |
| chr13:53911422-53911462 | Ch13 region 2 |
| chr13:53911611-53911651 | Ch13 region 2 |
| chr13:53911694-53911734 | Ch13 region 2 |
| chr13:53911742-53911782 | Ch13 region 2 |
| chr13:53911822-53911862 | Ch13 region 2 |
| chr13:53911897-53911937 | Ch13 region 2 |
| chr13:53911942-53911982 | Ch13 region 2 |
| chr13:53911988-53912028 | Ch13 region 2 |
| chr13:53912063-53912103 | Ch13 region 2 |
| chr13:53912108-53912148 | Ch13 region 2 |
| chr13:53912461-53912501 | Ch13 region 2 |
| chr13:53912548-53912588 | Ch13 region 2 |
| chr13:53912628-53912668 | Ch13 region 2 |
| chr13:53912673-53912713 | Ch13 region 2 |
| chr13:53912726-53912766 | Ch13 region 2 |
| chr13:53912794-53912834 | Ch13 region 2 |
| chr13:53912879-53912919 | Ch13 region 2 |
| chr13:53912924-53912964 | Ch13 region 2 |
| chr13:53912978-53913018 | Ch13 region 2 |
| chr13:53913024-53913064 | Ch13 region 2 |
| chr13:53913079-53913119 | Ch13 region 2 |
| chr13:53913126-53913166 | Ch13 region 2 |
| chr13:53913201-53913241 | Ch13 region 2 |
| chr13:53913257-53913297 | Ch13 region 2 |
| chr13:53913313-53913353 | Ch13 region 2 |
| chr13:53913364-53913404 | Ch13 region 2 |
| chr13:53913456-53913496 | Ch13 region 2 |

| | |
|---|---|
| chr13:53913548-53913588 | Ch13 region 2 |
| chr13:53913636-53913676 | Ch13 region 2 |
| chr13:53913681-53913721 | Ch13 region 2 |
| chr13:53913726-53913766 | Ch13 region 2 |
| chr13:53913781-53913821 | Ch13 region 2 |
| chr13:53913862-53913902 | Ch13 region 2 |
| chr13:53913970-53914010 | Ch13 region 2 |
| chr13:53914025-53914065 | Ch13 region 2 |
| chr13:53914084-53914124 | Ch13 region 2 |
| chr13:53914142-53914182 | Ch13 region 2 |
| chr13:53914216-53914256 | Ch13 region 2 |
| chr13:53914336-53914376 | Ch13 region 2 |
| chr13:53914381-53914421 | Ch13 region 2 |
| chr13:53914459-53914499 | Ch13 region 2 |
| chr13:53914513-53914553 | Ch13 region 2 |
| chr13:53914563-53914603 | Ch13 region 2 |
| chr13:53914608-53914648 | Ch13 region 2 |
| chr13:53914688-53914728 | Ch13 region 2 |
| chr13:53914734-53914774 | Ch13 region 2 |
| chr13:53914807-53914847 | Ch13 region 2 |
| chr13:53914852-53914892 | Ch13 region 2 |
| chr13:53914897-53914937 | Ch13 region 2 |
| chr13:53914978-53915018 | Ch13 region 2 |
| chr13:53915051-53915091 | Ch13 region 2 |
| chr13:53915096-53915136 | Ch13 region 2 |
| chr13:53915154-53915194 | Ch13 region 2 |
| chr13:53915203-53915243 | Ch13 region 2 |
| chr13:53915270-53915310 | Ch13 region 2 |
| chr13:53915391-53915431 | Ch13 region 2 |
| chr13:53915465-53915505 | Ch13 region 2 |
| chr13:53915512-53915552 | Ch13 region 2 |
| chr13:53915585-53915625 | Ch13 region 2 |
| chr13:53915631-53915671 | Ch13 region 2 |
| chr13:53915719-53915759 | Ch13 region 2 |
| chr13:53915777-53915817 | Ch13 region 2 |
| chr13:53915860-53915900 | Ch13 region 2 |
| chr13:53915924-53915964 | Ch13 region 2 |
| chr13:53915970-53916010 | Ch13 region 2 |
| chr13:53916031-53916071 | Ch13 region 2 |
| chr13:53916186-53916226 | Ch13 region 2 |
| chr13:53916233-53916273 | Ch13 region 2 |
| chr13:53916299-53916339 | Ch13 region 2 |
| chr13:53916409-53916449 | Ch13 region 2 |
| chr13:53916455-53916495 | Ch13 region 2 |
| chr13:53916523-53916563 | Ch13 region 2 |
| chr13:53916572-53916612 | Ch13 region 2 |
| chr13:53916655-53916695 | Ch13 region 2 |
| chr13:53916725-53916765 | Ch13 region 2 |
| chr13:53916784-53916824 | Ch13 region 2 |
| chr13:53916864-53916904 | Ch13 region 2 |
| chr13:53916911-53916951 | Ch13 region 2 |
| chr13:53916962-53917002 | Ch13 region 2 |

| | |
|---|---|
| chr13:53917023-53917063 | Ch13 region 2 |
| chr13:53917074-53917114 | Ch13 region 2 |
| chr13:53917156-53917196 | Ch13 region 2 |
| chr13:53917225-53917265 | Ch13 region 2 |
| chr13:53917291-53917331 | Ch13 region 2 |
| chr13:53917339-53917379 | Ch13 region 2 |
| chr13:53917403-53917443 | Ch13 region 2 |
| chr13:53917463-53917503 | Ch13 region 2 |
| chr13:53917529-53917569 | Ch13 region 2 |
| chr13:53917606-53917646 | Ch13 region 2 |
| chr13:53917677-53917717 | Ch13 region 2 |
| chr13:53917738-53917778 | Ch13 region 2 |
| chr13:53917796-53917836 | Ch13 region 2 |
| chr13:53917853-53917893 | Ch13 region 2 |
| chr13:53917916-53917956 | Ch13 region 2 |
| chr13:53918075-53918115 | Ch13 region 2 |
| chr13:53918139-53918179 | Ch13 region 2 |
| chr13:53918227-53918267 | Ch13 region 2 |
| chr13:53918346-53918386 | Ch13 region 2 |
| chr13:53918429-53918469 | Ch13 region 2 |
| chr13:53918536-53918576 | Ch13 region 2 |
| chr13:53918581-53918621 | Ch13 region 2 |
| chr13:53918647-53918687 | Ch13 region 2 |
| chr13:53918757-53918797 | Ch13 region 2 |
| chr13:53918830-53918870 | Ch13 region 2 |
| chr13:53918919-53918959 | Ch13 region 2 |
| chr13:53918964-53919004 | Ch13 region 2 |
| chr13:53919028-53919068 | Ch13 region 2 |
| chr13:53919092-53919132 | Ch13 region 2 |
| chr13:53919167-53919207 | Ch13 region 2 |
| chr13:53919247-53919287 | Ch13 region 2 |
| chr13:53919325-53919365 | Ch13 region 2 |
| chr13:53919377-53919417 | Ch13 region 2 |
| chr13:53919452-53919492 | Ch13 region 2 |
| chr13:53919539-53919579 | Ch13 region 2 |
| chr13:53919593-53919633 | Ch13 region 2 |
| chr13:53919643-53919683 | Ch13 region 2 |
| chr13:53919724-53919764 | Ch13 region 2 |
| chr13:53919782-53919822 | Ch13 region 2 |
| chr13:53919833-53919873 | Ch13 region 2 |
| chr13:53919914-53919954 | Ch13 region 2 |
| chr13:53919982-53920022 | Ch13 region 2 |
| chr13:53920062-53920102 | Ch13 region 2 |
| chr13:53920122-53920162 | Ch13 region 2 |
| chr13:53920177-53920217 | Ch13 region 2 |
| chr13:53920286-53920326 | Ch13 region 2 |
| chr13:53920356-53920396 | Ch13 region 2 |
| chr13:53920417-53920457 | Ch13 region 2 |
| chr13:53920475-53920515 | Ch13 region 2 |
| chr13:53920552-53920592 | Ch13 region 2 |
| chr13:53920610-53920650 | Ch13 region 2 |
| chr13:53920700-53920740 | Ch13 region 2 |

| | |
|---|---|
| chr13:53920755-53920795 | Ch13 region 2 |
| chr13:53920829-53920869 | Ch13 region 2 |
| chr13:53920875-53920915 | Ch13 region 2 |
| chr13:53920920-53920960 | Ch13 region 2 |
| chr13:53920968-53921008 | Ch13 region 2 |
| chr13:53921019-53921059 | Ch13 region 2 |
| chr13:53921095-53921135 | Ch13 region 2 |
| chr13:53921146-53921186 | Ch13 region 2 |
| chr13:53921217-53921257 | Ch13 region 2 |
| chr13:53921320-53921360 | Ch13 region 2 |
| chr13:53921372-53921412 | Ch13 region 2 |
| chr13:53921443-53921483 | Ch13 region 2 |
| chr13:53921499-53921539 | Ch13 region 2 |
| chr13:53921561-53921601 | Ch13 region 2 |
| chr13:53921628-53921668 | Ch13 region 2 |
| chr13:53921674-53921714 | Ch13 region 2 |
| chr13:53921730-53921770 | Ch13 region 2 |
| chr13:53921796-53921836 | Ch13 region 2 |
| chr13:53921853-53921893 | Ch13 region 2 |
| chr13:53921932-53921972 | Ch13 region 2 |
| chr13:53922001-53922041 | Ch13 region 2 |
| chr13:53922069-53922109 | Ch13 region 2 |
| chr13:53922165-53922205 | Ch13 region 2 |
| chr13:53922233-53922273 | Ch13 region 2 |
| chr13:53922314-53922354 | Ch13 region 2 |
| chr13:53922362-53922402 | Ch13 region 2 |
| chr13:53922410-53922450 | Ch13 region 2 |
| chr13:53922486-53922526 | Ch13 region 2 |
| chr13:53922533-53922573 | Ch13 region 2 |
| chr13:53922611-53922651 | Ch13 region 2 |
| chr13:53922698-53922738 | Ch13 region 2 |
| chr13:53922797-53922837 | Ch13 region 2 |
| chr13:53922843-53922883 | Ch13 region 2 |
| chr13:53922890-53922930 | Ch13 region 2 |
| chr13:53922942-53922982 | Ch13 region 2 |
| chr13:53923002-53923042 | Ch13 region 2 |
| chr13:53923100-53923140 | Ch13 region 2 |
| chr13:53923145-53923185 | Ch13 region 2 |
| chr13:53923220-53923260 | Ch13 region 2 |
| chr13:53923274-53923314 | Ch13 region 2 |
| chr13:53923319-53923359 | Ch13 region 2 |
| chr13:53923415-53923455 | Ch13 region 2 |
| chr13:53923508-53923548 | Ch13 region 2 |
| chr13:53923584-53923624 | Ch13 region 2 |
| chr13:53923643-53923683 | Ch13 region 2 |
| chr13:53923701-53923741 | Ch13 region 2 |
| chr13:53923797-53923837 | Ch13 region 2 |
| chr13:53923842-53923882 | Ch13 region 2 |
| chr13:53923908-53923948 | Ch13 region 2 |
| chr13:53923953-53923993 | Ch13 region 2 |
| chr13:53924007-53924047 | Ch13 region 2 |
| chr13:53924054-53924094 | Ch13 region 2 |

| | |
|---|---|
| chr13:53924153-53924193 | Ch13 region 2 |
| chr13:53924199-53924239 | Ch13 region 2 |
| chr13:53924323-53924363 | Ch13 region 2 |
| chr13:53924368-53924408 | Ch13 region 2 |
| chr13:53924428-53924468 | Ch13 region 2 |
| chr13:53924489-53924529 | Ch13 region 2 |
| chr13:53924534-53924574 | Ch13 region 2 |
| chr13:53924589-53924629 | Ch13 region 2 |
| chr13:53924643-53924683 | Ch13 region 2 |
| chr13:53924691-53924731 | Ch13 region 2 |
| chr13:53924745-53924785 | Ch13 region 2 |
| chr13:53924801-53924841 | Ch13 region 2 |
| chr13:53924895-53924935 | Ch13 region 2 |
| chr13:53925078-53925118 | Ch13 region 2 |
| chr13:53925123-53925163 | Ch13 region 2 |
| chr13:53925183-53925223 | Ch13 region 2 |
| chr13:53925232-53925272 | Ch13 region 2 |
| chr13:53925316-53925356 | Ch13 region 2 |
| chr13:53925401-53925441 | Ch13 region 2 |
| chr13:53925470-53925510 | Ch13 region 2 |
| chr13:53925521-53925561 | Ch13 region 2 |
| chr13:53925567-53925607 | Ch13 region 2 |
| chr13:53925635-53925675 | Ch13 region 2 |
| chr13:53925689-53925729 | Ch13 region 2 |
| chr13:53925751-53925791 | Ch13 region 2 |
| chr13:53925827-53925867 | Ch13 region 2 |
| chr13:53925917-53925957 | Ch13 region 2 |
| chr13:53926070-53926110 | Ch13 region 2 |
| chr13:53926150-53926190 | Ch13 region 2 |
| chr13:53926196-53926236 | Ch13 region 2 |
| chr13:53926261-53926301 | Ch13 region 2 |
| chr13:53926331-53926371 | Ch13 region 2 |
| chr13:53926376-53926416 | Ch13 region 2 |
| chr13:53926421-53926461 | Ch13 region 2 |
| chr13:53926512-53926552 | Ch13 region 2 |
| chr13:53926574-53926614 | Ch13 region 2 |
| chr13:53926654-53926694 | Ch13 region 2 |
| chr13:53926699-53926739 | Ch13 region 2 |
| chr13:53926926-53926966 | Ch13 region 2 |
| chr13:53926979-53927019 | Ch13 region 2 |
| chr13:53927050-53927090 | Ch13 region 2 |
| chr13:53927156-53927196 | Ch13 region 2 |
| chr13:53927206-53927246 | Ch13 region 2 |
| chr13:53927271-53927311 | Ch13 region 2 |
| chr13:53927320-53927360 | Ch13 region 2 |
| chr13:53927374-53927414 | Ch13 region 2 |
| chr13:53927473-53927513 | Ch13 region 2 |
| chr13:53927519-53927559 | Ch13 region 2 |
| chr13:53927612-53927652 | Ch13 region 2 |
| chr13:53927685-53927725 | Ch13 region 2 |
| chr13:53927814-53927854 | Ch13 region 2 |
| chr13:53927900-53927940 | Ch13 region 2 |

| | |
|---|---|
| chr13:53927989-53928029 | Ch13 region 2 |
| chr13:53928065-53928105 | Ch13 region 2 |
| chr13:53928161-53928201 | Ch13 region 2 |
| chr13:53928233-53928273 | Ch13 region 2 |
| chr13:53928304-53928344 | Ch13 region 2 |
| chr13:53928371-53928411 | Ch13 region 2 |
| chr13:53928423-53928463 | Ch13 region 2 |
| chr13:53928469-53928509 | Ch13 region 2 |
| chr13:53928570-53928610 | Ch13 region 2 |
| chr13:53928654-53928694 | Ch13 region 2 |
| chr13:53928717-53928757 | Ch13 region 2 |
| chr13:53928772-53928812 | Ch13 region 2 |
| chr13:53928817-53928857 | Ch13 region 2 |
| chr13:53928914-53928954 | Ch13 region 2 |
| chr13:53928975-53929015 | Ch13 region 2 |
| chr13:53929026-53929066 | Ch13 region 2 |
| chr13:53929097-53929137 | Ch13 region 2 |
| chr13:53929158-53929198 | Ch13 region 2 |
| chr13:53929203-53929243 | Ch13 region 2 |
| chr13:53929268-53929308 | Ch13 region 2 |
| chr13:53929333-53929373 | Ch13 region 2 |
| chr13:53929413-53929453 | Ch13 region 2 |
| chr13:53929459-53929499 | Ch13 region 2 |
| chr13:53929508-53929548 | Ch13 region 2 |
| chr13:53929579-53929619 | Ch13 region 2 |
| chr13:53929634-53929674 | Ch13 region 2 |
| chr13:53929705-53929745 | Ch13 region 2 |
| chr13:53929761-53929801 | Ch13 region 2 |
| chr13:53929828-53929868 | Ch13 region 2 |
| chr13:53929901-53929941 | Ch13 region 2 |
| chr13:53929961-53930001 | Ch13 region 2 |
| chr13:53930007-53930047 | Ch13 region 2 |
| chr13:53930220-53930260 | Ch13 region 2 |
| chr13:53930332-53930372 | Ch13 region 2 |
| chr13:53930379-53930419 | Ch13 region 2 |
| chr13:53930455-53930495 | Ch13 region 2 |
| chr13:53930504-53930544 | Ch13 region 2 |
| chr13:53930603-53930643 | Ch13 region 2 |
| chr13:53930676-53930716 | Ch13 region 2 |
| chr13:53930725-53930765 | Ch13 region 2 |
| chr13:53930816-53930856 | Ch13 region 2 |
| chr13:53930874-53930914 | Ch13 region 2 |
| chr13:53930961-53931001 | Ch13 region 2 |
| chr13:53931022-53931062 | Ch13 region 2 |
| chr13:53931076-53931116 | Ch13 region 2 |
| chr13:53931121-53931161 | Ch13 region 2 |
| chr13:53931190-53931230 | Ch13 region 2 |
| chr13:53931264-53931304 | Ch13 region 2 |
| chr13:53931384-53931424 | Ch13 region 2 |
| chr13:53931467-53931507 | Ch13 region 2 |
| chr13:53931515-53931555 | Ch13 region 2 |
| chr13:53931563-53931603 | Ch13 region 2 |

| | |
|---|---|
| chr13:53931636-53931676 | Ch13 region 2 |
| chr13:53931681-53931721 | Ch13 region 2 |
| chr13:53931793-53931833 | Ch13 region 2 |
| chr13:53931889-53931929 | Ch13 region 2 |
| chr13:53931976-53932016 | Ch13 region 2 |
| chr13:53932039-53932079 | Ch13 region 2 |
| chr13:53932098-53932138 | Ch13 region 2 |
| chr13:53932186-53932226 | Ch13 region 2 |
| chr13:53932255-53932295 | Ch13 region 2 |
| chr13:53932300-53932340 | Ch13 region 2 |
| chr13:53932355-53932395 | Ch13 region 2 |
| chr13:53932400-53932440 | Ch13 region 2 |
| chr13:53932447-53932487 | Ch13 region 2 |
| chr13:53932513-53932553 | Ch13 region 2 |
| chr13:53932558-53932598 | Ch13 region 2 |
| chr13:53932604-53932644 | Ch13 region 2 |
| chr13:53932691-53932731 | Ch13 region 2 |
| chr13:53932751-53932791 | Ch13 region 2 |
| chr13:53932848-53932888 | Ch13 region 2 |
| chr13:53932924-53932964 | Ch13 region 2 |
| chr13:53933114-53933154 | Ch13 region 2 |
| chr13:53933170-53933210 | Ch13 region 2 |
| chr13:53933223-53933263 | Ch13 region 2 |
| chr13:53933279-53933319 | Ch13 region 2 |
| chr13:53933328-53933368 | Ch13 region 2 |
| chr13:53933384-53933424 | Ch13 region 2 |
| chr13:53933433-53933473 | Ch13 region 2 |
| chr13:53933560-53933600 | Ch13 region 2 |
| chr13:53933623-53933663 | Ch13 region 2 |
| chr13:53933726-53933766 | Ch13 region 2 |
| chr13:53933771-53933811 | Ch13 region 2 |
| chr13:53933877-53933917 | Ch13 region 2 |
| chr13:53933927-53933967 | Ch13 region 2 |
| chr13:53934019-53934059 | Ch13 region 2 |
| chr13:53934085-53934125 | Ch13 region 2 |
| chr13:53934135-53934175 | Ch13 region 2 |
| chr13:53934185-53934225 | Ch13 region 2 |
| chr13:53934238-53934278 | Ch13 region 2 |
| chr13:53934283-53934323 | Ch13 region 2 |
| chr13:53934339-53934379 | Ch13 region 2 |
| chr13:53934410-53934450 | Ch13 region 2 |
| chr13:53934503-53934543 | Ch13 region 2 |
| chr13:53934566-53934606 | Ch13 region 2 |
| chr13:53934622-53934662 | Ch13 region 2 |
| chr13:53934667-53934707 | Ch13 region 2 |
| chr13:53934814-53934854 | Ch13 region 2 |
| chr13:53934863-53934903 | Ch13 region 2 |
| chr13:53934950-53934990 | Ch13 region 2 |
| chr13:53935037-53935077 | Ch13 region 2 |
| chr13:53935085-53935125 | Ch13 region 2 |
| chr13:53935181-53935221 | Ch13 region 2 |
| chr13:53935240-53935280 | Ch13 region 2 |

| | |
|---|---|
| chr13:53935435-53935475 | Ch13 region 2 |
| chr13:53935502-53935542 | Ch13 region 2 |
| chr13:53935547-53935587 | Ch13 region 2 |
| chr13:53935632-53935672 | Ch13 region 2 |
| chr13:53935707-53935747 | Ch13 region 2 |
| chr13:53935770-53935810 | Ch13 region 2 |
| chr13:53935838-53935878 | Ch13 region 2 |
| chr13:53935885-53935925 | Ch13 region 2 |
| chr13:53935930-53935970 | Ch13 region 2 |
| chr13:53935975-53936015 | Ch13 region 2 |
| chr13:53936028-53936068 | Ch13 region 2 |
| chr13:53936080-53936120 | Ch13 region 2 |
| chr13:53936170-53936210 | Ch13 region 2 |
| chr13:53936246-53936286 | Ch13 region 2 |
| chr13:53936291-53936331 | Ch13 region 2 |
| chr13:53936336-53936376 | Ch13 region 2 |
| chr13:53936448-53936488 | Ch13 region 2 |
| chr13:53936496-53936536 | Ch13 region 2 |
| chr13:53936565-53936605 | Ch13 region 2 |
| chr13:53936653-53936693 | Ch13 region 2 |
| chr13:53936715-53936755 | Ch13 region 2 |
| chr13:53936777-53936817 | Ch13 region 2 |
| chr13:53936851-53936891 | Ch13 region 2 |
| chr13:53936929-53936969 | Ch13 region 2 |
| chr13:53936981-53937021 | Ch13 region 2 |
| chr13:53937057-53937097 | Ch13 region 2 |
| chr13:53937186-53937226 | Ch13 region 2 |
| chr13:53937289-53937329 | Ch13 region 2 |
| chr13:53937334-53937374 | Ch13 region 2 |
| chr13:53937379-53937419 | Ch13 region 2 |
| chr13:53937464-53937504 | Ch13 region 2 |
| chr13:53937550-53937590 | Ch13 region 2 |
| chr13:53937642-53937682 | Ch13 region 2 |
| chr13:53937721-53937761 | Ch13 region 2 |
| chr13:53937781-53937821 | Ch13 region 2 |
| chr13:53937864-53937904 | Ch13 region 2 |
| chr13:53937920-53937960 | Ch13 region 2 |
| chr13:53938006-53938046 | Ch13 region 2 |
| chr13:53938078-53938118 | Ch13 region 2 |
| chr13:53938190-53938230 | Ch13 region 2 |
| chr13:53938242-53938282 | Ch13 region 2 |
| chr13:53938392-53938432 | Ch13 region 2 |
| chr13:53938467-53938507 | Ch13 region 2 |
| chr13:53938639-53938679 | Ch13 region 2 |
| chr13:53938711-53938751 | Ch13 region 2 |
| chr13:53938787-53938827 | Ch13 region 2 |
| chr13:53938859-53938899 | Ch13 region 2 |
| chr13:53938938-53938978 | Ch13 region 2 |
| chr13:53938986-53939026 | Ch13 region 2 |
| chr13:53939040-53939080 | Ch13 region 2 |
| chr13:53939087-53939127 | Ch13 region 2 |
| chr13:53939149-53939189 | Ch13 region 2 |

| | |
|---|---|
| chr13:53939204-53939244 | Ch13 region 2 |
| chr13:53939288-53939328 | Ch13 region 2 |
| chr13:53939354-53939394 | Ch13 region 2 |
| chr13:53939400-53939440 | Ch13 region 2 |
| chr13:53939449-53939489 | Ch13 region 2 |
| chr13:53939606-53939646 | Ch13 region 2 |
| chr13:53939683-53939723 | Ch13 region 2 |
| chr13:53939732-53939772 | Ch13 region 2 |
| chr13:53939903-53939943 | Ch13 region 2 |
| chr13:53940069-53940109 | Ch13 region 2 |
| chr13:53940123-53940163 | Ch13 region 2 |
| chr13:53940196-53940236 | Ch13 region 2 |
| chr13:53940294-53940334 | Ch13 region 2 |
| chr13:53940379-53940419 | Ch13 region 2 |
| chr13:53940467-53940507 | Ch13 region 2 |
| chr13:53940539-53940579 | Ch13 region 2 |
| chr13:53940626-53940666 | Ch13 region 2 |
| chr13:53940683-53940723 | Ch13 region 2 |
| chr13:53940740-53940780 | Ch13 region 2 |
| chr13:53940818-53940858 | Ch13 region 2 |
| chr13:53940907-53940947 | Ch13 region 2 |
| chr13:53940959-53940999 | Ch13 region 2 |
| chr13:53941013-53941053 | Ch13 region 2 |
| chr13:53941100-53941140 | Ch13 region 2 |
| chr13:53941146-53941186 | Ch13 region 2 |
| chr13:53941191-53941231 | Ch13 region 2 |
| chr13:53941313-53941353 | Ch13 region 2 |
| chr13:53941402-53941442 | Ch13 region 2 |
| chr13:53941447-53941487 | Ch13 region 2 |
| chr13:53941539-53941579 | Ch13 region 2 |
| chr13:53941584-53941624 | Ch13 region 2 |
| chr13:53941643-53941683 | Ch13 region 2 |
| chr13:53941703-53941743 | Ch13 region 2 |
| chr13:53941754-53941794 | Ch13 region 2 |
| chr13:53941851-53941891 | Ch13 region 2 |
| chr13:53941901-53941941 | Ch13 region 2 |
| chr13:53942063-53942103 | Ch13 region 2 |
| chr13:53942151-53942191 | Ch13 region 2 |
| chr13:53942313-53942353 | Ch13 region 2 |
| chr13:53942395-53942435 | Ch13 region 2 |
| chr13:53942498-53942538 | Ch13 region 2 |
| chr13:53942612-53942652 | Ch13 region 2 |
| chr13:53942659-53942699 | Ch13 region 2 |
| chr13:53942773-53942813 | Ch13 region 2 |
| chr13:53942831-53942871 | Ch13 region 2 |
| chr13:53942891-53942931 | Ch13 region 2 |
| chr13:53942957-53942997 | Ch13 region 2 |
| chr13:53943084-53943124 | Ch13 region 2 |
| chr13:53943159-53943199 | Ch13 region 2 |
| chr13:53943212-53943252 | Ch13 region 2 |
| chr13:53943278-53943318 | Ch13 region 2 |
| chr13:53943362-53943402 | Ch13 region 2 |

| | |
|---|---|
| chr13:53943442-53943482 | Ch13 region 2 |
| chr13:53943499-53943539 | Ch13 region 2 |
| chr13:53943556-53943596 | Ch13 region 2 |
| chr13:53943624-53943664 | Ch13 region 2 |
| chr13:53943669-53943709 | Ch13 region 2 |
| chr13:53943776-53943816 | Ch13 region 2 |
| chr13:53943857-53943897 | Ch13 region 2 |
| chr13:53943913-53943953 | Ch13 region 2 |
| chr13:53943960-53944000 | Ch13 region 2 |
| chr13:53944005-53944045 | Ch13 region 2 |
| chr13:53944065-53944105 | Ch13 region 2 |
| chr13:53944111-53944151 | Ch13 region 2 |
| chr13:53944162-53944202 | Ch13 region 2 |
| chr13:53944236-53944276 | Ch13 region 2 |
| chr13:53944297-53944337 | Ch13 region 2 |
| chr13:53944342-53944382 | Ch13 region 2 |
| chr13:53944430-53944470 | Ch13 region 2 |
| chr13:53944487-53944527 | Ch13 region 2 |
| chr13:53944564-53944604 | Ch13 region 2 |
| chr13:53944627-53944667 | Ch13 region 2 |
| chr13:53944682-53944722 | Ch13 region 2 |
| chr13:53944779-53944819 | Ch13 region 2 |
| chr13:53944845-53944885 | Ch13 region 2 |
| chr13:53944906-53944946 | Ch13 region 2 |
| chr13:53944972-53945012 | Ch13 region 2 |
| chr13:53945020-53945060 | Ch13 region 2 |
| chr13:53945074-53945114 | Ch13 region 2 |
| chr13:53945146-53945186 | Ch13 region 2 |
| chr13:53945197-53945237 | Ch13 region 2 |
| chr13:53945251-53945291 | Ch13 region 2 |
| chr13:53945301-53945341 | Ch13 region 2 |
| chr13:53945346-53945386 | Ch13 region 2 |
| chr13:53945416-53945456 | Ch13 region 2 |
| chr13:53945543-53945583 | Ch13 region 2 |
| chr13:53945601-53945641 | Ch13 region 2 |
| chr13:53945654-53945694 | Ch13 region 2 |
| chr13:53945722-53945762 | Ch13 region 2 |
| chr13:53945818-53945858 | Ch13 region 2 |
| chr13:53945863-53945903 | Ch13 region 2 |
| chr13:53945995-53946035 | Ch13 region 2 |
| chr13:53946073-53946113 | Ch13 region 2 |
| chr13:53946136-53946176 | Ch13 region 2 |
| chr13:53946192-53946232 | Ch13 region 2 |
| chr13:53946274-53946314 | Ch13 region 2 |
| chr13:53946383-53946423 | Ch13 region 2 |
| chr13:53946545-53946585 | Ch13 region 2 |
| chr13:53946675-53946715 | Ch13 region 2 |
| chr13:53946848-53946888 | Ch13 region 2 |
| chr13:53947098-53947138 | Ch13 region 2 |
| chr13:53947165-53947205 | Ch13 region 2 |
| chr13:53947230-53947270 | Ch13 region 2 |
| chr13:53947339-53947379 | Ch13 region 2 |

| | |
|---|---|
| chr13:53947478-53947518 | Ch13 region 2 |
| chr13:53947550-53947590 | Ch13 region 2 |
| chr13:53947596-53947636 | Ch13 region 2 |
| chr13:53947647-53947687 | Ch13 region 2 |
| chr13:53947708-53947748 | Ch13 region 2 |
| chr13:53947768-53947808 | Ch13 region 2 |
| chr13:53947849-53947889 | Ch13 region 2 |
| chr13:53947913-53947953 | Ch13 region 2 |
| chr13:53947984-53948024 | Ch13 region 2 |
| chr13:53948046-53948086 | Ch13 region 2 |
| chr13:53948112-53948152 | Ch13 region 2 |
| chr13:53948174-53948214 | Ch13 region 2 |
| chr13:53948244-53948284 | Ch13 region 2 |
| chr13:53948389-53948429 | Ch13 region 2 |
| chr13:53948482-53948522 | Ch13 region 2 |
| chr13:53948529-53948569 | Ch13 region 2 |
| chr13:53948577-53948617 | Ch13 region 2 |
| chr13:53948627-53948667 | Ch13 region 2 |
| chr13:53948688-53948728 | Ch13 region 2 |
| chr13:53948739-53948779 | Ch13 region 2 |
| chr13:53948796-53948836 | Ch13 region 2 |
| chr13:53948899-53948939 | Ch13 region 2 |
| chr13:53948969-53949009 | Ch13 region 2 |
| chr13:53949035-53949075 | Ch13 region 2 |
| chr13:53949121-53949161 | Ch13 region 2 |
| chr13:53949192-53949232 | Ch13 region 2 |
| chr13:53949282-53949322 | Ch13 region 2 |
| chr13:53949357-53949397 | Ch13 region 2 |
| chr13:53949478-53949518 | Ch13 region 2 |
| chr13:53949525-53949565 | Ch13 region 2 |
| chr13:53949609-53949649 | Ch13 region 2 |
| chr13:53949656-53949696 | Ch13 region 2 |
| chr13:53949735-53949775 | Ch13 region 2 |
| chr13:53949807-53949847 | Ch13 region 2 |
| chr13:53949945-53949985 | Ch13 region 2 |
| chr13:53950157-53950197 | Ch13 region 2 |
| chr13:53950250-53950290 | Ch13 region 2 |
| chr13:53950301-53950341 | Ch13 region 2 |
| chr13:53950362-53950402 | Ch13 region 2 |
| chr13:53950425-53950465 | Ch13 region 2 |
| chr13:53950521-53950561 | Ch13 region 2 |
| chr13:53950578-53950618 | Ch13 region 2 |
| chr13:53950638-53950678 | Ch13 region 2 |
| chr13:53950686-53950726 | Ch13 region 2 |
| chr13:53950759-53950799 | Ch13 region 2 |
| chr13:53950831-53950871 | Ch13 region 2 |
| chr13:53950877-53950917 | Ch13 region 2 |
| chr13:53950927-53950967 | Ch13 region 2 |
| chr13:53950984-53951024 | Ch13 region 2 |
| chr13:53951056-53951096 | Ch13 region 2 |
| chr13:53951103-53951143 | Ch13 region 2 |
| chr13:53951162-53951202 | Ch13 region 2 |

| | |
|---|---|
| chr13:53951252-53951292 | Ch13 region 2 |
| chr13:53951352-53951392 | Ch13 region 2 |
| chr13:53951401-53951441 | Ch13 region 2 |
| chr13:53951511-53951551 | Ch13 region 2 |
| chr13:53951571-53951611 | Ch13 region 2 |
| chr13:53951638-53951678 | Ch13 region 2 |
| chr13:53951737-53951777 | Ch13 region 2 |
| chr13:53951801-53951841 | Ch13 region 2 |
| chr13:53951859-53951899 | Ch13 region 2 |
| chr13:53951957-53951997 | Ch13 region 2 |
| chr13:53952033-53952073 | Ch13 region 2 |
| chr13:53952082-53952122 | Ch13 region 2 |
| chr13:53952215-53952255 | Ch13 region 2 |
| chr13:53952325-53952365 | Ch13 region 2 |
| chr13:53952542-53952582 | Ch13 region 2 |
| chr13:53952624-53952664 | Ch13 region 2 |
| chr13:53952728-53952768 | Ch13 region 2 |
| chr13:53952812-53952852 | Ch13 region 2 |
| chr13:53952864-53952904 | Ch13 region 2 |
| chr13:53952909-53952949 | Ch13 region 2 |
| chr13:53952989-53953029 | Ch13 region 2 |
| chr13:53953157-53953197 | Ch13 region 2 |
| chr13:53953206-53953246 | Ch13 region 2 |
| chr13:53953431-53953471 | Ch13 region 2 |
| chr13:53953514-53953554 | Ch13 region 2 |
| chr13:53953590-53953630 | Ch13 region 2 |
| chr13:53953662-53953702 | Ch13 region 2 |
| chr13:53953721-53953761 | Ch13 region 2 |
| chr13:53953778-53953818 | Ch13 region 2 |
| chr13:53953831-53953871 | Ch13 region 2 |
| chr13:53953935-53953975 | Ch13 region 2 |
| chr13:53954007-53954047 | Ch13 region 2 |
| chr13:53954109-53954149 | Ch13 region 2 |
| chr13:53954167-53954207 | Ch13 region 2 |
| chr13:53954326-53954366 | Ch13 region 2 |
| chr13:53954424-53954464 | Ch13 region 2 |
| chr13:53954486-53954526 | Ch13 region 2 |
| chr13:53954567-53954607 | Ch13 region 2 |
| chr13:53954665-53954705 | Ch13 region 2 |
| chr13:53954741-53954781 | Ch13 region 2 |
| chr13:53954793-53954833 | Ch13 region 2 |
| chr13:53954841-53954881 | Ch13 region 2 |
| chr13:53954910-53954950 | Ch13 region 2 |
| chr13:53954967-53955007 | Ch13 region 2 |
| chr13:53955054-53955094 | Ch13 region 2 |
| chr13:53955128-53955168 | Ch13 region 2 |
| chr13:53955237-53955277 | Ch13 region 2 |
| chr13:53955282-53955322 | Ch13 region 2 |
| chr13:53955353-53955393 | Ch13 region 2 |
| chr13:53955403-53955443 | Ch13 region 2 |
| chr13:53955502-53955542 | Ch13 region 2 |
| chr13:53955664-53955704 | Ch13 region 2 |

| | |
|---|---|
| chr13:53955803-53955843 | Ch13 region 2 |
| chr13:53956007-53956047 | Ch13 region 2 |
| chr13:53956077-53956117 | Ch13 region 2 |
| chr13:53956130-53956170 | Ch13 region 2 |
| chr13:53956182-53956222 | Ch13 region 2 |
| chr13:53956232-53956272 | Ch13 region 2 |
| chr13:53956279-53956319 | Ch13 region 2 |
| chr13:53956351-53956391 | Ch13 region 2 |
| chr13:53956416-53956456 | Ch13 region 2 |
| chr13:53956479-53956519 | Ch13 region 2 |
| chr13:53956532-53956572 | Ch13 region 2 |
| chr13:53956581-53956621 | Ch13 region 2 |
| chr13:53956678-53956718 | Ch13 region 2 |
| chr13:53956766-53956806 | Ch13 region 2 |
| chr13:53956873-53956913 | Ch13 region 2 |
| chr13:53956922-53956962 | Ch13 region 2 |
| chr13:53956985-53957025 | Ch13 region 2 |
| chr13:53957049-53957089 | Ch13 region 2 |
| chr13:53957127-53957167 | Ch13 region 2 |
| chr13:53957178-53957218 | Ch13 region 2 |
| chr13:53957232-53957272 | Ch13 region 2 |
| chr13:53957612-53957652 | Ch13 region 2 |
| chr13:53957670-53957710 | Ch13 region 2 |
| chr13:53957747-53957787 | Ch13 region 2 |
| chr13:53957869-53957909 | Ch13 region 2 |
| chr13:53957952-53957992 | Ch13 region 2 |
| chr13:53958139-53958179 | Ch13 region 2 |
| chr13:53958214-53958254 | Ch13 region 2 |
| chr13:53958262-53958302 | Ch13 region 2 |
| chr13:53958311-53958351 | Ch13 region 2 |
| chr13:53958371-53958411 | Ch13 region 2 |
| chr13:53958444-53958484 | Ch13 region 2 |
| chr13:53958535-53958575 | Ch13 region 2 |
| chr13:53958581-53958621 | Ch13 region 2 |
| chr13:53958639-53958679 | Ch13 region 2 |
| chr13:53958712-53958752 | Ch13 region 2 |
| chr13:53958767-53958807 | Ch13 region 2 |
| chr13:53958994-53959034 | Ch13 region 2 |
| chr13:53959067-53959107 | Ch13 region 2 |
| chr13:53959117-53959157 | Ch13 region 2 |
| chr13:53959256-53959296 | Ch13 region 2 |
| chr13:53959354-53959394 | Ch13 region 2 |
| chr13:53959454-53959494 | Ch13 region 2 |
| chr13:53959506-53959546 | Ch13 region 2 |
| chr13:53959561-53959601 | Ch13 region 2 |
| chr13:53959634-53959674 | Ch13 region 2 |
| chr13:53959697-53959737 | Ch13 region 2 |
| chr13:53959754-53959794 | Ch13 region 2 |
| chr13:53959824-53959864 | Ch13 region 2 |
| chr13:53959912-53959952 | Ch13 region 2 |
| chr13:53959958-53959998 | Ch13 region 2 |
| chr13:53960004-53960044 | Ch13 region 2 |

| | |
|---|---|
| chr13:53960059-53960099 | Ch13 region 2 |
| chr13:53960114-53960154 | Ch13 region 2 |
| chr13:53960163-53960203 | Ch13 region 2 |
| chr13:53960232-53960272 | Ch13 region 2 |
| chr13:53960300-53960340 | Ch13 region 2 |
| chr13:53960360-53960400 | Ch13 region 2 |
| chr13:53960429-53960469 | Ch13 region 2 |
| chr13:53960545-53960585 | Ch13 region 2 |
| chr13:53960593-53960633 | Ch13 region 2 |
| chr13:53960682-53960722 | Ch13 region 2 |
| chr13:53960741-53960781 | Ch13 region 2 |
| chr13:53960824-53960864 | Ch13 region 2 |
| chr13:53960872-53960912 | Ch13 region 2 |
| chr13:53960928-53960968 | Ch13 region 2 |
| chr13:53961013-53961053 | Ch13 region 2 |
| chr13:53961118-53961158 | Ch13 region 2 |
| chr13:53961203-53961243 | Ch13 region 2 |
| chr13:53961248-53961288 | Ch13 region 2 |
| chr13:53961326-53961366 | Ch13 region 2 |
| chr13:53961479-53961519 | Ch13 region 2 |
| chr13:53961537-53961577 | Ch13 region 2 |
| chr13:53961601-53961641 | Ch13 region 2 |
| chr13:53961663-53961703 | Ch13 region 2 |
| chr13:53961708-53961748 | Ch13 region 2 |
| chr13:53961819-53961859 | Ch13 region 2 |
| chr13:53961880-53961920 | Ch13 region 2 |
| chr13:53961956-53961996 | Ch13 region 2 |
| chr13:53962001-53962041 | Ch13 region 2 |
| chr13:53962069-53962109 | Ch13 region 2 |
| chr13:53962136-53962176 | Ch13 region 2 |
| chr13:53962242-53962282 | Ch13 region 2 |
| chr13:53962304-53962344 | Ch13 region 2 |
| chr13:53962378-53962418 | Ch13 region 2 |
| chr13:53962468-53962508 | Ch13 region 2 |
| chr13:53962543-53962583 | Ch13 region 2 |
| chr13:53962605-53962645 | Ch13 region 2 |
| chr13:53962651-53962691 | Ch13 region 2 |
| chr13:53962706-53962746 | Ch13 region 2 |
| chr13:53962768-53962808 | Ch13 region 2 |
| chr13:53962865-53962905 | Ch13 region 2 |
| chr13:53962912-53962952 | Ch13 region 2 |
| chr13:53962983-53963023 | Ch13 region 2 |
| chr13:53963038-53963078 | Ch13 region 2 |
| chr13:53963170-53963210 | Ch13 region 2 |
| chr13:53963241-53963281 | Ch13 region 2 |
| chr13:53963295-53963335 | Ch13 region 2 |
| chr13:53963351-53963391 | Ch13 region 2 |
| chr13:53963417-53963457 | Ch13 region 2 |
| chr13:53963487-53963527 | Ch13 region 2 |
| chr13:53963598-53963638 | Ch13 region 2 |
| chr13:53963700-53963740 | Ch13 region 2 |
| chr13:53963889-53963929 | Ch13 region 2 |

| | |
|---|---|
| chr13:53963934-53963974 | Ch13 region 2 |
| chr13:53963979-53964019 | Ch13 region 2 |
| chr13:53964036-53964076 | Ch13 region 2 |
| chr13:53964090-53964130 | Ch13 region 2 |
| chr13:53964144-53964184 | Ch13 region 2 |
| chr13:53964255-53964295 | Ch13 region 2 |
| chr13:53964331-53964371 | Ch13 region 2 |
| chr13:53964399-53964439 | Ch13 region 2 |
| chr13:53964468-53964508 | Ch13 region 2 |
| chr13:53964524-53964564 | Ch13 region 2 |
| chr13:53964586-53964626 | Ch13 region 2 |
| chr13:53964645-53964685 | Ch13 region 2 |
| chr13:53964704-53964744 | Ch13 region 2 |
| chr13:53964819-53964859 | Ch13 region 2 |
| chr13:53964867-53964907 | Ch13 region 2 |
| chr13:53964974-53965014 | Ch13 region 2 |
| chr13:53965112-53965152 | Ch13 region 2 |
| chr13:53965213-53965253 | Ch13 region 2 |
| chr13:53965369-53965409 | Ch13 region 2 |
| chr13:53965472-53965512 | Ch13 region 2 |
| chr13:53965527-53965567 | Ch13 region 2 |
| chr13:53965637-53965677 | Ch13 region 2 |
| chr13:53965826-53965866 | Ch13 region 2 |
| chr13:53965959-53965999 | Ch13 region 2 |
| chr13:53966048-53966088 | Ch13 region 2 |
| chr13:53966172-53966212 | Ch13 region 2 |
| chr13:53966303-53966343 | Ch13 region 2 |
| chr13:53966476-53966516 | Ch13 region 2 |
| chr13:53966625-53966665 | Ch13 region 2 |
| chr13:53966717-53966757 | Ch13 region 2 |
| chr13:53966784-53966824 | Ch13 region 2 |
| chr13:53966866-53966906 | Ch13 region 2 |
| chr13:53967002-53967042 | Ch13 region 2 |
| chr13:53967114-53967154 | Ch13 region 2 |
| chr13:53967183-53967223 | Ch13 region 2 |
| chr13:53967385-53967425 | Ch13 region 2 |
| chr13:53967549-53967589 | Ch13 region 2 |
| chr13:53967633-53967673 | Ch13 region 2 |
| chr13:53967694-53967734 | Ch13 region 2 |
| chr13:53967776-53967816 | Ch13 region 2 |
| chr13:53967937-53967977 | Ch13 region 2 |
| chr13:53968018-53968058 | Ch13 region 2 |
| chr13:53968069-53968109 | Ch13 region 2 |
| chr13:53968126-53968166 | Ch13 region 2 |
| chr13:53968171-53968211 | Ch13 region 2 |
| chr13:53968228-53968268 | Ch13 region 2 |
| chr13:53968338-53968378 | Ch13 region 2 |
| chr13:53968483-53968523 | Ch13 region 2 |
| chr13:53968550-53968590 | Ch13 region 2 |
| chr13:53968677-53968717 | Ch13 region 2 |
| chr13:53968801-53968841 | Ch13 region 2 |
| chr13:53968863-53968903 | Ch13 region 2 |

| | |
|---|---|
| chr13:53968965-53969005 | Ch13 region 2 |
| chr13:53969043-53969083 | Ch13 region 2 |
| chr13:53969131-53969171 | Ch13 region 2 |
| chr13:53969218-53969258 | Ch13 region 2 |
| chr13:53969315-53969355 | Ch13 region 2 |
| chr13:53969405-53969445 | Ch13 region 2 |
| chr13:53969467-53969507 | Ch13 region 2 |
| chr13:53969564-53969604 | Ch13 region 2 |
| chr13:53969633-53969673 | Ch13 region 2 |
| chr13:53969695-53969735 | Ch13 region 2 |
| chr13:53969757-53969797 | Ch13 region 2 |
| chr13:53969821-53969861 | Ch13 region 2 |
| chr13:53969883-53969923 | Ch13 region 2 |
| chr13:53969931-53969971 | Ch13 region 2 |
| chr13:53969993-53970033 | Ch13 region 2 |
| chr13:53970117-53970157 | Ch13 region 2 |
| chr13:53970175-53970215 | Ch13 region 2 |
| chr13:53970272-53970312 | Ch13 region 2 |
| chr13:53970322-53970362 | Ch13 region 2 |
| chr13:53970383-53970423 | Ch13 region 2 |
| chr13:53970449-53970489 | Ch13 region 2 |
| chr13:53970522-53970562 | Ch13 region 2 |
| chr13:53970594-53970634 | Ch13 region 2 |
| chr13:53970648-53970688 | Ch13 region 2 |
| chr13:53970737-53970777 | Ch13 region 2 |
| chr13:53970782-53970822 | Ch13 region 2 |
| chr13:53970866-53970906 | Ch13 region 2 |
| chr13:53970914-53970954 | Ch13 region 2 |
| chr13:53971012-53971052 | Ch13 region 2 |
| chr13:53971260-53971300 | Ch13 region 2 |
| chr13:53971406-53971446 | Ch13 region 2 |
| chr13:53971460-53971500 | Ch13 region 2 |
| chr13:53971509-53971549 | Ch13 region 2 |
| chr13:53971557-53971597 | Ch13 region 2 |
| chr13:53971646-53971686 | Ch13 region 2 |
| chr13:53971753-53971793 | Ch13 region 2 |
| chr13:53971807-53971847 | Ch13 region 2 |
| chr13:53971870-53971910 | Ch13 region 2 |
| chr13:53971931-53971971 | Ch13 region 2 |
| chr13:53971981-53972021 | Ch13 region 2 |
| chr13:53972040-53972080 | Ch13 region 2 |
| chr13:53972133-53972173 | Ch13 region 2 |
| chr13:53972197-53972237 | Ch13 region 2 |
| chr13:53972243-53972283 | Ch13 region 2 |
| chr13:53972304-53972344 | Ch13 region 2 |
| chr13:53972387-53972427 | Ch13 region 2 |
| chr13:53972451-53972491 | Ch13 region 2 |
| chr13:53972496-53972536 | Ch13 region 2 |
| chr13:53972565-53972605 | Ch13 region 2 |
| chr13:53972640-53972680 | Ch13 region 2 |
| chr13:53972688-53972728 | Ch13 region 2 |
| chr13:53972760-53972800 | Ch13 region 2 |

| | |
|---|---|
| chr13:53972808-53972848 | Ch13 region 2 |
| chr13:53972914-53972954 | Ch13 region 2 |
| chr13:53972959-53972999 | Ch13 region 2 |
| chr13:53973017-53973057 | Ch13 region 2 |
| chr13:53973066-53973106 | Ch13 region 2 |
| chr13:53973124-53973164 | Ch13 region 2 |
| chr13:53973185-53973225 | Ch13 region 2 |
| chr13:53973268-53973308 | Ch13 region 2 |
| chr13:53973362-53973402 | Ch13 region 2 |
| chr13:53973456-53973496 | Ch13 region 2 |
| chr13:53973504-53973544 | Ch13 region 2 |
| chr13:53973601-53973641 | Ch13 region 2 |
| chr13:53973652-53973692 | Ch13 region 2 |
| chr13:53973702-53973742 | Ch13 region 2 |
| chr13:53973747-53973787 | Ch13 region 2 |
| chr13:53973813-53973853 | Ch13 region 2 |
| chr13:53973882-53973922 | Ch13 region 2 |
| chr13:53973955-53973995 | Ch13 region 2 |
| chr13:53974025-53974065 | Ch13 region 2 |
| chr13:53974115-53974155 | Ch13 region 2 |
| chr13:53974229-53974269 | Ch13 region 2 |
| chr13:53974295-53974335 | Ch13 region 2 |
| chr13:53974369-53974409 | Ch13 region 2 |
| chr13:53974422-53974462 | Ch13 region 2 |
| chr13:53974568-53974608 | Ch13 region 2 |
| chr13:53974629-53974669 | Ch13 region 2 |
| chr13:53974677-53974717 | Ch13 region 2 |
| chr13:53974756-53974796 | Ch13 region 2 |
| chr13:53974824-53974864 | Ch13 region 2 |
| chr13:53974895-53974935 | Ch13 region 2 |
| chr13:53974956-53974996 | Ch13 region 2 |
| chr13:53975001-53975041 | Ch13 region 2 |
| chr13:53975071-53975111 | Ch13 region 2 |
| chr13:53975124-53975164 | Ch13 region 2 |
| chr13:53975193-53975233 | Ch13 region 2 |
| chr13:53975252-53975292 | Ch13 region 2 |
| chr13:53975364-53975404 | Ch13 region 2 |
| chr13:53975427-53975467 | Ch13 region 2 |
| chr13:53975556-53975596 | Ch13 region 2 |
| chr13:53975625-53975665 | Ch13 region 2 |
| chr13:53975680-53975720 | Ch13 region 2 |
| chr13:53975776-53975816 | Ch13 region 2 |
| chr13:53975821-53975861 | Ch13 region 2 |
| chr13:53975885-53975925 | Ch13 region 2 |
| chr13:53975945-53975985 | Ch13 region 2 |
| chr13:53976006-53976046 | Ch13 region 2 |
| chr13:53976051-53976091 | Ch13 region 2 |
| chr13:53976127-53976167 | Ch13 region 2 |
| chr13:53976187-53976227 | Ch13 region 2 |
| chr13:53976232-53976272 | Ch13 region 2 |
| chr13:53976291-53976331 | Ch13 region 2 |
| chr13:53976349-53976389 | Ch13 region 2 |

| | |
|---|---|
| chr13:53976499-53976539 | Ch13 region 2 |
| chr13:53976617-53976657 | Ch13 region 2 |
| chr13:53976670-53976710 | Ch13 region 2 |
| chr13:53976749-53976789 | Ch13 region 2 |
| chr13:53976872-53976912 | Ch13 region 2 |
| chr13:53976948-53976988 | Ch13 region 2 |
| chr13:53977018-53977058 | Ch13 region 2 |
| chr13:53977063-53977103 | Ch13 region 2 |
| chr13:53977144-53977184 | Ch13 region 2 |
| chr13:53977212-53977252 | Ch13 region 2 |
| chr13:53977300-53977340 | Ch13 region 2 |
| chr13:53977352-53977392 | Ch13 region 2 |
| chr13:53977421-53977461 | Ch13 region 2 |
| chr13:53977669-53977709 | Ch13 region 2 |
| chr13:53977717-53977757 | Ch13 region 2 |
| chr13:53977788-53977828 | Ch13 region 2 |
| chr13:53977894-53977934 | Ch13 region 2 |
| chr13:53977952-53977992 | Ch13 region 2 |
| chr13:53978037-53978077 | Ch13 region 2 |
| chr13:53978106-53978146 | Ch13 region 2 |
| chr13:53978183-53978223 | Ch13 region 2 |
| chr13:53978255-53978295 | Ch13 region 2 |
| chr13:53978300-53978340 | Ch13 region 2 |
| chr13:53978345-53978385 | Ch13 region 2 |
| chr13:53978410-53978450 | Ch13 region 2 |
| chr13:53978477-53978517 | Ch13 region 2 |
| chr13:53978531-53978571 | Ch13 region 2 |
| chr13:53978602-53978642 | Ch13 region 2 |
| chr13:53978650-53978690 | Ch13 region 2 |
| chr13:53978708-53978748 | Ch13 region 2 |
| chr13:53978782-53978822 | Ch13 region 2 |
| chr13:53978871-53978911 | Ch13 region 2 |
| chr13:53978931-53978971 | Ch13 region 2 |
| chr13:53979023-53979063 | Ch13 region 2 |
| chr13:53979073-53979113 | Ch13 region 2 |
| chr13:53979119-53979159 | Ch13 region 2 |
| chr13:53979209-53979249 | Ch13 region 2 |
| chr13:53979277-53979317 | Ch13 region 2 |
| chr13:53979338-53979378 | Ch13 region 2 |
| chr13:53979428-53979468 | Ch13 region 2 |
| chr13:53979532-53979572 | Ch13 region 2 |
| chr13:53979589-53979629 | Ch13 region 2 |
| chr13:53979676-53979716 | Ch13 region 2 |
| chr13:53979772-53979812 | Ch13 region 2 |
| chr13:53979829-53979869 | Ch13 region 2 |
| chr13:53979896-53979936 | Ch13 region 2 |
| chr13:53979951-53979991 | Ch13 region 2 |
| chr13:53980008-53980048 | Ch13 region 2 |
| chr13:53980105-53980145 | Ch13 region 2 |
| chr13:53980170-53980210 | Ch13 region 2 |
| chr13:53980304-53980344 | Ch13 region 2 |
| chr13:53980386-53980426 | Ch13 region 2 |

| | |
|---|---|
| chr13:53980452-53980492 | Ch13 region 2 |
| chr13:53980497-53980537 | Ch13 region 2 |
| chr13:53980559-53980599 | Ch13 region 2 |
| chr13:53980636-53980676 | Ch13 region 2 |
| chr13:53980701-53980741 | Ch13 region 2 |
| chr13:53980752-53980792 | Ch13 region 2 |
| chr13:53980823-53980863 | Ch13 region 2 |
| chr13:53980868-53980908 | Ch13 region 2 |
| chr13:53980919-53980959 | Ch13 region 2 |
| chr13:53981026-53981066 | Ch13 region 2 |
| chr13:53981231-53981271 | Ch13 region 2 |
| chr13:53981276-53981316 | Ch13 region 2 |
| chr13:53981383-53981423 | Ch13 region 2 |
| chr13:53981453-53981493 | Ch13 region 2 |
| chr13:53981541-53981581 | Ch13 region 2 |
| chr13:53981626-53981666 | Ch13 region 2 |
| chr13:53981681-53981721 | Ch13 region 2 |
| chr13:53981802-53981842 | Ch13 region 2 |
| chr13:53981959-53981999 | Ch13 region 2 |
| chr13:53982011-53982051 | Ch13 region 2 |
| chr13:53982104-53982144 | Ch13 region 2 |
| chr13:53982325-53982365 | Ch13 region 2 |
| chr13:53982386-53982426 | Ch13 region 2 |
| chr13:53982445-53982485 | Ch13 region 2 |
| chr13:53982562-53982602 | Ch13 region 2 |
| chr13:53982615-53982655 | Ch13 region 2 |
| chr13:53982672-53982712 | Ch13 region 2 |
| chr13:53982717-53982757 | Ch13 region 2 |
| chr13:53982779-53982819 | Ch13 region 2 |
| chr13:53982834-53982874 | Ch13 region 2 |
| chr13:53982881-53982921 | Ch13 region 2 |
| chr13:53982942-53982982 | Ch13 region 2 |
| chr13:53983017-53983057 | Ch13 region 2 |
| chr13:53983073-53983113 | Ch13 region 2 |
| chr13:53983182-53983222 | Ch13 region 2 |
| chr13:53983233-53983273 | Ch13 region 2 |
| chr13:53983286-53983326 | Ch13 region 2 |
| chr13:53983389-53983429 | Ch13 region 2 |
| chr13:53983442-53983482 | Ch13 region 2 |
| chr13:53983511-53983551 | Ch13 region 2 |
| chr13:53983575-53983615 | Ch13 region 2 |
| chr13:53983633-53983673 | Ch13 region 2 |
| chr13:53983713-53983753 | Ch13 region 2 |
| chr13:53983763-53983803 | Ch13 region 2 |
| chr13:53983837-53983877 | Ch13 region 2 |
| chr13:53983887-53983927 | Ch13 region 2 |
| chr13:53983954-53983994 | Ch13 region 2 |
| chr13:53984033-53984073 | Ch13 region 2 |
| chr13:53984112-53984152 | Ch13 region 2 |
| chr13:53984181-53984221 | Ch13 region 2 |
| chr13:53984226-53984266 | Ch13 region 2 |
| chr13:53984283-53984323 | Ch13 region 2 |

| | |
|---|---|
| chr13:53984366-53984406 | Ch13 region 2 |
| chr13:53984469-53984509 | Ch13 region 2 |
| chr13:53984622-53984662 | Ch13 region 2 |
| chr13:53984667-53984707 | Ch13 region 2 |
| chr13:53984745-53984785 | Ch13 region 2 |
| chr13:53984808-53984848 | Ch13 region 2 |
| chr13:53984901-53984941 | Ch13 region 2 |
| chr13:53985001-53985041 | Ch13 region 2 |
| chr13:53985055-53985095 | Ch13 region 2 |
| chr13:53985107-53985147 | Ch13 region 2 |
| chr13:53985160-53985200 | Ch13 region 2 |
| chr13:53985236-53985276 | Ch13 region 2 |
| chr13:53985315-53985355 | Ch13 region 2 |
| chr13:53985368-53985408 | Ch13 region 2 |
| chr13:53985414-53985454 | Ch13 region 2 |
| chr13:53985477-53985517 | Ch13 region 2 |
| chr13:53985578-53985618 | Ch13 region 2 |
| chr13:53985670-53985710 | Ch13 region 2 |
| chr13:53985782-53985822 | Ch13 region 2 |
| chr13:53985846-53985886 | Ch13 region 2 |
| chr13:53985896-53985936 | Ch13 region 2 |
| chr13:53985993-53986033 | Ch13 region 2 |
| chr13:53986044-53986084 | Ch13 region 2 |
| chr13:53986094-53986134 | Ch13 region 2 |
| chr13:53986152-53986192 | Ch13 region 2 |
| chr13:53986215-53986255 | Ch13 region 2 |
| chr13:53986273-53986313 | Ch13 region 2 |
| chr13:53986351-53986391 | Ch13 region 2 |
| chr13:53986435-53986475 | Ch13 region 2 |
| chr13:53986511-53986551 | Ch13 region 2 |
| chr13:53986556-53986596 | Ch13 region 2 |
| chr13:53986631-53986671 | Ch13 region 2 |
| chr13:53986676-53986716 | Ch13 region 2 |
| chr13:53986721-53986761 | Ch13 region 2 |
| chr13:53986800-53986840 | Ch13 region 2 |
| chr13:53986889-53986929 | Ch13 region 2 |
| chr13:53986955-53986995 | Ch13 region 2 |
| chr13:53987000-53987040 | Ch13 region 2 |
| chr13:53987063-53987103 | Ch13 region 2 |
| chr13:53987118-53987158 | Ch13 region 2 |
| chr13:53987165-53987205 | Ch13 region 2 |
| chr13:53987216-53987256 | Ch13 region 2 |
| chr13:53987266-53987306 | Ch13 region 2 |
| chr13:53987344-53987384 | Ch13 region 2 |
| chr13:53987410-53987450 | Ch13 region 2 |
| chr13:53987498-53987538 | Ch13 region 2 |
| chr13:53987582-53987622 | Ch13 region 2 |
| chr13:53987700-53987740 | Ch13 region 2 |
| chr13:53987763-53987803 | Ch13 region 2 |
| chr13:53987859-53987899 | Ch13 region 2 |
| chr13:53987904-53987944 | Ch13 region 2 |
| chr13:53987953-53987993 | Ch13 region 2 |

| | |
|---|---|
| chr13:53988009-53988049 | Ch13 region 2 |
| chr13:53988074-53988114 | Ch13 region 2 |
| chr13:53988144-53988184 | Ch13 region 2 |
| chr13:53988281-53988321 | Ch13 region 2 |
| chr13:53988466-53988506 | Ch13 region 2 |
| chr13:53988544-53988584 | Ch13 region 2 |
| chr13:53988596-53988636 | Ch13 region 2 |
| chr13:53988770-53988810 | Ch13 region 2 |
| chr13:53988951-53988991 | Ch13 region 2 |
| chr13:53989110-53989150 | Ch13 region 2 |
| chr13:53989164-53989204 | Ch13 region 2 |
| chr13:53989220-53989260 | Ch13 region 2 |
| chr13:53989271-53989311 | Ch13 region 2 |
| chr13:53989333-53989373 | Ch13 region 2 |
| chr13:53989402-53989442 | Ch13 region 2 |
| chr13:53989463-53989503 | Ch13 region 2 |
| chr13:53989536-53989576 | Ch13 region 2 |
| chr13:53989606-53989646 | Ch13 region 2 |
| chr13:53989764-53989804 | Ch13 region 2 |
| chr13:53989810-53989850 | Ch13 region 2 |
| chr13:53989889-53989929 | Ch13 region 2 |
| chr13:53989939-53989979 | Ch13 region 2 |
| chr13:53990031-53990071 | Ch13 region 2 |
| chr13:53990076-53990116 | Ch13 region 2 |
| chr13:53990177-53990217 | Ch13 region 2 |
| chr13:53990268-53990308 | Ch13 region 2 |
| chr13:53990322-53990362 | Ch13 region 2 |
| chr13:53990368-53990408 | Ch13 region 2 |
| chr13:53990416-53990456 | Ch13 region 2 |
| chr13:53990483-53990523 | Ch13 region 2 |
| chr13:53990539-53990579 | Ch13 region 2 |
| chr13:53990585-53990625 | Ch13 region 2 |
| chr13:53990655-53990695 | Ch13 region 2 |
| chr13:53990730-53990770 | Ch13 region 2 |
| chr13:53990786-53990826 | Ch13 region 2 |
| chr13:53990860-53990900 | Ch13 region 2 |
| chr13:53990939-53990979 | Ch13 region 2 |
| chr13:53990988-53991028 | Ch13 region 2 |
| chr13:53991033-53991073 | Ch13 region 2 |
| chr13:53991095-53991135 | Ch13 region 2 |
| chr13:53991141-53991181 | Ch13 region 2 |
| chr13:53991186-53991226 | Ch13 region 2 |
| chr13:53991282-53991322 | Ch13 region 2 |
| chr13:53991335-53991375 | Ch13 region 2 |
| chr13:53991383-53991423 | Ch13 region 2 |
| chr13:53991450-53991490 | Ch13 region 2 |
| chr13:53991516-53991556 | Ch13 region 2 |
| chr13:53991577-53991617 | Ch13 region 2 |
| chr13:53991656-53991696 | Ch13 region 2 |
| chr13:53991762-53991802 | Ch13 region 2 |
| chr13:53991812-53991852 | Ch13 region 2 |
| chr13:53991884-53991924 | Ch13 region 2 |

| | |
|---|---|
| chr13:53991936-53991976 | Ch13 region 2 |
| chr13:53992020-53992060 | Ch13 region 2 |
| chr13:53992117-53992157 | Ch13 region 2 |
| chr13:53992208-53992248 | Ch13 region 2 |
| chr13:53992271-53992311 | Ch13 region 2 |
| chr13:53992383-53992423 | Ch13 region 2 |
| chr13:53992428-53992468 | Ch13 region 2 |
| chr13:53992552-53992592 | Ch13 region 2 |
| chr13:53992739-53992779 | Ch13 region 2 |
| chr13:53992796-53992836 | Ch13 region 2 |
| chr13:53992887-53992927 | Ch13 region 2 |
| chr13:53992955-53992995 | Ch13 region 2 |
| chr13:53993009-53993049 | Ch13 region 2 |
| chr13:53993078-53993118 | Ch13 region 2 |
| chr13:53993175-53993215 | Ch13 region 2 |
| chr13:53993237-53993277 | Ch13 region 2 |
| chr13:53993282-53993322 | Ch13 region 2 |
| chr13:53993334-53993374 | Ch13 region 2 |
| chr13:53993379-53993419 | Ch13 region 2 |
| chr13:53993438-53993478 | Ch13 region 2 |
| chr13:53993486-53993526 | Ch13 region 2 |
| chr13:53993565-53993605 | Ch13 region 2 |
| chr13:53993635-53993675 | Ch13 region 2 |
| chr13:53993687-53993727 | Ch13 region 2 |
| chr13:53993778-53993818 | Ch13 region 2 |
| chr13:53993845-53993885 | Ch13 region 2 |
| chr13:53993895-53993935 | Ch13 region 2 |
| chr13:53993979-53994019 | Ch13 region 2 |
| chr13:53994087-53994127 | Ch13 region 2 |
| chr13:53994135-53994175 | Ch13 region 2 |
| chr13:53994232-53994272 | Ch13 region 2 |
| chr13:53994288-53994328 | Ch13 region 2 |
| chr13:53994419-53994459 | Ch13 region 2 |
| chr13:53994551-53994591 | Ch13 region 2 |
| chr13:53994598-53994638 | Ch13 region 2 |
| chr13:53994682-53994722 | Ch13 region 2 |
| chr13:53994751-53994791 | Ch13 region 2 |
| chr13:53994817-53994857 | Ch13 region 2 |
| chr13:53994924-53994964 | Ch13 region 2 |
| chr13:53994979-53995019 | Ch13 region 2 |
| chr13:53995090-53995130 | Ch13 region 2 |
| chr13:53995136-53995176 | Ch13 region 2 |
| chr13:53995183-53995223 | Ch13 region 2 |
| chr13:53995237-53995277 | Ch13 region 2 |
| chr13:53995329-53995369 | Ch13 region 2 |
| chr13:53995411-53995451 | Ch13 region 2 |
| chr13:53995456-53995496 | Ch13 region 2 |
| chr13:53995511-53995551 | Ch13 region 2 |
| chr13:53995566-53995606 | Ch13 region 2 |
| chr13:53995649-53995689 | Ch13 region 2 |
| chr13:53995695-53995735 | Ch13 region 2 |
| chr13:53995740-53995780 | Ch13 region 2 |

| | |
|---|---|
| chr13:53995817-53995857 | Ch13 region 2 |
| chr13:53995863-53995903 | Ch13 region 2 |
| chr13:53995984-53996024 | Ch13 region 2 |
| chr13:53996029-53996069 | Ch13 region 2 |
| chr13:53996077-53996117 | Ch13 region 2 |
| chr13:53996204-53996244 | Ch13 region 2 |
| chr13:53996272-53996312 | Ch13 region 2 |
| chr13:53996340-53996380 | Ch13 region 2 |
| chr13:53996443-53996483 | Ch13 region 2 |
| chr13:53996492-53996532 | Ch13 region 2 |
| chr13:53996588-53996628 | Ch13 region 2 |
| chr13:53996877-53996917 | Ch13 region 2 |
| chr13:53996950-53996990 | Ch13 region 2 |
| chr13:53996996-53997036 | Ch13 region 2 |
| chr13:53997316-53997356 | Ch13 region 2 |
| chr13:53997394-53997434 | Ch13 region 2 |
| chr13:53997474-53997514 | Ch13 region 2 |
| chr13:53997543-53997583 | Ch13 region 2 |
| chr13:53997588-53997628 | Ch13 region 2 |
| chr13:53997652-53997692 | Ch13 region 2 |
| chr13:53997703-53997743 | Ch13 region 2 |
| chr13:53997787-53997827 | Ch13 region 2 |
| chr13:53997915-53997955 | Ch13 region 2 |
| chr13:53998010-53998050 | Ch13 region 2 |
| chr13:53998077-53998117 | Ch13 region 2 |
| chr13:53998149-53998189 | Ch13 region 2 |
| chr13:53998314-53998354 | Ch13 region 2 |
| chr13:53998463-53998503 | Ch13 region 2 |
| chr13:53998523-53998563 | Ch13 region 2 |
| chr13:53998620-53998660 | Ch13 region 2 |
| chr13:53998670-53998710 | Ch13 region 2 |
| chr13:53998738-53998778 | Ch13 region 2 |
| chr13:53998822-53998862 | Ch13 region 2 |
| chr13:53998905-53998945 | Ch13 region 2 |
| chr13:53998952-53998992 | Ch13 region 2 |
| chr13:53999056-53999096 | Ch13 region 2 |
| chr13:53999128-53999168 | Ch13 region 2 |
| chr13:53999215-53999255 | Ch13 region 2 |
| chr13:53999262-53999302 | Ch13 region 2 |
| chr13:53999365-53999405 | Ch13 region 2 |
| chr13:53999432-53999472 | Ch13 region 2 |
| chr13:53999494-53999534 | Ch13 region 2 |
| chr13:53999544-53999584 | Ch13 region 2 |
| chr13:53999601-53999641 | Ch13 region 2 |
| chr13:53999683-53999723 | Ch13 region 2 |
| chr13:53999753-53999793 | Ch13 region 2 |
| chr13:53999834-53999874 | Ch13 region 2 |
| chr13:53999906-53999946 | Ch13 region 2 |
| chr13:54000007-54000047 | Ch13 region 2 |
| chr13:54000114-54000154 | Ch13 region 2 |
| chr13:54000176-54000216 | Ch13 region 2 |
| chr13:54000227-54000267 | Ch13 region 2 |

| | |
|---|---|
| chr13:54000297-54000337 | Ch13 region 2 |
| chr13:54000391-54000431 | Ch13 region 2 |
| chr13:54000604-54000644 | Ch13 region 2 |
| chr13:54000711-54000751 | Ch13 region 2 |
| chr13:54000784-54000824 | Ch13 region 2 |
| chr13:54000951-54000991 | Ch13 region 2 |
| chr13:54001066-54001106 | Ch13 region 2 |
| chr13:54001143-54001183 | Ch13 region 2 |
| chr13:54001214-54001254 | Ch13 region 2 |
| chr13:54001357-54001397 | Ch13 region 2 |
| chr13:54001500-54001540 | Ch13 region 2 |
| chr13:54001738-54001778 | Ch13 region 2 |
| chr13:54001810-54001850 | Ch13 region 2 |
| chr13:54002009-54002049 | Ch13 region 2 |
| chr13:54002119-54002159 | Ch13 region 2 |
| chr13:54002338-54002378 | Ch13 region 2 |
| chr13:54002383-54002423 | Ch13 region 2 |
| chr13:54002511-54002551 | Ch13 region 2 |
| chr13:54002584-54002624 | Ch13 region 2 |
| chr13:54002647-54002687 | Ch13 region 2 |
| chr13:54002725-54002765 | Ch13 region 2 |
| chr13:54002787-54002827 | Ch13 region 2 |
| chr13:54002854-54002894 | Ch13 region 2 |
| chr13:54002902-54002942 | Ch13 region 2 |
| chr13:54002948-54002988 | Ch13 region 2 |
| chr13:54003043-54003083 | Ch13 region 2 |
| chr13:54003096-54003136 | Ch13 region 2 |
| chr13:54003171-54003211 | Ch13 region 2 |
| chr13:54003273-54003313 | Ch13 region 2 |
| chr13:54003324-54003364 | Ch13 region 2 |
| chr13:54003391-54003431 | Ch13 region 2 |
| chr13:54003446-54003486 | Ch13 region 2 |
| chr13:54003504-54003544 | Ch13 region 2 |
| chr13:54003586-54003626 | Ch13 region 2 |
| chr13:54003918-54003958 | Ch13 region 2 |
| chr13:54003984-54004024 | Ch13 region 2 |
| chr13:54004073-54004113 | Ch13 region 2 |
| chr13:54004173-54004213 | Ch13 region 2 |
| chr13:54004219-54004259 | Ch13 region 2 |
| chr13:54004277-54004317 | Ch13 region 2 |
| chr13:54004351-54004391 | Ch13 region 2 |
| chr13:54004430-54004470 | Ch13 region 2 |
| chr13:54004494-54004534 | Ch13 region 2 |
| chr13:54004553-54004593 | Ch13 region 2 |
| chr13:54004621-54004661 | Ch13 region 2 |
| chr13:54004666-54004706 | Ch13 region 2 |
| chr13:54004921-54004961 | Ch13 region 2 |
| chr13:54004966-54005006 | Ch13 region 2 |
| chr13:54005039-54005079 | Ch13 region 2 |
| chr13:54005085-54005125 | Ch13 region 2 |
| chr13:54005154-54005194 | Ch13 region 2 |
| chr13:54005234-54005274 | Ch13 region 2 |

| | |
|---|---|
| chr13:54005304-54005344 | Ch13 region 2 |
| chr13:54005366-54005406 | Ch13 region 2 |
| chr13:54005415-54005455 | Ch13 region 2 |
| chr13:54005477-54005517 | Ch13 region 2 |
| chr13:54005522-54005562 | Ch13 region 2 |
| chr13:54005588-54005628 | Ch13 region 2 |
| chr13:54005636-54005676 | Ch13 region 2 |
| chr13:54005683-54005723 | Ch13 region 2 |
| chr13:54005746-54005786 | Ch13 region 2 |
| chr13:54005802-54005842 | Ch13 region 2 |
| chr13:54005847-54005887 | Ch13 region 2 |
| chr13:54005990-54006030 | Ch13 region 2 |
| chr13:54006039-54006079 | Ch13 region 2 |
| chr13:54006116-54006156 | Ch13 region 2 |
| chr13:54006173-54006213 | Ch13 region 2 |
| chr13:54006228-54006268 | Ch13 region 2 |
| chr13:54006281-54006321 | Ch13 region 2 |
| chr13:54006340-54006380 | Ch13 region 2 |
| chr13:54006389-54006429 | Ch13 region 2 |
| chr13:54006485-54006525 | Ch13 region 2 |
| chr13:54006551-54006591 | Ch13 region 2 |
| chr13:54006619-54006659 | Ch13 region 2 |
| chr13:54006705-54006745 | Ch13 region 2 |
| chr13:54006758-54006798 | Ch13 region 2 |
| chr13:54006843-54006883 | Ch13 region 2 |
| chr13:54006933-54006973 | Ch13 region 2 |
| chr13:54006978-54007018 | Ch13 region 2 |
| chr13:54007026-54007066 | Ch13 region 2 |
| chr13:54007092-54007132 | Ch13 region 2 |
| chr13:54007148-54007188 | Ch13 region 2 |
| chr13:54007272-54007312 | Ch13 region 2 |
| chr13:54007344-54007384 | Ch13 region 2 |
| chr13:54007423-54007463 | Ch13 region 2 |
| chr13:54007476-54007516 | Ch13 region 2 |
| chr13:54007571-54007611 | Ch13 region 2 |
| chr13:54007656-54007696 | Ch13 region 2 |
| chr13:54007701-54007741 | Ch13 region 2 |
| chr13:54007780-54007820 | Ch13 region 2 |
| chr13:54007835-54007875 | Ch13 region 2 |
| chr13:54007880-54007920 | Ch13 region 2 |
| chr13:54007949-54007989 | Ch13 region 2 |
| chr13:54008064-54008104 | Ch13 region 2 |
| chr13:54008160-54008200 | Ch13 region 2 |
| chr13:54008213-54008253 | Ch13 region 2 |
| chr13:54008281-54008321 | Ch13 region 2 |
| chr13:54008332-54008372 | Ch13 region 2 |
| chr13:54008382-54008422 | Ch13 region 2 |
| chr13:54008459-54008499 | Ch13 region 2 |
| chr13:54008531-54008571 | Ch13 region 2 |
| chr13:54008581-54008621 | Ch13 region 2 |
| chr13:54008649-54008689 | Ch13 region 2 |
| chr13:54008739-54008779 | Ch13 region 2 |

| | |
|---|---|
| chr13:54008809-54008849 | Ch13 region 2 |
| chr13:54008923-54008963 | Ch13 region 2 |
| chr13:54008989-54009029 | Ch13 region 2 |
| chr13:54009057-54009097 | Ch13 region 2 |
| chr13:54009189-54009229 | Ch13 region 2 |
| chr13:54009251-54009291 | Ch13 region 2 |
| chr13:54009355-54009395 | Ch13 region 2 |
| chr13:54009426-54009466 | Ch13 region 2 |
| chr13:54009471-54009511 | Ch13 region 2 |
| chr13:54009645-54009685 | Ch13 region 2 |
| chr13:54009759-54009799 | Ch13 region 2 |
| chr13:54009807-54009847 | Ch13 region 2 |
| chr13:54009868-54009908 | Ch13 region 2 |
| chr13:54009915-54009955 | Ch13 region 2 |
| chr13:54009971-54010011 | Ch13 region 2 |
| chr13:54010097-54010137 | Ch13 region 2 |
| chr13:54010186-54010226 | Ch13 region 2 |
| chr13:54010231-54010271 | Ch13 region 2 |
| chr13:54010277-54010317 | Ch13 region 2 |
| chr13:54010356-54010396 | Ch13 region 2 |
| chr13:54010401-54010441 | Ch13 region 2 |
| chr13:54010478-54010518 | Ch13 region 2 |
| chr13:54010542-54010582 | Ch13 region 2 |
| chr13:54010597-54010637 | Ch13 region 2 |
| chr13:54010652-54010692 | Ch13 region 2 |
| chr13:54010697-54010737 | Ch13 region 2 |
| chr13:54010755-54010795 | Ch13 region 2 |
| chr13:54010824-54010864 | Ch13 region 2 |
| chr13:54010874-54010914 | Ch13 region 2 |
| chr13:54010947-54010987 | Ch13 region 2 |
| chr13:54011022-54011062 | Ch13 region 2 |
| chr13:54011069-54011109 | Ch13 region 2 |
| chr13:54011125-54011165 | Ch13 region 2 |
| chr13:54011205-54011245 | Ch13 region 2 |
| chr13:54011281-54011321 | Ch13 region 2 |
| chr13:54012289-54012329 | Ch13 region 2 |
| chr13:54012657-54012697 | Ch13 region 2 |
| chr13:54012788-54012828 | Ch13 region 2 |
| chr13:54012885-54012925 | Ch13 region 2 |
| chr13:54012948-54012988 | Ch13 region 2 |
| chr13:54013023-54013063 | Ch13 region 2 |
| chr13:54013132-54013172 | Ch13 region 2 |
| chr13:54013201-54013241 | Ch13 region 2 |
| chr13:54013294-54013334 | Ch13 region 2 |
| chr13:54013383-54013423 | Ch13 region 2 |
| chr13:54013433-54013473 | Ch13 region 2 |
| chr13:54013589-54013629 | Ch13 region 2 |
| chr13:54013740-54013780 | Ch13 region 2 |
| chr13:54013835-54013875 | Ch13 region 2 |
| chr13:54013912-54013952 | Ch13 region 2 |
| chr13:54013957-54013997 | Ch13 region 2 |
| chr13:54014017-54014057 | Ch13 region 2 |

| | |
|---|---|
| chr13:54014107-54014147 | Ch13 region 2 |
| chr13:54014176-54014216 | Ch13 region 2 |
| chr13:54014237-54014277 | Ch13 region 2 |
| chr13:54014442-54014482 | Ch13 region 2 |
| chr13:54014596-54014636 | Ch13 region 2 |
| chr13:54014651-54014691 | Ch13 region 2 |
| chr13:54014700-54014740 | Ch13 region 2 |
| chr13:54014929-54014969 | Ch13 region 2 |
| chr13:54014983-54015023 | Ch13 region 2 |
| chr13:54015082-54015122 | Ch13 region 2 |
| chr13:54015147-54015187 | Ch13 region 2 |
| chr13:54015226-54015266 | Ch13 region 2 |
| chr13:54015283-54015323 | Ch13 region 2 |
| chr13:54015335-54015375 | Ch13 region 2 |
| chr13:54015400-54015440 | Ch13 region 2 |
| chr13:54015455-54015495 | Ch13 region 2 |
| chr13:54015513-54015553 | Ch13 region 2 |
| chr13:54015573-54015613 | Ch13 region 2 |
| chr13:54015657-54015697 | Ch13 region 2 |
| chr13:54015718-54015758 | Ch13 region 2 |
| chr13:54015764-54015804 | Ch13 region 2 |
| chr13:54015817-54015857 | Ch13 region 2 |
| chr13:54015862-54015902 | Ch13 region 2 |
| chr13:54015912-54015952 | Ch13 region 2 |
| chr13:54015969-54016009 | Ch13 region 2 |
| chr13:54016048-54016088 | Ch13 region 2 |
| chr13:54016134-54016174 | Ch13 region 2 |
| chr13:54016182-54016222 | Ch13 region 2 |
| chr13:54016266-54016306 | Ch13 region 2 |
| chr13:54016472-54016512 | Ch13 region 2 |
| chr13:54016531-54016571 | Ch13 region 2 |
| chr13:54016589-54016629 | Ch13 region 2 |
| chr13:54016634-54016674 | Ch13 region 2 |
| chr13:54016750-54016790 | Ch13 region 2 |
| chr13:54016959-54016999 | Ch13 region 2 |
| chr13:54017036-54017076 | Ch13 region 2 |
| chr13:54017113-54017153 | Ch13 region 2 |
| chr13:54017159-54017199 | Ch13 region 2 |
| chr13:54017207-54017247 | Ch13 region 2 |
| chr13:54017257-54017297 | Ch13 region 2 |
| chr13:54017342-54017382 | Ch13 region 2 |
| chr13:54017612-54017652 | Ch13 region 2 |
| chr13:54017725-54017765 | Ch13 region 2 |
| chr13:54017771-54017811 | Ch13 region 2 |
| chr13:54017853-54017893 | Ch13 region 2 |
| chr13:54017950-54017990 | Ch13 region 2 |
| chr13:54018205-54018245 | Ch13 region 2 |
| chr13:54018261-54018301 | Ch13 region 2 |
| chr13:54018359-54018399 | Ch13 region 2 |
| chr13:54018426-54018466 | Ch13 region 2 |
| chr13:54018542-54018582 | Ch13 region 2 |
| chr13:54018597-54018637 | Ch13 region 2 |

| | |
|---|---|
| chr13:54018651-54018691 | Ch13 region 2 |
| chr13:54018736-54018776 | Ch13 region 2 |
| chr13:54018782-54018822 | Ch13 region 2 |
| chr13:54018835-54018875 | Ch13 region 2 |
| chr13:54018909-54018949 | Ch13 region 2 |
| chr13:54018954-54018994 | Ch13 region 2 |
| chr13:54019032-54019072 | Ch13 region 2 |
| chr13:54019087-54019127 | Ch13 region 2 |
| chr13:54019157-54019197 | Ch13 region 2 |
| chr13:54019204-54019244 | Ch13 region 2 |
| chr13:54019281-54019321 | Ch13 region 2 |
| chr13:54019372-54019412 | Ch13 region 2 |
| chr13:54019417-54019457 | Ch13 region 2 |
| chr13:54019499-54019539 | Ch13 region 2 |
| chr13:54019605-54019645 | Ch13 region 2 |
| chr13:54019681-54019721 | Ch13 region 2 |
| chr13:54019761-54019801 | Ch13 region 2 |
| chr13:54019832-54019872 | Ch13 region 2 |
| chr13:54019900-54019940 | Ch13 region 2 |
| chr13:54019948-54019988 | Ch13 region 2 |
| chr13:54020047-54020087 | Ch13 region 2 |
| chr13:54020102-54020142 | Ch13 region 2 |
| chr13:54020151-54020191 | Ch13 region 2 |
| chr13:54020225-54020265 | Ch13 region 2 |
| chr13:54020306-54020346 | Ch13 region 2 |
| chr13:54020430-54020470 | Ch13 region 2 |
| chr13:54020496-54020536 | Ch13 region 2 |
| chr13:54020556-54020596 | Ch13 region 2 |
| chr13:54020614-54020654 | Ch13 region 2 |
| chr13:54020669-54020709 | Ch13 region 2 |
| chr13:54020736-54020776 | Ch13 region 2 |
| chr13:54020858-54020898 | Ch13 region 2 |
| chr13:54020910-54020950 | Ch13 region 2 |
| chr13:54020972-54021012 | Ch13 region 2 |
| chr13:54021046-54021086 | Ch13 region 2 |
| chr13:54021107-54021147 | Ch13 region 2 |
| chr13:54021187-54021227 | Ch13 region 2 |
| chr13:54021258-54021298 | Ch13 region 2 |
| chr13:54021335-54021375 | Ch13 region 2 |
| chr13:54021398-54021438 | Ch13 region 2 |
| chr13:54021468-54021508 | Ch13 region 2 |
| chr13:54021526-54021566 | Ch13 region 2 |
| chr13:54021592-54021632 | Ch13 region 2 |
| chr13:54021653-54021693 | Ch13 region 2 |
| chr13:54021759-54021799 | Ch13 region 2 |
| chr13:54021839-54021879 | Ch13 region 2 |
| chr13:54021887-54021927 | Ch13 region 2 |
| chr13:54021975-54022015 | Ch13 region 2 |
| chr13:54022081-54022121 | Ch13 region 2 |
| chr13:54022144-54022184 | Ch13 region 2 |
| chr13:54022199-54022239 | Ch13 region 2 |
| chr13:54022279-54022319 | Ch13 region 2 |

| | |
|---|---|
| chr13:54022347-54022387 | Ch13 region 2 |
| chr13:54022399-54022439 | Ch13 region 2 |
| chr13:54022445-54022485 | Ch13 region 2 |
| chr13:54022503-54022543 | Ch13 region 2 |
| chr13:54022566-54022606 | Ch13 region 2 |
| chr13:54022657-54022697 | Ch13 region 2 |
| chr13:54022705-54022745 | Ch13 region 2 |
| chr13:54022766-54022806 | Ch13 region 2 |
| chr13:54022811-54022851 | Ch13 region 2 |
| chr13:54022947-54022987 | Ch13 region 2 |
| chr13:54023052-54023092 | Ch13 region 2 |
| chr13:54023114-54023154 | Ch13 region 2 |
| chr13:54023195-54023235 | Ch13 region 2 |
| chr13:54023247-54023287 | Ch13 region 2 |
| chr13:54023324-54023364 | Ch13 region 2 |
| chr13:54023369-54023409 | Ch13 region 2 |
| chr13:54023448-54023488 | Ch13 region 2 |
| chr13:54023507-54023547 | Ch13 region 2 |
| chr13:54023581-54023621 | Ch13 region 2 |
| chr13:54023638-54023678 | Ch13 region 2 |
| chr13:54023756-54023796 | Ch13 region 2 |
| chr13:54023831-54023871 | Ch13 region 2 |
| chr13:54023917-54023957 | Ch13 region 2 |
| chr13:54024018-54024058 | Ch13 region 2 |
| chr13:54024072-54024112 | Ch13 region 2 |
| chr13:54024287-54024327 | Ch13 region 2 |
| chr13:54024388-54024428 | Ch13 region 2 |
| chr13:54024433-54024473 | Ch13 region 2 |
| chr13:54024557-54024597 | Ch13 region 2 |
| chr13:54024659-54024699 | Ch13 region 2 |
| chr13:54024716-54024756 | Ch13 region 2 |
| chr13:54024802-54024842 | Ch13 region 2 |
| chr13:54024916-54024956 | Ch13 region 2 |
| chr13:54024997-54025037 | Ch13 region 2 |
| chr13:54025091-54025131 | Ch13 region 2 |
| chr13:54025179-54025219 | Ch13 region 2 |
| chr13:54025231-54025271 | Ch13 region 2 |
| chr13:54025311-54025351 | Ch13 region 2 |
| chr13:54025388-54025428 | Ch13 region 2 |
| chr13:54025466-54025506 | Ch13 region 2 |
| chr13:54025556-54025596 | Ch13 region 2 |
| chr13:54025636-54025676 | Ch13 region 2 |
| chr13:54025681-54025721 | Ch13 region 2 |
| chr13:54025806-54025846 | Ch13 region 2 |
| chr13:54025901-54025941 | Ch13 region 2 |
| chr13:54026046-54026086 | Ch13 region 2 |
| chr13:54026147-54026187 | Ch13 region 2 |
| chr13:54026503-54026543 | Ch13 region 2 |
| chr13:54026571-54026611 | Ch13 region 2 |
| chr13:54026683-54026723 | Ch13 region 2 |
| chr13:54026819-54026859 | Ch13 region 2 |
| chr13:54027079-54027119 | Ch13 region 2 |

| | |
|---|---|
| chr13:54027202-54027242 | Ch13 region 2 |
| chr13:54027301-54027341 | Ch13 region 2 |
| chr13:54027384-54027424 | Ch13 region 2 |
| chr13:54027473-54027513 | Ch13 region 2 |
| chr13:54027522-54027562 | Ch13 region 2 |
| chr13:54027684-54027724 | Ch13 region 2 |
| chr13:54027729-54027769 | Ch13 region 2 |
| chr13:54027861-54027901 | Ch13 region 2 |
| chr13:54027966-54028006 | Ch13 region 2 |
| chr13:54028029-54028069 | Ch13 region 2 |
| chr13:54028086-54028126 | Ch13 region 2 |
| chr13:54028242-54028282 | Ch13 region 2 |
| chr13:54028300-54028340 | Ch13 region 2 |
| chr13:54028357-54028397 | Ch13 region 2 |
| chr13:54028456-54028496 | Ch13 region 2 |
| chr13:54028501-54028541 | Ch13 region 2 |
| chr13:54028563-54028603 | Ch13 region 2 |
| chr13:54028609-54028649 | Ch13 region 2 |
| chr13:54028660-54028700 | Ch13 region 2 |
| chr13:54028728-54028768 | Ch13 region 2 |
| chr13:54028801-54028841 | Ch13 region 2 |
| chr13:54028873-54028913 | Ch13 region 2 |
| chr13:54028973-54029013 | Ch13 region 2 |
| chr13:54029019-54029059 | Ch13 region 2 |
| chr13:54029071-54029111 | Ch13 region 2 |
| chr13:54029135-54029175 | Ch13 region 2 |
| chr13:54029206-54029246 | Ch13 region 2 |
| chr13:54029363-54029403 | Ch13 region 2 |
| chr13:54029436-54029476 | Ch13 region 2 |
| chr13:54029484-54029524 | Ch13 region 2 |
| chr13:54029561-54029601 | Ch13 region 2 |
| chr13:54029618-54029658 | Ch13 region 2 |
| chr13:54029665-54029705 | Ch13 region 2 |
| chr13:54029739-54029779 | Ch13 region 2 |
| chr13:54029823-54029863 | Ch13 region 2 |
| chr13:54029876-54029916 | Ch13 region 2 |
| chr13:54029933-54029973 | Ch13 region 2 |
| chr13:54029982-54030022 | Ch13 region 2 |
| chr13:54030034-54030074 | Ch13 region 2 |
| chr13:54030080-54030120 | Ch13 region 2 |
| chr13:54030131-54030171 | Ch13 region 2 |
| chr13:54030177-54030217 | Ch13 region 2 |
| chr13:54030293-54030333 | Ch13 region 2 |
| chr13:54030523-54030563 | Ch13 region 2 |
| chr13:54030580-54030620 | Ch13 region 2 |
| chr13:54030625-54030665 | Ch13 region 2 |
| chr13:54030706-54030746 | Ch13 region 2 |
| chr13:54030757-54030797 | Ch13 region 2 |
| chr13:54030844-54030884 | Ch13 region 2 |
| chr13:54030921-54030961 | Ch13 region 2 |
| chr13:54030966-54031006 | Ch13 region 2 |
| chr13:54031016-54031056 | Ch13 region 2 |

| | |
|---|---|
| chr13:54031072-54031112 | Ch13 region 2 |
| chr13:54031125-54031165 | Ch13 region 2 |
| chr13:54031223-54031263 | Ch13 region 2 |
| chr13:54031305-54031345 | Ch13 region 2 |
| chr13:54031380-54031420 | Ch13 region 2 |
| chr13:54031438-54031478 | Ch13 region 2 |
| chr13:54031505-54031545 | Ch13 region 2 |
| chr13:54031554-54031594 | Ch13 region 2 |
| chr13:54031655-54031695 | Ch13 region 2 |
| chr13:54031816-54031856 | Ch13 region 2 |
| chr13:54031898-54031938 | Ch13 region 2 |
| chr13:54031983-54032023 | Ch13 region 2 |
| chr13:54032028-54032068 | Ch13 region 2 |
| chr13:54032161-54032201 | Ch13 region 2 |
| chr13:54032236-54032276 | Ch13 region 2 |
| chr13:54032286-54032326 | Ch13 region 2 |
| chr13:54032333-54032373 | Ch13 region 2 |
| chr13:54032416-54032456 | Ch13 region 2 |
| chr13:54032491-54032531 | Ch13 region 2 |
| chr13:54032570-54032610 | Ch13 region 2 |
| chr13:54032643-54032683 | Ch13 region 2 |
| chr13:54032722-54032762 | Ch13 region 2 |
| chr13:54032848-54032888 | Ch13 region 2 |
| chr13:54032901-54032941 | Ch13 region 2 |
| chr13:54032951-54032991 | Ch13 region 2 |
| chr13:54033015-54033055 | Ch13 region 2 |
| chr13:54033093-54033133 | Ch13 region 2 |
| chr13:54033175-54033215 | Ch13 region 2 |
| chr13:54033222-54033262 | Ch13 region 2 |
| chr13:54033294-54033334 | Ch13 region 2 |
| chr13:54033442-54033482 | Ch13 region 2 |
| chr13:54033501-54033541 | Ch13 region 2 |
| chr13:54033612-54033652 | Ch13 region 2 |
| chr13:54033778-54033818 | Ch13 region 2 |
| chr13:54033847-54033887 | Ch13 region 2 |
| chr13:54033918-54033958 | Ch13 region 2 |
| chr13:54033973-54034013 | Ch13 region 2 |
| chr13:54034058-54034098 | Ch13 region 2 |
| chr13:54034108-54034148 | Ch13 region 2 |
| chr13:54034153-54034193 | Ch13 region 2 |
| chr13:54034223-54034263 | Ch13 region 2 |
| chr13:54034282-54034322 | Ch13 region 2 |
| chr13:54034338-54034378 | Ch13 region 2 |
| chr13:54034458-54034498 | Ch13 region 2 |
| chr13:54034515-54034555 | Ch13 region 2 |
| chr13:54034577-54034617 | Ch13 region 2 |
| chr13:54034622-54034662 | Ch13 region 2 |
| chr13:54034722-54034762 | Ch13 region 2 |
| chr13:54034790-54034830 | Ch13 region 2 |
| chr13:54034836-54034876 | Ch13 region 2 |
| chr13:54034920-54034960 | Ch13 region 2 |
| chr13:54035023-54035063 | Ch13 region 2 |

| | |
|---|---|
| chr13:54035097-54035137 | Ch13 region 2 |
| chr13:54035172-54035212 | Ch13 region 2 |
| chr13:54035227-54035267 | Ch13 region 2 |
| chr13:54035310-54035350 | Ch13 region 2 |
| chr13:54035397-54035437 | Ch13 region 2 |
| chr13:54035477-54035517 | Ch13 region 2 |
| chr13:54035527-54035567 | Ch13 region 2 |
| chr13:54035574-54035614 | Ch13 region 2 |
| chr13:54035623-54035663 | Ch13 region 2 |
| chr13:54035688-54035728 | Ch13 region 2 |
| chr13:54035846-54035886 | Ch13 region 2 |
| chr13:54035905-54035945 | Ch13 region 2 |
| chr13:54035982-54036022 | Ch13 region 2 |
| chr13:54036049-54036089 | Ch13 region 2 |
| chr13:54036110-54036150 | Ch13 region 2 |
| chr13:54036176-54036216 | Ch13 region 2 |
| chr13:54036283-54036323 | Ch13 region 2 |
| chr13:54036366-54036406 | Ch13 region 2 |
| chr13:54036450-54036490 | Ch13 region 2 |
| chr13:54036497-54036537 | Ch13 region 2 |
| chr13:54036593-54036633 | Ch13 region 2 |
| chr13:54036719-54036759 | Ch13 region 2 |
| chr13:54036927-54036967 | Ch13 region 2 |
| chr13:54037025-54037065 | Ch13 region 2 |
| chr13:54037090-54037130 | Ch13 region 2 |
| chr13:54037148-54037188 | Ch13 region 2 |
| chr13:54037254-54037294 | Ch13 region 2 |
| chr13:54037299-54037339 | Ch13 region 2 |
| chr13:54037371-54037411 | Ch13 region 2 |
| chr13:54037451-54037491 | Ch13 region 2 |
| chr13:54037496-54037536 | Ch13 region 2 |
| chr13:54037553-54037593 | Ch13 region 2 |
| chr13:54037625-54037665 | Ch13 region 2 |
| chr13:54037678-54037718 | Ch13 region 2 |
| chr13:54037730-54037770 | Ch13 region 2 |
| chr13:54037777-54037817 | Ch13 region 2 |
| chr13:54037841-54037881 | Ch13 region 2 |
| chr13:54037886-54037926 | Ch13 region 2 |
| chr13:54037935-54037975 | Ch13 region 2 |
| chr13:54038281-54038321 | Ch13 region 2 |
| chr13:54038360-54038400 | Ch13 region 2 |
| chr13:54038426-54038466 | Ch13 region 2 |
| chr13:54038472-54038512 | Ch13 region 2 |
| chr13:54038803-54038843 | Ch13 region 2 |
| chr13:54038890-54038930 | Ch13 region 2 |
| chr13:54038953-54038993 | Ch13 region 2 |
| chr13:54039034-54039074 | Ch13 region 2 |
| chr13:54039079-54039119 | Ch13 region 2 |
| chr13:54039175-54039215 | Ch13 region 2 |
| chr13:54039234-54039274 | Ch13 region 2 |
| chr13:54039291-54039331 | Ch13 region 2 |
| chr13:54039402-54039442 | Ch13 region 2 |

| | |
|---|---|
| chr13:54039610-54039650 | Ch13 region 2 |
| chr13:54039657-54039697 | Ch13 region 2 |
| chr13:54040001-54040041 | Ch13 region 2 |
| chr13:54040126-54040166 | Ch13 region 2 |
| chr13:54040284-54040324 | Ch13 region 2 |
| chr13:54040330-54040370 | Ch13 region 2 |
| chr13:54040418-54040458 | Ch13 region 2 |
| chr13:54040486-54040526 | Ch13 region 2 |
| chr13:54040545-54040585 | Ch13 region 2 |
| chr13:54040627-54040667 | Ch13 region 2 |
| chr13:54040681-54040721 | Ch13 region 2 |
| chr13:54040749-54040789 | Ch13 region 2 |
| chr13:54040837-54040877 | Ch13 region 2 |
| chr13:54040882-54040922 | Ch13 region 2 |
| chr13:54040936-54040976 | Ch13 region 2 |
| chr13:54041025-54041065 | Ch13 region 2 |
| chr13:54041162-54041202 | Ch13 region 2 |
| chr13:54041228-54041268 | Ch13 region 2 |
| chr13:54041286-54041326 | Ch13 region 2 |
| chr13:54041333-54041373 | Ch13 region 2 |
| chr13:54041479-54041519 | Ch13 region 2 |
| chr13:54041547-54041587 | Ch13 region 2 |
| chr13:54041614-54041654 | Ch13 region 2 |
| chr13:54041676-54041716 | Ch13 region 2 |
| chr13:54041723-54041763 | Ch13 region 2 |
| chr13:54041811-54041851 | Ch13 region 2 |
| chr13:54041901-54041941 | Ch13 region 2 |
| chr13:54041955-54041995 | Ch13 region 2 |
| chr13:54042022-54042062 | Ch13 region 2 |
| chr13:54042099-54042139 | Ch13 region 2 |
| chr13:54042148-54042188 | Ch13 region 2 |
| chr13:54042229-54042269 | Ch13 region 2 |
| chr13:54042336-54042376 | Ch13 region 2 |
| chr13:54042427-54042467 | Ch13 region 2 |
| chr13:54042520-54042560 | Ch13 region 2 |
| chr13:54042632-54042672 | Ch13 region 2 |
| chr13:54042744-54042784 | Ch13 region 2 |
| chr13:54042794-54042834 | Ch13 region 2 |
| chr13:54042879-54042919 | Ch13 region 2 |
| chr13:54042943-54042983 | Ch13 region 2 |
| chr13:54042990-54043030 | Ch13 region 2 |
| chr13:54043043-54043083 | Ch13 region 2 |
| chr13:54043139-54043179 | Ch13 region 2 |
| chr13:54043191-54043231 | Ch13 region 2 |
| chr13:54043261-54043301 | Ch13 region 2 |
| chr13:54043355-54043395 | Ch13 region 2 |
| chr13:54043411-54043451 | Ch13 region 2 |
| chr13:54043477-54043517 | Ch13 region 2 |
| chr13:54043524-54043564 | Ch13 region 2 |
| chr13:54043584-54043624 | Ch13 region 2 |
| chr13:54043696-54043736 | Ch13 region 2 |
| chr13:54043741-54043781 | Ch13 region 2 |

| | |
|---|---|
| chr13:54043845-54043885 | Ch13 region 2 |
| chr13:54043914-54043954 | Ch13 region 2 |
| chr13:54043990-54044030 | Ch13 region 2 |
| chr13:54044035-54044075 | Ch13 region 2 |
| chr13:54044080-54044120 | Ch13 region 2 |
| chr13:54044151-54044191 | Ch13 region 2 |
| chr13:54044333-54044373 | Ch13 region 2 |
| chr13:54044386-54044426 | Ch13 region 2 |
| chr13:54044454-54044494 | Ch13 region 2 |
| chr13:54044624-54044664 | Ch13 region 2 |
| chr13:54044669-54044709 | Ch13 region 2 |
| chr13:54044727-54044767 | Ch13 region 2 |
| chr13:54044777-54044817 | Ch13 region 2 |
| chr13:54044834-54044874 | Ch13 region 2 |
| chr13:54044882-54044922 | Ch13 region 2 |
| chr13:54044943-54044983 | Ch13 region 2 |
| chr13:54045038-54045078 | Ch13 region 2 |
| chr13:54045152-54045192 | Ch13 region 2 |
| chr13:54045221-54045261 | Ch13 region 2 |
| chr13:54045343-54045383 | Ch13 region 2 |
| chr13:54045394-54045434 | Ch13 region 2 |
| chr13:54045485-54045525 | Ch13 region 2 |
| chr13:54045567-54045607 | Ch13 region 2 |
| chr13:54045618-54045658 | Ch13 region 2 |
| chr13:54045686-54045726 | Ch13 region 2 |
| chr13:54045769-54045809 | Ch13 region 2 |
| chr13:54045822-54045862 | Ch13 region 2 |
| chr13:54045918-54045958 | Ch13 region 2 |
| chr13:54046021-54046061 | Ch13 region 2 |
| chr13:54046081-54046121 | Ch13 region 2 |
| chr13:54046145-54046185 | Ch13 region 2 |
| chr13:54046248-54046288 | Ch13 region 2 |
| chr13:54046310-54046350 | Ch13 region 2 |
| chr13:54046381-54046421 | Ch13 region 2 |
| chr13:54046445-54046485 | Ch13 region 2 |
| chr13:54046521-54046561 | Ch13 region 2 |
| chr13:54046603-54046643 | Ch13 region 2 |
| chr13:54046665-54046705 | Ch13 region 2 |
| chr13:54046779-54046819 | Ch13 region 2 |
| chr13:54046848-54046888 | Ch13 region 2 |
| chr13:54046924-54046964 | Ch13 region 2 |
| chr13:54046986-54047026 | Ch13 region 2 |
| chr13:54047089-54047129 | Ch13 region 2 |
| chr13:54047149-54047189 | Ch13 region 2 |
| chr13:54047227-54047267 | Ch13 region 2 |
| chr13:54047272-54047312 | Ch13 region 2 |
| chr13:54047330-54047370 | Ch13 region 2 |
| chr13:54047402-54047442 | Ch13 region 2 |
| chr13:54047501-54047541 | Ch13 region 2 |
| chr13:54047585-54047625 | Ch13 region 2 |
| chr13:54047641-54047681 | Ch13 region 2 |
| chr13:54047691-54047731 | Ch13 region 2 |

| | |
|---|---|
| chr13:54047754-54047794 | Ch13 region 2 |
| chr13:54047828-54047868 | Ch13 region 2 |
| chr13:54047909-54047949 | Ch13 region 2 |
| chr13:54047977-54048017 | Ch13 region 2 |
| chr13:54048045-54048085 | Ch13 region 2 |
| chr13:54048106-54048146 | Ch13 region 2 |
| chr13:54048166-54048206 | Ch13 region 2 |
| chr13:54048217-54048257 | Ch13 region 2 |
| chr13:54048283-54048323 | Ch13 region 2 |
| chr13:54048341-54048381 | Ch13 region 2 |
| chr13:54048400-54048440 | Ch13 region 2 |
| chr13:54048447-54048487 | Ch13 region 2 |
| chr13:54048559-54048599 | Ch13 region 2 |
| chr13:54048627-54048667 | Ch13 region 2 |
| chr13:54048695-54048735 | Ch13 region 2 |
| chr13:54048741-54048781 | Ch13 region 2 |
| chr13:54048789-54048829 | Ch13 region 2 |
| chr13:54048854-54048894 | Ch13 region 2 |
| chr13:54048930-54048970 | Ch13 region 2 |
| chr13:54048976-54049016 | Ch13 region 2 |
| chr13:54049049-54049089 | Ch13 region 2 |
| chr13:54049111-54049151 | Ch13 region 2 |
| chr13:54049176-54049216 | Ch13 region 2 |
| chr13:54049246-54049286 | Ch13 region 2 |
| chr13:54049296-54049336 | Ch13 region 2 |
| chr13:54049357-54049397 | Ch13 region 2 |
| chr13:54049402-54049442 | Ch13 region 2 |
| chr13:54049479-54049519 | Ch13 region 2 |
| chr13:54049535-54049575 | Ch13 region 2 |
| chr13:54049617-54049657 | Ch13 region 2 |
| chr13:54049689-54049729 | Ch13 region 2 |
| chr13:54049749-54049789 | Ch13 region 2 |
| chr13:54049794-54049834 | Ch13 region 2 |
| chr13:54049857-54049897 | Ch13 region 2 |
| chr13:54049953-54049993 | Ch13 region 2 |
| chr13:54050037-54050077 | Ch13 region 2 |
| chr13:54050082-54050122 | Ch13 region 2 |
| chr13:54050149-54050189 | Ch13 region 2 |
| chr13:54050298-54050338 | Ch13 region 2 |
| chr13:54050343-54050383 | Ch13 region 2 |
| chr13:54050389-54050429 | Ch13 region 2 |
| chr13:54050469-54050509 | Ch13 region 2 |
| chr13:54050529-54050569 | Ch13 region 2 |
| chr13:54050621-54050661 | Ch13 region 2 |
| chr13:54050672-54050712 | Ch13 region 2 |
| chr13:54050771-54050811 | Ch13 region 2 |
| chr13:54050865-54050905 | Ch13 region 2 |
| chr13:54050911-54050951 | Ch13 region 2 |
| chr13:54051070-54051110 | Ch13 region 2 |
| chr13:54051116-54051156 | Ch13 region 2 |
| chr13:54051230-54051270 | Ch13 region 2 |
| chr13:54051302-54051342 | Ch13 region 2 |

| | |
|---|---|
| chr13:54051359-54051399 | Ch13 region 2 |
| chr13:54051439-54051479 | Ch13 region 2 |
| chr13:54051515-54051555 | Ch13 region 2 |
| chr13:54051616-54051656 | Ch13 region 2 |
| chr13:54051690-54051730 | Ch13 region 2 |
| chr13:54051741-54051781 | Ch13 region 2 |
| chr13:54051801-54051841 | Ch13 region 2 |
| chr13:54051887-54051927 | Ch13 region 2 |
| chr13:54051996-54052036 | Ch13 region 2 |
| chr13:54052058-54052098 | Ch13 region 2 |
| chr13:54052122-54052162 | Ch13 region 2 |
| chr13:54052222-54052262 | Ch13 region 2 |
| chr13:54052275-54052315 | Ch13 region 2 |
| chr13:54052351-54052391 | Ch13 region 2 |
| chr13:54052401-54052441 | Ch13 region 2 |
| chr13:54052462-54052502 | Ch13 region 2 |
| chr13:54052537-54052577 | Ch13 region 2 |
| chr13:54052596-54052636 | Ch13 region 2 |
| chr13:54052660-54052700 | Ch13 region 2 |
| chr13:54052711-54052751 | Ch13 region 2 |
| chr13:54052803-54052843 | Ch13 region 2 |
| chr13:54052849-54052889 | Ch13 region 2 |
| chr13:54052901-54052941 | Ch13 region 2 |
| chr13:54052967-54053007 | Ch13 region 2 |
| chr13:54053033-54053073 | Ch13 region 2 |
| chr13:54053104-54053144 | Ch13 region 2 |
| chr13:54053170-54053210 | Ch13 region 2 |
| chr13:54053247-54053287 | Ch13 region 2 |
| chr13:54053339-54053379 | Ch13 region 2 |
| chr13:54053422-54053462 | Ch13 region 2 |
| chr13:54053502-54053542 | Ch13 region 2 |
| chr13:54053621-54053661 | Ch13 region 2 |
| chr13:54053756-54053796 | Ch13 region 2 |
| chr13:54053850-54053890 | Ch13 region 2 |
| chr13:54053895-54053935 | Ch13 region 2 |
| chr13:54053989-54054029 | Ch13 region 2 |
| chr13:54054090-54054130 | Ch13 region 2 |
| chr13:54054143-54054183 | Ch13 region 2 |
| chr13:54054191-54054231 | Ch13 region 2 |
| chr13:54054253-54054293 | Ch13 region 2 |
| chr13:54054298-54054338 | Ch13 region 2 |
| chr13:54054345-54054385 | Ch13 region 2 |
| chr13:54054404-54054444 | Ch13 region 2 |
| chr13:54054479-54054519 | Ch13 region 2 |
| chr13:54054575-54054615 | Ch13 region 2 |
| chr13:54054668-54054708 | Ch13 region 2 |
| chr13:54054726-54054766 | Ch13 region 2 |
| chr13:54054777-54054817 | Ch13 region 2 |
| chr13:54054850-54054890 | Ch13 region 2 |
| chr13:54054930-54054970 | Ch13 region 2 |
| chr13:54054977-54055017 | Ch13 region 2 |
| chr13:54055046-54055086 | Ch13 region 2 |

| | |
|---|---|
| chr13:54055091-54055131 | Ch13 region 2 |
| chr13:54055153-54055193 | Ch13 region 2 |
| chr13:54055216-54055256 | Ch13 region 2 |
| chr13:54055295-54055335 | Ch13 region 2 |
| chr13:54055378-54055418 | Ch13 region 2 |
| chr13:54055423-54055463 | Ch13 region 2 |
| chr13:54055479-54055519 | Ch13 region 2 |
| chr13:54055553-54055593 | Ch13 region 2 |
| chr13:54055605-54055645 | Ch13 region 2 |
| chr13:54055654-54055694 | Ch13 region 2 |
| chr13:54055772-54055812 | Ch13 region 2 |
| chr13:54055828-54055868 | Ch13 region 2 |
| chr13:54055979-54056019 | Ch13 region 2 |
| chr13:54056060-54056100 | Ch13 region 2 |
| chr13:54056123-54056163 | Ch13 region 2 |
| chr13:54056168-54056208 | Ch13 region 2 |
| chr13:54056242-54056282 | Ch13 region 2 |
| chr13:54056301-54056341 | Ch13 region 2 |
| chr13:54056349-54056389 | Ch13 region 2 |
| chr13:54056394-54056434 | Ch13 region 2 |
| chr13:54056483-54056523 | Ch13 region 2 |
| chr13:54056699-54056739 | Ch13 region 2 |
| chr13:54056784-54056824 | Ch13 region 2 |
| chr13:54056829-54056869 | Ch13 region 2 |
| chr13:54056919-54056959 | Ch13 region 2 |
| chr13:54056995-54057035 | Ch13 region 2 |
| chr13:54057042-54057082 | Ch13 region 2 |
| chr13:54057114-54057154 | Ch13 region 2 |
| chr13:54057173-54057213 | Ch13 region 2 |
| chr13:54057234-54057274 | Ch13 region 2 |
| chr13:54057281-54057321 | Ch13 region 2 |
| chr13:54057442-54057482 | Ch13 region 2 |
| chr13:54057509-54057549 | Ch13 region 2 |
| chr13:54057563-54057603 | Ch13 region 2 |
| chr13:54057619-54057659 | Ch13 region 2 |
| chr13:54057711-54057751 | Ch13 region 2 |
| chr13:54057757-54057797 | Ch13 region 2 |
| chr13:54057806-54057846 | Ch13 region 2 |
| chr13:54057884-54057924 | Ch13 region 2 |
| chr13:54057969-54058009 | Ch13 region 2 |
| chr13:54058014-54058054 | Ch13 region 2 |
| chr13:54058059-54058099 | Ch13 region 2 |
| chr13:54058244-54058284 | Ch13 region 2 |
| chr13:54058355-54058395 | Ch13 region 2 |
| chr13:54058573-54058613 | Ch13 region 2 |
| chr13:54058736-54058776 | Ch13 region 2 |
| chr13:54058781-54058821 | Ch13 region 2 |
| chr13:54058850-54058890 | Ch13 region 2 |
| chr13:54058912-54058952 | Ch13 region 2 |
| chr13:54058987-54059027 | Ch13 region 2 |
| chr13:54059038-54059078 | Ch13 region 2 |
| chr13:54059130-54059170 | Ch13 region 2 |

| | |
|---|---|
| chr13:54059232-54059272 | Ch13 region 2 |
| chr13:54059294-54059334 | Ch13 region 2 |
| chr13:54059412-54059452 | Ch13 region 2 |
| chr13:54059536-54059576 | Ch13 region 2 |
| chr13:54059619-54059659 | Ch13 region 2 |
| chr13:54059706-54059746 | Ch13 region 2 |
| chr13:54059751-54059791 | Ch13 region 2 |
| chr13:54059894-54059934 | Ch13 region 2 |
| chr13:54059941-54059981 | Ch13 region 2 |
| chr13:54060043-54060083 | Ch13 region 2 |
| chr13:54060089-54060129 | Ch13 region 2 |
| chr13:54060154-54060194 | Ch13 region 2 |
| chr13:54060203-54060243 | Ch13 region 2 |
| chr13:54060311-54060351 | Ch13 region 2 |
| chr13:54060356-54060396 | Ch13 region 2 |
| chr13:54060411-54060451 | Ch13 region 2 |
| chr13:54060491-54060531 | Ch13 region 2 |
| chr13:54060750-54060790 | Ch13 region 2 |
| chr13:54060805-54060845 | Ch13 region 2 |
| chr13:54060895-54060935 | Ch13 region 2 |
| chr13:54060965-54061005 | Ch13 region 2 |
| chr13:54061037-54061077 | Ch13 region 2 |
| chr13:54061084-54061124 | Ch13 region 2 |
| chr13:54061169-54061209 | Ch13 region 2 |
| chr13:54061215-54061255 | Ch13 region 2 |
| chr13:54061287-54061327 | Ch13 region 2 |
| chr13:54061386-54061426 | Ch13 region 2 |
| chr13:54061478-54061518 | Ch13 region 2 |
| chr13:54061538-54061578 | Ch13 region 2 |
| chr13:54061625-54061665 | Ch13 region 2 |
| chr13:54061734-54061774 | Ch13 region 2 |
| chr13:54061799-54061839 | Ch13 region 2 |
| chr13:54061864-54061904 | Ch13 region 2 |
| chr13:54062101-54062141 | Ch13 region 2 |
| chr13:54062184-54062224 | Ch13 region 2 |
| chr13:54062246-54062286 | Ch13 region 2 |
| chr13:54062360-54062400 | Ch13 region 2 |
| chr13:54062411-54062451 | Ch13 region 2 |
| chr13:54062461-54062501 | Ch13 region 2 |
| chr13:54062507-54062547 | Ch13 region 2 |
| chr13:54062602-54062642 | Ch13 region 2 |
| chr13:54062709-54062749 | Ch13 region 2 |
| chr13:54062848-54062888 | Ch13 region 2 |
| chr13:54062916-54062956 | Ch13 region 2 |
| chr13:54062963-54063003 | Ch13 region 2 |
| chr13:54063058-54063098 | Ch13 region 2 |
| chr13:54063140-54063180 | Ch13 region 2 |
| chr13:54063219-54063259 | Ch13 region 2 |
| chr13:54063287-54063327 | Ch13 region 2 |
| chr13:54063379-54063419 | Ch13 region 2 |
| chr13:54063424-54063464 | Ch13 region 2 |
| chr13:54063473-54063513 | Ch13 region 2 |

| | |
|---|---|
| chr13:54063590-54063630 | Ch13 region 2 |
| chr13:54063676-54063716 | Ch13 region 2 |
| chr13:54063758-54063798 | Ch13 region 2 |
| chr13:54063831-54063871 | Ch13 region 2 |
| chr13:54063878-54063918 | Ch13 region 2 |
| chr13:54063949-54063989 | Ch13 region 2 |
| chr13:54064035-54064075 | Ch13 region 2 |
| chr13:54064080-54064120 | Ch13 region 2 |
| chr13:54064125-54064165 | Ch13 region 2 |
| chr13:54064172-54064212 | Ch13 region 2 |
| chr13:54064268-54064308 | Ch13 region 2 |
| chr13:54064327-54064367 | Ch13 region 2 |
| chr13:54064413-54064453 | Ch13 region 2 |
| chr13:54064526-54064566 | Ch13 region 2 |
| chr13:54064571-54064611 | Ch13 region 2 |
| chr13:54064637-54064677 | Ch13 region 2 |
| chr13:54064711-54064751 | Ch13 region 2 |
| chr13:54064790-54064830 | Ch13 region 2 |
| chr13:54064872-54064912 | Ch13 region 2 |
| chr13:54064936-54064976 | Ch13 region 2 |
| chr13:54065009-54065049 | Ch13 region 2 |
| chr13:54065054-54065094 | Ch13 region 2 |
| chr13:54065101-54065141 | Ch13 region 2 |
| chr13:54065177-54065217 | Ch13 region 2 |
| chr13:54065240-54065280 | Ch13 region 2 |
| chr13:54065289-54065329 | Ch13 region 2 |
| chr13:54065392-54065432 | Ch13 region 2 |
| chr13:54065487-54065527 | Ch13 region 2 |
| chr13:54065616-54065656 | Ch13 region 2 |
| chr13:54065691-54065731 | Ch13 region 2 |
| chr13:54065773-54065813 | Ch13 region 2 |
| chr13:54065838-54065878 | Ch13 region 2 |
| chr13:54065887-54065927 | Ch13 region 2 |
| chr13:54065998-54066038 | Ch13 region 2 |
| chr13:54066081-54066121 | Ch13 region 2 |
| chr13:54066126-54066166 | Ch13 region 2 |
| chr13:54066205-54066245 | Ch13 region 2 |
| chr13:54066250-54066290 | Ch13 region 2 |
| chr13:54066298-54066338 | Ch13 region 2 |
| chr13:54066344-54066384 | Ch13 region 2 |
| chr13:54066401-54066441 | Ch13 region 2 |
| chr13:54066447-54066487 | Ch13 region 2 |
| chr13:54066576-54066616 | Ch13 region 2 |
| chr13:54066673-54066713 | Ch13 region 2 |
| chr13:54066737-54066777 | Ch13 region 2 |
| chr13:54066801-54066841 | Ch13 region 2 |
| chr13:54066868-54066908 | Ch13 region 2 |
| chr13:54066939-54066979 | Ch13 region 2 |
| chr13:54067032-54067072 | Ch13 region 2 |
| chr13:54067126-54067166 | Ch13 region 2 |
| chr13:54067179-54067219 | Ch13 region 2 |
| chr13:54067227-54067267 | Ch13 region 2 |

| | |
|---|---|
| chr13:54067302-54067342 | Ch13 region 2 |
| chr13:54067446-54067486 | Ch13 region 2 |
| chr13:54067506-54067546 | Ch13 region 2 |
| chr13:54067644-54067684 | Ch13 region 2 |
| chr13:54067815-54067855 | Ch13 region 2 |
| chr13:54067874-54067914 | Ch13 region 2 |
| chr13:54067923-54067963 | Ch13 region 2 |
| chr13:54067982-54068022 | Ch13 region 2 |
| chr13:54068040-54068080 | Ch13 region 2 |
| chr13:54068124-54068164 | Ch13 region 2 |
| chr13:54068188-54068228 | Ch13 region 2 |
| chr13:54068243-54068283 | Ch13 region 2 |
| chr13:54068288-54068328 | Ch13 region 2 |
| chr13:54068373-54068413 | Ch13 region 2 |
| chr13:54068468-54068508 | Ch13 region 2 |
| chr13:54068544-54068584 | Ch13 region 2 |
| chr13:54068592-54068632 | Ch13 region 2 |
| chr13:54068684-54068724 | Ch13 region 2 |
| chr13:54068775-54068815 | Ch13 region 2 |
| chr13:54068855-54068895 | Ch13 region 2 |
| chr13:54068900-54068940 | Ch13 region 2 |
| chr13:54068945-54068985 | Ch13 region 2 |
| chr13:54069036-54069076 | Ch13 region 2 |
| chr13:54069156-54069196 | Ch13 region 2 |
| chr13:54069230-54069270 | Ch13 region 2 |
| chr13:54069275-54069315 | Ch13 region 2 |
| chr13:54069519-54069559 | Ch13 region 2 |
| chr13:54069568-54069608 | Ch13 region 2 |
| chr13:54069624-54069664 | Ch13 region 2 |
| chr13:54069680-54069720 | Ch13 region 2 |
| chr13:54069729-54069769 | Ch13 region 2 |
| chr13:54069783-54069823 | Ch13 region 2 |
| chr13:54069850-54069890 | Ch13 region 2 |
| chr13:54069950-54069990 | Ch13 region 2 |
| chr13:54069995-54070035 | Ch13 region 2 |
| chr13:54070058-54070098 | Ch13 region 2 |
| chr13:54070122-54070162 | Ch13 region 2 |
| chr13:54070190-54070230 | Ch13 region 2 |
| chr13:54070318-54070358 | Ch13 region 2 |
| chr13:54070381-54070421 | Ch13 region 2 |
| chr13:54070475-54070515 | Ch13 region 2 |
| chr13:54070554-54070594 | Ch13 region 2 |
| chr13:54070839-54070879 | Ch13 region 2 |
| chr13:54070895-54070935 | Ch13 region 2 |
| chr13:54071016-54071056 | Ch13 region 2 |
| chr13:54071098-54071138 | Ch13 region 2 |
| chr13:54071190-54071230 | Ch13 region 2 |
| chr13:54071251-54071291 | Ch13 region 2 |
| chr13:54071354-54071394 | Ch13 region 2 |
| chr13:54071434-54071474 | Ch13 region 2 |
| chr13:54071509-54071549 | Ch13 region 2 |
| chr13:54071592-54071632 | Ch13 region 2 |

| | |
|---|---|
| chr13:54071684-54071724 | Ch13 region 2 |
| chr13:54071941-54071981 | Ch13 region 2 |
| chr13:54072047-54072087 | Ch13 region 2 |
| chr13:54072160-54072200 | Ch13 region 2 |
| chr13:54072244-54072284 | Ch13 region 2 |
| chr13:54072321-54072361 | Ch13 region 2 |
| chr13:54072393-54072433 | Ch13 region 2 |
| chr13:54072455-54072495 | Ch13 region 2 |
| chr13:54072501-54072541 | Ch13 region 2 |
| chr13:54072592-54072632 | Ch13 region 2 |
| chr13:54072699-54072739 | Ch13 region 2 |
| chr13:54072902-54072942 | Ch13 region 2 |
| chr13:54072959-54072999 | Ch13 region 2 |
| chr13:54073012-54073052 | Ch13 region 2 |
| chr13:54073096-54073136 | Ch13 region 2 |
| chr13:54073146-54073186 | Ch13 region 2 |
| chr13:54073197-54073237 | Ch13 region 2 |
| chr13:54073401-54073441 | Ch13 region 2 |
| chr13:54073468-54073508 | Ch13 region 2 |
| chr13:54073579-54073619 | Ch13 region 2 |
| chr13:54073624-54073664 | Ch13 region 2 |
| chr13:54073723-54073763 | Ch13 region 2 |
| chr13:54073769-54073809 | Ch13 region 2 |
| chr13:54073824-54073864 | Ch13 region 2 |
| chr13:54073886-54073926 | Ch13 region 2 |
| chr13:54073931-54073971 | Ch13 region 2 |
| chr13:54074000-54074040 | Ch13 region 2 |
| chr13:54074062-54074102 | Ch13 region 2 |
| chr13:54074117-54074157 | Ch13 region 2 |
| chr13:54074180-54074220 | Ch13 region 2 |
| chr13:54074227-54074267 | Ch13 region 2 |
| chr13:54074360-54074400 | Ch13 region 2 |
| chr13:54074409-54074449 | Ch13 region 2 |
| chr13:54074560-54074600 | Ch13 region 2 |
| chr13:54074605-54074645 | Ch13 region 2 |
| chr13:54074681-54074721 | Ch13 region 2 |
| chr13:54074737-54074777 | Ch13 region 2 |
| chr13:54074991-54075031 | Ch13 region 2 |
| chr13:54075079-54075119 | Ch13 region 2 |
| chr13:54075272-54075312 | Ch13 region 2 |
| chr13:54075456-54075496 | Ch13 region 2 |
| chr13:54075608-54075648 | Ch13 region 2 |
| chr13:54075672-54075712 | Ch13 region 2 |
| chr13:54075769-54075809 | Ch13 region 2 |
| chr13:54075814-54075854 | Ch13 region 2 |
| chr13:54075901-54075941 | Ch13 region 2 |
| chr13:54076102-54076142 | Ch13 region 2 |
| chr13:54076166-54076206 | Ch13 region 2 |
| chr13:54076260-54076300 | Ch13 region 2 |
| chr13:54076391-54076431 | Ch13 region 2 |
| chr13:54076476-54076516 | Ch13 region 2 |
| chr13:54076521-54076561 | Ch13 region 2 |

| | |
|---|---|
| chr13:54076593-54076633 | Ch13 region 2 |
| chr13:54076680-54076720 | Ch13 region 2 |
| chr13:54076838-54076878 | Ch13 region 2 |
| chr13:54076890-54076930 | Ch13 region 2 |
| chr13:54077038-54077078 | Ch13 region 2 |
| chr13:54077143-54077183 | Ch13 region 2 |
| chr13:54077336-54077376 | Ch13 region 2 |
| chr13:54077412-54077452 | Ch13 region 2 |
| chr13:54077521-54077561 | Ch13 region 2 |
| chr13:54077624-54077664 | Ch13 region 2 |
| chr13:54077673-54077713 | Ch13 region 2 |
| chr13:54077879-54077919 | Ch13 region 2 |
| chr13:54077998-54078038 | Ch13 region 2 |
| chr13:54078110-54078150 | Ch13 region 2 |
| chr13:54078163-54078203 | Ch13 region 2 |
| chr13:54078242-54078282 | Ch13 region 2 |
| chr13:54078307-54078347 | Ch13 region 2 |
| chr13:54078385-54078425 | Ch13 region 2 |
| chr13:54078434-54078474 | Ch13 region 2 |
| chr13:54078521-54078561 | Ch13 region 2 |
| chr13:54078613-54078653 | Ch13 region 2 |
| chr13:54078677-54078717 | Ch13 region 2 |
| chr13:54078722-54078762 | Ch13 region 2 |
| chr13:54078767-54078807 | Ch13 region 2 |
| chr13:54078812-54078852 | Ch13 region 2 |
| chr13:54078868-54078908 | Ch13 region 2 |
| chr13:54078914-54078954 | Ch13 region 2 |
| chr13:54079000-54079040 | Ch13 region 2 |
| chr13:54079065-54079105 | Ch13 region 2 |
| chr13:54079114-54079154 | Ch13 region 2 |
| chr13:54079219-54079259 | Ch13 region 2 |
| chr13:54079301-54079341 | Ch13 region 2 |
| chr13:54079368-54079408 | Ch13 region 2 |
| chr13:54079416-54079456 | Ch13 region 2 |
| chr13:54079463-54079503 | Ch13 region 2 |
| chr13:54079549-54079589 | Ch13 region 2 |
| chr13:54079619-54079659 | Ch13 region 2 |
| chr13:54079679-54079719 | Ch13 region 2 |
| chr13:54079730-54079770 | Ch13 region 2 |
| chr13:54079816-54079856 | Ch13 region 2 |
| chr13:54079902-54079942 | Ch13 region 2 |
| chr13:54080115-54080155 | Ch13 region 2 |
| chr13:54080166-54080206 | Ch13 region 2 |
| chr13:54080223-54080263 | Ch13 region 2 |
| chr13:54080319-54080359 | Ch13 region 2 |
| chr13:54080407-54080447 | Ch13 region 2 |
| chr13:54080577-54080617 | Ch13 region 2 |
| chr13:54080627-54080667 | Ch13 region 2 |
| chr13:54080701-54080741 | Ch13 region 2 |
| chr13:54080799-54080839 | Ch13 region 2 |
| chr13:54080999-54081039 | Ch13 region 2 |
| chr13:54081155-54081195 | Ch13 region 2 |

| | |
|---|---|
| chr13:54081336-54081376 | Ch13 region 2 |
| chr13:54081409-54081449 | Ch13 region 2 |
| chr13:54081487-54081527 | Ch13 region 2 |
| chr13:54081557-54081597 | Ch13 region 2 |
| chr13:54081634-54081674 | Ch13 region 2 |
| chr13:54081701-54081741 | Ch13 region 2 |
| chr13:54081762-54081802 | Ch13 region 2 |
| chr13:54081827-54081867 | Ch13 region 2 |
| chr13:54081872-54081912 | Ch13 region 2 |
| chr13:54081956-54081996 | Ch13 region 2 |
| chr13:54082054-54082094 | Ch13 region 2 |
| chr13:54082109-54082149 | Ch13 region 2 |
| chr13:54082195-54082235 | Ch13 region 2 |
| chr13:54082247-54082287 | Ch13 region 2 |
| chr13:54082322-54082362 | Ch13 region 2 |
| chr13:54082411-54082451 | Ch13 region 2 |
| chr13:54082473-54082513 | Ch13 region 2 |
| chr13:54082525-54082565 | Ch13 region 2 |
| chr13:54082597-54082637 | Ch13 region 2 |
| chr13:54082643-54082683 | Ch13 region 2 |
| chr13:54082708-54082748 | Ch13 region 2 |
| chr13:54082849-54082889 | Ch13 region 2 |
| chr13:54082917-54082957 | Ch13 region 2 |
| chr13:54082971-54083011 | Ch13 region 2 |
| chr13:54083066-54083106 | Ch13 region 2 |
| chr13:54083120-54083160 | Ch13 region 2 |
| chr13:54083215-54083255 | Ch13 region 2 |
| chr13:54083264-54083304 | Ch13 region 2 |
| chr13:54083311-54083351 | Ch13 region 2 |
| chr13:54083356-54083396 | Ch13 region 2 |
| chr13:54083405-54083445 | Ch13 region 2 |
| chr13:54083473-54083513 | Ch13 region 2 |
| chr13:54083523-54083563 | Ch13 region 2 |
| chr13:54083569-54083609 | Ch13 region 2 |
| chr13:54083657-54083697 | Ch13 region 2 |
| chr13:54083736-54083776 | Ch13 region 2 |
| chr13:54083824-54083864 | Ch13 region 2 |
| chr13:54083977-54084017 | Ch13 region 2 |
| chr13:54084023-54084063 | Ch13 region 2 |
| chr13:54084068-54084108 | Ch13 region 2 |
| chr13:54084121-54084161 | Ch13 region 2 |
| chr13:54084191-54084231 | Ch13 region 2 |
| chr13:54084256-54084296 | Ch13 region 2 |
| chr13:54084306-54084346 | Ch13 region 2 |
| chr13:54084380-54084420 | Ch13 region 2 |
| chr13:54084483-54084523 | Ch13 region 2 |
| chr13:54084560-54084600 | Ch13 region 2 |
| chr13:54084620-54084660 | Ch13 region 2 |
| chr13:54084665-54084705 | Ch13 region 2 |
| chr13:54084711-54084751 | Ch13 region 2 |
| chr13:54084805-54084845 | Ch13 region 2 |
| chr13:54084897-54084937 | Ch13 region 2 |

| | |
|---|---|
| chr13:54084948-54084988 | Ch13 region 2 |
| chr13:54085019-54085059 | Ch13 region 2 |
| chr13:54085074-54085114 | Ch13 region 2 |
| chr13:54085153-54085193 | Ch13 region 2 |
| chr13:54085201-54085241 | Ch13 region 2 |
| chr13:54085259-54085299 | Ch13 region 2 |
| chr13:54085356-54085396 | Ch13 region 2 |
| chr13:54085401-54085441 | Ch13 region 2 |
| chr13:54085494-54085534 | Ch13 region 2 |
| chr13:54085597-54085637 | Ch13 region 2 |
| chr13:54085736-54085776 | Ch13 region 2 |
| chr13:54085842-54085882 | Ch13 region 2 |
| chr13:54085914-54085954 | Ch13 region 2 |
| chr13:54085979-54086019 | Ch13 region 2 |
| chr13:54086048-54086088 | Ch13 region 2 |
| chr13:54086134-54086174 | Ch13 region 2 |
| chr13:54086218-54086258 | Ch13 region 2 |
| chr13:54086311-54086351 | Ch13 region 2 |
| chr13:54086392-54086432 | Ch13 region 2 |
| chr13:54086460-54086500 | Ch13 region 2 |
| chr13:54086572-54086612 | Ch13 region 2 |
| chr13:54086667-54086707 | Ch13 region 2 |
| chr13:54086719-54086759 | Ch13 region 2 |
| chr13:54086798-54086838 | Ch13 region 2 |
| chr13:54086853-54086893 | Ch13 region 2 |
| chr13:54086912-54086952 | Ch13 region 2 |
| chr13:54086975-54087015 | Ch13 region 2 |
| chr13:54087027-54087067 | Ch13 region 2 |
| chr13:54087087-54087127 | Ch13 region 2 |
| chr13:54087172-54087212 | Ch13 region 2 |
| chr13:54087228-54087268 | Ch13 region 2 |
| chr13:54087327-54087367 | Ch13 region 2 |
| chr13:54087391-54087431 | Ch13 region 2 |
| chr13:54087444-54087484 | Ch13 region 2 |
| chr13:54087510-54087550 | Ch13 region 2 |
| chr13:54087578-54087618 | Ch13 region 2 |
| chr13:54087664-54087704 | Ch13 region 2 |
| chr13:54087759-54087799 | Ch13 region 2 |
| chr13:54087814-54087854 | Ch13 region 2 |
| chr13:54087862-54087902 | Ch13 region 2 |
| chr13:54087913-54087953 | Ch13 region 2 |
| chr13:54087958-54087998 | Ch13 region 2 |
| chr13:54088037-54088077 | Ch13 region 2 |
| chr13:54088144-54088184 | Ch13 region 2 |
| chr13:54088198-54088238 | Ch13 region 2 |
| chr13:54088251-54088291 | Ch13 region 2 |
| chr13:54088323-54088363 | Ch13 region 2 |
| chr13:54088397-54088437 | Ch13 region 2 |
| chr13:54088442-54088482 | Ch13 region 2 |
| chr13:54088537-54088577 | Ch13 region 2 |
| chr13:54088582-54088622 | Ch13 region 2 |
| chr13:54088628-54088668 | Ch13 region 2 |

| | |
|---|---|
| chr13:54088673-54088713 | Ch13 region 2 |
| chr13:54088733-54088773 | Ch13 region 2 |
| chr13:54088788-54088828 | Ch13 region 2 |
| chr13:54088865-54088905 | Ch13 region 2 |
| chr13:54089008-54089048 | Ch13 region 2 |
| chr13:54089060-54089100 | Ch13 region 2 |
| chr13:54089238-54089278 | Ch13 region 2 |
| chr13:54089440-54089480 | Ch13 region 2 |
| chr13:54089615-54089655 | Ch13 region 2 |
| chr13:54089952-54089992 | Ch13 region 2 |
| chr13:54090015-54090055 | Ch13 region 2 |
| chr13:54090159-54090199 | Ch13 region 2 |
| chr13:54090207-54090247 | Ch13 region 2 |
| chr13:54090272-54090312 | Ch13 region 2 |
| chr13:54090419-54090459 | Ch13 region 2 |
| chr13:54090489-54090529 | Ch13 region 2 |
| chr13:54090664-54090704 | Ch13 region 2 |
| chr13:54090761-54090801 | Ch13 region 2 |
| chr13:54090924-54090964 | Ch13 region 2 |
| chr13:54090996-54091036 | Ch13 region 2 |
| chr13:54091043-54091083 | Ch13 region 2 |
| chr13:54091144-54091184 | Ch13 region 2 |
| chr13:54091216-54091256 | Ch13 region 2 |
| chr13:54091368-54091408 | Ch13 region 2 |
| chr13:54091422-54091462 | Ch13 region 2 |
| chr13:54091467-54091507 | Ch13 region 2 |
| chr13:54091564-54091604 | Ch13 region 2 |
| chr13:54091610-54091650 | Ch13 region 2 |
| chr13:54091743-54091783 | Ch13 region 2 |
| chr13:54091861-54091901 | Ch13 region 2 |
| chr13:54091940-54091980 | Ch13 region 2 |
| chr13:54092009-54092049 | Ch13 region 2 |
| chr13:54092070-54092110 | Ch13 region 2 |
| chr13:54092145-54092185 | Ch13 region 2 |
| chr13:54092221-54092261 | Ch13 region 2 |
| chr13:54092319-54092359 | Ch13 region 2 |
| chr13:54092371-54092411 | Ch13 region 2 |
| chr13:54092462-54092502 | Ch13 region 2 |
| chr13:54092614-54092654 | Ch13 region 2 |
| chr13:54092687-54092727 | Ch13 region 2 |
| chr13:54092750-54092790 | Ch13 region 2 |
| chr13:54092815-54092855 | Ch13 region 2 |
| chr13:54092873-54092913 | Ch13 region 2 |
| chr13:54092920-54092960 | Ch13 region 2 |
| chr13:54092965-54093005 | Ch13 region 2 |
| chr13:54093012-54093052 | Ch13 region 2 |
| chr13:54093064-54093104 | Ch13 region 2 |
| chr13:54093127-54093167 | Ch13 region 2 |
| chr13:54093172-54093212 | Ch13 region 2 |
| chr13:54093247-54093287 | Ch13 region 2 |
| chr13:54093375-54093415 | Ch13 region 2 |
| chr13:54093420-54093460 | Ch13 region 2 |

| | |
|---|---|
| chr13:54093468-54093508 | Ch13 region 2 |
| chr13:54093518-54093558 | Ch13 region 2 |
| chr13:54093582-54093622 | Ch13 region 2 |
| chr13:54093657-54093697 | Ch13 region 2 |
| chr13:54093711-54093751 | Ch13 region 2 |
| chr13:54093857-54093897 | Ch13 region 2 |
| chr13:54093903-54093943 | Ch13 region 2 |
| chr13:54093999-54094039 | Ch13 region 2 |
| chr13:54094044-54094084 | Ch13 region 2 |
| chr13:54094116-54094156 | Ch13 region 2 |
| chr13:54094178-54094218 | Ch13 region 2 |
| chr13:54094245-54094285 | Ch13 region 2 |
| chr13:54094293-54094333 | Ch13 region 2 |
| chr13:54094353-54094393 | Ch13 region 2 |
| chr13:54094400-54094440 | Ch13 region 2 |
| chr13:54094466-54094506 | Ch13 region 2 |
| chr13:54094512-54094552 | Ch13 region 2 |
| chr13:54094558-54094598 | Ch13 region 2 |
| chr13:54094606-54094646 | Ch13 region 2 |
| chr13:54094679-54094719 | Ch13 region 2 |
| chr13:54094736-54094776 | Ch13 region 2 |
| chr13:54094816-54094856 | Ch13 region 2 |
| chr13:54094886-54094926 | Ch13 region 2 |
| chr13:54094939-54094979 | Ch13 region 2 |
| chr13:54094987-54095027 | Ch13 region 2 |
| chr13:54095032-54095072 | Ch13 region 2 |
| chr13:54095089-54095129 | Ch13 region 2 |
| chr13:54095162-54095202 | Ch13 region 2 |
| chr13:54095211-54095251 | Ch13 region 2 |
| chr13:54095265-54095305 | Ch13 region 2 |
| chr13:54095351-54095391 | Ch13 region 2 |
| chr13:54095418-54095458 | Ch13 region 2 |
| chr13:54095463-54095503 | Ch13 region 2 |
| chr13:54095539-54095579 | Ch13 region 2 |
| chr13:54095586-54095626 | Ch13 region 2 |
| chr13:54095649-54095689 | Ch13 region 2 |
| chr13:54095705-54095745 | Ch13 region 2 |
| chr13:54095777-54095817 | Ch13 region 2 |
| chr13:54095857-54095897 | Ch13 region 2 |
| chr13:54095905-54095945 | Ch13 region 2 |
| chr13:54096010-54096050 | Ch13 region 2 |
| chr13:54096071-54096111 | Ch13 region 2 |
| chr13:54096142-54096182 | Ch13 region 2 |
| chr13:54096187-54096227 | Ch13 region 2 |
| chr13:54096288-54096328 | Ch13 region 2 |
| chr13:54096367-54096407 | Ch13 region 2 |
| chr13:54096429-54096469 | Ch13 region 2 |
| chr13:54096525-54096565 | Ch13 region 2 |
| chr13:54096582-54096622 | Ch13 region 2 |
| chr13:54096636-54096676 | Ch13 region 2 |
| chr13:54096717-54096757 | Ch13 region 2 |
| chr13:54096762-54096802 | Ch13 region 2 |

| | |
|---|---|
| chr13:54096848-54096888 | Ch13 region 2 |
| chr13:54096909-54096949 | Ch13 region 2 |
| chr13:54096954-54096994 | Ch13 region 2 |
| chr13:54097018-54097058 | Ch13 region 2 |
| chr13:54097066-54097106 | Ch13 region 2 |
| chr13:54097132-54097172 | Ch13 region 2 |
| chr13:54097178-54097218 | Ch13 region 2 |
| chr13:54097223-54097263 | Ch13 region 2 |
| chr13:54097278-54097318 | Ch13 region 2 |
| chr13:54097354-54097394 | Ch13 region 2 |
| chr13:54097419-54097459 | Ch13 region 2 |
| chr13:54097505-54097545 | Ch13 region 2 |
| chr13:54097552-54097592 | Ch13 region 2 |
| chr13:54097615-54097655 | Ch13 region 2 |
| chr13:54097662-54097702 | Ch13 region 2 |
| chr13:54097785-54097825 | Ch13 region 2 |
| chr13:54097851-54097891 | Ch13 region 2 |
| chr13:54097947-54097987 | Ch13 region 2 |
| chr13:54098017-54098057 | Ch13 region 2 |
| chr13:54098116-54098156 | Ch13 region 2 |
| chr13:54098171-54098211 | Ch13 region 2 |
| chr13:54098216-54098256 | Ch13 region 2 |
| chr13:54098301-54098341 | Ch13 region 2 |
| chr13:54098348-54098388 | Ch13 region 2 |
| chr13:54098414-54098454 | Ch13 region 2 |
| chr13:54098467-54098507 | Ch13 region 2 |
| chr13:54098617-54098657 | Ch13 region 2 |
| chr13:54098681-54098721 | Ch13 region 2 |
| chr13:54098735-54098775 | Ch13 region 2 |
| chr13:54098783-54098823 | Ch13 region 2 |
| chr13:54099004-54099044 | Ch13 region 2 |
| chr13:54099051-54099091 | Ch13 region 2 |
| chr13:54099152-54099192 | Ch13 region 2 |
| chr13:54099203-54099243 | Ch13 region 2 |
| chr13:54099254-54099294 | Ch13 region 2 |
| chr13:54099302-54099342 | Ch13 region 2 |
| chr13:54099389-54099429 | Ch13 region 2 |
| chr13:54099452-54099492 | Ch13 region 2 |
| chr13:54099497-54099537 | Ch13 region 2 |
| chr13:54099575-54099615 | Ch13 region 2 |
| chr13:54099620-54099660 | Ch13 region 2 |
| chr13:54099715-54099755 | Ch13 region 2 |
| chr13:54099761-54099801 | Ch13 region 2 |
| chr13:54099807-54099847 | Ch13 region 2 |
| chr13:54099874-54099914 | Ch13 region 2 |
| chr13:54099945-54099985 | Ch13 region 2 |
| chr13:54100077-54100117 | Ch13 region 2 |
| chr13:54100148-54100188 | Ch13 region 2 |
| chr13:54100206-54100246 | Ch13 region 2 |
| chr13:54100290-54100330 | Ch13 region 2 |
| chr13:54100371-54100411 | Ch13 region 2 |
| chr13:54100433-54100473 | Ch13 region 2 |

| | |
|---|---|
| chr13:54100516-54100556 | Ch13 region 2 |
| chr13:54100563-54100603 | Ch13 region 2 |
| chr13:54100615-54100655 | Ch13 region 2 |
| chr13:54100662-54100702 | Ch13 region 2 |
| chr13:54100744-54100784 | Ch13 region 2 |
| chr13:54100789-54100829 | Ch13 region 2 |
| chr13:54100835-54100875 | Ch13 region 2 |
| chr13:54100896-54100936 | Ch13 region 2 |
| chr13:54101011-54101051 | Ch13 region 2 |
| chr13:54101098-54101138 | Ch13 region 2 |
| chr13:54101190-54101230 | Ch13 region 2 |
| chr13:54101259-54101299 | Ch13 region 2 |
| chr13:54101333-54101373 | Ch13 region 2 |
| chr13:54101400-54101440 | Ch13 region 2 |
| chr13:54101608-54101648 | Ch13 region 2 |
| chr13:54101687-54101727 | Ch13 region 2 |
| chr13:54101753-54101793 | Ch13 region 2 |
| chr13:54101852-54101892 | Ch13 region 2 |
| chr13:54101906-54101946 | Ch13 region 2 |
| chr13:54101951-54101991 | Ch13 region 2 |
| chr13:54101999-54102039 | Ch13 region 2 |
| chr13:54102064-54102104 | Ch13 region 2 |
| chr13:54102147-54102187 | Ch13 region 2 |
| chr13:54102210-54102250 | Ch13 region 2 |
| chr13:54102268-54102308 | Ch13 region 2 |
| chr13:54102314-54102354 | Ch13 region 2 |
| chr13:54102371-54102411 | Ch13 region 2 |
| chr13:54102421-54102461 | Ch13 region 2 |
| chr13:54102500-54102540 | Ch13 region 2 |
| chr13:54102577-54102617 | Ch13 region 2 |
| chr13:54102672-54102712 | Ch13 region 2 |
| chr13:54102864-54102904 | Ch13 region 2 |
| chr13:54103000-54103040 | Ch13 region 2 |
| chr13:54103099-54103139 | Ch13 region 2 |
| chr13:54103184-54103224 | Ch13 region 2 |
| chr13:54103252-54103292 | Ch13 region 2 |
| chr13:54103304-54103344 | Ch13 region 2 |
| chr13:54103370-54103410 | Ch13 region 2 |
| chr13:54103442-54103482 | Ch13 region 2 |
| chr13:54103496-54103536 | Ch13 region 2 |
| chr13:54103593-54103633 | Ch13 region 2 |
| chr13:54103658-54103698 | Ch13 region 2 |
| chr13:54103728-54103768 | Ch13 region 2 |
| chr13:54103782-54103822 | Ch13 region 2 |
| chr13:54103842-54103882 | Ch13 region 2 |
| chr13:54103948-54103988 | Ch13 region 2 |
| chr13:54104022-54104062 | Ch13 region 2 |
| chr13:54104080-54104120 | Ch13 region 2 |
| chr13:54104142-54104182 | Ch13 region 2 |
| chr13:54104194-54104234 | Ch13 region 2 |
| chr13:54104252-54104292 | Ch13 region 2 |
| chr13:54104333-54104373 | Ch13 region 2 |

| | |
|---|---|
| chr13:54104390-54104430 | Ch13 region 2 |
| chr13:54104495-54104535 | Ch13 region 2 |
| chr13:54104562-54104602 | Ch13 region 2 |
| chr13:54104664-54104704 | Ch13 region 2 |
| chr13:54104761-54104801 | Ch13 region 2 |
| chr13:54104871-54104911 | Ch13 region 2 |
| chr13:54104956-54104996 | Ch13 region 2 |
| chr13:54105038-54105078 | Ch13 region 2 |
| chr13:54105148-54105188 | Ch13 region 2 |
| chr13:54105217-54105257 | Ch13 region 2 |
| chr13:54105426-54105466 | Ch13 region 2 |
| chr13:54105474-54105514 | Ch13 region 2 |
| chr13:54105627-54105667 | Ch13 region 2 |
| chr13:54105852-54105892 | Ch13 region 2 |
| chr13:54105897-54105937 | Ch13 region 2 |
| chr13:54105944-54105984 | Ch13 region 2 |
| chr13:54106024-54106064 | Ch13 region 2 |
| chr13:54106112-54106152 | Ch13 region 2 |
| chr13:54106192-54106232 | Ch13 region 2 |
| chr13:54106268-54106308 | Ch13 region 2 |
| chr13:54106313-54106353 | Ch13 region 2 |
| chr13:54106363-54106403 | Ch13 region 2 |
| chr13:54106472-54106512 | Ch13 region 2 |
| chr13:54106565-54106605 | Ch13 region 2 |
| chr13:54106620-54106660 | Ch13 region 2 |
| chr13:54106675-54106715 | Ch13 region 2 |
| chr13:54106720-54106760 | Ch13 region 2 |
| chr13:54106780-54106820 | Ch13 region 2 |
| chr13:54106847-54106887 | Ch13 region 2 |
| chr13:54106909-54106949 | Ch13 region 2 |
| chr13:54107007-54107047 | Ch13 region 2 |
| chr13:54107157-54107197 | Ch13 region 2 |
| chr13:54107325-54107365 | Ch13 region 2 |
| chr13:54107379-54107419 | Ch13 region 2 |
| chr13:54107435-54107475 | Ch13 region 2 |
| chr13:54107543-54107583 | Ch13 region 2 |
| chr13:54107629-54107669 | Ch13 region 2 |
| chr13:54107804-54107844 | Ch13 region 2 |
| chr13:54108011-54108051 | Ch13 region 2 |
| chr13:54108350-54108390 | Ch13 region 2 |
| chr13:54108452-54108492 | Ch13 region 2 |
| chr13:54108540-54108580 | Ch13 region 2 |
| chr13:54108594-54108634 | Ch13 region 2 |
| chr13:54108676-54108716 | Ch13 region 2 |
| chr13:54108799-54108839 | Ch13 region 2 |
| chr13:54108949-54108989 | Ch13 region 2 |
| chr13:54109109-54109149 | Ch13 region 2 |
| chr13:54109154-54109194 | Ch13 region 2 |
| chr13:54109352-54109392 | Ch13 region 2 |
| chr13:54109454-54109494 | Ch13 region 2 |
| chr13:54109509-54109549 | Ch13 region 2 |
| chr13:54109588-54109628 | Ch13 region 2 |

| | |
|---|---|
| chr13:54109669-54109709 | Ch13 region 2 |
| chr13:54109880-54109920 | Ch13 region 2 |
| chr13:54110150-54110190 | Ch13 region 2 |
| chr13:54110196-54110236 | Ch13 region 2 |
| chr13:54110300-54110340 | Ch13 region 2 |
| chr13:54110382-54110422 | Ch13 region 2 |
| chr13:54110439-54110479 | Ch13 region 2 |
| chr13:54110534-54110574 | Ch13 region 2 |
| chr13:54110586-54110626 | Ch13 region 2 |
| chr13:54110869-54110909 | Ch13 region 2 |
| chr13:54110942-54110982 | Ch13 region 2 |
| chr13:54111060-54111100 | Ch13 region 2 |
| chr13:54111128-54111168 | Ch13 region 2 |
| chr13:54111191-54111231 | Ch13 region 2 |
| chr13:54111385-54111425 | Ch13 region 2 |
| chr13:54111449-54111489 | Ch13 region 2 |
| chr13:54111615-54111655 | Ch13 region 2 |
| chr13:54111707-54111747 | Ch13 region 2 |
| chr13:54111771-54111811 | Ch13 region 2 |
| chr13:54111860-54111900 | Ch13 region 2 |
| chr13:54111924-54111964 | Ch13 region 2 |
| chr13:54111969-54112009 | Ch13 region 2 |
| chr13:54112019-54112059 | Ch13 region 2 |
| chr13:54112087-54112127 | Ch13 region 2 |
| chr13:54112160-54112200 | Ch13 region 2 |
| chr13:54112298-54112338 | Ch13 region 2 |
| chr13:54112356-54112396 | Ch13 region 2 |
| chr13:54112542-54112582 | Ch13 region 2 |
| chr13:54112597-54112637 | Ch13 region 2 |
| chr13:54112642-54112682 | Ch13 region 2 |
| chr13:54112701-54112741 | Ch13 region 2 |
| chr13:54112788-54112828 | Ch13 region 2 |
| chr13:54112881-54112921 | Ch13 region 2 |
| chr13:54113002-54113042 | Ch13 region 2 |
| chr13:54113100-54113140 | Ch13 region 2 |
| chr13:54113159-54113199 | Ch13 region 2 |
| chr13:54113221-54113261 | Ch13 region 2 |
| chr13:54113275-54113315 | Ch13 region 2 |
| chr13:54113339-54113379 | Ch13 region 2 |
| chr13:54113425-54113465 | Ch13 region 2 |
| chr13:54113490-54113530 | Ch13 region 2 |
| chr13:54113574-54113614 | Ch13 region 2 |
| chr13:54113819-54113859 | Ch13 region 2 |
| chr13:54113965-54114005 | Ch13 region 2 |
| chr13:54114084-54114124 | Ch13 region 2 |
| chr13:54114153-54114193 | Ch13 region 2 |
| chr13:54114230-54114270 | Ch13 region 2 |
| chr13:54114296-54114336 | Ch13 region 2 |
| chr13:54114341-54114381 | Ch13 region 2 |
| chr13:54114433-54114473 | Ch13 region 2 |
| chr13:54114502-54114542 | Ch13 region 2 |
| chr13:54114572-54114612 | Ch13 region 2 |

| | |
|---|---|
| chr13:54114705-54114745 | Ch13 region 2 |
| chr13:54114763-54114803 | Ch13 region 2 |
| chr13:54114834-54114874 | Ch13 region 2 |
| chr13:54114916-54114956 | Ch13 region 2 |
| chr13:54115002-54115042 | Ch13 region 2 |
| chr13:54115267-54115307 | Ch13 region 2 |
| chr13:54115316-54115356 | Ch13 region 2 |
| chr13:54115380-54115420 | Ch13 region 2 |
| chr13:54115459-54115499 | Ch13 region 2 |
| chr13:54115597-54115637 | Ch13 region 2 |
| chr13:54115647-54115687 | Ch13 region 2 |
| chr13:54115692-54115732 | Ch13 region 2 |
| chr13:54115778-54115818 | Ch13 region 2 |
| chr13:54115823-54115863 | Ch13 region 2 |
| chr13:54115882-54115922 | Ch13 region 2 |
| chr13:54115964-54116004 | Ch13 region 2 |
| chr13:54116016-54116056 | Ch13 region 2 |
| chr13:54116182-54116222 | Ch13 region 2 |
| chr13:54116248-54116288 | Ch13 region 2 |
| chr13:54116333-54116373 | Ch13 region 2 |
| chr13:54116393-54116433 | Ch13 region 2 |
| chr13:54116473-54116513 | Ch13 region 2 |
| chr13:54116548-54116588 | Ch13 region 2 |
| chr13:54116648-54116688 | Ch13 region 2 |
| chr13:54116693-54116733 | Ch13 region 2 |
| chr13:54116755-54116795 | Ch13 region 2 |
| chr13:54116831-54116871 | Ch13 region 2 |
| chr13:54116894-54116934 | Ch13 region 2 |
| chr13:54116943-54116983 | Ch13 region 2 |
| chr13:54117107-54117147 | Ch13 region 2 |
| chr13:54117185-54117225 | Ch13 region 2 |
| chr13:54117266-54117306 | Ch13 region 2 |
| chr13:54117344-54117384 | Ch13 region 2 |
| chr13:54117389-54117429 | Ch13 region 2 |
| chr13:54117437-54117477 | Ch13 region 2 |
| chr13:54117504-54117544 | Ch13 region 2 |
| chr13:54117565-54117605 | Ch13 region 2 |
| chr13:54117612-54117652 | Ch13 region 2 |
| chr13:54117696-54117736 | Ch13 region 2 |
| chr13:54117763-54117803 | Ch13 region 2 |
| chr13:54117820-54117860 | Ch13 region 2 |
| chr13:54117883-54117923 | Ch13 region 2 |
| chr13:54117929-54117969 | Ch13 region 2 |
| chr13:54118077-54118117 | Ch13 region 2 |
| chr13:54118177-54118217 | Ch13 region 2 |
| chr13:54118246-54118286 | Ch13 region 2 |
| chr13:54118329-54118369 | Ch13 region 2 |
| chr13:54118427-54118467 | Ch13 region 2 |
| chr13:54118516-54118556 | Ch13 region 2 |
| chr13:54118567-54118607 | Ch13 region 2 |
| chr13:54118623-54118663 | Ch13 region 2 |
| chr13:54118668-54118708 | Ch13 region 2 |

| | |
|---|---|
| chr13:54118785-54118825 | Ch13 region 2 |
| chr13:54118831-54118871 | Ch13 region 2 |
| chr13:54118910-54118950 | Ch13 region 2 |
| chr13:54118960-54119000 | Ch13 region 2 |
| chr13:54119083-54119123 | Ch13 region 2 |
| chr13:54119267-54119307 | Ch13 region 2 |
| chr13:54119348-54119388 | Ch13 region 2 |
| chr13:54119398-54119438 | Ch13 region 2 |
| chr13:54119472-54119512 | Ch13 region 2 |
| chr13:54119557-54119597 | Ch13 region 2 |
| chr13:54119633-54119673 | Ch13 region 2 |
| chr13:54119708-54119748 | Ch13 region 2 |
| chr13:54119829-54119869 | Ch13 region 2 |
| chr13:54119924-54119964 | Ch13 region 2 |
| chr13:54119982-54120022 | Ch13 region 2 |
| chr13:54120037-54120077 | Ch13 region 2 |
| chr13:54120099-54120139 | Ch13 region 2 |
| chr13:54120172-54120212 | Ch13 region 2 |
| chr13:54120238-54120278 | Ch13 region 2 |
| chr13:54120298-54120338 | Ch13 region 2 |
| chr13:54120381-54120421 | Ch13 region 2 |
| chr13:54120426-54120466 | Ch13 region 2 |
| chr13:54120519-54120559 | Ch13 region 2 |
| chr13:54120599-54120639 | Ch13 region 2 |
| chr13:54120704-54120744 | Ch13 region 2 |
| chr13:54120754-54120794 | Ch13 region 2 |
| chr13:54120972-54121012 | Ch13 region 2 |
| chr13:54121063-54121103 | Ch13 region 2 |
| chr13:54121168-54121208 | Ch13 region 2 |
| chr13:54121277-54121317 | Ch13 region 2 |
| chr13:54121348-54121388 | Ch13 region 2 |
| chr13:54121487-54121527 | Ch13 region 2 |
| chr13:54121541-54121581 | Ch13 region 2 |
| chr13:54121590-54121630 | Ch13 region 2 |
| chr13:54121741-54121781 | Ch13 region 2 |
| chr13:54121905-54121945 | Ch13 region 2 |
| chr13:54121950-54121990 | Ch13 region 2 |
| chr13:54122170-54122210 | Ch13 region 2 |
| chr13:54122225-54122265 | Ch13 region 2 |
| chr13:54122332-54122372 | Ch13 region 2 |
| chr13:54122399-54122439 | Ch13 region 2 |
| chr13:54122614-54122654 | Ch13 region 2 |
| chr13:54122675-54122715 | Ch13 region 2 |
| chr13:54122758-54122798 | Ch13 region 2 |
| chr13:54122937-54122977 | Ch13 region 2 |
| chr13:54123058-54123098 | Ch13 region 2 |
| chr13:54123166-54123206 | Ch13 region 2 |
| chr13:54123386-54123426 | Ch13 region 2 |
| chr13:54123464-54123504 | Ch13 region 2 |
| chr13:54123623-54123663 | Ch13 region 2 |
| chr13:54123668-54123708 | Ch13 region 2 |
| chr13:54123721-54123761 | Ch13 region 2 |

| | |
|---|---|
| chr13:54123772-54123812 | Ch13 region 2 |
| chr13:54123881-54123921 | Ch13 region 2 |
| chr13:54123988-54124028 | Ch13 region 2 |
| chr13:54124052-54124092 | Ch13 region 2 |
| chr13:54124105-54124145 | Ch13 region 2 |
| chr13:54124160-54124200 | Ch13 region 2 |
| chr13:54124238-54124278 | Ch13 region 2 |
| chr13:54124288-54124328 | Ch13 region 2 |
| chr13:54124352-54124392 | Ch13 region 2 |
| chr13:54124433-54124473 | Ch13 region 2 |
| chr13:54124522-54124562 | Ch13 region 2 |
| chr13:54124567-54124607 | Ch13 region 2 |
| chr13:54124646-54124686 | Ch13 region 2 |
| chr13:54124766-54124806 | Ch13 region 2 |
| chr13:54124826-54124866 | Ch13 region 2 |
| chr13:54124918-54124958 | Ch13 region 2 |
| chr13:54124967-54125007 | Ch13 region 2 |
| chr13:54125052-54125092 | Ch13 region 2 |
| chr13:54125101-54125141 | Ch13 region 2 |
| chr13:54125154-54125194 | Ch13 region 2 |
| chr13:54125220-54125260 | Ch13 region 2 |
| chr13:54125286-54125326 | Ch13 region 2 |
| chr13:54125385-54125425 | Ch13 region 2 |
| chr13:54125436-54125476 | Ch13 region 2 |
| chr13:54125495-54125535 | Ch13 region 2 |
| chr13:54125557-54125597 | Ch13 region 2 |
| chr13:54125604-54125644 | Ch13 region 2 |
| chr13:54125664-54125704 | Ch13 region 2 |
| chr13:54125716-54125756 | Ch13 region 2 |
| chr13:54125763-54125803 | Ch13 region 2 |
| chr13:54125870-54125910 | Ch13 region 2 |
| chr13:54125919-54125959 | Ch13 region 2 |
| chr13:54125987-54126027 | Ch13 region 2 |
| chr13:54126035-54126075 | Ch13 region 2 |
| chr13:54126083-54126123 | Ch13 region 2 |
| chr13:54126128-54126168 | Ch13 region 2 |
| chr13:54126193-54126233 | Ch13 region 2 |
| chr13:54126248-54126288 | Ch13 region 2 |
| chr13:54126330-54126370 | Ch13 region 2 |
| chr13:54126430-54126470 | Ch13 region 2 |
| chr13:54126545-54126585 | Ch13 region 2 |
| chr13:54126692-54126732 | Ch13 region 2 |
| chr13:54126785-54126825 | Ch13 region 2 |
| chr13:54126851-54126891 | Ch13 region 2 |
| chr13:54126907-54126947 | Ch13 region 2 |
| chr13:54126960-54127000 | Ch13 region 2 |
| chr13:54127269-54127309 | Ch13 region 2 |
| chr13:54127431-54127471 | Ch13 region 2 |
| chr13:54127674-54127714 | Ch13 region 2 |
| chr13:54127726-54127766 | Ch13 region 2 |
| chr13:54127790-54127830 | Ch13 region 2 |
| chr13:54127857-54127897 | Ch13 region 2 |

| | |
|---|---|
| chr13:54127912-54127952 | Ch13 region 2 |
| chr13:54127975-54128015 | Ch13 region 2 |
| chr13:54128034-54128074 | Ch13 region 2 |
| chr13:54128231-54128271 | Ch13 region 2 |
| chr13:54128407-54128447 | Ch13 region 2 |
| chr13:54128452-54128492 | Ch13 region 2 |
| chr13:54128554-54128594 | Ch13 region 2 |
| chr13:54128751-54128791 | Ch13 region 2 |
| chr13:54128823-54128863 | Ch13 region 2 |
| chr13:54128939-54128979 | Ch13 region 2 |
| chr13:54129162-54129202 | Ch13 region 2 |
| chr13:54129296-54129336 | Ch13 region 2 |
| chr13:54129341-54129381 | Ch13 region 2 |
| chr13:54129413-54129453 | Ch13 region 2 |
| chr13:54129534-54129574 | Ch13 region 2 |
| chr13:54129589-54129629 | Ch13 region 2 |
| chr13:54129661-54129701 | Ch13 region 2 |
| chr13:54129799-54129839 | Ch13 region 2 |
| chr13:54129858-54129898 | Ch13 region 2 |
| chr13:54129966-54130006 | Ch13 region 2 |
| chr13:54130036-54130076 | Ch13 region 2 |
| chr13:54130082-54130122 | Ch13 region 2 |
| chr13:54130139-54130179 | Ch13 region 2 |
| chr13:54130288-54130328 | Ch13 region 2 |
| chr13:54130361-54130401 | Ch13 region 2 |
| chr13:54130436-54130476 | Ch13 region 2 |
| chr13:54130487-54130527 | Ch13 region 2 |
| chr13:54130534-54130574 | Ch13 region 2 |
| chr13:54130613-54130653 | Ch13 region 2 |
| chr13:54130667-54130707 | Ch13 region 2 |
| chr13:54130736-54130776 | Ch13 region 2 |
| chr13:54130813-54130853 | Ch13 region 2 |
| chr13:54130859-54130899 | Ch13 region 2 |
| chr13:54130917-54130957 | Ch13 region 2 |
| chr13:54130992-54131032 | Ch13 region 2 |
| chr13:54131040-54131080 | Ch13 region 2 |
| chr13:54131088-54131128 | Ch13 region 2 |
| chr13:54131179-54131219 | Ch13 region 2 |
| chr13:54131250-54131290 | Ch13 region 2 |
| chr13:54131328-54131368 | Ch13 region 2 |
| chr13:54131374-54131414 | Ch13 region 2 |
| chr13:54131455-54131495 | Ch13 region 2 |
| chr13:54131519-54131559 | Ch13 region 2 |
| chr13:54131564-54131604 | Ch13 region 2 |
| chr13:54131641-54131681 | Ch13 region 2 |
| chr13:54131707-54131747 | Ch13 region 2 |
| chr13:54131753-54131793 | Ch13 region 2 |
| chr13:54131805-54131845 | Ch13 region 2 |
| chr13:54131896-54131936 | Ch13 region 2 |
| chr13:54131941-54131981 | Ch13 region 2 |
| chr13:54132026-54132066 | Ch13 region 2 |
| chr13:54132087-54132127 | Ch13 region 2 |

| | |
|---|---|
| chr13:54132157-54132197 | Ch13 region 2 |
| chr13:54132229-54132269 | Ch13 region 2 |
| chr13:54132304-54132344 | Ch13 region 2 |
| chr13:54132350-54132390 | Ch13 region 2 |
| chr13:54132426-54132466 | Ch13 region 2 |
| chr13:54132489-54132529 | Ch13 region 2 |
| chr13:54132595-54132635 | Ch13 region 2 |
| chr13:54132662-54132702 | Ch13 region 2 |
| chr13:54132790-54132830 | Ch13 region 2 |
| chr13:54132921-54132961 | Ch13 region 2 |
| chr13:54133061-54133101 | Ch13 region 2 |
| chr13:54133147-54133187 | Ch13 region 2 |
| chr13:54133192-54133232 | Ch13 region 2 |
| chr13:54133253-54133293 | Ch13 region 2 |
| chr13:54133363-54133403 | Ch13 region 2 |
| chr13:54133477-54133517 | Ch13 region 2 |
| chr13:54133691-54133731 | Ch13 region 2 |
| chr13:54133768-54133808 | Ch13 region 2 |
| chr13:54133819-54133859 | Ch13 region 2 |
| chr13:54133911-54133951 | Ch13 region 2 |
| chr13:54134304-54134344 | Ch13 region 2 |
| chr13:54134381-54134421 | Ch13 region 2 |
| chr13:54134465-54134505 | Ch13 region 2 |
| chr13:54134577-54134617 | Ch13 region 2 |
| chr13:54134757-54134797 | Ch13 region 2 |
| chr13:54134819-54134859 | Ch13 region 2 |
| chr13:54134864-54134904 | Ch13 region 2 |
| chr13:54135002-54135042 | Ch13 region 2 |
| chr13:54135061-54135101 | Ch13 region 2 |
| chr13:54135106-54135146 | Ch13 region 2 |
| chr13:54135151-54135191 | Ch13 region 2 |
| chr13:54135292-54135332 | Ch13 region 2 |
| chr13:54135455-54135495 | Ch13 region 2 |
| chr13:54135537-54135577 | Ch13 region 2 |
| chr13:54135596-54135636 | Ch13 region 2 |
| chr13:54135726-54135766 | Ch13 region 2 |
| chr13:54135912-54135952 | Ch13 region 2 |
| chr13:54135971-54136011 | Ch13 region 2 |
| chr13:54136116-54136156 | Ch13 region 2 |
| chr13:54136225-54136265 | Ch13 region 2 |
| chr13:54136348-54136388 | Ch13 region 2 |
| chr13:54136393-54136433 | Ch13 region 2 |
| chr13:54136589-54136629 | Ch13 region 2 |
| chr13:54136668-54136708 | Ch13 region 2 |
| chr13:54136756-54136796 | Ch13 region 2 |
| chr13:54136912-54136952 | Ch13 region 2 |
| chr13:54137013-54137053 | Ch13 region 2 |
| chr13:54137092-54137132 | Ch13 region 2 |
| chr13:54137164-54137204 | Ch13 region 2 |
| chr13:54137381-54137421 | Ch13 region 2 |
| chr13:54137512-54137552 | Ch13 region 2 |
| chr13:54137658-54137698 | Ch13 region 2 |

| | |
|---|---|
| chr13:54137787-54137827 | Ch13 region 2 |
| chr13:54137876-54137916 | Ch13 region 2 |
| chr13:54138095-54138135 | Ch13 region 2 |
| chr13:54138183-54138223 | Ch13 region 2 |
| chr13:54138258-54138298 | Ch13 region 2 |
| chr13:54138303-54138343 | Ch13 region 2 |
| chr13:54138350-54138390 | Ch13 region 2 |
| chr13:54138413-54138453 | Ch13 region 2 |
| chr13:54138514-54138554 | Ch13 region 2 |
| chr13:54138559-54138599 | Ch13 region 2 |
| chr13:54138627-54138667 | Ch13 region 2 |
| chr13:54138807-54138847 | Ch13 region 2 |
| chr13:54138901-54138941 | Ch13 region 2 |
| chr13:54139221-54139261 | Ch13 region 2 |
| chr13:54139402-54139442 | Ch13 region 2 |
| chr13:54139462-54139502 | Ch13 region 2 |
| chr13:54139508-54139548 | Ch13 region 2 |
| chr13:54139657-54139697 | Ch13 region 2 |
| chr13:54139703-54139743 | Ch13 region 2 |
| chr13:54139760-54139800 | Ch13 region 2 |
| chr13:54139852-54139892 | Ch13 region 2 |
| chr13:54139944-54139984 | Ch13 region 2 |
| chr13:54140034-54140074 | Ch13 region 2 |
| chr13:54140107-54140147 | Ch13 region 2 |
| chr13:54140316-54140356 | Ch13 region 2 |
| chr13:54140361-54140401 | Ch13 region 2 |
| chr13:54140546-54140586 | Ch13 region 2 |
| chr13:54140591-54140631 | Ch13 region 2 |
| chr13:54140644-54140684 | Ch13 region 2 |
| chr13:54140705-54140745 | Ch13 region 2 |
| chr13:54140787-54140827 | Ch13 region 2 |
| chr13:54140940-54140980 | Ch13 region 2 |
| chr13:54141007-54141047 | Ch13 region 2 |
| chr13:54141121-54141161 | Ch13 region 2 |
| chr13:54141173-54141213 | Ch13 region 2 |
| chr13:54141248-54141288 | Ch13 region 2 |
| chr13:54141293-54141333 | Ch13 region 2 |
| chr13:54141351-54141391 | Ch13 region 2 |
| chr13:54141396-54141436 | Ch13 region 2 |
| chr13:54141452-54141492 | Ch13 region 2 |
| chr13:54141622-54141662 | Ch13 region 2 |
| chr13:54141691-54141731 | Ch13 region 2 |
| chr13:54141760-54141800 | Ch13 region 2 |
| chr13:54141835-54141875 | Ch13 region 2 |
| chr13:54141914-54141954 | Ch13 region 2 |
| chr13:54141981-54142021 | Ch13 region 2 |
| chr13:54142029-54142069 | Ch13 region 2 |
| chr13:54142111-54142151 | Ch13 region 2 |
| chr13:54142190-54142230 | Ch13 region 2 |
| chr13:54142263-54142303 | Ch13 region 2 |
| chr13:54142308-54142348 | Ch13 region 2 |
| chr13:54142366-54142406 | Ch13 region 2 |

| | |
|---|---|
| chr13:54142411-54142451 | Ch13 region 2 |
| chr13:54142468-54142508 | Ch13 region 2 |
| chr13:54142515-54142555 | Ch13 region 2 |
| chr13:54142582-54142622 | Ch13 region 2 |
| chr13:54142630-54142670 | Ch13 region 2 |
| chr13:54142710-54142750 | Ch13 region 2 |
| chr13:54142800-54142840 | Ch13 region 2 |
| chr13:54142870-54142910 | Ch13 region 2 |
| chr13:54142921-54142961 | Ch13 region 2 |
| chr13:54142968-54143008 | Ch13 region 2 |
| chr13:54143024-54143064 | Ch13 region 2 |
| chr13:54143069-54143109 | Ch13 region 2 |
| chr13:54143115-54143155 | Ch13 region 2 |
| chr13:54143194-54143234 | Ch13 region 2 |
| chr13:54143247-54143287 | Ch13 region 2 |
| chr13:54143314-54143354 | Ch13 region 2 |
| chr13:54143488-54143528 | Ch13 region 2 |
| chr13:54143593-54143633 | Ch13 region 2 |
| chr13:54143655-54143695 | Ch13 region 2 |
| chr13:54143720-54143760 | Ch13 region 2 |
| chr13:54143783-54143823 | Ch13 region 2 |
| chr13:54143883-54143923 | Ch13 region 2 |
| chr13:54143946-54143986 | Ch13 region 2 |
| chr13:54144002-54144042 | Ch13 region 2 |
| chr13:54144047-54144087 | Ch13 region 2 |
| chr13:54144092-54144132 | Ch13 region 2 |
| chr13:54144151-54144191 | Ch13 region 2 |
| chr13:54144230-54144270 | Ch13 region 2 |
| chr13:54144287-54144327 | Ch13 region 2 |
| chr13:54144345-54144385 | Ch13 region 2 |
| chr13:54144394-54144434 | Ch13 region 2 |
| chr13:54144444-54144484 | Ch13 region 2 |
| chr13:54144526-54144566 | Ch13 region 2 |
| chr13:54144611-54144651 | Ch13 region 2 |
| chr13:54144672-54144712 | Ch13 region 2 |
| chr13:54144736-54144776 | Ch13 region 2 |
| chr13:54144860-54144900 | Ch13 region 2 |
| chr13:54144927-54144967 | Ch13 region 2 |
| chr13:54144975-54145015 | Ch13 region 2 |
| chr13:54145064-54145104 | Ch13 region 2 |
| chr13:54145135-54145175 | Ch13 region 2 |
| chr13:54145216-54145256 | Ch13 region 2 |
| chr13:54145315-54145355 | Ch13 region 2 |
| chr13:54145371-54145411 | Ch13 region 2 |
| chr13:54145476-54145516 | Ch13 region 2 |
| chr13:54145576-54145616 | Ch13 region 2 |
| chr13:54145637-54145677 | Ch13 region 2 |
| chr13:54145721-54145761 | Ch13 region 2 |
| chr13:54145786-54145826 | Ch13 region 2 |
| chr13:54145831-54145871 | Ch13 region 2 |
| chr13:54145918-54145958 | Ch13 region 2 |
| chr13:54146111-54146151 | Ch13 region 2 |

| | |
|---|---|
| chr13:54146158-54146198 | Ch13 region 2 |
| chr13:54146245-54146285 | Ch13 region 2 |
| chr13:54146295-54146335 | Ch13 region 2 |
| chr13:54146447-54146487 | Ch13 region 2 |
| chr13:54146614-54146654 | Ch13 region 2 |
| chr13:54146705-54146745 | Ch13 region 2 |
| chr13:54146777-54146817 | Ch13 region 2 |
| chr13:54146860-54146900 | Ch13 region 2 |
| chr13:54146931-54146971 | Ch13 region 2 |
| chr13:54146986-54147026 | Ch13 region 2 |
| chr13:54147058-54147098 | Ch13 region 2 |
| chr13:54147111-54147151 | Ch13 region 2 |
| chr13:54147214-54147254 | Ch13 region 2 |
| chr13:54147289-54147329 | Ch13 region 2 |
| chr13:54147352-54147392 | Ch13 region 2 |
| chr13:54147412-54147452 | Ch13 region 2 |
| chr13:54147504-54147544 | Ch13 region 2 |
| chr13:54147556-54147596 | Ch13 region 2 |
| chr13:54147613-54147653 | Ch13 region 2 |
| chr13:54147720-54147760 | Ch13 region 2 |
| chr13:54147774-54147814 | Ch13 region 2 |
| chr13:54147820-54147860 | Ch13 region 2 |
| chr13:54147921-54147961 | Ch13 region 2 |
| chr13:54147967-54148007 | Ch13 region 2 |
| chr13:54148026-54148066 | Ch13 region 2 |
| chr13:54148182-54148222 | Ch13 region 2 |
| chr13:54148253-54148293 | Ch13 region 2 |
| chr13:54148367-54148407 | Ch13 region 2 |
| chr13:54148444-54148484 | Ch13 region 2 |
| chr13:54148529-54148569 | Ch13 region 2 |
| chr13:54148603-54148643 | Ch13 region 2 |
| chr13:54148662-54148702 | Ch13 region 2 |
| chr13:54148761-54148801 | Ch13 region 2 |
| chr13:54148981-54149021 | Ch13 region 2 |
| chr13:54149074-54149114 | Ch13 region 2 |
| chr13:54149180-54149220 | Ch13 region 2 |
| chr13:54149244-54149284 | Ch13 region 2 |
| chr13:54149300-54149340 | Ch13 region 2 |
| chr13:54149346-54149386 | Ch13 region 2 |
| chr13:54149393-54149433 | Ch13 region 2 |
| chr13:54149459-54149499 | Ch13 region 2 |
| chr13:54149574-54149614 | Ch13 region 2 |
| chr13:54149638-54149678 | Ch13 region 2 |
| chr13:54149704-54149744 | Ch13 region 2 |
| chr13:54149760-54149800 | Ch13 region 2 |
| chr13:54149825-54149865 | Ch13 region 2 |
| chr13:54149894-54149934 | Ch13 region 2 |
| chr13:54149939-54149979 | Ch13 region 2 |
| chr13:54150015-54150055 | Ch13 region 2 |
| chr13:54150131-54150171 | Ch13 region 2 |
| chr13:54150198-54150238 | Ch13 region 2 |
| chr13:54150287-54150327 | Ch13 region 2 |

| | |
|---|---|
| chr13:54150355-54150395 | Ch13 region 2 |
| chr13:54150405-54150445 | Ch13 region 2 |
| chr13:54150520-54150560 | Ch13 region 2 |
| chr13:54150567-54150607 | Ch13 region 2 |
| chr13:54150637-54150677 | Ch13 region 2 |
| chr13:54150685-54150725 | Ch13 region 2 |
| chr13:54150756-54150796 | Ch13 region 2 |
| chr13:54150849-54150889 | Ch13 region 2 |
| chr13:54150917-54150957 | Ch13 region 2 |
| chr13:54150988-54151028 | Ch13 region 2 |
| chr13:54151193-54151233 | Ch13 region 2 |
| chr13:54151294-54151334 | Ch13 region 2 |
| chr13:54151391-54151431 | Ch13 region 2 |
| chr13:54151455-54151495 | Ch13 region 2 |
| chr13:54151503-54151543 | Ch13 region 2 |
| chr13:54151565-54151605 | Ch13 region 2 |
| chr13:54151638-54151678 | Ch13 region 2 |
| chr13:54151707-54151747 | Ch13 region 2 |
| chr13:54151825-54151865 | Ch13 region 2 |
| chr13:54151924-54151964 | Ch13 region 2 |
| chr13:54151970-54152010 | Ch13 region 2 |
| chr13:54152042-54152082 | Ch13 region 2 |
| chr13:54152087-54152127 | Ch13 region 2 |
| chr13:54152134-54152174 | Ch13 region 2 |
| chr13:54152193-54152233 | Ch13 region 2 |
| chr13:54152258-54152298 | Ch13 region 2 |
| chr13:54152317-54152357 | Ch13 region 2 |
| chr13:54152374-54152414 | Ch13 region 2 |
| chr13:54152492-54152532 | Ch13 region 2 |
| chr13:54152551-54152591 | Ch13 region 2 |
| chr13:54152601-54152641 | Ch13 region 2 |
| chr13:54152687-54152727 | Ch13 region 2 |
| chr13:54152772-54152812 | Ch13 region 2 |
| chr13:54152817-54152857 | Ch13 region 2 |
| chr13:54152915-54152955 | Ch13 region 2 |
| chr13:54153013-54153053 | Ch13 region 2 |
| chr13:54153066-54153106 | Ch13 region 2 |
| chr13:54153172-54153212 | Ch13 region 2 |
| chr13:54153232-54153272 | Ch13 region 2 |
| chr13:54153300-54153340 | Ch13 region 2 |
| chr13:54153374-54153414 | Ch13 region 2 |
| chr13:54153427-54153467 | Ch13 region 2 |
| chr13:54153506-54153546 | Ch13 region 2 |
| chr13:54153576-54153616 | Ch13 region 2 |
| chr13:54153621-54153661 | Ch13 region 2 |
| chr13:54153722-54153762 | Ch13 region 2 |
| chr13:54153789-54153829 | Ch13 region 2 |
| chr13:54153881-54153921 | Ch13 region 2 |
| chr13:54153959-54153999 | Ch13 region 2 |
| chr13:54154029-54154069 | Ch13 region 2 |
| chr13:54154092-54154132 | Ch13 region 2 |
| chr13:54154170-54154210 | Ch13 region 2 |

| | |
|---|---|
| chr13:54154228-54154268 | Ch13 region 2 |
| chr13:54154330-54154370 | Ch13 region 2 |
| chr13:54154389-54154429 | Ch13 region 2 |
| chr13:54154449-54154489 | Ch13 region 2 |
| chr13:54154520-54154560 | Ch13 region 2 |
| chr13:54154671-54154711 | Ch13 region 2 |
| chr13:54154815-54154855 | Ch13 region 2 |
| chr13:54154901-54154941 | Ch13 region 2 |
| chr13:54155033-54155073 | Ch13 region 2 |
| chr13:54155114-54155154 | Ch13 region 2 |
| chr13:54155228-54155268 | Ch13 region 2 |
| chr13:54155324-54155364 | Ch13 region 2 |
| chr13:54155424-54155464 | Ch13 region 2 |
| chr13:54155520-54155560 | Ch13 region 2 |
| chr13:54155601-54155641 | Ch13 region 2 |
| chr13:54155852-54155892 | Ch13 region 2 |
| chr13:54155921-54155961 | Ch13 region 2 |
| chr13:54155967-54156007 | Ch13 region 2 |
| chr13:54156117-54156157 | Ch13 region 2 |
| chr13:54156198-54156238 | Ch13 region 2 |
| chr13:54156263-54156303 | Ch13 region 2 |
| chr13:54156308-54156348 | Ch13 region 2 |
| chr13:54156377-54156417 | Ch13 region 2 |
| chr13:54156522-54156562 | Ch13 region 2 |
| chr13:54156686-54156726 | Ch13 region 2 |
| chr13:54156753-54156793 | Ch13 region 2 |
| chr13:54156822-54156862 | Ch13 region 2 |
| chr13:54156938-54156978 | Ch13 region 2 |
| chr13:54156997-54157037 | Ch13 region 2 |
| chr13:54157097-54157137 | Ch13 region 2 |
| chr13:54157317-54157357 | Ch13 region 2 |
| chr13:54157448-54157488 | Ch13 region 2 |
| chr13:54157536-54157576 | Ch13 region 2 |
| chr13:54157599-54157639 | Ch13 region 2 |
| chr13:54157659-54157699 | Ch13 region 2 |
| chr13:54157704-54157744 | Ch13 region 2 |
| chr13:54157930-54157970 | Ch13 region 2 |
| chr13:54157989-54158029 | Ch13 region 2 |
| chr13:54158056-54158096 | Ch13 region 2 |
| chr13:54158260-54158300 | Ch13 region 2 |
| chr13:54158305-54158345 | Ch13 region 2 |
| chr13:54158359-54158399 | Ch13 region 2 |
| chr13:54158613-54158653 | Ch13 region 2 |
| chr13:54158658-54158698 | Ch13 region 2 |
| chr13:54158755-54158795 | Ch13 region 2 |
| chr13:54158806-54158846 | Ch13 region 2 |
| chr13:54158873-54158913 | Ch13 region 2 |
| chr13:54158940-54158980 | Ch13 region 2 |
| chr13:54159083-54159123 | Ch13 region 2 |
| chr13:54159243-54159283 | Ch13 region 2 |
| chr13:54159373-54159413 | Ch13 region 2 |
| chr13:54159519-54159559 | Ch13 region 2 |

| | |
|---|---|
| chr13:54159574-54159614 | Ch13 region 2 |
| chr13:54159637-54159677 | Ch13 region 2 |
| chr13:54159695-54159735 | Ch13 region 2 |
| chr13:54159797-54159837 | Ch13 region 2 |
| chr13:54159855-54159895 | Ch13 region 2 |
| chr13:54159901-54159941 | Ch13 region 2 |
| chr13:54159980-54160020 | Ch13 region 2 |
| chr13:54160033-54160073 | Ch13 region 2 |
| chr13:54160086-54160126 | Ch13 region 2 |
| chr13:54160131-54160171 | Ch13 region 2 |
| chr13:54160208-54160248 | Ch13 region 2 |
| chr13:54160260-54160300 | Ch13 region 2 |
| chr13:54160390-54160430 | Ch13 region 2 |
| chr13:54160435-54160475 | Ch13 region 2 |
| chr13:54160590-54160630 | Ch13 region 2 |
| chr13:54160662-54160702 | Ch13 region 2 |
| chr13:54160775-54160815 | Ch13 region 2 |
| chr13:54160834-54160874 | Ch13 region 2 |
| chr13:54160883-54160923 | Ch13 region 2 |
| chr13:54160947-54160987 | Ch13 region 2 |
| chr13:54161012-54161052 | Ch13 region 2 |
| chr13:54161065-54161105 | Ch13 region 2 |
| chr13:54161328-54161368 | Ch13 region 2 |
| chr13:54161561-54161601 | Ch13 region 2 |
| chr13:54161682-54161722 | Ch13 region 2 |
| chr13:54161727-54161767 | Ch13 region 2 |
| chr13:54161835-54161875 | Ch13 region 2 |
| chr13:54161900-54161940 | Ch13 region 2 |
| chr13:54161949-54161989 | Ch13 region 2 |
| chr13:54162042-54162082 | Ch13 region 2 |
| chr13:54162087-54162127 | Ch13 region 2 |
| chr13:54162145-54162185 | Ch13 region 2 |
| chr13:54162234-54162274 | Ch13 region 2 |
| chr13:54162284-54162324 | Ch13 region 2 |
| chr13:54162341-54162381 | Ch13 region 2 |
| chr13:54162388-54162428 | Ch13 region 2 |
| chr13:54162460-54162500 | Ch13 region 2 |
| chr13:54162535-54162575 | Ch13 region 2 |
| chr13:54162592-54162632 | Ch13 region 2 |
| chr13:54162661-54162701 | Ch13 region 2 |
| chr13:54162708-54162748 | Ch13 region 2 |
| chr13:54162753-54162793 | Ch13 region 2 |
| chr13:54162824-54162864 | Ch13 region 2 |
| chr13:54162869-54162909 | Ch13 region 3 |
| chr13:77193633-77193673 | Ch13 region 3 |
| chr13:77193678-77193718 | Ch13 region 3 |
| chr13:77193742-77193782 | Ch13 region 3 |
| chr13:77193811-77193851 | Ch13 region 3 |
| chr13:77193870-77193910 | Ch13 region 3 |
| chr13:77193929-77193969 | Ch13 region 3 |
| chr13:77193984-77194024 | Ch13 region 3 |
| chr13:77194059-77194099 | Ch13 region 3 |

| | |
|---|---|
| chr13:77194138-77194178 | Ch13 region 3 |
| chr13:77194187-77194227 | Ch13 region 3 |
| chr13:77194276-77194316 | Ch13 region 3 |
| chr13:77194341-77194381 | Ch13 region 3 |
| chr13:77194425-77194465 | Ch13 region 3 |
| chr13:77194556-77194596 | Ch13 region 3 |
| chr13:77194601-77194641 | Ch13 region 3 |
| chr13:77194655-77194695 | Ch13 region 3 |
| chr13:77194709-77194749 | Ch13 region 3 |
| chr13:77194800-77194840 | Ch13 region 3 |
| chr13:77194847-77194887 | Ch13 region 3 |
| chr13:77194914-77194954 | Ch13 region 3 |
| chr13:77194959-77194999 | Ch13 region 3 |
| chr13:77195027-77195067 | Ch13 region 3 |
| chr13:77195094-77195134 | Ch13 region 3 |
| chr13:77195143-77195183 | Ch13 region 3 |
| chr13:77195216-77195256 | Ch13 region 3 |
| chr13:77195266-77195306 | Ch13 region 3 |
| chr13:77195319-77195359 | Ch13 region 3 |
| chr13:77195394-77195434 | Ch13 region 3 |
| chr13:77195451-77195491 | Ch13 region 3 |
| chr13:77195514-77195554 | Ch13 region 3 |
| chr13:77195559-77195599 | Ch13 region 3 |
| chr13:77195618-77195658 | Ch13 region 3 |
| chr13:77195826-77195866 | Ch13 region 3 |
| chr13:77195907-77195947 | Ch13 region 3 |
| chr13:77195965-77196005 | Ch13 region 3 |
| chr13:77196044-77196084 | Ch13 region 3 |
| chr13:77196148-77196188 | Ch13 region 3 |
| chr13:77196193-77196233 | Ch13 region 3 |
| chr13:77196437-77196477 | Ch13 region 3 |
| chr13:77196520-77196560 | Ch13 region 3 |
| chr13:77196596-77196636 | Ch13 region 3 |
| chr13:77196648-77196688 | Ch13 region 3 |
| chr13:77196710-77196750 | Ch13 region 3 |
| chr13:77196766-77196806 | Ch13 region 3 |
| chr13:77196852-77196892 | Ch13 region 3 |
| chr13:77196910-77196950 | Ch13 region 3 |
| chr13:77197010-77197050 | Ch13 region 3 |
| chr13:77197077-77197117 | Ch13 region 3 |
| chr13:77197133-77197173 | Ch13 region 3 |
| chr13:77197218-77197258 | Ch13 region 3 |
| chr13:77197276-77197316 | Ch13 region 3 |
| chr13:77197325-77197365 | Ch13 region 3 |
| chr13:77197387-77197427 | Ch13 region 3 |
| chr13:77197444-77197484 | Ch13 region 3 |
| chr13:77197493-77197533 | Ch13 region 3 |
| chr13:77197560-77197600 | Ch13 region 3 |
| chr13:77197620-77197660 | Ch13 region 3 |
| chr13:77197681-77197721 | Ch13 region 3 |
| chr13:77197741-77197781 | Ch13 region 3 |
| chr13:77197828-77197868 | Ch13 region 3 |

| | |
|---|---|
| chr13:77197889-77197929 | Ch13 region 3 |
| chr13:77197952-77197992 | Ch13 region 3 |
| chr13:77198005-77198045 | Ch13 region 3 |
| chr13:77198071-77198111 | Ch13 region 3 |
| chr13:77198182-77198222 | Ch13 region 3 |
| chr13:77198230-77198270 | Ch13 region 3 |
| chr13:77198293-77198333 | Ch13 region 3 |
| chr13:77198388-77198428 | Ch13 region 3 |
| chr13:77198433-77198473 | Ch13 region 3 |
| chr13:77198505-77198545 | Ch13 region 3 |
| chr13:77198615-77198655 | Ch13 region 3 |
| chr13:77198667-77198707 | Ch13 region 3 |
| chr13:77198807-77198847 | Ch13 region 3 |
| chr13:77199033-77199073 | Ch13 region 3 |
| chr13:77199078-77199118 | Ch13 region 3 |
| chr13:77199175-77199215 | Ch13 region 3 |
| chr13:77199292-77199332 | Ch13 region 3 |
| chr13:77199367-77199407 | Ch13 region 3 |
| chr13:77199550-77199590 | Ch13 region 3 |
| chr13:77199600-77199640 | Ch13 region 3 |
| chr13:77199683-77199723 | Ch13 region 3 |
| chr13:77199773-77199813 | Ch13 region 3 |
| chr13:77199853-77199893 | Ch13 region 3 |
| chr13:77199919-77199959 | Ch13 region 3 |
| chr13:77199965-77200005 | Ch13 region 3 |
| chr13:77200027-77200067 | Ch13 region 3 |
| chr13:77200083-77200123 | Ch13 region 3 |
| chr13:77200202-77200242 | Ch13 region 3 |
| chr13:77200363-77200403 | Ch13 region 3 |
| chr13:77200414-77200454 | Ch13 region 3 |
| chr13:77200475-77200515 | Ch13 region 3 |
| chr13:77200538-77200578 | Ch13 region 3 |
| chr13:77200588-77200628 | Ch13 region 3 |
| chr13:77200637-77200677 | Ch13 region 3 |
| chr13:77200682-77200722 | Ch13 region 3 |
| chr13:77200732-77200772 | Ch13 region 3 |
| chr13:77200822-77200862 | Ch13 region 3 |
| chr13:77200869-77200909 | Ch13 region 3 |
| chr13:77200924-77200964 | Ch13 region 3 |
| chr13:77201009-77201049 | Ch13 region 3 |
| chr13:77201055-77201095 | Ch13 region 3 |
| chr13:77201125-77201165 | Ch13 region 3 |
| chr13:77201170-77201210 | Ch13 region 3 |
| chr13:77201231-77201271 | Ch13 region 3 |
| chr13:77201311-77201351 | Ch13 region 3 |
| chr13:77201371-77201411 | Ch13 region 3 |
| chr13:77201440-77201480 | Ch13 region 3 |
| chr13:77201491-77201531 | Ch13 region 3 |
| chr13:77201561-77201601 | Ch13 region 3 |
| chr13:77201634-77201674 | Ch13 region 3 |
| chr13:77201706-77201746 | Ch13 region 3 |
| chr13:77201754-77201794 | Ch13 region 3 |

| | |
|---|---|
| chr13:77201799-77201839 | Ch13 region 3 |
| chr13:77201856-77201896 | Ch13 region 3 |
| chr13:77201925-77201965 | Ch13 region 3 |
| chr13:77202007-77202047 | Ch13 region 3 |
| chr13:77202053-77202093 | Ch13 region 3 |
| chr13:77202110-77202150 | Ch13 region 3 |
| chr13:77202178-77202218 | Ch13 region 3 |
| chr13:77202271-77202311 | Ch13 region 3 |
| chr13:77202460-77202500 | Ch13 region 3 |
| chr13:77202566-77202606 | Ch13 region 3 |
| chr13:77202616-77202656 | Ch13 region 3 |
| chr13:77202814-77202854 | Ch13 region 3 |
| chr13:77202920-77202960 | Ch13 region 3 |
| chr13:77202993-77203033 | Ch13 region 3 |
| chr13:77203066-77203106 | Ch13 region 3 |
| chr13:77203258-77203298 | Ch13 region 3 |
| chr13:77203310-77203350 | Ch13 region 3 |
| chr13:77203367-77203407 | Ch13 region 3 |
| chr13:77203538-77203578 | Ch13 region 3 |
| chr13:77203648-77203688 | Ch13 region 3 |
| chr13:77203701-77203741 | Ch13 region 3 |
| chr13:77203756-77203796 | Ch13 region 3 |
| chr13:77203802-77203842 | Ch13 region 3 |
| chr13:77203865-77203905 | Ch13 region 3 |
| chr13:77203919-77203959 | Ch13 region 3 |
| chr13:77203999-77204039 | Ch13 region 3 |
| chr13:77204132-77204172 | Ch13 region 3 |
| chr13:77204228-77204268 | Ch13 region 3 |
| chr13:77204289-77204329 | Ch13 region 3 |
| chr13:77204370-77204410 | Ch13 region 3 |
| chr13:77204471-77204511 | Ch13 region 3 |
| chr13:77204535-77204575 | Ch13 region 3 |
| chr13:77204602-77204642 | Ch13 region 3 |
| chr13:77204690-77204730 | Ch13 region 3 |
| chr13:77204741-77204781 | Ch13 region 3 |
| chr13:77204854-77204894 | Ch13 region 3 |
| chr13:77204911-77204951 | Ch13 region 3 |
| chr13:77204978-77205018 | Ch13 region 3 |
| chr13:77205047-77205087 | Ch13 region 3 |
| chr13:77205155-77205195 | Ch13 region 3 |
| chr13:77205205-77205245 | Ch13 region 3 |
| chr13:77205259-77205299 | Ch13 region 3 |
| chr13:77205307-77205347 | Ch13 region 3 |
| chr13:77205361-77205401 | Ch13 region 3 |
| chr13:77205420-77205460 | Ch13 region 3 |
| chr13:77205493-77205533 | Ch13 region 3 |
| chr13:77205585-77205625 | Ch13 region 3 |
| chr13:77205630-77205670 | Ch13 region 3 |
| chr13:77205683-77205723 | Ch13 region 3 |
| chr13:77205758-77205798 | Ch13 region 3 |
| chr13:77205847-77205887 | Ch13 region 3 |
| chr13:77205892-77205932 | Ch13 region 3 |

| | |
|---|---|
| chr13:77205998-77206038 | Ch13 region 3 |
| chr13:77206049-77206089 | Ch13 region 3 |
| chr13:77206135-77206175 | Ch13 region 3 |
| chr13:77206180-77206220 | Ch13 region 3 |
| chr13:77206252-77206292 | Ch13 region 3 |
| chr13:77206297-77206337 | Ch13 region 3 |
| chr13:77206379-77206419 | Ch13 region 3 |
| chr13:77206477-77206517 | Ch13 region 3 |
| chr13:77206644-77206684 | Ch13 region 3 |
| chr13:77206699-77206739 | Ch13 region 3 |
| chr13:77206775-77206815 | Ch13 region 3 |
| chr13:77206878-77206918 | Ch13 region 3 |
| chr13:77206925-77206965 | Ch13 region 3 |
| chr13:77206971-77207011 | Ch13 region 3 |
| chr13:77207049-77207089 | Ch13 region 3 |
| chr13:77207103-77207143 | Ch13 region 3 |
| chr13:77207209-77207249 | Ch13 region 3 |
| chr13:77207335-77207375 | Ch13 region 3 |
| chr13:77207434-77207474 | Ch13 region 3 |
| chr13:77207535-77207575 | Ch13 region 3 |
| chr13:77207718-77207758 | Ch13 region 3 |
| chr13:77207776-77207816 | Ch13 region 3 |
| chr13:77207821-77207861 | Ch13 region 3 |
| chr13:77207937-77207977 | Ch13 region 3 |
| chr13:77207986-77208026 | Ch13 region 3 |
| chr13:77208031-77208071 | Ch13 region 3 |
| chr13:77208086-77208126 | Ch13 region 3 |
| chr13:77208169-77208209 | Ch13 region 3 |
| chr13:77208225-77208265 | Ch13 region 3 |
| chr13:77208282-77208322 | Ch13 region 3 |
| chr13:77208390-77208430 | Ch13 region 3 |
| chr13:77208438-77208478 | Ch13 region 3 |
| chr13:77208488-77208528 | Ch13 region 3 |
| chr13:77208538-77208578 | Ch13 region 3 |
| chr13:77208617-77208657 | Ch13 region 3 |
| chr13:77208662-77208702 | Ch13 region 3 |
| chr13:77208716-77208756 | Ch13 region 3 |
| chr13:77208778-77208818 | Ch13 region 3 |
| chr13:77208831-77208871 | Ch13 region 3 |
| chr13:77208894-77208934 | Ch13 region 3 |
| chr13:77208941-77208981 | Ch13 region 3 |
| chr13:77209028-77209068 | Ch13 region 3 |
| chr13:77209085-77209125 | Ch13 region 3 |
| chr13:77209140-77209180 | Ch13 region 3 |
| chr13:77209196-77209236 | Ch13 region 3 |
| chr13:77209257-77209297 | Ch13 region 3 |
| chr13:77209325-77209365 | Ch13 region 3 |
| chr13:77209410-77209450 | Ch13 region 3 |
| chr13:77209488-77209528 | Ch13 region 3 |
| chr13:77209572-77209612 | Ch13 region 3 |
| chr13:77209659-77209699 | Ch13 region 3 |
| chr13:77209710-77209750 | Ch13 region 3 |

| | |
|---|---|
| chr13:77209763-77209803 | Ch13 region 3 |
| chr13:77209825-77209865 | Ch13 region 3 |
| chr13:77209892-77209932 | Ch13 region 3 |
| chr13:77209937-77209977 | Ch13 region 3 |
| chr13:77210008-77210048 | Ch13 region 3 |
| chr13:77210084-77210124 | Ch13 region 3 |
| chr13:77210131-77210171 | Ch13 region 3 |
| chr13:77210200-77210240 | Ch13 region 3 |
| chr13:77210289-77210329 | Ch13 region 3 |
| chr13:77210353-77210393 | Ch13 region 3 |
| chr13:77210442-77210482 | Ch13 region 3 |
| chr13:77210492-77210532 | Ch13 region 3 |
| chr13:77210537-77210577 | Ch13 region 3 |
| chr13:77210582-77210622 | Ch13 region 3 |
| chr13:77210627-77210667 | Ch13 region 3 |
| chr13:77210672-77210712 | Ch13 region 3 |
| chr13:77210740-77210780 | Ch13 region 3 |
| chr13:77210826-77210866 | Ch13 region 3 |
| chr13:77210871-77210911 | Ch13 region 3 |
| chr13:77210946-77210986 | Ch13 region 3 |
| chr13:77211016-77211056 | Ch13 region 3 |
| chr13:77211061-77211101 | Ch13 region 3 |
| chr13:77211143-77211183 | Ch13 region 3 |
| chr13:77211192-77211232 | Ch13 region 3 |
| chr13:77211321-77211361 | Ch13 region 3 |
| chr13:77211405-77211445 | Ch13 region 3 |
| chr13:77211521-77211561 | Ch13 region 3 |
| chr13:77211570-77211610 | Ch13 region 3 |
| chr13:77211661-77211701 | Ch13 region 3 |
| chr13:77211709-77211749 | Ch13 region 3 |
| chr13:77211769-77211809 | Ch13 region 3 |
| chr13:77211815-77211855 | Ch13 region 3 |
| chr13:77211875-77211915 | Ch13 region 3 |
| chr13:77211954-77211994 | Ch13 region 3 |
| chr13:77212000-77212040 | Ch13 region 3 |
| chr13:77212150-77212190 | Ch13 region 3 |
| chr13:77212231-77212271 | Ch13 region 3 |
| chr13:77212329-77212369 | Ch13 region 3 |
| chr13:77212403-77212443 | Ch13 region 3 |
| chr13:77212479-77212519 | Ch13 region 3 |
| chr13:77212535-77212575 | Ch13 region 3 |
| chr13:77212616-77212656 | Ch13 region 3 |
| chr13:77212664-77212704 | Ch13 region 3 |
| chr13:77212709-77212749 | Ch13 region 3 |
| chr13:77212796-77212836 | Ch13 region 3 |
| chr13:77212855-77212895 | Ch13 region 3 |
| chr13:77212912-77212952 | Ch13 region 3 |
| chr13:77212984-77213024 | Ch13 region 3 |
| chr13:77213072-77213112 | Ch13 region 3 |
| chr13:77213126-77213166 | Ch13 region 3 |
| chr13:77213184-77213224 | Ch13 region 3 |
| chr13:77213235-77213275 | Ch13 region 3 |

| | |
|---|---|
| chr13:77213312-77213352 | Ch13 region 3 |
| chr13:77213357-77213397 | Ch13 region 3 |
| chr13:77213425-77213465 | Ch13 region 3 |
| chr13:77213520-77213560 | Ch13 region 3 |
| chr13:77213607-77213647 | Ch13 region 3 |
| chr13:77213662-77213702 | Ch13 region 3 |
| chr13:77213753-77213793 | Ch13 region 3 |
| chr13:77213810-77213850 | Ch13 region 3 |
| chr13:77213871-77213911 | Ch13 region 3 |
| chr13:77213919-77213959 | Ch13 region 3 |
| chr13:77214000-77214040 | Ch13 region 3 |
| chr13:77214048-77214088 | Ch13 region 3 |
| chr13:77214146-77214186 | Ch13 region 3 |
| chr13:77214198-77214238 | Ch13 region 3 |
| chr13:77214270-77214310 | Ch13 region 3 |
| chr13:77214322-77214362 | Ch13 region 3 |
| chr13:77214381-77214421 | Ch13 region 3 |
| chr13:77214438-77214478 | Ch13 region 3 |
| chr13:77214595-77214635 | Ch13 region 3 |
| chr13:77214642-77214682 | Ch13 region 3 |
| chr13:77214713-77214753 | Ch13 region 3 |
| chr13:77214762-77214802 | Ch13 region 3 |
| chr13:77214924-77214964 | Ch13 region 3 |
| chr13:77215025-77215065 | Ch13 region 3 |
| chr13:77215075-77215115 | Ch13 region 3 |
| chr13:77215237-77215277 | Ch13 region 3 |
| chr13:77215302-77215342 | Ch13 region 3 |
| chr13:77215396-77215436 | Ch13 region 3 |
| chr13:77215441-77215481 | Ch13 region 3 |
| chr13:77215577-77215617 | Ch13 region 3 |
| chr13:77215623-77215663 | Ch13 region 3 |
| chr13:77215675-77215715 | Ch13 region 3 |
| chr13:77215721-77215761 | Ch13 region 3 |
| chr13:77215814-77215854 | Ch13 region 3 |
| chr13:77215878-77215918 | Ch13 region 3 |
| chr13:77215927-77215967 | Ch13 region 3 |
| chr13:77215972-77216012 | Ch13 region 3 |
| chr13:77216062-77216102 | Ch13 region 3 |
| chr13:77216166-77216206 | Ch13 region 3 |
| chr13:77216328-77216368 | Ch13 region 3 |
| chr13:77216420-77216460 | Ch13 region 3 |
| chr13:77216511-77216551 | Ch13 region 3 |
| chr13:77216586-77216626 | Ch13 region 3 |
| chr13:77216656-77216696 | Ch13 region 3 |
| chr13:77216736-77216776 | Ch13 region 3 |
| chr13:77216791-77216831 | Ch13 region 3 |
| chr13:77216930-77216970 | Ch13 region 3 |
| chr13:77216975-77217015 | Ch13 region 3 |
| chr13:77217160-77217200 | Ch13 region 3 |
| chr13:77217226-77217266 | Ch13 region 3 |
| chr13:77217287-77217327 | Ch13 region 3 |
| chr13:77217332-77217372 | Ch13 region 3 |

| | |
|---|---|
| chr13:77217395-77217435 | Ch13 region 3 |
| chr13:77217456-77217496 | Ch13 region 3 |
| chr13:77217732-77217772 | Ch13 region 3 |
| chr13:77217777-77217817 | Ch13 region 3 |
| chr13:77217829-77217869 | Ch13 region 3 |
| chr13:77217889-77217929 | Ch13 region 3 |
| chr13:77217975-77218015 | Ch13 region 3 |
| chr13:77218020-77218060 | Ch13 region 3 |
| chr13:77218092-77218132 | Ch13 region 3 |
| chr13:77218219-77218259 | Ch13 region 3 |
| chr13:77218276-77218316 | Ch13 region 3 |
| chr13:77218330-77218370 | Ch13 region 3 |
| chr13:77218472-77218512 | Ch13 region 3 |
| chr13:77218542-77218582 | Ch13 region 3 |
| chr13:77218603-77218643 | Ch13 region 3 |
| chr13:77218688-77218728 | Ch13 region 3 |
| chr13:77218771-77218811 | Ch13 region 3 |
| chr13:77218838-77218878 | Ch13 region 3 |
| chr13:77218911-77218951 | Ch13 region 3 |
| chr13:77219124-77219164 | Ch13 region 3 |
| chr13:77219175-77219215 | Ch13 region 3 |
| chr13:77219225-77219265 | Ch13 region 3 |
| chr13:77219292-77219332 | Ch13 region 3 |
| chr13:77219364-77219404 | Ch13 region 3 |
| chr13:77219451-77219491 | Ch13 region 3 |
| chr13:77219501-77219541 | Ch13 region 3 |
| chr13:77219711-77219751 | Ch13 region 3 |
| chr13:77219756-77219796 | Ch13 region 3 |
| chr13:77219835-77219875 | Ch13 region 3 |
| chr13:77219886-77219926 | Ch13 region 3 |
| chr13:77219943-77219983 | Ch13 region 3 |
| chr13:77220007-77220047 | Ch13 region 3 |
| chr13:77220077-77220117 | Ch13 region 3 |
| chr13:77220122-77220162 | Ch13 region 3 |
| chr13:77220187-77220227 | Ch13 region 3 |
| chr13:77220266-77220306 | Ch13 region 3 |
| chr13:77220313-77220353 | Ch13 region 3 |
| chr13:77220397-77220437 | Ch13 region 3 |
| chr13:77220467-77220507 | Ch13 region 3 |
| chr13:77220530-77220570 | Ch13 region 3 |
| chr13:77220607-77220647 | Ch13 region 3 |
| chr13:77220662-77220702 | Ch13 region 3 |
| chr13:77220731-77220771 | Ch13 region 3 |
| chr13:77220782-77220822 | Ch13 region 3 |
| chr13:77220859-77220899 | Ch13 region 3 |
| chr13:77220904-77220944 | Ch13 region 3 |
| chr13:77220956-77220996 | Ch13 region 3 |
| chr13:77221074-77221114 | Ch13 region 3 |
| chr13:77221125-77221165 | Ch13 region 3 |
| chr13:77221232-77221272 | Ch13 region 3 |
| chr13:77221294-77221334 | Ch13 region 3 |
| chr13:77221404-77221444 | Ch13 region 3 |

| | |
|---|---|
| chr13:77221490-77221530 | Ch13 region 3 |
| chr13:77221537-77221577 | Ch13 region 3 |
| chr13:77221595-77221635 | Ch13 region 3 |
| chr13:77221673-77221713 | Ch13 region 3 |
| chr13:77221738-77221778 | Ch13 region 3 |
| chr13:77221783-77221823 | Ch13 region 3 |
| chr13:77221828-77221868 | Ch13 region 3 |
| chr13:77221892-77221932 | Ch13 region 3 |
| chr13:77221942-77221982 | Ch13 region 3 |
| chr13:77221994-77222034 | Ch13 region 3 |
| chr13:77222099-77222139 | Ch13 region 3 |
| chr13:77222150-77222190 | Ch13 region 3 |
| chr13:77222209-77222249 | Ch13 region 3 |
| chr13:77222280-77222320 | Ch13 region 3 |
| chr13:77222354-77222394 | Ch13 region 3 |
| chr13:77222439-77222479 | Ch13 region 3 |
| chr13:77222484-77222524 | Ch13 region 3 |
| chr13:77222549-77222589 | Ch13 region 3 |
| chr13:77222627-77222667 | Ch13 region 3 |
| chr13:77222679-77222719 | Ch13 region 3 |
| chr13:77222792-77222832 | Ch13 region 3 |
| chr13:77222856-77222896 | Ch13 region 3 |
| chr13:77223009-77223049 | Ch13 region 3 |
| chr13:77223161-77223201 | Ch13 region 3 |
| chr13:77223401-77223441 | Ch13 region 3 |
| chr13:77223468-77223508 | Ch13 region 3 |
| chr13:77223577-77223617 | Ch13 region 3 |
| chr13:77223675-77223715 | Ch13 region 3 |
| chr13:77223746-77223786 | Ch13 region 3 |
| chr13:77223824-77223864 | Ch13 region 3 |
| chr13:77223920-77223960 | Ch13 region 3 |
| chr13:77223990-77224030 | Ch13 region 3 |
| chr13:77224077-77224117 | Ch13 region 3 |
| chr13:77224181-77224221 | Ch13 region 3 |
| chr13:77224257-77224297 | Ch13 region 3 |
| chr13:77224316-77224356 | Ch13 region 3 |
| chr13:77224411-77224451 | Ch13 region 3 |
| chr13:77224588-77224628 | Ch13 region 3 |
| chr13:77224696-77224736 | Ch13 region 3 |
| chr13:77224774-77224814 | Ch13 region 3 |
| chr13:77224884-77224924 | Ch13 region 3 |
| chr13:77225021-77225061 | Ch13 region 3 |
| chr13:77225093-77225133 | Ch13 region 3 |
| chr13:77225151-77225191 | Ch13 region 3 |
| chr13:77225385-77225425 | Ch13 region 3 |
| chr13:77225431-77225471 | Ch13 region 3 |
| chr13:77225494-77225534 | Ch13 region 3 |
| chr13:77225566-77225606 | Ch13 region 3 |
| chr13:77225691-77225731 | Ch13 region 3 |
| chr13:77225761-77225801 | Ch13 region 3 |
| chr13:77225845-77225885 | Ch13 region 3 |
| chr13:77225946-77225986 | Ch13 region 3 |

| | |
|---|---|
| chr13:77226091-77226131 | Ch13 region 3 |
| chr13:77226157-77226197 | Ch13 region 3 |
| chr13:77226492-77226532 | Ch13 region 3 |
| chr13:77226537-77226577 | Ch13 region 3 |
| chr13:77226688-77226728 | Ch13 region 3 |
| chr13:77226751-77226791 | Ch13 region 3 |
| chr13:77226989-77227029 | Ch13 region 3 |
| chr13:77227071-77227111 | Ch13 region 3 |
| chr13:77227301-77227341 | Ch13 region 3 |
| chr13:77227456-77227496 | Ch13 region 3 |
| chr13:77227563-77227603 | Ch13 region 3 |
| chr13:77227710-77227750 | Ch13 region 3 |
| chr13:77227899-77227939 | Ch13 region 3 |
| chr13:77228080-77228120 | Ch13 region 3 |
| chr13:77228131-77228171 | Ch13 region 3 |
| chr13:77228284-77228324 | Ch13 region 3 |
| chr13:77228493-77228533 | Ch13 region 3 |
| chr13:77228558-77228598 | Ch13 region 3 |
| chr13:77228645-77228685 | Ch13 region 3 |
| chr13:77228691-77228731 | Ch13 region 3 |
| chr13:77228755-77228795 | Ch13 region 3 |
| chr13:77228803-77228843 | Ch13 region 3 |
| chr13:77228848-77228888 | Ch13 region 3 |
| chr13:77228893-77228933 | Ch13 region 3 |
| chr13:77228983-77229023 | Ch13 region 3 |
| chr13:77229081-77229121 | Ch13 region 3 |
| chr13:77229275-77229315 | Ch13 region 3 |
| chr13:77229325-77229365 | Ch13 region 3 |
| chr13:77229380-77229420 | Ch13 region 3 |
| chr13:77229427-77229467 | Ch13 region 3 |
| chr13:77229487-77229527 | Ch13 region 3 |
| chr13:77229547-77229587 | Ch13 region 3 |
| chr13:77229639-77229679 | Ch13 region 3 |
| chr13:77229684-77229724 | Ch13 region 3 |
| chr13:77229814-77229854 | Ch13 region 3 |
| chr13:77229887-77229927 | Ch13 region 3 |
| chr13:77229991-77230031 | Ch13 region 3 |
| chr13:77230055-77230095 | Ch13 region 3 |
| chr13:77230127-77230167 | Ch13 region 3 |
| chr13:77230205-77230245 | Ch13 region 3 |
| chr13:77230260-77230300 | Ch13 region 3 |
| chr13:77230330-77230370 | Ch13 region 3 |
| chr13:77230376-77230416 | Ch13 region 3 |
| chr13:77230478-77230518 | Ch13 region 3 |
| chr13:77230543-77230583 | Ch13 region 3 |
| chr13:77230615-77230655 | Ch13 region 3 |
| chr13:77230669-77230709 | Ch13 region 3 |
| chr13:77230749-77230789 | Ch13 region 3 |
| chr13:77230800-77230840 | Ch13 region 3 |
| chr13:77230897-77230937 | Ch13 region 3 |
| chr13:77230976-77231016 | Ch13 region 3 |
| chr13:77231048-77231088 | Ch13 region 3 |

| | |
|---|---|
| chr13:77231100-77231140 | Ch13 region 3 |
| chr13:77231175-77231215 | Ch13 region 3 |
| chr13:77231231-77231271 | Ch13 region 3 |
| chr13:77231355-77231395 | Ch13 region 3 |
| chr13:77231475-77231515 | Ch13 region 3 |
| chr13:77231520-77231560 | Ch13 region 3 |
| chr13:77231608-77231648 | Ch13 region 3 |
| chr13:77231673-77231713 | Ch13 region 3 |
| chr13:77231729-77231769 | Ch13 region 3 |
| chr13:77231774-77231814 | Ch13 region 3 |
| chr13:77231854-77231894 | Ch13 region 3 |
| chr13:77231947-77231987 | Ch13 region 3 |
| chr13:77231998-77232038 | Ch13 region 3 |
| chr13:77232054-77232094 | Ch13 region 3 |
| chr13:77232113-77232153 | Ch13 region 3 |
| chr13:77232232-77232272 | Ch13 region 3 |
| chr13:77232282-77232322 | Ch13 region 3 |
| chr13:77232345-77232385 | Ch13 region 3 |
| chr13:77232690-77232730 | Ch13 region 3 |
| chr13:77232874-77232914 | Ch13 region 3 |
| chr13:77232927-77232967 | Ch13 region 3 |
| chr13:77233084-77233124 | Ch13 region 3 |
| chr13:77233219-77233259 | Ch13 region 3 |
| chr13:77233309-77233349 | Ch13 region 3 |
| chr13:77233461-77233501 | Ch13 region 3 |
| chr13:77233506-77233546 | Ch13 region 3 |
| chr13:77233570-77233610 | Ch13 region 3 |
| chr13:77233615-77233655 | Ch13 region 3 |
| chr13:77233697-77233737 | Ch13 region 3 |
| chr13:77233747-77233787 | Ch13 region 3 |
| chr13:77233840-77233880 | Ch13 region 3 |
| chr13:77233917-77233957 | Ch13 region 3 |
| chr13:77234002-77234042 | Ch13 region 3 |
| chr13:77234066-77234106 | Ch13 region 3 |
| chr13:77234221-77234261 | Ch13 region 3 |
| chr13:77234348-77234388 | Ch13 region 3 |
| chr13:77234405-77234445 | Ch13 region 3 |
| chr13:77234458-77234498 | Ch13 region 3 |
| chr13:77234513-77234553 | Ch13 region 3 |
| chr13:77234583-77234623 | Ch13 region 3 |
| chr13:77234632-77234672 | Ch13 region 3 |
| chr13:77234797-77234837 | Ch13 region 3 |
| chr13:77235017-77235057 | Ch13 region 3 |
| chr13:77235110-77235150 | Ch13 region 3 |
| chr13:77235286-77235326 | Ch13 region 3 |
| chr13:77235426-77235466 | Ch13 region 3 |
| chr13:77235529-77235569 | Ch13 region 3 |
| chr13:77235659-77235699 | Ch13 region 3 |
| chr13:77235793-77235833 | Ch13 region 3 |
| chr13:77235851-77235891 | Ch13 region 3 |
| chr13:77235957-77235997 | Ch13 region 3 |
| chr13:77236007-77236047 | Ch13 region 3 |

| | |
|---|---|
| chr13:77236129-77236169 | Ch13 region 3 |
| chr13:77236298-77236338 | Ch13 region 3 |
| chr13:77236507-77236547 | Ch13 region 3 |
| chr13:77236560-77236600 | Ch13 region 3 |
| chr13:77236639-77236679 | Ch13 region 3 |
| chr13:77236686-77236726 | Ch13 region 3 |
| chr13:77236807-77236847 | Ch13 region 3 |
| chr13:77236894-77236934 | Ch13 region 3 |
| chr13:77236939-77236979 | Ch13 region 3 |
| chr13:77237042-77237082 | Ch13 region 3 |
| chr13:77237138-77237178 | Ch13 region 3 |
| chr13:77237218-77237258 | Ch13 region 3 |
| chr13:77237327-77237367 | Ch13 region 3 |
| chr13:77237372-77237412 | Ch13 region 3 |
| chr13:77237447-77237487 | Ch13 region 3 |
| chr13:77237501-77237541 | Ch13 region 3 |
| chr13:77237561-77237601 | Ch13 region 3 |
| chr13:77237626-77237666 | Ch13 region 3 |
| chr13:77237695-77237735 | Ch13 region 3 |
| chr13:77237788-77237828 | Ch13 region 3 |
| chr13:77237834-77237874 | Ch13 region 3 |
| chr13:77237951-77237991 | Ch13 region 3 |
| chr13:77238017-77238057 | Ch13 region 3 |
| chr13:77238083-77238123 | Ch13 region 3 |
| chr13:77238154-77238194 | Ch13 region 3 |
| chr13:77238199-77238239 | Ch13 region 3 |
| chr13:77238259-77238299 | Ch13 region 3 |
| chr13:77238313-77238353 | Ch13 region 3 |
| chr13:77238391-77238431 | Ch13 region 3 |
| chr13:77238493-77238533 | Ch13 region 3 |
| chr13:77238583-77238623 | Ch13 region 3 |
| chr13:77238634-77238674 | Ch13 region 3 |
| chr13:77238698-77238738 | Ch13 region 3 |
| chr13:77238886-77238926 | Ch13 region 3 |
| chr13:77238967-77239007 | Ch13 region 3 |
| chr13:77239017-77239057 | Ch13 region 3 |
| chr13:77239069-77239109 | Ch13 region 3 |
| chr13:77239135-77239175 | Ch13 region 3 |
| chr13:77239184-77239224 | Ch13 region 3 |
| chr13:77239229-77239269 | Ch13 region 3 |
| chr13:77239308-77239348 | Ch13 region 3 |
| chr13:77239386-77239426 | Ch13 region 3 |
| chr13:77239438-77239478 | Ch13 region 3 |
| chr13:77239529-77239569 | Ch13 region 3 |
| chr13:77239581-77239621 | Ch13 region 3 |
| chr13:77239653-77239693 | Ch13 region 3 |
| chr13:77239731-77239771 | Ch13 region 3 |
| chr13:77239797-77239837 | Ch13 region 3 |
| chr13:77239873-77239913 | Ch13 region 3 |
| chr13:77239938-77239978 | Ch13 region 3 |
| chr13:77240018-77240058 | Ch13 region 3 |
| chr13:77240086-77240126 | Ch13 region 3 |

| | |
|---|---|
| chr13:77240147-77240187 | Ch13 region 3 |
| chr13:77240192-77240232 | Ch13 region 3 |
| chr13:77240243-77240283 | Ch13 region 3 |
| chr13:77240314-77240354 | Ch13 region 3 |
| chr13:77240359-77240399 | Ch13 region 3 |
| chr13:77240473-77240513 | Ch13 region 3 |
| chr13:77240543-77240583 | Ch13 region 3 |
| chr13:77240622-77240662 | Ch13 region 3 |
| chr13:77240709-77240749 | Ch13 region 3 |
| chr13:77240754-77240794 | Ch13 region 3 |
| chr13:77240814-77240854 | Ch13 region 3 |
| chr13:77240991-77241031 | Ch13 region 3 |
| chr13:77241119-77241159 | Ch13 region 3 |
| chr13:77241170-77241210 | Ch13 region 3 |
| chr13:77241221-77241261 | Ch13 region 3 |
| chr13:77241292-77241332 | Ch13 region 3 |
| chr13:77241348-77241388 | Ch13 region 3 |
| chr13:77241439-77241479 | Ch13 region 3 |
| chr13:77241629-77241669 | Ch13 region 3 |
| chr13:77241690-77241730 | Ch13 region 3 |
| chr13:77241749-77241789 | Ch13 region 3 |
| chr13:77241816-77241856 | Ch13 region 3 |
| chr13:77241867-77241907 | Ch13 region 3 |
| chr13:77241913-77241953 | Ch13 region 3 |
| chr13:77241994-77242034 | Ch13 region 3 |
| chr13:77242057-77242097 | Ch13 region 3 |
| chr13:77242110-77242150 | Ch13 region 3 |
| chr13:77242156-77242196 | Ch13 region 3 |
| chr13:77242201-77242241 | Ch13 region 3 |
| chr13:77242303-77242343 | Ch13 region 3 |
| chr13:77242355-77242395 | Ch13 region 3 |
| chr13:77242431-77242471 | Ch13 region 3 |
| chr13:77242499-77242539 | Ch13 region 3 |
| chr13:77242566-77242606 | Ch13 region 3 |
| chr13:77242631-77242671 | Ch13 region 3 |
| chr13:77242678-77242718 | Ch13 region 3 |
| chr13:77242741-77242781 | Ch13 region 3 |
| chr13:77242822-77242862 | Ch13 region 3 |
| chr13:77242867-77242907 | Ch13 region 3 |
| chr13:77242931-77242971 | Ch13 region 3 |
| chr13:77242984-77243024 | Ch13 region 3 |
| chr13:77243047-77243087 | Ch13 region 3 |
| chr13:77243103-77243143 | Ch13 region 3 |
| chr13:77243202-77243242 | Ch13 region 3 |
| chr13:77243248-77243288 | Ch13 region 3 |
| chr13:77243301-77243341 | Ch13 region 3 |
| chr13:77243399-77243439 | Ch13 region 3 |
| chr13:77243466-77243506 | Ch13 region 3 |
| chr13:77243521-77243561 | Ch13 region 3 |
| chr13:77243566-77243606 | Ch13 region 3 |
| chr13:77243638-77243678 | Ch13 region 3 |
| chr13:77243685-77243725 | Ch13 region 3 |

| | |
|---|---|
| chr13:77243758-77243798 | Ch13 region 3 |
| chr13:77243813-77243853 | Ch13 region 3 |
| chr13:77244019-77244059 | Ch13 region 3 |
| chr13:77244169-77244209 | Ch13 region 3 |
| chr13:77244217-77244257 | Ch13 region 3 |
| chr13:77244264-77244304 | Ch13 region 3 |
| chr13:77244422-77244462 | Ch13 region 3 |
| chr13:77244494-77244534 | Ch13 region 3 |
| chr13:77244540-77244580 | Ch13 region 3 |
| chr13:77244587-77244627 | Ch13 region 3 |
| chr13:77244649-77244689 | Ch13 region 3 |
| chr13:77244726-77244766 | Ch13 region 3 |
| chr13:77244797-77244837 | Ch13 region 3 |
| chr13:77244845-77244885 | Ch13 region 3 |
| chr13:77244919-77244959 | Ch13 region 3 |
| chr13:77244988-77245028 | Ch13 region 3 |
| chr13:77245067-77245107 | Ch13 region 3 |
| chr13:77245114-77245154 | Ch13 region 3 |
| chr13:77245173-77245213 | Ch13 region 3 |
| chr13:77245226-77245266 | Ch13 region 3 |
| chr13:77245315-77245355 | Ch13 region 3 |
| chr13:77245413-77245453 | Ch13 region 3 |
| chr13:77245485-77245525 | Ch13 region 3 |
| chr13:77245679-77245719 | Ch13 region 3 |
| chr13:77245767-77245807 | Ch13 region 3 |
| chr13:77245820-77245860 | Ch13 region 3 |
| chr13:77245873-77245913 | Ch13 region 3 |
| chr13:77245971-77246011 | Ch13 region 3 |
| chr13:77246069-77246109 | Ch13 region 3 |
| chr13:77246121-77246161 | Ch13 region 3 |
| chr13:77246166-77246206 | Ch13 region 3 |
| chr13:77246212-77246252 | Ch13 region 3 |
| chr13:77246429-77246469 | Ch13 region 3 |
| chr13:77246523-77246563 | Ch13 region 3 |
| chr13:77246652-77246692 | Ch13 region 3 |
| chr13:77246737-77246777 | Ch13 region 3 |
| chr13:77246799-77246839 | Ch13 region 3 |
| chr13:77246898-77246938 | Ch13 region 3 |
| chr13:77246952-77246992 | Ch13 region 3 |
| chr13:77247009-77247049 | Ch13 region 3 |
| chr13:77247056-77247096 | Ch13 region 3 |
| chr13:77247109-77247149 | Ch13 region 3 |
| chr13:77247176-77247216 | Ch13 region 3 |
| chr13:77247240-77247280 | Ch13 region 3 |
| chr13:77247331-77247371 | Ch13 region 3 |
| chr13:77247403-77247443 | Ch13 region 3 |
| chr13:77247519-77247559 | Ch13 region 3 |
| chr13:77247587-77247627 | Ch13 region 3 |
| chr13:77247695-77247735 | Ch13 region 3 |
| chr13:77247804-77247844 | Ch13 region 3 |
| chr13:77247853-77247893 | Ch13 region 3 |
| chr13:77247902-77247942 | Ch13 region 3 |

| | |
|---|---|
| chr13:77247950-77247990 | Ch13 region 3 |
| chr13:77247997-77248037 | Ch13 region 3 |
| chr13:77248066-77248106 | Ch13 region 3 |
| chr13:77248111-77248151 | Ch13 region 3 |
| chr13:77248223-77248263 | Ch13 region 3 |
| chr13:77248293-77248333 | Ch13 region 3 |
| chr13:77248391-77248431 | Ch13 region 3 |
| chr13:77248456-77248496 | Ch13 region 3 |
| chr13:77248571-77248611 | Ch13 region 3 |
| chr13:77248626-77248666 | Ch13 region 3 |
| chr13:77248714-77248754 | Ch13 region 3 |
| chr13:77248759-77248799 | Ch13 region 3 |
| chr13:77248976-77249016 | Ch13 region 3 |
| chr13:77249040-77249080 | Ch13 region 3 |
| chr13:77249143-77249183 | Ch13 region 3 |
| chr13:77249264-77249304 | Ch13 region 3 |
| chr13:77249385-77249425 | Ch13 region 3 |
| chr13:77249434-77249474 | Ch13 region 3 |
| chr13:77249492-77249532 | Ch13 region 3 |
| chr13:77249544-77249584 | Ch13 region 3 |
| chr13:77249593-77249633 | Ch13 region 3 |
| chr13:77249758-77249798 | Ch13 region 3 |
| chr13:77249804-77249844 | Ch13 region 3 |
| chr13:77249944-77249984 | Ch13 region 3 |
| chr13:77250084-77250124 | Ch13 region 3 |
| chr13:77250161-77250201 | Ch13 region 3 |
| chr13:77250238-77250278 | Ch13 region 3 |
| chr13:77250283-77250323 | Ch13 region 3 |
| chr13:77250389-77250429 | Ch13 region 3 |
| chr13:77250462-77250502 | Ch13 region 3 |
| chr13:77250538-77250578 | Ch13 region 3 |
| chr13:77250797-77250837 | Ch13 region 3 |
| chr13:77251057-77251097 | Ch13 region 3 |
| chr13:77251155-77251195 | Ch13 region 3 |
| chr13:77251219-77251259 | Ch13 region 3 |
| chr13:77251267-77251307 | Ch13 region 3 |
| chr13:77251343-77251383 | Ch13 region 3 |
| chr13:77251459-77251499 | Ch13 region 3 |
| chr13:77251504-77251544 | Ch13 region 3 |
| chr13:77251580-77251620 | Ch13 region 3 |
| chr13:77251637-77251677 | Ch13 region 3 |
| chr13:77251732-77251772 | Ch13 region 3 |
| chr13:77251779-77251819 | Ch13 region 3 |
| chr13:77251871-77251911 | Ch13 region 3 |
| chr13:77251917-77251957 | Ch13 region 3 |
| chr13:77251986-77252026 | Ch13 region 3 |
| chr13:77252045-77252085 | Ch13 region 3 |
| chr13:77252090-77252130 | Ch13 region 3 |
| chr13:77252137-77252177 | Ch13 region 3 |
| chr13:77252211-77252251 | Ch13 region 3 |
| chr13:77252256-77252296 | Ch13 region 3 |
| chr13:77252329-77252369 | Ch13 region 3 |

| | |
|---|---|
| chr13:77252413-77252453 | Ch13 region 3 |
| chr13:77252483-77252523 | Ch13 region 3 |
| chr13:77252544-77252584 | Ch13 region 3 |
| chr13:77252608-77252648 | Ch13 region 3 |
| chr13:77252664-77252704 | Ch13 region 3 |
| chr13:77252726-77252766 | Ch13 region 3 |
| chr13:77252778-77252818 | Ch13 region 3 |
| chr13:77252865-77252905 | Ch13 region 3 |
| chr13:77252932-77252972 | Ch13 region 3 |
| chr13:77253049-77253089 | Ch13 region 3 |
| chr13:77253125-77253165 | Ch13 region 3 |
| chr13:77253205-77253245 | Ch13 region 3 |
| chr13:77253403-77253443 | Ch13 region 3 |
| chr13:77253564-77253604 | Ch13 region 3 |
| chr13:77253635-77253675 | Ch13 region 3 |
| chr13:77253754-77253794 | Ch13 region 3 |
| chr13:77253815-77253855 | Ch13 region 3 |
| chr13:77253881-77253921 | Ch13 region 3 |
| chr13:77253946-77253986 | Ch13 region 3 |
| chr13:77253991-77254031 | Ch13 region 3 |
| chr13:77254070-77254110 | Ch13 region 3 |
| chr13:77254152-77254192 | Ch13 region 3 |
| chr13:77254206-77254246 | Ch13 region 3 |
| chr13:77254288-77254328 | Ch13 region 3 |
| chr13:77254345-77254385 | Ch13 region 3 |
| chr13:77254390-77254430 | Ch13 region 3 |
| chr13:77254445-77254485 | Ch13 region 3 |
| chr13:77254497-77254537 | Ch13 region 3 |
| chr13:77254545-77254585 | Ch13 region 3 |
| chr13:77254607-77254647 | Ch13 region 3 |
| chr13:77254659-77254699 | Ch13 region 3 |
| chr13:77254718-77254758 | Ch13 region 3 |
| chr13:77254787-77254827 | Ch13 region 3 |
| chr13:77255274-77255314 | Ch13 region 3 |
| chr13:77255343-77255383 | Ch13 region 3 |
| chr13:77255391-77255431 | Ch13 region 3 |
| chr13:77255453-77255493 | Ch13 region 3 |
| chr13:77255515-77255555 | Ch13 region 3 |
| chr13:77255616-77255656 | Ch13 region 3 |
| chr13:77255666-77255706 | Ch13 region 3 |
| chr13:77255756-77255796 | Ch13 region 3 |
| chr13:77255803-77255843 | Ch13 region 3 |
| chr13:77255884-77255924 | Ch13 region 3 |
| chr13:77255932-77255972 | Ch13 region 3 |
| chr13:77255997-77256037 | Ch13 region 3 |
| chr13:77256081-77256121 | Ch13 region 3 |
| chr13:77256172-77256212 | Ch13 region 3 |
| chr13:77256240-77256280 | Ch13 region 3 |
| chr13:77256285-77256325 | Ch13 region 3 |
| chr13:77256330-77256370 | Ch13 region 3 |
| chr13:77256375-77256415 | Ch13 region 3 |
| chr13:77256443-77256483 | Ch13 region 3 |

| | |
|---|---|
| chr13:77256521-77256561 | Ch13 region 3 |
| chr13:77256574-77256614 | Ch13 region 3 |
| chr13:77256639-77256679 | Ch13 region 3 |
| chr13:77256692-77256732 | Ch13 region 3 |
| chr13:77256737-77256777 | Ch13 region 3 |
| chr13:77256806-77256846 | Ch13 region 3 |
| chr13:77256870-77256910 | Ch13 region 3 |
| chr13:77256915-77256955 | Ch13 region 3 |
| chr13:77256963-77257003 | Ch13 region 3 |
| chr13:77257034-77257074 | Ch13 region 3 |
| chr13:77257110-77257150 | Ch13 region 3 |
| chr13:77257269-77257309 | Ch13 region 3 |
| chr13:77257314-77257354 | Ch13 region 3 |
| chr13:77257389-77257429 | Ch13 region 3 |
| chr13:77257445-77257485 | Ch13 region 3 |
| chr13:77257490-77257530 | Ch13 region 3 |
| chr13:77257555-77257595 | Ch13 region 3 |
| chr13:77257667-77257707 | Ch13 region 3 |
| chr13:77257712-77257752 | Ch13 region 3 |
| chr13:77257774-77257814 | Ch13 region 3 |
| chr13:77257829-77257869 | Ch13 region 3 |
| chr13:77257948-77257988 | Ch13 region 3 |
| chr13:77258006-77258046 | Ch13 region 3 |
| chr13:77258052-77258092 | Ch13 region 3 |
| chr13:77258133-77258173 | Ch13 region 3 |
| chr13:77258194-77258234 | Ch13 region 3 |
| chr13:77258274-77258314 | Ch13 region 3 |
| chr13:77258361-77258401 | Ch13 region 3 |
| chr13:77258435-77258475 | Ch13 region 3 |
| chr13:77258503-77258543 | Ch13 region 3 |
| chr13:77258549-77258589 | Ch13 region 3 |
| chr13:77258619-77258659 | Ch13 region 3 |
| chr13:77258701-77258741 | Ch13 region 3 |
| chr13:77258749-77258789 | Ch13 region 3 |
| chr13:77258839-77258879 | Ch13 region 3 |
| chr13:77258934-77258974 | Ch13 region 3 |
| chr13:77259019-77259059 | Ch13 region 3 |
| chr13:77259104-77259144 | Ch13 region 3 |
| chr13:77259150-77259190 | Ch13 region 3 |
| chr13:77259217-77259257 | Ch13 region 3 |
| chr13:77259287-77259327 | Ch13 region 3 |
| chr13:77259336-77259376 | Ch13 region 3 |
| chr13:77259381-77259421 | Ch13 region 3 |
| chr13:77259441-77259481 | Ch13 region 3 |
| chr13:77259507-77259547 | Ch13 region 3 |
| chr13:77259555-77259595 | Ch13 region 3 |
| chr13:77259625-77259665 | Ch13 region 3 |
| chr13:77259678-77259718 | Ch13 region 3 |
| chr13:77259812-77259852 | Ch13 region 3 |
| chr13:77259913-77259953 | Ch13 region 3 |
| chr13:77259975-77260015 | Ch13 region 3 |
| chr13:77260046-77260086 | Ch13 region 3 |

| | |
|---|---|
| chr13:77260133-77260173 | Ch13 region 3 |
| chr13:77260245-77260285 | Ch13 region 3 |
| chr13:77260374-77260414 | Ch13 region 3 |
| chr13:77260451-77260491 | Ch13 region 3 |
| chr13:77260504-77260544 | Ch13 region 3 |
| chr13:77260581-77260621 | Ch13 region 3 |
| chr13:77260799-77260839 | Ch13 region 3 |
| chr13:77260885-77260925 | Ch13 region 3 |
| chr13:77261039-77261079 | Ch13 region 3 |
| chr13:77261109-77261149 | Ch13 region 3 |
| chr13:77261162-77261202 | Ch13 region 3 |
| chr13:77261229-77261269 | Ch13 region 3 |
| chr13:77261295-77261335 | Ch13 region 3 |
| chr13:77261344-77261384 | Ch13 region 3 |
| chr13:77261405-77261445 | Ch13 region 3 |
| chr13:77261450-77261490 | Ch13 region 3 |
| chr13:77261680-77261720 | Ch13 region 3 |
| chr13:77261729-77261769 | Ch13 region 3 |
| chr13:77261803-77261843 | Ch13 region 3 |
| chr13:77261913-77261953 | Ch13 region 3 |
| chr13:77261958-77261998 | Ch13 region 3 |
| chr13:77262003-77262043 | Ch13 region 3 |
| chr13:77262049-77262089 | Ch13 region 3 |
| chr13:77262138-77262178 | Ch13 region 3 |
| chr13:77262189-77262229 | Ch13 region 3 |
| chr13:77262264-77262304 | Ch13 region 3 |
| chr13:77262310-77262350 | Ch13 region 3 |
| chr13:77262355-77262395 | Ch13 region 3 |
| chr13:77262413-77262453 | Ch13 region 3 |
| chr13:77262511-77262551 | Ch13 region 3 |
| chr13:77262573-77262613 | Ch13 region 3 |
| chr13:77262619-77262659 | Ch13 region 3 |
| chr13:77262671-77262711 | Ch13 region 3 |
| chr13:77262738-77262778 | Ch13 region 3 |
| chr13:77262918-77262958 | Ch13 region 3 |
| chr13:77263012-77263052 | Ch13 region 3 |
| chr13:77263057-77263097 | Ch13 region 3 |
| chr13:77263102-77263142 | Ch13 region 3 |
| chr13:77263184-77263224 | Ch13 region 3 |
| chr13:77263242-77263282 | Ch13 region 3 |
| chr13:77263392-77263432 | Ch13 region 3 |
| chr13:77263459-77263499 | Ch13 region 3 |
| chr13:77263543-77263583 | Ch13 region 3 |
| chr13:77263597-77263637 | Ch13 region 3 |
| chr13:77263665-77263705 | Ch13 region 3 |
| chr13:77263723-77263763 | Ch13 region 3 |
| chr13:77263827-77263867 | Ch13 region 3 |
| chr13:77263951-77263991 | Ch13 region 3 |
| chr13:77264012-77264052 | Ch13 region 3 |
| chr13:77264059-77264099 | Ch13 region 3 |
| chr13:77264111-77264151 | Ch13 region 3 |
| chr13:77264202-77264242 | Ch13 region 3 |

| | |
|---|---|
| chr13:77264359-77264399 | Ch13 region 3 |
| chr13:77264577-77264617 | Ch13 region 3 |
| chr13:77264631-77264671 | Ch13 region 3 |
| chr13:77264779-77264819 | Ch13 region 3 |
| chr13:77264824-77264864 | Ch13 region 3 |
| chr13:77264871-77264911 | Ch13 region 3 |
| chr13:77264949-77264989 | Ch13 region 3 |
| chr13:77264996-77265036 | Ch13 region 3 |
| chr13:77265057-77265097 | Ch13 region 3 |
| chr13:77265132-77265172 | Ch13 region 3 |
| chr13:77265188-77265228 | Ch13 region 3 |
| chr13:77265292-77265332 | Ch13 region 3 |
| chr13:77265392-77265432 | Ch13 region 3 |
| chr13:77265457-77265497 | Ch13 region 3 |
| chr13:77265505-77265545 | Ch13 region 3 |
| chr13:77265554-77265594 | Ch13 region 3 |
| chr13:77265601-77265641 | Ch13 region 3 |
| chr13:77265659-77265699 | Ch13 region 3 |
| chr13:77265709-77265749 | Ch13 region 3 |
| chr13:77265769-77265809 | Ch13 region 3 |
| chr13:77265833-77265873 | Ch13 region 3 |
| chr13:77265919-77265959 | Ch13 region 3 |
| chr13:77266065-77266105 | Ch13 region 3 |
| chr13:77266110-77266150 | Ch13 region 3 |
| chr13:77266195-77266235 | Ch13 region 3 |
| chr13:77266243-77266283 | Ch13 region 3 |
| chr13:77266337-77266377 | Ch13 region 3 |
| chr13:77266486-77266526 | Ch13 region 3 |
| chr13:77266540-77266580 | Ch13 region 3 |
| chr13:77266595-77266635 | Ch13 region 3 |
| chr13:77266686-77266726 | Ch13 region 3 |
| chr13:77266747-77266787 | Ch13 region 3 |
| chr13:77266810-77266850 | Ch13 region 3 |
| chr13:77266880-77266920 | Ch13 region 3 |
| chr13:77266970-77267010 | Ch13 region 3 |
| chr13:77267154-77267194 | Ch13 region 3 |
| chr13:77267291-77267331 | Ch13 region 3 |
| chr13:77267402-77267442 | Ch13 region 3 |
| chr13:77267461-77267501 | Ch13 region 3 |
| chr13:77267579-77267619 | Ch13 region 3 |
| chr13:77267636-77267676 | Ch13 region 3 |
| chr13:77267687-77267727 | Ch13 region 3 |
| chr13:77267832-77267872 | Ch13 region 3 |
| chr13:77267887-77267927 | Ch13 region 3 |
| chr13:77267972-77268012 | Ch13 region 3 |
| chr13:77268108-77268148 | Ch13 region 3 |
| chr13:77268248-77268288 | Ch13 region 3 |
| chr13:77268407-77268447 | Ch13 region 3 |
| chr13:77268575-77268615 | Ch13 region 3 |
| chr13:77268628-77268668 | Ch13 region 3 |
| chr13:77268703-77268743 | Ch13 region 3 |
| chr13:77268751-77268791 | Ch13 region 3 |

| | |
|---|---|
| chr13:77268801-77268841 | Ch13 region 3 |
| chr13:77268866-77268906 | Ch13 region 3 |
| chr13:77268946-77268986 | Ch13 region 3 |
| chr13:77269031-77269071 | Ch13 region 3 |
| chr13:77269085-77269125 | Ch13 region 3 |
| chr13:77269159-77269199 | Ch13 region 3 |
| chr13:77269204-77269244 | Ch13 region 3 |
| chr13:77269257-77269297 | Ch13 region 3 |
| chr13:77269305-77269345 | Ch13 region 3 |
| chr13:77269378-77269418 | Ch13 region 3 |
| chr13:77269446-77269486 | Ch13 region 3 |
| chr13:77269703-77269743 | Ch13 region 3 |
| chr13:77269751-77269791 | Ch13 region 3 |
| chr13:77269933-77269973 | Ch13 region 3 |
| chr13:77270069-77270109 | Ch13 region 3 |
| chr13:77270115-77270155 | Ch13 region 3 |
| chr13:77270170-77270210 | Ch13 region 3 |
| chr13:77270249-77270289 | Ch13 region 3 |
| chr13:77270332-77270372 | Ch13 region 3 |
| chr13:77270385-77270425 | Ch13 region 3 |
| chr13:77270473-77270513 | Ch13 region 3 |
| chr13:77270541-77270581 | Ch13 region 3 |
| chr13:77270619-77270659 | Ch13 region 3 |
| chr13:77270683-77270723 | Ch13 region 3 |
| chr13:77270761-77270801 | Ch13 region 3 |
| chr13:77270810-77270850 | Ch13 region 3 |
| chr13:77270899-77270939 | Ch13 region 3 |
| chr13:77270945-77270985 | Ch13 region 3 |
| chr13:77271005-77271045 | Ch13 region 3 |
| chr13:77271074-77271114 | Ch13 region 3 |
| chr13:77271139-77271179 | Ch13 region 3 |
| chr13:77271235-77271275 | Ch13 region 3 |
| chr13:77271430-77271470 | Ch13 region 3 |
| chr13:77271515-77271555 | Ch13 region 3 |
| chr13:77271562-77271602 | Ch13 region 3 |
| chr13:77271637-77271677 | Ch13 region 3 |
| chr13:77271702-77271742 | Ch13 region 3 |
| chr13:77271780-77271820 | Ch13 region 3 |
| chr13:77271884-77271924 | Ch13 region 3 |
| chr13:77271955-77271995 | Ch13 region 3 |
| chr13:77272028-77272068 | Ch13 region 3 |
| chr13:77272073-77272113 | Ch13 region 3 |
| chr13:77272122-77272162 | Ch13 region 3 |
| chr13:77272177-77272217 | Ch13 region 3 |
| chr13:77272290-77272330 | Ch13 region 3 |
| chr13:77272386-77272426 | Ch13 region 3 |
| chr13:77272431-77272471 | Ch13 region 3 |
| chr13:77272483-77272523 | Ch13 region 3 |
| chr13:77272533-77272573 | Ch13 region 3 |
| chr13:77272626-77272666 | Ch13 region 3 |
| chr13:77272690-77272730 | Ch13 region 3 |
| chr13:77272760-77272800 | Ch13 region 3 |

| | |
|---|---|
| chr13:77272819-77272859 | Ch13 region 3 |
| chr13:77272868-77272908 | Ch13 region 3 |
| chr13:77272936-77272976 | Ch13 region 3 |
| chr13:77272992-77273032 | Ch13 region 3 |
| chr13:77273086-77273126 | Ch13 region 3 |
| chr13:77273131-77273171 | Ch13 region 3 |
| chr13:77273179-77273219 | Ch13 region 3 |
| chr13:77273237-77273277 | Ch13 region 3 |
| chr13:77273331-77273371 | Ch13 region 3 |
| chr13:77273377-77273417 | Ch13 region 3 |
| chr13:77273444-77273484 | Ch13 region 3 |
| chr13:77273557-77273597 | Ch13 region 3 |
| chr13:77273611-77273651 | Ch13 region 3 |
| chr13:77273678-77273718 | Ch13 region 3 |
| chr13:77273734-77273774 | Ch13 region 3 |
| chr13:77273779-77273819 | Ch13 region 3 |
| chr13:77273850-77273890 | Ch13 region 3 |
| chr13:77273907-77273947 | Ch13 region 3 |
| chr13:77273997-77274037 | Ch13 region 3 |
| chr13:77274070-77274110 | Ch13 region 3 |
| chr13:77274135-77274175 | Ch13 region 3 |
| chr13:77274187-77274227 | Ch13 region 3 |
| chr13:77274284-77274324 | Ch13 region 3 |
| chr13:77274373-77274413 | Ch13 region 3 |
| chr13:77274455-77274495 | Ch13 region 3 |
| chr13:77274540-77274580 | Ch13 region 3 |
| chr13:77274585-77274625 | Ch13 region 3 |
| chr13:77274684-77274724 | Ch13 region 3 |
| chr13:77274762-77274802 | Ch13 region 3 |
| chr13:77274903-77274943 | Ch13 region 3 |
| chr13:77274966-77275006 | Ch13 region 3 |
| chr13:77275016-77275056 | Ch13 region 3 |
| chr13:77275119-77275159 | Ch13 region 3 |
| chr13:77275212-77275252 | Ch13 region 3 |
| chr13:77275279-77275319 | Ch13 region 3 |
| chr13:77275326-77275366 | Ch13 region 3 |
| chr13:77275430-77275470 | Ch13 region 3 |
| chr13:77275520-77275560 | Ch13 region 3 |
| chr13:77275569-77275609 | Ch13 region 3 |
| chr13:77275639-77275679 | Ch13 region 3 |
| chr13:77275686-77275726 | Ch13 region 3 |
| chr13:77275763-77275803 | Ch13 region 3 |
| chr13:77275838-77275878 | Ch13 region 3 |
| chr13:77275891-77275931 | Ch13 region 3 |
| chr13:77275937-77275977 | Ch13 region 3 |
| chr13:77275992-77276032 | Ch13 region 3 |
| chr13:77276040-77276080 | Ch13 region 3 |
| chr13:77276087-77276127 | Ch13 region 3 |
| chr13:77276155-77276195 | Ch13 region 3 |
| chr13:77276200-77276240 | Ch13 region 3 |
| chr13:77276265-77276305 | Ch13 region 3 |
| chr13:77276334-77276374 | Ch13 region 3 |

| | |
|---|---|
| chr13:77276488-77276528 | Ch13 region 3 |
| chr13:77276561-77276601 | Ch13 region 3 |
| chr13:77276779-77276819 | Ch13 region 3 |
| chr13:77276840-77276880 | Ch13 region 3 |
| chr13:77276893-77276933 | Ch13 region 3 |
| chr13:77276970-77277010 | Ch13 region 3 |
| chr13:77277020-77277060 | Ch13 region 3 |
| chr13:77277259-77277299 | Ch13 region 3 |
| chr13:77277369-77277409 | Ch13 region 3 |
| chr13:77277419-77277459 | Ch13 region 3 |
| chr13:77277489-77277529 | Ch13 region 3 |
| chr13:77277544-77277584 | Ch13 region 3 |
| chr13:77277602-77277642 | Ch13 region 3 |
| chr13:77277647-77277687 | Ch13 region 3 |
| chr13:77277753-77277793 | Ch13 region 3 |
| chr13:77277827-77277867 | Ch13 region 3 |
| chr13:77277900-77277940 | Ch13 region 3 |
| chr13:77278080-77278120 | Ch13 region 3 |
| chr13:77278169-77278209 | Ch13 region 3 |
| chr13:77278215-77278255 | Ch13 region 3 |
| chr13:77278266-77278306 | Ch13 region 3 |
| chr13:77278312-77278352 | Ch13 region 3 |
| chr13:77278427-77278467 | Ch13 region 3 |
| chr13:77278521-77278561 | Ch13 region 3 |
| chr13:77278642-77278682 | Ch13 region 3 |
| chr13:77278732-77278772 | Ch13 region 3 |
| chr13:77278808-77278848 | Ch13 region 3 |
| chr13:77278857-77278897 | Ch13 region 3 |
| chr13:77278934-77278974 | Ch13 region 3 |
| chr13:77279049-77279089 | Ch13 region 3 |
| chr13:77279102-77279142 | Ch13 region 3 |
| chr13:77279193-77279233 | Ch13 region 3 |
| chr13:77279238-77279278 | Ch13 region 3 |
| chr13:77279324-77279364 | Ch13 region 3 |
| chr13:77279395-77279435 | Ch13 region 3 |
| chr13:77279445-77279485 | Ch13 region 3 |
| chr13:77279497-77279537 | Ch13 region 3 |
| chr13:77279555-77279595 | Ch13 region 3 |
| chr13:77279607-77279647 | Ch13 region 3 |
| chr13:77279672-77279712 | Ch13 region 3 |
| chr13:77279774-77279814 | Ch13 region 3 |
| chr13:77279837-77279877 | Ch13 region 3 |
| chr13:77279979-77280019 | Ch13 region 3 |
| chr13:77280117-77280157 | Ch13 region 3 |
| chr13:77280188-77280228 | Ch13 region 3 |
| chr13:77280267-77280307 | Ch13 region 3 |
| chr13:77280337-77280377 | Ch13 region 3 |
| chr13:77280396-77280436 | Ch13 region 3 |
| chr13:77280493-77280533 | Ch13 region 3 |
| chr13:77280595-77280635 | Ch13 region 3 |
| chr13:77280641-77280681 | Ch13 region 3 |
| chr13:77280715-77280755 | Ch13 region 3 |

| | |
|---|---|
| chr13:77280785-77280825 | Ch13 region 3 |
| chr13:77280869-77280909 | Ch13 region 3 |
| chr13:77280976-77281016 | Ch13 region 3 |
| chr13:77281059-77281099 | Ch13 region 3 |
| chr13:77281107-77281147 | Ch13 region 3 |
| chr13:77281164-77281204 | Ch13 region 3 |
| chr13:77281236-77281276 | Ch13 region 3 |
| chr13:77281290-77281330 | Ch13 region 3 |
| chr13:77281343-77281383 | Ch13 region 3 |
| chr13:77281429-77281469 | Ch13 region 3 |
| chr13:77281474-77281514 | Ch13 region 3 |
| chr13:77281525-77281565 | Ch13 region 3 |
| chr13:77281572-77281612 | Ch13 region 3 |
| chr13:77281649-77281689 | Ch13 region 3 |
| chr13:77281730-77281770 | Ch13 region 3 |
| chr13:77281784-77281824 | Ch13 region 3 |
| chr13:77281831-77281871 | Ch13 region 3 |
| chr13:77281898-77281938 | Ch13 region 3 |
| chr13:77281966-77282006 | Ch13 region 3 |
| chr13:77282064-77282104 | Ch13 region 3 |
| chr13:77282158-77282198 | Ch13 region 3 |
| chr13:77282203-77282243 | Ch13 region 3 |
| chr13:77282275-77282315 | Ch13 region 3 |
| chr13:77282357-77282397 | Ch13 region 3 |
| chr13:77282447-77282487 | Ch13 region 3 |
| chr13:77282533-77282573 | Ch13 region 3 |
| chr13:77282617-77282657 | Ch13 region 3 |
| chr13:77282665-77282705 | Ch13 region 3 |
| chr13:77282755-77282795 | Ch13 region 3 |
| chr13:77282804-77282844 | Ch13 region 3 |
| chr13:77282849-77282889 | Ch13 region 3 |
| chr13:77282898-77282938 | Ch13 region 3 |
| chr13:77282954-77282994 | Ch13 region 3 |
| chr13:77283055-77283095 | Ch13 region 3 |
| chr13:77283152-77283192 | Ch13 region 3 |
| chr13:77283205-77283245 | Ch13 region 3 |
| chr13:77283257-77283297 | Ch13 region 3 |
| chr13:77283343-77283383 | Ch13 region 3 |
| chr13:77283402-77283442 | Ch13 region 3 |
| chr13:77283478-77283518 | Ch13 region 3 |
| chr13:77283551-77283591 | Ch13 region 3 |
| chr13:77283622-77283662 | Ch13 region 3 |
| chr13:77283679-77283719 | Ch13 region 3 |
| chr13:77283741-77283781 | Ch13 region 3 |
| chr13:77283811-77283851 | Ch13 region 3 |
| chr13:77283865-77283905 | Ch13 region 3 |
| chr13:77283916-77283956 | Ch13 region 3 |
| chr13:77284009-77284049 | Ch13 region 3 |
| chr13:77284091-77284131 | Ch13 region 3 |
| chr13:77284177-77284217 | Ch13 region 3 |
| chr13:77284224-77284264 | Ch13 region 3 |
| chr13:77284287-77284327 | Ch13 region 3 |

| | |
|---|---|
| chr13:77284348-77284388 | Ch13 region 3 |
| chr13:77284403-77284443 | Ch13 region 3 |
| chr13:77284481-77284521 | Ch13 region 3 |
| chr13:77284536-77284576 | Ch13 region 3 |
| chr13:77284591-77284631 | Ch13 region 3 |
| chr13:77284653-77284693 | Ch13 region 3 |
| chr13:77284761-77284801 | Ch13 region 3 |
| chr13:77284814-77284854 | Ch13 region 3 |
| chr13:77284899-77284939 | Ch13 region 3 |
| chr13:77284948-77284988 | Ch13 region 3 |
| chr13:77285058-77285098 | Ch13 region 3 |
| chr13:77285113-77285153 | Ch13 region 3 |
| chr13:77285158-77285198 | Ch13 region 3 |
| chr13:77285208-77285248 | Ch13 region 3 |
| chr13:77285253-77285293 | Ch13 region 3 |
| chr13:77285306-77285346 | Ch13 region 3 |
| chr13:77285351-77285391 | Ch13 region 3 |
| chr13:77285469-77285509 | Ch13 region 3 |
| chr13:77285587-77285627 | Ch13 region 3 |
| chr13:77285682-77285722 | Ch13 region 3 |
| chr13:77285745-77285785 | Ch13 region 3 |
| chr13:77285815-77285855 | Ch13 region 3 |
| chr13:77285868-77285908 | Ch13 region 3 |
| chr13:77285921-77285961 | Ch13 region 3 |
| chr13:77285969-77286009 | Ch13 region 3 |
| chr13:77286065-77286105 | Ch13 region 3 |
| chr13:77286110-77286150 | Ch13 region 3 |
| chr13:77286163-77286203 | Ch13 region 3 |
| chr13:77286254-77286294 | Ch13 region 3 |
| chr13:77286305-77286345 | Ch13 region 3 |
| chr13:77286353-77286393 | Ch13 region 3 |
| chr13:77286400-77286440 | Ch13 region 3 |
| chr13:77286479-77286519 | Ch13 region 3 |
| chr13:77286528-77286568 | Ch13 region 3 |
| chr13:77286588-77286628 | Ch13 region 3 |
| chr13:77286683-77286723 | Ch13 region 3 |
| chr13:77286729-77286769 | Ch13 region 3 |
| chr13:77286945-77286985 | Ch13 region 3 |
| chr13:77287077-77287117 | Ch13 region 3 |
| chr13:77287160-77287200 | Ch13 region 3 |
| chr13:77287246-77287286 | Ch13 region 3 |
| chr13:77287318-77287358 | Ch13 region 3 |
| chr13:77287410-77287450 | Ch13 region 3 |
| chr13:77287466-77287506 | Ch13 region 3 |
| chr13:77287541-77287581 | Ch13 region 3 |
| chr13:77287609-77287649 | Ch13 region 3 |
| chr13:77287672-77287712 | Ch13 region 3 |
| chr13:77287793-77287833 | Ch13 region 3 |
| chr13:77287847-77287887 | Ch13 region 3 |
| chr13:77287901-77287941 | Ch13 region 3 |
| chr13:77287967-77288007 | Ch13 region 3 |
| chr13:77288019-77288059 | Ch13 region 3 |

| | |
|---|---|
| chr13:77288079-77288119 | Ch13 region 3 |
| chr13:77288131-77288171 | Ch13 region 3 |
| chr13:77288205-77288245 | Ch13 region 3 |
| chr13:77288278-77288318 | Ch13 region 3 |
| chr13:77288328-77288368 | Ch13 region 3 |
| chr13:77288386-77288426 | Ch13 region 3 |
| chr13:77288431-77288471 | Ch13 region 3 |
| chr13:77288496-77288536 | Ch13 region 3 |
| chr13:77288545-77288585 | Ch13 region 3 |
| chr13:77288596-77288636 | Ch13 region 3 |
| chr13:77288653-77288693 | Ch13 region 3 |
| chr13:77288709-77288749 | Ch13 region 3 |
| chr13:77288760-77288800 | Ch13 region 3 |
| chr13:77288840-77288880 | Ch13 region 3 |
| chr13:77289098-77289138 | Ch13 region 3 |
| chr13:77289143-77289183 | Ch13 region 3 |
| chr13:77289205-77289245 | Ch13 region 3 |
| chr13:77289280-77289320 | Ch13 region 3 |
| chr13:77289332-77289372 | Ch13 region 3 |
| chr13:77289401-77289441 | Ch13 region 3 |
| chr13:77289463-77289503 | Ch13 region 3 |
| chr13:77289532-77289572 | Ch13 region 3 |
| chr13:77289594-77289634 | Ch13 region 3 |
| chr13:77289655-77289695 | Ch13 region 3 |
| chr13:77289710-77289750 | Ch13 region 3 |
| chr13:77289761-77289801 | Ch13 region 3 |
| chr13:77289842-77289882 | Ch13 region 3 |
| chr13:77289887-77289927 | Ch13 region 3 |
| chr13:77289980-77290020 | Ch13 region 3 |
| chr13:77290055-77290095 | Ch13 region 3 |
| chr13:77290228-77290268 | Ch13 region 3 |
| chr13:77290309-77290349 | Ch13 region 3 |
| chr13:77290354-77290394 | Ch13 region 3 |
| chr13:77290402-77290442 | Ch13 region 3 |
| chr13:77290498-77290538 | Ch13 region 3 |
| chr13:77290650-77290690 | Ch13 region 3 |
| chr13:77290695-77290735 | Ch13 region 3 |
| chr13:77290868-77290908 | Ch13 region 3 |
| chr13:77290991-77291031 | Ch13 region 3 |
| chr13:77291041-77291081 | Ch13 region 3 |
| chr13:77291091-77291131 | Ch13 region 3 |
| chr13:77291155-77291195 | Ch13 region 3 |
| chr13:77291306-77291346 | Ch13 region 3 |
| chr13:77291354-77291394 | Ch13 region 3 |
| chr13:77291399-77291439 | Ch13 region 3 |
| chr13:77291457-77291497 | Ch13 region 3 |
| chr13:77291503-77291543 | Ch13 region 3 |
| chr13:77291549-77291589 | Ch13 region 3 |
| chr13:77291641-77291681 | Ch13 region 3 |
| chr13:77291820-77291860 | Ch13 region 3 |
| chr13:77291975-77292015 | Ch13 region 3 |
| chr13:77292022-77292062 | Ch13 region 3 |

| | |
|---|---|
| chr13:77292131-77292171 | Ch13 region 3 |
| chr13:77292211-77292251 | Ch13 region 3 |
| chr13:77292292-77292332 | Ch13 region 3 |
| chr13:77292351-77292391 | Ch13 region 3 |
| chr13:77292426-77292466 | Ch13 region 3 |
| chr13:77292505-77292545 | Ch13 region 3 |
| chr13:77292568-77292608 | Ch13 region 3 |
| chr13:77292616-77292656 | Ch13 region 3 |
| chr13:77292673-77292713 | Ch13 region 3 |
| chr13:77292718-77292758 | Ch13 region 3 |
| chr13:77292765-77292805 | Ch13 region 3 |
| chr13:77292830-77292870 | Ch13 region 3 |
| chr13:77292878-77292918 | Ch13 region 3 |
| chr13:77292950-77292990 | Ch13 region 3 |
| chr13:77293019-77293059 | Ch13 region 3 |
| chr13:77293071-77293111 | Ch13 region 3 |
| chr13:77293145-77293185 | Ch13 region 3 |
| chr13:77293193-77293233 | Ch13 region 3 |
| chr13:77293265-77293305 | Ch13 region 3 |
| chr13:77293365-77293405 | Ch13 region 3 |
| chr13:77293466-77293506 | Ch13 region 3 |
| chr13:77293546-77293586 | Ch13 region 3 |
| chr13:77293601-77293641 | Ch13 region 3 |
| chr13:77293717-77293757 | Ch13 region 3 |
| chr13:77293822-77293862 | Ch13 region 3 |
| chr13:77293942-77293982 | Ch13 region 3 |
| chr13:77294005-77294045 | Ch13 region 3 |
| chr13:77294053-77294093 | Ch13 region 3 |
| chr13:77294105-77294145 | Ch13 region 3 |
| chr13:77294197-77294237 | Ch13 region 3 |
| chr13:77294258-77294298 | Ch13 region 3 |
| chr13:77294324-77294364 | Ch13 region 3 |
| chr13:77294396-77294436 | Ch13 region 3 |
| chr13:77294485-77294525 | Ch13 region 3 |
| chr13:77294558-77294598 | Ch13 region 3 |
| chr13:77294638-77294678 | Ch13 region 3 |
| chr13:77294697-77294737 | Ch13 region 3 |
| chr13:77294759-77294799 | Ch13 region 3 |
| chr13:77294824-77294864 | Ch13 region 3 |
| chr13:77294882-77294922 | Ch13 region 3 |
| chr13:77294947-77294987 | Ch13 region 3 |
| chr13:77295044-77295084 | Ch13 region 3 |
| chr13:77295101-77295141 | Ch13 region 3 |
| chr13:77295174-77295214 | Ch13 region 3 |
| chr13:77295243-77295283 | Ch13 region 3 |
| chr13:77295318-77295358 | Ch13 region 3 |
| chr13:77295364-77295404 | Ch13 region 3 |
| chr13:77295434-77295474 | Ch13 region 3 |
| chr13:77295497-77295537 | Ch13 region 3 |
| chr13:77295598-77295638 | Ch13 region 3 |
| chr13:77295661-77295701 | Ch13 region 3 |
| chr13:77295754-77295794 | Ch13 region 3 |

| | |
|---|---|
| chr13:77295815-77295855 | Ch13 region 3 |
| chr13:77295909-77295949 | Ch13 region 3 |
| chr13:77295983-77296023 | Ch13 region 3 |
| chr13:77296041-77296081 | Ch13 region 3 |
| chr13:77296092-77296132 | Ch13 region 3 |
| chr13:77296205-77296245 | Ch13 region 3 |
| chr13:77296299-77296339 | Ch13 region 3 |
| chr13:77296392-77296432 | Ch13 region 3 |
| chr13:77296563-77296603 | Ch13 region 3 |
| chr13:77296632-77296672 | Ch13 region 3 |
| chr13:77296692-77296732 | Ch13 region 3 |
| chr13:77296761-77296801 | Ch13 region 3 |
| chr13:77296807-77296847 | Ch13 region 3 |
| chr13:77296873-77296913 | Ch13 region 3 |
| chr13:77296935-77296975 | Ch13 region 3 |
| chr13:77296980-77297020 | Ch13 region 3 |
| chr13:77297035-77297075 | Ch13 region 3 |
| chr13:77297088-77297128 | Ch13 region 3 |
| chr13:77297160-77297200 | Ch13 region 3 |
| chr13:77297253-77297293 | Ch13 region 3 |
| chr13:77297313-77297353 | Ch13 region 3 |
| chr13:77297391-77297431 | Ch13 region 3 |
| chr13:77297592-77297632 | Ch13 region 3 |
| chr13:77297655-77297695 | Ch13 region 3 |
| chr13:77297726-77297766 | Ch13 region 3 |
| chr13:77297773-77297813 | Ch13 region 3 |
| chr13:77297818-77297858 | Ch13 region 3 |
| chr13:77297868-77297908 | Ch13 region 3 |
| chr13:77297953-77297993 | Ch13 region 3 |
| chr13:77297999-77298039 | Ch13 region 3 |
| chr13:77298086-77298126 | Ch13 region 3 |
| chr13:77298134-77298174 | Ch13 region 3 |
| chr13:77298180-77298220 | Ch13 region 3 |
| chr13:77298281-77298321 | Ch13 region 3 |
| chr13:77298363-77298403 | Ch13 region 3 |
| chr13:77298554-77298594 | Ch13 region 3 |
| chr13:77298607-77298647 | Ch13 region 3 |
| chr13:77298656-77298696 | Ch13 region 3 |
| chr13:77298755-77298795 | Ch13 region 3 |
| chr13:77298811-77298851 | Ch13 region 3 |
| chr13:77298918-77298958 | Ch13 region 3 |
| chr13:77298972-77299012 | Ch13 region 3 |
| chr13:77299048-77299088 | Ch13 region 3 |
| chr13:77299097-77299137 | Ch13 region 3 |
| chr13:77299185-77299225 | Ch13 region 3 |
| chr13:77299230-77299270 | Ch13 region 3 |
| chr13:77299275-77299315 | Ch13 region 3 |
| chr13:77299338-77299378 | Ch13 region 3 |
| chr13:77299389-77299429 | Ch13 region 3 |
| chr13:77299440-77299480 | Ch13 region 3 |
| chr13:77299512-77299552 | Ch13 region 3 |
| chr13:77299604-77299644 | Ch13 region 3 |

| | |
|---|---|
| chr13:77299679-77299719 | Ch13 region 3 |
| chr13:77299725-77299765 | Ch13 region 3 |
| chr13:77299782-77299822 | Ch13 region 3 |
| chr13:77299852-77299892 | Ch13 region 3 |
| chr13:77299983-77300023 | Ch13 region 3 |
| chr13:77300051-77300091 | Ch13 region 3 |
| chr13:77300100-77300140 | Ch13 region 3 |
| chr13:77300177-77300217 | Ch13 region 3 |
| chr13:77300232-77300272 | Ch13 region 3 |
| chr13:77300277-77300317 | Ch13 region 3 |
| chr13:77300348-77300388 | Ch13 region 3 |
| chr13:77300508-77300548 | Ch13 region 3 |
| chr13:77300645-77300685 | Ch13 region 3 |
| chr13:77300707-77300747 | Ch13 region 3 |
| chr13:77300830-77300870 | Ch13 region 3 |
| chr13:77300877-77300917 | Ch13 region 3 |
| chr13:77300923-77300963 | Ch13 region 3 |
| chr13:77301139-77301179 | Ch13 region 3 |
| chr13:77301248-77301288 | Ch13 region 3 |
| chr13:77301363-77301403 | Ch13 region 3 |
| chr13:77301408-77301448 | Ch13 region 3 |
| chr13:77301524-77301564 | Ch13 region 3 |
| chr13:77301737-77301777 | Ch13 region 3 |
| chr13:77301864-77301904 | Ch13 region 3 |
| chr13:77302036-77302076 | Ch13 region 3 |
| chr13:77302085-77302125 | Ch13 region 3 |
| chr13:77302132-77302172 | Ch13 region 3 |
| chr13:77302371-77302411 | Ch13 region 3 |
| chr13:77302491-77302531 | Ch13 region 3 |
| chr13:77302671-77302711 | Ch13 region 3 |
| chr13:77302927-77302967 | Ch13 region 3 |
| chr13:77302979-77303019 | Ch13 region 3 |
| chr13:77303119-77303159 | Ch13 region 3 |
| chr13:77303233-77303273 | Ch13 region 3 |
| chr13:77303287-77303327 | Ch13 region 3 |
| chr13:77303387-77303427 | Ch13 region 3 |
| chr13:77303447-77303487 | Ch13 region 3 |
| chr13:77303497-77303537 | Ch13 region 3 |
| chr13:77303558-77303598 | Ch13 region 3 |
| chr13:77303603-77303643 | Ch13 region 3 |
| chr13:77303696-77303736 | Ch13 region 3 |
| chr13:77303741-77303781 | Ch13 region 3 |
| chr13:77303827-77303867 | Ch13 region 3 |
| chr13:77303917-77303957 | Ch13 region 3 |
| chr13:77303972-77304012 | Ch13 region 3 |
| chr13:77304025-77304065 | Ch13 region 3 |
| chr13:77304090-77304130 | Ch13 region 3 |
| chr13:77304155-77304195 | Ch13 region 3 |
| chr13:77304207-77304247 | Ch13 region 3 |
| chr13:77304273-77304313 | Ch13 region 3 |
| chr13:77304329-77304369 | Ch13 region 3 |
| chr13:77304455-77304495 | Ch13 region 3 |

| | |
|---|---|
| chr13:77304507-77304547 | Ch13 region 3 |
| chr13:77304577-77304617 | Ch13 region 3 |
| chr13:77304630-77304670 | Ch13 region 3 |
| chr13:77304743-77304783 | Ch13 region 3 |
| chr13:77304815-77304855 | Ch13 region 3 |
| chr13:77304867-77304907 | Ch13 region 3 |
| chr13:77304932-77304972 | Ch13 region 3 |
| chr13:77304986-77305026 | Ch13 region 3 |
| chr13:77305034-77305074 | Ch13 region 3 |
| chr13:77305119-77305159 | Ch13 region 3 |
| chr13:77305208-77305248 | Ch13 region 3 |
| chr13:77305329-77305369 | Ch13 region 3 |
| chr13:77305425-77305465 | Ch13 region 3 |
| chr13:77305474-77305514 | Ch13 region 3 |
| chr13:77305531-77305571 | Ch13 region 3 |
| chr13:77305624-77305664 | Ch13 region 3 |
| chr13:77305670-77305710 | Ch13 region 3 |
| chr13:77305786-77305826 | Ch13 region 3 |
| chr13:77305840-77305880 | Ch13 region 3 |
| chr13:77306289-77306329 | Ch13 region 3 |
| chr13:77306393-77306433 | Ch13 region 3 |
| chr13:77306474-77306514 | Ch13 region 3 |
| chr13:77306535-77306575 | Ch13 region 3 |
| chr13:77306595-77306635 | Ch13 region 3 |
| chr13:77306640-77306680 | Ch13 region 3 |
| chr13:77306716-77306756 | Ch13 region 3 |
| chr13:77306786-77306826 | Ch13 region 3 |
| chr13:77306831-77306871 | Ch13 region 3 |
| chr13:77306876-77306916 | Ch13 region 3 |
| chr13:77306945-77306985 | Ch13 region 3 |
| chr13:77306990-77307030 | Ch13 region 3 |
| chr13:77307044-77307084 | Ch13 region 3 |
| chr13:77307156-77307196 | Ch13 region 3 |
| chr13:77307211-77307251 | Ch13 region 3 |
| chr13:77307403-77307443 | Ch13 region 3 |
| chr13:77307511-77307551 | Ch13 region 3 |
| chr13:77307597-77307637 | Ch13 region 3 |
| chr13:77307693-77307733 | Ch13 region 3 |
| chr13:77307750-77307790 | Ch13 region 3 |
| chr13:77307810-77307850 | Ch13 region 3 |
| chr13:77307878-77307918 | Ch13 region 3 |
| chr13:77307945-77307985 | Ch13 region 3 |
| chr13:77308014-77308054 | Ch13 region 3 |
| chr13:77308062-77308102 | Ch13 region 3 |
| chr13:77308115-77308155 | Ch13 region 3 |
| chr13:77308224-77308264 | Ch13 region 3 |
| chr13:77308272-77308312 | Ch13 region 3 |
| chr13:77308332-77308372 | Ch13 region 3 |
| chr13:77308387-77308427 | Ch13 region 3 |
| chr13:77308432-77308472 | Ch13 region 3 |
| chr13:77308497-77308537 | Ch13 region 3 |
| chr13:77308557-77308597 | Ch13 region 3 |

| | |
|---|---|
| chr13:77308602-77308642 | Ch13 region 3 |
| chr13:77308649-77308689 | Ch13 region 3 |
| chr13:77308778-77308818 | Ch13 region 3 |
| chr13:77308923-77308963 | Ch13 region 3 |
| chr13:77308987-77309027 | Ch13 region 3 |
| chr13:77309140-77309180 | Ch13 region 3 |
| chr13:77309198-77309238 | Ch13 region 3 |
| chr13:77309245-77309285 | Ch13 region 3 |
| chr13:77309347-77309387 | Ch13 region 3 |
| chr13:77309417-77309457 | Ch13 region 3 |
| chr13:77309474-77309514 | Ch13 region 3 |
| chr13:77309556-77309596 | Ch13 region 3 |
| chr13:77309608-77309648 | Ch13 region 3 |
| chr13:77309662-77309702 | Ch13 region 3 |
| chr13:77309709-77309749 | Ch13 region 3 |
| chr13:77309758-77309798 | Ch13 region 3 |
| chr13:77309822-77309862 | Ch13 region 3 |
| chr13:77309895-77309935 | Ch13 region 3 |
| chr13:77309951-77309991 | Ch13 region 3 |
| chr13:77310004-77310044 | Ch13 region 3 |
| chr13:77310050-77310090 | Ch13 region 3 |
| chr13:77310148-77310188 | Ch13 region 3 |
| chr13:77310214-77310254 | Ch13 region 3 |
| chr13:77310291-77310331 | Ch13 region 3 |
| chr13:77310365-77310405 | Ch13 region 3 |
| chr13:77310411-77310451 | Ch13 region 3 |
| chr13:77310469-77310509 | Ch13 region 3 |
| chr13:77310641-77310681 | Ch13 region 3 |
| chr13:77310720-77310760 | Ch13 region 3 |
| chr13:77310778-77310818 | Ch13 region 3 |
| chr13:77310879-77310919 | Ch13 region 3 |
| chr13:77310926-77310966 | Ch13 region 3 |
| chr13:77311022-77311062 | Ch13 region 3 |
| chr13:77311079-77311119 | Ch13 region 3 |
| chr13:77311124-77311164 | Ch13 region 3 |
| chr13:77311171-77311211 | Ch13 region 3 |
| chr13:77311311-77311351 | Ch13 region 3 |
| chr13:77311543-77311583 | Ch13 region 3 |
| chr13:77311595-77311635 | Ch13 region 3 |
| chr13:77311661-77311701 | Ch13 region 3 |
| chr13:77311713-77311753 | Ch13 region 3 |
| chr13:77311811-77311851 | Ch13 region 3 |
| chr13:77311905-77311945 | Ch13 region 3 |
| chr13:77311987-77312027 | Ch13 region 3 |
| chr13:77312041-77312081 | Ch13 region 3 |
| chr13:77312159-77312199 | Ch13 region 3 |
| chr13:77312216-77312256 | Ch13 region 3 |
| chr13:77312291-77312331 | Ch13 region 3 |
| chr13:77312354-77312394 | Ch13 region 3 |
| chr13:77312441-77312481 | Ch13 region 3 |
| chr13:77312503-77312543 | Ch13 region 3 |
| chr13:77312573-77312613 | Ch13 region 3 |

| | |
|---|---|
| chr13:77312627-77312667 | Ch13 region 3 |
| chr13:77312694-77312734 | Ch13 region 3 |
| chr13:77312739-77312779 | Ch13 region 3 |
| chr13:77312835-77312875 | Ch13 region 3 |
| chr13:77312918-77312958 | Ch13 region 3 |
| chr13:77312995-77313035 | Ch13 region 3 |
| chr13:77313041-77313081 | Ch13 region 3 |
| chr13:77313137-77313177 | Ch13 region 3 |
| chr13:77313216-77313256 | Ch13 region 3 |
| chr13:77313311-77313351 | Ch13 region 3 |
| chr13:77313356-77313396 | Ch13 region 3 |
| chr13:77313414-77313454 | Ch13 region 3 |
| chr13:77313493-77313533 | Ch13 region 3 |
| chr13:77313566-77313606 | Ch13 region 3 |
| chr13:77313612-77313652 | Ch13 region 3 |
| chr13:77313673-77313713 | Ch13 region 3 |
| chr13:77313718-77313758 | Ch13 region 3 |
| chr13:77313781-77313821 | Ch13 region 3 |
| chr13:77313826-77313866 | Ch13 region 3 |
| chr13:77313875-77313915 | Ch13 region 3 |
| chr13:77313991-77314031 | Ch13 region 3 |
| chr13:77314038-77314078 | Ch13 region 3 |
| chr13:77314115-77314155 | Ch13 region 3 |
| chr13:77314160-77314200 | Ch13 region 3 |
| chr13:77314229-77314269 | Ch13 region 3 |
| chr13:77314285-77314325 | Ch13 region 3 |
| chr13:77314346-77314386 | Ch13 region 3 |
| chr13:77314403-77314443 | Ch13 region 3 |
| chr13:77314513-77314553 | Ch13 region 3 |
| chr13:77314592-77314632 | Ch13 region 3 |
| chr13:77314653-77314693 | Ch13 region 3 |
| chr13:77314719-77314759 | Ch13 region 3 |
| chr13:77314770-77314810 | Ch13 region 3 |
| chr13:77314817-77314857 | Ch13 region 3 |
| chr13:77314873-77314913 | Ch13 region 3 |
| chr13:77314918-77314958 | Ch13 region 3 |
| chr13:77315001-77315041 | Ch13 region 3 |
| chr13:77315077-77315117 | Ch13 region 3 |
| chr13:77315143-77315183 | Ch13 region 3 |
| chr13:77315243-77315283 | Ch13 region 3 |
| chr13:77315288-77315328 | Ch13 region 3 |
| chr13:77315355-77315395 | Ch13 region 3 |
| chr13:77315425-77315465 | Ch13 region 3 |
| chr13:77315533-77315573 | Ch13 region 3 |
| chr13:77315615-77315655 | Ch13 region 3 |
| chr13:77315713-77315753 | Ch13 region 3 |
| chr13:77315805-77315845 | Ch13 region 3 |
| chr13:77315850-77315890 | Ch13 region 3 |
| chr13:77315924-77315964 | Ch13 region 3 |
| chr13:77316006-77316046 | Ch13 region 3 |
| chr13:77316062-77316102 | Ch13 region 3 |
| chr13:77316155-77316195 | Ch13 region 3 |

| | |
|---|---|
| chr13:77316299-77316339 | Ch13 region 3 |
| chr13:77316360-77316400 | Ch13 region 3 |
| chr13:77316421-77316461 | Ch13 region 3 |
| chr13:77316495-77316535 | Ch13 region 3 |
| chr13:77316549-77316589 | Ch13 region 3 |
| chr13:77316625-77316665 | Ch13 region 3 |
| chr13:77316683-77316723 | Ch13 region 3 |
| chr13:77316730-77316770 | Ch13 region 3 |
| chr13:77316778-77316818 | Ch13 region 3 |
| chr13:77316864-77316904 | Ch13 region 3 |
| chr13:77316938-77316978 | Ch13 region 3 |
| chr13:77316995-77317035 | Ch13 region 3 |
| chr13:77317059-77317099 | Ch13 region 3 |
| chr13:77317136-77317176 | Ch13 region 3 |
| chr13:77317218-77317258 | Ch13 region 3 |
| chr13:77317280-77317320 | Ch13 region 3 |
| chr13:77317443-77317483 | Ch13 region 3 |
| chr13:77317503-77317543 | Ch13 region 3 |
| chr13:77317559-77317599 | Ch13 region 3 |
| chr13:77317640-77317680 | Ch13 region 3 |
| chr13:77317689-77317729 | Ch13 region 3 |
| chr13:77317784-77317824 | Ch13 region 3 |
| chr13:77317859-77317899 | Ch13 region 3 |
| chr13:77317925-77317965 | Ch13 region 3 |
| chr13:77317977-77318017 | Ch13 region 3 |
| chr13:77318085-77318125 | Ch13 region 3 |
| chr13:77318168-77318208 | Ch13 region 3 |
| chr13:77318391-77318431 | Ch13 region 3 |
| chr13:77318455-77318495 | Ch13 region 3 |
| chr13:77318506-77318546 | Ch13 region 3 |
| chr13:77318551-77318591 | Ch13 region 3 |
| chr13:77318611-77318651 | Ch13 region 3 |
| chr13:77318656-77318696 | Ch13 region 3 |
| chr13:77318723-77318763 | Ch13 region 3 |
| chr13:77318779-77318819 | Ch13 region 3 |
| chr13:77318824-77318864 | Ch13 region 3 |
| chr13:77318913-77318953 | Ch13 region 3 |
| chr13:77318974-77319014 | Ch13 region 3 |
| chr13:77319195-77319235 | Ch13 region 3 |
| chr13:77319334-77319374 | Ch13 region 3 |
| chr13:77319430-77319470 | Ch13 region 3 |
| chr13:77319497-77319537 | Ch13 region 3 |
| chr13:77319558-77319598 | Ch13 region 3 |
| chr13:77319634-77319674 | Ch13 region 3 |
| chr13:77319747-77319787 | Ch13 region 3 |
| chr13:77319818-77319858 | Ch13 region 3 |
| chr13:77319890-77319930 | Ch13 region 3 |
| chr13:77319964-77320004 | Ch13 region 3 |
| chr13:77320027-77320067 | Ch13 region 3 |
| chr13:77320072-77320112 | Ch13 region 3 |
| chr13:77320119-77320159 | Ch13 region 3 |
| chr13:77320171-77320211 | Ch13 region 3 |

| | |
|---|---|
| chr13:77320218-77320258 | Ch13 region 3 |
| chr13:77320290-77320330 | Ch13 region 3 |
| chr13:77320356-77320396 | Ch13 region 3 |
| chr13:77320480-77320520 | Ch13 region 3 |
| chr13:77320694-77320734 | Ch13 region 3 |
| chr13:77320742-77320782 | Ch13 region 3 |
| chr13:77320844-77320884 | Ch13 region 3 |
| chr13:77320896-77320936 | Ch13 region 3 |
| chr13:77320953-77320993 | Ch13 region 3 |
| chr13:77321008-77321048 | Ch13 region 3 |
| chr13:77321107-77321147 | Ch13 region 3 |
| chr13:77321193-77321233 | Ch13 region 3 |
| chr13:77321272-77321312 | Ch13 region 3 |
| chr13:77321479-77321519 | Ch13 region 3 |
| chr13:77321524-77321564 | Ch13 region 3 |
| chr13:77321596-77321636 | Ch13 region 3 |
| chr13:77321659-77321699 | Ch13 region 3 |
| chr13:77321713-77321753 | Ch13 region 3 |
| chr13:77321763-77321803 | Ch13 region 3 |
| chr13:77321842-77321882 | Ch13 region 3 |
| chr13:77322055-77322095 | Ch13 region 3 |
| chr13:77322146-77322186 | Ch13 region 3 |
| chr13:77322191-77322231 | Ch13 region 3 |
| chr13:77322292-77322332 | Ch13 region 3 |
| chr13:77322337-77322377 | Ch13 region 3 |
| chr13:77322442-77322482 | Ch13 region 3 |
| chr13:77322507-77322547 | Ch13 region 3 |
| chr13:77322552-77322592 | Ch13 region 3 |
| chr13:77322654-77322694 | Ch13 region 3 |
| chr13:77322705-77322745 | Ch13 region 3 |
| chr13:77322792-77322832 | Ch13 region 3 |
| chr13:77322854-77322894 | Ch13 region 3 |
| chr13:77322910-77322950 | Ch13 region 3 |
| chr13:77322968-77323008 | Ch13 region 3 |
| chr13:77323038-77323078 | Ch13 region 3 |
| chr13:77323089-77323129 | Ch13 region 3 |
| chr13:77323148-77323188 | Ch13 region 3 |
| chr13:77323218-77323258 | Ch13 region 3 |
| chr13:77323275-77323315 | Ch13 region 3 |
| chr13:77323322-77323362 | Ch13 region 3 |
| chr13:77323376-77323416 | Ch13 region 3 |
| chr13:77323421-77323461 | Ch13 region 3 |
| chr13:77323526-77323566 | Ch13 region 3 |
| chr13:77323571-77323611 | Ch13 region 3 |
| chr13:77323724-77323764 | Ch13 region 3 |
| chr13:77323801-77323841 | Ch13 region 3 |
| chr13:77323860-77323900 | Ch13 region 3 |
| chr13:77323905-77323945 | Ch13 region 3 |
| chr13:77324008-77324048 | Ch13 region 3 |
| chr13:77324074-77324114 | Ch13 region 3 |
| chr13:77324204-77324244 | Ch13 region 3 |
| chr13:77324295-77324335 | Ch13 region 3 |

| | |
|---|---|
| chr13:77324340-77324380 | Ch13 region 3 |
| chr13:77324421-77324461 | Ch13 region 3 |
| chr13:77324500-77324540 | Ch13 region 3 |
| chr13:77324581-77324621 | Ch13 region 3 |
| chr13:77324626-77324666 | Ch13 region 3 |
| chr13:77324685-77324725 | Ch13 region 3 |
| chr13:77324742-77324782 | Ch13 region 3 |
| chr13:77324802-77324842 | Ch13 region 3 |
| chr13:77324847-77324887 | Ch13 region 3 |
| chr13:77324901-77324941 | Ch13 region 3 |
| chr13:77324975-77325015 | Ch13 region 3 |
| chr13:77325052-77325092 | Ch13 region 3 |
| chr13:77325127-77325167 | Ch13 region 3 |
| chr13:77325210-77325250 | Ch13 region 3 |
| chr13:77325266-77325306 | Ch13 region 3 |
| chr13:77325336-77325376 | Ch13 region 3 |
| chr13:77325398-77325438 | Ch13 region 3 |
| chr13:77325458-77325498 | Ch13 region 3 |
| chr13:77325559-77325599 | Ch13 region 3 |
| chr13:77325621-77325661 | Ch13 region 3 |
| chr13:77325688-77325728 | Ch13 region 3 |
| chr13:77325736-77325776 | Ch13 region 3 |
| chr13:77325783-77325823 | Ch13 region 3 |
| chr13:77325869-77325909 | Ch13 region 3 |
| chr13:77325914-77325954 | Ch13 region 3 |
| chr13:77325959-77325999 | Ch13 region 3 |
| chr13:77326016-77326056 | Ch13 region 3 |
| chr13:77326061-77326101 | Ch13 region 3 |
| chr13:77326206-77326246 | Ch13 region 3 |
| chr13:77326334-77326374 | Ch13 region 3 |
| chr13:77326380-77326420 | Ch13 region 3 |
| chr13:77326490-77326530 | Ch13 region 3 |
| chr13:77326562-77326602 | Ch13 region 3 |
| chr13:77326623-77326663 | Ch13 region 3 |
| chr13:77326722-77326762 | Ch13 region 3 |
| chr13:77326767-77326807 | Ch13 region 3 |
| chr13:77326846-77326886 | Ch13 region 3 |
| chr13:77326892-77326932 | Ch13 region 3 |
| chr13:77326964-77327004 | Ch13 region 3 |
| chr13:77327009-77327049 | Ch13 region 3 |
| chr13:77327064-77327104 | Ch13 region 3 |
| chr13:77327122-77327162 | Ch13 region 3 |
| chr13:77327237-77327277 | Ch13 region 3 |
| chr13:77327296-77327336 | Ch13 region 3 |
| chr13:77327362-77327402 | Ch13 region 3 |
| chr13:77327580-77327620 | Ch13 region 3 |
| chr13:77327713-77327753 | Ch13 region 3 |
| chr13:77327761-77327801 | Ch13 region 3 |
| chr13:77327840-77327880 | Ch13 region 3 |
| chr13:77327969-77328009 | Ch13 region 3 |
| chr13:77328065-77328105 | Ch13 region 3 |
| chr13:77328136-77328176 | Ch13 region 3 |

| | |
|---|---|
| chr13:77328213-77328253 | Ch13 region 3 |
| chr13:77328342-77328382 | Ch13 region 3 |
| chr13:77328404-77328444 | Ch13 region 3 |
| chr13:77328456-77328496 | Ch13 region 3 |
| chr13:77328604-77328644 | Ch13 region 3 |
| chr13:77328651-77328691 | Ch13 region 3 |
| chr13:77328696-77328736 | Ch13 region 3 |
| chr13:77328753-77328793 | Ch13 region 3 |
| chr13:77328801-77328841 | Ch13 region 3 |
| chr13:77328881-77328921 | Ch13 region 3 |
| chr13:77329034-77329074 | Ch13 region 3 |
| chr13:77329115-77329155 | Ch13 region 3 |
| chr13:77329174-77329214 | Ch13 region 3 |
| chr13:77329225-77329265 | Ch13 region 3 |
| chr13:77329312-77329352 | Ch13 region 3 |
| chr13:77329383-77329423 | Ch13 region 3 |
| chr13:77329487-77329527 | Ch13 region 3 |
| chr13:77329551-77329591 | Ch13 region 3 |
| chr13:77329615-77329655 | Ch13 region 3 |
| chr13:77329677-77329717 | Ch13 region 3 |
| chr13:77329734-77329774 | Ch13 region 3 |
| chr13:77329783-77329823 | Ch13 region 3 |
| chr13:77329870-77329910 | Ch13 region 3 |
| chr13:77329940-77329980 | Ch13 region 3 |
| chr13:77330027-77330067 | Ch13 region 3 |
| chr13:77330121-77330161 | Ch13 region 3 |
| chr13:77330183-77330223 | Ch13 region 3 |
| chr13:77330241-77330281 | Ch13 region 3 |
| chr13:77330331-77330371 | Ch13 region 3 |
| chr13:77330398-77330438 | Ch13 region 3 |
| chr13:77330451-77330491 | Ch13 region 3 |
| chr13:77330517-77330557 | Ch13 region 3 |
| chr13:77330562-77330602 | Ch13 region 3 |
| chr13:77330638-77330678 | Ch13 region 3 |
| chr13:77330712-77330752 | Ch13 region 3 |
| chr13:77330768-77330808 | Ch13 region 3 |
| chr13:77330844-77330884 | Ch13 region 3 |
| chr13:77330908-77330948 | Ch13 region 3 |
| chr13:77330990-77331030 | Ch13 region 3 |
| chr13:77331037-77331077 | Ch13 region 3 |
| chr13:77331114-77331154 | Ch13 region 3 |
| chr13:77331169-77331209 | Ch13 region 3 |
| chr13:77331219-77331259 | Ch13 region 3 |
| chr13:77331296-77331336 | Ch13 region 3 |
| chr13:77331362-77331402 | Ch13 region 3 |
| chr13:77331483-77331523 | Ch13 region 3 |
| chr13:77331600-77331640 | Ch13 region 3 |
| chr13:77331649-77331689 | Ch13 region 3 |
| chr13:77331765-77331805 | Ch13 region 3 |
| chr13:77331932-77331972 | Ch13 region 3 |
| chr13:77332126-77332166 | Ch13 region 3 |
| chr13:77332306-77332346 | Ch13 region 3 |

| | |
|---|---|
| chr13:77332355-77332395 | Ch13 region 3 |
| chr13:77332439-77332479 | Ch13 region 3 |
| chr13:77332540-77332580 | Ch13 region 3 |
| chr13:77332587-77332627 | Ch13 region 3 |
| chr13:77332632-77332672 | Ch13 region 3 |
| chr13:77332724-77332764 | Ch13 region 3 |
| chr13:77332840-77332880 | Ch13 region 3 |
| chr13:77332946-77332986 | Ch13 region 3 |
| chr13:77332993-77333033 | Ch13 region 3 |
| chr13:77333072-77333112 | Ch13 region 3 |
| chr13:77333135-77333175 | Ch13 region 3 |
| chr13:77333184-77333224 | Ch13 region 3 |
| chr13:77333241-77333281 | Ch13 region 3 |
| chr13:77333286-77333326 | Ch13 region 3 |
| chr13:77333359-77333399 | Ch13 region 3 |
| chr13:77333418-77333458 | Ch13 region 3 |
| chr13:77333464-77333504 | Ch13 region 3 |
| chr13:77333538-77333578 | Ch13 region 3 |
| chr13:77333587-77333627 | Ch13 region 3 |
| chr13:77333657-77333697 | Ch13 region 3 |
| chr13:77333702-77333742 | Ch13 region 3 |
| chr13:77333751-77333791 | Ch13 region 3 |
| chr13:77333811-77333851 | Ch13 region 3 |
| chr13:77333857-77333897 | Ch13 region 3 |
| chr13:77333975-77334015 | Ch13 region 3 |
| chr13:77334056-77334096 | Ch13 region 3 |
| chr13:77334130-77334170 | Ch13 region 3 |
| chr13:77334204-77334244 | Ch13 region 3 |
| chr13:77334262-77334302 | Ch13 region 3 |
| chr13:77334307-77334347 | Ch13 region 3 |
| chr13:77334397-77334437 | Ch13 region 3 |
| chr13:77334444-77334484 | Ch13 region 3 |
| chr13:77334494-77334534 | Ch13 region 3 |
| chr13:77334588-77334628 | Ch13 region 3 |
| chr13:77334644-77334684 | Ch13 region 3 |
| chr13:77334716-77334756 | Ch13 region 3 |
| chr13:77334787-77334827 | Ch13 region 3 |
| chr13:77334844-77334884 | Ch13 region 3 |
| chr13:77334892-77334932 | Ch13 region 3 |
| chr13:77334972-77335012 | Ch13 region 3 |
| chr13:77335028-77335068 | Ch13 region 3 |
| chr13:77335133-77335173 | Ch13 region 3 |
| chr13:77335230-77335270 | Ch13 region 3 |
| chr13:77335312-77335352 | Ch13 region 3 |
| chr13:77335426-77335466 | Ch13 region 3 |
| chr13:77335515-77335555 | Ch13 region 3 |
| chr13:77335597-77335637 | Ch13 region 3 |
| chr13:77335992-77336032 | Ch13 region 3 |
| chr13:77336037-77336077 | Ch13 region 3 |
| chr13:77336111-77336151 | Ch13 region 3 |
| chr13:77336204-77336244 | Ch13 region 3 |
| chr13:77336289-77336329 | Ch13 region 3 |

| | |
|---|---|
| chr13:77336334-77336374 | Ch13 region 3 |
| chr13:77336415-77336455 | Ch13 region 3 |
| chr13:77336579-77336619 | Ch13 region 3 |
| chr13:77336625-77336665 | Ch13 region 3 |
| chr13:77336685-77336725 | Ch13 region 3 |
| chr13:77336730-77336770 | Ch13 region 3 |
| chr13:77336782-77336822 | Ch13 region 3 |
| chr13:77336840-77336880 | Ch13 region 3 |
| chr13:77336890-77336930 | Ch13 region 3 |
| chr13:77336952-77336992 | Ch13 region 3 |
| chr13:77337012-77337052 | Ch13 region 3 |
| chr13:77337059-77337099 | Ch13 region 3 |
| chr13:77337109-77337149 | Ch13 region 3 |
| chr13:77337179-77337219 | Ch13 region 3 |
| chr13:77337232-77337272 | Ch13 region 3 |
| chr13:77337279-77337319 | Ch13 region 3 |
| chr13:77337337-77337377 | Ch13 region 3 |
| chr13:77337382-77337422 | Ch13 region 3 |
| chr13:77337556-77337596 | Ch13 region 3 |
| chr13:77337618-77337658 | Ch13 region 3 |
| chr13:77337754-77337794 | Ch13 region 3 |
| chr13:77337832-77337872 | Ch13 region 3 |
| chr13:77337912-77337952 | Ch13 region 3 |
| chr13:77337965-77338005 | Ch13 region 3 |
| chr13:77338030-77338070 | Ch13 region 3 |
| chr13:77338076-77338116 | Ch13 region 3 |
| chr13:77338207-77338247 | Ch13 region 3 |
| chr13:77338256-77338296 | Ch13 region 3 |
| chr13:77338301-77338341 | Ch13 region 3 |
| chr13:77338351-77338391 | Ch13 region 3 |
| chr13:77338504-77338544 | Ch13 region 3 |
| chr13:77338560-77338600 | Ch13 region 3 |
| chr13:77338634-77338674 | Ch13 region 3 |
| chr13:77338688-77338728 | Ch13 region 3 |
| chr13:77338798-77338838 | Ch13 region 3 |
| chr13:77338850-77338890 | Ch13 region 3 |
| chr13:77338916-77338956 | Ch13 region 3 |
| chr13:77338974-77339014 | Ch13 region 3 |
| chr13:77339040-77339080 | Ch13 region 3 |
| chr13:77339119-77339159 | Ch13 region 3 |
| chr13:77339190-77339230 | Ch13 region 3 |
| chr13:77339257-77339297 | Ch13 region 3 |
| chr13:77339307-77339347 | Ch13 region 3 |
| chr13:77339363-77339403 | Ch13 region 3 |
| chr13:77339440-77339480 | Ch13 region 3 |
| chr13:77339486-77339526 | Ch13 region 3 |
| chr13:77339549-77339589 | Ch13 region 3 |
| chr13:77339603-77339643 | Ch13 region 3 |
| chr13:77339783-77339823 | Ch13 region 3 |
| chr13:77339837-77339877 | Ch13 region 3 |
| chr13:77339905-77339945 | Ch13 region 3 |
| chr13:77339964-77340004 | Ch13 region 3 |

| | |
|---|---|
| chr13:77340020-77340060 | Ch13 region 3 |
| chr13:77340124-77340164 | Ch13 region 3 |
| chr13:77340197-77340237 | Ch13 region 3 |
| chr13:77340252-77340292 | Ch13 region 3 |
| chr13:77340297-77340337 | Ch13 region 3 |
| chr13:77340368-77340408 | Ch13 region 3 |
| chr13:77340422-77340462 | Ch13 region 3 |
| chr13:77340476-77340516 | Ch13 region 3 |
| chr13:77340544-77340584 | Ch13 region 3 |
| chr13:77340593-77340633 | Ch13 region 3 |
| chr13:77340713-77340753 | Ch13 region 3 |
| chr13:77340758-77340798 | Ch13 region 3 |
| chr13:77340835-77340875 | Ch13 region 3 |
| chr13:77340889-77340929 | Ch13 region 3 |
| chr13:77340950-77340990 | Ch13 region 3 |
| chr13:77341060-77341100 | Ch13 region 3 |
| chr13:77341124-77341164 | Ch13 region 3 |
| chr13:77341179-77341219 | Ch13 region 3 |
| chr13:77341237-77341277 | Ch13 region 3 |
| chr13:77341333-77341373 | Ch13 region 3 |
| chr13:77341385-77341425 | Ch13 region 3 |
| chr13:77341430-77341470 | Ch13 region 3 |
| chr13:77341489-77341529 | Ch13 region 3 |
| chr13:77341544-77341584 | Ch13 region 3 |
| chr13:77341592-77341632 | Ch13 region 3 |
| chr13:77341659-77341699 | Ch13 region 3 |
| chr13:77341799-77341839 | Ch13 region 3 |
| chr13:77341866-77341906 | Ch13 region 3 |
| chr13:77341985-77342025 | Ch13 region 3 |
| chr13:77342048-77342088 | Ch13 region 3 |
| chr13:77342097-77342137 | Ch13 region 3 |
| chr13:77342211-77342251 | Ch13 region 3 |
| chr13:77342274-77342314 | Ch13 region 3 |
| chr13:77342362-77342402 | Ch13 region 3 |
| chr13:77342429-77342469 | Ch13 region 3 |
| chr13:77342516-77342556 | Ch13 region 3 |
| chr13:77342610-77342650 | Ch13 region 3 |
| chr13:77342688-77342728 | Ch13 region 3 |
| chr13:77342749-77342789 | Ch13 region 3 |
| chr13:77342825-77342865 | Ch13 region 3 |
| chr13:77342874-77342914 | Ch13 region 3 |
| chr13:77342928-77342968 | Ch13 region 3 |
| chr13:77342998-77343038 | Ch13 region 3 |
| chr13:77343097-77343137 | Ch13 region 3 |
| chr13:77343142-77343182 | Ch13 region 3 |
| chr13:77343191-77343231 | Ch13 region 3 |
| chr13:77343238-77343278 | Ch13 region 3 |
| chr13:77343292-77343332 | Ch13 region 3 |
| chr13:77343364-77343404 | Ch13 region 3 |
| chr13:77343412-77343452 | Ch13 region 3 |
| chr13:77343468-77343508 | Ch13 region 3 |
| chr13:77343565-77343605 | Ch13 region 3 |

| | |
|---|---|
| chr13:77343657-77343697 | Ch13 region 3 |
| chr13:77343748-77343788 | Ch13 region 3 |
| chr13:77343800-77343840 | Ch13 region 3 |
| chr13:77343875-77343915 | Ch13 region 3 |
| chr13:77343927-77343967 | Ch13 region 3 |
| chr13:77343989-77344029 | Ch13 region 3 |
| chr13:77344043-77344083 | Ch13 region 3 |
| chr13:77344149-77344189 | Ch13 region 3 |
| chr13:77344199-77344239 | Ch13 region 3 |
| chr13:77344245-77344285 | Ch13 region 3 |
| chr13:77344302-77344342 | Ch13 region 3 |
| chr13:77344407-77344447 | Ch13 region 3 |
| chr13:77344463-77344503 | Ch13 region 3 |
| chr13:77344533-77344573 | Ch13 region 3 |
| chr13:77344587-77344627 | Ch13 region 3 |
| chr13:77344635-77344675 | Ch13 region 3 |
| chr13:77344720-77344760 | Ch13 region 3 |
| chr13:77344765-77344805 | Ch13 region 3 |
| chr13:77344833-77344873 | Ch13 region 3 |
| chr13:77344908-77344948 | Ch13 region 3 |
| chr13:77344987-77345027 | Ch13 region 3 |
| chr13:77345137-77345177 | Ch13 region 3 |
| chr13:77345197-77345237 | Ch13 region 3 |
| chr13:77345359-77345399 | Ch13 region 3 |
| chr13:77345552-77345592 | Ch13 region 3 |
| chr13:77345705-77345745 | Ch13 region 3 |
| chr13:77345777-77345817 | Ch13 region 3 |
| chr13:77345860-77345900 | Ch13 region 3 |
| chr13:77346040-77346080 | Ch13 region 3 |
| chr13:77346151-77346191 | Ch13 region 3 |
| chr13:77346277-77346317 | Ch13 region 3 |
| chr13:77346323-77346363 | Ch13 region 3 |
| chr13:77346408-77346448 | Ch13 region 3 |
| chr13:77346455-77346495 | Ch13 region 3 |
| chr13:77346517-77346557 | Ch13 region 3 |
| chr13:77346584-77346624 | Ch13 region 3 |
| chr13:77346629-77346669 | Ch13 region 3 |
| chr13:77346696-77346736 | Ch13 region 3 |
| chr13:77346797-77346837 | Ch13 region 3 |
| chr13:77346925-77346965 | Ch13 region 3 |
| chr13:77346972-77347012 | Ch13 region 3 |
| chr13:77347017-77347057 | Ch13 region 3 |
| chr13:77347115-77347155 | Ch13 region 3 |
| chr13:77347165-77347205 | Ch13 region 3 |
| chr13:77347229-77347269 | Ch13 region 3 |
| chr13:77347356-77347396 | Ch13 region 3 |
| chr13:77347452-77347492 | Ch13 region 3 |
| chr13:77347544-77347584 | Ch13 region 3 |
| chr13:77347604-77347644 | Ch13 region 3 |
| chr13:77347649-77347689 | Ch13 region 3 |
| chr13:77347743-77347783 | Ch13 region 3 |
| chr13:77347857-77347897 | Ch13 region 3 |

| | |
|---|---|
| chr13:77347908-77347948 | Ch13 region 3 |
| chr13:77347963-77348003 | Ch13 region 3 |
| chr13:77348008-77348048 | Ch13 region 3 |
| chr13:77348076-77348116 | Ch13 region 3 |
| chr13:77348209-77348249 | Ch13 region 3 |
| chr13:77348255-77348295 | Ch13 region 3 |
| chr13:77348324-77348364 | Ch13 region 3 |
| chr13:77348391-77348431 | Ch13 region 3 |
| chr13:77348497-77348537 | Ch13 region 3 |
| chr13:77348654-77348694 | Ch13 region 3 |
| chr13:77348721-77348761 | Ch13 region 3 |
| chr13:77348766-77348806 | Ch13 region 3 |
| chr13:77348815-77348855 | Ch13 region 3 |
| chr13:77348887-77348927 | Ch13 region 3 |
| chr13:77348936-77348976 | Ch13 region 3 |
| chr13:77348985-77349025 | Ch13 region 3 |
| chr13:77349043-77349083 | Ch13 region 3 |
| chr13:77349157-77349197 | Ch13 region 3 |
| chr13:77349220-77349260 | Ch13 region 3 |
| chr13:77349299-77349339 | Ch13 region 3 |
| chr13:77349367-77349407 | Ch13 region 3 |
| chr13:77349460-77349500 | Ch13 region 3 |
| chr13:77349514-77349554 | Ch13 region 3 |
| chr13:77349603-77349643 | Ch13 region 3 |
| chr13:77349664-77349704 | Ch13 region 3 |
| chr13:77349723-77349763 | Ch13 region 3 |
| chr13:77349784-77349824 | Ch13 region 3 |
| chr13:77349842-77349882 | Ch13 region 3 |
| chr13:77349888-77349928 | Ch13 region 3 |
| chr13:77349933-77349973 | Ch13 region 3 |
| chr13:77350049-77350089 | Ch13 region 3 |
| chr13:77350096-77350136 | Ch13 region 3 |
| chr13:77350165-77350205 | Ch13 region 3 |
| chr13:77350217-77350257 | Ch13 region 3 |
| chr13:77350312-77350352 | Ch13 region 3 |
| chr13:77350384-77350424 | Ch13 region 3 |
| chr13:77350497-77350537 | Ch13 region 3 |
| chr13:77350552-77350592 | Ch13 region 3 |
| chr13:77350598-77350638 | Ch13 region 3 |
| chr13:77350668-77350708 | Ch13 region 3 |
| chr13:77350744-77350784 | Ch13 region 3 |
| chr13:77350789-77350829 | Ch13 region 3 |
| chr13:77350871-77350911 | Ch13 region 3 |
| chr13:77350930-77350970 | Ch13 region 3 |
| chr13:77351017-77351057 | Ch13 region 3 |
| chr13:77351223-77351263 | Ch13 region 3 |
| chr13:77351390-77351430 | Ch13 region 3 |
| chr13:77351448-77351488 | Ch13 region 3 |
| chr13:77351498-77351538 | Ch13 region 3 |
| chr13:77351543-77351583 | Ch13 region 3 |
| chr13:77351647-77351687 | Ch13 region 3 |
| chr13:77351697-77351737 | Ch13 region 3 |

| | |
|---|---|
| chr13:77351749-77351789 | Ch13 region 3 |
| chr13:77351872-77351912 | Ch13 region 3 |
| chr13:77351924-77351964 | Ch13 region 3 |
| chr13:77352033-77352073 | Ch13 region 3 |
| chr13:77352094-77352134 | Ch13 region 3 |
| chr13:77352141-77352181 | Ch13 region 3 |
| chr13:77352222-77352262 | Ch13 region 3 |
| chr13:77352270-77352310 | Ch13 region 3 |
| chr13:77352334-77352374 | Ch13 region 3 |
| chr13:77352397-77352437 | Ch13 region 3 |
| chr13:77352453-77352493 | Ch13 region 3 |
| chr13:77352498-77352538 | Ch13 region 3 |
| chr13:77352561-77352601 | Ch13 region 3 |
| chr13:77352614-77352654 | Ch13 region 3 |
| chr13:77352678-77352718 | Ch13 region 3 |
| chr13:77352768-77352808 | Ch13 region 3 |
| chr13:77352881-77352921 | Ch13 region 3 |
| chr13:77352926-77352966 | Ch13 region 3 |
| chr13:77352988-77353028 | Ch13 region 3 |
| chr13:77353057-77353097 | Ch13 region 3 |
| chr13:77353102-77353142 | Ch13 region 3 |
| chr13:77353147-77353187 | Ch13 region 3 |
| chr13:77353202-77353242 | Ch13 region 3 |
| chr13:77353252-77353292 | Ch13 region 3 |
| chr13:77353341-77353381 | Ch13 region 3 |
| chr13:77353389-77353429 | Ch13 region 3 |
| chr13:77353435-77353475 | Ch13 region 3 |
| chr13:77353496-77353536 | Ch13 region 3 |
| chr13:77353541-77353581 | Ch13 region 3 |
| chr13:77353590-77353630 | Ch13 region 3 |
| chr13:77353663-77353703 | Ch13 region 3 |
| chr13:77353708-77353748 | Ch13 region 3 |
| chr13:77353793-77353833 | Ch13 region 3 |
| chr13:77353838-77353878 | Ch13 region 3 |
| chr13:77353887-77353927 | Ch13 region 3 |
| chr13:77353965-77354005 | Ch13 region 3 |
| chr13:77354044-77354084 | Ch13 region 3 |
| chr13:77354113-77354153 | Ch13 region 3 |
| chr13:77354175-77354215 | Ch13 region 3 |
| chr13:77354220-77354260 | Ch13 region 3 |
| chr13:77354307-77354347 | Ch13 region 3 |
| chr13:77354403-77354443 | Ch13 region 3 |
| chr13:77354465-77354505 | Ch13 region 3 |
| chr13:77354564-77354604 | Ch13 region 3 |
| chr13:77354609-77354649 | Ch13 region 3 |
| chr13:77354732-77354772 | Ch13 region 3 |
| chr13:77354819-77354859 | Ch13 region 3 |
| chr13:77354900-77354940 | Ch13 region 3 |
| chr13:77354959-77354999 | Ch13 region 3 |
| chr13:77355033-77355073 | Ch13 region 3 |
| chr13:77355124-77355164 | Ch13 region 3 |
| chr13:77355211-77355251 | Ch13 region 3 |

| | |
|---|---|
| chr13:77355282-77355322 | Ch13 region 3 |
| chr13:77355335-77355375 | Ch13 region 3 |
| chr13:77355440-77355480 | Ch13 region 3 |
| chr13:77355502-77355542 | Ch13 region 3 |
| chr13:77355576-77355616 | Ch13 region 3 |
| chr13:77355661-77355701 | Ch13 region 3 |
| chr13:77355725-77355765 | Ch13 region 3 |
| chr13:77355780-77355820 | Ch13 region 3 |
| chr13:77355846-77355886 | Ch13 region 3 |
| chr13:77355893-77355933 | Ch13 region 3 |
| chr13:77355942-77355982 | Ch13 region 3 |
| chr13:77356007-77356047 | Ch13 region 3 |
| chr13:77356060-77356100 | Ch13 region 3 |
| chr13:77356132-77356172 | Ch13 region 3 |
| chr13:77356204-77356244 | Ch13 region 3 |
| chr13:77356347-77356387 | Ch13 region 3 |
| chr13:77356417-77356457 | Ch13 region 3 |
| chr13:77356536-77356576 | Ch13 region 3 |
| chr13:77356644-77356684 | Ch13 region 3 |
| chr13:77356777-77356817 | Ch13 region 3 |
| chr13:77357002-77357042 | Ch13 region 3 |
| chr13:77357289-77357329 | Ch13 region 3 |
| chr13:77357334-77357374 | Ch13 region 3 |
| chr13:77357650-77357690 | Ch13 region 3 |
| chr13:77357713-77357753 | Ch13 region 3 |
| chr13:77357772-77357812 | Ch13 region 3 |
| chr13:77357914-77357954 | Ch13 region 3 |
| chr13:77357990-77358030 | Ch13 region 3 |
| chr13:77358035-77358075 | Ch13 region 3 |
| chr13:77358081-77358121 | Ch13 region 3 |
| chr13:77358143-77358183 | Ch13 region 3 |
| chr13:77358201-77358241 | Ch13 region 3 |
| chr13:77358257-77358297 | Ch13 region 3 |
| chr13:77358320-77358360 | Ch13 region 3 |
| chr13:77358557-77358597 | Ch13 region 3 |
| chr13:77358622-77358662 | Ch13 region 3 |
| chr13:77358676-77358716 | Ch13 region 3 |
| chr13:77358766-77358806 | Ch13 region 3 |
| chr13:77358848-77358888 | Ch13 region 3 |
| chr13:77358998-77359038 | Ch13 region 3 |
| chr13:77359093-77359133 | Ch13 region 3 |
| chr13:77359189-77359229 | Ch13 region 3 |
| chr13:77359296-77359336 | Ch13 region 3 |
| chr13:77359465-77359505 | Ch13 region 3 |
| chr13:77359566-77359606 | Ch13 region 3 |
| chr13:77359666-77359706 | Ch13 region 3 |
| chr13:77359830-77359870 | Ch13 region 3 |
| chr13:77359881-77359921 | Ch13 region 3 |
| chr13:77360095-77360135 | Ch13 region 3 |
| chr13:77360140-77360180 | Ch13 region 3 |
| chr13:77360219-77360259 | Ch13 region 3 |
| chr13:77360321-77360361 | Ch13 region 3 |

| | |
|---|---|
| chr13:77360483-77360523 | Ch13 region 3 |
| chr13:77360695-77360735 | Ch13 region 3 |
| chr13:77360765-77360805 | Ch13 region 3 |
| chr13:77360900-77360940 | Ch13 region 3 |
| chr13:77360975-77361015 | Ch13 region 3 |
| chr13:77361183-77361223 | Ch13 region 3 |
| chr13:77361346-77361386 | Ch13 region 3 |
| chr13:77361438-77361478 | Ch13 region 3 |
| chr13:77361496-77361536 | Ch13 region 3 |
| chr13:77361640-77361680 | Ch13 region 3 |
| chr13:77361738-77361778 | Ch13 region 3 |
| chr13:77361858-77361898 | Ch13 region 3 |
| chr13:77361904-77361944 | Ch13 region 3 |
| chr13:77361979-77362019 | Ch13 region 3 |
| chr13:77362042-77362082 | Ch13 region 3 |
| chr13:77362222-77362262 | Ch13 region 3 |
| chr13:77362276-77362316 | Ch13 region 3 |
| chr13:77362325-77362365 | Ch13 region 3 |
| chr13:77362423-77362463 | Ch13 region 3 |
| chr13:77362508-77362548 | Ch13 region 3 |
| chr13:77362555-77362595 | Ch13 region 3 |
| chr13:77362647-77362687 | Ch13 region 3 |
| chr13:77362834-77362874 | Ch13 region 3 |
| chr13:77363075-77363115 | Ch13 region 3 |
| chr13:77363163-77363203 | Ch13 region 3 |
| chr13:77363220-77363260 | Ch13 region 3 |
| chr13:77363283-77363323 | Ch13 region 3 |
| chr13:77363396-77363436 | Ch13 region 3 |
| chr13:77363493-77363533 | Ch13 region 3 |
| chr13:77363550-77363590 | Ch13 region 3 |
| chr13:77363598-77363638 | Ch13 region 3 |
| chr13:77363683-77363723 | Ch13 region 3 |
| chr13:77363791-77363831 | Ch13 region 3 |
| chr13:77363845-77363885 | Ch13 region 3 |
| chr13:77363916-77363956 | Ch13 region 3 |
| chr13:77363979-77364019 | Ch13 region 3 |
| chr13:77364024-77364064 | Ch13 region 3 |
| chr13:77364096-77364136 | Ch13 region 3 |
| chr13:77364171-77364211 | Ch13 region 3 |
| chr13:77364221-77364261 | Ch13 region 3 |
| chr13:77364308-77364348 | Ch13 region 3 |
| chr13:77364357-77364397 | Ch13 region 3 |
| chr13:77364430-77364470 | Ch13 region 3 |
| chr13:77364486-77364526 | Ch13 region 3 |
| chr13:77364532-77364572 | Ch13 region 3 |
| chr13:77364577-77364617 | Ch13 region 3 |
| chr13:77364626-77364666 | Ch13 region 3 |
| chr13:77364688-77364728 | Ch13 region 3 |
| chr13:77364761-77364801 | Ch13 region 3 |
| chr13:77364806-77364846 | Ch13 region 3 |
| chr13:77364911-77364951 | Ch13 region 3 |
| chr13:77364979-77365019 | Ch13 region 3 |

| | |
|---|---|
| chr13:77365073-77365113 | Ch13 region 3 |
| chr13:77365150-77365190 | Ch13 region 3 |
| chr13:77365217-77365257 | Ch13 region 3 |
| chr13:77365286-77365326 | Ch13 region 3 |
| chr13:77365331-77365371 | Ch13 region 3 |
| chr13:77365409-77365449 | Ch13 region 3 |
| chr13:77365491-77365531 | Ch13 region 3 |
| chr13:77365556-77365596 | Ch13 region 3 |
| chr13:77365646-77365686 | Ch13 region 3 |
| chr13:77365711-77365751 | Ch13 region 3 |
| chr13:77365785-77365825 | Ch13 region 3 |
| chr13:77365830-77365870 | Ch13 region 3 |
| chr13:77365875-77365915 | Ch13 region 3 |
| chr13:77365947-77365987 | Ch13 region 3 |
| chr13:77366011-77366051 | Ch13 region 3 |
| chr13:77366135-77366175 | Ch13 region 3 |
| chr13:77366283-77366323 | Ch13 region 3 |
| chr13:77366427-77366467 | Ch13 region 3 |
| chr13:77366531-77366571 | Ch13 region 3 |
| chr13:77366698-77366738 | Ch13 region 3 |
| chr13:77366831-77366871 | Ch13 region 3 |
| chr13:77366879-77366919 | Ch13 region 3 |
| chr13:77366947-77366987 | Ch13 region 3 |
| chr13:77367030-77367070 | Ch13 region 3 |
| chr13:77367087-77367127 | Ch13 region 3 |
| chr13:77367132-77367172 | Ch13 region 3 |
| chr13:77367193-77367233 | Ch13 region 3 |
| chr13:77367239-77367279 | Ch13 region 3 |
| chr13:77367311-77367351 | Ch13 region 3 |
| chr13:77367402-77367442 | Ch13 region 3 |
| chr13:77367447-77367487 | Ch13 region 3 |
| chr13:77367507-77367547 | Ch13 region 3 |
| chr13:77367575-77367615 | Ch13 region 3 |
| chr13:77367637-77367677 | Ch13 region 3 |
| chr13:77367738-77367778 | Ch13 region 3 |
| chr13:77367783-77367823 | Ch13 region 3 |
| chr13:77367862-77367902 | Ch13 region 3 |
| chr13:77367925-77367965 | Ch13 region 3 |
| chr13:77368101-77368141 | Ch13 region 3 |
| chr13:77368163-77368203 | Ch13 region 3 |
| chr13:77368226-77368266 | Ch13 region 3 |
| chr13:77368271-77368311 | Ch13 region 3 |
| chr13:77368341-77368381 | Ch13 region 3 |
| chr13:77368394-77368434 | Ch13 region 3 |
| chr13:77368462-77368502 | Ch13 region 3 |
| chr13:77368525-77368565 | Ch13 region 3 |
| chr13:77368598-77368638 | Ch13 region 3 |
| chr13:77368673-77368713 | Ch13 region 3 |
| chr13:77368766-77368806 | Ch13 region 3 |
| chr13:77368834-77368874 | Ch13 region 3 |
| chr13:77368879-77368919 | Ch13 region 3 |
| chr13:77368957-77368997 | Ch13 region 3 |

| | |
|---|---|
| chr13:77369010-77369050 | Ch13 region 3 |
| chr13:77369069-77369109 | Ch13 region 3 |
| chr13:77369123-77369163 | Ch13 region 3 |
| chr13:77369219-77369259 | Ch13 region 3 |
| chr13:77369265-77369305 | Ch13 region 3 |
| chr13:77369312-77369352 | Ch13 region 3 |
| chr13:77369357-77369397 | Ch13 region 3 |
| chr13:77369449-77369489 | Ch13 region 3 |
| chr13:77369514-77369554 | Ch13 region 3 |
| chr13:77369614-77369654 | Ch13 region 3 |
| chr13:77369664-77369704 | Ch13 region 3 |
| chr13:77369731-77369771 | Ch13 region 3 |
| chr13:77369808-77369848 | Ch13 region 3 |
| chr13:77369893-77369933 | Ch13 region 3 |
| chr13:77369941-77369981 | Ch13 region 3 |
| chr13:77370031-77370071 | Ch13 region 3 |
| chr13:77370100-77370140 | Ch13 region 3 |
| chr13:77370150-77370190 | Ch13 region 3 |
| chr13:77370205-77370245 | Ch13 region 3 |
| chr13:77370309-77370349 | Ch13 region 3 |
| chr13:77370364-77370404 | Ch13 region 3 |
| chr13:77370417-77370457 | Ch13 region 3 |
| chr13:77370462-77370502 | Ch13 region 3 |
| chr13:77370507-77370547 | Ch13 region 3 |
| chr13:77370580-77370620 | Ch13 region 3 |
| chr13:77370625-77370665 | Ch13 region 3 |
| chr13:77370710-77370750 | Ch13 region 3 |
| chr13:77370766-77370806 | Ch13 region 3 |
| chr13:77370833-77370873 | Ch13 region 3 |
| chr13:77370889-77370929 | Ch13 region 3 |
| chr13:77370991-77371031 | Ch13 region 3 |
| chr13:77371038-77371078 | Ch13 region 3 |
| chr13:77371112-77371152 | Ch13 region 3 |
| chr13:77371179-77371219 | Ch13 region 3 |
| chr13:77371272-77371312 | Ch13 region 3 |
| chr13:77371326-77371366 | Ch13 region 3 |
| chr13:77371377-77371417 | Ch13 region 3 |
| chr13:77371422-77371462 | Ch13 region 3 |
| chr13:77371469-77371509 | Ch13 region 3 |
| chr13:77371540-77371580 | Ch13 region 3 |
| chr13:77371585-77371625 | Ch13 region 3 |
| chr13:77371635-77371675 | Ch13 region 3 |
| chr13:77371719-77371759 | Ch13 region 3 |
| chr13:77371791-77371831 | Ch13 region 3 |
| chr13:77371865-77371905 | Ch13 region 3 |
| chr13:77371933-77371973 | Ch13 region 3 |
| chr13:77372065-77372105 | Ch13 region 3 |
| chr13:77372193-77372233 | Ch13 region 3 |
| chr13:77372238-77372278 | Ch13 region 3 |
| chr13:77372285-77372325 | Ch13 region 3 |
| chr13:77372371-77372411 | Ch13 region 3 |
| chr13:77372447-77372487 | Ch13 region 3 |

| | |
|---|---|
| chr13:77372512-77372552 | Ch13 region 3 |
| chr13:77372627-77372667 | Ch13 region 3 |
| chr13:77372679-77372719 | Ch13 region 3 |
| chr13:77372739-77372779 | Ch13 region 3 |
| chr13:77372828-77372868 | Ch13 region 3 |
| chr13:77372893-77372933 | Ch13 region 3 |
| chr13:77372958-77372998 | Ch13 region 3 |
| chr13:77373011-77373051 | Ch13 region 3 |
| chr13:77373057-77373097 | Ch13 region 3 |
| chr13:77373102-77373142 | Ch13 region 3 |
| chr13:77373150-77373190 | Ch13 region 3 |
| chr13:77373228-77373268 | Ch13 region 3 |
| chr13:77373273-77373313 | Ch13 region 3 |
| chr13:77373431-77373471 | Ch13 region 3 |
| chr13:77373534-77373574 | Ch13 region 3 |
| chr13:77373583-77373623 | Ch13 region 3 |
| chr13:77373875-77373915 | Ch13 region 3 |
| chr13:77373972-77374012 | Ch13 region 3 |
| chr13:77374029-77374069 | Ch13 region 3 |
| chr13:77374109-77374149 | Ch13 region 3 |
| chr13:77374154-77374194 | Ch13 region 3 |
| chr13:77374288-77374328 | Ch13 region 3 |
| chr13:77374340-77374380 | Ch13 region 3 |
| chr13:77374498-77374538 | Ch13 region 3 |
| chr13:77374626-77374666 | Ch13 region 3 |
| chr13:77374896-77374936 | Ch13 region 3 |
| chr13:77375052-77375092 | Ch13 region 3 |
| chr13:77375097-77375137 | Ch13 region 3 |
| chr13:77375227-77375267 | Ch13 region 3 |
| chr13:77375274-77375314 | Ch13 region 3 |
| chr13:77375319-77375359 | Ch13 region 3 |
| chr13:77375364-77375404 | Ch13 region 3 |
| chr13:77375416-77375456 | Ch13 region 3 |
| chr13:77375502-77375542 | Ch13 region 3 |
| chr13:77375553-77375593 | Ch13 region 3 |
| chr13:77375598-77375638 | Ch13 region 3 |
| chr13:77375659-77375699 | Ch13 region 3 |
| chr13:77375852-77375892 | Ch13 region 3 |
| chr13:77375944-77375984 | Ch13 region 3 |
| chr13:77376069-77376109 | Ch13 region 3 |
| chr13:77376153-77376193 | Ch13 region 3 |
| chr13:77376199-77376239 | Ch13 region 3 |
| chr13:77376379-77376419 | Ch13 region 3 |
| chr13:77376518-77376558 | Ch13 region 3 |
| chr13:77376585-77376625 | Ch13 region 3 |
| chr13:77376631-77376671 | Ch13 region 3 |
| chr13:77376724-77376764 | Ch13 region 3 |
| chr13:77376805-77376845 | Ch13 region 3 |
| chr13:77376861-77376901 | Ch13 region 3 |
| chr13:77376928-77376968 | Ch13 region 3 |
| chr13:77376986-77377026 | Ch13 region 3 |
| chr13:77377130-77377170 | Ch13 region 3 |

| | |
|---|---|
| chr13:77377191-77377231 | Ch13 region 3 |
| chr13:77377285-77377325 | Ch13 region 3 |
| chr13:77377416-77377456 | Ch13 region 3 |
| chr13:77377469-77377509 | Ch13 region 3 |
| chr13:77377524-77377564 | Ch13 region 3 |
| chr13:77377602-77377642 | Ch13 region 3 |
| chr13:77377663-77377703 | Ch13 region 3 |
| chr13:77377782-77377822 | Ch13 region 3 |
| chr13:77377836-77377876 | Ch13 region 3 |
| chr13:77377910-77377950 | Ch13 region 3 |
| chr13:77377958-77377998 | Ch13 region 3 |
| chr13:77378048-77378088 | Ch13 region 3 |
| chr13:77378093-77378133 | Ch13 region 3 |
| chr13:77378163-77378203 | Ch13 region 3 |
| chr13:77378208-77378248 | Ch13 region 3 |
| chr13:77378253-77378293 | Ch13 region 3 |
| chr13:77378301-77378341 | Ch13 region 3 |
| chr13:77378361-77378401 | Ch13 region 3 |
| chr13:77378410-77378450 | Ch13 region 3 |
| chr13:77378595-77378635 | Ch13 region 3 |
| chr13:77378658-77378698 | Ch13 region 3 |
| chr13:77378872-77378912 | Ch13 region 3 |
| chr13:77378949-77378989 | Ch13 region 3 |
| chr13:77379040-77379080 | Ch13 region 3 |
| chr13:77379090-77379130 | Ch13 region 3 |
| chr13:77379148-77379188 | Ch13 region 3 |
| chr13:77379223-77379263 | Ch13 region 3 |
| chr13:77379298-77379338 | Ch13 region 3 |
| chr13:77379380-77379420 | Ch13 region 3 |
| chr13:77379435-77379475 | Ch13 region 3 |
| chr13:77379584-77379624 | Ch13 region 3 |
| chr13:77379647-77379687 | Ch13 region 3 |
| chr13:77379692-77379732 | Ch13 region 3 |
| chr13:77379737-77379777 | Ch13 region 3 |
| chr13:77379814-77379854 | Ch13 region 3 |
| chr13:77379862-77379902 | Ch13 region 3 |
| chr13:77379907-77379947 | Ch13 region 3 |
| chr13:77379984-77380024 | Ch13 region 3 |
| chr13:77380047-77380087 | Ch13 region 3 |
| chr13:77380111-77380151 | Ch13 region 3 |
| chr13:77380160-77380200 | Ch13 region 3 |
| chr13:77380247-77380287 | Ch13 region 3 |
| chr13:77380369-77380409 | Ch13 region 3 |
| chr13:77380521-77380561 | Ch13 region 3 |
| chr13:77380568-77380608 | Ch13 region 3 |
| chr13:77380658-77380698 | Ch13 region 3 |
| chr13:77380760-77380800 | Ch13 region 3 |
| chr13:77380821-77380861 | Ch13 region 3 |
| chr13:77380888-77380928 | Ch13 region 3 |
| chr13:77380933-77380973 | Ch13 region 3 |
| chr13:77380986-77381026 | Ch13 region 3 |
| chr13:77381070-77381110 | Ch13 region 3 |

| | |
|---|---|
| chr13:77381115-77381155 | Ch13 region 3 |
| chr13:77381176-77381216 | Ch13 region 3 |
| chr13:77381227-77381267 | Ch13 region 3 |
| chr13:77381296-77381336 | Ch13 region 3 |
| chr13:77381390-77381430 | Ch13 region 3 |
| chr13:77381457-77381497 | Ch13 region 3 |
| chr13:77381507-77381547 | Ch13 region 3 |
| chr13:77381561-77381601 | Ch13 region 3 |
| chr13:77381614-77381654 | Ch13 region 3 |
| chr13:77381662-77381702 | Ch13 region 3 |
| chr13:77381752-77381792 | Ch13 region 3 |
| chr13:77381806-77381846 | Ch13 region 3 |
| chr13:77381869-77381909 | Ch13 region 3 |
| chr13:77381919-77381959 | Ch13 region 3 |
| chr13:77381968-77382008 | Ch13 region 3 |
| chr13:77382070-77382110 | Ch13 region 3 |
| chr13:77382188-77382228 | Ch13 region 3 |
| chr13:77382370-77382410 | Ch13 region 3 |
| chr13:77382415-77382455 | Ch13 region 3 |
| chr13:77382536-77382576 | Ch13 region 3 |
| chr13:77382681-77382721 | Ch13 region 3 |
| chr13:77382747-77382787 | Ch13 region 3 |
| chr13:77382816-77382856 | Ch13 region 3 |
| chr13:77382870-77382910 | Ch13 region 3 |
| chr13:77382940-77382980 | Ch13 region 3 |
| chr13:77382992-77383032 | Ch13 region 3 |
| chr13:77383102-77383142 | Ch13 region 3 |
| chr13:77383156-77383196 | Ch13 region 3 |
| chr13:77383275-77383315 | Ch13 region 3 |
| chr13:77383364-77383404 | Ch13 region 3 |
| chr13:77383452-77383492 | Ch13 region 3 |
| chr13:77383520-77383560 | Ch13 region 3 |
| chr13:77383574-77383614 | Ch13 region 3 |
| chr13:77383664-77383704 | Ch13 region 3 |
| chr13:77383713-77383753 | Ch13 region 3 |
| chr13:77383759-77383799 | Ch13 region 3 |
| chr13:77383804-77383844 | Ch13 region 3 |
| chr13:77383867-77383907 | Ch13 region 3 |
| chr13:77383920-77383960 | Ch13 region 3 |
| chr13:77383980-77384020 | Ch13 region 3 |
| chr13:77384076-77384116 | Ch13 region 3 |
| chr13:77384121-77384161 | Ch13 region 3 |
| chr13:77384185-77384225 | Ch13 region 3 |
| chr13:77384269-77384309 | Ch13 region 3 |
| chr13:77384489-77384529 | Ch13 region 3 |
| chr13:77384593-77384633 | Ch13 region 3 |
| chr13:77384660-77384700 | Ch13 region 3 |
| chr13:77384717-77384757 | Ch13 region 3 |
| chr13:77384806-77384846 | Ch13 region 3 |
| chr13:77384875-77384915 | Ch13 region 3 |
| chr13:77384932-77384972 | Ch13 region 3 |
| chr13:77385055-77385095 | Ch13 region 3 |

| | |
|---|---|
| chr13:77385100-77385140 | Ch13 region 3 |
| chr13:77385145-77385185 | Ch13 region 3 |
| chr13:77385206-77385246 | Ch13 region 3 |
| chr13:77385255-77385295 | Ch13 region 3 |
| chr13:77385305-77385345 | Ch13 region 3 |
| chr13:77385402-77385442 | Ch13 region 3 |
| chr13:77385488-77385528 | Ch13 region 3 |
| chr13:77385618-77385658 | Ch13 region 3 |
| chr13:77385699-77385739 | Ch13 region 3 |
| chr13:77385761-77385801 | Ch13 region 3 |
| chr13:77385835-77385875 | Ch13 region 3 |
| chr13:77385903-77385943 | Ch13 region 3 |
| chr13:77385961-77386001 | Ch13 region 3 |
| chr13:77386013-77386053 | Ch13 region 3 |
| chr13:77386110-77386150 | Ch13 region 3 |
| chr13:77386167-77386207 | Ch13 region 3 |
| chr13:77386218-77386258 | Ch13 region 3 |
| chr13:77386263-77386303 | Ch13 region 3 |
| chr13:77386369-77386409 | Ch13 region 3 |
| chr13:77386418-77386458 | Ch13 region 3 |
| chr13:77386521-77386561 | Ch13 region 3 |
| chr13:77386587-77386627 | Ch13 region 3 |
| chr13:77386640-77386680 | Ch13 region 3 |
| chr13:77386705-77386745 | Ch13 region 3 |
| chr13:77386766-77386806 | Ch13 region 3 |
| chr13:77386813-77386853 | Ch13 region 3 |
| chr13:77386872-77386912 | Ch13 region 3 |
| chr13:77386952-77386992 | Ch13 region 3 |
| chr13:77387011-77387051 | Ch13 region 3 |
| chr13:77387081-77387121 | Ch13 region 3 |
| chr13:77387139-77387179 | Ch13 region 3 |
| chr13:77387201-77387241 | Ch13 region 3 |
| chr13:77387257-77387297 | Ch13 region 3 |
| chr13:77387346-77387386 | Ch13 region 3 |
| chr13:77387393-77387433 | Ch13 region 3 |
| chr13:77387438-77387478 | Ch13 region 3 |
| chr13:77387534-77387574 | Ch13 region 3 |
| chr13:77387585-77387625 | Ch13 region 3 |
| chr13:77387692-77387732 | Ch13 region 3 |
| chr13:77387783-77387823 | Ch13 region 3 |
| chr13:77387857-77387897 | Ch13 region 3 |
| chr13:77387917-77387957 | Ch13 region 3 |
| chr13:77387969-77388009 | Ch13 region 3 |
| chr13:77388020-77388060 | Ch13 region 3 |
| chr13:77388102-77388142 | Ch13 region 3 |
| chr13:77388208-77388248 | Ch13 region 3 |
| chr13:77388280-77388320 | Ch13 region 3 |
| chr13:77388326-77388366 | Ch13 region 3 |
| chr13:77388373-77388413 | Ch13 region 3 |
| chr13:77388436-77388476 | Ch13 region 3 |
| chr13:77388526-77388566 | Ch13 region 3 |
| chr13:77388597-77388637 | Ch13 region 3 |

| | |
|---|---|
| chr13:77388701-77388741 | Ch13 region 3 |
| chr13:77388751-77388791 | Ch13 region 3 |
| chr13:77389107-77389147 | Ch13 region 3 |
| chr13:77389289-77389329 | Ch13 region 3 |
| chr13:77389371-77389411 | Ch13 region 3 |
| chr13:77389552-77389592 | Ch13 region 3 |
| chr13:77389635-77389675 | Ch13 region 3 |
| chr13:77389680-77389720 | Ch13 region 3 |
| chr13:77389734-77389774 | Ch13 region 3 |
| chr13:77389809-77389849 | Ch13 region 3 |
| chr13:77389868-77389908 | Ch13 region 3 |
| chr13:77389916-77389956 | Ch13 region 3 |
| chr13:77389967-77390007 | Ch13 region 3 |
| chr13:77390012-77390052 | Ch13 region 3 |
| chr13:77390059-77390099 | Ch13 region 3 |
| chr13:77390123-77390163 | Ch13 region 3 |
| chr13:77390182-77390222 | Ch13 region 3 |
| chr13:77390228-77390268 | Ch13 region 3 |
| chr13:77390351-77390391 | Ch13 region 3 |
| chr13:77390401-77390441 | Ch13 region 3 |
| chr13:77390501-77390541 | Ch13 region 3 |
| chr13:77390554-77390594 | Ch13 region 3 |
| chr13:77390614-77390654 | Ch13 region 3 |
| chr13:77390659-77390699 | Ch13 region 3 |
| chr13:77390728-77390768 | Ch13 region 3 |
| chr13:77390809-77390849 | Ch13 region 3 |
| chr13:77390906-77390946 | Ch13 region 3 |
| chr13:77390953-77390993 | Ch13 region 3 |
| chr13:77391005-77391045 | Ch13 region 3 |
| chr13:77391096-77391136 | Ch13 region 3 |
| chr13:77391230-77391270 | Ch13 region 3 |
| chr13:77391289-77391329 | Ch13 region 3 |
| chr13:77391342-77391382 | Ch13 region 3 |
| chr13:77391415-77391455 | Ch13 region 3 |
| chr13:77391520-77391560 | Ch13 region 3 |
| chr13:77391607-77391647 | Ch13 region 3 |
| chr13:77391699-77391739 | Ch13 region 3 |
| chr13:77391748-77391788 | Ch13 region 3 |
| chr13:77391819-77391859 | Ch13 region 3 |
| chr13:77391894-77391934 | Ch13 region 3 |
| chr13:77391946-77391986 | Ch13 region 3 |
| chr13:77391991-77392031 | Ch13 region 3 |
| chr13:77392038-77392078 | Ch13 region 3 |
| chr13:77392117-77392157 | Ch13 region 3 |
| chr13:77392219-77392259 | Ch13 region 3 |
| chr13:77392280-77392320 | Ch13 region 3 |
| chr13:77392334-77392374 | Ch13 region 3 |
| chr13:77392381-77392421 | Ch13 region 3 |
| chr13:77392459-77392499 | Ch13 region 3 |
| chr13:77392524-77392564 | Ch13 region 3 |
| chr13:77392581-77392621 | Ch13 region 3 |
| chr13:77392648-77392688 | Ch13 region 3 |

| | |
|---|---|
| chr13:77392713-77392753 | Ch13 region 3 |
| chr13:77392761-77392801 | Ch13 region 3 |
| chr13:77392829-77392869 | Ch13 region 3 |
| chr13:77392900-77392940 | Ch13 region 3 |
| chr13:77392958-77392998 | Ch13 region 3 |
| chr13:77393007-77393047 | Ch13 region 3 |
| chr13:77393098-77393138 | Ch13 region 3 |
| chr13:77393155-77393195 | Ch13 region 3 |
| chr13:77393201-77393241 | Ch13 region 3 |
| chr13:77393287-77393327 | Ch13 region 3 |
| chr13:77393360-77393400 | Ch13 region 3 |
| chr13:77393412-77393452 | Ch13 region 3 |
| chr13:77393482-77393522 | Ch13 region 3 |
| chr13:77393550-77393590 | Ch13 region 3 |
| chr13:77393619-77393659 | Ch13 region 3 |
| chr13:77393730-77393770 | Ch13 region 3 |
| chr13:77393914-77393954 | Ch13 region 3 |
| chr13:77393959-77393999 | Ch13 region 3 |
| chr13:77394119-77394159 | Ch13 region 3 |
| chr13:77394164-77394204 | Ch13 region 3 |
| chr13:77394291-77394331 | Ch13 region 3 |
| chr13:77394373-77394413 | Ch13 region 3 |
| chr13:77394500-77394540 | Ch13 region 3 |
| chr13:77394581-77394621 | Ch13 region 3 |
| chr13:77394662-77394702 | Ch13 region 3 |
| chr13:77394707-77394747 | Ch13 region 3 |
| chr13:77394879-77394919 | Ch13 region 3 |
| chr13:77394952-77394992 | Ch13 region 3 |
| chr13:77395014-77395054 | Ch13 region 3 |
| chr13:77395070-77395110 | Ch13 region 3 |
| chr13:77395121-77395161 | Ch13 region 3 |
| chr13:77395167-77395207 | Ch13 region 3 |
| chr13:77395242-77395282 | Ch13 region 3 |
| chr13:77395289-77395329 | Ch13 region 3 |
| chr13:77395346-77395386 | Ch13 region 3 |
| chr13:77395548-77395588 | Ch13 region 3 |
| chr13:77395619-77395659 | Ch13 region 3 |
| chr13:77395736-77395776 | Ch13 region 3 |
| chr13:77395882-77395922 | Ch13 region 3 |
| chr13:77396035-77396075 | Ch13 region 3 |
| chr13:77396109-77396149 | Ch13 region 3 |
| chr13:77396161-77396201 | Ch13 region 3 |
| chr13:77396225-77396265 | Ch13 region 3 |
| chr13:77396308-77396348 | Ch13 region 3 |
| chr13:77396358-77396398 | Ch13 region 3 |
| chr13:77396433-77396473 | Ch13 region 3 |
| chr13:77396490-77396530 | Ch13 region 3 |
| chr13:77396545-77396585 | Ch13 region 3 |
| chr13:77396618-77396658 | Ch13 region 3 |
| chr13:77396773-77396813 | Ch13 region 3 |
| chr13:77396935-77396975 | Ch13 region 3 |
| chr13:77396996-77397036 | Ch13 region 3 |

| | |
|---|---|
| chr13:77397073-77397113 | Ch13 region 3 |
| chr13:77397131-77397171 | Ch13 region 3 |
| chr13:77397197-77397237 | Ch13 region 3 |
| chr13:77397268-77397308 | Ch13 region 3 |
| chr13:77397335-77397375 | Ch13 region 3 |
| chr13:77397381-77397421 | Ch13 region 3 |
| chr13:77397426-77397466 | Ch13 region 3 |
| chr13:77397476-77397516 | Ch13 region 3 |
| chr13:77397591-77397631 | Ch13 region 3 |
| chr13:77397636-77397676 | Ch13 region 3 |
| chr13:77397686-77397726 | Ch13 region 3 |
| chr13:77397747-77397787 | Ch13 region 3 |
| chr13:77397807-77397847 | Ch13 region 3 |
| chr13:77397915-77397955 | Ch13 region 3 |
| chr13:77397996-77398036 | Ch13 region 3 |
| chr13:77398042-77398082 | Ch13 region 3 |
| chr13:77398092-77398132 | Ch13 region 3 |
| chr13:77398137-77398177 | Ch13 region 3 |
| chr13:77398186-77398226 | Ch13 region 3 |
| chr13:77398266-77398306 | Ch13 region 3 |
| chr13:77398329-77398369 | Ch13 region 3 |
| chr13:77398385-77398425 | Ch13 region 3 |
| chr13:77398432-77398472 | Ch13 region 3 |
| chr13:77398496-77398536 | Ch13 region 3 |
| chr13:77398566-77398606 | Ch13 region 3 |
| chr13:77398750-77398790 | Ch13 region 3 |
| chr13:77398805-77398845 | Ch13 region 3 |
| chr13:77398919-77398959 | Ch13 region 3 |
| chr13:77399026-77399066 | Ch13 region 3 |
| chr13:77399116-77399156 | Ch13 region 3 |
| chr13:77399195-77399235 | Ch13 region 3 |
| chr13:77399265-77399305 | Ch13 region 3 |
| chr13:77399362-77399402 | Ch13 region 3 |
| chr13:77399410-77399450 | Ch13 region 3 |
| chr13:77399467-77399507 | Ch13 region 3 |
| chr13:77399526-77399566 | Ch13 region 3 |
| chr13:77399611-77399651 | Ch13 region 3 |
| chr13:77399680-77399720 | Ch13 region 3 |
| chr13:77399740-77399780 | Ch13 region 3 |
| chr13:77399787-77399827 | Ch13 region 3 |
| chr13:77399834-77399874 | Ch13 region 3 |
| chr13:77399904-77399944 | Ch13 region 3 |
| chr13:77399979-77400019 | Ch13 region 3 |
| chr13:77400053-77400093 | Ch13 region 3 |
| chr13:77400098-77400138 | Ch13 region 3 |
| chr13:77400143-77400183 | Ch13 region 3 |
| chr13:77400209-77400249 | Ch13 region 3 |
| chr13:77400264-77400304 | Ch13 region 3 |
| chr13:77400360-77400400 | Ch13 region 3 |
| chr13:77400441-77400481 | Ch13 region 3 |
| chr13:77400500-77400540 | Ch13 region 3 |
| chr13:77400545-77400585 | Ch13 region 3 |

| | |
|---|---|
| chr13:77400636-77400676 | Ch13 region 3 |
| chr13:77400723-77400763 | Ch13 region 3 |
| chr13:77400773-77400813 | Ch13 region 3 |
| chr13:77400850-77400890 | Ch13 region 3 |
| chr13:77400896-77400936 | Ch13 region 3 |
| chr13:77400941-77400981 | Ch13 region 3 |
| chr13:77401018-77401058 | Ch13 region 3 |
| chr13:77401063-77401103 | Ch13 region 3 |
| chr13:77401137-77401177 | Ch13 region 3 |
| chr13:77401189-77401229 | Ch13 region 3 |
| chr13:77401300-77401340 | Ch13 region 3 |
| chr13:77401347-77401387 | Ch13 region 3 |
| chr13:77401421-77401461 | Ch13 region 3 |
| chr13:77401493-77401533 | Ch13 region 3 |
| chr13:77401541-77401581 | Ch13 region 3 |
| chr13:77401593-77401633 | Ch13 region 3 |
| chr13:77401639-77401679 | Ch13 region 3 |
| chr13:77401694-77401734 | Ch13 region 3 |
| chr13:77401806-77401846 | Ch13 region 3 |
| chr13:77401862-77401902 | Ch13 region 3 |
| chr13:77401910-77401950 | Ch13 region 3 |
| chr13:77401976-77402016 | Ch13 region 3 |
| chr13:77402038-77402078 | Ch13 region 3 |
| chr13:77402084-77402124 | Ch13 region 3 |
| chr13:77402214-77402254 | Ch13 region 3 |
| chr13:77402427-77402467 | Ch13 region 3 |
| chr13:77402504-77402544 | Ch13 region 3 |
| chr13:77402557-77402597 | Ch13 region 3 |
| chr13:77402624-77402664 | Ch13 region 3 |
| chr13:77402686-77402726 | Ch13 region 3 |
| chr13:77402742-77402782 | Ch13 region 3 |
| chr13:77402800-77402840 | Ch13 region 3 |
| chr13:77402863-77402903 | Ch13 region 3 |
| chr13:77402930-77402970 | Ch13 region 3 |
| chr13:77403028-77403068 | Ch13 region 3 |
| chr13:77403106-77403146 | Ch13 region 3 |
| chr13:77403156-77403196 | Ch13 region 3 |
| chr13:77403234-77403274 | Ch13 region 3 |
| chr13:77403288-77403328 | Ch13 region 3 |
| chr13:77403356-77403396 | Ch13 region 3 |
| chr13:77403401-77403441 | Ch13 region 3 |
| chr13:77403446-77403486 | Ch13 region 3 |
| chr13:77403502-77403542 | Ch13 region 3 |
| chr13:77403552-77403592 | Ch13 region 3 |
| chr13:77403647-77403687 | Ch13 region 3 |
| chr13:77403736-77403776 | Ch13 region 3 |
| chr13:77403784-77403824 | Ch13 region 3 |
| chr13:77403847-77403887 | Ch13 region 3 |
| chr13:77403907-77403947 | Ch13 region 3 |
| chr13:77404001-77404041 | Ch13 region 3 |
| chr13:77404050-77404090 | Ch13 region 3 |
| chr13:77404112-77404152 | Ch13 region 3 |

| | |
|---|---|
| chr13:77404157-77404197 | Ch13 region 3 |
| chr13:77404242-77404282 | Ch13 region 3 |
| chr13:77404287-77404327 | Ch13 region 3 |
| chr13:77404335-77404375 | Ch13 region 3 |
| chr13:77404400-77404440 | Ch13 region 3 |
| chr13:77404445-77404485 | Ch13 region 3 |
| chr13:77404517-77404557 | Ch13 region 3 |
| chr13:77404571-77404611 | Ch13 region 3 |
| chr13:77404635-77404675 | Ch13 region 3 |
| chr13:77404691-77404731 | Ch13 region 3 |
| chr13:77404765-77404805 | Ch13 region 3 |
| chr13:77404810-77404850 | Ch13 region 3 |
| chr13:77404856-77404896 | Ch13 region 3 |
| chr13:77404904-77404944 | Ch13 region 3 |
| chr13:77404970-77405010 | Ch13 region 3 |
| chr13:77405063-77405103 | Ch13 region 3 |
| chr13:77405111-77405151 | Ch13 region 3 |
| chr13:77405156-77405196 | Ch13 region 3 |
| chr13:77405210-77405250 | Ch13 region 3 |
| chr13:77405263-77405303 | Ch13 region 3 |
| chr13:77405312-77405352 | Ch13 region 3 |
| chr13:77405367-77405407 | Ch13 region 3 |
| chr13:77405437-77405477 | Ch13 region 3 |
| chr13:77405488-77405528 | Ch13 region 3 |
| chr13:77405561-77405601 | Ch13 region 3 |
| chr13:77405606-77405646 | Ch13 region 3 |
| chr13:77405691-77405731 | Ch13 region 3 |
| chr13:77405737-77405777 | Ch13 region 3 |
| chr13:77405798-77405838 | Ch13 region 3 |
| chr13:77405873-77405913 | Ch13 region 3 |
| chr13:77405962-77406002 | Ch13 region 3 |
| chr13:77406021-77406061 | Ch13 region 3 |
| chr13:77406080-77406120 | Ch13 region 3 |
| chr13:77406132-77406172 | Ch13 region 3 |
| chr13:77406184-77406224 | Ch13 region 3 |
| chr13:77406250-77406290 | Ch13 region 3 |
| chr13:77406324-77406364 | Ch13 region 3 |
| chr13:77406370-77406410 | Ch13 region 3 |
| chr13:77406425-77406465 | Ch13 region 3 |
| chr13:77406470-77406510 | Ch13 region 3 |
| chr13:77406524-77406564 | Ch13 region 3 |
| chr13:77406583-77406623 | Ch13 region 3 |
| chr13:77406669-77406709 | Ch13 region 3 |
| chr13:77406778-77406818 | Ch13 region 3 |
| chr13:77406856-77406896 | Ch13 region 3 |
| chr13:77406903-77406943 | Ch13 region 3 |
| chr13:77406953-77406993 | Ch13 region 3 |
| chr13:77407016-77407056 | Ch13 region 3 |
| chr13:77407144-77407184 | Ch13 region 3 |
| chr13:77407190-77407230 | Ch13 region 3 |
| chr13:77407277-77407317 | Ch13 region 3 |
| chr13:77407343-77407383 | Ch13 region 3 |

| | |
|---|---|
| chr13:77407414-77407454 | Ch13 region 3 |
| chr13:77407459-77407499 | Ch13 region 3 |
| chr13:77407531-77407571 | Ch13 region 3 |
| chr13:77407581-77407621 | Ch13 region 3 |
| chr13:77407630-77407670 | Ch13 region 3 |
| chr13:77407698-77407738 | Ch13 region 3 |
| chr13:77407754-77407794 | Ch13 region 3 |
| chr13:77407823-77407863 | Ch13 region 3 |
| chr13:77407909-77407949 | Ch13 region 3 |
| chr13:77408012-77408052 | Ch13 region 3 |
| chr13:77408057-77408097 | Ch13 region 3 |
| chr13:77408165-77408205 | Ch13 region 3 |
| chr13:77408330-77408370 | Ch13 region 3 |
| chr13:77408412-77408452 | Ch13 region 3 |
| chr13:77408484-77408524 | Ch13 region 3 |
| chr13:77408544-77408584 | Ch13 region 3 |
| chr13:77408611-77408651 | Ch13 region 3 |
| chr13:77408805-77408845 | Ch13 region 3 |
| chr13:77408896-77408936 | Ch13 region 3 |
| chr13:77408943-77408983 | Ch13 region 3 |
| chr13:77409027-77409067 | Ch13 region 3 |
| chr13:77409193-77409233 | Ch13 region 3 |
| chr13:77409311-77409351 | Ch13 region 3 |
| chr13:77409478-77409518 | Ch13 region 3 |
| chr13:77409545-77409585 | Ch13 region 3 |
| chr13:77409685-77409725 | Ch13 region 3 |
| chr13:77409814-77409854 | Ch13 region 3 |
| chr13:77409894-77409934 | Ch13 region 3 |
| chr13:77410027-77410067 | Ch13 region 3 |
| chr13:77410235-77410275 | Ch13 region 3 |
| chr13:77410336-77410376 | Ch13 region 3 |
| chr13:77410461-77410501 | Ch13 region 3 |
| chr13:77410568-77410608 | Ch13 region 3 |
| chr13:77410629-77410669 | Ch13 region 3 |
| chr13:77410680-77410720 | Ch13 region 3 |
| chr13:77410739-77410779 | Ch13 region 3 |
| chr13:77410829-77410869 | Ch13 region 3 |
| chr13:77410876-77410916 | Ch13 region 3 |
| chr13:77410932-77410972 | Ch13 region 3 |
| chr13:77410977-77411017 | Ch13 region 3 |
| chr13:77411042-77411082 | Ch13 region 3 |
| chr13:77411087-77411127 | Ch13 region 3 |
| chr13:77411178-77411218 | Ch13 region 3 |
| chr13:77411230-77411270 | Ch13 region 3 |
| chr13:77411314-77411354 | Ch13 region 3 |
| chr13:77411382-77411422 | Ch13 region 3 |
| chr13:77411447-77411487 | Ch13 region 3 |
| chr13:77411495-77411535 | Ch13 region 3 |
| chr13:77411557-77411597 | Ch13 region 3 |
| chr13:77411828-77411868 | Ch13 region 3 |
| chr13:77411905-77411945 | Ch13 region 3 |
| chr13:77411984-77412024 | Ch13 region 3 |

| | |
|---|---|
| chr13:77412076-77412116 | Ch13 region 3 |
| chr13:77412139-77412179 | Ch13 region 3 |
| chr13:77412212-77412252 | Ch13 region 3 |
| chr13:77412311-77412351 | Ch13 region 3 |
| chr13:77412382-77412422 | Ch13 region 3 |
| chr13:77412453-77412493 | Ch13 region 3 |
| chr13:77412525-77412565 | Ch13 region 3 |
| chr13:77412636-77412676 | Ch13 region 3 |
| chr13:77412754-77412794 | Ch13 region 3 |
| chr13:77412847-77412887 | Ch13 region 3 |
| chr13:77412939-77412979 | Ch13 region 3 |
| chr13:77412984-77413024 | Ch13 region 3 |
| chr13:77413030-77413070 | Ch13 region 3 |
| chr13:77413081-77413121 | Ch13 region 3 |
| chr13:77413148-77413188 | Ch13 region 3 |
| chr13:77413224-77413264 | Ch13 region 3 |
| chr13:77413295-77413335 | Ch13 region 3 |
| chr13:77413340-77413380 | Ch13 region 3 |
| chr13:77413487-77413527 | Ch13 region 3 |
| chr13:77413749-77413789 | Ch13 region 3 |
| chr13:77413800-77413840 | Ch13 region 3 |
| chr13:77413851-77413891 | Ch13 region 3 |
| chr13:77413943-77413983 | Ch13 region 3 |
| chr13:77413990-77414030 | Ch13 region 3 |
| chr13:77414052-77414092 | Ch13 region 3 |
| chr13:77414113-77414153 | Ch13 region 3 |
| chr13:77414161-77414201 | Ch13 region 3 |
| chr13:77414211-77414251 | Ch13 region 3 |
| chr13:77414306-77414346 | Ch13 region 3 |
| chr13:77414399-77414439 | Ch13 region 3 |
| chr13:77414447-77414487 | Ch13 region 3 |
| chr13:77414492-77414532 | Ch13 region 3 |
| chr13:77414552-77414592 | Ch13 region 3 |
| chr13:77414641-77414681 | Ch13 region 3 |
| chr13:77414743-77414783 | Ch13 region 3 |
| chr13:77414790-77414830 | Ch13 region 3 |
| chr13:77414906-77414946 | Ch13 region 3 |
| chr13:77414981-77415021 | Ch13 region 3 |
| chr13:77415119-77415159 | Ch13 region 3 |
| chr13:77415171-77415211 | Ch13 region 3 |
| chr13:77415284-77415324 | Ch13 region 3 |
| chr13:77415352-77415392 | Ch13 region 3 |
| chr13:77415503-77415543 | Ch13 region 3 |
| chr13:77415579-77415619 | Ch13 region 3 |
| chr13:77415788-77415828 | Ch13 region 3 |
| chr13:77415864-77415904 | Ch13 region 3 |
| chr13:77416064-77416104 | Ch13 region 3 |
| chr13:77416171-77416211 | Ch13 region 3 |
| chr13:77416238-77416278 | Ch13 region 3 |
| chr13:77416287-77416327 | Ch13 region 3 |
| chr13:77416334-77416374 | Ch13 region 3 |
| chr13:77416399-77416439 | Ch13 region 3 |

| | |
|---|---|
| chr13:77416449-77416489 | Ch13 region 3 |
| chr13:77416515-77416555 | Ch13 region 3 |
| chr13:77416577-77416617 | Ch13 region 3 |
| chr13:77416648-77416688 | Ch13 region 3 |
| chr13:77416696-77416736 | Ch13 region 3 |
| chr13:77416754-77416794 | Ch13 region 3 |
| chr13:77416818-77416858 | Ch13 region 3 |
| chr13:77416930-77416970 | Ch13 region 3 |
| chr13:77416980-77417020 | Ch13 region 3 |
| chr13:77417064-77417104 | Ch13 region 3 |
| chr13:77417232-77417272 | Ch13 region 3 |
| chr13:77417408-77417448 | Ch13 region 3 |
| chr13:77417592-77417632 | Ch13 region 3 |
| chr13:77417638-77417678 | Ch13 region 3 |
| chr13:77417707-77417747 | Ch13 region 3 |
| chr13:77417766-77417806 | Ch13 region 3 |
| chr13:77417846-77417886 | Ch13 region 3 |
| chr13:77417939-77417979 | Ch13 region 3 |
| chr13:77417994-77418034 | Ch13 region 3 |
| chr13:77418082-77418122 | Ch13 region 3 |
| chr13:77418173-77418213 | Ch13 region 3 |
| chr13:77418231-77418271 | Ch13 region 3 |
| chr13:77418319-77418359 | Ch13 region 3 |
| chr13:77418372-77418412 | Ch13 region 3 |
| chr13:77418422-77418462 | Ch13 region 3 |
| chr13:77418495-77418535 | Ch13 region 3 |
| chr13:77418627-77418667 | Ch13 region 3 |
| chr13:77418822-77418862 | Ch13 region 3 |
| chr13:77418887-77418927 | Ch13 region 3 |
| chr13:77418969-77419009 | Ch13 region 3 |
| chr13:77419054-77419094 | Ch13 region 3 |
| chr13:77419145-77419185 | Ch13 region 3 |
| chr13:77419246-77419286 | Ch13 region 3 |
| chr13:77419339-77419379 | Ch13 region 3 |
| chr13:77419460-77419500 | Ch13 region 3 |
| chr13:77419602-77419642 | Ch13 region 3 |
| chr13:77419785-77419825 | Ch13 region 3 |
| chr13:77419838-77419878 | Ch13 region 3 |
| chr13:77419912-77419952 | Ch13 region 3 |
| chr13:77420063-77420103 | Ch13 region 3 |
| chr13:77420130-77420170 | Ch13 region 3 |
| chr13:77420212-77420252 | Ch13 region 3 |
| chr13:77420318-77420358 | Ch13 region 3 |
| chr13:77420367-77420407 | Ch13 region 3 |
| chr13:77420470-77420510 | Ch13 region 3 |
| chr13:77420551-77420591 | Ch13 region 3 |
| chr13:77420611-77420651 | Ch13 region 3 |
| chr13:77420706-77420746 | Ch13 region 3 |
| chr13:77420849-77420889 | Ch13 region 3 |
| chr13:77420898-77420938 | Ch13 region 3 |
| chr13:77420991-77421031 | Ch13 region 3 |
| chr13:77421060-77421100 | Ch13 region 3 |

| | |
|---|---|
| chr13:77421123-77421163 | Ch13 region 3 |
| chr13:77421171-77421211 | Ch13 region 3 |
| chr13:77421216-77421256 | Ch13 region 3 |
| chr13:77421264-77421304 | Ch13 region 3 |
| chr13:77421317-77421357 | Ch13 region 3 |
| chr13:77421370-77421410 | Ch13 region 3 |
| chr13:77421419-77421459 | Ch13 region 3 |
| chr13:77421464-77421504 | Ch13 region 3 |
| chr13:77421512-77421552 | Ch13 region 3 |
| chr13:77421646-77421686 | Ch13 region 3 |
| chr13:77421718-77421758 | Ch13 region 3 |
| chr13:77421763-77421803 | Ch13 region 3 |
| chr13:77421844-77421884 | Ch13 region 3 |
| chr13:77421943-77421983 | Ch13 region 3 |
| chr13:77422003-77422043 | Ch13 region 3 |
| chr13:77422048-77422088 | Ch13 region 3 |
| chr13:77422093-77422133 | Ch13 region 3 |
| chr13:77422229-77422269 | Ch13 region 3 |
| chr13:77422277-77422317 | Ch13 region 3 |
| chr13:77422328-77422368 | Ch13 region 3 |
| chr13:77422408-77422448 | Ch13 region 3 |
| chr13:77422473-77422513 | Ch13 region 3 |
| chr13:77422552-77422592 | Ch13 region 3 |
| chr13:77422623-77422663 | Ch13 region 3 |
| chr13:77422682-77422722 | Ch13 region 3 |
| chr13:77422749-77422789 | Ch13 region 3 |
| chr13:77422812-77422852 | Ch13 region 3 |
| chr13:77422877-77422917 | Ch13 region 3 |
| chr13:77422923-77422963 | Ch13 region 3 |
| chr13:77423010-77423050 | Ch13 region 3 |
| chr13:77423116-77423156 | Ch13 region 3 |
| chr13:77423178-77423218 | Ch13 region 3 |
| chr13:77423260-77423300 | Ch13 region 3 |
| chr13:77423305-77423345 | Ch13 region 3 |
| chr13:77423521-77423561 | Ch13 region 3 |
| chr13:77423644-77423684 | Ch13 region 3 |
| chr13:77423693-77423733 | Ch13 region 3 |
| chr13:77423755-77423795 | Ch13 region 3 |
| chr13:77423835-77423875 | Ch13 region 3 |
| chr13:77423908-77423948 | Ch13 region 3 |
| chr13:77424052-77424092 | Ch13 region 3 |
| chr13:77424118-77424158 | Ch13 region 3 |
| chr13:77424195-77424235 | Ch13 region 3 |
| chr13:77424400-77424440 | Ch13 region 3 |
| chr13:77424468-77424508 | Ch13 region 3 |
| chr13:77424531-77424571 | Ch13 region 3 |
| chr13:77424607-77424647 | Ch13 region 3 |
| chr13:77424705-77424745 | Ch13 region 3 |
| chr13:77424794-77424834 | Ch13 region 3 |
| chr13:77424847-77424887 | Ch13 region 3 |
| chr13:77424903-77424943 | Ch13 region 3 |
| chr13:77424979-77425019 | Ch13 region 3 |

| | |
|---|---|
| chr13:77425217-77425257 | Ch13 region 3 |
| chr13:77425263-77425303 | Ch13 region 3 |
| chr13:77425318-77425358 | Ch13 region 3 |
| chr13:77425415-77425455 | Ch13 region 3 |
| chr13:77425466-77425506 | Ch13 region 3 |
| chr13:77425531-77425571 | Ch13 region 3 |
| chr13:77425611-77425651 | Ch13 region 3 |
| chr13:77425659-77425699 | Ch13 region 3 |
| chr13:77425751-77425791 | Ch13 region 3 |
| chr13:77425806-77425846 | Ch13 region 3 |
| chr13:77425861-77425901 | Ch13 region 3 |
| chr13:77425908-77425948 | Ch13 region 3 |
| chr13:77425983-77426023 | Ch13 region 3 |
| chr13:77426058-77426098 | Ch13 region 3 |
| chr13:77426125-77426165 | Ch13 region 3 |
| chr13:77426171-77426211 | Ch13 region 3 |
| chr13:77426261-77426301 | Ch13 region 3 |
| chr13:77426334-77426374 | Ch13 region 3 |
| chr13:77426383-77426423 | Ch13 region 3 |
| chr13:77426476-77426516 | Ch13 region 3 |
| chr13:77426607-77426647 | Ch13 region 3 |
| chr13:77426675-77426715 | Ch13 region 3 |
| chr13:77426737-77426777 | Ch13 region 3 |
| chr13:77426782-77426822 | Ch13 region 3 |
| chr13:77426829-77426869 | Ch13 region 3 |
| chr13:77426874-77426914 | Ch13 region 3 |
| chr13:77426922-77426962 | Ch13 region 3 |
| chr13:77426974-77427014 | Ch13 region 3 |
| chr13:77427051-77427091 | Ch13 region 3 |
| chr13:77427111-77427151 | Ch13 region 3 |
| chr13:77427161-77427201 | Ch13 region 3 |
| chr13:77427211-77427251 | Ch13 region 3 |
| chr13:77427256-77427296 | Ch13 region 3 |
| chr13:77427358-77427398 | Ch13 region 3 |
| chr13:77427426-77427466 | Ch13 region 3 |
| chr13:77427597-77427637 | Ch13 region 3 |
| chr13:77427648-77427688 | Ch13 region 3 |
| chr13:77427718-77427758 | Ch13 region 3 |
| chr13:77427777-77427817 | Ch13 region 3 |
| chr13:77427822-77427862 | Ch13 region 3 |
| chr13:77427913-77427953 | Ch13 region 3 |
| chr13:77428014-77428054 | Ch13 region 3 |
| chr13:77428096-77428136 | Ch13 region 3 |
| chr13:77428203-77428243 | Ch13 region 3 |
| chr13:77428275-77428315 | Ch13 region 3 |
| chr13:77428330-77428370 | Ch13 region 3 |
| chr13:77428410-77428450 | Ch13 region 3 |
| chr13:77428472-77428512 | Ch13 region 3 |
| chr13:77428534-77428574 | Ch13 region 3 |
| chr13:77428670-77428710 | Ch13 region 3 |
| chr13:77428742-77428782 | Ch13 region 3 |
| chr13:77428793-77428833 | Ch13 region 3 |

| | |
|---|---|
| chr13:77428915-77428955 | Ch13 region 3 |
| chr13:77429005-77429045 | Ch13 region 3 |
| chr13:77429064-77429104 | Ch13 region 3 |
| chr13:77429129-77429169 | Ch13 region 3 |
| chr13:77429235-77429275 | Ch13 region 3 |
| chr13:77429301-77429341 | Ch13 region 3 |
| chr13:77429378-77429418 | Ch13 region 3 |
| chr13:77429434-77429474 | Ch13 region 3 |
| chr13:77429609-77429649 | Ch13 region 3 |
| chr13:77429761-77429801 | Ch13 region 3 |
| chr13:77429807-77429847 | Ch13 region 3 |
| chr13:77429879-77429919 | Ch13 region 3 |
| chr13:77430140-77430180 | Ch13 region 3 |
| chr13:77430315-77430355 | Ch13 region 3 |
| chr13:77430434-77430474 | Ch13 region 3 |
| chr13:77430483-77430523 | Ch13 region 3 |
| chr13:77430542-77430582 | Ch13 region 3 |
| chr13:77430590-77430630 | Ch13 region 3 |
| chr13:77430671-77430711 | Ch13 region 3 |
| chr13:77430814-77430854 | Ch13 region 3 |
| chr13:77430906-77430946 | Ch13 region 3 |
| chr13:77430973-77431013 | Ch13 region 3 |
| chr13:77431110-77431150 | Ch13 region 3 |
| chr13:77431267-77431307 | Ch13 region 3 |
| chr13:77431464-77431504 | Ch13 region 3 |
| chr13:77431633-77431673 | Ch13 region 3 |
| chr13:77431789-77431829 | Ch13 region 3 |
| chr13:77431834-77431874 | Ch13 region 3 |
| chr13:77431898-77431938 | Ch13 region 3 |
| chr13:77431989-77432029 | Ch13 region 3 |
| chr13:77432039-77432079 | Ch13 region 3 |
| chr13:77432084-77432124 | Ch13 region 3 |
| chr13:77432138-77432178 | Ch13 region 3 |
| chr13:77432194-77432234 | Ch13 region 3 |
| chr13:77432242-77432282 | Ch13 region 3 |
| chr13:77432290-77432330 | Ch13 region 3 |
| chr13:77432378-77432418 | Ch13 region 3 |
| chr13:77432430-77432470 | Ch13 region 3 |
| chr13:77432513-77432553 | Ch13 region 3 |
| chr13:77432563-77432603 | Ch13 region 3 |
| chr13:77432653-77432693 | Ch13 region 3 |
| chr13:77432708-77432748 | Ch13 region 3 |
| chr13:77432790-77432830 | Ch13 region 3 |
| chr13:77432839-77432879 | Ch13 region 3 |
| chr13:77432910-77432950 | Ch13 region 3 |
| chr13:77433005-77433045 | Ch13 region 3 |
| chr13:77433055-77433095 | Ch13 region 3 |
| chr13:77433280-77433320 | Ch13 region 3 |
| chr13:77433426-77433466 | Ch13 region 3 |
| chr13:77433550-77433590 | Ch13 region 3 |
| chr13:77433660-77433700 | Ch13 region 3 |
| chr13:77433895-77433935 | Ch13 region 3 |

| | |
|---|---|
| chr13:77434030-77434070 | Ch13 region 3 |
| chr13:77434075-77434115 | Ch13 region 3 |
| chr13:77434177-77434217 | Ch13 region 3 |
| chr13:77434255-77434295 | Ch13 region 3 |
| chr13:77434310-77434350 | Ch13 region 3 |
| chr13:77434371-77434411 | Ch13 region 3 |
| chr13:77434449-77434489 | Ch13 region 3 |
| chr13:77434529-77434569 | Ch13 region 3 |
| chr13:77434580-77434620 | Ch13 region 3 |
| chr13:77434626-77434666 | Ch13 region 3 |
| chr13:77434708-77434748 | Ch13 region 3 |
| chr13:77434770-77434810 | Ch13 region 3 |
| chr13:77434817-77434857 | Ch13 region 3 |
| chr13:77434883-77434923 | Ch13 region 3 |
| chr13:77434931-77434971 | Ch13 region 3 |
| chr13:77435002-77435042 | Ch13 region 3 |
| chr13:77435070-77435110 | Ch13 region 3 |
| chr13:77435143-77435183 | Ch13 region 3 |
| chr13:77435224-77435264 | Ch13 region 3 |
| chr13:77435278-77435318 | Ch13 region 3 |
| chr13:77435329-77435369 | Ch13 region 3 |
| chr13:77435408-77435448 | Ch13 region 3 |
| chr13:77435453-77435493 | Ch13 region 3 |
| chr13:77435543-77435583 | Ch13 region 3 |
| chr13:77435601-77435641 | Ch13 region 3 |
| chr13:77435801-77435841 | Ch13 region 3 |
| chr13:77435854-77435894 | Ch13 region 3 |
| chr13:77435982-77436022 | Ch13 region 3 |
| chr13:77436039-77436079 | Ch13 region 3 |
| chr13:77436104-77436144 | Ch13 region 3 |
| chr13:77436176-77436216 | Ch13 region 3 |
| chr13:77436221-77436261 | Ch13 region 3 |
| chr13:77436300-77436340 | Ch13 region 3 |
| chr13:77436349-77436389 | Ch13 region 3 |
| chr13:77436412-77436452 | Ch13 region 3 |
| chr13:77436471-77436511 | Ch13 region 3 |
| chr13:77436614-77436654 | Ch13 region 3 |
| chr13:77436744-77436784 | Ch13 region 3 |
| chr13:77436913-77436953 | Ch13 region 3 |
| chr13:77436960-77437000 | Ch13 region 3 |
| chr13:77437005-77437045 | Ch13 region 3 |
| chr13:77437094-77437134 | Ch13 region 3 |
| chr13:77437175-77437215 | Ch13 region 3 |
| chr13:77437235-77437275 | Ch13 region 3 |
| chr13:77437329-77437369 | Ch13 region 3 |
| chr13:77437395-77437435 | Ch13 region 3 |
| chr13:77437494-77437534 | Ch13 region 3 |
| chr13:77437612-77437652 | Ch13 region 3 |
| chr13:77437833-77437873 | Ch13 region 3 |
| chr13:77437897-77437937 | Ch13 region 3 |
| chr13:77438058-77438098 | Ch13 region 3 |
| chr13:77438189-77438229 | Ch13 region 3 |

| | |
|---|---|
| chr13:77438276-77438316 | Ch13 region 3 |
| chr13:77438408-77438448 | Ch13 region 3 |
| chr13:77438460-77438500 | Ch13 region 3 |
| chr13:77438519-77438559 | Ch13 region 3 |
| chr13:77438589-77438629 | Ch13 region 3 |
| chr13:77438663-77438703 | Ch13 region 3 |
| chr13:77438708-77438748 | Ch13 region 3 |
| chr13:77438766-77438806 | Ch13 region 3 |
| chr13:77438823-77438863 | Ch13 region 3 |
| chr13:77438878-77438918 | Ch13 region 3 |
| chr13:77438949-77438989 | Ch13 region 3 |
| chr13:77438994-77439034 | Ch13 region 3 |
| chr13:77439168-77439208 | Ch13 region 3 |
| chr13:77439263-77439303 | Ch13 region 3 |
| chr13:77439378-77439418 | Ch13 region 3 |
| chr13:77439484-77439524 | Ch13 region 3 |
| chr13:77439542-77439582 | Ch13 region 3 |
| chr13:77439587-77439627 | Ch13 region 3 |
| chr13:77439665-77439705 | Ch13 region 3 |
| chr13:77439727-77439767 | Ch13 region 3 |
| chr13:77439774-77439814 | Ch13 region 3 |
| chr13:77439850-77439890 | Ch13 region 3 |
| chr13:77439911-77439951 | Ch13 region 3 |
| chr13:77439994-77440034 | Ch13 region 3 |
| chr13:77440062-77440102 | Ch13 region 3 |
| chr13:77440129-77440169 | Ch13 region 3 |
| chr13:77440187-77440227 | Ch13 region 3 |
| chr13:77440270-77440310 | Ch13 region 3 |
| chr13:77440420-77440460 | Ch13 region 3 |
| chr13:77440478-77440518 | Ch13 region 3 |
| chr13:77440708-77440748 | Ch13 region 3 |
| chr13:77440927-77440967 | Ch13 region 3 |
| chr13:77440991-77441031 | Ch13 region 3 |
| chr13:77441038-77441078 | Ch13 region 3 |
| chr13:77441245-77441285 | Ch13 region 3 |
| chr13:77441311-77441351 | Ch13 region 3 |
| chr13:77441411-77441451 | Ch13 region 3 |
| chr13:77441643-77441683 | Ch13 region 3 |
| chr13:77441808-77441848 | Ch13 region 3 |
| chr13:77441944-77441984 | Ch13 region 3 |
| chr13:77442005-77442045 | Ch13 region 3 |
| chr13:77442078-77442118 | Ch13 region 3 |
| chr13:77442183-77442223 | Ch13 region 3 |
| chr13:77442291-77442331 | Ch13 region 3 |
| chr13:77442337-77442377 | Ch13 region 3 |
| chr13:77442516-77442556 | Ch13 region 3 |
| chr13:77442574-77442614 | Ch13 region 3 |
| chr13:77442631-77442671 | Ch13 region 3 |
| chr13:77442694-77442734 | Ch13 region 3 |
| chr13:77442775-77442815 | Ch13 region 3 |
| chr13:77442846-77442886 | Ch13 region 3 |
| chr13:77442894-77442934 | Ch13 region 3 |

| | |
|---|---|
| chr13:77442959-77442999 | Ch13 region 3 |
| chr13:77443046-77443086 | Ch13 region 3 |
| chr13:77443091-77443131 | Ch13 region 3 |
| chr13:77443153-77443193 | Ch13 region 3 |
| chr13:77443467-77443507 | Ch13 region 3 |
| chr13:77443520-77443560 | Ch13 region 3 |
| chr13:77443575-77443615 | Ch13 region 3 |
| chr13:77443622-77443662 | Ch13 region 3 |
| chr13:77443685-77443725 | Ch13 region 3 |
| chr13:77443754-77443794 | Ch13 region 3 |
| chr13:77443805-77443845 | Ch13 region 3 |
| chr13:77443911-77443951 | Ch13 region 3 |
| chr13:77443980-77444020 | Ch13 region 3 |
| chr13:77444061-77444101 | Ch13 region 3 |
| chr13:77444121-77444161 | Ch13 region 3 |
| chr13:77444175-77444215 | Ch13 region 3 |
| chr13:77444225-77444265 | Ch13 region 3 |
| chr13:77444292-77444332 | Ch13 region 3 |
| chr13:77444365-77444405 | Ch13 region 3 |
| chr13:77444431-77444471 | Ch13 region 3 |
| chr13:77444476-77444516 | Ch13 region 3 |
| chr13:77444521-77444561 | Ch13 region 3 |
| chr13:77444587-77444627 | Ch13 region 3 |
| chr13:77444672-77444712 | Ch13 region 3 |
| chr13:77444718-77444758 | Ch13 region 3 |
| chr13:77444795-77444835 | Ch13 region 3 |
| chr13:77444872-77444912 | Ch13 region 3 |
| chr13:77444940-77444980 | Ch13 region 3 |
| chr13:77445020-77445060 | Ch13 region 3 |
| chr13:77445262-77445302 | Ch13 region 3 |
| chr13:77445312-77445352 | Ch13 region 3 |
| chr13:77445357-77445397 | Ch13 region 3 |
| chr13:77445440-77445480 | Ch13 region 3 |
| chr13:77445561-77445601 | Ch13 region 3 |
| chr13:77445634-77445674 | Ch13 region 3 |
| chr13:77445696-77445736 | Ch13 region 3 |
| chr13:77445830-77445870 | Ch13 region 3 |
| chr13:77445905-77445945 | Ch13 region 3 |
| chr13:77445965-77446005 | Ch13 region 3 |
| chr13:77446022-77446062 | Ch13 region 3 |
| chr13:77446217-77446257 | Ch13 region 3 |
| chr13:77446271-77446311 | Ch13 region 3 |
| chr13:77446386-77446426 | Ch13 region 3 |
| chr13:77446437-77446477 | Ch13 region 3 |
| chr13:77446520-77446560 | Ch13 region 3 |
| chr13:77446777-77446817 | Ch13 region 3 |
| chr13:77446828-77446868 | Ch13 region 3 |
| chr13:77446946-77446986 | Ch13 region 3 |
| chr13:77447045-77447085 | Ch13 region 3 |
| chr13:77447194-77447234 | Ch13 region 3 |
| chr13:77447255-77447295 | Ch13 region 3 |
| chr13:77447329-77447369 | Ch13 region 3 |

| | |
|---|---|
| chr13:77447462-77447502 | Ch13 region 3 |
| chr13:77447534-77447574 | Ch13 region 3 |
| chr13:77447695-77447735 | Ch13 region 3 |
| chr13:77447740-77447780 | Ch13 region 3 |
| chr13:77447820-77447860 | Ch13 region 3 |
| chr13:77448094-77448134 | Ch13 region 3 |
| chr13:77448139-77448179 | Ch13 region 3 |
| chr13:77448218-77448258 | Ch13 region 3 |
| chr13:77448314-77448354 | Ch13 region 3 |
| chr13:77448359-77448399 | Ch13 region 3 |
| chr13:77448409-77448449 | Ch13 region 3 |
| chr13:77448485-77448525 | Ch13 region 3 |
| chr13:77448538-77448578 | Ch13 region 3 |
| chr13:77448604-77448644 | Ch13 region 3 |
| chr13:77448702-77448742 | Ch13 region 3 |
| chr13:77448758-77448798 | Ch13 region 3 |
| chr13:77448803-77448843 | Ch13 region 3 |
| chr13:77448890-77448930 | Ch13 region 3 |
| chr13:77448999-77449039 | Ch13 region 3 |
| chr13:77449066-77449106 | Ch13 region 3 |
| chr13:77449113-77449153 | Ch13 region 3 |
| chr13:77449171-77449211 | Ch13 region 3 |
| chr13:77449225-77449265 | Ch13 region 3 |
| chr13:77449372-77449412 | Ch13 region 3 |
| chr13:77449432-77449472 | Ch13 region 3 |
| chr13:77449533-77449573 | Ch13 region 3 |
| chr13:77449613-77449653 | Ch13 region 3 |
| chr13:77449671-77449711 | Ch13 region 3 |
| chr13:77449755-77449795 | Ch13 region 3 |
| chr13:77449802-77449842 | Ch13 region 3 |
| chr13:77449853-77449893 | Ch13 region 3 |
| chr13:77449978-77450018 | Ch13 region 3 |
| chr13:77450264-77450304 | Ch13 region 3 |
| chr13:77450408-77450448 | Ch13 region 3 |
| chr13:77450481-77450521 | Ch13 region 3 |
| chr13:77450548-77450588 | Ch13 region 3 |
| chr13:77450593-77450633 | Ch13 region 3 |
| chr13:77450674-77450714 | Ch13 region 3 |
| chr13:77450721-77450761 | Ch13 region 3 |
| chr13:77450936-77450976 | Ch13 region 3 |
| chr13:77451094-77451134 | Ch13 region 3 |
| chr13:77451142-77451182 | Ch13 region 3 |
| chr13:77451229-77451269 | Ch13 region 3 |
| chr13:77451334-77451374 | Ch13 region 3 |
| chr13:77451402-77451442 | Ch13 region 3 |
| chr13:77451467-77451507 | Ch13 region 3 |
| chr13:77451530-77451570 | Ch13 region 3 |
| chr13:77451626-77451666 | Ch13 region 3 |
| chr13:77451718-77451758 | Ch13 region 3 |
| chr13:77451807-77451847 | Ch13 region 3 |
| chr13:77451859-77451899 | Ch13 region 3 |
| chr13:77451904-77451944 | Ch13 region 3 |

| | |
|---|---|
| chr13:77451966-77452006 | Ch13 region 3 |
| chr13:77452011-77452051 | Ch13 region 3 |
| chr13:77452061-77452101 | Ch13 region 3 |
| chr13:77452132-77452172 | Ch13 region 3 |
| chr13:77452197-77452237 | Ch13 region 3 |
| chr13:77452271-77452311 | Ch13 region 3 |
| chr13:77452325-77452365 | Ch13 region 3 |
| chr13:77452370-77452410 | Ch13 region 3 |
| chr13:77452435-77452475 | Ch13 region 3 |
| chr13:77452512-77452552 | Ch13 region 3 |
| chr13:77452602-77452642 | Ch13 region 3 |
| chr13:77452665-77452705 | Ch13 region 3 |
| chr13:77452714-77452754 | Ch13 region 3 |
| chr13:77452787-77452827 | Ch13 region 3 |
| chr13:77452884-77452924 | Ch13 region 3 |
| chr13:77452957-77452997 | Ch13 region 3 |
| chr13:77453011-77453051 | Ch13 region 3 |
| chr13:77453135-77453175 | Ch13 region 3 |
| chr13:77453201-77453241 | Ch13 region 3 |
| chr13:77453284-77453324 | Ch13 region 3 |
| chr13:77453345-77453385 | Ch13 region 3 |
| chr13:77453421-77453461 | Ch13 region 3 |
| chr13:77453503-77453543 | Ch13 region 3 |
| chr13:77453548-77453588 | Ch13 region 3 |
| chr13:77453642-77453682 | Ch13 region 3 |
| chr13:77453712-77453752 | Ch13 region 3 |
| chr13:77453768-77453808 | Ch13 region 3 |
| chr13:77453817-77453857 | Ch13 region 3 |
| chr13:77453909-77453949 | Ch13 region 3 |
| chr13:77453986-77454026 | Ch13 region 3 |
| chr13:77454063-77454103 | Ch13 region 3 |
| chr13:77454110-77454150 | Ch13 region 3 |
| chr13:77454155-77454195 | Ch13 region 3 |
| chr13:77454223-77454263 | Ch13 region 3 |
| chr13:77454284-77454324 | Ch13 region 3 |
| chr13:77454356-77454396 | Ch13 region 3 |
| chr13:77454427-77454467 | Ch13 region 3 |
| chr13:77454472-77454512 | Ch13 region 3 |
| chr13:77454653-77454693 | Ch13 region 3 |
| chr13:77454772-77454812 | Ch13 region 3 |
| chr13:77454857-77454897 | Ch13 region 3 |
| chr13:77454999-77455039 | Ch13 region 3 |
| chr13:77455077-77455117 | Ch13 region 3 |
| chr13:77455144-77455184 | Ch13 region 3 |
| chr13:77455194-77455234 | Ch13 region 3 |
| chr13:77455321-77455361 | Ch13 region 3 |
| chr13:77455429-77455469 | Ch13 region 3 |
| chr13:77455493-77455533 | Ch13 region 3 |
| chr13:77455571-77455611 | Ch13 region 3 |
| chr13:77455667-77455707 | Ch13 region 3 |
| chr13:77455718-77455758 | Ch13 region 3 |
| chr13:77455909-77455949 | Ch13 region 3 |

| | |
|---|---|
| chr13:77456063-77456103 | Ch13 region 3 |
| chr13:77456153-77456193 | Ch13 region 3 |
| chr13:77456199-77456239 | Ch13 region 3 |
| chr13:77456283-77456323 | Ch13 region 3 |
| chr13:77456337-77456377 | Ch13 region 3 |
| chr13:77456398-77456438 | Ch13 region 3 |
| chr13:77456463-77456503 | Ch13 region 3 |
| chr13:77456509-77456549 | Ch13 region 3 |
| chr13:77456597-77456637 | Ch13 region 3 |
| chr13:77456673-77456713 | Ch13 region 3 |
| chr13:77456718-77456758 | Ch13 region 3 |
| chr13:77456807-77456847 | Ch13 region 3 |
| chr13:77456854-77456894 | Ch13 region 3 |
| chr13:77456907-77456947 | Ch13 region 3 |
| chr13:77457039-77457079 | Ch13 region 3 |
| chr13:77457156-77457196 | Ch13 region 3 |
| chr13:77457201-77457241 | Ch13 region 3 |
| chr13:77457339-77457379 | Ch13 region 3 |
| chr13:77457460-77457500 | Ch13 region 3 |
| chr13:77457530-77457570 | Ch13 region 3 |
| chr13:77457585-77457625 | Ch13 region 3 |
| chr13:77457661-77457701 | Ch13 region 3 |
| chr13:77457751-77457791 | Ch13 region 3 |
| chr13:77457797-77457837 | Ch13 region 3 |
| chr13:77457861-77457901 | Ch13 region 3 |
| chr13:77457943-77457983 | Ch13 region 3 |
| chr13:77458036-77458076 | Ch13 region 3 |
| chr13:77458110-77458150 | Ch13 region 3 |
| chr13:77458196-77458236 | Ch13 region 3 |
| chr13:77458295-77458335 | Ch13 region 3 |
| chr13:77458369-77458409 | Ch13 region 3 |
| chr13:77458458-77458498 | Ch13 region 3 |
| chr13:77458503-77458543 | Ch13 region 3 |
| chr13:77458569-77458609 | Ch13 region 3 |
| chr13:77458637-77458677 | Ch13 region 3 |
| chr13:77458722-77458762 | Ch13 region 3 |
| chr13:77458790-77458830 | Ch13 region 3 |
| chr13:77458838-77458878 | Ch13 region 3 |
| chr13:77458915-77458955 | Ch13 region 3 |
| chr13:77458962-77459002 | Ch13 region 3 |
| chr13:77459014-77459054 | Ch13 region 3 |
| chr13:77459059-77459099 | Ch13 region 3 |
| chr13:77459134-77459174 | Ch13 region 3 |
| chr13:77459215-77459255 | Ch13 region 3 |
| chr13:77459302-77459342 | Ch13 region 3 |
| chr13:77459347-77459387 | Ch13 region 3 |
| chr13:77459447-77459487 | Ch13 region 3 |
| chr13:77459529-77459569 | Ch13 region 3 |
| chr13:77459620-77459660 | Ch13 region 3 |
| chr13:77459714-77459754 | Ch13 region 3 |
| chr13:77459820-77459860 | Ch13 region 3 |
| chr13:77459879-77459919 | Ch13 region 3 |

| | |
|---|---|
| chr13:77459977-77460017 | Ch13 region 3 |
| chr13:77460025-77460065 | Ch13 region 3 |
| chr13:77460118-77460158 | Ch13 region 3 |
| chr13:77460207-77460247 | Ch13 region 3 |
| chr13:77460274-77460314 | Ch13 region 3 |
| chr13:77460325-77460365 | Ch13 region 3 |
| chr13:77460421-77460461 | Ch13 region 3 |
| chr13:77460485-77460525 | Ch13 region 3 |
| chr13:77460557-77460597 | Ch13 region 3 |
| chr13:77460646-77460686 | Ch13 region 3 |
| chr13:77460730-77460770 | Ch13 region 3 |
| chr13:77460823-77460863 | Ch13 region 3 |
| chr13:77460868-77460908 | Ch13 region 3 |
| chr13:77460924-77460964 | Ch13 region 3 |
| chr13:77461002-77461042 | Ch13 region 3 |
| chr13:77461047-77461087 | Ch13 region 3 |
| chr13:77461146-77461186 | Ch13 region 3 |
| chr13:77461223-77461263 | Ch13 region 3 |
| chr13:77461303-77461343 | Ch13 region 3 |
| chr13:77461362-77461402 | Ch13 region 3 |
| chr13:77461428-77461468 | Ch13 region 3 |
| chr13:77461481-77461521 | Ch13 region 3 |
| chr13:77461528-77461568 | Ch13 region 3 |
| chr13:77461573-77461613 | Ch13 region 3 |
| chr13:77461671-77461711 | Ch13 region 3 |
| chr13:77461753-77461793 | Ch13 region 3 |
| chr13:77461799-77461839 | Ch13 region 3 |
| chr13:77461903-77461943 | Ch13 region 3 |
| chr13:77461951-77461991 | Ch13 region 3 |
| chr13:77462029-77462069 | Ch13 region 3 |
| chr13:77462107-77462147 | Ch13 region 3 |
| chr13:77462152-77462192 | Ch13 region 3 |
| chr13:77462258-77462298 | Ch13 region 3 |
| chr13:77462338-77462378 | Ch13 region 3 |
| chr13:77462440-77462480 | Ch13 region 3 |
| chr13:77462488-77462528 | Ch13 region 3 |
| chr13:77462534-77462574 | Ch13 region 3 |
| chr13:77462579-77462619 | Ch13 region 3 |
| chr13:77462662-77462702 | Ch13 region 3 |
| chr13:77462734-77462774 | Ch13 region 3 |
| chr13:77462793-77462833 | Ch13 region 3 |
| chr13:77462843-77462883 | Ch13 region 3 |
| chr13:77462928-77462968 | Ch13 region 3 |
| chr13:77462973-77463013 | Ch13 region 3 |
| chr13:77463043-77463083 | Ch13 region 3 |
| chr13:77463129-77463169 | Ch13 region 3 |
| chr13:77463175-77463215 | Ch13 region 3 |
| chr13:77463252-77463292 | Ch13 region 3 |
| chr13:77463303-77463343 | Ch13 region 3 |
| chr13:77463360-77463400 | Ch13 region 3 |
| chr13:77463436-77463476 | Ch13 region 3 |
| chr13:77463504-77463544 | Ch13 region 3 |

| | |
|---|---|
| chr13:77463565-77463605 | Ch13 region 3 |
| chr13:77463610-77463650 | Ch13 region 3 |
| chr13:77463699-77463739 | Ch13 region 3 |
| chr13:77463775-77463815 | Ch13 region 3 |
| chr13:77463826-77463866 | Ch13 region 3 |
| chr13:77463893-77463933 | Ch13 region 3 |
| chr13:77463951-77463991 | Ch13 region 3 |
| chr13:77464005-77464045 | Ch13 region 3 |
| chr13:77464074-77464114 | Ch13 region 3 |
| chr13:77464122-77464162 | Ch13 region 3 |
| chr13:77464167-77464207 | Ch13 region 3 |
| chr13:77464226-77464266 | Ch13 region 3 |
| chr13:77464357-77464397 | Ch13 region 3 |
| chr13:77464402-77464442 | Ch13 region 3 |
| chr13:77464448-77464488 | Ch13 region 3 |
| chr13:77464516-77464556 | Ch13 region 3 |
| chr13:77464573-77464613 | Ch13 region 3 |
| chr13:77464728-77464768 | Ch13 region 3 |
| chr13:77464795-77464835 | Ch13 region 3 |
| chr13:77464851-77464891 | Ch13 region 3 |
| chr13:77464926-77464966 | Ch13 region 3 |
| chr13:77464994-77465034 | Ch13 region 3 |
| chr13:77465068-77465108 | Ch13 region 3 |
| chr13:77465122-77465162 | Ch13 region 3 |
| chr13:77465185-77465225 | Ch13 region 3 |
| chr13:77465253-77465293 | Ch13 region 3 |
| chr13:77465330-77465370 | Ch13 region 3 |
| chr13:77465418-77465458 | Ch13 region 3 |
| chr13:77465621-77465661 | Ch13 region 3 |
| chr13:77465720-77465760 | Ch13 region 3 |
| chr13:77465785-77465825 | Ch13 region 3 |
| chr13:77465852-77465892 | Ch13 region 3 |
| chr13:77465914-77465954 | Ch13 region 3 |
| chr13:77465988-77466028 | Ch13 region 3 |
| chr13:77466088-77466128 | Ch13 region 3 |
| chr13:77466174-77466214 | Ch13 region 3 |
| chr13:77466235-77466275 | Ch13 region 3 |
| chr13:77466291-77466331 | Ch13 region 3 |
| chr13:77466345-77466385 | Ch13 region 3 |
| chr13:77466430-77466470 | Ch13 region 3 |
| chr13:77466497-77466537 | Ch13 region 3 |
| chr13:77466563-77466603 | Ch13 region 3 |
| chr13:77466615-77466655 | Ch13 region 3 |
| chr13:77466672-77466712 | Ch13 region 3 |
| chr13:77466740-77466780 | Ch13 region 3 |
| chr13:77466785-77466825 | Ch13 region 3 |
| chr13:77466832-77466872 | Ch13 region 3 |
| chr13:77466997-77467037 | Ch13 region 3 |
| chr13:77467080-77467120 | Ch13 region 3 |
| chr13:77467182-77467222 | Ch13 region 3 |
| chr13:77467245-77467285 | Ch13 region 3 |
| chr13:77467307-77467347 | Ch13 region 3 |

| | |
|---|---|
| chr13:77467405-77467445 | Ch13 region 3 |
| chr13:77467461-77467501 | Ch13 region 3 |
| chr13:77467527-77467567 | Ch13 region 3 |
| chr13:77467591-77467631 | Ch13 region 3 |
| chr13:77467652-77467692 | Ch13 region 3 |
| chr13:77467709-77467749 | Ch13 region 3 |
| chr13:77467776-77467816 | Ch13 region 3 |
| chr13:77467936-77467976 | Ch13 region 3 |
| chr13:77468046-77468086 | Ch13 region 3 |
| chr13:77468099-77468139 | Ch13 region 3 |
| chr13:77468167-77468207 | Ch13 region 3 |
| chr13:77468238-77468278 | Ch13 region 3 |
| chr13:77468290-77468330 | Ch13 region 3 |
| chr13:77468335-77468375 | Ch13 region 3 |
| chr13:77468387-77468427 | Ch13 region 3 |
| chr13:77468432-77468472 | Ch13 region 3 |
| chr13:77468478-77468518 | Ch13 region 3 |
| chr13:77468528-77468568 | Ch13 region 3 |
| chr13:77468575-77468615 | Ch13 region 3 |
| chr13:77468628-77468668 | Ch13 region 3 |
| chr13:77468675-77468715 | Ch13 region 3 |
| chr13:77468720-77468760 | Ch13 region 3 |
| chr13:77468816-77468856 | Ch13 region 3 |
| chr13:77468891-77468931 | Ch13 region 3 |
| chr13:77468939-77468979 | Ch13 region 3 |
| chr13:77468996-77469036 | Ch13 region 3 |
| chr13:77469056-77469096 | Ch13 region 3 |
| chr13:77469101-77469141 | Ch13 region 3 |
| chr13:77469177-77469217 | Ch13 region 3 |
| chr13:77469231-77469271 | Ch13 region 3 |
| chr13:77469299-77469339 | Ch13 region 3 |
| chr13:77469413-77469453 | Ch13 region 3 |
| chr13:77469507-77469547 | Ch13 region 3 |
| chr13:77469567-77469607 | Ch13 region 3 |
| chr13:77469655-77469695 | Ch13 region 3 |
| chr13:77469723-77469763 | Ch13 region 3 |
| chr13:77469786-77469826 | Ch13 region 3 |
| chr13:77469853-77469893 | Ch13 region 3 |
| chr13:77469937-77469977 | Ch13 region 3 |
| chr13:77469985-77470025 | Ch13 region 3 |
| chr13:77470078-77470118 | Ch13 region 3 |
| chr13:77470123-77470163 | Ch13 region 3 |
| chr13:77470220-77470260 | Ch13 region 3 |
| chr13:77470270-77470310 | Ch13 region 3 |
| chr13:77470326-77470366 | Ch13 region 3 |
| chr13:77470380-77470420 | Ch13 region 3 |
| chr13:77470441-77470481 | Ch13 region 3 |
| chr13:77470565-77470605 | Ch13 region 3 |
| chr13:77470643-77470683 | Ch13 region 3 |
| chr13:77470739-77470779 | Ch13 region 3 |
| chr13:77470898-77470938 | Ch13 region 3 |
| chr13:77470970-77471010 | Ch13 region 3 |

| | |
|---|---|
| chr13:77471086-77471126 | Ch13 region 3 |
| chr13:77471301-77471341 | Ch13 region 3 |
| chr13:77471398-77471438 | Ch13 region 3 |
| chr13:77471483-77471523 | Ch13 region 3 |
| chr13:77471560-77471600 | Ch13 region 3 |
| chr13:77471610-77471650 | Ch13 region 3 |
| chr13:77471680-77471720 | Ch13 region 3 |
| chr13:77471865-77471905 | Ch13 region 3 |
| chr13:77471946-77471986 | Ch13 region 3 |
| chr13:77472001-77472041 | Ch13 region 3 |
| chr13:77472078-77472118 | Ch13 region 3 |
| chr13:77472192-77472232 | Ch13 region 3 |
| chr13:77472434-77472474 | Ch13 region 3 |
| chr13:77472544-77472584 | Ch13 region 3 |
| chr13:77472628-77472668 | Ch13 region 3 |
| chr13:77472718-77472758 | Ch13 region 3 |
| chr13:77472874-77472914 | Ch13 region 3 |
| chr13:77472947-77472987 | Ch13 region 3 |
| chr13:77473140-77473180 | Ch13 region 3 |
| chr13:77473288-77473328 | Ch13 region 3 |
| chr13:77473334-77473374 | Ch13 region 3 |
| chr13:77473442-77473482 | Ch13 region 3 |
| chr13:77473595-77473635 | Ch13 region 3 |
| chr13:77473732-77473772 | Ch13 region 3 |
| chr13:77473806-77473846 | Ch13 region 3 |
| chr13:77473853-77473893 | Ch13 region 3 |
| chr13:77473900-77473940 | Ch13 region 3 |
| chr13:77473978-77474018 | Ch13 region 3 |
| chr13:77474033-77474073 | Ch13 region 3 |
| chr13:77474261-77474301 | Ch13 region 3 |
| chr13:77474320-77474360 | Ch13 region 3 |
| chr13:77474408-77474448 | Ch13 region 3 |
| chr13:77474525-77474565 | Ch13 region 3 |
| chr13:77474653-77474693 | Ch13 region 3 |
| chr13:77474698-77474738 | Ch13 region 3 |
| chr13:77474770-77474810 | Ch13 region 3 |
| chr13:77474847-77474887 | Ch13 region 3 |
| chr13:77474925-77474965 | Ch13 region 3 |
| chr13:77474980-77475020 | Ch13 region 3 |
| chr13:77475040-77475080 | Ch13 region 3 |
| chr13:77475112-77475152 | Ch13 region 3 |
| chr13:77475179-77475219 | Ch13 region 3 |
| chr13:77475287-77475327 | Ch13 region 3 |
| chr13:77475442-77475482 | Ch13 region 3 |
| chr13:77475497-77475537 | Ch13 region 3 |
| chr13:77475544-77475584 | Ch13 region 3 |
| chr13:77475593-77475633 | Ch13 region 3 |
| chr13:77475659-77475699 | Ch13 region 3 |
| chr13:77475733-77475773 | Ch13 region 3 |
| chr13:77475792-77475832 | Ch13 region 3 |
| chr13:77475885-77475925 | Ch13 region 3 |
| chr13:77475974-77476014 | Ch13 region 3 |

| | |
|---|---|
| chr13:77476024-77476064 | Ch13 region 3 |
| chr13:77476069-77476109 | Ch13 region 3 |
| chr13:77476136-77476176 | Ch13 region 3 |
| chr13:77476194-77476234 | Ch13 region 3 |
| chr13:77476258-77476298 | Ch13 region 3 |
| chr13:77476341-77476381 | Ch13 region 3 |
| chr13:77476407-77476447 | Ch13 region 3 |
| chr13:77476462-77476502 | Ch13 region 3 |
| chr13:77476535-77476575 | Ch13 region 3 |
| chr13:77476582-77476622 | Ch13 region 3 |
| chr13:77476635-77476675 | Ch13 region 3 |
| chr13:77476711-77476751 | Ch13 region 3 |
| chr13:77476756-77476796 | Ch13 region 3 |
| chr13:77476819-77476859 | Ch13 region 3 |
| chr13:77476920-77476960 | Ch13 region 3 |
| chr13:77476970-77477010 | Ch13 region 3 |
| chr13:77477090-77477130 | Ch13 region 3 |
| chr13:77477140-77477180 | Ch13 region 3 |
| chr13:77477219-77477259 | Ch13 region 3 |
| chr13:77477280-77477320 | Ch13 region 3 |
| chr13:77477346-77477386 | Ch13 region 3 |
| chr13:77477480-77477520 | Ch13 region 3 |
| chr13:77477550-77477590 | Ch13 region 3 |
| chr13:77477615-77477655 | Ch13 region 3 |
| chr13:77477697-77477737 | Ch13 region 3 |
| chr13:77477770-77477810 | Ch13 region 3 |
| chr13:77477826-77477866 | Ch13 region 3 |
| chr13:77477937-77477977 | Ch13 region 3 |
| chr13:77477996-77478036 | Ch13 region 3 |
| chr13:77478106-77478146 | Ch13 region 3 |
| chr13:77478157-77478197 | Ch13 region 3 |
| chr13:77478239-77478279 | Ch13 region 3 |
| chr13:77478306-77478346 | Ch13 region 3 |
| chr13:77478362-77478402 | Ch13 region 3 |
| chr13:77478457-77478497 | Ch13 region 3 |
| chr13:77478550-77478590 | Ch13 region 3 |
| chr13:77478595-77478635 | Ch13 region 3 |
| chr13:77478674-77478714 | Ch13 region 3 |
| chr13:77478738-77478778 | Ch13 region 3 |
| chr13:77478790-77478830 | Ch13 region 3 |
| chr13:77478860-77478900 | Ch13 region 3 |
| chr13:77478938-77478978 | Ch13 region 3 |
| chr13:77479025-77479065 | Ch13 region 3 |
| chr13:77479086-77479126 | Ch13 region 3 |
| chr13:77479160-77479200 | Ch13 region 3 |
| chr13:77479218-77479258 | Ch13 region 3 |
| chr13:77479268-77479308 | Ch13 region 3 |
| chr13:77479345-77479385 | Ch13 region 3 |
| chr13:77479445-77479485 | Ch13 region 3 |
| chr13:77479534-77479574 | Ch13 region 3 |
| chr13:77479604-77479644 | Ch13 region 3 |
| chr13:77479669-77479709 | Ch13 region 3 |

| | |
|---|---|
| chr13:77479740-77479780 | Ch13 region 3 |
| chr13:77479826-77479866 | Ch13 region 3 |
| chr13:77479906-77479946 | Ch13 region 3 |
| chr13:77479977-77480017 | Ch13 region 3 |
| chr13:77480027-77480067 | Ch13 region 3 |
| chr13:77480077-77480117 | Ch13 region 3 |
| chr13:77480133-77480173 | Ch13 region 3 |
| chr13:77480201-77480241 | Ch13 region 3 |
| chr13:77480253-77480293 | Ch13 region 3 |
| chr13:77480341-77480381 | Ch13 region 3 |
| chr13:77480418-77480458 | Ch13 region 3 |
| chr13:77480499-77480539 | Ch13 region 3 |
| chr13:77480546-77480586 | Ch13 region 3 |
| chr13:77480649-77480689 | Ch13 region 3 |
| chr13:77480695-77480735 | Ch13 region 3 |
| chr13:77480751-77480791 | Ch13 region 3 |
| chr13:77480805-77480845 | Ch13 region 3 |
| chr13:77480853-77480893 | Ch13 region 3 |
| chr13:77480944-77480984 | Ch13 region 3 |
| chr13:77481032-77481072 | Ch13 region 3 |
| chr13:77481375-77481415 | Ch13 region 3 |
| chr13:77481444-77481484 | Ch13 region 3 |
| chr13:77481539-77481579 | Ch13 region 3 |
| chr13:77481635-77481675 | Ch13 region 3 |
| chr13:77481705-77481745 | Ch13 region 3 |
| chr13:77481768-77481808 | Ch13 region 3 |
| chr13:77481833-77481873 | Ch13 region 3 |
| chr13:77481917-77481957 | Ch13 region 3 |
| chr13:77481968-77482008 | Ch13 region 3 |
| chr13:77482042-77482082 | Ch13 region 3 |
| chr13:77482106-77482146 | Ch13 region 3 |
| chr13:77482182-77482222 | Ch13 region 3 |
| chr13:77482365-77482405 | Ch13 region 3 |
| chr13:77482530-77482570 | Ch13 region 3 |
| chr13:77482589-77482629 | Ch13 region 3 |
| chr13:77482649-77482689 | Ch13 region 3 |
| chr13:77482727-77482767 | Ch13 region 3 |
| chr13:77482783-77482823 | Ch13 region 3 |
| chr13:77482860-77482900 | Ch13 region 3 |
| chr13:77482929-77482969 | Ch13 region 3 |
| chr13:77482989-77483029 | Ch13 region 3 |
| chr13:77483040-77483080 | Ch13 region 3 |
| chr13:77483184-77483224 | Ch13 region 3 |
| chr13:77483249-77483289 | Ch13 region 3 |
| chr13:77483331-77483371 | Ch13 region 3 |
| chr13:77483383-77483423 | Ch13 region 3 |
| chr13:77483428-77483468 | Ch13 region 3 |
| chr13:77483502-77483542 | Ch13 region 3 |
| chr13:77483548-77483588 | Ch13 region 3 |
| chr13:77483593-77483633 | Ch13 region 3 |
| chr13:77483648-77483688 | Ch13 region 3 |
| chr13:77483969-77484009 | Ch13 region 3 |

| | |
|---|---|
| chr13:77484152-77484192 | Ch13 region 3 |
| chr13:77484223-77484263 | Ch13 region 3 |
| chr13:77484287-77484327 | Ch13 region 3 |
| chr13:77484353-77484393 | Ch13 region 3 |
| chr13:77484399-77484439 | Ch13 region 3 |
| chr13:77484453-77484493 | Ch13 region 3 |
| chr13:77484519-77484559 | Ch13 region 3 |
| chr13:77484574-77484614 | Ch13 region 3 |
| chr13:77484647-77484687 | Ch13 region 3 |
| chr13:77484704-77484744 | Ch13 region 3 |
| chr13:77484752-77484792 | Ch13 region 3 |
| chr13:77484819-77484859 | Ch13 region 3 |
| chr13:77484864-77484904 | Ch13 region 3 |
| chr13:77484968-77485008 | Ch13 region 3 |
| chr13:77485019-77485059 | Ch13 region 3 |
| chr13:77485090-77485130 | Ch13 region 3 |
| chr13:77485146-77485186 | Ch13 region 3 |
| chr13:77485197-77485237 | Ch13 region 3 |
| chr13:77485260-77485300 | Ch13 region 3 |
| chr13:77485327-77485367 | Ch13 region 3 |
| chr13:77485383-77485423 | Ch13 region 3 |
| chr13:77485446-77485486 | Ch13 region 3 |
| chr13:77485525-77485565 | Ch13 region 3 |
| chr13:77485638-77485678 | Ch13 region 3 |
| chr13:77485731-77485771 | Ch13 region 3 |
| chr13:77485779-77485819 | Ch13 region 3 |
| chr13:77485909-77485949 | Ch13 region 3 |
| chr13:77485976-77486016 | Ch13 region 3 |
| chr13:77486031-77486071 | Ch13 region 3 |
| chr13:77486078-77486118 | Ch13 region 3 |
| chr13:77486126-77486166 | Ch13 region 3 |
| chr13:77486193-77486233 | Ch13 region 3 |
| chr13:77486279-77486319 | Ch13 region 3 |
| chr13:77486354-77486394 | Ch13 region 3 |
| chr13:77486418-77486458 | Ch13 region 3 |
| chr13:77486640-77486680 | Ch13 region 3 |
| chr13:77486751-77486791 | Ch13 region 3 |
| chr13:77486796-77486836 | Ch13 region 3 |
| chr13:77486842-77486882 | Ch13 region 3 |
| chr13:77486887-77486927 | Ch13 region 3 |
| chr13:77486965-77487005 | Ch13 region 3 |
| chr13:77487010-77487050 | Ch13 region 3 |
| chr13:77487062-77487102 | Ch13 region 3 |
| chr13:77487158-77487198 | Ch13 region 3 |
| chr13:77487251-77487291 | Ch13 region 3 |
| chr13:77487338-77487378 | Ch13 region 3 |
| chr13:77487410-77487450 | Ch13 region 3 |
| chr13:77487512-77487552 | Ch13 region 3 |
| chr13:77487578-77487618 | Ch13 region 3 |
| chr13:77487728-77487768 | Ch13 region 3 |
| chr13:77487838-77487878 | Ch13 region 3 |
| chr13:77487920-77487960 | Ch13 region 3 |

| | |
|---|---|
| chr13:77488005-77488045 | Ch13 region 3 |
| chr13:77488065-77488105 | Ch13 region 3 |
| chr13:77488112-77488152 | Ch13 region 3 |
| chr13:77488166-77488206 | Ch13 region 3 |
| chr13:77488263-77488303 | Ch13 region 3 |
| chr13:77488313-77488353 | Ch13 region 3 |
| chr13:77488374-77488414 | Ch13 region 3 |
| chr13:77488470-77488510 | Ch13 region 3 |
| chr13:77488524-77488564 | Ch13 region 3 |
| chr13:77488594-77488634 | Ch13 region 3 |
| chr13:77488641-77488681 | Ch13 region 3 |
| chr13:77488686-77488726 | Ch13 region 3 |
| chr13:77488737-77488777 | Ch13 region 3 |
| chr13:77488845-77488885 | Ch13 region 3 |
| chr13:77488913-77488953 | Ch13 region 3 |
| chr13:77489097-77489137 | Ch13 region 3 |
| chr13:77489157-77489197 | Ch13 region 3 |
| chr13:77489246-77489286 | Ch13 region 3 |
| chr13:77489310-77489350 | Ch13 region 3 |
| chr13:77489355-77489395 | Ch13 region 3 |
| chr13:77489426-77489466 | Ch13 region 3 |
| chr13:77489474-77489514 | Ch13 region 3 |
| chr13:77489530-77489570 | Ch13 region 3 |
| chr13:77489605-77489645 | Ch13 region 3 |
| chr13:77489659-77489699 | Ch13 region 3 |
| chr13:77489730-77489770 | Ch13 region 3 |
| chr13:77489873-77489913 | Ch13 region 3 |
| chr13:77490014-77490054 | Ch13 region 3 |
| chr13:77490203-77490243 | Ch13 region 3 |
| chr13:77490292-77490332 | Ch13 region 3 |
| chr13:77490338-77490378 | Ch13 region 3 |
| chr13:77490416-77490456 | Ch13 region 3 |
| chr13:77490474-77490514 | Ch13 region 3 |
| chr13:77490544-77490584 | Ch13 region 3 |
| chr13:77490608-77490648 | Ch13 region 3 |
| chr13:77490658-77490698 | Ch13 region 3 |
| chr13:77490706-77490746 | Ch13 region 3 |
| chr13:77490764-77490804 | Ch13 region 3 |
| chr13:77490832-77490872 | Ch13 region 3 |
| chr13:77490882-77490922 | Ch13 region 3 |
| chr13:77490980-77491020 | Ch13 region 3 |
| chr13:77491037-77491077 | Ch13 region 3 |
| chr13:77491107-77491147 | Ch13 region 3 |
| chr13:77491179-77491219 | Ch13 region 3 |
| chr13:77491258-77491298 | Ch13 region 3 |
| chr13:77491420-77491460 | Ch13 region 3 |
| chr13:77491470-77491510 | Ch13 region 3 |
| chr13:77491535-77491575 | Ch13 region 3 |
| chr13:77491740-77491780 | Ch13 region 3 |
| chr13:77491911-77491951 | Ch13 region 3 |
| chr13:77491985-77492025 | Ch13 region 3 |
| chr13:77492031-77492071 | Ch13 region 3 |

| | |
|---|---|
| chr13:77492153-77492193 | Ch13 region 3 |
| chr13:77492293-77492333 | Ch13 region 3 |
| chr13:77492348-77492388 | Ch13 region 3 |
| chr13:77492475-77492515 | Ch13 region 3 |
| chr13:77492695-77492735 | Ch13 region 3 |
| chr13:77492740-77492780 | Ch13 region 3 |
| chr13:77492958-77492998 | Ch13 region 3 |
| chr13:77493055-77493095 | Ch13 region 3 |
| chr13:77493254-77493294 | Ch13 region 3 |
| chr13:77493367-77493407 | Ch13 region 3 |
| chr13:77493607-77493647 | Ch13 region 3 |
| chr13:77493696-77493736 | Ch13 region 3 |
| chr13:77493887-77493927 | Ch13 region 3 |
| chr13:77493967-77494007 | Ch13 region 3 |
| chr13:77494140-77494180 | Ch13 region 3 |
| chr13:77494235-77494275 | Ch13 region 3 |
| chr13:77494294-77494334 | Ch13 region 3 |
| chr13:77494488-77494528 | Ch13 region 3 |
| chr13:77494571-77494611 | Ch13 region 3 |
| chr13:77494793-77494833 | Ch13 region 3 |
| chr13:77495007-77495047 | Ch13 region 3 |
| chr13:77495067-77495107 | Ch13 region 3 |
| chr13:77495174-77495214 | Ch13 region 3 |
| chr13:77495375-77495415 | Ch13 region 3 |
| chr13:77495425-77495465 | Ch13 region 3 |
| chr13:77495694-77495734 | Ch13 region 3 |
| chr13:77495905-77495945 | Ch13 region 3 |
| chr13:77495989-77496029 | Ch13 region 3 |
| chr13:77496047-77496087 | Ch13 region 3 |
| chr13:77496136-77496176 | Ch13 region 3 |
| chr13:77496202-77496242 | Ch13 region 3 |
| chr13:77496248-77496288 | Ch13 region 3 |
| chr13:77496317-77496357 | Ch13 region 3 |
| chr13:77496367-77496407 | Ch13 region 3 |
| chr13:77496439-77496479 | Ch13 region 3 |
| chr13:77496515-77496555 | Ch13 region 3 |
| chr13:77496591-77496631 | Ch13 region 3 |
| chr13:77496642-77496682 | Ch13 region 3 |
| chr13:77496724-77496764 | Ch13 region 3 |
| chr13:77496852-77496892 | Ch13 region 3 |
| chr13:77496924-77496964 | Ch13 region 3 |
| chr13:77497009-77497049 | Ch13 region 3 |
| chr13:77497057-77497097 | Ch13 region 3 |
| chr13:77497102-77497142 | Ch13 region 3 |
| chr13:77497203-77497243 | Ch13 region 3 |
| chr13:77497248-77497288 | Ch13 region 3 |
| chr13:77497293-77497333 | Ch13 region 3 |
| chr13:77497369-77497409 | Ch13 region 3 |
| chr13:77497415-77497455 | Ch13 region 3 |
| chr13:77497472-77497512 | Ch13 region 3 |
| chr13:77497545-77497585 | Ch13 region 3 |
| chr13:77497608-77497648 | Ch13 region 3 |

| | |
|---|---|
| chr13:77497660-77497700 | Ch13 region 3 |
| chr13:77497724-77497764 | Ch13 region 3 |
| chr13:77497790-77497830 | Ch13 region 3 |
| chr13:77497841-77497881 | Ch13 region 3 |
| chr13:77497886-77497926 | Ch13 region 3 |
| chr13:77497932-77497972 | Ch13 region 3 |
| chr13:77497994-77498034 | Ch13 region 3 |
| chr13:77498047-77498087 | Ch13 region 3 |
| chr13:77498092-77498132 | Ch13 region 3 |
| chr13:77498137-77498177 | Ch13 region 3 |
| chr13:77498208-77498248 | Ch13 region 3 |
| chr13:77498267-77498307 | Ch13 region 3 |
| chr13:77498333-77498373 | Ch13 region 3 |
| chr13:77498406-77498446 | Ch13 region 3 |
| chr13:77498480-77498520 | Ch13 region 3 |
| chr13:77498525-77498565 | Ch13 region 3 |
| chr13:77498579-77498619 | Ch13 region 3 |
| chr13:77498624-77498664 | Ch13 region 3 |
| chr13:77498689-77498729 | Ch13 region 3 |
| chr13:77498771-77498811 | Ch13 region 3 |
| chr13:77498851-77498891 | Ch13 region 3 |
| chr13:77498896-77498936 | Ch13 region 3 |
| chr13:77498986-77499026 | Ch13 region 3 |
| chr13:77499087-77499127 | Ch13 region 3 |
| chr13:77499150-77499190 | Ch13 region 3 |
| chr13:77499266-77499306 | Ch13 region 3 |
| chr13:77499358-77499398 | Ch13 region 3 |
| chr13:77499443-77499483 | Ch13 region 3 |
| chr13:77499488-77499528 | Ch13 region 3 |
| chr13:77499604-77499644 | Ch13 region 3 |
| chr13:77499680-77499720 | Ch13 region 3 |
| chr13:77499732-77499772 | Ch13 region 3 |
| chr13:77499800-77499840 | Ch13 region 3 |
| chr13:77499867-77499907 | Ch13 region 3 |
| chr13:77499927-77499967 | Ch13 region 3 |
| chr13:77500029-77500069 | Ch13 region 3 |
| chr13:77500093-77500133 | Ch13 region 3 |
| chr13:77500172-77500212 | Ch13 region 3 |
| chr13:77500226-77500266 | Ch13 region 3 |
| chr13:77500298-77500338 | Ch13 region 3 |
| chr13:77500372-77500412 | Ch13 region 3 |
| chr13:77500429-77500469 | Ch13 region 3 |
| chr13:77500475-77500515 | Ch13 region 3 |
| chr13:77500572-77500612 | Ch13 region 3 |
| chr13:77500679-77500719 | Ch13 region 3 |
| chr13:77500729-77500769 | Ch13 region 3 |
| chr13:77500817-77500857 | Ch13 region 3 |
| chr13:77500892-77500932 | Ch13 region 3 |
| chr13:77500974-77501014 | Ch13 region 3 |
| chr13:77501045-77501085 | Ch13 region 3 |
| chr13:77501097-77501137 | Ch13 region 3 |
| chr13:77501167-77501207 | Ch13 region 3 |

| | |
|---|---|
| chr13:77501389-77501429 | Ch13 region 3 |
| chr13:77501535-77501575 | Ch13 region 3 |
| chr13:77501620-77501660 | Ch13 region 3 |
| chr13:77501706-77501746 | Ch13 region 3 |
| chr13:77501751-77501791 | Ch13 region 3 |
| chr13:77501851-77501891 | Ch13 region 3 |
| chr13:77501924-77501964 | Ch13 region 3 |
| chr13:77502074-77502114 | Ch13 region 3 |
| chr13:77502146-77502186 | Ch13 region 3 |
| chr13:77502210-77502250 | Ch13 region 3 |
| chr13:77502271-77502311 | Ch13 region 3 |
| chr13:77502355-77502395 | Ch13 region 3 |
| chr13:77502428-77502468 | Ch13 region 3 |
| chr13:77502537-77502577 | Ch13 region 3 |
| chr13:77502597-77502637 | Ch13 region 3 |
| chr13:77502678-77502718 | Ch13 region 3 |
| chr13:77502724-77502764 | Ch13 region 3 |
| chr13:77502801-77502841 | Ch13 region 3 |
| chr13:77502896-77502936 | Ch13 region 3 |
| chr13:77502948-77502988 | Ch13 region 3 |
| chr13:77502997-77503037 | Ch13 region 3 |
| chr13:77503199-77503239 | Ch13 region 3 |
| chr13:77503396-77503436 | Ch13 region 3 |
| chr13:77503493-77503533 | Ch13 region 3 |
| chr13:77503638-77503678 | Ch13 region 3 |
| chr13:77503742-77503782 | Ch13 region 3 |
| chr13:77503837-77503877 | Ch13 region 3 |
| chr13:77503958-77503998 | Ch13 region 3 |
| chr13:77504023-77504063 | Ch13 region 3 |
| chr13:77504068-77504108 | Ch13 region 3 |
| chr13:77504129-77504169 | Ch13 region 3 |
| chr13:77504190-77504230 | Ch13 region 3 |
| chr13:77504304-77504344 | Ch13 region 3 |
| chr13:77504359-77504399 | Ch13 region 3 |
| chr13:77504434-77504474 | Ch13 region 3 |
| chr13:77504491-77504531 | Ch13 region 3 |
| chr13:77504626-77504666 | Ch13 region 3 |
| chr13:77504698-77504738 | Ch13 region 3 |
| chr13:77504760-77504800 | Ch13 region 3 |
| chr13:77504868-77504908 | Ch13 region 3 |
| chr13:77504924-77504964 | Ch13 region 3 |
| chr13:77504971-77505011 | Ch13 region 3 |
| chr13:77505022-77505062 | Ch13 region 3 |
| chr13:77505096-77505136 | Ch13 region 3 |
| chr13:77505143-77505183 | Ch13 region 3 |
| chr13:77505221-77505261 | Ch13 region 3 |
| chr13:77505331-77505371 | Ch13 region 3 |
| chr13:77505395-77505435 | Ch13 region 3 |
| chr13:77505514-77505554 | Ch13 region 3 |
| chr13:77505567-77505607 | Ch13 region 3 |
| chr13:77505675-77505715 | Ch13 region 3 |
| chr13:77505787-77505827 | Ch13 region 3 |

| | |
|---|---|
| chr13:77505953-77505993 | Ch13 region 3 |
| chr13:77506166-77506206 | Ch13 region 3 |
| chr13:77506269-77506309 | Ch13 region 3 |
| chr13:77506388-77506428 | Ch13 region 3 |
| chr13:77506438-77506478 | Ch13 region 3 |
| chr13:77506509-77506549 | Ch13 region 3 |
| chr13:77506587-77506627 | Ch13 region 3 |
| chr13:77506705-77506745 | Ch13 region 3 |
| chr13:77506775-77506815 | Ch13 region 3 |
| chr13:77506851-77506891 | Ch13 region 3 |
| chr13:77506923-77506963 | Ch13 region 3 |
| chr13:77507026-77507066 | Ch13 region 3 |
| chr13:77507078-77507118 | Ch13 region 3 |
| chr13:77507129-77507169 | Ch13 region 3 |
| chr13:77507263-77507303 | Ch13 region 3 |
| chr13:77507313-77507353 | Ch13 region 3 |
| chr13:77507375-77507415 | Ch13 region 3 |
| chr13:77507420-77507460 | Ch13 region 3 |
| chr13:77507513-77507553 | Ch13 region 3 |
| chr13:77507559-77507599 | Ch13 region 3 |
| chr13:77507766-77507806 | Ch13 region 3 |
| chr13:77507913-77507953 | Ch13 region 3 |
| chr13:77508057-77508097 | Ch13 region 3 |
| chr13:77508120-77508160 | Ch13 region 3 |
| chr13:77508201-77508241 | Ch13 region 3 |
| chr13:77508297-77508337 | Ch13 region 3 |
| chr13:77508399-77508439 | Ch13 region 3 |
| chr13:77508495-77508535 | Ch13 region 3 |
| chr13:77508541-77508581 | Ch13 region 3 |
| chr13:77508612-77508652 | Ch13 region 3 |
| chr13:77508711-77508751 | Ch13 region 3 |
| chr13:77508757-77508797 | Ch13 region 3 |
| chr13:77508832-77508872 | Ch13 region 3 |
| chr13:77508939-77508979 | Ch13 region 3 |
| chr13:77509002-77509042 | Ch13 region 3 |
| chr13:77509055-77509095 | Ch13 region 3 |
| chr13:77509201-77509241 | Ch13 region 3 |
| chr13:77509271-77509311 | Ch13 region 3 |
| chr13:77509439-77509479 | Ch13 region 3 |
| chr13:77509549-77509589 | Ch13 region 3 |
| chr13:77509620-77509660 | Ch13 region 3 |
| chr13:77509671-77509711 | Ch13 region 3 |
| chr13:77509766-77509806 | Ch13 region 3 |
| chr13:77509811-77509851 | Ch13 region 3 |
| chr13:77509904-77509944 | Ch13 region 3 |
| chr13:77509949-77509989 | Ch13 region 3 |
| chr13:77510020-77510060 | Ch13 region 3 |
| chr13:77510251-77510291 | Ch13 region 3 |
| chr13:77510397-77510437 | Ch13 region 3 |
| chr13:77510454-77510494 | Ch13 region 3 |
| chr13:77510501-77510541 | Ch13 region 3 |
| chr13:77510573-77510613 | Ch13 region 3 |

| | |
|---|---|
| chr13:77510646-77510686 | Ch13 region 3 |
| chr13:77510699-77510739 | Ch13 region 3 |
| chr13:77510751-77510791 | Ch13 region 3 |
| chr13:77510800-77510840 | Ch13 region 3 |
| chr13:77510847-77510887 | Ch13 region 3 |
| chr13:77510921-77510961 | Ch13 region 3 |
| chr13:77510992-77511032 | Ch13 region 3 |
| chr13:77511067-77511107 | Ch13 region 3 |
| chr13:77511112-77511152 | Ch13 region 3 |
| chr13:77511226-77511266 | Ch13 region 3 |
| chr13:77511373-77511413 | Ch13 region 3 |
| chr13:77511473-77511513 | Ch13 region 3 |
| chr13:77511552-77511592 | Ch13 region 3 |
| chr13:77511695-77511735 | Ch13 region 3 |
| chr13:77511740-77511780 | Ch13 region 3 |
| chr13:77511796-77511836 | Ch13 region 3 |
| chr13:77511841-77511881 | Ch13 region 3 |
| chr13:77511886-77511926 | Ch13 region 3 |
| chr13:77511962-77512002 | Ch13 region 3 |
| chr13:77512008-77512048 | Ch13 region 3 |
| chr13:77512069-77512109 | Ch13 region 3 |
| chr13:77512114-77512154 | Ch13 region 3 |
| chr13:77512185-77512225 | Ch13 region 3 |
| chr13:77512273-77512313 | Ch13 region 3 |
| chr13:77512336-77512376 | Ch13 region 3 |
| chr13:77512383-77512423 | Ch13 region 3 |
| chr13:77512434-77512474 | Ch13 region 3 |
| chr13:77512536-77512576 | Ch13 region 3 |
| chr13:77512595-77512635 | Ch13 region 3 |
| chr13:77512653-77512693 | Ch13 region 3 |
| chr13:77512727-77512767 | Ch13 region 3 |
| chr13:77512774-77512814 | Ch13 region 3 |
| chr13:77512829-77512869 | Ch13 region 3 |
| chr13:77512884-77512924 | Ch13 region 3 |
| chr13:77512955-77512995 | Ch13 region 3 |
| chr13:77513027-77513067 | Ch13 region 3 |
| chr13:77513072-77513112 | Ch13 region 3 |
| chr13:77513143-77513183 | Ch13 region 3 |
| chr13:77513286-77513326 | Ch13 region 3 |
| chr13:77513338-77513378 | Ch13 region 3 |
| chr13:77513400-77513440 | Ch13 region 3 |
| chr13:77513493-77513533 | Ch13 region 3 |
| chr13:77513604-77513644 | Ch13 region 3 |
| chr13:77513701-77513741 | Ch13 region 3 |
| chr13:77513752-77513792 | Ch13 region 3 |
| chr13:77513833-77513873 | Ch13 region 3 |
| chr13:77513897-77513937 | Ch13 region 3 |
| chr13:77513942-77513982 | Ch13 region 3 |
| chr13:77514011-77514051 | Ch13 region 3 |
| chr13:77514092-77514132 | Ch13 region 3 |
| chr13:77514171-77514211 | Ch13 region 3 |
| chr13:77514230-77514270 | Ch13 region 3 |

| | |
|---|---|
| chr13:77514312-77514352 | Ch13 region 3 |
| chr13:77514379-77514419 | Ch13 region 3 |
| chr13:77514473-77514513 | Ch13 region 3 |
| chr13:77514541-77514581 | Ch13 region 3 |
| chr13:77514591-77514631 | Ch13 region 3 |
| chr13:77514662-77514702 | Ch13 region 3 |
| chr13:77514851-77514891 | Ch13 region 3 |
| chr13:77515038-77515078 | Ch13 region 3 |
| chr13:77515105-77515145 | Ch13 region 3 |
| chr13:77515152-77515192 | Ch13 region 3 |
| chr13:77515204-77515244 | Ch13 region 3 |
| chr13:77515309-77515349 | Ch13 region 3 |
| chr13:77515405-77515445 | Ch13 region 3 |
| chr13:77515510-77515550 | Ch13 region 3 |
| chr13:77515705-77515745 | Ch13 region 3 |
| chr13:77515781-77515821 | Ch13 region 3 |
| chr13:77515840-77515880 | Ch13 region 3 |
| chr13:77515898-77515938 | Ch13 region 3 |
| chr13:77515982-77516022 | Ch13 region 3 |
| chr13:77516086-77516126 | Ch13 region 3 |
| chr13:77516144-77516184 | Ch13 region 3 |
| chr13:77516246-77516286 | Ch13 region 3 |
| chr13:77516291-77516331 | Ch13 region 3 |
| chr13:77516364-77516404 | Ch13 region 3 |
| chr13:77516438-77516478 | Ch13 region 3 |
| chr13:77516560-77516600 | Ch13 region 3 |
| chr13:77516609-77516649 | Ch13 region 3 |
| chr13:77516693-77516733 | Ch13 region 3 |
| chr13:77516741-77516781 | Ch13 region 3 |
| chr13:77516818-77516858 | Ch13 region 3 |
| chr13:77516865-77516905 | Ch13 region 3 |
| chr13:77516944-77516984 | Ch13 region 3 |
| chr13:77517019-77517059 | Ch13 region 3 |
| chr13:77517086-77517126 | Ch13 region 3 |
| chr13:77517184-77517224 | Ch13 region 3 |
| chr13:77517246-77517286 | Ch13 region 3 |
| chr13:77517293-77517333 | Ch13 region 3 |
| chr13:77517362-77517402 | Ch13 region 3 |
| chr13:77517478-77517518 | Ch13 region 3 |
| chr13:77517600-77517640 | Ch13 region 3 |
| chr13:77517697-77517737 | Ch13 region 3 |
| chr13:77517758-77517798 | Ch13 region 3 |
| chr13:77517847-77517887 | Ch13 region 3 |
| chr13:77517931-77517971 | Ch13 region 3 |
| chr13:77517976-77518016 | Ch13 region 3 |
| chr13:77518084-77518124 | Ch13 region 3 |
| chr13:77518156-77518196 | Ch13 region 3 |
| chr13:77518249-77518289 | Ch13 region 3 |
| chr13:77518450-77518490 | Ch13 region 3 |
| chr13:77518497-77518537 | Ch13 region 3 |
| chr13:77518598-77518638 | Ch13 region 3 |
| chr13:77518684-77518724 | Ch13 region 3 |

| | |
|---|---|
| chr13:77518791-77518831 | Ch13 region 3 |
| chr13:77518935-77518975 | Ch13 region 3 |
| chr13:77519017-77519057 | Ch13 region 3 |
| chr13:77519063-77519103 | Ch13 region 3 |
| chr13:77519265-77519305 | Ch13 region 3 |
| chr13:77519391-77519431 | Ch13 region 3 |
| chr13:77519436-77519476 | Ch13 region 3 |
| chr13:77519509-77519549 | Ch13 region 3 |
| chr13:77519558-77519598 | Ch13 region 3 |
| chr13:77519659-77519699 | Ch13 region 3 |
| chr13:77519785-77519825 | Ch13 region 3 |
| chr13:77519889-77519929 | Ch13 region 3 |
| chr13:77520043-77520083 | Ch13 region 3 |
| chr13:77520088-77520128 | Ch13 region 3 |
| chr13:77520179-77520219 | Ch13 region 3 |
| chr13:77520295-77520335 | Ch13 region 3 |
| chr13:77520472-77520512 | Ch13 region 3 |
| chr13:77520543-77520583 | Ch13 region 3 |
| chr13:77520624-77520664 | Ch13 region 3 |
| chr13:77520710-77520750 | Ch13 region 3 |
| chr13:77520755-77520795 | Ch13 region 3 |
| chr13:77520800-77520840 | Ch13 region 3 |
| chr13:77520877-77520917 | Ch13 region 3 |
| chr13:77520969-77521009 | Ch13 region 3 |
| chr13:77521060-77521100 | Ch13 region 3 |
| chr13:77521112-77521152 | Ch13 region 3 |
| chr13:77521184-77521224 | Ch13 region 3 |
| chr13:77521263-77521303 | Ch13 region 3 |
| chr13:77521330-77521370 | Ch13 region 3 |
| chr13:77521396-77521436 | Ch13 region 3 |
| chr13:77521489-77521529 | Ch13 region 3 |
| chr13:77521543-77521583 | Ch13 region 3 |
| chr13:77521608-77521648 | Ch13 region 3 |
| chr13:77521672-77521712 | Ch13 region 3 |
| chr13:77521717-77521757 | Ch13 region 3 |
| chr13:77521795-77521835 | Ch13 region 3 |
| chr13:77521858-77521898 | Ch13 region 3 |
| chr13:77522011-77522051 | Ch13 region 3 |
| chr13:77522188-77522228 | Ch13 region 3 |
| chr13:77522233-77522273 | Ch13 region 3 |
| chr13:77522351-77522391 | Ch13 region 3 |
| chr13:77522438-77522478 | Ch13 region 3 |
| chr13:77522509-77522549 | Ch13 region 3 |
| chr13:77522577-77522617 | Ch13 region 3 |
| chr13:77522656-77522696 | Ch13 region 3 |
| chr13:77522740-77522780 | Ch13 region 3 |
| chr13:77522805-77522845 | Ch13 region 3 |
| chr13:77522882-77522922 | Ch13 region 3 |
| chr13:77522938-77522978 | Ch13 region 3 |
| chr13:77522984-77523024 | Ch13 region 3 |
| chr13:77523034-77523074 | Ch13 region 3 |
| chr13:77523087-77523127 | Ch13 region 3 |

| | |
|---|---|
| chr13:77523185-77523225 | Ch13 region 3 |
| chr13:77523468-77523508 | Ch13 region 3 |
| chr13:77523517-77523557 | Ch13 region 3 |
| chr13:77523562-77523602 | Ch13 region 3 |
| chr13:77523671-77523711 | Ch13 region 3 |
| chr13:77523722-77523762 | Ch13 region 3 |
| chr13:77523779-77523819 | Ch13 region 3 |
| chr13:77523852-77523892 | Ch13 region 3 |
| chr13:77523897-77523937 | Ch13 region 3 |
| chr13:77523952-77523992 | Ch13 region 3 |
| chr13:77524019-77524059 | Ch13 region 3 |
| chr13:77524066-77524106 | Ch13 region 3 |
| chr13:77524126-77524166 | Ch13 region 3 |
| chr13:77524181-77524221 | Ch13 region 3 |
| chr13:77524259-77524299 | Ch13 region 3 |
| chr13:77524310-77524350 | Ch13 region 3 |
| chr13:77524411-77524451 | Ch13 region 3 |
| chr13:77524469-77524509 | Ch13 region 3 |
| chr13:77524603-77524643 | Ch13 region 3 |
| chr13:77524695-77524735 | Ch13 region 3 |
| chr13:77524741-77524781 | Ch13 region 3 |
| chr13:77524831-77524871 | Ch13 region 3 |
| chr13:77524910-77524950 | Ch13 region 3 |
| chr13:77524987-77525027 | Ch13 region 3 |
| chr13:77525075-77525115 | Ch13 region 3 |
| chr13:77525120-77525160 | Ch13 region 3 |
| chr13:77525178-77525218 | Ch13 region 3 |
| chr13:77525223-77525263 | Ch13 region 3 |
| chr13:77525373-77525413 | Ch13 region 3 |
| chr13:77525659-77525699 | Ch13 region 3 |
| chr13:77525813-77525853 | Ch13 region 3 |
| chr13:77525858-77525898 | Ch13 region 3 |
| chr13:77525929-77525969 | Ch13 region 3 |
| chr13:77525997-77526037 | Ch13 region 3 |
| chr13:77526073-77526113 | Ch13 region 3 |
| chr13:77526149-77526189 | Ch13 region 3 |
| chr13:77526228-77526268 | Ch13 region 3 |
| chr13:77526316-77526356 | Ch13 region 3 |
| chr13:77526361-77526401 | Ch13 region 3 |
| chr13:77526407-77526447 | Ch13 region 3 |
| chr13:77526488-77526528 | Ch13 region 3 |
| chr13:77526552-77526592 | Ch13 region 3 |
| chr13:77526598-77526638 | Ch13 region 3 |
| chr13:77526679-77526719 | Ch13 region 3 |
| chr13:77526756-77526796 | Ch13 region 3 |
| chr13:77526810-77526850 | Ch13 region 3 |
| chr13:77526856-77526896 | Ch13 region 3 |
| chr13:77526954-77526994 | Ch13 region 3 |
| chr13:77527009-77527049 | Ch13 region 3 |
| chr13:77527074-77527114 | Ch13 region 3 |
| chr13:77527198-77527238 | Ch13 region 3 |
| chr13:77527332-77527372 | Ch13 region 3 |

| | |
|---|---|
| chr13:77527405-77527445 | Ch13 region 3 |
| chr13:77527459-77527499 | Ch13 region 3 |
| chr13:77527518-77527558 | Ch13 region 3 |
| chr13:77527578-77527618 | Ch13 region 3 |
| chr13:77527664-77527704 | Ch13 region 3 |
| chr13:77527717-77527757 | Ch13 region 3 |
| chr13:77527802-77527842 | Ch13 region 3 |
| chr13:77527935-77527975 | Ch13 region 3 |
| chr13:77528020-77528060 | Ch13 region 3 |
| chr13:77528117-77528157 | Ch13 region 3 |
| chr13:77528162-77528202 | Ch13 region 3 |
| chr13:77528236-77528276 | Ch13 region 3 |
| chr13:77528296-77528336 | Ch13 region 3 |
| chr13:77528362-77528402 | Ch13 region 3 |
| chr13:77528412-77528452 | Ch13 region 3 |
| chr13:77528466-77528506 | Ch13 region 3 |
| chr13:77528553-77528593 | Ch13 region 3 |
| chr13:77528633-77528673 | Ch13 region 3 |
| chr13:77528700-77528740 | Ch13 region 3 |
| chr13:77528775-77528815 | Ch13 region 3 |
| chr13:77528823-77528863 | Ch13 region 3 |
| chr13:77528892-77528932 | Ch13 region 3 |
| chr13:77528937-77528977 | Ch13 region 3 |
| chr13:77529128-77529168 | Ch13 region 3 |
| chr13:77529356-77529396 | Ch13 region 3 |
| chr13:77529405-77529445 | Ch13 region 3 |
| chr13:77529452-77529492 | Ch13 region 3 |
| chr13:77529550-77529590 | Ch13 region 3 |
| chr13:77529653-77529693 | Ch13 region 3 |
| chr13:77529698-77529738 | Ch13 region 3 |
| chr13:77529779-77529819 | Ch13 region 3 |
| chr13:77529835-77529875 | Ch13 region 3 |
| chr13:77529898-77529938 | Ch13 region 3 |
| chr13:77530002-77530042 | Ch13 region 3 |
| chr13:77530069-77530109 | Ch13 region 3 |
| chr13:77530142-77530182 | Ch13 region 3 |
| chr13:77530235-77530275 | Ch13 region 3 |
| chr13:77530299-77530339 | Ch13 region 3 |
| chr13:77530613-77530653 | Ch13 region 3 |
| chr13:77530783-77530823 | Ch13 region 3 |
| chr13:77530926-77530966 | Ch13 region 3 |
| chr13:77530973-77531013 | Ch13 region 3 |
| chr13:77531065-77531105 | Ch13 region 3 |
| chr13:77531111-77531151 | Ch13 region 3 |
| chr13:77531188-77531228 | Ch13 region 3 |
| chr13:77531263-77531303 | Ch13 region 3 |
| chr13:77531322-77531362 | Ch13 region 3 |
| chr13:77531404-77531444 | Ch13 region 3 |
| chr13:77531478-77531518 | Ch13 region 3 |
| chr13:77531535-77531575 | Ch13 region 3 |
| chr13:77531582-77531622 | Ch13 region 3 |
| chr13:77531633-77531673 | Ch13 region 3 |

| | |
|---|---|
| chr13:77531692-77531732 | Ch13 region 3 |
| chr13:77531739-77531779 | Ch13 region 3 |
| chr13:77531793-77531833 | Ch13 region 3 |
| chr13:77531874-77531914 | Ch13 region 3 |
| chr13:77531929-77531969 | Ch13 region 3 |
| chr13:77531977-77532017 | Ch13 region 3 |
| chr13:77532077-77532117 | Ch13 region 3 |
| chr13:77532137-77532177 | Ch13 region 3 |
| chr13:77532183-77532223 | Ch13 region 3 |
| chr13:77532288-77532328 | Ch13 region 3 |
| chr13:77532390-77532430 | Ch13 region 3 |
| chr13:77532443-77532483 | Ch13 region 3 |
| chr13:77532491-77532531 | Ch13 region 3 |
| chr13:77532551-77532591 | Ch13 region 3 |
| chr13:77532613-77532653 | Ch13 region 3 |
| chr13:77532658-77532698 | Ch13 region 3 |
| chr13:77532705-77532745 | Ch13 region 3 |
| chr13:77532851-77532891 | Ch13 region 3 |
| chr13:77532927-77532967 | Ch13 region 3 |
| chr13:77533108-77533148 | Ch13 region 3 |
| chr13:77533185-77533225 | Ch13 region 3 |
| chr13:77533327-77533367 | Ch13 region 3 |
| chr13:77533438-77533478 | Ch13 region 3 |
| chr13:77533546-77533586 | Ch13 region 3 |
| chr13:77533712-77533752 | Ch13 region 3 |
| chr13:77533826-77533866 | Ch13 region 3 |
| chr13:77533932-77533972 | Ch13 region 3 |
| chr13:77534137-77534177 | Ch13 region 3 |
| chr13:77534214-77534254 | Ch13 region 3 |
| chr13:77534306-77534346 | Ch13 region 3 |
| chr13:77534374-77534414 | Ch13 region 3 |
| chr13:77534436-77534476 | Ch13 region 3 |
| chr13:77534490-77534530 | Ch13 region 3 |
| chr13:77534609-77534649 | Ch13 region 3 |
| chr13:77534685-77534725 | Ch13 region 3 |
| chr13:77534746-77534786 | Ch13 region 3 |
| chr13:77534791-77534831 | Ch13 region 3 |
| chr13:77534844-77534884 | Ch13 region 3 |
| chr13:77534896-77534936 | Ch13 region 3 |
| chr13:77535110-77535150 | Ch13 region 3 |
| chr13:77535169-77535209 | Ch13 region 3 |
| chr13:77535321-77535361 | Ch13 region 3 |
| chr13:77535367-77535407 | Ch13 region 3 |
| chr13:77535505-77535545 | Ch13 region 3 |
| chr13:77535611-77535651 | Ch13 region 3 |
| chr13:77535847-77535887 | Ch13 region 3 |
| chr13:77535987-77536027 | Ch13 region 3 |
| chr13:77536076-77536116 | Ch13 region 3 |
| chr13:77536153-77536193 | Ch13 region 3 |
| chr13:77536199-77536239 | Ch13 region 3 |
| chr13:77536263-77536303 | Ch13 region 3 |
| chr13:77536345-77536385 | Ch13 region 3 |

| | |
|---|---|
| chr13:77536390-77536430 | Ch13 region 3 |
| chr13:77536473-77536513 | Ch13 region 3 |
| chr13:77536520-77536560 | Ch13 region 3 |
| chr13:77536636-77536676 | Ch13 region 3 |
| chr13:77536683-77536723 | Ch13 region 3 |
| chr13:77536814-77536854 | Ch13 region 3 |
| chr13:77536883-77536923 | Ch13 region 3 |
| chr13:77536977-77537017 | Ch13 region 3 |
| chr13:77537036-77537076 | Ch13 region 3 |
| chr13:77537095-77537135 | Ch13 region 3 |
| chr13:77537191-77537231 | Ch13 region 3 |
| chr13:77537261-77537301 | Ch13 region 3 |
| chr13:77537307-77537347 | Ch13 region 3 |
| chr13:77537387-77537427 | Ch13 region 3 |
| chr13:77537483-77537523 | Ch13 region 3 |
| chr13:77537540-77537580 | Ch13 region 3 |
| chr13:77537750-77537790 | Ch13 region 3 |
| chr13:77538208-77538248 | Ch13 region 3 |
| chr13:77538454-77538494 | Ch13 region 3 |
| chr13:77538507-77538547 | Ch13 region 3 |
| chr13:77538586-77538626 | Ch13 region 3 |
| chr13:77538649-77538689 | Ch13 region 3 |
| chr13:77538695-77538735 | Ch13 region 3 |
| chr13:77538758-77538798 | Ch13 region 3 |
| chr13:77538824-77538864 | Ch13 region 3 |
| chr13:77538872-77538912 | Ch13 region 3 |
| chr13:77538924-77538964 | Ch13 region 3 |
| chr13:77538975-77539015 | Ch13 region 3 |
| chr13:77539077-77539117 | Ch13 region 3 |
| chr13:77539307-77539347 | Ch13 region 3 |
| chr13:77539536-77539576 | Ch13 region 3 |
| chr13:77539588-77539628 | Ch13 region 3 |
| chr13:77539637-77539677 | Ch13 region 3 |
| chr13:77539703-77539743 | Ch13 region 3 |
| chr13:77539777-77539817 | Ch13 region 3 |
| chr13:77539829-77539869 | Ch13 region 3 |
| chr13:77539896-77539936 | Ch13 region 3 |
| chr13:77539945-77539985 | Ch13 region 3 |
| chr13:77540005-77540045 | Ch13 region 3 |
| chr13:77540090-77540130 | Ch13 region 3 |
| chr13:77540139-77540179 | Ch13 region 3 |
| chr13:77540194-77540234 | Ch13 region 3 |
| chr13:77540243-77540283 | Ch13 region 3 |
| chr13:77540306-77540346 | Ch13 region 3 |
| chr13:77540358-77540398 | Ch13 region 3 |
| chr13:77540419-77540459 | Ch13 region 3 |
| chr13:77540469-77540509 | Ch13 region 3 |
| chr13:77540552-77540592 | Ch13 region 3 |
| chr13:77540625-77540665 | Ch13 region 3 |
| chr13:77540691-77540731 | Ch13 region 3 |
| chr13:77540737-77540777 | Ch13 region 3 |
| chr13:77540897-77540937 | Ch13 region 3 |

| | |
|---|---|
| chr13:77541030-77541070 | Ch13 region 3 |
| chr13:77541166-77541206 | Ch13 region 3 |
| chr13:77541223-77541263 | Ch13 region 3 |
| chr13:77541322-77541362 | Ch13 region 3 |
| chr13:77541505-77541545 | Ch13 region 3 |
| chr13:77541708-77541748 | Ch13 region 3 |
| chr13:77541765-77541805 | Ch13 region 3 |
| chr13:77541902-77541942 | Ch13 region 3 |
| chr13:77541971-77542011 | Ch13 region 3 |
| chr13:77542022-77542062 | Ch13 region 3 |
| chr13:77542079-77542119 | Ch13 region 3 |
| chr13:77542180-77542220 | Ch13 region 3 |
| chr13:77542244-77542284 | Ch13 region 3 |
| chr13:77542411-77542451 | Ch13 region 3 |
| chr13:77542489-77542529 | Ch13 region 3 |
| chr13:77542595-77542635 | Ch13 region 3 |
| chr13:77542699-77542739 | Ch13 region 3 |
| chr13:77542792-77542832 | Ch13 region 3 |
| chr13:77542871-77542911 | Ch13 region 3 |
| chr13:77543019-77543059 | Ch13 region 3 |
| chr13:77543121-77543161 | Ch13 region 3 |
| chr13:77543327-77543367 | Ch13 region 3 |
| chr13:77543425-77543465 | Ch13 region 3 |
| chr13:77543494-77543534 | Ch13 region 3 |
| chr13:77543543-77543583 | Ch13 region 3 |
| chr13:77543590-77543630 | Ch13 region 3 |
| chr13:77543648-77543688 | Ch13 region 3 |
| chr13:77543706-77543746 | Ch13 region 3 |
| chr13:77543778-77543818 | Ch13 region 3 |
| chr13:77543823-77543863 | Ch13 region 3 |
| chr13:77543874-77543914 | Ch13 region 3 |
| chr13:77543931-77543971 | Ch13 region 3 |
| chr13:77543978-77544018 | Ch13 region 3 |
| chr13:77544067-77544107 | Ch13 region 3 |
| chr13:77544149-77544189 | Ch13 region 3 |
| chr13:77544196-77544236 | Ch13 region 3 |
| chr13:77544281-77544321 | Ch13 region 3 |
| chr13:77544329-77544369 | Ch13 region 3 |
| chr13:77544445-77544485 | Ch13 region 3 |
| chr13:77544501-77544541 | Ch13 region 3 |
| chr13:77544632-77544672 | Ch13 region 3 |
| chr13:77544731-77544771 | Ch13 region 3 |
| chr13:77544810-77544850 | Ch13 region 3 |
| chr13:77545012-77545052 | Ch13 region 3 |
| chr13:77545105-77545145 | Ch13 region 3 |
| chr13:77545177-77545217 | Ch13 region 3 |
| chr13:77545228-77545268 | Ch13 region 3 |
| chr13:77545315-77545355 | Ch13 region 3 |
| chr13:77545375-77545415 | Ch13 region 3 |
| chr13:77545432-77545472 | Ch13 region 3 |
| chr13:77545478-77545518 | Ch13 region 3 |
| chr13:77545556-77545596 | Ch13 region 3 |

| | |
|---|---|
| chr13:77545639-77545679 | Ch13 region 3 |
| chr13:77545700-77545740 | Ch13 region 3 |
| chr13:77545748-77545788 | Ch13 region 3 |
| chr13:77545809-77545849 | Ch13 region 3 |
| chr13:77545860-77545900 | Ch13 region 3 |
| chr13:77545917-77545957 | Ch13 region 3 |
| chr13:77545963-77546003 | Ch13 region 3 |
| chr13:77546022-77546062 | Ch13 region 3 |
| chr13:77546067-77546107 | Ch13 region 3 |
| chr13:77546160-77546200 | Ch13 region 3 |
| chr13:77546205-77546245 | Ch13 region 3 |
| chr13:77546251-77546291 | Ch13 region 3 |
| chr13:77546475-77546515 | Ch13 region 3 |
| chr13:77546558-77546598 | Ch13 region 3 |
| chr13:77546642-77546682 | Ch13 region 3 |
| chr13:77546724-77546764 | Ch13 region 3 |
| chr13:77546770-77546810 | Ch13 region 3 |
| chr13:77546838-77546878 | Ch13 region 3 |
| chr13:77546883-77546923 | Ch13 region 3 |
| chr13:77546928-77546968 | Ch13 region 3 |
| chr13:77547062-77547102 | Ch13 region 3 |
| chr13:77547132-77547172 | Ch13 region 3 |
| chr13:77547360-77547400 | Ch13 region 3 |
| chr13:77547472-77547512 | Ch13 region 3 |
| chr13:77547569-77547609 | Ch13 region 3 |
| chr13:77547771-77547811 | Ch13 region 3 |
| chr13:77547843-77547883 | Ch13 region 3 |
| chr13:77547888-77547928 | Ch13 region 3 |
| chr13:77547987-77548027 | Ch13 region 3 |
| chr13:77548040-77548080 | Ch13 region 3 |
| chr13:77548092-77548132 | Ch13 region 3 |
| chr13:77548148-77548188 | Ch13 region 3 |
| chr13:77548240-77548280 | Ch13 region 3 |
| chr13:77548322-77548362 | Ch13 region 3 |
| chr13:77548434-77548474 | Ch13 region 3 |
| chr13:77548492-77548532 | Ch13 region 3 |
| chr13:77548550-77548590 | Ch13 region 3 |
| chr13:77548628-77548668 | Ch13 region 3 |
| chr13:77548680-77548720 | Ch13 region 3 |
| chr13:77548725-77548765 | Ch13 region 3 |
| chr13:77548815-77548855 | Ch13 region 3 |
| chr13:77548865-77548905 | Ch13 region 3 |
| chr13:77548917-77548957 | Ch13 region 3 |
| chr13:77548987-77549027 | Ch13 region 3 |
| chr13:77549042-77549082 | Ch13 region 3 |
| chr13:77549151-77549191 | Ch13 region 3 |
| chr13:77549212-77549252 | Ch13 region 3 |
| chr13:77549302-77549342 | Ch13 region 3 |
| chr13:77549356-77549396 | Ch13 region 3 |
| chr13:77549607-77549647 | Ch13 region 3 |
| chr13:77549713-77549753 | Ch13 region 3 |
| chr13:77549853-77549893 | Ch13 region 3 |

| | |
|---|---|
| chr13:77549969-77550009 | Ch13 region 3 |
| chr13:77550030-77550070 | Ch13 region 3 |
| chr13:77550087-77550127 | Ch13 region 3 |
| chr13:77550141-77550181 | Ch13 region 3 |
| chr13:77550230-77550270 | Ch13 region 3 |
| chr13:77550276-77550316 | Ch13 region 3 |
| chr13:77550360-77550400 | Ch13 region 3 |
| chr13:77550405-77550445 | Ch13 region 3 |
| chr13:77550480-77550520 | Ch13 region 3 |
| chr13:77550554-77550594 | Ch13 region 3 |
| chr13:77550603-77550643 | Ch13 region 3 |
| chr13:77550649-77550689 | Ch13 region 3 |
| chr13:77550738-77550778 | Ch13 region 3 |
| chr13:77550805-77550845 | Ch13 region 3 |
| chr13:77550850-77550890 | Ch13 region 3 |
| chr13:77550895-77550935 | Ch13 region 3 |
| chr13:77550960-77551000 | Ch13 region 3 |
| chr13:77551041-77551081 | Ch13 region 3 |
| chr13:77551124-77551164 | Ch13 region 3 |
| chr13:77551178-77551218 | Ch13 region 3 |
| chr13:77551257-77551297 | Ch13 region 3 |
| chr13:77551312-77551352 | Ch13 region 3 |
| chr13:77551360-77551400 | Ch13 region 3 |
| chr13:77551456-77551496 | Ch13 region 3 |
| chr13:77551540-77551580 | Ch13 region 3 |
| chr13:77551591-77551631 | Ch13 region 3 |
| chr13:77551673-77551713 | Ch13 region 3 |
| chr13:77551728-77551768 | Ch13 region 3 |
| chr13:77551813-77551853 | Ch13 region 3 |
| chr13:77552012-77552052 | Ch13 region 3 |
| chr13:77552074-77552114 | Ch13 region 3 |
| chr13:77552208-77552248 | Ch13 region 3 |
| chr13:77552370-77552410 | Ch13 region 3 |
| chr13:77552415-77552455 | Ch13 region 3 |
| chr13:77552491-77552531 | Ch13 region 3 |
| chr13:77552543-77552583 | Ch13 region 3 |
| chr13:77552593-77552633 | Ch13 region 3 |
| chr13:77552645-77552685 | Ch13 region 3 |
| chr13:77552691-77552731 | Ch13 region 3 |
| chr13:77552741-77552781 | Ch13 region 3 |
| chr13:77552800-77552840 | Ch13 region 3 |
| chr13:77552852-77552892 | Ch13 region 3 |
| chr13:77552914-77552954 | Ch13 region 3 |
| chr13:77553092-77553132 | Ch13 region 3 |
| chr13:77553196-77553236 | Ch13 region 3 |
| chr13:77553259-77553299 | Ch13 region 3 |
| chr13:77553324-77553364 | Ch13 region 3 |
| chr13:77553370-77553410 | Ch13 region 3 |
| chr13:77553441-77553481 | Ch13 region 3 |
| chr13:77553486-77553526 | Ch13 region 3 |
| chr13:77553534-77553574 | Ch13 region 3 |
| chr13:77553587-77553627 | Ch13 region 3 |

| | |
|---|---|
| chr13:77553656-77553696 | Ch13 region 3 |
| chr13:77553718-77553758 | Ch13 region 3 |
| chr13:77553801-77553841 | Ch13 region 3 |
| chr13:77554013-77554053 | Ch13 region 3 |
| chr13:77554206-77554246 | Ch13 region 3 |
| chr13:77554259-77554299 | Ch13 region 3 |
| chr13:77554323-77554363 | Ch13 region 3 |
| chr13:77554384-77554424 | Ch13 region 3 |
| chr13:77554495-77554535 | Ch13 region 3 |
| chr13:77554561-77554601 | Ch13 region 3 |
| chr13:77554668-77554708 | Ch13 region 3 |
| chr13:77554776-77554816 | Ch13 region 3 |
| chr13:77554861-77554901 | Ch13 region 3 |
| chr13:77554906-77554946 | Ch13 region 3 |
| chr13:77554991-77555031 | Ch13 region 3 |
| chr13:77555050-77555090 | Ch13 region 3 |
| chr13:77555145-77555185 | Ch13 region 3 |
| chr13:77555300-77555340 | Ch13 region 3 |
| chr13:77555405-77555445 | Ch13 region 3 |
| chr13:77555499-77555539 | Ch13 region 3 |
| chr13:77555609-77555649 | Ch13 region 3 |
| chr13:77555667-77555707 | Ch13 region 3 |
| chr13:77555798-77555838 | Ch13 region 3 |
| chr13:77555963-77556003 | Ch13 region 3 |
| chr13:77556019-77556059 | Ch13 region 3 |
| chr13:77556094-77556134 | Ch13 region 3 |
| chr13:77556146-77556186 | Ch13 region 3 |
| chr13:77556200-77556240 | Ch13 region 3 |
| chr13:77556249-77556289 | Ch13 region 3 |
| chr13:77556326-77556366 | Ch13 region 3 |
| chr13:77556414-77556454 | Ch13 region 3 |
| chr13:77556466-77556506 | Ch13 region 3 |
| chr13:77556511-77556551 | Ch13 region 3 |
| chr13:77556560-77556600 | Ch13 region 3 |
| chr13:77556656-77556696 | Ch13 region 3 |
| chr13:77556706-77556746 | Ch13 region 3 |
| chr13:77556878-77556918 | Ch13 region 3 |
| chr13:77557068-77557108 | Ch13 region 3 |
| chr13:77557143-77557183 | Ch13 region 3 |
| chr13:77557221-77557261 | Ch13 region 3 |
| chr13:77557312-77557352 | Ch13 region 3 |
| chr13:77557363-77557403 | Ch13 region 3 |
| chr13:77557444-77557484 | Ch13 region 3 |
| chr13:77557518-77557558 | Ch13 region 3 |
| chr13:77557665-77557705 | Ch13 region 3 |
| chr13:77557710-77557750 | Ch13 region 3 |
| chr13:77557902-77557942 | Ch13 region 3 |
| chr13:77558117-77558157 | Ch13 region 3 |
| chr13:77558162-77558202 | Ch13 region 3 |
| chr13:77558232-77558272 | Ch13 region 3 |
| chr13:77558282-77558322 | Ch13 region 3 |
| chr13:77558390-77558430 | Ch13 region 3 |

| | |
|---|---|
| chr13:77558445-77558485 | Ch13 region 3 |
| chr13:77558511-77558551 | Ch13 region 3 |
| chr13:77558597-77558637 | Ch13 region 3 |
| chr13:77558713-77558753 | Ch13 region 3 |
| chr13:77558770-77558810 | Ch13 region 3 |
| chr13:77558817-77558857 | Ch13 region 3 |
| chr13:77558889-77558929 | Ch13 region 3 |
| chr13:77558971-77559011 | Ch13 region 3 |
| chr13:77559061-77559101 | Ch13 region 3 |
| chr13:77559106-77559146 | Ch13 region 3 |
| chr13:77559156-77559196 | Ch13 region 3 |
| chr13:77559255-77559295 | Ch13 region 3 |
| chr13:77559311-77559351 | Ch13 region 3 |
| chr13:77559390-77559430 | Ch13 region 3 |
| chr13:77559450-77559490 | Ch13 region 3 |
| chr13:77559602-77559642 | Ch13 region 3 |
| chr13:77559706-77559746 | Ch13 region 3 |
| chr13:77559779-77559819 | Ch13 region 3 |
| chr13:77559849-77559889 | Ch13 region 3 |
| chr13:77559916-77559956 | Ch13 region 3 |
| chr13:77559972-77560012 | Ch13 region 3 |
| chr13:77560019-77560059 | Ch13 region 3 |
| chr13:77560069-77560109 | Ch13 region 3 |
| chr13:77560171-77560211 | Ch13 region 3 |
| chr13:77560265-77560305 | Ch13 region 3 |
| chr13:77560317-77560357 | Ch13 region 3 |
| chr13:77560411-77560451 | Ch13 region 3 |
| chr13:77560512-77560552 | Ch13 region 3 |
| chr13:77560591-77560631 | Ch13 region 3 |
| chr13:77560651-77560691 | Ch13 region 3 |
| chr13:77560742-77560782 | Ch13 region 3 |
| chr13:77560840-77560880 | Ch13 region 3 |
| chr13:77560906-77560946 | Ch13 region 3 |
| chr13:77561002-77561042 | Ch13 region 3 |
| chr13:77561134-77561174 | Ch13 region 3 |
| chr13:77561208-77561248 | Ch13 region 3 |
| chr13:77561342-77561382 | Ch13 region 3 |
| chr13:77561387-77561427 | Ch13 region 3 |
| chr13:77561510-77561550 | Ch13 region 3 |
| chr13:77561570-77561610 | Ch13 region 3 |
| chr13:77561653-77561693 | Ch13 region 3 |
| chr13:77561704-77561744 | Ch13 region 3 |
| chr13:77561766-77561806 | Ch13 region 3 |
| chr13:77561848-77561888 | Ch13 region 3 |
| chr13:77561937-77561977 | Ch13 region 3 |
| chr13:77561988-77562028 | Ch13 region 3 |
| chr13:77562090-77562130 | Ch13 region 3 |
| chr13:77562155-77562195 | Ch13 region 3 |
| chr13:77562280-77562320 | Ch13 region 3 |
| chr13:77562347-77562387 | Ch13 region 3 |
| chr13:77562394-77562434 | Ch13 region 3 |
| chr13:77562457-77562497 | Ch13 region 3 |

| | |
|---|---|
| chr13:77562504-77562544 | Ch13 region 3 |
| chr13:77562618-77562658 | Ch13 region 3 |
| chr13:77562669-77562709 | Ch13 region 3 |
| chr13:77562759-77562799 | Ch13 region 3 |
| chr13:77562839-77562879 | Ch13 region 3 |
| chr13:77562893-77562933 | Ch13 region 3 |
| chr13:77563114-77563154 | Ch13 region 3 |
| chr13:77563246-77563286 | Ch13 region 3 |
| chr13:77563338-77563378 | Ch13 region 3 |
| chr13:77563446-77563486 | Ch13 region 3 |
| chr13:77563492-77563532 | Ch13 region 3 |
| chr13:77563559-77563599 | Ch13 region 3 |
| chr13:77563621-77563661 | Ch13 region 3 |
| chr13:77563666-77563706 | Ch13 region 3 |
| chr13:77563748-77563788 | Ch13 region 3 |
| chr13:77563841-77563881 | Ch13 region 3 |
| chr13:77563888-77563928 | Ch13 region 3 |
| chr13:77563935-77563975 | Ch13 region 3 |
| chr13:77564096-77564136 | Ch13 region 3 |
| chr13:77564256-77564296 | Ch13 region 3 |
| chr13:77564306-77564346 | Ch13 region 3 |
| chr13:77564361-77564401 | Ch13 region 3 |
| chr13:77564443-77564483 | Ch13 region 3 |
| chr13:77564513-77564553 | Ch13 region 3 |
| chr13:77564687-77564727 | Ch13 region 3 |
| chr13:77564752-77564792 | Ch13 region 3 |
| chr13:77564859-77564899 | Ch13 region 3 |
| chr13:77564908-77564948 | Ch13 region 3 |
| chr13:77564972-77565012 | Ch13 region 3 |
| chr13:77565023-77565063 | Ch13 region 3 |
| chr13:77565082-77565122 | Ch13 region 3 |
| chr13:77565129-77565169 | Ch13 region 3 |
| chr13:77565190-77565230 | Ch13 region 3 |
| chr13:77565247-77565287 | Ch13 region 3 |
| chr13:77565322-77565362 | Ch13 region 3 |
| chr13:77565369-77565409 | Ch13 region 3 |
| chr13:77565457-77565497 | Ch13 region 3 |
| chr13:77565522-77565562 | Ch13 region 3 |
| chr13:77565567-77565607 | Ch13 region 3 |
| chr13:77565615-77565655 | Ch13 region 3 |
| chr13:77565660-77565700 | Ch13 region 3 |
| chr13:77565762-77565802 | Ch13 region 3 |
| chr13:77565818-77565858 | Ch13 region 3 |
| chr13:77565881-77565921 | Ch13 region 3 |
| chr13:77565930-77565970 | Ch13 region 3 |
| chr13:77565998-77566038 | Ch13 region 3 |
| chr13:77566073-77566113 | Ch13 region 3 |
| chr13:77566149-77566189 | Ch13 region 3 |
| chr13:77566212-77566252 | Ch13 region 3 |
| chr13:77566292-77566332 | Ch13 region 3 |
| chr13:77566420-77566460 | Ch13 region 3 |
| chr13:77566480-77566520 | Ch13 region 3 |

| | |
|---|---|
| chr13:77566543-77566583 | Ch13 region 3 |
| chr13:77566611-77566651 | Ch13 region 3 |
| chr13:77566661-77566701 | Ch13 region 3 |
| chr13:77566719-77566759 | Ch13 region 3 |
| chr13:77566764-77566804 | Ch13 region 3 |
| chr13:77566837-77566877 | Ch13 region 3 |
| chr13:77566936-77566976 | Ch13 region 3 |
| chr13:77567011-77567051 | Ch13 region 3 |
| chr13:77567068-77567108 | Ch13 region 3 |
| chr13:77567113-77567153 | Ch13 region 3 |
| chr13:77567229-77567269 | Ch13 region 3 |
| chr13:77567278-77567318 | Ch13 region 3 |
| chr13:77567350-77567390 | Ch13 region 3 |
| chr13:77567420-77567460 | Ch13 region 3 |
| chr13:77567504-77567544 | Ch13 region 3 |
| chr13:77567629-77567669 | Ch13 region 3 |
| chr13:77567689-77567729 | Ch13 region 3 |
| chr13:77567796-77567836 | Ch13 region 3 |
| chr13:77567850-77567890 | Ch13 region 3 |
| chr13:77567956-77567996 | Ch13 region 3 |
| chr13:77568035-77568075 | Ch13 region 3 |
| chr13:77568096-77568136 | Ch13 region 3 |
| chr13:77568162-77568202 | Ch13 region 3 |
| chr13:77568232-77568272 | Ch13 region 3 |
| chr13:77568277-77568317 | Ch13 region 3 |
| chr13:77568353-77568393 | Ch13 region 3 |
| chr13:77568410-77568450 | Ch13 region 3 |
| chr13:77568461-77568501 | Ch13 region 3 |
| chr13:77568522-77568562 | Ch13 region 3 |
| chr13:77568601-77568641 | Ch13 region 3 |
| chr13:77568646-77568686 | Ch13 region 3 |
| chr13:77568704-77568744 | Ch13 region 3 |
| chr13:77568881-77568921 | Ch13 region 3 |
| chr13:77568946-77568986 | Ch13 region 3 |
| chr13:77569071-77569111 | Ch13 region 3 |
| chr13:77569168-77569208 | Ch13 region 3 |
| chr13:77569219-77569259 | Ch13 region 3 |
| chr13:77569275-77569315 | Ch13 region 3 |
| chr13:77569349-77569389 | Ch13 region 3 |
| chr13:77569433-77569473 | Ch13 region 3 |
| chr13:77569556-77569596 | Ch13 region 3 |
| chr13:77569653-77569693 | Ch13 region 3 |
| chr13:77569734-77569774 | Ch13 region 3 |
| chr13:77569788-77569828 | Ch13 region 3 |
| chr13:77569883-77569923 | Ch13 region 3 |
| chr13:77569933-77569973 | Ch13 region 3 |
| chr13:77569983-77570023 | Ch13 region 3 |
| chr13:77570053-77570093 | Ch13 region 3 |
| chr13:77570114-77570154 | Ch13 region 3 |
| chr13:77570185-77570225 | Ch13 region 3 |
| chr13:77570230-77570270 | Ch13 region 3 |
| chr13:77570446-77570486 | Ch13 region 3 |

| | |
|---|---|
| chr13:77570570-77570610 | Ch13 region 3 |
| chr13:77570681-77570721 | Ch13 region 3 |
| chr13:77570774-77570814 | Ch13 region 3 |
| chr13:77570896-77570936 | Ch13 region 3 |
| chr13:77571037-77571077 | Ch13 region 3 |
| chr13:77571123-77571163 | Ch13 region 3 |
| chr13:77571168-77571208 | Ch13 region 3 |
| chr13:77571239-77571279 | Ch13 region 3 |
| chr13:77571291-77571331 | Ch13 region 3 |
| chr13:77571376-77571416 | Ch13 region 3 |
| chr13:77571561-77571601 | Ch13 region 3 |
| chr13:77571783-77571823 | Ch13 region 3 |
| chr13:77571862-77571902 | Ch13 region 3 |
| chr13:77572058-77572098 | Ch13 region 3 |
| chr13:77572105-77572145 | Ch13 region 3 |
| chr13:77572190-77572230 | Ch13 region 3 |
| chr13:77572244-77572284 | Ch13 region 3 |
| chr13:77572289-77572329 | Ch13 region 3 |
| chr13:77572360-77572400 | Ch13 region 3 |
| chr13:77572410-77572450 | Ch13 region 3 |
| chr13:77572478-77572518 | Ch13 region 3 |
| chr13:77572527-77572567 | Ch13 region 3 |
| chr13:77572606-77572646 | Ch13 region 3 |
| chr13:77572722-77572762 | Ch13 region 3 |
| chr13:77572768-77572808 | Ch13 region 3 |
| chr13:77572824-77572864 | Ch13 region 3 |
| chr13:77572902-77572942 | Ch13 region 3 |
| chr13:77572947-77572987 | Ch13 region 3 |
| chr13:77573013-77573053 | Ch13 region 3 |
| chr13:77573195-77573235 | Ch13 region 3 |
| chr13:77573240-77573280 | Ch13 region 3 |
| chr13:77573328-77573368 | Ch13 region 3 |
| chr13:77573417-77573457 | Ch13 region 3 |
| chr13:77573462-77573502 | Ch13 region 3 |
| chr13:77573508-77573548 | Ch13 region 3 |
| chr13:77573598-77573638 | Ch13 region 3 |
| chr13:77573657-77573697 | Ch13 region 3 |
| chr13:77573724-77573764 | Ch13 region 3 |
| chr13:77573812-77573852 | Ch13 region 3 |
| chr13:77573882-77573922 | Ch13 region 3 |
| chr13:77573951-77573991 | Ch13 region 3 |
| chr13:77574062-77574102 | Ch13 region 3 |
| chr13:77574145-77574185 | Ch13 region 3 |
| chr13:77574197-77574237 | Ch13 region 3 |
| chr13:77574305-77574345 | Ch13 region 3 |
| chr13:77574462-77574502 | Ch13 region 3 |
| chr13:77574582-77574622 | Ch13 region 3 |
| chr13:77574628-77574668 | Ch13 region 3 |
| chr13:77574698-77574738 | Ch13 region 3 |
| chr13:77574755-77574795 | Ch13 region 3 |
| chr13:77574806-77574846 | Ch13 region 3 |
| chr13:77574873-77574913 | Ch13 region 3 |

| | |
|---|---|
| chr13:77574933-77574973 | Ch13 region 3 |
| chr13:77575055-77575095 | Ch13 region 3 |
| chr13:77575103-77575143 | Ch13 region 3 |
| chr13:77575170-77575210 | Ch13 region 3 |
| chr13:77575229-77575269 | Ch13 region 3 |
| chr13:77575284-77575324 | Ch13 region 3 |
| chr13:77575333-77575373 | Ch13 region 3 |
| chr13:77575391-77575431 | Ch13 region 3 |
| chr13:77575456-77575496 | Ch13 region 3 |
| chr13:77575564-77575604 | Ch13 region 3 |
| chr13:77575629-77575669 | Ch13 region 3 |
| chr13:77575679-77575719 | Ch13 region 3 |
| chr13:77575770-77575810 | Ch13 region 3 |
| chr13:77575839-77575879 | Ch13 region 3 |
| chr13:77575884-77575924 | Ch13 region 3 |
| chr13:77575993-77576033 | Ch13 region 3 |
| chr13:77576058-77576098 | Ch13 region 3 |
| chr13:77576137-77576177 | Ch13 region 3 |
| chr13:77576182-77576222 | Ch13 region 3 |
| chr13:77576266-77576306 | Ch13 region 3 |
| chr13:77576358-77576398 | Ch13 region 3 |
| chr13:77576420-77576460 | Ch13 region 3 |
| chr13:77576466-77576506 | Ch13 region 3 |
| chr13:77576538-77576578 | Ch13 region 3 |
| chr13:77576661-77576701 | Ch13 region 3 |
| chr13:77576717-77576757 | Ch13 region 3 |
| chr13:77576780-77576820 | Ch13 region 3 |
| chr13:77576826-77576866 | Ch13 region 3 |
| chr13:77576875-77576915 | Ch13 region 3 |
| chr13:77576924-77576964 | Ch13 region 3 |
| chr13:77576978-77577018 | Ch13 region 3 |
| chr13:77577054-77577094 | Ch13 region 3 |
| chr13:77577138-77577178 | Ch13 region 3 |
| chr13:77577188-77577228 | Ch13 region 3 |
| chr13:77577282-77577322 | Ch13 region 3 |
| chr13:77577497-77577537 | Ch13 region 3 |
| chr13:77577566-77577606 | Ch13 region 3 |
| chr13:77577612-77577652 | Ch13 region 3 |
| chr13:77577699-77577739 | Ch13 region 3 |
| chr13:77577763-77577803 | Ch13 region 3 |
| chr13:77577812-77577852 | Ch13 region 3 |
| chr13:77577927-77577967 | Ch13 region 3 |
| chr13:77578112-77578152 | Ch13 region 3 |
| chr13:77578160-77578200 | Ch13 region 3 |
| chr13:77578207-77578247 | Ch13 region 3 |
| chr13:77578257-77578297 | Ch13 region 3 |
| chr13:77578308-77578348 | Ch13 region 3 |
| chr13:77578397-77578437 | Ch13 region 3 |
| chr13:77578482-77578522 | Ch13 region 3 |
| chr13:77578611-77578651 | Ch13 region 3 |
| chr13:77578661-77578701 | Ch13 region 3 |
| chr13:77578753-77578793 | Ch13 region 3 |

| | |
|---|---|
| chr13:77578827-77578867 | Ch13 region 3 |
| chr13:77578872-77578912 | Ch13 region 3 |
| chr13:77578933-77578973 | Ch13 region 3 |
| chr13:77579042-77579082 | Ch13 region 3 |
| chr13:77579213-77579253 | Ch13 region 3 |
| chr13:77579260-77579300 | Ch13 region 3 |
| chr13:77579542-77579582 | Ch13 region 3 |
| chr13:77579607-77579647 | Ch13 region 3 |
| chr13:77579709-77579749 | Ch13 region 3 |
| chr13:77579805-77579845 | Ch13 region 3 |
| chr13:77579862-77579902 | Ch13 region 3 |
| chr13:77579907-77579947 | Ch13 region 3 |
| chr13:77579969-77580009 | Ch13 region 3 |
| chr13:77580143-77580183 | Ch13 region 3 |
| chr13:77580223-77580263 | Ch13 region 3 |
| chr13:77580318-77580358 | Ch13 region 3 |
| chr13:77580481-77580521 | Ch13 region 3 |
| chr13:77580532-77580572 | Ch13 region 3 |
| chr13:77580611-77580651 | Ch13 region 3 |
| chr13:77580681-77580721 | Ch13 region 3 |
| chr13:77580762-77580802 | Ch13 region 3 |
| chr13:77580863-77580903 | Ch13 region 3 |
| chr13:77581011-77581051 | Ch13 region 3 |
| chr13:77581061-77581101 | Ch13 region 3 |
| chr13:77581120-77581160 | Ch13 region 3 |
| chr13:77581179-77581219 | Ch13 region 3 |
| chr13:77581258-77581298 | Ch13 region 3 |
| chr13:77581319-77581359 | Ch13 region 3 |
| chr13:77581391-77581431 | Ch13 region 3 |
| chr13:77581442-77581482 | Ch13 region 3 |
| chr13:77581528-77581568 | Ch13 region 3 |
| chr13:77581621-77581661 | Ch13 region 3 |
| chr13:77581707-77581747 | Ch13 region 3 |
| chr13:77581768-77581808 | Ch13 region 3 |
| chr13:77581822-77581862 | Ch13 region 3 |
| chr13:77581878-77581918 | Ch13 region 3 |
| chr13:77581978-77582018 | Ch13 region 3 |
| chr13:77582176-77582216 | Ch13 region 3 |
| chr13:77582253-77582293 | Ch13 region 3 |
| chr13:77582298-77582338 | Ch13 region 3 |
| chr13:77582539-77582579 | Ch13 region 3 |
| chr13:77582608-77582648 | Ch13 region 3 |
| chr13:77582691-77582731 | Ch13 region 3 |
| chr13:77582886-77582926 | Ch13 region 3 |
| chr13:77582975-77583015 | Ch13 region 3 |
| chr13:77583065-77583105 | Ch13 region 3 |
| chr13:77583124-77583164 | Ch13 region 3 |
| chr13:77583175-77583215 | Ch13 region 3 |
| chr13:77583233-77583273 | Ch13 region 3 |
| chr13:77583319-77583359 | Ch13 region 3 |
| chr13:77583376-77583416 | Ch13 region 3 |
| chr13:77583425-77583465 | Ch13 region 3 |

| | |
|---|---|
| chr13:77583644-77583684 | Ch13 region 3 |
| chr13:77583763-77583803 | Ch13 region 3 |
| chr13:77583992-77584032 | Ch13 region 3 |
| chr13:77584210-77584250 | Ch13 region 3 |
| chr13:77584399-77584439 | Ch13 region 3 |
| chr13:77584491-77584531 | Ch13 region 3 |
| chr13:77584579-77584619 | Ch13 region 3 |
| chr13:77584624-77584664 | Ch13 region 3 |
| chr13:77584677-77584717 | Ch13 region 3 |
| chr13:77584730-77584770 | Ch13 region 3 |
| chr13:77584813-77584853 | Ch13 region 3 |
| chr13:77584865-77584905 | Ch13 region 3 |
| chr13:77584943-77584983 | Ch13 region 3 |
| chr13:77585040-77585080 | Ch13 region 3 |
| chr13:77585110-77585150 | Ch13 region 3 |
| chr13:77585159-77585199 | Ch13 region 3 |
| chr13:77585233-77585273 | Ch13 region 3 |
| chr13:77585416-77585456 | Ch13 region 3 |
| chr13:77585538-77585578 | Ch13 region 3 |
| chr13:77585686-77585726 | Ch13 region 3 |
| chr13:77585910-77585950 | Ch13 region 3 |
| chr13:77586017-77586057 | Ch13 region 3 |
| chr13:77586064-77586104 | Ch13 region 3 |
| chr13:77586206-77586246 | Ch13 region 3 |
| chr13:77586309-77586349 | Ch13 region 3 |
| chr13:77586455-77586495 | Ch13 region 3 |
| chr13:77586507-77586547 | Ch13 region 3 |
| chr13:77586570-77586610 | Ch13 region 3 |
| chr13:77586616-77586656 | Ch13 region 3 |
| chr13:77586705-77586745 | Ch13 region 3 |
| chr13:77586788-77586828 | Ch13 region 3 |
| chr13:77586951-77586991 | Ch13 region 3 |
| chr13:77587193-77587233 | Ch13 region 3 |
| chr13:77587241-77587281 | Ch13 region 3 |
| chr13:77587359-77587399 | Ch13 region 3 |
| chr13:77587423-77587463 | Ch13 region 3 |
| chr13:77587504-77587544 | Ch13 region 3 |
| chr13:77587667-77587707 | Ch13 region 3 |
| chr13:77587717-77587757 | Ch13 region 3 |
| chr13:77587922-77587962 | Ch13 region 3 |
| chr13:77587984-77588024 | Ch13 region 3 |
| chr13:77588052-77588092 | Ch13 region 3 |
| chr13:77588105-77588145 | Ch13 region 3 |
| chr13:77588163-77588203 | Ch13 region 3 |
| chr13:77588229-77588269 | Ch13 region 3 |
| chr13:77588276-77588316 | Ch13 region 3 |
| chr13:77588358-77588398 | Ch13 region 3 |
| chr13:77588441-77588481 | Ch13 region 3 |
| chr13:77588516-77588556 | Ch13 region 3 |
| chr13:77588611-77588651 | Ch13 region 3 |
| chr13:77588660-77588700 | Ch13 region 3 |
| chr13:77588755-77588795 | Ch13 region 3 |

| | |
|---|---|
| chr13:77588810-77588850 | Ch13 region 3 |
| chr13:77588855-77588895 | Ch13 region 3 |
| chr13:77588913-77588953 | Ch13 region 3 |
| chr13:77588984-77589024 | Ch13 region 3 |
| chr13:77589050-77589090 | Ch13 region 3 |
| chr13:77589131-77589171 | Ch13 region 3 |
| chr13:77589177-77589217 | Ch13 region 3 |
| chr13:77589222-77589262 | Ch13 region 3 |
| chr13:77589269-77589309 | Ch13 region 3 |
| chr13:77589359-77589399 | Ch13 region 3 |
| chr13:77589436-77589476 | Ch13 region 3 |
| chr13:77589521-77589561 | Ch13 region 3 |
| chr13:77589579-77589619 | Ch13 region 3 |
| chr13:77589624-77589664 | Ch13 region 3 |
| chr13:77589678-77589718 | Ch13 region 3 |
| chr13:77589753-77589793 | Ch13 region 3 |
| chr13:77589849-77589889 | Ch13 region 3 |
| chr13:77589899-77589939 | Ch13 region 3 |
| chr13:77589962-77590002 | Ch13 region 3 |
| chr13:77590020-77590060 | Ch13 region 3 |
| chr13:77590305-77590345 | Ch13 region 3 |
| chr13:77590387-77590427 | Ch13 region 3 |
| chr13:77590432-77590472 | Ch13 region 3 |
| chr13:77590482-77590522 | Ch13 region 3 |
| chr13:77590534-77590574 | Ch13 region 3 |
| chr13:77590579-77590619 | Ch13 region 3 |
| chr13:77590624-77590664 | Ch13 region 3 |
| chr13:77590762-77590802 | Ch13 region 3 |
| chr13:77590882-77590922 | Ch13 region 3 |
| chr13:77590949-77590989 | Ch13 region 3 |
| chr13:77591147-77591187 | Ch13 region 3 |
| chr13:77591197-77591237 | Ch13 region 3 |
| chr13:77591242-77591282 | Ch13 region 3 |
| chr13:77591309-77591349 | Ch13 region 3 |
| chr13:77591361-77591401 | Ch13 region 3 |
| chr13:77591522-77591562 | Ch13 region 3 |
| chr13:77591567-77591607 | Ch13 region 3 |
| chr13:77591637-77591677 | Ch13 region 3 |
| chr13:77591693-77591733 | Ch13 region 3 |
| chr13:77591783-77591823 | Ch13 region 3 |
| chr13:77591876-77591916 | Ch13 region 3 |
| chr13:77591957-77591997 | Ch13 region 3 |
| chr13:77592006-77592046 | Ch13 region 3 |
| chr13:77592075-77592115 | Ch13 region 3 |
| chr13:77592148-77592188 | Ch13 region 3 |
| chr13:77592267-77592307 | Ch13 region 3 |
| chr13:77592319-77592359 | Ch13 region 3 |
| chr13:77592365-77592405 | Ch13 region 3 |
| chr13:77592416-77592456 | Ch13 region 3 |
| chr13:77592476-77592516 | Ch13 region 3 |
| chr13:77592718-77592758 | Ch13 region 3 |
| chr13:77592773-77592813 | Ch13 region 3 |

| | |
|---|---|
| chr13:77592835-77592875 | Ch13 region 3 |
| chr13:77592902-77592942 | Ch13 region 3 |
| chr13:77593021-77593061 | Ch13 region 3 |
| chr13:77593071-77593111 | Ch13 region 3 |
| chr13:77593240-77593280 | Ch13 region 3 |
| chr13:77593301-77593341 | Ch13 region 3 |
| chr13:77593352-77593392 | Ch13 region 3 |
| chr13:77593423-77593463 | Ch13 region 3 |
| chr13:77593508-77593548 | Ch13 region 3 |
| chr13:77593649-77593689 | Ch13 region 3 |
| chr13:77593729-77593769 | Ch13 region 3 |
| chr13:77593807-77593847 | Ch13 region 3 |
| chr13:77593861-77593901 | Ch13 region 3 |
| chr13:77593909-77593949 | Ch13 region 3 |
| chr13:77594010-77594050 | Ch13 region 3 |
| chr13:77594063-77594103 | Ch13 region 3 |
| chr13:77594186-77594226 | Ch13 region 3 |
| chr13:77594243-77594283 | Ch13 region 3 |
| chr13:77594291-77594331 | Ch13 region 3 |
| chr13:77594448-77594488 | Ch13 region 3 |
| chr13:77594502-77594542 | Ch13 region 3 |
| chr13:77594615-77594655 | Ch13 region 3 |
| chr13:77594667-77594707 | Ch13 region 3 |
| chr13:77594789-77594829 | Ch13 region 3 |
| chr13:77594834-77594874 | Ch13 region 3 |
| chr13:77594900-77594940 | Ch13 region 3 |
| chr13:77595009-77595049 | Ch13 region 3 |
| chr13:77595058-77595098 | Ch13 region 3 |
| chr13:77595103-77595143 | Ch13 region 3 |
| chr13:77595209-77595249 | Ch13 region 3 |
| chr13:77595255-77595295 | Ch13 region 3 |
| chr13:77595396-77595436 | Ch13 region 3 |
| chr13:77595554-77595594 | Ch13 region 3 |
| chr13:77595643-77595683 | Ch13 region 3 |
| chr13:77595719-77595759 | Ch13 region 3 |
| chr13:77595778-77595818 | Ch13 region 3 |
| chr13:77595827-77595867 | Ch13 region 3 |
| chr13:77595905-77595945 | Ch13 region 3 |
| chr13:77595950-77595990 | Ch13 region 3 |
| chr13:77595995-77596035 | Ch13 region 3 |
| chr13:77596061-77596101 | Ch13 region 3 |
| chr13:77596135-77596175 | Ch13 region 3 |
| chr13:77596329-77596369 | Ch13 region 3 |
| chr13:77596378-77596418 | Ch13 region 3 |
| chr13:77596433-77596473 | Ch13 region 3 |
| chr13:77596499-77596539 | Ch13 region 3 |
| chr13:77596580-77596620 | Ch13 region 3 |
| chr13:77596673-77596713 | Ch13 region 3 |
| chr13:77596732-77596772 | Ch13 region 3 |
| chr13:77596777-77596817 | Ch13 region 3 |
| chr13:77596841-77596881 | Ch13 region 3 |
| chr13:77596937-77596977 | Ch13 region 3 |

| | |
|---|---|
| chr13:77596998-77597038 | Ch13 region 3 |
| chr13:77597108-77597148 | Ch13 region 3 |
| chr13:77597196-77597236 | Ch13 region 3 |
| chr13:77597292-77597332 | Ch13 region 3 |
| chr13:77597430-77597470 | Ch13 region 3 |
| chr13:77597534-77597574 | Ch13 region 3 |
| chr13:77597583-77597623 | Ch13 region 3 |
| chr13:77597653-77597693 | Ch13 region 3 |
| chr13:77597745-77597785 | Ch13 region 3 |
| chr13:77597792-77597832 | Ch13 region 3 |
| chr13:77597860-77597900 | Ch13 region 3 |
| chr13:77597945-77597985 | Ch13 region 3 |
| chr13:77597990-77598030 | Ch13 region 3 |
| chr13:77598042-77598082 | Ch13 region 3 |
| chr13:77598088-77598128 | Ch13 region 3 |
| chr13:77598157-77598197 | Ch13 region 3 |
| chr13:77598216-77598256 | Ch13 region 3 |
| chr13:77598336-77598376 | Ch13 region 3 |
| chr13:77598420-77598460 | Ch13 region 3 |
| chr13:77598466-77598506 | Ch13 region 3 |
| chr13:77598543-77598583 | Ch13 region 3 |
| chr13:77598589-77598629 | Ch13 region 3 |
| chr13:77598823-77598863 | Ch13 region 3 |
| chr13:77598877-77598917 | Ch13 region 3 |
| chr13:77599003-77599043 | Ch13 region 3 |
| chr13:77599184-77599224 | Ch13 region 3 |
| chr13:77599268-77599308 | Ch13 region 3 |
| chr13:77599313-77599353 | Ch13 region 3 |
| chr13:77599397-77599437 | Ch13 region 3 |
| chr13:77599454-77599494 | Ch13 region 3 |
| chr13:77599542-77599582 | Ch13 region 3 |
| chr13:77599661-77599701 | Ch13 region 3 |
| chr13:77599849-77599889 | Ch13 region 3 |
| chr13:77599904-77599944 | Ch13 region 3 |
| chr13:77599952-77599992 | Ch13 region 3 |
| chr13:77600075-77600115 | Ch13 region 3 |
| chr13:77600223-77600263 | Ch13 region 3 |
| chr13:77600270-77600310 | Ch13 region 3 |
| chr13:77600349-77600389 | Ch13 region 3 |
| chr13:77600410-77600450 | Ch13 region 3 |
| chr13:77600482-77600522 | Ch13 region 3 |
| chr13:77600531-77600571 | Ch13 region 3 |
| chr13:77600618-77600658 | Ch13 region 3 |
| chr13:77600709-77600749 | Ch13 region 3 |
| chr13:77600762-77600802 | Ch13 region 3 |
| chr13:77600882-77600922 | Ch13 region 3 |
| chr13:77600934-77600974 | Ch13 region 3 |
| chr13:77601036-77601076 | Ch13 region 3 |
| chr13:77601084-77601124 | Ch13 region 3 |
| chr13:77601173-77601213 | Ch13 region 3 |
| chr13:77601262-77601302 | Ch13 region 3 |
| chr13:77601315-77601355 | Ch13 region 3 |

| | |
|---|---|
| chr13:77601431-77601471 | Ch13 region 3 |
| chr13:77601504-77601544 | Ch13 region 3 |
| chr13:77601575-77601615 | Ch13 region 3 |
| chr13:77601665-77601705 | Ch13 region 3 |
| chr13:77601732-77601772 | Ch13 region 3 |
| chr13:77601892-77601932 | Ch13 region 3 |
| chr13:77601956-77601996 | Ch13 region 3 |
| chr13:77602086-77602126 | Ch13 region 3 |
| chr13:77602148-77602188 | Ch13 region 3 |
| chr13:77602208-77602248 | Ch13 region 3 |
| chr13:77602259-77602299 | Ch13 region 3 |
| chr13:77602350-77602390 | Ch13 region 3 |
| chr13:77602440-77602480 | Ch13 region 3 |
| chr13:77602597-77602637 | Ch13 region 3 |
| chr13:77602669-77602709 | Ch13 region 3 |
| chr13:77602726-77602766 | Ch13 region 3 |
| chr13:77602821-77602861 | Ch13 region 3 |
| chr13:77602882-77602922 | Ch13 region 3 |
| chr13:77602982-77603022 | Ch13 region 3 |
| chr13:77603027-77603067 | Ch13 region 3 |
| chr13:77603147-77603187 | Ch13 region 3 |
| chr13:77603224-77603264 | Ch13 region 3 |
| chr13:77603311-77603351 | Ch13 region 3 |
| chr13:77603398-77603438 | Ch13 region 3 |
| chr13:77603481-77603521 | Ch13 region 3 |
| chr13:77603527-77603567 | Ch13 region 3 |
| chr13:77603581-77603621 | Ch13 region 3 |
| chr13:77603631-77603671 | Ch13 region 3 |
| chr13:77603679-77603719 | Ch13 region 3 |
| chr13:77603758-77603798 | Ch13 region 3 |
| chr13:77603830-77603870 | Ch13 region 3 |
| chr13:77603886-77603926 | Ch13 region 3 |
| chr13:77603953-77603993 | Ch13 region 3 |
| chr13:77604074-77604114 | Ch13 region 3 |
| chr13:77604159-77604199 | Ch13 region 3 |
| chr13:77604214-77604254 | Ch13 region 3 |
| chr13:77604262-77604302 | Ch13 region 3 |
| chr13:77604308-77604348 | Ch13 region 3 |
| chr13:77604385-77604425 | Ch13 region 3 |
| chr13:77604458-77604498 | Ch13 region 3 |
| chr13:77604651-77604691 | Ch13 region 3 |
| chr13:77604723-77604763 | Ch13 region 3 |
| chr13:77604775-77604815 | Ch13 region 3 |
| chr13:77604857-77604897 | Ch13 region 3 |
| chr13:77605089-77605129 | Ch13 region 3 |
| chr13:77605139-77605179 | Ch13 region 3 |
| chr13:77605211-77605251 | Ch13 region 3 |
| chr13:77605297-77605337 | Ch13 region 3 |
| chr13:77605350-77605390 | Ch13 region 3 |
| chr13:77605415-77605455 | Ch13 region 3 |
| chr13:77605485-77605525 | Ch13 region 3 |
| chr13:77605561-77605601 | Ch13 region 3 |

| | |
|---|---|
| chr13:77605606-77605646 | Ch13 region 3 |
| chr13:77605671-77605711 | Ch13 region 3 |
| chr13:77605755-77605795 | Ch13 region 3 |
| chr13:77605825-77605865 | Ch13 region 3 |
| chr13:77605877-77605917 | Ch13 region 3 |
| chr13:77605943-77605983 | Ch13 region 3 |
| chr13:77605993-77606033 | Ch13 region 3 |
| chr13:77606039-77606079 | Ch13 region 3 |
| chr13:77606119-77606159 | Ch13 region 3 |
| chr13:77606221-77606261 | Ch13 region 3 |
| chr13:77606268-77606308 | Ch13 region 3 |
| chr13:77606328-77606368 | Ch13 region 3 |
| chr13:77606387-77606427 | Ch13 region 3 |
| chr13:77606500-77606540 | Ch13 region 3 |
| chr13:77606610-77606650 | Ch13 region 3 |
| chr13:77606778-77606818 | Ch13 region 3 |
| chr13:77606833-77606873 | Ch13 region 3 |
| chr13:77606899-77606939 | Ch13 region 3 |
| chr13:77607021-77607061 | Ch13 region 3 |
| chr13:77607077-77607117 | Ch13 region 3 |
| chr13:77607144-77607184 | Ch13 region 3 |
| chr13:77607279-77607319 | Ch13 region 3 |
| chr13:77607357-77607397 | Ch13 region 3 |
| chr13:77607406-77607446 | Ch13 region 3 |
| chr13:77607498-77607538 | Ch13 region 3 |
| chr13:77607582-77607622 | Ch13 region 3 |
| chr13:77607627-77607667 | Ch13 region 3 |
| chr13:77607672-77607712 | Ch13 region 3 |
| chr13:77607717-77607757 | Ch13 region 3 |
| chr13:77607945-77607985 | Ch13 region 3 |
| chr13:77608037-77608077 | Ch13 region 3 |
| chr13:77608084-77608124 | Ch13 region 3 |
| chr13:77608149-77608189 | Ch13 region 3 |
| chr13:77608199-77608239 | Ch13 region 3 |
| chr13:77608244-77608284 | Ch13 region 3 |
| chr13:77608325-77608365 | Ch13 region 3 |
| chr13:77608373-77608413 | Ch13 region 3 |
| chr13:77608422-77608462 | Ch13 region 3 |
| chr13:77608476-77608516 | Ch13 region 3 |
| chr13:77608561-77608601 | Ch13 region 3 |
| chr13:77608627-77608667 | Ch13 region 3 |
| chr13:77608733-77608773 | Ch13 region 3 |
| chr13:77608808-77608848 | Ch13 region 3 |
| chr13:77608861-77608901 | Ch13 region 3 |
| chr13:77608931-77608971 | Ch13 region 3 |
| chr13:77608977-77609017 | Ch13 region 3 |
| chr13:77609050-77609090 | Ch13 region 3 |
| chr13:77609118-77609158 | Ch13 region 3 |
| chr13:77609178-77609218 | Ch13 region 3 |
| chr13:77609263-77609303 | Ch13 region 3 |
| chr13:77609308-77609348 | Ch13 region 3 |
| chr13:77609376-77609416 | Ch13 region 3 |

| | |
|---|---|
| chr13:77609424-77609464 | Ch13 region 3 |
| chr13:77609469-77609509 | Ch13 region 3 |
| chr13:77609518-77609558 | Ch13 region 3 |
| chr13:77609596-77609636 | Ch13 region 3 |
| chr13:77609729-77609769 | Ch13 region 3 |
| chr13:77609774-77609814 | Ch13 region 3 |
| chr13:77609961-77610001 | Ch13 region 3 |
| chr13:77610056-77610096 | Ch13 region 3 |
| chr13:77610204-77610244 | Ch13 region 3 |
| chr13:77610299-77610339 | Ch13 region 3 |
| chr13:77610382-77610422 | Ch13 region 3 |
| chr13:77610449-77610489 | Ch13 region 3 |
| chr13:77610497-77610537 | Ch13 region 3 |
| chr13:77610561-77610601 | Ch13 region 3 |
| chr13:77610633-77610673 | Ch13 region 3 |
| chr13:77610698-77610738 | Ch13 region 3 |
| chr13:77610761-77610801 | Ch13 region 3 |
| chr13:77610814-77610854 | Ch13 region 3 |
| chr13:77610859-77610899 | Ch13 region 3 |
| chr13:77610918-77610958 | Ch13 region 3 |
| chr13:77610994-77611034 | Ch13 region 3 |
| chr13:77611058-77611098 | Ch13 region 3 |
| chr13:77611117-77611157 | Ch13 region 3 |
| chr13:77611185-77611225 | Ch13 region 3 |
| chr13:77611253-77611293 | Ch13 region 3 |
| chr13:77611298-77611338 | Ch13 region 3 |
| chr13:77611391-77611431 | Ch13 region 3 |
| chr13:77611447-77611487 | Ch13 region 3 |
| chr13:77611498-77611538 | Ch13 region 3 |
| chr13:77611695-77611735 | Ch13 region 3 |
| chr13:77611748-77611788 | Ch13 region 3 |
| chr13:77611793-77611833 | Ch13 region 3 |
| chr13:77611838-77611878 | Ch13 region 3 |
| chr13:77611883-77611923 | Ch13 region 3 |
| chr13:77611932-77611972 | Ch13 region 3 |
| chr13:77612005-77612045 | Ch13 region 3 |
| chr13:77612073-77612113 | Ch13 region 3 |
| chr13:77612143-77612183 | Ch13 region 3 |
| chr13:77612228-77612268 | Ch13 region 3 |
| chr13:77612324-77612364 | Ch13 region 3 |
| chr13:77612392-77612432 | Ch13 region 3 |
| chr13:77612437-77612477 | Ch13 region 3 |
| chr13:77612489-77612529 | Ch13 region 3 |
| chr13:77612573-77612613 | Ch13 region 3 |
| chr13:77612630-77612670 | Ch13 region 3 |
| chr13:77612848-77612888 | Ch13 region 3 |
| chr13:77612949-77612989 | Ch13 region 3 |
| chr13:77613088-77613128 | Ch13 region 3 |
| chr13:77613150-77613190 | Ch13 region 3 |
| chr13:77613195-77613235 | Ch13 region 3 |
| chr13:77613248-77613288 | Ch13 region 3 |
| chr13:77613314-77613354 | Ch13 region 3 |

| | |
|---|---|
| chr13:77613392-77613432 | Ch13 region 3 |
| chr13:77613475-77613515 | Ch13 region 3 |
| chr13:77613584-77613624 | Ch13 region 3 |
| chr13:77613688-77613728 | Ch13 region 3 |
| chr13:77613783-77613823 | Ch13 region 3 |
| chr13:77613836-77613876 | Ch13 region 3 |
| chr13:77613909-77613949 | Ch13 region 3 |
| chr13:77613977-77614017 | Ch13 region 3 |
| chr13:77614055-77614095 | Ch13 region 3 |
| chr13:77614147-77614187 | Ch13 region 3 |
| chr13:77614219-77614259 | Ch13 region 3 |
| chr13:77614292-77614332 | Ch13 region 3 |
| chr13:77614337-77614377 | Ch13 region 3 |
| chr13:77614392-77614432 | Ch13 region 3 |
| chr13:77614488-77614528 | Ch13 region 3 |
| chr13:77614575-77614615 | Ch13 region 3 |
| chr13:77614653-77614693 | Ch13 region 3 |
| chr13:77614710-77614750 | Ch13 region 3 |
| chr13:77614755-77614795 | Ch13 region 3 |
| chr13:77614805-77614845 | Ch13 region 3 |
| chr13:77614853-77614893 | Ch13 region 3 |
| chr13:77614934-77614974 | Ch13 region 3 |
| chr13:77614993-77615033 | Ch13 region 3 |
| chr13:77615072-77615112 | Ch13 region 3 |
| chr13:77615137-77615177 | Ch13 region 3 |
| chr13:77615198-77615238 | Ch13 region 3 |
| chr13:77615256-77615296 | Ch13 region 3 |
| chr13:77615359-77615399 | Ch13 region 3 |
| chr13:77615404-77615444 | Ch13 region 3 |
| chr13:77615449-77615489 | Ch13 region 3 |
| chr13:77615539-77615579 | Ch13 region 3 |
| chr13:77615642-77615682 | Ch13 region 3 |
| chr13:77615739-77615779 | Ch13 region 3 |
| chr13:77615803-77615843 | Ch13 region 3 |
| chr13:77615870-77615910 | Ch13 region 3 |
| chr13:77615971-77616011 | Ch13 region 3 |
| chr13:77616043-77616083 | Ch13 region 3 |
| chr13:77616106-77616146 | Ch13 region 3 |
| chr13:77616170-77616210 | Ch13 region 3 |
| chr13:77616229-77616269 | Ch13 region 3 |
| chr13:77616274-77616314 | Ch13 region 3 |
| chr13:77616319-77616359 | Ch13 region 3 |
| chr13:77616382-77616422 | Ch13 region 3 |
| chr13:77616614-77616654 | Ch13 region 3 |
| chr13:77616705-77616745 | Ch13 region 3 |
| chr13:77616750-77616790 | Ch13 region 3 |
| chr13:77616821-77616861 | Ch13 region 3 |
| chr13:77616872-77616912 | Ch13 region 3 |
| chr13:77616976-77617016 | Ch13 region 3 |
| chr13:77617049-77617089 | Ch13 region 3 |
| chr13:77617118-77617158 | Ch13 region 3 |
| chr13:77617176-77617216 | Ch13 region 3 |

| | |
|---|---|
| chr13:77617258-77617298 | Ch13 region 3 |
| chr13:77617322-77617362 | Ch13 region 3 |
| chr13:77617400-77617440 | Ch13 region 3 |
| chr13:77617463-77617503 | Ch13 region 3 |
| chr13:77617562-77617602 | Ch13 region 3 |
| chr13:77617738-77617778 | Ch13 region 3 |
| chr13:77617784-77617824 | Ch13 region 3 |
| chr13:77617844-77617884 | Ch13 region 3 |
| chr13:77618086-77618126 | Ch13 region 3 |
| chr13:77618183-77618223 | Ch13 region 3 |
| chr13:77618250-77618290 | Ch13 region 3 |
| chr13:77618343-77618383 | Ch13 region 3 |
| chr13:77618401-77618441 | Ch13 region 3 |
| chr13:77618609-77618649 | Ch13 region 3 |
| chr13:77618696-77618736 | Ch13 region 3 |
| chr13:77618764-77618804 | Ch13 region 3 |
| chr13:77618823-77618863 | Ch13 region 3 |
| chr13:77618875-77618915 | Ch13 region 3 |
| chr13:77619068-77619108 | Ch13 region 3 |
| chr13:77619157-77619197 | Ch13 region 3 |
| chr13:77619219-77619259 | Ch13 region 3 |
| chr13:77619274-77619314 | Ch13 region 3 |
| chr13:77619347-77619387 | Ch13 region 3 |
| chr13:77619405-77619445 | Ch13 region 3 |
| chr13:77619460-77619500 | Ch13 region 3 |
| chr13:77619527-77619567 | Ch13 region 3 |
| chr13:77619601-77619641 | Ch13 region 3 |
| chr13:77619818-77619858 | Ch13 region 3 |
| chr13:77619869-77619909 | Ch13 region 3 |
| chr13:77619958-77619998 | Ch13 region 3 |
| chr13:77620069-77620109 | Ch13 region 3 |
| chr13:77620125-77620165 | Ch13 region 3 |
| chr13:77620330-77620370 | Ch13 region 3 |
| chr13:77620395-77620435 | Ch13 region 3 |
| chr13:77620455-77620495 | Ch13 region 3 |
| chr13:77620521-77620561 | Ch13 region 3 |
| chr13:77620567-77620607 | Ch13 region 3 |
| chr13:77620632-77620672 | Ch13 region 3 |
| chr13:77620721-77620761 | Ch13 region 3 |
| chr13:77620766-77620806 | Ch13 region 3 |
| chr13:77620815-77620855 | Ch13 region 3 |
| chr13:77620894-77620934 | Ch13 region 3 |
| chr13:77620943-77620983 | Ch13 region 3 |
| chr13:77621005-77621045 | Ch13 region 3 |
| chr13:77621057-77621097 | Ch13 region 3 |
| chr13:77621168-77621208 | Ch13 region 3 |
| chr13:77621213-77621253 | Ch13 region 3 |
| chr13:77621283-77621323 | Ch13 region 3 |
| chr13:77621351-77621391 | Ch13 region 3 |
| chr13:77621454-77621494 | Ch13 region 3 |
| chr13:77621503-77621543 | Ch13 region 3 |
| chr13:77621551-77621591 | Ch13 region 3 |

| | |
|---|---|
| chr13:77621596-77621636 | Ch13 region 3 |
| chr13:77621669-77621709 | Ch13 region 3 |
| chr13:77621728-77621768 | Ch13 region 3 |
| chr13:77621801-77621841 | Ch13 region 3 |
| chr13:77621868-77621908 | Ch13 region 3 |
| chr13:77621939-77621979 | Ch13 region 3 |
| chr13:77622017-77622057 | Ch13 region 3 |
| chr13:77622114-77622154 | Ch13 region 3 |
| chr13:77622189-77622229 | Ch13 region 3 |
| chr13:77622244-77622284 | Ch13 region 3 |
| chr13:77622306-77622346 | Ch13 region 3 |
| chr13:77622351-77622391 | Ch13 region 3 |
| chr13:77622403-77622443 | Ch13 region 3 |
| chr13:77622460-77622500 | Ch13 region 3 |
| chr13:77622552-77622592 | Ch13 region 3 |
| chr13:77622597-77622637 | Ch13 region 3 |
| chr13:77622644-77622684 | Ch13 region 3 |
| chr13:77622722-77622762 | Ch13 region 3 |
| chr13:77622779-77622819 | Ch13 region 3 |
| chr13:77622828-77622868 | Ch13 region 3 |
| chr13:77622892-77622932 | Ch13 region 3 |
| chr13:77622948-77622988 | Ch13 region 3 |
| chr13:77623063-77623103 | Ch13 region 3 |
| chr13:77623128-77623168 | Ch13 region 3 |
| chr13:77623173-77623213 | Ch13 region 3 |
| chr13:77623218-77623258 | Ch13 region 3 |
| chr13:77623265-77623305 | Ch13 region 3 |
| chr13:77623340-77623380 | Ch13 region 3 |
| chr13:77623397-77623437 | Ch13 region 3 |
| chr13:77623580-77623620 | Ch13 region 3 |
| chr13:77623656-77623696 | Ch13 region 3 |
| chr13:77623720-77623760 | Ch13 region 3 |
| chr13:77623776-77623816 | Ch13 region 3 |
| chr13:77623873-77623913 | Ch13 region 3 |
| chr13:77624050-77624090 | Ch13 region 3 |
| chr13:77624111-77624151 | Ch13 region 3 |
| chr13:77624161-77624201 | Ch13 region 3 |
| chr13:77624244-77624284 | Ch13 region 3 |
| chr13:77624314-77624354 | Ch13 region 3 |
| chr13:77624394-77624434 | Ch13 region 3 |
| chr13:77624476-77624516 | Ch13 region 3 |
| chr13:77624553-77624593 | Ch13 region 3 |
| chr13:77624598-77624638 | Ch13 region 3 |
| chr13:77624643-77624683 | Ch13 region 3 |
| chr13:77624689-77624729 | Ch13 region 3 |
| chr13:77624740-77624780 | Ch13 region 3 |
| chr13:77624861-77624901 | Ch13 region 3 |
| chr13:77624911-77624951 | Ch13 region 3 |
| chr13:77624959-77624999 | Ch13 region 3 |
| chr13:77625005-77625045 | Ch13 region 3 |
| chr13:77625112-77625152 | Ch13 region 3 |
| chr13:77625203-77625243 | Ch13 region 3 |

| | |
|---|---|
| chr13:77625285-77625325 | Ch13 region 3 |
| chr13:77625343-77625383 | Ch13 region 3 |
| chr13:77625450-77625490 | Ch13 region 3 |
| chr13:77625638-77625678 | Ch13 region 3 |
| chr13:77625817-77625857 | Ch13 region 3 |
| chr13:77625918-77625958 | Ch13 region 3 |
| chr13:77625993-77626033 | Ch13 region 3 |
| chr13:77626058-77626098 | Ch13 region 3 |
| chr13:77626115-77626155 | Ch13 region 3 |
| chr13:77626196-77626236 | Ch13 region 3 |
| chr13:77626313-77626353 | Ch13 region 3 |
| chr13:77626390-77626430 | Ch13 region 3 |
| chr13:77626619-77626659 | Ch13 region 3 |
| chr13:77626664-77626704 | Ch13 region 3 |
| chr13:77626812-77626852 | Ch13 region 3 |
| chr13:77626941-77626981 | Ch13 region 3 |
| chr13:77627020-77627060 | Ch13 region 3 |
| chr13:77627094-77627134 | Ch13 region 3 |
| chr13:77627305-77627345 | Ch13 region 3 |
| chr13:77627419-77627459 | Ch13 region 3 |
| chr13:77627490-77627530 | Ch13 region 3 |
| chr13:77627595-77627635 | Ch13 region 3 |
| chr13:77627686-77627726 | Ch13 region 3 |
| chr13:77627733-77627773 | Ch13 region 3 |
| chr13:77627819-77627859 | Ch13 region 3 |
| chr13:77627873-77627913 | Ch13 region 3 |
| chr13:77627918-77627958 | Ch13 region 3 |
| chr13:77627987-77628027 | Ch13 region 3 |
| chr13:77628073-77628113 | Ch13 region 3 |
| chr13:77628158-77628198 | Ch13 region 3 |
| chr13:77628217-77628257 | Ch13 region 3 |
| chr13:77628293-77628333 | Ch13 region 3 |
| chr13:77628361-77628401 | Ch13 region 3 |
| chr13:77628418-77628458 | Ch13 region 3 |
| chr13:77628464-77628504 | Ch13 region 3 |
| chr13:77628527-77628567 | Ch13 region 3 |
| chr13:77628601-77628641 | Ch13 region 3 |
| chr13:77628646-77628686 | Ch13 region 3 |
| chr13:77628871-77628911 | Ch13 region 3 |
| chr13:77629023-77629063 | Ch13 region 3 |
| chr13:77629068-77629108 | Ch13 region 3 |
| chr13:77629132-77629172 | Ch13 region 3 |
| chr13:77629195-77629235 | Ch13 region 3 |
| chr13:77629281-77629321 | Ch13 region 3 |
| chr13:77629357-77629397 | Ch13 region 3 |
| chr13:77629434-77629474 | Ch13 region 3 |
| chr13:77629554-77629594 | Ch13 region 3 |
| chr13:77629683-77629723 | Ch13 region 3 |
| chr13:77629731-77629771 | Ch13 region 3 |
| chr13:77629784-77629824 | Ch13 region 3 |
| chr13:77629831-77629871 | Ch13 region 3 |
| chr13:77629934-77629974 | Ch13 region 3 |

| | |
|---|---|
| chr13:77630049-77630089 | Ch13 region 3 |
| chr13:77630144-77630184 | Ch13 region 3 |
| chr13:77630190-77630230 | Ch13 region 3 |
| chr13:77630315-77630355 | Ch13 region 3 |
| chr13:77630369-77630409 | Ch13 region 3 |
| chr13:77630460-77630500 | Ch13 region 3 |
| chr13:77630509-77630549 | Ch13 region 3 |
| chr13:77630555-77630595 | Ch13 region 3 |
| chr13:77630612-77630652 | Ch13 region 3 |
| chr13:77630659-77630699 | Ch13 region 3 |
| chr13:77630721-77630761 | Ch13 region 3 |
| chr13:77630778-77630818 | Ch13 region 3 |
| chr13:77630828-77630868 | Ch13 region 3 |
| chr13:77630947-77630987 | Ch13 region 3 |
| chr13:77630993-77631033 | Ch13 region 3 |
| chr13:77631092-77631132 | Ch13 region 3 |
| chr13:77631163-77631203 | Ch13 region 3 |
| chr13:77631285-77631325 | Ch13 region 3 |
| chr13:77631405-77631445 | Ch13 region 3 |
| chr13:77631582-77631622 | Ch13 region 3 |
| chr13:77631733-77631773 | Ch13 region 3 |
| chr13:77631778-77631818 | Ch13 region 3 |
| chr13:77631824-77631864 | Ch13 region 3 |
| chr13:77631891-77631931 | Ch13 region 3 |
| chr13:77632035-77632075 | Ch13 region 3 |
| chr13:77632268-77632308 | Ch13 region 3 |
| chr13:77632316-77632356 | Ch13 region 3 |
| chr13:77632371-77632411 | Ch13 region 3 |
| chr13:77632461-77632501 | Ch13 region 3 |
| chr13:77632522-77632562 | Ch13 region 3 |
| chr13:77632635-77632675 | Ch13 region 3 |
| chr13:77632748-77632788 | Ch13 region 3 |
| chr13:77632803-77632843 | Ch13 region 3 |
| chr13:77633133-77633173 | Ch13 region 3 |
| chr13:77633213-77633253 | Ch13 region 3 |
| chr13:77633435-77633475 | Ch13 region 3 |
| chr13:77633481-77633521 | Ch13 region 3 |
| chr13:77633565-77633605 | Ch13 region 3 |
| chr13:77633640-77633680 | Ch13 region 3 |
| chr13:77633742-77633782 | Ch13 region 3 |
| chr13:77633842-77633882 | Ch13 region 3 |
| chr13:77633947-77633987 | Ch13 region 3 |
| chr13:77634050-77634090 | Ch13 region 3 |
| chr13:77634167-77634207 | Ch13 region 3 |
| chr13:77634216-77634256 | Ch13 region 3 |
| chr13:77634312-77634352 | Ch13 region 3 |
| chr13:77634473-77634513 | Ch13 region 3 |
| chr13:77634550-77634590 | Ch13 region 3 |
| chr13:77634595-77634635 | Ch13 region 3 |
| chr13:77634809-77634849 | Ch13 region 3 |
| chr13:77634917-77634957 | Ch13 region 3 |
| chr13:77634978-77635018 | Ch13 region 3 |

| | |
|---|---|
| chr13:77635033-77635073 | Ch13 region 3 |
| chr13:77635090-77635130 | Ch13 region 3 |
| chr13:77635210-77635250 | Ch13 region 3 |
| chr13:77635275-77635315 | Ch13 region 3 |
| chr13:77635321-77635361 | Ch13 region 3 |
| chr13:77635400-77635440 | Ch13 region 3 |
| chr13:77635452-77635492 | Ch13 region 3 |
| chr13:77635555-77635595 | Ch13 region 3 |
| chr13:77635645-77635685 | Ch13 region 3 |
| chr13:77635701-77635741 | Ch13 region 3 |
| chr13:77635788-77635828 | Ch13 region 3 |
| chr13:77635871-77635911 | Ch13 region 3 |
| chr13:77635932-77635972 | Ch13 region 3 |
| chr13:77636092-77636132 | Ch13 region 3 |
| chr13:77636188-77636228 | Ch13 region 3 |
| chr13:77636293-77636333 | Ch13 region 3 |
| chr13:77636355-77636395 | Ch13 region 3 |
| chr13:77636465-77636505 | Ch13 region 3 |
| chr13:77636520-77636560 | Ch13 region 3 |
| chr13:77636594-77636634 | Ch13 region 3 |
| chr13:77636718-77636758 | Ch13 region 3 |
| chr13:77636832-77636872 | Ch13 region 3 |
| chr13:77636918-77636958 | Ch13 region 3 |
| chr13:77636972-77637012 | Ch13 region 3 |
| chr13:77637048-77637088 | Ch13 region 3 |
| chr13:77637190-77637230 | Ch13 region 3 |
| chr13:77637240-77637280 | Ch13 region 3 |
| chr13:77637291-77637331 | Ch13 region 3 |
| chr13:77637340-77637380 | Ch13 region 3 |
| chr13:77637471-77637511 | Ch13 region 3 |
| chr13:77637551-77637591 | Ch13 region 3 |
| chr13:77637759-77637799 | Ch13 region 3 |
| chr13:77637808-77637848 | Ch13 region 3 |
| chr13:77637900-77637940 | Ch13 region 3 |
| chr13:77637961-77638001 | Ch13 region 3 |
| chr13:77638016-77638056 | Ch13 region 3 |
| chr13:77638102-77638142 | Ch13 region 3 |
| chr13:77638151-77638191 | Ch13 region 3 |
| chr13:77638244-77638284 | Ch13 region 3 |
| chr13:77638331-77638371 | Ch13 region 3 |
| chr13:77638435-77638475 | Ch13 region 3 |
| chr13:77638509-77638549 | Ch13 region 3 |
| chr13:77638564-77638604 | Ch13 region 3 |
| chr13:77638730-77638770 | Ch13 region 3 |
| chr13:77638807-77638847 | Ch13 region 3 |
| chr13:77638858-77638898 | Ch13 region 3 |
| chr13:77638948-77638988 | Ch13 region 3 |
| chr13:77638999-77639039 | Ch13 region 3 |
| chr13:77639093-77639133 | Ch13 region 3 |
| chr13:77639188-77639228 | Ch13 region 3 |
| chr13:77639263-77639303 | Ch13 region 3 |
| chr13:77639369-77639409 | Ch13 region 3 |

| | |
|---|---|
| chr13:77639414-77639454 | Ch13 region 3 |
| chr13:77639462-77639502 | Ch13 region 3 |
| chr13:77639510-77639550 | Ch13 region 3 |
| chr13:77639556-77639596 | Ch13 region 3 |
| chr13:77639610-77639650 | Ch13 region 3 |
| chr13:77639657-77639697 | Ch13 region 3 |
| chr13:77639764-77639804 | Ch13 region 3 |
| chr13:77639823-77639863 | Ch13 region 3 |
| chr13:77639894-77639934 | Ch13 region 3 |
| chr13:77639957-77639997 | Ch13 region 3 |
| chr13:77640026-77640066 | Ch13 region 3 |
| chr13:77640122-77640162 | Ch13 region 3 |
| chr13:77640212-77640252 | Ch13 region 3 |
| chr13:77640353-77640393 | Ch13 region 3 |
| chr13:77640410-77640450 | Ch13 region 3 |
| chr13:77640474-77640514 | Ch13 region 3 |
| chr13:77640540-77640580 | Ch13 region 3 |
| chr13:77640590-77640630 | Ch13 region 3 |
| chr13:77640637-77640677 | Ch13 region 3 |
| chr13:77640705-77640745 | Ch13 region 3 |
| chr13:77640824-77640864 | Ch13 region 3 |
| chr13:77640869-77640909 | Ch13 region 3 |
| chr13:77640931-77640971 | Ch13 region 3 |
| chr13:77640988-77641028 | Ch13 region 3 |
| chr13:77641240-77641280 | Ch13 region 3 |
| chr13:77641351-77641391 | Ch13 region 3 |
| chr13:77641490-77641530 | Ch13 region 3 |
| chr13:77641589-77641629 | Ch13 region 3 |
| chr13:77641651-77641691 | Ch13 region 3 |
| chr13:77641707-77641747 | Ch13 region 3 |
| chr13:77641790-77641830 | Ch13 region 3 |
| chr13:77641835-77641875 | Ch13 region 3 |
| chr13:77641933-77641973 | Ch13 region 3 |
| chr13:77642204-77642244 | Ch13 region 3 |
| chr13:77642349-77642389 | Ch13 region 3 |
| chr13:77642446-77642486 | Ch13 region 3 |
| chr13:77642529-77642569 | Ch13 region 3 |
| chr13:77642600-77642640 | Ch13 region 3 |
| chr13:77642653-77642693 | Ch13 region 3 |
| chr13:77642724-77642764 | Ch13 region 3 |
| chr13:77642784-77642824 | Ch13 region 3 |
| chr13:77642850-77642890 | Ch13 region 3 |
| chr13:77642901-77642941 | Ch13 region 3 |
| chr13:77642946-77642986 | Ch13 region 3 |
| chr13:77642998-77643038 | Ch13 region 3 |
| chr13:77643082-77643122 | Ch13 region 3 |
| chr13:77643133-77643173 | Ch13 region 3 |
| chr13:77643237-77643277 | Ch13 region 3 |
| chr13:77643288-77643328 | Ch13 region 3 |
| chr13:77643377-77643417 | Ch13 region 3 |
| chr13:77643443-77643483 | Ch13 region 3 |
| chr13:77643596-77643636 | Ch13 region 3 |

| | |
|---|---|
| chr13:77643713-77643753 | Ch13 region 3 |
| chr13:77643850-77643890 | Ch13 region 3 |
| chr13:77644074-77644114 | Ch13 region 3 |
| chr13:77644149-77644189 | Ch13 region 3 |
| chr13:77644252-77644292 | Ch13 region 3 |
| chr13:77644344-77644384 | Ch13 region 3 |
| chr13:77644421-77644461 | Ch13 region 3 |
| chr13:77644655-77644695 | Ch13 region 3 |
| chr13:77644747-77644787 | Ch13 region 3 |
| chr13:77644798-77644838 | Ch13 region 3 |
| chr13:77645033-77645073 | Ch13 region 3 |
| chr13:77645142-77645182 | Ch13 region 3 |
| chr13:77645204-77645244 | Ch13 region 3 |
| chr13:77645249-77645289 | Ch13 region 3 |
| chr13:77645311-77645351 | Ch13 region 3 |
| chr13:77645433-77645473 | Ch13 region 3 |
| chr13:77645551-77645591 | Ch13 region 3 |
| chr13:77645598-77645638 | Ch13 region 3 |
| chr13:77645668-77645708 | Ch13 region 3 |
| chr13:77645720-77645760 | Ch13 region 3 |
| chr13:77645774-77645814 | Ch13 region 3 |
| chr13:77645863-77645903 | Ch13 region 3 |
| chr13:77645926-77645966 | Ch13 region 3 |
| chr13:77645987-77646027 | Ch13 region 3 |
| chr13:77646084-77646124 | Ch13 region 3 |
| chr13:77646129-77646169 | Ch13 region 3 |
| chr13:77646508-77646548 | Ch13 region 3 |
| chr13:77646572-77646612 | Ch13 region 3 |
| chr13:77646635-77646675 | Ch13 region 3 |
| chr13:77646726-77646766 | Ch13 region 3 |
| chr13:77646798-77646838 | Ch13 region 3 |
| chr13:77646924-77646964 | Ch13 region 3 |
| chr13:77647181-77647221 | Ch13 region 3 |
| chr13:77647263-77647303 | Ch13 region 3 |
| chr13:77647314-77647354 | Ch13 region 3 |
| chr13:77647399-77647439 | Ch13 region 3 |
| chr13:77647444-77647484 | Ch13 region 3 |
| chr13:77647537-77647577 | Ch13 region 3 |
| chr13:77647585-77647625 | Ch13 region 3 |
| chr13:77647651-77647691 | Ch13 region 3 |
| chr13:77647745-77647785 | Ch13 region 3 |
| chr13:77647993-77648033 | Ch13 region 3 |
| chr13:77648071-77648111 | Ch13 region 3 |
| chr13:77648143-77648183 | Ch13 region 3 |
| chr13:77648268-77648308 | Ch13 region 3 |
| chr13:77648319-77648359 | Ch13 region 3 |
| chr13:77648403-77648443 | Ch13 region 3 |
| chr13:77648515-77648555 | Ch13 region 3 |
| chr13:77648602-77648642 | Ch13 region 3 |
| chr13:77648693-77648733 | Ch13 region 3 |
| chr13:77648786-77648826 | Ch13 region 3 |
| chr13:77648874-77648914 | Ch13 region 3 |

| | |
|---|---|
| chr13:77648930-77648970 | Ch13 region 3 |
| chr13:77649021-77649061 | Ch13 region 3 |
| chr13:77649163-77649203 | Ch13 region 3 |
| chr13:77649235-77649275 | Ch13 region 3 |
| chr13:77649317-77649357 | Ch13 region 3 |
| chr13:77649484-77649524 | Ch13 region 3 |
| chr13:77649529-77649569 | Ch13 region 3 |
| chr13:77649583-77649623 | Ch13 region 3 |
| chr13:77649631-77649671 | Ch13 region 3 |
| chr13:77649703-77649743 | Ch13 region 3 |
| chr13:77649781-77649821 | Ch13 region 3 |
| chr13:77649859-77649899 | Ch13 region 3 |
| chr13:77649910-77649950 | Ch13 region 3 |
| chr13:77649961-77650001 | Ch13 region 3 |
| chr13:77650022-77650062 | Ch13 region 3 |
| chr13:77650120-77650160 | Ch13 region 3 |
| chr13:77650203-77650243 | Ch13 region 3 |
| chr13:77650325-77650365 | Ch13 region 3 |
| chr13:77650374-77650414 | Ch13 region 3 |
| chr13:77650481-77650521 | Ch13 region 3 |
| chr13:77650584-77650624 | Ch13 region 3 |
| chr13:77650648-77650688 | Ch13 region 3 |
| chr13:77650885-77650925 | Ch13 region 3 |
| chr13:77650964-77651004 | Ch13 region 3 |
| chr13:77651023-77651063 | Ch13 region 3 |
| chr13:77651129-77651169 | Ch13 region 3 |
| chr13:77651239-77651279 | Ch13 region 3 |
| chr13:77651310-77651350 | Ch13 region 3 |
| chr13:77651375-77651415 | Ch13 region 3 |
| chr13:77651448-77651488 | Ch13 region 3 |
| chr13:77651523-77651563 | Ch13 region 3 |
| chr13:77651582-77651622 | Ch13 region 3 |
| chr13:77651642-77651682 | Ch13 region 3 |
| chr13:77651728-77651768 | Ch13 region 3 |
| chr13:77651775-77651815 | Ch13 region 3 |
| chr13:77651852-77651892 | Ch13 region 3 |
| chr13:77651919-77651959 | Ch13 region 3 |
| chr13:77652137-77652177 | Ch13 region 3 |
| chr13:77652351-77652391 | Ch13 region 3 |
| chr13:77652425-77652465 | Ch13 region 3 |
| chr13:77652477-77652517 | Ch13 region 3 |
| chr13:77652566-77652606 | Ch13 region 3 |
| chr13:77652620-77652660 | Ch13 region 3 |
| chr13:77652689-77652729 | Ch13 region 3 |
| chr13:77652738-77652778 | Ch13 region 3 |
| chr13:77652791-77652831 | Ch13 region 3 |
| chr13:77652850-77652890 | Ch13 region 3 |
| chr13:77652949-77652989 | Ch13 region 3 |
| chr13:77653051-77653091 | Ch13 region 3 |
| chr13:77653135-77653175 | Ch13 region 3 |
| chr13:77653187-77653227 | Ch13 region 3 |
| chr13:77653247-77653287 | Ch13 region 3 |

| | |
|---|---|
| chr13:77653314-77653354 | Ch13 region 3 |
| chr13:77653377-77653417 | Ch13 region 3 |
| chr13:77653442-77653482 | Ch13 region 3 |
| chr13:77653500-77653540 | Ch13 region 3 |
| chr13:77653574-77653614 | Ch13 region 3 |
| chr13:77653628-77653668 | Ch13 region 3 |
| chr13:77653673-77653713 | Ch13 region 3 |
| chr13:77653728-77653768 | Ch13 region 3 |
| chr13:77653850-77653890 | Ch13 region 3 |
| chr13:77653896-77653936 | Ch13 region 3 |
| chr13:77653986-77654026 | Ch13 region 3 |
| chr13:77654103-77654143 | Ch13 region 3 |
| chr13:77654149-77654189 | Ch13 region 3 |
| chr13:77654245-77654285 | Ch13 region 3 |
| chr13:77654304-77654344 | Ch13 region 3 |
| chr13:77654363-77654403 | Ch13 region 3 |
| chr13:77654411-77654451 | Ch13 region 3 |
| chr13:77654499-77654539 | Ch13 region 3 |
| chr13:77654547-77654587 | Ch13 region 3 |
| chr13:77654616-77654656 | Ch13 region 3 |
| chr13:77654677-77654717 | Ch13 region 3 |
| chr13:77654746-77654786 | Ch13 region 3 |
| chr13:77654856-77654896 | Ch13 region 3 |
| chr13:77655100-77655140 | Ch13 region 3 |
| chr13:77655146-77655186 | Ch13 region 3 |
| chr13:77655203-77655243 | Ch13 region 3 |
| chr13:77655256-77655296 | Ch13 region 3 |
| chr13:77655329-77655369 | Ch13 region 3 |
| chr13:77655437-77655477 | Ch13 region 3 |
| chr13:77655490-77655530 | Ch13 region 3 |
| chr13:77655587-77655627 | Ch13 region 3 |
| chr13:77655647-77655687 | Ch13 region 3 |
| chr13:77655711-77655751 | Ch13 region 3 |
| chr13:77655913-77655953 | Ch13 region 3 |
| chr13:77656010-77656050 | Ch13 region 3 |
| chr13:77656061-77656101 | Ch13 region 3 |
| chr13:77656106-77656146 | Ch13 region 3 |
| chr13:77656182-77656222 | Ch13 region 3 |
| chr13:77656314-77656354 | Ch13 region 3 |
| chr13:77656359-77656399 | Ch13 region 3 |
| chr13:77656433-77656473 | Ch13 region 3 |
| chr13:77656492-77656532 | Ch13 region 3 |
| chr13:77656541-77656581 | Ch13 region 3 |
| chr13:77656636-77656676 | Ch13 region 3 |
| chr13:77656708-77656748 | Ch13 region 3 |
| chr13:77656776-77656816 | Ch13 region 3 |
| chr13:77656862-77656902 | Ch13 region 3 |
| chr13:77656907-77656947 | Ch13 region 3 |
| chr13:77656965-77657005 | Ch13 region 3 |
| chr13:77657036-77657076 | Ch13 region 3 |
| chr13:77657105-77657145 | Ch13 region 3 |
| chr13:77657150-77657190 | Ch13 region 3 |

| | |
|---|---|
| chr13:77657231-77657271 | Ch13 region 3 |
| chr13:77657297-77657337 | Ch13 region 3 |
| chr13:77657346-77657386 | Ch13 region 3 |
| chr13:77657499-77657539 | Ch13 region 3 |
| chr13:77657553-77657593 | Ch13 region 3 |
| chr13:77657715-77657755 | Ch13 region 3 |
| chr13:77657787-77657827 | Ch13 region 3 |
| chr13:77657832-77657872 | Ch13 region 3 |
| chr13:77657944-77657984 | Ch13 region 3 |
| chr13:77658006-77658046 | Ch13 region 3 |
| chr13:77658059-77658099 | Ch13 region 3 |
| chr13:77658186-77658226 | Ch13 region 3 |
| chr13:77658265-77658305 | Ch13 region 3 |
| chr13:77658363-77658403 | Ch13 region 3 |
| chr13:77658438-77658478 | Ch13 region 3 |
| chr13:77658499-77658539 | Ch13 region 3 |
| chr13:77658603-77658643 | Ch13 region 3 |
| chr13:77658654-77658694 | Ch13 region 3 |
| chr13:77658735-77658775 | Ch13 region 3 |
| chr13:77658781-77658821 | Ch13 region 3 |
| chr13:77658854-77658894 | Ch13 region 3 |
| chr13:77658919-77658959 | Ch13 region 3 |
| chr13:77658966-77659006 | Ch13 region 3 |
| chr13:77659028-77659068 | Ch13 region 3 |
| chr13:77659094-77659134 | Ch13 region 3 |
| chr13:77659148-77659188 | Ch13 region 3 |
| chr13:77659201-77659241 | Ch13 region 3 |
| chr13:77659246-77659286 | Ch13 region 3 |
| chr13:77659313-77659353 | Ch13 region 3 |
| chr13:77659404-77659444 | Ch13 region 3 |
| chr13:77659471-77659511 | Ch13 region 3 |
| chr13:77659567-77659607 | Ch13 region 3 |
| chr13:77659615-77659655 | Ch13 region 3 |
| chr13:77659661-77659701 | Ch13 region 3 |
| chr13:77659722-77659762 | Ch13 region 3 |
| chr13:77659810-77659850 | Ch13 region 3 |
| chr13:77659899-77659939 | Ch13 region 3 |
| chr13:77659963-77660003 | Ch13 region 3 |
| chr13:77660068-77660108 | Ch13 region 3 |
| chr13:77660133-77660173 | Ch13 region 3 |
| chr13:77660240-77660280 | Ch13 region 3 |
| chr13:77660339-77660379 | Ch13 region 3 |
| chr13:77660434-77660474 | Ch13 region 3 |
| chr13:77660508-77660548 | Ch13 region 3 |
| chr13:77660572-77660612 | Ch13 region 3 |
| chr13:77660647-77660687 | Ch13 region 3 |
| chr13:77660698-77660738 | Ch13 region 3 |
| chr13:77660766-77660806 | Ch13 region 3 |
| chr13:77660815-77660855 | Ch13 region 3 |
| chr13:77661133-77661173 | Ch13 region 3 |
| chr13:77661178-77661218 | Ch13 region 3 |
| chr13:77661240-77661280 | Ch13 region 3 |

| | |
|---|---|
| chr13:77661341-77661381 | Ch13 region 3 |
| chr13:77661404-77661444 | Ch13 region 3 |
| chr13:77661456-77661496 | Ch13 region 3 |
| chr13:77661504-77661544 | Ch13 region 3 |
| chr13:77661634-77661674 | Ch13 region 3 |
| chr13:77661710-77661750 | Ch13 region 3 |
| chr13:77661758-77661798 | Ch13 region 3 |
| chr13:77661822-77661862 | Ch13 region 3 |
| chr13:77661987-77662027 | Ch13 region 3 |
| chr13:77662198-77662238 | Ch13 region 3 |
| chr13:77662244-77662284 | Ch13 region 3 |
| chr13:77662383-77662423 | Ch13 region 3 |
| chr13:77662428-77662468 | Ch13 region 3 |
| chr13:77662478-77662518 | Ch13 region 3 |
| chr13:77662549-77662589 | Ch13 region 3 |
| chr13:77662607-77662647 | Ch13 region 3 |
| chr13:77662864-77662904 | Ch13 region 3 |
| chr13:77662956-77662996 | Ch13 region 3 |
| chr13:77663001-77663041 | Ch13 region 3 |
| chr13:77663046-77663086 | Ch13 region 3 |
| chr13:77663137-77663177 | Ch13 region 3 |
| chr13:77663241-77663281 | Ch13 region 3 |
| chr13:77663351-77663391 | Ch13 region 3 |
| chr13:77663460-77663500 | Ch13 region 3 |
| chr13:77663658-77663698 | Ch13 region 3 |
| chr13:77663706-77663746 | Ch13 region 3 |
| chr13:77663751-77663791 | Ch13 region 3 |
| chr13:77663820-77663860 | Ch13 region 3 |
| chr13:77663881-77663921 | Ch13 region 3 |
| chr13:77663958-77663998 | Ch13 region 3 |
| chr13:77664033-77664073 | Ch13 region 3 |
| chr13:77664105-77664145 | Ch13 region 3 |
| chr13:77664211-77664251 | Ch13 region 3 |
| chr13:77664283-77664323 | Ch13 region 3 |
| chr13:77664348-77664388 | Ch13 region 3 |
| chr13:77664400-77664440 | Ch13 region 3 |
| chr13:77664478-77664518 | Ch13 region 3 |
| chr13:77664556-77664596 | Ch13 region 3 |
| chr13:77664648-77664688 | Ch13 region 3 |
| chr13:77664706-77664746 | Ch13 region 3 |
| chr13:77664813-77664853 | Ch13 region 3 |
| chr13:77664887-77664927 | Ch13 region 3 |
| chr13:77665133-77665173 | Ch13 region 3 |
| chr13:77665180-77665220 | Ch13 region 3 |
| chr13:77665232-77665272 | Ch13 region 3 |
| chr13:77665297-77665337 | Ch13 region 3 |
| chr13:77665372-77665412 | Ch13 region 3 |
| chr13:77665467-77665507 | Ch13 region 3 |
| chr13:77665596-77665636 | Ch13 region 3 |
| chr13:77665667-77665707 | Ch13 region 3 |
| chr13:77665750-77665790 | Ch13 region 3 |
| chr13:77665802-77665842 | Ch13 region 3 |

| | |
|---|---|
| chr13:77665999-77666039 | Ch13 region 3 |
| chr13:77666100-77666140 | Ch13 region 3 |
| chr13:77666176-77666216 | Ch13 region 3 |
| chr13:77666226-77666266 | Ch13 region 3 |
| chr13:77666297-77666337 | Ch13 region 3 |
| chr13:77666351-77666391 | Ch13 region 3 |
| chr13:77666485-77666525 | Ch13 region 3 |
| chr13:77666544-77666584 | Ch13 region 3 |
| chr13:77666693-77666733 | Ch13 region 3 |
| chr13:77666846-77666886 | Ch13 region 3 |
| chr13:77666906-77666946 | Ch13 region 3 |
| chr13:77666969-77667009 | Ch13 region 3 |
| chr13:77667022-77667062 | Ch13 region 3 |
| chr13:77667073-77667113 | Ch13 region 3 |
| chr13:77667136-77667176 | Ch13 region 3 |
| chr13:77667195-77667235 | Ch13 region 3 |
| chr13:77667245-77667285 | Ch13 region 3 |
| chr13:77667307-77667347 | Ch13 region 3 |
| chr13:77667376-77667416 | Ch13 region 3 |
| chr13:77667434-77667474 | Ch13 region 3 |
| chr13:77667504-77667544 | Ch13 region 3 |
| chr13:77667574-77667614 | Ch13 region 3 |
| chr13:77667620-77667660 | Ch13 region 3 |
| chr13:77667686-77667726 | Ch13 region 3 |
| chr13:77667744-77667784 | Ch13 region 3 |
| chr13:77667808-77667848 | Ch13 region 3 |
| chr13:77667853-77667893 | Ch13 region 3 |
| chr13:77667907-77667947 | Ch13 region 3 |
| chr13:77667975-77668015 | Ch13 region 3 |
| chr13:77668099-77668139 | Ch13 region 3 |
| chr13:77668160-77668200 | Ch13 region 3 |
| chr13:77668276-77668316 | Ch13 region 3 |
| chr13:77668342-77668382 | Ch13 region 3 |
| chr13:77668418-77668458 | Ch13 region 3 |
| chr13:77668465-77668505 | Ch13 region 3 |
| chr13:77668515-77668555 | Ch13 region 3 |
| chr13:77668586-77668626 | Ch13 region 3 |
| chr13:77668686-77668726 | Ch13 region 3 |
| chr13:77668731-77668771 | Ch13 region 3 |
| chr13:77668785-77668825 | Ch13 region 3 |
| chr13:77668830-77668870 | Ch13 region 3 |
| chr13:77668938-77668978 | Ch13 region 3 |
| chr13:77669037-77669077 | Ch13 region 3 |
| chr13:77669082-77669122 | Ch13 region 3 |
| chr13:77669163-77669203 | Ch13 region 3 |
| chr13:77669208-77669248 | Ch13 region 3 |
| chr13:77669311-77669351 | Ch13 region 3 |
| chr13:77669408-77669448 | Ch13 region 3 |
| chr13:77669471-77669511 | Ch13 region 3 |
| chr13:77669550-77669590 | Ch13 region 3 |
| chr13:77669627-77669667 | Ch13 region 3 |
| chr13:77669701-77669741 | Ch13 region 3 |

| | |
|---|---|
| chr13:77669869-77669909 | Ch13 region 3 |
| chr13:77670022-77670062 | Ch13 region 3 |
| chr13:77670073-77670113 | Ch13 region 3 |
| chr13:77670120-77670160 | Ch13 region 3 |
| chr13:77670206-77670246 | Ch13 region 3 |
| chr13:77670418-77670458 | Ch13 region 3 |
| chr13:77670480-77670520 | Ch13 region 3 |
| chr13:77670529-77670569 | Ch13 region 3 |
| chr13:77670579-77670619 | Ch13 region 3 |
| chr13:77670649-77670689 | Ch13 region 3 |
| chr13:77670776-77670816 | Ch13 region 3 |
| chr13:77671070-77671110 | Ch13 region 3 |
| chr13:77671151-77671191 | Ch13 region 3 |
| chr13:77671204-77671244 | Ch13 region 3 |
| chr13:77671295-77671335 | Ch13 region 3 |
| chr13:77671469-77671509 | Ch13 region 3 |
| chr13:77671548-77671588 | Ch13 region 3 |
| chr13:77671606-77671646 | Ch13 region 3 |
| chr13:77671667-77671707 | Ch13 region 3 |
| chr13:77671730-77671770 | Ch13 region 3 |
| chr13:77671776-77671816 | Ch13 region 3 |
| chr13:77671825-77671865 | Ch13 region 3 |
| chr13:77671908-77671948 | Ch13 region 3 |
| chr13:77671964-77672004 | Ch13 region 3 |
| chr13:77672033-77672073 | Ch13 region 3 |
| chr13:77672083-77672123 | Ch13 region 3 |
| chr13:77672129-77672169 | Ch13 region 3 |
| chr13:77672217-77672257 | Ch13 region 3 |
| chr13:77672272-77672312 | Ch13 region 3 |
| chr13:77672336-77672376 | Ch13 region 3 |
| chr13:77672389-77672429 | Ch13 region 3 |
| chr13:77672447-77672487 | Ch13 region 3 |
| chr13:77672519-77672559 | Ch13 region 3 |
| chr13:77672567-77672607 | Ch13 region 3 |
| chr13:77672614-77672654 | Ch13 region 3 |
| chr13:77672706-77672746 | Ch13 region 3 |
| chr13:77672751-77672791 | Ch13 region 3 |
| chr13:77672799-77672839 | Ch13 region 3 |
| chr13:77672889-77672929 | Ch13 region 3 |
| chr13:77672966-77673006 | Ch13 region 3 |
| chr13:77673031-77673071 | Ch13 region 3 |
| chr13:77673091-77673131 | Ch13 region 3 |
| chr13:77673136-77673176 | Ch13 region 3 |
| chr13:77673330-77673370 | Ch13 region 3 |
| chr13:77673395-77673435 | Ch13 region 3 |
| chr13:77673464-77673504 | Ch13 region 3 |
| chr13:77673534-77673574 | Ch13 region 3 |
| chr13:77673589-77673629 | Ch13 region 3 |
| chr13:77673655-77673695 | Ch13 region 3 |
| chr13:77673705-77673745 | Ch13 region 3 |
| chr13:77673766-77673806 | Ch13 region 3 |
| chr13:77673866-77673906 | Ch13 region 3 |

| | |
|---|---|
| chr13:77673935-77673975 | Ch13 region 3 |
| chr13:77674006-77674046 | Ch13 region 3 |
| chr13:77674090-77674130 | Ch13 region 3 |
| chr13:77674164-77674204 | Ch13 region 3 |
| chr13:77674266-77674306 | Ch13 region 3 |
| chr13:77674350-77674390 | Ch13 region 3 |
| chr13:77674401-77674441 | Ch13 region 3 |
| chr13:77674486-77674526 | Ch13 region 3 |
| chr13:77674584-77674624 | Ch13 region 3 |
| chr13:77674651-77674691 | Ch13 region 3 |
| chr13:77674711-77674751 | Ch13 region 3 |
| chr13:77674764-77674804 | Ch13 region 3 |
| chr8:127827010-127827050 | Myc |
| chr8:127827060-127827100 | Myc |
| chr8:127827232-127827272 | Myc |
| chr8:127827282-127827322 | Myc |
| chr8:127827632-127827672 | Myc |
| chr8:127827851-127827891 | Myc |
| chr8:127828064-127828104 | Myc |
| chr8:127828328-127828368 | Myc |
| chr8:127828419-127828459 | Myc |
| chr8:127828824-127828864 | Myc |
| chr8:127829091-127829131 | Myc |
| chr8:127829169-127829209 | Myc |
| chr8:127829229-127829269 | Myc |
| chr8:127829295-127829335 | Myc |
| chr8:127829367-127829407 | Myc |
| chr8:127829484-127829524 | Myc |
| chr8:127829551-127829591 | Myc |
| chr8:127829633-127829673 | Myc |
| chr8:127829695-127829735 | Myc |
| chr8:127829809-127829849 | Myc |
| chr8:127829921-127829961 | Myc |
| chr8:127830185-127830225 | Myc |
| chr8:127830270-127830310 | Myc |
| chr8:127830411-127830451 | Myc |
| chr8:127830652-127830692 | Myc |
| chr8:127830711-127830751 | Myc |
| chr8:127830766-127830806 | Myc |
| chr8:127831334-127831374 | Myc |
| chr8:127831398-127831438 | Myc |
| chr8:127831523-127831563 | Myc |
| chr8:127831573-127831613 | Myc |
| chr8:127831638-127831678 | Myc |
| chr8:127831756-127831796 | Myc |
| chr8:127831856-127831896 | Myc |
| chr8:127831924-127831964 | Myc |
| chr8:127831974-127832014 | Myc |
| chr8:127832024-127832064 | Myc |
| chr8:127832106-127832146 | Myc |
| chr8:127832156-127832196 | Myc |
| chr8:127832219-127832259 | Myc |

| | |
|---|---|
| chr8:127832331-127832371 | Myc |
| chr8:127832434-127832474 | Myc |
| chr8:127832484-127832524 | Myc |
| chr8:127832556-127832596 | Myc |
| chr8:127832623-127832663 | Myc |
| chr8:127832761-127832801 | Myc |
| chr8:127832815-127832855 | Myc |
| chr8:127832872-127832912 | Myc |
| chr8:127832954-127832994 | Myc |
| chr8:127833010-127833050 | Myc |
| chr8:127833100-127833140 | Myc |
| chr8:127833243-127833283 | Myc |
| chr8:127833293-127833333 | Myc |
| chr8:127833385-127833425 | Myc |
| chr8:127833445-127833485 | Myc |
| chr8:127833499-127833539 | Myc |
| chr8:127833549-127833589 | Myc |
| chr8:127833677-127833717 | Myc |
| chr8:127833769-127833809 | Myc |
| chr8:127833924-127833964 | Myc |
| chr8:127833974-127834014 | Myc |
| chr8:127834060-127834100 | Myc |
| chr8:127834132-127834172 | Myc |
| chr8:127834254-127834294 | Myc |
| chr8:127834310-127834350 | Myc |
| chr8:127834491-127834531 | Myc |
| chr8:127834696-127834736 | Myc |
| chr8:127834819-127834859 | Myc |
| chr8:127834870-127834910 | Myc |
| chr8:127834963-127835003 | Myc |
| chr8:127835067-127835107 | Myc |
| chr8:127835119-127835159 | Myc |
| chr8:127835181-127835221 | Myc |
| chr8:127835335-127835375 | Myc |
| chr8:127835421-127835461 | Myc |
| chr8:127835531-127835571 | Myc |
| chr8:127835604-127835644 | Myc |
| chr8:127835664-127835704 | Myc |
| chr8:127835782-127835822 | Myc |
| chr8:127835838-127835878 | Myc |
| chr8:127835890-127835930 | Myc |
| chr8:127835990-127836030 | Myc |
| chr8:127836141-127836181 | Myc |
| chr8:127836250-127836290 | Myc |
| chr8:127836326-127836366 | Myc |
| chr8:127836424-127836464 | Myc |
| chr8:127836480-127836520 | Myc |
| chr8:127836624-127836664 | Myc |
| chr8:127836691-127836731 | Myc |
| chr8:127836800-127836840 | Myc |
| chr8:127836925-127836965 | Myc |
| chr8:127836976-127837016 | Myc |

| | |
|---|---|
| chr8:127837036-127837076 | Myc |
| chr8:127837109-127837149 | Myc |
| chr8:127837192-127837232 | Myc |
| chr8:127837292-127837332 | Myc |
| chr8:127837364-127837404 | Myc |
| chr8:127837440-127837480 | Myc |
| chr8:127837504-127837544 | Myc |
| chr8:127837617-127837657 | Myc |
| chr8:127837753-127837793 | Myc |
| chr8:127837865-127837905 | Myc |
| chr8:127837940-127837980 | Myc |
| chr8:127838030-127838070 | Myc |
| chr8:127838193-127838233 | Myc |
| chr8:127838350-127838390 | Myc |
| chr8:127838436-127838476 | Myc |
| chr8:127838489-127838529 | Myc |
| chr8:127838567-127838607 | Myc |
| chr8:127838705-127838745 | Myc |
| chr8:127838760-127838800 | Myc |
| chr8:127838822-127838862 | Myc |
| chr8:127838879-127838919 | Myc |
| chr8:127838965-127839005 | Myc |
| chr8:127839018-127839058 | Myc |
| chr8:127839152-127839192 | Myc |
| chr8:127839217-127839257 | Myc |
| chr8:127839281-127839321 | Myc |
| chr8:127839342-127839382 | Myc |
| chr8:127839411-127839451 | Myc |
| chr8:127839528-127839568 | Myc |
| chr8:127839578-127839618 | Myc |
| chr8:127839735-127839775 | Myc |
| chr8:127839835-127839875 | Myc |
| chr8:127839941-127839981 | Myc |
| chr8:127840021-127840061 | Myc |
| chr8:127840132-127840172 | Myc |
| chr8:127840223-127840263 | Myc |
| chr8:127840310-127840350 | Myc |
| chr8:127840361-127840401 | Myc |
| chr8:127840482-127840522 | Myc |
| chr8:127840538-127840578 | Myc |
| chr8:127840642-127840682 | Myc |
| chr8:127840692-127840732 | Myc |
| chr8:127840775-127840815 | Myc |
| chr8:127840860-127840900 | Myc |
| chr8:127840921-127840961 | Myc |
| chr8:127841009-127841049 | Myc |
| chr8:127841138-127841178 | Myc |
| chr8:127841225-127841265 | Myc |
| chr8:127841275-127841315 | Myc |
| chr8:127841326-127841366 | Myc |
| chr8:127841422-127841462 | Myc |
| chr8:127841588-127841628 | Myc |

| | |
|---|---|
| chr8:127841640-127841680 | Myc |
| chr8:127841745-127841785 | Myc |
| chr8:127841840-127841880 | Myc |
| chr8:127841942-127841982 | Myc |
| chr8:127842012-127842052 | Myc |
| chr8:127842107-127842147 | Myc |
| chr8:127842199-127842239 | Myc |
| chr8:127842299-127842339 | Myc |
| chr8:127842387-127842427 | Myc |
| chr8:127842503-127842543 | Myc |
| chr8:127842561-127842601 | Myc |
| chr8:127842687-127842727 | Myc |
| chr8:127842778-127842818 | Myc |
| chr8:127842829-127842869 | Myc |
| chr8:127842881-127842921 | Myc |
| chr8:127842939-127842979 | Myc |
| chr8:127843111-127843151 | Myc |
| chr8:127843188-127843228 | Myc |
| chr8:127843262-127843302 | Myc |
| chr8:127843367-127843407 | Myc |
| chr8:127843426-127843466 | Myc |
| chr8:127843622-127843662 | Myc |
| chr8:127843690-127843730 | Myc |
| chr8:127843765-127843805 | Myc |
| chr8:127843823-127843863 | Myc |
| chr8:127843941-127843981 | Myc |
| chr8:127844033-127844073 | Myc |
| chr8:127844130-127844170 | Myc |
| chr8:127844230-127844270 | Myc |
| chr8:127844283-127844323 | Myc |
| chr8:127844364-127844404 | Myc |
| chr8:127844476-127844516 | Myc |
| chr8:127844562-127844602 | Myc |
| chr8:127844676-127844716 | Myc |
| chr8:127844827-127844867 | Myc |
| chr8:127844878-127844918 | Myc |
| chr8:127844975-127845015 | Myc |
| chr8:127845086-127845126 | Myc |
| chr8:127845139-127845179 | Myc |
| chr8:127845198-127845238 | Myc |
| chr8:127845248-127845288 | Myc |
| chr8:127845347-127845387 | Myc |
| chr8:127845397-127845437 | Myc |
| chr8:127845511-127845551 | Myc |
| chr8:127845610-127845650 | Myc |
| chr8:127845660-127845700 | Myc |
| chr8:127845710-127845750 | Myc |
| chr8:127845760-127845800 | Myc |
| chr8:127845868-127845908 | Myc |
| chr8:127845935-127845975 | Myc |
| chr8:127846003-127846043 | Myc |
| chr8:127846053-127846093 | Myc |

| | |
|---|---|
| chr8:127846126-127846166 | Myc |
| chr8:127846201-127846241 | Myc |
| chr8:127846269-127846309 | Myc |
| chr8:127846319-127846359 | Myc |
| chr8:127846372-127846412 | Myc |
| chr8:127846479-127846519 | Myc |
| chr8:127846529-127846569 | Myc |
| chr8:127846634-127846674 | Myc |
| chr8:127846684-127846724 | Myc |
| chr8:127846777-127846817 | Myc |
| chr8:127846862-127846902 | Myc |
| chr8:127847080-127847120 | Myc |
| chr8:127847235-127847275 | Myc |
| chr8:127847306-127847346 | Myc |
| chr8:127847374-127847414 | Myc |
| chr8:127847447-127847487 | Myc |
| chr8:127847497-127847537 | Myc |
| chr8:127847548-127847588 | Myc |
| chr8:127847626-127847666 | Myc |
| chr8:127847731-127847771 | Myc |
| chr8:127847781-127847821 | Myc |
| chr8:127847849-127847889 | Myc |
| chr8:127847908-127847948 | Myc |
| chr8:127848009-127848049 | Myc |
| chr8:127848060-127848100 | Myc |
| chr8:127848124-127848164 | Myc |
| chr8:127848197-127848237 | Myc |
| chr8:127848283-127848323 | Myc |
| chr8:127848511-127848551 | Myc |
| chr8:127848802-127848842 | Myc |
| chr8:127849057-127849097 | Myc |
| chr8:127849380-127849420 | Myc |
| chr8:127849465-127849505 | Myc |
| chr8:127849515-127849555 | Myc |
| chr8:127849600-127849640 | Myc |
| chr8:127849663-127849703 | Myc |
| chr8:127849771-127849811 | Myc |
| chr8:127849823-127849863 | Myc |
| chr8:127849876-127849916 | Myc |
| chr8:127849935-127849975 | Myc |
| chr8:127849994-127850034 | Myc |
| chr8:127850056-127850096 | Myc |
| chr8:127850144-127850184 | Myc |
| chr8:127850229-127850269 | Myc |
| chr8:127850350-127850390 | Myc |
| chr8:127850401-127850441 | Myc |
| chr8:127850493-127850533 | Myc |
| chr8:127850566-127850606 | Myc |
| chr8:127850673-127850713 | Myc |
| chr8:127850779-127850819 | Myc |
| chr8:127850912-127850952 | Myc |
| chr8:127851013-127851053 | Myc |

| | |
|---|---|
| chr8:127851072-127851112 | Myc |
| chr8:127851163-127851203 | Myc |
| chr8:127851214-127851254 | Myc |
| chr8:127851277-127851317 | Myc |
| chr8:127851366-127851406 | Myc |
| chr8:127851469-127851509 | Myc |
| chr8:127851553-127851593 | Myc |
| chr8:127851660-127851700 | Myc |
| chr8:127851759-127851799 | Myc |
| chr8:127851821-127851861 | Myc |
| chr8:127851875-127851915 | Myc |
| chr8:127852005-127852045 | Myc |
| chr8:127852063-127852103 | Myc |
| chr8:127852113-127852153 | Myc |
| chr8:127852217-127852257 | Myc |
| chr8:127852312-127852352 | Myc |
| chr8:127852403-127852443 | Myc |
| chr8:127852540-127852580 | Myc |
| chr8:127852619-127852659 | Myc |
| chr8:127852672-127852712 | Myc |
| chr8:127852752-127852792 | Myc |
| chr8:127852816-127852856 | Myc |
| chr8:127852866-127852906 | Myc |
| chr8:127852995-127853035 | Myc |
| chr8:127853089-127853129 | Myc |
| chr8:127853172-127853212 | Myc |
| chr8:127853222-127853262 | Myc |
| chr8:127853328-127853368 | Myc |
| chr8:127853407-127853447 | Myc |
| chr8:127853509-127853549 | Myc |
| chr8:127853593-127853633 | Myc |
| chr8:127853652-127853692 | Myc |
| chr8:127853745-127853785 | Myc |
| chr8:127853977-127854017 | Myc |
| chr8:127854055-127854095 | Myc |
| chr8:127854135-127854175 | Myc |
| chr8:127854207-127854247 | Myc |
| chr8:127854471-127854511 | Myc |
| chr8:127854578-127854618 | Myc |
| chr8:127854681-127854721 | Myc |
| chr8:127854766-127854806 | Myc |
| chr8:127854833-127854873 | Myc |
| chr8:127854938-127854978 | Myc |
| chr8:127855125-127855165 | Myc |
| chr8:127855179-127855219 | Myc |
| chr8:127855487-127855527 | Myc |
| chr8:127855554-127855594 | Myc |
| chr8:127855604-127855644 | Myc |
| chr8:127855757-127855797 | Myc |
| chr8:127855819-127855859 | Myc |
| chr8:127855959-127855999 | Myc |
| chr8:127856040-127856080 | Myc |

| | |
|---|---|
| chr8:127856108-127856148 | Myc |
| chr8:127856211-127856251 | Myc |
| chr8:127856297-127856337 | Myc |
| chr8:127856499-127856539 | Myc |
| chr8:127856576-127856616 | Myc |
| chr8:127856626-127856666 | Myc |
| chr8:127856697-127856737 | Myc |
| chr8:127856763-127856803 | Myc |
| chr8:127856888-127856928 | Myc |
| chr8:127856956-127856996 | Myc |
| chr8:127857010-127857050 | Myc |
| chr8:127857156-127857196 | Myc |
| chr8:127857233-127857273 | Myc |
| chr8:127857332-127857372 | Myc |
| chr8:127857384-127857424 | Myc |
| chr8:127857491-127857531 | Myc |
| chr8:127857593-127857633 | Myc |
| chr8:127857701-127857741 | Myc |
| chr8:127857770-127857810 | Myc |
| chr8:127857896-127857936 | Myc |
| chr8:127857977-127858017 | Myc |
| chr8:127858037-127858077 | Myc |
| chr8:127858110-127858150 | Myc |
| chr8:127858193-127858233 | Myc |
| chr8:127858278-127858318 | Myc |
| chr8:127858339-127858379 | Myc |
| chr8:127858395-127858435 | Myc |
| chr8:127858503-127858543 | Myc |
| chr8:127858567-127858607 | Myc |
| chr8:127858685-127858725 | Myc |
| chr8:127858800-127858840 | Myc |
| chr8:127858857-127858897 | Myc |
| chr8:127858909-127858949 | Myc |
| chr8:127859076-127859116 | Myc |
| chr8:127859209-127859249 | Myc |
| chr8:127859290-127859330 | Myc |
| chr8:127859387-127859427 | Myc |
| chr8:127859514-127859554 | Myc |
| chr8:127859567-127859607 | Myc |
| chr8:127859622-127859662 | Myc |
| chr8:127859698-127859738 | Myc |
| chr8:127859748-127859788 | Myc |
| chr8:127859899-127859939 | Myc |
| chr8:127859987-127860027 | Myc |
| chr8:127860043-127860083 | Myc |
| chr8:127860107-127860147 | Myc |
| chr8:127860160-127860200 | Myc |
| chr8:127860211-127860251 | Myc |
| chr8:127860264-127860304 | Myc |
| chr8:127860444-127860484 | Myc |
| chr8:127860516-127860556 | Myc |
| chr8:127860607-127860647 | Myc |

| | |
|---|---|
| chr8:127860692-127860732 | Myc |
| chr8:127860776-127860816 | Myc |
| chr8:127860907-127860947 | Myc |
| chr8:127860962-127861002 | Myc |
| chr8:127861058-127861098 | Myc |
| chr8:127861113-127861153 | Myc |
| chr8:127861189-127861229 | Myc |
| chr8:127861345-127861385 | Myc |
| chr8:127861431-127861471 | Myc |
| chr8:127861484-127861524 | Myc |
| chr8:127861612-127861652 | Myc |
| chr8:127861672-127861712 | Myc |
| chr8:127861755-127861795 | Myc |
| chr8:127861805-127861845 | Myc |
| chr8:127861909-127861949 | Myc |
| chr8:127862005-127862045 | Myc |
| chr8:127862058-127862098 | Myc |
| chr8:127862146-127862186 | Myc |
| chr8:127862196-127862236 | Myc |
| chr8:127862280-127862320 | Myc |
| chr8:127862699-127862739 | Myc |
| chr8:127862754-127862794 | Myc |
| chr8:127862846-127862886 | Myc |
| chr8:127862909-127862949 | Myc |
| chr8:127862986-127863026 | Myc |
| chr8:127863061-127863101 | Myc |
| chr8:127863160-127863200 | Myc |
| chr8:127863227-127863267 | Myc |
| chr8:127863278-127863318 | Myc |
| chr8:127863330-127863370 | Myc |
| chr8:127863380-127863420 | Myc |
| chr8:127863508-127863548 | Myc |
| chr8:127863566-127863606 | Myc |
| chr8:127863628-127863668 | Myc |
| chr8:127863687-127863727 | Myc |
| chr8:127863748-127863788 | Myc |
| chr8:127863798-127863838 | Myc |
| chr8:127863863-127863903 | Myc |
| chr8:127863947-127863987 | Myc |
| chr8:127864003-127864043 | Myc |
| chr8:127864175-127864215 | Myc |
| chr8:127864300-127864340 | Myc |
| chr8:127864351-127864391 | Myc |
| chr8:127864427-127864467 | Myc |
| chr8:127864479-127864519 | Myc |
| chr8:127864566-127864606 | Myc |
| chr8:127864661-127864701 | Myc |
| chr8:127864719-127864759 | Myc |
| chr8:127864818-127864858 | Myc |
| chr8:127864868-127864908 | Myc |
| chr8:127865029-127865069 | Myc |
| chr8:127865148-127865188 | Myc |

| | |
|---|---|
| chr8:127865198-127865238 | Myc |
| chr8:127865282-127865322 | Myc |
| chr8:127865354-127865394 | Myc |
| chr8:127865413-127865453 | Myc |
| chr8:127865526-127865566 | Myc |
| chr8:127865620-127865660 | Myc |
| chr8:127865717-127865757 | Myc |
| chr8:127865843-127865883 | Myc |
| chr8:127865925-127865965 | Myc |
| chr8:127866029-127866069 | Myc |
| chr8:127866081-127866121 | Myc |
| chr8:127866170-127866210 | Myc |
| chr8:127866308-127866348 | Myc |
| chr8:127866358-127866398 | Myc |
| chr8:127866486-127866526 | Myc |
| chr8:127866576-127866616 | Myc |
| chr8:127866646-127866686 | Myc |
| chr8:127866712-127866752 | Myc |
| chr8:127866808-127866848 | Myc |
| chr8:127866861-127866901 | Myc |
| chr8:127866953-127866993 | Myc |
| chr8:127867088-127867128 | Myc |
| chr8:127867235-127867275 | Myc |
| chr8:127867341-127867381 | Myc |
| chr8:127867434-127867474 | Myc |
| chr8:127867534-127867574 | Myc |
| chr8:127867615-127867655 | Myc |
| chr8:127867708-127867748 | Myc |
| chr8:127867798-127867838 | Myc |
| chr8:127867915-127867955 | Myc |
| chr8:127868013-127868053 | Myc |
| chr8:127868071-127868111 | Myc |
| chr8:127868124-127868164 | Myc |
| chr8:127868174-127868214 | Myc |
| chr8:127868294-127868334 | Myc |
| chr8:127868428-127868468 | Myc |
| chr8:127868517-127868557 | Myc |
| chr8:127868634-127868674 | Myc |
| chr8:127868785-127868825 | Myc |
| chr8:127868835-127868875 | Myc |
| chr8:127868892-127868932 | Myc |
| chr8:127869045-127869085 | Myc |
| chr8:127869122-127869162 | Myc |
| chr8:127869205-127869245 | Myc |
| chr8:127869270-127869310 | Myc |
| chr8:127869338-127869378 | Myc |
| chr8:127869401-127869441 | Myc |
| chr8:127869462-127869502 | Myc |
| chr8:127869539-127869579 | Myc |
| chr8:127869593-127869633 | Myc |
| chr8:127869659-127869699 | Myc |
| chr8:127869739-127869779 | Myc |

| | |
|---|---|
| chr8:127869819-127869859 | Myc |
| chr8:127869869-127869909 | Myc |
| chr8:127870010-127870050 | Myc |
| chr8:127870109-127870149 | Myc |
| chr8:127870245-127870285 | Myc |
| chr8:127870340-127870380 | Myc |
| chr8:127870444-127870484 | Myc |
| chr8:127870528-127870568 | Myc |
| chr8:127870592-127870632 | Myc |
| chr8:127870656-127870696 | Myc |
| chr8:127870754-127870794 | Myc |
| chr8:127870844-127870884 | Myc |
| chr8:127870935-127870975 | Myc |
| chr8:127871012-127871052 | Myc |
| chr8:127871126-127871166 | Myc |
| chr8:127871318-127871358 | Myc |
| chr8:127871460-127871500 | Myc |
| chr8:127871625-127871665 | Myc |
| chr8:127871762-127871802 | Myc |
| chr8:127871854-127871894 | Myc |
| chr8:127871906-127871946 | Myc |
| chr8:127872006-127872046 | Myc |
| chr8:127872065-127872105 | Myc |
| chr8:127872158-127872198 | Myc |
| chr8:127872229-127872269 | Myc |
| chr8:127872297-127872337 | Myc |
| chr8:127872373-127872413 | Myc |
| chr8:127872486-127872526 | Myc |
| chr8:127872583-127872623 | Myc |
| chr8:127872725-127872765 | Myc |
| chr8:127872778-127872818 | Myc |
| chr8:127872840-127872880 | Myc |
| chr8:127873160-127873200 | Myc |
| chr8:127873270-127873310 | Myc |
| chr8:127873337-127873377 | Myc |
| chr8:127873462-127873502 | Myc |
| chr8:127873555-127873595 | Myc |
| chr8:127873605-127873645 | Myc |
| chr8:127873678-127873718 | Myc |
| chr8:127873747-127873787 | Myc |
| chr8:127873847-127873887 | Myc |
| chr8:127873901-127873941 | Myc |
| chr8:127873985-127874025 | Myc |
| chr8:127874046-127874086 | Myc |
| chr8:127874104-127874144 | Myc |
| chr8:127874178-127874218 | Myc |
| chr8:127874228-127874268 | Myc |
| chr8:127874310-127874350 | Myc |
| chr8:127874372-127874412 | Myc |
| chr8:127874445-127874485 | Myc |
| chr8:127874495-127874535 | Myc |
| chr8:127874611-127874651 | Myc |

| | |
|---|---|
| chr8:127874684-127874724 | Myc |
| chr8:127874858-127874898 | Myc |
| chr8:127874954-127874994 | Myc |
| chr8:127875004-127875044 | Myc |
| chr8:127875060-127875100 | Myc |
| chr8:127875130-127875170 | Myc |
| chr8:127875186-127875226 | Myc |
| chr8:127875246-127875286 | Myc |
| chr8:127875357-127875397 | Myc |
| chr8:127875470-127875510 | Myc |
| chr8:127875556-127875596 | Myc |
| chr8:127875612-127875652 | Myc |
| chr8:127875691-127875731 | Myc |
| chr8:127875746-127875786 | Myc |
| chr8:127875829-127875869 | Myc |
| chr8:127875920-127875960 | Myc |
| chr8:127876022-127876062 | Myc |
| chr8:127876102-127876142 | Myc |
| chr8:127876280-127876320 | Myc |
| chr8:127876377-127876417 | Myc |
| chr8:127876435-127876475 | Myc |
| chr8:127876541-127876581 | Myc |
| chr8:127876617-127876657 | Myc |
| chr8:127876667-127876707 | Myc |
| chr8:127876775-127876815 | Myc |
| chr8:127876881-127876921 | Myc |
| chr8:127876998-127877038 | Myc |
| chr8:127877065-127877105 | Myc |
| chr8:127877118-127877158 | Myc |
| chr8:127877259-127877299 | Myc |
| chr8:127877376-127877416 | Myc |
| chr8:127877456-127877496 | Myc |
| chr8:127877544-127877584 | Myc |
| chr8:127877720-127877760 | Myc |
| chr8:127877798-127877838 | Myc |
| chr8:127877848-127877888 | Myc |
| chr8:127877926-127877966 | Myc |
| chr8:127877976-127878016 | Myc |
| chr8:127878228-127878268 | Myc |
| chr8:127878308-127878348 | Myc |
| chr8:127878367-127878407 | Myc |
| chr8:127878417-127878457 | Myc |
| chr8:127878503-127878543 | Myc |
| chr8:127878553-127878593 | Myc |
| chr8:127878606-127878646 | Myc |
| chr8:127878659-127878699 | Myc |
| chr8:127878738-127878778 | Myc |
| chr8:127878790-127878830 | Myc |
| chr8:127878860-127878900 | Myc |
| chr8:127878913-127878953 | Myc |
| chr8:127879031-127879071 | Myc |
| chr8:127879131-127879171 | Myc |

| | |
|---|---|
| chr8:127879224-127879264 | Myc |
| chr8:127879280-127879320 | Myc |
| chr8:127879440-127879480 | Myc |
| chr8:127879520-127879560 | Myc |
| chr8:127879581-127879621 | Myc |
| chr8:127879634-127879674 | Myc |
| chr8:127879715-127879755 | Myc |
| chr8:127879800-127879840 | Myc |
| chr8:127879912-127879952 | Myc |
| chr8:127880260-127880300 | Myc |
| chr8:127880321-127880361 | Myc |
| chr8:127880394-127880434 | Myc |
| chr8:127880464-127880504 | Myc |
| chr8:127880565-127880605 | Myc |
| chr8:127880616-127880656 | Myc |
| chr8:127880667-127880707 | Myc |
| chr8:127880732-127880772 | Myc |
| chr8:127880891-127880931 | Myc |
| chr8:127881211-127881251 | Myc |
| chr8:127881274-127881314 | Myc |
| chr8:127881401-127881441 | Myc |
| chr8:127881451-127881491 | Myc |
| chr8:127881505-127881545 | Myc |
| chr8:127881611-127881651 | Myc |
| chr8:127881667-127881707 | Myc |
| chr8:127881717-127881757 | Myc |
| chr8:127881809-127881849 | Myc |
| chr8:127881859-127881899 | Myc |
| chr8:127881914-127881954 | Myc |
| chr8:127881985-127882025 | Myc |
| chr8:127882115-127882155 | Myc |
| chr8:127882214-127882254 | Myc |
| chr8:127882302-127882342 | Myc |
| chr8:127882353-127882393 | Myc |
| chr8:127882491-127882531 | Myc |
| chr8:127882760-127882800 | Myc |
| chr8:127882857-127882897 | Myc |
| chr8:127882950-127882990 | Myc |
| chr8:127883041-127883081 | Myc |
| chr8:127883101-127883141 | Myc |
| chr8:127883199-127883239 | Myc |
| chr8:127883292-127883332 | Myc |
| chr8:127883373-127883413 | Myc |
| chr8:127883434-127883474 | Myc |
| chr8:127883586-127883626 | Myc |
| chr8:127883636-127883676 | Myc |
| chr8:127883710-127883750 | Myc |
| chr8:127883833-127883873 | Myc |
| chr8:127883922-127883962 | Myc |
| chr8:127884014-127884054 | Myc |
| chr8:127884066-127884106 | Myc |
| chr8:127884185-127884225 | Myc |

| | |
|---|---|
| chr8:127884296-127884336 | Myc |
| chr8:127884429-127884469 | Myc |
| chr8:127884486-127884526 | Myc |
| chr8:127884636-127884676 | Myc |
| chr8:127884686-127884726 | Myc |
| chr8:127884867-127884907 | Myc |
| chr8:127885015-127885055 | Myc |
| chr8:127885070-127885110 | Myc |
| chr8:127885211-127885251 | Myc |
| chr8:127885293-127885333 | Myc |
| chr8:127885355-127885395 | Myc |
| chr8:127885413-127885453 | Myc |
| chr8:127885512-127885552 | Myc |
| chr8:127885622-127885662 | Myc |
| chr8:127885672-127885712 | Myc |
| chr8:127885742-127885782 | Myc |
| chr8:127885861-127885901 | Myc |
| chr8:127885956-127885996 | Myc |
| chr8:127886007-127886047 | Myc |
| chr8:127886079-127886119 | Myc |
| chr8:127886171-127886211 | Myc |
| chr8:127886270-127886310 | Myc |
| chr8:127886339-127886379 | Myc |
| chr8:127886402-127886442 | Myc |
| chr8:127886452-127886492 | Myc |
| chr8:127886506-127886546 | Myc |
| chr8:127886590-127886630 | Myc |
| chr8:127886662-127886702 | Myc |
| chr8:127886821-127886861 | Myc |
| chr8:127886872-127886912 | Myc |
| chr8:127887012-127887052 | Myc |
| chr8:127887076-127887116 | Myc |
| chr8:127887126-127887166 | Myc |
| chr8:127887222-127887262 | Myc |
| chr8:127887385-127887425 | Myc |
| chr8:127887469-127887509 | Myc |
| chr8:127887544-127887584 | Myc |
| chr8:127887726-127887766 | Myc |
| chr8:127887909-127887949 | Myc |
| chr8:127887991-127888031 | Myc |
| chr8:127888041-127888081 | Myc |
| chr8:127888091-127888131 | Myc |
| chr8:127888142-127888182 | Myc |
| chr8:127888269-127888309 | Myc |
| chr8:127888429-127888469 | Myc |
| chr8:127888508-127888548 | Myc |
| chr8:127888605-127888645 | Myc |
| chr8:127888660-127888700 | Myc |
| chr8:127888736-127888776 | Myc |
| chr8:127888791-127888831 | Myc |
| chr8:127888919-127888959 | Myc |
| chr8:127888970-127889010 | Myc |

| | |
|---|---|
| chr8:127889073-127889113 | Myc |
| chr8:127889123-127889163 | Myc |
| chr8:127889180-127889220 | Myc |
| chr8:127889278-127889318 | Myc |
| chr8:127889417-127889457 | Myc |
| chr8:127889482-127889522 | Myc |
| chr8:127889571-127889611 | Myc |
| chr8:127889637-127889677 | Myc |
| chr8:127889694-127889734 | Myc |
| chr8:127889752-127889792 | Myc |
| chr8:127889919-127889959 | Myc |
| chr8:127890001-127890041 | Myc |
| chr8:127890051-127890091 | Myc |
| chr8:127890105-127890145 | Myc |
| chr8:127890187-127890227 | Myc |
| chr8:127890242-127890282 | Myc |
| chr8:127890406-127890446 | Myc |
| chr8:127890456-127890496 | Myc |
| chr8:127890518-127890558 | Myc |
| chr8:127890645-127890685 | Myc |
| chr8:127890787-127890827 | Myc |
| chr8:127890851-127890891 | Myc |
| chr8:127890929-127890969 | Myc |
| chr8:127890997-127891037 | Myc |
| chr8:127891138-127891178 | Myc |
| chr8:127891210-127891250 | Myc |
| chr8:127891290-127891330 | Myc |
| chr8:127891344-127891384 | Myc |
| chr8:127891423-127891463 | Myc |
| chr8:127891473-127891513 | Myc |
| chr8:127891543-127891583 | Myc |
| chr8:127891620-127891660 | Myc |
| chr8:127891670-127891710 | Myc |
| chr8:127891781-127891821 | Myc |
| chr8:127891861-127891901 | Myc |
| chr8:127891929-127891969 | Myc |
| chr8:127892021-127892061 | Myc |
| chr8:127892116-127892156 | Myc |
| chr8:127892354-127892394 | Myc |
| chr8:127892405-127892445 | Myc |
| chr8:127892490-127892530 | Myc |
| chr8:127892540-127892580 | Myc |
| chr8:127892688-127892728 | Myc |
| chr8:127892783-127892823 | Myc |
| chr8:127892850-127892890 | Myc |
| chr8:127892949-127892989 | Myc |
| chr8:127893006-127893046 | Myc |
| chr8:127893108-127893148 | Myc |
| chr8:127893161-127893201 | Myc |
| chr8:127893254-127893294 | Myc |
| chr8:127893381-127893421 | Myc |
| chr8:127893509-127893549 | Myc |

| | |
|---|---|
| chr8:127893604-127893644 | Myc |
| chr8:127893738-127893778 | Myc |
| chr8:127893889-127893929 | Myc |
| chr8:127893984-127894024 | Myc |
| chr8:127894069-127894109 | Myc |
| chr8:127894191-127894231 | Myc |
| chr8:127894586-127894626 | Myc |
| chr8:127894636-127894676 | Myc |
| chr8:127894745-127894785 | Myc |
| chr8:127894813-127894853 | Myc |
| chr8:127894894-127894934 | Myc |
| chr8:127894944-127894984 | Myc |
| chr8:127895026-127895066 | Myc |
| chr8:127895153-127895193 | Myc |
| chr8:127895270-127895310 | Myc |
| chr8:127895320-127895360 | Myc |
| chr8:127895443-127895483 | Myc |
| chr8:127895493-127895533 | Myc |
| chr8:127895604-127895644 | Myc |
| chr8:127895733-127895773 | Myc |
| chr8:127895803-127895843 | Myc |
| chr8:127895881-127895921 | Myc |
| chr8:127895950-127895990 | Myc |
| chr8:127896266-127896306 | Myc |
| chr8:127896371-127896411 | Myc |
| chr8:127896421-127896461 | Myc |
| chr8:127896567-127896607 | Myc |
| chr8:127896656-127896696 | Myc |
| chr8:127896762-127896802 | Myc |
| chr8:127896871-127896911 | Myc |
| chr8:127896921-127896961 | Myc |
| chr8:127897027-127897067 | Myc |
| chr8:127897109-127897149 | Myc |
| chr8:127897196-127897236 | Myc |
| chr8:127897273-127897313 | Myc |
| chr8:127897325-127897365 | Myc |
| chr8:127897375-127897415 | Myc |
| chr8:127897441-127897481 | Myc |
| chr8:127897517-127897557 | Myc |
| chr8:127897590-127897630 | Myc |
| chr8:127897649-127897689 | Myc |
| chr8:127897731-127897771 | Myc |
| chr8:127897818-127897858 | Myc |
| chr8:127897926-127897966 | Myc |
| chr8:127897976-127898016 | Myc |
| chr8:127898039-127898079 | Myc |
| chr8:127898196-127898236 | Myc |
| chr8:127898330-127898370 | Myc |
| chr8:127898407-127898447 | Myc |
| chr8:127898533-127898573 | Myc |
| chr8:127898627-127898667 | Myc |
| chr8:127898694-127898734 | Myc |

| | |
|---|---|
| chr8:127898760-127898800 | Myc |
| chr8:127898837-127898877 | Myc |
| chr8:127898938-127898978 | Myc |
| chr8:127899105-127899145 | Myc |
| chr8:127899245-127899285 | Myc |
| chr8:127899307-127899347 | Myc |
| chr8:127899382-127899422 | Myc |
| chr8:127899437-127899477 | Myc |
| chr8:127899496-127899536 | Myc |
| chr8:127899568-127899608 | Myc |
| chr8:127899629-127899669 | Myc |
| chr8:127899751-127899791 | Myc |
| chr8:127899801-127899841 | Myc |
| chr8:127899911-127899951 | Myc |
| chr8:127900020-127900060 | Myc |
| chr8:127900071-127900111 | Myc |
| chr8:127900121-127900161 | Myc |
| chr8:127900171-127900211 | Myc |
| chr8:127900290-127900330 | Myc |
| chr8:127900397-127900437 | Myc |
| chr8:127900450-127900490 | Myc |
| chr8:127900523-127900563 | Myc |
| chr8:127900709-127900749 | Myc |
| chr8:127901457-127901497 | Myc |
| chr8:127901548-127901588 | Myc |
| chr8:127901611-127901651 | Myc |
| chr8:127901774-127901814 | Myc |
| chr8:127901852-127901892 | Myc |
| chr8:127901986-127902026 | Myc |
| chr8:127902036-127902076 | Myc |
| chr8:127902142-127902182 | Myc |
| chr8:127902249-127902289 | Myc |
| chr8:127902304-127902344 | Myc |
| chr8:127902390-127902430 | Myc |
| chr8:127902475-127902515 | Myc |
| chr8:127902534-127902574 | Myc |
| chr8:127902673-127902713 | Myc |
| chr8:127902797-127902837 | Myc |
| chr8:127902907-127902947 | Myc |
| chr8:127902993-127903033 | Myc |
| chr8:127903107-127903147 | Myc |
| chr8:127903157-127903197 | Myc |
| chr8:127903236-127903276 | Myc |
| chr8:127903348-127903388 | Myc |
| chr8:127903443-127903483 | Myc |
| chr8:127903493-127903533 | Myc |
| chr8:127903649-127903689 | Myc |
| chr8:127903726-127903766 | Myc |
| chr8:127903810-127903850 | Myc |
| chr8:127903865-127903905 | Myc |
| chr8:127903946-127903986 | Myc |
| chr8:127904116-127904156 | Myc |

| | |
|---|---|
| chr8:127904320-127904360 | Myc |
| chr8:127904412-127904452 | Myc |
| chr8:127904542-127904582 | Myc |
| chr8:127904615-127904655 | Myc |
| chr8:127904721-127904761 | Myc |
| chr8:127904850-127904890 | Myc |
| chr8:127904901-127904941 | Myc |
| chr8:127904979-127905019 | Myc |
| chr8:127905059-127905099 | Myc |
| chr8:127905274-127905314 | Myc |
| chr8:127905350-127905390 | Myc |
| chr8:127905423-127905463 | Myc |
| chr8:127905735-127905775 | Myc |
| chr8:127905786-127905826 | Myc |
| chr8:127905852-127905892 | Myc |
| chr8:127905994-127906034 | Myc |
| chr8:127906046-127906086 | Myc |
| chr8:127906251-127906291 | Myc |
| chr8:127906415-127906455 | Myc |
| chr8:127906716-127906756 | Myc |
| chr8:127906777-127906817 | Myc |
| chr8:127906827-127906867 | Myc |
| chr8:127906890-127906930 | Myc |
| chr8:127907020-127907060 | Myc |
| chr8:127907135-127907175 | Myc |
| chr8:127907204-127907244 | Myc |
| chr8:127907309-127907349 | Myc |
| chr8:127907361-127907401 | Myc |
| chr8:127907419-127907459 | Myc |
| chr8:127907586-127907626 | Myc |
| chr8:127907636-127907676 | Myc |
| chr8:127907686-127907726 | Myc |
| chr8:127907738-127907778 | Myc |
| chr8:127907869-127907909 | Myc |
| chr8:127907967-127908007 | Myc |
| chr8:127908029-127908069 | Myc |
| chr8:127908079-127908119 | Myc |
| chr8:127908138-127908178 | Myc |
| chr8:127908188-127908228 | Myc |
| chr8:127908269-127908309 | Myc |
| chr8:127908327-127908367 | Myc |
| chr8:127908377-127908417 | Myc |
| chr8:127908485-127908525 | Myc |
| chr8:127908643-127908683 | Myc |
| chr8:127908707-127908747 | Myc |
| chr8:127908772-127908812 | Myc |
| chr8:127908822-127908862 | Myc |
| chr8:127908916-127908956 | Myc |
| chr8:127908971-127909011 | Myc |
| chr8:127909062-127909102 | Myc |
| chr8:127909177-127909217 | Myc |
| chr8:127909256-127909296 | Myc |

| | |
|---|---|
| chr8:127909357-127909397 | Myc |
| chr8:127909413-127909453 | Myc |
| chr8:127909547-127909587 | Myc |
| chr8:127909603-127909643 | Myc |
| chr8:127909710-127909750 | Myc |
| chr8:127909833-127909873 | Myc |
| chr8:127909908-127909948 | Myc |
| chr8:127909959-127909999 | Myc |
| chr8:127910023-127910063 | Myc |
| chr8:127910189-127910229 | Myc |
| chr8:127910292-127910332 | Myc |
| chr8:127910482-127910522 | Myc |
| chr8:127910563-127910603 | Myc |
| chr8:127910665-127910705 | Myc |
| chr8:127910746-127910786 | Myc |
| chr8:127910851-127910891 | Myc |
| chr8:127910943-127910983 | Myc |
| chr8:127911019-127911059 | Myc |
| chr8:127911084-127911124 | Myc |
| chr8:127911212-127911252 | Myc |
| chr8:127911388-127911428 | Myc |
| chr8:127911459-127911499 | Myc |
| chr8:127911584-127911624 | Myc |
| chr8:127911648-127911688 | Myc |
| chr8:127911772-127911812 | Myc |
| chr8:127911864-127911904 | Myc |
| chr8:127912047-127912087 | Myc |
| chr8:127912137-127912177 | Myc |
| chr8:127912230-127912270 | Myc |
| chr8:127912335-127912375 | Myc |
| chr8:127912396-127912436 | Myc |
| chr8:127912480-127912520 | Myc |
| chr8:127912530-127912570 | Myc |
| chr8:127912709-127912749 | Myc |
| chr8:127912802-127912842 | Myc |
| chr8:127912965-127913005 | Myc |
| chr8:127913051-127913091 | Myc |
| chr8:127913170-127913210 | Myc |
| chr8:127913234-127913274 | Myc |
| chr8:127913521-127913561 | Myc |
| chr8:127913623-127913663 | Myc |
| chr8:127913950-127913990 | Myc |
| chr8:127914066-127914106 | Myc |
| chr8:127914232-127914272 | Myc |
| chr8:127914304-127914344 | Myc |
| chr8:127914358-127914398 | Myc |
| chr8:127914468-127914508 | Myc |
| chr8:127914602-127914642 | Myc |
| chr8:127914735-127914775 | Myc |
| chr8:127914912-127914952 | Myc |
| chr8:127914980-127915020 | Myc |
| chr8:127915032-127915072 | Myc |

| | |
|---|---|
| chr8:127915156-127915196 | Myc |
| chr8:127915214-127915254 | Myc |
| chr8:127915341-127915381 | Myc |
| chr8:127915391-127915431 | Myc |
| chr8:127915514-127915554 | Myc |
| chr8:127915611-127915651 | Myc |
| chr8:127915710-127915750 | Myc |
| chr8:127915784-127915824 | Myc |
| chr8:127915904-127915944 | Myc |
| chr8:127915954-127915994 | Myc |
| chr8:127916053-127916093 | Myc |
| chr8:127916139-127916179 | Myc |
| chr8:127916215-127916255 | Myc |
| chr8:127916271-127916311 | Myc |
| chr8:127916343-127916383 | Myc |
| chr8:127916462-127916502 | Myc |
| chr8:127916516-127916556 | Myc |
| chr8:127916598-127916638 | Myc |
| chr8:127916658-127916698 | Myc |
| chr8:127916804-127916844 | Myc |
| chr8:127916929-127916969 | Myc |
| chr8:127916994-127917034 | Myc |
| chr8:127917084-127917124 | Myc |
| chr8:127917134-127917174 | Myc |
| chr8:127917257-127917297 | Myc |
| chr8:127917310-127917350 | Myc |
| chr8:127917422-127917462 | Myc |
| chr8:127917558-127917598 | Myc |
| chr8:127917644-127917684 | Myc |
| chr8:127917716-127917756 | Myc |
| chr8:127917883-127917923 | Myc |
| chr8:127918046-127918086 | Myc |
| chr8:127918139-127918179 | Myc |
| chr8:127918194-127918234 | Myc |
| chr8:127918250-127918290 | Myc |
| chr8:127918316-127918356 | Myc |
| chr8:127918386-127918426 | Myc |
| chr8:127918485-127918525 | Myc |
| chr8:127918574-127918614 | Myc |
| chr8:127918671-127918711 | Myc |
| chr8:127918781-127918821 | Myc |
| chr8:127918849-127918889 | Myc |
| chr8:127918900-127918940 | Myc |
| chr8:127919034-127919074 | Myc |
| chr8:127919165-127919205 | Myc |
| chr8:127919280-127919320 | Myc |
| chr8:127919348-127919388 | Myc |
| chr8:127919438-127919478 | Myc |
| chr8:127919612-127919652 | Myc |
| chr8:127919692-127919732 | Myc |
| chr8:127919946-127919986 | Myc |
| chr8:127920361-127920401 | Myc |

| | |
|---|---|
| chr8:127920415-127920455 | Myc |
| chr8:127920484-127920524 | Myc |
| chr8:127920593-127920633 | Myc |
| chr8:127920643-127920683 | Myc |
| chr8:127920753-127920793 | Myc |
| chr8:127920843-127920883 | Myc |
| chr8:127920959-127920999 | Myc |
| chr8:127921048-127921088 | Myc |
| chr8:127921279-127921319 | Myc |
| chr8:127921329-127921369 | Myc |
| chr8:127921457-127921497 | Myc |
| chr8:127921547-127921587 | Myc |
| chr8:127921665-127921705 | Myc |
| chr8:127921776-127921816 | Myc |
| chr8:127921863-127921903 | Myc |
| chr8:127921978-127922018 | Myc |
| chr8:127922029-127922069 | Myc |
| chr8:127922084-127922124 | Myc |
| chr8:127922239-127922279 | Myc |
| chr8:127922300-127922340 | Myc |
| chr8:127922464-127922504 | Myc |
| chr8:127922572-127922612 | Myc |
| chr8:127922646-127922686 | Myc |
| chr8:127922766-127922806 | Myc |
| chr8:127922836-127922876 | Myc |
| chr8:127922888-127922928 | Myc |
| chr8:127922956-127922996 | Myc |
| chr8:127923023-127923063 | Myc |
| chr8:127923122-127923162 | Myc |
| chr8:127923270-127923310 | Myc |
| chr8:127923521-127923561 | Myc |
| chr8:127923571-127923611 | Myc |
| chr8:127923624-127923664 | Myc |
| chr8:127923765-127923805 | Myc |
| chr8:127923824-127923864 | Myc |
| chr8:127924002-127924042 | Myc |
| chr8:127924072-127924112 | Myc |
| chr8:127924127-127924167 | Myc |
| chr8:127924204-127924244 | Myc |
| chr8:127924261-127924301 | Myc |
| chr8:127924360-127924400 | Myc |
| chr8:127924410-127924450 | Myc |
| chr8:127924465-127924505 | Myc |
| chr8:127924605-127924645 | Myc |
| chr8:127924702-127924742 | Myc |
| chr8:127924819-127924859 | Myc |
| chr8:127924870-127924910 | Myc |
| chr8:127924946-127924986 | Myc |
| chr8:127925023-127925063 | Myc |
| chr8:127925165-127925205 | Myc |
| chr8:127925337-127925377 | Myc |
| chr8:127925445-127925485 | Myc |

| | |
|---|---|
| chr8:127925551-127925591 | Myc |
| chr8:127925681-127925721 | Myc |
| chr8:127925743-127925783 | Myc |
| chr8:127925796-127925836 | Myc |
| chr8:127926135-127926175 | Myc |
| chr8:127926199-127926239 | Myc |
| chr8:127926292-127926332 | Myc |
| chr8:127926378-127926418 | Myc |
| chr8:127926470-127926510 | Myc |
| chr8:127926609-127926649 | Myc |
| chr8:127926661-127926701 | Myc |
| chr8:127926800-127926840 | Myc |
| chr8:127926917-127926957 | Myc |
| chr8:127927044-127927084 | Myc |
| chr8:127927097-127927137 | Myc |
| chr8:127927192-127927232 | Myc |
| chr8:127927257-127927297 | Myc |
| chr8:127927338-127927378 | Myc |
| chr8:127927395-127927435 | Myc |
| chr8:127927492-127927532 | Myc |
| chr8:127927651-127927691 | Myc |
| chr8:127927711-127927751 | Myc |
| chr8:127927796-127927836 | Myc |
| chr8:127927951-127927991 | Myc |
| chr8:127928204-127928244 | Myc |
| chr8:127928403-127928443 | Myc |
| chr8:127928549-127928589 | Myc |
| chr8:127928659-127928699 | Myc |
| chr8:127928718-127928758 | Myc |
| chr8:127928795-127928835 | Myc |
| chr8:127928878-127928918 | Myc |
| chr8:127928950-127928990 | Myc |
| chr8:127929114-127929154 | Myc |
| chr8:127929185-127929225 | Myc |
| chr8:127929287-127929327 | Myc |
| chr8:127929345-127929385 | Myc |
| chr8:127929434-127929474 | Myc |
| chr8:127929523-127929563 | Myc |
| chr8:127929611-127929651 | Myc |
| chr8:127929683-127929723 | Myc |
| chr8:127929749-127929789 | Myc |
| chr8:127929834-127929874 | Myc |
| chr8:127929918-127929958 | Myc |
| chr8:127929968-127930008 | Myc |
| chr8:127930043-127930083 | Myc |
| chr8:127930103-127930143 | Myc |
| chr8:127930212-127930252 | Myc |
| chr8:127930339-127930379 | Myc |
| chr8:127930389-127930429 | Myc |
| chr8:127930533-127930573 | Myc |
| chr8:127930620-127930660 | Myc |
| chr8:127930705-127930745 | Myc |

| | |
|---|---|
| chr8:127930784-127930824 | Myc |
| chr8:127930975-127931015 | Myc |
| chr8:127931095-127931135 | Myc |
| chr8:127931151-127931191 | Myc |
| chr8:127931226-127931266 | Myc |
| chr8:127931367-127931407 | Myc |
| chr8:127931467-127931507 | Myc |
| chr8:127931727-127931767 | Myc |
| chr8:127931853-127931893 | Myc |
| chr8:127932016-127932056 | Myc |
| chr8:127932095-127932135 | Myc |
| chr8:127932179-127932219 | Myc |
| chr8:127932346-127932386 | Myc |
| chr8:127932464-127932504 | Myc |
| chr8:127932533-127932573 | Myc |
| chr8:127932611-127932651 | Myc |
| chr8:127932749-127932789 | Myc |
| chr8:127932827-127932867 | Myc |
| chr8:127932927-127932967 | Myc |
| chr8:127933003-127933043 | Myc |
| chr8:127933189-127933229 | Myc |
| chr8:127933361-127933401 | Myc |
| chr8:127933441-127933481 | Myc |
| chr8:127933492-127933532 | Myc |
| chr8:127933662-127933702 | Myc |
| chr8:127933908-127933948 | Myc |
| chr8:127933990-127934030 | Myc |
| chr8:127934075-127934115 | Myc |
| chr8:127934142-127934182 | Myc |
| chr8:127934285-127934325 | Myc |
| chr8:127934407-127934447 | Myc |
| chr8:127934485-127934525 | Myc |
| chr8:127934610-127934650 | Myc |
| chr8:127934674-127934714 | Myc |
| chr8:127934794-127934834 | Myc |
| chr8:127934864-127934904 | Myc |
| chr8:127934935-127934975 | Myc |
| chr8:127935078-127935118 | Myc |
| chr8:127935175-127935215 | Myc |
| chr8:127935358-127935398 | Myc |
| chr8:127935476-127935516 | Myc |
| chr8:127935820-127935860 | Myc |
| chr8:127935989-127936029 | Myc |
| chr8:127936061-127936101 | Myc |
| chr8:127936218-127936258 | Myc |
| chr8:127936372-127936412 | Myc |
| chr8:127936691-127936731 | Myc |
| chr8:127936746-127936786 | Myc |
| chr8:127936850-127936890 | Myc |
| chr8:127936993-127937033 | Myc |
| chr8:127937161-127937201 | Myc |
| chr8:127937220-127937260 | Myc |

| | |
|---|---|
| chr8:127937289-127937329 | Myc |
| chr8:127937340-127937380 | Myc |
| chr8:127937423-127937463 | Myc |
| chr8:127937526-127937566 | Myc |
| chr8:127937612-127937652 | Myc |
| chr8:127937721-127937761 | Myc |
| chr8:127937772-127937812 | Myc |
| chr8:127937838-127937878 | Myc |
| chr8:127937927-127937967 | Myc |
| chr8:127938022-127938062 | Myc |
| chr8:127938072-127938112 | Myc |
| chr8:127938198-127938238 | Myc |
| chr8:127938306-127938346 | Myc |
| chr8:127938480-127938520 | Myc |
| chr8:127938543-127938583 | Myc |
| chr8:127938609-127938649 | Myc |
| chr8:127938685-127938725 | Myc |
| chr8:127938743-127938783 | Myc |
| chr8:127938817-127938857 | Myc |
| chr8:127938934-127938974 | Myc |
| chr8:127939049-127939089 | Myc |
| chr8:127939146-127939186 | Myc |
| chr8:127939200-127939240 | Myc |
| chr8:127939281-127939321 | Myc |
| chr8:127939344-127939384 | Myc |
| chr8:127939404-127939444 | Myc |
| chr8:127939466-127939506 | Myc |
| chr8:127939544-127939584 | Myc |
| chr8:127939742-127939782 | Myc |
| chr8:127939834-127939874 | Myc |
| chr8:127939962-127940002 | Myc |
| chr8:127940024-127940064 | Myc |
| chr8:127940111-127940151 | Myc |
| chr8:127940223-127940263 | Myc |
| chr8:127940280-127940320 | Myc |
| chr8:127940372-127940412 | Myc |
| chr8:127940454-127940494 | Myc |
| chr8:127940728-127940768 | Myc |
| chr8:127940854-127940894 | Myc |
| chr8:127940904-127940944 | Myc |
| chr8:127940954-127940994 | Myc |
| chr8:127941007-127941047 | Myc |
| chr8:127941343-127941383 | Myc |
| chr8:127941409-127941449 | Myc |
| chr8:127941643-127941683 | Myc |
| chr8:127941714-127941754 | Myc |
| chr8:127941810-127941850 | Myc |
| chr8:127941860-127941900 | Myc |
| chr8:127941911-127941951 | Myc |
| chr8:127941990-127942030 | Myc |
| chr8:127942103-127942143 | Myc |
| chr8:127942177-127942217 | Myc |

| | |
|---|---|
| chr8:127942290-127942330 | Myc |
| chr8:127942429-127942469 | Myc |
| chr8:127942488-127942528 | Myc |
| chr8:127942550-127942590 | Myc |
| chr8:127942613-127942653 | Myc |
| chr8:127942679-127942719 | Myc |
| chr8:127942734-127942774 | Myc |
| chr8:127942877-127942917 | Myc |
| chr8:127942961-127943001 | Myc |
| chr8:127943122-127943162 | Myc |
| chr8:127943215-127943255 | Myc |
| chr8:127943334-127943374 | Myc |
| chr8:127943421-127943461 | Myc |
| chr8:127943481-127943521 | Myc |
| chr8:127943597-127943637 | Myc |
| chr8:127943742-127943782 | Myc |
| chr8:127943950-127943990 | Myc |
| chr8:127944041-127944081 | Myc |
| chr8:127944091-127944131 | Myc |
| chr8:127944178-127944218 | Myc |
| chr8:127944293-127944333 | Myc |
| chr8:127944350-127944390 | Myc |
| chr8:127944708-127944748 | Myc |
| chr8:127944775-127944815 | Myc |
| chr8:127944834-127944874 | Myc |
| chr8:127945054-127945094 | Myc |
| chr8:127945171-127945211 | Myc |
| chr8:127945240-127945280 | Myc |
| chr8:127945313-127945353 | Myc |
| chr8:127945481-127945521 | Myc |
| chr8:127945759-127945799 | Myc |
| chr8:127945826-127945866 | Myc |
| chr8:127945918-127945958 | Myc |
| chr8:127946019-127946059 | Myc |
| chr8:127946125-127946165 | Myc |
| chr8:127946184-127946224 | Myc |
| chr8:127946247-127946287 | Myc |
| chr8:127946338-127946378 | Myc |
| chr8:127946402-127946442 | Myc |
| chr8:127946575-127946615 | Myc |
| chr8:127946625-127946665 | Myc |
| chr8:127946690-127946730 | Myc |
| chr8:127946846-127946886 | Myc |
| chr8:127946973-127947013 | Myc |
| chr8:127947090-127947130 | Myc |
| chr8:127947178-127947218 | Myc |
| chr8:127947275-127947315 | Myc |
| chr8:127947325-127947365 | Myc |
| chr8:127947440-127947480 | Myc |
| chr8:127947491-127947531 | Myc |
| chr8:127947546-127947586 | Myc |
| chr8:127947608-127947648 | Myc |

| | |
|---|---|
| chr8:127947682-127947722 | Myc |
| chr8:127947783-127947823 | Myc |
| chr8:127947935-127947975 | Myc |
| chr8:127948016-127948056 | Myc |
| chr8:127948155-127948195 | Myc |
| chr8:127948262-127948302 | Myc |
| chr8:127948317-127948357 | Myc |
| chr8:127948381-127948421 | Myc |
| chr8:127948434-127948474 | Myc |
| chr8:127948488-127948528 | Myc |
| chr8:127948574-127948614 | Myc |
| chr8:127948624-127948664 | Myc |
| chr8:127948746-127948786 | Myc |
| chr8:127948802-127948842 | Myc |
| chr8:127948961-127949001 | Myc |
| chr8:127949038-127949078 | Myc |
| chr8:127949148-127949188 | Myc |
| chr8:127949198-127949238 | Myc |
| chr8:127949295-127949335 | Myc |
| chr8:127949394-127949434 | Myc |
| chr8:127949507-127949547 | Myc |
| chr8:127949614-127949654 | Myc |
| chr8:127949720-127949760 | Myc |
| chr8:127949830-127949870 | Myc |
| chr8:127949881-127949921 | Myc |
| chr8:127949931-127949971 | Myc |
| chr8:127949981-127950021 | Myc |
| chr8:127950079-127950119 | Myc |
| chr8:127950161-127950201 | Myc |
| chr8:127950282-127950322 | Myc |
| chr8:127950393-127950433 | Myc |
| chr8:127950469-127950509 | Myc |
| chr8:127950576-127950616 | Myc |
| chr8:127950708-127950748 | Myc |
| chr8:127950877-127950917 | Myc |
| chr8:127951004-127951044 | Myc |
| chr8:127951098-127951138 | Myc |
| chr8:127951202-127951242 | Myc |
| chr8:127951252-127951292 | Myc |
| chr8:127951327-127951367 | Myc |
| chr8:127951490-127951530 | Myc |
| chr8:127951570-127951610 | Myc |
| chr8:127951674-127951714 | Myc |
| chr8:127951728-127951768 | Myc |
| chr8:127951778-127951818 | Myc |
| chr8:127951925-127951965 | Myc |
| chr8:127951983-127952023 | Myc |
| chr8:127952060-127952100 | Myc |
| chr8:127952190-127952230 | Myc |
| chr8:127952246-127952286 | Myc |
| chr8:127952370-127952410 | Myc |
| chr8:127952434-127952474 | Myc |

| | |
|---|---|
| chr8:127952545-127952585 | Myc |
| chr8:127952703-127952743 | Myc |
| chr8:127952848-127952888 | Myc |
| chr8:127952910-127952950 | Myc |
| chr8:127952978-127953018 | Myc |
| chr8:127953049-127953089 | Myc |
| chr8:127953127-127953167 | Myc |
| chr8:127953226-127953266 | Myc |
| chr8:127953358-127953398 | Myc |
| chr8:127953427-127953467 | Myc |
| chr8:127953531-127953571 | Myc |
| chr8:127953609-127953649 | Myc |
| chr8:127953689-127953729 | Myc |
| chr8:127953750-127953790 | Myc |
| chr8:127953809-127953849 | Myc |
| chr8:127953902-127953942 | Myc |
| chr8:127954001-127954041 | Myc |
| chr8:127954304-127954344 | Myc |
| chr8:127954379-127954419 | Myc |
| chr8:127954483-127954523 | Myc |
| chr8:127954575-127954615 | Myc |
| chr8:127954654-127954694 | Myc |
| chr8:127954915-127954955 | Myc |
| chr8:127954982-127955022 | Myc |
| chr8:127955055-127955095 | Myc |
| chr8:127955168-127955208 | Myc |
| chr8:127955297-127955337 | Myc |
| chr8:127955432-127955472 | Myc |
| chr8:127955579-127955619 | Myc |
| chr8:127955683-127955723 | Myc |
| chr8:127955733-127955773 | Myc |
| chr8:127955953-127955993 | Myc |
| chr8:127956118-127956158 | Myc |
| chr8:127956213-127956253 | Myc |
| chr8:127956332-127956372 | Myc |
| chr8:127956484-127956524 | Myc |
| chr8:127956594-127956634 | Myc |
| chr8:127956662-127956702 | Myc |
| chr8:127956730-127956770 | Myc |
| chr8:127956780-127956820 | Myc |
| chr8:127956910-127956950 | Myc |
| chr8:127957038-127957078 | Myc |
| chr8:127957105-127957145 | Myc |
| chr8:127957221-127957261 | Myc |
| chr8:127957316-127957356 | Myc |
| chr8:127957501-127957541 | Myc |
| chr8:127957597-127957637 | Myc |
| chr8:127957671-127957711 | Myc |
| chr8:127957723-127957763 | Myc |
| chr8:127957773-127957813 | Myc |
| chr8:127957872-127957912 | Myc |
| chr8:127957978-127958018 | Myc |

| | |
|---|---|
| chr8:127958063-127958103 | Myc |
| chr8:127958113-127958153 | Myc |
| chr8:127958190-127958230 | Myc |
| chr8:127958383-127958423 | Myc |
| chr8:127958520-127958560 | Myc |
| chr8:127958570-127958610 | Myc |
| chr8:127958640-127958680 | Myc |
| chr8:127958743-127958783 | Myc |
| chr8:127958810-127958850 | Myc |
| chr8:127958860-127958900 | Myc |
| chr8:127958916-127958956 | Myc |
| chr8:127959025-127959065 | Myc |
| chr8:127959132-127959172 | Myc |
| chr8:127959231-127959271 | Myc |
| chr8:127959354-127959394 | Myc |
| chr8:127959467-127959507 | Myc |
| chr8:127959547-127959587 | Myc |
| chr8:127959684-127959724 | Myc |
| chr8:127959784-127959824 | Myc |
| chr8:127959875-127959915 | Myc |
| chr8:127959926-127959966 | Myc |
| chr8:127959996-127960036 | Myc |
| chr8:127960089-127960129 | Myc |
| chr8:127960139-127960179 | Myc |
| chr8:127960208-127960248 | Myc |
| chr8:127960258-127960298 | Myc |
| chr8:127960311-127960351 | Myc |
| chr8:127960386-127960426 | Myc |
| chr8:127960453-127960493 | Myc |
| chr8:127960503-127960543 | Myc |
| chr8:127960563-127960603 | Myc |
| chr8:127960634-127960674 | Myc |
| chr8:127960843-127960883 | Myc |
| chr8:127960893-127960933 | Myc |
| chr8:127960993-127961033 | Myc |
| chr8:127961046-127961086 | Myc |
| chr8:127961099-127961139 | Myc |
| chr8:127961172-127961212 | Myc |
| chr8:127961276-127961316 | Myc |
| chr8:127961609-127961649 | Myc |
| chr8:127961779-127961819 | Myc |
| chr8:127961957-127961997 | Myc |
| chr8:127962007-127962047 | Myc |
| chr8:127962078-127962118 | Myc |
| chr8:127962220-127962260 | Myc |
| chr8:127962284-127962324 | Myc |
| chr8:127962423-127962463 | Myc |
| chr8:127962491-127962531 | Myc |
| chr8:127962543-127962583 | Myc |
| chr8:127962612-127962652 | Myc |
| chr8:127962726-127962766 | Myc |
| chr8:127962817-127962857 | Myc |

| | |
|---|---|
| chr8:127962922-127962962 | Myc |
| chr8:127963073-127963113 | Myc |
| chr8:127963158-127963198 | Myc |
| chr8:127963254-127963294 | Myc |
| chr8:127963358-127963398 | Myc |
| chr8:127963444-127963484 | Myc |
| chr8:127963606-127963646 | Myc |
| chr8:127963742-127963782 | Myc |
| chr8:127963800-127963840 | Myc |
| chr8:127963862-127963902 | Myc |
| chr8:127963929-127963969 | Myc |
| chr8:127964011-127964051 | Myc |
| chr8:127964116-127964156 | Myc |
| chr8:127964220-127964260 | Myc |
| chr8:127964270-127964310 | Myc |
| chr8:127964375-127964415 | Myc |
| chr8:127964459-127964499 | Myc |
| chr8:127964534-127964574 | Myc |
| chr8:127964603-127964643 | Myc |
| chr8:127964707-127964747 | Myc |
| chr8:127964774-127964814 | Myc |
| chr8:127964856-127964896 | Myc |
| chr8:127964932-127964972 | Myc |
| chr8:127965001-127965041 | Myc |
| chr8:127965064-127965104 | Myc |
| chr8:127965114-127965154 | Myc |
| chr8:127965195-127965235 | Myc |
| chr8:127965420-127965460 | Myc |
| chr8:127965489-127965529 | Myc |
| chr8:127965573-127965613 | Myc |
| chr8:127965626-127965666 | Myc |
| chr8:127965756-127965796 | Myc |
| chr8:127965834-127965874 | Myc |
| chr8:127965951-127965991 | Myc |
| chr8:127966066-127966106 | Myc |
| chr8:127966136-127966176 | Myc |
| chr8:127966199-127966239 | Myc |
| chr8:127966311-127966351 | Myc |
| chr8:127966396-127966436 | Myc |
| chr8:127966535-127966575 | Myc |
| chr8:127966616-127966656 | Myc |
| chr8:127966734-127966774 | Myc |
| chr8:127966860-127966900 | Myc |
| chr8:127966936-127966976 | Myc |
| chr8:127967066-127967106 | Myc |
| chr8:127967130-127967170 | Myc |
| chr8:127967225-127967265 | Myc |
| chr8:127967302-127967342 | Myc |
| chr8:127967353-127967393 | Myc |
| chr8:127967475-127967515 | Myc |
| chr8:127967573-127967613 | Myc |
| chr8:127967666-127967706 | Myc |

| | |
|---|---|
| chr8:127967836-127967876 | Myc |
| chr8:127967929-127967969 | Myc |
| chr8:127968067-127968107 | Myc |
| chr8:127968315-127968355 | Myc |
| chr8:127968386-127968426 | Myc |
| chr8:127968450-127968490 | Myc |
| chr8:127968513-127968553 | Myc |
| chr8:127968578-127968618 | Myc |
| chr8:127968658-127968698 | Myc |
| chr8:127968708-127968748 | Myc |
| chr8:127968881-127968921 | Myc |
| chr8:127968936-127968976 | Myc |
| chr8:127969032-127969072 | Myc |
| chr8:127969150-127969190 | Myc |
| chr8:127969201-127969241 | Myc |
| chr8:127969311-127969351 | Myc |
| chr8:127969372-127969412 | Myc |
| chr8:127969458-127969498 | Myc |
| chr8:127969544-127969584 | Myc |
| chr8:127969659-127969699 | Myc |
| chr8:127969731-127969771 | Myc |
| chr8:127970069-127970109 | Myc |
| chr8:127970126-127970166 | Myc |
| chr8:127970187-127970227 | Myc |
| chr8:127970508-127970548 | Myc |
| chr8:127970601-127970641 | Myc |
| chr8:127970679-127970719 | Myc |
| chr8:127970739-127970779 | Myc |
| chr8:127970928-127970968 | Myc |
| chr8:127970994-127971034 | Myc |
| chr8:127971096-127971136 | Myc |
| chr8:127971230-127971270 | Myc |
| chr8:127971304-127971344 | Myc |
| chr8:127971381-127971421 | Myc |
| chr8:127971436-127971476 | Myc |
| chr8:127971507-127971547 | Myc |
| chr8:127971734-127971774 | Myc |
| chr8:127971846-127971886 | Myc |
| chr8:127971978-127972018 | Myc |
| chr8:127972069-127972109 | Myc |
| chr8:127972120-127972160 | Myc |
| chr8:127972197-127972237 | Myc |
| chr8:127972251-127972291 | Myc |
| chr8:127972348-127972388 | Myc |
| chr8:127972461-127972501 | Myc |
| chr8:127972518-127972558 | Myc |
| chr8:127972623-127972663 | Myc |
| chr8:127972673-127972713 | Myc |
| chr8:127972723-127972763 | Myc |
| chr8:127972868-127972908 | Myc |
| chr8:127972919-127972959 | Myc |
| chr8:127972991-127973031 | Myc |

| | |
|---|---|
| chr8:127973080-127973120 | Myc |
| chr8:127973228-127973268 | Myc |
| chr8:127973296-127973336 | Myc |
| chr8:127973354-127973394 | Myc |
| chr8:127973407-127973447 | Myc |
| chr8:127973601-127973641 | Myc |
| chr8:127973651-127973691 | Myc |
| chr8:127973743-127973783 | Myc |
| chr8:127973803-127973843 | Myc |
| chr8:127973998-127974038 | Myc |
| chr8:127974080-127974120 | Myc |
| chr8:127974171-127974211 | Myc |
| chr8:127974224-127974264 | Myc |
| chr8:127974295-127974335 | Myc |
| chr8:127974401-127974441 | Myc |
| chr8:127974451-127974491 | Myc |
| chr8:127974514-127974554 | Myc |
| chr8:127974586-127974626 | Myc |
| chr8:127974655-127974695 | Myc |
| chr8:127974839-127974879 | Myc |
| chr8:127974900-127974940 | Myc |
| chr8:127974988-127975028 | Myc |
| chr8:127975057-127975097 | Myc |
| chr8:127975153-127975193 | Myc |
| chr8:127975235-127975275 | Myc |
| chr8:127975319-127975359 | Myc |
| chr8:127975438-127975478 | Myc |
| chr8:127975506-127975546 | Myc |
| chr8:127975556-127975596 | Myc |
| chr8:127975761-127975801 | Myc |
| chr8:127975813-127975853 | Myc |
| chr8:127975864-127975904 | Myc |
| chr8:127975970-127976010 | Myc |
| chr8:127976037-127976077 | Myc |
| chr8:127976154-127976194 | Myc |
| chr8:127976274-127976314 | Myc |
| chr8:127976360-127976400 | Myc |
| chr8:127976513-127976553 | Myc |
| chr8:127976613-127976653 | Myc |
| chr8:127976676-127976716 | Myc |
| chr8:127976862-127976902 | Myc |
| chr8:127976913-127976953 | Myc |
| chr8:127976980-127977020 | Myc |
| chr8:127977073-127977113 | Myc |
| chr8:127977251-127977291 | Myc |
| chr8:127977313-127977353 | Myc |
| chr8:127977363-127977403 | Myc |
| chr8:127977441-127977481 | Myc |
| chr8:127977558-127977598 | Myc |
| chr8:127977901-127977941 | Myc |
| chr8:127977964-127978004 | Myc |
| chr8:127978068-127978108 | Myc |

| | |
|---|---|
| chr8:127978221-127978261 | Myc |
| chr8:127978342-127978382 | Myc |
| chr8:127978481-127978521 | Myc |
| chr8:127978533-127978573 | Myc |
| chr8:127978781-127978821 | Myc |
| chr8:127978865-127978905 | Myc |
| chr8:127978926-127978966 | Myc |
| chr8:127978976-127979016 | Myc |
| chr8:127979055-127979095 | Myc |
| chr8:127979144-127979184 | Myc |
| chr8:127979214-127979254 | Myc |
| chr8:127979359-127979399 | Myc |
| chr8:127979439-127979479 | Myc |
| chr8:127979616-127979656 | Myc |
| chr8:127979687-127979727 | Myc |
| chr8:127979767-127979807 | Myc |
| chr8:127979876-127979916 | Myc |
| chr8:127979938-127979978 | Myc |
| chr8:127980088-127980128 | Myc |
| chr8:127980164-127980204 | Myc |
| chr8:127980282-127980322 | Myc |
| chr8:127980417-127980457 | Myc |
| chr8:127980494-127980534 | Myc |
| chr8:127980554-127980594 | Myc |
| chr8:127980757-127980797 | Myc |
| chr8:127980902-127980942 | Myc |
| chr8:127981011-127981051 | Myc |
| chr8:127981083-127981123 | Myc |
| chr8:127981133-127981173 | Myc |
| chr8:127981208-127981248 | Myc |
| chr8:127981283-127981323 | Myc |
| chr8:127981414-127981454 | Myc |
| chr8:127981503-127981543 | Myc |
| chr8:127981616-127981656 | Myc |
| chr8:127981780-127981820 | Myc |
| chr8:127981929-127981969 | Myc |
| chr8:127982096-127982136 | Myc |
| chr8:127982236-127982276 | Myc |
| chr8:127982288-127982328 | Myc |
| chr8:127982449-127982489 | Myc |
| chr8:127982503-127982543 | Myc |
| chr8:127982569-127982609 | Myc |
| chr8:127982634-127982674 | Myc |
| chr8:127982732-127982772 | Myc |
| chr8:127982824-127982864 | Myc |
| chr8:127982877-127982917 | Myc |
| chr8:127982934-127982974 | Myc |
| chr8:127982984-127983024 | Myc |
| chr8:127983045-127983085 | Myc |
| chr8:127983098-127983138 | Myc |
| chr8:127983222-127983262 | Myc |
| chr8:127983313-127983353 | Myc |

| | |
|---|---|
| chr8:127983417-127983457 | Myc |
| chr8:127983467-127983507 | Myc |
| chr8:127983553-127983593 | Myc |
| chr8:127983635-127983675 | Myc |
| chr8:127983736-127983776 | Myc |
| chr8:127983837-127983877 | Myc |
| chr8:127983894-127983934 | Myc |
| chr8:127984038-127984078 | Myc |
| chr8:127984089-127984129 | Myc |
| chr8:127984139-127984179 | Myc |
| chr8:127984282-127984322 | Myc |
| chr8:127984396-127984436 | Myc |
| chr8:127984446-127984486 | Myc |
| chr8:127984622-127984662 | Myc |
| chr8:127984711-127984751 | Myc |
| chr8:127984777-127984817 | Myc |
| chr8:127984845-127984885 | Myc |
| chr8:127984904-127984944 | Myc |
| chr8:127984966-127985006 | Myc |
| chr8:127985063-127985103 | Myc |
| chr8:127985189-127985229 | Myc |
| chr8:127985367-127985407 | Myc |
| chr8:127985489-127985529 | Myc |
| chr8:127985547-127985587 | Myc |
| chr8:127985630-127985670 | Myc |
| chr8:127985680-127985720 | Myc |
| chr8:127985744-127985784 | Myc |
| chr8:127985810-127985850 | Myc |
| chr8:127985917-127985957 | Myc |
| chr8:127985971-127986011 | Myc |
| chr8:127986036-127986076 | Myc |
| chr8:127986403-127986443 | Myc |
| chr8:127986474-127986514 | Myc |
| chr8:127986558-127986598 | Myc |
| chr8:127986682-127986722 | Myc |
| chr8:127986760-127986800 | Myc |
| chr8:127986836-127986876 | Myc |
| chr8:127986932-127986972 | Myc |
| chr8:127987012-127987052 | Myc |
| chr8:127987068-127987108 | Myc |
| chr8:127987157-127987197 | Myc |
| chr8:127987273-127987313 | Myc |
| chr8:127987372-127987412 | Myc |
| chr8:127987430-127987470 | Myc |
| chr8:127987521-127987561 | Myc |
| chr8:127987622-127987662 | Myc |
| chr8:127987800-127987840 | Myc |
| chr8:127987872-127987912 | Myc |
| chr8:127987958-127987998 | Myc |
| chr8:127988034-127988074 | Myc |
| chr8:127988114-127988154 | Myc |
| chr8:127988253-127988293 | Myc |

| | |
|---|---|
| chr8:127988324-127988364 | Myc |
| chr8:127988381-127988421 | Myc |
| chr8:127988477-127988517 | Myc |
| chr8:127988604-127988644 | Myc |
| chr8:127988655-127988695 | Myc |
| chr8:127988706-127988746 | Myc |
| chr8:127988834-127988874 | Myc |
| chr8:127988968-127989008 | Myc |
| chr8:127989018-127989058 | Myc |
| chr8:127989076-127989116 | Myc |
| chr8:127989154-127989194 | Myc |
| chr8:127989276-127989316 | Myc |
| chr8:127989347-127989387 | Myc |
| chr8:127989413-127989453 | Myc |
| chr8:127989520-127989560 | Myc |
| chr8:127989572-127989612 | Myc |
| chr8:127989636-127989676 | Myc |
| chr8:127989732-127989772 | Myc |
| chr8:127989782-127989822 | Myc |
| chr8:127989962-127990002 | Myc |
| chr8:127990015-127990055 | Myc |
| chr8:127990072-127990112 | Myc |
| chr8:127990128-127990168 | Myc |
| chr8:127990201-127990241 | Myc |
| chr8:127990291-127990331 | Myc |
| chr8:127990350-127990390 | Myc |
| chr8:127990400-127990440 | Myc |
| chr8:127990461-127990501 | Myc |
| chr8:127990629-127990669 | Myc |
| chr8:127990709-127990749 | Myc |
| chr8:127990759-127990799 | Myc |
| chr8:127990810-127990850 | Myc |
| chr8:127990902-127990942 | Myc |
| chr8:127991053-127991093 | Myc |
| chr8:127991127-127991167 | Myc |
| chr8:127991234-127991274 | Myc |
| chr8:127991284-127991324 | Myc |
| chr8:127991374-127991414 | Myc |
| chr8:127991426-127991466 | Myc |
| chr8:127991566-127991606 | Myc |
| chr8:127991641-127991681 | Myc |
| chr8:127991703-127991743 | Myc |
| chr8:127991783-127991823 | Myc |
| chr8:127992132-127992172 | Myc |
| chr8:127992271-127992311 | Myc |
| chr8:127992423-127992463 | Myc |
| chr8:127992496-127992536 | Myc |
| chr8:127992612-127992652 | Myc |
| chr8:127993967-127994007 | Myc |
| chr8:127994304-127994344 | Myc |
| chr8:127994537-127994577 | Myc |
| chr8:127995718-127995758 | Myc |

| | |
|---|---|
| chr8:127996398-127996438 | Myc |
| chr8:127996792-127996832 | Myc |
| chr8:127997211-127997251 | Myc |
| chr8:127998251-127998291 | Myc |
| chr8:127998303-127998343 | Myc |
| chr8:127998460-127998500 | Myc |
| chr8:127998528-127998568 | Myc |
| chr8:127998629-127998669 | Myc |
| chr8:127998740-127998780 | Myc |
| chr8:127998854-127998894 | Myc |
| chr8:127998947-127998987 | Myc |
| chr8:127999078-127999118 | Myc |
| chr8:127999270-127999310 | Myc |
| chr8:127999338-127999378 | Myc |
| chr8:127999404-127999444 | Myc |
| chr8:127999484-127999524 | Myc |
| chr8:127999561-127999601 | Myc |
| chr8:127999611-127999651 | Myc |
| chr8:127999689-127999729 | Myc |
| chr8:127999762-127999802 | Myc |
| chr8:127999840-127999880 | Myc |
| chr8:127999909-127999949 | Myc |
| chr8:127999964-128000004 | Myc |
| chr8:128000098-128000138 | Myc |
| chr8:128000195-128000235 | Myc |
| chr8:128000329-128000369 | Myc |
| chr8:128000406-128000446 | Myc |
| chr8:128000472-128000512 | Myc |
| chr8:128000597-128000637 | Myc |
| chr8:128000665-128000705 | Myc |
| chr8:128000751-128000791 | Myc |
| chr8:128000860-128000900 | Myc |
| chr8:128001046-128001086 | Myc |
| chr8:128001185-128001225 | Myc |
| chr8:128001265-128001305 | Myc |
| chr8:128001384-128001424 | Myc |
| chr8:128001439-128001479 | Myc |
| chr8:128001506-128001546 | Myc |
| chr8:128001574-128001614 | Myc |
| chr8:128001662-128001702 | Myc |
| chr8:128001798-128001838 | Myc |
| chr8:128001849-128001889 | Myc |
| chr8:128001942-128001982 | Myc |
| chr8:128001994-128002034 | Myc |
| chr8:128002044-128002084 | Myc |
| chr8:128002141-128002181 | Myc |
| chr8:128002281-128002321 | Myc |
| chr8:128002332-128002372 | Myc |
| chr8:128002407-128002447 | Myc |
| chr8:128002480-128002520 | Myc |
| chr8:128002545-128002585 | Myc |
| chr8:128002613-128002653 | Myc |

| | |
|---|---|
| chr8:128002663-128002703 | Myc |
| chr8:128002713-128002753 | Myc |
| chr8:128002809-128002849 | Myc |
| chr8:128002945-128002985 | Myc |
| chr8:128003088-128003128 | Myc |
| chr8:128003191-128003231 | Myc |
| chr8:128003273-128003313 | Myc |
| chr8:128003368-128003408 | Myc |
| chr8:128003479-128003519 | Myc |
| chr8:128003598-128003638 | Myc |
| chr8:128003673-128003713 | Myc |
| chr8:128003764-128003804 | Myc |
| chr8:128003849-128003889 | Myc |
| chr8:128003903-128003943 | Myc |
| chr8:128003954-128003994 | Myc |
| chr8:128004058-128004098 | Myc |
| chr8:128004119-128004159 | Myc |
| chr8:128004175-128004215 | Myc |
| chr8:128004296-128004336 | Myc |
| chr8:128004365-128004405 | Myc |
| chr8:128004436-128004476 | Myc |
| chr8:128004487-128004527 | Myc |
| chr8:128004578-128004618 | Myc |
| chr8:128004692-128004732 | Myc |
| chr8:128004788-128004828 | Myc |
| chr8:128004909-128004949 | Myc |
| chr8:128004962-128005002 | Myc |
| chr8:128005042-128005082 | Myc |
| chr8:128005152-128005192 | Myc |
| chr8:128005208-128005248 | Myc |
| chr8:128005287-128005327 | Myc |
| chr8:128005344-128005384 | Myc |
| chr8:128005445-128005485 | Myc |
| chr8:128005511-128005551 | Myc |
| chr8:128005591-128005631 | Myc |
| chr8:128005677-128005717 | Myc |
| chr8:128005793-128005833 | Myc |
| chr8:128005866-128005906 | Myc |
| chr8:128005925-128005965 | Myc |
| chr8:128005983-128006023 | Myc |
| chr8:128006036-128006076 | Myc |
| chr8:128006120-128006160 | Myc |
| chr8:128006255-128006295 | Myc |
| chr8:128006327-128006367 | Myc |
| chr8:128006405-128006445 | Myc |
| chr8:128006493-128006533 | Myc |
| chr8:128006591-128006631 | Myc |
| chr8:128006718-128006758 | Myc |
| chr8:128006881-128006921 | Myc |
| chr8:128006993-128007033 | Myc |
| chr8:128007093-128007133 | Myc |
| chr8:128007192-128007232 | Myc |

| | |
|---|---|
| chr8:128007243-128007283 | Myc |
| chr8:128007293-128007333 | Myc |
| chr8:128007360-128007400 | Myc |
| chr8:128007472-128007512 | Myc |
| chr8:128007667-128007707 | Myc |
| chr8:128007739-128007779 | Myc |
| chr8:128007810-128007850 | Myc |
| chr8:128007921-128007961 | Myc |
| chr8:128007977-128008017 | Myc |
| chr8:128008198-128008238 | Myc |
| chr8:128008360-128008400 | Myc |
| chr8:128008422-128008462 | Myc |
| chr8:128008559-128008599 | Myc |
| chr8:128008645-128008685 | Myc |
| chr8:128008696-128008736 | Myc |
| chr8:128008790-128008830 | Myc |
| chr8:128008880-128008920 | Myc |
| chr8:128008946-128008986 | Myc |
| chr8:128009038-128009078 | Myc |
| chr8:128009234-128009274 | Myc |
| chr8:128009333-128009373 | Myc |
| chr8:128009427-128009467 | Myc |
| chr8:128009608-128009648 | Myc |
| chr8:128009673-128009713 | Myc |
| chr8:128009755-128009795 | Myc |
| chr8:128009816-128009856 | Myc |
| chr8:128009894-128009934 | Myc |
| chr8:128009991-128010031 | Myc |
| chr8:128010134-128010174 | Myc |
| chr8:128010189-128010229 | Myc |
| chr8:128010245-128010285 | Myc |
| chr8:128010309-128010349 | Myc |
| chr8:128010363-128010403 | Myc |
| chr8:128010413-128010453 | Myc |
| chr8:128010541-128010581 | Myc |
| chr8:128010630-128010670 | Myc |
| chr8:128010680-128010720 | Myc |
| chr8:128010797-128010837 | Myc |
| chr8:128010869-128010909 | Myc |
| chr8:128011025-128011065 | Myc |
| chr8:128011116-128011156 | Myc |
| chr8:128011181-128011221 | Myc |
| chr8:128011297-128011337 | Myc |
| chr8:128011363-128011403 | Myc |
| chr8:128011458-128011498 | Myc |
| chr8:128011588-128011628 | Myc |
| chr8:128011677-128011717 | Myc |
| chr8:128011938-128011978 | Myc |
| chr8:128012021-128012061 | Myc |
| chr8:128012107-128012147 | Myc |
| chr8:128012219-128012259 | Myc |
| chr8:128012330-128012370 | Myc |

| | |
|---|---|
| chr8:128012592-128012632 | Myc |
| chr8:128012672-128012712 | Myc |
| chr8:128012725-128012765 | Myc |
| chr8:128012799-128012839 | Myc |
| chr8:128012849-128012889 | Myc |
| chr8:128012956-128012996 | Myc |
| chr8:128013016-128013056 | Myc |
| chr8:128013091-128013131 | Myc |
| chr8:128013191-128013231 | Myc |
| chr8:128013353-128013393 | Myc |
| chr8:128013431-128013471 | Myc |
| chr8:128013502-128013542 | Myc |
| chr8:128013666-128013706 | Myc |
| chr8:128013749-128013789 | Myc |
| chr8:128013832-128013872 | Myc |
| chr8:128013896-128013936 | Myc |
| chr8:128014256-128014296 | Myc |
| chr8:128014306-128014346 | Myc |
| chr8:128014517-128014557 | Myc |
| chr8:128014717-128014757 | Myc |
| chr8:128014841-128014881 | Myc |
| chr8:128014931-128014971 | Myc |
| chr8:128015057-128015097 | Myc |
| chr8:128015115-128015155 | Myc |
| chr8:128015241-128015281 | Myc |
| chr8:128015291-128015331 | Myc |
| chr8:128015427-128015467 | Myc |
| chr8:128015800-128015840 | Myc |
| chr8:128015877-128015917 | Myc |
| chr8:128015971-128016011 | Myc |
| chr8:128016068-128016108 | Myc |
| chr8:128016120-128016160 | Myc |
| chr8:128016180-128016220 | Myc |
| chr8:128016255-128016295 | Myc |
| chr8:128016309-128016349 | Myc |
| chr8:128016473-128016513 | Myc |
| chr8:128016550-128016590 | Myc |
| chr8:128016673-128016713 | Myc |
| chr8:128016781-128016821 | Myc |
| chr8:128016947-128016987 | Myc |
| chr8:128017007-128017047 | Myc |
| chr8:128017116-128017156 | Myc |
| chr8:128017166-128017206 | Myc |
| chr8:128017270-128017310 | Myc |
| chr8:128017346-128017386 | Myc |
| chr8:128017400-128017440 | Myc |
| chr8:128017506-128017546 | Myc |
| chr8:128017676-128017716 | Myc |
| chr8:128017729-128017769 | Myc |
| chr8:128017832-128017872 | Myc |
| chr8:128017893-128017933 | Myc |
| chr8:128018012-128018052 | Myc |

| | |
|---|---|
| chr8:128018127-128018167 | Myc |
| chr8:128018183-128018223 | Myc |
| chr8:128018375-128018415 | Myc |
| chr8:128018537-128018577 | Myc |
| chr8:128018594-128018634 | Myc |
| chr8:128018693-128018733 | Myc |
| chr8:128018747-128018787 | Myc |
| chr8:128018802-128018842 | Myc |
| chr8:128018852-128018892 | Myc |
| chr8:128018926-128018966 | Myc |
| chr8:128018976-128019016 | Myc |
| chr8:128019106-128019146 | Myc |
| chr8:128019197-128019237 | Myc |
| chr8:128019289-128019329 | Myc |
| chr8:128019398-128019438 | Myc |
| chr8:128019456-128019496 | Myc |
| chr8:128019625-128019665 | Myc |
| chr8:128019778-128019818 | Myc |
| chr8:128019828-128019868 | Myc |
| chr8:128019948-128019988 | Myc |
| chr8:128020004-128020044 | Myc |
| chr8:128020055-128020095 | Myc |
| chr8:128020181-128020221 | Myc |
| chr8:128020250-128020290 | Myc |
| chr8:128020325-128020365 | Myc |
| chr8:128020415-128020455 | Myc |
| chr8:128020475-128020515 | Myc |
| chr8:128020546-128020586 | Myc |
| chr8:128020692-128020732 | Myc |
| chr8:128020777-128020817 | Myc |
| chr8:128020829-128020869 | Myc |
| chr8:128020952-128020992 | Myc |
| chr8:128021059-128021099 | Myc |
| chr8:128021156-128021196 | Myc |
| chr8:128021220-128021260 | Myc |
| chr8:128021341-128021381 | Myc |
| chr8:128021395-128021435 | Myc |
| chr8:128021446-128021486 | Myc |
| chr8:128021630-128021670 | Myc |
| chr8:128021704-128021744 | Myc |
| chr8:128021769-128021809 | Myc |
| chr8:128021830-128021870 | Myc |
| chr8:128021916-128021956 | Myc |
| chr8:128021989-128022029 | Myc |
| chr8:128022039-128022079 | Myc |
| chr8:128022132-128022172 | Myc |
| chr8:128022263-128022303 | Myc |
| chr8:128022350-128022390 | Myc |
| chr8:128022407-128022447 | Myc |
| chr8:128022478-128022518 | Myc |
| chr8:128022540-128022580 | Myc |
| chr8:128022618-128022658 | Myc |

| | |
|---|---|
| chr8:128022701-128022741 | Myc |
| chr8:128022793-128022833 | Myc |
| chr8:128022929-128022969 | Myc |
| chr8:128022979-128023019 | Myc |
| chr8:128023101-128023141 | Myc |
| chr8:128023189-128023229 | Myc |
| chr8:128023258-128023298 | Myc |
| chr8:128023309-128023349 | Myc |
| chr8:128023360-128023400 | Myc |
| chr8:128023423-128023463 | Myc |
| chr8:128023513-128023553 | Myc |
| chr8:128023583-128023623 | Myc |
| chr8:128023648-128023688 | Myc |
| chr8:128023709-128023749 | Myc |
| chr8:128023804-128023844 | Myc |
| chr8:128023977-128024017 | Myc |
| chr8:128024031-128024071 | Myc |
| chr8:128024110-128024150 | Myc |
| chr8:128024167-128024207 | Myc |
| chr8:128024260-128024300 | Myc |
| chr8:128024588-128024628 | Myc |
| chr8:128024652-128024692 | Myc |
| chr8:128024702-128024742 | Myc |
| chr8:128024763-128024803 | Myc |
| chr8:128024876-128024916 | Myc |
| chr8:128024964-128025004 | Myc |
| chr8:128025070-128025110 | Myc |
| chr8:128025217-128025257 | Myc |
| chr8:128025425-128025465 | Myc |
| chr8:128025475-128025515 | Myc |
| chr8:128025579-128025619 | Myc |
| chr8:128025629-128025669 | Myc |
| chr8:128025716-128025756 | Myc |
| chr8:128025788-128025828 | Myc |
| chr8:128025850-128025890 | Myc |
| chr8:128025900-128025940 | Myc |
| chr8:128025969-128026009 | Myc |
| chr8:128026115-128026155 | Myc |
| chr8:128026232-128026272 | Myc |
| chr8:128026283-128026323 | Myc |
| chr8:128026347-128026387 | Myc |
| chr8:128026405-128026445 | Myc |
| chr8:128026501-128026541 | Myc |
| chr8:128026558-128026598 | Myc |
| chr8:128026612-128026652 | Myc |
| chr8:128026712-128026752 | Myc |
| chr8:128026832-128026872 | Myc |
| chr8:128027201-128027241 | Myc |
| chr8:128027276-128027316 | Myc |
| chr8:128027334-128027374 | Myc |
| chr8:128027415-128027455 | Myc |
| chr8:128027465-128027505 | Myc |

| | |
|---|---|
| chr8:128027612-128027652 | Myc |
| chr8:128027826-128027866 | Myc |
| chr8:128027902-128027942 | Myc |
| chr8:128027952-128027992 | Myc |
| chr8:128028126-128028166 | Myc |
| chr8:128028188-128028228 | Myc |
| chr8:128028249-128028289 | Myc |
| chr8:128028310-128028350 | Myc |
| chr8:128028413-128028453 | Myc |
| chr8:128028544-128028584 | Myc |
| chr8:128028601-128028641 | Myc |
| chr8:128028681-128028721 | Myc |
| chr8:128028736-128028776 | Myc |
| chr8:128028791-128028831 | Myc |
| chr8:128028924-128028964 | Myc |
| chr8:128029008-128029048 | Myc |
| chr8:128029058-128029098 | Myc |
| chr8:128029146-128029186 | Myc |
| chr8:128029197-128029237 | Myc |
| chr8:128029273-128029313 | Myc |
| chr8:128029361-128029401 | Myc |
| chr8:128029516-128029556 | Myc |
| chr8:128029584-128029624 | Myc |
| chr8:128029639-128029679 | Myc |
| chr8:128029734-128029774 | Myc |
| chr8:128029832-128029872 | Myc |
| chr8:128029889-128029929 | Myc |
| chr8:128029972-128030012 | Myc |
| chr8:128030037-128030077 | Myc |
| chr8:128030135-128030175 | Myc |
| chr8:128030304-128030344 | Myc |
| chr8:128030402-128030442 | Myc |
| chr8:128030467-128030507 | Myc |
| chr8:128030557-128030597 | Myc |
| chr8:128030611-128030651 | Myc |
| chr8:128030742-128030782 | Myc |
| chr8:128030818-128030858 | Myc |
| chr8:128030868-128030908 | Myc |
| chr8:128031063-128031103 | Myc |
| chr8:128031117-128031157 | Myc |
| chr8:128031192-128031232 | Myc |
| chr8:128031299-128031339 | Myc |
| chr8:128031382-128031422 | Myc |
| chr8:128031432-128031472 | Myc |
| chr8:128031504-128031544 | Myc |
| chr8:128031562-128031602 | Myc |
| chr8:128031651-128031691 | Myc |
| chr8:128031717-128031757 | Myc |
| chr8:128031887-128031927 | Myc |
| chr8:128031965-128032005 | Myc |
| chr8:128032075-128032115 | Myc |
| chr8:128032147-128032187 | Myc |

| | |
|---|---|
| chr8:128032223-128032263 | Myc |
| chr8:128032349-128032389 | Myc |
| chr8:128032677-128032717 | Myc |
| chr8:128032790-128032830 | Myc |
| chr8:128032893-128032933 | Myc |
| chr8:128033014-128033054 | Myc |
| chr8:128033110-128033150 | Myc |
| chr8:128033278-128033318 | Myc |
| chr8:128033382-128033422 | Myc |
| chr8:128033434-128033474 | Myc |
| chr8:128033505-128033545 | Myc |
| chr8:128033577-128033617 | Myc |
| chr8:128033666-128033706 | Myc |
| chr8:128033830-128033870 | Myc |
| chr8:128033921-128033961 | Myc |
| chr8:128033971-128034011 | Myc |
| chr8:128034090-128034130 | Myc |
| chr8:128034151-128034191 | Myc |
| chr8:128034231-128034271 | Myc |
| chr8:128034331-128034371 | Myc |
| chr8:128034413-128034453 | Myc |
| chr8:128034471-128034511 | Myc |
| chr8:128034608-128034648 | Myc |
| chr8:128034681-128034721 | Myc |
| chr8:128034755-128034795 | Myc |
| chr8:128034894-128034934 | Myc |
| chr8:128035063-128035103 | Myc |
| chr8:128035120-128035160 | Myc |
| chr8:128035195-128035235 | Myc |
| chr8:128035300-128035340 | Myc |
| chr8:128035351-128035391 | Myc |
| chr8:128035443-128035483 | Myc |
| chr8:128035507-128035547 | Myc |
| chr8:128035568-128035608 | Myc |
| chr8:128035725-128035765 | Myc |
| chr8:128035959-128035999 | Myc |
| chr8:128036045-128036085 | Myc |
| chr8:128036121-128036161 | Myc |
| chr8:128036246-128036286 | Myc |
| chr8:128036399-128036439 | Myc |
| chr8:128036551-128036591 | Myc |
| chr8:128036601-128036641 | Myc |
| chr8:128036821-128036861 | Myc |
| chr8:128036921-128036961 | Myc |
| chr8:128037031-128037071 | Myc |
| chr8:128037085-128037125 | Myc |
| chr8:128037252-128037292 | Myc |
| chr8:128037790-128037830 | Myc |
| chr8:128037840-128037880 | Myc |
| chr8:128037898-128037938 | Myc |
| chr8:128037992-128038032 | Myc |
| chr8:128038071-128038111 | Myc |

| | |
|---|---|
| chr8:128038163-128038203 | Myc |
| chr8:128038316-128038356 | Myc |
| chr8:128038424-128038464 | Myc |
| chr8:128038533-128038573 | Myc |
| chr8:128038583-128038623 | Myc |
| chr8:128038675-128038715 | Myc |
| chr8:128038771-128038811 | Myc |
| chr8:128038821-128038861 | Myc |
| chr8:128038906-128038946 | Myc |
| chr8:128038957-128038997 | Myc |
| chr8:128039046-128039086 | Myc |
| chr8:128039123-128039163 | Myc |
| chr8:128039185-128039225 | Myc |
| chr8:128039235-128039275 | Myc |
| chr8:128039297-128039337 | Myc |
| chr8:128039349-128039389 | Myc |
| chr8:128039431-128039471 | Myc |
| chr8:128039526-128039566 | Myc |
| chr8:128039609-128039649 | Myc |
| chr8:128039702-128039742 | Myc |
| chr8:128039803-128039843 | Myc |
| chr8:128039862-128039902 | Myc |
| chr8:128039971-128040011 | Myc |
| chr8:128040082-128040122 | Myc |
| chr8:128040171-128040211 | Myc |
| chr8:128040243-128040283 | Myc |
| chr8:128040377-128040417 | Myc |
| chr8:128040462-128040502 | Myc |
| chr8:128040551-128040591 | Myc |
| chr8:128040619-128040659 | Myc |
| chr8:128040747-128040787 | Myc |
| chr8:128040825-128040865 | Myc |
| chr8:128040919-128040959 | Myc |
| chr8:128040996-128041036 | Myc |
| chr8:128041060-128041100 | Myc |
| chr8:128041136-128041176 | Myc |
| chr8:128041297-128041337 | Myc |
| chr8:128041394-128041434 | Myc |
| chr8:128041460-128041500 | Myc |
| chr8:128041510-128041550 | Myc |
| chr8:128041561-128041601 | Myc |
| chr8:128041652-128041692 | Myc |
| chr8:128041728-128041768 | Myc |
| chr8:128041787-128041827 | Myc |
| chr8:128041837-128041877 | Myc |
| chr8:128041895-128041935 | Myc |
| chr8:128041978-128042018 | Myc |
| chr8:128042068-128042108 | Myc |
| chr8:128042158-128042198 | Myc |
| chr8:128042240-128042280 | Myc |
| chr8:128042300-128042340 | Myc |
| chr8:128042455-128042495 | Myc |

| | |
|---|---|
| chr8:128042534-128042574 | Myc |
| chr8:128042584-128042624 | Myc |
| chr8:128042814-128042854 | Myc |
| chr8:128043037-128043077 | Myc |
| chr8:128043239-128043279 | Myc |
| chr8:128043484-128043524 | Myc |
| chr8:128043537-128043577 | Myc |
| chr8:128043618-128043658 | Myc |
| chr8:128043744-128043784 | Myc |
| chr8:128043919-128043959 | Myc |
| chr8:128043989-128044029 | Myc |
| chr8:128044040-128044080 | Myc |
| chr8:128044097-128044137 | Myc |
| chr8:128044159-128044199 | Myc |
| chr8:128044209-128044249 | Myc |
| chr8:128044282-128044322 | Myc |
| chr8:128044381-128044421 | Myc |
| chr8:128044530-128044570 | Myc |
| chr8:128044636-128044676 | Myc |
| chr8:128044758-128044798 | Myc |
| chr8:128044822-128044862 | Myc |
| chr8:128044997-128045037 | Myc |
| chr8:128045108-128045148 | Myc |
| chr8:128045253-128045293 | Myc |
| chr8:128045438-128045478 | Myc |
| chr8:128045548-128045588 | Myc |
| chr8:128045618-128045658 | Myc |
| chr8:128045816-128045856 | Myc |
| chr8:128045881-128045921 | Myc |
| chr8:128045933-128045973 | Myc |
| chr8:128046029-128046069 | Myc |
| chr8:128046116-128046156 | Myc |
| chr8:128046212-128046252 | Myc |
| chr8:128046263-128046303 | Myc |
| chr8:128046370-128046410 | Myc |
| chr8:128046478-128046518 | Myc |
| chr8:128046542-128046582 | Myc |
| chr8:128046633-128046673 | Myc |
| chr8:128046772-128046812 | Myc |
| chr8:128046853-128046893 | Myc |
| chr8:128046903-128046943 | Myc |
| chr8:128046972-128047012 | Myc |
| chr8:128047114-128047154 | Myc |
| chr8:128047221-128047261 | Myc |
| chr8:128047366-128047406 | Myc |
| chr8:128047422-128047462 | Myc |
| chr8:128047480-128047520 | Myc |
| chr8:128047539-128047579 | Myc |
| chr8:128047593-128047633 | Myc |
| chr8:128047656-128047696 | Myc |
| chr8:128047733-128047773 | Myc |
| chr8:128047798-128047838 | Myc |

| | |
|---|---|
| chr8:128047862-128047902 | Myc |
| chr8:128047950-128047990 | Myc |
| chr8:128048031-128048071 | Myc |
| chr8:128048139-128048179 | Myc |
| chr8:128048257-128048297 | Myc |
| chr8:128048314-128048354 | Myc |
| chr8:128048374-128048414 | Myc |
| chr8:128048433-128048473 | Myc |
| chr8:128048592-128048632 | Myc |
| chr8:128048652-128048692 | Myc |
| chr8:128048707-128048747 | Myc |
| chr8:128048794-128048834 | Myc |
| chr8:128048886-128048926 | Myc |
| chr8:128048941-128048981 | Myc |
| chr8:128049042-128049082 | Myc |
| chr8:128049177-128049217 | Myc |
| chr8:128049294-128049334 | Myc |
| chr8:128049382-128049422 | Myc |
| chr8:128049438-128049478 | Myc |
| chr8:128049615-128049655 | Myc |
| chr8:128049708-128049748 | Myc |
| chr8:128049822-128049862 | Myc |
| chr8:128049907-128049947 | Myc |
| chr8:128049972-128050012 | Myc |
| chr8:128050056-128050096 | Myc |
| chr8:128050145-128050185 | Myc |
| chr8:128050205-128050245 | Myc |
| chr8:128050269-128050309 | Myc |
| chr8:128050376-128050416 | Myc |
| chr8:128050469-128050509 | Myc |
| chr8:128050526-128050566 | Myc |
| chr8:128050607-128050647 | Myc |
| chr8:128050666-128050706 | Myc |
| chr8:128050733-128050773 | Myc |
| chr8:128050785-128050825 | Myc |
| chr8:128050950-128050990 | Myc |
| chr8:128051043-128051083 | Myc |
| chr8:128051137-128051177 | Myc |
| chr8:128051193-128051233 | Myc |
| chr8:128051326-128051366 | Myc |
| chr8:128051480-128051520 | Myc |
| chr8:128051557-128051597 | Myc |
| chr8:128051715-128051755 | Myc |
| chr8:128051796-128051836 | Myc |
| chr8:128051856-128051896 | Myc |
| chr8:128051958-128051998 | Myc |
| chr8:128052057-128052097 | Myc |
| chr8:128052213-128052253 | Myc |
| chr8:128052317-128052357 | Myc |
| chr8:128052368-128052408 | Myc |
| chr8:128052419-128052459 | Myc |
| chr8:128052506-128052546 | Myc |

| | |
|---|---|
| chr8:128052576-128052616 | Myc |
| chr8:128052687-128052727 | Myc |
| chr8:128052737-128052777 | Myc |
| chr8:128052816-128052856 | Myc |
| chr8:128052930-128052970 | Myc |
| chr8:128052981-128053021 | Myc |
| chr8:128053040-128053080 | Myc |
| chr8:128053149-128053189 | Myc |
| chr8:128053252-128053292 | Myc |
| chr8:128053309-128053349 | Myc |
| chr8:128053375-128053415 | Myc |
| chr8:128053668-128053708 | Myc |
| chr8:128053779-128053819 | Myc |
| chr8:128053836-128053876 | Myc |
| chr8:128053925-128053965 | Myc |
| chr8:128054026-128054066 | Myc |
| chr8:128054129-128054169 | Myc |
| chr8:128054199-128054239 | Myc |
| chr8:128054249-128054289 | Myc |
| chr8:128054376-128054416 | Myc |
| chr8:128054426-128054466 | Myc |
| chr8:128054510-128054550 | Myc |
| chr8:128054578-128054618 | Myc |
| chr8:128054638-128054678 | Myc |
| chr8:128054730-128054770 | Myc |
| chr8:128054824-128054864 | Myc |
| chr8:128054881-128054921 | Myc |
| chr8:128055084-128055124 | Myc |
| chr8:128055188-128055228 | Myc |
| chr8:128055265-128055305 | Myc |
| chr8:128055377-128055417 | Myc |
| chr8:128055437-128055477 | Myc |
| chr8:128055575-128055615 | Myc |
| chr8:128055626-128055666 | Myc |
| chr8:128055715-128055755 | Myc |
| chr8:128055770-128055810 | Myc |
| chr8:128055834-128055874 | Myc |
| chr8:128055906-128055946 | Myc |
| chr8:128056020-128056060 | Myc |
| chr8:128056100-128056140 | Myc |
| chr8:128056196-128056236 | Myc |
| chr8:128056246-128056286 | Myc |
| chr8:128056406-128056446 | Myc |
| chr8:128056466-128056506 | Myc |
| chr8:128056555-128056595 | Myc |
| chr8:128056804-128056844 | Myc |
| chr8:128056938-128056978 | Myc |
| chr8:128057019-128057059 | Myc |
| chr8:128057251-128057291 | Myc |
| chr8:128057392-128057432 | Myc |
| chr8:128057503-128057543 | Myc |
| chr8:128057582-128057622 | Myc |

| | |
|---|---|
| chr8:128057655-128057695 | Myc |
| chr8:128057823-128057863 | Myc |
| chr8:128057992-128058032 | Myc |
| chr8:128058255-128058295 | Myc |
| chr8:128058559-128058599 | Myc |
| chr8:128058799-128058839 | Myc |
| chr8:128058888-128058928 | Myc |
| chr8:128059050-128059090 | Myc |
| chr8:128059102-128059142 | Myc |
| chr8:128059260-128059300 | Myc |
| chr8:128059421-128059461 | Myc |
| chr8:128059514-128059554 | Myc |
| chr8:128059641-128059681 | Myc |
| chr8:128060008-128060048 | Myc |
| chr8:128060157-128060197 | Myc |
| chr8:128060405-128060445 | Myc |
| chr8:128060528-128060568 | Myc |
| chr8:128060578-128060618 | Myc |
| chr8:128060675-128060715 | Myc |
| chr8:128060823-128060863 | Myc |
| chr8:128060896-128060936 | Myc |
| chr8:128061002-128061042 | Myc |
| chr8:128061145-128061185 | Myc |
| chr8:128061219-128061259 | Myc |
| chr8:128061271-128061311 | Myc |
| chr8:128061482-128061522 | Myc |
| chr8:128061540-128061580 | Myc |
| chr8:128061591-128061631 | Myc |
| chr8:128061687-128061727 | Myc |
| chr8:128061769-128061809 | Myc |
| chr8:128061834-128061874 | Myc |
| chr8:128061899-128061939 | Myc |
| chr8:128062009-128062049 | Myc |
| chr8:128062074-128062114 | Myc |
| chr8:128062148-128062188 | Myc |
| chr8:128062253-128062293 | Myc |
| chr8:128062395-128062435 | Myc |
| chr8:128062510-128062550 | Myc |
| chr8:128062574-128062614 | Myc |
| chr8:128062640-128062680 | Myc |
| chr8:128062695-128062735 | Myc |
| chr8:128062810-128062850 | Myc |
| chr8:128062876-128062916 | Myc |
| chr8:128062968-128063008 | Myc |
| chr8:128063065-128063105 | Myc |
| chr8:128063128-128063168 | Myc |
| chr8:128063227-128063267 | Myc |
| chr8:128063296-128063336 | Myc |
| chr8:128063405-128063445 | Myc |
| chr8:128063491-128063531 | Myc |
| chr8:128063610-128063650 | Myc |
| chr8:128063680-128063720 | Myc |

| | |
|---|---|
| chr8:128063853-128063893 | Myc |
| chr8:128063955-128063995 | Myc |
| chr8:128064041-128064081 | Myc |
| chr8:128064148-128064188 | Myc |
| chr8:128064316-128064356 | Myc |
| chr8:128064383-128064423 | Myc |
| chr8:128064448-128064488 | Myc |
| chr8:128064580-128064620 | Myc |
| chr8:128064630-128064670 | Myc |
| chr8:128064792-128064832 | Myc |
| chr8:128064891-128064931 | Myc |
| chr8:128064985-128065025 | Myc |
| chr8:128065089-128065129 | Myc |
| chr8:128065141-128065181 | Myc |
| chr8:128065210-128065250 | Myc |
| chr8:128065327-128065367 | Myc |
| chr8:128065483-128065523 | Myc |
| chr8:128065548-128065588 | Myc |
| chr8:128065626-128065666 | Myc |
| chr8:128065776-128065816 | Myc |
| chr8:128065900-128065940 | Myc |
| chr8:128065951-128065991 | Myc |
| chr8:128066031-128066071 | Myc |
| chr8:128066084-128066124 | Myc |
| chr8:128066142-128066182 | Myc |
| chr8:128066208-128066248 | Myc |
| chr8:128066309-128066349 | Myc |
| chr8:128066384-128066424 | Myc |
| chr8:128066471-128066511 | Myc |
| chr8:128066582-128066622 | Myc |
| chr8:128066655-128066695 | Myc |
| chr8:128066784-128066824 | Myc |
| chr8:128066915-128066955 | Myc |
| chr8:128067024-128067064 | Myc |
| chr8:128067077-128067117 | Myc |
| chr8:128067141-128067181 | Myc |
| chr8:128067199-128067239 | Myc |
| chr8:128067316-128067356 | Myc |
| chr8:128067480-128067520 | Myc |
| chr8:128067643-128067683 | Myc |
| chr8:128067733-128067773 | Myc |
| chr8:128067785-128067825 | Myc |
| chr8:128067907-128067947 | Myc |
| chr8:128067969-128068009 | Myc |
| chr8:128068121-128068161 | Myc |
| chr8:128068209-128068249 | Myc |
| chr8:128068265-128068305 | Myc |
| chr8:128068358-128068398 | Myc |
| chr8:128068457-128068497 | Myc |
| chr8:128068671-128068711 | Myc |
| chr8:128068930-128068970 | Myc |
| chr8:128069029-128069069 | Myc |

| | |
|---|---|
| chr8:128069170-128069210 | Myc |
| chr8:128069281-128069321 | Myc |
| chr8:128069335-128069375 | Myc |
| chr8:128069392-128069432 | Myc |
| chr8:128069526-128069566 | Myc |
| chr8:128069579-128069619 | Myc |
| chr8:128069669-128069709 | Myc |
| chr8:128069765-128069805 | Myc |
| chr8:128069815-128069855 | Myc |
| chr8:128069916-128069956 | Myc |
| chr8:128070044-128070084 | Myc |
| chr8:128070136-128070176 | Myc |
| chr8:128070238-128070278 | Myc |
| chr8:128070293-128070333 | Myc |
| chr8:128070363-128070403 | Myc |
| chr8:128070417-128070457 | Myc |
| chr8:128070504-128070544 | Myc |
| chr8:128070601-128070641 | Myc |
| chr8:128070662-128070702 | Myc |
| chr8:128070799-128070839 | Myc |
| chr8:128071069-128071109 | Myc |
| chr8:128071119-128071159 | Myc |
| chr8:128071191-128071231 | Myc |
| chr8:128071272-128071312 | Myc |
| chr8:128071351-128071391 | Myc |
| chr8:128071486-128071526 | Myc |
| chr8:128071595-128071635 | Myc |
| chr8:128071696-128071736 | Myc |
| chr8:128071765-128071805 | Myc |
| chr8:128071885-128071925 | Myc |
| chr8:128071953-128071993 | Myc |
| chr8:128072017-128072057 | Myc |
| chr8:128072099-128072139 | Myc |
| chr8:128072219-128072259 | Myc |
| chr8:128072270-128072310 | Myc |
| chr8:128072355-128072395 | Myc |
| chr8:128072540-128072580 | Myc |
| chr8:128072600-128072640 | Myc |
| chr8:128072679-128072719 | Myc |
| chr8:128072913-128072953 | Myc |
| chr8:128072963-128073003 | Myc |
| chr8:128073017-128073057 | Myc |
| chr8:128073160-128073200 | Myc |
| chr8:128073210-128073250 | Myc |
| chr8:128073296-128073336 | Myc |
| chr8:128073386-128073426 | Myc |
| chr8:128073436-128073476 | Myc |
| chr8:128073590-128073630 | Myc |
| chr8:128073689-128073729 | Myc |
| chr8:128073750-128073790 | Myc |
| chr8:128073831-128073871 | Myc |
| chr8:128073940-128073980 | Myc |

| | |
|---|---|
| chr8:128074126-128074166 | Myc |
| chr8:128074261-128074301 | Myc |
| chr8:128074317-128074357 | Myc |
| chr8:128074367-128074407 | Myc |
| chr8:128074429-128074469 | Myc |
| chr8:128074517-128074557 | Myc |
| chr8:128074582-128074622 | Myc |
| chr8:128074654-128074694 | Myc |
| chr8:128074750-128074790 | Myc |
| chr8:128074813-128074853 | Myc |
| chr8:128074939-128074979 | Myc |
| chr8:128074997-128075037 | Myc |
| chr8:128075198-128075238 | Myc |
| chr8:128075486-128075526 | Myc |
| chr8:128075562-128075602 | Myc |
| chr8:128075673-128075713 | Myc |
| chr8:128075759-128075799 | Myc |
| chr8:128075812-128075852 | Myc |
| chr8:128075862-128075902 | Myc |
| chr8:128075920-128075960 | Myc |
| chr8:128075976-128076016 | Myc |
| chr8:128076042-128076082 | Myc |
| chr8:128076111-128076151 | Myc |
| chr8:128076225-128076265 | Myc |
| chr8:128076361-128076401 | Myc |
| chr8:128076422-128076462 | Myc |
| chr8:128076579-128076619 | Myc |
| chr8:128076755-128076795 | Myc |
| chr8:128076849-128076889 | Myc |
| chr8:128076926-128076966 | Myc |
| chr8:128076976-128077016 | Myc |
| chr8:128077060-128077100 | Myc |
| chr8:128077211-128077251 | Myc |
| chr8:128077332-128077372 | Myc |
| chr8:128077401-128077441 | Myc |
| chr8:128077484-128077524 | Myc |
| chr8:128077579-128077619 | Myc |
| chr8:128077657-128077697 | Myc |
| chr8:128077823-128077863 | Myc |
| chr8:128077906-128077946 | Myc |
| chr8:128078027-128078067 | Myc |
| chr8:128078108-128078148 | Myc |
| chr8:128078264-128078304 | Myc |
| chr8:128078333-128078373 | Myc |
| chr8:128078400-128078440 | Myc |
| chr8:128078513-128078553 | Myc |
| chr8:128078566-128078606 | Myc |
| chr8:128078648-128078688 | Myc |
| chr8:128078741-128078781 | Myc |
| chr8:128078791-128078831 | Myc |
| chr8:128078890-128078930 | Myc |
| chr8:128078990-128079030 | Myc |

| | |
|---|---|
| chr8:128079074-128079114 | Myc |
| chr8:128079198-128079238 | Myc |
| chr8:128079285-128079325 | Myc |
| chr8:128079414-128079454 | Myc |
| chr8:128079472-128079512 | Myc |
| chr8:128079531-128079571 | Myc |
| chr8:128079678-128079718 | Myc |
| chr8:128079882-128079922 | Myc |
| chr8:128080022-128080062 | Myc |
| chr8:128080089-128080129 | Myc |
| chr8:128080161-128080201 | Myc |
| chr8:128080225-128080265 | Myc |
| chr8:128080284-128080324 | Myc |
| chr8:128080400-128080440 | Myc |
| chr8:128080467-128080507 | Myc |
| chr8:128080517-128080557 | Myc |
| chr8:128080621-128080661 | Myc |
| chr8:128080671-128080711 | Myc |
| chr8:128080785-128080825 | Myc |
| chr8:128080860-128080900 | Myc |
| chr8:128080948-128080988 | Myc |
| chr8:128080998-128081038 | Myc |
| chr8:128081062-128081102 | Myc |
| chr8:128081112-128081152 | Myc |
| chr8:128081242-128081282 | Myc |
| chr8:128081321-128081361 | Myc |
| chr8:128081415-128081455 | Myc |
| chr8:128081484-128081524 | Myc |
| chr8:128081540-128081580 | Myc |
| chr8:128081637-128081677 | Myc |
| chr8:128081708-128081748 | Myc |
| chr8:128081873-128081913 | Myc |
| chr8:128081931-128081971 | Myc |
| chr8:128082045-128082085 | Myc |
| chr8:128082095-128082135 | Myc |
| chr8:128082179-128082219 | Myc |
| chr8:128082242-128082282 | Myc |
| chr8:128082359-128082399 | Myc |
| chr8:128082443-128082483 | Myc |
| chr8:128082510-128082550 | Myc |
| chr8:128082665-128082705 | Myc |
| chr8:128082755-128082795 | Myc |
| chr8:128082828-128082868 | Myc |
| chr8:128082972-128083012 | Myc |
| chr8:128083032-128083072 | Myc |
| chr8:128083222-128083262 | Myc |
| chr8:128083304-128083344 | Myc |
| chr8:128083410-128083450 | Myc |
| chr8:128083503-128083543 | Myc |
| chr8:128083573-128083613 | Myc |
| chr8:128083693-128083733 | Myc |
| chr8:128083745-128083785 | Myc |

| | |
|---|---|
| chr8:128083944-128083984 | Myc |
| chr8:128084080-128084120 | Myc |
| chr8:128084214-128084254 | Myc |
| chr8:128084279-128084319 | Myc |
| chr8:128084479-128084519 | Myc |
| chr8:128084569-128084609 | Myc |
| chr8:128084650-128084690 | Myc |
| chr8:128084702-128084742 | Myc |
| chr8:128084825-128084865 | Myc |
| chr8:128084909-128084949 | Myc |
| chr8:128085018-128085058 | Myc |
| chr8:128085068-128085108 | Myc |
| chr8:128085119-128085159 | Myc |
| chr8:128085272-128085312 | Myc |
| chr8:128085390-128085430 | Myc |
| chr8:128085460-128085500 | Myc |
| chr8:128085514-128085554 | Myc |
| chr8:128085588-128085628 | Myc |
| chr8:128085687-128085727 | Myc |
| chr8:128085765-128085805 | Myc |
| chr8:128085845-128085885 | Myc |
| chr8:128085940-128085980 | Myc |
| chr8:128086004-128086044 | Myc |
| chr8:128086080-128086120 | Myc |
| chr8:128086182-128086222 | Myc |
| chr8:128086240-128086280 | Myc |
| chr8:128086290-128086330 | Myc |
| chr8:128086416-128086456 | Myc |
| chr8:128086616-128086656 | Myc |
| chr8:128086753-128086793 | Myc |
| chr8:128086803-128086843 | Myc |
| chr8:128086946-128086986 | Myc |
| chr8:128087053-128087093 | Myc |
| chr8:128087103-128087143 | Myc |
| chr8:128087171-128087211 | Myc |
| chr8:128087229-128087269 | Myc |
| chr8:128087282-128087322 | Myc |
| chr8:128087356-128087396 | Myc |
| chr8:128087539-128087579 | Myc |
| chr8:128087640-128087680 | Myc |
| chr8:128087692-128087732 | Myc |
| chr8:128087752-128087792 | Myc |
| chr8:128087842-128087882 | Myc |
| chr8:128087893-128087933 | Myc |
| chr8:128087974-128088014 | Myc |
| chr8:128088052-128088092 | Myc |
| chr8:128088123-128088163 | Myc |
| chr8:128088223-128088263 | Myc |
| chr8:128088291-128088331 | Myc |
| chr8:128088411-128088451 | Myc |
| chr8:128088562-128088602 | Myc |
| chr8:128088761-128088801 | Myc |

| | |
|---|---|
| chr8:128088860-128088900 | Myc |
| chr8:128088926-128088966 | Myc |
| chr8:128088988-128089028 | Myc |
| chr8:128089048-128089088 | Myc |
| chr8:128089184-128089224 | Myc |
| chr8:128089280-128089320 | Myc |
| chr8:128089380-128089420 | Myc |
| chr8:128089495-128089535 | Myc |
| chr8:128089562-128089602 | Myc |
| chr8:128089682-128089722 | Myc |
| chr8:128089739-128089779 | Myc |
| chr8:128089791-128089831 | Myc |
| chr8:128089909-128089949 | Myc |
| chr8:128089972-128090012 | Myc |
| chr8:128090040-128090080 | Myc |
| chr8:128090166-128090206 | Myc |
| chr8:128090260-128090300 | Myc |
| chr8:128090321-128090361 | Myc |
| chr8:128090444-128090484 | Myc |
| chr8:128090497-128090537 | Myc |
| chr8:128090565-128090605 | Myc |
| chr8:128090637-128090677 | Myc |
| chr8:128090761-128090801 | Myc |
| chr8:128090912-128090952 | Myc |
| chr8:128090984-128091024 | Myc |
| chr8:128091118-128091158 | Myc |
| chr8:128091202-128091242 | Myc |
| chr8:128091255-128091295 | Myc |
| chr8:128091343-128091383 | Myc |
| chr8:128091565-128091605 | Myc |
| chr8:128091699-128091739 | Myc |
| chr8:128091814-128091854 | Myc |
| chr8:128091875-128091915 | Myc |
| chr8:128091952-128091992 | Myc |
| chr8:128092123-128092163 | Myc |
| chr8:128092207-128092247 | Myc |
| chr8:128092356-128092396 | Myc |
| chr8:128092435-128092475 | Myc |
| chr8:128092485-128092525 | Myc |
| chr8:128092584-128092624 | Myc |
| chr8:128092661-128092701 | Myc |
| chr8:128092796-128092836 | Myc |
| chr8:128092850-128092890 | Myc |
| chr8:128092911-128092951 | Myc |
| chr8:128092976-128093016 | Myc |
| chr8:128093061-128093101 | Myc |
| chr8:128093198-128093238 | Myc |
| chr8:128093259-128093299 | Myc |
| chr8:128093323-128093363 | Myc |
| chr8:128093401-128093441 | Myc |
| chr8:128093452-128093492 | Myc |
| chr8:128093503-128093543 | Myc |

| | |
|---|---|
| chr8:128093647-128093687 | Myc |
| chr8:128093733-128093773 | Myc |
| chr8:128093929-128093969 | Myc |
| chr8:128094026-128094066 | Myc |
| chr8:128094076-128094116 | Myc |
| chr8:128094135-128094175 | Myc |
| chr8:128094247-128094287 | Myc |
| chr8:128094322-128094362 | Myc |
| chr8:128094476-128094516 | Myc |
| chr8:128094533-128094573 | Myc |
| chr8:128094639-128094679 | Myc |
| chr8:128094690-128094730 | Myc |
| chr8:128094744-128094784 | Myc |
| chr8:128094911-128094951 | Myc |
| chr8:128095100-128095140 | Myc |
| chr8:128095188-128095228 | Myc |
| chr8:128095265-128095305 | Myc |
| chr8:128095333-128095373 | Myc |
| chr8:128095462-128095502 | Myc |
| chr8:128095527-128095567 | Myc |
| chr8:128095618-128095658 | Myc |
| chr8:128095734-128095774 | Myc |
| chr8:128095798-128095838 | Myc |
| chr8:128095877-128095917 | Myc |
| chr8:128095930-128095970 | Myc |
| chr8:128096019-128096059 | Myc |
| chr8:128096110-128096150 | Myc |
| chr8:128096236-128096276 | Myc |
| chr8:128096289-128096329 | Myc |
| chr8:128096360-128096400 | Myc |
| chr8:128096416-128096456 | Myc |
| chr8:128096467-128096507 | Myc |
| chr8:128096557-128096597 | Myc |
| chr8:128096676-128096716 | Myc |
| chr8:128096726-128096766 | Myc |
| chr8:128096837-128096877 | Myc |
| chr8:128096920-128096960 | Myc |
| chr8:128096996-128097036 | Myc |
| chr8:128097134-128097174 | Myc |
| chr8:128097193-128097233 | Myc |
| chr8:128097256-128097296 | Myc |
| chr8:128097368-128097408 | Myc |
| chr8:128097479-128097519 | Myc |
| chr8:128097564-128097604 | Myc |
| chr8:128097645-128097685 | Myc |
| chr8:128097743-128097783 | Myc |
| chr8:128097842-128097882 | Myc |
| chr8:128097914-128097954 | Myc |
| chr8:128097984-128098024 | Myc |
| chr8:128098118-128098158 | Myc |
| chr8:128098245-128098285 | Myc |
| chr8:128098345-128098385 | Myc |

| | |
|---|---|
| chr8:128098448-128098488 | Myc |
| chr8:128098502-128098542 | Myc |
| chr8:128098567-128098607 | Myc |
| chr8:128098785-128098825 | Myc |
| chr8:128098870-128098910 | Myc |
| chr8:128098920-128098960 | Myc |
| chr8:128099113-128099153 | Myc |
| chr8:128099254-128099294 | Myc |
| chr8:128099466-128099506 | Myc |
| chr8:128099549-128099589 | Myc |
| chr8:128099610-128099650 | Myc |
| chr8:128099667-128099707 | Myc |
| chr8:128099733-128099773 | Myc |
| chr8:128100071-128100111 | Myc |
| chr8:128100288-128100328 | Myc |
| chr8:128100345-128100385 | Myc |
| chr8:128100418-128100458 | Myc |
| chr8:128100498-128100538 | Myc |
| chr8:128100585-128100625 | Myc |
| chr8:128100645-128100685 | Myc |
| chr8:128100842-128100882 | Myc |
| chr8:128100892-128100932 | Myc |
| chr8:128100969-128101009 | Myc |
| chr8:128101023-128101063 | Myc |
| chr8:128101073-128101113 | Myc |
| chr8:128101177-128101217 | Myc |
| chr8:128101245-128101285 | Myc |
| chr8:128101297-128101337 | Myc |
| chr8:128101350-128101390 | Myc |
| chr8:128101433-128101473 | Myc |
| chr8:128101483-128101523 | Myc |
| chr8:128101628-128101668 | Myc |
| chr8:128101715-128101755 | Myc |
| chr8:128101777-128101817 | Myc |
| chr8:128101875-128101915 | Myc |
| chr8:128101933-128101973 | Myc |
| chr8:128102065-128102105 | Myc |
| chr8:128102159-128102199 | Myc |
| chr8:128102248-128102288 | Myc |
| chr8:128102482-128102522 | Myc |
| chr8:128102569-128102609 | Myc |
| chr8:128102639-128102679 | Myc |
| chr8:128102742-128102782 | Myc |
| chr8:128102834-128102874 | Myc |
| chr8:128102888-128102928 | Myc |
| chr8:128102951-128102991 | Myc |
| chr8:128103023-128103063 | Myc |
| chr8:128103095-128103135 | Myc |
| chr8:128103229-128103269 | Myc |
| chr8:128103535-128103575 | Myc |
| chr8:128103652-128103692 | Myc |
| chr8:128103759-128103799 | Myc |

| | |
|---|---|
| chr8:128103886-128103926 | Myc |
| chr8:128104055-128104095 | Myc |
| chr8:128104107-128104147 | Myc |
| chr8:128104182-128104222 | Myc |
| chr8:128104254-128104294 | Myc |
| chr8:128104304-128104344 | Myc |
| chr8:128104355-128104395 | Myc |
| chr8:128104412-128104452 | Myc |
| chr8:128104478-128104518 | Myc |
| chr8:128104630-128104670 | Myc |
| chr8:128104680-128104720 | Myc |
| chr8:128104730-128104770 | Myc |
| chr8:128104818-128104858 | Myc |
| chr8:128104887-128104927 | Myc |
| chr8:128104993-128105033 | Myc |
| chr8:128105077-128105117 | Myc |
| chr8:128105272-128105312 | Myc |
| chr8:128105338-128105378 | Myc |
| chr8:128105426-128105466 | Myc |
| chr8:128105620-128105660 | Myc |
| chr8:128105693-128105733 | Myc |
| chr8:128105781-128105821 | Myc |
| chr8:128105898-128105938 | Myc |
| chr8:128105997-128106037 | Myc |
| chr8:128106060-128106100 | Myc |
| chr8:128106261-128106301 | Myc |
| chr8:128106317-128106357 | Myc |
| chr8:128106417-128106457 | Myc |
| chr8:128106513-128106553 | Myc |
| chr8:128106593-128106633 | Myc |
| chr8:128106762-128106802 | Myc |
| chr8:128106812-128106852 | Myc |
| chr8:128106993-128107033 | Myc |
| chr8:128107221-128107261 | Myc |
| chr8:128107271-128107311 | Myc |
| chr8:128107357-128107397 | Myc |
| chr8:128107407-128107447 | Myc |
| chr8:128107527-128107567 | Myc |
| chr8:128107586-128107626 | Myc |
| chr8:128108114-128108154 | Myc |
| chr8:128108241-128108281 | Myc |
| chr8:128108323-128108363 | Myc |
| chr8:128108394-128108434 | Myc |
| chr8:128108656-128108696 | Myc |
| chr8:128108708-128108748 | Myc |
| chr8:128108852-128108892 | Myc |
| chr8:128108925-128108965 | Myc |
| chr8:128108984-128109024 | Myc |
| chr8:128109049-128109089 | Myc |
| chr8:128109119-128109159 | Myc |
| chr8:128109205-128109245 | Myc |
| chr8:128109256-128109296 | Myc |

| | |
|---|---|
| chr8:128109342-128109382 | Myc |
| chr8:128109407-128109447 | Myc |
| chr8:128109464-128109504 | Myc |
| chr8:128109514-128109554 | Myc |
| chr8:128109565-128109605 | Myc |
| chr8:128109638-128109678 | Myc |
| chr8:128109738-128109778 | Myc |
| chr8:128109809-128109849 | Myc |
| chr8:128109863-128109903 | Myc |
| chr8:128109913-128109953 | Myc |
| chr8:128109964-128110004 | Myc |
| chr8:128110031-128110071 | Myc |
| chr8:128110122-128110162 | Myc |
| chr8:128110186-128110226 | Myc |
| chr8:128110248-128110288 | Myc |
| chr8:128110303-128110343 | Myc |
| chr8:128110429-128110469 | Myc |
| chr8:128110606-128110646 | Myc |
| chr8:128110662-128110702 | Myc |
| chr8:128110718-128110758 | Myc |
| chr8:128110768-128110808 | Myc |
| chr8:128110819-128110859 | Myc |
| chr8:128110955-128110995 | Myc |
| chr8:128111070-128111110 | Myc |
| chr8:128111332-128111372 | Myc |
| chr8:128111450-128111490 | Myc |
| chr8:128111524-128111564 | Myc |
| chr8:128111617-128111657 | Myc |
| chr8:128111723-128111763 | Myc |
| chr8:128111777-128111817 | Myc |
| chr8:128111945-128111985 | Myc |
| chr8:128111995-128112035 | Myc |
| chr8:128112076-128112116 | Myc |
| chr8:128112129-128112169 | Myc |
| chr8:128112268-128112308 | Myc |
| chr8:128112318-128112358 | Myc |
| chr8:128112502-128112542 | Myc |
| chr8:128112556-128112596 | Myc |
| chr8:128112641-128112681 | Myc |
| chr8:128112695-128112735 | Myc |
| chr8:128112799-128112839 | Myc |
| chr8:128112902-128112942 | Myc |
| chr8:128112952-128112992 | Myc |
| chr8:128113188-128113228 | Myc |
| chr8:128113455-128113495 | Myc |
| chr8:128113665-128113705 | Myc |
| chr8:128113845-128113885 | Myc |
| chr8:128113895-128113935 | Myc |
| chr8:128113949-128113989 | Myc |
| chr8:128114118-128114158 | Myc |
| chr8:128114220-128114260 | Myc |
| chr8:128114300-128114340 | Myc |

| | |
|---|---|
| chr8:128114380-128114420 | Myc |
| chr8:128114485-128114525 | Myc |
| chr8:128114538-128114578 | Myc |
| chr8:128114664-128114704 | Myc |
| chr8:128114774-128114814 | Myc |
| chr8:128114833-128114873 | Myc |
| chr8:128115096-128115136 | Myc |
| chr8:128115191-128115231 | Myc |
| chr8:128115250-128115290 | Myc |
| chr8:128115620-128115660 | Myc |
| chr8:128115712-128115752 | Myc |
| chr8:128115909-128115949 | Myc |
| chr8:128115972-128116012 | Myc |
| chr8:128116081-128116121 | Myc |
| chr8:128116211-128116251 | Myc |
| chr8:128116299-128116339 | Myc |
| chr8:128116363-128116403 | Myc |
| chr8:128116463-128116503 | Myc |
| chr8:128116520-128116560 | Myc |
| chr8:128116570-128116610 | Myc |
| chr8:128116646-128116686 | Myc |
| chr8:128116710-128116750 | Myc |
| chr8:128116760-128116800 | Myc |
| chr8:128116828-128116868 | Myc |
| chr8:128116930-128116970 | Myc |
| chr8:128117002-128117042 | Myc |
| chr8:128117115-128117155 | Myc |
| chr8:128117218-128117258 | Myc |
| chr8:128117313-128117353 | Myc |
| chr8:128117454-128117494 | Myc |
| chr8:128117778-128117818 | Myc |
| chr8:128117888-128117928 | Myc |
| chr8:128117982-128118022 | Myc |
| chr8:128118061-128118101 | Myc |
| chr8:128118111-128118151 | Myc |
| chr8:128118198-128118238 | Myc |
| chr8:128118259-128118299 | Myc |
| chr8:128118309-128118349 | Myc |
| chr8:128118414-128118454 | Myc |
| chr8:128118513-128118553 | Myc |
| chr8:128118571-128118611 | Myc |
| chr8:128118713-128118753 | Myc |
| chr8:128118763-128118803 | Myc |
| chr8:128118890-128118930 | Myc |
| chr8:128119017-128119057 | Myc |
| chr8:128119152-128119192 | Myc |
| chr8:128119493-128119533 | Myc |
| chr8:128119577-128119617 | Myc |
| chr8:128119673-128119713 | Myc |
| chr8:128119743-128119783 | Myc |
| chr8:128119802-128119842 | Myc |
| chr8:128119852-128119892 | Myc |

| | |
|---|---|
| chr8:128119930-128119970 | Myc |
| chr8:128120073-128120113 | Myc |
| chr8:128121877-128121917 | Myc |
| chr8:128122308-128122348 | Myc |
| chr8:128122401-128122441 | Myc |
| chr8:128122498-128122538 | Myc |
| chr8:128122581-128122621 | Myc |
| chr8:128122684-128122724 | Myc |
| chr8:128122740-128122780 | Myc |
| chr8:128122867-128122907 | Myc |
| chr8:128123058-128123098 | Myc |
| chr8:128123259-128123299 | Myc |
| chr8:128123339-128123379 | Myc |
| chr8:128123454-128123494 | Myc |
| chr8:128123511-128123551 | Myc |
| chr8:128123616-128123656 | Myc |
| chr8:128123736-128123776 | Myc |
| chr8:128123816-128123856 | Myc |
| chr8:128124074-128124114 | Myc |
| chr8:128124175-128124215 | Myc |
| chr8:128124278-128124318 | Myc |
| chr8:128124344-128124384 | Myc |
| chr8:128124425-128124465 | Myc |
| chr8:128124493-128124533 | Myc |
| chr8:128124581-128124621 | Myc |
| chr8:128124666-128124706 | Myc |
| chr8:128124716-128124756 | Myc |
| chr8:128124852-128124892 | Myc |
| chr8:128124950-128124990 | Myc |
| chr8:128125007-128125047 | Myc |
| chr8:128125091-128125131 | Myc |
| chr8:128125169-128125209 | Myc |
| chr8:128125219-128125259 | Myc |
| chr8:128125279-128125319 | Myc |
| chr8:128125365-128125405 | Myc |
| chr8:128125448-128125488 | Myc |
| chr8:128125515-128125555 | Myc |
| chr8:128125570-128125610 | Myc |
| chr8:128125657-128125697 | Myc |
| chr8:128125720-128125760 | Myc |
| chr8:128125804-128125844 | Myc |
| chr8:128125902-128125942 | Myc |
| chr8:128125965-128126005 | Myc |
| chr8:128126146-128126186 | Myc |
| chr8:128126212-128126252 | Myc |
| chr8:128126275-128126315 | Myc |
| chr8:128126351-128126391 | Myc |
| chr8:128126401-128126441 | Myc |
| chr8:128126457-128126497 | Myc |
| chr8:128126544-128126584 | Myc |
| chr8:128126596-128126636 | Myc |
| chr8:128126708-128126748 | Myc |

| | |
|---|---|
| chr8:128126758-128126798 | Myc |
| chr8:128126820-128126860 | Myc |
| chr8:128126921-128126961 | Myc |
| chr8:128126983-128127023 | Myc |
| chr8:128127034-128127074 | Myc |
| chr8:128127095-128127135 | Myc |
| chr8:128127193-128127233 | Myc |
| chr8:128127281-128127321 | Myc |
| chr8:128127331-128127371 | Myc |
| chr8:128127503-128127543 | Myc |
| chr8:128127660-128127700 | Myc |
| chr8:128127710-128127750 | Myc |
| chr8:128127806-128127846 | Myc |
| chr8:128127897-128127937 | Myc |
| chr8:128127968-128128008 | Myc |
| chr8:128128056-128128096 | Myc |
| chr8:128128106-128128146 | Myc |
| chr8:128128192-128128232 | Myc |
| chr8:128128263-128128303 | Myc |
| chr8:128128337-128128377 | Myc |
| chr8:128128420-128128460 | Myc |
| chr8:128128476-128128516 | Myc |
| chr8:128128648-128128688 | Myc |
| chr8:128128712-128128752 | Myc |
| chr8:128128808-128128848 | Myc |
| chr8:128128987-128129027 | Myc |
| chr8:128129120-128129160 | Myc |
| chr8:128129207-128129247 | Myc |
| chr8:128129289-128129329 | Myc |
| chr8:128129406-128129446 | Myc |
| chr8:128129605-128129645 | Myc |
| chr8:128129858-128129898 | Myc |
| chr8:128129962-128130002 | Myc |
| chr8:128130058-128130098 | Myc |
| chr8:128130150-128130190 | Myc |
| chr8:128130251-128130291 | Myc |
| chr8:128130330-128130370 | Myc |
| chr8:128130410-128130450 | Myc |
| chr8:128130496-128130536 | Myc |
| chr8:128130598-128130638 | Myc |
| chr8:128130788-128130828 | Myc |
| chr8:128130867-128130907 | Myc |
| chr8:128130917-128130957 | Myc |
| chr8:128131024-128131064 | Myc |
| chr8:128131119-128131159 | Myc |
| chr8:128131215-128131255 | Myc |
| chr8:128131394-128131434 | Myc |
| chr8:128131444-128131484 | Myc |
| chr8:128131501-128131541 | Myc |
| chr8:128131575-128131615 | Myc |
| chr8:128131673-128131713 | Myc |
| chr8:128131771-128131811 | Myc |

| | |
|---|---|
| chr8:128131821-128131861 | Myc |
| chr8:128131871-128131911 | Myc |
| chr8:128131940-128131980 | Myc |
| chr8:128132003-128132043 | Myc |
| chr8:128132081-128132121 | Myc |
| chr8:128132173-128132213 | Myc |
| chr8:128132276-128132316 | Myc |
| chr8:128132338-128132378 | Myc |
| chr8:128132403-128132443 | Myc |
| chr8:128132553-128132593 | Myc |
| chr8:128132616-128132656 | Myc |
| chr8:128132718-128132758 | Myc |
| chr8:128132859-128132899 | Myc |
| chr8:128132942-128132982 | Myc |
| chr8:128133023-128133063 | Myc |
| chr8:128133092-128133132 | Myc |
| chr8:128133332-128133372 | Myc |
| chr8:128133427-128133467 | Myc |
| chr8:128133538-128133578 | Myc |
| chr8:128133675-128133715 | Myc |
| chr8:128133738-128133778 | Myc |
| chr8:128133797-128133837 | Myc |
| chr8:128133914-128133954 | Myc |
| chr8:128133983-128134023 | Myc |
| chr8:128134078-128134118 | Myc |
| chr8:128134175-128134215 | Myc |
| chr8:128134242-128134282 | Myc |
| chr8:128134355-128134395 | Myc |
| chr8:128134501-128134541 | Myc |
| chr8:128134654-128134694 | Myc |
| chr8:128134704-128134744 | Myc |
| chr8:128134765-128134805 | Myc |
| chr8:128134826-128134866 | Myc |
| chr8:128134881-128134921 | Myc |
| chr8:128134964-128135004 | Myc |
| chr8:128135036-128135076 | Myc |
| chr8:128135089-128135129 | Myc |
| chr8:128135176-128135216 | Myc |
| chr8:128135252-128135292 | Myc |
| chr8:128135391-128135431 | Myc |
| chr8:128135500-128135540 | Myc |
| chr8:128135590-128135630 | Myc |
| chr8:128135671-128135711 | Myc |
| chr8:128135813-128135853 | Myc |
| chr8:128135888-128135928 | Myc |
| chr8:128136085-128136125 | Myc |
| chr8:128136149-128136189 | Myc |
| chr8:128136217-128136257 | Myc |
| chr8:128136313-128136353 | Myc |
| chr8:128136371-128136411 | Myc |
| chr8:128136481-128136521 | Myc |
| chr8:128136557-128136597 | Myc |

| | |
|---|---|
| chr8:128136607-128136647 | Myc |
| chr8:128136658-128136698 | Myc |
| chr8:128136724-128136764 | Myc |
| chr8:128136776-128136816 | Myc |
| chr8:128136833-128136873 | Myc |
| chr8:128136944-128136984 | Myc |
| chr8:128136994-128137034 | Myc |
| chr8:128137061-128137101 | Myc |
| chr8:128137137-128137177 | Myc |
| chr8:128137225-128137265 | Myc |
| chr8:128137279-128137319 | Myc |
| chr8:128137336-128137376 | Myc |
| chr8:128137425-128137465 | Myc |
| chr8:128137499-128137539 | Myc |
| chr8:128137625-128137665 | Myc |
| chr8:128137691-128137731 | Myc |
| chr8:128137805-128137845 | Myc |
| chr8:128137900-128137940 | Myc |
| chr8:128137962-128138002 | Myc |
| chr8:128138012-128138052 | Myc |
| chr8:128138132-128138172 | Myc |
| chr8:128138229-128138269 | Myc |
| chr8:128138305-128138345 | Myc |
| chr8:128138390-128138430 | Myc |
| chr8:128138499-128138539 | Myc |
| chr8:128138588-128138628 | Myc |
| chr8:128138659-128138699 | Myc |
| chr8:128138716-128138756 | Myc |
| chr8:128138766-128138806 | Myc |
| chr8:128138869-128138909 | Myc |
| chr8:128138951-128138991 | Myc |
| chr8:128139068-128139108 | Myc |
| chr8:128139209-128139249 | Myc |
| chr8:128139420-128139460 | Myc |
| chr8:128139475-128139515 | Myc |
| chr8:128139644-128139684 | Myc |
| chr8:128139726-128139766 | Myc |
| chr8:128139799-128139839 | Myc |
| chr8:128139934-128139974 | Myc |
| chr8:128139993-128140033 | Myc |
| chr8:128140073-128140113 | Myc |
| chr8:128140175-128140215 | Myc |
| chr8:128140235-128140275 | Myc |
| chr8:128140337-128140377 | Myc |
| chr8:128140388-128140428 | Myc |
| chr8:128140446-128140486 | Myc |
| chr8:128140496-128140536 | Myc |
| chr8:128140560-128140600 | Myc |
| chr8:128140679-128140719 | Myc |
| chr8:128140777-128140817 | Myc |
| chr8:128140898-128140938 | Myc |
| chr8:128140983-128141023 | Myc |

| | |
|---|---|
| chr8:128141041-128141081 | Myc |
| chr8:128141119-128141159 | Myc |
| chr8:128141238-128141278 | Myc |
| chr8:128141288-128141328 | Myc |
| chr8:128141353-128141393 | Myc |
| chr8:128141441-128141481 | Myc |
| chr8:128141497-128141537 | Myc |
| chr8:128141586-128141626 | Myc |
| chr8:128141657-128141697 | Myc |
| chr8:128141708-128141748 | Myc |
| chr8:128141788-128141828 | Myc |
| chr8:128141962-128142002 | Myc |
| chr8:128142046-128142086 | Myc |
| chr8:128142166-128142206 | Myc |
| chr8:128142255-128142295 | Myc |
| chr8:128142308-128142348 | Myc |
| chr8:128142410-128142450 | Myc |
| chr8:128142471-128142511 | Myc |
| chr8:128142538-128142578 | Myc |
| chr8:128142588-128142628 | Myc |
| chr8:128142700-128142740 | Myc |
| chr8:128142771-128142811 | Myc |
| chr8:128142856-128142896 | Myc |
| chr8:128142962-128143002 | Myc |
| chr8:128143046-128143086 | Myc |
| chr8:128143116-128143156 | Myc |
| chr8:128143311-128143351 | Myc |
| chr8:128143427-128143467 | Myc |
| chr8:128143514-128143554 | Myc |
| chr8:128143815-128143855 | Myc |
| chr8:128143898-128143938 | Myc |
| chr8:128144242-128144282 | Myc |
| chr8:128144399-128144439 | Myc |
| chr8:128144475-128144515 | Myc |
| chr8:128144580-128144620 | Myc |
| chr8:128144630-128144670 | Myc |
| chr8:128144682-128144722 | Myc |
| chr8:128144743-128144783 | Myc |
| chr8:128144806-128144846 | Myc |
| chr8:128144920-128144960 | Myc |
| chr8:128144975-128145015 | Myc |
| chr8:128145056-128145096 | Myc |
| chr8:128145133-128145173 | Myc |
| chr8:128145219-128145259 | Myc |
| chr8:128145383-128145423 | Myc |
| chr8:128145438-128145478 | Myc |
| chr8:128145547-128145587 | Myc |
| chr8:128145661-128145701 | Myc |
| chr8:128145756-128145796 | Myc |
| chr8:128145840-128145880 | Myc |
| chr8:128145890-128145930 | Myc |
| chr8:128145979-128146019 | Myc |

| | |
|---|---|
| chr8:128146078-128146118 | Myc |
| chr8:128146160-128146200 | Myc |
| chr8:128146460-128146500 | Myc |
| chr8:128146613-128146653 | Myc |
| chr8:128146663-128146703 | Myc |
| chr8:128147011-128147051 | Myc |
| chr8:128147166-128147206 | Myc |
| chr8:128147249-128147289 | Myc |
| chr8:128147325-128147365 | Myc |
| chr8:128147443-128147483 | Myc |
| chr8:128147564-128147604 | Myc |
| chr8:128147626-128147666 | Myc |
| chr8:128147686-128147726 | Myc |
| chr8:128147779-128147819 | Myc |
| chr8:128147941-128147981 | Myc |
| chr8:128147997-128148037 | Myc |
| chr8:128148069-128148109 | Myc |
| chr8:128148179-128148219 | Myc |
| chr8:128148346-128148386 | Myc |
| chr8:128148432-128148472 | Myc |
| chr8:128148521-128148561 | Myc |
| chr8:128148621-128148661 | Myc |
| chr8:128148732-128148772 | Myc |
| chr8:128148855-128148895 | Myc |
| chr8:128148913-128148953 | Myc |
| chr8:128148970-128149010 | Myc |
| chr8:128149079-128149119 | Myc |
| chr8:128149163-128149203 | Myc |
| chr8:128149237-128149277 | Myc |
| chr8:128149330-128149370 | Myc |
| chr8:128149403-128149443 | Myc |
| chr8:128149505-128149545 | Myc |
| chr8:128149555-128149595 | Myc |
| chr8:128149711-128149751 | Myc |
| chr8:128149764-128149804 | Myc |
| chr8:128149871-128149911 | Myc |
| chr8:128149958-128149998 | Myc |
| chr8:128150030-128150070 | Myc |
| chr8:128150111-128150151 | Myc |
| chr8:128150241-128150281 | Myc |
| chr8:128150302-128150342 | Myc |
| chr8:128150505-128150545 | Myc |
| chr8:128150628-128150668 | Myc |
| chr8:128150679-128150719 | Myc |
| chr8:128150732-128150772 | Myc |
| chr8:128150807-128150847 | Myc |
| chr8:128150889-128150929 | Myc |
| chr8:128150944-128150984 | Myc |
| chr8:128151203-128151243 | Myc |
| chr8:128151316-128151356 | Myc |
| chr8:128151418-128151458 | Myc |
| chr8:128151472-128151512 | Myc |

| | |
|---|---|
| chr8:128151534-128151574 | Myc |
| chr8:128151598-128151638 | Myc |
| chr8:128151734-128151774 | Myc |
| chr8:128151833-128151873 | Myc |
| chr8:128151889-128151929 | Myc |
| chr8:128152365-128152405 | Myc |
| chr8:128152418-128152458 | Myc |
| chr8:128152520-128152560 | Myc |
| chr8:128152633-128152673 | Myc |
| chr8:128152724-128152764 | Myc |
| chr8:128152788-128152828 | Myc |
| chr8:128152865-128152905 | Myc |
| chr8:128152923-128152963 | Myc |
| chr8:128153043-128153083 | Myc |
| chr8:128153183-128153223 | Myc |
| chr8:128153278-128153318 | Myc |
| chr8:128153340-128153380 | Myc |
| chr8:128153433-128153473 | Myc |
| chr8:128153515-128153555 | Myc |
| chr8:128153629-128153669 | Myc |
| chr8:128153710-128153750 | Myc |
| chr8:128153767-128153807 | Myc |
| chr8:128153865-128153905 | Myc |
| chr8:128153956-128153996 | Myc |
| chr8:128154052-128154092 | Myc |
| chr8:128154152-128154192 | Myc |
| chr8:128154252-128154292 | Myc |
| chr8:128154306-128154346 | Myc |
| chr8:128154357-128154397 | Myc |
| chr8:128154435-128154475 | Myc |
| chr8:128154521-128154561 | Myc |
| chr8:128154591-128154631 | Myc |
| chr8:128154665-128154705 | Myc |
| chr8:128154732-128154772 | Myc |
| chr8:128154823-128154863 | Myc |
| chr8:128154915-128154955 | Myc |
| chr8:128155009-128155049 | Myc |
| chr8:128155117-128155157 | Myc |
| chr8:128155167-128155207 | Myc |
| chr8:128155217-128155257 | Myc |
| chr8:128155299-128155339 | Myc |
| chr8:128155391-128155431 | Myc |
| chr8:128155551-128155591 | Myc |
| chr8:128155628-128155668 | Myc |
| chr8:128155688-128155728 | Myc |
| chr8:128155781-128155821 | Myc |
| chr8:128155852-128155892 | Myc |
| chr8:128155930-128155970 | Myc |
| chr8:128156019-128156059 | Myc |
| chr8:128156084-128156124 | Myc |
| chr8:128156151-128156191 | Myc |
| chr8:128156255-128156295 | Myc |

| | |
|---|---|
| chr8:128156310-128156350 | Myc |
| chr8:128156457-128156497 | Myc |
| chr8:128156546-128156586 | Myc |
| chr8:128156624-128156664 | Myc |
| chr8:128156699-128156739 | Myc |
| chr8:128156980-128157020 | Myc |
| chr8:128157095-128157135 | Myc |
| chr8:128157156-128157196 | Myc |
| chr8:128157223-128157263 | Myc |
| chr8:128157373-128157413 | Myc |
| chr8:128157485-128157525 | Myc |
| chr8:128157640-128157680 | Myc |
| chr8:128157705-128157745 | Myc |
| chr8:128157814-128157854 | Myc |
| chr8:128157873-128157913 | Myc |
| chr8:128157934-128157974 | Myc |
| chr8:128158010-128158050 | Myc |
| chr8:128158118-128158158 | Myc |
| chr8:128158222-128158262 | Myc |
| chr8:128158304-128158344 | Myc |
| chr8:128158473-128158513 | Myc |
| chr8:128158537-128158577 | Myc |
| chr8:128158588-128158628 | Myc |
| chr8:128158654-128158694 | Myc |
| chr8:128158704-128158744 | Myc |
| chr8:128158779-128158819 | Myc |
| chr8:128158928-128158968 | Myc |
| chr8:128159009-128159049 | Myc |
| chr8:128159106-128159146 | Myc |
| chr8:128159272-128159312 | Myc |
| chr8:128159358-128159398 | Myc |
| chr8:128159787-128159827 | Myc |
| chr8:128159837-128159877 | Myc |
| chr8:128160046-128160086 | Myc |
| chr8:128160232-128160272 | Myc |
| chr8:128160286-128160326 | Myc |
| chr8:128160505-128160545 | Myc |
| chr8:128160567-128160607 | Myc |
| chr8:128160642-128160682 | Myc |
| chr8:128160696-128160736 | Myc |
| chr8:128160777-128160817 | Myc |
| chr8:128160868-128160908 | Myc |
| chr8:128160961-128161001 | Myc |
| chr8:128161025-128161065 | Myc |
| chr8:128161106-128161146 | Myc |
| chr8:128161303-128161343 | Myc |
| chr8:128161381-128161421 | Myc |
| chr8:128161701-128161741 | Myc |
| chr8:128161794-128161834 | Myc |
| chr8:128161860-128161900 | Myc |
| chr8:128161913-128161953 | Myc |
| chr8:128162005-128162045 | Myc |

| | |
|---|---|
| chr8:128162055-128162095 | Myc |
| chr8:128162219-128162259 | Myc |
| chr8:128162314-128162354 | Myc |
| chr8:128162445-128162485 | Myc |
| chr8:128162518-128162558 | Myc |
| chr8:128162593-128162633 | Myc |
| chr8:128162704-128162744 | Myc |
| chr8:128162791-128162831 | Myc |
| chr8:128162842-128162882 | Myc |
| chr8:128162944-128162984 | Myc |
| chr8:128163040-128163080 | Myc |
| chr8:128163107-128163147 | Myc |
| chr8:128163187-128163227 | Myc |
| chr8:128163259-128163299 | Myc |
| chr8:128163328-128163368 | Myc |
| chr8:128163501-128163541 | Myc |
| chr8:128163585-128163625 | Myc |
| chr8:128163685-128163725 | Myc |
| chr8:128163776-128163816 | Myc |
| chr8:128163882-128163922 | Myc |
| chr8:128163934-128163974 | Myc |
| chr8:128164019-128164059 | Myc |
| chr8:128164143-128164183 | Myc |
| chr8:128164193-128164233 | Myc |
| chr8:128164294-128164334 | Myc |
| chr8:128164347-128164387 | Myc |
| chr8:128164426-128164466 | Myc |
| chr8:128164535-128164575 | Myc |
| chr8:128164637-128164677 | Myc |
| chr8:128164694-128164734 | Myc |
| chr8:128164759-128164799 | Myc |
| chr8:128164832-128164872 | Myc |
| chr8:128164953-128164993 | Myc |
| chr8:128165080-128165120 | Myc |
| chr8:128165160-128165200 | Myc |
| chr8:128165247-128165287 | Myc |
| chr8:128165347-128165387 | Myc |
| chr8:128165406-128165446 | Myc |
| chr8:128165456-128165496 | Myc |
| chr8:128165529-128165569 | Myc |
| chr8:128165662-128165702 | Myc |
| chr8:128165817-128165857 | Myc |
| chr8:128165926-128165966 | Myc |
| chr8:128165989-128166029 | Myc |
| chr8:128166110-128166150 | Myc |
| chr8:128166194-128166234 | Myc |
| chr8:128166285-128166325 | Myc |
| chr8:128166338-128166378 | Myc |
| chr8:128166429-128166469 | Myc |
| chr8:128166548-128166588 | Myc |
| chr8:128166600-128166640 | Myc |
| chr8:128166674-128166714 | Myc |

| | |
|---|---|
| chr8:128166760-128166800 | Myc |
| chr8:128166820-128166860 | Myc |
| chr8:128166870-128166910 | Myc |
| chr8:128166993-128167033 | Myc |
| chr8:128167052-128167092 | Myc |
| chr8:128167102-128167142 | Myc |
| chr8:128167184-128167224 | Myc |
| chr8:128167313-128167353 | Myc |
| chr8:128167432-128167472 | Myc |
| chr8:128167520-128167560 | Myc |
| chr8:128167643-128167683 | Myc |
| chr8:128167714-128167754 | Myc |
| chr8:128167854-128167894 | Myc |
| chr8:128167935-128167975 | Myc |
| chr8:128168077-128168117 | Myc |
| chr8:128168146-128168186 | Myc |
| chr8:128168237-128168277 | Myc |
| chr8:128168316-128168356 | Myc |
| chr8:128168366-128168406 | Myc |
| chr8:128168455-128168495 | Myc |
| chr8:128168554-128168594 | Myc |
| chr8:128168639-128168679 | Myc |
| chr8:128169040-128169080 | Myc |
| chr8:128169097-128169137 | Myc |
| chr8:128169149-128169189 | Myc |
| chr8:128169225-128169265 | Myc |
| chr8:128169424-128169464 | Myc |
| chr8:128169487-128169527 | Myc |
| chr8:128169537-128169577 | Myc |
| chr8:128169669-128169709 | Myc |
| chr8:128169753-128169793 | Myc |
| chr8:128169989-128170029 | Myc |
| chr8:128170234-128170274 | Myc |
| chr8:128170285-128170325 | Myc |
| chr8:128170372-128170412 | Myc |
| chr8:128170426-128170466 | Myc |
| chr8:128170535-128170575 | Myc |
| chr8:128170615-128170655 | Myc |
| chr8:128170725-128170765 | Myc |
| chr8:128170805-128170845 | Myc |
| chr8:128170855-128170895 | Myc |
| chr8:128170988-128171028 | Myc |
| chr8:128171048-128171088 | Myc |
| chr8:128171113-128171153 | Myc |
| chr8:128171228-128171268 | Myc |
| chr8:128171310-128171350 | Myc |
| chr8:128171366-128171406 | Myc |
| chr8:128171464-128171504 | Myc |
| chr8:128171531-128171571 | Myc |
| chr8:128171581-128171621 | Myc |
| chr8:128171671-128171711 | Myc |
| chr8:128171782-128171822 | Myc |

| | |
|---|---|
| chr8:128171856-128171896 | Myc |
| chr8:128171927-128171967 | Myc |
| chr8:128172035-128172075 | Myc |
| chr8:128172098-128172138 | Myc |
| chr8:128172208-128172248 | Myc |
| chr8:128172302-128172342 | Myc |
| chr8:128172394-128172434 | Myc |
| chr8:128172499-128172539 | Myc |
| chr8:128172640-128172680 | Myc |
| chr8:128172698-128172738 | Myc |
| chr8:128172827-128172867 | Myc |
| chr8:128172885-128172925 | Myc |
| chr8:128172972-128173012 | Myc |
| chr8:128173065-128173105 | Myc |
| chr8:128173119-128173159 | Myc |
| chr8:128173267-128173307 | Myc |
| chr8:128173320-128173360 | Myc |
| chr8:128173417-128173457 | Myc |
| chr8:128173477-128173517 | Myc |
| chr8:128173539-128173579 | Myc |
| chr8:128173668-128173708 | Myc |
| chr8:128173751-128173791 | Myc |
| chr8:128173822-128173862 | Myc |
| chr8:128173904-128173944 | Myc |
| chr8:128173956-128173996 | Myc |
| chr8:128174014-128174054 | Myc |
| chr8:128174161-128174201 | Myc |
| chr8:128174215-128174255 | Myc |
| chr8:128174266-128174306 | Myc |
| chr8:128174347-128174387 | Myc |
| chr8:128174522-128174562 | Myc |
| chr8:128174593-128174633 | Myc |
| chr8:128174688-128174728 | Myc |
| chr8:128174763-128174803 | Myc |
| chr8:128174849-128174889 | Myc |
| chr8:128174918-128174958 | Myc |
| chr8:128174971-128175011 | Myc |
| chr8:128175094-128175134 | Myc |
| chr8:128175170-128175210 | Myc |
| chr8:128175255-128175295 | Myc |
| chr8:128175368-128175408 | Myc |
| chr8:128175447-128175487 | Myc |
| chr8:128175525-128175565 | Myc |
| chr8:128175635-128175675 | Myc |
| chr8:128175829-128175869 | Myc |
| chr8:128175965-128176005 | Myc |
| chr8:128176053-128176093 | Myc |
| chr8:128176183-128176223 | Myc |
| chr8:128176249-128176289 | Myc |
| chr8:128176319-128176359 | Myc |
| chr8:128176387-128176427 | Myc |
| chr8:128176490-128176530 | Myc |

| | |
|---|---|
| chr8:128176576-128176616 | Myc |
| chr8:128176694-128176734 | Myc |
| chr8:128176808-128176848 | Myc |
| chr8:128176976-128177016 | Myc |
| chr8:128177052-128177092 | Myc |
| chr8:128177102-128177142 | Myc |
| chr8:128177222-128177262 | Myc |
| chr8:128177289-128177329 | Myc |
| chr8:128177339-128177379 | Myc |
| chr8:128177427-128177467 | Myc |
| chr8:128177482-128177522 | Myc |
| chr8:128177554-128177594 | Myc |
| chr8:128177604-128177644 | Myc |
| chr8:128177696-128177736 | Myc |
| chr8:128177808-128177848 | Myc |
| chr8:128178030-128178070 | Myc |
| chr8:128178174-128178214 | Myc |
| chr8:128178253-128178293 | Myc |
| chr8:128178319-128178359 | Myc |
| chr8:128178465-128178505 | Myc |
| chr8:128178620-128178660 | Myc |
| chr8:128178672-128178712 | Myc |
| chr8:128178748-128178788 | Myc |
| chr8:128178854-128178894 | Myc |
| chr8:128178959-128178999 | Myc |
| chr8:128179024-128179064 | Myc |
| chr8:128179074-128179114 | Myc |
| chr8:128179143-128179183 | Myc |
| chr8:128179226-128179266 | Myc |
| chr8:128179285-128179325 | Myc |
| chr8:128179368-128179408 | Myc |
| chr8:128179465-128179505 | Myc |
| chr8:128179520-128179560 | Myc |
| chr8:128179570-128179610 | Myc |
| chr8:128179663-128179703 | Myc |
| chr8:128179793-128179833 | Myc |
| chr8:128179892-128179932 | Myc |
| chr8:128179975-128180015 | Myc |
| chr8:128180029-128180069 | Myc |
| chr8:128180135-128180175 | Myc |
| chr8:128180248-128180288 | Myc |
| chr8:128180299-128180339 | Myc |
| chr8:128180482-128180522 | Myc |
| chr8:128180735-128180775 | Myc |
| chr8:128180858-128180898 | Myc |
| chr8:128180969-128181009 | Myc |
| chr8:128181137-128181177 | Myc |
| chr8:128181207-128181247 | Myc |
| chr8:128181257-128181297 | Myc |
| chr8:128181320-128181360 | Myc |
| chr8:128181411-128181451 | Myc |
| chr8:128181492-128181532 | Myc |

| | |
|---|---|
| chr8:128181594-128181634 | Myc |
| chr8:128181671-128181711 | Myc |
| chr8:128181806-128181846 | Myc |
| chr8:128181860-128181900 | Myc |
| chr8:128181937-128181977 | Myc |
| chr8:128182050-128182090 | Myc |
| chr8:128182161-128182201 | Myc |
| chr8:128182222-128182262 | Myc |
| chr8:128182308-128182348 | Myc |
| chr8:128182382-128182422 | Myc |
| chr8:128182445-128182485 | Myc |
| chr8:128182527-128182567 | Myc |
| chr8:128182592-128182632 | Myc |
| chr8:128182642-128182682 | Myc |
| chr8:128182757-128182797 | Myc |
| chr8:128182821-128182861 | Myc |
| chr8:128182902-128182942 | Myc |
| chr8:128182973-128183013 | Myc |
| chr8:128183061-128183101 | Myc |
| chr8:128183147-128183187 | Myc |
| chr8:128183206-128183246 | Myc |
| chr8:128183291-128183331 | Myc |
| chr8:128183356-128183396 | Myc |
| chr8:128183470-128183510 | Myc |
| chr8:128183551-128183591 | Myc |
| chr8:128183656-128183696 | Myc |
| chr8:128183752-128183792 | Myc |
| chr8:128184161-128184201 | Myc |
| chr8:128184288-128184328 | Myc |
| chr8:128184396-128184436 | Myc |
| chr8:128184446-128184486 | Myc |
| chr8:128184593-128184633 | Myc |
| chr8:128184692-128184732 | Myc |
| chr8:128184861-128184901 | Myc |
| chr8:128184924-128184964 | Myc |
| chr8:128184976-128185016 | Myc |
| chr8:128185061-128185101 | Myc |
| chr8:128185113-128185153 | Myc |
| chr8:128185360-128185400 | Myc |
| chr8:128185412-128185452 | Myc |
| chr8:128185552-128185592 | Myc |
| chr8:128185648-128185688 | Myc |
| chr8:128185707-128185747 | Myc |
| chr8:128185773-128185813 | Myc |
| chr8:128185851-128185891 | Myc |
| chr8:128185979-128186019 | Myc |
| chr8:128186037-128186077 | Myc |
| chr8:128186108-128186148 | Myc |
| chr8:128186205-128186245 | Myc |
| chr8:128186272-128186312 | Myc |
| chr8:128186367-128186407 | Myc |
| chr8:128186444-128186484 | Myc |

| | |
|---|---|
| chr8:128186548-128186588 | Myc |
| chr8:128186601-128186641 | Myc |
| chr8:128186737-128186777 | Myc |
| chr8:128186802-128186842 | Myc |
| chr8:128186961-128187001 | Myc |
| chr8:128187026-128187066 | Myc |
| chr8:128187102-128187142 | Myc |
| chr8:128187175-128187215 | Myc |
| chr8:128187237-128187277 | Myc |
| chr8:128187298-128187338 | Myc |
| chr8:128187465-128187505 | Myc |
| chr8:128187535-128187575 | Myc |
| chr8:128187603-128187643 | Myc |
| chr8:128187697-128187737 | Myc |
| chr8:128187799-128187839 | Myc |
| chr8:128187945-128187985 | Myc |
| chr8:128188101-128188141 | Myc |
| chr8:128188164-128188204 | Myc |
| chr8:128188219-128188259 | Myc |
| chr8:128188269-128188309 | Myc |
| chr8:128188385-128188425 | Myc |
| chr8:128188437-128188477 | Myc |
| chr8:128188498-128188538 | Myc |
| chr8:128188584-128188624 | Myc |
| chr8:128188718-128188758 | Myc |
| chr8:128188826-128188866 | Myc |
| chr8:128188961-128189001 | Myc |
| chr8:128189022-128189062 | Myc |
| chr8:128189212-128189252 | Myc |
| chr8:128189298-128189338 | Myc |
| chr8:128189348-128189388 | Myc |
| chr8:128189430-128189470 | Myc |
| chr8:128189549-128189589 | Myc |
| chr8:128189626-128189666 | Myc |
| chr8:128189707-128189747 | Myc |
| chr8:128189760-128189800 | Myc |
| chr8:128189858-128189898 | Myc |
| chr8:128189908-128189948 | Myc |
| chr8:128190107-128190147 | Myc |
| chr8:128190157-128190197 | Myc |
| chr8:128190216-128190256 | Myc |
| chr8:128190351-128190391 | Myc |
| chr8:128190507-128190547 | Myc |
| chr8:128190558-128190598 | Myc |
| chr8:128190693-128190733 | Myc |
| chr8:128190821-128190861 | Myc |
| chr8:128190904-128190944 | Myc |
| chr8:128191030-128191070 | Myc |
| chr8:128191130-128191170 | Myc |
| chr8:128191207-128191247 | Myc |
| chr8:128191324-128191364 | Myc |
| chr8:128191452-128191492 | Myc |

| | |
|---|---|
| chr8:128191542-128191582 | Myc |
| chr8:128191735-128191775 | Myc |
| chr8:128191842-128191882 | Myc |
| chr8:128191964-128192004 | Myc |
| chr8:128192018-128192058 | Myc |
| chr8:128192103-128192143 | Myc |
| chr8:128192153-128192193 | Myc |
| chr8:128192266-128192306 | Myc |
| chr8:128192319-128192359 | Myc |
| chr8:128192502-128192542 | Myc |
| chr8:128192567-128192607 | Myc |
| chr8:128192735-128192775 | Myc |
| chr8:128192893-128192933 | Myc |
| chr8:128192994-128193034 | Myc |
| chr8:128193075-128193115 | Myc |
| chr8:128193251-128193291 | Myc |
| chr8:128193324-128193364 | Myc |
| chr8:128193399-128193439 | Myc |
| chr8:128193488-128193528 | Myc |
| chr8:128193560-128193600 | Myc |
| chr8:128193636-128193676 | Myc |
| chr8:128193773-128193813 | Myc |
| chr8:128193880-128193920 | Myc |
| chr8:128193946-128193986 | Myc |
| chr8:128194031-128194071 | Myc |
| chr8:128194136-128194176 | Myc |
| chr8:128194219-128194259 | Myc |
| chr8:128194274-128194314 | Myc |
| chr8:128194367-128194407 | Myc |
| chr8:128194444-128194484 | Myc |
| chr8:128194563-128194603 | Myc |
| chr8:128194920-128194960 | Myc |
| chr8:128194999-128195039 | Myc |
| chr8:128195057-128195097 | Myc |
| chr8:128195267-128195307 | Myc |
| chr8:128195404-128195444 | Myc |
| chr8:128195463-128195503 | Myc |
| chr8:128195633-128195673 | Myc |
| chr8:128195773-128195813 | Myc |
| chr8:128196160-128196200 | Myc |
| chr8:128196212-128196252 | Myc |
| chr8:128196310-128196350 | Myc |
| chr8:128196415-128196455 | Myc |
| chr8:128196694-128196734 | Myc |
| chr8:128196829-128196869 | Myc |
| chr8:128196906-128196946 | Myc |
| chr8:128196994-128197034 | Myc |
| chr8:128197047-128197087 | Myc |
| chr8:128197120-128197160 | Myc |
| chr8:128197188-128197228 | Myc |
| chr8:128197318-128197358 | Myc |
| chr8:128197428-128197468 | Myc |

| | |
|---|---|
| chr8:128197485-128197525 | Myc |
| chr8:128197535-128197575 | Myc |
| chr8:128197597-128197637 | Myc |
| chr8:128197739-128197779 | Myc |
| chr8:128197797-128197837 | Myc |
| chr8:128197871-128197911 | Myc |
| chr8:128197976-128198016 | Myc |
| chr8:128198047-128198087 | Myc |
| chr8:128198143-128198183 | Myc |
| chr8:128198233-128198273 | Myc |
| chr8:128198566-128198606 | Myc |
| chr8:128198627-128198667 | Myc |
| chr8:128198722-128198762 | Myc |
| chr8:128198929-128198969 | Myc |
| chr8:128198990-128199030 | Myc |
| chr8:128199106-128199146 | Myc |
| chr8:128199156-128199196 | Myc |
| chr8:128199254-128199294 | Myc |
| chr8:128199326-128199366 | Myc |
| chr8:128199404-128199444 | Myc |
| chr8:128199482-128199522 | Myc |
| chr8:128199550-128199590 | Myc |
| chr8:128199600-128199640 | Myc |
| chr8:128199689-128199729 | Myc |
| chr8:128199764-128199804 | Myc |
| chr8:128199916-128199956 | Myc |
| chr8:128199995-128200035 | Myc |
| chr8:128200056-128200096 | Myc |
| chr8:128200106-128200146 | Myc |
| chr8:128200188-128200228 | Myc |
| chr8:128200283-128200323 | Myc |
| chr8:128200483-128200523 | Myc |
| chr8:128200542-128200582 | Myc |
| chr8:128200600-128200640 | Myc |
| chr8:128200729-128200769 | Myc |
| chr8:128200788-128200828 | Myc |
| chr8:128200936-128200976 | Myc |
| chr8:128201270-128201310 | Myc |
| chr8:128201353-128201393 | Myc |
| chr8:128201423-128201463 | Myc |
| chr8:128201474-128201514 | Myc |
| chr8:128201549-128201589 | Myc |
| chr8:128201627-128201667 | Myc |
| chr8:128201685-128201725 | Myc |
| chr8:128201742-128201782 | Myc |
| chr8:128201863-128201903 | Myc |
| chr8:128201932-128201972 | Myc |
| chr8:128201984-128202024 | Myc |
| chr8:128202062-128202102 | Myc |
| chr8:128202165-128202205 | Myc |
| chr8:128202215-128202255 | Myc |
| chr8:128202335-128202375 | Myc |

| | |
|---|---|
| chr8:128202385-128202425 | Myc |
| chr8:128202458-128202498 | Myc |
| chr8:128202562-128202602 | Myc |
| chr8:128202665-128202705 | Myc |
| chr8:128202732-128202772 | Myc |
| chr8:128202853-128202893 | Myc |
| chr8:128202907-128202947 | Myc |
| chr8:128203082-128203122 | Myc |
| chr8:128203250-128203290 | Myc |
| chr8:128203308-128203348 | Myc |
| chr8:128203388-128203428 | Myc |
| chr8:128203481-128203521 | Myc |
| chr8:128203543-128203583 | Myc |
| chr8:128203627-128203667 | Myc |
| chr8:128203679-128203719 | Myc |
| chr8:128203735-128203775 | Myc |
| chr8:128203788-128203828 | Myc |
| chr8:128203838-128203878 | Myc |
| chr8:128203891-128203931 | Myc |
| chr8:128203975-128204015 | Myc |
| chr8:128204166-128204206 | Myc |
| chr8:128204277-128204317 | Myc |
| chr8:128204407-128204447 | Myc |
| chr8:128206398-128206438 | Myc |
| chr8:128206866-128206906 | Myc |
| chr8:128206990-128207030 | Myc |
| chr8:128207092-128207132 | Myc |
| chr8:128207145-128207185 | Myc |
| chr8:128207359-128207399 | Myc |
| chr8:128207448-128207488 | Myc |
| chr8:128207522-128207562 | Myc |
| chr8:128207609-128207649 | Myc |
| chr8:128207676-128207716 | Myc |
| chr8:128207842-128207882 | Myc |
| chr8:128207921-128207961 | Myc |
| chr8:128208020-128208060 | Myc |
| chr8:128208070-128208110 | Myc |
| chr8:128208136-128208176 | Myc |
| chr8:128208201-128208241 | Myc |
| chr8:128208311-128208351 | Myc |
| chr8:128208433-128208473 | Myc |
| chr8:128208621-128208661 | Myc |
| chr8:128208727-128208767 | Myc |
| chr8:128208783-128208823 | Myc |
| chr8:128208887-128208927 | Myc |
| chr8:128209018-128209058 | Myc |
| chr8:128209089-128209129 | Myc |
| chr8:128209140-128209180 | Myc |
| chr8:128209252-128209292 | Myc |
| chr8:128209333-128209373 | Myc |
| chr8:128209404-128209444 | Myc |
| chr8:128209454-128209494 | Myc |

| | |
|---|---|
| chr8:128209515-128209555 | Myc |
| chr8:128209566-128209606 | Myc |
| chr8:128209667-128209707 | Myc |
| chr8:128209743-128209783 | Myc |
| chr8:128209793-128209833 | Myc |
| chr8:128209877-128209917 | Myc |
| chr8:128209928-128209968 | Myc |
| chr8:128209982-128210022 | Myc |
| chr8:128210034-128210074 | Myc |
| chr8:128210151-128210191 | Myc |
| chr8:128210276-128210316 | Myc |
| chr8:128210380-128210420 | Myc |
| chr8:128210430-128210470 | Myc |
| chr8:128210523-128210563 | Myc |
| chr8:128210602-128210642 | Myc |
| chr8:128210678-128210718 | Myc |
| chr8:128210859-128210899 | Myc |
| chr8:128210940-128210980 | Myc |
| chr8:128210995-128211035 | Myc |
| chr8:128211052-128211092 | Myc |
| chr8:128211152-128211192 | Myc |
| chr8:128211283-128211323 | Myc |
| chr8:128211389-128211429 | Myc |
| chr8:128211520-128211560 | Myc |
| chr8:128211801-128211841 | Myc |
| chr8:128211862-128211902 | Myc |
| chr8:128211985-128212025 | Myc |
| chr8:128212035-128212075 | Myc |
| chr8:128212092-128212132 | Myc |
| chr8:128212162-128212202 | Myc |
| chr8:128212212-128212252 | Myc |
| chr8:128212279-128212319 | Myc |
| chr8:128212340-128212380 | Myc |
| chr8:128212403-128212443 | Myc |
| chr8:128212517-128212557 | Myc |
| chr8:128212596-128212636 | Myc |
| chr8:128212685-128212725 | Myc |
| chr8:128212738-128212778 | Myc |
| chr8:128212795-128212835 | Myc |
| chr8:128212846-128212886 | Myc |
| chr8:128212932-128212972 | Myc |
| chr8:128213012-128213052 | Myc |
| chr8:128213065-128213105 | Myc |
| chr8:128213164-128213204 | Myc |
| chr8:128213245-128213285 | Myc |
| chr8:128213306-128213346 | Myc |
| chr8:128213389-128213429 | Myc |
| chr8:128213469-128213509 | Myc |
| chr8:128213539-128213579 | Myc |
| chr8:128213683-128213723 | Myc |
| chr8:128213812-128213852 | Myc |
| chr8:128213864-128213904 | Myc |

| | |
|---|---|
| chr8:128213933-128213973 | Myc |
| chr8:128214046-128214086 | Myc |
| chr8:128214112-128214152 | Myc |
| chr8:128214184-128214224 | Myc |
| chr8:128214236-128214276 | Myc |
| chr8:128214334-128214374 | Myc |
| chr8:128214428-128214468 | Myc |
| chr8:128214609-128214649 | Myc |
| chr8:128214780-128214820 | Myc |
| chr8:128214862-128214902 | Myc |
| chr8:128214966-128215006 | Myc |
| chr8:128215017-128215057 | Myc |
| chr8:128215070-128215110 | Myc |
| chr8:128215161-128215201 | Myc |
| chr8:128215221-128215261 | Myc |
| chr8:128215427-128215467 | Myc |
| chr8:128215504-128215544 | Myc |
| chr8:128216161-128216201 | Myc |
| chr8:128216496-128216536 | Myc |
| chr8:128216565-128216605 | Myc |
| chr8:128216743-128216783 | Myc |
| chr8:128216975-128217015 | Myc |
| chr8:128217083-128217123 | Myc |
| chr8:128217173-128217213 | Myc |
| chr8:128217343-128217383 | Myc |
| chr8:128217403-128217443 | Myc |
| chr8:128217469-128217509 | Myc |
| chr8:128217568-128217608 | Myc |
| chr8:128217627-128217667 | Myc |
| chr8:128217723-128217763 | Myc |
| chr8:128217848-128217888 | Myc |
| chr8:128217985-128218025 | Myc |
| chr8:128218035-128218075 | Myc |
| chr8:128218113-128218153 | Myc |
| chr8:128218173-128218213 | Myc |
| chr8:128218231-128218271 | Myc |
| chr8:128218344-128218384 | Myc |
| chr8:128218404-128218444 | Myc |
| chr8:128218473-128218513 | Myc |
| chr8:128218537-128218577 | Myc |
| chr8:128218599-128218639 | Myc |
| chr8:128218676-128218716 | Myc |
| chr8:128218729-128218769 | Myc |
| chr8:128218844-128218884 | Myc |
| chr8:128218894-128218934 | Myc |
| chr8:128219013-128219053 | Myc |
| chr8:128219268-128219308 | Myc |
| chr8:128219363-128219403 | Myc |
| chr8:128219413-128219453 | Myc |
| chr8:128219481-128219521 | Myc |
| chr8:128219561-128219601 | Myc |
| chr8:128219747-128219787 | Myc |

| | |
|---|---|
| chr8:128219804-128219844 | Myc |
| chr8:128219864-128219904 | Myc |
| chr8:128219957-128219997 | Myc |
| chr8:128220054-128220094 | Myc |
| chr8:128220105-128220145 | Myc |
| chr8:128220179-128220219 | Myc |
| chr8:128220276-128220316 | Myc |
| chr8:128220444-128220484 | Myc |
| chr8:128220514-128220554 | Myc |
| chr8:128220586-128220626 | Myc |
| chr8:128220690-128220730 | Myc |
| chr8:128220746-128220786 | Myc |
| chr8:128220871-128220911 | Myc |
| chr8:128220973-128221013 | Myc |
| chr8:128221062-128221102 | Myc |
| chr8:128221166-128221206 | Myc |
| chr8:128221218-128221258 | Myc |
| chr8:128221271-128221311 | Myc |
| chr8:128221330-128221370 | Myc |
| chr8:128221401-128221441 | Myc |
| chr8:128221502-128221542 | Myc |
| chr8:128221624-128221664 | Myc |
| chr8:128221718-128221758 | Myc |
| chr8:128221783-128221823 | Myc |
| chr8:128221838-128221878 | Myc |
| chr8:128221929-128221969 | Myc |
| chr8:128222098-128222138 | Myc |
| chr8:128222148-128222188 | Myc |
| chr8:128222201-128222241 | Myc |
| chr8:128222255-128222295 | Myc |
| chr8:128222374-128222414 | Myc |
| chr8:128222484-128222524 | Myc |
| chr8:128222577-128222617 | Myc |
| chr8:128222663-128222703 | Myc |
| chr8:128222732-128222772 | Myc |
| chr8:128222864-128222904 | Myc |
| chr8:128222919-128222959 | Myc |
| chr8:128223048-128223088 | Myc |
| chr8:128223099-128223139 | Myc |
| chr8:128223152-128223192 | Myc |
| chr8:128223344-128223384 | Myc |
| chr8:128223451-128223491 | Myc |
| chr8:128223555-128223595 | Myc |
| chr8:128223702-128223742 | Myc |
| chr8:128223782-128223822 | Myc |
| chr8:128223887-128223927 | Myc |
| chr8:128224089-128224129 | Myc |
| chr8:128224163-128224203 | Myc |
| chr8:128224213-128224253 | Myc |
| chr8:128224278-128224318 | Myc |
| chr8:128224360-128224400 | Myc |
| chr8:128224420-128224460 | Myc |

| | |
|---|---|
| chr8:128224488-128224528 | Myc |
| chr8:128224585-128224625 | Myc |
| chr8:128224640-128224680 | Myc |
| chr8:128224700-128224740 | Myc |
| chr8:128224750-128224790 | Myc |
| chr8:128224955-128224995 | Myc |
| chr8:128225009-128225049 | Myc |
| chr8:128225059-128225099 | Myc |
| chr8:128237794-128237834 | Myc |
| chr8:128237921-128237961 | Myc |
| chr8:128237975-128238015 | Myc |
| chr8:128238103-128238143 | Myc |
| chr8:128238191-128238231 | Myc |
| chr8:128238272-128238312 | Myc |
| chr8:128238342-128238382 | Myc |
| chr8:128238536-128238576 | Myc |
| chr8:128238596-128238636 | Myc |
| chr8:128238648-128238688 | Myc |
| chr8:128238778-128238818 | Myc |
| chr8:128238828-128238868 | Myc |
| chr8:128238941-128238981 | Myc |
| chr8:128238994-128239034 | Myc |
| chr8:128239162-128239202 | Myc |
| chr8:128239213-128239253 | Myc |
| chr8:128239263-128239303 | Myc |
| chr8:128239343-128239383 | Myc |
| chr8:128239443-128239483 | Myc |
| chr8:128239543-128239583 | Myc |
| chr8:128239630-128239670 | Myc |
| chr8:128239714-128239754 | Myc |
| chr8:128239878-128239918 | Myc |
| chr8:128239972-128240012 | Myc |
| chr8:128240023-128240063 | Myc |
| chr8:128240105-128240145 | Myc |
| chr8:128240177-128240217 | Myc |
| chr8:128240232-128240272 | Myc |
| chr8:128240361-128240401 | Myc |
| chr8:128240414-128240454 | Myc |
| chr8:128240587-128240627 | Myc |
| chr8:128240637-128240677 | Myc |
| chr8:128240716-128240756 | Myc |
| chr8:128240766-128240806 | Myc |
| chr8:128240841-128240881 | Myc |
| chr8:128241022-128241062 | Myc |
| chr8:128241139-128241179 | Myc |
| chr8:128241253-128241293 | Myc |
| chr8:128241325-128241365 | Myc |
| chr8:128241439-128241479 | Myc |
| chr8:128241574-128241614 | Myc |
| chr8:128241700-128241740 | Myc |
| chr8:128241750-128241790 | Myc |
| chr8:128241825-128241865 | Myc |

| | |
|---|---|
| chr8:128241946-128241986 | Myc |
| chr8:128242071-128242111 | Myc |
| chr8:128242198-128242238 | Myc |
| chr8:128242383-128242423 | Myc |
| chr8:128242574-128242614 | Myc |
| chr8:128242653-128242693 | Myc |
| chr8:128242709-128242749 | Myc |
| chr8:128242770-128242810 | Myc |
| chr8:128242857-128242897 | Myc |
| chr8:128243017-128243057 | Myc |
| chr8:128243143-128243183 | Myc |
| chr8:128243193-128243233 | Myc |
| chr8:128243268-128243308 | Myc |
| chr8:128243326-128243366 | Myc |
| chr8:128243377-128243417 | Myc |
| chr8:128243431-128243471 | Myc |
| chr8:128243481-128243521 | Myc |
| chr8:128243628-128243668 | Myc |
| chr8:128243678-128243718 | Myc |
| chr8:128243734-128243774 | Myc |
| chr8:128243787-128243827 | Myc |
| chr8:128243838-128243878 | Myc |
| chr8:128243905-128243945 | Myc |
| chr8:128243996-128244036 | Myc |
| chr8:128244105-128244145 | Myc |
| chr8:128244221-128244261 | Myc |
| chr8:128244274-128244314 | Myc |
| chr8:128244374-128244414 | Myc |
| chr8:128244425-128244465 | Myc |
| chr8:128244481-128244521 | Myc |
| chr8:128244686-128244726 | Myc |
| chr8:128244736-128244776 | Myc |
| chr8:128244849-128244889 | Myc |
| chr8:128244927-128244967 | Myc |
| chr8:128244998-128245038 | Myc |
| chr8:128245063-128245103 | Myc |
| chr8:128245115-128245155 | Myc |
| chr8:128245229-128245269 | Myc |
| chr8:128245331-128245371 | Myc |
| chr8:128245412-128245452 | Myc |
| chr8:128245487-128245527 | Myc |
| chr8:128245561-128245601 | Myc |
| chr8:128245833-128245873 | Myc |
| chr8:128245901-128245941 | Myc |
| chr8:128246088-128246128 | Myc |
| chr8:128246151-128246191 | Myc |
| chr8:128246214-128246254 | Myc |
| chr8:128246364-128246404 | Myc |
| chr8:128246448-128246488 | Myc |
| chr8:128246593-128246633 | Myc |
| chr8:128246673-128246713 | Myc |
| chr8:128246729-128246769 | Myc |

| | |
|---|---|
| chr8:128246800-128246840 | Myc |
| chr8:128246885-128246925 | Myc |
| chr8:128246956-128246996 | Myc |
| chr8:128247007-128247047 | Myc |
| chr8:128247057-128247097 | Myc |
| chr8:128247109-128247149 | Myc |
| chr8:128247227-128247267 | Myc |
| chr8:128247298-128247338 | Myc |
| chr8:128247348-128247388 | Myc |
| chr8:128247402-128247442 | Myc |
| chr8:128247484-128247524 | Myc |
| chr8:128247547-128247587 | Myc |
| chr8:128247662-128247702 | Myc |
| chr8:128247771-128247811 | Myc |
| chr8:128247874-128247914 | Myc |
| chr8:128247946-128247986 | Myc |
| chr8:128248284-128248324 | Myc |
| chr8:128248668-128248708 | Myc |
| chr8:128248724-128248764 | Myc |
| chr8:128248793-128248833 | Myc |
| chr8:128248913-128248953 | Myc |
| chr8:128248982-128249022 | Myc |
| chr8:128249078-128249118 | Myc |
| chr8:128249160-128249200 | Myc |
| chr8:128249212-128249252 | Myc |
| chr8:128249263-128249303 | Myc |
| chr8:128249315-128249355 | Myc |
| chr8:128249376-128249416 | Myc |
| chr8:128249428-128249468 | Myc |
| chr8:128249526-128249566 | Myc |
| chr8:128249582-128249622 | Myc |
| chr8:128249632-128249672 | Myc |
| chr8:128249682-128249722 | Myc |
| chr8:128249736-128249776 | Myc |
| chr8:128249797-128249837 | Myc |
| chr8:128250089-128250129 | Myc |
| chr8:128250139-128250179 | Myc |
| chr8:128250292-128250332 | Myc |
| chr8:128250369-128250409 | Myc |
| chr8:128250538-128250578 | Myc |
| chr8:128250589-128250629 | Myc |
| chr8:128250659-128250699 | Myc |
| chr8:128250803-128250843 | Myc |
| chr8:128250856-128250896 | Myc |
| chr8:128250942-128250982 | Myc |
| chr8:128251007-128251047 | Myc |
| chr8:128251108-128251148 | Myc |
| chr8:128251193-128251233 | Myc |
| chr8:128251273-128251313 | Myc |
| chr8:128251335-128251375 | Myc |
| chr8:128251462-128251502 | Myc |
| chr8:128251512-128251552 | Myc |

| | |
|---|---|
| chr8:128251584-128251624 | Myc |
| chr8:128251649-128251689 | Myc |
| chr8:128251764-128251804 | Myc |
| chr8:128251874-128251914 | Myc |
| chr8:128251973-128252013 | Myc |
| chr8:128252045-128252085 | Myc |
| chr8:128252100-128252140 | Myc |
| chr8:128252203-128252243 | Myc |
| chr8:128252352-128252392 | Myc |
| chr8:128252468-128252508 | Myc |
| chr8:128252518-128252558 | Myc |
| chr8:128252625-128252665 | Myc |
| chr8:128252733-128252773 | Myc |
| chr8:128252804-128252844 | Myc |
| chr8:128252910-128252950 | Myc |
| chr8:128252980-128253020 | Myc |
| chr8:128253049-128253089 | Myc |
| chr8:128253118-128253158 | Myc |
| chr8:128253173-128253213 | Myc |
| chr8:128253324-128253364 | Myc |
| chr8:128253390-128253430 | Myc |
| chr8:128253466-128253506 | Myc |
| chr8:128253536-128253576 | Myc |
| chr8:128253720-128253760 | Myc |
| chr8:128253770-128253810 | Myc |
| chr8:128253842-128253882 | Myc |
| chr8:128253892-128253932 | Myc |
| chr8:128253958-128253998 | Myc |
| chr8:128254038-128254078 | Myc |
| chr8:128254104-128254144 | Myc |
| chr8:128254159-128254199 | Myc |
| chr8:128254287-128254327 | Myc |
| chr8:128254372-128254412 | Myc |
| chr8:128254486-128254526 | Myc |
| chr8:128254540-128254580 | Myc |
| chr8:128254604-128254644 | Myc |
| chr8:128254700-128254740 | Myc |
| chr8:128254756-128254796 | Myc |
| chr8:128254826-128254866 | Myc |
| chr8:128254881-128254921 | Myc |
| chr8:128254931-128254971 | Myc |
| chr8:128255120-128255160 | Myc |
| chr8:128255254-128255294 | Myc |
| chr8:128255432-128255472 | Myc |
| chr8:128255488-128255528 | Myc |
| chr8:128255556-128255596 | Myc |
| chr8:128255612-128255652 | Myc |
| chr8:128255713-128255753 | Myc |
| chr8:128255817-128255857 | Myc |
| chr8:128255915-128255955 | Myc |
| chr8:128255992-128256032 | Myc |
| chr8:128256069-128256109 | Myc |

| | |
|---|---|
| chr8:128256151-128256191 | Myc |
| chr8:128256201-128256241 | Myc |
| chr8:128256341-128256381 | Myc |
| chr8:128256391-128256431 | Myc |
| chr8:128256447-128256487 | Myc |
| chr8:128256572-128256612 | Myc |
| chr8:128256622-128256662 | Myc |
| chr8:128256703-128256743 | Myc |
| chr8:128256837-128256877 | Myc |
| chr8:128256938-128256978 | Myc |
| chr8:128256995-128257035 | Myc |
| chr8:128257227-128257267 | Myc |
| chr8:128257286-128257326 | Myc |
| chr8:128257417-128257457 | Myc |
| chr8:128257467-128257507 | Myc |
| chr8:128257541-128257581 | Myc |
| chr8:128257630-128257670 | Myc |
| chr8:128257687-128257727 | Myc |
| chr8:128257751-128257791 | Myc |
| chr8:128257869-128257909 | Myc |
| chr8:128257931-128257971 | Myc |
| chr8:128258064-128258104 | Myc |
| chr8:128258158-128258198 | Myc |
| chr8:128258258-128258298 | Myc |
| chr8:128258359-128258399 | Myc |
| chr8:128258517-128258557 | Myc |
| chr8:128258601-128258641 | Myc |
| chr8:128258665-128258705 | Myc |
| chr8:128258794-128258834 | Myc |
| chr8:128258878-128258918 | Myc |
| chr8:128258941-128258981 | Myc |
| chr8:128258991-128259031 | Myc |
| chr8:128259086-128259126 | Myc |
| chr8:128259195-128259235 | Myc |
| chr8:128259245-128259285 | Myc |
| chr8:128259321-128259361 | Myc |
| chr8:128259403-128259443 | Myc |
| chr8:128259459-128259499 | Myc |
| chr8:128259522-128259562 | Myc |
| chr8:128259572-128259612 | Myc |
| chr8:128259622-128259662 | Myc |
| chr8:128259729-128259769 | Myc |
| chr8:128259806-128259846 | Myc |
| chr8:128259856-128259896 | Myc |
| chr8:128259919-128259959 | Myc |
| chr8:128260026-128260066 | Myc |
| chr8:128260131-128260171 | Myc |
| chr8:128260182-128260222 | Myc |
| chr8:128260248-128260288 | Myc |
| chr8:128260355-128260395 | Myc |
| chr8:128260459-128260499 | Myc |
| chr8:128260581-128260621 | Myc |

| | |
|---|---|
| chr8:128260653-128260693 | Myc |
| chr8:128260712-128260752 | Myc |
| chr8:128261048-128261088 | Myc |
| chr8:128261193-128261233 | Myc |
| chr8:128261258-128261298 | Myc |
| chr8:128261331-128261371 | Myc |
| chr8:128261692-128261732 | Myc |
| chr8:128261765-128261805 | Myc |
| chr8:128261827-128261867 | Myc |
| chr8:128261968-128262008 | Myc |
| chr8:128262042-128262082 | Myc |
| chr8:128262103-128262143 | Myc |
| chr8:128262153-128262193 | Myc |
| chr8:128262234-128262274 | Myc |
| chr8:128262325-128262365 | Myc |
| chr8:128262455-128262495 | Myc |
| chr8:128262505-128262545 | Myc |
| chr8:128262571-128262611 | Myc |
| chr8:128262659-128262699 | Myc |
| chr8:128262723-128262763 | Myc |
| chr8:128262825-128262865 | Myc |
| chr8:128262884-128262924 | Myc |
| chr8:128262999-128263039 | Myc |
| chr8:128263139-128263179 | Myc |
| chr8:128263189-128263229 | Myc |
| chr8:128263248-128263288 | Myc |
| chr8:128263341-128263381 | Myc |
| chr8:128263442-128263482 | Myc |
| chr8:128263516-128263556 | Myc |
| chr8:128263709-128263749 | Myc |
| chr8:128263763-128263803 | Myc |
| chr8:128263841-128263881 | Myc |
| chr8:128263963-128264003 | Myc |
| chr8:128264013-128264053 | Myc |
| chr8:128264202-128264242 | Myc |
| chr8:128264277-128264317 | Myc |
| chr8:128264334-128264374 | Myc |
| chr8:128264393-128264433 | Myc |
| chr8:128264469-128264509 | Myc |
| chr8:128264526-128264566 | Myc |
| chr8:128264614-128264654 | Myc |
| chr8:128264720-128264760 | Myc |
| chr8:128264777-128264817 | Myc |
| chr8:128264938-128264978 | Myc |
| chr8:128265013-128265053 | Myc |
| chr8:128265071-128265111 | Myc |
| chr8:128265125-128265165 | Myc |
| chr8:128265492-128265532 | Myc |
| chr8:128265545-128265585 | Myc |
| chr8:128265661-128265701 | Myc |
| chr8:128265737-128265777 | Myc |
| chr8:128265814-128265854 | Myc |

| | |
|---|---|
| chr8:128265888-128265928 | Myc |
| chr8:128265978-128266018 | Myc |
| chr8:128266089-128266129 | Myc |
| chr8:128266248-128266288 | Myc |
| chr8:128266298-128266338 | Myc |
| chr8:128266348-128266388 | Myc |
| chr8:128266401-128266441 | Myc |
| chr8:128266453-128266493 | Myc |
| chr8:128266534-128266574 | Myc |
| chr8:128266606-128266646 | Myc |
| chr8:128266671-128266711 | Myc |
| chr8:128266769-128266809 | Myc |
| chr8:128266825-128266865 | Myc |
| chr8:128266989-128267029 | Myc |
| chr8:128267042-128267082 | Myc |
| chr8:128267092-128267132 | Myc |
| chr8:128267149-128267189 | Myc |
| chr8:128267222-128267262 | Myc |
| chr8:128267299-128267339 | Myc |
| chr8:128267434-128267474 | Myc |
| chr8:128267536-128267576 | Myc |
| chr8:128267631-128267671 | Myc |
| chr8:128267698-128267738 | Myc |
| chr8:128267802-128267842 | Myc |
| chr8:128267918-128267958 | Myc |
| chr8:128268014-128268054 | Myc |
| chr8:128268098-128268138 | Myc |
| chr8:128268194-128268234 | Myc |
| chr8:128268320-128268360 | Myc |
| chr8:128268370-128268410 | Myc |
| chr8:128268494-128268534 | Myc |
| chr8:128268566-128268606 | Myc |
| chr8:128268722-128268762 | Myc |
| chr8:128268776-128268816 | Myc |
| chr8:128268833-128268873 | Myc |
| chr8:128268957-128268997 | Myc |
| chr8:128269073-128269113 | Myc |
| chr8:128269150-128269190 | Myc |
| chr8:128269226-128269266 | Myc |
| chr8:128269358-128269398 | Myc |
| chr8:128269431-128269471 | Myc |
| chr8:128269481-128269521 | Myc |
| chr8:128269561-128269601 | Myc |
| chr8:128269640-128269680 | Myc |
| chr8:128269805-128269845 | Myc |
| chr8:128269894-128269934 | Myc |
| chr8:128269971-128270011 | Myc |
| chr8:128270060-128270100 | Myc |
| chr8:128270110-128270150 | Myc |
| chr8:128270196-128270236 | Myc |
| chr8:128270246-128270286 | Myc |
| chr8:128270332-128270372 | Myc |

| | |
|---|---|
| chr8:128270445-128270485 | Myc |
| chr8:128270586-128270626 | Myc |
| chr8:128270653-128270693 | Myc |
| chr8:128270791-128270831 | Myc |
| chr8:128270847-128270887 | Myc |
| chr8:128270954-128270994 | Myc |
| chr8:128271037-128271077 | Myc |
| chr8:128271177-128271217 | Myc |
| chr8:128271262-128271302 | Myc |
| chr8:128271350-128271390 | Myc |
| chr8:128271427-128271467 | Myc |
| chr8:128271488-128271528 | Myc |
| chr8:128271600-128271640 | Myc |
| chr8:128271654-128271694 | Myc |
| chr8:128271733-128271773 | Myc |
| chr8:128271798-128271838 | Myc |
| chr8:128271894-128271934 | Myc |
| chr8:128272040-128272080 | Myc |
| chr8:128272142-128272182 | Myc |
| chr8:128272204-128272244 | Myc |
| chr8:128272303-128272343 | Myc |
| chr8:128272419-128272459 | Myc |
| chr8:128272477-128272517 | Myc |
| chr8:128272563-128272603 | Myc |
| chr8:128272751-128272791 | Myc |
| chr8:128272886-128272926 | Myc |
| chr8:128273005-128273045 | Myc |
| chr8:128273292-128273332 | Myc |
| chr8:128273371-128273411 | Myc |
| chr8:128273438-128273478 | Myc |
| chr8:128273559-128273599 | Myc |
| chr8:128273610-128273650 | Myc |
| chr8:128273736-128273776 | Myc |
| chr8:128273786-128273826 | Myc |
| chr8:128274040-128274080 | Myc |
| chr8:128274167-128274207 | Myc |
| chr8:128274218-128274258 | Myc |
| chr8:128274286-128274326 | Myc |
| chr8:128274356-128274396 | Myc |
| chr8:128274451-128274491 | Myc |
| chr8:128274501-128274541 | Myc |
| chr8:128274569-128274609 | Myc |
| chr8:128274707-128274747 | Myc |
| chr8:128274826-128274866 | Myc |
| chr8:128274961-128275001 | Myc |
| chr8:128275052-128275092 | Myc |
| chr8:128275289-128275329 | Myc |
| chr8:128275411-128275451 | Myc |
| chr8:128275534-128275574 | Myc |
| chr8:128275611-128275651 | Myc |
| chr8:128275710-128275750 | Myc |
| chr8:128275941-128275981 | Myc |

| | |
|---|---|
| chr8:128276011-128276051 | Myc |
| chr8:128276144-128276184 | Myc |
| chr8:128276198-128276238 | Myc |
| chr8:128276249-128276289 | Myc |
| chr8:128276352-128276392 | Myc |
| chr8:128276410-128276450 | Myc |
| chr8:128276480-128276520 | Myc |
| chr8:128276530-128276570 | Myc |
| chr8:128276747-128276787 | Myc |
| chr8:128276854-128276894 | Myc |
| chr8:128276904-128276944 | Myc |
| chr8:128276961-128277001 | Myc |
| chr8:128277012-128277052 | Myc |
| chr8:128277062-128277102 | Myc |
| chr8:128277127-128277167 | Myc |
| chr8:128277194-128277234 | Myc |
| chr8:128277268-128277308 | Myc |
| chr8:128277324-128277364 | Myc |
| chr8:128277391-128277431 | Myc |
| chr8:128277474-128277514 | Myc |
| chr8:128277584-128277624 | Myc |
| chr8:128277634-128277674 | Myc |
| chr8:128277688-128277728 | Myc |
| chr8:128277795-128277835 | Myc |
| chr8:128277850-128277890 | Myc |
| chr8:128277948-128277988 | Myc |
| chr8:128278049-128278089 | Myc |
| chr8:128278203-128278243 | Myc |
| chr8:128278261-128278301 | Myc |
| chr8:128278360-128278400 | Myc |
| chr8:128278426-128278466 | Myc |
| chr8:128278550-128278590 | Myc |
| chr8:128278670-128278710 | Myc |
| chr8:128278768-128278808 | Myc |
| chr8:128278842-128278882 | Myc |
| chr8:128278928-128278968 | Myc |
| chr8:128278979-128279019 | Myc |
| chr8:128279140-128279180 | Myc |
| chr8:128279239-128279279 | Myc |
| chr8:128279303-128279343 | Myc |
| chr8:128279410-128279450 | Myc |
| chr8:128279559-128279599 | Myc |
| chr8:128279632-128279672 | Myc |
| chr8:128279738-128279778 | Myc |
| chr8:128279933-128279973 | Myc |
| chr8:128279983-128280023 | Myc |
| chr8:128280033-128280073 | Myc |
| chr8:128280087-128280127 | Myc |
| chr8:128280149-128280189 | Myc |
| chr8:128280227-128280267 | Myc |
| chr8:128280281-128280321 | Myc |
| chr8:128280340-128280380 | Myc |

| | |
|---|---|
| chr8:128280450-128280490 | Myc |
| chr8:128280552-128280592 | Myc |
| chr8:128280630-128280670 | Myc |
| chr8:128280688-128280728 | Myc |
| chr8:128280749-128280789 | Myc |
| chr8:128280842-128280882 | Myc |
| chr8:128280892-128280932 | Myc |
| chr8:128280972-128281012 | Myc |
| chr8:128281054-128281094 | Myc |
| chr8:128281218-128281258 | Myc |
| chr8:128281298-128281338 | Myc |
| chr8:128281360-128281400 | Myc |
| chr8:128281518-128281558 | Myc |
| chr8:128281592-128281632 | Myc |
| chr8:128281700-128281740 | Myc |
| chr8:128281756-128281796 | Myc |
| chr8:128281857-128281897 | Myc |
| chr8:128281935-128281975 | Myc |
| chr8:128282057-128282097 | Myc |
| chr8:128282110-128282150 | Myc |
| chr8:128282195-128282235 | Myc |
| chr8:128282269-128282309 | Myc |
| chr8:128282321-128282361 | Myc |
| chr8:128282419-128282459 | Myc |
| chr8:128282520-128282560 | Myc |
| chr8:128282596-128282636 | Myc |
| chr8:128282666-128282706 | Myc |
| chr8:128282742-128282782 | Myc |
| chr8:128282792-128282832 | Myc |
| chr8:128282866-128282906 | Myc |
| chr8:128282953-128282993 | Myc |
| chr8:128283048-128283088 | Myc |
| chr8:128283121-128283161 | Myc |
| chr8:128283237-128283277 | Myc |
| chr8:128283288-128283328 | Myc |
| chr8:128283372-128283412 | Myc |
| chr8:128283446-128283486 | Myc |
| chr8:128283513-128283553 | Myc |
| chr8:128283579-128283619 | Myc |
| chr8:128283696-128283736 | Myc |
| chr8:128283751-128283791 | Myc |
| chr8:128283802-128283842 | Myc |
| chr8:128283873-128283913 | Myc |
| chr8:128283986-128284026 | Myc |
| chr8:128284053-128284093 | Myc |
| chr8:128284152-128284192 | Myc |
| chr8:128284240-128284280 | Myc |
| chr8:128284290-128284330 | Myc |
| chr8:128284340-128284380 | Myc |
| chr8:128284426-128284466 | Myc |
| chr8:128284559-128284599 | Myc |
| chr8:128284708-128284748 | Myc |

| | |
|---|---|
| chr8:128284762-128284802 | Myc |
| chr8:128284819-128284859 | Myc |
| chr8:128284914-128284954 | Myc |
| chr8:128285025-128285065 | Myc |
| chr8:128285103-128285143 | Myc |
| chr8:128285190-128285230 | Myc |
| chr8:128285303-128285343 | Myc |
| chr8:128285528-128285568 | Myc |
| chr8:128285602-128285642 | Myc |
| chr8:128285727-128285767 | Myc |
| chr8:128285805-128285845 | Myc |
| chr8:128285859-128285899 | Myc |
| chr8:128286009-128286049 | Myc |
| chr8:128286104-128286144 | Myc |
| chr8:128286185-128286225 | Myc |
| chr8:128286244-128286284 | Myc |
| chr8:128286296-128286336 | Myc |
| chr8:128286384-128286424 | Myc |
| chr8:128286449-128286489 | Myc |
| chr8:128286525-128286565 | Myc |
| chr8:128286605-128286645 | Myc |
| chr8:128286676-128286716 | Myc |
| chr8:128286726-128286766 | Myc |
| chr8:128286832-128286872 | Myc |
| chr8:128286914-128286954 | Myc |
| chr8:128286965-128287005 | Myc |
| chr8:128287016-128287056 | Myc |
| chr8:128287106-128287146 | Myc |
| chr8:128287229-128287269 | Myc |
| chr8:128287343-128287383 | Myc |
| chr8:128287678-128287718 | Myc |
| chr8:128287740-128287780 | Myc |
| chr8:128287790-128287830 | Myc |
| chr8:128288028-128288068 | Myc |
| chr8:128288143-128288183 | Myc |
| chr8:128288193-128288233 | Myc |
| chr8:128288278-128288318 | Myc |
| chr8:128288396-128288436 | Myc |
| chr8:128288446-128288486 | Myc |
| chr8:128288576-128288616 | Myc |
| chr8:128288653-128288693 | Myc |
| chr8:128288722-128288762 | Myc |
| chr8:128288884-128288924 | Myc |
| chr8:128288977-128289017 | Myc |
| chr8:128289064-128289104 | Myc |
| chr8:128289163-128289203 | Myc |
| chr8:128289264-128289304 | Myc |
| chr8:128289370-128289410 | Myc |
| chr8:128289428-128289468 | Myc |
| chr8:128289501-128289541 | Myc |
| chr8:128289684-128289724 | Myc |
| chr8:128289734-128289774 | Myc |

| | |
|---|---|
| chr8:128289812-128289852 | Myc |
| chr8:128289954-128289994 | Myc |
| chr8:128290006-128290046 | Myc |
| chr8:128290154-128290194 | Myc |
| chr8:128290204-128290244 | Myc |
| chr8:128290255-128290295 | Myc |
| chr8:128290449-128290489 | Myc |
| chr8:128290499-128290539 | Myc |
| chr8:128290689-128290729 | Myc |
| chr8:128290844-128290884 | Myc |
| chr8:128290969-128291009 | Myc |
| chr8:128291030-128291070 | Myc |
| chr8:128291192-128291232 | Myc |
| chr8:128291282-128291322 | Myc |
| chr8:128291360-128291400 | Myc |
| chr8:128291411-128291451 | Myc |
| chr8:128291551-128291591 | Myc |
| chr8:128291616-128291656 | Myc |
| chr8:128291941-128291981 | Myc |
| chr8:128291992-128292032 | Myc |
| chr8:128292050-128292090 | Myc |
| chr8:128292124-128292164 | Myc |
| chr8:128292196-128292236 | Myc |
| chr8:128292335-128292375 | Myc |
| chr8:128292404-128292444 | Myc |
| chr8:128292476-128292516 | Myc |
| chr8:128292555-128292595 | Myc |
| chr8:128292607-128292647 | Myc |
| chr8:128292657-128292697 | Myc |
| chr8:128292826-128292866 | Myc |
| chr8:128292921-128292961 | Myc |
| chr8:128292971-128293011 | Myc |
| chr8:128293172-128293212 | Myc |
| chr8:128293301-128293341 | Myc |
| chr8:128293365-128293405 | Myc |
| chr8:128293430-128293470 | Myc |
| chr8:128293496-128293536 | Myc |
| chr8:128293672-128293712 | Myc |
| chr8:128293734-128293774 | Myc |
| chr8:128293805-128293845 | Myc |
| chr8:128293902-128293942 | Myc |
| chr8:128293961-128294001 | Myc |
| chr8:128294063-128294103 | Myc |
| chr8:128294163-128294203 | Myc |
| chr8:128294231-128294271 | Myc |
| chr8:128294293-128294333 | Myc |
| chr8:128294444-128294484 | Myc |
| chr8:128294494-128294534 | Myc |
| chr8:128294595-128294635 | Myc |
| chr8:128294677-128294717 | Myc |
| chr8:128294729-128294769 | Myc |
| chr8:128294903-128294943 | Myc |

| | |
|---|---|
| chr8:128295008-128295048 | Myc |
| chr8:128295118-128295158 | Myc |
| chr8:128295168-128295208 | Myc |
| chr8:128295411-128295451 | Myc |
| chr8:128295536-128295576 | Myc |
| chr8:128295681-128295721 | Myc |
| chr8:128295750-128295790 | Myc |
| chr8:128295820-128295860 | Myc |
| chr8:128295885-128295925 | Myc |
| chr8:128295984-128296024 | Myc |
| chr8:128296055-128296095 | Myc |
| chr8:128296120-128296160 | Myc |
| chr8:128296190-128296230 | Myc |
| chr8:128296310-128296350 | Myc |
| chr8:128296366-128296406 | Myc |
| chr8:128296479-128296519 | Myc |
| chr8:128296702-128296742 | Myc |
| chr8:128296752-128296792 | Myc |
| chr8:128296836-128296876 | Myc |
| chr8:128296904-128296944 | Myc |
| chr8:128296954-128296994 | Myc |
| chr8:128297072-128297112 | Myc |
| chr8:128297185-128297225 | Myc |
| chr8:128297245-128297285 | Myc |
| chr8:128297305-128297345 | Myc |
| chr8:128297367-128297407 | Myc |
| chr8:128297481-128297521 | Myc |
| chr8:128297625-128297665 | Myc |
| chr8:128297867-128297907 | Myc |
| chr8:128297979-128298019 | Myc |
| chr8:128298065-128298105 | Myc |
| chr8:128298123-128298163 | Myc |
| chr8:128298263-128298303 | Myc |
| chr8:128298463-128298503 | Myc |
| chr8:128298535-128298575 | Myc |
| chr8:128298834-128298874 | Myc |
| chr8:128299269-128299309 | Myc |
| chr8:128299386-128299426 | Myc |
| chr8:128299445-128299485 | Myc |
| chr8:128299496-128299536 | Myc |
| chr8:128299685-128299725 | Myc |
| chr8:128299735-128299775 | Myc |
| chr8:128299785-128299825 | Myc |
| chr8:128299896-128299936 | Myc |
| chr8:128299952-128299992 | Myc |
| chr8:128300051-128300091 | Myc |
| chr8:128300226-128300266 | Myc |
| chr8:128300301-128300341 | Myc |
| chr8:128300365-128300405 | Myc |
| chr8:128300487-128300527 | Myc |
| chr8:128300559-128300599 | Myc |
| chr8:128300650-128300690 | Myc |

| | |
|---|---|
| chr8:128300727-128300767 | Myc |
| chr8:128300777-128300817 | Myc |
| chr8:128300951-128300991 | Myc |
| chr8:128301089-128301129 | Myc |
| chr8:128301144-128301184 | Myc |
| chr8:128301259-128301299 | Myc |
| chr8:128301349-128301389 | Myc |
| chr8:128301448-128301488 | Myc |
| chr8:128301517-128301557 | Myc |
| chr8:128301584-128301624 | Myc |
| chr8:128301646-128301686 | Myc |
| chr8:128301739-128301779 | Myc |
| chr8:128301807-128301847 | Myc |
| chr8:128302002-128302042 | Myc |
| chr8:128302133-128302173 | Myc |
| chr8:128302213-128302253 | Myc |
| chr8:128302263-128302303 | Myc |
| chr8:128302346-128302386 | Myc |
| chr8:128302428-128302468 | Myc |
| chr8:128302511-128302551 | Myc |
| chr8:128302569-128302609 | Myc |
| chr8:128302629-128302669 | Myc |
| chr8:128302682-128302722 | Myc |
| chr8:128302732-128302772 | Myc |
| chr8:128302787-128302827 | Myc |
| chr8:128302848-128302888 | Myc |
| chr8:128302947-128302987 | Myc |
| chr8:128303108-128303148 | Myc |
| chr8:128303223-128303263 | Myc |
| chr8:128303323-128303363 | Myc |
| chr8:128303406-128303446 | Myc |
| chr8:128303506-128303546 | Myc |
| chr8:128303606-128303646 | Myc |
| chr8:128303659-128303699 | Myc |
| chr8:128303794-128303834 | Myc |
| chr8:128303848-128303888 | Myc |
| chr8:128303971-128304011 | Myc |
| chr8:128304101-128304141 | Myc |
| chr8:128304155-128304195 | Myc |
| chr8:128304238-128304278 | Myc |
| chr8:128304404-128304444 | Myc |
| chr8:128304498-128304538 | Myc |
| chr8:128304585-128304625 | Myc |
| chr8:128304661-128304701 | Myc |
| chr8:128304711-128304751 | Myc |
| chr8:128304910-128304950 | Myc |
| chr8:128305056-128305096 | Myc |
| chr8:128305134-128305174 | Myc |
| chr8:128305198-128305238 | Myc |
| chr8:128305381-128305421 | Myc |
| chr8:128305432-128305472 | Myc |
| chr8:128305527-128305567 | Myc |

| | |
|---|---|
| chr8:128305594-128305634 | Myc |
| chr8:128305650-128305690 | Myc |
| chr8:128305762-128305802 | Myc |
| chr8:128305875-128305915 | Myc |
| chr8:128305941-128305981 | Myc |
| chr8:128305991-128306031 | Myc |
| chr8:128306080-128306120 | Myc |
| chr8:128306196-128306236 | Myc |
| chr8:128306302-128306342 | Myc |
| chr8:128306395-128306435 | Myc |
| chr8:128315477-128315517 | Myc |
| chr8:128315527-128315567 | Myc |
| chr8:128315616-128315656 | Myc |
| chr8:128315713-128315753 | Myc |
| chr8:128315763-128315803 | Myc |
| chr8:128315820-128315860 | Myc |
| chr8:128315905-128315945 | Myc |
| chr8:128315959-128315999 | Myc |
| chr8:128316092-128316132 | Myc |
| chr8:128316161-128316201 | Myc |
| chr8:128316211-128316251 | Myc |
| chr8:128316378-128316418 | Myc |
| chr8:128316428-128316468 | Myc |
| chr8:128316665-128316705 | Myc |
| chr8:128316765-128316805 | Myc |
| chr8:128316847-128316887 | Myc |
| chr8:128316898-128316938 | Myc |
| chr8:128317011-128317051 | Myc |
| chr8:128317127-128317167 | Myc |
| chr8:128317179-128317219 | Myc |
| chr8:128317229-128317269 | Myc |
| chr8:128317290-128317330 | Myc |
| chr8:128317359-128317399 | Myc |
| chr8:128317409-128317449 | Myc |
| chr8:128317472-128317512 | Myc |
| chr8:128317523-128317563 | Myc |
| chr8:128317650-128317690 | Myc |
| chr8:128317700-128317740 | Myc |
| chr8:128317779-128317819 | Myc |
| chr8:128317878-128317918 | Myc |
| chr8:128317950-128317990 | Myc |
| chr8:128318039-128318079 | Myc |
| chr8:128318124-128318164 | Myc |
| chr8:128318254-128318294 | Myc |
| chr8:128318304-128318344 | Myc |
| chr8:128318354-128318394 | Myc |
| chr8:128318420-128318460 | Myc |
| chr8:128318475-128318515 | Myc |
| chr8:128318555-128318595 | Myc |
| chr8:128318667-128318707 | Myc |
| chr8:128318717-128318757 | Myc |
| chr8:128318816-128318856 | Myc |

| | |
|---|---|
| chr8:128318892-128318932 | Myc |
| chr8:128319006-128319046 | Myc |
| chr8:128319117-128319157 | Myc |
| chr8:128319227-128319267 | Myc |
| chr8:128319341-128319381 | Myc |
| chr8:128319459-128319499 | Myc |
| chr8:128319510-128319550 | Myc |
| chr8:128319583-128319623 | Myc |
| chr8:128319659-128319699 | Myc |
| chr8:128319738-128319778 | Myc |
| chr8:128319824-128319864 | Myc |
| chr8:128319971-128320011 | Myc |
| chr8:128320023-128320063 | Myc |
| chr8:128320130-128320170 | Myc |
| chr8:128320181-128320221 | Myc |
| chr8:128320265-128320305 | Myc |
| chr8:128320375-128320415 | Myc |
| chr8:128320458-128320498 | Myc |
| chr8:128320624-128320664 | Myc |
| chr8:128320717-128320757 | Myc |
| chr8:128320889-128320929 | Myc |
| chr8:128321071-128321111 | Myc |
| chr8:128321169-128321209 | Myc |
| chr8:128321276-128321316 | Myc |
| chr8:128321327-128321367 | Myc |
| chr8:128321407-128321447 | Myc |
| chr8:128321496-128321536 | Myc |
| chr8:128321597-128321637 | Myc |
| chr8:128321665-128321705 | Myc |
| chr8:128321716-128321756 | Myc |
| chr8:128321766-128321806 | Myc |
| chr8:128321899-128321939 | Myc |
| chr8:128321972-128322012 | Myc |
| chr8:128322067-128322107 | Myc |
| chr8:128322154-128322194 | Myc |
| chr8:128322215-128322255 | Myc |
| chr8:128322308-128322348 | Myc |
| chr8:128322366-128322406 | Myc |
| chr8:128322523-128322563 | Myc |
| chr8:128322593-128322633 | Myc |
| chr8:128322900-128322940 | Myc |
| chr8:128322950-128322990 | Myc |
| chr8:128323036-128323076 | Myc |
| chr8:128323127-128323167 | Myc |
| chr8:128323263-128323303 | Myc |
| chr8:128323357-128323397 | Myc |
| chr8:128323525-128323565 | Myc |
| chr8:128323635-128323675 | Myc |
| chr8:128323685-128323725 | Myc |
| chr8:128323801-128323841 | Myc |
| chr8:128323873-128323913 | Myc |
| chr8:128324091-128324131 | Myc |

| | |
|---|---|
| chr8:128324756-128324796 | Myc |
| chr8:128326395-128326435 | Myc |
| chr8:128326816-128326856 | Myc |
| chr8:128327710-128327750 | Myc |
| chr8:128328575-128328615 | Myc |
| chr8:128330292-128330332 | Myc |
| chr8:128330345-128330385 | Myc |
| chr8:128330487-128330527 | Myc |
| chr8:128330587-128330627 | Myc |
| chr8:128331005-128331045 | Myc |
| chr8:128331555-128331595 | Myc |
| chr8:128331773-128331813 | Myc |
| chr8:128333144-128333184 | Myc |
| chr8:128333268-128333308 | Myc |
| chr8:128333345-128333385 | Myc |
| chr8:128333430-128333470 | Myc |
| chr8:128333512-128333552 | Myc |
| chr8:128333650-128333690 | Myc |
| chr8:128333743-128333783 | Myc |
| chr8:128333847-128333887 | Myc |
| chr8:128334047-128334087 | Myc |
| chr8:128334128-128334168 | Myc |
| chr8:128334178-128334218 | Myc |
| chr8:128334242-128334282 | Myc |
| chr8:128334341-128334381 | Myc |
| chr8:128334547-128334587 | Myc |
| chr8:128334597-128334637 | Myc |
| chr8:128334649-128334689 | Myc |
| chr8:128334749-128334789 | Myc |
| chr8:128334799-128334839 | Myc |
| chr8:128334904-128334944 | Myc |
| chr8:128335054-128335094 | Myc |
| chr8:128335179-128335219 | Myc |
| chr8:128335273-128335313 | Myc |
| chr8:128335323-128335363 | Myc |
| chr8:128335487-128335527 | Myc |
| chr8:128335541-128335581 | Myc |
| chr8:128335595-128335635 | Myc |
| chr8:128335694-128335734 | Myc |
| chr8:128335774-128335814 | Myc |
| chr8:128335838-128335878 | Myc |
| chr8:128335911-128335951 | Myc |
| chr8:128336020-128336060 | Myc |
| chr8:128336078-128336118 | Myc |
| chr8:128336180-128336220 | Myc |
| chr8:128336295-128336335 | Myc |
| chr8:128336383-128336423 | Myc |
| chr8:128336503-128336543 | Myc |
| chr8:128336579-128336619 | Myc |
| chr8:128336723-128336763 | Myc |
| chr8:128336775-128336815 | Myc |
| chr8:128336836-128336876 | Myc |

| | |
|---|---|
| chr8:128336923-128336963 | Myc |
| chr8:128337049-128337089 | Myc |
| chr8:128337112-128337152 | Myc |
| chr8:128337253-128337293 | Myc |
| chr8:128337329-128337369 | Myc |
| chr8:128337411-128337451 | Myc |
| chr8:128337584-128337624 | Myc |
| chr8:128337634-128337674 | Myc |
| chr8:128337820-128337860 | Myc |
| chr8:128337913-128337953 | Myc |
| chr8:128338102-128338142 | Myc |
| chr8:128338169-128338209 | Myc |
| chr8:128338229-128338269 | Myc |
| chr8:128338284-128338324 | Myc |
| chr8:128338456-128338496 | Myc |
| chr8:128338507-128338547 | Myc |
| chr8:128338639-128338679 | Myc |
| chr8:128338802-128338842 | Myc |
| chr8:128338913-128338953 | Myc |
| chr8:128338975-128339015 | Myc |
| chr8:128339030-128339070 | Myc |
| chr8:128339145-128339185 | Myc |
| chr8:128339200-128339240 | Myc |
| chr8:128339308-128339348 | Myc |
| chr8:128339368-128339408 | Myc |
| chr8:128339465-128339505 | Myc |
| chr8:128339556-128339596 | Myc |
| chr8:128339638-128339678 | Myc |
| chr8:128339826-128339866 | Myc |
| chr8:128339885-128339925 | Myc |
| chr8:128339991-128340031 | Myc |
| chr8:128340067-128340107 | Myc |
| chr8:128340117-128340157 | Myc |
| chr8:128340188-128340228 | Myc |
| chr8:128340289-128340329 | Myc |
| chr8:128340367-128340407 | Myc |
| chr8:128340450-128340490 | Myc |
| chr8:128340567-128340607 | Myc |
| chr8:128340626-128340666 | Myc |
| chr8:128340737-128340777 | Myc |
| chr8:128340802-128340842 | Myc |
| chr8:128340871-128340911 | Myc |
| chr8:128340963-128341003 | Myc |
| chr8:128341095-128341135 | Myc |
| chr8:128341177-128341217 | Myc |
| chr8:128341267-128341307 | Myc |
| chr8:128341317-128341357 | Myc |
| chr8:128341460-128341500 | Myc |
| chr8:128341541-128341581 | Myc |
| chr8:128341617-128341657 | Myc |
| chr8:128341667-128341707 | Myc |
| chr8:128342114-128342154 | Myc |

| | |
|---|---|
| chr8:128342182-128342222 | Myc |
| chr8:128342288-128342328 | Myc |
| chr8:128342351-128342391 | Myc |
| chr8:128342501-128342541 | Myc |
| chr8:128342621-128342661 | Myc |
| chr8:128342671-128342711 | Myc |
| chr8:128342744-128342784 | Myc |
| chr8:128342916-128342956 | Myc |
| chr8:128342966-128343006 | Myc |
| chr8:128343051-128343091 | Myc |
| chr8:128343155-128343195 | Myc |
| chr8:128343266-128343306 | Myc |
| chr8:128343367-128343407 | Myc |
| chr8:128343439-128343479 | Myc |
| chr8:128343539-128343579 | Myc |
| chr8:128343636-128343676 | Myc |
| chr8:128343702-128343742 | Myc |
| chr8:128343755-128343795 | Myc |
| chr8:128343806-128343846 | Myc |
| chr8:128343873-128343913 | Myc |
| chr8:128344015-128344055 | Myc |
| chr8:128344136-128344176 | Myc |
| chr8:128344211-128344251 | Myc |
| chr8:128344347-128344387 | Myc |
| chr8:128344399-128344439 | Myc |
| chr8:128344449-128344489 | Myc |
| chr8:128344542-128344582 | Myc |
| chr8:128344607-128344647 | Myc |
| chr8:128344748-128344788 | Myc |
| chr8:128344852-128344892 | Myc |
| chr8:128344991-128345031 | Myc |
| chr8:128345059-128345099 | Myc |
| chr8:128345176-128345216 | Myc |
| chr8:128345241-128345281 | Myc |
| chr8:128345390-128345430 | Myc |
| chr8:128345479-128345519 | Myc |
| chr8:128345529-128345569 | Myc |
| chr8:128345645-128345685 | Myc |
| chr8:128345717-128345757 | Myc |
| chr8:128345796-128345836 | Myc |
| chr8:128345849-128345889 | Myc |
| chr8:128345910-128345950 | Myc |
| chr8:128345981-128346021 | Myc |
| chr8:128346118-128346158 | Myc |
| chr8:128346171-128346211 | Myc |
| chr8:128346253-128346293 | Myc |
| chr8:128346311-128346351 | Myc |
| chr8:128346472-128346512 | Myc |
| chr8:128346549-128346589 | Myc |
| chr8:128346721-128346761 | Myc |
| chr8:128346822-128346862 | Myc |
| chr8:128346929-128346969 | Myc |

| | |
|---|---|
| chr8:128346989-128347029 | Myc |
| chr8:128347040-128347080 | Myc |
| chr8:128347112-128347152 | Myc |
| chr8:128347209-128347249 | Myc |
| chr8:128347396-128347436 | Myc |
| chr8:128347446-128347486 | Myc |
| chr8:128347498-128347538 | Myc |
| chr8:128347574-128347614 | Myc |
| chr8:128347658-128347698 | Myc |
| chr8:128347744-128347784 | Myc |
| chr8:128347816-128347856 | Myc |
| chr8:128347945-128347985 | Myc |
| chr8:128347995-128348035 | Myc |
| chr8:128348046-128348086 | Myc |
| chr8:128348154-128348194 | Myc |
| chr8:128348270-128348310 | Myc |
| chr8:128348383-128348423 | Myc |
| chr8:128348453-128348493 | Myc |
| chr8:128348538-128348578 | Myc |
| chr8:128348600-128348640 | Myc |
| chr8:128348650-128348690 | Myc |
| chr8:128348744-128348784 | Myc |
| chr8:128348798-128348838 | Myc |
| chr8:128349050-128349090 | Myc |
| chr8:128349103-128349143 | Myc |
| chr8:128349161-128349201 | Myc |
| chr8:128349235-128349275 | Myc |
| chr8:128349350-128349390 | Myc |
| chr8:128349489-128349529 | Myc |
| chr8:128349583-128349623 | Myc |
| chr8:128349633-128349673 | Myc |
| chr8:128349728-128349768 | Myc |
| chr8:128349804-128349844 | Myc |
| chr8:128349876-128349916 | Myc |
| chr8:128349926-128349966 | Myc |
| chr8:128349990-128350030 | Myc |
| chr8:128350040-128350080 | Myc |
| chr8:128350144-128350184 | Myc |
| chr8:128350194-128350234 | Myc |
| chr8:128350329-128350369 | Myc |
| chr8:128350393-128350433 | Myc |
| chr8:128350471-128350511 | Myc |
| chr8:128350569-128350609 | Myc |
| chr8:128350698-128350738 | Myc |
| chr8:128350748-128350788 | Myc |
| chr8:128350911-128350951 | Myc |
| chr8:128350965-128351005 | Myc |
| chr8:128351142-128351182 | Myc |
| chr8:128351211-128351251 | Myc |
| chr8:128351288-128351328 | Myc |
| chr8:128351402-128351442 | Myc |
| chr8:128351492-128351532 | Myc |

| | |
|---|---|
| chr8:128351614-128351654 | Myc |
| chr8:128351748-128351788 | Myc |
| chr8:128351810-128351850 | Myc |
| chr8:128351874-128351914 | Myc |
| chr8:128351975-128352015 | Myc |
| chr8:128352099-128352139 | Myc |
| chr8:128352155-128352195 | Myc |
| chr8:128352231-128352271 | Myc |
| chr8:128352297-128352337 | Myc |
| chr8:128352361-128352401 | Myc |
| chr8:128352415-128352455 | Myc |
| chr8:128352512-128352552 | Myc |
| chr8:128352573-128352613 | Myc |
| chr8:128352645-128352685 | Myc |
| chr8:128352745-128352785 | Myc |
| chr8:128352800-128352840 | Myc |
| chr8:128352881-128352921 | Myc |
| chr8:128352931-128352971 | Myc |
| chr8:128353014-128353054 | Myc |
| chr8:128353106-128353146 | Myc |
| chr8:128353157-128353197 | Myc |
| chr8:128353218-128353258 | Myc |
| chr8:128353325-128353365 | Myc |
| chr8:128353375-128353415 | Myc |
| chr8:128353429-128353469 | Myc |
| chr8:128353497-128353537 | Myc |
| chr8:128353570-128353610 | Myc |
| chr8:128353686-128353726 | Myc |
| chr8:128353777-128353817 | Myc |
| chr8:128353870-128353910 | Myc |
| chr8:128353973-128354013 | Myc |
| chr8:128354102-128354142 | Myc |
| chr8:128354152-128354192 | Myc |
| chr8:128354289-128354329 | Myc |
| chr8:128354354-128354394 | Myc |
| chr8:128354414-128354454 | Myc |
| chr8:128354527-128354567 | Myc |
| chr8:128354628-128354668 | Myc |
| chr8:128354686-128354726 | Myc |
| chr8:128354811-128354851 | Myc |
| chr8:128354898-128354938 | Myc |
| chr8:128354989-128355029 | Myc |
| chr8:128355055-128355095 | Myc |
| chr8:128355106-128355146 | Myc |
| chr8:128355283-128355323 | Myc |
| chr8:128355333-128355373 | Myc |
| chr8:128355385-128355425 | Myc |
| chr8:128355504-128355544 | Myc |
| chr8:128355615-128355655 | Myc |
| chr8:128355710-128355750 | Myc |
| chr8:128355760-128355800 | Myc |
| chr8:128355812-128355852 | Myc |

| | |
|---|---|
| chr8:128355862-128355902 | Myc |
| chr8:128355929-128355969 | Myc |
| chr8:128356041-128356081 | Myc |
| chr8:128356117-128356157 | Myc |
| chr8:128356222-128356262 | Myc |
| chr8:128356272-128356312 | Myc |
| chr8:128356386-128356426 | Myc |
| chr8:128356497-128356537 | Myc |
| chr8:128356558-128356598 | Myc |
| chr8:128356670-128356710 | Myc |
| chr8:128356778-128356818 | Myc |
| chr8:128356882-128356922 | Myc |
| chr8:128356958-128356998 | Myc |
| chr8:128357049-128357089 | Myc |
| chr8:128357106-128357146 | Myc |
| chr8:128357178-128357218 | Myc |
| chr8:128357235-128357275 | Myc |
| chr8:128357287-128357327 | Myc |
| chr8:128357399-128357439 | Myc |
| chr8:128357476-128357516 | Myc |
| chr8:128357565-128357605 | Myc |
| chr8:128357680-128357720 | Myc |
| chr8:128357797-128357837 | Myc |
| chr8:128357870-128357910 | Myc |
| chr8:128357955-128357995 | Myc |
| chr8:128358013-128358053 | Myc |
| chr8:128358087-128358127 | Myc |
| chr8:128358137-128358177 | Myc |
| chr8:128358245-128358285 | Myc |
| chr8:128358306-128358346 | Myc |
| chr8:128358542-128358582 | Myc |
| chr8:128358614-128358654 | Myc |
| chr8:128358726-128358766 | Myc |
| chr8:128358864-128358904 | Myc |
| chr8:128358973-128359013 | Myc |
| chr8:128359087-128359127 | Myc |
| chr8:128359196-128359236 | Myc |
| chr8:128359308-128359348 | Myc |
| chr8:128359415-128359455 | Myc |
| chr8:128359486-128359526 | Myc |
| chr8:128359582-128359622 | Myc |
| chr8:128359652-128359692 | Myc |
| chr8:128359712-128359752 | Myc |
| chr8:128359980-128360020 | Myc |
| chr8:128360081-128360121 | Myc |
| chr8:128360133-128360173 | Myc |
| chr8:128360243-128360283 | Myc |
| chr8:128360317-128360357 | Myc |
| chr8:128360367-128360407 | Myc |
| chr8:128360584-128360624 | Myc |
| chr8:128360634-128360674 | Myc |
| chr8:128360771-128360811 | Myc |

| | |
|---|---|
| chr8:128360828-128360868 | Myc |
| chr8:128360934-128360974 | Myc |
| chr8:128361083-128361123 | Myc |
| chr8:128361134-128361174 | Myc |
| chr8:128361214-128361254 | Myc |
| chr8:128361268-128361308 | Myc |
| chr8:128361349-128361389 | Myc |
| chr8:128361487-128361527 | Myc |
| chr8:128361603-128361643 | Myc |
| chr8:128361688-128361728 | Myc |
| chr8:128361760-128361800 | Myc |
| chr8:128361843-128361883 | Myc |
| chr8:128361964-128362004 | Myc |
| chr8:128362184-128362224 | Myc |
| chr8:128362243-128362283 | Myc |
| chr8:128362497-128362537 | Myc |
| chr8:128362580-128362620 | Myc |
| chr8:128362720-128362760 | Myc |
| chr8:128362854-128362894 | Myc |
| chr8:128362938-128362978 | Myc |
| chr8:128363011-128363051 | Myc |
| chr8:128363282-128363322 | Myc |
| chr8:128363412-128363452 | Myc |
| chr8:128363632-128363672 | Myc |
| chr8:128363695-128363735 | Myc |
| chr8:128363757-128363797 | Myc |
| chr8:128363824-128363864 | Myc |
| chr8:128363980-128364020 | Myc |
| chr8:128364045-128364085 | Myc |
| chr8:128364099-128364139 | Myc |
| chr8:128364227-128364267 | Myc |
| chr8:128364300-128364340 | Myc |
| chr8:128364401-128364441 | Myc |
| chr8:128364452-128364492 | Myc |
| chr8:128364518-128364558 | Myc |
| chr8:128364640-128364680 | Myc |
| chr8:128364709-128364749 | Myc |
| chr8:128364760-128364800 | Myc |
| chr8:128364810-128364850 | Myc |
| chr8:128364966-128365006 | Myc |
| chr8:128365034-128365074 | Myc |
| chr8:128365087-128365127 | Myc |
| chr8:128365144-128365184 | Myc |
| chr8:128365225-128365265 | Myc |
| chr8:128365397-128365437 | Myc |
| chr8:128365472-128365512 | Myc |
| chr8:128365535-128365575 | Myc |
| chr8:128365675-128365715 | Myc |
| chr8:128365786-128365826 | Myc |
| chr8:128365883-128365923 | Myc |
| chr8:128365964-128366004 | Myc |
| chr8:128366024-128366064 | Myc |

| | |
|---|---|
| chr8:128366084-128366124 | Myc |
| chr8:128366168-128366208 | Myc |
| chr8:128366218-128366258 | Myc |
| chr8:128366283-128366323 | Myc |
| chr8:128366364-128366404 | Myc |
| chr8:128366429-128366469 | Myc |
| chr8:128366493-128366533 | Myc |
| chr8:128366594-128366634 | Myc |
| chr8:128366761-128366801 | Myc |
| chr8:128366876-128366916 | Myc |
| chr8:128366937-128366977 | Myc |
| chr8:128367013-128367053 | Myc |
| chr8:128367084-128367124 | Myc |
| chr8:128367134-128367174 | Myc |
| chr8:128367298-128367338 | Myc |
| chr8:128367374-128367414 | Myc |
| chr8:128367578-128367618 | Myc |
| chr8:128367780-128367820 | Myc |
| chr8:128367830-128367870 | Myc |
| chr8:128367887-128367927 | Myc |
| chr8:128367963-128368003 | Myc |
| chr8:128368042-128368082 | Myc |
| chr8:128368111-128368151 | Myc |
| chr8:128368182-128368222 | Myc |
| chr8:128368275-128368315 | Myc |
| chr8:128368326-128368366 | Myc |
| chr8:128368392-128368432 | Myc |
| chr8:128368507-128368547 | Myc |
| chr8:128368627-128368667 | Myc |
| chr8:128368723-128368763 | Myc |
| chr8:128368859-128368899 | Myc |
| chr8:128368913-128368953 | Myc |
| chr8:128369030-128369070 | Myc |
| chr8:128369130-128369170 | Myc |
| chr8:128369215-128369255 | Myc |
| chr8:128369302-128369342 | Myc |
| chr8:128369352-128369392 | Myc |
| chr8:128369428-128369468 | Myc |
| chr8:128369569-128369609 | Myc |
| chr8:128369631-128369671 | Myc |
| chr8:128369741-128369781 | Myc |
| chr8:128369793-128369833 | Myc |
| chr8:128369911-128369951 | Myc |
| chr8:128369961-128370001 | Myc |
| chr8:128370011-128370051 | Myc |
| chr8:128370128-128370168 | Myc |
| chr8:128370178-128370218 | Myc |
| chr8:128370256-128370296 | Myc |
| chr8:128370358-128370398 | Myc |
| chr8:128370410-128370450 | Myc |
| chr8:128370473-128370513 | Myc |
| chr8:128370548-128370588 | Myc |

| | |
|---|---|
| chr8:128370704-128370744 | Myc |
| chr8:128370785-128370825 | Myc |
| chr8:128370839-128370879 | Myc |
| chr8:128370929-128370969 | Myc |
| chr8:128371089-128371129 | Myc |
| chr8:128371157-128371197 | Myc |
| chr8:128371209-128371249 | Myc |
| chr8:128371314-128371354 | Myc |
| chr8:128371364-128371404 | Myc |
| chr8:128371414-128371454 | Myc |
| chr8:128371590-128371630 | Myc |
| chr8:128371674-128371714 | Myc |
| chr8:128371743-128371783 | Myc |
| chr8:128371826-128371866 | Myc |
| chr8:128371900-128371940 | Myc |
| chr8:128371950-128371990 | Myc |
| chr8:128372117-128372157 | Myc |
| chr8:128372206-128372246 | Myc |
| chr8:128372328-128372368 | Myc |
| chr8:128372392-128372432 | Myc |
| chr8:128372588-128372628 | Myc |
| chr8:128372638-128372678 | Myc |
| chr8:128372711-128372751 | Myc |
| chr8:128372780-128372820 | Myc |
| chr8:128372835-128372875 | Myc |
| chr8:128372928-128372968 | Myc |
| chr8:128372982-128373022 | Myc |
| chr8:128373084-128373124 | Myc |
| chr8:128373290-128373330 | Myc |
| chr8:128373343-128373383 | Myc |
| chr8:128373476-128373516 | Myc |
| chr8:128373580-128373620 | Myc |
| chr8:128373770-128373810 | Myc |
| chr8:128373827-128373867 | Myc |
| chr8:128373940-128373980 | Myc |
| chr8:128373990-128374030 | Myc |
| chr8:128374086-128374126 | Myc |
| chr8:128374184-128374224 | Myc |
| chr8:128374269-128374309 | Myc |
| chr8:128374326-128374366 | Myc |
| chr8:128374390-128374430 | Myc |
| chr8:128374515-128374555 | Myc |
| chr8:128374599-128374639 | Myc |
| chr8:128374681-128374721 | Myc |
| chr8:128374842-128374882 | Myc |
| chr8:128374892-128374932 | Myc |
| chr8:128374943-128374983 | Myc |
| chr8:128375001-128375041 | Myc |
| chr8:128375144-128375184 | Myc |
| chr8:128375243-128375283 | Myc |
| chr8:128375296-128375336 | Myc |
| chr8:128375365-128375405 | Myc |

| | |
|---|---|
| chr8:128375456-128375496 | Myc |
| chr8:128375659-128375699 | Myc |
| chr8:128375724-128375764 | Myc |
| chr8:128375795-128375835 | Myc |
| chr8:128375858-128375898 | Myc |
| chr8:128375917-128375957 | Myc |
| chr8:128375978-128376018 | Myc |
| chr8:128376096-128376136 | Myc |
| chr8:128376197-128376237 | Myc |
| chr8:128376247-128376287 | Myc |
| chr8:128376341-128376381 | Myc |
| chr8:128376421-128376461 | Myc |
| chr8:128376479-128376519 | Myc |
| chr8:128376564-128376604 | Myc |
| chr8:128376704-128376744 | Myc |
| chr8:128376755-128376795 | Myc |
| chr8:128376829-128376869 | Myc |
| chr8:128376941-128376981 | Myc |
| chr8:128376991-128377031 | Myc |
| chr8:128377145-128377185 | Myc |
| chr8:128377323-128377363 | Myc |
| chr8:128377411-128377451 | Myc |
| chr8:128377473-128377513 | Myc |
| chr8:128377600-128377640 | Myc |
| chr8:128377663-128377703 | Myc |
| chr8:128377740-128377780 | Myc |
| chr8:128377799-128377839 | Myc |
| chr8:128377858-128377898 | Myc |
| chr8:128378006-128378046 | Myc |
| chr8:128378056-128378096 | Myc |
| chr8:128378140-128378180 | Myc |
| chr8:128378204-128378244 | Myc |
| chr8:128378283-128378323 | Myc |
| chr8:128378351-128378391 | Myc |
| chr8:128378446-128378486 | Myc |
| chr8:128378506-128378546 | Myc |
| chr8:128378556-128378596 | Myc |
| chr8:128378659-128378699 | Myc |
| chr8:128378802-128378842 | Myc |
| chr8:128378852-128378892 | Myc |
| chr8:128378942-128378982 | Myc |
| chr8:128379015-128379055 | Myc |
| chr8:128379065-128379105 | Myc |
| chr8:128379166-128379206 | Myc |
| chr8:128379280-128379320 | Myc |
| chr8:128379352-128379392 | Myc |
| chr8:128379413-128379453 | Myc |
| chr8:128379534-128379574 | Myc |
| chr8:128379584-128379624 | Myc |
| chr8:128379645-128379685 | Myc |
| chr8:128379916-128379956 | Myc |
| chr8:128380011-128380051 | Myc |

| | |
|---|---|
| chr8:128380073-128380113 | Myc |
| chr8:128380136-128380176 | Myc |
| chr8:128380262-128380302 | Myc |
| chr8:128380319-128380359 | Myc |
| chr8:128380413-128380453 | Myc |
| chr8:128380473-128380513 | Myc |
| chr8:128380528-128380568 | Myc |
| chr8:128380589-128380629 | Myc |
| chr8:128380647-128380687 | Myc |
| chr8:128380727-128380767 | Myc |
| chr8:128380896-128380936 | Myc |
| chr8:128381012-128381052 | Myc |
| chr8:128381102-128381142 | Myc |
| chr8:128381163-128381203 | Myc |
| chr8:128381389-128381429 | Myc |
| chr8:128381441-128381481 | Myc |
| chr8:128381494-128381534 | Myc |
| chr8:128381607-128381647 | Myc |
| chr8:128381681-128381721 | Myc |
| chr8:128381738-128381778 | Myc |
| chr8:128381815-128381855 | Myc |
| chr8:128381944-128381984 | Myc |
| chr8:128381997-128382037 | Myc |
| chr8:128382102-128382142 | Myc |
| chr8:128382152-128382192 | Myc |
| chr8:128382256-128382296 | Myc |
| chr8:128382333-128382373 | Myc |
| chr8:128382441-128382481 | Myc |
| chr8:128382497-128382537 | Myc |
| chr8:128382567-128382607 | Myc |
| chr8:128382651-128382691 | Myc |
| chr8:128382701-128382741 | Myc |
| chr8:128382764-128382804 | Myc |
| chr8:128382846-128382886 | Myc |
| chr8:128382922-128382962 | Myc |
| chr8:128382980-128383020 | Myc |
| chr8:128383087-128383127 | Myc |
| chr8:128383162-128383202 | Myc |
| chr8:128383231-128383271 | Myc |
| chr8:128383379-128383419 | Myc |
| chr8:128383440-128383480 | Myc |
| chr8:128383504-128383544 | Myc |
| chr8:128383570-128383610 | Myc |
| chr8:128383655-128383695 | Myc |
| chr8:128383705-128383745 | Myc |
| chr8:128383766-128383806 | Myc |
| chr8:128383828-128383868 | Myc |
| chr8:128383924-128383964 | Myc |
| chr8:128383987-128384027 | Myc |
| chr8:128384126-128384166 | Myc |
| chr8:128384189-128384229 | Myc |
| chr8:128384275-128384315 | Myc |

| | |
|---|---|
| chr8:128384357-128384397 | Myc |
| chr8:128384470-128384510 | Myc |
| chr8:128384705-128384745 | Myc |
| chr8:128384768-128384808 | Myc |
| chr8:128384874-128384914 | Myc |
| chr8:128384974-128385014 | Myc |
| chr8:128385032-128385072 | Myc |
| chr8:128385104-128385144 | Myc |
| chr8:128385188-128385228 | Myc |
| chr8:128385274-128385314 | Myc |
| chr8:128385349-128385389 | Myc |
| chr8:128385461-128385501 | Myc |
| chr8:128385766-128385806 | Myc |
| chr8:128386015-128386055 | Myc |
| chr8:128386146-128386186 | Myc |
| chr8:128386260-128386300 | Myc |
| chr8:128386321-128386361 | Myc |
| chr8:128386371-128386411 | Myc |
| chr8:128386432-128386472 | Myc |
| chr8:128386504-128386544 | Myc |
| chr8:128386593-128386633 | Myc |
| chr8:128386643-128386683 | Myc |
| chr8:128386737-128386777 | Myc |
| chr8:128386801-128386841 | Myc |
| chr8:128386851-128386891 | Myc |
| chr8:128386901-128386941 | Myc |
| chr8:128386962-128387002 | Myc |
| chr8:128387019-128387059 | Myc |
| chr8:128387099-128387139 | Myc |
| chr8:128387260-128387300 | Myc |
| chr8:128387339-128387379 | Myc |
| chr8:128387423-128387463 | Myc |
| chr8:128387473-128387513 | Myc |
| chr8:128387548-128387588 | Myc |
| chr8:128387599-128387639 | Myc |
| chr8:128387692-128387732 | Myc |
| chr8:128387742-128387782 | Myc |
| chr8:128387793-128387833 | Myc |
| chr8:128387850-128387890 | Myc |
| chr8:128387936-128387976 | Myc |
| chr8:128388018-128388058 | Myc |
| chr8:128388126-128388166 | Myc |
| chr8:128388230-128388270 | Myc |
| chr8:128388498-128388538 | Myc |
| chr8:128388580-128388620 | Myc |
| chr8:128388807-128388847 | Myc |
| chr8:128388924-128388964 | Myc |
| chr8:128389015-128389055 | Myc |
| chr8:128389121-128389161 | Myc |
| chr8:128389201-128389241 | Myc |
| chr8:128389328-128389368 | Myc |
| chr8:128389378-128389418 | Myc |

| | |
|---|---|
| chr8:128389450-128389490 | Myc |
| chr8:128389506-128389546 | Myc |
| chr8:128389581-128389621 | Myc |
| chr8:128389706-128389746 | Myc |
| chr8:128389776-128389816 | Myc |
| chr8:128389879-128389919 | Myc |
| chr8:128389936-128389976 | Myc |
| chr8:128390019-128390059 | Myc |
| chr8:128390075-128390115 | Myc |
| chr8:128390133-128390173 | Myc |
| chr8:128390189-128390229 | Myc |
| chr8:128390240-128390280 | Myc |
| chr8:128390314-128390354 | Myc |
| chr8:128390412-128390452 | Myc |
| chr8:128390494-128390534 | Myc |
| chr8:128390550-128390590 | Myc |
| chr8:128390632-128390672 | Myc |
| chr8:128390721-128390761 | Myc |
| chr8:128390787-128390827 | Myc |
| chr8:128390856-128390896 | Myc |
| chr8:128391081-128391121 | Myc |
| chr8:128391158-128391198 | Myc |
| chr8:128391250-128391290 | Myc |
| chr8:128391363-128391403 | Myc |
| chr8:128391430-128391470 | Myc |
| chr8:128391501-128391541 | Myc |
| chr8:128391610-128391650 | Myc |
| chr8:128391732-128391772 | Myc |
| chr8:128391782-128391822 | Myc |
| chr8:128391882-128391922 | Myc |
| chr8:128392003-128392043 | Myc |
| chr8:128394164-128394204 | Myc |
| chr8:128394399-128394439 | Myc |
| chr8:128394590-128394630 | Myc |
| chr8:128394653-128394693 | Myc |
| chr8:128394703-128394743 | Myc |
| chr8:128394859-128394899 | Myc |
| chr8:128394909-128394949 | Myc |
| chr8:128395035-128395075 | Myc |
| chr8:128395110-128395150 | Myc |
| chr8:128395164-128395204 | Myc |
| chr8:128395257-128395297 | Myc |
| chr8:128395399-128395439 | Myc |
| chr8:128395461-128395501 | Myc |
| chr8:128395573-128395613 | Myc |
| chr8:128395649-128395689 | Myc |
| chr8:128395699-128395739 | Myc |
| chr8:128395752-128395792 | Myc |
| chr8:128395867-128395907 | Myc |
| chr8:128395971-128396011 | Myc |
| chr8:128396053-128396093 | Myc |
| chr8:128396207-128396247 | Myc |

| | |
|---|---|
| chr8:128396269-128396309 | Myc |
| chr8:128396371-128396411 | Myc |
| chr8:128396439-128396479 | Myc |
| chr8:128396514-128396554 | Myc |
| chr8:128396608-128396648 | Myc |
| chr8:128396870-128396910 | Myc |
| chr8:128396950-128396990 | Myc |
| chr8:128397003-128397043 | Myc |
| chr8:128397180-128397220 | Myc |
| chr8:128397256-128397296 | Myc |
| chr8:128397386-128397426 | Myc |
| chr8:128397455-128397495 | Myc |
| chr8:128397527-128397567 | Myc |
| chr8:128397588-128397628 | Myc |
| chr8:128397996-128398036 | Myc |
| chr8:128398087-128398127 | Myc |
| chr8:128398137-128398177 | Myc |
| chr8:128398211-128398251 | Myc |
| chr8:128398301-128398341 | Myc |
| chr8:128398351-128398391 | Myc |
| chr8:128398463-128398503 | Myc |
| chr8:128398593-128398633 | Myc |
| chr8:128398643-128398683 | Myc |
| chr8:128398713-128398753 | Myc |
| chr8:128398779-128398819 | Myc |
| chr8:128398857-128398897 | Myc |
| chr8:128398922-128398962 | Myc |
| chr8:128398973-128399013 | Myc |
| chr8:128399025-128399065 | Myc |
| chr8:128399131-128399171 | Myc |
| chr8:128399233-128399273 | Myc |
| chr8:128399297-128399337 | Myc |
| chr8:128399400-128399440 | Myc |
| chr8:128399450-128399490 | Myc |
| chr8:128399549-128399589 | Myc |
| chr8:128399634-128399674 | Myc |
| chr8:128399684-128399724 | Myc |
| chr8:128399795-128399835 | Myc |
| chr8:128399879-128399919 | Myc |
| chr8:128399995-128400035 | Myc |
| chr8:128400045-128400085 | Myc |
| chr8:128400106-128400146 | Myc |
| chr8:128400203-128400243 | Myc |
| chr8:128400292-128400332 | Myc |
| chr8:128400381-128400421 | Myc |
| chr8:128400437-128400477 | Myc |
| chr8:128400503-128400543 | Myc |
| chr8:128400559-128400599 | Myc |
| chr8:128400617-128400657 | Myc |
| chr8:128400701-128400741 | Myc |
| chr8:128400778-128400818 | Myc |
| chr8:128400829-128400869 | Myc |

| | |
|---|---|
| chr8:128400921-128400961 | Myc |
| chr8:128401014-128401054 | Myc |
| chr8:128401093-128401133 | Myc |
| chr8:128401181-128401221 | Myc |
| chr8:128401259-128401299 | Myc |
| chr8:128401312-128401352 | Myc |
| chr8:128401396-128401436 | Myc |
| chr8:128401453-128401493 | Myc |
| chr8:128423383-128423423 | Myc |
| chr8:128423558-128423598 | Myc |
| chr8:128423622-128423662 | Myc |
| chr8:128423675-128423715 | Myc |
| chr8:128423772-128423812 | Myc |
| chr8:128423865-128423905 | Myc |
| chr8:128423934-128423974 | Myc |
| chr8:128423984-128424024 | Myc |
| chr8:128424034-128424074 | Myc |
| chr8:128424150-128424190 | Myc |
| chr8:128424262-128424302 | Myc |
| chr8:128424335-128424375 | Myc |
| chr8:128424452-128424492 | Myc |
| chr8:128424518-128424558 | Myc |
| chr8:128424696-128424736 | Myc |
| chr8:128424809-128424849 | Myc |
| chr8:128424860-128424900 | Myc |
| chr8:128424910-128424950 | Myc |
| chr8:128425011-128425051 | Myc |
| chr8:128425093-128425133 | Myc |
| chr8:128425143-128425183 | Myc |
| chr8:128425216-128425256 | Myc |
| chr8:128425287-128425327 | Myc |
| chr8:128425337-128425377 | Myc |
| chr8:128425425-128425465 | Myc |
| chr8:128425550-128425590 | Myc |
| chr8:128425653-128425693 | Myc |
| chr8:128425703-128425743 | Myc |
| chr8:128425824-128425864 | Myc |
| chr8:128425876-128425916 | Myc |
| chr8:128425930-128425970 | Myc |
| chr8:128425987-128426027 | Myc |
| chr8:128426105-128426145 | Myc |
| chr8:128426155-128426195 | Myc |
| chr8:128426302-128426342 | Myc |
| chr8:128426414-128426454 | Myc |
| chr8:128426517-128426557 | Myc |
| chr8:128426616-128426656 | Myc |
| chr8:128426681-128426721 | Myc |
| chr8:128426733-128426773 | Myc |
| chr8:128426843-128426883 | Myc |
| chr8:128426893-128426933 | Myc |
| chr8:128426979-128427019 | Myc |
| chr8:128427096-128427136 | Myc |

| | |
|---|---|
| chr8:128427196-128427236 | Myc |
| chr8:128427301-128427341 | Myc |
| chr8:128427366-128427406 | Myc |
| chr8:128427492-128427532 | Myc |
| chr8:128427559-128427599 | Myc |
| chr8:128427611-128427651 | Myc |
| chr8:128427711-128427751 | Myc |
| chr8:128427770-128427810 | Myc |
| chr8:128427820-128427860 | Myc |
| chr8:128428010-128428050 | Myc |
| chr8:128428189-128428229 | Myc |
| chr8:128428286-128428326 | Myc |
| chr8:128428472-128428512 | Myc |
| chr8:128428603-128428643 | Myc |
| chr8:128428773-128428813 | Myc |
| chr8:128428862-128428902 | Myc |
| chr8:128429133-128429173 | Myc |
| chr8:128429197-128429237 | Myc |
| chr8:128429301-128429341 | Myc |
| chr8:128429492-128429532 | Myc |
| chr8:128429551-128429591 | Myc |
| chr8:128429644-128429684 | Myc |
| chr8:128429694-128429734 | Myc |
| chr8:128429965-128430005 | Myc |
| chr8:128430224-128430264 | Myc |
| chr8:128430383-128430423 | Myc |
| chr8:128430462-128430502 | Myc |
| chr8:128430517-128430557 | Myc |
| chr8:128430567-128430607 | Myc |
| chr8:128430653-128430693 | Myc |
| chr8:128430713-128430753 | Myc |
| chr8:128430766-128430806 | Myc |
| chr8:128430833-128430873 | Myc |
| chr8:128430884-128430924 | Myc |
| chr8:128431083-128431123 | Myc |
| chr8:128431139-128431179 | Myc |
| chr8:128431200-128431240 | Myc |
| chr8:128431251-128431291 | Myc |
| chr8:128431322-128431362 | Myc |
| chr8:128431452-128431492 | Myc |
| chr8:128431509-128431549 | Myc |
| chr8:128431580-128431620 | Myc |
| chr8:128431639-128431679 | Myc |
| chr8:128431772-128431812 | Myc |
| chr8:128431835-128431875 | Myc |
| chr8:128431959-128431999 | Myc |
| chr8:128432011-128432051 | Myc |
| chr8:128432096-128432136 | Myc |
| chr8:128432147-128432187 | Myc |
| chr8:128432244-128432284 | Myc |
| chr8:128432352-128432392 | Myc |
| chr8:128432453-128432493 | Myc |

| | |
|---|---|
| chr8:128432510-128432550 | Myc |
| chr8:128432560-128432600 | Myc |
| chr8:128432702-128432742 | Myc |
| chr8:128432768-128432808 | Myc |
| chr8:128432886-128432926 | Myc |
| chr8:128432936-128432976 | Myc |
| chr8:128432996-128433036 | Myc |
| chr8:128433069-128433109 | Myc |
| chr8:128433120-128433160 | Myc |
| chr8:128433229-128433269 | Myc |
| chr8:128433300-128433340 | Myc |
| chr8:128433356-128433396 | Myc |
| chr8:128433419-128433459 | Myc |
| chr8:128433500-128433540 | Myc |
| chr8:128433618-128433658 | Myc |
| chr8:128433673-128433713 | Myc |
| chr8:128433818-128433858 | Myc |
| chr8:128433921-128433961 | Myc |
| chr8:128433989-128434029 | Myc |
| chr8:128434079-128434119 | Myc |
| chr8:128434145-128434185 | Myc |
| chr8:128434274-128434314 | Myc |
| chr8:128434327-128434367 | Myc |
| chr8:128434454-128434494 | Myc |
| chr8:128434531-128434571 | Myc |
| chr8:128434636-128434676 | Myc |
| chr8:128434736-128434776 | Myc |
| chr8:128434882-128434922 | Myc |
| chr8:128434967-128435007 | Myc |
| chr8:128435060-128435100 | Myc |
| chr8:128435119-128435159 | Myc |
| chr8:128435276-128435316 | Myc |
| chr8:128435344-128435384 | Myc |
| chr8:128435409-128435449 | Myc |
| chr8:128435477-128435517 | Myc |
| chr8:128435540-128435580 | Myc |
| chr8:128435629-128435669 | Myc |
| chr8:128435687-128435727 | Myc |
| chr8:128435777-128435817 | Myc |
| chr8:128435895-128435935 | Myc |
| chr8:128436002-128436042 | Myc |
| chr8:128436109-128436149 | Myc |
| chr8:128436225-128436265 | Myc |
| chr8:128436277-128436317 | Myc |
| chr8:128436459-128436499 | Myc |
| chr8:128436576-128436616 | Myc |
| chr8:128436626-128436666 | Myc |
| chr8:128436696-128436736 | Myc |
| chr8:128436822-128436862 | Myc |
| chr8:128436878-128436918 | Myc |
| chr8:128437011-128437051 | Myc |
| chr8:128437093-128437133 | Myc |

| | |
|---|---|
| chr8:128437184-128437224 | Myc |
| chr8:128437299-128437339 | Myc |
| chr8:128437350-128437390 | Myc |
| chr8:128437422-128437462 | Myc |
| chr8:128437503-128437543 | Myc |
| chr8:128437585-128437625 | Myc |
| chr8:128437721-128437761 | Myc |
| chr8:128437772-128437812 | Myc |
| chr8:128437847-128437887 | Myc |
| chr8:128437957-128437997 | Myc |
| chr8:128438015-128438055 | Myc |
| chr8:128438113-128438153 | Myc |
| chr8:128438170-128438210 | Myc |
| chr8:128438254-128438294 | Myc |
| chr8:128438304-128438344 | Myc |
| chr8:128438421-128438461 | Myc |
| chr8:128438486-128438526 | Myc |
| chr8:128438537-128438577 | Myc |
| chr8:128438587-128438627 | Myc |
| chr8:128438667-128438707 | Myc |
| chr8:128438726-128438766 | Myc |
| chr8:128438793-128438833 | Myc |
| chr8:128438844-128438884 | Myc |
| chr8:128438940-128438980 | Myc |
| chr8:128439020-128439060 | Myc |
| chr8:128439077-128439117 | Myc |
| chr8:128439141-128439181 | Myc |
| chr8:128439199-128439239 | Myc |
| chr8:128439249-128439289 | Myc |
| chr8:128439303-128439343 | Myc |
| chr8:128439355-128439395 | Myc |
| chr8:128439420-128439460 | Myc |
| chr8:128439498-128439538 | Myc |
| chr8:128439609-128439649 | Myc |
| chr8:128439671-128439711 | Myc |
| chr8:128439750-128439790 | Myc |
| chr8:128439812-128439852 | Myc |
| chr8:128439964-128440004 | Myc |
| chr8:128440021-128440061 | Myc |
| chr8:128440137-128440177 | Myc |
| chr8:128440192-128440232 | Myc |
| chr8:128440275-128440315 | Myc |
| chr8:128440409-128440449 | Myc |
| chr8:128440502-128440542 | Myc |
| chr8:128440555-128440595 | Myc |
| chr8:128440632-128440672 | Myc |
| chr8:128440779-128440819 | Myc |
| chr8:128440843-128440883 | Myc |
| chr8:128440899-128440939 | Myc |
| chr8:128440989-128441029 | Myc |
| chr8:128441117-128441157 | Myc |
| chr8:128441167-128441207 | Myc |

| | |
|---|---|
| chr8:128441344-128441384 | Myc |
| chr8:128441501-128441541 | Myc |
| chr8:128441555-128441595 | Myc |
| chr8:128441686-128441726 | Myc |
| chr8:128441763-128441803 | Myc |
| chr8:128441822-128441862 | Myc |
| chr8:128441910-128441950 | Myc |
| chr8:128441964-128442004 | Myc |
| chr8:128442021-128442061 | Myc |
| chr8:128442103-128442143 | Myc |
| chr8:128442209-128442249 | Myc |
| chr8:128442329-128442369 | Myc |
| chr8:128442445-128442485 | Myc |
| chr8:128442543-128442583 | Myc |
| chr8:128442603-128442643 | Myc |
| chr8:128442654-128442694 | Myc |
| chr8:128442724-128442764 | Myc |
| chr8:128442781-128442821 | Myc |
| chr8:128442839-128442879 | Myc |
| chr8:128442943-128442983 | Myc |
| chr8:128443112-128443152 | Myc |
| chr8:128443177-128443217 | Myc |
| chr8:128443281-128443321 | Myc |
| chr8:128443356-128443396 | Myc |
| chr8:128443406-128443446 | Myc |
| chr8:128443466-128443506 | Myc |
| chr8:128443577-128443617 | Myc |
| chr8:128443696-128443736 | Myc |
| chr8:128443768-128443808 | Myc |
| chr8:128443893-128443933 | Myc |
| chr8:128443973-128444013 | Myc |
| chr8:128444093-128444133 | Myc |
| chr8:128444398-128444438 | Myc |
| chr8:128444610-128444650 | Myc |
| chr8:128444782-128444822 | Myc |
| chr8:128444970-128445010 | Myc |
| chr8:128445127-128445167 | Myc |
| chr8:128445216-128445256 | Myc |
| chr8:128445454-128445494 | Myc |
| chr8:128445510-128445550 | Myc |
| chr8:128445630-128445670 | Myc |
| chr8:128445709-128445749 | Myc |
| chr8:128445902-128445942 | Myc |
| chr8:128445976-128446016 | Myc |
| chr8:128446358-128446398 | Myc |
| chr8:128446478-128446518 | Myc |
| chr8:128446812-128446852 | Myc |
| chr8:128446875-128446915 | Myc |
| chr8:128447508-128447548 | Myc |
| chr8:128447582-128447622 | Myc |
| chr8:128448246-128448286 | Myc |
| chr8:128448310-128448350 | Myc |

| | |
|---|---|
| chr8:128448551-128448591 | Myc |
| chr8:128448899-128448939 | Myc |
| chr8:128448981-128449021 | Myc |
| chr8:128449180-128449220 | Myc |
| chr8:128449734-128449774 | Myc |
| chr8:128449830-128449870 | Myc |
| chr8:128449895-128449935 | Myc |
| chr8:128449981-128450021 | Myc |
| chr8:128450062-128450102 | Myc |
| chr8:128450128-128450168 | Myc |
| chr8:128450450-128450490 | Myc |
| chr8:128450564-128450604 | Myc |
| chr8:128450644-128450684 | Myc |
| chr8:128450695-128450735 | Myc |
| chr8:128450786-128450826 | Myc |
| chr8:128450840-128450880 | Myc |
| chr8:128450940-128450980 | Myc |
| chr8:128451087-128451127 | Myc |
| chr8:128451190-128451230 | Myc |
| chr8:128451290-128451330 | Myc |
| chr8:128451374-128451414 | Myc |
| chr8:128451562-128451602 | Myc |
| chr8:128451733-128451773 | Myc |
| chr8:128451813-128451853 | Myc |
| chr8:128451895-128451935 | Myc |
| chr8:128452033-128452073 | Myc |
| chr8:128452092-128452132 | Myc |
| chr8:128452183-128452223 | Myc |
| chr8:128452341-128452381 | Myc |
| chr8:128452559-128452599 | Myc |
| chr8:128452671-128452711 | Myc |
| chr8:128452721-128452761 | Myc |
| chr8:128452859-128452899 | Myc |
| chr8:128452909-128452949 | Myc |
| chr8:128452978-128453018 | Myc |
| chr8:128453114-128453154 | Myc |
| chr8:128453171-128453211 | Myc |
| chr8:128453339-128453379 | Myc |
| chr8:128453401-128453441 | Myc |
| chr8:128453472-128453512 | Myc |
| chr8:128453522-128453562 | Myc |
| chr8:128453684-128453724 | Myc |
| chr8:128453801-128453841 | Myc |
| chr8:128453864-128453904 | Myc |
| chr8:128453951-128453991 | Myc |
| chr8:128454065-128454105 | Myc |
| chr8:128454220-128454260 | Myc |
| chr8:128454369-128454409 | Myc |
| chr8:128454468-128454508 | Myc |
| chr8:128454533-128454573 | Myc |
| chr8:128454598-128454638 | Myc |
| chr8:128454674-128454714 | Myc |

| | |
|---|---|
| chr8:128454756-128454796 | Myc |
| chr8:128454832-128454872 | Myc |
| chr8:128455016-128455056 | Myc |
| chr8:128455083-128455123 | Myc |
| chr8:128455165-128455205 | Myc |
| chr8:128455528-128455568 | Myc |
| chr8:128455695-128455735 | Myc |
| chr8:128455856-128455896 | Myc |
| chr8:128455931-128455971 | Myc |
| chr8:128455988-128456028 | Myc |
| chr8:128456046-128456086 | Myc |
| chr8:128456154-128456194 | Myc |
| chr8:128456279-128456319 | Myc |
| chr8:128456358-128456398 | Myc |
| chr8:128456408-128456448 | Myc |
| chr8:128456509-128456549 | Myc |
| chr8:128456584-128456624 | Myc |
| chr8:128456678-128456718 | Myc |
| chr8:128456840-128456880 | Myc |
| chr8:128456944-128456984 | Myc |
| chr8:128457023-128457063 | Myc |
| chr8:128457096-128457136 | Myc |
| chr8:128457147-128457187 | Myc |
| chr8:128457233-128457273 | Myc |
| chr8:128457308-128457348 | Myc |
| chr8:128457377-128457417 | Myc |
| chr8:128457509-128457549 | Myc |
| chr8:128457591-128457631 | Myc |
| chr8:128457661-128457701 | Myc |
| chr8:128457752-128457792 | Myc |
| chr8:128457818-128457858 | Myc |
| chr8:128457898-128457938 | Myc |
| chr8:128457994-128458034 | Myc |
| chr8:128458049-128458089 | Myc |
| chr8:128458137-128458177 | Myc |
| chr8:128458221-128458261 | Myc |
| chr8:128458366-128458406 | Myc |
| chr8:128458446-128458486 | Myc |
| chr8:128458552-128458592 | Myc |
| chr8:128458602-128458642 | Myc |
| chr8:128458735-128458775 | Myc |
| chr8:128458800-128458840 | Myc |
| chr8:128458862-128458902 | Myc |
| chr8:128458943-128458983 | Myc |
| chr8:128459143-128459183 | Myc |
| chr8:128459215-128459255 | Myc |
| chr8:128459373-128459413 | Myc |
| chr8:128459451-128459491 | Myc |
| chr8:128459503-128459543 | Myc |
| chr8:128459554-128459594 | Myc |
| chr8:128459628-128459668 | Myc |
| chr8:128459698-128459738 | Myc |

| | |
|---|---|
| chr8:128459757-128459797 | Myc |
| chr8:128459941-128459981 | Myc |
| chr8:128460030-128460070 | Myc |
| chr8:128460115-128460155 | Myc |
| chr8:128460199-128460239 | Myc |
| chr8:128460249-128460289 | Myc |
| chr8:128460370-128460410 | Myc |
| chr8:128460436-128460476 | Myc |
| chr8:128460549-128460589 | Myc |
| chr8:128460599-128460639 | Myc |
| chr8:128460654-128460694 | Myc |
| chr8:128460787-128460827 | Myc |
| chr8:128460904-128460944 | Myc |
| chr8:128460969-128461009 | Myc |
| chr8:128461026-128461066 | Myc |
| chr8:128461077-128461117 | Myc |
| chr8:128461210-128461250 | Myc |
| chr8:128461261-128461301 | Myc |
| chr8:128461312-128461352 | Myc |
| chr8:128461467-128461507 | Myc |
| chr8:128461520-128461560 | Myc |
| chr8:128461637-128461677 | Myc |
| chr8:128461702-128461742 | Myc |
| chr8:128461769-128461809 | Myc |
| chr8:128461856-128461896 | Myc |
| chr8:128461972-128462012 | Myc |
| chr8:128462035-128462075 | Myc |
| chr8:128462150-128462190 | Myc |
| chr8:128462215-128462255 | Myc |
| chr8:128462272-128462312 | Myc |
| chr8:128462381-128462421 | Myc |
| chr8:128462532-128462572 | Myc |
| chr8:128462592-128462632 | Myc |
| chr8:128462716-128462756 | Myc |
| chr8:128462788-128462828 | Myc |
| chr8:128462839-128462879 | Myc |
| chr8:128462955-128462995 | Myc |
| chr8:128463032-128463072 | Myc |
| chr8:128463208-128463248 | Myc |
| chr8:128463258-128463298 | Myc |
| chr8:128463362-128463402 | Myc |
| chr8:128463476-128463516 | Myc |
| chr8:128463544-128463584 | Myc |
| chr8:128463645-128463685 | Myc |
| chr8:128463695-128463735 | Myc |
| chr8:128463816-128463856 | Myc |
| chr8:128463977-128464017 | Myc |
| chr8:128464108-128464148 | Myc |
| chr8:128464213-128464253 | Myc |
| chr8:128464268-128464308 | Myc |
| chr8:128464344-128464384 | Myc |
| chr8:128464412-128464452 | Myc |

| | |
|---|---|
| chr8:128464530-128464570 | Myc |
| chr8:128464620-128464660 | Myc |
| chr8:128464680-128464720 | Myc |
| chr8:128464734-128464774 | Myc |
| chr8:128464784-128464824 | Myc |
| chr8:128464892-128464932 | Myc |
| chr8:128464947-128464987 | Myc |
| chr8:128465024-128465064 | Myc |
| chr8:128465115-128465155 | Myc |
| chr8:128465172-128465212 | Myc |
| chr8:128465246-128465286 | Myc |
| chr8:128465320-128465360 | Myc |
| chr8:128465392-128465432 | Myc |
| chr8:128465468-128465508 | Myc |
| chr8:128465518-128465558 | Myc |
| chr8:128465611-128465651 | Myc |
| chr8:128465721-128465761 | Myc |
| chr8:128465790-128465830 | Myc |
| chr8:128465916-128465956 | Myc |
| chr8:128465995-128466035 | Myc |
| chr8:128466112-128466152 | Myc |
| chr8:128466181-128466221 | Myc |
| chr8:128466287-128466327 | Myc |
| chr8:128466341-128466381 | Myc |
| chr8:128466418-128466458 | Myc |
| chr8:128466517-128466557 | Myc |
| chr8:128466646-128466686 | Myc |
| chr8:128466704-128466744 | Myc |
| chr8:128466780-128466820 | Myc |
| chr8:128466901-128466941 | Myc |
| chr8:128466993-128467033 | Myc |
| chr8:128467079-128467119 | Myc |
| chr8:128467163-128467203 | Myc |
| chr8:128467284-128467324 | Myc |
| chr8:128467379-128467419 | Myc |
| chr8:128467534-128467574 | Myc |
| chr8:128467584-128467624 | Myc |
| chr8:128467693-128467733 | Myc |
| chr8:128467775-128467815 | Myc |
| chr8:128467854-128467894 | Myc |
| chr8:128467907-128467947 | Myc |
| chr8:128467957-128467997 | Myc |
| chr8:128468007-128468047 | Myc |
| chr8:128468063-128468103 | Myc |
| chr8:128468169-128468209 | Myc |
| chr8:128468323-128468363 | Myc |
| chr8:128468376-128468416 | Myc |
| chr8:128468436-128468476 | Myc |
| chr8:128468498-128468538 | Myc |
| chr8:128468571-128468611 | Myc |
| chr8:128468663-128468703 | Myc |
| chr8:128468943-128468983 | Myc |

| | |
|---|---|
| chr8:128469027-128469067 | Myc |
| chr8:128469122-128469162 | Myc |
| chr8:128469174-128469214 | Myc |
| chr8:128469273-128469313 | Myc |
| chr8:128469411-128469451 | Myc |
| chr8:128469461-128469501 | Myc |
| chr8:128469520-128469560 | Myc |
| chr8:128469616-128469656 | Myc |
| chr8:128469697-128469737 | Myc |
| chr8:128469850-128469890 | Myc |
| chr8:128469984-128470024 | Myc |
| chr8:128470055-128470095 | Myc |
| chr8:128470108-128470148 | Myc |
| chr8:128470240-128470280 | Myc |
| chr8:128470355-128470395 | Myc |
| chr8:128470412-128470452 | Myc |
| chr8:128470481-128470521 | Myc |
| chr8:128470570-128470610 | Myc |
| chr8:128470621-128470661 | Myc |
| chr8:128470793-128470833 | Myc |
| chr8:128470845-128470885 | Myc |
| chr8:128470960-128471000 | Myc |
| chr8:128471021-128471061 | Myc |
| chr8:128471071-128471111 | Myc |
| chr8:128471182-128471222 | Myc |
| chr8:128471319-128471359 | Myc |
| chr8:128471386-128471426 | Myc |
| chr8:128471626-128471666 | Myc |
| chr8:128471676-128471716 | Myc |
| chr8:128471739-128471779 | Myc |
| chr8:128471866-128471906 | Myc |
| chr8:128471916-128471956 | Myc |
| chr8:128472116-128472156 | Myc |
| chr8:128472181-128472221 | Myc |
| chr8:128472250-128472290 | Myc |
| chr8:128472326-128472366 | Myc |
| chr8:128472422-128472462 | Myc |
| chr8:128472536-128472576 | Myc |
| chr8:128472599-128472639 | Myc |
| chr8:128472654-128472694 | Myc |
| chr8:128472802-128472842 | Myc |
| chr8:128472854-128472894 | Myc |
| chr8:128472924-128472964 | Myc |
| chr8:128472997-128473037 | Myc |
| chr8:128473068-128473108 | Myc |
| chr8:128473359-128473399 | Myc |
| chr8:128473409-128473449 | Myc |
| chr8:128473797-128473837 | Myc |
| chr8:128473945-128473985 | Myc |
| chr8:128473995-128474035 | Myc |
| chr8:128474079-128474119 | Myc |
| chr8:128474129-128474169 | Myc |

| | |
|---|---|
| chr8:128474261-128474301 | Myc |
| chr8:128474313-128474353 | Myc |
| chr8:128474409-128474449 | Myc |
| chr8:128474463-128474503 | Myc |
| chr8:128474559-128474599 | Myc |
| chr8:128474628-128474668 | Myc |
| chr8:128474717-128474757 | Myc |
| chr8:128474796-128474836 | Myc |
| chr8:128474877-128474917 | Myc |
| chr8:128474952-128474992 | Myc |
| chr8:128475125-128475165 | Myc |
| chr8:128475193-128475233 | Myc |
| chr8:128475335-128475375 | Myc |
| chr8:128475507-128475547 | Myc |
| chr8:128475609-128475649 | Myc |
| chr8:128475699-128475739 | Myc |
| chr8:128475754-128475794 | Myc |
| chr8:128475806-128475846 | Myc |
| chr8:128475895-128475935 | Myc |
| chr8:128476022-128476062 | Myc |
| chr8:128476072-128476112 | Myc |
| chr8:128476218-128476258 | Myc |
| chr8:128476296-128476336 | Myc |
| chr8:128476418-128476458 | Myc |
| chr8:128476485-128476525 | Myc |
| chr8:128476538-128476578 | Myc |
| chr8:128476631-128476671 | Myc |
| chr8:128476681-128476721 | Myc |
| chr8:128476784-128476824 | Myc |
| chr8:128476873-128476913 | Myc |
| chr8:128476968-128477008 | Myc |
| chr8:128477020-128477060 | Myc |
| chr8:128477118-128477158 | Myc |
| chr8:128477168-128477208 | Myc |
| chr8:128477295-128477335 | Myc |
| chr8:128477374-128477414 | Myc |
| chr8:128477472-128477512 | Myc |
| chr8:128477530-128477570 | Myc |
| chr8:128477621-128477661 | Myc |
| chr8:128477709-128477749 | Myc |
| chr8:128477767-128477807 | Myc |
| chr8:128477823-128477863 | Myc |
| chr8:128477883-128477923 | Myc |
| chr8:128477936-128477976 | Myc |
| chr8:128478003-128478043 | Myc |
| chr8:128478103-128478143 | Myc |
| chr8:128478200-128478240 | Myc |
| chr8:128478356-128478396 | Myc |
| chr8:128478428-128478468 | Myc |
| chr8:128478541-128478581 | Myc |
| chr8:128478621-128478661 | Myc |
| chr8:128478735-128478775 | Myc |

| | |
|---|---|
| chr8:128478860-128478900 | Myc |
| chr8:128478933-128478973 | Myc |
| chr8:128478991-128479031 | Myc |
| chr8:128479068-128479108 | Myc |
| chr8:128479174-128479214 | Myc |
| chr8:128479292-128479332 | Myc |
| chr8:128479384-128479424 | Myc |
| chr8:128479446-128479486 | Myc |
| chr8:128479567-128479607 | Myc |
| chr8:128479696-128479736 | Myc |
| chr8:128479774-128479814 | Myc |
| chr8:128479824-128479864 | Myc |
| chr8:128479942-128479982 | Myc |
| chr8:128480025-128480065 | Myc |
| chr8:128480076-128480116 | Myc |
| chr8:128480175-128480215 | Myc |
| chr8:128480294-128480334 | Myc |
| chr8:128480438-128480478 | Myc |
| chr8:128480579-128480619 | Myc |
| chr8:128480655-128480695 | Myc |
| chr8:128480723-128480763 | Myc |
| chr8:128480871-128480911 | Myc |
| chr8:128481035-128481075 | Myc |
| chr8:128481097-128481137 | Myc |
| chr8:128481152-128481192 | Myc |
| chr8:128481235-128481275 | Myc |
| chr8:128481293-128481333 | Myc |
| chr8:128481394-128481434 | Myc |
| chr8:128481469-128481509 | Myc |
| chr8:128481535-128481575 | Myc |
| chr8:128481645-128481685 | Myc |
| chr8:128481706-128481746 | Myc |
| chr8:128481776-128481816 | Myc |
| chr8:128481872-128481912 | Myc |
| chr8:128481968-128482008 | Myc |
| chr8:128482022-128482062 | Myc |
| chr8:128482096-128482136 | Myc |
| chr8:128482166-128482206 | Myc |
| chr8:128482234-128482274 | Myc |
| chr8:128482331-128482371 | Myc |
| chr8:128482420-128482460 | Myc |
| chr8:128482541-128482581 | Myc |
| chr8:128482623-128482663 | Myc |
| chr8:128482732-128482772 | Myc |
| chr8:128482795-128482835 | Myc |
| chr8:128482861-128482901 | Myc |
| chr8:128483060-128483100 | Myc |
| chr8:128483161-128483201 | Myc |
| chr8:128483219-128483259 | Myc |
| chr8:128483282-128483322 | Myc |
| chr8:128483381-128483421 | Myc |
| chr8:128483445-128483485 | Myc |

| | |
|---|---|
| chr8:128483534-128483574 | Myc |
| chr8:128483632-128483672 | Myc |
| chr8:128483682-128483722 | Myc |
| chr8:128483733-128483773 | Myc |
| chr8:128483822-128483862 | Myc |
| chr8:128483890-128483930 | Myc |
| chr8:128484239-128484279 | Myc |
| chr8:128484309-128484349 | Myc |
| chr8:128484383-128484423 | Myc |
| chr8:128484454-128484494 | Myc |
| chr8:128484519-128484559 | Myc |
| chr8:128484601-128484641 | Myc |
| chr8:128484658-128484698 | Myc |
| chr8:128484774-128484814 | Myc |
| chr8:128484893-128484933 | Myc |
| chr8:128484996-128485036 | Myc |
| chr8:128485089-128485129 | Myc |
| chr8:128485188-128485228 | Myc |
| chr8:128485247-128485287 | Myc |
| chr8:128485355-128485395 | Myc |
| chr8:128485450-128485490 | Myc |
| chr8:128485500-128485540 | Myc |
| chr8:128485604-128485644 | Myc |
| chr8:128485660-128485700 | Myc |
| chr8:128485717-128485757 | Myc |
| chr8:128485790-128485830 | Myc |
| chr8:128485913-128485953 | Myc |
| chr8:128485981-128486021 | Myc |
| chr8:128486065-128486105 | Myc |
| chr8:128486115-128486155 | Myc |
| chr8:128486223-128486263 | Myc |
| chr8:128486289-128486329 | Myc |
| chr8:128486349-128486389 | Myc |
| chr8:128486467-128486507 | Myc |
| chr8:128486584-128486624 | Myc |
| chr8:128486672-128486712 | Myc |
| chr8:128486812-128486852 | Myc |
| chr8:128486880-128486920 | Myc |
| chr8:128486930-128486970 | Myc |
| chr8:128487119-128487159 | Myc |
| chr8:128487232-128487272 | Myc |
| chr8:128487288-128487328 | Myc |
| chr8:128487405-128487445 | Myc |
| chr8:128487497-128487537 | Myc |
| chr8:128487599-128487639 | Myc |
| chr8:128487670-128487710 | Myc |
| chr8:128487767-128487807 | Myc |
| chr8:128487882-128487922 | Myc |
| chr8:128487971-128488011 | Myc |
| chr8:128488676-128488716 | Myc |
| chr8:128488749-128488789 | Myc |
| chr8:128488801-128488841 | Myc |

| | |
|---|---|
| chr8:128488877-128488917 | Myc |
| chr8:128488970-128489010 | Myc |
| chr8:128489055-128489095 | Myc |
| chr8:128489159-128489199 | Myc |
| chr8:128489221-128489261 | Myc |
| chr8:128489410-128489450 | Myc |
| chr8:128489576-128489616 | Myc |
| chr8:128489643-128489683 | Myc |
| chr8:128489927-128489967 | Myc |
| chr8:128489990-128490030 | Myc |
| chr8:128490040-128490080 | Myc |
| chr8:128490205-128490245 | Myc |
| chr8:128490394-128490434 | Myc |
| chr8:128490452-128490492 | Myc |
| chr8:128490513-128490553 | Myc |
| chr8:128490623-128490663 | Myc |
| chr8:128490735-128490775 | Myc |
| chr8:128490820-128490860 | Myc |
| chr8:128490940-128490980 | Myc |
| chr8:128490990-128491030 | Myc |
| chr8:128491102-128491142 | Myc |
| chr8:128491217-128491257 | Myc |
| chr8:128491271-128491311 | Myc |
| chr8:128491403-128491443 | Myc |
| chr8:128491549-128491589 | Myc |
| chr8:128491614-128491654 | Myc |
| chr8:128491672-128491712 | Myc |
| chr8:128491825-128491865 | Myc |
| chr8:128491935-128491975 | Myc |
| chr8:128491995-128492035 | Myc |
| chr8:128492087-128492127 | Myc |
| chr8:128492162-128492202 | Myc |
| chr8:128492252-128492292 | Myc |
| chr8:128492336-128492376 | Myc |
| chr8:128492422-128492462 | Myc |
| chr8:128492517-128492557 | Myc |
| chr8:128492593-128492633 | Myc |
| chr8:128492644-128492684 | Myc |
| chr8:128492726-128492766 | Myc |
| chr8:128492785-128492825 | Myc |
| chr8:128492842-128492882 | Myc |
| chr8:128492972-128493012 | Myc |
| chr8:128493022-128493062 | Myc |
| chr8:128493088-128493128 | Myc |
| chr8:128493178-128493218 | Myc |
| chr8:128493254-128493294 | Myc |
| chr8:128493393-128493433 | Myc |
| chr8:128493450-128493490 | Myc |
| chr8:128493514-128493554 | Myc |
| chr8:128493566-128493606 | Myc |
| chr8:128493625-128493665 | Myc |
| chr8:128493731-128493771 | Myc |

| | |
|---|---|
| chr8:128493786-128493826 | Myc |
| chr8:128493876-128493916 | Myc |
| chr8:128493947-128493987 | Myc |
| chr8:128494008-128494048 | Myc |
| chr8:128494079-128494119 | Myc |
| chr8:128494169-128494209 | Myc |
| chr8:128494340-128494380 | Myc |
| chr8:128494393-128494433 | Myc |
| chr8:128494443-128494483 | Myc |
| chr8:128494516-128494556 | Myc |
| chr8:128494605-128494645 | Myc |
| chr8:128494667-128494707 | Myc |
| chr8:128494771-128494811 | Myc |
| chr8:128494823-128494863 | Myc |
| chr8:128495001-128495041 | Myc |
| chr8:128495179-128495219 | Myc |
| chr8:128495241-128495281 | Myc |
| chr8:128495296-128495336 | Myc |
| chr8:128495353-128495393 | Myc |
| chr8:128495468-128495508 | Myc |
| chr8:128495570-128495610 | Myc |
| chr8:128495720-128495760 | Myc |
| chr8:128495886-128495926 | Myc |
| chr8:128495957-128495997 | Myc |
| chr8:128496084-128496124 | Myc |
| chr8:128496136-128496176 | Myc |
| chr8:128496189-128496229 | Myc |
| chr8:128496255-128496295 | Myc |
| chr8:128496422-128496462 | Myc |
| chr8:128496534-128496574 | Myc |
| chr8:128496599-128496639 | Myc |
| chr8:128496699-128496739 | Myc |
| chr8:128496784-128496824 | Myc |
| chr8:128496850-128496890 | Myc |
| chr8:128496941-128496981 | Myc |
| chr8:128496991-128497031 | Myc |
| chr8:128497090-128497130 | Myc |
| chr8:128497192-128497232 | Myc |
| chr8:128497294-128497334 | Myc |
| chr8:128497415-128497455 | Myc |
| chr8:128497524-128497564 | Myc |
| chr8:128497583-128497623 | Myc |
| chr8:128497713-128497753 | Myc |
| chr8:128497763-128497803 | Myc |
| chr8:128497886-128497926 | Myc |
| chr8:128498063-128498103 | Myc |
| chr8:128498116-128498156 | Myc |
| chr8:128498386-128498426 | Myc |
| chr8:128498473-128498513 | Myc |
| chr8:128498569-128498609 | Myc |
| chr8:128498671-128498711 | Myc |
| chr8:128498723-128498763 | Myc |

| | |
|---|---|
| chr8:128498864-128498904 | Myc |
| chr8:128498954-128498994 | Myc |
| chr8:128499005-128499045 | Myc |
| chr8:128499056-128499096 | Myc |
| chr8:128499123-128499163 | Myc |
| chr8:128499189-128499229 | Myc |
| chr8:128499309-128499349 | Myc |
| chr8:128499377-128499417 | Myc |
| chr8:128499436-128499476 | Myc |
| chr8:128499556-128499596 | Myc |
| chr8:128499941-128499981 | Myc |
| chr8:128500002-128500042 | Myc |
| chr8:128500052-128500092 | Myc |
| chr8:128500103-128500143 | Myc |
| chr8:128500179-128500219 | Myc |
| chr8:128500238-128500278 | Myc |
| chr8:128500336-128500376 | Myc |
| chr8:128500557-128500597 | Myc |
| chr8:128500718-128500758 | Myc |
| chr8:128500775-128500815 | Myc |
| chr8:128500872-128500912 | Myc |
| chr8:128500922-128500962 | Myc |
| chr8:128501027-128501067 | Myc |
| chr8:128501083-128501123 | Myc |
| chr8:128501159-128501199 | Myc |
| chr8:128501237-128501277 | Myc |
| chr8:128501307-128501347 | Myc |
| chr8:128501357-128501397 | Myc |
| chr8:128501460-128501500 | Myc |
| chr8:128501538-128501578 | Myc |
| chr8:128501632-128501672 | Myc |
| chr8:128501703-128501743 | Myc |
| chr8:128501855-128501895 | Myc |
| chr8:128501998-128502038 | Myc |
| chr8:128502087-128502127 | Myc |
| chr8:128502192-128502232 | Myc |
| chr8:128502303-128502343 | Myc |
| chr8:128502362-128502402 | Myc |
| chr8:128502471-128502511 | Myc |
| chr8:128502531-128502571 | Myc |
| chr8:128502583-128502623 | Myc |
| chr8:128502658-128502698 | Myc |
| chr8:128502796-128502836 | Myc |
| chr8:128502877-128502917 | Myc |
| chr8:128503114-128503154 | Myc |
| chr8:128503177-128503217 | Myc |
| chr8:128503239-128503279 | Myc |
| chr8:128503306-128503346 | Myc |
| chr8:128503363-128503403 | Myc |
| chr8:128503444-128503484 | Myc |
| chr8:128503517-128503557 | Myc |
| chr8:128503576-128503616 | Myc |

| | |
|---|---|
| chr8:128503826-128503866 | Myc |
| chr8:128503884-128503924 | Myc |
| chr8:128503978-128504018 | Myc |
| chr8:128504063-128504103 | Myc |
| chr8:128504209-128504249 | Myc |
| chr8:128504294-128504334 | Myc |
| chr8:128504356-128504396 | Myc |
| chr8:128504583-128504623 | Myc |
| chr8:128504766-128504806 | Myc |
| chr8:128504887-128504927 | Myc |
| chr8:128504963-128505003 | Myc |
| chr8:128505073-128505113 | Myc |
| chr8:128505215-128505255 | Myc |
| chr8:128505274-128505314 | Myc |
| chr8:128505334-128505374 | Myc |
| chr8:128505456-128505496 | Myc |
| chr8:128505529-128505569 | Myc |
| chr8:128505632-128505672 | Myc |
| chr8:128505772-128505812 | Myc |
| chr8:128505822-128505862 | Myc |
| chr8:128505952-128505992 | Myc |
| chr8:128506017-128506057 | Myc |
| chr8:128506142-128506182 | Myc |
| chr8:128506248-128506288 | Myc |
| chr8:128506378-128506418 | Myc |
| chr8:128506504-128506544 | Myc |
| chr8:128506591-128506631 | Myc |
| chr8:128506689-128506729 | Myc |
| chr8:128506739-128506779 | Myc |
| chr8:128506830-128506870 | Myc |
| chr8:128506922-128506962 | Myc |
| chr8:128506972-128507012 | Myc |
| chr8:128507079-128507119 | Myc |
| chr8:128507129-128507169 | Myc |
| chr8:128507221-128507261 | Myc |
| chr8:128507273-128507313 | Myc |
| chr8:128507329-128507369 | Myc |
| chr8:128507430-128507470 | Myc |
| chr8:128507480-128507520 | Myc |
| chr8:128507530-128507570 | Myc |
| chr8:128507696-128507736 | Myc |
| chr8:128507748-128507788 | Myc |
| chr8:128507827-128507867 | Myc |
| chr8:128507880-128507920 | Myc |
| chr8:128507940-128507980 | Myc |
| chr8:128508062-128508102 | Myc |
| chr8:128508134-128508174 | Myc |
| chr8:128508233-128508273 | Myc |
| chr8:128508297-128508337 | Myc |
| chr8:128508408-128508448 | Myc |
| chr8:128508460-128508500 | Myc |
| chr8:128508531-128508571 | Myc |

| | |
|---|---|
| chr8:128508640-128508680 | Myc |
| chr8:128508791-128508831 | Myc |
| chr8:128508849-128508889 | Myc |
| chr8:128508939-128508979 | Myc |
| chr8:128509053-128509093 | Myc |
| chr8:128509190-128509230 | Myc |
| chr8:128509240-128509280 | Myc |
| chr8:128509364-128509404 | Myc |
| chr8:128509453-128509493 | Myc |
| chr8:128509503-128509543 | Myc |
| chr8:128509556-128509596 | Myc |
| chr8:128509662-128509702 | Myc |
| chr8:128509745-128509785 | Myc |
| chr8:128509825-128509865 | Myc |
| chr8:128509928-128509968 | Myc |
| chr8:128509981-128510021 | Myc |
| chr8:128510088-128510128 | Myc |
| chr8:128510214-128510254 | Myc |
| chr8:128510265-128510305 | Myc |
| chr8:128510378-128510418 | Myc |
| chr8:128510455-128510495 | Myc |
| chr8:128510548-128510588 | Myc |
| chr8:128510673-128510713 | Myc |
| chr8:128510752-128510792 | Myc |
| chr8:128510816-128510856 | Myc |
| chr8:128510873-128510913 | Myc |
| chr8:128510933-128510973 | Myc |
| chr8:128510998-128511038 | Myc |
| chr8:128511097-128511137 | Myc |
| chr8:128511147-128511187 | Myc |
| chr8:128511273-128511313 | Myc |
| chr8:128511343-128511383 | Myc |
| chr8:128511405-128511445 | Myc |
| chr8:128511521-128511561 | Myc |
| chr8:128511629-128511669 | Myc |
| chr8:128511745-128511785 | Myc |
| chr8:128511845-128511885 | Myc |
| chr8:128511897-128511937 | Myc |
| chr8:128511973-128512013 | Myc |
| chr8:128512025-128512065 | Myc |
| chr8:128512086-128512126 | Myc |
| chr8:128512137-128512177 | Myc |
| chr8:128512199-128512239 | Myc |
| chr8:128512311-128512351 | Myc |
| chr8:128512400-128512440 | Myc |
| chr8:128512518-128512558 | Myc |
| chr8:128512655-128512695 | Myc |
| chr8:128512828-128512868 | Myc |
| chr8:128512891-128512931 | Myc |
| chr8:128513018-128513058 | Myc |
| chr8:128513120-128513160 | Myc |
| chr8:128513255-128513295 | Myc |

| | |
|---|---|
| chr8:128513356-128513396 | Myc |
| chr8:128513406-128513446 | Myc |
| chr8:128513526-128513566 | Myc |
| chr8:128513576-128513616 | Myc |
| chr8:128513657-128513697 | Myc |
| chr8:128513753-128513793 | Myc |
| chr8:128513804-128513844 | Myc |
| chr8:128513969-128514009 | Myc |
| chr8:128514074-128514114 | Myc |
| chr8:128514199-128514239 | Myc |
| chr8:128514288-128514328 | Myc |
| chr8:128514422-128514462 | Myc |
| chr8:128514490-128514530 | Myc |
| chr8:128514544-128514584 | Myc |
| chr8:128514594-128514634 | Myc |
| chr8:128514682-128514722 | Myc |
| chr8:128514732-128514772 | Myc |
| chr8:128514857-128514897 | Myc |
| chr8:128514908-128514948 | Myc |
| chr8:128515019-128515059 | Myc |
| chr8:128515108-128515148 | Myc |
| chr8:128515158-128515198 | Myc |
| chr8:128515361-128515401 | Myc |
| chr8:128515609-128515649 | Myc |
| chr8:128515727-128515767 | Myc |
| chr8:128515836-128515876 | Myc |
| chr8:128515899-128515939 | Myc |
| chr8:128515969-128516009 | Myc |
| chr8:128516019-128516059 | Myc |
| chr8:128516189-128516229 | Myc |
| chr8:128516326-128516366 | Myc |
| chr8:128516384-128516424 | Myc |
| chr8:128516435-128516475 | Myc |
| chr8:128516557-128516597 | Myc |
| chr8:128516723-128516763 | Myc |
| chr8:128516865-128516905 | Myc |
| chr8:128516922-128516962 | Myc |
| chr8:128516985-128517025 | Myc |
| chr8:128517053-128517093 | Myc |
| chr8:128517115-128517155 | Myc |
| chr8:128517167-128517207 | Myc |
| chr8:128517253-128517293 | Myc |
| chr8:128517303-128517343 | Myc |
| chr8:128517398-128517438 | Myc |
| chr8:128517453-128517493 | Myc |
| chr8:128517555-128517595 | Myc |
| chr8:128517634-128517674 | Myc |
| chr8:128517719-128517759 | Myc |
| chr8:128517791-128517831 | Myc |
| chr8:128517911-128517951 | Myc |
| chr8:128517989-128518029 | Myc |
| chr8:128518115-128518155 | Myc |

| | |
|---|---|
| chr8:128518238-128518278 | Myc |
| chr8:128518359-128518399 | Myc |
| chr8:128518443-128518483 | Myc |
| chr8:128518553-128518593 | Myc |
| chr8:128518619-128518659 | Myc |
| chr8:128518813-128518853 | Myc |
| chr8:128518889-128518929 | Myc |
| chr8:128518941-128518981 | Myc |
| chr8:128519015-128519055 | Myc |
| chr8:128519094-128519134 | Myc |
| chr8:128519158-128519198 | Myc |
| chr8:128519332-128519372 | Myc |
| chr8:128519382-128519422 | Myc |
| chr8:128519469-128519509 | Myc |
| chr8:128519554-128519594 | Myc |
| chr8:128519613-128519653 | Myc |
| chr8:128519681-128519721 | Myc |
| chr8:128519748-128519788 | Myc |
| chr8:128519809-128519849 | Myc |
| chr8:128519935-128519975 | Myc |
| chr8:128520062-128520102 | Myc |
| chr8:128520118-128520158 | Myc |
| chr8:128520191-128520231 | Myc |
| chr8:128520289-128520329 | Myc |
| chr8:128520427-128520467 | Myc |
| chr8:128520494-128520534 | Myc |
| chr8:128520683-128520723 | Myc |
| chr8:128520918-128520958 | Myc |
| chr8:128521002-128521042 | Myc |
| chr8:128521116-128521156 | Myc |
| chr8:128521224-128521264 | Myc |
| chr8:128521299-128521339 | Myc |
| chr8:128521429-128521469 | Myc |
| chr8:128521479-128521519 | Myc |
| chr8:128521579-128521619 | Myc |
| chr8:128521631-128521671 | Myc |
| chr8:128521714-128521754 | Myc |
| chr8:128521767-128521807 | Myc |
| chr8:128521830-128521870 | Myc |
| chr8:128521916-128521956 | Myc |
| chr8:128521967-128522007 | Myc |
| chr8:128522032-128522072 | Myc |
| chr8:128522130-128522170 | Myc |
| chr8:128522189-128522229 | Myc |
| chr8:128522322-128522362 | Myc |
| chr8:128522372-128522412 | Myc |
| chr8:128522440-128522480 | Myc |
| chr8:128522602-128522642 | Myc |
| chr8:128522670-128522710 | Myc |
| chr8:128522742-128522782 | Myc |
| chr8:128522837-128522877 | Myc |
| chr8:128522914-128522954 | Myc |

| | |
|---|---|
| chr8:128522977-128523017 | Myc |
| chr8:128523048-128523088 | Myc |
| chr8:128523147-128523187 | Myc |
| chr8:128523197-128523237 | Myc |
| chr8:128523366-128523406 | Myc |
| chr8:128523612-128523652 | Myc |
| chr8:128523715-128523755 | Myc |
| chr8:128523861-128523901 | Myc |
| chr8:128523911-128523951 | Myc |
| chr8:128523961-128524001 | Myc |
| chr8:128524434-128524474 | Myc |
| chr8:128524493-128524533 | Myc |
| chr8:128524589-128524629 | Myc |
| chr8:128524715-128524755 | Myc |
| chr8:128524831-128524871 | Myc |
| chr8:128524886-128524926 | Myc |
| chr8:128524989-128525029 | Myc |
| chr8:128525048-128525088 | Myc |
| chr8:128525100-128525140 | Myc |
| chr8:128525163-128525203 | Myc |
| chr8:128525248-128525288 | Myc |
| chr8:128525341-128525381 | Myc |
| chr8:128525397-128525437 | Myc |
| chr8:128525457-128525497 | Myc |
| chr8:128525555-128525595 | Myc |
| chr8:128525632-128525672 | Myc |
| chr8:128525735-128525775 | Myc |
| chr8:128525842-128525882 | Myc |
| chr8:128525942-128525982 | Myc |
| chr8:128526000-128526040 | Myc |
| chr8:128526172-128526212 | Myc |
| chr8:128526224-128526264 | Myc |
| chr8:128526284-128526324 | Myc |
| chr8:128526337-128526377 | Myc |
| chr8:128526429-128526469 | Myc |
| chr8:128526490-128526530 | Myc |
| chr8:128526566-128526606 | Myc |
| chr8:128526657-128526697 | Myc |
| chr8:128526751-128526791 | Myc |
| chr8:128526831-128526871 | Myc |
| chr8:128526910-128526950 | Myc |
| chr8:128526961-128527001 | Myc |
| chr8:128527026-128527066 | Myc |
| chr8:128527156-128527196 | Myc |
| chr8:128527275-128527315 | Myc |
| chr8:128527399-128527439 | Myc |
| chr8:128527564-128527604 | Myc |
| chr8:128527831-128527871 | Myc |
| chr8:128527917-128527957 | Myc |
| chr8:128527967-128528007 | Myc |
| chr8:128528018-128528058 | Myc |
| chr8:128528150-128528190 | Myc |

| | |
|---|---|
| chr8:128528200-128528240 | Myc |
| chr8:128528251-128528291 | Myc |
| chr8:128528329-128528369 | Myc |
| chr8:128528675-128528715 | Myc |
| chr8:128528768-128528808 | Myc |
| chr8:128528820-128528860 | Myc |
| chr8:128528951-128528991 | Myc |
| chr8:128529057-128529097 | Myc |
| chr8:128529114-128529154 | Myc |
| chr8:128529260-128529300 | Myc |
| chr8:128529420-128529460 | Myc |
| chr8:128529536-128529576 | Myc |
| chr8:128529601-128529641 | Myc |
| chr8:128529677-128529717 | Myc |
| chr8:128529753-128529793 | Myc |
| chr8:128529847-128529887 | Myc |
| chr8:128529897-128529937 | Myc |
| chr8:128529949-128529989 | Myc |
| chr8:128530051-128530091 | Myc |
| chr8:128530101-128530141 | Myc |
| chr8:128530154-128530194 | Myc |
| chr8:128530224-128530264 | Myc |
| chr8:128530347-128530387 | Myc |
| chr8:128530410-128530450 | Myc |
| chr8:128530545-128530585 | Myc |
| chr8:128530598-128530638 | Myc |
| chr8:128530822-128530862 | Myc |
| chr8:128530874-128530914 | Myc |
| chr8:128530986-128531026 | Myc |
| chr8:128531081-128531121 | Myc |
| chr8:128531166-128531206 | Myc |
| chr8:128531248-128531288 | Myc |
| chr8:128531303-128531343 | Myc |
| chr8:128531614-128531654 | Myc |
| chr8:128531808-128531848 | Myc |
| chr8:128531962-128532002 | Myc |
| chr8:128532034-128532074 | Myc |
| chr8:128532196-128532236 | Myc |
| chr8:128532252-128532292 | Myc |
| chr8:128532396-128532436 | Myc |
| chr8:128532482-128532522 | Myc |
| chr8:128532642-128532682 | Myc |
| chr8:128532759-128532799 | Myc |
| chr8:128532902-128532942 | Myc |
| chr8:128533022-128533062 | Myc |
| chr8:128533129-128533169 | Myc |
| chr8:128533184-128533224 | Myc |
| chr8:128533286-128533326 | Myc |
| chr8:128533417-128533457 | Myc |
| chr8:128533992-128534032 | Myc |
| chr8:128534053-128534093 | Myc |
| chr8:128534131-128534171 | Myc |

| | |
|---|---|
| chr8:128534256-128534296 | Myc |
| chr8:128534338-128534378 | Myc |
| chr8:128534450-128534490 | Myc |
| chr8:128534606-128534646 | Myc |
| chr8:128534697-128534737 | Myc |
| chr8:128534795-128534835 | Myc |
| chr8:128534845-128534885 | Myc |
| chr8:128534931-128534971 | Myc |
| chr8:128534981-128535021 | Myc |
| chr8:128535032-128535072 | Myc |
| chr8:128535115-128535155 | Myc |
| chr8:128535165-128535205 | Myc |
| chr8:128535284-128535324 | Myc |
| chr8:128535572-128535612 | Myc |
| chr8:128535849-128535889 | Myc |
| chr8:128535928-128535968 | Myc |
| chr8:128535981-128536021 | Myc |
| chr8:128536065-128536105 | Myc |
| chr8:128536202-128536242 | Myc |
| chr8:128536275-128536315 | Myc |
| chr8:128536358-128536398 | Myc |
| chr8:128536408-128536448 | Myc |
| chr8:128536496-128536536 | Myc |
| chr8:128536547-128536587 | Myc |
| chr8:128536652-128536692 | Myc |
| chr8:128536712-128536752 | Myc |
| chr8:128536835-128536875 | Myc |
| chr8:128536895-128536935 | Myc |
| chr8:128536988-128537028 | Myc |
| chr8:128537073-128537113 | Myc |
| chr8:128537125-128537165 | Myc |
| chr8:128537225-128537265 | Myc |
| chr8:128537296-128537336 | Myc |
| chr8:128537393-128537433 | Myc |
| chr8:128537518-128537558 | Myc |
| chr8:128537576-128537616 | Myc |
| chr8:128537663-128537703 | Myc |
| chr8:128537757-128537797 | Myc |
| chr8:128537825-128537865 | Myc |
| chr8:128537886-128537926 | Myc |
| chr8:128538026-128538066 | Myc |
| chr8:128538092-128538132 | Myc |
| chr8:128538142-128538182 | Myc |
| chr8:128538342-128538382 | Myc |
| chr8:128538436-128538476 | Myc |
| chr8:128538599-128538639 | Myc |
| chr8:128538659-128538699 | Myc |
| chr8:128538770-128538810 | Myc |
| chr8:128538841-128538881 | Myc |
| chr8:128538939-128538979 | Myc |
| chr8:128539050-128539090 | Myc |
| chr8:128539163-128539203 | Myc |

| | |
|---|---|
| chr8:128539284-128539324 | Myc |
| chr8:128539334-128539374 | Myc |
| chr8:128539399-128539439 | Myc |
| chr8:128539507-128539547 | Myc |
| chr8:128539563-128539603 | Myc |
| chr8:128539675-128539715 | Myc |
| chr8:128539736-128539776 | Myc |
| chr8:128539822-128539862 | Myc |
| chr8:128539873-128539913 | Myc |
| chr8:128539955-128539995 | Myc |
| chr8:128540012-128540052 | Myc |
| chr8:128540135-128540175 | Myc |
| chr8:128540193-128540233 | Myc |
| chr8:128540671-128540711 | Myc |
| chr8:128540742-128540782 | Myc |
| chr8:128540795-128540835 | Myc |
| chr8:128540958-128540998 | Myc |
| chr8:128541025-128541065 | Myc |
| chr8:128541076-128541116 | Myc |
| chr8:128541144-128541184 | Myc |
| chr8:128541217-128541257 | Myc |
| chr8:128541268-128541308 | Myc |
| chr8:128541373-128541413 | Myc |
| chr8:128541475-128541515 | Myc |
| chr8:128541603-128541643 | Myc |
| chr8:128541653-128541693 | Myc |
| chr8:128541742-128541782 | Myc |
| chr8:128541792-128541832 | Myc |
| chr8:128541918-128541958 | Myc |
| chr8:128542034-128542074 | Myc |
| chr8:128542104-128542144 | Myc |
| chr8:128542156-128542196 | Myc |
| chr8:128542275-128542315 | Myc |
| chr8:128542367-128542407 | Myc |
| chr8:128542447-128542487 | Myc |
| chr8:128542501-128542541 | Myc |
| chr8:128542580-128542620 | Myc |
| chr8:128542692-128542732 | Myc |
| chr8:128543010-128543050 | Myc |
| chr8:128543090-128543130 | Myc |
| chr8:128543207-128543247 | Myc |
| chr8:128543281-128543321 | Myc |
| chr8:128543333-128543373 | Myc |
| chr8:128543395-128543435 | Myc |
| chr8:128543478-128543518 | Myc |
| chr8:128543554-128543594 | Myc |
| chr8:128543608-128543648 | Myc |
| chr8:128543664-128543704 | Myc |
| chr8:128543826-128543866 | Myc |
| chr8:128544011-128544051 | Myc |
| chr8:128544081-128544121 | Myc |
| chr8:128544138-128544178 | Myc |

| | |
|---|---|
| chr8:128544267-128544307 | Myc |
| chr8:128544404-128544444 | Myc |
| chr8:128544476-128544516 | Myc |
| chr8:128544540-128544580 | Myc |
| chr8:128544673-128544713 | Myc |
| chr8:128544767-128544807 | Myc |
| chr8:128544821-128544861 | Myc |
| chr8:128544951-128544991 | Myc |
| chr8:128545007-128545047 | Myc |
| chr8:128545060-128545100 | Myc |
| chr8:128545146-128545186 | Myc |
| chr8:128545251-128545291 | Myc |
| chr8:128545325-128545365 | Myc |
| chr8:128545388-128545428 | Myc |
| chr8:128545470-128545510 | Myc |
| chr8:128545522-128545562 | Myc |
| chr8:128545626-128545666 | Myc |
| chr8:128545758-128545798 | Myc |
| chr8:128545814-128545854 | Myc |
| chr8:128545887-128545927 | Myc |
| chr8:128545950-128545990 | Myc |
| chr8:128546073-128546113 | Myc |
| chr8:128546132-128546172 | Myc |
| chr8:128546198-128546238 | Myc |
| chr8:128546254-128546294 | Myc |
| chr8:128546304-128546344 | Myc |
| chr8:128546407-128546447 | Myc |
| chr8:128546461-128546501 | Myc |
| chr8:128546511-128546551 | Myc |
| chr8:128546610-128546650 | Myc |
| chr8:128546671-128546711 | Myc |
| chr8:128546877-128546917 | Myc |
| chr8:128546976-128547016 | Myc |
| chr8:128547064-128547104 | Myc |
| chr8:128547115-128547155 | Myc |
| chr8:128547210-128547250 | Myc |
| chr8:128547352-128547392 | Myc |
| chr8:128547461-128547501 | Myc |
| chr8:128547580-128547620 | Myc |
| chr8:128547643-128547683 | Myc |
| chr8:128547772-128547812 | Myc |
| chr8:128547855-128547895 | Myc |
| chr8:128547944-128547984 | Myc |
| chr8:128547994-128548034 | Myc |
| chr8:128548061-128548101 | Myc |
| chr8:128548119-128548159 | Myc |
| chr8:128548251-128548291 | Myc |
| chr8:128548325-128548365 | Myc |
| chr8:128548379-128548419 | Myc |
| chr8:128548474-128548514 | Myc |
| chr8:128548682-128548722 | Myc |
| chr8:128548767-128548807 | Myc |

| | |
|---|---|
| chr8:128548855-128548895 | Myc |
| chr8:128549005-128549045 | Myc |
| chr8:128549102-128549142 | Myc |
| chr8:128549235-128549275 | Myc |
| chr8:128549289-128549329 | Myc |
| chr8:128549410-128549450 | Myc |
| chr8:128549528-128549568 | Myc |
| chr8:128549587-128549627 | Myc |
| chr8:128549638-128549678 | Myc |
| chr8:128549759-128549799 | Myc |
| chr8:128549909-128549949 | Myc |
| chr8:128549965-128550005 | Myc |
| chr8:128550066-128550106 | Myc |
| chr8:128550183-128550223 | Myc |
| chr8:128550233-128550273 | Myc |
| chr8:128550289-128550329 | Myc |
| chr8:128550362-128550402 | Myc |
| chr8:128550412-128550452 | Myc |
| chr8:128550469-128550509 | Myc |
| chr8:128550549-128550589 | Myc |
| chr8:128550701-128550741 | Myc |
| chr8:128550757-128550797 | Myc |
| chr8:128550852-128550892 | Myc |
| chr8:128550902-128550942 | Myc |
| chr8:128550962-128551002 | Myc |
| chr8:128551084-128551124 | Myc |
| chr8:128551139-128551179 | Myc |
| chr8:128551265-128551305 | Myc |
| chr8:128551316-128551356 | Myc |
| chr8:128551443-128551483 | Myc |
| chr8:128551494-128551534 | Myc |
| chr8:128551544-128551584 | Myc |
| chr8:128551629-128551669 | Myc |
| chr8:128551681-128551721 | Myc |
| chr8:128551739-128551779 | Myc |
| chr8:128551792-128551832 | Myc |
| chr8:128551884-128551924 | Myc |
| chr8:128551934-128551974 | Myc |
| chr8:128552109-128552149 | Myc |
| chr8:128552183-128552223 | Myc |
| chr8:128552261-128552301 | Myc |
| chr8:128552315-128552355 | Myc |
| chr8:128552393-128552433 | Myc |
| chr8:128552443-128552483 | Myc |
| chr8:128552742-128552782 | Myc |
| chr8:128552803-128552843 | Myc |
| chr8:128552896-128552936 | Myc |
| chr8:128552970-128553010 | Myc |
| chr8:128553026-128553066 | Myc |
| chr8:128553109-128553149 | Myc |
| chr8:128553314-128553354 | Myc |
| chr8:128553454-128553494 | Myc |

| | |
|---|---|
| chr8:128554114-128554154 | Myc |
| chr8:128554332-128554372 | Myc |
| chr8:128554418-128554458 | Myc |
| chr8:128554472-128554512 | Myc |
| chr8:128554527-128554567 | Myc |
| chr8:128554590-128554630 | Myc |
| chr8:128554683-128554723 | Myc |
| chr8:128554745-128554785 | Myc |
| chr8:128554870-128554910 | Myc |
| chr8:128554928-128554968 | Myc |
| chr8:128554978-128555018 | Myc |
| chr8:128555056-128555096 | Myc |
| chr8:128555152-128555192 | Myc |
| chr8:128555213-128555253 | Myc |
| chr8:128555372-128555412 | Myc |
| chr8:128555468-128555508 | Myc |
| chr8:128555577-128555617 | Myc |
| chr8:128555645-128555685 | Myc |
| chr8:128555788-128555828 | Myc |
| chr8:128555853-128555893 | Myc |
| chr8:128555956-128555996 | Myc |
| chr8:128556035-128556075 | Myc |
| chr8:128556085-128556125 | Myc |
| chr8:128556262-128556302 | Myc |
| chr8:128556312-128556352 | Myc |
| chr8:128556383-128556423 | Myc |
| chr8:128556433-128556473 | Myc |
| chr8:128556541-128556581 | Myc |
| chr8:128556620-128556660 | Myc |
| chr8:128556680-128556720 | Myc |
| chr8:128556765-128556805 | Myc |
| chr8:128556845-128556885 | Myc |
| chr8:128556900-128556940 | Myc |
| chr8:128556961-128557001 | Myc |
| chr8:128557083-128557123 | Myc |
| chr8:128557205-128557245 | Myc |
| chr8:128557288-128557328 | Myc |
| chr8:128557348-128557388 | Myc |
| chr8:128557440-128557480 | Myc |
| chr8:128557531-128557571 | Myc |
| chr8:128557620-128557660 | Myc |
| chr8:128557699-128557739 | Myc |
| chr8:128557813-128557853 | Myc |
| chr8:128557918-128557958 | Myc |
| chr8:128558001-128558041 | Myc |
| chr8:128558140-128558180 | Myc |
| chr8:128558245-128558285 | Myc |
| chr8:128558319-128558359 | Myc |
| chr8:128558377-128558417 | Myc |
| chr8:128558481-128558521 | Myc |
| chr8:128558561-128558601 | Myc |
| chr8:128558646-128558686 | Myc |

| | |
|---|---|
| chr8:128558724-128558764 | Myc |
| chr8:128558837-128558877 | Myc |
| chr8:128558921-128558961 | Myc |
| chr8:128559022-128559062 | Myc |
| chr8:128559084-128559124 | Myc |
| chr8:128559135-128559175 | Myc |
| chr8:128559220-128559260 | Myc |
| chr8:128559420-128559460 | Myc |
| chr8:128559527-128559567 | Myc |
| chr8:128559577-128559617 | Myc |
| chr8:128559707-128559747 | Myc |
| chr8:128559791-128559831 | Myc |
| chr8:128559852-128559892 | Myc |
| chr8:128559909-128559949 | Myc |
| chr8:128560027-128560067 | Myc |
| chr8:128560094-128560134 | Myc |
| chr8:128560144-128560184 | Myc |
| chr8:128560287-128560327 | Myc |
| chr8:128560337-128560377 | Myc |
| chr8:128560404-128560444 | Myc |
| chr8:128560478-128560518 | Myc |
| chr8:128560571-128560611 | Myc |
| chr8:128560628-128560668 | Myc |
| chr8:128560713-128560753 | Myc |
| chr8:128560780-128560820 | Myc |
| chr8:128560931-128560971 | Myc |
| chr8:128560994-128561034 | Myc |
| chr8:128561051-128561091 | Myc |
| chr8:128561172-128561212 | Myc |
| chr8:128561245-128561285 | Myc |
| chr8:128561322-128561362 | Myc |
| chr8:128561389-128561429 | Myc |
| chr8:128561498-128561538 | Myc |
| chr8:128561566-128561606 | Myc |
| chr8:128561620-128561660 | Myc |
| chr8:128561772-128561812 | Myc |
| chr8:128561852-128561892 | Myc |
| chr8:128561967-128562007 | Myc |
| chr8:128562051-128562091 | Myc |
| chr8:128562101-128562141 | Myc |
| chr8:128562203-128562243 | Myc |
| chr8:128562358-128562398 | Myc |
| chr8:128562420-128562460 | Myc |
| chr8:128562585-128562625 | Myc |
| chr8:128562677-128562717 | Myc |
| chr8:128562736-128562776 | Myc |
| chr8:128562850-128562890 | Myc |
| chr8:128562934-128562974 | Myc |
| chr8:128562997-128563037 | Myc |
| chr8:128563086-128563126 | Myc |
| chr8:128563144-128563184 | Myc |
| chr8:128563317-128563357 | Myc |

| | |
|---|---|
| chr8:128563497-128563537 | Myc |
| chr8:128563571-128563611 | Myc |
| chr8:128563646-128563686 | Myc |
| chr8:128563703-128563743 | Myc |
| chr8:128563820-128563860 | Myc |
| chr8:128563870-128563910 | Myc |
| chr8:128563920-128563960 | Myc |
| chr8:128563999-128564039 | Myc |
| chr8:128564078-128564118 | Myc |
| chr8:128564153-128564193 | Myc |
| chr8:128564215-128564255 | Myc |
| chr8:128564294-128564334 | Myc |
| chr8:128564348-128564388 | Myc |
| chr8:128564439-128564479 | Myc |
| chr8:128564528-128564568 | Myc |
| chr8:128564600-128564640 | Myc |
| chr8:128564662-128564702 | Myc |
| chr8:128564727-128564767 | Myc |
| chr8:128564839-128564879 | Myc |
| chr8:128564936-128564976 | Myc |
| chr8:128564999-128565039 | Myc |
| chr8:128565083-128565123 | Myc |
| chr8:128565136-128565176 | Myc |
| chr8:128565186-128565226 | Myc |
| chr8:128565277-128565317 | Myc |
| chr8:128565335-128565375 | Myc |
| chr8:128565416-128565456 | Myc |
| chr8:128565471-128565511 | Myc |
| chr8:128565547-128565587 | Myc |
| chr8:128565675-128565715 | Myc |
| chr8:128565756-128565796 | Myc |
| chr8:128565806-128565846 | Myc |
| chr8:128565953-128565993 | Myc |
| chr8:128566005-128566045 | Myc |
| chr8:128566125-128566165 | Myc |
| chr8:128566187-128566227 | Myc |
| chr8:128566282-128566322 | Myc |
| chr8:128566383-128566423 | Myc |
| chr8:128566447-128566487 | Myc |
| chr8:128566514-128566554 | Myc |
| chr8:128566599-128566639 | Myc |
| chr8:128566692-128566732 | Myc |
| chr8:128566750-128566790 | Myc |
| chr8:128566987-128567027 | Myc |
| chr8:128567065-128567105 | Myc |
| chr8:128567245-128567285 | Myc |
| chr8:128567431-128567471 | Myc |
| chr8:128567488-128567528 | Myc |
| chr8:128567550-128567590 | Myc |
| chr8:128567645-128567685 | Myc |
| chr8:128567801-128567841 | Myc |
| chr8:128567880-128567920 | Myc |

| | |
|---|---|
| chr8:128589517-128589557 | Myc |
| chr8:128589567-128589607 | Myc |
| chr8:128589617-128589657 | Myc |
| chr8:128589669-128589709 | Myc |
| chr8:128589732-128589772 | Myc |
| chr8:128589831-128589871 | Myc |
| chr8:128589885-128589925 | Myc |
| chr8:128589984-128590024 | Myc |
| chr8:128590101-128590141 | Myc |
| chr8:128590155-128590195 | Myc |
| chr8:128590225-128590265 | Myc |
| chr8:128590302-128590342 | Myc |
| chr8:128590353-128590393 | Myc |
| chr8:128590464-128590504 | Myc |
| chr8:128590524-128590564 | Myc |
| chr8:128590575-128590615 | Myc |
| chr8:128590628-128590668 | Myc |
| chr8:128590724-128590764 | Myc |
| chr8:128590840-128590880 | Myc |
| chr8:128590897-128590937 | Myc |
| chr8:128590986-128591026 | Myc |
| chr8:128591057-128591097 | Myc |
| chr8:128591119-128591159 | Myc |
| chr8:128591171-128591211 | Myc |
| chr8:128591245-128591285 | Myc |
| chr8:128591590-128591630 | Myc |
| chr8:128591659-128591699 | Myc |
| chr8:128591736-128591776 | Myc |
| chr8:128591831-128591871 | Myc |
| chr8:128592232-128592272 | Myc |
| chr8:128592305-128592345 | Myc |
| chr8:128592356-128592396 | Myc |
| chr8:128592458-128592498 | Myc |
| chr8:128592521-128592561 | Myc |
| chr8:128592572-128592612 | Myc |
| chr8:128592705-128592745 | Myc |
| chr8:128592806-128592846 | Myc |
| chr8:128592915-128592955 | Myc |
| chr8:128592965-128593005 | Myc |
| chr8:128593031-128593071 | Myc |
| chr8:128593166-128593206 | Myc |
| chr8:128593273-128593313 | Myc |
| chr8:128593344-128593384 | Myc |
| chr8:128593394-128593434 | Myc |
| chr8:128593490-128593530 | Myc |
| chr8:128593549-128593589 | Myc |
| chr8:128593677-128593717 | Myc |
| chr8:128593730-128593770 | Myc |
| chr8:128593789-128593829 | Myc |
| chr8:128593839-128593879 | Myc |
| chr8:128593990-128594030 | Myc |
| chr8:128594146-128594186 | Myc |

| | |
|---|---|
| chr8:128594220-128594260 | Myc |
| chr8:128594306-128594346 | Myc |
| chr8:128594411-128594451 | Myc |
| chr8:128594476-128594516 | Myc |
| chr8:128594533-128594573 | Myc |
| chr8:128594777-128594817 | Myc |
| chr8:128594902-128594942 | Myc |
| chr8:128595033-128595073 | Myc |
| chr8:128595175-128595215 | Myc |
| chr8:128595251-128595291 | Myc |
| chr8:128595380-128595420 | Myc |
| chr8:128595436-128595476 | Myc |
| chr8:128595510-128595550 | Myc |
| chr8:128595585-128595625 | Myc |
| chr8:128595728-128595768 | Myc |
| chr8:128595804-128595844 | Myc |
| chr8:128595873-128595913 | Myc |
| chr8:128595947-128595987 | Myc |
| chr8:128596002-128596042 | Myc |
| chr8:128596053-128596093 | Myc |
| chr8:128596109-128596149 | Myc |
| chr8:128596210-128596250 | Myc |
| chr8:128596281-128596321 | Myc |
| chr8:128596398-128596438 | Myc |
| chr8:128596509-128596549 | Myc |
| chr8:128596586-128596626 | Myc |
| chr8:128596721-128596761 | Myc |
| chr8:128596791-128596831 | Myc |
| chr8:128596861-128596901 | Myc |
| chr8:128596957-128596997 | Myc |
| chr8:128597034-128597074 | Myc |
| chr8:128597091-128597131 | Myc |
| chr8:128597238-128597278 | Myc |
| chr8:128597349-128597389 | Myc |
| chr8:128597413-128597453 | Myc |
| chr8:128597464-128597504 | Myc |
| chr8:128597514-128597554 | Myc |
| chr8:128597573-128597613 | Myc |
| chr8:128597658-128597698 | Myc |
| chr8:128597811-128597851 | Myc |
| chr8:128598091-128598131 | Myc |
| chr8:128598241-128598281 | Myc |
| chr8:128598328-128598368 | Myc |
| chr8:128598384-128598424 | Myc |
| chr8:128598462-128598502 | Myc |
| chr8:128598599-128598639 | Myc |
| chr8:128598681-128598721 | Myc |
| chr8:128598778-128598818 | Myc |
| chr8:128598910-128598950 | Myc |
| chr8:128598960-128599000 | Myc |
| chr8:128599083-128599123 | Myc |
| chr8:128599230-128599270 | Myc |

| | |
|---|---|
| chr8:128599282-128599322 | Myc |
| chr8:128599354-128599394 | Myc |
| chr8:128599415-128599455 | Myc |
| chr8:128599538-128599578 | Myc |
| chr8:128599608-128599648 | Myc |
| chr8:128599747-128599787 | Myc |
| chr8:128599817-128599857 | Myc |
| chr8:128599871-128599911 | Myc |
| chr8:128599944-128599984 | Myc |
| chr8:128600016-128600056 | Myc |
| chr8:128600075-128600115 | Myc |
| chr8:128600147-128600187 | Myc |
| chr8:128600238-128600278 | Myc |
| chr8:128600328-128600368 | Myc |
| chr8:128600392-128600432 | Myc |
| chr8:128600475-128600515 | Myc |
| chr8:128600531-128600571 | Myc |
| chr8:128600598-128600638 | Myc |
| chr8:128600719-128600759 | Myc |
| chr8:128601101-128601141 | Myc |
| chr8:128601166-128601206 | Myc |
| chr8:128601216-128601256 | Myc |
| chr8:128601329-128601369 | Myc |
| chr8:128601386-128601426 | Myc |
| chr8:128601436-128601476 | Myc |
| chr8:128601562-128601602 | Myc |
| chr8:128601672-128601712 | Myc |
| chr8:128601796-128601836 | Myc |
| chr8:128601963-128602003 | Myc |
| chr8:128602053-128602093 | Myc |
| chr8:128602118-128602158 | Myc |
| chr8:128602189-128602229 | Myc |
| chr8:128602266-128602306 | Myc |
| chr8:128602317-128602357 | Myc |
| chr8:128602417-128602457 | Myc |
| chr8:128602482-128602522 | Myc |
| chr8:128602542-128602582 | Myc |
| chr8:128602628-128602668 | Myc |
| chr8:128602701-128602741 | Myc |
| chr8:128602765-128602805 | Myc |
| chr8:128602883-128602923 | Myc |
| chr8:128603001-128603041 | Myc |
| chr8:128603051-128603091 | Myc |
| chr8:128603140-128603180 | Myc |
| chr8:128603250-128603290 | Myc |
| chr8:128603334-128603374 | Myc |
| chr8:128603431-128603471 | Myc |
| chr8:128603524-128603564 | Myc |
| chr8:128603576-128603616 | Myc |
| chr8:128603731-128603771 | Myc |
| chr8:128603784-128603824 | Myc |
| chr8:128603873-128603913 | Myc |

| | |
|---|---|
| chr8:128603939-128603979 | Myc |
| chr8:128603991-128604031 | Myc |
| chr8:128604058-128604098 | Myc |
| chr8:128604130-128604170 | Myc |
| chr8:128604180-128604220 | Myc |
| chr8:128604251-128604291 | Myc |
| chr8:128604501-128604541 | Myc |
| chr8:128604596-128604636 | Myc |
| chr8:128604744-128604784 | Myc |
| chr8:128604794-128604834 | Myc |
| chr8:128604869-128604909 | Myc |
| chr8:128604959-128604999 | Myc |
| chr8:128605106-128605146 | Myc |
| chr8:128605181-128605221 | Myc |
| chr8:128605320-128605360 | Myc |
| chr8:128605419-128605459 | Myc |
| chr8:128605541-128605581 | Myc |
| chr8:128605664-128605704 | Myc |
| chr8:128605714-128605754 | Myc |
| chr8:128605819-128605859 | Myc |
| chr8:128605872-128605912 | Myc |
| chr8:128605934-128605974 | Myc |
| chr8:128606019-128606059 | Myc |
| chr8:128606085-128606125 | Myc |
| chr8:128606198-128606238 | Myc |
| chr8:128606284-128606324 | Myc |
| chr8:128606334-128606374 | Myc |
| chr8:128606415-128606455 | Myc |
| chr8:128606471-128606511 | Myc |
| chr8:128606583-128606623 | Myc |
| chr8:128606692-128606732 | Myc |
| chr8:128606748-128606788 | Myc |
| chr8:128606838-128606878 | Myc |
| chr8:128606937-128606977 | Myc |
| chr8:128607051-128607091 | Myc |
| chr8:128607103-128607143 | Myc |
| chr8:128607166-128607206 | Myc |
| chr8:128607389-128607429 | Myc |
| chr8:128607445-128607485 | Myc |
| chr8:128607532-128607572 | Myc |
| chr8:128607621-128607661 | Myc |
| chr8:128607736-128607776 | Myc |
| chr8:128607818-128607858 | Myc |
| chr8:128607881-128607921 | Myc |
| chr8:128607935-128607975 | Myc |
| chr8:128608002-128608042 | Myc |
| chr8:128608089-128608129 | Myc |
| chr8:128608190-128608230 | Myc |
| chr8:128608244-128608284 | Myc |
| chr8:128608520-128608560 | Myc |
| chr8:128608596-128608636 | Myc |
| chr8:128608654-128608694 | Myc |

| | |
|---|---|
| chr8:128608735-128608775 | Myc |
| chr8:128608867-128608907 | Myc |
| chr8:128608960-128609000 | Myc |
| chr8:128609054-128609094 | Myc |
| chr8:128609169-128609209 | Myc |
| chr8:128609253-128609293 | Myc |
| chr8:128609377-128609417 | Myc |
| chr8:128609465-128609505 | Myc |
| chr8:128609567-128609607 | Myc |
| chr8:128609626-128609666 | Myc |
| chr8:128609676-128609716 | Myc |
| chr8:128609747-128609787 | Myc |
| chr8:128609816-128609856 | Myc |
| chr8:128609876-128609916 | Myc |
| chr8:128609944-128609984 | Myc |
| chr8:128610644-128610684 | Myc |
| chr8:128610727-128610767 | Myc |
| chr8:128610834-128610874 | Myc |
| chr8:128610936-128610976 | Myc |
| chr8:128610986-128611026 | Myc |
| chr8:128611149-128611189 | Myc |
| chr8:128611200-128611240 | Myc |
| chr8:128611287-128611327 | Myc |
| chr8:128611357-128611397 | Myc |
| chr8:128611454-128611494 | Myc |
| chr8:128611524-128611564 | Myc |
| chr8:128611607-128611647 | Myc |
| chr8:128611662-128611702 | Myc |
| chr8:128611716-128611756 | Myc |
| chr8:128611768-128611808 | Myc |
| chr8:128611859-128611899 | Myc |
| chr8:128611950-128611990 | Myc |
| chr8:128612008-128612048 | Myc |
| chr8:128612081-128612121 | Myc |
| chr8:128612178-128612218 | Myc |
| chr8:128612234-128612274 | Myc |
| chr8:128612310-128612350 | Myc |
| chr8:128612367-128612407 | Myc |
| chr8:128612427-128612467 | Myc |
| chr8:128612485-128612525 | Myc |
| chr8:128612552-128612592 | Myc |
| chr8:128612610-128612650 | Myc |
| chr8:128612723-128612763 | Myc |
| chr8:128612850-128612890 | Myc |
| chr8:128612917-128612957 | Myc |
| chr8:128613023-128613063 | Myc |
| chr8:128613087-128613127 | Myc |
| chr8:128613150-128613190 | Myc |
| chr8:128613230-128613270 | Myc |
| chr8:128613280-128613320 | Myc |
| chr8:128613331-128613371 | Myc |
| chr8:128613392-128613432 | Myc |

| | |
|---|---|
| chr8:128613543-128613583 | Myc |
| chr8:128613663-128613703 | Myc |
| chr8:128613758-128613798 | Myc |
| chr8:128613824-128613864 | Myc |
| chr8:128613876-128613916 | Myc |
| chr8:128614018-128614058 | Myc |
| chr8:128614094-128614134 | Myc |
| chr8:128614215-128614255 | Myc |
| chr8:128614331-128614371 | Myc |
| chr8:128614445-128614485 | Myc |
| chr8:128614529-128614569 | Myc |
| chr8:128614579-128614619 | Myc |
| chr8:128614693-128614733 | Myc |
| chr8:128614743-128614783 | Myc |
| chr8:128614793-128614833 | Myc |
| chr8:128614856-128614896 | Myc |
| chr8:128614923-128614963 | Myc |
| chr8:128615097-128615137 | Myc |
| chr8:128615149-128615189 | Myc |
| chr8:128615235-128615275 | Myc |
| chr8:128615289-128615329 | Myc |
| chr8:128615371-128615411 | Myc |
| chr8:128615443-128615483 | Myc |
| chr8:128615532-128615572 | Myc |
| chr8:128615601-128615641 | Myc |
| chr8:128615667-128615707 | Myc |
| chr8:128615758-128615798 | Myc |
| chr8:128615856-128615896 | Myc |
| chr8:128615983-128616023 | Myc |
| chr8:128616036-128616076 | Myc |
| chr8:128616130-128616170 | Myc |
| chr8:128616223-128616263 | Myc |
| chr8:128616301-128616341 | Myc |
| chr8:128616404-128616444 | Myc |
| chr8:128616463-128616503 | Myc |
| chr8:128616661-128616701 | Myc |
| chr8:128616716-128616756 | Myc |
| chr8:128616850-128616890 | Myc |
| chr8:128616922-128616962 | Myc |
| chr8:128616974-128617014 | Myc |
| chr8:128617028-128617068 | Myc |
| chr8:128617108-128617148 | Myc |
| chr8:128617226-128617266 | Myc |
| chr8:128617278-128617318 | Myc |
| chr8:128617432-128617472 | Myc |
| chr8:128617509-128617549 | Myc |
| chr8:128617570-128617610 | Myc |
| chr8:128617637-128617677 | Myc |
| chr8:128617688-128617728 | Myc |
| chr8:128617775-128617815 | Myc |
| chr8:128617832-128617872 | Myc |
| chr8:128617883-128617923 | Myc |

| | |
|---|---|
| chr8:128618055-128618095 | Myc |
| chr8:128618130-128618170 | Myc |
| chr8:128618229-128618269 | Myc |
| chr8:128618282-128618322 | Myc |
| chr8:128618360-128618400 | Myc |
| chr8:128618454-128618494 | Myc |
| chr8:128618560-128618600 | Myc |
| chr8:128618624-128618664 | Myc |
| chr8:128618700-128618740 | Myc |
| chr8:128618794-128618834 | Myc |
| chr8:128618886-128618926 | Myc |
| chr8:128618950-128618990 | Myc |
| chr8:128619015-128619055 | Myc |
| chr8:128619134-128619174 | Myc |
| chr8:128619219-128619259 | Myc |
| chr8:128619340-128619380 | Myc |
| chr8:128619393-128619433 | Myc |
| chr8:128619484-128619524 | Myc |
| chr8:128619683-128619723 | Myc |
| chr8:128619759-128619799 | Myc |
| chr8:128619854-128619894 | Myc |
| chr8:128619970-128620010 | Myc |
| chr8:128620025-128620065 | Myc |
| chr8:128620087-128620127 | Myc |
| chr8:128620185-128620225 | Myc |
| chr8:128620245-128620285 | Myc |
| chr8:128620377-128620417 | Myc |
| chr8:128620471-128620511 | Myc |
| chr8:128620540-128620580 | Myc |
| chr8:128620713-128620753 | Myc |
| chr8:128620936-128620976 | Myc |
| chr8:128621039-128621079 | Myc |
| chr8:128621206-128621246 | Myc |
| chr8:128621279-128621319 | Myc |
| chr8:128621329-128621369 | Myc |
| chr8:128621452-128621492 | Myc |
| chr8:128621553-128621593 | Myc |
| chr8:128621657-128621697 | Myc |
| chr8:128621719-128621759 | Myc |
| chr8:128621769-128621809 | Myc |
| chr8:128621835-128621875 | Myc |
| chr8:128621885-128621925 | Myc |
| chr8:128621935-128621975 | Myc |
| chr8:128622011-128622051 | Myc |
| chr8:128622096-128622136 | Myc |
| chr8:128622181-128622221 | Myc |
| chr8:128622237-128622277 | Myc |
| chr8:128622353-128622393 | Myc |
| chr8:128622415-128622455 | Myc |
| chr8:128622570-128622610 | Myc |
| chr8:128622648-128622688 | Myc |
| chr8:128622746-128622786 | Myc |

| | |
|---|---|
| chr8:128622850-128622890 | Myc |
| chr8:128622999-128623039 | Myc |
| chr8:128623123-128623163 | Myc |
| chr8:128623180-128623220 | Myc |
| chr8:128623238-128623278 | Myc |
| chr8:128623366-128623406 | Myc |
| chr8:128623526-128623566 | Myc |
| chr8:128623604-128623644 | Myc |
| chr8:128623685-128623725 | Myc |
| chr8:128623785-128623825 | Myc |
| chr8:128623852-128623892 | Myc |
| chr8:128623907-128623947 | Myc |
| chr8:128623963-128624003 | Myc |
| chr8:128624017-128624057 | Myc |
| chr8:128624076-128624116 | Myc |
| chr8:128624177-128624217 | Myc |
| chr8:128624296-128624336 | Myc |
| chr8:128624359-128624399 | Myc |
| chr8:128624409-128624449 | Myc |
| chr8:128624475-128624515 | Myc |
| chr8:128624569-128624609 | Myc |
| chr8:128624630-128624670 | Myc |
| chr8:128624766-128624806 | Myc |
| chr8:128624864-128624904 | Myc |
| chr8:128624932-128624972 | Myc |
| chr8:128624999-128625039 | Myc |
| chr8:128625062-128625102 | Myc |
| chr8:128625190-128625230 | Myc |
| chr8:128625240-128625280 | Myc |
| chr8:128625319-128625359 | Myc |
| chr8:128625385-128625425 | Myc |
| chr8:128625455-128625495 | Myc |
| chr8:128625538-128625578 | Myc |
| chr8:128625650-128625690 | Myc |
| chr8:128625713-128625753 | Myc |
| chr8:128625790-128625830 | Myc |
| chr8:128625913-128625953 | Myc |
| chr8:128626030-128626070 | Myc |
| chr8:128626218-128626258 | Myc |
| chr8:128626278-128626318 | Myc |
| chr8:128626416-128626456 | Myc |
| chr8:128626466-128626506 | Myc |
| chr8:128626516-128626556 | Myc |
| chr8:128626789-128626829 | Myc |
| chr8:128626859-128626899 | Myc |
| chr8:128626943-128626983 | Myc |
| chr8:128627045-128627085 | Myc |
| chr8:128627164-128627204 | Myc |
| chr8:128627216-128627256 | Myc |
| chr8:128627306-128627346 | Myc |
| chr8:128627363-128627403 | Myc |
| chr8:128627472-128627512 | Myc |

| | |
|---|---|
| chr8:128627550-128627590 | Myc |
| chr8:128627640-128627680 | Myc |
| chr8:128627764-128627804 | Myc |
| chr8:128627894-128627934 | Myc |
| chr8:128627969-128628009 | Myc |
| chr8:128628020-128628060 | Myc |
| chr8:128628096-128628136 | Myc |
| chr8:128628161-128628201 | Myc |
| chr8:128628257-128628297 | Myc |
| chr8:128628404-128628444 | Myc |
| chr8:128628510-128628550 | Myc |
| chr8:128628584-128628624 | Myc |
| chr8:128628641-128628681 | Myc |
| chr8:128628728-128628768 | Myc |
| chr8:128628778-128628818 | Myc |
| chr8:128628948-128628988 | Myc |
| chr8:128629070-128629110 | Myc |
| chr8:128629124-128629164 | Myc |
| chr8:128629199-128629239 | Myc |
| chr8:128629272-128629312 | Myc |
| chr8:128629349-128629389 | Myc |
| chr8:128629562-128629602 | Myc |
| chr8:128629970-128630010 | Myc |
| chr8:128630026-128630066 | Myc |
| chr8:128630076-128630116 | Myc |
| chr8:128630179-128630219 | Myc |
| chr8:128630245-128630285 | Myc |
| chr8:128630317-128630357 | Myc |
| chr8:128630391-128630431 | Myc |
| chr8:128630473-128630513 | Myc |
| chr8:128630558-128630598 | Myc |
| chr8:128630628-128630668 | Myc |
| chr8:128630684-128630724 | Myc |
| chr8:128630742-128630782 | Myc |
| chr8:128630804-128630844 | Myc |
| chr8:128630892-128630932 | Myc |
| chr8:128630960-128631000 | Myc |
| chr8:128631049-128631089 | Myc |
| chr8:128631163-128631203 | Myc |
| chr8:128631213-128631253 | Myc |
| chr8:128631268-128631308 | Myc |
| chr8:128631386-128631426 | Myc |
| chr8:128631472-128631512 | Myc |
| chr8:128631578-128631618 | Myc |
| chr8:128631633-128631673 | Myc |
| chr8:128631722-128631762 | Myc |
| chr8:128631774-128631814 | Myc |
| chr8:128631856-128631896 | Myc |
| chr8:128631906-128631946 | Myc |
| chr8:128631962-128632002 | Myc |
| chr8:128632041-128632081 | Myc |
| chr8:128632092-128632132 | Myc |

| | |
|---|---|
| chr8:128632157-128632197 | Myc |
| chr8:128632210-128632250 | Myc |
| chr8:128632281-128632321 | Myc |
| chr8:128632357-128632397 | Myc |
| chr8:128632447-128632487 | Myc |
| chr8:128632500-128632540 | Myc |
| chr8:128632637-128632677 | Myc |
| chr8:128632755-128632795 | Myc |
| chr8:128632839-128632879 | Myc |
| chr8:128632924-128632964 | Myc |
| chr8:128632999-128633039 | Myc |
| chr8:128633101-128633141 | Myc |
| chr8:128633183-128633223 | Myc |
| chr8:128633271-128633311 | Myc |
| chr8:128633321-128633361 | Myc |
| chr8:128633388-128633428 | Myc |
| chr8:128633495-128633535 | Myc |
| chr8:128633802-128633842 | Myc |
| chr8:128633853-128633893 | Myc |
| chr8:128634006-128634046 | Myc |
| chr8:128634056-128634096 | Myc |
| chr8:128634115-128634155 | Myc |
| chr8:128634238-128634278 | Myc |
| chr8:128634296-128634336 | Myc |
| chr8:128634525-128634565 | Myc |
| chr8:128634589-128634629 | Myc |
| chr8:128634713-128634753 | Myc |
| chr8:128634999-128635039 | Myc |
| chr8:128635050-128635090 | Myc |
| chr8:128635164-128635204 | Myc |
| chr8:128635230-128635270 | Myc |
| chr8:128635295-128635335 | Myc |
| chr8:128635362-128635402 | Myc |
| chr8:128635412-128635452 | Myc |
| chr8:128635518-128635558 | Myc |
| chr8:128635573-128635613 | Myc |
| chr8:128635644-128635684 | Myc |
| chr8:128635786-128635826 | Myc |
| chr8:128636026-128636066 | Myc |
| chr8:128636113-128636153 | Myc |
| chr8:128636194-128636234 | Myc |
| chr8:128636244-128636284 | Myc |
| chr8:128636316-128636356 | Myc |
| chr8:128636391-128636431 | Myc |
| chr8:128636449-128636489 | Myc |
| chr8:128636576-128636616 | Myc |
| chr8:128636644-128636684 | Myc |
| chr8:128636718-128636758 | Myc |
| chr8:128636833-128636873 | Myc |
| chr8:128636982-128637022 | Myc |
| chr8:128637038-128637078 | Myc |
| chr8:128637132-128637172 | Myc |

| | |
|---|---|
| chr8:128637195-128637235 | Myc |
| chr8:128637369-128637409 | Myc |
| chr8:128637419-128637459 | Myc |
| chr8:128637469-128637509 | Myc |
| chr8:128637519-128637559 | Myc |
| chr8:128637591-128637631 | Myc |
| chr8:128637664-128637704 | Myc |
| chr8:128637718-128637758 | Myc |
| chr8:128637858-128637898 | Myc |
| chr8:128637909-128637949 | Myc |
| chr8:128638011-128638051 | Myc |
| chr8:128638123-128638163 | Myc |
| chr8:128638173-128638213 | Myc |
| chr8:128638263-128638303 | Myc |
| chr8:128638382-128638422 | Myc |
| chr8:128638467-128638507 | Myc |
| chr8:128638565-128638605 | Myc |
| chr8:128638699-128638739 | Myc |
| chr8:128638913-128638953 | Myc |
| chr8:128639014-128639054 | Myc |
| chr8:128639076-128639116 | Myc |
| chr8:128639134-128639174 | Myc |
| chr8:128639211-128639251 | Myc |
| chr8:128639290-128639330 | Myc |
| chr8:128639352-128639392 | Myc |
| chr8:128639402-128639442 | Myc |
| chr8:128639482-128639522 | Myc |
| chr8:128639589-128639629 | Myc |
| chr8:128639655-128639695 | Myc |
| chr8:128639727-128639767 | Myc |
| chr8:128639853-128639893 | Myc |
| chr8:128639938-128639978 | Myc |
| chr8:128640066-128640106 | Myc |
| chr8:128640173-128640213 | Myc |
| chr8:128640244-128640284 | Myc |
| chr8:128640317-128640357 | Myc |
| chr8:128640367-128640407 | Myc |
| chr8:128640448-128640488 | Myc |
| chr8:128640616-128640656 | Myc |
| chr8:128640706-128640746 | Myc |
| chr8:128640756-128640796 | Myc |
| chr8:128640808-128640848 | Myc |
| chr8:128640887-128640927 | Myc |
| chr8:128640957-128640997 | Myc |
| chr8:128641025-128641065 | Myc |
| chr8:128641144-128641184 | Myc |
| chr8:128641242-128641282 | Myc |
| chr8:128641565-128641605 | Myc |
| chr8:128641660-128641700 | Myc |
| chr8:128641713-128641753 | Myc |
| chr8:128641815-128641855 | Myc |
| chr8:128641875-128641915 | Myc |

| | |
|---|---|
| chr8:128641944-128641984 | Myc |
| chr8:128642077-128642117 | Myc |
| chr8:128642183-128642223 | Myc |
| chr8:128642244-128642284 | Myc |
| chr8:128642347-128642387 | Myc |
| chr8:128642402-128642442 | Myc |
| chr8:128643179-128643219 | Myc |
| chr8:128643229-128643269 | Myc |
| chr8:128643281-128643321 | Myc |
| chr8:128643341-128643381 | Myc |
| chr8:128643451-128643491 | Myc |
| chr8:128643575-128643615 | Myc |
| chr8:128643842-128643882 | Myc |
| chr8:128643949-128643989 | Myc |
| chr8:128644000-128644040 | Myc |
| chr8:128644136-128644176 | Myc |
| chr8:128644244-128644284 | Myc |
| chr8:128644309-128644349 | Myc |
| chr8:128644456-128644496 | Myc |
| chr8:128644508-128644548 | Myc |
| chr8:128644579-128644619 | Myc |
| chr8:128644645-128644685 | Myc |
| chr8:128644761-128644801 | Myc |
| chr8:128644816-128644856 | Myc |
| chr8:128644870-128644910 | Myc |
| chr8:128644938-128644978 | Myc |
| chr8:128645061-128645101 | Myc |
| chr8:128645199-128645239 | Myc |
| chr8:128645310-128645350 | Myc |
| chr8:128645574-128645614 | Myc |
| chr8:128645698-128645738 | Myc |
| chr8:128645750-128645790 | Myc |
| chr8:128645898-128645938 | Myc |
| chr8:128645948-128645988 | Myc |
| chr8:128646011-128646051 | Myc |
| chr8:128646139-128646179 | Myc |
| chr8:128646190-128646230 | Myc |
| chr8:128646304-128646344 | Myc |
| chr8:128646363-128646403 | Myc |
| chr8:128646460-128646500 | Myc |
| chr8:128646528-128646568 | Myc |
| chr8:128646599-128646639 | Myc |
| chr8:128646763-128646803 | Myc |
| chr8:128646841-128646881 | Myc |
| chr8:128646928-128646968 | Myc |
| chr8:128647064-128647104 | Myc |
| chr8:128647131-128647171 | Myc |
| chr8:128647230-128647270 | Myc |
| chr8:128647350-128647390 | Myc |
| chr8:128647401-128647441 | Myc |
| chr8:128647485-128647525 | Myc |
| chr8:128647593-128647633 | Myc |

| | |
|---|---|
| chr8:128647661-128647701 | Myc |
| chr8:128647711-128647751 | Myc |
| chr8:128647845-128647885 | Myc |
| chr8:128647901-128647941 | Myc |
| chr8:128648038-128648078 | Myc |
| chr8:128648097-128648137 | Myc |
| chr8:128648187-128648227 | Myc |
| chr8:128648285-128648325 | Myc |
| chr8:128648354-128648394 | Myc |
| chr8:128648410-128648450 | Myc |
| chr8:128648478-128648518 | Myc |
| chr8:128648607-128648647 | Myc |
| chr8:128648709-128648749 | Myc |
| chr8:128648773-128648813 | Myc |
| chr8:128648848-128648888 | Myc |
| chr8:128648901-128648941 | Myc |
| chr8:128648973-128649013 | Myc |
| chr8:128649160-128649200 | Myc |
| chr8:128649260-128649300 | Myc |
| chr8:128649368-128649408 | Myc |
| chr8:128649472-128649512 | Myc |
| chr8:128649522-128649562 | Myc |
| chr8:128649587-128649627 | Myc |
| chr8:128649639-128649679 | Myc |
| chr8:128649700-128649740 | Myc |
| chr8:128649756-128649796 | Myc |
| chr8:128649818-128649858 | Myc |
| chr8:128649913-128649953 | Myc |
| chr8:128649981-128650021 | Myc |
| chr8:128650041-128650081 | Myc |
| chr8:128650133-128650173 | Myc |
| chr8:128650483-128650523 | Myc |
| chr8:128650587-128650627 | Myc |
| chr8:128650655-128650695 | Myc |
| chr8:128650725-128650765 | Myc |
| chr8:128650790-128650830 | Myc |
| chr8:128650847-128650887 | Myc |
| chr8:128650910-128650950 | Myc |
| chr8:128650979-128651019 | Myc |
| chr8:128651040-128651080 | Myc |
| chr8:128651126-128651166 | Myc |
| chr8:128651230-128651270 | Myc |
| chr8:128651335-128651375 | Myc |
| chr8:128651388-128651428 | Myc |
| chr8:128651500-128651540 | Myc |
| chr8:128651630-128651670 | Myc |
| chr8:128651691-128651731 | Myc |
| chr8:128651808-128651848 | Myc |
| chr8:128651881-128651921 | Myc |
| chr8:128652005-128652045 | Myc |
| chr8:128652134-128652174 | Myc |
| chr8:128652257-128652297 | Myc |

| | |
|---|---|
| chr8:128652414-128652454 | Myc |
| chr8:128652490-128652530 | Myc |
| chr8:128652540-128652580 | Myc |
| chr8:128652644-128652684 | Myc |
| chr8:128652754-128652794 | Myc |
| chr8:128652810-128652850 | Myc |
| chr8:128652892-128652932 | Myc |
| chr8:128652943-128652983 | Myc |
| chr8:128653039-128653079 | Myc |
| chr8:128653152-128653192 | Myc |
| chr8:128653248-128653288 | Myc |
| chr8:128653343-128653383 | Myc |
| chr8:128653399-128653439 | Myc |
| chr8:128653526-128653566 | Myc |
| chr8:128653677-128653717 | Myc |
| chr8:128653795-128653835 | Myc |
| chr8:128653862-128653902 | Myc |
| chr8:128654000-128654040 | Myc |
| chr8:128654053-128654093 | Myc |
| chr8:128654171-128654211 | Myc |
| chr8:128654272-128654312 | Myc |
| chr8:128654329-128654369 | Myc |
| chr8:128654469-128654509 | Myc |
| chr8:128654521-128654561 | Myc |
| chr8:128654576-128654616 | Myc |
| chr8:128654688-128654728 | Myc |
| chr8:128654903-128654943 | Myc |
| chr8:128654996-128655036 | Myc |
| chr8:128655046-128655086 | Myc |
| chr8:128655204-128655244 | Myc |
| chr8:128655258-128655298 | Myc |
| chr8:128655308-128655348 | Myc |
| chr8:128655398-128655438 | Myc |
| chr8:128655457-128655497 | Myc |
| chr8:128655555-128655595 | Myc |
| chr8:128655649-128655689 | Myc |
| chr8:128655744-128655784 | Myc |
| chr8:128655853-128655893 | Myc |
| chr8:128655948-128655988 | Myc |
| chr8:128656013-128656053 | Myc |
| chr8:128656066-128656106 | Myc |
| chr8:128656211-128656251 | Myc |
| chr8:128656307-128656347 | Myc |
| chr8:128656359-128656399 | Myc |
| chr8:128656447-128656487 | Myc |
| chr8:128656514-128656554 | Myc |
| chr8:128656564-128656604 | Myc |
| chr8:128656631-128656671 | Myc |
| chr8:128656693-128656733 | Myc |
| chr8:128656874-128656914 | Myc |
| chr8:128656951-128656991 | Myc |
| chr8:128657026-128657066 | Myc |

| | |
|---|---|
| chr8:128657114-128657154 | Myc |
| chr8:128657165-128657205 | Myc |
| chr8:128657264-128657304 | Myc |
| chr8:128657406-128657446 | Myc |
| chr8:128657582-128657622 | Myc |
| chr8:128657694-128657734 | Myc |
| chr8:128657779-128657819 | Myc |
| chr8:128657833-128657873 | Myc |
| chr8:128657964-128658004 | Myc |
| chr8:128658076-128658116 | Myc |
| chr8:128658157-128658197 | Myc |
| chr8:128658232-128658272 | Myc |
| chr8:128658289-128658329 | Myc |
| chr8:128658339-128658379 | Myc |
| chr8:128658516-128658556 | Myc |
| chr8:128658588-128658628 | Myc |
| chr8:128658711-128658751 | Myc |
| chr8:128658788-128658828 | Myc |
| chr8:128658932-128658972 | Myc |
| chr8:128659023-128659063 | Myc |
| chr8:128659174-128659214 | Myc |
| chr8:128659243-128659283 | Myc |
| chr8:128659338-128659378 | Myc |
| chr8:128659405-128659445 | Myc |
| chr8:128659503-128659543 | Myc |
| chr8:128659558-128659598 | Myc |
| chr8:128659669-128659709 | Myc |
| chr8:128659746-128659786 | Myc |
| chr8:128659813-128659853 | Myc |
| chr8:128660026-128660066 | Myc |
| chr8:128660091-128660131 | Myc |
| chr8:128660197-128660237 | Myc |
| chr8:128660268-128660308 | Myc |
| chr8:128660608-128660648 | Myc |
| chr8:128660664-128660704 | Myc |
| chr8:128660719-128660759 | Myc |
| chr8:128661174-128661214 | Myc |
| chr8:128661259-128661299 | Myc |
| chr8:128661336-128661376 | Myc |
| chr8:128661460-128661500 | Myc |
| chr8:128661537-128661577 | Myc |
| chr8:128661587-128661627 | Myc |
| chr8:128661666-128661706 | Myc |
| chr8:128661718-128661758 | Myc |
| chr8:128661772-128661812 | Myc |
| chr8:128661846-128661886 | Myc |
| chr8:128661910-128661950 | Myc |
| chr8:128661960-128662000 | Myc |
| chr8:128662091-128662131 | Myc |
| chr8:128662143-128662183 | Myc |
| chr8:128662216-128662256 | Myc |
| chr8:128662294-128662334 | Myc |

| | |
|---|---|
| chr8:128662344-128662384 | Myc |
| chr8:128662437-128662477 | Myc |
| chr8:128662520-128662560 | Myc |
| chr8:128662612-128662652 | Myc |
| chr8:128662707-128662747 | Myc |
| chr8:128662820-128662860 | Myc |
| chr8:128663000-128663040 | Myc |
| chr8:128663122-128663162 | Myc |
| chr8:128663172-128663212 | Myc |
| chr8:128663251-128663291 | Myc |
| chr8:128663349-128663389 | Myc |
| chr8:128663431-128663471 | Myc |
| chr8:128663497-128663537 | Myc |
| chr8:128663586-128663626 | Myc |
| chr8:128663674-128663714 | Myc |
| chr8:128663829-128663869 | Myc |
| chr8:128663896-128663936 | Myc |
| chr8:128663946-128663986 | Myc |
| chr8:128664017-128664057 | Myc |
| chr8:128664102-128664142 | Myc |
| chr8:128664162-128664202 | Myc |
| chr8:128664261-128664301 | Myc |
| chr8:128664318-128664358 | Myc |
| chr8:128664465-128664505 | Myc |
| chr8:128664544-128664584 | Myc |
| chr8:128664712-128664752 | Myc |
| chr8:128664778-128664818 | Myc |
| chr8:128664845-128664885 | Myc |
| chr8:128664897-128664937 | Myc |
| chr8:128664990-128665030 | Myc |
| chr8:128665172-128665212 | Myc |
| chr8:128665268-128665308 | Myc |
| chr8:128665358-128665398 | Myc |
| chr8:128665409-128665449 | Myc |
| chr8:128665564-128665604 | Myc |
| chr8:128665633-128665673 | Myc |
| chr8:128665716-128665756 | Myc |
| chr8:128665887-128665927 | Myc |
| chr8:128666013-128666053 | Myc |
| chr8:128666144-128666184 | Myc |
| chr8:128666199-128666239 | Myc |
| chr8:128666331-128666371 | Myc |
| chr8:128666422-128666462 | Myc |
| chr8:128666474-128666514 | Myc |
| chr8:128666565-128666605 | Myc |
| chr8:128666616-128666656 | Myc |
| chr8:128666734-128666774 | Myc |
| chr8:128666821-128666861 | Myc |
| chr8:128666899-128666939 | Myc |
| chr8:128667072-128667112 | Myc |
| chr8:128667215-128667255 | Myc |
| chr8:128667481-128667521 | Myc |

| | |
|---|---|
| chr8:128667683-128667723 | Myc |
| chr8:128667739-128667779 | Myc |
| chr8:128667897-128667937 | Myc |
| chr8:128667947-128667987 | Myc |
| chr8:128668022-128668062 | Myc |
| chr8:128668138-128668178 | Myc |
| chr8:128668202-128668242 | Myc |
| chr8:128668263-128668303 | Myc |
| chr8:128668335-128668375 | Myc |
| chr8:128668452-128668492 | Myc |
| chr8:128668587-128668627 | Myc |
| chr8:128668649-128668689 | Myc |
| chr8:128668744-128668784 | Myc |
| chr8:128668822-128668862 | Myc |
| chr8:128668886-128668926 | Myc |
| chr8:128669021-128669061 | Myc |
| chr8:128669151-128669191 | Myc |
| chr8:128669328-128669368 | Myc |
| chr8:128669414-128669454 | Myc |
| chr8:128669473-128669513 | Myc |
| chr8:128669564-128669604 | Myc |
| chr8:128669673-128669713 | Myc |
| chr8:128669746-128669786 | Myc |
| chr8:128669799-128669839 | Myc |
| chr8:128669888-128669928 | Myc |
| chr8:128669967-128670007 | Myc |
| chr8:128670048-128670088 | Myc |
| chr8:128670160-128670200 | Myc |
| chr8:128670232-128670272 | Myc |
| chr8:128670372-128670412 | Myc |
| chr8:128670439-128670479 | Myc |
| chr8:128670527-128670567 | Myc |
| chr8:128670588-128670628 | Myc |
| chr8:128670706-128670746 | Myc |
| chr8:128670757-128670797 | Myc |
| chr8:128670822-128670862 | Myc |
| chr8:128670931-128670971 | Myc |
| chr8:128671023-128671063 | Myc |
| chr8:128671165-128671205 | Myc |
| chr8:128671278-128671318 | Myc |
| chr8:128671328-128671368 | Myc |
| chr8:128671469-128671509 | Myc |
| chr8:128671596-128671636 | Myc |
| chr8:128671649-128671689 | Myc |
| chr8:128671732-128671772 | Myc |
| chr8:128671873-128671913 | Myc |
| chr8:128672033-128672073 | Myc |
| chr8:128672128-128672168 | Myc |
| chr8:128672210-128672250 | Myc |
| chr8:128672265-128672305 | Myc |
| chr8:128672391-128672431 | Myc |
| chr8:128672614-128672654 | Myc |

| | |
|---|---|
| chr8:128672748-128672788 | Myc |
| chr8:128672833-128672873 | Myc |
| chr8:128672905-128672945 | Myc |
| chr8:128672955-128672995 | Myc |
| chr8:128673046-128673086 | Myc |
| chr8:128673105-128673145 | Myc |
| chr8:128673179-128673219 | Myc |
| chr8:128673238-128673278 | Myc |
| chr8:128673340-128673380 | Myc |
| chr8:128673438-128673478 | Myc |
| chr8:128673537-128673577 | Myc |
| chr8:128673698-128673738 | Myc |
| chr8:128673753-128673793 | Myc |
| chr8:128673856-128673896 | Myc |
| chr8:128673906-128673946 | Myc |
| chr8:128673978-128674018 | Myc |
| chr8:128674045-128674085 | Myc |
| chr8:128674128-128674168 | Myc |
| chr8:128674200-128674240 | Myc |
| chr8:128674341-128674381 | Myc |
| chr8:128674440-128674480 | Myc |
| chr8:128674534-128674574 | Myc |
| chr8:128674649-128674689 | Myc |
| chr8:128674763-128674803 | Myc |
| chr8:128674888-128674928 | Myc |
| chr8:128674971-128675011 | Myc |
| chr8:128675053-128675093 | Myc |
| chr8:128675119-128675159 | Myc |
| chr8:128675193-128675233 | Myc |
| chr8:128675249-128675289 | Myc |
| chr8:128675341-128675381 | Myc |
| chr8:128675392-128675432 | Myc |
| chr8:128675450-128675490 | Myc |
| chr8:128675500-128675540 | Myc |
| chr8:128675566-128675606 | Myc |
| chr8:128675642-128675682 | Myc |
| chr8:128675720-128675760 | Myc |
| chr8:128675770-128675810 | Myc |
| chr8:128675841-128675881 | Myc |
| chr8:128675928-128675968 | Myc |
| chr8:128676004-128676044 | Myc |
| chr8:128676093-128676133 | Myc |
| chr8:128676151-128676191 | Myc |
| chr8:128676202-128676242 | Myc |
| chr8:128676321-128676361 | Myc |
| chr8:128676459-128676499 | Myc |
| chr8:128676516-128676556 | Myc |
| chr8:128676811-128676851 | Myc |
| chr8:128676930-128676970 | Myc |
| chr8:128676995-128677035 | Myc |
| chr8:128677149-128677189 | Myc |
| chr8:128677202-128677242 | Myc |

| | |
|---|---|
| chr8:128677317-128677357 | Myc |
| chr8:128677420-128677460 | Myc |
| chr8:128677470-128677510 | Myc |
| chr8:128677531-128677571 | Myc |
| chr8:128677587-128677627 | Myc |
| chr8:128677639-128677679 | Myc |
| chr8:128677694-128677734 | Myc |
| chr8:128677745-128677785 | Myc |
| chr8:128677832-128677872 | Myc |
| chr8:128678006-128678046 | Myc |
| chr8:128678086-128678126 | Myc |
| chr8:128678186-128678226 | Myc |
| chr8:128678239-128678279 | Myc |
| chr8:128678318-128678358 | Myc |
| chr8:128678400-128678440 | Myc |
| chr8:128678508-128678548 | Myc |
| chr8:128678624-128678664 | Myc |
| chr8:128678731-128678771 | Myc |
| chr8:128678798-128678838 | Myc |
| chr8:128678897-128678937 | Myc |
| chr8:128678957-128678997 | Myc |
| chr8:128679109-128679149 | Myc |
| chr8:128679186-128679226 | Myc |
| chr8:128679275-128679315 | Myc |
| chr8:128679395-128679435 | Myc |
| chr8:128679495-128679535 | Myc |
| chr8:128679589-128679629 | Myc |
| chr8:128679686-128679726 | Myc |
| chr8:128679820-128679860 | Myc |
| chr8:128679870-128679910 | Myc |
| chr8:128679943-128679983 | Myc |
| chr8:128680011-128680051 | Myc |
| chr8:128680080-128680120 | Myc |
| chr8:128680199-128680239 | Myc |
| chr8:128680299-128680339 | Myc |
| chr8:128680354-128680394 | Myc |
| chr8:128680422-128680462 | Myc |
| chr8:128680488-128680528 | Myc |
| chr8:128680542-128680582 | Myc |
| chr8:128680701-128680741 | Myc |
| chr8:128680751-128680791 | Myc |
| chr8:128680868-128680908 | Myc |
| chr8:128680964-128681004 | Myc |
| chr8:128681015-128681055 | Myc |
| chr8:128681067-128681107 | Myc |
| chr8:128681153-128681193 | Myc |
| chr8:128681267-128681307 | Myc |
| chr8:128681339-128681379 | Myc |
| chr8:128681390-128681430 | Myc |
| chr8:128681440-128681480 | Myc |
| chr8:128681550-128681590 | Myc |
| chr8:128681638-128681678 | Myc |

| | |
|---|---|
| chr8:128681716-128681756 | Myc |
| chr8:128681768-128681808 | Myc |
| chr8:128681818-128681858 | Myc |
| chr8:128681953-128681993 | Myc |
| chr8:128682112-128682152 | Myc |
| chr8:128682162-128682202 | Myc |
| chr8:128682217-128682257 | Myc |
| chr8:128682273-128682313 | Myc |
| chr8:128682329-128682369 | Myc |
| chr8:128682467-128682507 | Myc |
| chr8:128682538-128682578 | Myc |
| chr8:128682655-128682695 | Myc |
| chr8:128682706-128682746 | Myc |
| chr8:128682775-128682815 | Myc |
| chr8:128682843-128682883 | Myc |
| chr8:128683025-128683065 | Myc |
| chr8:128683075-128683115 | Myc |
| chr8:128683134-128683174 | Myc |
| chr8:128683200-128683240 | Myc |
| chr8:128683250-128683290 | Myc |
| chr8:128683301-128683341 | Myc |
| chr8:128683396-128683436 | Myc |
| chr8:128683492-128683532 | Myc |
| chr8:128683584-128683624 | Myc |
| chr8:128683634-128683674 | Myc |
| chr8:128683740-128683780 | Myc |
| chr8:128683851-128683891 | Myc |
| chr8:128684030-128684070 | Myc |
| chr8:128684186-128684226 | Myc |
| chr8:128684259-128684299 | Myc |
| chr8:128684380-128684420 | Myc |
| chr8:128684493-128684533 | Myc |
| chr8:128684548-128684588 | Myc |
| chr8:128684599-128684639 | Myc |
| chr8:128684649-128684689 | Myc |
| chr8:128684729-128684769 | Myc |
| chr8:128684787-128684827 | Myc |
| chr8:128684870-128684910 | Myc |
| chr8:128684980-128685020 | Myc |
| chr8:128685035-128685075 | Myc |
| chr8:128685107-128685147 | Myc |
| chr8:128685254-128685294 | Myc |
| chr8:128685304-128685344 | Myc |
| chr8:128685421-128685461 | Myc |
| chr8:128685515-128685555 | Myc |
| chr8:128685633-128685673 | Myc |
| chr8:128685698-128685738 | Myc |
| chr8:128685778-128685818 | Myc |
| chr8:128685896-128685936 | Myc |
| chr8:128686287-128686327 | Myc |
| chr8:128686357-128686397 | Myc |
| chr8:128686420-128686460 | Myc |

| | |
|---|---|
| chr8:128686542-128686582 | Myc |
| chr8:128686646-128686686 | Myc |
| chr8:128686741-128686781 | Myc |
| chr8:128686799-128686839 | Myc |
| chr8:128686863-128686903 | Myc |
| chr8:128686981-128687021 | Myc |
| chr8:128687034-128687074 | Myc |
| chr8:128687095-128687135 | Myc |
| chr8:128687242-128687282 | Myc |
| chr8:128687359-128687399 | Myc |
| chr8:128687436-128687476 | Myc |
| chr8:128687520-128687560 | Myc |
| chr8:128687640-128687680 | Myc |
| chr8:128687773-128687813 | Myc |
| chr8:128687883-128687923 | Myc |
| chr8:128687962-128688002 | Myc |
| chr8:128688086-128688126 | Myc |
| chr8:128688173-128688213 | Myc |
| chr8:128688243-128688283 | Myc |
| chr8:128688424-128688464 | Myc |
| chr8:128688545-128688585 | Myc |
| chr8:128688617-128688657 | Myc |
| chr8:128688711-128688751 | Myc |
| chr8:128688783-128688823 | Myc |
| chr8:128688861-128688901 | Myc |
| chr8:128688911-128688951 | Myc |
| chr8:128688970-128689010 | Myc |
| chr8:128689044-128689084 | Myc |
| chr8:128689094-128689134 | Myc |
| chr8:128689206-128689246 | Myc |
| chr8:128689318-128689358 | Myc |
| chr8:128689421-128689461 | Myc |
| chr8:128689476-128689516 | Myc |
| chr8:128689582-128689622 | Myc |
| chr8:128689694-128689734 | Myc |
| chr8:128689752-128689792 | Myc |
| chr8:128689863-128689903 | Myc |
| chr8:128689947-128689987 | Myc |
| chr8:128689997-128690037 | Myc |
| chr8:128690103-128690143 | Myc |
| chr8:128690216-128690256 | Myc |
| chr8:128690266-128690306 | Myc |
| chr8:128690352-128690392 | Myc |
| chr8:128690432-128690472 | Myc |
| chr8:128690500-128690540 | Myc |
| chr8:128690553-128690593 | Myc |
| chr8:128690634-128690674 | Myc |
| chr8:128690684-128690724 | Myc |
| chr8:128690788-128690828 | Myc |
| chr8:128690842-128690882 | Myc |
| chr8:128690932-128690972 | Myc |
| chr8:128691023-128691063 | Myc |

| | |
|---|---|
| chr8:128691078-128691118 | Myc |
| chr8:128691160-128691200 | Myc |
| chr8:128691315-128691355 | Myc |
| chr8:128691365-128691405 | Myc |
| chr8:128691480-128691520 | Myc |
| chr8:128691577-128691617 | Myc |
| chr8:128691627-128691667 | Myc |
| chr8:128691717-128691757 | Myc |
| chr8:128691795-128691835 | Myc |
| chr8:128691883-128691923 | Myc |
| chr8:128691940-128691980 | Myc |
| chr8:128692040-128692080 | Myc |
| chr8:128692090-128692130 | Myc |
| chr8:128692199-128692239 | Myc |
| chr8:128692269-128692309 | Myc |
| chr8:128692395-128692435 | Myc |
| chr8:128692487-128692527 | Myc |
| chr8:128692569-128692609 | Myc |
| chr8:128692660-128692700 | Myc |
| chr8:128692716-128692756 | Myc |
| chr8:128692816-128692856 | Myc |
| chr8:128692929-128692969 | Myc |
| chr8:128692982-128693022 | Myc |
| chr8:128693084-128693124 | Myc |
| chr8:128693160-128693200 | Myc |
| chr8:128693240-128693280 | Myc |
| chr8:128693329-128693369 | Myc |
| chr8:128693440-128693480 | Myc |
| chr8:128693490-128693530 | Myc |
| chr8:128693595-128693635 | Myc |
| chr8:128693716-128693756 | Myc |
| chr8:128693768-128693808 | Myc |
| chr8:128693837-128693877 | Myc |
| chr8:128693932-128693972 | Myc |
| chr8:128693984-128694024 | Myc |
| chr8:128694034-128694074 | Myc |
| chr8:128694139-128694179 | Myc |
| chr8:128694260-128694300 | Myc |
| chr8:128694310-128694350 | Myc |
| chr8:128694402-128694442 | Myc |
| chr8:128694454-128694494 | Myc |
| chr8:128694514-128694554 | Myc |
| chr8:128694648-128694688 | Myc |
| chr8:128694797-128694837 | Myc |
| chr8:128694880-128694920 | Myc |
| chr8:128694962-128695002 | Myc |
| chr8:128695109-128695149 | Myc |
| chr8:128695159-128695199 | Myc |
| chr8:128695215-128695255 | Myc |
| chr8:128695345-128695385 | Myc |
| chr8:128695398-128695438 | Myc |
| chr8:128695480-128695520 | Myc |

| | |
|---|---|
| chr8:128695591-128695631 | Myc |
| chr8:128695681-128695721 | Myc |
| chr8:128695761-128695801 | Myc |
| chr8:128696192-128696232 | Myc |
| chr8:128696264-128696304 | Myc |
| chr8:128696344-128696384 | Myc |
| chr8:128696478-128696518 | Myc |
| chr8:128696545-128696585 | Myc |
| chr8:128696600-128696640 | Myc |
| chr8:128696715-128696755 | Myc |
| chr8:128696818-128696858 | Myc |
| chr8:128696930-128696970 | Myc |
| chr8:128697020-128697060 | Myc |
| chr8:128697166-128697206 | Myc |
| chr8:128697221-128697261 | Myc |
| chr8:128697298-128697338 | Myc |
| chr8:128697434-128697474 | Myc |
| chr8:128697484-128697524 | Myc |
| chr8:128697588-128697628 | Myc |
| chr8:128697685-128697725 | Myc |
| chr8:128697831-128697871 | Myc |
| chr8:128697881-128697921 | Myc |
| chr8:128697968-128698008 | Myc |
| chr8:128698086-128698126 | Myc |
| chr8:128698182-128698222 | Myc |
| chr8:128698254-128698294 | Myc |
| chr8:128698338-128698378 | Myc |
| chr8:128698400-128698440 | Myc |
| chr8:128698488-128698528 | Myc |
| chr8:128698539-128698579 | Myc |
| chr8:128698616-128698656 | Myc |
| chr8:128698667-128698707 | Myc |
| chr8:128698753-128698793 | Myc |
| chr8:128698857-128698897 | Myc |
| chr8:128698959-128698999 | Myc |
| chr8:128699167-128699207 | Myc |
| chr8:128699265-128699305 | Myc |
| chr8:128699352-128699392 | Myc |
| chr8:128699419-128699459 | Myc |
| chr8:128699515-128699555 | Myc |
| chr8:128699630-128699670 | Myc |
| chr8:128699688-128699728 | Myc |
| chr8:128699790-128699830 | Myc |
| chr8:128699908-128699948 | Myc |
| chr8:128699962-128700002 | Myc |
| chr8:128700074-128700114 | Myc |
| chr8:128700126-128700166 | Myc |
| chr8:128700210-128700250 | Myc |
| chr8:128700323-128700363 | Myc |
| chr8:128700373-128700413 | Myc |
| chr8:128700453-128700493 | Myc |
| chr8:128700519-128700559 | Myc |

| | |
|---|---|
| chr8:128700592-128700632 | Myc |
| chr8:128700644-128700684 | Myc |
| chr8:128700706-128700746 | Myc |
| chr8:128700892-128700932 | Myc |
| chr8:128701032-128701072 | Myc |
| chr8:128701135-128701175 | Myc |
| chr8:128701190-128701230 | Myc |
| chr8:128701300-128701340 | Myc |
| chr8:128701456-128701496 | Myc |
| chr8:128701507-128701547 | Myc |
| chr8:128701573-128701613 | Myc |
| chr8:128701623-128701663 | Myc |
| chr8:128701721-128701761 | Myc |
| chr8:128701809-128701849 | Myc |
| chr8:128701871-128701911 | Myc |
| chr8:128702002-128702042 | Myc |
| chr8:128702077-128702117 | Myc |
| chr8:128702164-128702204 | Myc |
| chr8:128702426-128702466 | Myc |
| chr8:128702489-128702529 | Myc |
| chr8:128702560-128702600 | Myc |
| chr8:128702660-128702700 | Myc |
| chr8:128702768-128702808 | Myc |
| chr8:128702825-128702865 | Myc |
| chr8:128703024-128703064 | Myc |
| chr8:128703122-128703162 | Myc |
| chr8:128703425-128703465 | Myc |
| chr8:128703532-128703572 | Myc |
| chr8:128703606-128703646 | Myc |
| chr8:128703778-128703818 | Myc |
| chr8:128703853-128703893 | Myc |
| chr8:128703941-128703981 | Myc |
| chr8:128704022-128704062 | Myc |
| chr8:128704094-128704134 | Myc |
| chr8:128704202-128704242 | Myc |
| chr8:128704291-128704331 | Myc |
| chr8:128704463-128704503 | Myc |
| chr8:128704519-128704559 | Myc |
| chr8:128704630-128704670 | Myc |
| chr8:128704705-128704745 | Myc |
| chr8:128704831-128704871 | Myc |
| chr8:128704899-128704939 | Myc |
| chr8:128705090-128705130 | Myc |
| chr8:128705160-128705200 | Myc |
| chr8:128705233-128705273 | Myc |
| chr8:128705339-128705379 | Myc |
| chr8:128705397-128705437 | Myc |
| chr8:128705449-128705489 | Myc |
| chr8:128705505-128705545 | Myc |
| chr8:128705571-128705611 | Myc |
| chr8:128705734-128705774 | Myc |
| chr8:128705791-128705831 | Myc |

| | |
|---|---|
| chr8:128705872-128705912 | Myc |
| chr8:128705924-128705964 | Myc |
| chr8:128705990-128706030 | Myc |
| chr8:128706082-128706122 | Myc |
| chr8:128706176-128706216 | Myc |
| chr8:128706279-128706319 | Myc |
| chr8:128706366-128706406 | Myc |
| chr8:128706528-128706568 | Myc |
| chr8:128706618-128706658 | Myc |
| chr8:128706690-128706730 | Myc |
| chr8:128706769-128706809 | Myc |
| chr8:128706820-128706860 | Myc |
| chr8:128706894-128706934 | Myc |
| chr8:128707005-128707045 | Myc |
| chr8:128707064-128707104 | Myc |
| chr8:128707125-128707165 | Myc |
| chr8:128707214-128707254 | Myc |
| chr8:128707316-128707356 | Myc |
| chr8:128707393-128707433 | Myc |
| chr8:128707443-128707483 | Myc |
| chr8:128707567-128707607 | Myc |
| chr8:128707658-128707698 | Myc |
| chr8:128707708-128707748 | Myc |
| chr8:128707840-128707880 | Myc |
| chr8:128707936-128707976 | Myc |
| chr8:128708155-128708195 | Myc |
| chr8:128708235-128708275 | Myc |
| chr8:128708377-128708417 | Myc |
| chr8:128708447-128708487 | Myc |
| chr8:128708509-128708549 | Myc |
| chr8:128708621-128708661 | Myc |
| chr8:128708773-128708813 | Myc |
| chr8:128708905-128708945 | Myc |
| chr8:128709072-128709112 | Myc |
| chr8:128709162-128709202 | Myc |
| chr8:128709212-128709252 | Myc |
| chr8:128709270-128709310 | Myc |
| chr8:128709334-128709374 | Myc |
| chr8:128709419-128709459 | Myc |
| chr8:128709504-128709544 | Myc |
| chr8:128709554-128709594 | Myc |
| chr8:128709606-128709646 | Myc |
| chr8:128709785-128709825 | Myc |
| chr8:128709844-128709884 | Myc |
| chr8:128709976-128710016 | Myc |
| chr8:128710099-128710139 | Myc |
| chr8:128710258-128710298 | Myc |
| chr8:128710364-128710404 | Myc |
| chr8:128710502-128710542 | Myc |
| chr8:128710552-128710592 | Myc |
| chr8:128710602-128710642 | Myc |
| chr8:128710693-128710733 | Myc |

| | |
|---|---|
| chr8:128710791-128710831 | Myc |
| chr8:128710841-128710881 | Myc |
| chr8:128710950-128710990 | Myc |
| chr8:128711011-128711051 | Myc |
| chr8:128711061-128711101 | Myc |
| chr8:128711139-128711179 | Myc |
| chr8:128711219-128711259 | Myc |
| chr8:128711343-128711383 | Myc |
| chr8:128711393-128711433 | Myc |
| chr8:128711504-128711544 | Myc |
| chr8:128711584-128711624 | Myc |
| chr8:128711658-128711698 | Myc |
| chr8:128711821-128711861 | Myc |
| chr8:128711874-128711914 | Myc |
| chr8:128711988-128712028 | Myc |
| chr8:128712112-128712152 | Myc |
| chr8:128712180-128712220 | Myc |
| chr8:128712239-128712279 | Myc |
| chr8:128712302-128712342 | Myc |
| chr8:128712361-128712401 | Myc |
| chr8:128712562-128712602 | Myc |
| chr8:128712617-128712657 | Myc |
| chr8:128712706-128712746 | Myc |
| chr8:128712764-128712804 | Myc |
| chr8:128712841-128712881 | Myc |
| chr8:128712946-128712986 | Myc |
| chr8:128713031-128713071 | Myc |
| chr8:128713119-128713159 | Myc |
| chr8:128713257-128713297 | Myc |
| chr8:128713378-128713418 | Myc |
| chr8:128713486-128713526 | Myc |
| chr8:128713536-128713576 | Myc |
| chr8:128713622-128713662 | Myc |
| chr8:128713748-128713788 | Myc |
| chr8:128713831-128713871 | Myc |
| chr8:128713885-128713925 | Myc |
| chr8:128713948-128713988 | Myc |
| chr8:128713999-128714039 | Myc |
| chr8:128714117-128714157 | Myc |
| chr8:128714206-128714246 | Myc |
| chr8:128714292-128714332 | Myc |
| chr8:128714425-128714465 | Myc |
| chr8:128714505-128714545 | Myc |
| chr8:128714569-128714609 | Myc |
| chr8:128714625-128714665 | Myc |
| chr8:128714723-128714763 | Myc |
| chr8:128714773-128714813 | Myc |
| chr8:128714827-128714867 | Myc |
| chr8:128714895-128714935 | Myc |
| chr8:128714947-128714987 | Myc |
| chr8:128715055-128715095 | Myc |
| chr8:128715256-128715296 | Myc |

| | |
|---|---|
| chr8:128715306-128715346 | Myc |
| chr8:128715384-128715424 | Myc |
| chr8:128715450-128715490 | Myc |
| chr8:128715678-128715718 | Myc |
| chr8:128715732-128715772 | Myc |
| chr8:128715809-128715849 | Myc |
| chr8:128716124-128716164 | Myc |
| chr8:128716201-128716241 | Myc |
| chr8:128716251-128716291 | Myc |
| chr8:128716350-128716390 | Myc |
| chr8:128716437-128716477 | Myc |
| chr8:128716532-128716572 | Myc |
| chr8:128716582-128716622 | Myc |
| chr8:128716670-128716710 | Myc |
| chr8:128716737-128716777 | Myc |
| chr8:128716856-128716896 | Myc |
| chr8:128716982-128717022 | Myc |
| chr8:128717045-128717085 | Myc |
| chr8:128717256-128717296 | Myc |
| chr8:128717337-128717377 | Myc |
| chr8:128717387-128717427 | Myc |
| chr8:128717506-128717546 | Myc |
| chr8:128717802-128717842 | Myc |
| chr8:128717877-128717917 | Myc |
| chr8:128718040-128718080 | Myc |
| chr8:128718102-128718142 | Myc |
| chr8:128718307-128718347 | Myc |
| chr8:128718439-128718479 | Myc |
| chr8:128718523-128718563 | Myc |
| chr8:128718578-128718618 | Myc |
| chr8:128718672-128718712 | Myc |
| chr8:128718726-128718766 | Myc |
| chr8:128719089-128719129 | Myc |
| chr8:128719139-128719179 | Myc |
| chr8:128719250-128719290 | Myc |
| chr8:128719301-128719341 | Myc |
| chr8:128719398-128719438 | Myc |
| chr8:128719516-128719556 | Myc |
| chr8:128719661-128719701 | Myc |
| chr8:128719712-128719752 | Myc |
| chr8:128719772-128719812 | Myc |
| chr8:128719823-128719863 | Myc |
| chr8:128719885-128719925 | Myc |
| chr8:128719985-128720025 | Myc |
| chr8:128720119-128720159 | Myc |
| chr8:128720266-128720306 | Myc |
| chr8:128720318-128720358 | Myc |
| chr8:128720392-128720432 | Myc |
| chr8:128720530-128720570 | Myc |
| chr8:128720597-128720637 | Myc |
| chr8:128720655-128720695 | Myc |
| chr8:128720705-128720745 | Myc |

| | |
|---|---|
| chr8:128720822-128720862 | Myc |
| chr8:128720877-128720917 | Myc |
| chr8:128721010-128721050 | Myc |
| chr8:128721060-128721100 | Myc |
| chr8:128721110-128721150 | Myc |
| chr8:128721254-128721294 | Myc |
| chr8:128721515-128721555 | Myc |
| chr8:128721621-128721661 | Myc |
| chr8:128721682-128721722 | Myc |
| chr8:128721732-128721772 | Myc |
| chr8:128721816-128721856 | Myc |
| chr8:128721914-128721954 | Myc |
| chr8:128721986-128722026 | Myc |
| chr8:128722070-128722110 | Myc |
| chr8:128722183-128722223 | Myc |
| chr8:128722238-128722278 | Myc |
| chr8:128722372-128722412 | Myc |
| chr8:128722431-128722471 | Myc |
| chr8:128722551-128722591 | Myc |
| chr8:128722610-128722650 | Myc |
| chr8:128722699-128722739 | Myc |
| chr8:128722792-128722832 | Myc |
| chr8:128722890-128722930 | Myc |
| chr8:128722951-128722991 | Myc |
| chr8:128723009-128723049 | Myc |
| chr8:128723094-128723134 | Myc |
| chr8:128723233-128723273 | Myc |
| chr8:128723332-128723372 | Myc |
| chr8:128723388-128723428 | Myc |
| chr8:128723669-128723709 | Myc |
| chr8:128723737-128723777 | Myc |
| chr8:128723859-128723899 | Myc |
| chr8:128724054-128724094 | Myc |
| chr8:128724128-128724168 | Myc |
| chr8:128724216-128724256 | Myc |
| chr8:128724273-128724313 | Myc |
| chr8:128724457-128724497 | Myc |
| chr8:128724557-128724597 | Myc |
| chr8:128724697-128724737 | Myc |
| chr8:128724769-128724809 | Myc |
| chr8:128724919-128724959 | Myc |
| chr8:128725020-128725060 | Myc |
| chr8:128725125-128725165 | Myc |
| chr8:128725184-128725224 | Myc |
| chr8:128725309-128725349 | Myc |
| chr8:128725368-128725408 | Myc |
| chr8:128725546-128725586 | Myc |
| chr8:128725664-128725704 | Myc |
| chr8:128725734-128725774 | Myc |
| chr8:128725786-128725826 | Myc |
| chr8:128725897-128725937 | Myc |
| chr8:128725984-128726024 | Myc |

| | |
|---|---|
| chr8:128726042-128726082 | Myc |
| chr8:128726121-128726161 | Myc |
| chr8:128726212-128726252 | Myc |
| chr8:128726309-128726349 | Myc |
| chr8:128726537-128726577 | Myc |
| chr8:128726587-128726627 | Myc |
| chr8:128726720-128726760 | Myc |
| chr8:128726843-128726883 | Myc |
| chr8:128726988-128727028 | Myc |
| chr8:128727038-128727078 | Myc |
| chr8:128727130-128727170 | Myc |
| chr8:128727236-128727276 | Myc |
| chr8:128727323-128727363 | Myc |
| chr8:128727511-128727551 | Myc |
| chr8:128727598-128727638 | Myc |
| chr8:128727667-128727707 | Myc |
| chr8:128727801-128727841 | Myc |
| chr8:128727896-128727936 | Myc |
| chr8:128727983-128728023 | Myc |
| chr8:128728053-128728093 | Myc |
| chr8:128728132-128728172 | Myc |
| chr8:128728241-128728281 | Myc |
| chr8:128728342-128728382 | Myc |
| chr8:128728392-128728432 | Myc |
| chr8:128728469-128728509 | Myc |
| chr8:128728523-128728563 | Myc |
| chr8:128728647-128728687 | Myc |
| chr8:128728740-128728780 | Myc |
| chr8:128728912-128728952 | Myc |
| chr8:128728965-128729005 | Myc |
| chr8:128729050-128729090 | Myc |
| chr8:128729100-128729140 | Myc |
| chr8:128729199-128729239 | Myc |
| chr8:128729250-128729290 | Myc |
| chr8:128729321-128729361 | Myc |
| chr8:128729482-128729522 | Myc |
| chr8:128729544-128729584 | Myc |
| chr8:128729630-128729670 | Myc |
| chr8:128729731-128729771 | Myc |
| chr8:128729915-128729955 | Myc |
| chr8:128730002-128730042 | Myc |
| chr8:128730222-128730262 | Myc |
| chr8:128730272-128730312 | Myc |
| chr8:128730420-128730460 | Myc |
| chr8:128762795-128762835 | Myc |
| chr8:128762908-128762948 | Myc |
| chr8:128762962-128763002 | Myc |
| chr8:128763013-128763053 | Myc |
| chr8:128763098-128763138 | Myc |
| chr8:128763240-128763280 | Myc |
| chr8:128763312-128763352 | Myc |
| chr8:128763379-128763419 | Myc |

| | |
|---|---|
| chr8:128763490-128763530 | Myc |
| chr8:128763556-128763596 | Myc |
| chr8:128763621-128763661 | Myc |
| chr8:128763698-128763738 | Myc |
| chr8:128763755-128763795 | Myc |
| chr8:128763815-128763855 | Myc |
| chr8:128763911-128763951 | Myc |
| chr8:128763973-128764013 | Myc |
| chr8:128764116-128764156 | Myc |
| chr8:128764177-128764217 | Myc |
| chr8:128764302-128764342 | Myc |
| chr8:128764409-128764449 | Myc |
| chr8:128764470-128764510 | Myc |
| chr8:128764525-128764565 | Myc |
| chr8:128764612-128764652 | Myc |
| chr8:128764706-128764746 | Myc |
| chr8:128764908-128764948 | Myc |
| chr8:128764962-128765002 | Myc |
| chr8:128765033-128765073 | Myc |
| chr8:128765128-128765168 | Myc |
| chr8:128765207-128765247 | Myc |
| chr8:128765324-128765364 | Myc |
| chr8:128765384-128765424 | Myc |
| chr8:128765563-128765603 | Myc |
| chr8:128765726-128765766 | Myc |
| chr8:128765819-128765859 | Myc |
| chr8:128765870-128765910 | Myc |
| chr8:128765938-128765978 | Myc |
| chr8:128765989-128766029 | Myc |
| chr8:128766053-128766093 | Myc |
| chr8:128766103-128766143 | Myc |
| chr8:128766230-128766270 | Myc |
| chr8:128766307-128766347 | Myc |
| chr8:128766393-128766433 | Myc |
| chr8:128766526-128766566 | Myc |
| chr8:128766592-128766632 | Myc |
| chr8:128766643-128766683 | Myc |
| chr8:128766707-128766747 | Myc |
| chr8:128766803-128766843 | Myc |
| chr8:128766869-128766909 | Myc |
| chr8:128766955-128766995 | Myc |
| chr8:128767011-128767051 | Myc |
| chr8:128767061-128767101 | Myc |
| chr8:128767207-128767247 | Myc |
| chr8:128767415-128767455 | Myc |
| chr8:128767527-128767567 | Myc |
| chr8:128767611-128767651 | Myc |
| chr8:128767664-128767704 | Myc |
| chr8:128767717-128767757 | Myc |
| chr8:128768028-128768068 | Myc |
| chr8:128768088-128768128 | Myc |
| chr8:128768163-128768203 | Myc |

| | |
|---|---|
| chr8:128768219-128768259 | Myc |
| chr8:128768308-128768348 | Myc |
| chr8:128768372-128768412 | Myc |
| chr8:128768473-128768513 | Myc |
| chr8:128768562-128768602 | Myc |
| chr8:128768612-128768652 | Myc |
| chr8:128768734-128768774 | Myc |
| chr8:128768981-128769021 | Myc |
| chr8:128769063-128769103 | Myc |
| chr8:128769122-128769162 | Myc |
| chr8:128769185-128769225 | Myc |
| chr8:128769263-128769303 | Myc |
| chr8:128769316-128769356 | Myc |
| chr8:128769396-128769436 | Myc |
| chr8:128769513-128769553 | Myc |
| chr8:128769629-128769669 | Myc |
| chr8:128770040-128770080 | Myc |
| chr8:128770105-128770145 | Myc |
| chr8:128770183-128770223 | Myc |
| chr8:128770254-128770294 | Myc |
| chr8:128770327-128770367 | Myc |
| chr8:128770377-128770417 | Myc |
| chr8:128770430-128770470 | Myc |
| chr8:128770528-128770568 | Myc |
| chr8:128770582-128770622 | Myc |
| chr8:128770774-128770814 | Myc |
| chr8:128770922-128770962 | Myc |
| chr8:128771001-128771041 | Myc |
| chr8:128771099-128771139 | Myc |
| chr8:128771214-128771254 | Myc |
| chr8:128771366-128771406 | Myc |
| chr8:128771462-128771502 | Myc |
| chr8:128771558-128771598 | Myc |
| chr8:128771734-128771774 | Myc |
| chr8:128771792-128771832 | Myc |
| chr8:128771863-128771903 | Myc |
| chr8:128771919-128771959 | Myc |
| chr8:128771978-128772018 | Myc |
| chr8:128772031-128772071 | Myc |
| chr8:128772111-128772151 | Myc |
| chr8:128772169-128772209 | Myc |
| chr8:128772230-128772270 | Myc |
| chr8:128772284-128772324 | Myc |
| chr8:128772402-128772442 | Myc |
| chr8:128772470-128772510 | Myc |
| chr8:128772560-128772600 | Myc |
| chr8:128772644-128772684 | Myc |
| chr8:128772742-128772782 | Myc |
| chr8:128772827-128772867 | Myc |
| chr8:128772968-128773008 | Myc |
| chr8:128773095-128773135 | Myc |
| chr8:128773157-128773197 | Myc |

| | |
|---|---|
| chr8:128773333-128773373 | Myc |
| chr8:128773483-128773523 | Myc |
| chr8:128773892-128773932 | Myc |
| chr8:128774020-128774060 | Myc |
| chr8:128774093-128774133 | Myc |
| chr8:128774181-128774221 | Myc |
| chr8:128774313-128774353 | Myc |
| chr8:128774376-128774416 | Myc |
| chr8:128774528-128774568 | Myc |
| chr8:128774661-128774701 | Myc |
| chr8:128774711-128774751 | Myc |
| chr8:128774794-128774834 | Myc |
| chr8:128774844-128774884 | Myc |
| chr8:128774905-128774945 | Myc |
| chr8:128775007-128775047 | Myc |
| chr8:128775127-128775167 | Myc |
| chr8:128775233-128775273 | Myc |
| chr8:128775458-128775498 | Myc |
| chr8:128775633-128775673 | Myc |
| chr8:128775690-128775730 | Myc |
| chr8:128775763-128775803 | Myc |
| chr8:128775815-128775855 | Myc |
| chr8:128775979-128776019 | Myc |
| chr8:128776035-128776075 | Myc |
| chr8:128776115-128776155 | Myc |
| chr8:128776229-128776269 | Myc |
| chr8:128776286-128776326 | Myc |
| chr8:128776348-128776388 | Myc |
| chr8:128776451-128776491 | Myc |
| chr8:128776535-128776575 | Myc |
| chr8:128776669-128776709 | Myc |
| chr8:128776895-128776935 | Myc |
| chr8:128776985-128777025 | Myc |
| chr8:128777061-128777101 | Myc |
| chr8:128777111-128777151 | Myc |
| chr8:128777170-128777210 | Myc |
| chr8:128777317-128777357 | Myc |
| chr8:128777483-128777523 | Myc |
| chr8:128777555-128777595 | Myc |
| chr8:128777791-128777831 | Myc |
| chr8:128777901-128777941 | Myc |
| chr8:128778009-128778049 | Myc |
| chr8:128778064-128778104 | Myc |
| chr8:128778151-128778191 | Myc |
| chr8:128778245-128778285 | Myc |
| chr8:128778360-128778400 | Myc |
| chr8:128778425-128778465 | Myc |
| chr8:128778511-128778551 | Myc |
| chr8:128778616-128778656 | Myc |
| chr8:128778680-128778720 | Myc |
| chr8:128778791-128778831 | Myc |
| chr8:128778841-128778881 | Myc |

| | |
|---|---|
| chr8:128778903-128778943 | Myc |
| chr8:128778962-128779002 | Myc |
| chr8:128779040-128779080 | Myc |
| chr8:128779167-128779207 | Myc |
| chr8:128779218-128779258 | Myc |
| chr8:128779326-128779366 | Myc |
| chr8:128779466-128779506 | Myc |
| chr8:128779589-128779629 | Myc |
| chr8:128779641-128779681 | Myc |
| chr8:128779747-128779787 | Myc |
| chr8:128779901-128779941 | Myc |
| chr8:128779959-128779999 | Myc |
| chr8:128780039-128780079 | Myc |
| chr8:128780166-128780206 | Myc |
| chr8:128780275-128780315 | Myc |
| chr8:128780370-128780410 | Myc |
| chr8:128780447-128780487 | Myc |
| chr8:128780513-128780553 | Myc |
| chr8:128780587-128780627 | Myc |
| chr8:128780680-128780720 | Myc |
| chr8:128780743-128780783 | Myc |
| chr8:128781048-128781088 | Myc |
| chr8:128781124-128781164 | Myc |
| chr8:128781238-128781278 | Myc |
| chr8:128781314-128781354 | Myc |
| chr8:128781716-128781756 | Myc |
| chr8:128782085-128782125 | Myc |
| chr8:128782974-128783014 | Myc |
| chr8:128783090-128783130 | Myc |
| chr8:128783174-128783214 | Myc |
| chr8:128783683-128783723 | Myc |
| chr8:128784276-128784316 | Myc |
| chr8:128784343-128784383 | Myc |
| chr8:128784395-128784435 | Myc |
| chr8:128784482-128784522 | Myc |
| chr8:128784588-128784628 | Myc |
| chr8:128784649-128784689 | Myc |
| chr8:128784778-128784818 | Myc |
| chr8:128784857-128784897 | Myc |
| chr8:128784942-128784982 | Myc |
| chr8:128785094-128785134 | Myc |
| chr8:128785148-128785188 | Myc |
| chr8:128785218-128785258 | Myc |
| chr8:128785276-128785316 | Myc |
| chr8:128785411-128785451 | Myc |
| chr8:128785471-128785511 | Myc |
| chr8:128785554-128785594 | Myc |
| chr8:128785615-128785655 | Myc |
| chr8:128785672-128785712 | Myc |
| chr8:128785957-128785997 | Myc |
| chr8:128786306-128786346 | Myc |
| chr8:128786392-128786432 | Myc |

| | |
|---|---|
| chr8:128786458-128786498 | Myc |
| chr8:128786509-128786549 | Myc |
| chr8:128786576-128786616 | Myc |
| chr8:128786669-128786709 | Myc |
| chr8:128786751-128786791 | Myc |
| chr8:128786801-128786841 | Myc |
| chr8:128786954-128786994 | Myc |
| chr8:128787011-128787051 | Myc |
| chr8:128787101-128787141 | Myc |
| chr8:128787215-128787255 | Myc |
| chr8:128787287-128787327 | Myc |
| chr8:128787397-128787437 | Myc |
| chr8:128787472-128787512 | Myc |
| chr8:128787553-128787593 | Myc |
| chr8:128787635-128787675 | Myc |
| chr8:128787704-128787744 | Myc |
| chr8:128787934-128787974 | Myc |
| chr8:128788092-128788132 | Myc |
| chr8:128788142-128788182 | Myc |
| chr8:128788204-128788244 | Myc |
| chr8:128788390-128788430 | Myc |
| chr8:128788440-128788480 | Myc |
| chr8:128788535-128788575 | Myc |
| chr8:128788599-128788639 | Myc |
| chr8:128788688-128788728 | Myc |
| chr8:128788766-128788806 | Myc |
| chr8:128788878-128788918 | Myc |
| chr8:128788947-128788987 | Myc |
| chr8:128789016-128789056 | Myc |
| chr8:128789136-128789176 | Myc |
| chr8:128789250-128789290 | Myc |
| chr8:128789300-128789340 | Myc |
| chr8:128789429-128789469 | Myc |
| chr8:128790011-128790051 | Myc |
| chr8:128790070-128790110 | Myc |
| chr8:128790174-128790214 | Myc |
| chr8:128790227-128790267 | Myc |
| chr8:128790311-128790351 | Myc |
| chr8:128790391-128790431 | Myc |
| chr8:128790452-128790492 | Myc |
| chr8:128790598-128790638 | Myc |
| chr8:128790660-128790700 | Myc |
| chr8:128790847-128790887 | Myc |
| chr8:128790915-128790955 | Myc |
| chr8:128790966-128791006 | Myc |
| chr8:128791165-128791205 | Myc |
| chr8:128791274-128791314 | Myc |
| chr8:128791360-128791400 | Myc |
| chr8:128791417-128791457 | Myc |
| chr8:128791543-128791583 | Myc |
| chr8:128791746-128791786 | Myc |
| chr8:128791807-128791847 | Myc |

| | |
|---|---|
| chr8:128791933-128791973 | Myc |
| chr8:128792000-128792040 | Myc |
| chr8:128792091-128792131 | Myc |
| chr8:128792196-128792236 | Myc |
| chr8:128792246-128792286 | Myc |
| chr8:128792333-128792373 | Myc |
| chr8:128792386-128792426 | Myc |
| chr8:128792497-128792537 | Myc |
| chr8:128792566-128792606 | Myc |
| chr8:128792707-128792747 | Myc |
| chr8:128792821-128792861 | Myc |
| chr8:128792872-128792912 | Myc |
| chr8:128792960-128793000 | Myc |
| chr8:128793113-128793153 | Myc |
| chr8:128793521-128793561 | Myc |
| chr8:128793719-128793759 | Myc |
| chr8:128793913-128793953 | Myc |
| chr8:128794080-128794120 | Myc |
| chr8:128794190-128794230 | Myc |
| chr8:128794244-128794284 | Myc |
| chr8:128794294-128794334 | Myc |
| chr8:128794417-128794457 | Myc |
| chr8:128794467-128794507 | Myc |
| chr8:128794555-128794595 | Myc |
| chr8:128794705-128794745 | Myc |
| chr8:128794760-128794800 | Myc |
| chr8:128794882-128794922 | Myc |
| chr8:128794955-128794995 | Myc |
| chr8:128795070-128795110 | Myc |
| chr8:128795123-128795163 | Myc |
| chr8:128795173-128795213 | Myc |
| chr8:128795228-128795268 | Myc |
| chr8:128795322-128795362 | Myc |
| chr8:128795465-128795505 | Myc |
| chr8:128795574-128795614 | Myc |
| chr8:128795677-128795717 | Myc |
| chr8:128795800-128795840 | Myc |
| chr8:128795917-128795957 | Myc |
| chr8:128795972-128796012 | Myc |
| chr8:128796022-128796062 | Myc |
| chr8:128796091-128796131 | Myc |
| chr8:128796198-128796238 | Myc |
| chr8:128796365-128796405 | Myc |
| chr8:128796438-128796478 | Myc |
| chr8:128796557-128796597 | Myc |
| chr8:128796666-128796706 | Myc |
| chr8:128796733-128796773 | Myc |
| chr8:128796802-128796842 | Myc |
| chr8:128796975-128797015 | Myc |
| chr8:128797055-128797095 | Myc |
| chr8:128797152-128797192 | Myc |
| chr8:128797333-128797373 | Myc |

| | |
|---|---|
| chr8:128797419-128797459 | Myc |
| chr8:128797507-128797547 | Myc |
| chr8:128797563-128797603 | Myc |
| chr8:128797631-128797671 | Myc |
| chr8:128797723-128797763 | Myc |
| chr8:128797773-128797813 | Myc |
| chr8:128797859-128797899 | Myc |
| chr8:128797978-128798018 | Myc |
| chr8:128798075-128798115 | Myc |
| chr8:128798195-128798235 | Myc |
| chr8:128798247-128798287 | Myc |
| chr8:128798334-128798374 | Myc |
| chr8:128798495-128798535 | Myc |
| chr8:128798560-128798600 | Myc |
| chr8:128798611-128798651 | Myc |
| chr8:128798760-128798800 | Myc |
| chr8:128798950-128798990 | Myc |
| chr8:128799010-128799050 | Myc |
| chr8:128799060-128799100 | Myc |
| chr8:128799197-128799237 | Myc |
| chr8:128799359-128799399 | Myc |
| chr8:128799548-128799588 | Myc |
| chr8:128799659-128799699 | Myc |
| chr8:128799723-128799763 | Myc |
| chr8:128799840-128799880 | Myc |
| chr8:128799892-128799932 | Myc |
| chr8:128800012-128800052 | Myc |
| chr8:128800273-128800313 | Myc |
| chr8:128800411-128800451 | Myc |
| chr8:128800537-128800577 | Myc |
| chr8:128800673-128800713 | Myc |
| chr8:128800842-128800882 | Myc |
| chr8:128800928-128800968 | Myc |
| chr8:128800997-128801037 | Myc |
| chr8:128801083-128801123 | Myc |
| chr8:128801151-128801191 | Myc |
| chr8:128801203-128801243 | Myc |
| chr8:128801280-128801320 | Myc |
| chr8:128801376-128801416 | Myc |
| chr8:128801480-128801520 | Myc |
| chr8:128801677-128801717 | Myc |
| chr8:128801739-128801779 | Myc |
| chr8:128801811-128801851 | Myc |
| chr8:128801941-128801981 | Myc |
| chr8:128802003-128802043 | Myc |
| chr8:128802061-128802101 | Myc |
| chr8:128802126-128802166 | Myc |
| chr8:128802176-128802216 | Myc |
| chr8:128802249-128802289 | Myc |
| chr8:128802310-128802350 | Myc |
| chr8:128802360-128802400 | Myc |
| chr8:128802427-128802467 | Myc |

| | |
|---|---|
| chr8:128802533-128802573 | Myc |
| chr8:128802692-128802732 | Myc |
| chr8:128802758-128802798 | Myc |
| chr8:128802900-128802940 | Myc |
| chr8:128802985-128803025 | Myc |
| chr8:128803096-128803136 | Myc |
| chr8:128803218-128803258 | Myc |
| chr8:128803291-128803331 | Myc |
| chr8:128803408-128803448 | Myc |
| chr8:128803468-128803508 | Myc |
| chr8:128803566-128803606 | Myc |
| chr8:128803670-128803710 | Myc |
| chr8:128803760-128803800 | Myc |
| chr8:128803820-128803860 | Myc |
| chr8:128804034-128804074 | Myc |
| chr8:128804132-128804172 | Myc |
| chr8:128804222-128804262 | Myc |
| chr8:128804458-128804498 | Myc |
| chr8:128804723-128804763 | Myc |
| chr8:128804777-128804817 | Myc |
| chr8:128804861-128804901 | Myc |
| chr8:128804960-128805000 | Myc |
| chr8:128805054-128805094 | Myc |
| chr8:128805146-128805186 | Myc |
| chr8:128805204-128805244 | Myc |
| chr8:128805393-128805433 | Myc |
| chr8:128805516-128805556 | Myc |
| chr8:128805566-128805606 | Myc |
| chr8:128805628-128805668 | Myc |
| chr8:128805919-128805959 | Myc |
| chr8:128805989-128806029 | Myc |
| chr8:128806069-128806109 | Myc |
| chr8:128806146-128806186 | Myc |
| chr8:128806224-128806264 | Myc |
| chr8:128806287-128806327 | Myc |
| chr8:128806351-128806391 | Myc |
| chr8:128806474-128806514 | Myc |
| chr8:128806852-128806892 | Myc |
| chr8:128806936-128806976 | Myc |
| chr8:128807036-128807076 | Myc |
| chr8:128807136-128807176 | Myc |
| chr8:128807187-128807227 | Myc |
| chr8:128807278-128807318 | Myc |
| chr8:128807359-128807399 | Myc |
| chr8:128807458-128807498 | Myc |
| chr8:128807517-128807557 | Myc |
| chr8:128807606-128807646 | Myc |
| chr8:128807656-128807696 | Myc |
| chr8:128807717-128807757 | Myc |
| chr8:128807837-128807877 | Myc |
| chr8:128807904-128807944 | Myc |
| chr8:128808025-128808065 | Myc |

| | |
|---|---|
| chr8:128808085-128808125 | Myc |
| chr8:128808187-128808227 | Myc |
| chr8:128808257-128808297 | Myc |
| chr8:128808307-128808347 | Myc |
| chr8:128808357-128808397 | Myc |
| chr8:128808419-128808459 | Myc |
| chr8:128808542-128808582 | Myc |
| chr8:128808592-128808632 | Myc |
| chr8:128808655-128808695 | Myc |
| chr8:128808771-128808811 | Myc |
| chr8:128808864-128808904 | Myc |
| chr8:128808931-128808971 | Myc |
| chr8:128808981-128809021 | Myc |
| chr8:128809031-128809071 | Myc |
| chr8:128809084-128809124 | Myc |
| chr8:128809474-128809514 | Myc |
| chr8:128809535-128809575 | Myc |
| chr8:128809608-128809648 | Myc |
| chr8:128809658-128809698 | Myc |
| chr8:128809708-128809748 | Myc |
| chr8:128809807-128809847 | Myc |
| chr8:128809948-128809988 | Myc |
| chr8:128810002-128810042 | Myc |
| chr8:128810101-128810141 | Myc |
| chr8:128810165-128810205 | Myc |
| chr8:128810247-128810287 | Myc |
| chr8:128810480-128810520 | Myc |
| chr8:128810593-128810633 | Myc |
| chr8:128810673-128810713 | Myc |
| chr8:128810747-128810787 | Myc |
| chr8:128810797-128810837 | Myc |
| chr8:128811060-128811100 | Myc |
| chr8:128811168-128811208 | Myc |
| chr8:128811244-128811284 | Myc |
| chr8:128811345-128811385 | Myc |
| chr8:128811464-128811504 | Myc |
| chr8:128811524-128811564 | Myc |
| chr8:128811614-128811654 | Myc |
| chr8:128811739-128811779 | Myc |
| chr8:128811816-128811856 | Myc |
| chr8:128811952-128811992 | Myc |
| chr8:128812048-128812088 | Myc |
| chr8:128812126-128812166 | Myc |
| chr8:128812202-128812242 | Myc |
| chr8:128812280-128812320 | Myc |
| chr8:128812337-128812377 | Myc |
| chr8:128812516-128812556 | Myc |
| chr8:128812586-128812626 | Myc |
| chr8:128812646-128812686 | Myc |
| chr8:128812720-128812760 | Myc |
| chr8:128812814-128812854 | Myc |
| chr8:128812879-128812919 | Myc |

| | |
|---|---|
| chr8:128812960-128813000 | Myc |
| chr8:128813018-128813058 | Myc |
| chr8:128813135-128813175 | Myc |
| chr8:128813187-128813227 | Myc |
| chr8:128813237-128813277 | Myc |
| chr8:128813301-128813341 | Myc |
| chr8:128813357-128813397 | Myc |
| chr8:128813436-128813476 | Myc |
| chr8:128813490-128813530 | Myc |
| chr8:128813572-128813612 | Myc |
| chr8:128813764-128813804 | Myc |
| chr8:128813818-128813858 | Myc |
| chr8:128813927-128813967 | Myc |
| chr8:128814007-128814047 | Myc |
| chr8:128814082-128814122 | Myc |
| chr8:128814391-128814431 | Myc |
| chr8:128814441-128814481 | Myc |
| chr8:128814513-128814553 | Myc |
| chr8:128814578-128814618 | Myc |
| chr8:128814681-128814721 | Myc |
| chr8:128814742-128814782 | Myc |
| chr8:128814920-128814960 | Myc |
| chr8:128815009-128815049 | Myc |
| chr8:128815110-128815150 | Myc |
| chr8:128815203-128815243 | Myc |
| chr8:128815270-128815310 | Myc |
| chr8:128815320-128815360 | Myc |
| chr8:128815506-128815546 | Myc |
| chr8:128815666-128815706 | Myc |
| chr8:128815721-128815761 | Myc |
| chr8:128815834-128815874 | Myc |
| chr8:128815903-128815943 | Myc |
| chr8:128816083-128816123 | Myc |
| chr8:128816291-128816331 | Myc |
| chr8:128816352-128816392 | Myc |
| chr8:128816473-128816513 | Myc |
| chr8:128816607-128816647 | Myc |
| chr8:128816700-128816740 | Myc |
| chr8:128816801-128816841 | Myc |
| chr8:128816897-128816937 | Myc |
| chr8:128817030-128817070 | Myc |
| chr8:128817129-128817169 | Myc |
| chr8:128817451-128817491 | Myc |
| chr8:128817537-128817577 | Myc |
| chr8:128817621-128817661 | Myc |
| chr8:128817704-128817744 | Myc |
| chr8:128817880-128817920 | Myc |
| chr8:128817947-128817987 | Myc |
| chr8:128818102-128818142 | Myc |
| chr8:128818154-128818194 | Myc |
| chr8:128818219-128818259 | Myc |
| chr8:128818403-128818443 | Myc |

| | |
|---|---|
| chr8:128818664-128818704 | Myc |
| chr8:128818761-128818801 | Myc |
| chr8:128818921-128818961 | Myc |
| chr8:128818973-128819013 | Myc |
| chr8:128819081-128819121 | Myc |
| chr8:128819136-128819176 | Myc |
| chr8:128819280-128819320 | Myc |
| chr8:128819336-128819376 | Myc |
| chr8:128819388-128819428 | Myc |
| chr8:128819483-128819523 | Myc |
| chr8:128819619-128819659 | Myc |
| chr8:128819716-128819756 | Myc |
| chr8:128819797-128819837 | Myc |
| chr8:128819893-128819933 | Myc |
| chr8:128819960-128820000 | Myc |
| chr8:128820148-128820188 | Myc |
| chr8:128820297-128820337 | Myc |
| chr8:128820407-128820447 | Myc |
| chr8:128820572-128820612 | Myc |
| chr8:128820653-128820693 | Myc |
| chr8:128820708-128820748 | Myc |
| chr8:128820845-128820885 | Myc |
| chr8:128820911-128820951 | Myc |
| chr8:128820966-128821006 | Myc |
| chr8:128821218-128821258 | Myc |
| chr8:128821335-128821375 | Myc |
| chr8:128821385-128821425 | Myc |
| chr8:128821515-128821555 | Myc |
| chr8:128821581-128821621 | Myc |
| chr8:128821661-128821701 | Myc |
| chr8:128821724-128821764 | Myc |
| chr8:128821791-128821831 | Myc |
| chr8:128821860-128821900 | Myc |
| chr8:128821968-128822008 | Myc |
| chr8:128822062-128822102 | Myc |
| chr8:128822186-128822226 | Myc |
| chr8:128822237-128822277 | Myc |
| chr8:128822333-128822373 | Myc |
| chr8:128822414-128822454 | Myc |
| chr8:128822505-128822545 | Myc |
| chr8:128822563-128822603 | Myc |
| chr8:128822663-128822703 | Myc |
| chr8:128822733-128822773 | Myc |
| chr8:128822783-128822823 | Myc |
| chr8:128822845-128822885 | Myc |
| chr8:128822990-128823030 | Myc |
| chr8:128823116-128823156 | Myc |
| chr8:128823254-128823294 | Myc |
| chr8:128823308-128823348 | Myc |
| chr8:128823395-128823435 | Myc |
| chr8:128823445-128823485 | Myc |
| chr8:128823599-128823639 | Myc |

| | |
|---|---|
| chr8:128823696-128823736 | Myc |
| chr8:128823756-128823796 | Myc |
| chr8:128823851-128823891 | Myc |
| chr8:128823928-128823968 | Myc |
| chr8:128824012-128824052 | Myc |
| chr8:128824067-128824107 | Myc |
| chr8:128824119-128824159 | Myc |
| chr8:128824182-128824222 | Myc |
| chr8:128824263-128824303 | Myc |
| chr8:128824397-128824437 | Myc |
| chr8:128824546-128824586 | Myc |
| chr8:128824674-128824714 | Myc |
| chr8:128824739-128824779 | Myc |
| chr8:128824810-128824850 | Myc |
| chr8:128824908-128824948 | Myc |
| chr8:128824986-128825026 | Myc |
| chr8:128825099-128825139 | Myc |
| chr8:128825169-128825209 | Myc |
| chr8:128825312-128825352 | Myc |
| chr8:128825533-128825573 | Myc |
| chr8:128825607-128825647 | Myc |
| chr8:128825708-128825748 | Myc |
| chr8:128825773-128825813 | Myc |
| chr8:128825824-128825864 | Myc |
| chr8:128825896-128825936 | Myc |
| chr8:128825955-128825995 | Myc |
| chr8:128826032-128826072 | Myc |
| chr8:128826302-128826342 | Myc |
| chr8:128826426-128826466 | Myc |
| chr8:128826488-128826528 | Myc |
| chr8:128826551-128826591 | Myc |
| chr8:128826601-128826641 | Myc |
| chr8:128826662-128826702 | Myc |
| chr8:128826737-128826777 | Myc |
| chr8:128826795-128826835 | Myc |
| chr8:128826845-128826885 | Myc |
| chr8:128826970-128827010 | Myc |
| chr8:128827076-128827116 | Myc |
| chr8:128827133-128827173 | Myc |
| chr8:128827283-128827323 | Myc |
| chr8:128827490-128827530 | Myc |
| chr8:128827540-128827580 | Myc |
| chr8:128827715-128827755 | Myc |
| chr8:128827778-128827818 | Myc |
| chr8:128827860-128827900 | Myc |
| chr8:128827967-128828007 | Myc |
| chr8:128828068-128828108 | Myc |
| chr8:128828130-128828170 | Myc |
| chr8:128828431-128828471 | Myc |
| chr8:128828520-128828560 | Myc |
| chr8:128828601-128828641 | Myc |
| chr8:128828672-128828712 | Myc |

| | |
|---|---|
| chr8:128828818-128828858 | Myc |
| chr8:128829004-128829044 | Myc |
| chr8:128829088-128829128 | Myc |
| chr8:128829138-128829178 | Myc |
| chr8:128829229-128829269 | Myc |
| chr8:128829311-128829351 | Myc |
| chr8:128829470-128829510 | Myc |
| chr8:128829553-128829593 | Myc |
| chr8:128829829-128829869 | Myc |
| chr8:128830122-128830162 | Myc |
| chr8:128830177-128830217 | Myc |
| chr8:128830320-128830360 | Myc |
| chr8:128830392-128830432 | Myc |
| chr8:128830452-128830492 | Myc |
| chr8:128830566-128830606 | Myc |
| chr8:128830622-128830662 | Myc |
| chr8:128830689-128830729 | Myc |
| chr8:128830752-128830792 | Myc |
| chr8:128830813-128830853 | Myc |
| chr8:128830897-128830937 | Myc |
| chr8:128830956-128830996 | Myc |
| chr8:128831054-128831094 | Myc |
| chr8:128831181-128831221 | Myc |
| chr8:128831231-128831271 | Myc |
| chr8:128831337-128831377 | Myc |
| chr8:128831387-128831427 | Myc |
| chr8:128831476-128831516 | Myc |
| chr8:128831586-128831626 | Myc |
| chr8:128831733-128831773 | Myc |
| chr8:128831802-128831842 | Myc |
| chr8:128831901-128831941 | Myc |
| chr8:128831951-128831991 | Myc |
| chr8:128832001-128832041 | Myc |
| chr8:128832096-128832136 | Myc |
| chr8:128832146-128832186 | Myc |
| chr8:128832292-128832332 | Myc |
| chr8:128832358-128832398 | Myc |
| chr8:128832410-128832450 | Myc |
| chr8:128832491-128832531 | Myc |
| chr8:128832541-128832581 | Myc |
| chr8:128832630-128832670 | Myc |
| chr8:128832785-128832825 | Myc |
| chr8:128832854-128832894 | Myc |
| chr8:128832904-128832944 | Myc |
| chr8:128833224-128833264 | Myc |
| chr8:128833274-128833314 | Myc |
| chr8:128833415-128833455 | Myc |
| chr8:128833484-128833524 | Myc |
| chr8:128833756-128833796 | Myc |
| chr8:128834110-128834150 | Myc |
| chr8:128834190-128834230 | Myc |
| chr8:128834316-128834356 | Myc |

| | |
|---|---|
| chr8:128834381-128834421 | Myc |
| chr8:128834466-128834506 | Myc |
| chr8:128834518-128834558 | Myc |
| chr8:128834576-128834616 | Myc |
| chr8:128834671-128834711 | Myc |
| chr8:128834866-128834906 | Myc |
| chr8:128834977-128835017 | Myc |
| chr8:128835063-128835103 | Myc |
| chr8:128835147-128835187 | Myc |
| chr8:128835217-128835257 | Myc |
| chr8:128835271-128835311 | Myc |
| chr8:128835357-128835397 | Myc |
| chr8:128835486-128835526 | Myc |
| chr8:128835544-128835584 | Myc |
| chr8:128835594-128835634 | Myc |
| chr8:128835719-128835759 | Myc |
| chr8:128835856-128835896 | Myc |
| chr8:128835924-128835964 | Myc |
| chr8:128836017-128836057 | Myc |
| chr8:128836137-128836177 | Myc |
| chr8:128836227-128836267 | Myc |
| chr8:128836299-128836339 | Myc |
| chr8:128836443-128836483 | Myc |
| chr8:128836528-128836568 | Myc |
| chr8:128836588-128836628 | Myc |
| chr8:128836677-128836717 | Myc |
| chr8:128836731-128836771 | Myc |
| chr8:128836816-128836856 | Myc |
| chr8:128836945-128836985 | Myc |
| chr8:128837004-128837044 | Myc |
| chr8:128837067-128837107 | Myc |
| chr8:128837167-128837207 | Myc |
| chr8:128837256-128837296 | Myc |
| chr8:128837306-128837346 | Myc |
| chr8:128837420-128837460 | Myc |
| chr8:128837532-128837572 | Myc |
| chr8:128837649-128837689 | Myc |
| chr8:128837754-128837794 | Myc |
| chr8:128837820-128837860 | Myc |
| chr8:128837949-128837989 | Myc |
| chr8:128838009-128838049 | Myc |
| chr8:128838209-128838249 | Myc |
| chr8:128838325-128838365 | Myc |
| chr8:128838382-128838422 | Myc |
| chr8:128838442-128838482 | Myc |
| chr8:128838549-128838589 | Myc |
| chr8:128838614-128838654 | Myc |
| chr8:128838668-128838708 | Myc |
| chr8:128838730-128838770 | Myc |
| chr8:128838781-128838821 | Myc |
| chr8:128838870-128838910 | Myc |
| chr8:128838928-128838968 | Myc |

| | |
|---|---|
| chr8:128838978-128839018 | Myc |
| chr8:128839103-128839143 | Myc |
| chr8:128839174-128839214 | Myc |
| chr8:128839474-128839514 | Myc |
| chr8:128839570-128839610 | Myc |
| chr8:128839659-128839699 | Myc |
| chr8:128839752-128839792 | Myc |
| chr8:128839876-128839916 | Myc |
| chr8:128839956-128839996 | Myc |
| chr8:128840026-128840066 | Myc |
| chr8:128840104-128840144 | Myc |
| chr8:128840164-128840204 | Myc |
| chr8:128840228-128840268 | Myc |
| chr8:128840317-128840357 | Myc |
| chr8:128840393-128840433 | Myc |
| chr8:128840507-128840547 | Myc |
| chr8:128840560-128840600 | Myc |
| chr8:128840688-128840728 | Myc |
| chr8:128840827-128840867 | Myc |
| chr8:128840878-128840918 | Myc |
| chr8:128840947-128840987 | Myc |
| chr8:128840997-128841037 | Myc |
| chr8:128841085-128841125 | Myc |
| chr8:128841146-128841186 | Myc |
| chr8:128841234-128841274 | Myc |
| chr8:128841595-128841635 | Myc |
| chr8:128841687-128841727 | Myc |
| chr8:128841739-128841779 | Myc |
| chr8:128841803-128841843 | Myc |
| chr8:128841867-128841907 | Myc |
| chr8:128841939-128841979 | Myc |
| chr8:128842005-128842045 | Myc |
| chr8:128842098-128842138 | Myc |
| chr8:128842178-128842218 | Myc |
| chr8:128842229-128842269 | Myc |
| chr8:128842293-128842333 | Myc |
| chr8:128842393-128842433 | Myc |
| chr8:128842444-128842484 | Myc |
| chr8:128842578-128842618 | Myc |
| chr8:128842628-128842668 | Myc |
| chr8:128842686-128842726 | Myc |
| chr8:128842773-128842813 | Myc |
| chr8:128842854-128842894 | Myc |
| chr8:128842932-128842972 | Myc |
| chr8:128843085-128843125 | Myc |
| chr8:128843440-128843480 | Myc |
| chr8:128843490-128843530 | Myc |
| chr8:128843610-128843650 | Myc |
| chr8:128843673-128843713 | Myc |
| chr8:128843766-128843806 | Myc |
| chr8:128843839-128843879 | Myc |
| chr8:128843892-128843932 | Myc |

| | |
|---|---|
| chr8:128843958-128843998 | Myc |
| chr8:128844056-128844096 | Myc |
| chr8:128844106-128844146 | Myc |
| chr8:128844157-128844197 | Myc |
| chr8:128844246-128844286 | Myc |
| chr8:128844376-128844416 | Myc |
| chr8:128844467-128844507 | Myc |
| chr8:128844547-128844587 | Myc |
| chr8:128844621-128844661 | Myc |
| chr8:128844696-128844736 | Myc |
| chr8:128844799-128844839 | Myc |
| chr8:128845146-128845186 | Myc |
| chr8:128845262-128845302 | Myc |
| chr8:128845341-128845381 | Myc |
| chr8:128845488-128845528 | Myc |
| chr8:128845603-128845643 | Myc |
| chr8:128845759-128845799 | Myc |
| chr8:128845826-128845866 | Myc |
| chr8:128846010-128846050 | Myc |
| chr8:128846155-128846195 | Myc |
| chr8:128846215-128846255 | Myc |
| chr8:128846319-128846359 | Myc |
| chr8:128846374-128846414 | Myc |
| chr8:128846571-128846611 | Myc |
| chr8:128846706-128846746 | Myc |
| chr8:128846854-128846894 | Myc |
| chr8:128846904-128846944 | Myc |
| chr8:128847397-128847437 | Myc |
| chr8:128847600-128847640 | Myc |
| chr8:128847710-128847750 | Myc |
| chr8:128847784-128847824 | Myc |
| chr8:128847874-128847914 | Myc |
| chr8:128847929-128847969 | Myc |
| chr8:128847982-128848022 | Myc |
| chr8:128848131-128848171 | Myc |
| chr8:128848211-128848251 | Myc |
| chr8:128848304-128848344 | Myc |
| chr8:128848355-128848395 | Myc |
| chr8:128848478-128848518 | Myc |
| chr8:128848551-128848591 | Myc |
| chr8:128848715-128848755 | Myc |
| chr8:128848765-128848805 | Myc |
| chr8:128848831-128848871 | Myc |
| chr8:128848889-128848929 | Myc |
| chr8:128848961-128849001 | Myc |
| chr8:128849072-128849112 | Myc |
| chr8:128849122-128849162 | Myc |
| chr8:128849173-128849213 | Myc |
| chr8:128849269-128849309 | Myc |
| chr8:128849396-128849436 | Myc |
| chr8:128849493-128849533 | Myc |
| chr8:128849580-128849620 | Myc |

| | |
|---|---|
| chr8:128849679-128849719 | Myc |
| chr8:128849731-128849771 | Myc |
| chr8:128849862-128849902 | Myc |
| chr8:128849958-128849998 | Myc |
| chr8:128850042-128850082 | Myc |
| chr8:128850094-128850134 | Myc |
| chr8:128850417-128850457 | Myc |
| chr8:128850547-128850587 | Myc |
| chr8:128850644-128850684 | Myc |
| chr8:128850905-128850945 | Myc |
| chr8:128851001-128851041 | Myc |
| chr8:128851085-128851125 | Myc |
| chr8:128851227-128851267 | Myc |
| chr8:128851278-128851318 | Myc |
| chr8:128851339-128851379 | Myc |
| chr8:128851399-128851439 | Myc |
| chr8:128851481-128851521 | Myc |
| chr8:128851541-128851581 | Myc |
| chr8:128851649-128851689 | Myc |
| chr8:128851862-128851902 | Myc |
| chr8:128851927-128851967 | Myc |
| chr8:128852040-128852080 | Myc |
| chr8:128852100-128852140 | Myc |
| chr8:128852150-128852190 | Myc |
| chr8:128852207-128852247 | Myc |
| chr8:128852257-128852297 | Myc |
| chr8:128852357-128852397 | Myc |
| chr8:128852424-128852464 | Myc |
| chr8:128852505-128852545 | Myc |
| chr8:128852582-128852622 | Myc |
| chr8:128852639-128852679 | Myc |
| chr8:128852736-128852776 | Myc |
| chr8:128852849-128852889 | Myc |
| chr8:128852899-128852939 | Myc |
| chr8:128853002-128853042 | Myc |
| chr8:128853109-128853149 | Myc |
| chr8:128853206-128853246 | Myc |
| chr8:128853260-128853300 | Myc |
| chr8:128853366-128853406 | Myc |
| chr8:128853452-128853492 | Myc |
| chr8:128853525-128853565 | Myc |
| chr8:128853748-128853788 | Myc |
| chr8:128853861-128853901 | Myc |
| chr8:128853922-128853962 | Myc |
| chr8:128854102-128854142 | Myc |
| chr8:128854253-128854293 | Myc |
| chr8:128854462-128854502 | Myc |
| chr8:128854576-128854616 | Myc |
| chr8:128854688-128854728 | Myc |
| chr8:128854753-128854793 | Myc |
| chr8:128854854-128854894 | Myc |
| chr8:128854904-128854944 | Myc |

| | |
|---|---|
| chr8:128854955-128854995 | Myc |
| chr8:128855048-128855088 | Myc |
| chr8:128855169-128855209 | Myc |
| chr8:128855240-128855280 | Myc |
| chr8:128855299-128855339 | Myc |
| chr8:128855653-128855693 | Myc |
| chr8:128855918-128855958 | Myc |
| chr8:128856011-128856051 | Myc |
| chr8:128856120-128856160 | Myc |
| chr8:128856170-128856210 | Myc |
| chr8:128856508-128856548 | Myc |
| chr8:128856587-128856627 | Myc |
| chr8:128856648-128856688 | Myc |
| chr8:128856698-128856738 | Myc |
| chr8:128856769-128856809 | Myc |
| chr8:128856827-128856867 | Myc |
| chr8:128856891-128856931 | Myc |
| chr8:128857238-128857278 | Myc |
| chr8:128857290-128857330 | Myc |
| chr8:128857418-128857458 | Myc |
| chr8:128857468-128857508 | Myc |
| chr8:128857521-128857561 | Myc |
| chr8:128857673-128857713 | Myc |
| chr8:128857739-128857779 | Myc |
| chr8:128857793-128857833 | Myc |
| chr8:128857867-128857907 | Myc |
| chr8:128857948-128857988 | Myc |
| chr8:128858024-128858064 | Myc |
| chr8:128858089-128858129 | Myc |
| chr8:128858215-128858255 | Myc |
| chr8:128858301-128858341 | Myc |
| chr8:128858351-128858391 | Myc |
| chr8:128858505-128858545 | Myc |
| chr8:128858591-128858631 | Myc |
| chr8:128858702-128858742 | Myc |
| chr8:128858814-128858854 | Myc |
| chr8:128858885-128858925 | Myc |
| chr8:128859116-128859156 | Myc |
| chr8:128859252-128859292 | Myc |
| chr8:128859366-128859406 | Myc |
| chr8:128859442-128859482 | Myc |
| chr8:128859595-128859635 | Myc |
| chr8:128859680-128859720 | Myc |
| chr8:128859751-128859791 | Myc |
| chr8:128859842-128859882 | Myc |
| chr8:128859939-128859979 | Myc |
| chr8:128860075-128860115 | Myc |
| chr8:128860160-128860200 | Myc |
| chr8:128860211-128860251 | Myc |
| chr8:128860290-128860330 | Myc |
| chr8:128860377-128860417 | Myc |
| chr8:128860482-128860522 | Myc |

| | |
|---|---|
| chr8:128860548-128860588 | Myc |
| chr8:128860620-128860660 | Myc |
| chr8:128860904-128860944 | Myc |
| chr8:128861001-128861041 | Myc |
| chr8:128861109-128861149 | Myc |
| chr8:128861172-128861212 | Myc |
| chr8:128861237-128861277 | Myc |
| chr8:128861308-128861348 | Myc |
| chr8:128861433-128861473 | Myc |
| chr8:128861578-128861618 | Myc |
| chr8:128861644-128861684 | Myc |
| chr8:128861713-128861753 | Myc |
| chr8:128861772-128861812 | Myc |
| chr8:128861835-128861875 | Myc |
| chr8:128862226-128862266 | Myc |
| chr8:128862277-128862317 | Myc |
| chr8:128862357-128862397 | Myc |
| chr8:128862465-128862505 | Myc |
| chr8:128862520-128862560 | Myc |
| chr8:128862612-128862652 | Myc |
| chr8:128862662-128862702 | Myc |
| chr8:128862775-128862815 | Myc |
| chr8:128862826-128862866 | Myc |
| chr8:128862959-128862999 | Myc |
| chr8:128863101-128863141 | Myc |
| chr8:128863207-128863247 | Myc |
| chr8:128863413-128863453 | Myc |
| chr8:128863506-128863546 | Myc |
| chr8:128863559-128863599 | Myc |
| chr8:128863662-128863702 | Myc |
| chr8:128863740-128863780 | Myc |
| chr8:128863810-128863850 | Myc |
| chr8:128863913-128863953 | Myc |
| chr8:128863965-128864005 | Myc |
| chr8:128864066-128864106 | Myc |
| chr8:128864133-128864173 | Myc |
| chr8:128864234-128864274 | Myc |
| chr8:128864332-128864372 | Myc |
| chr8:128864420-128864460 | Myc |
| chr8:128864485-128864525 | Myc |
| chr8:128864535-128864575 | Myc |
| chr8:128864684-128864724 | Myc |
| chr8:128864757-128864797 | Myc |
| chr8:128864869-128864909 | Myc |
| chr8:128864984-128865024 | Myc |
| chr8:128865121-128865161 | Myc |
| chr8:128865185-128865225 | Myc |
| chr8:128865299-128865339 | Myc |
| chr8:128865357-128865397 | Myc |
| chr8:128865441-128865481 | Myc |
| chr8:128865491-128865531 | Myc |
| chr8:128865581-128865621 | Myc |

| | |
|---|---|
| chr8:128865631-128865671 | Myc |
| chr8:128865681-128865721 | Myc |
| chr8:128865935-128865975 | Myc |
| chr8:128865985-128866025 | Myc |
| chr8:128866078-128866118 | Myc |
| chr8:128866152-128866192 | Myc |
| chr8:128866216-128866256 | Myc |
| chr8:128866296-128866336 | Myc |
| chr8:128866384-128866424 | Myc |
| chr8:128866460-128866500 | Myc |
| chr8:128866533-128866573 | Myc |
| chr8:128866622-128866662 | Myc |
| chr8:128866676-128866716 | Myc |
| chr8:128866749-128866789 | Myc |
| chr8:128866815-128866855 | Myc |
| chr8:128866882-128866922 | Myc |
| chr8:128866962-128867002 | Myc |
| chr8:128867087-128867127 | Myc |
| chr8:128867160-128867200 | Myc |
| chr8:128867242-128867282 | Myc |
| chr8:128867347-128867387 | Myc |
| chr8:128867509-128867549 | Myc |
| chr8:128867564-128867604 | Myc |
| chr8:128867736-128867776 | Myc |
| chr8:128867912-128867952 | Myc |
| chr8:128867962-128868002 | Myc |
| chr8:128868073-128868113 | Myc |
| chr8:128868129-128868169 | Myc |
| chr8:128868181-128868221 | Myc |
| chr8:128868325-128868365 | Myc |
| chr8:128868479-128868519 | Myc |
| chr8:128868540-128868580 | Myc |
| chr8:128868621-128868661 | Myc |
| chr8:128868866-128868906 | Myc |
| chr8:128869013-128869053 | Myc |
| chr8:128869077-128869117 | Myc |
| chr8:128869168-128869208 | Myc |
| chr8:128869429-128869469 | Myc |
| chr8:128869504-128869544 | Myc |
| chr8:128869593-128869633 | Myc |
| chr8:128869678-128869718 | Myc |
| chr8:128869791-128869831 | Myc |
| chr8:128869881-128869921 | Myc |
| chr8:128870011-128870051 | Myc |
| chr8:128870136-128870176 | Myc |
| chr8:128870233-128870273 | Myc |
| chr8:128870300-128870340 | Myc |
| chr8:128870420-128870460 | Myc |
| chr8:128870700-128870740 | Myc |
| chr8:128870750-128870790 | Myc |
| chr8:128870817-128870857 | Myc |
| chr8:128870896-128870936 | Myc |

| | |
|---|---|
| chr8:128870967-128871007 | Myc |
| chr8:128871077-128871117 | Myc |
| chr8:128871172-128871212 | Myc |
| chr8:128871261-128871301 | Myc |
| chr8:128871321-128871361 | Myc |
| chr8:128871407-128871447 | Myc |
| chr8:128871514-128871554 | Myc |
| chr8:128871580-128871620 | Myc |
| chr8:128871695-128871735 | Myc |
| chr8:128871854-128871894 | Myc |
| chr8:128871959-128871999 | Myc |
| chr8:128872009-128872049 | Myc |
| chr8:128872181-128872221 | Myc |
| chr8:128872284-128872324 | Myc |
| chr8:128872348-128872388 | Myc |
| chr8:128872482-128872522 | Myc |
| chr8:128872532-128872572 | Myc |
| chr8:128872591-128872631 | Myc |
| chr8:128872646-128872686 | Myc |
| chr8:128872704-128872744 | Myc |
| chr8:128873079-128873119 | Myc |
| chr8:128873205-128873245 | Myc |
| chr8:128873269-128873309 | Myc |
| chr8:128873324-128873364 | Myc |
| chr8:128873401-128873441 | Myc |
| chr8:128873488-128873528 | Myc |
| chr8:128873710-128873750 | Myc |
| chr8:128873764-128873804 | Myc |
| chr8:128873891-128873931 | Myc |
| chr8:128874025-128874065 | Myc |
| chr8:128874099-128874139 | Myc |
| chr8:128874171-128874211 | Myc |
| chr8:128874270-128874310 | Myc |
| chr8:128874354-128874394 | Myc |
| chr8:128874479-128874519 | Myc |
| chr8:128874696-128874736 | Myc |
| chr8:128874791-128874831 | Myc |
| chr8:128874883-128874923 | Myc |
| chr8:128874936-128874976 | Myc |
| chr8:128875008-128875048 | Myc |
| chr8:128875058-128875098 | Myc |
| chr8:128875141-128875181 | Myc |
| chr8:128875263-128875303 | Myc |
| chr8:128875363-128875403 | Myc |
| chr8:128875491-128875531 | Myc |
| chr8:128875575-128875615 | Myc |
| chr8:128875652-128875692 | Myc |
| chr8:128875783-128875823 | Myc |
| chr8:128875841-128875881 | Myc |
| chr8:128875897-128875937 | Myc |
| chr8:128876015-128876055 | Myc |
| chr8:128876073-128876113 | Myc |

| | |
|---|---|
| chr8:128876124-128876164 | Myc |
| chr8:128876213-128876253 | Myc |
| chr8:128876266-128876306 | Myc |
| chr8:128876381-128876421 | Myc |
| chr8:128876495-128876535 | Myc |
| chr8:128876545-128876585 | Myc |
| chr8:128876632-128876672 | Myc |
| chr8:128876719-128876759 | Myc |
| chr8:128877121-128877161 | Myc |
| chr8:128877177-128877217 | Myc |
| chr8:128877243-128877283 | Myc |
| chr8:128877306-128877346 | Myc |
| chr8:128877416-128877456 | Myc |
| chr8:128877529-128877569 | Myc |
| chr8:128877581-128877621 | Myc |
| chr8:128877707-128877747 | Myc |
| chr8:128877797-128877837 | Myc |
| chr8:128877875-128877915 | Myc |
| chr8:128878008-128878048 | Myc |
| chr8:128878064-128878104 | Myc |
| chr8:128878159-128878199 | Myc |
| chr8:128878265-128878305 | Myc |
| chr8:128878320-128878360 | Myc |
| chr8:128878383-128878423 | Myc |
| chr8:128878452-128878492 | Myc |
| chr8:128878559-128878599 | Myc |
| chr8:128878637-128878677 | Myc |
| chr8:128878702-128878742 | Myc |
| chr8:128878806-128878846 | Myc |
| chr8:128878873-128878913 | Myc |
| chr8:128878958-128878998 | Myc |
| chr8:128879030-128879070 | Myc |
| chr8:128879080-128879120 | Myc |
| chr8:128879153-128879193 | Myc |
| chr8:128879248-128879288 | Myc |
| chr8:128879361-128879401 | Myc |
| chr8:128879493-128879533 | Myc |
| chr8:128879571-128879611 | Myc |
| chr8:128879679-128879719 | Myc |
| chr8:128879733-128879773 | Myc |
| chr8:128879798-128879838 | Myc |
| chr8:128879954-128879994 | Myc |
| chr8:128880054-128880094 | Myc |
| chr8:128880110-128880150 | Myc |
| chr8:128880207-128880247 | Myc |
| chr8:128880334-128880374 | Myc |
| chr8:128880403-128880443 | Myc |
| chr8:128880465-128880505 | Myc |
| chr8:128880536-128880576 | Myc |
| chr8:128880586-128880626 | Myc |
| chr8:128880746-128880786 | Myc |
| chr8:128880877-128880917 | Myc |

| | |
|---|---|
| chr8:128880969-128881009 | Myc |
| chr8:128881161-128881201 | Myc |
| chr8:128881250-128881290 | Myc |
| chr8:128881416-128881456 | Myc |
| chr8:128881657-128881697 | Myc |
| chr8:128881749-128881789 | Myc |
| chr8:128881852-128881892 | Myc |
| chr8:128881902-128881942 | Myc |
| chr8:128881969-128882009 | Myc |
| chr8:128882072-128882112 | Myc |
| chr8:128882192-128882232 | Myc |
| chr8:128882328-128882368 | Myc |
| chr8:128882402-128882442 | Myc |
| chr8:128882468-128882508 | Myc |
| chr8:128882533-128882573 | Myc |
| chr8:128882588-128882628 | Myc |
| chr8:128882642-128882682 | Myc |
| chr8:128882774-128882814 | Myc |
| chr8:128882871-128882911 | Myc |
| chr8:128882934-128882974 | Myc |
| chr8:128882985-128883025 | Myc |
| chr8:128883068-128883108 | Myc |
| chr8:128883177-128883217 | Myc |
| chr8:128883227-128883267 | Myc |
| chr8:128883309-128883349 | Myc |
| chr8:128883409-128883449 | Myc |
| chr8:128883522-128883562 | Myc |
| chr8:128883656-128883696 | Myc |
| chr8:128883721-128883761 | Myc |
| chr8:128883780-128883820 | Myc |
| chr8:128883830-128883870 | Myc |
| chr8:128883911-128883951 | Myc |
| chr8:128884035-128884075 | Myc |
| chr8:128884357-128884397 | Myc |
| chr8:128884512-128884552 | Myc |
| chr8:128884630-128884670 | Myc |
| chr8:128884748-128884788 | Myc |
| chr8:128884945-128884985 | Myc |
| chr8:128885003-128885043 | Myc |
| chr8:128885121-128885161 | Myc |
| chr8:128885228-128885268 | Myc |
| chr8:128885278-128885318 | Myc |
| chr8:128885384-128885424 | Myc |
| chr8:128885444-128885484 | Myc |
| chr8:128885528-128885568 | Myc |
| chr8:128885618-128885658 | Myc |
| chr8:128885668-128885708 | Myc |
| chr8:128885772-128885812 | Myc |
| chr8:128885826-128885866 | Myc |
| chr8:128885906-128885946 | Myc |
| chr8:128885975-128886015 | Myc |
| chr8:128886025-128886065 | Myc |

| | |
|---|---|
| chr8:128886103-128886143 | Myc |
| chr8:128886208-128886248 | Myc |
| chr8:128886307-128886347 | Myc |
| chr8:128886386-128886426 | Myc |
| chr8:128886473-128886513 | Myc |
| chr8:128886533-128886573 | Myc |
| chr8:128886600-128886640 | Myc |
| chr8:128886658-128886698 | Myc |
| chr8:128886735-128886775 | Myc |
| chr8:128886806-128886846 | Myc |
| chr8:128886856-128886896 | Myc |
| chr8:128886921-128886961 | Myc |
| chr8:128887017-128887057 | Myc |
| chr8:128887099-128887139 | Myc |
| chr8:128887180-128887220 | Myc |
| chr8:128887230-128887270 | Myc |
| chr8:128887341-128887381 | Myc |
| chr8:128887435-128887475 | Myc |
| chr8:128887499-128887539 | Myc |
| chr8:128887561-128887601 | Myc |
| chr8:128887615-128887655 | Myc |
| chr8:128887677-128887717 | Myc |
| chr8:128887742-128887782 | Myc |
| chr8:128887843-128887883 | Myc |
| chr8:128887930-128887970 | Myc |
| chr8:128888029-128888069 | Myc |
| chr8:128888085-128888125 | Myc |
| chr8:128888212-128888252 | Myc |
| chr8:128888313-128888353 | Myc |
| chr8:128888365-128888405 | Myc |
| chr8:128888422-128888462 | Myc |
| chr8:128888558-128888598 | Myc |
| chr8:128888608-128888648 | Myc |
| chr8:128888774-128888814 | Myc |
| chr8:128888857-128888897 | Myc |
| chr8:128888920-128888960 | Myc |
| chr8:128888990-128889030 | Myc |
| chr8:128889051-128889091 | Myc |
| chr8:128889146-128889186 | Myc |
| chr8:128889215-128889255 | Myc |
| chr8:128889302-128889342 | Myc |
| chr8:128889358-128889398 | Myc |
| chr8:128889432-128889472 | Myc |
| chr8:128889497-128889537 | Myc |
| chr8:128889572-128889612 | Myc |
| chr8:128889709-128889749 | Myc |
| chr8:128889767-128889807 | Myc |
| chr8:128889818-128889858 | Myc |
| chr8:128889943-128889983 | Myc |
| chr8:128890184-128890224 | Myc |
| chr8:128890294-128890334 | Myc |
| chr8:128890367-128890407 | Myc |

| | |
|---|---|
| chr8:128890439-128890479 | Myc |
| chr8:128890515-128890555 | Myc |
| chr8:128890594-128890634 | Myc |
| chr8:128890690-128890730 | Myc |
| chr8:128890807-128890847 | Myc |
| chr8:128890912-128890952 | Myc |
| chr8:128891015-128891055 | Myc |
| chr8:128891082-128891122 | Myc |
| chr8:128891148-128891188 | Myc |
| chr8:128891205-128891245 | Myc |
| chr8:128891288-128891328 | Myc |
| chr8:128891355-128891395 | Myc |
| chr8:128891456-128891496 | Myc |
| chr8:128891524-128891564 | Myc |
| chr8:128891828-128891868 | Myc |
| chr8:128891878-128891918 | Myc |
| chr8:128892032-128892072 | Myc |
| chr8:128892082-128892122 | Myc |
| chr8:128892137-128892177 | Myc |
| chr8:128892187-128892227 | Myc |
| chr8:128892299-128892339 | Myc |
| chr8:128892357-128892397 | Myc |
| chr8:128892440-128892480 | Myc |
| chr8:128892494-128892534 | Myc |
| chr8:128892903-128892943 | Myc |
| chr8:128893180-128893220 | Myc |
| chr8:128893279-128893319 | Myc |
| chr8:128893329-128893369 | Myc |
| chr8:128893426-128893466 | Myc |
| chr8:128893517-128893557 | Myc |
| chr8:128893567-128893607 | Myc |
| chr8:128893883-128893923 | Myc |
| chr8:128894040-128894080 | Myc |
| chr8:128894116-128894156 | Myc |
| chr8:128894379-128894419 | Myc |
| chr8:128894429-128894469 | Myc |
| chr8:128894523-128894563 | Myc |
| chr8:128894579-128894619 | Myc |
| chr8:128894643-128894683 | Myc |
| chr8:128894694-128894734 | Myc |
| chr8:128895008-128895048 | Myc |
| chr8:128895111-128895151 | Myc |
| chr8:128895245-128895285 | Myc |
| chr8:128895339-128895379 | Myc |
| chr8:128895432-128895472 | Myc |
| chr8:128895487-128895527 | Myc |
| chr8:128895563-128895603 | Myc |
| chr8:128895664-128895704 | Myc |
| chr8:128895768-128895808 | Myc |
| chr8:128896008-128896048 | Myc |
| chr8:128896087-128896127 | Myc |
| chr8:128896145-128896185 | Myc |

| | |
|---|---|
| chr8:128896195-128896235 | Myc |
| chr8:128896314-128896354 | Myc |
| chr8:128896371-128896411 | Myc |
| chr8:128896476-128896516 | Myc |
| chr8:128896540-128896580 | Myc |
| chr8:128896611-128896651 | Myc |
| chr8:128896661-128896701 | Myc |
| chr8:128896729-128896769 | Myc |
| chr8:128896822-128896862 | Myc |
| chr8:128896943-128896983 | Myc |
| chr8:128897026-128897066 | Myc |
| chr8:128897090-128897130 | Myc |
| chr8:128897191-128897231 | Myc |
| chr8:128897249-128897289 | Myc |
| chr8:128897306-128897346 | Myc |
| chr8:128897381-128897421 | Myc |
| chr8:128897431-128897471 | Myc |
| chr8:128897481-128897521 | Myc |
| chr8:128897570-128897610 | Myc |
| chr8:128897654-128897694 | Myc |
| chr8:128897704-128897744 | Myc |
| chr8:128897811-128897851 | Myc |
| chr8:128897872-128897912 | Myc |
| chr8:128897944-128897984 | Myc |
| chr8:128898017-128898057 | Myc |
| chr8:128898186-128898226 | Myc |
| chr8:128898379-128898419 | Myc |
| chr8:128898452-128898492 | Myc |
| chr8:128898503-128898543 | Myc |
| chr8:128898623-128898663 | Myc |
| chr8:128898699-128898739 | Myc |
| chr8:128898791-128898831 | Myc |
| chr8:128898880-128898920 | Myc |
| chr8:128899007-128899047 | Myc |
| chr8:128899138-128899178 | Myc |
| chr8:128899233-128899273 | Myc |
| chr8:128899299-128899339 | Myc |
| chr8:128899353-128899393 | Myc |
| chr8:128899487-128899527 | Myc |
| chr8:128899538-128899578 | Myc |
| chr8:128899588-128899628 | Myc |
| chr8:128899689-128899729 | Myc |
| chr8:128899778-128899818 | Myc |
| chr8:128900051-128900091 | Myc |
| chr8:128900129-128900169 | Myc |
| chr8:128900544-128900584 | Myc |
| chr8:128900697-128900737 | Myc |
| chr8:128900763-128900803 | Myc |
| chr8:128900836-128900876 | Myc |
| chr8:128900887-128900927 | Myc |
| chr8:128900940-128900980 | Myc |
| chr8:128901012-128901052 | Myc |

| | |
|---|---|
| chr8:128901062-128901102 | Myc |
| chr8:128901224-128901264 | Myc |
| chr8:128901457-128901497 | Myc |
| chr8:128901601-128901641 | Myc |
| chr8:128901656-128901696 | Myc |
| chr8:128901716-128901756 | Myc |
| chr8:128901906-128901946 | Myc |
| chr8:128902000-128902040 | Myc |
| chr8:128902065-128902105 | Myc |
| chr8:128902124-128902164 | Myc |
| chr8:128902211-128902251 | Myc |
| chr8:128902310-128902350 | Myc |
| chr8:128902404-128902444 | Myc |
| chr8:128902494-128902534 | Myc |
| chr8:128902555-128902595 | Myc |
| chr8:128902651-128902691 | Myc |
| chr8:128902736-128902776 | Myc |
| chr8:128902867-128902907 | Myc |
| chr8:128902974-128903014 | Myc |
| chr8:128903072-128903112 | Myc |
| chr8:128903223-128903263 | Myc |
| chr8:128903320-128903360 | Myc |
| chr8:128903378-128903418 | Myc |
| chr8:128903462-128903502 | Myc |
| chr8:128903527-128903567 | Myc |
| chr8:128903594-128903634 | Myc |
| chr8:128903697-128903737 | Myc |
| chr8:128903770-128903810 | Myc |
| chr8:128903824-128903864 | Myc |
| chr8:128903883-128903923 | Myc |
| chr8:128903974-128904014 | Myc |
| chr8:128904047-128904087 | Myc |
| chr8:128904201-128904241 | Myc |
| chr8:128904309-128904349 | Myc |
| chr8:128904382-128904422 | Myc |
| chr8:128904451-128904491 | Myc |
| chr8:128904558-128904598 | Myc |
| chr8:128904630-128904670 | Myc |
| chr8:128904791-128904831 | Myc |
| chr8:128904929-128904969 | Myc |
| chr8:128904982-128905022 | Myc |
| chr8:128905044-128905084 | Myc |
| chr8:128905126-128905166 | Myc |
| chr8:128905181-128905221 | Myc |
| chr8:128905262-128905302 | Myc |
| chr8:128905340-128905380 | Myc |
| chr8:128905406-128905446 | Myc |
| chr8:128905459-128905499 | Myc |
| chr8:128905736-128905776 | Myc |
| chr8:128905855-128905895 | Myc |
| chr8:128905923-128905963 | Myc |
| chr8:128906021-128906061 | Myc |

| | |
|---|---|
| chr8:128906078-128906118 | Myc |
| chr8:128906146-128906186 | Myc |
| chr8:128906212-128906252 | Myc |
| chr8:128906284-128906324 | Myc |
| chr8:128906392-128906432 | Myc |
| chr8:128906469-128906509 | Myc |
| chr8:128906528-128906568 | Myc |
| chr8:128906580-128906620 | Myc |
| chr8:128906736-128906776 | Myc |
| chr8:128906786-128906826 | Myc |
| chr8:128906876-128906916 | Myc |
| chr8:128907011-128907051 | Myc |
| chr8:128907074-128907114 | Myc |
| chr8:128907131-128907171 | Myc |
| chr8:128907181-128907221 | Myc |
| chr8:128907237-128907277 | Myc |
| chr8:128907287-128907327 | Myc |
| chr8:128907413-128907453 | Myc |
| chr8:128907675-128907715 | Myc |
| chr8:128907801-128907841 | Myc |
| chr8:128907868-128907908 | Myc |
| chr8:128907948-128907988 | Myc |
| chr8:128908068-128908108 | Myc |
| chr8:128908185-128908225 | Myc |
| chr8:128908235-128908275 | Myc |
| chr8:128908315-128908355 | Myc |
| chr8:128908384-128908424 | Myc |
| chr8:128908498-128908538 | Myc |
| chr8:128908609-128908649 | Myc |
| chr8:128908665-128908705 | Myc |
| chr8:128908770-128908810 | Myc |
| chr8:128908926-128908966 | Myc |
| chr8:128909026-128909066 | Myc |
| chr8:128909076-128909116 | Myc |
| chr8:128909165-128909205 | Myc |
| chr8:128909244-128909284 | Myc |
| chr8:128909389-128909429 | Myc |
| chr8:128909451-128909491 | Myc |
| chr8:128909504-128909544 | Myc |
| chr8:128909572-128909612 | Myc |
| chr8:128909629-128909669 | Myc |
| chr8:128909681-128909721 | Myc |
| chr8:128909814-128909854 | Myc |
| chr8:128909870-128909910 | Myc |
| chr8:128909933-128909973 | Myc |
| chr8:128910009-128910049 | Myc |
| chr8:128910082-128910122 | Myc |
| chr8:128910288-128910328 | Myc |
| chr8:128910428-128910468 | Myc |
| chr8:128910478-128910518 | Myc |
| chr8:128910641-128910681 | Myc |
| chr8:128910725-128910765 | Myc |

| | |
|---|---|
| chr8:128910783-128910823 | Myc |
| chr8:128910949-128910989 | Myc |
| chr8:128911051-128911091 | Myc |
| chr8:128911175-128911215 | Myc |
| chr8:128911228-128911268 | Myc |
| chr8:128911278-128911318 | Myc |
| chr8:128911337-128911377 | Myc |
| chr8:128911468-128911508 | Myc |
| chr8:128911552-128911592 | Myc |
| chr8:128911712-128911752 | Myc |
| chr8:128911788-128911828 | Myc |
| chr8:128911847-128911887 | Myc |
| chr8:128911911-128911951 | Myc |
| chr8:128911980-128912020 | Myc |
| chr8:128912030-128912070 | Myc |
| chr8:128912081-128912121 | Myc |
| chr8:128912199-128912239 | Myc |
| chr8:128912565-128912605 | Myc |
| chr8:128912631-128912671 | Myc |
| chr8:128912681-128912721 | Myc |
| chr8:128912734-128912774 | Myc |
| chr8:128912809-128912849 | Myc |
| chr8:128912866-128912906 | Myc |
| chr8:128912947-128912987 | Myc |
| chr8:128912997-128913037 | Myc |
| chr8:128913068-128913108 | Myc |
| chr8:128913132-128913172 | Myc |
| chr8:128913202-128913242 | Myc |
| chr8:128913302-128913342 | Myc |
| chr8:128913396-128913436 | Myc |
| chr8:128913476-128913516 | Myc |
| chr8:128913543-128913583 | Myc |
| chr8:128913596-128913636 | Myc |
| chr8:128913668-128913708 | Myc |
| chr8:128913750-128913790 | Myc |
| chr8:128913868-128913908 | Myc |
| chr8:128913926-128913966 | Myc |
| chr8:128914316-128914356 | Myc |
| chr8:128914439-128914479 | Myc |
| chr8:128914489-128914529 | Myc |
| chr8:128914586-128914626 | Myc |
| chr8:128914747-128914787 | Myc |
| chr8:128914817-128914857 | Myc |
| chr8:128914922-128914962 | Myc |
| chr8:128914977-128915017 | Myc |
| chr8:128915069-128915109 | Myc |
| chr8:128915176-128915216 | Myc |
| chr8:128915241-128915281 | Myc |
| chr8:128915292-128915332 | Myc |
| chr8:128915365-128915405 | Myc |
| chr8:128915473-128915513 | Myc |
| chr8:128915553-128915593 | Myc |

| | |
|---|---|
| chr8:128915609-128915649 | Myc |
| chr8:128915742-128915782 | Myc |
| chr8:128915845-128915885 | Myc |
| chr8:128915918-128915958 | Myc |
| chr8:128916013-128916053 | Myc |
| chr8:128916089-128916129 | Myc |
| chr8:128916327-128916367 | Myc |
| chr8:128916498-128916538 | Myc |
| chr8:128916622-128916662 | Myc |
| chr8:128916672-128916712 | Myc |
| chr8:128916755-128916795 | Myc |
| chr8:128916831-128916871 | Myc |
| chr8:128916887-128916927 | Myc |
| chr8:128916959-128916999 | Myc |
| chr8:128917015-128917055 | Myc |
| chr8:128917124-128917164 | Myc |
| chr8:128917183-128917223 | Myc |
| chr8:128917241-128917281 | Myc |
| chr8:128917331-128917371 | Myc |
| chr8:128917498-128917538 | Myc |
| chr8:128917624-128917664 | Myc |
| chr8:128917675-128917715 | Myc |
| chr8:128917755-128917795 | Myc |
| chr8:128917818-128917858 | Myc |
| chr8:128917957-128917997 | Myc |
| chr8:128918067-128918107 | Myc |
| chr8:128918117-128918157 | Myc |
| chr8:128918195-128918235 | Myc |
| chr8:128918253-128918293 | Myc |
| chr8:128918354-128918394 | Myc |
| chr8:128918411-128918451 | Myc |
| chr8:128918537-128918577 | Myc |
| chr8:128918599-128918639 | Myc |
| chr8:128918652-128918692 | Myc |
| chr8:128918728-128918768 | Myc |
| chr8:128918828-128918868 | Myc |
| chr8:128918879-128918919 | Myc |
| chr8:128918958-128918998 | Myc |
| chr8:128919016-128919056 | Myc |
| chr8:128919101-128919141 | Myc |
| chr8:128919263-128919303 | Myc |
| chr8:128919329-128919369 | Myc |
| chr8:128919507-128919547 | Myc |
| chr8:128919596-128919636 | Myc |
| chr8:128919702-128919742 | Myc |
| chr8:128919784-128919824 | Myc |
| chr8:128919834-128919874 | Myc |
| chr8:128919902-128919942 | Myc |
| chr8:128919976-128920016 | Myc |
| chr8:128920074-128920114 | Myc |
| chr8:128920133-128920173 | Myc |
| chr8:128920218-128920258 | Myc |

| | |
|---|---|
| chr8:128920279-128920319 | Myc |
| chr8:128920376-128920416 | Myc |
| chr8:128920458-128920498 | Myc |
| chr8:128920527-128920567 | Myc |
| chr8:128920642-128920682 | Myc |
| chr8:128920905-128920945 | Myc |
| chr8:128921008-128921048 | Myc |
| chr8:128921113-128921153 | Myc |
| chr8:128921172-128921212 | Myc |
| chr8:128921271-128921311 | Myc |
| chr8:128921321-128921361 | Myc |
| chr8:128921387-128921427 | Myc |
| chr8:128921456-128921496 | Myc |
| chr8:128921538-128921578 | Myc |
| chr8:128921634-128921674 | Myc |
| chr8:128921691-128921731 | Myc |
| chr8:128921805-128921845 | Myc |
| chr8:128921939-128921979 | Myc |
| chr8:128922011-128922051 | Myc |
| chr8:128922078-128922118 | Myc |
| chr8:128922164-128922204 | Myc |
| chr8:128922290-128922330 | Myc |
| chr8:128922426-128922466 | Myc |
| chr8:128922485-128922525 | Myc |
| chr8:128922612-128922652 | Myc |
| chr8:128922669-128922709 | Myc |
| chr8:128922740-128922780 | Myc |
| chr8:128922884-128922924 | Myc |
| chr8:128922944-128922984 | Myc |
| chr8:128923030-128923070 | Myc |
| chr8:128923084-128923124 | Myc |
| chr8:128923157-128923197 | Myc |
| chr8:128923217-128923257 | Myc |
| chr8:128923379-128923419 | Myc |
| chr8:128923523-128923563 | Myc |
| chr8:128923588-128923628 | Myc |
| chr8:128923702-128923742 | Myc |
| chr8:128923830-128923870 | Myc |
| chr8:128923889-128923929 | Myc |
| chr8:128923951-128923991 | Myc |
| chr8:128924049-128924089 | Myc |
| chr8:128924210-128924250 | Myc |
| chr8:128924266-128924306 | Myc |
| chr8:128924381-128924421 | Myc |
| chr8:128924440-128924480 | Myc |
| chr8:128924538-128924578 | Myc |
| chr8:128924660-128924700 | Myc |
| chr8:128924713-128924753 | Myc |
| chr8:128924763-128924803 | Myc |
| chr8:128924825-128924865 | Myc |
| chr8:128924876-128924916 | Myc |
| chr8:128925076-128925116 | Myc |

| | |
|---|---|
| chr8:128925223-128925263 | Myc |
| chr8:128925373-128925413 | Myc |
| chr8:128925511-128925551 | Myc |
| chr8:128925606-128925646 | Myc |
| chr8:128925656-128925696 | Myc |
| chr8:128925775-128925815 | Myc |
| chr8:128925875-128925915 | Myc |
| chr8:128925975-128926015 | Myc |
| chr8:128926111-128926151 | Myc |
| chr8:128926168-128926208 | Myc |
| chr8:128926237-128926277 | Myc |
| chr8:128926317-128926357 | Myc |
| chr8:128926391-128926431 | Myc |
| chr8:128926464-128926504 | Myc |
| chr8:128926619-128926659 | Myc |
| chr8:128926697-128926737 | Myc |
| chr8:128926754-128926794 | Myc |
| chr8:128926805-128926845 | Myc |
| chr8:128926886-128926926 | Myc |
| chr8:128927008-128927048 | Myc |
| chr8:128927184-128927224 | Myc |
| chr8:128927349-128927389 | Myc |
| chr8:128927412-128927452 | Myc |
| chr8:128927548-128927588 | Myc |
| chr8:128927675-128927715 | Myc |
| chr8:128927915-128927955 | Myc |
| chr8:128927978-128928018 | Myc |
| chr8:128928043-128928083 | Myc |
| chr8:128928129-128928169 | Myc |
| chr8:128928214-128928254 | Myc |
| chr8:128928346-128928386 | Myc |
| chr8:128928404-128928444 | Myc |
| chr8:128928459-128928499 | Myc |
| chr8:128928582-128928622 | Myc |
| chr8:128928729-128928769 | Myc |
| chr8:128928779-128928819 | Myc |
| chr8:128928949-128928989 | Myc |
| chr8:128929008-128929048 | Myc |
| chr8:128929114-128929154 | Myc |
| chr8:128929168-128929208 | Myc |
| chr8:128929226-128929266 | Myc |
| chr8:128929299-128929339 | Myc |
| chr8:128929445-128929485 | Myc |
| chr8:128929539-128929579 | Myc |
| chr8:128929616-128929656 | Myc |
| chr8:128929711-128929751 | Myc |
| chr8:128929761-128929801 | Myc |
| chr8:128929829-128929869 | Myc |
| chr8:128929907-128929947 | Myc |
| chr8:128930008-128930048 | Myc |
| chr8:128930066-128930106 | Myc |
| chr8:128930131-128930171 | Myc |

| | |
|---|---|
| chr8:128930223-128930263 | Myc |
| chr8:128930361-128930401 | Myc |
| chr8:128930454-128930494 | Myc |
| chr8:128930505-128930545 | Myc |
| chr8:128930607-128930647 | Myc |
| chr8:128930714-128930754 | Myc |
| chr8:128930843-128930883 | Myc |
| chr8:128930894-128930934 | Myc |
| chr8:128930945-128930985 | Myc |
| chr8:128931015-128931055 | Myc |
| chr8:128931124-128931164 | Myc |
| chr8:128931199-128931239 | Myc |
| chr8:128931278-128931318 | Myc |
| chr8:128931336-128931376 | Myc |
| chr8:128931386-128931426 | Myc |
| chr8:128931477-128931517 | Myc |
| chr8:128931528-128931568 | Myc |
| chr8:128931683-128931723 | Myc |
| chr8:128931803-128931843 | Myc |
| chr8:128931899-128931939 | Myc |
| chr8:128931970-128932010 | Myc |
| chr8:128932021-128932061 | Myc |
| chr8:128932119-128932159 | Myc |
| chr8:128932226-128932266 | Myc |
| chr8:128932309-128932349 | Myc |
| chr8:128932386-128932426 | Myc |
| chr8:128932486-128932526 | Myc |
| chr8:128932575-128932615 | Myc |
| chr8:128932672-128932712 | Myc |
| chr8:128932722-128932762 | Myc |
| chr8:128932825-128932865 | Myc |
| chr8:128932880-128932920 | Myc |
| chr8:128933070-128933110 | Myc |
| chr8:128933120-128933160 | Myc |
| chr8:128933300-128933340 | Myc |
| chr8:128933372-128933412 | Myc |
| chr8:128933457-128933497 | Myc |
| chr8:128933508-128933548 | Myc |
| chr8:128933602-128933642 | Myc |
| chr8:128933679-128933719 | Myc |
| chr8:128933924-128933964 | Myc |
| chr8:128934025-128934065 | Myc |
| chr8:128934197-128934237 | Myc |
| chr8:128934398-128934438 | Myc |
| chr8:128934528-128934568 | Myc |
| chr8:128934578-128934618 | Myc |
| chr8:128934628-128934668 | Myc |
| chr8:128934721-128934761 | Myc |
| chr8:128934782-128934822 | Myc |
| chr8:128934851-128934891 | Myc |
| chr8:128934947-128934987 | Myc |
| chr8:128935007-128935047 | Myc |

| | |
|---|---|
| chr8:128935077-128935117 | Myc |
| chr8:128935133-128935173 | Myc |
| chr8:128935210-128935250 | Myc |
| chr8:128935349-128935389 | Myc |
| chr8:128935440-128935480 | Myc |
| chr8:128935522-128935562 | Myc |
| chr8:128935670-128935710 | Myc |
| chr8:128935745-128935785 | Myc |
| chr8:128935806-128935846 | Myc |
| chr8:128935982-128936022 | Myc |
| chr8:128936053-128936093 | Myc |
| chr8:128936111-128936151 | Myc |
| chr8:128936266-128936306 | Myc |
| chr8:128936316-128936356 | Myc |
| chr8:128936384-128936424 | Myc |
| chr8:128936434-128936474 | Myc |
| chr8:128936530-128936570 | Myc |
| chr8:128937034-128937074 | Myc |
| chr8:128937171-128937211 | Myc |
| chr8:128937231-128937271 | Myc |
| chr8:128937292-128937332 | Myc |
| chr8:128937360-128937400 | Myc |
| chr8:128937447-128937487 | Myc |
| chr8:128937517-128937557 | Myc |
| chr8:128937629-128937669 | Myc |
| chr8:128937713-128937753 | Myc |
| chr8:128937763-128937803 | Myc |
| chr8:128937831-128937871 | Myc |
| chr8:128938168-128938208 | Myc |
| chr8:128938327-128938367 | Myc |
| chr8:128938389-128938429 | Myc |
| chr8:128938455-128938495 | Myc |
| chr8:128938582-128938622 | Myc |
| chr8:128938705-128938745 | Myc |
| chr8:128938847-128938887 | Myc |
| chr8:128938942-128938982 | Myc |
| chr8:128939011-128939051 | Myc |
| chr8:128939065-128939105 | Myc |
| chr8:128939185-128939225 | Myc |
| chr8:128939252-128939292 | Myc |
| chr8:128939324-128939364 | Myc |
| chr8:128939374-128939414 | Myc |
| chr8:128939468-128939508 | Myc |
| chr8:128939552-128939592 | Myc |
| chr8:128939602-128939642 | Myc |
| chr8:128939654-128939694 | Myc |
| chr8:128939715-128939755 | Myc |
| chr8:128939772-128939812 | Myc |
| chr8:128939823-128939863 | Myc |
| chr8:128939902-128939942 | Myc |
| chr8:128939985-128940025 | Myc |
| chr8:128940035-128940075 | Myc |

| | |
|---|---|
| chr8:128940104-128940144 | Myc |
| chr8:128940190-128940230 | Myc |
| chr8:128940245-128940285 | Myc |
| chr8:128940312-128940352 | Myc |
| chr8:128940378-128940418 | Myc |
| chr8:128940453-128940493 | Myc |
| chr8:128940569-128940609 | Myc |
| chr8:128940643-128940683 | Myc |
| chr8:128940727-128940767 | Myc |
| chr8:128940777-128940817 | Myc |
| chr8:128940873-128940913 | Myc |
| chr8:128940976-128941016 | Myc |
| chr8:128941034-128941074 | Myc |
| chr8:128941091-128941131 | Myc |
| chr8:128941143-128941183 | Myc |
| chr8:128941234-128941274 | Myc |
| chr8:128941347-128941387 | Myc |
| chr8:128941397-128941437 | Myc |
| chr8:128941525-128941565 | Myc |
| chr8:128941582-128941622 | Myc |
| chr8:128941661-128941701 | Myc |
| chr8:128942029-128942069 | Myc |
| chr8:128942117-128942157 | Myc |
| chr8:128942180-128942220 | Myc |
| chr8:128942270-128942310 | Myc |
| chr8:128942320-128942360 | Myc |
| chr8:128942376-128942416 | Myc |
| chr8:128942459-128942499 | Myc |
| chr8:128942683-128942723 | Myc |
| chr8:128942767-128942807 | Myc |
| chr8:128942890-128942930 | Myc |
| chr8:128942942-128942982 | Myc |
| chr8:128942992-128943032 | Myc |
| chr8:128943042-128943082 | Myc |
| chr8:128943098-128943138 | Myc |
| chr8:128943204-128943244 | Myc |
| chr8:128943377-128943417 | Myc |
| chr8:128943570-128943610 | Myc |
| chr8:128943651-128943691 | Myc |
| chr8:128943742-128943782 | Myc |
| chr8:128943890-128943930 | Myc |
| chr8:128943986-128944026 | Myc |
| chr8:128944136-128944176 | Myc |
| chr8:128944234-128944274 | Myc |
| chr8:128944319-128944359 | Myc |
| chr8:128944399-128944439 | Myc |
| chr8:128944490-128944530 | Myc |
| chr8:128944547-128944587 | Myc |
| chr8:128944629-128944669 | Myc |
| chr8:128944753-128944793 | Myc |
| chr8:128944815-128944855 | Myc |
| chr8:128944866-128944906 | Myc |

| | |
|---|---|
| chr8:128945023-128945063 | Myc |
| chr8:128945087-128945127 | Myc |
| chr8:128945159-128945199 | Myc |
| chr8:128945239-128945279 | Myc |
| chr8:128945466-128945506 | Myc |
| chr8:128945599-128945639 | Myc |
| chr8:128945710-128945750 | Myc |
| chr8:128945815-128945855 | Myc |
| chr8:128945866-128945906 | Myc |
| chr8:128945916-128945956 | Myc |
| chr8:128946011-128946051 | Myc |
| chr8:128946093-128946133 | Myc |
| chr8:128946185-128946225 | Myc |
| chr8:128946277-128946317 | Myc |
| chr8:128946329-128946369 | Myc |
| chr8:128946449-128946489 | Myc |
| chr8:128946597-128946637 | Myc |
| chr8:128946651-128946691 | Myc |
| chr8:128946701-128946741 | Myc |
| chr8:128946761-128946801 | Myc |
| chr8:128946850-128946890 | Myc |
| chr8:128946969-128947009 | Myc |
| chr8:128947031-128947071 | Myc |
| chr8:128947091-128947131 | Myc |
| chr8:128947181-128947221 | Myc |
| chr8:128947322-128947362 | Myc |
| chr8:128947490-128947530 | Myc |
| chr8:128947575-128947615 | Myc |
| chr8:128947638-128947678 | Myc |
| chr8:128947691-128947731 | Myc |
| chr8:128947742-128947782 | Myc |
| chr8:128947836-128947876 | Myc |
| chr8:128947893-128947933 | Myc |
| chr8:128948025-128948065 | Myc |
| chr8:128948133-128948173 | Myc |
| chr8:128948191-128948231 | Myc |
| chr8:128948241-128948281 | Myc |
| chr8:128948429-128948469 | Myc |
| chr8:128948576-128948616 | Myc |
| chr8:128948626-128948666 | Myc |
| chr8:128948727-128948767 | Myc |
| chr8:128948804-128948844 | Myc |
| chr8:128948929-128948969 | Myc |
| chr8:128949054-128949094 | Myc |
| chr8:128949135-128949175 | Myc |
| chr8:128949193-128949233 | Myc |
| chr8:128949285-128949325 | Myc |
| chr8:128949347-128949387 | Myc |
| chr8:128949417-128949457 | Myc |
| chr8:128949584-128949624 | Myc |
| chr8:128949685-128949725 | Myc |
| chr8:128949749-128949789 | Myc |

| | |
|---|---|
| chr8:128949852-128949892 | Myc |
| chr8:128949914-128949954 | Myc |
| chr8:128949977-128950017 | Myc |
| chr8:128950065-128950105 | Myc |
| chr8:128950179-128950219 | Myc |
| chr8:128950259-128950299 | Myc |
| chr8:128950309-128950349 | Myc |
| chr8:128950384-128950424 | Myc |
| chr8:128950439-128950479 | Myc |
| chr8:128950518-128950558 | Myc |
| chr8:128950640-128950680 | Myc |
| chr8:128950706-128950746 | Myc |
| chr8:128950798-128950838 | Myc |
| chr8:128950910-128950950 | Myc |
| chr8:128951018-128951058 | Myc |
| chr8:128951071-128951111 | Myc |
| chr8:128951128-128951168 | Myc |
| chr8:128951195-128951235 | Myc |
| chr8:128951298-128951338 | Myc |
| chr8:128951377-128951417 | Myc |
| chr8:128951478-128951518 | Myc |
| chr8:128951555-128951595 | Myc |
| chr8:128951625-128951665 | Myc |
| chr8:128951683-128951723 | Myc |
| chr8:128951763-128951803 | Myc |
| chr8:128951850-128951890 | Myc |
| chr8:128951922-128951962 | Myc |
| chr8:128952015-128952055 | Myc |
| chr8:128952085-128952125 | Myc |
| chr8:128952205-128952245 | Myc |
| chr8:128952255-128952295 | Myc |
| chr8:128952327-128952367 | Myc |
| chr8:128952379-128952419 | Myc |
| chr8:128952436-128952476 | Myc |
| chr8:128952545-128952585 | Myc |
| chr8:128952625-128952665 | Myc |
| chr8:128952676-128952716 | Myc |
| chr8:128952727-128952767 | Myc |
| chr8:128952843-128952883 | Myc |
| chr8:128952914-128952954 | Myc |
| chr8:128952966-128953006 | Myc |
| chr8:128953132-128953172 | Myc |
| chr8:128953204-128953244 | Myc |
| chr8:128953288-128953328 | Myc |
| chr8:128953366-128953406 | Myc |
| chr8:128953433-128953473 | Myc |
| chr8:128953496-128953536 | Myc |
| chr8:128953627-128953667 | Myc |
| chr8:128953678-128953718 | Myc |
| chr8:128953744-128953784 | Myc |
| chr8:128953832-128953872 | Myc |
| chr8:128954021-128954061 | Myc |

| | |
|---|---|
| chr8:128954089-128954129 | Myc |
| chr8:128954158-128954198 | Myc |
| chr8:128954263-128954303 | Myc |
| chr8:128954313-128954353 | Myc |
| chr8:128954390-128954430 | Myc |
| chr8:128954477-128954517 | Myc |
| chr8:128954531-128954571 | Myc |
| chr8:128954626-128954666 | Myc |
| chr8:128954728-128954768 | Myc |
| chr8:128954810-128954850 | Myc |
| chr8:128954862-128954902 | Myc |
| chr8:128954916-128954956 | Myc |
| chr8:128955021-128955061 | Myc |
| chr8:128955071-128955111 | Myc |
| chr8:128955209-128955249 | Myc |
| chr8:128955322-128955362 | Myc |
| chr8:128955395-128955435 | Myc |
| chr8:128955445-128955485 | Myc |
| chr8:128955537-128955577 | Myc |
| chr8:128955698-128955738 | Myc |
| chr8:128955791-128955831 | Myc |
| chr8:128955893-128955933 | Myc |
| chr8:128955983-128956023 | Myc |
| chr8:128956106-128956146 | Myc |
| chr8:128956219-128956259 | Myc |
| chr8:128956308-128956348 | Myc |
| chr8:128956375-128956415 | Myc |
| chr8:128956425-128956465 | Myc |
| chr8:128956476-128956516 | Myc |
| chr8:128956628-128956668 | Myc |
| chr8:128956725-128956765 | Myc |
| chr8:128956887-128956927 | Myc |
| chr8:128956979-128957019 | Myc |
| chr8:128957056-128957096 | Myc |
| chr8:128957127-128957167 | Myc |
| chr8:128957182-128957222 | Myc |
| chr8:128957243-128957283 | Myc |
| chr8:128957300-128957340 | Myc |
| chr8:128957367-128957407 | Myc |
| chr8:128957474-128957514 | Myc |
| chr8:128957530-128957570 | Myc |
| chr8:128957635-128957675 | Myc |
| chr8:128957760-128957800 | Myc |
| chr8:128957837-128957877 | Myc |
| chr8:128957921-128957961 | Myc |
| chr8:128957983-128958023 | Myc |
| chr8:128958091-128958131 | Myc |
| chr8:128958141-128958181 | Myc |
| chr8:128958192-128958232 | Myc |
| chr8:128958295-128958335 | Myc |
| chr8:128958359-128958399 | Myc |
| chr8:128958411-128958451 | Myc |

| | |
|---|---|
| chr8:128958557-128958597 | Myc |
| chr8:128958609-128958649 | Myc |
| chr8:128958678-128958718 | Myc |
| chr8:128958734-128958774 | Myc |
| chr8:128958829-128958869 | Myc |
| chr8:128958879-128958919 | Myc |
| chr8:128958940-128958980 | Myc |
| chr8:128959009-128959049 | Myc |
| chr8:128959097-128959137 | Myc |
| chr8:128959161-128959201 | Myc |
| chr8:128959323-128959363 | Myc |
| chr8:128959404-128959444 | Myc |
| chr8:128959454-128959494 | Myc |
| chr8:128959651-128959691 | Myc |
| chr8:128959711-128959751 | Myc |
| chr8:128959775-128959815 | Myc |
| chr8:128959867-128959907 | Myc |
| chr8:128959928-128959968 | Myc |
| chr8:128959989-128960029 | Myc |
| chr8:128960104-128960144 | Myc |
| chr8:128960181-128960221 | Myc |
| chr8:128960324-128960364 | Myc |
| chr8:128960407-128960447 | Myc |
| chr8:128960525-128960565 | Myc |
| chr8:128960615-128960655 | Myc |
| chr8:128960721-128960761 | Myc |
| chr8:128960849-128960889 | Myc |
| chr8:128960950-128960990 | Myc |
| chr8:128961000-128961040 | Myc |
| chr8:128961072-128961112 | Myc |
| chr8:128961124-128961164 | Myc |
| chr8:128961196-128961236 | Myc |
| chr8:128961247-128961287 | Myc |
| chr8:128961308-128961348 | Myc |
| chr8:128961389-128961429 | Myc |
| chr8:128961454-128961494 | Myc |
| chr8:128961512-128961552 | Myc |
| chr8:128961562-128961602 | Myc |
| chr8:128961672-128961712 | Myc |
| chr8:128961764-128961804 | Myc |
| chr8:128961829-128961869 | Myc |
| chr8:128961945-128961985 | Myc |
| chr8:128962031-128962071 | Myc |
| chr8:128962089-128962129 | Myc |
| chr8:128962211-128962251 | Myc |
| chr8:128962263-128962303 | Myc |
| chr8:128962313-128962353 | Myc |
| chr8:128962491-128962531 | Myc |
| chr8:128962586-128962626 | Myc |
| chr8:128962652-128962692 | Myc |
| chr8:128962702-128962742 | Myc |
| chr8:128962778-128962818 | Myc |

| | |
|---|---|
| chr8:128962832-128962872 | Myc |
| chr8:128962914-128962954 | Myc |
| chr8:128963006-128963046 | Myc |
| chr8:128963103-128963143 | Myc |
| chr8:128963154-128963194 | Myc |
| chr8:128963435-128963475 | Myc |
| chr8:128963492-128963532 | Myc |
| chr8:128963556-128963596 | Myc |
| chr8:128963665-128963705 | Myc |
| chr8:128963731-128963771 | Myc |
| chr8:128963836-128963876 | Myc |
| chr8:128963901-128963941 | Myc |
| chr8:128963960-128964000 | Myc |
| chr8:128964011-128964051 | Myc |
| chr8:128964195-128964235 | Myc |
| chr8:128964339-128964379 | Myc |
| chr8:128964485-128964525 | Myc |
| chr8:128964547-128964587 | Myc |
| chr8:128964640-128964680 | Myc |
| chr8:128964710-128964750 | Myc |
| chr8:128964807-128964847 | Myc |
| chr8:128964923-128964963 | Myc |
| chr8:128964976-128965016 | Myc |
| chr8:128965288-128965328 | Myc |
| chr8:128965338-128965378 | Myc |
| chr8:128965493-128965533 | Myc |
| chr8:128965601-128965641 | Myc |
| chr8:128965658-128965698 | Myc |
| chr8:128965799-128965839 | Myc |
| chr8:128965917-128965957 | Myc |
| chr8:128965990-128966030 | Myc |
| chr8:128966098-128966138 | Myc |
| chr8:128966173-128966213 | Myc |
| chr8:128966299-128966339 | Myc |
| chr8:128966349-128966389 | Myc |
| chr8:128966414-128966454 | Myc |
| chr8:128966499-128966539 | Myc |
| chr8:128966857-128966897 | Myc |
| chr8:128966914-128966954 | Myc |
| chr8:128967024-128967064 | Myc |
| chr8:128967115-128967155 | Myc |
| chr8:128967300-128967340 | Myc |
| chr8:128967427-128967467 | Myc |
| chr8:128967502-128967542 | Myc |
| chr8:128967625-128967665 | Myc |
| chr8:128967826-128967866 | Myc |
| chr8:128967885-128967925 | Myc |
| chr8:128968009-128968049 | Myc |
| chr8:128968105-128968145 | Myc |
| chr8:128968160-128968200 | Myc |
| chr8:128968211-128968251 | Myc |
| chr8:128968290-128968330 | Myc |

| | |
|---|---|
| chr8:128968377-128968417 | Myc |
| chr8:128968455-128968495 | Myc |
| chr8:128968525-128968565 | Myc |
| chr8:128968622-128968662 | Myc |
| chr8:128968672-128968712 | Myc |
| chr8:128969070-128969110 | Myc |
| chr8:128969162-128969202 | Myc |
| chr8:128969257-128969297 | Myc |
| chr8:128969408-128969448 | Myc |
| chr8:128969561-128969601 | Myc |
| chr8:128969674-128969714 | Myc |
| chr8:128969937-128969977 | Myc |
| chr8:128970063-128970103 | Myc |
| chr8:128970120-128970160 | Myc |
| chr8:128970200-128970240 | Myc |
| chr8:128970285-128970325 | Myc |
| chr8:128970338-128970378 | Myc |
| chr8:128970412-128970452 | Myc |
| chr8:128970593-128970633 | Myc |
| chr8:128970657-128970697 | Myc |
| chr8:128970816-128970856 | Myc |
| chr8:128970866-128970906 | Myc |
| chr8:128970975-128971015 | Myc |
| chr8:128971067-128971107 | Myc |
| chr8:128971206-128971246 | Myc |
| chr8:128971289-128971329 | Myc |
| chr8:128971352-128971392 | Myc |
| chr8:128971416-128971456 | Myc |
| chr8:128971526-128971566 | Myc |
| chr8:128971711-128971751 | Myc |
| chr8:128971909-128971949 | Myc |
| chr8:128972032-128972072 | Myc |
| chr8:128972082-128972122 | Myc |
| chr8:128972157-128972197 | Myc |
| chr8:128972223-128972263 | Myc |
| chr8:128972273-128972313 | Myc |
| chr8:128972346-128972386 | Myc |
| chr8:128972416-128972456 | Myc |
| chr8:128972474-128972514 | Myc |
| chr8:128972573-128972613 | Myc |
| chr8:128972651-128972691 | Myc |
| chr8:128972713-128972753 | Myc |
| chr8:128972797-128972837 | Myc |
| chr8:128972882-128972922 | Myc |
| chr8:128972933-128972973 | Myc |
| chr8:128972993-128973033 | Myc |
| chr8:128973074-128973114 | Myc |
| chr8:128973192-128973232 | Myc |
| chr8:128973247-128973287 | Myc |
| chr8:128973297-128973337 | Myc |
| chr8:128973353-128973393 | Myc |
| chr8:128973414-128973454 | Myc |

| | |
|---|---|
| chr8:128973516-128973556 | Myc |
| chr8:128973592-128973632 | Myc |
| chr8:128973720-128973760 | Myc |
| chr8:128973771-128973811 | Myc |
| chr8:128973832-128973872 | Myc |
| chr8:128973884-128973924 | Myc |
| chr8:128973934-128973974 | Myc |
| chr8:128974044-128974084 | Myc |
| chr8:128974094-128974134 | Myc |
| chr8:128974171-128974211 | Myc |
| chr8:128974352-128974392 | Myc |
| chr8:128974493-128974533 | Myc |
| chr8:128974543-128974583 | Myc |
| chr8:128974594-128974634 | Myc |
| chr8:128974645-128974685 | Myc |
| chr8:128974754-128974794 | Myc |
| chr8:128974804-128974844 | Myc |
| chr8:128974882-128974922 | Myc |
| chr8:128975123-128975163 | Myc |
| chr8:128975258-128975298 | Myc |
| chr8:128975315-128975355 | Myc |
| chr8:128975426-128975466 | Myc |
| chr8:128975515-128975555 | Myc |
| chr8:128975576-128975616 | Myc |
| chr8:128975626-128975666 | Myc |
| chr8:128975693-128975733 | Myc |
| chr8:128975790-128975830 | Myc |
| chr8:128975861-128975901 | Myc |
| chr8:128975933-128975973 | Myc |
| chr8:128976006-128976046 | Myc |
| chr8:128976065-128976105 | Myc |
| chr8:128976197-128976237 | Myc |
| chr8:128976318-128976358 | Myc |
| chr8:128976378-128976418 | Myc |
| chr8:128976489-128976529 | Myc |
| chr8:128976595-128976635 | Myc |
| chr8:128976671-128976711 | Myc |
| chr8:128976748-128976788 | Myc |
| chr8:128976848-128976888 | Myc |
| chr8:128976902-128976942 | Myc |
| chr8:128976956-128976996 | Myc |
| chr8:128977132-128977172 | Myc |
| chr8:128977184-128977224 | Myc |
| chr8:128977295-128977335 | Myc |
| chr8:128977358-128977398 | Myc |
| chr8:128977474-128977514 | Myc |
| chr8:128977527-128977567 | Myc |
| chr8:128977612-128977652 | Myc |
| chr8:128977688-128977728 | Myc |
| chr8:128977748-128977788 | Myc |
| chr8:128977801-128977841 | Myc |
| chr8:128977851-128977891 | Myc |

| | |
|---|---|
| chr8:128977923-128977963 | Myc |
| chr8:128978033-128978073 | Myc |
| chr8:128978120-128978160 | Myc |
| chr8:128978194-128978234 | Myc |
| chr8:128978265-128978305 | Myc |
| chr8:128978333-128978373 | Myc |
| chr8:128978449-128978489 | Myc |

**ing the DNA FISH probes used in this study.**

er sequences.

| Sequence (5'->3')* |
|---|
| gcagtcatatagagcagacgGAAAATGACAGTCAAATGAGAAATCAGATGTGTGGAAAACaagttatactgccct |
| gcagtcatatagagcagacgTCCCAGGAATTCCATAGCTCACCAGTAACTTTCCACTCTGaagttatactgccc |
| gcagtcatatagagcagacgCAAAACGAGTTGGAATTGTGGGAAGAATCTAGAAAATCAAaagttatactgccct |
| gcagtcatatagagcagacgCTGCAAGTGGAGTTAAAAGTAATAAGGAAAATAGTTTGGCaagttatactgccct |
| gcagtcatatagagcagacgTATCTTATTGTTGCACCTATCAAGTCTATTCCTGTTCTTCaagttatactgccctgt |
| gcagtcatatagagcagacgACTATCCTTAGTATCATCCCTCTAGACTAAACAAAAAGAGaagttatactgccct |
| gcagtcatatagagcagacgGCACAATTAGCAGAGGGTAGGGATGTATTCATTTAGCAAAaagttatactgccct |
| gcagtcatatagagcagacgGTATCTACTGTGTGCCAGCCTTGGCCCAGCCTAAAGACACaagttatactgcc |
| gcagtcatatagagcagacgTCTTAGCTTAGAGAGAGACAAACCTTTGCGCATCGAGTGAaagttatactgccc |
| gcagtcatatagagcagacgAACAAAGTGCTGTTCATACTGAGGGGGAAGAAGTGATTCTaagttatactgccct |
| gcagtcatatagagcagacgCAGAGAAGGTGACGTAAGAGCTGAGCCATGGGGTTTGATAaagttatactgcc |
| gcagtcatatagagcagacgCATTTCAGACAGATGGAAGAGCAAAGCCCAAAAGTAGGAAaagttatactgcc |
| gcagtcatatagagcagacgGTGATCACTGGTGTTGGGAAGAACTTGGATCATAACCAAAaagttatactgccct |
| gcagtcatatagagcagacgGACATAGTGGTATTGCCAAAGTCTAGGGATCTTCAGTTGAaagttatactgccct |
| gcagtcatatagagcagacgATGTCACAGCTCATGGGAGGACCTTAGTTTTGTGCTTGTGaagttatactgccct |
| gcagtcatatagagcagacgAGCCAGTGATAGGGTATATTCAGTAAGAAAGAAGAGTAGCaagttatactgccc |
| gcagtcatatagagcagacgGCCTTTGCCTCAAATATACAGTACATCCCACATGCAGATTaagttatactgccct |
| gcagtcatatagagcagacgCTGCGTCCATTAAAAAATCTTAATTCCAGGTGTCAAAAAGaagttatactgccct |
| gcagtcatatagagcagacgAGTAAAAAAACTGACTGTCCCTACCTTGCTTCGGAATGATaagttatactgccct |
| gcagtcatatagagcagacgCCACTTGATTGTAGGTTGAGGGATACCATATGCGGTACAAaagttatactgccct |
| gcagtcatatagagcagacgGGGTCTGGAAACGATGACACGGCCTTTTCGTAAATCTGGGaagttatactgccc |
| gcagtcatatagagcagacgGAGCAGTTCTAATGACTTTCTCCTGGGCCACACATGAAGGaagttatactgccc |
| gcagtcatatagagcagacgCACTGGCTTTGATGTCAGGGAAACAAACAAGAAACAAAAAaagttatactgccc |
| gcagtcatatagagcagacgATTGTGATTACAGTGATAAAAGATCATGCCGCCTATGGAAaagttatactgccct |
| gcagtcatatagagcagacgCACGAGATTGTTAAGATGGGAACTTAACGTTAAGTGTTTTaagttatactgccctg |
| gcagtcatatagagcagacgTTCACTAATTTGTTGCCTACCAGAACCAGAAGAAAGTATGaagttatactgccct |
| gcagtcatatagagcagacgTTGACAATTAGAGTGGCAGTGAGGTTTTTAAACACATTTGaagttatactgccctg |
| gcagtcatatagagcagacgCAAGATTGTATAATTCCCTGCATCCTCTTCAGTTACGTCCaagttatactgccct |
| gcagtcatatagagcagacgCGAGCAGATTTCTCAGTCGCAGGTAACCCATCTTTTAACCaagttatactgccc |
| gcagtcatatagagcagacgTGATCAGTAAGTCATTTCACACGGCCTCTCAATATCACTTaagttatactgccct |
| gcagtcatatagagcagacgTCATTGAAGTTTTTGAGCTCAAGTTGACCAATTTCAAACCaagttatactgccctg |
| gcagtcatatagagcagacgGATTTTTAGACTGGGTAACACTTCATATTTATTGGCAAGGaagttatactgccctg |
| gcagtcatatagagcagacgCTGGTGGCCACTGTGACTTAAGTATTAACCCAGAAGCAGGaagttatactgccc |
| gcagtcatatagagcagacgATCCGAGATGTTTTTCAGAGTTTTCCACGAGGCTCCATCAaagttatactgccct |
| gcagtcatatagagcagacgAGACTTTTCTATTGTGGTCTCATGTGGAGTTCTTATTAGTaagttatactgccctgt |
| gcagtcatatagagcagacgCACCAGTTGTCCCTTTCATAAACTAGAGTGAAATCAAGACaagttatactgccct |
| gcagtcatatagagcagacgCTCTGAGGAAGAACAGTTAAGGCACAGAAAAAAAAAAAACaagttatactgccc |
| gcagtcatatagagcagacgTTAAATACTGAATCGATGCATTGATATCACTCTGTTGCACaagttatactgccctg |
| gcagtcatatagagcagacgAGAGGCTTGGGTGCTTGACCTGGCTCTGCTACTCACTGCAaagttatactgccc |
| gcagtcatatagagcagacgCTGCAAATGTGACTAACCAGTTTGGGCTCCAATTTTCTCAaagttatactgccct |
| gcagtcatatagagcagacgGCATGGTTCTTTAAATAGGCAGTTTTAATCCATTAGGAGTaagttatactgccctg |
| gcagtcatatagagcagacgACAAACAAATTCCCATTACTAGTGTGTCTCAGATCCAAAGaagttatactgccct |
| gcagtcatatagagcagacgTGAGGATGCTACACTTTGTGATTACAAATATTGTCTTGGTaagttatactgccctg |
| gcagtcatatagagcagacgTTTGACGGTGGATTGAATTTTTTTTTAATGCCCTTGCTACaagttatactgccctgt |
| gcagtcatatagagcagacgGAAGAAATCAACACTAACAGATGAGCGTGCCCTCAGACACaagttatactgcc |
| gcagtcatatagagcagacgGAGTGAATGTACCTCCTGGAGTCATCAGATTACTATTCAAaagttatactgccct |
| gcagtcatatagagcagacgAAAAGATCATTTTTCTACTGCAATGGCATGAAGACATAGCaagttatactgccct |
| gcagtcatatagagcagacgGGGAACTGCAAACAAACACTGCCCTCATATGACATTGAAAaagttatactgccc |

gcagtcatatagagcagacgTCAGCTCTGGTTAGGGAGATTCTAAAATGGCGCCACATCCaagttatactgccc
gcagtcatatagagcagacgAACAGATTACATTGTTTTACATCAGGTGTTGTACCAGTGTaagttatactgccctg
gcagtcatatagagcagacgAACAACTGCTGAAGGAGGTTTTGAATATGTATCTTTCTAGaagttatactgccctg
gcagtcatatagagcagacgACTTAATTAGTTCAAATAGGCCCCATCGTCCCCCTCTTCAaagttatactgccct
gcagtcatatagagcagacgCAGGACTGGAACTTTCAACCCAGCTGTATATAAAAGAATTaagttatactgccct
gcagtcatatagagcagacgCCACTTCTTCAGGGTTAATAGAGCATGTTACATATTCAAAaagttatactgccctg
gcagtcatatagagcagacgATCATAGGAAGGTTTAGAGATGCAGAAAGTTAGCCACAGTaagttatactgccct
gcagtcatatagagcagacgAGCACTCAAACAAACCTCATCTCTCGCTTACCTTGCTACAaagttatactgccc
gcagtcatatagagcagacgTCTCACCTCCACCACCCAAGAATCTACAGAGTTCTGAGCTaagttatactgccc
gcagtcatatagagcagacgAGGAACGTCTCTTGGTATTTTGTCAGGAATTCGAGTCATTaagttatactgccctg
gcagtcatatagagcagacgGCAGCTTCCCACTTACAAAGTATGTCTTAAACTGTTATGGaagttatactgccct
gcagtcatatagagcagacgGTCCTACCATACAACTTCCGGCGAGGGAAATGCCTTCACTaagttatactgccc
gcagtcatatagagcagacgCAGCTACGGTTTCAAGCACCTGCTGTTTTCGATGTTTCACaagttatactgccct
gcagtcatatagagcagacgAGAATAATTTCCATAAACAAAACCTTAGGGCCTCATTCAGaagttatactgccct
gcagtcatatagagcagacgATAGAGCCTATCAATGAGCCCAAATGTTGATCATGAATAGaagttatactgccct
gcagtcatatagagcagacgAACGCCTTTCAAATCAAATGGTATCACGTCCTAGCCACATaagttatactgccct
gcagtcatatagagcagacgGCAGAGCAACAGAAAGTGAAACACCTTAAATATCTCTATCaagttatactgccct
gcagtcatatagagcagacgGAAGATATTTCTAAAAATCTACCTGGCCACTATAAGCAAGaagttatactgccct
gcagtcatatagagcagacgAAATCAACTGGAGAAAGACAAAAGGAAATAAAGGCTACTGaagttatactgccc
gcagtcatatagagcagacgATTGTTCCATAGTTTCTGCCCTCCACACTGTTTCCTTTTCaagttatactgccctg
gcagtcatatagagcagacgCTGAAATTAAGCAACAGCATTAGGATTCCTTGAGAGTAAAaagttatactgccct
gcagtcatatagagcagacgATTAGCCATAATAAGCATCTAAAGAAAGGACTCTCTGTTCaagttatactgccct
gcagtcatatagagcagacgACTTAATGTGGTTTGCTTTTTGACAGGGGGAGGAGGGAACaagttatactgccct
gcagtcatatagagcagacgAATGTAACTCTTGGCCAACAGAAAACTGACTTATTTAGAGaagttatactgccct
gcagtcatatagagcagacgTCTACAATGATTAGAAAGCATAGCATGAGTTGGCAACGCTaagttatactgccct
gcagtcatatagagcagacgAGGTGTTAACAGATTTGAATGATGGTCCACTCCTTACACGaagttatactgccct
gcagtcatatagagcagacgGCATTTTGTCAATAGTAAGAACACTGTAGAATATGTTGGCaagttatactgccctg
gcagtcatatagagcagacgCTTTGGTCAATTCGTCGCCTTACGGAAGCTCTCTTATTTTaagttatactgccctg
gcagtcatatagagcagacgGAAAATGTATAACAAATGTAGAAGGAGTGAGTGTTCATGCaagttatactgccct
gcagtcatatagagcagacgTCTGTTTTATACTAAGAATCATGTCAAATCCACCTAGAGCaagttatactgccctg
gcagtcatatagagcagacgTTGGAAACTACATCTGTGTCTAGAGGTGCTTCAAATATGTaagttatactgccctg
gcagtcatatagagcagacgCACTTTCACATATACTATCTCAATTGTTTGTCCCAACAACaagttatactgccctg
gcagtcatatagagcagacgAGTGATGAAAAAACTGAAGGAAGTGATACCCTGGCCCAAGaagttatactgccc
gcagtcatatagagcagacgACTGAAGTTGTTTGAGCATAAATCTAGTGTCCTTCCTCTGaagttatactgccctg
gcagtcatatagagcagacgCAGTAACCTGTTCCCTAGGCATAGATATCGCACACAAACGaagttatactgccc
gcagtcatatagagcagacgCTGGCTTAATGACAGTGCTTCTCTCACCAAATCACCATGTaagttatactgccct
gcagtcatatagagcagacgCTACGAGATATGAAGGAAAATATAATGAGTCTTGGCTGTAaagttatactgccct
gcagtcatatagagcagacgCACTATTTGAACAGTGCTTGCCCATGTCAGCTAAATCACAaagttatactgccct
gcagtcatatagagcagacgCTGTCCCAGGTTTGTCTCAGAGAAGATCAAGAATTAGCCTaagttatactgccct
gcagtcatatagagcagacgTCTGCAATCTGACCTGGCCACAATGACAGTCCTGCTCACTaagttatactgccc
gcagtcatatagagcagacgGACAAGATCACATTGCCGTCTCCTTTTTTTTTTAAGCACTaagttatactgccctg
gcagtcatatagagcagacgAATAAGTTATTTCCACCTCTCAATCCTTCCCCATAGTCATaagttatactgccctg
gcagtcatatagagcagacgGCGGAGGGGAGGAGCATGGAAAAATGTCTTTGGGAGCTGTaagttatactgcc
gcagtcatatagagcagacgGATTTTTGGATCTTGAGCTTAGTTCTCCTCTGTGCCACGTaagttatactgccctg
gcagtcatatagagcagacgACAGACAGACACATGTTGAGGCCATTTTCTTCATTCTCAGaagttatactgccct
gcagtcatatagagcagacgACCAGAGAGAAAAGTCAATTGCCACTATCAAAAGTCAGAGaagttatactgccc
gcagtcatatagagcagacgCTTTCTGCAGCTCTATTTATACAGGTACTACTTCCTTTGCaagttatactgccctg
gcagtcatatagagcagacgACTCATCCTGTGTTTTAGAATTGCTTCGCCCAGTACTGCAaagttatactgccct
gcagtcatatagagcagacgCACATTTCAAGTGCTCATCTGCCATATGTGGCTAGTGGCAaagttatactgccct
gcagtcatatagagcagacgCTGGAGCACAAATACGGGATATTTCCATCACAGAAAGTTCaagttatactgccct
gcagtcatatagagcagacgAATAGATGTGTTGTAGATACTCTTACTTACAGCAGCAATGaagttatactgccctg
gcagtcatatagagcagacgGGATAAAAGTTTCTTCCCTCTTGGCACTTACGGAGAATATaagttatactgccct

gcagtcatatagagcagacgCTCCTAGAGAATAGTTTATGACTGATAAGTCCTTAATGTGaagttatactgccctg
gcagtcatatagagcagacgACAGGGCACTCGTTACACGTTTTTGTTGCTAAGTTATTCCaagttatactgccct
gcagtcatatagagcagacgATGGCGAGAAATGTCGATGAAAAACAACACTAGCTTGAGGaagttatactgccc
gcagtcatatagagcagacgTTGACTTTGTAATTGGGCAATCTGAACGACATAACAAATGaagttatactgccct
gcagtcatatagagcagacgCATAAATCATGCTGACACACAAGAAATGGGGGCTAGAACTaagttatactgccc
gcagtcatatagagcagacgATTATTGTTTGCAAGGATGTTCGATTTGAATAAGGGAACCaagttatactgccctg
gcagtcatatagagcagacgAGAATTGCTCCCCTGACACTTAACAAATGGAATATTGGATaagttatactgccct
gcagtcatatagagcagacgGGTTGTAATATCTTTTCCAAACATGCTCCAATCAGGTGCCaagttatactgccct
gcagtcatatagagcagacgCTGACAGAGACAGCCTCCTGGAAAGTTGTTGTTGTTGTTAaagttatactgccct
gcagtcatatagagcagacgTTGGTGATAAATTTCTCTCTCATCTCTAAATCCTACAGAGaagttatactgccctg
gcagtcatatagagcagacgGAAACAATCCCTTTTCTTCCACCTAGAATCATGCAATAGAaagttatactgccct
gcagtcatatagagcagacgGTCTACACTCTTCAAAACAAAACCCCCAGACATCAAAAGAaagttatactgccc
gcagtcatatagagcagacgTGAAATGCCAGGCAAAGAAAGCACTCAAACCAAACCCAAAaagttatactgcc
gcagtcatatagagcagacgGGAGTCAGGATTTTCTGATTTTTCCCATCTCAAAACCCCTaagttatactgccct
gcagtcatatagagcagacgAAGAAATAGAAAAATGGGTCACTCACTTCATCAGGGTAACaagttatactgccc
gcagtcatatagagcagacgCTCTCGTGTTCAAGGGAGTGGTAGCAGTACAATTGAGGACaagttatactgccc
gcagtcatatagagcagacgCGTGGTGTGCTTATTTGGACATCTATGATTGTATTGGCTGaagttatactgccctg
gcagtcatatagagcagacgAAGATTTCATTACACAGATAAAAATACAGAGCACTTCGGCaagttatactgccct
gcagtcatatagagcagacgTCACTTACTTTGTCGATGTGTGAGATAGTTTGTCTTATACaagttatactgccctgt
gcagtcatatagagcagacgTTCACAGGTCAGAAGCCCTATTTCTTTGTACGTTGCATTTaagttatactgccctg
gcagtcatatagagcagacgAGCCCTTTCTACTGTCCCAGATTATGCGTTTTCCATCAGGaagttatactgccct
gcagtcatatagagcagacgAGGAAGAGGCATGCATTAACAAGGTCCTATTAGCACTGGTaagttatactgccc
gcagtcatatagagcagacgACTGTTTCACCAGCAGTTTGTGAAAACATGATTTTTCATCaagttatactgccctg
gcagtcatatagagcagacgGATAATTTACTATCAGCCTTGGCAACAAGTGGCATCTACCaagttatactgccct
gcagtcatatagagcagacgCCTACTTGTCATGTCAGACTTTCAGTTGGAGCCCATAACTaagttatactgccct
gcagtcatatagagcagacgCTCCGGGAATCTTCATAATAAGCTGGTCAGACACTCAATAaagttatactgccc
gcagtcatatagagcagacgTGGGCGGATCACCTGAGGTTAGCAGTTAGAGACCAGCCTGaagttatactgcc
gcagtcatatagagcagacgAAACAGAGAGAGAAACCAATTAGCATTTTTTTTCTACCGGaagttatactgccct
gcagtcatatagagcagacgCTTTTAACAAGCCATAGTTTTTCCTCCCATTTTGCATAGTaagttatactgccctg
gcagtcatatagagcagacgAAGTGGCTGTCTTCAAAAGAGTGTAAATAATTCCAGGTCCaagttatactgccct
gcagtcatatagagcagacgCACTGGGGCTGTCATTCTGACGATCCTGCTGTGTCACAAGaagttatactgccc
gcagtcatatagagcagacgCCTCGGAGGCCAGAGACTCTGCAATGAATGAAGTGCTCTCaagttatactgcc
gcagtcatatagagcagacgAAGTATTCCCAAGCACGGTGAGTCCAAACAGAAGAAAATAaagttatactgccc
gcagtcatatagagcagacgTGATCTGACTTTCTCCTTATGCCTTCTTCTGACATCAAGAaagttatactgccctg
gcagtcatatagagcagacgACATTTAAAGACTACAAGATAAACACTCTGCCTTGGGATTaagttatactgccct
gcagtcatatagagcagacgGGTCATTGCTCCATAAAGTGGCCCAGCCAAATTCCCCTGAaagttatactgccc
gcagtcatatagagcagacgGTTGCCAGCCCAATTTCAGTGGTGGTAGACTCCCAACTACaagttatactgccc
gcagtcatatagagcagacgCCTTCCCCCAACAGATTATAAAGGAAATACCAAACAGCCAaagttatactgccc
gcagtcatatagagcagacgGGCATCCCTCGCTCTATCCAGTTATCATTTATTTCTATTGaagttatactgccctg
gcagtcatatagagcagacgGATTTGTGATCCTATGGCAGAGATTTCTAGTCCAGAAGCCaagttatactgccct
gcagtcatatagagcagacgACCGGTGTTCCTTGGGTTTGAAATTAATTACATCCTGTTTaagttatactgccctg
gcagtcatatagagcagacgAAGGATTCGCTTTCATGTTTCAGTAAATCAAGGTGCTACCaagttatactgccct
gcagtcatatagagcagacgCAGGAATGCAGTCGGTGGGAAAAAAAGAACCTTCTCTTCTaagttatactgccc
gcagtcatatagagcagacgAGTGCAAGCAAAGGAAAAACCCAGAAAAATTTAAGTCAGGaagttatactgccc
gcagtcatatagagcagacgGCATGTGCTGAATCTAAAAGAAAGGATTGCTGACAGACGTaagttatactgccct
gcagtcatatagagcagacgACTAAAATTTCCTCCGAAGTGTTTCTCTTACTCTCATCTAaagttatactgccctg
gcagtcatatagagcagacgGTAGCATTCCATTGACAAAGTAACTTTACATTTAGCTGACaagttatactgccctg
gcagtcatatagagcagacgATGTGTAAGAATGAAGGCAGATGGGCAGGTGGATCTCAGCaagttatactgccc
gcagtcatatagagcagacgAGCAAGGCCTGTCTGCAGCTGACCATGGCTCCGACAAAGCaagttatactgcc
gcagtcatatagagcagacgGGAGAGGGAGGGGACAGAAAATTAGAAGAAAAGGGACAATaagttatactgcc
gcagtcatatagagcagacgGATATTAACTTCATGTCAGCAGAAATACGTCATAGGTTAGaagttatactgccct
gcagtcatatagagcagacgTTCCAGTATATATGTGCCAAACCAAAATCCCTGTGAAAATaagttatactgccct

gcagtcatatagagcagacgTCTCTTTCTCTCTTTTTAAATTGTGCACTCCAGATCGTAGaagttatactgccctgt
gcagtcatatagagcagacgAAAAACAGTAAATAGACTCCCACCAGCCCCAGATTTTTATaagttatactgccct
gcagtcatatagagcagacgGATGACCAAACAACCCATCTGCATTAGCGCTCCCATGAAAaagttatactgccc
gcagtcatatagagcagacgCGTTTCCCTCTCCACGTGGTGTTTTCCATATTAATCATACaagttatactgccctg
gcagtcatatagagcagacgATCTCAGACTAATGGGAGTAATGCCATGCTATTGAAAACAaagttatactgccctg
gcagtcatatagagcagacgGGGTGTTAAACTGTTAAATAATGCTAAAGCTAACCCTTGAaagttatactgccctg
gcagtcatatagagcagacgAGTGCTTTCCATATATCACCTGACTTAAGTTTTCTAACAGaagttatactgccctg
gcagtcatatagagcagacgTCAGATGGGGAGAGTGAGGTGGAGAGAGGTTTGATAACTCaagttatactgccc
gcagtcatatagagcagacgCCCAGTGACAAGTGCAGTATGTTCAGAGTGCAGGCAATGTaagttatactgccc
gcagtcatatagagcagacgGCCTCTGCTCTTAACCACACTTGTCTTCCTCCTATATGAAaagttatactgccctg
gcagtcatatagagcagacgGCTCAAATGCCCAGTATGACCTGCGAAGTTCCTGACACTCaagttatactgccc
gcagtcatatagagcagacgACCAGGGTGTTTTTATCTTGACCAACTGAGGAGTTCAACTaagttatactgccctg
gcagtcatatagagcagacgAGCTCTGCTGTATAATGACTTTCTATTTCCTGGCGATGGCaagttatactgccctg
gcagtcatatagagcagacgCGTGAGTAACAACCCAGGAGAGTGAATGCAAGGAAAGAGGaagttatactgcc
gcagtcatatagagcagacgGAATCCTCCACCACATTTCAGGACTCAGGGTTGATGGGTGaagttatactgccc
gcagtcatatagagcagacgTCATTCTCTGAACTTTTCCAGCCTAACTGCCACAGCTACCaagttatactgccct
gcagtcatatagagcagacgCACTGTGCGAGGTGCTGCTCTGGGGCGGACTTGACTTTCTaagttatactgccc
gcagtcatatagagcagacgACTTAGCACAGTTTTCCGTTTGGGGAGCTTTTATCCAGTTaagttatactgccctg
gcagtcatatagagcagacgTCCTGGTATACTCCTAACTCTTACTGCTGATTAATTAGCTaagttatactgccctg
gcagtcatatagagcagacgAAAAGTTTCACGTATGGCCTTCTTTCCTCCCTTTTTCCCTaagttatactgccctg
gcagtcatatagagcagacgTGAAGTTTGGGTGCTCAGCAGGAGACAGGCCTTTTATCACaagttatactgccc
gcagtcatatagagcagacgCTTGCCAATTCAAACCAAACGTGCTGGCTCCTTATGCATGaagttatactgccct
gcagtcatatagagcagacgACATTCTGTCTTCTCTGCAGTAATCTGGATAATCCACTCTaagttatactgccctg
gcagtcatatagagcagacgAGAATTGATTTTTCTCACATTTGCACACAGTTGGGTTTGTaagttatactgccctgt
gcagtcatatagagcagacgAGACACCATCATTTTATTCTATTTAACCACATGCATGGACaagttatactgccctg
gcagtcatatagagcagacgGGAAAGTCCACTGAGAAGCCCAGGTGTTTGTAAGGAAATTaagttatactgccc
gcagtcatatagagcagacgCAGTGAAAGTATGCTGAGAATAGCGGTGTTCAAATTTACCaagttatactgccctg
gcagtcatatagagcagacgGACGAGCTCCCTTCCTTCAGTCATGTGTATAATTTCGGGGaagttatactgccct
gcagtcatatagagcagacgACTGTACATCTCTACGAAAGGTCTACCTGTATCTGAATGAaagttatactgccctg
gcagtcatatagagcagacgGCATTGCCACAGTCACGTCATTCTCCCTTAATGTAATCCAaagttatactgccct
gcagtcatatagagcagacgACATCTTTGTCTCTGAATTTGGCCCTCTGTGTTTCTACCCaagttatactgccctg
gcagtcatatagagcagacgAGCTTCCCTAATAAAACCTAATTTCATATCTGTTGTTGGCaagttatactgccctg
gcagtcatatagagcagacgTGGTATGGGGGGACCACTGTGGGAAAAAGCCTAAAAGTGCaagttatactgccc
gcagtcatatagagcagacgCATAGACTTCTCACCACAGCTATGCTAAGTGCCTCAGGAAaagttatactgccc
gcagtcatatagagcagacgGCAGTGGCGAAGGTCAACTCTCTTTGGAGAATTCAAGGTGaagttatactgccc
gcagtcatatagagcagacgAGCAAGTTGAGGAGAGGATTATCATTTGAATGACAGAAGCaagttatactgccct
gcagtcatatagagcagacgTCGGAGGCTAATAATAATGAATGAGAAGGTCTGAGCTGCTaagttatactgccct
gcagtcatatagagcagacgCAGGCCGGAGAAGCTGTGTTGAATGCTGCATTTATTTCATaagttatactgccctg
gcagtcatatagagcagacgAGGCTTATTTCTTTAGTATGCGCTACTTACCTGCTAATAGaagttatactgccctg
gcagtcatatagagcagacgCTGGAAAAGACTTTAAGCACCAATACTTCCTTAAAAAGCGaagttatactgccct
gcagtcatatagagcagacgCCAGTCTCAGAGGCAAGGAGGATTCAAACACCTGTTCAGCaagttatactgccc
gcagtcatatagagcagacgAATGAGTCCAAGATCCTTTTAGAAACATTCTCAGTCTACCaagttatactgccctg
gcagtcatatagagcagacgCCTCGGGAATCTATTGACATCCTCAGGCAACGATCTCTTCaagttatactgccc
gcagtcatatagagcagacgTCATTGTGTTCTGATCGTCCCGCATTTGTTCCAGCCTTATaagttatactgccctg
gcagtcatatagagcagacgGGAAAGAAGTCCTGTGTGAACTCTCTCCATTTGGACCTGAaagttatactgccct
gcagtcatatagagcagacgCCTGCTTGCTGGTCCTGCCTCAGGTTTGTGGATTACGAAGaagttatactgccc
gcagtcatatagagcagacgAGGATGCTGAATGGAGAAGGAATTGGAGAGAAACAAAAGCaagttatactgccc
gcagtcatatagagcagacgTTGCCCAGACTAGATGGTGGCTACAGTGATCATTATGGTAaagttatactgccct
gcagtcatatagagcagacgAAGTCCTAACCCCCCTGTACTTCTGAATGTAACCTCATTTaagttatactgccctg
gcagtcatatagagcagacgCTTAAGCTAAGTAAGGGAGGGCTCCTCATCCAATGTGATTaagttatactgccct
gcagtcatatagagcagacgCTTGTAAGAAGGCAGTCATGTGAAGACAGACAGGGTGTGCaagttatactgccc
gcagtcatatagagcagacgGCAGATGGAGGCAGAGACAGGTGTGATGCAGCGGCAAGCCaagttatactgc

gcagtcatatagagcagacgGGGCAACCACCAGCATGTAGGAAGAGGCGAGGGAGGGTTCaagttatactgc
gcagtcatatagagcagacgCATGGCCCAGATGATTTGATTTCAGACTTAGAGTTTCCAGaagttatactgccctg
gcagtcatatagagcagacgGGCCATTGCTGTAGCCTGCTGAAGATGGGATAGTTTTCTGaagttatactgccct
gcagtcatatagagcagacgCAGATGTGAGCTCTAGAGAGACCCTTTCACCACAGCCCATaagttatactgccc
gcagtcatatagagcagacgGGTCCCTGGCACCTATATGTTAATTATTCCCACTCATCCTaagttatactgccctg
gcagtcatatagagcagacgGCCTTGCTAGCCCCCTTCCTTCCTGCACCCTTCTCCACCCaagttatactgcc
gcagtcatatagagcagacgGGACTTTCCCAGTGTTCCCATTGATTCTGTTACAACACGGaagttatactgccct
gcagtcatatagagcagacgGCAGTTTTTTTCTCTGGGTACTCCTTGGATGCTACATTCCaagttatactgccctg
gcagtcatatagagcagacgCATGTCTGTCTCATGTGTACTGAATGAATGAGTGAACGGGaagttatactgccct
gcagtcatatagagcagacgACAGATGAGAAAAGAGGCACAAACAGGTGAAATGAGTTGTaagttatactgccc
gcagtcatatagagcagacgGCCAGTTACTTCTGCCTCCAATCTAGAGCCCCTTCCCTGGaagttatactgccc
gcagtcatatagagcagacgGCTGAACCTCTGGCTACGTTCTCTTGAAGGCCCCCATAGAaagttatactgccc
gcagtcatatagagcagacgGCAGGTGACAATGGTTTTTCAGAGATGAAGGCTGATTTCCaagttatactgccct
gcagtcatatagagcagacgTGTTGGAGAAATTCTGAAAAAGGGCTGAACACCAAGGCATaagttatactgccc
gcagtcatatagagcagacgCTTTGGGGCAGGCTTTGAGTCGGAGGGAGGTGGTTCAGAGaagttatactgcc
gcagtcatatagagcagacgCTGGGTTTAATCCTGGCTGTGGTCCTGTGTAACTGTGAGAaagttatactgccct
gcagtcatatagagcagacgTGTAAAATCTAAGTGAAAGGGCTATGATGAGGATTACATGaagttatactgccctg
gcagtcatatagagcagacgATGACTCATTTGAACTTATGAACAAAGTGTTAGAGTTGGCaagttatactgccctg
gcagtcatatagagcagacgCAGACATGAGTGGGACAGGAGAGTGCCCCTCCTCCCAAGAaagttatactgcc
gcagtcatatagagcagacgTTAGTCACAGCTGGCACCAGAGACCGTCTCCCAACAGATAaagttatactgcc
gcagtcatatagagcagacgCAGGCTCAAGCATGCACACTCAAAGGCAAAATGGCACAGTaagttatactgcc
gcagtcatatagagcagacgCGGTATATGACTTTCCTCTGTGAATGCTCAACTGGTAAGGaagttatactgccctg
gcagtcatatagagcagacgAACTTCAGTAAACACAGTGCGCATGTGGCTCCTCCCAGGTaagttatactgccc
gcagtcatatagagcagacgGGACAGCCCATCCCAAAGGAAAAATCAGGGGAAAATAAACaagttatactgcc
gcagtcatatagagcagacgGCCCTCTGCCAAGTGTAGTTTGCTTCCTCTCATTCCTGCTaagttatactgccct
gcagtcatatagagcagacgTCTCCCTCTGCCTTACACCTACTAATGCCCTTGGCTGAATaagttatactgccct
gcagtcatatagagcagacgCAGATTCGCTGCTGGTACCAGTAGGTAGATATGAAAATCAaagttatactgccct
gcagtcatatagagcagacgGATATAGTGGCATAGGAAAAAGGATTGGAAACAAACATACaagttatactgccct
gcagtcatatagagcagacgGCAAGATCCCTTCTCTGCCCAAAAAAAGTTAAAAATGATTaagttatactgccctg
gcagtcatatagagcagacgTGGGTGTTGGCTTTATGGATTATTGTTGTTCTTATTTACCaagttatactgccctgtg
gcagtcatatagagcagacgACTTATGTGTTTATTAGGAAATCTCAGTGGGTGTAGTAACaagttatactgccctg
gcagtcatatagagcagacgCAGAGAGAAAAAGTTAATCAGATAGACACAAGGGTTTCTTaagttatactgccctg
gcagtcatatagagcagacgCTGAGCTAGACATTCAGCCATGGACTATTGGGATTTGATCaagttatactgccct
gcagtcatatagagcagacgCCTTCCTCTTGTGAGTCTATATTTTTGTCTCTTTTGTGCAaagttatactgccctgtg
gcagtcatatagagcagacgCTTAAGCAACTGTTTTCAACAAAGGACACTGGCACAAAAAaagttatactgccct
gcagtcatatagagcagacgATCTTTGAATTTCTTCTTTCCCTTTGGTACTGTGGATCCCaagttatactgccctgt
gcagtcatatagagcagacgGCCTCTAAAAGCCAATACCAATTTCAGCACTGGGTCCACTaagttatactgccc
gcagtcatatagagcagacgGAAGATATCAGAACTTCAGTGATTTGTGTCCCTCCAAACAaagttatactgccctg
gcagtcatatagagcagacgTGGCATTATAAATCCACTGTGTTTGCAAATGAGCATATGAaagttatactgccctg
gcagtcatatagagcagacgCGAACACACGAACAATACTCTCTGGGCTTATTAATAGTATaagttatactgccctg
gcagtcatatagagcagacgCTCTGCAATTTACTTCAAGGAAGGGCTAAGAATGAATTTGaagttatactgccctg
gcagtcatatagagcagacgATATAGAAGAGGTCACGGTTCCATTAATATCACCAGTAAAaagttatactgccctg
gcagtcatatagagcagacgTCACAAAATTAAAAATGACTTCCAATCCCAGCCAGCACTGaagttatactgccct
gcagtcatatagagcagacgCCAAATTGGCTGGTTGTTGAAAGAGTGAGGAAGAGCTGCTaagttatactgccc
gcagtcatatagagcagacgACACACTGTCACCAGGGACTCATGCCAACACTCAGCTAGGaagttatactgcc
gcagtcatatagagcagacgCACACTGCCCTCACCTCTCTGAACTCTTAGAGCTCAAAGCaagttatactgccc
gcagtcatatagagcagacgCACTGGGAGTATCTGGGCTGTCACCTATCCTTCAGGACAAaagttatactgccc
gcagtcatatagagcagacgAGAAAGATCAAACAGGGACCCTTGCTCAGAATGGAACGATaagttatactgccc
gcagtcatatagagcagacgTGTGTTTTTTCCTTTTGAAATCTGTAATCATGCTGGCACAaagttatactgccctgt
gcagtcatatagagcagacgGCATACAGACATAGTGGCCCCTGCATGTTATCCTCAATGGaagttatactgccc
gcagtcatatagagcagacgAACAATGTTCAAAATTTAGGACTTGTGACTTGAAGAGGAGaagttatactgccctg
gcagtcatatagagcagacgTTTCCTGGTTGCCTTTCTTTTAATTGCATTTTTATCCTGGaagttatactgccctgtg

gcagtcatatagagcagacgTACTGAAGATGAAGGGGAGAGGATAAAAAATGAATTAACCaagttatactgccct
gcagtcatatagagcagacgAAAAGGGTTCTATCTAGACATAAGAGTTTTCACATCACACaagttatactgccctg
gcagtcatatagagcagacgGTGGACCGTAAGTAATACTCAAGGAGAATGTACTGATTGGaagttatactgccct
gcagtcatatagagcagacgGTTCTTCCATGACTTACACTGCCTATCAAAGCCACTGACAaagttatactgccct
gcagtcatatagagcagacgGAAATTATTAGTGAGGAAATGGTTAGTGAAGATAGAGAGGaagttatactgccctg
gcagtcatatagagcagacgTATGTACCTAATAACAGAGCACTAACACACAGGAAGCAAAaagttatactgccc
gcagtcatatagagcagacgTAGACAGTTCAACAATAACAGGTGAAGATTTCAATACCCCaagttatactgccct
gcagtcatatagagcagacgCAATAGTAGATAGTGGGCTAGGCATAGTGGCTCATGTCTGaagttatactgccct
gcagtcatatagagcagacgCACAAGTTAGCCAGTGTGGTGATGCACGCCTGAAGTCTCAaagttatactgccc
gcagtcatatagagcagacgAGTGGATAGAAGAACTGGACAGAAGATTTTAAAAAGCTAGaagttatactgccct
gcagtcatatagagcagacgCACCAGAACAACACTACTAACCAACTATACCCAACGACATaagttatactgccc
gcagtcatatagagcagacgCACCCATCACAGCAGGGTATATATTTTCCTCAAGTGCACAaagttatactgccct
gcagtcatatagagcagacgATATGTGGAAATTGACACATCCCTAAATAATCAATGGGTCaagttatactgccctg
gcagtcatatagagcagacgAACCTAAACTTCCACCTTAGAACATTGAGTAAAGAACAGAaagttatactgccct
gcagtcatatagagcagacgAAGACAAACACAACTAATAAACCTTAAGCCAGACAAGCCAaagttatactgccc
gcagtcatatagagcagacgAGACTCAAATTACTAGATTCAGAAATGAAACAGGGCCAGGaagttatactgccc
gcagtcatatagagcagacgGACGGGTGGATTATCTGAGGCAAGGGTTCAAGACCAGCCTaagttatactgcc
gcagtcatatagagcagacgGGGTGACAGAGCTAGACTAAAAAGAAATGAAACAGTGGACaagttatactgccc
gcagtcatatagagcagacgTCACAGATGAAATGGATAAATGCCGGGAAAGACACAAACTaagttatactgccc
gcagtcatatagagcagacgCTCAACAAGAATTAGACAACCCAAGCAGACTTTTAACAAGaagttatactgccct
gcagtcatatagagcagacgACAACAACAACAACAAGCTGCTCACGAGGGAAAGTCAGGGaagttatactgcc
gcagtcatatagagcagacgACTCATTCTAGGAGGCTAGTTTTACTCTGATATCAAAATCaagttatactgccctg
gcagtcatatagagcagacgATTATATACAATGCCCAAGTGGGATTTTTTTCAGGAATGCaagttatactgccctg
gcagtcatatagagcagacgAAACATCAAAACCCACATGACTATCTTAAAATGGCAGAAGaagttatactgccct
gcagtcatatagagcagacgGACAAAAATCCAAGCTCCTTTCGTGATGAAAACACTCAGGaagttatactgccc
gcagtcatatagagcagacgAACCTAATAAATGGCATCTAGAAAAAACCCTACGTCTAACaagttatactgccct
gcagtcatatagagcagacgACATCGTTGTCCACCCTCACCGCTTCTATTCAATAGTGTAaagttatactgccctg
gcagtcatatagagcagacgTGGGCAAGAAAAATAAATCCAGATTGGGAAGATCAAAATGaagttatactgccct
gcagtcatatagagcagacgTTTCAGATGACATGATCTTACAAAATGCTAAGGAATCCACaagttatactgccctg
gcagtcatatagagcagacgGAGTTCAGCAAGATTGCAGGATACAATGTACAAAATGAATaagttatactgccctg
gcagtcatatagagcagacgTAAAGAAGATCTACTAAGTGGGAAACATCTCATGTTCATGaagttatactgccctg
gcagtcatatagagcagacgCAGATTAAATGCAATAACTGTAAGAATCCCAGATGACTTCaagttatactgccctg
gcagtcatatagagcagacgCAATCTTGAAAAGCAAAAACAATCTTGGAAGACCGCACTCaagttatactgccc
gcagtcatatagagcagacgGTGCTGGCACAATGGATTAGTGGAATAGAATTGAGACTCCaagttatactgccct
gcagtcatatagagcagacgCCATGTGTCTATGGCCACTTGATTTTCAAGGAGACCATTCaagttatactgccctg
gcagtcatatagagcagacgCATGCAAAAGAATGGAGTTGTACTCTTATATCACACCATAaagttatactgccctg
gcagtcatatagagcagacgATATATATTTGCAAAAGGGTCTCACTTTTCTAGGCTGAAGaagttatactgccctg
gcagtcatatagagcagacgGCAGCTTCAACATCCCAGGCGCAAGTGATGCCCCCACCTCaagttatactgcc
gcagtcatatagagcagacgCCACACCTGGCCTAATACAATATTCTCAGAAGAAACCAGAaagttatactgccc
gcagtcatatagagcagacgTTTGAAACTTGCATGCTTCAAAGAACACTGTGTAGAAAGTaagttatactgccctg
gcagtcatatagagcagacgAAATGGGAAAAGGATCTGAACTGACATTTACCCAAAGAAGaagttatactgccct
gcagtcatatagagcagacgCTTAACATCATAAGCCATCATGGAAATACCAATCAAAACCaagttatactgccct
gcagtcatatagagcagacgTAAAAAAGACATAATTACAAGTGTTGGCAAGGATGTGGAGaagttatactgccctg
gcagtcatatagagcagacgTGGGAATGTGAAGTAGTGCACCCACTTTGGAAAGCAGTCTaagttatactgccc
gcagtcatatagagcagacgCATCAGTTGATTGATGGATAGCTAAAATGTGATATGTCCAaagttatactgccctg
gcagtcatatagagcagacgACTCAGAAATAGAAAGAAATCAAGTGCTGATACATGCATGaagttatactgccct
gcagtcatatagagcagacgTATCCATGAGACAAATCTATGGGAACAGAAAGTAGATAAGaagttatactgccct
gcagtcatatagagcagacgGTTTAGGGCTGGGGGGATAAGGGGTGATAGCTAAAGGCTAaagttatactgccc
gcagtcatatagagcagacgAATGACATATTCTAAAATTGACCGTGATGATGATGGTTGCaagttatactgccctg
gcagtcatatagagcagacgATGCTACAAACCACTGAATCATCATTTTAAATGGGTGTGGaagttatactgccctg
gcagtcatatagagcagacgAGGTTTAGTCTCTTTCTTAGAAATGATGATCCACTGTTTCaagttatactgccctgt
gcagtcatatagagcagacgCACCTTATGAGTCTTTGCAGCATTTCTCTCCCATTCCCCTaagttatactgccctg

gcagtcatatagagcagacgTCCCCTGTGATAGAACCATCATGATTTAAATATGTCTTTCaagttatactgccctg
gcagtcatatagagcagacgTTGGGAGCCCAAGACAGAAGCATTGTTTGAGGCTAGGAGTaagttatactgccc
gcagtcatatagagcagacgCTAAGATAGGAGGATCAATCGAAGCCCAGGAATTCAAGGCaagttatactgcc
gcagtcatatagagcagacgTGTCTCTAAGAAAAAAAAATGTCCTTCAGTTCCTTGTTGGaagttatactgccctg
gcagtcatatagagcagacgATCTCTGCCTTCACATAGATGGGAATACTTGCCTGGTGTGaagttatactgccct
gcagtcatatagagcagacgTGCGCAATTTAATATCTCCTAGTAAGCTATCCTGATTATCaagttatactgccctg
gcagtcatatagagcagacgTTATCAGCTAAAAATATAACAACTGTCGCCACTATTGGTGaagttatactgccct
gcagtcatatagagcagacgAATTCAAAGGTGATAAAAAATCTCACTGAGACTGGCTTAGaagttatactgccct
gcagtcatatagagcagacgTGCCATCCAGACAGACACTTACATGTTACACAAATATTTCaagttatactgccct
gcagtcatatagagcagacgGAGCAGGGTAAACTTAGAGAAAAGTCATTGTGTTTCAGACaagttatactgccct
gcagtcatatagagcagacgCTTTCTAAAAAAGAGCTCTTTGCCTACTACCTGCCTTTCCaagttatactgccct
gcagtcatatagagcagacgTTGTTTGTTTTGCATTGCTTCACTTATAACAGCAGATCGAaagttatactgccctg
gcagtcatatagagcagacgGCAGAGCAAAGGAGAAGAATTAGAAAGGAGGAACAGGAGTaagttatactgcc
gcagtcatatagagcagacgCTGGGATGCTCTTTGGAATGTGGGTGTCTAGCGGCAGGGGaagttatactgcc
gcagtcatatagagcagacgCAGATGTTTATGGAGAAGATATAAGATTGAAGCAAGCAAGaagttatactgccct
gcagtcatatagagcagacgGAAATCAGGACGTATTATGTACCCAGCACTGTGCACACGGaagttatactgcc
gcagtcatatagagcagacgACACACACCTCACACACAAGAATGCTCATGTTGATTAGAGaagttatactgccc
gcagtcatatagagcagacgAGGGAAGTAGGGACCTCATTTTAAAAATTTACAGCTAAGAaagttatactgccct
gcagtcatatagagcagacgTCAGTGAGAGTGGAATGCTGGCTAGATTTAAGCTTGATGGaagttatactgccct
gcagtcatatagagcagacgGTGCTCTCTGGGTTTTCACACTAAGCTGGCTGGCGGTTTCaagttatactgccc
gcagtcatatagagcagacgACCTCAGTCAGAGATCTTCATCTTGATCAGTTGAGACATCaagttatactgccct
gcagtcatatagagcagacgATCTTTCTTCTACTGCTAAGATTTTTGGCCATGTCCCAAGaagttatactgccctg
gcagtcatatagagcagacgAGCTCTGCCTCTGAATCATAACATTATAAAACTGTCAAAGaagttatactgccct
gcagtcatatagagcagacgAAGATGCCCTAATTCTGCCTATCATGTTACAGATGAGAAAaagttatactgccct
gcagtcatatagagcagacgCAGAGAGGAGAGACTGTCCCACAAGCACAGGAATACAATTaagttatactgcc
gcagtcatatagagcagacgGCAAAAACTGAATTTACGCTTCCTGATCTGAGGCTAGTCTaagttatactgccct
gcagtcatatagagcagacgCACTCCTCAATGACTCAAGCGTGACAATTACATCTTCAAAaagttatactgccct
gcagtcatatagagcagacgGCATGGGGCCTTCTGACGGGGACTAGAAATGGGAAGCAACaagttatactgcc
gcagtcatatagagcagacgTGCTGTAAGGTGCTGAGGACTTTTTAGGAGATGGTGAGGAaagttatactgccc
gcagtcatatagagcagacgCCAGGCACGGTGGCTGACGCCTGTCATCCTAGCACTTTGGaagttatactgcc
gcagtcatatagagcagacgATACAAAAAATTAGCTAGACGTGGCGGCAGGCACCTGTAGaagttatactgcc
gcagtcatatagagcagacgACAAAACAAAAACCAACAGTCACACCAGAGCAAGCCCTGTaagttatactgcc
gcagtcatatagagcagacgAGGATGCCTGGTCAGACTCCTCTCCAGCAGGGACACTGGGaagttatactgc
gcagtcatatagagcagacgGGGGCGGCGGGGGAGGGGAGCGGGTCGGGGTGGAGGAGGGaagttatact
gcagtcatatagagcagacgCAGGAGTCAGGGGAGGAGTCGGGCTGTGTCGAAGTGCTGCaagttatactgc
gcagtcatatagagcagacgTGACCTGCGCCACCACGGGACTCTCTAGGGCTCAGCTTCCaagttatactgc
gcagtcatatagagcagacgGGCACTTTCCTGTTCCTAAACTCGCGATTTGTTATGCATTaagttatactgccctg
gcagtcatatagagcagacgCGTAGCACACTGACATCCCTCCGGGGGCCAGACACACGAGaagttatactgc
gcagtcatatagagcagacgGGTGGGACAGGAGGTGTTGTGGCAGCTGGGAACGTGAGCCaagttatactgc
gcagtcatatagagcagacgTGCAGTCCATTGTTGCCAAGTGGAAATATGCGCCAGTGTGaagttatactgccc
gcagtcatatagagcagacgGCTAAAAGTGCTGTGTCTGCCAATCAAAACATGTGTGCCAaagttatactgccc
gcagtcatatagagcagacgCAACCCTGCATGCACTCCTGTCCCTTCTCTCCCATTTTATaagttatactgccct
gcagtcatatagagcagacgCCAGGCAAGGACACGTTAGAAAGACCGGGGGCTTTGGAGTaagttatactgcc
gcagtcatatagagcagacgAAAGCTTTGACAGGATAATTTACGAAGTATGCAGCCTTAAaagttatactgccct
gcagtcatatagagcagacgCTTTTCTGTAAAGGAATTTCTCTTATTGGGTTGTTTCGAGaagttatactgccctgt
gcagtcatatagagcagacgTGAGATGATGTATGTATTAATAACAAGCCTGAGCCGGACAaagttatactgccct
gcagtcatatagagcagacgCCTACACCAGGTGCTTAATGAATCCCAGCAGAGGAAAGGTaagttatactgcc
gcagtcatatagagcagacgTCATCTGATCTGGTCCCATCAAGTGAGTGTCTGCTTCCTAaagttatactgccct
gcagtcatatagagcagacgGGACTAATCACTCCCAAGACATTCCTTCCTTCCACTTTGGaagttatactgccc
gcagtcatatagagcagacgCTTTGCATGTCAGAAAGTGTTTCCTTAAGTCAAGCTGAAAaagttatactgccct
gcagtcatatagagcagacgAGATAGTTGTTCTGCCATCTAGGATTATAAAGACGAAGCTaagttatactgccct
gcagtcatatagagcagacgGACTGTCACTCATGTCTCCTATTTTCCCACCCAGTTCTTTaagttatactgccct

gcagtcatatagagcagacgAAAATCATCAAATTACTTCATGCTCACAACCCTATGAGGCaagttatactgccct
gcagtcatatagagcagacgTTGCTTGAAATTCCTATTTTTAATGTCCCTCTTGAGGTGTaagttatactgccctgt
gcagtcatatagagcagacgCCTTTATTCCTGGTGCTATATATTTCTTCCTTTCCTCCCCaagttatactgccctg
gcagtcatatagagcagacgCTTTCCTTTCCTTTCCTTTCCTTTCTGACAGAGTTTCACTaagttatactgccctgt
gcagtcatatagagcagacgCAGGCTGGAGTGCAATGCTGCCATCTCAGCTCACTGCAAGaagttatactgcc
gcagtcatatagagcagacgTAAGCCACCACGCCTGGCAGGGTGCTACATATTTCTTAATaagttatactgccc
gcagtcatatagagcagacgAGGTTGGATACATTCTCTTGGTAAATATGGAGTCAACTGAaagttatactgccct
gcagtcatatagagcagacgTGGCTTCCCCCATCCAGTACTAGGGTAGTTAGTGGGTTTTaagttatactgccct
gcagtcatatagagcagacgCATGTATTCCTGTTAAATCTCATTTTGCTAATCCCTGTTCaagttatactgccctgt
gcagtcatatagagcagacgGACATGATTTTGACCCCCACATTTGTCTCATTTTCTCAATaagttatactgccctg
gcagtcatatagagcagacgGTCAAGGAAGTCAGGTAAGACTAATATACTTAAAAATCCCaagttatactgccct
gcagtcatatagagcagacgACCTTCTGTGTACTATACCCTGTGCTGGGTCCTGGGGCTAaagttatactgccc
gcagtcatatagagcagacgCTCAGTCCTGTCCGTAGAGCTTGTGAGAGGTGAGTCCTGTaagttatactgccc
gcagtcatatagagcagacgCCAGAGGGGGGATATTTAACTTGAGACCTGAAAAGTGAGGaagttatactgccc
gcagtcatatagagcagacgGAATGTCCCAGGCAGAGCACCCCCAGTCCACACCCCTCTTaagttatactgc
gcagtcatatagagcagacgTTAATTGTAACTCATTTTCTTTGACCTACCAAACTGCAGGaagttatactgccctg
gcagtcatatagagcagacgAAAGTTGTTGGTATCATCTTTGTACTTCCTACTGTGTCCAaagttatactgccctg
gcagtcatatagagcagacgCTAAGTGTTAGGCAGTCTGTATTGTCTCATGTAGTCTCCTaagttatactgccctg
gcagtcatatagagcagacgAATAAACTCGTATATAGTGCATACTATACACCAAACACCGaagttatactgccct
gcagtcatatagagcagacgCACATATGCACTCTTGATTATGGGAGGTGGTTATAGGCAAaagttatactgccct
gcagtcatatagagcagacgTTGTTGCCATCTCTGCCATACACTAGTTATATAACCTTGAaagttatactgccctg
gcagtcatatagagcagacgCTACTTAACCTTTCTGAGAAATAAATGGAGATGACTGTGAaagttatactgccct
gcagtcatatagagcagacgATCAGGATCAAGGCAAGGATCTTCCCTCTCACTACTACTTaagttatactgccc
gcagtcatatagagcagacgACACTGGAAGTCTTAACTCATGCAATAAGACAAGAAAGGGaagttatactgccc
gcagtcatatagagcagacgACAAAAATACCCCTGGAACTAGTAAAAGATTATAGCAAGCaagttatactgccc
gcagtcatatagagcagacgCACAATACCATTTGCATTAGCACCCCCAAAAGTGAAACACaagttatactgccc
gcagtcatatagagcagacgAATCAAAATCCCAGAAGTTACTTTGTGGATATGGACAAACaagttatactgccct
gcagtcatatagagcagacgATAATATTGAAGAAGAGCAAAGTTGGAAGACTGACACTACaagttatactgccct
gcagtcatatagagcagacgTTGACAAAGGAGCAAAAATCATCTTGTCAATAAATGGTGCaagttatactgccct
gcagtcatatagagcagacgCAAACAAACAAAAAATGAGGCTAGACACAGACCTTACACTaagttatactgccc
gcagtcatatagagcagacgACAGGAGAAAACCTAGATGAACTTCGGTATGACAATGACTaagttatactgccc
gcagtcatatagagcagacgACACTAAAGGCATGATCTACAAAACAATTGATAAGCTGGAaagttatactgccct
gcagtcatatagagcagacgACAACTCTGTTTAAAATGGGTCATTTCACCAAAGAAGACCaagttatactgccct
gcagtcatatagagcagacgATGAAAAGGTGCTCCACGTCATGTCATTGGGAAAATGCAAaagttatactgccc
gcagtcatatagagcagacgATGAGAGACCATTACATACTTATTAGAGCAGCCAAAATCCaagttatactgccct
gcagtcatatagagcagacgATGTTGGCAAGGAGGCAGAGTGATAGGAATTGTCATTTGCaagttatactgccc
gcagtcatatagagcagacgTTCTTACAAAACTAGCCTTTCTATTGCCCTAAAATTGAGCaagttatactgccctg
gcagtcatatagagcagacgCCCAAAGGACCTGAAGAATTACATCCACCCAAAACCTGCAaagttatactgcc
gcagtcatatagagcagacgGATGTCCTTTGGTAGGTGAATGGGTAAATAAACTATGAATaagttatactgccctg
gcagtcatatagagcagacgAACTATGGAAACAGTAAGAAAGATAAGTGTTTGCCGGCCGaagttatactgccc
gcagtcatatagagcagacgAAAAGAAAGAAAAATAAGGATAGGTGTTTGCCAGGGGTTGaagttatactgccc
gcagtcatatagagcagacgGGAAAGGATGACTAAGTAGAGCACAGTAGATTTTAGGGCAaagttatactgccc
gcagtcatatagagcagacgACATGTCATTGTATGTTTCTCCAATCCATAGAATGTCCAAaagttatactgccctg
gcagtcatatagagcagacgACCCTCATGTACACTTTGCACTTGGGTGATTATGATGTCTaagttatactgccct
gcagtcatatagagcagacgGGGGAATGTTGATAATGAAGACTATGCATGTGGGGACAAGaagttatactgccc
gcagtcatatagagcagacgGACATGCACACACATCTAATTTTTAAAATGAGAGCACTACaagttatactgccct
gcagtcatatagagcagacgGGATAAAATCTGTTAATGAATGCAAAGTGCTTAGGATGGCaagttatactgccct
gcagtcatatagagcagacgTGGATTAAGACCCACCTAAATGTTAACCATTCTCCTTATTaagttatactgccctg
gcagtcatatagagcagacgATTTGTCCAAAGTCAAAGACCTAGTGAGTATCCGAACAGGaagttatactgccc
gcagtcatatagagcagacgAAGCCTGGATCATGTTGTTTCAACTGTAGCTTATTAATTGaagttatactgccctg
gcagtcatatagagcagacgGCTACCTATTTTGAACAACACAATCAGCTAGCCAACATAAaagttatactgccct
gcagtcatatagagcagacgGTTAAAGGAAAAACTGTAGAATATTAGAACTGGAAGGACCaagttatactgccct

gcagtcatatagagcagacgATAAAAACAGATCACCTCCTCATCTCCAAGCAAGGCAAGCaagttatactgccc
gcagtcatatagagcagacgTGAAATGCCAGAAACTTAAAGCCTTTTCAGCTTAAGGGGAaagttatactgccct
gcagtcatatagagcagacgCCTTGCTTATTTTTCTTAAAAGATCACTTGAGGGAGAGTTaagttatactgccctg
gcagtcatatagagcagacgCACTTCTTGATTGTAGGGGAAAAATGAGAGAATATGGTCAaagttatactgccctg
gcagtcatatagagcagacgCCTTTGGACTTCATTGCCATCTTTAGATTTCTTTTCCTTTaagttatactgccctgtg
gcagtcatatagagcagacgCCGAGTAGCTGGGACTATACACATGCGCCATCACACCTGGaagttatactgcc
gcagtcatatagagcagacgACCACACCCGGCCCATCTTTAGATTTCATAATTCCACCACaagttatactgccc
gcagtcatatagagcagacgCCAAGTGGCTTATTCAGGACTATGTTCATTCTGATTATTTaagttatactgccctg
gcagtcatatagagcagacgGTTCTACTAAATGCCTGATGATGTATAACTGCGTCGTTGCaagttatactgccctg
gcagtcatatagagcagacgATGCCAAATTTTTTCTGGTATGGATTTTAGAGAGCCCCAGaagttatactgccctg
gcagtcatatagagcagacgAGGGAATGGCCTGAACATTTTACATGTCCACACACCAAGTaagttatactgccc
gcagtcatatagagcagacgAGGGGCCAAAGAGAGAAGATGCATTTGTTCCTGACTGCCAaagttatactgccc
gcagtcatatagagcagacgAGTGGCTTCTGTCCCAATCATATATCAGGGTCTTCTATAGaagttatactgccctg
gcagtcatatagagcagacgGAAATGTTTTTAGAAGGGAAATGTGACATGTCAGGGCTTAaagttatactgccctg
gcagtcatatagagcagacgCCACGTAGCTGGAAAGGAAAATGGTTAAGAGTTAAGTTACaagttatactgccct
gcagtcatatagagcagacgTCTAAACTCTCACCATGTGGTCCTAATCAGCTGTCTGCCAaagttatactgccct
gcagtcatatagagcagacgCTAGAAGGAGTGCACAGCCAGGATGTAGAGGGGTCTATTTaagttatactgccc
gcagtcatatagagcagacgCGATTGCAAACAACATCATCAGTGTGTATTTTAAACCCTAaagttatactgccctg
gcagtcatatagagcagacgCACTAGGTTTTTGATGAAGATGTGTGCAGATGGGCCAGAAaagttatactgccct
gcagtcatatagagcagacgAATTTGAAGGATAATTCTGCAGATGCACGTCAGTCTCAGGaagttatactgccct
gcagtcatatagagcagacgACTGAGAGTCCTACCTTTGTTCACCCCAACATGTTCACCTaagttatactgccct
gcagtcatatagagcagacgGGATCCATCACCCAACCTGTCTGCACTTGTGGCCTCCTGGaagttatactgccc
gcagtcatatagagcagacgGAACCCTAGACAATTCCTGATCAAAACTTTGGAAACCAGTaagttatactgccct
gcagtcatatagagcagacgATGGAAGGAGCCACATGTTAGGGATAGAGGTGAGATGAGTaagttatactgccc
gcagtcatatagagcagacgGGAAGCTCTTTGGATGGGCAGGTGGAGGAGAGAATGAGGCaagttatactgcc
gcagtcatatagagcagacgGAAGACTCTAGACATTAACCAAGCAATATTAGATAACGTCaagttatactgccctg
gcagtcatatagagcagacgAACTAGTGTGGCTATGAGTTTGGGTTTTATTTTGGGCGCTaagttatactgccctg
gcagtcatatagagcagacgAAAATGTAAACTGTTTTTGCCTATTTTCCCTTGGAGGAGAaagttatactgccctg
gcagtcatatagagcagacgAGTGATTCCATTGACATATTTTTGAAGCTTTGGCAATCTGaagttatactgccctg
gcagtcatatagagcagacgGCTCTCTGCCCAACACAGTGCTGACCTGCATGAAAGCGTGaagttatactgcc
gcagtcatatagagcagacgGCCCACATTCCTGGTGCATTTCCATCTTACAAGGGTCCTTaagttatactgccct
gcagtcatatagagcagacgACAGGGGGAAAGTGGAGGCGGCAAGGCTCGGCGATGAAGCaagttatactgc
gcagtcatatagagcagacgCTACTCTGTTGATGTCCCTGCTGGTTCCATGGAGATGTTTaagttatactgccctg
gcagtcatatagagcagacgGCCATACCAGTCTGGCAGATTGGGCAACAAACACCATCTGaagttatactgccc
gcagtcatatagagcagacgCTGAGCCTTCTGTGTGTGCCAAATTGTGTCAGATCTGGGAaagttatactgccct
gcagtcatatagagcagacgCTGGAGCAACAGCCACGCCTCACAGAACATGAATAGGGCTaagttatactgcc
gcagtcatatagagcagacgGAGCCCTCTTCACTCCTTAGAGACCCACTTCTGAACAATAaagttatactgccc
gcagtcatatagagcagacgACAAATCTGTGAGCTATGGTCCTAGAGAGTAACTGGGGGAaagttatactgccc
gcagtcatatagagcagacgGTGACTTTGCTTTTTATGTCTCTCACAGATCAAACAACAGaagttatactgccctg
gcagtcatatagagcagacgTGTGGAGTAGGTATGTAATCAACATCTGCAGACAGATTCCaagttatactgccct
gcagtcatatagagcagacgGCCAAGCCTCCGAGTGGGAAAACTCAGTTCAGGAGTGAAAaagttatactgcc
gcagtcatatagagcagacgCAAAAGCAGTGGACCCAAATCAGCTAAAGGAGAGGGCTTCaagttatactgcc
gcagtcatatagagcagacgCCAGACCTCTGTTTCAGTACTGGTGTGCCCCTCCCCCCACaagttatactgcc
gcagtcatatagagcagacgGCTTACAATTGGCACTGCTTACCTAAATAATAGATATGGTaagttatactgccctg
gcagtcatatagagcagacgCACAAAAAGAAATACATTTTACGTCAGACCCATTACACACaagttatactgccct
gcagtcatatagagcagacgACACACACACGCACGTATGTACTTTACATTTATGAGGGTAaagttatactgccctg
gcagtcatatagagcagacgACTAAACAACACTTACCCTTACTCTTTGCAATTTACTTGGaagttatactgccctg
gcagtcatatagagcagacgTGAGACTACAGGTGGATGCCCCAGCTCCTAGCTTTTTTTTaagttatactgccctg
gcagtcatatagagcagacgGGTTGAAACCCATTCATTTGATTCCATAGCCCATTAATGAaagttatactgccctg
gcagtcatatagagcagacgGAGGCACATTTTTAAAAGCCCTGCTCAACTTAGTAAATTTaagttatactgccctg
gcagtcatatagagcagacgCAATTAAAACAACAACTAGCTCGTAGTTTGATGTCAAGAGaagttatactgccctg
gcagtcatatagagcagacgGGGGAAATAGACCTAACCTAGCTGGATCCTCATCCTGGTTaagttatactgccc

gcagtcatatagagcagacgCTCATGCAAGTCACTGAGGTTGTTTTCTCATATATAACATaagttatactgccctg
gcagtcatatagagcagacgACCAACTCCCTTTCTCTCAGAGATAATTAAGTAAAAGCTCaagttatactgccct
gcagtcatatagagcagacgGAGTTCATTGAATCCTCCTCAGAATCGCCACTGCACTCCAaagttatactgccc
gcagtcatatagagcagacgAGAATAAAAAAAGAATCCTCAGAATATCTGCACTGGTAGGaagttatactgccct
gcagtcatatagagcagacgGGTAGAGCCTTTTTTTCCCCCGCTAAACACATTTATATAAaagttatactgccctg
gcagtcatatagagcagacgTGCTTTCAGTTATATACCCAATGTGTCCGATAATGTTTCTaagttatactgccctg
gcagtcatatagagcagacgTTATGCCGCTGTGCAAATCTCATGAGTTTTATGCATATGGaagttatactgccctg
gcagtcatatagagcagacgTCAACTGCGGAAAAATTTCTTATAAAAATGCTCCAACCCAaagttatactgccct
gcagtcatatagagcagacgGAGCCAAAGAGGAAAATAATCAAGTAAAGAGAAACAGAAGaagttatactgccc
gcagtcatatagagcagacgGCCCCATCATTTTGCTTTTCGATTGTAATTTTTCAAGAGGaagttatactgccctg
gcagtcatatagagcagacgTTACGGCTGGTGAGAGAGAGATATAAGCTAACTTCACAGTaagttatactgccc
gcagtcatatagagcagacgAGCTCAACACCCTAAGTTTAAGGTCTTGCTAACTAAATTAaagttatactgccct
gcagtcatatagagcagacgGTGTTAACTACTTCAGAGATGCCATTAAAAAACTCGTCTGaagttatactgccct
gcagtcatatagagcagacgATTTTCAATGAAATCCTCTAACTGTGAGAAAAGGCTGACCaagttatactgccct
gcagtcatatagagcagacgACCAGAGTGTCAGAAAAGAAGTGCAAAAGTGCTCCTTTCTaagttatactgccc
gcagtcatatagagcagacgAGTTCACTTAAATGGCCCCAACATTACTGTTCTTCTAGAAaagttatactgccct
gcagtcatatagagcagacgGGTGCTTTGTAATTTGTTTTTGTTTATGACCTGGTGTTTCaagttatactgccctgt
gcagtcatatagagcagacgCATATTTTCCACTATTATGATTCAAAAGCCATACCAAGGGaagttatactgccct
gcagtcatatagagcagacgGTGCCTTTGCATCAAGGTAATTTAGTGTGGCAAACCTCCAaagttatactgccct
gcagtcatatagagcagacgAGTGTCACATTGATTCCCCTCTTTCCCTCAGCCACTCAACaagttatactgccc
gcagtcatatagagcagacgAATCCCCCAATTCTGTCAACTCTCCTTCCCCACCACGACCaagttatactgcc
gcagtcatatagagcagacgGCCTTCCTGCCTCAGTCTTGCCCCTTCCGACCTCCTCCTGaagttatactgcc
gcagtcatatagagcagacgTGCGAGCACTTTCCCATGACTCCCCACAGAGCCCTAATGTaagttatactgcc
gcagtcatatagagcagacgTTCTTGGTCCTTTGCTTGTAAAATGAAGAGAACGATGCTTaagttatactgccctg
gcagtcatatagagcagacgAATACATTATGCTAGAGGTTTGGTGGATTTATACAAGCTGaagttatactgccctg
gcagtcatatagagcagacgCATATCAGATGTTACATCCTAATCAGTGCTCAGGTGGGAAaagttatactgccct
gcagtcatatagagcagacgAATGCCTAAGTAAGAGTTTGTTCATAATTTACATGCTGGGaagttatactgccctg
gcagtcatatagagcagacgTGACATGTAGACAGAGCTATACTCCTGATTCTTTTTCTTAaagttatactgccctg
gcagtcatatagagcagacgCTACCTTTTCAAAAGATGTGTTTACATAGGGTTACTCATCaagttatactgccctg
gcagtcatatagagcagacgGGGCTGAAATCCCCCAAACATATTCTAAAGTAAATATTCAaagttatactgccct
gcagtcatatagagcagacgCAGCTGTATATTAGGAAATGGTTGGAGTTTTATTAATGCCaagttatactgccctg
gcagtcatatagagcagacgGTTTTGTGCTCCTGAACTAAAATAAACCTGTCATGGAAGTaagttatactgccct
gcagtcatatagagcagacgGTAACAATGGAAGCAGAAGATCAAATTTTCTCAGAAAGAGaagttatactgccct
gcagtcatatagagcagacgAATGAGAGAGAGACATAAAATATTAGACTCCTCGGAAAAGaagttatactgccc
gcagtcatatagagcagacgCCTTGGACTTTAATAAGGTGTGATTATGAGACCAATATCAaagttatactgccctg
gcagtcatatagagcagacgGTTACAGGCTCATAGTAACTGTTTCCTAAATATTTGCTGAaagttatactgccctg
gcagtcatatagagcagacgTTAGCAAACAAGATTCAGAAGGCTCTTCTATGGTTACCATaagttatactgccct
gcagtcatatagagcagacgTACAACGCATTTCACAATCCTTCCTTATCCAAAAACAATCaagttatactgccct
gcagtcatatagagcagacgTATGATATTGGAACTTGTAATTTTCTGCGAATGGTTGTGGaagttatactgccctg
gcagtcatatagagcagacgGCCACTGCTCCCCGACAATCCCACAAGAGACATCAATAATaagttatactgcc
gcagtcatatagagcagacgTCCTCAATGTCAATTTTAACATAGTATTCCACAAAGCACCaagttatactgccct
gcagtcatatagagcagacgCTGTACCAGGGAGGCACATTGCAGATAAAAGTAACCAGGAaagttatactgcc
gcagtcatatagagcagacgCCATCACAGGGCTTAGATTAACAATGGATTTGACACTCAAaagttatactgccct
gcagtcatatagagcagacgGAACTAATTCTAAATAGCAGTGTCTCTCCTTTGTGATAGTaagttatactgccctg
gcagtcatatagagcagacgAAGCCATGATGTCTATTGAATAGTTTAGGAGTTAAAGGTGaagttatactgccctg
gcagtcatatagagcagacgTAAGTGTTCAATTTGCAAGTCAGACTGATAAACCTATAGGaagttatactgccct
gcagtcatatagagcagacgAGAATGAATAAATTTTTCTCCCACTCCCCTATAGAGACAGaagttatactgccct
gcagtcatatagagcagacgTTTCTCCACAGATGTCGGAAAGAACACAATATGCTACGTGaagttatactgccc
gcagtcatatagagcagacgAGATGGCTGAGTACATGGCTTTTTCCACCACTATCAAGCGaagttatactgccc
gcagtcatatagagcagacgCAGCTGCAAAATTACCTCTCTGAAAGTCTCTGCTCTTAATaagttatactgccct
gcagtcatatagagcagacgCTACCCTTCCTCATTCCTGCCTACCATCCTTTCATCCCTGaagttatactgccc
gcagtcatatagagcagacgCTTCCCCAAGCAAAATTCCCCGGCCTACCTCCCACTCATAaagttatactgcc

gcagtcatatagagcagacgCTGCTTGGATGGTTAGTCCCTTACAGTTATCTACCCCCTGaagttatactgccct
gcagtcatatagagcagacgATGTCAAAACAGAATTTGGAGGATCGGGGACATGAAAGTGaagttatactgccc
gcagtcatatagagcagacgTGATAGCAGCTACAAAGAGGTTACTGTCAATAACAGGCAAaagttatactgccct
gcagtcatatagagcagacgCTGAATTTGGATATGTACATGTTATGCTCAGGGAAAAACAaagttatactgccctg
gcagtcatatagagcagacgATTTCACAAATTAAAAAAAATCTCATGGCCAGGCACGGTAaagttatactgccctg
gcagtcatatagagcagacgTGAGGCAGGAGTAATCGCTTGCCCCTGGGAGGCAGAAGTTaagttatactgcc
gcagtcatatagagcagacgCAAACAACAACGAAAAAGAACTGCCTCACTAAAATGTTAGaagttatactgccct
gcagtcatatagagcagacgACTACCTCCAATGCAAAGACCCAGTTCCTGACCTCACCAGaagttatactgccc
gcagtcatatagagcagacgGACCCACAATGTATATTTAGCTATTGATGCTGATGGTTGGaagttatactgccctg
gcagtcatatagagcagacgTTTTTAAACCTGATTCTGTGCCTGAAGTGCCTACTAATATaagttatactgccctgt
gcagtcatatagagcagacgTTGTGTGTGTGCGTTTTTTTCTTTAATCCCCTCAAGATTTaagttatactgccctgtg
gcagtcatatagagcagacgTTGGAACTAAACAAGGAAGAAGAAAGTGACAGGCCATTTTaagttatactgccct
gcagtcatatagagcagacgCAGGGTTCCAATTTTACTGTGAGTTTCTACTTCTTAAAAGaagttatactgccctgt
gcagtcatatagagcagacgGCATTTCTCCCCTTTGAATGTTACTTCATTTGTTCATTCTaagttatactgccctgtg
gcagtcatatagagcagacgCTGTATCTTGGTCAGTGCCAGTTCTCATGTCATTATAGTAaagttatactgccctg
gcagtcatatagagcagacgCAACATTGTCCATTGGAACCAGTTATATAACACTCGAGATaagttatactgccctg
gcagtcatatagagcagacgACAGTATTTTACAGTTCCAACATATAAGAGAATGCAGCTGaagttatactgccctg
gcagtcatatagagcagacgTTATTCACTTATTCATCCAACTATGGGCCACATACTGTGCaagttatactgccctg
gcagtcatatagagcagacgAGGTGACCTATACAGACCAGTCTCTCTCTTTGTGGAGCACaagttatactgccc
gcagtcatatagagcagacgACTGGGCCTTGACAATTCTGGTTAAAATTGACATCCACGAaagttatactgccct
gcagtcatatagagcagacgGAATCTCTTTCCTAACCCGTTTCTTTCATACTCACTAGGAaagttatactgccctg
gcagtcatatagagcagacgCTTTCCAATGGATCAGGGTAGATTTATGACTCATTTGCAAaagttatactgccctg
gcagtcatatagagcagacgTATAGATTGCAGATTCTGTTTCCTTCTTCTTTGAAGTAGGaagttatactgccctgt
gcagtcatatagagcagacgGCTGTGTTCAAGGTTAAAGTACTGCAGAATTGTTTGCCATaagttatactgccctg
gcagtcatatagagcagacgGCCTTTCGCTTTCCTTACTCACCATTTCAGGCAAAGACCAaagttatactgccct
gcagtcatatagagcagacgGGCTGCTTCCCCCCTGCAATCACAGATAAGAAAACAAAACaagttatactgccc
gcagtcatatagagcagacgCCCTAGGCTCAGCACCAGTTCCTCAGCATTCATTTTATTAaagttatactgccctg
gcagtcatatagagcagacgAGGTCCTCTCTTAAGTGTAGTCATCTTCCTCTTTACGGGAaagttatactgccctg
gcagtcatatagagcagacgGTAGAACAATACTCAGAATAAAGATGCAGAACTATTTGGGaagttatactgccctg
gcagtcatatagagcagacgTCCCCCATTTCCCTGAGTGAGTAATGCTCTCCCTTCCCCAaagttatactgccc
gcagtcatatagagcagacgCCCTTTCATGGGAAGCTGGATGCCTATGGTTTGCACTTTGaagttatactgccct
gcagtcatatagagcagacgGCAACTAGATTATTCCCAAACCAACACAGCCACTTACCTGaagttatactgccc
gcagtcatatagagcagacgTGGGTGCCTTTTAAACTCAGTTCAGGATCTTTTAATTTTGaagttatactgccctgt
gcagtcatatagagcagacgTCTCTTAATTCACAAATTTTTTCTAGCTCTACCCTTGCTCaagttatactgccctgt
gcagtcatatagagcagacgTATTTTCAATACTGAACTTTTTCCAGTGTGGGTCATCTAGaagttatactgccctgt
gcagtcatatagagcagacgAAATTTTCCTGGGCTCTGAATTTCATGCAATCATTAGTTGaagttatactgccctgt
gcagtcatatagagcagacgTGATATTTTTGGCCAATCATTTCCCTTTCTGAACCAGTTTaagttatactgccctgt
gcagtcatatagagcagacgACATTTGGTTTGTCTTACGTAATCTATGAGTTCCCTCTACaagttatactgccctgt
gcagtcatatagagcagacgCGCTTACGTGACATTCGACAAACATCTCGGAGGATTGTTGaagttatactgccct
gcagtcatatagagcagacgCAATAAAGTGAGGTGGTGAAAGTGCTTTAAAATCTACACAaagttatactgccctg
gcagtcatatagagcagacgTAGCTGTATCACCCTAGACAAGTCATCTCTCTTTCCAGATaagttatactgccctg
gcagtcatatagagcagacgGATTGGACTAGACGACCTTGAAATCTCTGACCTTCCATTTaagttatactgccctg
gcagtcatatagagcagacgGCCTGATATAGCTTGGATGCTGACCCTCCAATTCTGATGGaagttatactgccct
gcagtcatatagagcagacgCATGGTGCTGTTGTTTAGATCTGATTATCTGAAAGTGTGTaagttatactgccctgt
gcagtcatatagagcagacgCACCATGTGATACACCTGCTCCTGTTTTGCCTTTGACATGaagttatactgccctg
gcagtcatatagagcagacgGTGAGTCAATTATACCTCTTAATAAATTACCAAGCCTCAGaagttatactgccctg
gcagtcatatagagcagacgTAAGAACAGCCTAACAGACAGCCTCATGATTTTTTGAACCaagttatactgccct
gcagtcatatagagcagacgTGACCAATCAGAATACTATCACTAAGGATCTCTCTCGCCTaagttatactgccct
gcagtcatatagagcagacgTCTCCTTCTCCCTCTTCTTCACCAGCCTCCTCCTCCTCCCaagttatactgccc
gcagtcatatagagcagacgCCTTAGGCGGGAATAGAAGTTAGCAGGAGAGAGGAAAGAAaagttatactgccc
gcagtcatatagagcagacgTGGAGGGAGGAAAACTTAGTATTTGGCACCAATCTCAGGGaagttatactgccc
gcagtcatatagagcagacgAAAAATCAGTTTAAACAGTGGAAGCAAAGGTAGAAGCTCTaagttatactgccct

gcagtcatatagagcagacgTCTGTGAGATGATGTGTGTGTCAAATACCCTGGTTATGCCaagttatactgccctg
gcagtcatatagagcagacgTTCAGTTAGAAGATCTGCGTTTGAGGCTTAGCTTTTCCACaagttatactgccctg
gcagtcatatagagcagacgGACTGCATGACCTTGGCATCACTTAATTGGCTGCTCCTCAaagttatactgccct
gcagtcatatagagcagacgACAGGCAGGGTGTGTGCATTCGGTGCTGTGCTGTGGAAGAaagttatactgccc
gcagtcatatagagcagacgGAATCACTTCCCCTCCCCCTGCTTCTGTCAGAATATTTTGaagttatactgccctg
gcagtcatatagagcagacgAGTGCCTTGGTGCGATCTTGACTCACTCTGCAACCTCTGCaagttatactgccc
gcagtcatatagagcagacgTGAGATTAGAGGCATGAGCCCAGCCTAAATTAATATATTAaagttatactgccctg
gcagtcatatagagcagacgACAGACAGTAGATCCCGGTATCAAAGTACTCCTAAATAAAaagttatactgccct
gcagtcatatagagcagacgGGTCTATCAAACAACCTCCTAGATCAAGTCAACCCCTTCAaagttatactgccc
gcagtcatatagagcagacgTCATACTTATCTTTATGAGTTTCTTGAGGCAACATGTGGCaagttatactgccctg
gcagtcatatagagcagacgTGGATAACACCTGCCCTGAGAAACTTCCATGCCAGAATAGaagttatactgccc
gcagtcatatagagcagacgCCTTCTTTATGGCATGGAACTCTGGGGATTTAATTGAGCAaagttatactgccctg
gcagtcatatagagcagacgAACTTCTGGCATTACTTGTAGTTATCTCTGTGGTCTGTCTaagttatactgccctg
gcagtcatatagagcagacgAGTTCCCCTGTTTGTGACATCCCACTGCCTACATCTTGGTaagttatactgccct
gcagtcatatagagcagacgGACCACATGAATCTTTCTTTATCAGCAACTTAGAACACCAaagttatactgccctg
gcagtcatatagagcagacgGAGGAAACCGCTAACTTCAAAGAACCACCCAAAGCCCAGCaagttatactgcc
gcagtcatatagagcagacgCGTGAGCCACCGCGCCGCGCAAGAAGTGCTATTTAGAGAAaagttatactgcc
gcagtcatatagagcagacgTGTCCACACTCCCCATGCTGTCCCCCAACCAGAAGACAACaagttatactgcc
gcagtcatatagagcagacgGACACTGACAAAAATGGGGTTATTCTAAAAAAAAGCGTTGaagttatactgccctg
gcagtcatatagagcagacgCTGGAATGAGGAAGCTAGATAAAAAACAGTTAACCAAAACaagttatactgccct
gcagtcatatagagcagacgGAACAAATTTCCAAGTAACTGGTGACTTTGAGTGCATCACaagttatactgccctg
gcagtcatatagagcagacgGGAAACTAGAATCTCATAAAGGTTATTGGAGGTGTCAAGAaagttatactgccctg
gcagtcatatagagcagacgCCTTAACACATTTCTTTGAGGCAGTTTCATGTCCATGTACaagttatactgccctg
gcagtcatatagagcagacgCATCTACTGCATGCTAGACACTTGCCCATCTTATTAGTGAaagttatactgccctg
gcagtcatatagagcagacgGTGTATGTACGTACCCATAAACACATGCATAGTCATAGCAaagttatactgccct
gcagtcatatagagcagacgCTTGATCGTCTCAGTAGCTCAGAGTCATGTTGCCCATTTAaagttatactgccctg
gcagtcatatagagcagacgCCTACAGGCTATTTCATTTTCAAGGGTGGAGTGCTATAATaagttatactgccctg
gcagtcatatagagcagacgGCTTCACATGGCAGCCTAGAAATCTGATTTAAAAGAATGAaagttatactgccctg
gcagtcatatagagcagacgGTCCTCCCAATAAATGATCTTGTCCCTTCTGTGAACTACTaagttatactgccctg
gcagtcatatagagcagacgCAATATAATCATAAGAGGCATACAGATCTCCTTACATCACaagttatactgccctg
gcagtcatatagagcagacgCAAAAAAACTCAACATAGAGCATGAAAATGGGCTTTATCCaagttatactgccct
gcagtcatatagagcagacgCTGTTAAGGTCAAGCAGTGTAGGCTAATAGCATTAAAAAAaagttatactgccctg
gcagtcatatagagcagacgGAACACTGACACTTTGTATTGATCTTATCTGAGTCTTTTCaagttatactgccctgt
gcagtcatatagagcagacgGTGGAACAGAACTTTATGACTCTTTGACACAAATCCCTTCaagttatactgccctg
gcagtcatatagagcagacgAGAGTGTTCTAAAATGCCACCCCTAATTCCCCACCTTAGTaagttatactgccct
gcagtcatatagagcagacgTTCATTGGGAAGAGTGAATACATATCCAAGTAAATGATGGaagttatactgccctg
gcagtcatatagagcagacgCCCAGATGATTGAGGCTCTTTCTTCAAATGCTCTTCAAAGaagttatactgccctg
gcagtcatatagagcagacgGTCATTATGAACCAGACAGCCTTGAGGTCACTTGTAAAACaagttatactgccct
gcagtcatatagagcagacgCTCTCAATGTAAAGTCTGTAAAGTCTGACTAAGAGTATGTaagttatactgccctg
gcagtcatatagagcagacgAACCCAACTTCAAGACAAATCCCCTGGTTTCTCTCCCACCaagttatactgccc
gcagtcatatagagcagacgTCTGTTGAAGTCTTAGGGTTATTCTCTGTAAGAAAGGAGCaagttatactgccctg
gcagtcatatagagcagacgTCCATTTTTTATTATGAGTCATTAATCTCCCTAGCTCTGCaagttatactgccctgt
gcagtcatatagagcagacgTGTCAACACAGGAGATTAGGGTTCGGTTTTTAGGATCCTTaagttatactgccctg
gcagtcatatagagcagacgCTAAAAATTCTGATCCTGGGGTCACAGTAATTGATAACAAaagttatactgccctg
gcagtcatatagagcagacgGTAAAAAGACAACAGCCTAATTTTAAAGATCAACCAAGGCaagttatactgccct
gcagtcatatagagcagacgCAGGAACATGCAAATAAAAAACACAGGGAGACACCACTTCaagttatactgccc
gcagtcatatagagcagacgAAACTGGAACCCTCTTACATCGCAGGTAGGACTTTTAAACaagttatactgccct
gcagtcatatagagcagacgGCAGTATGGCAATTCCTCGATGGTTAAACATAAGAGTTGAaagttatactgccctg
gcagtcatatagagcagacgCTACCCAAGACAGATGAAAACATCCATCCACATAAAAACTaagttatactgccc
gcagtcatatagagcagacgACAGCAGTATTACCTATAACAGCCAAAAAGAAACAACCCAaagttatactgccc
gcagtcatatagagcagacgGTGACAGAGCAAGACTATATCTCAAAAGAAAGAAAGAAAGaagttatactgccc
gcagtcatatagagcagacgAGAAAGAAAGAAAGAAAGAACCCAATGTCCATCATCTGATaagttatactgccct

gcagtcatatagagcagacgCAATAAAAAGGAATGCTGATATGTGCTGCCACATAGATGAaagttatactgccct
gcagtcatatagagcagacgCAGAGAAAACGTCATGTAAGCACACAAGGAGAAAGCAGCTaagttatactgcc
gcagtcatatagagcagacgCCTGCCAGCACCTTGATCTCCTTGATCTCAAGCTTCCAGCaagttatactgccc
gcagtcatatagagcagacgTCCTCTAGTTTTATTGTTAAAAAACAAAAAACAGGCCGGGaagttatactgccct
gcagtcatatagagcagacgTCCCAGCATTTTAGGAGGCTGAGGCAGCTGTATCACCTGAaagttatactgccc
gcagtcatatagagcagacgGAGTTCGAGACCAGTTTGAACAACATGGTGAAATCCTGTCaagttatactgccc
gcagtcatatagagcagacgAGAATCACTTGAACCTGGAGGAGGAGATTTCAGTGAGCTGaagttatactgccc
gcagtcatatagagcagacgACAAAACAAAACAAAACAACCACAGACACACACACACACAaagttatactgcc
gcagtcatatagagcagacgAATGTTCAGAGCAGGCAAATCCTGGAGACAGGAAGGTAGAaagttatactgcc
gcagtcatatagagcagacgCCAGGGCTTGCGGGGAGGGGATGGAGGGTGACTGATAATCaagttatactgc
gcagtcatatagagcagacgGATGGTTGCACAACTCTTGAATGGACCAAAAACCACTGAAaagttatactgccc
gcagtcatatagagcagacgTTCTGGTCTCCTATAGTATGTGAGATTTACCTCAATAAAGaagttatactgccctg
gcagtcatatagagcagacgATGACTAAAGTAATCACAGAGTTATGTTGAAGGCCTAGAAaagttatactgccct
gcagtcatatagagcagacgTATTTTGAAATTTGTTGCCACATATTACCCCATCAGTGTGaagttatactgccctg
gcagtcatatagagcagacgAGGCTGATTTTAGAGACCTAATTTTCCCTTAATGTTCTGTaagttatactgccctg
gcagtcatatagagcagacgACTTGCTGTTCCCTCCCCCGTTACAATCTTTTCAATTTAAaagttatactgccctg
gcagtcatatagagcagacgCACTAGGACTCTGAATAAATCTGGTGTTTCCTTTTATATCaagttatactgccctg
gcagtcatatagagcagacgCATCTGTCTTTGATTTTGCATACTGTGTTTTCAGTAGTACaagttatactgccctg
gcagtcatatagagcagacgATGGCCTCTTTATCTAAATAGAGATCCCTAAGTATCATGTaagttatactgccctg
gcagtcatatagagcagacgGAATTTAAGATCATGAGCCTTTTACAAAGGTTGTTCGAGGaagttatactgccct
gcagtcatatagagcagacgCCTTAAGATGTAGCTTATTAAAGGCCGAAATCTCATAGACaagttatactgccct
gcagtcatatagagcagacgCTTTGGGGTTGGTGGCAAGGATGGGAGAAGGGTTAAGAATaagttatactgccc
gcagtcatatagagcagacgGCTACCAACATTTTTATTTGCGATGCTTATTCTTATAGGGaagttatactgccctg
gcagtcatatagagcagacgACTCTTAGTTCTTAACCTCTGTGTCATATCTCTATCCAAGaagttatactgccctg
gcagtcatatagagcagacgAGTGAATGCCTAGATTAAAGTAACCACATTAAATGGAGACaagttatactgccct
gcagtcatatagagcagacgATGAAGAGCATTATAATGTCCCATCTCCTCTCACTTCCAGaagttatactgccct
gcagtcatatagagcagacgTGGAGAAAGGCCAGAGAGTTCATATTCCATCCACCAACAGaagttatactgccc
gcagtcatatagagcagacgGAAGATGTGAAAGTCATGGGGATTCGTATGAACTGAAAGCaagttatactgccc
gcagtcatatagagcagacgAAATGTCAGGTTTGTCTTCCAAAACAATTGGCCTTAAATCaagttatactgccctg
gcagtcatatagagcagacgCACACACACACACGCTTGCTGACACGCATGCATACACTTTaagttatactgccc
gcagtcatatagagcagacgAAGTTTGTTTGAGTGGTGGTGGGGAGGTTGATGTAGAACTaagttatactgccct
gcagtcatatagagcagacgTTGGTGTGTGCTTTAATTACATTTTTCATTCAGCTTAGGCaagttatactgccctg
gcagtcatatagagcagacgTTTCTAGGCGGGGATTTAGAAATTGACTTGAATGTGGCAAaagttatactgccct
gcagtcatatagagcagacgCTGCTGACTAATCCCTTATGCAGATAAGATTATTTGTGACaagttatactgccctg
gcagtcatatagagcagacgAGAGTAATTGTGTTATTGGTCTTTAAGGTCCACAATAGGAaagttatactgccctg
gcagtcatatagagcagacgAGTTTGTAGAAAGTTTCTAATCCTAGGAACCTGCTTTTTCaagttatactgccctg
gcagtcatatagagcagacgTAAACGCACTATTTTACCACACTGATTTCTTGCTTTAGAGaagttatactgccctg
gcagtcatatagagcagacgCCTCAAACTATTAGATTCAATATGTGAAGTCTCTGATCTCaagttatactgccctg
gcagtcatatagagcagacgAATAGTCACGTATATAGCATTTATGTCTTGAATCCAGGTGaagttatactgccctg
gcagtcatatagagcagacgTTGGCACTTCTCCGCAATTATGTTCATTTCAATAATTCTGaagttatactgccctg
gcagtcatatagagcagacgCTGTCAATGAAACTCATAGATAGGTTTTGTAGCTCTTGGCaagttatactgccct
gcagtcatatagagcagacgCACTGAAGTGGCCAAAGTCTAAAATCATCAAAACTTGGACaagttatactgccc
gcagtcatatagagcagacgTGGGTCATTGACTTTACTCATTTTTCCTTAAGCTTCAGAAaagttatactgccctg
gcagtcatatagagcagacgTGTTTACCAGGAAAAAATGCTCTAATAACGTCAACATTCCaagttatactgccct
gcagtcatatagagcagacgTGTCTTCCGCATGCCTCTCTGAGCTTTTTACTTGACACTGaagttatactgccct
gcagtcatatagagcagacgTTTGAGGGTCACAGTTTAACAGGGCACATTCATTGATCATaagttatactgccct
gcagtcatatagagcagacgGGACCTACCCTTCCAATGACCTATAGCCATGCACAACTCCaagttatactgcc
gcagtcatatagagcagacgATGTGACATCAGAGTTAATTAAGGTTGGTTTGGGAGAAATaagttatactgccctg
gcagtcatatagagcagacgGAGTTACCTTTTCCCAAAAAAATGCTCTACTCCTACACCGCaagttatactgccc
gcagtcatatagagcagacgGTTCAAAAACCACCTCCATAGAGCTGTCTAACAATATGTCaagttatactgccct
gcagtcatatagagcagacgAGTTTACCATACGATGTCATGAAAGTTATCTGAGCTGCCTaagttatactgccct
gcagtcatatagagcagacgGAGACATAACTCGTAAAATACTCGGTAGAGAAGTCTGGTAaagttatactgccc

gcagtcatatagagcagacgAGAAAACACTCTTTACACCCTAAAAATGCATACAGGAACGaagttatactgccct
gcagtcatatagagcagacgCTGTTAGGAAGGGCTTGGACACATCTCTGTTTTCTCTATCaagttatactgccctg
gcagtcatatagagcagacgCCCACGGGCTTATGTTCTTATCCAATGTCACTGTTTAGAGaagttatactgccctg
gcagtcatatagagcagacgCGTCACGTCTTATAGAATGGATCATAATTGCCAGTAGTGTaagttatactgccctg
gcagtcatatagagcagacgCATTTTCAGATTGTGAACACCTGGAAGAGACATGACAACTaagttatactgccct
gcagtcatatagagcagacgTCAATGTATGTGAGACTAAATATACAGGTGGCTTAGTGAAaagttatactgccctg
gcagtcatatagagcagacgATCAATTTGTGCATCCTCTTTAGAATCATAGTGTCTGATCaagttatactgccctgt
gcagtcatatagagcagacgAAGACTAGTAGTACTAATAACATGCTTGAACTGCCTTAACaagttatactgccctg
gcagtcatatagagcagacgAGGCATCCTCCGTGTTCTCTTTAAATAAACAATTACCAGTaagttatactgccctg
gcagtcatatagagcagacgAGTTTTCTGTTTTACATAGCATCAGGATTACAGTTTGGTCaagttatactgccctgt
gcagtcatatagagcagacgACTCTGCTTTCATTACTGTTCAGTGTCAAAGTTCCAGGCAaagttatactgccctg
gcagtcatatagagcagacgGCATGTTTCAATGTTATGGATTTGTCATTTATGTGGATCCaagttatactgccctgt
gcagtcatatagagcagacgCGTATAGGAGCTTGCCATGTAACAGTTTAAGTATAAAAGCaagttatactgccctg
gcagtcatatagagcagacgTCATGTGACATAATCTTCAGATCCACACACAGTTCCTTCTaagttatactgccctg
gcagtcatatagagcagacgATATGTCATGGAGCTGTAATGTGATGCTGTTAAACTACCAaagttatactgccctg
gcagtcatatagagcagacgTGTTCCAGTTTCTATTTTGATTCCCAAATAGTGCCTTCACaagttatactgccctgt
gcagtcatatagagcagacgTGAATAAGAAAGGCTGGCAAACAGGCAGACAGACAAGTGTaagttatactgccc
gcagtcatatagagcagacgGCCAGAAAAGCCAAAAATGTTATTATAGCAATACAAAGCCaagttatactgccct
gcagtcatatagagcagacgATTCTCCTCATGCAGACTTTCAGCTTCAGAATCGATTACAaagttatactgccctg
gcagtcatatagagcagacgAAAAATGGAGGAGAGATGAAATGAGATTGGAAGGCCGAATaagttatactgccc
gcagtcatatagagcagacgAAAACATCTTTCTGGAGTCCTTTAGACTGTTCATAAAGGGaagttatactgccctg
gcagtcatatagagcagacgCTCTATCTTTGCAACCAATAATATATACAAACCCTCTGACaagttatactgccctg
gcagtcatatagagcagacgGATTTAAATGGCCAAGAGTGTGCAATAAAGTTATCCATCAaagttatactgccctg
gcagtcatatagagcagacgGCCTTTCATTACACATATTTTGGAGATTAATCATGCTTCGaagttatactgccctgt
gcagtcatatagagcagacgGGGGTGGACATGTTGATTAAACTGTTATTTTTTATTCTGGaagttatactgccctgt
gcagtcatatagagcagacgAAGATATCCTTCACGTAGTAAGTATTTGTAAGAGCTGTGCaagttatactgccctg
gcagtcatatagagcagacgCATGTTATACAAGTTTTGCTGAGAGTCAAATTTCTGGAAGaagttatactgccctg
gcagtcatatagagcagacgATGTTTTTCACTCATTTCGGCATGTGATATTAAGGCCTTCaagttatactgccctgt
gcagtcatatagagcagacgCGTTGTTTGTATTCAATGCCTGGGGAATGCTGGTTTTCATaagttatactgccctg
gcagtcatatagagcagacgGGATATATACCCATACACCTTCATTACTCTATCTAGAACTaagttatactgccctg
gcagtcatatagagcagacgTGGCCACCAACCCAACAGCTTTAAAATAACTTATGAATCAaagttatactgccct
gcagtcatatagagcagacgAACCAAAGTAGAAGTTACATAAAACCAAAAGGAACAGCTCaagttatactgccc
gcagtcatatagagcagacgGCCGAAGCTGCTCCCATGAAATTAAGCCACCACCTCCTTTaagttatactgccc
gcagtcatatagagcagacgAGACTGATGAACTGAGTTAAGGGTTGGACTCTTAGGTGCAaagttatactgccct
gcagtcatatagagcagacgAGCTATGCTTCTTATCTCTCTTTTACTGCCACCAATCCTGaagttatactgccctg
gcagtcatatagagcagacgCATTCTCTATATGACCCCTTTCTCTCACTCCCTTACCTTGaagttatactgccctg
gcagtcatatagagcagacgGTATACCGATAGAGATACAACAATAAACAAGACAAACGAGaagttatactgccc
gcagtcatatagagcagacgTTCCTTTACAGTGTTTATAGCTTAGCATGGACAACAGATAaagttatactgccctg
gcagtcatatagagcagacgACATTATATTTGTGTGTTATCATGGGGAAGTGCCAGGTGTaagttatactgccctg
gcagtcatatagagcagacgTCAGCCCAGGGAGTGAGAGTCTTCTATTAGAATAAAGTTGaagttatactgccct
gcagtcatatagagcagacgAGAAATAGCAGGGTCTCTTTCTCCTTCAAAGACTTCCACAaagttatactgccct
gcagtcatatagagcagacgCTCTTGTTTTCCTCAGCCTCTGTGCTGGGAGGAGAAAGCTaagttatactgccct
gcagtcatatagagcagacgTCTTTCAATGCTGTGGTTCGCATATTTTTTGTCTGGGGTCaagttatactgccctgt
gcagtcatatagagcagacgGTAATCATACTTTAGATACATACCCAGGTCACGTTGTGCTaagttatactgccctg
gcagtcatatagagcagacgAGCAATGAGGGAATGAATGAGGACTGTCAGATGTGATGCGaagttatactgccc
gcagtcatatagagcagacgGTCACCTAGCCAGTACCACACATGACCCAGAGCCAAGAGGaagttatactgcc
gcagtcatatagagcagacgATGTTCTCATTCCAGGTGACAAAACAATGGCAGCAATAGTaagttatactgccct
gcagtcatatagagcagacgCCTAGAAATTAAAGGTTCCAATCAACTATCTTCTGTGCTGaagttatactgccctg
gcagtcatatagagcagacgGCCAGTTAACAGTCTCCCAGCAGGAAGGCTCATCTCAAGGaagttatactgcc
gcagtcatatagagcagacgTTCTCTAGAAGATGGGCAGTGTAGATCCTCATTCTCAATCaagttatactgccctg
gcagtcatatagagcagacgTCTACAGTCGAGTCAGGAGAACATGGCCAGCTTAGCCAAAaagttatactgccc
gcagtcatatagagcagacgTGGACCTAGAAGATAACCCAAATGGAAGGAACAGAATATTaagttatactgccct

gcagtcatatagagcagacgCTCCTGGCCATATTTCTCCTAACCTCCTCCGCTTATACAcaagttatactgccct
gcagtcatatagagcagacgCACGTACTTAGAAGGTGAACCAATGAGGAACCATGACTCCaagttatactgccc
gcagtcatatagagcagacgGCTCAGGATATGATTTCAGTCAGTTCAGCCAGTATTTATTaagttatactgccctg
gcagtcatatagagcagacgCACTGTACTGGATTCACAATTGCAAAGATGGGTCAGAAACaagttatactgccct
gcagtcatatagagcagacgCCTCTTAGGGGCTGTCATTTAGAAGTAGAATAGGTGAATAaagttatactgccct
gcagtcatatagagcagacgATCTAAGTCAACAAATAGAGATCTCCTGCAAACTATGTACaagttatactgccct
gcagtcatatagagcagacgGAACACATCCTTTAATATAGGAAGTCACACATTCTTCACCaagttatactgccct
gcagtcatatagagcagacgAAAAATCTTCTACACATTCCTGAACTCAGCTTGGAAATAGaagttatactgccct
gcagtcatatagagcagacgGAATCTAGAATAATGACTGGCTATAAAATCGGGATCAGGAaagttatactgccct
gcagtcatatagagcagacgAATATGGCTCATGTTTTTCCTCATGCAGTTGTGGGAGAAAaagttatactgccct
gcagtcatatagagcagacgCACTCTGGGAGGCCAAGGCGGGTGGGAGTTTGAGACCAGCaagttatactgc
gcagtcatatagagcagacgAAAAATACAATATTAGCCCAGCGTGGCGGTGCATGCCTGTaagttatactgccc
gcagtcatatagagcagacgAAAAGAAAAGAAAATACAGCCCTCCAAATCATTGGAAGAGaagttatactgccc
gcagtcatatagagcagacgAAACCTCCCCTCCTAAGTTTGTATTTAACATTATCCCAAAaagttatactgccctg
gcagtcatatagagcagacgGGTTTGGACTATAAAGCGCATTTAAGAAATAACAACAACCaagttatactgccct
gcagtcatatagagcagacgCAGTTTAGGATTTTACTCCCTCTAAGAAAAGATAATCTGGaagttatactgccctg
gcagtcatatagagcagacgGTCAGTCTCAAAAGGCTAAAGTGAAGTAGAAATTCCTGAAaagttatactgccct
gcagtcatatagagcagacgACTCCTGTTAGTTTCCTGAATTTCCCATTTGCCTTCTAAGaagttatactgccctg
gcagtcatatagagcagacgGGTATTTTTCATTGACCAGTTTTGAAGACAGAGGATGGGAaagttatactgccct
gcagtcatatagagcagacgCAATGTGTGTCAAATATTTCCTAACCCTCAAGAAGTTACAaagttatactgccct
gcagtcatatagagcagacgAGGGCACTGCAATTACTCTATTTCCACACATATATTTTTCaagttatactgccctg
gcagtcatatagagcagacgGGAGAGGATTCTGAAGCCATAAAAATTTTCAAAGAACAGCaagttatactgccct
gcagtcatatagagcagacgCTGATCTAGCAGGAGACATTAAGTGGAAAAAGAGAGTTAAaagttatactgccct
gcagtcatatagagcagacgACAATACCATTAATCATAGATGCAGTGTTGTTGCCAGAGTaagttatactgccct
gcagtcatatagagcagacgCATATCCCTGCCTCCCCATCTGTTTACATACATTTTTGGTaagttatactgccctg
gcagtcatatagagcagacgCCAACAGTGGATTTTTACATACATAGACACCATGCATATTaagttatactgccct
gcagtcatatagagcagacgATACACCATTCCGTATAATTTCCATCTAAAGCATGTATCCaagttatactgccctg
gcagtcatatagagcagacgGAGTGAAAGATAAAACTCCTCAATAGGTGGCAGGATAGGAaagttatactgccc
gcagtcatatagagcagacgTTCTGGTTGAAAAAGTGTCCTTCTCCGAGTAACAGGTCAAaagttatactgccct
gcagtcatatagagcagacgAAACGCTATTTTCTCCTTCTTGACTTCTTGCAGGTCTATGaagttatactgccctg
gcagtcatatagagcagacgGGAATGATCTAAGAAGGAGGGAGAGACACAGGAGCTACGGaagttatactgcc
gcagtcatatagagcagacgCTGTACAAGGACTGCCAATCAAACAGGGAAATCAAAGTGAaagttatactgccc
gcagtcatatagagcagacgAAGGCTCTTGGATGGTTGTCTTTTCTCATAAATATGACAGaagttatactgccctg
gcagtcatatagagcagacgTCATTCAAGATACTTTGAGTAAATGTTGGTGGAAAGAGTCaagttatactgccctg
gcagtcatatagagcagacgTAGGTTTGAATCTTGGCTTGGCTGTTCCATAAATGGATGGaagttatactgccct
gcagtcatatagagcagacgCCTTGGGGCTTCAATCCATAAAACAAAGTTACAGGTGTGCaagttatactgccct
gcagtcatatagagcagacgAGGTGGCTAGGAAGAGACTTGATGGAGTTAACACAGAAAGaagttatactgccc
gcagtcatatagagcagacgACAGTTAATGTCCTCCCCTATCCAATTCATGTAGCATCTGaagttatactgccct
gcagtcatatagagcagacgCAGGTCGTTCTGTGAGGTATTTTAGTTTCATTCTGGAAAAaagttatactgccctg
gcagtcatatagagcagacgAATCTGAAAGGTGCAGAGGAGCCCACGTTGGATTCCTGGAaagttatactgcc
gcagtcatatagagcagacgGCTTTTTCCTCGGACCTGCCTGATATGAGCTTTTCTAGAAaagttatactgccct
gcagtcatatagagcagacgTGCTGTCCTCATGCTGTTGTTTGTCATCTGAACTGGTTTAaagttatactgccctg
gcagtcatatagagcagacgTATCTAGGAGAGCAGTTTGGAAGCCCACATAAAATCATGTaagttatactgccct
gcagtcatatagagcagacgTCCTTTTCTGACTAAGTTTTTTGGGAGGTCCTTGAATGCCaagttatactgccctg
gcagtcatatagagcagacgTTAAAATTCTGTCTTATTGGAGATGTGAACAGTGTCTGCCaagttatactgccctg
gcagtcatatagagcagacgGTGGAATCTCCCAAGGCCACTTTAAGCTGGAGTAGCAGTGaagttatactgccc
gcagtcatatagagcagacgCACAGTCCCCTTGATGCTTTTTCCTTCTCCTCTTTTCATTaagttatactgccctg
gcagtcatatagagcagacgCATGGTAGGAGGATCCCTAAATGTCGGGTTTGCAAGTCTGaagttatactgccc
gcagtcatatagagcagacgAATTCCAAGTTTACTTACCACTTCTGTGATCTTAGGGAAAaagttatactgccctg
gcagtcatatagagcagacgTCGGGACTGAGTAAAATAATATATGTAACTCCTGTTGTGGaagttatactgccct
gcagtcatatagagcagacgTTACATATATTGATTGTATAGTCAGTAGGGTTTGGTTGCCaagttatactgccctg
gcagtcatatagagcagacgTTCCCCAAAGCATGTCTGTCCCTGCAGTCTCGGCACCTCCaagttatactgcc

gcagtcatatagagcagacgTCTATGATAAGAGCCCCAAGAAGCTGCAGAGCTGCCTGCAaagttatactgcc
gcagtcatatagagcagacgGCTGTCCTCCCTTGCTGGTGTCATGTTAGTCATCAGAGGCaagttatactgccc
gcagtcatatagagcagacgAATTTGAATTAAGAATAGAAAGCTCTTCTCTGGGGCCTGTaagttatactgccct
gcagtcatatagagcagacgCCAGATACTGTTTAGAAGGGTCAAGACATTATGTAGCATTaagttatactgccct
gcagtcatatagagcagacgTATTTACTGTATTTTCTTGGCCAAACTCCAGCTTAATAGCaagttatactgccctg
gcagtcatatagagcagacgGTGTGGTCCATTGTGAGGCTGTGCTCCTCTCCACGCAGAGaagttatactgcc
gcagtcatatagagcagacgCCAATAGAATGTGGTAGAGCGGTGTTGTAGGCGTTCCAAAaagttatactgccc
gcagtcatatagagcagacgGCCTCAATATGCCTTGCAGCTTCCACCATCACACTCTTGCaagttatactgccc
gcagtcatatagagcagacgAGGAAAGATCTCATGAAGAGGGAGGCCCAGCACCAAGGACaagttatactgc
gcagtcatatagagcagacgTGCCAAGGACTATAGATGCATGAGAAACCCAGGTGAGACCaagttatactgcc
gcagtcatatagagcagacgCCCAAAGGTCTAGAAGAATCATGAAGCATAATGCATTGTTaagttatactgccct
gcagtcatatagagcagacgGGGTGACTAAGTTTTGGCACATCTCGTTATGTAGAAATAGaagttatactgccct
gcagtcatatagagcagacgACTGTGTGGTAGTTCAAGTGTATGCTCTTTCTGTATTTGAaagttatactgccctg
gcagtcatatagagcagacgGCTCACGCAATATTAAGAATTACATGTTGTACGGGTTACCaagttatactgccct
gcagtcatatagagcagacgCTTGGTGTTTCACAAGAAGTGTTCCCAGGACCGTGGTTTCaagttatactgccc
gcagtcatatagagcagacgGGCCAGCTCTGTCTCCAACCAGCAGTAGCTTTTCACAACAaagttatactgcc
gcagtcatatagagcagacgGACAGAACCCTGGCACATCATGAAAGTTAGGAGGCCACCCaagttatactgc
gcagtcatatagagcagacgCCAACAGATAGAATTCAGCCACCCAAGTGGGAGCCATACTaagttatactgcc
gcagtcatatagagcagacgCGGATGGAGAGGAAATTTTGCAGTCATACTGGAGGAAACCaagttatactgccc
gcagtcatatagagcagacgGTGTTTGCCAATATTTACATAGAGAAAAGACTGAGCACTCaagttatactgccct
gcagtcatatagagcagacgCATAAGGCAGAGAAGGAGGACCATTGTTAGTCTAGACGGGaagttatactgcc
gcagtcatatagagcagacgCGTACCTCACTCGGCAAAACAGTCCCCCTCTCCTCAGAAGaagttatactgc
gcagtcatatagagcagacgCCCTGGCATTTGATCTAAGTATTCTTTCTTGCACACTCTAaagttatactgccctg
gcagtcatatagagcagacgGAATTCTCGTAAATAGTTTCTTCCACCTTGTCGTATGTCTaagttatactgccctg
gcagtcatatagagcagacgTGATTTAGTGCCTTTTACAGATTGGCATTTGGGCAGGTGCaagttatactgccct
gcagtcatatagagcagacgAAGTGTGATCTCTGGACCAGGCACAACAGCATCCCCTGGGaagttatactgc
gcagtcatatagagcagacgACCAGGTGATTCTAATGTGTGATGAAATTTGAGGTTCTTAaagttatactgccctg
gcagtcatatagagcagacgCTCTCAAGAGCTCATGATTAAATTGGTCAGAGGTACAGCCaagttatactgccc
gcagtcatatagagcagacgCCCTAGAGAGAGGCAAAGGCTTTCATAGACATCACTTGGCaagttatactgcc
gcagtcatatagagcagacgGCTACAGGTGGCCTTGAGTTTAAGTGGTTTTATATTTGTGaagttatactgccctg
gcagtcatatagagcagacgAGGCTTTTCATTTATGCTAGGTTCACTTCAGGCCTCTTTTaagttatactgccctg
gcagtcatatagagcagacgAAGAATCTGATAATTGTTGTAGCTCTGCCTGTGTAGTCCAaagttatactgccct
gcagtcatatagagcagacgGTGCTCCACCCATTAAATAGAAGGATAGAAGACCAGGCTGaagttatactgcc
gcagtcatatagagcagacgACCGTGATAATCTGTGAAAAGGTAAAAGCATTTGTCAAGAaagttatactgccct
gcagtcatatagagcagacgTGTTAAGTGCAAATACATTCCATTTCAAGACAGTAAGAGGaagttatactgccct
gcagtcatatagagcagacgAGCGAGACAATGACACCTCCCTGACAACCAAAAATTCTGCaagttatactgcc
gcagtcatatagagcagacgACCGCTGCTGGTTTTTGCCACCAAGTCATTTTCTTTCAAAaagttatactgccct
gcagtcatatagagcagacgAAGGCTTGTAGAACTTTATGGTGAAAACCTGGCTATTTAAaagttatactgccctg
gcagtcatatagagcagacgAATCCATATTTTCCAAGACCTTGATCTCGCCACCCATGATaagttatactgccct
gcagtcatatagagcagacgGTATTCTTCAAGACCACAACCTCTGTTTACTTTGATTACAaagttatactgccctg
gcagtcatatagagcagacgACACCGATGTCCCAGCAGAAGGCGGCATGAGACCAACGTGaagttatactgc
gcagtcatatagagcagacgCTGGCCCAGGTCTAGTCCCTTAGTCACACACGTCAAAGTTaagttatactgccc
gcagtcatatagagcagacgTAGGGTGTTAGTGAAGAGGAAATGGGACCAGGCGCGGTGGaagttatactgcc
gcagtcatatagagcagacgCTCAAAAAAAAGGAAATGGACCCTGACACCAGAACTGACTaagttatactgccc
gcagtcatatagagcagacgCCACGTGGGGATTTTATTAAAGTATAGGCTCTGAGTCATCaagttatactgccct
gcagtcatatagagcagacgCGGAGCACACAGAGAGAAGCACGTTTTGGAGTATGATGTGaagttatactgccc
gcagtcatatagagcagacgGTGCAGTGTGGGCCGGGATGGGGGTGAGGATAGGGAGGCAaagttatactgc
gcagtcatatagagcagacgGGGATTCCAATTTTTAAGAACATGTCTCGCACCTAGCAGGaagttatactgccc
gcagtcatatagagcagacgCTCATCCTAAAGAAAGAGGAAGTTAGACAGACCAGTGTGGaagttatactgccc
gcagtcatatagagcagacgCCAAGGCCCCTATTAAAACAGAGTTGATTCTGACTGTGGCaagttatactgccc
gcagtcatatagagcagacgCTGTCATTCAGGTATGCATCTCAGCCCAGCACCCAAGGACaagttatactgcc
gcagtcatatagagcagacgAGGACTCTGTTCTCTCTAAAAATTGTATAACACGCTGCAGaagttatactgccct

gcagtcatatagagcagacgCCAACAGAGGCTAACACAGGACTTCTGCAGTGACCAGAGCaagttatactgcc
gcagtcatatagagcagacgCGAAGCATCATGCCCACACAGGAGGGGAGGACATGAGACAaagttatactgc
gcagtcatatagagcagacgATCATAGCTGATCTCATACATCTCAATCGCTCACTGAAGAaagttatactgccct
gcagtcatatagagcagacgACTTTAAGTAATCCCTTGTAAATGAGGCCAAATGAACAGGaagttatactgccct
gcagtcatatagagcagacgCTCATGGAATAAATTTGTGTCTGAGATTGCTTCTGCGAAAaagttatactgccctg
gcagtcatatagagcagacgCTCATTCGTTCCAACACTTACCGAGAGTCTCTGTACTAGGaagttatactgccct
gcagtcatatagagcagacgCTACCAAAGAGAAAGGACACAGACACATAAAATCCTGACAaagttatactgcc
gcagtcatatagagcagacgAGTACACCTCAGGGCTACAGGAGAGCAGAAGAAATGGCAAaagttatactgcc
gcagtcatatagagcagacgCAGCAGGAGAGGAGGAGACACAGAGGGAGGGAAAAGGTCTaagttatactgc
gcagtcatatagagcagacgAGACGAGCAGCATGAACAGAGTGATGAAAGTGAAGGCGCCaagttatactgcc
gcagtcatatagagcagacgGAAGGGGGAATAATTTGTTTGACTAGGGTAGAGTTTGTGTaagttatactgccct
gcagtcatatagagcagacgGTGTGTGTGGAGAGAAGTGGCGAAAGACAAAGCTAGTACCaagttatactgcc
gcagtcatatagagcagacgGCCACAGCTGTTCAGGAAAAGGATACCAAGTGCTCGTAAGaagttatactgcc
gcagtcatatagagcagacgCTAAGCAGCCATCATTTTGATTCTTAGTTTAAACCCCTTAaagttatactgccctg
gcagtcatatagagcagacgGTTAGTCAATAAATATTTGAACATCTGTCTTCCGTCTGGGaagttatactgccctg
gcagtcatatagagcagacgGGAAATACACCAGTGAAGAAGAGATGTTGTCTGTAGCAGGaagttatactgccc
gcagtcatatagagcagacgAGGTTCTGTTCCTAGGCCTGCCGCTAGCCAGCTGTTGACCaagttatactgcc
gcagtcatatagagcagacgCTCTGGCATGAGTAGATTGGGTCAATGAGGTTCCATCTAGaagttatactgccct
gcagtcatatagagcagacgACGTGTGTGTGTATCTGTGAGACAAGATCAAGTTCCTCAAaagttatactgccct
gcagtcatatagagcagacgTCAAAATTGAGGAAAGGAGCAGCAATCAGAAACAGCTTTCaagttatactgccc
gcagtcatatagagcagacgTCCCCAGGAATTCTTGACATCTCCCCACATCCGCCCCAGCaagttatactgcc
gcagtcatatagagcagacgTGCAACTGTTTCCTTAATGAACCCCCCTGCAGCCCCGTGAaagttatactgcc
gcagtcatatagagcagacgCTTTCTTTTCAAGCCCTCCCTCAAATACCCCATTCCTCCAaagttatactgccct
gcagtcatatagagcagacgCTTCCCCTCCCTCAATCTGGTATAAAATCCAGCAATTTAAaagttatactgccct
gcagtcatatagagcagacgAGCATGCAATTTCCACAGCACCCAAGAGTGTGAGGAACCAaagttatactgcc
gcagtcatatagagcagacgCTGGAAAGCATCTGGCAGGAGATCTTGTCATTTCTTACAAaagttatactgccct
gcagtcatatagagcagacgGAATGGACTGTTCGGTTCTACCAAATTGCTTAAACCAAATaagttatactgccct
gcagtcatatagagcagacgCTTAAAAAAAAGTCTGGTATGATTTCAAATGCCCTTCCTCaagttatactgccctg
gcagtcatatagagcagacgAAATGATGATATTTACTTGATACCCAACACACCCCTAACTaagttatactgccct
gcagtcatatagagcagacgTCCAAACCAAGAATTAGACACGCCTGGCCTCTACTAACACaagttatactgcc
gcagtcatatagagcagacgAGATAGCTTCTCTCACTTTTTCTCCATTCTTATCCCCACCaagttatactgccctg
gcagtcatatagagcagacgAGAAGCTACTTACTCTGTTATTTGTTCTTGTCTTGCCCAAaagttatactgccctg
gcagtcatatagagcagacgGCCATAACTCTGCTGTGTAGATCTCAATAGGCTGCATAACaagttatactgccct
gcagtcatatagagcagacgGGTCAATATGAAACCATTTTTTGGTAACACATCACCTTCAaagttatactgccctg
gcagtcatatagagcagacgGACTGCTATCCTTAGAAGAGTTAATGCAACTAGAGTTTTGaagttatactgccct
gcagtcatatagagcagacgGAGTTATACATTCTGCTAGTGAAAAAAGCCCAAGATAAAGaagttatactgccct
gcagtcatatagagcagacgGCATCCTCTACCAGCCCATCTCTTCCAAAACAGCCTATTCaagttatactgccc
gcagtcatatagagcagacgCAAATTGACTGGCTGTTTAGGTACATGCTACTAAAGAAAAaagttatactgccct
gcagtcatatagagcagacgGAAACCCAGTTCATCCTAACAGCCCCAGCATCATATGGTTaagttatactgccc
gcagtcatatagagcagacgGGGAAGAGAAAAGAGAAAGACAGACACGGTGTTCAGGCCAaagttatactgcc
gcagtcatatagagcagacgATTCATTGAATTTTGGAGTCAGAAAAGGGCCTTAGAAATCaagttatactgccct
gcagtcatatagagcagacgTCGTTAGAAAAGAAACCAGATAGCCTATTATAGAATGTCCaagttatactgccct
gcagtcatatagagcagacgATAGTAAGCAACAATATTTGAGGAACTGGAGTAAGTAGTCaagttatactgccct
gcagtcatatagagcagacgTTGGAAAACTGTAGGCTGCCAAAGCTCCTAAATAATTAATaagttatactgccct
gcagtcatatagagcagacgGTGCAGCTTCATAAAACCTTGGAGAAAACAACAAAAATTGaagttatactgccct
gcagtcatatagagcagacgGGACTTCCTTCACAAAATTCACAATTTCAGTATTTCCACTaagttatactgccctg
gcagtcatatagagcagacgGTAAATGTTAGCTGTTCAAGAACTCAAGAAAAATGAGGGCaagttatactgccct
gcagtcatatagagcagacgAGGTCTGTAGCTTTCATCCAGGTTGTGATACCTTAAAAGTaagttatactgccct
gcagtcatatagagcagacgGGACTGGTATCCTTTATAGAGAAAGAGCAATAGAGAGGACaagttatactgccc
gcagtcatatagagcagacgAGAATGTCTGGAAGAGGTAAATAAGCGCTTTAGAGGGGAAaagttatactgccc
gcagtcatatagagcagacgTCTTTTTGTAATACTGAGCCTCTGGTTATTTTACCTCTCAaagttatactgccctgt
gcagtcatatagagcagacgCACACCAGGCATGCACACACACAGAATATTAATAGCTGATaagttatactgccc

gcagtcatatagagcagacgTTCAAAGATTACATAGTACCTGAAGGAAGGCAGACCCAGGaagttatactgccc
gcagtcatatagagcagacgGCTTCTTACTAGACATTGCTTCCCGGATATGCAGCCTCTCaagttatactgccc
gcagtcatatagagcagacgCATGAAAGCCCTGATGCGGTTTGTAAAGTAGGCGGTAGCTaagttatactgccc
gcagtcatatagagcagacgCTGTGAGATGCTTGTTAACCAGACAGAGAAACTATGAGTTaagttatactgccct
gcagtcatatagagcagacgGCACCCATGATGCCGACTGAGCTAACATGAACTAACAGAGaagttatactgcc
gcagtcatatagagcagacgGAGGTTCTCAGCTTGCCAATGTTAGTCCTTTCTTCCCAAAaagttatactgccct
gcagtcatatagagcagacgTAATCTAAGAAAAATCACAAGAGGATCCCACTCACATTTCaagttatactgccct
gcagtcatatagagcagacgTTTCCACTGATCTAAGATGCTTCCAATTACAAAGAAAACGaagttatactgccct
gcagtcatatagagcagacgTGTGAAGCAGAAAATCTAATAAATCACGTACTTTTCCCTCaagttatactgccct
gcagtcatatagagcagacgTTTAATATAGTTATGTTAGCACCTTTCCCAACCTACAGTGaagttatactgccctg
gcagtcatatagagcagacgCTTCGCCCTGTCTCCGGGTTTAGTAGTTTCTCCAATCTAAaagttatactgccct
gcagtcatatagagcagacgGACGGTCTAGTTAGTCTTCCCACTGAGAAAGAATATTTAAaagttatactgccct
gcagtcatatagagcagacgGTTGTTGTTGTTGTTGTACTACGCTCCATCCTATTGACTTaagttatactgccctg
gcagtcatatagagcagacgTTTTCTCCACGATGCTTAAAAGAAAGGCGCTCCGAAATCTaagttatactgccct
gcagtcatatagagcagacgGCGCCCAGGGGAACGAGGCACAAATCTCCGCCACTCAGCCaagttatactg
gcagtcatatagagcagacgCCCTCAAGTTCCTCGGGGCTTTGACCCCTCAATGGCGGCGaagttatactgcc
gcagtcatatagagcagacgGGCCCCTGACTGTTCCACCGCCCGTCTCCCAAACGCCCAGaagttatactg
gcagtcatatagagcagacgCCACATCGTTTGGTCAAAATCGCCCGCCCTCTCTTCGCTCaagttatactgcc
gcagtcatatagagcagacgGGCTTGTAAGTCAGTAAGTTAACGGTATTTGGGCCCTGGGaagttatactgccc
gcagtcatatagagcagacgCGCGACACGGACTGGACCGCGCACCACGCCTGCTCCCGGCaagttatactg
gcagtcatatagagcagacgCCGGGTGGCTGAGCGCAGCGCACTGGGGCCCCGCGGTCCGaagttatactg
gcagtcatatagagcagacgGCCGAGCGTTCCCCGGGACAGTCCGGCTTGGGGACTCAAGaagttatactg
gcagtcatatagagcagacgCGCAGGCACGAGTCCCCTCCAAATCGGCTCCCCTCCTACAaagttatactg
gcagtcatatagagcagacgCGAGGTCGCGGCCAGTGCCCAATATCACCAAAGTCCCGGCaagttatactgc
gcagtcatatagagcagacgCCAGCCCCGGGCAGCCTGGCAGGCTTTCAGGAGCCTCTGCaagttatactg
gcagtcatatagagcagacgCAGGTCCTCTGGGAATGGGCTCTGGGTGCCAGGCAGGTCCaagttatactgc
gcagtcatatagagcagacgTCGGCAGTGAGACCCCAACCCGACAGCTGGATAAGGGATCaagttatactgc
gcagtcatatagagcagacgGTCGCAAACGGACAGCCCCAGCGCGGACCCGGTCTGAGCGaagttatactg
gcagtcatatagagcagacgGCTCGCGGGTCCAGGCCAGCCACTCCTCGAGGCGCTCCCTaagttatactg
gcagtcatatagagcagacgTGCAACTAGTGTCCCAGGTTCTCGCCGTAGCCAGCTTCTCaagttatactgcc
gcagtcatatagagcagacgTACAGCCTCGTCTCCCCGCGTGCACCCCGCCCTGGCCTCGaagttatactg
gcagtcatatagagcagacgGCCGCCCCATCGCAGCCCGCCTCAGGCCCCGGCCCCCAGCaagttatact
gcagtcatatagagcagacgGACCCCGGTGTCCCAGTAGCTGACCATGGTGAGCGCGACGaagttatactgc
gcagtcatatagagcagacgGCCAACGACCCGGCCGCCAGAGTCCGTCCTCTCGTTCGCCaagttatactg
gcagtcatatagagcagacgGCCAGGAGACAACCACTTCCCCGGGTAATCCTCGCCGCCAaagttatactgc
gcagtcatatagagcagacgGCACCCGAGCCCCGGAGCCCGCTCCGAGCCGCCGCCGCTGaagttatact
gcagtcatatagagcagacgGGCGATTTATAACCTTTCCCCAGCCCACTTCCTACCCCGGaagttatactgcc
gcagtcatatagagcagacgGTCCCCCACCCCTCATCCGCCTCCCCACCTACCCTCTTCTaagttatactgc
gcagtcatatagagcagacgGGCTACGGAGAGGGGCGGGGGCCGACGAGGGCGGGTGGCCaagttatact
gcagtcatatagagcagacgGGTCTTTGCTCCGCGCCCTCTTGCCCTTGCCCCTCCTCCTaagttatactgcc
gcagtcatatagagcagacgCTCAGGAAATCCTTGCAGACTCCCTCCACCCGGCTTACTCaagttatactgcc
gcagtcatatagagcagacgCCGAAACTGGGGAAGGGACAAGGCTGAAGCTCCGCCGCCTaagttatactg
gcagtcatatagagcagacgTCCGCGACGGTGTTCTTGGCACGGAGGACTGAGGCGCAGTaagttatactgc
gcagtcatatagagcagacgTCTCCCCTTGCCTGCTCTTCCCGCGCTGTCCCGCCGCTCCaagttatactgc
gcagtcatatagagcagacgTCAGACTGTCCCGAGAAGTCTAGGAAGGCACCGGAGACCCaagttatactgc
gcagtcatatagagcagacgCTGAGGGGAACTGACGCTTGCTTGGCACCTTTGGGAAGACaagttatactgcc
gcagtcatatagagcagacgCCAGCTCAGGTCCCAGGGGAGCGTGTCCGTGTCTTTTTCGaagttatactgcc
gcagtcatatagagcagacgGCACCCGCCCAAGTCATTTCCTCAGTCTTCTGCTTCCTAGaagttatactgccc
gcagtcatatagagcagacgAGGAATAACATTATGATACCTTTAAAAAGCTGAGGCGTGAaagttatactgccct
gcagtcatatagagcagacgGGGCACTTGAACTTTATTCCCTAAATGCTCGAAAAAGAAAaagttatactgccct
gcagtcatatagagcagacgTCAAGATGAGTGACTCTGACCACAACCGGCTGAAATTAACaagttatactgccc
gcagtcatatagagcagacgATTTGCTATGACACTAATTCCACGTTGTTCCTCAATTATGaagttatactgccctg

gcagtcatatagagcagacgCGGAGCAGCCCAGTCTTGTCTCTGTAAATACCGCAGATTTaagttatactgccc
gcagtcatatagagcagacgCGTCAGAGGGATGTAATACAAGGGACTGTGATGGCTCTTGaagttatactgccc
gcagtcatatagagcagacgGAGACTCTTCCTGAAGGACTGGACTACCTCCCCGCCTAGGaagttatactgcc
gcagtcatatagagcagacgACCCAAAGTTGCCACATTTCTGGGCTGCACACCAAGGAGAaagttatactgcc
gcagtcatatagagcagacgCGCAGGACCTGAGCCACTAACTGTCCCTGGATCCCTTCTGaagttatactgcc
gcagtcatatagagcagacgCTTCACCAGCGCTTTCTGTTTTTCCAGTTCTCCTCACCACaagttatactgccct
gcagtcatatagagcagacgCACCCCTGCCTCCTCTAAAACCCACAGCATCATCCCTCAGaagttatactgcc
gcagtcatatagagcagacgTGGAAGCGGACAGCTGATCCTTCTCCTTGCTGGTCCCCCAaagttatactgcc
gcagtcatatagagcagacgTCACCTGAGAGCTGCTTGTCACTCCCACACCCTCTTTCAAaagttatactgccc
gcagtcatatagagcagacgGCATTGTCCTTCTGCAGCATTCAACAGGGGCCAGGCATTGaagttatactgccc
gcagtcatatagagcagacgTTCTAAAGGGCTTGGACTGTGATAAGAAGGTGGACACTTGaagttatactgccct
gcagtcatatagagcagacgGGGGTGTTTAGTCCTCCATCAGCTCAGAGAAGTAAAGGCAaagttatactgccc
gcagtcatatagagcagacgGAGACACACATTTGACTCTGGGGGAAAGTGGTCCATAGTAaagttatactgccc
gcagtcatatagagcagacgAAGAGACAACACTTTCCAAACCTGCTGCCAGCAATGAATCaagttatactgccc
gcagtcatatagagcagacgTGTAAAGCCAGAACAGGTATTTGATGGACAGCTGAAACAAaagttatactgccct
gcagtcatatagagcagacgTGTTGCTCAATTATTTAAGGTTAGTAGAGTCACAGTCGAAaagttatactgccctg
gcagtcatatagagcagacgGATTTATATTAAAACTATGCTGCAGTAAGGGTGGGCTTCCaagttatactgccct
gcagtcatatagagcagacgTATTGACTATGACTTCTTTGTTAGTTGCATTAGCGTTCTCaagttatactgccctgt
gcagtcatatagagcagacgGGAATAAATGTATAGCTCCCAATCCTCTCCCCAAAAGAGAaagttatactgccc
gcagtcatatagagcagacgCTATAGCACTCTCCTAACTAAATCACTAATGAAAGAGGTTaagttatactgccct
gcagtcatatagagcagacgGATAATACTGATGGAGAAGATGGAATTCAACACGCCTGGTaagttatactgccct
gcagtcatatagagcagacgCAATCTGCTTGCTTCCTGCTCACATACAAAAGAACATTTCaagttatactgccct
gcagtcatatagagcagacgAACCAGACACTCTCTCTTGTTATCAGCTACAAACATTCTCaagttatactgccct
gcagtcatatagagcagacgGCACACTGTTGCTAAATCCATCAGCTTTAATATTCGAGACaagttatactgccct
gcagtcatatagagcagacgAGAGTGTTGGAGCACCAGTGTGAGAGACAGGAGGGAGCCCaagttatactgcc
gcagtcatatagagcagacgGAATTTCTTTTCCACTTGGGTTAGGTAGCTTGATACAAAAaagttatactgccctg
gcagtcatatagagcagacgGTTGAATACAAGAACCAAATGATCTCATAAAACCCTTCTGaagttatactgccct
gcagtcatatagagcagacgACATATCCTCTGGACAGTTCAGGCTCCTTGGTCTAAAAAGaagttatactgccct
gcagtcatatagagcagacgTGCGGAGTTCACCAGGCAGGCACATGTATGAGTTTACTTAaagttatactgccct
gcagtcatatagagcagacgCCATTGGCTCAAAAGTCTTGCTATGGGAAGCTACCACTGTaagttatactgccct
gcagtcatatagagcagacgATGCCCCCATTTTCAAAGAACTTAAACATCTTGTACTGGGaagttatactgccct
gcagtcatatagagcagacgAGCCATTCAACCCATCTACCGAGTCAATTCGTGGAATTCCaagttatactgccc
gcagtcatatagagcagacgCATGAAAATTTATTGAACACTTAAAAACACCTCCTGTGCCaagttatactgccct
gcagtcatatagagcagacgCATCTTGTCTCCAGCCTCAACATCCACATCCCTTTGGTCAaagttatactgccct
gcagtcatatagagcagacgACACGCTCCAGAAATTCCATCCAGCATATCAGTGTCCCTGaagttatactgccc
gcagtcatatagagcagacgAATGCAGCACAATTATCATTTCCCTACTCCTAGAGGTAGTaagttatactgccct
gcagtcatatagagcagacgAAAATCGTTCAGGTGGGAAGAGCAGCGTGGAGAAGGAAGCaagttatactgcc
gcagtcatatagagcagacgGCCCCCTCCTGGAGATAAGGCCAGTGCCACCGAGAGCAGGaagttatactgc
gcagtcatatagagcagacgGGGCTCCCTGGCTCCTCCTGTTCCTGCTACTCATGCTCCCaagttatactgcc
gcagtcatatagagcagacgCCTGACTACCTTCTCCCGGGCATGGGTGGGCGCTGCTGGTaagttatactgcc
gcagtcatatagagcagacgCCATTGGCTTTACTTGGAATTCTCCTCCTGCTACTCCCAGaagttatactgccct
gcagtcatatagagcagacgAACTTGCTTCTGTTCCTGGGACCTAAGATCAAAGGAGCCTaagttatactgccct
gcagtcatatagagcagacgGTCAGCGCCTTCATACCTTGGACTCTTCCAGAACTTATTCaagttatactgccct
gcagtcatatagagcagacgCCCCAAGGCTTTCTCATGAGAGCTGCTATCTGGCTGCCCTaagttatactgccc
gcagtcatatagagcagacgGCCCTGAGCGTGTTGTTTAAGCCTTTTGTTTGTTTTAGTCaagttatactgccctg
gcagtcatatagagcagacgCAGCATGTCGAAGTCTTTCTGGTCTCTGATTCTGCCATCCaagttatactgccct
gcagtcatatagagcagacgCGGTAAACGTGCCTTCTGGGTCATCATCAAAATCACTGACaagttatactgccc
gcagtcatatagagcagacgAATGAATGGATGGATGAAAGTAGAGATCTGTGGAGGTTGAaagttatactgccct
gcagtcatatagagcagacgATTATTCCTTTGTGTCATTATCTCTAGCTGTATGTTCACCaagttatactgccctgt
gcagtcatatagagcagacgGGAATAATTGATGGTCCTAGCAAGAGAAAATAAGACAGGTaagttatactgccct
gcagtcatatagagcagacgAGTTGAAAGGAAAAAAAACAAAAAGACCAGACTGGGATAGaagttatactgccc
gcagtcatatagagcagacgCATGGGGCAGGGGAAGCGGCAAATCAGAGAAGCAAATAAGaagttatactgcc

gcagtcatatagagcagacgGGAACCTCCTCTAATCAGTATCACTACTATCTTGCCTAAGaagttatactgccct
gcagtcatatagagcagacgCACTTAACTGCCAGCCATTGACTCTTCTCCTTACTCTTTTaagttatactgccctg
gcagtcatatagagcagacgTTTAGAAAGATTAAAAGAAGGGAAGGTCTGGAGGATGAGTaagttatactgccct
gcagtcatatagagcagacgCACTGTGGTGTATTTCATTCGAGCCTTTTCCATACAGACGaagttatactgccct
gcagtcatatagagcagacgTCTTTCCATGTTTCTATAGAGCCTTCATAACTGTTGTAGCaagttatactgccctg
gcagtcatatagagcagacgAAACTATTATCCTCTTGCTAGACATTCCAGTTTCTGCATTaagttatactgccctg
gcagtcatatagagcagacgATTTGACACTGGAATGTTTTTAGTCAGATGTACACACGCAaagttatactgccct
gcagtcatatagagcagacgACGCTCATCAATATCCCTGCTTTACTTAAAACCCTACCAGaagttatactgccct
gcagtcatatagagcagacgCGAAGGAAACAGCTCGTAGCATGGTGCATGATCCAGGAAAaagttatactgcc
gcagtcatatagagcagacgGTAACATAGGAGATACAATTATTACTCTAGAAGACCTTGCaagttatactgccct
gcagtcatatagagcagacgGAAACATTTCTAAGGCGATTAAAGGCAAGATACAGTTATGaagttatactgccct
gcagtcatatagagcagacgTTGGGAAATGGCTTTGAAGGTAGTAACAGTTTATATCAGCaagttatactgccct
gcagtcatatagagcagacgCTGGAAAGGCGGAGAAGGTAATAAAACCAGTCAACAGTGGaagttatactgcc
gcagtcatatagagcagacgTGAGGGGTGGAGGTGTTGGGAACAGGGAGGCTGAGTTTGCaagttatactgcc
gcagtcatatagagcagacgCACTTGCGAGATACTCACGTGTGTTTCATTCTGTCAGTACaagttatactgccct
gcagtcatatagagcagacgCAATAGTGGGTTGCAAGGGAATGAGACTGGAGGGAAATGAaagttatactgcc
gcagtcatatagagcagacgTTGAGAGGCAATAGAATAGGACTGGTACCATGAGGGGAAGaagttatactgcc
gcagtcatatagagcagacgTGGTTAGCAGTTCAGGGAAGGGATGTAGAGATAATTAAGTaagttatactgccct
gcagtcatatagagcagacgCATTGGCTGATGGGGATGTTGTGAGAGGTGGCTGGAGAGGaagttatactgcc
gcagtcatatagagcagacgCATGTTGGGATGAGTGGGCAATTGAAAGCAACTGACTTTAaagttatactgccct
gcagtcatatagagcagacgCCAAGGCAGAGTTTAAAGGGATGGCGTGCTGGGCGTGGTGaagttatactgcc
gcagtcatatagagcagacgTAAATATTTTAAAAGACAACAGGATAGTGTGGCTCAGGTCaagttatactgccct
gcagtcatatagagcagacgCTGGGACTACAGGTGGCTGCCACCGTGCCCGGCTAATTTTaagttatactgcc
gcagtcatatagagcagacgGCGCCCGGCCAGCTCAGGTCTAAACATTTTAAGGAAGTAGaagttatactgcc
gcagtcatatagagcagacgAGGTAGTGCTATATCCTGACTTCACATGTCGTATATATGTaagttatactgccctg
gcagtcatatagagcagacgCATATATTTGAGATAGATAGCCTGAGTTTTCTGTAATGGCaagttatactgccctg
gcagtcatatagagcagacgGCCCAGCAGAGTACTCCAGTTTTAGCCTTCCTTCTTCAAAaagttatactgccct
gcagtcatatagagcagacgCCAGACACACCCTCAGGGCAGATTTGCTGGCGACTTCTGTaagttatactgcc
gcagtcatatagagcagacgACCAGCCCTTTCCTTCCTGCGTCTCTTCCAGGCTCCTCCTaagttatactgccc
gcagtcatatagagcagacgGCAGCAGTCACGGAGCAGCCAGCCACACTCTGCCCTGGACaagttatactgc
gcagtcatatagagcagacgCCTAGAGGCAAGGCTGGCATTTGTAAGAAAGTTAGATCCCaagttatactgccc
gcagtcatatagagcagacgAGTGAAACCTACTGTGGAGGGAAGAATTGTGTATCATAATaagttatactgccct
gcagtcatatagagcagacgTAATCCATAAAAATAAGCTCCAGTTATACCACTAGTGTGGaagttatactgccct
gcagtcatatagagcagacgTTTGCTTCAAAATTATCCAAGACCCCAATTACACCGATTTaagttatactgccctg
gcagtcatatagagcagacgTTTTCACACGTACCACAAAAATATGTACATTTCCTATGCCaagttatactgccctg
gcagtcatatagagcagacgCAAGAGGAGTGAAGATGAAACATGATAGGCCATGTATTGAaagttatactgccct
gcagtcatatagagcagacgGAGCAAAAGTTAATTTTCTCTTCTAGTCTGAAATTGGTCCaagttatactgccctg
gcagtcatatagagcagacgGGCTACAGAGAAAAGGCGGGATAATGAATGTAACCTGTTGaagttatactgccc
gcagtcatatagagcagacgGACATATGTCCTCCCCGTGAGTTTCTAGCAAATTGTGTGTaagttatactgccct
gcagtcatatagagcagacgATGAAGAGGCATGAAGAGGTGTAAAGCAATCAGAGAGGAAaagttatactgcc
gcagtcatatagagcagacgGCAGAAAGAAGTAGCACGGAGAAGTAAAAAACATATAAACaagttatactgccc
gcagtcatatagagcagacgAGAAAAAGAACAGGAAAACCCTTTGAAAGGAAAAATCCAGaagttatactgccc
gcagtcatatagagcagacgGCTGGAGATCATGAAGTAGAGAAAATGAAATTTGGGGTGAaagttatactgccct
gcagtcatatagagcagacgACCCATGCTGCATCGTCTAATGACATTATTAAAAAGGACTaagttatactgccct
gcagtcatatagagcagacgGCAAGCATTCTAGGCAATTTGGGAACAGGAAGATAATATTaagttatactgccct
gcagtcatatagagcagacgCCATTAGCCATTTTTTTCTAGAACTTATGCACAAGTCCCCaagttatactgccct
gcagtcatatagagcagacgAGGAAAGAAATTCAGCAGGTTGATTGGTAAAGATAAACACaagttatactgccct
gcagtcatatagagcagacgCATACAATCTTAGTGTGCAAGTGGACCTTATAGATCTAGTaagttatactgccct
gcagtcatatagagcagacgCTGATCTTGCTTTGAGGAAAGAGGCAGCGAGGTGATGTGCaagttatactgcc
gcagtcatatagagcagacgCAGACTATTGCTGAGAGCGTCTCCCCACACGATCCCCATGaagttatactgcc
gcagtcatatagagcagacgGAGCCTAAAGATGCATTGCTCTGTTCTCTGGATGCTTGTGaagttatactgccct
gcagtcatatagagcagacgGGATTTGCCTGCAGGTCAGCTCAGAATTTATTTAGGTAAGaagttatactgccct

gcagtcatatagagcagacgGCACCCAGAAAGCAACATAGAGCAGAGGGCAAAGAACTTCaagttatactgcc
gcagtcatatagagcagacgTGTGCACAGCTCCTAGGCCAGCCTACCGGGGGTTTGAATCaagttatactgcc
gcagtcatatagagcagacgCCACCTAATGTCTCTGGGCTCTAGTCTCTTAATCTCCAAAaagttatactgccct
gcagtcatatagagcagacgCAGAAAAGCATTTATAATAGTGTCTGGACCAAACTAAGCAaagttatactgccct
gcagtcatatagagcagacgCGTGAGTAATGTTTGCCATATGCCTCCTGCCTGCTGCCTGaagttatactgccc
gcagtcatatagagcagacgTCCTTCAGTACTTATCTCCATGGGTTAGCTATACTTCTGCaagttatactgccctg
gcagtcatatagagcagacgACACCGTGTACATTTGCTAGCCTTTTTACTCAAGACTGAAaagttatactgccct
gcagtcatatagagcagacgAGACCCCGCCGCCTCTCTGACTTCAAGTGTGCCCTGATGGaagttatactgcc
gcagtcatatagagcagacgGCCCCATCCTCTGGACTGGAAGCCCTCAGATGCCCATTCGaagttatactgcc
gcagtcatatagagcagacgCCTGCCCCTGGGTAACTGTTGGATTGCGTGGCTGTTCCTGaagttatactgccc
gcagtcatatagagcagacgAGTTCATTGCTGTCTTGTTCCCTGCTCCCTTGCTTCTCTGaagttatactgccct
gcagtcatatagagcagacgCCCCAGTCCAGTTGCTCATCTTCTCTCTCTACCAGCCTCTaagttatactgccc
gcagtcatatagagcagacgGTCCTAGCACAAAATCCTTATCTCTATGTAAGAACCCTCCaagttatactgccct
gcagtcatatagagcagacgCTTGATCTGCCAAGGTTGATCTCAGAGGGAGGAGCTCATCaagttatactgccc
gcagtcatatagagcagacgGACCTCTCTGTCCTCCTCTTGTCCTGACATCTCCTCTCCTaagttatactgccct
gcagtcatatagagcagacgTGGTCTTCAGGGCTCCATGGTGCCTTATGGGGTTTCACACaagttatactgccc
gcagtcatatagagcagacgGACAACTGCTTCCCTGAGCTTCCATCAGCATCCCCGGGATaagttatactgcc
gcagtcatatagagcagacgCAGGACCCAGCTAAAGGCTCAGGCAACACAACAGCGGAGGaagttatactgc
gcagtcatatagagcagacgCTTCTGTCTTGAGCACCACCCTTCCCTTTTCCATCAGACTaagttatactgccct
gcagtcatatagagcagacgGAAGAATCCACAGGTATGGGCTTCAGTAAAAAAGCTGGAGaagttatactgccc
gcagtcatatagagcagacgCATGGCTCACTTTTGTCTTGGCTTATTTTGTACTTTGCACaagttatactgccctg
gcagtcatatagagcagacgAAAAATGATAGTGGAAAAGGAAAAGGTAGGAATCCTGTGAaagttatactgccct
gcagtcatatagagcagacgCAGTTTCAGTTTAGAAAAGCCCCCAAAACCTGTTCCTTATaagttatactgccct
gcagtcatatagagcagacgAGTAATTGCTTTTCTTCTATAGTCACAGTGCCAGGTCTTGaagttatactgccctg
gcagtcatatagagcagacgTCATAGCTCACTGCATCCTTGCACTTCAGGGCTCAAACAAaagttatactgccc
gcagtcatatagagcagacgACCATGCCCAGCCAAGAGTCACTTAAACATGTATTTTTTGaagttatactgccct
gcagtcatatagagcagacgAATCGTTAATAGTGTAAGATGATTTGAAGAGCCTGACCACaagttatactgccct
gcagtcatatagagcagacgAGGAGTACCTTTAGCTTAAAAGACTTCTCTGGGTCCTGGAaagttatactgccct
gcagtcatatagagcagacgTCTGATCCCCTGACACTAAAATTAAGACTTGGGCCACAAAaagttatactgccc
gcagtcatatagagcagacgCAGGAGGAAGGTATATTAAGAAGTCAGGAAATTCACCCAGaagttatactgccc
gcagtcatatagagcagacgCCTGAAGGTGGGGTAAGAGAGGGGGGCTGTTATTGTGCTCaagttatactgcc
gcagtcatatagagcagacgGAGCTTCCTGTCTCAACACATGCCTGACTTGATGATCCTCaagttatactgccc
gcagtcatatagagcagacgCTTCCTGAAGCACCCACGAACGTTAGAACACGCAGGCAAGaagttatactgcc
gcagtcatatagagcagacgGGTCCACTAGGGTATCCAAAGGAGCTGTCATCGTTTGCTCaagttatactgccc
gcagtcatatagagcagacgCAATTTCTGTCAGACTGTTCAACTCTCACCTACCATGTATaagttatactgccct
gcagtcatatagagcagacgGAAGTCTTTCCTGAGCCCTCTCGTCAACCAGACCCTTCTCaagttatactgccc
gcagtcatatagagcagacgTTTATCTTGTACTTCTTTGTATAGAAACTGTGGGGAACAGaagttatactgccctg
gcagtcatatagagcagacgTTAGCCAGGAGCAGTGGCACCCACCTCAGCTACTCAGGAGaagttatactgcc
gcagtcatatagagcagacgAATATATGTGAAAAAGAAAACTGTAGGGAACTGCAAGCAGaagttatactgccct
gcagtcatatagagcagacgTATAATGGTGAATATACCAGTTAGGAATGCTTTTGGTTGCaagttatactgccctg
gcagtcatatagagcagacgAAAGACAAAATTCAATAATCTCACATAGTAAGGGGTCTGCaagttatactgccct
gcagtcatatagagcagacgGCAGTTTCAGGTTGGATAAGGCAATTTCACAATGTGATGAaagttatactgccct
gcagtcatatagagcagacgTTCTGCCCTGCCAACTCTGATGTCGCTTTATGGTCACAAGaagttatactgccct
gcagtcatatagagcagacgTGTACAACTACATTCAAGGCAGGAGGTGGGGCAAAGAAGTaagttatactgccc
gcagtcatatagagcagacgAGGAACTACCCAGGATACTCTTTATATCTCATTGGCCAGAaagttatactgccct
gcagtcatatagagcagacgCCAATCATTGGCAAAGGGAAGTGGAATGAGCATGACTGGCaagttatactgcc
gcagtcatatagagcagacgACACACACACATATACACGCTGGGCCGGGCACTCCCTCTCaagttatactgcc
gcagtcatatagagcagacgCATCAACCTGAACAATATGGGGGGGCCTGTTAGCAGCCAAaagttatactgcc
gcagtcatatagagcagacgCCATCAAAATTGGCCTTCCTACAACAAAGCATACATCATGaagttatactgccct
gcagtcatatagagcagacgGCTTTCCATGTTAAAAACTGCAGAGTATCCATATTGTGCCaagttatactgccct
gcagtcatatagagcagacgCCTTTGCTTGGAATCAGAAGAGTGGATTACTTACTAGCTAaagttatactgccct
gcagtcatatagagcagacgGGTTTGCTTCATCCCTTTTGCTTCCAACACCCTCTGGCTTaagttatactgccct

gcagtcatatagagcagacgGCTTCCTCAAATCTGGTCAGACGTCAATATTTAATCATATaagttatactgccctg
gcagtcatatagagcagacgATTTAATCCTATCAAGAGTCTCACTTCCTTTCATCTAGTCaagttatactgccctg
gcagtcatatagagcagacgTCATTTATTCCACAGACATTCACTGAGTGACTAGACATGCaagttatactgccctg
gcagtcatatagagcagacgTTCTTTGGGAAATGGAGCAGCAACAATTTAGAGGGACGTGaagttatactgccct
gcagtcatatagagcagacgAAGCTCCATACTACCTACTTCTAGATTGCTTCATGAGAGAaagttatactgccctg
gcagtcatatagagcagacgTCTCCTGACTACTTGTTGAGAGCTTAGTTGTGTCCCCTCAaagttatactgccctg
gcagtcatatagagcagacgTATGTTGAAGCCCTGATCCCCAGTACCTCCGAATGTGACTaagttatactgccc
gcagtcatatagagcagacgGCTGGTCTCGAACTCCGTACTTCAAATGATCTGCCTGCCTaagttatactgccc
gcagtcatatagagcagacgGACTGTATTTGGAGATAAGAGGTCTTAGTTTGCTTGTGCTaagttatactgccctg
gcagtcatatagagcagacgAAGATCAAGGTGCTAGCATTTGCTGTTGAGTGAGGGCTGCaagttatactgccc
gcagtcatatagagcagacgCTCATGTGGCAGAGGGTGGACAAAATACATTAATCCTCTTaagttatactgccct
gcagtcatatagagcagacgGCGCGATCTCAGCTTACTACAATTTCCGTCTCCCAGGTTCaagttatactgccc
gcagtcatatagagcagacgGCTTTAATCCCATTCATGAAAGGAGATGCCTTCATGACCTaagttatactgccctg
gcagtcatatagagcagacgTTAAAGAGGTGATAATGAAGTGATTAAAGTGGCCCTAATCaagttatactgccctg
gcagtcatatagagcagacgATGGCCATCTAAAAGCTAAGGAGAAGAAAAAAACATACCTaagttatactgccct
gcagtcatatagagcagacgTTAGCCTTTAGAACCATGAATGAATAGATTTCTGTGAAGCaagttatactgccctg
gcagtcatatagagcagacgGCCCTAGTAAACACCATTCCCGCCTATCTGCTTTCTTAATaagttatactgccctg
gcagtcatatagagcagacgACACTTTTGTTCAAGGCTTACTAAGGGTAGAAAACATGTCaagttatactgccctg
gcagtcatatagagcagacgAAGAAAAAAGACAGAGGAGACAAAAATTAACAAAGGACCaagttatactgccc
gcagtcatatagagcagacgACAGTCTGGAGTTTGTAAAAATCATTCAAAAACCTTCTGGaagttatactgccctg
gcagtcatatagagcagacgAATTGGACTGAATAGATTCTAGATGAAAATGACAGATGGGaagttatactgccctg
gcagtcatatagagcagacgAAATCACAAAAACCAACCCCCAAATATTCATTAACAGCACaagttatactgccct
gcagtcatatagagcagacgATACCTTAATGAAACACACCTCAAATATTAGTAGCCAAGGaagttatactgccctg
gcagtcatatagagcagacgGGCTAGGCTTCTAATCCTACCCTCTGCTCCAAAACCATGCaagttatactgccc
gcagtcatatagagcagacgAGAAAAATCCTAAGAATTCTCATTAACGCCTCCCACTAATaagttatactgccctg
gcagtcatatagagcagacgCTGGAGAGGTATAAGCTATATAGGTGTCTTAATCTTTTTCaagttatactgccctg
gcagtcatatagagcagacgCCAAATCTGAGACGGTCCTTACATGGGGATTGGGAGTCCTaagttatactgccc
gcagtcatatagagcagacgCTCAGAGGAGGGAAAAGGTTCACATATTCTAAAACATACCaagttatactgccct
gcagtcatatagagcagacgGAACCACAGATAAACTCTTCCTTTCCAACTATCCATTTTTaagttatactgccctg
gcagtcatatagagcagacgGAAATGTTCAATCCACAAACTATAACTCGGAGGGGAAAATaagttatactgccct
gcagtcatatagagcagacgAAAGTCATCCACATTGTATTACAGAAAATAGAAGGTCTGGaagttatactgccctg
gcagtcatatagagcagacgCAAGATGAAAAGAAACAGCATTATAACTAGGCAAAGATGCaagttatactgccct
gcagtcatatagagcagacgAAAGAAAAAAGAAAAGCAGACAGGCAAATAAGCACAAGACaagttatactgccc
gcagtcatatagagcagacgATAAGGGCTCAAGAATGATATGATACAGTAAATGTGACAGaagttatactgccctg
gcagtcatatagagcagacgATTCACCAAATCTTGTGTCGTTTTCTTCTTCCTGGGCACAaagttatactgccctg
gcagtcatatagagcagacgTTCCCTTGAATCCTACCATCTGACCCAAAAGTGTATATTCaagttatactgccctg
gcagtcatatagagcagacgAATTGATAGGTGAACATTCCAGGCCAATTCAAGACAATGCaagttatactgccct
gcagtcatatagagcagacgGCAAGTGTGTGTGAATTCTTTATGCTGTCATTTCTCCTGCaagttatactgccctg
gcagtcatatagagcagacgACCATCAACTCCATGTGGTATAGCGTTAAGGTTCCATGTGaagttatactgccct
gcagtcatatagagcagacgGCTTTACAATGCAAGTACCCTGGACTCTGTGTTATCTCTTaagttatactgccctg
gcagtcatatagagcagacgGCTGTCAGCCCTCACCCAGTTTTGCATGAAAAAAAAAAAAaagttatactgccct
gcagtcatatagagcagacgGTTAGATTTGTTACTGTATCATAACCTAGTTCATCCTGACaagttatactgccctg
gcagtcatatagagcagacgCAGACCTAAGGAAAACAACTGAATGTCCAAACCATATCATaagttatactgccct
gcagtcatatagagcagacgACCTGAGTCTGGTGCTTTCTGTTTCGGAAGGTAATCAATTaagttatactgccctg
gcagtcatatagagcagacgTAAAAGATATAGGCCTGTTCAGATCATCCATTTCTTCTTGaagttatactgccctg
gcagtcatatagagcagacgTTGGCAGACTGTGTCTTTCAGGAACTGGTCCATTTTATCTaagttatactgccctg
gcagtcatatagagcagacgTATCCTTTAAATGTCCATGGAATCTGTAGCAATGTCCCCGaagttatactgccctg
gcagtcatatagagcagacgCAATCGTATTGAGCTTTTCAAAAAACCAGGTTTTGGTTTCaagttatactgccctg
gcagtcatatagagcagacgTTTCTAAGCACTGTTTTTGCCTGTTCCCACAAATTTTGGTaagttatactgccctg
gcagtcatatagagcagacgCCATGTATTTTGGTATTTTCTGATTATCTTCCCACTAGTGaagttatactgccctg
gcagtcatatagagcagacgGTCTGAGAGCAGACGTTATATAATTTCTACTCTTCTGAATaagttatactgccctg
gcagtcatatagagcagacgTTTATGTCCCAGGATATGGTGTCTTGGTGAATGTTCCACAaagttatactgccctg

gcagtcatatagagcagacgGGCTTCAACCACATCGTTACTGACTTTTTGCCTGCTGAATaagttatactgccct
gcagtcatatagagcagacgGGATGTTGAAGTCTCCAACCGTAATAGTGGATTTATCTATaagttatactgccctg
gcagtcatatagagcagacgTTTTCTTTTTGAGACAGAGTTGAGTGCAGTGGTGCCATCAaagttatactgccct
gcagtcatatagagcagacgATAGGTGTGAGCCACTGCCCGGGGCCTGCCTCATGTAATTaagttatactgcc
gcagtcatatagagcagacgTGCATACATGTTAAGGATTACTATGTCTTTTTGGAGAACCaagttatactgccctg
gcagtcatatagagcagacgATTATGTAATGCCTGTCTTGATCCCTGATAGTTTTCCTTGaagttatactgccctg
gcagtcatatagagcagacgCATAACTACTCCTGCATTCTGTTGATTAGTGTTAATATGGaagttatactgccctg
gcagtcatatagagcagacgTATTTTTGTTTTCTGCTCTTTTCTGCCTGTTGCAGTTTCAaagttatactgccctgt
gcagtcatatagagcagacgTTCTCTTCTTTCTTAGCACACCAGTTGCATTTCTTTCTTTaagttatactgccctgt
gcagtcatatagagcagacgCTCCTGAGTAGCTGGAACGACAAGCACATGCTACCACACCaagttatactgcc
gcagtcatatagagcagacgAAATAATATTAGTTCCTCCTTCTCACCCCTTGTATCATTGaagttatactgccctg
gcagtcatatagagcagacgGCATACACAACCGAATACATTGTTAGTGCTTTGAACAAGCaagttatactgccct
gcagtcatatagagcagacgTTCTCTAAACATTTCTTGTAAGGGAAGTTTACTGGCAACAaagttatactgccctg
gcagtcatatagagcagacgTTTTGAAGGATAATTTTGCCCAGTACAGACTTCTAGGTTGaagttatactgccctg
gcagtcatatagagcagacgTTTCACTCCACCTTTTTCTTATTTGCATGGCTTCTGATGAaagttatactgccctgt
gcagtcatatagagcagacgAAAATATTTGCTCCTCCATCAGTAAGGCATTTTTTTCCTCaagttatactgccctg
gcagtcatatagagcagacgTTATCCTGCTTGGTGTTCCCTGAGTTTCCTGGACCTGTGGaagttatactgccct
gcagtcatatagagcagacgTCAAATACTTCTGCTCCTTTCTATCTCCCTTGTCCTCTTGaagttatactgccctg
gcagtcatatagagcagacgAGTCCTTGGATATACTGTTCTGGTTTGTTTGTTTTCTTTCaagttatactgccctgt
gcagtcatatagagcagacgTCTCTTTGCTTTTCAGTTTCGGAGGATTCTATTGACATATaagttatactgccctgt
gcagtcatatagagcagacgTTAGCTGTGTCTAGACTACTAAGAAGTCCCTCTGAGGCCTaagttatactgccct
gcagtcatatagagcagacgGGAGGCTAAGGTGGGGTGATCGTCTGATGTCAGGAGTTTGaagttatactgccc
gcagtcatatagagcagacgTAAAAATACAAAAAAAAATTAGTCGAATGTGGTGGCGGGCAaagttatactgccct
gcagtcatatagagcagacgTTTCCATCTCTCTGCTCATGCTGCTTGTCTGTTCTTGCATaagttatactgccctg
gcagtcatatagagcagacgGCTTCATCCATTAGCACCCACAGCATATTAATCATAGTCAaagttatactgccct
gcagtcatatagagcagacgCCTCCCAAGTAACCAGGATTACAAGCACCTGCCACCATGCaagttatactgcc
gcagtcatatagagcagacgGCGTAAGCCACCGTGCTGGGATATTAATCACAGTTTTAAAaagttatactgccct
gcagtcatatagagcagacgTTCCAATATCCCTGCCACATCTGCTTCTGAAGTTTGCTCTaagttatactgccct
gcagtcatatagagcagacgAGTATGCCTTGTAACTTTTTGCTAAAAGCCATACATGATGaagttatactgccctg
gcagtcatatagagcagacgCTTTAGTGATGTGGAGGTAAGGTGCAGGGGGAGGAAAAGCaagttatactgcc
gcagtcatatagagcagacgCTCAGTCTTTTTTTTTTTTTTTTCCGACGGAATCTCACTCaagttatactgccctgt
gcagtcatatagagcagacgGCCACCATGCCCTGCCAGGCCTGTCTTTTAGTGAGCCTGTaagttatactgcc
gcagtcatatagagcagacgTGCCTTTCACAAGTGCTTCTTAGTTTTTTCCTCCCCTTTTaagttatactgccctgt
gcagtcatatagagcagacgGATGTTTCCGTTCTCCCAGGTCACTTAGGCTGTAATTAAAaagttatactgccct
gcagtcatatagagcagacgGTTGACCAGTTTCTCCTGAAAGCAAACCTTGTTAAGAAGAaagttatactgccct
gcagtcatatagagcagacgATTTCAAAATACAGGTCCTCTCTCCCTGCTGGAAGCGCAGaagttatactgccc
gcagtcatatagagcagacgCTAATATTTACTATGAGAACTTGGTGGAGTTCCTGGAGGTaagttatactgccct
gcagtcatatagagcagacgCCACTAGAACTGGGTCCTCTGGAGTCTTTGACTTGTCCACaagttatactgccc
gcagtcatatagagcagacgGTCCAGCAATTTGTAATTACATTTCAGGTTTTCCTACTCCaagttatactgccctg
gcagtcatatagagcagacgCAGTATATTGTGATTCTCTGCATCTACCTCTCTGTCTGTCaagttatactgccctg
gcagtcatatagagcagacgGGCAGCAGTTTTCCCTGTGGATTTTTCTCTTATGGATCTAaagttatactgccctg
gcagtcatatagagcagacgCTTCGAAGCTCCTTACCTGTCCAACTAGCCATTATAGACAaagttatactgccct
gcagtcatatagagcagacgAGCACTGAAAATCCAATTATTGACATAAAGGTAAGTCCTGaagttatactgccct
gcagtcatatagagcagacgTGAAGTAATTCATATGAAAAAGACAAAGGGACCAGGCATGaagttatactgccc
gcagtcatatagagcagacgCTGAGGCGTGAGGACTGCCTAAGCCCAGCAGGTTGAGGCTaagttatactgcc
gcagtcatatagagcagacgTTGTCCCACTGCACTCCAGCATGGATGTGGTGAAAAGGGAaagttatactgcc
gcagtcatatagagcagacgTCCTTAAAGAACTAAAGTAGAACTACCGTTTGATCCAGCAaagttatactgccct
gcagtcatatagagcagacgTATGAAAAAGATACTTGTGTACGCATGTTTATAGCAGCACaagttatactgccct
gcagtcatatagagcagacgACTCAGCCATAAAATGGAATCAAATAATGGCTTTCATAGCaagttatactgccct
gcagtcatatagagcagacgGGAAAACTAAACATTGTACGTTCTCATTTATAAGTGGGAGaagttatactgccct
gcagtcatatagagcagacgCGGGCATGGTGATGTGTGCCGGTAGTACCAGCTACTTGGGaagttatactgcc
gcagtcatatagagcagacgGATACAATGCACTTTGGGGATTTGGGGATCTGGGGGAAAGaagttatactgccc

gcagtcatatagagcagacgTAAAAGACTACACTGGGTACGGCATACACTGCTTGGGTGAaagttatactgccc
gcagtcatatagagcagacgATATTCCAACTTAAAGACAATTGATGCCCCTAAAGCAAAGaagttatactgccct
gcagtcatatagagcagacgCTCTGCAAATCAAGAGAACTAATGACATTCCAGGAAAATTaagttatactgccct
gcagtcatatagagcagacgATTCCAGGATAAAGAACGAAGTTTTTAGCTAATCAAGCAGaagttatactgccct
gcagtcatatagagcagacgTTCTCCATCGCAACCTCCAGGGGCAGAAGACAAGAACACAaagttatactgcc
gcagtcatatagagcagacgCCAAGATGTTATTCAAGTAGAAAGGTAGAGGCAGGAATTTaagttatactgccct
gcagtcatatagagcagacgGGAATGCAATGCCCATCACTATTCATGCCCATCACTCATGaagttatactgccc
gcagtcatatagagcagacgAGGAATGAGGAGCAATGGGGAAAGGACCAAAGGTGAACATaagttatactgcc
gcagtcatatagagcagacgCCATTTGAAGACATAGAGACATACACCTACATACATGCTAaagttatactgccct
gcagtcatatagagcagacgTGCTTAGAATGTAAATGTTATAAAACGGGCAAAGTTCAGGaagttatactgccct
gcagtcatatagagcagacgCTGTTTTTCTCTTCTTTTAACGGCAGGGAGTCAATTGATAaagttatactgccctg
gcagtcatatagagcagacgGTGGGGTTTGAAGTAACTTTAAAAAAAAATGTTGGCTAGGaagttatactgccct
gcagtcatatagagcagacgCCAGCCTAGGCAACATACTAAGACACTGTCTCTACAAAAAaagttatactgccc
gcagtcatatagagcagacgGCAGGGAGCTGTGATAGTGCAACTGCATTCCAGCCTAAGAaagttatactgcc
gcagtcatatagagcagacgACGAAAACAAAAAACTTCCACCTTCATACTTTTCTGCAAGaagttatactgccct
gcagtcatatagagcagacgACTAAGATAAAAAAATTTCTTTCCTATCCATCACCCCTCCaagttatactgccct
gcagtcatatagagcagacgAATCCTAAGCCAAAAGAATACAGCAAGAAGCATCACATTAaagttatactgccc
gcagtcatatagagcagacgTGGTCCTAGGAAAATTGGAGGGCCATTTGCAGAAGAATGAaagttatactgccc
gcagtcatatagagcagacgGACCCATATCTCAACATATACAAACATTAATGCAAGATGGaagttatactgccct
gcagtcatatagagcagacgAAAACCTAGGAAAAGCTCTTTAGACATTAGCCTAGACAAAaagttatactgccct
gcagtcatatagagcagacgGGAAACACATAAAAAAAAAGCCAAAAAATAGGCAAATGGGACaagttatactgccc
gcagtcatatagagcagacgCCAACAGGGTAAATAGAAAACCTCCAGAACATGAGAAAATaagttatactgccc
gcagtcatatagagcagacgTAAACAATGCATCCAACAAACGCCTAATATCCAGAATCATaagttatactgccct
gcagtcatatagagcagacgCATAAAAATGTTCCAGATAAGCAGAGTGAAAGCTACCCAAaagttatactgccc
gcagtcatatagagcagacgGGTTACATAACACTGATCCCAAAACCAAATAAGAAAATGGaagttatactgccc
gcagtcatatagagcagacgCCAGGTTCGAGCAATTCTTCCCGCTTCATCCTCCCAAGTAaagttatactgccc
gcagtcatatagagcagacgTATAGGCGTGAGCCATCTTGTCTGGCCAAAAATCACAATTaagttatactgccct
gcagtcatatagagcagacgAACATACCCATCACTTCAAACACTTACTTTTTGTGGTAAGaagttatactgccctg
gcagtcatatagagcagacgGTACACTACATTATTTACCACATTCACTACTCTGTGCAATaagttatactgccctg
gcagtcatatagagcagacgAAAACAACTTGCTCTTTTCATCTACTGTGACCCTCTACCCaagttatactgccct
gcagtcatatagagcagacgTCTCTCCATCCTGCCTCCCGGCTGCATTCAATAACCACCAaagttatactgcc
gcagtcatatagagcagacgCTCTCTGTCTCTTTGAGTTCAATTGTTTTTAGATCTCACAaagttatactgccctgt
gcagtcatatagagcagacgCATTTAGCACAATGTTCTCCAATTCGGTCCATCCATGTTGaagttatactgccct
gcagtcatatagagcagacgATTGAGAATAGTGTTGCAATGTACATGGGGAGGAGATATCaagttatactgccct
gcagtcatatagagcagacgCTGATTTCACACTTTTTGAGAAAATATCCAGCAACACGATaagttatactgccct
gcagtcatatagagcagacgGCTGGATCATATGTAGTTCTTTAAGGAGACTCTATATTGTaagttatactgccctg
gcagtcatatagagcagacgCCATTTTAACAGGTGTGGGGGTGCTATCCCATTGTGGTTTaagttatactgccct
gcagtcatatagagcagacgGATTAGAGATGATGAACATTGTTTCACGTATCTGTTGGCCaagttatactgccct
gcagtcatatagagcagacgTATTTTCTCCCAATTCGTAGACGGTCTCTTTACTCTGTTAaagttatactgccctg
gcagtcatatagagcagacgAGATTGACTGATGATTCTACTGTCTTGAACCCTTGTTAGGaagttatactgccctg
gcagtcatatagagcagacgAAAGAAGTTGCAATTGATTGGCTCAGAAAAACAAAATGCCaagttatactgccct
gcagtcatatagagcagacgTTTGAATGTTTACAAAAGCAGCTGGGCATGATGTCTCACGaagttatactgccct
gcagtcatatagagcagacgGAGCCGAGATTTCACCATTGCACTCCAGCTTGGGCAACAGaagttatactgcc
gcagtcatatagagcagacgCTAGAAATATTATCATCAACAATGCACTTGATAAGTCCCCaagttatactgccct
gcagtcatatagagcagacgAGAAAGACATTAAGGGAAAATCCATAGGTGCACAGCCAGCaagttatactgcc
gcagtcatatagagcagacgTTCAGTTGCACAGAACCCCGGGCTCAGAAGGGCTCCATACaagttatactgcc
gcagtcatatagagcagacgTTTTTAGAAAGGTTGGTGGGGTGGGCTCCACATGTTCATTaagttatactgccct
gcagtcatatagagcagacgCCTTGCAAATCATGTAGCCGGAACTTCACAGATGGGTAACaagttatactgccc
gcagtcatatagagcagacgTGGTCAACCGGCTAGCTCTATTGGTCCGTGAAGATGGCAGaagttatactgccc
gcagtcatatagagcagacgTGGGGATTCCTCTGGAAAGAAGATAATACGGACATGGTTCaagttatactgccc
gcagtcatatagagcagacgTGAAGAGTGTCTGATCATAGATTTTTTTCTCATCATGTCCaagttatactgccctg
gcagtcatatagagcagacgAAACTTGAAGGTGGGAATTTTATGACCAAGGCTACTCTGGaagttatactgccct

gcagtcatatagagcagacgTATTTCTCTCAAAGTATAGGCCACTGAGCAACTCAAACTAaagttatactgccctg
gcagtcatatagagcagacgGCTACGAAAACTATGAAGCATTATCTTGAAGGGGAATATAaagttatactgccctg
gcagtcatatagagcagacgGCATGCCAAACCAATAGTATATATTCCTACAACACCAACAaagttatactgccct
gcagtcatatagagcagacgCACCCCAAAACCCTTCTCAGCCTGGTCCTCAATGATGGGAaagttatactgccc
gcagtcatatagagcagacgACGGATAGCTCTGTCAATTAACTCATCAATGCTCTGTATTaagttatactgccctg
gcagtcatatagagcagacgGAACATCAGAGCCTTGGTCTGGAAGGTTGTATAGGCATGTaagttatactgccct
gcagtcatatagagcagacgGAATATTCTGGATAACAATGTGAACAACCGTTTATGTCCTaagttatactgccctg
gcagtcatatagagcagacgCCAAACCAGGAATTCTATAAGCACAACTTAGCAGACAATGaagttatactgccc
gcagtcatatagagcagacgTGAATCTCAGAGAAGTGAAGCAATTTGTACAGGATCGTGCaagttatactgccct
gcagtcatatagagcagacgAACTCAAGTGCAAGTTCTTTGTCCTCTGCCACACTGTTGGaagttatactgccct
gcagtcatatagagcagacgGTGTGAGACCTTCGAGCTGGGCAGACTGAGAATTAGGCACaagttatactgccc
gcagtcatatagagcagacgCATTCGCATCCAGAAAATTATTGCCAGGAAGACACAGATGaagttatactgccc
gcagtcatatagagcagacgCTGATTAGGCTGATTATCCATTCAAGAAATATAGGAGTGCaagttatactgccctg
gcagtcatatagagcagacgGTGAGTCATTTGTCCTCCAAGATCCTTAAAACCCAGTAGGaagttatactgccct
gcagtcatatagagcagacgGTGCCTAGGAATTACACCAAGATTTTTAGCAGAAATTTACaagttatactgccctg
gcagtcatatagagcagacgGACCTTTCCACTATTTTAAAATTATACGCATGTGTTGGGAaagttatactgccctg
gcagtcatatagagcagacgCTGCTCCTAAAAAATAAAGTCGGCTGGACGCAGTGCTCCTaagttatactgccc
gcagtcatatagagcagacgGAGGCCGAGGCGAGCAGACCATGAGGTCAAGAAATCGAGAaagttatactgcc
gcagtcatatagagcagacgAAAAATTATCTGGCCATGGTGGTGTGCCTGTAGCCCCAGCaagttatactgccc
gcagtcatatagagcagacgCTACACTCCAGCCTAGCAACAGAGCAGGACTCGGTCTCAAaagttatactgcc
gcagtcatatagagcagacgTGAGGCACAATTTCTTCAGTATTTCCCAGGGTCTGTCAAAaagttatactgccctg
gcagtcatatagagcagacgCGTCATTATGTGTTAAAAGAAAATTAGAGGGCATATGTGGaagttatactgccctg
gcagtcatatagagcagacgCACAAATGAAAAGAGGAAGACATAAACCTAAAAGTGGTGAaagttatactgccc
gcagtcatatagagcagacgATTTATAGAATGCACGTACCGTAACACTCAATTCTCATTCaagttatactgccctg
gcagtcatatagagcagacgAGCAACAGTTACTCACCTTCTCACTCATCTTGGGAATTTTaagttatactgccctg
gcagtcatatagagcagacgCATTTATGATGTACGACATGAGTTTTTCAAATACGTACCCaagttatactgccctg
gcagtcatatagagcagacgCATATGTATTCCCTCACATACTTACTATTTGTTTGTGGTGaagttatactgccctgt
gcagtcatatagagcagacgCTGTCTCAGCAATTGTCAAGCATACCCTGCGTTGTTACTAaagttatactgccctg
gcagtcatatagagcagacgACATTAGGGCTCTTGAACTTGGTCCTCTGTCAAACCTCTTaagttatactgccctg
gcagtcatatagagcagacgGAAGAATGTAAAAGAGTTTTTAGTTCCTGAGACAAAGCAGaagttatactgccctg
gcagtcatatagagcagacgTCTGAGGAGTCTTTGATGCCAAGTGGAGACCAAAATAACCaagttatactgccc
gcagtcatatagagcagacgGCGAGATGGCAGAGGAAGTGGGGGTGACTGGGATTTAGATaagttatactgccc
gcagtcatatagagcagacgGGGATCCCAGAAAACCTGCACATGTGGCTTCCCAAAGGGGaagttatactgcc
gcagtcatatagagcagacgCCATGAGGCAAGGAAGCTTAGGATCCCATCTCCATCACTGaagttatactgccc
gcagtcatatagagcagacgGAGCTTCAGTCTCCTCATTTGCAACACAGGGATAAAAACAaagttatactgccct
gcagtcatatagagcagacgACAGGGTGACAGGAAGGAAAAGGCATGCAGGTGCTCAATGaagttatactgcc
gcagtcatatagagcagacgTTGTAAACGGATGGTACTTATAGTCATTAGGATGCTCCCGaagttatactgccctg
gcagtcatatagagcagacgCATCTGTCAACTCACTGCTTGTATCTCTTCATGGTTTAAAaagttatactgccctg
gcagtcatatagagcagacgTTGCTGGTTCAGTCAAGTCTGTGAATTGGTTTGCCCTAGGaagttatactgccctg
gcagtcatatagagcagacgTGCACGCACACACTCAGAGGGGTATACCAGAATATCAGATaagttatactgccc
gcagtcatatagagcagacgGCCAAAATATAAAATCTTCTGATTGGCAAGAAGACAACTGaagttatactgccctg
gcagtcatatagagcagacgTCCATGTTGCTGCCTGCATGTGCTGTTTCTCCCTCTTTGTaagttatactgccctg
gcagtcatatagagcagacgTCATATGTTCAAACCCTAATCTCAATGTGATGGTAGTAAGaagttatactgccctg
gcagtcatatagagcagacgAGGGTGGAGCTCTCACGTATAGGACTGTGCATTTGTAAGAaagttatactgccct
gcagtcatatagagcagacgGCTCCCTTGCTGTCTTTTCTCCTGGTGAGGACATACCCAGaagttatactgccc
gcagtcatatagagcagacgAGAACCTGACCATGCTGACCCCCTGATCTGAGACATCCACaagttatactgcc
gcagtcatatagagcagacgTTTGTTGTTTAATCCTTAGCCTATGGTGTTTAGTCTATGGaagttatactgccctgtg
gcagtcatatagagcagacgCAAGCGGACTAAGACAGGATGGAAACAAAGAGGGCAAGAGaagttatactgcc
gcagtcatatagagcagacgTGTGTATATATATATAGCTCCCGCTACACTGGTGGAGGACaagttatactgccct
gcagtcatatagagcagacgTTCTGTTATAATGTTCCAGAGCTTATACAGAGGAGAGTCTaagttatactgccctg
gcagtcatatagagcagacgCAAGAGGGGTTTGAGATGTGTGAGAAAGGTGTTTTCAACTaagttatactgccctg
gcagtcatatagagcagacgACATGGAAACAGAGACTGAAAACAGCTACAGATTGGTGACaagttatactgccc

gcagtcatatagagcagacgAACTAATTAGGACAGGGTAGAGCTATTCAGAAGTTTAAACaagttatactgccctg
gcagtcatatagagcagacgTATTGCTCTTTAATATTCAAGTTTAAGATGGGCATGGTGGaagttatactgccctgt
gcagtcatatagagcagacgTGAGCCGAGATCAGGCCAGTGTGCTCCAGCCTGGGAGACAaagttatactgcc
gcagtcatatagagcagacgGCCTGTAATCCCAGTTATTTAAGAGGCTGTGGTGGGAGGAaagttatactgccc
gcagtcatatagagcagacgGGCAACAGAGTGAGACCCTGGCTAAAAAAATAAAATAAAAaagttatactgccc
gcagtcatatagagcagacgGGGGGATCACTTGAGGTCAACTGTTCAAGATCAGCCTAGCaagttatactgccc
gcagtcatatagagcagacgCATTGAGTTTCTATATTAAGAACATCATTGGCCCAGCGTGaagttatactgccctg
gcagtcatatagagcagacgAGTTCGAGACCAGCCTGGCGCAACCTCATCTCTACTAAAAaagttatactgccc
gcagtcatatagagcagacgAGTGTAATTTATTAAAGGCATGCTATAGTCTTGGACTCAGaagttatactgccctg
gcagtcatatagagcagacgTTTACTTACGGGAGAATCTAGGCATTGAAAAATCTAAGGTaagttatactgccctg
gcagtcatatagagcagacgCCAATGTGGCAGCTTGCTTAAACTGTGAGCAGGATTCCAGaagttatactgccc
gcagtcatatagagcagacgCAGACCCAAACCTGCAAGGCGATCGCAAACAAACCCGCAAaagttatactgcc
gcagtcatatagagcagacgCGCAAACATCCCTAGGTATTTCAAACCTCATTAGGATTCAaagttatactgccctg
gcagtcatatagagcagacgTTTAACGTAGATCTACCTTTGTTCAGAGAGATTTTTGAGCaagttatactgccctgt
gcagtcatatagagcagacgCAGCAAGTGAAAAAGAGGAAGGCCCCAGCAAAGGAGCGGAaagttatactgc
gcagtcatatagagcagacgGGGCGTCCACAGTTGATCATAATCTTTTCAAGTCCAGGAAaagttatactgccct
gcagtcatatagagcagacgAGCAATGGGAAAAGGAGTCTAAGATGAATAAAAGGATGTCaagttatactgccct
gcagtcatatagagcagacgATGTGGGCAGAACAGCATGCTAAGAAATCTGGGTGGGCATaagttatactgccc
gcagtcatatagagcagacgGCTAGCAACCAGTCCCCAGCCTCAGGATGAAAGATCTAGAaagttatactgcc
gcagtcatatagagcagacgTAGGAATAGAGCGGGGAAATGGGGAGGGAGGAGGACACCTaagttatactgcc
gcagtcatatagagcagacgCGTTAAGTCCTCGAACTGATTTATTTCATATAGACCTCTAaagttatactgccctg
gcagtcatatagagcagacgAGTTTCAGCAATGCGTGGGAAAATAAAAGGTCCATAGTAAaagttatactgccct
gcagtcatatagagcagacgACCAGAAGGCAAGGTGAAGGCGAGCTGAAACCATTCTTCAaagttatactgccc
gcagtcatatagagcagacgCCTAGCTGAAGGTTATCAAAGAAACACAACTCAATAAGCCaagttatactgccc
gcagtcatatagagcagacgAAGTTTAAGTTCTGCTAATCTTTTTGTGAGAAGACTGTGGaagttatactgccctgt
gcagtcatatagagcagacgGCAGCGATAAAGTGAATTGGAAATGCCAGTAGGGTGTGTGaagttatactgccc
gcagtcatatagagcagacgACAGCTATTTCTGTTGAAGAGAACAAAAGATGCAAAGAGTaagttatactgccctg
gcagtcatatagagcagacgCCCACTCCCCTCCAACTCTGAGCATTTCTTCTCTGAAACGaagttatactgccc
gcagtcatatagagcagacgCAGAAAAAAAATAAGAGTTGATGATAGGGGAAGTTCACTCaagttatactgccct
gcagtcatatagagcagacgTCTTCCTCCACTATTCATTGGTTTGGGATTGCAACCTGCCaagttatactgccctg
gcagtcatatagagcagacgCCATTGCTGTCATTCTTCAGAGGTAGATGTTTGCAAAAATaagttatactgccctg
gcagtcatatagagcagacgAGAAGACTCTCCTCCACTGATCCCTTGAAGCACATCCATGaagttatactgccc
gcagtcatatagagcagacgCATTTAATTTGCTCCATGATGACTTTTGACCTCTGAATTCaagttatactgccctgt
gcagtcatatagagcagacgGACCAGGCTAGTCCCATCGTCACCCTCAGACACTTATTTGaagttatactgccc
gcagtcatatagagcagacgTCTAGGGGCAGAAAACAGTGCAAGCCTTGCAGATATAAGGaagttatactgccc
gcagtcatatagagcagacgGGGACACATATATAAAAAGATAACAACTACAGTAGGTTCCaagttatactgccct
gcagtcatatagagcagacgCATTATATGTAAAAGCCACAGCAACACAGCTCTCAGGGGCaagttatactgccc
gcagtcatatagagcagacgGGTCTTATACACTCTGCTGCCCTAACTCTCACTTCATCTTaagttatactgccctg
gcagtcatatagagcagacgTAGGATATGCCCAATTTATACTATTTAGGGATTGTTCCACaagttatactgccctg
gcagtcatatagagcagacgTTTCCCCAAATCTATTTCTACAGACCAGGAGCTATAATTTaagttatactgccctg
gcagtcatatagagcagacgCCTAAGCTTATAAATAAAATAACGCCGGGTGCGGTTGCTCaagttatactgccct
gcagtcatatagagcagacgCAGAAGGTGGAGGTTGCGATGAGCAGAGATGGTGCCATTGaagttatactgccc
gcagtcatatagagcagacgTTCATTCATTCATCCGTTCAAGAAGCACTAAGTGGCCTTGaagttatactgccctg
gcagtcatatagagcagacgGCAAAGGGCAGAGGACTAAGCAACTGGGAGTGAAGCAATTaagttatactgcc
gcagtcatatagagcagacgAAAGAGTCCCAGAGGGTTCCTTTTAGTGTGCTTCCCCAGGaagttatactgccc
gcagtcatatagagcagacgTTAAATATTGTTATACAGATCCTTAAGTCTTGCAGCTGGGaagttatactgccctg
gcagtcatatagagcagacgGACAGCTGAGGGTGGTCAGGCAAAAAGCAAAGATTTAAATaagttatactgccc
gcagtcatatagagcagacgAACTGATCTCTAATTCCAATACTAGTTATGCCCTACCAATaagttatactgccctg
gcagtcatatagagcagacgGAAAGTCCGCAAAGCACTTCTTCGACCTTCTCTCTTTTGGaagttatactgccct
gcagtcatatagagcagacgAATGTTTGAAGAGTTTGTCTGTGGCAAAAAGAGATTAACGaagttatactgccctg
gcagtcatatagagcagacgATGACAGATTTCACACATAGCTTCCATTTTCTTTCTCATCaagttatactgccctgt
gcagtcatatagagcagacgTGGGGTGGGGAGCGTGGGATGGTGGTGGTATTGCAGTTACaagttatactgccc

gcagtcatatagagcagacgCACACATGTGGTACATATAAAACAGTGTGCATGTCATATAaagttatactgccct
gcagtcatatagagcagacgTGTGTGTATGCTCAACAGTCAGATGATAGTGTTCCCAGCGaagttatactgccc
gcagtcatatagagcagacgCAAGATATGGTGGTAGGGGTGTCCAGGAGAGGCAAGAATGaagttatactgcc
gcagtcatatagagcagacgGCCTCTAAGCCCAGAAAATGCAGAAACAAAATAGACCTTTaagttatactgccc
gcagtcatatagagcagacgCTCTTCCACTAGAGCACCCAGGACACTATATGCAGCAATTaagttatactgccc
gcagtcatatagagcagacgACTAGTAGTTTTGTGGGGCCAAATGTTGTTCAGACTTGTAaagttatactgccct
gcagtcatatagagcagacgGACCTAGACAAGTCAGCCCATTGCCCAAGCTTGGCATGCTaagttatactgcc
gcagtcatatagagcagacgGGAGCCCCTTCTGCCAAGTTCCCCTTCCCAACAGATCGAGaagttatactgcc
gcagtcatatagagcagacgGGGAGGGTCACGGTTGAGACAGCAGCCCCATCTCATTGTTaagttatactgcc
gcagtcatatagagcagacgCCGGGGCTTCCTCTTTTGGCGGAGCGGGAGCGGGAGCGGGaagttatactg
gcagtcatatagagcagacgCCTTCGGGATGTCTATTGCTGCCGCTCCCTCCCCTCGCCGaagttatactgcc
gcagtcatatagagcagacgTGCGGATTCTCAATAAAAAGGGAAAAGCACGATCCAAGTTaagttatactgccc
gcagtcatatagagcagacgCGTACAACGCTGGTGCCAACAAGACTGACACAGGAGCCCAaagttatactgc
gcagtcatatagagcagacgGGGAAGGACGCCTGAGAAAACAGACCCCGGCCCCGTGAGCaagttatactg
gcagtcatatagagcagacgGTGTCTGTGTGGGTGGGTGTATTTGTGCATGTGTGGTGTAaagttatactgccct
gcagtcatatagagcagacgGAGTGTGTCTATGTGTGTCTCCGTAGTATGTGTATCTGTGaagttatactgccctg
gcagtcatatagagcagacgTGTTTGTGTCTGTGTGAGTAGGTGTGTTTGTGCATGAGTGaagttatactgccctg
gcagtcatatagagcagacgAGTGCGTGAGTGTATGAGTGCGTGAGTGTGAAAGTGTGTGaagttatactgccc
gcagtcatatagagcagacgCCTCTGCTCCCCTGGCGGGCTCCCAAAGTGGCAAAAGAGAaagttatactgc
gcagtcatatagagcagacgAGAAAAAGAAAGGGGGAAACGGGGGAAAAGAAAAGCATGGaagttatactgcc
gcagtcatatagagcagacgGGAAAAGAAAATAAAATCTCCCCAAAGTCTCCCCAGAGGCaagttatactgcc
gcagtcatatagagcagacgGCCTCTCGTCCACCCTGCGCCGCGTCCAGTCCACCCCCTTaagttatactgc
gcagtcatatagagcagacgCGCCCCATGTCCACCCGCCCGCTGTCCACCCTGCGCCCCAaagttatactg
gcagtcatatagagcagacgCGCCCCCTGTCCACCCGCTCACTGTCCAACCCGCGCCCCAaagttatactg
gcagtcatatagagcagacgCTGCCCACCCCGCGCCCCCAGTGCACCCGCCCAGCCCCGAaagttatact
gcagtcatatagagcagacgGCTGGGCTGGCCGGCGCGAACCCCCGGGGGCGTCTGGGCGaagttatact
gcagtcatatagagcagacgGCTGCGCACGTAGGAGTACCCCCTGAGCCCAGCCGCTCGGaagttatactg
gcagtcatatagagcagacgGGTGACCGGATACGAGTAACGCGTGCGCCCCGATCTGTGCaagttatactgc
gcagtcatatagagcagacgCGGGTTGACTGTCGGGGGCGATTTCCGCGGGGCAAGGCGGaagttatactgc
gcagtcatatagagcagacgGCAGCTGCGGGGCCATTCCACCTGAGCGCCCTGAGGTCAGaagttatactgc
gcagtcatatagagcagacgGGGCCTGAGGACACGGCGCTGGGCTCCGGGGGACGCGGGTaagttatactg
gcagtcatatagagcagacgCGAGCGCCACCTCCCCGCCCTCCGCCGGCTCACGTAGTCCaagttatactg
gcagtcatatagagcagacgAGGATACACAACACCCTGGACTGTCACAAGCCCCTCGCAGaagttatactgc
gcagtcatatagagcagacgCCGGGCTGAACAAAAACTAGGAGGTGGGGGGCGGTGGGGCaagttatactgc
gcagtcatatagagcagacgACAACAGAACCGAAGAATTGCCGCACGGAGCCTGGGCCGGaagttatactgc
gcagtcatatagagcagacgTCTCCTTCCCTTTGGTGACTATCCGTGGGGGTTTCTCCCGaagttatactgccc
gcagtcatatagagcagacgCACACCACACCACACACTACGTTCAAACTTTGTCCTTGAAaagttatactgccc
gcagtcatatagagcagacgCTTTTGGAGACCAAGTGGCTTTTCTGGACGAGGAGGCTGCaagttatactgccc
gcagtcatatagagcagacgGGATTAGAGCAGGCAGCGACCCAAACCACCCTTACACCCGaagttatactgc
gcagtcatatagagcagacgCCTGTTTAAAAAGAAGGGGAGGAGCACCAAAATCCAAACCaagttatactgcc
gcagtcatatagagcagacgCAGATACTCCATGGGCACAATAAAATGTCTGTTAACAATAaagttatactgccct
gcagtcatatagagcagacgACGCTTGTAATCCCATGCTTTAGAAGGCAGAGGCAAGAGGaagttatactgccc
gcagtcatatagagcagacgACACAAAAAAATTAAAAATTAGCGGAGCGTGGTGGCACATaagttatactgccc
gcagtcatatagagcagacgACCCTGTCTCTTAGAGAAGAACCAAAAACCTACAGGGTTTaagttatactgccc
gcagtcatatagagcagacgTGGTAGCCCTTGATTTCTCCCACCTGGATTTGGAGTGACCaagttatactgccc
gcagtcatatagagcagacgGACTGGCCTTTGTGATTCACCCCTGTCTTACTTTGTAGACaagttatactgccct
gcagtcatatagagcagacgAGGCTGCTGGTCTCCACTGCCCAAACAGGATCACTTAAGAaagttatactgcc
gcagtcatatagagcagacgGCACTGCCCAAACTGCACTACATGAGATTTATAACTTACAaagttatactgccct
gcagtcatatagagcagacgCTTAGGCACCCCCCACATCAATTTCCCCTGCATAGGGTATaagttatactgccc
gcagtcatatagagcagacgCTCCAGGAGCTCATACTAGTCTGTCTTTTCAATAATCTTTaagttatactgccctg
gcagtcatatagagcagacgAGCAGGGGTTCAGAATGGGGGGATGCGGTGAGAGTGTGGGaagttatactgcc
gcagtcatatagagcagacgACTCAATACAATTCATAGCAGGCTGATATTCTCATTTCTCaagttatactgccctg

gcagtcatatagagcagacgTAGATTTCTAGATGTTACTCAAGCTTTCTCATGCCCGCCAaagttatactgccct
gcagtcatatagagcagacgGTGAGACATCTGGGTTGGGTGCTGGAACAAGTTGGTGGCGaagttatactgcc
gcagtcatatagagcagacgGGAGGCAAGCAGCACCCCCACCACTTCCCCCACAACCCGCaagttatactg
gcagtcatatagagcagacgCCAGGTTGTATGCCAGGCAAGGGGAAGCTCCAGACGCCTCaagttatactgc
gcagtcatatagagcagacgGGATACATTGACCTTGCCATAGATGCCAGCCAAGACCCAGaagttatactgcc
gcagtcatatagagcagacgCCTAGGCTGTGAGGGAGAGCGATTGGGTGCTGTGCTCCACaagttatactgcc
gcagtcatatagagcagacgGCCAGGGACAAGCGGGAGCCATGCTCCTTCCAAGTTGGGAaagttatactgc
gcagtcatatagagcagacgCTGTGTCTGCCCAAGCTGCTGCTGCAACACGGGCACCCCTaagttatactgc
gcagtcatatagagcagacgGCCCCAGCCACCTCCCCATCCCAACCCAACCCCCCCATCCaagttatactg
gcagtcatatagagcagacgCTCTTGGGGATCCTGGGAAGCCCTGCCCTGTTGCCCTTGCaagttatactgcc
gcagtcatatagagcagacgCAGAGTTGGGGCCAAGCCCTGACACTGTCACTGCCTGGCAaagttatactgc
gcagtcatatagagcagacgCTGACGTGCCAGCCCCCTGCTTCCTTGGCCCCCTCTGGACaagttatactgc
gcagtcatatagagcagacgGAATGGAGGAGCTCGGGAGGAGAAGCTGAGGGGGACTGAGaagttatactgc
gcagtcatatagagcagacgGCCTGGGTACCATGGCTGGCCGTGGGAGGCAGACAGGCTCaagttatactgc
gcagtcatatagagcagacgTCCTGTGGACTGGAGGGAAAACTGATGGTGTTTTTTCCTGaagttatactgccct
gcagtcatatagagcagacgTGGACCAATCAGCACGCAGTTCCCCCCTCTGAGGCCCATGaagttatactgc
gcagtcatatagagcagacgGGACAGAGAGGATGGAGAGACATGGCCGGGATGACCAGCTaagttatactgc
gcagtcatatagagcagacgCCAGCTGCAGAAAGGAGCAATCACTCTCCAGGGCCTCCTTaagttatactgcc
gcagtcatatagagcagacgGAGCAGCAGATGTGAGGAGCAAGCCTTTTCCAGGACCTCCaagttatactgcc
gcagtcatatagagcagacgCTGAGAGCTGAACACTGGACTGGACTGACTTGTCTACGGAaagttatactgcc
gcagtcatatagagcagacgCTCCTCTGAGCTGTTACTGGACAAACGAGTCCTAATTTTAaagttatactgccct
gcagtcatatagagcagacgGGTAAGAGATTGCCTTGTATGACAGAAGTAGTTAGTAAACaagttatactgccct
gcagtcatatagagcagacgAGGCTGTCTTCTGAAAAGGAGTTTTGGAATTGAAATGGTAaagttatactgccct
gcagtcatatagagcagacgCAGGCTGGTGGTGGAGAGGAATTGAAAGGAATGAAGGATTaagttatactgcc
gcagtcatatagagcagacgTCTTTCCAGATGTTTAGAAGGGCACAAAGTATGTTAAAAGaagttatactgccct
gcagtcatatagagcagacgGATTCATGTGCAATATTGTCTCTTGGAAATTGATCAAACCaagttatactgccctg
gcagtcatatagagcagacgATTATATTTGTGCTGGTTCAAAAAGGAAGGAGGAGCAGAAaagttatactgccct
gcagtcatatagagcagacgCCATTGTCCCTTATGTCTGCGTTTTCTCTTACTGTGCTGTaagttatactgccctg
gcagtcatatagagcagacgGGAGTCCTAATTTGCAATGTCTAGTCTTTCTCGATGTTAAaagttatactgccctg
gcagtcatatagagcagacgATACATTGAGTAACTTTGTGTTAAGATTCATAGGGCAGACaagttatactgccctg
gcagtcatatagagcagacgCAAAAACACAGAAGTGAAGTTGTTAAAATCCCCTCTAAGCaagttatactgccc
gcagtcatatagagcagacgACAGAGGTCAGAAGGGAAAGGGTTCTAGGTCAGAATATTCaagttatactgcc
gcagtcatatagagcagacgGCACCATTTATTTATTCAATGCTACTGTGAAGATAGTGGAaagttatactgccctg
gcagtcatatagagcagacgAAAGTGCACAACATAGGTTAAGAGGGCTAAAATAACACTCaagttatactgccc
gcagtcatatagagcagacgTTTTGGAGAGTATGCTGACTATTCACTGGGTGGCTGTGGGaagttatactgccct
gcagtcatatagagcagacgAGGGTTCTGTGTCCTTGAGTTGGATTAGTTCAGGAGGTCTaagttatactgccct
gcagtcatatagagcagacgACAAGTCCTAATGCTCAAAGAATGTGAAAAGTAGAGGTAGaagttatactgccc
gcagtcatatagagcagacgCTTCTTTGTTAGCTTTTAGGAAGGCCTGTCTGGGTGTGACaagttatactgccct
gcagtcatatagagcagacgTATACTTAGTCACTGGGATGATGAAAGAGGGAAAGAGGAAaagttatactgccc
gcagtcatatagagcagacgCGGAAGGGTTTTTCGCTAGTATCTGCATATCTAGAAGATGaagttatactgccct
gcagtcatatagagcagacgTAGAAAAGATGCATCTTGGATATGGAATTGTTAAACCACCaagttatactgccct
gcagtcatatagagcagacgGCAGTGTATGTCAGGACTTATTCATTGGGTTGGCAGCAGAaagttatactgccc
gcagtcatatagagcagacgTGTGCATCTCTTTTGGCTGTGCTGTAGAAATACATTAAGTaagttatactgccctg
gcagtcatatagagcagacgCACTATACTTAGTGGATTCTTTTGCTGTGTGTTTCTTTCTaagttatactgccctgt
gcagtcatatagagcagacgCGCCTCCCGGGTTCACTCCATTCTCCTCCCTCAGCCTCCCaagttatactgcc
gcagtcatatagagcagacgAACTCTGCCAATGTTTTTATATAGAGGTGTGTTGCCATTTaagttatactgccctg
gcagtcatatagagcagacgAGGCAGGATTACATTTTTTTCTAACAGAGTGAGTTGGTGTaagttatactgccctg
gcagtcatatagagcagacgCCTATAGCTGTGGGATTTTGAGTTGGTAAATATTCATGATaagttatactgccctg
gcagtcatatagagcagacgAGCATGATATATACTGTACAGTCTCAGTCCCATAAAATTGaagttatactgccct
gcagtcatatagagcagacgTGTCTACACACACACAGGACCTGGAAAGACTCGTCAAACTaagttatactgcc
gcagtcatatagagcagacgCTTTGCTTCTTGCGTTTTTCGGTTTTCTATAATGCACATAaagttatactgccctgt
gcagtcatatagagcagacgAGAGACATAAGTGATGTTTAGTTTGGAAAAGAGGAGATACaagttatactgccct

gcagtcatatagagcagacgCATGAGCGCTGCTTTCATGTTTGAAATGTATGAAAAGATGaagttatactgccctg
gcagtcatatagagcagacgCCGAGGAAAGACCCATGGGCAAGAACTTCAGAGAAACCGAaagttatactgcc
gcagtcatatagagcagacgGTGCCCCACCAACAATAATAAACCTATCACATAGTCAAAGaagttatactgccc
gcagtcatatagagcagacgGCCTGCCAGAGAAGCCTGAAAATGAGAGTGATTATCTACAaagttatactgccc
gcagtcatatagagcagacgGGAAGTAAGATCCCCCAAAAGAAGTCAGATTACAGTGGAAaagttatactgccc
gcagtcatatagagcagacgACTCATTACATAGTAAGATAGGAGGCAAGCATCCCTTCGGaagttatactgccc
gcagtcatatagagcagacgGACCTCTTCTATCACCACCCATGTAAATTCCATTATCTGCaagttatactgccctg
gcagtcatatagagcagacgGCAAAACTCTGCCTTGCTGTAATTTCTTAAACCTAATAAGaagttatactgccctg
gcagtcatatagagcagacgCGCCTCCTGGGTTCACGCTGTTCTCGTGCCTCAGCCTCCCaagttatactgcc
gcagtcatatagagcagacgGCGCCCGGCCTCATTAAGATACTTTTATATTACTGTTTTAaagttatactgccctg
gcagtcatatagagcagacgAAAGTATGCATCATGTAAACAACTGAGAAAGGCACAATTCaagttatactgccct
gcagtcatatagagcagacgTTACAGATTGGAAAGTTTCAACACATAACCACTCATTGCTaagttatactgccctg
gcagtcatatagagcagacgTGTTCATGAATACTGTGGGTTTGAATAATTAAGACACTGCaagttatactgccctg
gcagtcatatagagcagacgAATTTAAATATGTGCTTTCCAAATGACTAAGCTACCCCACaagttatactgccctg
gcagtcatatagagcagacgGACCTATATGCAAGTGTTTGTAGCAGCTTCATTTATAATGaagttatactgccctg
gcagtcatatagagcagacgAATCGGAAAGAATTTGAATGTTTGTCAGCTAGTGAATGGAaagttatactgccctg
gcagtcatatagagcagacgCTCAGTGCTAAAAAGGAGGTGAACTGGTGACACATGCAAAaagttatactgccc
gcagtcatatagagcagacgGAAGCCAGACACAAAAGCTATGTACCTCTCATACTTTATGaagttatactgccct
gcagtcatatagagcagacgTGGAAAACACAGAACTACAGGTACAAAAATCAGATTAGTGaagttatactgccct
gcagtcatatagagcagacgCAAAGGAGCACAAGAGAACTATTTGGGGTGATGGAAATATaagttatactgccct
gcagtcatatagagcagacgTCATGGTGCTGGTTATACAATGATATACATCTGTCAAACTaagttatactgccctg
gcagtcatatagagcagacgATTTTACATTTTTAGAGGTGGGGATTTCAGTATATTGCCCaagttatactgccctgt
gcagtcatatagagcagacgTAGCTGAGATTACAGTTACCACCGTGCCCGGCTGTATGTAaagttatactgccc
gcagtcatatagagcagacgAAAAAGTTCTGTCCTCATTTTCAATAGCTATCATCTGGTGaagttatactgccctg
gcagtcatatagagcagacgCAGTTGATATCGAATAAATGTTGAATGAATCTGGGTGGAAaagttatactgccctg
gcagtcatatagagcagacgGTTTTTCAGCCAAGTCAAATTGTACAGAGATGAACTGGACaagttatactgccctg
gcagtcatatagagcagacgCACTCTGTTCCTCTCGAAGGTGCTCTGCTGTCCGTGTGAGaagttatactgccc
gcagtcatatagagcagacgTCTGTGATCCCTTGTGACTTAGAGGTGGTATGGGAGGGCGaagttatactgccc
gcagtcatatagagcagacgAAGCAGTGCCACTATATCAGGTCAGCTTTTTGATGAGTCTaagttatactgccctg
gcagtcatatagagcagacgGGAAACTCCCTGCAGACAACCCAGTATTGAAAAACGCAAAaagttatactgcc
gcagtcatatagagcagacgCTCATCTTGGGGCAGCAGAGGATTCTAAAAATGAAAGTGCaagttatactgccc
gcagtcatatagagcagacgCAAGTCGATTATAGAAAAAATAGATCATAAAACACCCCCCaagttatactgccct
gcagtcatatagagcagacgAATAATTTACTGCCAGAAATTGCTAGAATACTGTAGCTGGaagttatactgccctg
gcagtcatatagagcagacgCCGTTTTTAAGTGTACAGCTCCATAGTGTTGTTTATTCATaagttatactgccctgt
gcagtcatatagagcagacgCCAGTCTTCAGAACTCATCTGTCTCATGGAACTGAAACTCaagttatactgccct
gcagtcatatagagcagacgAACTCCCCATTGTTCCCTCCCTGCCAGCCCGTTTTTGGTAaagttatactgccc
gcagtcatatagagcagacgCTTCTCTAAGGACCTCATATGAGTGAACTCATACAGCATTaagttatactgccctg
gcagtcatatagagcagacgTATGTGTCAGAATTTCCTTCACCTTTAAGGCTGAATATCCaagttatactgccctg
gcagtcatatagagcagacgATTCATCTGTCTATGGATACTTGGGTCACTTCAGCCTTTTaagttatactgccctg
gcagtcatatagagcagacgCTCTTCAAGATCCTGCTTTTATTCTTTTGGGTATATAGCCaagttatactgccctgt
gcagtcatatagagcagacgCAAGGGTTTTAATTTTTGCGCATCCTTGTCAATGTTTGTTaagttatactgccctgt
gcagtcatatagagcagacgTGACTTATATTTTTGATAGCAGCTCTCCTAATGGCTGTGAaagttatactgccctg
gcagtcatatagagcagacgTTGCATTTCCCCATTGACTAGTCAGGTTGAACATCTTTCCaagttatactgccctg
gcagtcatatagagcagacgATATATCTTTTTTGGCAAAATGCCTATTCAAGGCCTTTGCaagttatactgccctg
gcagtcatatagagcagacgATCACGGGATTCTTTTAAGCAACAATTTCTTCTAGTGCAGaagttatactgccctg
gcagtcatatagagcagacgCTTTAGCAAGCTCCCCGGGTGATACTGATAAATCATATGGaagttatactgccct
gcagtcatatagagcagacgGTTCTGATGCACATATGATGGGATTGAACTTTCATTAACAaagttatactgccctg
gcagtcatatagagcagacgCCTAGCATCCTTGGGAAGTGAAGTCTGATGACAGCCTAAGaagttatactgccc
gcagtcatatagagcagacgTTTTAGATCAAAAGTTTCTCCTACCTGTTTCCATGATCCAaagttatactgccctg
gcagtcatatagagcagacgGGTTCTCCTTCTACTTCTCTATGTGTACCATGTTAGTCCCaagttatactgccctg
gcagtcatatagagcagacgGGACTGTCTCATACTCCTTCTATTCCCTAAATGTGAAATTaagttatactgccctg
gcagtcatatagagcagacgGCTCTTTTCTCAGCCCTATTCTTTCTCAACAGGTACCTTCaagttatactgccctg

gcagtcatatagagcagacgTGACTCCCAGATTGAGCACCCCAGCCTAGCCCAGACCCAGaagttatactgcc
gcagtcatatagagcagacgGAACTAAATGAATCCGAATCTTACTTGTTCTTTCCCCCTCaagttatactgccctg
gcagtcatatagagcagacgTTTGCATTTGTCAAAGCCAGAAACTTCACAGTCTATCTCTaagttatactgccctg
gcagtcatatagagcagacgAACCCCCATGTTCACTAAGTCATTTAATTCTGGTAATGCCaagttatactgccctg
gcagtcatatagagcagacgCCTCTGATCCTCCTCTTCATTCTGGCTTGACATAGTCTTGaagttatactgccctg
gcagtcatatagagcagacgAGGCTTCTCTCTACACACACGTGGACACACTCATTCAACTaagttatactgccc
gcagtcatatagagcagacgCTGGTGAACAGGCCGTGAAATTGCCAAGAGCTTTCTCCATaagttatactgccc
gcagtcatatagagcagacgCATAGAATGGTATGGTGGTTTGGGAGGGGAGGGCAATTTTaagttatactgccct
gcagtcatatagagcagacgCCGGGGGATGTTTGGTGATGTCTAGAGATTGGGGTGGTGAaagttatactgccc
gcagtcatatagagcagacgGACAGCTCCTCCCACCAACAAAAAATTGTCCAGCCCTCAAaagttatactgcc
gcagtcatatagagcagacgGATCTTTAACCTTTTGTTCTACACCCAGATCTCTCTTACCaagttatactgccctg
gcagtcatatagagcagacgATGCACCAAAGCCCAGCTGTTCAGTGTCAATGGCTGCCAGaagttatactgcc
gcagtcatatagagcagacgCTCAATGTCTGCGAGATTTCCTGTCCCTCCTTTTCAATCCaagttatactgccctg
gcagtcatatagagcagacgTGGATACAGGGATACAAAAGCCCAGACACCTATCTTCCAAaagttatactgccc
gcagtcatatagagcagacgGTGGTGCCATTTATGTTCCAGAGCAACTGCGGGATCAAGCaagttatactgccc
gcagtcatatagagcagacgCAACAGACTGAATGCTTGTGCCCTCCCAAAATTAATATGTaagttatactgccctg
gcagtcatatagagcagacgCAATGTGATGATGGTATTAGGGAGGTAATTGGGTCATAAAaagttatactgccctg
gcagtcatatagagcagacgAAGAGGCCAGAGAACTAGCTTGCTCCCTTTTCTCCATGTAaagttatactgccct
gcagtcatatagagcagacgATAAATGTGTATTGTGTGAGTCCTCAGTCTGTGGTAATTTaagttatactgccctgt
gcagtcatatagagcagacgGCTGCTCTGCTTCTATTTCCTACTTCTCTCTTCTCCCTTCaagttatactgccctg
gcagtcatatagagcagacgTTGTACAAGAATCTCCTGCTCAGCTCAGCTTCTAGGAAACaagttatactgccct
gcagtcatatagagcagacgCAACATTCACCAGTGTGGGAGAGCAGAGGATGGAAAGGTCaagttatactgcc
gcagtcatatagagcagacgAGCTTCTCTGGCAATGATTTCTGGCCCCTAGGTGCTCAGCaagttatactgccc
gcagtcatatagagcagacgCCTGGGGAAGCTGAGTTAACAATATCATGTGATCAGTTAAaagttatactgccctg
gcagtcatatagagcagacgAATTCTGTCAGTAAATGTTAGAGTTGGTAGAGGATGGGTTaagttatactgccctg
gcagtcatatagagcagacgTCAGTGAAAATGAGCCCCTGGAGACAACCCTTCTGTATTAaagttatactgccct
gcagtcatatagagcagacgAAAAGAAAGAGGTTTAATGACCTATAGTTCCACATGGCTAaagttatactgccctg
gcagtcatatagagcagacgAGATCTCGTGAGACTTATTTTCTCTCATGAGAACAGCACAaagttatactgccctg
gcagtcatatagagcagacgCCAAACCATATCACCTTCCTCTTGTCTTCCAGATTTCTATaagttatactgccctg
gcagtcatatagagcagacgCCATTCGTTTTTCTATTAGTTCCAGTCAGATCAGAGCCTTaagttatactgccctg
gcagtcatatagagcagacgGCAAGTGATTCTTTGGCTCCCCATCATCCACAGGTGAAACaagttatactgccc
gcagtcatatagagcagacgTTGCATGACAGAGGTCCTTATGATCTAGCCCTTACCTGCCaagttatactgccct
gcagtcatatagagcagacgGCTCTCCTATAATATACTTTAAGCTCTTGAAAACATGGACaagttatactgccctg
gcagtcatatagagcagacgACCTTCTGAATATGCTGGTCCCCTTCTCTTTCCCCATTGTaagttatactgccctg
gcagtcatatagagcagacgCCCCATTGACACCTGGCAATCTCCCACTCACCCTCCAAGTaagttatactgcc
gcagtcatatagagcagacgTGAGGGCTGACTTCAGGAATCCTTTCTCTGAGTTCTTTTTaagttatactgccctg
gcagtcatatagagcagacgACCGCACCTGGCCCTTTCTCTGAGTTCTAACTGCACGTTGaagttatactgccc
gcagtcatatagagcagacgTCTACTATAATTGTTCATCTGCATTGTCTTTCCTGTAACCaagttatactgccctgt
gcagtcatatagagcagacgGCTCGGCAACTGTCTATTTTTATTTTGTATCCTAGGACCTaagttatactgccctg
gcagtcatatagagcagacgCTGGAGTCTATTTAGTGCTTCAGGAAGTAGATTCTGAGCTaagttatactgccctg
gcagtcatatagagcagacgCTATCCTGGGACTTTGGACAAGCTACTTCGCATTCTGTTTaagttatactgccctg
gcagtcatatagagcagacgAAATGGAGATAACAATGTTGACTTCTCAAGTTACTGTGAGaagttatactgccctg
gcagtcatatagagcagacgCAGGTGGGTAAGGGCTGGATCATAAGGCACAATTCCTGGAaagttatactgcc
gcagtcatatagagcagacgCATAGTGCTATCCTGACTGCTAATCCCTAACCTCAAAGTAaagttatactgccct
gcagtcatatagagcagacgATAAACAACAGATACAAAGAAGTGCTGCTCTACCAGACAGaagttatactgccc
gcagtcatatagagcagacgATATAAGAGACGTATTTACTTACTTTAGAGAGAGCTGACGaagttatactgccctg
gcagtcatatagagcagacgTTTACTAATCTACCCATCATTGCTCTATCAGGCCCTTTTCaagttatactgccctg
gcagtcatatagagcagacgAAGATGGTATTTGAGATAAGCCCTGAAAGATAAGATGAAGaagttatactgccctg
gcagtcatatagagcagacgAGGCAAAAACAAAAATCAAGCAAAACCCCTCACTCCCTCCaagttatactgcc
gcagtcatatagagcagacgTACTAACCCCCTACTAACCCCACCATTAACGAGAAAAATAaagttatactgccc
gcagtcatatagagcagacgTGTTCCAGGGACCAAGAGGAAGCTGCTGTATCTGATGCTTaagttatactgccc
gcagtcatatagagcagacgCAGAGATGTGTCAATGGACCTGGTGCTAAATAATAATAATaagttatactgccctg

gcagtcatatagagcagacgGGATGAGAAGAAGTTGACAAAGAAAAATTAGATCATGCAGaagttatactgccct
gcagtcatatagagcagacgCAAGGTGTGATGGGAGGTTTTTAGGCCGTGTAATAAGACGaagttatactgccct
gcagtcatatagagcagacgAGATTGTCTGACTCAGGAGGCTTTTGTGTTAATAGATAAAaagttatactgccctg
gcagtcatatagagcagacgAGCCATTTTAATTTCCAGAATGTATAGGAAGGGTTAAGGGaagttatactgccct
gcagtcatatagagcagacgGCCTCCAGAGTAGCTGGGCTTCAGGTATGCACCACCACATaagttatactgcc
gcagtcatatagagcagacgGTAGATTTACAAAAGTCCACAGGCATGTAAGTGTAACGTTaagttatactgccct
gcagtcatatagagcagacgTGCCACAGCAAAGGATGAATTCTTGTTGGTAATTTTTTGAaagttatactgccctg
gcagtcatatagagcagacgGCAGATCTTACATTCAGAAGTTTTATTGCCATAGCTACTAaagttatactgccctg
gcagtcatatagagcagacgGTAAAATTCTGCTTGCGTCTATTGCTGTTTTAAAAATGGCaagttatactgccctg
gcagtcatatagagcagacgGGAAATCATTTCTTTAAGTGAAAGGGGATTAAATGGTACCaagttatactgccct
gcagtcatatagagcagacgAAGTGACTTTAATTTCCGGCATTTTCTTTTGCAAGAGTTGaagttatactgccctg
gcagtcatatagagcagacgGGTTGTTTCGTGGTAAATATATGAAGTGGAGGCAGCAGTGaagttatactgccct
gcagtcatatagagcagacgGTAAGGAGGCCGTTGCGAGAGACAGGCGAGCTGAGGGCGGaagttatactgc
gcagtcatatagagcagacgCCGTGGGGGAAAAGGAGGGAGCTGAGATGCAAGCATACCCaagttatactgc
gcagtcatatagagcagacgCATTGGATGTGGAGGCAAGAACGAGGCAGGAATCACAGTGaagttatactgcc
gcagtcatatagagcagacgCCGTAGATGGCTGAATGGTGTCACCAGTGAGTGAGATGGGaagttatactgcc
gcagtcatatagagcagacgTGGTTTGCCAGATGGTGAGGGAATCAAGGGCTCCACTTTGaagttatactgccc
gcagtcatatagagcagacgTTACATTTAAAGCAAACCATGCCAGAAAGTGGACCTCAGGaagttatactgccc
gcagtcatatagagcagacgGAGGCAGGGAAGGGAAGGGGTGGATGAAGAGAAGTTGGCTaagttatactgcc
gcagtcatatagagcagacgAGTTAGGTCAAAGGAATAAGTCCTAGCATTTGATAGTACAaagttatactgccct
gcagtcatatagagcagacgTTTCAAAATGGCTGTGAGAGGAGATTTGTAATGTTCCCAAaagttatactgccct
gcagtcatatagagcagacgTGATGGATATGGCAGTTGCCTGTTTCGATCACACACATTGaagttatactgccct
gcagtcatatagagcagacgCAAAGCATGCCTCCAAGTGGATTTGTCCTATAGGAAGCTGaagttatactgccc
gcagtcatatagagcagacgGGAGATCCTTAGCATAGAAATGGCATCCTCATTAGAAAGGaagttatactgccc
gcagtcatatagagcagacgGTCACCCAGAAGAGAGCATAGAGGGGAGAAGAGAGGCTAGaagttatactgcc
gcagtcatatagagcagacgAAAGCCAACCTTTAGAGTTGTGCATCCATGCAGGGGGAGGaagttatactgcc
gcagtcatatagagcagacgCTGAAAAAGAACAGCCAAAGAGGAAGGAGACAGAGCAGTGaagttatactgcc
gcagtcatatagagcagacgCAGGAGGAAAGTGAGGAAGTAGTTAGGTAAATCCAGTGCAaagttatactgccc
gcagtcatatagagcagacgAAGCCGGTTATTTGGATTTGCAATATGGAGGTGTTAAGCGaagttatactgccct
gcagtcatatagagcagacgCTGAGGACAGAAGCTGGGTTAGAGTGGGTTTGATGAGCACaagttatactgcc
gcagtcatatagagcagacgACAGTTAGTTTTGAGGTGACAGGGTGGCAGAGAAAGGCATaagttatactgccc
gcagtcatatagagcagacgAGTAGGAGGCTGAGGAATAAAACGGGTATGCTGGTAAATGaagttatactgccc
gcagtcatatagagcagacgCATTTGTGGTAATATGAATACTCTCACCATGGCTAACATCaagttatactgccct
gcagtcatatagagcagacgGCTCATAAAATTCCTGAAAAATCTAGCATTGGCCCCTGGGaagttatactgccc
gcagtcatatagagcagacgCTATCACCCAGCTCCAGCACACCACATCATCGGGGAAGGGaagttatactgcc
gcagtcatatagagcagacgGGCACTAAAACACACTGGCAAGCCCTTGGAAAGCCCTCCTaagttatactgcc
gcagtcatatagagcagacgGCATGAGCAGGGAAGGTGAGACAGGAGGGAACAAGGTGCCaagttatactgc
gcagtcatatagagcagacgCTAATAGGTTTGTTGATTTGAGGGTGGGAAGAGAGGAGAGaagttatactgccc
gcagtcatatagagcagacgGTAAGTGAAAAAAACAAAAATAAAGGCCTTCAGCTGACAGaagttatactgccc
gcagtcatatagagcagacgGGAAGTGGTCTGGATGGAGTGGGAGGATGGTTATGGTGGAaagttatactgcc
gcagtcatatagagcagacgACAGGAAGGTGTGGAGGGTAAGGGGGATGGGATCTGATATaagttatactgcc
gcagtcatatagagcagacgGGAATTTGATCTCCAGTCTCGGAAGTGGGGCGTAATGGGAaagttatactgccc
gcagtcatatagagcagacgGTTCAAGTGAGAGCTGGTTGCTGTAAAGAGCCTGGCATGGaagttatactgccc
gcagtcatatagagcagacgCAACACTTTGGGAGATCAAAGCAGGATTGCTTGAGGCCAAaagttatactgccc
gcagtcatatagagcagacgCCAGCCTGGACAACACAGCAAGATCTCATCTCTACAAAAAaagttatactgcc
gcagtcatatagagcagacgTCTTTCTTTCTCCCTTTCTCGCTGTGTGACATGCTGGTTTaagttatactgccctg
gcagtcatatagagcagacgATGAAATGGACTAAAACAGATTGAGAGACTGGGGAGAATAaagttatactgccc
gcagtcatatagagcagacgGCCTGTGGGTGGCTGCTGTCAAGACTATTGAGTAAAGAGCaagttatactgccc
gcagtcatatagagcagacgCACGAATGCTGCAAGAGTTGGGGTGGAAAGGAAGACAGGCaagttatactgcc
gcagtcatatagagcagacgTGTGTGTGGGGGGTGGGTAGCGGGAAAAATCCTGGAAGTCaagttatactgcc
gcagtcatatagagcagacgCATGTGAGCATGTGAGTGGTGTGAGGGAGGGAGAGACATGaagttatactgcc
gcagtcatatagagcagacgGAACACACAAATTTGTTTCTGGTCCTGAGATCCAAGCGGGaagttatactgccc

gcagtcatatagagcagacgAGTCTCTCATGGGTCGGACTGCAGTGTGGAACATTCTCAGaagttatactgccc
gcagtcatatagagcagacgCGAAGGAATGAGGTAGAGCTGGAGGGGTTTGCTGATTGTAaagttatactgccc
gcagtcatatagagcagacgCTCCTGGGGGGTCAATGGAGGGTTGGTGGGGATTGGGTCCaagttatactgcc
gcagtcatatagagcagacgCTGGATTTCTGGCAGTGATTATGGTAAGAATGGATGTGAGaagttatactgccctg
gcagtcatatagagcagacgGCACATTCCACACCAGTATGCCTCCAACATTATCAACAAGaagttatactgccc
gcagtcatatagagcagacgGAGGAGCTATGAGACATTACCTGATTTAACCTCAGTGAACaagttatactgccct
gcagtcatatagagcagacgGCATCATCTGGCATCCCCCAATCACCGAGTTCCTTCATTTaagttatactgccct
gcagtcatatagagcagacgAGTTTTCTGGCAACAAATTTTGATTTCTTGGCCTAGGATAaagttatactgccctgt
gcagtcatatagagcagacgGTCTCAGAGAGGTTGTCTTTCCGAAATTTAACTTCACATTaagttatactgccctg
gcagtcatatagagcagacgTGTGTTCCCTTCGGTTCTCTTCTCCTGGCTGGGTTTTTGTaagttatactgccctg
gcagtcatatagagcagacgGGATTTGTTTTCCCTTAGTCGTCAGCTTGTCAACTCAAAAaagttatactgccctg
gcagtcatatagagcagacgTTTGCGAGGGTGTTTTCTTTAATGACTCAAACTTTTCTCCaagttatactgccctgt
gcagtcatatagagcagacgGGCCCCCCAAAATTCTCCCTAGTCCAGTTTATCTCCTCGAaagttatactgccc
gcagtcatatagagcagacgAGTATTTCTCCCTCCTTGTATAAGTCATTTGAAAAAGTCCaagttatactgccctg
gcagtcatatagagcagacgGAAGATTCTTAGTTGTCAACCAGATATGAGAATGCAGTCTaagttatactgccctg
gcagtcatatagagcagacgGTTGGCAACTCCTATTTCAAAGGAAAAACATCACATTATGaagttatactgccctg
gcagtcatatagagcagacgGAAGTCTTCATCTTTAGAACCCAGCCTCCAGAAACAAGAGaagttatactgccc
gcagtcatatagagcagacgTTCTGTTTTTTAGAGTGGTGGCTTTCAGTGTAGTAATAGTaagttatactgccctgt
gcagtcatatagagcagacgCTGATCAGAAATATGAGTCAAAGTTAATTTGGTGTGGGTGaagttatactgccctg
gcagtcatatagagcagacgAACAAAAACAAAAAAGCAATAGAACATCAGACCCCCCCACaagttatactgcc
gcagtcatatagagcagacgAATCTTGGCTCAATGCAGCTTCCAACCTTCTGGGCTCAAGaagttatactgccc
gcagtcatatagagcagacgTGAGCCACTGTGTCCAACATAGCCTCATAATAATAATTTAaagttatactgccctg
gcagtcatatagagcagacgTGTATTCAGAAGCCCATAAGAGCCTGCAAAATAAGGTTTTaagttatactgccctg
gcagtcatatagagcagacgTCCATAATCTAAGCCTGTTTTCATGATCCACTTATTCTACaagttatactgccctgt
gcagtcatatagagcagacgTGAGTGTCTGATATATGCAAAGCACCATTAAGTCTTGTGGaagttatactgccctg
gcagtcatatagagcagacgCACATGTGATAAGAAATAAAAGGAAACCTCAAACGCTCTGaagttatactgccct
gcagtcatatagagcagacgAGATGGAATCTTAGAGGAAGTGGAATTTGATGGTGTGTCAaagttatactgccctg
gcagtcatatagagcagacgAAAGGAGACATGATGGGAAGAACAATCAAGGCAGGGAAATaagttatactgccc
gcagtcatatagagcagacgGAAATGCCAAGTAGCCTGGTTTGAGGGGAACACAGGACACaagttatactgcc
gcagtcatatagagcagacgCCCAGACTCGTATGGCTAACAAGCAAAAGAAATGTGATCTaagttatactgccc
gcagtcatatagagcagacgGGTTAATCGCCCAGCACAAGTAGGATGGTTTGGAGAAAGGaagttatactgccc
gcagtcatatagagcagacgCTTCGGCTAAAACTCTATGTGTCTCATGGTCAAGATCGGTaagttatactgccctg
gcagtcatatagagcagacgATTATCTTTCGCAATATTTATTACATGCCTACCACATGCCaagttatactgccctg
gcagtcatatagagcagacgGATCAAATCAGACTCCAATGGATTGCATATTCTCATCGGAaagttatactgccctg
gcagtcatatagagcagacgCCAAGAGTAAGGGGGACATTCCAGGTTTGATACGTGTTATaagttatactgccct
gcagtcatatagagcagacgGAATACTTTCTTTTCCTGGAATTGAATAGGGGAGGTTACTaagttatactgccctg
gcagtcatatagagcagacgTCTTGAAAGCTCAGATACAATAGTAAGAAATTCGCTAGAGaagttatactgccctg
gcagtcatatagagcagacgAAAAAGAGAGACGAGGAACCCAGAGAGTTGCGAAAGTCCTaagttatactgcc
gcagtcatatagagcagacgGGAGAACCTGAAGCTCGGATTTGGTGACCTAGAGGAGTAGaagttatactgccc
gcagtcatatagagcagacgCCTCCTTAAATTGAGGGGCTGGAAATATGACAGGCTCATCaagttatactgccc
gcagtcatatagagcagacgGAAGAAAACCTCCATCCCCATACTTTGCAATATAATCTCTaagttatactgccctg
gcagtcatatagagcagacgAGGAGTTAGGCCCTGTTCAAATACACAATTAACTTTTTCTaagttatactgccctg
gcagtcatatagagcagacgCCTTGAAGGTGGCTGAGATGATCGGAAATGAACTACCAAAaagttatactgccc
gcagtcatatagagcagacgAGTTATTCGGGGTGCTCTGTGGATAGACAGGATAAATCCAaagttatactgccct
gcagtcatatagagcagacgCTTGGCATGAAGAGGAGATCAAAGGATACTGCACCCTTCTaagttatactgccc
gcagtcatatagagcagacgCTGAAGTTCACAGCAGTCATAGAATTAGGTAGTTGGGAGAaagttatactgccct
gcagtcatatagagcagacgACTTGGTATTTTCATGATCTGCATATGCCAGAAAGCGAAGaagttatactgccctg
gcagtcatatagagcagacgGCAGTAACTAATATCGTCCTCTAAAACAGCCGGCACCATCaagttatactgccc
gcagtcatatagagcagacgGCTCCTTCCTTCTCAGCTTGATTACCCAAAAGGCAGTGTGaagttatactgccct
gcagtcatatagagcagacgGGCCAGATGGAGTCAGGAGCCTTCATTCTGGTCACAGGATaagttatactgccc
gcagtcatatagagcagacgAGAATGTCATTACACCTGCAGCATGGGAACAGAAAATGAAaagttatactgccc
gcagtcatatagagcagacgGATAATTTTCCACTGGCTGGTTCCAATTACCTAATAAAGCaagttatactgccctg

gcagtcatatagagcagacgAGGATCACGCTCTGTCACCCTGACTGGTGTGCAGTGGCATaagttatactgccc
gcagtcatatagagcagacgAAATTTTTTTAGAGATTGGGGTGGGTGGGTGGTCTCACTAaagttatactgccct
gcagtcatatagagcagacgTGCCTCGCCTTCCCAAGTAGTTACACTATTTTCTTAGTCTaagttatactgccctg
gcagtcatatagagcagacgTGGCAGTTATTAGACATCTTACATTTCCTAAGCTTTACAGaagttatactgccctg
gcagtcatatagagcagacgCAGTGAGAGAAAGTAGGTTCAGTGTGATTCTTGTATAACAaagttatactgccct
gcagtcatatagagcagacgTGTTTAAGGCAGGTCTCCCCTCCAGGCTATGAGCTCTGAGaagttatactgccc
gcagtcatatagagcagacgCCTGTATACGCAAGGTTAAGCAAGGGATATATATTAGGTGaagttatactgccct
gcagtcatatagagcagacgCGGTTGAATGGAAGAACAAATGAAGGAAAGAAAACTGTGTaagttatactgccc
gcagtcatatagagcagacgAGAGATATTTTGGTGAAGTCCTTATTTGAGAGGAAATGGCaagttatactgccct
gcagtcatatagagcagacgTCCCTAACATTCTTACATTAGTGTGTTTCGGAAGCTGGAAaagttatactgccct
gcagtcatatagagcagacgAGCAGCTGGTTATTCTAACTCATCGCTCAGTCCATAGAACaagttatactgccc
gcagtcatatagagcagacgTTGCTTTAAGATTGAAGATTGAAGTGGTCAGGTGTGTTCTaagttatactgccctg
gcagtcatatagagcagacgGAGCGTCACTCTCTCTCACATCTGATAAAATACCTTATGGaagttatactgccct
gcagtcatatagagcagacgCCATGTCTCTAGAACATTGTTGTATTTTCTGTTGTTTTCCaagttatactgccctgt
gcagtcatatagagcagacgTAATGTTTCTGTAGTCCGTCCCCCTGCGCCTACCTTCCGCaagttatactgccc
gcagtcatatagagcagacgCTTGAGTTCCCAAATGAGCATATACGACCAAACTGATATAaagttatactgccct
gcagtcatatagagcagacgCTCTGGGATTGGAGGGGTTGACTGTGTCCTAATAAAGCAGaagttatactgccc
gcagtcatatagagcagacgACAAATATTTACACCATTTTTTTAAGGGGCAGGATCATGCaagttatactgccctg
gcagtcatatagagcagacgATTTTTTTTAGAGATTTGATTGTGGGGGGGTCTTACTATGaagttatactgccctgt
gcagtcatatagagcagacgACCAGATGCGGTAGCTAACTTGTGTTTGGGATCTATGAGAaagttatactgccct
gcagtcatatagagcagacgGGTGCCAGCCAGCAGCCTTGGGTTTCTCCTCTCTTTCTCAaagttatactgccc
gcagtcatatagagcagacgGGATTCTGTCTCTCCTTACCATCCCTCTCTCTCTCACCTCaagttatactgccct
gcagtcatatagagcagacgTCAATCCACTGCACTTGGCCCAGGAGACACAAAGAAAAGGaagttatactgcc
gcagtcatatagagcagacgTTCACAGCCATGAACCAAAAATTACACTACAAAGACAGGGaagttatactgccc
gcagtcatatagagcagacgTTGGGAGGCCAAAGCGGGCAGATCACTTGATGTCAGGAGTaagttatactgcc
gcagtcatatagagcagacgATACAAAATTTAGCCAGGCGTGCCAGCTACTTGGGAGGCGaagttatactgcc
gcagtcatatagagcagacgAAATGAAATATAAAGAAAGGTGTGTACAAATGGTGTGGGGaagttatactgccct
gcagtcatatagagcagacgAGCTAGCTGAGTGTGGGAGCTGGGAAGGCTGCAGGGAAGAaagttatactgcc
gcagtcatatagagcagacgACCCGGGTCTCAAGGGCGGAGTGTGAGTTCGCTGGGCCTAaagttatactgcc
gcagtcatatagagcagacgCCAGTTATAGCAAGAGGGGAAGGGAAGAATGAGGTGGGCAaagttatactgcc
gcagtcatatagagcagacgGACTGACCTTTTCCTTTGAACTGACGTATTTTATGAGAAAaagttatactgccctg
gcagtcatatagagcagacgTGTAAGACTTAAACATTTCATTTCACCCTGCGCACCAGCCaagttatactgccc
gcagtcatatagagcagacgCACAAGGCTCGCATAAGTAAAGGAAAAAGACCAAACTAAAaagttatactgccc
gcagtcatatagagcagacgCCTTCCCAGGCCCAGACCACTTCTTTTCAGCAACCTTAGTaagttatactgccc
gcagtcatatagagcagacgGGACACCCCTTCACACCCAGACCCCAGCAACTGCCCCTGTaagttatactgc
gcagtcatatagagcagacgTGAAAAGCTCACACTCAGTATAGTTCATGACACAGTAAAGaagttatactgccct
gcagtcatatagagcagacgGGCTAATATGTAGTCTTGAACTTGTAGTGTTTCCTTCCTAaagttatactgccctg
gcagtcatatagagcagacgAAAGAAAATTTATGTTCCAAGTTAAGGGTAATCGGTCCAGaagttatactgccct
gcagtcatatagagcagacgCAACATAGGGAGACCCCTGGCTCTATAAAAATTTAAAAACaagttatactgccc
gcagtcatatagagcagacgTGTGGTGATATATGCCTGCAGCCCAAGCTATTTAGGAGGCaagttatactgccc
gcagtcatatagagcagacgATTCCAGGTTGCAGTGAGCTCTGCTCACACCACTGCACTCaagttatactgccc
gcagtcatatagagcagacgTGTTCCCGTATTATTTTGCAGTTCCCTATTATATGAGTAGaagttatactgccctgt
gcagtcatatagagcagacgAGCACAAAGAAGGCTTGTTCCATTTATAAGAAAGGTGAAAaagttatactgccct
gcagtcatatagagcagacgGATTTGACAGAGACAAGAGGGTAATCAACTAGGAAGCTATaagttatactgccc
gcagtcatatagagcagacgGTTACAGAAAACTCCAGAAATACCACTACCCACAAAATTAaagttatactgccc
gcagtcatatagagcagacgGAAAGGTCAGTTGGTATACACAATTTTCTCTTGGAGTCAGaagttatactgccct
gcagtcatatagagcagacgGAAATCTTTGGGAATACACAGAAAAAGTAGAGAAAGAGTGaagttatactgccc
gcagtcatatagagcagacgAATGCCACCCTCTCCTAAAATATTCTAAATAAAACTCAGGaagttatactgccct
gcagtcatatagagcagacgATATGCAAGGAATTCTAAAAGCCAAGAGGAAGTACATTTCaagttatactgccct
gcagtcatatagagcagacgTGATGTCATAATCACGGCATGAAGAGAAATACCAAAGGCTaagttatactgccc
gcagtcatatagagcagacgAGATTTTCTCCTCGTGTTGTTTTACTGTTCCTGTGGAAAGaagttatactgccctg
gcagtcatatagagcagacgGATTTTCTCAAACAGAATTAATTAGGAGTCAAGGAGTTCCaagttatactgccct

gcagtcatatagagcagacgCTAAGTGGTATAAAGACAAAAGGCGGTCGAAAAGATACATaagttatactgccct
gcagtcatatagagcagacgAAATGTTTTGCTGCTCTCTCTTTCCATTGTCTGACCTTTCaagttatactgccctgt
gcagtcatatagagcagacgCCTGTTTGTTTTTCTTTGGTCCATTTTCCACTCTGGGAGTaagttatactgccctgt
gcagtcatatagagcagacgTTGATTAGGAGAACCCGAAAAAGTAAAAACATGGCCACCGaagttatactgccc
gcagtcatatagagcagacgAGCTCACTCACCTATTTCTCCTTGGCCAGCTTTTAAAGTCaagttatactgccctg
gcagtcatatagagcagacgACATTAGAATCCCTGGAGTTGGGCCAAGGTGTCAGTCTTTaagttatactgccct
gcagtcatatagagcagacgCCACCGTCCTCAACTGAACAACGTCATGGCTGGGAGACTCaagttatactgcc
gcagtcatatagagcagacgCCAAGTGCTTCCCAGGCCTTTGGTGTCACAGACCAGCAACaagttatactgcc
gcagtcatatagagcagacgCGAAAAGGGCTTGGGAGGCCAATATATATTTTGGTTTATCaagttatactgccctg
gcagtcatatagagcagacgATGTTTACATATGTACACACACCTAGGTATGTGTTATGTGaagttatactgccctg
gcagtcatatagagcagacgCTTCTTTTTTTTCTGGGAAACAAGAGTTTCACTTTGTTGCaagttatactgccctgt
gcagtcatatagagcagacgTCACTGTGCCCGGCTGTTCTAATTATTTTCAAGTGTGCAAaagttatactgccctg
gcagtcatatagagcagacgGCATTACGCACCTTTATATACTCATGCAACCATCACTACCaagttatactgccct
gcagtcatatagagcagacgGAGCAGTCATTCCCCATTCTTGCCTCCCCCAGCCCCTGGCaagttatactgcc
gcagtcatatagagcagacgTATGAACTTGACTACTCTAGCTTCCTCACATAAGTGGAATaagttatactgccctg
gcagtcatatagagcagacgTAGCATAACATCTTCAAGATCCATCTGCGTGTGTCAGTGTaagttatactgccctg
gcagtcatatagagcagacgGATAGCCCAGCCCATATTGTGTTCATCCACCCATCTGTTGaagttatactgccct
gcagtcatatagagcagacgGGCCATTTGTGTACCTTCTTCCGAGAGATGTCTATTGATGaagttatactgccctg
gcagtcatatagagcagacgTTTTGAGTTGTATAAAGCACAGATTCTTGGATAACAGACCaagttatactgccctg
gcagtcatatagagcagacgCCTTCCCTCATTTCAAGTCCAAGGATTTTAAGTATTATGTaagttatactgccctgt
gcagtcatatagagcagacgGAATATTTCTATCCAAAGACATTCAGGCAAAAGGTGAGGCaagttatactgccct
gcagtcatatagagcagacgCTGAGAGGAGAAAGAGTTGGAAATCTGGCAGCTGGTATTAaagttatactgccc
gcagtcatatagagcagacgGTCTCTAATGTTGATTTGGATGGAAGGCCTCAGATGGTGGaagttatactgccct
gcagtcatatagagcagacgGCTCCATTTGGCTGACATTATTGACACTTCTGCTCCCCATaagttatactgccctg
gcagtcatatagagcagacgTGAGGAAAAATAGCACGGTTCCCTGCATGAAACTTTTTGGaagttatactgccct
gcagtcatatagagcagacgGGCATGCCTCCAGTTCTTCCAAATAGTTCTGAGAATCCCTaagttatactgccct
gcagtcatatagagcagacgGAACTTATCCTCTTGGAGTTGTGTGGCTTCAGCTCTGCTCaagttatactgccctg
gcagtcatatagagcagacgCACCATGTCTTCTGAACAGTGCCTGCCAAAGCTCCCTCCAaagttatactgccc
gcagtcatatagagcagacgCTGCTGGAACTGGGCTGTAAACACACCATGTCTGATTGTTaagttatactgccct
gcagtcatatagagcagacgGCCTTGCACCAAACTTCTCTCATTCTTCTAACTTATATTTaagttatactgccctgt
gcagtcatatagagcagacgGCCCCCTTACCACGAATCAGGCTGGATCCCCATGTGCAGGaagttatactgcc
gcagtcatatagagcagacgGCTGTCTGACCCACCGCATGAGCCATCCCCACCCTACCACaagttatactgc
gcagtcatatagagcagacgGCACGTTGATCTCTGGCCAGACATGTACAAAGGCGCTCCCaagttatactgcc
gcagtcatatagagcagacgAGTTGTCCCCTAGCAAGATACTCTAATTTCTCCCCCTTATaagttatactgccctg
gcagtcatatagagcagacgCAAAAATAAAAAAGCCCAAACCAAACAGATATGGTTTGGCaagttatactgccct
gcagtcatatagagcagacgGTAATTCCCATGTGTTATGAAAGGGAGCTGGTGGGAGATAaagttatactgccct
gcagtcatatagagcagacgTGGTGGTATAAGGGGTTTCCTCTTTGCTTGGTTCTCATTGaagttatactgccctg
gcagtcatatagagcagacgTTCCTGCTGACATGTGAGACGTGGCTTTCGCCTTCTGCCAaagttatactgccc
gcagtcatatagagcagacgCTAATATGCAAACCAAAACAAATATATCCTATTAGGGCCCaagttatactgccctg
gcagtcatatagagcagacgCCAAATGTGAATAAATCTCAGAGTGGGCACAGATCAGGCTaagttatactgccc
gcagtcatatagagcagacgACCATATTTCAGCCACAGTTATTCAATCCACTCATTTTTCaagttatactgccctg
gcagtcatatagagcagacgGACTCTGCAACAAAGATGCTTCCATGTTTCAAACTGCAAAaagttatactgccct
gcagtcatatagagcagacgATTCCTCTAATGAGAATAGTCCCACTATTTTGGCATTTTGaagttatactgccctgt
gcagtcatatagagcagacgCATATACATATTAATACGTTCATTTTTCAACCCTCCCAGGaagttatactgccctg
gcagtcatatagagcagacgGAATTTGAGTGTGTAGTCTTCCTGGAATATCTAAACGAACaagttatactgccctg
gcagtcatatagagcagacgCCAAAGGGTAATGTGATGCATTGATTATTCTGCTAGGGTTaagttatactgccctg
gcagtcatatagagcagacgGAAAATTTCTTCTTGGACTGCTCGACTTTGGGAACAGCAAaagttatactgccctg
gcagtcatatagagcagacgCTCCTTCCAAGAGACTGTCACTAACTGGGCATCTACTGTAaagttatactgccct
gcagtcatatagagcagacgAAATCACTTAATGCTCAAAACCCTAGAAAATAGGTGCTGTaagttatactgccctg
gcagtcatatagagcagacgTATCATCATTTTTCAGATAAATAAACACATCGCAGGGAGGaagttatactgccctg
gcagtcatatagagcagacgCCAGGTCACACAGTAAAGGATGGAGGCTGACTTAGAACCCaagttatactgcc
gcagtcatatagagcagacgAGAATCCCCCATGTTCTTAACTGTTTCGTTGCACAAGTAGaagttatactgccctg

gcagtcatatagagcagacgGCTAGGCACCATGATGAGACCCAGCAATAAAGAGATTAAGaagttatactgccc
gcagtcatatagagcagacgTGGACACACTTATTTCCCACTTCTGATATTTCCACAGATGaagttatactgccctg
gcagtcatatagagcagacgAGGCCATCCAGCTTCTTTCTGTCCCCCTGCTCACTCCTGGaagttatactgccc
gcagtcatatagagcagacgGTGGCTTATGATGGTTTGTGATGATCAGTGATGGTTTATGaagttatactgccctg
gcagtcatatagagcagacgTTATGGCTTATAATGATCAGTGACTGGGGAGATGAAACCTaagttatactgccctg
gcagtcatatagagcagacgCCTGGGAGGGGATGATCTGCATGGCCTGTGGCTGCCCATGaagttatactgcc
gcagtcatatagagcagacgAGGTCTAGCTGATTCACTCCTATCCTGGTATCCCTACTGGaagttatactgccct
gcagtcatatagagcagacgTGAGGAGCCTGCCCAAAAGCCGAGTACCTGCTACCACTGGaagttatactgcc
gcagtcatatagagcagacgAGAATTTGTCTCCCTGTGAAAAATGGTCAGCAGGTGGAACaagttatactgccct
gcagtcatatagagcagacgCACGGGGATCTGCACTGAGAACATGAGTCGGGGAAAGCTGaagttatactgcc
gcagtcatatagagcagacgCTGTGCCAGCATTCTTATACCCACAAAGGCTACCCAGGGCaagttatactgcc
gcagtcatatagagcagacgTGGAGAATGTGTAAGGAAGGGCCCACAGAGGCAAACCAGGaagttatactgcc
gcagtcatatagagcagacgCCAGGTGAGGAGCAGGAGATGGGTCAGGGACTTCTCTCTGaagttatactgcc
gcagtcatatagagcagacgGCAGGACCACCAGGCATTTGAGATTGTGCAGGGGCTGTGGaagttatactgcc
gcagtcatatagagcagacgTGGGCAGGGGTAGTAGCAAATGCAGAGGCGGGAGAAGGCGaagttatactgc
gcagtcatatagagcagacgCCAAGGGAGAAGAAACGAGGCTAGGAGGATGGGCAGGGATaagttatactgc
gcagtcatatagagcagacgCTGATGGCAATGAGGAGCTCTCGAAGGATTTTATAAACAAaagttatactgccct
gcagtcatatagagcagacgTGATCAATTTTGCTTCCCCCGTGATAATATAGAAAACTGGaagttatactgccctg
gcagtcatatagagcagacgCTTGAAATAAGGAAGTGGTCATGGGGATGTAGAGGAGAGAaagttatactgccc
gcagtcatatagagcagacgAAAAGGACTGATTTCTGGCTTAGTGGCTGAGTGCGTGTTGaagttatactgccct
gcagtcatatagagcagacgCCAAGACAGAGTGTAGGGTGAACAGGGAGAAGAGGATGGTaagttatactgcc
gcagtcatatagagcagacgCTCTTGCCTTGAGCTCTGGTCAGCCCCTCCCATGGTCCACaagttatactgcc
gcagtcatatagagcagacgATTCATGGCCTTGAATCCATTTTGTTTCTTCCCCCTGCTCaagttatactgccctg
gcagtcatatagagcagacgCCACTGAGGCTCTAAGGCGCGGTTCTGGGCTCCTTTCCAGaagttatactgcc
gcagtcatatagagcagacgCTCCTGCATCCTCCACACTAACCGGTTTCACAAGTTTTTTaagttatactgccctg
gcagtcatatagagcagacgAGCCAGTTTAGAGCTTTCTTGATCAAATCGCCTAACCTCCaagttatactgccct
gcagtcatatagagcagacgGCACCCCATCCCAGCCAAGGAGCCACCATCCCACTATCTTaagttatactgcc
gcagtcatatagagcagacgCCCGTCTCATGTGGGCCTGTTTGTAGTCAGTTCTGCCTAGaagttatactgccct
gcagtcatatagagcagacgGCAGATTTCGATTAAACCTCCCACTAGGTCCACTCTCTTCaagttatactgccct
gcagtcatatagagcagacgAGGCACACTTGGGTCTTTGAACAGGAATGGAAGGTGATTAaagttatactgccct
gcagtcatatagagcagacgCCGGGGTTGGCAGAAGCGCTTGGGGCTACACCAGTGTGGGaagttatactgc
gcagtcatatagagcagacgCTCTAAGCACCTCCCACTGCCCCACCACAGATCCAAGTTTaagttatactgcc
gcagtcatatagagcagacgCAGAGACTGGACCTGGCTTGTCAGAAGATGCCTGAATTCTaagttatactgccc
gcagtcatatagagcagacgGAAGATCACAGAAACAGTTAATTTTAGTAGACAATCAGGGaagttatactgccct
gcagtcatatagagcagacgCTGAGGTAGATCTTGAATTTCCTTCTGGACCCCGCAACTGaagttatactgccct
gcagtcatatagagcagacgCTTGCCTTCCAAATGTGGTTGATTGGTGTTGGCCGGCACGaagttatactgccc
gcagtcatatagagcagacgGGCCGCCCAGCTCGTTCATCTACAATGTCCATGCTTTCCCaagttatactgccc
gcagtcatatagagcagacgGGCTTCATTCTGATGTAAATTGGAGTGCAAACATGGCTCGaagttatactgccct
gcagtcatatagagcagacgATTGCTAAATGAAAACTGTCACAAAGAACCCAGACAAATGaagttatactgccct
gcagtcatatagagcagacgACAGCCAAAATACAAAAATATCACACTGTTGACACTGCTTaagttatactgccct
gcagtcatatagagcagacgCTAAGCCAACAGAATAGCCCCAGGGTCATATATGTCAACCaagttatactgccc
gcagtcatatagagcagacgCATCTTTTCTCCTGCCCCTTCCAGTTTTGGACTGCCATGGaagttatactgccct
gcagtcatatagagcagacgAGAGTATCTGGATCCTTGTTATTATCCTGTCCCTTACTGAaagttatactgccctg
gcagtcatatagagcagacgACACAAGCTGGTCATCAAAATTCTTTGGGTTTCACTTTTCaagttatactgccctg
gcagtcatatagagcagacgGAAATGGAAGGACATTAAGTGAGAAAATGTATGTGGAAGCaagttatactgccct
gcagtcatatagagcagacgCCAGGCTGGAGTGTGGTGTTATGATCACGGCTCACTGTAAaagttatactgccc
gcagtcatatagagcagacgTTTTTAAGGAGATGGGGTCCAGTTGTATTGCCCGGGCTGGaagttatactgccct
gcagtcatatagagcagacgGTGCCTGGCCTAAATGCCTTCTTTTAAACTAACACTTTCTaagttatactgccctg
gcagtcatatagagcagacgAATCTTATTAAATGGTTTGTCTCTGGTATTTTCTGCCACCaagttatactgccctgt
gcagtcatatagagcagacgCTCTATTTTGACTGTCCTCAAGGTCGAGAACTTAGAGCAGaagttatactgccct
gcagtcatatagagcagacgTCACAGCAGAGATAAGAGTTTAAATGAAAAAGTTTGTCGCaagttatactgccctg
gcagtcatatagagcagacgGCCTTGAAGAACTTAGGACATACAGTTGGCATCATAAATGaagttatactgccct

gcagtcatatagagcagacgGTTAGAAGAAGGAATGATCAGATGAGAATGAGATTTTGCTaagttatactgccctg
gcagtcatatagagcagacgTGAAGTAATCCAGACCATAGGGCAGGCAGGGAAGACGTTGaagttatactgcc
gcagtcatatagagcagacgCCTAGGTGGGTTAATGGACTGTAGAGGAGAGCTGTGCTTCaagttatactgccc
gcagtcatatagagcagacgGCCTGGCTTGTTGGCAAAGCTGTGTCATGCTTGTGGGGCAaagttatactgccc
gcagtcatatagagcagacgGCCAATCCACACAGGGAAGGTCTGAGACAGCCCCGCCAAGaagttatactgc
gcagtcatatagagcagacgGGGCATCCAAAAATTTACCATCTCTCCTTCTCTAAAAAAGaagttatactgccctg
gcagtcatatagagcagacgGCCCACTCCCAGAGACCAGAAGAAATGCGGCCATCTCAGCaagttatactgc
gcagtcatatagagcagacgCCAGTGGAAGAGGGAGCATGTTGGCAGATGTCAGCCTGGAaagttatactgcc
gcagtcatatagagcagacgGCACTATGTTAGCACCCTAGAACTCAGACGCAACTCCAGGaagttatactgcc
gcagtcatatagagcagacgGAGTGGGGAGGGACACGTCTGAATCAACAGATAAAATCCTaagttatactgccc
gcagtcatatagagcagacgCAGAATGGGGGCTCAAGAAACAAAATTAAATTCCGTACTTaagttatactgccct
gcagtcatatagagcagacgAACCACCAAAGTATGTTACGTTTTATCACCTGTTGAGGCTaagttatactgccctg
gcagtcatatagagcagacgGCACGTTCACACATCATTTATTTCAGGGCTGATCTGCTGAaagttatactgccctg
gcagtcatatagagcagacgCCATACCAGTACCTGTTCCAATTGGCTGATGATTGCACCAaagttatactgccct
gcagtcatatagagcagacgATTTTGAAATCGCCCCTGAAATTGAATGCTTATGATGTGCaagttatactgccctg
gcagtcatatagagcagacgCATTGTGTTTGGGTCACTGTGTCTCACTTAGATTGTAAACaagttatactgccctg
gcagtcatatagagcagacgATTTAAATTCCAGGAACCATGCAAGCAACTTAATGGTGAAaagttatactgccctg
gcagtcatatagagcagacgAATCACAGTTTCTGCTGCCATGGGCCTTACGGTCTAGTGGaagttatactgccc
gcagtcatatagagcagacgTCACAAAAATGAATGGGAAATTACAATGATGATGCCTGGTaagttatactgccctg
gcagtcatatagagcagacgCGACGACCGGGCTGAATATAAAGAAAGAGGAAGAGTTAAGaagttatactgccc
gcagtcatatagagcagacgGATGGAGGAAAGATGGTAGGTGGGAAAGAGGTCCTGTGCGaagttatactgcc
gcagtcatatagagcagacgTGTGGTCAGAGGCAGCATGGTACATTTGAAGGACTGAAAGaagttatactgccc
gcagtcatatagagcagacgACTGACATGCTGCGTATAACAGGGAGTGTGGTCCAGGGCGaagttatactgcc
gcagtcatatagagcagacgCCTGGTAGGTCATGATGATTTAGATCTTTGTCCTCTGGGGaagttatactgccctg
gcagtcatatagagcagacgCCCTCTGGAGTTAATTTTTAGTACAGGCATACAAACCATAaagttatactgccctg
gcagtcatatagagcagacgTGATTCATTTCAGCACCACCCCCTCATGGCCACCTTTTCCaagttatactgccc
gcagtcatatagagcagacgGAAACTTGTGGCTTCGTAACAGTTCACCCATACAGCTTGGaagttatactgccct
gcagtcatatagagcagacgGTGCAAGCTATTTCTGTCTGTGGCAGTCCCTGAATGTTTCaagttatactgccctg
gcagtcatatagagcagacgTGGACATCTCTTGCCACTGCCTGCCCCACATTCAATTCCCaagttatactgccc
gcagtcatatagagcagacgCCTGATGGTCCTTTTGAGAATCACATCGTGGCAACTCTAGaagttatactgccct
gcagtcatatagagcagacgCTGGCCAACTCCACCCCACCCCTGCTCCCTCCTTCCCCAGaagttatactgc
gcagtcatatagagcagacgCATGATCCACTCAGGCTAATGAACATACTCCATCCTATCAaagttatactgccct
gcagtcatatagagcagacgGGATCAACCTCATTCCGAGTCTCATTCCTAGAGCTTCTTCaagttatactgccct
gcagtcatatagagcagacgAGGGCTGCTCTGTTGTTCCTCTGGGGTTGCTGTGCTGAGGaagttatactgccc
gcagtcatatagagcagacgTGAAGCCAGTGTATGAGGAACTAGAGCTGAGAGAAAGAGAaagttatactgccc
gcagtcatatagagcagacgAGAGAAGCCAAGGCCTGATAGTATCTCTTAGAGCTTTCAGaagttatactgccct
gcagtcatatagagcagacgAGCTAATTGGAGCTGGCTTCTCTTACTGGTGACCCAAAGAaagttatactgccct
gcagtcatatagagcagacgCCCGGCACTCAGGCTTTGCTAACGCAATGGGCATCCTGTCaagttatactgcc
gcagtcatatagagcagacgCAGTTGGGTCCTCACTGATACCGCCCTCACCATCCCCCGAaagttatactgcc
gcagtcatatagagcagacgGCTGAACCTTGCTTTTACTCTGACAAAACTAGAGCTTTCTaagttatactgccctg
gcagtcatatagagcagacgGCCTGGTTCTCTTGTGCTCTTGACTGGCCATTCCCTCTTTaagttatactgccctg
gcagtcatatagagcagacgCATAAAGTGCACATCCCTAGGTTATGCCACCCAAGAATTCaagttatactgccc
gcagtcatatagagcagacgCAGCTCTCCTCAACTGGACTCATTCTACAGGAACCAAGAGaagttatactgccc
gcagtcatatagagcagacgGCACTGCTCTCCTTCTCACGTTCTCAAGCCATAACCTGCAaagttatactgccc
gcagtcatatagagcagacgACTGACGTCTTCCCTCCCGGCCCTGCAGTGGCTTTCCTCCaagttatactgcc
gcagtcatatagagcagacgGCTTTTCCTCCTCACCTGTCATCACCTTAAGCCCCTCCATaagttatactgccct
gcagtcatatagagcagacgCAACATCTGCTTCCTACGTTCTAGCGATTCTCGAGCCTCAaagttatactgccct
gcagtcatatagagcagacgTGATCTTCATCTACCTATCAGGCTGTGGGTCTCTGTCGCCaagttatactgccct
gcagtcatatagagcagacgTTCTCATTGTCTTCAGAATCAGGGCTCTCTAACTCACGTTaagttatactgccctg
gcagtcatatagagcagacgGCCCTCGCCCTTTGGTTTACAACAGAACACACTATACACAaagttatactgccc
gcagtcatatagagcagacgGTAGACAGCGTGTATTCTGCCCTCTCACTTTATACGTAAGaagttatactgccctg
gcagtcatatagagcagacgCCAGATAGGTTGTAGTTTGTCCTGATTACACAGATGGTTAaagttatactgccctg

gcagtcatatagagcagacgAAGATCACCCAGCTACTCTTGTTGTTTGACTTTAATGAAAaagttatactgccctg
gcagtcatatagagcagacgTTAGTGAAACAATAACCAGTCCCCTCCCCCTACTCCCATTaagttatactgccc
gcagtcatatagagcagacgCTGGGAATGCTTTGACTAATTTAGATTTGGGAAAAAAGACaagttatactgccctg
gcagtcatatagagcagacgATTAAGCTGAAAGAGTAATATGAACAGTGGAAGGGACAAAaagttatactgccct
gcagtcatatagagcagacgGTTTCTGCACGTCTGTACATAACTTCTTCACTCCAATTGAaagttatactgccctg
gcagtcatatagagcagacgTCATTCAATGGCTGCTTGATGTCCCTGGAACTACTAATTAaagttatactgccctg
gcagtcatatagagcagacgGATAAAAGCTCAGATGAGCCCCAGGCTGACCTTGAGGCAGaagttatactgcc
gcagtcatatagagcagacgACTGCAGAGTCACAAAGCCAGGATTTGGATTCAGTTTTCCaagttatactgccct
gcagtcatatagagcagacgACATTTTCAACCTCTTCATTCTGTCCCAGGTAAAGGATCTaagttatactgccctg
gcagtcatatagagcagacgCAGCCAGGGCAACATGGTCTTAAAAAAAATTTAGCTGGATaagttatactgccct
gcagtcatatagagcagacgAAAAATGACAACTTCCTGGATTGAGTAATGACTCTGTGCTaagttatactgccctg
gcagtcatatagagcagacgTTTTTGAGACAGAGTTTCACTCTGTTGCGCAGGCTGGAGTaagttatactgccctg
gcagtcatatagagcagacgTGTATTTTTGGTAGAGATCATGTTGCCGGGGCTGGTCTGGaagttatactgccctg
gcagtcatatagagcagacgCGTGCCCAGCCAGATTCTTTATGTTTTAAGAGACAATGTCaagttatactgccctg
gcagtcatatagagcagacgTCTTGGTGGGGTAAAGAAGTATTTTATCATAGGCGCACGCaagttatactgccct
gcagtcatatagagcagacgAGTGAAGGACTTGTTTGGACAATTGGAGAGTGAGTTTGTGaagttatactgccctg
gcagtcatatagagcagacgTCCTGAGGAAATATGCAAAACCTGACTGGGGGCGAGCGGAaagttatactgcc
gcagtcatatagagcagacgAGCCATTTCTCTTTAGCTATTTACTGCAACCTTTGGAACAaagttatactgccctg
gcagtcatatagagcagacgAATTGTGAGATTCTGCTCCTTTCTGTTAGTTACCTTTCTTaagttatactgccctgtg
gcagtcatatagagcagacgTTAGAGACAAACTCTCGCGCTGATGCCCAGACTGCAGTGGaagttatactgccc
gcagtcatatagagcagacgTCCTCCCACCCCAGCCTCCTCTGAGTCACTGGGACTACAGaagttatactgcc
gcagtcatatagagcagacgCATGCCTGGATGTTACCTTTCTATGTGCCTGTGTCTACATaagttatactgccctg
gcagtcatatagagcagacgGTAGAGTGCTAGCTCCTTTTCTTTGTCTTTGTGTACTTCAaagttatactgccctgt
gcagtcatatagagcagacgCAGGTCTCAGTCAATGCTGCTTGGTTGGTTAATTGTCATCaagttatactgccctg
gcagtcatatagagcagacgAAGAGGATGGCTTGGCTTGGGAGCTTATACCTTGAGCATAaagttatactgccct
gcagtcatatagagcagacgAGACAAATTTAAAATGTGGTCTTGGTCTGGCTGTCTGAGCaagttatactgccctg
gcagtcatatagagcagacgCAGATATGCCTGAGAGAACCCAGCAGAGCGGTCAGACAGCaagttatactgcc
gcagtcatatagagcagacgGCTTGTCTCACCTTGGCTCACCCAGGCCTGTCCTCTGTCCaagttatactgccc
gcagtcatatagagcagacgATGGCTTGGAATTTAGACGTGATTTTTCAGCTGGATCAGTaagttatactgccctg
gcagtcatatagagcagacgCTGAGTCCTGTGTCACTTGCTTCTCTCTGCGTCTCCGTCTaagttatactgccct
gcagtcatatagagcagacgCTTTGTGGCATGTAAAAAAAAGGAAATGGATCAGGTGTGGaagttatactgccct
gcagtcatatagagcagacgGGGAGGCTGAGGCAGGCCTCTCATTTGAGGTCAGGTGTTCaagttatactgcc
gcagtcatatagagcagacgGGGCAACATGGCGAAATGCCGTCATTACTAAAAATACAAAaagttatactgccc
gcagtcatatagagcagacgAAACAAAAAGGAGGAAAAGGTATGGCAGTTCCTATAACAAaagttatactgccct
gcagtcatatagagcagacgGAGTGATCCTTGTTTTTAAAGAAGAGTGATTTCTACCTGTaagttatactgccctgt
gcagtcatatagagcagacgCAGAAATGTCTCTTCTCCCACACCCCCAATTCCTATAGTGaagttatactgccct
gcagtcatatagagcagacgAATATTAACACTCTTCAGGAGCTCATTCAAAACATGGTAGaagttatactgccctg
gcagtcatatagagcagacgCTAGAAAAGAAGCAGATAAAAAGCATGGGTCCTGGGAAAGaagttatactgccc
gcagtcatatagagcagacgGAGATGTAGCTTTAGGAGAAGAGGAGTTGGGGGGAAAACGaagttatactgccc
gcagtcatatagagcagacgAAACTTTTCTTCTTCGTCATTTTCCCCAATATCTGCCCTGaagttatactgccctg
gcagtcatatagagcagacgATTCCTATGTGTCTCCTTAGGCATGCCAACACCATTCCACaagttatactgccct
gcagtcatatagagcagacgTGAAATCACTTCAACATTTGGCATTGAAACCATGGAGAATaagttatactgccctg
gcagtcatatagagcagacgTAGAAGATATGACAGATAAGTTGGTTTCACTTGTATAGGGaagttatactgccctg
gcagtcatatagagcagacgGTAAGAAAAAAATGTATACCTGTTTCACTTAGTGAGGGAGaagttatactgccctg
gcagtcatatagagcagacgGTGAGACATTGAAAGTACGAGGAAATTGGTGAGTTTTTGGaagttatactgccctg
gcagtcatatagagcagacgGTGAATGTCTTAATTTTTCATGTCGCAATTCAAGCCATTGaagttatactgccctgt
gcagtcatatagagcagacgGAACAAATTCCTGGAAAGTCATTTCATACCATCAAGCCACaagttatactgccct
gcagtcatatagagcagacgGCAATCTTGCTAATATTGCTTTGGGGGGCTTTTCTGCAAAaagttatactgccctg
gcagtcatatagagcagacgGGAATTCTTTAAAATGAGTCCATCCAAATCTACGCTAAAGaagttatactgccctg
gcagtcatatagagcagacgCAGTCACCCTCCACTGCCAGAAGCCCTCGACATGATACGAaagttatactgcc
gcagtcatatagagcagacgCTTCCTTCTCAAGCATCCTCCTCATTGAATATGTACAAGTaagttatactgccctg
gcagtcatatagagcagacgTCCTCGAACGGTAGTAGCCAAAAATTTCTCTTTCCTTGCTaagttatactgccctg

gcagtcatatagagcagacgAGTCTGTCACAAGTCCCAGGCTGTTCATCTGGCATGTGCAaagttatactgccc
gcagtcatatagagcagacgAGTGCAAAATGAAAATGGCAAGCCCCTGTTCAAAAATTAAaagttatactgccct
gcagtcatatagagcagacgCAAAATGATGACAAGAGAGCATTGAACCAAGCATGGGCTCaagttatactgccc
gcagtcatatagagcagacgTGTAGTACATGACGCCGGCCTTGCGCTCCTCTGGATGGACaagttatactgccc
gcagtcatatagagcagacgGCTTTGAGACACTGGCCATGCTACCCTCCTAACCCACATCaagttatactgccc
gcagtcatatagagcagacgGCAATTCTCTGCCTATCTTGAGCAATGTCCCTCTCTTGGCaagttatactgccct
gcagtcatatagagcagacgCCCACCGCTTCTCTCTTAAGTTCCCTTTCTTCCTACCTCCaagttatactgccct
gcagtcatatagagcagacgGGATCTCTGGATTCAAACGCAAGGCCTGTTCCAAAATCAGaagttatactgccc
gcagtcatatagagcagacgACATTTCCTTGTTATCTCCATGCTCTCCTCTTTGGGGTGTaagttatactgccctg
gcagtcatatagagcagacgTGGATCTTAACTACCAAGTCCTCAAGTGTCACACAGGTTGaagttatactgccct
gcagtcatatagagcagacgAGTGGGCACATGGGAGGTGGGATGCCAGGGGTTAATCTGGaagttatactgcc
gcagtcatatagagcagacgATGATGTTTCCAACCATTTCCTTCACTTCACCTCTGTATCaagttatactgccctg
gcagtcatatagagcagacgAATTCCACAATTTTCCACCTCCTGCCATTTCTCTTGGTCTaagttatactgccctg
gcagtcatatagagcagacgCTGTGAAGCCCTTGTCAAGGCCATCTTGGACCCACAACTAaagttatactgccc
gcagtcatatagagcagacgCCATGCACACTAGGCCACGAACTTCCCGTCACACATGCTCaagttatactgcc
gcagtcatatagagcagacgTCGCTCAGATTGCCTTGGTGGAAACTGCTTTCGATGGTTCaagttatactgccct
gcagtcatatagagcagacgCGATGACTCACAAAACAGCCTTTATTCATTGACGTGTTCTaagttatactgccct
gcagtcatatagagcagacgCTCTATTTACTAGGTTTTATTGGGAATGTGAGGTGAAATCaagttatactgccctg
gcagtcatatagagcagacgCAGGAAGTTTTCAGTGTTTCTTAGAAGAGGACGTGGGGACaagttatactgccc
gcagtcatatagagcagacgACACACATGTTGTTGATGGGCGGACTTTCTCTATCAGGGAaagttatactgccc
gcagtcatatagagcagacgAGCATTTCTGGAGGAGGGCATATCAGTTTTGACACCTGTTaagttatactgccct
gcagtcatatagagcagacgCAGAAGGTAAGGCTGGACGAAATGTTTCCTGTTGTGAATAaagttatactgccct
gcagtcatatagagcagacgGAGAATCAGAAAGTATTTTACGGAGCATTGATGAAAGGTTaagttatactgccct
gcagtcatatagagcagacgCCTATTGGGAATATGAGGGTATCCACAGTCAACGGGAAACaagttatactgccc
gcagtcatatagagcagacgAGTCAACCCCAGCACCCCAAGGATGATTGCTTGCTCACGGaagttatactgcc
gcagtcatatagagcagacgCCATAGACTTTGGACTCCCAGACAGGTGTGACAAGCAACAaagttatactgccc
gcagtcatatagagcagacgGACTTTACACCAGAGGCCAGCAGATACCACAAAGCAGATGaagttatactgcc
gcagtcatatagagcagacgGCCACAGAACGAAACAGCCACACTCCCCATCCCCCAACATaagttatactgc
gcagtcatatagagcagacgGCAAGGCCATCTGAGAGAGTGAGCGATTATACAAGAGAAGaagttatactgccc
gcagtcatatagagcagacgCACAAGTGATTATAAAAGAGAAATAATTGAGGCCGGGCATaagttatactgccc
gcagtcatatagagcagacgAATACAAAATTAGCCTGGTGCAGTGCTGCATGCCTGTAATaagttatactgccct
gcagtcatatagagcagacgAATTTAAATTTTGGAGTGGGGTAAGCTTTAACCTGGATGAaagttatactgccctg
gcagtcatatagagcagacgCAATGGATGGGGACAGAGAAAAATAAAAATGATCCAGTTAaagttatactgccc
gcagtcatatagagcagacgTAAATCATGGTGGGTTAATCTATATGTCAAATGGTACCTGaagttatactgccctg
gcagtcatatagagcagacgATTGGCAACCCTGCTGCACAATTGGTTACGTAACACTTGTaagttatactgccct
gcagtcatatagagcagacgCAGTTTAAGACAGAGGCAGCTGGGAAACAATTACTATAGCaagttatactgccc
gcagtcatatagagcagacgCCCAGGGGCCCAGAAGAAGGTTATTTAATTCAGAATACAGaagttatactgccc
gcagtcatatagagcagacgCTTCCTGGGAGTTGGGACCTGAGCTGAATTTGGAAGGACAaagttatactgccc
gcagtcatatagagcagacgGGTGTTACCTAGAACTTAGATTCTAAATGAACAAACAGAGaagttatactgccct
gcagtcatatagagcagacgCAGTGGGGAGGCACAGAGGTGAGAAGCCCGTGATGAGGTCaagttatactgc
gcagtcatatagagcagacgCACCTGGGTGTGCAGAAGAACCAAGAGATGAGGCCAGAGGaagttatactgc
gcagtcatatagagcagacgCTTGGGAGGTGAGCTAAGATGTTTGAATTTTGTTTTGAAGaagttatactgccctg
gcagtcatatagagcagacgAATAATTATCAAAGATTCTAACTAGGGGTAGCACAGAAGCaagttatactgccct
gcagtcatatagagcagacgCTCATGGGAAAAAGGTAGTTAAAGTGGGAGGGTGGAGGATaagttatactgccc
gcagtcatatagagcagacgCAGTGAGGCAACGATGGTAGAAATGATGACAAACCGTGAAaagttatactgccc
gcagtcatatagagcagacgCAGATGGTGGTGTTGCAAGTAAGAAGCAGGATTTGGGTTGaagttatactgccc
gcagtcatatagagcagacgATAACTAATCAGATGTGGGAGGTGAATGAGAGGGGCAGAGaagttatactgccc
gcagtcatatagagcagacgCTTGCTCCAAGGGGAATACCAGGGCCACAGTGTTCGGGGGaagttatactgcc
gcagtcatatagagcagacgTTGGGCTTGGGGTGCGATTGATGATGTGGTACCTGACAGCaagttatactgccc
gcagtcatatagagcagacgCCAAAAAGTAAAATGAAAACTTGGACTTCTCCTTGCACTTaagttatactgccct
gcagtcatatagagcagacgCTTCATAAGAGCAGGTTCAGTGGGGTGACAGGGATAAAAGaagttatactgccc
gcagtcatatagagcagacgAAGTTACTCTCTTCTCCTGATGCCTTATTCTTCCTGGGGGaagttatactgccct

gcagtcatatagagcagacgATCACGGACTCCTCCCTCTTTCCACATGATGTACTTGGCCaagttatactgccc
gcagtcatatagagcagacgTCTCTTGATGCCATTCCCTGTTCTCCACACTCCCTCCCGCaagttatactgccc
gcagtcatatagagcagacgCCAGTCTTTCCAAGATTTGCTGCCTAACTTTAAACAAAACaagttatactgccctg
gcagtcatatagagcagacgCTTATAAAGTTCTTCATGACCAAGTCCCCAAGATTCCTCTaagttatactgccctg
gcagtcatatagagcagacgGCTCCCCAAAGCCCTCTGCTCTGCCTGGACTGAGACCCCAaagttatactgcc
gcagtcatatagagcagacgGCACTGCCACCTGTTCGTCCTTGCTCTCCCCACCCCACCGaagttatactgcc
gcagtcatatagagcagacgGTGTTCATCTTTTGTGTCCACAGCCGCCAGTAAAGTAGCCaagttatactgccct
gcagtcatatagagcagacgAAGAGACTTGATTATGAGGGGGTTTATGATTCAAAGTGATaagttatactgccctg
gcagtcatatagagcagacgCTGAGAGGAGTGAGCCAGCAGAGACAGAGAGAGGTTGAAAaagttatactgcc
gcagtcatatagagcagacgCCTGAGCATGCGGGGTGGGATGAGAAGGGGGTGGGAGCTGaagttatactgc
gcagtcatatagagcagacgCAGGAAGCAGGGACCTGGGGGTGGGGAGATGCCAATTCCAaagttatactgcc
gcagtcatatagagcagacgATGTTCCCATGTGCTTGAAAACGAAGTTGAGGGCTTTTTCaagttatactgccctg
gcagtcatatagagcagacgAGCAATGAGGAGGCAAAGTGTGAGAGTGAATGGGATTGGGaagttatactgccc
gcagtcatatagagcagacgGCAGAGGGTTAGCAGAAAAGTGAAAGTTTGGAAGAACCACaagttatactgccc
gcagtcatatagagcagacgTGACATGACACACGACCCACACTCATTCTCACTCTACTTAaagttatactgccct
gcagtcatatagagcagacgCATCAATGATCTGATTCTTAGAATTTCTCAAGGGCCCAGGaagttatactgccctg
gcagtcatatagagcagacgTTTGGGAAAATGCAGTGCCCCACCTTGCGGCTGGGATGCCaagttatactgcc
gcagtcatatagagcagacgCCCTTCCCGCAGGTTCTGCTGCACCCACCCATAGCCGATGaagttatactgcc
gcagtcatatagagcagacgCCTCCGAGGGGCTGCCATCTCCACACTGGAATGCACTTGGaagttatactgcc
gcagtcatatagagcagacgCATAAGGCCCTGTTTTAGCTTCCCACTGAGTGTGTCGGGGaagttatactgccc
gcagtcatatagagcagacgTCTGCCCTGAGATGCTGCAAGCCGTATGCCCAACTGCTGTaagttatactgccc
gcagtcatatagagcagacgGAGGCCTTTTGGGAGGTGCTTTGGTCACTGGGGCTCTGCCaagttatactgccc
gcagtcatatagagcagacgTCACTCTCTCTTGCCCTTTCGTCTTCTGCCTTCTGCCACAaagttatactgccctg
gcagtcatatagagcagacgAGAAGTCCTGCACCAGATGCCAGCACCTTGGATCCCGGACaagttatactgcc
gcagtcatatagagcagacgAGTCTCCAGGACCGTGAGATAGTCAATTTCTGTTCTTTATaagttatactgccctg
gcagtcatatagagcagacgCAGACTAAGACACCAGCCTGCCACATGTGTCCTCCTGGGCaagttatactgcc
gcagtcatatagagcagacgCGGTGGAACGTGGTCTCCTCACTTCGTGTGACCCAGCTACaagttatactgccc
gcagtcatatagagcagacgCTGTAGTAGGGTGGGGGCAGGGAGGGGAAATTGGGCAGGGaagttatactgcc
gcagtcatatagagcagacgCGGGAGAAGAACAGATGTCTGAGAAAACATCCTGGGAGTCaagttatactgccc
gcagtcatatagagcagacgCCTGGCAGGCTCCATTGTCTCGTTACACTTCACTTAGGGAaagttatactgccc
gcagtcatatagagcagacgCTGAAGAATTTCAAGAGAGCAATTATAGAGTGTTAAACCCaagttatactgccctg
gcagtcatatagagcagacgGCCCTGCCTCTTCCCTCCTTCCCCTCCTTAACATCCTGCCaagttatactgccc
gcagtcatatagagcagacgTCCTTCCTCTCCTTAACATCTTGCAGTGGCTCACATTCTTaagttatactgccctg
gcagtcatatagagcagacgTTGCAAAAAAATCCAACTGCTTTAACCTGGCTTTCAGTTTaagttatactgccctg
gcagtcatatagagcagacgTTCTCACTGCCTGAACATGTTTACGCCCTCCTCAAATGTCaagttatactgccct
gcagtcatatagagcagacgGCCTAAGTCTTATTCATCATTGAGATCTCGATTTCAGCACaagttatactgccctg
gcagtcatatagagcagacgACATTCCGAGACTCTTTTGTTTAAATGCCTCACTTATTTGaagttatactgccctgt
gcagtcatatagagcagacgGGAACTGCACGTGGAAAGGCCAGCCACATACATCACAAAAaagttatactgcc
gcagtcatatagagcagacgTTCTTCAAGAATGGCTGGAGTGACCCTGGGGACTTGGTAGaagttatactgccc
gcagtcatatagagcagacgGGATGGAAGCCTTTGTGATTGTGCTGTTATTAACACATACaagttatactgccctg
gcagtcatatagagcagacgTAATTGCCAATGGTCCACACTTTTGTTTACTTGCCACGTGaagttatactgccctg
gcagtcatatagagcagacgTCTAAGTATCCTGGGATTTTATCGTCATTACCTTGGCACAaagttatactgccctg
gcagtcatatagagcagacgAATACTCAGTTCCTATTTGCGGCCGAATCATTGAATAAAAaagttatactgccctg
gcagtcatatagagcagacgATAGTTATATGCTCAGGTTATTGTAGCACCCTAAGAATGTaagttatactgccctg
gcagtcatatagagcagacgATTCTCTGAAACAGAGGATCTTTACCGGAGTTCCCCAGGGaagttatactgccc
gcagtcatatagagcagacgTAGGGAATCCATGAATTGGTTTGGGGTGGAAGGACTACAAaagttatactgccct
gcagtcatatagagcagacgCGTGGGTACATGTAGTTTGGGGAGGCAGATGATCAAAGTTaagttatactgccct
gcagtcatatagagcagacgTTTTTAGACAGAGTCTGGCCCTGACACCCAGGCTAGAGTGaagttatactgccc
gcagtcatatagagcagacgATGAGCCACCACACCCAACAAACAAATCCTTTTTTTTTTTaagttatactgccctg
gcagtcatatagagcagacgGCAGGTTTGTTACATATGCAGATTGAGTGTCTCTGGGGTTaagttatactgccctg
gcagtcatatagagcagacgCAAGTAGGCCCAGTGTTTATCGTTTCCTTCTTTGTGTCGAaagttatactgccctg
gcagtcatatagagcagacgAGCTCCCACTTACAAATAAGAACATTCAGTGTTTGGTTTTaagttatactgccctg

gcagtcatatagagcagacgGTCTTTGCTTTTGTGAACAGGGCTGTGATGAACATACGCAaagttatactgccct
gcagtcatatagagcagacgGAACAATTTATATTCCTTAAGGCATACACCCAGTAATGAGaagttatactgccct
gcagtcatatagagcagacgTTCCACAATAGCTGAACTGATTTACATTCCCACCGGCAGTaagttatactgccct
gcagtcatatagagcagacgGTCTTTGGAAAAGTTGTCTGTTCACATCCTTTAAAACAACaagttatactgccctg
gcagtcatatagagcagacgACAGTAGTTTTTAAAGTGTTCTCCCAGAGGAATAGCATCAaagttatactgccct
gcagtcatatagagcagacgGGCCCAATGGTCTATGTGAAGTCTCCAGTCTTCTGTTGCAaagttatactgccct
gcagtcatatagagcagacgGAAATAGGCTGTTATGTTTCCAAATGTAGAGGTAAGAAGTaagttatactgccct
gcagtcatatagagcagacgAATCAGTACTCAAAATACTTGTGAACATACTAAGGTCACCaagttatactgccct
gcagtcatatagagcagacgGTTAAAAGCAACCCAAAGTATCACATTGTACCTGGTAAATaagttatactgccct
gcagtcatatagagcagacgGGATTATGCCTCAATAAACCTGGTGGGGGGGAAAGTTTACaagttatactgccc
gcagtcatatagagcagacgGAAAACTTACTAACATTTCCAAAGCTCCCACTTCATCGTCaagttatactgccct
gcagtcatatagagcagacgCTAGAGTTTTACAGTTCTGCGAGGTTGCGAGGTCACGGGCaagttatactgccc
gcagtcatatagagcagacgAAGGTCATGAGCTGGTGGGGATCAGAGCAGGAGGAGGCCCaagttatactgc
gcagtcatatagagcagacgCTAAGCAGGATCAGGTCTCTCTCCACTGTGCGGTGTGTGCaagttatactgccc
gcagtcatatagagcagacgGTTCCAACCTACGTGCCTGACTATCATCTCTTATCCTTATaagttatactgccct
gcagtcatatagagcagacgAAGGCATTTGTTTTACTCTAGTAGGTTTTCATAAGGCTTCaagttatactgccctg
gcagtcatatagagcagacgTTTTGTTTCAGCTGCAAGAAGCAAGAGGATTGGGCAAAGGaagttatactgccc
gcagtcatatagagcagacgTGTAACAGGATATGCACAGTGAGACATGGCGGGGGAGAGCaagttatactgcc
gcagtcatatagagcagacgGACATCATAGTCACAGTCCTACATCTTCCTCTGTTTCCCCaagttatactgccct
gcagtcatatagagcagacgCCAAGTAGGGCTGGAAAGAGACACTCGGCAGGTAGATCTTaagttatactgcc
gcagtcatatagagcagacgGGAGAAGAAACTCATGGATTTCACCTAAGACCAGAGAAGGaagttatactgccc
gcagtcatatagagcagacgAGACTTGGAGGTTCCATCTCTCAAGCAACCTGAAATTAGCaagttatactgccc
gcagtcatatagagcagacgCATGATGTGGGGAAAAGGGAAGTGGTTTTGACGCAAGGGAaagttatactgccc
gcagtcatatagagcagacgGGAGTGTTGGCGGTGGCAGGGGGAGCAGGGGCAGCAGCTCaagttatactgc
gcagtcatatagagcagacgTGAAGTTCGACATCAGAATATACATTCGAGTAAGGGTGGTaagttatactgccct
gcagtcatatagagcagacgTGAGACTCACTTTCCTTGTCTTTATACATGAGGTTTAACAaagttatactgccctg
gcagtcatatagagcagacgCCATAAATATTGGTTCCCTTGGCTATATTCGTCCCCTCTTaagttatactgccctg
gcagtcatatagagcagacgCCTTGCATTGAGGCTATCTAGGTAATCTATTTAACAGCATaagttatactgccctg
gcagtcatatagagcagacgGAAACGTCTATATGTGCGCCTCTCTCTGTAATGTCCTGATaagttatactgccct
gcagtcatatagagcagacgCATCAGTGGTTCAGATATTCTCTCTATATTTGTGTTTGCAaagttatactgccctgt
gcagtcatatagagcagacgAATTTTCTCCCAGGAATTCCCGACCGAGGCCATCTGCTCTaagttatactgccc
gcagtcatatagagcagacgCCAGGAAGGCAGAGGCTGAGAGGAGGGAGTGTGGCATCAGaagttatactgc
gcagtcatatagagcagacgCGGGGCTCTGGCTTCCTCCTCCGGGATCAAGGGAAATGTGaagttatactgcc
gcagtcatatagagcagacgTGGGGGAACAGATAAAACACGAGTGAGGGGGAGATTTTGTaagttatactgccc
gcagtcatatagagcagacgCAGAAATGCTGTGTGCACATGGGCGTGGTGATTGATACTTaagttatactgccct
gcagtcatatagagcagacgTGAAGCAAACACCGTGAGACTACTCAGCATTGGGAGAAGCaagttatactgcc
gcagtcatatagagcagacgAGAATCCCAACTACTGACGAGAATGAGTAAAAGAATGCACaagttatactgccc
gcagtcatatagagcagacgGTTAAATACCCTGGCAGCTCTTGAAAACAAACCCCTTTTTaagttatactgccct
gcagtcatatagagcagacgTGCTCAGAAACCTCCGTTCTCATGGTCACTCGGCAACATGaagttatactgccc
gcagtcatatagagcagacgGAACAGCCATGCCCGGGTTCGATCCCAGTCCTAGCTTGTGaagttatactgcc
gcagtcatatagagcagacgCCTAATCTGTAAAACGTAGGGGCTGAAGGAGGAAGCGGGAaagttatactgcc
gcagtcatatagagcagacgTGGAGACCATTAGAATTAATGTATGAGGCCAGGCGCAGTGaagttatactgccc
gcagtcatatagagcagacgGGCAGGAGAATTGTTTGAATATGGGGGGCCAAGGTTGCAGaagttatactgccc
gcagtcatatagagcagacgATGTATAAAAAGTGAAGTCCCTGGTACCTGTTAAATGTTCaagttatactgccctg
gcagtcatatagagcagacgAAAGTGAGTTTTGCCCTCTCCCTTGTCCGTGGGTGAATAAaagttatactgccct
gcagtcatatagagcagacgTTCCTATGGACCATATCCTCAAATAGCACATACCTACCGGaagttatactgccct
gcagtcatatagagcagacgTCCATGTCACCCTCTGAGCAGGACACCAAAACCACATTTTaagttatactgccc
gcagtcatatagagcagacgCTGATGCAACATTCCAGGGCTATTACAAGTCAAATTCCAAaagttatactgccct
gcagtcatatagagcagacgGAGAATCTGGCTGGCTAAGGTTTTAAAATCTCTTGGTTTGaagttatactgccctg
gcagtcatatagagcagacgGATTTTACTTGTGGCATTACTAGCAAACTTTGTAAATCCCaagttatactgccctg
gcagtcatatagagcagacgTACTTTGGCAGGGTAGATGGATGGAATTGTTGGGAAAGCCaagttatactgccct
gcagtcatatagagcagacgAAGAAAAAATGTACAAAGGAAATGAGGAAAGGATAGGCAGaagttatactgccc

gcagtcatatagagcagacgGGTCCTGTGCCTGATATCTTCTTCCTCTTGGTGTGTTTCTaagttatactgccctg
gcagtcatatagagcagacgCCCTCAAGATAACAGGGGCATCTGAGAAATGTAGGAAGCAaagttatactgccc
gcagtcatatagagcagacgGGTTGCTTGGGGAAATATTTGTGGACTGCCTTCTTCTAAGaagttatactgccct
gcagtcatatagagcagacgGGGTTAACATCACGAAAGAGTTTTCCTTTACTCCCAATCTaagttatactgccct
gcagtcatatagagcagacgAATACCTACTATTGATTCCCTTCTTTCCAGTCCCTAGTAAaagttatactgccctg
gcagtcatatagagcagacgAAGATTACTTATGGCTTCAGTGCTAATTCTTGCCACATAAaagttatactgccctg
gcagtcatatagagcagacgTTTCACTTTCAAGATTGTGACACAAGACAAAGGGCAGCTCaagttatactgccc
gcagtcatatagagcagacgGACGTGATCATGTCAGGGGATCAGACAACCAGCTTTATGAaagttatactgccc
gcagtcatatagagcagacgCCAACAGATCATTGTGAATCTCTTTCAGACACTCTGCCACaagttatactgccc
gcagtcatatagagcagacgCTTCATTTTACCTAAACAAGAAAAGAAAAGAGGCCACCATaagttatactgccc
gcagtcatatagagcagacgAAGTCAGTGAATTATGGGGATTGAGTTTTTGCTGGGATGCaagttatactgccct
gcagtcatatagagcagacgTTCTATAGGAAAGAAGATGAAATGGACCATAGACAGCGAAaagttatactgccc
gcagtcatatagagcagacgGTTGTGATTAAGAAAATGTAGATCTCCAGCTTGGGCAGCAaagttatactgccc
gcagtcatatagagcagacgATTAAATTAGCCAGGCACCATGATGTGTGCTTGTAGTCCCaagttatactgccc
gcagtcatatagagcagacgCAGGAGTTGGAGACTGCAGAGAGCAAAGATCACAGCACTGaagttatactgcc
gcagtcatatagagcagacgGATCAACTTCAAAGACTTACTTTCTGGAGATCAGTTAAAGaagttatactgccct
gcagtcatatagagcagacgGGCACACATACACAAATCCTACGACACGCCTAAAACAGTGaagttatactgcc
gcagtcatatagagcagacgGATTTAGTTCGGAATATTTGCACAAATTAACTTCGGCTTCaagttatactgccctg
gcagtcatatagagcagacgAGGATTCTTATTCAATGCTTCATGAAGGGCACTATATTTGaagttatactgccctg
gcagtcatatagagcagacgAAACCATGTTTCCTCAATGTACAGGCTCCATATTGCTGAGaagttatactgccct
gcagtcatatagagcagacgGGGACAGTGAGAGCCCCAGGGAGCAGGAGACAATTAGAAAaagttatactgc
gcagtcatatagagcagacgCTGTTTATTTTCTCACATTCCTCCTGCCAGCCACCTCATAaagttatactgccct
gcagtcatatagagcagacgCAGAACTGAATGGAAGGTTGCAAATGAAAGAGGCTGGAGGaagttatactgcc
gcagtcatatagagcagacgGGCAGAAAGGTGGGATGGCTGAGAAGGGATTTAATGACAAaagttatactgcc
gcagtcatatagagcagacgTGGTGTCCACAGGGAAGGAAATGAGAGGAACAGGTTTACAaagttatactgcc
gcagtcatatagagcagacgCTTCTGGAAGTTTTTGACAAAAGATGGGTGACAGGGCCTGaagttatactgccc
gcagtcatatagagcagacgAAGGCATGTGACATACTTCCTCCCTCCCTACACTAAATGTaagttatactgccc
gcagtcatatagagcagacgGCTCACCTCCTATTGTATTTTGGCCCTTGTTTCTTTCTTCaagttatactgccctg
gcagtcatatagagcagacgTTCTTCTTCTTCTTCCTCATCGTCGTCTTCTTCTTCTTCTaagttatactgccctgt
gcagtcatatagagcagacgTAGCTGGGAGTAGCTGAGTAGCTACAGGTGCATGCTACCAaagttatactgcc
gcagtcatatagagcagacgCCACCACATCTGGCTGCATTACTGGCTATGAGAGAGAAGGaagttatactgcc
gcagtcatatagagcagacgCTAAAATTTAAATACCCACACATCAGAGGCAGTGTCCATTaagttatactgccct
gcagtcatatagagcagacgTGGTTGCCATAGAAATCTTTAGTTGAAATTATCTGAACGCaagttatactgccctg
gcagtcatatagagcagacgCTGGCTAGGCTGGGACTTAGATTTCACCGAGAGGAACTTTaagttatactgccc
gcagtcatatagagcagacgGGCTTAGCTTCCAGAACATGAGCCTGGCTAGGGATTTATAaagttatactgccc
gcagtcatatagagcagacgGGAAAATATTTGCAATCTGTGACAAAGGCAAATCACCTTAaagttatactgccct
gcagtcatatagagcagacgCAACATGATAAAATCGCAAATGAAGAAAAACACTCCACACaagttatactgccc
gcagtcatatagagcagacgTTGTTTCCTTTTTTTAGTGATGAAATTCAGTATTGGCCGGaagttatactgccctg
gcagtcatatagagcagacgTGTAAACTGGCAAACCTTTTCTGGAGAGACATTTGGAATAaagttatactgccct
gcagtcatatagagcagacgGCTCTTCAAAGCAGTATTTCTCATTATATGCACCACAGAGaagttatactgccct
gcagtcatatagagcagacgAGGTAAGCTACGGTACATCCACATGGCGCCATTAAAATGTaagttatactgccc
gcagtcatatagagcagacgTAAAAAATGTATAGAAGGACCTGAAGAACCGGATTCTTAGaagttatactgccct
gcagtcatatagagcagacgTTCAAGTGATTCTTCTGCCTCGGCCTCTGGAGTAGCTGGGaagttatactgccc
gcagtcatatagagcagacgCCGTGCTGGCTTAAATATTTAATAGGAAGAATTTCCCTAGaagttatactgccct
gcagtcatatagagcagacgTTTTTGATGGAATCGCATTCTGTCGCCAGGCTGGAGTGCAaagttatactgccc
gcagtcatatagagcagacgCTGGGATCACAGGTGCTGGACCACGCCCAGCTAATTTTTTaagttatactgccc
gcagtcatatagagcagacgGCCCACCACCATGCCCGGCTAATTTCTTCTAGTAGAGACGaagttatactgcc
gcagtcatatagagcagacgCCACCCCGCCCGGCCTTATGAATATTTATTAAACTCCTCCaagttatactgccc
gcagtcatatagagcagacgAGAAGTGAATAAGGCAAACGTCTAGCTTTTATATCCCTTTaagttatactgccctg
gcagtcatatagagcagacgTAAGCAAAACCAACAGAAATAAGAGGGTGATAGTTTCATGaagttatactgccc
gcagtcatatagagcagacgCAGAGATTGTAAGGAGGTCAGGGGGCAACATTACGTACAAaagttatactgcc
gcagtcatatagagcagacgAAATGATATTTGAGTTGGGACCAGAGTGACGAAAAGGATTaagttatactgccct

gcagtcatatagagcagacgCCTGGGGTGGGAACAGCACTGAAATACTTATGGAACAGAAaagttatactgccc
gcagtcatatagagcagacgACAGCAAGAGAGTAAGAAATGAAGTCAGAGTTAGGCAGGCaagttatactgccc
gcagtcatatagagcagacgTCACTTACACTCTTAGGTCAATGGAAGGCAATGGGAAGCCaagttatactgccc
gcagtcatatagagcagacgTCTTCTTTAATAAAACAATTCTGTGGGACAGATCTGCTACaagttatactgccctg
gcagtcatatagagcagacgGTCACACAGCTGATAGGTAGCAGGATTGGGACTAGAGCCTaagttatactgccc
gcagtcatatagagcagacgTTACTAGGCATGAATAGAAACACCGCACTGCAACACTGAAaagttatactgccc
gcagtcatatagagcagacgATTACCAAATGAGTGAGCCGATTGAGCAAGTGAGTTCTCCaagttatactgccct
gcagtcatatagagcagacgCATTGGCTTTGAAGACGGTAACCGGGAAAACTCAGACTGGaagttatactgccc
gcagtcatatagagcagacgAGGACTGGAGACTCTGAGAAGCTGGTTGCAACATTTGGCGaagttatactgccc
gcagtcatatagagcagacgCTATAAGCCTTTTGAGAGATGAGCCTGAGTTTGTGGCGGCaagttatactgccct
gcagtcatatagagcagacgCTGGAGGAGGATTGAGCATTATTCCAAAAAAGAAGGGTGGaagttatactgccc
gcagtcatatagagcagacgCAATGTTTGTTTTGGACTTTAAGCTTCTCTTTCACCCTTCaagttatactgccctgt
gcagtcatatagagcagacgCAGAGCCTTTTCCAGCAGCTGCAGAAGCCGCCTCACAACAaagttatactgcc
gcagtcatatagagcagacgCTGGTGCTTCTTCCTCTCTGTGTGGGATGTTGCCTCTTACaagttatactgccctg
gcagtcatatagagcagacgTGGGGAAGCATAGGGTACCCTCAGTGGTCAGTTTTATCTCaagttatactgccct
gcagtcatatagagcagacgAGAATGCTGCTCTTGATTAGCTGGACCTTTCCCAGTCTGGaagttatactgccct
gcagtcatatagagcagacgACAGAGTCCTTCAAACACATGGCTCTTTGAGATTTGAGTTaagttatactgccctg
gcagtcatatagagcagacgTTTTTCTTTAGAGATGGGGTTTTGTACTGTCACCCAGGCTaagttatactgccctg
gcagtcatatagagcagacgGCTCGCTGCAGCCTCCAACTCATGGATTCAAGCAATCCTCaagttatactgccc
gcagtcatatagagcagacgTGAGCCACTGCAGCCGGCTTCCAGACTTCTTAAATAAGGAaagttatactgccc
gcagtcatatagagcagacgGGTACCTGGAGTAGTGAAATGTATCAAGACAGAAAGTAGAaagttatactgccct
gcagtcatatagagcagacgAACTTAAAAAATAGTTAAACCAACCTGGCTGACACAGTGAaagttatactgccct
gcagtcatatagagcagacgAGATGGCCAGTTTTCTGTTATGTGTTTTTTAAAACCATGGaagttatactgccctgt
gcagtcatatagagcagacgGAGTGGACCTGATCTCCCCATAGTTTGCTGCCCTCTTGTCaagttatactgccc
gcagtcatatagagcagacgATCTTTGGTAATGAGATGCCTTGACCTGCCACTTGATGTCaagttatactgccctg
gcagtcatatagagcagacgCAGTCCACCGCCTCAAAACACTCCACTGATGCCTCCAACCaagttatactgcc
gcagtcatatagagcagacgCATTTTCTGGTCTTCCTCTCAGCTTCTCTCTTGGACACAAaagttatactgccctg
gcagtcatatagagcagacgCCACCTCGATGTCCTGGTTTCTGCCTAACTTAATAAACATaagttatactgccctg
gcagtcatatagagcagacgCACCTCTTCTCAAATGACTAACTGTTGGTGTAGCCCAGGGaagttatactgccc
gcagtcatatagagcagacgTCCTTGGCCTGCTTCTCTTTTCCCTCAGCATTTTCTCTCTaagttatactgccctg
gcagtcatatagagcagacgCAGATAAGTCAACAAGATAATTCCCAGATCTGTGCTTTCAaagttatactgccctg
gcagtcatatagagcagacgAGTTTTAAATTATGCCTCCAACTGCCTTCTTGACCTCTCCaagttatactgccctg
gcagtcatatagagcagacgAATTTTCAATTCATTGGGGGTATAAACATGAAGTCGGTTCaagttatactgccctg
gcagtcatatagagcagacgATAACTGCCCTGATTATGCTAGTCTCAATACTCAAGAATCaagttatactgccctg
gcagtcatatagagcagacgGTGTGATCTCGACTCACTGCAACCACCATCTCCCAGGTTCaagttatactgccc
gcagtcatatagagcagacgACTGCGCCCGGCCGATGTTGAGATTACTAAAGTAGTCTCAaagttatactgccc
gcagtcatatagagcagacgTAACAGAAAGCAAGGAATCGATAAAAAAAATCTTAGGCCCaagttatactgccct
gcagtcatatagagcagacgAGGAGCCAGCATGAAGTCTCCTTCTGTCTAAAGATGGAAAaagttatactgccc
gcagtcatatagagcagacgGTTCACAATGAAACTTCAAAAAACTCTCTCATTTGTCCTCaagttatactgccctg
gcagtcatatagagcagacgAGGATGACTAGGCAACCAACACATTATTTTGAAAACTTGTaagttatactgccctg
gcagtcatatagagcagacgAAATAAATGTTTCCAGCATAAAATGAGCCTCAGGACAACCaagttatactgccct
gcagtcatatagagcagacgGGATTAAATTGGAATATCACATTTTTTGGAACCTCTGGTGaagttatactgccctg
gcagtcatatagagcagacgGCTAAAAAGACAACTGTAGAAAAAAGCATAAGGGAAAAGCaagttatactgccc
gcagtcatatagagcagacgGGATTTCCTCAGTATGATACCAAAAGAACAGTCAACAAAAaagttatactgccct
gcagtcatatagagcagacgAAAAGCTTCTGTGTATCAAAAGACACAATCGAAAGAGTGAaagttatactgccct
gcagtcatatagagcagacgGGTAGAACATATTTGCAATTCATATAAGGGGTTATAAGGGaagttatactgccctg
gcagtcatatagagcagacgGCATATGAAAGATGCTCAGCGTCACCGTCACTAATCATTAaagttatactgccct
gcagtcatatagagcagacgCCTGTGAGGATGGCTAATCAAAAAACAAAAAAAGTACGCAaagttatactgccc
gcagtcatatagagcagacgAAAACAAGGGTTGGAGCAGAGGCAGAGAAATTGGAACGCTaagttatactgccc
gcagtcatatagagcagacgAAATAGAATTGCCTTAGGATCCAGTCATTTGATTTCTAGGaagttatactgccctg
gcagtcatatagagcagacgCATGTTTATAGCAGCATTATAGTCAAAAGGTGTAAACGGCaagttatactgccctg
gcagtcatatagagcagacgGATAAACAAAATGAGGTACACACATACAATGGAGACCCTGaagttatactgccc

gcagtcatatagagcagacgACACATGCTACAACATGAATCCTGAGAACACTATGATAAGaagttatactgccc
gcagtcatatagagcagacgCATGTGCTGGAATGGGGTCAATGAGGGGCTTTTGTTTAATaagttatactgccct
gcagtcatatagagcagacgTAGTTCTGCAAGATGGAAAGGTTCTGGAGAGGGATGGTGGaagttatactgccc
gcagtcatatagagcagacgTGCAAAACAGTGTGAAGATACTAACGCTACTGAACCGTACaagttatactgccc
gcagtcatatagagcagacgAAATCTGATCAAGCCTCTCCATCTAACTACCAATTTATAGaagttatactgccct
gcagtcatatagagcagacgGTGGGTCAGAGGAACATCTTAACAAGGTCGCTTGGAAGCAaagttatactgcc
gcagtcatatagagcagacgCTGTGGGAAACTCTACAGGAAAAACAACCAAATTTCTTTAaagttatactgccct
gcagtcatatagagcagacgTTTTTGAGACACAGTTTCACTCTTGTTTCCCAGGCTGGAGaagttatactgccct
gcagtcatatagagcagacgATGATCTCAGGTCACCACAACCTGTGCCTCCCGGATTCAAaagttatactgcc
gcagtcatatagagcagacgCCATGCCCAGCCAAGGACTCTTAAAAGATATAAACATCAAaagttatactgccc
gcagtcatatagagcagacgAAATTATGGCACAATCTGGGAACATCTGATAATATTCAGGaagttatactgccct
gcagtcatatagagcagacgATTTATGGTAAAATAGCCTGACGTCTGAGACCTGCTTAAAaagttatactgccct
gcagtcatatagagcagacgTTGAGGTGGGAAAGTTGGAGGGTAAGTATGGATGAGCAAAaagttatactgccc
gcagtcatatagagcagacgGGCGGGAGTCAGCACACTATTCTTTCTGTTTTGTGAAATAaagttatactgccct
gcagtcatatagagcagacgTCCCCCTTTTGGCTGTCTCTCATATCTTTCTCCACATCTGaagttatactgccct
gcagtcatatagagcagacgACTCTGCAGTTGCTTATCTTTGAGTCTGTCCTTTCATTTAaagttatactgccctgt
gcagtcatatagagcagacgCCTTTTAAAATAGTAACCTTAGATTCTGCTTGAAGTCAGCaagttatactgccctg
gcagtcatatagagcagacgTGACATAGGAGACAAAGTCATGGCGGGTACACATTTTTCCaagttatactgccc
gcagtcatatagagcagacgCTGGGTCGCAGCAGAAGGTGGAACAGAGCAGCAGGGACAGaagttatactgc
gcagtcatatagagcagacgGAGGCAGTCTCCTCTCCCTGGATCGCTGTAGCCTTGTGGGaagttatactgcc
gcagtcatatagagcagacgCAGGGGTCACATGCTGGATCTCTCCTGGAACCAATCTTTGaagttatactgccc
gcagtcatatagagcagacgCAGCTCTGCGGCGTGAAGATCGGGTTCTGGCTGGCGTTCAaagttatactgcc
gcagtcatatagagcagacgTCACCACAGCTCCTTGGTTGGCCCTGCATGAAGGACAGTGaagttatactgcc
gcagtcatatagagcagacgAGCTTAAGTCTAGCCACCTCTCAGCTGTCTAGAAAACCCAaagttatactgccc
gcagtcatatagagcagacgATTTCTGTGTCAAATGGAGGAAGTGCAGGCTGCCCAAGCGaagttatactgcc
gcagtcatatagagcagacgCCCTCTTTTGGCTGTCTCTCCTACACCTTTCTCCACAACTaagttatactgccct
gcagtcatatagagcagacgACTCTGCAGTTGCTTATCTTGGAGTCCGTCCTTTCATTTAaagttatactgccctg
gcagtcatatagagcagacgGCTGCACGTAAAATATGATCAAACTGAGCATACAATTAAGaagttatactgccct
gcagtcatatagagcagacgCCAAAGAGGGAGAGTCAGGACTTCATGAGATACACAGAATaagttatactgcc
gcagtcatatagagcagacgCAATAAGCTTGTTTCTAATTTCTCTCTGTGTTCTCTGTTCaagttatactgccctgt
gcagtcatatagagcagacgCTTTTCTTTTAAAGACAGGGTCTTGCCCCTTTGCCCAGGCaagttatactgccct
gcagtcatatagagcagacgTCCTGCCTGGGCTTCCCAATTTGTTGTGATTACAGGCGTGaagttatactgccc
gcagtcatatagagcagacgCTGTGCCTGGCCTAATGTTATTTTTTAAATTACTCAGTGTaagttatactgccctgt
gcagtcatatagagcagacgTTTATGAATCTTCCTACTATGAAAGTGTGTTTCCTAGCACaagttatactgccctg
gcagtcatatagagcagacgCATAAAACGCATATGAGACAGTGCCACAAAGAAGATAACAaagttatactgccc
gcagtcatatagagcagacgTCTTGCTACCAGCACAAGTCTGTTGTATATGTGATATATAaagttatactgccctg
gcagtcatatagagcagacgTACCACATGTAAGTCCCCTAACAATCCCACACGATTTGCAaagttatactgccc
gcagtcatatagagcagacgAGAGATGAGGAAAGCAAAGCTCTGAGAAATTACTTTAGGAaagttatactgccc
gcagtcatatagagcagacgAGCCAGGATTCAACCCAACACTGTATTGCTTAAATCTCAAaagttatactgccct
gcagtcatatagagcagacgACAACCATATTTAATAAGAGCACACCATGTACCAAGCAGTaagttatactgccc
gcagtcatatagagcagacgCCCAAGGGAATCATCACGAGTTATATTTTCTAAATATAGGaagttatactgccct
gcagtcatatagagcagacgACATGCTGCTCAGTCAAATTCCTTATCTCCCTATATCTTAaagttatactgccctg
gcagtcatatagagcagacgAACCAGGGAATGATTTGTTGCTTTTGTTTGCTAGAGGAAAaagttatactgccctg
gcagtcatatagagcagacgGGCAAAACCTATTTCTCCTTAAGGGAAAAAATAGATTTGGaagttatactgccct
gcagtcatatagagcagacgGAAGTTAATTCAGACCTTCTATGTCTTGCCTAAATAAAGCaagttatactgccctg
gcagtcatatagagcagacgATCGTATTAGTTACCTGCTTCTGCATAACAAGTTATTCCAaagttatactgccctg
gcagtcatatagagcagacgTCAGGAATTTGGGAATGACTTAGCTGGTGGTTGTGGCTCAaagttatactgccct
gcagtcatatagagcagacgGGGGGATCTGCTTCTAAGATGGCACACTCATGTGTTAGGCaagttatactgccc
gcagtcatatagagcagacgACGCCTCAATTTCTCATCACGTGGACCTCCCTATACAGCTaagttatactgccc
gcagtcatatagagcagacgAGTGTTCTCATGACAAGGTAGCTAATTTTCCCAAAGTAAGaagttatactgccct
gcagtcatatagagcagacgGGTCTTTTTATAACCTAACCTCTGAAGTAACACAGTCATTaagttatactgccctg
gcagtcatatagagcagacgCCATCTTAGAGGCTGGTTATCACAGTAATCAATGTGCATTaagttatactgccct

gcagtcatatagagcagacgAATTTGTTGCTAATTTCTAAGGATCATGTCGCAAGTCAGGaagttatactgccctg
gcagtcatatagagcagacgTCGAGAGTAACTGGAATGAAATTGTTGACAGCTTTGATGAaagttatactgccctg
gcagtcatatagagcagacgCATGGCATCTACGCCTATGGTTCTGAGAAGCCCTCTGCCAaagttatactgccc
gcagtcatatagagcagacgGGGTTATGTTGTGATCGTTCAAGCCCAATCTGGCACTGGGaagttatactgccc
gcagtcatatagagcagacgAATTCTGCATCAGATTGAATTAGATGTAAAAGCCACCCAGaagttatactgccct
gcagtcatatagagcagacgGGTCCTAGCACCCACTTGAGAATCGGCTCAGCAGATACAGaagttatactgcc
gcagtcatatagagcagacgGGAGACCACATGGATGCCTCCTGTCACACCTGTATTGGGGaagttatactgcc
gcagtcatatagagcagacgCAGATGGAAGCTCCCCATATAATTGTGGGTACCCCTGGCCaagttatactgcc
gcagtcatatagagcagacgTTGTGCTGGACAAAACTGACGAAATGTTAAGCCATGGATTaagttatactgccct
gcagtcatatagagcagacgCAACACCCAGGTAGTTTTGCCATCAGCTACAATGCCTTCTaagttatactgccc
gcagtcatatagagcagacgTTCATGAGGGAACACATTTGGATTCTGGTCAAGAAGGAAGaagttatactgccct
gcagtcatatagagcagacgGGTATCTGCCAATTCTACATCAACATGGAACGAGAAGAGTaagttatactgccc
gcagtcatatagagcagacgTTGTATGAAACCCTGACCATTGCCCAGGCAGTCATCTTCCaagttatactgccc
gcagtcatatagagcagacgCTTACCAAGAAGATGCATGCTCAAGATTTCACTGTCTCTGaagttatactgccct
gcagtcatatagagcagacgGGAGATATGGACCAGAAGGAATGAGACGGGATCATGAGGGaagttatactgcc
gcagtcatatagagcagacgTCTAGCAGAGTTTTGATTATCACTGACCTGCTGGCCAGAGaagttatactgccct
gcagtcatatagagcagacgCACAGAATCGGTCGACGTGGACAGTTTGGCCGTAAAGGTGaagttatactgcc
gcagtcatatagagcagacgCTCTTTGAGACATCGAGACCTTCTATAATACCTCCATTGAaagttatactgccctg
gcagtcatatagagcagacgGCTGTCCTGCTTCCTAGCCCCAGCCAGAGTTCAATCTTGGaagttatactgccc
gcagtcatatagagcagacgGGGAGCCAAGGGATGGACATTTTGTCATTTTTTTCTGTGAaagttatactgccctg
gcagtcatatagagcagacgCTCATATGGAGAGCATAAAACACACTGTGGCATATATTCTaagttatactgccctg
gcagtcatatagagcagacgATGGTCCTCAATACATCACATCTCCCTGAACTCATGTCCTaagttatactgccct
gcagtcatatagagcagacgATGCTAGGTTTGGCGGTGTGACTTGCTTCAATCAATGGGAaagttatactgccct
gcagtcatatagagcagacgAAAGCAAATTTACAGTGGGACCTACTCTCTCTAGCTGGTGaagttatactgccct
gcagtcatatagagcagacgTCTTCCTTCAATATGTAAACAGCCCAAGCTAGCCTCCTAGaagttatactgccct
gcagtcatatagagcagacgGGAGACTCAGCCAGCCCAGCCTTTCTAGCCAACAAGACCTaagttatactgcc
gcagtcatatagagcagacgTGATTGAGCACCGCCAACACCACTCAGAGCACAGATTAGCaagttatactgcc
gcagtcatatagagcagacgAGCACAGTCCAAATTACTAACCCATAAATTAGTGAGCAAAaagttatactgccct
gcagtcatatagagcagacgCTGTGTTTTGAGATCAGGGGTAGGTAAATGGCATTACTGTaagttatactgccctg
gcagtcatatagagcagacgCAGGTAGTGGTTGTGAGGGAGGAGGATCTGGGAGAAGACCaagttatactgcc
gcagtcatatagagcagacgTGACAGAAATAGCCTCATTGGGAATCAACTGCACTTCATCaagttatactgccct
gcagtcatatagagcagacgCAAAAGCCTTTCACTGTGTACTTAATGGAATCTTCTGTAAaagttatactgccctg
gcagtcatatagagcagacgTTTCCATGTACCTGAACACTACCTGGGTCACTGCTACAGGaagttatactgccc
gcagtcatatagagcagacgGACCCTTTAAATATTTTTTAGTCCATACATGCTAGCCTGTaagttatactgccctg
gcagtcatatagagcagacgTTCCTCAAAGCAATCCATTCTCACCAACATATAGGAAAGAaagttatactgccct
gcagtcatatagagcagacgATTTTTCTCTATGGAAATATTACTCAGCTCCTTCATGGGGaagttatactgccctg
gcagtcatatagagcagacgGGATGTTTCTACCCGTTTCATTGCACTTGCTATCTTTCTTaagttatactgccctg
gcagtcatatagagcagacgATTGAGCTAATGGTCTAATAGAAGAGATGGATTTTTGCTGaagttatactgccctg
gcagtcatatagagcagacgAGAAAGCTCTGAGGCTCTTTATTCTAAGCAAGTGTGGTGAaagttatactgccctg
gcagtcatatagagcagacgGCTTAGGGAGGCATCAAGAAACAGAAAATCAGACAGACATaagttatactgccc
gcagtcatatagagcagacgCCTCTCTGAGTCTCATCTAGAAAATAAGGACGAAGCTATGaagttatactgccct
gcagtcatatagagcagacgAGGTAAGAATCCCTAAAGTGCTGTGTTTAAATCGGGGCCCaagttatactgccc
gcagtcatatagagcagacgGGCTCACTCCACTTTCCCTTCTTTCCCTTTAGTCACCATAaagttatactgccctg
gcagtcatatagagcagacgGTGACCACAGGCCAGAACCCAAGAAAAAACAGAACATGATaagttatactgcc
gcagtcatatagagcagacgAGGATCCCTTATTCTAGAAGTTTGCAGTGTCAGATGGAGCaagttatactgccct
gcagtcatatagagcagacgGCAAACCAGCCCGAGACAAACAACAAATGTATTAATTAGTaagttatactgccct
gcagtcatatagagcagacgAATGAATAAAGAAATGGTGCAAGAGTTTTTGGTGTGGTTGaagttatactgccctg
gcagtcatatagagcagacgCCGTGCATGGCAGGTCCAGACAGGATGGGTTTTGGGGGTGaagttatactgcc
gcagtcatatagagcagacgGAAGCGTGATTTTAGAATTATTTGAATATTGGTAGGAGGGaagttatactgccctg
gcagtcatatagagcagacgGACACATTTCAAGGGACAGTCAGAGTAGAAGGAAGCAAACaagttatactgccc
gcagtcatatagagcagacgGATGGGGTCTGTGATTTTCTGAAAAGTTGGCAAATTGAGTaagttatactgccctg
gcagtcatatagagcagacgGGTATCAGTTCAGTTTAGGGCCCAGACTTAGAGTGAGGTGaagttatactgccc

gcagtcatatagagcagacgCATGAGGAAAAGATAAAAGGATGGGGCTAGTTTACTGGGGaagttatactgccc
gcagtcatatagagcagacgATTTTCTTAATGATGGCTCTGGAAAACATGTAGTTGCTGCaagttatactgccctg
gcagtcatatagagcagacgTCAGCAGAAGAATATAGAAGAGACAAGGAAAAATCAGTGTaagttatactgccct
gcagtcatatagagcagacgGCAACAAAAGCACAGTTTAATCAGAGGCCCACAAAATTACaagttatactgccc
gcagtcatatagagcagacgGAGAAACCCGCTCTGTCTTCATGCCTCATTTGTTCATAATaagttatactgccctg
gcagtcatatagagcagacgGTCATAATACAGACTATAACCTTAGCATACTCAACACCCAaagttatactgccct
gcagtcatatagagcagacgCCATGATATATTGTTCAGGTCCCATCAAGCTTAAAGTTTCaagttatactgccctg
gcagtcatatagagcagacgTCCTGGAATTTTCTACTGGATTTCCCATAGACTCTGCTGCaagttatactgccctg
gcagtcatatagagcagacgTGTTCCGCTCAGACAGATCTTTGCCTTTCTGCAAACCAGCaagttatactgccct
gcagtcatatagagcagacgGCAACTGAGCTGCTGGAGAGATATTCCCCTGTCACAGAATaagttatactgccc
gcagtcatatagagcagacgCTAGGAGCAAACATATTCACACTCATAATTTTTGGGGGTCaagttatactgccctg
gcagtcatatagagcagacgCACCAAATCCTGTCTCTTCTTCCTCCCAGCATACAGTGAGaagttatactgccc
gcagtcatatagagcagacgAGTTGGAGCTGTATGATGAATCCTGGGGAATTAGGAACGAaagttatactgccct
gcagtcatatagagcagacgCAAGCCTGGCCCTTAAAAAAACTCCTGCAAGATCCTCCATaagttatactgccc
gcagtcatatagagcagacgATCCAGAAGATCTAGTGAAGGACAATAAGCCCTTGGGGACaagttatactgccc
gcagtcatatagagcagacgCTGGTCAGCAGCTAAGGGAAAAAAGAGGACTATAAAGCCCaagttatactgcc
gcagtcatatagagcagacgCCTCTTGCTGGAAGGAGACTGAGTCCTGAGTACGTGCATGaagttatactgccc
gcagtcatatagagcagacgCGATTCACATGGGACTACGATAAGAGAAAAATAAACTTTGaagttatactgccctg
gcagtcatatagagcagacgTGTTTGTTACAGTACTTGGTCTACCCTGGCTAATGCACATaagttatactgccctg
gcagtcatatagagcagacgCTGGGCTCTATACACCAGTCTGTGTTAAAATGGGTGCCTCaagttatactgccct
gcagtcatatagagcagacgCTGAGCTATGAGGCCAGAAATGATTAATGGAAATAAATAGaagttatactgccctg
gcagtcatatagagcagacgAAGAAATTAGGGGTAGCATCAGGTGAAAAAGTAAGGTAATaagttatactgccct
gcagtcatatagagcagacgAGAATAAAGTTGGATCTTTCCCTTACACTGTACACAAAAGaagttatactgccctg
gcagtcatatagagcagacgTCACAGACCTAAATTTAAGGGCTAAAGCTACAAAACTCCTaagttatactgccct
gcagtcatatagagcagacgAGCCTTCTTAGATATATAATGCCAAAAGCACAACAACAACaagttatactgccct
gcagtcatatagagcagacgTTTCATAAGCAACTTGAATCCAGAAGTGTTACAACTCAACaagttatactgccctg
gcagtcatatagagcagacgCACACCCACTAGGATGCCTAAAATAAAGAAATCAGATAGCaagttatactgccc
gcagtcatatagagcagacgGGTGCAGCCTCTTTGGAAAATATTTTGCAGTTCCTCAAATaagttatactgccctg
gcagtcatatagagcagacgACACAGAGTTAGCATATTACCCCCACAATTTCACTCCTCGaagttatactgccct
gcagtcatatagagcagacgTGCCCACATAAAATCTTGTACACGAATACTCATAGCAGCAaagttatactgccct
gcagtcatatagagcagacgAAAAAGGTAGTATATCCAACCAAGGGAATATTATGCAGCAaagttatactgccct
gcagtcatatagagcagacgCATTATATTATGTCTAAGAAGTGGGTCAGAAAAAGCCATCaagttatactgccctg
gcagtcatatagagcagacgCAGATAGACAAATCTCTGGAGACCAAAAGAAGACTAGTGGaagttatactgccc
gcagtcatatagagcagacgGAGGGGAGAGGAGAGGGAAGACTAAGGGGAAATGGAGAGTaagttatactgcc
gcagtcatatagagcagacgATCTCCTAACGCTATCCCTCCCCCACACCCCCACCACACAaagttatactgcc
gcagtcatatagagcagacgTATTCCACGGTGCATATGTGACACATTCTCTTAATCCAATaagttatactgccctg
gcagtcatatagagcagacgATTACAGGCATGCACTACCACGCCTGGCAATTTTTGTATTaagttatactgccctg
gcagtcatatagagcagacgGAGCAACTGCGCCCAGCCCCAGGGTTTCTTTTTAGGGTGTaagttatactgccc
gcagtcatatagagcagacgTCTAAAATTGATTGTGGTGTCAGCTGCACAACTTTGTGAAaagttatactgccctg
gcagtcatatagagcagacgGAAATATTTCCAACCACAGAATGTCATAATGCCCCAATACaagttatactgccct
gcagtcatatagagcagacgCACACATTGCTCAGGATAGCTGTCCTCAGAGGCCACCAAGaagttatactgcc
gcagtcatatagagcagacgCCCCAAGGCCCATGAAAACTTAAGTAAATCTATTCCTAAGaagttatactgccct
gcagtcatatagagcagacgCAGTGATGTGCTAGTGAATGTTTAACAGCTGACTCTGGTGaagttatactgccctg
gcagtcatatagagcagacgGGAAAAACATGAGGCCACTAAGCATAGGGTTGAGGAAGAGaagttatactgccc
gcagtcatatagagcagacgACACCATACAGATTTAACAGACATAAGTAACCTCAAAAGCaagttatactgccct
gcagtcatatagagcagacgGTATAATTTTTAGTAACAGATGTGTTTATCAGCCGCTTGGaagttatactgccctgt
gcagtcatatagagcagacgCAGCGACCACTGGGCCTCTTAAGTTTATTTCAGTCTGTGGaagttatactgccct
gcagtcatatagagcagacgAGGAACAAATGAGCAAAGTTTCTCTCTCTTCTTACACTCCaagttatactgccctg
gcagtcatatagagcagacgGCATGCCTTTCCCTTGAATCAACAGAACTCCCCTGCCTCCaagttatactgccc
gcagtcatatagagcagacgAGACGGTAAATTCATTGGGAGTTTCCTTTTCCTTCTTATTaagttatactgccctgt
gcagtcatatagagcagacgGTTCTACTCACCATATTCACCTTCATCCCCTAATACTCTGaagttatactgccctg
gcagtcatatagagcagacgCCAAGTAAAAATAACAAGGGAACAAGAAGAGAGCAGGCAGaagttatactgcc

gcagtcatatagagcagacgAAGGATGATGCTCAGGTCTCTTCAGTTCACAATCAATACAaagttatactgccct
gcagtcatatagagcagacgAAGTAATTTGGCTTGAATCTATCACCTCCCCACCCAGAGTaagttatactgccct
gcagtcatatagagcagacgTTTCAGGTTGTTGGCCCTAAAAGCTTGAGGTAATATGTTTaagttatactgccctg
gcagtcatatagagcagacgTTGTCAAAACAAATTGAACCTTTTAGTACAGAGCGAGGGCaagttatactgccct
gcagtcatatagagcagacgGGCCTCTCCCTTCTTCTCCTAGTGAGCTGGTGCCCATCCCaagttatactgccc
gcagtcatatagagcagacgCAGTCCTACCTCAATGCCCCCACAGCTCTTTGTTTACTTTaagttatactgccct
gcagtcatatagagcagacgAGGTAGCTAGATATACAGTCATGAAATGAATAAGTGTCTGaagttatactgccct
gcagtcatatagagcagacgGAACTCATCACTTTGACTTCTGGCTCCTCATGTACCTCATaagttatactgccct
gcagtcatatagagcagacgTGAGGAGTCCGTACCACACACCATGATGGGTTTCTAGAGCaagttatactgccc
gcagtcatatagagcagacgTGTTGAATGAATGAAATGGATCTATAGGGGAGGCTATTCCaagttatactgccct
gcagtcatatagagcagacgTGAAAGTCTTTAATCTTCCAGAATCCAGCCAGAAATATCCaagttatactgccct
gcagtcatatagagcagacgGAAATCTTCCCCATGACTTACTCACGTATCCTTAAGCCACaagttatactgccct
gcagtcatatagagcagacgCAGATCCGTGAATTAATACGTAAATCAGTAACTTAAGAGCaagttatactgccct
gcagtcatatagagcagacgAGAGACACCCTGAAGAGAGAGCAAAATGATTGTTTAAAAGaagttatactgccct
gcagtcatatagagcagacgGGTCTATTTTCAACCAGCCATTGCATAAGAACTTAAGAAAaagttatactgccct
gcagtcatatagagcagacgGGTATTTTCCAAACCCTTTCAGCCATGCCTATGTATCATCaagttatactgccct
gcagtcatatagagcagacgGGGACCCAAAATCAAACACCCGCCAAGAGATGCATAAACAaagttatactgcc
gcagtcatatagagcagacgCACACTTACACATGTGCAGTCACCGATGACCTGGAGGACCaagttatactgcc
gcagtcatatagagcagacgAAGTATTTCATAGGCAACACTGAATAGCAATTTTACAGGCaagttatactgccct
gcagtcatatagagcagacgCCCACTGGACTTTTATTAATTAGGTTCATTCTCTCATAACaagttatactgccctg
gcagtcatatagagcagacgAGGTTATCTGATGTAGTGAATGCCTCTTTTTGTCTATAGAaagttatactgccctg
gcagtcatatagagcagacgGGTCGTTGTATATGTCCTGTCCAAAGTAGCTTTAAATCAAaagttatactgccctg
gcagtcatatagagcagacgCATCACATGACTCTACGAACCACTAGATGTCTATAGCAATaagttatactgccct
gcagtcatatagagcagacgCTATCTTCATGGTCATTCAAGAGATGGAGCTTTAGGTACTaagttatactgccct
gcagtcatatagagcagacgCTTTTAGAGATAGGAGGGCTTAGAATTAGAGGAGAGTTGAaagttatactgccct
gcagtcatatagagcagacgCAAAAAGTAAGCAAGGAAACTGAGTATTGTATAGGCACGGaagttatactgccc
gcagtcatatagagcagacgAAAGATTTTGATACTCATATGGGGTCCATCAGCATCCTTAaagttatactgccct
gcagtcatatagagcagacgCCTTTGCAAATCTACTCTCCATCTTTCTTCTGCTGTTTGTaagttatactgccctg
gcagtcatatagagcagacgGAGGCTGATCTGCATGGGCTACGTCAGTGGTTCTCAAAGTaagttatactgccc
gcagtcatatagagcagacgGAACCTACTGAATCAGAAACGGTGTGGTGGGGTCCAGCAAaagttatactgcc
gcagtcatatagagcagacgGTTGTACCAGGCCTTCCAGGGAGTGCTGATGTTAGAGAGCaagttatactgccc
gcagtcatatagagcagacgCAGATAGGAGTGAGTTCAGAGTTGTTTTTTTCCCAGGTTCaagttatactgccct
gcagtcatatagagcagacgGCTGCAGCAGGGCACCAATGTCACTGTACTTCTGAAGGCAaagttatactgcc
gcagtcatatagagcagacgGCAACGTGGCCCCTTCCTTCCAGAGCCAGTACTCACTCCCaagttatactgcc
gcagtcatatagagcagacgAGTACACTGCTCTCTTCCTCATGGTTCCCCTACACTCCACaagttatactgccc
gcagtcatatagagcagacgCTTTGTAAAGTGTCCCTATATTTAGCCCTCCTCTAATCCTaagttatactgccctg
gcagtcatatagagcagacgAGATAGAAAGTTGCTTGTCTGGGTCATGGGAATGGAGCATaagttatactgccct
gcagtcatatagagcagacgGGCCCCACAAGACTTTTCTCAGATGGGATAAGAATCTGTCaagttatactgccc
gcagtcatatagagcagacgGCACTGGGATATTCCTAAGTACCATATTTATAGCAAAAACaagttatactgccct
gcagtcatatagagcagacgGGTCATTCCCAAGACAAAATGTATTTCTCCCCATAGATGGaagttatactgccct
gcagtcatatagagcagacgGTTTATTTTTTTCTTCCTCTCCCCTCTCATTGGTAGGTACaagttatactgccctgt
gcagtcatatagagcagacgGCAAAAAACTTGGCTTGCCCTGGCTCGAGACCCTCCCCAAaagttatactgcc
gcagtcatatagagcagacgGGTCTTTATTTCCTAAACCCTTGGCTCACAGACTCTTTTCaagttatactgccctg
gcagtcatatagagcagacgACATTGTTGTGTTATCATCCACCACCATCCTCCCACAGAAaagttatactgccct
gcagtcatatagagcagacgCTGTCTCTATAAATATGATGACTAAGTACCTCGATATGAGaagttatactgccctg
gcagtcatatagagcagacgTGGCTAATTTAACTTAGCATCATATTCTCAAGGTTCATCCaagttatactgccctg
gcagtcatatagagcagacgACTTGGGTTGGTTGCTTCCACGTTTCCGCTATTATGAATAaagttatactgccctg
gcagtcatatagagcagacgTGAATATGAGTGTACAAACATCTCCTTGAGATTCTGCTTTaagttatactgccctg
gcagtcatatagagcagacgATGGTAGCTCTATTTTCAATTTTTCGAGGAACTGTCATACaagttatactgccctg
gcagtcatatagagcagacgGGTATCTCACTGTGGTTCTGTTTTTATAAATCTGCCAGGTaagttatactgccctg
gcagtcatatagagcagacgTGCTATTTCCTCCACCTAGAATGTAATCCCTCTATGCCTAaagttatactgccct
gcagtcatatagagcagacgTTGAAATCAATCTCCTCCAGCTTCCTATTATCTTAACATGaagttatactgccctg

gcagtcatatagagcagacgCTCTTATTTTAACACCTACCACCCCCTCCCTTATGCTTTCaagttatactgccctg
gcagtcatatagagcagacgTTCTTTTTTTTTCATTTGCCTGTCTGCTCAATTTGGATGCaagttatactgccctgtg
gcagtcatatagagcagacgGATTGTTCATCTTTATATCTCCCAGTGTGTTTAGCCCAAGaagttatactgccctg
gcagtcatatagagcagacgACAAACTGCAGGCTCTAAAATCCCAACTGACTTGAAATACaagttatactgccct
gcagtcatatagagcagacgGGTTTTGGGCCTCGCTCTTAGTTCAATACTTTTTCTGCTGaagttatactgccctg
gcagtcatatagagcagacgTGTCTGAAAAAGAGATGTGACTTCCTGACATTGATTACTCaagttatactgccctg
gcagtcatatagagcagacgCACACACACAAACACCAACATTGTGCTTTCAAGCCCACAAaagttatactgccc
gcagtcatatagagcagacgGTGGTCCACGAAGATCAGAGTAACGGGCTTTCTGGCAATCaagttatactgccc
gcagtcatatagagcagacgGTTTTGTAGTTTTCCTCATTGTCGCTTGCCCTGGTGTGTTaagttatactgccctgt
gcagtcatatagagcagacgTTCATCCTGATGTGTTTCAGTGCGCTTCCCCTCCTTTGAGaagttatactgccctg
gcagtcatatagagcagacgTGTGTTCTCGCAGAAAATTCAGAAGCTAAATTAAAGGGCAaagttatactgccctg
gcagtcatatagagcagacgCCCAAGCATCTTCCATTCATCCCCCATTTTCCTTTAATCTaagttatactgccctg
gcagtcatatagagcagacgGCATTCATGTCAACTCTTCACGGTTTTATCCACAGGGCTTaagttatactgccctg
gcagtcatatagagcagacgCTGACAAGACTGCCGTAAGCGCCAAGGACCACTCAGATCTaagttatactgcc
gcagtcatatagagcagacgTCACATCAGGGAAGAGGCCTCTGTTTGGCAGTAAAGCACCaagttatactgccc
gcagtcatatagagcagacgCAGGATAAAGGCTACTTCCTGTCTTCAGTCTCTTTGATTTaagttatactgccctg
gcagtcatatagagcagacgCTTTTAAAGGACTTCTCAGGGAACTCACTGCTCTTAGCCAaagttatactgccct
gcagtcatatagagcagacgAGTCATCTTACTGTTAGTTCCATACACCCTTTGAATAGGCaagttatactgccctg
gcagtcatatagagcagacgGGACAACCTAAAGAACTTGCATTCAACCGTTTCCTTCCTAaagttatactgccct
gcagtcatatagagcagacgCCTCAGTGGCAGGGCCATGTGTCAACAGCCAAAACATCTCaagttatactgcc
gcagtcatatagagcagacgGGCTGTTGATTTTGGAGGCATATCTCGATCCCTTAGGTGGaagttatactgccct
gcagtcatatagagcagacgCTGCCCTCCCAGCCACCCTACTTTCCTGTCAGTGCTTGTCaagttatactgccc
gcagtcatatagagcagacgTCTTGGAGAGATAGGGCATGACAGGGATTTTCCAGGACTGaagttatactgccc
gcagtcatatagagcagacgAGGCATTACTTCCTTTTATCACCGGGCTCCCTCCTAGTCAaagttatactgccct
gcagtcatatagagcagacgCTTTTCTCAGTCAATTTCAAGCCTCAAGGTCATAGATTCCaagttatactgccctg
gcagtcatatagagcagacgTCATTCCTGAGGACTAGACCCCAAGCCAGAAGACAAACACaagttatactgcc
gcagtcatatagagcagacgCACTGAAGTGGGAGTCCAATCTGAGATTTAAGGTTTATTCaagttatactgccctg
gcagtcatatagagcagacgTGGGACAAGAAGAGGAGGGAGGCAATTACGGAACAGGGTTaagttatactgcc
gcagtcatatagagcagacgGCTCCTTACTTGTCTCTGCCATTGTTTGTGTGACCTCAGGaagttatactgccctg
gcagtcatatagagcagacgAATCTCCACAACAATCATGTAAAGCTAATTCCGCTAAAACaagttatactgccctg
gcagtcatatagagcagacgTGAAGTATTAGGCACAGAGTAAGTAGAGGATGTTGTTTTTaagttatactgccctg
gcagtcatatagagcagacgAGTCATTAGTCATCTGAAAGAGTGCATATATATAACCTGGaagttatactgccctg
gcagtcatatagagcagacgAGAAGTCATTTCCAGAGCCTTCTTCCATAGTCAGCTTTGAaagttatactgccctg
gcagtcatatagagcagacgCCGTCAGGTTTATTCTCAAGCCCCCAAGTGTTTGCTATTCaagttatactgccctg
gcagtcatatagagcagacgAAATCCTGTTCCTCTGTTGCCCAAATCTACCTTTGTCCAGaagttatactgccctg
gcagtcatatagagcagacgATGGGTGAAACTACATCAGTGGGGAAGCTATTGTAAGATGaagttatactgccct
gcagtcatatagagcagacgCCCAGTGTGAGTCATTAGGGGCAAAACCTTGAGCCAGGCAaagttatactgcc
gcagtcatatagagcagacgTCAAGGAGCTTACAGTTTGGATTAGAGAGTAATATAGTTCaagttatactgccctg
gcagtcatatagagcagacgGCCACAACAGCTTCTTTACCGTCTCCTGAATATGTCAAGCaagttatactgccct
gcagtcatatagagcagacgTGGCTGCTTCCACACTTGAACATTACCCTCCCACACACCTaagttatactgccc
gcagtcatatagagcagacgACGTGTTCCCTCCATTCCTTCAAATCTCTAATCAAATGTTaagttatactgccctg
gcagtcatatagagcagacgGATCTCCATGTGATGAGCCCCTGGTCAGCTGCAACAGTGTaagttatactgccc
gcagtcatatagagcagacgCACCTGTGTCTGCATCATCCTTCCTCCCCCATACTTCACAaagttatactgccc
gcagtcatatagagcagacgCTCCTTGGCCATTGAAACCACCTCCACAATCTTCCTGTCTaagttatactgccct
gcagtcatatagagcagacgTGACCCCACCCTGACTAAGTATGTTCCACCCACAATGCATaagttatactgccc
gcagtcatatagagcagacgAAGTGAATTGTCTGCTCTTTGCCCTTGGTTCCTCCGCATGaagttatactgccctg
gcagtcatatagagcagacgGATTTATCAGTATGCTTCCAGAGGTCAAGAACTATCTGTTaagttatactgccctg
gcagtcatatagagcagacgTTACGAGGTAAATGCATAAATAAGCCAAACGAGAAAACACaagttatactgccct
gcagtcatatagagcagacgCAGAGAAGTGACAATGGAGAGTAGTTACTGGTAGTAAAGGaagttatactgccc
gcagtcatatagagcagacgGTGAATCCTCTCCCCAGCCACATGTATCCCCATTTGTAGGaagttatactgccc
gcagtcatatagagcagacgCCAGCTTAAGTAGGGGTTTCACTATGGAAGACAAATTACTaagttatactgccctg
gcagtcatatagagcagacgGAAATGACTCTCTTTCCTTCTCTCCTCATCCCAGCCAATGaagttatactgccct

gcagtcatatagagcagacgCTTCACTTTCCCACTGAGTTACTTCAATCCAAAGGAAAAAaagttatactgccct
gcagtcatatagagcagacgGAGTTTAAATGAATGTACACTCCCCACTATAGACAACTTTaagttatactgccct
gcagtcatatagagcagacgGGGTCCTCCTAGCTGCTGAGATGTTAACCTGAAGCCAATTaagttatactgccc
gcagtcatatagagcagacgAATTACGTATTTTAAAGTGTATTATAGGCTGGGCATGGTGaagttatactgccctg
gcagtcatatagagcagacgGTGGATCACCAAAGGTCAGGAGTTCGAGATCAGCCAGGCCaagttatactgcc
gcagtcatatagagcagacgGCCTGGGTGACAGAGCGAGAGACCTGTCTCAAAAATAAATaagttatactgcc
gcagtcatatagagcagacgATATTTTAAGGTGGTAGGACTATAATTTCTTTTTGCGGGGaagttatactgccctg
gcagtcatatagagcagacgTGTACTTTTAGTAGAGGTGGGATTGGCCAGGCTGGTCTCAaagttatactgccc
gcagtcatatagagcagacgTTTGTTTGATTTAATTTATCTGGCTCTCACTCTGCATGTGaagttatactgccctg
gcagtcatatagagcagacgGAAATCCATTCCCTTCCTGTCTATATATTTATTTCTTGGGaagttatactgccctg
gcagtcatatagagcagacgCATACAAAGTGGTGATTCTCTTGCTTGCCTGAATACTGGGaagttatactgccct
gcagtcatatagagcagacgACTAATGTTTAAGTCCCAAGAGCAGAGATTTTGATGTAGTaagttatactgccctg
gcagtcatatagagcagacgTTAAAGCCTCCCAAGTAATTGTAATTTGCATTGAAGGTCGaagttatactgccct
gcagtcatatagagcagacgAGGAATGAACACCACTGAGAATTCATCAGAATAAAGCATCaagttatactgccc
gcagtcatatagagcagacgAGAATACAACCTACCTTGTTTTGCAGTTCAAGAACGAAAAaagttatactgccct
gcagtcatatagagcagacgTATTAAAAATAATAATGACTCCCGCTACTTCATACGTGCCaagttatactgccct
gcagtcatatagagcagacgCATCTGTCATCTCTAATCATCATAAGGACACTGAAAGGTAaagttatactgccct
gcagtcatatagagcagacgGATAGTGATCATAGTACCTGGTTAGGTAGTTTTTCAATCCaagttatactgccctg
gcagtcatatagagcagacgGCCCCAGTGCCTATGGTTCCCTTCTTCATGTCCATATGTAaagttatactgccct
gcagtcatatagagcagacgTCCAGCTGCAACCACATTGCTGCAAAGAACATGATCTCGTaagttatactgccc
gcagtcatatagagcagacgGTGTAGTATTCTATGGTGTACATGTACCCCATTTTCTTTAaagttatactgccctg
gcagtcatatagagcagacgCACTGTTGATGGGCATCTAGGTTGATTTCCTGTCTTTGCTaagttatactgccctg
gcagtcatatagagcagacgCACATGCGTGCATGTGTCTTTATGATGAGTGATTTTTTTTaagttatactgccctg
gcagtcatatagagcagacgACTCCTGACCTCGTGAGTGATGAGTGATTTATATTCCTTTaagttatactgccctg
gcagtcatatagagcagacgCAAATGGCAGTTCTGTTTTAATTCTTTGAGACATCTCCAAaagttatactgccctg
gcagtcatatagagcagacgCTAATGATCAGTTATGTGGAGGAGTTTTTCATATGCTTGTaagttatactgccctg
gcagtcatatagagcagacgTCTTCTTGCATGAGACCCATTTTAAAGATGAGACTTAAAGaagttatactgccctg
gcagtcatatagagcagacgAGTAATTAGCCCAAGGACACGCAGCTCTACTAGGAGGGGGaagttatactgcc
gcagtcatatagagcagacgGCCTGTGCTTAGCCACTACCCTACATTGCTTTCATCTAAAaagttatactgccct
gcagtcatatagagcagacgCCTTCAGTGTACAGGCATCAAAAAAAAAATCTGAATGGATaagttatactgccct
gcagtcatatagagcagacgCGGGGGACGGGCCTACCACTTTTAACTTTAAATACTTACAaagttatactgccc
gcagtcatatagagcagacgCAAGTATTACCAATTGGAGATAACCATCACTACTAGTTACaagttatactgccct
gcagtcatatagagcagacgAACTATCAATACTAGTGTGGTGGTTAAGGGCAAACATCTAaagttatactgccct
gcagtcatatagagcagacgCATTCCTATCTCCCTTTGCTGTCACTTCCCTAAACTTTGAaagttatactgccctg
gcagtcatatagagcagacgCTGTCAGCCAGGGAGAATCATAGTACTCACCTCAGAAATTaagttatactgccc
gcagtcatatagagcagacgAATTTCTTATTTAAACATGGCAGCCATCCTGTGGTAACTCaagttatactgccctg
gcagtcatatagagcagacgGGAAACCAAGATCTAAATCATCCAATACATGAATGGCTATaagttatactgccct
gcagtcatatagagcagacgCAGCTCAGAAGATTCTTAATTCCAAACCTTTCCTTGCACCaagttatactgccct
gcagtcatatagagcagacgCTCTTTGGGTTCTCACGAGGTTCCTGATTTGGACAATGACaagttatactgccct
gcagtcatatagagcagacgTACAGGTGCCTGCAACCACGACGACTAGTTTTTGTATTTTaagttatactgccct
gcagtcatatagagcagacgTTTTGATCAAACCTCATTTATAAAACCCTCCTCAGCCACCaagttatactgccct
gcagtcatatagagcagacgCGCAGGAGACCAACAACCCACCTTGTGATTTCTAAACGTGaagttatactgcc
gcagtcatatagagcagacgGTTTTGAGCATGAGACAGAAAAAACAAGCTTTGCAGTCGGaagttatactgccc
gcagtcatatagagcagacgCTGGACGGTAAGGATGGGGAAGCCACGGAGACACGTGTTAaagttatactgcc
gcagtcatatagagcagacgGGAGCTGATGTGAGCCAGATGTGATTGTATGGGCAAGACCaagttatactgcc
gcagtcatatagagcagacgCACTCCTTAAAACCCTGCAACTTCTCATGTAAAAGACCCGaagttatactgccc
gcagtcatatagagcagacgAACCGCTACTTTCTGACATGTCCTGAGGAACGCCCGCTTCaagttatactgcc
gcagtcatatagagcagacgGGAAGTCGGTTGCAAATTAAAGGAGGAAGGAGAAATTGATaagttatactgccc
gcagtcatatagagcagacgGAAGCAATGAGATCCTCGATGTCTCCATGGCCGTTTCATTaagttatactgccc
gcagtcatatagagcagacgCGGGTTTTAGGTCGAGTTAATCAGGAGCATCTACGTAGGAaagttatactgccc
gcagtcatatagagcagacgGCTGGCCGAGCGCACGTAGGGGTTCAGCTGTCCTGCCGGGaagttatactgc
gcagtcatatagagcagacgCCAGATTAAAACGTGGTGCCAGTATCCCAGACAGAAAAACaagttatactgcc

gcagtcatatagagcagacgCTCACACACACAGAGGATGGGAACACGAACCACAAGACCCaagttatactgcc
gcagtcatatagagcagacgCACGTCCTTGTAAGAGTGAGATCAGAGAACATTGAAAAGTaagttatactgccct
gcagtcatatagagcagacgCCATTTAGGGCATTTGGGACAAGCATGAAATTGAAGGGCCaagttatactgccc
gcagtcatatagagcagacgTTTTAATAAAGGGAGAAACATAGTTAAGGGAGAGTGTGTGaagttatactgccctg
gcagtcatatagagcagacgAGATGAAATCTTTGTGAGCATGAAGGACGTTATCAGAGATaagttatactgccctg
gcagtcatatagagcagacgAGGAGTGTCTGTGAATGAAAGAGCTCATCAAGGAAAGAGGaagttatactgccc
gcagtcatatagagcagacgCAGGTTCCTGGGATGAGAAAGAATGAAACAAGAATCAGCAaagttatactgccc
gcagtcatatagagcagacgGCAATGAGAGAAAGGGTGACACAGGGAAGACGTGAGAGGCaagttatactgcc
gcagtcatatagagcagacgGGTGATCTTCAGCTATGTCCTCTCTCTGCTCCCCCTGAACaagttatactgccc
gcagtcatatagagcagacgCTTTTCTCTTTTAATTTCACTTTAGATTCGGGGGTACATGaagttatactgccctgtg
gcagtcatatagagcagacgCGTGAGTATATTGCATGACGCCGAGGTTTAGGCTTCTAATaagttatactgccctg
gcagtcatatagagcagacgAACATAGTACCCAATAGATCGTTTTTCAACCCTTAATCCCaagttatactgccctg
gcagtcatatagagcagacgGAATCCCCAGTGTCTCTTGATCTCATCTTTGTGTCTGTGTaagttatactgccctg
gcagtcatatagagcagacgCTTAGAAGTGAGAAGATGCGATTTTTGGTTTACTGTTTCTaagttatactgccctgt
gcagtcatatagagcagacgTTGAGTAACATGAGTAGCTTGCACAGTCATTAGAGGGGATaagttatactgccct
gcagtcatatagagcagacgAGTTCTGCCTGATGCCAAAGCCCCGCCGCCATGTTGTTAAaagttatactgccc
gcagtcatatagagcagacgCAGCTTAAGTAAACAGCCTAAGAAAGTTGAGAGCAATGTGaagttatactgccct
gcagtcatatagagcagacgCAGCCTGTGACTGACCCTGTCAATGACCCTGATATTTTAAaagttatactgccct
gcagtcatatagagcagacgCAGTGGCTGGGTAGGATATTTTAATGTTTTGGCATATCGTaagttatactgccctg
gcagtcatatagagcagacgTGCACTATTGTTACACATGCCGCACACCCAGACAACATTGaagttatactgccc
gcagtcatatagagcagacgGGGCACCAGGATTTGAACTGCAATCATGAAGGGGTCCACCaagttatactgcc
gcagtcatatagagcagacgTCCTTAACCCAGGACTTTATGAAGATAACAAAAGGAGGTAaagttatactgccct
gcagtcatatagagcagacgGAACCCATGCTATAAAATAAGACCAATCCAGTTCATTTCCaagttatactgccctg
gcagtcatatagagcagacgCACACATGGTATAGAAAGGCATTTTGGCATAAACCACCAAaagttatactgccct
gcagtcatatagagcagacgAGAAAAACTCAGCGTGGAATGGCTAAGGAAATGTTCTCTGaagttatactgccct
gcagtcatatagagcagacgCAAGTTCTAGTGATTCTCCTAAGCATATTCCACAATTATCaagttatactgccctg
gcagtcatatagagcagacgCTACCTCTCTAGTAAGGCCTCTGTCTCTACACTGGAACTTaagttatactgccct
gcagtcatatagagcagacgAGGTTTGTTTCTCCTTCCCTTGGGGTTAACCAAAATGCTTaagttatactgccctg
gcagtcatatagagcagacgCTCCATATTTTTGAGGCCTGTCATATCTGAGAACCCAAAGaagttatactgccctg
gcagtcatatagagcagacgGCTAGTTGTATTTGATTTCTTCCTAATTTGTGTGGTCAGGaagttatactgccctgt
gcagtcatatagagcagacgTTTTGACAAGAAGAGTGACCTTTTTAAATGGACCCCTCTaagttatactgccctg
gcagtcatatagagcagacgGCATGTACCTGTAGTCCCTGACTCCTGGGAGCCTGGGGCTaagttatactgcc
gcagtcatatagagcagacgCATGGTGAGACCCTGTCTTGGAAACAACCACAACAACAAAaagttatactgccc
gcagtcatatagagcagacgTCTCGTGCAGCCCTATTACAAATCCTTTGGGTGTAGCAGCaagttatactgccct
gcagtcatatagagcagacgTACTGTTAACCATTTTATCCTCAGTACTGTGTGGGCTGCAaagttatactgccctg
gcagtcatatagagcagacgGTTCTTAAAAGGGAACCATTTAGGTTTTGATGATAGTGGAaagttatactgccctg
gcagtcatatagagcagacgGCGCTGATGGTACGTTGGAGGCTTCACATACATGCTTTCAaagttatactgccct
gcagtcatatagagcagacgGAATATCTTTATGAGTGGGTATTCTACGCCCATCTTACAGaagttatactgccctg
gcagtcatatagagcagacgACAAAAAAAATTAGGTGACTTGTGCAAAGTCAAGACCACAaagttatactgccct
gcagtcatatagagcagacgTGTTTTCAAGGGTACAATACAACAGTGTTAACTACGTGCAaagttatactgccctg
gcagtcatatagagcagacgTCTTGCAAAACTGAAACTGACCCATTAAGCCACAACTCCTaagttatactgccct
gcagtcatatagagcagacgATTCTACTTTATGTTTCTACGAATCGGATCGCCCTAGGTAaagttatactgccctg
gcagtcatatagagcagacgTTCACTTAGCATAACATCCACAAGATCCAACCATGTTATAaagttatactgccctg
gcagtcatatagagcagacgCAGAATTTCCTGGTTTTTTTAAGGCTGAATTACACTCCATaagttatactgccctgt
gcagtcatatagagcagacgGATAGACCACATTTTCTTTCCAGATCTAGGATTTTAATCCaagttatactgccctg
gcagtcatatagagcagacgTGTGAGTCTCCAAAACCCAAACTCTTTTCAGGTGAGTACAaagttatactgccct
gcagtcatatagagcagacgTTATACATAAAACAAATGTGAGCTGAGACAGTGGTGTCCCaagttatactgccct
gcagtcatatagagcagacgTGTTCTTAACTTTCCAAGCTCAGAGACGGAGGGACAGGGGaagttatactgccc
gcagtcatatagagcagacgCTAAGGGGAGCACTTGTCTTTCTTGTTGGATAGAGAACAGaagttatactgccct
gcagtcatatagagcagacgGCCTAGAACTGCCTCCCACCACCCTCCTAGCCCAAGACACaagttatactgcc
gcagtcatatagagcagacgGGCCTTCCTTCTTCTATAAGTCACGTACCAAGGGCTCCAGaagttatactgccc
gcagtcatatagagcagacgGAGACCTCAAACTCCCCACCCTTCCCTCACCCAGCCTTGCaagttatactgcc

gcagtcatatagagcagacgAGCTTTGCCAGTCCCCATCGCTTCCGCAAGCCTGCAGCCTaagttatactgcc
gcagtcatatagagcagacgATCCATGCCGCCTCTCTAGTTTTCATCATACCCCGTCAGCaagttatactgccc
gcagtcatatagagcagacgGCCTGACAGCTGAGCGTATCCTTCCTTTCCCCAGAATCCCaagttatactgcc
gcagtcatatagagcagacgGTTTGCAACATTGTTCTTTATATCCACCCCCAATCTCCACaagttatactgccct
gcagtcatatagagcagacgAGCCATTTCTCCTGCTTGACCATGTTACTCCACTCTGCTGaagttatactgccct
gcagtcatatagagcagacgGTCAGTGCTGTCCTGCTTGGATGTTTCTTCTGGCTGCTAAaagttatactgccct
gcagtcatatagagcagacgTGCCTCCCTCTTCACCTACACCCTCCTTGGGCAGCTTGGTaagttatactgcc
gcagtcatatagagcagacgCTGGGCAACATACACAGACCACCTCCCTACTGCAAACTCTaagttatactgcc
gcagtcatatagagcagacgGGATCGCTTAAATCTGGGAGGTCAAAGCAGCAGGGAGCCAaagttatactgcc
gcagtcatatagagcagacgAAAAAGATTGTTCAGTGCCTCCCACCCAGTTCTACAGCCTaagttatactgccc
gcagtcatatagagcagacgACTTCCAGCAACCCCTGGACATTCACCAAAACCCACTTCCaagttatactgcc
gcagtcatatagagcagacgTTCCACTGCTGGTGTGAATCTCTCCTTTCCTCTCCTCCCTaagttatactgccct
gcagtcatatagagcagacgCTCATTTGTATCTCTGAGCACTTGGATTTCAATGTAACTAaagttatactgccctg
gcagtcatatagagcagacgCTTTTTATGTAAATTCTTTGGAGAATGAACTAGGCCTACCaagttatactgccctg
gcagtcatatagagcagacgGGCCCCTCACCCTCATTGCTTATATTATTCCATAAATAAAaagttatactgccct
gcagtcatatagagcagacgTTGTAAGCGAATCAGGAGGGAGAGGATTATTTTTCTTTTaagttatactgccctg
gcagtcatatagagcagacgCTGGCCAGGGGGGATTCAGGATAACATAACAGCTGCTTTCaagttatactgcc
gcagtcatatagagcagacgAGTAATGGGGAAGAGGAGAATTGAAGGATATGGGGATAATaagttatactgccc
gcagtcatatagagcagacgTTTGAAGAAAAGGCCTTAATACAGAACCTAGATGAGTCTGaagttatactgccct
gcagtcatatagagcagacgCCTTTAAATGATACTGAAGAGGGAAAGGGAAGTGCTGGGTaagttatactgccc
gcagtcatatagagcagacgCCTAGCAGGCAGACACAGACACCTCCTGACATGGAGAGAAaagttatactgcc
gcagtcatatagagcagacgATGGAGAGGAACGCATGGACAGGCACCAGCAGGCCACCAAaagttatactgc
gcagtcatatagagcagacgCTTGCACGCATTCTCAAACACCAAGGCAGGAACACTGGGAaagttatactgcc
gcagtcatatagagcagacgTGTAACACACGCCCACTGGGTCTGTAAACATTCATCCCTAaagttatactgccc
gcagtcatatagagcagacgCTCACCTAGAGGTTTGAGCAACAGGGCACTGAAGAAGCAAaagttatactgcc
gcagtcatatagagcagacgAGGGAACTTCGCTCGTTTCAACGTCTGACATCTACCTTCAaagttatactgccc
gcagtcatatagagcagacgGCACCGTAATTGGGCCTGCATGTCAGCGTGATTCCAAGGCaagttatactgcc
gcagtcatatagagcagacgGCACCACCTGCCTCTGCCTAGGATGCATATAAGTGGCTGAaagttatactgcc
gcagtcatatagagcagacgAGCCACGGGGATGCCACTTGCAGGACACTACAACCAGGGGaagttatactgc
gcagtcatatagagcagacgCTCTTTGAAGACAACAGCAACCACAGAACAGAAACAGCCTaagttatactgcc
gcagtcatatagagcagacgGAAGAAGGAGAAAGGGAGGGGCCTGAATTGACCGGGGGCTaagttatactgc
gcagtcatatagagcagacgCAGAATTCAAACTGAAAGGGACTAAGTTGCTTGTATTGGCaagttatactgccct
gcagtcatatagagcagacgCCAAGTCTGTGGAATTCCTCCCCTCCTACAAACTATGCACaagttatactgccc
gcagtcatatagagcagacgCAGCCCCAGGCCTGGATGACTTGTGAATCTGGCCCTTTGGaagttatactgcc
gcagtcatatagagcagacgGTGGAGCCCATCACCTTTAGTGGCAAGGACCTAAGCCCCAaagttatactgcc
gcagtcatatagagcagacgCCTCCCATCACCTTCACAGCTGTGAGGCTCTGTTCTCTTCaagttatactgccc
gcagtcatatagagcagacgTTAACCTGATCTAATATGACAGCATACGTTTCAATGAAGGaagttatactgccct
gcagtcatatagagcagacgTCTGTGTATGTTAAGAGGGCTAAGGTGATATGAGATGAGAaagttatactgccct
gcagtcatatagagcagacgAATCCATTGTTTATTTTTACGTCAAAATCAAGGTGGGACCaagttatactgccctg
gcagtcatatagagcagacgCTTGCTAGGCTTAGGTCCAGGTGACTTGGAGGTGAAGGTGaagttatactgccc
gcagtcatatagagcagacgGAAATAGTGGGTCCTTGCTGAGCATTTCTAGAGGGTTAAAaagttatactgccct
gcagtcatatagagcagacgGAGTTCGAGACAAGCCTGACTAACATGGTAAAACCCCGTCaagttatactgcc
gcagtcatatagagcagacgACCTTCAGTCATTCTTTACTCTAAAAACATTTGCTTTGCCaagttatactgccctg
gcagtcatatagagcagacgAAAAAGTTGACACCCATTTGCACCCTTATCTAACACCTTCaagttatactgccct
gcagtcatatagagcagacgAGAAGCAATCCCAACTTGTTCTTAACTTGTCTGAGAGTACaagttatactgccct
gcagtcatatagagcagacgTGTTGAATTTAGGAGTCACGGATCATCATTCTGCAGTGGTaagttatactgccct
gcagtcatatagagcagacgTTCACCTTCCATATGTTAATGCTTCATTTCCCCCATAGAAaagttatactgccctg
gcagtcatatagagcagacgGCTTATTATGAGTATGTTTGTGCCAAGTGATAAGTGAAAGaagttatactgccctg
gcagtcatatagagcagacgACCTCTTTATACAGTGTCTGCATAGGTCCCCATGATGTTTaagttatactgccct
gcagtcatatagagcagacgTGAATAAAAAAGCGTCTCAGTGATTATAGCCAAATGTCTCaagttatactgccct
gcagtcatatagagcagacgATAATTTGGGGACCATTATGCTAATCACATCTGCCATAATaagttatactgccctg
gcagtcatatagagcagacgACTTTTCCAGTTTGTGCATGCGCCCAGTCATACTATTCCTaagttatactgccct

gcagtcatatagagcagacgCTTTTAAGGGTGGGATTGTCGGTGGGAGAGAAGGGAGGGTaagttatactgccc
gcagtcatatagagcagacgCCTCTGGCTGTGGGGTGATCATGTTCTTGTGCTTTTGTCCaagttatactgccct
gcagtcatatagagcagacgAGCTAGGAAGGGGTATGTGCTCAATGGTTTTATGTGTGCAaagttatactgccct
gcagtcatatagagcagacgAAGGCTCATAGCTTTCTGCTCACAATCCTCCAGGCCATGTaagttatactgccc
gcagtcatatagagcagacgGCACCATTTCCTTGCATTGAATATTGAAGAATAGCTAATCaagttatactgccctg
gcagtcatatagagcagacgAACAGCAATAATAATGTCCTTGTGAATGTCATGTTATGCCaagttatactgccctg
gcagtcatatagagcagacgCTTCACACACACTTTCTCATTCAGTGCTTCTCCACACCGTaagttatactgccct
gcagtcatatagagcagacgCAGATTAGGAGTGTCTGTGGGCAGTGTATAAAAATGTAAGaagttatactgccct
gcagtcatatagagcagacgTTACCCCTTACGTATATGCATGTTTGCTCTCTTTATAACAaagttatactgccctg
gcagtcatatagagcagacgTTTGTTTACTCTTCATCATGGCCCTGAGACTGGTTACTTGaagttatactgccctg
gcagtcatatagagcagacgGTGCCTGTCTGTGCCAAACCCTCATTCTACCCCCACCATGaagttatactgcc
gcagtcatatagagcagacgACATTCAGCAGCATGTCCCTCTCCTTCATAGCCATAACCAaagttatactgccc
gcagtcatatagagcagacgTGGTCCACTCCTTCTCCAGCCATGTACTCCACAACAGCAAaagttatactgcc
gcagtcatatagagcagacgAGGACTCAACACAAAACCTGACAGATGGCAGGCCCTCAAAaagttatactgcc
gcagtcatatagagcagacgTGAAGCCAGGAGTTCGAAACCAACCTGAGCAACAAAATGAaagttatactgcc
gcagtcatatagagcagacgATAGAAATATTAGCCCGGTGTGGTAGCATGAACCTGTAGTaagttatactgccct
gcagtcatatagagcagacgGAATTCAACGATCTGCTTTAGTCATCTGTAATATGTGTAGaagttatactgccctg
gcagtcatatagagcagacgACACGCTAAGTATGAGACATTGCAAAGATCTGAAATAATCaagttatactgccct
gcagtcatatagagcagacgCTAGTCAACTGAGAGCAATTGAGATAACATCTATATTCAGaagttatactgccct
gcagtcatatagagcagacgGGTATAATCTTGCCAATTTCACCTTTTTCCCTCAACTTCAaagttatactgccctg
gcagtcatatagagcagacgGTTCAGGCCACACTATCATTGTTCCAAGCATGGTCCTCTCaagttatactgccc
gcagtcatatagagcagacgTAAATAATAAAATGAACCGACTGCCCAACCCAAAAACTAGaagttatactgccc
gcagtcatatagagcagacgGGCTCCTCCCACTCCACTCCCTTCCTCTTCCCAAGTGCAAaagttatactgcc
gcagtcatatagagcagacgTTAGTTATTTTGGACTTACTTATTTGTCTGTTGACTCGGCaagttatactgccctg
gcagtcatatagagcagacgATTCCTTACACCCATCTATTTGTCACATGTAGCTGAAGTTaagttatactgccctg
gcagtcatatagagcagacgATTTCCTCTGCTGTGAAATCTTTCCATACCGTCTGTTGATaagttatactgccctg
gcagtcatatagagcagacgGTTGCTTTGCATCCTTTGATATTGTGAAAAACTGTTCATCaagttatactgccctg
gcagtcatatagagcagacgGAGAAGTTAAGCAAGTCACGTAAGATCTCACAGCCCATGAaagttatactgccc
gcagtcatatagagcagacgAGCCTTGATTTCTCTAGCTCTAAAAACATGACTCTTTCACaagttatactgccctg
gcagtcatatagagcagacgGTCCTGCTGTGATGAAAAGATCATAGAAAGTAAAGAGCACaagttatactgccc
gcagtcatatagagcagacgCTGGAAAGTTACTTCACTCACTCTTTGTCTACTAGAGAAGaagttatactgccct
gcagtcatatagagcagacgGGATGCATTTACTCTCCACTCCTCCCCATTTTAAAAAGTAaagttatactgccct
gcagtcatatagagcagacgGCTGAGTACAATACTTTATCTCCAAAACCATATGTCTAGTaagttatactgccctg
gcagtcatatagagcagacgTGGTTTTGTGCTTAAATTCTCTTTGCAATAGAACTAGCTGaagttatactgccctg
gcagtcatatagagcagacgAAAGTGATATGTTAGCTTCTACAGCCTGTTATAATGTTCCaagttatactgccctg
gcagtcatatagagcagacgCTAAACTCTCCTCTTCAGATTATAAATAGGCTTTATGTGGaagttatactgccctg
gcagtcatatagagcagacgGAAATGATTTCACTTATGTGCAAAGGAATTTTCCAGAGGAaagttatactgccct
gcagtcatatagagcagacgCTGAAAAAAAAATAGGTATTTCTGTACTTCAGGGCGCTAAaagttatactgccct
gcagtcatatagagcagacgAGATTGAGGTCTGGTTTCTACAAGGTTATGTCCCCTACTCaagttatactgccct
gcagtcatatagagcagacgTGGCTCCAGAAACTTCCCTGTCCCTCTTCTTCCCCCCGCCaagttatactgcc
gcagtcatatagagcagacgGCTCATTTCTTCCACAGCTATTTCTCATTTCTCTTTCCAGaagttatactgccctg
gcagtcatatagagcagacgGTTTTCTTAAAAATATCAATCAGGCTGGTCCAAGTGCAGTaagttatactgccct
gcagtcatatagagcagacgCCACTCCCACTGCTTCACCTTCACCTGACTAGCCTAATTTaagttatactgccc
gcagtcatatagagcagacgTAGAGACGGGGTTTCGTCATATTGATCAGGCTGGTCTCAAaagttatactgccc
gcagtcatatagagcagacgCTAACAACTTTAGGACATTTAGGAAATAGGGGAATCTTACaagttatactgccct
gcagtcatatagagcagacgCTCACCATCCTCATTCAACTCCCACAATTGTTCATTCCTTaagttatactgccct
gcagtcatatagagcagacgCAGCAGCAGAAGACAGAGATACACCCACATTCCCTAACCTaagttatactgcc
gcagtcatatagagcagacgAAAAGGATTCATGAGAGGATTATTGATTTGGGAAGTTCAGaagttatactgccct
gcagtcatatagagcagacgGGAAGCAAAACACAGAAGGAAAGGCAACCAATAAAGAGGCaagttatactgcc
gcagtcatatagagcagacgAATAAATAAAAAAAGAGGGGCTTAAGTGATGGAGGCTTACaagttatactgccct
gcagtcatatagagcagacgCTCTAGGAGTGATGGAGCTGGGGTATCTATGCATTAACTGaagttatactgccc
gcagtcatatagagcagacgGGTGATTGCTTGAGGACTGTTCATGGGGAGGAAAGGGGGAaagttatactgcc

gcagtcatatagagcagacgGGTGGTTAGCTGGAACTTGGCTGCAACACCCTGAGAGCCCaagttatactgcc
gcagtcatatagagcagacgCAGAGTGTCTGCTATCCTGCTTTATTCACTTAACATTAATaagttatactgccctg
gcagtcatatagagcagacgTGTTGCTACACAGTCTATACTTAGGAGATTAATTTCTGCAaagttatactgccctg
gcagtcatatagagcagacgATACTGGATATTTAGGTCGTTTCAAAATGGAATGAGTGAGaagttatactgccctg
gcagtcatatagagcagacgGAAATAGCTAACTCTTTTCGATCCAGACTCTAAAAGCAGGaagttatactgccct
gcagtcatatagagcagacgGAAGTCTTCCAGTCCTACGAGCCAAGCAGAATACCACAGGaagttatactgcc
gcagtcatatagagcagacgTGCTCTGTTATTTAGGAGGACACTTTAATTGCTGGTCTGTaagttatactgccctg
gcagtcatatagagcagacgTTGTATATCAAAACTTTGCATCAGGGCCGTGATAGTTGAGaagttatactgccctg
gcagtcatatagagcagacgAGATAGCTATCCCATCATTTTGTGCCGAGAAGTAATAATGaagttatactgccctg
gcagtcatatagagcagacgAGCCCGGGATTTTGCTATGGTTAACCACTAAACAGCTCTGaagttatactgccct
gcagtcatatagagcagacgGGAGGCTTCCGTGTGTCTCTGTGCCCCACCTATAAATCAAaagttatactgccc
gcagtcatatagagcagacgGACCAGTCTGCAGCTTCTTCTACAACCATTTCCACCCTTCaagttatactgccct
gcagtcatatagagcagacgCAAGTTTGGTGCAGATGCCAGGTGGAGAGACCCTGTTCCTaagttatactgccc
gcagtcatatagagcagacgGGATGCTGGGCTTGAGTCACTTGCTGTCATTCCATTCTTCaagttatactgccctg
gcagtcatatagagcagacgAAATAATAATTCTTACACAACATAACAGTTTCCACGGGCCaagttatactgccctg
gcagtcatatagagcagacgGTGTTAACTCATATCATTCTCACAACAACCCCAAGAGCTAaagttatactgccct
gcagtcatatagagcagacgTATTATCATCCTGTTTGACTGAGAAGGAAGCCCTGGAACAaagttatactgccct
gcagtcatatagagcagacgTCAAGGTCACATGGCTAGTAGATGTTGGAACAAGGACTCTaagttatactgccct
gcagtcatatagagcagacgTGGCTCCAAGATCCATGCTGCTGTGTTATAGTCAGACTATaagttatactgccctg
gcagtcatatagagcagacgTCTTCATGTAACCCAAAGCATATCTAAGCATTATCCCTGCaagttatactgccctg
gcagtcatatagagcagacgGCTTTTTCACATAGCCTTACTCATGAGCAGCAGGAATGACaagttatactgccct
gcagtcatatagagcagacgCCTTAAGGGCCTCATCGTTCATTCTAAACATTAGTCAAAGaagttatactgccctg
gcagtcatatagagcagacgGTTCAAGCAATTATCCTGCCTCAGCCTTGCGAGCAGCTGGaagttatactgccc
gcagtcatatagagcagacgCCTTTCCTCCTCTTTTAAGGCTAGATTGTATAGTGTTAGTaagttatactgccctgt
gcagtcatatagagcagacgCAAATTGCTTTCCCTCCATTTCCACTCTGTGTTGTGCACGaagttatactgccctg
gcagtcatatagagcagacgATGCATACGCCCCCCATACACCCTTCACCCTCTGCCGGGTaagttatactgcc
gcagtcatatagagcagacgGTTTTCCTTCCACGTACTCAGTCCCTAGCTTCTTACTTGAaagttatactgccctg
gcagtcatatagagcagacgCATTTCCTCAAACCTCACCCCAACCTGCTCTCAACTCTTCaagttatactgccc
gcagtcatatagagcagacgTCGTTTCTTTAGTCTTTACCTTTGATCCTTATTTTTGGGCaagttatactgccctgtg
gcagtcatatagagcagacgCTGTAAACTTCCAGGCTAGAAATACGGGTTCTCTTGACAGaagttatactgccct
gcagtcatatagagcagacgCCAATTCCACGGGACCTTTAGCTGTCACATACTTTGGCCTaagttatactgccct
gcagtcatatagagcagacgCAGAGATTCCAGGCAAGGACTCCTTAACTCTGTTATTTTTaagttatactgccctg
gcagtcatatagagcagacgTGTATTTATTAGAGACTGGGTCGTTAGGTTCCTCACACTTaagttatactgccctg
gcagtcatatagagcagacgGCACGAGCCACCACGCCCTTTTTGTTATTAGGGACACAGTaagttatactgccc
gcagtcatatagagcagacgCTCATAGAGTACCAGCCAGCACCACTACCGTAGAGTAAGTaagttatactgccc
gcagtcatatagagcagacgAATACATATTTCATATATACAGTTTCACCTGAGGCCCTAGaagttatactgccctg
gcagtcatatagagcagacgGAACGCACCCTCAGTTCTCTTTGAGAAATCATACAGGATGaagttatactgccct
gcagtcatatagagcagacgTTCCAGAACACTGCTTATAGGAAAACATTCTCCCAGGTTTaagttatactgccctg
gcagtcatatagagcagacgTGTATTACAGAAACTATAAACTCTTCAGCTCAACCTCTCCaagttatactgccctg
gcagtcatatagagcagacgCTGAAGAGTCAACGTGAATTCACTAAGTGCAAATCAGGCCaagttatactgccc
gcagtcatatagagcagacgACCTTGTTTCCGTTTTGGGGTGGGTTGGTAAGTTAGAGAAaagttatactgccctg
gcagtcatatagagcagacgACCCTCAGCTCTCAATACCCTGGTGTGTAACAATTAGTGGaagttatactgccct
gcagtcatatagagcagacgAGGAAGGAAGGGTTATAACCAGTCTCCAGAAAATTATACAaagttatactgccct
gcagtcatatagagcagacgTCATCCCTTTAAAATGTAAAAGAGAAAGCTCCAAACTGAGaagttatactgccctg
gcagtcatatagagcagacgTCTTTAGAGGGCTTTGTATAGAAGAAGGATTAGATTTCAGaagttatactgccctg
gcagtcatatagagcagacgCCAGGATAAGTGTGTGCAAATTACAGGGAGGCAGTATTGTaagttatactgccct
gcagtcatatagagcagacgCTAACGCTTAGGACACTGTTTCTTAAACCTGAGTCATGTAaagttatactgccctg
gcagtcatatagagcagacgCGCAAATTTCCAGGGTTTACGAATTGTTTTTCTATAACAGaagttatactgccctg
gcagtcatatagagcagacgAAACAACCTTCTTTTAATGGGAGAGATGATGTTTTGCTGAaagttatactgccctg
gcagtcatatagagcagacgCTTTTGCCTTGGGCAGTTGTAGACCGCCGTGTGAATTGTGaagttatactgccct
gcagtcatatagagcagacgAGGTTACTTTCCTTCACTAAAATTGACATCATGGTTACCTaagttatactgccctg
gcagtcatatagagcagacgCACTCCCAATTAAAGTGGATCTACTCTCCTAACACATAAAaagttatactgccctg

gcagtcatatagagcagacgGGTAGCCCTCAGATAAAAAGTGATTACTACATTCGCCAAAaagttatactgccct
gcagtcatatagagcagacgCCTAATGGGAAGGCTGAACTTTAGTTAATGATGTTGTATCaagttatactgccctg
gcagtcatatagagcagacgCACTAATGTCAGATGCTAAAAGTAGAAGAAATTGGAAGCCaagttatactgccct
gcagtcatatagagcagacgAGGAATTTGAGGCTGGAGTGTGAACTGCCACTGCACTCCAaagttatactgccc
gcagtcatatagagcagacgAAAAAGTGTGGGGGAGAGGAAATTAGGGATGGAGGGCCAGaagttatactgcc
gcagtcatatagagcagacgGAATTCTCTGTGCTTTTTGTTCAATGTCTCTGTAAACCTAaagttatactgccctgt
gcagtcatatagagcagacgACAACTATTTCTAGATCTTGTATGAAACTCTTGTACCTCCaagttatactgccctg
gcagtcatatagagcagacgGGTCTGTACCGTAGTTTTTAAAAATGCTGAAATTGAGAGTaagttatactgccctg
gcagtcatatagagcagacgCTCAAAGTGTTTCAGAAAATGCCTAGAAGGTCATCTTTTAaagttatactgccctg
gcagtcatatagagcagacgGTTCTTCTAGGGATTTTTGCTTTTGGAGAGAGGTCAGAGCaagttatactgccct
gcagtcatatagagcagacgGCTTTGAATTCATTTGCTTTTACTGCTACTTTTCACAGACaagttatactgccctgt
gcagtcatatagagcagacgCCATGAGGTGATGAAATTTGTAGCTTTCTAAGGTTTTTCTaagttatactgccctg
gcagtcatatagagcagacgCAATCCTGTTCAAATAGCAACAATTTCATTTAGGCCAAACaagttatactgccct
gcagtcatatagagcagacgGAAAGGATAACTAATAAACCTCACATCATTCCAGTGAACCaagttatactgccct
gcagtcatatagagcagacgTGCATTCCTTTCTGATGGCTTATCACATTGCACTCTGGGTaagttatactgccct
gcagtcatatagagcagacgGCCATAGTTATAGCTCCATTTACATATTTGGGTAATCCATaagttatactgccctg
gcagtcatatagagcagacgCTTGCAGGAATCTAAGGCTAGAATTGAATCTGTAAAAGGAaagttatactgccct
gcagtcatatagagcagacgATTCACATACGTGCATTTTCGTTTAGTTTTAGCAGACAACaagttatactgccctg
gcagtcatatagagcagacgCATGATAGGAGTTGTAACGTCATAGGCATTGAGTAGGTCAaagttatactgccct
gcagtcatatagagcagacgTGGCACTGGGTCTAAAGATTTGATTCCTCCAATTGGGGTGaagttatactgccct
gcagtcatatagagcagacgTTCCATTAGAAAGTAAAGAAGCTGGGTCCTCGTCATAAATaagttatactgccct
gcagtcatatagagcagacgTGAGGGAGGAGTACAGTCACGTTTTCTGATTTTCAGCCTGaagttatactgccct
gcagtcatatagagcagacgGCATTCGCCCCCTGTCCCATTACATCAGACCACTCACGATaagttatactgcc
gcagtcatatagagcagacgTGGCTTCTTATCACCACCCACTGTCTGCTCCAAGAACTGCaagttatactgccc
gcagtcatatagagcagacgGCAGGTGACTCTCCCAACCGCAGAAAGAGGAAATGTACTAaagttatactgcc
gcagtcatatagagcagacgAAAGAAAGAATAAAACGGAAGAACAAATGAGCATCTGAGGaagttatactgccc
gcagtcatatagagcagacgCCAAAAGGAAACCCGTATTTATAAAAGATCACAACCAAAGaagttatactgccc
gcagtcatatagagcagacgCATATTAAAAACAGTTGTTTGATAGCATTCCCATGGAGACaagttatactgccct
gcagtcatatagagcagacgTTTTTCTAATTGTTATCCCAGTGTGCTCCTGTTTGAAAGCaagttatactgccctgt
gcagtcatatagagcagacgCTGGGCAACATGGCAAAATCATGTCTTTACAAAAAATACAaagttatactgccct
gcagtcatatagagcagacgTTGAGCCTAGGAATTCAAGGTTGCAGTAAGCCATGATTGTaagttatactgccct
gcagtcatatagagcagacgAAAACACGCCTCCTTGATTTGAGATTCTGCTTCTTTTTCCaagttatactgccctg
gcagtcatatagagcagacgCCAGTTTCAGGAAAGGGTGGGTTAGAGTCAGGGGCTTCCTaagttatactgccc
gcagtcatatagagcagacgAACAAGAATTCTAATGAAGCAGTGTGTTCAGAGGGGGGTCaagttatactgccc
gcagtcatatagagcagacgCCTCATGTGGTATTATCTTTGCTACACTTTGCAGTTTCGTaagttatactgccctg
gcagtcatatagagcagacgGGTCTTTAACAAAGCATGTCCATGTTTATCTTAAATGCTGaagttatactgccctg
gcagtcatatagagcagacgCAATGAATGCTGAAACATAAAAGTCAGACAGAATGTTACCaagttatactgccct
gcagtcatatagagcagacgAAGAAAGGGCTTCCTAAAGATCAAAAGATTGCTGTTTGTTaagttatactgccct
gcagtcatatagagcagacgAGTACTTTCACTTTAGGGGATTCAGATTGCTATTGAAGGAaagttatactgccctg
gcagtcatatagagcagacgGCCCCACCCCCACCTAGATAAAGGCTTCCTCTACCTCATTaagttatactgcc
gcagtcatatagagcagacgGTGGGAAACTTGCTTATTGATACTAAAAACATTGACTGCGaagttatactgccct
gcagtcatatagagcagacgGACATTTGATTAAAGATAGTATAGGATTCAAGCCAATGGCaagttatactgccct
gcagtcatatagagcagacgGACCATGTCTTTCAGTCTTACTTAAAATTAGAGGGGAAATaagttatactgccctg
gcagtcatatagagcagacgACCTTGGGTTTTGCCTTATTCTGTGTACGGGTAGTAAGTAaagttatactgccctg
gcagtcatatagagcagacgCTGAGTGCCAGTGCTACCAAGTTCCACCCCCCTCTCATGTTaagttatactgccc
gcagtcatatagagcagacgGCCACCACACCCAGGCTTAAGCTGATTTTAATGCCACCTTaagttatactgccc
gcagtcatatagagcagacgAAGCTCTCTTTCCCTCTAAATATTTCCAGATTTGGGTTTTaagttatactgccctgt
gcagtcatatagagcagacgACAAGATGAAAATGTACCCTCAGTCCAATCTAATCTGCAGaagttatactgccct
gcagtcatatagagcagacgAGATGGGTTTGTATAATGGTGGTCCCTTTCTTTACGGGGCaagttatactgccct
gcagtcatatagagcagacgGATGTAATATCTGGTCATTTCCATCTACAGCTATCATATGaagttatactgccctg
gcagtcatatagagcagacgGAGAAATAACAGTTCTTGTCTTGTGCTACCTCGAAATAGCaagttatactgccct
gcagtcatatagagcagacgTTTGCACATGGCTAGATCATGATTTCTGGTTTTTGTTTTGaagttatactgccctgt

gcagtcatatagagcagacgGTTCTCTACCCAGGCAGTATAGTAAAGTTTTTAAGAGCTTaagttatactgccctg
gcagtcatatagagcagacgACTCTCACTTACTTATGTGATCTGAGCAGATTAATGAAGCaagttatactgccctg
gcagtcatatagagcagacgACTTGAAAGGTTGTTGTGAGGATTGAACATGTGTAAGGAGaagttatactgccctg
gcagtcatatagagcagacgCTTAAATCAATTACAAGGAACCTAAAAGACTTTCTGCAGCaagttatactgccctg
gcagtcatatagagcagacgGAAATCAAACACCACTTCTCTTTTCAACTCGCTTCCTTTCaagttatactgccctg
gcagtcatatagagcagacgCTGTTCTCACCAGCATGCATGTTTGTAAGCAGTCTACAGCaagttatactgccct
gcagtcatatagagcagacgAGTGACCCTCATCCATATAGAAATGACAAAATATGCATGCaagttatactgccct
gcagtcatatagagcagacgCTTTAAGACTCTAATGAAATCCGGCTAAAGGAAATTACAGaagttatactgccctg
gcagtcatatagagcagacgCAGGAAATCTTACCAGAAAAACCCAAAAACCTCTCATGGTaagttatactgccc
gcagtcatatagagcagacgATGCAAAGATTCTAAAATACCAGATACTCCCCAGATCCCCaagttatactgccc
gcagtcatatagagcagacgTTTGTTTTGCTTAATTTACTTTAATGGGAGGCAGTATGGCaagttatactgccctgt
gcagtcatatagagcagacgATTGCTTGTGATTCTTGAGTAGCATTCCATAATTACATGCaagttatactgccctg
gcagtcatatagagcagacgAGAATGGTGGGAAAGGTAGTGCATGTGTATTTGAGATAATaagttatactgccctg
gcagtcatatagagcagacgATTTTAAAGTATCAGAGGGCTGGAGGCAGTGGCTCATGCCaagttatactgccc
gcagtcatatagagcagacgGCAGAAGAATGGCTTGAGCCCGGAGGTGGGGGTTGCAGTGaagttatactgcc
gcagtcatatagagcagacgTTCTAAGGGAGGAGTCAGAAAACAGGAATAGTGATTTATTaagttatactgccctg
gcagtcatatagagcagacgGAAGAAAGATTTGGGTTTCTATGAGAATGTATGAAAGCTCaagttatactgccctg
gcagtcatatagagcagacgAGATAATTTTAGAAAATCCTTCATTTCCTCCACAGCATGGaagttatactgccctg
gcagtcatatagagcagacgTTTAGACTGAGTTTATGAGTGTTTTCCCAGCATTGATTTCaagttatactgccctgt
gcagtcatatagagcagacgGTTCTGGACGTAAAGGACAAGCTTTGATCTGAAGTGCGGCaagttatactgccc
gcagtcatatagagcagacgCAGTATGAGGGCAGATTGTGCTATGGAGATGGTTTTATGAaagttatactgccctg
gcagtcatatagagcagacgCAGTTACATCCTGACCTGCCGAATTATTAGTAAGTAGACAaagttatactgccctg
gcagtcatatagagcagacgAGCTCCAATCCCATAACTCGACCACAAAAATAGAATTCTAaagttatactgccct
gcagtcatatagagcagacgGGTCTTTCCTGACAAGTTATATGATTTAGTGCTTTGTTTCaagttatactgccctgt
gcagtcatatagagcagacgTTTTTACGCATCTAAGTGTACAAGGAAATCAGACTGCTTCaagttatactgccctg
gcagtcatatagagcagacgAAGACATGCTCGTTGTTGAGTATCCTTTTTAGGTCAGCAAaagttatactgccctg
gcagtcatatagagcagacgAGCGGAAAGGCCTGAAGCTCTGATTTATCACAACTCCCTGaagttatactgccc
gcagtcatatagagcagacgCATGGCTGGAGAAAGGAAAACATTTGTTGTACTAATACTGaagttatactgccctg
gcagtcatatagagcagacgTCCTGTGTAAAATAACCTTTAACTCTGTGGAGATCCCCCAaagttatactgccctg
gcagtcatatagagcagacgGAAATCCAAATTCCCCACAGGCAGCCTCTCTCCTCAACAGaagttatactgcc
gcagtcatatagagcagacgCTCACGACAGAAACGCTTACTGAGGTAATGCCCCGCTAGGaagttatactgcc
gcagtcatatagagcagacgTTTCACCTGTTTGAAGACACGACTCCATGAATTGTCATCAaagttatactgccctg
gcagtcatatagagcagacgGTGTGACATTTTGACCTATGAAGAACACTGGGCGACAGCAaagttatactgccc
gcagtcatatagagcagacgGACCGTCCTCTCATGCCAGCTTCAGTTATAGGCACTGCTGaagttatactgccc
gcagtcatatagagcagacgTGACCAATAGCAGAACCAGCCCTCAAACCCCAAGCCAGACaagttatactgcc
gcagtcatatagagcagacgTCACCATATCTTTACTGTAGAAACACTGTTAAGCCTGCAGaagttatactgccctg
gcagtcatatagagcagacgCATGAGAATGCCTCTTTCTTTGCGAAACTTGAAGGGTATTaagttatactgccctg
gcagtcatatagagcagacgTGGTCAACACGATTAACAAACCTTTCAAGTCTCTTTGGATaagttatactgccctg
gcagtcatatagagcagacgTGGGCTCAAGCAATCCTATCGCCTCAGCCTCTCAAGTAGCaagttatactgccc
gcagtcatatagagcagacgACTGTGCCTGGCCTCTTTGGGTTTTTGAAAGCCTCTATAGaagttatactgccctg
gcagtcatatagagcagacgCTCTCTGAGTTCTGTGGAAAGTGCATCCATGAGATGGGGAaagttatactgccc
gcagtcatatagagcagacgGCTTTCAGAGCTTACCTAGCACGCTGGCACTACAATGTCTaagttatactgccc
gcagtcatatagagcagacgCATGTTTTATCTCAGCACCTTTATAGTGATCTTGTATTCCaagttatactgccctgt
gcagtcatatagagcagacgCCACAGACACAAGAGCTTTCCAGAGGCAATCAATGGTCCAaagttatactgccc
gcagtcatatagagcagacgTTTTAAAAGACAACTGAGCTGTTTGAGAGAGAGAAGGAGGaagttatactgccct
gcagtcatatagagcagacgCCTCACCTTCCTCCCTCTCTGCACTTACATCTGCAACTCCaagttatactgccc
gcagtcatatagagcagacgGCCAGATCTCCCAATTCATCTTGATTTTCATGTCTTGCCTaagttatactgccctg
gcagtcatatagagcagacgCTGTCTCAGTGAGTGCATACAAACCTTAACAATACAAGAAaagttatactgccct
gcagtcatatagagcagacgCTGTATCAGTGGGTCATGAAACTCCATTTAAGTCTGACATaagttatactgccctg
gcagtcatatagagcagacgGGTACCTTAGCAGGTATTTGAGAAATATTGATTATCAACCaagttatactgccctg
gcagtcatatagagcagacgAACACTGGTTAGGAATCATGAAACTCAGGCTCTGTCAAGAaagttatactgccct
gcagtcatatagagcagacgCAATTTTTGTCAGTCTCTTAGCCTCTCTGGTTTACGATTTaagttatactgccctgt

gcagtcatatagagcagacgAATTCAGGATTGTGTCAGGGATGTATATAGCCAGAAGATAaagttatactgccct
gcagtcatatagagcagacgAGAGTTTCATTCTTGTTGCAATGGCTGGATCTGCAATGGCaagttatactgccct
gcagtcatatagagcagacgCTCGGCTCACCCGCAACCTCCACCTCCCGGGTCCTGGTTCaagttatactgc
gcagtcatatagagcagacgTGGCTGAAGTTTTGTTTTCTGTTGTTGATGAAACCAAACAaagttatactgccctg
gcagtcatatagagcagacgGTGTAAATTTTGTGCTAGCAGTCAACAGTTTTGAATCATGaagttatactgccctg
gcagtcatatagagcagacgCTACTGCCCCTCCTTCCTCGCCCTCTGTTAATATCCACCTaagttatactgccc
gcagtcatatagagcagacgCTACTTTCCTTTTAGCTCCTTCCTGTTTCTTTCCTATTCTaagttatactgccctgt
gcagtcatatagagcagacgGCTTTGGTTAATGCCTCTATGCTGATGTCTCATTTATGTAaagttatactgccctg
gcagtcatatagagcagacgTGTCTTTCAGCAGTCTACTTGAGATTTCCTTTTGAATACCaagttatactgccctg
gcagtcatatagagcagacgAACATGTCTAAAGTAAGAACTCATATTGACTCCCCCAAACaagttatactgccct
gcagtcatatagagcagacgTGCCACTACACCCATTAATGCACAAATTAGAAACTTCAGTaagttatactgccct
gcagtcatatagagcagacgTAAAAATTAGCTGGACATGATACCAGCTACTTGGGAAGCTaagttatactgccct
gcagtcatatagagcagacgTGAGCCCAGGAGTTCGAAGTTAAAGTGAGCCGTGATTGCAaagttatactgccc
gcagtcatatagagcagacgCCCTGTTTCTAAAAAAAAAACCAAGGCCGGGCGCGGTGGCaagttatactgcc
gcagtcatatagagcagacgTCAGGAGATCAAGACCATACTGGCTAACGCGGTGAAACCCaagttatactgcc
gcagtcatatagagcagacgACCGTACTCCAGCCTGGGAGTCAGCGAGACTCCGTCTCAAaagttatactgcc
gcagtcatatagagcagacgCATCCTTTGACATCATTCTCATAATTCCCTATGGCCAATTaagttatactgccctg
gcagtcatatagagcagacgTTGCTGTCAAATCCCCATCCTCAAGCTACTGTCCTCTGTTaagttatactgccct
gcagtcatatagagcagacgCTCACTGTTCTTCTGGCCCCTCTACCTGCAAGACCCTGCGaagttatactgcc
gcagtcatatagagcagacgTGGCTTTCTTCTAGTCCCGCATAGTCACCAGGCACCCTCAaagttatactgccc
gcagtcatatagagcagacgGGACCTTTGCCTGTATCTATTCTCACTCCTTTTTCACTCAaagttatactgccctg
gcagtcatatagagcagacgACTCATCAGATCAGAGAAGCCTTCTGACCTCACTGACTCCaagttatactgccc
gcagtcatatagagcagacgATCATTTGATTAAGGTGTGCTCCTCCAAGGCTGAAACTCCaagttatactgccct
gcagtcatatagagcagacgCATTGTGCTCCAAGAAACTAACACAGTAAACACTTCAGTTaagttatactgccct
gcagtcatatagagcagacgAGTTATTCACTACCAAAAAAAGGGGAACTATTGACAAGTGaagttatactgccct
gcagtcatatagagcagacgGCATTTTGTGAATCTAAACTGCTGTTCAACGCCAGGCACAaagttatactgccct
gcagtcatatagagcagacgGGCGAAACCTGTCTCTACAAGAAACAAAAACTTAGCCGGTaagttatactgccc
gcagtcatatagagcagacgAGGCTACAGTGAGCTACAACAGAGCAACTGCACTCCAGTCaagttatactgcc
gcagtcatatagagcagacgGGCTGAGGCGGGAGGATACTTGAGCAGGAATTCGAGGTTAaagttatactgcc
gcagtcatatagagcagacgTCAAGTATTAGGTTGTGTTTGAAGTGACTAGTTAGGATGTaagttatactgccctg
gcagtcatatagagcagacgGTGTAGACTACTTGAATACAGTGTCCAGATTTATGAAGACaagttatactgccct
gcagtcatatagagcagacgGAGTCAAGCCCTCCTCCAAAGTCTTTAACAGAACTGCTGAaagttatactgccc
gcagtcatatagagcagacgGCATTTGTATACGTGCACATAAACTCTCCATATCTGTTAAaagttatactgccctg
gcagtcatatagagcagacgAATACATTTTTCACAGTGTCTCCCTCCAAAAACAAATCTCaagttatactgccct
gcagtcatatagagcagacgGGGAGTGCCAGCAATAAAAATAACAAAAGCCTATAATCATaagttatactgccct
gcagtcatatagagcagacgACGGTCAGTAGTTGATTCCCACAAAGAAAACGAGCCAGGCaagttatactgcc
gcagtcatatagagcagacgAGGCCGAGGCAGGCAGCTCACCTGCAGTCAGGAGTTGGAGaagttatactgc
gcagtcatatagagcagacgGCACTCCAGCACAGGCGAAAAGAGCGAGACTCAGAAACAAaagttatactgc
gcagtcatatagagcagacgCAACAAAAAAAAACAAACACCAGGTCCAGGTCCAAGTAGTaagttatactgccc
gcagtcatatagagcagacgGGCAAGTTTACCGGGATCCAAAGGCAGATAATTCCAATCCaagttatactgccc
gcagtcatatagagcagacgAACCACAGAAGAAAAAAATTACAGTCTAACTCCACTCTTGaagttatactgccct
gcagtcatatagagcagacgGATAATGATCAATAGGGTTTATTCCCGGAATGTAAGGTTAaagttatactgccctg
gcagtcatatagagcagacgGAAAACAAACAACTCTCATCCAAAAAGGGTAGAAAAGAACaagttatactgccc
gcagtcatatagagcagacgCGGTATCTGCAAAAATAAATCTCTGGCAAATGTAATTCTCaagttatactgccct
gcagtcatatagagcagacgACTTTGTGTTGGAGTTCTAGCCATTGCAGAATATCAAAAAaagttatactgccctg
gcagtcatatagagcagacgAATTCAATAATTCTCTACATGGGTAACTCAAAGAGTCTGCaagttatactgccct
gcagtcatatagagcagacgGCAGGATTGCTAGACATAAAATCAACAGGCAAAAGTCGATaagttatactgccc
gcagtcatatagagcagacgAGTCTGGTCTTTAACTTTTGGACTCAATTGATCCTTCTGCaagttatactgccctg
gcagtcatatagagcagacgGATTACAGACTAAGCCACTACACCTGGTTCAATGTTTTTAaagttatactgccct
gcagtcatatagagcagacgCTAGACCATAGAAAAAGACAGAAAGCTCCACATTTAATTGaagttatactgccct
gcagtcatatagagcagacgAGAATGACCTCACTTAAGACTATAACTGCAAAAATTCTGGaagttatactgccct
gcagtcatatagagcagacgAAATATAAACAAGTGGGGTTAGTCTAGGAATACAGGTATGaagttatactgccct

gcagtcatatagagcagacgAAGAAACCTGTCAACATAATTCATCATGCCAGCAAATTAGaagttatactgccctg
gcagtcatatagagcagacgAGGCATTTGAAAAAGTTGAGCAGTCATCCCAATTAAAAACaagttatactgccctg
gcagtcatatagagcagacgGCAGAAGCTAACTTGTCCTACAAAGAGCCCCAAACTTTATaagttatactgccct
gcagtcatatagagcagacgACACAGTAAGTTATTTTTCCTCATGTCATAATCCAGTGTGaagttatactgccctg
gcagtcatatagagcagacgAAGACCTGGGCTCTTTCCATCAGTAGTTCTTCCATCTTCTaagttatactgccctg
gcagtcatatagagcagacgCTGGTAGAGAAAGGAAGGCAGTGTGGAGAGTGTGGGAGGCaagttatactgccc
gcagtcatatagagcagacgGAAACATAATCTCTTTTTTCCACCCACATTCTTCTGATCAaagttatactgccctg
gcagtcatatagagcagacgCCATAAGCAAGACTGGAAAATGTAGTTGATCTCTGTTCTCaagttatactgccctg
gcagtcatatagagcagacgTGCCACAGAAAGTTACCACCTAAAATATTAAAGACCTTACaagttatactgccctg
gcagtcatatagagcagacgTGAAATCAGGAAAAAAACTGGGACATTCACTATCACCATTaagttatactgccctg
gcagtcatatagagcagacgACATTATTTGGAGGTTTTAGCAGGTACAGTAAGATAAGGaagttatactgccctg
gcagtcatatagagcagacgGCTGATAATATGCCAACATACTTAGAAGCTCTAAGATATGaagttatactgccctg
gcagtcatatagagcagacgCATACGTGTGTACATTGTTCTCCATATGTTCTCTTCGTTTaagttatactgccctgt
gcagtcatatagagcagacgATAACTCTTGGGGTTTACACAAGCTCTCAATGTCATCTGTaagttatactgccctg
gcagtcatatagagcagacgACAGACTATTGCTTATGGCAGAAAAAATGCACTCAACTTGaagttatactgccctg
gcagtcatatagagcagacgCTCCACTTTTACCAGATGAGTTGTTTGTTTACCAATTTCTaagttatactgccctgt
gcagtcatatagagcagacgTAAACCAAGTAACTTGAACAAATACTACTGAAGCCAACACaagttatactgccct
gcagtcatatagagcagacgTTTTTGTTTTCATGGAAGCTGTTGTCTGCTTTAGAAAGGTaagttatactgccctgt
gcagtcatatagagcagacgCTGTTGGTTTGGGCATAACAATTGTAAAGCATAAGGAGAAaagttatactgccctg
gcagtcatatagagcagacgCCACATTAGAGAAATCCAAACTGAAAACTGTATATGATGCaagttatactgccctg
gcagtcatatagagcagacgGTACTCCAAATCAGTAGACCCATACTTAAGAGAAGGACCTaagttatactgccc
gcagtcatatagagcagacgAGCCTATCCTGATTAAAGCTAGGCTAGGTTATATTACAGTaagttatactgccctg
gcagtcatatagagcagacgTCTCAGTGACTTAAAACCACAGACTCTTTATCTGGATGACaagttatactgccctg
gcagtcatatagagcagacgTCTTCTATATGATGACATTCTAGCTACTTTACCCTTTCGGaagttatactgccctgt
gcagtcatatagagcagacgCTACATCAGCATATACTTCTATTATCACTATGACAGGGAAaagttatactgccctg
gcagtcatatagagcagacgGAATCATGCTTCAACTTCTCAATGCTTCTGTGCAGGAATGaagttatactgccctg
gcagtcatatagagcagacgTTTTTGGAGATAGAGTCTCGGTGTATCCTCTAGGCTGGAGaagttatactgccctg
gcagtcatatagagcagacgTGGTGAGATCTCGGTTCAATGCAACTTCCGCCTCCCGGGTaagttatactgccc
gcagtcatatagagcagacgATGACCCACTGCACCTGGCTGGCTATATTTAAGAGACAAGaagttatactgccc
gcagtcatatagagcagacgGCGTCACCTTCTTGCATACTTGAAAGTAAGAGAAAAACCAaagttatactgccct
gcagtcatatagagcagacgATTGCATAGGTGTCATTTTCTAGGATTTCTCAATCTCGCCaagttatactgccctg
gcagtcatatagagcagacgAAAATATTTGCTGCCCTTCAAGTTGAAAGAGTACTGATCCaagttatactgccctg
gcagtcatatagagcagacgTTCTTAGGTGTCAGGCTTGGCATTGTGACTTGAAGGGCCCaagttatactgccct
gcagtcatatagagcagacgCGGATGCTTCAATGTCGGATATGGGCACTTCTTGCTTTGGaagttatactgccct
gcagtcatatagagcagacgGCTTTCAGAACCAGTAGATGCTATACTTTGACTCCTCTCTaagttatactgccctg
gcagtcatatagagcagacgACTGCAATGTGCCCGACAGCAATGGTTTCATTAGCCTGGGaagttatactgccc
gcagtcatatagagcagacgCAGCTTGGGATGGGCATGCAGTATGAGCAGGAAATTTTTTaagttatactgccct
gcagtcatatagagcagacgTTTCAGGTTGCTTGTTATTGCAGTGTAATTTAGACAAACCaagttatactgccctgt
gcagtcatatagagcagacgCCAAAAAGAAGTGAGAATGGAAAAACACAGTCATATGCCAaagttatactgccc
gcagtcatatagagcagacgTGTATTACAGTGGCCTACAGGATTCCATACAGTAACATACaagttatactgccctg
gcagtcatatagagcagacgCCACCTAGGTTTGTGTAACGACACGATGACATTTTTCACAaagttatactgccctg
gcagtcatatagagcagacgAAAAGTACAGAATGCTTAAGGCTAAATTTGACAAGACAGGaagttatactgccct
gcagtcatatagagcagacgTGGAACAGAATTTACCCACAATGATGCTGCCAGGATTAAAaagttatactgccct
gcagtcatatagagcagacgAGTATGTAAAGCACCCAGGACAGGATTTGACATCTATAGTaagttatactgccct
gcagtcatatagagcagacgGGGAGTGCCAAAATGTATCTTAGTATACATGGGATTTTAAaagttatactgccctg
gcagtcatatagagcagacgATACACTACAGGTTAGGAAAACATTCTCAACTGAAATAGCaagttatactgccctg
gcagtcatatagagcagacgCAGTAGACTTCAAGAAATATCTGGGATGTATCAGAAAAAGaagttatactgccctg
gcagtcatatagagcagacgAAAACTCCACAGAAAAACGAGTAAAGAATGTGTGAATAGGaagttatactgccct
gcagtcatatagagcagacgGAACAGTAATCAGAAAGATCCAAACTAAAGTAATCCTGAGaagttatactgccct
gcagtcatatagagcagacgCTTCTCACCCATCAGAACCACAAAAGTTAAATTTCTGTTGaagttatactgccctg
gcagtcatatagagcagacgACTTGGAATCATGTTTACCATTTGGGGAAGACGTTTGAAAaagttatactgccctg
gcagtcatatagagcagacgCTTTAATATAGTGGTCAAGGAGGAAATTATGTGGCATTGCaagttatactgccctg

gcagtcatatagagcagacgAAATGGATTTTGACAAGACTCTGACTTAGGCTGCTCGTTGaagttatactgccct
gcagtcatatagagcagacgCAGTGTATGTTCCTCTCGGCTCCAAACTCAGAATTTTGCAaagttatactgccct
gcagtcatatagagcagacgTTGTCACTGCACAGCGTTTTCTCATTTCATGACCACTCCAaagttatactgccct
gcagtcatatagagcagacgGAAGCTCCTAACGGCACTGCCTGGCACACCCTAACCACACaagttatactgcc
gcagtcatatagagcagacgCTGGCTAACTCGACCTTTTCAGATCTGTGACTCGTATGGCaagttatactgccct
gcagtcatatagagcagacgCTGATTCTTCACCTAGTTTCTCAGCCCCTACTTTCTCCACaagttatactgccct
gcagtcatatagagcagacgCCTCTCAGCACATGGATACCTACTGTAGTAAATAAATTAGaagttatactgccct
gcagtcatatagagcagacgGAGCTAATTTCACCCAGTTCCCACCACCACACCTTTCACAaagttatactgccc
gcagtcatatagagcagacgCTCTGGAGATGGGGAGAATCACTTCAAAAGCAGGTAACAGaagttatactgccc
gcagtcatatagagcagacgAGGCTATTACGATGTTCCTAATGAAAGATGACAGCTCCAGaagttatactgccct
gcagtcatatagagcagacgTGGTGAATGGAGTTTATAGTCTAATAAGGAGAAGCACATGaagttatactgccct
gcagtcatatagagcagacgATAAACAATGAGGCATAAAGAGTCTATTTTCCCCAAGCAGaagttatactgccct
gcagtcatatagagcagacgGTAGGGCAGCGGAGATGGTAAGGAAAGGTTTTGTAGAGGAaagttatactgccc
gcagtcatatagagcagacgTCCAAAAAGAGGGGAGAACAGGAAAAAATGTATAGGGGTAaagttatactgccc
gcagtcatatagagcagacgACTGGAGTAAAGAATTTAAGAGTGGGGTTGACAGATTACAaagttatactgccct
gcagtcatatagagcagacgTAAAGAGTCATAAATTAAATCTGTAGGGCGGGTAAAGAGGaagttatactgccct
gcagtcatatagagcagacgTTTGAGAAAACAACTTTGGCAGCGGTGTGGAGGACGGTTTaagttatactgccc
gcagtcatatagagcagacgCTTGTGGCTCGGGTGTCAGGTAATTAATTCTGTTATCCATaagttatactgccctg
gcagtcatatagagcagacgACTTCTCAACCAGGCAAGTTCATCAAGGAATTTAAGAAACaagttatactgccct
gcagtcatatagagcagacgAATAAATGAGTGAGGCTGAATAGTTTTTGGCATTAATCCCaagttatactgccctg
gcagtcatatagagcagacgCAGTGAGAAAACAACAAAACATGGAACATAGTATCTATCCaagttatactgccct
gcagtcatatagagcagacgGGGTTACTGGAGAATTTTAAGAGTTATGATTTCCCTGGCAaagttatactgccct
gcagtcatatagagcagacgCCAAAAAACATACCTTCAAAAATTCCAAACAGCCATTTGGaagttatactgccct
gcagtcatatagagcagacgAATTGCCTACGTGTAATTTCTCATTCCCGAAGAAGTAAAGaagttatactgccct
gcagtcatatagagcagacgTGGTCATAAAAAAGCCAAAAGAAAATTTCCGAGCTCACTTaagttatactgccct
gcagtcatatagagcagacgAGATATATTCTGTCACATAGAAGAACGTCTAAGGATATCCaagttatactgccct
gcagtcatatagagcagacgCGTTGGTTATCTCTGGGTAATGCGTTTAAGTGTTCTCTATaagttatactgccctg
gcagtcatatagagcagacgGTGAACATGCATTGCTTCTGGAAGAAAAAACAAAAAAGTCaagttatactgccct
gcagtcatatagagcagacgCAACCATACAATCAGAATGTCGCAGGCCATCTTTTTCAGCaagttatactgccc
gcagtcatatagagcagacgAATGTAAACTGCTTTCAAGGTAACATTCTGGTGTGTTTTGaagttatactgccctg
gcagtcatatagagcagacgTCAAAATACAGATATTCCCACTAATTATTCATGCCAGTGGaagttatactgccct
gcagtcatatagagcagacgTGTATACATGTATATATTAGGCTTTGTGAGGCTCTGCACAaagttatactgccctg
gcagtcatatagagcagacgTTTGGCTAATATTAAAATTACTCCATGCTCCCACATGTGGaagttatactgccctg
gcagtcatatagagcagacgCAAATGATGGGAGAAGGCTATAGTAAAAGAAATAGGTGTAaagttatactgccct
gcagtcatatagagcagacgAGCTGTATGCTGTGCAGGCAAATGAATAGATTATGGTGAAaagttatactgccct
gcagtcatatagagcagacgGCATGAAAAATGGAATGCTAACTTGCGCCATGTCTGATGCaagttatactgccc
gcagtcatatagagcagacgAAACTCCTGGATTCCAAATAGTCTACTCTAGTACTTATTCaagttatactgccctg
gcagtcatatagagcagacgCTTTTCATAATCAGCAGACACTCTTTTAAAAGCAGCTGCAaagttatactgccct
gcagtcatatagagcagacgACACTCATCTCTTAGGGTTAAAGGGTCAAATGAACTTGATaagttatactgccct
gcagtcatatagagcagacgACATAGGAAGGGGCTTGCAGAGAAGATTTGTAAATATGACaagttatactgccct
gcagtcatatagagcagacgCTAAATTATGCATATTGAACCCTTCCCAACACTGTAAGAAaagttatactgccct
gcagtcatatagagcagacgTTATAACTTATAGATTCTTTTCAGGCCCTCGAAAACCTAGaagttatactgccctg
gcagtcatatagagcagacgGATTTGTATAATACCACTACCCTTATATGAACTACTCCTCaagttatactgccctg
gcagtcatatagagcagacgTTACAGGAGTGAGTCTCTGCGCCTGGGCCTGCCTCCTTTTaagttatactgccc
gcagtcatatagagcagacgCAGCTGCATCTTTGTCCTTGATTCCCTCTCACAACTGTTCaagttatactgccct
gcagtcatatagagcagacgTCCTTCTTGCTGCTTCTCTTCCCTGCTGTCTTCACTCTTGaagttatactgccctg
gcagtcatatagagcagacgACCAACTCTTTCTAACATCTGCCTAACAAACTTTGGTCTGaagttatactgccct
gcagtcatatagagcagacgCTTTCTGAAAAGAATGATCTATAAAATTTGGCTGCCTCCAaagttatactgccctg
gcagtcatatagagcagacgATGACATCCCATTTGCCAAACCCCATCGATCTTCCCATAAaagttatactgccc
gcagtcatatagagcagacgGGTCATTCCCTTATCTCAACATCGTATCTTCCTTAGCTCTaagttatactgccctg
gcagtcatatagagcagacgATTTTCAGACCTCTCTTCTCTATGCCTCTTCCAGCTTCTTaagttatactgccctg
gcagtcatatagagcagacgATTTCAAAATCCTGCTGAGTACCTGCCTAAAGTATGCAAGaagttatactgccct

gcagtcatatagagcagacgTGTGGAGACAAAGCAGCTTTTAAGAGGGGGCTAACCAAGCaagttatactgcc
gcagtcatatagagcagacgCCAAGCTTAGGCAAATAACAGAAAAGAGCAAAGTTAAGTGaagttatactgccc
gcagtcatatagagcagacgAGCTGACAAGATGAAGTACAAGAATAGTACCCAAACAGGAaagttatactgcc
gcagtcatatagagcagacgATGAGGTAACAAATGGGGAAAGAGCTAGGAGGAGGGAGGAaagttatactgcc
gcagtcatatagagcagacgAGCTGAATTTGTCCTCAGTTTACCTAGCTATGTCACCAGAaagttatactgccct
gcagtcatatagagcagacgAAAGTAAAACAAAATAGGGTGCTTGTTCTCATACAATGCCaagttatactgccct
gcagtcatatagagcagacgTTGAAACAATATATGCAAAAAACTAAACCCTGGCCTGACAaagttatactgccct
gcagtcatatagagcagacgCTCAAAAGTACCCCGTATTAGTTAATACTGTAACAAACCAaagttatactgccct
gcagtcatatagagcagacgGGAATTTTCTCCTTGCTGAGCCACTAAAACAACTCTTCACaagttatactgccct
gcagtcatatagagcagacgGGCTTTATGATCCCTTTTTAGCTACTTTTGAACTACTTTCaagttatactgccctgt
gcagtcatatagagcagacgGGCTCCCCTTCTTCTGGCTGTCTTTTGAATTCTGGTATTTaagttatactgccctg
gcagtcatatagagcagacgCTTTTTCCTTCCTTTCAAAGCCTTTCCAGTATGTGACACTaagttatactgccctg
gcagtcatatagagcagacgATTAGTGGCTTAAATGGTATCCTGGTTCCTACTTTCTCCCaagttatactgccctg
gcagtcatatagagcagacgTTTAATGCCAAATGTTTGTGAAAAGACCTCCATTATCTCCaagttatactgccctg
gcagtcatatagagcagacgCTATAAGAAAAAGCCCTAAACCTCAGCATACCATAGAAGGaagttatactgccc
gcagtcatatagagcagacgTGGCCCCAACCTGTCCCTCCAACCTCATCTCATTTTGCTTaagttatactgccc
gcagtcatatagagcagacgGTTTTCCATTACTCAGATAACAACACGACTTTTCACAGATaagttatactgccctg
gcagtcatatagagcagacgCATGTTCCTCTTCCGTGAATGCTGTTTTCTCCTGCTCCCAaagttatactgccct
gcagtcatatagagcagacgTTAGCCCCTTTCTCTATCAAGCATTTATCACACTGCATTTaagttatactgccctg
gcagtcatatagagcagacgCTGCAACCTCTACCTCTCAGGTTCAAGCGACTCTCGTGCCaagttatactgcc
gcagtcatatagagcagacgCTGCACCCAGCCCATGCTGCATTTTAAATATCTGTTTGCAaagttatactgccct
gcagtcatatagagcagacgTCCCCAGGAAAATATATAAGCATCTCAGGTACAGGGAATTaagttatactgccc
gcagtcatatagagcagacgTTCTTCTTTATAGCTCCAGAATCTAGTACCAAGCAAATGTaagttatactgccctg
gcagtcatatagagcagacgTTCTCAGGTGGAATTGACGGAAAGAAAAAAAGATGCAAACaagttatactgccc
gcagtcatatagagcagacgATCTCAGGATTAAACATGGCTATTGTGAAGGACCAGAAGAaagttatactgccc
gcagtcatatagagcagacgAAAGGACAGCATGGCAGAACACATAACATCTGGAACTTCAaagttatactgcc
gcagtcatatagagcagacgAACATATATGGCCTTATCATTTTCCTCTCTCAAATCCTTCaagttatactgccctg
gcagtcatatagagcagacgCTTTTTAAAGACAAAGCCTCATCCTTAATCTTGCCTACAAaagttatactgccctg
gcagtcatatagagcagacgAGGTCCCATGATGCACGCCTGTAGTTCCCATTACTCAGCCaagttatactgcc
gcagtcatatagagcagacgGGGCAACATAGCTAAACCCTATCATTATTTAGAACAAACAaagttatactgccct
gcagtcatatagagcagacgAACAACAACAAAAACCACTTGCCTCCATCTACTTACAACCaagttatactgccc
gcagtcatatagagcagacgCCAGACTTAGGCCTTCTTTCCAGTTCCGTTTTTGCACAAGaagttatactgccct
gcagtcatatagagcagacgATGTTGTTTTAAATATCCTACTACCCCAAACTAGTTCGCTaagttatactgccctg
gcagtcatatagagcagacgTTTTAGCTGTCAGTCCAAAGATTCCCTAAGATGTCTCCATaagttatactgccct
gcagtcatatagagcagacgGGCAGCAGACTATAAGGCCGTTTTATTACATTATATTTGAaagttatactgccctg
gcagtcatatagagcagacgGCATATTTATTTAAGCCCTTCATGTCTCGGTTTCTTCACCaagttatactgccctg
gcagtcatatagagcagacgAGAGATAAAGGATGTAAGCACTTAGAACTGTGTTGATAACaagttatactgccct
gcagtcatatagagcagacgTGATCACATTTACTTAATTTGTAGTCACTATCGTCCCCTCaagttatactgccctg
gcagtcatatagagcagacgCCACACCGGCCTGGTCCTTTTTTGTATTCCTAGCCTCAGCaagttatactgccc
gcagtcatatagagcagacgCTTTATTAGCCGAACGATCTCAAGCAAATCACTCAAACCAaagttatactgccc
gcagtcatatagagcagacgGCCTCGTTTACCCTTCTGTGAAATGTAATAGCTACCTCAGaagttatactgccct
gcagtcatatagagcagacgTTTGTGTTAAATAAGGCGATGTACCTACAAAAGCACACTGaagttatactgccct
gcagtcatatagagcagacgTTTATTAATAAACCGGAGTCCGTGTCTTCCCTCCCCTCAGaagttatactgccc
gcagtcatatagagcagacgCGTCCCGACCCCTCCCTCCTCCCAGCCCACACCCTTACTGaagttatactgc
gcagtcatatagagcagacgGGGTCCCTTGGGATTGCAAGCGTGGGAGCCTCGGCCCCGCaagttatactgc
gcagtcatatagagcagacgGCCCTCCTTCCTTCCTCGCCAGGACGGACGCACTTCCGGCaagttatactgc
gcagtcatatagagcagacgCAGCCCTCGGAAACGGAAGTGAGCGGCGGGGTCGACTGACaagttatactgc
gcagtcatatagagcagacgCTGCGGAAGATGGTGAGTGGGTACCCGGGCGGCTGGAGTTaagttatactgc
gcagtcatatagagcagacgGCTCGGCAGAAACAGGCAGTCGCCTCCCAACCCGTCCCCGaagttatactg
gcagtcatatagagcagacgCCTCAGGCGCCATAATCTGTCGAGTCACGGGGTTTGGGTGaagttatactgcc
gcagtcatatagagcagacgGCTCCTCTTAGCTCGGTTTGCGCCCTTCCATCCCTTAGACaagttatactgccc
gcagtcatatagagcagacgGGCCCAGACCTCCTCCCCTTACTTGGCGCAGAAGTGGAAAaagttatactgcc

gcagtcatatagagcagacgGCCCAGAGCTCTCAGACTCCTCATCTCCCCCAGCCCCCGCaagttatactgc
gcagtcatatagagcagacgACTGCCTAGTAAACAGCCCACAACTATAGGTCGCATTTGTaagttatactgccc
gcagtcatatagagcagacgAGCTTGTAGTGTGCTAGATGTTAGAGGGGAAAAAATCAGAaagttatactgccct
gcagtcatatagagcagacgTGTTTGCCAGTATTGTTAGTATATCTCCTCCCTCCTTCTCaagttatactgccctg
gcagtcatatagagcagacgCTTAGAAGTTGTGGTATTTGAGAAGAACACTGAGTCACCAaagttatactgccct
gcagtcatatagagcagacgAGACTCTGATATCGGTACTGCTGGTAGGCCATCCATCCAAaagttatactgccc
gcagtcatatagagcagacgATGTAACTAGTAGTTAGGTTTTCGTAGATTTGCCTTTCTCaagttatactgccctgt
gcagtcatatagagcagacgACAGAAGGAAATTGGGAAAAATGACAATAACAGAATCCAGaagttatactgccc
gcagtcatatagagcagacgAGAATTCACCTTTTCTTTGTGCGTTGAAACCTTGTTACTTaagttatactgccctgt
gcagtcatatagagcagacgCAGAAGGAGGAGAATGTTCATATAGGGAAGTAATAGAAGAaagttatactgccc
gcagtcatatagagcagacgCAACTTCTGGTGGTTCAACAATCTTCTGACTTTATGATGGaagttatactgccctg
gcagtcatatagagcagacgAATCCATACACAGTCAGTAGAGGCCATATTTGAGTACCAAaagttatactgccc
gcagtcatatagagcagacgCTTTGTATTAGATGATTTTGGCCGACTGTAGGCTAGTGTAaagttatactgccctg
gcagtcatatagagcagacgAGGTAGGCTAGGCTAAGGCCTGATGTTTAGTAGGTTAGGTaagttatactgccc
gcagtcatatagagcagacgAGTCAAGGAGCATTTGTAGTCCTGGTAGGAGTTCAGCTCCaagttatactgccc
gcagtcatatagagcagacgGCAGATGAGTAAACAAGGCCAGAGAGGTTGGTGCTTTGCTaagttatactgccc
gcagtcatatagagcagacgTGACAGTCCACACTAGAGTCCAGATTCTTTTCCTCGTCCAaagttatactgccc
gcagtcatatagagcagacgACTTCTTGTATGTAGGGGAGTTGGAACCTATTATGCAGGTaagttatactgccct
gcagtcatatagagcagacgTTGTGCTTCAGACATCGATGAACCAACAAAGATTTTATAGaagttatactgccct
gcagtcatatagagcagacgGTTTCATGCTTGAGGAAAACCTTGATATACAGAACTGGAAaagttatactgccct
gcagtcatatagagcagacgAGTAGAATGCTGTTGAAGTTATCCATGTGTGATAGGCTGGaagttatactgccct
gcagtcatatagagcagacgGTGGTATAGTGAAGAAACCCTGCAAAGGATAAAATACCTGaagttatactgccc
gcagtcatatagagcagacgTGTAGCTGTTTTACATTTTTCTGCAGAATCACAGGGATATaagttatactgccctg
gcagtcatatagagcagacgCAAGTGATTTAAAGTTAGCGTCTTGGGCCGTGCATGCTGGaagttatactgccc
gcagtcatatagagcagacgACTAGCCTGGCCATCATAGTTGAGACCTTATCTCCACAAAaagttatactgccc
gcagtcatatagagcagacgCAGTTAGCCGTGATTGCTCGACTGCATTCCAGCCTGGATGaagttatactgccc
gcagtcatatagagcagacgCATCTCATGCTTTGCCTGTTGTAACACCATGTTTTCTTTGaagttatactgccctg
gcagtcatatagagcagacgCGGGAGGTAGACATGTAGCTTCAGTATCTAGCAGAGTACCaagttatactgccc
gcagtcatatagagcagacgTAGTAGATGGCTAATAAGTATCTGGTGATTGCGCATTTCAaagttatactgccctg
gcagtcatatagagcagacgGGTAATCATACACCAATACCAAATGCACTGTCTTTTTTACaagttatactgccctg
gcagtcatatagagcagacgACTATCCTGATTAGATACACATAGTTTTTATAGGTAGCGCaagttatactgccctg
gcagtcatatagagcagacgAATTCCCTTTTTTTCCCAGCTTCTCTACTTTTCTTTATCCaagttatactgccctgt
gcagtcatatagagcagacgGTCTGTTAATCCCATTTCTGTGTTGTTTTTCCTTGTCCTCaagttatactgccctgt
gcagtcatatagagcagacgGTATTCCCGAGGATGCTGTCTTCAACTTTTTTATTCTGTGaagttatactgccctg
gcagtcatatagagcagacgTTCCAGGGGAAACAAATTCAATCATGCCATAACCTTTCAGaagttatactgccct
gcagtcatatagagcagacgATGTTTCACTTTTGTATTTCATCAGCCTTTACTTCTGCCTaagttatactgccctgt
gcagtcatatagagcagacgAACATATTTAGCCAGTTATCTCAAACTACCTGAAGCTTGTaagttatactgccctg
gcagtcatatagagcagacgATGCCTTTCAAACTGACAGCATTCAGAGATCACCTCAGGAaagttatactgccc
gcagtcatatagagcagacgTGTACCATGTGTGTCTCTATCATATCATAATGTTTAAGGCaagttatactgccctg
gcagtcatatagagcagacgGTCTGGACTTTTTGAGGGCCTCTTATTTTTCTGTCTCCTAaagttatactgccctg
gcagtcatatagagcagacgAATGTTTTTTATTTGTGTTGTAGAATGCCAGAGGACTTGGaagttatactgccctg
gcagtcatatagagcagacgTCAGCAAGTGGACCTTTTGAAAGTCATGATCTTCTTCGGAaagttatactgccct
gcagtcatatagagcagacgATTTTGATAGGCTATACATGTGTATTTAGTTGGCAGATGGaagttatactgccctg
gcagtcatatagagcagacgCCTTACATGGGGTGCAGAGGGGAAGAGGCTTAGATATTTTaagttatactgccc
gcagtcatatagagcagacgAGGTGTCTCATCAAAGTAGTAGGTTCCCAATTAGAGCATCaagttatactgccct
gcagtcatatagagcagacgCAGTGGTGCCCATTAGCATGATTTGGTAGGATTCTTTTTAaagttatactgccctg
gcagtcatatagagcagacgGCCATTTGCCTTCTTGTCTTAACTATTAGGGTAATTCACTaagttatactgccctg
gcagtcatatagagcagacgTTGGGAGGCACTAGAACATAGCAGTTGAGAGATTAAACTCaagttatactgccc
gcagtcatatagagcagacgCCTGGGTTTGGATCGTGGCTCTGCTCTTTACTAGATGTGTaagttatactgccct
gcagtcatatagagcagacgCATAATTAATGTCTCAGTGTCCTTATGGATAGAATAGGCCaagttatactgccct
gcagtcatatagagcagacgAATGGTACCTGACATTAGAAGCACTCAGTATGTATTGGCTaagttatactgccct
gcagtcatatagagcagacgATGCAGTTACGCCATGTTTTTGATTTCCTTTTTGTTTCTGaagttatactgccctgt

gcagtcatatagagcagacgCAGTTTCTGGTACTAGACCATACTCTTTGCAGGTCACTTTaagttatactgccct
gcagtcatatagagcagacgAAAATAAGGATAACGACCTTTCCCCCATAAACATTTTGTGaagttatactgccct
gcagtcatatagagcagacgAGTGCAATGGCGCGATCTCCCCTCATCGCAGCCTCCGCCTaagttatactgcc
gcagtcatatagagcagacgCCACTGCGTCTGGCCGTAATTTACTCTTACTATAATAGAAaagttatactgccct
gcagtcatatagagcagacgTTCTAATGCTGGTGTCTGAGTACAAATATAATCCCTCATTaagttatactgccctg
gcagtcatatagagcagacgTGTAGTAAAAAAATGCAGCATATAGCCTGTGAAATTGGAGaagttatactgccct
gcagtcatatagagcagacgAGTAACTAAGGTCTGAAGATTTTTTTCCCCCCACATGAAAaagttatactgccct
gcagtcatatagagcagacgCTGATTGGACTGGGGCAGGACCATTTTCAAAACCCACTTCaagttatactgccc
gcagtcatatagagcagacgAAGGTAGTCATCTTTATATAGCTAGAGTGATAAAGACCAGaagttatactgccct
gcagtcatatagagcagacgGGTTCTAGTTGTGCCTTACCTAACATACTTTATACATAGTaagttatactgccctg
gcagtcatatagagcagacgCTGAAGGGAATAGGCTTAGTTTATATAGTGCAAGAAGTACaagttatactgccct
gcagtcatatagagcagacgCATTCATTAAATTTCTTCTAGATGTGGCACCAGATTTTGGaagttatactgccctg
gcagtcatatagagcagacgCCCTAGTGATTGAAGTCTTGACCACCAATGTCTATAATGGaagttatactgccct
gcagtcatatagagcagacgCCTTTTGGTGTGTAGGTCTATGAATTTTTAGCACATGTATaagttatactgccctgt
gcagtcatatagagcagacgTTGTCACCCCAAAAAACTCTCTCATGCCATCCCTTTATAGaagttatactgccct
gcagtcatatagagcagacgATTTTGAGCTCTGGGTATTGAGTGATATTTAATCTCCTCTaagttatactgccctgt
gcagtcatatagagcagacgATGAACTTTTGCCTAGTCATCCCCTTGAATTATCAGAAAAaagttatactgccctg
gcagtcatatagagcagacgAAACAATTTTGGTACTGAGGATTTGCCCATGAAGAATATGaagttatactgccct
gcagtcatatagagcagacgCTTGTAGGATAGACTTAAAAATTACACTGCTCCTCTAAAGaagttatactgccct
gcagtcatatagagcagacgTTTCTATGCGTTCTTACCTGCCTTTTTAATGTACGTCTCCaagttatactgccctg
gcagtcatatagagcagacgCCTCTCACAGGAATTTTCTCTTTCAATAATGCTGTTTCTTaagttatactgccctgt
gcagtcatatagagcagacgCTTCCAAGTTCATATCTCTAGTTTTGACCTCTTTAAGTCCaagttatactgccctg
gcagtcatatagagcagacgCCACTAGGCACAGTAGGTACTTCGATGAATATGACATCATaagttatactgccc
gcagtcatatagagcagacgATAACTTAAAATTTAATGAGGAAGACAAGCAGTGGGTAGGaagttatactgccct
gcagtcatatagagcagacgTCGTAAAATGCTAGAGAAGACCAAGGATGTGACAGCCACCaagttatactgcc
gcagtcatatagagcagacgGAGGGACCTTTGAACAGTTTTAGACCAGAAAGTTTACTCCaagttatactgccct
gcagtcatatagagcagacgAAAAAGGTTTCTGACTACTTGGGGAAGATGATGCAGTTGTaagttatactgccct
gcagtcatatagagcagacgTGTTGTTTGTTTGTTTCGCTTTCTAATAAGAGGATAGGTCaagttatactgccctgt
gcagtcatatagagcagacgATTCCTCATCGATTTAAGTGAATTCAGGTTTCAGGTGCAGaagttatactgccct
gcagtcatatagagcagacgGACAGCCCAGAGTACACTTATAAGTATGTGAGTATGATTAaagttatactgccct
gcagtcatatagagcagacgGCACAGTTGATCATCTTTGAGAACTAATGGGAGAAATGAAaagttatactgccct
gcagtcatatagagcagacgAAAATAAGATTGTCTGTGGTAACATTGATCCCTGAAACTGaagttatactgccct
gcagtcatatagagcagacgGTGAGCATTTAATAGGGTAAGCAGTGATCATCAGCATCTGaagttatactgccct
gcagtcatatagagcagacgCTCAGTTGCCAAATGGGTCAGTCAAGGGATCAAATTATCTaagttatactgccct
gcagtcatatagagcagacgGGGATAAGATTGAGGATTACATGGTGGATTATTAACCGAGaagttatactgccct
gcagtcatatagagcagacgCCAGAGAACGGTGATTAGGGAATAGTCAATTTGGAGGGAAaagttatactgccc
gcagtcatatagagcagacgTCAGGTGGACACATAGGAAGGATAAGTATTTTTCAAGGACaagttatactgccct
gcagtcatatagagcagacgAGGTTTTTGGAAAAAAGGTTGAGAAGGATAGTTCATTCTGaagttatactgccctg
gcagtcatatagagcagacgTTTAAGGAAAACATTTATAGAGGTGAGTGTGAAGTTCTGCaagttatactgccctg
gcagtcatatagagcagacgCAAAAAATGAATTCTACACCTTTGGGCAGGGTAAAGTGACaagttatactgccc
gcagtcatatagagcagacgGACTATTTTAATTGACTGTATACTCACTATAAGCCAGCAGaagttatactgccctg
gcagtcatatagagcagacgGTTAGTTACTAAAAAGAAAGGTTAATGCAGTCTTGGGCTTaagttatactgccctg
gcagtcatatagagcagacgCAAGGGAGATTATATTATTTTCCTGTGTATTAGTCTGGCTaagttatactgccctg
gcagtcatatagagcagacgGCAATACATATTAAGAGAAACTGACAAATCTGAGTGAGACaagttatactgccct
gcagtcatatagagcagacgAACTTCAGAGATCTTTTTGAGACTTACATACATCTACTGCaagttatactgccctg
gcagtcatatagagcagacgTATGTGACTCTTACTGTCTATTATAAATCCTTCAGCATGGaagttatactgccctg
gcagtcatatagagcagacgCCCACTTTGCTAAGTAAACTTGGGGCATATTGCTTAATTTaagttatactgccctg
gcagtcatatagagcagacgAGCCTTAGTCTTCTCATCTGGAGTTAGTAATAGCCCATCCaagttatactgccct
gcagtcatatagagcagacgCTCTGCATGATTGGGAGCTTCAGTGGTGATAGTGATGGAAaagttatactgccc
gcagtcatatagagcagacgCTACAAGGGAACAGTTGGCAAAGTCTGAGGACATTTTAGGaagttatactgccc
gcagtcatatagagcagacgACAAGTGGTGCGGTGGGGGATGGGGGGATGTAATATTGGCaagttatactgcc
gcagtcatatagagcagacgGAATTATCTGGCCCAAGAAATCAGTAGTGCGGAGGGTTGAaagttatactgccc

gcagtcatatagagcagacgGATTATAATCTCCAAGGTAGGCTGTTCAGTTTTGTTGTGTaagttatactgccctg
gcagtcatatagagcagacgGCACACGGTTGCTAACTTACATGTTGATTGATAATTACAGaagttatactgccctg
gcagtcatatagagcagacgTATTATACATAAAATTACATTTACTGCTGGGTGTGGTGGGaagttatactgccctg
gcagtcatatagagcagacgGCCTGGTAATTCGAGGCTGTAGTGCTCTATAATCGTGCCTaagttatactgccct
gcagtcatatagagcagacgGCAAGACCCGTCTCTTAGAATAAAATTGTATTATCATGAGaagttatactgccctg
gcagtcatatagagcagacgGCCAATATTTAAAGCAAACCCTTCACTTAGAACTTCAGATaagttatactgccctg
gcagtcatatagagcagacgACCCATAATAAAATACTGCAGGTTTTCCCTAAGGACCATCaagttatactgccct
gcagtcatatagagcagacgGTTAGTCACCTTACTTTCAAACTTCATAGTCATATCCCAAaagttatactgccctg
gcagtcatatagagcagacgGATTACTATTAGAAGACAATGTTATCAAGAGAGTAGCGTGaagttatactgccctg
gcagtcatatagagcagacgTGACTTTACCTCATTATAATCACCTTAGCTCAGACAAATCaagttatactgccctg
gcagtcatatagagcagacgACAGTCTGTTTGCATATTACATTCACTATCCTTAGGAGCCaagttatactgccctg
gcagtcatatagagcagacgCAAGTCGATTCATTTTGCCTGAAATACATTAAGTTATGGCaagttatactgccctg
gcagtcatatagagcagacgAGTGAAAGTATGATCCAGGAAAAATTGCTGATTAGTACCCaagttatactgccctg
gcagtcatatagagcagacgCTTTTGAGAGATGTTTGTTTTTCCCCTGAGGCAAGTACCAaagttatactgccctg
gcagtcatatagagcagacgTAACTATTTTTGTCAGTGGATGTGACATTTTCTGGATCTCaagttatactgccctgt
gcagtcatatagagcagacgACACCTTTGCTAGTTTTGCAGTACTTTTAATCAAAAGCGCaagttatactgccctg
gcagtcatatagagcagacgACTCTGGTAAAAGTACACCTGCATGCTAAATGAAATATAGaagttatactgccctg
gcagtcatatagagcagacgTGTGTTATATATATGCCACTGCTAACTGTTTTGAAAGCAGaagttatactgccctg
gcagtcatatagagcagacgCATTGTCTTTTGTAGTTCCAGCTCAACCAAGATAAAATGAaagttatactgccctg
gcagtcatatagagcagacgGAAACATTCAGGGTCTATTTGCTCCGCTAAAATTACAGATaagttatactgccctg
gcagtcatatagagcagacgGAGTTGATGGATCTCTGTTGCATATGTGTCGATATCATTCaagttatactgccctg
gcagtcatatagagcagacgTTTATACCTTATCTACCTTACAACCTATTCTACTTCCCTCaagttatactgccctgt
gcagtcatatagagcagacgAGTCATTAAATGGAAAACTTCAAATTACAGGCTCAGTGTCaagttatactgccctg
gcagtcatatagagcagacgTTACATTGAATTTGAGATCCCTCTAAGAATATCCAAGTGGaagttatactgccctg
gcagtcatatagagcagacgGAAAATGTCCTTCCAATTATGTAAGAACCACCCTCCTACCaagttatactgccct
gcagtcatatagagcagacgGGAAAATCCCTTTATCAATGATGCCTTCTCAAGCTGATATaagttatactgccctg
gcagtcatatagagcagacgGCTCTGTCTCAGCAGTAGCATTTGAACCGCTGTTGTCTTTaagttatactgccctg
gcagtcatatagagcagacgTCAAATTCCTTTATAGTAACTTGTCTCTCATTGAGTGTGCaagttatactgccctgt
gcagtcatatagagcagacgAAGTTGTCTACATTTGCTTTCTATACTCCTTTCCCTGCTAaagttatactgccctgt
gcagtcatatagagcagacgTCTCCCATGACACCAATGAAACTGCTCCTGAACAATGTGGaagttatactgccc
gcagtcatatagagcagacgACTTCATCTCCCTGTAGTATTTGACACTAATAATGGCTTCaagttatactgccctg
gcagtcatatagagcagacgGCAGGGACTACAGTCACTTGCTACCTCGCCTGGCTAGTTTaagttatactgccc
gcagtcatatagagcagacgAGCTATCATCCCACCTCCTTGGCTTTTAGAACACCACACTaagttatactgccct
gcagtcatatagagcagacgCTGAATCCTTGTTCTCAGGTGTCTCTGTGTAGGTTCCATTaagttatactgccctg
gcagtcatatagagcagacgAAATGCCAGTTTTGTTAGTGCCCCATCTCTAGTCCTTTCCaagttatactgccctg
gcagtcatatagagcagacgTAGTTATCAATTATATGTTGATGATTCCAACTGGGCGTGGaagttatactgccctg
gcagtcatatagagcagacgACAAAAAAAAATACAACAAATGTGCCAGCCGTGGTGGTGCaagttatactgccc
gcagtcatatagagcagacgTAAAATATGTTGATGACCCCCAAGTTACAATTCTCATGAGaagttatactgccctg
gcagtcatatagagcagacgCAGTTTTACATGGACAAGTACAGAATAACTCATAATCCCAaagttatactgccctg
gcagtcatatagagcagacgCATACTCCTTATCCCATATGAATAGGTTTCTGCCTAAGGCaagttatactgccctg
gcagtcatatagagcagacgGCACCCATCCTAACCTCTTCCCCTGCTCCCCCAAATCATTaagttatactgccc
gcagtcatatagagcagacgCCATTGCTACTATGCTGGTCCTTGCCCCTGTCAGACTTCAaagttatactgccct
gcagtcatatagagcagacgCAACACTCATCCCTGCTGCCCCTCCAATCCTTGTTAAGACaagttatactgccc
gcagtcatatagagcagacgGATTTGATTATGTCATTCCCCTCACTTTACCCTCTAAATAaagttatactgccctg
gcagtcatatagagcagacgCTCCTTGCTTCCCACCACACTTCTTATCCTCTTAAACTACaagttatactgccctg
gcagtcatatagagcagacgTTTCCTAATTCATTAAGTTCTTTCACTGTTAGGCGTTTGCaagttatactgccctgt
gcagtcatatagagcagacgCAGATTCCTTTCTCTGACCTAGCTAAATTGTCTTTTTCAAaagttatactgccctgt
gcagtcatatagagcagacgTTCTATTGGCCGAGCTGGGTGGCTCATGCCTGTAATCCCAaagttatactgccc
gcagtcatatagagcagacgTACTATCAATTGTATTTGGCTAAGTCATTCTGGAAGCTACaagttatactgccctg
gcagtcatatagagcagacgGATAAAGCTCTTAAAAAACATGGTTTTGAGTAACCTAGGGaagttatactgccctg
gcagtcatatagagcagacgCTTAGCCATCTACTAGTCTTAAGAACTTGAGTAAGCTGGAaagttatactgccctg
gcagtcatatagagcagacgGTGAAATAAGGAGAACAGTAAGAGTAGCATAGGATTATTCaagttatactgccctg

gcagtcatatagagcagacgGCTTCCCTTTTTATTTTCCTCTTACTCAGATTTCAACTACaagttatactgccctgt
gcagtcatatagagcagacgTCTTATCTCTATACTTTTGCAGTTTTGGGTTGTCAAGATGaagttatactgccctgt
gcagtcatatagagcagacgGCTTTTGATGCATTCTTAGGACTGTACTCTGATCTAACTAaagttatactgccctg
gcagtcatatagagcagacgTAATTTACAGGTTCCTATTTTTGTTTCTTCCTCTTCCTCCaagttatactgccctgt
gcagtcatatagagcagacgAATAATATATTTAGTCTGGATACATTTTGTTGGGGGTGGGaagttatactgccctg
gcagtcatatagagcagacgGAAACTGCCCTCTTAGATTCTAACGGTGTGATTATCTTTAaagttatactgccctg
gcagtcatatagagcagacgGGTAGTCTGTGGGAATATCAAAAATTATCTGCATTAGCTGaagttatactgccct
gcagtcatatagagcagacgGGACCTGTAATCCCAGTTTTAGTGGGAGGATCACTAGAGCaagttatactgccc
gcagtcatatagagcagacgAACCTATTGAGTTGGACTTTCTTCAGGAATTTGATCATCCaagttatactgccctg
gcagtcatatagagcagacgAGGAATGAGGCGTGATGTTAAGCTCTGGGATTTTGCATTTaagttatactgccct
gcagtcatatagagcagacgGGTATATGCCGTAGTTTTCCTGAGCAGAAGCAAAGCCTCAaagttatactgccc
gcagtcatatagagcagacgTATTCTGTCATTTTCCCCTGAGTTTTTTCTAATCTTGTCCaagttatactgccctgt
gcagtcatatagagcagacgCCCAAAGGTTCAGCGTCTTCCATTTCTTTCAAGCTCAAATaagttatactgccct
gcagtcatatagagcagacgACTGGATGTTTTGAGGGGTAATGATGAGACTATTGGATTTaagttatactgccctg
gcagtcatatagagcagacgAATGGTAAGTGTCATTCAGCACCTTTTTATGGAGCCCTTGaagttatactgccct
gcagtcatatagagcagacgACTTACAGCTGATTGGGTTCACTTTATTCATTGCATTAAGaagttatactgccctg
gcagtcatatagagcagacgCTGTTCTTTAACTGTTTTTAAAAGTGAAGATTGGCTGGGTaagttatactgccctg
gcagtcatatagagcagacgATTCTAAGAGTACATAGTCATTTCAGTCTAGTAATGGTGGaagttatactgccctg
gcagtcatatagagcagacgACTTCATGTCCTGAGCTTCAAGTGTAATATGTCAGTCTGAaagttatactgccct
gcagtcatatagagcagacgCATTGTTGGGATGTGACTGGTGTTGTTTTTGTGTGTTTAGaagttatactgccctgt
gcagtcatatagagcagacgTTTGTACTTCTACAGGTGATAAATAGGAAAGAGGAATCTGaagttatactgccct
gcagtcatatagagcagacgAGATGGAGTCTTGTGCTGTTACCCGGGCTGGAGTGCAGTGaagttatactgcc
gcagtcatatagagcagacgCCACTGCACCCGGCCGAATCTGTTATTTTTAATAAAATTCaagttatactgccct
gcagtcatatagagcagacgAGAACAATATATGCAGCATCCTCTATTTTTTCTGGTCCTGaagttatactgccctg
gcagtcatatagagcagacgACAATCCTGTCCAATGCTGCCTCTGCTCGTTATGCCCCAGaagttatactgcc
gcagtcatatagagcagacgTCCAACAGGATTAAGTTTTATTAGTTTTGCTTGGCTGCTGaagttatactgccctg
gcagtcatatagagcagacgGACATAACAGAAGTCTAGTGAGCCTCCTTTCAGAATCTTCaagttatactgccct
gcagtcatatagagcagacgGCTGGGATTCTTAAGCTTTTACATTGAACTTAGACCTTATaagttatactgccctg
gcagtcatatagagcagacgCTAAGGTGATAGTTCAGGTCTTTAGTTACAAAAGCATTCAaagttatactgccctg
gcagtcatatagagcagacgCTGTTTAAAGAAAGGGAGAGGGACTTAGGAGCATTTATCTaagttatactgccct
gcagtcatatagagcagacgCAGCTGTGAACAAAACAAATATCTGTGCTTTCAAATGGGGaagttatactgccct
gcagtcatatagagcagacgAATTCTAGGAATAAAAAGGATGTTTTTGGGTTGGGAGGGCaagttatactgccct
gcagtcatatagagcagacgCTAGAAGTAGAAGGAGAGTATTTCAGACAGAAGAGATGAGaagttatactgccc
gcagtcatatagagcagacgATGTAACCTGAGAACAGCTGAAGAACTCTGAACGGTTTGAaagttatactgccc
gcagtcatatagagcagacgGAGACTCCTAAAGGACAGTTTCTTATTCATAGCAATAGCGaagttatactgccct
gcagtcatatagagcagacgGAGTATCAGCATTTCTTTTGGTTCCCTTAGCCCCACTTCCaagttatactgccct
gcagtcatatagagcagacgGGTGAAGCCTTGTACATTGAGTGTGTTACGTTAGAGGAGAaagttatactgccct
gcagtcatatagagcagacgCAGTAAACCTGCCTGTCCTGTGCTGCTTTGTCTTATGAGGaagttatactgccct
gcagtcatatagagcagacgCTCCCAGCTTGCAGAATGGAGAGACCAATAGAGAACTGTCaagttatactgcc
gcagtcatatagagcagacgCATGTTTTGAGCAAAGTAGAGAGTAGAATAAGGTTGAGATaagttatactgccct
gcagtcatatagagcagacgAGCTCAGCAAGGAACAGATGAATCAGGGCCTACAGACCATaagttatactgcc
gcagtcatatagagcagacgCCATTGAGCAGTTTTAAGTAAGGATGTAATGTGATGGAATaagttatactgccctg
gcagtcatatagagcagacgGTTTAAAAAGGTCACTAGCTGGCTGGATGTGGTGGCTCAGaagttatactgccc
gcagtcatatagagcagacgGGCAGAGGTGGGAGGATCAGTTTAGCCAAGGAGTTTGAGAaagttatactgcc
gcagtcatatagagcagacgTTTTTAAATTAATTGGGCATAGTGGTGTGCAGCTGTAGTCaagttatactgccctg
gcagtcatatagagcagacgCACTTGCTATCTTGTGGGCATCTTTTAAAATTCATTCCTTaagttatactgccctgt
gcagtcatatagagcagacgCTAAGGTAGCTATTAGGAAAAATCACCTGTTTAGGTAGATaagttatactgccct
gcagtcatatagagcagacgTTGTTTTGGAGACACGAGTCTAGCTCTGTCACCCAGGCTGaagttatactgccc
gcagtcatatagagcagacgTTACAGGTGCCCACGACCACACCTGCCTAATTTTGTATTTaagttatactgccct
gcagtcatatagagcagacgGTAGATGATTTTTATCCACCTGGTAGATGATTACAGTGAGaagttatactgccctg
gcagtcatatagagcagacgGTATTTGGGGTAGTTTTTTTCTTGTGGGAACTGTTATTGCaagttatactgccctgt
gcagtcatatagagcagacgGGGACCTGAATTAGCTGTTCCTTTCTGCCTCAAATAGTTTaagttatactgccct

gcagtcatatagagcagacgTTCAGATCCCAACCTAAATATTACTCAGAGAGTCCTCCGTaagttatactgccct
gcagtcatatagagcagacgGTGGCAGAAGACAAAGGGGGGCCAGTGAATTATATGGCGAaagttatactgcc
gcagtcatatagagcagacgACTCAGAACAAGAACTCACTTATCCGGGGGGTGGTGCTAGaagttatactgccc
gcagtcatatagagcagacgCCGCACTGGGAATCACATGGCTTTGTTTTTTTTCTCTCTCaagttatactgccctg
gcagtcatatagagcagacgCCGCGCCTGGCCGGGAATCACATTTCCAACATGAGATTTGaagttatactgccc
gcagtcatatagagcagacgCATCCAAACCATATCAACTACCTAATCTAAAGTAATCCCCaagttatactgccct
gcagtcatatagagcagacgTATTATTTGTCCAACTTAAATATACGTGTCAGGAGAGCTGaagttatactgccctg
gcagtcatatagagcagacgTGTTGATCACATTCTTATCACCTTTGACACAAAACACAGTaagttatactgccctg
gcagtcatatagagcagacgGAGTAGCTTGTTAATCTAAATACTGTGTAAAGACTGTGACaagttatactgccctg
gcagtcatatagagcagacgTGGAGATGATGACTTTTTTGCCTGCTTATCAGTGAAGTACaagttatactgccctg
gcagtcatatagagcagacgTACAAGATTAAGAAGGACGTATAAGGACACTAGTAGAATCaagttatactgccct
gcagtcatatagagcagacgAGACTTTTTCTTAATGCTGGAAGGGGACTTTCATGTACTCaagttatactgccctg
gcagtcatatagagcagacgTCACTCAGAGTCTCACGGCGAGCTTGTGGTACAACAGATGaagttatactgccc
gcagtcatatagagcagacgCCCGTCTCCACCAGTGCCTTGCTTACATGATATGCTGTCTaagttatactgccct
gcagtcatatagagcagacgAAAAGCAGGGTGTGGAGAAAAGTTTAGTTACATAAATGAGaagttatactgccct
gcagtcatatagagcagacgAGATTTGCTGTAGTGACCCTCAATCCCATTTTTCTCGTTCaagttatactgccctg
gcagtcatatagagcagacgGACTGTTCTGCGTGTCTTTCTATGTAGTCAAAAATATTGCaagttatactgccctg
gcagtcatatagagcagacgGTGCCTTTTATATACACAGCAGTTTATTAATCTCCAGTCTaagttatactgccctg
gcagtcatatagagcagacgGAAACATCCACATTGTACTGAACTGTTCCTCTTGGTAGCCaagttatactgccct
gcagtcatatagagcagacgTAGGTTTTTATTCTGTTTCTAGTCCTATGTTGAGTAGAGGaagttatactgccctgt
gcagtcatatagagcagacgCAGTTAAAATAAGGCTTTTGTCTGACTCATCTGTAAACACaagttatactgccctg
gcagtcatatagagcagacgAAGAATTCTTACGTTTGTGTGGATACAACAGGTCAGAAGAaagttatactgccct
gcagtcatatagagcagacgTCCCGTCCTGTGGGAAGCCACTAGAAAACTTAAGAACTTTaagttatactgccct
gcagtcatatagagcagacgTTCTGTGGGTTTTAATTAACATGGAACCTGGTCTCCTAAAaagttatactgccctg
gcagtcatatagagcagacgAAGTTCCCTTGGTCCCTTTTCAGTCAGTCACCCCACCCCAaagttatactgccc
gcagtcatatagagcagacgCTTTAGGTTAGTTTTGACTCTTCTGGATTCTTATGTAAGCaagttatactgccctgt
gcagtcatatagagcagacgTTTCAGATCCATCACAAAGCGAAGTCATGGGAGAGCCACAaagttatactgccc
gcagtcatatagagcagacgAAACCGAGGGCTGCATCTTGTTTATAGTCATCTTTGTACTaagttatactgccctg
gcagtcatatagagcagacgTACACAACATTAAAAGTACTGACACCTGAGAATTTCTGCTaagttatactgccct
gcagtcatatagagcagacgAGTATCAGTGATCATTTAAAATTTGGAGGGGTCTTTGGTTaagttatactgccctg
gcagtcatatagagcagacgGTGACAATTAAAAGCACTAAAGGGAGATCATGTTAAAGCTaagttatactgccct
gcagtcatatagagcagacgTTCAGGTTGAAAAGTAGAAGTTCCAAGGTTTTGATTTTGGaagttatactgccctg
gcagtcatatagagcagacgAGCAACCAGTAGATGTAGGTTAAACTTTTACTTCATAGACaagttatactgccct
gcagtcatatagagcagacgGCAATGTCTAGAGACTTTTTGGAGTTACAACTTTGGGGTTaagttatactgccctg
gcagtcatatagagcagacgGGAATACAGCTAAAGATCCTAAAATGCACGGAACATCAGCaagttatactgccc
gcagtcatatagagcagacgATGGTCTTGTAACTAGTTGAAGACACTAAAGGGAAATGAAaagttatactgccct
gcagtcatatagagcagacgATGAACATTCTAAAACCATCTACATACAGTTTGCACTCTGaagttatactgccct
gcagtcatatagagcagacgGTCAGTAGTTGCTAAAGTTGACAAACCTTGAGAAAAGTCAaagttatactgccct
gcagtcatatagagcagacgGGCGAGATCATGGTTCATTGCAGCTGAGACTTCCCAGGCTaagttatactgccc
gcagtcatatagagcagacgCCACTGCACCTGGCCCGTTACACATTTTTAAAATAGATATaagttatactgccct
gcagtcatatagagcagacgGATTAAGTTTAGAATCGTCTTTGCTAATACAGTGTTCTCCaagttatactgccctg
gcagtcatatagagcagacgTTAAGGAAGTTGGAGATTTCTTCTGTACCTTAAACCTAGGaagttatactgccctg
gcagtcatatagagcagacgGTAGAGGGTACTGTGTTTAAGAGTGTAGTTTAAATGGAAAaagttatactgccctg
gcagtcatatagagcagacgATCACAAATAACACTTTCTGATGGTCTTTTTGCCTCCTCTaagttatactgccctg
gcagtcatatagagcagacgCTTATAGGAATCAGCTGGGAATCTTGAAGAAATTTTAGTGaagttatactgccctg
gcagtcatatagagcagacgCTAGTCTGAAATGGATCCCAGGAAGTATATTTTTTGAAAGaagttatactgccctg
gcagtcatatagagcagacgTGAGAACCTTTTATATGCTTTATTCCCTTTCTTTGCTTGCaagttatactgccctgt
gcagtcatatagagcagacgCTGTGCTTCTGACCGACTGGTTGCAAATTGGGGTTCCCACaagttatactgccc
gcagtcatatagagcagacgCTCTCCTTGGGTTTCATAATTTGCTAGGGCAGCTCATAGAaagttatactgccct
gcagtcatatagagcagacgCAGATGGAGGTGATGCATAGGGCAAGGTAAGGGGGAAGGTaagttatactgcc
gcagtcatatagagcagacgGCACACCACCCTCCGGACACCTCCACGTGTTTACCAACCTaagttatactgcc
gcagtcatatagagcagacgCAATTATGAACTTACTCTCCAATCCCTCTCCCCTTCCTAGaagttatactgccct

gcagtcatatagagcagacgGGTTGGTTGAATGGGGCTGTAGATTCCAAGCTTCTAGTCAaagttatactgccct
gcagtcatatagagcagacgAATATATATGAGAACAAAGGAGTTGGTGGCTATTACCATCaagttatactgccctg
gcagtcatatagagcagacgCCACGGGATGGCTCTTGCACTAAACACAACTTTTCACTCTaagttatactgccct
gcagtcatatagagcagacgCAGGCTTATTACTTTTTCTTAATTTTGCCCTTTACCATGGaagttatactgccctgt
gcagtcatatagagcagacgGAATACATCTCAGTGCTCTTCTACATTGTGTTATTCCTTTaagttatactgccctgt
gcagtcatatagagcagacgGTGTACGCCACCATGCCCCCAGCTTGTTATTTCTTTGAATaagttatactgccctg
gcagtcatatagagcagacgGAGAAAAACGATCTGAACAAGTGCTTTTGAAAAATGGAAGaagttatactgccct
gcagtcatatagagcagacgATATAAGTTAAAATCAACTCATCTCTTCAGAAGTGTGGCCaagttatactgccctg
gcagtcatatagagcagacgCAAAAAATGCTGTGACACTTTGTTGGACTTACTTTCCTCAaagttatactgccctg
gcagtcatatagagcagacgGCCTCAGCCTCCCAAGTGGCTGTCTTTGTTTGTTTGTTTTaagttatactgccctg
gcagtcatatagagcagacgGGCTGGTCTCTAACTCCTAACCTCATGATCCGCCCGCCTCaagttatactgccc
gcagtcatatagagcagacgAGCCACAGTGCCCAGCCGACTTTAAAAACTTTAAAGATGAaagttatactgccc
gcagtcatatagagcagacgCTTCTGGAGCCAGTAGACCTGTTTTGTCAGCTGTACTTTGaagttatactgccctg
gcagtcatatagagcagacgCTTGGCTTAATGGAGACAGGAAAAAGTTATATTGTTCAGCaagttatactgccctg
gcagtcatatagagcagacgGACAAAATAGAGGTCACGGAGGATATGGCTTTGGGTCAGTaagttatactgccc
gcagtcatatagagcagacgGCTTTGACTTTTGCCACACTTTTGGAGAGATGAATTGATTaagttatactgccctg
gcagtcatatagagcagacgACAAAAGCACCCATATAACCTTCATGTCTCTAACATATTGaagttatactgccctg
gcagtcatatagagcagacgCGTGAGTAAATTCAGGAGAAACATTTTTGGCAGGACATCCaagttatactgccct
gcagtcatatagagcagacgGCTTCTTACTGCAGCTCATCAGGGGTCATACAATATCAGGaagttatactgccct
gcagtcatatagagcagacgCCTGGTTAATGGGTGATGGCTATCAGAGATCTTTACTGTAaagttatactgccctg
gcagtcatatagagcagacgTCAATAAGTAGTCTGTGAAGCAATAATTTGAGACCTTGTGaagttatactgccctg
gcagtcatatagagcagacgTTTCACTGAGGTTTTAGTATCCATGGAAAATCTTTGCCCAaagttatactgccctg
gcagtcatatagagcagacgCATGGTCATGGGGGTCAAAAGAGGGCTACAAAATCGTGGCaagttatactgccc
gcagtcatatagagcagacgACAAGTTGTAAAGCAGAGTTTCCCCACATCTCTCTTCCTCaagttatactgccct
gcagtcatatagagcagacgTCTTCTTAAATGCTTTAGAGTCTATTTCTACACAGTACCCaagttatactgccctgt
gcagtcatatagagcagacgTGGCCTAAACCTAACCATTTTATTGAGCAGGACTTCAATTaagttatactgccctg
gcagtcatatagagcagacgCATAATCATAGATATTACGGCCAAAGAAATACCATGTTGGaagttatactgccctg
gcagtcatatagagcagacgCAACTGCTGACCCTAACACTACCAATGTTCTGCCCATCACaagttatactgccc
gcagtcatatagagcagacgCATTCTGTGTGGTAGCTTTGAAGTAGGCACTAGGGTATGAaagttatactgccctg
gcagtcatatagagcagacgAGAATTGTTTGAACCTGGAGCAGGCTAAGAAAGAAACCAGaagttatactgccc
gcagtcatatagagcagacgAATTTAAGAGAAGATAAGATGGAGGAAACTGATAAGGCACaagttatactgccct
gcagtcatatagagcagacgATATCCAGAAGTAGAAAAATACAACATAATCCCATGTCCCaagttatactgccct
gcagtcatatagagcagacgAATCATGTTTCATCTATATTTACACCCATTGTTCCCCTCaagttatactgccctgt
gcagtcatatagagcagacgAGTAATTCCTGACAAATACCCATCGTTCAAATTCCTAATGaagttatactgccctg
gcagtcatatagagcagacgGTTCCACCTCCATCTCATTTTTCTTTGTCTTGCTAGTTGTaagttatactgccctgt
gcagtcatatagagcagacgGATTTCACTGTCTGCCTGTTTGTGATAGTCTAACATGTTGaagttatactgccctg
gcagtcatatagagcagacgCCTTAGTATTGCCTATAAACTGCTAGTTGAATCTACAATCaagttatactgccctg
gcagtcatatagagcagacgACTTCATAGGTGATGTGTTCTTCCATAAATCTGCCTGCAGaagttatactgccctg
gcagtcatatagagcagacgCCTGGCCTCTTGCTGCTCCCGGCACCTGCTCCAATTTCTGaagttatactgccc
gcagtcatatagagcagacgCAACTCGGCCGGGTGTGCATGCTCAGGGTGGCAATGACATaagttatactgcc
gcagtcatatagagcagacgACTTTGGGTGCCGACAAACGTGAGAGGGAGGCCAAGGTGGaagttatactgcc
gcagtcatatagagcagacgTGGATGACATGACTTACAGCAGCAGGACGCAGACAGGCTCaagttatactgcc
gcagtcatatagagcagacgGGCGTGTCTCCAGTGAAGCCACACCTTCAAGTCAGGGATGaagttatactgcc
gcagtcatatagagcagacgAGTTCTGTGAGGAGTCTGCAGCCTGGATGGAGTAAGAACTaagttatactgccc
gcagtcatatagagcagacgCCCAGAAAAGCACTGGGTTCAGCCAGACTTGCAGAGGTGTaagttatactgcc
gcagtcatatagagcagacgGCCTGCGGATAGGAACTACCAACTCTATATCTCCTCTCTGaagttatactgccc
gcagtcatatagagcagacgTCCATTGAGAGCTGCACTCATCCAGATGACCTGCCCGTGGaagttatactgccc
gcagtcatatagagcagacgAAGAACTTGGGACCTGACAAATGGCAGAACTGAAAGATCTaagttatactgccc
gcagtcatatagagcagacgCACACCCCCCAACCCACTGGCCATGTTGTGAGAAGGAGAAaagttatactgcc
gcagtcatatagagcagacgCAAGCTTCCAGATGCCACCACATTTCTTGGATCCTGCAGTaagttatactgccc
gcagtcatatagagcagacgAGTGGCCAGACCCCGTGCTCGCTCACTCACGCACCCCTTGaagttatactgc
gcagtcatatagagcagacgCCTTGGCAGGAATGGGATCTGGGCTGGTGGCACAAGCCAAaagttatactgcc

gcagtcatatagagcagacgGGGCCCAAACAAAACTCCCGCAAAGGCGCCACAGCCACAGaagttatactgc
gcagtcatatagagcagacgTATAATTTGAAGTCCGGTAACTTTATGCCTCCAGATTTGTaagttatactgccctgt
gcagtcatatagagcagacgTCCCATCCATGAGCATGGGTCATATTTCCATTTGTTTGTGaagttatactgccctg
gcagtcatatagagcagacgCACCTCCTTGGTTAAGTAGAGTCCTAGGTTGTTTGTTTGTaagttatactgccctg
gcagtcatatagagcagacgCTGAATTCATTTATCAGACGTAGGAGCCTTTTGGATGAGTaagttatactgccctg
gcagtcatatagagcagacgCTTCTAGAACTATGTTGAGGAGAAATGGTGAAAGTGGGCAaagttatactgccct
gcagtcatatagagcagacgCAACTTTTCCCCAATCAGAATGATGTTGGCTAGTTGGCTAaagttatactgccctg
gcagtcatatagagcagacgTATGTTAAACCATCCCTGGAGTGAAACTCACTTGATTATGaagttatactgccctg
gcagtcatatagagcagacgTTTTTAAATGTCCTTCCCTGCTATTGGTATTAGGGTGATAaagttatactgccctgt
gcagtcatatagagcagacgGTCATAGAATGATTTACGAAGGATTCCCTCTTTCTCATCAaagttatactgccctg
gcagtcatatagagcagacgAGGGCTTCTGTTTCTTTCTGATTTGATCTAAGAGGGTTATaagttatactgccctgt
gcagtcatatagagcagacgATTGAGCTTATTTGGACCTCTCTCTTCTTAATCTCACTAAaagttatactgccctgt
gcagtcatatagagcagacgAGGTGTGACATTAGATTGTCGGTTTGTTCTCTTTCAGACTaagttatactgccctg
gcagtcatatagagcagacgGTAGGTATTTAATGCTATGAGTTTTCCTCTTAGCACTGCTaagttatactgccctgt
gcagtcatatagagcagacgTTCCAGTTTTATTCCACTCTGATCTGAGAAGATACTTGATaagttatactgccctgt
gcagtcatatagagcagacgTTGGAGGATGTTCCGTGTGCCTATGAGAAGAATGTATACTaagttatactgccctg
gcagtcatatagagcagacgTTGATGATCTCTCTAGTGCTTTCGGTGGAGTATTAAGTCCaagttatactgccctg
gcagtcatatagagcagacgGAGTTCCAGTGTTAGGTAAATATAAATCCAGGATTGTAGTaagttatactgccctg
gcagtcatatagagcagacgACTCCTGCTCACTTTTGATTGCCATTTGTGTGGAATATCTaagttatactgccctg
gcagtcatatagagcagacgCACACCTTTACCTTGAGTTTATGAGTCCTTATCTGTCAGGaagttatactgccctg
gcagtcatatagagcagacgGGTATACATGTGCCATGTTGGTCATTTACATTAGGTATTTaagttatactgccctgt
gcagtcatatagagcagacgCACCCCCCAGCAGGCCCCAGCATGTAATGTTCCCCTCCCTaagttatactgcc
gcagtcatatagagcagacgGGTTTTCTGTTCTTGTGTAACTTTGCTGAAAATGATGGTTaagttatactgccctgt
gcagtcatatagagcagacgCACTCCCAACAGTGTAAAATTGTTCCTATTACTCCACATCaagttatactgccctg
gcagtcatatagagcagacgTCTCATTGTGGTATTGATTTGCATTCTCTAATGACCAGTGaagttatactgccctgt
gcagtcatatagagcagacgGCATTTTTTCATAAGTTTGCTGGCTGCATAAACATCTTCTaagttatactgccctgt
gcagtcatatagagcagacgTCCGTTGCCCATTTTTTGATGAGGCTGTTTGTTTTTGTCTaagttatactgccctgt
gcagtcatatagagcagacgTGTGTGGTGTTATTTCTGTGGCATCTGTTCTGTTCCATTGaagttatactgccctgt
gcagtcatatagagcagacgGTGGGCTCTTTTTTGGTTCCATTTGAAATTTCAAGTAGTTaagttatactgccctgt
gcagtcatatagagcagacgAATACTATGTTGAACAGGAGAGAGAGGGCCTCCTTGTCTTaagttatactgccct
gcagtcatatagagcagacgGAGAGTTTTTAGTATGAAGAGGTGTTGAATTTTGTCGAAGaagttatactgccctg
gcagtcatatagagcagacgTAATCATGTGGTTTTGTTGTTGGTTCTGTTCATGTGATGGaagttatactgccctgt
gcagtcatatagagcagacgTTTATTGATTGGTGTATGTTGAACCAGCGTTGCATCCCAGaagttatactgccctg
gcagtcatatagagcagacgTGATACCAGCTCCTCTTTGTCACTTTGGTAGAGTTCAGCTaagttatactgccctg
gcagtcatatagagcagacgCTGGTTTAGTCTTGGGATGATATATGTGTCCAGAAATTTAaagttatactgccctgt
gcagtcatatagagcagacgTCCTTCAATTCCGCTCTGATGTTAGTTATTGCTTGTCTTCaagttatactgccctgt
gcagtcatatagagcagacgTTAGTGCTAAAAATTTCTCTCTCCACCCTGCTTTAAATGTaagttatactgccctgt
gcagtcatatagagcagacgAGAGATTCTGGTAAGCTGTGTCATTGTTCTCACTGGTTTCaagttatactgccctg
gcagtcatatagagcagacgGCAGGTTGTTCAGTTTCCGCGTAGTTGTGTGGTTTTGAGTaagttatactgccctg
gcagtcatatagagcagacgGAATGTATATTCTGTTAATCTGGAGTAGAGAGTTCTGTAGaagttatactgccctg
gcagtcatatagagcagacgCCTGGATATCCTTGGTAATCTTCTGTCTCATCAATCTAATaagttatactgccctg
gcagtcatatagagcagacgTAGTTAGCTCTTCCTGTTGCGTTGATCACTTTACCATTGTaagttatactgccctgt
gcagtcatatagagcagacgTTTGAGCCTATGTGTGTTTTAGCATGTGAGATGGTCTCCTaagttatactgccctg
gcagtcatatagagcagacgTCTGTGTCTTTTAGTTGGGGCATTTAACCCATTGACATTTaagttatactgccctgt
gcagtcatatagagcagacgCCTTTCCATGTTTAGGACTTCCCTCAGGAGCTCTTGAAATaagttatactgccctg
gcagtcatatagagcagacgGGCCCCCACTCTCTGGCTTGGAGGGTATCTTCCAAGAGATaagttatactgcc
gcagtcatatagagcagacgGTCTGATGGGCTTCACTTTGCGGGAAACCCAGCCTTTGTTaagttatactgccct
gcagtcatatagagcagacgCCTTGGTGAATCTGATCATTTTGTGTATTGGGGTTGCTATaagttatactgccctgt
gcagtcatatagagcagacgTGTCACTTTCAGGTACAGCCATCAAACGTAGATTTGATCTaagttatactgccctg
gcagtcatatagagcagacgTAGTCCCATATTTCTCGGAGCCTTTGTTTCTTTTCACTCCaagttatactgccctg
gcagtcatatagagcagacgTTCTTCTGCATGATTGATTCGGCTACTGAAGCTTGTGTATaagttatactgccctg
gcagtcatatagagcagacgCAGGTCATTTATGTTCTCTACATTGGTTATGTAGCAATTCaagttatactgccctgt

gcagtcatatagagcagacgAGTTTTTTTCTCTTGCTGGCGAGAAGTTATGATTCTTTGGaagttatactgccctgt
gcagtcatatagagcagacgGTTGATGCTATTCTTTTCATCTGTTAGTTTCCCTTCTAACaagttatactgccctgt
gcagtcatatagagcagacgTCCAGACCCTGTTTGCCGGGATATCACCAGTGGAGTCTGCaagttatactgcc
gcagtcatatagagcagacgCTCTATGAGGTGTCTGTTGGACCTTACTGGGAGGTGTCTCaagttatactgccc
gcagtcatatagagcagacgACTTGAGGTGGCAGTCTGTCTGTTATCAGAGCTCAAACACaagttatactgccc
gcagtcatatagagcagacgGGTTTTATCTATAGGTCCCTGACTGGGGCTGCTGTCTTTTaagttatactgccct
gcagtcatatagagcagacgTTACACTGTGAGCATAATACTGCTTGCTCAAGCCTCAGCAaagttatactgccc
gcagtcatatagagcagacgTCTCCTGGTCTGCCGTTTGCAAAGAATGTGGGAAAAGTACaagttatactgccc
gcagtcatatagagcagacgGCTCACCCTCCGTGGGCTGTTCCCACTGTCCAACCAGTCCaagttatactgcc
gcagtcatatagagcagacgGTTCCTATTCAGCCATCTGGCTGGAAATCCCACTGTGTATaagttatactgccct
gcagtcatatagagcagacgACTTTGTTTTTTTCACTGTGTTACTGCTTTATAGACCCTGaagttatactgccctgt
gcagtcatatagagcagacgTCAGCAGGTTCCATTTTGGTACATACCAAGTTTTGTTTCAaagttatactgccctg
gcagtcatatagagcagacgGACTCAACACCACTTAATGAAGGAATAGATTTTGAATGTGaagttatactgccct
gcagtcatatagagcagacgAGATAATTCTAGACTCAAATATTTCAAGGAGCTGGATGTGaagttatactgccct
gcagtcatatagagcagacgATACTTTGTAAAATTGGATTCACCACAGGCACACACTAAGaagttatactgccct
gcagtcatatagagcagacgGCATCAGTTCTCAAGCTTTTGTGTGCACCAGAATCATAGGaagttatactgccct
gcagtcatatagagcagacgCTCCTAGAGTTTGTGATTCAGAAAGGTTGAGTTGAAGCCTaagttatactgccct
gcagtcatatagagcagacgGGTAATGCTACTAAACTAAGGACCATATTTTGAAAGTCACaagttatactgccct
gcagtcatatagagcagacgTGTTGGAGTGCTTTGCCTACGTAAATCCCCTCTTCATTTGaagttatactgccct
gcagtcatatagagcagacgCAAAATAAAAAATATGTCCATCATCCCAGAACGTTCTCTCaagttatactgccct
gcagtcatatagagcagacgGGCAACCCCTCATCTCATTTCTCACACTGTAGATTAGTTTaagttatactgccct
gcagtcatatagagcagacgCCAGTTTGGGGCTATTATAGATGAAGCATCTATAAACATTaagttatactgccctg
gcagtcatatagagcagacgGTCTTGGGGCAAATGACATAGGAGAGAGCTACTGCATCATaagttatactgcc
gcagtcatatagagcagacgTGTATGCGCGAATTTCCACTTGCTTAACGGTGTCATCAGCaagttatactgccct
gcagtcatatagagcagacgCACTTTTTAAACTTTAGCCGCTGTAGTGAATATGTAGTAGaagttatactgccctg
gcagtcatatagagcagacgTTTGCGTTTCCTTGATGACTGATGGTGTTGAACATTACTGaagttatactgccctg
gcagtcatatagagcagacgAATACTTTGTCCACTTCCCCACACTCTTTTTAAATTTGTCaagttatactgccctg
gcagtcatatagagcagacgATGTATTGCAAATATCTTCTGGAGTCTTGGCTTGCTTTTTaagttatactgccctgt
gcagtcatatagagcagacgGGTTCTCATTCTGTTGCCCAGGCTGGCACATTCACGGCTCaagttatactgcc
gcagtcatatagagcagacgGGCTCAAACGATCTGTCTGCATTAGCCTCCCAGGATGCTGaagttatactgcc
gcagtcatatagagcagacgTGTGTTTTACATCTTTGGGGTCCTGTCTAAAATCTTTGCCaagttatactgccctg
gcagtcatatagagcagacgTTTTCCTTCCCAATCTATATCCCTTTTATCGCCTTCTCTTaagttatactgccctgt
gcagtcatatagagcagacgTATCTAGGGCTCCCAGTATGGTGTTGAATAGGAGTTGTGAaagttatactgccct
gcagtcatatagagcagacgGGAAAGCAGCCAGGCCTTTTACCATAAAGTATCATTAGCAaagttatactgccc
gcagtcatatagagcagacgGATGCCCTTTATCAGTTTGATCCCTAGTTTCTTGAGAATTaagttatactgccctg
gcagtcatatagagcagacgATGTTGAGTCAGCCTTGCATTCACAGGACAAACCTCATTTaagttatactgccct
gcagtcatatagagcagacgTTCTTGTAACTTCTTTGGTTTCACATGAAGGTAATGCTGAaagttatactgccctg
gcagtcatatagagcagacgAAGTGTTCCTTGCTTCTATTTTCTAGAAAAGCTGTAGGTAaagttatactgccctg
gcagtcatatagagcagacgACCATTTGCGCCAGGTCCCATCTTGATCACACACAGCTGTaagttatactgcc
gcagtcatatagagcagacgTATTTATGTATGGTCTTTGGTTACTTTGTGCTACAATGGCaagttatactgccctgt
gcagtcatatagagcagacgGTAATGTGACAGACTGTATAATCCAGAAAGCCTAAACTGTaagttatactgccct
gcagtcatatagagcagacgGAAAATGGTTTCTAACCCCTGACCTAGAGTAGTGATTCTCaagttatactgccct
gcagtcatatagagcagacgGAATGTTTGAAAAAAAATACTGTCTGGCTCTCATACCTAGaagttatactgccct
gcagtcatatagagcagacgTTTAATGTGTTTGAGCTGAGGCTAGGCCATTGGAAGTTTCaagttatactgccct
gcagtcatatagagcagacgAGTGATTTGACTTGGCAGGTTGAGAACTACTGGATGAGAAaagttatactgccct
gcagtcatatagagcagacgATGTATTCATTTTACTTCCCACCCCAAAGCTGATTATTCTaagttatactgccctg
gcagtcatatagagcagacgGCATTTAGGGTTGAAAATCAGTCATCTAGGGCAGTAGTTTaagttatactgccct
gcagtcatatagagcagacgGTTTTAGAAATACATTCCAAGACCTCACTGCAGAGCTACTaagttatactgccct
gcagtcatatagagcagacgTTCTTCAGATGGTTCTCATATGCAGGGATTTCTTTAAAAGaagttatactgccctg
gcagtcatatagagcagacgTTCTTGGGCATTGAATGAGTAATTAGAAGATTATGAGAGGaagttatactgccctg
gcagtcatatagagcagacgTGCTCTGTGAAAACGCACCATCTGTGCTCTGCAGCTAGCTaagttatactgcc
gcagtcatatagagcagacgGCCACCCCCAAGCCAGCAGCCACAACCCCCTCAGGTCACTaagttatactg

gcagtcatatagagcagacgCCAACTCTGGACACAATTATCTAGTGAGTATGGAAGTGGAaagttatactgccct
gcagtcatatagagcagacgCAATGCAGGCTGTGGACAATCCCATTGACTAGTCCTCTGTaagttatactgccc
gcagtcatatagagcagacgGGTAATCTAACCCCCTTTGTACACTGCTGGATAACATGGAaagttatactgccct
gcagtcatatagagcagacgAAGACCAGTCCTTCCATTAACATTCTTTCTTCAGATAATCaagttatactgccctg
gcagtcatatagagcagacgTTTTTCATTTCTGTTTTGAGTCTATAGTTCCCCATCCCATaagttatactgccctgtg
gcagtcatatagagcagacgGGTTCTCTAGTCTCTTCAGCACCACATTTACTAATTGTCTaagttatactgccctg
gcagtcatatagagcagacgAGACACTTTGGAGATGGCTAAGTTCATTTGTGCTACATTTaagttatactgccctg
gcagtcatatagagcagacgCTTGGCCACCTCAATCTAATCTCCTGATATACTTGCTACAaagttatactgccctg
gcagtcatatagagcagacgAGTTTTTGCTCAAGTAATCATCTGTTAACCCTTCAACTGTaagttatactgccctgt
gcagtcatatagagcagacgTCTTCAGTTCAGATGAGATGTGTAAAATCATGGAATGTTGaagttatactgccctg
gcagtcatatagagcagacgGAAGAGAGTTTATAGACCATCTTGTCTAAGCTCAAAAAAGaagttatactgccctg
gcagtcatatagagcagacgCCCTGTACAATTCTTGCTTTGGGGTTTATAATCTATTACTaagttatactgccctgt
gcagtcatatagagcagacgAGTAGGGTGAAGTCCATTCATGTGTCATACTGATAGGCACaagttatactgccct
gcagtcatatagagcagacgGTCCAGCCAAATCTCTGTGAGACTTATCTTCCTCTTCTGTaagttatactgccctg
gcagtcatatagagcagacgTGCTTTTTGTTTTGGTTTGGGAAGAACCTTGGACAGGAAGaagttatactgccctg
gcagtcatatagagcagacgTGTATCAGTCAATCTGAGAATTAGTTATTACACGGGCTCCaagttatactgccctg
gcagtcatatagagcagacgCAGAAGATACTAGCTAAACCTAAAAGGAATGATCTTGGAAaagttatactgccct
gcagtcatatagagcagacgGCTATTCACTAGTCAAGCAATCATTGATTTTCTTAAAGCCaagttatactgccctg
gcagtcatatagagcagacgAGAGATTACAATGAAGTTACTCATTTAGTTGCACGTTAGGaagttatactgccctg
gcagtcatatagagcagacgTATTTATGGTAACTTCGAAAATGGGAGGTATTAAAGGACCaagttatactgccctg
gcagtcatatagagcagacgGTATAAACTGGCATATTTTTCAATTTGTTTGCAAGGCTGGaagttatactgccctgt
gcagtcatatagagcagacgTGATTGGCATGATAACAGTGGTTACTCTCATAAGGCATTAaagttatactgccctg
gcagtcatatagagcagacgAATCAAAGGTTGTCCCAGTACTCAAGGCGATGATTGTCAGaagttatactgccct
gcagtcatatagagcagacgGATGGAAAATGCTGGGATCGAAGTGCACACAAAGAGCTATaagttatactgccc
gcagtcatatagagcagacgGGGACTGGGGAGTGAGGAGGGATGTCAAGAAGGCTGCTATaagttatactgcc
gcagtcatatagagcagacgGAAAGTTTCAATGTAATGTGGGAGGAAGAGCAAATGGAACaagttatactgccct
gcagtcatatagagcagacgCGTAAGGCAGTAAGAAATGAGATGGGAAGGAGTCAAATCAaagttatactgccc
gcagtcatatagagcagacgGGCTTGCTTGTTTGATCATTTTCCTATGAGTGACCGTGTTaagttatactgccctg
gcagtcatatagagcagacgAGCAGTATTTATAATTGAACAAGGATAGACAACAGCTGGGaagttatactgccct
gcagtcatatagagcagacgCAGGGATCATGGAATGTGAATATCCTTTTGCTGACACAAGaagttatactgccct
gcagtcatatagagcagacgTGTTGGCTCCAAACATTAGTTACTTTATACCAGTATTAGCaagttatactgccctg
gcagtcatatagagcagacgTTACCCATTACATATAGAAATATTATCATGCCTGTCACCGaagttatactgccctg
gcagtcatatagagcagacgATTCATGTATAGTCTTCAGAAGTCACAGTATATAAGCTCCaagttatactgccctg
gcagtcatatagagcagacgAGCAGCTTAACAGGCTCTATAAAATAAAGCATTACCAAGTaagttatactgccctg
gcagtcatatagagcagacgATTTTCATTATGCCTTCAAAATTATCCCTGAGAGAGGCCAaagttatactgccctg
gcagtcatatagagcagacgTCTTTGGTCTCAGTGTAAGAGCCTGGAATATGTCAGAGAGaagttatactgccct
gcagtcatatagagcagacgAAGTTTCTTGATCCCTCCTTACACCCTTAAGTTTTAATGTaagttatactgccctgt
gcagtcatatagagcagacgCTAAGTCTAAAAGTGAGGCGTATTTGAAATTTGTTCTGTGaagttatactgccctg
gcagtcatatagagcagacgCACAAAGTGCTCAAAGTGCGTGGTACCACAAGAAGCTAAAaagttatactgccc
gcagtcatatagagcagacgGCTGTCTCTCTGACTGTTCAGTTTAGAAGAAAAGATAGGTaagttatactgccctg
gcagtcatatagagcagacgCACTGATTGTGGAATTCATTTATAGTCTTCAGAAGTCATGaagttatactgccctg
gcagtcatatagagcagacgTTTTTGTTTGTTTAAATCACAGTCACCTGTCTGAAGCATCaagttatactgccctgt
gcagtcatatagagcagacgGCTTCCCTCATTCCAGCTGGTACACTTGACAACACTATAAaagttatactgccct
gcagtcatatagagcagacgAAATTATGCAGATTATTAGGAGGAGTATCCTGGCATTTTGaagttatactgccctg
gcagtcatatagagcagacgTAAAACTGCATGATTTTTTTTCTCTTCTAGACCCCTAAGGaagttatactgccctgt
gcagtcatatagagcagacgCATCTGTCCCATGGTTAACTCTGTGTAGTCCACATAAACTaagttatactgccctg
gcagtcatatagagcagacgATTACTGAAGTTCCTGTTCAATAGCTATAGCTGAAGCTCAaagttatactgccctg
gcagtcatatagagcagacgAGCAGTGTAAGGTTAAAAAAAAAGCTCAGAAGCCCATCTAaagttatactgccct
gcagtcatatagagcagacgAAATCTGTTTTTTTTTGGTAAAGATGGGGGTCCCACTATGaagttatactgccctg
gcagtcatatagagcagacgTGTGAGCCACGGTGCCTGCTACAATTCTTGTTAGTTAAGTaagttatactgccctg
gcagtcatatagagcagacgGGCTGTACCATCATTTACTTAAACATTCTCCCACTGTTGGaagttatactgccctg
gcagtcatatagagcagacgGAGTATGTTTCTAAAAGGAATTGCTGGATCAATCACTATGaagttatactgccctg

gcagtcatatagagcagacgACTATTATCAACAATGTGTAAGATCAGTTGTCTCATGTGCaagttatactgccctg
gcagtcatatagagcagacgTCATCAAAATTAGAGGAACTGCAAATACTGTTGTTCTCTGaagttatactgccctg
gcagtcatatagagcagacgAGGTATCAGAATTCAATAGAAACAATAAGTACTGAGCTCCaagttatactgccct
gcagtcatatagagcagacgTAAGACATGAGCCAGCCCTTTAGGGAGTTTGCCATCTTGGaagttatactgccc
gcagtcatatagagcagacgCACTTTCCTCCCTAACTCAGAGCCTTACCCCAAAGTGCTTaagttatactgccc
gcagtcatatagagcagacgGCCCCATTGCCTTGATTCCCGTCACTGTGCTATGCTATCAaagttatactgccct
gcagtcatatagagcagacgTGAGCTCTTCATTAATGTGTCAAATAAATTTGCTTGGACGaagttatactgccctg
gcagtcatatagagcagacgCATTAAAACAAGACAAAGTCATTAAACAAGCCAACTCTGGaagttatactgccct
gcagtcatatagagcagacgCACATCCTATCTGCTTTTGTCATTTATACAAGAATCCATGaagttatactgccctg
gcagtcatatagagcagacgCAATCTCATTACAGCAATCTTTGACCATCAGGGGATATTGaagttatactgccct
gcagtcatatagagcagacgATCAATGGTTTAAAGGTGGGGAGTTATAATGAACACAATCaagttatactgccct
gcagtcatatagagcagacgTTTAGTTCTTCACCCATGTCACTCTCTTTTTCTTCCTAAGaagttatactgccctgt
gcagtcatatagagcagacgCTCAAACACCTGGGCTCAAAATCATCTTCCCGTCTCAGTCaagttatactgccc
gcagtcatatagagcagacgTGAGACTTCAGACATACACCACCATGACCGGTTCTCTTTTaagttatactgccct
gcagtcatatagagcagacgAGGGAGCATGCCAATCTTCACTGTGTCATTCCAATTTTGTaagttatactgccct
gcagtcatatagagcagacgCGAGCACTTCATGTCTAAAAGTTCTTGAGAAAAGCACGTTaagttatactgccct
gcagtcatatagagcagacgACAGACCTATGTAATTTACTGGTGAGGTTAGGGAATTCACaagttatactgccct
gcagtcatatagagcagacgGATTAAAAATAATGACATACCATAGACTGATGGCTGGAAGaagttatactgccct
gcagtcatatagagcagacgGGTACCAAGCAACAGTGGCTGAGTAAATTCCATTAAAAGTaagttatactgccct
gcagtcatatagagcagacgCTGCAGTGACTCCTCCAAGTGTTTCTAAGAAAGCAATACAaagttatactgccct
gcagtcatatagagcagacgGGGTACCTGTTTGGAGTAGGGATAGAGAGAGAGGAAAAAAaagttatactgccc
gcagtcatatagagcagacgCAAAACAACAAACACAAACTAATGGAGACAACTGTTACTCaagttatactgccct
gcagtcatatagagcagacgACTGCTGGGTGAATGAAATGAAAGTTATTCCCTCAAAGTCaagttatactgccct
gcagtcatatagagcagacgGGTTCTCCTTGGACTTGATGTGAGAGATGTTGGAAACTCCaagttatactgccct
gcagtcatatagagcagacgCAGCTGACTCTTTTGATAACTGCATGTCTCTCCTCTCTGCaagttatactgccct
gcagtcatatagagcagacgGGATGCTGTCTCCCACCCTCCAACCCCATGCTAATCTGTGaagttatactgccc
gcagtcatatagagcagacgCTCCTAACAAGCAATGCTTCCTGCAAGAGCCTTTTGAAAGaagttatactgccct
gcagtcatatagagcagacgAAAACCTTCTTATCTGTTCCTCAGTCTGGCAAGTGACAGCaagttatactgccct
gcagtcatatagagcagacgGAATTTTATCCCTCTTCAAAGTCACCAAAAAATCCCATCAaagttatactgccct
gcagtcatatagagcagacgACCATCTTGTGCTCTAGTGAAGGTTTATGCCACAGTTTAAaagttatactgccct
gcagtcatatagagcagacgACATATGTTTCCACCTTATTGCTTGCAACTGCCTCTGCCCaagttatactgccct
gcagtcatatagagcagacgCCTTACAAGATGAAGAGTGGGGATTCTGCTATGGTCACACaagttatactgccc
gcagtcatatagagcagacgCAAAGTTGGTCAAGGGTAGGCCACTCAGCATTGTTCATGAaagttatactgccc
gcagtcatatagagcagacgGACAGTATTCTACCTATGTTTAGTTTTGTCTGGAATCCAAaagttatactgccctg
gcagtcatatagagcagacgAGTTAAACTTCCAAAAGACAGGCTGGCCATCTTCATGTGCaagttatactgccct
gcagtcatatagagcagacgCCTGGCTTTCTTGGTCTCAGTTCCTATATATGTAGCTTCTaagttatactgccctg
gcagtcatatagagcagacgAAGTCACTATGCTTGTTCCCTACTTTGCCAAGTTTACTTTaagttatactgccctg
gcagtcatatagagcagacgCATGTAAATAATTGTGTTTGCTTGGACTGATAGGAAGGAAaagttatactgccctg
gcagtcatatagagcagacgATGGCTTATGTGTATATTCTAATTTTCCCCATGGTCTTCAaagttatactgccctgt
gcagtcatatagagcagacgGATAGGACAAACATTCATAAAACATACGTCTATCTCTGCAaagttatactgccct
gcagtcatatagagcagacgCTTTTGACTAAAATGAAAGCTTTGTCGATCTGCCTTGTACaagttatactgccctg
gcagtcatatagagcagacgCTGTATCCCTAGGGCTCAGCTGTCCGGGTGCATGGCAGACaagttatactgcc
gcagtcatatagagcagacgAATCAAAGCTTAATTCAGGATCCCATTTACCAGTCATCAAaagttatactgccct
gcagtcatatagagcagacgCATTATTTTCTTTTATTTCTGATCTCTTTCTGCAGCCAGGaagttatactgccctgt
gcagtcatatagagcagacgTGCTATGGAAAACAGACTGCACAGTACTAGTTCTATTATGaagttatactgccct
gcagtcatatagagcagacgTTAATCTATTCTTGATGGGAAATAGGTCAGACACTGCACCaagttatactgccct
gcagtcatatagagcagacgCTTATTGCCCAACTTCTCTGAGCAGTTCGACTTGTTTAATaagttatactgccctg
gcagtcatatagagcagacgCCAGAGATAAAACAGGATTAAAGGATAAGTACAACTGAGTaagttatactgccct
gcagtcatatagagcagacgGACCTCAAAAATCAAGTCTGTTTTACTGTCTCCCAATTTAaagttatactgccctg
gcagtcatatagagcagacgCCCAGGTAAAATAGTCCAAGGCAGCTTCAGCTATGAGTTCaagttatactgccc
gcagtcatatagagcagacgGGTGCAGAGCGCAGAGATTGCAAAATAATTACCAGAATTCaagttatactgccc
gcagtcatatagagcagacgGAAAACTTCGGCAGTATTTCCCCACTCCTCTGATTATTTAaagttatactgccct

gcagtcatatagagcagacgAAGTATCCTAGAATAATGAAGTCACAATTCACAGTTACCGaagttatactgccctg
gcagtcatatagagcagacgTCCCTCTGTTGCCCACACTGCTTTTGAACTCCTGGTCTCAaagttatactgccct
gcagtcatatagagcagacgTGCACCTGGCGGAGATGACTTTTTAAAAGCCTTTATAAGGaagttatactgccctg
gcagtcatatagagcagacgTCATTCTCAGCAAACCATCGCAAGAGCAAAAAACCAAACAaagttatactgccc
gcagtcatatagagcagacgCCTGTCGTGGAGTGGGGGGAGGGAGGAGGGGGGAGGGATAaagttatactgc
gcagtcatatagagcagacgTAAACTATACCTCTTGACAGCTCTATTTTAGCTCTAAGTCaagttatactgccctg
gcagtcatatagagcagacgGATATAATTAACATCCCACAAACCTCACCCTTTTAAAGTCaagttatactgccctg
gcagtcatatagagcagacgTCATCACCCCGAAAAGAAACTGTATACCCATAAGTAGTCAaagttatactgccct
gcagtcatatagagcagacgTCCCCAGCCCCAGTCAAGCCCTAATCAAATTTCTGTCTCTaagttatactgccc
gcagtcatatagagcagacgTTCTTTTTATTGCAGAAGAGCAGTTTGCTGGATGGATAAAaagttatactgccctg
gcagtcatatagagcagacgACCCATTTGTCAGTTCATGAACATTTGCATAGGTTTCACTaagttatactgccctg
gcagtcatatagagcagacgCTTCTCATGGTATGTAAGTAGGGGAGGGATTGCTGAGTCAaagttatactgccct
gcagtcatatagagcagacgGGAACTGCCAAACTGCTTTGTACAGCTGCTGCACCATTTTaagttatactgccct
gcagtcatatagagcagacgCCTATTTTATTTTGGCCACCCCAGCCCACTCTGTGGTTTTaagttatactgccctg
gcagtcatatagagcagacgCCCTGATGGCTAATGATATCCAGCATTTTCATGCGCATATaagttatactgccctg
gcagtcatatagagcagacgAGAGTTCTTAAATATGCTGGATACGAGTTCCTGCTCAGACaagttatactgccctg
gcagtcatatagagcagacgCATTCTGTGGGTTGTCGGTTGTCATTTCACTTTCTTTCTTaagttatactgccctgtg
gcagtcatatagagcagacgTCTCAAACTCCTAACCTCGGGTCATCCACCCGCTTTGGCCaagttatactgccc
gcagtcatatagagcagacgAGCCACTGCGCCTGGCCTCACTTTCTTGATGTTGTCCTTTaagttatactgccct
gcagtcatatagagcagacgCAAAGCAATCATCATCACATCATCTATAAACAGATGTGTGaagttatactgccctg
gcagtcatatagagcagacgTAGCTACATCTTATGAGTGAAAAATAAGAGGCCGGGTGCGaagttatactgccc
gcagtcatatagagcagacgTATTGAATTATGTAGCCTTCTCCAGTATTGACTAAACTGGaagttatactgccctg
gcagtcatatagagcagacgATTATTCATAAATCAAATAAGTACATTCACAGCACCGGGCaagttatactgccctg
gcagtcatatagagcagacgGCCAATGTCTAATAAGATTCATATGATACAAAAGGGAAGGaagttatactgccctg
gcagtcatatagagcagacgATTTAATTTTTGTCTTTGCTGCATCCTGGTTATTCTCTGCaagttatactgccctgtg
gcagtcatatagagcagacgACTGTTTGACTTTGTGTACCCTGCTCTGTGCTTTGGAGGCaagttatactgccctg
gcagtcatatagagcagacgCGAGTTCCTGCTGCACTTGGTTAAAAGGATGATGGGGGAGaagttatactgccc
gcagtcatatagagcagacgGTGTTTCTTCTCTGCTTCCTATCTGGGGCATCTCTCTGGCaagttatactgccctg
gcagtcatatagagcagacgCGACAATGCCCACTCTCTATAATGGCCTTCATGAAAGTTTaagttatactgccctg
gcagtcatatagagcagacgCTGTGACAGTTGTAGGGGCCAGTGTAAGAACATTATCAGAaagttatactgccct
gcagtcatatagagcagacgCTGGCCAACAAACTGTTAACTAAAATATATTATGCCTTCCaagttatactgccctg
gcagtcatatagagcagacgCCTTTGTAATGAATGATCTAGGAGACCCGCACACACTTAGaagttatactgccct
gcagtcatatagagcagacgTCAGACGTTTTCAGAATTCCACGTAGGAACAGGGTAGCCCaagttatactgccc
gcagtcatatagagcagacgAGCCCACCCCCATTCTGCTCTACCCCTTGCACCACCCTGCaagttatactgcc
gcagtcatatagagcagacgCAGCCCCACCCTGTGATTCCAAAGTCCAGCCACACCCTTTaagttatactgcc
gcagtcatatagagcagacgCCTTGGCTCCTTCCTTGGTCTACAGGTGAGCACACTGGGCaagttatactgccc
gcagtcatatagagcagacgGAGGAAGAGGATCATCCAGGCCCTTACTGGACATAAGCCCaagttatactgcc
gcagtcatatagagcagacgCCAGGTGGGCAGAGTTCAGTCTAGAAGGAAGGAGATGTGGaagttatactgcc
gcagtcatatagagcagacgGGCATCTCCCACTTGGCCCTGAGAGCTCCTTTCTTCTTAGaagttatactgccc
gcagtcatatagagcagacgGGCTCCCTTGCAAGAGTGTAAGCCTTTCCCTCACAGAGCTaagttatactgccc
gcagtcatatagagcagacgTTATTCAAAAGGAGATCCGTGCATAAAAAGGGCTGTAGAAaagttatactgccct
gcagtcatatagagcagacgATTTCTTAAGCAAATTAAACTCACCTTGTTCAGTCGAACGaagttatactgccctg
gcagtcatatagagcagacgCTTTTGACAACATGGACCGTAGAACAGAGAATGGCACAATaagttatactgccc
gcagtcatatagagcagacgCACCCTGGCTGTGAGAGAAAGGATTCAAAATCACTGGCTGaagttatactgccc
gcagtcatatagagcagacgAACACTTTCACTCTGTCTAAATACTTCCTTTTCTACAAGCaagttatactgccctg
gcagtcatatagagcagacgGTAGAACTCTGTTTGATGAATTAAAGTAGACTATGGACAGaagttatactgccctg
gcagtcatatagagcagacgCTTTACTCTAGAATTCTTATTCCAGTACAGAGAACAAGAGaagttatactgccctg
gcagtcatatagagcagacgTTTATATAATCTGCCGATTAAGACTGATCTGCTGGCCGGGaagttatactgccctg
gcagtcatatagagcagacgGAGGCCGAGGAGGGCAGATCACCTGAGTTCGGGAGTTCGAaagttatactgcc
gcagtcatatagagcagacgTCATTTTTGTTTGTGCTTTTTTCAGACTTGGCACTCTTAGaagttatactgccctgtg
gcagtcatatagagcagacgTTATTGGTAAACTGGTTCTTTATCATCTTGCCTGTTCATCaagttatactgccctgtg
gcagtcatatagagcagacgGGCAGAATAACTTCATAAACACAAATTACCTCTGGTCTTCaagttatactgccctg

gcagtcatatagagcagacgTTTTCACATAGATTAACTTACCTGACCTTGGCAGCTCTTAaagttatactgccctg
gcagtcatatagagcagacgACTGGCTATTATTGAATAGTAGGGATTGTGCCAAGCGAGTaagttatactgccct
gcagtcatatagagcagacgTGTTCACTGTAATCCTCTAAATTAGGTAGTACCACGCCCAaagttatactgccct
gcagtcatatagagcagacgGATGAGAAAATGAGGGCTGAGAGTAGTTCAGTAACTTGCCaagttatactgccc
gcagtcatatagagcagacgCAATTCTATTTGGTCACACTGCTTCCCTGCAAGATAGTAAaagttatactgccct
gcagtcatatagagcagacgGACCTTTCCCAGAGTCTCATTTTATAGTTCCACAGCCTTTaagttatactgccct
gcagtcatatagagcagacgTGCAAAAACAGTTATAAGGTAGGTAGTTAAGGAAGCAGTCaagttatactgccct
gcagtcatatagagcagacgGCCAACCAGCAATTCATCAAAAATCAATTTATGAAATGGCaagttatactgccct
gcagtcatatagagcagacgGGGAAGTAGTCTTATTAGCAGCATTTTAAGAAATCTTCACaagttatactgccctg
gcagtcatatagagcagacgTGAGGCAAATATACACATTTGGGAAATTGTGGTTATTCAGaagttatactgccctg
gcagtcatatagagcagacgCTTTTGGCTTATTGACTCCTGGCAAACTAGTTATTACCCTaagttatactgccctg
gcagtcatatagagcagacgATGGCTTAAATAGAGGGCTTTAGATTAAAGTTTCTCCCCCaagttatactgccct
gcagtcatatagagcagacgTTCTAAAGCTACACTTGGAACTCACCTAAAAACATCATCAaagttatactgccct
gcagtcatatagagcagacgGCAAACGCGGTCATAGCCATTCCTGTCATCGTGGTAAAAAaagttatactgccc
gcagtcatatagagcagacgGGAAACTAGTCAAAAAAGAGGCACTACCCAAAGCTGATCCaagttatactgcc
gcagtcatatagagcagacgAACTTCATTCCAGCAGAGTGGTGAATCCAATTCATAAAGGaagttatactgccct
gcagtcatatagagcagacgGTAAAAAGTAACAAACAGAGCAAAAATCGTCTCTTACCAGaagttatactgccct
gcagtcatatagagcagacgGGTTTTTGAAGCCTTCACTACATGACATAGTAACATTCTGaagttatactgccctg
gcagtcatatagagcagacgAATTAACAGCAAGAGACACAGCAATAATTAGAAACGACCAaagttatactgccc
gcagtcatatagagcagacgTTAGTCCAGTAACAAGTCCAAGTAGAAAGTCAATGATAGCaagttatactgccct
gcagtcatatagagcagacgTGTTCTTTACACTGATCAACTATATAGGCAAAGCAAGCTCaagttatactgccct
gcagtcatatagagcagacgCCAATGAAGGTAGATGCAATCAAAAGCTGGAATGGAAAGGaagttatactgccc
gcagtcatatagagcagacgCAAGAGCACCAACGGAAGACAAAATCATAGGGAATTTTCGaagttatactgccc
gcagtcatatagagcagacgGATCACTAATAGACAAAAGTATGAATGTAGACACTAGACCaagttatactgccct
gcagtcatatagagcagacgAATCCACTTATGTCCATCTGCAGATTAAAACGTGACACTTaagttatactgccct
gcagtcatatagagcagacgCGTGTATAAAAATATCATGGTGCATATGCTTTTTTCCCTGaagttatactgccctg
gcagtcatatagagcagacgCTCTGGTGGATGGGCGGGACTAATAGAGCTTACATTTTCAaagttatactgccct
gcagtcatatagagcagacgTGCATGTATATAATGAGATACCTTGTGGATGGGACTCACAaagttatactgccct
gcagtcatatagagcagacgACAAATTTGTGCATAAAACAGTTGCCATTGAGTACTTGTGaagttatactgccctg
gcagtcatatagagcagacgTTTACACTTGTGGTAAAGTGTGGTGATCAAAAAGTTTCAGaagttatactgccctg
gcagtcatatagagcagacgCCTTTATTTAGGGAGAGCTTGGCTGTTGCCCTTATTTTTTaagttatactgccctg
gcagtcatatagagcagacgACCTATGTCTTCACAGATAGATTAGTGAGAAACATCTTCCaagttatactgccct
gcagtcatatagagcagacgGATAAAATTGAAAGAGGAGACTTACACTGTCTACAGATGGaagttatactgccct
gcagtcatatagagcagacgTACCGTGGCTGAGTGTGGTGGCTTACGCTTATAATCCCAGaagttatactgccc
gcagtcatatagagcagacgTCTAAAAATAAATTTAAAAAAACTAGGCCAGGCACGGTGGaagttatactgccct
gcagtcatatagagcagacgGCGTGGTGGCGTACGCCTGTAGTCTCCGCTACTTGGGAGGaagttatactgcc
gcagtcatatagagcagacgTAAAAAGTAAATAAAGGTGGTAGTATGTCATGACCTGAGGaagttatactgccct
gcagtcatatagagcagacgGATAAACTCAACTTCCAAGGACCAAAATGACAATGACAGAaagttatactgccc
gcagtcatatagagcagacgTTGGAAGATGTGTCAATCATTACACTAGTTACTTGATGGAaagttatactgccctg
gcagtcatatagagcagacgTGCTCTGTATTTTCCAAAGTCAATATAAAAAGTGTGTGCCaagttatactgccctg
gcagtcatatagagcagacgTGTAAACCGTTTTTAGTCATTGAATGAATCACTGAAGGCGaagttatactgccct
gcagtcatatagagcagacgCTTTGTCCTAAGGAGCAAACAGAGAGTAGGCCACCACTCCaagttatactgcc
gcagtcatatagagcagacgGGTCTAAGAGCACCAACTTTAAGGGAAGGGATCTAGAGTAaagttatactgccc
gcagtcatatagagcagacgTTTGAAAATTCGGCAAAAATCTTTGACATGAGACCACTGCaagttatactgccct
gcagtcatatagagcagacgGTAGGGGGCTCTCAGCAGCTGGGCCAGGCTGGAGGTGGGCaagttatactg
gcagtcatatagagcagacgCCACCAAGCTGGCTGGACCAAGGCCCGTCTTCCCTCAGTCaagttatactgc
gcagtcatatagagcagacgACATTCTTGTCCTCTCCTCCCTGGTCCCTCCCACTGCAACaagttatactgccc
gcagtcatatagagcagacgGCCAAAAGAAAGCACAGAATCAAATACAGAAATTGATGAGaagttatactgccc
gcagtcatatagagcagacgAACACAAAGAAATGAGAGATGTCTGAGTTGATGGATATGCaagttatactgccct
gcagtcatatagagcagacgCACTATACATTGTATGTATCAAAACACCACTGTATACTCCaagttatactgccct
gcagtcatatagagcagacgATCCCAACACTTTGAGAGGACCAGACAGGAGGCTCACTTGaagttatactgcc
gcagtcatatagagcagacgAGTTCAAAGTTGCAGCGAGCTATTACCACGCCACTCCACTaagttatactgccc

gcagtcatatagagcagacgACTTCAGGTGATTCCCATGTCCAGTCAAGGTTAAAAACAaagttatactgccct
gcagtcatatagagcagacgCAGGATGGGATTCAAGTATGTATATTAACAAACGTCTCCAaagttatactgccctg
gcagtcatatagagcagacgCTGCTGTAGGACCCCAACTTTGAAACTAAGAGGATTTGTTaagttatactgccctg
gcagtcatatagagcagacgATTCTGTTTATGTGTGTTTTTCTCAACCTAGAGAGACTCTaagttatactgccctgt
gcagtcatatagagcagacgTCGGGTTCAGAGCCGTTTTGCTTATCCTTGTCACCACAGGaagttatactgccct
gcagtcatatagagcagacgAATGCATGAATAAACACTTACAAAAAGAAACTGTGAGCCGaagttatactgccct
gcagtcatatagagcagacgAATGACCTAGGAGACATTCTTGAAGGGAGATAAGACCGCTaagttatactgccc
gcagtcatatagagcagacgAGACTCAGGGCCCTTTTTTTTATGACCTTACTCTACGCCaagttatactgccctg
gcagtcatatagagcagacgCTTGATGTCTTCCAGGCTGCTCAGAAATAATCACTCCCAAaagttatactgccct
gcagtcatatagagcagacgATGGGAATGAGGAAATGAGTAAGATTAGGGATTGTGCCAAaagttatactgccct
gcagtcatatagagcagacgCCAGCAAGGTTCCACCTGACACCTCGGGAAGTGCTAGAACaagttatactgcc
gcagtcatatagagcagacgGCCATTTCCTCCTTCTCAGGCTCTTCTCCCAGTCACAGAGaagttatactgccc
gcagtcatatagagcagacgGCAGCACCACTCCTTTACAATCACAAAATATCCCCCAGAAaagttatactgccc
gcagtcatatagagcagacgGTTTAGAGTTTACAGGTGTGATTTGTTTTTGAGGCATTGTaagttatactgccctgtg
gcagtcatatagagcagacgCAAACCTGGATACTATCTCTTTGCATTTTGAAATGGTAGAaagttatactgccctg
gcagtcatatagagcagacgGAGAATGGTGAAGTGCCTGCCTCCTGTTTTACACACAAAAaagttatactgccct
gcagtcatatagagcagacgCCTCCTTAATATCTGGCTAAACATCTTTGGTGTCAACGGTaagttatactgccctg
gcagtcatatagagcagacgTTTGATACCACCACCAAATAATAAATGCTCCTCCACCCCAaagttatactgccct
gcagtcatatagagcagacgACAATCTTCAGTGATGGGCCACTTTCTAGAGTCCCTGAAAaagttatactgccct
gcagtcatatagagcagacgGACATGCAGAAACAGTAAAAGGATTGGGGAAGAATAGAGCaagttatactgccc
gcagtcatatagagcagacgACACCATGTTTCAGCCACCCCTGTCCACCAAGTAGCTCATaagttatactgccc
gcagtcatatagagcagacgTTTTTAGGACAAAGCAAGGAGACAGCCACAGAAGAGATGAaagttatactgccc
gcagtcatatagagcagacgCACAGAGCAGGAGGGTGTTTTAAGAAGAAGAGGCAGATTTaagttatactgccc
gcagtcatatagagcagacgCTTCAAATGAGGTGAGTTCCCAGTAAAGGCCATTAGGTTTaagttatactgccctg
gcagtcatatagagcagacgAGGACTTTGATGGCCGCCCTAGAGAGACCAATGGATGATGaagttatactgccc
gcagtcatatagagcagacgCAAAGCCCGGTATACATTCCCATTTTATTAAATACTATGGaagttatactgccctg
gcagtcatatagagcagacgCACTCAGAAATATTGCTATCAGATGAAAAAGTGTAGTTGCaagttatactgccctg
gcagtcatatagagcagacgTCTTCCCATATTCTCCGATAAACATATTGCGTTGTCAGTGaagttatactgccctg
gcagtcatatagagcagacgTGCACTAAGGAAAATGGCAGGTTCTACAAATAAAATCTTCaagttatactgccctg
gcagtcatatagagcagacgCTGGGTAGGTAGCTGTATTCTATAAAAAGTGAAAACGGAGaagttatactgccct
gcagtcatatagagcagacgCAAAGATGGGTAAGTTTGAGCAATTTTGTGGAGTAAATGTaagttatactgccctg
gcagtcatatagagcagacgGCAGTAAGCACTTTCTCAGAGAATAACTTTTACTTTGGAGaagttatactgccctg
gcagtcatatagagcagacgCTCTGTAACAACCTGAGGCTGCCATGCTTTCCATTTTTATaagttatactgccctg
gcagtcatatagagcagacgAAGCCGCCTTTGAGTCTTTGAAATCATAAACTGTCACCCAaagttatactgccct
gcagtcatatagagcagacgAGTTCCCACACTGTATATTGCCAACTGATTTTTCGTAAGTaagttatactgccctg
gcagtcatatagagcagacgTTGTTATTGTTATTTTGCTGCGTATTAAAAACGGACAGGGaagttatactgccctgt
gcagtcatatagagcagacgGGGCCTTATTTTTAACTCACCCCACGATTACAGTCTTCCTaagttatactgccctg
gcagtcatatagagcagacgACAGGTCGGGCCGTGCCAGGGGCTGTCGCCTCGGATGCCCaagttatactgc
gcagtcatatagagcagacgTCGTGCGCGCCCGGGTTCAAGGGCAGCCTCTGGACTGGGGaagttatactgc
gcagtcatatagagcagacgCCGGCGCTGGGGTGGGGAGAGTACTTTCCCGTCGAACTCGaagttatactgcc
gcagtcatatagagcagacgCGGGGAGTCTCAAGCGTATTGTAAAGCAAACCTCGGCAAGaagttatactgcc
gcagtcatatagagcagacgGTGACACCTGCAGGGTTCTCCGGCAGCCCTCGGCTTCCCCaagttatactgc
gcagtcatatagagcagacgAGCAGCTGCTCCTTCTGCGCAGTGGACAATCGGCCAATCGaagttatactgcc
gcagtcatatagagcagacgTCGGTCAGCGGCCCTGCCAGCGCTAGGCGATGACTCGGCCaagttatactgc
gcagtcatatagagcagacgCGCGGCGCTGTCACTGTTCTACCCGGCTCTCCTGCGATTGaagttatactgcc
gcagtcatatagagcagacgCCCCAGCGGCGATTGGTCCACCCGGACGTCACTCTCCCCTaagttatactgc
gcagtcatatagagcagacgGGAGGCCCGGAGTGGATCCCGCGCGACCGAGCAGCCACTCaagttatactg
gcagtcatatagagcagacgCTCTGGAGTTGCAAAGCATTTCTGAATCCGAATATCCTTTaagttatactgccctg
gcagtcatatagagcagacgCACATAAAACAACAGTTGGAAAAGTAACATGTTTGGGCTGaagttatactgccct
gcagtcatatagagcagacgGCAGAAGAGGGGAATGGGCGCAGGTGGAGGGCATAGGAAAaagttatactgc
gcagtcatatagagcagacgGGAATGTGGTCGCTGGCTTTTCCTGGGGAGGGGAGGGGGGaagttatactgcc
gcagtcatatagagcagacgCGTTAAATTTTCATGAACCTCCATGTCTTTATCTGCCACGaagttatactgccctg

gcagtcatatagagcagacgTTGTCTAATTAAACACTCAGCATGTATACACAACACAGCCaagttatactgccct
gcagtcatatagagcagacgGTGCTCTCCAACACACCCACCAGGAACAACTTGTAGTAAAaagttatactgccc
gcagtcatatagagcagacgTTCTTGAAGCCAGGAGAAATGCATGCATGAGGAACCGTGGaagttatactgccc
gcagtcatatagagcagacgGTAAGAAGGTGTTAGAATTTACTATGGGCTTGCGTTATGTaagttatactgccctg
gcagtcatatagagcagacgGCAAGGCCTTACTCTGGATTGGTTGCTGTCAGGAAATGGCaagttatactgccc
gcagtcatatagagcagacgGCTAAAGCCATGAATGATAAAGAAGCAACAGTCACACATAaagttatactgccc
gcagtcatatagagcagacgTGTGGTTTAGACAGTGCCTAAATTTTGTCCGTGTTCAGACaagttatactgccct
gcagtcatatagagcagacgGTCTTGTTTTTGTCTTGATCATGGTTACAGAATGCTCTTGaagttatactgccctgt
gcagtcatatagagcagacgAAATTGTTTTCATTCGACAGAACACCATGGTCCAGAAAAGaagttatactgccct
gcagtcatatagagcagacgCTGCTTTCTCTTCCTCAGCACTCCTTAAATGACAGTTGTGaagttatactgccct
gcagtcatatagagcagacgGTCCTAACTGATACTTAGTAGCTGGTAACCTTGGACTTGCaagttatactgccct
gcagtcatatagagcagacgCATTTATGAAGCACTTGTTATGAGTTTCAAACAATGGAGCaagttatactgccctg
gcagtcatatagagcagacgGATAAACAAACATGAGTAAGATACAGTTTGCAAGGAAGACaagttatactgccct
gcagtcatatagagcagacgTGTCACCCAGGCCGGAGTGCAGTTGCAGGATCTTGGCTCAaagttatactgcc
gcagtcatatagagcagacgGCGCCACCGTGCCCTGCTGAAAATGTCCTTTCTTATGGGGaagttatactgccc
gcagtcatatagagcagacgTAATTAATGTCCAATATACTTCCACGCTTTGCATTCCTACaagttatactgccctg
gcagtcatatagagcagacgAGAAAGCCAGGTCTTCTTAATGAGTTCTGCTGCTGATCGCaagttatactgccct
gcagtcatatagagcagacgATGTGAAAAATTAGCAGAGTTTCTGCCTTCACAATTAGAGaagttatactgccct
gcagtcatatagagcagacgTCATCTAGTGAAAACAGACAATGTGGTATTTTTTCAGCCAaagttatactgccctg
gcagtcatatagagcagacgATAGTTAGGCCCTGAGTTATAATTTGAGATAAGTGATTCCaagttatactgccctg
gcagtcatatagagcagacgAGGACCTTGTTTCAGATTATATTAAAACCAGAGTTCACTGaagttatactgccctg
gcagtcatatagagcagacgGATTCTAGAGAGGAAGCAGGGAGGGCCAGGCATGTGGCTCaagttatactgcc
gcagtcatatagagcagacgAGTTCAAGACCAGCCTAGCCAAAAGAGTGAAACCCTGTCTaagttatactgccc
gcagtcatatagagcagacgTACTCAGGAGGCTGATGTAGACAATTGCTTAAACCCGGGAaagttatactgccc
gcagtcatatagagcagacgAAAAAGAGAGAGAGAGGAAGCAGGGAACAACTGAGCCAAAaagttatactgcc
gcagtcatatagagcagacgCTTCCACATAAATGTGACCTTGAAGACCTAGATACATATTaagttatactgccct
gcagtcatatagagcagacgATCAAATATGGTCCCGGATCTCTTAATTTCCTATATTCAGaagttatactgccctg
gcagtcatatagagcagacgGTATTCTCTCATTCAAGGCTTTAGATGTTCAATTATTCCCaagttatactgccctg
gcagtcatatagagcagacgACCTGAGGTTTCCAAGTATCTCACCGGGATTTTTATTCTTaagttatactgccctg
gcagtcatatagagcagacgTCTGCCAATTACCTAGATCCTCCTGAACAATTGTTTTACTaagttatactgccctg
gcagtcatatagagcagacgCAGTGATGATCACATATATTGGTTCCTTGCAACCTTTTTCaagttatactgccctg
gcagtcatatagagcagacgTCTTCGTTTGTAATGGATAAGTGGGTTTTCAGGGGTGTGAaagttatactgccctg
gcagtcatatagagcagacgTAGGCAAAATTTTCCCCAATTCAACGATGACACCCAGCTCaagttatactgccc
gcagtcatatagagcagacgGGGTTATCCTGTAAGTAGCAATCAGGATTAAAGTCCAGGCaagttatactgccc
gcagtcatatagagcagacgCTGGAGGAATGGTATACCCTACCACAGTCTGAGGAGTTCTaagttatactgccc
gcagtcatatagagcagacgGTAAATTTAGATCCTTGAGTTGGTCATAAGTTAAAGGAGGaagttatactgccctg
gcagtcatatagagcagacgTGTCTCTGGCCAAAAAGAAGTCCATCCAAGCACTAGTAGTaagttatactgccc
gcagtcatatagagcagacgGAAAATATCCTTGATTTCCCAATGAGCTCTGTCCCTGCATaagttatactgccct
gcagtcatatagagcagacgATTTCCTTTCCATGCAAACAGTGGCTAGCTGTTAAAATTAaagttatactgccctg
gcagtcatatagagcagacgCTTTCTCCCCAACTCTGAAAGTACTCCTTTGTTTCTTTTTaagttatactgccctgt
gcagtcatatagagcagacgCATGCTGTTTAAAGTGGGCTTTGTTAAGGCCACTTTTTAAaagttatactgccctg
gcagtcatatagagcagacgCCCTTCCCAAACACAGGTGTCGTCAGGTGACTGTAGACATaagttatactgccc
gcagtcatatagagcagacgTTTTACTATTAAAAAGCAGCGCAGCAGCATCACTCCCCAGaagttatactgccc
gcagtcatatagagcagacgACAGGTCCATACTTCTCATATGCATTTCTAGAAATTCAACaagttatactgccctg
gcagtcatatagagcagacgCAGGGAATGGAACTGGGGAGAAAACGTATGGAGGACTTTTaagttatactgccc
gcagtcatatagagcagacgCCAAGTGGCCGGCGGCCAGACGGAACAGGTAGACCAGGCTaagttatactgc
gcagtcatatagagcagacgGCATGGACAAGAGGTTGCCTCCTGTCACCTTCTCCATCGCaagttatactgccc
gcagtcatatagagcagacgAGCAACATCATCCCCGCCGCTGCCGCCATTTCACTTCATTaagttatactgccc
gcagtcatatagagcagacgCTCCTCCTAATCGACGGAGCGAGAGAAGCTGGCGGATGGTaagttatactgcc
gcagtcatatagagcagacgGGTCTCCTACTAAACGACAACAGTCATTATGGAAGCCTTTaagttatactgccct
gcagtcatatagagcagacgGCTTGAGTATGTATCAAAGAAGCCTTGACTGTTAATTGTCaagttatactgccctg
gcagtcatatagagcagacgTGGGCACTGTGGAGTAAAAGCAATTCTAAAACCAACTACAaagttatactgccc

gcagtcatatagagcagacgACCATCAAGACCTGTCACTCCCGTTTCTCCACAGCTATTGaagttatactgccc
gcagtcatatagagcagacgCCTCACAGGTTCCCACTCCAGCCTTCTTCCTTCAAAGACCaagttatactgccc
gcagtcatatagagcagacgATCTCCCTGGCCGTGTTTCTGAAACGTAGATCAAGAGGTGaagttatactgccc
gcagtcatatagagcagacgTTCGGGAAAGGGGTCTCCAGGATTTTGAAATTTTATATTTaagttatactgccctg
gcagtcatatagagcagacgTATATTTGTATTTCTGCTTATGATCACATTGGCTGCAAGCaagttatactgccctg
gcagtcatatagagcagacgATGAGAAAAGAAGTGGACTAAATATGTAAATGATACGCCCaagttatactgccct
gcagtcatatagagcagacgACAATGTCATATCATGCTGTTCAAGAGAAGTTTCCATCGTaagttatactgccct
gcagtcatatagagcagacgACTGAAGTCATTGCCTTACACACAAATAGACATACTGAACaagttatactgccct
gcagtcatatagagcagacgCAAAGTGGAATAGAGTAGCTATGAGATTTTGTTGGTTTCAaagttatactgccctg
gcagtcatatagagcagacgGACCTCAATTATACCAATTTCTGCTTAGTAGATGATTGTCaagttatactgccctg
gcagtcatatagagcagacgGTAATGTGATACAACTCTAAGCACAAATTGTGGAAGGGAAaagttatactgccct
gcagtcatatagagcagacgTCATATGACCTTGGTTAATAAGTTTCCAAAATAGGCCTCAaagttatactgccctg
gcagtcatatagagcagacgGTCAGCCTTGTCTTTCTTGACCTCTGTTGCTTTGGATACTaagttatactgccctg
gcagtcatatagagcagacgCCTCCTTCTGGAAATCTTTTCTCCCTTTGGGTTCCATAACaagttatactgccct
gcagtcatatagagcagacgCGTCTCGGTTTCCTCTGTGAGCTTGCTTCTCAGATGGTGCaagttatactgccct
gcagtcatatagagcagacgCTCAGGCCCTATCTCAGCTCCGACCTTTTCCCAGTCAATTaagttatactgccc
gcagtcatatagagcagacgATGCTGACCATTTCCAAATCTCTTATCTCAAGCCCACCCTaagttatactgccct
gcagtcatatagagcagacgCCCAAACCCCATAAAACTGTATACTTTAAATGTACAAAGCaagttatactgccct
gcagtcatatagagcagacgTCATCTTCCTCATACATCTGATACCCTCTTATAATCTTTGaagttatactgccctg
gcagtcatatagagcagacgGTGACATCACCTATCCAGTTAACAGATACGGAGACTTAGGaagttatactgccc
gcagtcatatagagcagacgCCTTTTTCAGGTGTGTGGCTGTAAGGAAAGTACATCCCTGaagttatactgccct
gcagtcatatagagcagacgTAGCATTCAGACCGAACATCTCTTTTTTGGCTTGGCTAAGaagttatactgccct
gcagtcatatagagcagacgGCTCTGTCTTGCTATTGGTCTATTGCTGAGTGGCATTTGAaagttatactgccctg
gcagtcatatagagcagacgGTTGAAGAGGTGAACAAAATGCAAAGCGATCAAGGAGTTCaagttatactgccc
gcagtcatatagagcagacgCCTCTAAGGTCTCAAGTTATTACAACACTTCTGATGGCCTaagttatactgccct
gcagtcatatagagcagacgCCTAAGAAGTCTCAGTTGTCAGAAATGTACAATAACAGCCaagttatactgccct
gcagtcatatagagcagacgACAGTGGTGGTGGAGTTGGTGGTGGTGTCTTATCCTGGGGaagttatactgccc
gcagtcatatagagcagacgGTTTGTCAAGAGAAATGGCTGATTAACTGTAAATGCAACTaagttatactgccct
gcagtcatatagagcagacgCCATTGCCACAACCTTTATGCCTGGGAGCTGAACATCTTGaagttatactgccc
gcagtcatatagagcagacgGAGTGACTAGACTTCTCAGTTTCTTTTCTGCGTAACGATAaagttatactgccctg
gcagtcatatagagcagacgCTCATTTGGACTAAGTAAGCAGCTGGAGGATTTTACATTTaagttatactgccctg
gcagtcatatagagcagacgGCAGTAGGATGGAAGCTCTAATTTGATTTATCTGTGTACTaagttatactgccctg
gcagtcatatagagcagacgCTTATTTCCTATAGCAGCAGGTAAAAGCTCCTAGAACAGGaagttatactgccct
gcagtcatatagagcagacgAATCTCATTCTTGCTTAATCAGACACACCTCATACTATCTaagttatactgccctg
gcagtcatatagagcagacgAAATTATAAATGACAAGCCGGCCACCACTCCAAACTGACAaagttatactgccc
gcagtcatatagagcagacgCCTTAACAGGCTTGGGAATAGCAGGAAGAAAGGGTTTAAAaagttatactgccc
gcagtcatatagagcagacgGGTTAGCCCACCTGTTTCCACCTCCAGCTCCTTACCCTTAaagttatactgccc
gcagtcatatagagcagacgAGCCACCACTGTTTCTCACTTGCATTCCTCACTGTTTCCTaagttatactgccct
gcagtcatatagagcagacgAGTAACACTCAACCTCCTCACTATGACCTGAAAGACCCCTaagttatactgccc
gcagtcatatagagcagacgGGCCACATCTCTCCTGCAACCTCAGCCCGCCAAAATCCCTaagttatactgcc
gcagtcatatagagcagacgACACTGCTCCCTCCCTTCCTTTAGGTGTAGCTGAAATGTCaagttatactgccct
gcagtcatatagagcagacgAAAGTATTCTCTGACTGTCACACCCAAAGCAGGAAAGTGCaagttatactgccc
gcagtcatatagagcagacgTATATTTTTGTATGTTTGCCTTTTTTGGTCATCTGTGCCCaagttatactgccctgt
gcagtcatatagagcagacgTCTGTGCATGCAGGGACCATGGCTGACTCTTGTATGACTGaagttatactgccc
gcagtcatatagagcagacgATGCAGGAAATCTTAGTCAAATGGATGAAATGAGGTGGTTaagttatactgccct
gcagtcatatagagcagacgTGGAATTATACACCTAATCTACCCTTGGGAACCACAGAGAaagttatactgccct
gcagtcatatagagcagacgGCTCTGTCTGGCTGAATCACTTAAAATACATGTGTACCAGaagttatactgccct
gcagtcatatagagcagacgACCAATCTAAAATGTGTCACTCCCAGACTAATTGTAATTCaagttatactgccct
gcagtcatatagagcagacgTCTCAACCAGAAAGAGGGACAAAAATGTCCAACCTCACTAaagttatactgccc
gcagtcatatagagcagacgAGCTTAAATTACAGTAGTTTGTGCCTATAATCCCAACTACaagttatactgccctg
gcagtcatatagagcagacgCATTTGGCCAGCCTGAAGTAAAATGAACAAAATGTCATATaagttatactgccct
gcagtcatatagagcagacgTATACACTAAAATCGTTATACATTGTCTGCTCAGCAAGTCaagttatactgccctg

gcagtcatatagagcagacgTAGGGGAACAGTGCATACATATAGTTTTATGTTGAGAAAGaagttatactgccctg
gcagtcatatagagcagacgTGAGTGAAAAAAAGGCAACAAAACATATTTGGCATAGCACaagttatactgccct
gcagtcatatagagcagacgATGGTACTAACAATTGTCTCTGAGGGGTAGGATTATTAACaagttatactgccctg
gcagtcatatagagcagacgGAGAGGAAAACAATAGAACAATTACATGCAAAGTCATTGCaagttatactgccct
gcagtcatatagagcagacgGGTTGATATTTTCCAAGTGGTTCTACAAGCTTTAACGTTGaagttatactgccctg
gcagtcatatagagcagacgCCATAGGTAGCCCAAGATAAGAGGATAGTGGATGTTAACTaagttatactgccct
gcagtcatatagagcagacgGAAAAGTAGAGTCAAGAATGGAGAAATAACTGTTCTGAAGaagttatactgccct
gcagtcatatagagcagacgATCTATTTAAAACCAGACTCAGAAATCAGTAAGACTCACGaagttatactgccctg
gcagtcatatagagcagacgCTAGTCCAGCCCTCTCACCCTACCCATAAGAAAACACAAGaagttatactgccc
gcagtcatatagagcagacgTGTTATCCTCCACGGAACACATTGCCCTGACCTAGGCCTGaagttatactgccc
gcagtcatatagagcagacgGGAACCCAGGCCCTGCTACCACCCATCTGTCCACCTTTGTaagttatactgcc
gcagtcatatagagcagacgCTGGTCTCAAACTCCCAGCCGCCTCAGGTGATCTGCCCACaagttatactgcc
gcagtcatatagagcagacgAAGTGGAGTTTTTAAGGAGCAGTTTAAGGAGGCAGTTAAGaagttatactgccctg
gcagtcatatagagcagacgGGTAAACTGCCCTCTCTTATCTTTCCACTCCTACAGCTGCaagttatactgccct
gcagtcatatagagcagacgCCTCGAGGTCACTGCACTGCCCAGGGTTATGAAAACAGGAaagttatactgcc
gcagtcatatagagcagacgAAATTAGGGGAGATGGAATTTGTCTGGTCAGCATTCATTCaagttatactgccctg
gcagtcatatagagcagacgTGATCAATTATGGTTCAGTTTATTTCATTCTCCCTGCAGCaagttatactgccctg
gcagtcatatagagcagacgCAGGAAAGCTGTAGGTAATCAGAGTGGAATCTATAAGTATaagttatactgccctg
gcagtcatatagagcagacgAAAGAATCTGCAAGTATTCAGAACACAACTAAGCAAAACGaagttatactgccct
gcagtcatatagagcagacgTTAAAAATTATTTTATTTGGCCAGGCATGATGGCTCATGCaagttatactgccctg
gcagtcatatagagcagacgAGCCAGGTGTGGTGGCGGCTACCTATAGTCCCAGCTACTCaagttatactgcc
gcagtcatatagagcagacgGGAGGCGGAGCTGGCAATGAGCTGAGATGGCGCCACTGCAaagttatactgc
gcagtcatatagagcagacgAAAAATTTCCTTCCCCAAATAATCTGCTATCCCACAGTATaagttatactgccctg
gcagtcatatagagcagacgTTTTCCTGAAGACTCGAGAGTCAGTTTCCTAGGGCTGCGGaagttatactgccc
gcagtcatatagagcagacgCTGCAGGCCAGAAGACAGGAATCAAGTGTCACCAGGGTTGaagttatactgcc
gcagtcatatagagcagacgGCTTCTGGTGGCTGCTGGCGATCCTTGACATCCTCTGGCTaagttatactgccc
gcagtcatatagagcagacgCTCTCTGTGTGTCTGGGTCTCGGTGCCCCTCTTATAAAGAaagttatactgccct
gcagtcatatagagcagacgATGACCTCATCTTCACTCATTATATCCTGTAATGACACTAaagttatactgccctg
gcagtcatatagagcagacgCTTTTTTAAGGACCATGATTCAACCATAATACCCTACATCaagttatactgccctg
gcagtcatatagagcagacgTGCTTTCTTTCTTATTTGATGGTTTACTAAGGGCAGGGAAaagttatactgccctg
gcagtcatatagagcagacgACAACAAGAGGCAATTGTCTGTAAAGGATGTATAAACGTAaagttatactgccctg
gcagtcatatagagcagacgTGCTTTTTATTATCTACTTGGGCCAGCAGTTCTAAACAACaagttatactgccctg
gcagtcatatagagcagacgGAAATTTTTCCATACTCTTCCCTTCCAATTCCACTCTCTTaagttatactgccctg
gcagtcatatagagcagacgAATTACTGAAAATAGTTTTGTTGTTTACAGCCAAGGAGGCaagttatactgccctg
gcagtcatatagagcagacgTTTGGTATGGTTGTCAAATATGCTACGCATTATGCCACTGaagttatactgccctg
gcagtcatatagagcagacgCTCTGAGACAGTATAAAATTTCTAGTATCTAAGGAAGACCaagttatactgccctg
gcagtcatatagagcagacgGAGAGTTAAGATCATCTAAAGACTGAAATTATATCCCTCCaagttatactgccctg
gcagtcatatagagcagacgACCAAAGATTTCTTGAGTACCAACTATGCGCTTAGAACACaagttatactgccct
gcagtcatatagagcagacgTTCTAGGGAACGGAGACAGCACACCAAAAGGTTAGAAGTAaagttatactgccc
gcagtcatatagagcagacgAAAAAGGTTAGGGGAAGAAAAAGTCAAAAAAGGACCTGAAaagttatactgccc
gcagtcatatagagcagacgCATGCTGAGGTGGGTTGGAGTGGGCGGGCTGTAGGATTACaagttatactgccc
gcagtcatatagagcagacgCTCACTGAGGAGAGAAGCTTGAGCAGAGACTTGCCGGAAGaagttatactgcc
gcagtcatatagagcagacgCAGATAGCTGGGAAAGCATTCCAGGAAACAGACCAGTGAAaagttatactgccc
gcagtcatatagagcagacgGCTGGAGGAAAGGCAAGGAAACCAATGAGGGCAGGAAGGAaagttatactgcc
gcagtcatatagagcagacgCTCTCTTCCATGGGGGAGTGCCTGTAGACCAACTCAGCTGaagttatactgccc
gcagtcatatagagcagacgAGCAGCTGGGAAATGAGGCTTCAGCCCTGAAAAAAGTCTCaagttatactgccc
gcagtcatatagagcagacgCAGAGCTGGATTTATAGGAGTCATTCCATTTGGATAGGGAaagttatactgccctg
gcagtcatatagagcagacgAAATTATATTTAAGAGGGGAGCAGGAGCGGTAGGAGCGGTaagttatactgccc
gcagtcatatagagcagacgGAGTTCGAGACCCGCCTGGAGTAACATGGTGAAACCCAATaagttatactgccc
gcagtcatatagagcagacgCTGTAGTCCCAGTTACTCGCAAGGCTAAGATGGGAGGACCaagttatactgccc
gcagtcatatagagcagacgGGTCACCCTCCTCCTCATCTATATCCTATAATTTCTTGCAaagttatactgccctg
gcagtcatatagagcagacgCCTTAACGCAACTATTTAATTATTCTACACATTTCCTGGGaagttatactgccctg

gcagtcatatagagcagacgTGTTGGGTGTGGAGTTAAGATGCCCTTGGGGTCCCTGCGCaagttatactgccc
gcagtcatatagagcagacgCATTTAATGTGCCATCAGTTTCTACTCTCGTTGTCATTGAaagttatactgccctg
gcagtcatatagagcagacgAGCCTCCTTTAATTTAAACAAACAAACAAAAAAGCCAGGCaagttatactgccc
gcagtcatatagagcagacgATTTTTTTTTTTGAGACAAGGTCTTACGCTGTCACCCAGGaagttatactgccctg
gcagtcatatagagcagacgCCCACCTCAGCCTCCCCAGGGACTGGGACTATAGGCACATaagttatactgc
gcagtcatatagagcagacgCGCACCCGGCCTGTAACCTCCTTTTATTACTACAGATCAAaagttatactgccc
gcagtcatatagagcagacgAATTCAGTATTCTTGCTTAACCCTGCCTCCACCCACCTCTaagttatactgccct
gcagtcatatagagcagacgCTGCTATACATGATCAATCCCATATACCAGACTTACCAACaagttatactgccct
gcagtcatatagagcagacgTGTTTGCCAGGGATTCTAGGTTTGAGGAAAAAGTAAATTTaagttatactgccctg
gcagtcatatagagcagacgAACTCATCCAACTCCAAAATGTTATGGTATTAAGTCCTGAaagttatactgccct
gcagtcatatagagcagacgTATGGACTGAATTGTTTCCCAAAATGCACTGGTTTAAGAGaagttatactgccct
gcagtcatatagagcagacgCTCAAATGTGACAGAAATACTTGGAGAAGGGCCTTTAAAGaagttatactgccc
gcagtcatatagagcagacgTGAGGTCTTTAGAGTGGGTCCTAATTCGAGAGGACTGGGTaagttatactgccc
gcagtcatatagagcagacgAAGAAGAGGAAGGCAGGCTGGCATGTTGGCTCACACCCGTaagttatactgc
gcagtcatatagagcagacgTGAGCACAGCCTGGCCAACATATATGAAACCCCATCTCTAaagttatactgcc
gcagtcatatagagcagacgAAATTAGCTAGGTGTAGTAGTATACGCCTGTAGTCCCAGCaagttatactgccc
gcagtcatatagagcagacgGAAAAAAAAGGGAAGAGACACCAGGGGCTCGTGTGCACAGaagttatactgc
gcagtcatatagagcagacgGAGGACAGTGAACAGACAAGACATCCCTGCAAGTCAAAGTaagttatactgc
gcagtcatatagagcagacgTTTGCAAGCCTCAGCAGAAATCAAACCTACTGACACCTTGaagttatactgccc
gcagtcatatagagcagacgCCTCCAGGAGTATGAGAAAATAACTCCAAAGGGCCATCATaagttatactgcc
gcagtcatatagagcagacgGCCTTGGTTGTTGCTGCTTCTCCTGTGGCCTTCACTCCTCaagttatactgccc
gcagtcatatagagcagacgGGAACCAGACCTATGACACGCCCTCTACCTTTCCATATTTaagttatactgccc
gcagtcatatagagcagacgCCGGGGAAGAACATTTTGACAGCATATCAAGAAGGTGCCTaagttatactgcc
gcagtcatatagagcagacgTACTGGAGGGTATTTGGGTGGAGGTTAAGGGGTCTGAATAaagttatactgccc
gcagtcatatagagcagacgGGCCCTTTTTATAGGACTTGCCTGTTCTAACTCACTGTCAaagttatactgccct
gcagtcatatagagcagacgAGTTCTTCCTTAAATAGTCAGAGGGCTTAGAGGCTCAAATaagttatactgccct
gcagtcatatagagcagacgCAATTCTGTTGTTCTCAGGCCAAATTTAGCTTCTTTCCCAaagttatactgccctg
gcagtcatatagagcagacgAGCCCCAAACAGTTTCACTTTCAACCTGCTTACCTTTCTTaagttatactgccct
gcagtcatatagagcagacgGAAAAGATTTGGGGTCATCACAATTTTCATTTCATTGCAGaagttatactgccctg
gcagtcatatagagcagacgTTACAGGCATGAGCCAATGGAGTGAATTTCATTGATAATTaagttatactgccctg
gcagtcatatagagcagacgCATCTATTCTCTGCTGGATACTCTACCATGAAATACAAAAaagttatactgccct
gcagtcatatagagcagacgCGAAAAATCCTTGCTCTAAAGTGATTGCAAAACAGACAGGaagttatactgccc
gcagtcatatagagcagacgACAGAAACTTGTCATTGTCACTGGAGACATGCGCAGAGGGaagttatactgcc
gcagtcatatagagcagacgCCATTTTAGAGAGGACCTGATGATGGCAGCCTTGCTTCTGaagttatactgccc
gcagtcatatagagcagacgTGGGAATCTCTCATCATTGACATCAGATAAGGACCTGTAAaagttatactgccct
gcagtcatatagagcagacgGTTCTGACTCTGCAAGACTTAGCCTTGAGTGAAACGCAGGaagttatactgccc
gcagtcatatagagcagacgAGTGTGGACTGTGGGGCGGAAACCTCTTACAAATTGAAAAaagttatactgccc
gcagtcatatagagcagacgTGAGCTCTTCAGAAACAGATATGGCGTAAACATAAAGACTaagttatactgccct
gcagtcatatagagcagacgGTTGTTCCATTTGTACCAGCAACTGCTATTATTCACGGAAaagttatactgccct
gcagtcatatagagcagacgAATGTTAGTATATTTATGTCTGTCTCGTTTCCTCTAGGAGaagttatactgccctgt
gcagtcatatagagcagacgCATTGAGCCAAATATGTAACTTAGGTAATGAGTACAGGAAaagttatactgccct
gcagtcatatagagcagacgCTCAAAACCACATAGAGGAAAAACACCAAGATTTAGCTGCaagttatactgccc
gcagtcatatagagcagacgTCCTGCCTTCTGTCAGCTCCTAGGGGGACCACAGCTCAGCaagttatactgcc
gcagtcatatagagcagacgCTCTGGGAGTGTCATCTACAGTGAAATGTCAAAGGAGAATaagttatactgccc
gcagtcatatagagcagacgTATCTTGTACTTGGTATGAATTTTGAGACAGTTCTGAAGCaagttatactgccctg
gcagtcatatagagcagacgGTTTGCATAAACCTACCATCACTTTCTCTTGTTTTTGTCAaagttatactgccctgt
gcagtcatatagagcagacgCGGCTTTAGGGTATGGGGATGTTCTAGAAGTCCACTGGGGaagttatactgccc
gcagtcatatagagcagacgCTAAGTGTGATTTGGGCTGAGTTGGGCTACATGCTTGGAAaagttatactgccct
gcagtcatatagagcagacgCTTAAAGAGATCCCACAGAGACTATAAATTAGCCTTTCACaagttatactgccct
gcagtcatatagagcagacgCAATTCTGGCATGGGATTTGAAGTATGAACTAGATTTGTCaagttatactgccctg
gcagtcatatagagcagacgACTCAAGACTGGCTGATTGGTTATGTATGAGCAGGTTCAGaagttatactgccct
gcagtcatatagagcagacgCATTGACTTAGATGCCTTTAATCCACTTGCACTGCTTTACaagttatactgccctg

gcagtcatatagagcagacgTAGTATTCCCTAGGTAGTGTCTGGATCTTTGTAATGTACTaagttatactgccctg
gcagtcatatagagcagacgGTCTGTATGTCCTTTGTTATTAAAAGTTATTTCAGCCGGaagttatactgccctgt
gcagtcatatagagcagacgTACAAAAATTATCAGGGCATGGTGGCGGGTGCCTGTAGTCaagttatactgccc
gcagtcatatagagcagacgTTTTTGAGACAATTTTGCTCTTGTTGCCAGGCTGGAGTGCaagttatactgccctg
gcagtcatatagagcagacgCCTCCCGAGTAGCTAAAATTACAGATGCCTGCCACCGCACaagttatactgcc
gcagtcatatagagcagacgCAGACATGAGCCACTGCCTAGCCTATTTTTTTCTTTTAAaagttatactgccctg
gcagtcatatagagcagacgAATTTTTTTTGGAAGGTGGGTACTTTTTAACATCTAGGGGaagttatactgccctgt
gcagtcatatagagcagacgACTTTGTAGTAAATTCTGTGAGAAGGTATGAGTGCAAGCCaagttatactgccctg
gcagtcatatagagcagacgGATGCCTAATGAAAATAGAGGCTAGGCTAGGCACAGTGGCaagttatactgccc
gcagtcatatagagcagacgAGGAGTTCTGAGACCAGCTGGACAACATAACAAGATCTTGaagttatactgccc
gcagtcatatagagcagacgCCACTGCACTCCACCTCAGGTGAAGGAGCTCATCAATTGAaagttatactgccc
gcagtcatatagagcagacgGTACATACACATATGCTCATCAATTTATACATGTCTCCACaagttatactgccctg
gcagtcatatagagcagacgGCTGGGATTACAGATGCGTCCCTCCATGCCAGCTAATTTTaagttatactgccct
gcagtcatatagagcagacgTGCCCGGTCACAGCTACATCTTTTATATATGCATATTTATaagttatactgccctgt
gcagtcatatagagcagacgGTGAACAATTGCACCCAGCCAGACATCGTTTAATGTTGAAaagttatactgccct
gcagtcatatagagcagacgCAAACTGGGTAACTACTTGTGTGAGCTCCTTGGGACAATCaagttatactgccct
gcagtcatatagagcagacgCTACTTCTCCCCTTAGACCTTGGGATTTCTTAACCATCCaagttatactgccctg
gcagtcatatagagcagacgTGAGAGGAGAAGCTACTTTTCACTTTCTAAAGCCTTGTGCaagttatactgccctg
gcagtcatatagagcagacgGAGAAAATCTTGAGGTGCTTGGGATCTCTGACAACTATGCaagttatactgccct
gcagtcatatagagcagacgCTGGGAATCTGCAAACTGAGTGGAAAGTAAAAGGGAGAAGaagttatactgccc
gcagtcatatagagcagacgTGGGCTCCAGCTTGGGCATGCAGTGGTGTACTCAGAGTATaagttatactgccc
gcagtcatatagagcagacgTGTTTTCTACACACAAAATTACTTCCACTACTGGTCATGTaagttatactgccctg
gcagtcatatagagcagacgAATTGCAATTGCCAAACCAAAAAATCACACTTCATAGGTGaagttatactgccctg
gcagtcatatagagcagacgCACTCTTTTACATTTTAAGCACACAAAAAGTTCCAGCTGGaagttatactgccctg
gcagtcatatagagcagacgGCCAGGAATTCAAGGCCAGACTTGACAGCGTAGTGAGATCaagttatactgccc
gcagtcatatagagcagacgTTTTCCTAAAGCCAAGCATAATGGTACATACCTGTAGTCCaagttatactgccctg
gcagtcatatagagcagacgGGTGGGAGAATCACTTCAGCTCAGGAGGTCAAGGCTGCAGaagttatactgcc
gcagtcatatagagcagacgCAATAGGCAGTCTGCAAGCTAAGGAGCATGAAGCCAGTCCaagttatactgcc
gcagtcatatagagcagacgCTAGTCCTTCCACGTTCTTTCGCCTGCTTTTATTCTGGCTaagttatactgccctg
gcagtcatatagagcagacgATCAAAGTATTGCAAGTTCTGCTTCTTCATCTTCACAATCaagttatactgccctg
gcagtcatatagagcagacgACTCTCCTCATTTATTTCAAATCTGCCTTAGTCTCACTCTaagttatactgccctgt
gcagtcatatagagcagacgCCTCCCAGGTTCAAGGGATTGTCGTGTCTCAGCCTCCCGAaagttatactgccc
gcagtcatatagagcagacgCACCCAGCATTTTAAACAGAGTAGGAGATGGTACCACAGGaagttatactgccc
gcagtcatatagagcagacgGCAAACTTGAGTGATTCCAAAGTGTTACCAATTTATTCTGaagttatactgccctg
gcagtcatatagagcagacgGATTCTCCAGAGTAACTTCTGTAAGTGTAAGACGATGATTaagttatactgccctg
gcagtcatatagagcagacgACTTACATATGTACTAGTAAAACTCACCAATAACTGCAGCaagttatactgccctg
gcagtcatatagagcagacgAAAAAGATTTCTCAGGAAGGTGCATTTTGTGTAGAATCGCaagttatactgccctg
gcagtcatatagagcagacgGGTTTTACGATTTTTTTGTATGTGTGTCTACTTGTGATGCaagttatactgccctgtg
gcagtcatatagagcagacgCTCAGGGAGTGGGAATCTGGACTTGGAAGATATGGACGTAaagttatactgccc
gcagtcatatagagcagacgCAGTATTGTGGTTGCTGACTTCTTAGTAAAATGGAAATGAaagttatactgccctg
gcagtcatatagagcagacgTGCTGACCTCACATAGCTGTCACAGAGATTCCAGAACTATaagttatactgccct
gcagtcatatagagcagacgTGTACTCAGAGCCCAAGACATTGCAGTTTGCCACAGTAAGaagttatactgccc
gcagtcatatagagcagacgCCTGCTGCTTCTGTCTTCCAGGATGGCTCCTCTCTCTGTCaagttatactgccct
gcagtcatatagagcagacgGCTTTGGCTCTGGGCTCCTCTCACTTCCCCCTTTTCTGCCaagttatactgccc
gcagtcatatagagcagacgTCCTCCTTTCTTTCCCTGCTGAGCTTACCCACTTATTGCTaagttatactgccctg
gcagtcatatagagcagacgAGCAACAGCTCCCAAATCCACCCCTCCCATGAATGCCAGGaagttatactgcc
gcagtcatatagagcagacgGAACCTTCTATCTCTTCCCCTTCCACTTCACAGCAGCATCaagttatactgccct
gcagtcatatagagcagacgCTGAATGTCCCACTTTTGTAAACTGAAACCCTGTCTTTCAaagttatactgccctg
gcagtcatatagagcagacgAGTCTGCATCACACTCTCTTTTTATTAGTCCTTACACCCAaagttatactgccctg
gcagtcatatagagcagacgTGTGGAGGAGCCATGGAGTCAGCTGCCTGGATTCAGATCCaagttatactgccc
gcagtcatatagagcagacgACCATGTCTGACCTTGAGAAAGTTTTTTTAACGTCTATGAaagttatactgccctg
gcagtcatatagagcagacgTTGTTTGTTTGTTTTAAATAGGCAGGGTCTCACTCTGTTGaagttatactgccctgt

gcagtcatatagagcagacgGTATTTTTTATTGAGATGGAATTTCACCGTGTTGTCCAGaagttatactgccctgt
gcagtcatatagagcagacgTAACTAGTATTTTTCCATCTCCTACCTGAAAAACAGTCACaagttatactgccctg
gcagtcatatagagcagacgTTTGTGTAATGTCCTTTCCCTTTAAGGGGTATTGAATAGTaagttatactgccctgt
gcagtcatatagagcagacgACATTCTTGTGCAACCACCACCAGCCGTCTCCAGAACTCTaagttatactgccc
gcagtcatatagagcagacgTGTACCCATTGAACAGCTCCCTATTCTCTCCTCCCCAAAGaagttatactgccc
gcagtcatatagagcagacgCTACCATTCTACTTTCTCTATGTGTGAAGTTGACCATTCTaagttatactgccctg
gcagtcatatagagcagacgCCTTTTGTGACTGGCTTACTAAACGTAGCATAATGTCCGCaagttatactgccct
gcagtcatatagagcagacgAGAGTGTCACCACGTCTCTGAATCAGGCTGGTCTCAAACTaagttatactgccc
gcagtcatatagagcagacgACCACATTTTGTTTCTCCACTCATCCTGTCAGTGGGCCCTaagttatactgccct
gcagtcatatagagcagacgGCTGCTAGGAACATGAGTGTGCAACTGTCTCTTCGAGTCCaagttatactgccc
gcagtcatatagagcagacgGGTTCCCCTTTCTCCTCATCTTACACTTAATTTCTGTTTTaagttatactgccctgt
gcagtcatatagagcagacgCTGAAAATCACTACTCAGAATCATCTGGAGGGGCAGGTGCaagttatactgccc
gcagtcatatagagcagacgTGCAGACTCCTAACCCCACCTCAGACCAAGTAGGACCCCTaagttatactgcc
gcagtcatatagagcagacgGAGTGCAGTGGTGCCATGACTGCTCACTGTAGCCTCCAACaagttatactgcc
gcagtcatatagagcagacgCACTGTGCACGGCCTGAAGTGTGGCTTTTAAACAAGCTCCaagttatactgccc
gcagtcatatagagcagacgAGGTGGGGTTCTGGACATTATTAGCTGAGGGGAAGAGGGAaagttatactgccc
gcagtcatatagagcagacgTCCAGCACATAGGAGAACTCAAGCCACAGTCATCAGGATAaagttatactgccc
gcagtcatatagagcagacgTCATCCATATTGTTTCATCAAGTCTTCAGTCTTTTCAGTCaagttatactgccctgt
gcagtcatatagagcagacgGATCGAGGTCTGGAAGCATGACATTTTTAGCTGAGGAAGTaagttatactgccct
gcagtcatatagagcagacgATTTTGACTGGCGATACTCAAGTGAAATACCCTATTATCAaagttatactgccctg
gcagtcatatagagcagacgACAGATCTCACCAGAGCCATTCTCAGGCATTAGGGAAGCAaagttatactgccc
gcagtcatatagagcagacgCGCCTACTGTGAGATGAGTCATGCCTTTTGTCTTGTGCCTaagttatactgccct
gcagtcatatagagcagacgCGTACAAACGTCATATCACTTCCAGCTCCATTGCCACCTGaagttatactgccc
gcagtcatatagagcagacgCTTTCTACTGTTTTACTAGGGATTATTTAAAGGCTGTTGGaagttatactgccctgt
gcagtcatatagagcagacgAGTAATATTGAAACACACATTCACGTCAACTGTACTGTAGaagttatactgccct
gcagtcatatagagcagacgGCTGAGGCCCTTACAAGAAAAAAAGACAGATCAATAAGAGaagttatactgccc
gcagtcatatagagcagacgAGTTTTACATGATGCAGGAGACTTTATAAGGAAACGAAGAaagttatactgccct
gcagtcatatagagcagacgGTCATGGAGAAGTACAACTGAATAAAAAACAAGTATGGTCaagttatactgccct
gcagtcatatagagcagacgCTCGGCAGAGTCTGTTCAGATTCCTCTCTGTGGTCCTGTGaagttatactgccct
gcagtcatatagagcagacgAGATAAGGAAGTTCCTTTGCCTGGGTAGAGGGAGGGCACCaagttatactgcc
gcagtcatatagagcagacgGGCCTGAGACGCTTTTGCTGGTGCTATGGCCTGCTTCAGGaagttatactgccc
gcagtcatatagagcagacgCATGTTTTGGGGTAGTATGTTTTGGGGTTGTGTGTCTTGAaagttatactgccctg
gcagtcatatagagcagacgCAATTTTATCTTTTGAGGGCCGAGTGGAGATTCTTGAGGTaagttatactgccct
gcagtcatatagagcagacgAGTACAGTGGCACGATCACAGCTCGCTACAGCCTCAATCTaagttatactgcc
gcagtcatatagagcagacgTCCCAAGTGTCTAGGAATACAGGCACATGCTGCTATGCCCaagttatactgccc
gcagtcatatagagcagacgAAGTATCTTTTTAAATGCAGCCTCCTTGAGTTTGAGTACAaagttatactgccctg
gcagtcatatagagcagacgCACTCTGCTATTGAACATTAGCGTTTTAAGGTAAAGCATGaagttatactgccct
gcagtcatatagagcagacgAGAATGGGTCACATCAGCTTTCTCCCCCGGTGGCATCTCTaagttatactgccc
gcagtcatatagagcagacgTGAATCCCTCAGACACAGGCACGAGAGTAGCCCAGATGAAaagttatactgcc
gcagtcatatagagcagacgGCTAGTCTCAAACAAAACAGCAGAATAAAAACCACAACATaagttatactgccc
gcagtcatatagagcagacgAACAGACATAATGTTAAGCAAATGGCCATCTAACCTATGTaagttatactgccct
gcagtcatatagagcagacgAGGGTGAGTCCATAGAGTAAAGTAAAGCAAGTTTATTAGGaagttatactgccct
gcagtcatatagagcagacgTGGCATTTGTAAACCGTCATAGCATTGGTGGGAGTATAGTaagttatactgccct
gcagtcatatagagcagacgTATGACTTGTATTTTGTGCTTACCTCCTATCTCTTCCTGTaagttatactgccctgt
gcagtcatatagagcagacgCAATGGTGATGGAGAAAGGAAAGGGAGTTAGAAATAAAATaagttatactgccct
gcagtcatatagagcagacgTTTTTATAGAGAAACCCAGTCCCTACAAACAGCATCCTGTaagttatactgccct
gcagtcatatagagcagacgTGCCATAAACATTTCCCTCTCATTTTTCTTCCCATATTTCaagttatactgccctgt
gcagtcatatagagcagacgAATTACATTTTGTCAGGAGTGAAAGGTCATTTTGTGCTGGaagttatactgccctg
gcagtcatatagagcagacgCTGTGGAAATTCAGTGAGTCAAAGATGAAGGCGAAATTGAaagttatactgccct
gcagtcatatagagcagacgAAATTGTACTTTTAGGAGCTATGGGGACATACTGAAATAGaagttatactgccct
gcagtcatatagagcagacgCTTGGAAGCAGATGAATTAATACTTGGGCATTTGGCGCTAaagttatactgccct
gcagtcatatagagcagacgGAGCCTGGATTCTCCTTTGTTCTGTTCATTAGAGGGCAGTaagttatactgccct

gcagtcatatagagcagacgAGAACAGGGTCTCTCAGATTTTAATGTGGCTGCATATCACaagttatactgccct
gcagtcatatagagcagacgATGTAGGTTTTCATTTAGCAGGTCAGGATGGTGCCTCAGAaagttatactgccct
gcagtcatatagagcagacgCATCGCTGCTATTGCTTGATGGAAGGACCACACTTTGCACaagttatactgccc
gcagtcatatagagcagacgTGTTGTAGCATTACTGAACAAGTTGAATTTGCACCTCATTaagttatactgccctg
gcagtcatatagagcagacgGCAAACAGGTGCTTTGTATCCTCTTAGAGCTGCAGAGCAGaagttatactgccc
gcagtcatatagagcagacgCCTTCTGGGCTGTTAACCTGCTTGGCTCTCTTATCAATTAaagttatactgccct
gcagtcatatagagcagacgGTCCTTGTTTGGTTTGAGACTTAAAAGTGTGCTTTATACTaagttatactgccctgt
gcagtcatatagagcagacgGTGAGGTTGCAGAGTGGAAACAAGGCTTGGTATGGATGGTaagttatactgccc
gcagtcatatagagcagacgTTCAACAGCTTCTCAGACACATTAGTGTTCCCCTTTTATTaagttatactgccctg
gcagtcatatagagcagacgGTTCAAGCAATTATCCTGTCTCAGCCTCCCAGGTAGCTGGaagttatactgccc
gcagtcatatagagcagacgCACCTGGCCAGTGCTCCCATTTTATAGAGAAACACAGTCCaagttatactgccc
gcagtcatatagagcagacgTTTTTGGGAAAAGCTCTGGGGGACAAAGCCATAAACATTTaagttatactgccct
gcagtcatatagagcagacgCCATATTTCTGGTCCTCCCTGCTGCTTTATTTCACAAGAAaagttatactgccct
gcagtcatatagagcagacgAGTGGCTTGGCATGGAGAACACCAGGGACTGTGGGCATTTaagttatactgccc
gcagtcatatagagcagacgGCCAAAGCCATGAAGTCCCATGTAGAGAGATTGCTACCAAaagttatactgccc
gcagtcatatagagcagacgCAGGGCTGTGGACAAAATAGTCTCAAGTATTTTCTCTTCCaagttatactgccct
gcagtcatatagagcagacgTTCAAATATACCACCCTCTCTGTGACCTCGAGACCCATAGaagttatactgccc
gcagtcatatagagcagacgAGGTTTCTGTTGTCCTTGTCTCTGTAGTCATTGGCCACTCaagttatactgccct
gcagtcatatagagcagacgCCATCTGGCCCTTTCCCCATGTGCACATATACCTACTAAAaagttatactgccc
gcagtcatatagagcagacgATCTTAAAGGGTGTATTTGTCTTATTCATAAACAGGGAGCaagttatactgccctg
gcagtcatatagagcagacgTGGCAAAGGGGTAGGGGTCAGAGGTGGGGGAGAAGTCTTTaagttatactgcc
gcagtcatatagagcagacgTTGTTGAAGTACAGTAATAGCGTCATTTGGATTATTCCAGaagttatactgccctg
gcagtcatatagagcagacgCCATTGACATTGAGTTGGGTTCTGGATTGCTCACCTAGGAaagttatactgccct
gcagtcatatagagcagacgGGCATGGGAGCAACCCCAACTTAATGGCCTCCTAATTAATaagttatactgccc
gcagtcatatagagcagacgCAGCTAATTTTTGCATTTATAGTAGAAACAGGGTTTCACCaagttatactgccctg
gcagtcatatagagcagacgATTTTTAACACCTGTCATCATTCTTGAGTCCACTTTATGCaagttatactgccctg
gcagtcatatagagcagacgTTTAGTCTGAGGTGCACAAAGTTAAAAGGGGCCATTTTTGaagttatactgccct
gcagtcatatagagcagacgTGCTTCCATATTCTAACATCCAATTTCCTTTACTCAATGCaagttatactgccctg
gcagtcatatagagcagacgTGTCATAGTCATTAGGGCCACACTTAAGTGAATAGTGCAGaagttatactgccct
gcagtcatatagagcagacgCTGGGCACATACTGCTCTGGCTTAGATGTGGTTTATCCCCaagttatactgccc
gcagtcatatagagcagacgGGTGGGGCCTAATAAGATCTCTCTTGAATAGATTAATGCCaagttatactgccct
gcagtcatatagagcagacgGGAGCGAATGAGTTTTCTCTCTCCACGGAATGAATTAGTTaagttatactgccct
gcagtcatatagagcagacgCAGGTGTACACCCTGTGAGAATGGGTTGTTCAAAAAAGTCaagttatactgccc
gcagtcatatagagcagacgATGTTTTTGCACATGCTTGCCCCCTTCCACTTTCTGCCATaagttatactgccct
gcagtcatatagagcagacgAAATTACCCAGCCTCATGAGCTTAGACATAAACTAAGTTTaagttatactgccct
gcagtcatatagagcagacgAATGTGCGTAGAAATTATATGAGACTTCCAGGAGGCAACAaagttatactgccc
gcagtcatatagagcagacgACAATTCTGTTCTCATTCTTCCAGCATCGTGGGAGCATTTaagttatactgccct
gcagtcatatagagcagacgATCAGAGTCCTTGGATGGTTGCAATGAGAAGAGTGCTCCCaagttatactgccc
gcagtcatatagagcagacgACTGCATGTGGTCTGTGAGCCTTGAATAAGTGTTGTGTTAaagttatactgccct
gcagtcatatagagcagacgGGTGTTACAGCAGCCTAACCTATTTTATTCCAGCAAATCCaagttatactgccct
gcagtcatatagagcagacgTGTAATCAACCATCTGTGAGTATACTGCCAGGAAACTACCaagttatactgccc
gcagtcatatagagcagacgATAGTCTTTATTTTCTATTGTAGTGTGGGGAGTGTTTAGGaagttatactgccctgt
gcagtcatatagagcagacgTGATTTCCTCATCTTCTCAAACAGCTGGCAGTTCAGTCAAaagttatactgccct
gcagtcatatagagcagacgGGAATGAATTCTTGTGCATGGTAAGGACTCCATAAATATTaagttatactgccctg
gcagtcatatagagcagacgCGTTAACTGTCCACAACGCATTACTGCTCATTCCTCCAACaagttatactgccc
gcagtcatatagagcagacgCAATCTTCAATGAGAGCTGGTAAAAGACTGCCAAGTATAGaagttatactgccct
gcagtcatatagagcagacgGTTTGTTGGTTTGTTTTGTGGGAAGAATCAGGTACTTGACaagttatactgccctg
gcagtcatatagagcagacgGGTGTTTTATTGTTTCTGTTCCACTCGGTCATTAATCTGGaagttatactgccctg
gcagtcatatagagcagacgGCCCTGCCAGTTCTTCCAGTTCTTGACTTTCTGACCCGAAaagttatactgccc
gcagtcatatagagcagacgATTTTCTACTCACCACATCTCACTTCTACTGCTATTACCAaagttatactgccctg
gcagtcatatagagcagacgTGGTAATGACAAATCTGGGCTTCACGTGGGCATTTTCTGCaagttatactgccct
gcagtcatatagagcagacgCCCTATTGTTTAACATTGTTGACTTATTTGCATAGAGTGCaagttatactgccctg

gcagtcatatagagcagacgGAAACGACACACGGCTGGGTGCAGTTTGGGACCAGCCTGGaagttatactgcc
gcagtcatatagagcagacgACCCTGTCTCTACAATCAATATAAAAATTATCTGGGCATGaagttatactgccctg
gcagtcatatagagcagacgAAGGAAGAAAGGAAACGACATAACGATATATGTAAGACTGaagttatactgccct
gcagtcatatagagcagacgTTCTTGCTGCATGAGATATGGTATTGTGAAATGATTCAGAaagttatactgccctg
gcagtcatatagagcagacgTGAGCTCAGATTGCCTAGGTCCAAATCCCACTCTGGCACAaagttatactgccc
gcagtcatatagagcagacgGGTGATCTTGGACAGCTCCCTCAGCCTATTAGTTTCTTCAaagttatactgccctg
gcagtcatatagagcagacgAAAATGGGATGCTGATCGTGTATCTTTCATTGGCTATTGCaagttatactgccctg
gcagtcatatagagcagacgATTGCAAGTGTTAAAAGCATTGTACCCTGCATAGGCCAGAaagttatactgccct
gcagtcatatagagcagacgCAGAGCCGACGGTACAGGCCATGTTGTCTGACCACATCTGaagttatactgcc
gcagtcatatagagcagacgCTTCAGAGGAGTTTTGTGAGGCTAAAATGAAATTGCATGTaagttatactgccctg
gcagtcatatagagcagacgTGTTGGCACATACAAAATGCTCCGTTGATACCAGGTATTTaagttatactgccctg
gcagtcatatagagcagacgAAAGAAATCTGCCACCTCCTTTCCAAACACCCTATACTTTaagttatactgccctg
gcagtcatatagagcagacgTTAATTCTTGATGTTGTCCAGTTTTTCTTTGAAGGGCTAGaagttatactgccctgt
gcagtcatatagagcagacgGCAACGCCCAGGAATTATTATTTTTCTAATGACCTAACGTaagttatactgccctg
gcagtcatatagagcagacgTCCAGAAATTCTCTAAAAAACAAACTTCTTCGGGGACAAAaagttatactgccctg
gcagtcatatagagcagacgTTGAGACTTCCATTAGTTCTTGCAAACACTTTATAACCCAaagttatactgccctg
gcagtcatatagagcagacgCAGATGCTAACATTTTCCTGCGCATTCTGACCGGGTGCCAaagttatactgccc
gcagtcatatagagcagacgTGGCTCTGACATCTAAAATGAGGATTGTTTGGCTGTTCTCaagttatactgccctg
gcagtcatatagagcagacgGGAATTACCTTGTATCTTATATTTAGCCTACCGTAACTTGaagttatactgccctgt
gcagtcatatagagcagacgTTTACAGTTATCACAAGGACACTACAAAGTCTCTTTTGTGaagttatactgccctg
gcagtcatatagagcagacgTCTTTCACCCTCCAAGAAACTCTACATTCATCTCAGTTCCaagttatactgccctg
gcagtcatatagagcagacgCATATTTGTTGGTTCAAGGTAAGTACTAATTGTGCTTCTCaagttatactgccctgt
gcagtcatatagagcagacgTTTCTTCAGAAATAATGCTAAGCCTAGACAACAGGGCAAGaagttatactgccct
gcagtcatatagagcagacgGCTCTGTCACCCAAGCTGGAGTACGATAGTGCGATCTCAGaagttatactgccc
gcagtcatatagagcagacgCTCCTGCCCCAGCTTCCCTAGCAGCTGGCATTACAGGTGCaagttatactgcc
gcagtcatatagagcagacgGAGTTTGAGACTTCAGTGAGTGACGATGGTGCCACTGCACaagttatactgccc
gcagtcatatagagcagacgAGAGAGAGAGAGACCCTATCTGAAAAAAGAAAGAAAAAAGaagttatactgccc
gcagtcatatagagcagacgAAGAAAGAAAGAAAATATTGATGGCCAGAGAGAGAGAGAGaagttatactgccc
gcagtcatatagagcagacgGACGAAAGAAAGAAATGCTGAATGTTACAGATATTGACACaagttatactgccct
gcagtcatatagagcagacgCCTCTGGGCTTTTACAGTTAGGGAAGTGCTGAGTCACTGCaagttatactgccc
gcagtcatatagagcagacgCATGCTGGCAGAAACAGTGTAGAACCAGTGGGCCATGTGGaagttatactgcc
gcagtcatatagagcagacgACAAGCAAGGCGGGTGACAGGGCAGAGAGCTCCTTAGCAGaagttatactgc
gcagtcatatagagcagacgTGGCTCGCTTGGTTGAAATAATTTTTTGAAGCTTTTGACAaagttatactgccctgt
gcagtcatatagagcagacgAAAGATTACTAAGAAAGCCTGGCGGTTTGCAGCAAAAATTaagttatactgccct
gcagtcatatagagcagacgGCTCTGGAATCTTCTCCTAGGGCCTTCTATGCACTTCCTTaagttatactgccctg
gcagtcatatagagcagacgCTAAGTCAGTTAAGCAAGAATCCCCCACCTACAATATCTGaagttatactgccct
gcagtcatatagagcagacgTCATGTCTGATTGGGTTCTTCTCCCTCCACCAGCCCCCAGaagttatactgccc
gcagtcatatagagcagacgAAGAATCCTGTTGTTGGTTTAGCTAAAGTCTTTCTCAGTCaagttatactgccctg
gcagtcatatagagcagacgTCCCTCTTAGTGATTTTCCCTCCACCGACCCCCACCCTGCaagttatactgccc
gcagtcatatagagcagacgGTTGAGCCCAATTTCTGTCTCCCACTGCAAAATCTAATTGaagttatactgccctg
gcagtcatatagagcagacgGCTTTAACAAGTATCACTGGGTCATTTTTTTCTGTAACAGaagttatactgccctgt
gcagtcatatagagcagacgCTTAGAAGGGAGCTCAGGAGAAAGGGGAAACACAGGGCAGaagttatactgcc
gcagtcatatagagcagacgGGGATTAGGAATAGAAGCAGCGTGCTGGGCATCTAAGGGCaagttatactgcc
gcagtcatatagagcagacgCAGGAGCCTCTGTGTAAAATATGAAAGAAGAAAGACATGCaagttatactgccc
gcagtcatatagagcagacgATGGGAAGTTGGACATGATGGCGTGGGAGAAAGTAACTTTaagttatactgccct
gcagtcatatagagcagacgCGTATTTTGATAAGCAGTCTGATATTTGGGAAGATAGAGTaagttatactgccctg
gcagtcatatagagcagacgCGTCCATTCGAGATAGACTACCCGGTTCCTCAGGCCCACGaagttatactgcc
gcagtcatatagagcagacgGGTGCTTCGAGTACGCCTCCACCAGTGGGCAGTGTGGTCCaagttatactgcc
gcagtcatatagagcagacgGCCATTCTAGCCAACGGTAGGTTCTTCCTGTGGAGCCCTGaagttatactgccc
gcagtcatatagagcagacgATTTCTGGGTCTCTGTTCTGCGGTGTTCGGTGCCTTAACTaagttatactgccctg
gcagtcatatagagcagacgCCCCCGCATCCCAGAGCCAGCCGGCCATCTTCTGCAGTGAaagttatactgc
gcagtcatatagagcagacgGCCCAGCCCTGAAGCTCAGCCGCAGGAACAAACGCCCTGGaagttatactgc

gcagtcatatagagcagacgGCCGCCTGCACTCCGCCTCTTCTTCCTGCCGCCTCCTCTCaagttatactgcc
gcagtcatatagagcagacgAAAGTCCTCCTCCCCTTCCTTTGCGCTGATAACATCGAGCaagttatactgccc
gcagtcatatagagcagacgGCTTTTTACTGCCTAACTGATCCTCTACACTGTGCTAAGGaagttatactgccct
gcagtcatatagagcagacgTTCTCAGGAAACCTCAATATCTAGGTGGGAAAGGGCAGGTaagttatactgccc
gcagtcatatagagcagacgTTCTAGAAAGAGCACCCCCTCCTAAGAAACAAAACAAACTaagttatactgccc
gcagtcatatagagcagacgGGTACTTCCTGTTTCCTCCGTTTAAAGCACTCCCCCATCTaagttatactgccct
gcagtcatatagagcagacgTGTAAAAGTACTTCAAAGACTCAGCTGGCCGGGTTTGGACaagttatactgccc
gcagtcatatagagcagacgAACTCTTTGTGGTTTACTTTAATTGGGCTTTTAGTTGTCCaagttatactgccctgt
gcagtcatatagagcagacgAGGTTATGAAATCCTCGTCAATTCCGCTAAAACTCTTGTAaagttatactgccct
gcagtcatatagagcagacgCTTGACGCATATATTATGTCATGCAGAGGGGAACACGTGGaagttatactgccc
gcagtcatatagagcagacgAAGGCACACAGTGGATGCTGGGGTCACTTGTAAATGTAAAaagttatactgccc
gcagtcatatagagcagacgTTCTATTTCTCCTTGTAGTTATTCTCCTTGGGAGGCACAAaagttatactgccctg
gcagtcatatagagcagacgTGGCACATAAGAGTCTCAAGACGACCAGGCTGGCCAACATaagttatactgcc
gcagtcatatagagcagacgTACAAAAGAAAATTAGCTGGTGTAGAGGCGCAATCCCAGCaagttatactgccc
gcagtcatatagagcagacgAGAGTCTCAAGATTAGGGAATCACTTCTTAGATACAAATGaagttatactgccct
gcagtcatatagagcagacgAGTGAATCTTTCTTGGAGGGAACATGTGACGGAACTAGCCaagttatactgccc
gcagtcatatagagcagacgTCTATTGGAGATTGGAAAGATCTTGAAAAGAGACATGAGAaagttatactgccct
gcagtcatatagagcagacgTTCCTGTAGAAAAGTGGGAAGGATGACTATCTCACAAAGGaagttatactgccc
gcagtcatatagagcagacgATCAAAATCCAAAGGAGATGATGAATGGGACAGTGCTTCAaagttatactgccc
gcagtcatatagagcagacgGCTATTATGGTTTTAACAGCAGCAGAGGAAAGAGGAGGCTaagttatactgccc
gcagtcatatagagcagacgGAGTTCCACACCAGCCTGAGCAACATAACATAGAGACCTTaagttatactgccc
gcagtcatatagagcagacgAAGAATAACCAAGTTTAACAATTTCTGTGGGGTCTTCATGaagttatactgccctg
gcagtcatatagagcagacgCTCCCTTCCTTCTCTCCTCTACTCTCACTGATTTTTTGGTaagttatactgccctg
gcagtcatatagagcagacgATAACATGAAAGATACCATCCCAACCATTTAAGTGTACAGaagttatactgccct
gcagtcatatagagcagacgAACTCTACACCCACTAAACAACAATGCTTTTCCCCCTCACaagttatactgccc
gcagtcatatagagcagacgTTGACTACTTTAGATACACATATAAGGAGAATCATGCAGCaagttatactgccct
gcagtcatatagagcagacgGCTTATTTCTCTTACCATAACATCCCGACAAGTTTCATCCaagttatactgccct
gcagtcatatagagcagacgCCTTTTTTGAGGCTCAATAATACTCCTTTGTAGGTATGTAaagttatactgccctg
gcagtcatatagagcagacgTGTTCATTTATAGACATTCAGGTTACTTCTGTATCTTGGCaagttatactgccctgt
gcagtcatatagagcagacgGCTGCAATGAATAAGAGTGTGAAACTATTTCTTTGAGATCaagttatactgccctg
gcagtcatatagagcagacgACCATTTTACATGCCTACTAACAGTCCACAAGGGCTCGAAaagttatactgccc
gcagtcatatagagcagacgTCTGTTTTCCTGATAGTGGCTATTCTAATAGGCATAAGGTaagttatactgccctg
gcagtcatatagagcagacgAAGTATATTTTCATGTGCTTGTGGGCCATTTGTAGTTCTTaagttatactgccctgt
gcagtcatatagagcagacgCTGTAAGTTGCCTTTTCATGCTGTCAATTATTCCCTTGCTaagttatactgccctg
gcagtcatatagagcagacgTGCTTTGGTGTCATGTCTAAGATCCTTGTAAAATCTAACAaagttatactgccctg
gcagtcatatagagcagacgCCAACTTCATTTTTTTACATCTGGATATCCGGTTTTCCCAaagttatactgccctg
gcagtcatatagagcagacgAGACAATCTGTTCTCCATTGAGTGACTTTGGCACCCTTGTaagttatactgccct
gcagtcatatagagcagacgATGTGAAGGTTTATTTCTGCATTTTTCTGTTCCTTTGGTCaagttatactgccctgt
gcagtcatatagagcagacgGTAGCTTTGTATTCTGTTTTGAAATTAAGAAGTGTGAGGCaagttatactgccctg
gcagtcatatagagcagacgTGGCTATTCAGGGTGCTTTGATATTTCATATGAATTTGAGaagttatactgccctg
gcagtcatatagagcagacgTTTTCCATTTTTGCAAAAGAGGCCATTGGGATTTTGATAGaagttatactgccctg
gcagtcatatagagcagacgGCATTGAATCTGTGGATCAGTTTGGGTAGTATGAACATTTaagttatactgccctg
gcagtcatatagagcagacgTGTTTAGTGTAGAGAAACACAATTCCTTTTTGTATCCTGCaagttatactgccctg
gcagtcatatagagcagacgCTCTGGCTAGGAATTCTAGTGCTACGTTGAATAGAAGTGAaagttatactgccct
gcagtcatatagagcagacgCTGATCTTAGAGGCAGTTTTTCACAATTTCTATGGTGTTAaagttatactgccctg
gcagtcatatagagcagacgGTTGGGTGTTTTATCATGAAGGGTCTGAATTTTGTGAAATaagttatactgccctg
gcagtcatatagagcagacgTTGAGCCATCCTTGCATTCCACAAATAAATCTCACTTGGTaagttatactgccct
gcagtcatatagagcagacgCTTTTCTTTTCTTTTCTGAGTCTGAGTTTTGCTCTTGTTGaagttatactgccctgtg
gcagtcatatagagcagacgTAGCTGGCATTATAGGCATGCAACATCATGCCCTGCTAATaagttatactgccc
gcagtcatatagagcagacgATTTTGTTATCAGAGTAGTGGGGTCCTGAAAAAAATGAGTTaagttatactgccctg
gcagtcatatagagcagacgGAAGCCATCTAGTCCTGGGCCAATGTAATTTTTTTGATATTaagttatactgccctg
gcagtcatatagagcagacgAGGTCTCTCTATACTATTGAACTAAATTTTCCAACTGACCaagttatactgccctg

gcagtcatatagagcagacgAAAATCATGCATGGGCAAAAGAGTCCTTCAAAGTGCAAGAaagttatactgccc
gcagtcatatagagcagacgACAAATTATTCTTTCACAATCACCTATTTGTGTGAGGCTGaagttatactgccctg
gcagtcatatagagcagacgCATGTACTTTAACCTGAACAACAGATTATAACGGATTGTGaagttatactgccctg
gcagtcatatagagcagacgTGTAATGTTGAAGAAGGCAACATTACATGCCTGTAATCCCaagttatactgccctg
gcagtcatatagagcagacgTTCAAGACCACTCTGGCCAACATAATGAAACCCCGTCTCTaagttatactgccc
gcagtcatatagagcagacgGGAGACTGAGATAGGAGAATGACTTGAACCGGGAGGTAAAaagttatactgccc
gcagtcatatagagcagacgACTAACAATTTATTCTCATCTTTCTGTAATGCCACATGGCaagttatactgccctg
gcagtcatatagagcagacgGTGTATGTGTGTGTGTCCATTCATTCTTTTGCCCTTGTTAaagttatactgccctgt
gcagtcatatagagcagacgTAGTGAAAAATTGAAGCAACCCAAGTAACCAATGAATAGCaagttatactgccct
gcagtcatatagagcagacgATGACAGACAACCAGTGCAAGAATATTAAATTCTGGATACaagttatactgccctg
gcagtcatatagagcagacgCACAAACAGGCAAGGTGAACCATTTTAGTGTTGGCGGAGGaagttatactgccc
gcagtcatatagagcagacgCAGGGGGAGTAGGAGGGCAGTGCAGTTGTGGGGGTCGACAaagttatactgcc
gcagtcatatagagcagacgCTGACTGGGGTCCATGAAGAGGAGCAGCAGAAGTTGGGGCaagttatactgcc
gcagtcatatagagcagacgCAGGGATGCTTGGTTGACTTGGCAGTACATCAGACTTCATaagttatactgccct
gcagtcatatagagcagacgGTTTGCAATCCAACCCAACCAGATTAATCACACATTTATTaagttatactgccctg
gcagtcatatagagcagacgTACTGTGTACCAGGTACTCTACTAGGCCTTGGAACCAAAGaagttatactgccc
gcagtcatatagagcagacgGCCCATAAGCAGCTCTCAATCTAGACGAACATAGCCAGATaagttatactgccc
gcagtcatatagagcagacgTTGAAGACAAGACATCCTGAAAACCCCCACTGATGAATAAaagttatactgccc
gcagtcatatagagcagacgTTTTTAGAATAATTGACTCAAGCTGCTCACAGGTGGCCCTaagttatactgccctg
gcagtcatatagagcagacgCTTTGAAGAGGCTCAGAAGGGGGAAAAGACCAATGCAAACaagttatactgcc
gcagtcatatagagcagacgAGCAACAACATAAACAGGGCAGCTTCGTCACAAATTGTATaagttatactgccct
gcagtcatatagagcagacgTGGAGGAAGATGAATGAAAATAATTCTGTTTCTGAAGGGAaagttatactgccctg
gcagtcatatagagcagacgATTTCAAGGTCATCTTTGGGAGCCTAAAAATTGATGCACAaagttatactgccctg
gcagtcatatagagcagacgGGAAGCACACAGAGCAAGGAGAGAAGTCTGGGTTTTAAGCaagttatactgcc
gcagtcatatagagcagacgGAGCCACTTCCTCAATTCCCTGTTGTCAAATATTATCTTCaagttatactgccctg
gcagtcatatagagcagacgTGGAGAATTGTGAGAATACATAATATGGGATTGTGAACTGaagttatactgccctg
gcagtcatatagagcagacgAGAGCCTATAAAGTGTTCTGTAGTGATGATAAAAGCAATGaagttatactgccctg
gcagtcatatagagcagacgATGCTGTTTTTCCTAAGACTCCTTGCATGCTGTTACATAAaagttatactgccctg
gcagtcatatagagcagacgACTCTGCCCTAAATTCTAATTAATCCTGAGTCTTGGCACTaagttatactgccctg
gcagtcatatagagcagacgTTCTGGGTGAAAAGTAGAACAAACTCATTAGCAAGTGCCAaagttatactgccct
gcagtcatatagagcagacgCGTCTGGTGTGGTTTGCTAGAGGGCGCTGATGATATTTCTaagttatactgccctg
gcagtcatatagagcagacgTGAAGAGAGTTGTCATTAGGAACTCCTGCCCAGGTAACAGaagttatactgccc
gcagtcatatagagcagacgCAGATGCAACAGGAAAGATGAAGGCGGGATGAGGTGGTGTaagttatactgcc
gcagtcatatagagcagacgTGACTCACTCACTCTGACATCCTTAGCAACCTGGGACATCaagttatactgccc
gcagtcatatagagcagacgGGTCTCCCAGCCCCTCAGTGTTGTAAATAATGCCACAGGTaagttatactgccc
gcagtcatatagagcagacgGAGGAGCGGGGAGGGGAGGGTAAGGACTGTCGAAGCAAACaagttatactgc
gcagtcatatagagcagacgGAGCTTGTCCCCGTGCCGACTCCTCTGTCCATGCTGGCCTaagttatactgcc
gcagtcatatagagcagacgGTTCTAGAACCAGGAGTGTGGCAGCTTTGAGCTCAAACCTaagttatactgccc
gcagtcatatagagcagacgGTGTGTGTAGGGGCCTGGTACTCACGGGAATCCAGTTAGGaagttatactgccc
gcagtcatatagagcagacgGCTGAGAGGAGCTCCAGAGTGAGGTCAGCATGCCCAGTCTaagttatactgcc
gcagtcatatagagcagacgCCTGAGAGCCTGGCAACAAGTCCTGACGGGGAACAAGTCCaagttatactgcc
gcagtcatatagagcagacgCTGAGACCTGGGCCAGGCTGAGAGGGGAGACGGGAGAGGTaagttatactgc
gcagtcatatagagcagacgAAATTTCCAAATTCTTGACCGCATCATTCCAGTTGTAATGaagttatactgccctg
gcagtcatatagagcagacgAAGGTGCTTAAGGTGGTGTTCTTCAAGGGACAGCCCCCAGaagttatactgccc
gcagtcatatagagcagacgAAAATGTCTACTTTGATTTTGAGGGTGTAGGGTGGGGCTCaagttatactgccctg
gcagtcatatagagcagacgTGCGCACTTTAAACAGGGACTCCAAATGCTTCTTTTTGCAaagttatactgccctg
gcagtcatatagagcagacgGGCAGATATTTTTTGACCTAGCTTTTCAGTATTCCTCAAGaagttatactgccctg
gcagtcatatagagcagacgAAACCTTGAGAGAAAAATATGTTTGAGCATGGTAGAGAGTaagttatactgccctg
gcagtcatatagagcagacgCATTTCTCATTTCCTTTTCCCTTCTCATATTCAGTGCAATaagttatactgccctgt
gcagtcatatagagcagacgGAATTCCTAGCTTGGAGTCATGGATATCACCTGCCCCACCaagttatactgccc
gcagtcatatagagcagacgCAGCAGGAAACAGAAACTTCATCATCAAGCATGTATTAGTaagttatactgccct
gcagtcatatagagcagacgGCTTAATGCTCCCCGCCATTTATCATCCCACCATTTTCATaagttatactgccctg

gcagtcatatagagcagacgCAGGCTGCAATCAAGGTATTAGCCAGACTGTGTTCTCATCaagttatactgccc
gcagtcatatagagcagacgGAATTCTCTTCCAAGCTTTCATGATTGTGGGCAGATCTCAaagttatactgccct
gcagtcatatagagcagacgTTGTGGTTATAGAATGGAGGGTCTCAGCTTCTTACTTGCTaagttatactgccctg
gcagtcatatagagcagacgCACCATCAGATTCTAGAGGTGCCTGTAGTTACCAGTCACAaagttatactgccc
gcagtcatatagagcagacgAGAATTTCTCACTATACAGGCAGCTAAGACGAGGTTTTATaagttatactgccct
gcagtcatatagagcagacgGGCACATCAGCTTGGCCGTATTGGTTATGTATTAGTTGGCaagttatactgccct
gcagtcatatagagcagacgATCAAGGTGTCGGCAGAGTTAGTTTCTCCTGAGGTTCTCTaagttatactgccct
gcagtcatatagagcagacgGTGTGTGTCTGTGTCTGAATCCCTTTTTCTTTTGCTTTTTaagttatactgccctgt
gcagtcatatagagcagacgCAGCTCACTGCAACCTCTGACCACTGGGTTCAAGCAATTCaagttatactgccc
gcagtcatatagagcagacgTGGGATTACAGGTGCCCACCACTAAGTCCAGCTAATTTTTaagttatactgccct
gcagtcatatagagcagacgAGTCACATTGGATTAGGGCAAACCCATATGGCTTCATTTAaagttatactgccct
gcagtcatatagagcagacgACAGAGTAATGTGGGATATATGAGAAAAGAAAAAGTGCGGaagttatactgccc
gcagtcatatagagcagacgGCGACAGTAAGGAAAATACAAGATGCCTTGGGGAAATTCCaagttatactgccc
gcagtcatatagagcagacgGATCCAGACTTAGGCCAGGGTCAGATCCAGAATTTAGCCTaagttatactgccc
gcagtcatatagagcagacgGCATCAGAAACAGCTAACAGGATAAAGAGGGAAGGGACCAaagttatactgcc
gcagtcatatagagcagacgGTAAGAGTATATCGCATTGATTAAAAGTGACTTGAAGCAGaagttatactgccct
gcagtcatatagagcagacgCTAAGGAGGTGGCATAGGAAACTTCTGGCCTTGGTTCAGCaagttatactgccc
gcagtcatatagagcagacgTGTTCCTTACAGTCACAAGATAGCTATTCCTGCCTGATTAaagttatactgccct
gcagtcatatagagcagacgTGTTCAAGGCACAAAAAATCAGGGAAAGTTATTAGCAATGaagttatactgccct
gcagtcatatagagcagacgCCTTTTAATCAGGAAAGCAAAACCTGTCCTAGAAGCTTCCaagttatactgccc
gcagtcatatagagcagacgTGGCTTAACTTCCATAGGTTAGACAGTGTCACAAATGGCTaagttatactgccct
gcagtcatatagagcagacgCTGTTGTGGAAGTGGAAGGGAGATAGAAGCTGAGAGTGAAaagttatactgccc
gcagtcatatagagcagacgGGGTGGCCATAAGCATTGAAAGGGCTGAGTGTGAAGGAAAaagttatactgcc
gcagtcatatagagcagacgTCAAGTGGGAGAAATGGCAAGTGTAGGGATGGGGTCATGAaagttatactgcc
gcagtcatatagagcagacgCCTTGAAGACCTCAGAGGATGTCCCAGTTATCTATTGCCAaagttatactgccc
gcagtcatatagagcagacgACCAAACAGTGACCATTTTGTTATGTGCAGATTCTGTGAGaagttatactgccct
gcagtcatatagagcagacgGGATGGCTTGTCCCTGCTATATGGTATGTGGAGCCTTAGCaagttatactgccc
gcagtcatatagagcagacgCCTGGACATGGCTCAAGTGGCTGGGGACAGGGTCATTGAAaagttatactgcc
gcagtcatatagagcagacgCCTGTCTGGCGCCTGGGCTGGGATGGCCTGAAGGTTGGAAaagttatactgcc
gcagtcatatagagcagacgCTCTCCATGAGGTTGACTTGGGCTCCTTCCAGTGTGCTGCaagttatactgccc
gcagtcatatagagcagacgCTGAGAGTAAGTGTCTGGAGACTGGGCATATCAAGAGACCaagttatactgccc
gcagtcatatagagcagacgCACCCTAGCCTTGGAAGCCACAGAGCACCATTTTGGTTCTaagttatactgccc
gcagtcatatagagcagacgCCATTCAGAGGCAGCCCAGATCCAAGAGAAGGTGAATTAGaagttatactgcc
gcagtcatatagagcagacgTTGAAGGGGGAGAGGCAAGATGACATTGTAGAGCAGCATGaagttatactgcc
gcagtcatatagagcagacgGGGAGATGTGGCAGCCATATTTTGAAAAGCCCATATGCCCaagttatactgccc
gcagtcatatagagcagacgCTGAAAGGTTTTAGGCAAGGGTAATAAGTTTAGACTGTGCaagttatactgccct
gcagtcatatagagcagacgTTGAGAACTAGTTGGAATGGTCAAAGACTGTAGGTGGGGAaagttatactgccc
gcagtcatatagagcagacgGGTTGGCCTCAGTAAATATGGTAGGTGTTGCATCTGGACAaagttatactgccc
gcagtcatatagagcagacgTTAGGACTAAGTAAATGAGAGGTTTGAGGGAGAGGAGGAGaagttatactgccc
gcagtcatatagagcagacgCAGGATTAAGCAGCTGGTAACCGGTGCCTTTAACCAACACaagttatactgccc
gcagtcatatagagcagacgAATTGTGAACATTTGACTTTTGAGTTTCTGTGAGACATCCaagttatactgccctg
gcagtcatatagagcagacgGAGTCAGTAATTGAGATGTTGAGAGAAAATGAGATTTACCaagttatactgccct
gcagtcatatagagcagacgTGAGAAAAGAAATGAGGGTAAACAGAATTCTGGGGGGCACaagttatactgccc
gcagtcatatagagcagacgCAGGGAAAGAAGTCTATAAAGGAGGTAGGTGAGTGGTCAAaagttatactgccc
gcagtcatatagagcagacgAGAATGTAGTAGGTTGATAATAGCTTCCAAAGATAGGTCCaagttatactgccct
gcagtcatatagagcagacgAGCATGTTACCTTATTTGAAAATGGGTGTTTGTAGATGTGaagttatactgccctg
gcagtcatatagagcagacgGGATAGACCCTAAATGCCATGACAAATATCCTTATAAAAGaagttatactgccct
gcagtcatatagagcagacgAAGGTGATGTGAAGACAGGGACAGATTAGAGTGATGTAGCaagttatactgccc
gcagtcatatagagcagacgCTACTAGAAGCAGGAAGAATCAAAGAACAGGTTGTCCCTAaagttatactgccc
gcagtcatatagagcagacgCAGGGGGAAGATGGTCCTGTTGACACCTCGATTTCAAACTaagttatactgccc
gcagtcatatagagcagacgCCCCTAGGAAATTAATGAAGATGGAAATCAGGACCGTGTAaagttatactgccc
gcagtcatatagagcagacgCCAAGGAAGAGAGGATTTTAAGAGGACGAGTGGTAAATTCaagttatactgccc

gcagtcatatagagcagacgCAGACCCTGTCCAAGATGCTGAAGATGCTGTCCTCATGAAaagttatactgccc
gcagtcatatagagcagacgAAGAATGGCTACTCCACAGGAAGAGCAGCCTTGGGGGCTGaagttatactgcc
gcagtcatatagagcagacgTGGATTATTCATGAGTGTTCACAGAAAAGGGTGGACAATTaagttatactgccct
gcagtcatatagagcagacgCCATATCGGGTAACTTATGGCATTTGTAAATTGTTATGGCaagttatactgccctg
gcagtcatatagagcagacgGTGAGGACGACCAGAGGTCTTTCATTACCATCTTAGTTTTaagttatactgccct
gcagtcatatagagcagacgCAAGGCCTTTGTGAGCTTACCTTGTGCCAGCCTATCTCATaagttatactgccc
gcagtcatatagagcagacgCCAGCCGCTATTCAAAATGGCATCACTGTGGTTCAAACACaagttatactgccc
gcagtcatatagagcagacgCTGTAACTTCTTCAGGCTAGAGAGGGGTGATGATATTCCTaagttatactgccct
gcagtcatatagagcagacgAGTCAGAAAGCATGGGTGTGTCGAGGACTGCTCATAACTCaagttatactgccc
gcagtcatatagagcagacgCCAAATAAACTAAATAAGAACGCTTTCCATGAACTGGGTAaagttatactgccct
gcagtcatatagagcagacgGAACACTAATCCAAATTGTTACATTACCAATTCCTCTGACaagttatactgccct
gcagtcatatagagcagacgGACACATCATGGTAAGATCAGTTTGCTTTATTATTCTTGGaagttatactgccctg
gcagtcatatagagcagacgGAGGAATCACAAATAAAGACTTTTTTTTGAGAGGTGCAGTaagttatactgccct
gcagtcatatagagcagacgAGTGATTCTTCTGCCTCAGCCTCCCGTAGGTGGGATTACAaagttatactgccc
gcagtcatatagagcagacgTTTTTAAAGCTGAGCCCACCATGGGTTTGTACCATAAATTaagttatactgccctg
gcagtcatatagagcagacgACATTCTTTAATTATCACAGTTCAGAAAGCCTGTACAGAGaagttatactgccct
gcagtcatatagagcagacgTTCCTTAAAGAAAAGTATACCATTACAGGAAAAGCCTTGCaagttatactgccct
gcagtcatatagagcagacgAACCAGTTTCTCAATTGTGTCCTATTGCAAAAGAAAACAGaagttatactgccct
gcagtcatatagagcagacgTGCTCCAAATTTTGTTCACTCAATTGTTAAAAGCTGTCAGaagttatactgccctg
gcagtcatatagagcagacgTGTTCTGCTTGACATTCATAAGTATTACAGTTCTCCATGAaagttatactgccctg
gcagtcatatagagcagacgAGTTATCAGAAACCTGCTTTAAGAACACCTGTTAAAGTCCaagttatactgccct
gcagtcatatagagcagacgATGACACAACATCTTAGTTAGGACAGCCAATATTAAAGCCaagttatactgccc
gcagtcatatagagcagacgTGGAACACACACCAATAACATATGTATACAAATAAACCCCaagttatactgccc
gcagtcatatagagcagacgGCCAAATATGTCATTTTTGTACTTTAGGGGACTTAATACCaagttatactgccctg
gcagtcatatagagcagacgCATCATCTAGGTTTTAAGACTGGCATGCAGTAGGTATTTGaagttatactgccct
gcagtcatatagagcagacgATGCTCTCCCTCCCCTTGCCCCACAGCCCCTGACAGGCCCaagttatactgc
gcagtcatatagagcagacgTGATGGGCATTTGTGTCGGTTCCAAGTCTTTGCTATTGTAaagttatactgccctg
gcagtcatatagagcagacgTCCACAATGGTTGAACTGATTTACGCTCCCACCAACAGTGaagttatactgccc
gcagtcatatagagcagacgGAGAAGCATCTATTCATATCCTTCACCCTCTTTTTGATGGaagttatactgccctg
gcagtcatatagagcagacgAGGTTGCCTTTTCACTCTGATGATAGTTCCTTTTGCTGAGaagttatactgccctg
gcagtcatatagagcagacgGGTTTTAGGTTTTACATTTCAGTCTTTACTCCATCTTGAGaagttatactgccctgt
gcagtcatatagagcagacgCGTGTAAGGAAGGGGTCCTGTTTCTGTTTTCTGCATATGGaagttatactgccct
gcagtcatatagagcagacgGGAATCCTTTGCCCACTTCTTGTTTTTGTCAGACTTGTTGaagttatactgccctg
gcagtcatatagagcagacgCTGTTCTGTTCAATTGGTCTGTATATCTGCTTTGGTACCAaagttatactgccctg
gcagtcatatagagcagacgATACGGGCTCCTTTTTGGTTCCATATGAAACTTAAAGTAGaagttatactgccctg
gcagtcatatagagcagacgTTCTCCTTGAAGAAGTCTGTCACAGCTCTTGTAAGTTGTTaagttatactgccctg
gcagtcatatagagcagacgTATTTCTTTCTCTTTCCTGATTGCCCTGGTCAGAACTTCCaagttatactgccctg
gcagtcatatagagcagacgTTGCCCATTCAGTGTGATATTGGCTAGGGGTTTGTCATAAaagttatactgccct
gcagtcatatagagcagacgTATGGTTTGGTCATCAATTCTGTTTATGTGATGGATTACGaagttatactgccctgt
gcagtcatatagagcagacgATTTTTGCATTGGTGTTCACCAGGAATATTGTCCTGAAATaagttatactgccctg
gcagtcatatagagcagacgAGCTCCTCTTTGTACCTTTCATAGAATTCAACTGTGAATCaagttatactgccctg
gcagtcatatagagcagacgTCCTGTTTTAGTCTTGGGAGGGCATATGTATCCAGGAATTaagttatactgccct
gcagtcatatagagcagacgCAGTGGTGATAACCCCCTTATCAGTTTTTATTGTGTATATaagttatactgccctg
gcagtcatatagagcagacgGATATCTCTTTTCTCTTTCATTAGTCTGGCTAGTAGTCTCaagttatactgccctgt
gcagtcatatagagcagacgGATTTTTTGAAGGGTTTTTATCTCCTTCAGTTTTGCTCTGaagttatactgccctgt
gcagtcatatagagcagacgGCTCTTGCTTCTCTAGTTTGTTTAATTTTGATGTTAGGGTaagttatactgccctgt
gcagtcatatagagcagacgTTTAGTGCTATAAATTTTCCTCAAAACACGGCTTTAGCTGaagttatactgccctg
gcagtcatatagagcagacgTTTTGCCTTAACTTCATTATTTACCAAGTAGTCACTCAGGaagttatactgccctg
gcagtcatatagagcagacgTTTGCTGAGGAGTATTTTAGTTCCAATTGTGTGGTCGATTaagttatactgccctg
gcagtcatatagagcagacgAGGTCCACTTTGTCCAGAGCCGAGTTCAAGTTCTGATTATaagttatactgccct
gcagtcatatagagcagacgGTGTTAAAGTTCCTCACTATTATTGTGTAGGAGTCTAAGTaagttatactgccctg
gcagtcatatagagcagacgACTTGGGTGCTTCTGTGTTGGGTGCATATATGTTTAGGATaagttatactgccctg

gcagtcatatagagcagacgTTTATCAGAGACTAAGATTGCAACCCCTCCTTTTTCTTGaagttatactgccctg
gcagtcatatagagcagacgCTGGGACTACACATGTGTGCCACCACGCCTGGCTAATTTTaagttatactgccc
gcagtcatatagagcagacgCTGTGTTAGCCAGGATGGGCTTCATCTCCTGACCTCGTGAaagttatactgccc
gcagtcatatagagcagacgGAGCCACTGTGCCCAGCCCAACCCCTGCTTTTTTTTTTTTaagttatactgccct
gcagtcatatagagcagacgTCCTCCATCCCTTTTTTTTGAGCCTGTGTGTGTCTTTGCAaagttatactgccctg
gcagtcatatagagcagacgTGAGTACAGCACACCCATGGGTCTTGACTTTTTATCCAATaagttatactgccct
gcagtcatatagagcagacgGGGGCATTTAGCCCATTTGCATCTAATGTTAATATAGTTAaagttatactgccctg
gcagtcatatagagcagacgGGCCTGGTGGTGACAAAGCCTCCCTGCATTTGCTTGTCTGaagttatactgccc
gcagtcatatagagcagacgTTCTTTAAGAATGTTGAGGCTGGGCACGATGGCTCACACCaagttatactgccc
gcagtcatatagagcagacgGAGGTTGAGGTGGGCAGATCCCGAAGTCAAGAGATTGAGAaagttatactgcc
gcagtcatatagagcagacgTCCCAGCTATGTGAGAGGCTGAGGCAGGAAAATCGCTTGAaagttatactgcc
gcagtcatatagagcagacgAATGTTGAATATTGGCCCCCTTGGCCCCCACTCTCTTCTGaagttatactgccc
gcagtcatatagagcagacgTTCCCTTTGTGCATAACCTGACGTTTCTTTCTGGCTGCCCaagttatactgccct
gcagtcatatagagcagacgGAGTTGCTCTTCTCGAGGAGTATATTAGTGGTGTTCTCTGaagttatactgccct
gcagtcatatagagcagacgGTGTTTTCCAGCTTGGTTCCACTTTCAGGTACACCAGTCAaagttatactgccct
gcagtcatatagagcagacgCTTTTCACATAGTCTCACATTTCCTGGAGGGTTTGTTCATaagttatactgccctg
gcagtcatatagagcagacgTACATGTGTATTTGTTCTCGTACTGTGTTTTTCAGGTCCAaagttatactgccctgt
gcagtcatatagagcagacgTGCATTGGGTCAGAACATGCTTCTTTAGCTTGGAGGAGTTaagttatactgccct
gcagtcatatagagcagacgCTTCTGTCAATTCATCAAAATCATTCTCCATCCAGTTTTGaagttatactgccctg
gcagtcatatagagcagacgATTCTGGTTTTTGAAATTTTCAGCCTTGTTGCACTGGTTTaagttatactgccctgt
gcagtcatatagagcagacgGTTGGTGACCTTCAGATGGGGTTTTGTGTGGACGTCCTTTaagttatactgccct
gcagtcatatagagcagacgGCACCCCTGAGATGCCAGCCTAAGCTCTCCTGTATGAGGTaagttatactgcc
gcagtcatatagagcagacgCGTTTAATTCTGCAGAAGCTACACCCATAGCTGCCCCTTCaagttatactgccc
gcagtcatatagagcagacgTGTCCCAGGAAGATGGGAGTCTTGTCTATAAGCCCCTGACaagttatactgccc
gcagtcatatagagcagacgTTCAGAGATGTCCTGGCCAAAGAGGAGGAATGTAGAGAGGaagttatactgcc
gcagtcatatagagcagacgCAGTGGGCTCCGTCCAGTTGGTACTTCCTGGCAGCTTTGTaagttatactgccc
gcagtcatatagagcagacgCTCAAGCCTCAGTAATGGCCAACGCCTCTCCTCCAACCAAaagttatactgcc
gcagtcatatagagcagacgACAGTTCTGTTTCACTGGTGTTTTTGGCGCCACTGGGGTAaagttatactgccct
gcagtcatatagagcagacgAGTATCTGGGCCGGAATGCATTGTCCCCCATGGCACAGTCaagttatactgcc
gcagtcatatagagcagacgGATGCCCTACCCTGCTTCATCTAGCCCTCCATGGTGTGCAaagttatactgccc
gcagtcatatagagcagacgCCATCTTGCCAGCCACTCTGCCTTTTCTTTCTATAATTCTaagttatactgccctg
gcagtcatatagagcagacgTGATAGGCAATTTCCCCCACAGTCTAGGGCTTGTTTCACAaagttatactgccc
gcagtcatatagagcagacgATGATCTACTTGCGTTAGTTTTTGTATATAGTGTGAGGTGaagttatactgccctgt
gcagtcatatagagcagacgAGGGAGAAGGCAGAAGTGGGCGGGCAGGGGAAAGGTAAGGaagttatactg
gcagtcatatagagcagacgTCTCTTCAGTGTGGGCTCCACAGGCTAAGGAGAGGCGGGGaagttatactgcc
gcagtcatatagagcagacgGGGAAGTGCCCTGGTGGGGTCAGTGGCTCCTGTGGGGGTAaagttatactgcc
gcagtcatatagagcagacgAGGCTGGGTGTGCTCAGAGGTGCATGGTCTCCCCTGGGCGaagttatactgcc
gcagtcatatagagcagacgGCCCAAGACACCGAGACATTGCTGCACCTACTCCTAGAGCaagttatactgcc
gcagtcatatagagcagacgGTGGCTCTGCGACTGGGCTGTGGCCGGGGTACTTCCTGCAaagttatactgcc
gcagtcatatagagcagacgCTTCTCCTGGAAGACACTGTTGCTGGAGAGTGGAGGCTTCaagttatactgccc
gcagtcatatagagcagacgACCTCAAACCAGAGTGGCTCGTCGAATTTCCTTTTTCCCTaagttatactgccct
gcagtcatatagagcagacgAGCTGGGGCCTCGGGGGGAGGTGACGCTTGTATAGGCTCCaagttatactgc
gcagtcatatagagcagacgCCGGCGGGGCGATTTTCACGGTCACCAGGCTGCGGACGGGaagttatactg
gcagtcatatagagcagacgCCTCGGAAAAGTCTGGCCCAGTGAGGCTTAGGAGTGGTGGaagttatactgcc
gcagtcatatagagcagacgGGACCATCTCTCTCCGGGTGCGGGCATAGTCCCAATCGAGaagttatactgcc
gcagtcatatagagcagacgGTGTGGACCCCGAGCTTCCCATGGACGTAGACAACTTTGCaagttatactgcc
gcagtcatatagagcagacgCCTGGACCTGGGAGCGGAGGACCGGCCTGGACGGCGGGGAaagttatactg
gcagtcatatagagcagacgGGCCAGCCCAGGAACCAGCTCAGGTAGTTGCCCATGACAGaagttatactgc
gcagtcatatagagcagacgCAGAAGTTTCCGAAGTCTGCCGCCAGGGCCAAGTGTGCTGaagttatactgcc
gcagtcatatagagcagacgCCGCTGGCCTCAGGACGCTACAAAGCGCGCAAAACATCGCaagttatactgc
gcagtcatatagagcagacgCCATAGGGTCCCGCACGGGCCAGGACGGCACCTCGTGGAAaagttatactg
gcagtcatatagagcagacgGTCACAGGGCGCAGCTGCAGTGGGGGAGGGCAGCACCCAGaagttatactg

gcagtcatatagagcagacgGTCCCAGTCACTTGGCTCTGAGCAGAGGGCCCAGCAGCTaagttatactgc
gcagtcatatagagcagacgACTTTTGTGCCTGACCTTTCAAGGTAGAAGATAGAATTTCaagttatactgccctg
gcagtcatatagagcagacgTGAACACAATACACTTGGATGCTCTTTCAAGGAAAAACAAaagttatactgccct
gcagtcatatagagcagacgTTTATGTGGATTTCCTATAATTTCCAGCCTTCGTATATCCaagttatactgccctgt
gcagtcatatagagcagacgTTTTGGGGAAAAGGTGATGTTTGGTTACACATGTAAGTTAaagttatactgccctg
gcagtcatatagagcagacgTGTGTAGTCTTTCATCCCTCTCTCACCTCCCATCCATTCCaagttatactgccct
gcagtcatatagagcagacgGGCCATTTAGTATTCCATAGTACGTTTGTACCAAATTTTCaagttatactgccctg
gcagtcatatagagcagacgGCTGATGCCATAGTTTTGCAATTGCTATTTCTGCTGCTATaagttatactgccctg
gcagtcatatagagcagacgCTTTCCTCTGGGTAGGTACCGAGTAGTGGCATTGCTGGATaagttatactgccc
gcagtcatatagagcagacgCATTCCCACCAGCAGTATACCAGTGTTTCCTTTTCACCACaagttatactgccc
gcagtcatatagagcagacgTGCAGGAGTAATGTGGTATCACATTATGGTTTTTATTTGCaagttatactgccctg
gcagtcatatagagcagacgTTGAGTTCCTCATAGATTCTGGATATTATTCCTTTGTCAGaagttatactgccctgt
gcagtcatatagagcagacgTCTGTGGGTTGTCTGTTTATCTGCTGATTATTGCTTTTACaagttatactgccctgt
gcagtcatatagagcagacgTGCTTTTGGGTTCGTGGTTATAAAATCTTTGCCTAAGCCAaagttatactgccctg
gcagtcatatagagcagacgCACTTTACGTTTTTGTTTGCGTTGTTGAAGATCAGTTTGCaagttatactgccctgt
gcagtcatatagagcagacgGTCAGGTAATGTGCTGCCTCAAGATGTGTTCTTTTGCTTAaagttatactgccctg
gcagtcatatagagcagacgTTGTAGATTGCTTTTAGCAGTGTGGTCATTTTCACAATACaagttatactgccctg
gcagtcatatagagcagacgCAATATTTTGCAGTTTTCTTTGTAGGGGTCTTTCACTTCCaagttatactgccctgt
gcagtcatatagagcagacgGAGTTTGTATCCTGAATTCATTTGTCAGTTTGCTGAATTCaagttatactgccctgt
gcagtcatatagagcagacgCTAGGTATACCATTATATCAATGGTGAACAGCAACAGTTTaagttatactgccct
gcagtcatatagagcagacgATGCTTTCAACTTGTCCCCATTCAGTGTAATGTTGGCTGTaagttatactgccctg
gcagtcatatagagcagacgTCTATGCTGATTTTGCTGAGGGATGCTGGATTTTGTCAAAaagttatactgccctg
gcagtcatatagagcagacgGATTTTTGCAACCATGTTCATTGAGGATATTCGTCTGTAAaagttatactgccctg
gcagtcatatagagcagacgAGAATGATGTAGGGAGGGTTGCCTCTTTCTCTATTTTTTGaagttatactgccctg
gcagtcatatagagcagacgTCAACTGTGAACTTGTCTGGTCCTGGAATTTTTCTTGTTGaagttatactgccctg
gcagtcatatagagcagacgGCTGCTTATTACTGGTCTATTCAGAGTTTTAATTTCTTCCaagttatactgccctgt
gcagtcatatagagcagacgATCCATCTCCTCTAGGTTTTCTTTCCATGTTTAGTGCTTCaagttatactgccctg
gcagtcatatagagcagacgAATTATTTCCTTTAAGAATGTTGGATATTGGTCCCCACTCaagttatactgccctg
gcagtcatatagagcagacgGGTGTTCTCTGTATTCCCTGTACTTGAATGTTGTCCTGTCaagttatactgccctg
gcagtcatatagagcagacgAGTTGATCTTCAATCTCCGACATCCTTTTTTTCCGCTTGAaagttatactgccctg
gcagtcatatagagcagacgCTTCTCTAAACTGGTTTTCTAGTTAGCAATTCATCTAACCaagttatactgccctg
gcagtcatatagagcagacgCCTTTTTGCTCTGGTTTCTCAACATCTTTGTGGATTCATCaagttatactgccctgt
gcagtcatatagagcagacgTGGTCTTTGATGTTGGTTATCTTTGAATGGGGTCTTTGAGaagttatactgccctgt
gcagtcatatagagcagacgGAGGTGTCTCCCATTCAGGATACACGGGGGCCAGGGACCCaagttatactgc
gcagtcatatagagcagacgTGTGCTGGGAGGTCCGCCGCCCTCTTCAGAGCCATCATGCaagttatactgcc
gcagtcatatagagcagacgGCCCTGTCCAGAGAGGAGAAGTCTAGCAGTCTGGCCACAGaagttatactgcc
gcagtcatatagagcagacgGTGGATCTCAGACTGCTGCTCCACTGGGAGCAAGAATTTCaagttatactgccc
gcagtcatatagagcagacgGGGGTGGGACCTGCCAAGCCGCATCACTTGGCTCCCTGGCaagttatactgc
gcagtcatatagagcagacgGGTATGGCAAAAAGAAAACCCTGAAGCTAGTTTGATGTCTaagttatactgccct
gcagtcatatagagcagacgATCTCCTGGTCTGCAGGTTACGAAGACCACTGGAAAAGTGaagttatactgccc
gcagtcatatagagcagacgGGCAGAATGCATGGTTCCTCTGGCTCAGTCCCTCATGGCTaagttatactgccc
gcagtcatatagagcagacgGGGGAGAAAATTCCCTGACCGCTTCACTTCCCGAGTGAGGaagttatactgcc
gcagtcatatagagcagacgCTATTCGGCTATCTTGCCAGCCAAATGCTTCCAGCTTTTCaagttatactgccct
gcagtcatatagagcagacgTGGCTGTGGGCTTGTCGTAAATAACTCTTCTTATTTTGAGaagttatactgccctg
gcagtcatatagagcagacgGAGATAATCACGTGGTTTTTGTCATTGGTTGTGTTTATGTaagttatactgccctgt
gcagtcatatagagcagacgAGCCGACTTGATTGTGATGGATAAACTTTCTGATGTGCTGaagttatactgccct
gcagtcatatagagcagacgGCTCCTCTTTTTGCCTCTGGTAGAATTCAACTGTGAATCCaagttatactgccct
gcagtcatatagagcagacgGTGTCCAGGAATGTACCCATTTCTTTTAGAATTTCTAGTTaagttatactgccctg
gcagtcatatagagcagacgGTCTCTATTTCCCTCAGTTCTGCTCTGAGCTTAGTTATTTaagttatactgccctg
gcagtcatatagagcagacgCTCTTGCTTCTCTAGTTATTTGAATTGTGATGTTAGGGTGaagttatactgccctgt
gcagtcatatagagcagacgTCAGTTTCCATGTAGTTGCAAAGTTTTGAGTCAGTTTCTTaagttatactgccctgt
gcagtcatatagagcagacgTATGTGGTCGATTTTACAATAAGAGTGATGTGGTGCTGAGaagttatactgccct

gcagtcatatagagcagacgAGCTGAGTTCAAGTTCTGAATATCCCTGTTAATTTTCTGTaagttatactgccctgt
gcagtcatatagagcagacgTGTTTTATGAATCTGGGTCCTGCTGTATTGGGTGCATATAaagttatactgccctg
gcagtcatatagagcagacgGCACTTCTTTGTCTCTTTTGTCTTTTTTGATCTTTGTTGGaagttatactgccctgtg
gcagtcatatagagcagacgTAGTAGAGATGGGGTTTCTCTGTGTTAGCCTGGATGGTCTaagttatactgccctg
gcagtcatatagagcagacgGCCCAATGCCTGCTTTTTTCCTTTTGCTTTCCATTTGCTTaagttatactgccctgt
gcagtcatatagagcagacgTTGAGCCTATATGTGTCTTCGCACCTGAGATGGGTCTCCTaagttatactgccct
gcagtcatatagagcagacgAGCACACTGATGGGTCTTGACTCTATTTAATTTGCCAGTCaagttatactgccctg
gcagtcatatagagcagacgTGGTGCCAGTTTTTTTTTCCCCCCCACATTTAGTGCTTCaagttatactgccctg
gcagtcatatagagcagacgGGTGACAAAATCTCTCTGCATTTACTTGTCTGTAAAGGATaagttatactgccctg
gcagtcatatagagcagacgATGAGGTCAAGAGATCAAGGGTATCCTGGCCAGGATGGTGaagttatactgccc
gcagtcatatagagcagacgTCCCAGCTACGTGAGAGGCTGAGGCAGAAGAATCGCTTGAaagttatactgcc
gcagtcatatagagcagacgGCTTGTAGGGTTTCAGCAGAGAGATCCACTGTTAGTCTGAaagttatactgccct
gcagtcatatagagcagacgTTCCCTTTGTGCATAACCCGATCTTTCTTTCTGGCTGCCCaagttatactgccctg
gcagtcatatagagcagacgCAACTTGGTTCCATTCCCTCTGTTACTTTCAGGTAAACCAaagttatactgccctg
gcagtcatatagagcagacgTTCATCAAGCTGATCTTCAATCTGATATCCTTTTTTCCACaagttatactgccctgt
gcagtcatatagagcagacgAAGTTCTCATGCTCTGTTTTGCAGCTTCGTCAGGTCATTTaagttatactgccctg
gcagtcatatagagcagacgCATCTAACCTTTTTTAAGATTCTTAGCTTCCTTGCATTGGaagttatactgccctgt
gcagtcatatagagcagacgAGTTCATCAAATTCATTCTCTCTCCAGTTTTGTGCCTTTGaagttatactgccctgt
gcagtcatatagagcagacgCCTTTGGATGGGTTTTCTGTGTGTGGAGGTCTTTTTTGTTaagttatactgccctgt
gcagtcatatagagcagacgCAAAGATTGCTACCTACTTCTGCCCCTGGAAGCTTTGTCCaagttatactgccct
gcagtcatatagagcagacgGGCAGAAATGTTTAAGTCTGGTGAAGTTGCGCCCACAGCCaagttatactgccc
gcagtcatatagagcagacgGGACGTCCCTCTCCCCACCACGCTCAATTGTCTCAGGTTGaagttatactgcc
gcagtcatatagagcagacgGGGGGTGGGACCCGCCAAACGAGACCAGTTGGCTCCCTGGaagttatactgc
gcagtcatatagagcagacgTCCTGGTCTGCAGATTGCAGATATCATGGGAGAAGCATAGaagttatactgccc
gcagtcatatagagcagacgGGCTGGGAAAGGGAGTCCCCCATCCTTGCCCTTTCCGGGTaagttatactgcc
gcagtcatatagagcagacgGCCATCTTGCCAGATCCCCCAAAGACATAATTTATAATTTaagttatactgccctg
gcagtcatatagagcagacgAATAGAGGGAAGACAGCTTTCAAGTTTTATGTCTCTTTTCaagttatactgccctg
gcagtcatatagagcagacgTTCAAGAGAGAAAGCCAAATCTCATCCTTGCATTAGTGTGaagttatactgccctg
gcagtcatatagagcagacgGACACCATAAACCTTTTGTACCCCTTTTCAAATTTTTGTGaagttatactgccctgt
gcagtcatatagagcagacgATACCCCTTTAACTTCAGCCTGTATGTTCACACACAGAATaagttatactgccctg
gcagtcatatagagcagacgTTTTCGCAAACCTTCCACAATGTGCTCAAACCTTCAGCTTaagttatactgccctg
gcagtcatatagagcagacgTTTTACCAAAAACACATTTCGCTTCCCTTATACACTCTGCaagttatactgccctg
gcagtcatatagagcagacgCAATTAATCAAAGCTCTTTCTTACTACACACACAACACACaagttatactgccctg
gcagtcatatagagcagacgCACAGACAGAAGATCCAGTGCTTGTAAGATTTTTCATTTGaagttatactgccctg
gcagtcatatagagcagacgCATGCAGTTTCTATGGTCTAATTAGCAGGCACAATTGGAAaagttatactgccctg
gcagtcatatagagcagacgGGTTCCATTTTAATACCGAATCCTGGATCCCTCAAAGAGAaagttatactgccct
gcagtcatatagagcagacgACAGAACAACAATCCCATGATTTGACCGTGCATTTCATTGaagttatactgccctg
gcagtcatatagagcagacgGGGGGTGGGGGGAGTGGAGACCTTTCCATAACTTCCAGGTaagttatactgcc
gcagtcatatagagcagacgCTATATTTCCTACATAGTTATTAGACACCAAAGCTCTCTCaagttatactgccctg
gcagtcatatagagcagacgCCAAAAGTCAAAACCATCAGTTAATGCAGTGCAGAACAGAaagttatactgccc
gcagtcatatagagcagacgACCTTAGATTTTGAGAGGGCTCGATCCACTTTTAATTAAGaagttatactgccctg
gcagtcatatagagcagacgTTAGATCTCTTATTACCCAATTTTAACCATGCCAAACTGCaagttatactgccctg
gcagtcatatagagcagacgGGTTAACTCCCAGGTGAAACCAGTAAATCTGAACGAAGGCaagttatactgccc
gcagtcatatagagcagacgACTTCATCCAGTTTTTTTCGTTTGTTTCGGGGACCTCCTAaagttatactgccctg
gcagtcatatagagcagacgGAGAAAGAGAGAGAAAGCATTACCTGTGGTGAGGTGGGGAaagttatactgcc
gcagtcatatagagcagacgAGTTCAGGTGGCCGCTTGTCAGTCGTGAAGGGATCTTTTCaagttatactgccct
gcagtcatatagagcagacgCCTTTTAGGTAATTCTCTCTGCAGGTGAGCATCTCTTGTTaagttatactgccctg
gcagtcatatagagcagacgGCAAGTTGCCTAACCCTCTAAGCCTCACCTTCCTAATTGGaagttatactgccc
gcagtcatatagagcagacgCAAATCACTTAGCACAGCCCATGCCTGATATGCAGTAGACaagttatactgccc
gcagtcatatagagcagacgGCTATTATTAATGAGCATCTCTGTTGGGGCTTGTGATGTAaagttatactgccctg
gcagtcatatagagcagacgCTTCTGTAATACAAGTCTTTACTCAAGATCATCTCTGCAGaagttatactgccctg
gcagtcatatagagcagacgGCCTCCATAGCCCCAATAATTAACATTTTGTTTATCTCATaagttatactgccctg

gcagtcatatagagcagacgGAAGGGGAAGAGTCTGTGCGCATCTTATTGTTCTTATGTCaagttatactgccct
gcagtcatatagagcagacgAGCAAACACTGGTCATGAGCATCCTATCAGCACAACTCTTaagttatactgccc
gcagtcatatagagcagacgTAATGGAGATTAGTTTTAGGTCCACATTAAATCCACCCTGaagttatactgccct
gcagtcatatagagcagacgGATCATTTCCACTCAGTTACTTTTGTCCCTGGCTTGCTGGaagttatactgccct
gcagtcatatagagcagacgGTGACCACAGCAGCTATATCCCCTCTCCAGAGATTTACTTaagttatactgccc
gcagtcatatagagcagacgGAACCATCAACTCTGTCACAAGGGCCTCTCCACCCATGGCaagttatactgcc
gcagtcatatagagcagacgCAACTTTGAGGCTAATTGGGCAGGAAATATTACTTGCACTaagttatactgccct
gcagtcatatagagcagacgAGATTTTAAATACCTCTGTAAGACTTTTCCCCCTTTTCTGaagttatactgccctg
gcagtcatatagagcagacgTTTAATTTATCATCCCTAGGTACTTGGCATTGACAACTGGaagttatactgccctg
gcagtcatatagagcagacgTTTTTCATAGAGTCAGGGTCTTGACTTGTTACCCAGGCTGaagttatactgccct
gcagtcatatagagcagacgTGGGCTCAGGCAATCCCCCTCCTCAGCCTCTGAGTGTTGGaagttatactgcc
gcagtcatatagagcagacgCCCAGATGATAACTGTATTATTTAGGGAATTGGGTATCAGaagttatactgccct
gcagtcatatagagcagacgCAACATATATTTCTTGAGAACTGCCCATGTTACAGACACAaagttatactgccct
gcagtcatatagagcagacgCCTGATAGAAAAGGGTGAGAGGCAGAAAACAGACAAGAAAaagttatactgcc
gcagtcatatagagcagacgATCAGGCCAGTTGTACCAGGTATAGATTTTACTTAATAACaagttatactgccctg
gcagtcatatagagcagacgTTTAACAAAAGGCAAAAGAAGGATTTTACTGGGAGGATACaagttatactgccct
gcagtcatatagagcagacgGTTCACAGAATCAAAGAAAAGGCTGAAGAACCAAAATTTGaagttatactgccc
gcagtcatatagagcagacgGGAAACTAGAAGTGACTGACTTTCTTTTTAGGACCGCACTaagttatactgccct
gcagtcatatagagcagacgGGTCAGGATGCCAGAAAGAGAAAGAGTGAGCAGTTGGTTGaagttatactgcc
gcagtcatatagagcagacgTCTTGGCCAGAGTAAGACATACCATGTGCATGAGCAGTTTaagttatactgccc
gcagtcatatagagcagacgTCCAGTGGAGAGAGTGGTAATTTTCCAAAGCCAAACTGTTaagttatactgccct
gcagtcatatagagcagacgGTTGCAGAAGAAGAAATGGATACTGAGCAGTCAAAAAGGAaagttatactgccc
gcagtcatatagagcagacgTGATTTTAAGGCAGTTTACATAGAAGAGGCTTAGGGGTTTaagttatactgccctg
gcagtcatatagagcagacgGATTTGGTTTGCATTAATTAAATTCACCTGGGTTTCAAGCaagttatactgccctg
gcagtcatatagagcagacgAACCCATAGTTCCCTAAAGTGGTGAGAATAGAATTGAGAAaagttatactgccct
gcagtcatatagagcagacgTTGAAATATATCAAGGGCAATCAGGTAATCAGAAGGATCTaagttatactgccct
gcagtcatatagagcagacgAGCCTCTGAAAGAAATATTAGCTGTCTCACTATTGTCATGaagttatactgccct
gcagtcatatagagcagacgTTGTTTATGCTTCTGTAAAATGAATTTGAACCAGATCGCCaagttatactgccctg
gcagtcatatagagcagacgGTCTCTCCTAGTTTCAAAATTCTGTGTTCTACAAATCCTCaagttatactgccctg
gcagtcatatagagcagacgTTTAGAGATTAGGAAATGAGAAACAGTGGTTCATGACAAGaagttatactgccct
gcagtcatatagagcagacgTCTAGTGACCCTGAAGTGTATTATTCTTGTCCCTTTTGTAaagttatactgccctg
gcagtcatatagagcagacgGATAATCATGTATCCAAAAGTTATCTGTGGTGCAAAGTCTaagttatactgccctg
gcagtcatatagagcagacgACTTGAGCATAGGAACTTGACGATAAAATCTAAGATCTTCaagttatactgccct
gcagtcatatagagcagacgATTAGGAAAACACAATTAAAGGGATGCAGTGTCTATTCTCaagttatactgccct
gcagtcatatagagcagacgACAAAGGCAACCCTTACGTGAAATGTAGTTGTGGCTAATTaagttatactgccct
gcagtcatatagagcagacgCTCTTCAGCGGCCAGAGATCAACGATCATTTAGGAAGCAAaagttatactgccc
gcagtcatatagagcagacgCTTAGAATATGCTTGTCTGGTGCAATGTTTCACAATACATaagttatactgccctg
gcagtcatatagagcagacgACTCCCTATGTTTTCTCTCCATATTCGATATGCAACTGCTaagttatactgccctg
gcagtcatatagagcagacgAAGAATTATTGATATGCTAACCCTGAACTGAAACCAAAGCaagttatactgccct
gcagtcatatagagcagacgGGACTGATCTCTGGAAACAAGCTGAGTAAACTCCCAAGGTaagttatactgccc
gcagtcatatagagcagacgGTCTCTTCTGCTGCCTCCTGACCTTGAATATAAAACTACAaagttatactgccct
gcagtcatatagagcagacgAGAAAAGAATGATCTCTGAAAACAGCACAGGATAAAACTCaagttatactgccc
gcagtcatatagagcagacgTAAGATCTTTACTTCTCTGGGCTTTCTGTATTTTACCTGTaagttatactgccctg
gcagtcatatagagcagacgAGCTTAGGAGATGGGAAATATAAAGATACTATGTGAAGCTaagttatactgccct
gcagtcatatagagcagacgAGAGTTCTTTCCTGTACCAGCCACTCAGTGACTTACTTACaagttatactgccct
gcagtcatatagagcagacgAACCCTGTCCTGGTTCTTCCTTGGTCTCATTAGTAGGCACaagttatactgccct
gcagtcatatagagcagacgCAGCCAGTAGGTTATCAAGACATTTATTCTCACTTCAACAaagttatactgccct
gcagtcatatagagcagacgGGAGCTGCAGATAGTCATGAACAAAAAAACTCCCCCACAAaagttatactgcc
gcagtcatatagagcagacgATTACAGGCATGAGCTAAGGTGCCTTGCCAGACTGTCCAGaagttatactgcc
gcagtcatatagagcagacgGGATGATATGCGATATGAGGAAAAATAAAGCTGGTTAAAGaagttatactgccct
gcagtcatatagagcagacgCTATTTTAGCTGGAGAAAGCGTCCCTGATCTGAATGAAGGaagttatactgccc
gcagtcatatagagcagacgCTGGGAAGGAGCAGTCCAGGGAAGCACACAGTTAAGGGCCaagttatactgc

gcagtcatatagagcagacgTGGGTGTTCCAGGAACGTGGTGGAGATCAATATAGCCAGAaagttatactgccc
gcagtcatatagagcagacgAGGAGATTCTGTTCCTGATGGTAGTGAGCATCCCGGGGTGaagttatactgccc
gcagtcatatagagcagacgCCTTCTCTTCCAGGAACTAGAAAGCAGGCAGGTGACAGCAaagttatactgcc
gcagtcatatagagcagacgTGGGCTTAGGTGTACCAGCTGGTTGGCGTTGGTTAGGCTCaagttatactgccc
gcagtcatatagagcagacgCCCTCTGGACAGGTGTGTGGCGAGGTACGCCGGCAGGGCCaagttatactgc
gcagtcatatagagcagacgCCATCTTGAAATTCTCAACACAGTTATCCTTCCTTTTGTTaagttatactgccctg
gcagtcatatagagcagacgTCCCCAGGGCCTGGAGGCTCGGCTCCTGTGCGGTCCTGCCaagttatactgc
gcagtcatatagagcagacgCCCACCTCGCGGACTTGGGTCTGGGTCACCTGTTCCTGCTaagttatactgcc
gcagtcatatagagcagacgCAGAACTGCTGCTGCCCTTCATCCTGAGAGGCCCGACACGaagttatactgcc
gcagtcatatagagcagacgGGCAGGGCCAGGCATGACCCTCTGCACGCCTAACCACAGGaagttatactgc
gcagtcatatagagcagacgTGCCAGCATCTCTGTGCGTGAGGGAGTGTGACATTAAATAaagttatactgccc
gcagtcatatagagcagacgGAAACGCTTTTTTTCTCTTTTTTAACAAGGGACCCGCATTaagttatactgccctg
gcagtcatatagagcagacgAGGAATGATGTCACCCACCCCTTGGCCGCCGCTTTCTCTCaagttatactgcc
gcagtcatatagagcagacgCCGAGTAGCTGGGAAGATGGCTTTTAAAGCGGGTGAACACaagttatactgcc
gcagtcatatagagcagacgGGGCAAACCAGGAATTAGCTCTACCTCAAAATCTGTGAAGaagttatactgccc
gcagtcatatagagcagacgCAATGGTTTGAGCATCAGTTTTCTCCATCTGCACTTTTGAaagttatactgccctg
gcagtcatatagagcagacgGATTTGAATAGCCAGTTCTAAAGTGAATTACCTTGTAAGCaagttatactgccctg
gcagtcatatagagcagacgCTTAAAACCAAGCAGTCACAAACTTTAGTATCCAGATCTAaagttatactgccct
gcagtcatatagagcagacgAGTTGGTTCCACTGTTGAATTGTGCCATCATATAACCTTGaagttatactgccctg
gcagtcatatagagcagacgTCTTCATAATCTTTAAAAGTTATTGGAATGGGCCTACACCaagttatactgccctg
gcagtcatatagagcagacgTTTTGTTCAGGGCTTAAATAATGTAATCAGGCCAGGTGCGaagttatactgccct
gcagtcatatagagcagacgAATGTAATCACTCCCATTTCCAAGTTCTCAACATGTGGTGaagttatactgccct
gcagtcatatagagcagacgTGGACATAGGGAGAATTACCTGAGAAATAGTACTTTAATGaagttatactgccct
gcagtcatatagagcagacgTCACGGACTCTATGATAACCTTTCTTTTCTGGCAATGATGaagttatactgccctg
gcagtcatatagagcagacgCCTGGTGGACTTTTGCTGTCTTTTCCCTATTTCACTCTGTaagttatactgccctg
gcagtcatatagagcagacgGAAATTGTAAAGCATAAGTAAAAACTGAAAGATGGCCAGGaagttatactgccc
gcagtcatatagagcagacgGAGGCAAGTGGATGACTTGAACCTCTGGAGTTTGAGACCAaagttatactgccc
gcagtcatatagagcagacgAGACCCTGCCTAAAAAAAACTGATGAAAGATGAGCTTCCTaagttatactgccc
gcagtcatatagagcagacgTGATTTACACCACGGAAGTCCCTGTCTAGACTCTACTGTGaagttatactgccc
gcagtcatatagagcagacgAGTGGCATGCGCTTCATACATTTTGTCTCATTTAGCAACAaagttatactgccct
gcagtcatatagagcagacgCAATCAGTGAATTTAAATAATGTGCCCTAGGTTGTTAAGCaagttatactgccctg
gcagtcatatagagcagacgACTGATATTTTGCCCTCAATGCCAGATCCACCTGACTCCAaagttatactgccc
gcagtcatatagagcagacgCCCTGTTTTTCCACTGTGCAATGTGGTTGCCTAATGAATGaagttatactgccct
gcagtcatatagagcagacgCCTAAGCAGGTGTTCAGGGGAAGAGCTGGACTTGGACTCCaagttatactgcc
gcagtcatatagagcagacgGATGATCCTTTAAAATTCCCCCTTTCTTATCTCCTATGTGaagttatactgccctg
gcagtcatatagagcagacgTCTAGATTGAGAGATATAAAAGAATGACCATTTTGGCCAGaagttatactgccct
gcagtcatatagagcagacgAAAAATACAAAAGTCAGCCAAGTGTGGTGGTGGGTGCCTGaagttatactgccc
gcagtcatatagagcagacgTGAAGTGGATTATATATTGATCTTGGTCTAACTGTTCTGGaagttatactgccctg
gcagtcatatagagcagacgGAAGCTAAGCAAATAGGCCACCCACTTTAACATCAACCAGaagttatactgccc
gcagtcatatagagcagacgCCAAGATAGGTATTAGACAACTGGAAATAAATAAGGCATGaagttatactgccct
gcagtcatatagagcagacgGCAATGCAAGTTCTACACACACACAGCTACGGGACTACAGaagttatactgcc
gcagtcatatagagcagacgCCTGACGGACACTCACCCATGCCAGCATACCTTCAATATGaagttatactgcc
gcagtcatatagagcagacgAACAACAAAGTGTATATTCTTCCATTCCAGATTGTACAGGaagttatactgccct
gcagtcatatagagcagacgAGCAGCTTCATTATTTTGGCTAATAAGTAACAGAGATGTGaagttatactgccctg
gcagtcatatagagcagacgGGTTCTAGTAAATTTAGATTCTCTAGAGAAATTGGGCAGCaagttatactgccct
gcagtcatatagagcagacgTTGGTTAAACACAATGTAATTGGAATAGTTGCACTTTGGGaagttatactgccctg
gcagtcatatagagcagacgGGAGAATGGCGTGAACCCGGGCGGCAGAGATTGCAGTGAGaagttatactgc
gcagtcatatagagcagacgGGCATGCTAATTTAATTAAAAACTTAAGCCAGGTGCAGCGaagttatactgccct
gcagtcatatagagcagacgTAATAAATTAGCTGGGCATGAAGTCTGCTGGTTACACTGGaagttatactgccct
gcagtcatatagagcagacgTTCAAGGCTACAGTGAGCTGTGATTATTCCACTGCACTCCaagttatactgccc
gcagtcatatagagcagacgTTGAAGACTTCTTTAAAAATGACATAGGCTGGGCACGGTGaagttatactgccct
gcagtcatatagagcagacgGAGGCAGAGGCAAGTGGATCACGAGGTTAGGAGTTCAAGAaagttatactgcc

gcagtcatatagagcagacgCCAACATGGTGAAGCTGTGTCTCTACTAAAAATACAAAAAaagttatactgccct
gcagtcatatagagcagacgACATAAAGAAAATTAAAGTCAAGCCACACTCAGGAGAACAaagttatactgccc
gcagtcatatagagcagacgAATACATATCACCAACAAAGGATTAGTATAGTGCATACGCaagttatactgccct
gcagtcatatagagcagacgTGACCACACAATTGAAAAATAGGCAAAAGAAATGAACTGGaagttatactgccc
gcagtcatatagagcagacgATATGTATGGCATATAACCATAAGAAGAGATGCTGTGTCTaagttatactgccct
gcagtcatatagagcagacgTACCAAGTAGAACCACAGTTAGATATTTTATACCCACTAGaagttatactgccct
gcagtcatatagagcagacgGTTTTGAAGAGGATGAGAACAAATGGGAAATCTATATTGCaagttatactgccct
gcagtcatatagagcagacgAGACGGGACATTGTGCAGCCACATTAGAAAAATGTTTGGCaagttatactgccc
gcagtcatatagagcagacgATGGATTCATATTAACCAAAACTAGAAACAACCCATCAGGaagttatactgccct
gcagtcatatagagcagacgATTGTATAGCAGAGAAAATGAGTTGACATCAATTGTGTGCaagttatactgccct
gcagtcatatagagcagacgCTGAGTAAAAATTGCAAGTTGCAGAACCTTATTAAGGATGaagttatactgccct
gcagtcatatagagcagacgTAAAAGACAGAAAGAATCGACCTAAAATTCAGGATGAAGGaagttatactgccc
gcagtcatatagagcagacgTTGGGGGGTGGCATGAGGAGGAGGAGGCGAAGGGAATGGGaagttatactgc
gcagtcatatagagcagacgCCTTGGCTTGGGTCATGAGTTCATGTGTTATTTTGTTTTAaagttatactgccctgt
gcagtcatatagagcagacgAAAGGTGAAACAATAGGCTTTCTTGTACATTGATGTTAGGaagttatactgccctg
gcagtcatatagagcagacgAATTGGTGCAGTCTTTTTAGTTGAATCTCTCTCTTTACACaagttatactgccctgt
gcagtcatatagagcagacgCTTCTGTATCCCAATAATTCAATTGTGAGGAAATAAAGGCaagttatactgccct
gcagtcatatagagcagacgAAGATACAATTAGTGAGTTCTTTTTCACGTCAGGGAATTGaagttatactgccctg
gcagtcatatagagcagacgCTGGAGACTCACAGCCAAAGTACATACTCAACCCTTCCAAaagttatactgccc
gcagtcatatagagcagacgGGATCTTATTTATTTTTTCCATGGTCCTTGCACAATCTGAaagttatactgccctgt
gcagtcatatagagcagacgGGATGTGAATTTCCTAAGTACAGAGACTATGTGTGATGTGaagttatactgccct
gcagtcatatagagcagacgCCTGTTGAGAAATGAATGCCTAGGAAAAAAAAAGAAAAGGaagttatactgccc
gcagtcatatagagcagacgAATATGGAAGGCTCAAAGGAATATAAGTAATGTGTGTACCaagttatactgccct
gcagtcatatagagcagacgCATATTCAGATTTTGGGGGTTAATATTTACTACTTGGCAGaagttatactgccctg
gcagtcatatagagcagacgCTTTTTGAGATCATAGGTAAAATGCTAGGGGATTGTCATTaagttatactgccctg
gcagtcatatagagcagacgGATGAGGACTTTAGATCTTTAACATAAATAACAGGTAGGGaagttatactgccct
gcagtcatatagagcagacgAAGGATTCTTTGTGTGCTCTACGGCAAATTCTGTGGTAAAaagttatactgccct
gcagtcatatagagcagacgGTGTACACTTTCCAATAGGCTTTGCCATAATAACTAAAGTaagttatactgccctg
gcagtcatatagagcagacgTTAGTTATAGCCCTAAATTAATGTCATATTCCCAAGGGACaagttatactgccctg
gcagtcatatagagcagacgCAAAGAGGGTTGCAGGCATTTTATTTTACTAATGTTTTCGaagttatactgccctg
gcagtcatatagagcagacgGGAATTTACCACAGGTCTAGGGACAGAATTTATGATTTTGaagttatactgccct
gcagtcatatagagcagacgAGCAATCTCAAATCTCAGTAAAAACAGACTTTCAAGGCAAaagttatactgccct
gcagtcatatagagcagacgCAAAAGGAGTGTGCATTATTGAGACAAAAACAGCCAACGCaagttatactgccc
gcagtcatatagagcagacgTGCCTTTTTTTTCTTTAGGGTGTTAGCTATAGTACTCCAAaagttatactgccctgt
gcagtcatatagagcagacgCCAAAAGTAAATCATAGGCCATTTTCAAGAACGGTAACCAaagttatactgccc
gcagtcatatagagcagacgCGGTGTTGATGGTATTTAGTAACTGCAGGCAATGTGTCATaagttatactgccct
gcagtcatatagagcagacgCTCTAAGCAACACGGCCAGAAGCTAGAGTGGTCTGCCTTTaagttatactgccc
gcagtcatatagagcagacgTTGGTTGTTCTGTTCCATTTTCCCTGCTCAGGATGAAAGGaagttatactgccct
gcagtcatatagagcagacgGGTGGTGGCAAAGCAGCAGCGGAAGGATGAGGAGAAGCTGaagttatactgc
gcagtcatatagagcagacgCAGGAGCTGGGCAGTGTGTTTGAAATGGGGAGGTATGGCTaagttatactgcc
gcagtcatatagagcagacgTGCACACACACCCCCAGCCTTACGCAGATCCAGGACAAAAaagttatactgc
gcagtcatatagagcagacgCCATTGATCCTTGATGCCGTGGGTGCTCTTGTGAAAATCGaagttatactgccc
gcagtcatatagagcagacgATCCACCTCCAGAACTCTTTCCATTTTGTGAAACTAAAACaagttatactgccct
gcagtcatatagagcagacgACCCATTAAACATCAACTCCACATTCCCCCTGCCCCCAGCaagttatactgcc
gcagtcatatagagcagacgACTTCTTGTCTCCATCAACTTGACTAGTCTAGGTACCTTGaagttatactgccct
gcagtcatatagagcagacgTCCATTGTAGGGATAGATCATATTTGGTTTATCCATTCACaagttatactgccctg
gcagtcatatagagcagacgCCTTTTGGCTGGTGTAAATAATGCTACTATTAACATGGGTaagttatactgccctg
gcagtcatatagagcagacgTGGGAATATACCCAGAAGAAAAATTGCTGGAGCATATGGTaagttatactgccc
gcagtcatatagagcagacgTGTCTTTAGTTTTCTGGGGAGCCTCCACACTGTTTTCTTCaagttatactgccctg
gcagtcatatagagcagacgTGTACCATCTCATATTCACATCACAGTGCACAAGGGTTCCaagttatactgccc
gcagtcatatagagcagacgACTGGAACACGTCTGCTTCATTGTAAAGAATACTATCACAaagttatactgccct
gcagtcatatagagcagacgCTTTTTTCTCTCTCCTCGAAAGCATCCATCTCATGTCAATaagttatactgccctg

gcagtcatatagagcagacgCCAAGTCTCAGTGTGGGTTTTGTTCTGGGAGTGCAGAGGAaagttatactgccc
gcagtcatatagagcagacgAACACACTTGCATTTATTACATCTTTGATTCCTGTCCTACaagttatactgccctg
gcagtcatatagagcagacgTTTTCTTATTCCCCTTCCTTTTTGATGGAGAAACTGTGCCaagttatactgccctg
gcagtcatatagagcagacgGCTCAGGTTATATAGCCAATGTAAGACCAGAAAAAATATCaagttatactgccct
gcagtcatatagagcagacgGCTTTGACATATTTGGTGCTTTTCCCATCCCACTGTGAGAaagttatactgccct
gcagtcatatagagcagacgGAGACTAGTGTGGTTATTCTCCTTCCAGATTTAGAGAGACaagttatactgccct
gcagtcatatagagcagacgGCTGTGATGGATGCTGCTTGTGAGATCTCTGCAAATTCCCaagttatactgccct
gcagtcatatagagcagacgGCAGTACCCTCACCTTGATTATGCCCTGTGGATCTCTCGCaagttatactgccc
gcagtcatatagagcagacgGGTACCTGTAGACACTTCTCATCTCTCACACATGCACACTaagttatactgccct
gcagtcatatagagcagacgAGTTAAAACTCCTGCACCGCCCTTGACCAATAGAAAATGGaagttatactgccc
gcagtcatatagagcagacgGAGGATATTTCTGAGACGCATTTTTTAATCTTCCCCAATTaagttatactgccctg
gcagtcatatagagcagacgTCCTCACTTCCCAGGCGGTGTGGCCACCAGAGGGCGCTCCaagttatactgc
gcagtcatatagagcagacgGAGGTGCTCTGAGATGCGTTTTATAACATGTTCTTTCAAAaagttatactgccctg
gcagtcatatagagcagacgGTGGCCAACTCAATAAAGCATCCAATATTTTACATGACCTaagttatactgccct
gcagtcatatagagcagacgGTTCTACAACTGCCTTCTATATAGCTGCTTTTTGGGATCAaagttatactgccctg
gcagtcatatagagcagacgGTAATCTACTTGAACATGAACTTCCTCTTACACCCCCTCGaagttatactgccct
gcagtcatatagagcagacgAGACAGTTCTAGAAGTAGCTCTGGAAAACAGCCCCTCAGGaagttatactgcc
gcagtcatatagagcagacgTGGATCACTTTCCAGTCAGACAGCAATAAGGACTCCATTGaagttatactgccc
gcagtcatatagagcagacgCAGTGGCACCAAAATTAATAAGATTCTCTTGGCCGGGCGCaagttatactgcc
gcagtcatatagagcagacgAAAATTAGCCGGGAGCGGTGGTGGGCACCTGTAGTCCCAGaagttatactgcc
gcagtcatatagagcagacgAAAAAGATTCTCTCATCTGATGGATTAGACTGAGGCAAAAaagttatactgccct
gcagtcatatagagcagacgACAGTTTGGGGAAAATAATTATAAGGATTGTGGCAGAGCTaagttatactgccct
gcagtcatatagagcagacgGGCACAGCCTCAACTGGTCTCCCGTGTCAGAATCAGGGCTaagttatactgcc
gcagtcatatagagcagacgAGAAGGTCCTGAGGTCTTGGATAGGGATGTTAGGGCTGACaagttatactgcc
gcagtcatatagagcagacgTTTGAATCCCTAGATTTCCTTGAACATCTGGGCTGGTAGAaagttatactgccct
gcagtcatatagagcagacgAGAGAAAACTCCACTTGCTTGATTTCCTGTGTAAATATCAaagttatactgccctg
gcagtcatatagagcagacgGTGGTGTTGTGCTCTTCAAGATCTGCCCACGTGGTCCCGCaagttatactgcc
gcagtcatatagagcagacgTGAGCAGAAAAATACAGACCCTCCTCCAGGAATAAGAAGCaagttatactgcc
gcagtcatatagagcagacgCATACACTCACAAACCTGCATGCCCACACACGTATATACAaagttatactgcc
gcagtcatatagagcagacgAGGTATGTGTGCTAACCTATTGGCTAATGTGGTGCTGTGTaagttatactgccct
gcagtcatatagagcagacgTATTGGTCCAAGAACCACAGAACAGAAGAAGGAGTAGGTCaagttatactgcc
gcagtcatatagagcagacgACTCATAATGTGTGCTTCCTCTGAAACCTTGGACTTTGCTaagttatactgccct
gcagtcatatagagcagacgGGTACAACTACCATCTGGTCACTTTGAGCTTCCCATGCCAaagttatactgccc
gcagtcatatagagcagacgCAAAGAAAGTAGTTATTATTCTGGCAGGAGATGTTGATCCaagttatactgccct
gcagtcatatagagcagacgGCAGGAGTATGCCAGAATGCAGGGGAGTGCTTCCATGACCaagttatactgcc
gcagtcatatagagcagacgGCAATGAATGGGCAATTGAAGAGGTCATGGCTAAACAGAGaagttatactgcc
gcagtcatatagagcagacgCCAAGCCCCACCTGACAAGTAAACTAGCCTGGCCAAAGTCaagttatactgcc
gcagtcatatagagcagacgAGAGGTAAAGAAAATCTGGAATGGAAAATAGAGGAGGAAAaagttatactgccc
gcagtcatatagagcagacgTCAGTTACCTTCTAGGCACCAGGCACAACAGAAGAGGCTGaagttatactgcc
gcagtcatatagagcagacgCCCACTGGTATTAAGGCTTTTTAGAGATTACTGCTGGCTGaagttatactgccct
gcagtcatatagagcagacgTTTACGAATTGAGTTTAGTAGGAAGCACAAGAGATATGAGaagttatactgccct
gcagtcatatagagcagacgATCTGTGATGCATTTTATAAGAGCCCAGCAAGATCAAGCAaagttatactgccct
gcagtcatatagagcagacgTTTAATATTGTTTTTCCCCCCTTCACTCATGTCTTTTTGCaagttatactgccctgt
gcagtcatatagagcagacgCATAAGATTTTGTCTCATGGTCTGCATTTAGGAGAACTCAaagttatactgccctg
gcagtcatatagagcagacgTTTAAATGCATAGCTAAATTCACAAGAAAGTAAGCCAGGCaagttatactgccct
gcagtcatatagagcagacgGAAAAAGAGAAAGATGAGGCCAGAGTGGTAAGTAAGTGGGaagttatactgcc
gcagtcatatagagcagacgATAGATGACCTATAAACTAGGGTTTTGATGTTTTGTGGGTaagttatactgccctg
gcagtcatatagagcagacgGAGATGTCGGAGCTAAAATCTCTGCATAAAGCCTTTAAGGaagttatactgccct
gcagtcatatagagcagacgGGATTATAAAAAGACCAACCCATCAGTAAAATGAGTCCAGaagttatactgccct
gcagtcatatagagcagacgACAGTGTGAGAGTCAGCAGAAAAAGTGTTACAAATTGATAaagttatactgccct
gcagtcatatagagcagacgTCTGTGTCCCATCAAATCTCACGTTGAATCCCCAATGTTGaagttatactgccct
gcagtcatatagagcagacgTATTTAAAACTGTGTAGCACGTCCCCGCTCTCTTTCTTGCaagttatactgccct

gcagtcatatagagcagacgCTCTGGCCATGTGAGATGCTTTTCTCCCCCTTTGCCTTCTaagttatactgccct
gcagtcatatagagcagacgGTAGGGCATTACTATAAAGACACTGGAACATGTGGAAGTGaagttatactgccct
gcagtcatatagagcagacgGAACTTCCTAGAGACTGGCTAAATGGCAGTGATCAAAATGaagttatactgccct
gcagtcatatagagcagacgGACAGAGAAGGCCAGGATGAGGAGGTGTCACATAAAAACGaagttatactgcc
gcagtcatatagagcagacgAAGGTCAGTTTTGTTGTCTTAGCTGTGAACTTGGCTGCATaagttatactgccctg
gcagtcatatagagcagacgCATAGGGATCTGTAAAACCTCGACCTGGAGAGTGATGATTaagttatactgccct
gcagtcatatagagcagacgAGCAGTAAAGTATTCAATATTTGTCCTGGCTGCTTCTAACaagttatactgccctg
gcagtcatatagagcagacgAGTTTGCAGACTGACCATATAGAGAAAAGAAAAGCCCATTaagttatactgccct
gcagtcatatagagcagacgCATTTGCTAGAGAAATTTGCGAGACTAAAAAGGCAAGTGCaagttatactgccct
gcagtcatatagagcagacgTCAAAGAGGACTTTTGAGACCTCACATTTCAGAGACCTTGaagttatactgccct
gcagtcatatagagcagacgAGAGAATGGTTTAGTGTGCCAGGCCCAGGGACGTGCTACCaagttatactgcc
gcagtcatatagagcagacgCACTGCTCCCCATATCCCAGGCACTCTAGCTCTAACTCCAaagttatactgccc
gcagtcatatagagcagacgCACGGAATGCAAGAGTGGAAGTTTGGAGCTTCTGCCTAGAaagttatactgccc
gcagtcatatagagcagacgCCTCTATTAGGGCAGTGGGAAGGGGAAATGTGGGGTAGGAaagttatactgcc
gcagtcatatagagcagacgGGACCACCATCCACTAGACCCTAGAATAGTTGATCCATCAaagttatactgccc
gcagtcatatagagcagacgGGGGGTTGAACCCTGCAAAGACACAAGGGCAGAGCTGCCCaagttatactgc
gcagtcatatagagcagacgGCCCACCTCTCATACCAGTGTGCACTGGGTGTGAAACATGaagttatactgcc
gcagtcatatagagcagacgCCTATAGCCCCTTTCTTTTGTCAGAGTTCTCCTTTTTGGAaagttatactgccctg
gcagtcatatagagcagacgAATATTTACTCAACCCTTGTACCCTTATTGCATCTTGGAAaagttatactgccctg
gcagtcatatagagcagacgAAAGACATGAGATTTGGGGAGACTGGCTTTGGATCTGTGTaagttatactgccct
gcagtcatatagagcagacgCATGGGGGTGGATTTTTCCATGTATGGCTCAGCACCATCCaagttatactgccc
gcagtcatatagagcagacgAGCACCTCCCCCCATACCTCTCTTTCTTCTGCTCCTACCAaagttatactgccc
gcagtcatatagagcagacgCCCCAGAAGCAGAAGCCACCGCATTTCCTATACAGCCTGCaagttatactgcc
gcagtcatatagagcagacgAAACCTCTTTTCTCATAAAATACCCAGTCTCTGGTATTTCaagttatactgccctg
gcagtcatatagagcagacgCAAAATTTGTATGTTGACATATAACATCCAAGGTGACGGCaagttatactgccct
gcagtcatatagagcagacgTGCCCTTATAGGAAGAGACATGAAAACTAGATCACTATTTaagttatactgccct
gcagtcatatagagcagacgCCATTTGAGGATATAAGAAAAGGCTTTAGCCACCTGCAAAaagttatactgccct
gcagtcatatagagcagacgTGCTGGCACCCTGATCTTAGGTTTATAGCTTCCAGAGCTGaagttatactgccct
gcagtcatatagagcagacgCCTGAGCTGATTAAAACAAAAGGCAGGCCGGATACGGTGGaagttatactgcc
gcagtcatatagagcagacgACTAAACAAAATACAAAAAAAATTCTCTGGGCATGGTGGCaagttatactgccct
gcagtcatatagagcagacgATAATATTATTAGACCCTTGAGGGCTTCAGGTATTAGACTaagttatactgccctg
gcagtcatatagagcagacgAACGAATGTCAAGACAAGTTTAAAATGGGGTAAACTGATGaagttatactgccct
gcagtcatatagagcagacgGCATTACAGATTTAAAAATTAGTGAAATGGGCTGGGCGCAaagttatactgccct
gcagtcatatagagcagacgTACCAACCTGACCAACATGGCGAAATCCCTTCTCTACTAAaagttatactgccc
gcagtcatatagagcagacgTGGAATAAAGATGTGAAGAAGTTATCCAGAATGTAGAGGGaagttatactgccct
gcagtcatatagagcagacgGAGAATGAGAGACTAAGACAGTGGGAGACAGAAGGGAACAaagttatactgcc
gcagtcatatagagcagacgTGATTAAGGGACATGGAATAGAATAAGAAAGTACAACGTGaagttatactgccct
gcagtcatatagagcagacgAGACAAGAATTCTTAGATGTAGGAAGCACAATAAACCATGaagttatactgccct
gcagtcatatagagcagacgCCAAGACAATGGGAAAATCTTATAAGCAAACATAGAAGAGaagttatactgccct
gcagtcatatagagcagacgTGGCATAATTGGATTGGTAGCTGACTTCCTAGCAGCATCAaagttatactgccct
gcagtcatatagagcagacgTCACAAAGGAATTTTTGATGATGCACTTCCAAAAAGAAGGaagttatactgccct
gcagtcatatagagcagacgGAAGATCTACGATTTAAGAAGGAATGATGAGAAAACACTCaagttatactgccct
gcagtcatatagagcagacgCTATAAAATAGCCTAATTCTCAAGGTAAAAGAACCCCACTaagttatactgccct
gcagtcatatagagcagacgAATGTAGAACAGGTGGAGAATCATTGGTGTATAAGTGCTCaagttatactgccct
gcagtcatatagagcagacgTGGGGGGAGACACAAGACGTAAAAAAATTAGAATTTGGCTaagttatactgccct
gcagtcatatagagcagacgTCGCACCATTGCACTCTAGCCCTGGGCAACAAGAGCGAAAaagttatactgcc
gcagtcatatagagcagacgAACAATGGAAAGTTTCCCTTAGTCAATCCAAGGAAGGTAGaagttatactgccct
gcagtcatatagagcagacgAGAAAAGATGCATTGAAAAGCAGGACCAATAGAAAGCACAaagttatactgccc
gcagtcatatagagcagacgTATATAAGTCGTTGAAATGTTTTAAATGGGGCAGATCCACaagttatactgccctg
gcagtcatatagagcagacgCGTCAGACTGAATAAAAATAACCAGATATTGCAAGCTTACaagttatactgccct
gcagtcatatagagcagacgAGATATATGCCTCTTATTTACCAAAAGGAAGCTTCTGTAGaagttatactgccctg
gcagtcatatagagcagacgTAGCATTTCAAAACTTATTTGCCTCTAGTAACGTAGCTTCaagttatactgccctg

gcagtcatatagagcagacgCTCACAGGATTGCAAAGAGAAATTAACAAATTCAGCATCAaagttatactgccct
gcagtcatatagagcagacgCTCTCAGTAATTGATAGCTCATGAAGACAAACTCAGAAAAaagttatactgccctg
gcagtcatatagagcagacgAAAAGCTCAATCTAAAAAGTCCAATGTAGAGCCCTACACCaagttatactgccc
gcagtcatatagagcagacgAAGAATGTGCAGTATTTTTGAGTAGGCCGCAAAGCAAGTGaagttatactgccct
gcagtcatatagagcagacgCAAAGAATTTGTATCAAAAAGACCATGTTTTCTGGCCACAaagttatactgccctg
gcagtcatatagagcagacgTTCATGGGTCAAAGAAGAAATCACAACGGAAATTAAGAACaagttatactgccct
gcagtcatatagagcagacgGGAAGTGAGAAAAAGAATAGAATAAGCCCAAAGAACATAGaagttatactgccc
gcagtcatatagagcagacgTGTAAAGAAAATACAGTAGGGTCAACAAAGTCAAAAGCTGaagttatactgccct
gcagtcatatagagcagacgGGCTGTAACTATAGAAGTAGCAGGTATTTAAAACGAGTAAaagttatactgccctg
gcagtcatatagagcagacgTATAAACAACTCCATGCTAGTTGTGAAGGATACAGAACTGaagttatactgccctg
gcagtcatatagagcagacgTGTGAGCAGATTGTCTTGGGCCGTCTACTCCTTGTGTTTGaagttatactgccctg
gcagtcatatagagcagacgCCTGCAGAGAGCCCCCTCGCCCAAGAATACACCTTCCCTGaagttatactgcc
gcagtcatatagagcagacgGGTAGGGACATAGAGACCTTAACCATTCAGCCCACCCTGGaagttatactgcc
gcagtcatatagagcagacgAGCAGTGTGCATGTAAATGTTTAATAACGGGCTCCTTTAAaagttatactgccctg
gcagtcatatagagcagacgGTTTGCTAATTTCTGTGGTGAAAATACATCCTCCATAGATaagttatactgccctg
gcagtcatatagagcagacgATGTGAAGTTGGGAATAGATGCAAAATAGTACACTACTACaagttatactgccctg
gcagtcatatagagcagacgTCCATCATACAGCTATGCCAGTGGTAAATAAACTCAACACaagttatactgccct
gcagtcatatagagcagacgTTGAGATCAGGAGTTCGGGAGCAGACTGAGCAATATAGCAaagttatactgccc
gcagtcatatagagcagacgTCATGTAGTAATTAGGGAAGGATGAGTTTTGAGTATTCACaagttatactgccctg
gcagtcatatagagcagacgACAACCAATTCACAAAAATTTAACAAGTGATGCTCAGGAGaagttatactgccct
gcagtcatatagagcagacgTGCTCCAGGACTTCTTGTGGTTAGGACAGCGGCTTTGTTGaagttatactgccct
gcagtcatatagagcagacgTCATGGTTCAAACTCTCCCTCTGTCCAATCCTGGCTCCCAaagttatactgccc
gcagtcatatagagcagacgAGGCCCTGACCCCTAATACCTACCTTGCTCTCTGGTACATaagttatactgccc
gcagtcatatagagcagacgGTGTTATTCATGACTTGCAAACGCCACTCTATTCTTTCTTaagttatactgccctgt
gcagtcatatagagcagacgCCACTGGCATTTCATCATTTTGGAATTTCACAAGGCAGATaagttatactgccctg
gcagtcatatagagcagacgTGTTCTACCCACATGCTCCTCAGTCTAGTAAGGCCCTGGGaagttatactgccc
gcagtcatatagagcagacgGCCCTGCACATCTCCTTGGCTCCCTCTCAGCCATAGTTCTaagttatactgccc
gcagtcatatagagcagacgATCTCAAGGATGCGGCTATGGGTCACTCTGATTCTCTGCCaagttatactgccc
gcagtcatatagagcagacgCCTTTTCTAAAGCACTAGGCGCCTTTGGGAGTTAAAGCTCaagttatactgccct
gcagtcatatagagcagacgGGTAAATGCTCGGCCCTGTGTCTTTGAAATGACCATTGTAaagttatactgccctg
gcagtcatatagagcagacgCTCAGAGATCCAAGTGGTATTATGTCCTTGTGTGCCTGTTaagttatactgccctg
gcagtcatatagagcagacgAGTTGCCTTTCATACTTCTCTGTCTTAACATCTCAATTTCaagttatactgccctgt
gcagtcatatagagcagacgCTTCCTGAGATCACCTTCAAAAATAAACTATCTAGCTGGGaagttatactgccctg
gcagtcatatagagcagacgCACCCGTAATCCCAGCAGGATTGAGAGGCTGAGGTGTGTGaagttatactgcc
gcagtcatatagagcagacgACAAATATTAGCCGGGCACGGCGGCGGGCGCCTATAATCCaagttatactgcc
gcagtcatatagagcagacgAACAATTAACTATCTGTATCCAAGTCATTCTACCAGTCTCaagttatactgccctg
gcagtcatatagagcagacgAGGAATCTACACTACGATACTTGATACACTATGATACTTGaagttatactgccctg
gcagtcatatagagcagacgCTTTTTTTTTTTTTAATTTAATTTGGCTGGGCGCGGTGGTaagttatactgccctgtg
gcagtcatatagagcagacgAAAATACAAAAATTAGCCTGGATATGGTGGCATGCACCTGaagttatactgccct
gcagtcatatagagcagacgCTGTTTGTTACATGAGCATACTGGAGATTGGGCTTCTAGTaagttatactgccctg
gcagtcatatagagcagacgTAGGTAATTTTTCAGCTCTCGCCCCCTGCCCACCCTCCCCaagttatactgccc
gcagtcatatagagcagacgCCATGATGCATTAATATATGGACCACATCTTCTTTATCCAaagttatactgccctg
gcagtcatatagagcagacgTGCCATTGTTCTTTTCACAGTTTCTAAATACAGCATAACCaagttatactgccctg
gcagtcatatagagcagacgGACTTATCTTGGAATGCTGCCAGGTGGGCCAATAGATAACaagttatactgccc
gcagtcatatagagcagacgTAGATGTGCCAAAATATTTTAAAGAAAACACCTGTGCAGGaagttatactgccctg
gcagtcatatagagcagacgGCAGGAGGGAGAAGGAGCCTGGCATGCAGGTGAGGAGTGCaagttatactgc
gcagtcatatagagcagacgAGAAGATTGGGTAGGAAGAGCTCCAGACTGCAGCACATTCaagttatactgccc
gcagtcatatagagcagacgGGTGGTAGGAATTCCAGAGCAAAGACTGCCCACTAGAGGCaagttatactgcc
gcagtcatatagagcagacgTGGAAACAGCTGGGGAATGTGTGACCTTGGTATGAGCTCTaagttatactgccc
gcagtcatatagagcagacgGAAGGCTGTCAGTTGCTCAAGCTTCCAAGATCTCAGGGGAaagttatactgccc
gcagtcatatagagcagacgACCACACAGATAGGTATCCCCACACATAAATCCATATAAAaagttatactgccct
gcagtcatatagagcagacgTCTTTGTAGGATTTGCTAAGTTGGGGTTCTGCCACCTTCAaagttatactgccctg

gcagtcatatagagcagacgAGCAATCAAGGGAGAAGATGAGAACGACACGGTGCTGACAaagttatactgcc
gcagtcatatagagcagacgCTTCCAAAGAACTACTTGGAGAGCCTGGCATGTGTCACCTaagttatactgccc
gcagtcatatagagcagacgTGAGAAGCTTCAAGGAAAGAGGCCGGTCAGAGTTACCTGTaagttatactgccc
gcagtcatatagagcagacgGAGAAGAAGAAAAGAGAACTGCAGTGACAGTGGCCTGCACaagttatactgcc
gcagtcatatagagcagacgGCGTGATGGGCCAAATGCCAGTTGGTGTCCGTGGGTCAGCaagttatactgcc
gcagtcatatagagcagacgCGTGGTGGAAGGTAGCTGGGACTGACTGCCAGGGCATGGTaagttatactgcc
gcagtcatatagagcagacgGGTCAGAACCCCACCCTGCCCAACTTGAGAAACCAGTGTTaagttatactgcc
gcagtcatatagagcagacgTGCAGTGGAGCGATCTCCGTGAACTGCAACCTCCACCTCCaagttatactgcc
gcagtcatatagagcagacgCCTCAGCCTCCGAGTAGAGGAGCCTAGAGGCATGTGGCACaagttatactgc
gcagtcatatagagcagacgGTCTCGATCTCTTGACCTCGTGATTCATCCACCTCGGCCTaagttatactgccc
gcagtcatatagagcagacgCAAGCAATTCTCCTGAGGAGATGGAATTGCAGGTGCACGCaagttatactgcc
gcagtcatatagagcagacgCTCCGTGCCCGGCGGTGGAACCCGTCTTCACAGAGCTAGCaagttatactgc
gcagtcatatagagcagacgTGAGCTTGGGTCAGGTTGGGGTCTGGTGGGTGGTGGGAGGaagttatactgcc
gcagtcatatagagcagacgCTTGAATGGATTGCGATTAAAGTTTTGATTTGGATTGGGCaagttatactgccctg
gcagtcatatagagcagacgCTGAAAGTGACAGGAAAAGCTATGAAATTTGCCCCAGATTaagttatactgccc
gcagtcatatagagcagacgAAAAAGTGCCATTTCATGATTTCCCTTCACCAAGTAGTCCaagttatactgccct
gcagtcatatagagcagacgCTGTCATAAAGTATTGGTTAACACAGTGGTTTGGAAAGACaagttatactgccct
gcagtcatatagagcagacgCTAAAAACAAAAAACGCCAACCTAAATAGTGCAAAGACCAaagttatactgccc
gcagtcatatagagcagacgTGGTAAATGTTTATTCCATTGCTGTTTGAATCAGTCTGCAaagttatactgccctg
gcagtcatatagagcagacgTCTGGTTTCGCTTGCCTTTTCAAACCTCCAGGTGCACCGCaagttatactgccc
gcagtcatatagagcagacgTGAGCATTTCAGGTAGGACTGGAAAAAGGTCTTTGAGAAAaagttatactgccct
gcagtcatatagagcagacgAATCTATTTACCTTACCATAAGAGCAGTCAAGGCTCGCTGaagttatactgccct
gcagtcatatagagcagacgGGTGCTCATTAAACATAGAAATAAGCTTGCTTCTAGAAAGaagttatactgccct
gcagtcatatagagcagacgCCTGCCTTTTGCGCAAATCTTTTCAGAGACTGGCTTTGGGaagttatactgccct
gcagtcatatagagcagacgAATCAGGCACCCTAGCGTCCTCCAAAGTTCTTCCCGATGCaagttatactgcc
gcagtcatatagagcagacgCGCCTAATTCGAAGCGGCTCCCTCTTCTGTTCTGAGCGGAaagttatactgcc
gcagtcatatagagcagacgGCCCTGGAACGGCCAACTCATTCTTACTCCTTTCTCCTGGaagttatactgccc
gcagtcatatagagcagacgCGCGCCGCGCCCACCGCCTGCGGAGGCTGGGTCTGGGTCTaagttatactg
gcagtcatatagagcagacgGCTGGGGCTGAGGCGGGAGCTGGGGCGGGGGCGGGGGCTGaagttatact
gcagtcatatagagcagacgGCAGGAGCGGGCGCAGGGGCCGTCGGCAGAGGGCGCTACGaagttatact
gcagtcatatagagcagacgGGAGCGCGGGCCGGGAGGAGGCGCCTGGGTCTCGGCGAGTaagttatactg
gcagtcatatagagcagacgCGCGCATCAGCTGCCCCGCGCCGCCTACCGCTCGGGCCCGaagttatactg
gcagtcatatagagcagacgCGCCGGCTGCGTGGCGCGCTGCGAAGGTTGGGCTGGGCGGaagttatactg
gcagtcatatagagcagacgCCCCGCGGCTTCTCTTTGTCCAGGCGAGGCGGCTGGAGGAaagttatactgc
gcagtcatatagagcagacgGCCTGGGCGTCGTGCGGGCGGCTCCGGTCCCCGGCTGCGCaagttatactg
gcagtcatatagagcagacgCGGCGGCCTGGGCGGTGCCTGCGCCGAGGTGAGTGCGGGGaagttatactg
gcagtcatatagagcagacgGGCGGGCAGAGGTGATAGCGAGCGCCGGCGGCTCCAGGTGaagttatactg
gcagtcatatagagcagacgCCGCCGGCTCTCAGGGCAAGTTTATCTCCCCCCTTCTGCGaagttatactgcc
gcagtcatatagagcagacgCGGAGTACAGACCTGTCGGTAAATAGAAAACTCGAGCTGGaagttatactgcc
gcagtcatatagagcagacgGGAGGAAGCAGCGTGGCGGAACCCGGCCCGAGATGAGCGTaagttatactg
gcagtcatatagagcagacgCTCTTGGACATTACATCTTTCTTGGCTCTTTTTGACCCAGaagttatactgccctg
gcagtcatatagagcagacgCCTCTGCTGCGCGGTCCGGTGGGGTCGGCTGGCTCTGCTGaagttatactgc
gcagtcatatagagcagacgCGGAGACAGGTGAGCGGAGTTGGGGAGCGCGCAGCCGGGCaagttatactg
gcagtcatatagagcagacgGGCTTCTCGTCTCCAGAGCGCTGATGCTTCCTGCCCACATaagttatactgcc
gcagtcatatagagcagacgTCTGGCTTAGGGACTGCAGCTCTTTTGGCACCTTCGCTTGaagttatactgccc
gcagtcatatagagcagacgGCTTTAGAGGCGTTGGTAGGTTAGTTAAGGCGAGTTCTCCaagttatactgccc
gcagtcatatagagcagacgCAGCTTTGGATAATTTGGTCTGAGTCTTCGTCTTAATTTAaagttatactgccctgt
gcagtcatatagagcagacgAGCAGTGAAGCGGCTTGCTACATCCATTTTTTTTTTTCATaagttatactgccctg
gcagtcatatagagcagacgTGTGCACAGAAGGAATTACAGGCACACAAGTGATGTTACTaagttatactgccc
gcagtcatatagagcagacgAGGATACCAAAGGCGCCTGTACTCTCTGTTATATAAATGCaagttatactgccct
gcagtcatatagagcagacgCACCTAAGAAGAGAGTGTCATGCTTTTAGTTTACTTGGAGaagttatactgccct
gcagtcatatagagcagacgAGTGGTCTTAGCTTGAGGTACCCTTTTAAGATAATATAGGaagttatactgccctg

gcagtcatatagagcagacgCTTAGTAATTGTGAGTTGCCAGTTATTAAAGATGTGTCTCaagttatactgccctg
gcagtcatatagagcagacgCCCTTCCCTTTCCCCAGGCCCCAAAACCAGTACTCACATTaagttatactgccc
gcagtcatatagagcagacgCCTTGTCCCTCCTCCACCCAGTATTCTGTTCTAATTGGTCaagttatactgccct
gcagtcatatagagcagacgTGATACCTTGATACCTGTAATGTTAAAAAGCTTTCCTGGTaagttatactgccctg
gcagtcatatagagcagacgGTGGAGCATTGTATTGTCACTTTCTGTAGCCTATGGTGTTaagttatactgccctg
gcagtcatatagagcagacgTAGAAACATGGGAAGAAAAAAATGGTATTGGAAAGGGGACaagttatactgccc
gcagtcatatagagcagacgGTGTTTTGTCTAAATGCTCTTTCTTGATTTTCCTGCGAAGaagttatactgccctgt
gcagtcatatagagcagacgTTTATTAGGGACCCATAATTCCATGTAGAATACTTACGGGaagttatactgccct
gcagtcatatagagcagacgAGTTGAGTAGTAGTGGCAGGATATTCATTTATGTTTACACaagttatactgccctg
gcagtcatatagagcagacgAGTTTTTATTGTTTTTAAGGTCAGAAAGAAACCTGGCCTCaagttatactgccctg
gcagtcatatagagcagacgCAGTGTTTTCTGTCTCCCTTATGTAATCCCAAATATCCAAaagttatactgccctg
gcagtcatatagagcagacgGGAGCAGATCTTATGGATGGTTTATGAAGGCTTATGTAAGaagttatactgccct
gcagtcatatagagcagacgCCTGTTTGGGCTATTACATAATTGACACGGAACTAGACTGaagttatactgccct
gcagtcatatagagcagacgCCGAAACCCACTCCGGCATACGCACTTTTATAATGAATTGaagttatactgccc
gcagtcatatagagcagacgCAGGGATGGTGGTTTGTGAAAGAGAGGTAAGAATAGCATCaagttatactgccc
gcagtcatatagagcagacgGAAGACTGTTATGCTTAGCGATTTTCTAGAGATTTGAATGaagttatactgccctg
gcagtcatatagagcagacgCCTCTCTAGTCCTTTCTATCTCCAGCAGAGGCAGATCGACaagttatactgccc
gcagtcatatagagcagacgAGCTAATGAACCTAAGCCTCAGGTCCCTCCGTCACAGAGGaagttatactgcc
gcagtcatatagagcagacgACTGGCAGTGTGTTCATGTGGTCATCCCTGAGTTTGTATTaagttatactgccct
gcagtcatatagagcagacgGAGTACTCTTTCTATATTCCCACCCCCCAATTATATAAACaagttatactgccct
gcagtcatatagagcagacgAGACCCTGAATCTATCCCTGTCTTCAAACCTGTGTTCAAGaagttatactgccct
gcagtcatatagagcagacgCAATATCATAGTTGACTTGCTGAGAAAATGCAAACTGCTAaagttatactgccct
gcagtcatatagagcagacgCACATACCTGGAGAACTAGCTCTTTGCCTCGTGATCGCACaagttatactgccc
gcagtcatatagagcagacgTGGAGGTGGAGAAGAGTGGAAACGGGTTGGGGAAGGGAGGaagttatactgc
gcagtcatatagagcagacgCAGGATGCCAAGGGGAAAGGAAGGAAAACTATTGAAAGATaagttatactgcc
gcagtcatatagagcagacgCTGTAAGCTCAGGATTGGTCTCTAATACGAATTTTCAACAaagttatactgccct
gcagtcatatagagcagacgACTCATTCCTCGAAAATATCCTACAGGAAACAAATGCAGGaagttatactgccc
gcagtcatatagagcagacgGGGAGGCTTATAATATCTACCACACACATGTTTGAAACGAaagttatactgccct
gcagtcatatagagcagacgGGAGTTTCCCCTTTTCTCTGAAATGATTTCTACTTCCTATaagttatactgccctg
gcagtcatatagagcagacgTGCCTTTAGATCTTATCATCTTGACCCTTCCCACCAACGGaagttatactgccct
gcagtcatatagagcagacgCATCCACCCACCCATCCACAGAGACTCTCTTTTTGACCAAaagttatactgccc
gcagtcatatagagcagacgCCTCAACCTTGGCCCGTAAGAACTGCAGCTTTCAACACAAaagttatactgcc
gcagtcatatagagcagacgTGCTCCCACCCTCCAACGGATTTGAAAACACTGTAGCATGaagttatactgccc
gcagtcatatagagcagacgAAAATGAGCAAAAGGGGGAAAAACACCGTATAAAACCAGCaagttatactgcc
gcagtcatatagagcagacgCATGAGAACTTACTGTCGCGAGAACAACATGAGGGTAGTTaagttatactgccc
gcagtcatatagagcagacgACAGCCAAACCATATAACATGGCATCTAGCAGCCTAAGGCaagttatactgcc
gcagtcatatagagcagacgACCCAGCCTCACTTGCATTCCTACCTAGAAAAGCTCAAGGaagttatactgcc
gcagtcatatagagcagacgATGATAATAGGCCCCTGACTTTCATTTTGTTAGATCGTTTaagttatactgccctg
gcagtcatatagagcagacgGATATATGTGTATCTCCTACCACCTGGAGTGTCTTCCTCAaagttatactgccct
gcagtcatatagagcagacgCCATTCCTTTGAAGTGTAATCATCAGGGAGGATAGGGCTCaagttatactgccc
gcagtcatatagagcagacgCCATTTAGCAGTCACAGATGGCCTAGTCTCATTGGCAAGCaagttatactgccc
gcagtcatatagagcagacgGCCTGATTGCTGTAATGTTCTGTGTCCTTGACCCTGCTTGaagttatactgccct
gcagtcatatagagcagacgCCACACATTCTCATTTCTCCTCCCTACTTCCTTATTCTCCaagttatactgccct
gcagtcatatagagcagacgACACCCAGTCACCTCTACCTGAATTGGAGTTGAGCTCAGTaagttatactgccc
gcagtcatatagagcagacgCTGAAGTCTCTTTCTCTCACTGCAGTTGTACTGGGTAAAAaagttatactgccct
gcagtcatatagagcagacgTGTCTCCATAGGCCAGGCACTTTTACATATTAGGTATAGAaagttatactgccct
gcagtcatatagagcagacgACTTACAGTGTAATGGAATGAATCAGATGAGGTTAAGTACaagttatactgccct
gcagtcatatagagcagacgTATCAGGTATGGATACTGCTATGGAGAAAATAAATCAGAGaagttatactgccct
gcagtcatatagagcagacgGCAGGAGTGGGTTATGGGGAATTGCTATTTCAAGTAGTGTaagttatactgccct
gcagtcatatagagcagacgGGGAATGCCTCACTGAAAGGAGATGTTGAGCAGAGCCCTAaagttatactgcc
gcagtcatatagagcagacgCCAGAGTGCAGCTGGCATTTGTGGGGAGCACAGGGAGGCAaagttatactgcc
gcagtcatatagagcagacgCTTTTCCTCATGACAGCAGCCCCATCCCCCACTCCAGCCAaagttatactgcc

gcagtcatatagagcagacgCACTGGTGTTGTTCTTGGGGCCTTAGTGGTATGGTGGCTTaagttatactgccctg
gcagtcatatagagcagacgCTGAATAGGAGTAAGGAATGAGAGTGTATGTTTTCAATGAaagttatactgccctg
gcagtcatatagagcagacgCCACTGGTTTGTCATAGGAAAGAGGCTGGGGGCAAGCTCAaagttatactgccc
gcagtcatatagagcagacgGGGAACAAATGGAACTATGATGTATAAAAGATGTGGACAGaagttatactgccct
gcagtcatatagagcagacgCTCTTTCTCGAGGAATGTCTTAATATCAGGAACCTAGAACaagttatactgccctg
gcagtcatatagagcagacgTGATTTGAACTGAGCCTAGAGAGAATCCCCAGCTTAAAGGaagttatactgccc
gcagtcatatagagcagacgGGGGTTGGTTAGGGTTTGAGGAGGGGAAATTACTCAGTCAaagttatactgccc
gcagtcatatagagcagacgGATGGCTGCTCAACATCTGCTAACCAGAACAACACACACAaagttatactgccc
gcagtcatatagagcagacgCAACAGGCTGTCATGAATTTCTCCATGGAGAAGGTGAAGAaagttatactgccc
gcagtcatatagagcagacgGCAGCCAGGGCCTGTCTGCTGGTTTTTAGTTACAGCCTTTaagttatactgccct
gcagtcatatagagcagacgGGGGACAGGGGTATGGAACCAGGATGGGATAGCTGTTTCAaagttatactgccc
gcagtcatatagagcagacgGCTGTTAGCTTGTAACCATTTTAAACTCTCAACAACCAAAaagttatactgccctg
gcagtcatatagagcagacgTCATATAGAGGTTCCTTGGATTGTTTGCTGGTGGGTAGGGaagttatactgccctg
gcagtcatatagagcagacgTGCAGCTGGCATTGGGGAGGTGGCATTTATAGGAGTCTGGaagttatactgccc
gcagtcatatagagcagacgGACTTCATGTGGTCTCTTACTTTGCAAGCTTCTCTCACGTaagttatactgccctg
gcagtcatatagagcagacgGGTCATTCCTGGGTATTTGATACTCCTCTTTAGCTTTTGTaagttatactgccctgt
gcagtcatatagagcagacgATTTTCCTTGGATTTTCCAGATAGATGGTCATATCATCTGaagttatactgccctgt
gcagtcatatagagcagacgATTATCAGACTAGATAGAAGACGAGACTGTTTATCTGTTGaagttatactgccctg
gcagtcatatagagcagacgCAAGATTATTGTATTTCAGCCAAATGTACTTCCCTTGTCCaagttatactgccctg
gcagtcatatagagcagacgGGTCCTCTCCTTATTAGTCTTTTTACTCTGTTTACTGTAGaagttatactgccctgt
gcagtcatatagagcagacgACCAGTATGTTCAGTTCCCATAGAAAAACAAAAGAGTGTTaagttatactgccctg
gcagtcatatagagcagacgACAGTCATTATAGATAAGCCAGCAGTGTGTGTGATTAGGGaagttatactgccct
gcagtcatatagagcagacgCTTTAGAAAGTTTGACTCACCTTGCAGAAATGAAAAGCCAaagttatactgccctg
gcagtcatatagagcagacgAAATACATTTGTCCAGAAACAGGCTGATATTGATACACTGaagttatactgccctg
gcagtcatatagagcagacgTTTCCCCAAAAGTACAATAGAGAAAGCCAGGAATATATAGaagttatactgccct
gcagtcatatagagcagacgAAGTTTATTTTCCACACTTCATCAGATTGCTACTGCAGAGaagttatactgccctg
gcagtcatatagagcagacgAATATGAACACTAGCTCTGTAAATACGGGAGAAGTAATCTaagttatactgccctg
gcagtcatatagagcagacgCTGTTTGACATTACTCATTGAAAATAGCTTCTCAAGACAGaagttatactgccctg
gcagtcatatagagcagacgCAAATAATTGTGTTCTCAGGTTGGAAATGTTGCCACATATaagttatactgccctg
gcagtcatatagagcagacgTTGATGTTTCTAAGCCTGATGCATTTTCCATTAGTGTTACaagttatactgccctgt
gcagtcatatagagcagacgGTTTGTACAGTGAACATGCCAAACATTAGGTTTAGTTTTCaagttatactgccctg
gcagtcatatagagcagacgTAGTTTTATATTGAGTGAGCTAATTTGCCACTACTAGCAGaagttatactgccctg
gcagtcatatagagcagacgACATTGTCCTCTACCCCCAAACTTTATACTGAAATTTTTCaagttatactgccctg
gcagtcatatagagcagacgCTCAGTTTAGAGAATAACAATAAAAGCAGGAATGTGACCTaagttatactgccctg
gcagtcatatagagcagacgAAATTCAGCCCCCAATATTTGACGTGGGTCCTTTTCTATTaagttatactgccctg
gcagtcatatagagcagacgAAGAGAGAAATTTTAGAGCTGGGTGTCTGGGGGAGACATCaagttatactgccc
gcagtcatatagagcagacgTTCACAAGGTAATAAAATATCACATGGCAAATGGAGGCAGaagttatactgccct
gcagtcatatagagcagacgAGACTGGGGCTTATTTCACCGCTTATCCACAACCGTAAAAaagttatactgccct
gcagtcatatagagcagacgAATATGGCTCTGTTCTGCCAAGCCCACAGGCAGCCAGACTaagttatactgccc
gcagtcatatagagcagacgATGAAGATGACAGGATTAAGCAATTAAAGTAAAGACAGGCaagttatactgccct
gcagtcatatagagcagacgTGCAACACTCCAGGGTCAGCAGTAGAGAGAGGCTAGTTGAaagttatactgccc
gcagtcatatagagcagacgTGAGTTTTCCTTTTGTGTCAGGCTCCAGTTTGGTTTGTAAaagttatactgccctgt
gcagtcatatagagcagacgTCTTGGAACACATCTGAATTATTGGAGTAAAGACATTCTGaagttatactgccctg
gcagtcatatagagcagacgAGAATGGTGGAAATTAACGAAGGGTTTATAACAATCTGAGaagttatactgccctg
gcagtcatatagagcagacgTGTCATTTTTAAACTTACCTTATTCCCATTCCTTCATCCGaagttatactgccctgt
gcagtcatatagagcagacgAGCCTTCAAAACTAATAGCCTAAGTCAGGCACAGTGGCTCaagttatactgccc
gcagtcatatagagcagacgGTAATCCTAGCACTTAGAGAGGCTGAGACAGGTGGGTCACaagttatactgccc
gcagtcatatagagcagacgCCAACAAGGTGAAATGCTGTCTCTCCTAAAAGTACAAAAAaagttatactgccct
gcagtcatatagagcagacgAAAACAAACAATAACTAATAGCCTAGAAGCCACTGAAAGGaagttatactgccc
gcagtcatatagagcagacgTTAGAGCCCTCTAAGATCCCCAGTCTCAGAAAATTGTCACaagttatactgccct
gcagtcatatagagcagacgAGCTTCTTAAAAAACTCCATTCTCAAAGCTTGTTTTGCAGaagttatactgccctg
gcagtcatatagagcagacgCAATCAGCAGTAATTGTTTAACCTCTCAGCTGCCTGAGGCaagttatactgccct

gcagtcatatagagcagacgCCTATGGGGGCTTTAAAAACCTTTGATGTAGTCTTTGGAAaagttatactgccctg
gcagtcatatagagcagacgTATGTCCAGGAAATACTTAATAAGGCCTTAATCTGTCATCaagttatactgccctg
gcagtcatatagagcagacgGCATAAGTAAAGTAGGAAGAGAAGACTAAGAGTTGTAAACaagttatactgccct
gcagtcatatagagcagacgGCATGTACCCCACCCCCCAACACGCGTAATCCCTCAGCAAaagttatactgcc
gcagtcatatagagcagacgGCTAGACTTCAGCAGCTGCACATGATAAACAAAGTAGACTaagttatactgccc
gcagtcatatagagcagacgGCAACAAATATAACAACAAACCTTTATGCAGAGTGGAATCaagttatactgccct
gcagtcatatagagcagacgAATCCAGTTTTCAACATAAAATTTCCACACACACAAGCAGaagttatactgccctg
gcagtcatatagagcagacgTGGGCCATAACAGGAAAAACAAAAGTATCCCTGAGAAAGTaagttatactgccc
gcagtcatatagagcagacgTTCGAAGAGCTGAAGGAAACCTTGTCTGAAGAATTAAAGTaagttatactgccctg
gcagtcatatagagcagacgAAAATTTACTAGAGAGGCTCATCAACAGATTTGAACTAGCaagttatactgccctg
gcagtcatatagagcagacgCTATGAGGAACAGAAAGTCAAAGGAATGAAGAGAAATGAAaagttatactgccc
gcagtcatatagagcagacgTCAAAAGGAAAAGAATGAGAGTAAGGGGACAGAAAGATTAaagttatactgccc
gcagtcatatagagcagacgGGAAATAATGGACAAAAACTCAAGCAGCAAGAGAGAAGTGaagttatactgccc
gcagtcatatagagcagacgGGAGGCTAGAAGATAGTGGAATGATGTATTGAAAATGCTGaagttatactgccct
gcagtcatatagagcagacgGAATTTGTCACTAGCAGACTGTCCCTACAAGAAATATTAAaagttatactgccctg
gcagtcatatagagcagacgGACACAAGTCAGTAACACACATCCACAGGAAGAAATAAAGaagttatactgccc
gcagtcatatagagcagacgTCTATTTAACACAAAAGAGGGCAGTAGTGGAGGAACAAAAaagttatactgccct
gcagtcatatagagcagacgACTAATCACTGTAGTCAAAAGGCATAGGTTGGCAGAATTAaagttatactgccctg
gcagtcatatagagcagacgAACATGACTCAACTATATGCCATTCACCACACTCACTTTAaagttatactgccctg
gcagtcatatagagcagacgGATATAACCTGAACATAGTAATAAAAGGAGAGCTGGAATGaagttatactgccct
gcagtcatatagagcagacgCTGTTGTTAAAGGGATCAATTTCTCAGGAAGTCAAATATGaagttatactgccctg
gcagtcatatagagcagacgACAATAGATCCCCTAAATATGTGAAGCAGAAACAGAATTGaagttatactgccct
gcagtcatatagagcagacgCAACAATAGTATTTGGAGATTTCAATACACATTCCCTCCCaagttatactgccctg
gcagtcatatagagcagacgTGTTGGCCAGGCTTGTCTCAAACTTCTGAACTCAGGTGATaagttatactgccct
gcagtcatatagagcagacgCACTGTGCCTGGCATCAATACCTCATTTTCAAAAATGGGTaagttatactgccctg
gcagtcatatagagcagacgGATCAGCAAGGAAATAAAAGGCTTGAGCAACTTTATAAGCaagttatactgccct
gcagtcatatagagcagacgAACAGACACCTAAAGAACACTTCACCCAGTGGTAGCAGAAaagttatactgccc
gcagtcatatagagcagacgATATACTCCTAGTGGATTAAAGAAGAAATCAGAAGGCAAGaagttatactgccct
gcagtcatatagagcagacgAGCTGTTAATGACTGTATGAACAAAAGAAAGATCTCAGACaagttatactgccctg
gcagtcatatagagcagacgTTATATAACCTGAATGAAATGGGTAAATTCCCAGGAAGACaagttatactgccctg
gcagtcatatagagcagacgAGGGCAGATCTGAAGTGACCTATAATGAGAAAGGAGATTGaagttatactgccc
gcagtcatatagagcagacgCCAGGCTGGGATGGCTTCAACGGTAAATTCTACCAAATGTaagttatactgccc
gcagtcatatagagcagacgCAGTTTTGCATAATGTTTTTCAAGAAGGTAGACAAGGAGGaagttatactgccctg
gcagtcatatagagcagacgGCTCATTCTTTAAGGCCAGTAGTACTTTGATACAAAAGCCaagttatactgccctg
gcagtcatatagagcagacgCAGCAACATATAAAAAGGTTTGTATACCGTGACTAAGTGGaagttatactgccctg
gcagtcatatagagcagacgCAGCACTTCATGATAAAAATACAAAACTAGGAATGGAAGGaagttatactgccct
gcagtcatatagagcagacgCCACAGCTAACATCGTACTTGTGAAAGACTGTTTTCCTCTaagttatactgccctg
gcagtcatatagagcagacgGGATGTCTACTGTTTTCACATTTATTCAGCCTAATACTGGaagttatactgccctg
gcagtcatatagagcagacgTGTAGAAAATCCTAAGGACTGTATACACAGACTGTGAGAAaagttatactgccct
gcagtcatatagagcagacgAGTTCAGTAAGGTGGCCATATGCAAGATTAATAAAAGTCAaagttatactgccctg
gcagtcatatagagcagacgTAATATTATTAAGGTGACAGTGCTTCCCATATTGATCTGCaagttatactgccctg
gcagtcatatagagcagacgTGTAATCGCTATCAAAAACACAGAGGTCTTTTCCTCCCAAaagttatactgccctg
gcagtcatatagagcagacgACCCAGAATAGTCAATGTAATCCTGAAAAAGCAGAACAAAaagttatactgccct
gcagtcatatagagcagacgCAAAACTTAACACAAAACTACCGTAATAAAGACAGCATGGaagttatactgccct
gcagtcatatagagcagacgATCAGTGGTATAAAATTGATCCATAAGGCCCAGAAGCAAAaagttatactgccct
gcagtcatatagagcagacgCTGGAATAACTAGATGCCTATGTGTAAAAGGATGAAGTTGaagttatactgccctg
gcagtcatatagagcagacgGGCAATATTTTCTTATATGTAACAAAAAGCAGGAGTGACGaagttatactgccctg
gcagtcatatagagcagacgTTGTTTCCTTCAGAGGATATCATCAAGAAAGAGAAAAGACaagttatactgccctg
gcagtcatatagagcagacgTGATTTCATTCAGCAGTATAACTTTTTGGGGGTGATAAAGaagttatactgccctg
gcagtcatatagagcagacgCAGTAAGCATATGGAAAGGCAGTCTACATTAGTCATTAGGaagttatactgccct
gcagtcatatagagcagacgTGTTGGTGAAGATATGGAAATGTTGGAAGCCTGGTACTTGaagttatactgccctg
gcagtcatatagagcagacgTGGGAAACAGTTGATAGTTACTCAAATTATAGGCATAGAGaagttatactgccctg

gcagtcatatagagcagacgAAAAACTTGCACATGAATGTGTACATAGGAGCCTCAAACTaagttatactgccct
gcagtcatatagagcagacgCAGCCATAAAAAGCAGTGAAGTACCGATTTCTGGTACAGAaagttatactgccc
gcagtcatatagagcagacgAGCCAGACACAAAATACCACCTGTTATATGATTCTGTTTAaagttatactgccct
gcagtcatatagagcagacgAATAGGCAAATTCAAGTGACAGAATGGATTACTGGTTGTCaagttatactgccct
gcagtcatatagagcagacgGTGGAGTATCTGCTAATGGTCTGGGTTTCTTTGGGGGGTGaagttatactgccct
gcagtcatatagagcagacgTGCTTGCATGACCTTGCGAATGTGATAAAAACCATTGAATaagttatactgccct
gcagtcatatagagcagacgTCATGAATTTTATGGTATCTGGGCTACATATCAGTAAAGCaagttatactgccctg
gcagtcatatagagcagacgAATTGTTTATCTTCATTTCTGAGAGTTTTAGTTCCCCCCAaagttatactgccctgt
gcagtcatatagagcagacgGTGCTCTGCCTTGCCCTGGTCCTGCTCAGATTTCAGTAAAaagttatactgccc
gcagtcatatagagcagacgTTGGAAGATTAGAAACCTGAGAAGTTACATGTGAAGTGCCaagttatactgccct
gcagtcatatagagcagacgGAAAAAATATTCTGTGAGCTTTTCGTGTAGGGACAAACTCaagttatactgccct
gcagtcatatagagcagacgCTCTGGCTCTATGTCATTCGGAAAGGCTAAATTATGCTTAaagttatactgccct
gcagtcatatagagcagacgGTGTTAATGGGTTTATTAATAAGGAGGAGTTAAGGTACACaagttatactgccctg
gcagtcatatagagcagacgTGTGAGCTGTATATGTAATTTTCTGTTTTCTAGTACCTGCaagttatactgccctgt
gcagtcatatagagcagacgTCCTCAAAATAAAGTATTTCATACTTACAGCACGTCTCAGaagttatactgccct
gcagtcatatagagcagacgGACAGTGTTAAGAGTGATTTTCATGAAAGTAGATGCAAAGaagttatactgccct
gcagtcatatagagcagacgTGGTTTTGAGTTGGATCCAGATTCAAATTCTAGCTTTGGCaagttatactgccctg
gcagtcatatagagcagacgCATCTGTTCTCAAAAAATCATAGTCTGTTTCTACATAGGGaagttatactgccctg
gcagtcatatagagcagacgGTATCATGTTTTATTGTGCTAATTATTGCAAGACCCAAGGaagttatactgccctg
gcagtcatatagagcagacgCTGGGAAAGAGTTGGTAGTGGTCAGGAAGGGCTTTGCAAAaagttatactgccc
gcagtcatatagagcagacgCTTGGAAGAGGAGGTAGATGAGGACAGAGGCATTCTAGGAaagttatactgccc
gcagtcatatagagcagacgGCAACAGCACTGCCCAGGAGATTGCTTATAGCTTGGTTCAaagttatactgccc
gcagtcatatagagcagacgAGGAATGGGTGGTTGAAAGGGTGGTGTGGCTGAGCTGGAGaagttatactgcc
gcagtcatatagagcagacgCCTACAAGGGGCTTTAGAATGCATTTCATATTAGGCAGTGaagttatactgccct
gcagtcatatagagcagacgAAGTAAGTGATGAGGTGGGATTTGATCTTTCTTGTGTTGTaagttatactgccctg
gcagtcatatagagcagacgAAGTTATTGACCTTTCAGATCTGCCCCCCTTAGTCTCCTAaagttatactgccct
gcagtcatatagagcagacgCAACATTGGGTGATTGAAATCTACTCAGGCTGTTGCTATGaagttatactgccct
gcagtcatatagagcagacgTATGTGAAGCATACCTGATGGTCCAGCTCAGACTCCTGCTaagttatactgccc
gcagtcatatagagcagacgGGTAGTGTCCATCTGTTGCCCTAGCTCCTGTTCTTCATTGaagttatactgccct
gcagtcatatagagcagacgCTCCAGCAGTACCCAGGGAAAGCTTTCAAGAATATAAATTaagttatactgccc
gcagtcatatagagcagacgCCTCAGGGTGGTGTCCAGGAATCTATGTTTAACAAATCTAaagttatactgccct
gcagtcatatagagcagacgCTGTTTCGGGTTTGCATACCAATGTGCAGAGTTCACGTCCaagttatactgccc
gcagtcatatagagcagacgTCTTATCTAGGATTCTGGGCCATTCGGTAACATCAGCTCTaagttatactgccct
gcagtcatatagagcagacgCCTCTTACATCAGCTTCCGTGTACTGTTTCCTGGCCGGTGaagttatactgccc
gcagtcatatagagcagacgTCTGTTTGAGGTTGACCAGTTGGTTCTTAGGTGGCTCTTCaagttatactgccct
gcagtcatatagagcagacgTTGGCCTCATTGAAATGGTACCTGACATCTTCTACACATGaagttatactgccct
gcagtcatatagagcagacgATCTACTTCACAGACTTTAGCCAGTACTCCTTAATTTTGTaagttatactgccctg
gcagtcatatagagcagacgATCTGTATTAACATAAAGTCACGTTCTAACTCCTTTCCACaagttatactgccctg
gcagtcatatagagcagacgCTGTTCTATTTGCAGCTCTTTATCTATGACCATACCCATTaagttatactgccctg
gcagtcatatagagcagacgTTTCTGTTACAATTAGTACTATCACTGCCATCATTGGTACaagttatactgccctg
gcagtcatatagagcagacgGATCCAAATATCTCTGTTTAAAGGGGAAAGTCGCTATTGGaagttatactgccct
gcagtcatatagagcagacgGAATGTCAGATTTACTTTGCTTGCATTTCATTTCATGGTCaagttatactgccctgt
gcagtcatatagagcagacgAAGATTCTGATTGGATGTGAACAAAATGACAGGTTTCATCaagttatactgccct
gcagtcatatagagcagacgCCATGTCACAGAATGCTGATTTTAATCTTACATCTTATCCaagttatactgccctg
gcagtcatatagagcagacgCTAAAAAATAGGGAAAGAGAAATGGGTGGCACTAAAAGGAaagttatactgccc
gcagtcatatagagcagacgGCTCTTTCATGCAAAAACAAAACAATTTATAGTAGGCTGCaagttatactgccct
gcagtcatatagagcagacgACCCTTTGTTATTCCTTCTCAGAAGAAAATGCACTTTGGGaagttatactgccct
gcagtcatatagagcagacgGGTACCCAAATGGCCTTGATTCCACTAGCTTTTACTTCCAaagttatactgccct
gcagtcatatagagcagacgACATTGTGTTGGGGTGGTTGTGATGCTGTTGTGCTGGTGGaagttatactgccct
gcagtcatatagagcagacgGTTGTGGTAGGTCCTAAAATAGATGGGTTACTACATTTATaagttatactgccctg
gcagtcatatagagcagacgGACAGAAGTAAATGTTCCCTTCATTGCCCCATGGACTTGGaagttatactgccc
gcagtcatatagagcagacgGATGCTGCTCCCTGGCCATGCCTGTCCTGAGCTAGTGGGCaagttatactgcc

gcagtcatatagagcagacgCCCACGCTTCCCTCTGCATCTTATTCCTGCATCCATCTGCaagttatactgccc
gcagtcatatagagcagacgGCTCTTGCCTGCTTTCCCTTTCACCTGCCCCTGTTAGAGAaagttatactgccct
gcagtcatatagagcagacgCCCTCAGCTTTTCTTTCTGTCATATCTATGCTCTTGGAAAaagttatactgccctg
gcagtcatatagagcagacgAGTAATTGGGGACTGGGTGGGAAGAAAGAATTGAAAAGATaagttatactgccct
gcagtcatatagagcagacgCAAATGCTGGAGCTAATTCCTTTCCTTTCTTATCTTTTAGaagttatactgccctgt
gcagtcatatagagcagacgATTGATGGTAAGTTAGAATGGAATCAGTTACTCATACCCAaagttatactgccctg
gcagtcatatagagcagacgGCTCAGAAAAATGATGTATGTTGAGCAATAGGTAGTTTAGaagttatactgccctg
gcagtcatatagagcagacgATTGCGATATGTGTAGGTAAAAAAATGGTTGAGTATCAGCaagttatactgccctg
gcagtcatatagagcagacgAGTCAACCTAATAATAGCAGGCATTCTGTGGTGTCTATGCaagttatactgccct
gcagtcatatagagcagacgAAATTTGTCTCTTTTCTACCACTAGCAACTAATCTTCCTGaagttatactgccctg
gcagtcatatagagcagacgTACTGTAGGATTCATAGCTACTCTGGTTTTTTATTGCTTCaagttatactgccctgt
gcagtcatatagagcagacgAAATGGACGTGATTATTCATATATCAGAGGTATTCTTCCCaagttatactgccctg
gcagtcatatagagcagacgGAAAGAAAACTGAAGTCATGTAATCCACTGAGGATTAATCaagttatactgccct
gcagtcatatagagcagacgGTATCTCTAAAATCCTAGACTCAGCTAATGGAAAAAATGGaagttatactgccct
gcagtcatatagagcagacgTATGAAGAGTTTGTGGTATAATTGTTGCATAGATTCAGGGaagttatactgccctg
gcagtcatatagagcagacgTTTCTTTTTAAGGTTTCCACAGCTGGATTGTTTACTAGGGaagttatactgccctg
gcagtcatatagagcagacgTCATTTTCTGGGGAATAGGGTGAGTTCATGTGGATTTCTAaagttatactgccctg
gcagtcatatagagcagacgTTTATTATTCCATTCTATTTGGGCCTGTGATTGTCTCTCAaagttatactgccctgt
gcagtcatatagagcagacgTGTTTACTTCCTTATACTCAAGGTAAATAGCTGGTGGTAGaagttatactgccctg
gcagtcatatagagcagacgTTAAAGCTGTGAGGAATCTAGTTAGATATTGACTGAGTTCaagttatactgccctg
gcagtcatatagagcagacgAGTGATTTTAAATTTGTAGGGGGTGTTTAGAATTGACACCaagttatactgccctg
gcagtcatatagagcagacgGCATGTGTTTAGTAATATTAATGGACAGACATAGGACATGaagttatactgccct
gcagtcatatagagcagacgGGATGTGCCTGATAACTGAAGTTACTGAGAGAAGACGACCaagttatactgccc
gcagtcatatagagcagacgGGGAGGAAAAACGTGAATGCAGTGCAAACAGTCCATTAAAaagttatactgccc
gcagtcatatagagcagacgCTTTAGTAAACAGCCAAGACCACAGAAAGCTTAAAACATTaagttatactgccct
gcagtcatatagagcagacgCTGCTGCTAGCCCATATGACGCTTTGTGCAAATTAGACAAaagttatactgccc
gcagtcatatagagcagacgTGTAAGACATTCACTTCCATCTGATGGGAAATTATTCAGCaagttatactgccct
gcagtcatatagagcagacgTACAAACAACTTTTAAACGAGTAGAGAGATGCTCATGATGaagttatactgccct
gcagtcatatagagcagacgTTGGGATTCAACTCATGATAGACTCTAAGGAAGGGACACTaagttatactgccc
gcagtcatatagagcagacgAGTTTGAGGTTTGGCTTTGATTAGAAACTTGGGTGGCTCAaagttatactgccct
gcagtcatatagagcagacgAACTTACCAGAAGCACAGTAGCTGTAGGTTTGGGGTCCCAaagttatactgccc
gcagtcatatagagcagacgTTAGGGCTAAAACTGGGAAAGAAACACCTAAACTGTGCTTaagttatactgccc
gcagtcatatagagcagacgATGACTGAGTTTCTGCCATGTGGTGATTTATAGTATGTGCaagttatactgccctg
gcagtcatatagagcagacgGAAAGCTTCATTCTATAGACCACCACCTGTGTGATGTCCTaagttatactgccct
gcagtcatatagagcagacgAAGAGATTTGCTTTCAGAATGTCGAGTGTATTCATAATGGaagttatactgccctg
gcagtcatatagagcagacgTTTGATAGATATTGGTTGACAAATGCAGGCTAAATGGGATaagttatactgccctg
gcagtcatatagagcagacgGTATGTGACTGGTGATTGGAAAGTTATTGAGGAAATTAGAaagttatactgccctg
gcagtcatatagagcagacgTAGGGGAAAAAGTCATAGGGTAAGTAGGTGGATGTGGCAGaagttatactgccc
gcagtcatatagagcagacgCATGGTGAAAGTGGTGCAGGTTGTGGGGTGGTGAGGATGGaagttatactgccc
gcagtcatatagagcagacgCCACTGTGGCGCTGAAGACACATGAGAAGGAAGAGATGACaagttatactgcc
gcagtcatatagagcagacgAAGTAAAAGCTTTTGGTTTCTTGGGTGCTGGAGACAAATTaagttatactgccctg
gcagtcatatagagcagacgGAGGGATATTCAAGGGGCCTCCTCTTTGTGGTGATGCAAAaagttatactgccc
gcagtcatatagagcagacgCGTGCCACCACGCCTGGCTAGATTTTGTATTTTTAGTAGAaagttatactgccct
gcagtcatatagagcagacgCGCCCGGCCAAAGTTTGTTTTCTTATTGTATCATTTACCCaagttatactgccctg
gcagtcatatagagcagacgTGAATACAGAAGCACAGAAACATGATCGATTACAAACACCaagttatactgccc
gcagtcatatagagcagacgGTTTACATCTAAAACGACCTCTCCTCTGCTCCCCCAACACaagttatactgccc
gcagtcatatagagcagacgGTGGATTTTTTTTTTAATGAGTGGAGGAAGCAGAAGGAAAaagttatactgccctg
gcagtcatatagagcagacgAGAAGACTTAGATCAGTTAATTCCTCCCACCCCTGCAGTTaagttatactgccc
gcagtcatatagagcagacgGTGAGAATATGAAGACACTTTGGATCTGATGTTTGGCAGCaagttatactgccct
gcagtcatatagagcagacgCAGTAGAGAGAATGATATTCCTTATTAGAAAACACAGACGaagttatactgccct
gcagtcatatagagcagacgCAGAATGATAGGTTTGATGATGATCTGGTATTCTCACTGTaagttatactgccctg
gcagtcatatagagcagacgTGGATTGAAATAATTTATCGCTGTATGGTCCACATAGTTGaagttatactgccctg

gcagtcatatagagcagacgCCTTTATGATGATTCTGTAAAGATGTATATACTCTGCAGCaagttatactgccctg
gcagtcatatagagcagacgTTTTGAAAAAGGGGAGTTAGTGGGTTTTCCTGCAACGAATaagttatactgccctg
gcagtcatatagagcagacgCATGTGCTTGATGTGTGCTTAATGGGATGGAATGACTTTAaagttatactgccctg
gcagtcatatagagcagacgGGTGCTCCTAGTTCAGTACTGTGCTGAGCCCAACACTCATaagttatactgccc
gcagtcatatagagcagacgCCCCTCTGGGACAGGTGCTGCCATGGCTCTTATTTTACAGaagttatactgccc
gcagtcatatagagcagacgAGAGGTGAAGGATGGAGCTAATGTAGCCAGACATGCCTGTaagttatactgccc
gcagtcatatagagcagacgGTCTCCTTGGATCCGGACTGGCTCTACTCTTCAATGTCTTaagttatactgccctg
gcagtcatatagagcagacgACTATGCATTTTAGACCAGCCTGTCTCATCCCCCAGAGCTaagttatactgccc
gcagtcatatagagcagacgAAAATTGAGCAGAGAATTCTATCATTAGAGGACACCAGCTaagttatactgccct
gcagtcatatagagcagacgGGGGAATCCTAGAACTAAATTTGTGCCACATAATGAGATGaagttatactgccct
gcagtcatatagagcagacgTTGGTTGCAATTGGTTTCCCAGAGACCTGAGAGATTCCTGaagttatactgccct
gcagtcatatagagcagacgATTTCGAGAATGTGAAGGAACCCAGCGGTGTGTACATCTTaagttatactgccct
gcagtcatatagagcagacgAAATTATCCTCAAGAATAGATTGGAAGCACCGGGGTCACAaagttatactgccc
gcagtcatatagagcagacgCCTTAGGCAGAAAACACAGGAGGAGAAAAAAGACCAAACAaagttatactgcc
gcagtcatatagagcagacgATTCATAGATCCAGAGCCCAGAATTTTAGTATGTTGGTTAaagttatactgccctg
gcagtcatatagagcagacgTGGATTCAGTAAATGTCACATGGGCAATTCTCTTAAGATGaagttatactgccctg
gcagtcatatagagcagacgCAATTTACCCTGCACCAATCTCTTGTATGTACTGTTATGTaagttatactgccctg
gcagtcatatagagcagacgTTTGATATGAGGCCAGATTTTATTAAACTAGGTCATCAGGaagttatactgccctg
gcagtcatatagagcagacgGGATGGGTCATGCTGGCACTGTGTCGTCTTTGGAAATTAAaagttatactgccct
gcagtcatatagagcagacgGCTGTGTAATATTTGAACATGCATCCAGTTAATGAGGAATaagttatactgccctg
gcagtcatatagagcagacgCCTGGAAAAGTTCCTGTGATACTCATATTATGTCAAAGGAaagttatactgccctg
gcagtcatatagagcagacgCATTCTTAATACAATGTGGATTTCTGCCCTCTCCTTGTAGaagttatactgccctg
gcagtcatatagagcagacgCTGAGCTCACATCAGCCATTTGGTCGTGATTTCCTGGGGCaagttatactgccc
gcagtcatatagagcagacgCTGGGGAGTTTGCATAGGGAGATCATTTATAGTTCTTTTGaagttatactgccctg
gcagtcatatagagcagacgAAATGGCTTTTTAATGTCTGCTTTTGCCTGAAAGGGATTTaagttatactgccctgt
gcagtcatatagagcagacgTTCATTGTACTTAGCGTCCAGATTTTTAAAATGCTGTGTGaagttatactgccctgt
gcagtcatatagagcagacgGTGCGTGTGTGTGTATTTAATGTTCAGTTGCTAATATGGGaagttatactgccctg
gcagtcatatagagcagacgAAAGGCCTTTTCAGAAGTACTCGTCCACCCCACATGGCTTaagttatactgccc
gcagtcatatagagcagacgTTTGGCACATAGGGAGGGAGGAAGGGGTCTGCAGCATTTGaagttatactgccc
gcagtcatatagagcagacgGAGAGAGAGAGACGGAGTCTAACTCTGTAGCCCAAGCTGGaagttatactgcc
gcagtcatatagagcagacgCCAGGGTGGTCTCGAACTCGAGCTCAGGAGATCTGTCTGCaagttatactgcc
gcagtcatatagagcagacgAAGCAGTCCTCCCATCTCAGGCTTTTGAAGCTAAGACTATaagttatactgccct
gcagtcatatagagcagacgCTCAAGCTGATCTCAAACAACCATCCTTCTGCCTCCGCCTaagttatactgccc
gcagtcatatagagcagacgCCACCGCTCCTGGCTAGCAACATTTTCTAATCTTTCTGCTaagttatactgccctg
gcagtcatatagagcagacgGTGGGGATTTTGTCTGTTTATTCACTACCATTTTCCTGACaagttatactgccctgt
gcagtcatatagagcagacgAACATGCATTGGACACTAAGGAAATGTTTCTGGAATGAATaagttatactgccctg
gcagtcatatagagcagacgCATTTGAAGCTTGTGTCTCTGGGAAGTAGAATTCTTTTTCaagttatactgccctgt
gcagtcatatagagcagacgACAAATTTACCTAGTCTTTTGCTTATTCTTTCAGCAGCACaagttatactgccctgt
gcagtcatatagagcagacgAGAGATCAAGGAGAGACACTTCATTTAAGAAATTGGAAGGaagttatactgccct
gcagtcatatagagcagacgCCTTTATGGTGTAATACTTATTTAATGGAAGGACTGGCTTaagttatactgccctgt
gcagtcatatagagcagacgGAGGTCCCTGTTATGTTCCCTCTCTGTTTTGTTTGTTTTTaagttatactgccctgtg
gcagtcatatagagcagacgACTTGCTATACATGTTGCTCAGAATAAGAGGGTACGTAAGaagttatactgccctg
gcagtcatatagagcagacgAAAGAAGGACGATGTAGACTTTGGTGACACTCTGGGTCATaagttatactgccct
gcagtcatatagagcagacgCTCTGGCATTTTTTAGTGTGTGACTTTACTTATGTTACAGaagttatactgccctgtg
gcagtcatatagagcagacgTTACCTTAAAAGGGGATGGTACCTGGCTTGCAGCTCTGTAaagttatactgccct
gcagtcatatagagcagacgCACTCAATAATATAACAGTGCCTGCCACAAAGTAAATGTTaagttatactgccctg
gcagtcatatagagcagacgTGAAGCCTCATTTAAGTCTCCTTTAAGAACTTGACATTACaagttatactgccctg
gcagtcatatagagcagacgATATATTCCTCCTTCAGAATAGGGCATATTAAACAGAGTCaagttatactgccctg
gcagtcatatagagcagacgGCTGCTTGAGTTATAGCCTGGCACTGTGGATCGTGTTTAGaagttatactgccct
gcagtcatatagagcagacgACATAACTGCACTGATCTGAACTAGTTTGTTAATGAACGAaagttatactgccctg
gcagtcatatagagcagacgGGGAAATCTAAACTAAGTTTTTGAGGTCTCCCAATCCAGGaagttatactgccct
gcagtcatatagagcagacgTTTGGAGTTTAAAGGAATCGTCAGGTCTCTGTAATAACTAaagttatactgccctg

gcagtcatatagagcagacgGGCTAATTATAAGTTCTAAGAGGCATTTCAGTTCACTCAAaagttatactgccctg
gcagtcatatagagcagacgCACTTCTCAGTCCATCAAGTCAAAAGCTGTGTGTGTCTTTaagttatactgccctg
gcagtcatatagagcagacgGAAATTATCAGGGCAGGTTAATCTCCACAGACTATTATAGaagttatactgccctg
gcagtcatatagagcagacgTAATATGCCACTTACAGTGCGTTTTTAATAGACTGCTTGTaagttatactgccctg
gcagtcatatagagcagacgGAAGCAGTTCTTGTATATGCATTCAGAAAACAAAGTAGTCaagttatactgccctg
gcagtcatatagagcagacgAAACAATGAATCTTAATAGAAGAGGGTATCAGGACTCATCaagttatactgccct
gcagtcatatagagcagacgAATTGATAGTCTATGAAACTCACTAATTTGTCCTAGCTCCaagttatactgccctg
gcagtcatatagagcagacgTTTTTCATACCTCATGCATCCCTGTACCTTGTATTATAGTaagttatactgccctgt
gcagtcatatagagcagacgCTTCTTCAGTGGTGTAATCCATGTCTTATTCATTCCCAATaagttatactgccctg
gcagtcatatagagcagacgAATACTTGGTGAATCAATGAGGATTTTCTGAGCAAAGTACaagttatactgccctg
gcagtcatatagagcagacgGCTGGGTTCATTGTTTGTGGTTCTCATAGTTTTAATTGGGaagttatactgccctg
gcagtcatatagagcagacgGGTTCCTGCTGAGCCAGTGTGCTGTATTGTATTTAGGATGaagttatactgccctg
gcagtcatatagagcagacgGAAGCCTGCCAGAAAATCACCTAAGGTAATTATTTTTGACaagttatactgccctg
gcagtcatatagagcagacgGTCAATATCACTTGAGGTTTTTTTTTGGTTCCTTGGATTCaagttatactgccctgt
gcagtcatatagagcagacgGGTGGAGTATAAGTTTTTGCCTCCCAGATAGAGGGACACAaagttatactgccc
gcagtcatatagagcagacgTTGGATTTCTCAGTGCAAAGGAAGAGGGAGGGGAGTAGTCaagttatactgccc
gcagtcatatagagcagacgGCAAAGGCAAGAGGATCTCTAGAAATGGGGCTCAGTGGGAaagttatactgcc
gcagtcatatagagcagacgAGGATAACTTATTGTGATTTTGTACCTCTCCCCCTTGCCTaagttatactgccctg
gcagtcatatagagcagacgCCGCTGCATCCTCTGTCTTGTTGTGGCTCTGGCAAATTGAaagttatactgccct
gcagtcatatagagcagacgCCTCATTTCCTTTCCTGTGTGGCCTAGTTCTTCCATCTGTaagttatactgccctg
gcagtcatatagagcagacgTCTTGATAGCATCTTCCTAGACTTTGTTTCATCTTGGCTTaagttatactgccctgt
gcagtcatatagagcagacgCTGGCTGGTCTCAAGTCTTGCTAGGGCACTGTCCAAGGAAaagttatactgccc
gcagtcatatagagcagacgTGATGCTGTCTCAGAGTTATACTTCCTGATGCTGGATGGAaagttatactgccctg
gcagtcatatagagcagacgCTCAGACTTGGGAGGAGAGCGGGAAGGAGGGGAGTAGCTAaagttatactgcc
gcagtcatatagagcagacgGCTCAGCCCTCCTGTCATCTGACCTCACTATACACTTATCaagttatactgccct
gcagtcatatagagcagacgGGAATATAAGACCTGTCCCACACTCTATGTTTTATCCACAaagttatactgccctg
gcagtcatatagagcagacgTATATATATTGAATGCCTGCTTTGTGATAATCACTGTGCCaagttatactgccctg
gcagtcatatagagcagacgTGAGAACGTCAAAAGGAATAAACTTTGGTTTCTGCTTTCCaagttatactgccctg
gcagtcatatagagcagacgAAATAGTTTAAAGGAATGTTAATGGGGAAACACAGAGGTGaagttatactgccctg
gcagtcatatagagcagacgCACAAGCCCCTTTTCTCCATCCTTCCTAAGCCACTCATGGaagttatactgccc
gcagtcatatagagcagacgTTGGGGTCAGCCAATATTGGTTTGATTGTGATGTTTTATAaagttatactgccctgt
gcagtcatatagagcagacgCAAACTTGTTGGAATGTTAGTGGTACCTGGTGTGTGAGTGaagttatactgccctg
gcagtcatatagagcagacgGAGAGATGAATATTTCAGAAGTATTTTCAGTAACAAGGGGaagttatactgccctg
gcagtcatatagagcagacgTAACTTAGAAGAGTCTTTCAGTAAACATTCCCGTTATCACaagttatactgccctg
gcagtcatatagagcagacgATAATTGATATTTCTAGCCCCACCCTATTCCCTGCCAATCaagttatactgccctg
gcagtcatatagagcagacgGAGGAGATATTTTTGGCTCGAATGGTCTCGATGTCAACCTaagttatactgccctg
gcagtcatatagagcagacgGCTGTCATAACTGGGGACACTGTGCCACTAGCATCTAACAaagttatactgccc
gcagtcatatagagcagacgGATAGTTCCCCACAAAGGATCTTACCACCCCAAATGCCAGaagttatactgccc
gcagtcatatagagcagacgCAGTCTGGATGAATAGCCTATATTGCTGCATGGTCTTCTTaagttatactgccctg
gcagtcatatagagcagacgAGATGAATAGTTTCGATGATCACTTAGCATGTTTTGGAGAaagttatactgccctg
gcagtcatatagagcagacgCAGACCCTGTAACCTAGTCCACAATTCATATGTATTTGTTaagttatactgccctg
gcagtcatatagagcagacgCCTGAAGATAGTAGCATTTTATTTGTTTTCCAAACACTCCaagttatactgccctg
gcagtcatatagagcagacgTGGAAAATATGTGCCGGAGTTTATGAGAGTTATTTCAAAGaagttatactgccctg
gcagtcatatagagcagacgAGCAAGTGTTAGGTCAGTAAAAATAAGACACCTGGATCTCaagttatactgccct
gcagtcatatagagcagacgAACAGGCAAAAACACAGCAGAAGTTCATGAAAGCCCAGTTaagttatactgccc
gcagtcatatagagcagacgGCACCAAAAAGCAAACTCAACACAAACCTCTAATAGGAAAaagttatactgccc
gcagtcatatagagcagacgGGGCATCATACAGAGTCCTTAGAGGAACTAGAAACTGAGCaagttatactgccc
gcagtcatatagagcagacgGCACAGACAGATATAACTACAGCCAGCTCAGAAATTACCTaagttatactgccc
gcagtcatatagagcagacgGTGCTTGACTTCCCAATTTCTCTCTCACCGTTTTCCTGCTaagttatactgccctg
gcagtcatatagagcagacgGCTCCAGCCAGACTTTCCTGCTTCTCCTCTAAACATGCCTaagttatactgccc
gcagtcatatagagcagacgTCTGTCTAGAATGCTGTTTTCCCTCTCCTCTCTACCTCTGaagttatactgccctg
gcagtcatatagagcagacgTCCAAGACTCAGCTCAAGTTCAGCAGCCTGTATGAAACCCaagttatactgccc

gcagtcatatagagcagacgAAAAAGTCTTTAACAATCTTATTTGGCCCTCTCTTACCCCaagttatactgccctg
gcagtcatatagagcagacgGGTAGTGTCATGTTTTATCAGATTCCATTCTATAGAAACCaagttatactgccctg
gcagtcatatagagcagacgGCTTCGAAACGATGAATGGGACAAATGCAGCTTTTCATGGaagttatactgccc
gcagtcatatagagcagacgGAGAGCATCTCCCTTTGGTTCATCAGGGCCCACTGGATGGaagttatactgcc
gcagtcatatagagcagacgAACAGTCTGCTCGTTGGTATAAGGTCCTTTGAAGGTTCTGaagttatactgccct
gcagtcatatagagcagacgCTTACCTGGACAAAGAAAGAGACCTGAAGGAGTACTTTTAaagttatactgccc
gcagtcatatagagcagacgTGACATTAGACAGATACATTCCCCTCTGTATAGGGCCCCAaagttatactgccc
gcagtcatatagagcagacgCATGGAAAGTGTGTGTGTCTCTTGTTGTGCTCTGATCTTGaagttatactgccctg
gcagtcatatagagcagacgTGGGCTAGGCATTCTGTCAGTCTCACCAGGCACCCTCTCTaagttatactgcc
gcagtcatatagagcagacgTTCAACCCAGAACTTTGCAATATTGTGTAGAATTCCCAGCaagttatactgccct
gcagtcatatagagcagacgAAAAACATAAATTGCCCAGAGGGTATGCTTAACATTAAGGaagttatactgccct
gcagtcatatagagcagacgAACAGTAGAGGGAGAAGGGACTTGTGAAAATGAAGTTTAAaagttatactgccc
gcagtcatatagagcagacgCATTCTTTGTGCAGGATGTCTTAAGGATGTCTCTAAACCAaagttatactgccct
gcagtcatatagagcagacgCTTTAGACAGTTGATTCATTTTCTAGCAAAACTCAGACAGaagttatactgccctg
gcagtcatatagagcagacgTACCTTGATTTTTGCATTCAGAATTCAGTTGCCTGTTTACaagttatactgccctgt
gcagtcatatagagcagacgTTTTAGATGAAAATTTCCCCTTTTTGTGTGCAGCTGAGTGaagttatactgccctg
gcagtcatatagagcagacgCGTTTGCCAGAAGGGAGGGGAATATCGCTGTTAAATGAGGaagttatactgccc
gcagtcatatagagcagacgGTTAAAACATATTCTGAAAAGGATGGAAGCTTGTTCTTGCaagttatactgccctg
gcagtcatatagagcagacgGCTGTTTGTTCTGTGAGGCTTTGAGGTGTCTCTGAGGGCTaagttatactgccct
gcagtcatatagagcagacgTGCTCTATTTTGTCTTGTTTCTTAGCAGTAAAGCCTTTACaagttatactgccctgt
gcagtcatatagagcagacgGTGATAAAAATATCTGGAGTGGACCTAAGAGAAATAAGAGaagttatactgccct
gcagtcatatagagcagacgAACATTCACATAATTCAAATGCAGGCCTGGTGATTCTGAAaagttatactgccct
gcagtcatatagagcagacgGTAAGCACATACATTTGTCCATTGACCGTGATGGAATTCAaagttatactgccct
gcagtcatatagagcagacgCCATGGCATTGGCTGATCTACAAAGAAAAGGAATCAGAGTaagttatactgccc
gcagtcatatagagcagacgATTAAGAAAACATGTACTGAGTGACTACAACATAGTGGGCaagttatactgccct
gcagtcatatagagcagacgTGTCAGAGATACGAAGATATATATGATATGGTTCTTGCCCaagttatactgccct
gcagtcatatagagcagacgTTAGAGTCTAAGAGGAAATATAGATGAGTATAGATCCGGTaagttatactgccct
gcagtcatatagagcagacgTCTGACCAGGGTCCTGAAGCACGCACAGTTCCAGAGGCCCaagttatactgc
gcagtcatatagagcagacgGGAGCACCTGGTAGAGAGATGAATGTGTATGTTGAAGGGGaagttatactgcc
gcagtcatatagagcagacgAGTGAGGATCCCCGAAGTGTCAGTTTCCGCCCCGGCCCTGaagttatactgc
gcagtcatatagagcagacgTCTGAGTGGGTAGGAGCACTTGAGCAGGACTTAGCGCCCAaagttatactgcc
gcagtcatatagagcagacgGTGGCATTGTAGGTGTGTGCTGTTTCCACTCTTCCCTCCTaagttatactgccct
gcagtcatatagagcagacgACAGTAGCCAGCGTGAGTCTTTAAAACCTAAATGAGATGAaagttatactgccc
gcagtcatatagagcagacgCTCTCATGTTCACACTTGCTTTCCATCACACCGGGACTTGaagttatactgccc
gcagtcatatagagcagacgCATCTATCTCCCTCCCCCTCGCTCCTGATGCCTCCTTGCTaagttatactgcc
gcagtcatatagagcagacgCTTGCTGTGCCCTCTGCTTGCAGGATTACTCCCTCCTATTaagttatactgccc
gcagtcatatagagcagacgCTGGCTCTTCTCCACCTGCTCAAATGTCCGTCCTCAGAGGaagttatactgcc
gcagtcatatagagcagacgACTATCTGGTGCCCCTCTGTGTTCCCTGCCCTCTTTATTCaagttatactgccct
gcagtcatatagagcagacgAGCATTCTCCGTACATTACATACTCCTTTGTTCATCTGTCaagttatactgccctg
gcagtcatatagagcagacgAGGATGACTGTATGGCTCTTATTCTCCGTTATTTCCCCACaagttatactgccct
gcagtcatatagagcagacgTAGAAAAAAAAGTGAGTGAAACTAATACAGACATGGTGGGaagttatactgccc
gcagtcatatagagcagacgTGTCTTCAACAGTTTGTGAAAAGGGAGGCTAATGGTCTTCaagttatactgccct
gcagtcatatagagcagacgAACTTACAGAATTTTAAAATTCCTTGGAAAAAGGCTGGGCaagttatactgccct
gcagtcatatagagcagacgGGGTGACAGAGCGAGCCTCTATCTTAAATACGTACATACAaagttatactgccc
gcagtcatatagagcagacgTTCATTCCTTGGAAAAAACTTTCATGCAAGATTCCTATGGaagttatactgccctg
gcagtcatatagagcagacgTGGCATCTGATGATCTGAAAAAGGTCTGAGATTGTATTCTaagttatactgccctg
gcagtcatatagagcagacgCTTAAATGTTCTGTTGTCTTAAATGCCTTCATGTTGGATCaagttatactgccctgt
gcagtcatatagagcagacgATTTAAATCTCGAAAAGTGGGAAAGGGAGGGGACAGTTCAaagttatactgccc
gcagtcatatagagcagacgTGAATGTGTGGTAGGCGGGGAGTTTAATTTTCAAGAGGTAaagttatactgccct
gcagtcatatagagcagacgGTTAACTTTATGAAAATGTATTGATCCTGAAAGCAGCAGCaagttatactgccct
gcagtcatatagagcagacgAATTGTTAAAGTGATCTCTTGTTCTCCATGTTCCTACTGGaagttatactgccctg
gcagtcatatagagcagacgGAGGCTCCTAAAATTCTGCTGGGGGTGGCTGATGAGACAGaagttatactgcc

gcagtcatatagagcagacgTATTTTGCTATAAGGACTGACACAGATGGCCTCACTTGGGaagttatactgccct
gcagtcatatagagcagacgGGTGTCTTGAGGCAGCTGGTGGAAGTTCCATTGTGTGGAGaagttatactgccc
gcagtcatatagagcagacgACACCTTCATACCCTTGTCCTTTTTGCTGCATATGTGTATaagttatactgccctg
gcagtcatatagagcagacgCACAAAATACAGAAGTGGCTTAAAAACCAGAGATTGACCAaagttatactgccc
gcagtcatatagagcagacgAAAAAGTAAAAGTGCAAATGAGCATCAAAGAGGTGGCAGAaagttatactgccc
gcagtcatatagagcagacgTGTTTGTGTGAGAATTGCCTTAGCCAAATAAAAACAGATGaagttatactgccctg
gcagtcatatagagcagacgGGCCAAGGAGTTCAAGATCATCAAGAAGATTATTTGAAATaagttatactgccctg
gcagtcatatagagcagacgGCTAGTATTATAACAGTGTATCTTGTGCTATCTGTATAGGaagttatactgccctg
gcagtcatatagagcagacgAGTAGTTACTGGTCAGAATACATGTAATCTACAATGCATGaagttatactgccctg
gcagtcatatagagcagacgACACTTGTATTGGGGGATGCATGCTCAAAGCTTCTTACAGaagttatactgccct
gcagtcatatagagcagacgACTTGCTATTCAAAGCGTGGTCTGCAAATGAGGCATCAGCaagttatactgccc
gcagtcatatagagcagacgAGAATCTCAAGACTTACCCTAGACCTCTGGAATCATAATCaagttatactgccct
gcagtcatatagagcagacgTGCACATTACAGTCTGGGAAACACTCCTTTAGGGAATATTaagttatactgccctg
gcagtcatatagagcagacgCTTCTGTGCATCTGAAACCAGGAAGTCATTTCTCTCCACAaagttatactgccct
gcagtcatatagagcagacgCTGATTTTTACTTCTAGTTATGTTGCTTACAAACTTGGCCaagttatactgccctgt
gcagtcatatagagcagacgGCTTAATGACCTGCTGAGATTCTGTTCCCACATCTGTAAAaagttatactgccctg
gcagtcatatagagcagacgTCCTTGCCTCCCAGAAGATTTATAAGGGTTGTCTGAATGTaagttatactgccctg
gcagtcatatagagcagacgAACCACCGTTCAGTTTTGCCTTTTCCCCTCTGCGTTGTTAaagttatactgccctg
gcagtcatatagagcagacgGCACAGAGATCTTTAAACTTTTGCTCATGTACCTCTTAAAaagttatactgccctg
gcagtcatatagagcagacgTAAGAAATAAAACCAGCTATATTCCGCTTTTCAGTCTAGCaagttatactgccctg
gcagtcatatagagcagacgGAGAATGTCTTCCCTCCCACTCTCTGACTTCCGTGGCGTGaagttatactgccc
gcagtcatatagagcagacgTAGCAGCTTAGTATCCTGGCAGTTCTAGCTCCCACTGGATaagttatactgccct
gcagtcatatagagcagacgGATGCTCTGGTGACACATCCCCTCCTTTTGTCCCTTTAGCaagttatactgccct
gcagtcatatagagcagacgGGTGGTGCTAGCTTCCAGCCCTTACTTATCTTGGTGTTTCaagttatactgccctg
gcagtcatatagagcagacgCCAGCTCTTCCTTCATTGGTTTAGTCAGTTCCCTGCAGTAaagttatactgccctg
gcagtcatatagagcagacgACCAGTTTTTATTGATATGCCGATACGAATTGTGCTTTTGaagttatactgccctgt
gcagtcatatagagcagacgATTCTTTAGTAGCATAATTCTATCATCTCACAGGCACTAGaagttatactgccctg
gcagtcatatagagcagacgCCTGACGCTCTCCCTCCCCCTAGACTGCCCCACTACTGACaagttatactgcc
gcagtcatatagagcagacgGTGTGTCGTTCCCCTCCCTGTGTCTGTATGTTCTCATTGTaagttatactgccctg
gcagtcatatagagcagacgTGCTATTGTGAATAGTCCCGTACTGAACATACACATGCATaagttatactgccctg
gcagtcatatagagcagacgTTGCTGGGTCAAATGGTATGTTTGGTTCTAGGTCTTTGAGaagttatactgccctg
gcagtcatatagagcagacgTTTGCGTTTCTCTGATGATTGTCACTGGAAGGTTTTATACaagttatactgccctgt
gcagtcatatagagcagacgGAAAGATTCAGGATGTGTGAGGGGTATTCTTCTCTTTAAGaagttatactgccctg
gcagtcatatagagcagacgGAAGAAGGGTAGTTGTGAGAGAAAAGATGTTCCCAGGCAGaagttatactgccc
gcagtcatatagagcagacgAACTAATTCCCTTAAAATGAGCACCTTTGAAATGTGTCTGaagttatactgccctg
gcagtcatatagagcagacgGCTGGACACATTGCACATTTAGGCACACTAAGTATTGTTTaagttatactgccctg
gcagtcatatagagcagacgGAAGTCCCAGATTGTGCATAAAATCTTTTTTAGCAGTGGTaagttatactgccctg
gcagtcatatagagcagacgCAGCCAACAATGTGGGATGCATGCTCAGTCTGACACATCCaagttatactgccc
gcagtcatatagagcagacgTCATAGCTGGTCAGACTGGGTGTCTCCGAGGGCCTGGCGCaagttatactgcc
gcagtcatatagagcagacgGATAGAGCCACTGGGAAAGCAGGCAGGATCTTGGAAGCCCaagttatactgcc
gcagtcatatagagcagacgTGGAAACAGGGCTTCTGTTGTCACTTTGCCAGCAACAACAaagttatactgccc
gcagtcatatagagcagacgCAGAACTCTGCCGCTTATCTGGGGAAATCTCTCTGACAGAaagttatactgccc
gcagtcatatagagcagacgGCCAACAAAACTGAAAATGGCCTAAGGAGGATTTAAAAAAaagttatactgccct
gcagtcatatagagcagacgGTAGAACAGAAGCCTCCCTACCCCGCCAAAATAAAGAGTCaagttatactgcc
gcagtcatatagagcagacgGCTTTTTGCTGTGATTTTGTTAAGTTGCAGTATATCCGTAaagttatactgccctgtg
gcagtcatatagagcagacgTCTCCCACAAGAAACTCCACTAAAACTCCAAACCAAAAAAaagttatactgccc
gcagtcatatagagcagacgCTAACACAACTTCGTAGCCACTGAGTCAAAACATCTGAGCaagttatactgccc
gcagtcatatagagcagacgCGCACATTCACCCCAGACACTCTAAGGAAGTGAGGAGCTGaagttatactgcc
gcagtcatatagagcagacgAGGGAAGACCTCGGCATTGAGTGTATTAAGACAGGTTAAAaagttatactgccct
gcagtcatatagagcagacgGCTATTAATACTGTTTCAAAGAGCCATCTCGAGTATAGAAaagttatactgccctg
gcagtcatatagagcagacgGGCTTCTTACAGGGAATGTTGTACCAACTGAGAAAATTTAaagttatactgccctg
gcagtcatatagagcagacgAGCCTTCTTAAATGTAGAAACTTGACCATCCAACTAGTGAaagttatactgccctg

gcagtcatatagagcagacgCTTTTTAAAAATCTGAAAGGCTAAAGTCATAGGGTCCGTGaagttatactgccctg
gcagtcatatagagcagacgGATCAGTCTTTAGGAAGATATCATGTGGGAAATGGAGTAGaagttatactgccctg
gcagtcatatagagcagacgCAAAATTGGCTCCTGAAAGAAAAGGTAAAGATACTGCACAaagttatactgccct
gcagtcatatagagcagacgGGGTATCCAGGAAGAAAGAGGGTTTTCATATCAAGAGGCAaagttatactgccc
gcagtcatatagagcagacgTGAAAGTCAAACTACAGAACTGCCCTTTTCAAGTGTTTTAaagttatactgccctg
gcagtcatatagagcagacgATGGAAATACATTTGACATAATATGGAAGCAGTGAAGGACaagttatactgccctg
gcagtcatatagagcagacgATGGAACTTTGATGATCACTATGTAAGAAAGTACTTAGGGaagttatactgccctg
gcagtcatatagagcagacgAACAAATCAAGGTCAGGAGAGATGCCTACTTATGTTCAAGaagttatactgccct
gcagtcatatagagcagacgGGTTCACTTCGGTATTTCCAGCATTACCCATCATCATCTAaagttatactgccctg
gcagtcatatagagcagacgCCATCTACCTTTCCATTCTTTCCAGAGATCCTTTTGAGTCaagttatactgccctg
gcagtcatatagagcagacgTCCTGGACTGGGCACTAAGAACACTGTAGTGTATAAAGCCaagttatactgccc
gcagtcatatagagcagacgTTAAATCTCATTAAAAGCACAAGTGTACGATTGCAACAGCaagttatactgccctg
gcagtcatatagagcagacgCAGTGTGGTGCTCTGGATATAAAGTAGAGGGATTTGACCTaagttatactgccct
gcagtcatatagagcagacgCCTTAGAGGCTAGGTCTTCATATGCTTGGGAGTTTCAGGAaagttatactgccct
gcagtcatatagagcagacgCTTATAGGAAGGATTCTTGAATGAAGGTATCTAGGATTGAaagttatactgccctg
gcagtcatatagagcagacgTTAGTTTATGTAAGTGTATTATACTGGGGTTATGGCTCAGaagttatactgccctgt
gcagtcatatagagcagacgAAAGCATGGAATTGCAAATTTGTCTTTTTCACTGACCTGCaagttatactgccctg
gcagtcatatagagcagacgTTATTTAAACTTGTATCTCTACTCTGTCCCTGGCTCTCCTaagttatactgccctgt
gcagtcatatagagcagacgCCAGGCAGGTAGGACAGGCAGAACATTGTTGTCATTATCTaagttatactgccc
gcagtcatatagagcagacgTTGGAAGTTGACAATAAAATTACAGCACGTGCCCACCATCaagttatactgccct
gcagtcatatagagcagacgCTCAAACTTAGCACTTAGAGAGGGGACATATACTTTAAAAaagttatactgccctg
gcagtcatatagagcagacgTGCATTAATTTCTCCAGAGGCATGTACAAGGCTTGTTGGAaagttatactgccctg
gcagtcatatagagcagacgGTGTGGAGAGGGGACAGGGAGTAGGGTGTATGTTTGTCGGaagttatactgccc
gcagtcatatagagcagacgTGGATGAGTGACAGAGTGGGTGGGGATGAAGTTCAGATAAaagttatactgccc
gcagtcatatagagcagacgGAGTTGAGGCCAGATCTGTCAGTTCCTTCTCCAGTGCACAaagttatactgccc
gcagtcatatagagcagacgTCAGGTGGAGAGAGGAGTGGGAAACGGTAAAGACAGCCAAaagttatactgcc
gcagtcatatagagcagacgCCAGGAAGGGAAGCAAGAATAATTTGGTAGCAGGAGACGGaagttatactgcc
gcagtcatatagagcagacgCAGTCATCACATTTGCTCTCTCCATAACATCCTGAATTAGaagttatactgccctg
gcagtcatatagagcagacgCTAGCTGCATTTTACAGAAGGAGAAGTTGAGGCCTAGTGGaagttatactgccc
gcagtcatatagagcagacgAACCAGGTTGGACTTGACCCTGTGTTGCCCTGTTCAGTGGaagttatactgccc
gcagtcatatagagcagacgAACTGTGGGCTGCTTGTGGCATTGTCCCTGGTTGGATGTCaagttatactgccct
gcagtcatatagagcagacgCCATAGTGGGCAGGCATGGCTGGGTTCTTCAGAGATGTAAaagttatactgccc
gcagtcatatagagcagacgCCTCATCCTTGCGGTCCAGTGACTTCTGTGGAAAAATGTGaagttatactgccct
gcagtcatatagagcagacgCCATTCGTAAGCTGGCCCTATCTGAATCCTCCTTTACTTTaagttatactgccctg
gcagtcatatagagcagacgTGGGGAATTAGGCATGTGCTGGCTGCTTAGGCCCTGTGTGaagttatactgccc
gcagtcatatagagcagacgGTAATGAAGGCTCTCATGACTTTGCAGAACTCAGTGTCGTaagttatactgccct
gcagtcatatagagcagacgGTAGAAGTATCCATTATTAGACTTAAGGACAGAGGTACATaagttatactgccctg
gcagtcatatagagcagacgGCAAAACATATTTAACTTATGCAGGATGATGGGGTAGACCaagttatactgccct
gcagtcatatagagcagacgGTGTTTATTTAAGGAGCTGCACAGAGAAACATACAGGAGAaagttatactgccct
gcagtcatatagagcagacgGAATATGGGAGATTTGTATGTTGTTCAGGGACAAAGTTTGaagttatactgccctg
gcagtcatatagagcagacgGCCCTTCCTGTAGCATCTTCCCCTGAGGCCTGACGAGAGAaagttatactgcc
gcagtcatatagagcagacgAGAATTAATCTTACTGATCAGTACACTCCATCCTGCCTACaagttatactgccctg
gcagtcatatagagcagacgGCTTGGGTTGAGGCAGTGTTGAAAGGAGGGATTGATTTATaagttatactgccctg
gcagtcatatagagcagacgAAGTGGTAGATAATCTTTGGCAGGATGGATTTAGAATTTGaagttatactgccctg
gcagtcatatagagcagacgCTGCAGGGAGAGCTTCCATTTGGAGATAAAATCAAAATACaagttatactgccct
gcagtcatatagagcagacgGGTCGTCCTCCCTTAAACAAATACTGTTGTTTCTTTTTTCaagttatactgccctgt
gcagtcatatagagcagacgCAGCATTTCTAATAGTCACTCTTTTGCCAGCTTTTTGTAGaagttatactgccctgt
gcagtcatatagagcagacgAGGTTGTTTGACTTTCTTCCCCCTAAGAGGTTAGTATCTGaagttatactgccctg
gcagtcatatagagcagacgGGTACCATCACCAGGCCGGAATGTGGGATTTGGAAAGGAGaagttatactgcc
gcagtcatatagagcagacgCAGTGTCATGTGGAAAATGCGGCTAGGAAGGGGGCTATCTaagttatactgccc
gcagtcatatagagcagacgGCGGCAGTGGAGTGCACCTTAGCAAAACATCGCCACATGCaagttatactgcc
gcagtcatatagagcagacgTCTGACTGACCGCACCCTCCTTGCTCCCATCCACCTCAGAaagttatactgcc

gcagtcatatagagcagacgCTGTGTGTGCTGGATATTGGAGGAACTAGGGCAAGTCATTaagttatactgccct
gcagtcatatagagcagacgCATATATATGTCAACTCTCCTCCTGTCACGCTCAGTTTTTaagttatactgccctg
gcagtcatatagagcagacgACAGTGTATGTCCCTCGTTTTGCATTGTACCTGTTGTAGGaagttatactgccct
gcagtcatatagagcagacgTCAGGCAGACAGTCCCTTGCTAGTTCCATCAGTAACAAATaagttatactgccc
gcagtcatatagagcagacgATCATGATGGTAACCAGTCTGTTAGGGAATCTTATCAGCCaagttatactgccct
gcagtcatatagagcagacgTGCAGGGCAGGGACCAGGCTGGTAGAGGGCAGGGCAGCTTaagttatactgc
gcagtcatatagagcagacgGGCAGTTTCCCAGTGTTGTGGCATTAAGTAGAGAAATTAGaagttatactgccct
gcagtcatatagagcagacgTGGAGTTAGGATGGTACGTACAATGGGAAGTCCTATGTGAaagttatactgccc
gcagtcatatagagcagacgATTTAACAATGATTGCTGGAGTTAGAGAATGAATCACAGGaagttatactgccct
gcagtcatatagagcagacgCAAAATTATTGCTCTGCATGTAGTTGAAGGCTGTGGGGCCaagttatactgccc
gcagtcatatagagcagacgGCATGGGTACAGTTCTGTCATTTAATGACTTTGTGTAGGCaagttatactgccct
gcagtcatatagagcagacgCCTTAATGGTGTTAACATAGCCCAGGGATTGTCTTGAAGAaagttatactgccct
gcagtcatatagagcagacgCAATAGTTGCAGTTACTTTAAGGCCACAATGTAATCTTTGaagttatactgccctg
gcagtcatatagagcagacgCAATCAGGAGTCTAGGGTCTTTGTTCAGTGGTGTTTTTACaagttatactgccctg
gcagtcatatagagcagacgGGCACATGGTAGGCACTTGGTTGGTACTTACTGGCTGTGTaagttatactgccc
gcagtcatatagagcagacgGATATTCCCCGTATCCAGCAGCCTGAGGAGTGCCTGTGTGaagttatactgcc
gcagtcatatagagcagacgTCTGGAGGCGTGAGTGGAGGTGGCGACCCTGTGGAGTGTTaagttatactgcc
gcagtcatatagagcagacgGAAATGCCCTCCTGTGCCTGGTTTCCCTGACCCGTCCCTAaagttatactgcc
gcagtcatatagagcagacgGAACCCCCATACGCTCCTTCCCTTCAATTTTGCTGCAATaagttatactgccct
gcagtcatatagagcagacgAACCCTTTTTAGTTTTCACCCTTAACATAGGAGAGAGAGAaagttatactgccct
gcagtcatatagagcagacgATGCATGTGTGTATACCCCTTGTCTATAGATACCCCCTTGaagttatactgccct
gcagtcatatagagcagacgCACATGAATAGAAAATGTGTGCATGACAAAGGAAATAGCAaagttatactgccc
gcagtcatatagagcagacgGAATGGGCTTATTCAACAATATTGTCACATGGTTGGTTTGaagttatactgccctg
gcagtcatatagagcagacgCCTGATAGGATATACCCAAATTCTGCATGATTACATAGTAaagttatactgccct
gcagtcatatagagcagacgGCAGGAACGCAATCAAACATTAGAAAAGCCAGAAGAAAACaagttatactgcc
gcagtcatatagagcagacgCAAACCAAACCAAACCCTCTACACACAAGTTGTTGATTGTaagttatactgccc
gcagtcatatagagcagacgGTGGAAAATTGGAAACATCTTATGTGCTCAGAGATTGGGGaagttatactgccct
gcagtcatatagagcagacgCCCTTGGTGAATTCTCCAGTCATTGAAAGTGATCATTTTGaagttatactgccctg
gcagtcatatagagcagacgCATCATAATAGATATTTCCCAGTAGAGTCATGGTTTTGTGaagttatactgccctg
gcagtcatatagagcagacgGACATTATATACTTAAGTGATTACCTTTAGTCTGCTGAGGaagttatactgccctg
gcagtcatatagagcagacgGAAAATAACTCACATGCTGTCTCATCTTTTGTTTCGAAACaagttatactgccctg
gcagtcatatagagcagacgTTAAGCTTCATCTTTATCCAGAGAAAGTTATAACAGAGCCaagttatactgccct
gcagtcatatagagcagacgGAACTTTCTTCTGCATTCTTTCCTACTTCTAGCTTTCCTTaagttatactgccctgt
gcagtcatatagagcagacgCCAGGGCTCTCAGAAGCTGGTCTACAGGCTTTGTCCCCAGaagttatactgcc
gcagtcatatagagcagacgTGTTCCTTATTAAAATCCAGACGCTTTTCTGTTTCAAACCaagttatactgccctg
gcagtcatatagagcagacgGGTCCTCTGCTAGCCTCCCACTCGGTTACAATTTGTTGGTaagttatactgccc
gcagtcatatagagcagacgGGTGTGTCAAGCCTTGTCAACGTTCTCATATAGCCCGAGGaagttatactgcc
gcagtcatatagagcagacgAGACGTAAGAGGGTATTAGTAAACACCGTTTCTTTCTGCTaagttatactgccct
gcagtcatatagagcagacgCCTGCGAGTGAGCTTTGTATGCCTGTCTTCCTAGGTTTGCaagttatactgccct
gcagtcatatagagcagacgGGAGGCTTACCAGTGAGGAAAAAACACATCTTAGCAGAAGaagttatactgcc
gcagtcatatagagcagacgGTGCTGCAAAGTGCTTCCGCATTCTTCTGAAAATAGATGTaagttatactgccct
gcagtcatatagagcagacgGTGTCACCTTTTCTTACTGCAGCTTCAAACCCCTGAGCCCaagttatactgccc
gcagtcatatagagcagacgCGCTCAGCCCCAGATGTGTTTTAATATAAGTTGATTTTCAaagttatactgccctg
gcagtcatatagagcagacgGCTGTGGGCTAGGAATCACTAATTACATCCCTGCAAATTGaagttatactgccc
gcagtcatatagagcagacgTCCTCTTCTACCCCTTTGCCTACTTGAAAATCTATGAAGAaagttatactgccct
gcagtcatatagagcagacgGTGAATTTCTAACCATCTGTACCTCCTCCCCTACACATTAaagttatactgccct
gcagtcatatagagcagacgGGATGCTCCCTTAGAAACATACCAATTGTGAGTTCATACTaagttatactgccct
gcagtcatatagagcagacgGTATGTTATAAGGGTCCATGAAAAATAAAAGGAAAGTGGGaagttatactgccct
gcagtcatatagagcagacgTGATTTCAGGATATCATGTTAGGCCATAGTGTGGGAATGTaagttatactgccct
gcagtcatatagagcagacgAGAGACTGACACCTGCAGCTCTTGAAATTCTCAGGCCTCCaagttatactgcc
gcagtcatatagagcagacgCATGTCGGTGAGGGAGGAGCAAATGAACAAATTTCAAACAaagttatactgccc
gcagtcatatagagcagacgTAGCTGAAGACCATGAATTTAAAAACCCTATGATGATAGCaagttatactgccct

gcagtcatatagagcagacgAGTGGCCAAGCTAAATTAAGAAGGTTAAAAAGGTGTGGGCaagttatactgccc
gcagtcatatagagcagacgTTTACTAAACAAAAATACCCCACTCAGATTCTTCACACCTaagttatactgccctg
gcagtcatatagagcagacgTAGGTCATTACTTAGTTCTGTTGGGCATTTTCACTTCAGTaagttatactgccctgt
gcagtcatatagagcagacgGGAGACCCATACATTGGTGACTGGTAAATGTGAACCTTGGaagttatactgccc
gcagtcatatagagcagacgAGTCTGTGATCACTAGCTTATTTTCATGAGTATTGTCCTTaagttatactgccctgt
gcagtcatatagagcagacgATAGTTGATGTTTTTTGTTTAGTAGAGACATAGCAGCTCGaagttatactgccctgt
gcagtcatatagagcagacgACATCTAAAATTCTTTCAAGTGACTGCAAATGCATGTGTGaagttatactgccctg
gcagtcatatagagcagacgACGTGTGTGTGTCTATTCAGCTGATCTACCTGTCTGGGAAaagttatactgccctg
gcagtcatatagagcagacgGGCTCCTACGACCCTTGAGATTCTGGTTGAGTAGCTCTGGaagttatactgccc
gcagtcatatagagcagacgAGGATTCTGTGTCTCTAATAAGCACTGTTAAAGATTGGGCaagttatactgccctg
gcagtcatatagagcagacgGTGGTACAGAGGTCCATGATTTAAAGAAACATCGCTTCAGaagttatactgccct
gcagtcatatagagcagacgCATGGTGCTAATGGAGTCCTTTTTATTTTCATGGTGGTTCaagttatactgccctgt
gcagtcatatagagcagacgCAGTTCTTTGGGTCAGGCCTGCACTCATATGTCAGAAGCAaagttatactgccc
gcagtcatatagagcagacgAAATTGCCATGCTTTTCTCAGCCATTTGTGCTTCCAAGCGaagttatactgccctg
gcagtcatatagagcagacgCTTGATAATGAGAGGCCGAGCTTAGTTATGCAAATTGTTTaagttatactgccctg
gcagtcatatagagcagacgTGCACATTTTAAGCAGGGAGAAAATACAACTTGTCCTGTGaagttatactgccctg
gcagtcatatagagcagacgCATGGATGGATTATGGCTTGTCTGGGGGTATTTCTCTTGAaagttatactgccctg
gcagtcatatagagcagacgGCCCATTGATTGAACCAGCAGAGAAGGAGCAGTATTATTGaagttatactgccc
gcagtcatatagagcagacgGGTTTTCCCTATGATGACATTCTGGGTTGACTGGCCACACaagttatactgccct
gcagtcatatagagcagacgCTGTTGCAGGTCTTTGTAATTGCTTTTCTCTTATGATTTCaagttatactgccctgtg
gcagtcatatagagcagacgAGACAAACTGTTCTCTGGAAAACAATACTAGAATCAAACCaagttatactgccct
gcagtcatatagagcagacgAGAATGTTTTTTAGGTTTCTCACTTGTCTGTAAAAGGCCAaagttatactgccctgt
gcagtcatatagagcagacgTCATACTTGGGGGAGGGCTGTTCACTGATGCTGTATTTAAaagttatactgccctg
gcagtcatatagagcagacgCAGGATTTTCGTGCCTCCCTGGGTAGGTGGTTCTGATGCTaagttatactgccct
gcagtcatatagagcagacgGCGAGAAGGTGTTTGCTCTATGCATGGGAGCACATGTGAGaagttatactgccc
gcagtcatatagagcagacgTGCTGATTACTGCACTATCTCGTTGTTTTGACCCTGTGAGaagttatactgccctg
gcagtcatatagagcagacgGGATCACCATAAGCTGTTTGACTTTTTCTTCTCTCTTGTGaagttatactgccctgt
gcagtcatatagagcagacgTGCTTATACAGGTGTCCACGGAGAGCATGATAGTAGCGAAaagttatactgccc
gcagtcatatagagcagacgAGCTTCTTGTTGTGTGGTCTTTGTGCTTTTTATTATTGGTaagttatactgccctgtg
gcagtcatatagagcagacgGCATCTGTGCTTCTTGGGTTAGAAATATTCTGTATCTAAAaagttatactgccctgt
gcagtcatatagagcagacgCTGATTCTTGGTTTTAAGCCAGGTTTCAGAAGTTATTCTGaagttatactgccctgt
gcagtcatatagagcagacgATCTTAAAGATGCCTTTGTATAAGATTTAGAGCACGGTTGaagttatactgccctg
gcagtcatatagagcagacgTGAATTCAGCCTTTATTTTTGGTTGAGTACTAGACAGGTTaagttatactgccctgt
gcagtcatatagagcagacgTGTCTACCATACCTGCATTTTAAAGTCACTACATCCTCCTaagttatactgccctg
gcagtcatatagagcagacgAAATTCTGATAAAGTTTTTAAGGACAGCAGTAGGATCGTGaagttatactgccctg
gcagtcatatagagcagacgGTTTAAGTTGCATAAATAATTCCTAAATCCCTGCTGAACCaagttatactgccctg
gcagtcatatagagcagacgAGTTTTAAATCATACTTACCCTATTTCACTGGCAGTTGGTaagttatactgccctgt
gcagtcatatagagcagacgTGGAAGGGTGGAATCATGGTGAGAGAATGTAAGCTGGTCTaagttatactgccc
gcagtcatatagagcagacgAAGGACCTGTGCTCTTTAATGTTAAGCAAATACTTCTATCaagttatactgccctg
gcagtcatatagagcagacgCCTTGGGAATATGTAACCAAAGCTCTTTTTTTCTTCTCTGaagttatactgccctgt
gcagtcatatagagcagacgAATACAAAGTGAAATTTCTAGACCCACTGAATGTACTAGGaagttatactgccctg
gcagtcatatagagcagacgCCGGAGAAGAATGTAATTTAGAACCTCAGTAAATCTCTTGaagttatactgccctg
gcagtcatatagagcagacgCTGGTTTTCTTTTCTGCCAAATATCAGCTTGTATTGCTTAaagttatactgccctgtg
gcagtcatatagagcagacgCTCACATGTTTAAACTCTATTACAACTCTTAATCTGTGGGaagttatactgccctg
gcagtcatatagagcagacgCAGAGAGTGCAGCATTTTATTGTAGGTCATGAGTCTAGGCaagttatactgccct
gcagtcatatagagcagacgCAGCTTTCACACCTGCATCCCGTGCCTACTGTTTGCCAAGaagttatactgccc
gcagtcatatagagcagacgCAGCTCCTGTTCTTTTTTCTGAAATACTGTACAAGGATCCaagttatactgccctg
gcagtcatatagagcagacgAAGATGTGTTGCTTGCTCAGACTCTAGGGCCCTGATGGTTaagttatactgccct
gcagtcatatagagcagacgGGGAATCCTGAAAACACCATCACAAATAATTAGACAATTGaagttatactgccct
gcagtcatatagagcagacgTTTATCTGCTTATAACAAAGATGAAGGGATTACAGGGCTGaagttatactgccctg
gcagtcatatagagcagacgAAAGATGTCAGCAGCACTTTCTCACCAATTCAGAACCTAGaagttatactgccct
gcagtcatatagagcagacgAGTGTTATTCTTCGTGATAGGTTTATCCATCTGTTTCTTGaagttatactgccctgtg

gcagtcatatagagcagacgACCTTTATTTCTGCAGTGGTTTAGCTCTCACCCTGAGTTAaagttatactgccctg
gcagtcatatagagcagacgCACTGGTGTTTTTTCTTCGTTAGTAAGCTCTTTCTCAGCCaagttatactgccctg
gcagtcatatagagcagacgGCCACAGCCTCCATGTTATATTGCAGATTCTGGTCACATCaagttatactgccct
gcagtcatatagagcagacgGGTGTTCCTTTTGTCCAGTTTGTGTCTGTTTATGGTGCTCaagttatactgccctg
gcagtcatatagagcagacgGACGGACTTTCATCAGCAGGAGAATTCAGGCCTTACAAAGaagttatactgccc
gcagtcatatagagcagacgTGAATGCATCCTTAGATTTTTGGATTTCAGGAACCAGATCaagttatactgccctg
gcagtcatatagagcagacgTTTTGACAATCCAGTGGTTTTAAAATAGCCTCAGTATCACaagttatactgccctg
gcagtcatatagagcagacgACCATCAGTGTCATTCTCTCCACGTTTTGTTGCCCATAGTaagttatactgccct
gcagtcatatagagcagacgCCAGAAAAGCAAGAGAACTATAATTTTAGACTATCAGTCCaagttatactgccct
gcagtcatatagagcagacgTATATAAGGTCCCTATCTGCCATCTTTTAAACATTTCACCaagttatactgccctg
gcagtcatatagagcagacgGAGTGATAAAAAGTTCATGTCAGACTAGACTTTGGGAAGCaagttatactgccct
gcagtcatatagagcagacgTGAGTCAGATCCCTTGTGCTCCTCTGAGTGATCCCATTCCaagttatactgccc
gcagtcatatagagcagacgAGGTTCTCAGTCTCCCTTTACTCTTAGGTGCAAGCACGAAaagttatactgccct
gcagtcatatagagcagacgGGCGGACATTTAACTTGTATCCAGAGCATTGCCAGAAACTaagttatactgccc
gcagtcatatagagcagacgATTTGATAATCTGCACCCCATTCACAAAGAAGAACCATTTaagttatactgccct
gcagtcatatagagcagacgAGTTCTTTGAAATCTTGACCCTGGCTTCCACTCCTACACTaagttatactgccct
gcagtcatatagagcagacgCCTGCTGTGGGTTTCTGAGTTCTTAAAGCTAAGTAATTAAaagttatactgccctg
gcagtcatatagagcagacgGCTCTCTCAGGTTCCACTGAATTTAACTAGATAACTCATTaagttatactgccctg
gcagtcatatagagcagacgATGACTTCACATTGACCTCGATCATAGGGCCTTTTCAAAGaagttatactgccct
gcagtcatatagagcagacgTATTCGTAAAGCTTTGTTAAAAGTAGGTAGTAAAGGCCAGaagttatactgccct
gcagtcatatagagcagacgGGTGGATCACGGGGTCAAGAAATAGAGACCATTCTGGCCAaagttatactgcc
gcagtcatatagagcagacgCAAACAAAAAAAGTAGTAAAACCAAAAGCGAAGTGAGCAGaagttatactgccc
gcagtcatatagagcagacgTAATTGCACATATTCATGGGTATGGGTCATACAGTGATATaagttatactgccctg
gcagtcatatagagcagacgACACATTGTTGGTAACTATCAGACTCTGATTTCCTCATACaagttatactgccctg
gcagtcatatagagcagacgAGAAGGAATCCCAATGAAATATCTCTTACTGTTTCAGTAGaagttatactgccct
gcagtcatatagagcagacgTTGATATTATATTCATTATCCACCTGGCTGGACAAACGGCaagttatactgccct
gcagtcatatagagcagacgTGGTAAGGCAGGCAGATGACAGAGTATGGGAGGGGGAAGGaagttatactgcc
gcagtcatatagagcagacgACTCCATGTCACAGCCTCTCCTATCATTTTTTCAGTGAAGaagttatactgccct
gcagtcatatagagcagacgGGTTGGCCCATTCCCCATCTCCTTACATTGACTGCCCAGGaagttatactgcc
gcagtcatatagagcagacgTGTTTGATTAAATTTGCAGTTGATCAGGTGTGGTAGTTAGaagttatactgccctg
gcagtcatatagagcagacgAGGTCCTGAGAGAGATAGTAAGTTGTTTTGAAGCCAGAGAaagttatactgccc
gcagtcatatagagcagacgAGTGGAAGAACTGCAGAGCTATAATTTGTTTAAGTATGGGaagttatactgccct
gcagtcatatagagcagacgGATTGCTTTGAAATCCCCCTCTCTTCCTCCTCCCCATGAAaagttatactgccc
gcagtcatatagagcagacgCCCAGTTGTTCAGTTTTAGTTCTTGCCACAGCTTGCACATaagttatactgccct
gcagtcatatagagcagacgCAAAGGAGGGAATTTTGGTGCATCATTATCAAATATCTACaagttatactgccct
gcagtcatatagagcagacgACTGGGCTAAGTGTTGGGAATAGAAAGGTATGAGAGACATaagttatactgccc
gcagtcatatagagcagacgCAACTACTGTCATGTGCTTTATCTTGTGTAATCCTTATAGaagttatactgccctg
gcagtcatatagagcagacgTTTCTTAACTTCTGGAAGAAAGGACACAGTGTTGGTAGCAaagttatactgccct
gcagtcatatagagcagacgCCACAAGCCCTTCTATCTCCTTATGTTATTTCAGCTGGAGaagttatactgccct
gcagtcatatagagcagacgAGCTGGAAGGTTTTGACGTGTCCTAGTTATGAGGAGCTTTaagttatactgccct
gcagtcatatagagcagacgGGTGTGGGCATCTGGACTTTATTCTGGGATTATAGTGGCTaagttatactgccct
gcagtcatatagagcagacgTGGATAGGAGAGAAATAGAAGAATGAGAGAGAAATTACAGaagttatactgccc
gcagtcatatagagcagacgAGTGCTTACTATATACCAACTACTCTATTAATCACTCAGGaagttatactgccctg
gcagtcatatagagcagacgCTTGTGGAGCTTTTATTTTAGTGGTAGGAGTCAATCGTTAaagttatactgccctg
gcagtcatatagagcagacgCAGGATAAGGAGATAGGATTGCTGAAGTGGGGGGATTACTaagttatactgccc
gcagtcatatagagcagacgACCTGTGTCATGCAAAAGAGCCCACCATTCCAATATCTCAaagttatactgccc
gcagtcatatagagcagacgCAAGTGCAAAGGCCCGGTGGAGGGATCAAGGATCTACAGAaagttatactgcc
gcagtcatatagagcagacgGGCAGATCACAATGGCTTGGGACCTCATGCTAAAGGATGGaagttatactgcc
gcagtcatatagagcagacgTGTGGCAGGACATTGGAGCTTTGTAGGAAGAGAGGGAAGCaagttatactgcc
gcagtcatatagagcagacgGCTGACTCATGTTTTAAAATGTTGACCATGGTTATAAGTGaagttatactgccctg
gcagtcatatagagcagacgTCTGGGTAAGAGACAGTGGTATGGACCTTGGTGCTGGTGGaagttatactgcc
gcagtcatatagagcagacgTGGAGGAGGAGTGGATAACTTAGAGGATGTTTTAAAAGATaagttatactgccct

gcagtcatatagagcagacgAATTCATGGGGGGTGTAAGGGAAAGAGAGGAATTGAACAGaagttatactgccc
gcagtcatatagagcagacgAGGTTTTTGTCCTGAGCATTTGAGTAGATACGATGGCTTTaagttatactgccctg
gcagtcatatagagcagacgCATTGGGGGAATACTTGGAGAATAATAGGTTTTTGGGAAAaagttatactgccctg
gcagtcatatagagcagacgCTGAGATCCGTATTAGGTGTCTTGGTTCATTTTGAGTTGCaagttatactgccctg
gcagtcatatagagcagacgAATATTTATTTGGTGCATTATTCTGCTGGATGGAAGACTGaagttatactgccctg
gcagtcatatagagcagacgCATGGCAAGAGAGGAAGTAACAGAGTGGGTGGGAGGGAGCaagttatactgcc
gcagtcatatagagcagacgGCAATCAGCTCTTGTGGGAATGAGTACAGTGAGAACTCACaagttatactgccc
gcagtcatatagagcagacgCTAACATCTCCCATTAGGCCATACCTCCAACATTTGGGATaagttatactgccct
gcagtcatatagagcagacgACAAATATTCCAACTATAGCCATCCAAGGAGAGAACTCTGaagttatactgccct
gcagtcatatagagcagacgTACAAACCTCAAATACACAATTCCTTCAACAGACTAACAGaagttatactgccct
gcagtcatatagagcagacgCTGACAGCTGACTTCTCAGGAGAAACAATGAAAGTTAGAAaagttatactgccct
gcagtcatatagagcagacgTTAGACAAATGAAAACTGAAAAGCTGTATTTCCCACAGACaagttatactgccctg
gcagtcatatagagcagacgTGGAAGTGTGGAGATGTGGGATTGAATAATATATTTCGGTaagttatactgccctg
gcagtcatatagagcagacgTGTTTGAGGCACATTCTGTTTACTCTGTCTGATGTTTTTCaagttatactgccctgt
gcagtcatatagagcagacgGAGAAGTTAACCCACTCTCATCATTTAGATAGAAGCACAAaagttatactgccct
gcagtcatatagagcagacgTTCCCTGAAGCCTTCTCTGAACCCTGTCTAGTTTAGAACTaagttatactgccctg
gcagtcatatagagcagacgCCAAATATTACTAGAGTGTTACACACAACATTGAATAGGGaagttatactgccctg
gcagtcatatagagcagacgAAAATTAGTTCCCACTCAGACTTTTTGCCTTAACACTAGTaagttatactgccctg
gcagtcatatagagcagacgTGGGTTTCAAAGTAAAATCACACATATTGCCAGCTATGCAaagttatactgccctg
gcagtcatatagagcagacgTGATGTTGAACTCCTAGAATCACATGCATTGGTAATGACTaagttatactgccctg
gcagtcatatagagcagacgTGGCAGAGTTATTAATAATTGAAGAAGCCAGAGTGGACATaagttatactgccctg
gcagtcatatagagcagacgACTATCATTTGGTAATGTTATCCTGCTCTCTAATCAACACaagttatactgccctg
gcagtcatatagagcagacgCTAATTTTTGTATTTTTAGTACATCTTGGCCAGGCTGGTCaagttatactgccctgt
gcagtcatatagagcagacgTGAACCACCATGCCCCGCCACTTAATTTCATTTTTAGCCAaagttatactgccct
gcagtcatatagagcagacgATAGCTTAAAACTATGGGAATCTTTTGTTGTCTCTCATGGaagttatactgccctgt
gcagtcatatagagcagacgCTAGGTGGCTCGCTTTTAGGATCTCTCAGACAGTTGTAAAaagttatactgccctg
gcagtcatatagagcagacgTAGGTGTGGCTAGGCTGGAATGGCTCACTTATATGGCTGGaagttatactgccc
gcagtcatatagagcagacgCTTGGGGCCTCAGTTGCTTCCCATGTGTGTCTTTCTGTGGaagttatactgccct
gcagtcatatagagcagacgTCCCCAGAGAGCAAACCGTCCAAGAGGCCAAGGTAGAAGCaagttatactgcc
gcagtcatatagagcagacgGACGTAACATCAGAAGTCACACTGTCACTTTTACCAACTTaagttatactgccctg
gcagtcatatagagcagacgCTGTGGGATGGAGCTATGTAAGGACAGGAATACCAGGAGTaagttatactgccc
gcagtcatatagagcagacgTCATTGTGAATAATCTTGAGGGTTGGCTGCCACATAAGTTaagttatactgccctg
gcagtcatatagagcagacgTTTTGTTTTGTTTGAAGTAGATTCCCTCTAAGAGGTCTTGaagttatactgccctgt
gcagtcatatagagcagacgGGATCTCCTACACCAGTCTTTTGTTCATATTCTTACCACAaagttatactgccctg
gcagtcatatagagcagacgGAGGTATTTTGTTGTCTCCCCTTTAGAAAAATGGTTTGTGaagttatactgccctgt
gcagtcatatagagcagacgAAACTTAGATAAAGCACCAGATAGCACAGTACTTGGTACAaagttatactgccct
gcagtcatatagagcagacgGGTTCTTGATTAGAGAAGCTCCAAACTTAGATAAAGTGGAaagttatactgccctg
gcagtcatatagagcagacgTTTGTTTCGTTTTGTTTATTATCCTGGCTATGCCCTTACAaagttatactgccctgt
gcagtcatatagagcagacgTTGTAGTCAACCTATGTCTTCAGTGTTCTGTATTCATGTTaagttatactgccctgt
gcagtcatatagagcagacgTTTTAAAAGAACTGTTAGTATATGTGCTGTTGTGGAAGGGaagttatactgccctg
gcagtcatatagagcagacgCAGCCCACATAAGTAAATCTGCACTGTTGAATTGTATGGAaagttatactgccctg
gcagtcatatagagcagacgTCAGCTGAATTCTGTAATTGATCATTTGGTGTTTCTCAAGaagttatactgccctgt
gcagtcatatagagcagacgCCATTTTCAACAGCTTGGGATGAGTAAGGAAGATAAAATTaagttatactgccctg
gcagtcatatagagcagacgCTTTTAATTTAAACATTGATGAGACTTGGCTGGGTAGCGGaagttatactgccctg
gcagtcatatagagcagacgGCTGCCTGCACACCATTATTCTGAAATGCTGCTATTTAAAaagttatactgccctg
gcagtcatatagagcagacgGGTAGCTCCTCTAATTTTCTGCCTTTGCCTCTGCCTTTCTaagttatactgccctg
gcagtcatatagagcagacgTGTTAGTCAGTGCACAATTTCCTTATGGATTCCCAGTTGGaagttatactgccctg
gcagtcatatagagcagacgGTGGACACAGCATCACAGAGAGCCTAATACTGAATATTGGaagttatactgccc
gcagtcatatagagcagacgCTCCCTAGATTAGAGATTCTGAAGCAAAGAGACCATGTATaagttatactgccct
gcagtcatatagagcagacgTCAGAGGTTGGTTTCAGATGTTTACACAGCATATCCAGTAaagttatactgccctg
gcagtcatatagagcagacgTTGGCACACATAAATATGGACACTGAGCAGTACGGCAGGGaagttatactgccc
gcagtcatatagagcagacgCAGATTCATTTCCATGTGACTGTCCTGGCTCTCTTCTCCAaagttatactgccctg

gcagtcatatagagcagacgATGGCTGCAGCAGTTCCAAAACTCTCTTCTCAACACACCGaagttatactgccc
gcagtcatatagagcagacgAAGAGAGAGTTGTTTCTTGAAAGCTGCAGAAGATCGAGGAaagttatactgccc
gcagtcatatagagcagacgTTTTCCAAAAGTGTCCATGAAAGTTATCACCCTCTTTGTGaagttatactgccctg
gcagtcatatagagcagacgCACTAGTGATGAATATGGAATTACCCTTGGATCGGTCAGAaagttatactgccct
gcagtcatatagagcagacgTGAACAGAATCAGGGTTTCATTAGGAAGGAGGGAAAACACaagttatactgccc
gcagtcatatagagcagacgACAATGTTCCTTATCCTCTTTGTTATGTCCCCCATACTTAaagttatactgccctg
gcagtcatatagagcagacgCTTCTGATTTAATTTCTGAGTTCTATGATTGCTCGTTTCCaagttatactgccctgt
gcagtcatatagagcagacgGAACTTTAAAAAATACTTATTTAAGGCTGGGTATGGTGGCaagttatactgccctg
gcagtcatatagagcagacgAGATCAAGACCACCCTGGCTATGATGAAACCCCGTCTCTAaagttatactgccc
gcagtcatatagagcagacgGAAAAACAGAACTTATTTAAGTAGCCCGTTGTTATTTCGCaagttatactgccctg
gcagtcatatagagcagacgTATATTCAGATAACTTTGCTTGCATATTACAGTCCAAGGGaagttatactgccctg
gcagtcatatagagcagacgCAGTAGAAGTATAACCGAGAATGTCATATTATAGTCATGGaagttatactgccctg
gcagtcatatagagcagacgAGGAGCCATAAAACCTCACATTACTCATCGAAACTTGTTCaagttatactgccct
gcagtcatatagagcagacgAGAAGTGAATTTGGTGTGAGATTACTATAAATCCTGTGACaagttatactgccctg
gcagtcatatagagcagacgATGAATCCATAAGAAGATACTCAGAAGCTTTCATGCTGATaagttatactgccctg
gcagtcatatagagcagacgGATGCCATAATCTGTAATTAAATTCTTCTATTGCTGGTGCaagttatactgccctg
gcagtcatatagagcagacgATAAATTTAGCATCTGGATTCCTTAGCTGATTCTCTATGCaagttatactgccctg
gcagtcatatagagcagacgAGAGGTGCTGGCTTGTTTGTTTTATCATATCCGAAAAATGaagttatactgccctg
gcagtcatatagagcagacgTGAGTTCAGGGCATGGGCTGAGGTATTTTCTTCCATTATGaagttatactgccctg
gcagtcatatagagcagacgGCGTTTGAGTAGTGGCTGGTGGATTAGGGTGCATGCTTTGaagttatactgccct
gcagtcatatagagcagacgAACTTTTTGGTGACGTTTAGTCTAATTTGGTGACATTTCGaagttatactgccctgt
gcagtcatatagagcagacgCCCCAAATACTTTCAAACATTGAGCTTTGTGGCTTAAAAAaagttatactgccctg
gcagtcatatagagcagacgCATTTTTTGTTTTTTCTTCAACTCTTAAGTCCTGGGGTACaagttatactgccctgt
gcagtcatatagagcagacgATTAAGTCTTGCATGCATTAACTGTTGTTCCTGAGGCTCTaagttatactgccctg
gcagtcatatagagcagacgACTGTACTTATAGGCCCTAGTATGTGTTGTTCTTCTCCCTaagttatactgccctg
gcagtcatatagagcagacgCATGTGTTCTCCTTGTTCAGCTCTCACTTACAAGTAAGAAaagttatactgccctg
gcagtcatatagagcagacgCCAGGTCTTTGCTATTGTGATAAATGCTGCAGAAAACATAaagttatactgccctg
gcagtcatatagagcagacgTATAGTAGAATGATTGCTGTTCCTCTGAGTATATACACAGaagttatactgccctg
gcagtcatatagagcagacgCTTCTGCAATGGTTGAACTACTTACACTCTCACTAACAGTaagttatactgccctg
gcagtcatatagagcagacgAGCATTTCTTTTTATTTGCAACCTCACCAGCATCTGTTGTaagttatactgccctg
gcagtcatatagagcagacgTCTGACTGGCATGAGATGTATCTTGTTGTGGTTTTGATTTaagttatactgccctgt
gcagtcatatagagcagacgTTCCTTAAGTGCCTGCTGGTGGATAGGAAAACTCAAGTGAaagttatactgccct
gcagtcatatagagcagacgACCTTTTCCTCAATCATTTATGAAACCAGTGGGAAGCAATaagttatactgccctg
gcagtcatatagagcagacgGAACCTGAACGTAAGGTCATAGTATGTGGTAGGGCAGGGCaagttatactgccc
gcagtcatatagagcagacgGGGGCTCCATTTTACTTTTTAGAAGGTGTATTTCATTGGTaagttatactgccctgt
gcagtcatatagagcagacgACTTCTGACCTATTATTTTGTGAGCTGGTTATGTGAACGGaagttatactgccctg
gcagtcatatagagcagacgGTTGAAATCTTGACCTGGGGCCTCTCCTTGTTTGAAAATAaagttatactgccctg
gcagtcatatagagcagacgACATCATTGCTGTAAATTTGAGAGAGTGAATTTCTCCTGAaagttatactgccctg
gcagtcatatagagcagacgATCAAATAATAAAGCCCCTTGATGATCTAGCAGAGGTTGCaagttatactgccct
gcagtcatatagagcagacgGATGGGAGGGAAAACTCTCTAAAGTGAGGTCAAAGAATTTaagttatactgccct
gcagtcatatagagcagacgGTGTGCAACTGATACTGATATTAGAGATGTATCTAAGTGTaagttatactgccctg
gcagtcatatagagcagacgACCCAGGTAGAAGTGGTTGAATTGATCCAGGTCAGAGAGAaagttatactgccc
gcagtcatatagagcagacgAAGAAGTTGAGGGTATTTGTGCTAATAATATCTGAAGTCGaagttatactgccctg
gcagtcatatagagcagacgTGAGGCAAAGAAATAAAGCTAGAAAGGACCTGATATATTGaagttatactgccct
gcagtcatatagagcagacgTCAGCTACTGAAATACTCCATTCACCATTTCATTACGTATaagttatactgccctg
gcagtcatatagagcagacgGTTGATACTTCTTTTCACCAACATGTTGTCTCACTGTTCTaagttatactgccctg
gcagtcatatagagcagacgAAGGAAGAAATCTGTATTCATTATGCTGAACTCACACCAAaagttatactgccctg
gcagtcatatagagcagacgCTCATTTCCTTCCTGTCTACTGGTTATTGTTTCTTATTTGaagttatactgccctgt
gcagtcatatagagcagacgTTTATTAAGATCAGAACAAATCAAGCAAGGTCGATTCCTGaagttatactgccctg
gcagtcatatagagcagacgAGTTTGTTGTTCTATCTTGGATTTTGTGAAGTGGAAAAGGaagttatactgccctg
gcagtcatatagagcagacgCCAGGACTCGTGGTAAACAGTGGTGTATTCATGCTAACAAaagttatactgccct
gcagtcatatagagcagacgGGAAAATTTGGGAAACTGTAAAAAGGAAAACCTAGTCATCaagttatactgccct

gcagtcatatagagcagacgCTCAGCTGTAACCACAGGTAATATTTTGAGTATACCAGATaagttatactgccct
gcagtcatatagagcagacgAGTGTTAGGGATTATTTTAGTAGGTTTCAGCTATTCCTGaagttatactgccctgt
gcagtcatatagagcagacgTGAAAAGATGAGGCAGACACGATTTTACCCATGGTTAACAaagttatactgccc
gcagtcatatagagcagacgGCATGGTCTCTTTTTGAGAGAAATCTTATAAAAGGTGGAGaagttatactgccct
gcagtcatatagagcagacgTGGAAACACAGAGGACCTAGAATCACCAAGATACTTTTGTaagttatactgccc
gcagtcatatagagcagacgTTTAATTTTTAGACTTGGGGGGTTTGTTACGTGGATATACaagttatactgccctgt
gcagtcatatagagcagacgACCCAATAGATAATTTTTTAAGCCCTCACTGCCCCTGCCTaagttatactgccct
gcagtcatatagagcagacgCCAGTGTCTATCGTTGCCATTGTTATGTCCATGTGTACTTaagttatactgccctg
gcagtcatatagagcagacgTGTTGCTGCAAAGGACATTAGTCCGTTCTTTTTAGTGGCTaagttatactgccctg
gcagtcatatagagcagacgTGGACACTTAGGTTGATTCTATGATTTTGCTGTTGTGAATaagttatactgccctgt
gcagtcatatagagcagacgCCCTTTTCTCTGCATCCTTCAATATCTGCTGTTTTTTTTAaagttatactgccctgt
gcagtcatatagagcagacgGAAATGGCCAAGACAGTTTTGAAAAAAGATGACAGTTTTGaagttatactgccct
gcagtcatatagagcagacgAGAATAGAAAGTCTAGAAATGAGCCTACATATTTATGGCCaagttatactgccct
gcagtcatatagagcagacgTTTTTGACAAAGGTATCAAGGTATTGCAATGGAGAAAGGAaagttatactgccct
gcagtcatatagagcagacgCAAATGGTGCTAGGACAATAGGATATCCATGTGCAGAACAaagttatactgccc
gcagtcatatagagcagacgTCAATCCTTCCCTCACACCATGTGTACAAAAATTATTTCAaagttatactgccctg
gcagtcatatagagcagacgGATTACAGGTGCATGCCATCCACCTGGCAAATTTTTATATaagttatactgccct
gcagtcatatagagcagacgTAGATATAACACCGAAAGCATGGCCCATAAAGGAAAACATaagttatactgccc
gcagtcatatagagcagacgAAACAGACTTTTGGAGCTTGGTGCAATGGCTCACACCTGTaagttatactgccc
gcagtcatatagagcagacgCTCTACAAACAAAACAAAACATTACCTGGGTGTGGTGACAaagttatactgccc
gcagtcatatagagcagacgAATGAAAAGGCAAGCCACAGTTAGAAACTTTTGCTCTAAAaagttatactgccct
gcagtcatatagagcagacgAAGACATGAGCAGACTTTACAGACATTTCAGTGAGTAGGAaagttatactgccc
gcagtcatatagagcagacgTGGCAAATAAGACACGAAAAGATGTTCATATCTTTACCCAaagttatactgccct
gcagtcatatagagcagacgCAGAATGGCTAGAATAAAAATAGCGAGAATACCACATACTaagttatactgccc
gcagtcatatagagcagacgCCTGGATCACTGATGTATTTCTGGAGTAATGTAAAATAGTaagttatactgccctg
gcagtcatatagagcagacgGTAAAGCTAAAAATGTACTAACTACACAACTCGGCAATTGaagttatactgccct
gcagtcatatagagcagacgATTTTCATACATGAACCTGCACGAAAGTGTTCATAGCAGCaagttatactgccct
gcagtcatatagagcagacgGGAAACCACCAAAGTATTCTTTAATGAGTGGATAAACAACaagttatactgccct
gcagtcatatagagcagacgCATGGAATACTATGTAACAATGAGAAGGAATGAGCTGTTGaagttatactgccct
gcagtcatatagagcagacgGATCTCCAGGGAATTATGTGGATTGAAAAAAAGCCAATATaagttatactgccct
gcagtcatatagagcagacgCAGGGGTATGGAATGGGGGAGGGTCTGGCTATAAAGGGGTaagttatactgcc
gcagtcatatagagcagacgGAGGAGGCAGGAGATTTGTGGTGATGGAGCAGTTCAGTATaagttatactgccc
gcagtcatatagagcagacgTTCTAACTAAACATGTTATCAGATTGCACTGAAGAGATGCaagttatactgccct
gcagtcatatagagcagacgGGAAAATGCAAACCAAAATTGTAGTGAGATACTTCTTCACaagttatactgccct
gcagtcatatagagcagacgTAGAAAGTTAGGGAAGTTAGGGAAATAAAAAGTGTTGGTGaagttatactgccct
gcagtcatatagagcagacgCATTGGCTGGGGGGAATGTGAAATGGTGTAACTGCTATGGaagttatactgccc
gcagtcatatagagcagacgCACTTCTGGATATATAGTCTGAATGGAAAGCAGGGATTCAaagttatactgccct
gcagtcatatagagcagacgAGCAGCATTATTCACTTTAGACAAAGGTAAAAACAACCCAaagttatactgccct
gcagtcatatagagcagacgAGGAAATTCTGACCTATGCTGAGGTATGAACGAACCTTGAaagttatactgccc
gcagtcatatagagcagacgTGGGAGGAGAGAGGAATGGGTATAGACTTTGGCAAGATGAaagttatactgccc
gcagtcatatagagcagacgATGGATGGTGGTGATGTCTGTACAGCAGTGTAAATGTACTaagttatactgccct
gcagtcatatagagcagacgATAAAAATTGCATAGATCTACACACATGAGTACACACACGaagttatactgccct
gcagtcatatagagcagacgCGGGAGTGAGTGCATTTGAAGCTAGTGAAATCTGAATGAAaagttatactgccc
gcagtcatatagagcagacgAGTTTTCTGGTTTGATATTTTACTGTAGTTAAGCAGGGTGaagttatactgccctgt
gcagtcatatagagcagacgAATGTACATAGGACCTTGCTGCATTTTTTGTTACTTCCTGaagttatactgccctg
gcagtcatatagagcagacgTGGTATACAGGAAACAGAAAAAACTAACTGCAGTTATTGGaagttatactgccct
gcagtcatatagagcagacgCAATTGTTGGGAGAGTTTATCATTTTGTCAGCGTCCTCACaagttatactgccct
gcagtcatatagagcagacgATTTATCATCAATCATTTTGACTCCTGGTCATTAATCCCCaagttatactgccctg
gcagtcatatagagcagacgTCTGAATGGTAGCTTTATCTTAACAGATTTGATTTGGGTGaagttatactgccctg
gcagtcatatagagcagacgTTATACATCATCTGGTAAACATTAGTGAGAAAGTTTGGCGaagttatactgccctg
gcagtcatatagagcagacgTATGTGTTCAACTAGATTTAAAAACGTTCTTGGCAACTCGaagttatactgccctg
gcagtcatatagagcagacgAGGAGTGGCTGGGCAGGGGTTACTTCTGTGGATAAAGGGAaagttatactgcc

gcagtcatatagagcagacgACACTGATTGTCAACATTTATGGAAACCTGTTCTGCAACTaagttatactgccctg
gcagtcatatagagcagacgTGTTCTTTACAAGGGAAAATTGAAGCCTGGAAACACTGAGaagttatactgccct
gcagtcatatagagcagacgCCCAAGGTTAGACAGCTGGTGAATGCCCAAGTTAGGATCCaagttatactgccc
gcagtcatatagagcagacgGGAAGACGCCAATTTTACTACCTTTGTTGATCTTTAGCTCaagttatactgccctg
gcagtcatatagagcagacgCATGCTTGTATGGACCAATCTTAAAGGTGAATATTCAACAaagttatactgccctg
gcagtcatatagagcagacgATTGAGTTTTTAATTATCCCAGGCAATGTTTTAGGGACTGaagttatactgccctgt
gcagtcatatagagcagacgAAGGTCTATTGGAAAACCTTTAGAAGGGAAGGCTGTAGGAaagttatactgccct
gcagtcatatagagcagacgGGGTTGATATTTCATCAGGGAGAGCCTCACAGGGTGGGTGaagttatactgccc
gcagtcatatagagcagacgTGGAGGCAGTGAGGCTTGTGGAGCAGTTGGGGAAGGGCATaagttatactgcc
gcagtcatatagagcagacgGTGAGTCTCCAAGCCCCAGATTGGAATGTGCCTGGTGTGGaagttatactgccc
gcagtcatatagagcagacgGGGGAGAGGGGTGGTCAGGAGTGAGGTCAGAGGGATGTAGaagttatactgcc
gcagtcatatagagcagacgCCATTGTTAGGACTTTTGCTTTTACTTTGCGAGATGGGAAaagttatactgccctg
gcagtcatatagagcagacgTCATCTGACTTAATTTTGAAAGGATCACTCAAGGGGTTTGaagttatactgccctg
gcagtcatatagagcagacgAGAGATGGAAGGGGGCCAGCAGGGGAGCAGGGAGGCAGGTaagttatactgc
gcagtcatatagagcagacgGACCTGGGTCTAGGGTTAGTGGTGGAGGTGGGGAGAAGTGaagttatactgcc
gcagtcatatagagcagacgCTGATGGATTAGCTATGAGGTATGAGAGGAAGAGAGGATTaagttatactgccct
gcagtcatatagagcagacgAAGTTGTTTGGTCTAAGTAGGGAAAGAGATGAAGTTGTGAaagttatactgccctg
gcagtcatatagagcagacgTGTGTTGAGATGGGGAAGAGTGTGGCAGGGCAGATTAGGGaagttatactgccc
gcagtcatatagagcagacgCGGATTAAGTATGAGAGTGCTAAGGGGAAATGTGTGTTGCaagttatactgccct
gcagtcatatagagcagacgAGTGTAAAATACTCTCCAGGAATGTGCTCTCCAGAAACTCaagttatactgccct
gcagtcatatagagcagacgAGGGGGGATAAAGTATGCACAGACTTACAAAGTACACGACaagttatactgccc
gcagtcatatagagcagacgAAACCATTCATTCATTCTCACCATTGAGACCAAAGCTGCTaagttatactgccctg
gcagtcatatagagcagacgAATGATGGCCATGGGCACAGCAGTTTAGGATTAGGTGAACaagttatactgccc
gcagtcatatagagcagacgGGGCTTGGCAGGTTGGAGCGCTAGGAGGCCAGTGTCTCTGaagttatactgcc
gcagtcatatagagcagacgCGTGAGGTTCGAGTTGGAGATCAGCGGAGGTTAAGGTTGGaagttatactgccc
gcagtcatatagagcagacgTCATTTTGTGGATAATTGGCCATGATTGAAAGGGGTTGATaagttatactgccctg
gcagtcatatagagcagacgGAAGTTTGATCATTCCCTCAGGTTTCTTAGAAGTAGGCCTaagttatactgccctg
gcagtcatatagagcagacgTGTATAATCATGCTTATTCTCTGGGCAGCTTTGACATGGCaagttatactgccctg
gcagtcatatagagcagacgCCAGATACCCTCGCCCCTTTCGGCTCCCTGGACGCTACTCaagttatactgcc
gcagtcatatagagcagacgTGACCACAGCTGCCCCATCATTGACTCAGGAGATACTTAGaagttatactgccc
gcagtcatatagagcagacgGCGGGTGAGGCAGGCAGGGTCTAGTGGAGAGCAGACATTCaagttatactgcc
gcagtcatatagagcagacgCTGGGAAGGGAAACTACAGAGAACTTAAATAAGGGCTTTGaagttatactgccc
gcagtcatatagagcagacgCTGGACTTCAGTGGGCCAAAGGGAGGGAGGAAGAGCATTCaagttatactgcc
gcagtcatatagagcagacgATATGGCTGCACATGTGAGGAGGGTCCCTTGAGGTCTGCTaagttatactgccc
gcagtcatatagagcagacgATCATCTCTTCCCTTGATCTCAGCTATCCTCTGGGCAGTTaagttatactgccctg
gcagtcatatagagcagacgCTCTTCTTACCTCCTTCATGACATCTAGACTGCCTCTCTGaagttatactgccctg
gcagtcatatagagcagacgCCAAAGGCATGGTTAACTTCCTCTGTCCTAAACTCTAAGAaagttatactgccct
gcagtcatatagagcagacgCTTCCCACTTTGCCATTTCTACTTTCCTTCAAGCTTTTCTaagttatactgccctgt
gcagtcatatagagcagacgCTCAGACACCTTTTCTACCTGCTGTTGGCCATCCAGTCGCaagttatactgccc
gcagtcatatagagcagacgCACTCTTCTGTGAAACCTTTCCTACCTGTTTATTCCTGCCaagttatactgccctg
gcagtcatatagagcagacgGTTTAATCCTGGGACCCTATAAAACACTGTTCATTCCTCTaagttatactgccctg
gcagtcatatagagcagacgTATTTGTCTGAGTTCTTAATGGATTGAATGCTTCTGAACCaagttatactgccctgt
gcagtcatatagagcagacgTGGGGCTTTGCATGATATTTGGGTCTCAGAGTTTTACTAAaagttatactgccctg
gcagtcatatagagcagacgGTTTTAAATAGGACAGTGAAAGTAGAGAGAACAGAAAGAGaagttatactgccct
gcagtcatatagagcagacgGACCCCATCTATCTGGTTTTCCTCTTGGCATCTTTTTTTTaagttatactgccctgt
gcagtcatatagagcagacgTGCAATGGCATTATCTTGGCTCACCGCAAACTTTGCCTCCaagttatactgccct
gcagtcatatagagcagacgCATGAGACACCGCGCTCGGCTCCTCTTGGCATCTTAATCAaagttatactgccc
gcagtcatatagagcagacgTGCTGTTGATTATTATTCTGCCTGCAAACCCCATATTGATaagttatactgccctg
gcagtcatatagagcagacgGAAAAACAAGAGATATTTCGAGAGAGAGAATTGGTCCCTTaagttatactgccct
gcagtcatatagagcagacgGCATCCTTTACATTTAGAATTGCATTGTCAAGTGTCCAGGaagttatactgccctg
gcagtcatatagagcagacgAGGATAAGAAATAAAATTGTATGCTAGCATGAACCATCCCaagttatactgccctg
gcagtcatatagagcagacgCTCTCCCAGGAAAAGTGTAATTGATATTGAAGTTCCGACAaagttatactgccctg

gcagtcatatagagcagacgACCGAGGGTGCTTCATAGAATCCATTTGTTTTAATTTCTTaagttatactgccctgt
gcagtcatatagagcagacgCAGGTTCATTGTTCATTTGTTCTCAGCTGTTCCCTGGAGGaagttatactgccctg
gcagtcatatagagcagacgTGTGTGGCAGGCTTCAGAAAAAAAAGAAAGGATCTCATGTaagttatactgccct
gcagtcatatagagcagacgGGAAAAAAAGTTGGAGGCATTTCGGGTTCACTTACTTGCAaagttatactgccct
gcagtcatatagagcagacgGAATTAAGATAAGCATTGCCTACCCCAGAGTAATTAAACTaagttatactgccctg
gcagtcatatagagcagacgGTGTCACGCTTGCAGGTACATGTTCATTATTTCAGACTTTaagttatactgccctg
gcagtcatatagagcagacgCCTTGTCATTAGGTAATAACTCTCTGTGTCGTTGGGCTTGaagttatactgccctg
gcagtcatatagagcagacgGAGCATGGGTGTGTCTCTTTTTCAAGGATTCCACGTGTTCaagttatactgccctg
gcagtcatatagagcagacgACTTATAGAGTGGTGCTTTATGACGCATGCTGAAATATGCaagttatactgccctg
gcagtcatatagagcagacgATGCCTGGGGAAACATGGACAAAACACTAAACTCTGAGATaagttatactgccc
gcagtcatatagagcagacgGTGATTCCATAATAATCAATAGCAATAAGGGTAAGAGTCCaagttatactgccctg
gcagtcatatagagcagacgAGCAATTAGAGACCCAGGCATGACCCCCATAAGCTCCCACaagttatactgcc
gcagtcatatagagcagacgTCTTAAGTGGAGTTTGGGCATGAGAGGGCAGATCTGTTCCaagttatactgccct
gcagtcatatagagcagacgCCCAGTGATACTCGGTTAAGGTCTTGGCTACATTTGTCCAaagttatactgccct
gcagtcatatagagcagacgCATTCCAGTGTTTGTTTTGGAGGGCATAGCAAGAAAGTTTaagttatactgccctg
gcagtcatatagagcagacgTGTAAAGTGCAGTGTTGTCCCTGGAACATCCTTATTTTGGaagttatactgccctg
gcagtcatatagagcagacgTGGTTTGTACTCCATGATCACAGAACTGCATGAGTTATTAaagttatactgccctg
gcagtcatatagagcagacgATCATAATTCTATAAGATGGCTCGGAAGGGACCTTACTGAaagttatactgccctg
gcagtcatatagagcagacgCACCAGTGATTTGAGTGTTATTATAGCCATGCTTTTTGCTaagttatactgccctgt
gcagtcatatagagcagacgAAGAACTACCATTATTAGTTACGTCTCTCTGGATTCTATCaagttatactgccctg
gcagtcatatagagcagacgGCATACTGTTTGGATTTGATAATGTTGCAGCTAGGACTCTaagttatactgccctg
gcagtcatatagagcagacgCTGGTGTGGCAACTGGGAGCACGTCACTACATTTCTTTGTaagttatactgccct
gcagtcatatagagcagacgGAGACAATAATAATCTCTGTCTCCTAGCGTTTTTATGAGAaagttatactgccctg
gcagtcatatagagcagacgCACATAGTAAGTGCTCACAGAATGCTAGTCTTCTTAGTTAaagttatactgccctg
gcagtcatatagagcagacgTAATGGAGTATGTATTTTTCTGTGAGAAGCATTGATCCTGaagttatactgccctgt
gcagtcatatagagcagacgGCTTTTTGGTGTTTAATTGTTGCCTACATTGCCCTAGACCaagttatactgccctg
gcagtcatatagagcagacgGGGATTCCACTTCATTTGAGCCAGCTGTGTCTGTTAGAATaagttatactgccctg
gcagtcatatagagcagacgTGGCTTCCTTCCATCCATTGCATTTTACCCTTTGACATCAaagttatactgccctg
gcagtcatatagagcagacgCAACTCTAATCTCTCTTCTCCATTACAGCCCTTTGCATAAaagttatactgccctg
gcagtcatatagagcagacgAGTCTCCTCTTCATCAGCCCTACAAAGTGTCAGTTTTCCCaagttatactgccct
gcagtcatatagagcagacgCGAGCCCCTGCCTGCCTTGCTTGTATAAGTCAGCCCTTTGaagttatactgccc
gcagtcatatagagcagacgAGGACATGTGGGATACGTTTTGCTTGTACTAATTAGTGCTaagttatactgccctg
gcagtcatatagagcagacgCACCTGGAAGCTGGAAGCAGCCAATGTCTTCTCAGATGCAaagttatactgccc
gcagtcatatagagcagacgCCAGAAACGGTAATTTAAAGAAGGCTTGGAAAACTGGTGCaagttatactgccc
gcagtcatatagagcagacgCTCTTGAAAGGGAACTGGGTGGCAGGTGAAACTTACTTTTaagttatactgccct
gcagtcatatagagcagacgCCGTTGAACTGAAAGGCATTAAAAAATAGTATGAGGGAGGaagttatactgccct
gcagtcatatagagcagacgCTAAGGGAACAAAAAGATAATAGGGGCAATGTGCTGAAAGaagttatactgccc
gcagtcatatagagcagacgGATCTTGAGATCAGGAGGTGGAGACTACCCTGACCAACATaagttatactgccc
gcagtcatatagagcagacgGAATAACAACAAAGGACAAAACCTTATCTCTAGGCTTTTCaagttatactgccctg
gcagtcatatagagcagacgTGATATACTCTATTTGGTCTAGCTTCTCCCCTAAGATCCGaagttatactgccctg
gcagtcatatagagcagacgAGCCCTCCTTAACCCTCTAACCCCTTCACCAGCCTGGCTCaagttatactgcc
gcagtcatatagagcagacgCTTCCCTTCCTGTTTCTCAGCAACTCCCACAGCCTTCTAGaagttatactgccct
gcagtcatatagagcagacgCAGATACCATTATTCATTTACTTCTCAGCCTTTTTCACTCaagttatactgccctgt
gcagtcatatagagcagacgGTATTTATATACCCTGTGTTATGTCAGTGAGATGCCCGGCaagttatactgccctg
gcagtcatatagagcagacgAATGCGTGGGATGTATCTGTACTGAATGATCACATGTGTGaagttatactgccctg
gcagtcatatagagcagacgAGCTGGTGCTGGAGCCATCCTGAGGTCTTGGTAAGCATTTaagttatactgccc
gcagtcatatagagcagacgTGAGAAGAAAGAGACAACCCCCTGGGCAAGGTGAACAGCAaagttatactgcc
gcagtcatatagagcagacgCAGAGGCATGACTGAGGTTATCCTTTTAGAACAACCCATTaagttatactgccct
gcagtcatatagagcagacgTAACCCAAAACACACCTAACATTTTATAGCAAGACATACCaagttatactgccct
gcagtcatatagagcagacgTCAGATTGAAGATGACTCTGGGTTCATTGAGTGCATTTTGaagttatactgccctg
gcagtcatatagagcagacgGAGGTTATCTGGATTGGGCATTTTTTCCTTCTTGACACTTaagttatactgccctgt
gcagtcatatagagcagacgTCTCTCCTTCTCATTCTTCTCTTTCCCTTATCTCCTGATTaagttatactgccctgt

gcagtcatatagagcagacgACCTTGGTGAAGAGCTTATGGCCTTGCAATGAACCTCAGGaagttatactgccc
gcagtcatatagagcagacgAAAACAAGGCAGAAAGTGATGCAAGCCACAGGAGAAAACCaagttatactgcc
gcagtcatatagagcagacgCCAAAGAAGTCTTCATGGTGGGAATGATTTTTTTTTTAGCAaagttatactgccctg
gcagtcatatagagcagacgAATTTAGATGTGAGGAGTCTTGGGGTGGTAGGCATTCATTaagttatactgccct
gcagtcatatagagcagacgCTTCCACATGGTTGGAGAGGAGGAGCAGCAAGAAATAACCaagttatactgcc
gcagtcatatagagcagacgGGGTTCATTTTCAGGTGTATATGGAGGTTCTTACTGCAAGaagttatactgccct
gcagtcatatagagcagacgGTTTGGACTTTCTTCTTTAGGTAGACGGGAGCCAGTGAGAaagttatactgccct
gcagtcatatagagcagacgGCAGTTGATGTATCAGAGCTGTAGGCAGAGTCAGGTGTCCaagttatactgcc
gcagtcatatagagcagacgGCTATTTATTAAGCCAGCTGGGAAGAGGTATTCATGTAGGaagttatactgccct
gcagtcatatagagcagacgCACATGGTGCTTTGACGAGATCATTAAGGATTCATTCTGTaagttatactgccct
gcagtcatatagagcagacgGCAAACATAGACGGGAAGTGGCCAAGGAAACAATAAGAATaagttatactgcc
gcagtcatatagagcagacgGTAAAGCCATTGGTCAACTGATGTGATTTGTGGTTAGAAGaagttatactgccct
gcagtcatatagagcagacgTGTACGTTTCTATTGGGATTGGAATGTCTAGTAATTATCCaagttatactgccctg
gcagtcatatagagcagacgCATCCCATGAAGATACTGTGCTTTGCTGGTTCCCTGGAGGaagttatactgccc
gcagtcatatagagcagacgGCAACATCAGAAGCAACATTGAGATACCCTCCACGTGGTTaagttatactgcc
gcagtcatatagagcagacgCTGACTAAATGAGAAGACAGGTTAAACAGACATTCCATATaagttatactgccct
gcagtcatatagagcagacgAATTATTCTTATCCTTGAGTTGTGAGTCTTGGTTTCTGTCaagttatactgccctg
gcagtcatatagagcagacgTTTATGACTCAAAGAAATATGTTGTAAAGCTAGGCACCAGaagttatactgccct
gcagtcatatagagcagacgTGGTTTTGCTAAAGATTTGAGATTTTGCATGTACAGCAAGaagttatactgccctg
gcagtcatatagagcagacgCATAGAGTGAGTTTGTTGTTGAGAACTTAGAGTTGTATCAaagttatactgccctg
gcagtcatatagagcagacgTTTAACTTCCCTCTTCATCTCTTTTTCTGCTATGGACTGAaagttatactgccctgt
gcagtcatatagagcagacgTCATATATTGAAGTTCCAGCTCCTAACATGATGGTATTGGaagttatactgccctg
gcagtcatatagagcagacgTTATAAGAAGAAGGAGAGTTGGTGGGGGGGGGGGAGAGAGaagttatactgcc
gcagtcatatagagcagacgAGAGAGAGAACAGAGCTCACTGTCTCTGCTATGTGGGGACaagttatactgcc
gcagtcatatagagcagacgGGGTACCCTCACCACAAACAAAATCTACTGGCACCTTGATaagttatactgcc
gcagtcatatagagcagacgTGTTTGAGCCAGCCAGTCTGTCCACTCCGTGGTGTTTTGCaagttatactgccc
gcagtcatatagagcagacgAGCTGACCAATACAGTGCTACCATGTGCAAAACTTAGGACaagttatactgccc
gcagtcatatagagcagacgTCTGATAGAGACAGGCCAAGATGAAAAACTGGGGTGGGTTaagttatactgcc
gcagtcatatagagcagacgAAGAATCAGTAAAATCTTTGCAGCCAGGGAAAATGTATGCaagttatactgccct
gcagtcatatagagcagacgGGAGCCTTTTGTGTGTTTTACTAGCTTCATGTAGAAATATaagttatactgccctg
gcagtcatatagagcagacgGGGACCACAGTAGCAATAAAAGAAGTACGTAAGATGAAACaagttatactgcc
gcagtcatatagagcagacgGAAGATCTCGAAATCCATCTGTCAGTTTTAGCTCTTCTAAaagttatactgccctg
gcagtcatatagagcagacgCCTCATATATGCCAGGTTATATTTAGAAAAACAAAAGGGGaagttatactgccct
gcagtcatatagagcagacgGAAGTAGTCCCAGCTATGTTTTTATACCTCCCAGATCCGAaagttatactgccct
gcagtcatatagagcagacgGGGACTCCCTTAGAATGCAGTTTCATGGGCCTCTGAGACAaagttatactgcc
gcagtcatatagagcagacgGAGCATGTCCTCTAAGCATTTTATTGTACTAGGCTGATATaagttatactgccctg
gcagtcatatagagcagacgTCATGCTAATTTGTGTAATAATGAACTGTGCTTGTGATGGaagttatactgccctg
gcagtcatatagagcagacgGAATTTGGAACCACATAACTCCCCAATATTAATTCAGACAaagttatactgccct
gcagtcatatagagcagacgCAGGCATAGTTCAAGCAGTGGGGAGATTTAGTATATAATCaagttatactgccct
gcagtcatatagagcagacgAAAACTTGTGCCCTTAAAATATGGTTGACCGTCTGTAGGTaagttatactgccct
gcagtcatatagagcagacgCAATAACTGAGTTGTTCATTGTACACACATCTGCACCAGTaagttatactgccct
gcagtcatatagagcagacgGCAGAGAAGCAGTAGCCCTCCACTGTTGGATATGGTAGCCaagttatactgcc
gcagtcatatagagcagacgGAATTCCATTTCTTAGTTGCACAGCCACATTTCACATACTaagttatactgccctg
gcagtcatatagagcagacgACAGCTAGTGGTAATTCTGTGAGATGGTGCAGATGTAGGAaagttatactgccct
gcagtcatatagagcagacgCCGGCTCAGGAAGTTCTATTGGATAATGTGGTGTGGGAACaagttatactgccc
gcagtcatatagagcagacgCTGGAATAGAAAGATGTGCAGAAGGAGAGGATAAAAGAAAaagttatactgccc
gcagtcatatagagcagacgAGTGTCTCTGGGCAGCATGTGATTGATTATTGGAGTAATTaagttatactgccctg
gcagtcatatagagcagacgACAAGATGTTAGACTGAGATGATTAGGGAAGAGTTGTAACaagttatactgccct
gcagtcatatagagcagacgCTGGATTTTCAAATAACCAGTGGGATTCCAACAGCAGTGAaagttatactgccct
gcagtcatatagagcagacgTGGGAGGCACTCTATAATCAATGTGCAAAAATCACAAGCAaagttatactgccc
gcagtcatatagagcagacgGAATCAATATTGTGAAAATGGCCCTACTGTCCAAAGTAATaagttatactgccct
gcagtcatatagagcagacgCCAAAAGAACAAAGCTGGTGGCGTCACGCTACCTGACTTCaagttatactgcc

gcagtcatatagagcagacgAATGGTGCTGGGAAGACTGGCTAGCCTTATGTAGAAAGCTaagttatactgccc
gcagtcatatagagcagacgCAAGTGGGTGAAGGATATGAACACATGCTTCTCAAAAGAAaagttatactgccct
gcagtcatatagagcagacgTATAAAGACACACACACACACATGTATGTTTACTGTGGCAaagttatactgccctg
gcagtcatatagagcagacgCTGGAAACCATCGTTCTCAGCATACTATCACAAGGACAAAaagttatactgccc
gcagtcatatagagcagacgAATTGAACAATGAGAACATGGACACAAGAAGGGGAACATCaagttatactgccc
gcagtcatatagagcagacgGTATAAAAAAAAAGAACAGGTTGAAGTTTTGTTTGGGGGGaagttatactgccctg
gcagtcatatagagcagacgGAGTGGGCAGTATAAGATGTGTTTGATGCAGGAACAGCTGaagttatactgccc
gcagtcatatagagcagacgGGGTTTCCTGTTAAAAAATTCGTAGGTATTTGTAGTCCATaagttatactgccctg
gcagtcatatagagcagacgATGGGGTAGCATGGCATGGTTTAAAGGCTGGTGAGTGAGCaagttatactgccc
gcagtcatatagagcagacgAAGAGATACATGGCCAAAAACACTATCACAACCTAAAAACaagttatactgccc
gcagtcatatagagcagacgAGCATGGTATGTTCCTCGTGGGTTGTTGTTTGGGGCCAGAaagttatactgccct
gcagtcatatagagcagacgGTGGTTTGTCTTTCCATCCTGAACATCACACCGAAGATAGaagttatactgccct
gcagtcatatagagcagacgGTACATGATGAACTACCGTATCCCTGCTGATTGTGAACATaagttatactgccctg
gcagtcatatagagcagacgAAAATAAATGATGTGGTTTAGAAAGTACCATCTGAGTGCCaagttatactgccctg
gcagtcatatagagcagacgGCTTGGCATTCATTAGTTTGTTAGCTGACAATTACACCAGaagttatactgccctg
gcagtcatatagagcagacgCTCTTTCATTTTGAATGTCTCAATGGGAATCATTACTGCAaagttatactgccctg
gcagtcatatagagcagacgTTAGACATACTTAGCAAATAATATCATGTAATGGGCTGCGaagttatactgccctg
gcagtcatatagagcagacgTTAAATGTGTGATTGAACTGTGCATAGTGTTTGCCTGCCAaagttatactgccctg
gcagtcatatagagcagacgGGAAAATAAAATAACACAAAGACTCGATGAAGATGCGTACaagttatactgccct
gcagtcatatagagcagacgTTTGGCAAATAATTCAAATTCCCTTTCCTCCCACCTTTTTaagttatactgccctg
gcagtcatatagagcagacgGTCTCAGTTTTTCTTCTTCAGGCAATAAATACATTTACCCaagttatactgccctg
gcagtcatatagagcagacgGTTGCAAGAAATAATAGAGATCTCACGTACCGTTTTCCTCaagttatactgccctg
gcagtcatatagagcagacgCTGTAACCTAAATGAACTTGTAAACAGAGTGTAGCCAACTaagttatactgccctg
gcagtcatatagagcagacgGTTCAAATAAGGCAAATGCGAAGCTGTAAGAAATCGAGCTaagttatactgccct
gcagtcatatagagcagacgGCCACCATGCCCAGCACAATCCAGCTGTTTTTATATCTGAaagttatactgccct
gcagtcatatagagcagacgGTCTCTGAACCTGCTAACGTTTTGGAGGCTGCCCGATTCAaagttatactgccc
gcagtcatatagagcagacgTTTTGAGATGGAGTAAAAAAGTACTCCATCACCCAGGCTGaagttatactgccct
gcagtcatatagagcagacgACCTCTGCCTCCAGGGTACAATCCATTCTCCTGCCTCAGCaagttatactgccc
gcagtcatatagagcagacgCGTGAGCCATTGCGTGCAGCCCCAGTTAAACTCTTTTGGCaagttatactgccc
gcagtcatatagagcagacgCAATCCACTTACCTTGTTCAGATTCTCCCACCAGTTTTACaagttatactgccctg
gcagtcatatagagcagacgTGTATGTGTGTGTGTTCCATGCAGTCTAATCAGTGTGGGTaagttatactgccctg
gcagtcatatagagcagacgTCACTACCACAGGCGAGATTCTGAATTCCTTCACCACCAGaagttatactgccc
gcagtcatatagagcagacgGTTCCCCCAACCTGCTCTCTGCCATCATCCTTAATCCCTTaagttatactgccct
gcagtcatatagagcagacgACAGTGTGTAACCATTAAGATTGACTTTTTCCCCTCAGAAaagttatactgccctg
gcagtcatatagagcagacgGCTGAGTAGTAGTAATGATATAGAGGTACCACAGTCCATCaagttatactgccct
gcagtcatatagagcagacgATCTAGGTGGTTTCTCGTTCTAGGCTGTCACAAATAAAGCaagttatactgccctg
gcagtcatatagagcagacgACATACATTGTATTAGTTGACTAGGCTGGCATAACAAAGCaagttatactgccctg
gcagtcatatagagcagacgAACTAGGTTAGACAACAGACATTTAGGAGTTCAGCATATGaagttatactgccctg
gcagtcatatagagcagacgGTGTGAGACTCAATTCAGCGCTTAACATTCATGTGCAGGTaagttatactgccct
gcagtcatatagagcagacgGCAGTTTCTCAGTAGTATTGTGATTGCATGTCTAGGCTTTaagttatactgccctg
gcagtcatatagagcagacgCCAGTTTACATTCCCACAGTACAAGTGAACCTGTTTCTCCaagttatactgccct
gcagtcatatagagcagacgCATTTGGTGGTGTTGCTATTTCCTATTTTTGCCATTTTGAaagttatactgccctgtg
gcagtcatatagagcagacgCTTTTCTTGTGCTTATTGCATCTAAATATCCTCTTTGGAGaagttatactgccctgtg
gcagtcatatagagcagacgTGCAAAGATTTTCTCCCAATGCGTAGCTTCTCTTTTTATCaagttatactgccctg
gcagtcatatagagcagacgGCATTAAGCCTAGGAACTCTTTTTGTCTAGCCTTAGATCCaagttatactgccctg
gcagtcatatagagcagacgGAGTTAATTTTTGCAATAGGATATAAGGCTTAGGTTGAGGaagttatactgccctg
gcagtcatatagagcagacgCACAACAGCTCCAGCACTGTGTGTTAGGAAGGCTCTCTGTaagttatactgccc
gcagtcatatagagcagacgTTAGTTAGGCATATTTTTGTGAATCTATCTCTGGGTTCTCaagttatactgccctgtg
gcagtcatatagagcagacgGGTCTGTGTGTCTATGCCTCTACCAATACCATACTATTTTaagttatactgccctg
gcagtcatatagagcagacgTCTTCCTATTGTGTTTTTTCCCAAGATTATTATGGTAGGGaagttatactgccctgtg
gcagtcatatagagcagacgCTTTTCAGCTTGAAGTTTCCGTATGTGTTTTGTTAAGTACaagttatactgccctgtg
gcagtcatatagagcagacgTGATCATGTGTCCTGTGAATTTGCTGAACTTATTATTTCCaagttatactgccctgt

gcagtcatatagagcagacgTCTTGGGAAATTCTATGTAGCAAATTATATGCAAACCAGGaagttatactgccctg
gcagtcatatagagcagacgGTGAGAGTGGATCACCTTAACTTGCTCCTGATCTTAGGGGaagttatactgccc
gcagtcatatagagcagacgCACTTAAATATAATGTCCCCTGTAGGTTTTTTGTAGGTACaagttatactgccctg
gcagtcatatagagcagacgCTCCTAACTTGCTGAGAGTATTTGTCATGAGTGGGTGTTGaagttatactgccctg
gcagtcatatagagcagacgCCAGCCTTGCATACCTGGATTAATCCTAGTGTGTGTAATTaagttatactgccctg
gcagtcatatagagcagacgTTATCTTTGTCTGGTTTGGTCAGGGTAATAGTGGCTTCTTaagttatactgccctg
gcagtcatatagagcagacgTAGTGCTAATTCTACTTTAAACATTGGTGGAATTCTCCAGaagttatactgccctg
gcagtcatatagagcagacgCTCTTCAGTAGTTTATAGGGCTGTTCAAATGATTTCTTTCaagttatactgccctg
gcagtcatatagagcagacgATTGAGTTGTGGTAATTTGCGGTTTTTGAGAAATTGGACCaagttatactgccctg
gcagtcatatagagcagacgAACTTCCTTTAGCCACTCTGTGAGGATAGGTCTTTTAGAAaagttatactgccctg
gcagtcatatagagcagacgTTTTAGTTGTTCTTCATGTGAGAGTGTCTTTATTTCCCTCaagttatactgccctg
gcagtcatatagagcagacgTCCTGAATGAGAGTTTTGCTGGATAAATAATTTGCAGTTGaagttatactgccctg
gcagtcatatagagcagacgATGCTGTGTTTGGTTTTTTGTGCTTGGGATAGTTTGCTGAaagttatactgccctg
gcagtcatatagagcagacgCTCCACATCCTCTCCAACACCTGGTGTTTCCTGACTTTTTaagttatactgccctg
gcagtcatatagagcagacgTCTTTTGAGAGGTGTCTGTTTATATCCTTCGCCCACTTGTaagttatactgccctg
gcagtcatatagagcagacgTTAGATTCCATTTGTCAATTTTGTTGCCCTTGCTTTTGGTaagttatactgccctg
gcagtcatatagagcagacgTTGGCTCTCTGTTTGCCTGTTATTGGTTTATAAGAATGCTaagttatactgccctg
gcagtcatatagagcagacgGGCATCCCTGTCTTGTGCCTCTTTTCGAAGGGAATGCTTCaagttatactgccc
gcagtcatatagagcagacgGTTTCAGAAGGAATGCTAGCAGCTCCTCCTTGTACCTCTGaagttatactgccc
gcagtcatatagagcagacgGGTCTGCTTGGCGCAGAGCTGAGGTGATGGGGAGAATGGAaagttatactgcc
gcagtcatatagagcagacgGTTGGAAAACACTTTGCACGGTGTTATCCAGGAGAACTTCaagttatactgccc
gcagtcatatagagcagacgAGCCACTGCAAAAACATGCAGAATTATAAAGACCATTAAGaagttatactgccc
gcagtcatatagagcagacgCAGCTAACATCATAATGAGAGGCTCAAATTCACACATAACaagttatactgccc
gcagtcatatagagcagacgGGAGCACCCAAATTCATAACGCAAGTACTGAGTGACCTACaagttatactgcc
gcagtcatatagagcagacgTCTTTCAGCACTTAAAAAACCATTGTGTCACTTCCTTCTGaagttatactgccctg
gcagtcatatagagcagacgAGTAATCTGAATTGTGTTTCCCTGTGGGCCGTCTGTTATTaagttatactgccctg
gcagtcatatagagcagacgTTTAGTTTTGAGATGTTTAATCGTGTTGTGTCTTGGTGTGaagttatactgccctg
gcagtcatatagagcagacgTCCACTCAGTTTCTTGATTCTCTAGATTTATGTCTTTCACaagttatactgccctg
gcagtcatatagagcagacgACATTAGGTATATCTCCAAACGCTATCCCTTCCACCTCCCaagttatactgccc
gcagtcatatagagcagacgACTTCCACAATGGTTGAATTCGTTTTAGACATGAGTCCTTaagttatactgccctg
gcagtcatatagagcagacgATTTTTGTATAAGATATAAGGAAGGGACCCAGTTTCAGCTaagttatactgccctg
gcagtcatatagagcagacgGATATGCGGCATTGTTTCTGAGGGCTGTGTTCTGTTCCATaagttatactgccctg
gcagtcatatagagcagacgGTTTTTTCCAATTCTGTGTAGCAAGTCATTGGTAGCTTAAaagttatactgccctg
gcagtcatatagagcagacgGTTCACTCATGATTTGGCTCTCTGTTTAAAACTCTCAATAaagttatactgccctg
gcagtcatatagagcagacgCCGCCAATATCATACTGAATCACAAAAACTGGAAGCATTCaagttatactgccc
gcagtcatatagagcagacgCAGTGTGCAAAAATCACAGCCGTTCTTCAGCAATTGTTTCaagttatactgccctg
gcagtcatatagagcagacgTCACATAACAGAAAATGGGGTATCCGTTCCTTCAAGCATTaagttatactgccctg
gcagtcatatagagcagacgAACTATTACTGACTGTAGTCTGTCTGTTGTGCTATCAAATaagttatactgccctg
gcagtcatatagagcagacgTTATTTTATATCTATTAACCACTCCCATTTCCCTCCATCCaagttatactgccctg
gcagtcatatagagcagacgTTTGTTCTTATTTTTTTCCCCACCTTTTAAGTTCCAGGGAaagttatactgccctg
gcagtcatatagagcagacgATGCCATGGTGGTTTGCCGCGTAGATCAACCTATCACCTAaagttatactgccc
gcagtcatatagagcagacgCAGTCATTACCTGTTCTTTCTTTTCCTGATGCTCTTCCTCaagttatactgccctg
gcagtcatatagagcagacgTTCCCCTCCATGTGTTCTCATCGTTAAGCTCTCACTTATAaagttatactgccctg
gcagtcatatagagcagacgTTGTGAATAGTGGCTGCAATGAACACACATATACATGGCTaagttatactgccctg
gcagtcatatagagcagacgGAGTCTTGCTCTGTTGTGCAGACTGGAGTACAGTGGCGTGaagttatactgccc
gcagtcatatagagcagacgGCAACGTCCACCTCCTGGTTTCAAGTGATTACCCTGCCTCaagttatactgccc
gcagtcatatagagcagacgGTGAGCCACCTTGCCCGGTCATATTTTTATAATATAATAAaagttatactgccctg
gcagtcatatagagcagacgAACTAATCTACACTCCCATCTACAGAGTAAAAGCTTTCCTaagttatactgccctg
gcagtcatatagagcagacgGTGGTTTTGATTTGTATTTCGTGATGATCGGTGATGTTGAaagttatactgccctg
gcagtcatatagagcagacgTAAATTTGTTTACATTTTTTGTAGGCCCTGGGTAGTAGACaagttatactgccctg
gcagtcatatagagcagacgCATTGCAAAAATTGTCTTCTATTCTGTAGGTTGTCTCTTCaagttatactgccctg
gcagtcatatagagcagacgCTTTTGTTGCAATTGCTTCTGGAGTTTTTGTCATGAAATCaagttatactgccctg

gcagtcatatagagcagacgCCATGCCTGTGTCCTCAATGGTATTTTCTAGGTTTTCTTCaagttatactgccctg
gcagtcatatagagcagacgAGGAAAGGTTCCAATTTCAAATTTTTTGCATATGGCTAGCaagttatactgccctg
gcagtcatatagagcagacgCCTTTCCCCATTACTTGTTTTTGCCAGGTTTGTCGAAGATaagttatactgccctg
gcagtcatatagagcagacgTGTATCTATTCTTGTAGCAATATCATGCTGTTTTGGTCACaagttatactgccctgt
gcagtcatatagagcagacgGTGAAGAATGTCAATGGTAGTTTAATGGGAATCACATTGAaagttatactgccctg
gcagtcatatagagcagacgCAGTGGTTTGTTGTTCTTCTTAAAGAGGTCCTTTACTTCCaagttatactgccctgt
gcagtcatatagagcagacgTCTGTTTGCCTATTGTTGGTTTATTGGAATGCTAGCGATTaagttatactgccctgt
gcagtcatatagagcagacgTTTTAGGGTACATGTACACAATACCTAATGCTAAATGACGaagttatactgccctg
gcagtcatatagagcagacgAAGATAATTTGACTTCCTGTCTCCCTGTTTGAATACCCTTaagttatactgccctg
gcagtcatatagagcagacgGCTTTTGCCCATTTAATATGATATTGTCTGTGGGTTTGTCaagttatactgccctgt
gcagtcatatagagcagacgATTTGGATATGTTGAACCAATCTTGCGTCACGGGGATGAAaagttatactgccctg
gcagtcatatagagcagacgCTTTTGTATCTCTGATATCTAGTTTGGTATCAGGATGATGaagttatactgccctgt
gcagtcatatagagcagacgGAGTCCTTCCTTTTCAATTGTTGGGAATAGTTTCAGTAGAaagttatactgccctg
gcagtcatatagagcagacgTTCTTTGTACCTCTGGTAGTATTCAGCTGTAAATCCATCTaagttatactgccctgt
gcagtcatatagagcagacgGTTTCTTCTTGATTCAGTCTTGTGAGGGTGTGTGTGTCTAaagttatactgccctgt
gcagtcatatagagcagacgCTTGTCTTCTCCTAGCTTTGCGGTTCTCTAGTTCTTTTAGaagttatactgccctgt
gcagtcatatagagcagacgAGGAGTTTAATTTGAGATGTTTCAGCTTTTCGATATGGGCaagttatactgccctg
gcagtcatatagagcagacgTTACCCAGGAGTCATTCAGAGGAAGGTAGTTTATTTTCCAaagttatactgccctg
gcagtcatatagagcagacgTAGAGCAAATGTGGCAATGATAAGGATACATATTCTGTTGaagttatactgccctg
gcagtcatatagagcagacgCTCGATGATCTAATATTTTCAGTTGGGTGTTAAAGTTTCCaagttatactgccctgt
gcagtcatatagagcagacgCTCCTGTATTAGGTGCAAGCACATTTTGGGTAGTTAGTTCaagttatactgccctg
gcagtcatatagagcagacgAATCTTTGCTTAAAGCCTGCTTTGTCAGAAACTAGTATTGaagttatactgccctg
gcagtcatatagagcagacgGAAGACAGGATACCAATGGGGCTCGACTCTATCCAGCTTGaagttatactgccc
gcagtcatatagagcagacgGAATTGAGGATCTCATTCTTTTTTAATGGCAGAACAGTACaagttatactgccctg
gcagtcatatagagcagacgCCTTTGTTTTGAATATATACTAAGCAGTGGAATTGGTGGAaagttatactgccctg
gcagtcatatagagcagacgAATGTATGAGGGTTCTCTTTGCTCCATATCCTCACTAGGAaagttatactgccctg
gcagtcatatagagcagacgTTGGATAAAAGTCATTTTACCTGGGGTGAGGTAATATCTCaagttatactgccctg
gcagtcatatagagcagacgCCTTTTCACAAGCCTGTTTGTTGTTTGTAAGTCTTCTTTTaagttatactgccctgtg
gcagtcatatagagcagacgCACATATTTTTCTCCGATTTTGTGGGTTCTCTCTTCATTTaagttatactgccctgtg
gcagtcatatagagcagacgATTTGTCCATTTTGCTTTGGATGCCTGTCCTTGTGGGTTAaagttatactgccctgt
gcagtcatatagagcagacgAGGGGTTTAGTTTCATTCTCCTGTGTATGGATATCCAGTTaagttatactgccctg
gcagtcatatagagcagacgAATGAGTTCGCTGTAGGTGTTTGGGTTGTTTCTGGGTTCTaagttatactgccctg
gcagtcatatagagcagacgAGTATAATTTTAAGTCGGGCAAAGTAACTCCTTCAGATTCaagttatactgccctg
gcagtcatatagagcagacgGTGATTCCACAGACATATTAGCGTTGTTTTTTCTGTTTCTaagttatactgccctgtg
gcagtcatatagagcagacgTGTTTATCTGTTTTAATCGGATTTTTGTGGCGTCTTTAGGaagttatactgccctgtg
gcagtcatatagagcagacgGGACTTCCAGTACTATCCTGAGTGACAATGGTGAAAGTGGaagttatactgccc
gcagtcatatagagcagacgGATACTAGCTGTGGGTGTATGACATACAGCCTTTATTATAaagttatactgccctg
gcagtcatatagagcagacgTCACATTGATTCACTTAGGTCGAACCATCCTTGTATCTCAaagttatactgccctg
gcagtcatatagagcagacgTAGAATGAATTTGGAAGTGTCCTCTTATCCTCTCTTGTTTaagttatactgccctgt
gcagtcatatagagcagacgCTGGGGGACTTTATTGCAGTTTCTGTCTCATTATTTGTTAaagttatactgccctgt
gcagtcatatagagcagacgTCTGATTTTATTTGTTCGGGTAGTCTCTCTGTTTTTCTTCaagttatactgccctgtg
gcagtcatatagagcagacgCTTCTTTTTTGAGGTAGTTACTTAAAGCTGTAAACTTCCCaagttatactgccctgt
gcagtcatatagagcagacgGGTTTTGGTATGTTGTGTTTCTATTACCATTTGTGTCAAGaagttatactgccctgtg
gcagtcatatagagcagacgTTTCCTTCTTAATTTCTTCACTGGTAATTCAGGAGCATGTaagttatactgccctgt
gcagtcatatagagcagacgTTTCGAACGCTGTAAGACTTGTTTTGTGACATAACATATGaagttatactgccctg
gcagtcatatagagcagacgTGAGCCATATGTTTAGGAAAAGAATATGTATTCTGAAGCCaagttatactgccctg
gcagtcatatagagcagacgATATCTGGGTGCTCTATCACTGGGATATTTGTAAATGTTAaagttatactgccctgt
gcagtcatatagagcagacgGTTCTTTTTTGGTTTCTGTGGACATGGGATATTTTTTTCCaagttatactgccctgtg
gcagtcatatagagcagacgGTGTGTGTGGGGAACAGATAATTGATTCTTATTTTTCTTGaagttatactgccctgt
gcagtcatatagagcagacgGTAAGAGCTTACTCTTGCTATTTTGCTGTTTGGTTTTTGTaagttatactgccctgtg
gcagtcatatagagcagacgGTGAAGGTGATGTTCTCTGGTAGTATGATTTAATTTCTTGaagttatactgccctgt
gcagtcatatagagcagacgAGCCTATCATTTTAAACTGATGACAGCTTAACACTGATTGaagttatactgccctg

gcagtcatatagagcagacgGTGCTATTACCAGTGAATTTCATACCTTCAGATGATTCATaagttatactgccctg
gcagtcatatagagcagacgGCATTTCTTATAGGACAAGTGCAGTGTTGCTGAAATCACAaagttatactgccct
gcagtcatatagagcagacgATATGTTATGCCACTCTCTCTCGCCCTGCAAGGTTTCCACaagttatactgccc
gcagtcatatagagcagacgCCTGCTGGTAGACATATTGGAGCTCCATTTTATATTATTTaagttatactgccctg
gcagtcatatagagcagacgATGCCTTGAGGTAGTCCTCTTTGGATTAAATCTGCTTGGTaagttatactgccctg
gcagtcatatagagcagacgTTCTACCCCTATCTCTGTCTCTACTTTCTCTTTAAGGCCAaagttatactgccctg
gcagtcatatagagcagacgTTTCCTCTGACTGTATTTTCGAATCGCTTGTCCTCAAGCTaagttatactgccctg
gcagtcatatagagcagacgATTCTTTCTTCTACATCAATTCTGCCCTAAATAGACTCTGaagttatactgccctg
gcagtcatatagagcagacgTGTCAATTGCATTTTCAGCTCTAGAATTTCAGCTTTGTTCaagttatactgccctgt
gcagtcatatagagcagacgCCTTCTCTGTGTTATCTTGATTTCTTTGAGTTTCCTCAACaagttatactgccctgt
gcagtcatatagagcagacgGCTTATAGATGTTCATCAGTAACATTGATGAGTTAGGCATaagttatactgccctg
gcagtcatatagagcagacgGCTTGTTTTTGCCAGTTTTCCTTGGGAAAGCTTTCTAGATaagttatactgccctg
gcagtcatatagagcagacgCCTTGGTGGTCTTTGATAAAATCGGAAGAATTCTCTGGATaagttatactgccct
gcagtcatatagagcagacgTACTTTCTCCCAAACAATCAGAGTCTCTGTGATGAGTCACaagttatactgccct
gcagtcatatagagcagacgCCACCACTGGAATTGTGTTAATATGGACCTGAAGCTGGCAaagttatactgccc
gcagtcatatagagcagacgCTATGTTTACTCAAGGTCCTATAGCTCTGTGATCAGCAGTaagttatactgccct
gcagtcatatagagcagacgCCGAGCTGGGTTTGTGTCCCTTTCTTCAGGGTGGTGAGTTaagttatactgccc
gcagtcatatagagcagacgTAGAGTCAAAAAGCCTAGATATTTAGTTCTGTTCTACTGCaagttatactgccctg
gcagtcatatagagcagacgCCACCAGCAACACCACCACAAGGGAGTTTCTCTCAGGCCAaagttatactgc
gcagtcatatagagcagacgAGAAATGCTGACCAAGGGCCTAGGCCTGCACTCAGGGACCaagttatactgc
gcagtcatatagagcagacgCAAGACAGAGTCCCTTTTACATTTCCCCTTGTTTTTCTCAaagttatactgccctg
gcagtcatatagagcagacgCTCCCACAGCTGAGAATTTGCTTGGTTACACCTGAAGCCAaagttatactgccc
gcagtcatatagagcagacgATGGCCTATCACCTGGGTATGGCTGCTGGGTATTTAGGGCaagttatactgccc
gcagtcatatagagcagacgGCAGTGGTTCCCTTTTGGCCTATTCTGTCTCTAGAAATGTaagttatactgccct
gacttatcagtcttgtgtacTAGATGGTTTTTGTGCCTCAGAGTAAAGGAAATTGGTTTCtatagttacttgtcctaaac
gacttatcagtcttgtgtacCAAGAAGTTGTAGCCCAGTAAGTTGTAAATGGTACTTGAAtatagttacttgtcctaaac
gacttatcagtcttgtgtacAACACAGTATCAGAGCAGAATCCATTTCTTATAGGACTCCtatagttacttgtcctaaa
gacttatcagtcttgtgtacTCTATTTAGAAGGCCTTTTACATCTGAGAGGGTGCACTTTtatagttacttgtcctaaac
gacttatcagtcttgtgtacGCAGGTGGTAAGCCACAGTTTTACATTTTAGATAGTTAGAtatagttacttgtcctaaac
gacttatcagtcttgtgtacAGGTACTTTATATTTAGGCTTTCAAGGAACAAGGGGACAAtatagttacttgtcctaaac
gacttatcagtcttgtgtacAGTCTACAAGCAACTTTATAAAATAGCATCCACATGGCCTtatagttacttgtcctaaa
gacttatcagtcttgtgtacCTGTAGGAGGGAGAAAATAATGATTAGGCTTAAGCAAAGTtatagttacttgtcctaaa
gacttatcagtcttgtgtacAGTTGTTAATACTTCAGCTGATAGAGACTTGGAGAATAACtatagttacttgtcctaaac
gacttatcagtcttgtgtacGAATAATTCACAGGAAGAGAAATCGCAGGAGAGAGATAACtatagttacttgtcctaaa
gacttatcagtcttgtgtacGATCTGAGGCTGATCGTTTTCCTGCTTATGGGGATGTTGCtatagttacttgtcctaaa
gacttatcagtcttgtgtacCCTAAGTCTTGGGCTGACAGCTCTTCTGGACATAGTCTCCtatagttacttgtcctaaa
gacttatcagtcttgtgtacAAGTCAGTGGAGGGCTTTACATTTTCATAGAAGACATAATtatagttacttgtcctaaac
gacttatcagtcttgtgtacGAGACATAAGCACAGCGTTCAGAATTCTAAACCTCCTCTGtatagttacttgtcctaaa
gacttatcagtcttgtgtacAATCCAACTAGTGAGAAACAGTCACTAATTAGAAGTAGAGtatagttacttgtcctaaa
gacttatcagtcttgtgtacTTTCTTCATATGGCATGTGCAGTTTTATAATCTTCTTGGGtatagttacttgtcctaaac
gacttatcagtcttgtgtacGTTCTGACGGGTAATAGTTAAATGTCTATACTCTGGAATCtatagttacttgtcctaaac
gacttatcagtcttgtgtacACTTACTATCTAGGTGGCATAAGTCTCCATTTCCTCTTCTtatagttacttgtcctaaac
gacttatcagtcttgtgtacAAGATAATGTTACTTGGATTGAAAGGTGTTTGTAGGGACAtatagttacttgtcctaaac
gacttatcagtcttgtgtacTCTTTCTGTCTTCCTTTTTCTATCTCTACCACATTGGTGCtatagttacttgtcctaaac
gacttatcagtcttgtgtacTTAGGATTAAGATACCTTGACAGACGAAGAAAAGAGTAAGtatagttacttgtcctaaa
gacttatcagtcttgtgtacCTGAACATGTTTAGGTTTGGGTGCTGCTTATATTTTGCCTtatagttacttgtcctaaac
gacttatcagtcttgtgtacTGTCTTGATTATCAGGCTTCACTATATCTGTCTCTTTAAGtatagttacttgtcctaaac
gacttatcagtcttgtgtacATTGAGCCAATAATAATCAGGATCTCACTTCTCTGACAGAtatagttacttgtcctaaac
gacttatcagtcttgtgtacGATGTCCTTACTTCTACTTCTGTTTTAACAATGTTTGAGGtatagttacttgtcctaaac
gacttatcagtcttgtgtacGCATGAAAAAGTTAAGCCCCCAGTCAAGGATGAGAGCTGCtatagttacttgtcctaa
gacttatcagtcttgtgtacGAATGGACTCTTCTGACTTTCCCCGGGACAAGCAAATGGAtatagttacttgtcctaaa
gacttatcagtcttgtgtacGTCATGGACTAACAGTGATCATGTATAGTGGCGTATCTACtatagttacttgtcctaaa

gacttatcagtcttgtgtacCTTCCTTGGGTGCAGCAGTGGGGCAACAGGTGAAGCATGCtatagttacttgtcctaa

gacttatcagtcttgtgtacCCGAGTAAGTGGAGCAATTTCTTTGTCAAAGTGCCTGGTGtatagttacttgtcctaaac

gacttatcagtcttgtgtacTGCACTGTTTAGTTGGTCAACATGAGTGGGCCTGGGAAGCtatagttacttgtcctaaa

gacttatcagtcttgtgtacTGGGCATGGTGGTGCATGTTGCCAGCAGTCTCAGCTACTCtatagttacttgtcctaaa

gacttatcagtcttgtgtacGAGAGAATAGTACAAATGAACCCATGAACCCACCACACAAtatagttacttgtcctaaa

gacttatcagtcttgtgtacATTTCATCTTTTGCCATATCACTTTCTTTCCTTGCTTCACtatagttacttgtcctaaac

gacttatcagtcttgtgtacAAGAAATTCTCTAAGTGCTTCTGCAGGTATCTCTAAAAGAtatagttacttgtcctaaac

gacttatcagtcttgtgtacAACCCATAATCACAAACCCATTCATACCTAACAAAACTAGtatagttacttgtcctaaac

gacttatcagtcttgtgtacAGCCTAATATCTATTTCAAGTTCAGATTTGCCTGATTACCtatagttacttgtcctaaac

gacttatcagtcttgtgtacGTTTCTTCAAATCAGAATCCAATCAAAGGCTATACATTGCtatagttacttgtcctaaac

gacttatcagtcttgtgtacTGTTTTAATTACAACAGTTCTTCCTTCCCCTCCCCCTATTtatagttacttgtcctaaac

gacttatcagtcttgtgtacAGGGATGACTTCGGATGAGCTATTTTTTTCAGGCCATAAGtatagttacttgtcctaaac

gacttatcagtcttgtgtacGGCTAGCTTGGAGGTCAGGTGGAACAATGTAATAGAAGTGtatagttacttgtcctaaa

gacttatcagtcttgtgtacATCAGATGATTGAGTGATTGATGGGATGTGGGAGAACAGCtatagttacttgtcctaaac

gacttatcagtcttgtgtacGCACGAGATATGGAGTTGCGACTCACTGCTTAAAGGGAACtatagttacttgtcctaaa

gacttatcagtcttgtgtacGCAAGACAGCTGCCTTGTGAATTATCTCATTGATCACACTtatagttacttgtcctaaac

gacttatcagtcttgtgtacACCCCTCCCTGGCCTACTGGTGGTATGGTTTGTGAGCCTGtatagttacttgtcctaaa

gacttatcagtcttgtgtacCAATTTTGTGTGTACTGACTCTACTATCCCCCAGCAGCTGtatagttacttgtcctaaac

gacttatcagtcttgtgtacGCACCGGCTTTCCCATGTAAGACTCTCTTGGCATTTCTAGtatagttacttgtcctaaac

gacttatcagtcttgtgtacATACCTACTATGTGCTCTGTGGAAGAACAGGTGTGTGGTGtatagttacttgtcctaaac

gacttatcagtcttgtgtacAAGTCTCTGCTTTCTATTCCAAATTCAAACAGACTAAGTCtatagttacttgtcctaaac

gacttatcagtcttgtgtacGGTTTTTAAAAGCTACTTAGCTCATTCTTTGGGCTTTGAAtatagttacttgtcctaaac

gacttatcagtcttgtgtacGAAATGAATTCTTTCTTAACCTGACATCTGGGTTACTCCCtatagttacttgtcctaaac

gacttatcagtcttgtgtacCTAAATAGTTAGGAAAGGGGTTAGAAATGTGGCATAAAGCtatagttacttgtcctaaac

gacttatcagtcttgtgtacACGTGGACTGTGCAATATCTCCTGAACCTACTTCATGATTtatagttacttgtcctaaac

gacttatcagtcttgtgtacAGAAATGTGCTCCTTCTTTGTCTCTTTGAAATCTCCTCAAtatagttacttgtcctaaac

gacttatcagtcttgtgtacAGCTCCCCAACCAGGTGTGCTTTTCTCTCCTCTGAACTTTtatagttacttgtcctaaac

gacttatcagtcttgtgtacAGAGCCCCCTGTCACTTGTCCTCTTTCCTCTAAGCCCCTCtatagttacttgtcctaaa

gacttatcagtcttgtgtacATGAAAAAGGGACATGTCACCAACTAGCCAATAAAGAAGCtatagttacttgtcctaaa

gacttatcagtcttgtgtacGGTGAATTTTTCTTACCTATGCCTTGTTTGTGAGTGGAAAtatagttacttgtcctaaac

gacttatcagtcttgtgtacGTGCAGTGGTGTGATCTCGGCTCAGCCCAACCTCCGCCTCtatagttacttgtcctaa

gacttatcagtcttgtgtacCTGGCTCATTTTTGTATTTGTGTTTTAGTAGAGACACGGCtatagttacttgtcctaaac

gacttatcagtcttgtgtacCTATGAGCTAAAGATGATGTGAGATACTGAGAATACAAAGtatagttacttgtcctaaac

gacttatcagtcttgtgtacCATCACACCATGTTCTAAGCACTTTACAGCTATTATTACTtatagttacttgtcctaaac

gacttatcagtcttgtgtacCCTCAGAACAATGCTATGAAGTTGATGCTATTATTATCTCtatagttacttgtcctaaac

gacttatcagtcttgtgtacAAAAATAGAGGCATTGAGAGGTTAAACAAGTTACCCAGGTtatagttacttgtcctaaac

gacttatcagtcttgtgtacTGTAAGTAACACAGCTGAGATTTGAACCAAAGCAGTCTTGtatagttacttgtcctaaac

gacttatcagtcttgtgtacCGTAGTCCTTTAGCCTTCAACTGCTTATAGTCTGAGGGTTtatagttacttgtcctaaac

gacttatcagtcttgtgtacTGGTTCTTTTTCAGTCTCAGAATCCCAGACACATTTCACAtatagttacttgtcctaaac

gacttatcagtcttgtgtacAGTGGAGTGCAGTGTAAGCTTTGGGTTAACAGACTATTGGtatagttacttgtcctaaac

gacttatcagtcttgtgtacCAAGTGTTGAGAAGTGGTTGTCTTAGCATGCCTCAGAAAGtatagttacttgtcctaaac

gacttatcagtcttgtgtacGCTGGCCTGAGCTCCCCATTAAACTGTGAATCCTTATGCAtatagttacttgtcctaaa

gacttatcagtcttgtgtacGAATTTTCCCTTTGTTCTGCTTCATCATCATCAGTGAATGtatagttacttgtcctaaac

gacttatcagtcttgtgtacCAATTTTGTGGTATAGACTCACTGCACTGCCTGTTAAGAAtatagttacttgtcctaaac

gacttatcagtcttgtgtacATTTCCTTAGAATTTCCTTCCAAAGTAACTGATCATGCTGtatagttacttgtcctaaac

gacttatcagtcttgtgtacCTAGGACGGCATAGCAATCTCCACAAACATTCCTCTGGGGtatagttacttgtcctaaa

gacttatcagtcttgtgtacTAGCCAAATGAGTTTTCTGAGAAAGAAGAGGTTTTAAGAGtatagttacttgtcctaaac

gacttatcagtcttgtgtacGAAATGGTTCCAAAACTCTGGCATAAATCTGGGAAGTTGTtatagttacttgtcctaaac

gacttatcagtcttgtgtacGGTCTCTATAATTTGCTCACTTCTACCCACAGTTTCTAAAtatagttacttgtcctaaac

gacttatcagtcttgtgtacGTAACAAACAACCATTCTTATCACCCTAGAGATTAAGCACtatagttacttgtcctaaac

gacttatcagtcttgtgtacCTGATCCTCTTCTTCTTATGTGAGATTCTGTTGAGATGGCtatagttacttgtcctaaac

gacttatcagtcttgtgtacCACTTGGGGTTCCGTGATATTTGTCCTGCTTTGCCCATGAtatagttacttgtcctaaac

gacttatcagtcttgtgtacCCAGAACGGTTGCTTCCCCTAGTAGGGACATGGACAGCTCtatagttacttgtcctaa
gacttatcagtcttgtgtacAGAGGACTCTGAGCTCCATGACTTCTGGACTCTAGACAGTtatagttacttgtcctaaa
gacttatcagtcttgtgtacCAGTCCCTGTTGAATTGTCCAGTGATTGTTCTTTCAAAGAtatagttacttgtcctaaac
gacttatcagtcttgtgtacTGTTGGAAATATTACACACAGCCAGATTCAGTTCAGCTCAtatagttacttgtcctaaac
gacttatcagtcttgtgtacTGACATTTCTATAAACACATGGCACTGAATTAGAGGCTGGtatagttacttgtcctaaac
gacttatcagtcttgtgtacTCACTAGGGGCTCTGAGTCTAATGGGAAAGATGAAAATGTtatagttacttgtcctaaa
gacttatcagtcttgtgtacAGTATGAGGGATGCTGTAATAGAAGTGGTAGACAAGGTGTtatagttacttgtcctaaa
gacttatcagtcttgtgtacAGGAAGTTAATTACCTGAGGTCAAGAAAGGCATTACAGAAtatagttacttgtcctaaa
gacttatcagtcttgtgtacAGAGTTTGAAAATGAATTGGTGCTTGCCCACTGGACAGAGtatagttacttgtcctaaa
gacttatcagtcttgtgtacTCTGGCAAAGCTAAGAACACAGAACAAAAGCACAGAGACAtatagttacttgtcctaa
gacttatcagtcttgtgtacGAAACTTTCTGGTATATTTAGGGAACAACAGGTATGGTCTtatagttacttgtcctaaac
gacttatcagtcttgtgtacCAGAGTTTGAGATAGAAAGAATGGTAGGAGAGGAGGATGGtatagttacttgtcctaaa
gacttatcagtcttgtgtacAGGGGCATGAGAGAGAGGTTGTGAATGGCATTGTATGAAAtatagttacttgtcctaaa
gacttatcagtcttgtgtacTGGCCACCTACCCAACACTTTTTCCCACTTTGTTAATTCTtatagttacttgtcctaaac
gacttatcagtcttgtgtacCTTCTTTAAGCCAGTCACAGTGATCTAATTTCTCTTGTCAtatagttacttgtcctaaac
gacttatcagtcttgtgtacCAAAAGAGATGCAAAGAGATGTGTGTGTGTTGAGAGTCCTtatagttacttgtcctaaa
gacttatcagtcttgtgtacAAAGATGCTCTACTCTCCTGCTAGAAGTTGTCATGTTTGCtatagttacttgtcctaaac
gacttatcagtcttgtgtacTTGAAGACAAGCTGACTTGCTGAGAGTGATAGAGCAGAAAtatagttacttgtcctaaa
gacttatcagtcttgtgtacAAAGCTGCTGGGGCCTTGGAGTCACTTTCCCTTAGGACAGtatagttacttgtcctaa
gacttatcagtcttgtgtacACACTTAATTAGATGGCATGATGTGTTGCAAAGTTGAGACtatagttacttgtcctaaac
gacttatcagtcttgtgtacGTGACAGGAGAAAAGGAGTAATGTTGCATGAGTTTTGTTAtatagttacttgtcctaaac
gacttatcagtcttgtgtacGCACCATCTTAGATGCTGAAGATTCACTGTCAAATAAGATtatagttacttgtcctaaac
gacttatcagtcttgtgtacCTTGCAGTTTCTTAGCGGAGAGAGGTAAGTAAATCAATTTtatagttacttgtcctaaac
gacttatcagtcttgtgtacAGGTGAAGCTGTGGCAGGTGCTTTGATAAAGTTACCCCCAtatagttacttgtcctaaa
gacttatcagtcttgtgtacCATGCTATAGGGGCTCCTAAGTCAGTTGTATAAATTGTGGtatagttacttgtcctaaac
gacttatcagtcttgtgtacGTGTGATTGGACTGGACCAGTTGAAGACTCCTAAAGCAGTtatagttacttgtcctaaa
gacttatcagtcttgtgtacCCTGTGCACCCTCCCATCCTTGACTTACTTGTTTCCACCGtatagttacttgtcctaaa
gacttatcagtcttgtgtacTGACTTTAACTCATTTCTGGATGCTGGAGACCTGTTTCGGtatagttacttgtcctaaac
gacttatcagtcttgtgtacCTAGATTTAGTTTCTTGCTTTGGTCTCTTGGATTGACTAGtatagttacttgtcctaaac
gacttatcagtcttgtgtacCTATTCTGACTTGCCTTTTCATTGGTTCTAGCTTCTGGAAtatagttacttgtcctaaac
gacttatcagtcttgtgtacGATTTCACAGATTTGGGCTTTTTGCTGTCTATTTTTAGACtatagttacttgtcctaaac
gacttatcagtcttgtgtacGAACACTTCTAGCTTGTCTGGGTTCTCCACACCTTGCCTTtatagttacttgtcctaaa
gacttatcagtcttgtgtacGGTCTATACATGGCATTAGTTGTCAGGGCACATAGGTTCAtatagttacttgtcctaaa
gacttatcagtcttgtgtacTGGGTCACAGTCATATAGTCGTTTTGGGATTGATGTAGAGtatagttacttgtcctaaac
gacttatcagtcttgtgtacCCATCTTCCTTCTTGGCATCTTTGTTTTTAGGGGGTTTTGtatagttacttgtcctaaac
gacttatcagtcttgtgtacACATCCAAAGAAGCTATATATGCATAACAGGATAACTTCCtatagttacttgtcctaaa
gacttatcagtcttgtgtacATTATGAATGATGTTGTTTAGTGTGGGGACAGTATTAACGtatagttacttgtcctaaac
gacttatcagtcttgtgtacCTTTTTCTAGAAGGTAAAAGCTAAAGAATTGTCTGGGGCTtatagttacttgtcctaaac
gacttatcagtcttgtgtacAGCCTTCCTGAGAATTGATTATGTGAAGTATCCCCTGAGGtatagttacttgtcctaaa
gacttatcagtcttgtgtacGCTTTTCTGAGGGCCCCTTACAGTTTTAACATAGCTTAATtatagttacttgtcctaaac
gacttatcagtcttgtgtacCCTCTAGGATCACAGAGCTTGGAATTCAGTAGGTAGCACCtatagttacttgtcctaaa
gacttatcagtcttgtgtacCTTGGGAGGCAAGAGGATCTACCTGAGGCAAGAGGATCTCtatagttacttgtcctaa
gacttatcagtcttgtgtacAGAAAGTCAGTAGGTAGAAAGGTTTAAGGAAAAAGAAGACtatagttacttgtcctaaa
gacttatcagtcttgtgtacGCTCATTATAAAGAAGTCAAGCAGCAAAGAAAGGTACAAAtatagttacttgtcctaaa
gacttatcagtcttgtgtacAGCAAAAAAGTCACTCCGTATCCCACTATACAAAAATAACtatagttacttgtcctaaa
gacttatcagtcttgtgtacCAAAGGGCAATATCATCCTATGTATCTCTCTCTATAGAAAtatagttacttgtcctaaac
gacttatcagtcttgtgtacGTGTCATTATACTCTTTAACTTCATGCACCAAGTGTAGGCtatagttacttgtcctaaac
gacttatcagtcttgtgtacCTGGGAAAACTGCTATATTACTATTTTTATGCGATTCACCtatagttacttgtcctaaac
gacttatcagtcttgtgtacATGTAAATGAATTCAGTGCTCATAACTAGCCCTTTTCCCAtatagttacttgtcctaaac
gacttatcagtcttgtgtacGAATGGCTGTGATTCTTTGTTTGCTTCTTTCCCTCAGAACtatagttacttgtcctaaac
gacttatcagtcttgtgtacGTCTGGAGACTCTTCATCTGCATTCCTTCTTCATTTTTGAtatagttacttgtcctaaac
gacttatcagtcttgtgtacTGAACTATTAATCATATAGAACCTACCTGATGTTAGTGGGtatagttacttgtcctaaac

gacttatcagtcttgtgtacCAGGTAGAAGTATTAGGTAGACTGGCGACTGATTTGGAGCtatagttacttgtcctaaa
gacttatcagtcttgtgtacTCAATTGTTAGTTCTCTTGCCTGGGTGGTGTGGGAGGATCtatagttacttgtcctaaac
gacttatcagtcttgtgtacGAGCAGATGTTACTCACTCAAAAGTCCCACGTCACAGTCTtatagttacttgtcctaaa
gacttatcagtcttgtgtacTATTGAAATCTGACTGTTACTGCCTTCTCTCCCTTCTGTAtatagttacttgtcctaaac
gacttatcagtcttgtgtacGAAGTCCCACTCAACATGATTTCTCCTGGCTCCACTTGTAtatagttacttgtcctaaac
gacttatcagtcttgtgtacGGTTTACCACATCTATCCATCTTATGCTTGAAAACATAGTtatagttacttgtcctaaac
gacttatcagtcttgtgtacCTTGGCCTCTATGATGGACCTGCACAATATTTCAACAAACtatagttacttgtcctaaac
gacttatcagtcttgtgtacAGAATCAGGAGATATTGCACACAAAAATATGAGTTTCTGGtatagttacttgtcctaaac
gacttatcagtcttgtgtacCAATGACTTGATGATACTGGGACCACATTCCTACAGCCATtatagttacttgtcctaaac
gacttatcagtcttgtgtacGGATGCCCTGTTCTGAAATGCCACAGTCCCCACATTCCCTtatagttacttgtcctaaa
gacttatcagtcttgtgtacTTACGTTAAATGCCTGGCAGCTGTAAGATTTGAATGTGTAtatagttacttgtcctaaac
gacttatcagtcttgtgtacAACATCTTGCAATTGAACTTTTTAGGTATAGGGTGATAGCtatagttacttgtcctaaac
gacttatcagtcttgtgtacATAAACCTTTTGAGTGATTATTTATGGAGCACCTGTTTGGtatagttacttgtcctaaac
gacttatcagtcttgtgtacGTGTCAGATGCTAGAGATATAAGATAACAAGATGTAGTTGtatagttacttgtcctaaac
gacttatcagtcttgtgtacAACAGATAGAAATCGTATCTTTTGACATGTGCCTCTGATAtatagttacttgtcctaaac
gacttatcagtcttgtgtacTAAAGCATGTTTTAGGAAGAAAAGAGTGATCCCATCTTGGtatagttacttgtcctaaac
gacttatcagtcttgtgtacTTGTTCTTAACGGCTTTGCAAGATTTAATTGGCAGGGGGtatagttacttgtcctaaac
gacttatcagtcttgtgtacAGAAATGAACAAGAAACGAGATAACCCGGGCACTGCAAGTtatagttacttgtcctaaa
gacttatcagtcttgtgtacGCTTCATTTTTGGCAGTTGGGAAAGATATGGCTTAGAAAGtatagttacttgtcctaaac
gacttatcagtcttgtgtacCAATGCTAAGATTTGAGTTTTATCCTTCGTACAGTGAGAAtatagttacttgtcctaaac
gacttatcagtcttgtgtacTCAGGATGCAATGAGTGGAATGACATGATCAAAGAAAGGTtatagttacttgtcctaaac
gacttatcagtcttgtgtacATTCTACTCAAGCAGGATGAATTAAAAGGAGAGAAATGCAtatagttacttgtcctaaac
gacttatcagtcttgtgtacGTTAAAAAAAAAAACCTGGATGCTTATGGGATTTTCTCCCtatagttacttgtcctaaac
gacttatcagtcttgtgtacTATCGATGTATTCATTGTATGATACTTGGTGAACCCTTTCtatagttacttgtcctaaac
gacttatcagtcttgtgtacGATCTCTCTTTCTGGAATTCTTGTTACACATAAGTACTAGtatagttacttgtcctaaac
gacttatcagtcttgtgtacCTGGATCATTCCCCCTGTGTTTCTCATTTATTTAATACTTtatagttacttgtcctaaac
gacttatcagtcttgtgtacCCTTTATCTCTTACACTCTAAGTTATGGGGAAATTTCTTGtatagttacttgtcctaaac
gacttatcagtcttgtgtacATACATAATCCATTTGCTATGCACTGCACTTATTGCATTCtatagttacttgtcctaaac
gacttatcagtcttgtgtacAAAAATGCTCTATGAAGTTATCTCACTGAAATCCTCAGGAtatagttacttgtcctaaac
gacttatcagtcttgtgtacTTTCCCAAAGTCTCAAATCTAGTATGTGAAAGTGCCTGGAtatagttacttgtcctaaac
gacttatcagtcttgtgtacTCGTCTATATCATGAGGAAGGACTGGCATCACAGAGAGCTtatagttacttgtcctaaa
gacttatcagtcttgtgtacATATCACATGTGCCAAGATACTAAAGAGATAAAAATGCGCtatagttacttgtcctaaac
gacttatcagtcttgtgtacCCTTCTGAATTTTGGAGAGAATTTGATAGGGGTGGAGAAGtatagttacttgtcctaaac
gacttatcagtcttgtgtacAAGCATATCCTTTAATCCTCAACTGATCTATTAGCTCCCGtatagttacttgtcctaaac
gacttatcagtcttgtgtacCTTTTAAGCTCTCAAGGGAAGGGAGAAAAGGTGGGCAAGTtatagttacttgtcctaaa
gacttatcagtcttgtgtacAGGGTAATGAGACTCTCCTGCTCCAAATTATTCAGAACACtatagttacttgtcctaaac
gacttatcagtcttgtgtacCCCAGAATCACCATGCCGGCAGAGACCAGTAGAGAACAAAtatagttacttgtcctaa
gacttatcagtcttgtgtacGGTTTTTGGAGTAGTCTGGCACAGTGGGTTTTCTTCTCACtatagttacttgtcctaaac
gacttatcagtcttgtgtacCATATACTAGTGCCACACTGACACAGCTGTTAGAATAAGCtatagttacttgtcctaaac
gacttatcagtcttgtgtacAGGATTTTTTTCTCTCTAGTTCCAGGTAAGCTTTCCTAACtatagttacttgtcctaaac
gacttatcagtcttgtgtacTCATAGAAAAACCGCGAGAAAGAGAATTTAAAAAGCTAGCtatagttacttgtcctaaac
gacttatcagtcttgtgtacACAAGGTTTATGTATCTAGGAAGGAGAGAGGAAAAAGCCCtatagttacttgtcctaaa
gacttatcagtcttgtgtacATGCTAGCTAGTAAATTGTGGCACAGGTCTAGGATTTCCCtatagttacttgtcctaaac
gacttatcagtcttgtgtacCCACTTATGCACAACCCCCATTCCACTTCTTCGGAGGCTCtatagttacttgtcctaaa
gacttatcagtcttgtgtacGAGCCATTTCCTATTCCTTTGTGTTTCACCTTATTTAGTGtatagttacttgtcctaaac
gacttatcagtcttgtgtacAGCAACAATGTATGGGAGGTAAGGCTGAAGACGTCTTTCCtatagttacttgtcctaaa
gacttatcagtcttgtgtacCTTCTTCAGTGTACCTGGACTTGTCAACATCTAGCTTTGCtatagttacttgtcctaaac
gacttatcagtcttgtgtacCAATAATGATGATTAAGGCTCAATGTGAATGACAGCTGGGtatagttacttgtcctaaac
gacttatcagtcttgtgtacAGCAATCTGACAAAATCCTGAAACAGCCTTAGGGATAATGtatagttacttgtcctaaac
gacttatcagtcttgtgtacCAGAGTGTGGCATCTTCCAACTACATCCAGGGTGTTAAGTtatagttacttgtcctaaac
gacttatcagtcttgtgtacTGACAGGAACACATTTGAGCACATTCAGAGGAGGGCAATGtatagttacttgtcctaaa
gacttatcagtcttgtgtacCAGGGTAGACTGGTTGTGTGCTGTTTCAGGCAAAGATAAAtatagttacttgtcctaaac

gacttatcagtcttgtgtacAAGGAATAACTTTTCTAACAACGACAAAATGGGCTACTGTtatagttacttgtcctaaac
gacttatcagtcttgtgtacAGCTCCCTAACATCAGAAATGCTCAAATGTATGCACCAGAtatagttacttgtcctaaa
gacttatcagtcttgtgtacAGATTCTGTGATCTTCTGATTTACTGGAGTCTTGCAAGTGtatagttacttgtcctaaac
gacttatcagtcttgtgtacGTTGATGCCTCTTTCCTTTCCTAACAGTCTCTTCAACCCAtatagttacttgtcctaaac
gacttatcagtcttgtgtacACCAGCAACTCTCAGTCACCAAACCCCAATCTTCTCTGTCtatagttacttgtcctaaa
gacttatcagtcttgtgtacCATTTAGCACAATGGCAACTCCTACTTTCTGGAAATAGATtatagttacttgtcctaaac
gacttatcagtcttgtgtacTGACCTCATATTCACCCAGTGGTTTCCTTCCTACCTCACTtatagttacttgtcctaaac
gacttatcagtcttgtgtacTCCTATCAGTCTCGTTTGCTAATTATTCATCTTCTCCTCAtatagttacttgtcctaaac
gacttatcagtcttgtgtacAGAATTCTCTTTTCTTCTCTGTTTGTGTTGACTCTATGTGtatagttacttgtcctaaac
gacttatcagtcttgtgtacAAACATGTGTGCAAACTCCTTGACACTCCTCTCATCCAGAtatagttacttgtcctaaa
gacttatcagtcttgtgtacTGTCTTCTCCCCTTGAATTTGGGTTGAACATAGTAATTCAtatagttacttgtcctaaac
gacttatcagtcttgtgtacCATTAGAATGCAGCAGGAGAGATAATTTGTAACACCAAGGtatagttacttgtcctaaa
gacttatcagtcttgtgtacGTCCTAAAAGACCATGAAGTGTCCACCGGGTTCCTTGGGAtatagttacttgtcctaaa
gacttatcagtcttgtgtacCTTAGGAATCCTGAGCTTCCGGCACTGAGGTGGTCACAGTtatagttacttgtcctaaa
gacttatcagtcttgtgtacCCCAAGCCTTTCAGGGGGCCATGTGTAGTGCTCCAGCTGAtatagttacttgtcctaa
gacttatcagtcttgtgtacCAGGCATCACGGAGCAGAGACAAACCTTCCCTGTTGTGTTtatagttacttgtcctaaa
gacttatcagtcttgtgtacAAAATGGTTGCTTTAGGTCACTAAATTTTGGGGTGGCTTGtatagttacttgtcctaaac
gacttatcagtcttgtgtacCTGTAGCATGACTTTAAGTACCGCCTATAAAAGCTTATGAtatagttacttgtcctaaac
gacttatcagtcttgtgtacTGAACTCCAGACTCTTAAATCCAATTATCCACATGTTTACtatagttacttgtcctaaac
gacttatcagtcttgtgtacATATGACTAAAACAGAAAGCTCAACATTAACCTTTGTCCCtatagttacttgtcctaaac
gacttatcagtcttgtgtacCTTATTTCTTCTCTCTCCCCCATTTCTGTATACATCTAGTtatagttacttgtcctaaac
gacttatcagtcttgtgtacGTCAAAACTGTTAGTCTTATTTCTGCCTTTTACCCCTAAAtatagttacttgtcctaaac
gacttatcagtcttgtgtacGAATCTGTCCCGTTCTTTCCATCTCATCTGCTTCTGTCCTtatagttacttgtcctaaac
gacttatcagtcttgtgtacAGTTATCATAATCTCTCTTCTCATCTACTGTCATATCCACtatagttacttgtcctaaac
gacttatcagtcttgtgtacCAGTTTTCTTTCTTTCACTCTGGCTCTCTAAAATTCCATTtatagttacttgtcctaaac
gacttatcagtcttgtgtacGCAAATTACCCCCAAATGCAGTAACTCAAAACAGTAATATtatagttacttgtcctaaa
gacttatcagtcttgtgtacGCTTATCTGGGTTCTCTGGCAATGATCTCAAAGCTTGACTtatagttacttgtcctaaac
gacttatcagtcttgtgtacTGGCAGGATTCAGTTTTACCAAGAGCCCAGCTTCCTCATGtatagttacttgtcctaaa
gacttatcagtcttgtgtacTGTCAGGTGGGACTCTCCCTCAGGCAGCTTACAACATGGCtatagttacttgtcctaa
gacttatcagtcttgtgtacAGAGGGAAAGAGAAGGAGGACTGGCAAGTCATAGCGTTTTtatagttacttgtcctaaa
gacttatcagtcttgtgtacCCATGTTCTGTTAGAAGCAAGTCCCTCAGTCTGGCCCACTtatagttacttgtcctaaa
gacttatcagtcttgtgtacACAAGAGTGTGCATACCAGAAGGTGGGAATCACTGAGGGCtatagttacttgtcctaa
gacttatcagtcttgtgtacTAAAATCATAAATCACCCCTCCTTAAATCATATCACACCCtatagttacttgtcctaaac
gacttatcagtcttgtgtacAATATCTCATTGTTTTGCAGAGCCCCACATGTGTAGCCTGtatagttacttgtcctaaac
gacttatcagtcttgtgtacCCTCTAATGTTACGATGCTTTATTTAATCAGTTTCACTCCtatagttacttgtcctaaac
gacttatcagtcttgtgtacTTCTTTCTGTTCCTTGAATTCTCCAAGTATGCTCCTTCCTtatagttacttgtcctaaac
gacttatcagtcttgtgtacCCTTTTTGGATCACTTTCCCCTGTTTATATAGAGATAGCTtatagttacttgtcctaaac
gacttatcagtcttgtgtacGTGGATCTCTAGCCTCTGTCACATTAACCTCTTTCATTCTtatagttacttgtcctaaac
gacttatcagtcttgtgtacATTCATTAATTTACTTACCATCAGTCTCTTTCCTCCCTTCtatagttacttgtcctaaac
gacttatcagtcttgtgtacCTTTGGGTCATTCACTCTACACCTTTAGTAATATCAGTCTtatagttacttgtcctaaac
gacttatcagtcttgtgtacCAGGGACTTTCTGGCAATTTTCATTACTACACAATTTTCAtatagttacttgtcctaaac
gacttatcagtcttgtgtacAGCTGACAGTTATATGATACTTATAAGGTGGCAAATACTGtatagttacttgtcctaaac
gacttatcagtcttgtgtacTAATCATCACAACATCCACATTCTACAGATAAAGAAAGCCtatagttacttgtcctaaac
gacttatcagtcttgtgtacAGATCACATAGGTAGTAAGTAGGATGGCTGGAATATGACCtatagttacttgtcctaaa
gacttatcagtcttgtgtacTGAGCACACTTTCCTCCCATTCGTGACATGTACAATTCTGtatagttacttgtcctaaac
gacttatcagtcttgtgtacGCAGGTTGCCAAATTCATGCAATTCCCACTAGTCACTGGAtatagttacttgtcctaaa
gacttatcagtcttgtgtacTCCCCCCCAAAGTATGCAGAACCTGTAGTTTTTAAATTTGtatagttacttgtcctaaac
gacttatcagtcttgtgtacATATTTCTACCATTTTAGTGCCTGTGTTTCAACTATCTGGtatagttacttgtcctaaac
gacttatcagtcttgtgtacGCTCTTTGAAGCCATCGTTTTACCCAGGGAACAGATTATGtatagttacttgtcctaaac
gacttatcagtcttgtgtacATATCTCTCAGATCTGCCCCGGCCTGGAATCGCAAGAGGGtatagttacttgtcctaa
gacttatcagtcttgtgtacCAGGAAGAGAACAACTGCTAGCTTGGTTATGACCTAAAAAtatagttacttgtcctaaa
gacttatcagtcttgtgtacTCCCTTGCTCTAAAGATGTTATTGACTCAATGATATTCTCtatagttacttgtcctaaac

gacttatcagtcttgtgtacAGAGTTTTACTGTGTAATTGCTTCATTTTCCATCCATGTGtatagttacttgtcctaaac
gacttatcagtcttgtgtacCCGTACGGTTCCACTATCCCATCTAGTCCCCCTGCAAACTtatagttacttgtcctaaa
gacttatcagtcttgtgtacCCCAGATAATGATGATCACATGGCAGACAACAGGGGCAGGtatagttacttgtcctaa
gacttatcagtcttgtgtacCGAGACGGAATAAACATCACCCAGTTTCCTTTTCCCTCCCtatagttacttgtcctaaa
gacttatcagtcttgtgtacAAGGAACGGTGATGTCTTGAGAGTCAGAAAGAAAACAGTGtatagttacttgtcctaaa
gacttatcagtcttgtgtacACATGATTTACTGAAAAATATGCAGGGCTTTGTTGGGAGGtatagttacttgtcctaaac
gacttatcagtcttgtgtacATGTTGTCTCCTCAGGGAGCCCGCCTCTTCCCACTGCTGCtatagttacttgtcctaa
gacttatcagtcttgtgtacCGGACGCCATTTTCTTTGGTACACGCTGTCTCTGGGTAGAtatagttacttgtcctaaa
gacttatcagtcttgtgtacGTTTAAGCTGAATATTCACCTGAACAAATAGCAGATGTTGtatagttacttgtcctaaac
gacttatcagtcttgtgtacTTAATTTCTGTTTTACTATTCAGGCCCTGATTGCAATGAGtatagttacttgtcctaaac
gacttatcagtcttgtgtacCTGGCTGGGGGCGGGGCTGGAGGTCTTCAGGGAGTGGAGCtatagttacttgtccta
gacttatcagtcttgtgtacCAATGCAGAGATAAAATTGTCTCCAGTGGAGTCCAGATTCtatagttacttgtcctaaa
gacttatcagtcttgtgtacTGGCACAGTGGAGAGCTGGCAGCTCCTGAGAGACAGAGAGtatagttacttgtccta
gacttatcagtcttgtgtacAGAGGGGAGAGACAGCAAGAGAGCGAGAGCTGAGCAAGTGtatagttacttgtccta
gacttatcagtcttgtgtacGGATGATCCCCTCGTGGTCTCCTTTTAACATATCTGAGTCtatagttacttgtcctaaa
gacttatcagtcttgtgtacGAAGTTTAATTCTAATTTGTATCCTCCTAAACCCACATGGtatagttacttgtcctaaac
gacttatcagtcttgtgtacTGAGTCTCAGTTTCCTCCCTCGTAATACAGAGTTAGTGAGtatagttacttgtcctaaa
gacttatcagtcttgtgtacCCAAGTGGCACTCCGTAAATTAGATCCCTATCCTCCTTCCtatagttacttgtcctaaa
gacttatcagtcttgtgtacATCTCTCTGTTGTGTATGTTTATTTCTTTACGTTGCTCAGtatagttacttgtcctaaac
gacttatcagtcttgtgtacCTATTTTGAGTTTTTCTGCCAGAATAATGTGAATGCCATGtatagttacttgtcctaaac
gacttatcagtcttgtgtacTGGCACCATATTCCATATTTGTCTTGTTCGTCCTTGACCTtatagttacttgtcctaaac
gacttatcagtcttgtgtacTGAATTTCTGTTTCATTGAGAATTGCCCTTTGAAGTCTGGtatagttacttgtcctaaac
gacttatcagtcttgtgtacTCCATCTCAATAAAAAGAGAATTGCCCTTTGCAGACCTAAtatagttacttgtcctaaa
gacttatcagtcttgtgtacCATTTTGCCTTCTTGTTCTTTCCACCTCGCACACTCTCATtatagttacttgtcctaaac
gacttatcagtcttgtgtacACTGCAATCAAAAGAATCCTGAGTAATGCCAGTGTGTAGGtatagttacttgtcctaaa
gacttatcagtcttgtgtacGGATTCCCACCTCCCTCTAGTTCTCTTCCTGTTATGTTTGtatagttacttgtcctaaac
gacttatcagtcttgtgtacTTTATTATTTCATGGTTGTGGAGACGAAGTCCAAAATCACtatagttacttgtcctaaac
gacttatcagtcttgtgtacCCAGCTTCTAGTGGTGACCTTCGATCTTTAGTGTTCCTTGtatagttacttgtcctaaac
gacttatcagtcttgtgtacCTGTCTTTCCATGGTGTTATTCTTGTGTTTTCACATTATGtatagttacttgtcctaaac
gacttatcagtcttgtgtacTATGAGGATAGTAGTTATGTTGGATTTGGTTTCATCCTAGtatagttacttgtcctaaac
gacttatcagtcttgtgtacTTGTGAGATATTGGTTATATATGAGCTCAGGACTTCTTACtatagttacttgtcctaaac
gacttatcagtcttgtgtacCAAACTTGTATTGATTCAGCAACGACCTTCATTTTCTCTTtatagttacttgtcctaaac
gacttatcagtcttgtgtacTTAACCCTAAGAAATCCTGTGTGTCCTTGCTATCCCCTCCtatagttacttgtcctaaa
gacttatcagtcttgtgtacCCTCTGTCTTTATCTTTGGGATTTCTGTGACTCTGGCCTCtatagttacttgtcctaaac
gacttatcagtcttgtgtacGGACATTTCTTTTCCTTGCCCTTCTCCCTTCCATCTCACTtatagttacttgtcctaaac
gacttatcagtcttgtgtacGGTCCCTGATATCATTTACCTATATTGAAAACTGGCTGGTtatagttacttgtcctaaac
gacttatcagtcttgtgtacCCATCTTCTTTGTTTATACATCAGCACAATTCTTTACCACtatagttacttgtcctaaac
gacttatcagtcttgtgtacCTTGACACTCCTCTGTTTTTTTCAAAGCAACACACATAGCtatagttacttgtcctaaac
gacttatcagtcttgtgtacGCCCAGTTTCCTACCTCTGCTGCCTCTTCCAAACACGACGtatagttacttgtcctaaa
gacttatcagtcttgtgtacCCCCTGGGGGATATTGCCTGTTCTTTCCCAAGCGCTGAGAtatagttacttgtcctaaa
gacttatcagtcttgtgtacCCTCTTGCTAATGTTATGGCATGGTGCTTTTTCAACAAAAtatagttacttgtcctaaac
gacttatcagtcttgtgtacGTCTAGTTGGCTGCTCTTTTAACAAGCTTAGTGCCCTTCCtatagttacttgtcctaaac
gacttatcagtcttgtgtacGTTACTTGAACTCATTTGAAAGATTACCTGGCTGATACTTtatagttacttgtcctaaac
gacttatcagtcttgtgtacTGAAAGAATAACTATCCTAAAGTACCCGAGGTAGGTGAAAtatagttacttgtcctaaa
gacttatcagtcttgtgtacGTTTAACTCCCTTGATTAAAAATAATCCCCAATTTCAGGGtatagttacttgtcctaaac
gacttatcagtcttgtgtacGATCTTCTTTATCACTTTTTAGGTACTATATGTGTGCTGGtatagttacttgtcctaaac
gacttatcagtcttgtgtacCAGATCGATATTTTTAGTTCTCTGTCATCTCCCTGGGTGTtatagttacttgtcctaaac
gacttatcagtcttgtgtacCATTCCTCCGTCCCTGCTGCCTTCTTCCTCTTGTAGTCTAtatagttacttgtcctaaa
gacttatcagtcttgtgtacGTTACCAGCCTCAGTCACATCCCCCATCTCCCAAGTTATAtatagttacttgtcctaaa
gacttatcagtcttgtgtacGTCGTGTCTCTAATCAGTTAGTGAACCCTACCATTCCAGCtatagttacttgtcctaaa
gacttatcagtcttgtgtacTGAACTCCAGCATCTCTACCAGTCTTCCTCTCCAGTCCCTtatagttacttgtcctaaa
gacttatcagtcttgtgtacAGTTGACCATGTAACTGTTTTTTCTTTGAAACTCTGTTGGtatagttacttgtcctaaac

gacttatcagtcttgtgtacGTCATTCTGGGCTTCTCAAATCTGACTCTGACCTTCTCATtatagttacttgtcctaaac
gacttatcagtcttgtgtacGCTCTGGCCCTCTACTCTCTCTCTCCACCCCCAAGCGTACtatagttacttgtcctaa
gacttatcagtcttgtgtacGCCTCTCATTGTGTCTTGGAAACAACATACCGTTCATGCCtatagttacttgtcctaaa
gacttatcagtcttgtgtacTGCCCTTTCCTCCCTTCATTACCAGACAAATTCCTTCTCGtatagttacttgtcctaaac
gacttatcagtcttgtgtacTTCTTGGTAATAGTAGTAATAGAATCCCCCTTGCTAGAGTtatagttacttgtcctaaac
gacttatcagtcttgtgtacCCTCTGTACGGCATCCCATGGTACTTTATTCACAGTTGCAtatagttacttgtcctaaa
gacttatcagtcttgtgtacTATAATTTCCTTTGCCTCCTTCATTAGACTATTCACTCACtatagttacttgtcctaaac
gacttatcagtcttgtgtacGTGTTCAACAGGAAAGCCATGAATGTAACTCAACTCTATGtatagttacttgtcctaaa
gacttatcagtcttgtgtacTCTCCCCTTAGAGAAAAATCTGAGATAGTGTTTCAAAATCtatagttacttgtcctaaac
gacttatcagtcttgtgtacACTAGCTGAGGTTTTAAAAAAGGTGGTGGCAGTGGATACTtatagttacttgtcctaaa
gacttatcagtcttgtgtacGGGAACTAAATAATGTGTACCTATGGACGTGGAGTGTGGAtatagttacttgtcctaaa
gacttatcagtcttgtgtacAGGAGTGGGGGTGAGGTGGGAAGGGGCTGGATGATGGGAAtatagttacttgtcctaa
gacttatcagtcttgtgtacAAGGCTCTGACTTCATCATAATTCAATGTATCAATGTAGCtatagttacttgtcctaaac
gacttatcagtcttgtgtacAATTAAAAAAAAAAAAGGTAGTGGGATCAAGTGAGGGATGGtatagttacttgtcctaaa
gacttatcagtcttgtgtacGTGTATCTCCTGCCTGTCTTGAGTAGCCCAGCACTAGAGAtatagttacttgtcctaaa
gacttatcagtcttgtgtacCTTCCCAGCAGCACAAACAGAGGGTACAAGATCAGTTTATtatagttacttgtcctaaa
gacttatcagtcttgtgtacATAACTCATTTCTCTGGGAAACAGCAACCACTCTAAATCCtatagttacttgtcctaaa
gacttatcagtcttgtgtacAGAAGTTTCTCATGAGGGCATATGTGATTTAGTGAACACTtatagttacttgtcctaaac
gacttatcagtcttgtgtacTTCCTGGCTGAGCTATGGAAAAATCGTATAATATTTGTCCtatagttacttgtcctaaac
gacttatcagtcttgtgtacTGATAGGCAGGGACAGAAAAAAAGAGTAGAGAGGAGAGAGtatagttacttgtcctaa
gacttatcagtcttgtgtacTTTTTGGACTAAAAGCCCTGACAGATTCATTCATTCACTCtatagttacttgtcctaaac
gacttatcagtcttgtgtacGAAATTTCCCTTATACCCCTATTGCGCAATCTCTCCTATCtatagttacttgtcctaaac
gacttatcagtcttgtgtacCTGGAAATTATCTTACCACATATCAGCTCTTTCAAATCAGtatagttacttgtcctaaac
gacttatcagtcttgtgtacGGATGCATCGCCTAAATTGCTTTTTAAGGATTCAGCACTCtatagttacttgtcctaaac
gacttatcagtcttgtgtacTGGCAGCTAATGGTTGTCATAGGGAGGAACAAAAATGGCTtatagttacttgtcctaaa
gacttatcagtcttgtgtacCACAGAGCAATTTTAGCCATTATCAACTGAGAATAAAACCtatagttacttgtcctaaa
gacttatcagtcttgtgtacAATTAAATTAGCTAGTCTTCCCTCCCAGATTCATTTTTCCtatagttacttgtcctaaac
gacttatcagtcttgtgtacATGAAACCCATCTCTATTTAGCAGGCCATGTAACCTCTCCtatagttacttgtcctaaa
gacttatcagtcttgtgtacCAAATATAAACACCAGTGCAATTAAGATCTCGACTCCATTtatagttacttgtcctaaac
gacttatcagtcttgtgtacATGCGCATCAGGGTTTTAATGTTTTCTCCGGTTTAGTGCTtatagttacttgtcctaaac
gacttatcagtcttgtgtacAACAATTGCCTCTTGGGTTGGAAAATAGTTTGAATAGGATtatagttacttgtcctaaac
gacttatcagtcttgtgtacGAACCAAACAGTTGTTCTTCAAATTGACAGTGTATTCAGTtatagttacttgtcctaaac
gacttatcagtcttgtgtacTTGCTGAAAACATGGTGAGCCTTTATTTTTTTTCCTATGCtatagttacttgtcctaaac
gacttatcagtcttgtgtacAGAGGCACAAACAAGAATGAGGGAAATTGAAAAAAAGGACtatagttacttgtcctaa
gacttatcagtcttgtgtacGCCTTTATTTTCTCTCCATTACCCTCAAATTATTTCTCTCtatagttacttgtcctaaac
gacttatcagtcttgtgtacTTTATCAGACAATTCTTCTTCACAGTCAGCTCTAAATGTGtatagttacttgtcctaaac
gacttatcagtcttgtgtacGGCCGTCCATCTTTCTCCCTGCATAACAGCCTCCCTTCCTtatagttacttgtcctaaa
gacttatcagtcttgtgtacTGAAAAGCAGCTCCTCTCCCCTTTCAGTTCCTCTCCTGACtatagttacttgtcctaaa
gacttatcagtcttgtgtacCCTTCAGGAAATAACCATTGTCAGCCCTTAGAATATGAGGtatagttacttgtcctaaa
gacttatcagtcttgtgtacAGAACTGTAACTAACGTGTCTGAAAGGAGAATAATGAGCCtatagttacttgtcctaaa
gacttatcagtcttgtgtacCTCTCCTTCTCTCCGTCTCTCTATTTTGTCTTTGCCTTGTtatagttacttgtcctaaac
gacttatcagtcttgtgtacTCTCTCCTTATGCCTTTCTTTGTGCCTGTGTTTTTCTGTGtatagttacttgtcctaaac
gacttatcagtcttgtgtacGGTCCATCCGTTTCCTCTCCCTGAGCACCACCTCATCCCGtatagttacttgtcctaa
gacttatcagtcttgtgtacCTCCCTTCTCCTCTCCTCTTAAATTCTAAGTCTCTGGGTTtatagttacttgtcctaaac
gacttatcagtcttgtgtacCAAGATCCTTGAAGCCAACCAGGCGTGGTATCTCACGCCTtatagttacttgtcctaa
gacttatcagtcttgtgtacTGGGAGGCCCAGGCAGGTGGATCAGCTGAGGTCAGGAGTTtatagttacttgtcctaa
gacttatcagtcttgtgtacAGGCAGAAGAATCAATTGAACTCAGAAGGCGGAGGTTGCAtatagttacttgtcctaa
gacttatcagtcttgtgtacTTGAAGCCAAATAAAGAGTGATGGAGCATGTCTCTGGAGAtatagttacttgtcctaaa
gacttatcagtcttgtgtacCTGGCAAGACTCAGGAACAGCTATTAATGAATCAAGGCAGtatagttacttgtcctaaa
gacttatcagtcttgtgtacCTGGGGCAGCACATTCAAATACTTTTGGTCGTCATTGAGAtatagttacttgtcctaaa
gacttatcagtcttgtgtacAGTTTGGGCCTTAAGTGCTGATTTCTCTTCTGATGGAGTGtatagttacttgtcctaaac
gacttatcagtcttgtgtacTCTGTGTGATACTTAAAGTTACCTCTCTTGGTGTTGTCCAtatagttacttgtcctaaac

gacttatcagtcttgtgtacACTGTCACCACCAAGGCATAGCCCACAGTGAGCTCTATGAtatagttacttgtcctaa
gacttatcagtcttgtgtacCTGAGGTAGGTGGGGACAGGATTTAAACTTTCCTTAGCCAtatagttacttgtcctaaa
gacttatcagtcttgtgtacCCTGAGCCACATGCGGCCTGCAGGCTGCACATTGGACAAGtatagttacttgtcctaa
gacttatcagtcttgtgtacGTGTACCAAAGAAATATGTCCCCTTAAACACCTTCTACATtatagttacttgtcctaaac
gacttatcagtcttgtgtacGCCAGCTTCTCACAGGGGACTGGTAGCTTTCTGATAAACAtatagttacttgtcctaaa
gacttatcagtcttgtgtacAGCATACTTGTTTGCTCGTGATGTGCAGGCTTGAGAAACAtatagttacttgtcctaaa
gacttatcagtcttgtgtacATAGAAACTTTATTGATTTGGCACGCTTGACTTTCCCTTTtatagttacttgtcctaaac
gacttatcagtcttgtgtacCTTTTGATGAATGTCTAGAACGAATGTATAACAAACACCCtatagttacttgtcctaaac
gacttatcagtcttgtgtacACTGCTGTCTACTGATTTCTGATTCTTTTTCCCTCTGTGAtatagttacttgtcctaaac
gacttatcagtcttgtgtacCAGACCTTGTTTTACAGACTCTTCAAGACAGTTGAAATAAtatagttacttgtcctaaac
gacttatcagtcttgtgtacGGTTTTAGCCTTGTTCACCTAGCAAGAATAGAGGAAAAGAtatagttacttgtcctaaa
gacttatcagtcttgtgtacTTACTTACTCTTACTTCTTACAATCTTTCAGTCCTTGGTGtatagttacttgtcctaaac
gacttatcagtcttgtgtacGACTTAGAAAATCCTCAGGTCAGTTCAGTCTAAAACATATtatagttacttgtcctaaac
gacttatcagtcttgtgtacGGTACTGATGAAGGGCAGGGATCCTGGGGTAAGTAGGACAtatagttacttgtcctaa
gacttatcagtcttgtgtacCCTTCCAGAGATTGTATGAGGAAAGTGCTGAGTAAATGTCtatagttacttgtcctaaa
gacttatcagtcttgtgtacCTTGATATAAGGACTTTCTAAGGACAGTGGGCAAACACTCtatagttacttgtcctaaa
gacttatcagtcttgtgtacGGTTTCTTCCTATTTGTTTTGTGTTCTTGGTAAAGTGGTAtatagttacttgtcctaaac
gacttatcagtcttgtgtacATACCACTTTGCATATACTTTTACTTCCAAACTGTGACCAtatagttacttgtcctaaac
gacttatcagtcttgtgtacCATGCATGAGGGCCTGAATTGTTTTGCAATTACTTTTTAAtatagttacttgtcctaaac
gacttatcagtcttgtgtacATGTCTTTCTCATGTTTGTGTCCCCAAATAATATGTGCTCtatagttacttgtcctaaac
gacttatcagtcttgtgtacTCTTGGAGTTACATTTAATTTCATGCCTCCTTTTTGGGTGtatagttacttgtcctaaac
gacttatcagtcttgtgtacGATGCTTTTCTCCTGGCCCTCATCCCGAACACACTGGGTGtatagttacttgtcctaa
gacttatcagtcttgtgtacCTTGGAAAGCGTGGTTTTGGGGGTGAGTGGTGGCAGCTATtatagttacttgtcctaaa
gacttatcagtcttgtgtacCAGTGTGTTTCTGGGTGTTCTGCTCTGAAGCGTTTCTGCAtatagttacttgtcctaaac
gacttatcagtcttgtgtacGCAACAATTAAGTAAGCATTTTTTTCTTCCTTCTGCCAGCtatagttacttgtcctaaac
gacttatcagtcttgtgtacCCTTCTGGTTTCCTCTAAATTAGCATACCCCTCTCTTGGTtatagttacttgtcctaaac
gacttatcagtcttgtgtacATATTGTATATAAATCCTCTGTGGGTGCTTGTTCATTAGCtatagttacttgtcctaaac
gacttatcagtcttgtgtacCTCTCCTCCCTTCCTTCTCTCAGAATCTCCCTGACATAAGtatagttacttgtcctaaa
gacttatcagtcttgtgtacGGCTTATCACTTTGTCTCATTGTCCTCTTGCCTGGTTTAGtatagttacttgtcctaaac
gacttatcagtcttgtgtacCCTGATTTTGGAATTCTTAAGATACTACTGCAATGAAGAGtatagttacttgtcctaaac
gacttatcagtcttgtgtacCCAGTTTCATCTTTGGTCCAGGGAATAGAACAGCCTAGGGtatagttacttgtcctaaa
gacttatcagtcttgtgtacCACCTTGTTCTGTATCTGTGGAGAAAGCTGTGATTCCCCTtatagttacttgtcctaaac
gacttatcagtcttgtgtacTCAACTGGAGGTGTAGGAGTAGGGGAAGATTATGGGTGGAtatagttacttgtcctaaa
gacttatcagtcttgtgtacGCATCTTTCCTTCTTACTCACAGGACACACAGCTAGCCTTtatagttacttgtcctaaa
gacttatcagtcttgtgtacCCAGGTATACAGCAGAAGAGGTTTGTGGATGTCATTTGCAtatagttacttgtcctaaa
gacttatcagtcttgtgtacCAAGTTCATATGCTGGCTCCTAGAAGGGAAAAGTTGGGAAtatagttacttgtcctaaa
gacttatcagtcttgtgtacAGAACTTTTTTGGGATTAGAAACTATTGAGAATGCTTGCCtatagttacttgtcctaaac
gacttatcagtcttgtgtacAAAAGATTCTAGAACAGGACAGATCTAAGGAGATACATCTtatagttacttgtcctaaa
gacttatcagtcttgtgtacTTTTCTTGTTTTGGAACACTCGGAGTCTGGCCCTGTCACCtatagttacttgtcctaaac
gacttatcagtcttgtgtacGATCTTGGCTCACTACAACCCCTGCTTCCCGGGTTCAAGCtatagttacttgtcctaa
gacttatcagtcttgtgtacCCACCATGCCAGGCCTGATCCTCGACTTTTAAAGTCATTCtatagttacttgtcctaaa
gacttatcagtcttgtgtacGACAGTGTCAAGGGCAAAGGCAGCAAAAATCTAAGGGTTTtatagttacttgtcctaaa
gacttatcagtcttgtgtacAGGGCCTGGCTCTGTAACTCTTTCTCATTTTTCAGGTGTCtatagttacttgtcctaaac
gacttatcagtcttgtgtacGTGCCCCCTTAACACACTGTGCCTACAATTCTGCCAGCCCtatagttacttgtcctaa
gacttatcagtcttgtgtacCCATCCAAGGCCCATGTTTGTGAGACCGTAACAGTACTGAtatagttacttgtcctaaa
gacttatcagtcttgtgtacTCACACAGACATTACGGGCATTCACTGGGGACTTGCTGAAtatagttacttgtcctaaa
gacttatcagtcttgtgtacGCAGATCCTTTATAGCACAGGTGGTCCACTCTAGGAAAAGtatagttacttgtcctaaa
gacttatcagtcttgtgtacGGTAACTTGTGCCCAGAATGTGCCCATGGAGTATTAACTTtatagttacttgtcctaaa
gacttatcagtcttgtgtacTGAACACAATACCTTCTCTTCATGAATTTATAAAGGCCACtatagttacttgtcctaaac
gacttatcagtcttgtgtacATCTTCGATGAGTACTATTTCCCTTTTGATGATATGCAGGtatagttacttgtcctaaac
gacttatcagtcttgtgtacGTCCAACAGGAATTAGATAAAAATCTGGAGAAATTTGTGGtatagttacttgtcctaaac
gacttatcagtcttgtgtacCATTATGCAGACTAGAACATTATTGCTTAAACTACCGAAGtatagttacttgtcctaaac

gacttatcagtcttgtgtacGAAGTCTCCAAAATATAACATGGAAGTACTCTTGGCTTGAtatagttacttgtcctaaac
gacttatcagtcttgtgtacATTCTATTTAACACCTTTCAATAGGGCAACATCTGGATGTtatagttacttgtcctaaac
gacttatcagtcttgtgtacGCACTCTTGTCTTTGCCTGTAATTGCCTTAAAGATATAAAtatagttacttgtcctaaac
gacttatcagtcttgtgtacAACTCCCCTAGTGTAACTATTCTTCATTTCTATTCCACAAtatagttacttgtcctaaac
gacttatcagtcttgtgtacGAGCTAATCCTTAAATATGTGTGCCCTGCATCTCAAATAAtatagttacttgtcctaaac
gacttatcagtcttgtgtacCCTATTAACTGGAATAATTGTTGCACTAAGATAAGACGCCtatagttacttgtcctaaac
gacttatcagtcttgtgtacTATTATTTATTTCCTAAAGTCCTGTTCTCCAGTGATAGGGtatagttacttgtcctaaac
gacttatcagtcttgtgtacATGGTTGTTTCATTTTTAATCTGGATCCTTTGCCCTGACAtatagttacttgtcctaaac
gacttatcagtcttgtgtacATCTGAAGAGTGCTGTTAAGAATGTGATGAATTGGGCTTCtatagttacttgtcctaaac
gacttatcagtcttgtgtacGGGAGCAAGATACAGCAGCCTGGAACTACTTTCATTGCACtatagttacttgtcctaaa
gacttatcagtcttgtgtacGTAGAGCCAACCATTGCCGAGAAGACTTAAGCAAGGAGAAtatagttacttgtcctaaa
gacttatcagtcttgtgtacTGCCTTGAATTCTGTCAGTGCACTTCATCTCTCATTTCAGtatagttacttgtcctaaac
gacttatcagtcttgtgtacGGTCTTTGGATTGAGCTGGAATGATGGAGAACACTTGAGTtatagttacttgtcctaaac
gacttatcagtcttgtgtacGTGTCTGAGTTAGTTTTGCCCCTGGGTACCATGACACTTTtatagttacttgtcctaaac
gacttatcagtcttgtgtacCAGAGAAATGGGTGCTTCTGTGTTACAGTTCAACAGCTCTtatagttacttgtcctaaac
gacttatcagtcttgtgtacAGAGGGTAAGGCATGTCTGAGTCACTCTTGGTACCACGTGtatagttacttgtcctaaa
gacttatcagtcttgtgtacGCCAAGCATAAAGATAAGCAGCAGTTTAATCCTTATTTGAtatagttacttgtcctaaac
gacttatcagtcttgtgtacAGCCAAACTGTGTGATTACCATTCCCAACTAGACTTTTGAtatagttacttgtcctaaac
gacttatcagtcttgtgtacGGTCACGAGTTCGAGGCCAGCCTGCCTAACATGGTGAAACtatagttacttgtcctaa
gacttatcagtcttgtgtacAGAAAATAAAGAAAGAACAAAAGTGGTAAAGCAGTCAGGGtatagttacttgtcctaaa
gacttatcagtcttgtgtacGAGACAGAAATCACAATAGTATAGTCATGAGCCGCATAATtatagttacttgtcctaaac
gacttatcagtcttgtgtacGTGGTCCCATAGATTACAACACTGTTTTTACTGTACCTTTtatagttacttgtcctaaac
gacttatcagtcttgtgtacAGCAATAGACTATGCCATATGGCCTAGGGTTGTAATAGGCtatagttacttgtcctaaac
gacttatcagtcttgtgtacTCATTAAGTGAGACATGACTAATAGTTTGGACAGGGAAAAtatagttacttgtcctaaac
gacttatcagtcttgtgtacCTAAGAAGTATAAAGGGAATTCTCAAGAAAACTGGGGTAAtatagttacttgtcctaaac
gacttatcagtcttgtgtacGGCATAGGGCTGAAAGACAGTATTCAAAGAAAGTATAACCtatagttacttgtcctaaa
gacttatcagtcttgtgtacAGTTTCAAGGCTGTGATTCATTTAATGTGGTCTAGTAGATtatagttacttgtcctaaac
gacttatcagtcttgtgtacCTCTTTGGAGGGTTTGTGTCATCAGGTCTTCTGCATGCCCtatagttacttgtcctaaac
gacttatcagtcttgtgtacTTTTCTAAGAACAGAGCTGGAACCAGAAGAGAAGCCCCTTtatagttacttgtcctaaa
gacttatcagtcttgtgtacCTAACGAAAGAAATTAGGTCCAGCTCCAGTTATCATAATGtatagttacttgtcctaaac
gacttatcagtcttgtgtacCAGAAAAGGGCATATGTGGGGTTCAGAGACAGTAAATTGGtatagttacttgtcctaaa
gacttatcagtcttgtgtacAGGTTGTCTTAGAGAGGGGAGAGGAGGTTGGAGTGGGTGGtatagttacttgtcctaaa
gacttatcagtcttgtgtacCGGATAAATTAGGATGAGACTAAGCGATGCACAGACCATGtatagttacttgtcctaaa
gacttatcagtcttgtgtacGAAGTGTGGCTGAAAGAGTGGGGATGGAAAGGAGGGGGAAtatagttacttgtcctaa
gacttatcagtcttgtgtacAGAATACTCAGGGGATGTAATAAATAGGTCTCTGTGGCGGtatagttacttgtcctaaac
gacttatcagtcttgtgtacGGCATGCTCCCTATGTTTGGTGGCTTTCCTATCTGAGAAAtatagttacttgtcctaaac
gacttatcagtcttgtgtacTTGTTTGGACTTCTCAATTTATGGTACCTGTAGACTATCCtatagttacttgtcctaaac
gacttatcagtcttgtgtacGTTCCACAACTGATACAAAACTCACATGCAAAGAAAAGGAtatagttacttgtcctaaa
gacttatcagtcttgtgtacCTTGGAGCTGCAGTACTTTTTTATAATATGGGAAGTTGTAtatagttacttgtcctaaac
gacttatcagtcttgtgtacACAGGTGTGCACAGAAACTGTAATTTGAATAAAGATCACCtatagttacttgtcctaaac
gacttatcagtcttgtgtacTATTTTTACAGGCTGTTACATTTTATTGCAGGCCAAAAGCtatagttacttgtcctaaac
gacttatcagtcttgtgtacAAAGGAAGGAGGTGGAGCTAATGGGCTGAGAAGTCAAAAAtatagttacttgtcctaaa
gacttatcagtcttgtgtacACAAAACAAGTGTAATTTTTCCTTAGCGCTCATTTCTCGAtatagttacttgtcctaaac
gacttatcagtcttgtgtacTTCTATTTTGTGTCCAGTTGCTTTCTGTATATAAGTCCTGtatagttacttgtcctaaac
gacttatcagtcttgtgtacGCAGGACAGATACCACATGCTATTTATTCTTCACAATTTAtatagttacttgtcctaaac
gacttatcagtcttgtgtacCTTACTAAGCATCATTCTGAGCCAACACATGCCTAGGTGAtatagttacttgtcctaaac
gacttatcagtcttgtgtacCCTGTCCTTGAGGAACTCATGATTTAAATGAAGAAAAATGtatagttacttgtcctaaac
gacttatcagtcttgtgtacGGTCAGACTCTTGAATTTTGAATGGATCTTTGAAATAAGGtatagttacttgtcctaaac
gacttatcagtcttgtgtacGTACCATCCATTCTTTCCACCAAAGTTTGTGAGTTTACTGtatagttacttgtcctaaac
gacttatcagtcttgtgtacCACAGTGTGGGATAATCAGAACTCCAGTAGTATTTATATGtatagttacttgtcctaaac
gacttatcagtcttgtgtacCTGGAGAGGACATCAGAACAGTCTAAAGCTTTATAAACTTtatagttacttgtcctaaac
gacttatcagtcttgtgtacCAAAATGCTTTGAACACAGAAAGTACCAACATAAGTGCTAtatagttacttgtcctaaac

gacttatcagtcttgtgtacTATTATTAGCCCCAAGACCTTTTTCCCTGTACTGTTAGATtatagttacttgtcctaaac
gacttatcagtcttgtgtacTGCTTAACTTCAAGGGAAAGTATTGTTCTTTACATGAGCAtatagttacttgtcctaaac
gacttatcagtcttgtgtacACATTGTGTCATTTATCTACTCTGTAAGGTCCCTAGCAAAtatagttacttgtcctaaac
gacttatcagtcttgtgtacGCAATGGCTGGCTTGCCTTAAGAAGAAGCAAAATGTATATtatagttacttgtcctaaa
gacttatcagtcttgtgtacGGATCACCTGAGCCCCTGGGAAATCGAGGCTGCGGTGAGCtatagttacttgtccta
gacttatcagtcttgtgtacAACAACAACAGAAAAAACCCCAAAGAATCTTAGCTAGGACtatagttacttgtcctaaa
gacttatcagtcttgtgtacAAATATTTGCATTTTAAAGTGGTTACCGGGGAGACTATAGtatagttacttgtcctaaac
gacttatcagtcttgtgtacTTCTTGTCATCTGGAATTGCTTTCAAAACCACTTGTGAAAtatagttacttgtcctaaac
gacttatcagtcttgtgtacAAAAGCTAGTGCCATTACTTCATTTCACATCTGATACACAtatagttacttgtcctaaac
gacttatcagtcttgtgtacGAACATGAGAGACCAAAAGTGAAAAATATATCCAAAGGGAtatagttacttgtcctaaa
gacttatcagtcttgtgtacCTGTGTTGGTGTGTGCAAGGCTCAGAAAGTCAATGAGTTAtatagttacttgtcctaaa
gacttatcagtcttgtgtacTGAAGCAAATCATTCTATACACACCCACATAAACTCAAGAtatagttacttgtcctaaa
gacttatcagtcttgtgtacCAAATTTCGGTTATACTGGATCTCGTGGGAAAAAAAAATCtatagttacttgtcctaaac
gacttatcagtcttgtgtacAAAAGACATTCACCAAAGAAGGAAAGAAGGCAAACAGGGAtatagttacttgtcctaa
gacttatcagtcttgtgtacTCTTTCCAGGAGTCTAATCAAATGTTCTGAAAAACCAACTtatagttacttgtcctaaac
gacttatcagtcttgtgtacACCTCAGGTGACAATAGTCAGAAATTTTTGGTTACTTTCCtatagttacttgtcctaaac
gacttatcagtcttgtgtacCTGGAAGAGGAAAGAGAACTAGATAATGGGCAGCTGGAAGtatagttacttgtcctaa
gacttatcagtcttgtgtacAGGCTCTTATTCCTTTCCCTGTCTTATACATCTTTTTAAGtatagttacttgtcctaaac
gacttatcagtcttgtgtacTGGCAGGGTTGTTTTTTTCTTAAGACGTGAGGAAGAATCTtatagttacttgtcctaaac
gacttatcagtcttgtgtacTCACCTTGATCTCTGCGTTGATGTTCACTTGCTGTTTTTTtatagttacttgtcctaaac
gacttatcagtcttgtgtacTGTATGTCTGCCTCTGGATTTCCCCTTTTATCAGGACACCtatagttacttgtcctaaac
gacttatcagtcttgtgtacGACTTCAACATGTAAATTTGTGGGGAACACAATTTGACCCtatagttacttgtcctaaa
gacttatcagtcttgtgtacCACGGAGAAATGAAAATGTGGGGAAAAAGAAAGCCTAACAtatagttacttgtcctaa
gacttatcagtcttgtgtacGCACTATGGCATATATAAACAGGACGTCGTTCTTCTCACTtatagttacttgtcctaaa
gacttatcagtcttgtgtacAGGCACCTTTTCATATCCATCTCAGATAAATTCCTATAGCtatagttacttgtcctaaac
gacttatcagtcttgtgtacACCTATCTATGACACTCATAATTTTCTACCTTGTACTGCAtatagttacttgtcctaaac
gacttatcagtcttgtgtacGCAGTATCTAATCCATCTTTGTTTCTATTTCTTGACCTAGtatagttacttgtcctaaac
gacttatcagtcttgtgtacATCATTGAATCCATAAATGGCTGTCCAGACTTTATAGGCCtatagttacttgtcctaaa
gacttatcagtcttgtgtacCTGTTAAGCACTATATTGGTCAGGATAAGTTAGGTTGTACtatagttacttgtcctaaac
gacttatcagtcttgtgtacGCTGAGGCAAAAATCTTGTAGAAGTCCAGGCAGTTCCTTAtatagttacttgtcctaaa
gacttatcagtcttgtgtacGAAGCAAAGTACCCTGGACAATTTCATCTTAGCATTAAGCtatagttacttgtcctaaa
gacttatcagtcttgtgtacAAATGTGAAATGTCAATCTCAGGGACCTTTTCTTTTTCCCtatagttacttgtcctaaac
gacttatcagtcttgtgtacTTTACTGAGCCCTTAACTATGTACCAGGCAGTATTGTAGAtatagttacttgtcctaaac
gacttatcagtcttgtgtacCATTCCTGTGTTGAACAAGGATTGAAAAAAGGAGTGGCCAtatagttacttgtcctaaa
gacttatcagtcttgtgtacCTTGTGATTTCAGTGCCAGTATGTTCTAAAAATCTTAGTCtatagttacttgtcctaaac
gacttatcagtcttgtgtacTGTATATTTGCTCAGAATTCTGTGATTTGGGTAAAGCTTGtatagttacttgtcctaaac
gacttatcagtcttgtgtacAACCTCATCTGTGCTCCATATCTGTTGGGCTGGAATGTCCtatagttacttgtcctaaa
gacttatcagtcttgtgtacCACATGACTATTGACTGAGTTGAATGTCTAAATTATGGAGtatagttacttgtcctaaac
gacttatcagtcttgtgtacCTCCAAATGGCTAGCTTGGGGTTCCTCACAGTATTGCAGCtatagttacttgtcctaaa
gacttatcagtcttgtgtacCTCAGGGCCCTATCAATCCTTTGCTTGCATCACACTTACTtatagttacttgtcctaaa
gacttatcagtcttgtgtacCCCTAGAGTCAACCTGGCAGAGAACTACACAAGGGCATGAtatagttacttgtcctaa
gacttatcagtcttgtgtacGGTTCATTGAGAGCTACCAAAATATCTGTTTATCACTGTCtatagttacttgtcctaaac
gacttatcagtcttgtgtacTCTGGCCTCTAATAATTCTCATTCCTCCTGCAATACCCTCtatagttacttgtcctaaac
gacttatcagtcttgtgtacTTCTGTTACAGCATGAGATTTGGACTTGAAGTCTACAATCtatagttacttgtcctaaac
gacttatcagtcttgtgtacGCCCAGGTGAAAGATGAGGCTTCTTAGGTATAATCACTTTtatagttacttgtcctaaa
gacttatcagtcttgtgtacGTTATATTCTCCCTCATGCCCATAACCATCAACATATAATtatagttacttgtcctaaac
gacttatcagtcttgtgtacATGGCAATAAAAGGTCCATTCAAAAAGGGGGAAATGTGAAtatagttacttgtcctaaa
gacttatcagtcttgtgtacCTGCAATCTAGCTGGGTACATATTTTCAGATCTTAGGGCCtatagttacttgtcctaaa
gacttatcagtcttgtgtacAAGTGATTCTCCATGGCTCATGCTTACAACTGAGTGGCTTtatagttacttgtcctaaa
gacttatcagtcttgtgtacGAAATGCTTCTGTCTCTTTTAGTTCAATCTGGCAATGTTTtatagttacttgtcctaaac
gacttatcagtcttgtgtacACTTTGTGGGTTTCTTAGGTATCTTATTAGGGGTTAACTTtatagttacttgtcctaaac
gacttatcagtcttgtgtacACAAAAGTTATACCCACAAAGCTCTTTAAAACTGATCCCTtatagttacttgtcctaaa

gacttatcagtcttgtgtacGCAAGTCAGGGTGCTATGGATAATACCTTTACTATTTTTtatagttacttgtcctaaac
gacttatcagtcttgtgtacTTTTTGGTGAGAGAGTTTGAGGCAATTTGTGAGGATTCCTtatagttacttgtcctaaac
gacttatcagtcttgtgtacAGAGCCCTAAATCTTTTCTGAGATCTCACAAGCAGTATCCtatagttacttgtcctaaac
gacttatcagtcttgtgtacGTCCCTATTTTATTGGTATTGTTCTGGATTTGATATGAGGtatagttacttgtcctaaac
gacttatcagtcttgtgtacCTTTGAGAATATTTTTCCAAGGAACACTGGGGATGAGAATtatagttacttgtcctaaac
gacttatcagtcttgtgtacTTGAACCCCAAAGCCTTTGCATTTTCTTCAAATTCTGCTTtatagttacttgtcctaaac
gacttatcagtcttgtgtacTCTTTAGTTCATCTTACACTTCCCATAACTTAACTTAGGCtatagttacttgtcctaaac
gacttatcagtcttgtgtacGATATCTTCATAGCTGGATCCACAAGTTAATTAGGTATCTtatagttacttgtcctaaac
gacttatcagtcttgtgtacACTGTTGGTGATAGTTTTGCTAATGAATCTACACATCCATtatagttacttgtcctaaac
gacttatcagtcttgtgtacAGCAATTTCCTCTCCGCTCTTTTAACTAACATTAAGTGTCtatagttacttgtcctaaac
gacttatcagtcttgtgtacGTCTCCACCTATTGCACAATTCCAAATCCCACATGTCTTAtatagttacttgtcctaaac
gacttatcagtcttgtgtacCACTTTGGGTATCAATTTCTAACTTCCAGTTTACTTCAGGtatagttacttgtcctaaac
gacttatcagtcttgtgtacTTTTACCAATTGTTCTGCCAGTACATGATTCAAGCTTTCTtatagttacttgtcctaaac
gacttatcagtcttgtgtacGATAGCTATTTTATCACTGCCCTTCCATTCTTCACTAATAtatagttacttgtcctaaac
gacttatcagtcttgtgtacGGAATAAGTACACATTTAATTTGGTCCCCTATATCTTCTGtatagttacttgtcctaaac
gacttatcagtcttgtgtacGATTCAGTTTTCTTAGAAAGATTGCAAACAATTGCCTGTCtatagttacttgtcctaaac
gacttatcagtcttgtgtacTTCCCTTCCTTCCTCCCTGCCCTTCCTTCTTCTTCTTTCCtatagttacttgtcctaaac
gacttatcagtcttgtgtacTCCTTCCTTCCTTTTATCCAGGTACATAGTATTTAGCTTTtatagttacttgtcctaaac
gacttatcagtcttgtgtacTTCTTGCATGATACTTACTGTCTAGTGAGAAACTTACTTGtatagttacttgtcctaaac
gacttatcagtcttgtgtacCAAAACTTGTCTTTGGCTTACCTCACTTCACTTGTTGGCAtatagttacttgtcctaaac
gacttatcagtcttgtgtacAGTCACTGAGGTCTACTGTTTTTATGTTTTTCTTAGGAAGtatagttacttgtcctaaac
gacttatcagtcttgtgtacGTCCTTTGCTTGATTTCTTCCAGAACGGGCCTCAGAATCTtatagttacttgtcctaaac
gacttatcagtcttgtgtacATACATATATATCTCTACCTTCTTCTTTGATTTCCCCAGGtatagttacttgtcctaaac
gacttatcagtcttgtgtacAATGTGATATTGGGGCATCATTGCTTTCTCTGATAAAATGtatagttacttgtcctaaac
gacttatcagtcttgtgtacAGACAGATACTTTTCTTTCTAGAGATGAAACATGTGAACCtatagttacttgtcctaaac
gacttatcagtcttgtgtacATGACTAGGCATGTGAGGAAGATATGTAGGGGTAAACTGGtatagttacttgtcctaaa
gacttatcagtcttgtgtacCATGTGGCTTCTGTCTCTCCTCCTGGGATTCTCCATGTGAtatagttacttgtcctaaac
gacttatcagtcttgtgtacTGGAGTAGGCATGCTTTCAAGAGTGGATAGAACCAGTGGCtatagttacttgtcctaaa
gacttatcagtcttgtgtacAAAGAGGTTTGCAATGTAGGTCTGGCTTTACACCCTTACTtatagttacttgtcctaaac
gacttatcagtcttgtgtacCGAAGGAACCTAATTAGCCACATAGTAAAATCTGTACTTTtatagttacttgtcctaaac
gacttatcagtcttgtgtacCTTTGACAGCAAGTGTAAGTAACTTTGTGACCAGCATTTGtatagttacttgtcctaaac
gacttatcagtcttgtgtacTCTGTGTATTTAAGATCTGAAGAAAAAGATTTCTGACCCCtatagttacttgtcctaaac
gacttatcagtcttgtgtacGGGTATGACCCTCTGGCAGGGTTTTCAGCTCTTTACCCGCtatagttacttgtcctaaa
gacttatcagtcttgtgtacTCTCGCTCTGCCACCCAGGTGGGGGTGCAGAGGCAGGATCtatagttacttgtcctaa
gacttatcagtcttgtgtacGCGATTCTCCTGTGTCAGCCTCCCCAGCAGCTGGGAGTACtatagttacttgtcctaa
gacttatcagtcttgtgtacCCGGCCTGGGATTTTTATCTGTGAGGATTAAGTGAGGTGTtatagttacttgtcctaaac
gacttatcagtcttgtgtacAGCCAGACTCTGCCTCTTGCTGCTTGTGGGACCTATTGGGtatagttacttgtcctaaa
gacttatcagtcttgtgtacCTATAAATGTGGAATGTTCTCAAGTGAAGAGAAGAATGGGtatagttacttgtcctaaac
gacttatcagtcttgtgtacTCACGGTTCACTACAACCTCGACGTCCTGTGCTAAAGCAAtatagttacttgtcctaaa
gacttatcagtcttgtgtacCCTGGGCTCAAGCGATCCGGGTCCTGTTTTGGCCTCCCAAtatagttacttgtcctaa
gacttatcagtcttgtgtacGGAGTGAACCACCGCCCGCAGTCTTCAATGATAGTTTATCtatagttacttgtcctaaa
gacttatcagtcttgtgtacATCTGCCCAGCCTTCCCAAAGATGAATTATATTTACAGTTtatagttacttgtcctaaac
gacttatcagtcttgtgtacTCCTGGGGAAGAAGATGTTCATAATTTCTGAGAAAGTTTCtatagttacttgtcctaaac
gacttatcagtcttgtgtacCTGTAGGGAAGAGCAGGTCTCTCTGACTTTCTGACTCAGCtatagttacttgtcctaaa
gacttatcagtcttgtgtacTGCTTTTCTGTGCTTCTTTGAGAGCAAGATGAAACCCAGAtatagttacttgtcctaaac
gacttatcagtcttgtgtacAGGGACATTCCTTTGTACATTTACTGCTACGTGGAAGAATtatagttacttgtcctaaac
gacttatcagtcttgtgtacGAATCTAAAAATGTTCCCTTTTTTTGGGTGTAGTTAGGTGtatagttacttgtcctaaac
gacttatcagtcttgtgtacAGTTATGACAAAGCTACTTCTTTTTTTCAGGGAGAAAAGGtatagttacttgtcctaaac
gacttatcagtcttgtgtacTTAGTGCAGCTAATCACCATGGGCTCCTCTAGTACTTAGCtatagttacttgtcctaaac
gacttatcagtcttgtgtacTACTCCTGTAGGCACATTTTCTAAGTAAAGTTCTGATTTGtatagttacttgtcctaaac
gacttatcagtcttgtgtacCATTGCAAATGGATGATTTATAAAGAGGAACAACAACGACtatagttacttgtcctaaac
gacttatcagtcttgtgtacCTTATCTAATCTAAAAGGAACTAAGTCCTAGAGAGAAAGGtatagttacttgtcctaaac

gacttatcagtcttgtgtacCACTCGCTGTCCAGTCAAAGACATACTTCTCATCCACTTAtatagttacttgtcctaaac
gacttatcagtcttgtgtacCCATTTGACCCTTTAACAATGATTCGTTTGATTAGGATAGtatagttacttgtcctaaac
gacttatcagtcttgtgtacTTACTAATAACTTGGACCTCTGAGCTTTTTAATGCAAAGGtatagttacttgtcctaaac
gacttatcagtcttgtgtacGCAAGTCCATTATAATCACCTGGCTGATCTCCATGCAAAAtatagttacttgtcctaaac
gacttatcagtcttgtgtacATTAATTAATTGTTTTGAAGCAGAGTCTCACTCTGTCACCtatagttacttgtcctaaac
gacttatcagtcttgtgtacGTTGGACAGGCTGGTCATGAACCTCTGACCTCAAGTAATCtatagttacttgtcctaaa
gacttatcagtcttgtgtacCAGTAGGCATTTTAAACTGGAACTTGGCTGAGGGGCAATCtatagttacttgtcctaaa
gacttatcagtcttgtgtacAGAAATTCTCTGTTAGCAACACCCACTACCTGTCAATTCAtatagttacttgtcctaaac
gacttatcagtcttgtgtacTCCTGTCCTAGTGAGATCCTGGAGTTGATTTAGATGAGGGtatagttacttgtcctaaac
gacttatcagtcttgtgtacGCTTGATCACCTGCCCACTGTTACCTATGGCCCCATGTTGtatagttacttgtcctaaa
gacttatcagtcttgtgtacTCAGAGGAAGAGACAGGAGGGTTCACAACTTAGTCAAACTtatagttacttgtcctaaa
gacttatcagtcttgtgtacCATTCTGATGGTTGGTCATTATTGTCCCCTCAATGTTTGAtatagttacttgtcctaaac
gacttatcagtcttgtgtacTTTCATGCTACTGATAAACAATACCCAAAACTGGGAACAAtatagttacttgtcctaaac
gacttatcagtcttgtgtacCACTTCTTACACAGTGGTAGAAAGAGAAAAATGATGAAGAtatagttacttgtcctaaac
gacttatcagtcttgtgtacACCCATCAGATCTCACGAGACTTAATCACTATCACAAGAAtatagttacttgtcctaaa
gacttatcagtcttgtgtacCCCCATGATTCAATTACCTCCTCCTGGGCCCCTCCCACAAtatagttacttgtcctaa
gacttatcagtcttgtgtacAAGTTGAGATTTGGGTGGGGCACAGTCAAACCATATCATTtatagttacttgtcctaaac
gacttatcagtcttgtgtacGGTGAATACAGCCCTTTCAAATGAGAGAAACTAGCCAAAAtatagttacttgtcctaaa
gacttatcagtcttgtgtacTGCAAGTCCAAAATCCAGCCGGGCAGTCAAATTTTAAAGCtatagttacttgtcctaaa
gacttatcagtcttgtgtacGTCTTGGGCAGCTTTGCCCCAGTGGTTTGCAGGGTACAGCtatagttacttgtcctaa
gacttatcagtcttgtgtacGGATTTTCCAGGTGCATGATGTAAGTTATCAGTGGATCTAtatagttacttgtcctaaac
gacttatcagtcttgtgtacGCCATGGCTGGAGTCAGGGGGACACAGGGCACCAAGGTCCtatagttacttgtccta
gacttatcagtcttgtgtacAAACCACTTTTTCATTCTAGGTCCCTGGGCCTGTGATGGGtatagttacttgtcctaaac
gacttatcagtcttgtgtacTCTCCCACATGGCCTGGAGACGTTTTTCCCATGGTCTTGGtatagttacttgtcctaaa
gacttatcagtcttgtgtacGGTTCCTTATACAAATTTCTGTAGCCAGCTTTAATTTCTCtatagttacttgtcctaaac
gacttatcagtcttgtgtacGCTGCAAATTTTCCAAATTTTTAATGCTCTGCTTCCCTTAtatagttacttgtcctaaac
gacttatcagtcttgtgtacCAGGGGGAAAATGCCACCATTCCCTTTGCTAAAACATAAAtatagttacttgtcctaaa
gacttatcagtcttgtgtacTTCGGGTTTCTTTTCAGCAACATCCCAATTCTGGTACCAAtatagttacttgtcctaaac
gacttatcagtcttgtgtacCTCAGAATCATGGCGAGAGAAAAAAGGCACTTCCTAAATGtatagttacttgtcctaaa
gacttatcagtcttgtgtacCCATATCATTGGGTCTGCATTTCTTTTAAGGCTGTTTCTCtatagttacttgtcctaaac
gacttatcagtcttgtgtacTTTGTTCTCCTCCCTTGAAAAAAGTCCATATTTGCAGAGAtatagttacttgtcctaaac
gacttatcagtcttgtgtacAGAGGATAATTAGTCTCAAAGAGTTGCAAATATCTGTGGTtatagttacttgtcctaaac
gacttatcagtcttgtgtacGCTGGGTGCTGGTCTGGAAACAGAGGATGCTTATTTTTAAtatagttacttgtcctaaac
gacttatcagtcttgtgtacTTTGGGAATATGAAAGAGTAGAATGTTTACTGGGGAAGGGtatagttacttgtcctaaac
gacttatcagtcttgtgtacTGTAGTCATTTTGAGTTTTGCAGATAAAGGCAATCCACACtatagttacttgtcctaaac
gacttatcagtcttgtgtacCTCTCACAGGACTAAAGAGCCGCTCCCCAATGTGTAGCCAtatagttacttgtcctaa
gacttatcagtcttgtgtacGCTGCTGTTTCAAAGGCTTTCAATTCTGTGTGTCATTATCtatagttacttgtcctaaac
gacttatcagtcttgtgtacCTTTTTTGTACTCGGGTGTTTATTATTGGATTGCAACAACtatagttacttgtcctaaac
gacttatcagtcttgtgtacGCTAATTAGCCAGTTTCCTTTTGATCTTTTTCTCACTGAAtatagttacttgtcctaaac
gacttatcagtcttgtgtacGAATCTTAGTGCTGTACATTCATCTCCTCATCTCCATAATtatagttacttgtcctaaac
gacttatcagtcttgtgtacCAATTTTACTGTTGAATGAGAGCTACTAGAGGCTTTAGTTtatagttacttgtcctaaac
gacttatcagtcttgtgtacCAAGGAGGAAGACGCAAAAGGGGAAAAAGGCACAAATATTtatagttacttgtcctaa
gacttatcagtcttgtgtacGCTCAGTTGCTTTTATTTTTCCTTGAAGCATTCAATCCATtatagttacttgtcctaaac
gacttatcagtcttgtgtacACTGAATATTTAGTTGGAAGAAAATGGCCTGAATAGTGTGtatagttacttgtcctaaac
gacttatcagtcttgtgtacCCGTTGGAGGATAAGCTGCCTTTCAAAAGACACAGTAAATtatagttacttgtcctaaa
gacttatcagtcttgtgtacCTGGAGGCACATGTATTAGATTATGTTTGGGAAAGGATGGtatagttacttgtcctaaac
gacttatcagtcttgtgtacAGTTAGTAGATTCTTCGTTTGATTGTTTATTGGTCCACTGtatagttacttgtcctaaac
gacttatcagtcttgtgtacCTGGGAAGTTGAGTGGGAAATTTCATCATTAATATCATCAtatagttacttgtcctaaac
gacttatcagtcttgtgtacCATGGCGTTACAAGGACCAAGCAAGTTGATATTTAAATACtatagttacttgtcctaaac
gacttatcagtcttgtgtacGACCTATATCCAGCCCCTACAGATACACATTCACAAACATtatagttacttgtcctaaa
gacttatcagtcttgtgtacCATTTTTGGTTGAGCTACAGGAAATCACTGTTTTTGCAAGtatagttacttgtcctaaac
gacttatcagtcttgtgtacCTGTTTGGGAGGTGTGATCAGGGTAATCAAGTTTTATTTAtatagttacttgtcctaaac

gacttatcagtcttgtgtacTTTAAAAAGCGCTCATACATCCAATAGAGTATTGCCTCCTtatagttacttgtcctaaac
gacttatcagtcttgtgtacCATTTTTAGAATAGTCCTAAACCTGAGGTGTTTCATTTCCtatagttacttgtcctaaac
gacttatcagtcttgtgtacCTTCGAAGATGGGTTTGAATTTTGGAAATAAAGGAAAGTCtatagttacttgtcctaaac
gacttatcagtcttgtgtacACTGTATAAAACTGCTGAATGAGAGCTACTATTTTGAGTGtatagttacttgtcctaaac
gacttatcagtcttgtgtacAAATAAGCAAAATTTGAATGCAATGGTTTCCTTGGGTGCCtatagttacttgtcctaaac
gacttatcagtcttgtgtacTGAGGTCATCTGGTTGAACAATCCTTAAACTTTATCAAACtatagttacttgtcctaaac
gacttatcagtcttgtgtacGCCATAAAGATTCTGATTTATTAGACTTAGAAATGACCCGtatagttacttgtcctaaac
gacttatcagtcttgtgtacCTTTTACTAAGTTTGTCAGTGATTCCAATGCAGGAGGTGGtatagttacttgtcctaaac
gacttatcagtcttgtgtacACTTGGAATGGCTGCTCTTACCTATAGAGATAGATGCTGAtatagttacttgtcctaaac
gacttatcagtcttgtgtacGAAATTATTCCACTACACAAAAGCTGGTCCAGAGCAGAATtatagttacttgtcctaaac
gacttatcagtcttgtgtacAGGGACTTCTTGGGTTTGAATCACAGTATGGTACTTAACAtatagttacttgtcctaaac
gacttatcagtcttgtgtacGCATCCAACCATTTTTAAAGCCCATGTTTTCTCATATGTAtatagttacttgtcctaaac
gacttatcagtcttgtgtacGGATAATTCTGATACCAAGCTCACAGTGTTGTGTCCTGCAtatagttacttgtcctaaac
gacttatcagtcttgtgtacCATCACTTACCCTACGTTTAGCCAAAGAATTTTATTATGGtatagttacttgtcctaaac
gacttatcagtcttgtgtacTTTCCTTTTCTGTACTTGTCACTTTTTATGATTTCTCCCCtatagttacttgtcctaaac
gacttatcagtcttgtgtacCCTTTTCACAGGCATCTCATCTTGACATTCCTTCCCCATGtatagttacttgtcctaaac
gacttatcagtcttgtgtacCTGCCACCAGACTAGAAAAAGTCACTGAAAGATAACAATCtatagttacttgtcctaaa
gacttatcagtcttgtgtacCTGAGAGGAACAGCTGGGTCTTTTCACATTTTGTCTTTATtatagttacttgtcctaaac
gacttatcagtcttgtgtacAGCTCTACTGGGTTGTTATTTAAAAAATTCCTCAGCTAGTtatagttacttgtcctaaac
gacttatcagtcttgtgtacAGATTGAAGTTATCAGCACAGAAATTCTCTCTTTATGCCAtatagttacttgtcctaaac
gacttatcagtcttgtgtacGTGGAGGGCAAAGAGCAGGATATAGAGAAAAGCAGCCAGCtatagttacttgtcctaa
gacttatcagtcttgtgtacGGCATCTGCACTTGCCCACGGGCTCTTCAGACCTGGCTCTtatagttacttgtcctaaa
gacttatcagtcttgtgtacCACTAATGATGGTGCCAGAGGGAGAAAGTCAGCATGAGGCtatagttacttgtcctaaa
gacttatcagtcttgtgtacACTTGTGTCTCTTCCCCATCTGTTCCCATCACCCGTACCCtatagttacttgtcctaaa
gacttatcagtcttgtgtacCAACCAAATGCTGCCTGAGCTACCTGATGCTGTTAATTACtatagttacttgtcctaaac
gacttatcagtcttgtgtacCTTAGAGCCAATTATCTCCACAAAAGCAAAACATCTTCATtatagttacttgtcctaaac
gacttatcagtcttgtgtacGAAACTGTTTCATCTGGGGCAGAGACAAATTGAACACTTTtatagttacttgtcctaaac
gacttatcagtcttgtgtacAGTTCAAGCTAAAAAGGCTCTTCTTTGGCATGGATTTCAAtatagttacttgtcctaaac
gacttatcagtcttgtgtacGTAGCTTTGCAGATGCCACAACTACTAAAAATATAGCAGGtatagttacttgtcctaaac
gacttatcagtcttgtgtacTGGGGTGTGATACTCTGTAAAGATAGATTTGTATATGTCTtatagttacttgtcctaaac
gacttatcagtcttgtgtacCTATAAGTATTTCTCTATTCATTCAGTCTCTCTCAGTCTCtatagttacttgtcctaaac
gacttatcagtcttgtgtacGGCTAAAGAGTGTCATGTGAACTTATGGAGAGGTTTGCATtatagttacttgtcctaaac
gacttatcagtcttgtgtacCATTCAAACTTTGTTGGAAATCTAGTCTAGGACTTCATTCtatagttacttgtcctaaac
gacttatcagtcttgtgtacCTGGTCAAAACATTTCAACTAAGGAAAAAGTTAGGCTATGtatagttacttgtcctaaac
gacttatcagtcttgtgtacGGACAATTGCTCAGCTCTCTTTGTTCATAGGTACAAGTTGtatagttacttgtcctaaac
gacttatcagtcttgtgtacTTCTCTCCCATGTCACTGTTGGAAGAAACAAAGATAATTTtatagttacttgtcctaaac
gacttatcagtcttgtgtacAGTAATATGCATGATTTCAGTGTCCTCTAAATGAACGAAGtatagttacttgtcctaaac
gacttatcagtcttgtgtacACAATGAAGTCAAGCACCTGGAGGAGAAGGATGCAGATAAtatagttacttgtcctaaa
gacttatcagtcttgtgtacCCTTGGGCATGCTTCTAATTTCCCCTTGGTTTCTCTAAAGtatagttacttgtcctaaac
gacttatcagtcttgtgtacGGAAAACATCCACATATTGACTCATCCTGGATAAAGTTCAtatagttacttgtcctaaac
gacttatcagtcttgtgtacATCCAGTCATTCATTCATTTGCCATGGGGATGCAGTGGAGtatagttacttgtcctaaac
gacttatcagtcttgtgtacATTGTAGCTGAGGTGAAATCCATGGGAAAAACAGTTTTTAtatagttacttgtcctaaac
gacttatcagtcttgtgtacGGAGTTAGAGAAGCCATACCGAGAAACTTCCAGGCCTACGtatagttacttgtcctaaa
gacttatcagtcttgtgtacCAGTCCCAATGTTGTGAGGATAGCATACTTTTTTGAGATTtatagttacttgtcctaaac
gacttatcagtcttgtgtacTCAAGGTTGTTTTTGATAATGCCTGCCGGTTATCCTTAGTtatagttacttgtcctaaac
gacttatcagtcttgtgtacCATTTGGATGTGTGTGGTAGTATGTGCTAGTATCAAACGAtatagttacttgtcctaaac
gacttatcagtcttgtgtacCTGAAGCTTACTAGGTACCCCCATTTCATCATGTGAGTACtatagttacttgtcctaaac
gacttatcagtcttgtgtacGGATCTCTCTTCACTGAGCTCCCTCCCAGCTTGTTAATTGtatagttacttgtcctaaac
gacttatcagtcttgtgtacAGAGAGAGAGAATATGTATGGGCAAACATGTATGTGAACAtatagttacttgtcctaaac
gacttatcagtcttgtgtacGTTTGTGCATAATGGGTGTGGACATGCATATATGTGAGTGtatagttacttgtcctaaac
gacttatcagtcttgtgtacTGTCTTCAGAATGCTTGTAATATAGATCATCATAGGCCTCtatagttacttgtcctaaac
gacttatcagtcttgtgtacTTATAGAGCAGTTGAGTAATTTTAGTAACCCAGCACATTCtatagttacttgtcctaaac

gacttatcagtcttgtgtacACCCTTTTCAATAAGCTTTAGGTTCTATCATACTATGAGGtatagttacttgtcctaaac
gacttatcagtcttgtgtacGCAGTCCATGCTGCATTCATTTTAAATTGGTCTTTTTAAGtatagttacttgtcctaaac
gacttatcagtcttgtgtacGGAGAAGATTCCTCAGTTCTTTACCCATGTGATTTGTAACtatagttacttgtcctaaac
gacttatcagtcttgtgtacGGTGTAGAACATTTCTGCCCAGAATATCATTATACATTACtatagttacttgtcctaaac
gacttatcagtcttgtgtacTTCATTTTTCTTACAAAATTATGCTCTGAATTGTCCCCGCtatagttacttgtcctaaac
gacttatcagtcttgtgtacTCAAAGGCTGATTTCATAATGGCTGTGTTCTGGTGTGCTTtatagttacttgtcctaaac
gacttatcagtcttgtgtacATTTGAGACTGGGTCTTCATGACATACAGTGAGTGCAACTtatagttacttgtcctaaac
gacttatcagtcttgtgtacCATCTATAATGACAGGTTCTGGGGAGAAAATGATGAGTATtatagttacttgtcctaaac
gacttatcagtcttgtgtacCTCAAGGTGCTCTCGGAGTGGTGGGGGAGGCATTCTCTAAtatagttacttgtcctaaa
gacttatcagtcttgtgtacAAGGGAGTTCTTTTCTTGGCATTATGGAAGCTTTACAGAAtatagttacttgtcctaaac
gacttatcagtcttgtgtacCAGATATTGTAGGGGGGTAGTGAGAAAACATTCCAGGTTAtatagttacttgtcctaaac
gacttatcagtcttgtgtacTGTCGAATTTCCATTTTCAGAGCTAAGTAGTCACAATTCAtatagttacttgtcctaaac
gacttatcagtcttgtgtacCTATTCTAGTCTTGCCATTTTCTCTCCTCTCCTTTGCCTGtatagttacttgtcctaaac
gacttatcagtcttgtgtacAAGCACTTGACTCACTATAAAAATATAAGACACCTAGCCTtatagttacttgtcctaaac
gacttatcagtcttgtgtacCTCTCCTTCCTTGAGTTTATTATGCTGCCTTCTTTGTCTAtatagttacttgtcctaaac
gacttatcagtcttgtgtacACTGATTTCTTGGTTTAGAACTTTTTTGGCAGTAGGGTGAtatagttacttgtcctaaac
gacttatcagtcttgtgtacCTACTCCAGCCTCTATGTAACAGCCCAGAAAAAGGACAAAtatagttacttgtcctaaa
gacttatcagtcttgtgtacAGCTTATCCTGGGTTGTGTAACTTCAAGAGGCTCACTTTCtatagttacttgtcctaaac
gacttatcagtcttgtgtacGGTCTCCATTTGCTTTTTATTTCATGCTTTGAATTGTGAGtatagttacttgtcctaaac
gacttatcagtcttgtgtacGGACAAGCCATAAAACCAGACCAAAACAAAAAAACAAAAGtatagttacttgtcctaaa
gacttatcagtcttgtgtacAAACCCCCCAAAAGTGCCTTGTCTGATTTGCAACACATTTtatagttacttgtcctaaac
gacttatcagtcttgtgtacTTCGTTAGTAAACAGCAAGTGAAGATTTATTGGAGTTCGGtatagttacttgtcctaaac
gacttatcagtcttgtgtacCAGCAACCATATTCTGTTCTGTTTCATACTAGGTGGGACTtatagttacttgtcctaaac
gacttatcagtcttgtgtacTGCAAAGGTCAGAAAGCAGCTTCCCTTCAGGCTCATATGAtatagttacttgtcctaaa
gacttatcagtcttgtgtacAGTACCAGTTTTAGTTACAATGTTACATAGCAAACAAGGCtatagttacttgtcctaaac
gacttatcagtcttgtgtacTGTGGTTTAATTCCTGCATGTGTGTTCTTGATTTGTGTTGtatagttacttgtcctaaac
gacttatcagtcttgtgtacCAGCTTATGACCCCTGAAATGATCTATTCTTGTACTAACAtatagttacttgtcctaaac
gacttatcagtcttgtgtacCCTTTGTGTTGCATAAATGATAATCTCCTTTTTGTCTAGGtatagttacttgtcctaaac
gacttatcagtcttgtgtacTCTATATGGTGTGAGAATAGAGTTCTTTCTATCCTTTTGGtatagttacttgtcctaaac
gacttatcagtcttgtgtacTCTTTAATTCCATAGTGTAATATAAGCACAGGAAGTGTGGtatagttacttgtcctaaac
gacttatcagtcttgtgtacTACTATTTACTGTGATACTTCTTCTTGTCAATTGCTGCAGtatagttacttgtcctaaac
gacttatcagtcttgtgtacACTGGCTTCCATTGTCTTCCACGCTTTTGATTTCTTATTTtatagttacttgtcctaaac
gacttatcagtcttgtgtacTGTACAAAATTCTGTGATTGGGTAGGCAATTCAAGTTAGTtatagttacttgtcctaaac
gacttatcagtcttgtgtacTATCAAATGTATGTGTAAGACTCTGTATTAGGTGTGTAGGtatagttacttgtcctaaac
gacttatcagtcttgtgtacCAATAGCATTATGGGCAAAGATAAACAAGAATCTGTCCTTtatagttacttgtcctaaac
gacttatcagtcttgtgtacTTTTGAAGGGGAAATGCTGACTTATAAACAAGAAGGAATGtatagttacttgtcctaaac
gacttatcagtcttgtgtacCTTTTATAGTAGAAGCATAAGCACAGTGCTCTGGCAACCTtatagttacttgtcctaaac
gacttatcagtcttgtgtacCTGGAACTCATTACTTCTAGCTTGATATATATTCTCTGGGtatagttacttgtcctaaac
gacttatcagtcttgtgtacGATCCAGCTTCATTTCACATGTATATTTCAAAATTGCCTCtatagttacttgtcctaaac
gacttatcagtcttgtgtacTGGTTCCAAAGGCAGGTAGTATGTAGTTAAATAGTTATGAtatagttacttgtcctaaac
gacttatcagtcttgtgtacGTCTCCTTTGTTTGGATGACTTTATCTTGTATATTGTCTCtatagttacttgtcctaaac
gacttatcagtcttgtgtacACTTTCAATTGTGGCTCAGAAGTTTTGCTCATCCATTAATtatagttacttgtcctaaac
gacttatcagtcttgtgtacTATTTTCTTAACCTGGAATACATGGATGAGCTTTAGGGAAtatagttacttgtcctaaac
gacttatcagtcttgtgtacCTTTTTTTTTCTTTTGACATAGAGTTTTGCTCTGTTGCCCtatagttacttgtcctaaac
gacttatcagtcttgtgtacGGGATTACAGGGACCTGCCACCATGCCCTGTTAATTTTTTtatagttacttgtcctaaac
gacttatcagtcttgtgtacGAACTCCTGACCTCGAGTGGTCTGCCTGCCTTGGCCTCCCtatagttacttgtcctaa
gacttatcagtcttgtgtacGCCACCGCGCCCGGTCAGCTGTTATTATTTATTATTATTtatagttacttgtcctaaac
gacttatcagtcttgtgtacTGTATTTTTAGTAGTGATTGGGTTTCACCATGTTGGCAAGtatagttacttgtcctaaac
gacttatcagtcttgtgtacCGTGAGCCACCATGTCCGGATCTCTTCTCTGGAGAGCACAtatagttacttgtcctaaa
gacttatcagtcttgtgtacCTTATGAACCATACGATGTAAAACTACCATATTTCTCGCtatagttacttgtcctaaac
gacttatcagtcttgtgtacATGTTGATGACTTTAAACCTGTATTTCTAGCCTAAACCTCtatagttacttgtcctaaac
gacttatcagtcttgtgtacCCAAATGTTTACCAGCCATCTTTGCTTTTTTGGGCCAATAtatagttacttgtcctaaac

gacttatcagtcttgtgtacTCATGATCTTTCCTTCTAAATCTTGCTCACACATTTTCTGtatagttacttgtcctaaac
gacttatcagtcttgtgtacATGCCACCAACATCTACTCAATTACCCAAGATAGGAAACTtatagttacttgtcctaaac
gacttatcagtcttgtgtacCTTTTAAGACAACTCAAATATCACTTGCATAAGGAAGCACtatagttacttgtcctaaac
gacttatcagtcttgtgtacCTTTTGGGTTCCCATAGCACTCTGTACGTATCTCCATCATtatagttacttgtcctaaac
gacttatcagtcttgtgtacTCTATTTTTCTGTTTCCCTTATTAGTCTGTAAGCTCCTTGtatagttacttgtcctaaac
gacttatcagtcttgtgtacTGACCCATCTCTTATTTGGCTTTGTAAGTCAGCACTGAGCtatagttacttgtcctaaac
gacttatcagtcttgtgtacAATGAATTAATTAAGAGGACGTACATTGTACATGGCTTCCtatagttacttgtcctaaac
gacttatcagtcttgtgtacTGTAGTGATAGTGACTTAAAATATTCTGATTTCATGGCCCtatagttacttgtcctaaac
gacttatcagtcttgtgtacCAGTGTGGTCTGCTAACATGAAATTATCTTAAAAGTCAGAtatagttacttgtcctaaac
gacttatcagtcttgtgtacCCAACTGTGTGGTAATAGCCTATATTCTACATGTCCTTTCtatagttacttgtcctaaac
gacttatcagtcttgtgtacTTGTGTTAGATAGAAGTATAAAATTCCTGAGTTCTGGTGGtatagttacttgtcctaaac
gacttatcagtcttgtgtacAGAATATGTCTCTCTGTCCTTCACCTACCAAACCTCTTTTtatagttacttgtcctaaac
gacttatcagtcttgtgtacCAAATGAATCAGAATTTTGTATCCCCTCCCCTCCTCCTGGtatagttacttgtcctaaac
gacttatcagtcttgtgtacTGCTTCCTCCTACTCTATGCCATGTAAGAAGGACTTGAGAtatagttacttgtcctaaac
gacttatcagtcttgtgtacCTCACCCTGGCTGACTGTTAGCTGAGCTATAGAATCCCATtatagttacttgtcctaaa
gacttatcagtcttgtgtacTAGGGAGAAGTCAAACTACATCCTCTAGAGTCTGGACCTAtatagttacttgtcctaaa
gacttatcagtcttgtgtacCTGTAGCTTGGAAAATCTTAAAAGCAAGAGGCTTGTGGAGtatagttacttgtcctaaac
gacttatcagtcttgtgtacCAATGATAGAAGGCCACAGTGGGAGCGTCGTGTACATCCAtatagttacttgtcctaaa
gacttatcagtcttgtgtacCAGAGGTGCAAGTTTTCTCATGGGCTAACTGGACATGGTGtatagttacttgtcctaaa
gacttatcagtcttgtgtacGGTATCTTGAAAGTCCACTGCCTAAGAGATCTGATGGTCAtatagttacttgtcctaaac
gacttatcagtcttgtgtacACTGGGGTAGGGAAGTGAGGTGCATAGTCTGGGAATGGTGtatagttacttgtcctaaa
gacttatcagtcttgtgtacTTGTGTATGTTTTGGGTTGAGGTCTTTCCCCATAGTCCTGtatagttacttgtcctaaac
gacttatcagtcttgtgtacTTGCTGACTTCAAGAATGAAGATTGAGAGGTGAAGCCAGCtatagttacttgtcctaaa
gacttatcagtcttgtgtacTTCCTGGGTTGAGTGGGGACTCGGAGAACTTTTCTGTCTAtatagttacttgtcctaaac
gacttatcagtcttgtgtacCCCAGCCAACAGTGGCAACCAGCTGTGGTCCCCTTCCACGtatagttacttgtcctaa
gacttatcagtcttgtgtacTGCTGCTGCTCACTGGGTCCGCACTACCTCTATGAGCTGTtatagttacttgtcctaaa
gacttatcagtcttgtgtacGTCAGTGAGACCACGAACCGACTGAGAGGAACAAACAACTtatagttacttgtcctaa
gacttatcagtcttgtgtacAATAAATTCTAGACACGGTCCCATTCTCAGACTTACTCAAtatagttacttgtcctaaac
gacttatcagtcttgtgtacCTTGTTGAGAAGCTAGAAAAAGAAGTTAGTGGGCCGGGCTtatagttacttgtcctaaa
gacttatcagtcttgtgtacGTAATCCTAGCTACTCAAGGGGCTGAAGCGGGAGAATTGCtatagttacttgtcctaaa
gacttatcagtcttgtgtacAAAGAAAGAAAAAGAAGTTAGCCATGGGACAGGCAAAGGAtatagttacttgtcctaaa
gacttatcagtcttgtgtacAAATTTATTTATCATATTTTCAGATGCAGGGAGGCCCGAGtatagttacttgtcctaaac
gacttatcagtcttgtgtacCTGTCTCTGTTTCCAGGATGATGCTTTGAATGCTGTGTCCtatagttacttgtcctaaac
gacttatcagtcttgtgtacGAATAGTTCCCCCATACCACTTTTTCTAAGAGCACCCATCtatagttacttgtcctaaac
gacttatcagtcttgtgtacTGAATTTTGGAGGACACATATACCCCATAGCAGCTGTGTTtatagttacttgtcctaaac
gacttatcagtcttgtgtacAGGCCCTTCTAGACCTCCTTTTCACCTAATACAGCAGTCTtatagttacttgtcctaaac
gacttatcagtcttgtgtacGCTTTCCTGTGCTGCTGAGAATTGTGCTTGGCCAAATAGGtatagttacttgtcctaaac
gacttatcagtcttgtgtacTCTCACCCAGGAAGCTATGAATGCCGCCTTAGTAAACCTAtatagttacttgtcctaaa
gacttatcagtcttgtgtacCTGCTGGCATAGGATGTTAAAAATGTTAGAAGTATTTTGGtatagttacttgtcctaaac
gacttatcagtcttgtgtacAAAGTCTACCAAATCAACTAGAACTTTAGTGGCCAAGGTGtatagttacttgtcctaaac
gacttatcagtcttgtgtacGACAAGAGAAGTACACAGAGGAAATGATTTTCAGGGCTGCtatagttacttgtcctaaa
gacttatcagtcttgtgtacAGAGAGGCTGAGAAATTGAGCAGAAATTGATGGCTGAAAGtatagttacttgtcctaaa
gacttatcagtcttgtgtacCCTGGTGGTCTCATGGGCTAGTGGACAAAAATTGGAATTCtatagttacttgtcctaaac
gacttatcagtcttgtgtacCCCTCCAGGGTTTTAATTAGGACTTTGGATCGGCTACTCCtatagttacttgtcctaaac
gacttatcagtcttgtgtacAAATCAGTTCATTCTCAAGTCAGAGTAAGATGATCTTTCCtatagttacttgtcctaaac
gacttatcagtcttgtgtacGCAAAAGTAAATGCTGTCTTGAAGAAGCTGACGTTATCCAtatagttacttgtcctaaac
gacttatcagtcttgtgtacGTGTGGCATTTAATAAAAAGTTACCAGGGCACTAGTAGAAtatagttacttgtcctaaac
gacttatcagtcttgtgtacGTCTGTTTGAAGAAATTAGAGGAAAGCTTAATGTTTCTGGtatagttacttgtcctaaac
gacttatcagtcttgtgtacAATAGACTTAGAGGAGATGCAGGTATTGAAATTATCTGACtatagttacttgtcctaaac
gacttatcagtcttgtgtacCAACTGACATTAATAACTCCCTAGATAGGTTTAATAGCAGtatagttacttgtcctaaac
gacttatcagtcttgtgtacACAACAACAAAAACCTGCTAAAGAGAGAATTGGCACTCTGtatagttacttgtcctaaa
gacttatcagtcttgtgtacCATAAGACAAATATGGAATATGGTGGATAGGCCAAACAGTtatagttacttgtcctaaac

gacttatcagtcttgtgtacCATGATTGAAGGGATAATGGCTAAGAAGTCTCCAAAAATAtatagttacttgtcctaaac
gacttatcagtcttgtgtacAGTCCAGAAGCACTACAACTCTCTACTAGGATACATATAAtatagttacttgtcctaaac
gacttatcagtcttgtgtacACCATACCTAGCCATATCATAGTACAATTGCCACAAACCAtatagttacttgtcctaaac
gacttatcagtcttgtgtacTCTTAAAAGCAGACAGAAGCCAAAATAAAACTAATGCGACtatagttacttgtcctaaac
gacttatcagtcttgtgtacCTTTTGAAGAAAATAAACTGTTAACCTAGGATTCCATCCCtatagttacttgtcctaaac
gacttatcagtcttgtgtacTTATCCAGCTGACTTGTATGATAGTAAATACTAGAGCAAGtatagttacttgtcctaaac
gacttatcagtcttgtgtacGCCAGAAACAGGGAGAAAATGAAAAGCAATGCAAAAAATAtatagttacttgtcctaaa
gacttatcagtcttgtgtacATGTTTTGTGGGGTTTACATATCTATGGGATAGACACATGtatagttacttgtcctaaac
gacttatcagtcttgtgtacGGAATGGAATGGAATTATCCTCAGGTCTTTTCATTATGACtatagttacttgtcctaaac
gacttatcagtcttgtgtacATGAAAAAAATAAGGCAACAATGAGAGCAAAGGAAACACGtatagttacttgtcctaaa
gacttatcagtcttgtgtacTGTTGGGGGAAAATAGACAATGTACAGCATGATGGTAGATtatagttacttgtcctaaac
gacttatcagtcttgtgtacTCAAAAGTGAAAAGGCAAAAAGGTTGACAGTAAAAGGATGtatagttacttgtcctaaac
gacttatcagtcttgtgtacGACGAATATTTCAGAACAATAAAGTGGTTAATTTGCCAGGtatagttacttgtcctaaac
gacttatcagtcttgtgtacACTGACATGACAATATTACGTGGATGTAATAACAAGCCTCtatagttacttgtcctaaac
gacttatcagtcttgtgtacTCTAATATACCTCTTTCAATGACTGACTGAATAAGCAGAGtatagttacttgtcctaaac
gacttatcagtcttgtgtacACTACATCCAACAACTGCAGTGGAACCTAAGACAAGTCCCtatagttacttgtcctaaa
gacttatcagtcttgtgtacCCACACATTAAATTTATACAGGGCATGTTCTTTGATCACAtatagttacttgtcctaaac
gacttatcagtcttgtgtacAGGTAACTAGAAAATCCCTAGGTGTTTGGAAATTAAGCAAtatagttacttgtcctaaac
gacttatcagtcttgtgtacGAAGATGACGAATATTGTGCTAAAATGAATACTGTAGGAGtatagttacttgtcctaaac
gacttatcagtcttgtgtacTAGCTAGCCCAACGAAATGAAGTAGATTCTTTCTTTTTCAtatagttacttgtcctaaac
gacttatcagtcttgtgtacCAGAAATGTCATGGTAGGGCTTTTGAAGTGTTTGTTTTGTtatagttacttgtcctaaac
gacttatcagtcttgtgtacTATAGATCAAATGGCCAATTTAGAGTAGCTTTCTGGGACAtatagttacttgtcctaaac
gacttatcagtcttgtgtacTGATTATTTCTATCACCCCAAAGATCTGTAAAATGGCCTGtatagttacttgtcctaaac
gacttatcagtcttgtgtacGGGATTATAGACACCCACCGTCGTGCCTGGTTAATTTTTGtatagttacttgtcctaaac
gacttatcagtcttgtgtacACTCTTGACCTCAGGCGACCCCCCCTAACCCCATGGCCTCtatagttacttgtcctaa
gacttatcagtcttgtgtacAATGTTACTTTTCTGTGGATACAATTTGGCCGAGTTTTAGtatagttacttgtcctaaac
gacttatcagtcttgtgtacCACACTGTGCATAATAATTAAGAATCCTGCTATCTGTTTCtatagttacttgtcctaaac
gacttatcagtcttgtgtacGATACTAGATGTTTTTGTGTGCAGAATAAGGACAAAGATGtatagttacttgtcctaaac
gacttatcagtcttgtgtacATAATAGTAATCATGCCAGTAAGTAATGAAATGGAGGGTGtatagttacttgtcctaaac
gacttatcagtcttgtgtacATCCCTCTGATGAAGATCATAAGTTTCCAATGCCAAAAACtatagttacttgtcctaaac
gacttatcagtcttgtgtacACCTTAAGAAGCCCATGAGGTCAAGACTTATGTGATGGCAtatagttacttgtcctaaa
gacttatcagtcttgtgtacCTCTGAGGACCATAGTCTAACATAGCAGGTTAACACAAATtatagttacttgtcctaaac
gacttatcagtcttgtgtacTAGATTAATCACAAGAGCTAAATTTAGGGAGTTAAGGAGCtatagttacttgtcctaaac
gacttatcagtcttgtgtacTATTCCTCCCATATCCCTACCATAGCTTTAACCTCCAAGGtatagttacttgtcctaaac
gacttatcagtcttgtgtacAACACTGCTGGCTTCTATTAGAAATGCAAAATGTTTCCTTtatagttacttgtcctaaac
gacttatcagtcttgtgtacGGATGAAGGGGCTGCATTTGTAATATTGTGCTCATAAAACtatagttacttgtcctaaac
gacttatcagtcttgtgtacGGTGTGTTAGTACCTTTCTTATGTATTTCTCTCCATGCTCtatagttacttgtcctaaac
gacttatcagtcttgtgtacTAGCACACTCTTCTTCACTTGCTAAAGATGAATTTGGATTtatagttacttgtcctaaac
gacttatcagtcttgtgtacAATTTGTGGTTAGGCTAGAACACATTAAATCATAAGGTCCtatagttacttgtcctaaac
gacttatcagtcttgtgtacGTCCCTAGGATTTTGACACCAGAGCAAGTTTTCTTTTTCTtatagttacttgtcctaaac
gacttatcagtcttgtgtacTGGTTTGTTCTGAAAGACTCACTTGGATTTTATATGATGCtatagttacttgtcctaaac
gacttatcagtcttgtgtacAGTGTCCTGCCTTTTAACCTACTATAGAACCCAATTTGAAtatagttacttgtcctaaac
gacttatcagtcttgtgtacTTTCGGAGGACAGATCAAACTTACTTTAAAGTGATTGCTTtatagttacttgtcctaaac
gacttatcagtcttgtgtacTTGTAAAATGCTGACACTGAAGTGGTTCTGAGACACATATtatagttacttgtcctaaac
gacttatcagtcttgtgtacAGAATGAAAACATTTCTGAAGATGACCGCCATGTTAAAGCtatagttacttgtcctaaac
gacttatcagtcttgtgtacCTTGGTATCAGGGTCACTTGGTGGAATCAGGGCTCAGGGAtatagttacttgtcctaaa
gacttatcagtcttgtgtacATGAGTGAGAGAATTCAGAAATAAGTTTTTCCATCCATCGtatagttacttgtcctaaac
gacttatcagtcttgtgtacATAACACTCCCTCCTCATTCCCACACATTCTTAAGCCATGtatagttacttgtcctaaac
gacttatcagtcttgtgtacAGGAACGAAATCTCAAATATCTTCTGCCTCCTATTTATCAtatagttacttgtcctaaac
gacttatcagtcttgtgtacAATATATAAAGTAAAACAGCAGTGTGAGAGACTCAAGTGCtatagttacttgtcctaaac
gacttatcagtcttgtgtacCAGGACTGAATCTCATGAAAATCAATTGGGATGCTTTACTtatagttacttgtcctaaac
gacttatcagtcttgtgtacAACAAACAAAAAAGAGCATATTGGCCTAACAGTAGAAAGGtatagttacttgtcctaaa

gacttatcagtcttgtgtacAGCTTGCTTCTCCTGTCTTCTTGTCCTATGATGTCACTAAtatagttacttgtcctaaac
gacttatcagtcttgtgtacACTTGATCTTTCTGGGCCTGCTTCTTCATGTGTAATCTGAtatagttacttgtcctaaac
gacttatcagtcttgtgtacATGTTAGAGAGCCCATGAGTCTGTGAATTACGCCTAAAGTtatagttacttgtcctaaa
gacttatcagtcttgtgtacATTTACCAACCTTATAAGTACTCAACAGTCAAAATGGAGGtatagttacttgtcctaaac
gacttatcagtcttgtgtacCCTGTGCGTGCTAGGTAAATATCAACTACACTTTTGGGAAtatagttacttgtcctaaac
gacttatcagtcttgtgtacAGTCCATCTGCCTGATTTTACACACTGTTGCAACCTCTGTtatagttacttgtcctaaac
gacttatcagtcttgtgtacACATTGTGATACTGAAAAATTGCAGCAGAAGGATGACACAtatagttacttgtcctaaa
gacttatcagtcttgtgtacAATAATGGCATTAATGAATCACACCTCCTTGTATCTACACtatagttacttgtcctaaac
gacttatcagtcttgtgtacATTAATCCTTTCCCACAGTAACTCTGGGCTCAGCCATGTGtatagttacttgtcctaaa
gacttatcagtcttgtgtacCTCATTAGCGTTTACCCTATCTTTTGTTACTTTTGGATCAtatagttacttgtcctaaac
gacttatcagtcttgtgtacGGCCCAACTGACAGCCAGCACCACCCACCAGACTTGCGAGtatagttacttgtccta
gacttatcagtcttgtgtacCGTGAGTGACCCCAAGTGAGACCAGAAAAGATACCACCCAtatagttacttgtcctaa
gacttatcagtcttgtgtacCCCACAGAATCAGAGCAAATGAAATGTTGGTTACTTAAGCtatagttacttgtcctaaa
gacttatcagtcttgtgtacTTGACACAGCAAGAAAAAGATCTATAAGTTGGGGAGGAACtatagttacttgtcctaaa
gacttatcagtcttgtgtacGAAGGGGAAGTTGATGGGAGGAAAGACAGTGGTGATATCCtatagttacttgtcctaa
gacttatcagtcttgtgtacTTCCTAATTGAAGTTGGAACCAAGGATTTGTGTCTAGTTCtatagttacttgtcctaaac
gacttatcagtcttgtgtacAGCTCCTTCATACTCATCTTTCTGGTCAAGCACACTTAGTtatagttacttgtcctaaac
gacttatcagtcttgtgtacACTCTCAGTGAAGAAACATGGGTGGATTTCCTGCTTCCATtatagttacttgtcctaaa
gacttatcagtcttgtgtacGAATGTGACAGGGGACAACTGTTCTGGCCTTTTCTTGTTGtatagttacttgtcctaaac
gacttatcagtcttgtgtacGGTGGCTACCAGCAAAAAAATTCATGGAATACTATGAGAAtatagttacttgtcctaaa
gacttatcagtcttgtgtacGAAGATTTTTTCCCAAATGTCCATATAAGACCTGCAACCTtatagttacttgtcctaaac
gacttatcagtcttgtgtacATCAATTGTATAATCGTACTAAAGCTTCTGAAGCCTTCCCtatagttacttgtcctaaac
gacttatcagtcttgtgtacCCTGAGGACTTCCACTTTTTGTCATAAATCATTCTAAAACtatagttacttgtcctaaac
gacttatcagtcttgtgtacTTTTCAGGTATCAAACAAGCAGCACGATTCCCAAGAGAAGtatagttacttgtcctaaa
gacttatcagtcttgtgtacACACTCTCTCCATGTTCACTTCACTCTCTGCCTAATTTTCtatagttacttgtcctaaac
gacttatcagtcttgtgtacTGGCACAGGAAGCTGGAGTTCATGCAAAACACGATGGTCCtatagttacttgtcctaa
gacttatcagtcttgtgtacGCATGTAGGGCAGTAAGGACGAAAGAACTATGCTGGGTCAtatagttacttgtcctaa
gacttatcagtcttgtgtacCAAAAAATTACCCGAGAAAGAAATCTTTATAGGAGTTCCCtatagttacttgtcctaaac
gacttatcagtcttgtgtacTTGACTGAGGACTGAGATGTATAGGTGTGGGGCAAGATCAtatagttacttgtcctaaa
gacttatcagtcttgtgtacCTGTGGGGCTGAGAGTTAGAACAGAAATACCAGATAGAACtatagttacttgtcctaaa
gacttatcagtcttgtgtacGCTCTGACCCAGCCTGAGGAGAGAGTTCTCATGAATACCTtatagttacttgtcctaa
gacttatcagtcttgtgtacAAGCAAACAGGTTCTTCTTCACATGGCAGCCAGAAGGAGAtatagttacttgtcctaa
gacttatcagtcttgtgtacGGAGACTGCTGCCATAATCTAATTACTTCCCACAAGGTCCtatagttacttgtcctaaa
gacttatcagtcttgtgtacCCAGACCATATCAACACCCCAGGCATTCATCTGAGGCAACtatagttacttgtcctaa
gacttatcagtcttgtgtacTTAGAATAAAGGATCATGTCTCGCGGTAGGCCCTACTCTGtatagttacttgtcctaaa
gacttatcagtcttgtgtacCAAGATTTGACAAGAGATGGAATTTGGAGGTTGAGTACACtatagttacttgtcctaaa
gacttatcagtcttgtgtacGGCTTTTCACAGGCTAATAGGTTCAAATTTACTATCCAGTtatagttacttgtcctaaac
gacttatcagtcttgtgtacTCTAAAAAAAAGGTAGGATATCATGCACAAATCAAGCTGGtatagttacttgtcctaaa
gacttatcagtcttgtgtacGTGAAACACTGAATGCTTTTCTCTTAAACTGGGAACATGGtatagttacttgtcctaaac
gacttatcagtcttgtgtacATGTCTGCTCTCACTGCTTTTATTTGATGATATACTGGAAtatagttacttgtcctaaac
gacttatcagtcttgtgtacAAGAAATAAAGATGAGAAGGAAGAAGTGAGCTTGTCTTAGtatagttacttgtcctaaa
gacttatcagtcttgtgtacGAATTAATGAATAAATTTAGTAATGCCCCAGGATGCATCCtatagttacttgtcctaaac
gacttatcagtcttgtgtacTAGATTCAATGCAACCCCAACCATTTTTTTTTATCCCAACtatagttacttgtcctaaac
gacttatcagtcttgtgtacGTAGAAATTGAGAAGCTGATGCTAACGTTTGCACACAAATtatagttacttgtcctaaac
gacttatcagtcttgtgtacCTATTGGAATTAAGTATGAAATTGTCAGAAGGGCAGACATtatagttacttgtcctaaac
gacttatcagtcttgtgtacACAGAATTCAGAAATAGAGCCCTACATATATGATCAATCCtatagttacttgtcctaaac
gacttatcagtcttgtgtacGACAAAGGCAACAAGATGATTCAATAGAAAAGGACACTTtatagttacttgtcctaaa
gacttatcagtcttgtgtacAAAATGGCAAAAGACTTAGCACTTCACAAAAGACACAAAGtatagttacttgtcctaaa
gacttatcagtcttgtgtacTTTGAACAAACACTGCACCAAGAAAATATAGAGATGGTTCtatagttacttgtcctaaac
gacttatcagtcttgtgtacGAAAATTAAAACCATGGTGGATATCACTTTACACATACCtatagttacttgtcctaaac
gacttatcagtcttgtgtacTTTAAAAGACTGTAAATAGAAAATGCCAGTGAGGATGTGGtatagttacttgtcctaaac
gacttatcagtcttgtgtacGGAATGTAAAATTGCACACCATGATGGAAAATAGTTTCACtatagttacttgtcctaaac

gacttatcagtcttgtgtacCTTGTAGGTATTTACCCCCCGTCTTCAAGATAAAGAACATtatagttacttgtcctaaac
gacttatcagtcttgtgtacCTTGTACACAAACGTTCATAACTGCTTTATTCCTAAGAGCtatagttacttgtcctaaac
gacttatcagtcttgtgtacACAACAAGCAAACAAAAATGGAATACATGGAACAACATGGtatagttacttgtcctaaa
gacttatcagtcttgtgtacCTGGTAGAAGTAACACAGTGATAGGATTCAGATCAGGAGTtatagttacttgtcctaaa
gacttatcagtcttgtgtacATGGAAGTGTTCTATCTACATCTTAACCGTGGTGGCAATAtatagttacttgtcctaaac
gacttatcagtcttgtgtacAACAAAACAGCAGTAAGAAAAATAAAGCCTTCAGGAATGCtatagttacttgtcctaaa
gacttatcagtcttgtgtacCTTTACACATTAACAAATTTAAGCTTCACAAAGGCAGGAGtatagttacttgtcctaaac
gacttatcagtcttgtgtacGGCATTAACTACCCCATCTTCGTATTAAAGAAAAAGAGCCtatagttacttgtcctaaa
gacttatcagtcttgtgtacGAGGTTTTAAATATCTTGTCTAGAGTCATAGTAAGGTGGAtatagttacttgtcctaaac
gacttatcagtcttgtgtacTGGCACTAAATCCAATAACATTTCCACTATACTATGCTGTtatagttacttgtcctaaac
gacttatcagtcttgtgtacACAATGTCCTTTCTGTTGCTAGGCCTTAAATTGCATTTCCtatagttacttgtcctaaac
gacttatcagtcttgtgtacAGTTGGCACTGAGTTTTTTTCTGAAATCTTCTCAATCACTtatagttacttgtcctaaac
gacttatcagtcttgtgtacTGTGGGGATCTTTATCTTCCTTGCAGAAAGTTTGGAATCTtatagttacttgtcctaaac
gacttatcagtcttgtgtacATTAAATTAAATCTTATAACAGCATTTTCTCCCCCCAGCCtatagttacttgtcctaaac
gacttatcagtcttgtgtacATGTATCAAAACTCTTGGCATATTCTTGGGCTGAGCAATTtatagttacttgtcctaaac
gacttatcagtcttgtgtacCCAAGGCAATTATTAGAGACGTGTGTATGGATGTGTGCACtatagttacttgtcctaaa
gacttatcagtcttgtgtacCTGTTGTTTTCCTTCCATTCTCATCTCTTCTTCCTTAGCAtatagttacttgtcctaaac
gacttatcagtcttgtgtacATGCATTTCATACCTTGGCCCTCCTTTACAGCTCATTTGTtatagttacttgtcctaaac
gacttatcagtcttgtgtacAAAATATTTTGTGTGACTTCTAAGAACTTGCCTTACAGGGtatagttacttgtcctaaac
gacttatcagtcttgtgtacCAGCCTACTGGATTGTGAATGTGATGGCTGGACTTGAAGCtatagttacttgtcctaaa
gacttatcagtcttgtgtacGTTGGAACATGGGGCAACACTCTAAGGATGATGGAACAGCtatagttacttgtcctaaa
gacttatcagtcttgtgtacACTCTGGGTCCTGACACTGTGGGAATCAACCATACAAGCCtatagttacttgtcctaaa
gacttatcagtcttgtgtacTGTCACAAACAGCCAAATCGAATCCCAACAAATGCAGTGTtatagttacttgtcctaaa
gacttatcagtcttgtgtacGCAAAGGGAGAGTAAAATTGTGATATAGGCACAGGGTAGAtatagttacttgtcctaaa
gacttatcagtcttgtgtacGCAACTGCTCAAATGACATGAAAAAATTTGCTAAGAGGAAtatagttacttgtcctaaa
gacttatcagtcttgtgtacGCATAAGTGGTTATAATTACTCTTAAGACTGTATGTGAGCtatagttacttgtcctaaac
gacttatcagtcttgtgtacAGATGGGCGAATGTATATACTTTACAGCTCCTACTGCTCAtatagttacttgtcctaaa
gacttatcagtcttgtgtacGAGGGACATTCTAAAGATGACATCTAGTGGTTGCTGTGGTtatagttacttgtcctaaa
gacttatcagtcttgtgtacACTTGCAGTTATTAACCCTTACAGGCTGTATTTGTGATAAtatagttacttgtcctaaac
gacttatcagtcttgtgtacCAGTGGAAGTGTTCAAGAGCAGCCCTACAATGATTAAAGGtatagttacttgtcctaaa
gacttatcagtcttgtgtacACCTGTGTTAGAATTTTCATGACAACCTTAAAGAACTGTCtatagttacttgtcctaaac
gacttatcagtcttgtgtacAAGTTCATGTTTATTACAGAGCACTTACTATTCAACAGGCtatagttacttgtcctaaac
gacttatcagtcttgtgtacTAAGAGAGAACAAAACTGAGACTTAGAGAATTTAGGTGAGtatagttacttgtcctaaa
gacttatcagtcttgtgtacGGATTCAAACCCAAGTCATCCGTCCACACTCATAAGCACTtatagttacttgtcctaaa
gacttatcagtcttgtgtacCCGGTCTGGGTCTATCATAAACATGCACCTTTAAGGGAAAtatagttacttgtcctaaa
gacttatcagtcttgtgtacGTTCATCCAGTCTAGCCTTAATCATCTACACTCATTAAACtatagttacttgtcctaaac
gacttatcagtcttgtgtacCTGTTGTGATTTCTCAAAGTATGATGCACCCTTGTTATGAtatagttacttgtcctaaac
gacttatcagtcttgtgtacGTGCCACATCATAATTTTATTTTGAAGCCTAGCTCTAAAGtatagttacttgtcctaaac
gacttatcagtcttgtgtacAATGAGATAAGCCTAATAATCATGGTAGGGGAGATGCTGCtatagttacttgtcctaaa
gacttatcagtcttgtgtacAGGCAGTGAGAGGATGCTTCCCTAAAGAGCTGACCTTGGAtatagttacttgtcctaaa
gacttatcagtcttgtgtacTGCCTTGTTTGAAATGCAGCATCTTAGACCCTACCCTAGAtatagttacttgtcctaaa
gacttatcagtcttgtgtacGCATTTTAACAAGATCCTCTTGAGATTTGCAGGCACATTGtatagttacttgtcctaaac
gacttatcagtcttgtgtacCAGATAGCATTTCAGCAGCATATACAGAGGGAACAGCACCtatagttacttgtcctaaa
gacttatcagtcttgtgtacACAGTGGTGGGAATGAATAAAGCCATTTGGGGGGAATATCtatagttacttgtcctaaa
gacttatcagtcttgtgtacACAAACTGGAATGAAAGATGGTGAGTAGTGGGAAGATAAGtatagttacttgtcctaaa
gacttatcagtcttgtgtacGAACTACATCTTGTTCTTAAGGTAATGGGAATACCAAAGTtatagttacttgtcctaaac
gacttatcagtcttgtgtacATACTTAAGCTTTGTGATACACTCAGAACATAACCTTTGGtatagttacttgtcctaaac
gacttatcagtcttgtgtacATGACTCCACATATTTCCTTTAGCCTTCATATCACCAAACtatagttacttgtcctaaac
gacttatcagtcttgtgtacTAAAAGAGAATGAAGTGAATCCTGATGGCAAAAAGAACACtatagttacttgtcctaaa
gacttatcagtcttgtgtacATCATGCATTTACTGTTAGGGATTTGATGGTTAAACTGACtatagttacttgtcctaaac
gacttatcagtcttgtgtacTGTCCTTCATGAGTAGATAGAACAAGCAGACAATGTATATtatagttacttgtcctaaac
gacttatcagtcttgtgtacTCGTTCTGAGAAGGGCTCCCAAATCCCCCCAGGCTCCTTTtatagttacttgtcctaaa

gacttatcagtcttgtgtacCTGAATAAGGCAGCCAGGAATAAGAGAAACAGATAAGAGGtatagttacttgtcctaaa
gacttatcagtcttgtgtacCTTGGATTAAAAGGAGAAGGAAAGAGGATTTTGGAGGGTGtatagttacttgtcctaaa
gacttatcagtcttgtgtacTGCAGGCTGACAGACACGTTACAGACACAAAATGAGTCAAtatagttacttgtcctaaa
gacttatcagtcttgtgtacTGAGGAACAAAGACAAGGAAGGGATAAGTGCAGCTAGACAtatagttacttgtcctaaa
gacttatcagtcttgtgtacTCAGGGAGGGTGAGTTAGAGGGAAAAGGGGAAAGGTAAAAtatagttacttgtcctaaa
gacttatcagtcttgtgtacAATCTCTGTTCCTGTACCTTGGGCCTTGCTAACTAGGACAtatagttacttgtcctaaac
gacttatcagtcttgtgtacAGCTCAGAGACTGATATAATTGCTGGGTCCTACAAGTCAGtatagttacttgtcctaaa
gacttatcagtcttgtgtacCAGGACTGGGCATAGCTGCTCAAAATCTGACTATATGTATtatagttacttgtcctaaac
gacttatcagtcttgtgtacAATGCAATTAATCTCATCCGGAAAAGGGAAACCCACACATtatagttacttgtcctaaa
gacttatcagtcttgtgtacTGCCATTATCCATAGCCCATGAGAATATCCTGTTGGCCAGtatagttacttgtcctaaa
gacttatcagtcttgtgtacCTCCTTACCCTGCCGGAGGTGTGTGTTCCTAGTGTGGCCTtatagttacttgtcctaaa
gacttatcagtcttgtgtacTGGTCTTTCTTAGCTTTTGTCCTTTGTGACCACAACTAAGtatagttacttgtcctaaac
gacttatcagtcttgtgtacCAACATAACAGGCCATTTATAATTGGGAGTTCAGCTGTCAtatagttacttgtcctaaac
gacttatcagtcttgtgtacTAATTCAAATTTACTCAGAGCCTAGCGATTTTCATCTTCGtatagttacttgtcctaaac
gacttatcagtcttgtgtacTTGTTCATGCCCTTAGCCCTCAGCTTAAATGACTTCTCCCtatagttacttgtcctaaac
gacttatcagtcttgtgtacTGGAGGTCACAGCTAATATTCCCCTTTTGCCCCCAGGTTGtatagttacttgtcctaaa
gacttatcagtcttgtgtacCTGCCATTTTTCTTCAATTAGATATGCCAGACTTAAGGAGtatagttacttgtcctaaac
gacttatcagtcttgtgtacAGGCTTCACTCTTATCTCCTTGATTATGTTTTCCTTTCATtatagttacttgtcctaaac
gacttatcagtcttgtgtacGATGGAGAAAAGTTTGCATATTCTGTCCTTAAAATTTGCCtatagttacttgtcctaaac
gacttatcagtcttgtgtacAGCTCTCCTGGTTTGTTGGCCTCAGTCAGAAAGGGTCTATtatagttacttgtcctaaa
gacttatcagtcttgtgtacTATTTTCTTCTCTTTCTGCTTTCTGAGGGACTATGTTCAGtatagttacttgtcctaaac
gacttatcagtcttgtgtacTATATGATCCCTTCCAATTCCTGACTCTTTGGTCTCTGGCtatagttacttgtcctaaac
gacttatcagtcttgtgtacAGAATGCAGATCAGCCTCTCAACTTAGTATTGGTAGGTCTtatagttacttgtcctaaac
gacttatcagtcttgtgtacAACCTCCTCCTAGATCCTCTCAGCTAATCACTCTTGTTAGtatagttacttgtcctaaac
gacttatcagtcttgtgtacATTTGGAATTTGTTTTGATTGTTTAGGGAGTCAGTGACAGtatagttacttgtcctaaac
gacttatcagtcttgtgtacCACTAATTGGCTTAAACATATTGGGGGAACTTATTTCACTtatagttacttgtcctaaac
gacttatcagtcttgtgtacAAATTCATTTGGAGATCAGGTATGGTTGGATATAAAAGCCtatagttacttgtcctaaac
gacttatcagtcttgtgtacGCTCTTCTTTGCAAAACACTGATTTTATTCTCAGGCAAGCtatagttacttgtcctaaac
gacttatcagtcttgtgtacGTTTCACATCCCAGCCTTTATCTTCTAAACTCAGCAACTCtatagttacttgtcctaaac
gacttatcagtcttgtgtacGAATTATATCTCTTTCATAGTCGTTGCAGCAATAATCCTGtatagttacttgtcctaaac
gacttatcagtcttgtgtacCCTGTCTGGATAAGTCACTGCAGCTAGTTGGATGGAATGCtatagttacttgtcctaaa
gacttatcagtcttgtgtacCCTACTTGCATGATTAATTTGGTGAAGGGATGGATTCTTAtatagttacttgtcctaaac
gacttatcagtcttgtgtacTGATAGGTCAGTTTTGAAGTTGATCCAAGAAACCAGTTTTtatagttacttgtcctaaac
gacttatcagtcttgtgtacAAAGATTCTAAAAAACCAATTTTATCACATGTGGCTGGGGtatagttacttgtcctaaac
gacttatcagtcttgtgtacGGTGAAGCCTCATATAATATACTAGTTCCACATACTAAGCtatagttacttgtcctaaac
gacttatcagtcttgtgtacTCCTTTAGTCAGTCTTCTGGTTTCAAATTCAGATGATGTTtatagttacttgtcctaaac
gacttatcagtcttgtgtacGTGTTCCACAAATACTAATGATCCCCTTTTATGCTATAGGtatagttacttgtcctaaac
gacttatcagtcttgtgtacCCTAGAAGGACTCAGAATGGGGGGTAAGACATATACTTTAtatagttacttgtcctaaa
gacttatcagtcttgtgtacACACAGTTGTCTTCTAAAGTCCCACAGATTGCTAGGCACTtatagttacttgtcctaaa
gacttatcagtcttgtgtacCTCCTCTCACATTTTCTTCCTTAAATTTCGAAGCAGCTCCtatagttacttgtcctaaac
gacttatcagtcttgtgtacAGATATTGAGGTTTGAAGAGATGAAGTCTTTTTGGTCAACtatagttacttgtcctaaac
gacttatcagtcttgtgtacAAGCATGACCATTCTTTTAGCCATCCACAAGTTTTGGGACtatagttacttgtcctaaac
gacttatcagtcttgtgtacAAGTCCTACCAATTCTTCCTAAGTAATGTTTCTCAGGTCTtatagttacttgtcctaaac
gacttatcagtcttgtgtacTATACTTCTTGCTACACATGGGTCAGGCTTTCAACATCCTtatagttacttgtcctaaac
gacttatcagtcttgtgtacACCTTCTTTTTACTTTCGGTCTGTCTTTAATTCATTCTGCtatagttacttgtcctaaac
gacttatcagtcttgtgtacAGTGGGTTTTTGAGGTTGGCAAAAGTTGTACACCATACAAtatagttacttgtcctaaac
gacttatcagtcttgtgtacATAAATATTTATTACAAGGCAGTCATTGTGCTGGATGCTGtatagttacttgtcctaaac
gacttatcagtcttgtgtacTTTTCCTAATAAAACACACATCAACAATCCTTTGCTCCACtatagttacttgtcctaaac
gacttatcagtcttgtgtacAAGTGTTGAAAATTTTTAGCTTAGCATCAAGGTCTTCCTCtatagttacttgtcctaaac
gacttatcagtcttgtgtacTGCCATCCATTCATACTTGTTCTGCATATTGTTCTTGACCtatagttacttgtcctaaac
gacttatcagtcttgtgtacGTGCCTGCCCTTCTCCACATCTTGTTCACATCATTCCTAGtatagttacttgtcctaaa
gacttatcagtcttgtgtacCTCCATTACTGGAAATTTTGGCTGATTGTGATCTTTTTCAtatagttacttgtcctaaac

gacttatcagtcttgtgtacTTTTGAAGATGAGTCTCCCTCTGTCACCCAGGCTGGAGTGtatagttacttgtcctaaa
gacttatcagtcttgtgtacGTGTGAGCCACCATGTGCGGCCTGATGGTAAACTTATCTGtatagttacttgtcctaaa
gacttatcagtcttgtgtacAGATGGCTTCTCATTTGTACTTCCTATAACTTGGTCTGTGtatagttacttgtcctaaac
gacttatcagtcttgtgtacGGCACTTGTGACCTTCTCTAAACTATGGTGAGTTAATTTTtatagttacttgtcctaaac
gacttatcagtcttgtgtacTGTAAGTATTTTCTTCTAACAAGGCTGTTTGCTTCATTGCtatagttacttgtcctaaac
gacttatcagtcttgtgtacCATCTTTTGATAAAGGGCTTGAAAGTGTTCTAATTGCTGGtatagttacttgtcctaaac
gacttatcagtcttgtgtacCAGCTAAGAGGAAAAGAGATGAGGACAAAATAAGGCACAGtatagttacttgtcctaa
gacttatcagtcttgtgtacCTGGGCTCAAGAGATTCTCCCCCTCAGCTTCTTGAGTAGCtatagttacttgtcctaaa
gacttatcagtcttgtgtacCTTCATCTTGCTTTGATACATTCTCTGGTGAAGGAGAAAGtatagttacttgtcctaaac
gacttatcagtcttgtgtacGCTTGCAGGGTGGGAAGGACACAAAGCATGAACCAGTAACtatagttacttgtcctaa
gacttatcagtcttgtgtacCAAAGTCAATTTCACTAACTCACTTTGGATTAATGAGCCAtatagttacttgtcctaaac
gacttatcagtcttgtgtacCCACTGTGCCCAGCCAGTAAAATGTTTTTTTATTATTCATtatagttacttgtcctaaac
gacttatcagtcttgtgtacGAAGACTACATCATTTAAATCCTTGTTTTATGTTGAGGGGtatagttacttgtcctaaac
gacttatcagtcttgtgtacAGAGGTTTTCTGTGATTTTATCTGGGTTAAAGCACTTAAGtatagttacttgtcctaaac
gacttatcagtcttgtgtacTCTTGTAGAGGGTGTTTTGGCTGACTTCAGGGCTGAGTTAtatagttacttgtcctaaac
gacttatcagtcttgtgtacTGCTGACAACTTTTGTTATAAATAACCGCATAGTCAAGACtatagttacttgtcctaaac
gacttatcagtcttgtgtacAACTTAAAACAATTTCTCTCTGCTTTGGAGACTCATACTGtatagttacttgtcctaaac
gacttatcagtcttgtgtacTCAATTTCTGGAACTTACTTCTTGAAGACAGTAACATTGCtatagttacttgtcctaaac
gacttatcagtcttgtgtacTCGTCTCCAGAATGTTTGTTCAGTTGAGAAATGAAAATGGtatagttacttgtcctaaac
gacttatcagtcttgtgtacCTAAACTGTTTCTGGCCTAGGGAAGTATTAGTTCAGGGTCtatagttacttgtcctaaac
gacttatcagtcttgtgtacCTTGATGCTTACTCTGACACATGGATGCAATAATCCAGCCtatagttacttgtcctaaa
gacttatcagtcttgtgtacTATGTTGTTGGTTTTCTGTAATAGGAAAGGGGAAATATGCtatagttacttgtcctaaac
gacttatcagtcttgtgtacCCCATGTTTTTGATGAAAGTTGCGTTTCTTTACATACAGTtatagttacttgtcctaaac
gacttatcagtcttgtgtacTCTGATTTCTTGACATTTGCGGGGAACACTAGCATATCTGtatagttacttgtcctaaac
gacttatcagtcttgtgtacGAGTTCACCAAAATCCTCACCCTTTAAAACCAAACTAAGAtatagttacttgtcctaaa
gacttatcagtcttgtgtacCTGTGGTATGGTATCCCCCTTCAGTCTATTGTTTGTTTTGtatagttacttgtcctaaac
gacttatcagtcttgtgtacGCAATATGCTGGCAAATGGGAAAACACATAACATAAAGCAtatagttacttgtcctaaa
gacttatcagtcttgtgtacTTTATCTAACAGAGTCAACAGAATGTAACATGAGTGTACCtatagttacttgtcctaaac
gacttatcagtcttgtgtacCTAGGGGGACAGATCATACAGTTTCTATATTATTCAGCTTtatagttacttgtcctaaac
gacttatcagtcttgtgtacTTTCTGAAACTCCAGCTGTCTCTCATTCATAAAACGGTGTtatagttacttgtcctaaac
gacttatcagtcttgtgtacCAATGATAGCGCTAATAATGAATCACTAACAATATGGCACtatagttacttgtcctaaac
gacttatcagtcttgtgtacCAAACAGCTGGAGGAAACGTGGGGGAAAATGCAATCAAGCtatagttacttgtcctaa
gacttatcagtcttgtgtacCCAGATGCTTAGTTTTGTGGAGGTAATATGTTTGATTGCTtatagttacttgtcctaaac
gacttatcagtcttgtgtacTCTTTTGATGTTCTGTAGGTGGAGTCATGGACTGTTCTTCtatagttacttgtcctaaac
gacttatcagtcttgtgtacATCATGGATAATCAGAAAAACAGTAATGCTGAAGACTAGCtatagttacttgtcctaaa
gacttatcagtcttgtgtacGGAACTGTGTGAGCTAAAGGAGAAGAGGAAGACATGAGACtatagttacttgtcctaa
gacttatcagtcttgtgtacCTGTGGTTAAATGGGTCCCTCTCCTGGCTAATACATATGGtatagttacttgtcctaaa
gacttatcagtcttgtgtacCTAAAGCAGATTCAGTTTTCAGTTAAGGGACATATAGCAAtatagttacttgtcctaaac
gacttatcagtcttgtgtacTGAGACCCAGAGGCTGAATAACCAGGAACTTCCAGAGATAtatagttacttgtcctaa
gacttatcagtcttgtgtacTTCTCAGAAAGAAAGAAAGCTGTCCAACATGCCATTGAGGtatagttacttgtcctaaa
gacttatcagtcttgtgtacCTGTTCTTACCAGGCTGAAGTAGAGCTCTTCTCTTCCTCAtatagttacttgtcctaaac
gacttatcagtcttgtgtacAAATAGAGTACTTTTTCTTTTTAGGATCTCAGAGTACCCCtatagttacttgtcctaaac
gacttatcagtcttgtgtacTGAACTCATGCTTCCCAGTATACTTCCCCAATACCGGTCAtatagttacttgtcctaaa
gacttatcagtcttgtgtacCATTGCCAAATGCTTCACTAAATGAAAACCCCTGAAAATTtatagttacttgtcctaaac
gacttatcagtcttgtgtacTCAAAGATGGCTCACTTTCAGAACAGGATTCATATGTCCTtatagttacttgtcctaaac
gacttatcagtcttgtgtacATGCCAACTGCCTCCTCGGAATGAAAGGAAAAAAAAGTCAtatagttacttgtcctaaa
gacttatcagtcttgtgtacCTGCAGGAATCAATAACATTACAGATCATAGAGGAGACAAtatagttacttgtcctaaa
gacttatcagtcttgtgtacGCCTGGGCATATTAATAACTCCGCCTACTCACTGATTTGAtatagttacttgtcctaaac
gacttatcagtcttgtgtacTTAAATAACATTCTGCACTGATGAGGACTCCATGGCACGAtatagttacttgtcctaaa
gacttatcagtcttgtgtacGTCATCATTAACATGTGAGACTGTTTAGCATTTTGAACACtatagttacttgtcctaaac
gacttatcagtcttgtgtacAAAGAACTTAACACAGGACTATTTATTGGGCTTTAGACAGtatagttacttgtcctaaac
gacttatcagtcttgtgtacTTAATGTTTTTTTAGCCTTCAGCCTATCAGACCACCCGTGtatagttacttgtcctaaac

gacttatcagtcttgtgtacATTGGTCTACAGCAATTTTCTAACAAGACTGACTGCAATAtatagttacttgtcctaaac
gacttatcagtcttgtgtacATCTTTTTCTTACCCTTAAGTATCCCTTACAGTGGTTTGTtatagttacttgtcctaaac
gacttatcagtcttgtgtacGACTTCTCTGTCTGATACTGGTTCCTCACATATAAGTAATtatagttacttgtcctaaac
gacttatcagtcttgtgtacTGAAGGGGCTGAAATTTCATCATTAATGGTTTCTTTGCTGtatagttacttgtcctaaac
gacttatcagtcttgtgtacGAAAACAGTTCTTCCCTAATTAGACATTGCCTGACACTCTtatagttacttgtcctaaac
gacttatcagtcttgtgtacAATCTTTCAATATAGGAAATGACAACCCAACATAGATGGGtatagttacttgtcctaaac
gacttatcagtcttgtgtacATACGTTTATCTTAAAATGGTAAAGAGGGTCAAATGTGCCtatagttacttgtcctaaac
gacttatcagtcttgtgtacGGAGTAAAGTGGCTGTGGATTCCTGCTCTTATTGTCGAAGtatagttacttgtcctaaac
gacttatcagtcttgtgtacCAGGATGTGTTTTTATAGTGCAGAGATCTTAATAACCACCtatagttacttgtcctaaac
gacttatcagtcttgtgtacGCCCTTCTGTGGGTCCTTTAGTGAAACAGCATGGCCCAGAtatagttacttgtcctaaa
gacttatcagtcttgtgtacAATGGATGATTATTTGCATATTCTTCTGTATGTAGCTCCCtatagttacttgtcctaaac
gacttatcagtcttgtgtacCTTGGGGCAGAGAAAAGCAAGTATATAGCAGAGAGGACTTtatagttacttgtcctaaa
gacttatcagtcttgtgtacTGGGATGTTCTAAGGGGAAAAGTGTTTTCTGACTGAGCACtatagttacttgtcctaaac
gacttatcagtcttgtgtacCGATTGGCTTTTCTCTTTGAGGAATAAATGTCTCGCTGGAtatagttacttgtcctaaac
gacttatcagtcttgtgtacCAGGGTTGGATTGGAGTCCTTGCTGATTCTGTGGCTGCTGtatagttacttgtcctaaa
gacttatcagtcttgtgtacCAGCGTCGGTGTTTTGTCTGATCACGTCCAGTTTAATTCGtatagttacttgtcctaaac
gacttatcagtcttgtgtacGAAACAATTCAACAGCTCTCTGCTCCGGGTGCGCGAGGGCtatagttacttgtcctaa
gacttatcagtcttgtgtacGCAGCTGACCGTCGGGGACACCGGCCTGGACTGCGAGGCGtatagttacttgtccta
gacttatcagtcttgtgtacCTGGTCTTCGACGTGGTCAGCTTCTCGCAGGAACAGTTCCtatagttacttgtcctaaa
gacttatcagtcttgtgtacTGGAGGTGGAGGTGAGGAATGTCAACGACCACGCGCCGCGtatagttacttgtcctaa
gacttatcagtcttgtgtacGCCCAGATCCCGGTGGAGATGTTTGAGGGTGCGGCCGTGTtatagttacttgtcctaa
gacttatcagtcttgtgtacGGTGGACTAGGACGTGGGCACCAATGGGCTGCAGAGCGTGtatagttacttgtcctaa
gacttatcagtcttgtgtacCAACCGCACAGCCCCTTTTCGCGTGGAGCTGCAGAAGCGCtatagttacttgtcctaa
gacttatcagtcttgtgtacCTGCTGCAGGAGCTGGACCGCGAGAGCCAGGCCGCCGACAtatagttacttgtccta
gacttatcagtcttgtgtacCGGCTGTCTTCAACGTGTGTGTCCTGGACGCCAATGACCAtatagttacttgtcctaaa
gacttatcagtcttgtgtacCGCAGGGCACGGTGGCCGAAGTGGAGCTGGCGGAGGCGCGtatagttacttgtccta
gacttatcagtcttgtgtacGGCTCCCTGCTTCTTGACCTGGACGCAGCCGACCCCGACGtatagttacttgtcctaa
gacttatcagtcttgtgtacGCGACGTGGTGTTCGCTTTTGGCGCCTGCACCCAGAGGGAtatagttacttgtcctaa
gacttatcagtcttgtgtacCGGGCCCGTGGATTACCAGAGTCAGGACACCTACGAGCTGtatagttacttgtcctaa
gacttatcagtcttgtgtacCGACGTCAATGACAACGCGCCCGACATCGCCATCACCCCGtatagttacttgtcctaa
gacttatcagtcttgtgtacCCCCAGGCGCGCCGGCTGCCTCGCCCTTCGCCGCTGCCGCtatagttacttgtccta
gacttatcagtcttgtgtacCTGCCCTCGGGGGAGCGGACGCTAGCTAGTCGGCGGGAGCtatagttacttgtccta
gacttatcagtcttgtgtacGCTCTGGTCAGCACTTCGGACAGGGACTCGGGCGCCAACCtatagttacttgtcctaa
gacttatcagtcttgtgtacCTGCGCCCTCTACTGGCAGGTGCACTTCCGGCTGCAGCGGtatagttacttgtcctaa
gacttatcagtcttgtgtacGCCCATCGCCTAGTACAACCTGACGCTGGTGACCGCGGACtatagttacttgtcctaa
gacttatcagtcttgtgtacGGTGCGCGTGGGCTACGAGAAAGACAATGCTCCGCTCTTCtatagttacttgtcctaa
gacttatcagtcttgtgtacGCCGGTCTATGAGGTATCTGTGAGTGAGAACACCCCGCCCtatagttacttgtcctaa
gacttatcagtcttgtgtacCCAACGGCCAGGTCACCTACCGGCTGCTGGAGGCCGAGGTtatagttacttgtccta
gacttatcagtcttgtgtacGGACCCAACCACCAGGAGCTGCGCGCTCGACGGCTCTTTGtatagttacttgtcctaa
gacttatcagtcttgtgtacGCTGGCGGAGATGCAGCTGCGGTTGGAGGCGCTAGACGGCtatagttacttgtcctaa
gacttatcagtcttgtgtacGGCTGCAGGTGGAGGACCAGAACTACCACGCGCCCCGCCTtatagttacttgtccta
gacttatcagtcttgtgtacCCCCACTGCTCTCCCACGGCTCAGCCCAGCTGCCTCTGCCtatagttacttgtcctaa
gacttatcagtcttgtgtacGTCTGCAGGCAAAAGACGAGGACTAGGGCGCCGACGTCGAtatagttacttgtcctaa
gacttatcagtcttgtgtacCTTACTTAGTCCTCGACAGCGACGGCGGCCCCGGGGTGCTtatagttacttgtcctaa
gacttatcagtcttgtgtacCTGTCTCTGCAGCGCCGCCTGTCCCCGAAGCCCGCCGGCCtatagttacttgtccta
gacttatcagtcttgtgtacCGGGACTGAGGCCGGCCCACTCTGAGTGCACAGCCACACTtatagttacttgtcctaa
gacttatcagtcttgtgtacCAGAAGTGGTGCTGATGCCGCCGCCGCACGAGAGAGGAGAtatagttacttgtcctaa
gacttatcagtcttgtgtacGGTGGGTGTTTTGGCCAGCGGCTGCATCCTTCTGCTAGTGtatagttacttgtcctaaa
gacttatcagtcttgtgtacCCAAGGCAAGGCTAGGACGCTGGTGAAGAAGATCCGTGAGtatagttacttgtcctaa
gacttatcagtcttgtgtacGGGCTTCTGCTCCAGGTTCTTTATAGACACACGGACAGTTtatagttacttgtcctaaac
gacttatcagtcttgtgtacCAGGCGTTCGAGTCTGTGGAGGATTCCTAGTCTGAGGTCAtatagttacttgtcctaaa
gacttatcagtcttgtgtacCACCCCTGCCGTTCTTAGTGTCCTCATTCTCTTGGAGTAGtatagttacttgtcctaaac

gacttatcagtcttgtgtacTTGGGACTTCAAATGGCTGATATTTCCGGGCTTGTTTTTtatagttacttgtcctaaac
gacttatcagtcttgtgtacTGGAAAGGAAAAATTATCCACAGGCAAGGTTAGGGGGAGAtatagttacttgtcctaaa
gacttatcagtcttgtgtacGGCCCATGTCTGTTTCTGCTGTTTTTGCCTATGATAACTTtatagttacttgtcctaaac
gacttatcagtcttgtgtacGCGTACAATTTGGATACAGTCGACCCTCAGGTTAATGCTGtatagttacttgtcctaaa
gacttatcagtcttgtgtacCTGTCGGGTTTGATATTAGACGGTGAAATATTATTAACTGtatagttacttgtcctaaac
gacttatcagtcttgtgtacAGGGTTCAATCTCAGTTCCTTTTCATTCCTCATCCCCTTAtatagttacttgtcctaaac
gacttatcagtcttgtgtacCATTCAAATCTAGATGAATTCAGTATGAAGGACAGTGGAAtatagttacttgtcctaaac
gacttatcagtcttgtgtacAATGACAGCCATTTGGATATCAGCAGAGAGGGTTTCAAGAtatagttacttgtcctaaa
gacttatcagtcttgtgtacTCCTTTCAAATGATTGCAAAACAAGGGGGTAAGTTCAAACtatagttacttgtcctaaac
gacttatcagtcttgtgtacCTCTTCCTCTCAAAGCATCATAAGACTCATTCCTGACTTAtatagttacttgtcctaaac
gacttatcagtcttgtgtacTATATATTCATAGAGGGAGTGTCTCGGTCATTTCAGGCTGtatagttacttgtcctaaac
gacttatcagtcttgtgtacCATTTGCTGGAAACCTATGAGTATGTGATTTTGTTTGGCTtatagttacttgtcctaaac
gacttatcagtcttgtgtacAATGCTGCACCAACGCAACCTCATTCTATCCAGTCCATCTtatagttacttgtcctaaa
gacttatcagtcttgtgtacGCTTTAGGCCCAATTCTCTGTAATGGTGTCTGGTAATGGAtatagttacttgtcctaaac
gacttatcagtcttgtgtacGCTCCTGTGGTGAAGACATTATCAATTATGAGGTATTTTGtatagttacttgtcctaaac
gacttatcagtcttgtgtacGAAAATATCTTACTTAAAGAGGAGTCCTAATGGTATGTCCtatagttacttgtcctaaac
gacttatcagtcttgtgtacACCATAACAGGTGTTTTTGCACAATCTATGTTCATTTCCTtatagttacttgtcctaaac
gacttatcagtcttgtgtacCCTCCTGGTTACCTTCCTCTGAATATTCTCTAGTGTTCAAtatagttacttgtcctaaac
gacttatcagtcttgtgtacACATGAAGTTGTATACAGTGATTCGGTACCAGCCTCTCTTtatagttacttgtcctaaac
gacttatcagtcttgtgtacCATTTGGATTCTGAATGGACACGCTATATGTTCAAAACTAtatagttacttgtcctaaac
gacttatcagtcttgtgtacAACCTGTTTTTCCTCTAGTGTCGATATCTCAGTAAAAATGtatagttacttgtcctaaac
gacttatcagtcttgtgtacATTCACTTAGTTGCTTGCTCTGGCTTAAATCCTTGGTGTTtatagttacttgtcctaaac
gacttatcagtcttgtgtacTGATTTATTTCTTCCTACACACAACTTTCAATCCATCAGCtatagttacttgtcctaaac
gacttatcagtcttgtgtacTATTCTCAATCAATTCACTTCTCACCACATTCATTGCTACtatagttacttgtcctaaac
gacttatcagtcttgtgtacCCTCAAACTTGGCTCCTCTGCTTCTACTCTTGCCCTCTTGtatagttacttgtcctaaa
gacttatcagtcttgtgtacGACTGGTCTCTTACTTCATAAATCAGGTCACATTACTCAAtatagttacttgtcctaaac
gacttatcagtcttgtgtacAATTACTATTGCTGTGTAACAGGCCAGGCTAAAATGTAGTtatagttacttgtcctaaac
gacttatcagtcttgtgtacCCATTTTTTTATGCTCGTGGATTCTGTGGTCAATTTAGACtatagttacttgtcctaaac
gacttatcagtcttgtgtacCCTTAGCTGGGATGATATGATGGCTGGAGGCTGGAACCGTtatagttacttgtcctaaa
gacttatcagtcttgtgtacAAGTTTAGGAGAGCTGACTTGGTGGAGTGGGAAAAAGGGCtatagttacttgtcctaaa
gacttatcagtcttgtgtacTCTCTTTACACGTGGCTTGACTTCTCACAGCGTGGCAGCCtatagttacttgtcctaaa
gacttatcagtcttgtgtacCTCGGGCTGCAAACATGAATGATTCATCCAACAGGACAGAtatagttacttgtcctaaa
gacttatcagtcttgtgtacCATCTTCTTTTATGACCCAGCTTCTGAAATTACATGATCAtatagttacttgtcctaaac
gacttatcagtcttgtgtacGTCATATTCTATAGTCCAAGCTGTCTTAAGCCCATCCAGAtatagttacttgtcctaaac
gacttatcagtcttgtgtacACAAAGACCCTGTGCCTACCCTAGAAAATTGTCAAAGAGTtatagttacttgtcctaaa
gacttatcagtcttgtgtacCCCTTGGCTCTTTCTTCTACCTGATTTCTTACCCCTCTTCtatagttacttgtcctaaac
gacttatcagtcttgtgtacTGTTTTTTGAATACACAACGCATCTCTCCATCTCAGACCTtatagttacttgtcctaaac
gacttatcagtcttgtgtacGTAAACTTTCTCTAGATACCCATATTGCTTCCCACCTTCAtatagttacttgtcctaaac
gacttatcagtcttgtgtacGCCTTTGTTCATATGTCCATCTAATCAAAAGAGGTCTTCCtatagttacttgtcctaaac
gacttatcagtcttgtgtacTCAGATTGCCCCTCCTCCCTGTCACTCTCCATCTTCTTTAtatagttacttgtcctaaac
gacttatcagtcttgtgtacTTTTCTTTCCTGGCATTTATCACTCCCTGGCATTATTTGTtatagttacttgtcctaaac
gacttatcagtcttgtgtacTGGCCACCATCAGATATATACCGAAACAAAAAAACAGAGCtatagttacttgtcctaaa
gacttatcagtcttgtgtacTCAGCAAATTTACTGGGTGGATAAATGAAAGGTGTGGTCTtatagttacttgtcctaaac
gacttatcagtcttgtgtacTGTATTGGTATTTTTTGGAAGTTCCATTGTACTCCCTAACtatagttacttgtcctaaac
gacttatcagtcttgtgtacATTTATGTTTCCGGGTCACTTAGCGTTAAATTAGCCTTTTtatagttacttgtcctaaac
gacttatcagtcttgtgtacGGAAATTACCAGCTTATTCTTGTCTTAGTTTCTTCTTCCTtatagttacttgtcctaaac
gacttatcagtcttgtgtacCTATGGCAAACGATAATGTGATCAAGGCAATGCATTCAACtatagttacttgtcctaaa
gacttatcagtcttgtgtacCTTCTTAGTAATAATAGTCACAACGTTGCTTCTTACCTGTtatagttacttgtcctaaac
gacttatcagtcttgtgtacTAACATTTACAGAGGACTTCTTATGTACCAGAAACTGTTCtatagttacttgtcctaaac
gacttatcagtcttgtgtacGCTATTAGGCAGGTACTATCATTTGACCCCATTTCACACAtatagttacttgtcctaaac
gacttatcagtcttgtgtacTCAGGCACAGAGCTGTTCGAGTAATTTGGTTGAGGCCGCTtatagttacttgtcctaaa
gacttatcagtcttgtgtacAGTCTGGTTCTAGAGTTCATGCTTTTAATGTTCTGCTGTAtatagttacttgtcctaaac

gacttatcagtcttgtgtacTATGAGTCTTTTAAAGATTTAGGGCCAGTTTACATTCCTGtatagttacttgtcctaaac
gacttatcagtcttgtgtacTTCTGTCTTTGAGTTTGAATGAGATGCTCTGATTTATGGTtatagttacttgtcctaaac
gacttatcagtcttgtgtacCTTTTAGTGGGATTATTGAAAGGATGACACATAGGTAATGtatagttacttgtcctaaac
gacttatcagtcttgtgtacACTACCAGCCCTCAGAAACTTCATTTTAGATTGTAATTTCtatagttacttgtcctaaac
gacttatcagtcttgtgtacGACTTGCTTGAATTTTATTTCCACTTCATTTCCTCAGTACtatagttacttgtcctaaac
gacttatcagtcttgtgtacCAAGTCATTGGCAGAAAATCCAGAGGAAGAAAAAAGCAAAtatagttacttgtcctaaa
gacttatcagtcttgtgtacCCAGCCCGGGAACAACTGAATATTGTTTACATTTTAAAGTtatagttacttgtcctaaac
gacttatcagtcttgtgtacTTAAACCAGGAATCTCAAACCAGCAAAAATTGTTCATCACtatagttacttgtcctaaac
gacttatcagtcttgtgtacCTTTCTTCAAGTACCCTTTACCCTCACATTTCCCAGATGAtatagttacttgtcctaaac
gacttatcagtcttgtgtacACAAATTTCCAACCAGAACACTCAGAACTCAATTGATGTTtatagttacttgtcctaaac
gacttatcagtcttgtgtacCCCAAGTGAAGGGCTGCCAAGTGTCTATGACAGAGTTCAAtatagttacttgtcctaaa
gacttatcagtcttgtgtacAATGCCACTTCCCCATTGTTGGACAGAATGATGAATTAAAtatagttacttgtcctaaac
gacttatcagtcttgtgtacGGATCACCTTCACATACTTCAGAAAATGCCACTTCATTCTtatagttacttgtcctaaac
gacttatcagtcttgtgtacCATAGATGGTCATTTCTGTCAAGTGTTTACAAGCCAGGTTtatagttacttgtcctaaac
gacttatcagtcttgtgtacACATTTTCTTAAATTAAGTTGACAATCCGTGACACAGCACtatagttacttgtcctaaac
gacttatcagtcttgtgtacGCTACTCATTTACATTTGAATGTGTATGTGATTCTACTGCtatagttacttgtcctaaac
gacttatcagtcttgtgtacTTAAGAACCTGGACCCGGACCTTAATCATGAGACCCAGTGtatagttacttgtcctaaa
gacttatcagtcttgtgtacAGTGGGGGTTACTCAAATTGTCTCTCTAGCCTGGCTTCTGtatagttacttgtcctaaac
gacttatcagtcttgtgtacTGCTGAGTGTTTCATGGGTATAACAGTGGCAGATTATGGCtatagttacttgtcctaaac
gacttatcagtcttgtgtacAAAATCAAATGGCAGTGGCTGATGATGTGTTTCTGGGCACtatagttacttgtcctaaac
gacttatcagtcttgtgtacTCAATCAGAGGCTGCGGCTACAGGGAGGAGCATGGTGAGAtatagttacttgtcctaa
gacttatcagtcttgtgtacAGAGAGCAAGAGAGAGTGAGTGTCTTATGGCCTCAGCCATtatagttacttgtcctaaa
gacttatcagtcttgtgtacAGAAAACTGAGGCATAAAAGCAAGGATTAAAGGAAGAGTTtatagttacttgtcctaaac
gacttatcagtcttgtgtacAAAAGCTACTTGAGCAGCTGGGCTTCATCTGGGTCCAACCtatagttacttgtcctaaa
gacttatcagtcttgtgtacATGGAAGCATCTGATTGTGTCTTGTTTTCTGTAAGATTGGtatagttacttgtcctaaac
gacttatcagtcttgtgtacAAACTGCAGGATGTTTTGTGTCTCCAACTAGAAGAGCCTGtatagttacttgtcctaaac
gacttatcagtcttgtgtacGGAACTGTGCTGCCCAATACTATAACCACTAGCCACATGTtatagttacttgtcctaaa
gacttatcagtcttgtgtacGAAAAATGCTTCAGTCTCAGACAGAGTCACTATAAAATGTtatagttacttgtcctaaac
gacttatcagtcttgtgtacGTTCTGATGGACGGCATTGGTCTGGAAGAAGGGATGAGCCtatagttacttgtcctaaa
gacttatcagtcttgtgtacGAAATGTGTGTTTTTGTCTGTTTTCCACCCTTCCAAATGTtatagttacttgtcctaaac
gacttatcagtcttgtgtacCTAACTAGCTCATGACAGTTAGCCTTTTAATGTTTCATGTtatagttacttgtcctaaac
gacttatcagtcttgtgtacGAAAAAATGTCTAAAGGCAAGAATTCTCAATGGTAGAAGGtatagttacttgtcctaaac
gacttatcagtcttgtgtacGCTCCTCTGCTATAGTGCCTTAAATTTATAGTCCTCATATtatagttacttgtcctaaac
gacttatcagtcttgtgtacCTTCAGAGATTTTGTAGCAATAGTCAGATAAAACCTACTGtatagttacttgtcctaaac
gacttatcagtcttgtgtacTCAGCAGTTTTTAAAACCAGTAATTTATTCCTTAGCCACGtatagttacttgtcctaaac
gacttatcagtcttgtgtacGAAGCTAGGAAGAACTTCAATCCTTTGGCCACTGGTCTGCtatagttacttgtcctaaa
gacttatcagtcttgtgtacTGCTGTTGGAAATTAGTTTCTGAGTCCCATGTTATCTTCGtatagttacttgtcctaaac
gacttatcagtcttgtgtacGTGTGAGCTTTCCAGATCGGGTGGTCTAGGCCCCAGAGAGtatagttacttgtcctaaa
gacttatcagtcttgtgtacGGCAAATGTGAATGTTCCTTGCTTTATCTGATACATATCTtatagttacttgtcctaaac
gacttatcagtcttgtgtacAGTGCATTCTGTGGGAAACATTTTTTAAAAGCTGAGAATCtatagttacttgtcctaaac
gacttatcagtcttgtgtacAGGCTCTCATGTGTTTGTGATATTAGCTTCTACCTTAATTtatagttacttgtcctaaac
gacttatcagtcttgtgtacTAACTCCTGAATCTTGTCTGTTTATGATTGCATTCCTAAGtatagttacttgtcctaaac
gacttatcagtcttgtgtacTAGGCAATCTTTATCAAACACTATATGCAAGGTTCTGTGCtatagttacttgtcctaaac
gacttatcagtcttgtgtacGAGGAAATGAGTTTAAAGAGGGTAATATGCTTGAGGTTACtatagttacttgtcctaaac
gacttatcagtcttgtgtacATGAGCTTAGAGTTTAAAAAATTATGGTATAGGCTGGGTGtatagttacttgtcctaaac
gacttatcagtcttgtgtacCTATCATTTTGGGAGGCCGAGCAGGTGGGTCTCTTGAGCCtatagttacttgtcctaaa
gacttatcagtcttgtgtacATAAAAATTAGCCATGCATGGCAGTATGTGCCTGTAGTCCtatagttacttgtcctaaac
gacttatcagtcttgtgtacACTAAAGTGCATGAGTTTCGAGTTTACAGCCAGATGAAGTtatagttacttgtcctaaac
gacttatcagtcttgtgtacGAATATTTCCAGATCCCTAGACGGCTCCCACATATACCTTtatagttacttgtcctaaac
gacttatcagtcttgtgtacTCCAGCAACCCCACTTCTAGATGTAAACCCAAATGAAACAtatagttacttgtcctaaa
gacttatcagtcttgtgtacCCTAAGTGTCTATAGATGGATAAGTAGATAAAAATGGCAGtatagttacttgtcctaaac
gacttatcagtcttgtgtacGCCTTAAAGAAGAAAATGTTAACCATTTGCGACAATATGGtatagttacttgtcctaaac

gacttatcagtcttgtgtacATGTGTGCAATCTAAAAAAGTCAGATTCATAGAAGAGTGGtatagttacttgtcctaaac
gacttatcagtcttgtgtacGGTGTGCTAGGTGAGGGGGAGAGGAAACTGAGAGATGTTGtatagttacttgtcctaaa
gacttatcagtcttgtgtacAGGGTACAAAGTTTCAGTGAGGATACATAAATTCTAGAGAtatagttacttgtcctaaac
gacttatcagtcttgtgtacGTTCAGCATAGTGACTATATTGCTGAATTTGATTCTTGAGtatagttacttgtcctaaac
gacttatcagtcttgtgtacTCTCACCACATGTGGAAAAAACAACTATCTGATTTTTAGCtatagttacttgtcctaaac
gacttatcagtcttgtgtacTTATACTTTTGGGGCCTGGTTCAAAATTATGTCTATGAACtatagttacttgtcctaaac
gacttatcagtcttgtgtacCTGTGTATATCATAGTCTGTGCATTCTCACTGCTGTATAGtatagttacttgtcctaaac
gacttatcagtcttgtgtacTTGTATAGTTGGTGGAGATTAGGGCAATTTTCAGTTTTTGtatagttacttgtcctaaac
gacttatcagtcttgtgtacTAAACACTTATTTCTGTTGGCCATATACCCAGGAGTAGAAtatagttacttgtcctaaac
gacttatcagtcttgtgtacGTGTCAAAGAGTGATGTATCTAGGGTTTGAACCTTGAATTtatagttacttgtcctaaac
gacttatcagtcttgtgtacGCACCCCTGCACCCAGCTTGAACCTTGAGTTTCTGACTCtatagttacttgtcctaaa
gacttatcagtcttgtgtacACTCATTACATCAATAGTGGCTAATAAAGACTGAGAGCTCtatagttacttgtcctaaac
gacttatcagtcttgtgtacTAAGCTTTCATTTTCTAAGTAGGGTGTTGATTCAGTAGAGtatagttacttgtcctaaac
gacttatcagtcttgtgtacAACTAATAACAGCAGGGGGAAGTCTTCTATTTGTTCAACAtatagttacttgtcctaaac
gacttatcagtcttgtgtacTGGAACACTTAGTAGCTTCTCTTTGAAAACAGCATGATAAtatagttacttgtcctaaac
gacttatcagtcttgtgtacATCAATTTAAAACCAATTTCCATAGGCTACACATGCACAGtatagttacttgtcctaaac
gacttatcagtcttgtgtacAAATGAAGCTGGCTTATCTCAATTTGTATTTCATCGAGTGtatagttacttgtcctaaac
gacttatcagtcttgtgtacAGGCAATCTGTCTGTCATTCTTTAAAGGTAGCACAGCACGtatagttacttgtcctaaac
gacttatcagtcttgtgtacTGCCGTTTAATTAGGTGTGGTCTAGTCATTTCACTGTTTGtatagttacttgtcctaaac
gacttatcagtcttgtgtacGAAAAACCTCAGCCAGATGCCTGCTCTATCACCCTCTTTCtatagttacttgtcctaaa
gacttatcagtcttgtgtacATGCCCATTGATTCTTATTGTTAGATGTTGTACAGAGTAGtatagttacttgtcctaaac
gacttatcagtcttgtgtacGTGTTGGAAGACAAAGCTAAAATGGACTTTACAGGCTAACtatagttacttgtcctaaac
gacttatcagtcttgtgtacGCTTTCTTGAACTCTTGACAATGCAGCATTACATTCTAGGtatagttacttgtcctaaac
gacttatcagtcttgtgtacGAATAGGTTTCCACAGTGTGCCAGGTTTTCTTAGAGAGTAtatagttacttgtcctaaac
gacttatcagtcttgtgtacTGCAATGTCTTTTTCCCTGGCGCAACACAGAAGATGTAATtatagttacttgtcctaaac
gacttatcagtcttgtgtacAGAAGTATCACAACCTCTTTATCTCTTCGTTTTCGGTTTCtatagttacttgtcctaaac
gacttatcagtcttgtgtacTGGAGGAATAGTACCTACTTCATTGCTATTTTATGCTAGCtatagttacttgtcctaaac
gacttatcagtcttgtgtacCAGTATGTTTAAAATGCATACAGCAATGCCTTTCACCTAGtatagttacttgtcctaaac
gacttatcagtcttgtgtacCTCAGCTCATTGCAACCTGTGGCTCCTGGATTCAAGCGATtatagttacttgtcctaaa
gacttatcagtcttgtgtacCCACCATGCCTGGCCAAAAATGTTTTTAATGTCATTAATAtatagttacttgtcctaaac
gacttatcagtcttgtgtacCTCTATTTCTTTCTGCTTGGTGCAATTAATACTTTCCATCtatagttacttgtcctaaac
gacttatcagtcttgtgtacCTTTTCAAATAGGTGACCAAAAATGATTTAAATCCCTGGGtatagttacttgtcctaaac
gacttatcagtcttgtgtacTAGATCAGTTGGTCATTTTGTCTACAGCTATATTTCAAGCtatagttacttgtcctaaac
gacttatcagtcttgtgtacCTTGACAGATGTATGAGGTATTTAAATGGAAAGATCAGAGtatagttacttgtcctaaac
gacttatcagtcttgtgtacGTAGGTATCCCCCTTTTACTTACTTTGACACGTGTTAATGtatagttacttgtcctaaac
gacttatcagtcttgtgtacTTTAATTGAACCTGTAAGGCTACCTGCTCTGTATCCCAAAtatagttacttgtcctaaac
gacttatcagtcttgtgtacGCAAGAGGCATTAAGTAATAAAAGGGTAAATGATCAGCTTtatagttacttgtcctaaac
gacttatcagtcttgtgtacACTGTAACTGGGGAGATTATTGATAGTGAGGCATTGCTACtatagttacttgtcctaaac
gacttatcagtcttgtgtacCAGGTAGGACTCAATCTGAAAGGTATTTTTCTTCCATCTAtatagttacttgtcctaaac
gacttatcagtcttgtgtacGGAAAACCTATAGAACATCTCCTACATAACCCAGAAGACAtatagttacttgtcctaaa
gacttatcagtcttgtgtacGACATTTTTGCAGACATGTTTATAAAAGGCATACTGGTCCtatagttacttgtcctaaac
gacttatcagtcttgtgtacCAGCTTCATCTCTGGTCACCTTCTTACTTGCACTTCCTGTtatagttacttgtcctaaac
gacttatcagtcttgtgtacTCACCGTAGAAAGGGCAGCCACAGAGACAATAAGTCAAGCtatagttacttgtcctaa
gacttatcagtcttgtgtacAAGAGTCTTTAGAGAAATAGGTACCTGAAATGCATGGATAtatagttacttgtcctaaac
gacttatcagtcttgtgtacTTCATTTGAGTCCATGTCCCTGAGCAACCTTCCTCCTTCTtatagttacttgtcctaaac
gacttatcagtcttgtgtacCGTAGTGTTCTGTGTAGGTATGTCTTTAGGGGGAGAATGGtatagttacttgtcctaaac
gacttatcagtcttgtgtacGTGTTTTGGTTTTATACATTAGTTCCATGAGGTTGAGTTGtatagttacttgtcctaaac
gacttatcagtcttgtgtacTTCATATTGCTGAATAACCAGGGTTCTGAGTCTGCATATCtatagttacttgtcctaaac
gacttatcagtcttgtgtacCTGACAAAGACAATAAATGCAAGTCCCTAAGGCTTCCTAGtatagttacttgtcctaaa
gacttatcagtcttgtgtacCAGGTCCTTCTTTAGGCAGTATTCTACCCTGATCCCAACCtatagttacttgtcctaaac
gacttatcagtcttgtgtacTCAGTCACGATGATGGAGTAATAACTAATATCAGTAGAGCtatagttacttgtcctaaac
gacttatcagtcttgtgtacCTGTGAAGCAGGTACTGTGATTATCTTTCTCCTCCATATAtatagttacttgtcctaaac

gacttatcagtcttgtgtacCCTGGGCATGTCTTCTGGGTTCAGCGCATATTCCACACTGtatagttacttgtcctaaa
gacttatcagtcttgtgtacATTATGTGTATAGGCTTGGAAATCAAAGTATCGAAAACCCtatagttacttgtcctaaac
gacttatcagtcttgtgtacGCTTAAATCACCAGGAGCAAATGAACAGTCTAAATGCACAtatagttacttgtcctaaa
gacttatcagtcttgtgtacCACTTGAAGCTGTGATCGGGTCCCTGGTTGAGTGCCGCTCtatagttacttgtcctaa
gacttatcagtcttgtgtacGGAGCTTTTTTCCCTCTCTCGGTGGTACTTCCTTTGAGGCtatagttacttgtcctaaa
gacttatcagtcttgtgtacCAGATATTGTGCAACTTCCATTAATGTTCTATTTCTCCTGtatagttacttgtcctaaac
gacttatcagtcttgtgtacCTTGGAACTTTCTCAAGGGAATTTAATTTTCCATGTCTGTtatagttacttgtcctaaac
gacttatcagtcttgtgtacCTGGTCCGGGGTTTATACTTGTAATTCAATGATTCTTTTAtatagttacttgtcctaaac
gacttatcagtcttgtgtacAACCTATTTTGTTGTCGTTGTTCTGTAATTCACAAACCACtatagttacttgtcctaaac
gacttatcagtcttgtgtacGTATTTAACTTTCTTTTGCCCTCCTCTATATGGTTATGGTtatagttacttgtcctaaac
gacttatcagtcttgtgtacGGAAGAAAACCCTAAGCTGAGAAACTATGAAAATCCATTTtatagttacttgtcctaaa
gacttatcagtcttgtgtacATTATACACCTGAACAGTTCTATAATGCCTCCAATGATACtatagttacttgtcctaaac
gacttatcagtcttgtgtacATATAAATCAGTGCCCCAGTCTTTACCTAGCTCCATGTATtatagttacttgtcctaaac
gacttatcagtcttgtgtacGACTATGTCAGCTTTGTCAATAACCTGAATATTTTCCTTCtatagttacttgtcctaaac
gacttatcagtcttgtgtacCCTCTGTGGCTTTAATTTTTTTACCTCCTGGATACTGTTTtatagttacttgtcctaaac
gacttatcagtcttgtgtacCTAGTATTAGAGCTAAAGTAGAAGAGTGAGGAAAGTCAAAtatagttacttgtcctaaa
gacttatcagtcttgtgtacATTTGACTTTTTTGTGATTTGGCCCATCAGGTGTTTTAGAtatagttacttgtcctaaac
gacttatcagtcttgtgtacAAGCAACCAGTACAAAGTTTAGAGCTTCAGTTTCTACTAAtatagttacttgtcctaaac
gacttatcagtcttgtgtacTTCTAGATACATGGAATAAATTGGAGTAGGGAGTGGATACtatagttacttgtcctaaac
gacttatcagtcttgtgtacGGGTACTTTTCTTTTTCTGGCATCCCACACAGTCATACTTtatagttacttgtcctaaac
gacttatcagtcttgtgtacCAATTCATTGCCCACATTTCCTCAGAAAGACTAAGCTAAAtatagttacttgtcctaaa
gacttatcagtcttgtgtacGACTACAAGAGTCACATTTATAGACTTCACTTAGCCATAAtatagttacttgtcctaaac
gacttatcagtcttgtgtacTTTAAACAGTTGCCATAGCTGAAATCCATGCTTTCTAAAGtatagttacttgtcctaaac
gacttatcagtcttgtgtacCCTTAGGAGAAAAAGTCATACATGCATGTCCTACCAGCTTtatagttacttgtcctaaa
gacttatcagtcttgtgtacGCTGGACTTGTGAGCCCCTTTTATAGAGTTTTCCTTCTTGtatagttacttgtcctaaac
gacttatcagtcttgtgtacTCTAGAAGGGAAGGAGACAGCTGAGCTTGGGGACAGACTGtatagttacttgtcctaa
gacttatcagtcttgtgtacCAGCTGTCTGGAAGGAGGGTTCTTTTCTATAGCCATCTGCtatagttacttgtcctaaa
gacttatcagtcttgtgtacGGAAAGGTGATGGATTTACTGCTGTCCCTGATGGTTGAGAtatagttacttgtcctaaa
gacttatcagtcttgtgtacGTCCTTGATGATTCAGATAACCTCCCATCTTCAGTTTTGTtatagttacttgtcctaaac
gacttatcagtcttgtgtacCCTGGGCTGGGATGTTTTGGGAAATGGAAAATATATACTAtatagttacttgtcctaaac
gacttatcagtcttgtgtacTCCCTCTCCATTTTCTCTTTATGCTTAAGATCTAATCAAGtatagttacttgtcctaaac
gacttatcagtcttgtgtacAGTATTAAAAGTCTGATCTCTATGCAGCCAGTCCAGAGTGtatagttacttgtcctaaa
gacttatcagtcttgtgtacCTTTTGCCTTCTCCCTGTATAGGGTTGTTATGGACTGAATtatagttacttgtcctaaac
gacttatcagtcttgtgtacAAGGTGGGACCTTTGGGTGATAATTGGGTCATGAGAGTGGtatagttacttgtcctaaa
gacttatcagtcttgtgtacATCTTTTTCATCCTTTTTCTGCTATCTGAGGGTACAACAAtatagttacttgtcctaaac
gacttatcagtcttgtgtacCAGTACAGTTTTCAATCTGGAAGAGGGCCCTAAGCAGGACtatagttacttgtcctaa
gacttatcagtcttgtgtacAGCCATCCAGTTAATGGTATTCTGTTATAGCAAGCCTGAAtatagttacttgtcctaaac
gacttatcagtcttgtgtacAGGCCTTAGTAATTATCTTGTTTGACATCAGTAGACTTTCtatagttacttgtcctaaac
gacttatcagtcttgtgtacTGTTAAATAATGAGATGACTAAGCCGAGAAAAGTGTTAGGtatagttacttgtcctaaac
gacttatcagtcttgtgtacCTTCCTTTTTTAGCCCTGGATGAAATAGAGTCAAGCTTTAtatagttacttgtcctaaac
gacttatcagtcttgtgtacTGAAAATCTACATGGCATTATATATAGTTCAGCCACTCTGtatagttacttgtcctaaac
gacttatcagtcttgtgtacGTTGCAAAAACTTAGGCATAAATTCATCTCCTTGCAGAAGtatagttacttgtcctaaac
gacttatcagtcttgtgtacGTTCTGAAGACCTATAGAGTTTGTGTTGGGAAATGATTAGtatagttacttgtcctaaac
gacttatcagtcttgtgtacGAGATTGGGTTGCATGAATGTGGCTTGGTAGCCTGGAGGCtatagttacttgtcctaaa
gacttatcagtcttgtgtacACTTCAGGCTATGAATGTCAGTATGTGTGCCAGGCATAGGtatagttacttgtcctaaa
gacttatcagtcttgtgtacGCTTTCTGTTGCATACCTCTAACCTTTGCAGACTTCTGAAtatagttacttgtcctaaac
gacttatcagtcttgtgtacGTTAAGTGACTTACCTAAGGGCACAAGTTATAAAATTGTCtatagttacttgtcctaaac
gacttatcagtcttgtgtacTGTTTTACTATATTTTGCTGCCTTTTTTGGTAAACCTGGCtatagttacttgtcctaaac
gacttatcagtcttgtgtacGTTTCAGACATGTATCAATAAACACAGTGGCAAATGTATCtatagttacttgtcctaaac
gacttatcagtcttgtgtacTGTTTCACACGTGTCACTAAACACAGTTAAATGGAGACTGtatagttacttgtcctaaa
gacttatcagtcttgtgtacTTCACACATATATTATTAAACACAATATGCCCCCAGTGACtatagttacttgtcctaaac
gacttatcagtcttgtgtacCCACAGCCAGTTGTGTTTTTAGTGGTTAGGTGTCAAAAAAtatagttacttgtcctaaac

gacttatcagtcttgtgtacAACAAAATCTTATATATCAATAATCTCCCTGGCAAGCTGCtatagttacttgtcctaaac
gacttatcagtcttgtgtacTTAATATGTACATTTCTCCTTTCAGGGGATTATCTGATGGtatagttacttgtcctaaac
gacttatcagtcttgtgtacCCCATGCATTCTTCTAGATACCCTAAAATGTACCTATAGAtatagttacttgtcctaaac
gacttatcagtcttgtgtacCAAGCGTTTTTTGATCTATATACCTGGAATGAATCACTCTtatagttacttgtcctaaac
gacttatcagtcttgtgtacCCAACTTGGGGGTAACAGTGTAAGAGGAAGAGTTATGAAAtatagttacttgtcctaaa
gacttatcagtcttgtgtacGGATGTTTTTGCCTATTTCCCCCGTTAGTTCTTTCAGTTTtatagttacttgtcctaaac
gacttatcagtcttgtgtacAGATTGTTGTATCTTTCTGCTAGATTGACCCTTTCGTTATtatagttacttgtcctaaac
gacttatcagtcttgtgtacTCCCTATTTATCTCTAGTGATGCTTCTTTCTTTCCTTACAtatagttacttgtcctaaac
gacttatcagtcttgtgtacACTTTCAGTATTTGTGTGTCCACACATTAAAGGTATGTCTtatagttacttgtcctaaac
gacttatcagtcttgtgtacATCTGTATAGGTTGTTTCATTTTTCTTCTAGTGGTTAGCCtatagttacttgtcctaaac
gacttatcagtcttgtgtacATGAAGAACAGTTTGCATTGACCCCTCTCTTATCTTTTGTtatagttacttgtcctaaac
gacttatcagtcttgtgtacTTCTGTTATTCTTAATACATTTCTATGCTACCATCTGGGGtatagttacttgtcctaaac
gacttatcagtcttgtgtacAGGAACTCATTTTAATAATTCTTTTCCTCTGAACCTGCTGtatagttacttgtcctaaac
gacttatcagtcttgtgtacCAAAACGTCATCAAACTACAACATACATGCAGAAAATTGCtatagttacttgtcctaaa
gacttatcagtcttgtgtacTGAAAATCAGCGCTGTGTAAAAGCTCAAGAAAGAGAACATtatagttacttgtcctaaa
gacttatcagtcttgtgtacGGTGTCCATTCTTCTGACCCCTAACATCATAGATTATATTtatagttacttgtcctaaac
gacttatcagtcttgtgtacTTCACTGATATTGTTGCATGTAGTTTCATAAAGTTGGTGCtatagttacttgtcctaaac
gacttatcagtcttgtgtacCATTCACTATTGATATGGATTTGGGTAGTTGCCAATTTTGtatagttacttgtcctaaac
gacttatcagtcttgtgtacAAATACATTTGCCCCATCCCCACAGCCCTCAAAGTGTTAAtatagttacttgtcctaaa
gacttatcagtcttgtgtacGATGTATGATTCATCCTGAATTAAAATTCCTCTCCAGCTGtatagttacttgtcctaaac
gacttatcagtcttgtgtacGCGGGACTGGCATAGAATCGACATTCTTATTCTAAAAAGGtatagttacttgtcctaaa
gacttatcagtcttgtgtacGGTCCCAGCTAAGTTCAAAACCCAGGAGGAAAATTCCATTtatagttacttgtcctaaa
gacttatcagtcttgtgtacCTTGATGCTCTGCTTTCTGGGCCCACTGGAGTGAATGTTTtatagttacttgtcctaaa
gacttatcagtcttgtgtacTCTGGTTGGCAGCCTTGAGCTTGAGGCACTGGGTGGTGGCtatagttacttgtcctaa
gacttatcagtcttgtgtacCAAGGAGGAGCACACTTGCCCCTGGGGCCGGCCTGTGTATtatagttacttgtcctaa
gacttatcagtcttgtgtacGCCTGTGGTGGGAGTGGCACCCCTGCTAATCTCTGAATAGtatagttacttgtcctaa
gacttatcagtcttgtgtacAAGGATGATGCATGTTTCTAGCCAAATAGCTTCACTGTTCtatagttacttgtcctaaac
gacttatcagtcttgtgtacATCCCAGAAATCCAACAGTCTTCCTGCATTTTGTCCATGCtatagttacttgtcctaaa
gacttatcagtcttgtgtacCTGACATGGCTGCTTGGATCCACTGACAATCTCTTTTTGGtatagttacttgtcctaaa
gacttatcagtcttgtgtacCCCTTGGTGTTTACAACTTGCATTATACTTTTTTATCCTGtatagttacttgtcctaaac
gacttatcagtcttgtgtacCCTTTCACATTTAAGTAGTAAGGAGAAACCAGGCCACACCtatagttacttgtcctaaa
gacttatcagtcttgtgtacTTTTACCATCTATAAAACGCTAGAAAACAATTCAGCCCAGtatagttacttgtcctaaa
gacttatcagtcttgtgtacACTTACTCCAAGTTCTCACTAGAATCACCTTTAATGTCCAtatagttacttgtcctaaac
gacttatcagtcttgtgtacCCACCAACCTGTCTCAAAAACTCTTTTGGCCTGTGCCCATtatagttacttgtcctaaa
gacttatcagtcttgtgtacCTTTAGCAGCACAGTACAGCACAGTACCAAAATCTGTTTTtatagttacttgtcctaaa
gacttatcagtcttgtgtacGCCATAATGAAATACTACAGCCTGGGCTTAAACAATGGAAtatagttacttgtcctaaa
gacttatcagtcttgtgtacCAAAATCAAGGTGCCAGTAGTGTTGCTTTCTCCTGAGGCCtatagttacttgtcctaaa
gacttatcagtcttgtgtacTGTCCTCATATAATCTCTTCCCTGGGCACATGTACTCCTAtatagttacttgtcctaaac
gacttatcagtcttgtgtacCTATTGGATTAGGGCCTCTTCCTTATAATCTCATTTAACCtatagttacttgtcctaaac
gacttatcagtcttgtgtacGGACCCTGTTTCCAAATGTGGTTACACTGAAGGTTGACACtatagttacttgtcctaaa
gacttatcagtcttgtgtacATGTAGTCCTTAACAGAGCTCAAGTTACATTTTTTCCCCAtatagttacttgtcctaaac
gacttatcagtcttgtgtacATCCAACAGACCCAACACCATTTAGTATGAAGATAAACTTtatagttacttgtcctaaa
gacttatcagtcttgtgtacCCATGTATGTGAGGCTCTGTTTCTTAGGTCTGTGTTCTCTtatagttacttgtcctaaac
gacttatcagtcttgtgtacAGGTTTGTCTATTACTGCATCATTACCTTGCTGACCCTACtatagttacttgtcctaaac
gacttatcagtcttgtgtacCTATAGGTTTTAAAAAATGTCTCTCATCAGTGTAGAGGTCtatagttacttgtcctaaac
gacttatcagtcttgtgtacTGATGTTCTTTGATGCTGTTGCGAAATATCTTATTACACGtatagttacttgtcctaaac
gacttatcagtcttgtgtacTTCAGCACTTAAAAATGATACTCCATTATCTTCTGGATGCtatagttacttgtcctaaac
gacttatcagtcttgtgtacAAGTTACTGTCAGTCCTATTATTGCAACTTTGAAGGTAGTtatagttacttgtcctaaac
gacttatcagtcttgtgtacCTGCTTTTCAGAGTTTCCTCTTCATCTTTGTTTTTCAATGtatagttacttgtcctaaac
gacttatcagtcttgtgtacGATAGTTAGTTTCCAACGATGTTCCTCTAATGAATTATGCtatagttacttgtcctaaac
gacttatcagtcttgtgtacCCTTTAATCTAGGATAGCTGTGACACTTCTGACTAGTAAAtatagttacttgtcctaaac
gacttatcagtcttgtgtacGCTTGCTTGGCTTTTTGGAACACTCACTATTAGGATGCTCtatagttacttgtcctaaac

gacttatcagtcttgtgtacCTCTGAGACCACTGTGCTGCAAGAGAGTCAAGCCACATAGtatagttacttgtcctaaa
gacttatcagtcttgtgtacAGTTTCTACTGGGGAGCGAGAGAGGGGAAGATAGGCAGAGtatagttacttgtcctaa
gacttatcagtcttgtgtacCATTTTGGAAATCTAGCCCAGTCCAGCTTTCAGACAATGAtatagttacttgtcctaaac
gacttatcagtcttgtgtacGCTGCATGAGAGACCCAAGCAAGAACTGTGTAGCTGAAGTtatagttacttgtcctaaa
gacttatcagtcttgtgtacCAGAACTAGGAGAGCTGTTATAATAATTGTTTTAAGGCCGtatagttacttgtcctaaac
gacttatcagtcttgtgtacACATCTTAAAAAATGTGGCGTCTGCTTTGGGACCAGTAGCtatagttacttgtcctaaac
gacttatcagtcttgtgtacATGCTCAGGGATTGTTCATGGAAACTTGAAGGACAGTGAAtatagttacttgtcctaaac
gacttatcagtcttgtgtacCATGCTATGTAGTGTCAGAAAATTTAGGAGAACTGTTGTCtatagttacttgtcctaaac
gacttatcagtcttgtgtacTTTAGCCATGTATAATAAGGTAAAGAGGAAGAGAGAGAAGtatagttacttgtcctaaac
gacttatcagtcttgtgtacAAGCAGAAATTAGAGAAGATATAAAGAACTCAGGGCTTACtatagttacttgtcctaaac
gacttatcagtcttgtgtacGAAACTGTATATTAGACTTAGCCACTCTGTATTTCAAAGGtatagttacttgtcctaaac
gacttatcagtcttgtgtacGTGCAACTGTAAGACACATTGTTAAGATTTCAGGAAGACTtatagttacttgtcctaaac
gacttatcagtcttgtgtacGTGTGCCTTACAGAGCATTTGAATTAGACAAAAAGTTTTCtatagttacttgtcctaaac
gacttatcagtcttgtgtacTGTTATGTCTCAACACTCTCACCTTCAGCACGAGGCGAACtatagttacttgtcctaaa
gacttatcagtcttgtgtacGAGATTTCTGAGTGAAGTTGTTGTTTAGTGAAGTGAACCCtatagttacttgtcctaaac
gacttatcagtcttgtgtacCTTTAAGGGAATTGTGTATACAGAAACATCATAAGCTTGGtatagttacttgtcctaaac
gacttatcagtcttgtgtacGAGTCAGTTTGAAATGAAAAGATGCCTCTGGGTTCCTAATtatagttacttgtcctaaac
gacttatcagtcttgtgtacCTGAGACATTTACTTAGCTGCAAACATGAACCATTTCTGTtatagttacttgtcctaaac
gacttatcagtcttgtgtacAGGAAGATTGGGATTAATAAACAAGAGGGAGACAACAGTAtatagttacttgtcctaaa
gacttatcagtcttgtgtacGAGGATAGAAATAAAAGCCATGGAAAACTACTCTCAGAACtatagttacttgtcctaaa
gacttatcagtcttgtgtacGCAACATAAGCCCAATTAGATTTCAGAAATGCTTTGGACCtatagttacttgtcctaaac
gacttatcagtcttgtgtacACTCTTGTCCTCTTATCTCCTCCACTCCCATCCCCAGCTTtatagttacttgtcctaaac
gacttatcagtcttgtgtacGTACTACCATTGTTATGTTGAGTGTGTGGGGGCGGATAACtatagttacttgtcctaaac
gacttatcagtcttgtgtacTCAAGAGGAGTATAAGCTGCATCTGGGAAATTGCACTTCAtatagttacttgtcctaaac
gacttatcagtcttgtgtacGCCTGATGTAAATGCCTGGCTTCTCATTACGGGGAATTCAtatagttacttgtcctaaac
gacttatcagtcttgtgtacAAAATGTTGACTCACTTCCCTACAATTCTTTTTTGTCCAGtatagttacttgtcctaaac
gacttatcagtcttgtgtacTTGCATTGTACTCATTTTGTAATTGCTCTTAGTAGGAAGCtatagttacttgtcctaaac
gacttatcagtcttgtgtacCTACATAGTTCAAGCAATCGTCCTCCATACTTAAGCAATCtatagttacttgtcctaaac
gacttatcagtcttgtgtacGAAGGATCTAAAAAAGAATTTCTCTCTAGTCTCAGTACCAtatagttacttgtcctaaac
gacttatcagtcttgtgtacGTATTTTTATTAAAAAGATAAAATAGACTGGGCGCGGTGGtatagttacttgtcctaaac
gacttatcagtcttgtgtacATATTTTTTTCCTGGGGGAATATTAGCTTCATATATGGCCtatagttacttgtcctaaac
gacttatcagtcttgtgtacTGGTTTGGCCAAATCTCATCTTTGTAGCTCCCATAATTCCtatagttacttgtcctaaac
gacttatcagtcttgtgtacCTGATGGTTTTATAAACAGGGGGTCCCCTGGACATGCTCTtatagttacttgtcctaaac
gacttatcagtcttgtgtacTTGCTTCTCCTTTGCCTTTGACCGTGATTGTGAGGCCTCCtatagttacttgtcctaaac
gacttatcagtcttgtgtacAAGAAACAATGATAAATGTTCAGATTTGGAAGGTGGGAAGtatagttacttgtcctaaac
gacttatcagtcttgtgtacGTTAACCAATGTGTCAAAGATGATGTTTATAGGCAGGTATtatagttacttgtcctaaac
gacttatcagtcttgtgtacGGGCAACATAGGGAGAATCTCGTTTCTAAAATTAGCTGGGtatagttacttgtcctaaac
gacttatcagtcttgtgtacATTGCTCGAGCCTGAGAGGTTTTGGCTGCAGTGCCATTGCtatagttacttgtcctaaa
gacttatcagtcttgtgtacAAATCTTTCAAACAAAATGAGGGTAACATTGGTCAGATCCtatagttacttgtcctaaac
gacttatcagtcttgtgtacGATCTATACTACTCCCTAAATACAAACTATTCATCCATGGtatagttacttgtcctaaac
gacttatcagtcttgtgtacCTGGATTTTCAGGTAATAAATGATCTCCTCTAAGACTCTAtatagttacttgtcctaaac
gacttatcagtcttgtgtacTTAAATATAAGGGCTAAACGCTTAATTCACTGACAGGAGGtatagttacttgtcctaaac
gacttatcagtcttgtgtacGACAACCAGCCTGTACTAATTCAAATATATTAAACTGCAGtatagttacttgtcctaaac
gacttatcagtcttgtgtacGTTTATTTGTCTGAGGAATAGAGTCCTATGATTAGCTAACtatagttacttgtcctaaac
gacttatcagtcttgtgtacCATGTCTAAGGAATAGAGTTCTATGCATAGTAGCTTACAAtatagttacttgtcctaaac
gacttatcagtcttgtgtacATGTGATAAGTAATCTTTATGCGATAAGCTACTATGCCGGtatagttacttgtcctaaac
gacttatcagtcttgtgtacATTCCTTTGAGCCTTCCCTTCAGTTTTTCTACTTCTCTTCtatagttacttgtcctaaac
gacttatcagtcttgtgtacTCATGGGGTATCACTCTGATGCTTGGGCCAAATAAACAGGtatagttacttgtcctaaa
gacttatcagtcttgtgtacGCACAGTATTTCTATTGTTGAATTATGGGCATTGGGCAATtatagttacttgtcctaaac
gacttatcagtcttgtgtacTGGGTGCCAATCAATCAGAACAGCTGCCTCATCCATCAGAtatagttacttgtcctaaa
gacttatcagtcttgtgtacTAGAACCTCCTTCCTCTCTCTTCTACACCTACACCCACCAtatagttacttgtcctaaa
gacttatcagtcttgtgtacTGATAGAAAGCTTCCTGAGGACTCCCCCAAAGCAGAAGCTtatagttacttgtcctaaa

gacttatcagtcttgtgtacAGAGAATGGGCTAATACATGAACCAATCAGAAAGGCTGCTtatagttacttgtcctaaa
gacttatcagtcttgtgtacGCTGTACCTGTGCATAGTAGCCAAATACCAGCTCTCATTAtatagttacttgtcctaaa
gacttatcagtcttgtgtacTTTAACAGAGGAAAAGTAGGGAAGTAAGAAGTTAAAACCCtatagttacttgtcctaaa
gacttatcagtcttgtgtacCTATTCCCATAGTCTGTTGCCTTCTAGATCATATCATTATtatagttacttgtcctaaac
gacttatcagtcttgtgtacTTTATTTTTCTGTTCCAAGATGTTTAAAGCAGAAGCTGCCtatagttacttgtcctaaac
gacttatcagtcttgtgtacCACTCTTTGTTTTTCTTGAAGGAAGTATTCAGGGTAATGAtatagttacttgtcctaaac
gacttatcagtcttgtgtacCTGGTGAGTTTTGAGGAAGAGGAATAGGAGCAAGAAGATAtatagttacttgtcctaaa
gacttatcagtcttgtgtacTGGAAACAAAGTGTGTGGTTGTATTAATGAAGAGAATGTGtatagttacttgtcctaaac
gacttatcagtcttgtgtacAAGATAAGTTAAATTATATTATGTGTTGTGGTGGGGCTGGtatagttacttgtcctaaac
gacttatcagtcttgtgtacAAATGGCATGGAATCAGCGTTTTATGGTTGCGAGGAAAATtatagttacttgtcctaaa
gacttatcagtcttgtgtacGCCATTAAATCCTGCTACCTCTCATTTTACTAGAAGAAATtatagttacttgtcctaaac
gacttatcagtcttgtgtacATGGCTTGCTCAGGATCACATATCTAATTATTTACTGAGAtatagttacttgtcctaaac
gacttatcagtcttgtgtacCAACAACACTTAAGTGGGTGGTGCCCCTGTAGTCCCCACCtatagttacttgtcctaa
gacttatcagtcttgtgtacTACTGTCATGATTGCCTTGTGAGCAGCCACTGTACTGTAGtatagttacttgtcctaaa
gacttatcagtcttgtgtacGAAAATTGTGTCCCTATTCCCTGAGGTCTTCCTGCTGCTCtatagttacttgtcctaaa
gacttatcagtcttgtgtacGCAAGTCTCTCTGCCCATTAAACCTCCCCCGCCCTGTATCtatagttacttgtcctaa
gacttatcagtcttgtgtacCACAATTGTCAACTGAGACCTAACGCATCTGTGACATATTtatagttacttgtcctaaa
gacttatcagtcttgtgtacAGAAGTGGTTTTTATTTCTCTCAACGTTTTTTGGGGGTAAtatagttacttgtcctaaac
gacttatcagtcttgtgtacTCTGTATCCATTAAACAATTCTGCATTCTCCTTCCCCTACtatagttacttgtcctaaac
gacttatcagtcttgtgtacGAATTTGAGTACTTTAGATGCCTCCTATATGAAGAATCACtatagttacttgtcctaaac
gacttatcagtcttgtgtacAATGTTCTTAAGATTTACCCACATCGTAGCATGTGGTGGAtatagttacttgtcctaaac
gacttatcagtcttgtgtacGACACTTGGGTTGCTTCCTACTATTCCCTGTTTAGCCATTtatagttacttgtcctaaac
gacttatcagtcttgtgtacAGTAGAATCATTAGTGTTGCCTACGTGATCCTAGTGTTACtatagttacttgtcctaaac
gacttatcagtcttgtgtacTGTCTTCACAAACTTACACAGCTGTGGGGTGAGATGCAGCtatagttacttgtcctaaa
gacttatcagtcttgtgtacGGTGAGAAAAGGGACTTACAACTGGTCTGGCGCTTGATGTtatagttacttgtcctaaa
gacttatcagtcttgtgtacACTTTAAAGGTGGTAAGGGTAGAACTGGGCATTTGGATTTtatagttacttgtcctaaac
gacttatcagtcttgtgtacCTGTGTGACCGGGAGCTGATTATTTAAATTCTGGAATACAtatagttacttgtcctaaac
gacttatcagtcttgtgtacCAACTAAAGTGGCTAACAACATCCTATTACAGAGTGAGTAtatagttacttgtcctaaa
gacttatcagtcttgtgtacTGACTTTCTTATGATCACACAGACAAAACAGACAAAACCAtatagttacttgtcctaaa
gacttatcagtcttgtgtacAATAAGGAGAGGGGTCTATTTTCAACAATCATCTTTCTTGtatagttacttgtcctaaac
gacttatcagtcttgtgtacATTTATGAATGCTCACAGACACTCATGCCTAGCAGAAACAtatagttacttgtcctaaa
gacttatcagtcttgtgtacATTCAATAACCATTGAGTGCCTATCATATGACAACCATTGtatagttacttgtcctaaac
gacttatcagtcttgtgtacTCATGTGGAGGTGACAGGCAAGTATGCTAAGTGTTCCAGTtatagttacttgtcctaaa
gacttatcagtcttgtgtacAGAGAGCATGACTGACAAGAGGAGAGGTGCCTAGCCCACAtatagttacttgtcctaa
gacttatcagtcttgtgtacGTTCAGGAAGGGTGTAGGGAACTGGAGTGGGCATGTGTTCtatagttacttgtcctaaa
gacttatcagtcttgtgtacGGTTACTGTCTGCATAAAACTGAGTAAGCCTGGCATTTGCtatagttacttgtcctaaa
gacttatcagtcttgtgtacACCTAGTGAGATAATTACCCCATAAACAGCAACTGAGAACtatagttacttgtcctaaa
gacttatcagtcttgtgtacCTGAAGTGCATTTGGGAGGGGTGATGATATGGCTGGAGAGtatagttacttgtcctaaa
gacttatcagtcttgtgtacCTGACAAAGGAGGTAAGTCTTGCAGGATGAGGAGATTATCtatagttacttgtcctaaa
gacttatcagtcttgtgtacGGGACAGTTAATCTAGGCAAACACCCAAGAAGCATGCGGGtatagttacttgtcctaa
gacttatcagtcttgtgtacTGGGATACAGGTTGTTCACGAGACACTGGTGATAGAGTGCtatagttacttgtcctaaa
gacttatcagtcttgtgtacGACTTACCTGCTGATGAGTGTAATGGTGTTATGTTAACATtatagttacttgtcctaaac
gacttatcagtcttgtgtacGTCCTGATTCTGATGGTTGTTCACAAAGAAGTTCTAATCCtatagttacttgtcctaaac
gacttatcagtcttgtgtacCATCTGTTTTGCCTAGACCCAGGATCAACTAGTCTCAGTCtatagttacttgtcctaaa
gacttatcagtcttgtgtacTGGCCTCATTGAGTTTCCTTGAGTTCACCACTGCTGACTGtatagttacttgtcctaaa
gacttatcagtcttgtgtacAATGTTTTCTGCCTAGCCCAGGGTACTTTCTTTGGTGGCCtatagttacttgtcctaaa
gacttatcagtcttgtgtacATCTGTGACTGCAAATGGGGTCGTGGGGATAGATCAAGCTtatagttacttgtcctaaa
gacttatcagtcttgtgtacAGAGAGAGAGAGGGAGAGTCCTGTGAGGTTGTATTTCTCGtatagttacttgtcctaaa
gacttatcagtcttgtgtacTTCTCTCCTGAATACACTTCTCTTTCCTTATTAGGACTTAtatagttacttgtcctaaac
gacttatcagtcttgtgtacGACATTTACCATTCCACTGTCTTCTCTGACTTTATCTCTCtatagttacttgtcctaaac
gacttatcagtcttgtgtacACACAGGCATCATTATTAATGCACCATTATCTTCCTCTTGtatagttacttgtcctaaac
gacttatcagtcttgtgtacCATATGAAGAAATGAAAACAGATACTTGGTCAGGGAATGTtatagttacttgtcctaaa

gacttatcagtcttgtgtacTCCTATCCTTCTGGTTCTCAATTTTGGCCCAGCCAGCAGCtatagttacttgtcctaaac
gacttatcagtcttgtgtacTCATTGCCTAGCTGAGAAGTTCCAACATTCAAGAAACAATtatagttacttgtcctaaac
gacttatcagtcttgtgtacGGGTAGCCTCATTTGTTTGATGTCAGGTGAATTGTGTAATtatagttacttgtcctaaac
gacttatcagtcttgtgtacCTGTGGATTTTAAAAGAGAGGTATGGCAGGAAGACCAAGTtatagttacttgtcctaaac
gacttatcagtcttgtgtacAGATGCTTAAGCTAGGAAAGTGATTGGTAGGGATGCAGTGtatagttacttgtcctaaac
gacttatcagtcttgtgtacAAGTTGGAGACAGATTCGGTTCAGGAGAGCACAGGATAGAtatagttacttgtcctaaa
gacttatcagtcttgtgtacAGTTTGTCTTGAGAGAGTTGGGTGGTTGTTGATGAGATGGtatagttacttgtcctaaac
gacttatcagtcttgtgtacAAGTGATTTGTCCAAGGTCAGTGAACATACTGATAGCTGCtatagttacttgtcctaaac
gacttatcagtcttgtgtacAACTCCAAACAAGAATAGTTGCTACTTCATTACCCCCATGtatagttacttgtcctaaac
gacttatcagtcttgtgtacGAGATGAGAAAATATTAAAATACCTGACCTGCGTGGGTATtatagttacttgtcctaaac
gacttatcagtcttgtgtacATGATCTCATCCATTTAGTTCTAGTGTTTGGTGTCCTTATtatagttacttgtcctaaac
gacttatcagtcttgtgtacAGGCGAACTGAAGGAGGATATCTTAATATACAGTGAAATAtatagttacttgtcctaaac
gacttatcagtcttgtgtacAGTAGTGAAATACATTAACAGGTGCATAGGTTCAACTTTGtatagttacttgtcctaaac
gacttatcagtcttgtgtacCCAAAATTAATCTGAGAGGTGTCAAATCTGTGCTGGGTTAtatagttacttgtcctaaac
gacttatcagtcttgtgtacCCAAATTGTAAAGGGATCAGAATTAATAATGCTCTGTCACtatagttacttgtcctaaac
gacttatcagtcttgtgtacCCCGAACACACAGGAGTTTAAGGAATTCACATCTTGGAGAtatagttacttgtcctaaa
gacttatcagtcttgtgtacCTGGTCATCAGTTGGATTGATCTGTCTTATGAATAACTTCtatagttacttgtcctaaac
gacttatcagtcttgtgtacTCTTTTATGTTAGTGGGGCTTATGTATTTCAACAAACCAGtatagttacttgtcctaaac
gacttatcagtcttgtgtacTACAGTATTTTTCTCCCCCATGAATCATTTAAACCCATACtatagttacttgtcctaaac
gacttatcagtcttgtgtacCTCTACCTGTTTATTTAATTCAGAGCTTCTTGGAGATGATtatagttacttgtcctaaac
gacttatcagtcttgtgtacCCAATCTTCATTCATCAGTGGCTTTCATTATGGGCTCTTCtatagttacttgtcctaaac
gacttatcagtcttgtgtacTATGAAAATGAGCATAAAGAGAAGAAGGAGGCGGGGAATGtatagttacttgtcctaaa
gacttatcagtcttgtgtacCACTCCAAATATCTCTCTCTCTCCAAATACCTAAAATAACtatagttacttgtcctaaac
gacttatcagtcttgtgtacCAAAAACATATCCTTTGGTCAATTCGCAGGCAGATTTTCAtatagttacttgtcctaaac
gacttatcagtcttgtgtacAGGGTGATCTATTAATGGGACACGTGCACATCTCTTCCTCtatagttacttgtcctaaac
gacttatcagtcttgtgtacCAGTGGGGAGGGATATGGCTGATGTGTCTTCTGAGCTGGGtatagttacttgtcctaaa
gacttatcagtcttgtgtacGAAGCTGACATTTGAGCCTGCCAACCAACTTCGCCTCATGtatagttacttgtcctaaa
gacttatcagtcttgtgtacGCCAAAGGTTTAGCAACTCTGTAACTGACATTTTCCCCCAtatagttacttgtcctaaac
gacttatcagtcttgtgtacTGGAATTTCTGAAGAACTGGAATAGTTTTAGGAAGACTCAtatagttacttgtcctaaac
gacttatcagtcttgtgtacCTGTATCTATAGGAATAATTCTTGCTTGCGGTATCAGAGTtatagttacttgtcctaaac
gacttatcagtcttgtgtacTTTTATTCCCCACCCAAAACCCTTTCCTATAAGCATGATTtatagttacttgtcctaaac
gacttatcagtcttgtgtacAGGGAAAAAAGGAAGAAGGGACAGGTTAAAAAAAACTAGGtatagttacttgtcctaaa
gacttatcagtcttgtgtacGAACTTTCCATCTCTTGCCCACCCTCCTTCCATACTCTTTtatagttacttgtcctaaac
gacttatcagtcttgtgtacTTTTGATTCTCATCCGTTGAACCTGTCATTTCTTCTTATCtatagttacttgtcctaaac
gacttatcagtcttgtgtacGAACCATCATCTTCCCAATTACTCAACCCAAAAGTCTAGAtatagttacttgtcctaaac
gacttatcagtcttgtgtacTTGGTTTTAATTTCAATCCCTTTGCCTGCTTGGATCTGGGtatagttacttgtcctaaac
gacttatcagtcttgtgtacTTAGATTCTTCTAAGCTTATTCTCACCTTAGCAGTTTGTGtatagttacttgtcctaaac
gacttatcagtcttgtgtacGCAAGGTTGGCATCTTTTTCTTATTCAAATATCACCGCTTtatagttacttgtcctaaac
gacttatcagtcttgtgtacGTTTATCACAGCACTCCACTATAATTTTGTTACGACATAGtatagttacttgtcctaaac
gacttatcagtcttgtgtacAGAATAGGGACCTTATCATTCTTAATCTTTGTCATAACCCtatagttacttgtcctaaac
gacttatcagtcttgtgtacTGTTTTCAATTACTCTTAAGTTGTGGCAGTCATGGAGATGtatagttacttgtcctaaac
gacttatcagtcttgtgtacCTGACAGCTTCCATTGTTACTGCCATCAAGGTTAACGTCAtatagttacttgtcctaaac
gacttatcagtcttgtgtacGAATACCAGGTTTAGTTCTGCCTAAGACAGATCACCACTCtatagttacttgtcctaaac
gacttatcagtcttgtgtacAATCCTTAATCTGGAGTTTCTGTGGGCTAAGACTTTGTTAtatagttacttgtcctaaac
gacttatcagtcttgtgtacCTTGGTGGAGACTTTATTAGGACTGCATAAGGGATGTTGCtatagttacttgtcctaaac
gacttatcagtcttgtgtacGCTTATTTGCGGCCACAGAACTTTTGTATGACTATTAGTTtatagttacttgtcctaaac
gacttatcagtcttgtgtacTAACTGATAAAAACAAGGGTGGAAGAAAACAAGCAAACAGtatagttacttgtcctaaa
gacttatcagtcttgtgtacCTGTGCCAGGCTGTTTTAAAAACTATTGATAAGATCTCACtatagttacttgtcctaaac
gacttatcagtcttgtgtacTCCTGGGTTCAAGCAACTCTCAAGCCTCAACCTCCTGAGTtatagttacttgtcctaaa
gacttatcagtcttgtgtacTTACAGGTGTGAGACAGTGTACCCGGCCTAGTGATCACACtatagttacttgtcctaaa
gacttatcagtcttgtgtacTGAAAGGTGGGTAGCAGCATGTTTGCTTTTACAGAAAGAGtatagttacttgtcctaaac
gacttatcagtcttgtgtacAACTGTTGTTTGGTTATGTAAGTACTGAAGCTAGGATTCAtatagttacttgtcctaaac

gacttatcagtcttgtgtacAAAGCCCTTATTTTTTCAAACTCACCATGCTGTCTCCATCtatagttacttgtcctaaac
gacttatcagtcttgtgtacCGACTCTCAATTCAGGATCTTATAAGGGCAGCAAAAGGACtatagttacttgtcctaaa
gacttatcagtcttgtgtacCAGCAAATGAGATACAATCCATGGTATGACTTTCTGTTAAtatagttacttgtcctaaac
gacttatcagtcttgtgtacTACCCATCTTTTGTATTTTTGCAATGGTTCAGTTGAGAACtatagttacttgtcctaaac
gacttatcagtcttgtgtacAGAACTGTCTCTTTGTCCTTAAGATAATGGAAACCGAACAtatagttacttgtcctaaac
gacttatcagtcttgtgtacCTCACTTGCAGCTGGGACTGTGTACATGATGAGGTTCCACtatagttacttgtcctaaa
gacttatcagtcttgtgtacGGGACACTTTCATTTGGAACACATCTATTTGCTGTCACAGtatagttacttgtcctaaac
gacttatcagtcttgtgtacGAGCTTCCTTGGTGGTCAGTTCTGGGCTGTGATTCTTGAGtatagttacttgtcctaaac
gacttatcagtcttgtgtacATCATCTGTTTACCTGTTTCTTCAGCCCTTTAATGATTCTtatagttacttgtcctaaac
gacttatcagtcttgtgtacACTCATACACACCCACACACCTCCTGTATATAATATTCCAtatagttacttgtcctaaac
gacttatcagtcttgtgtacTCAATTAAGAACTTTGATCAAGCAGCTCATAATCTTAGGCtatagttacttgtcctaaac
gacttatcagtcttgtgtacTGTGAGGAATGAGATGCTCTCTTTAAAAGTAAGAGCTAAGtatagttacttgtcctaaac
gacttatcagtcttgtgtacTTGCACAAAATCTCCATCTCAATGATAAGTTTCCATGAACtatagttacttgtcctaaac
gacttatcagtcttgtgtacTGCTATAACTCTTGCCACTTTGCTACTCAGATTGATTATTtatagttacttgtcctaaac
gacttatcagtcttgtgtacAGGTCATCACTTTCCTAAGGGGTGCACAATGCAAAATTCTtatagttacttgtcctaaac
gacttatcagtcttgtgtacCCTTCTACTTTCCATCTTTTTTTCCTCTAGGCTGTTTCCCtatagttacttgtcctaaac
gacttatcagtcttgtgtacGCCTCAGAACTGGTAGGTAATTTAAAGGCTATATTTCCTTtatagttacttgtcctaaac
gacttatcagtcttgtgtacCTGAAAATATGTCCCTCTTTCTGGTTAGAAGTTGAAGGATtatagttacttgtcctaaac
gacttatcagtcttgtgtacTATTTCTTGTGTCTCATTGCCAACCCCTTCCCCCAACCCCtatagttacttgtcctaaac
gacttatcagtcttgtgtacCAATTTTGTGCTTATCTTATTTGGGCTATCATGTGTATGGtatagttacttgtcctaaac
gacttatcagtcttgtgtacGTGTATATTAGTCTGAACTCACATTATAACACACTCTGCAtatagttacttgtcctaaac
gacttatcagtcttgtgtacATGCAAATGTTGGTTTTTCTTCTTGGCTTAATTCCTGATGtatagttacttgtcctaaac
gacttatcagtcttgtgtacATAAAGAGTTTCATAGAGTCCCAACACATCATAAGAATGCtatagttacttgtcctaaac
gacttatcagtcttgtgtacCCTTCTTGGGGGGATCAGAGCTTCTCACCTCACTTCGGCCtatagttacttgtcctaaa
gacttatcagtcttgtgtacAGTTTGAAAAATCACTGATCTCATTGCATCCTTAGCCCATtatagttacttgtcctaaac
gacttatcagtcttgtgtacTCCCTTTATAATCTCTGATAGGGTTTAGATGTGTGTCCCCtatagttacttgtcctaaac
gacttatcagtcttgtgtacTCTCTTGCTCCCTCTTTTGCTCTGTGACATGCCTACTCCTtatagttacttgtcctaaac
gacttatcagtcttgtgtacCTTCCACCATGAGTAAAGGCAAGCAGATACCGGCACCATCtatagttacttgtcctaa
gacttatcagtcttgtgtacAAGAATGGACTAACACAACCTCTCTCATAGCTAGTTGTCCtatagttacttgtcctaaac
gacttatcagtcttgtgtacACTTAATGATGCAGCTCACTGTATTTGAAGACTGTTTTGTtatagttacttgtcctaaac
gacttatcagtcttgtgtacAACTGATTTCATTGTTGGGGTCTGGTTAATTAAGGCTTTCtatagttacttgtcctaaac
gacttatcagtcttgtgtacGCACAGATAATCATTCTAAGGCATTTTAGTAATTCACACCtatagttacttgtcctaaac
gacttatcagtcttgtgtacTTATGCCCTTTCTGGGTAGGGATGAGGGTGGATATATTACtatagttacttgtcctaaac
gacttatcagtcttgtgtacCAGTGAGTTACCATAGGCAATTAAACAGATAATATTGTGGtatagttacttgtcctaaac
gacttatcagtcttgtgtacTGCAGAACTGAGGTGGGTAGGCAGAGAAAAGAAGGGGCAGtatagttacttgtcctaa
gacttatcagtcttgtgtacGGGCCTACTTTTTGCTAAACCCATCATATATTTTTTCCATtatagttacttgtcctaaac
gacttatcagtcttgtgtacAAAATCAAACTTAGGTGCAATTATCTTGAAGTCCAGATGCtatagttacttgtcctaaac
gacttatcagtcttgtgtacGAAAATAGTGTAAAACTTAACAGCTAGCAAGTAGTGGAGTtatagttacttgtcctaaac
gacttatcagtcttgtgtacATTCTGAAATCCGAGTTCTTTCTTCTTTCTTCCACTTCAGtatagttacttgtcctaaac
gacttatcagtcttgtgtacGACTGGCGACCACTGACATCATTATTATGGCATTGTTGTAtatagttacttgtcctaaac
gacttatcagtcttgtgtacAGTCAATAATGTCTGCTTAAAAATTTCCTCTTCCTGCCTCtatagttacttgtcctaaac
gacttatcagtcttgtgtacGAAGAGTAAAAAAGCAATGATAAGTTATGTCCTCAGTGTCtatagttacttgtcctaaac
gacttatcagtcttgtgtacAGCCTGGGATTTCTGTTAACTTTGTATGATTATTTATGCCtatagttacttgtcctaaac
gacttatcagtcttgtgtacTGTCCCTCACTGAGTAAAATTAGAAAGCACAAAATCACATtatagttacttgtcctaaac
gacttatcagtcttgtgtacTTCATGTGTTAAAATGTCTTAAATCTCTTTTCTGCCTCGGtatagttacttgtcctaaac
gacttatcagtcttgtgtacCAGGGGCTGGGCATGGGGGTGGGAGGCAGTGGAAGTGGAGtatagttacttgtccta
gacttatcagtcttgtgtacCACCATCTAGTTGCTTCTAGGCTAAGGAGGACCCTGGAGGtatagttacttgtcctaaa
gacttatcagtcttgtgtacATATGTGGATATCAGAGAAAAAGCGCCCAGTTGATAGGCAtatagttacttgtcctaaa
gacttatcagtcttgtgtacGCATCACCGTGGCACCAGTTTTATCCTCTTTCTATTCATGtatagttacttgtcctaaac
gacttatcagtcttgtgtacAGCCCAGCTTGTGATGTGGTTGTCAGGCAGCAGTTTCCCTtatagttacttgtcctaaa
gacttatcagtcttgtgtacGTGCAGAATCCTAACCTCAGGCTGTCCTACCACCCGGGCAtatagttacttgtcctaa
gacttatcagtcttgtgtacCACTTCCCTAATGTGTCACAATTACGGAGACAACTGTGCTtatagttacttgtcctaaac

gacttatcagtcttgtgtacCTCTGAAAACTTGCAGTGTGTTCAACAGCCCTCAAAATTTtatagttacttgtcctaaac
gacttatcagtcttgtgtacCCTGTGGTGAGGTGGTCTGGTGGCGGTTATACTGAGCTGTtatagttacttgtcctaaa
gacttatcagtcttgtgtacAAATACAGTCACCTCCAGAGATGTTTTTCCTCCCCTTTGGtatagttacttgtcctaaac
gacttatcagtcttgtgtacGAATTTGGGTTCCTTTTCAGCATCATTTGTATCTCAGTTTtatagttacttgtcctaaac
gacttatcagtcttgtgtacTGAAAGAGGAGGAAGAGAGGTTGCAAAGATAGCACAAGTGtatagttacttgtcctaaa
gacttatcagtcttgtgtacGCTGCAGAACCGGATCTTGTACATTGCTAAGAATCAGGCGtatagttacttgtcctaaa
gacttatcagtcttgtgtacAAGGCTGCTTACACGAGTTATCCCATTAGCGCTGTCAGTGtatagttacttgtcctaaa
gacttatcagtcttgtgtacGTGGCAAACACAGCAGGTAATGGAGCAGGGAATGTGACTTtatagttacttgtcctaaa
gacttatcagtcttgtgtacAGTCCCTCACTTGTTGTTTCTGGAGAGTAGCGATTTAGAGtatagttacttgtcctaaac
gacttatcagtcttgtgtacAGCGCTACAGAGAAGCACTCATTATTTACCCAGGATACAAtatagttacttgtcctaaa
gacttatcagtcttgtgtacCAGGAGCACCACTTCTCAAGTCCCAGCTTTGCACTTCAGAtatagttacttgtcctaaa
gacttatcagtcttgtgtacGTGACAATGAGATCAAAGAAAGGTGGAGGCTGCAATTCATtatagttacttgtcctaaa
gacttatcagtcttgtgtacAGGGGAATGCATATCAACACTTCTCTTTCAGGTGACAGGGtatagttacttgtcctaaa
gacttatcagtcttgtgtacGGAGACCCACATTCCAGTTAATAAGGTATAGCTAGGCCAAtatagttacttgtcctaaa
gacttatcagtcttgtgtacGGAGAAGGCTGTAAATGAAGTGGGCTGAGCACGAATGGGCtatagttacttgtcctaa
gacttatcagtcttgtgtacCAGGAGTTCAAGAATAGGCTGAGTTTCTCTAGGTTACTGTtatagttacttgtcctaaac
gacttatcagtcttgtgtacAAATGATTTCTTTAGTTTCAGGACCTGCTTTGTCTTTTGGtatagttacttgtcctaaac
gacttatcagtcttgtgtacCCCTGGTTGGTTTATCACAGCGGGTAGAGGGAACTTCACTtatagttacttgtcctaaa
gacttatcagtcttgtgtacCCTTTCTTGAGGGCAGGGATAGAAGTGTTGCTGATGTGCAtatagttacttgtcctaaac
gacttatcagtcttgtgtacGCGTTAGACCCTCAGGAATTAAAAGGCCACCAATGTCATCtatagttacttgtcctaaa
gacttatcagtcttgtgtacGCTTACTTTCCAGATGCTTCCTCCATTCAGTGACCTGTACtatagttacttgtcctaaac
gacttatcagtcttgtgtacGGGTGCCGTGGTCTTCAAACACTCTGTGCTTCCTCATGTCtatagttacttgtcctaaa
gacttatcagtcttgtgtacGGTCCCCCTCCCTTCCCTACCTAACCATCTCTGTTAGTCTtatagttacttgtcctaaa
gacttatcagtcttgtgtacGTCGTGGAAACATCCCCGTGACACTTTGAGCTTACCTGGCtatagttacttgtcctaaa
gacttatcagtcttgtgtacCTGCCTTTCACAACGTGCGACAATTATTTATTTTCTTCTCtatagttacttgtcctaaac
gacttatcagtcttgtgtacAGAACCCATGTCTTATTTATTTTTACAACCTCAGGGCCTCtatagttacttgtcctaaac
gacttatcagtcttgtgtacCATTTACCGAAAGTTTCCCAAAAGAAATGAATGGAGTCGCtatagttacttgtcctaaac
gacttatcagtcttgtgtacGCGGTCCTATTGTGCTGTGTATCTTTAATGAAGCTATAACtatagttacttgtcctaaac
gacttatcagtcttgtgtacGTTAAAAAGGAAACAGGTATGAGAATAGTACAGATCTACCtatagttacttgtcctaaac
gacttatcagtcttgtgtacGACCTTGTATGAATTAGCAAACTAGAGAATTATGCAGACGtatagttacttgtcctaaac
gacttatcagtcttgtgtacTGGCTGACAAACCTGAAGATGTAGTTGAGCTTATGATTGGtatagttacttgtcctaaac
gacttatcagtcttgtgtacAGCCAGGCTGCTTCAACCCACTTAAGCATAGATAATCAAAtatagttacttgtcctaaa
gacttatcagtcttgtgtacGCTTCTCTGACAGCTGCTGATCACAGACACTATTAAAGAAtatagttacttgtcctaaac
gacttatcagtcttgtgtacAATACAGGTGACAAAGAGAGAGGCAAGCCTTCTGAGTGGCtatagttacttgtcctaaa
gacttatcagtcttgtgtacACTAGGAAAGTGACAAAGCTTCAGAGGCCTCTCCCCAAAGtatagttacttgtcctaaa
gacttatcagtcttgtgtacCTGTTGGAAACGAGCGGGCTTATGTAGACAATCATTTACAtatagttacttgtcctaaac
gacttatcagtcttgtgtacGCAAACATGCAACTGTCAGGCCTGGATGGAAAAGAAACATtatagttacttgtcctaaa
gacttatcagtcttgtgtacAATTGTGAGAAATTCTAACGTCGTAGCAAAAAATCAGGAGtatagttacttgtcctaaac
gacttatcagtcttgtgtacGTGGAGAGCTGTCATCTTGGCATTATATTAAAATAACTTGtatagttacttgtcctaaac
gacttatcagtcttgtgtacCAGTATTTCTTCTTTCAAATCTGTCAGTTCTCGGGAGGCCtatagttacttgtcctaaac
gacttatcagtcttgtgtacAGGGACACTGGGAGGCTTTCTGGGGGCAGGAGCGGGGGCGtatagttacttgtccta
gacttatcagtcttgtgtacCAAATCATTTGCCGACCTATTTCTTCTTGACTTAGTGGGGtatagttacttgtcctaaac
gacttatcagtcttgtgtacGCAGATAAATTATTTTCAGTAGTGCTCCTCAGACAGCGATtatagttacttgtcctaaac
gacttatcagtcttgtgtacTGAACTGCATTTGTTTATGAAATCCGTTGCTTAATAGGACtatagttacttgtcctaaac
gacttatcagtcttgtgtacGTGTCTCATTTCAGCCTCTAGTCATGATAGTTTTAATGATtatagttacttgtcctaaac
gacttatcagtcttgtgtacCAAGTGGGAGCAGAACAATGGGAAAAAAAACCTAAATCGGtatagttacttgtcctaaa
gacttatcagtcttgtgtacCGATGCTGTGTCAGGGACCACGTCTCCTGCTCTGTCGAATtatagttacttgtcctaaa
gacttatcagtcttgtgtacTGTCTCTGCACTGTCTGTTTCCTCTCGCTTTGGCTCAAGAtatagttacttgtcctaaac
gacttatcagtcttgtgtacACCTCTTTCCCTAGTCAAATGCTTCCCAAGGATCCCAGGTtatagttacttgtcctaaa
gacttatcagtcttgtgtacCCTGTTTCTCTGCTTTCCTGATATGTGCTGTATAATATAGtatagttacttgtcctaaac
gacttatcagtcttgtgtacGTAACACCACAATTTGCATTATCTTTTTGGCTTTTGAAGGtatagttacttgtcctaaac
gacttatcagtcttgtgtacTCTCACAAATTCCTAACAAAAGAGAGGCAGAGTGGACTTTtatagttacttgtcctaaac

gacttatcagtcttgtgtacTGGTGTGTAAGGCCTGTCTGTCCATATTGTGTAATTAGTTtatagttacttgtcctaaac
gacttatcagtcttgtgtacGGTTATCCCCATTACATAGGCAAGCAAAGTAACGTAGAGGtatagttacttgtcctaaa
gacttatcagtcttgtgtacGCTTATTTTTATGTGATCTTGGGCCAGCTACTTAACCTGCtatagttacttgtcctaaac
gacttatcagtcttgtgtacTTAGTGGTAGTATTGCAGAAATAAAGATTATGAGCCTTGGtatagttacttgtcctaaac
gacttatcagtcttgtgtacCCATGGTAAGGGCTGGAGTTATACATACCTGGTTACTGGAtatagttacttgtcctaaa
gacttatcagtcttgtgtacGGTCTCCTGAGTTTTCATCACATTCTGCCCTGCCCTTGCGtatagttacttgtcctaaa
gacttatcagtcttgtgtacGTGTGCAGATTTTCCACCTTTCAGCAATTCCAGATACAGAtatagttacttgtcctaaac
gacttatcagtcttgtgtacTTTGCCAAAAGTGCAGTATTTCTTCAAAAGCCCTTAAATGtatagttacttgtcctaaac
gacttatcagtcttgtgtacCTCCTTGCTCAGTCTGGCTGGAAACGCCCCCACCATTGCCtatagttacttgtcctaa
gacttatcagtcttgtgtacGCCTGAGCCATGTGTGTCTGAAAGAGGGATTCACCTTGACtatagttacttgtcctaaa
gacttatcagtcttgtgtacAGGTCATCAAAAACATCACAGTTCCTCCCACTCAGCACCTtatagttacttgtcctaaa
gacttatcagtcttgtgtacTGACCCCCCATCGATCAGCCATAAAACATAAAATCAACAGtatagttacttgtcctaaa
gacttatcagtcttgtgtacATGCAGAAAAATCAACATTGACCAGCATTGTCAGCCAAGCtatagttacttgtcctaaa
gacttatcagtcttgtgtacGTGCCCACAGACTTCCCTCTCAGATCCTCTTAAGCCTCTCtatagttacttgtcctaaa
gacttatcagtcttgtgtacGCTGCTAAGTACTAAGTGGGAGAAGCGTTCATCTTTGTCCtatagttacttgtcctaaa
gacttatcagtcttgtgtacGTTGGTGACTGACAGCCCGAATAAGCAGTTATTTCTCCAGtatagttacttgtcctaaa
gacttatcagtcttgtgtacGCGAGAGACACATGTAACCCAGTGCCTGACCTTATGCAGCtatagttacttgtcctaa
gacttatcagtcttgtgtacGCATTCTGGAGCCAGCTTGCTGCCACTATTACCTGTTAAGtatagttacttgtcctaaa
gacttatcagtcttgtgtacGCTCTGATCATGCTACTCTCCCCCTGAAATAATTTTTGAGtatagttacttgtcctaaac
gacttatcagtcttgtgtacCCAAATTCCTTATTCTGGTGTTCATGGCTTATTATGATCTtatagttacttgtcctaaac
gacttatcagtcttgtgtacCCTTCCTAGCTGTGTCTCTTTTTACCAGGGTTCATGACTCtatagttacttgtcctaaac
gacttatcagtcttgtgtacCTGAATCTGCCCCCACAGTCTCTCACCTTTCAGTTGACACtatagttacttgtcctaaa
gacttatcagtcttgtgtacTTTTTCCTTCCCCTGGTATGCTCTTTTCCTTGACTGTTCAtatagttacttgtcctaaac
gacttatcagtcttgtgtacCAAGCACCATTGATATACTACCTCCTCCGTGAAGAGCTACtatagttacttgtcctaaa
gacttatcagtcttgtgtacCTATGGCAAGTTTTCCCTTCTCTCAAATCCCTTAGCTCCTtatagttacttgtcctaaac
gacttatcagtcttgtgtacCCTTCTCAGGACAGTTATGCCTTCTTCCTAGTATTATGTAtatagttacttgtcctaaac
gacttatcagtcttgtgtacGCACCTGGCCTATATTTTTTAATAGACTCTGAGCTCCTTGtatagttacttgtcctaaac
gacttatcagtcttgtgtacGATCCCTGATCCCCACAGAACGAACTTAGAGCTTGTTATTtatagttacttgtcctaaa
gacttatcagtcttgtgtacCTCCTTTCTATTCATTTAATGAGAGCTTCACATGGGTTGAtatagttacttgtcctaaac
gacttatcagtcttgtgtacCTCATGTTTGTGGCGATGCATGGGTAGAACCAAGGTCTGCtatagttacttgtcctaaa
gacttatcagtcttgtgtacAGGGATTCGCTCAATATGATTTCCTAGACTGTTTTTAAGGtatagttacttgtcctaaac
gacttatcagtcttgtgtacCCATATCAGCAAGGAATCTAGTTGTCATACACACACAGACtatagttacttgtcctaaa
gacttatcagtcttgtgtacTTTGGCCAAATTCTCTTTAAAATAATCTGCTCCTATCTCCtatagttacttgtcctaaac
gacttatcagtcttgtgtacACAACTGTGGCTGTTAGTGAGAAAAGAATTCATGAGCAACtatagttacttgtcctaaa
gacttatcagtcttgtgtacAGAATAATGTATTGTGGGCCAAAAAGATCAGAGTAAGTCAtatagttacttgtcctaaac
gacttatcagtcttgtgtacGGAACCTACTTGTTATAAACAGTTTTCTCAGGAATTGGCCtatagttacttgtcctaaac
gacttatcagtcttgtgtacTGGTTAATGATTGTCACCTTCTCCTCTACCACCACAAACAtatagttacttgtcctaaac
gacttatcagtcttgtgtacTATTAACTATTCACGGCATCCATTAAATATTCTCCTGGACtatagttacttgtcctaaac
gacttatcagtcttgtgtacTTCTAATTTTTACTGCCAGTAAAGCTTTACCTCTGTGTTGtatagttacttgtcctaaac
gacttatcagtcttgtgtacCTTTGTTTAAGGAATATGAACTCAGTCGTCTCTAGGAAGCtatagttacttgtcctaaac
gacttatcagtcttgtgtacTTTGTGCCACGCCAGGGCTGTTCTCTACTTGCATATTTGCtatagttacttgtcctaaac
gacttatcagtcttgtgtacGGAAGTTTGCCTGCATATCGAAATGATATATATTTCCACTtatagttacttgtcctaaac
gacttatcagtcttgtgtacTAGGTTATGCTGACTACTGATTCCCTTGCACCAAAAAGCAtatagttacttgtcctaaa
gacttatcagtcttgtgtacCAGAGATGACCTTTAGCTAAAGAGTTGCAAGTTATATTACtatagttacttgtcctaaac
gacttatcagtcttgtgtacTAGAATACAGAAAAGAATCCTTCCCACCATCAAGCCACTAtatagttacttgtcctaaa
gacttatcagtcttgtgtacGGAGAGATCACCTCCTAAGCGCTGGTGTTAGAACATCCGTtatagttacttgtcctaaa
gacttatcagtcttgtgtacATTTTCATTTCCATGCTAACAAGGTAGACCTATCCTAGCGtatagttacttgtcctaaac
gacttatcagtcttgtgtacCAGGGCCTGGGGAGCTTTAACATAGTTAGATGTACTCTAAtatagttacttgtcctaaa
gacttatcagtcttgtgtacGGATTAAAATAAAAGAGAAATGCACTTGCAGTAGGATGGCtatagttacttgtcctaaa
gacttatcagtcttgtgtacTTATTTAACACATTTCAAGTGCTTGGTGGGAAGATGCAGCtatagttacttgtcctaaac
gacttatcagtcttgtgtacTCAAGGTTTTATGTTATGATCATGGATTTTTGGAGCTGAGtatagttacttgtcctaaac
gacttatcagtcttgtgtacCACAACAAGGTGCTGGGAGAGGGCTTTCTTGCTATCCAAGtatagttacttgtcctaaa

gacttatcagtcttgtgtacGGATTTGTACTCACTGTCCAGCCACTCCGCCTAGAATGTGtatagttacttgtcctaaa
gacttatcagtcttgtgtacGGTTCTGCCTTGTTTATTTCTCAATTCCCAGCATCTAGAAtatagttacttgtcctaaac
gacttatcagtcttgtgtacCAGTGAATATGTGTTATCTGAAAAAAGTGTCCCAAGCATGtatagttacttgtcctaaac
gacttatcagtcttgtgtacGAACAAGTTTCTTAAGAATCCTAAGTGCCAGTTTTTCATCtatagttacttgtcctaaac
gacttatcagtcttgtgtacACTACAGTTGACTTTCCAGTGTTCTTGGGAGGGCGTCTAAtatagttacttgtcctaaac
gacttatcagtcttgtgtacAAGCTATTTTTGTATCTGGCAGGCTGCGTTTCATGGACAGtatagttacttgtcctaaac
gacttatcagtcttgtgtacGACGCTTCTGGAGGGCTCTGCTTCATGTCTCACTAGAAAGtatagttacttgtcctaaa
gacttatcagtcttgtgtacAGCAGGTGGTGGTGGTATCAGGAGGAAAGAGAGATCCTTAtatagttacttgtcctaaa
gacttatcagtcttgtgtacTGGAATGAGCCAGCTGAACCAAGTCCCTGAAACTCACAGCtatagttacttgtcctaa
gacttatcagtcttgtgtacAGGGAGATCTTAATGTAGCACTTATTTCTGGCACCTAGAGtatagttacttgtcctaaac
gacttatcagtcttgtgtacTGGCTTGGTGTCATCAAAATCTACTTGTCTTGAAATACTAtatagttacttgtcctaaac
gacttatcagtcttgtgtacTAAGCCTATATTCTGCCATCTCATAGTCAGGTCCCCTAACtatagttacttgtcctaaac
gacttatcagtcttgtgtacGCTTTCCACTATTTTTATGCTGTTTTACTTGGGTCAATGTtatagttacttgtcctaaac
gacttatcagtcttgtgtacCAATTCTCCATTTGCCCCACCCCACCAGATCACTTTTTGTtatagttacttgtcctaaac
gacttatcagtcttgtgtacCAATGCATTGCCTCACTCTCTTGCCCTTTGGTTTCCAGTTtatagttacttgtcctaaac
gacttatcagtcttgtgtacCATGGGCAAGAGATCAGAAGTTAGAAGAGATAGGTCAGGAtatagttacttgtcctaaa
gacttatcagtcttgtgtacCCTACTGACTCCCTGTTTTGACATCATGGTCTGGCAATGGtatagttacttgtcctaaac
gacttatcagtcttgtgtacAAGGTTTTGCTTCAAGTTATCATCTCTCATTGGGCACTGGtatagttacttgtcctaaac
gacttatcagtcttgtgtacCCAAGGGATAATAAGGGCTTCCTACTGCCATCGGTCTCTGtatagttacttgtcctaaa
gacttatcagtcttgtgtacGTGTCCAGATCACTAAAAACCACCTCATCATCAAGTCTCTtatagttacttgtcctaaac
gacttatcagtcttgtgtacCAATTGTGCCCATCTTAAAGAAACTATGAGCTACCATCCTtatagttacttgtcctaaac
gacttatcagtcttgtgtacGACTTTTATGATGTCCTCCTCAATTCCACTGTATTAGCCAtatagttacttgtcctaaac
gacttatcagtcttgtgtacTAGGACATATAAAAGAGAGTTGATTAGGAAGAATTGGCTCtatagttacttgtcctaaac
gacttatcagtcttgtgtacCAAGTCTAAAAGCCTTGAAGCCAGGGAAGCCAATCGTGCAtatagttacttgtcctaaa
gacttatcagtcttgtgtacGTCGAAGGCCTGAGAGCTCCCTGGAGGGCACTGGTGTAAGtatagttacttgtcctaa
gacttatcagtcttgtgtacGCAGGAGGAGAGGAAGCAAGAGTCTGACACAGGGAGAGAGtatagttacttgtcctaa
gacttatcagtcttgtgtacCTCAGTAAGTCGCTTATCCCCTTTCTTCTGCCTGCTTTGTtatagttacttgtcctaaac
gacttatcagtcttgtgtacTATGCTGGCAGCCTGTGGGATGGTACACTCTCACTGAGGGtatagttacttgtcctaaa
gacttatcagtcttgtgtacCAATACTTTACCAGCCATCCAGCCATGCCTCAATCCAGTCtatagttacttgtcctaaa
gacttatcagtcttgtgtacCAAACCCTGGCTCTCCACTCTCTGAGAAATCCTACCCACCtatagttacttgtcctaaa
gacttatcagtcttgtgtacTGTCTCCCACAGGCTTATGATCATCAGATTCATTCATTTGtatagttacttgtcctaaac
gacttatcagtcttgtgtacTTCTAGGCTTGGGATGTGCTGGTGAACAAAACAGTTATGGtatagttacttgtcctaaac
gacttatcagtcttgtgtacGGCTTTTATGGGATCAAATCTGTGCTATAAGCCTCTCACAtatagttacttgtcctaaac
gacttatcagtcttgtgtacCAGTTGCCTGCCAGACATCCCCAAAGCTCCATCAGTGCAGtatagttacttgtcctaa
gacttatcagtcttgtgtacCTGAACCCAAATTCATTATTACGTGAACCCAAAACTGTTCtatagttacttgtcctaaac
gacttatcagtcttgtgtacATGGCACCATCACCCCCTAGTTTTATGTAAGTTAACCATCtatagttacttgtcctaaac
gacttatcagtcttgtgtacCCAATAAGATTTTTTCGATTTTGCTTTGGACTTTGCTCCTtatagttacttgtcctaaac
gacttatcagtcttgtgtacCTTTGTCTAAGGTTAAAGATCTTATCATTTCTCACCTGGAtatagttacttgtcctaaac
gacttatcagtcttgtgtacAGCCTCTTACTGGTCTACTGGCCCCCAGCTTGTCACTCTGtatagttacttgtcctaaa
gacttatcagtcttgtgtacCATAATTACTGTGCATTGTGGGTGGGACTAGTGGGAGATAtatagttacttgtcctaaac
gacttatcagtcttgtgtacGGTTTTAAAAATTGGAGTTTCCCTGCATATGCTCTCTCTCtatagttacttgtcctaaac
gacttatcagtcttgtgtacGGACTAATACACACTCCAAGCCTTTCTCATTACTACTGCCtatagttacttgtcctaaac
gacttatcagtcttgtgtacAAGAATTCTGCCAGCTCCCTATTGCCACACTTTTTTGTAAtatagttacttgtcctaaac
gacttatcagtcttgtgtacACCACTGGTTTCTGCAGGCCTCACCCATCACATCAACTCAtatagttacttgtcctaaa
gacttatcagtcttgtgtacCCTTTGCACATGTACTTTTCCTGAGTTGTTAGCACCTCCAtatagttacttgtcctaaac
gacttatcagtcttgtgtacCCCTTTTTAAAATTCAAGGCCTTATTGATGACACCTTCTCtatagttacttgtcctaaac
gacttatcagtcttgtgtacAGTTAGTTACTCCATCCTTTTCACTCCCATACTTCTTTGTtatagttacttgtcctaaac
gacttatcagtcttgtgtacCAGTTAGTTATAAAAGTGTTGATTCTCCTTCTAGCCTGAGtatagttacttgtcctaaac
gacttatcagtcttgtgtacGCACTGTTTGGCATAGAAGAATTATCGGGATATACTTTTGtatagttacttgtcctaaac
gacttatcagtcttgtgtacCAGGAAAAGCATCTTCTGACCCCCCAACTCTTCTTGCCACtatagttacttgtcctaaa
gacttatcagtcttgtgtacGGTAGCTGCTGTTGTGATACCTTGTCCAAAGAACCTACTCtatagttacttgtcctaaac
gacttatcagtcttgtgtacGTCCCTAACATGTTACAATACCTAAGTTACCCAACAGCTGtatagttacttgtcctaaac

gacttatcagtcttgtgtacGCCAGTGCAAATAATCTTCCTGACCTCTACTTTATTTTTCtatagttacttgtcctaaac
gacttatcagtcttgtgtacGAAGGGGGCTGGTGATGAGATACGGAGTTTTGTGAGGACAtatagttacttgtcctaaa
gacttatcagtcttgtgtacGTCATCTCATGGAGGGCACTAACAGGCAGAAACTCAAAATtatagttacttgtcctaaa
gacttatcagtcttgtgtacTCTATGTGCAGTCAAAATGTCACCAAAGCCCTGGAGCTGCtatagttacttgtcctaaa
gacttatcagtcttgtgtacCCACAGCATCCCCTTCTCAGCCCCTCACCCCAATCTGCGTtatagttacttgtcctaa
gacttatcagtcttgtgtacGGGAGAGAAAGGAGAGAGGAATTAAAACATGTAGTAGTTCtatagttacttgtcctaaa
gacttatcagtcttgtgtacTGCCTAGAGATGGTGTTGCTAATAATACCCTGAGCAGAAGtatagttacttgtcctaaa
gacttatcagtcttgtgtacCAAATTTTAAAATACTGTGAGGTTTGGGGCTGGCTTCAATtatagttacttgtcctaaac
gacttatcagtcttgtgtacACTCAACTTATTTTATGTGTTTAACCCTAACCCCTTCTCCtatagttacttgtcctaaac
gacttatcagtcttgtgtacCCAACTAACAGCTAAAGAAGGAGAGATGAGATGGCACCAGtatagttacttgtcctaa
gacttatcagtcttgtgtacCATGTTGATGGGGTTGACAAGTGACCTGAGGCTACCTGTAtatagttacttgtcctaaa
gacttatcagtcttgtgtacCATGTGACCACCATGAAGAACAAAGGCTCCAGGGAATTGTtatagttacttgtcctaaa
gacttatcagtcttgtgtacCATATATGGGAAAGAAGACATGAAGAAGGATGAGGATGGTtatagttacttgtcctaaa
gacttatcagtcttgtgtacGCTTTACTGTGTTTCAAGCGCAAAGTAGATTTACAGCAGCtatagttacttgtcctaaac
gacttatcagtcttgtgtacTAAATTCCTGACAACTCACCAAACCAGAGTGCACAGCTGCtatagttacttgtcctaaa
gacttatcagtcttgtgtacTCCCAGGAGATTTCAAAACATATGAGAGGAGAAAGCAATGtatagttacttgtcctaaa
gacttatcagtcttgtgtacGTCCCATCTGATCCTACATTCTGCTCTCACTGTGACCTGAtatagttacttgtcctaaa
gacttatcagtcttgtgtacAGACATTGCTGGCCTCTGATGCTCTTTGAAATGGTTTCTCtatagttacttgtcctaaac
gacttatcagtcttgtgtacGAATGGGTCTAAAGTGCAGTGCCTAGAAATTGTTTTAAAAtatagttacttgtcctaaac
gacttatcagtcttgtgtacCACTGAAAAATAGCTCAATTTCTCCATCTTTTTTAGGTGGtatagttacttgtcctaaac
gacttatcagtcttgtgtacGAAGTTACCATGGTTCTAATGAGGGTATTCTTTATTGTCAtatagttacttgtcctaaac
gacttatcagtcttgtgtacTGCTTCCTCAAAAAAACCACGTTCTTTTATACATTTCTGCtatagttacttgtcctaaac
gacttatcagtcttgtgtacGTTAAAGACTATCTGGGCAAGTGATAAAGAATAGGAGGCTtatagttacttgtcctaaa
gacttatcagtcttgtgtacGATTTTAGCTATTTATGAAAAGTCAAGTACAGAGGCGAAGtatagttacttgtcctaaac
gacttatcagtcttgtgtacTGTTTCACATTTTTCGAGCATGTAATACAAGCTGAAGGGCtatagttacttgtcctaaac
gacttatcagtcttgtgtacCAAGGTGAAATTGACCAATAAATTGAGAACTCCATTACAGtatagttacttgtcctaaa
gacttatcagtcttgtgtacCAGCATTTGAGTGATAACCTAAACCTGTCTCAACTTTGTGtatagttacttgtcctaaac
gacttatcagtcttgtgtacCAGAAGGAAAGGATGAGGAACACTGGAGGAGGCGGAGGGGtatagttacttgtccta
gacttatcagtcttgtgtacGTGGGCACTGATGTTTATATATTTTCACCAAGTTTTCTAGtatagttacttgtcctaaac
gacttatcagtcttgtgtacGCAGACCTGTGCTTCAGTGTTCAGTTTTTTTACAAGGTTAtatagttacttgtcctaaac
gacttatcagtcttgtgtacGAAAAATGACAGCTTGAATACAATAGTCCAATTTAGAGGGtatagttacttgtcctaaa
gacttatcagtcttgtgtacTGCTGGCTATTGGAGACATTTGCAATGAATACAAATCCGCtatagttacttgtcctaaa
gacttatcagtcttgtgtacACCTTGGGTTTGTGTGGGTAAGTGTGTGCTAGCTTGTTCTtatagttacttgtcctaaac
gacttatcagtcttgtgtacACCTGTTTCTTAGTTGATTTTGGCCCCAAGCAGTATGAATtatagttacttgtcctaaac
gacttatcagtcttgtgtacACATTGGTGTGAGGGTGCTGGGGGTGGAAGGACTGCAGGAtatagttacttgtcctaa
gacttatcagtcttgtgtacTGACAGCTCTACATGGCTCTTGGCACTGCAATATTTCAGGtatagttacttgtcctaaa
gacttatcagtcttgtgtacTCTCATTTAAATATTCCAAAGAGAGTGGCCTACATACTGAtatagttacttgtcctaaac
gacttatcagtcttgtgtacTCTCTCAAAAGGAGTGGGGTGGGGGAGAAGGAAAGGAAGGtatagttacttgtcctaa
gacttatcagtcttgtgtacAGAAAGAAACGATGGTAAGGAAATGCAGCAAGTGAGTGATtatagttacttgtcctaaa
gacttatcagtcttgtgtacGTGAGCACAGAATTCAATATCCAGGACGAGGACATATATGtatagttacttgtcctaaa
gacttatcagtcttgtgtacCTCAGAACTGATGGAGAAATGGGCAGGAAGAGAGATAATCtatagttacttgtcctaaa
gacttatcagtcttgtgtacGAAATAGCTAAGGCTAAATCACCCTCAGAAGAAAGCAGGAtatagttacttgtcctaaa
gacttatcagtcttgtgtacCTGCTCCAAGTCACCCATGAGATTTCAGAGTGCAGGCCACtatagttacttgtcctaaa
gacttatcagtcttgtgtacTGAGTTCAGGGTCAGTCAGTATATGGAAACCTCAGTCAGTtatagttacttgtcctaaa
gacttatcagtcttgtgtacGTCTAGGCTAAGCTTCTGATTTCTGTTTTATTTAGGAAACtatagttacttgtcctaaac
gacttatcagtcttgtgtacAGAATCTGAAATCTATGCAAAAAGCAAAAAGGAGAGGAGAtatagttacttgtcctaaa
gacttatcagtcttgtgtacTTCCACAGCTTCATTTGTTCCTGGGAAAACCATGTATCTAtatagttacttgtcctaaac
gacttatcagtcttgtgtacTTCTAACATTTCACATGCCCACTCACATATTTGAAAGTCTtatagttacttgtcctaaac
gacttatcagtcttgtgtacAAGCTAATCATTACTGCAGCAGGATTTTAATAGTTTGGAGtatagttacttgtcctaaac
gacttatcagtcttgtgtacGGATGATGAGAACAAACCCTGGTGCTTGAAAGAAAGTGGCtatagttacttgtcctaaa
gacttatcagtcttgtgtacCATATCCTTCCTTTATGGTATTTGGACTTGCTTTCCCACAtatagttacttgtcctaaac
gacttatcagtcttgtgtacATCTGTGATCAAAGAAGTGAAAAAAAGGCTCAGAAAATGCtatagttacttgtcctaaa

gacttatcagtcttgtgtacACTAACTTGTGAAGCATGAAGTCCATTATCTGCACTGGGCtatagttacttgtcctaaac
gacttatcagtcttgtgtacCATCATACAACCAAGCCATTGAAATTGATGTTCTTTGAGTtatagttacttgtcctaaac
gacttatcagtcttgtgtacTGGAAGATATTTCAGTCCTCCAATCAGCTAGCTTAGTTAGtatagttacttgtcctaaac
gacttatcagtcttgtgtacCTTTAGGGAATCATATTGGTGTGGTTTAGTTATCCTTGCAtatagttacttgtcctaaac
gacttatcagtcttgtgtacCCATTTCAGACTTTGCAATTATAAGCAGACAGGAAAACTCtatagttacttgtcctaaac
gacttatcagtcttgtgtacGTTGTGAACCTACAAAGAATCAACTTTTCTAACCGCATTCtatagttacttgtcctaaac
gacttatcagtcttgtgtacAGTGGCACTGTAGAACTTCATCTCATTAGCATGGTTCAGAtatagttacttgtcctaaac
gacttatcagtcttgtgtacCAGGATGGCTTGCTCTAGTTACCTTCGTCATTGCTGGGACtatagttacttgtcctaaac
gacttatcagtcttgtgtacTTGCTTTATTTGGTTCTGTTTACTTCTTCTAGGTAGTGGTtatagttacttgtcctaaac
gacttatcagtcttgtgtacTTGAAATGGGTAAATCAACAGGTAGAGGCCAAGGAGTTTCtatagttacttgtcctaaac
gacttatcagtcttgtgtacTCCACTTCTGAATGATACTGGCAAGGATTCTTCCATCTCTtatagttacttgtcctaaac
gacttatcagtcttgtgtacTCATTTTCCTTGTTAATCAAAGGGGCACAGTAATACTTGCtatagttacttgtcctaaac
gacttatcagtcttgtgtacTGTGGTTCAAGTAAATCTGTGAGAATAAGTTGACAAGAACtatagttacttgtcctaaac
gacttatcagtcttgtgtacACATAAGTTTATAATTAACCATGAGGAAGGAGGAAGGGGGtatagttacttgtcctaaac
gacttatcagtcttgtgtacCAATTCACTGCATATTTCTGGTTCTTTTTCCTTCCTGCCCtatagttacttgtcctaaac
gacttatcagtcttgtgtacGCTTTGGCCAGAGAATGTGAGTGAAAGGAATTATGTGTCAtatagttacttgtcctaaac
gacttatcagtcttgtgtacAGAATCTGGTCTGAATGGTTGTATTCTGTTCTCCCCACCTtatagttacttgtcctaaac
gacttatcagtcttgtgtacCTGGAAGATTTTTAGAGATGGAGCTATTGTCAGCCTGAGTtatagttacttgtcctaaac
gacttatcagtcttgtgtacAAGCCACCTTTCCAACTCATATTGTATATGTAACTTCATGtatagttacttgtcctaaac
gacttatcagtcttgtgtacGTAGAAGTAAGAAGTGAACTTTTGTTGTGCCAAGCTGTTGtatagttacttgtcctaaac
gacttatcagtcttgtgtacTGTTATTACTGCAAAACCTAGACTCTTCTGATTGATACACtatagttacttgtcctaaac
gacttatcagtcttgtgtacTGCTTTCTGATTTATTTTCTGTAGACTCTCAAGGCAATGCtatagttacttgtcctaaac
gacttatcagtcttgtgtacATCACACAGTCTTTATGATTTCTTCAGACCTATTTCTCTGtatagttacttgtcctaaac
gacttatcagtcttgtgtacCATATATGGTCTATCTGCCTGCCACAACATTCTCAACTTCtatagttacttgtcctaaac
gacttatcagtcttgtgtacTGCTGGCTGAAACATCACTTTTACTGGGGCATCTTCCTGGtatagttacttgtcctaaac
gacttatcagtcttgtgtacGGATTGTCTCTTCTTTACGCTCCCAAGCAGCTTTTGCCACtatagttacttgtcctaaac
gacttatcagtcttgtgtacAATTGCTTTTTCTGGTGTGTCTTTATGTATGACGTTTTCCtatagttacttgtcctaaac
gacttatcagtcttgtgtacCTTAGTGAAGTGAATGGAATCTGGAATTTTTAAACCTACGtatagttacttgtcctaaac
gacttatcagtcttgtgtacCTCTAAATTGACCATAATGAGAGAGTATTGGTGGGAGAAGtatagttacttgtcctaaac
gacttatcagtcttgtgtacCACCCAAGATATACATTGTTGATTTACAGCCTGAGGGGGGtatagttacttgtcctaaa
gacttatcagtcttgtgtacCCTTTGACTTTCCACCTGCAGCTCTCTGCTATGGGGCCCCtatagttacttgtcctaaa
gacttatcagtcttgtgtacACCACTCGCTAGGTTACTGCTTCTTCCACTGCCTCCGGATtatagttacttgtcctaaa
gacttatcagtcttgtgtacTCCCCTCTGTTGATTGTGTTGTTGTTACTGAAGGAGGCCAtatagttacttgtcctaaac
gacttatcagtcttgtgtacAGATGAAAGTGGATAAAAAGAGGACCTAGGGCAGAAAAAAtatagttacttgtcctaaa
gacttatcagtcttgtgtacCCCTGTCCCAGAAATGATCCAGAAAGTTTAGGCCTTGAGCtatagttacttgtcctaaa
gacttatcagtcttgtgtacAGGTGGGAAATATCTGCAGATAGTACTCTTCCTTCTCCGGtatagttacttgtcctaaac
gacttatcagtcttgtgtacTCCCGTCTGCCAACCAAGGTGCAAGGTAGCTGTCCATGGTtatagttacttgtcctaaa
gacttatcagtcttgtgtacTTCATTGAAGGATGATGGGCAAGTAAAAAGCTGTACATATtatagttacttgtcctaaac
gacttatcagtcttgtgtacGTATACATCTGTTTAACCCTATCCCTTATTCTTAATCTCCtatagttacttgtcctaaac
gacttatcagtcttgtgtacAGAGTTAACATTCATAGGTTGCTTAGCAAATTCCATGAACtatagttacttgtcctaaac
gacttatcagtcttgtgtacGAATTTTATAAATGCGATTTCAGCTACCACTTGCAACAGCtatagttacttgtcctaaac
gacttatcagtcttgtgtacCAATTTGCATGTGAGAAATTAGAGCTCAGTGTAGTGAAGCtatagttacttgtcctaaac
gacttatcagtcttgtgtacTTGATTCCCACCCTAGGCTTTTTCCTCTTATGTTCTATCCtatagttacttgtcctaaac
gacttatcagtcttgtgtacATCAAATTAAATGAAGGGAAGTGATGTTTTGGTTGGGAGCtatagttacttgtcctaaac
gacttatcagtcttgtgtacCTTACTCTTAAACCCCATCTCAGGCAAAAGAAAAAGTCAGtatagttacttgtcctaaac
gacttatcagtcttgtgtacAAGAAATTTTCCAGGAGATATGATTTTGGCTGGCACACAAtatagttacttgtcctaaac
gacttatcagtcttgtgtacAGCATTATGGTTCTCTGGCATTTTCATTCTGGGGTGCTGAtatagttacttgtcctaaac
gacttatcagtcttgtgtacAATGAAACAAGGAAAGCATTGAGAGCCAAGGTCATTATGAtatagttacttgtcctaaac
gacttatcagtcttgtgtacCTATTTGCAGAAAAATATTCCTTAGGCCTGGTTGAACTAGtatagttacttgtcctaaac
gacttatcagtcttgtgtacCAATTGCTTGGCCCTGTACTCCTTTTGCCTGTGTGGACATtatagttacttgtcctaaac
gacttatcagtcttgtgtacGGCCACACATTAAAACGGAATATTGTTATTGCCCTCATTAtatagttacttgtcctaaac
gacttatcagtcttgtgtacCTCTAACAGGAGCCTTTGGAAAAACAAGAGAAATTTCAGAtatagttacttgtcctaaac

gacttatcagtcttgtgtacTGAGCAATTAACAAAGTGACATAGTTAACACTACTGCTAGtatagttacttgtcctaaac
gacttatcagtcttgtgtacCTTGGGAAGGAGGCTATTAGAATTAGGTGCCACTGCATATtatagttacttgtcctaaa
gacttatcagtcttgtgtacTGTGAGGCTTACAGAGGTTGAAAAATCTACTTTTGGTGGTtatagttacttgtcctaaac
gacttatcagtcttgtgtacATGTTATTTCCATAGTGCATGTTTCTCTTAGAGGCAAAAGtatagttacttgtcctaaac
gacttatcagtcttgtgtacCTGGTGCTGCTCCTATCTGTGTGTCCTGGATTAGGCTGGGtatagttacttgtcctaaa
gacttatcagtcttgtgtacTCCTGAGGCTTGAGGAGACCCTTGTAGGACCTTCACACCTtatagttacttgtcctaa
gacttatcagtcttgtgtacCTTCTTGCTGGGTTTACATGGTGACAATCACTGAAATGAAtatagttacttgtcctaaac
gacttatcagtcttgtgtacTTAGATTAGAAGCCTATCCAATATTCAACTAGAGTGACTCtatagttacttgtcctaaac
gacttatcagtcttgtgtacCTGGTTCCAGGGTCACTTTGAATAGCTCTAATTAGTAATGtatagttacttgtcctaaac
gacttatcagtcttgtgtacGGTGGCAGTGAGGGGAAAAGGCAGCTCAAGTAGGGTAGGGtatagttacttgtccta
gacttatcagtcttgtgtacTGTCCCCAGATGGAAACTGTTGTAAATTGGACAAGCTCTGtatagttacttgtcctaaa
gacttatcagtcttgtgtacTGGGGCAGTTTGGATAGCAACTCACAGCCTGAGGGTTGGGtatagttacttgtcctaa
gacttatcagtcttgtgtacGAACCTTTCATTTAGGCTGGGAATGGGGGATTAGCAGGGCtatagttacttgtcctaaa
gacttatcagtcttgtgtacTTATGCAACAACTAGTGGGTCTGGGTTAGCTCAATCCTAGtatagttacttgtcctaaa
gacttatcagtcttgtgtacTGGGAGATGGGGGCTGGAGGGTCTGTTCACACATGGCAAGtatagttacttgtcctaa
gacttatcagtcttgtgtacCTGGCAAAGAAGGCTGGTGGAGCAAGGTGGGCCACTGGGAtatagttacttgtccta
gacttatcagtcttgtgtacTCATACCAGGATATCGAGCAGGTGTCTCCAGACCTAGCCTtatagttacttgtcctaa
gacttatcagtcttgtgtacTGTCTTATCAACAGGGTCAATGTGGGAGGGGCTTACAGGAtatagttacttgtcctaaa
gacttatcagtcttgtgtacTCCTGAGACTGTTGGATTTCAAAGACACTTGTTGGGAGGGtatagttacttgtcctaaa
gacttatcagtcttgtgtacGGAACCTAACAATGATAAAGACTCTTCAGATCTCAGGTCTtatagttacttgtcctaaa
gacttatcagtcttgtgtacCTATGCTTTGATTGGAGGCCACTAAGAGCTGATGGAGTGGtatagttacttgtcctaaa
gacttatcagtcttgtgtacGGCCAGGGGTGGACCTTTCCTAAAACTCAAGCTTCTGTGAtatagttacttgtcctaaa
gacttatcagtcttgtgtacTAGTAATAGTCCTTTCATGTCCCATAAAGCCTGCCCCTTCtatagttacttgtcctaaac
gacttatcagtcttgtgtacATCAAGGTACCAAGCAAACAGCTGCCAGATGAAGATGCCAtatagttacttgtcctaa
gacttatcagtcttgtgtacTCCTTTATATTTCTGATGTCTTTGGATGATGTGTGATAGCtatagttacttgtcctaaac
gacttatcagtcttgtgtacCATGCTCCAATTCCAGATGTAAACTTTGTAACTCTGAATAtatagttacttgtcctaaac
gacttatcagtcttgtgtacCAAGTCTCTATGTAATGTAGTATCCGGAGTTTGATAGAATtatagttacttgtcctaaac
gacttatcagtcttgtgtacGCTCTGTCACTCATTAGCTAATGACATTGGTCCAAAGAGTtatagttacttgtcctaaac
gacttatcagtcttgtgtacCAATGGTCATAAGATTGTCCTTTTCAGGCATATAGGAGATtatagttacttgtcctaaac
gacttatcagtcttgtgtacCTGGCACAGAGTTACTGTTTAATAAATGGAAATGGCATATtatagttacttgtcctaaac
gacttatcagtcttgtgtacGCCCAAAGCCTTGTCATGGAGCACTGCACTAAACCTGTAGtatagttacttgtcctaa
gacttatcagtcttgtgtacGGGGATATCAGAAACAGTGGTCACTAGGCATGGTGGTGTGtatagttacttgtcctaa
gacttatcagtcttgtgtacAGTCCAGGAGTTCTGGATTGTGGTGTGCTATGCTGATTGGtatagttacttgtcctaaa
gacttatcagtcttgtgtacTGCACTAAGTTTGGCAACAATATGTTGATCTCCCATGAGCtatagttacttgtcctaaac
gacttatcagtcttgtgtacCTGATTAGTAGTGAGGTTGCACCTGTGAATCGGCACTGCAtatagttacttgtcctaaa
gacttatcagtcttgtgtacTTTGATACTGACAAAAATCATATGGACCCTCTCTGTTGTAtatagttacttgtcctaaac
gacttatcagtcttgtgtacACTAGTGATTCCCATCCCTCAACTTGTCCCCTCACCCCCAtatagttacttgtcctaa
gacttatcagtcttgtgtacGATCACTGGTTTAAGGAGAACTATAAAGGACAGAGTGTTTtatagttacttgtcctaaac
gacttatcagtcttgtgtacTGATAGAATGAAAGGTTGAGAATCACTGATATGGCTTGCAtatagttacttgtcctaaac
gacttatcagtcttgtgtacGAAACACCAGGACATACTATAGCCCCACTTACTTTTCTTTtatagttacttgtcctaaac
gacttatcagtcttgtgtacGCATTTTGCTCAAGTTTATGCCAACACAAAAGCCAGAAGGtatagttacttgtcctaaa
gacttatcagtcttgtgtacAAAATGAAACATCTCCTTGGTCCAATAGTAAAAAGACTGCtatagttacttgtcctaaac
gacttatcagtcttgtgtacCTTGCAGAGGTTAATTTCTAACACTTGTGGCTCTTATGACtatagttacttgtcctaaac
gacttatcagtcttgtgtacCAGCTGCTTCTGGTCACCACACCCTCCTACGCTGAAGAGGtatagttacttgtcctaa
gacttatcagtcttgtgtacGTTTTGGAGGGTGCAGGAAGAAAATGATGGATGAGATGGTtatagttacttgtcctaaa
gacttatcagtcttgtgtacCACTAAGAAAATAAATAGGTATATGAGGTCTCAGGAAACCtatagttacttgtcctaaa
gacttatcagtcttgtgtacGGGAGAAAGGAGATGGGGGAGGGGGGAATTCAGCTGGCCTtatagttacttgtccta
gacttatcagtcttgtgtacGGTTCTGTCTGAATCCGCTATCTTGGTCCTGGGTTTCTCTtatagttacttgtcctaaac
gacttatcagtcttgtgtacGGAACAAAAAGAAATAGAACTTCACACGTAACTGTTAGTCtatagttacttgtcctaaa
gacttatcagtcttgtgtacTCTGAGTAACAATGCCCATAGGTATGAAGAGCGAAGGGGTtatagttacttgtcctaaa
gacttatcagtcttgtgtacTGCATTATGTCCAGCAGAGCACCAATGAAACACCTCTTACtatagttacttgtcctaaa
gacttatcagtcttgtgtacAAGGACTTGAAGTTCAATACAATTCTATAGGAAACTGAGGtatagttacttgtcctaaac

gacttatcagtcttgtgtacCTCCAAGTACATGAGGGTAAATCAAAGACTAATTGGTGACtatagttacttgtcctaaa
gacttatcagtcttgtgtacATTTCAATTTTAGACTTAGGGCTGCAATTCCATTATGAGGtatagttacttgtcctaaac
gacttatcagtcttgtgtacCAGTGCTTGGTGTTCATCTGCTGGTGTCTCCTCATGCCCAtatagttacttgtcctaaa
gacttatcagtcttgtgtacAACTATATTGGGTTCTGTGGATGGGATTAAGCAAAATTCTtatagttacttgtcctaaac
gacttatcagtcttgtgtacGCAGAAAGCGAGGGGGGTTGATGAGTAGTCCATAGAAGATtatagttacttgtcctaa
gacttatcagtcttgtgtacGGATGCTATTTTCCCTGCTTGATGACGGTTTCAGCCCACTtatagttacttgtcctaaa
gacttatcagtcttgtgtacCCAGAGCTATTTATATTCAGAGGCACAGATTCCACAGCTGtatagttacttgtcctaaa
gacttatcagtcttgtgtacAGTGCCTTCGAGTGCCTCTGAAATCAGTTTCTTCTCCCCTtatagttacttgtcctaaa
gacttatcagtcttgtgtacTTTGAATATAATCATGAAGAAAAGGCAGAGTTGGCTGGGGtatagttacttgtcctaaa
gacttatcagtcttgtgtacGAGGTTCCACTGTGCTCCCTATGTAGCTTCCTCTTCTCTCtatagttacttgtcctaaa
gacttatcagtcttgtgtacGTAACATTACTTCCTCTCATTCCCAGGCACCACATCTCATtatagttacttgtcctaaa
gacttatcagtcttgtgtacAAAGGTGTTTTTCTCTTGGCTAACTCTACAAAGATTCTACtatagttacttgtcctaaac
gacttatcagtcttgtgtacTTCAGATGAGGGTCATTCTGTAAGCAAAGCTATTAACTTTtatagttacttgtcctaaac
gacttatcagtcttgtgtacCCATGTGAGGGGGGTGGTGTTAGGAAGGGCATTGGGACATtatagttacttgtcctaa
gacttatcagtcttgtgtacGGTATGAAGAGAAATGGAATCTCAGCCAGGTTCTGCCCCGtatagttacttgtcctaa
gacttatcagtcttgtgtacGGGTTTACCTTTCTCTCCTCTCTTCTCATCTGTTCCTGATtatagttacttgtcctaaac
gacttatcagtcttgtgtacCTGCAAGGCTTATTTTCCTCCTTCCTGTGCAATGATAACAtatagttacttgtcctaaac
gacttatcagtcttgtgtacTGGAAAGTGATCTAACAGTGTTGCGACATGAATGAAAATGtatagttacttgtcctaaac
gacttatcagtcttgtgtacCAGAGAAAAACACCCACCCCCTGCCGCCTCCCTGCCGCTCtatagttacttgtccta
gacttatcagtcttgtgtacCCACCACTTTGTATTTTTACCTTTCGTGATAAAACTGTAGtatagttacttgtcctaaac
gacttatcagtcttgtgtacGTTTGACAAGTGGCACATTTAAAACCTGATTTAGCAAAAGtatagttacttgtcctaaac
gacttatcagtcttgtgtacACTGTCACTTGAACTAAAAATATAGGCTAGCAAGTTGAACtatagttacttgtcctaaac
gacttatcagtcttgtgtacTTTGCAATTCATTTTGTTCAGCTTTGATTAAAGGAGTGGCtatagttacttgtcctaaac
gacttatcagtcttgtgtacGACTTGGCTAATTAGTCTTTGGGAAGATGAGAGTGTGACCtatagttacttgtcctaaa
gacttatcagtcttgtgtacACACCAGCCCTTGTTAGTTTTAGCAGCTTAGCCATATATCtatagttacttgtcctaaac
gacttatcagtcttgtgtacAAGAACACTTGACTTAAAGAGGGCAAAGAAAACTGGAGGAtatagttacttgtcctaaa
gacttatcagtcttgtgtacTTTCCCCCTGACCATAATCTCAAGGTTTTTACATGTTTGTtatagttacttgtcctaaac
gacttatcagtcttgtgtacGCTATAGCCCCAGATTAAATGGTAATGATGGCAAGTCAGAtatagttacttgtcctaaa
gacttatcagtcttgtgtacGAAAGGATCTGGAACAAAAAAATGATGAACCCGGGTGTCCtatagttacttgtcctaa
gacttatcagtcttgtgtacTGTGGAATATTAAAAACGGCTACTCAGGATTCACAAAATGtatagttacttgtcctaaac
gacttatcagtcttgtgtacTGACCTTAGGCAAGTCACTGCATTTTTCTGAATTTTAGTGtatagttacttgtcctaaac
gacttatcagtcttgtgtacGCAAATATTCTTGTATGCCCTAAAATCTCTATTTGTGCCTtatagttacttgtcctaaac
gacttatcagtcttgtgtacGCAGAGCTTGTTCATGAAGGTATTAATATTTGTTGGATTCtatagttacttgtcctaaac
gacttatcagtcttgtgtacTCGGAAGAGATAAGTCAGCTCGGGGTTCTGGGATTGGTCCtatagttacttgtcctaa
gacttatcagtcttgtgtacAGCTTCTTTGTTTGAAAATGGGGATTATAATCTTCACTCCtatagttacttgtcctaaac
gacttatcagtcttgtgtacGACTGTATAAAATGGAAGGTATTAGTATAAAGGCCCTTTCtatagttacttgtcctaaac
gacttatcagtcttgtgtacCATCTTTATGCACACACGGACTTCATTATTGAACTGTTTAtatagttacttgtcctaaac
gacttatcagtcttgtgtacGCTTCTAATTCCAGACCATTGATCTATCATAAGGGCAAAAtatagttacttgtcctaaac
gacttatcagtcttgtgtacTACTGATGTATCTATCTGGGGCTGATATCCTGGAAATGACtatagttacttgtcctaaac
gacttatcagtcttgtgtacAGTGATGGAAGGGGTGGTTGAGGTCAGGGAGGGAAACTTAtatagttacttgtcctaa
gacttatcagtcttgtgtacCTTTAGGTCCCTTAGTGCATTTGATTTTTATCACCATTCTtatagttacttgtcctaaac
gacttatcagtcttgtgtacTAGAGTTAGGATAAAGATCTTATTCCGTCTACATGCCATAtatagttacttgtcctaaac
gacttatcagtcttgtgtacTCTCTTTTGACAGATTCTCCTCCCCATCTTTTGACAGTTCtatagttacttgtcctaaac
gacttatcagtcttgtgtacAACAAATCAACCAACAAACACACCCCAAACTATTCCTTATtatagttacttgtcctaaa
gacttatcagtcttgtgtacGGCTACTCAAAACATAGTTCCTGGGACCAGCATCATAGGCtatagttacttgtcctaa
gacttatcagtcttgtgtacTGCATTTCAGTGAGATCCAGAGTGATCCATGAGCATGTTAtatagttacttgtcctaaa
gacttatcagtcttgtgtacCTCTAGAGCACACAGAGCAGATCACATTTCTCCTTTTCATtatagttacttgtcctaaa
gacttatcagtcttgtgtacCTTCGTCCCAAAAAGAACCATATGCCTGAGACACCTTTAAtatagttacttgtcctaaa
gacttatcagtcttgtgtacAGAATCCTGAAAGCCTCTAGAAAGTTTCCTTGACTAACTAtatagttacttgtcctaaac
gacttatcagtcttgtgtacGGTGGCCCTTTCTGTGACCCTCCCCCAAATATATATTTCAtatagttacttgtcctaaa
gacttatcagtcttgtgtacGGTGAAAAATGCCTGGATATTTGTGATTCTGTGAATTGTAtatagttacttgtcctaaac
gacttatcagtcttgtgtacAAGTCAGGTGGAGGAATTTAGATTTTAAGAGGCTAGGTTTtatagttacttgtcctaaac

gacttatcagtcttgtgtacCAGGAGAGTGGCGTATAGAAAATGATGAACTAAGCTGAAAtatagttacttgtcctaaa
gacttatcagtcttgtgtacGATTTATCTTGTGGTGTTCTGCAGTGTGGTTAAAGAGGGTtatagttacttgtcctaaac
gacttatcagtcttgtgtacCTATTTGAAAGCTTTTATTGGCTGTCATTTGACGCATTGAtatagttacttgtcctaaac
gacttatcagtcttgtgtacCCCTTTATAAATGCACCAAATTTTGTCAAAATTAGTCGGCtatagttacttgtcctaaac
gacttatcagtcttgtgtacTTAAGTTGAACCATAGTCCTCACTTTGTTCAGATAATGGTtatagttacttgtcctaaac
gacttatcagtcttgtgtacTGTCAATCATTCTTTATAAGGGCAGGGTATAAGTCTGTTCtatagttacttgtcctaaac
gacttatcagtcttgtgtacTCATTGATCACATGAAGCTCTTTCCCCTTTCTGGATGTTTtatagttacttgtcctaaac
gacttatcagtcttgtgtacATAAAAGACACTGTTTCTCCCAGGGAGTTACGAAGCCCCTtatagttacttgtcctaaa
gacttatcagtcttgtgtacCTGACAACAAAGACAGAAGAGAACAGACTATTCCAGGTGGtatagttacttgtcctaaa
gacttatcagtcttgtgtacAAATCTTGTTTCTTCTTGCGTCCTCCATCTTCTGCCTGCTtatagttacttgtcctaaac
gacttatcagtcttgtgtacTTGTATAACACCTCCCTCCAGTTCACTACTCTAATATATTtatagttacttgtcctaaac
gacttatcagtcttgtgtacTCCGATGCTGATGATCCCCAATAAAAAACTTCAAAATCTCtatagttacttgtcctaaac
gacttatcagtcttgtgtacGGTTGTTTAAATACCTCCAAATTCATACAAATCCAGCCCTtatagttacttgtcctaaac
gacttatcagtcttgtgtacAAATTTATTTTTCAACCCAAGTAACATCTGGAGAGAGCTGtatagttacttgtcctaaac
gacttatcagtcttgtgtacCAACCCAGTTCCAGGACGACTGCTGTAATCTCTGACATGAtatagttacttgtcctaaa
gacttatcagtcttgtgtacAAAAATGAGTTAGAATTATCAGTCTGTTTTTCTGGCCACCtatagttacttgtcctaaac
gacttatcagtcttgtgtacGAATCATATTTTCCTCTTTACTTCCTCCCTACTACTACGTtatagttacttgtcctaaac
gacttatcagtcttgtgtacGAACCAATCACGTAGCAGTAGGGAGAAAGTAAAAAGGAGGtatagttacttgtcctaaa
gacttatcagtcttgtgtacCCATTGCTTAAGGATCCTTCAAATGGCCATGGGTGAAGATtatagttacttgtcctaaac
gacttatcagtcttgtgtacTCTATCAAAGCCTTAATAGCCATGAGGAGGACCCTACAGGtatagttacttgtcctaaa
gacttatcagtcttgtgtacGGACTAGATCATCAACAGAGAGATAGGTGCAGAAAGGGATtatagttacttgtcctaaa
gacttatcagtcttgtgtacGATGAATAATCTTGAGTTTCCATTGTAGATCAGGCTCTGGtatagttacttgtcctaaac
gacttatcagtcttgtgtacTGGAAAACCAGCTTTGCTCTAACACTTATCAACACTGCAGtatagttacttgtcctaaac
gacttatcagtcttgtgtacCAGGATTTAGTGAGATTATGTTATTTGCATTGCCTAGTACtatagttacttgtcctaaac
gacttatcagtcttgtgtacTTAATAAATGTTAGCCTTTTCTCCTTTGTCTACTTAGGGGtatagttacttgtcctaaac
gacttatcagtcttgtgtacGGGGCTGTTGAGAGAAGTAAGATGTGTTTGGGATCTGAAAtatagttacttgtcctaaac
gacttatcagtcttgtgtacCCAATTGGGGATGAAAGACATGCACACAACTAACTGTAATtatagttacttgtcctaaac
gacttatcagtcttgtgtacAGAAAAAAAGCCATGATTAATTCTACTGGGGAGTGAGTTGtatagttacttgtcctaaac
gacttatcagtcttgtgtacTGGGGGACAGCCAACATCAGCACTGGTCCTGAAGATTTGAtatagttacttgtcctaaa
gacttatcagtcttgtgtacCCTGCTGGAGAAACTCAGGAATGTGCACTCAGAGCCATCTtatagttacttgtcctaaa
gacttatcagtcttgtgtacCAGATGTGTGTAGAGACTCAGGAAGAAAAGCAAGAAAACTtatagttacttgtcctaaa
gacttatcagtcttgtgtacGGCAAAGTCTAATTACCCAAATTGAAATCTGGCCTCTGTGtatagttacttgtcctaaac
gacttatcagtcttgtgtacGCTTTATACTGCAGCTCTTGTTTGGGAATTCTTGGTTTGTtatagttacttgtcctaaac
gacttatcagtcttgtgtacAGGAAGCCTGAGAGGTATTTTGAGAATGGTTAGCAAATTGtatagttacttgtcctaaac
gacttatcagtcttgtgtacGCAGGGCAGGAAAATGTAAAATCAATAAATAATCAGGCTGtatagttacttgtcctaaac
gacttatcagtcttgtgtacTCCTAAGGCCATGCTGACATGTAAATTTAAAAAACGACTCtatagttacttgtcctaaac
gacttatcagtcttgtgtacCCTCTGTGGGTACTTCCATTCCAGGAGATGGCACAAAGATtatagttacttgtcctaaa
gacttatcagtcttgtgtacTTTGCAATTCCTTCCCACATTTTCTGTTCTCTTTCCTGAAtatagttacttgtcctaaac
gacttatcagtcttgtgtacCAGTGTTAAACCATCTTCCTACGTTTCTCTTTTCTGCAAAtatagttacttgtcctaaac
gacttatcagtcttgtgtacTTCATGTAGAAGCAAACAACAGAGAGGAAGGCAGAGTCCTtatagttacttgtcctaaa
gacttatcagtcttgtgtacCTTTTGGGTAGAGATGTAGTTACTTTTCAGAGGGCATTGGtatagttacttgtcctaaac
gacttatcagtcttgtgtacCACATCTGAAGGTCAATAATGTCATAGATATGTGTCCTAGtatagttacttgtcctaaac
gacttatcagtcttgtgtacTTTGGTGAAGTTATTTCTGAAATTTTGAGTGAGCTGCATGtatagttacttgtcctaaac
gacttatcagtcttgtgtacGCTAAGGAGAGCTACTTTAGACCAAACTGGGGCTCAGGAAtatagttacttgtcctaaa
gacttatcagtcttgtgtacCAGGATGGGTGGGGTGGAAGGAATCAAAGTGAGAAGTCCCtatagttacttgtcctaa
gacttatcagtcttgtgtacCTTTCAGGCATTTCAATTTGTATTTGATTTTGTAGCACCGtatagttacttgtcctaaac
gacttatcagtcttgtgtacGTCTTCTTGTCACCTTAACCTCTGCACAGACACTGTTGCTtatagttacttgtcctaaac
gacttatcagtcttgtgtacCAAATTTGTTTCTACATGAGTTTAATTAGGCACCTACCAGtatagttacttgtcctaaac
gacttatcagtcttgtgtacCAGGGATATGGAAGATTTATGAGACACAGGCTTTGACTTTtatagttacttgtcctaaac
gacttatcagtcttgtgtacAGAAAGCTGGTAAATAAATTTTAGCGTATACGAGCCTCCAtatagttacttgtcctaaac
gacttatcagtcttgtgtacGAGATTCTGATTCTGTAGCCCTGGGAGGAAGCTCTTGAATtatagttacttgtcctaaac
gacttatcagtcttgtgtacCTGGTTCAGGTAATTCATGACTCAGACTTTAAAAGACAGTtatagttacttgtcctaaac

gacttatcagtcttgtgtacTTTCATTAGTGAGAAAGACCTGCTGTTTAAAGACCCCCTCtatagttacttgtcctaaac
gacttatcagtcttgtgtacCAAGATACTCACATTCTGCAACCTTTACTCGTCTATAAAAtatagttacttgtcctaaac
gacttatcagtcttgtgtacTGTAAGAATTAAATGAGATACAAACTAGCACAGTTCCTGGtatagttacttgtcctaaac
gacttatcagtcttgtgtacTCCTAATGCTATTCTTCTCCTGGCCCCCCACCCCGCGACAtatagttacttgtcctaa
gacttatcagtcttgtgtacTCATTGTTCAACTCCTAACCATGAGCAAAAACATGTGGTGtatagttacttgtcctaaac
gacttatcagtcttgtgtacTCTATGTCCCTGCAAACGGCATGAACTTATCCTTTTTTATtatagttacttgtcctaaac
gacttatcagtcttgtgtacCAGTAATGAGATTGCTGAGGCAAATAGTATTTCTGGTTCTtatagttacttgtcctaaac
gacttatcagtcttgtgtacCTTGAGGAACCAGACTGTCTTCCACCACACTGTCTTCCACtatagttacttgtcctaaa
gacttatcagtcttgtgtacCTCTAATGACCATTGATGAGCTTTTTTCATGTTTTTTGGCtatagttacttgtcctaaac
gacttatcagtcttgtgtacGTGTCTGTTCATATCTTTCACCCACCTTTTGATGGGGTTGtatagttacttgtcctaaac
gacttatcagtcttgtgtacGGTAGATTGCAAGAATTTTCTCCCATTCTATAGGTTGCCTtatagttacttgtcctaaac
gacttatcagtcttgtgtacAAACCATTTTTTTAACCCTCCTTCTCAATGATCTAGTTGGtatagttacttgtcctaaac
gacttatcagtcttgtgtacCACAGGCTACTTCAGCTTTCAAGTTATTTCTTATTCGCTGtatagttacttgtcctaaac
gacttatcagtcttgtgtacAGTGAAGTTTATCATCTATGTGACACAACTGGCTGTTTATtatagttacttgtcctaaac
gacttatcagtcttgtgtacATCGTATTGTTAGTTTTCCATTTGTGTGTGTGTGTGTTTGtatagttacttgtcctaaac
gacttatcagtcttgtgtacTTTGGCTCACAGCAACCTCCACTTCCCAGGCTCAAGTGATtatagttacttgtcctaaa
gacttatcagtcttgtgtacGTGAGCCACTGCGTTCGGACTATTTGTGTTTTTAACTGAAtatagttacttgtcctaaac
gacttatcagtcttgtgtacCCCAGGAATTATGAGATGTCACCTTAGGCCACCAGAAATTtatagttacttgtcctaaa
gacttatcagtcttgtgtacTGGGTTTGGGTCTTACACCAAGATAGGAGAGAAAAGCTGCtatagttacttgtcctaaa
gacttatcagtcttgtgtacGTTAATAAAAGAGAGAACTAAGGCCGTAAACTCCATTGTAtatagttacttgtcctaaac
gacttatcagtcttgtgtacCCAATTGAGGAGAAATAATAGCACAACTTAGAGGACAGTTtatagttacttgtcctaaa
gacttatcagtcttgtgtacGATCTAGATCTTGTCCATATCCTTGCTGATACTGGAGACAtatagttacttgtcctaaac
gacttatcagtcttgtgtacGCTACCAAAAAGATCTATGAACAGTTGCCTCTGGGACCTAtatagttacttgtcctaaa
gacttatcagtcttgtgtacGTTTATAACTCGAGGCCAAAAGAGCTGGGATGATCAAGAGtatagttacttgtcctaaa
gacttatcagtcttgtgtacTGGAGCTGGCTGTGGTGACAAGGAAGTGGAGGGAAGTTGGtatagttacttgtcctaa
gacttatcagtcttgtgtacGAAGAACTATCAAGGATTATGCTAAGGACTCTAGATCATCtatagttacttgtcctaaac
gacttatcagtcttgtgtacCAGTAGGAATTCAGGCATAGGTGAGGGGCTGGTAATGTCAtatagttacttgtcctaaa
gacttatcagtcttgtgtacATAGAAAGGACTATGGGTAATTAATTCTCTCTGACCAAGAtatagttacttgtcctaaac
gacttatcagtcttgtgtacGCCTGAAGCTAAGAGATCCAGCCAAGCCAAATGACTTTTGtatagttacttgtcctaaa
gacttatcagtcttgtgtacCTCTTGGTTCTAAGTGCATACTCCCTACCCACCCTATTCTtatagttacttgtcctaaac
gacttatcagtcttgtgtacGGGCCTGGGGAATGAGTACTGTGTTTATGGGAATGTGCTCtatagttacttgtcctaaa
gacttatcagtcttgtgtacTGAATGCTTCTCCATACAACCCTACTCAATACATATTTCAtatagttacttgtcctaaac
gacttatcagtcttgtgtacTCACAGAGTGTTAGAAAGGAAAGTGGCCTTAGGGATCACCtatagttacttgtcctaaa
gacttatcagtcttgtgtacTCTGGTAAGGAAGCAAACTCAGAAAAAGTATACGACCTTCtatagttacttgtcctaaa
gacttatcagtcttgtgtacCCACTCACATTACTAGGCTGCACTTTCTTATGATGATTTAtatagttacttgtcctaaac
gacttatcagtcttgtgtacGCATACCATTGTTCAGCGTTTAAAATCCAACAGGCTCCTTtatagttacttgtcctaaac
gacttatcagtcttgtgtacGCTCTTCACATGCTAACCACGGAGTAGGAGATGGAGCTGCtatagttacttgtcctaa
gacttatcagtcttgtgtacACAATTTCATTAACTCTGTGTCGTTGGGCTTTGGCAATCCtatagttacttgtcctaaac
gacttatcagtcttgtgtacCTCATAATTGTGCTAAGCTCATTCTGGTGAAAAGTACATGtatagttacttgtcctaaac
gacttatcagtcttgtgtacTCATAAATACTTAAAAACACCAGCATCTGAGATGTCAGAGtatagttacttgtcctaaac
gacttatcagtcttgtgtacGGTCCTGGCCAGTGTCCTCCTACATTTGCAAACTTAACTTtatagttacttgtcctaaa
gacttatcagtcttgtgtacGTCTGGTTAGAGCTGTCTTTGAAGCTGCCTGGCTGTAAAAtatagttacttgtcctaaa
gacttatcagtcttgtgtacGCATCCTTCACTTAGCAAACTGCCATCATGAAATGTGAGCtatagttacttgtcctaaa
gacttatcagtcttgtgtacCTTCTTTGCTCTTTTCTGATCCAAAATTAGTGTTTCTAGCtatagttacttgtcctaaac
gacttatcagtcttgtgtacCTATCAAGTTAGAGGAATACGTTTTGGTACCACAGCCAGCtatagttacttgtcctaaa
gacttatcagtcttgtgtacGATTTCTCTGTTTAGATGGAAAGCCAGGGCAATGTAAAACtatagttacttgtcctaaac
gacttatcagtcttgtgtacGTCAATAGTTCTGTGTTGTATGCCATTTAAGTTTCAGCCAtatagttacttgtcctaaac
gacttatcagtcttgtgtacTTCAGATTGAGATTTTTTAGACTCATAGAAGGGGGGGGGGtatagttacttgtcctaaa
gacttatcagtcttgtgtacCTCTGTCTTGTCCTTCCGCAGTGTCCACCCCTGTTGTAGTtatagttacttgtcctaaa
gacttatcagtcttgtgtacCTGCCAATAGTCAGTTAGATCTCCCACTTGTGAAAATATGtatagttacttgtcctaaac
gacttatcagtcttgtgtacCCCTTTACTTCCTTCTAATTTTTTGGGGGTCTATTTCCTAtatagttacttgtcctaaac
gacttatcagtcttgtgtacAAAGTGCATTTGTAGTTGTCAGAATCTAAGTGGCTTAAACtatagttacttgtcctaaac

gacttatcagtcttgtgtacGGGTTTATTTCATGCTGACACTACAAAGATTGGATGGGGTtatagttacttgtcctaaac
gacttatcagtcttgtgtacCCGAAGTGTGGCCAGTCCCTGAAGGAGAGAGAAAGAAGGAtatagttacttgtcctaa
gacttatcagtcttgtgtacCTCTTTTAGCTTCCACTCAGAAGGGCTACAAATCCTTTCCtatagttacttgtcctaaac
gacttatcagtcttgtgtacTTAAAGGTAACAAAAATACAATCCTATCATGTGTCCAGGGtatagttacttgtcctaaac
gacttatcagtcttgtgtacGACATTTGTTTAGCAGCAATAATGACTATCAACACTTTCGtatagttacttgtcctaaac
gacttatcagtcttgtgtacCTGTCATGAGCATAGCTGAGAGAGAATTATAGATTGTATTtatagttacttgtcctaaac
gacttatcagtcttgtgtacTTTTTAATTTGAGGAGTGTGTAGTTCACATAGGGTTGCATtatagttacttgtcctaaac
gacttatcagtcttgtgtacTTCAGGGATTGTACAATTTTGAAAACTCAGAAAACATGGGtatagttacttgtcctaaac
gacttatcagtcttgtgtacTTCCCCTTACTTTTTCTCTATCTTCCTGATAACTTTTGTGtatagttacttgtcctaaac
gacttatcagtcttgtgtacCTCTGAGAGGTTATAATGCCAAACTGCCAAGAAACATGAGtatagttacttgtcctaaa
gacttatcagtcttgtgtacGTAGGGTATACACACAAGGTATAAGAGTAGATGCAAGGTTtatagttacttgtcctaaa
gacttatcagtcttgtgtacTGCAAATAGAAGGTAGAGTGGCAGGTTGAAGGTTAAAAATtatagttacttgtcctaaa
gacttatcagtcttgtgtacCTTGAAAGGCTAATTTGGAGAAGGATAGATTGAACAAACCtatagttacttgtcctaaa
gacttatcagtcttgtgtacGAATTGAGGAGGAAAGGAGGGGTGAGTATTAACTTGGGCAtatagttacttgtcctaaa
gacttatcagtcttgtgtacAAGCCTTGACTCTGGGGTTTTCAAAACAATGCATGGAGAAtatagttacttgtcctaaa
gacttatcagtcttgtgtacGGGAAGTAGCAGTGAAATAAGAAGTGGGAGAGATCATGGCtatagttacttgtcctaa
gacttatcagtcttgtgtacATGGATGCCTTTGGTCCCTTGGAGCTTCTATACCTGTACAtatagttacttgtcctaaac
gacttatcagtcttgtgtacGATCTTGGTATCTGCATGGAGAAAAGACATTTGGGATCATtatagttacttgtcctaaac
gacttatcagtcttgtgtacCCATTTAGAAGATTGCCTATTCTGTGGACCAGTGAAGAAGtatagttacttgtcctaaa
gacttatcagtcttgtgtacAGAAAGAGCTACACAGGAAGGATATTTTGCTTTGGGATAAtatagttacttgtcctaaac
gacttatcagtcttgtgtacCTAACCCAGTTCAAAAGTAAAGGAAAGAAAAGAGTTGAGAtatagttacttgtcctaaa
gacttatcagtcttgtgtacTTGCATACACCAGGAGAACAAGAAGTCTATTAAGAGAATCtatagttacttgtcctaaa
gacttatcagtcttgtgtacATAGTACATATTATCCATTTTTTAGCCCCTTTCTTCTCCGtatagttacttgtcctaaac
gacttatcagtcttgtgtacAAATACTAAACATTAAAGGACAACTATGAGAACCTCCCGTtatagttacttgtcctaaa
gacttatcagtcttgtgtacTGAAGGTCTCCATATTCTGCAGAGAACACCATCACAATCAtatagttacttgtcctaaa
gacttatcagtcttgtgtacCAGATCGGTAGTGGAGCAGGGTTGTAATTCTAGAAAATGCtatagttacttgtcctaaa
gacttatcagtcttgtgtacCTGAAATCACACCCAGATAGAATAGAATTTATATGAAGGGtatagttacttgtcctaaac
gacttatcagtcttgtgtacAAGTTTTCCTTTATTTAGCATGGATTTTGTTAGCCCTTCGtatagttacttgtcctaaac
gacttatcagtcttgtgtacGAAGTTAGTGAATTCTGTCCAGAGGAACTTGCATTTCCAGtatagttacttgtcctaaac
gacttatcagtcttgtgtacCAGTAGGACGTCTGGGGTCAGGTCTGGCTCTGTTCACTGAtatagttacttgtcctaa
gacttatcagtcttgtgtacCCTAATCTCTCTAGGTTTCCACTTCCTCATGTATAAAATCtatagttacttgtcctaaac
gacttatcagtcttgtgtacGTTTGTAATTCTCAAAAAGAAGATAGAAAATGCCTCCTGGtatagttacttgtcctaaac
gacttatcagtcttgtgtacTGGTTGGACTTTGTGCCAATCCAAAATCTTTTCAGAATGGtatagttacttgtcctaaac
gacttatcagtcttgtgtacCCTTTTCTGGCTTTTATTTTTCCATTCTATGACTCAGGATtatagttacttgtcctaaac
gacttatcagtcttgtgtacCCTTTTTAAAATTAAGCCTTCCCCTGCTGTCTGATTCATTtatagttacttgtcctaaac
gacttatcagtcttgtgtacATGTTACAACCCACAGCTTGAGATGCGACATTTCTTTTCAtatagttacttgtcctaaac
gacttatcagtcttgtgtacTGTCACAAAACTTGAATTGATGCATCCAACTGAATTAAGCtatagttacttgtcctaaac
gacttatcagtcttgtgtacACATATGTGTCTAGGTAATCAGCAATGAAGGGAAAGGAAAtatagttacttgtcctaaa
gacttatcagtcttgtgtacTGCATTTTTAGACGCAGCATCCCGTTTTAAAGATTGTAGCtatagttacttgtcctaaac
gacttatcagtcttgtgtacCTTGAATCTATACCAGGAACGAAGCATTTTACAATTACCTtatagttacttgtcctaaac
gacttatcagtcttgtgtacAGTCATATCACTTGAACTTTGGCCAAATGTATTAACCTGAtatagttacttgtcctaaac
gacttatcagtcttgtgtacACTTGCACATCATAAGTCTTACAGACAGCTTTTAAAAAGCtatagttacttgtcctaaac
gacttatcagtcttgtgtacGCATCAATTCAGAAAATGTTAACAGTATTGCAGCTTCATGtatagttacttgtcctaaac
gacttatcagtcttgtgtacCTTTTGTTTTTGATATGGACTCTTGATGACATTGCTGAACtatagttacttgtcctaaac
gacttatcagtcttgtgtacGTTCAATAACGTCACTTACTGAGTTAAAACATGAGCAATCtatagttacttgtcctaaac
gacttatcagtcttgtgtacAAGATCTCTTCAATTTTCACCCATACAGAGAGGGCTGCCCtatagttacttgtcctaaa
gacttatcagtcttgtgtacTATAGAATTTTGCAGCTTATACAATGCTTTATAGCCCCACtatagttacttgtcctaaac
gacttatcagtcttgtgtacTCACTGACATGCTAGATTAGTCTTACCCACATCTCAAATTtatagttacttgtcctaaac
gacttatcagtcttgtgtacGTTCTACGTGCACATCCACTATTGTATCTTAGAATTAAGTtatagttacttgtcctaaac
gacttatcagtcttgtgtacTTCGGCATTTCTGAAGATAGCACAACTTCCCTTCCTCACTtatagttacttgtcctaaac
gacttatcagtcttgtgtacTCTTCTTTCCTCTTTTCTTTGATCATTTGACAACTCCTAGtatagttacttgtcctaaac
gacttatcagtcttgtgtacTGCCTCTCTAAATTATATTTGTGTCTGTTCTTGCTACCTCtatagttacttgtcctaaac

gacttatcagtcttgtgtacTCATATTGACTACACATATACTGGGGCCATGACCTCTGCAtatagttacttgtcctaaac
gacttatcagtcttgtgtacCAGAGTTTTCCTGATCAAATCCATTTTGCACATTGCTGATtatagttacttgtcctaaac
gacttatcagtcttgtgtacATGTCATTCTTCTGCTTAAAAATCTCCAATGGGCCGGGTGtatagttacttgtcctaaac
gacttatcagtcttgtgtacCTATTGAGCTTGGCATTATTAGATCTCTAAGCTTTTTAGCtatagttacttgtcctaaac
gacttatcagtcttgtgtacGCTCTAGCCAGTATAGAAACCAGAAAGTAGAATTTTCTTGtatagttacttgtcctaaac
gacttatcagtcttgtgtacAATGTAATATGGTGTATTGGAAGGTGTTATGGACTGAAAGtatagttacttgtcctaaac
gacttatcagtcttgtgtacTGTGATGGACATTAGCAGGTGGGATCTCTGGGCAGAAATTtatagttacttgtcctaaac
gacttatcagtcttgtgtacGAGGGATGCTCTGTCTTTGTCTGCTGTCATGCAAGGGCAAtatagttacttgtcctaaa
gacttatcagtcttgtgtacGCCCCTTGATTTGGAATATACCAGCCTCCATAACTGTTAAtatagttacttgtcctaaac
gacttatcagtcttgtgtacCCAAACTGACTAAGAGAGATGGGATCCTAAAACAAACAAAtatagttacttgtcctaaa
gacttatcagtcttgtgtacGTAACATGTCAATATTGGCTCATTACTGGACAAATATACCtatagttacttgtcctaaac
gacttatcagtcttgtgtacATGTGTTAGCAATTGAAGACACTGTGTGTAGGGAGAGAGTtatagttacttgtcctaaac
gacttatcagtcttgtgtacCTGTAAGAATAGAATAAGTAGGGGAACTAGGATGTGAAAGtatagttacttgtcctaaac
gacttatcagtcttgtgtacGATGATAAAGTATGATTTTAAAGGAAGTGTTGTGGGCAGAtatagttacttgtcctaaac
gacttatcagtcttgtgtacAGATTGTGTGTCTGTTTTTAGGGGAGAAAGCTTACTGGAAtatagttacttgtcctaaac
gacttatcagtcttgtgtacGGCTTCCCTACAAGTGACCAAGACAACCTTAACATTCCCCtatagttacttgtcctaaa
gacttatcagtcttgtgtacTTGACTCAATTTTAGATAAGCTTTTTCCTGACTATGGGCCtatagttacttgtcctaaac
gacttatcagtcttgtgtacACTTTCAGTTGTAAATTCTTTCTGTGCCTCTTTGAGATGTtatagttacttgtcctaaac
gacttatcagtcttgtgtacACAACCTAGGAGTGCCTTTCTCAAAGACCTGGAAGCCATCtatagttacttgtcctaaa
gacttatcagtcttgtgtacTCCTGTCTCTATGGGAAAGTAGGAACCTGACTTCGTTAACtatagttacttgtcctaaac
gacttatcagtcttgtgtacTGACTCAATCACATAGACCACCCTCCCTGCTAATGTTCCAtatagttacttgtcctaaa
gacttatcagtcttgtgtacGCTCAAAGCTGTGCTTAAAAATCTTCCCACCTTTTGTTTCtatagttacttgtcctaaac
gacttatcagtcttgtgtacCAATAGTTTGACTCCTCCCCTGTTGTAATAGTCTTAGTGAtatagttacttgtcctaaac
gacttatcagtcttgtgtacTCCCTGCCTTTCGTAATAAGTATCCAGTGAAACTTTTCCTtatagttacttgtcctaaac
gacttatcagtcttgtgtacAGATCCTGAATGAGGCTAAGAAAATGTGGCATAGCGTGGCtatagttacttgtcctaaa
gacttatcagtcttgtgtacGTGGTGAACAGCTTTGAGGTAAAGGCTTTCGAGTTTGAGAtatagttacttgtcctaaac
gacttatcagtcttgtgtacACTGTAGGTAAACGTCGGAAAACTCATTTAAGATACTAAGtatagttacttgtcctaaac
gacttatcagtcttgtgtacTGTATGTTTTCTCTAAGACATTTGGATCAGCATGCTTATGtatagttacttgtcctaaac
gacttatcagtcttgtgtacAGCTATCTGATGAGATTTCTCTAGGATTTGGCATGAAAGTtatagttacttgtcctaaac
gacttatcagtcttgtgtacGATGTTTAGGTAAAATCCGGGACTTTCAATTCACAGACATtatagttacttgtcctaaac
gacttatcagtcttgtgtacTTCTGAGCTTTGTAATTTAGGGACAACACAAACATAACTGtatagttacttgtcctaaac
gacttatcagtcttgtgtacAGTCGCAAGTTGTGTATCTCCCTTTGAAATTGTTTATGTCtatagttacttgtcctaaac
gacttatcagtcttgtgtacGCATTCAGCCTTGCGTATTCATTTTCACACTGTCTATGGTtatagttacttgtcctaaac
gacttatcagtcttgtgtacCAGAGTTGAGTAGCTGCAAGGGAGATGGTATGGATCTCAAtatagttacttgtcctaaa
gacttatcagtcttgtgtacTATTATACGATCCTTTATTAAAGAAGCATGCCAACCCCTGtatagttacttgtcctaaac
gacttatcagtcttgtgtacCAATAAATACTCAGAAGCAGCATATTATGGTGGTGAATAGtatagttacttgtcctaaac
gacttatcagtcttgtgtacCAGATATTGGCTGGCTTTGAAATTTGACTTTACCATGTACtatagttacttgtcctaaac
gacttatcagtcttgtgtacGAATGACAAGTCACCTTGCTAGTAACCTCTCTGTGCCTCAtatagttacttgtcctaaac
gacttatcagtcttgtgtacAATGAATTAGTAATGCAAGTACTCAGAACATTGGCACATGtatagttacttgtcctaaac
gacttatcagtcttgtgtacCACCATGAAGATAGAAAAGATCTCTATTTTGTAAAGTGGGtatagttacttgtcctaaac
gacttatcagtcttgtgtacTTGCACAAGCATATACCCTGTTACATATAAGAGCAAAGGCtatagttacttgtcctaaac
gacttatcagtcttgtgtacTATCCATTTTATTTTCAGCACTCTCCTACCTCTTTTTGACtatagttacttgtcctaaac
gacttatcagtcttgtgtacCCGTTTTAGCCGGGATTTTCTCAATTTTTTGATAACCAATtatagttacttgtcctaaac
gacttatcagtcttgtgtacCAGAAGGTGAAGGGGAGGCACGCATGTCTTACATGCTGGAtatagttacttgtcctaaa
gacttatcagtcttgtgtacATTCTCTAGAGGGACAGAACACATAGGATAGATGTGTATAtatagttacttgtcctaaac
gacttatcagtcttgtgtacGACTCACATGATTACAAGGCGAGGTCCCATAATAGGCCATtatagttacttgtcctaaa
gacttatcagtcttgtgtacGAGCAAGGAAGCCAGCCAGTCCAAGTCTCCAAACCTCAAAtatagttacttgtcctaa
gacttatcagtcttgtgtacAGCCTTCAGTCTGTGGTTGGAGGTTCAGGAGTCCCAAAGCtatagttacttgtcctaaa
gacttatcagtcttgtgtacTGTAGGCTGGAAGACTAAGCTAGTGTAGTTCTTCCACGTTtatagttacttgtcctaaac
gacttatcagtcttgtgtacACTATCATGAGAACAGCACCTAGGGAATGGTGCTAACCCAtatagttacttgtcctaaa
gacttatcagtcttgtgtacGTGAGGACGCAGATCCAAACTATGTCACCAATGTCAGATCtatagttacttgtcctaaa
gacttatcagtcttgtgtacTAATGGTGGAATTTCATGCATAATGGTGAGAGGTGAAGCCtatagttacttgtcctaaac

gacttatcagtcttgtgtacGACTTTCTGGGTTGAGTGGTGACTTGAAGGACTTTTGTGTtatagttacttgtcctaaac
gacttatcagtcttgtgtacGCACTCTCTGTCTAGCTAAAAGATTATAAACCCACCAATCtatagttacttgtcctaaac
gacttatcagtcttgtgtacGCTGTGGAAGCTTTGTGCTGTTGCTCTTCATAATAAATCTtatagttacttgtcctaaac
gacttatcagtcttgtgtacAGCGAGACCAAGAACCCACGAGAAGGAATAAATTCCAGACtatagttacttgtcctaa
gacttatcagtcttgtgtacCCTAAAGTTTATATTTTGGTTCAAGCTCATTTGGTGGCATtatagttacttgtcctaaac
gacttatcagtcttgtgtacATCGCCATCTCTTCTTTTTGCTCTGTCTGCTAACTCCCCTtatagttacttgtcctaaac
gacttatcagtcttgtgtacAAGTCGAACTTGTTTCCTCAGCTATAAATAGGGATGAAGCtatagttacttgtcctaaac
gacttatcagtcttgtgtacGCTTTTCCATAGCTTACAGGCCTGATGTAACATACTCTCTtatagttacttgtcctaaac
gacttatcagtcttgtgtacCATTTCTGACCCTCTCCTTGCTTGATTCTCATCACAGTGCtatagttacttgtcctaaac
gacttatcagtcttgtgtacCCTCAGGGCTTTTGGACTTGCTATTGTTTCTGTCTGGAAGtatagttacttgtcctaaac
gacttatcagtcttgtgtacCAAATTCTTTCTTCTTTTGGGTCTCTGCTTTGTTGTCCATtatagttacttgtcctaaac
gacttatcagtcttgtgtacTGATCACTGTTTATCATATTGCAACCTCTTATCCTGCTTTtatagttacttgtcctaaac
gacttatcagtcttgtgtacCTGATCTTTTATATTTCTTCATTGTTGTCTGCTTCCTCTCtatagttacttgtcctaaac
gacttatcagtcttgtgtacGTCATTGCTGTATTCCTAACACTGAGAACTGACTGACACAtatagttacttgtcctaaac
gacttatcagtcttgtgtacATGAAAGGACATGAGAGTTTGCATATTAACTTGCTCGAAAtatagttacttgtcctaaac
gacttatcagtcttgtgtacGGCAATTTAGGAGATGTGGCTGAAGAACTTTTGCTTTTTCtatagttacttgtcctaaac
gacttatcagtcttgtgtacGCTCCTGTGGCTGCCAAAACATTCCTTCTGTCTTTCCCTAtatagttacttgtcctaaac
gacttatcagtcttgtgtacGTGCCCGGCTTCCTTTCATCAACTTTCTCTCACCAGCTTTtatagttacttgtcctaaac
gacttatcagtcttgtgtacGAAATCTCATTGTTTCCCTCTTACTTCTTAATCTCCTCCCtatagttacttgtcctaaac
gacttatcagtcttgtgtacTCTATCACCGAGAGTACATCAATGCATCAGAGAGAAGAGCtatagttacttgtcctaaa
gacttatcagtcttgtgtacGTATGAGACAGTAAGACAATTAAGCAATGGATAAAGGACTtatagttacttgtcctaaac
gacttatcagtcttgtgtacCACAGCATCTTGTGAGCTATTATTAATTTAGGAACGATCAtatagttacttgtcctaaac
gacttatcagtcttgtgtacGAGTATCTGGCAAACACTAGAAAAGACTGTACATGATTTGtatagttacttgtcctaaac
gacttatcagtcttgtgtacTTTCTGACAAATATATCAAGACAAGATGAGGAGAGAAACGtatagttacttgtcctaaac
gacttatcagtcttgtgtacGATCCACAATCTCTATTCCAATTCTCCCTGATCCCTGGTTtatagttacttgtcctaaac
gacttatcagtcttgtgtacATATGGAGCAGGGATGTCTTCTGTCATTCAGACCTTGGCCtatagttacttgtcctaaa
gacttatcagtcttgtgtacGACCACGTTGATCCTCCTAATGTGAATTCTTCTGATTTTAtatagttacttgtcctaaac
gacttatcagtcttgtgtacAGGAACTCAGGAAACAAACGTCCATGACAGCAGCAATAATtatagttacttgtcctaaa
gacttatcagtcttgtgtacAGAAACAAATGAACTAAGACAGGTCTGGAAGTTGGATGGAtatagttacttgtcctaaa
gacttatcagtcttgtgtacCAGCTGAGGTTTTTTTGCTATCAAGTAGGAAGAACATAATtatagttacttgtcctaaac
gacttatcagtcttgtgtacCCTAGTCTCTCAAGTGATGTATCAGAGGAAGAAGGAATTTtatagttacttgtcctaaac
gacttatcagtcttgtgtacGTGGAGAGAGAATTAAAAACATGAGATTGGTCTGCAGTTGtatagttacttgtcctaaac
gacttatcagtcttgtgtacGCAGTCCCGGCAACTTGAAGATACATGTCAGGCTCCTAAGtatagttacttgtcctaaa
gacttatcagtcttgtgtacCTCTAGAGCTTTGCAGCAGAGGATAATGCATTTTCGTGGCtatagttacttgtcctaaac
gacttatcagtcttgtgtacGCAATTAGGAGAAGGCTAACACTAGAAACATCAGAAAATGtatagttacttgtcctaaa
gacttatcagtcttgtgtacCAGAGATCACTGCAGTGCTACTTAGCCCACACAGGTAAGGtatagttacttgtcctaaa
gacttatcagtcttgtgtacGGTTTACAGGAGCTGGACCTTTTCACTTGCCCCCAATAAAtatagttacttgtcctaaa
gacttatcagtcttgtgtacCTGGGGATCATTTACATTTTTCAGGTTTGTCTTGGGGAAAtatagttacttgtcctaaac
gacttatcagtcttgtgtacGCTATTTGGTTTACAGACAAGCCTCATTTCATCACCTTCAtatagttacttgtcctaaac
gacttatcagtcttgtgtacCCATCCCTAGAACCCCAGGCATTGCTTAAGAACTTGTTACtatagttacttgtcctaaa
gacttatcagtcttgtgtacCAAGGCTGGAGTGCTAGAGTGTTGTGGCATGGTCATGGCTtatagttacttgtcctaaa
gacttatcagtcttgtgtacAGTTGAGACTACAGATGTTCTCCACCATACCTGACTACTTtatagttacttgtcctaaac
gacttatcagtcttgtgtacACTATGTTGTCCAAGCTAGTTCCAAACTCCTGGGCTCAGGtatagttacttgtcctaaa
gacttatcagtcttgtgtacCCCCTGTACCTGGCCTGCTTTTCTAATTTACTTAATCCTTtatagttacttgtcctaaac
gacttatcagtcttgtgtacATGATCCCAATTTCATTTGCCCAAGTTCACAGAGATTGCAtatagttacttgtcctaaac
gacttatcagtcttgtgtacTCTGTCCGACTATAAAACCCAAGCATTTAACTTCTATCCTtatagttacttgtcctaaac
gacttatcagtcttgtgtacGGTTTAGCCCATGTAGATACTCAAAAATGTTAAGGGGACTtatagttacttgtcctaaac
gacttatcagtcttgtgtacTATATACATCTTTTATGAGACTGGCCCACATGATTAAAGCtatagttacttgtcctaaac
gacttatcagtcttgtgtacGCTATCGGCGTGGTTTGAATTCATGTTCTTGAGTTAAAAGtatagttacttgtcctaaac
gacttatcagtcttgtgtacCTGTGTCAGCAAGTCCAGAAACCAGAGATATTGGCAGAGTtatagttacttgtcctaaa
gacttatcagtcttgtgtacAAAAATCTCTATGTAAGCTAACTGTAAGCCAACCAGATCTtatagttacttgtcctaaac
gacttatcagtcttgtgtacGCTTCTATGCCCACCTCTCTTTGCCCTCCCATAAGCTCTTtatagttacttgtcctaaac

gacttatcagtcttgtgtacGCTACTCTCTCCTACAAGCCTGCCTTCCTGACTTCTGCCTtatagttacttgtcctaaa
gacttatcagtcttgtgtacCACCACCTTGCCTACTCAAGTCTTTCAGGTAGAGTTTTTGtatagttacttgtcctaaac
gacttatcagtcttgtgtacGCCAAAGTTTATAAAAGACTTGCTTGATTTTCAGGATGTGtatagttacttgtcctaaac
gacttatcagtcttgtgtacCTCTTTCCTGAGTGAGAAAAGTTGTGAAGGTTGCGAGGAAtatagttacttgtcctaaac
gacttatcagtcttgtgtacGCTACCCAAAAGTCAGTTAAAGCTAGCAAAGGCAATTACTtatagttacttgtcctaaac
gacttatcagtcttgtgtacAGGTTCCTTAAGTTCCTTGAGTCTTTTGGGGTGGAGGCAGtatagttacttgtcctaaac
gacttatcagtcttgtgtacCTGTGCTGGCTTCTGTCCTTCTTTAGGTACCCGTTCTGCCtatagttacttgtcctaaac
gacttatcagtcttgtgtacTGAATAAGCAAGATGAGATTATTACATTCTTTCTCCTCGGtatagttacttgtcctaaac
gacttatcagtcttgtgtacGCTACTGATTCCCCAGCCCCTTTAATTTTAGATTCCCTGGtatagttacttgtcctaaac
gacttatcagtcttgtgtacAGCAATTACAAGTTAAAAAAAACACCCTGGGCTCCTGTGCtatagttacttgtcctaaa
gacttatcagtcttgtgtacATTACACAGTGCTCACCTAATTAATAAGTTAACAGTGCCCtatagttacttgtcctaaac
gacttatcagtcttgtgtacTTCTTGAAGAACAGCAAATGGAACTGAACTCAGCACAATAtatagttacttgtcctaaac
gacttatcagtcttgtgtacCCTGACAGGGGTGATGGAGTGCAGAGGCTAATGTTTCCAGtatagttacttgtcctaaa
gacttatcagtcttgtgtacCCAAGGATCACAGTGGTTTGGATAATATTGGGCAAATCATtatagttacttgtcctaaac
gacttatcagtcttgtgtacAGTCCTACTAAATGCTGATAATTTTGGGGGAATGTTGGGAtatagttacttgtcctaaac
gacttatcagtcttgtgtacTGCCATGATACATTTAGCAGAACAGTGACCAATACGTAAGtatagttacttgtcctaaac
gacttatcagtcttgtgtacTGAGGCTATCTCCTTAAAATGTTTACCCCAGTAGGTCCTGtatagttacttgtcctaaac
gacttatcagtcttgtgtacCTGTGGGAATATTGTTTTGCTGTGAAGCCCTCACACTGATtatagttacttgtcctaaac
gacttatcagtcttgtgtacGCTTTCTGTGTTCTTAAAACTTCCAGTGGCTCTCCATCAGtatagttacttgtcctaaac
gacttatcagtcttgtgtacTGTGAAGATTGTTGTTTGATTGTGACATTTCTGTGTGGAGtatagttacttgtcctaaac
gacttatcagtcttgtgtacTGTAGATCACAAAGTCTGCTCCAGAGGCTGCATGTCCTGGtatagttacttgtcctaaa
gacttatcagtcttgtgtacAAACATTCATGCATTGAGAAGCCCAACTCCATTTTAGATTtatagttacttgtcctaaac
gacttatcagtcttgtgtacCTCCAGGCTGAGGTTTCACTATTTTTAAGGAGAATTCAGTtatagttacttgtcctaaac
gacttatcagtcttgtgtacCAATGACTGCACTATGTTTCTCATTCCATATAGGTTACTTtatagttacttgtcctaaac
gacttatcagtcttgtgtacTATACATATTATTTTCTAACTTCCCCTGCTGACCCACATTtatagttacttgtcctaaac
gacttatcagtcttgtgtacCTTTGTAGTTTATGTTGGATCTTAAGGAACTCTCAAGGCAtatagttacttgtcctaaac
gacttatcagtcttgtgtacGGGACACACTCTATTCATAACCAGATCTCCCATGAACAACtatagttacttgtcctaaa
gacttatcagtcttgtgtacATGCTAGGGAGAATGGATGAGGAGTTGATGTTTGCTGAAAtatagttacttgtcctaaac
gacttatcagtcttgtgtacACTCCAGGAATCTTTGACCAGCTCTGCATACCTCTTTCAGtatagttacttgtcctaaac
gacttatcagtcttgtgtacAGGGGACAGTTATCTTGTGGCTACTTCAAAAAAATCACAGtatagttacttgtcctaaac
gacttatcagtcttgtgtacGGACTCCCCTCACCTAGTACTAGTGGGTCTAAAGTATAAGtatagttacttgtcctaaa
gacttatcagtcttgtgtacACTGGCAAGTCATAGGGAAAGCACATTAAGGAAAAGAAAGtatagttacttgtcctaaa
gacttatcagtcttgtgtacTGTCTCCTAAAAATAAACATGCTGAAGTCCTACACTAACTtatagttacttgtcctaaac
gacttatcagtcttgtgtacGGGTCACTGCAGATGTAATCATTTAAGATAAAGTCATTAGtatagttacttgtcctaaac
gacttatcagtcttgtgtacAGCCGGAGATCAGGGTGATGCTTTTACAATCCAAGGAACAtatagttacttgtcctaaa
gacttatcagtcttgtgtacGAACTGTGAGACCATTAATTTCTGTTCCTTAAGCTACCCAtatagttacttgtcctaaac
gacttatcagtcttgtgtacAGCAGCCCTAGAAAACTAAAGCAGGTACCAATGCCAAACCtatagttacttgtcctaa
gacttatcagtcttgtgtacGGACAATAAAGACCCCCTACTCCAGAGTTGTTTTGAAGGAtatagttacttgtcctaaa
gacttatcagtcttgtgtacTCAAATACTTAGCACAGTTCATGACACACATTTGAAGCTGtatagttacttgtcctaaac
gacttatcagtcttgtgtacTACCTATCATTATATCTGTCTGAATATACTCCACCTTCTCtatagttacttgtcctaaac
gacttatcagtcttgtgtacCAGTTAGATTGTGAAGTTCTGAAGCTGGTTGAAAGCTAACtatagttacttgtcctaaac
gacttatcagtcttgtgtacAGAGCTGAGTTAAGGAGATCTGAGCCTGCCTTTGTGCCACtatagttacttgtcctaaa
gacttatcagtcttgtgtacCGTGAGAGTCGCCTCCCATAACTTAATCTTTAAAGCTACTtatagttacttgtcctaaac
gacttatcagtcttgtgtacAGGTTTCATTATGGCGTGATTCTATTATTAAAATCGGCAGtatagttacttgtcctaaac
gacttatcagtcttgtgtacCTGGCTTGCATTACTGACACCAGCATTCTTTTTAATGGGTtatagttacttgtcctaaac
gacttatcagtcttgtgtacCCTTCAGGAAGCTCTAAAATAACCCTTTCTCAAAAATATGtatagttacttgtcctaaac
gacttatcagtcttgtgtacAACTCGTTGAAAGTGCAGGATATCCACTGAGAAGACCGATtatagttacttgtcctaaa
gacttatcagtcttgtgtacGAGGGAGGGAGAGAGGCAGAAGCTGGGAGTTCAAAGCTGCtatagttacttgtcctaa
gacttatcagtcttgtgtacACGCTTTGATGCTGTCCTAACACCGTAAATGATGGCTGGTtatagttacttgtcctaaac
gacttatcagtcttgtgtacCCTGTCATCTCTATAAACCTCGATCAGCCAGCTTCCCTATtatagttacttgtcctaaac
gacttatcagtcttgtgtacCAAAAGATTGCTCAGACTCCAGTGGAACTTAGTAGACCCAtatagttacttgtcctaaa
gacttatcagtcttgtgtacGGGCTTGTCATAATTCTCACTGTGCAACCCTTTTTATGGTtatagttacttgtcctaaac

gacttatcagtcttgtgtacCTTCAGTTGCCAATTTAAGACAGTCTGGACTTAGAGATGCtatagttacttgtcctaaac
gacttatcagtcttgtgtacCTTATTGGACTAAACTGACCTTTCCTTATGGGTTTCTATTtatagttacttgtcctaaac
gacttatcagtcttgtgtacTTCGTCTACTACAATCCACTGGAAAAAACCTTCCAAAACTtatagttacttgtcctaaac
gacttatcagtcttgtgtacTTCCAACCAATGTGCCAGAGGAAATAATCAATAGCTCTAGtatagttacttgtcctaaa
gacttatcagtcttgtgtacCTATTGCTTTTCTCTTCCCACTAACATTTTAGATTGTCCCtatagttacttgtcctaaac
gacttatcagtcttgtgtacTTGTTCAAGGTTAGCAATCTTATTTCAGATGTGTCTTCGTtatagttacttgtcctaaac
gacttatcagtcttgtgtacGCACTCATATTTTATAAGCTGCTAATTGTTGTCTCTTGCAtatagttacttgtcctaaac
gacttatcagtcttgtgtacTACTCATGAAACAAACAGCCAAAAAGCACTAGCAAATATCtatagttacttgtcctaaa
gacttatcagtcttgtgtacGAACGACTCATGATACTAGCCCACAACCATCAGCAATGCTtatagttacttgtcctaaa
gacttatcagtcttgtgtacATTGTGAGATTCTCAGCTATCGCAGTGGTTACAATCCTTGtatagttacttgtcctaaac
gacttatcagtcttgtgtacATGGGCTTAGCATTTTCTTCACTCATGGCAATATTATCACtatagttacttgtcctaaac
gacttatcagtcttgtgtacTTGATGATGCTTGGCACAGCCTATGTCAGTGATATGTTCAtatagttacttgtcctaaac
gacttatcagtcttgtgtacGGCTCACATTTTTTTGATCCAGTGGTAAGAGTCTTTCAATtatagttacttgtcctaaac
gacttatcagtcttgtgtacATTATAAAATATGATCTTGTGTTCTCCCACCCTATGCCTGtatagttacttgtcctaaac
gacttatcagtcttgtgtacGGTTTCCCCATAAGTTGAGGCAGAAGAACTAACCGTAGTGtatagttacttgtcctaaa
gacttatcagtcttgtgtacAGTCATTGGAAAGAATTAAAGCTGAGATTCTAACGGTTTCtatagttacttgtcctaaac
gacttatcagtcttgtgtacCTATTAATCATTCCTCTGGCTATTTCTAGTTCTTACGTGGtatagttacttgtcctaaac
gacttatcagtcttgtgtacTCTTTGATTTACTGATTTTGGACATGTATCTATTGCCACCtatagttacttgtcctaaac
gacttatcagtcttgtgtacGGAATATAGCTTACTTATGCCTTTCTCTCCCTTTACTCCTtatagttacttgtcctaaac
gacttatcagtcttgtgtacGATTACTTGTTTGTTAGTTTTAGACCTTCTACTGACTACCtatagttacttgtcctaaac
gacttatcagtcttgtgtacCAATGTCTCTTCACTCCTTATTATGTAAGATGAGATTTGCtatagttacttgtcctaaac
gacttatcagtcttgtgtacCTTTTCCAAACATCTATCAGCTCTAGACTATTGCTTTTACtatagttacttgtcctaaac
gacttatcagtcttgtgtacACTTCCTGAGGTTGTGTTTAGGTGTAGGGTTTTTTTTTCtatagttacttgtcctaaac
gacttatcagtcttgtgtacTATACTGGAGTGAAGTCTTCATCTTTTCATAATCTTGCTCtatagttacttgtcctaaac
gacttatcagtcttgtgtacTTTTTGGCCTCCCTTTTATCTTCTGTCTTCCACTGTATTCtatagttacttgtcctaaac
gacttatcagtcttgtgtacAGCATCCTCTTCTAATTTATGGCTGGGTTATCTTTCAAATtatagttacttgtcctaaac
gacttatcagtcttgtgtacGTTATCTTTTGTTTCCTAAACTATCTCTATATCCTCCAGGtatagttacttgtcctaaac
gacttatcagtcttgtgtacACTTTTGTCTTTTCCTACAGCATTTGGAGTCCCTTGATTGtatagttacttgtcctaaac
gacttatcagtcttgtgtacCATGGGCTTCCTCTGGCATTGTGTGGGTGTGCTTTTCCAAtatagttacttgtcctaaa
gacttatcagtcttgtgtacCAGGTGACTGCAAATTCAGTGTGTGTGGCCAGGCATGTCAtatagttacttgtcctaaa
gacttatcagtcttgtgtacCTTGTTCAGCTTCTTCAGAGAGCTCATCGCCTGCTTCCCCtatagttacttgtcctaaa
gacttatcagtcttgtgtacTGTTAGCATCTTTGCACAGAGTCGTAGGGAGACAGGTGTGtatagttacttgtcctaaa
gacttatcagtcttgtgtacTGGTGTTCCGCTCCTTCTCAGTACAGATGGTTCAGAGTCGtatagttacttgtcctaaa
gacttatcagtcttgtgtacTGGCTCCCACTTCCACTGAAGCTTTCTTCTGAGTCTTCCTtatagttacttgtcctaaac
gacttatcagtcttgtgtacGGTAAACCTCAGTATGATTCTGTTGATTCCTATCTTTCAGtatagttacttgtcctaaac
gacttatcagtcttgtgtacTCTCTTAACAAATGAAGATTGACCACCTGTTCACCACGCGtatagttacttgtcctaaa
gacttatcagtcttgtgtacCATTCATGTTGTTTCATTGTGATTCCTCTTGAGAATGTGGtatagttacttgtcctaaac
gacttatcagtcttgtgtacAAGATACATGCTTAGTTTATTCTCAGAAGCCCCCTGTGAGtatagttacttgtcctaaac
gacttatcagtcttgtgtacGAACACTGGCTTTTCTATAGTTTATTCCTGCTTCTTTCAGtatagttacttgtcctaaac
gacttatcagtcttgtgtacCAGCATGAAAGCCTAGGTCTACTTCTAGGAATATATTTAAtatagttacttgtcctaaac
gacttatcagtcttgtgtacGTAGAGAGAAGAGTATGATGCACCTCAATGTGGTTGTCACtatagttacttgtcctaaa
gacttatcagtcttgtgtacCCTTGTTTCCTCTTTATCTTTCCTCACTCTACTCCTTTCAtatagttacttgtcctaaac
gacttatcagtcttgtgtacTGAAATAAAAGCTAGCAATGTGGGAATAAAATCTCAAGCCtatagttacttgtcctaaa
gacttatcagtcttgtgtacAGGTTAGGTTTGCCTCTTCTGTGTCATGTGCCTGCTATCAtatagttacttgtcctaaac
gacttatcagtcttgtgtacTGTTCCCCAGGGCCTAGATAGATTAAGCAATAGGTGACAAtatagttacttgtcctaaa
gacttatcagtcttgtgtacCCCCCAGCTGTAATGTATTTTTTCTATTAAAAGGTAAGTGtatagttacttgtcctaaac
gacttatcagtcttgtgtacTAGAATATAGTGGCTTTAAAAAGGTATCTACAGGCTGGGTtatagttacttgtcctaaac
gacttatcagtcttgtgtacGAAACCCTGTTTCTACTAACGGCTGAGGCAGGAAAATCGCtatagttacttgtcctaaa
gacttatcagtcttgtgtacGAAGATATATATAAAGTTGCTTCTCATACTCTGGCAGTAGtatagttacttgtcctaaac
gacttatcagtcttgtgtacTATTTGATTCTCTTCCTTTTTTCCTCTGGGCGGTTAGTACtatagttacttgtcctaaac
gacttatcagtcttgtgtacGCAGGAGGAGGGAATAAGAAAGAGTCAGGAGTATGCAAGCtatagttacttgtcctaa
gacttatcagtcttgtgtacCTGAAAGCTCCTAAAACCAGTGTCTCAGGGATTCCCATGAtatagttacttgtcctaaa

gacttatcagtcttgtgtacACATCCCCGCACCCTTAGTTTTTGTCTCTGGTCTAGAAAGtatagttacttgtcctaaac
gacttatcagtcttgtgtacATGTGTCTCAGTGTGATCATCTTTTCTTATAGCTTCAACCtatagttacttgtcctaaac
gacttatcagtcttgtgtacGTCTATATTAATAATTCCTGGCAGATCTCTTTTCAGCTCAtatagttacttgtcctaaac
gacttatcagtcttgtgtacCAAATTGCAGGGTTACTCGTTGGAAGCACCTATAGCAGTAtatagttacttgtcctaaac
gacttatcagtcttgtgtacTTTGTTTTAGTCTCAAATCCACCTTCTCCTGGTTCTACACtatagttacttgtcctaaac
gacttatcagtcttgtgtacAGCTTCCTCCCTGTTCTTTGCTTGGCACATCTAATTGTCCtatagttacttgtcctaaac
gacttatcagtcttgtgtacGACCTGTGTTGACTCTCTTTGCATTTTCCCACAAATACACtatagttacttgtcctaaac
gacttatcagtcttgtgtacAATTTCCACATCTGCTACCTTAGTTCTCCTTTTTGAATCAtatagttacttgtcctaaac
gacttatcagtcttgtgtacTAGTGTTTTTGCTCTAGTGTTCCCTCCTACATGCCATCCTtatagttacttgtcctaaac
gacttatcagtcttgtgtacAATGTCTTAAAAATCTAGTTTGACTGTCACTCCCCTCTCAtatagttacttgtcctaaac
gacttatcagtcttgtgtacGGACCAAAACCCCTCAGTTATTTCCTGTTGCCTTTAGAAGtatagttacttgtcctaaac
gacttatcagtcttgtgtacCAGGCTGTCCACATGCAATGCTCAGTGTACCTTTCCAGTCtatagttacttgtcctaaa
gacttatcagtcttgtgtacTTTGTCTTTACTGTTTGTCACTCACACTGGAAATAAACTCtatagttacttgtcctaaac
gacttatcagtcttgtgtacATCCATGTCAACTGATGTATCTATCCACAATGCCTAACGTtatagttacttgtcctaaac
gacttatcagtcttgtgtacGTTCTTTACTCCTCTCTGTAATTCTATGCTGAATCAATTGtatagttacttgtcctaaac
gacttatcagtcttgtgtacAAATGATATTTCCTACCCTCTCCTTTTGTCTTCTGCTCTTtatagttacttgtcctaaac
gacttatcagtcttgtgtacAGTTACATTCTCTCCACCTTTACTGCTAAGACCCAACTCTtatagttacttgtcctaaac
gacttatcagtcttgtgtacTTCTGCATCTTGCCTTCCCTGCACTGGGAGGGTCCTTGGAtatagttacttgtcctaaa
gacttatcagtcttgtgtacATTATCTTGTCTTTCTCTGAGAAATTGTAAGCTCCTAGAGtatagttacttgtcctaaac
gacttatcagtcttgtgtacCATATTTTGGAGGCATAATTCCTATGTTTTTGTACCTTCCtatagttacttgtcctaaac
gacttatcagtcttgtgtacCACTAAGGATGTTTGAATCACTCAGACTTGTATTCACTCGtatagttacttgtcctaaac
gacttatcagtcttgtgtacGCTATAAATATTTCACCCTTTTCTTCTACACTTACTCTCCtatagttacttgtcctaaac
gacttatcagtcttgtgtacTCATTGACTGACCATTGAGTTTTCTTCTATTTTCTGCGCGtatagttacttgtcctaaac
gacttatcagtcttgtgtacTGATATTGTGGAACTCCTCACTGGCATTGGGATTTGGTGAtatagttacttgtcctaaac
gacttatcagtcttgtgtacCACGATGTGTTTTTAGACTCAAACTCCTATAATTGACCATtatagttacttgtcctaaac
gacttatcagtcttgtgtacTTATTATTAATCTGGATAGAAATGCTGTGCTGAAACCCAGtatagttacttgtcctaaac
gacttatcagtcttgtgtacCACTGTATGGAGTAGCTTCTGAATTGCATGTCCTTAATTTtatagttacttgtcctaaac
gacttatcagtcttgtgtacGTGTAAAACATGGGAATTAAGGAGCCAGCTCTGCCACTTGtatagttacttgtcctaaa
gacttatcagtcttgtgtacAAATTACTTAACCTCTTTTTGACTCAGCTTCCTCACCTATtatagttacttgtcctaaac
gacttatcagtcttgtgtacGGGGTAATAATAGAGGCTAATTTCCATGTTTGTTGTGAAGtatagttacttgtcctaaac
gacttatcagtcttgtgtacACAAACAAATTAATTAGAATAGTGTCTGGCATGCTGGAAGtatagttacttgtcctaaac
gacttatcagtcttgtgtacTTCTCTGCTGTTATTGAGGGAATGCCATTTTGAAAAGTTTtatagttacttgtcctaaac
gacttatcagtcttgtgtacCTCATTCCATTCTGAGCAAGGACTTCATATCCCTGAGTTAtatagttacttgtcctaaac
gacttatcagtcttgtgtacGTCTTTCTTCCTTCAATGACTTTGGGAAACTCTTTAAATCtatagttacttgtcctaaac
gacttatcagtcttgtgtacATTTCTGGCCAGACATGCTAATTTTTTTCCTTCCCTTGTAtatagttacttgtcctaaac
gacttatcagtcttgtgtacCAAGCAATCCCTTGGTATACATTTACCCAACTTTAGTATCtatagttacttgtcctaaac
gacttatcagtcttgtgtacGTGTCTTCCCTGATAATCACTTTCCTCACTCCTAAAATTAtatagttacttgtcctaaac
gacttatcagtcttgtgtacATGAGTCTAGCATTGGCATAATCAATATGGTCTTACAACTtatagttacttgtcctaaac
gacttatcagtcttgtgtacGGAACCTCTGGCATCTTTCCTTTAGTTGCTGAGTTGAGAGtatagttacttgtcctaaac
gacttatcagtcttgtgtacCTGAGGGAGCTAGGGTGATGAAGAGAGGCATTCTGGGAGCtatagttacttgtcctaa
gacttatcagtcttgtgtacGCAACAGCTACACACCAGATGGTACAGGCCTACTATAATGtatagttacttgtcctaaa
gacttatcagtcttgtgtacGAAGACCAGCCTTGATTGAGACCCTTTATGCTTCCTCTAGtatagttacttgtcctaaac
gacttatcagtcttgtgtacCCACCAAACTTTGAGTCTTAATCAATATACCCTTTTAAGGtatagttacttgtcctaaac
gacttatcagtcttgtgtacCTTAAGTGTGTTCCCCTGAAAGTAGAGTCTGAGTCAAGAAtatagttacttgtcctaaac
gacttatcagtcttgtgtacCAGACTAAAGACTGGGGAGAGTGAGAGGAGAAAGAAGAAAtatagttacttgtcctaaa
gacttatcagtcttgtgtacGGGGTGTGGTTTCTAAGTCACTGCTATAGGGAATAAGGGTtatagttacttgtcctaaac
gacttatcagtcttgtgtacTCATCTTCTTGGTTGTGAGAGAATTCCTCCCAAAATTTTCtatagttacttgtcctaaac
gacttatcagtcttgtgtacGGAGGTTGGAGCACTTTTCCCCAGGCACCCATTCCCCTTGtatagttacttgtcctaaa
gacttatcagtcttgtgtacACTTCTGGATTGCATGTGTGTATAAATAGAGGTGACTTGTtatagttacttgtcctaaac
gacttatcagtcttgtgtacGCATGTCTGAGGGGGTGCATAAATCAGTGCAGTGTGAATCtatagttacttgtcctaaa
gacttatcagtcttgtgtacCTGAAGGTGCAGCTAACATATGGTAGGAGAAAATTTGACAtatagttacttgtcctaaac
gacttatcagtcttgtgtacGTATTTTTCACATGATATAGTACCACCAAGAACCCAGCTTtatagttacttgtcctaaac

gacttatcagtcttgtgtacTCATGTTCTTCTCCAATTCTTTCTCAATAGCTTCTCTTCCtatagttacttgtcctaaac
gacttatcagtcttgtgtacATTGACTACCTAGTACACATGAGATTTTGTTGATAGGTCCtatagttacttgtcctaaac
gacttatcagtcttgtgtacCAGTGATAAGACCCAGTTGCTGCCTTTCATAGAGCTTCACtatagttacttgtcctaaac
gacttatcagtcttgtgtacTTTTGATTTAGTAGAATATTGGGTACAGATGACAGAGACCtatagttacttgtcctaaac
gacttatcagtcttgtgtacCCTAGACCCTGATATTTCTTATTCCTTCTATACCTAGGTTtatagttacttgtcctaaac
gacttatcagtcttgtgtacATGGGAATTGTGTCACCTATCTTTTCTTACTCTGCACTAGtatagttacttgtcctaaac
gacttatcagtcttgtgtacTGCCTACATAAGCAAGATATGAAACAAGCTCAAACACACCtatagttacttgtcctaaa
gacttatcagtcttgtgtacTGAAATGTATTACAGTATCTAAAATCTCCCCTCTCCTCCTtatagttacttgtcctaaac
gacttatcagtcttgtgtacTTGAGACTTTCAGGGTAAAGCTTATAAAGTGCACGTCAGTtatagttacttgtcctaaac
gacttatcagtcttgtgtacATTATAATGTAAACCAATGCCTTAAACTAGGCCCTTCCCCtatagttacttgtcctaaac
gacttatcagtcttgtgtacCATATGCAGATCTTGTAACCATAATATGTTTGTCACTCCAtatagttacttgtcctaaac
gacttatcagtcttgtgtacAAGTCAATACACAGAGACACCAAATTGCAGCAGAGAAAGAtatagttacttgtcctaaa
gacttatcagtcttgtgtacTGGGAGGAAACCTCAAATTAATCTCTCCAAGGAGTCTGGGtatagttacttgtcctaaa
gacttatcagtcttgtgtacGAGTGCAGGGTGAGATCATGAGACAGGAAGTTGAAGAAGCtatagttacttgtcctaaa
gacttatcagtcttgtgtacCCATTTCTCTGTGGGGGGTCATCAAACTAGTGGGTGTCAGtatagttacttgtcctaaa
gacttatcagtcttgtgtacAATGGTCAGTATCTAGTGTTGTGGTGACTTTTAGCAACAAtatagttacttgtcctaaac
gacttatcagtcttgtgtacCGGGGCAAAAGTACAGCCTTGCTAATGCTTAATTATAATGtatagttacttgtcctaaac
gacttatcagtcttgtgtacCAATTTTTGCCAACCTTGTGGGGATAGTTTCAATCTCTTGtatagttacttgtcctaaac
gacttatcagtcttgtgtacAGGTGATGCATCTAGTCTTTGGTTCCAGCCTCCTCTTGTGtatagttacttgtcctaaac
gacttatcagtcttgtgtacTGGTTATTTCCTCTTTGTCCTTTCTCACCACTCTCACAACtatagttacttgtcctaaac
gacttatcagtcttgtgtacTCAACAGATTCAATATTTCTCACCTCACTCCTTTCCATTTtatagttacttgtcctaaac
gacttatcagtcttgtgtacCATTCCTTGCCTTAGTATCTTGCTATGTGTCAGGTTCATGtatagttacttgtcctaaac
gacttatcagtcttgtgtacAAGGAATTAGTGAAGGATACTGACATGAGGACATTGCATTtatagttacttgtcctaaac
gacttatcagtcttgtgtacTGAGATTTTCAGGTAATACAGCATTCAGTGGGAGAGAAATtatagttacttgtcctaaac
gacttatcagtcttgtgtacGGACGAAGTGGATATTTTGCTCATTCTCAAGTGACAGCACtatagttacttgtcctaaac
gacttatcagtcttgtgtacGCTTATATCATAATGTCAAGCAACCTGTGGCATCGTTTGAtatagttacttgtcctaaac
gacttatcagtcttgtgtacTTTCCATCATCATTATCAATTCTCATGTATCTGACTGCAGtatagttacttgtcctaaac
gacttatcagtcttgtgtacTGGGACATTAGCAAAAGTAAGAAGAATACAGGGGAGAAAAtatagttacttgtcctaaa
gacttatcagtcttgtgtacCTCTACTATTGGCTTTGAGTTGTAACCTATGGAGGCAGCCtatagttacttgtcctaaac
gacttatcagtcttgtgtacCCTCAAATATTGGCAGAGCTATCCTTGTCTCAGATTCATTtatagttacttgtcctaaac
gacttatcagtcttgtgtacTTATGACACCTTTGCAGTTGTTGTAATGATCATCACTCAAtatagttacttgtcctaaac
gacttatcagtcttgtgtacAGCACGAGTGTCCATTTTTCTTAAAAACCATTGTAAAGAGtatagttacttgtcctaaac
gacttatcagtcttgtgtacGAACACCACAAGCACCATTCCTGTACTTTTAAATTCTGGTtatagttacttgtcctaaac
gacttatcagtcttgtgtacTTTTAACAATGGTGCCAAAGTATCAGAGGTGTTTAAACCAtatagttacttgtcctaaac
gacttatcagtcttgtgtacGCATTCCCAGGAGGTTACACATTCTTTTTTTTGTTTTGTTtatagttacttgtcctaaac
gacttatcagtcttgtgtacTTCAAGCAATTCTTATGCCTCAACCTTCTGAGTAGCTGGGtatagttacttgtcctaaac
gacttatcagtcttgtgtacCCAGGTTACACATTCTTAGTCACTGGGTGAGATAGCAGGTtatagttacttgtcctaaac
gacttatcagtcttgtgtacCTGCTGATAAAACAGGACATAGTAAAGATATGGAACCAAAtatagttacttgtcctaaac
gacttatcagtcttgtgtacGTGCTGGGAAAACTGGCTCGCCGTATGTAGAAAGCTGAAAtatagttacttgtcctaaa
gacttatcagtcttgtgtacGCCAAAAGACCCATGAAAAAATGCTCATCAGCACTGGCCAtatagttacttgtcctaaa
gacttatcagtcttgtgtacCACCATGGAATACTATTGCCATAAAAAAATGGTGAGTTCAtatagttacttgtcctaaac
gacttatcagtcttgtgtacGCTGGCCAAAACCCGCCAAATCCAAAATGGCAATGAAAGTtatagttacttgtcctaaa
gacttatcagtcttgtgtacTAAAGCATTGGCATGCTAAATGAAACTCCCACCAGTGCCAtatagttacttgtcctaaa
gacttatcagtcttgtgtacAGTTACCCTATATGGACTAAGAAAGGGAGGAACCCTCAGTtatagttacttgtcctaaa
gacttatcagtcttgtgtacTTTCTGGAAAACTCATGAATACCCACCCTTTGTTTAGCATtatagttacttgtcctaaac
gacttatcagtcttgtgtacCAAGAACACTCTCTTGGTACCCCGTTCTGGTAACAAAAGTtatagttacttgtcctaaac
gacttatcagtcttgtgtacTTTACTTTTGTTCTTTTAGAACGCTTCCACTGGCTCAAACtatagttacttgtcctaaac
gacttatcagtcttgtgtacATTTAACATTTCTGATGAGGAGAAAACCTTAGGCAAACACtatagttacttgtcctaaac
gacttatcagtcttgtgtacTTACAAAACAGGCTCAGAATCAGAAAAGACATAGAATCTGtatagttacttgtcctaaac
gacttatcagtcttgtgtacAACTGTAATTAGGTTAGTTGAAATTTGCATGACCCTGTGGtatagttacttgtcctaaac
gacttatcagtcttgtgtacTACAGAAAGCTATTATATCTAGGATGGGAGGCTCTGTCTCtatagttacttgtcctaaac
gacttatcagtcttgtgtacCAAACACTGCACATTTTCTCTCCTCTTTTTCCTCACTAGCtatagttacttgtcctaaac

gacttatcagtcttgtgtacGACTTCGCAGATATCAGCAGGGAACAATGACCAGGTAGATtatagttacttgtcctaaa
gacttatcagtcttgtgtacCAACCTGCCCCATCTTCTACATCTTCATATGCTAAGCCCAtatagttacttgtcctaaac
gacttatcagtcttgtgtacCATGCTTTTCACTTTGGCCATGCTCATTAGACTTAAAATAtatagttacttgtcctaaac
gacttatcagtcttgtgtacCTTGGAAAAGCAATAAAACAGGTTTTGAAAATGGAGAAGCtatagttacttgtcctaaac
gacttatcagtcttgtgtacTGTTATCTTGAACACTAGACTCATACTATCTGGAAAGCAAtatagttacttgtcctaaac
gacttatcagtcttgtgtacTGTGATTAAGTATGTGTGGCAAATGCTGTATCTGCTGCTAtatagttacttgtcctaaac
gacttatcagtcttgtgtacGTGGTCCATTAACTGCTGCCTGTCCCCAGACTGTTTCCTGtatagttacttgtcctaaa
gacttatcagtcttgtgtacATGTGATGCATACAAAAATGAGACTGTGCAGAGACCTGAAtatagttacttgtcctaaac
gacttatcagtcttgtgtacAAAAACCCAAAACTGATCCTTCACCACAGATAGTCAGAGAtatagttacttgtcctaaa
gacttatcagtcttgtgtacGCTATATAATAAAGATTCTGAGAAGCCACACAGTAAAGAGtatagttacttgtcctaaac
gacttatcagtcttgtgtacATGATAAGCAATGTTCATAAAGTAGTTTGGGCCGGGCATGtatagttacttgtcctaaac
gacttatcagtcttgtgtacAGTTCCATACCAGCCTGACCAGCATGGCGAAACTCCATCTtatagttacttgtcctaaa
gacttatcagtcttgtgtacCTGCACTCCAGTCTGGGCATACTTTATATTTTTGAATGCAtatagttacttgtcctaaac
gacttatcagtcttgtgtacAATACCACACACCAAACATGATGCCATATTACATATTCAGtatagttacttgtcctaaac
gacttatcagtcttgtgtacTAAAAAGTTGTAGTACCTAGTCACATACCTTCAAGAGTTGtatagttacttgtcctaaac
gacttatcagtcttgtgtacGTTAAAAATAGAAGACACTGAATCCAGTTTTCATGCAGACtatagttacttgtcctaaac
gacttatcagtcttgtgtacGTAAATTCCTATCCTGTTCATGACTATGTCTTTAATGGTGtatagttacttgtcctaaac
gacttatcagtcttgtgtacTCAAGAAGAGCCTGGAGATTTGGTTTACATAAACTGGAGGtatagttacttgtcctaaac
gacttatcagtcttgtgtacCAGCTTTTGGATTCCCAGTGATGTAGAAGACTGCACTTTCtatagttacttgtcctaaac
gacttatcagtcttgtgtacTCTGTACTCAGTCAAATTGAAACTGCAACAAAGAAAACAGtatagttacttgtcctaaac
gacttatcagtcttgtgtacCAAGTGTTGAATTTAAAAATCAAGGTGCACAGGGATGGCAtatagttacttgtcctaaac
gacttatcagtcttgtgtacCCTTGGGGAATTCTCTATGGTCCTGGGTCTAGCTGTACTCtatagttacttgtcctaaac
gacttatcagtcttgtgtacTATGTTTGGATCATATGTTTGGAGCAAGTGATGTTGAAGCtatagttacttgtcctaaac
gacttatcagtcttgtgtacGAATACATGTTCTAGATGCAAGTTCCCTAGGTTCAAATCCtatagttacttgtcctaaac
gacttatcagtcttgtgtacAAAATGAGTTTAAAGTGCTTAGAATGTTACTGGTGTGGAGtatagttacttgtcctaaac
gacttatcagtcttgtgtacCACTGATAAATGCAATGTCAAGTGGAGTATACTTTCAAACtatagttacttgtcctaaac
gacttatcagtcttgtgtacCCAAATTATCAGCAGCTGCCTGGCAATCACGGTGTTGGAAtatagttacttgtcctaaa
gacttatcagtcttgtgtacCTTTATAGCACTAGGTTATGACCACATGATGCATTCTGTCtatagttacttgtcctaaac
gacttatcagtcttgtgtacAGAGTTGCTGTCCTGAGAGATATCCTGCCTGTTTATGGTGtatagttacttgtcctaaac
gacttatcagtcttgtgtacAGGGTAGATCTCAGAGGGCAACTTTATTTCTATACTCACAtatagttacttgtcctaaac
gacttatcagtcttgtgtacCTTCTTCCTGTTGTTCTTACCACATACCTACCTCCTATGAtatagttacttgtcctaaac
gacttatcagtcttgtgtacAAGAGACAAGACTGAGACTTCTATTAGCAAACAACCTACAtatagttacttgtcctaaac
gacttatcagtcttgtgtacCAGAAATGATGTGGATCTTATTGAGTGCAATTACATTTCCtatagttacttgtcctaaac
gacttatcagtcttgtgtacCAATTATCAAGTGTCTGCTGTATTGGTTTGGTCAAATTTCtatagttacttgtcctaaac
gacttatcagtcttgtgtacCCCACTATACAAGGCATATCAGAATAATCCTGGAAAGTTTtatagttacttgtcctaaac
gacttatcagtcttgtgtacAGATTATCGTCTATGAAGTTGTCTAAATTGGATGCTGGAAtatagttacttgtcctaaac
gacttatcagtcttgtgtacAGTGATGTGAACCTGTGGTGAGATTAGAGTTATGCATAGCtatagttacttgtcctaaac
gacttatcagtcttgtgtacCATGGTCTTGTTAAACACTGTTGAATGGTTACAACTAAACtatagttacttgtcctaaac
gacttatcagtcttgtgtacACGTTCCATTTTTAAAATGAGCAGAAATGTTGAACAGGTGtatagttacttgtcctaaac
gacttatcagtcttgtgtacGAAGACATACAAATATTCCTCAAATTCCCAAAAGGTATGGtatagttacttgtcctaaac
gacttatcagtcttgtgtacCTATATACCTAGAATAGTTAAAGACTGAGAGTACCAAGTGtatagttacttgtcctaaac
gacttatcagtcttgtgtacCTGGTACAATCATTAGGGAAAACTCTTTGTCAGTATCTACtatagttacttgtcctaaac
gacttatcagtcttgtgtacATAATTTCACGACTCAGTAATTTCACTCTCAGGAATATCCtatagttacttgtcctaaac
gacttatcagtcttgtgtacTTGCTCTTTTTATGAATAAAGTGGATCAAGCTGGTTTTGGtatagttacttgtcctaaac
gacttatcagtcttgtgtacACTGTTAAAAGTTGACGTCCATAATTGGCAGTGATTTAGCtatagttacttgtcctaaac
gacttatcagtcttgtgtacCATTTGTGTGTGGATCATTGTAGAGCTGATTATTAAGAAGtatagttacttgtcctaaac
gacttatcagtcttgtgtacGTCCCCATTTAACACAGCATATAGATTTTGTGTATCATTCtatagttacttgtcctaaac
gacttatcagtcttgtgtacGCAATCTAGTGAAGAAAATAGACTCATATCCCAATATTGCtatagttacttgtcctaaac
gacttatcagtcttgtgtacAAATAATCTCCATCATTTCAACTTGGCAATGAACCTACAGtatagttacttgtcctaaac
gacttatcagtcttgtgtacAGACTGTATCTTTACCCAAACTCAAAGCACAAGCAATTTGtatagttacttgtcctaaac
gacttatcagtcttgtgtacCTGCTTATCCAAAACTTCCTCCTAAAGAGAAAAAGTGAAAtatagttacttgtcctaaac
gacttatcagtcttgtgtacGAGGAGGTAAGTACAGTGTTCAATAACTTCATCAAATAGGtatagttacttgtcctaaac

gacttatcagtcttgtgtacAGGTCAGTGAGTCTTTTCAAATGGTACACGGGCTGCCTATtatagttacttgtcctaaa
gacttatcagtcttgtgtacAAAAGAATTGATTTTGGTGGTTGGAATAAGGCCATTTCAGtatagttacttgtcctaaac
gacttatcagtcttgtgtacCTGCAAGCTTTCCAGAAATAATAGTGCTATACTAGGAGATtatagttacttgtcctaaac
gacttatcagtcttgtgtacCTAAATAGCAAGTGTCAACCTGATTTTTAATGTTCTCCAGtatagttacttgtcctaaac
gacttatcagtcttgtgtacGCAGTTGTGTAGATTCTAGGGAAAGCTGTGCTGTCTGGCCtatagttacttgtcctaaa
gacttatcagtcttgtgtacATTCTCACCTTTTGTTTTCTTCCTTTTAGGTCTGTTATGCtatagttacttgtcctaaac
gacttatcagtcttgtgtacAGGACCTTTGATGCCTGGCAAGCTGTACCACTGGGAGGTAtatagttacttgtcctaa
gacttatcagtcttgtgtacTGCTTTTGTGTGAGATATTAATTGCAGAAAGATTGGTAGGtatagttacttgtcctaaac
gacttatcagtcttgtgtacTGAACTCTTATTTTGGGATCACTTCTCAGCATGGATACTTtatagttacttgtcctaaac
gacttatcagtcttgtgtacAATTTTTCACTTTACCCATTCTGAGAGGTCACAAGGGCTTtatagttacttgtcctaaac
gacttatcagtcttgtgtacATAGCTATATGTCTTCAGTGGTTAGGAATTGAATATCTGGtatagttacttgtcctaaac
gacttatcagtcttgtgtacTCTGTAGAGAAAGCCTTATCCAATTCCTTTAAGCATACTTtatagttacttgtcctaaac
gacttatcagtcttgtgtacCATCTGCGTATACCTGTTGCCCTTTCAATGTTGCTCTGTTtatagttacttgtcctaaac
gacttatcagtcttgtgtacTCCTTGATTAGGGCAGGCATCCCTACCATCTTTGCTTCCCtatagttacttgtcctaaa
gacttatcagtcttgtgtacATCGTAGGTGATTTTGTTTGTTAATTAACTGCCACCCTATtatagttacttgtcctaaac
gacttatcagtcttgtgtacCTAATGTGAAAACGAGATTTAGAACTGAGTCTCTATGATGtatagttacttgtcctaaac
gacttatcagtcttgtgtacCAAAAAAAAGATACATGTTTCCACAGTGTAAATGCTTCCCtatagttacttgtcctaaac
gacttatcagtcttgtgtacTGGATGGATCTCAATGGTCCTTTCAAGCTTTAAATTTTTCtatagttacttgtcctaaac
gacttatcagtcttgtgtacTATTTCAAAGATCTTATGTTCTTGCTGCCTTTCAGAATGCtatagttacttgtcctaaac
gacttatcagtcttgtgtacAGATATTACAAAGAAAACTTCCAAACCTTGCGCATTAGTCtatagttacttgtcctaaac
gacttatcagtcttgtgtacCATCTGCTTCTGATGAGTATGCAGAAAGCTTACAATCATGtatagttacttgtcctaaac
gacttatcagtcttgtgtacATGGTGAGACAGGGAGCAAGGGGGTGGTGGGGAGGTGCGTtatagttacttgtccta
gacttatcagtcttgtgtacACATTATGGTGAGGATGGCAGCCAGGTATTTGTATTAGTCtatagttacttgtcctaaac
gacttatcagtcttgtgtacTGGCAGCAAGGAGAAGTGTAAGTGAAGGGTTGGGGGGAAGtatagttacttgtcctaa
gacttatcagtcttgtgtacCTCACTATCACGAAAACAGTTTGGGGGTAACTGCCCCCATtatagttacttgtcctaaa
gacttatcagtcttgtgtacCCACCAGGCCCTACCTCAAATGCAGGGGATTACATTTCAAtatagttacttgtcctaa
gacttatcagtcttgtgtacTGCCTCAATAATATTTGCAACTGGCATAACAAATGACATGtatagttacttgtcctaaac
gacttatcagtcttgtgtacTAATACCTGCCAACTGTGCTTTAAAATGTAAATGAAGGGCtatagttacttgtcctaaac
gacttatcagtcttgtgtacGAGGCGGCCAGATCATGAGGTCAGGACATCGAGAGCATCCtatagttacttgtccta
gacttatcagtcttgtgtacCTAATTTTAATAGAATCATCTCAAATTACCTTCCCTCCCGtatagttacttgtcctaaac
gacttatcagtcttgtgtacATTTCATTGTTAAGTAAAAATCCTCTTCTTTCTGTGGGCCtatagttacttgtcctaaac
gacttatcagtcttgtgtacGTTGTTTAACGGCACACAATCAAGGACAGTCTAACCAGTCtatagttacttgtcctaaa
gacttatcagtcttgtgtacGTGGCTCATTTAATCACTTGGAATAATAGCGGTTCTCAGAtatagttacttgtcctaaac
gacttatcagtcttgtgtacCCAAGGTGTTCAGGGTTACTACAGGCAAGGCAGTCTCTTAtatagttacttgtcctaaa
gacttatcagtcttgtgtacGCCTCCTTTCCATTGCCTACTCTTCATGTTTGACTCCCTGtatagttacttgtcctaaac
gacttatcagtcttgtgtacCTGGGGGAGCGTGTGTTCTTATGTGTGTTCTTTACCATGGtatagttacttgtcctaaac
gacttatcagtcttgtgtacCCTCTAAGGTCATCTGTACGTTTCATAGATGAGGAAGCTGtatagttacttgtcctaaa
gacttatcagtcttgtgtacGGTCACACAGCTAGTGCGGGGAGTGCTGGACCTGGACTCTtatagttacttgtcctaa
gacttatcagtcttgtgtacCTCTTAGGCTTAGAAAGGGACAGTAGAGGCATCAGAGATTtatagttacttgtcctaaa
gacttatcagtcttgtgtacCATTTTCTTATATGGAAGGATAATCTGACATTCTGTGCAGtatagttacttgtcctaaac
gacttatcagtcttgtgtacCAAGTAGCACAAGTATTTAGGGATATCAGGGGTCCTCAGTtatagttacttgtcctaaa
gacttatcagtcttgtgtacGCTTCCAACAGCTCAGTAAACATCCCCTCCCTGATTGTCAtatagttacttgtcctaaa
gacttatcagtcttgtgtacAGTGTTCAGAGGACTGGGGATTACAGGAGGAGCAGTGTTGtatagttacttgtcctaa
gacttatcagtcttgtgtacAGTAAGTGAGTGAGTGAGTAGTATTTTTGGCATTGCAATAtatagttacttgtcctaaac
gacttatcagtcttgtgtacCTCAGAAAGCCTAGCAGTCAACAAGACTCATTTCTAAGGCtatagttacttgtcctaaa
gacttatcagtcttgtgtacAAGATACTGCATGTCACATCGGAGATCCTAGAAGAGAAGCtatagttacttgtcctaaa
gacttatcagtcttgtgtacGTAGATCTACAGGAAGTGTGGTATCCATCTCACAGTGGGAtatagttacttgtcctaaa
gacttatcagtcttgtgtacGCAACAGTAATACCAACACGCAGGGGGAATGACTAGATGCtatagttacttgtcctaa
gacttatcagtcttgtgtacAAGCAGGAGTGGTAAAGAATACTGAGCGGAGCGGGGGACGtatagttacttgtccta
gacttatcagtcttgtgtacTGGTGGGGATCCATGCAGAGGAAGTTAGAAAAGACTGAAAtatagttacttgtcctaaa
gacttatcagtcttgtgtacAGACTTCCCCGATTCTCTTACCTTTGACCCTTGTTTCTTTtatagttacttgtcctaaac
gacttatcagtcttgtgtacTTTTCTCCCTTTGCCCTTCATGCTACCACCCCATCCCCAAtatagttacttgtcctaaa

gacttatcagtcttgtgtacTGTTGTGACAGTTTTGGGGATGAGATTGCATGAATGCCTGtatagttacttgtcctaaac
gacttatcagtcttgtgtacCAAATTTCAGAGCAGTTAAAGATGAAACTGGGCCTGGTGCtatagttacttgtcctaaa
gacttatcagtcttgtgtacACAAAAAATTAGCCAGGTACTGGGGCATGTGCCTGTAGTCtatagttacttgtcctaaa
gacttatcagtcttgtgtacAAAAAGATGAAACTGCTACCTGAGCTGGACCACAAGGACAtatagttacttgtcctaaa
gacttatcagtcttgtgtacTTCTCTTATCTCCGCACACTATAAAACTCCAATCCTGCTTtatagttacttgtcctaaac
gacttatcagtcttgtgtacGGTTTCCTGATGTACATTGGCTGGATGAAGTCATTAATCTtatagttacttgtcctaaac
gacttatcagtcttgtgtacTTTTCTTTTGACAGAGGGAGTAGTGATCATAAATTAAGGGtatagttacttgtcctaaac
gacttatcagtcttgtgtacATGGAGTGTTACAATGAGACTGAGTTAGAAAATAAGGAAGtatagttacttgtcctaaac
gacttatcagtcttgtgtacGGTTTGACTCAGGGCCAGTAGAAAAAGCAGGTGGAACTAGtatagttacttgtcctaaa
gacttatcagtcttgtgtacCTTTCAGAAAGTAGGTTTGCACCCAGAATCAGCCAGAATCtatagttacttgtcctaaa
gacttatcagtcttgtgtacACAATGTCCTCCATAGTGTCAAGCCTGTCAAAGACTTCTCtatagttacttgtcctaaac
gacttatcagtcttgtgtacTCACCAGGACTATTTTCACTTTAGACCAGTTGGCTACAAAtatagttacttgtcctaaac
gacttatcagtcttgtgtacCCCTCAGGTTTGATAATTTGGTGGAGTAACTCATAGAACTtatagttacttgtcctaaac
gacttatcagtcttgtgtacCAATGGAAGAGAAACATAGGGTAGAATCTGGGACTGGGAAtatagttacttgtcctaaa
gacttatcagtcttgtgtacAACACCAAACTTTCATCTTCTCAGGGACACAGCACCTTCCtatagttacttgtcctaaa
gacttatcagtcttgtgtacCAGAACATTGCCAACTTAGAAAACACATCCTAGCTTTAGTtatagttacttgtcctaaac
gacttatcagtcttgtgtacGAAGTTTCATTTGATGGGCATAATTGATTGAGTTGTTCCCtatagttacttgtcctaaac
gacttatcagtcttgtgtacAGCTCAGTCTCCAGCCCTTTCGTCTCCAGAGGTCAGGTTGtatagttacttgtcctaaa
gacttatcagtcttgtgtacATGTTTGCTCTTTGTGGCATGACCCCATCCTAAACCTATCtatagttacttgtcctaaac
gacttatcagtcttgtgtacGCTCCACCACATATCATCTCTTTAGCATAAACTGTCACGTtatagttacttgtcctaaac
gacttatcagtcttgtgtacTGTGAGTCACTTTGTTAGCACAGATGGTCATGTATGGTCTtatagttacttgtcctaaac
gacttatcagtcttgtgtacACCATGATGAATAACAAAGGCGCCACTATCAGTTGGGCAAtatagttacttgtcctaaa
gacttatcagtcttgtgtacAGAGGTTACGTCTAAGGAGCCAGGAAAAAGGCCAGACTTCtatagttacttgtcctaaa
gacttatcagtcttgtgtacGACCCAAAGCTCATGTATGTGTGAACTTGTCAGGTTGCAAtatagttacttgtcctaaac
gacttatcagtcttgtgtacTATGGCAAGCAGGTGGGCAGGGAGCATGAGTGGTGGGACAtatagttacttgtcctaa
gacttatcagtcttgtgtacCATGGTCCAGAACTGCATGGGCTGAACTCTGCTTCACCAAtatagttacttgtcctaaa
gacttatcagtcttgtgtacCTATTGCACATATTTGGTCAGAACAGTCCAATCTCTAGGCtatagttacttgtcctaaac
gacttatcagtcttgtgtacCCAGTTTTTACATTCAAGTAGACCACTAGTGGCTATGAATtatagttacttgtcctaaac
gacttatcagtcttgtgtacGTTGCTTAAAACAAATTCTCAATCCTTTAGAGTGAGAGAGtatagttacttgtcctaaac
gacttatcagtcttgtgtacGAGAGAGAGAGAGAGTAAGCGAGCGAGGTAGCAGGCCGGAtatagttacttgtccta
gacttatcagtcttgtgtacGGGGAAATTGTCAGGTTATCTCTAAATAATTAGCATTTGGtatagttacttgtcctaaac
gacttatcagtcttgtgtacAGACACTCTGGTTTAGGAAATAGTAATCTTGTTTTGCTCTtatagttacttgtcctaaac
gacttatcagtcttgtgtacCTAATGCAATGCAAGTGGCTAAGCTGATTTGTGTTTTCTCtatagttacttgtcctaaac
gacttatcagtcttgtgtacAGGTGCTAGGGACTGGGTAGCTTATTGGACTTCAAAGGGCtatagttacttgtcctaaa
gacttatcagtcttgtgtacAGATTCTTCTCTCCTAGGACATGCTTCTCTCAGATCACCTtatagttacttgtcctaaac
gacttatcagtcttgtgtacAGTGGCACTGAGGATAATGGGATGAAGGTAAGGAAAGATTtatagttacttgtcctaaa
gacttatcagtcttgtgtacTGGGCATTTTGTATTCCTCGGTTATATCTCAGGACTTTTAtatagttacttgtcctaaac
gacttatcagtcttgtgtacAAACTTCTTTTAAACTATCTCCAAGGTGGTTCAGTGCAAGtatagttacttgtcctaaac
gacttatcagtcttgtgtacTTCAGAAGAGCTAGGTTTGAATAAGACTGCCTCACTTACTtatagttacttgtcctaaac
gacttatcagtcttgtgtacGTCAATCCACTACACTTTCTACCTCTGTTTTCTCATCTGTtatagttacttgtcctaaac
gacttatcagtcttgtgtacGCATATAAAAATGGAACACACTGTCTTTTCATAATGGGTGtatagttacttgtcctaaac
gacttatcagtcttgtgtacGCATTTCTTTTCCTCACCTTGCCTGCTTCCTTTTACTCCAtatagttacttgtcctaaac
gacttatcagtcttgtgtacAGAAATCTGCATTGTCTCTAAAAGTCTAATTCACCCTGCAtatagttacttgtcctaaac
gacttatcagtcttgtgtacAAATTCTGCTTAATTGCTTCCTTCTACCATTGCTCTTGGCtatagttacttgtcctaaac
gacttatcagtcttgtgtacCTGTATAGCTCAGTTTGAAATTTAACCCTATTTGCCCCTTtatagttacttgtcctaaac
gacttatcagtcttgtgtacTTGGGGGGTAAGCTGTTTCTGAAGAATTGTGCAATTTAGGtatagttacttgtcctaaac
gacttatcagtcttgtgtacGCTTAGACTGCTTAGTACTTAGCAATAATGGAGAAAATGCtatagttacttgtcctaaac
gacttatcagtcttgtgtacCCCACCAGGTAATTGTGTATTGAATATGTCTTGGCTCAAAtatagttacttgtcctaaac
gacttatcagtcttgtgtacTCGCTTAAATGAACTGAGGACAAAACTGCTTGTTGTAAGTtatagttacttgtcctaaac
gacttatcagtcttgtgtacAGTGATCTGGGTAATTAAAGATGCAAACTAGCAGCACAAGtatagttacttgtcctaaac
gacttatcagtcttgtgtacCGAGGTGCGGGGGAGGGCAAACCATTCCTGTTTAATGAACtatagttacttgtcctaaa
gacttatcagtcttgtgtacGGCACCAGGAAAGAGAGGCAGTCATTCAAACCTAGCTGTAtatagttacttgtcctaaa

gacttatcagtcttgtgtacAAGATTAGAGATATTGGTTGTCCTGGGTTCACAACTGCTCtatagttacttgtcctaaac
gacttatcagtcttgtgtacGGTAAAGAATTTGAAGGAGATAAGTACAAAACCACAGAGTtatagttacttgtcctaaac
gacttatcagtcttgtgtacGTGCTTTAGTCACTTGATCCTGCTAATCTTTGTTCTCTAAtatagttacttgtcctaaac
gacttatcagtcttgtgtacATGACTGTCATTGTGTAATCCCCACATATTTGCACCCTATtatagttacttgtcctaaac
gacttatcagtcttgtgtacCAGTAATCAGAAATAGGCAGCTGGGGAAGGCAGGAGAGAGtatagttacttgtcctaa
gacttatcagtcttgtgtacCTCTCAATTTCCCGGCACATTGTGTTTTTGTGTTGCAGCAtatagttacttgtcctaaac
gacttatcagtcttgtgtacATTCATTGAATAGCACCCTAGTGGCTTGAAAGAGTTAAAGtatagttacttgtcctaaac
gacttatcagtcttgtgtacTCCAGAAGTGCCTAACCTTCTCCCAGGGAGTAGCTTAAGTtatagttacttgtcctaaa
gacttatcagtcttgtgtacGAGAATCAACTGATAAGGGTCAAACTCCACAAAACAAAATtatagttacttgtcctaaac
gacttatcagtcttgtgtacAGACCATTTAATCCAGTACCTTCTCCATAAATCTCTTTTCtatagttacttgtcctaaac
gacttatcagtcttgtgtacCCTTTGAGGTCAGGTTAATATCCATTTATTCCTTTCATTCtatagttacttgtcctaaac
gacttatcagtcttgtgtacACCTACTGTGCACGGCTCTGTAATAAATAAGAAACAGTCTtatagttacttgtcctaaac
gacttatcagtcttgtgtacCTGGAAGAGAGAGATGGGATGAGATGGAAATATATAACCAtatagttacttgtcctaaa
gacttatcagtcttgtgtacGAGAATCTCCTCCAGAAGTCCTCACTAATATAACCATGTAtatagttacttgtcctaaac
gacttatcagtcttgtgtacTGAATATTTTAAAGAGGTAATAGCAGGAGCTGGCTACAGAtatagttacttgtcctaaac
gacttatcagtcttgtgtacATTACGCAGGCAAAATTTAAAATTTGTGTGTATGGGACACtatagttacttgtcctaaac
gacttatcagtcttgtgtacATCTATCTCTATCTCTATCAACATATTTGGGCATCCTAGAtatagttacttgtcctaaac
gacttatcagtcttgtgtacAAATCTGTACATTTTTCTCTATCCTTACTGCCACTACCTTtatagttacttgtcctaaac
gacttatcagtcttgtgtacCACTTGCTCCTGTCTTGCTACATTCCAGTCTTACTCCTTTtatagttacttgtcctaaac
gacttatcagtcttgtgtacTCTATTCTTCAAATTGAAACCAGAGTGTGCCTTGCTTAAGtatagttacttgtcctaaac
gacttatcagtcttgtgtacCCTAGTTGTGCACAGGATAAATAAAGTCTTAACTCCTTAGtatagttacttgtcctaaac
gacttatcagtcttgtgtacTCTCCTGATGCTCCTCTATTGAATTCCCTGCCCTAAGACTtatagttacttgtcctaaac
gacttatcagtcttgtgtacTGCACCATGTTCTATCTTGCCTCCTAGGTTTTGTGCGTCTtatagttacttgtcctaaac
gacttatcagtcttgtgtacGAGTATCCTGTTCCAGCCAAGCCCAAGCCCCACGCCCTTTtatagttacttgtcctaa
gacttatcagtcttgtgtacCTCAACTAAGTTCCATGAGCTAAAACATCTCTTCATTCAAtatagttacttgtcctaaac
gacttatcagtcttgtgtacTTCTTTCTTCCAAAACACAGTGTGCTTCGCATCATTTCTTtatagttacttgtcctaaac
gacttatcagtcttgtgtacCATCTCGGCTCTCTACAACCCCCGCTGCCCGGGTTGAAGCtatagttacttgtcctaa
gacttatcagtcttgtgtacCACCCAGCCTATTTCTTTGTCCTCTCATAGCATCCGTTGTtatagttacttgtcctaaac
gacttatcagtcttgtgtacCATCTATCTTTCTAGTTGTTTGATGGCAAGTTCTAAGTCTtatagttacttgtcctaaac
gacttatcagtcttgtgtacCTGAAAGTAAGATAAATGCTCAATAAAGGGTGAGTTCAGTtatagttacttgtcctaaac
gacttatcagtcttgtgtacAATTTTATTGATGGGTGAATTGGAAATAGAAGGGAAGAGCtatagttacttgtcctaaac
gacttatcagtcttgtgtacCCTTTGGTGAGTCAAGTCCTTGTTTCTAAAAACACTAAACtatagttacttgtcctaaac
gacttatcagtcttgtgtacAAAATTAGGGTGATCAACTGTCCGGATTTTCCTGGGACTGtatagttacttgtcctaaac
gacttatcagtcttgtgtacCAGGAAAGCTCTCACTCTTGATTTAGCTCTGAAAGTCTCAtatagttacttgtcctaaac
gacttatcagtcttgtgtacGCAGTTGTTTGCCCTAGAAGTCTTATGTCTTAGAGGTTATtatagttacttgtcctaaac
gacttatcagtcttgtgtacAGGGGATGTAGCTGAATGAGAATCTAAAGGGAGTGTGAGCtatagttacttgtcctaaa
gacttatcagtcttgtgtacAGGTAGAAGGCTTGGCAATATCCTCACAGTAGTCCAGTGTtatagttacttgtcctaaa
gacttatcagtcttgtgtacGCTGATACTCAAATCCACGATGGAACATCTGCAAAGCTCCtatagttacttgtcctaaa
gacttatcagtcttgtgtacAGTCCTGGGCTGGAGTTGGAACACCTTGGGAAATCTAGTTtatagttacttgtcctaaa
gacttatcagtcttgtgtacTCTCCAAAGCTAGTTGACTTTAGCCTCAGTGTCTTCATCTtatagttacttgtcctaaac
gacttatcagtcttgtgtacCCTGTATGCATCCCAGGTTTGTTGGAAATTTGAAAGGTCAtatagttacttgtcctaaac
gacttatcagtcttgtgtacGTGTGTGTGAGAGTAGTTTGAAAGCCATAAGTGCTCTGCAtatagttacttgtcctaaac
gacttatcagtcttgtgtacAACACACGCTGAGTTTTTCCATTAAAAGTTATAGTCAGGGtatagttacttgtcctaaac
gacttatcagtcttgtgtacGAAGTGTCTGTTCTCAGGGATATAAAAAGGCCTGGCCGGGtatagttacttgtcctaaa
gacttatcagtcttgtgtacGATGAAAAAAATATGGTACATTTCCACCATGGAGTACTACtatagttacttgtcctaaac
gacttatcagtcttgtgtacATAATGTCTTTTGCAGAACTTGGGTGGAGCTAGAGGCCATtatagttacttgtcctaaac
gacttatcagtcttgtgtacTGTTCTCATTTATAAGTGAGGTAACCTGTGGGCCTACAATtatagttacttgtcctaaac
gacttatcagtcttgtgtacACAGAGTGATATAACAGATGTTGGAGACTCAGAAGAGGGAtatagttacttgtcctaaa
gacttatcagtcttgtgtacGGAAGGGGGTGAGGCATAAGAAACTACATATTGGGTACAAtatagttacttgtcctaaa
gacttatcagtcttgtgtacAGCCTGTGCAAAAACATAAAGCAAAATCTATAAGCACATGtatagttacttgtcctaaac
gacttatcagtcttgtgtacTTCGAATGCTGGAAGTTCTTGGCTGTGACATAGGGATAGCtatagttacttgtcctaaac
gacttatcagtcttgtgtacAAGCAGTGTCTTTCTAACGTTACAGCCTAGGATCATCTCTtatagttacttgtcctaaac

gacttatcagtcttgtgtacAAGTTGGCTTCCTACCATATACAAGATGTGTTGAATTTTGtatagttacttgtcctaaac
gacttatcagtcttgtgtacGTTGCATTGCTATTTTATTCTCTTTCTGCAAAATCACAGGtatagttacttgtcctaaac
gacttatcagtcttgtgtacCTATTGCAGGAAATCTATGTCATCCCCCCTCACTAGTTCAtatagttacttgtcctaaac
gacttatcagtcttgtgtacTAATGTGTTTCAACCAAAGAAAATTACTGACCCCATGATCtatagttacttgtcctaaac
gacttatcagtcttgtgtacGCTTTTCTTTGCTTAGCATGTCTGGAGTTTCAGGAGCACTtatagttacttgtcctaaac
gacttatcagtcttgtgtacGCTGAATTGCATACTCTCACTGGTTTATGGTAATAAGAAGtatagttacttgtcctaaac
gacttatcagtcttgtgtacGGATAGCCTTATTTCTTCTACTACTCTGCTTGTCACACGCtatagttacttgtcctaaac
gacttatcagtcttgtgtacTGATTCTGACTGTCTTGTTGGCCAGATTAGACTAGCTAGAtatagttacttgtcctaaac
gacttatcagtcttgtgtacACACACACACACATTTGCTTCCATTAAGGAAGTCATATCAtatagttacttgtcctaaac
gacttatcagtcttgtgtacGGTGGATTTATGGTATGTGCTTTCTCACCTTAGTGAAATAtatagttacttgtcctaaac
gacttatcagtcttgtgtacCCTTGTGTAATGTTCATGGATCCTCAGAAGCACCAAATTGtatagttacttgtcctaaac
gacttatcagtcttgtgtacTCATCTTTTTACCTCCATTCCCCACTCTGTAGAGTCAGTGtatagttacttgtcctaaac
gacttatcagtcttgtgtacCAGTTTGATCCTGTATGACTCTATAAGGGTTATATGACTTtatagttacttgtcctaaac
gacttatcagtcttgtgtacAGTTCCTAGATGAAGTTACATTTGGCTTGGGATGATCTGAtatagttacttgtcctaaac
gacttatcagtcttgtgtacCAGGAGGATGTATGAGACCAATGCCATATTTACGGAGCTGtatagttacttgtcctaaa
gacttatcagtcttgtgtacCTTTTTGAAGTTTCTGGTTAGGTTTTGTAGAAGGATTGGTtatagttacttgtcctaaac
gacttatcagtcttgtgtacTGCTGGGTACTGTGCACATCTGGAAATGAGATCAAACGGCtatagttacttgtcctaaa
gacttatcagtcttgtgtacCTTTGGAAAGTTTAACATCTAGGGGGCTACTAGCCTTGCAtatagttacttgtcctaaac
gacttatcagtcttgtgtacATAGAAAGAGGATAGATGTAGAAGAATCACAGTTTGCTTGtatagttacttgtcctaaac
gacttatcagtcttgtgtacGCGGTGTTAGGGAAATTTTTTTTGGTCAGAATAGAATTTGtatagttacttgtcctaaac
gacttatcagtcttgtgtacACAAACAAAAAAGAACAGAACACTTCAGGAGGAGGAAAATtatagttacttgtcctaaa
gacttatcagtcttgtgtacGGTCTGGAGGCAAAAGAGAGCATGGTTGTTTAAAGCACGGtatagttacttgtcctaaa
gacttatcagtcttgtgtacTGGGGGCACTTGGAGGGGAGATGTTCTGGGCTAGATGGGAtatagttacttgtcctaa
gacttatcagtcttgtgtacCAGCCAGATTATGAAGAACATTGGTTGCCAAGCAAAGGAAtatagttacttgtcctaaa
gacttatcagtcttgtgtacATTTTTATCATAGAAGTTGTGACAAATGGCAGTTGGGGAGtatagttacttgtcctaaac
gacttatcagtcttgtgtacATTACACATCTCCAATACCCTATCTGTAACCTCCTCTGAGtatagttacttgtcctaaac
gacttatcagtcttgtgtacGGTGTGCCCTTGAGAATATGTCAAATGTAAGTGGAAAAGAtatagttacttgtcctaaac
gacttatcagtcttgtgtacTGAACTATTGAACACATTTGAGCAGGGGCACACTGGATCAtatagttacttgtcctaaa
gacttatcagtcttgtgtacATCTAGTGATGTTTGAGGTAGGATTTCATTCAGGTTGACTtatagttacttgtcctaaac
gacttatcagtcttgtgtacTGAAGGGGAGACTAGAAGCCACAGAGTAGATAAGGATTTAtatagttacttgtcctaaa
gacttatcagtcttgtgtacCATAGAATGGAAAGGGAACAAGATCTTAAGAGGAAATGTAtatagttacttgtcctaaac
gacttatcagtcttgtgtacTGGGCATCTAGTAGAAGTTACACAATTTTTGGTTAAAGAGtatagttacttgtcctaaac
gacttatcagtcttgtgtacTGGAGAGTTATAGCAGAGTTCAGAGGGAGTGATAGTAGGAtatagttacttgtcctaaa
gacttatcagtcttgtgtacTCATGTTGATTTTGACCTGAATGATCTAGTTGGAGGGTTGtatagttacttgtcctaaac
gacttatcagtcttgtgtacCCCATTGGAGCTGAATTTGGTGTAACATAGTTCCCACTTGtatagttacttgtcctaaac
gacttatcagtcttgtgtacGTGGCATTAGTTTTTAGAAACATCAGAAGTTCCAAGGGCAtatagttacttgtcctaaac
gacttatcagtcttgtgtacCATTCTTAAAATTTACAATCTTTTGGGTCTAGGAGAGCTCtatagttacttgtcctaaac
gacttatcagtcttgtgtacGCATTAAATACACAAAGGGTATGGCCATCAAACGGGAAGCtatagttacttgtcctaaa
gacttatcagtcttgtgtacCAACTCCTTGTCCAGCGGGCTTTATCTTAGACCTGCCACAtatagttacttgtcctaaa
gacttatcagtcttgtgtacTCTTGACCTAGCAGTTCTGTGATTTCATTACTTCAAACCTtatagttacttgtcctaaac
gacttatcagtcttgtgtacACATTTTGTCAAAACTCAACTCCCACTGTGGCCTTCAATTtatagttacttgtcctaaac
gacttatcagtcttgtgtacGAATCTTCTTCTTTCTATTTCTTGATACCTTGGAAGTGTCtatagttacttgtcctaaac
gacttatcagtcttgtgtacAACAATAACTGGTGCACTGGAGATTCAACATGTAACTCCAtatagttacttgtcctaaac
gacttatcagtcttgtgtacCTTTTTCAGTGTTTTGCATACCAAGAATATGGACCCTAAAtatagttacttgtcctaaac
gacttatcagtcttgtgtacTTTATATTTTCCTTGGACACATACCATGCTCTACACATCGtatagttacttgtcctaaac
gacttatcagtcttgtgtacAATTCTTGATTAAACTTCTTGAAGTCCTTGAACCAGCCTTtatagttacttgtcctaaac
gacttatcagtcttgtgtacTAAAATGACCTAAGTGGAAGGACCTAGTCTTGTGCCTGCCtatagttacttgtcctaaa
gacttatcagtcttgtgtacGCCACCTCCCCCAGACCACCAAAACCCTCATCCCAAAATTtatagttacttgtcctaa
gacttatcagtcttgtgtacAAATTTTGAAAGTGGGCAAATAATACTCTGCTCTGTCTTGtatagttacttgtcctaaac
gacttatcagtcttgtgtacGTTTACTTTCTGAAAAACAATCTGGCTGTTGAGTTCTGATtatagttacttgtcctaaac
gacttatcagtcttgtgtacCAAGGATGAGATATTAGAGGAAAAGATAACGCTCGAAACCtatagttacttgtcctaaa
gacttatcagtcttgtgtacCAACTATAAAACTGGGGAAAGGTGAGTAATAGATTTGACAtatagttacttgtcctaaac

gacttatcagtcttgtgtacTGAGATATAAATTATAACCAGGCATTTCCTATCAGCACCCtatagttacttgtcctaaac
gacttatcagtcttgtgtacGAGTCTTTGATGACAGTTTACTCTCAAACATTATGCTTAGtatagttacttgtcctaaac
gacttatcagtcttgtgtacATCTGTTCTGATTTTCCCTTTTCATGAGACTCACATTTTCtatagttacttgtcctaaac
gacttatcagtcttgtgtacTTCTTTTTTCCTTCCCTAGACTAAACCCTTAAACACTACAtatagttacttgtcctaaac
gacttatcagtcttgtgtacGTGATAACTTGCGGGGAAAGAGTTCAGTGAATTTCAAGAAtatagttacttgtcctaaac
gacttatcagtcttgtgtacATGAGGAGTCTAAAGAGGAGGCATGTTGTGTTAAAGTATTtatagttacttgtcctaaac
gacttatcagtcttgtgtacCAGGTTGGTGGCTGTAGGAAAGTTATGACATACAATAGCAtatagttacttgtcctaaac
gacttatcagtcttgtgtacAGGGCGGGGAGAAGGGCCTTTAACCATGTGAAGGGGTGGCtatagttacttgtcctaa
gacttatcagtcttgtgtacAGGTCACATCTCATTCTCTCCTGGAAAGAACCCCACATTGtatagttacttgtcctaaa
gacttatcagtcttgtgtacCTGTTGACAGATTGACTGCAGCTGGAGAGGGCACGCCAAGtatagttacttgtcctaa
gacttatcagtcttgtgtacCATGCAGTTCAAGTGTACACAGCTGTTGGATCTGTTGGCTtatagttacttgtcctaaac
gacttatcagtcttgtgtacACGGCAGGGAATGTGACTTTGGGTAGCAGGTGTGGGATTTtatagttacttgtcctaaa
gacttatcagtcttgtgtacTAGAACTGCTGGTCTATAGAAAATGTGGTCTTTCAGGAAGtatagttacttgtcctaaac
gacttatcagtcttgtgtacCTTTGAAGACAGGTATTTTCCAAAGGTGGTAGTATTAGAAtatagttacttgtcctaaac
gacttatcagtcttgtgtacAAGCATTCTTATCCCTTCTATTCTATGTTCTCTGTCCCATtatagttacttgtcctaaac
gacttatcagtcttgtgtacACCCCTTACACCTGCCCTACCTATCCAAGAGCCACATGTTtatagttacttgtcctaaa
gacttatcagtcttgtgtacGGCTGCCACCTGTGCCCCATTTGTTCTTGTTCTGACTTCAtatagttacttgtcctaaac
gacttatcagtcttgtgtacCCAACTCTGAAATCTTGGCCCTAACATTTCCTGCACTGGGtatagttacttgtcctaaa
gacttatcagtcttgtgtacGCTTGCCTCTCTGCCTCTTTGTACACGTGCTACTCAGAATtatagttacttgtcctaaac
gacttatcagtcttgtgtacTGGATTAGGAGCCTAACCTTGTCAATGACTATTTCACTAGtatagttacttgtcctaaac
gacttatcagtcttgtgtacGTGGACAATCAGGCACATACCTCGTAAATAAAACATTCAAtatagttacttgtcctaaac
gacttatcagtcttgtgtacACAATAACATCACTATCTATTTAGGCTCAACTATGCACCAtatagttacttgtcctaaac
gacttatcagtcttgtgtacACATATATTTCTTCCTAGGGCTTACTATATAGCCTTGTGAtatagttacttgtcctaaac
gacttatcagtcttgtgtacTCCTCATTTTACAACTAAGGGAAACTGAGTTTTTGAGATGtatagttacttgtcctaaac
gacttatcagtcttgtgtacGAGCAGAGATTCAAATCTAAACCTCGATATTCTTTCAGGAtatagttacttgtcctaaac
gacttatcagtcttgtgtacAGGAACTCTGATTTCTCATGCCCCCATCCATCTTACTGTCtatagttacttgtcctaaac
gacttatcagtcttgtgtacGAGGTCTCAGCTAGTACATTTATCTTTCTTTTTCATGAGGtatagttacttgtcctaaac
gacttatcagtcttgtgtacTGATCATTCTTGCCTGAAAGCTGATTGATTACAAGTGTTAtatagttacttgtcctaaac
gacttatcagtcttgtgtacAATGGAAAGAAAGCATTGGAATTCTCTGAGGCCTGAGTCAtatagttacttgtcctaaac
gacttatcagtcttgtgtacGGTTATCCTTGAGAAATTCATGAGTTACTAGTTTTGGTGTtatagttacttgtcctaaac
gacttatcagtcttgtgtacACTGACAGTGGAAGTCTGTGGCATGCACTTAAATTATTGCtatagttacttgtcctaaac
gacttatcagtcttgtgtacAGGAGTGGTTGTCAGGGAGAATTATGAAAGCAAGAGAGCGtatagttacttgtcctaaa
gacttatcagtcttgtgtacGACATTAAAGGGTCTTAAAAGGGTCATGGACAGAAGGCAAtatagttacttgtcctaaa
gacttatcagtcttgtgtacCTGAGTGAGGGCTTGGAACCAGCAAATGAGAGGCACATATtatagttacttgtcctaaa
gacttatcagtcttgtgtacGATCAAGTGGTTATTGCTCAGACTGGGTTAGAAAATCTTTtatagttacttgtcctaaac
gacttatcagtcttgtgtacCTGTCTGTCAGTCTTTGGGGCTTTGGAAATACAATATTACtatagttacttgtcctaaac
gacttatcagtcttgtgtacATAGGAGATGAAAAATAATGGATGGGCTACCAAATGCCTTtatagttacttgtcctaaac
gacttatcagtcttgtgtacTGTATAGCCTTTCTCCAAGGTTTCAAAAAACAACTTGTCCtatagttacttgtcctaaac
gacttatcagtcttgtgtacACCAGGAGAAAGCACGTGGAATGGTTCTAAATGAAGATATtatagttacttgtcctaaac
gacttatcagtcttgtgtacAGCTAAACCTTGCTAATTTGCACTAATGACTGGGAAGCCTtatagttacttgtcctaaac
gacttatcagtcttgtgtacGATTTCAAAAAACCCAAAGACAGCACCAGAAAATGAACCTtatagttacttgtcctaaa
gacttatcagtcttgtgtacATGTATGTGCATTTTACAGAGCAATGGTGAGCCTTTAAAAtatagttacttgtcctaaac
gacttatcagtcttgtgtacCTGTGTTCATTTCCTTTCGTAGTTATCAAACAGGATATACtatagttacttgtcctaaac
gacttatcagtcttgtgtacATTGCACAAAAGACCCATAAGCATTAGTGAAATCTCTCAAtatagttacttgtcctaaac
gacttatcagtcttgtgtacCAATTATGTGAAAAATGGAAATCGACAGAGTGGGAGTGAGtatagttacttgtcctaaac
gacttatcagtcttgtgtacGCCTCGTAGTTTTGGTAGCAGTGGCAGCCCCGGTCTAAATtatagttacttgtcctaaa
gacttatcagtcttgtgtacAGCAATGACCTACCCACCAATGTGTTGTGTTTTCTTTTCTtatagttacttgtcctaaac
gacttatcagtcttgtgtacACCTTCACTTTCATACCTAATTTCGTATTTGCAACTTTGCtatagttacttgtcctaaac
gacttatcagtcttgtgtacAAAAATTGAATCCTCCCTGATCATTGGAGCTAAGTGTCCAtatagttacttgtcctaaac
gacttatcagtcttgtgtacATTTGAGGAGAGTTATTGGAGTGGGAAAAAAGAATCGAGGtatagttacttgtcctaaac
gacttatcagtcttgtgtacGGACTAATACTCTAAAAATGGAGAAGGAATGAACTCTGGGtatagttacttgtcctaaac
gacttatcagtcttgtgtacGGATGTACAAATCTGTCCTTGTCCTCCTCCCTCTTCCCTAtatagttacttgtcctaaac

gacttatcagtcttgtgtacCTTTTCTGGGAGAGAAGAGGGCCCCTGCCTTCCTTGACCCtatagttacttgtcctaaa
gacttatcagtcttgtgtacCAGCTCTGCTGAAGATGAGGGCTAGGCAGACAACAGGAGTtatagttacttgtcctaa
gacttatcagtcttgtgtacAAGTTTTTAAAATTACCCCTCTCACCTTTGTTCTCTCCTTtatagttacttgtcctaaac
gacttatcagtcttgtgtacAAGTTTGGTTTTCTTACCTCACATAGTAAGTCAAGTTAGGtatagttacttgtcctaaac
gacttatcagtcttgtgtacCCAGATGTTCTTGGTCCGCTGTTGGTCAATTCCAGATAGAtatagttacttgtcctaaac
gacttatcagtcttgtgtacCTGGTAAAGTCCAGCTCTGAAGAAGGCAGGCTTGAGGCTGtatagttacttgtcctaaa
gacttatcagtcttgtgtacTGTGCACATCCTCTTGCCTATGGTAGTAAAGATTTAGGGCtatagttacttgtcctaaac
gacttatcagtcttgtgtacCAGATTGAGCTGGAATGAGGACCACTAAAAAGACTAGAAAtatagttacttgtcctaaa
gacttatcagtcttgtgtacTCCACCAGCTAACACCACCTTTTGCCTTTAAACCACTTATtatagttacttgtcctaaac
gacttatcagtcttgtgtacTGCATTAAGAGCCCATGAATTTCAAAGACAGCCCTGGACAtatagttacttgtcctaaa
gacttatcagtcttgtgtacGAAAATGTGCATCCCAGGTGTAGAGCTTAGAGTTGAGAATtatagttacttgtcctaaac
gacttatcagtcttgtgtacCCCTCTGGAAGCAAAAATAGAAACATGAAAGAAGCACAGAtatagttacttgtcctaaa
gacttatcagtcttgtgtacTGATGGCTAACTTTAACTTTACCTAAACTTTCAACTGCACtatagttacttgtcctaaac
gacttatcagtcttgtgtacCTTCAAAGCTTCAAGCATTTTGGAGTGCTGGGCTTGTTGGtatagttacttgtcctaaac
gacttatcagtcttgtgtacACTTTGTGAGTTGTTTCCCTGGAGACTTGATGGTAATGGTtatagttacttgtcctaaac
gacttatcagtcttgtgtacTCGCTCTTACAAGGATGCAGTTGGTGTGATTGGGTAAAAAtatagttacttgtcctaaac
gacttatcagtcttgtgtacTGTAGAACTTAAAGCATCAGCAGAATACACTGGAAACACTtatagttacttgtcctaaac
gacttatcagtcttgtgtacCAGCTCAGATATTATTACTTCTTAAACTTTCTTCTGCAGGtatagttacttgtcctaaac
gacttatcagtcttgtgtacAATGTCCTTTTTCAATCTCTCTAAAACATTCTCCAAGGAGtatagttacttgtcctaaac
gacttatcagtcttgtgtacGTCTCAAATTTTATTGTGTTTTTAAGAGCCCACTAGATGCtatagttacttgtcctaaac
gacttatcagtcttgtgtacCCTTAGGAAGAACAATCAGAGTAATCCTTCTTTTATTCAGtatagttacttgtcctaaac
gacttatcagtcttgtgtacCCAATGCCCTTAGCTGGTTATAATGAAATTTTCACAAAAGtatagttacttgtcctaaac
gacttatcagtcttgtgtacCATGAACAAATAGACATGGTGAATTCCCTTACACAGCTGAtatagttacttgtcctaaac
gacttatcagtcttgtgtacCAGCTATTGGAATCCCTCTGGACCATGACAATTTGATCCCtatagttacttgtcctaaac
gacttatcagtcttgtgtacTTGTTGAATTAAAATGTCCAGCTAAAAAGCAACACTCTGGtatagttacttgtcctaaac
gacttatcagtcttgtgtacCAGTCACATATTGAAAGGAAAAAGAGGCTAATGGATGTTTtatagttacttgtcctaaac
gacttatcagtcttgtgtacAGGGGACTTTGTTGTTTCATTATACTGTACTCCATATTCAtatagttacttgtcctaaac
gacttatcagtcttgtgtacTGTTAGAGTCTGACACCTCCTCGATCTGTTTCTTCATTTTtatagttacttgtcctaaac
gacttatcagtcttgtgtacCCCATTTCTAGAAGTGTTTTTAAAAAGAAGGGAAGAGTTGtatagttacttgtcctaaac
gacttatcagtcttgtgtacTGCAGGTATCTTCCATTGAAAGATTTCTTTATTCAGCTTCtatagttacttgtcctaaac
gacttatcagtcttgtgtacACTTGGAGTTGCCATTTTTTTGATCACTGAATTTCTGAACtatagttacttgtcctaaac
gacttatcagtcttgtgtacGATCTCAACAAGGGCAAATTTTAGCTAAGATAATAACGACtatagttacttgtcctaaac
gacttatcagtcttgtgtacCATCAAGATGTAGAATTTTGGTGTGTATGTTACGGTTAACtatagttacttgtcctaaac
gacttatcagtcttgtgtacTGCCCACATAAAATCTGGAGAAACATTGAAGGCCATTTAAtatagttacttgtcctaaac
gacttatcagtcttgtgtacACTGGCCTGCTTTCAAGTGGTTTATAACTGACATAATTATtatagttacttgtcctaaac
gacttatcagtcttgtgtacCATGAGGTAGTAAGAGAATGTGAAGTCTGACAGTATGGACtatagttacttgtcctaaac
gacttatcagtcttgtgtacGGCTTCCTGGGTTGTGCTAGAAAAGTGAGGTATAAATAAGtatagttacttgtcctaaac
gacttatcagtcttgtgtacGAGTAGAGGTATGTATTGGTCTATAAGAGATATATAAGGGtatagttacttgtcctaaac
gacttatcagtcttgtgtacAGGGAGAGACAGCACTGCAGGTAGAGGGACTGACACAATTtatagttacttgtcctaa
gacttatcagtcttgtgtacGCTGGGGTTAGCTGGGAAGGTCTGGGAACTGCTCGGTCAAtatagttacttgtcctaa
gacttatcagtcttgtgtacAGTAGTGGGAAATAACAACAGAGGGTGTTTGCTCTTGCTTtatagttacttgtcctaaac
gacttatcagtcttgtgtacCTTTTAATGGTGATGTTAGGGCATGCTGCTATAAAGACACtatagttacttgtcctaaac
gacttatcagtcttgtgtacCGAGTTAGTGGGTGCAGTGCTCCAGCACGGCACATGTATAtatagttacttgtcctaaa
gacttatcagtcttgtgtacAAAAACAACAACAGAGGGGAGACTAGTAAAATGCCCAATGtatagttacttgtcctaaa
gacttatcagtcttgtgtacCAAACTAGGATTTGCAACTGTGGTTTCAACAAGTCAGGAAtatagttacttgtcctaaac
gacttatcagtcttgtgtacAGGAATTAACCCTGTTGTCTCAGTAACTGACTTGAGTTTAtatagttacttgtcctaaac
gacttatcagtcttgtgtacAAAGACCGGATGACTAAAGAGTACTACCTAATGATCTTTAtatagttacttgtcctaaac
gacttatcagtcttgtgtacACATGTTTTCTGCTCAAAGTGTGTCTGTACATCAGCCATTtatagttacttgtcctaaac
gacttatcagtcttgtgtacTGGTTCTCAGTGGCATGTCTTTTTGTCTGGGGGTAACACAtatagttacttgtcctaaac
gacttatcagtcttgtgtacATGCAAATCCATTTATATACATATACAGTCGCCCTTCAGTtatagttacttgtcctaaac
gacttatcagtcttgtgtacTAGTATTTGCATATAACTTGCTCACATCCTCCCATGCACTtatagttacttgtcctaaac
gacttatcagtcttgtgtacTTTTCCTGAATATTTTTAATACGTAGTTGGTTGAGGCCACtatagttacttgtcctaaac

gacttatcagtcttgtgtacCTGCATATGTATTTTGGACAGAGAGTTTAAATAGTGGTTGtatagttacttgtcctaaac
gacttatcagtcttgtgtacGATATACATCTCCTCTTGTTAGTTATAGGTGCTTTTGATGtatagttacttgtcctaaac
gacttatcagtcttgtgtacAATGGACCAAATATGCCACAGTCTGTTCATTTCTTACAAAtatagttacttgtcctaaac
gacttatcagtcttgtgtacATAATTAACCTATATTCAAAGACTGCCTGCAAAAACCACCtatagttacttgtcctaaac
gacttatcagtcttgtgtacTTTTTATTTTTTGCAGGTGAATTCTGTGAGCAGGTGTCAAtatagttacttgtcctaaac
gacttatcagtcttgtgtacGGCCAAAGCCAAGAACACACATTAAAATAGTACTATAGTAtatagttacttgtcctaaac
gacttatcagtcttgtgtacGAACAACTACATCTAATCTTGCTTTCACAATGTACTACTGtatagttacttgtcctaaac
gacttatcagtcttgtgtacCTGTGAGACACATTGATGACAGGTTCGGATTTTAGAAAACtatagttacttgtcctaaac
gacttatcagtcttgtgtacGTATTTTACTTATTCTTCCCTCTGCTTTACCTTCCAACATtatagttacttgtcctaaac
gacttatcagtcttgtgtacTTCATATCCTTGCTAACCTTTGCAAATGATTTCCCCAAAGtatagttacttgtcctaaac
gacttatcagtcttgtgtacAACTAACCCATTGTCATCATCTTTTAAGAGGTAATTAGCCtatagttacttgtcctaaac
gacttatcagtcttgtgtacTTGCTCCATCCTGACATCTTTCTAACTGAATTGATTTTCTtatagttacttgtcctaaac
gacttatcagtcttgtgtacAATGCTATCACCCTTTCATTTTCCCATTTCTTACTGTGCTtatagttacttgtcctaaac
gacttatcagtcttgtgtacGGATTACCCTAGGCTTGTCGAGTTTAATGGCATGTGAACAtatagttacttgtcctaaac
gacttatcagtcttgtgtacATTCTATTCCAAAGCAGAAGAATGGGTTTTTCTATCCAGCtatagttacttgtcctaaac
gacttatcagtcttgtgtacTGGAATATTTTACTTAATGGGTTATTTTCATGGCAGTCCCtatagttacttgtcctaaac
gacttatcagtcttgtgtacATTTTTCCCCAGGATTGTTCTTGAAATTATCTTTTGGCTGtatagttacttgtcctaaac
gacttatcagtcttgtgtacATATACACATACCTTTTTTTTCTGGTCATGAGTTGGTTGCtatagttacttgtcctaaac
gacttatcagtcttgtgtacCAAATACAATAATTGGAGGACGGCCATGAAAAAGAAGAAGtatagttacttgtcctaaa
gacttatcagtcttgtgtacGACCATGTCTGTATAGTTCCTGTTGATAACACCCAGATGGtatagttacttgtcctaaac
gacttatcagtcttgtgtacTCTACTAAGGAGCTGCCCTGGAAAATTATTCAAAAGCTTTtatagttacttgtcctaaac
gacttatcagtcttgtgtacCTTTTGCTCGATAGTCTCCATTAGTCACTGATTTGCTTTTtatagttacttgtcctaaac
gacttatcagtcttgtgtacCTGGTTATCTTGGCAGTCATGTTTCTACCCGCTTTAAAGGtatagttacttgtcctaaac
gacttatcagtcttgtgtacGAGACTCCTTGATAGAATTATATATGATGAGAAGCCAGTCtatagttacttgtcctaaac
gacttatcagtcttgtgtacAACCTTAATGTTGTGTGACTAGATTTTGGCAGTAAACAGGtatagttacttgtcctaaac
gacttatcagtcttgtgtacTGTTCTGCCAAGATTTTACTATCAGATACACTCTCCTGTTtatagttacttgtcctaaac
gacttatcagtcttgtgtacTTAATGAGACAAGGTCTTTTTTCCCCTTTTGCTATATTGCtatagttacttgtcctaaac
gacttatcagtcttgtgtacCAGTTTCTTCCTGGTATTTAAAATCTGAACTATGTCTGGTtatagttacttgtcctaaac
gacttatcagtcttgtgtacTTTGAGAAGACTTTGTAGCTTGGATACTGATCTTATATGGtatagttacttgtcctaaac
gacttatcagtcttgtgtacCACAGTTCATTAGTAGAGTGTTGAAAGCCTGGGATATACTtatagttacttgtcctaaac
gacttatcagtcttgtgtacTTTGAAACCAACTCAGCCATAAAATCACTCTCTTTCTTGGtatagttacttgtcctaaac
gacttatcagtcttgtgtacGTAATCTCGTCCTACTTCAATTTCCTGAGTTAGAATATAGtatagttacttgtcctaaac
gacttatcagtcttgtgtacTTGCCTTCTATGGAGAATGATAAAGATTAATCACTGGAAGtatagttacttgtcctaaac
gacttatcagtcttgtgtacGGTCTCTCCCTTAGTAATCAGAAAAATCTCCTCGGTAATCtatagttacttgtcctaaac
gacttatcagtcttgtgtacGCAGCTCAATTTGACCTTCTACTATTTTGATCATCTAGTTtatagttacttgtcctaaac
gacttatcagtcttgtgtacCCTCTCCTTGTGCTGGCAACTTTTAAGAAATTGAAACAAAtatagttacttgtcctaaac
gacttatcagtcttgtgtacGCAAGCTGTACATATTGTCTGAAAATTTATAGGCCATTTGtatagttacttgtcctaaac
gacttatcagtcttgtgtacTCTTGGGGATGTTTAACACGTAGCTTCTGGACCTAAAATCtatagttacttgtcctaaac
gacttatcagtcttgtgtacGAGAAGGTCACCATCACCCAGCCAACTCTGCCTGCTAAAAtatagttacttgtcctaaa
gacttatcagtcttgtgtacGCTGTCCATAGGTATTGCTTTGTGAATTTGGTAAGTTTATtatagttacttgtcctaaac
gacttatcagtcttgtgtacCAGGGCAGGCACCGTGCTAGATGAACAAGGCAGATAGGTCtatagttacttgtcctaa
gacttatcagtcttgtgtacCTTCATGACAAACAAAAGGTAGTAAACGAGAAACAGACAAtatagttacttgtcctaaa
gacttatcagtcttgtgtacCTGGGAAAGGAAAGGATGATGCATTGAGGTAGGTAATAGTtatagttacttgtcctaaac
gacttatcagtcttgtgtacCAGGAAGTTTGCTGGGAGGAGGGGATATTTAAGCTGAGGCtatagttacttgtcctaaa
gacttatcagtcttgtgtacACAAGGAGGAGCCCTCAGTGTGGAAGTTGTCGGAAGAGGGtatagttacttgtcctaa
gacttatcagtcttgtgtacGGAACTAGCATGCGAAAAACATGGAAACAGAGAGCCAGGGtatagttacttgtcctaa
gacttatcagtcttgtgtacGGACAGCAGAGAGAGGCCCGGAGACAGTCTTGTTGGAAATtatagttacttgtcctaa
gacttatcagtcttgtgtacCAGTTTTTATCTGACTTGCCTTCCAAATCTTGCTGCTGTGtatagttacttgtcctaaac
gacttatcagtcttgtgtacAGGGCTCTTGAATGTGCTCATTTCTCTCCTTTTCACTGCCtatagttacttgtcctaaac
gacttatcagtcttgtgtacCCACCATCATCTCTCACCCCCCTCACTGCAATAGCCTCTTtatagttacttgtcctaaa
gacttatcagtcttgtgtacCTCTCCTCACATTCACCCTTCCTTCTCTTTAATCCATTCTtatagttacttgtcctaaac
gacttatcagtcttgtgtacAACTTGATTTTGTTCCTCTCCTGCTTAGAAACCTTGCAGTtatagttacttgtcctaaac

gacttatcagtcttgtgtacCATTTGTACATATCTCTCTAGGGACATTGTGAGGCTGGCGtatagttacttgtcctaaac
gacttatcagtcttgtgtacCTAAATGTTCTAAAGTCTATAATTTGCTCCTTCTGTTGCCtatagttacttgtcctaaac
gacttatcagtcttgtgtacGAAGTCTAGAGTTAGGATTTGGGGATCAGGAGCAGCATCCtatagttacttgtcctaaa
gacttatcagtcttgtgtacAGAGAAATGATGTGTGCTCTGACATGTGGGAGGTTGGTGCtatagttacttgtcctaaa
gacttatcagtcttgtgtacCATCTCTTCACGCCTCACAAAATGGATAATCAGCACACCTtatagttacttgtcctaaa
gacttatcagtcttgtgtacAATAGTTAATACGTGTACATTCTATAAACAGTGCCTGACGtatagttacttgtcctaaac
gacttatcagtcttgtgtacTGCTGATCATACAAAGAATTCAAATGACTGCTAATAGTGGtatagttacttgtcctaaac
gacttatcagtcttgtgtacTTAATCAGTGAAGTGGAGCATGAAGAATTTGTGGATTTTCtatagttacttgtcctaaac
gacttatcagtcttgtgtacACTATGTGTCTGGCCTGTCTGTAGTGTGTGGAGCCCAAAAtatagttacttgtcctaaac
gacttatcagtcttgtgtacCAGAATCTGGGATTTCGGCTCCAGCTGGCTCTACCTGGCTtatagttacttgtcctaaa
gacttatcagtcttgtgtacGGGTTACACTCCTGATGGTCATCCCCTTTTTCATTTCTGTtatagttacttgtcctaaac
gacttatcagtcttgtgtacCCTCTGCTTGGAATAAACCCATTTTTGCTTTGATATCATCtatagttacttgtcctaaac
gacttatcagtcttgtgtacCGTCACGATTCAAAATAATTGTCACCAATTCCAGAACGCTtatagttacttgtcctaaac
gacttatcagtcttgtgtacGATGCCCCATGTATTCCAGTTGCAACTTGAGCTTCCTTTTtatagttacttgtcctaaac
gacttatcagtcttgtgtacATTATTGTAGGACTGTAATTGTAGAGAGGAGGTGTTGTTTtatagttacttgtcctaaac
gacttatcagtcttgtgtacGGCAACTTGACAGAAGCAATCCATTGTGTTCCCATTGTTTtatagttacttgtcctaaac
gacttatcagtcttgtgtacCTTGTGACTGGAGGGATGGGGCATCAGGAGGCACTAGTTTtatagttacttgtcctaaa
gacttatcagtcttgtgtacTGACTTTATTTACATACTTTTCCAAGGAGGCATTTCCATGtatagttacttgtcctaaac
gacttatcagtcttgtgtacTCAACAAATATTATACTGAGTCCTTAAGAGGGGGGACCACtatagttacttgtcctaaac
gacttatcagtcttgtgtacAAACTCTTTCTCTCCACAACTCACCCTTCACTTATGTAACtatagttacttgtcctaaac
gacttatcagtcttgtgtacCAATGACATACAAGAGAATAACAGCAAAAAGTAGCATACCtatagttacttgtcctaaa
gacttatcagtcttgtgtacGTGTCTACCTAGGAGAGTAGAACACTCTGTTGAGGCCCATtatagttacttgtcctaaa
gacttatcagtcttgtgtacCCTCTGTTTGGCAGGTACCATGAGGCTGCTATTTGAAAAAtatagttacttgtcctaaac
gacttatcagtcttgtgtacTTCACTGAGAACTTAGGAAACAGATGTTGATGCTGAAGGCtatagttacttgtcctaaac
gacttatcagtcttgtgtacGTGAGGCTTTTCTTCTTACCGACCATGTTTTAAGTCAACTtatagttacttgtcctaaac
gacttatcagtcttgtgtacAGACTCAGGCAGCTGTTCTTTCTTTGGCCAGACCAGTTGGtatagttacttgtcctaaa
gacttatcagtcttgtgtacGCATCTCAGCACAGAGTGTCTTCTGCACACATTTGATATTtatagttacttgtcctaaac
gacttatcagtcttgtgtacAAATAAATTTGGGGTAATTCACACTTAAACTTGCCTGGTGtatagttacttgtcctaaac
gacttatcagtcttgtgtacTCAAATTTTTCTGAATAATGAACCCAAGTCCAACCTCCAAtatagttacttgtcctaaac
gacttatcagtcttgtgtacTTCTCTTTGAAACAGTGGACTTCTAAATCATTGGCCTAAGtatagttacttgtcctaaac
gacttatcagtcttgtgtacAGCTTTCCCTTCTCTGAAGTCCAAGAGCCATCGTCCAGGGtatagttacttgtcctaaa
gacttatcagtcttgtgtacTCTTTTCAGTTGAGGAGATCTTTAGGTCAAGCGCTGTGTTtatagttacttgtcctaaac
gacttatcagtcttgtgtacTGCATTGCACATTGAGGTTAGTCTGCAGATCATTTGAAATtatagttacttgtcctaaac
gacttatcagtcttgtgtacTGTACACAAAAGCGGTATACATTTCATTCCTGGCCTCTGTtatagttacttgtcctaaac
gacttatcagtcttgtgtacTATTTCTGATACTTACCTCTGACCACCACCCCACATGTAAtatagttacttgtcctaaac
gacttatcagtcttgtgtacACATTTAAAGTCTCAGAAGGCAACAAAATTGAAAGACTGCtatagttacttgtcctaaac
gacttatcagtcttgtgtacCTATATATCTATCCATCTATGAGCTGCTTATGTGCATACTtatagttacttgtcctaaac
gacttatcagtcttgtgtacGCCACACGTACATACAAGATCAAGGACAATGTCAGGCTCCtatagttacttgtcctaaa
gacttatcagtcttgtgtacTGATAATAAATGATCCCCTAGTTCAAGGCATCTATCAACTtatagttacttgtcctaaac
gacttatcagtcttgtgtacCTTTGGGAAGCTAAGGCGGGTGGATCCTGAGATCAGGAGTtatagttacttgtcctaaa
gacttatcagtcttgtgtacGCGCGCCTGTAATCCTGGCTACTTGATAGGCTGAGGAAGGtatagttacttgtcctaaa
gacttatcagtcttgtgtacAAGGGATACTTTCATTTGGGAACTTGTTGATGGAACTCACtatagttacttgtcctaaac
gacttatcagtcttgtgtacGATAAAGGACACATACATTACAAAAGAAGGAAAGAACAGGtatagttacttgtcctaaa
gacttatcagtcttgtgtacAAATAAAACTCTCTGTAAGTATCTGAGAAGAGTAGCTCTCtatagttacttgtcctaaac
gacttatcagtcttgtgtacTTTACAAAATCTCCTGTCAAGGCAAAACATGGATTTACAGtatagttacttgtcctaaac
gacttatcagtcttgtgtacTGAGGCTAGACAACATCAGTGAAGAGAAGCAAGGAAGGGGtatagttacttgtcctaa
gacttatcagtcttgtgtacTCTGGGATGGAATGAGGATGGTAGAGAATGTTTGAAAGTGtatagttacttgtcctaaac
gacttatcagtcttgtgtacAAGAATGGAGAGAAATTCAATAATAGGATGAAGGTCAGTGtatagttacttgtcctaaac
gacttatcagtcttgtgtacATTCTGCTATGATTTGCCAGAGTGTATATTGGTGTTAAGGtatagttacttgtcctaaac
gacttatcagtcttgtgtacCAAGATTGTGTCAAAGTAGATGTGTCAAAATAGCTGTCTAtatagttacttgtcctaaac
gacttatcagtcttgtgtacGGATAAAACAATCGCATCAGAATGGCTGTGTCAAATGGGCtatagttacttgtcctaaa
gacttatcagtcttgtgtacAACAACATGGTCAAGCTCCCTTTAGTATTACAATGGTCAGtatagttacttgtcctaaac

gacttatcagtcttgtgtacCAGCATCGACCACACAACACTGGCAGGGTAAGAATCCATTtatagttacttgtcctaaa
gacttatcagtcttgtgtacTTATTATAGTCGTTCATTGATTGATGTGTAAGTAGGGGACtatagttacttgtcctaaac
gacttatcagtcttgtgtacGCCTCAATACTGGAAATAGGTAGCTTTCTAGGCATGTGGTtatagttacttgtcctaaac
gacttatcagtcttgtgtacCATGTCATGGGGAGGGACCCCGTAGAAGGTAATTGAATCAtatagttacttgtcctaaa
gacttatcagtcttgtgtacACATGCCTTTGCTCCTCTTTTGTTTTCTGCCATGATTGTGtatagttacttgtcctaaac
gacttatcagtcttgtgtacATGGACTAATACATCTATCCTCCTTTTGAATATAGGCTTGtatagttacttgtcctaaac
gacttatcagtcttgtgtacAAAGATTTGTGAACATTGAACAGAAGTGATATGTGTGGCTtatagttacttgtcctaaac
gacttatcagtcttgtgtacCATGTATTGAGATGGTGCCTCTGTTCACTTTGGTCCCTGCtatagttacttgtcctaaac
gacttatcagtcttgtgtacTGAATACACTGGGCAAGGCCTCCACTGCTACCTTATATTAtatagttacttgtcctaaac
gacttatcagtcttgtgtacTTAAGCCACTGAGATTTGGTGCCATAGCCCATCTTAACTGtatagttacttgtcctaaac
gacttatcagtcttgtgtacCAAAGTCATTCAGCCTGTTGATGCATATGTAAAACTGTATtatagttacttgtcctaaac
gacttatcagtcttgtgtacCTGGAAAATTACTGCCACTTCATTAAACATCCTATAAAGCtatagttacttgtcctaaac
gacttatcagtcttgtgtacAATGTGGCTATCTTATTGGTAAGTATTGGGATTGTATTGCtatagttacttgtcctaaac
gacttatcagtcttgtgtacTTGAGAGCAGGTGACAAAGGATGATTAAAAAGAAATCTAGtatagttacttgtcctaaac
gacttatcagtcttgtgtacTGATAACAATCTGTAAGGGGAGTGGAAATGTACTGGGATTtatagttacttgtcctaaac
gacttatcagtcttgtgtacGTGATCCAAAACACCTCAGGACTGCACACTGTTCTTAAGTtatagttacttgtcctaaa
gacttatcagtcttgtgtacGAGAAATGAGGTGTGACATTGCCAAGTTTCCATGTGAAAAtatagttacttgtcctaaac
gacttatcagtcttgtgtacTTCTATTGTATATTCTGGTTGACTGACACACAGATTTGCCtatagttacttgtcctaaac
gacttatcagtcttgtgtacGAGGACGTGTGCTGCCAATAATTGAGTGTTTGCTATTTTTtatagttacttgtcctaaac
gacttatcagtcttgtgtacAAGTTTTTTTTTTTCCCCAGCAAGAATCATTCACATGCCCtatagttacttgtcctaaac
gacttatcagtcttgtgtacCATTTATATTCTTCTGTTATGGTTTGGTTGTAATCCAGGGtatagttacttgtcctaaac
gacttatcagtcttgtgtacCAGCTTTCTTCTGGCCTTGTTGTTTCTTTTTCATGTGTAAtatagttacttgtcctaaac
gacttatcagtcttgtgtacGAGGTTTCCTAACTTCGCCTGGAAATGTGGTTATTACTGCtatagttacttgtcctaaac
gacttatcagtcttgtgtacATAAAAATATTACCTCGCAACAAGTCTACCATCATATGCCtatagttacttgtcctaaac
gacttatcagtcttgtgtacATCATCTCTCTCCTCTAGCCGAGAGAAAAAAAGAAAAACTtatagttacttgtcctaaac
gacttatcagtcttgtgtacCCTTCCTAAATGTCTCAAGGTATTGTTACTTAATGTCAGAtatagttacttgtcctaaac
gacttatcagtcttgtgtacATTATATATTAGAAAGAAGAATGATTTGCCCCCCTCCCCAtatagttacttgtcctaaac
gacttatcagtcttgtgtacCTCTAAATTCAGGTGATACTATCTCTCTTTAAACTTCTCCtatagttacttgtcctaaac
gacttatcagtcttgtgtacCTTTGCATTGCTCCTTATTTGTTCCTTGAAAAGCTTCTGCtatagttacttgtcctaaac
gacttatcagtcttgtgtacACTTAATAGGAAAAATACTTGTAAGCAGCACAGTGGTATCtatagttacttgtcctaaac
gacttatcagtcttgtgtacTCTGATCCTTTTTCTTTTCCTTAGACTCTGCCTTCTGATAtatagttacttgtcctaaac
gacttatcagtcttgtgtacTTGCATATCAGTGACTTAAATGGACAAAAAGCAAGTGCCTtatagttacttgtcctaaac
gacttatcagtcttgtgtacCCCTGACTCAGAGGGTCTCAGCTTTCCGTTTTTCCATGTGtatagttacttgtcctaaa
gacttatcagtcttgtgtacGCAAGGAGGTGGGGCAGGCAGGTGTTTCAGCAATTCCGGGtatagttacttgtcctaa
gacttatcagtcttgtgtacGATGTGTGGTTTGAATGGTCTCCAGGCCATCTTTTTTTTTtatagttacttgtcctaaac
gacttatcagtcttgtgtacGGCCATCTTGATGCAGCCTGGTGGTGAGGAGAGGGTGTGGtatagttacttgtcctaa
gacttatcagtcttgtgtacGTTTCAACAGTGCATTTTGGAAAGAGAATTAGATAACCCCtatagttacttgtcctaaac
gacttatcagtcttgtgtacGGTATTGAGAGTTTCAGATAGCTTATTCTTGATAGCCTTTtatagttacttgtcctaaac
gacttatcagtcttgtgtacGTTAATGATGGCTACTAAGATTCTGGATCCCACACAATGGtatagttacttgtcctaaac
gacttatcagtcttgtgtacGGAGTCTACTCTGCCACTGTTCAATAAGTAGGTGCCCATCtatagttacttgtcctaaa
gacttatcagtcttgtgtacGAAGTGGGTCCATAAAAATCACCTCCAAGGGTCCTGACAAtatagttacttgtcctaaa
gacttatcagtcttgtgtacCCTGTGTGGTTTTTGAGTGGCCAACTTGGGTCCCTTTGGTtatagttacttgtcctaaac
gacttatcagtcttgtgtacGGTTTACTAAGGCAGCAATAAGTCTGTGCATAAGGATCTTtatagttacttgtcctaaac
gacttatcagtcttgtgtacTTTTTGAATTATTTGTGGCTTCCTTGTCTAATGCTTACCCtatagttacttgtcctaaac
gacttatcagtcttgtgtacACCTAACCACAGCATCCATGGCCTTCAATATTATACAGCCtatagttacttgtcctaaa
gacttatcagtcttgtgtacGGCCTCTTCAGTCTAGTGCCCCCTAGCTTCCTGACATGTAtatagttacttgtcctaaa
gacttatcagtcttgtgtacTTCCAGCCAAATGTTCACTGACTTTCCTGAAAGCTTTGCTtatagttacttgtcctaaac
gacttatcagtcttgtgtacCATCTTGTATAAAATGTGCTTCTATATGTTGCTGAATCCGtatagttacttgtcctaaac
gacttatcagtcttgtgtacTCCCAATTCAGTGAATGGCGTCTTCAAGCACCATGTTGCCtatagttacttgtcctaaa
gacttatcagtcttgtgtacGGTCTGTTTGATTTCTTCTTTTCCCTTCTTCCACTATTTGtatagttacttgtcctaaac
gacttatcagtcttgtgtacCGATTATGTTACCAAAAAATGTTACCCAGATTCCAAGAACtatagttacttgtcctaaac
gacttatcagtcttgtgtacGCCAAAGCAGACAGCCTTCCAAGTGGTCTCTCTAGTTCCAtatagttacttgtcctaaa

gacttatcagtcttgtgtacAAGTATAAATCAGATCATTTCAGTCCTTGCTTAAAACCCCtatagttacttgtcctaaac
gacttatcagtcttgtgtacAGCTTCACTGCCTACTCTTCTTTCCTGGATTATTATATTAtatagttacttgtcctaaac
gacttatcagtcttgtgtacTCCTGCTTTGTGGGCCTTCTTTCTGTTCCTTGGACAACTGtatagttacttgtcctaaac
gacttatcagtcttgtgtacTATATATCCCTCAGAGAGTTTACATGTATGGCTCAATCTGtatagttacttgtcctaaac
gacttatcagtcttgtgtacGCCCCAAATGTCTGCTTCTTAAAATTCAGGACTGAGCTCAtatagttacttgtcctaaac
gacttatcagtcttgtgtacGAAAATAATTCCTTCCCTGAGAAATTCCATATCCGTTGGTtatagttacttgtcctaaac
gacttatcagtcttgtgtacGAGGGATATTAATTTTTTTGTTAAGCACTCACTGGATTGCtatagttacttgtcctaaac
gacttatcagtcttgtgtacGGCCAGCCTATAGATATCGATCCCCTGAGAGAAAGAGTCTtatagttacttgtcctaaa
gacttatcagtcttgtgtacGAATGTTCATAGCTTTAACTTCTCTTATGCACCTACCCTAtatagttacttgtcctaaac
gacttatcagtcttgtgtacCCTTCTTATAGTTAAAGTTTTTCAAGGGACTGTGTTCTACtatagttacttgtcctaaac
gacttatcagtcttgtgtacGTAAACACTGAGCCAATATTTACTGAATTCAATTATCCCGtatagttacttgtcctaaac
gacttatcagtcttgtgtacCTCAAACAAGGACTACACTCAGCGTACTCATTTCCAGGTGtatagttacttgtcctaaa
gacttatcagtcttgtgtacTCTCAGATATGCTTTCATTTCAGACCTCTTATTCAAAAGCtatagttacttgtcctaaac
gacttatcagtcttgtgtacAGGAAATGAAGTAGGCCTGGAAAATAGTTGAGCAAAAGAAtatagttacttgtcctaaa
gacttatcagtcttgtgtacCCAAGGCTTAATTAATACTTCCTAGGTGTCATGAATTCCTtatagttacttgtcctaaac
gacttatcagtcttgtgtacAGGAAAAGGCATGGATTGGTGGTTTAGGAAGCAAGGTGGCtatagttacttgtcctaaa
gacttatcagtcttgtgtacCTAAGCAAGTAGAGGAGTGATAAGAAGGAGATTGATGTTCtatagttacttgtcctaaac
gacttatcagtcttgtgtacTAACTTTCTGACTGTCTATACTGATATTGGTTACTGTGCTtatagttacttgtcctaaac
gacttatcagtcttgtgtacGGCAGATAAAGAAAAGACACTCTCAGCCCTCTGTAATGTGtatagttacttgtcctaaa
gacttatcagtcttgtgtacAAGATGGTGTCAGCTTGGAAGTTGTTAGGCTGCCAATCCTtatagttacttgtcctaaac
gacttatcagtcttgtgtacCAGGCAACCAGGATATTTGAATTCAGATGTGTTGAAATTCtatagttacttgtcctaaac
gacttatcagtcttgtgtacGTGAGTTTTCCACTACAAATCCCTAACATACAGAGAGACCtatagttacttgtcctaaac
gacttatcagtcttgtgtacTTCAAATATATTTTAAATGGCCCTGCTCAAAGAATCCTGGtatagttacttgtcctaaac
gacttatcagtcttgtgtacTGAGGACAGGTAAGAGGGAGAATACAGTTATTTCAGGAAGtatagttacttgtcctaaa
gacttatcagtcttgtgtacCAGTCATCCATTTTGTCATGATGGAGTCATTCTTTCATTTtatagttacttgtcctaaac
gacttatcagtcttgtgtacCTGGTTGAAACACAGAGTATCACAATGCTTAAGAATTTAGtatagttacttgtcctaaac
gacttatcagtcttgtgtacCAGAAGGTTGGATTCAAATCTACTGTGATGGTTAGTTTTAtatagttacttgtcctaaac
gacttatcagtcttgtgtacTCAATTAAGCATGAATCTAGGTGTTGTGAAGGCATTTTTGtatagttacttgtcctaaac
gacttatcagtcttgtgtacGTTGAAAGGCCTTAGGAGCAAAACAGGTTTCGCCAAGGAAtatagttacttgtcctaaa
gacttatcagtcttgtgtacCCTGCCTGGGAGTTTCTAGCCTGCTGCTCGTCTGCCCCACtatagttacttgtcctaa
gacttatcagtcttgtgtacAGACTTGCCCAGGCTGACTATACAATCACATGAGACAATTtatagttacttgtcctaaac
gacttatcagtcttgtgtacCTCTGTCTCTATGTCTGTCTGTCTCTCAGTTTGTTTTGTTtatagttacttgtcctaaac
gacttatcagtcttgtgtacGTAGAACCCTGGCTGACAAATCTATTAACTGAATTGACCTtatagttacttgtcctaaac
gacttatcagtcttgtgtacATAAAGATATCAAGAGTGTCCATCTGTATCCATCTCACAAtatagttacttgtcctaaac
gacttatcagtcttgtgtacGTGAGTTAATAAATATAAAGTCCTTAGACTGGGAACTGACtatagttacttgtcctaaac
gacttatcagtcttgtgtacCTTAGTGTAGCTGCTGGGGAGAAATAAGAATAAAGATGAGtatagttacttgtcctaaac
gacttatcagtcttgtgtacCCATTCCTCGGAGGCTTCTAAGTCATAATAACAAGTAAACtatagttacttgtcctaaac
gacttatcagtcttgtgtacAGAATATGACCCCCAGTGCATGCTCATAATTAGGAAAAGGtatagttacttgtcctaaa
gacttatcagtcttgtgtacCCTACTAATGAGGGTAGATAATATTTCTCTATTTTGTGCCtatagttacttgtcctaaac
gacttatcagtcttgtgtacGCTGAGAATATAGTAGAAAATAGACAGAAAAGATCTCCTGtatagttacttgtcctaaac
gacttatcagtcttgtgtacATGTATATCGCAGTAGGAGCTACACAAACAAAAAGAATGTtatagttacttgtcctaaac
gacttatcagtcttgtgtacTTTTATGAAGAAGATAAAGCATGGTAAGAACAGCCTGGTGtatagttacttgtcctaaac
gacttatcagtcttgtgtacTTGAGAAACAGTGTGAAGGCCCTTAGATTACAAGTAGCTTtatagttacttgtcctaaac
gacttatcagtcttgtgtacGTGGAGGGAAGGAAAGAAGACCGTGTAGCTGGAGCAGAGTtatagttacttgtcctaa
gacttatcagtcttgtgtacATGTTGGGAAAGAAAGGAAGTGGAGCCCAGATCTCATAGGtatagttacttgtcctaaa
gacttatcagtcttgtgtacGCCATGCCAAGGGTTTTTATTTTATTCAAAGTGTGATGTGtatagttacttgtcctaaac
gacttatcagtcttgtgtacTAGATGTGGGTAAAGTCTATGGGCTATGGTGGTGTTGGGCtatagttacttgtcctaaac
gacttatcagtcttgtgtacCACAGGAGGCCATTGGGGTAGAACAGGTGAGGGAGGTGAGtatagttacttgtcctaa
gacttatcagtcttgtgtacCATGGAGGGCGTGAGAAGTGGATGAAATTCAGGATGGATCtatagttacttgtcctaaa
gacttatcagtcttgtgtacGCTGCCTTGACATCTTTAAGCTTCACAGTGCCCCAGAGTCtatagttacttgtcctaaa
gacttatcagtcttgtgtacCAAGTTTCCCTGCTATTACACTCCCCACCCACTAGTCCCTtatagttacttgtcctaaa
gacttatcagtcttgtgtacACCAGCTACACTCTAACAGGTCCTCAACCCAACAGGTTTGtatagttacttgtcctaaa

gacttatcagtcttgtgtacATTGTTCAAACCAGTTAATCATATTCTCCTATGGGAAACGtatagttacttgtcctaaac
gacttatcagtcttgtgtacACCCTCTTGTTACTACAGAGTCTCTCCCCAGCAGCCCCTGtatagttacttgtcctaaa
gacttatcagtcttgtgtacTGGTATGCATGGTACTTTATCCTGGGCTGTGAGAATGTATtatagttacttgtcctaaac
gacttatcagtcttgtgtacCAGTGTTGGGTTGCTTGTATTTGTCCATGTTGTGCTAATTtatagttacttgtcctaaac
gacttatcagtcttgtgtacTGGAGTGAATAGGAGGCAATCAGAACAATGGATGGCAAAAtatagttacttgtcctaaa
gacttatcagtcttgtgtacGCCCAACCACCACCAAGAACTTGTAGCTTTAGTTATCTGCtatagttacttgtcctaaa
gacttatcagtcttgtgtacATAGGGCCAGAATGTCTGCACTATAAGTGACTGGATGGAGtatagttacttgtcctaaa
gacttatcagtcttgtgtacCTTAATGCAGCCAGAAACCAAAATACAGTGAGGAAATTTGtatagttacttgtcctaaa
gacttatcagtcttgtgtacGGAGCCCACCCTGCAGGAAAAGGCAGTGTTGTAGCAGAGGtatagttacttgtcctaa
gacttatcagtcttgtgtacCACTGTTTTGCTGCATCACTGGGCTGCCTGTGGCTTTGTCtatagttacttgtcctaaa
gacttatcagtcttgtgtacATACCCCAAACAACAAAACACTCATGGTATCCCTGTGGCAtatagttacttgtcctaaa
gacttatcagtcttgtgtacCCTGACTTAAACCTAGGGCCCTTAAACCATAACAATTGCCtatagttacttgtcctaaa
gacttatcagtcttgtgtacCCTGACAATACTGAGGCTCTCTGGAAGAAGACTTTTTATAtatagttacttgtcctaaac
gacttatcagtcttgtgtacGGCGACCCTCCCTATTGAAGTATATCCAGTGACAAACAGAtatagttacttgtcctaaa
gacttatcagtcttgtgtacCTGGATACAGGAGGAGAGAAAATTTAAACATATGTGCATAtatagttacttgtcctaaac
gacttatcagtcttgtgtacGAAGTGTTAATTGGCTTTAGTGTCTCTATGATATAGGTTCtatagttacttgtcctaaac
gacttatcagtcttgtgtacAATTAACAATTCATGGACCTTCATATACTTCAACACTGGGtatagttacttgtcctaaac
gacttatcagtcttgtgtacAAGGTACCGCTATAATCTTGGGAGAGTTACTGAATATAAAtatagttacttgtcctaaac
gacttatcagtcttgtgtacCATTGCTCTAAAATGTCTCACAGTAAGCATGGATAATCTGtatagttacttgtcctaaac
gacttatcagtcttgtgtacTGGCACCTAAAAATTGGCTTGACAAAATGGAACTACATAAtatagttacttgtcctaaac
gacttatcagtcttgtgtacTATAAGACCTAATTAGTAACAGGATGATTATCTCCACTGCtatagttacttgtcctaaac
gacttatcagtcttgtgtacCATTTGACAGCCCAATTTTGTGCTTTCTTATACCTAGAAAtatagttacttgtcctaaac
gacttatcagtcttgtgtacCAGTCTTAATCCTGTGGTTGCATCTGCTAAGGTCCCCATAtatagttacttgtcctaaac
gacttatcagtcttgtgtacTAGGTTTAGTTAATGCAGTCTATTTAGTGCCTATTTTGGCtatagttacttgtcctaaac
gacttatcagtcttgtgtacAGGGACCCAACACACCTTTACCAGGCTGCCCATGGAGTACtatagttacttgtcctaa
gacttatcagtcttgtgtacGCATCTACATTTCCCAGAAGCACTGCTATGAAACTATATTtatagttacttgtcctaaac
gacttatcagtcttgtgtacCATCCTCCTGAGGTCATTTGACACATTAATTAAGGACATGtatagttacttgtcctaaac
gacttatcagtcttgtgtacGGGAATAGGCAATTGCCCCACATAGTATAAAGCCCTGCCAtatagttacttgtcctaaa
gacttatcagtcttgtgtacTTGGTAAATCAAGAGATGTTTCATTCCTGACAGTATCAAGtatagttacttgtcctaaac
gacttatcagtcttgtgtacCAAGTCCATTGTTTTTAGTAATTTTGGGATTCTGAAGGCAtatagttacttgtcctaaac
gacttatcagtcttgtgtacCTGTCACTTGAAATCAGCCCACCTTAAATGAGCCCCTCTTtatagttacttgtcctaaa
gacttatcagtcttgtgtacGGAATTCCCCTTAGTGCCCTTAGAGAGTCCTTTGTTTTAGtatagttacttgtcctaaac
gacttatcagtcttgtgtacCATTCCTTATGCCACTGGAGTTGCTGACTTACTTATGATGtatagttacttgtcctaaac
gacttatcagtcttgtgtacAAGTTGCCTTTCCTCAGACCTGTGCTATATATCATTAGAGtatagttacttgtcctaaac
gacttatcagtcttgtgtacCAGACTGTGAACCTGTTACCCTCCATATCCAGCTGTCCATtatagttacttgtcctaaa
gacttatcagtcttgtgtacCCCTCAAGGTTCAGTATGGCTACCAAGGTCTCTTTGCTACtatagttacttgtcctaaa
gacttatcagtcttgtgtacCAAATCTGGGCCATCTGTATATTCTACTTGCAGGAGAGGAtatagttacttgtcctaaa
gacttatcagtcttgtgtacGGAGAAAGACACCACTTTTCCAGACCCCTTGGCTACCCAGtatagttacttgtcctaa
gacttatcagtcttgtgtacTGGAGTTATTTGCAATGAGATTTTGTACACTTCACAGATGtatagttacttgtcctaaac
gacttatcagtcttgtgtacATCCTATGTGATGGAGCCCAGTGGAATGTTGCTGGTTTCCtatagttacttgtcctaaa
gacttatcagtcttgtgtacGGATAAGGACAATTGGCCAAACTTTACACAGTCATCTTACtatagttacttgtcctaaa
gacttatcagtcttgtgtacCCCCATCTGCACATTTTTACGTACTCTTGGACTACTGCCTtatagttacttgtcctaaa
gacttatcagtcttgtgtacTTGCCTCACAGATGCCCAAAATATAAATCAAGGTAAGAGAtatagttacttgtcctaaa
gacttatcagtcttgtgtacCCACAACACGAGGCTTCACCAGTACACTCCTTAGCTCCTGtatagttacttgtcctaa
gacttatcagtcttgtgtacCATCACCCATGTTGACAAGACACTCATTCACTTCTCAGAAtatagttacttgtcctaaa
gacttatcagtcttgtgtacGTTTCACTGTGAGGCCTAATGAGCTCCCCAAATAACTCCTtatagttacttgtcctaaa
gacttatcagtcttgtgtacCTACTGGGTATTGCTATTGCCATAGGTGTTGATTAATCTCtatagttacttgtcctaaac
gacttatcagtcttgtgtacGCACTTCCTCCACTTATGCCTCATGGGTCTATCAATTATCtatagttacttgtcctaaac
gacttatcagtcttgtgtacAGCTACCTCTAAGTTCATGTTCTTGATGTCTACTCCACCAtatagttacttgtcctaaac
gacttatcagtcttgtgtacTCCAAGGGGAATGCTTTCACTCTCTAAAAATCTTAGTTATtatagttacttgtcctaaac
gacttatcagtcttgtgtacCCTCAAGTTATCAAGGTACTGAATGTGGTATAAGGGTTAAtatagttacttgtcctaaac
gacttatcagtcttgtgtacGCCCCAACCCCAAGCCCAAGTCCCCTGCTCTCCTCAGATAtatagttacttgtcctaa

gacttatcagtcttgtgtacAGCTCTCACATCAGTTGGACCAGCTCTACCCTATCCAGTCtatagttacttgtcctaaa
gacttatcagtcttgtgtacGAAAACTTATTCAAACAAGCCAATTATATCCTTCCACTGGtatagttacttgtcctaaac
gacttatcagtcttgtgtacGGTTCACCCCAGACTCTTGTTACTACAAAGCATGCTTCCCtatagttacttgtcctaaa
gacttatcagtcttgtgtacCTGTTCTTGTTTACTCTGTTCCTGAATGCAACCCCAGTGTtatagttacttgtcctaaac
gacttatcagtcttgtgtacAGAGCATTTAGTGTCTTCCTCCTTGGGCTGTGAATATATGtatagttacttgtcctaaac
gacttatcagtcttgtgtacCTGCTGTCATTCTCATTTGCCCAGTGTTGTGTGTTTGCCAtatagttacttgtcctaaac
gacttatcagtcttgtgtacCCAGTGGGTGAATAGGAGGCTATCAGAACAAGAGCCAGCAtatagttacttgtcctaa
gacttatcagtcttgtgtacTGGTATGCATGTGAGATTAAAGGGAAGAATCAAGAGTGTTtatagttacttgtcctaaac
gacttatcagtcttgtgtacAAATGTCCATGCTATTTATTGAGAAGGGAGAGAGGAGAAAtatagttacttgtcctaaac
gacttatcagtcttgtgtacGATGAAATGGGATAAATAGGGTTTTGAGAAATGATTAGGGtatagttacttgtcctaaac
gacttatcagtcttgtgtacCAGTTGGAGGAATAAAGAGGCTCACATGAAAGGAGATATTtatagttacttgtcctaaac
gacttatcagtcttgtgtacTTGTCACTAGTGGTGTTGGAATTATACATCATAACAGAAGtatagttacttgtcctaaac
gacttatcagtcttgtgtacTCAGCAAGGTAGAGAAGCCCAGAATGACAAGTGATTTCAGtatagttacttgtcctaaa
gacttatcagtcttgtgtacCCAAGGCTATAATGTTATTCTGGGATCTTCCTTATTGAAGtatagttacttgtcctaaac
gacttatcagtcttgtgtacAGAGTTTACTTTGATCAGGAAGGCATTAGAATATGTTGAGtatagttacttgtcctaaac
gacttatcagtcttgtgtacCCAGTAGTGATTAGGGTTGTGTGTTCTGAAGATTAGTCTCtatagttacttgtcctaaac
gacttatcagtcttgtgtacCATCTTGGGAAGGAGTGTGGTGGACATTGTGAGTGTAGGTtatagttacttgtcctaaac
gacttatcagtcttgtgtacAGAAATGATGAAAGTCTTCCTAGAATGGAATCAGGAACAGtatagttacttgtcctaaac
gacttatcagtcttgtgtacGGAAGAGGATAAAAATGAAAACAATTTGAATGGGTCTTGGtatagttacttgtcctaaac
gacttatcagtcttgtgtacAACTAGGAAATTATGTAGTTGCCAGCAAATATGAAGATGCtatagttacttgtcctaaac
gacttatcagtcttgtgtacAGATTTTAGTTTCAACATGTTAAGTCTGAGATGCTACTGGtatagttacttgtcctaaac
gacttatcagtcttgtgtacGGGGCTACTCCTTTGGCAAGAGATTAGGTTCCCAGTTATCtatagttacttgtcctaaac
gacttatcagtcttgtgtacAAGCTCTTCTGACATCATTCTCTCTTACTTTCTCTTATTCtatagttacttgtcctaaac
gacttatcagtcttgtgtacCCATTCCAGGAACTCATTGAGCACGTTCTCACATCGGGGCtatagttacttgtcctaaa
gacttatcagtcttgtgtacTCTGCACAGATCACTGCCTCTCAATTTATTCAGATCTTTTtatagttacttgtcctaaac
gacttatcagtcttgtgtacTTCCCCATCTAAGAGGCTTCCGCTGACCATTCTATATAGAtatagttacttgtcctaaac
gacttatcagtcttgtgtacCTTTCTTCCTTCTGCTGTCATTCTTAGCCCCTTGCTCTGCtatagttacttgtcctaaac
gacttatcagtcttgtgtacTATTTTGTGTGTCACTACCTTCATTAGAACATAAGCTCTCtatagttacttgtcctaaac
gacttatcagtcttgtgtacAAGGGCTTTGTTCTAGGTAATGTTCCATCTGTGTGTCTGGtatagttacttgtcctaaac
gacttatcagtcttgtgtacGGGAAGAAAAGAGCAGGAGAGAAGAAGATGGGTCGGAATGtatagttacttgtcctaa
gacttatcagtcttgtgtacACATCTTGTTTATCTGTTCACTCCTTTTCCTTTGCTGGTGtatagttacttgtcctaaac
gacttatcagtcttgtgtacCAGTGAGTCTGCTCCCAGATCTGTGTGGCATGTGATTCAGtatagttacttgtcctaaa
gacttatcagtcttgtgtacCTAGTCAAAGCATCAAGTTCAAGGGGATTGGCGCAGGAATtatagttacttgtcctaaa
gacttatcagtcttgtgtacTGATCCACTTCCCCCCACTGACATCAGGCTGTAGCATGGTtatagttacttgtcctaaa
gacttatcagtcttgtgtacAGACTCTGGATCCTGACAGCTGGGCTCAAATGCTACCTCTtatagttacttgtcctaaa
gacttatcagtcttgtgtacCTTCGGTAAGTTATCTCACTCCTCTCTCTCTCAGTGTAATtatagttacttgtcctaaac
gacttatcagtcttgtgtacTGTCACAGAGTAAGAGCTATAAAAGTTTTATCATCTGGTGtatagttacttgtcctaaac
gacttatcagtcttgtgtacAGAGAAGTGCTCCTTTCAGTGGAACTCGAAGCCAAGATGCtatagttacttgtcctaaa
gacttatcagtcttgtgtacCACGTGGACCCTGAAGATGAAGCCCCATGGAGGAGAACAGtatagttacttgtcctaa
gacttatcagtcttgtgtacGAGATGCTGACTCCTGATGACACCTCTTGACTCACTTAAGtatagttacttgtcctaaac
gacttatcagtcttgtgtacTTGAGTTGGCTTGTAAGTAAGTTGTAATGGGAGATTCGCAtatagttacttgtcctaaac
gacttatcagtcttgtgtacCTTGCTAGCTTCAGGTGATTCTGTGACTTCTTTAATTCTCtatagttacttgtcctaaac
gacttatcagtcttgtgtacTTTTGCCTTTGGTAATTGTTTTTGTTATTCCCTGAACTCCtatagttacttgtcctaaac
gacttatcagtcttgtgtacGACTCCAGATTCACATTTTATCCTTTTTCTCAGAGGCCAGtatagttacttgtcctaaac
gacttatcagtcttgtgtacCCCTTTGTCTTAGGCCTGACTCTTGTTCTTTTTTGTTTTGtatagttacttgtcctaaac
gacttatcagtcttgtgtacACACACACCTCCTTCCATGTTATTTTTCTGTCCCCTCACTtatagttacttgtcctaaac
gacttatcagtcttgtgtacAGATATGTGTCCAGAGCCTTGCCCTTTTCAAATTCAGTGTtatagttacttgtcctaaac
gacttatcagtcttgtgtacTCCCCTTATTCACTTATTCCTCCCAAGACTCTCCTTCCTGtatagttacttgtcctaaac
gacttatcagtcttgtgtacCAGGGGAGAGAGGGCTGTGGGGGTTATTATTCTTAGCTTTtatagttacttgtcctaaac
gacttatcagtcttgtgtacTTGGCATTCTCTCTCACAACTGTATCCACTCAATTCCCAGtatagttacttgtcctaaac
gacttatcagtcttgtgtacCCCCATAGATGTTCCATGCTGGCTTCTCATATTGACTGGCtatagttacttgtcctaaac
gacttatcagtcttgtgtacCTTCTGAAATGGGAGCAAGGAGAAGTTGAATGAACACACAtatagttacttgtcctaaa

gacttatcagtcttgtgtacCTCTGAGAGTAAGAAGAAAGTGAGGGAATGTGGCAATGATtatagttacttgtcctaaa
gacttatcagtcttgtgtacTCCTTGGGATGGAAAGTAGGGAGAGTTTCAACCAGAGATCtatagttacttgtcctaaa
gacttatcagtcttgtgtacGGCCTCTCGTTTAGTCCATTCTCTCGCCCTCCGTGTCTAAtatagttacttgtcctaaa
gacttatcagtcttgtgtacGTTTCTTTGGGTCCTCAGATATATCAAGCTCTATTTTACCtatagttacttgtcctaaac
gacttatcagtcttgtgtacTTTCGCTATGCCAAGAAAACTCTAACTTCACGTCCTTCTTtatagttacttgtcctaaac
gacttatcagtcttgtgtacGCATCCCTCCCTTCAGGCCTTGGTTAGCTCCTCTTGCTATtatagttacttgtcctaaa
gacttatcagtcttgtgtacCCCGTAGCTCCTTAAACTTCCCCATAGCAACATACATTGCtatagttacttgtcctaaa
gacttatcagtcttgtgtacGTTATAGAACTGTCTTTCTCCACTGCGAGATCATGGGCTCtatagttacttgtcctaaa
gacttatcagtcttgtgtacGGCATTTCCTCTACATTTAATCACCACTGTAACTCCAGGAtatagttacttgtcctaaa
gacttatcagtcttgtgtacATTTCTTGGCCATATTTGGTACTCAATCAATGTTTGCTGAtatagttacttgtcctaaac
gacttatcagtcttgtgtacGCGTGAATCTTAAAATATTATCAGGGCAAGAGGTAATTGTtatagttacttgtcctaaac
gacttatcagtcttgtgtacAATGGAATTATAGCTGCTTCATCTGTGGCCAATTTTCTCCtatagttacttgtcctaaac
gacttatcagtcttgtgtacGCATGTCTGAACAAGAGCAATTAACATAGAATCACCCAAAtatagttacttgtcctaaa
gacttatcagtcttgtgtacGGCAACTATTTTATCCAGAAAAGTACTTCATCTCCAGACTtatagttacttgtcctaaac
gacttatcagtcttgtgtacTTAGATGTGGATGTGTAGACTCCTCTATTGGGCTGGAATAtatagttacttgtcctaaac
gacttatcagtcttgtgtacGCTACTTCTACAGCCCATTACCTACATTCCAAGCTACCTAtatagttacttgtcctaaa
gacttatcagtcttgtgtacCATTCACTTTCTGCTATGATAAACTCTGCTTTCTCTGTCTtatagttacttgtcctaaac
gacttatcagtcttgtgtacCTGAAAGTCTATGTAAGTGGAGGGGGCAGGGAAGAGAGGCtatagttacttgtcctaa
gacttatcagtcttgtgtacCTCAATAGCTTTTTTTCTAGTCTCCTTCTGACACTTGGGAtatagttacttgtcctaaac
gacttatcagtcttgtgtacTTAGCCAAGAAATTACAAGTGAAGCGGAGAGAAAATATGAtatagttacttgtcctaaa
gacttatcagtcttgtgtacAGAGAGTTTCTGAGTTTGACAGGCAGAGGTAAAGCAAAAAtatagttacttgtcctaaa
gacttatcagtcttgtgtacTGACTAGTGCCGGTATAACAGTTTTCAAAGTTGACATTTGtatagttacttgtcctaaac
gacttatcagtcttgtgtacGTTTGAGTGGAGAAACAATAGTGCTTAAGTTAAGTTCAACtatagttacttgtcctaaac
gacttatcagtcttgtgtacACCTTTATACCTTTTCGTTTTTATCTTGTCTTCTAGGGACtatagttacttgtcctaaac
gacttatcagtcttgtgtacGGTGGCAGTATCTATGACAACAAACTCATTTCATTTTCTTtatagttacttgtcctaaac
gacttatcagtcttgtgtacTTGATTTCTTTTCTACATACATAATTAAGCACCCGGAGACtatagttacttgtcctaaac
gacttatcagtcttgtgtacTGAAGGGCAATACTGTGGGTGGGGACTGGACATAATTAATtatagttacttgtcctaaa
gacttatcagtcttgtgtacCAAGAGAAATAATTCAAAGAGAAGCAAGAAAGGATGAGACtatagttacttgtcctaaa
gacttatcagtcttgtgtacGGCATGACCTGAGGCCCCAGGGGGGAAGCAAAAGCCTGAAtatagttacttgtccta
gacttatcagtcttgtgtacATGCTTGTTCTCCCTGAAGGTTTGCACAGACAACCCACGTtatagttacttgtcctaaa
gacttatcagtcttgtgtacGGGTCTTGCCTTTATTGACTGTCTTAAAGTTAATCCCAGTtatagttacttgtcctaaac
gacttatcagtcttgtgtacCCAGGGGAAACGAATTTAGGGGAAAAACACCTATCTGGCAtatagttacttgtcctaa
gacttatcagtcttgtgtacGTAGCTTTTACTGTTCAAGGGACAAGAGGTTAACTATTTTtatagttacttgtcctaaac
gacttatcagtcttgtgtacCCCATGAGCCAGTTTCCAGTGACACTCTGATTATCACATAtatagttacttgtcctaaa
gacttatcagtcttgtgtacGGATTCCTGGCTACTTTCCCCCCCATTTTTGTTCTGAATTtatagttacttgtcctaaac
gacttatcagtcttgtgtacAATGTGAATATTTTTCAGCACTTTCTCTCATTGGTCTGTGtatagttacttgtcctaaac
gacttatcagtcttgtgtacCCTGACTGGTTGTTAAAGTTGGCTTGGTAGCTTTATAAAAtatagttacttgtcctaaac
gacttatcagtcttgtgtacGATTTGTAAGTCAGTCCAGATCTACCTAGTTAGAATCTCTtatagttacttgtcctaaac
gacttatcagtcttgtgtacGTATGTGAGATAAGAGTGTTCCTTTTGATATTGGTTAGAGtatagttacttgtcctaaac
gacttatcagtcttgtgtacAATAGTAGATTCGCCACAATTCAGCTGCAACAGGACTTTAtatagttacttgtcctaaa
gacttatcagtcttgtgtacCGAGTTAATACTATTACCTAATGAATAGAATTGCTGTGGGtatagttacttgtcctaaac
gacttatcagtcttgtgtacCACACACACACAGAGTACTCCCCCGCTTATCCTCAGTTTTtatagttacttgtcctaaa
gacttatcagtcttgtgtacGTTTCAGTTGCCTGCAGTCATCCATGGTCTAAAAATATTAtatagttacttgtcctaaac
gacttatcagtcttgtgtacTTCATAAGTTTTAAGTTGCCACCATTCTGTCACGAAGAAAtatagttacttgtcctaaac
gacttatcagtcttgtgtacATGCCACTTTGTCCTGCCTGTAACATGAATCGTCTCTCTGtatagttacttgtcctaaa
gacttatcagtcttgtgtacACTGTCTACATCGCCCGCCAGTTAGTCACTAAGTTGCCATtatagttacttgtcctaaa
gacttatcagtcttgtgtacTGCAAGAGTAATGATGCTGGAAATTCAGATACGCCAAAGAtatagttacttgtcctaaa
gacttatcagtcttgtgtacTTAAAGTGCTTGCTTTAAGTGCAAAGTGAAAATTCTCCACtatagttacttgtcctaaac
gacttatcagtcttgtgtacATCTTCTGTCCATGAAATTGCAAGGAAGGAAACAGAAATTtatagttacttgtcctaaa
gacttatcagtcttgtgtacGGAGTTCCTGCCACAGTGCCTGATAGCTTTTATTACAGTAtatagttacttgtcctaaa
gacttatcagtcttgtgtacTTTTTATAGAAGAAAACCCCACAGCCTATACAGGGATTGAtatagttacttgtcctaaa
gacttatcagtcttgtgtacCCCCATGGATAAGGAGTGACTATGGTGTATCACATTTTAAtatagttacttgtcctaaa

gacttatcagtcttgtgtacTTCTGTAAACAGTTCTTAAGATGGAAACTATATGGTCTCCtatagttacttgtcctaaac
gacttatcagtcttgtgtacGCACATACTGGCTGCTGAAGAAAAAAATCAGATTGATCTAtatagttacttgtcctaaac
gacttatcagtcttgtgtacCCAGTGAAAGTTAAAGAAAACAGTTCAGAGATTTGGTCACtatagttacttgtcctaaac
gacttatcagtcttgtgtacGATACACATCCAATGTCCCCACTTAGCATGCCTGATTTTGtatagttacttgtcctaaac
gacttatcagtcttgtgtacCCAGGAAAGTAAGGAGGCAAAACACTGTGATATTAGAGAAtatagttacttgtcctaaa
gacttatcagtcttgtgtacCATTGGTTCCAGTGCAAGGTCATTGTTCGTTAATACTTGTtatagttacttgtcctaaac
gacttatcagtcttgtgtacTTAACCACCCAGAAAATTCACTTCAAGTTTTCAAGCTGCAtatagttacttgtcctaaac
gacttatcagtcttgtgtacTCAAAATCATTACCTGAATGTAGATTAGGAGATGCGGGCTtatagttacttgtcctaaac
gacttatcagtcttgtgtacGAGATTATTCCATGTATGACTTATCCCACCCTACTCACCCtatagttacttgtcctaaac
gacttatcagtcttgtgtacGAAGGATTCAGCACTGCACACCAGAAGGACAAGTTGGAGTtatagttacttgtcctaa
gacttatcagtcttgtgtacTGGAAACTTTTGACTTGAGTGCCCACTAAACAGTGACAATtatagttacttgtcctaaac
gacttatcagtcttgtgtacCTTCTGGATCTGCTTCTCTAGGCATCTTTTAATCCCTTGCtatagttacttgtcctaaac
gacttatcagtcttgtgtacCTAGTGTGGCACCTCCTTGCACAATAAAATGTCAATTATTtatagttacttgtcctaaac
gacttatcagtcttgtgtacATCTAATGGGATGGGGTTAGGAATTTACTCAGTGCACTTCtatagttacttgtcctaaac
gacttatcagtcttgtgtacTGACTATCTTATAGCCAAATTTACCTTACCTGGGAGGCCCtatagttacttgtcctaaac
gacttatcagtcttgtgtacTGGCTGCAAAAAGTTCCATTTACCAAGTACCTGTAGCTCAtatagttacttgtcctaaac
gacttatcagtcttgtgtacGCGGTAAACATTTAATCCATTCAAAGTTCAAGCTTTTCTCtatagttacttgtcctaaac
gacttatcagtcttgtgtacTCGCCTGTCTCTGTTCCATGTGAGATGCCCTCTGGGTGCAtatagttacttgtcctaaa
gacttatcagtcttgtgtacATGCTTAAGTGTTCACATGGGAGTAAAGACCTTTGTTTACtatagttacttgtcctaaac
gacttatcagtcttgtgtacCATTCTCTGCAATATCTCACCCTATCTGGTCACTGAGCAAtatagttacttgtcctaaac
gacttatcagtcttgtgtacGCTGGTCCTGCCTTGTGCTCTGAGCATTGTACTGGTATCAtatagttacttgtcctaaac
gacttatcagtcttgtgtacTCATCCTGACCCTGACTTGATCCTGTGGCCTCTGTTCTTAtatagttacttgtcctaaac
gacttatcagtcttgtgtacCTTCTCTTAGCCTACAACTTCCCTCAGCACTCACTTATCCtatagttacttgtcctaaac
gacttatcagtcttgtgtacGCACATGGGGCTTCTATATCCTTGGATTGGGGAGAAACATtatagttacttgtcctaaa
gacttatcagtcttgtgtacTCCCCCCAGACAATTGTACTTCTACCCCCATGCAAGTTGGtatagttacttgtcctaaa
gacttatcagtcttgtgtacGCATCTTGACCAACTCAGAATGTCTCTAAGGGGCTTGCAAtatagttacttgtcctaaa
gacttatcagtcttgtgtacAGGGGTGTTTCTACTGTACCTCTTCTCAGAACATCAACTCtatagttacttgtcctaaac
gacttatcagtcttgtgtacAGAAATTTAAGACACTTCTGAGCCTTCTCTCTTTTCGTATtatagttacttgtcctaaac
gacttatcagtcttgtgtacTCTCCCTTTCTTTCTCCCCCAACTCGGCTCTTTCCCTTGAtatagttacttgtcctaaac
gacttatcagtcttgtgtacCAAGAGCTGCTCCTATATCCTATCTTCTTTATATCCTCTGtatagttacttgtcctaaac
gacttatcagtcttgtgtacCCTTGGCTGCAGGGATTATGAAAAGGCTGTAAGGAAAAAAtatagttacttgtcctaaa
gacttatcagtcttgtgtacGGTTGCTGCACAGTTGGCCCTTGCTAACTGTGGGTTCTGCtatagttacttgtcctaaa
gacttatcagtcttgtgtacATATAAGAGGATGTGCATATCCTATATGCAAATGCTATGCtatagttacttgtcctaaac
gacttatcagtcttgtgtacAGATACTAAGGGATGACTGTGTTTTTTTTTTTAGGGCTGAtatagttacttgtcctaaac
gacttatcagtcttgtgtacAAGTACCACAATCCAGAAGGCCAGAGTATAAAATCAAGGTtatagttacttgtcctaaa
gacttatcagtcttgtgtacGAATCTGTTCCAGACCTCTTTCCATGGTTTGTAGATGGCCtatagttacttgtcctaaac
gacttatcagtcttgtgtacCTGTATCTTTATATCATCTTCCCTGTCTATGTGTCTGTGTtatagttacttgtcctaaac
gacttatcagtcttgtgtacCCTCATCTTAACTTCATTAGCTCTATAAAGACCTAATCTGtatagttacttgtcctaaac
gacttatcagtcttgtgtacCCAGGAAGGGAAAATACATTGGTAAGCATTTCACAATATTtatagttacttgtcctaaac
gacttatcagtcttgtgtacCCATCACATTAAATCTGTACATTAGGATGGAGCAAGAATGtatagttacttgtcctaaac
gacttatcagtcttgtgtacAGTTGTCTTAGCATTTTCTTATACTGTCACTGACACTCTAtatagttacttgtcctaaac
gacttatcagtcttgtgtacTGAAGATTTGGACTGGGGTGTAACTGGAAGGGGCAAACCCtatagttacttgtcctaa
gacttatcagtcttgtgtacCAAGAGATGAAGCCGATGTTATATTTTTCCCAGGTAAACTtatagttacttgtcctaaac
gacttatcagtcttgtgtacTACATCTTAGTAGGAGAGCTGAAGGGAGAGTATTTGAGCCtatagttacttgtcctaaa
gacttatcagtcttgtgtacAGTGAGTAGAGCTGATGGTGGAGGATTAGGCAAGGAACATtatagttacttgtcctaaa
gacttatcagtcttgtgtacGTGGGAGTCTGGTGGTGACAGTAATCAGAGTGGTTGCTTCtatagttacttgtcctaaa
gacttatcagtcttgtgtacTCATGTATTCATTCATTTTCTGTTCTGTAGGGGCAGTGCCtatagttacttgtcctaaac
gacttatcagtcttgtgtacCCTTGCACGGCTCCAAATCGTACTTCTTCACTGACTTATTtatagttacttgtcctaaac
gacttatcagtcttgtgtacAGGGTTATTGCAATGATTGAGTGAATTCATGTATGTAGAGtatagttacttgtcctaaac
gacttatcagtcttgtgtacTCTACTGTTATTAAAAATATTAGGGGCTAGGTGATGGGGAtatagttacttgtcctaaac
gacttatcagtcttgtgtacAAGATATCTATTGCACCCTGAAGAAACTCACAATCCATGGtatagttacttgtcctaaac
gacttatcagtcttgtgtacAAGCATCCAAACACACAATTGCTATTGAATGATGAGCACTtatagttacttgtcctaaac

gacttatcagtcttgtgtacCATCCAGTGCATCTCATTCATGTTGTGAGCCTTAGAGAGGtatagttacttgtcctaaa
gacttatcagtcttgtgtacCTGTGGAGAGGATGCCTAATTTAGATTTCAGATGATAAGTtatagttacttgtcctaaac
gacttatcagtcttgtgtacAGATTCCTGAGCCCCATACCTGAGATGTTGATTTGTCTGTtatagttacttgtcctaaac
gacttatcagtcttgtgtacGATGAAGTGGCAACGATCTAGGGGAGGAAGAGAAGTGGGTtatagttacttgtcctaa
gacttatcagtcttgtgtacAGTGCATGTGGGTTGAGCAGTGTTAGCAGAGAAGAATAAAtatagttacttgtcctaaa
gacttatcagtcttgtgtacCTAAGTGTCTTATTCTGTTTTAGGAGCTTGGACTTGGTGAtatagttacttgtcctaaac
gacttatcagtcttgtgtacCCAGTGGAAGGATTTAGGTGGGGGAGCAAGAGTATCAGATtatagttacttgtcctaa
gacttatcagtcttgtgtacAGGGAAAAAGCTCAGAAAGTAGGAAGGCCAGTCATTCTTGtatagttacttgtcctaaa
gacttatcagtcttgtgtacTCCAGCTAGAATAAAAGGAGACAGAGACCAGATTTGTTAAtatagttacttgtcctaaa
gacttatcagtcttgtgtacCAGTTTTGACAGCATTTGGAGGTACAACTAGATATGGAAAtatagttacttgtcctaaac
gacttatcagtcttgtgtacCAAGTCAGTGATGCCTCCTATGATTTTTGATTTGAGTCATtatagttacttgtcctaaac
gacttatcagtcttgtgtacTGGGACAGGCATGATGGTGGAGGGAGAGAAATTCATAAGTtatagttacttgtcctaaa
gacttatcagtcttgtgtacTGGGATGTGTTGGGTTTAAAGTGCCTGGAAGATATTCAAGtatagttacttgtcctaaac
gacttatcagtcttgtgtacGCCTTGTGCTTATCCATGCTTTGTGCTTTATACACGCTGTtatagttacttgtcctaaac
gacttatcagtcttgtgtacGCACTCAACTAACAATGGAAGAGGCATTACGGTCCCAATTtatagttacttgtcctaaa
gacttatcagtcttgtgtacAGGAGGGGCTGGAATTCACAGAAGTTTTGTGGCTTACTCAtatagttacttgtcctaaa
gacttatcagtcttgtgtacAAATTTTGCAGACATTTTGGTACAGATTCAAGCAGCCAGGtatagttacttgtcctaaac
gacttatcagtcttgtgtacGACACATATTTCATACTTTTGTCCTAGACCTAGAACCTTTtatagttacttgtcctaaac
gacttatcagtcttgtgtacGCCCAGGGTTGCTATTTGTTAATCACATGTTAGAAAATACtatagttacttgtcctaaac
gacttatcagtcttgtgtacGACTATAATATCTACATTAGAATTCACCAGAGTCTTGGAGtatagttacttgtcctaaac
gacttatcagtcttgtgtacGAAGGGTAAAGGGAAAACTAGTGGCTTCTAACTTAAACACtatagttacttgtcctaaa
gacttatcagtcttgtgtacAAGAAATTTTGTGGGCCATTAGTTCCTTAGCTATTTTCTGtatagttacttgtcctaaac
gacttatcagtcttgtgtacGTACTAAACTGGTAAGATACTGAGATTTAAAGAACAGCACtatagttacttgtcctaaac
gacttatcagtcttgtgtacCATGCAAAAGAGCTGTCTATAATGAATGACTTGATAACAGtatagttacttgtcctaaac
gacttatcagtcttgtgtacTCTAAGGGAATGCTGAAAGTCATTGGGTGTTTCTTGGTGGtatagttacttgtcctaaac
gacttatcagtcttgtgtacCCTCTGACGGTGGGATACATAACATGGCCAAAAATCTGACtatagttacttgtcctaaa
gacttatcagtcttgtgtacTGGCTGGAGTGCTACCTTAGGAGACTGCTACCAGAGACTGtatagttacttgtcctaa
gacttatcagtcttgtgtacAGTGAAATGCCTTCTCTGCAGACACAGATGTACGTGGGTGtatagttacttgtcctaaa
gacttatcagtcttgtgtacAAGAATTGAAATAACATCCTATCTGCAGCTTTAAGTGAGCtatagttacttgtcctaaac
gacttatcagtcttgtgtacCTATGATGTAGTTTTAAAGAACCTCAACTCTGTATTCAGGtatagttacttgtcctaaac
gacttatcagtcttgtgtacTACCCTAAAATTCCACCAATTAGTATCTATGTGAACTTGGtatagttacttgtcctaaac
gacttatcagtcttgtgtacCACTTCAATGTGTTGCTGTGCACAATAAATGTTTTCAAAGtatagttacttgtcctaaac
gacttatcagtcttgtgtacAATGTCTGGTCTGCTCATACTCAGCACAAGCATTTACTCTtatagttacttgtcctaaac
gacttatcagtcttgtgtacAGATATCTTGGTTCTTTTTGCCCTGCCTTCTATTAGAAAAtatagttacttgtcctaaac
gacttatcagtcttgtgtacGAGCCCTACCTCAAGAATCCTCTTCTCCACCAACAAATTGtatagttacttgtcctaaa
gacttatcagtcttgtgtacAAGCAGAGAAAGGGGCGTTATGGTATGGGATAGAGTCAGAtatagttacttgtcctaa
gacttatcagtcttgtgtacCCTGTGGATCTGGGACATCTTTAAGTCTAAGTGGGGGGCCtatagttacttgtcctaaa
gacttatcagtcttgtgtacCAATGGGTGGGGCTTTCTTTTCGGTCATTCTTCTTTTATCtatagttacttgtcctaaac
gacttatcagtcttgtgtacCCTACATTCTCAGTGCCTTCTCCATTCTCTCTTATTTAGTtatagttacttgtcctaaac
gacttatcagtcttgtgtacTTCTTTTCTAGTAATAGCCTGAGCTAGTGAGAGTTGGGCTtatagttacttgtcctaaac
gacttatcagtcttgtgtacCCCTGTGACACGGCCATTCATTTATTTAGCTCCATAAACCtatagttacttgtcctaaa
gacttatcagtcttgtgtacCAGTGCTGGGCTCTCATTCCTCCATCATTTTTCTGAGTTCtatagttacttgtcctaaac
gacttatcagtcttgtgtacCCATTTTTCTTTTGGTTCTCATCTTCATACATGGTTCTCTtatagttacttgtcctaaac
gacttatcagtcttgtgtacCAACTTCGTCTTGCTGAGAGCCTTCTCCAACTGCTTCCATtatagttacttgtcctaaa
gacttatcagtcttgtgtacCTGCTTCAGAAACTGAACTCTTCCTTAGCTAGTTTAAAAAtatagttacttgtcctaaac
gacttatcagtcttgtgtacTTGATAAATACTGTTTCCACTTTTGGCAACCCAGCTTCCCtatagttacttgtcctaaac
gacttatcagtcttgtgtacTCTGGATAATCCTGAGTTCTAACAGAGGCTTCAAACGAACtatagttacttgtcctaaa
gacttatcagtcttgtgtacAGGAACATCTTCGCTGCTCACACCTGTAGTCTTTCGTAAGtatagttacttgtcctaaa
gacttatcagtcttgtgtacAGACAGTCTTATCTTTCTTCTTTCTTTCAGAGGCAGAACCtatagttacttgtcctaaac
gacttatcagtcttgtgtacAGCAATCCTGAAGCTTAGGCAATATTTGGGACCTCTTTAAtatagttacttgtcctaaac
gacttatcagtcttgtgtacACGACAGCAACTTACAATTTTTAAAAGCTGACAAAACCACtatagttacttgtcctaaa
gacttatcagtcttgtgtacAATTGTATCCCTTCATTAATTTACCACTTGACACACCAACtatagttacttgtcctaaac

gacttatcagtcttgtgtacGTGTGTTCTTGCTACACTTTTTATTATTGGTAGTGTCATGtatagttacttgtcctaaac
gacttatcagtcttgtgtacTGGATTCACTCACAGACGTATTTGCTGTTTGTAGACTGTTtatagttacttgtcctaaac
gacttatcagtcttgtgtacTTTGCACCCTACAAACACAGGGATACTGATAAATTCTATTtatagttacttgtcctaaac
gacttatcagtcttgtgtacAATGGAATAATCTGCATTAGCTAGAATGCATTAGCCTGAGtatagttacttgtcctaaac
gacttatcagtcttgtgtacGACTCCCTGCTTACAATTTTACATATCCAATACTTGAAAGtatagttacttgtcctaaac
gacttatcagtcttgtgtacGCTCCAGGTGTTTCAAGCTTTGTTTCTTATGCACTCCCATtatagttacttgtcctaaac
gacttatcagtcttgtgtacACATTCATTTGGTGAGGTTATCTTTGGGATATATGGTACCtatagttacttgtcctaaac
gacttatcagtcttgtgtacAACGAGGCTGTTGAGTTGACACAAAGGGCAATAGGAATTTtatagttacttgtcctaaac
gacttatcagtcttgtgtacGAAGCCATTTCTATACTAGGATGATTGAAAATAGCAGCTAtatagttacttgtcctaaac
gacttatcagtcttgtgtacCAAGTGACTGCAAACTACATACATATATTCTGCCAGATCCtatagttacttgtcctaaac
gacttatcagtcttgtgtacACTCAACTTACTTTAATTGGATCCCAAATTACCCATGGCCtatagttacttgtcctaaac
gacttatcagtcttgtgtacCTGAACACGAGGGCAAGTTTAACAGAGGTAGAGTCAGAATtatagttacttgtcctaaa
gacttatcagtcttgtgtacTTTCACAAAATTACAAAAGCACATTGCTAGAGCCCCTCTCtatagttacttgtcctaaac
gacttatcagtcttgtgtacGACACTTAAGTTTCATTAGTCTTACGGTAAGTCCATCTCTtatagttacttgtcctaaac
gacttatcagtcttgtgtacTTACCTGCCTGCTTGCCTTTTCCGTTCTTGCTACCCTCCAtatagttacttgtcctaaac
gacttatcagtcttgtgtacTTCTTTTTCCTTCCTTATTTCTTCCATCCCTTTCTCCATCtatagttacttgtcctaaac
gacttatcagtcttgtgtacATTGAAGAATGAGTAGAAGCATAGTGATTAAGATGGGTTGtatagttacttgtcctaaac
gacttatcagtcttgtgtacGACATTAAAGCTGCATTACTTAACTGTGTGTCTTTGAGTGtatagttacttgtcctaaac
gacttatcagtcttgtgtacCCTCATCTATTCAACAAGGATAATGATAGTACCAGTCACAtatagttacttgtcctaaac
gacttatcagtcttgtgtacCATACTGACTGTTCAATCAATGTTAGCTGCTGTTCTATATtatagttacttgtcctaaac
gacttatcagtcttgtgtacAAAACAGAAGAGAGACCACACTCCTTGGGGATGTGAGACAtatagttacttgtcctaaa
gacttatcagtcttgtgtacATGAATCTTTATAATTGGTAAATGCTCCCATAGCAGGTACtatagttacttgtcctaaac
gacttatcagtcttgtgtacCTGAATAAGCGATTGAATCATGGGTTAAATGGATGCAGAAtatagttacttgtcctaaac
gacttatcagtcttgtgtacAGTGATTGGATTTAGATAAACAAAAAGGAGGAGGATGAAGtatagttacttgtcctaaac
gacttatcagtcttgtgtacCTGGGGGCTGATACAAGGAAAGGAAAGGCATGGAGGCTAGtatagttacttgtcctaa
gacttatcagtcttgtgtacTGAACTATATTTCCTTAGAACACAATAAACTAGACCCTGGtatagttacttgtcctaaac
gacttatcagtcttgtgtacTAGAAGTTTTATCCCAATTTAAAAATATTAGGGGGCTGGCtatagttacttgtcctaaac
gacttatcagtcttgtgtacATCCAAAATAAAATTCTGTGAATGACTTTGGTGGATACCGtatagttacttgtcctaaac
gacttatcagtcttgtgtacTGACCAGCACAACTGCCTTTCTTGATCTATTATGAACAATtatagttacttgtcctaaac
gacttatcagtcttgtgtacGTTAAATGTTCATATGATCAACCTCTCTTGACAAAGCTCTtatagttacttgtcctaaac
gacttatcagtcttgtgtacCTGATGGGGATCTACAGGGTTGGTAACACAGCTTCTTTCTtatagttacttgtcctaaac
gacttatcagtcttgtgtacTTGATATCAAGAAGGGACCACCAAGAGAACTGCAGAGACAtatagttacttgtcctaaa
gacttatcagtcttgtgtacGAGCAAAACACCATCTCTCTAGTTATTTATGCCCACTAACtatagttacttgtcctaaac
gacttatcagtcttgtgtacGCCACTTAAGGGTTTTCATTTATATACTAAACAGGAAGTCtatagttacttgtcctaaac
gacttatcagtcttgtgtacCCAAATTATCTATGTAACCTTGGCTGTTATTTAACCTCCAtatagttacttgtcctaaac
gacttatcagtcttgtgtacGGCATATCATGAGTTCTAGTAAATAACTCTAAAGTGCTTCtatagttacttgtcctaaac
gacttatcagtcttgtgtacGATAATGATCATGCCAGCAGAACCACTTTTGTACGACACAtatagttacttgtcctaaac
gacttatcagtcttgtgtacATAGTATTTCTTATAGGGAGAGAGCATGCATATTCCTAAGtatagttacttgtcctaaac
gacttatcagtcttgtgtacTTGAAAACTAAATGGAGAAGATAATGGTGTCATGATTGGGtatagttacttgtcctaaac
gacttatcagtcttgtgtacGCATTATATGTCATGACCAGGGAAATGTTAGCTTCTTATTtatagttacttgtcctaaac
gacttatcagtcttgtgtacGCTCACCTTTTAGCCTCTCTCTGAATTTTGCTCATCTTTTtatagttacttgtcctaaac
gacttatcagtcttgtgtacCAGTTATGCCTGCTGAGAGCCTGCTTTATTTTTGGTGTGAtatagttacttgtcctaaac
gacttatcagtcttgtgtacGCTTTGCCATGTAAATTCTTCAAAAAATTGCCAAACCCCAtatagttacttgtcctaaac
gacttatcagtcttgtgtacTTCTTGAATCTATGCCTGAGTAGAGAAGTAAAGGTGTATGtatagttacttgtcctaaac
gacttatcagtcttgtgtacGCTACATGCATTCTAATGTTTTCTTGGGCTGGATAGTTCAtatagttacttgtcctaaac
gacttatcagtcttgtgtacGGTTTCGATCCTACCTAATGAACTAGCCAGATTTTCACAGtatagttacttgtcctaaac
gacttatcagtcttgtgtacGCTTCAATTTGCTAATATGTTCTGTTGATTCCAAGTGAGAtatagttacttgtcctaaac
gacttatcagtcttgtgtacCTATGGAAGCAAGGATAGAATTAGCATCTACAACATCATGtatagttacttgtcctaaac
gacttatcagtcttgtgtacAAAAAGGAGGGGACAGTGAATAATACTCTAGTGTATGTTTtatagttacttgtcctaaac
gacttatcagtcttgtgtacGGTAGCTCGGATTTGGACCTTCTGTTCTCTTTTTGTTGTGtatagttacttgtcctaaac
gacttatcagtcttgtgtacGTCCTGGATTGATCAGCACTAATAGTTTATCTGAGACTCAtatagttacttgtcctaaac
gacttatcagtcttgtgtacCAAATCCTTTGTCACAAGTTAAGTTTGAAAAGCAGATGGCtatagttacttgtcctaaac

gacttatcagtcttgtgtacCCCACCCTCACTTTTTTAAACGCAGGTGCAGTTTTAAGGtatagttacttgtcctaaac
gacttatcagtcttgtgtacCTTAACGCTACTTTTCTCCCTTCTCTCTGAATACTAAAAAtatagttacttgtcctaaac
gacttatcagtcttgtgtacCAATTGATGACTCTCCTGTAGTTATTAGTGACCAGTAGTTtatagttacttgtcctaaac
gacttatcagtcttgtgtacCTCATCAGGGTAGAACAACTAGGAAGAGAGACTGGGACCTtatagttacttgtcctaa
gacttatcagtcttgtgtacTCTCAGACATTCCTTTAGCAGACTTAGTAGGTAGGTATATtatagttacttgtcctaaac
gacttatcagtcttgtgtacTTTTTCAGTGCACAGCTAGTACTTTAGGTACTTGAAGTAGtatagttacttgtcctaaac
gacttatcagtcttgtgtacAGATGGACACACTTGCTCACTCCTTTCTCTTAACATTCATtatagttacttgtcctaaac
gacttatcagtcttgtgtacCCTTTAGGGTGGTTTATTATATGTAATTGAGAGCCAGTGTtatagttacttgtcctaaac
gacttatcagtcttgtgtacGTGTACTTCTGTAAACCCTTTAAATTTACCCCTTAATGAGtatagttacttgtcctaaac
gacttatcagtcttgtgtacTATTAGAAAAAAATAAAAGGGTGGGGGAAGCAGTGCATCAtatagttacttgtcctaaa
gacttatcagtcttgtgtacAGGAGCTGATAATGTCCTTTTGCAAGAAACACTCTTAAAAtatagttacttgtcctaaac
gacttatcagtcttgtgtacGTTGGAAGTATCTTTCTCTACTTGATGAATTTGATAGTGCtatagttacttgtcctaaac
gacttatcagtcttgtgtacAATAAAGAGAATAGAATGGTTATTTTTGAGGGGGCTGACTtatagttacttgtcctaaac
gacttatcagtcttgtgtacAAGAATCAAGCAACATTGTTTTAGACATCTTTGACTGGCAtatagttacttgtcctaaac
gacttatcagtcttgtgtacGGAAATTGGATATCTCTGGACTGAAAAATTAGTGTTTGTGtatagttacttgtcctaaac
gacttatcagtcttgtgtacAGATTTTTATTTCTTGAGAGGAAGTGAGCCAGACATTTAGtatagttacttgtcctaaac
gacttatcagtcttgtgtacAAAAATTGGAGTTCAAAAAAGATTTTCCAGGGGCAGATGTtatagttacttgtcctaaac
gacttatcagtcttgtgtacAGAGTTTTTATTTTAACCTCCCACGTTCCTCTCCTCAGTGtatagttacttgtcctaaac
gacttatcagtcttgtgtacTATCTCAGTGGTGATAAATAAAATCTGGAAACCTGTCATCtatagttacttgtcctaaac
gacttatcagtcttgtgtacCTTTACTCAGAAAGGGCCTGGACCAAATTATCTCTTGAGGtatagttacttgtcctaaa
gacttatcagtcttgtgtacGATTAGTATCCAAATGATCCAAAGAATTTCACTGGCTTTCtatagttacttgtcctaaac
gacttatcagtcttgtgtacCACTGCTATAGGGTCATAATCTCTTGTTACATGTCAAAAAtatagttacttgtcctaaac
gacttatcagtcttgtgtacTATTGGCAATATTTCCCTTACTAACTGTGTTCTGGACAAAtatagttacttgtcctaaac
gacttatcagtcttgtgtacTTTAAATGGTATGAAAGAGTGCTTAGAGAAGACAGTTTGGtatagttacttgtcctaaac
gacttatcagtcttgtgtacTCCAGCTGATAGCCTTAGAACCAAGTGAAATGTCTGATTGtatagttacttgtcctaaa
gacttatcagtcttgtgtacCAAGGTTAATGATATTCAAGACTAATATCCTCTGCACTCAtatagttacttgtcctaaac
gacttatcagtcttgtgtacTTCAAAAGATTTGAGAGACCTCTTATGTCATTACCAGTAGtatagttacttgtcctaaac
gacttatcagtcttgtgtacTGTTCTTACACTACTGCCAGACTAATCAGAATCTGTGGCTtatagttacttgtcctaaac
gacttatcagtcttgtgtacGCGGAGAGTTTCTCATCCATTTTGTTTACTTTTTTAGACTtatagttacttgtcctaaac
gacttatcagtcttgtgtacTGCATGCTTCTGTTAATTATACTAAAAGAATGGGATGGTCtatagttacttgtcctaaac
gacttatcagtcttgtgtacGAAGAATCGCAATAACATGGCATAAGAACTCAACCTTTGTtatagttacttgtcctaaa
gacttatcagtcttgtgtacACGAAAGTAAACATTTGTGGCTGTTGAGTGGCTCTTTCCAtatagttacttgtcctaaac
gacttatcagtcttgtgtacGCCTGACACATCTGCCATTTATAATACATGCTTTCTAAGTtatagttacttgtcctaaac
gacttatcagtcttgtgtacACCAAGCAGAGAGCAGAAAGAAAAAGTCAAGGATCACGCCtatagttacttgtcctaa
gacttatcagtcttgtgtacCATAGACATGCATGGATACGAGAGAGGGCGGGAAACAATAtatagttacttgtcctaa
gacttatcagtcttgtgtacTGTAGGCAGGAAGGAAGAAAGGACTTTGGCAGACAGCTTGtatagttacttgtcctaa
gacttatcagtcttgtgtacACCGCAATTACTTTGAACCCACCTAATACAAAGAAAAAGtatagttacttgtcctaaa
gacttatcagtcttgtgtacCTGTTCATTCCATATGTCTTTTCTAGACATATAGAGAGTGtatagttacttgtcctaaac
gacttatcagtcttgtgtacTAGAAGAAAAAGAAGCTGAAGTGACATTTAAAGGATTGCCtatagttacttgtcctaaa
gacttatcagtcttgtgtacCCACTAAGGACATAAAGACTTTTTACAAGATTGAAGCCAAtatagttacttgtcctaaa
gacttatcagtcttgtgtacAGGACTCTTCTACTTTTCACTTTTCCTCCCCTCCCTCTACtatagttacttgtcctaaac
gacttatcagtcttgtgtacCCCTCTGTTGATCATTATGAGTTCTCTATTTAAATGGAAGtatagttacttgtcctaaac
gacttatcagtcttgtgtacCAGTGCTAGTCCTTTTACCATAACTTTTCCATTACTGAGTtatagttacttgtcctaaac
gacttatcagtcttgtgtacTTTTAGCAAGGGGATTATAAGTTCTTCCTTCTTAGTGTTCtatagttacttgtcctaaac
gacttatcagtcttgtgtacCAGCCTATTGCCTTTATGCTTGCAACACTTGACTGGGTGTtatagttacttgtcctaaa
gacttatcagtcttgtgtacCTGCCCTTTAGTATGTTGCAGACATTGTTCTGTTGACTTCtatagttacttgtcctaaac
gacttatcagtcttgtgtacCATGAGGCCAATCTGAGTTTTCCTACTATGGGTAATTTTTtatagttacttgtcctaaac
gacttatcagtcttgtgtacGCCAAATTGCTATGTCTAGGTCCTATTCTAAGAATTTCACtatagttacttgtcctaaac
gacttatcagtcttgtgtacTTGAGGTAGCAGGTAATGTTATTCATATTTCTCTGACAAGtatagttacttgtcctaaac
gacttatcagtcttgtgtacATGTAAGTGAAGAACCTTAATTGTGGTGGCAGGACCAGGAtatagttacttgtcctaaa
gacttatcagtcttgtgtacAGTGTGGCCATAATCACTATGCCTCCTTTCTCAGTAGTGTtatagttacttgtcctaaac
gacttatcagtcttgtgtacCAGCATCCTATTGTTACCTGGAGATTCAGTTCTTTATATAtatagttacttgtcctaaac

gacttatcagtcttgtgtacTTCTATTTCATATATATTCAGCTCTCTATAGCAGGGGTTGtatagttacttgtcctaaac
gacttatcagtcttgtgtacGGCCAAATCTAGCCTTCTGCATGTTTTTGTTAACTAAAGTtatagttacttgtcctaaac
gacttatcagtcttgtgtacATGCTAATTCGTTTACATGTTGCTTCCTGGATACATCAGCtatagttacttgtcctaaac
gacttatcagtcttgtgtacCTCATGGCCTGCACAGTTTAAAATATTTACTATTAGATCCtatagttacttgtcctaaac
gacttatcagtcttgtgtacAGAAAAGTCTGAAGACCCCCGTTTTAAGTGATTGATTGTTtatagttacttgtcctaaac
gacttatcagtcttgtgtacGTTGCTGTTTATCACTTTAGTTTGGGCATTTGTTTTACAGtatagttacttgtcctaaac
gacttatcagtcttgtgtacCTGCTTCCATTTGTAGTACTATGATTTTTCTTCTCTACACtatagttacttgtcctaaac
gacttatcagtcttgtgtacTACTTTCTCTTTGCATACTATGGTCTCTTCTTTTTCTCTCtatagttacttgtcctaaac
gacttatcagtcttgtgtacCTTTCTCTCCTTTTGATTATAGTCTGTTTGCAAAGTTTCCtatagttacttgtcctaaac
gacttatcagtcttgtgtacGTTCCTTTCTATCTTGCTCCTTTTTAACCTGAGTGGCTGTtatagttacttgtcctaaac
gacttatcagtcttgtgtacTGTTTTCTCTTTCTCAGAACCACCTAGCCTTCCTTCCCTCtatagttacttgtcctaaac
gacttatcagtcttgtgtacAGTTTGTGCCTGAGAGTTGCCTTCTACACATTCCTTGGGGtatagttacttgtcctaaac
gacttatcagtcttgtgtacGGAGGTGGGACAGATGGAGAGCTCTTCTAGCACAAAATGCtatagttacttgtcctaa
gacttatcagtcttgtgtacTTATTTAAATGTTCTGTTTGAAATCCCTCTCTGTCCAGAGtatagttacttgtcctaaac
gacttatcagtcttgtgtacGGCTTCACCGTGTTTCCTCCAGCTTAGCTCATCAGTTCTGtatagttacttgtcctaaac
gacttatcagtcttgtgtacATGCTCTCCCTAACTTTGCTCTCCCTAATAATGCTCTCCTtatagttacttgtcctaaac
gacttatcagtcttgtgtacAAGCTTTTCTTCAAACTAGAGGACATTAATGGGACTTTTCtatagttacttgtcctaaac
gacttatcagtcttgtgtacCCTTTATTCAGAACTACTTGGAGGTATGGGTGAGATTAAGtatagttacttgtcctaaac
gacttatcagtcttgtgtacCAATGCTTTAATGATCAAGTAGTGCTCTGAAATCTAACCAtatagttacttgtcctaaac
gacttatcagtcttgtgtacCAAATGGCTCTGACCTGTAATTTTGAATTCTATCTTTGGTtatagttacttgtcctaaac
gacttatcagtcttgtgtacTCTAGAGTTCATTGCTTTCTACTTCACAGAGATCTTGGTTtatagttacttgtcctaaac
gacttatcagtcttgtgtacATGCCTCATGTAGCATTTGAGACAGCTGCTTACCCTATTTtatagttacttgtcctaaac
gacttatcagtcttgtgtacGGGGAATAGCCGCCTATCAAAAAATAACAAACACTGAAAAtatagttacttgtcctaaa
gacttatcagtcttgtgtacTTACCACAGCTGCTCACTTCATTTTAAAAAAATTACCGTGtatagttacttgtcctaaac
gacttatcagtcttgtgtacCTGTTTGAGGAGTTGCAGTGTTTATGTGGCTGTTCTTTCTtatagttacttgtcctaaac
gacttatcagtcttgtgtacGCCCCACCCCAAGGAGAAGAGATATTGTATAATAAATTCCtatagttacttgtcctaaa
gacttatcagtcttgtgtacCTCCTGGGAACTAATGATTATACTATCTGTTATTGTCAACtatagttacttgtcctaaac
gacttatcagtcttgtgtacAGTTACTTGCTCATTCCCAAACTCCTTATAGTCAGCTGTTtatagttacttgtcctaaac
gacttatcagtcttgtgtacGCTGGGACAATTGGGAGTGCTGCTTTCAACAAAATCAATTtatagttacttgtcctaaac
gacttatcagtcttgtgtacAGGTGAATATTCTGGTCTTCTGATTAAGTATTTGGAGATCtatagttacttgtcctaaac
gacttatcagtcttgtgtacCTTGTGAGCCAATGTCAACTTCTTGTGTGCACCATTCAACtatagttacttgtcctaaac
gacttatcagtcttgtgtacTGGTGTTCACTTCTGCCAGTTCAGCTCAGAGCCAGCCTTGtatagttacttgtcctaaa
gacttatcagtcttgtgtacCTGCTGCCAGTCCCTAAGTGTCAGTGCATAAGTTGAAACTtatagttacttgtcctaaa
gacttatcagtcttgtgtacATTCTTAAACCCTTATAGAATCATGGAATGTCAGGTGTTCtatagttacttgtcctaaac
gacttatcagtcttgtgtacGATTGCATCGTAGAAATACAGGAGTAATGTGAATATTCAGtatagttacttgtcctaaac
gacttatcagtcttgtgtacCATTCATTTTATCCAGAAATATGCCTTGAGTGTCTACCAAtatagttacttgtcctaaac
gacttatcagtcttgtgtacAGACACCATCTCTCTTTTAAGGTTCTGAGTCCAGTGAGGGtatagttacttgtcctaaac
gacttatcagtcttgtgtacGGAAGCCATTGATTGCGCCTAGGAGTGTTGAGGGTAGAAGtatagttacttgtcctaa
gacttatcagtcttgtgtacCACAGACACAGATTCCCTTGAACTAGGTCCTGAAGCAGGTtatagttacttgtcctaa
gacttatcagtcttgtgtacGTAATAGCAAGAGAGTAGTTGAAGATAATTGTCTTTTGGGtatagttacttgtcctaaac
gacttatcagtcttgtgtacGAGAACTTGCCTCCTCTACCTACAGCTGCCTTACTCACTCtatagttacttgtcctaaa
gacttatcagtcttgtgtacCAGACCTCTCTCTCATCTCTATTCTGTGGATTTTTCATGTtatagttacttgtcctaaac
gacttatcagtcttgtgtacGTCCAATGAACGCAAGCAGACACTTCTTCAGACTAGCTCAtatagttacttgtcctaaa
gacttatcagtcttgtgtacTATATTTTAAATTTCACAATCTTACTCCCTCCCACCTCTCtatagttacttgtcctaaac
gacttatcagtcttgtgtacACCTATGCGCATCTTCTCATATACTTTAAATCATCTCTAGtatagttacttgtcctaaac
gacttatcagtcttgtgtacACATATAATGTGTGAAGATGAAATCAGGGCACTTTGGATAtatagttacttgtcctaaac
gacttatcagtcttgtgtacTCAACTATAGTGACTCTGCTGATTTATCAAACGTTAGGTCtatagttacttgtcctaaac
gacttatcagtcttgtgtacCATGGATGTGGAACTTGTGGGTATGAAGAGCTAACTGTATtatagttacttgtcctaaac
gacttatcagtcttgtgtacTTTAGGAAAGCAAATTCAGTGGAGAATGTAAATCACAAGGtatagttacttgtcctaaac
gacttatcagtcttgtgtacATTTGAGAATTACCAGTCTGCTTACCAGGAATTGTTTTGCtatagttacttgtcctaaac
gacttatcagtcttgtgtacTTTTGTTATGTTTCTCTCCTTGTGTTACTACAAAGCTCAGtatagttacttgtcctaaac
gacttatcagtcttgtgtacTCTTTTTTTCCTGGAGTTCTGAAGTTGCTAAAAAAGGAACtatagttacttgtcctaaac

gacttatcagtcttgtgtacCAGACAAGTGAGCACCGAAAATCTCTGAATTCTGTAATGTtatagttacttgtcctaaac
gacttatcagtcttgtgtacCACTCCTTGCTTTTAATGTGTTTGTGCCATCATTCCATTGtatagttacttgtcctaaac
gacttatcagtcttgtgtacGTTCAGGCTCATGCACTTGAAATTCTTTAGTTACAGTGTGtatagttacttgtcctaaac
gacttatcagtcttgtgtacATGAAAATGGCATCGAGAGCTTGGAAATATTTGCACCTGTtatagttacttgtcctaaac
gacttatcagtcttgtgtacGAGATAGCTCCATATTTCAACATGCTCCTATGGAACGTTGtatagttacttgtcctaaac
gacttatcagtcttgtgtacGTAATTCACAGGTTGTTATGGTAGAGATTTTGACAAGAACtatagttacttgtcctaaac
gacttatcagtcttgtgtacTAAAAACCCTAAGCATGCTAATTACTTCAACGATGACATGtatagttacttgtcctaaac
gacttatcagtcttgtgtacCTTGGTGCTTTGGAACACGGGTGATGCTGTGACTACCCAAtatagttacttgtcctaaa
gacttatcagtcttgtgtacACCTATGGTAATTGTGTCCCGAGTGAAAACAGGCTTTATGtatagttacttgtcctaaac
gacttatcagtcttgtgtacAAGTCAGAGGAAGCAAATGAACAGCAAGATACTGGCTCAGtatagttacttgtcctaaa
gacttatcagtcttgtgtacCAGCCTGTCTGGGAGTCTATAGGTCTTCTGTGTTTCACCTtatagttacttgtcctaaac
gacttatcagtcttgtgtacCAGATGTTTCCCAAATGAACAGGTACCTGAAACCAAAATAtatagttacttgtcctaaac
gacttatcagtcttgtgtacATGGTTCTCATCATATGAGTGAACATCTGTACTGGTTTCAtatagttacttgtcctaaac
gacttatcagtcttgtgtacTTTTTCGAAAATACATCGGTGGTTCACAGATTTCCCAGCAtatagttacttgtcctaaac
gacttatcagtcttgtgtacCAGAAAACCTGGCAGATTTATTAATACCTGTGCATGCTCTtatagttacttgtcctaaac
gacttatcagtcttgtgtacTGCCACTGTTACTATTGTTACTGTCACGAGCTTTTCTTCAtatagttacttgtcctaaac
gacttatcagtcttgtgtacGGAGAAAAGTCCCAGCAACAGTAAGAGAGAGATGAAGATTtatagttacttgtcctaaa
gacttatcagtcttgtgtacCATTCTCTTTCTGCTCCTTTGTTAAAAGCCTCCTGGTTCTtatagttacttgtcctaaac
gacttatcagtcttgtgtacCTTCATTTTCTCTGCTTCTTCCGTGTCGAGAAAGAAGTTAtatagttacttgtcctaaac
gacttatcagtcttgtgtacATCTTAACAAATGTTTATTGGCTACTTTCTCTGTGCTAGCtatagttacttgtcctaaac
gacttatcagtcttgtgtacCTTGGGATGGAGCAATGAACAAGTCACCCTGTCATGTCCTtatagttacttgtcctaaa
gacttatcagtcttgtgtacGTGCCAGAAATTTATCATCTAAACTTTATTCTTCAACCCCtatagttacttgtcctaaac
gacttatcagtcttgtgtacTGTAAGTAGATATTTTCTGGGGGTTTCAATTTCTGATGAGtatagttacttgtcctaaac
gacttatcagtcttgtgtacAAATGTTTTCAACAGATGCTCAAGTAGTCTCCTGAAGAATtatagttacttgtcctaaac
gacttatcagtcttgtgtacTCCTTCCAAATTCCTTAGGAACTTTAGTCAGATAGCTTTTtatagttacttgtcctaaac
gacttatcagtcttgtgtacGTTAATTAGAATTCCACATAAAACCCCCACAGAGACCATGtatagttacttgtcctaaac
gacttatcagtcttgtgtacAAAGTTTCTTCAACAAAGTTTCTCATATGTCCTGGATGCCtatagttacttgtcctaaac
gacttatcagtcttgtgtacGAATACAACACCATAGGAGTAGGAGTAGACATTAAAATACtatagttacttgtcctaaac
gacttatcagtcttgtgtacCAGGTAAGGATACACAATGAGCTGATTTACTTTTAATGAGtatagttacttgtcctaaac
gacttatcagtcttgtgtacCTGTCATTGTCCGTTCTTACACTTATATTTTCCAAATCAGtatagttacttgtcctaaac
gacttatcagtcttgtgtacTAGAGACTTATTTTTCAGAGGGCTTCTTTTGTTTTGGAACtatagttacttgtcctaaac
gacttatcagtcttgtgtacTGTCCTTTTTTTCACACATGTAGTTACTTTTCAGGAGTCTtatagttacttgtcctaaac
gacttatcagtcttgtgtacAGGAAACTCAGGGCCTCAAAACAATATTTTCTTGGTGAGCtatagttacttgtcctaaac
gacttatcagtcttgtgtacACATTTGGTACATTACGAACAGTTTCTTATACACCCAGGTtatagttacttgtcctaaac
gacttatcagtcttgtgtacGCAAGAAAGTAATAATGCCATAACATCCTCTTTGCCTTTAtatagttacttgtcctaaac
gacttatcagtcttgtgtacAACAACAAGGCATTTTAATCATGTATGAAATCTCAGGCACtatagttacttgtcctaaac
gacttatcagtcttgtgtacGATGGTATTAAATAAAGAGGGGCAAAACATCAATAGTTGGtatagttacttgtcctaaac
gacttatcagtcttgtgtacTGGACAACATTTGGTTCAGAGATAGATGACAACAGAAGTTtatagttacttgtcctaaac
gacttatcagtcttgtgtacTCAAATCATGCCTTTTATTTATACATGTTTCAGGCAGCGTtatagttacttgtcctaaac
gacttatcagtcttgtgtacAAGTAATAGAAAGAAACTTACGGGGGCATAAGGTCAAACTtatagttacttgtcctaaac
gacttatcagtcttgtgtacAATGGAAGATAAAGTCTGAAAAGGTAGGAATAGTAGGAACtatagttacttgtcctaaac
gacttatcagtcttgtgtacGCAATATGAGCTGTAAATTCTAATGTTAACTCTGTCATCCtatagttacttgtcctaaac
gacttatcagtcttgtgtacCCCCTTGGCCTTTCTCCTCTGTAACTTAAGAACAATTGAGtatagttacttgtcctaaac
gacttatcagtcttgtgtacCACTTTTCTTCAGGTTATTTTAGACACAATTGCCTTGCCCtatagttacttgtcctaaac
gacttatcagtcttgtgtacCACTAGAGAGCCATTGAAGACAGCACAGTAGAAGCAAGATtatagttacttgtcctaaa
gacttatcagtcttgtgtacCAAGATTCGATATGTCTGTTCTTGGCCTTGAGAAGTCAGGtatagttacttgtcctaaac
gacttatcagtcttgtgtacTTGTCCAAAAAAGCAAACAGACACAAAGCACTTTTCTAAGtatagttacttgtcctaaac
gacttatcagtcttgtgtacACACTTGGGTCTTCTGTAGGGCACTTTAGATTTCTCAGCTtatagttacttgtcctaaac
gacttatcagtcttgtgtacGGTTAAGTGCTGACTACACCAGGTGTAATTGCCTAGTTCAtatagttacttgtcctaaac
gacttatcagtcttgtgtacCTGAAATCCGGCAGTTTGAGGTCAAGTTTGGCATTGGTCTtatagttacttgtcctaaac
gacttatcagtcttgtgtacCATACTTGCCTCGGGAGACCTGTGCGTAGAGACAGTACTGtatagttacttgtcctaaa
gacttatcagtcttgtgtacCCGTGGAGGGTGTGTGTGGGTGGGAGCTGGGGATTTTGTTtatagttacttgtcctaaa

gacttatcagtcttgtgtacGGTTTTAGAATCACCATCAGAGGCAACCTGAAACTAATAAtatagttacttgtcctaaac
gacttatcagtcttgtgtacGAAATGAAAAATCCAAAAGTCTTCCAACTTCACCTTATGGtatagttacttgtcctaaac
gacttatcagtcttgtgtacTCTTTGACCACACGGATATTTCTCAGAGGTTGATTATACAtatagttacttgtcctaaac
gacttatcagtcttgtgtacACCAGAGCTTAAATAAATCAAAATATGGTGACTCCCTGAGtatagttacttgtcctaaac
gacttatcagtcttgtgtacGGTACTGGCCTTATTGAGAAAACTGGCTTTTTCTGTTTTGtatagttacttgtcctaaac
gacttatcagtcttgtgtacAATCTTTAAACTCCCCTTTCTATTAATAAGCAGACTCCTCtatagttacttgtcctaaac
gacttatcagtcttgtgtacATTGTATGTAGATAAGAAAATAGTCACTCAGGGGATATGCtatagttacttgtcctaaac
gacttatcagtcttgtgtacACTTTTGTTCTCTCTAGTTTTGTACCTGCTCCTCCCTGAAtatagttacttgtcctaaac
gacttatcagtcttgtgtacAGAGGAAATCTTTTGGGATCAGAAGCAAAGTGAGAATTTGtatagttacttgtcctaaac
gacttatcagtcttgtgtacGAGCTGTGGTTCTTGGTGAGAAACTCTGAGTTGGCTAAAGtatagttacttgtcctaaac
gacttatcagtcttgtgtacAGCAGATTTTATTTCCAGAAAGTGTGGTTTGGGATGGAGGtatagttacttgtcctaaac
gacttatcagtcttgtgtacATAAATGGATTTTCCAGAATTACTAAGGGGTGGGGAAACAtatagttacttgtcctaaac
gacttatcagtcttgtgtacCATGGATCAGGGCGAGAGATGGTTTTGGGATTTTTCAAGTtatagttacttgtcctaaac
gacttatcagtcttgtgtacCTCACCATAATGTAGAATCCCCGGGAGCCCTGAGCTATTTtatagttacttgtcctaaa
gacttatcagtcttgtgtacAGTAATGTGAGAGATGGGAAGTGTCCGTAAATACAGATGAtatagttacttgtcctaaac
gacttatcagtcttgtgtacGGAGACCCCTGGCCTAGAAGATAGTTTGTCCTTTATTGGGtatagttacttgtcctaaa
gacttatcagtcttgtgtacTCATCAAATTAAAATATGCACTCTAATTGGCCAAGCTCTGtatagttacttgtcctaaac
gacttatcagtcttgtgtacAGTGTGGGGTAAACTGAAATGGTTCACTAAGCAATCCTCCtatagttacttgtcctaaa
gacttatcagtcttgtgtacTCTTATAGCCCTCAATCAGTTTAATTCAATGTAAAACCCCtatagttacttgtcctaaac
gacttatcagtcttgtgtacTTTCCTCGATGTGATTTCAAATTGTCCTGAATTCTTCAGGtatagttacttgtcctaaac
gacttatcagtcttgtgtacAATCAGGAAACTACAAAAAAGGGGAAAGAATAGTCCAATGtatagttacttgtcctaaa
gacttatcagtcttgtgtacATGACATTTCACTCATAAATCCTTAGCGCTGATCTTCTAAtatagttacttgtcctaaac
gacttatcagtcttgtgtacAAGATGTTTTCCTGTTTACCACCATATCATTATTACACCCtatagttacttgtcctaaac
gacttatcagtcttgtgtacTTCTGTGTCAAAGACTTTTTTCAAAAATCTAGGCCAGTTGtatagttacttgtcctaaac
gacttatcagtcttgtgtacATTAGAATCAAGTTAGACCTTTGTTTGGCAGGAATATTGCtatagttacttgtcctaaac
gacttatcagtcttgtgtacAGTTTCACTACTCTTCTCCCCTTTTCTTCCCCAGCATTCCtatagttacttgtcctaaac
gacttatcagtcttgtgtacAGTTTGACTTGCTAAGGCCAATGGGAACTCAGTTGAACTTtatagttacttgtcctaaac
gacttatcagtcttgtgtacTCCTGTGTGCTCCTGCCATTACATAAGAGGAACATTTTCTtatagttacttgtcctaaac
gacttatcagtcttgtgtacGTGATCTCCAGACTCAACCTGAATTGAAAACTTAGCTTGCtatagttacttgtcctaaac
gacttatcagtcttgtgtacCTAGATCAGCAGAACCCCAGCCAACCTACAGATGTGCAAGtatagttacttgtcctaa
gacttatcagtcttgtgtacTAGCAATCCTGAGATGTGAGATAGCAATCATGTCTATTTCtatagttacttgtcctaaac
gacttatcagtcttgtgtacGCTCTACGCTCTTGACTTTTTAAGCACTTAGTACAAATATtatagttacttgtcctaaac
gacttatcagtcttgtgtacACTCTATATCCATTGAATAGTTCCCCTTCCCCCATCCCCTtatagttacttgtcctaaac
gacttatcagtcttgtgtacTTCTGCTTCTATGAATCTGGCTACAGATACCTTATATAAGtatagttacttgtcctaaac
gacttatcagtcttgtgtacTTCCATTGCATGTATATATCGCATTTGCTTTCTCCACTCTtatagttacttgtcctaaac
gacttatcagtcttgtgtacTGAACATGACAGTGCATATGCCTCTTCAAGATCCTGATTTtatagttacttgtcctaaac
gacttatcagtcttgtgtacAGAAGTGGGATTGTTGGAACACATGGTAGTTGTATTTTTAtatagttacttgtcctaaac
gacttatcagtcttgtgtacACCATTTTACATTCCCGTGAATAGTGCTCAAGGATTCCAAtatagttacttgtcctaaac
gacttatcagtcttgtgtacAATAACTATCCTAACAGGCTGAAGGTGATATTTCATTGTGtatagttacttgtcctaaac
gacttatcagtcttgtgtacGAGCATCTTTTTGTATACTTTTTGGCCGCTTGTGTGTCTTtatagttacttgtcctaaac
gacttatcagtcttgtgtacGCCTATTCAAATACTTCCGCTCATTTTTTAACCAAGTTACtatagttacttgtcctaaac
gacttatcagtcttgtgtacTTTTTGATATTGAGTTGGAGGGGTTCTGTATATATTTGGGtatagttacttgtcctaaac
gacttatcagtcttgtgtacTGCAAATATTTTCCATCAATCCATATGTTGACTTTCCACCtatagttacttgtcctaaac
gacttatcagtcttgtgtacGCACAGAAGCCTTTAATTTCATGTAGGTCCCACTTATCTAtatagttacttgtcctaaac
gacttatcagtcttgtgtacTTGCCAAGTCCAATATCATGAAGCGTTTCCCCTATATTTTtatagttacttgtcctaaac
gacttatcagtcttgtgtacTAAGCCTCTAACACATTTTGTGTTGATTTCTGTGTATGGTtatagttacttgtcctaaac
gacttatcagtcttgtgtacAGACTATCTATTCCCCATTGTGTATCCTGGTATTTTCATTtatagttacttgtcctaaac
gacttatcagtcttgtgtacTCAGTTGACTATGTATATGTGGCTTTATGTCTGTACTCTCtatagttacttgtcctaaac
gacttatcagtcttgtgtacATGCCAGTACTACACTGTACCATTGATTACTGTAGCTTTGtatagttacttgtcctaaac
gacttatcagtcttgtgtacTGAAGTCAAAAAGTATGAGGCAGGTGGCTTTGTTCTTTCTtatagttacttgtcctaaac
gacttatcagtcttgtgtacTCTATTTCTGCAAAAAAAGCCATTACGATTTTGATGGGGAtatagttacttgtcctaaac
gacttatcagtcttgtgtacGTTTTCTAGTTTTCAGTATACAAGCATATCACCTCATTGGtatagttacttgtcctaaac

gacttatcagtcttgtgtacCTGACTTTGCATGTTGATTTTTTATTCTGCAAGTTTGCTGtatagttacttgtcctaaac
gacttatcagtcttgtgtacGTACTATGTTGAATATAAATTGTGAGAGCAAGCATCCTTGtatagttacttgtcctaaac
gacttatcagtcttgtgtacCAGTTTTTCACCATTGAGCAGCATGTTAACTGTAGGCTTTtatagttacttgtcctaaac
gacttatcagtcttgtgtacGGTAATTTCCTTCTATTTTTTTGATTCTCTTGGGCAGTCTtatagttacttgtcctaaac
gacttatcagtcttgtgtacGAATATTGTATGCATGTTCCAACTCTTTCTCTTCCTAGCTtatagttacttgtcctaaac
gacttatcagtcttgtgtacAGATGTGGATTTGTTTCCTCCTATTTCATGCTGAGCTGGGtatagttacttgtcctaaac
gacttatcagtcttgtgtacTAGGGCCAGTAAGTGTGTGCTAGTCTAAATTATTGCTTTTtatagttacttgtcctaaac
gacttatcagtcttgtgtacCTGTCAGCACTTAGATTCAGGCAAGAGAAAAACTGGTTCCtatagttacttgtcctaaa
gacttatcagtcttgtgtacAACTTGTGTTTCAGACTTTTTTCCCCCCGCTGAGAGATAAtatagttacttgtcctaaac
gacttatcagtcttgtgtacCATTTTGCATCAAACTGAGCCAGAGGGAGGGACTACGGTAtatagttacttgtcctaaa
gacttatcagtcttgtgtacGATTCATCTATTGGCTTGATATGGCTGATTTTACACTTGTtatagttacttgtcctaaac
gacttatcagtcttgtgtacAGAAGTCTTTTAGCTCATTTCTGGGTTTCTGACAAAGGGAtatagttacttgtcctaaac
gacttatcagtcttgtgtacGGCTTTCCATTCTGCTATCTTACTGATGTCACTCCTCCTTtatagttacttgtcctaaac
gacttatcagtcttgtgtacGTGTCACATTTAATCATATGAAACAGGCAGTGATCAATAGtatagttacttgtcctaaac
gacttatcagtcttgtgtacGGATATCATTTTTCTTAACAGCCTTTCATTCAGTGTTAGCtatagttacttgtcctaaac
gacttatcagtcttgtgtacTGGTCCTTACCAGATCAATTATTACGTTAGGGCTCTCACCtatagttacttgtcctaaac
gacttatcagtcttgtgtacTCTTCTGATTCCCCCAAGAAAGAAAGATTAGACTTATGTAtatagttacttgtcctaaac
gacttatcagtcttgtgtacGAAATTATCTCTCTCAGGAACATGAAATAACCCCAAACATtatagttacttgtcctaaac
gacttatcagtcttgtgtacATCTGAGTTTTTGTTATAAGATGGAGAAGGGGGCTATAGAtatagttacttgtcctaaac
gacttatcagtcttgtgtacTCCTTCTGAATTATTGTGGGTATTTTCACTTTAATCGAGCtatagttacttgtcctaaac
gacttatcagtcttgtgtacGGCATGTTAAGAGGGGTAATTGCTTAATGACAACTTAGGAtatagttacttgtcctaaac
gacttatcagtcttgtgtacCCATTGGGTTAGACTAAAGGAAAGAATTAAAGGAAAGGCCtatagttacttgtcctaaa
gacttatcagtcttgtgtacAATTCAAACCATTCTGTCTTTAAACTAGCTATCAGATCCCtatagttacttgtcctaaac
gacttatcagtcttgtgtacAAATGTTCCTGCTTTTCTTCAAGAATGATTCCCCTCTTTCtatagttacttgtcctaaac
gacttatcagtcttgtgtacTTGAGAAAAGCAATAAATAGGTCCTGAAATCTGTACGCAAtatagttacttgtcctaaac
gacttatcagtcttgtgtacGGTGTGTGAGAATATGTATACTAAACCCTGTGAAAAAGTAtatagttacttgtcctaaac
gacttatcagtcttgtgtacCAAATGATTGTGATTCTCCCTCCTCCATCTTTCTCCTCCTtatagttacttgtcctaaac
gacttatcagtcttgtgtacTTTTTAAAAAACAAATGACTGCGATTCTCCCTCCTGCATCtatagttacttgtcctaaac
gacttatcagtcttgtgtacCCTCCTCCTTCTTCTTCCTTAATTAGAGCTTCAGTCCTGAtatagttacttgtcctaaac
gacttatcagtcttgtgtacGGCAGTAAATAGGCCTTGGGCTTAAAATAATTTTTTTTCCtatagttacttgtcctaaac
gacttatcagtcttgtgtacATTCATAGATTAGATATTTGTGGGTTGTATGCACTCTGCCtatagttacttgtcctaaac
gacttatcagtcttgtgtacCATGATCCAGGGTCCTAGGTGATGGCTCTCTTCTTTAGGGtatagttacttgtcctaaac
gacttatcagtcttgtgtacTTAGTCCCAAACAATTGTACAAATAACCCCAAATATGGTGtatagttacttgtcctaaac
gacttatcagtcttgtgtacTGTCCTCTTCAAACTGAGTATAAGTTAACACCTTCACTGTtatagttacttgtcctaaac
gacttatcagtcttgtgtacTGAGCTCAGATCAAAGTTTCCCAAGGAGTGTCAAAATGCAtatagttacttgtcctaaac
gacttatcagtcttgtgtacCAGTTGATAGTTGTTAAAATAAAGACCCTAGTTCTTGGTCtatagttacttgtcctaaac
gacttatcagtcttgtgtacCTCTTTACATTGAAGATCCTATTTTTATTCTACCTCAGCCtatagttacttgtcctaaac
gacttatcagtcttgtgtacGATAACATTTATTGGGCACTACTTTGCATCTAACACTATGtatagttacttgtcctaaac
gacttatcagtcttgtgtacCACACATTAATCCTGTGTGATAGTAACTTTTTTATCCCCAtatagttacttgtcctaaac
gacttatcagtcttgtgtacAAATAAACGGAGACTTATAGAATGTCTTAGTCCATTCCTGtatagttacttgtcctaaac
gacttatcagtcttgtgtacAGAGTCAATATCCGGTGAGGGCCTGCTCTGTTTCAAACATtatagttacttgtcctaaac
gacttatcagtcttgtgtacTTTATAAGGGCACTAACCTCGTACTAATCACCTCCTAAAGtatagttacttgtcctaaac
gacttatcagtcttgtgtacCTCCAACATTCAGACCACAGCGGAGAGTTTAAGTGACTTGtatagttacttgtcctaaa
gacttatcagtcttgtgtacGCTGATAAAATATAGGGTCCAAATTCTGAAGTTCATACTCtatagttacttgtcctaaac
gacttatcagtcttgtgtacGAGACAAAATCTTTTATTTCTAGAGAACAAAGCCTTGAGGtatagttacttgtcctaaac
gacttatcagtcttgtgtacTAATAAAGAGATCAAGCAATGAGAAGTTTCTTGGAGGTACtatagttacttgtcctaaac
gacttatcagtcttgtgtacCAAGAAGGTGGGAAAAAAGAGGGCTCAGAAAAGGAACTCAtatagttacttgtcctaaa
gacttatcagtcttgtgtacCTGGGAATGGGGGGAAAGCAGCATGTTGAGGGTAGAGGACtatagttacttgtcctaa
gacttatcagtcttgtgtacTGGCTATGAAACAGAAGTGAAGGGCATCTAGAGTGGTATTtatagttacttgtcctaaac
gacttatcagtcttgtgtacCTGGTGGTTACGAGCGAATTTATTGATAGAAGTAAAATCTtatagttacttgtcctaaac
gacttatcagtcttgtgtacGCATGCTTAATGTCAATCTAGTTTACAGCTTTAGGTAATGtatagttacttgtcctaaac
gacttatcagtcttgtgtacAAGGGTGACAAATTCAGAACAGATGGCAAGAAGCAAGTCTtatagttacttgtcctaaa

gacttatcagtcttgtgtacACAAATGCCTGTGAAATGGACATCCTGAAGGGACAGCACTtatagttacttgtcctaaa
gacttatcagtcttgtgtacGGAACTTCACAATCCCTATCAGTGATTTACATGTGACAGTtatagttacttgtcctaaac
gacttatcagtcttgtgtacAGGCACCCACTAGCAAAGCTTCCTCCAGTTTTCATCAAGGtatagttacttgtcctaaa
gacttatcagtcttgtgtacCTGACTGCATTTTAGGGGTAGCAAAATAACCTCCTGTTAGtatagttacttgtcctaaac
gacttatcagtcttgtgtacAGATGTACTGTGATAATCAAATGGCAAGTCCCTTCCCTCAtatagttacttgtcctaaa
gacttatcagtcttgtgtacCTTCTGCTTTGGAAAATTGTTTGGAAATGTTTGGGCAAGTtatagttacttgtcctaaac
gacttatcagtcttgtgtacACGGAACTGTTTGACTAAAATGAGAATTGGAGGAAAATCAtatagttacttgtcctaaac
gacttatcagtcttgtgtacGATTCTTTTGATAGTACGATACAGCACAATGACAAATTGGtatagttacttgtcctaaac
gacttatcagtcttgtgtacAAGGCAGAACACTTTACTACTTACAGCTCCAAAGGAGAGAtatagttacttgtcctaaa
gacttatcagtcttgtgtacCAGCAAGTTGAAAATGTAGCAGGTGACTTATGTGTGGCAAtatagttacttgtcctaaa
gacttatcagtcttgtgtacAGGGGCTTCCTGGGGATCAGCTAATTTGCATAATTCCCGGtatagttacttgtcctaaa
gacttatcagtcttgtgtacCCTGGGGAGGTTGGGATATGTGCCTAGTTGCTCTGGAGAGtatagttacttgtcctaa
gacttatcagtcttgtgtacTGATAAGAAAAGTGGTTGGGTTTTGGGCACAGCAAACTGCtatagttacttgtcctaaa
gacttatcagtcttgtgtacAGAGTTTTCAACCATGACTTCAGCACTGAGTCCAGACGGCtatagttacttgtcctaaa
gacttatcagtcttgtgtacCAGCTTGAACCTGCAGAAACAGTTGTTTTTCTTGTTCTTTtatagttacttgtcctaaac
gacttatcagtcttgtgtacTATATTCTCTCTTAGGATTCCACACTTTTTCAAACTGCTCtatagttacttgtcctaaac
gacttatcagtcttgtgtacAAATCATACTTGAATTGTTCACTGATCCTTCTTTCTGCAGtatagttacttgtcctaaac
gacttatcagtcttgtgtacTGCCTGCTCTTCCCTAATTTATTTATGCCTTTCTATTCCTtatagttacttgtcctaaac
gacttatcagtcttgtgtacATTATTTTAATAGCCTCTTTCAGCCACTTTCCTTTCCTCTtatagttacttgtcctaaac
gacttatcagtcttgtgtacTGCTTTCATGTTACAAGGGCAGAATACAGTCATTGCAATAtatagttacttgtcctaaac
gacttatcagtcttgtgtacACCTGAAATATTTATGGTCTGCCTTTACAGAAAAAGTGTGtatagttacttgtcctaaac
gacttatcagtcttgtgtacTACTAAATATCCATTCAAAATCCTGATCATGTCCATCCAGtatagttacttgtcctaaac
gacttatcagtcttgtgtacGAACTACATTCATTGAGAACTCCATGTCAACTCTGGGCTGtatagttacttgtcctaaa
gacttatcagtcttgtgtacTTTGTGTGAGTAAAGCCCAGGTCTATAAACTCAGGGGAGTtatagttacttgtcctaaa
gacttatcagtcttgtgtacGGATGTCAGAAACTTAAATCCCTGTTAGTTCAGTTAATTCtatagttacttgtcctaaac
gacttatcagtcttgtgtacCCTGGGATTTTATCTGTTCTCTGGGTGAGGGGAAGTGCACtatagttacttgtcctaaa
gacttatcagtcttgtgtacAGCCTCACTGTTGTCATGTAATTGTCACTCACCTATGAAGtatagttacttgtcctaaac
gacttatcagtcttgtgtacCTATATTGTTTCTTGTTCCCATCCTGAAAGTCCCATTGGAtatagttacttgtcctaaac
gacttatcagtcttgtgtacCCTTGGGATTTACAGAAAATGTGATATATTTCCCACTTGGtatagttacttgtcctaaac
gacttatcagtcttgtgtacGAGGCCTCATCCGCTTCTGTGTCACAGTAGGTTCATTTGAtatagttacttgtcctaaa
gacttatcagtcttgtgtacTCTCTAGATACGTTTCAATAATTTCTTCTTAGGGGTTTGCtatagttacttgtcctaaac
gacttatcagtcttgtgtacAGGCCTCCTTTACTCTCAAATCCCATAAACGTACACAGTGtatagttacttgtcctaaa
gacttatcagtcttgtgtacGGTACCCATAAAAGGTTCTACTTTCCTTGCCCTGCAGCATtatagttacttgtcctaaa
gacttatcagtcttgtgtacGTTGAAGACTGGAGTGGGGGTAAGAAAAAGGAAAGTTAGCtatagttacttgtcctaaa
gacttatcagtcttgtgtacGATAGACATGATTTTTTTTTCCATTCCCTCTCAAGTTAGGtatagttacttgtcctaaac
gacttatcagtcttgtgtacGACTGGCTCCTGTTTACTCTTCTTCTGCTGTTTGATGCCTtatagttacttgtcctaaac
gacttatcagtcttgtgtacAGGCAATAGAATCCTACCTGACCAGAGTTTATAGTAATTCtatagttacttgtcctaaac
gacttatcagtcttgtgtacGTTAATCTCCTAATAGTAACCTTTCTCTCATCTCCACTCCtatagttacttgtcctaaac
gacttatcagtcttgtgtacCTCACTTGGTCCTTACAATAGCACTGACAGATAAAGCTTTtatagttacttgtcctaaac
gacttatcagtcttgtgtacAATATGAATTCAGGTGTAATGGTCCCCAAATGAGTGTTCTtatagttacttgtcctaaac
gacttatcagtcttgtgtacTGGCATTACAGCATTAATTTTACACTGAAGTTTGGTTCTCtatagttacttgtcctaaac
gacttatcagtcttgtgtacCACCTCATGTTCCTATCCTATTCTTCTTCTCTGTTTAAAGtatagttacttgtcctaaac
gacttatcagtcttgtgtacACCTGTCTTCCTATTTCCTTTGGTTGTAATGGTTTTATAGtatagttacttgtcctaaac
gacttatcagtcttgtgtacAATATATATACATTGTCCCTTTGAATGCAAAGCAGTGTGCtatagttacttgtcctaaac
gacttatcagtcttgtgtacCCATAGGGGTATATAATATAGGAAGGGAAAAAGACATTCAtatagttacttgtcctaaa
gacttatcagtcttgtgtacACACAAGTTGTCATTGTTAATGGGGCTAAATCAATGTGCAtatagttacttgtcctaaac
gacttatcagtcttgtgtacTCCATGGAAGAATATTTAATTTGTCTGGAGGAATCAGGGCtatagttacttgtcctaaac
gacttatcagtcttgtgtacAACTGGGTCTTAGAGGATGATTAGAAGTTTCAATGGGGGAtatagttacttgtcctaaa
gacttatcagtcttgtgtacGTACACCAAGAAGAAGAGACACCAAAGAAAGACACCAAGTtatagttacttgtcctaa
gacttatcagtcttgtgtacATAAATAAAAAGAGATTTAGTAGACTCACAGTTCCGCGTGtatagttacttgtcctaaac
gacttatcagtcttgtgtacGAGAGAATGAGAGACAAGCAAAATGGGAAACTCCTTATCAtatagttacttgtcctaaa
gacttatcagtcttgtgtacCAGTCAAACAATATCAGTGTTCTATTCCTGGGTTGGTAAAtatagttacttgtcctaaac

gacttatcagtcttgtgtacTGGCAAGTAAAAACTACAAGGTCAATGGAGGGATGTCATAtatagttacttgtcctaaa
gacttatcagtcttgtgtacCAGGCCTCTAAATATAAATCAGCACCTGGACATGAAGGACtatagttacttgtcctaaa
gacttatcagtcttgtgtacGGAGCATGAACATTATTCCTAGGTAAAAGAAATCCACTGAtatagttacttgtcctaaa
gacttatcagtcttgtgtacGAGTGTCTCTGGCCAATCTAAAATCAACATATTCAGTAATtatagttacttgtcctaaac
gacttatcagtcttgtgtacTGCAGGGAGAACAAAGGAAGGGCAAATCTTGGGTTAAAATtatagttacttgtcctaaa
gacttatcagtcttgtgtacACTTGGTTTGTGGTTTTGATACCATGATGAAGTAGAGATTtatagttacttgtcctaaac
gacttatcagtcttgtgtacCTGTGTTCATTAGTCCAGAGGTGTTAGTTCTTCAATTCCTtatagttacttgtcctaaac
gacttatcagtcttgtgtacGGCTAATAATGGATGTGTCAAAGTTAATATGTCTCAACTGtatagttacttgtcctaaac
gacttatcagtcttgtgtacTGGTGACATAATTTTCGAGATGATATATGCCCTTCTTGTGtatagttacttgtcctaaac
gacttatcagtcttgtgtacCAAGTTTTGGCCTTTGACTACATCTTTTTTTTTCCAGGTAtatagttacttgtcctaaac
gacttatcagtcttgtgtacGCTTTTGATTTGCTCTTCCTGCCACATTAAGTATACAGCTtatagttacttgtcctaaac
gacttatcagtcttgtgtacGCATAGTTAGCAAATGGTCAATCTATTCTCCTTGTAGAAGtatagttacttgtcctaaac
gacttatcagtcttgtgtacAGAAGATTAAAAGCAAAATGATCAGTTTCCACCTCCAAACtatagttacttgtcctaaa
gacttatcagtcttgtgtacTTTGATTGACTGATTTATCAGGAAACATTGATCACATGCCtatagttacttgtcctaaac
gacttatcagtcttgtgtacAATGTTGGATTGGAAAGTAAGGTGAGGTGGGAGTATGAAGtatagttacttgtcctaaa
gacttatcagtcttgtgtacCCTGGCTGGAATTCAGTGGGAAGACCACAGCTCACTGCAGtatagttacttgtcctaa
gacttatcagtcttgtgtacATTTGTAGAGATGCAGTCTCGTCATTTTGTCCAGGCTGGTtatagttacttgtcctaaac
gacttatcagtcttgtgtacGCATGAACCAACATACCTGGCCACATAATCTTAAAATTAAtatagttacttgtcctaaac
gacttatcagtcttgtgtacTGTGCTATGGTGCTTTGCTGAACCCTTCAACCTGTCATCTtatagttacttgtcctaaac
gacttatcagtcttgtgtacGTGTGATGTTCCCCTCCTTGTGCTCATATGTTCTCATTGTtatagttacttgtcctaaac
gacttatcagtcttgtgtacGAGTGAGAACATGTCGTGTTTGGTTTCCTGTTCCTGTGTTtatagttacttgtcctaaac
gacttatcagtcttgtgtacTGGCTGCATAATGTTCCATGGTGTATGTGTGCCACATTTTtatagttacttgtcctaaac
gacttatcagtcttgtgtacGGTCAAATGGCATTTCTGGAAATGTTTTAGATCCTTGAGGtatagttacttgtcctaaac
gacttatcagtcttgtgtacCTGTTGTTTCCTGACTTGTTAATTATCACCATTCTAACTGtatagttacttgtcctaaac
gacttatcagtcttgtgtacCAAAGATGATGAGCTTTTTTTTTTACATCTGTTGGCTGCAtatagttacttgtcctaaac
gacttatcagtcttgtgtacTTAAGCTCCTTGTAGATGCCGGGCAGTATGGCCATTTTCAtatagttacttgtcctaaa
gacttatcagtcttgtgtacTTGAAGAGGTCTTTCACAACCCTTATGAGTTGTATTCCTAtatagttacttgtcctaaac
gacttatcagtcttgtgtacGTTGCTTATCAGCTTAAGGAGAATTTGGGCCAAGACAATGtatagttacttgtcctaaa
gacttatcagtcttgtgtacACTTCCAATACTATGTTGACTAGGAGTAGTGAGAGAGGGCtatagttacttgtcctaaa
gacttatcagtcttgtgtacCAGTATGATATTGGCTATGGGTTTGTATTAAACAGCTCTTtatagttacttgtcctaaac
gacttatcagtcttgtgtacCGTAGTTTATTGAGAGTTTTTACCATTAAGGGGTGTTGAAtatagttacttgtcctaaac
gacttatcagtcttgtgtacGGTTTTTGTCTTTGGTTCTGCTTTTGTGATGGATTATGTTtatagttacttgtcctaaac
gacttatcagtcttgtgtacTGCCAGTAGTTTATTGAGGATTTTTGTGTCGATGTCCATCtatagttacttgtcctaaac
gacttatcagtcttgtgtacATAAAATGAGTTAGGGACAAGTCCCCCTTTTTCTGTTGTTtatagttacttgtcctaaac
gacttatcagtcttgtgtacCTCCTCTTTGTACCTCCGGTAGAATTCTGCTGTGAATCTGtatagttacttgtcctaaac
gacttatcagtcttgtgtacGTGTAGAAGTATTTATAGTATCCTCTGGTGGTAGTTTGTAtatagttacttgtcctaaac
gacttatcagtcttgtgtacTTACATTTAATCTGTGTAAGACTTGTACTGGGTGCTATGGtatagttacttgtcctaaac
gacttatcagtcttgtgtacTGTTGGCTAATGACATAAGTAGAAATTAAGGATCGACTCAtatagttacttgtcctaaac
gacttatcagtcttgtgtacAAGAGGGGGAACTTCCTAGGGACTTGATGATTAGGGAATGtatagttacttgtcctaaa
gacttatcagtcttgtgtacCAGCTACCACTTCACTCCCATGTCTGAATAAACAATTTCAtatagttacttgtcctaaac
gacttatcagtcttgtgtacCTTTTAGAGATGAGAAAATTAGTTATGGGTGAGCATAGTCtatagttacttgtcctaaac
gacttatcagtcttgtgtacATGGCTTCACAACATGGCCCCAGAGGGTTAAATTTTGCATtatagttacttgtcctaaa
gacttatcagtcttgtgtacCATTTGCTGAAGATGAGTGAAATTGGATGACTCCCTGTTGtatagttacttgtcctaaac
gacttatcagtcttgtgtacGCTACTGTGTGTTGTTTATTTCCCTAGAGAGCATGAATTCtatagttacttgtcctaaac
gacttatcagtcttgtgtacCATCCTGCTTTTAGTAAAATTATGCCTCTTTCTCTCCAAAtatagttacttgtcctaaac
gacttatcagtcttgtgtacAGGCTAGATAAACTCTAGAAGTTCACAGTGACAGACCTCTtatagttacttgtcctaaa
gacttatcagtcttgtgtacCAAGTATACGAAATAATGTAATAAAGACCATGCACCCTGTtatagttacttgtcctaaac
gacttatcagtcttgtgtacAATAGAATAGTTATCAATACAGCTAATGCCTCTCCTTTGCtatagttacttgtcctaaac
gacttatcagtcttgtgtacGTAGTTGCTATTCTGAAGTTGGTTGATCTATGTTATTTCCtatagttacttgtcctaaac
gacttatcagtcttgtgtacCATTAGAATGTATATATCACTGCACTGCACCTTGCATTTTtatagttacttgtcctaaac
gacttatcagtcttgtgtacATTGTCTGAGTCAATCATGGTTTTTTTAAGCTCTGTTTGCtatagttacttgtcctaaac
gacttatcagtcttgtgtacTGTGTGTATGTTTCTTTGGGAGGAGCACATTAAAAGTGGAtatagttacttgtcctaaac

gacttatcagtcttgtgtacAGTTAATTTCTTTTCTCATAATATCCCAGAACAGGTGCTCtatagttacttgtcctaaac
gacttatcagtcttgtgtacCTTGACTGTATCACATGATTCCAAGATTATCTGGTTAAACtatagttacttgtcctaaac
gacttatcagtcttgtgtacAGCATTTAAATTAGTGAACTGAACAAAAGTGATGACCCTCtatagttacttgtcctaaac
gacttatcagtcttgtgtacCAAAAGGTGGAGGAAGGGAGGATTGGTTGGTTCTCTGACTtatagttacttgtcctaaa
gacttatcagtcttgtgtacCCTTGAGACTGGAACTGGAATCTATAACATGGGCTTTTCAtatagttacttgtcctaaac
gacttatcagtcttgtgtacCACACAGCAGATCATAGCCTCAGCCTCCTTTTCATAATTAtatagttacttgtcctaaac
gacttatcagtcttgtgtacCATCTGCATCTCTCTGTATATGTCTGTCTGCCTATCTATCtatagttacttgtcctaaac
gacttatcagtcttgtgtacTCATTTATCTATCTAAACTATCTGCCTATCTACCTGCCTAtatagttacttgtcctaaac
gacttatcagtcttgtgtacTTTTGTTTCTCTGGAGAACGATGAGTAATACAAGGTGGTTtatagttacttgtcctaaac
gacttatcagtcttgtgtacGACTTATTTGTCTTCCAACTGTGTCTCCATCTTTTCCTGGtatagttacttgtcctaaac
gacttatcagtcttgtgtacCTGCAGGAAGTGGAAAGAAAGCATGGAGGAGACACATTCGtatagttacttgtcctaa
gacttatcagtcttgtgtacGGCACAAACCACTTCTACTGTTCCCAGAAATATAACTGATtatagttacttgtcctaaac
gacttatcagtcttgtgtacGAAATTATCCTTAGTATGGAGTCATTCACAGTGGAGGTTGtatagttacttgtcctaaac
gacttatcagtcttgtgtacCTGTTATTTTCAGGTCAGCTTTCACAAACTGCCAGAATTAtatagttacttgtcctaaac
gacttatcagtcttgtgtacACATTTTCTCCTTGTAGGCCATGATAGTGTACATGCAATCtatagttacttgtcctaaac
gacttatcagtcttgtgtacGTTTCTTAAGGTAGATCACAATGGTTAATGGAGTTTGTTCtatagttacttgtcctaaac
gacttatcagtcttgtgtacCCTTCATGAAATATTGTACAAGGCAAATTGACAGATTTGGtatagttacttgtcctaaac
gacttatcagtcttgtgtacAAATTTACCAAAAGTGAACACAATGTAGAAAAGAGGGTGCtatagttacttgtcctaaac
gacttatcagtcttgtgtacAAGTAAATTCTCTACGTGTCTGCAACATTTCTGCACAGCAtatagttacttgtcctaaac
gacttatcagtcttgtgtacCCTTTGCTCTGGACTGTCTTGATTCTGGTTTCCTAATGATtatagttacttgtcctaaac
gacttatcagtcttgtgtacCAGTCATGATTAGTGTCCATTGTAGAAGAATTAGATCTCCtatagttacttgtcctaaac
gacttatcagtcttgtgtacACTGTGTGTGCGGGTACCACCTTGTCCTCTTTAAAGCTCCtatagttacttgtcctaaac
gacttatcagtcttgtgtacAGAAAGTGCTGATATTCTGGCTGCTATTGTTGCTGTGAGTtatagttacttgtcctaaac
gacttatcagtcttgtgtacTCTCTGACCCAGGAGACTTGCATTAAGACTTCTTTTAGCAtatagttacttgtcctaaac
gacttatcagtcttgtgtacTGGGACATCTTTCAAAGTTCTTGGCAATGAGAAAATAGAAtatagttacttgtcctaaac
gacttatcagtcttgtgtacCTTGGATGATCTTGGGAAACGAACCTAACCGCCACGAGTCtatagttacttgtcctaaa
gacttatcagtcttgtgtacAGATTACTGGTAAGGGCCCTCCGAGTCTGAAGAACTATGGtatagttacttgtcctaaa
gacttatcagtcttgtgtacTTCCTTGGGGGTAGTGCTTAGGCCTTTGCTCTGCGGGACCtatagttacttgtcctaaa
gacttatcagtcttgtgtacCACTTCTTGCTTTACTCTCCTATATCTTGACACCGTATGCtatagttacttgtcctaaac
gacttatcagtcttgtgtacACAGGTGCTCACCACCCTGCTTGGCTAGTTTTGGTATTTTtatagttacttgtcctaaac
gacttatcagtcttgtgtacCCACCATGTCCGGCTGTTATATCATCATTTTCAAAAGGACtatagttacttgtcctaaac
gacttatcagtcttgtgtacATCCATAGTTAAAGACACAGAGGGCATTTTAAAGACAATCtatagttacttgtcctaaac
gacttatcagtcttgtgtacGCTGGTGATTGAAAGAAATCCGTGTGTGTAGCCTCTGTTGtatagttacttgtcctaaac
gacttatcagtcttgtgtacAGCATATCAAGGCAGTTCTGCCATCCAGGCTTGTTTGAGGtatagttacttgtcctaaa
gacttatcagtcttgtgtacAGAGATCAGAAGCTAAGCAAAGGACAGTGACTCAGGACACtatagttacttgtcctaa
gacttatcagtcttgtgtacTCACACTGACAAACTTGGTTGCTAATGATATTTTTCCATGtatagttacttgtcctaaac
gacttatcagtcttgtgtacTCTTGACTGGAGGGTAGAGGGCAGAGAGAGTCCTGAGTGGtatagttacttgtcctaa
gacttatcagtcttgtgtacTTTATTTTTTTGATTTTTTTGCGGGGTCTCGAAGCGTGGTtatagttacttgtcctaaac
gacttatcagtcttgtgtacTTGCACCTCAAATGATTTTTTAACTCAAATGCTGTTCCTCtatagttacttgtcctaaac
gacttatcagtcttgtgtacCAGGAGGTGATGGGTCAAATGTTGGTGAGGGCTTGATTGAtatagttacttgtcctaaa
gacttatcagtcttgtgtacTGGAGGGAAGGCAGAGTGGGGAGGAAAACCGTGAAAGGCAtatagttacttgtcctaa
gacttatcagtcttgtgtacCTTGCTCTCCTGGCTTGCTTCCCTAATCTTTATATTCAAGtatagttacttgtcctaaac
gacttatcagtcttgtgtacCAACACACTTGCTTATTTACTGCTCTTTCTCTGGAACACTtatagttacttgtcctaaac
gacttatcagtcttgtgtacCTCTGGCTTGACGGGGTGTACAGATACCTATTCTATCTCTtatagttacttgtcctaaac
gacttatcagtcttgtgtacCTCTTGTCCTTCCCGATCACCTAGAGAGAGGCTTTCACAAtatagttacttgtcctaaa
gacttatcagtcttgtgtacTGTCCTCACATTGTCAGATGCTGCATAATCCTCATAAACTtatagttacttgtcctaaac
gacttatcagtcttgtgtacAAGTGCAGCTGCTACCTTTTCACATGGCATCTTCTCCTATtatagttacttgtcctaaac
gacttatcagtcttgtgtacGCATTTTCCTCTGCATTTTTCAAGCTTGTAATTCTCTAACtatagttacttgtcctaaac
gacttatcagtcttgtgtacCTTGCTTTTTTCTCATTAAGACCCAGAATCACAAGAATGTtatagttacttgtcctaaac
gacttatcagtcttgtgtacAGTGCTCTGTTTAATCTTCAGTTTTCTTATTCATACTGGGtatagttacttgtcctaaac
gacttatcagtcttgtgtacACACTTTTCTCCTTCTTGTTACTTTGGACTCCCTAGGTACtatagttacttgtcctaaac
gacttatcagtcttgtgtacACATGTGTCTCGTAAACAGTAGTCGGAATAGATCAGCGTTtatagttacttgtcctaaac

gacttatcagtcttgtgtacTCGAGTACATTTGTTAGCTATAAAGTAAGATATGAGAGGtatagttacttgtcctaaac
gacttatcagtcttgtgtacTGTATCCAAATATTCAACACACAAAAAGGCAGCATCCTAAtatagttacttgtcctaaac
gacttatcagtcttgtgtacTTTTCCTTCTCTCCTCACCTGACTGTGTTTTTCCTTACCTtatagttacttgtcctaaac
gacttatcagtcttgtgtacAGTTCTGCACTTCACCTTGTCGTCCCCCCTTATCTCCTTCtatagttacttgtcctaaac
gacttatcagtcttgtgtacTTATTAATCAAGTAATCCATCTTTACCGTGTAGGCAGTGCtatagttacttgtcctaaac
gacttatcagtcttgtgtacAGCTGCTCACACTGACTATAAAATAAACCTCTTTGAGAAGtatagttacttgtcctaaac
gacttatcagtcttgtgtacCATAAAGTAGATTATGCCTGTCTGGAGCCCCCTGGAAATTtatagttacttgtcctaaac
gacttatcagtcttgtgtacGATAGTTTATCTTTGAAATGCTCCTTTCCAGACTATGAAGtatagttacttgtcctaaac
gacttatcagtcttgtgtacAGATACTCTCTTCACAATACTATACCTCAGAAATGTTCTCtatagttacttgtcctaaac
gacttatcagtcttgtgtacAGTGTACTTGGTGTTGTTGAGGGGCAGGGAAGGAGAGAGAtatagttacttgtcctaaa
gacttatcagtcttgtgtacGAAAGATGGAGAGAAAGAGAATTACCATTTTAATGGTGCTtatagttacttgtcctaaac
gacttatcagtcttgtgtacCTGTCTTTATCTTTCAAATACATACCACCTTAGTGAAGAGtatagttacttgtcctaaac
gacttatcagtcttgtgtacATTACTTTGACAATTCTACTTTATACCCTCTAATGCCAGGtatagttacttgtcctaaac
gacttatcagtcttgtgtacAGATACATTTAAGGCTACTCACTCACTTATTTCTTTCCTCtatagttacttgtcctaaac
gacttatcagtcttgtgtacTGAGTGGTCAAAATTAGCAAGCAGAAGTTGAGTCTGTTTCtatagttacttgtcctaaac
gacttatcagtcttgtgtacGAGGTTCTCATCTGGGACTGATTCTTCCCCACTCCCTCCAtatagttacttgtcctaaa
gacttatcagtcttgtgtacTTTGGTTGTCACAACTGTGTATGGGAGGGTACTGCTGGCAtatagttacttgtcctaaac
gacttatcagtcttgtgtacGTATCCTATAATACCTAGTACAGCCCCAACAACAAAAAATtatagttacttgtcctaaac
gacttatcagtcttgtgtacGGCTGAGAAGTCCTGATTTATGCTAACCAGATTTAGTAGCtatagttacttgtcctaaac
gacttatcagtcttgtgtacTAAATCAAGAAGTTACTGCAGGCCTGATATATACACATTCtatagttacttgtcctaaac
gacttatcagtcttgtgtacTTCTGTCAAAGGAAGAATCTGAGACCCAGAGAAGGTAAGTtatagttacttgtcctaaa
gacttatcagtcttgtgtacCAAGTGTAATTCTGAAGCCCTTGCCTTTCCTGTTTCTCTTtatagttacttgtcctaaac
gacttatcagtcttgtgtacCTCAGAGTGGAGAGAATGGAGGCTGGATTGGTACACTTTAtatagttacttgtcctaaa
gacttatcagtcttgtgtacACAGGTAGATGCTCATATTGTATCAAATAACCAATGTCCAtatagttacttgtcctaaac
gacttatcagtcttgtgtacAAAATCTCCTTAAGAAACTTCGCCAATCACTTAGATCTTGtatagttacttgtcctaaac
gacttatcagtcttgtgtacACAGAAAAATGTGTAGGAGAGGTCAATGTGCCGTGGGTCAtatagttacttgtcctaaa
gacttatcagtcttgtgtacGAACACATAAACTACAGATGCTCAACTAAAAGCTTAAGCCtatagttacttgtcctaaac
gacttatcagtcttgtgtacAGAATGTCTCAAAAGAACTGAATCATCTTTTTCTCTGGCTtatagttacttgtcctaaac
gacttatcagtcttgtgtacCCACTGAAACAGTTTAGTACTTTTTTCCTTGAAGATTCCCtatagttacttgtcctaaac
gacttatcagtcttgtgtacCAATTTTTGATCAACTGCAATACAGAGTGGGATAAAAACGtatagttacttgtcctaaac
gacttatcagtcttgtgtacCTTTGGGAGTTTCGAAGTATAAATGACAAGTGCCAGTGGAtatagttacttgtcctaaac
gacttatcagtcttgtgtacCCATTTCTTTCCATTGACCTTGGCCTTCTGGAATCAGCGTtatagttacttgtcctaaac
gacttatcagtcttgtgtacTTCCATAAGTTCCACAGGAGATTTTATTTTAACAAGCACCtatagttacttgtcctaaac
gacttatcagtcttgtgtacAGTGCCACTCAATAATTCTTGTTGATGATAGTTCTCTTGGtatagttacttgtcctaaac
gacttatcagtcttgtgtacGCCTGGAACCCAAGCCAAATCTACGTTTGTTAGACTTTTAtatagttacttgtcctaaac
gacttatcagtcttgtgtacTGTTATAAGGAGATTTTTTAAACGTGGCTGTGAATTACCCtatagttacttgtcctaaac
gacttatcagtcttgtgtacTAATTTCTCTGATTTTCTCTGAGGTTCTCAATCTGGAAAGtatagttacttgtcctaaac
gacttatcagtcttgtgtacCCTCGTTTATGGGTTTAAGGATAGACTGAAGTATTATCTAtatagttacttgtcctaaac
gacttatcagtcttgtgtacTCTTTAAACTTGACCTTTGGTTTTAGAAGGACACTGAATCtatagttacttgtcctaaac
gacttatcagtcttgtgtacAGGTGCAATCAGGCCCACAGATGTGTATTGTTTAGCTCACtatagttacttgtcctaaac
gacttatcagtcttgtgtacAAAACCAGCTGTCAATGGTTAAAAACCAAGCAATTTCACAtatagttacttgtcctaaac
gacttatcagtcttgtgtacCCCAGTGCAGCAATTAGTAACCTTTTCCAGATCCTACAGGtatagttacttgtcctaaa
gacttatcagtcttgtgtacGAATTTGTCACCCATACTCTACAACTGAAGTTCTCAATGTtatagttacttgtcctaaac
gacttatcagtcttgtgtacCATTGCTGAAAACGTGTTGGAAATGGTAATTCTAGGGCCTtatagttacttgtcctaaac
gacttatcagtcttgtgtacAGAAGCTCTGGAGATCTGGCCTAGTACACTGTTTTGTCAGtatagttacttgtcctaaac
gacttatcagtcttgtgtacCCCACTGGAGCCTGAGAACCACTTCTCCACAACATTGCAGtatagttacttgtcctaa
gacttatcagtcttgtgtacTGTTCCGTGACTTAAGTCTAGTGCTGGGCAAGTCAGTCTTtatagttacttgtcctaaac
gacttatcagtcttgtgtacCAACATTTTATTTCCTTGCCCCTCTCACCATTGCTGCGTTtatagttacttgtcctaaac
gacttatcagtcttgtgtacGGACTTGAGATGAATAAAAAAACAAAATCCTTGCGGTCCTtatagttacttgtcctaaac
gacttatcagtcttgtgtacTTTTCAACCAACAGCAGCAATTTGTGAAGATAAGGCCTCTtatagttacttgtcctaaac
gacttatcagtcttgtgtacGATGTCTTGCCACTCTTTGAGGCCATCCTTACAACTGTCTtatagttacttgtcctaaac
gacttatcagtcttgtgtacAAACAAAAACCTCACTTTCTCCTCTTTCAAAACTAAGCAGtatagttacttgtcctaaac

gacttatcagtcttgtgtacTCTCTAGAGATTATACTCCATTGGTGTTCCCCCCTTGAGGtatagttacttgtcctaaac
gacttatcagtcttgtgtacCCAACTTTTGTTTAATATATTCAGTGTCACCTTGTAAGGCtatagttacttgtcctaaac
gacttatcagtcttgtgtacTTGTGTACCCTATCAATTTGTTTCTTGTTTGACATCTGTGtatagttacttgtcctaaac
gacttatcagtcttgtgtacCAATTAATGTTGATGGTTTCGAGTTAGAGAGAAGAGAGTTtatagttacttgtcctaaac
gacttatcagtcttgtgtacTGAATACTTCAATTGGGTAATGGAATCACAGGCTTTCAGAtatagttacttgtcctaaac
gacttatcagtcttgtgtacATTCCTCTGCTATTGAACAATGGGACAACAGAAAATACATtatagttacttgtcctaaac
gacttatcagtcttgtgtacCAAGTTTATGTTTTCTTGTGGAGGTGAAATCTTGGGCCAAtatagttacttgtcctaaac
gacttatcagtcttgtgtacCTTCTCAAACAACTGTAACATAGCCATTTGTAAGACATTGtatagttacttgtcctaaac
gacttatcagtcttgtgtacGGTTGGAGAATCACCTAGGGACCATGTTAAAAATATTACTtatagttacttgtcctaaac
gacttatcagtcttgtgtacGCTGTATTTCATGCTCCTTCCCAAGTGATTTTTAATATGTtatagttacttgtcctaaac
gacttatcagtcttgtgtacAAAGAGCCAAAAGAGAACATTTTTCAGCCCATAGGACCAGtatagttacttgtcctaaa
gacttatcagtcttgtgtacCTTTCCAAAGCATCTAGTACCTTTTGATCTTATAGGACAAtatagttacttgtcctaaac
gacttatcagtcttgtgtacCTTTTAGATTTGACTATAAGCTCAGAGGCAATTTTGACCCtatagttacttgtcctaaac
gacttatcagtcttgtgtacAAATAAATATGATAGCTGTAGCAAAAAGACTGCCACACAGtatagttacttgtcctaaac
gacttatcagtcttgtgtacTGGGGCTATTTTATGGAGTGTTACTTAGTAATCCTGAACTtatagttacttgtcctaaac
gacttatcagtcttgtgtacATGCAATGATTCTGTGACAGCTAAGTTAAAAAAGAGAACCtatagttacttgtcctaaac
gacttatcagtcttgtgtacCAGTGTAAAGACTCTGAAAAGACCTGCAATAAAGACATTTtatagttacttgtcctaaac
gacttatcagtcttgtgtacCATTTCCCAAGCTTTTCAAAAACCAATGACTATCTCCCTCtatagttacttgtcctaaac
gacttatcagtcttgtgtacGGAATGCCATATTATTGCTATGCTTGAAAGTGTTGAATGGtatagttacttgtcctaaac
gacttatcagtcttgtgtacTGTGGGACAAACAGAAATTATCAATCATAGTGGAGGGAATtatagttacttgtcctaaac
gacttatcagtcttgtgtacGCTAAATTGGCACTTGTGAAGAAGCATATATAATAAGGAGtatagttacttgtcctaaac
gacttatcagtcttgtgtacCAGTTGCAGAAGGAACAGTACTTATAGCACACATTTCAAGtatagttacttgtcctaaac
gacttatcagtcttgtgtacTGAAGTAAATCTAAGGTCTTAGCTGATCTCTACCATGGATtatagttacttgtcctaaac
gacttatcagtcttgtgtacTTCTGGGTGGTATCATGTTAGGATGTTATGTAAACGAAGTtatagttacttgtcctaaac
gacttatcagtcttgtgtacCTGGTAGAGAGAATAGTGGGAGGTCTAGGGAGAAATCTGGtatagttacttgtcctaaa
gacttatcagtcttgtgtacATGGGGGAAGAATTTGTGCATGTATCAGTTAAACGCTCCCtatagttacttgtcctaaac
gacttatcagtcttgtgtacCTCCCTAAGTGTTGCAGATATGTTGTACCTCCTGACAGCCtatagttacttgtcctaaac
gacttatcagtcttgtgtacATGAATCCATACTTACTCACTTTTCCTCTCTGTATCCCCAtatagttacttgtcctaaac
gacttatcagtcttgtgtacGGGGAGAAATGGTTTTATTTTAGGGAAATTCTGCTCTGTTtatagttacttgtcctaaac
gacttatcagtcttgtgtacGACATCCTATTAATTGGTCACTTAGTTGCCTAGTCTTTACtatagttacttgtcctaaac
gacttatcagtcttgtgtacACTCTTGCTTTTGAATGAACAGAAATTTAATAGAGGTGGGtatagttacttgtcctaaac
gacttatcagtcttgtgtacTAACCTAAATTATCCATGAAAGTAAACTTGGAGTCCAGTGtatagttacttgtcctaaac
gacttatcagtcttgtgtacCTTAACAAGAGACAGGCATGAAATGAGAACACCTAACACAtatagttacttgtcctaaa
gacttatcagtcttgtgtacGGTGCTCTATTCCTATTCTGTGAAGGTGCCATACGTCATAtatagttacttgtcctaaac
gacttatcagtcttgtgtacAGTGTCCTATCAGTAACAGCAAAATCGGTCAGTGAAAATTtatagttacttgtcctaaac
gacttatcagtcttgtgtacCCCAAAGAAGTGAGTGGGCTGAAACAGTGTTGGTCATCACtatagttacttgtcctaaa
gacttatcagtcttgtgtacTTTGTTTCATCATTAAGTGGATCACTTGAATCCATTCCCCtatagttacttgtcctaaac
gacttatcagtcttgtgtacGACATCTTCTCTGTCTTTTTTAGCCAAGAAAATATACCTCtatagttacttgtcctaaac
gacttatcagtcttgtgtacAGACAAATGCCTAAGCAGATGGGGGGAGTTCCTGGTGAAAtatagttacttgtcctaaa
gacttatcagtcttgtgtacGGTCCCTAATGAAACTTGTGACCCAGAGTGAGAAATTCTGtatagttacttgtcctaaac
gacttatcagtcttgtgtacTTGCCTTTTCCAATACTACCAATGGCCTGCCCCTCCCCTAtatagttacttgtcctaaa
gacttatcagtcttgtgtacCCCTGGACTCAGTCACACGGGGGGACTTTGCTGCCTTCAGtatagttacttgtcctaa
gacttatcagtcttgtgtacCACTCAATAAAACTCCCTGCATTGCTCATTCTTTAATTGTtatagttacttgtcctaaac
gacttatcagtcttgtgtacGGGTACCCAGAAATGCTGTGACAGTGTCCTTTTGTTCTTGtatagttacttgtcctaaac
gacttatcagtcttgtgtacCCTCATACAACGGGGCCAGGGGCTGACTGAGCAGCTAACAtatagttacttgtcctaa
gacttatcagtcttgtgtacTGGAACTAAAAAAGCTAATTAGCTCACTAACGCCACCTCTtatagttacttgtcctaaac
gacttatcagtcttgtgtacCAGGAACTCTTGCCTGGGTGCTTCTGTGTTCCCCTCGAGGtatagttacttgtcctaaa
gacttatcagtcttgtgtacCCACAGGCCTTGCACAGTGCTTGCTTCTGTGTCAGTGCTTtatagttacttgtcctaaa
gacttatcagtcttgtgtacTGAGCACAGTGGGCCGAGTAGACGGGGCACCCCTGCCGGGtatagttacttgtccta
gacttatcagtcttgtgtacAATCCTGCTTCACTTTCATATCTCTTTATGAGCTTATCTCtatagttacttgtcctaaac
gacttatcagtcttgtgtacGGAAATCTTGGTTACCCTACTAGCCTACCTCAAGCCATTAtatagttacttgtcctaaac
gacttatcagtcttgtgtacGCAACTGGAATTACATATAGTTCTAGCTCTGAAATTTGAGtatagttacttgtcctaaac

gacttatcagtcttgtgtacAATTGAGTTTACTAATTGAAGAATGATTTGAGCAGGGCAGtatagttacttgtcctaaac
gacttatcagtcttgtgtacGTGGTGTGGTGGTGTCAGGTTAACAGTCTTAGCAATATGAtatagttacttgtcctaaac
gacttatcagtcttgtgtacACAATTCAGAACAGAAGCAGGAGTTGTAACATAGGGAATGtatagttacttgtcctaaa
gacttatcagtcttgtgtacGGAACCTTAAGAACTGGTTTCTAATGGCAATAGTAAGAAAtatagttacttgtcctaaac
gacttatcagtcttgtgtacGCTTCAGGATAGTATGGTATGCTTGCTATATCTTCATAAAtatagttacttgtcctaaac
gacttatcagtcttgtgtacATGTTATTATTATCACCAATTTCTGGGAAACCAAGGCCCAtatagttacttgtcctaaac
gacttatcagtcttgtgtacAGTAAGTGATAGAACTGTAGTTTAGATCCAAAAGGCAGCAtatagttacttgtcctaaac
gacttatcagtcttgtgtacTCAAACTCTCTACTCTTATTGGTCCTACTTCCTTTCTTTGtatagttacttgtcctaaac
gacttatcagtcttgtgtacTAAGAAATAGAGTCAGAAACTATGGGTTGAGTGTGGTGGCtatagttacttgtcctaaac
gacttatcagtcttgtgtacACCAGCCTGGCTAACTTGGCGAAACCCCATCTCTACTAAAtatagttacttgtcctaaa
gacttatcagtcttgtgtacACAAAACAAAACATTGTGTAGAGGCATGATAATTGTGTGGtatagttacttgtcctaaac
gacttatcagtcttgtgtacCACCTCTTAAGGACAAGGATATGTCCTGAAAAATGCATCAtatagttacttgtcctaaac
gacttatcagtcttgtgtacAGCCTACCATACACCTAGGCCGTATGGCATAGCCTATGGCtatagttacttgtcctaaa
gacttatcagtcttgtgtacCTGTACTGAATACTGTGGCCAATTGTAACATAACGGTAATtatagttacttgtcctaaac
gacttatcagtcttgtgtacTATGGCATAAAAGATAAAAGATGGTACATCTGCATAGGGTtatagttacttgtcctaaac
gacttatcagtcttgtgtacAGTCAGTGAGTGAATAGTGAGTGAACCTGAAGGCCTAGGAtatagttacttgtcctaaa
gacttatcagtcttgtgtacCTGTAGACTTTATAAACACTGTACATAGGCTACACTAAACtatagttacttgtcctaaac
gacttatcagtcttgtgtacACACTTAGCTTAAAACACGAACACATTGTACAGCTGTAAAtatagttacttgtcctaaac
gacttatcagtcttgtgtacCTACTTAGGGTCAGAATCATCAACAGCGCTGTCTTCCACCtatagttacttgtcctaaa
gacttatcagtcttgtgtacCCCATTGAAAGGTCTTCAGGGGGCACTAACACGCATGGAGtatagttacttgtcctaa
gacttatcagtcttgtgtacGCCTGAGGCTGTTTTGTAGTGAACTTTGTAATAATTAGAAtatagttacttgtcctaaac
gacttatcagtcttgtgtacAATGCATGAGTAATGTATTGCGCTAAAATGTTAATGAGGGtatagttacttgtcctaaac
gacttatcagtcttgtgtacAGAATTTTGAGGTCCATTATATTCTTATGGGACTACCTTGtatagttacttgtcctaaac
gacttatcagtcttgtgtacGTTGTTTTGTTTTGAAAATTTCTTTTTTGGGAGGGCAGCAtatagttacttgtcctaaac
gacttatcagtcttgtgtacAGGCTTCCTTCCACAGGCAACTTTCCCACCCATAGCACTCtatagttacttgtcctaa
gacttatcagtcttgtgtacCCTCGTGTTGCAATCTGGCTGGTGAGGGTGGCCTGGCATCtatagttacttgtcctaa
gacttatcagtcttgtgtacTGGGAAGAGTACGGTGACAAAGGCTGGGGGAAGGAGGTCCtatagttacttgtcctaa
gacttatcagtcttgtgtacGGACAGGGACGCGAAGAGCACACCACCCTCTTCTTGGGAAtatagttacttgtcctaa
gacttatcagtcttgtgtacTCACAAGGGGAGGGGAGAAGAGATGGGGAGCCCAAACTGCtatagttacttgtccta
gacttatcagtcttgtgtacGAGATCAAGCCCCTCCCTGCACGCGCCCCCTCCCCCAGATtatagttacttgtccta
gacttatcagtcttgtgtacTTTGGAAACTTGAAGTTCTGAGGGAGAAGCTGAGGTCTGCtatagttacttgtcctaaa
gacttatcagtcttgtgtacCAGAAGTGGGGTCGTGGGAGTGGGTGGAGGGGGCAATGACtatagttacttgtcctaa
gacttatcagtcttgtgtacTCCAGAGCCTGCCACTGTCCTCTCCTCCTGGAGCTGCTGCtatagttacttgtcctaa
gacttatcagtcttgtgtacCGATTCTGTGGCAGAAGTGGCAGGGACCATGGGGGCAGGTtatagttacttgtcctaa
gacttatcagtcttgtgtacGCCTTGCCACAGATGCTGCAGTGGTGGTGCTTGCGGTTCGtatagttacttgtcctaa
gacttatcagtcttgtgtacCTGTGGTTCTTAAAAGACAGCAGGTTGGGGGCAGCGGGAGtatagttacttgtcctaa
gacttatcagtcttgtgtacCTGTGGTGTGGGTCTTCTGGTGTGTCAGCAGGCTGCCGGCtatagttacttgtcctaa
gacttatcagtcttgtgtacGGACGTCACCCCTTATCCTCTTCTAGCTCTCTCCGCACTCtatagttacttgtcctaaa
gacttatcagtcttgtgtacCTGGTGTTTCTAATGGAGCCTCCCAGGGTTTCCCTTGACCtatagttacttgtcctaaa
gacttatcagtcttgtgtacCGGGCTTGGGACATCTGGCCTCCTCGATCTGGGCCGGCACtatagttacttgtcctaa
gacttatcagtcttgtgtacCTGCAGAGGTGGCTGGCCAGTTTGTAGTGGGACTCAAAGGtatagttacttgtcctaa
gacttatcagtcttgtgtacAGGCAAAGGTCTGGGCATCTGTGTGGTTGTGGTTGTGGCTtatagttacttgtcctaaa
gacttatcagtcttgtgtacAGGCAATAGTGTTGGTCTGTCTTGTGAGTGTTGTGGTTCAtatagttacttgtcctaaac
gacttatcagtcttgtgtacTGGCTGGCAGTAAGTCTCGCCACATTGGCTGCAGGAGGAGtatagttacttgtcctaa
gacttatcagtcttgtgtacCATTGATCCAGCTATCCATGTGACTCAGACTAGGTCCATTtatagttacttgtcctaaac
gacttatcagtcttgtgtacTCACTGGCATGCCAAGGACTCCTGGGGACACTGTCATCTGtatagttacttgtcctaa
gacttatcagtcttgtgtacGAACCCAGCCTACATTGTGATTCCCCAGTCCCCCACCTCCtatagttacttgtcctaa
gacttatcagtcttgtgtacGCACATGCCTATACTGGACAGAGTTAAATTGTAATGCAATtatagttacttgtcctaaac
gacttatcagtcttgtgtacAGGCATTTCTTTTTACCCTACAGATCCAACATGAGTCAGCtatagttacttgtcctaaac
gacttatcagtcttgtgtacCCTTCTTGGGGACTCAGGTTTAGGGGTTGCATGTCACTTGtatagttacttgtcctaaac
gacttatcagtcttgtgtacCAGTGGGAAGTCAATAGGAAGAACAAGTATTATGGTATATtatagttacttgtcctaaac
gacttatcagtcttgtgtacCCAGGCGATGGGCTAAGTATCACTTCCACTCCCAGCTCAAtatagttacttgtcctaa

gacttatcagtcttgtgtacCAAGAAGCACGATCCTTACAAGAGCATAGAACAGCGGTCCtatagttacttgtcctaa
gacttatcagtcttgtgtacGGAGCACATAACCTAGACCCCTCTTAAGAGCAGTTCATGGtatagttacttgtcctaa
gacttatcagtcttgtgtacCCAGTACAGGTGTGTGGCGCAGGCATTGGGAACCCTTGGCtatagttacttgtcctaa
gacttatcagtcttgtgtacAGGAGAAGAGAGACTGATAAGGTGGAACAGGTGAATGACAtatagttacttgtcctaa
gacttatcagtcttgtgtacGGATGTCACTTTCCTTTTTACTGTTTTCATTTCCTCCTCCtatagttacttgtcctaaac
gacttatcagtcttgtgtacCTTGAGCTATGTGTAACTTTGTTTCCTAAAATTGTTGGTCtatagttacttgtcctaaac
gacttatcagtcttgtgtacCCTCTCCAGGCCTAATTTTATCAACGGTAAATAGATGGATtatagttacttgtcctaaa
gacttatcagtcttgtgtacAATCCTTCTCTTTTGATTATCTTGCACTGACCAAGTAGGCtatagttacttgtcctaaac
gacttatcagtcttgtgtacATGCCCAGGGAAACATAATCCAGCAGTGGAGAGATAAGCAtatagttacttgtcctaa
gacttatcagtcttgtgtacTGGTGCTGAGCCAGGAGAGATAGCAAGGATAAACTACTGAtatagttacttgtcctaa
gacttatcagtcttgtgtacCTCCAATACGAGTTCAAAGGCAGGTAAGAAATAGTGAGGAtatagttacttgtcctaaa
gacttatcagtcttgtgtacTTAGTGGTGATTTAGTGATTACTTAAAGTAAGCGCGCGCGtatagttacttgtcctaaa
gacttatcagtcttgtgtacACACACGTATATATATAAACTCCCCCCAACAAGAGAAGACtatagttacttgtcctaaa
gacttatcagtcttgtgtacCTTCAAAGTTCAGGCAGTGAGTCTCTTTTTAAGTTGTCAGtatagttacttgtcctaaac
gacttatcagtcttgtgtacACACCTAGGGCTTATGTTTTATGGTTTGCACAGTATGGTCtatagttacttgtcctaaac
gacttatcagtcttgtgtacCCTAGTGGCAAGGTAACCAAATGACAGCATGAAATTTTTGtatagttacttgtcctaaa
gacttatcagtcttgtgtacGTCTTGGTAAGGCTTGTTTTAAATAATTCAGACGATTTGCtatagttacttgtcctaaac
gacttatcagtcttgtgtacTCTAAAGTAAACATATTGGTTCATACACTAGGGTATGTGGtatagttacttgtcctaaac
gacttatcagtcttgtgtacCACAAAGGCTGATAATTGGTAATAGAAGCTGAGTGGAAAAtatagttacttgtcctaaa
gacttatcagtcttgtgtacCTACAGACACAAGCAAAGGAAAGGCTCCACTTACCCACATtatagttacttgtcctaa
gacttatcagtcttgtgtacAGATGTCTACTCTGATCTCCCTCCTCTGTCTTCTTACATAtatagttacttgtcctaaac
gacttatcagtcttgtgtacTAAAGAAATGAGATAGATTGGGAACCTAAATCCAGATCTCtatagttacttgtcctaaa
gacttatcagtcttgtgtacGAAGCTGTTTTGTACTTCTGGAGTTTTCTAGCACATAAGCtatagttacttgtcctaaac
gacttatcagtcttgtgtacGCTTTTATGTTTAGGATAGTTTTGGCTCAATTTCTTCCACtatagttacttgtcctaaac
gacttatcagtcttgtgtacCTGACTTATATACAGAATACGGATTACTGGGCTAAGGTTAtatagttacttgtcctaaac
gacttatcagtcttgtgtacGCTTGAGCAAAGGCATCTAGGCAGAAAATCTCAGAATGTAtatagttacttgtcctaaa
gacttatcagtcttgtgtacTTAAATGGAAAAAATGGGAGGTAGGTCTAGATAAGAAGTGtatagttacttgtcctaaa
gacttatcagtcttgtgtacTAAAAAAAATGAGGAACTGTGGAAAGTTTTTGAGTAGGGCtatagttacttgtcctaaac
gacttatcagtcttgtgtacCAAAGGAGAAATATTTGGAAGGGGAGATTTAGGAGTTTATtatagttacttgtcctaaac
gacttatcagtcttgtgtacACTTTGGCAGTGAATAGGAAGTCTTATTATATGTGGTGTAtatagttacttgtcctaaac
gacttatcagtcttgtgtacGACAGGAGGCAAACGTCAAGATTGATTCTAAAATTTTAAGtatagttacttgtcctaaac
gacttatcagtcttgtgtacAAATGGTTTTGAAAGGCTGGTTAACTAGATAAATAGGGGAtatagttacttgtcctaaac
gacttatcagtcttgtgtacGCTGGCAGATTGTTGGATATAGAAGAAGAGAATGTGTTTTtatagttacttgtcctaaac
gacttatcagtcttgtgtacGCAGCCAAGGATATGCTAAATCAGTTAATTTTGAAACTGTtatagttacttgtcctaaac
gacttatcagtcttgtgtacACTCAAATGACATTTATCTGTGTATCTGAAACCTGAATGGtatagttacttgtcctaaac
gacttatcagtcttgtgtacAAAAGAAACTCTGATTTTTCCTGTTAATGGAGGCAACATGtatagttacttgtcctaaac
gacttatcagtcttgtgtacGGCAACCCACAGTAGACAGGCCAAGGCAGCAGGAGATGATtatagttacttgtccta
gacttatcagtcttgtgtacGTGCTTAATGTTACCATTCTAACCATCAAACAAAATGCCAtatagttacttgtcctaaac
gacttatcagtcttgtgtacCCTCCTCCAGGTTCCATCTTCTCACATGACATCTCCAGTCtatagttacttgtcctaaa
gacttatcagtcttgtgtacATTGCCTTCCTGGTGATTCAAATTATTCAGAGTACAACCCtatagttacttgtcctaaac
gacttatcagtcttgtgtacCCCACCCCCAAAACTCTCAGTTCCATATGTCCCATTGCACtatagttacttgtcctaa
gacttatcagtcttgtgtacAATTTACCTCTCTCAAGACCTACCCATTTAAAAAAATCGCtatagttacttgtcctaaac
gacttatcagtcttgtgtacACTCCTCTTCAATCCATTTTCTTTTTTACTTGCAAGTCCAtatagttacttgtcctaaac
gacttatcagtcttgtgtacCTGGTGTCAGGAGAAAAATACTCATATGTGTCAACTTGGCtatagttacttgtcctaaa
gacttatcagtcttgtgtacCAAAGTCTAATTTAGGTGTTACCGGGAAGCTATTTTATAGtatagttacttgtcctaaac
gacttatcagtcttgtgtacGTAGATAATACAGGCAGGCCCCATCTAATCAGTTGAAAGTtatagttacttgtcctaaa
gacttatcagtcttgtgtacTGCCTCCAGACCACAACATCAACTCCTGCTGTCAAGTTTCtatagttacttgtcctaaa
gacttatcagtcttgtgtacTCTTTCTCTCTCCCCCTCTCAATTCTACTGGTTCTGTGTTtatagttacttgtcctaaac
gacttatcagtcttgtgtacAGTTCTTTGAAGTTATTTGCATGGTGGTGTGACTGACCAAtatagttacttgtcctaaac
gacttatcagtcttgtgtacGAAAAACAAAATTTGCCTTCATAGTATAGCTCAAAGGGTGtatagttacttgtcctaaac
gacttatcagtcttgtgtacGCCTATCCCCCCTACCTGTTTAAACTGGCTTTCACTTGGAtatagttacttgtcctaaa
gacttatcagtcttgtgtacGTGTAATTAGAATTAGTGGCACCTACTGATGGAGTAATACtatagttacttgtcctaaac

gacttatcagtcttgtgtacCTGAATGTGCCAGAAGCCATCAGCTATCTCTTAAATGTAGtatagttacttgtcctaaac
gacttatcagtcttgtgtacAAAAATTTTGAACCTTAGCACTGGGGATTTGCAAGAGATCtatagttacttgtcctaaac
gacttatcagtcttgtgtacTCAAATATAATGAAATAGAGGCCCAGGGAAAACTGTGTTGtatagttacttgtcctaaac
gacttatcagtcttgtgtacAAAGCATGTTGTTTTGGCCTGGACAATATTCCATCATTTGtatagttacttgtcctaaac
gacttatcagtcttgtgtacGAAATTTCTAGGTCAAACAGTATGCACACTATGAAGGATTtatagttacttgtcctaaac
gacttatcagtcttgtgtacCCAATTTCTAATACTCAGTGATGGATGGGAGCCTGTTTCCtatagttacttgtcctaaac
gacttatcagtcttgtgtacTATTTTCATTAAAAAACAAACAAATGGCCAGGCATGGTGGtatagttacttgtcctaaac
gacttatcagtcttgtgtacATGGGTGCCTATAATCTCAGCTACTCCAGGAGGCTGAGGCtatagttacttgtcctaaa
gacttatcagtcttgtgtacGGAGTCCTGTTTTAATTCACCTTATTTATTAGCAAGGTTGtatagttacttgtcctaaac
gacttatcagtcttgtgtacAGAATCCACAAAACCATTCCTCAATAAATGATGGGTATTGtatagttacttgtcctaaac
gacttatcagtcttgtgtacGTTGTTTGTTTTGTTGTAGTTGTTGATATTATTGTGCCGCtatagttacttgtcctaaac
gacttatcagtcttgtgtacCTATTTAGCAAGAAACTACAAAGAGATCACAGCCTGGAGTtatagttacttgtcctaaac
gacttatcagtcttgtgtacTGCTAGGGTCTAATAGCCTATGCTAGGTTGATCAGAGGGTtatagttacttgtcctaaac
gacttatcagtcttgtgtacGGAAAAATGCACCAAAGCATATCCCAATTATGCTGTTGATtatagttacttgtcctaaac
gacttatcagtcttgtgtacCAGGCTCTATTCTCCCTGGAATAAAGTAAAATGTGGTTTAtatagttacttgtcctaaac
gacttatcagtcttgtgtacGGCATGCAAACCACTCCACCTCTTTCGTTTCCTCTCTAACtatagttacttgtcctaaac
gacttatcagtcttgtgtacATGGGAATTGACCTCAGATAATACATAAATTGCAAAGTGCtatagttacttgtcctaaac
gacttatcagtcttgtgtacTCCTATTTTTTCCCCTCTAGAAAGTGTGAAATTGCATATCtatagttacttgtcctaaac
gacttatcagtcttgtgtacTCTTTTTCATTTTTATAGTAGGCTTCAAACTCCTGTAGGCtatagttacttgtcctaaac
gacttatcagtcttgtgtacTCATTCATTTGCCTTCTGCTGTGTTTGAACTCCTCTCCTAtatagttacttgtcctaaac
gacttatcagtcttgtgtacAGTCTTTGGTGAGGAAGGATGAGAGATGGCTAAGGGATATtatagttacttgtcctaaac
gacttatcagtcttgtgtacCTTTTATGACAACTGAAGAATACACATTAGACACTTTCGCtatagttacttgtcctaaac
gacttatcagtcttgtgtacGCTTAGCATGATTCCTGACACATAGCAGATATATATCAAAtatagttacttgtcctaaac
gacttatcagtcttgtgtacTATTTCCTTTCTGTTTATCAATCAGGGAACAAGACCAACAtatagttacttgtcctaaac
gacttatcagtcttgtgtacCAGTGAATGTGGAGGGAGGTGTTTGAAATTTGTAGGCCAAtatagttacttgtcctaaac
gacttatcagtcttgtgtacCATACAGTGGGCTTAGATTGGAAAGTCAGGCAGGATTTCTtatagttacttgtcctaaac
gacttatcagtcttgtgtacAGGCAGAATTCCTTTTACTCTAGGGAACCTTGGTTTTTGCtatagttacttgtcctaaac
gacttatcagtcttgtgtacGGCCTTCAATGGATTATATGAAGCACCTCTATCTTATAAAtatagttacttgtcctaaac
gacttatcagtcttgtgtacTCCTTTACCTAAAGTCAAATGACTGTGGATGTTAATCACAtatagttacttgtcctaaac
gacttatcagtcttgtgtacCAGACTAATGCTGGACCAAACAAGTGAGCACCATAGCTCGtatagttacttgtcctaaa
gacttatcagtcttgtgtacAAATTAACCATCACACCTTTATTTTAACTCCTCCTTTCCCtatagttacttgtcctaaac
gacttatcagtcttgtgtacAAAGGGAAGTAGGGATCTATCAAGAAACCAGAGGTGTTTTtatagttacttgtcctaaac
gacttatcagtcttgtgtacGAACTCACCGAATTCTTTTAATGAACATAAACAGCAAGACtatagttacttgtcctaaac
gacttatcagtcttgtgtacGAGAAAGAAAAAGAAGGTTTGTGCTGATTTTGGAACTAAGtatagttacttgtcctaaac
gacttatcagtcttgtgtacGTGAGAGCACCCCAAAGATGATGCAACTGTCTTCCTGGTTtatagttacttgtcctaaa
gacttatcagtcttgtgtacAGTAGTACTGAGAGTTGTGGGTTGGAGGGAAAACATAGAAtatagttacttgtcctaaac
gacttatcagtcttgtgtacCTGCTACAAAACAGGGGTGTCATGGGCTGAGTGAGGCAAGtatagttacttgtcctaaa
gacttatcagtcttgtgtacGATTCTCTGCAAGCATCCTAGATAAGTCCAAAGAAATTCCtatagttacttgtcctaaac
gacttatcagtcttgtgtacCCTTCTTCAGTATAGCAGTATAGACACAGACCAGAGTCGTtatagttacttgtcctaaac
gacttatcagtcttgtgtacGGCATTGAAGAAACAATCTCCAGAGCAGAAGGAAAATGAGtatagttacttgtcctaaa
gacttatcagtcttgtgtacTGAAACAAGAGTTCATCATATGCTGGTGGTAAAGTTGCTTtatagttacttgtcctaaac
gacttatcagtcttgtgtacTTCAGCTGAGCTAAGTATTTAGACATGTAGGTGGATAATAtatagttacttgtcctaaac
gacttatcagtcttgtgtacCACATCAATCAGCAAATTCAAAGCACAGCACTATAATTAGtatagttacttgtcctaaac
gacttatcagtcttgtgtacCCCGTAGAATATCCATGTGGGCAAAATTAAAGCATACTGAtatagttacttgtcctaaac
gacttatcagtcttgtgtacAGAAGTTGTTTTAATTTCGGGTATAGCCTGGTTCAAATGAtatagttacttgtcctaaac
gacttatcagtcttgtgtacGGATACCTACATATAACATAGACTTCCTTTAGAGACGGAAtatagttacttgtcctaaac
gacttatcagtcttgtgtacCCGATCTTTCTTAGGCCATGGTCAACAACTTAAAACAGAAtatagttacttgtcctaaac
gacttatcagtcttgtgtacGAGGTCTCCAAGGTGAAATTTGTTAATAGTGAACACAAAAtatagttacttgtcctaaac
gacttatcagtcttgtgtacGTAGATCACATCATTTTACTGCATATTTTTGTAAGGCCTGtatagttacttgtcctaaac
gacttatcagtcttgtgtacTATTCAATGGCTGATTTTCAAGGATTCTCTTAGGATTTGCtatagttacttgtcctaaac
gacttatcagtcttgtgtacGCAATATATGTGCAAATACACTTGATTCACTGCCTTTTGAtatagttacttgtcctaaac
gacttatcagtcttgtgtacCAAGACAGAACAAATGCACAGTAATCCCCTTTTTAACAGAtatagttacttgtcctaaac

gacttatcagtcttgtgtacGCTGAGAATCTTGAGCCTCCCATAAAATTGGTATAAATGAtatagttacttgtcctaaac
gacttatcagtcttgtgtacACATAAAAGGCAATGTTACACTATGGTAATAATGGGGGCAtatagttacttgtcctaaac
gacttatcagtcttgtgtacCAGGTCTTTGCATTGTTAGGCTGGTATTTGTTTAATGTGAtatagttacttgtcctaaac
gacttatcagtcttgtgtacCTCCAGGAAGTAGTTAAGAGAAATGTAATGGCACTATTTGtatagttacttgtcctaaac
gacttatcagtcttgtgtacGAAGGGATCCAATTTTTTCCAGTCTTTATTCACTAGGGAGtatagttacttgtcctaaac
gacttatcagtcttgtgtacTGTCACCAGTTTCTTCCACAACATTAAAATCTTTTCAGCCtatagttacttgtcctaaac
gacttatcagtcttgtgtacGCAGGGAGGATGTTACATACCCGGGTTTAGTGATCCAAGGtatagttacttgtcctaaa
gacttatcagtcttgtgtacCTGGATTCATAACAGCATTTTTCTCAAAAACCTTTTGCGTtatagttacttgtcctaaac
gacttatcagtcttgtgtacAGACATCTCCAATTTCTATCAGAATACTGCCATTGTATAGtatagttacttgtcctaaac
gacttatcagtcttgtgtacCACTGAGTGTTATAGTCCCCTTTCCACTTGAGAGGCAAATtatagttacttgtcctaaac
gacttatcagtcttgtgtacCTGAGAAGATGTGATGGTTCCCACAGAAGAGGAGAGAGAAtatagttacttgtcctaaa
gacttatcagtcttgtgtacTTGGTTATATATTGTAAAGAACTGGCCAAGAACTAAGACCtatagttacttgtcctaaac
gacttatcagtcttgtgtacATGATGTTTGTCTCTCACACAGATGATCTAGTTGCCGTGGtatagttacttgtcctaaac
gacttatcagtcttgtgtacCCCTTTGCATCCATTTCTAATTGTTCCTTAGCTGAGAATCtatagttacttgtcctaaac
gacttatcagtcttgtgtacCCAAGATGTGTAGTTAGAATTTGGTTAGACTGCAAGCCATtatagttacttgtcctaaac
gacttatcagtcttgtgtacAAGAGTACACCTTGAGACCAGGACTTTAGCGATTCTCAGAtatagttacttgtcctaaa
gacttatcagtcttgtgtacAGTGAGGACGAGTGAGGTGATGAAGAAAGTGAGACAAGAAtatagttacttgtcctaaa
gacttatcagtcttgtgtacCACTAAGTGTATTTTGAAGGGTAAATTACCTCTTTGGGAAtatagttacttgtcctaaac
gacttatcagtcttgtgtacGGACTCAAGTACAGTGAGCACCCTTTAAAAAATCAAGAGGtatagttacttgtcctaaa
gacttatcagtcttgtgtacCAGAAGGACAGGGATTATTTTTCAGTAGCGGAAGGTTGCCtatagttacttgtcctaaa
gacttatcagtcttgtgtacTGCCAACTGCCCCGTAGTGCATAGTACCAAGTTGCAGCTGtatagttacttgtcctaaa
gacttatcagtcttgtgtacCTCAAGCAGGGAATGGAGAGACACAGGATCTTAGGCTAGAtatagttacttgtcctaaa
gacttatcagtcttgtgtacAGAGGTGAGCTGAGAAACTATGAAGATGGATATCAAAAATtatagttacttgtcctaaac
gacttatcagtcttgtgtacCAATAAACTCAGAAATAATGGTGCCTTTGAAGAGTAGAGGtatagttacttgtcctaaac
gacttatcagtcttgtgtacTTGTCATCTTTACAATCTTCAGATCCCAAAACTCTTCCCAtatagttacttgtcctaaac
gacttatcagtcttgtgtacGGTTCACCTTCAAAATTTAAAATGGCTGCCATCATATGGTtatagttacttgtcctaaac
gacttatcagtcttgtgtacATATCCAAGGTGGTAGAACTTAGGAAGGGGCACAGAAGGCtatagttacttgtcctaaa
gacttatcagtcttgtgtacATTTCTGCCTATATCACATCAGTTAGAATTCAATGACAGGtatagttacttgtcctaaac
gacttatcagtcttgtgtacAAAGGTGGTTTTATTAGTATGAATGATAGGGAGAATGGGAtatagttacttgtcctaaac
gacttatcagtcttgtgtacCCTAACCATCTTTGCATTGTAGGGTCTTCATCAACCAGAGtatagttacttgtcctaaac
gacttatcagtcttgtgtacATTAATACATTGACTACTTGAGGATCCTGAAATTACTGGGtatagttacttgtcctaaac
gacttatcagtcttgtgtacAGGCTGACCATCTTCACAAAATTTAAGATAATGACCTTGCtatagttacttgtcctaaac
gacttatcagtcttgtgtacAATGTCAATGGCAGTAATGATTATATTAGTCTGCTAAGGCtatagttacttgtcctaaac
gacttatcagtcttgtgtacGCAGGTTTGGTTTCTCTTTTGCTTGCAGACAGCCACATTCtatagttacttgtcctaaac
gacttatcagtcttgtgtacGTCTTCACATGACCTTTCCTTGCATGAGCATACCCCTGGGtatagttacttgtcctaaa
gacttatcagtcttgtgtacGGCCCATCCTTATAACCATATTTAACCTTAATCATGTCTTtatagttacttgtcctaaac
gacttatcagtcttgtgtacTTGGAGGTCAAGATTTTAACATGTAAATTTGGTATGGGGAtatagttacttgtcctaaac
gacttatcagtcttgtgtacAAATACCAATTTAGTGCTTCCTATGTGTTGAGCCTGATGCtatagttacttgtcctaaac
gacttatcagtcttgtgtacATTTTACAAGGAAGGAAACAAATGGAGGGAAACTGACACAtatagttacttgtcctaaa
gacttatcagtcttgtgtacAGAACAAGTGAGGGCAGCAGACCTTTGCCAGAGCCTGGGTtatagttacttgtcctaa
gacttatcagtcttgtgtacGTGTGATGGTGCTTGTGAGATGATGCAGAGGACACGGAGTtatagttacttgtcctaaa
gacttatcagtcttgtgtacCATGAGAGATGGAGAAGTGCTATAGGTGGTCCCAGATGGAtatagttacttgtcctaaa
gacttatcagtcttgtgtacCTATTGGTTGTAGGTGATGTTGGAGGAGGATGCTTCGAAAtatagttacttgtcctaaac
gacttatcagtcttgtgtacGAGAATCTTCATCTTCCCCTGAACAAACAGATGTATATCCtatagttacttgtcctaaac
gacttatcagtcttgtgtacGGATGTGGCCTTAGGTCAGTTAGTCAAACGAATTAGCTAGtatagttacttgtcctaaac
gacttatcagtcttgtgtacGCCCAATTGAGCCTCCCAACTTCTACAGGATGGAGTGTCTtatagttacttgtcctaaa
gacttatcagtcttgtgtacAGCTGTGGAAAGTCAACTCCCTCCCAGTTTTAAGCTCCCAtatagttacttgtcctaaa
gacttatcagtcttgtgtacGAACCAAGCTGAACAAACTTATTACTTTTTCCCAGAGGTAtatagttacttgtcctaaac
gacttatcagtcttgtgtacCTTCTCATTAGTGGGCAGAAAGAGTAAAAGGAGCATACAGtatagttacttgtcctaaa
gacttatcagtcttgtgtacCTAGATAACAGGATTTAATGGAAGTTTCTTTGAGCTTGTCtatagttacttgtcctaaac
gacttatcagtcttgtgtacGCTTGGGAGTCTTTTCACCATTATAAATTGGCATCCACTCtatagttacttgtcctaaac
gacttatcagtcttgtgtacCTGAGACAAGGGGAAAGAAGATCTTTTTTTTCCTCATTAtatagttacttgtcctaaac

gacttatcagtcttgtgtacGTTGGGAGCAATACAAATCATTCAAATAATGTTAAGAGGCtatagttacttgtcctaaac
gacttatcagtcttgtgtacGCTTGGGATTGTGTGTAGTAGCTATGGTTTGGGAATAAATtatagttacttgtcctaaac
gacttatcagtcttgtgtacATTCAGGAAATTTCTCTCTAAATAATGGCTCTGCTTTCAGtatagttacttgtcctaaac
gacttatcagtcttgtgtacAAACAGTAGATCTAAACAAAGGAATCAGAGCCAATAGAAGtatagttacttgtcctaaa
gacttatcagtcttgtgtacATTTTAATCATTTTCTTGGCACGTTCAGCCTGTGATGAACtatagttacttgtcctaaac
gacttatcagtcttgtgtacTTCTCCAGAAAAATAAAGCCTACTGCCAACAAATTTAGCCtatagttacttgtcctaaac
gacttatcagtcttgtgtacTGGTTTCATGCACCTTTATTTAAGAATTCCTCCCTATACCtatagttacttgtcctaaac
gacttatcagtcttgtgtacCAGAATGGAGACTTGAGAACTTATGGAGAAAAATGGAATAtatagttacttgtcctaaac
gacttatcagtcttgtgtacAGTGGACTGATTGAGGATTCACTTCATGGATATAAAGTTGtatagttacttgtcctaaac
gacttatcagtcttgtgtacTCTCCTGAAGAATATAATACCTACATTTTCATGCTTGTGGtatagttacttgtcctaaac
gacttatcagtcttgtgtacCTTTTTTTTCAATTAATCCCTCTCCACTTGCAAATCTCACtatagttacttgtcctaaac
gacttatcagtcttgtgtacTTAGGTAGCTAACAAACTTTACTTTTTTCTCTGTACAGCGtatagttacttgtcctaaac
gacttatcagtcttgtgtacGGCTTCAAACAGATCCTTCCCCTTCTCTTTTCCTCCCCCTtatagttacttgtcctaaac
gacttatcagtcttgtgtacTTCCTTCATCATCTTCATCTCCTTTTTTGCTATGCTCTTCtatagttacttgtcctaaac
gacttatcagtcttgtgtacTCACTTATACAAGCTTGAGATATTCTATACACGTTTGTGCtatagttacttgtcctaaac
gacttatcagtcttgtgtacGAAAACAGTGACTTCTTCAGCAACAATATTACCAAAACAGtatagttacttgtcctaaac
gacttatcagtcttgtgtacGGACTGGTATCCGAGGGTTTAGTAAATTTGGAGCTTTATAtatagttacttgtcctaaac
gacttatcagtcttgtgtacCAATTCTCTGTTCTTTGGATTACTGATTTTGGAAGCTTGGtatagttacttgtcctaaac
gacttatcagtcttgtgtacCATGTCCTCAAAAGTAGGGTTCTTTCAACTGGAAATGTTGtatagttacttgtcctaaac
gacttatcagtcttgtgtacCTTCTCTAATGATTTTTAGCAGATGGGTGTCAGAGCAAGCtatagttacttgtcctaaac
gacttatcagtcttgtgtacAGTGGCTTTTACTTGTATAGTTAGGATTGGAGAGTCTTCCtatagttacttgtcctaaac
gacttatcagtcttgtgtacCTGATGAAAGGGTCAGGCAGATAACTGTTGGCCTTGTAAAtatagttacttgtcctaaac
gacttatcagtcttgtgtacTTGAGGACCAAAGAAGCAGTTTTCTGTGAGTGACCTGAGAtatagttacttgtcctaaac
gacttatcagtcttgtgtacCAACAGCCTGCTAAACTTCAGAAAGTAGACATCCTCAGAAtatagttacttgtcctaaa
gacttatcagtcttgtgtacACTTCCTTTAAAATGCACACTAAACGTATGCCAAATATGCtatagttacttgtcctaaac
gacttatcagtcttgtgtacCTCTTGCTATTTTAAAAATTAATTTTGGCTGGGCACAGTGtatagttacttgtcctaaac
gacttatcagtcttgtgtacAGGCATGGTGGCAAGCAACTGTGGCCCCAGCAACTCAGGAtatagttacttgtcctaa
gacttatcagtcttgtgtacTGAATGAGCGTATCATAGTAAAATATCTCTGATGTGGAACtatagttacttgtcctaaac
gacttatcagtcttgtgtacTGTTTGGAACACAGTCTGAAAATCAGGATGACTTTAGTTTtatagttacttgtcctaaac
gacttatcagtcttgtgtacAGGGCAAGTTGATCTGAGTACCTAGAATTGATGTTCCTAAtatagttacttgtcctaaac
gacttatcagtcttgtgtacGGTGTTTCCAATCCTATGCCTATTTTTATTTCTTTCTGTGtatagttacttgtcctaaac
gacttatcagtcttgtgtacGAACTTATGGTCCCAAATGGTTCATTATGTTCTTCTGTTAtatagttacttgtcctaaac
gacttatcagtcttgtgtacTTTCTGAAAACAAAGGGACACTTACATCTGATGAGTGGGGtatagttacttgtcctaaac
gacttatcagtcttgtgtacACGGTGTCTGATTGAGTCTGTAGGTATTATTCTTTGTAAGtatagttacttgtcctaaac
gacttatcagtcttgtgtacCAGCTCCACAGGTGGATTGAACTCCCTGAAAGAAAGTAGGtatagttacttgtcctaaa
gacttatcagtcttgtgtacTGGTTTCTGGCTAGTAGTTTCTCCTAATTCTTATACAAACtatagttacttgtcctaaac
gacttatcagtcttgtgtacGAATTAAGACACACATTTGCTAAAACTTAGGGAAAGTTCCtatagttacttgtcctaaac
gacttatcagtcttgtgtacAGAATGGCTGAGTGGTTAAGAAGATAGACAGACTTGGTTCtatagttacttgtcctaaac
gacttatcagtcttgtgtacGGTTCTGTCACTGTCTTGCTGTGTAACTTGGGCAGTTTGCtatagttacttgtcctaaac
gacttatcagtcttgtgtacGGTTGTTTTTTTCCAAGCTTCCAAACTCAGTAATCCGAAGtatagttacttgtcctaaac
gacttatcagtcttgtgtacTGAATTAATACTGTGTTTATACATGTTGAACAATGCCGGGtatagttacttgtcctaaac
gacttatcagtcttgtgtacGTTAGCTATTATTATGCAACGGCCCAACTATACTGTGTTTtatagttacttgtcctaaac
gacttatcagtcttgtgtacAGATCAAACTGTCCTTCATCCCCACTCACTCTCTGAATCGtatagttacttgtcctaaac
gacttatcagtcttgtgtacGCTGAATCAATAAGAAAACGGAGAAGCTAATTTATGTGCTtatagttacttgtcctaaac
gacttatcagtcttgtgtacCTGGCACTCTTATGAATGTCTGTTCCTATTACCTTTTTATtatagttacttgtcctaaac
gacttatcagtcttgtgtacACTAAGCTGTTGTAAGTTCTTATATTACCTGTGGGGAACTtatagttacttgtcctaaac
gacttatcagtcttgtgtacAAAGAACTTGGACTTTTCATCAGCAGGCCTATTGTAGAGAtatagttacttgtcctaaac
gacttatcagtcttgtgtacGTAGAATTTACCAGTGCTTAACTGTTTTAAACATGCACTGtatagttacttgtcctaaac
gacttatcagtcttgtgtacCTTAGGCAATGTTTGTGGGTACTTCATCTTTGGGAAACACtatagttacttgtcctaaac
gacttatcagtcttgtgtacAATATTTACTGCATTTCTTCTAGGTATCAGGCATCATCTCtatagttacttgtcctaaac
gacttatcagtcttgtgtacCAAACCATATTGCTATTTTACAGTTGAAAAGATTGGGGCTtatagttacttgtcctaaac
gacttatcagtcttgtgtacACTTCCCACTGGTCACAAACTGATCTTTGATGTAAGGGTCtatagttacttgtcctaaac

gacttatcagtcttgtgtacATTTTTTTAGACGGAGTTTCGCTCTCTTGCCCAGGCTGGtatagttacttgtcctaaac
gacttatcagtcttgtgtacATTCTCCTGCCTCAGCCTGCAGAGTATCTGGGACTACAGGtatagttacttgtcctaaa
gacttatcagtcttgtgtacTTTTAGATTCAGAAGGTACGTGGTCAGATTTATTATGTGGtatagttacttgtcctaaac
gacttatcagtcttgtgtacCAATAGGTAGGTTTTTAGCTTTTGCCCCCTCCTCTCTCTTtatagttacttgtcctaaac
gacttatcagtcttgtgtacCATCATTATGTTCATAAGTACTCAGTGCTTAGCTCTCACTtatagttacttgtcctaaac
gacttatcagtcttgtgtacTTCTATGGTGTCTATGTAGTACATTCTCTTTATCCAATCCtatagttacttgtcctaaac
gacttatcagtcttgtgtacTTTGGTAGACTTATTTATTTTCCTTTCGCTATATACCCCGtatagttacttgtcctaaac
gacttatcagtcttgtgtacAGTTGTTTAAGAAATCTGCCAACTTATTTTCACAGTGGCTtatagttacttgtcctaaac
gacttatcagtcttgtgtacAACAGTATATAAACGTTGTCTTTTCTCTGCAGCCACTTCAtatagttacttgtcctaaac
gacttatcagtcttgtgtacACCTCCGCCTCTGGTTTCAAGCAGTTCCCTGCCTCAGCCTtatagttacttgtcctaaa
gacttatcagtcttgtgtacCCTGAGCTCAGGCAGTCCGCCCCTCTCTGCCTCCCAAAGTtatagttacttgtcctaa
gacttatcagtcttgtgtacCCACTTTTAATGGGCTTATCTGCTTTTTGCTTGTTGAATTtatagttacttgtcctaaac
gacttatcagtcttgtgtacCCTTTGTTGGATACAGAGTTTGTGAATATTTCCTCTCATTtatagttacttgtcctaaac
gacttatcagtcttgtgtacGTAGGTTTTCTTCTAGGACTGATAGTTTGAAGTCTCACATtatagttacttgtcctaaac
gacttatcagtcttgtgtacGTTCTTCTGGAAACTCTATGATAGAACTAAAAAGTATCCCtatagttacttgtcctaaac
gacttatcagtcttgtgtacTCTGCTTCTGTGTTTTGACTGTTTTGAGTTTGATTTGTACtatagttacttgtcctaaac
gacttatcagtcttgtgtacCTGGATCTAAACACTTGTACTCTTACTTGTATCTCATTGGtatagttacttgtcctaaac
gacttatcagtcttgtgtacAAGGTGAAGCTTAGTAACATGATTAAAGTTAGGGTGGAAAtatagttacttgtcctaaac
gacttatcagtcttgtgtacAACTTGTTTTTGCCACCTATTTCTTATTGCGCCATCATATtatagttacttgtcctaaac
gacttatcagtcttgtgtacCCAATAGATAAAGTTTTCCATTTCTTCCTAAGCTCTTCTCtatagttacttgtcctaaac
gacttatcagtcttgtgtacGATTTGTTAGTCCTCTCATGAAGGTAATAAAGCTATCTCAtatagttacttgtcctaaac
gacttatcagtcttgtgtacTTTGTAACAAATTGGACCAAAGTAGGCATCGGCATAGATTtatagttacttgtcctaaac
gacttatcagtcttgtgtacTGGTTCTCAGTTAAGTCAGCATGTCCCCGCTCCTCTTGTGtatagttacttgtcctaaac
gacttatcagtcttgtgtacTTTGTATTGCAGGGGTTCAAAGTCTACATTTGTTTTCAGAtatagttacttgtcctaaac
gacttatcagtcttgtgtacTGCTGATAGAATTCAAGGTTAGATTTTGCCAAAAAGAAGCtatagttacttgtcctaaac
gacttatcagtcttgtgtacGGCTGGCAGATAGGCTTCATTTTCTATGAGTTGCTTACCCtatagttacttgtcctaaac
gacttatcagtcttgtgtacAGCTGTTCTCTTGCTAGGAGTTTCTCACAGTTCTGCAGTTtatagttacttgtcctaaac
gacttatcagtcttgtgtacCAGAAGCAGCTTTCCCAGATACTTACTCTTTCACCCTTCTtatagttacttgtcctaaac
gacttatcagtcttgtgtacTGCTGACAAAACTCTCACTGTGACACTGATATTAACCAAGtatagttacttgtcctaaac
gacttatcagtcttgtgtacATGACCTTTGCCAAGACATCTCAACTTTCTGTCCTATCATtatagttacttgtcctaaac
gacttatcagtcttgtgtacAACCTTTTTATAGAATTTTGATTCTGGCCAGAATGTGCTCtatagttacttgtcctaaac
gacttatcagtcttgtgtacTTGGAAGTAAGAGACAAGAGATAGTTCTGTACCAGTGGATtatagttacttgtcctaaac
gacttatcagtcttgtgtacTTATGTGTAAGTGTTGTTTGGCCTACTCAGTCTTAGTATCtatagttacttgtcctaaac
gacttatcagtcttgtgtacCAGTGTGGAGTTGTGAACTGTCTTTAAAATTCAGAAGATCtatagttacttgtcctaaac
gacttatcagtcttgtgtacAAAAAGGCAGAAGATCTGATAACAATGGCCCCATATTCCTtatagttacttgtcctaaac
gacttatcagtcttgtgtacGACACAACTACTGCATTCTCATCTTGTCCTATTTGGCCTAtatagttacttgtcctaaac
gacttatcagtcttgtgtacCCTTACAGGAAAGAGTTATTGTATTGAACCCATTCTCCTAtatagttacttgtcctaaac
gacttatcagtcttgtgtacCTGTTTTGTGCCAAATGTGTCTACAGATGTCTACATACTTtatagttacttgtcctaaac
gacttatcagtcttgtgtacCAATTCCTGACATACTCCACACATTTATTTCATTTGGCTGtatagttacttgtcctaaac
gacttatcagtcttgtgtacGTTTATGAGAACTGGTCTAGACTCTAGTGCTGGCTTTTTGtatagttacttgtcctaaac
gacttatcagtcttgtgtacATGAAAACTACTGGACACTACTCTAGAAATGTAAGAACACtatagttacttgtcctaaac
gacttatcagtcttgtgtacTGCTTGCAACTAAGTTTAATAAGATCCTTTACTCCTGAGCtatagttacttgtcctaaac
gacttatcagtcttgtgtacCAGATTCAGTAGGGCTGGGAGGTTTCTGAGATTGTGTATTtatagttacttgtcctaaac
gacttatcagtcttgtgtacATGTTAACATTGTTGGTTCGGGGGTCACACTTTGAGTAGCtatagttacttgtcctaaac
gacttatcagtcttgtgtacTGGATATTAGATCTCTAGAGACAGTTTTTCTTTGAGATGGtatagttacttgtcctaaac
gacttatcagtcttgtgtacTGCTCTTGTCGCCAAGTCTGGAGTGCAATGGTGCACTCTTtatagttacttgtcctaaac
gacttatcagtcttgtgtacGCCACCACGCCTGGCACAGTTTTTTATAAAGGTTAATGCCtatagttacttgtcctaaac
gacttatcagtcttgtgtacAATGGTACTTTTAAATAGCTCCCATCTGAGTGGTTGAAAAtatagttacttgtcctaaac
gacttatcagtcttgtgtacATAAGAAGAGACAGGAAGAAAAGCCAGTAAGAGGGGTCTAtatagttacttgtcctaaa
gacttatcagtcttgtgtacAGGTGGATGTTGTAATGGAGAGAAGTGGTGCCAACTAAGAtatagttacttgtcctaaa
gacttatcagtcttgtgtacTTGTTTTTCCAAGACTGACCAGTTTCTTTTTCCTATAGTGtatagttacttgtcctaaac
gacttatcagtcttgtgtacAGTGATCTGCTGTCATTGAAATCATAGAGATGGCTTAGCAtatagttacttgtcctaaac

gacttatcagtcttgtgtacGTCAGTTCCACAAGTAGACCATGAGTTGAAGTAAGAAAGAtatagttacttgtcctaaa
gacttatcagtcttgtgtacTTCTAGTCATTGTTTATTAGTTCCAAGGCAGTATTTTCCCtatagttacttgtcctaaac
gacttatcagtcttgtgtacTGTCCTGTCCCAAGCTACTAAAAAGGCACAAAAGTTTCTGtatagttacttgtcctaaa
gacttatcagtcttgtgtacAAAATTAGTTTAAAACAGGGTTCATTGACAGTTGTAGGGCtatagttacttgtcctaaac
gacttatcagtcttgtgtacCCTTCTTTCACAGGTAGTCCATAACAATTTAGGAGAAAACtatagttacttgtcctaaac
gacttatcagtcttgtgtacGTCCATGACACTGCTTGATCCAGGCTTCAAAGACAGAATTtatagttacttgtcctaaa
gacttatcagtcttgtgtacAACTTAAATTGGTGTCAGATCAGTGGAAAGAAGTTCTCACtatagttacttgtcctaaac
gacttatcagtcttgtgtacATCACAGTCTGGGTCATTAATGTTCCTTTCCCCTCTCCTTtatagttacttgtcctaaac
gacttatcagtcttgtgtacCCTCTAATTAGTGCAGCTGGAACCAAGGAGGCTTCTGATGtatagttacttgtcctaaa
gacttatcagtcttgtgtacTTTCCAAAGACGCATTTCCATTTTAAAGAGAACCTAAAGCtatagttacttgtcctaaac
gacttatcagtcttgtgtacCTGAATCAGCTCCCCCTTCAAAAATTCTATTCCTGCAGGGtatagttacttgtcctaaa
gacttatcagtcttgtgtacTAATAGTCCTAAAACTCTAAGGTTCAAAGAAACCCTTCAGtatagttacttgtcctaaac
gacttatcagtcttgtgtacGCTCCTGATGCCTATATATATACTTACTAAGTGATGTCCTtatagttacttgtcctaaac
gacttatcagtcttgtgtacCAATTAAACTTCTCTATCATGCACCATCAAGTCATTAGTCtatagttacttgtcctaaac
gacttatcagtcttgtgtacCTTTGACTTCTGAACATGTGGTAGAGTTATGTAAAAATGGtatagttacttgtcctaaac
gacttatcagtcttgtgtacCATCTCAAAATACTTATGGAATATTGCTCTTCTCTGTCTGtatagttacttgtcctaaac
gacttatcagtcttgtgtacAGAGAAGAAAGCAGACGATGGAATCTCTCCGAGTGATGACtatagttacttgtcctaaa
gacttatcagtcttgtgtacCAAGTGGTGAAGCATCCAGGAGCCAGCAGCTGCCAGTGGTtatagttacttgtcctaa
gacttatcagtcttgtgtacCCATGAGGTCAGCATCTCGAACACATATCCTTCTCTTCACtatagttacttgtcctaaa
gacttatcagtcttgtgtacTCTCTTCTACTCTGTTTCTGGTGATTTCCTCTTTGATCCtatagttacttgtcctaaac
gacttatcagtcttgtgtacCCACTAAGGTTGGTTTTACATTGTTTTGGCTTTCAAGGAGtatagttacttgtcctaaac
gacttatcagtcttgtgtacGCATAAATTGTTCTTGTAATTAGCGTGGTTAATGAACTCCtatagttacttgtcctaaac
gacttatcagtcttgtgtacGGTACTGTATAAAGATGAGAGGAAGGCTGGTGATTATGGAtatagttacttgtcctaaa
gacttatcagtcttgtgtacTCTCTAAGACCAAATCCCAAACGGTTTAATCATCATCAATtatagttacttgtcctaaac
gacttatcagtcttgtgtacGTTGCATTTACATTTCATCTTGACTTTGATATTTGCTCCCtatagttacttgtcctaaac
gacttatcagtcttgtgtacTGTTTTAAAAATATGCATGTGCCATTACCAATTTCCCACGtatagttacttgtcctaaac
gacttatcagtcttgtgtacTTCTCTCTTTTACTGTGTGTGTTTACCAAAGGCTTTTTGTtatagttacttgtcctaaac
gacttatcagtcttgtgtacGTTAGATGTAGTTATGGCTAGAAAAGATTGAGTCTGTGGCtatagttacttgtcctaaac
gacttatcagtcttgtgtacTGACAATTACCAAGAGCTCCATAGTACGATTTTTTTAGTCtatagttacttgtcctaaac
gacttatcagtcttgtgtacGAAGAAGGAGGTCTCTTCATGTAGAATGTAAACTTATTGAtatagttacttgtcctaaac
gacttatcagtcttgtgtacCAACCTATAATTATTGTCATCCTCTTCCTACACTCCTTTCtatagttacttgtcctaaac
gacttatcagtcttgtgtacATAATATTGGGCAATCATAGATGTCAAGAATCTTCTGGACtatagttacttgtcctaaac
gacttatcagtcttgtgtacAGTGGTTAGATGTCTGCACTCAGAGCTATTCCTGCCAAACtatagttacttgtcctaaa
gacttatcagtcttgtgtacGGTGCCACATCTCTCTACCACTTCTCTCAGGATTTGCTGCtatagttacttgtcctaaa
gacttatcagtcttgtgtacAATAGATAATTCCATTTTGCGGCTCTGTACCCTAATTATGtatagttacttgtcctaaac
gacttatcagtcttgtgtacTTTATGACTATAGACGTTTACTTAAAGAAGAGTGGGCATGtatagttacttgtcctaaac
gacttatcagtcttgtgtacACCAGAAAGATAGTGCAGAAGAAAAGTGTCAGTAATGAGAtatagttacttgtcctaaa
gacttatcagtcttgtgtacTTGTTATTTATGGAGCATATACCTATAGAATGTCGACTGGtatagttacttgtcctaaac
gacttatcagtcttgtgtacCAAAGTTGGGGAATTGGAGTCTATTGTCAAGGAACACACAtatagttacttgtcctaaa
gacttatcagtcttgtgtacGCGTTCTCTCCATTTAGGGGTGCCTTTGAATGCTGCTGAGtatagttacttgtcctaaa
gacttatcagtcttgtgtacGGTTCAAATATATACATTTCCCAGCTGTGTGACTTGTAGGtatagttacttgtcctaaac
gacttatcagtcttgtgtacGAAGTGGTAAGTCTAAAAATTATTTGAAAACTGGAGAGGGtatagttacttgtcctaaac
gacttatcagtcttgtgtacGTAGGACACCATAAACTTAAATTCACAAGGTCTGAAAATGtatagttacttgtcctaaac
gacttatcagtcttgtgtacACCATTTCAGAACTTTATTACATGTTCTCCAATATCTCCGtatagttacttgtcctaaac
gacttatcagtcttgtgtacAGTTGGTGGGTTCTACTAATTGACTTTGTAAGCTGAATGAtatagttacttgtcctaaac
gacttatcagtcttgtgtacGATTTTTCTATTTTTCACGGTTGTTCAATGGAAGGCAGTTtatagttacttgtcctaaac
gacttatcagtcttgtgtacCACTGTCTTCTTCAACAAAGTCACAAGAGTAAATATTTGGtatagttacttgtcctaaac
gacttatcagtcttgtgtacTGTTATGCTGACCTAATTCAAACACTACTCCTGGAAAAAGtatagttacttgtcctaaac
gacttatcagtcttgtgtacCTCTTGGTGTGATTCTTGTGAGACTGTACTAAAATTATCTtatagttacttgtcctaaac
gacttatcagtcttgtgtacCTTCTTGTTTTTACTTGGTATAGGTTGTGGGAAGAAAGCtatagttacttgtcctaaac
gacttatcagtcttgtgtacAATTCTTTGGACTGACATGGTTATTTTTATTTGCAGCAGGtatagttacttgtcctaaac
gacttatcagtcttgtgtacGTTGTGCCAGAGCAGGTGCGCCTTAAATGACAGGTTTAGAtatagttacttgtcctaaa

gacttatcagtcttgtgtacCAAGCTGGTTGTCAAGCGACACAGTGGCTGCCGTCCGTTAtatagttacttgtcctaaa
gacttatcagtcttgtgtacGGTGTCTCCTTAAAATGGTCAGATGATTGGACTCTTTGTAtatagttacttgtcctaaac
gacttatcagtcttgtgtacCCCATTACTGACTATCCAAGGGTGGCAATGCTTATTTGGTtatagttacttgtcctaaac
gacttatcagtcttgtgtacGGACAGTGAGTTCCTCCTTTGTAAAGACCTGAGAAGCTCCtatagttacttgtcctaaa
gacttatcagtcttgtgtacCTTCGTTTCATCCATTAATTTGTCAAGACAATGTTGAGAGtatagttacttgtcctaaac
gacttatcagtcttgtgtacCCTGTCAGAAAAACCCTTTCAGGGCCTTGAGGAAAAAAATtatagttacttgtcctaaa
gacttatcagtcttgtgtacATGAGTCTGAACATGAATTCTTATTCCAGATAGCTGCAAGtatagttacttgtcctaaac
gacttatcagtcttgtgtacCAATTAGTAATTCCTATATCCCACGATTATTTGTGCAGTGtatagttacttgtcctaaac
gacttatcagtcttgtgtacAAATTCAACCCTGTGGATTTCAGTCTGCTCATCTATAAAAtatagttacttgtcctaaac
gacttatcagtcttgtgtacGTCTAACCATCATATCTGAATTTAGTACTTAAAGCTTGGCtatagttacttgtcctaaac
gacttatcagtcttgtgtacTATTTGGGAGTAAACTGTAATAATATGCATCTATGCTGGGtatagttacttgtcctaaac
gacttatcagtcttgtgtacAACCCCAAACAGCCCTTTTGTCTTTTAGATTTTGAATGGCtatagttacttgtcctaaac
gacttatcagtcttgtgtacGTCCCAAACAATGCATGGACAGTCTTTGTGTCAGTAATAGtatagttacttgtcctaaac
gacttatcagtcttgtgtacTGCATGTACTGTGTGTATGTAGTAGCAAATATTAAGGCAAtatagttacttgtcctaaac
gacttatcagtcttgtgtacCTGATGGCCTTTGATAAAGTCAAATAAGAACCAACTGAGCtatagttacttgtcctaaac
gacttatcagtcttgtgtacGCACAATTCTCTGCCATCACCATTGGTTTTCACTAGCACCtatagttacttgtcctaaac
gacttatcagtcttgtgtacAACTTGATATCTGCATTTCTATAACTACTGGTACCTCTTCtatagttacttgtcctaaac
gacttatcagtcttgtgtacCGGTCAAACTCGATACAAGTCTCTGTGCTTAAACTTCTCAtatagttacttgtcctaaac
gacttatcagtcttgtgtacCAGCTTGTGATGTGATGTGGGGTCTGAAGGGTGAGAGATGtatagttacttgtcctaaa
gacttatcagtcttgtgtacCAGGTCATGTATTGGTAGGAAATCCTTCTTGCCGTGGAATtatagttacttgtcctaaac
gacttatcagtcttgtgtacCAGAGTTGAAGCGTAGATGACAAAATCAAAACAGAACCCTtatagttacttgtcctaaa
gacttatcagtcttgtgtacCTTTTATCACTTTGATCATGCTACTTTGTGGAGTTGTTTGtatagttacttgtcctaaac
gacttatcagtcttgtgtacCCCACTGACTTCCATCTAATAGGGTCATATTTATGTTGCTtatagttacttgtcctaaac
gacttatcagtcttgtgtacTGGCTTCATTTCCCCAGGAAGTAACTCTTGGCTAGCTCTGtatagttacttgtcctaaac
gacttatcagtcttgtgtacGCTTGCAGGCTTTATCAAATCTGTGAAACCAAGTGTGATTtatagttacttgtcctaaac
gacttatcagtcttgtgtacAGAGAGTTGGGGCAAAGAAACATACAGCAGTTGGATTCTAtatagttacttgtcctaaa
gacttatcagtcttgtgtacCTGAAGAGGTAGAGGGAATTTGAGTCAGAAAGCCAACAAAtatagttacttgtcctaaa
gacttatcagtcttgtgtacTTCTAGACTTATTCATAAATTTGACAGGGTGCTATCAGAGtatagttacttgtcctaaac
gacttatcagtcttgtgtacTGAAATGATAAAAAGTAAACACTCGACATGAGACCCAGAGtatagttacttgtcctaaa
gacttatcagtcttgtgtacCAGAGAGCAACAGTGACAACAGAATATGATGCCACATTCTtatagttacttgtcctaaa
gacttatcagtcttgtgtacTGCAAATAGGGTGGGAAATATTCAGTCAAAAATGCACTTTtatagttacttgtcctaaac
gacttatcagtcttgtgtacAAAATTAAAACAAAAAACAAGGGAGGGTAGAGTGGGCAGTtatagttacttgtcctaaa
gacttatcagtcttgtgtacCCATAGGAAAATGTCAGGAGAACAAAATTGGCATAAGTTCtatagttacttgtcctaaac
gacttatcagtcttgtgtacAGGAGAGATTTCTTTTCCTTTTGGTTGTTCACAGGGCATTtatagttacttgtcctaaac
gacttatcagtcttgtgtacGTGGAACACAAGCAGAACTTTCCATGATTATTCTATTTGAtatagttacttgtcctaaac
gacttatcagtcttgtgtacTTGTAGGTTGCCTGTTCACTCTGATGGTGCTGGGAAAACTtatagttacttgtcctaaac
gacttatcagtcttgtgtacGTGCCACAAAAATGTATTATTTTTCACTCTCCAGTAGGCCtatagttacttgtcctaaac
gacttatcagtcttgtgtacATATGTGTATATATCTATTCAGTTCTCCATGTCCACGGGGtatagttacttgtcctaaac
gacttatcagtcttgtgtacTAAAATGACATAGTATGGTTGGCCCTCGGTATCTGGGGGTtatagttacttgtcctaaac
gacttatcagtcttgtgtacTGTAGATGCAGAAACCCTCAGATAGATGGGTAATTGTACAtatagttacttgtcctaaac
gacttatcagtcttgtgtacATGGATATATACCTTTTGATTGAGTTTCTGACCAGCTTGCtatagttacttgtcctaaac
gacttatcagtcttgtgtacCTAACCTGGGCAGCATTATCAAGAGTAAGTGTGTGTGTGTtatagttacttgtcctaaac
gacttatcagtcttgtgtacCAAAAATAATTGGAGGAGTTCCATCGTAATCTTTGAGGATtatagttacttgtcctaaac
gacttatcagtcttgtgtacAGGAATAAGTCATGTATTTCAAAGTAGTCTCACACAAAGCtatagttacttgtcctaaac
gacttatcagtcttgtgtacTACTGTGGAGGAAAACTTTATCTTTTTTAGTAGGGGATAGtatagttacttgtcctaaac
gacttatcagtcttgtgtacTAAGGAGCCAGGGCAAAGGGAAAAATAGCCTTCCAGCAGTtatagttacttgtcctaaa
gacttatcagtcttgtgtacTGGAACATTTGGAAAAGGAGAGGTGAGCTCTTTGATTGTGtatagttacttgtcctaaac
gacttatcagtcttgtgtacCCCTCTCTGGGCTGACTGAGACAGGAGCCGGCTGCCTCTGtatagttacttgtcctaa
gacttatcagtcttgtgtacGCACTGCCGGTGTGGGGCTGCGCTTGAATTCTTGCTGGGCtatagttacttgtcctaa
gacttatcagtcttgtgtacCAGCCTGCCATGCCTGAGTCCCCAAACCCGCCTCGCCCCTtatagttacttgtcctaa
gacttatcagtcttgtgtacCCCCCATCCCCCTCCCCCCTTATCCCCCTCCCCCATCCCCtatagttacttgtccta
gacttatcagtcttgtgtacCCTCCCCCTCCCCATCCCTCCCCCTCCCTTGCCCCTCCCCtatagttacttgtccta

gacttatcagtcttgtgtacGTGCCACCCCCTGCTCCGCTGCGACCAGTCCCATCGACCAtatagttacttgtccta
gacttatcagtcttgtgtacTGTGGGACTGGTAGGCAGCTTTGCCCATGGCCCCTGTGCAtatagttacttgtcctaa
gacttatcagtcttgtgtacCTGGGCTCCTGAGTCAGGTGGGGACTTGCAGAACTTTTATtatagttacttgtcctaaa
gacttatcagtcttgtgtacGCTCGGGGATTGTAAACTCACCAAGCAGCACCCTGTCAAAtatagttacttgtcctaa
gacttatcagtcttgtgtacATAAAAACAGGCTGCCAGAGGCTGCAGTGGTAACCCGCTTtatagttacttgtcctaa
gacttatcagtcttgtgtacCTCTGGAAGCTTGGTTCTTTCACCATTCGCAATAAATCTTtatagttacttgtcctaaac
gacttatcagtcttgtgtacCACTCTTTGGGTCCGCGCTGCCTTTGTGAGCTGTAACGCTtatagttacttgtcctaaa
gacttatcagtcttgtgtacACAATTGTATCCCTGAAGCTCATGTGTTCTCAATGCCCAAtatagttacttgtcctaaa
gacttatcagtcttgtgtacTTTTGCAGGGCATCTGGGGACGAGAGAGAGAGAGTTGGGGtatagttacttgtcctaa
gacttatcagtcttgtgtacCATTTGCAAAACTGACTTTTATCCCATGGCACCCTTTGAGtatagttacttgtcctaaa
gacttatcagtcttgtgtacTCTGACATTTTAATTTTCACTTCTTCCAAGGAATCCTACCtatagttacttgtcctaaac
gacttatcagtcttgtgtacGCATTTAAATAAGCAGATCTGACAAAACAATGAAAGCTGCtatagttacttgtcctaaa
gacttatcagtcttgtgtacAAATTGCCACTTTGTGAACTTTGAAATTTGCACGCACGGTtatagttacttgtcctaaac
gacttatcagtcttgtgtacTGCTAATTAACTCTATGCACATGACCAGAAATAGGCTCACtatagttacttgtcctaaa
gacttatcagtcttgtgtacTGTCTTTACAGTATGCCAAACAAAATGCACCTTCAGATGAtatagttacttgtcctaaac
gacttatcagtcttgtgtacATGTTGTGTATCATAATATTTTTAGCTTCCTGGTCTCACCtatagttacttgtcctaaac
gacttatcagtcttgtgtacGGCGCAATTGCCTGAGAACATTTTCTATTGCTGGTTGTCAtatagttacttgtcctaaac
gacttatcagtcttgtgtacAGAATACTTGCTTTAAGCCATACAGTATACTTTGGGGAAAtatagttacttgtcctaaac
gacttatcagtcttgtgtacCCTGTCAGGTATATATGAAATTATGGAATCTTGGAATAGGtatagttacttgtcctaaac
gacttatcagtcttgtgtacTCTACTTAACCCCACATATACAGAGACTCAATCAAGGCAGtatagttacttgtcctaaa
gacttatcagtcttgtgtacATTTTGTTGACATGTAGACCAAGGGCGAGCAGTTCAGGATtatagttacttgtcctaaa
gacttatcagtcttgtgtacCTGCTTAAATAGTCATTCAGAGTCCCAAATTCCTTCAATCtatagttacttgtcctaaac
gacttatcagtcttgtgtacGCTCCATCCTTCTTCAGAATGTTGTCCGTGTCTATTTAGTtatagttacttgtcctaaac
gacttatcagtcttgtgtacTCATTGACAAGTCCAGATGCCATCCAGTAGAAACAAAAAAtatagttacttgtcctaaa
gacttatcagtcttgtgtacCAGGCAGCTTTATTTTTAAGATGATGATGCAGAAATTGGCtatagttacttgtcctaaac
gacttatcagtcttgtgtacCTGTCACGTGGATAGAGAGGTCTGTGGATAATGAAAGCTGtatagttacttgtcctaaa
gacttatcagtcttgtgtacTGGCTGGCAAATCCACAGAAATGGGTAAATCACATTGGATtatagttacttgtcctaaa
gacttatcagtcttgtgtacGAAACTCCTTGGGCTTTTGCGTGCCTTCATTTCTATTGATtatagttacttgtcctaaac
gacttatcagtcttgtgtacGAAACCTAAAATAGCTTTCTCTTGATCTATGAGTTGTGTGtatagttacttgtcctaaac
gacttatcagtcttgtgtacATGCTATTCATTGTACACCTCCTATGAGCCTGTGGTTGTTtatagttacttgtcctaaac
gacttatcagtcttgtgtacCAAAGCTGAATAATTTGCATAGGTCTCAAAAGTCATTGCAtatagttacttgtcctaaac
gacttatcagtcttgtgtacAGATATTCTGACACCTTACAAAATGTGGGTATGAAAGACTtatagttacttgtcctaaac
gacttatcagtcttgtgtacCTCTGATCCTTTTTGTATTGCCTCTGCCTTTGTTTTGTTTtatagttacttgtcctaaac
gacttatcagtcttgtgtacCAACCTCTGCCTCCTCAGCTCACGCCATTCTCCTGCCTCAtatagttacttgtcctaa
gacttatcagtcttgtgtacACTGTGCCTGGCCTGTATTGCCTCTGGATGTGGCATTCATtatagttacttgtcctaaa
gacttatcagtcttgtgtacCAATAGATAGTTGCAGGTCCCCCAATCCCCTAGTTAGTTCtatagttacttgtcctaaa
gacttatcagtcttgtgtacCCTTGTAAGCATCTGAAAACAATGGTGGAGAAGATAGTTTtatagttacttgtcctaaac
gacttatcagtcttgtgtacTCATGAAGCTTATGTTCTACATCAGTCATTCCCAATCACTtatagttacttgtcctaaac
gacttatcagtcttgtgtacGCACAGTCCAGATATGCTTGCAAAGTTTTTCATTCAGTAGtatagttacttgtcctaaac
gacttatcagtcttgtgtacCACTACAACTATAAACATGGTGCCCCTTCCTTTTTTTTTTtatagttacttgtcctaaac
gacttatcagtcttgtgtacCAGAGGCTTTGAACTACAGGAAATATTCCGGGGTCCAAACtatagttacttgtcctaaa
gacttatcagtcttgtgtacAGGGTTTCGCCGTGTTAGCCAGGATGTTCTTGATCTCCAGtatagttacttgtcctaaa
gacttatcagtcttgtgtacGAGTGGATCTTAACGTCTCTTCTCTCAGAAACTTCCCATGtatagttacttgtcctaaac
gacttatcagtcttgtgtacTCATGAGCAGTTGATTTGGAGAATTACATTTTTGCATGCAtatagttacttgtcctaaac
gacttatcagtcttgtgtacCATGTATAATTGGTGATGGGTCTGCTAGGTTTCATTGATAtatagttacttgtcctaaac
gacttatcagtcttgtgtacCACAATGTGCCTCAGTCTGACACCATCTTTTAGGTTTAACtatagttacttgtcctaaac
gacttatcagtcttgtgtacCAAAAAAAACAGCATCAGACATCCATCAATCATTTGAAGCtatagttacttgtcctaaa
gacttatcagtcttgtgtacGAACTGATCAATTTAAGGACATTCACTTATTTCTCTGTGCtatagttacttgtcctaaac
gacttatcagtcttgtgtacACATTACCTGTCAAATATGAACTTTGTATCCAAAGGTTCCtatagttacttgtcctaaac
gacttatcagtcttgtgtacTTGTTAACTATTTTTCTTTCTTCTCACTTTCTTTCCCCCCtatagttacttgtcctaaac
gacttatcagtcttgtgtacAGATTCTTCTTTCTGATGCCTCTTCTAGAGACCATTGAAAtatagttacttgtcctaaac
gacttatcagtcttgtgtacCTGCTTGGCTTTAAACTGTACAAATTTGAATGTCATTCTGtatagttacttgtcctaaac

gacttatcagtcttgtgtacCCTGAGGACTGATGACATTTGATCTGAGTAACAGCTATGGtatagttacttgtcctaaa
gacttatcagtcttgtgtacCTCAGAAAATCCATTAAAGACAGAGTTAGAGCCGAAATAAtatagttacttgtcctaaa
gacttatcagtcttgtgtacTGACATAGCCATTTGAGGTGAGAAAGGTGAGCTTCTTCAGtatagttacttgtcctaaa
gacttatcagtcttgtgtacAGTGCATAATGAAGACAATTTTGGAGCCATGCCTCTTTGGtatagttacttgtcctaaac
gacttatcagtcttgtgtacAGCCCTAGGATGAAGTGCAATTGTGACCTGATTAAACGCTtatagttacttgtcctaaa
gacttatcagtcttgtgtacTGCTCCCACTGTCCCGCCTCTATCCCTCATCGAAACACCTtatagttacttgtcctaa
gacttatcagtcttgtgtacCTTGCTAATATAAGGACTGAGTTTTATGATGCCGCTCTGAtatagttacttgtcctaaac
gacttatcagtcttgtgtacCAACATCAACTGCTTTTTTTCCCTCTCCAGAGCACACACGtatagttacttgtcctaaa
gacttatcagtcttgtgtacCATATAATGGAGAGGGATTACATCTACTTCTAGCATCACTtatagttacttgtcctaaac
gacttatcagtcttgtgtacTTACCTGTTAACTTCTATTAGAACTCATTAAGCAGTGCCTtatagttacttgtcctaaac
gacttatcagtcttgtgtacGATGAACAAGAAATCCAACCATCCTCAACAACCTACATGCtatagttacttgtcctaaa
gacttatcagtcttgtgtacGTATTTTGTTTCTTATTCTAGACAACCAGTATCATCCGTCtatagttacttgtcctaaac
gacttatcagtcttgtgtacGGGTTACTGTTGGTAGTCTTGTCACTCAGACTCAGCCACAtatagttacttgtcctaaa
gacttatcagtcttgtgtacAACTGTTCTGTTACCTCATTTGCCTGCTAAAAACCCATAGtatagttacttgtcctaaac
gacttatcagtcttgtgtacGCTCATTAACAGCTCCCCCCAACAATCGATTAAAAAGTTTtatagttacttgtcctaaa
gacttatcagtcttgtgtacGAACTCCTTTCTCATCTTTGGAATCTAATTGAAGTCACTTtatagttacttgtcctaaac
gacttatcagtcttgtgtacCATGGAAGAAGCATGCAGTGGGAACAGAAGGATGTGGCATtatagttacttgtcctaa
gacttatcagtcttgtgtacGAAGAACCAGCTGAAGGTGTGGTGTGCACAGTCAAAGGCAtatagttacttgtcctaa
gacttatcagtcttgtgtacATCAGGATGTACAAGGGTGGAGGTTTCTAATGTGGGCAGAtatagttacttgtcctaaa
gacttatcagtcttgtgtacATGGTGAATACAAGAGCACATGACCCTCAAAAGTACAAGGtatagttacttgtcctaaa
gacttatcagtcttgtgtacGCTAGATTTCAGGTGGAAGATGGCCAGATTGCAGCAGGATtatagttacttgtcctaaa
gacttatcagtcttgtgtacGCAGTAAACTAGGCAGAAGACTTCAAAAGGAGTAATAAAAtatagttacttgtcctaaa
gacttatcagtcttgtgtacAAACACTGCCCTCTAATCCATGTTTGCCATAACATCTGAAtatagttacttgtcctaaac
gacttatcagtcttgtgtacACTATGCTGTGTAAAGTTTTCAGTGACTTCCTAAGCCAGTtatagttacttgtcctaaac
gacttatcagtcttgtgtacTTGTTAAAGTGCAAATTCCGACTTGGTGGTCTGGGGTGGCtatagttacttgtcctaaa
gacttatcagtcttgtgtacTGGCCTCATCTCACCCCATTCTCACCCATCTCTCTCCAAAtatagttacttgtcctaaa
gacttatcagtcttgtgtacCCTCATAGGCTTTGTAATTTCTGTTTGCTGTGCATGGAATtatagttacttgtcctaaac
gacttatcagtcttgtgtacGGCTCTTCCTTCTCTGGTCTCAGCTTAAATGGCATCTTCTtatagttacttgtcctaaac
gacttatcagtcttgtgtacCTGCCCTATCCAAAACAGCCCCCTCACTCCCAGTTATTCTtatagttacttgtcctaaa
gacttatcagtcttgtgtacTGATTTTCAACTGTGATCATAGCTAGCTCTTAGTCTACAAtatagttacttgtcctaaac
gacttatcagtcttgtgtacGTATATATGTCCCCAATTAGATACCACCTTTGAAAAACACtatagttacttgtcctaaac
gacttatcagtcttgtgtacTTTATCTCATTTATTCCTAGCATCCAGTCTGTAAGTTAGGtatagttacttgtcctaaac
gacttatcagtcttgtgtacCTAGGTTACAGACAAGATAATAGACACCCTTTAAAAACACtatagttacttgtcctaaa
gacttatcagtcttgtgtacGAGCAAAATAGAGATCCAGAAAAATAGCTATTGCATGTGCtatagttacttgtcctaaa
gacttatcagtcttgtgtacTGGAGATATTCAAACTGCAATGACCACTACCATTTCCACAtatagttacttgtcctaaa
gacttatcagtcttgtgtacCTAAAGAAGGACAACAATTCCATTGCATGGCTGAGCTAGTtatagttacttgtcctaaa
gacttatcagtcttgtgtacGAAATTTAATAAGGAAAGCAACTAGAGTCATCAAGGCTACtatagttacttgtcctaaa
gacttatcagtcttgtgtacTGCTCCAATCACAGGCCCCTCACTTTAAGAATATCAGTCAtatagttacttgtcctaaa
gacttatcagtcttgtgtacGTATTATATACCTTGAATAGACATATGCTCCTCTGAACCTtatagttacttgtcctaaac
gacttatcagtcttgtgtacCAGGAATACTCTGGCCAGGTTTGTTTTCTCCTTTCTGTAGtatagttacttgtcctaaac
gacttatcagtcttgtgtacAGATTTAGCATCTGGGGTCCATACATAGCTTTTCCATGCAtatagttacttgtcctaaac
gacttatcagtcttgtgtacCCTTTGCTTTATCAGGGAGGATAATCCAGGGTTGAGAATTtatagttacttgtcctaaac
gacttatcagtcttgtgtacTGATCTTGAACAGAGGCATATGAGGTCACGTACATAATTCtatagttacttgtcctaaac
gacttatcagtcttgtgtacTTTCTTATTGTGGCGGAAAGGAGGAGTAGGAGAGTATGGAtatagttacttgtcctaaa
gacttatcagtcttgtgtacAGGGGCCTGCCGTGTGTGCTTATTATTTTTGATGGATTATtatagttacttgtcctaaac
gacttatcagtcttgtgtacCATTGTTGTAGCTTGCAAAGATGTTGAGGCTTGAAATATTtatagttacttgtcctaaac
gacttatcagtcttgtgtacGCCAGGCTCAATGAGAATTTCAGGTACTATCTTCAGAGTCtatagttacttgtcctaaa
gacttatcagtcttgtgtacGCCAAATTGCATTCCTCTCCCACACCGTTCTACACATACAtatagttacttgtcctaaa
gacttatcagtcttgtgtacGCTAAAGCTGTATTCATTAATCTGAACAGGGAAGAGCAGGtatagttacttgtcctaaa
gacttatcagtcttgtgtacTTCAGTAGTCATGCAGAACAGGTGAGAGAACAGGCGGCACtatagttacttgtcctaa
gacttatcagtcttgtgtacGCCGATCTCCATCAGCCCCATCACCCAGAGACAAAGAGAGtatagttacttgtcctaa
gacttatcagtcttgtgtacGGCATACCTCAAAGTCCACCATCCCAGGGCATCCCAGTGAtatagttacttgtcctaa

gacttatcagtcttgtgtacCTTCAGAGCTTTTAAATGTCATTTTAGAATGGAATGCCACtatagttacttgtcctaaac
gacttatcagtcttgtgtacTTGGTTAACATTTATCAAACGTGGATGGGGTGGGGTTGGTtatagttacttgtcctaaac
gacttatcagtcttgtgtacTGGAGTCATTCTTTGGGGAAATAAGAACAAAAGTTACAGCtatagttacttgtcctaaac
gacttatcagtcttgtgtacCGTGAGTAAAATTTGACAGTATTTCCATGTGTTCTCTTCGtatagttacttgtcctaaac
gacttatcagtcttgtgtacACCTTTTTGTGTTGAGATTTGGTGGGGTATGGTGAAATGAtatagttacttgtcctaaac
gacttatcagtcttgtgtacACTTATGATGATGTATTTTAGAGAAGTCACAACCTACCTGtatagttacttgtcctaaac
gacttatcagtcttgtgtacGGAGATGTAGGTAGGAAAGAGTGTGCTAGAATCACCCGAGtatagttacttgtcctaaa
gacttatcagtcttgtgtacGTATAGATGCTTACCTTAGAGATTCTGATCCAGTTTGTCTtatagttacttgtcctaaac
gacttatcagtcttgtgtacGGGGTATATATAGCTGTGAAATCTCCCCAGGTGACAGATAtatagttacttgtcctaaac
gacttatcagtcttgtgtacAAGAACAACATATGGTGTTGAAGGACAAAAATATAGGGTGtatagttacttgtcctaaac
gacttatcagtcttgtgtacTGACAATTCTTATCAAATTCAAAGTTTTAAGTAGGGGGCCtatagttacttgtcctaaac
gacttatcagtcttgtgtacGGCAGGGACCATGAATCCGGAAGTGTGTTATAGTGATAGCtatagttacttgtcctaaa
gacttatcagtcttgtgtacCAGAAAATTGCTGATTAAACTTATCAACTCTCCTTCCTCCtatagttacttgtcctaaac
gacttatcagtcttgtgtacTCTTTTATTTCTCTGTTCTGATTACAGTTAAGTGGTGGCAtatagttacttgtcctaaac
gacttatcagtcttgtgtacCCTTCCTGTATTTTCTTTCCCAACTCCATTCCTGCATTCAtatagttacttgtcctaaac
gacttatcagtcttgtgtacATAGTTGGCTTTGTAATTTTCACCTCAGGGGAAGGATGAGtatagttacttgtcctaaac
gacttatcagtcttgtgtacTTGTCTGTTTAAGGAGATTGTTCTTATATAGGCAATGCCCtatagttacttgtcctaaac
gacttatcagtcttgtgtacCAAGGTGGAATGAAAGGTAATAAAGGGGAATGAAAAGGGAtatagttacttgtcctaaa
gacttatcagtcttgtgtacCCAGGAGATATGTTGTGGGCTTTGTTTGCAGGAGGGGTGAtatagttacttgtcctaaa
gacttatcagtcttgtgtacCTAAGAAGGGCATATAAAACTAAGGAAAAGGGTGATTAAGtatagttacttgtcctaaac
gacttatcagtcttgtgtacTATAAAGAAAAATGGTTTAAGGACTAACAGTTCCACAGGCtatagttacttgtcctaaac
gacttatcagtcttgtgtacCTTACAATCGTGGCAGAAGGGTGAAGGAGAATCAAACATGtatagttacttgtcctaaa
gacttatcagtcttgtgtacTGAAGGGGAAGTGCTACACATGTTCTGTTTTTTTGTTTTGtatagttacttgtcctaaac
gacttatcagtcttgtgtacCTCACTGCAACCTCCACCCCCCAGGATCAAGTGATTCTCCtatagttacttgtcctaaa
gacttatcagtcttgtgtacGAGTAGCTGGGGTTACAGGTGCGCACCACAATGCCTGGCTtatagttacttgtcctaa
gacttatcagtcttgtgtacCAGCAAAGGGGAAATCTGCTCGCATGATTCAATCACCTCCtatagttacttgtcctaaa
gacttatcagtcttgtgtacAGAGCCAAAACATATCACTCCTACAATGGAGAAGCCCAGTtatagttacttgtcctaaa
gacttatcagtcttgtgtacGAAACCAACCTAATTGTCCCATACAACTGATATTTAATGGtatagttacttgtcctaaac
gacttatcagtcttgtgtacAGGAAATCAACCATCAGGCCATCCAGATAGTATAAAGAAAtatagttacttgtcctaaa
gacttatcagtcttgtgtacTGCCTAAGTGACCACCTGCTCTTCCTGTTTACCAACTCTTtatagttacttgtcctaaac
gacttatcagtcttgtgtacAATTCCTGTTTTCCTCCATGTCATTACATATCTTCCCTGCtatagttacttgtcctaaac
gacttatcagtcttgtgtacGATCTCCCATCTCCTTCGCTGCAGCACACACAATGAAAGCtatagttacttgtcctaaa
gacttatcagtcttgtgtacGGTGTTTTGGTAACAGTAAGATGGCTGTGAGGTCTATTTAtatagttacttgtcctaaac
gacttatcagtcttgtgtacTGTAAGCAACAGCAAGAACCACCAGACCCAAAGACGCCTTtatagttacttgtcctaa
gacttatcagtcttgtgtacCAGAGGATGGACAAGTCAGAGGATGCTGGTAAAAATGAGTtatagttacttgtcctaaa
gacttatcagtcttgtgtacCCAAGAAGAAAATGAAATGGAGTTGTGTGGTGGAACAGGGtatagttacttgtcctaaa
gacttatcagtcttgtgtacAATTGTTGTCCTTTAAAAACCTCTGGAGATATAAACCCTCtatagttacttgtcctaaac
gacttatcagtcttgtgtacCATGAAGTGTAGTTTGGCAAGTACATATAGTCATGTACGAtatagttacttgtcctaaac
gacttatcagtcttgtgtacCTATAGAGTATTAAAATGAATCCCCACTTAGAACACGTTCtatagttacttgtcctaaac
gacttatcagtcttgtgtacCAGAGAAACAGACTGGTAAGAGATAATTCCATTTGCAAGAtatagttacttgtcctaaac
gacttatcagtcttgtgtacATTCATCACTTCATTAATCTAATTTCAAAAGGGAAGGCCCtatagttacttgtcctaaac
gacttatcagtcttgtgtacTCAGGCATGAGTGAGCAGAGCATTCAACTGTTAGACTTTGtatagttacttgtcctaaac
gacttatcagtcttgtgtacTCTTTACCAGCTGGAGATAAGCACAATGAGGGTCACGCTAtatagttacttgtcctaaa
gacttatcagtcttgtgtacGGTGGCGCGTCTGGAGTTGTGCGTTCCTCCCGGTGGGCTCtatagttacttgtcctaa
gacttatcagtcttgtgtacCTCATAAAAGCAGCGTGGACGCAAAGAGTGAGCACTTGCAtatagttacttgtcctaaa
gacttatcagtcttgtgtacGCCCCACCCACATCTTGCTGACTGGTAGAGCCCAGTGGCCtatagttacttgtcctaa
gacttatcagtcttgtgtacACGTCCCCATCAGATTAGATACAGAGTTTTGACACACAGGtatagttacttgtcctaaa
gacttatcagtcttgtgtacGCCTGCCAGTCCCGCGCTGCGCGCTCGCATTCCTCAGCCCtatagttacttgtccta
gacttatcagtcttgtgtacGGAGCAGGGGGTGGTGCTGGCTGGGGAGGCTTCGGTCGCAtatagttacttgtccta
gacttatcagtcttgtgtacGCAGCTAAGGCTCGTTAGAAATCGAGAGCAGCGCCAGTGGtatagttacttgtcctaa
gacttatcagtcttgtgtacCGAGTGCGGGGCCCGCCAAGTCCATGCCCACCCGGAACTCtatagttacttgtccta
gacttatcagtcttgtgtacGCTGTCACCTCTCACTATCTAATCTTAAATATGGATAGATtatagttacttgtcctaaac

gacttatcagtcttgtgtacGGATAACATGTTCCACGGTATCTATTTCCATCTTGTGTTAtatagttacttgtcctaaac
gacttatcagtcttgtgtacATTGTATATATCAAACACTATTCTGCAACATGACAGAGGGtatagttacttgtcctaaac
gacttatcagtcttgtgtacACATCTGGTAAGTCAACAACATTATCACAAACATTATGGCtatagttacttgtcctaaac
gacttatcagtcttgtgtacCTGAGAAAGCCACACAGAACATTCCCACCATCTAGCAGATtatagttacttgtcctaaa
gacttatcagtcttgtgtacGGAAGCCAATATTTTCTGTGGAAACCAAAGTTTCTTCTTCtatagttacttgtcctaaac
gacttatcagtcttgtgtacTTTTTCCCCTTAAGATTGTATGAAGAGATTGCTAGACCCAtatagttacttgtcctaaac
gacttatcagtcttgtgtacGTTTGGTGGTGGTGAGAATGGCTGCTACTTTCCTATGTGGtatagttacttgtcctaaac
gacttatcagtcttgtgtacTGGAAATGAGTTTGCTCTTAGCTATGTCTAGATTTACAACtatagttacttgtcctaaac
gacttatcagtcttgtgtacCAAGGCTCAATTACCACACATTTATTTAGAAAGAGTAGGtatagttacttgtcctaaac
gacttatcagtcttgtgtacAACGTTGTTTGAGTAGCATTGAGTTAACACCTCTAGTTTCtatagttacttgtcctaaac
gacttatcagtcttgtgtacGTGCCAGGAAAATGGTCAATTCACCTTAAAATATTTTACCtatagttacttgtcctaaac
gacttatcagtcttgtgtacTTTGTTATTTAAGAAGTTTAGAGGACCATGTTGAAGTGGGtatagttacttgtcctaaac
gacttatcagtcttgtgtacAGTAATAACTGCAAATGAGTGTTTTAATGTCCCTGCAATCtatagttacttgtcctaaac
gacttatcagtcttgtgtacAAAAATTAATTGCGATACTGCCATGCAAAGATATTGGCGTtatagttacttgtcctaaac
gacttatcagtcttgtgtacAGTTACTCTGTACCTACTATGTACAAGACAATATGCTAAGtatagttacttgtcctaaac
gacttatcagtcttgtgtacCTGGAGACGCAAGAAATTTACACTGAAGTTTAATGGTGCAtatagttacttgtcctaaac
gacttatcagtcttgtgtacGTTATTAATCATATCTGCCTTCCTCCTTTAAGAAACTAGGtatagttacttgtcctaaac
gacttatcagtcttgtgtacAGCTGAAACATCCATTTGGTGGTTGCTTGGAGTCTGGCTTtatagttacttgtcctaaac
gacttatcagtcttgtgtacTGCAAACTGGAGTCAAATGTGTTATTCAGGGATGGCTAATtatagttacttgtcctaaac
gacttatcagtcttgtgtacTCTTCTTGGAGTAGAGTAAGGTTGAAAATCAAGAGAATCAtatagttacttgtcctaaac
gacttatcagtcttgtgtacGAATAATTTCCCCTCATTCTCCAAAGATTCCCATGTCCTAtatagttacttgtcctaaac
gacttatcagtcttgtgtacTGGCAAAAGGACTTTTGCTGATGTTAGTAAATTAAGGATCtatagttacttgtcctaaac
gacttatcagtcttgtgtacATAAGAAGAAAACAGGAGGAACTGAATCAGAAAAGGTATGtatagttacttgtcctaaa
gacttatcagtcttgtgtacCAAGCAGAGATTTGAGTGATATGTTTTGAGGATGGAGGACtatagttacttgtcctaaac
gacttatcagtcttgtgtacCAAGGAATGCAGGTAGCTTGGTAGATAAAGCAAGGAAACTtatagttacttgtcctaaa
gacttatcagtcttgtgtacGGGAACACAGCCCTGGCAGCACCTTGACAAATACACTTTTtatagttacttgtcctaaa
gacttatcagtcttgtgtacCAGAAATATAAGATAATAAGCTTGTGTTGTGATAAGCCCCtatagttacttgtcctaaac
gacttatcagtcttgtgtacAATAGCAATAATTATTCTTTCCCAACTCCGGCAGGTCTATtatagttacttgtcctaaac
gacttatcagtcttgtgtacCTCAAGAGGAACAAGGAACAAGAGGAAAAATGTGGAAAGAtatagttacttgtcctaaa
gacttatcagtcttgtgtacCTGCTCTCTGCAGCCACCTAAGTTTTGCTCAGATCTATATtatagttacttgtcctaaac
gacttatcagtcttgtgtacGCACATATAATCTCCTCTTGCCAACTTGTAGACAGACTTCtatagttacttgtcctaaac
gacttatcagtcttgtgtacTCCTAGAAAAGTCTCCATTGAATATTACTGGAGTACTAACtatagttacttgtcctaaac
gacttatcagtcttgtgtacTTTAATATGACTTGTTTCCATAGCAAACACACCATTCTGGtatagttacttgtcctaaac
gacttatcagtcttgtgtacCCTGAAGTTCCATGAATTCCTTTGCCATCTATTATATTTGtatagttacttgtcctaaac
gacttatcagtcttgtgtacCTTAGCGGAAAGATATCAGAGAGAACAGAAACTGGGAAAAtatagttacttgtcctaaa
gacttatcagtcttgtgtacAAAGCATTTAACGTGTAAAAGGGAAGAATCATCACAGGGAtatagttacttgtcctaaac
gacttatcagtcttgtgtacGCTGAATATTTTCCATTACTTCTGCACATTCCTGTACAATtatagttacttgtcctaaac
gacttatcagtcttgtgtacCAAGAGCAGCAGTCCTCCATCTGGGTCCTGTTTGTACCAAtatagttacttgtcctaaa
gacttatcagtcttgtgtacGTCCCACTTTTGATTCACCTTTATTCTGTAGGACAAGATGtatagttacttgtcctaaac
gacttatcagtcttgtgtacGATTGTATTTCTCTTCAGTCTAAGCCTCCTGAATGCCGGAtatagttacttgtcctaaac
gacttatcagtcttgtgtacCCTCCGAGTGACTCTAAAGAAACCCACCATCTTTCCCTGGtatagttacttgtcctaaa
gacttatcagtcttgtgtacCCATAGGAGCTTGGTCATCTGGATTATAGTCCTTATCAAAtatagttacttgtcctaaac
gacttatcagtcttgtgtacGGCTCACTACAACCTCAACCTCCCAGGTGCAAGCAATTCTtatagttacttgtcctaaa
gacttatcagtcttgtgtacTATTTTAATTTTTTATCTGCCCCCTTCCCCAATAGGTTGTtatagttacttgtcctaaac
gacttatcagtcttgtgtacGCATGGCATCTCATTCGTCTTTGGATTTCTGGTACCTAGGtatagttacttgtcctaaac
gacttatcagtcttgtgtacGACTTCACTCACTGAGCACTTCATACATACTGATTTCAGTtatagttacttgtcctaaac
gacttatcagtcttgtgtacCATTTTTGTAATCCTTTCCTCCCACTAAGACTTGAATCCTtatagttacttgtcctaaac
gacttatcagtcttgtgtacGGACCCTATTCTTTACTTTTCCATCCACTCAAATATTCATtatagttacttgtcctaaac
gacttatcagtcttgtgtacCTAAGAAACTCTGCATATTATCCCCTTGGCTATATTGATTtatagttacttgtcctaaac
gacttatcagtcttgtgtacGCTGCACCATAATTACCCAAGGAGTTTATTACAAAACAGGtatagttacttgtcctaaac
gacttatcagtcttgtgtacACCCCATAGTTTCTGATTGAATAGGTATCACATTGGATTCtatagttacttgtcctaaac
gacttatcagtcttgtgtacACAGTTTACCAGGTGTTATAGGATGAATTGTATCTCTCCAtatagttacttgtcctaaac

gacttatcagtcttgtgtacTCCTAACCCCTAGTACCTTATAATGTGAACTCATCTGGAAtatagttacttgtcctaaac
gacttatcagtcttgtgtacAGATGTAATTGGGTAGATTTAGATGAGTTCATGCTAGAGTtatagttacttgtcctaaac
gacttatcagtcttgtgtacCATACATGTGCATAGGGAGAATACTATGTGAAGGCAGAGAtatagttacttgtcctaaa
gacttatcagtcttgtgtacCAGAGATCAACAGAAGCTATGGAGGGGCATGTAAGAAATtatagttacttgtcctaaa
gacttatcagtcttgtgtacCAACAACTTGGCCTTGCATATCTAATCTCTAGAACTGTGAtatagttacttgtcctaaac
gacttatcagtcttgtgtacGACATCTAGTTTGTCTCACTGTGATACAATAGCCCTAGAAtatagttacttgtcctaaac
gacttatcagtcttgtgtacATTGACATGATTTTTGACCTGGACTCTTACCACTCCTTTAtatagttacttgtcctaaac
gacttatcagtcttgtgtacCTTCAATATCTCAAAGGGCCATTATGAGGACTAAATGAATtatagttacttgtcctaaac
gacttatcagtcttgtgtacTTTTTCAAAATCCTTACTGTGGAAAGAAGCAACATTGGTGtatagttacttgtcctaaac
gacttatcagtcttgtgtacACACCAGGGATTCAGTCTAGATCTGTCTGCTCACTGCACAtatagttacttgtcctaaa
gacttatcagtcttgtgtacGCTTTAGTTGGGTGCTGCAGTTGAGAGGATGGGAGCTCAGtatagttacttgtcctaaa
gacttatcagtcttgtgtacGCAGGGAAGAAATGTAACAAAGTGTAAGAAATCAAGAACTtatagttacttgtcctaaa
gacttatcagtcttgtgtacGGGAAAAGGAGACATCTGGTGCCCTGATCTAGTGAGCTTCtatagttacttgtcctaaa
gacttatcagtcttgtgtacGAAGGGCCAATTTCTATGTTGATCTAAAAGAACAGTCTATtatagttacttgtcctaaac
gacttatcagtcttgtgtacTGGTAATAATCCACTTTAGAATACATTGCATCTTAGAGGCtatagttacttgtcctaaac
gacttatcagtcttgtgtacGCTCATGAATCTTCAAACTTTAAGTCACTGGAGCATCTGTtatagttacttgtcctaaac
gacttatcagtcttgtgtacTGCTGGGACATAGGCAGATTTCTGGGATGGAGTGTTGTCAtatagttacttgtcctaaa
gacttatcagtcttgtgtacTGGCACATTCTTTGGTTTATGGATAGACAACATAGTGAAGtatagttacttgtcctaaac
gacttatcagtcttgtgtacATCCCACAAAATTAATATGAACAAATAACCTACCAGGCACtatagttacttgtcctaaa
gacttatcagtcttgtgtacCATGAAAACCCATGGAGGTAGTGGTATATTATCCCCATTTtatagttacttgtcctaaac
gacttatcagtcttgtgtacGCTCAGATGAATTAAGTAGTAAGCTATAGAACACAGAACTtatagttacttgtcctaaac
gacttatcagtcttgtgtacACACCAATGCCTGTCCTTCTTATTTTGTTTAGTGCCTTTtatagttacttgtcctaaac
gacttatcagtcttgtgtacCTTGTAAGACAGACACGTGTATAAGCACTAATCCAATCAAtatagttacttgtcctaaac
gacttatcagtcttgtgtacGCCAGTGCTCTAGAAACTTCCTGTGCAGCCTGCTATGATTtatagttacttgtcctaaa
gacttatcagtcttgtgtacGTGACATCAGAAGAAGGCAACACTTACACTTTTTGATAATtatagttacttgtcctaaac
gacttatcagtcttgtgtacCTGTCATTTCATTATCTATGTTCATAACCCTCAACAATGCtatagttacttgtcctaaac
gacttatcagtcttgtgtacCTATAGGCTCTGATCTTTTTTCCCATTATTTTGCTACTTGtatagttacttgtcctaaac
gacttatcagtcttgtgtacCATGCTGTATGTATTGCTTCTTCCTTCCCATTTATTCCAGtatagttacttgtcctaaac
gacttatcagtcttgtgtacCTTTGTCCAAGCCTTCTCTCCCCACATTTACCCTTTCTTTtatagttacttgtcctaaac
gacttatcagtcttgtgtacAGGTTTCCTTCTCCCCTGAACTGCTCTAGTCCTTGTATCTtatagttacttgtcctaaac
gacttatcagtcttgtgtacTTATGCATTTGACAAACATCTATTAAACACACAGTGTGGCtatagttacttgtcctaaac
gacttatcagtcttgtgtacCTGTAATCCCAGCTCTTGGGGAGTTTGAGGTGGGAAGATTtatagttacttgtcctaaa
gacttatcagtcttgtgtacATTAGCTAGGTATGGTGGTGTGTGCGTGTAGTTCCAGCTAtatagttacttgtcctaaac
gacttatcagtcttgtgtacACAAACACAGTGTGGCAGATATAGGTTCTACATATATAGCtatagttacttgtcctaaac
gacttatcagtcttgtgtacTCATGGAGCTTGCATTCTGGAGCAAAGAGACAAATAATAAtatagttacttgtcctaaa
gacttatcagtcttgtgtacCAGACTATGATAAACACTATGAAGGAAATGAAACAGAGGTtatagttacttgtcctaaa
gacttatcagtcttgtgtacAAGAGGACACTGTGTGAGCTTAGGTCTTTTCTTCCATCTGtatagttacttgtcctaaac
gacttatcagtcttgtgtacTGGGGCAGAAACAAGCTTGGCATGGAGTGGATGAAGAGAGtatagttacttgtcctaaa
gacttatcagtcttgtgtacCAAGAGGTTGGAGAAGAGGTGTTGAATAATCACGTGGGGGtatagttacttgtcctaaa
gacttatcagtcttgtgtacTAGGCCCTGATAATGAACTTAGTTGCTTTTTTAGGTGTAAtatagttacttgtcctaaac
gacttatcagtcttgtgtacAAGGTACATACTTTCACTTGCTTATGAGCTCTTACCTTATtatagttacttgtcctaaac
gacttatcagtcttgtgtacTGTACACATCTTTTGCCTAAACTAGAGCGTTCAATTTTTGtatagttacttgtcctaaac
gacttatcagtcttgtgtacTTACTGTTGGTGCTGCTAATATTTTTGTAAAACCTGTGTGtatagttacttgtcctaaac
gacttatcagtcttgtgtacGACTCAGAAAGGCATCGATTCAGTAAAGTTTCAATTGGACtatagttacttgtcctaaac
gacttatcagtcttgtgtacGTGGTTATTACACTGTCAACAAACATGACCTGCCTTGCGAtatagttacttgtcctaaa
gacttatcagtcttgtgtacACTTGTTAGTTGTTCAAAATCTAGAGTGTGCCTTCTTTAGtatagttacttgtcctaaac
gacttatcagtcttgtgtacGGGGTAACTTGATGAGACTTGAATTCCAGCTGACCTTTGGtatagttacttgtcctaaa
gacttatcagtcttgtgtacCTGACAGACATAGCCATTGCCTTTTCAGTTAAATTTTCTAtatagttacttgtcctaaac
gacttatcagtcttgtgtacGTATGAACTTCTGAGGATTTTAAACAAGGAAAGCAACTGTtatagttacttgtcctaaac
gacttatcagtcttgtgtacCTGAAAATGAGCCCTAACCAAATGTATCAACGTACAGAACtatagttacttgtcctaaa
gacttatcagtcttgtgtacAGAGATCTCTTTTTTGACAGCTTGAGAGCATGCATCTGGCtatagttacttgtcctaaac
gacttatcagtcttgtgtacCTTTGGAGGCTGAAAGAAAGGTCTGTAATTCTGTGAAACAtatagttacttgtcctaaac

gacttatcagtcttgtgtacCTCAGGCAGGGGTTTTCAGTTCCTAAAGTGGGGGCGGGTCtatagttacttgtcctaaa
gacttatcagtcttgtgtacCATTCCCAAGGTGCCACTGACTCCAAGGAAAGCATTGAAGtatagttacttgtcctaaa
gacttatcagtcttgtgtacATTACTCCTCTACTTACTAATGAGCCAGATTTCCAAAAGGtatagttacttgtcctaaac
gacttatcagtcttgtgtacCTGCAATTGTACTTACAGATGGATTATACTTAAATGGCTCtatagttacttgtcctaaac
gacttatcagtcttgtgtacATGACTCATTGCTCCAAGAATAGTGTGAAAAACCTAGGTCtatagttacttgtcctaaac
gacttatcagtcttgtgtacAGAGATGGAGTTTAGAAGGAAACAGTAGAGTTCCAGTGGCtatagttacttgtcctaaa
gacttatcagtcttgtgtacCTCTTCAATTTTAAAATCAACAGACATTTTACAGGCCCTGtatagttacttgtcctaaac
gacttatcagtcttgtgtacTGCTAAGGATAGAGAGTAGTGAGCAAAACAGAAAGGGGTGtatagttacttgtcctaaa
gacttatcagtcttgtgtacTGGAGCAGACATTCTATGAGAGACAGAAACACACACGCACtatagttacttgtcctaaa
gacttatcagtcttgtgtacATAGATGGATGGAAAAGATGCATTTAAGGCCCAGGGAGGAtatagttacttgtcctaaa
gacttatcagtcttgtgtacAAAATAAAAGGAGTTGGTTATGAAAATATTGGCCTCAGGGtatagttacttgtcctaaac
gacttatcagtcttgtgtacCAATGGGCACAATAACCTGACAGTGGAGAAAACCTGGATAtatagttacttgtcctaaa
gacttatcagtcttgtgtacTGTGTACGGGCAGAAGGAAAGAGTGATGTGTAATGGACAGtatagttacttgtcctaaa
gacttatcagtcttgtgtacGTCATGGGAAGACATTTGGATTTATTCTAAGTGCAATGTTtatagttacttgtcctaaac
gacttatcagtcttgtgtacCTGGGGGAGATAGGATCAGACTAGTGTTTTGGACGGCATGtatagttacttgtcctaaa
gacttatcagtcttgtgtacGCTTGGCATATTGATATAACTTGGAGTTCTTCAGAAAACCtatagttacttgtcctaaac
gacttatcagtcttgtgtacTGTCTAATGTTTGCTATGTAGTATCTTTGGACACATGGAGtatagttacttgtcctaaac
gacttatcagtcttgtgtacATAAAAAATCTTCATACATATGACCCATGTGTCCCCAACCtatagttacttgtcctaaac
gacttatcagtcttgtgtacCCCAGAGGTTATGAAAATTTCTTTGAGCAACGATCAGGAAtatagttacttgtcctaaac
gacttatcagtcttgtgtacGGAACCCTAACCAAATGTATCAACTTATAGACAACTGAGAtatagttacttgtcctaaac
gacttatcagtcttgtgtacGACAACAAAAAAACTCCTTGAGGAGGTTGCTCTGTCTGGCtatagttacttgtcctaaa
gacttatcagtcttgtgtacTTTTTAAAAAATAGCCTGTCATGGTGGCATGCCCCTGTGGtatagttacttgtcctaaac
gacttatcagtcttgtgtacATGATGTCATTATGTGTAGTAGGAAGATACATGGGAATCCtatagttacttgtcctaaac
gacttatcagtcttgtgtacGATTTAAAAAAGGAATCTAGGAGAACACAGGTCTTACCCAtatagttacttgtcctaaac
gacttatcagtcttgtgtacTTACTTAAAAATCTGCTTCCATTACGTCAACTAGAAACCCtatagttacttgtcctaaac
gacttatcagtcttgtgtacAGCTGGGATTCTTGCTATCATCGAATAAAACATAGGTGCTtatagttacttgtcctaaac
gacttatcagtcttgtgtacCCTTTGTCTGGTGGAGGTTTAGTGCTGCTGTTGTCGGTATtatagttacttgtcctaaac
gacttatcagtcttgtgtacCCAGAAAGAGAGAGGGGCGCTGGGAGATACTTTAACTGCCtatagttacttgtcctaa
gacttatcagtcttgtgtacAAATTGCCTGATTTCTTCTCTTGATAAAACTAGACATGGGtatagttacttgtcctaaac
gacttatcagtcttgtgtacAGTCCTTACTTCTCTTCCTTTATCATTCGCCAGTGCCTTTtatagttacttgtcctaaac
gacttatcagtcttgtgtacTTGGAGGGGAGTAAGGTAACTACTTGTAATGAAATGCACTtatagttacttgtcctaaac
gacttatcagtcttgtgtacTAGATTTCACTTTTGAGGCACAGTTGATAGAAGCATTATCtatagttacttgtcctaaac
gacttatcagtcttgtgtacTGCTGAAGCAAGCAGAGATTAAGAGATGCGAAAAGGGTTTtatagttacttgtcctaaa
gacttatcagtcttgtgtacGCTGCGGCGAGAACTTCTGTTGGCTTACCTTTGCTCCGGCtatagttacttgtcctaaa
gacttatcagtcttgtgtacTGTGCTTCTGTGGGGATCACGCTCATTTTCTCTGGTGAAGtatagttacttgtcctaaac
gacttatcagtcttgtgtacTCCTCACATGCCCTTAAAGTTCAGTCGGGACACCTCAGCAtatagttacttgtcctaaa
gacttatcagtcttgtgtacGCCACAGCCCACTTTGTCTTTCCCACCTCACTTCCATCACtatagttacttgtcctaaa
gacttatcagtcttgtgtacCCCTATGTCGTACCCATCCCAAACTTTCCATCATTCTGCAtatagttacttgtcctaaac
gacttatcagtcttgtgtacGAACTTCTTGTCCCTTTCTATGTCTTTGTTCCTGATTTTTtatagttacttgtcctaaac
gacttatcagtcttgtgtacCCACTGCACCTGGCCGTTCTGATATTCTTATCACATTAAAtatagttacttgtcctaaac
gacttatcagtcttgtgtacCTTCGTAATAACCCTGTTTAGATCATTACTATCCAGTTTCtatagttacttgtcctaaac
gacttatcagtcttgtgtacCTGTAGTTTTAACATTTTTACATTCCTCTCTGCATTCTGGtatagttacttgtcctaaac
gacttatcagtcttgtgtacTTCTATATCCCCTTGCCTTCAGACTGTAAGTTCCTTGAGAtatagttacttgtcctaaac
gacttatcagtcttgtgtacCACTGCACCCGGCCAGAGAATAATAAATCATTTTCATCATtatagttacttgtcctaaac
gacttatcagtcttgtgtacGCTAATCGCTAATGAGTGCCAGGAAAACAAAGAGAAGAAGtatagttacttgtcctaaa
gacttatcagtcttgtgtacGGAGTTGATATTCCACTGGCAGGAGAATAATAATAGATGGtatagttacttgtcctaaac
gacttatcagtcttgtgtacCTATAAAGCAAGCACACAGGGCAATGGGATAAGAGAGTCTtatagttacttgtcctaaa
gacttatcagtcttgtgtacAGGGGGATATTTCGTTAGATAGGGTATAGGAGCTGGCTTCtatagttacttgtcctaaac
gacttatcagtcttgtgtacTCTGGAGAGTAGCAACAATGTCCTAAGGCAGGCAAAAGCTtatagttacttgtcctaaa
gacttatcagtcttgtgtacTCAGACGTCATTTTGTGGTAAGGGGTAGACAGGTAGAAGGtatagttacttgtcctaaa
gacttatcagtcttgtgtacGCCGCTGCCAGATCATTCAGGACAAGGAGTTTGGACTGTGtatagttacttgtcctaaa
gacttatcagtcttgtgtacCACTGCAACCTTCACTTCCCGGGTTCAAACAATTCTCCTCtatagttacttgtcctaaa

gacttatcagtcttgtgtacTGAATCACCATGCCCAGCCCGAACTGTGTTTTAAGTTTAGtatagttacttgtcctaaac
gacttatcagtcttgtgtacCCATTGCATGGTTTTAAGCAGGAATGTGATATTGTATAACtatagttacttgtcctaaac
gacttatcagtcttgtgtacGCATGAGCTGCCATGGCCAGCCAAATGATATGTATTTTTAtatagttacttgtcctaaac
gacttatcagtcttgtgtacTTTCTTCATATGAAACATTTGCAGAGCCTTCAAAGTCTCCtatagttacttgtcctaaac
gacttatcagtcttgtgtacCAATAATAAACAATCTTCTAGTATAAACAGAGCCCTGGAGtatagttacttgtcctaaac
gacttatcagtcttgtgtacGGGTGGTAACTGTTTGCGTGAGTATGATGCCCCTTTAATTtatagttacttgtcctaaac
gacttatcagtcttgtgtacATAAAACCCCCTCCCTTCACTTTATTTCTCTCTTCATTCAtatagttacttgtcctaaac
gacttatcagtcttgtgtacCTTTTATTGTTTTAAGCAGGAGTTTACACATACCTGGGAGtatagttacttgtcctaaac
gacttatcagtcttgtgtacCAGATTTCTTTGTAGTGTGGGGGTAACTTTCTGACCTTTCtatagttacttgtcctaaac
gacttatcagtcttgtgtacAGAATAGTCTAAAGAAGGGGAAAAGAGATGTGATGGGAATtatagttacttgtcctaaac
gacttatcagtcttgtgtacCCAAATGGGGAAGAATCTAAAGAACAGACTTTCTACTTGAtatagttacttgtcctaaac
gacttatcagtcttgtgtacAGTTTTCTGGTAATAACTCAACTTGGAATAATGCAGCAAGtatagttacttgtcctaaac
gacttatcagtcttgtgtacCACACAATGAAACCCAGCTATGAGGCAAACTGACATAAACtatagttacttgtcctaaa
gacttatcagtcttgtgtacGTTCCTGATAGATCCACCTACTCTTTGCAAACAGAATGGGtatagttacttgtcctaaac
gacttatcagtcttgtgtacGAGTCTGGTCCATCTCTAATTTTATTCTGTCTTATTGGTAtatagttacttgtcctaaac
gacttatcagtcttgtgtacCATATAACTACTCCAGGATGTATTTCTAAGTTCAAGGTACtatagttacttgtcctaaac
gacttatcagtcttgtgtacTCAGGATGGTAATTGCTGTAAGGGGTAAAATTATTATGTGtatagttacttgtcctaaac
gacttatcagtcttgtgtacTGAAGTAATCCAGTAAAATTGACTTCTAGTCAGTCCCCGCtatagttacttgtcctaaac
gacttatcagtcttgtgtacAACATTTCATCTGTGGGTTGCTGTTGAGACATTTTTACTGtatagttacttgtcctaaac
gacttatcagtcttgtgtacGATGGGGTAAGATTTTACATAACTTAAAACATCTGCTGTCtatagttacttgtcctaaac
gacttatcagtcttgtgtacGTGAAAGACATTTAACTTTTCCCTCTCTTTTTTTCTCGTGtatagttacttgtcctaaac
gacttatcagtcttgtgtacGGGTCATGAATATAGTGAAATATTTAGATGCATCTCCTACtatagttacttgtcctaaac
gacttatcagtcttgtgtacACTGTTGTTAATGTCACCTCTGCTTTCCCGTACACACCTGtatagttacttgtcctaaac
gacttatcagtcttgtgtacTATAAAATACAGAAAGCCATTTTAGATCATGCCAAGGGTCtatagttacttgtcctaaac
gacttatcagtcttgtgtacCAAAATGTCTTCAAAAATGCTGTAAGGATAATGCTGCTCCtatagttacttgtcctaaac
gacttatcagtcttgtgtacAGTTATTTACTTGGGCTTTGAACTAGCCAATAGTGACAATtatagttacttgtcctaaac
gacttatcagtcttgtgtacGTTATAGATGTAACTATTGCTGTCCCATTTTATAGACGAGtatagttacttgtcctaaac
gacttatcagtcttgtgtacAGAGCATTTAACAACTTACCTAAGGTTAGGGTGGTGAAGCtatagttacttgtcctaaac
gacttatcagtcttgtgtacTTCTCCAAGAACCTCATTTCATTCCAACATAGCCAAGTCTtatagttacttgtcctaaac
gacttatcagtcttgtgtacCATGTTCATGATGCTGAGAGGTTAGAATGAGAAGTTTCCTtatagttacttgtcctaaac
gacttatcagtcttgtgtacGGTTGGAATATGGTGCGTAGAGCCTGTTGCTTCTGTGGGCtatagttacttgtcctaaa
gacttatcagtcttgtgtacGTTGAATTGTGTCTACCCAAACATATCCATAGTCCTAATGtatagttacttgtcctaaac
gacttatcagtcttgtgtacTGAGATTATCCTGGACTTAGCTTGGGCCCTGACTCCAATGtatagttacttgtcctaaac
gacttatcagtcttgtgtacCCAAATAAAGATGGAGACAAAGATTGGAGTTATGCAGCTAtatagttacttgtcctaaac
gacttatcagtcttgtgtacCATGGAATGGATTCTCTCGCTGAGCCTCTAGAAGGAACCAtatagttacttgtcctaaa
gacttatcagtcttgtgtacTTAAGTGATGAAGTGTGTGGTAGTTTTTAAACGGTAGACtatagttacttgtcctaaac
gacttatcagtcttgtgtacGCCGTATTTGTTTGATATCGAAAATGAGAAAAAAGGCAAGtatagttacttgtcctaaac
gacttatcagtcttgtgtacGCTACTGACAGAAAAAAAATTAAGAAGCAGCACCATTAGTtatagttacttgtcctaaac
gacttatcagtcttgtgtacGTAATATTTGCCTGTTTTATCTGACCTGCCATTTGAGGAAtatagttacttgtcctaaac
gacttatcagtcttgtgtacTTAGGAGTTTAGGACATCACTAATTAATTATGCCACCCTGtatagttacttgtcctaaac
gacttatcagtcttgtgtacACTTAAATTCTCAGAGCCTCCTTCATAAAATCAACTCACCtatagttacttgtcctaaac
gacttatcagtcttgtgtacGCCTGACACTGCCAGTCACCCAGCTTAGTAGAAACAGGGCtatagttacttgtcctaa
gacttatcagtcttgtgtacCAGGTTTGCAAATCTCAACGTTGTTTCTAGACCATAATGAtatagttacttgtcctaaac
gacttatcagtcttgtgtacCTTTAAGAGGTATTGAGTGGGTTAAATGGAATAATCTGTGtatagttacttgtcctaaac
gacttatcagtcttgtgtacCCCAAATCTTTTAGGGCAAGTTTATCATCATGCTAACTCAtatagttacttgtcctaaac
gacttatcagtcttgtgtacTCATGTTTTAGATTTGTATCCTTTAAGGGCTTTCTGGTTCtatagttacttgtcctaaac
gacttatcagtcttgtgtacCAATGGATGAGGGGTGTGTGTTATATTCTCTTTAGTTTGTtatagttacttgtcctaaac
gacttatcagtcttgtgtacTGTATTTTCTTTCAAACTATACGCTCATCACACCTCACACtatagttacttgtcctaaac
gacttatcagtcttgtgtacAGTTTGAAAAATCTCTGGACTAATATTCTGGCTGCAACACtatagttacttgtcctaaac
gacttatcagtcttgtgtacATTGTCCGGAAAATATCAATGTCAAGCATTTCTCACTGGCtatagttacttgtcctaaac
gacttatcagtcttgtgtacCCTGCTTAAAAGTAATAGTATAAAATCAAACCAGGGACCCtatagttacttgtcctaaac
gacttatcagtcttgtgtacTGATAAAAACGCAAAATACACCCTTACAAGTCCGGATGAAtatagttacttgtcctaaac

gacttatcagtcttgtgtacGTTTCAAAGTAACAAATTTTAATTGGCATCTCTCAGCCTGtatagttacttgtcctaaac
gacttatcagtcttgtgtacAGCACTTACAAAAATTCTCTTTCTTTTCTCCTCTCGTTTCtatagttacttgtcctaaac
gacttatcagtcttgtgtacCTGCAGCTCCCAGACTTACTTCATATTCCCAAACCCAGTGtatagttacttgtcctaaa
gacttatcagtcttgtgtacCCTCCTATGCTCTGTATCTCCACACACTCCATTCCTTAGCtatagttacttgtcctaaac
gacttatcagtcttgtgtacTTGTTGACCTATCCCATCTCTAGCTGCACACACTTCCTTCtatagttacttgtcctaaac
gacttatcagtcttgtgtacACCTGGGCAATGTGAGAAATCTCTTTTGTGGGTCATAGGAtatagttacttgtcctaaac
gacttatcagtcttgtgtacTATTATTATTTTCAGAGAAGCCTACTAACTAGATGGGAGGtatagttacttgtcctaaac
gacttatcagtcttgtgtacGCTGTCTCTTTTAGCACCTGACTTTCCTGGCTTGGCTAAGtatagttacttgtcctaaac
gacttatcagtcttgtgtacGGTCTTTGAACTCCTAGTCCCATGTTTTCTGTCTGCCTGTtatagttacttgtcctaaac
gacttatcagtcttgtgtacTCTATCTACCTACCTACATCATCTATCTACATGTATCTGCtatagttacttgtcctaaac
gacttatcagtcttgtgtacTTCATCCATCTAATTTCATTGCCTCACTGTACCTGTTTGAtatagttacttgtcctaaac
gacttatcagtcttgtgtacAAGTCTGTGAGTTGAATCCAGGAGGAATAGTCATGTGGTAtatagttacttgtcctaaac
gacttatcagtcttgtgtacCAACTGCACAGCAATGGGTGATAGGAGCCAGCCCTTATGGtatagttacttgtcctaa
gacttatcagtcttgtgtacCTTCCTAAAGTGTCATACCAACCAGAGGGGAACTATCAGAtatagttacttgtcctaaa
gacttatcagtcttgtgtacCCAAATGGGGGGAAATAAGGCACTGAGACAATTTTATGGGtatagttacttgtcctaaa
gacttatcagtcttgtgtacAGTGTCACTTATTCCTAATTCCACCTTTCCTGAGCAATGCtatagttacttgtcctaaac
gacttatcagtcttgtgtacGAAATAATTAATTCCCCAGTCCATAAGTGGAAATCCAAAGtatagttacttgtcctaaac
gacttatcagtcttgtgtacATTTATCATTCCATTCAAAGACCTCTTGGGTAGAAAAAGCtatagttacttgtcctaaac
gacttatcagtcttgtgtacACAGTCAGTAGGGTCAGATTCTCATTCCATGCTCATTGTAtatagttacttgtcctaaac
gacttatcagtcttgtgtacTCTTTTCCCAGATTAGAGGTCACCAAACTTTTCTTGTAAAtatagttacttgtcctaaac
gacttatcagtcttgtgtacCAAATACTTTAGGTTTGAGGGATATATGGTTTCTGTCTCAtatagttacttgtcctaaac
gacttatcagtcttgtgtacTATGTGCCAAATTAATATATGTGCCTGTGTTCCAAACAACtatagttacttgtcctaaac
gacttatcagtcttgtgtacGCTGTGGTTTGTTGGCCCCTTTTCTAGATCAAATGGAGAAtatagttacttgtcctaaac
gacttatcagtcttgtgtacGGGTGTAATCCTGGGAAAATCTGTATTTTTGCTGGACTGAtatagttacttgtcctaaac
gacttatcagtcttgtgtacCTGAACATAGCCACACGAACCTATCCCTATTCATTAATATtatagttacttgtcctaaac
gacttatcagtcttgtgtacCTTACATTTTCATAGCAAAGAAACAAAGGCACTGTTCTCTtatagttacttgtcctaaac
gacttatcagtcttgtgtacTGCTCTGTAGGGGAATGGAAACAATTGCAATTGGAAAAGCtatagttacttgtcctaaac
gacttatcagtcttgtgtacAGTGAGAACTGCGCCTTGGATGCTCATTGGTTTTGAAAGCtatagttacttgtcctaaac
gacttatcagtcttgtgtacAAGTGAACTTTCCATTTCAACCCCATTCTCCCTTCATTCAtatagttacttgtcctaaac
gacttatcagtcttgtgtacAGTCCTCCCTGAGTTAAACTACATTATGGTACCAAGCAAAtatagttacttgtcctaaac
gacttatcagtcttgtgtacCAAAATGACAACCGCACTGCAAATCTTAGAAAATGTAACAtatagttacttgtcctaaac
gacttatcagtcttgtgtacGCACCTGACCTCCTGAGATGCTTAATAAAATTGCCTACAAtatagttacttgtcctaaac
gacttatcagtcttgtgtacCAAGAGGCATTTGTGGAGAGATAAATAAAGCTATTTTCCAtatagttacttgtcctaaac
gacttatcagtcttgtgtacAAATGACAGTTTCTTTATGGGATGGGAGATGGGCAGGAAAtatagttacttgtcctaaac
gacttatcagtcttgtgtacTGCCCACCACTAGTAGAAGTTCTATAATGAGATTCCAAATtatagttacttgtcctaaac
gacttatcagtcttgtgtacCTGTGAGATGTTCAGCTTGACTTTATAATGAGTTCACCGAtatagttacttgtcctaaac
gacttatcagtcttgtgtacATGAAATAAGTGGGGATCAATCATAGTCAGGGAAGTGGCAtatagttacttgtcctaaa
gacttatcagtcttgtgtacACTAAAGGGGGAAAAATGAGAGTGTAAAATGGAGCAGATGtatagttacttgtcctaaa
gacttatcagtcttgtgtacGATGATATCTAAAGAGGGTGGGATGGTGAGAAGTATAAATtatagttacttgtcctaaac
gacttatcagtcttgtgtacCTACTCAGGAAGCAGCCATAATCATCGCCATCTTAGGAAGtatagttacttgtcctaaa
gacttatcagtcttgtgtacTGGTGACAGACTTTGGAAAAGCAAACAAATTAGTCATGGTtatagttacttgtcctaaac
gacttatcagtcttgtgtacTGCCCACATTCTAGATTAAGCCCACATAACCCACCTACTGtatagttacttgtcctaaa
gacttatcagtcttgtgtacCTGCAAAAGCTCTCAGGAACCAGTATCTCTCCCCACACAGtatagttacttgtcctaa
gacttatcagtcttgtgtacCTGAGAGAAGTATCGGCACAAAGGTCTGCAGGCTGGCATGtatagttacttgtcctaa
gacttatcagtcttgtgtacACTCTGCTGTGATTAGTATACCTGCCATAGCTAGGGTTCCtatagttacttgtcctaaac
gacttatcagtcttgtgtacGGTAGGGGTCAGCAGATGGAAATAATCATGGATGCCCGAGtatagttacttgtcctaa
gacttatcagtcttgtgtacGTTTCAGAACCAATATCATTACCTGTTTCAATGTCTAGCGtatagttacttgtcctaaac
gacttatcagtcttgtgtacGACAGTATAGAGAGTCATAGTCTAACTTCTGGCCTTTCTGtatagttacttgtcctaaac
gacttatcagtcttgtgtacTTTCTTTTTTCATACCACAACTGTCACTCATAGGGTTCCAtatagttacttgtcctaaac
gacttatcagtcttgtgtacCCTGGGAGGTCATGATGGTTTATTGCAGCCTTTGTTTTTAtatagttacttgtcctaaac
gacttatcagtcttgtgtacGCCAGTGTCTCTCTCGCCTACCTTCCCTTTTAAGATACTCtatagttacttgtcctaaac
gacttatcagtcttgtgtacACAGTCTTTTGGCTGTCAAAGCCCCATCCAAATCCCAGCAtatagttacttgtcctaaa

gacttatcagtcttgtgtacGACATGGTCCTGCTGCTGGCCTCAGGCTGCCTTCTGGAATtatagttacttgtcctaaa
gacttatcagtcttgtgtacCAAGTTTTTGTGTTTCATCCTTTTTGTTCCTAGCTGAGTCtatagttacttgtcctaaac
gacttatcagtcttgtgtacCCCCATCTGGTGCTCTCCTGCCCCCTGGTAAGATTATAGTtatagttacttgtcctaaa
gacttatcagtcttgtgtacAGTCTCTAGCTGCTACTATAGTTCTTTCTCTAACCTCACCtatagttacttgtcctaaac
gacttatcagtcttgtgtacAATGAAATAGAACTAGGAATGACATGATGGAAGGGTTTGGtatagttacttgtcctaaac
gacttatcagtcttgtgtacCACAGTATAATGCAAAGTTAAACAATGAGGATGTGTGGGAtatagttacttgtcctaaac
gacttatcagtcttgtgtacTTTCCAAAACTATCTGATCTAATAACTTGTCCTCCAGCGTtatagttacttgtcctaaac
gacttatcagtcttgtgtacGTTATTGTAGGGAGAACATTAAGGGTTATTACATAGAGGGtatagttacttgtcctaaac
gacttatcagtcttgtgtacACATATAGAGGTACGAGACTTAAAACACACATGTGCTTACtatagttacttgtcctaaac
gacttatcagtcttgtgtacACACACACACACACTGGACATTGCTCTCAAGGAGACAAGGtatagttacttgtcctaaa
gacttatcagtcttgtgtacAGACAAGTACAAATTGATCAGGGGTCTTCAAGGGCAAACCtatagttacttgtcctaaa
gacttatcagtcttgtgtacGAGGTGCTATTGTTCTGCTATTTGTCTCCCTCTCTCTTTCtatagttacttgtcctaaac
gacttatcagtcttgtgtacGCATTTTTCTAAGTTGAATCTCATTTTGCTATTCTTGCCCtatagttacttgtcctaaac
gacttatcagtcttgtgtacGCTGGAATTCTTGTGTGTATGACTGATAAAGCCTTTAGAAtatagttacttgtcctaaac
gacttatcagtcttgtgtacAGATCAAAGCATCTCCAGCATAGATATAACACAACAGCTAtatagttacttgtcctaaac
gacttatcagtcttgtgtacCTCAAAAGAAGTCCTGTCATTAATATGCTGGGGATGGGAGtatagttacttgtcctaaac
gacttatcagtcttgtgtacTGAGTGGTCAGGCAGGCGGGCAGCATTTACTAAGAACCTGtatagttacttgtcctaaa
gacttatcagtcttgtgtacCACAGCAGGGAATTATGTGGGAAAACGGCACTCTAATAAAtatagttacttgtcctaaa
gacttatcagtcttgtgtacCACAAAAAACCCTTATATTCAGAGACATGTGTAATCCTGTtatagttacttgtcctaaac
gacttatcagtcttgtgtacCTATCTGAGTGTTAAGGATGTTGCTATGATGAAAGAAGGAtatagttacttgtcctaaac
gacttatcagtcttgtgtacTTGAATACCAAGACACACTCGCAACTAGTAGAAAAAAGACtatagttacttgtcctaaa
gacttatcagtcttgtgtacTGGAACAGTGAGGGTGCAAAGCAACAGGAAGACATTACCAtatagttacttgtcctaaa
gacttatcagtcttgtgtacCTCTTGGGAGTTCCTATATATGTTCCATCTTAGCTTGGATtatagttacttgtcctaaac
gacttatcagtcttgtgtacTGTCCATGAATCACAGAAAACTTGAAGCCTTAGGAAACATtatagttacttgtcctaaac
gacttatcagtcttgtgtacCATGTAACCACAGGTGCATTACCATGTCTTCCATCTCAGGtatagttacttgtcctaaac
gacttatcagtcttgtgtacTTTGACTTCTCTTAGCTAACATATGTCCCAATAAGATCAGtatagttacttgtcctaaac
gacttatcagtcttgtgtacTATGTCACTGTCTTCTCTGGCCCTTTCAACGTTGCTAGGCtatagttacttgtcctaaac
gacttatcagtcttgtgtacATAGCCAATGAAGTCTCTCAAAGCAACTGTCTTAGATGTTtatagttacttgtcctaaac
gacttatcagtcttgtgtacATGTCTTGTGCTTGCCGTCATCTTCCTAGCTCTGCAGTCCtatagttacttgtcctaaac
gacttatcagtcttgtgtacTTGGGACTACTTCAAGATATTAGAGAACAGATCCTCAGTTtatagttacttgtcctaaac
gacttatcagtcttgtgtacCAGTGTATGTGGGGGCTGAGAGTGGCATTGTTGGCTGAATtatagttacttgtcctaaa
gacttatcagtcttgtgtacTCACCCTCTGCCTCCTTCACCTCCCCACTTTGCCCTCTACtatagttacttgtcctaaa
gacttatcagtcttgtgtacTTACGTCCCCTCCTTCCTTGCCATTCTTCCTGCCTGCGTAtatagttacttgtcctaaac
gacttatcagtcttgtgtacTGGTACAGATGTCTCAGTGATTAAAGTTCTCATACTGATTtatagttacttgtcctaaac
gacttatcagtcttgtgtacGTTAAAATAGTGATAGAATGGAAGTGGACAAATCTAAGGGtatagttacttgtcctaaac
gacttatcagtcttgtgtacAGTTTTGCTCCACTCCCCAGGGTACATTTGGCAATATCTGtatagttacttgtcctaaac
gacttatcagtcttgtgtacACTGATTTTCCTAATTGTGCTGGGTGAAGTTAAGGGTTTGtatagttacttgtcctaaac
gacttatcagtcttgtgtacGGATGTCCCTAAATATACTTCAATACACAGGCCTTCCAAAtatagttacttgtcctaaac
gacttatcagtcttgtgtacCGTTTTCCGTAGCTAGGCAAATGGATGCAATTTTCTATGCtatagttacttgtcctaaac
gacttatcagtcttgtgtacATTTTTTGGCTCCTTCTCTTACTCTGTTTCCATTAAGTGAtatagttacttgtcctaaac
gacttatcagtcttgtgtacCCAGAGATCACCAAACTTTTTCTGTTCCAGACGGTAAATGtatagttacttgtcctaaac
gacttatcagtcttgtgtacTTATGTCTTAAGAGAGAAAACTCCTGCTGTTTCCTGCTATtatagttacttgtcctaaac
gacttatcagtcttgtgtacCCCTTGGCGTTGGGCAAACTTTATATGCAGTTGTATTTCAtatagttacttgtcctaaac
gacttatcagtcttgtgtacTGGTGAGATGACTCCTGGACTGAGATTCTACCACTCGATGtatagttacttgtcctaaa
gacttatcagtcttgtgtacGTCCCCAGTGTTCAGGTGTCAAAGAGTGCTGGTCCTGTGCtatagttacttgtcctaaa
gacttatcagtcttgtgtacCAGATGCCCTGTGCACCTCATAATGCTAGCTGGTTCATGCtatagttacttgtcctaaa
gacttatcagtcttgtgtacTTGTATCCCCCAACCCCTGATTCTGTATCGTCTGTTTCTCtatagttacttgtcctaaac
gacttatcagtcttgtgtacCCTTTTTCTCCGAGAGGATTATTTAATCTCAGTCTGAAGGtatagttacttgtcctaaac
gacttatcagtcttgtgtacCTTTCTCTAAGTGGAATTAGCCTTGTTCAGATCTGCCATGtatagttacttgtcctaaac
gacttatcagtcttgtgtacCAGCTTTGACTAACCTTTGCGGGGCACTTTGAATCTGTTTtatagttacttgtcctaaac
gacttatcagtcttgtgtacGCTATTAGACATGAGCTTTTCTTCATCCACTAAACCTACAtatagttacttgtcctaaac
gacttatcagtcttgtgtacTTGGCAGAATGTTATAAGAGTTAGTCTGACTAAGTTAAGCtatagttacttgtcctaaac

gacttatcagtcttgtgtacTAAAAGAAAGGGAGGTAATTACTAGAAAACAGCCAAAACCtatagttacttgtcctaaa
gacttatcagtcttgtgtacATTCTCTTTTGTGTACTTCTTTGTTCAAGCCTATTCCTTCtatagttacttgtcctaaac
gacttatcagtcttgtgtacAGAAATAAGCTCTTTAACAATCTTCCCATTGGTATCTACCtatagttacttgtcctaaac
gacttatcagtcttgtgtacGAGAAAAACACACAAATGCTTGATATCACTGCTATATAGCtatagttacttgtcctaaac
gacttatcagtcttgtgtacAAGGAAAAAAATCTACTTGCTGGTTGTATTAGACAGGGTTtatagttacttgtcctaaac
gacttatcagtcttgtgtacATCAGTAGGACAGATAGATATAGAAAGGGGGATTTATTAGtatagttacttgtcctaaac
gacttatcagtcttgtgtacATGGTGTCTGAGGGGTCCCACAAAAGTCCATCTGCAAGCTtatagttacttgtcctaaa
gacttatcagtcttgtgtacCCTCAGAACCAGGAAAGCTTATGGCATAACTCTCAGTCAGtatagttacttgtcctaaa
gacttatcagtcttgtgtacGAGTTGTGATGTCCAAGGTCTAGAGAAGAAGAGTGTCCCAtatagttacttgtcctaaa
gacttatcagtcttgtgtacATCCTTCCTCTGCCTTTTTGATTCCATCCACCCCCCAGCTtatagttacttgtcctaaac
gacttatcagtcttgtgtacGTCTACTGACTCACCTGGCTATCTCCTCTAGAAACACCCTtatagttacttgtcctaaa
gacttatcagtcttgtgtacGTTGATGCCTAAAATTAACTTTCACACTGGTGTATCTTCTtatagttacttgtcctaaac
gacttatcagtcttgtgtacTGGTGTGCTCTCCAAACGTATATCCTGAATGCATTTACATtatagttacttgtcctaaac
gacttatcagtcttgtgtacCCTCCTGTGGCTTATAGTATATGAATCAAGAGTCCAAATCtatagttacttgtcctaaac
gacttatcagtcttgtgtacATTCTGACTTTTCTCCTGAATTGCAAACTCTTATCCTCCAtatagttacttgtcctaaac
gacttatcagtcttgtgtacTTTGCCCCACAGACATCTCAAAGCCAACATATTGAACATAtatagttacttgtcctaaac
gacttatcagtcttgtgtacAACTCTTTCTTTATTTACAGGGCTTATTCACAGCAACATGtatagttacttgtcctaaac
gacttatcagtcttgtgtacGAAATCTCCACTCTTTCTATATTTCCAATCATGGGAACAGtatagttacttgtcctaaac
gacttatcagtcttgtgtacGAATCTTGGGAGTCATTCCACGCTACTCATTTCATGCATAtatagttacttgtcctaaac
gacttatcagtcttgtgtacAAATTCCATTTCCTTACTATGTCACAATTTCTGTCCCCTTtatagttacttgtcctaaac
gacttatcagtcttgtgtacCATAACTATTATAATAATATATTGATCGGCCTCCCTGCTGtatagttacttgtcctaaac
gacttatcagtcttgtgtacTTATACCCCTTCATAATATTGCAATTAGAGAGACCTGTTGtatagttacttgtcctaaac
gacttatcagtcttgtgtacCTAATGATGTTATTTCCCGACTTAAACTGTTTCTGGAATCtatagttacttgtcctaaac
gacttatcagtcttgtgtacACTGTTTAGCATAATACATAAGGTCTTCCATTTTCTAGCCtatagttacttgtcctaaac
gacttatcagtcttgtgtacTTATTCTTCGGATTTATCACCACATTTACTGAGCACTATGtatagttacttgtcctaaac
gacttatcagtcttgtgtacGTAATCTCAACCAACCGTAAAATGAAGGTACTGTTTTAACtatagttacttgtcctaaac
gacttatcagtcttgtgtacGCATAAAGCCTTTAAGGAATTTGTTCAAGATCACATAGCTtatagttacttgtcctaaac
gacttatcagtcttgtgtacTGATACCAGAGCACAGAATACCCTTTGTCCCTACCCCCACtatagttacttgtcctaaa
gacttatcagtcttgtgtacTCTATGAGCTGCAATTACACAAAATTGGTTTAGCGTTTTGtatagttacttgtcctaaac
gacttatcagtcttgtgtacTATTTTTAATAATGTGATAAATACCCCCCAGTGCTACCCCtatagttacttgtcctaaac
gacttatcagtcttgtgtacATGTGGTACTTTTATTTATTTCCTTTCCCCTATTTCCCTGtatagttacttgtcctaaac
gacttatcagtcttgtgtacTATGTTAGATGCTTTTCACTTCAAAAAACCTATCGTGGTGtatagttacttgtcctaaac
gacttatcagtcttgtgtacTTATTGATCTACTTTTTAGTTGATTGGCATTGGGTTGTGGtatagttacttgtcctaaac
gacttatcagtcttgtgtacTCTATATTTTGTGTATACAGGTTTTCTTTGGACAGAGCTCtatagttacttgtcctaaac
gacttatcagtcttgtgtacCTTCATCCCTATACTGTACTCTCTCCCACCCTGGCTACCTtatagttacttgtcctaaac
gacttatcagtcttgtgtacTGTTTTAGGTGCAGAAGAAAAAAATAGATGAAACTCCCACtatagttacttgtcctaaac
gacttatcagtcttgtgtacGCACAGCTTACTTTCTTCCATTTGTCACTTTGCAGCTAATtatagttacttgtcctaaac
gacttatcagtcttgtgtacATTCCTTGCACATGAAGTTTAATCCTCTAAATCTTTGCTCtatagttacttgtcctaaac
gacttatcagtcttgtgtacTCCTTTGAATAAAATATCCTCTGAGCCCTCATTTTCCAGCtatagttacttgtcctaaac
gacttatcagtcttgtgtacATTTAACTCAGATATTGTATCCCTCATGAAACCATAGGTGtatagttacttgtcctaaac
gacttatcagtcttgtgtacTCTCTGACTGGAATAAGGACCTATTTTGCACTCACATAGCtatagttacttgtcctaaac
gacttatcagtcttgtgtacCATAACTTTATCATATCTCTTTTCTAGTTTTCCCCTCCTCtatagttacttgtcctaaac
gacttatcagtcttgtgtacCTACTTATCTTTAAATAGCTCTAGTGACCAGTGTGATGACtatagttacttgtcctaaac
gacttatcagtcttgtgtacTCGAAAAACAAAACTGAACTGATTCTGAAAATTCTGGGACtatagttacttgtcctaaac
gacttatcagtcttgtgtacTCTGTATATGAGGATTACATAGTTATGATGAGAGAGTCTCtatagttacttgtcctaaac
gacttatcagtcttgtgtacGACAGCCTTCTCATTAATTCACTCAAGAGACATATTATTCtatagttacttgtcctaaac
gacttatcagtcttgtgtacCTGTTCCTTTTGTACAGGAGATCAAAATGGATTTAGAAACtatagttacttgtcctaaac
gacttatcagtcttgtgtacGGTTTGAAAAGCAATGGGATTTAATGGAAAGAGCTCTGCCtatagttacttgtcctaaac
gacttatcagtcttgtgtacTATGCAGAGGGGGATGGGCTTACGTAATGATGAAGAAAAAtatagttacttgtcctaaa
gacttatcagtcttgtgtacGATGTTCTACAGCTACCCCTTTCATTAACAGAGACTTTTCtatagttacttgtcctaaac
gacttatcagtcttgtgtacGGATAAAACCCTCCCTAAATATTTAAACTTTAACCTGGCCtatagttacttgtcctaaac
gacttatcagtcttgtgtacCTTTTGATCTCTATTCATAACAGAAGATACAGGCTATACCtatagttacttgtcctaaac

gacttatcagtcttgtgtacTGACTATGTTTGTTTTCCTTTGCCTCAGTGATCTTTGGATtatagttacttgtcctaaac
gacttatcagtcttgtgtacGTCCCCCTCTCCCCATTTTCTGTTCTGGACTGCAATAAAAtatagttacttgtcctaaac
gacttatcagtcttgtgtacTGATTTGTAAGCAAGGTAATAGGAATTAGGGGACATTCTCtatagttacttgtcctaaac
gacttatcagtcttgtgtacTGAGGAAAAGAGAAGAAAGATGGGGAAGAAATCTGTACAAtatagttacttgtcctaaa
gacttatcagtcttgtgtacGGTGTCTATGACCAGAAATTTGACCTGAACCACATATATCtatagttacttgtcctaaac
gacttatcagtcttgtgtacTGTGTGTGGTTTTTTAAGGTAAGAATAGGTTTCGTGTCCAtatagttacttgtcctaaac
gacttatcagtcttgtgtacAGAAGAAAGGAAAGAGTAGATTCAGAAAGCAAGAAGAGAAtatagttacttgtcctaaa
gacttatcagtcttgtgtacGAGGGTTTCAGATTAAGAAGAAGTTAAGCACTCCAAGGACtatagttacttgtcctaaa
gacttatcagtcttgtgtacCTGCTGTTTCTGGGTTGCATAGAACTGGGCTGTTTTATTTtatagttacttgtcctaaac
gacttatcagtcttgtgtacAGGAAAAGGTGACACAGGATGACCAGAGTTCTTGAAGACAtatagttacttgtcctaaa
gacttatcagtcttgtgtacTATTAAGGATATGAAATAACACCAGGATGTTTCAGTGTGGtatagttacttgtcctaaac
gacttatcagtcttgtgtacGATATAGGTTTAATGTTCACACAGATTTCGTTTTCACCAGtatagttacttgtcctaaac
gacttatcagtcttgtgtacTTAAAGTGTTAGAGTTCAGTATTAGAGGTGACCAGAAAAGtatagttacttgtcctaaac
gacttatcagtcttgtgtacTTTTCTTAAGAGCACCCATTATTCGGATTTCTGAAAGTCCtatagttacttgtcctaaac
gacttatcagtcttgtgtacCCACTGCACTTTTACATTTGTAGGTAGGAACAATCAGCCGtatagttacttgtcctaaac
gacttatcagtcttgtgtacTGGTATAGTGCCCCCATGGAAGACAGAAAGACTCAGTTTCtatagttacttgtcctaaa
gacttatcagtcttgtgtacGTCTTCTACACTGAGATCCAAGATGGCTGCTTAAGATCCAtatagttacttgtcctaaac
gacttatcagtcttgtgtacTCAGAGTGTATGAAGAAGAAATGGTTTGAAAAATGGCATGtatagttacttgtcctaaac
gacttatcagtcttgtgtacGGAACATCCTGTAGTTTTTACCTTTGACTTATGCTTACATtatagttacttgtcctaaac
gacttatcagtcttgtgtacCTAAAAGAACGTAGGCTGGGAAGTCGGGCGCGGTGGCTCAtatagttacttgtcctaa
gacttatcagtcttgtgtacTCAGGAGATAGAGACCATTCTGACTAACACGGTGAAACCCtatagttacttgtcctaaa
gacttatcagtcttgtgtacAAAAAGAAAGTAGGCTGGGAAATGTTGTTTATTCTGGGCAtatagttacttgtcctaaac
gacttatcagtcttgtgtacTCTCTGTGACCATGTTCTGTATAACCTTTTTTGGGCTATAtatagttacttgtcctaaac
gacttatcagtcttgtgtacTGGTAGAGATGACATTGATAAGGAACAATGAAATGGTGAGtatagttacttgtcctaaac
gacttatcagtcttgtgtacAAATAGTAGCCAAACAGATGTTCGTTATCACTATGTTGTGtatagttacttgtcctaaac
gacttatcagtcttgtgtacCTGGGGGGAGGGAGAAAGAGTGGCAGAGAGAGTGAGGGAGtatagttacttgtccta
gacttatcagtcttgtgtacTGTGACTTATCTATTAAGGGAATATTTAAAGAGCAAGGCGtatagttacttgtcctaaac
gacttatcagtcttgtgtacTTCAGTTGGAGCCAAATATCAATATAGGTAAACTTGGTAGtatagttacttgtcctaaac
gacttatcagtcttgtgtacTTCACAGAGCTTAACAAAATGAATGCATGGCTTGATCTTAtatagttacttgtcctaaac
gacttatcagtcttgtgtacAAGAGAATATTGATCAGCTCGCTTTAGGACACCATCTGTAtatagttacttgtcctaaac
gacttatcagtcttgtgtacATTGTTTGAAATAGCCTTATTGCCATGAAGTTTACGTAGGtatagttacttgtcctaaac
gacttatcagtcttgtgtacAACTTTGCTGGTTTTGAGAAATGGCCTAATATTAGGAGTGtatagttacttgtcctaaac
gacttatcagtcttgtgtacATGTAAATTCTGGGTTCTTGTCTTACCTCAGTTAGTGAAAtatagttacttgtcctaaac
gacttatcagtcttgtgtacTGAAACAGTACAATATCTTTGCCTCAGTTACATTCTTAGGtatagttacttgtcctaaac
gacttatcagtcttgtgtacTGCATTGTACATTTTTTGAGCATAAGGGGAGAGGATGTCAtatagttacttgtcctaaac
gacttatcagtcttgtgtacACCTTCCTGAAAGTAGAATGATTCCTATGTTATTTTCACCtatagttacttgtcctaaac
gacttatcagtcttgtgtacCTTCCTTTGACTCACTTTAAAGCCAGCTCTTATTATTCTGtatagttacttgtcctaaac
gacttatcagtcttgtgtacTCATAAAAACAGCCTGTAGCTACAACAATGTCTAAAAGCAtatagttacttgtcctaaac
gacttatcagtcttgtgtacTACCAAGTTGACACACTTCATTTATCTAAATTTACCCAGCtatagttacttgtcctaaac
gacttatcagtcttgtgtacACACAATTATATCATAACTCCACTTATTTTGGAATGGCCCtatagttacttgtcctaaac
gacttatcagtcttgtgtacGAAAACTACATATAGGCTCTACATACCTTGAAATCACAGAtatagttacttgtcctaaac
gacttatcagtcttgtgtacAGAACGCTCTTATATCTATTGGAGGTATTGAAGATTAGGAtatagttacttgtcctaaac
gacttatcagtcttgtgtacTTCTAGGTATCTAAATGCTTAAAATAAGAGTTTGGCTGGGtatagttacttgtcctaaac
gacttatcagtcttgtgtacTCAAGATCAGCCTGGTCAACTTAGTAAGTCCCCATCTCTAtatagttacttgtcctaaac
gacttatcagtcttgtgtacAAAGTGATCGAGAAAACAGAAAATAAGAGCCCAATATTCCtatagttacttgtcctaaac
gacttatcagtcttgtgtacATCAGTAGGCAAGTACCAAATGAGTGTTTAAGATACAAACtatagttacttgtcctaaac
gacttatcagtcttgtgtacGGGAATTAAGGTTTTCATAGATAGAAAGGTTGGGGAAGCCtatagttacttgtcctaaac
gacttatcagtcttgtgtacACAGATAGATCAATCAGATAAAGATTAATGGAGATGGCAGtatagttacttgtcctaaac
gacttatcagtcttgtgtacCAATGAAGGAGGAAGTGCTTGAAAGTAGGAACAACTAAGAtatagttacttgtcctaaa
gacttatcagtcttgtgtacCCTAGGGGGTGATAAAAATGAGAATGATTAAGTTGGTCTGtatagttacttgtcctaaac
gacttatcagtcttgtgtacTCCCGGACTTGATCCACTAGTCTGTAGATACAATTTCAAGtatagttacttgtcctaaac
gacttatcagtcttgtgtacCAACTACTGCATGAGCTGGTGATATGATTTGGATCCGTAGtatagttacttgtcctaaac

gacttatcagtcttgtgtacAGGTGATTGGATCATAAGGATGGTTTCTCATGGTTTCCCAtatagttacttgtcctaaac
gacttatcagtcttgtgtacAGTGAGTTCTCATGAAAATTGGTTGCTTAAATGTATGCAGtatagttacttgtcctaaac
gacttatcagtcttgtgtacAGTTTCCTGAGGTCTTCACAGAAGCTGAACATATAGCATCtatagttacttgtcctaaac
gacttatcagtcttgtgtacGTGATGCGAGAATGGAATAATACAGATAGGTAGTTAATAGtatagttacttgtcctaaac
gacttatcagtcttgtgtacCTCAAGTTAAAATCACCTGGGCCAATTGTTAAACACATAGtatagttacttgtcctaaac
gacttatcagtcttgtgtacGTTGAATTAAACTGAACTTTCTCCAATCTTGAGTCTATGCtatagttacttgtcctaaac
gacttatcagtcttgtgtacCAGGTGTTATTTGTTCAGGGCTTATGAGTGATAATCATTAtatagttacttgtcctaaac
gacttatcagtcttgtgtacGTTAATAAATATTGAGCATCCACAGTGTGCTTCAACTCATtatagttacttgtcctaaac
gacttatcagtcttgtgtacCCAGGGTATGGGGCAAATGAAATAGACAAAAATAGATAAAtatagttacttgtcctaaac
gacttatcagtcttgtgtacCATATTGTTGTATAATGGGGAATATATCTGGTCTATGTCCtatagttacttgtcctaaac
gacttatcagtcttgtgtacCCTCATGCAGAGCTTCAGAAACACTTGAAATTTTTTCAGTtatagttacttgtcctaaac
gacttatcagtcttgtgtacGAAGAGACTAATAGTAGACTTCTAGATAGTTTTAGGATGGtatagttacttgtcctaaac
gacttatcagtcttgtgtacACCAGAGAGACCCATCACATGATTAGAGTTGGAACTTTGGtatagttacttgtcctaaa
gacttatcagtcttgtgtacATAAAACTTGCTAACATAATGAACCCCCCAAAAAACTCTGtatagttacttgtcctaaac
gacttatcagtcttgtgtacGAGATTGGTGAGCACATTGACATGCAGGGAGGATGACATGtatagttacttgtcctaaa
gacttatcagtcttgtgtacGGCATGGAGGAAGCTCTGTGTTCCCTTTCAGATCTTTTTCtatagttacttgtcctaaac
gacttatcagtcttgtgtacTTGTTGCCCAGGCTGTAGTGACTGCAGTGTCGTGATCTTGtatagttacttgtcctaaac
gacttatcagtcttgtgtacCCATGCCCAGCCTCCCTTTCAGATCTTACTTAGGTATATCtatagttacttgtcctaaac
gacttatcagtcttgtgtacGGAATCATAAGCATAGCACATTTCCAAGTTGTTCTAGTGAtatagttacttgtcctaaac
gacttatcagtcttgtgtacTCAGAAGACATGGTGCGCTGGGGATCCTCAAAGTGTGGCTtatagttacttgtcctaaa
gacttatcagtcttgtgtacGACTGAGCCCTTAACATGTGAGTTCTGCATTACTCTAGTTtatagttacttgtcctaaac
gacttatcagtcttgtgtacCAGTTGGGAATGGAAACAGGAAAACCAGAGTATTTATAAAtatagttacttgtcctaaac
gacttatcagtcttgtgtacATTGGATGCACTGAAGTATTAGGTGTGTTGCATTTTAAGAtatagttacttgtcctaaac
gacttatcagtcttgtgtacACTGACCTGATTGCCTCTTAAAATATGTTCATAAAAAGGGtatagttacttgtcctaaac
gacttatcagtcttgtgtacAATACAACTTACTATTGGCTCTTTGGTGGTTGCTTTAAAGtatagttacttgtcctaaac
gacttatcagtcttgtgtacCTGGTGATATTTGGATAGACAGAGAATGGAATGTAAAGAAtatagttacttgtcctaaac
gacttatcagtcttgtgtacTCAGGGCCATTATTGTTTTATTTGTCTTCTTTGTGTGATGtatagttacttgtcctaaac
gacttatcagtcttgtgtacAAGCATTACTCCTGTGTATGTCTTTGAGGGTGTTTTTAGAtatagttacttgtcctaaac
gacttatcagtcttgtgtacGTGGACTGAGTAAGAAAGATTCACCCTCACCCAGTATGGGtatagttacttgtcctaaa
gacttatcagtcttgtgtacAGAAAAGGTGAATTCTTCTCTCTCTGTCTTGAAGCTGGGAtatagttacttgtcctaaac
gacttatcagtcttgtgtacCCAGCAGACCCTCAGTTTCTCAGGCCTTTGGTTTCAGACTtatagttacttgtcctaaa
gacttatcagtcttgtgtacTTCTTAGGCCTTTGAACTTGAACTAAGCTACACTACTGGCtatagttacttgtcctaaac
gacttatcagtcttgtgtacGGCATATCATGGGACTTCTGAGCCTCCTAATCACATAAGCtatagttacttgtcctaaa
gacttatcagtcttgtgtacGAACCCTGACTAATAAAGTTCAGTTTGTTTTACTCCTCCTtatagttacttgtcctaaac
gacttatcagtcttgtgtacCAAAATTAATTGGTATTGAAGGAGGATGGTGATGGAATGTtatagttacttgtcctaaac
gacttatcagtcttgtgtacGTCCTGTTTATCATTGACCATTTAAGTTCATTCCATAAGGtatagttacttgtcctaaac
gacttatcagtcttgtgtacGTGCACTCAAAAAACAAAATCTGTAAACTGTAAACTGAGGtatagttacttgtcctaaac
gacttatcagtcttgtgtacGCAGAATTGGGGCCCGTTTAAATGAATGATTCCTCTAGAAtatagttacttgtcctaaac
gacttatcagtcttgtgtacTGTTTCTACCTTGTGAAATGACCTTCAGTGAACATTTAAGtatagttacttgtcctaaac
gacttatcagtcttgtgtacTCAGTGCACTAAACCTATACATTTTCAGCCATTTTGAAACtatagttacttgtcctaaac
gacttatcagtcttgtgtacATGAATGACTCCTCCATGTAACAAGATAGAAGACACCAGTtatagttacttgtcctaaac
gacttatcagtcttgtgtacCTTGGTGGTGAATTATTTATATCTTTCAAGTCTTTGCTGCtatagttacttgtcctaaac
gacttatcagtcttgtgtacACAGAAGCAAATGAAATGTGCTAATGTGCTCCCAATTAGCtatagttacttgtcctaaac
gacttatcagtcttgtgtacGTAATGAACTTTTTGCTTCTGATTGAGGAGAGAATTCACAtatagttacttgtcctaaac
gacttatcagtcttgtgtacATTAGTGAGTGTATCTGCTGGAAGTTGCCAAATTATAGGGtatagttacttgtcctaaac
gacttatcagtcttgtgtacGAATCTGAAAACGTGAAAAACATCCTTCACATAAGTGTAGtatagttacttgtcctaaac
gacttatcagtcttgtgtacATAAATGTGGCTTCAACTTGTTGATTTATTTGCCCAAGAGtatagttacttgtcctaaac
gacttatcagtcttgtgtacGGTGACTCCCATTAAACCTAGAGTGAAATGAAAAAATCTTtatagttacttgtcctaaac
gacttatcagtcttgtgtacACACTGCAGGAATCACAAAGCCTTAGGCATGTTAACCAGCtatagttacttgtcctaaa
gacttatcagtcttgtgtacTGGAACACTGGAAACTGCTACCTGATACGTTCGACCCTATtatagttacttgtcctaaa
gacttatcagtcttgtgtacGTGATAGGATAGTAGGTAGTAGAAGTAGGTGTGAAGATGTtatagttacttgtcctaaac
gacttatcagtcttgtgtacTGGTGGTGATGATGATAGGAACGGTTTCAATTTGTTGAGTtatagttacttgtcctaaac

gacttatcagtcttgtgtacGAGAAAGAAAATGTTCTCTTTCTCCAGAGGAAGAAACAAAtatagttacttgtcctaaac
gacttatcagtcttgtgtacAGACCTTCTCTACATTCCTACAAGATCTTTTTTGCCTGGAtatagttacttgtcctaaac
gacttatcagtcttgtgtacCTAATTGAGAAGTGTTGACTATCCAGCTAAAGAGAGTTCAtatagttacttgtcctaaac
gacttatcagtcttgtgtacTTTCTAGTTGTGTTTTCCATCTTCCTACTCAAAGTTCTTCtatagttacttgtcctaaac
gacttatcagtcttgtgtacGAGAATGCCCTCTATTTATTTTGATCCTCCCAAGACTGTAtatagttacttgtcctaaac
gacttatcagtcttgtgtacATAAGCTTTATTGTTCAGTGAGATGAGACTCTACCATGTTtatagttacttgtcctaaac
gacttatcagtcttgtgtacCATCCCATGGTTTATCAGCATTAACTATCAAGAAATTTGCtatagttacttgtcctaaac
gacttatcagtcttgtgtacGTTCTTGTCTGAATACCTCCTGCCATTTCCTCTCTTCACTtatagttacttgtcctaaac
gacttatcagtcttgtgtacCAAACATTCTCTGTATGATAGCCCTTTATATACTTGGAAGtatagttacttgtcctaaac
gacttatcagtcttgtgtacTCTTCCTTCAGCTGCTTCTTAAGACAAAATCATGGGCCTTtatagttacttgtcctaaac
gacttatcagtcttgtgtacAAAGAACACCAGAGGAAACATGCAAACACTAAATAAAACCtatagttacttgtcctaaa
gacttatcagtcttgtgtacGTTCTCTTGTATTTTTGCTCATAGCAAAGTCATGACATACtatagttacttgtcctaaac
gacttatcagtcttgtgtacTATTAGACTTATTTGCAGTTCTCCATGAGCTTGAGTTTTCtatagttacttgtcctaaac
gacttatcagtcttgtgtacATTAAAGGTGGTGGTCAGTGGGGTGGAGGGGATAAGTCAGtatagttacttgtcctaaa
gacttatcagtcttgtgtacTGCTGACTACTTCTAGCCAGAAATGTTGAACTTTTTATTCtatagttacttgtcctaaac
gacttatcagtcttgtgtacAGATAGAAGTTATGTTTATAAAGCTGGATTTTCTCACGGCtatagttacttgtcctaaac
gacttatcagtcttgtgtacGTTTCAGTATTCTCCCCAAACAGACCCAGCCCCTAATCCCtatagttacttgtcctaaa
gacttatcagtcttgtgtacAATCTTTCCATTTGCTTCACTCCACAACGTTTAATACTTCtatagttacttgtcctaaac
gacttatcagtcttgtgtacGCTGGATAGGGCTAAGTTGTGTGCATATTGTGTTTGGGATtatagttacttgtcctaaac
gacttatcagtcttgtgtacTGGAGTCGGATTTAACTGCTCTATGATGCTAAACTGTGAGtatagttacttgtcctaaac
gacttatcagtcttgtgtacGCTATAAAAGAAAAGTGCGATGTATTTCAGATGCCTAATCtatagttacttgtcctaaac
gacttatcagtcttgtgtacAGCAAAAGGTTAGCTCTTCAAGGAGGGGTAGTGTGTAGACtatagttacttgtcctaaa
gacttatcagtcttgtgtacCGTCATCTCTCTCTGGGGAACTCTGGGAGTTAAGGCTGTCtatagttacttgtcctaaa
gacttatcagtcttgtgtacTTTCCTAGGAGACATGATAAAGAAACTTCTGCAACAATTGtatagttacttgtcctaaac
gacttatcagtcttgtgtacGGGAAAACAAAACTTTAAAGCAGTAAGGAAATGAATGTGGtatagttacttgtcctaaac
gacttatcagtcttgtgtacAGAGTGTCTCCTGCATACTTAATAGAAAAAAGCATGTCAGtatagttacttgtcctaaac
gacttatcagtcttgtgtacCTTGCTCTTATTTTTGATGACCAGGAAGTTTTCAGCTCAAtatagttacttgtcctaaac
gacttatcagtcttgtgtacTCTGTGCTCCAACTTAGTATTTTTCCATCTAGTTTAACTGtatagttacttgtcctaaac
gacttatcagtcttgtgtacTATCTACCCATCTGTCTATTCTGAAGGATGACACAAAGGGtatagttacttgtcctaaac
gacttatcagtcttgtgtacCATTTAGTCCAGTGTCTCAACTTGATGTGTAGTGCTTTTAtatagttacttgtcctaaac
gacttatcagtcttgtgtacCATTAATGAGCCATAGACCACTTGTAAATTTCTGACTTACtatagttacttgtcctaaac
gacttatcagtcttgtgtacTTCTTCTGGAAATGTCTTCACCTGACAAGCTGGTAATAAAtatagttacttgtcctaaac
gacttatcagtcttgtgtacTTCTTTTATCCACTTAACCCTTCTACCATATCAATTTGGGtatagttacttgtcctaaac
gacttatcagtcttgtgtacCATTTGTATGATGCTTACTGTGTGCTGAGTTTGAGTTTTAtatagttacttgtcctaaac
gacttatcagtcttgtgtacAGAGCATTCTTCATCTATAAACTCATGTAATCCACCTGACtatagttacttgtcctaaac
gacttatcagtcttgtgtacGGTGATGTGACTACTCTTATTATGCCCATTTTATTCATGAtatagttacttgtcctaaac
gacttatcagtcttgtgtacGATCTTCTAAGTATTGGATCTGGAATTTGAATTGGGAAAGtatagttacttgtcctaaac
gacttatcagtcttgtgtacCTCTAGAGTATGTCTTCTAAAGTGTGGTCTTATGAACTAAtatagttacttgtcctaaac
gacttatcagtcttgtgtacTCATCCTCAGGAAGTACAAAAGGGCTTTACAAAGAAATTGtatagttacttgtcctaaac
gacttatcagtcttgtgtacTCACCTATAATATCCAAAGCAGTTGCACCACAATCAAGCTtatagttacttgtcctaaac
gacttatcagtcttgtgtacCTTTTTGCTTTTTTGATAGAACCCTTGGTGTCTGTACCCAtatagttacttgtcctaaac
gacttatcagtcttgtgtacTTCAGTTAGCTCTAAGGCAGCATTCTGGGTACTATGTTTTtatagttacttgtcctaaac
gacttatcagtcttgtgtacCACAGTCCACATACTTTGAAAGCAATAAATGATGATATGCtatagttacttgtcctaaac
gacttatcagtcttgtgtacAATGTATGTAAGGCATTTGAGGACTTGACTTTGATAAGCAtatagttacttgtcctaaac
gacttatcagtcttgtgtacCCTGAAGCTAGAAGAAAAAGTTGTCATATCTGGAATGTGCtatagttacttgtcctaaac
gacttatcagtcttgtgtacATCATTCCCATTCTTCAAAGTCCATGTCTTCAGAGAGCTTtatagttacttgtcctaaac
gacttatcagtcttgtgtacCTGTGTGCTTCTTTGAGATTTCCTCTAAAACAGTGCCTTGtatagttacttgtcctaaac
gacttatcagtcttgtgtacATTACTTTGTAAATTCTAGGAGGGCAAGGACAATATCTTCtatagttacttgtcctaaac
gacttatcagtcttgtgtacCTGGCAGCCTTAGGTGGTCTCTAGTGGTTTTCTACATTTGtatagttacttgtcctaaac
gacttatcagtcttgtgtacACTTGTAATGAGTGGATAGAACCTAAAGTGATCAGAGAAAtatagttacttgtcctaaac
gacttatcagtcttgtgtacAGCAATTACATATGGACAGTCTAATAAGGAGAGCTGCCTCtatagttacttgtcctaaac
gacttatcagtcttgtgtacAAGTAAGTTGAATGAACAAGTTCTCAGAAGAGGATCAGGAtatagttacttgtcctaaac

gacttatcagtcttgtgtacCCTGATGGGCTAAAGATTCTTTTATCTATGTAAGCAGAAAtatagttacttgtcctaaac
gacttatcagtcttgtgtacGTTGTCTTAATGGGATATATTATAATGAGAACTGGAGGGTtatagttacttgtcctaaac
gacttatcagtcttgtgtacCCCTGAATTAGTCAAAAGTTGGGGAGACAATATTTAATAGtatagttacttgtcctaaac
gacttatcagtcttgtgtacGATTTAACAAAATACTTTCAGATTACCAAGTTGGCGTTCCtatagttacttgtcctaaac
gacttatcagtcttgtgtacATGGGTAATGTGAATCAATTGACAAAATGTGTTCATTCCCtatagttacttgtcctaaac
gacttatcagtcttgtgtacGGCCAACCCTCTTGCTACAATGCTTTCACGGTGCGGGGCAtatagttacttgtcctaaa
gacttatcagtcttgtgtacTGTATATAATCTAGGACAAAAAACGTATGCACAAACACCCtatagttacttgtcctaaac
gacttatcagtcttgtgtacTTTTATGCATATTCAGCCAGACCATAACAGTTACTCTCCTtatagttacttgtcctaaac
gacttatcagtcttgtgtacAGATATCACCATTATGTCTGGGCAGATTGTTTGGGTCCTCtatagttacttgtcctaaac
gacttatcagtcttgtgtacATAATTTGCTTCATCCTGCATTTCTGCTTTTCCAAAAGTCtatagttacttgtcctaaac
gacttatcagtcttgtgtacAAAATGGAACTTTTCTCCCAAGGACAGAGTACATCTTGAAtatagttacttgtcctaaac
gacttatcagtcttgtgtacCTTAGCCAAAGTTTATCTTTTAAATAAAACACGGCGCCTGtatagttacttgtcctaaac
gacttatcagtcttgtgtacACCCTTGGCTCATATATTTGCCTTTCTTGTATAATCCCTTtatagttacttgtcctaaac
gacttatcagtcttgtgtacTAGTACCCCATTAAAATTATTGGTTGCATTTGGTAACAGGtatagttacttgtcctaaac
gacttatcagtcttgtgtacCTTCAGAAGTTATCATGAAAGATAGGAGTCCTCAAAAATCtatagttacttgtcctaaac
gacttatcagtcttgtgtacCAATGACTGAGTGCTGAGTTCCAGAAGAAGTGGGGATCTCtatagttacttgtcctaaa
gacttatcagtcttgtgtacACCCTTGCACTTTTATTTTGCAGTAGACATTGTAAAAGGCtatagttacttgtcctaaac
gacttatcagtcttgtgtacCACACTGTGTTTTGAGTTGAAAATCTTTTGGGAATAAGTCtatagttacttgtcctaaac
gacttatcagtcttgtgtacTCGGGTGGAGGGAAGGGAATAGAGAATGTTTTTAAAGTGtatagttacttgtcctaaac
gacttatcagtcttgtgtacAGCACTCCAATTTTCTAGTCTGGTTCATTAATCACATTAGtatagttacttgtcctaaac
gacttatcagtcttgtgtacTTTAGGGAAAACACAAGACAGAATGAAACACAGTTCCAACtatagttacttgtcctaaa
gacttatcagtcttgtgtacTATTGCAGCATTTTGGGTTTCTAGAATACAGCATAGGACTtatagttacttgtcctaaac
gacttatcagtcttgtgtacCAGTGTTTCGAGGTTCTGAGTTCATTTTAAAGAACATAAGtatagttacttgtcctaaac
gacttatcagtcttgtgtacAATTAGAGATAGTTACCTAAAAGGAAATAGGCATTGCAGGtatagttacttgtcctaaac
gacttatcagtcttgtgtacCTTTGGGAGGATGAGATGGGTGGATCACGAAGTCAGGAGAtatagttacttgtcctaaa
gacttatcagtcttgtgtacACCATCCTAGCTAACACAATGAAACCCCATCTCTACTAAAtatagttacttgtcctaaac
gacttatcagtcttgtgtacTAGGCATTGCATTTGTTCTGAGTATGATAATTTACTCCTCtatagttacttgtcctaaac
gacttatcagtcttgtgtacGCTATGATTATCAGCTTTATGGGGTTCACTACAAAATCCAtatagttacttgtcctaaac
gacttatcagtcttgtgtacTTTCTTCAAGATTCAGGTTAATGTTTCAGCTATTCCTACCtatagttacttgtcctaaac
gacttatcagtcttgtgtacTCTCTCATCTCAAGCACAACTCATTATATTTTTGACATCCtatagttacttgtcctaaac
gacttatcagtcttgtgtacATCCTTCCCTAATTTCCTGTTGCTAGAAATGACCAAACCAtatagttacttgtcctaaac
gacttatcagtcttgtgtacCGGGCTTGGAATCCTGAAGTTATCTTTGACTCATGAGGTGtatagttacttgtcctaaac
gacttatcagtcttgtgtacACTAAGTTTTTCTAGTTCTACCTGCAAGTATCTCTTCCCtatagttacttgtcctaaac
gacttatcagtcttgtgtacGTTCTTTTCAGAGCTTCCAAACTGATTTACCTGGATCAAGtatagttacttgtcctaaac
gacttatcagtcttgtgtacCCTGTCCTTGTCTACTCTCTCACTATCTTTAATAAATGTCtatagttacttgtcctaaac
gacttatcagtcttgtgtacAATTTCCTCCCCTCAAAAACATTCAGACATTTTACTTTGCtatagttacttgtcctaaac
gacttatcagtcttgtgtacCTTTACTGTCTTCAAAAGCCTTAACAACACGACTCTATTGtatagttacttgtcctaaac
gacttatcagtcttgtgtacATTAAATGTCAAGTTTTCCATGTCTTCTCAAGATACCTGGtatagttacttgtcctaaac
gacttatcagtcttgtgtacATGTAAAGAAGTATAAGGGCCAAATGAAGGATTCAGATAGtatagttacttgtcctaaac
gacttatcagtcttgtgtacGAAAGAATTCAAACAAAAAACTCTTCATGTCCTTTGCAGGtatagttacttgtcctaaac
gacttatcagtcttgtgtacAACCAAACACTGCATGTTCTCGCTCATAGATGGGAATTGAtatagttacttgtcctaaac
gacttatcagtcttgtgtacCACCAGGGACTGTTGTGAGGTGTGGGGAGGGGGGAGGGAAtatagttacttgtcctaa
gacttatcagtcttgtgtacACATGTGTGGTTTGAATCTCAGTGATATGTAAGATCTTAGtatagttacttgtcctaaac
gacttatcagtcttgtgtacCTAACTAGAGATGCTCCTTGGTATAGGATGGGGTTATGCTtatagttacttgtcctaaac
gacttatcagtcttgtgtacGCCTAGCCTGCCTTAAACATTCTCAGTACTCTGTCAGTACtatagttacttgtcctaaac
gacttatcagtcttgtgtacAAATAGGTTTTGTGTGGCATGTTCTGATCCAGCATAATGAtatagttacttgtcctaaac
gacttatcagtcttgtgtacTGGAAGTCTGTTTTCTACGAATACATATTACTGTTGTACCtatagttacttgtcctaaac
gacttatcagtcttgtgtacAAGTTGGGGATTATCGGTACTTACTAAGCAGATTATGAATtatagttacttgtcctaaac
gacttatcagtcttgtgtacCATAATACTTATTCATGAAAATTGCCACGAGTCTTCAGCAtatagttacttgtcctaaac
gacttatcagtcttgtgtacCGAATTATACGTAACACTATAAACAAGCCCAACAGCATACtatagttacttgtcctaaac
gacttatcagtcttgtgtacTCAGATTGTATATGAGGGGAGTGAATATTTGGGAAGCACTtatagttacttgtcctaaac
gacttatcagtcttgtgtacGTGTCAAAACCATTCAGATACCATGTTTCTACTTCTTTGGtatagttacttgtcctaaac

gacttatcagtcttgtgtacGTATTATTAGTGTATGTTTGCATGGGCAATGAATACACAGtatagttacttgtcctaaac
gacttatcagtcttgtgtacCATTATATGAGGTTTATTGCTTTGCTAGGCAGATACTGTTtatagttacttgtcctaaac
gacttatcagtcttgtgtacGATTGGTTACCAGCTATGTATAAAATATTCTGAACAGTGCtatagttacttgtcctaaac
gacttatcagtcttgtgtacCTTGTTTCTATGAGAATGTATATAAGCGGGACCTCTGTAGtatagttacttgtcctaaac
gacttatcagtcttgtgtacGCCCAAAACAAACGATAAGAAGCATTTATTAAGAGATAGGtatagttacttgtcctaaac
gacttatcagtcttgtgtacCATAATATATGCAGAGTGGCCTAGATGAGAGAAGACTTAGtatagttacttgtcctaaac
gacttatcagtcttgtgtacCTGGAGGGGAAGCAGCAATAGGGAGGATAGAGTAAAAGAGtatagttacttgtcctaa
gacttatcagtcttgtgtacAGTTGGTAAAGAACTCCTCATCTAACTCAAAGTTATTGGAtatagttacttgtcctaaac
gacttatcagtcttgtgtacTCTAAGAGCAACTAGAAGACTATGAGGGGTTTTAAGCAGAtatagttacttgtcctaaac
gacttatcagtcttgtgtacTGAATCACTATGTTACTTTGGAAAAGCCTGTACATTTCAGtatagttacttgtcctaaac
gacttatcagtcttgtgtacGAGGCTGAGGTGGGAGGAACTGAGGCCAAGAGTTAGAGGCtatagttacttgtcctaa
gacttatcagtcttgtgtacCTCCAGCCTGAGTAATAGAATGAGACCCCATCAATAAAAAtatagttacttgtcctaaa
gacttatcagtcttgtgtacTTGAGGTAATAACAATGAATCATGGAGTTCTTCTAGGGAGtatagttacttgtcctaaac
gacttatcagtcttgtgtacTAGTGGAAGCGTTTGTAAACCAATAAGTTGAGAGAGAGAAtatagttacttgtcctaaac
gacttatcagtcttgtgtacGAATGTGTAAAATGCACTGTTCAAAAAACCTGGCCCATGGtatagttacttgtcctaaac
gacttatcagtcttgtgtacTTTAAAACTCTTACCTTCCACATTATTCCACATTGACACCtatagttacttgtcctaaac
gacttatcagtcttgtgtacCTTGTGAGGTAAGGAGTGGGAAGACGAGCACAAAAAATTGtatagttacttgtcctaaa
gacttatcagtcttgtgtacTGGAGTGGCATTAAGATACTTCTTAGTATATAAACCAGAGtatagttacttgtcctaaac
gacttatcagtcttgtgtacCCATTGAAGGGTCCACTTAGAAAAAGCTTTTTTCTAGAGCtatagttacttgtcctaaac
gacttatcagtcttgtgtacGTATCTGACCTGCAGTAAAGACAGCACCTAACTATTTTGGtatagttacttgtcctaaac
gacttatcagtcttgtgtacTGGGTTTTTCTGTCCAGTTATAAAAGATCTCCAGGGCAAAtatagttacttgtcctaaac
gacttatcagtcttgtgtacAAGCACTAAGCCAAAGTTCAGAGTGATTAAGGAAAACAATtatagttacttgtcctaaac
gacttatcagtcttgtgtacTCTAAAAGCAAGAAATACCCCAAGAGAACAACGTCGAGGAtatagttacttgtcctaa
gacttatcagtcttgtgtacGACTTGTCCTAAAAAGACTGTAAAGGATTTGACTGGAATAtatagttacttgtcctaaac
gacttatcagtcttgtgtacACCCATTAGGTGACTGGCACTGTTCCAAACAAACCTTCTGtatagttacttgtcctaaa
gacttatcagtcttgtgtacTGGAAGGCTGGTCATTTTCTGCAGAGAGGAAAATTGATTTtatagttacttgtcctaaac
gacttatcagtcttgtgtacAGCTATTTGTTTTATTTATGGCTTGAACTCAGCTCCCTGCtatagttacttgtcctaaac
gacttatcagtcttgtgtacGTACAAACCTTCTTATTTAAATATTGACTGGCAAGGGAGTtatagttacttgtcctaaac
gacttatcagtcttgtgtacCCCACAGTTAGTTGCAGGGGTTTGATTTATGCTAATGAATtatagttacttgtcctaaac
gacttatcagtcttgtgtacGCTTTTATTATTAAATCCTCAGGACACTACTTTCCAAGACtatagttacttgtcctaaac
gacttatcagtcttgtgtacTGGGCTTTACAATAAGAACAGAAGTCTATATGCTCTATCCtatagttacttgtcctaaac
gacttatcagtcttgtgtacCAGAAAGTAATAAAAGATCAGGAAGCATATTCATGGAGAGtatagttacttgtcctaaac
gacttatcagtcttgtgtacTGATAGCGTCTTGAGAGTCAGAAGTTCCAACTTAACCTCAtatagttacttgtcctaaac
gacttatcagtcttgtgtacGTGAAAAACTTACTTCAGCCTTTACCTTAATCATTCCTGGtatagttacttgtcctaaac
gacttatcagtcttgtgtacATATGTTCTCTTTACTGACAGCAGGAAGCCAAAGACCTTTtatagttacttgtcctaaac
gacttatcagtcttgtgtacTGAACTACAGTAAGGTTATGGACAAATCTACTTGGACATAtatagttacttgtcctaaac
gacttatcagtcttgtgtacAGAAACTATACAACTAGATGTGATATGATTGGCATGCCCCtatagttacttgtcctaaac
gacttatcagtcttgtgtacTTTTTCATTTTGAAATAGAGTCTCCCTCTGTCACCCAGGCtatagttacttgtcctaaac
gacttatcagtcttgtgtacGTTCAAGCCATTCTAGTGCCTCAGCCATCCAAGTAGCTAGtatagttacttgtcctaaa
gacttatcagtcttgtgtacCCATGCCTGGCCTCACCTCAGTTTTAAAATCCTTTCTTGGtatagttacttgtcctaaac
gacttatcagtcttgtgtacTATGTTTTCTTCTAACTCTGTCCTTCATATAATCAGACCCtatagttacttgtcctaaac
gacttatcagtcttgtgtacAATTTTCTCACTGCCTTTGGATTGACTAACAATCTTCAACtatagttacttgtcctaaac
gacttatcagtcttgtgtacCAGAAAGGCTGAGGAATGAGTCATCACAAACATTTATTTTtatagttacttgtcctaaac
gacttatcagtcttgtgtacGCATTACCACTGCCCCTAGAGTTTTACCACATAAGATAGAtatagttacttgtcctaaac
gacttatcagtcttgtgtacACACCAATATCCAGCTTCTAAGAAGGCATAGACAGTAGAAtatagttacttgtcctaaa
gacttatcagtcttgtgtacATAATTTACACCTAATGTGATATAACAGTGGTCAGGGAACtatagttacttgtcctaaac
gacttatcagtcttgtgtacTTCTCACCCCATGCCCACCCTCTCACCCTGTTGAATCTCCtatagttacttgtcctaaa
gacttatcagtcttgtgtacCATGAGTACACATTATTTAGATCCCCCTTTTAAGTGACAAtatagttacttgtcctaaac
gacttatcagtcttgtgtacCACTTCCATCTATGTTGCTGTAGAATATATGGCTTCATTAtatagttacttgtcctaaac
gacttatcagtcttgtgtacATCAGTTGACAGACACTTAGATCAATTCCAAACCTTTGCTtatagttacttgtcctaaac
gacttatcagtcttgtgtacTTGATATTGTGGATTTCTTGTCCTTTGTGTAGATACTCAGtatagttacttgtcctaaac
gacttatcagtcttgtgtacGGATTGCCGGATCAAATGATAATTGCATTTTCAGTTCTTTtatagttacttgtcctaaac

gacttatcagtcttgtgtacGTACTAATTTACCTTTCTGCCAACAGTATATAATGTTCCCtatagttacttgtcctaaac
gacttatcagtcttgtgtacAAAAGCCATTTTGAGTGATGTAAGATGTTGTCTCATTGTGtatagttacttgtcctaaac
gacttatcagtcttgtgtacGGGTAACCTGTTTTTTGTATTATTGTTGATTTGGCTGAGTtatagttacttgtcctaaac
gacttatcagtcttgtgtacCTGGATATTAGTCCCTTGTAAGATGCAAATCTTTTCTGTTtatagttacttgtcctaaac
gacttatcagtcttgtgtacGGCATGAGCCACCGCACTGGCCTATTTTTTTTTTTTTTTTtatagttacttgtcctaaac
gacttatcagtcttgtgtacCACTGCAAGTTCTGCCTCCCAAGTTGGAGTGCAGTGGCACtatagttacttgtcctaaa
gacttatcagtcttgtgtacAGCCAGGATGGTCTCCATTTACTGACCTCATGATCCGCTGtatagttacttgtcctaaa
gacttatcagtcttgtgtacAGCTACTGTGCCTGGCCCAAGTACATTATCATTAACTGTAtatagttacttgtcctaaac
gacttatcagtcttgtgtacCCAACATCTCCCCAATCTCCTCCCTTTCTCCTTCCACCTAtatagttacttgtcctaaa
gacttatcagtcttgtgtacGTGAGATCATGCATTATTTATCTCTCTGTGACTGGATTATtatagttacttgtcctaaac
gacttatcagtcttgtgtacAATGATTTCCTCCAGGTTTATTTATGTTGTCTCAAGTGACtatagttacttgtcctaaac
gacttatcagtcttgtgtacATTTTATTTACTTTGGATATGCACCCAGATGTGGGATTACtatagttacttgtcctaaac
gacttatcagtcttgtgtacTTCTCCTGTCTTCCTTCTCTGATCCCTAAAATAGTGACTGtatagttacttgtcctaaac
gacttatcagtcttgtgtacAACATTATAAAAGGTTGACTGAGAATGATGACTGTCCTACtatagttacttgtcctaaac
gacttatcagtcttgtgtacCAGAGGGAGTGTACTAAATGTTTAACAAACAAGAAAGTGTtatagttacttgtcctaaac
gacttatcagtcttgtgtacAATTGGCAATTCTTTGTCACTTGGTGTAGTGAACAGTTACtatagttacttgtcctaaac
gacttatcagtcttgtgtacCCTTCCATAGTTTGTAGGCTTCTGTCTTGATCTGTATACTtatagttacttgtcctaaac
gacttatcagtcttgtgtacGACTGTCCTCAAAATACTAGAGCTGCTTTGTCTGAAGTTGtatagttacttgtcctaaac
gacttatcagtcttgtgtacGCAAATGCCTAGGAGCTTATGACCCCCTTCATGTTGCCCCtatagttacttgtcctaaa
gacttatcagtcttgtgtacCCCTGAGTAGAGATGAACCCTAAGGCATAATTTATACTACtatagttacttgtcctaaac
gacttatcagtcttgtgtacCCTTGCTTGGATTGCTGCCTTTCACTGTCCTGTTACCCAAtatagttacttgtcctaaac
gacttatcagtcttgtgtacTTGTACACAAGTCCTGCCTCAGTCCTATTACCCAACTCCCtatagttacttgtcctaaa
gacttatcagtcttgtgtacCTCTTATAAATCACTCATACACAAATCCTGCCTCGGAGACtatagttacttgtcctaaa
gacttatcagtcttgtgtacTGAGAAACGCTAAGACATGAGTGAATATTCAGCTAAAGAAtatagttacttgtcctaaac
gacttatcagtcttgtgtacATTTCTACTTGGTCTTGATCTTATTTCTCCTTTTTCCCCCtatagttacttgtcctaaac
gacttatcagtcttgtgtacCTTTTTTTCTTTCTTACACTGCTCTGCCTCCTTCATTGATtatagttacttgtcctaaac
gacttatcagtcttgtgtacTGGGCCACACACCATTCATGACCTATGCCATTTTGGGTTGtatagttacttgtcctaaa
gacttatcagtcttgtgtacTGTTTCTCTGGCAGGAAGAGTGCAGCTTCTGTGTTGAGTTtatagttacttgtcctaaac
gacttatcagtcttgtgtacGGTAGAAGCCATGAATTGTATAGCACTCCCTGTTTTATACtatagttacttgtcctaaac
gacttatcagtcttgtgtacGCTGCTTGGGCTGAGTAGTATCTCTGGGTTAAAAATTTTTtatagttacttgtcctaaac
gacttatcagtcttgtgtacGCACAGCTCAATAAATGTAGTAAAATAAACGGAGTTGTACtatagttacttgtcctaaac
gacttatcagtcttgtgtacGAAATGAGATCCTGCTCCTCTATGGTTATTACACGCCTTTtatagttacttgtcctaaac
gacttatcagtcttgtgtacATTGCATAGTGGAGGATTAATATTATCCAAAGAGAGGTCTtatagttacttgtcctaaac
gacttatcagtcttgtgtacCCCTTGGTTACTTGAAGCTGCTCTCTAGGCCTCTGACGTGtatagttacttgtcctaaa
gacttatcagtcttgtgtacTGAGAGCAGGGTCTTTGCTTGCTTGGGGATTTTGGCCACCtatagttacttgtcctaaa
gacttatcagtcttgtgtacTCAATTCTGATCTCTGGAAAAGCTGGAAACTAAAGGTATTtatagttacttgtcctaaac
gacttatcagtcttgtgtacTTGCAAACAGTAACTAATCAAGCCCCAGTAAAAATTTTGGtatagttacttgtcctaaac
gacttatcagtcttgtgtacCCTCTTTGGTTGGCAATTCTCTACGTATGTTGTGACACATtatagttacttgtcctaaac
gacttatcagtcttgtgtacGAATCCACCTCATAACAATTGTCTAAATTTCTGGAAGATGtatagttacttgtcctaaac
gacttatcagtcttgtgtacTAGTCAGAGTTGAAGACCACCGCTCTAAATTGGTTCAGGAtatagttacttgtcctaaa
gacttatcagtcttgtgtacCCCAGTTAAGTTTGGATGGTTTAAATTTCACTAAAGAACCtatagttacttgtcctaaac
gacttatcagtcttgtgtacGTGAGATGGAGGAGGTGCACACTTAAGGTCAATTCACTAAtatagttacttgtcctaaa
gacttatcagtcttgtgtacGATTTCTGATCACATGGAAGTTTGGAAAAGAAAGCTTATCtatagttacttgtcctaaac
gacttatcagtcttgtgtacTGAATGGATAGAGTTTTAATTTTCACTTTGGGGAGTAGCAtatagttacttgtcctaaac
gacttatcagtcttgtgtacTGAGTCATAACAAGACATCTTAAATACACTTACCTTTGGCtatagttacttgtcctaaac
gacttatcagtcttgtgtacTCTGGTTGGTCTCTACTTGAAATATTGACTGTAATTAAGCtatagttacttgtcctaaac
gacttatcagtcttgtgtacAAGTAACTCTTTCTCTCACCCTTGTTCCACAGACCCATGTtatagttacttgtcctaaac
gacttatcagtcttgtgtacAACAATCACTCTTACTTGTGTCTGGTACATTCTTCAAGAGtatagttacttgtcctaaac
gacttatcagtcttgtgtacTACAAACTTAAGTATGCAATCACATTGTTCATCACCTTGCtatagttacttgtcctaaac
gacttatcagtcttgtgtacGGTATAAATATGTGTGTATTGCCTTATTTTCAACTGCTGCtatagttacttgtcctaaac
gacttatcagtcttgtgtacCTGTAAGATAAATTTCTATAGGTAACGTTGCTGGGTCAAAtatagttacttgtcctaaac
gacttatcagtcttgtgtacTTCAAAAAGCTTATAATCAGGAAGTCTGTCTTATTGAGGCtatagttacttgtcctaaac

gacttatcagtcttgtgtacTTTAAGTTGAAATTGTGTAGGCTAGTTGTATCCAACAAGGtatagttacttgtcctaaac
gacttatcagtcttgtgtacACTCGATGTTTTTCAGATCTAGTTCCACATAACCAAGACTtatagttacttgtcctaaac
gacttatcagtcttgtgtacATTGGAGAAAATAATTAGTTAGTGATTGGAAACGTGGCTGtatagttacttgtcctaaac
gacttatcagtcttgtgtacCTCACCAAGAATTGAGGACCATTTATTTCCAAGCTCTGTGtatagttacttgtcctaaac
gacttatcagtcttgtgtacAGAACCCTCTGATAGACTTTTAGCTTTCAATTGCACAAGAtatagttacttgtcctaaac
gacttatcagtcttgtgtacTTTTATGGTGTTAAAATGCTATGGTTCTCTCGCTAGTAGAtatagttacttgtcctaaac
gacttatcagtcttgtgtacCAAGAATGGAGTTGACTGTGCACTAACCAGCATTTTACTAtatagttacttgtcctaaac
gacttatcagtcttgtgtacTCCTATCTCTTCACTGGAATAATCATGAGAGCCACCATGCtatagttacttgtcctaaac
gacttatcagtcttgtgtacTGTCCTCATAGTCACACAGTGGCTTTCATTATTATTTTCAtatagttacttgtcctaaac
gacttatcagtcttgtgtacATTTTCTTGTGTGGTGATGCTTTGTCAGACTGCCAGTTGAtatagttacttgtcctaaac
gacttatcagtcttgtgtacTCTAAAGGAAGTCAGGGAGTTAAACAAAGAAATACGATACtatagttacttgtcctaaac
gacttatcagtcttgtgtacATGGCTGAAGAATTCGAGAGTGCTATAGTTATGTATAAGAtatagttacttgtcctaaac
gacttatcagtcttgtgtacTCTAGTTAGTAAAATTAAAAGGTACAAACCCTATCCCTGCtatagttacttgtcctaaac
gacttatcagtcttgtgtacCAAATGCCTGCAAACCTTGAGACTTCCAAGATCTGCTGTTtatagttacttgtcctaaac
gacttatcagtcttgtgtacACAATTAGCAGATGTTGTGGACTGATTGAAAAATTACCACtatagttacttgtcctaaac
gacttatcagtcttgtgtacGTTCTTCTCACCAAAAGACGGACTATTTTCCCCCTAATTCtatagttacttgtcctaaac
gacttatcagtcttgtgtacCATCGTGGTAGAAGAAAAGATGTGCCAGTCCCAAGCCCATtatagttacttgtcctaaa
gacttatcagtcttgtgtacTTGAATCTGTGCCTATACTATGAGATCAAGCCTGGCCTCCtatagttacttgtcctaaac
gacttatcagtcttgtgtacGGTCAACTCAGTTGTCCCAGACAAGGCTCTTAATTATTTTtatagttacttgtcctaaac
gacttatcagtcttgtgtacGAGCAGTGTGGTCTTATACTCATGGGAAATAATAAATGATtatagttacttgtcctaaac
gacttatcagtcttgtgtacAAAATGGTTTGTTATGCAGCCACATGTAACATCAGAGGATtatagttacttgtcctaaac
gacttatcagtcttgtgtacTTCTTTAATTTATAGTTCCCCTCCTTTATACACATGCACGtatagttacttgtcctaaac
gacttatcagtcttgtgtacACTTGGTTGGTTGTCTGGTAGAGTTTCCCATAGTCTGTATtatagttacttgtcctaaac
gacttatcagtcttgtgtacCCTGTGGAATTGATAACATGTTTCTCTGCACCCTCTATTTtatagttacttgtcctaaac
gacttatcagtcttgtgtacGTTAGTTTTCTTCTCTTGGCTTTAGTGTGCACAAAAATACtatagttacttgtcctaaac
gacttatcagtcttgtgtacTTACTGGATAAATTAGTGTTTTCCATCTGAAAGTCTGCTGtatagttacttgtcctaaac
gacttatcagtcttgtgtacAGTATAATTATTATCCTTAGCTTGAGCTCCATGATGAGGGtatagttacttgtcctaaac
gacttatcagtcttgtgtacCTCAGGGCTTGAATGTTCACTTATTTTGTATACTCTTCTTtatagttacttgtcctaaac
gacttatcagtcttgtgtacAGAGTTTGCTACCATTACTACCCTTTGTGTCTATCCCCAAtatagttacttgtcctaaac
gacttatcagtcttgtgtacAGGCTTGACACTGTTAAGGTAGCTGGTTGTGAACGTAATCtatagttacttgtcctaaac
gacttatcagtcttgtgtacGCTTGAAAAATCCTTGTAACTCCACAGCTTCCCCCTCATTtatagttacttgtcctaaac
gacttatcagtcttgtgtacACTGCCATGAGTGCGCTTAAATTCTATAATGTAAACTTACtatagttacttgtcctaaac
gacttatcagtcttgtgtacGCATTTTTATATGGTTGGCTATCCTGAAATATTGGAAGACtatagttacttgtcctaaac
gacttatcagtcttgtgtacAGGGGCCAGTTGCTACCAAGTTTGGAGTACTAACAGGAGTtatagttacttgtcctaaa
gacttatcagtcttgtgtacCTTTAAGCTCCTTCAAATAATTTATGGATACCATCAGCTGtatagttacttgtcctaaac
gacttatcagtcttgtgtacATGACTTGTGCTTTGAATCTAGTTATCTTTTACTTTGCCCtatagttacttgtcctaaac
gacttatcagtcttgtgtacCAGTTATTTTGTGCTAGAACAATTTTTAGAGTTCTCCCAGtatagttacttgtcctaaac
gacttatcagtcttgtgtacTATGAAAGATATTGTAGAGAGTCTTAGTTTGGGTTCTTCCtatagttacttgtcctaaac
gacttatcagtcttgtgtacGTACCCCGGGATGAAGATTCAAATGCAAGTGGTTTATTCCtatagttacttgtcctaaac
gacttatcagtcttgtgtacGGTGGGTGGAGCTTAATTCTATGGGAAAATCTGAGAAAAGtatagttacttgtcctaaac
gacttatcagtcttgtgtacATCTCAGAAGTAATCCATTTAAAGGGCTAAGAAACTGAAGtatagttacttgtcctaaac
gacttatcagtcttgtgtacAATGGCTGCTGTGGAAGCGAGGCAGGTGTTAATTTTGCATtatagttacttgtcctaaac
gacttatcagtcttgtgtacGGAGAAAATGTCCTGAGGCACAAAATGAAGTTACTGGCAAtatagttacttgtcctaaa
gacttatcagtcttgtgtacGTCAGGGAACATTGGCAGCTACAAGGCATACGGGTGGAGGtatagttacttgtcctaa
gacttatcagtcttgtgtacCATATTATTTCATTAGATCCTTACAGTTCAAGGAGGAAGGtatagttacttgtcctaaac
gacttatcagtcttgtgtacCCAAGGTTATATACTAATAAGGGGCAGAGTCACAATTTAGtatagttacttgtcctaaac
gacttatcagtcttgtgtacAGTAATGTCTTCCCAAATGTCAAACAGCTGCTATTTAACAtatagttacttgtcctaaac
gacttatcagtcttgtgtacGTCCCCTGACGTGCAGCCTGGTGACCTTTCTGTAGACCGAtatagttacttgtcctaaa
gacttatcagtcttgtgtacTCATTCTCATTCCCTGTACAGCCCCACTTCCATCTCAGTGtatagttacttgtcctaaac
gacttatcagtcttgtgtacGAGTTTGATCCTTGAGCTTAGTTCTTACATGTCTTCCTTTtatagttacttgtcctaaac
gacttatcagtcttgtgtacCTCTGTAAGTTCCCCCTTCCAACTTGTGAGATGCTTATTAtatagttacttgtcctaaac
gacttatcagtcttgtgtacGTTTATTGATAGCTCTGTTGCCTTGAAATTGCTTTGTTTGtatagttacttgtcctaaac

gacttatcagtcttgtgtacGCATCTTTTTACATGTTTCTCCCTCTTTTTGAATGCACTTtatagttacttgtcctaaac
gacttatcagtcttgtgtacCCTTCTTATCATAGGTATCATTTCCCCAGGAATATCTCCCtatagttacttgtcctaaac
gacttatcagtcttgtgtacACATAAAATTGAAGCACCAAGAAGATCTTCTCTATAGCACtatagttacttgtcctaaac
gacttatcagtcttgtgtacTTCATTAAAGCCTGAAAACTATGATAAGAGGGACCATGTCtatagttacttgtcctaaac
gacttatcagtcttgtgtacTCACTGCATCCTCCTCACCTAAAACATAGAAAATCCTCTGtatagttacttgtcctaaac
gacttatcagtcttgtgtacTGGATGAATAAACAAATGCATTTTAGTCAAAGCACCGTCAtatagttacttgtcctaaac
gacttatcagtcttgtgtacTTATATAGATAAATTACGTGTTGCAGGGTTTGGGTGTACAtatagttacttgtcctaaac
gacttatcagtcttgtgtacTTTAGTCGCCCAGGTAACATAAGCATAGTACAGTACCCGAtatagttacttgtcctaaa
gacttatcagtcttgtgtacCTACTGTTCTCTTCTTTGTGCTCGTATGTACTCAATGTTTtatagttacttgtcctaaac
gacttatcagtcttgtgtacGGATAATGGCCTCCAGCTTGTTCCATGTTGCTGCAAAGGAtatagttacttgtcctaaa
gacttatcagtcttgtgtacCCATGTTTTTGCTGTTGTGAGTAATGTTGCGATGAATATTtatagttacttgtcctaaac
gacttatcagtcttgtgtacGCTGGGTCAAATGGTAGTCCCAAGTTCTTTCAGGAATTGCtatagttacttgtcctaaac
gacttatcagtcttgtgtacGCCTTCCCGCCAGCAGTGTATTCTTTTTTCTCCGCAACCTtatagttacttgtcctaaac
gacttatcagtcttgtgtacTTTAATAACCGCCATTCTGATTGATGTGGGATGATATCTCtatagttacttgtcctaaac
gacttatcagtcttgtgtacACTTAACTCTTGGTCACAGCCTCCGTCTAGCACTTAAAAAtatagttacttgtcctaaac
gacttatcagtcttgtgtacCTAAAAGCAACAAACCCAGGTGAGTCTGTAAACATTTAAAtatagttacttgtcctaaac
gacttatcagtcttgtgtacCATACACACGAGGAAAGAATATGATCATGCCATACACATCtatagttacttgtcctaaa
gacttatcagtcttgtgtacAGAGGTTAGAAAGTTACTCTCTAGTACATCTCCGCACTCTtatagttacttgtcctaaac
gacttatcagtcttgtgtacGGACTCTTCCCAAGCCTGTTTTTGCCACATGCACTTGCTAtatagttacttgtcctaaac
gacttatcagtcttgtgtacATGTAAGCCATTGAAATATGAGCTCAAAAGGAGACTGTGGtatagttacttgtcctaaac
gacttatcagtcttgtgtacGCTGAGTTAATCATGAATGAGATGATTGTAAAAGATGGAGtatagttacttgtcctaaac
gacttatcagtcttgtgtacAGATACTGAGATGATGGAATGATGCACTTTGTTGTAGTTTtatagttacttgtcctaaac
gacttatcagtcttgtgtacTGAAGCCAGATTTCATTGAAGAGATGCACATTTTGTTTTCtatagttacttgtcctaaac
gacttatcagtcttgtgtacTTATGGATTAGGTATAGTAATGCTTTTCAAGTGTCTTGCCtatagttacttgtcctaaac
gacttatcagtcttgtgtacAGTAAGTGATTAGTCTATTTTGCCTCCCATTCTGTGTGTGtatagttacttgtcctaaac
gacttatcagtcttgtgtacGGTGCATAAATGTCAGTTGTTGTTCCAAGGGGCCAGGGAGtatagttacttgtcctaaa
gacttatcagtcttgtgtacGTTTGGAAAGTCTGGAGATGAAACTACACAAGGGAAGAAGtatagttacttgtcctaaa
gacttatcagtcttgtgtacCAGGGCTGCAGAAGGTTGATTTTTCCTTTAATTGAGGGGCtatagttacttgtcctaaac
gacttatcagtcttgtgtacTGATAGAAGATGGGCATGATCAGGTTCATCAGAAGAAGCAtatagttacttgtcctaaa
gacttatcagtcttgtgtacCCACAACGGTTTTTATTTGGAGGGTAAAGTACAACACACAtatagttacttgtcctaaac
gacttatcagtcttgtgtacCAAAAAACAAAAAACAGGGGAAAAAGCTAGTGATGCATCAtatagttacttgtcctaaa
gacttatcagtcttgtgtacAGTAAACTTAACAAATTCCTATGGGTCAAAGGCTCGTTCCtatagttacttgtcctaaac
gacttatcagtcttgtgtacGTTTCACTTTTTGCAAACTACTTATTATTTCCGGAGTCTCtatagttacttgtcctaaac
gacttatcagtcttgtgtacTCCTATCCTCTCTTCCTGCTAAGAGCATTTAAGCACTCAGtatagttacttgtcctaaac
gacttatcagtcttgtgtacTTCACAAAGTGTAACCTCTGGAGTATGGCAAGCAGGCTGCtatagttacttgtcctaaa
gacttatcagtcttgtgtacGCCTATTATGCAACATCAACCATTTAAAAAACTGCCAACAtatagttacttgtcctaaac
gacttatcagtcttgtgtacTTCAATGGTCATGCCTGGCTTGTTGGTGCAGTGAAGAGGTtatagttacttgtcctaaac
gacttatcagtcttgtgtacCATGGGGTGGGAGGAGTAAGATAGGGGAGATGTATTAGGCtatagttacttgtcctaaa
gacttatcagtcttgtgtacGTTCCTTTTGTGTCTGTTGTCTTAGAAATTTCTTGAGTACtatagttacttgtcctaaac
gacttatcagtcttgtgtacTGAAATCAGACTTATAGTGGAAAGCAATGAGAAAAACTGGtatagttacttgtcctaaac
gacttatcagtcttgtgtacGAAAAAAACACACCCAACTGAATTCTTCTTCTTTACCAACtatagttacttgtcctaaac
gacttatcagtcttgtgtacCATTTTCCTCTCTCTTTCATTTGATAATTTTTGCCTCAGCtatagttacttgtcctaaac
gacttatcagtcttgtgtacCAGGAGCTTGTTTTACTTACCCAGGTATAGCCCTCATCCAtatagttacttgtcctaaac
gacttatcagtcttgtgtacGTAATTCTAAAGAGGAAAAACTTGACCCACACAGGAGTTTtatagttacttgtcctaaac
gacttatcagtcttgtgtacGTGTGTAAATTAGTTGTTTAGAATACATACCACTCTGGACtatagttacttgtcctaaac
gacttatcagtcttgtgtacCAAATATAGGCTTTGAGAATATCTGCTTCATGTCTGTGGCtatagttacttgtcctaaac
gacttatcagtcttgtgtacTGGTATAGGTTAATTTAGTTCTATGTTTGGCAGCAAAAGCtatagttacttgtcctaaac
gacttatcagtcttgtgtacCAGAGCCATAAATTCCGTCACCTGCTATCAAGAGCCACGCtatagttacttgtcctaaa
gacttatcagtcttgtgtacAGGAGGAGTCAGGAGAAGAGAGAATGAGATAATTTTATTTtatagttacttgtcctaaac
gacttatcagtcttgtgtacAAGTATCTTTGGCACCTTCTCCTTTCCTTTTGTTTTTGCAtatagttacttgtcctaaac
gacttatcagtcttgtgtacTTGCCTAATTTCCCATTGCTTTCGTTTATCTGTGCATTCAtatagttacttgtcctaaac
gacttatcagtcttgtgtacTCTTTTCACATTCCAAAATTATTACTCCCCTCACCATTTCtatagttacttgtcctaaac

gacttatcagtcttgtgtacCGAGCCCTCCACGTATAGCAGAGAACTTCCATCAACCCTGtatagttacttgtcctaa
gacttatcagtcttgtgtacGCTTAACCCATATGCATCTCTGACTGGTGGAATTTCAAACtatagttacttgtcctaaa
gacttatcagtcttgtgtacAAATAGCGTTTGTGTGTTTGTGTCTGTATTATTCTGGGCAtatagttacttgtcctaaac
gacttatcagtcttgtgtacAATGTTGTCCCCAAATTCATTTTCTTAAGGTGCGTCTAGTtatagttacttgtcctaaac
gacttatcagtcttgtgtacATTATAGATTCTCCAAATTCTCCAAGGCTTTTTTCACGAGtatagttacttgtcctaaac
gacttatcagtcttgtgtacGAGAGAATTGACTGTCTGAGGATGGGAGAACTGGCATTGGtatagttacttgtcctaaa
gacttatcagtcttgtgtacCGCCTCTCAGGTTCAAGCTATTACCCTGCCTTAGCCTCCCtatagttacttgtcctaa
gacttatcagtcttgtgtacCACCGCGCCTGGCCATGTGTGAATATTTCTATGCCCAATGtatagttacttgtcctaaa
gacttatcagtcttgtgtacCTCTTTGACAAACTTTTCTTGATGCTTTTTTCCTGTTAGCtatagttacttgtcctaaac
gacttatcagtcttgtgtacTCATTTTGTTTCACTGTATATTCTGAGAAGCCTTATTCCCtatagttacttgtcctaaac
gacttatcagtcttgtgtacCCTACTCACCACTTCTAGGTGACCTTTTTGTTGTTATTTTtatagttacttgtcctaaac
gacttatcagtcttgtgtacCTTCTGTGCTTTTTTCATGTAACGACATATCCTCTAAATCtatagttacttgtcctaaac
gacttatcagtcttgtgtacTGATTTCATAGTGATCTTTCTCAATCTGTTTTAGCAGCTGtatagttacttgtcctaaac
gacttatcagtcttgtgtacTCAAGTTTATTTAATTGGCTTTCTATGGATGGGCATTGAGtatagttacttgtcctaaac
gacttatcagtcttgtgtacACTGCTGAGTCAAATAATAAATGTATACCTTACCGTTTGCtatagttacttgtcctaaac
gacttatcagtcttgtgtacCCTAAGGTTGCATTGTGTCTTTTATATCTGAGGATTCCTGtatagttacttgtcctaaac
gacttatcagtcttgtgtacTTGTACTGCATTTTGGGTAAGTTCATTAGGACTATCTTTCtatagttacttgtcctaaac
gacttatcagtcttgtgtacTTCCATATCATCTATTGCAAAAGTTTTTCAGTGGCTATCCtatagttacttgtcctaaac
gacttatcagtcttgtgtacTCGGATTCATAAAATATCTATTTTAAGCCTGAGCATGGAGtatagttacttgtcctaaac
gacttatcagtcttgtgtacCCAAGGTGGGATGATCATTCGAGGGCAGGAATTCAAGGCCtatagttacttgtcctaa
gacttatcagtcttgtgtacGCAATATGATGAGGCCCCATCTACCTGCACACACACACACtatagttacttgtcctaa
gacttatcagtcttgtgtacTGCACCTGTAATCCTTGCTATCTTGGAGGCTGAGGTGGGAtatagttacttgtcctaaa
gacttatcagtcttgtgtacGTGAACTATAATTGTGCCACCAGCTCTCCAGCTTCTGCTCtatagttacttgtcctaaa
gacttatcagtcttgtgtacGACATAGTCTTTCATTTCACGAGTAATTTGGCTGTTTTCTtatagttacttgtcctaaac
gacttatcagtcttgtgtacTTTCTGTGTTTTGAAATATCTGTGGTAGCTATTCCTGAGTtatagttacttgtcctaaac
gacttatcagtcttgtgtacTTTTACTGTTATTTGCTTTATTTCCTCATTCCCACCCATCtatagttacttgtcctaaac
gacttatcagtcttgtgtacCATGTCTTATGTTGCAAATTGGGGCTTTATTATGAGAGGAtatagttacttgtcctaaac
gacttatcagtcttgtgtacGGAACCCATCTTTGAGCCAACATGCAGCAAATTACTCAACtatagttacttgtcctaaa
gacttatcagtcttgtgtacTACCTCATACCTCATACATGAATATACATGTACCTCATAGtatagttacttgtcctaaac
gacttatcagtcttgtgtacTCATCATCATCGTCACAATCACTGGTATATTGCTGGCATAtatagttacttgtcctaaac
gacttatcagtcttgtgtacCAATTTTTTGTACTTCTGGTGTCTTTCTAACTTTCAGGCCtatagttacttgtcctaaac
gacttatcagtcttgtgtacCAATATACTCTCTTTTTACCTTTGTGAATCTTTCCCAGTGtatagttacttgtcctaaac
gacttatcagtcttgtgtacAGGGCTCATCTCAGAACACTATTCCCTTCCCTTCCATAAGtatagttacttgtcctaaa
gacttatcagtcttgtgtacGATTCTACTGAGTCAAAACTTCCTTACCTTCTGAGCCAGCtatagttacttgtcctaaa
gacttatcagtcttgtgtacCCAATGATGTTAGTTAGAGATGGGTTTTTGGTTGGCTTCAtatagttacttgtcctaaac
gacttatcagtcttgtgtacCATCAAGGCAATTAGGAATCAGAAGCTCAAAAATTAAGAGtatagttacttgtcctaaa
gacttatcagtcttgtgtacCCTTACTAATAATGCTAGGCTGAGGAAACAAGGTACTGTTtatagttacttgtcctaaa
gacttatcagtcttgtgtacAGTCTTCTGGAGTTCACATTAAACGCTTCCACCTTTCTATtatagttacttgtcctaaac
gacttatcagtcttgtgtacTTGACTATTATATTAACATGGATCTTGGACCAAAGAAGGGtatagttacttgtcctaaac
gacttatcagtcttgtgtacCCAAGGATAATGATTTCATTGACAAGAAAGATAACCTGTCtatagttacttgtcctaaa
gacttatcagtcttgtgtacCTGTGAAGTCTTAGAATGAGTGCATAAGATGAGAAGCTGGtatagttacttgtcctaaa
gacttatcagtcttgtgtacATAGTTTATGATACAGATCTATCCCAGTCACAGAGAAGTTtatagttacttgtcctaaac
gacttatcagtcttgtgtacTGCTGCTCACATCTGGTCAGTCTCTGGAATTTTTGTCTTCtatagttacttgtcctaaac
gacttatcagtcttgtgtacCTTTCTTTTCCTGGCATTGGCTCTCAAAGACTAAAGTTGGtatagttacttgtcctaaac
gacttatcagtcttgtgtacTCTTGTGTGTTGAGGCAGTGTTTCTTCTTCTTAAGTCATAtatagttacttgtcctaaac
gacttatcagtcttgtgtacTCGTGGCCAGATTATACCAGATGGTGTGAGAAAAAGAGGAtatagttacttgtcctaaa
gacttatcagtcttgtgtacAGTGGCAAATGGGGAGACTGAAGAAGATACATCCAGGGGAtatagttacttgtcctaa
gacttatcagtcttgtgtacCTGAGTCACACAGCTTTTCCCTTGGTTACAATGTCATTGGtatagttacttgtcctaaa
gacttatcagtcttgtgtacAACAGCGTTTTGCCTATTCTTTAAATTTATGGCATAGGGTtatagttacttgtcctaaac
gacttatcagtcttgtgtacGCAGCTGAAATGCTTTTATAGAAAATATCTGATAGGCTGTtatagttacttgtcctaaac
gacttatcagtcttgtgtacTCAGAATTTAATTTGGCTTGTTCAACAGTTTGCTTTCCAGtatagttacttgtcctaaac
gacttatcagtcttgtgtacAGGCATTAAGGGAGGAGATTATGTATAAATCAGTATGACAtatagttacttgtcctaaa

gacttatcagtcttgtgtacTGTGAAGAAGTATCATGAAGAATGCTTGCAAAAGTTAGGTtatagttacttgtcctaaac
gacttatcagtcttgtgtacCCCTAATTTGTAGCTAATTTTAAGGTTTCCAAGACCAAGGtatagttacttgtcctaaac
gacttatcagtcttgtgtacATGATACAAGTCCAACATCCATATTTAAATGTGAGTACGCtatagttacttgtcctaaac
gacttatcagtcttgtgtacTGATCACTCAAACAGAATTGACCAACTTTTGCAAACTTTCtatagttacttgtcctaaac
gacttatcagtcttgtgtacCTATTGAAATTAAGTAACCCTCCTCTAAATACTCGGAATCtatagttacttgtcctaaac
gacttatcagtcttgtgtacCTCATTCAAGAGAGAGAACCAAAACAATTTCCTCTGGCAGtatagttacttgtcctaaa
gacttatcagtcttgtgtacCTTTAGCTTGTCAACTGGAGAACCTGATGTCTTTGCCAAAtatagttacttgtcctaaac
gacttatcagtcttgtgtacGGCTTTTCTTTCCATTTCTAACTTCATTCATTTCATGCTCtatagttacttgtcctaaac
gacttatcagtcttgtgtacACCAAGCCTAACTAAGACTCCATGACTTTCAATAGGACAGtatagttacttgtcctaaa
gacttatcagtcttgtgtacCTGTGAGCTCCCTATAAGTTCAAGGTAGTGTGGATGAGTGtatagttacttgtcctaaac
gacttatcagtcttgtgtacCATGTGTGGGGGTTTTGGAGGTGGGGTTTGGGCATGGGTAtatagttacttgtcctaaa
gacttatcagtcttgtgtacACAACAACAAAAAAACTCATCCTGTCAAAAGTGCATGTGCtatagttacttgtcctaaac
gacttatcagtcttgtgtacTATCACCAGGATTACACTGAGGTTTTATTTTAGAGTATCCtatagttacttgtcctaaac
gacttatcagtcttgtgtacAAATATAATTTGCCACAAGAAGCATGAAGTGAGTGACTAGtatagttacttgtcctaaac
gacttatcagtcttgtgtacCTAGCAAGGCCCAGCATGACAAATCAGTGTCTCTATGATAtatagttacttgtcctaaa
gacttatcagtcttgtgtacTGTGCTGCAGTTGAGTGGTAGAAGTAGGCTGGATGGCTCTtatagttacttgtcctaaa
gacttatcagtcttgtgtacATGCAAAGGCCACATATTCTAGATTGGTTCAACTGGTGTGtatagttacttgtcctaaac
gacttatcagtcttgtgtacAAAACCAGGCAGACACCATTCACTTCTTTCATTGCTCAGGtatagttacttgtcctaaac
gacttatcagtcttgtgtacCACTATTTTAACTCCTATGATTCTGTGCAAACTTTGGGCAtatagttacttgtcctaaac
gacttatcagtcttgtgtacAGAAAGTCTGCCTGGAGAAACAGCACTGGAGGGATATGAAtatagttacttgtcctaaa
gacttatcagtcttgtgtacGCTAGACTTGTGCCTTGAACTGTGATTCATACTTTACATGtatagttacttgtcctaaac
gacttatcagtcttgtgtacTTTGCAAATGAAGAAACAAACTTTGAGGAACTGAGTGTCTtatagttacttgtcctaaac
gacttatcagtcttgtgtacAGGAAAATTAACCTTCACTAAGCCTCTACTACGTGACTCCtatagttacttgtcctaaac
gacttatcagtcttgtgtacTGCCTGGTGTACACATACATGATTTCTCGGGTTAGTTTTTtatagttacttgtcctaaac
gacttatcagtcttgtgtacCCTTGGAAGTTGGTATTAGGATCATGTTTTATAGAGGAAGtatagttacttgtcctaaac
gacttatcagtcttgtgtacGAGTCTCACAGATGATAAACTAACCCAAGCTGAAAGAAGCtatagttacttgtcctaaa
gacttatcagtcttgtgtacAACTTTTTAATCCAACTCTGCCTGAAAGCAAAGCCCATGGtatagttacttgtcctaaac
gacttatcagtcttgtgtacTCATAAGCTGAAGGATGAACTGATTTTCAGAGAGAGACCTtatagttacttgtcctaaac
gacttatcagtcttgtgtacAGGGTGTCCTGGGAATATCTGTTTTCCCTTAGAAAATAACtatagttacttgtcctaaac
gacttatcagtcttgtgtacTTTCTAGTCTTTTCTTTAGGACTGCCTCACTTGTATCAGGtatagttacttgtcctaaac
gacttatcagtcttgtgtacTTACAAAAACCAACTCTGTTAAATTTCCCTTCAGGTCACCtatagttacttgtcctaaac
gacttatcagtcttgtgtacGCTAATTTTTGCACACTCCCTGACCACCTTCAATTCTTTCtatagttacttgtcctaaac
gacttatcagtcttgtgtacACTCACAAATTTAGATAGCTTATATGCACACCGTTCATCTtatagttacttgtcctaaac
gacttatcagtcttgtgtacCAAAGCTGAAGAGATGGAGTGTGTAGTTACCCTGGTTCCGtatagttacttgtcctaaa
gacttatcagtcttgtgtacCATTTGAAAGGCCTGGATATGATTCAGTAAGGATGAGTGCtatagttacttgtcctaaac
gacttatcagtcttgtgtacAGAAGGCTATCTGGGAACATTGCACATAAACACAATTCTCtatagttacttgtcctaaac
gacttatcagtcttgtgtacTCTTAGTGAGATTTGTTCTGGTATGGAGAGCTACAGGTTTtatagttacttgtcctaaac
gacttatcagtcttgtgtacGGCTTTTTTCCCCCTAGGCATCTTAAACAAATGTATCTATtatagttacttgtcctaaac
gacttatcagtcttgtgtacTGAACTTCATCATCAGCTGTCCCTTCACTCTGTTTTTACCtatagttacttgtcctaaac
gacttatcagtcttgtgtacCTTCAGCATTTAAAGACTAAGAACATCCCCTTTCAGCCCCtatagttacttgtcctaaac
gacttatcagtcttgtgtacGGACTTCTGAACTTCCTGGATTGGTTGAATTTGAATAACAtatagttacttgtcctaaac
gacttatcagtcttgtgtacATTTCCTCATCTGTAAAGAGATATCCTTACCCCCTTCTCAtatagttacttgtcctaaac
gacttatcagtcttgtgtacTGGAAACAGCTAGCAAAAGACTTAACATATATTACAGGCTtatagttacttgtcctaaac
gacttatcagtcttgtgtacAAACCTTCTCTTGTTGGATGAAAGCTGTGTGACCTTAGGCtatagttacttgtcctaaac
gacttatcagtcttgtgtacACTTGATAGGGTAGCTTTGAGGATTAAATGGCACAACATAtatagttacttgtcctaaac
gacttatcagtcttgtgtacCTTTTATTGTTACAAAGCCTGTGGTTGTGAAAGTGAAAGGtatagttacttgtcctaaac
gacttatcagtcttgtgtacCATGCATTCTTGGTTGCTCTGGGCAGAATCTGTTTATTCCtatagttacttgtcctaaac
gacttatcagtcttgtgtacATTCTGTGGTCACTCTAAATATAACTAATATGTAGGGAGGtatagttacttgtcctaaac
gacttatcagtcttgtgtacAGGAGACAGAGAGACCATAATAACGATTCCTTTCCCAGAAtatagttacttgtcctaaa
gacttatcagtcttgtgtacGTTCTTCCAACAGCGTAGTTTAGTTCTTACTAGTTAGCTAtatagttacttgtcctaaac
gacttatcagtcttgtgtacGGCAATTCAGAAACATCTAAATACACAGAAAATACATGGGtatagttacttgtcctaaac
gacttatcagtcttgtgtacGTTTTGATGAAGTGGTTTGTTCCCTTTTTCTGGTCATTTTtatagttacttgtcctaaac

gacttatcagtcttgtgtacCCCTCTGGCTGCCCCCAATCACTTTTCACTGTTAACTCTAtatagttacttgtcctaaac
gacttatcagtcttgtgtacCTGGCTGCCTGGACACCCTCTTGCACATACTGGTAATTTTtatagttacttgtcctaaac
gacttatcagtcttgtgtacGGCTCAGTTTCTCTTCACTCAGTCCTTTGGCCCACAAGCAtatagttacttgtcctaaac
gacttatcagtcttgtgtacGAGACTTCCTGATATGCCAGACAACCTTTGGATAGGGATAtatagttacttgtcctaaac
gacttatcagtcttgtgtacGGCTAACTTATTCCTCATGATCTCATTGAACTCTGCTGGCtatagttacttgtcctaaac
gacttatcagtcttgtgtacAGAAATGCATGGATCTGTGCCTTGAACTGGGAGTAAGGGCtatagttacttgtcctaaac
gacttatcagtcttgtgtacGAATATATTCCTGAGTCCTGCGTCTTCCTTTGGCACTGTCtatagttacttgtcctaaac
gacttatcagtcttgtgtacTCTCCTCTTCCTACTGCTCTTTTTCCTCCCACTACACCCCtatagttacttgtcctaaac
gacttatcagtcttgtgtacCGTCTATGGAGGAGTAAATGAGAAACAAAATGGTGGTCTTtatagttacttgtcctaaac
gacttatcagtcttgtgtacCTGAAAGACACATCCTAGGAGAGCTGGAATCATCAAGAGAtatagttacttgtcctaaac
gacttatcagtcttgtgtacCAAATCCTGACCCTCTCCCTTTCCTTTTTAGTTCACTTCAtatagttacttgtcctaaac
gacttatcagtcttgtgtacGCTTTTATTTCTGATGATTATTGTCTCTTACCTGGTATCCtatagttacttgtcctaaac
gacttatcagtcttgtgtacTGGAATGTAGATCATCTTTCTTAGCAGTCACTATGTGGAGtatagttacttgtcctaaac
gacttatcagtcttgtgtacCTGTTATTTCAGTGTTTGCCCCAAGTAAAGATTCTCAGATtatagttacttgtcctaaac
gacttatcagtcttgtgtacACATGGGTAACTGACTTTTCCTTTCATTTCCTCCAGCAGAtatagttacttgtcctaaac
gacttatcagtcttgtgtacGAAGGTTATCAAATGTTTTTCAGAAAAGGTCCAGACAATGtatagttacttgtcctaaac
gacttatcagtcttgtgtacACACCTCTGTTGCATAGATTCTTCCTCCATTCCCTCCCTCtatagttacttgtcctaaac
gacttatcagtcttgtgtacAAAAACATTCTTAGCTCACGTGCCAAACATAAGCATTCTAtatagttacttgtcctaaac
gacttatcagtcttgtgtacGCAGGACATATGCCTTCTCATTACCCTTCTTTTTCAAGGTtatagttacttgtcctaaac
gacttatcagtcttgtgtacTCCAAAACCCGTCATCATTTGTATTTCATGGAATCTATACtatagttacttgtcctaaac
gacttatcagtcttgtgtacGTAATTTATACTTTCTCTCCTCCTTTTTCCCACATGACGGtatagttacttgtcctaaac
gacttatcagtcttgtgtacCCTTCTCTCTCTTGACAATTAAGTTTCTGTATCAGGAGTTtatagttacttgtcctaaac
gacttatcagtcttgtgtacATCACTCTGTTCAGAAGGTCTTATGGGAAATGATCAATGCtatagttacttgtcctaaac
gacttatcagtcttgtgtacCAACATCAATAAACTCTGATTCCTCTGTTTTACTGGACTCtatagttacttgtcctaaac
gacttatcagtcttgtgtacCTATCCAATGGCTATTTTTTTTTTCTAGATGACTGGGTAGtatagttacttgtcctaaac
gacttatcagtcttgtgtacCCTGGGTGACTTTGGCATCTAGATACTTGCATACATCTGGtatagttacttgtcctaaac
gacttatcagtcttgtgtacATTAGATCCAAATTATTATCATTCCACCTCCTCTCACATCtatagttacttgtcctaaac
gacttatcagtcttgtgtacCCTAAGAAGTCAGGGAAGAATGTGAAGATTTGAGATCCAAtatagttacttgtcctaaac
gacttatcagtcttgtgtacTGACTACATACTCAGGAAAACACTTTTCTCTAAGCACTGGtatagttacttgtcctaaac
gacttatcagtcttgtgtacGATCCTACTCACCTCAGCCCATGCTTATGCTTGCCTTTCAtatagttacttgtcctaaac
gacttatcagtcttgtgtacTTGGCAATAATTCAAACACCCAAGCTACTGCAGCCACCACtatagttacttgtcctaaac
gacttatcagtcttgtgtacGCCAGTGCCTGATATTATAGCAACTACATATGAAATCCTTtatagttacttgtcctaaac
gacttatcagtcttgtgtacAGTCAACGAACTCACTGTCTGTCATTCCCAACCTTTTCCCtatagttacttgtcctaaac
gacttatcagtcttgtgtacTTACAGAAGTCCTTAAATATCAGCAGCACACTAAAAGTTCtatagttacttgtcctaaac
gacttatcagtcttgtgtacCACCAAATGAAATCCCATCTTTAACTCTAGCCCTCTCAACtatagttacttgtcctaaac
gacttatcagtcttgtgtacTCTGTGTTCAAATAACGTTGAGGGACAAATTTTCAGTGAAtatagttacttgtcctaaac
gacttatcagtcttgtgtacATCAGTACCCAACTAAAATGCATCAAAATTTGAGTACTCCtatagttacttgtcctaaac
gacttatcagtcttgtgtacGCATTCTACTAATTTAACTCTGCTGTAGGTTGAGCCCTGAtatagttacttgtcctaaac
gacttatcagtcttgtgtacTGTTTTCACAGCAAGATGAGCCATGTTCCAGCACTTTCCAtatagttacttgtcctaaac
gacttatcagtcttgtgtacAACATGTCATTTTAAAATCTTTGATTGCCTGCTCTATGGGtatagttacttgtcctaaac
gacttatcagtcttgtgtacGCTTATGAGACTCTATGTAGTGGGTCATCACACTCTGACAtatagttacttgtcctaaac
gacttatcagtcttgtgtacAGAAATGCTGGGGAGGTAGAAGAAAAATAAACAGGGAAAAtatagttacttgtcctaaac
gacttatcagtcttgtgtacGTTGTTTTAAGAAGCAGATTGTCAAAGTGGAGATTTGCCAtatagttacttgtcctaaac
gacttatcagtcttgtgtacGAAGCTGAATTTAGTTTGGTCTTCCCGGAGATAGTTATGGtatagttacttgtcctaaac
gacttatcagtcttgtgtacCTGGGGACTGTAGCTGCCATTGGAACTAGACACAACACTGtatagttacttgtcctaaac
gacttatcagtcttgtgtacCCTTGCAGTCTTTTTATTCTGAGGAGTTAAATTGGCACTTtatagttacttgtcctaaac
gacttatcagtcttgtgtacTCAAATGGATTATACATAATAAACAACCTTGGGACAGAGGtatagttacttgtcctaaac
gacttatcagtcttgtgtacCATGTATTCTGTCCCCAAATTCCCATGCTCTTTCTACCATtatagttacttgtcctaaac
gacttatcagtcttgtgtacCTCCAAATTTTAGTGAGAAGAAACCCAAGAGTACTAATGAtatagttacttgtcctaaac
gacttatcagtcttgtgtacAAGTAGGTCAAAATGAAGCTTCTATGATCACTGAATTGTGtatagttacttgtcctaaac
gacttatcagtcttgtgtacTTGAACTAGGTTTAATGAGGCTAAAGTGACCATGTAAATCtatagttacttgtcctaaac
gacttatcagtcttgtgtacCTGATCTGGGAAAGGTGCATTACTCTCTCGTAGCTGTCGTtatagttacttgtcctaaac

gacttatcagtcttgtgtacGAGAAGGCAAGGCTTTTTAGAGTATTAACAAAATCAATCGtatagttacttgtcctaaac
gacttatcagtcttgtgtacAGAATAGTTGGATTGAGTTTCTTAAGCACCTCAGGGTTGCtatagttacttgtcctaaac
gacttatcagtcttgtgtacCTGGGGAAGGAAGATGTAGACCACGTCAGAGGCAAAGTCTtatagttacttgtcctaa
gacttatcagtcttgtgtacCTGAGAGAAGAAACAAACTTTGCACCTTGGTCAGTTGTATtatagttacttgtcctaaac
gacttatcagtcttgtgtacAGAATCTCAGGAGCAGTGAATTTAATTAATATGAGCACACtatagttacttgtcctaaac
gacttatcagtcttgtgtacTCCTTTCCTTGACCCCACTCTTCCTTCTCAGAAAAGATTTtatagttacttgtcctaaac
gacttatcagtcttgtgtacATCTGAGAACCAAGAGACCTGCATCCTATAGCTCTCTATAtatagttacttgtcctaaa
gacttatcagtcttgtgtacCTTCTCTGTGGCTCTGAATTTCATTATTTAATAAATGGCCtatagttacttgtcctaaac
gacttatcagtcttgtgtacCTACAGAGAACAATGCAGGCTCCTAGCACAGTATGTAGTAtatagttacttgtcctaaa
gacttatcagtcttgtgtacTTTTATTGGTTCTGATTTAGTCACCATTTGTGAGTTTCCCtatagttacttgtcctaaac
gacttatcagtcttgtgtacGGATGATTCCTGATTTCAAAGTACTTGTGTAAGTTTCCATtatagttacttgtcctaaac
gacttatcagtcttgtgtacGACTTACCTGTCTGCACTAACTTGACTGTATAGGCCCTTAtatagttacttgtcctaaac
gacttatcagtcttgtgtacGTATCTAACAATGCATGCCCATCTGCCTCAAAAGACAGGTtatagttacttgtcctaaa
gacttatcagtcttgtgtacTAGTAGTTCTATCCCAAAGCCCATCAACAGTTGAATAAACtatagttacttgtcctaaac
gacttatcagtcttgtgtacCACAACTAAACCAAGCCTTTATAAATGCATGTTTACAAGGtatagttacttgtcctaaac
gacttatcagtcttgtgtacCTTTATGGGAGGGGTAGGTGGTTCTAAAGCATGAATCTGCtatagttacttgtcctaaa
gacttatcagtcttgtgtacCTCTATTTGATGATTTGTGCAGAGTGACAGAGGAGCTTGCtatagttacttgtcctaaac
gacttatcagtcttgtgtacTTTTAAGGAGTATATGTAAAATTCCAGGTTCCTAGTAGGCtatagttacttgtcctaaac
gacttatcagtcttgtgtacGGTTACCTAATAAGCATTCCACCTTTCTTCTTCCCCTGATtatagttacttgtcctaaac
gacttatcagtcttgtgtacCTGCACAGCTAACCTTAAATAGTAATGAATAATCTTGGTGtatagttacttgtcctaaac
gacttatcagtcttgtgtacCAACCCAGACATCCATTCCCAGAAGAAAAGTTGCCCAGAGtatagttacttgtcctaa
gacttatcagtcttgtgtacGTGTTCATTCCTGCCTCATCCGATTGCCTCTTACACCAGCtatagttacttgtcctaaa
gacttatcagtcttgtgtacCACTCCGACGCTTCATAATTACTGAAACACATGTGCTAAGtatagttacttgtcctaaa
gacttatcagtcttgtgtacAGAATTCTCAGTGTTCAATTGCCCTCAAAATTTCTCATCTtatagttacttgtcctaaac
gacttatcagtcttgtgtacCTTTAGTTTAGCAGCTGCACATGGGGATTGCATTTAAATCtatagttacttgtcctaaac
gacttatcagtcttgtgtacAGTTTGGAGTTCAATATGTCCTAAAAGCAGCCTCTCTTCTtatagttacttgtcctaaac
gacttatcagtcttgtgtacGGTACACCTAGATTTTTTCTACTTCAAAGTGAGGAATATCtatagttacttgtcctaaac
gacttatcagtcttgtgtacGGAGGAAACCTTCAAAAATACATTAACTCTTTGAATGCACtatagttacttgtcctaaac
gacttatcagtcttgtgtacACTGTTAGTCACTCCTTGGTAGTAGCTTTTTAAAGTCATTtatagttacttgtcctaaac
gacttatcagtcttgtgtacCACAGTGAATCAGGTGTATGGCTAGCTAAAGGTCAAAAGAtatagttacttgtcctaaa
gacttatcagtcttgtgtacATAGTCTCTATGATATAACAAACAGGACCAGCCATGGGGCtatagttacttgtcctaaa
gacttatcagtcttgtgtacACTCGGAAGGCTGAGGCTGGGGAATAACTTGAACCTGGGAtatagttacttgtcctaa
gacttatcagtcttgtgtacAACAAACAAAAAACAGAACCTCTCCCTGAAATCTCCCAACtatagttacttgtcctaaa
gacttatcagtcttgtgtacTGTAAATTTCCTCAGAATCCATCACATTGACACACTTATGtatagttacttgtcctaaac
gacttatcagtcttgtgtacTTTTGCAGAGAAGTAAAAAAGAGCAGCCAGTGGTATTTTGtatagttacttgtcctaaac
gacttatcagtcttgtgtacGCCCCAAACAAATTTCACCATTTAATGACTGGGCCAAGCCtatagttacttgtcctaaa
gacttatcagtcttgtgtacTTGTTAATTTTAGAACTTCTCATTACCCCAGCTGTGTTAGtatagttacttgtcctaaac
gacttatcagtcttgtgtacCTCAAAATTAAAGCAGGAATTTGGGACAATGGAAGAAGGCtatagttacttgtcctaaa
gacttatcagtcttgtgtacGCTTGGCCTCTTATGCTTAAATAGCTTTATTTTCAGAAGAtatagttacttgtcctaaac
gacttatcagtcttgtgtacGAAAGAACAAGCTCCATCTGTAGTTTTCTATGATAGACGTtatagttacttgtcctaaac
gacttatcagtcttgtgtacGCCAGGTACAAACTGCCATTTTATAGGGACTTCTAGAATGtatagttacttgtcctaaac
gacttatcagtcttgtgtacCGCATCATTAATCATTGTGAATTGAGTACATGGGGGCTGTtatagttacttgtcctaaac
gacttatcagtcttgtgtacTGTGGGGAGTTTGGATGGAGATTGTCAGAAAACACAGAGGtatagttacttgtcctaaa
gacttatcagtcttgtgtacCCAAAGTGAACCTCCAGGCCTAGCTCCCAGTTAGACAGAGtatagttacttgtcctaa
gacttatcagtcttgtgtacCACTGTTGGGTGTGATTTGCCAGTCACTGAGGTGTCCTCTtatagttacttgtcctaaa
gacttatcagtcttgtgtacGCTAGTGAACATTTCCCACCTTTTATCCTCTGGCTTCATAtatagttacttgtcctaaac
gacttatcagtcttgtgtacGCAGCTCAACTGCCAAAGGGGAGCGTTAGCCATCGTGATGtatagttacttgtcctaa
gacttatcagtcttgtgtacGGGTTCGGACTCTGTTTGTTCTAACGCTCCTCGCTCAACTtatagttacttgtcctaaa
gacttatcagtcttgtgtacACTTGGCAATAAAAGTTGTTGACTAAAGGACTGGAGAATCtatagttacttgtcctaaac
gacttatcagtcttgtgtacATGAATCTGAAAGGAGCATGAGGGTTTGTCCTTCTAAGTTtatagttacttgtcctaaac
gacttatcagtcttgtgtacAGAGGTAATCTAATGTGCAGAAATTTAAGGTCATACTCCTtatagttacttgtcctaaac
gacttatcagtcttgtgtacAGTCAGTGCTATTTCCATTCTGGGTGCATTTCTTACTTTTtatagttacttgtcctaaac

gacttatcagtcttgtgtacTTGGTTATCTATGTAATATGAGGAGGCAAAAGGTATCTACtatagttacttgtcctaaac
gacttatcagtcttgtgtacTCATTGACCTTTGACTCATTGTATCAAATATTTCCTAGGCtatagttacttgtcctaaac
gacttatcagtcttgtgtacCTGTTTATTTCTGACCAGACTACCTTACCCCAGAGTGAGTtatagttacttgtcctaaac
gacttatcagtcttgtgtacGAATTCCAGTTTCCCATCTGCCAGTTATATGATCTGTACCtatagttacttgtcctaaac
gacttatcagtcttgtgtacAGTAGTCTTTTGTGTCTTGTCTAAGATTTCTTTGCGTAACtatagttacttgtcctaaac
gacttatcagtcttgtgtacAGACTTTTTCTTCATTGAATTACCTCAGTGCCTTTGTTGAtatagttacttgtcctaaac
gacttatcagtcttgtgtacCTTATACCAATGCCACACTGCCATAAGTACTATAGCTTTGtatagttacttgtcctaaac
gacttatcagtcttgtgtacCTCTTGAGTTCATGATTGTTTTTGGCTATCCTAGGTTAGTtatagttacttgtcctaaac
gacttatcagtcttgtgtacATTACATTGAACCTATATATCCAATTTGTGGAGAGTTGCCtatagttacttgtcctaaac
gacttatcagtcttgtgtacGAGCATAGCATGGCTCTCCATTTTTTTTAGGTCGTTAATTtatagttacttgtcctaaac
gacttatcagtcttgtgtacTCTGCCTCCTGGGTTCAAGCGATTCTACCATCTCAGCCTCtatagttacttgtcctaaa
gacttatcagtcttgtgtacCTGGGACTACAGGCATTAGCAACCACACCCTGCTAATTTTtatagttacttgtcctaaa
gacttatcagtcttgtgtacAGCTACTGCGCCCAGCCTAATAGTTACATTTTTAAAGATTtatagttacttgtcctaaac
gacttatcagtcttgtgtacTTCTTTTTATTACTTTTTCTGGCCTTACTGAGTTGGCTAGtatagttacttgtcctaaac
gacttatcagtcttgtgtacCCAGTACAATATTAAAGTGAAGTGGGGAAGAACAGACATCtatagttacttgtcctaaa
gacttatcagtcttgtgtacTTTTCCTGATCTTGGCTAAAAGTTTTTGAACATGCCACAGtatagttacttgtcctaaac
gacttatcagtcttgtgtacAAACTATGCCTCCTATTCTTCGTTTTCAGACACTTATTACtatagttacttgtcctaaac
gacttatcagtcttgtgtacTTCAATCACAAATTCATGGGCATTCTCAAAATAACAGGTCtatagttacttgtcctaaac
gacttatcagtcttgtgtacGGAACTGTACTGGACTGGAAGGAACTAAGAAGGCATGGCAtatagttacttgtcctaaa
gacttatcagtcttgtgtacCCAGAGAAGGGACATTAAGGGGGTAAATGGGTGGAACTCCtatagttacttgtcctaa
gacttatcagtcttgtgtacGGTTTGCAGAGTAGTGAGTTAAAGTGTGTCAGTGTCAGTGtatagttacttgtcctaaac
gacttatcagtcttgtgtacATGTGCAATGTTAACACTAGGAGAAACTGGGTGACGTCGTtatagttacttgtcctaaa
gacttatcagtcttgtgtacACCAGGGTCAAACATCTCATAACCTCTAGTTGAATACGTCtatagttacttgtcctaaa
gacttatcagtcttgtgtacGGCCCGTCAAACCTCACCAAACCCAGATATTCTAAGATGGtatagttacttgtcctaaa
gacttatcagtcttgtgtacCTCAGCAGCATCTTCTTCATAGCCTGTCACCCCAGCCCCAtatagttacttgtcctaaa
gacttatcagtcttgtgtacTTCTTTGGAGCCCTTTGAGAGAAGTATGGTGGTAAACATAtatagttacttgtcctaaac
gacttatcagtcttgtgtacGACCTCCTAAAATGCCCATGCCTGATCCCTAGATCCTGTGtatagttacttgtcctaaa
gacttatcagtcttgtgtacCAGGAATGTGCAATATGAATGATTAAGGATCCCGAGATGCtatagttacttgtcctaaa
gacttatcagtcttgtgtacCTTGTAAGAAGGAGGCAGGGTTCAGAGTCAGAAAGAGACAtatagttacttgtcctaaa
gacttatcagtcttgtgtacCTTTGAAGATGGGGGGAAGGGGGCAGGAGCCAAGGAGTGCtatagttacttgtccta
gacttatcagtcttgtgtacAAGCAATTAGACCTGGTGACATCTTCATTTGAGCCTAGTGtatagttacttgtcctaaac
gacttatcagtcttgtgtacACATTGATTTTCAGATGTTACACCAACTTAGCCATTCCTGtatagttacttgtcctaaac
gacttatcagtcttgtgtacTATAATATTTTCTGATTTAGGGATCTGGGTAATACCGTCCtatagttacttgtcctaaac
gacttatcagtcttgtgtacTGGAAGTGTTCTTTACTATTTTCTGACAGAGTTTGCATAGtatagttacttgtcctaaac
gacttatcagtcttgtgtacTGTGATAGAATTCAACAGCAGAGCTACTTGATCCTGGAGTtatagttacttgtcctaaa
gacttatcagtcttgtgtacGTGTTTCCTTTGGTTGAATTAGGTACTTTATGACCTTAAGtatagttacttgtcctaaac
gacttatcagtcttgtgtacTAGTATTCCGTTATTGTGCTTTCATTGTATGTAGGATCTGtatagttacttgtcctaaac
gacttatcagtcttgtgtacTCACTGCAACCTCAATCTCCCAGGTTTAAGCAATTCTCCTtatagttacttgtcctaaa
gacttatcagtcttgtgtacCGTGAGCTACTGTGCCTGGACCCTCTTTTCTTTATTAGCAtatagttacttgtcctaaac
gacttatcagtcttgtgtacATGATGTTATAATTTCTCTTGGGTGTCTTCTTTAACCCACtatagttacttgtcctaaac
gacttatcagtcttgtgtacGCTATTGATTTCCAATTTAGTCTCATTGAGGTTCAGAAACtatagttacttgtcctaaac
gacttatcagtcttgtgtacTGGCACATCATATGATATATCCTGGGAAAAGCACCTTAGGtatagttacttgtcctaaa
gacttatcagtcttgtgtacAGTATTTTGCTGTTGGGTATAGTTCCTTTTTTCCATCCTTtatagttacttgtcctaaac
gacttatcagtcttgtgtacTTTTTGAAATGGAGTCCGGTTCTGTCGCCGAGGCTAGAGTtatagttacttgtcctaaa
gacttatcagtcttgtgtacACTACAGGTGACCGCCGCCGCGCTTGGCTAATTTTTCGTAtatagttacttgtcctaaa
gacttatcagtcttgtgtacACTGTGCCCGGCCCCTCCCTTCTTTTATATCTATCTATCTtatagttacttgtcctaaac
gacttatcagtcttgtgtacTTGTCTAGTTTTATCAAGTGAGAGGAGGGTCTTTAAATAGtatagttacttgtcctaaac
gacttatcagtcttgtgtacGATGATTGTGAGGTTGTCTATTTCTCCCTTAAATACTGTAtatagttacttgtcctaaac
gacttatcagtcttgtgtacATCATAAATTTTTCTCTTTTGAAGTGCAGAGCTCAGAGAGtatagttacttgtcctaaac
gacttatcagtcttgtgtacCTCTCAAAATAAATTCTGTACCTATTAGCAGTGATTCTCCtatagttacttgtcctaaac
gacttatcagtcttgtgtacAAGCACTTGGAAACCACTCATTTACTCTCTGTCTCTTTAGtatagttacttgtcctaaac
gacttatcagtcttgtgtacTATTATATGGCCTTCTGTGTCAGGTTTTTTTTTTCCTCTGtatagttacttgtcctaaac

gacttatcagtcttgtgtacCAGTCACTCCACTCTTTCTATGTTTACTGTTTGTATGTCTtatagttacttgtcctaaac
gacttatcagtcttgtgtacTTCCACCGTTTGTTACTATTTCTTTTTGTCTCTGTTTCTGtatagttacttgtcctaaac
gacttatcagtcttgtgtacTTTTAGTGTACTTTTATGTGATAAGCGGAACTATATGCCCtatagttacttgtcctaaac
gacttatcagtcttgtgtacATCCACATGCCCTAATTTCCAGAGCTTGTAAATGTGTTACtatagttacttgtcctaaac
gacttatcagtcttgtgtacGTGGACCAAATCTAATCATAATGTCTTTAAAGAAGGAGAGtatagttacttgtcctaaac
gacttatcagtcttgtgtacAAGGATTGAAAAAGGGATTTGACCTGCTATTTGTTTGAAGtatagttacttgtcctaaac
gacttatcagtcttgtgtacTGAGACAAGGAATAAGAGTGTCCCATAGAAATTAGGAATTtatagttacttgtcctaaac
gacttatcagtcttgtgtacCTTCTGCCAATAACCCAATGAGCAAAGAAGCAAAATCTTCtatagttacttgtcctaaac
gacttatcagtcttgtgtacAGTGAAAAACACATCTGCCTCCTTGATTTTAGCCTGGTGAtatagttacttgtcctaaac
gacttatcagtcttgtgtacCATTTTTGATGTTCTTTCTTCTGCTCTGAGAGCCTAAGTTtatagttacttgtcctaaac
gacttatcagtcttgtgtacACTTCCTTTAGCATCTATTTTAGTGCAGTAATGTTGACAGtatagttacttgtcctaaac
gacttatcagtcttgtgtacTTCTGGTTGGCAAGATTTTGTTTGTTTCAGCTCTTTAAAGtatagttacttgtcctaaac
gacttatcagtcttgtgtacTGGTTTTCATGAGAAGTCCCTAAGGTAATTTCTATTGATGtatagttacttgtcctaaac
gacttatcagtcttgtgtacTCCTTTGTAATATTTATTTCCCTGTTAAGTCCATCCAGTGtatagttacttgtcctaaac
gacttatcagtcttgtgtacCTGCTATAGCACTGAAATGTGATGTGATAGTATTGTTATCtatagttacttgtcctaaac
gacttatcagtcttgtgtacCATTTGGTAGTTCTAGCATTGTGTTCATTTAGGGATTGAGtatagttacttgtcctaaac
gacttatcagtcttgtgtacTTTTTCCAACTAGTATTGTTGTTTCTCTTCTTCCTCTTCCtatagttacttgtcctaaac
gacttatcagtcttgtgtacCCAGATTTTCGGGTGGTAGCTTTGCTTTCAGCTCAGAGTTtatagttacttgtcctaaac
gacttatcagtcttgtgtacCTGAAATTCTGCTTTGCTTTGTTCTAAATATTTACAGCCCtatagttacttgtcctaaac
gacttatcagtcttgtgtacCTCTTTTCTTTCTGAGATTTCATTGCTGTTTTCAACAACCtatagttacttgtcctaaac
gacttatcagtcttgtgtacTGGTTACCTAGGCAACAGAGACTATATTTTTTCAATCAGCtatagttacttgtcctaaac
gacttatcagtcttgtgtacACTGTACCAACTCTACTTTCCTAAGCTAAAAGCCATAAAGtatagttacttgtcctaaac
gacttatcagtcttgtgtacATTTGTAAATCACCCAGCTTTCTTCCTCTGAGTATGTACTtatagttacttgtcctaaac
gacttatcagtcttgtgtacCTTCTAGTACCTTTATGGAATTGCTTTTTGTATTGCTTCCtatagttacttgtcctaaac
gacttatcagtcttgtgtacCTGATAAGTTCTTAGACTATCATTTCTGCAAGCTGACTGGtatagttacttgtcctaaac
gacttatcagtcttgtgtacTAAATATTTTTATGCTCCCATTATGCTTCAGCTATGCTGCtatagttacttgtcctaaac
gacttatcagtcttgtgtacTTGTGAACAATTTGACCCTGGTATCTTCTCTGATGAAAATtatagttacttgtcctaaac
gacttatcagtcttgtgtacTGAAACCTGAAGAAATCATAGCTTGGCATCAGATAAATGGtatagttacttgtcctaaac
gacttatcagtcttgtgtacTTTGGAAAGTAAATTTGGGCATTGCTAGGAGAATTAGTTCtatagttacttgtcctaaac
gacttatcagtcttgtgtacAATGATAAGGTAGAGGGCAGGAGTAGAAAAAGAAAAATAGtatagttacttgtcctaaa
gacttatcagtcttgtgtacGGAGAAGATAATGTCCTGGAAGAGAATGATGATAGTGGGAtatagttacttgtcctaaa
gacttatcagtcttgtgtacAAGTAAGATCTCGTGTTTACTTCAATAAAGTCAGACAGAGtatagttacttgtcctaaac
gacttatcagtcttgtgtacTCTTTTATGGAGATGGGGCACCTTAGAGGGGGAATAAATTtatagttacttgtcctaaac
gacttatcagtcttgtgtacCAATTAGGTATGCATAAGAGAGTTCTTAGCTGAAAACATCtatagttacttgtcctaaac
gacttatcagtcttgtgtacGGAACACTTTCAGCTCTTCCTCCTCACTGACTCCTCTTTTtatagttacttgtcctaaac
gacttatcagtcttgtgtacTCTTACTGATAAATCTTGTAGAAGGAGCTTAGCATGCATGtatagttacttgtcctaaac
gacttatcagtcttgtgtacTCTTGGCAGAGCTCACATGTCTTCAGGACACAGGAGTAGAtatagttacttgtcctaaa
gacttatcagtcttgtgtacAGATGCCAGCGGCTAGACCTTATTTACTCTACCTGTAAGtatagttacttgtcctaaac
gacttatcagtcttgtgtacGGAGACATTCAAGGAACCCAACCTAGCTGCAGAACTAAAGtatagttacttgtcctaaa
gacttatcagtcttgtgtacGGATTTGCCCTTTTCGTTTCTCTTATTTCTTAGCTGGTTCtatagttacttgtcctaaac
gacttatcagtcttgtgtacGGCATTTACTGCCTCTCTCTAATCCCAAAAATTATAACAAtatagttacttgtcctaaac
gacttatcagtcttgtgtacAAAAATATATATGTTCAAGCCCTCATTCAAAGAGTGGTGCtatagttacttgtcctaaac
gacttatcagtcttgtgtacCCTGCCTAATTAAGTTTTTTTTTTGTAGGGACTAGATCTCtatagttacttgtcctaaac
gacttatcagtcttgtgtacAAAGATAGACATTAGCACTTTCAAACTTTGTTCCACCTCCtatagttacttgtcctaaac
gacttatcagtcttgtgtacTCTTATATACTTACAGGTGAATTAACTAAAACTGGGGCTGtatagttacttgtcctaaac
gacttatcagtcttgtgtacGTGAATTACTATCAGTGGACTTGAATTAGTTGACTGTTGCtatagttacttgtcctaaac
gacttatcagtcttgtgtacTGACTCTAAAAAATGTGAATAGTGAAACTTTGCCCTTTGCtatagttacttgtcctaaac
gacttatcagtcttgtgtacCTATCCCCTAGCAGAGAAGTCAGTTTGTATGTTCTGTCACtatagttacttgtcctaaac
gacttatcagtcttgtgtacGAGCAGACTAGAGAAATTGAAGAAAATACTTTCAGCTGTTtatagttacttgtcctaaac
gacttatcagtcttgtgtacCTGTCTCAGTGTAACGTTTGCCTGGTCCATCTAACAGTTCtatagttacttgtcctaaac
gacttatcagtcttgtgtacCATTTGAATGTCTCAGGAGGGTAATCTGAATTTGAGGAGAtatagttacttgtcctaaac
gacttatcagtcttgtgtacGGATTAAGTGTAAGAGTACCTTGTTAGAGTATGTTATGACtatagttacttgtcctaaac

gacttatcagtcttgtgtacCATATGTGATGGTGGTCAAGATATTTGAATGTCCTGAGCTtatagttacttgtcctaaac
gacttatcagtcttgtgtacAAATGTGGGCAATGCTTTCCATACTGTTACATAAAAGTTGtatagttacttgtcctaaac
gacttatcagtcttgtgtacAATGCAAATCAAAACCACACAATGAGATACCATCTCACACtatagttacttgtcctaaa
gacttatcagtcttgtgtacGCTGGTGAGGCTGTGGAGAAAAAGGAACATTTTTACACTGtatagttacttgtcctaaa
gacttatcagtcttgtgtacCGATTCCTCAAAGACCTAGGACAGAAATACCATTTGATCCtatagttacttgtcctaaa
gacttatcagtcttgtgtacGACAGAACTGGAGGCCATCACCCTCGGTAAACTAATACAGtatagttacttgtcctaaa
gacttatcagtcttgtgtacACCAAATATCACATGTTCTGGCTTAGAAGTGGGAGCAAAAtatagttacttgtcctaaa
gacttatcagtcttgtgtacTGTGGAGAAAACACACACTGGGGTCATTGGAGGGTGAGGGtatagttacttgtcctaa
gacttatcagtcttgtgtacGATGCTGGGCTTAATACATGAGTGATGGTATGATCTGTGCtatagttacttgtcctaaac
gacttatcagtcttgtgtacGTTTTGATATAGTTTACATGTTGGTATGCAGGTGTCAAAGtatagttacttgtcctaaac
gacttatcagtcttgtgtacGTTGTGGACATTCAAACTCAGTTTTTATACTAGCCTGCTTtatagttacttgtcctaaac
gacttatcagtcttgtgtacTCATACCTGCTGAATTTCTCACTGAGTATCCCTTTAGGGTtatagttacttgtcctaaac
gacttatcagtcttgtgtacACAGAGTGTTTGGTTACATGAAATAGTGATTTGTGGAAGCtatagttacttgtcctaaac
gacttatcagtcttgtgtacAGAATTGACATTGTAGGAAAGCTGCATGTTGCTGGGGTTTtatagttacttgtcctaaac
gacttatcagtcttgtgtacCAGTTTTGATGCTGCCTTTATGATTTACTGTTTCTGTGACtatagttacttgtcctaaac
gacttatcagtcttgtgtacTCTGTAAAATAAGGACAATACTATTACCTCCCCTGTTAACtatagttacttgtcctaaac
gacttatcagtcttgtgtacAAAGAATTCATATGTGTAAAGCGGTTATATTAGTGCCTGGtatagttacttgtcctaaac
gacttatcagtcttgtgtacTTCAATAAACTAGACCATGTGAGAAGGAAATACGTAAGAGtatagttacttgtcctaaa
gacttatcagtcttgtgtacCCTTAAGTGTATTGGCAGGAGGGATTTGAGAATGAATAAAtatagttacttgtcctaaac
gacttatcagtcttgtgtacTGGGAAAGAATAGCCGCAAACAACATGATCAGAAATAAAAtatagttacttgtcctaaa
gacttatcagtcttgtgtacTTTGAGATGGCTTTAATATTAGTCAAGAAACTCTACAGGGtatagttacttgtcctaaac
gacttatcagtcttgtgtacCCAAAGAGAAGAGCTGGGAAATAAGGTAAAAAGGCTTAAAtatagttacttgtcctaaa
gacttatcagtcttgtgtacCTCAGCCTTAGAGTCTTGGGTCTAGTGCAATGCCTGGCCTtatagttacttgtcctaaa
gacttatcagtcttgtgtacGTGACTCCTGCACAGCTATAAGGACCCTCACGGATTAGATtatagttacttgtcctaaa
gacttatcagtcttgtgtacCAAGAGCTTTACAGAGCCACAAATCCTCTCCGTGACTTGGtatagttacttgtcctaaa
gacttatcagtcttgtgtacGGTATTAATGAACCAAGACAAACTTCACACTGGCAGGCTAtatagttacttgtcctaaa
gacttatcagtcttgtgtacTCTGGGAGAGAATAAACATGTTATGGCAATTATTGGTGAAtatagttacttgtcctaaac
gacttatcagtcttgtgtacAGACATTACCTAACCTAATTCAAGCACAATCCATGTTAGCtatagttacttgtcctaaac
gacttatcagtcttgtgtacGAAGACTTTCATTGATTCAGGTCTTAGTCAAAAGATAAGGtatagttacttgtcctaaac
gacttatcagtcttgtgtacGTTGTTGTTGTTGGATGAACACTAAGCTAAATTAAACAGCtatagttacttgtcctaaac
gacttatcagtcttgtgtacTCTATGGAGAAAGAACTTGGCTACTTAATAAGTCCATCTCtatagttacttgtcctaaac
gacttatcagtcttgtgtacCTTTAGGATAAGATGAGAATAGAAGGGATAAGAAAAGCAGtatagttacttgtcctaaa
gacttatcagtcttgtgtacTTACTTTATCTTTGTTACTCTGGAGTTTCATGAGAACACCtatagttacttgtcctaaac
gacttatcagtcttgtgtacTGAAGAGTTGTAAAAACGTTCTCTGTCAGTGTTGTGGGCAtatagttacttgtcctaaac
gacttatcagtcttgtgtacGGGGCCACTTACTTTACAGTCCTACATGTAGGTCACTTGGtatagttacttgtcctaaa
gacttatcagtcttgtgtacCTATCTGTGTGAGGTGGAGTAAAGAGAGGCATATTCAAAAtatagttacttgtcctaaa
gacttatcagtcttgtgtacGTCTATGGCTGGCCCTTGACTTCACTCACCCTTCCCTTGGtatagttacttgtcctaaa
gacttatcagtcttgtgtacTGCCAGGCTGTTTTACATAAGGTGTTGGGCAGAAAAGTTCtatagttacttgtcctaaa
gacttatcagtcttgtgtacCAAGAACACATGCACACAAATCACTTTATAACAGGAGTCCtatagttacttgtcctaaa
gacttatcagtcttgtgtacAGAGATTTAGTAACTATAAAATGTAGTGTCACACGCCACAtatagttacttgtcctaaac
gacttatcagtcttgtgtacTGTTGTGTTAAGGGCCATATTATTTTAATCATCGGATCTCtatagttacttgtcctaaac
gacttatcagtcttgtgtacCATATGATGCTAATAATGACTGACAGTTTCTGTGAGGATTtatagttacttgtcctaaac
gacttatcagtcttgtgtacCATAATACATGTTAAAGTGTTAGCCTTGTGCATGGCAATTtatagttacttgtcctaaac
gacttatcagtcttgtgtacCATGAGCATATCCTCACAATATTCTAAAGTGAAAATGGCAtatagttacttgtcctaaac
gacttatcagtcttgtgtacCAATGGTACTTGTGGTATACGGTACAGGTTGAGCATCTCTtatagttacttgtcctaaac
gacttatcagtcttgtgtacCAAAATTTGAAACTTCTGAATGCCTACATGATGCTCAATGtatagttacttgtcctaaac
gacttatcagtcttgtgtacGATTTTCTGATTAGGGATGCAAAACCAGTAAGTATAATGCtatagttacttgtcctaaac
gacttatcagtcttgtgtacAAGCATTGTGGATAAGATATACTCTACATGTAAAGTCACCtatagttacttgtcctaaac
gacttatcagtcttgtgtacTATTTGCTATCCATGTGTATCCTTACATTTGAAGTGAGTCtatagttacttgtcctaaac
gacttatcagtcttgtgtacCCATACAAGTGGGTCTAATATTCAGTATTCTTGATTAGCAtatagttacttgtcctaaac
gacttatcagtcttgtgtacATTTTGAATATATTGTCCCACTATCTACTGTCCTGTAAGGtatagttacttgtcctaaac
gacttatcagtcttgtgtacGCTGAGAGATTTCATGCTAACTTTATTGAAACTCCCTTATtatagttacttgtcctaaac

gacttatcagtcttgtgtacTCTTTTGATGTAATCTTCTTTGGATTGAATCCGATTGGGtatagttacttgtcctaaac
gacttatcagtcttgtgtacGGCCTTCCTGTACTCTGTATCTGGATATTTATATTTTTTCtatagttacttgtcctaaac
gacttatcagtcttgtgtacAGCTTTCTATCCCTTTGTCTCTCTATTTTTTCACAAACTCtatagttacttgtcctaaac
gacttatcagtcttgtgtacAATATTTGCTCTTCTGATGCCCTGCTATTAATACCATAAGtatagttacttgtcctaaac
gacttatcagtcttgtgtacCCTGCTTGATCAATTCTCCTGGTGACATTCTTATTGCATTtatagttacttgtcctaaac
gacttatcagtcttgtgtacGAGCTTCCCTGAAAAAATTATCTTTAAATCTTTGTCAGGCtatagttacttgtcctaaac
gacttatcagtcttgtgtacATTGTGTTCCTTTGGTGGTGCTGTGTCTCCTTGGTTTTTCtatagttacttgtcctaaac
gacttatcagtcttgtgtacTTGCCTTATGTTGATCTGTGTACATTTGAAGAAATAGGGAtatagttacttgtcctaaac
gacttatcagtcttgtgtacGAAAGCCCTTCACCAGTCACCCCATCCAGAGATTCAAGGCtatagttacttgtcctaa
gacttatcagtcttgtgtacCAGCAAGGCTGTCTTAGTGCCTGGGTCAGAATGTGGGTGGtatagttacttgtcctaa
gacttatcagtcttgtgtacTGTATCAACAGAGGTTGGTCTGAAGTCTGGGTCCAAAGGAtatagttacttgtcctaaa
gacttatcagtcttgtgtacCCACTAGGTGGGTCTAGAGACTGAATTCATGGGGGCTGACtatagttacttgtcctaa
gacttatcagtcttgtgtacCATCCATGGGGGTTGGCCTGGCACTGGGGTAGAATGAACAtatagttacttgtcctaa
gacttatcagtcttgtgtacGCTGTCTGAGGCTAGTGTTGTTGACTGGGGGGAAGGGTACtatagttacttgtcctaa
gacttatcagtcttgtgtacGGCTATGCTCTGGGATAGTCTGAAATCTGGGACAGGTCTGtatagttacttgtcctaaa
gacttatcagtcttgtgtacGGGATTTGCCTGGGGCTGGTGGACCTGGAGCCTGTATCCAtatagttacttgtcctaa
gacttatcagtcttgtgtacATCTGCAAGTGCTGGCCAGATGACTAGAGTTTCTAAGCTGtatagttacttgtcctaaa
gacttatcagtcttgtgtacCTCGTGGATCAGACAGGTGTTGAGAATATTCTGGAGGCTGtatagttacttgtcctaaa
gacttatcagtcttgtgtacGGTAGGCCTAGAAACCCAGAGTCTGAGGAAAAGTCTAGTGtatagttacttgtcctaa
gacttatcagtcttgtgtacCCATTCTGTATTGCCTGGGATTCGGAGAGAGAAAACTTGGtatagttacttgtcctaaa
gacttatcagtcttgtgtacAAATTTTCTCTGTTACTCACAGGTGTTGTATTCTTTCACCtatagttacttgtcctaaac
gacttatcagtcttgtgtacTTGTTAGCTCTTGTGAAGTTGTATTTGTATGGGGATAGTTtatagttacttgtcctaaac
gacttatcagtcttgtgtacATCTTGCTGATGTCGGTTTTTGGGATGGGTTTTGAGAAACtatagttacttgtcctaaac
gacttatcagtcttgtgtacCTCAGACAGAAATGAGACATGTATTATTGGAAACTGGAGGtatagttacttgtcctaaa
gacttatcagtcttgtgtacGGTAGAACTTGTGAGCAACAAAATTGAGTATCTAGCTGAGtatagttacttgtcctaaa
gacttatcagtcttgtgtacAGCAAATTGTTAAAGGAATAGTTTGGTTCCTCTTGATCTCtatagttacttgtcctaaac
gacttatcagtcttgtgtacCATATTCCATTTTTATGTGATCTGGTAGAAGAGGCAGGGCtatagttacttgtcctaaac
gacttatcagtcttgtgtacCAATACTATGTTTATTAATTGCAACTAATGCCCAAGCAGCtatagttacttgtcctaaac
gacttatcagtcttgtgtacGGTTTTTCATCCCTTCTCATACCGTTTCACTTTGAACTAAtatagttacttgtcctaaac
gacttatcagtcttgtgtacCAGTTACTGTGCTTACTCTTCCACCACAACTCACAAAAAGtatagttacttgtcctaaa
gacttatcagtcttgtgtacTGTACTCCAGTATATTCAAAGTGGTTGTTTTGTCTTTTGGtatagttacttgtcctaaac
gacttatcagtcttgtgtacTGAATCCTTTTAAAGACTTACGAATATGTATCAGTGAGGCtatagttacttgtcctaaac
gacttatcagtcttgtgtacTCTTGGGCTCAACTATAATAATCAACTAAGTGCTTTGAAGtatagttacttgtcctaaac
gacttatcagtcttgtgtacTATTGTGGTCTGTGTATGTGTTTTTATTTTTTCCCCTTCCtatagttacttgtcctaaac
gacttatcagtcttgtgtacTAGTTTTCTGATCCTCACCAACCTCCTCCCACCCTCCATCtatagttacttgtcctaaa
gacttatcagtcttgtgtacATAATGGCTTCTAGCTCCAAGTGTGTTGCTGCAAAGGGCAtatagttacttgtcctaaa
gacttatcagtcttgtgtacTTGTTTGTTTCTATAGCCGTGTAATATTCCATGGTGTGTAtatagttacttgtcctaaac
gacttatcagtcttgtgtacTTTTAAGTTCTTTGAGAAGCTTCCCAACTGCTTTCAAAAGtatagttacttgtcctaaac
gacttatcagtcttgtgtacACATTCCCGACAACAGTGTAGCAGCATTCCCTCTTCTCTGtatagttacttgtcctaaa
gacttatcagtcttgtgtacGTTTTAACCTGCATTTCCCTAATGATGAGTGATATTAAGCtatagttacttgtcctaaac
gacttatcagtcttgtgtacATATGTCTTCTTTTGAGGATTATCTGTTCATGTCCTTTGCtatagttacttgtcctaaac
gacttatcagtcttgtgtacGTCAATTTTTGCTTTAGTTTCAAGTGCTTTTGGTGTCTTCtatagttacttgtcctaaac
gacttatcagtcttgtgtacCCTATGTCCAGAATAGTATTTCTTAGGTTATCTTCTAGAGtatagttacttgtcctaaac
gacttatcagtcttgtgtacTGAGTTTATTTTTGTATGTAGTGAAAGGAAGGGGTCCAGTtatagttacttgtcctaaac
gacttatcagtcttgtgtacACCTGTTCCATTGGTCTGTGCCTCTGTTTTTGTACCATTAtatagttacttgtcctaaac
gacttatcagtcttgtgtacTTTTCTTAGGATTCCTTTGGTTATTTGGGCCTTTTTTGGTtatagttacttgtcctaaac
gacttatcagtcttgtgtacAGTACTGAATCTGTAAATTTGCATTGGGAACTATGGCCACtatagttacttgtcctaaac
gacttatcagtcttgtgtacTCTTTGTTTGATTCTCAGCTCGTACTTTACTGGTGTATAGtatagttacttgtcctaaac
gacttatcagtcttgtgtacAAACTGGTGAATTTGTTTGTCAACTCTATGAGCCTTTGGGtatagttacttgtcctaaac
gacttatcagtcttgtgtacAGGACTTCAAGTACTATATTCAACAGAAGTGGTGAGAGTGtatagttacttgtcctaaa
gacttatcagtcttgtgtacCCTTTTCTTGCTTCATTCTCAAGTGGAATAGTTCTAGCTTtatagttacttgtcctaaac
gacttatcagtcttgtgtacTGAGTAAGATAGAACACTGTTACTGAGATATGTTCCCTCAtatagttacttgtcctaaac

gacttatcagtcttgtgtacCTTTAAATCAACCGTGAACAAAAAGGACAAACAAGGGCATtatagttacttgtcctaaa
gacttatcagtcttgtgtacGGAACACCCAGAGCCATAAAACAAGTTTTCAGAAACCTACtatagttacttgtcctaaa
gacttatcagtcttgtgtacATCATTGAGGCAGAAAACCAGCAAAAGTATTCAGGATCTAtatagttacttgtcctaaac
gacttatcagtcttgtgtacCTAACAGACATCTACAGGACCCTCCACCCAATAAAAACAGtatagttacttgtcctaaa
gacttatcagtcttgtgtacTTTCGTCTGAGCATGAACCTTGATTTGAAAAAGAGATCAGtatagttacttgtcctaaac
gacttatcagtcttgtgtacGCTCAGCCATCTCAATAGAAGACAGGAATAAACATGGAATtatagttacttgtcctaaa
gacttatcagtcttgtgtacCAGAGAATGTTAGATGAGATAAAAGATAGCTGCTGGACTTtatagttacttgtcctaaac
gacttatcagtcttgtgtacGCACCAGACCATAGAACTCCTCAGATATGAACATGCCTGTtatagttacttgtcctaaa
gacttatcagtcttgtgtacCATAACATGTCTGGTTTCACCAATTCACAGCTGGAGAGAAtatagttacttgtcctaaac
gacttatcagtcttgtgtacACCTGCAAATTAAGATAATATTTAGTTGAGCCTTCAGACCtatagttacttgtcctaaac
gacttatcagtcttgtgtacGCTGTTGAGATAGCCTCACATGATGTATTTTGTATATAAGtatagttacttgtcctaaac
gacttatcagtcttgtgtacTGCAGATTTATTTGAAGGCCAAGGATGGAGTGTTATAGACtatagttacttgtcctaaac
gacttatcagtcttgtgtacCACAGGTGTAATCTTTGAGAAGTCATGAGTGTGAGCCTTCtatagttacttgtcctaaac
gacttatcagtcttgtgtacCACAGTGAGAAGACAGCCACATGCAACACAAAGAGACAGCtatagttacttgtcctaa
gacttatcagtcttgtgtacCATGGACCTCGATTATGAACTTCTAGCCCCAGAACTGTAAtatagttacttgtcctaaa
gacttatcagtcttgtgtacGCTTAAGCTGACTAAAACAGATAGTCAAGACAGGATTCCTtatagttacttgtcctaaac
gacttatcagtcttgtgtacTGCTTAGAATATATTGGCATGCTGTGACACATACAGAAAGtatagttacttgtcctaaac
gacttatcagtcttgtgtacAGAGAGATAGAAAGAAAAAACAGTGGTATGAATGCATGAGtatagttacttgtcctaaa
gacttatcagtcttgtgtacAGCTTATTCTGTCTACTGAGCTAGAAGGTTTAAATGCTTCtatagttacttgtcctaaac
gacttatcagtcttgtgtacGATTCGTGCTGGAATTATTTGCAATTACTAATATCATGGGtatagttacttgtcctaaac
gacttatcagtcttgtgtacGTACTTTTCTTATTTATACTTCCTTGTATTAGGCTGTCCCtatagttacttgtcctaaac
gacttatcagtcttgtgtacCTCAGGAAGGATTTAGGCAGAAATGTTAGGGCTAGAAAACtatagttacttgtcctaaa
gacttatcagtcttgtgtacTGAGAAAGTACGCCAGGATTCACTAACAGAAAAAGATTTTtatagttacttgtcctaaac
gacttatcagtcttgtgtacGAAATTTCCCCCTTTTATTTCCAGAGGCAACCCAAGGAGAtatagttacttgtcctaaa
gacttatcagtcttgtgtacTTTAAGCTATAGTATAGAAAGCGTATGACCAATGCCAAGCtatagttacttgtcctaaac
gacttatcagtcttgtgtacCATGTGGCAAAGAGGATAATAGAAACTAGATGAGGAGTGGtatagttacttgtcctaaa
gacttatcagtcttgtgtacAGAGGTGCACCAGAAGAGGACGGACTAGAGTGAGGTGACTtatagttacttgtcctaa
gacttatcagtcttgtgtacCTAGAATCTACACTCCTAGGTCTCAGCCACAAGATAATATtatagttacttgtcctaaac
gacttatcagtcttgtgtacATCCCAGATTATTCTCTCTTTTGCTTCCAATGTTGCCCAAtatagttacttgtcctaaac
gacttatcagtcttgtgtacAGAGAGAGCAAGAGCATAGAGAAAGGGATGTCTAGATAGAtatagttacttgtcctaaa
gacttatcagtcttgtgtacATGTGCATGGCTTTGAGAATAGAGTAGGGGGTTTAGGGAGtatagttacttgtcctaaa
gacttatcagtcttgtgtacGGACCAGGGACCAGCAGAGAGCCTAAGTGGCACATGTGTGtatagttacttgtcctaa
gacttatcagtcttgtgtacCAAAGTGCCTACCAAGACTAGGTAAGATTAATTGCTATGTtatagttacttgtcctaaac
gacttatcagtcttgtgtacATCAGGACAGGGAAGGCAGGGTTCATACTGGATCTATACAtatagttacttgtcctaaa
gacttatcagtcttgtgtacCATCAGTGCAAGCTTCTCACAATAAAAGTGGACCAAACTCtatagttacttgtcctaaa
gacttatcagtcttgtgtacAGGGGAGAATAGTCAAGACTCAAACCAAAGCTACATCACTtatagttacttgtcctaaa
gacttatcagtcttgtgtacCCAATGTTATGGGTAAAGATATAATGAAGAGAAAACCCTGtatagttacttgtcctaaac
gacttatcagtcttgtgtacTGGCTCTCATGGGGATTTTAAGTTTCAAATTGGTAGATGAtatagttacttgtcctaaac
gacttatcagtcttgtgtacGGATTCACCTGGGAAGCATTTAAAATTCCTTAAAGCTAAGtatagttacttgtcctaaac
gacttatcagtcttgtgtacCATATGATTAAAAAGTATAGCCAGATTCTAGAACCACTGCtatagttacttgtcctaaac
gacttatcagtcttgtgtacTTTCCCCTCATTAATGAAATTCTTAGATATTCCAGAGCAGtatagttacttgtcctaaac
gacttatcagtcttgtgtacACTTTCAATTTGCAGGATGTTTGTCTAAGTGTTTTGTCCAtatagttacttgtcctaaac
gacttatcagtcttgtgtacTAATAATCCTCCTTTATATCAGAAAACTGGCACAAAGAGGtatagttacttgtcctaaac
gacttatcagtcttgtgtacAGCAAGTATTCACACAGAAACAGGCTGCCTTCAAAGCTAGtatagttacttgtcctaaa
gacttatcagtcttgtgtacTGGCTTCCTGGCATCACTCAAACATTGCCTTTCCTCTCCCtatagttacttgtcctaaa
gacttatcagtcttgtgtacCATGGTGGAAAGATGCTTTACCCTCAATTTATAAAAAGGTtatagttacttgtcctaaac
gacttatcagtcttgtgtacGTCTCAAACCCCTGACCTTGTGATCTGCCAGCCTCAGCCTtatagttacttgtcctaaa
gacttatcagtcttgtgtacCTCATAATCCCCACATGCTGCAGGAGGAACTTGGTGGGAGtatagttacttgtcctaaa
gacttatcagtcttgtgtacATCTCTTTCCTTTGAAAATTACCTAGTCTTTGGGTATGTCtatagttacttgtcctaaac
gacttatcagtcttgtgtacGTGAAAACAGACTAATACTCTTGCCTTTGGTTATAACAACtatagttacttgtcctaaac
gacttatcagtcttgtgtacGAACAAACTTTTGCATACCCTGTTCTGGTCATCAATTTTAtatagttacttgtcctaaac
gacttatcagtcttgtgtacGAATGTTTCTCACTATTCTTAGCCTGTTGACTTTACCTTTtatagttacttgtcctaaac

gacttatcagtcttgtgtacTCAAACACTACCTCTTTCTTGAAGCCTTTTCTAATTCTTCtatagttacttgtcctaaac
gacttatcagtcttgtgtacTTGAATCTCACAACAAACTCTGCTCCATTTTCCTTTACCAtatagttacttgtcctaaac
gacttatcagtcttgtgtacATTATGGAACCAATCGTGTGCTGCATAATGATGATTTGGTtatagttacttgtcctaaac
gacttatcagtcttgtgtacAGCAATAGGCTATACCATTCAGGCTAGTTGTGTAGGAGGCtatagttacttgtcctaaa
gacttatcagtcttgtgtacTCTCCCCATTTCTTTTTCTAAGTTCTTCTAGTATCTGTTCtatagttacttgtcctaaac
gacttatcagtcttgtgtacGGCAAGAACTGTCTTATTTACCACTGTATGTCTTGAATTAtatagttacttgtcctaaac
gacttatcagtcttgtgtacCACAACTCAAATCTAACATATACATAGGAAAATGGTTCCCtatagttacttgtcctaaac
gacttatcagtcttgtgtacACTTGTGTATCTCTCAAACTAACATGGCTAAGACTATTTGtatagttacttgtcctaaac
gacttatcagtcttgtgtacACTTTAAGTTGCTTGTCCTTCCATATTCTCATGACTTGTTtatagttacttgtcctaaac
gacttatcagtcttgtgtacGAAGTATTTGTTTTTAAAAAATTAAAGGGTAGGGCCAGGCtatagttacttgtcctaaac
gacttatcagtcttgtgtacGAGAATGGTGTGAACCTGGTGGGCAGAGGTGGCAGTGAGtatagttacttgtcctaa
gacttatcagtcttgtgtacGTTATTTTACTTACTACTGGCATAGATTTGTTTGTCCACGtatagttacttgtcctaaac
gacttatcagtcttgtgtacTTTGTATCAGCTTTGCTTTGGGCACAAATACTACACCTTGtatagttacttgtcctaaac
gacttatcagtcttgtgtacGGATCCAATTATTCCCTTTTGATTTTACGTTTTCTGGTTCtatagttacttgtcctaaac
gacttatcagtcttgtgtacATGTGAAAAAGATATCAGGCTTAACAGTAATTTCTCTGGCtatagttacttgtcctaaac
gacttatcagtcttgtgtacATGGATAAAGTTATATTGAACCAATTGGCACTGAACTGGCtatagttacttgtcctaaac
gacttatcagtcttgtgtacCATGAACTGACAACTATCTAGTAAAAAGACATATCGAAGGtatagttacttgtcctaaac
gacttatcagtcttgtgtacTTTAATCTTCCTCCAAATTCTTCTTGTACTACTGATGTGGtatagttacttgtcctaaac
gacttatcagtcttgtgtacTAAAAAGAAGTTTGAGACATATTTGCCAACTCTCTGAGGAtatagttacttgtcctaaac
gacttatcagtcttgtgtacCACAAATGAATGCTGCTAGTTGAAGGTAGAGTTCAGCTGTtatagttacttgtcctaaac
gacttatcagtcttgtgtacAACAAACTCATTCTAAAACTCCGTCGTAACTATGGAAACCtatagttacttgtcctaaac
gacttatcagtcttgtgtacAAATGAATTCTATTAGAAATACAAAAAGCCACCCCAGCAGtatagttacttgtcctaaac
gacttatcagtcttgtgtacCTCTTATTGATTTATGAAGGTTCCAGATGATGAAAAGTGGtatagttacttgtcctaaac
gacttatcagtcttgtgtacTCAGTCACTTTGCCTTCATTACAAGAAGCTATCAGCTTTTtatagttacttgtcctaaac
gacttatcagtcttgtgtacAGGAGAGCTTGACTAAAGAACCATTTCTGATTATGTTTGGtatagttacttgtcctaaac
gacttatcagtcttgtgtacATGGGCTGCACATTTATTTTGAGGGAGAATTAGTTGATTTtatagttacttgtcctaaac
gacttatcagtcttgtgtacACACTAGCATTTGGACAAAACTGTACAATTCACTTGGCTTtatagttacttgtcctaaac
gacttatcagtcttgtgtacCTGGTGCTTGTAGTGGGATTTGTTGAAAACAGGAAAATTAtatagttacttgtcctaaac
gacttatcagtcttgtgtacATTCGATTCAAGTAATAATGCGAGGCTAAAATGAAAGAGCtatagttacttgtcctaaac
gacttatcagtcttgtgtacTTAGAGTTGTTAATGTCCCAGAAGGTTTGGTGTGCAGGTGtatagttacttgtcctaaac
gacttatcagtcttgtgtacAGCATGTACTGTACCCACTTCTACCCTTTCCGTAAAATCAtatagttacttgtcctaaac
gacttatcagtcttgtgtacACTCTGACTTTTCTCCTTCAAAATATTGACAGTAGTCCTGtatagttacttgtcctaaac
gacttatcagtcttgtgtacGAGTATCTACAAACCAGGATGAAGAAGATAAGACATACACtatagttacttgtcctaaa
gacttatcagtcttgtgtacTCTTTAGATCTTTGGGTTTCATCATATGCATTTCCTCATGtatagttacttgtcctaaac
gacttatcagtcttgtgtacCTGTTGCAGTGAGTTAGGAAATGTGTTTTTCTAAGACGACtatagttacttgtcctaaac
gacttatcagtcttgtgtacGTCACAGAAACAAAACAGAAAAGTATGAAAGATCAAGGGCtatagttacttgtcctaaa
gacttatcagtcttgtgtacCCTTAAATAATTTCCTGTTCTCGTGTTTCCTTACACTCCAtatagttacttgtcctaaac
gacttatcagtcttgtgtacGAGGTTTATTTAGGTTGTATATTCGCAGTTTTATTTCGGGtatagttacttgtcctaaac
gacttatcagtcttgtgtacATTTCCGAAATAAACTATAGTCAACCCAAGCTCTTGCCTAtatagttacttgtcctaaac
gacttatcagtcttgtgtacTGTGGTTCAGGTGTGTTGTTTCTTGAAAGTCTCCGTGTTTtatagttacttgtcctaaac
gacttatcagtcttgtgtacGGCCTGGGTCCAGTGTTGAAGTAGCCTCCTAAATGCCATTtatagttacttgtcctaaa
gacttatcagtcttgtgtacGCTTCAAGGTTTTATAGGCATTTTTATTACATCTCACAGCtatagttacttgtcctaaac
gacttatcagtcttgtgtacGCTTTGAATAACAACATATGAGAATTGGAGGGGATTTCAGtatagttacttgtcctaaac
gacttatcagtcttgtgtacACCTCATCTAAGCGGCTCATTTTGCAGATAAGAAATTTAGtatagttacttgtcctaaac
gacttatcagtcttgtgtacGCTCATTAACAACACAGCTGGAATAAGACCTTGGTTTCCCtatagttacttgtcctaaac
gacttatcagtcttgtgtacTGTACTGCACAACCTCATGTTTTCTCTTATGTCTGAAGGAtatagttacttgtcctaaac
gacttatcagtcttgtgtacTTTCAGCTCTTCCTTAACATATAAAGCCACAGACAGAAGGtatagttacttgtcctaaac
gacttatcagtcttgtgtacTTCTCAACCGAATGCTCCATCCATTACTGCCCATTATACAtatagttacttgtcctaaac
gacttatcagtcttgtgtacTTCATTATCTACTCTGTGCTGTTAGTTGCTTCGTTTTTTGtatagttacttgtcctaaac
gacttatcagtcttgtgtacGCTTTTACTGATTAACTTGGTCTCTGCGATATAAAATGGTtatagttacttgtcctaaac
gacttatcagtcttgtgtacTTCTAGTTATGGGTCATTATTTCCGATAGATGTAATCGTCtatagttacttgtcctaaac
gacttatcagtcttgtgtacTGGTCTCACTAGAAGTTATAACGGTTTTCTTAATGGTCTGtatagttacttgtcctaaac

gacttatcagtcttgtgtacCAGAGAGAAATTGGTCCCAGAGCTAGCTTGCTTCTGCACTtatagttacttgtcctaaa
gacttatcagtcttgtgtacCATATTTTGTTACCAATCAGCTGGGTATTTTTTTCTGCCAtatagttacttgtcctaaac
gacttatcagtcttgtgtacTATTTATTTTCTATTTACAGCCTGCCCCGCAATAATACTGtatagttacttgtcctaaac
gacttatcagtcttgtgtacACCCTACAATGCTTTGCTTTTCTAATTCTTAGAGGAAACAtatagttacttgtcctaaac
gacttatcagtcttgtgtacCAAGCAATGAGATATGATGAAATCCAGTTGACCAGGCAAAtatagttacttgtcctaaa
gacttatcagtcttgtgtacGAAATCAACAGAGAATTTTTCCATTAGAGGTAGCTCATTGtatagttacttgtcctaaac
gacttatcagtcttgtgtacACTTTAGTATAGGTAAGAGGCATGCACACCAGAAATTATTtatagttacttgtcctaaac
gacttatcagtcttgtgtacTGATTGTTCTTTCAAATTTTTCCTTGCACAGAGCATAACCtatagttacttgtcctaaac
gacttatcagtcttgtgtacTATTTCTTGCACATATCCTAATTCTTGGTCTACTTAAGCCtatagttacttgtcctaaac
gacttatcagtcttgtgtacAGAGAGAGAGAAGGAGGTAGACTGTCCATGGGCATATGTCtatagttacttgtcctaaa
gacttatcagtcttgtgtacGTTCTAAGATGAATGTGTTTTCCCTCATATTTAACTCCTCtatagttacttgtcctaaac
gacttatcagtcttgtgtacGCCATGATTGTCATTGCCTATCCATATTTGAATGTATTGGtatagttacttgtcctaaac
gacttatcagtcttgtgtacCTGAAAAAGTGTAACTCCTTGACATGTTAATCAACACATCtatagttacttgtcctaaac
gacttatcagtcttgtgtacGGAATGTGAGATTGCTATTCTGAAACTCTTCTAAGAATGCtatagttacttgtcctaaac
gacttatcagtcttgtgtacCTTTTAGGAAGTGCTGCATCCATGTTCTTGGCTGAGTAAAtatagttacttgtcctaaac
gacttatcagtcttgtgtacACTTCCCTTTTCAATGTATAAACAAGAACACTACATGGAGtatagttacttgtcctaaac
gacttatcagtcttgtgtacTAGTCATCATATCATAGAACTAAATACACCTAAAGCCAGGtatagttacttgtcctaaac
gacttatcagtcttgtgtacATTTGAGGAAGTAAGACGCAAAGAAATTAAATGAGTAGCCtatagttacttgtcctaaac
gacttatcagtcttgtgtacAGCTCCCTGATGTATGATGTTGACACCGAGGTTATTTCAGtatagttacttgtcctaaac
gacttatcagtcttgtgtacGCTATCTTTTCCCATACAGTGAGCATACTTCTTCCTTAATtatagttacttgtcctaaac
gacttatcagtcttgtgtacGAAAGGGATAAAGTATTTTTCATATTTGGGAACTTCCCAGtatagttacttgtcctaaac
gacttatcagtcttgtgtacAAGTACTTTGGAGTTTAGTGGTGGTTTCCTAATATTTCAGtatagttacttgtcctaaac
gacttatcagtcttgtgtacTAGATAATTCTAGATAAACAACAGGGCCAATAATCAGTGGtatagttacttgtcctaaac
gacttatcagtcttgtgtacGCACCACCAAAATTATGTGAGTTTTAACTTTTACCTTGCAtatagttacttgtcctaaac
gacttatcagtcttgtgtacAGCCTTATAATGAAAAATAACAGCCTCTTTTTCTGTACCCtatagttacttgtcctaaac
gacttatcagtcttgtgtacTTTCACCATGTGTAATGTATTATCTTCTTGAGTCCAACACtatagttacttgtcctaaac
gacttatcagtcttgtgtacACTTCCAGACATCTCCCTTGACACTTCTAAATGATATATAtatagttacttgtcctaaac
gacttatcagtcttgtgtacATTTAAGTGTTCAATTCCATAAGTTTTGGCAGATGTGCTGtatagttacttgtcctaaac
gacttatcagtcttgtgtacCTGTGAAGACAAGGTTCATATTATTCCCACTAAAAGTTTCtatagttacttgtcctaaac
gacttatcagtcttgtgtacGCCCCTCTGTCATCTCCCTCCCACTTCTCCCTGCAACTTCtatagttacttgtcctaaa
gacttatcagtcttgtgtacCCACCCCAAGCAAACTGATTTACTTTCTATCACTATAGGTtatagttacttgtcctaaac
gacttatcagtcttgtgtacTGCACATAATAGTACTTTATTCCTATTTGTTGCTGAGCAGtatagttacttgtcctaaac
gacttatcagtcttgtgtacATATGCTTTCACTTATCTGGGGTAAATATATGAGTGGAACtatagttacttgtcctaaac
gacttatcagtcttgtgtacCTGACAAACTGCTTTTCAAAATGGGGCTTATAGTACTTTAtatagttacttgtcctaaac
gacttatcagtcttgtgtacTGAGAGTCCAAGTCACTTTTTGCTCTTACTAACATTACAAtatagttacttgtcctaaac
gacttatcagtcttgtgtacATAATTTTAAGCAGAGACAATATGATGAGGTGCATGAGACtatagttacttgtcctaaac
gacttatcagtcttgtgtacCTGATTTGTTCATTTCAGGTACCAATATGGCTTGAACTACtatagttacttgtcctaaac
gacttatcagtcttgtgtacTCATGTACTTCTTTGAGCTCCATATACTCTCTTTATGAAGtatagttacttgtcctaaac
gacttatcagtcttgtgtacGATTTATGTTTTCTTTCTCTGAACAGTGCCTTTCAAAGAGtatagttacttgtcctaaac
gacttatcagtcttgtgtacCTTTTGGTGTTGTTGCAAAGGAAGCCGTGTTTAACAAAAAtatagttacttgtcctaaac
gacttatcagtcttgtgtacGAGTTAATATTTATGTGGCTCAAAGTTCTTCCTTCCTCTCtatagttacttgtcctaaac
gacttatcagtcttgtgtacCTCTCCTCCTCTTTGTCTTCTTCTTTTTGGTATATGGCTAtatagttacttgtcctaaac
gacttatcagtcttgtgtacTTCAATTTCTACAGATAAAGCTGCTCACTTTTGATTGAGCtatagttacttgtcctaaac
gacttatcagtcttgtgtacTGTCTATAACGAGTTTTCTGATTCATGAGCATGGTTTATCtatagttacttgtcctaaac
gacttatcagtcttgtgtacCTTTTCAGTGTACAAGTCTTCAACATCTTTTGCCAGAGTTtatagttacttgtcctaaac
gacttatcagtcttgtgtacTAGGAATGCAATTGACTGTTTTATATTATCTCGGATCCTGtatagttacttgtcctaaac
gacttatcagtcttgtgtacCCATAGATAGTCATGTTGTCTTCAAATAAGAAACACTTGGtatagttacttgtcctaaac
gacttatcagtcttgtgtacGGAGGCTGAGGTGGACAGATCACCTATGGTCAGGAGTTTGtatagttacttgtcctaaa
gacttatcagtcttgtgtacTCTACTAAAAATATAAAATTAGCCAGGCAAGGTGGTGCATtatagttacttgtcctaaac
gacttatcagtcttgtgtacAAACTTCTTTTTGCTTGACCCATTGTGGTGGCTAGAACCTtatagttacttgtcctaaac
gacttatcagtcttgtgtacTATGCTGTTAGTTATAGGTTTTTGGTGATTCCTACTGTTCtatagttacttgtcctaaac
gacttatcagtcttgtgtacTATGGTCGATTACGTTGACTGATCTTTTTTTTGATGTAGGtatagttacttgtcctaaac

gacttatcagtcttgtgtacTGCACCCGTTAACTTGTCATGGAACATTGGGTATATCTCCtatagttacttgtcctaaac
gacttatcagtcttgtgtacGTGATGTTCCCTTTCCTGTAAGACATTAAGCTTATCTTCCtatagttacttgtcctaaac
gacttatcagtcttgtgtacCATGTTGAAATTTGATCTTCCATGTGGCAGCATTGGGAAGtatagttacttgtcctaaac
gacttatcagtcttgtgtacGAGGTTGGGGGAGGTGGTGGTGAGTGAGATTTTGCTGTCAtatagttacttgtcctaaa
gacttatcagtcttgtgtacGGAGTCTGACTTCCTTGCTTAATTTCTCTTTCTTGCTTCCtatagttacttgtcctaaac
gacttatcagtcttgtgtacTTACCACGTGATCTCTTTGCAAATATCCACTTCCACTCTGtatagttacttgtcctaaac
gacttatcagtcttgtgtacCTGCCATGTTTTGATACAGCACACGACCCTTACTAGAAGCtatagttacttgtcctaaa
gacttatcagtcttgtgtacCTTGAACTTCCCAACCTGCATAACGATGATCTAAATAAACtatagttacttgtcctaaac
gacttatcagtcttgtgtacCAATGTCTTTGTCCAATTTCGATACCATAATATGCTGGCCtatagttacttgtcctaaac
gacttatcagtcttgtgtacGGATTAATTGGGAAGTATTCTCTCTTCCTTTTCTAGAAGAtatagttacttgtcctaaac
gacttatcagtcttgtgtacCTCAAAATATTTTCTAGAATTCACTAGTGAGGCCTTATGGtatagttacttgtcctaaac
gacttatcagtcttgtgtacTGATTGTCAAAATTACTGGCACAGAAGGAAGTGGTCGATTtatagttacttgtcctaaac
gacttatcagtcttgtgtacTCTCTTGAAATATCTTTAGAAGATACAGTGTCACCATCTCtatagttacttgtcctaaac
gacttatcagtcttgtgtacTGTTATGATTACCATTTCAATACTACTTTGGCTGCATTCCtatagttacttgtcctaaac
gacttatcagtcttgtgtacCTTCTAACTTAAAGAACATGTGTGGTTTGCACACTTTTCAtatagttacttgtcctaaac
gacttatcagtcttgtgtacTGGTACAGAAGTTTGTCTAGCTTGCTAAATATTCCAGATGtatagttacttgtcctaaac
gacttatcagtcttgtgtacAGGAATGTATTTTTTGCTGTTACTGGGTAGGATTGTCATGtatagttacttgtcctaaac
gacttatcagtcttgtgtacATAGCAGTGTTGAAATCTCCGATCATATTTGAGAAATCTGtatagttacttgtcctaaac
gacttatcagtcttgtgtacTTCCATCCAAAGATCCTGTTATTTTCATATAAGACATCCCtatagttacttgtcctaaac
gacttatcagtcttgtgtacGTTAGAAGAACTCAGCCACTCTTAAATCTCTTGTACCAGCtatagttacttgtcctaaac
gacttatcagtcttgtgtacCCATCTTAAATACAAGGGAATCTGGGAAACAAAAAGGAAGtatagttacttgtcctaaa
gacttatcagtcttgtgtacTTGACCCCCAATTCAGAATGAGGTTCATCCCAGATCTATTtatagttacttgtcctaaac
gacttatcagtcttgtgtacTTTAAATTTGTCTTGTGCCACACAATTTACAGGTTGAACCtatagttacttgtcctaaac
gacttatcagtcttgtgtacAACTAAAAGCCTAATTTCTCCCATGAAATCTTTGGGACAGtatagttacttgtcctaaac
gacttatcagtcttgtgtacGCATATTAACATAGCAGCTAGCTTCAATTATTTAGTGGACtatagttacttgtcctaaac
gacttatcagtcttgtgtacTGGAGGAGAAGGTATTATGTTGACAATATATGGCCTTCTGtatagttacttgtcctaaac
gacttatcagtcttgtgtacTGTGTGTGTTTATGTGTTTCCATTTTCCATGCTCAGATAAtatagttacttgtcctaaac
gacttatcagtcttgtgtacGGTTTCTCAATCCCCAAATTGTTATCTGAGGAAAAGAATAtatagttacttgtcctaaac
gacttatcagtcttgtgtacTCAGTACCTTGAATGATTTGGCTGACTCTTCCAACCTCCAtatagttacttgtcctaaac
gacttatcagtcttgtgtacGAAACTGGAACTCTTATCTTCCCATTCACAATCGCAGAATtatagttacttgtcctaaac
gacttatcagtcttgtgtacGTGAAAGACCTATAACTCAAAAAGTCTCAAGCAGGAGGGAtatagttacttgtcctaaa
gacttatcagtcttgtgtacGAGACACAAATTAACCTTGCCAACATGAATGAAAAATCAGtatagttacttgtcctaaac
gacttatcagtcttgtgtacTGGAAGTGAAATCTCTTTTCAATGTTTGGTCAAGTTTTGCtatagttacttgtcctaaac
gacttatcagtcttgtgtacAAGTTTTTATAACGTTTGAGGCCATGGCAGAACAAGGAAAtatagttacttgtcctaaac
gacttatcagtcttgtgtacCAGGAAAAATGATCTGCCATCTTGTACATTTTAGTTTCACtatagttacttgtcctaaac
gacttatcagtcttgtgtacACATTCCTGTTTTCCTTCTGACTCGACATTTTGCTCTCTTtatagttacttgtcctaaac
gacttatcagtcttgtgtacAGCTTGTTCACATGTCCCCCAAACCCCGACCTGCTGTTTCtatagttacttgtcctaaa
gacttatcagtcttgtgtacCTTGAGCCAAGCTGTGTATGGTATTCTGTAGTTTTTATATtatagttacttgtcctaaac
gacttatcagtcttgtgtacAATAGAATGAAAATCTGTGTCGCTCTTTCAAGCTCTAAGAtatagttacttgtcctaaac
gacttatcagtcttgtgtacAGGCTTGACAATCACCTTTTGACAATTCTATTTTCTGAAGtatagttacttgtcctaaac
gacttatcagtcttgtgtacTTCATGATCTTTTTAAAAAGTCCTTTGCTGTGCAGATGTGtatagttacttgtcctaaac
gacttatcagtcttgtgtacATCTATAATAACATCCATTTCCACTTTCTATCTCCAGCTCtatagttacttgtcctaaac
gacttatcagtcttgtgtacTGTGCTAGTACATAATGCCCTTGGACGTGAATTCAGACTAtatagttacttgtcctaaac
gacttatcagtcttgtgtacAAGAAATTTGACTATCTCTACAACCTATGTAGGACTGACAtatagttacttgtcctaaac
gacttatcagtcttgtgtacTGTAAAGAAAGATGAACACATTTGTAGCCACAGGGTTTGTtatagttacttgtcctaaac
gacttatcagtcttgtgtacATTCAGTTAGAGTCATGGTCACATTTTCTGTAATATCTGGtatagttacttgtcctaaac
gacttatcagtcttgtgtacTCAACATTAGTAAATAGGATGGCAGACAATTGCTAAAGTGtatagttacttgtcctaaac
gacttatcagtcttgtgtacTTCTAAGAATTGAAAACTCCTAAAGACTCCAAAGCTTCAGtatagttacttgtcctaaac
gacttatcagtcttgtgtacGACTGAATTGGTGATTAATAGGAGTCATGAATGTAGGTATtatagttacttgtcctaaac
gacttatcagtcttgtgtacCATCTTTAAAGAGGTATGTCTAACTAGTAGCTTGCTTTTCtatagttacttgtcctaaac
gacttatcagtcttgtgtacTACCATGATACCTTTCCTTTGTGCACTTACCTGTGTGATTtatagttacttgtcctaaac
gacttatcagtcttgtgtacGCATTCTTTGGTTTACTTTACTTGCTCCCTTTACTACCCCtatagttacttgtcctaaac

gacttatcagtcttgtgtacAGGCCCTGCAATTACTGTAATACTCCCTGCCTCTTGCACAtatagttacttgtcctaaa
gacttatcagtcttgtgtacTTGTAGTCCAATTTGTTTCTTCACCCCCTCCTGCCCATGCtatagttacttgtcctaaac
gacttatcagtcttgtgtacTTGAACCATGTTTTCTCCTCTCCCAGCCAAGGCCTTGACTtatagttacttgtcctaaac
gacttatcagtcttgtgtacCATTTTTGGCAATGGCTCCACATACAGTTCAGATTTGGGGtatagttacttgtcctaaac
gacttatcagtcttgtgtacGGAGCGGCTGACCCTCTCTTTATACAAGAGCATTATCATGtatagttacttgtcctaaac
gacttatcagtcttgtgtacCAATACTGACATCCTGAGGAAAATAAAAATTCTGTGTGTGtatagttacttgtcctaaac
gacttatcagtcttgtgtacCTCTAAAGGGAATATAGGCAATAAGAACTGAAGCTCTGACtatagttacttgtcctaaac
gacttatcagtcttgtgtacAGCTAACTCACAAAAGGATCTCAACTAGAAGCCAGGCTAAtatagttacttgtcctaaa
gacttatcagtcttgtgtacCCCCAGGAGTACTCACAAATTTATGGTGCCTATCTTATCAtatagttacttgtcctaaac
gacttatcagtcttgtgtacTTGGACTCAGAATTAAGGAGCTTCTGATAAATCACTTTGAtatagttacttgtcctaaac
gacttatcagtcttgtgtacTGGGAATAAAACTTGGACAGAGAAATACAAGATGTTGGCAtatagttacttgtcctaaac
gacttatcagtcttgtgtacCCTTCTGAGTAAGATTAGAGATGGAATTTTCTAGGGCAGAtatagttacttgtcctaaac
gacttatcagtcttgtgtacCCTTTTAGTTCTTGAGGGAATACCATGCATCAATCTAGTTtatagttacttgtcctaaac
gacttatcagtcttgtgtacGGAACGTAGTGTTTTCCTTTAGTTATTTTGTTAGCTTGACtatagttacttgtcctaaac
gacttatcagtcttgtgtacAGTAGAGACCCGTGGTGCTATCAGAGCTCATTTTAACACAtatagttacttgtcctaaac
gacttatcagtcttgtgtacGCTTCCATGTAGCTATATTTTGTTTTGACCTGCAGATTTAtatagttacttgtcctaaac
gacttatcagtcttgtgtacAGCATCCTATGATCAAAGCAGAATAATAAGGTCATCATTCtatagttacttgtcctaaac
gacttatcagtcttgtgtacTCTTCTAACTGAAATGAAAGTATTCAAGGTGCAAATCCTGtatagttacttgtcctaaac
gacttatcagtcttgtgtacCAGGCAAGTCTGCTCTGTGATTTTCTCTACCTCTGTGAGCtatagttacttgtcctaaac
gacttatcagtcttgtgtacGCTGCTCTGCATCAGTTCAATGAGGTTTTCTTCCTGACTTtatagttacttgtcctaaac
gacttatcagtcttgtgtacTGCTTGAATTAAAGTGGGTTGGAAGTGTAGAAAGGCAGAAtatagttacttgtcctaaac
gacttatcagtcttgtgtacATGTTTCTGGCATTGGTATACCTCTTTCCATTCAAGTAATtatagttacttgtcctaaac
gacttatcagtcttgtgtacAGCTGGAGCTCTGGCTTTACTTTCCTTCTGCTAGTCTTGGtatagttacttgtcctaaac
gacttatcagtcttgtgtacCCTTTTGGCCCCCTCAATGTTCTTTTTCTGAATTTTTTGGtatagttacttgtcctaaac
gacttatcagtcttgtgtacTCAAGAATAGACTAAGTAAGTAAAACAAGCCTCTCCTCTTtatagttacttgtcctaaac
gacttatcagtcttgtgtacCACCTGTGCAGACACTTTGCTACATGCGTCTTTGAGATGTtatagttacttgtcctaaac
gacttatcagtcttgtgtacTTCCTTTCACCCTGAGAATCCCCTGGTACAGCAATCTTAGtatagttacttgtcctaaac
gacttatcagtcttgtgtacCACAGAAGTGCACTAGACCCTATGCTTTTGGAAGAGAAGGtatagttacttgtcctaaa
gacttatcagtcttgtgtacGGACAGATGCTTTCCTTCCCCTTTCTGCATATCCATGGAGtatagttacttgtcctaaac
gacttatcagtcttgtgtacCTGATGGGAGGCTTAGGTGAGGTAGAGCATATGGAGATAGtatagttacttgtcctaaa
gacttatcagtcttgtgtacCTGTGATTTCTCAGTAAACTTGCATTTCATCCCCTTTGTGtatagttacttgtcctaaac
gacttatcagtcttgtgtacAAGGCATTTCCAACTGCTCTTTTCAACCCTGAATGGATGGtatagttacttgtcctaaac
gacttatcagtcttgtgtacCTCTGGAACTGAGGCACAATAGATCAATGGGAAGGAGAGGtatagttacttgtcctaaa
gacttatcagtcttgtgtacCAATAACTGGGAGAGAAAGAAAAGCTGTGGTTTTTAGCAAtatagttacttgtcctaaac
gacttatcagtcttgtgtacCAAAGATTTGCTATAGATGTAACTATTTCAGGCCACTAAGtatagttacttgtcctaaac
gacttatcagtcttgtgtacGGTTGATGCCCACAGATTTATGATATTGACTAACATTACAtatagttacttgtcctaaac
gacttatcagtcttgtgtacTTATAAGTATGAACATATATGCATGACAATGGCTCCTAGGtatagttacttgtcctaaac
gacttatcagtcttgtgtacCCTGTCAGGATTCTTAGACATTATGGTGATTTTTCAAAGAtatagttacttgtcctaaac
gacttatcagtcttgtgtacAGGCTGACTGGTGGAATCTATCTGTGATGCAAAATGGTTCtatagttacttgtcctaaac
gacttatcagtcttgtgtacAGAGGTAAGAATTTGGAGAAAAAATTTTACTCAGAGGGAGtatagttacttgtcctaaac
gacttatcagtcttgtgtacCCATGGAGAGTTGGGAGTGGGTTGTTGAGGGAAACTTCACtatagttacttgtcctaaa
gacttatcagtcttgtgtacTGTTTTCCTCCTTGGCAAGACAAATAAGACAGAAAGAAAAtatagttacttgtcctaaac
gacttatcagtcttgtgtacGAGTTTAATTATTGTTTTTCTCTGCATGATTCGCTCCTTGtatagttacttgtcctaaac
gacttatcagtcttgtgtacCCTACCACTTAGTTTTGATTCTTCTTTTATCCTTCATCTGtatagttacttgtcctaaac
gacttatcagtcttgtgtacATTAAATATCAATTCAGGATCACAGCTTCAAGCTAGTTGCtatagttacttgtcctaaac
gacttatcagtcttgtgtacACTCAAATATTTTTTCATTTCCCTCTCTTCTTCCTTCTCCtatagttacttgtcctaaac
gacttatcagtcttgtgtacCCGCTAGGCTTGTTCACTCATTCCCCTTCCCTGTTCCTCCtatagttacttgtcctaaa
gacttatcagtcttgtgtacTGTTTTTTCTATAGCGTGCTTTAAACACTGAAAACCGGAGtatagttacttgtcctaaac
gacttatcagtcttgtgtacTGCTAATTCGCAAATCTACTTCCAGTTGCCAGTTCCTTTGtatagttacttgtcctaaac
gacttatcagtcttgtgtacGGCATCTTTGAAATGGTTTCCATTATATGCTTGATTCAGAtatagttacttgtcctaaac
gacttatcagtcttgtgtacAATTCACCCTGAGCTTTTATTTGGAAAAATAATTGCCTGCtatagttacttgtcctaaac
gacttatcagtcttgtgtacTCAGTATTGTGAATTTGAAAGGGCTAGGCTGAATTAGACAtatagttacttgtcctaaac

gacttatcagtcttgtgtacCCTGGAATGAGAGTTTTGCCTATACACTATAGTAATGAAGtatagttacttgtcctaaac
gacttatcagtcttgtgtacGCATTTATATGTTTTTGCAAGGTTACCTGAACACAAAAGGtatagttacttgtcctaaac
gacttatcagtcttgtgtacCAGTTCAAAATTACATACTGGTAGGTGTAGTCCATTATTCtatagttacttgtcctaaac
gacttatcagtcttgtgtacTATCCTCCTCTCCACATATAAACATGAAAATATACACAGGtatagttacttgtcctaaac
gacttatcagtcttgtgtacTTCTTCTTTATGCTCTTCTTCCTCCTTTGTTTACTTACGAtatagttacttgtcctaaac
gacttatcagtcttgtgtacTGACAGATTCAGTATACTTCTGCCTAAGTTCATTCTCTTAtatagttacttgtcctaaac
gacttatcagtcttgtgtacGAACTCATCATTTTGTACATCCTTTCCCCTTCAGCTGTGTtatagttacttgtcctaaac
gacttatcagtcttgtgtacCTTGATTCATCACGTTCACAGGTTCTATTCTTCACTAGTCtatagttacttgtcctaaac
gacttatcagtcttgtgtacAGTGTACTGATTCTCTTCTCCCTCCCATCTCCTTTCTTTTtatagttacttgtcctaaac
gacttatcagtcttgtgtacTTGGGAAACATGTTATTATAGACTTTTCTAGGAGGTGGGGtatagttacttgtcctaaac
gacttatcagtcttgtgtacTGGACTGTGGCTGATGTGCATGGATAGAAACTATTTGTGAtatagttacttgtcctaaac
gacttatcagtcttgtgtacGTATTATACATGAGTCTTGCATCCCTACTGGTCAATATAGtatagttacttgtcctaaac
gacttatcagtcttgtgtacTCTGAAGGATTCAGTTGTGATATAAAATAAGTGCCTTAGCtatagttacttgtcctaaac
gacttatcagtcttgtgtacGACAAGATAACAAGTGGAAAGGAGTCATGTGAGAGGAAGCtatagttacttgtcctaaa
gacttatcagtcttgtgtacCTGAAAGAATAGAAGACTAGGAAAGAAAGGCATATGGATTtatagttacttgtcctaaac
gacttatcagtcttgtgtacTGAATAGAACATGCCATAGAAGATAACTTTGAAAGTCCTGtatagttacttgtcctaaac
gacttatcagtcttgtgtacCAATATTTGCAAATCAGTAAGTAGTACAAGCCAGACAGAGtatagttacttgtcctaaac
gacttatcagtcttgtgtacGGCAGAAGAAACGATGGCAGAAAAATGGAATAGAACTTTCtatagttacttgtcctaaa
gacttatcagtcttgtgtacCAGGTGATCAAACAAAATAAATGGTCTAATTGCTACCTAGtatagttacttgtcctaaac
gacttatcagtcttgtgtacCCCTAAGAGTCTGGCCTTCTGAATTCCTCAACTTTCTCTGtatagttacttgtcctaaac
gacttatcagtcttgtgtacCTAGTTATTGAATAGTTTCCCAGTCATCTGCCTCCTCTATtatagttacttgtcctaaac
gacttatcagtcttgtgtacCTGAGGATGCCAAAGAGGAGGTACATGACTCTTATTTTCCtatagttacttgtcctaaac
gacttatcagtcttgtgtacCAAATTATATCATTTGCTGGGGGGATGGAGTAAGCATTGCtatagttacttgtcctaaac
gacttatcagtcttgtgtacGAGTGGTAAAATGCTTCTGAAATACTTAGAGAAAATGCTCtatagttacttgtcctaaac
gacttatcagtcttgtgtacCAATATTGTTTTACCTTTCAGTGTTCATTTGTTGTGGGTCtatagttacttgtcctaaac
gacttatcagtcttgtgtacAGTAGCAGGTTAACATTGCCACTTAGGAAAAAGCAGAAATtatagttacttgtcctaaac
gacttatcagtcttgtgtacTATTTAAAATTCTCTACTCACCCCACATTTCACAAACGTCtatagttacttgtcctaaac
gacttatcagtcttgtgtacCTTTCTAGCAAAGTGATGGCTTGTATTGGTTCACATTTAAtatagttacttgtcctaaac
gacttatcagtcttgtgtacGCTGGTTGAGCCATTTCACTTGCCTACTTAGAATATTAATtatagttacttgtcctaaac
gacttatcagtcttgtgtacGGTACATTTCCTCCACCTCTTCTCTGCAGTCCCTAAATTTtatagttacttgtcctaaac
gacttatcagtcttgtgtacTTATCTATGGATACTAGCAAACAGGTAAATCATGGAATCCtatagttacttgtcctaaac
gacttatcagtcttgtgtacCATTAACTCCATGTAAGTTGATGGCCAAATTCTTCTCATAtatagttacttgtcctaaac
gacttatcagtcttgtgtacCTGCTCAATTTATTTTCAGGGGACGCATGCACGCACGCACtatagttacttgtcctaaa
gacttatcagtcttgtgtacTTGCAGATAAATTGTCATGCCAAATCTTTCCAAGATTAGGtatagttacttgtcctaaac
gacttatcagtcttgtgtacTTTCAGAGAAAATCTTGCTATAAACTACACAGGATCACTGtatagttacttgtcctaaac
gacttatcagtcttgtgtacGTGAGTTAGCTTGCACTATTCATGGGGTAATCCTGCCTCAtatagttacttgtcctaaac
gacttatcagtcttgtgtacTTCTACTTGGAATGCTTATTAGAGTCACTTTTTTGTGTGCtatagttacttgtcctaaac
gacttatcagtcttgtgtacTCCTCTTTAAATGCAAATATGTGACCCAGGAGAATACATTtatagttacttgtcctaaac
gacttatcagtcttgtgtacTTCATTGTACTTTCATGAGGAACACTACACCAAAGGATGAtatagttacttgtcctaaac
gacttatcagtcttgtgtacATCTAGGTGCCAGAGAGGAAAATCGTGTCAGTAGTGTGGCtatagttacttgtcctaaa
gacttatcagtcttgtgtacTTAACACAACTCTGTGGGTTAATATTCCCATCATACAGATtatagttacttgtcctaaac
gacttatcagtcttgtgtacCTGTGCATTGAGTGCTATTCTATAAAACTATTGATATGGCtatagttacttgtcctaaac
gacttatcagtcttgtgtacATCATATTGCCATACTAGTATCTTCTGATGAAAGTTCACCtatagttacttgtcctaaac
gacttatcagtcttgtgtacAGTAGCCTCATATTTGATACTGTACTCCTGTCATGGTTGCtatagttacttgtcctaaac
gacttatcagtcttgtgtacCATGATGTTTTCTTGTTTTAGAGTATGAGGAGGTGGAGGTtatagttacttgtcctaaac
gacttatcagtcttgtgtacAAGAGTCACATAAGTTTACCTGAGAACTGCAAAGTGCGCGtatagttacttgtcctaaa
gacttatcagtcttgtgtacTGGTGATATTATGTTGTAAGAAGTAGAAAGCCAGGCGGTGtatagttacttgtcctaaac
gacttatcagtcttgtgtacTCAAAATCATCAAGGCAATGTGCTGCTTCTCTTGCTTCTAtatagttacttgtcctaaac
gacttatcagtcttgtgtacGAAAGTCTATGTAATAACAATTTTTCACGTCCATGGCTGTtatagttacttgtcctaaac
gacttatcagtcttgtgtacACTATCTTCAAGTATATGAGGGATTGTTACAGCATTTGGAtatagttacttgtcctaaac
gacttatcagtcttgtgtacCAAAGAGGAAAACAAGAGAAAATGAATTTGCAAAGTGGCAtatagttacttgtcctaaa
gacttatcagtcttgtgtacCATGAGGACTATTCAATGGTGAAAGTGGCAACTTCTAGGGtatagttacttgtcctaaac

gacttatcagtcttgtgtacTGTTTGGGGTGAACATCACTTAGTACCCTCCCTCCCTCTGtatagttacttgtcctaaac
gacttatcagtcttgtgtacATACAATATTGAGAACAAAATTACTGGGGCACATCTGAGAtatagttacttgtcctaaac
gacttatcagtcttgtgtacAATTCCAGGTAAAATTAGTTTTTCCAAACCCGATGATGACtatagttacttgtcctaaac
gacttatcagtcttgtgtacGCAGAATATTTACCTTTTTCTCCACTGTGACTCAATTTTGtatagttacttgtcctaaac
gacttatcagtcttgtgtacGAGTCTTTAAAGTGTTGGTGAAACTCCAGGCATGTTTATCtatagttacttgtcctaaac
gacttatcagtcttgtgtacTGGTCATCACTATACATATTTCATCAGTGAGATGGAGTTTtatagttacttgtcctaaac
gacttatcagtcttgtgtacAGGTCTCTTTTTGTTTCACTAGGATTCTCTATGGTTTTAGtatagttacttgtcctaaac
gacttatcagtcttgtgtacGAGAGGAAAAGATTCTGGAAGATGTTTGATCACTTTGGAAtatagttacttgtcctaaac
gacttatcagtcttgtgtacGTTCTAACTGATAAACTGATGAAGCTGATGCAAAATCACCtatagttacttgtcctaaac
gacttatcagtcttgtgtacTGGCACTCCTGTCAAGGAACCAACATGAATCTTTTTAAAAtatagttacttgtcctaaac
gacttatcagtcttgtgtacACATAATACAATCATGGGTAAACTTTCTCGATCACCTTTGtatagttacttgtcctaaac
gacttatcagtcttgtgtacGTAATTCCTACAGAGTGGCCCTTGTGACTTATCAACTTTTtatagttacttgtcctaaac
gacttatcagtcttgtgtacTGTTGTCTTTACCTGTATTTCAATGTATATGAAGTGGAGGtatagttacttgtcctaaac
gacttatcagtcttgtgtacTGATGTTACTTCTTTGGCAAGAAAACCCATATTTCTCTTGtatagttacttgtcctaaac
gacttatcagtcttgtgtacAAATAGCCCTCCATGTACTTTTTTATACCAGTATGAGTTGtatagttacttgtcctaaac
gacttatcagtcttgtgtacATAGATTTCTTGTTAATGGCTGGTTTATAACTGAATGCGGtatagttacttgtcctaaac
gacttatcagtcttgtgtacGAACAATTTCTTAGGCTCATGGCACCTTGGTCTGGAGTTTtatagttacttgtcctaaac
gacttatcagtcttgtgtacTTTCATGTAAATTTGCTCCTAGAGTAGAAAGGTAGTCTGCtatagttacttgtcctaaac
gacttatcagtcttgtgtacTAGAGACATCAATTCTATGGAACGTACTGAAAACAAGAGGtatagttacttgtcctaaac
gacttatcagtcttgtgtacGATACCATCTATCTTAGCACAGAAACTCAATAAACAGAGTtatagttacttgtcctaaac
gacttatcagtcttgtgtacCATTGCCAACACTTCCTACCACCCTTTCCTAGCAAATTTGtatagttacttgtcctaaac
gacttatcagtcttgtgtacACTCCTCAATTTCTAACAAACTATTTCCTCTCTCCTTATGtatagttacttgtcctaaac
gacttatcagtcttgtgtacCTTAAAGAGAAAATCAAGGCAATAGAAAGGAAACCTCATCtatagttacttgtcctaaac
gacttatcagtcttgtgtacTCAGGTCCAATGGGAGAACAGAGGAGGCAGGTTGAGGGGAtatagttacttgtcctaa
gacttatcagtcttgtgtacTGGGAAGACGACCAGAGACAGAGCCTTTATTTCAGTTTGCtatagttacttgtcctaaa
gacttatcagtcttgtgtacCCATTGTCTTGGTTCATACAGAAAACTCTTATTTGGGAAAtatagttacttgtcctaaac
gacttatcagtcttgtgtacCTTTTTAGAAACTGTATCCTGGCTTGCAAACACCTGAATCtatagttacttgtcctaaac
gacttatcagtcttgtgtacATGCAAATTCCACATTTGGTGTGTACAAGGTTAGAGTTAAtatagttacttgtcctaaac
gacttatcagtcttgtgtacCTTCATAGAGATTTAGAATTTCTAAGGGCATTGGGTTATGtatagttacttgtcctaaac
gacttatcagtcttgtgtacAAATTTCTCCTGATCCAGTCAATAAAGCAATGAGTTATCCtatagttacttgtcctaaac
gacttatcagtcttgtgtacCTGATTGCTAGGCAGAATAATCCTTTAAAACTTTTTTGCCtatagttacttgtcctaaac
gacttatcagtcttgtgtacCAGAAAATGAGTGCTGCTCCGTGTGCTCTATCCTGGAACAtatagttacttgtcctaaa
gacttatcagtcttgtgtacGTTTGGAAGTTACTGTTTAATCTGAGTTATTTTGGGGAAGtatagttacttgtcctaaac
gacttatcagtcttgtgtacGACTCGTAGAAGCTAAGTGTTCATACAGATTGTGATTTTGtatagttacttgtcctaaac
gacttatcagtcttgtgtacCAATTAGAATGTTGCTGCTGTGATGAATCCATACTGTACAtatagttacttgtcctaaac
gacttatcagtcttgtgtacATTTTCATTTTGGGTAATTCGATAAATGCTTGTGCTGTGCtatagttacttgtcctaaac
gacttatcagtcttgtgtacACCCCTTGAATTGTCCCTCAAAGCTAGAGAAGAGTTTTCGtatagttacttgtcctaaac
gacttatcagtcttgtgtacACACCTCCATAATAACTTGATATTTGCACATAGTTTAGGGtatagttacttgtcctaaac
gacttatcagtcttgtgtacAGAGCTAGATATTAAATATTTAGGCTTGCAGGACATATGGtatagttacttgtcctaaac
gacttatcagtcttgtgtacTGGTAGCTCAAATGCAGCCATACACAACTTACAGACAACTtatagttacttgtcctaaa
gacttatcagtcttgtgtacAAAATAGCTGGCTTATGAGACATAGTTTTCTGACCCCCTCtatagttacttgtcctaaac
gacttatcagtcttgtgtacCTGGAGCATGGGGATGTGGAAAACATGTTATCACTTTCAAtatagttacttgtcctaaac
gacttatcagtcttgtgtacATCTTATAGGTCACTTTAAATGCTGCTATTTTCTGGTGTCtatagttacttgtcctaaac
gacttatcagtcttgtgtacAATACATATATTCTTTTATCCCTGCTTCACACTGGCACAAtatagttacttgtcctaaac
gacttatcagtcttgtgtacCCAACTGGGCATATTGTTTTTCTCCTTCCTCTCTCATAATtatagttacttgtcctaaac
gacttatcagtcttgtgtacCATTGCTCCCTCTTCTAATAATGCCCTACTAGCGTGATAGtatagttacttgtcctaaac
gacttatcagtcttgtgtacCATCCAGATATTCAAATGATCAGCTAATACTTACTATGCCtatagttacttgtcctaaac
gacttatcagtcttgtgtacGACACATAGAAATAGACTCTCAAGGAAAGCAAATGCCAGAtatagttacttgtcctaaa
gacttatcagtcttgtgtacTTTTTGAGAAAGCTAGATTTGCAGTTGGTGAATGGTTAGTtatagttacttgtcctaaac
gacttatcagtcttgtgtacGCTTATTGAAGTGTAGCTGCTAAGTACTATAACAATGCAAtatagttacttgtcctaaac
gacttatcagtcttgtgtacCAGGATGCACATGTAGTAGGAGAGGAATCAAATGTCTTTGtatagttacttgtcctaaac
gacttatcagtcttgtgtacCTTTTGGAGTTTCAATAGAGCTTGACTATAGGGTGATGCAtatagttacttgtcctaaac

gacttatcagtcttgtgtacTTTACTCACTCATCCAACAGTTCATTCAAGGACTTCTCCCtatagttacttgtcctaaac
gacttatcagtcttgtgtacACAGACGCTCACAAGGGAAGATGTATGTTTGGTGTTAAGGtatagttacttgtcctaaa
gacttatcagtcttgtgtacTGGATAGAGTCTGCAGAAGTCAAGAAAGATTTACCAGAGGtatagttacttgtcctaaa
gacttatcagtcttgtgtacGCCAATAATGCAAAGAGAGATCATAAGAAATAGTCTAGAGtatagttacttgtcctaaa
gacttatcagtcttgtgtacGTCTGTTCTCATTTTGCTGAAAGAGACATACCTGAGACTGtatagttacttgtcctaaac
gacttatcagtcttgtgtacGGCAGGCAAGAGAGTGTGTGGCAGGGGAAGTGCCCTTTATtatagttacttgtcctaa
gacttatcagtcttgtgtacGTGGGGATTATGGGAGCTATGTTTCAAGATGAGATTTGGGtatagttacttgtcctaaac
gacttatcagtcttgtgtacAAACTGTATTTGGGCTTACCTTGTGATATGGTTTGTCTGTtatagttacttgtcctaaac
gacttatcagtcttgtgtacCCATAAACTTGGCTAGAGACAATAAGGGTGTTAGAGTGAAtatagttacttgtcctaaa
gacttatcagtcttgtgtacTTTTATCACTTACTTAAAATGTCACCTCAGGGAAGTTGTCtatagttacttgtcctaaac
gacttatcagtcttgtgtacGTATGAATAACCACACAATAGCTTCAAGATGGCACTTCGAtatagttacttgtcctaaa
gacttatcagtcttgtgtacGATAATTTCATGTGGCTTGCTTTACAGTGAAGACAATTTCtatagttacttgtcctaaac
gacttatcagtcttgtgtacATTCCTGAATGGATCATTTTCAAGGAGAATCAAGTAGAACtatagttacttgtcctaaac
gacttatcagtcttgtgtacGTGGGAAAGGGCTGGGAACTTACACAAAAACAAAATTCTGtatagttacttgtcctaaa
gacttatcagtcttgtgtacGGATTAAAATGGATCATGTTGAAAAGAGCATTAAACAGCCtatagttacttgtcctaaac
gacttatcagtcttgtgtacTGAAGTGTAACTTATACACAGCAACATGTCAGATCTTAAGtatagttacttgtcctaaac
gacttatcagtcttgtgtacATAAATTTTTGAACCTGCCATCTAGTGGAACGTGTCTGCCtatagttacttgtcctaaac
gacttatcagtcttgtgtacAGGGATTTCTTCATTCTCCTTTCTAGTAACCACTCCAAAGtatagttacttgtcctaaac
gacttatcagtcttgtgtacATGATTTTGAGATTCTTTCAAGCCGTTGGTTATTTTGGTGtatagttacttgtcctaaac
gacttatcagtcttgtgtacTGTCAAGTGTTAATGAAGCAGTGTGTATATTTGTGTTCAGtatagttacttgtcctaaac
gacttatcagtcttgtgtacTTTATCTTTGATATATACCTGAAAGTACGGTTACTGGCTGtatagttacttgtcctaaac
gacttatcagtcttgtgtacTGAAGCTTTGCAAGAAATGGCCAAACATGGCTGGGCGCGGtatagttacttgtcctaa
gacttatcagtcttgtgtacGGAGGCTGAGGTGGGAGGGTGATGAGGTCAAGAGATTGAGtatagttacttgtcctaa
gacttatcagtcttgtgtacGGCAGGAGAATTACTTGAATCTGAGAGGCAGAGGTTGCAGtatagttacttgtcctaaa
gacttatcagtcttgtgtacGGTTGTGCCATTTTACATTACTACTAGCAATGAGTAAGAAtatagttacttgtcctaaac
gacttatcagtcttgtgtacCTTCATATTGTCTTTTTTTCCCTGTTATTCTAGTGGTGTGtatagttacttgtcctaaac
gacttatcagtcttgtgtacGCATTTTTCTGTTGCCTTATGATGCTGACCATCTTTTATAtatagttacttgtcctaaac
gacttatcagtcttgtgtacTATCATTGATCTAATTGTGTCCTCTGTGTATTCTGACTACtatagttacttgtcctaaac
gacttatcagtcttgtgtacTCTATTTCCATTTTCTTACTAGTGTCTTCTGATGAACTGCtatagttacttgtcctaaac
gacttatcagtcttgtgtacTGTCTTATCACAAGGATGCTCTGCCGATCTCCTGTATTTTtatagttacttgtcctaaac
gacttatcagtcttgtgtacAAAAATTTTTTATTGCCATAGGTTCTTGGGGAACACATGGtatagttacttgtcctaaac
gacttatcagtcttgtgtacGTCTTTTATCCCTAACCCCCTCCCATGCTTTCCCCTTAGTtatagttacttgtcctaaac
gacttatcagtcttgtgtacAGCTCACATTTATGAGTGAAAAACTTACGTTTGGTTTTCCtatagttacttgtcctaaac
gacttatcagtcttgtgtacATAATAGTCTCCAGTTCCATACAGGTTGCTGTGAGTGCCAtatagttacttgtcctaaa
gacttatcagtcttgtgtacGCTGAGCAGTATTCCATATATCACAATTTCTTTAGCCACTtatagttacttgtcctaaac
gacttatcagtcttgtgtacATTGATGAACATTTGGGATGGTTCCATGTTTTTGCAATTGtatagttacttgtcctaaac
gacttatcagtcttgtgtacGAAGTGTTCCCGTTTTATCACGTCTATGCTAACATCTATTtatagttacttgtcctaaac
gacttatcagtcttgtgtacATGGTCATTTTTGCAGGGGTAAGTGCTATCACATTGTGGTtatagttacttgtcctaaac
gacttatcagtcttgtgtacTGATGTTGAGCACTTCTTCACATATTTGTTGGTCATTTGTtatagttacttgtcctaaac
gacttatcagtcttgtgtacTTTACTCTGCTGACTGTTCAATTGCTGTGCAGAAACCCTTtatagttacttgtcctaaac
gacttatcagtcttgtgtacTAATTAAGTCCCACCTATTCATCCCTGTTTTTGTTGCATTtatagttacttgtcctaaac
gacttatcagtcttgtgtacTGGTCATGAAGTCTTTGTCGAAGCCAATGTCTAGAAAAGTtatagttacttgtcctaaac
gacttatcagtcttgtgtacGTATTTTTTCTTCTTTCCTGCTCCCTACACAAATATACTCtatagttacttgtcctaaac
gacttatcagtcttgtgtacTGAACCACATAATCTATTTGTCTATCTACCGACCTACCAAtatagttacttgtcctaaac
gacttatcagtcttgtgtacTGAGTCTTTGTAGTTAAATGATATCTAGGTTTGACAGAGGtatagttacttgtcctaaac
gacttatcagtcttgtgtacGCAATGTCTTCTTTCTCAATTCATACCTGCTCTTCTACTTtatagttacttgtcctaaac
gacttatcagtcttgtgtacACAACATGATGCAATTGTGATATTTTCTTCCATTTCTCCCtatagttacttgtcctaaac
gacttatcagtcttgtgtacTGACATTTGTTTTCCCTTTTCTTCATTAGTGATGTGGAAGtatagttacttgtcctaaac
gacttatcagtcttgtgtacTCGATATAAGATGAGAACTACTTATGCTTTCTTCCTTCTCtatagttacttgtcctaaac
gacttatcagtcttgtgtacGAAATAAGTATTCTGTCTTGAATTCTGCTAGGAAAGTTCCtatagttacttgtcctaaac
gacttatcagtcttgtgtacAAATTCTTTCTAAATGTATACTGGATCTGTCCTCGAAGCCtatagttacttgtcctaaac
gacttatcagtcttgtgtacGTGTGTGTGCATGTGTACAAGCGTGTGTGTGTATACCTGTtatagttacttgtcctaaac

gacttatcagtcttgtgtacGTATGGAAATACCTTCATTGTATAGACAGCTTTAGATCACtatagttacttgtcctaaac
gacttatcagtcttgtgtacTGGGAACTTTCTCACTCCTTTGTATGTAGTAGTTCATATTtatagttacttgtcctaaac
gacttatcagtcttgtgtacACTTCTTGGTTGATGACCCAGTGTGTATTAACCTAATAATtatagttacttgtcctaaac
gacttatcagtcttgtgtacAGCACATCAACTACAATAAAAAGAGATTTAACTGCAAGGGtatagttacttgtcctaaa
gacttatcagtcttgtgtacTAGCATCAGTTTTCATATTATTCACATTTCCTTCTTGGCGtatagttacttgtcctaaac
gacttatcagtcttgtgtacACATCGGCTTCAGTTGTGAATCATTCTACGTGAATATTAAtatagttacttgtcctaaac
gacttatcagtcttgtgtacACTGAATTTTATGATCAAAACATGTTCTGTGGGGTGGGACtatagttacttgtcctaaac
gacttatcagtcttgtgtacCTTTTAATAGAGGAACAGAATATTCTTGAGAGGTTGGATGtatagttacttgtcctaaac
gacttatcagtcttgtgtacCATACATCAATCAAACAAGGTGTTCCATTTTGAACTAAGCtatagttacttgtcctaaac
gacttatcagtcttgtgtacTGACAGTACTGCTTGTGGTTCATGCTGCATTTTAATGTAGtatagttacttgtcctaaac
gacttatcagtcttgtgtacGCTGGGTTGCTTGTCTAGAGTCTCCATTCATTCAATCTTTtatagttacttgtcctaaac
gacttatcagtcttgtgtacGTCCTATCATTTCCTCTCTAAGTTGCAGCTTAAAATTATCtatagttacttgtcctaaac
gacttatcagtcttgtgtacGTCAGTAACATTTGGCAACATTTAGTCTTTGGGATATTGGtatagttacttgtcctaaac
gacttatcagtcttgtgtacTAGTTTGTTGTTTGCTTTGAATATTTGCTCCCACACATTGtatagttacttgtcctaaac
gacttatcagtcttgtgtacTTCTATTGTGAACCAATAATTTTAGTGCTAACTGCCACAGtatagttacttgtcctaaac
gacttatcagtcttgtgtacCAGAAAATCTGCATGCCACTCATCTTGGTAATGATATGTAtatagttacttgtcctaaac
gacttatcagtcttgtgtacATAGTGAATGAAAGAGTCCATAAATTAAAGACACGAAGCCtatagttacttgtcctaaa
gacttatcagtcttgtgtacACATCATCCAGAATAACAACACCAACATACAAAAAACAGCtatagttacttgtcctaaa
gacttatcagtcttgtgtacCATAAAGTATGGCTGTAGATTGTACTGATTGAATTTCAGCtatagttacttgtcctaaac
gacttatcagtcttgtgtacAATCATCTCTAAGTTGTTAGGTTCTATCATCAGGAGAATGtatagttacttgtcctaaac
gacttatcagtcttgtgtacACTATGGACTACTATGCAATGATTACTATAAAGGCACATGtatagttacttgtcctaaac
gacttatcagtcttgtgtacATTGCAGCACTAGTATATATAGCAAAGACTTGAAACCAACtatagttacttgtcctaaa
gacttatcagtcttgtgtacGGGGAACATCACACATCGGACCCTGTCTGGGGATACGGTGtatagttacttgtcctaa
gacttatcagtcttgtgtacCAACACCATGCTCAATTCTTAAAATGTAGATTCTAGGAAGtatagttacttgtcctaaac
gacttatcagtcttgtgtacTGTGCGAAGGAAATTTACATATCAAAAGTCACACATTGGTtatagttacttgtcctaaac
gacttatcagtcttgtgtacTTGAATGGATTTTTCTGTGGAATAGGGGAAATTGAATGGGtatagttacttgtcctaaac
gacttatcagtcttgtgtacATGACAGCATGCCAATAGTTGAGGATTATGAGGATTATAAtatagttacttgtcctaaac
gacttatcagtcttgtgtacGGTAAATCTGTATAACAATGTTGCCCTTCTGGAAAGACTAtatagttacttgtcctaaac
gacttatcagtcttgtgtacGAGTCAGAACACTTGCCTCTCATATTTTCTCTGCCATATTtatagttacttgtcctaaac
gacttatcagtcttgtgtacGACCAGAATTTCCTCTCTCCAAGTGCAGGTATTTGATAGTtatagttacttgtcctaaac
gacttatcagtcttgtgtacCCGCAAAGCATCTTCTGGCTTTAAACGTTGATATGAACATtatagttacttgtcctaaac
gacttatcagtcttgtgtacACAGCAAGAGTTACACACATATGTATGATTACTAGAAGCAtatagttacttgtcctaaa
gacttatcagtcttgtgtacTCTTTTGTCAATATTAGGCAAAGAGACTTCCTATCCTGCGtatagttacttgtcctaaac
gacttatcagtcttgtgtacCCCCTGCAGATATGGTTTGGGTGCATATACTCCTCATTTTtatagttacttgtcctaaac
gacttatcagtcttgtgtacCATTTTTTGGGTGCATGTACTTTGATGGTACTCTTGTGAAtatagttacttgtcctaaac
gacttatcagtcttgtgtacGAATATGTGTTGTTGAGGTAAGTTATGGCTATGTCCAAGGtatagttacttgtcctaaac
gacttatcagtcttgtgtacCAGAGGTTGGATTAGAACTTCAGGAATTGTTCACTATTGCtatagttacttgtcctaaac
gacttatcagtcttgtgtacCACTCCTTATGCCTCGCTGTTCCCTGCTTTGTCATTCATCtatagttacttgtcctaaac
gacttatcagtcttgtgtacGCTTTCTTATCTGCCTGACTTTTTGGTTTGTTGCTATTACtatagttacttgtcctaaac
gacttatcagtcttgtgtacCTTCTTAGGCATATTTGGTGGTGGTTGATTGCAGCTGCTAtatagttacttgtcctaaac
gacttatcagtcttgtgtacAAAGCTTCTATAAAGGAAAGAACTGAATCTGCATCTACTGtatagttacttgtcctaaac
gacttatcagtcttgtgtacTGATTCTTCACTGCCCTATATCCCTTTGCTTCCACCACACtatagttacttgtcctaaa
gacttatcagtcttgtgtacGTTGGGTTAAGAGATATAGAATAGGTACTTGGTAACATTGtatagttacttgtcctaaac
gacttatcagtcttgtgtacACTGTCTCACTGTCGTTCAAATTTTGGATTGGACTAAGTTtatagttacttgtcctaaac
gtagagtgtagcacctatccTAATACCAATTGCCCAGTGGGTGTTTTTCTCTCTTTTCCGggttacctcttatctagtg
gtagagtgtagcacctatccGCCCGCACAAAATTGACTCTTGTTTTATAGCCTTTGGAAAggttacctcttatctagt
gtagagtgtagcacctatccATTGACATATTCCCAGAGGCAATGCACAGCTCCAGTAGAAggttacctcttatctag
gtagagtgtagcacctatccAGAGATAAAGACCCATCATGAACCACTCTGTGCTCAGTGCggttacctcttatctag
gtagagtgtagcacctatccTTAATTACCCATTCCTTCAGCACATTTATGTCAGTTCTAGggttacctcttatctagtg
gtagagtgtagcacctatccGTCTCCTTTTATGTGGCCACTTGAAACTTTGACAGCAACCggttacctcttatctagt
gtagagtgtagcacctatccTCATTTGATTCTGACTGCTTTCTGTACCAAATTGTAATGGggttacctcttatctagtg
gtagagtgtagcacctatccCACTGCTCATCAATGTCATTCTCACAGCCAGGGGTCTAGCggttacctcttatctag

gtagagtgtagcacctatccGACATTTAATTGGGCTGTGCTAAAAGCTTAACTTGTTTTCggttacctcttatctagtg
gtagagtgtagcacctatccGTCTGAAATACACTGGGGTTCTAGAAAGGCCAAATATGCGggttacctcttatctag
gtagagtgtagcacctatccCATTAGTTCATTGCAATGGATGTGCTTTAAAACAGCAGGGggttacctcttatctagt
gtagagtgtagcacctatccTCAACACCTGAACTTGTAAAACAAATATGCTAGAGGAAGTggttacctcttatctagt
gtagagtgtagcacctatccAAATGAAATTCTTTTGTCTACCGAAACCACAGAACCACATggttacctcttatctagt
gtagagtgtagcacctatccTTGCCTACATTATCTCATTTGAACCTATTTAAAGCCCTGTggttacctcttatctagtg
gtagagtgtagcacctatccTGCAGAGTAGCTCTTATTTCCCATGTATAGATGTTGAAATggttacctcttatctagtg
gtagagtgtagcacctatccCAGGTTTTCTTCAAAGAAATGTTCATCCCTTGCATATACCggttacctcttatctagtg
gtagagtgtagcacctatccCACTTGAAAGCTAGAAATGGAGTGCCTCATTGACTACCTCggttacctcttatctagt
gtagagtgtagcacctatccTTGCTTTCAAAATCATACCTTTCTTAAGGGGTTTCAGAGAggttacctcttatctagtg
gtagagtgtagcacctatccCTGCAAGACTAACCCACATGGTAGAATACTGATGTAGCAAggttacctcttatctag
gtagagtgtagcacctatccGAGAAGAAAGATCTCTTGATTTATACTCTGGATGCAAGTCggttacctcttatctagt
gtagagtgtagcacctatccTTGGTTGGAATGTCAGCTTGGCTTGCTTTATCATTTACAAggttacctcttatctagtg
gtagagtgtagcacctatccAACAAATAAGGTAAAATCTAACTAGAGTGGACAACTTGGggttacctcttatctagt
gtagagtgtagcacctatccCTGAAGTACTGAAGCTATTAAATAGCTCTCCTATTTTGACggttacctcttatctagtg
gtagagtgtagcacctatccGGAACTTGAAGAATGCCAATAAAATGGCTTCAGTGGAGAAggttacctcttatctagt
gtagagtgtagcacctatccTATAATGGTTGGCTTTGTATACATAGTTTTGTGACAGTGGggttacctcttatctagtg
gtagagtgtagcacctatccGTTTAGTAATCATCTGTTTCCACGTATAACCATCTTTACCggttacctcttatctagtg
gtagagtgtagcacctatccGAAGGGTCTGGATCTTATTCATCTCTGCACTGCCACAGCCggttacctcttatctag
gtagagtgtagcacctatccTGGTAAGTACGTTAATTAATGAATAAACTGCAGCTGAGTGggttacctcttatctagt
gtagagtgtagcacctatccGGCAGAGAAAAAAGCACCGGCCTTAGTCAGCAACACTTTGggttacctcttatcta
gtagagtgtagcacctatccAGAGTAGTTGTGTGGTCTTAAATTTCTCAATCTTTCTGGGggttacctcttatctagtg
gtagagtgtagcacctatccGTTGTCTCAATTGTAAGAAGAAGGTAATCATGAGTGACAAggttacctcttatctagt
gtagagtgtagcacctatccCTCTGGCAAGAAAATGGGAGAATGTATGTCAAGTAACAAGggttacctcttatctagt
gtagagtgtagcacctatccCTTACAGTTCTGGAGCTCAGGAAGTCCCAAATCCAGGTGCggttacctcttatcta
gtagagtgtagcacctatccAGAAGGCAGAAGAGCAACAGAGTATGAGAGAGTAAGAGAGggttacctcttatcta
gtagagtgtagcacctatccCCTTTTACTAGGAAAACACTCCCATCTAATCCACTGTGGCggttacctcttatctagt
gtagagtgtagcacctatccTGAGTGAATCATCTCTTAAAAGAACCACATCTCAACACTGggttacctcttatctagt
gtagagtgtagcacctatccCTAAACTTCAAACCTCAAAATAGCACAGCCTTGAATTCTCggttacctcttatctagt
gtagagtgtagcacctatccCCAAAACAAAAGACTCTAGCTAAATGTGTCATCCTTTTTGggttacctcttatctagtg
gtagagtgtagcacctatccTCTGGAGTTATTGACAGCAAATTCTACATAGGAAAAGTACggttacctcttatctagtg
gtagagtgtagcacctatccCAATTTAAAGAACAAAAATGTGTGCAAAACCTCAGTGGGAggttacctcttatctagt
gtagagtgtagcacctatccGCCTGGTATGATTTCTAATTCCCTGCAAGCCTTTCCACAAggttacctcttatctagt
gtagagtgtagcacctatccGCTGAAAATTTTGGCAATTGCTTTAGGGAAAGGTAAGTTGggttacctcttatctagtg
gtagagtgtagcacctatccTGACATTTCAATCACCCTTCCATCCTGTTTTACAAAAGAAggttacctcttatctagtg
gtagagtgtagcacctatccGGGCTTTCATCTTATTTAGTATACTGACCCTGGAAATAAAggttacctcttatctagtg
gtagagtgtagcacctatccGCTCTGCAAGAAAAAGCAAGAACTCTAAGCCATCTCCCAAggttacctcttatcta
gtagagtgtagcacctatccAAATTGCTGTTTTTCATGATGTTTATCACCATCTCCCTACggttacctcttatctagtg
gtagagtgtagcacctatccAGGGAGGAGTCTGATGTTGGAAGGTTCTGGTCATGGTTTTggttacctcttatctagt
gtagagtgtagcacctatccTAAACCTCTTTATCTCTTAGACCCTCTGTTTTCTTCTCTGggttacctcttatctagtg
gtagagtgtagcacctatccCACCAGATGATTTTCAAAGTTCCTTCTACTTCCAATATACggttacctcttatctagtg
gtagagtgtagcacctatccACTACCTGTGTGATGATGTGATTCCCCCAATGCAGCCAACggttacctcttatctag
gtagagtgtagcacctatccGAAACCCCAGTGTTAGAATAAAATAACCAAAATGTGCTGAggttacctcttatctagt
gtagagtgtagcacctatccGGCAGACTCACAGTTATTCAGCTCCAGAGCCCAATATTTGggttacctcttatctag
gtagagtgtagcacctatccATACTGGATAAATAAAGGCACTTCTACTTCTTCTCTTGTGggttacctcttatctagtg
gtagagtgtagcacctatccCTTGTAGGCAGTAGAAATCACACCAAGGTAGCTTAAGGAAggttacctcttatctag
gtagagtgtagcacctatccCTGGTTATACAAATATAATCTTGACCCAAGAAGGAAGTGGggttacctcttatctagt
gtagagtgtagcacctatccAGTTTTTCTCTTACTGTATTCTGTCTCTGCTCCTTTCTGCggttacctcttatctagtg
gtagagtgtagcacctatccTCCTCTCCTCCTTTTCCTCCCTCCACATTTTCTTCCTCTAggttacctcttatctagtg
gtagagtgtagcacctatccTTCCTTGTGTCTTTCTCCTCCTCTTCATCATTCTGATCCTggttacctcttatctagtg
gtagagtgtagcacctatccTTCTTCTCTTTCTCTTTCCCCCACTGACAGGCTTTCTCTAggttacctcttatctagtg
gtagagtgtagcacctatccTATGAAGAAATGACCATTCCTGAATCCAATTCCAATGTTCggttacctcttatctagt

gtagagtgtagcacctatccACCAGGTGTCTATGCCTTGTCCCAATAATTGTGACAAAGAggttacctcttatctagt
gtagagtgtagcacctatccAGTTCTAAAGCCTCAACTGTAAGTCCTAACCTTATTAAGCggttacctcttatctagtg
gtagagtgtagcacctatccAAGCAAACTTCAGTAATCACCTGGTTCTATCTGCCCATTTggttacctcttatctagtg
gtagagtgtagcacctatccGCTCACTGACATTTATGATGTAGCTAATGGCCAAAATTTTggttacctcttatctagtg
gtagagtgtagcacctatccCTTCAAACACCTTTCAAGATCTTACTCAATTTAGTTGCTGggttacctcttatctagtg
gtagagtgtagcacctatccATGTCATAATTCAGAATATTGCACTTCAGAGACTACTTCCggttacctcttatctagtg
gtagagtgtagcacctatccTTCATTCCTAATCAACATAAAACACTGTTCTTGAGTTCCCggttacctcttatctagtg
gtagagtgtagcacctatccTGAGGACAGCTTGAGGAATTGCGAGAAGAGAGAAAGAAATggttacctcttatctag
gtagagtgtagcacctatccGAGATATTTGAGCACTTCTTTCCTAAGGTAGCATCAAATAggttacctcttatctagtg
gtagagtgtagcacctatccAGAGCCTTTCTATGCTTAGCTATCCAGGCATATCAATATGggttacctcttatctagtg
gtagagtgtagcacctatccTTCTTTCTTCCACAGGTAATTAGATTGGTCATTTGTTACCggttacctcttatctagtgg
gtagagtgtagcacctatccGTTCTCCACTATGCCCCACTAAGAGTCACTTACGAGTCGTggttacctcttatctag
gtagagtgtagcacctatccATAAAGCAGAACATAAAAGTCTCCTGTAATCTCAGCCTTTggttacctcttatctagtg
gtagagtgtagcacctatccAAATGTTGTTCTTACCCACCTCAATGTTTTAAAAGGCAGCggttacctcttatctagtg
gtagagtgtagcacctatccCCATGATTTGGATATAGCAAATGCCATTCAATCAATCTCCggttacctcttatctagtg
gtagagtgtagcacctatccCACTCTTAGATGTATACCTTTGTGCACTGGAGTCAGCATTggttacctcttatctagtg
gtagagtgtagcacctatccAACTGCCTTCCAAAAAGCTTTAACCAAATTATACTTCACCggttacctcttatctagtg
gtagagtgtagcacctatccAGCAGTAGGTGTTTAACAAATCATCATTAGCTGGCCAGATggttacctcttatctagtg
gtagagtgtagcacctatccCATTATCATTAACCTTTTGTTGCCTTTCATATGACCTTCCggttacctcttatctagtgg
gtagagtgtagcacctatccCTTGCCCTTCAGCAGGTTTGTGATTTGGAAATCATATCAAggttacctcttatctagtg
gtagagtgtagcacctatccTTTTCCGGTTTTCATATATTTTTTGCATCTTGCATGCTGCggttacctcttatctagtgg
gtagagtgtagcacctatccGCTAACAGTTGCAGAGGTCAGAAGACTTATAATCAAGGTGggttacctcttatctagt
gtagagtgtagcacctatccTCCATTTCCTTGCCTTTCCAAGCTTCTAGAAGCCACCTGCggttacctcttatctagt
gtagagtgtagcacctatccCATCTTCAAATCCAGCAGTGAGGCCTCCTTTCTCCCTCCTggttacctcttatctag
gtagagtgtagcacctatccCTTCTCCTTCTCCCTCTCCCACTCTCCTACTTTCCGTTGTggttacctcttatctagtg
gtagagtgtagcacctatccGTCCTCCTGCTTCCTTCTCATAAAGACCCTTGTGATTATAggttacctcttatctagtg
gtagagtgtagcacctatccAATCAAAAATAATCTTCCCATCCCAACATCCTTAACTTGCggttacctcttatctagtg
gtagagtgtagcacctatccAGGGGACATTATTTCTTTACCATTCTTCTCAACCAAAAGggttacctcttatctagtg
gtagagtgtagcacctatccAAATTTGACTCTCATATCTCCAGTAACTTTCTTCCTGTTGggttacctcttatctagtgg
gtagagtgtagcacctatccCCAACAAGCTTGCTAAAGTTCTATAGATGATTGATAGATGggttacctcttatctagtg
gtagagtgtagcacctatccTAGATATATGCAAAGCACTCGTGATCTTGTAGCTCTCTGTggttacctcttatctagtg
gtagagtgtagcacctatccTTGTGTCACAATTAAGCTTCTCATGGGAACAATCTGCAGCggttacctcttatctagtg
gtagagtgtagcacctatccAGTGCTCACCATCTCAAATCATCTCTCACTTAACTTCCCAggttacctcttatctagtg
gtagagtgtagcacctatccCGGCTTCCAATTTCATTCTCCATCCTCAATGCTGCACTCAggttacctcttatctagt
gtagagtgtagcacctatccTTCTTCCCCAAATTCAAACCAATCAAGTCACTTCCTGGCTggttacctcttatctagtg
gtagagtgtagcacctatccCAAAGCTTCTCTATTACCTGCAAGATGAAATGCAAACTCCggttacctcttatctagt
gtagagtgtagcacctatccAGGATTCCCTCTGCCACCCCCCTCTGTACTCTAGCCACACggttacctcttatctag
gtagagtgtagcacctatccCTTTTAGTTTCACCAATTTTCAATGCCTTCTTTGACCTTCggttacctcttatctagtgg
gtagagtgtagcacctatccGTGCTTCTCCACCTAGAATACTCTACCCTTCCCCATACATggttacctcttatctagt
gtagagtgtagcacctatccGAGACTTCCCTGACCAACCTCCCTACCTTCAAACCCACATggttacctcttatctag
gtagagtgtagcacctatccGACCAGCCACCCCTTTTCAGTACCTTTCTCATTTTTTTTTggttacctcttatctagtgg
gtagagtgtagcacctatccAAAGGGTTGGTTGGGGCAGCTTTTATACTTTCTCAAATTTggttacctcttatctagtg
gtagagtgtagcacctatccGCTTTTCCACTTTACACCAATCAAAATTAGCGTGCATTTGggttacctcttatctagtg
gtagagtgtagcacctatccCATACAAATGGTTTCTCAAGATGTTTCGTAGTACCTCTCAggttacctcttatctagtg
gtagagtgtagcacctatccCTGGCATATGGACAGGGAAAGGTACTATTCTACATGTATGggttacctcttatctagt
gtagagtgtagcacctatccCAAAAATTGAATGCTTAAAAGTTGTTGGAAGAGCTGGAGGggttacctcttatctagtg
gtagagtgtagcacctatccATTGGATTCAAGAATATTCTGCTATAGGGCTTAGAAAGACggttacctcttatctagtg
gtagagtgtagcacctatccTCCAATTGATGCAGCCACCACAACTGTCTTAGGACTCCCAggttacctcttatctag
gtagagtgtagcacctatccCTGTTTTAACTCGCTCTCCACATTCACAGTTCTGACCCTTggttacctcttatctagtg
gtagagtgtagcacctatccTGCCTTTCTTCTGCTTTCCAAGTCTCATGCAAATGCATCTggttacctcttatctagtg
gtagagtgtagcacctatccTAGCTGCAAGGGAGTCTGGAGAAACTAAATTTTTGCTTTCggttacctcttatctagtg
gtagagtgtagcacctatccTGCAGCACAAGACAGGACATAGAAAGAGGTGGAAATAGATggttacctcttatctag

gtagagtgtagcacctatccGAGCCAACTGACAACATAAACTTCTCCCTCCTATGCAGAAggttacctcttatctag
gtagagtgtagcacctatccTCTGTACACATGTATCTGAGGACCTATAAGATTTTATGGTggttacctcttatctagtg
gtagagtgtagcacctatccCTAAGGTTCTCGAGGATGGGGACAGTGTCAGGTTCATCTCggttacctcttatctag
gtagagtgtagcacctatccAACAATAAATTAAGAAATGTGCAAAGAGAGCCAGGGTATCggttacctcttatctagt
gtagagtgtagcacctatccGCATAGCCCTTCTTGCCTCACTACACATACACACCTCTTGggttacctcttatctag
gtagagtgtagcacctatccATACATGCACAAAGTAAGAAAAGAAATCAACTAGGAACGCggttacctcttatctag
gtagagtgtagcacctatccAGTTCAACCATTGTGACAGACAGTGTGGCAATTCCTCAAGggttacctcttatctag
gtagagtgtagcacctatccCCCAAAGGATTGTAAACCATTCTACTATAAAGACACATACggttacctcttatctagt
gtagagtgtagcacctatccTGCAGGGACATGGATGATGTAACAAAGCCATCATCCTCAGggttacctcttatctag
gtagagtgtagcacctatccGAGAACACATGGAGACAGGATGAGGAACAACACACACCAGggttacctcttatcta
gtagagtgtagcacctatccCTGGGGCCAGGGGTGGTGGAAGCAAGCAAGGGAGAGCATCggttacctcttatct
gtagagtgtagcacctatccCCTATCTTGAGGGCAGGTTTTGGAGTTAGAAAGGTCTACAggttacctcttatctagt
gtagagtgtagcacctatccTTCTTCAGAGGAGCTTATTATAAATATTGAATGAGGTGGCggttacctcttatctagtg
gtagagtgtagcacctatccCTTTGGGAGGCTGAAACGGGAGGATCACGAGGTCAGGAGAggttacctcttatcta
gtagagtgtagcacctatccTCTCTACTAAAAGTAAAAAAAATTAGTCGGGAGTGGTGGCggttacctcttatctagt
gtagagtgtagcacctatccAAAACTTGAATGAGGCAATACATGAAAACGCATAACATGCggttacctcttatctagt
gtagagtgtagcacctatccAGGTTTTTGGCTCCTTTTATCCCACTGTCACTGTGTCTTTggttacctcttatctagtg
gtagagtgtagcacctatccCTGCAACAGGGAGGATAAAAACTAAGGATAATGTAATATGggttacctcttatctagt
gtagagtgtagcacctatccACAACAGTAGCTAAAGTTTAGGGGTTTCATGATTTGTAAGggttacctcttatctagtg
gtagagtgtagcacctatccGCTTTTTTGCTTTTTCCTGTCTGTTTAAATTGTGGCTAAGggttacctcttatctagtgg
gtagagtgtagcacctatccCAATGACCCTTTTAAATATGAATTGTTGGCTAGTGATCTGggttacctcttatctagtg
gtagagtgtagcacctatccTTAAGGACAGATAAAGGTAAGAATTGAGGAAACTGGAGCCggttacctcttatctag
gtagagtgtagcacctatccTCCACCATTTCTGTTTCTAATCTAACACGATTTAGGGATTggttacctcttatctagtg
gtagagtgtagcacctatccCTATTTTTCTTCAGATTCCTCAAGCTTCACTTCCCACCACggttacctcttatctagtg
gtagagtgtagcacctatccACTGGGTTGAGTAAGTCTAAATCTCTGGCGTGGCAGGGGTggttacctcttatctag
gtagagtgtagcacctatccTCACTCAGGTTGGAGTGCACGGGCATAATCTTGGCTCACTggttacctcttatctag
gtagagtgtagcacctatccTTGTATTTATTGTCTCAAAGTCTAGTCTCAAAGGCTAGTCggttacctcttatctagtg
gtagagtgtagcacctatccCACGCTGGGCTAAATCTCATCTCTACATTGATTGAACTGGggttacctcttatctagt
gtagagtgtagcacctatccTGGTTCTGTGAGTTTCCATGTACAAATGGAGGCCTCTGGAggttacctcttatctagt
gtagagtgtagcacctatccAAGTCAAAAACATAGTCTTTTTACCTTCCTTTCTGAGTGGggttacctcttatctagtg
gtagagtgtagcacctatccCCATAGGGTTGAGGGTTGAGAGTTGCATGCAGGGGCCAAAggttacctcttatcta
gtagagtgtagcacctatccCAGCTTATAGCCAAGATGCTTAATGCTCCAGATGCACTCCggttacctcttatctag
gtagagtgtagcacctatccCTTGAGCTGTTGATGGTTCTGTTTACTGTGCAATCTAAGGggttacctcttatctagtg
gtagagtgtagcacctatccAGGAGGCTTCTGTGTAATGACTTGGTTCTTTCTATGGGTAggttacctcttatctagtg
gtagagtgtagcacctatccTGTAGGAATGCAAAGAGCTAAAGACAAAATATTGCTTCGTggttacctcttatctagt
gtagagtgtagcacctatccTTGATTTGGAGGAAAAACTGATAAGCAAGAGTCTAAAAGGggttacctcttatctagt
gtagagtgtagcacctatccTGATGATTCAGAAATAAACAAATGATGACTGGTGGAGCTTggttacctcttatctagt
gtagagtgtagcacctatccCTGTGTAGATTGAAGTCATTCACCCAGCAGTCCAGGCCATggttacctcttatctag
gtagagtgtagcacctatccGAGTTAGAGCTTATAAACCACTCAGTCTCTACTCATGTGAggttacctcttatctagt
gtagagtgtagcacctatccTGTTGTTCAGTGAGGATCTACTTTTGACAGCAGCTAGCATggttacctcttatctagt
gtagagtgtagcacctatccCTATTGACAGCCAGCATAAGTGACATCCATTGAAGCCCCGggttacctcttatctag
gtagagtgtagcacctatccCTCTATCAAACTCACTTCTATATAAAAATCCCTTTGTCCCggttacctcttatctagtg
gtagagtgtagcacctatccCCTAAACTCCAGAGCTGTATGAACACGGGAAGCAAAAGGCggttacctcttatcta
gtagagtgtagcacctatccGGAGTACAGCAGAACCCCTCTCCATATCTGTAATTAGCTGggttacctcttatctag
gtagagtgtagcacctatccTGAGATCTATCCCAGAGGTAAATGTGTGTGTAAAAATGAGggttacctcttatctagt
gtagagtgtagcacctatccATCTCAGAGTTCCAAGATAGCCCTAAGATTGAAGATGTTTggttacctcttatctagt
gtagagtgtagcacctatccAGTCCAGAGAGTTGTCATTTGAAAAGATAGAGCATAATGGggttacctcttatctagt
gtagagtgtagcacctatccCCTTTCAATTGGGCTTTGTGGGAAATTAATTTTGTATGGGggttacctcttatctagtg
gtagagtgtagcacctatccGATTGACCATCCAGTATTGATTCTACCACTTCTCATTGTTggttacctcttatctagtg
gtagagtgtagcacctatccACACTAAGTAGTCCATGCCAATCACGGAAATTTCTTTACCggttacctcttatctagt
gtagagtgtagcacctatccCCAGGAAACAGACATCAGTTCAGGAGTAAGCATGTGATTCggttacctcttatctag
gtagagtgtagcacctatccAGGTCAATAAGACATCAGGGGACAATTCCAGAGGCTTCTCggttacctcttatctag

gtagagtgtagcacctatccGCTTCATTTGTTACTGGCAGCATTTTATTATTTTCAGGGggttacctcttatctagtg
gtagagtgtagcacctatccGTATGAAGTTTACAATGTGAATGGCAGACAAAGAGATGGAggttacctcttatctagt
gtagagtgtagcacctatccCAATCCTGGTCCCTTTCCTGCCTCCACTAAGAGAGACTATggttacctcttatctag
gtagagtgtagcacctatccTCCTCTGTGTTTTATCCAGTTAAATCAGGTTTCTGTTACTggttacctcttatctagtg
gtagagtgtagcacctatccGCTATGATGCTAGTTGAAGGGAATATTGTATTTTGTGGCTggttacctcttatctagtg
gtagagtgtagcacctatccGACCAAACCACTTCTTATACATAGAGTCAATTTTCACCAAggttacctcttatctagt
gtagagtgtagcacctatccTCCTCAGATTAAAAGTTAGAAAGAATTAGATGACCTCAGGggttacctcttatctagt
gtagagtgtagcacctatccCAAGAACTGAGATGCAATTGGAAAGCTAGGAAATGAGAAAggttacctcttatctag
gtagagtgtagcacctatccGCATTTGAAATTGTTTTTGAAAGGTTTAGCATACCTCTGCggttacctcttatctagtg
gtagagtgtagcacctatccCCAAGAGTATATGCGTCATGGTAATAGAAACGAAAGTGAGggttacctcttatctag
gtagagtgtagcacctatccGATGCTGAATCCTTCTCTTTGGAGATTATACTACCCAGGCggttacctcttatctagt
gtagagtgtagcacctatccTGTATTAAAATGGACAAAACTCAACATCTAGATGGCCTGAggttacctcttatctagt
gtagagtgtagcacctatccCACTGGTGACATATCCATTGCCCCTGGATGGTGTGCCCTCggttacctcttatcta
gtagagtgtagcacctatccGGGATGCCTAGGTTAGCAGAGTCGCTGTCCAGAGTGCTTAggttacctcttatcta
gtagagtgtagcacctatccGGACCTACTTCCTTCCATGTAACACTTGCAGCTTGAAAACggttacctcttatctag
gtagagtgtagcacctatccATCAGTACCAGCCCTATTTATAGATATTAAGACCCTGCCCggttacctcttatctagt
gtagagtgtagcacctatccGTATCAGCACAATCAGCAATGCACAAAAGCAGAGACAAGGggttacctcttatcta
gtagagtgtagcacctatccAAATACAGTTTTAGTGTGTCAACCTTTACTGGCTGATTGTggttacctcttatctagtg
gtagagtgtagcacctatccCTAAAAAGGCTTTCAAACCATTTCATTGGGAACACAACCAggttacctcttatctagt
gtagagtgtagcacctatccTAACATTTTAAAATTTGGTTGGGTGTTTCTGCATATCCCCggttacctcttatctagtg
gtagagtgtagcacctatccATTATGATATGGTATAAAATCTTCATTCTGTGCCCTGGAGggttacctcttatctagtg
gtagagtgtagcacctatccTTTGTTGTTATTTTGAAGTTAGAAGTCAGTCACGCAAGAGggttacctcttatctagtg
gtagagtgtagcacctatccCAGCGGCACCCCCCTTTGAAAATCTCAACTATGTACTGTAggttacctcttatctag
gtagagtgtagcacctatccTGCACCACATGACATCCCTTTATTTTAAAATAGAAAGCCAggttacctcttatctagt
gtagagtgtagcacctatccGCAGGAATTTGTAGTTATATTGGATCTATGCTTCTGGTTAggttacctcttatctagtg
gtagagtgtagcacctatccACTTTCATTATTCCTCAGTAATTTGGTTCATAGGGCAGTAggttacctcttatctagtg
gtagagtgtagcacctatccTGAGACTCATTTTCAGCACAGCAAGCAGTTATAGGGTTTCggttacctcttatctagt
gtagagtgtagcacctatccTTTCCCCCACGATGATGAAATTGATGCAAGTAGGACCACAggttacctcttatctag
gtagagtgtagcacctatccACTGAAATAGGTAGGCTGGAGCATCTGTGGTTAGGTAGGTggttacctcttatctag
gtagagtgtagcacctatccAAGAATAAGGATCTCACAGAGGTCAGTTAATTTGAACAGTggttacctcttatctagt
gtagagtgtagcacctatccAATCATATAATGTTAGATGTTGTTTTCAACTGCAGGCAGCggttacctcttatctagtg
gtagagtgtagcacctatccAGGTGCGTATGATGTAATTCAGATATCTGGTTTTTTTCTGggttacctcttatctagtg
gtagagtgtagcacctatccCTAAGACGATTTGTGTAGAGTGGAGGCGGTAGTTTACTTCggttacctcttatctagt
gtagagtgtagcacctatccCTAAAGCCATTATGACAATTAACAAATTCAGCCAACAGTCggttacctcttatctagt
gtagagtgtagcacctatccAGATAGGATTCACACATCTACTCATTACTCCACAAACATGggttacctcttatctagt
gtagagtgtagcacctatccGGACCCTGCTATGTTAAGGCACTGTGCTAGGAATTGAGGTggttacctcttatctag
gtagagtgtagcacctatccGCCGCCATTGAGCCTGCCGTCCTCTGGTTGACACCCGAAGggttacctcttatcta
gtagagtgtagcacctatccGCTGATCCTATTTCTATATGGCCTTGTACAGTGATTACAGggttacctcttatctagtg
gtagagtgtagcacctatccGGCAGACAGACCTAGATCACAACCCAGGCTCAAATCTCAGggttacctcttatcta
gtagagtgtagcacctatccGGGAAAATCGTTTCTCTGAGCCTTAGTTCTTTCATCTGTAggttacctcttatctagtg
gtagagtgtagcacctatccAATACTAGCACCTACCTTGCAGATATTTTTAGGATTGAGAggttacctcttatctagtg
gtagagtgtagcacctatccTTAGATGACGATAAGTTTTAAATGGGCAAGGAAACCAATCggttacctcttatctagt
gtagagtgtagcacctatccGTAACATTGACCACACTCTTCTATATCTGACTTGAAAGATggttacctcttatctagtg
gtagagtgtagcacctatccGTGACAACCACCAGTGGGGCTATATCCTGTCTTGAAGCTCggttacctcttatctag
gtagagtgtagcacctatccGACCTTGTGTTCTGTTAAAAGAACATGCATAGACATTTTCggttacctcttatctagtg
gtagagtgtagcacctatccGAATTACTTCATTGTGTCCTACATGGAGGGCCCAGGTGACggttacctcttatctag
gtagagtgtagcacctatccGGTAGGTCCAGAGAATACTGCATTGTTATCATGAATATATggttacctcttatctagtg
gtagagtgtagcacctatccGAAGGGAAAAAGGGAAGAAGAGATAGCAGTAAGAGGGAAGggttacctcttatcta
gtagagtgtagcacctatccCTATTCTAAAAATCTTACACACGTCTCACATGGTACTTTCggttacctcttatctagtg
gtagagtgtagcacctatccCTGAGGCATCTGAAACAAAGAGAAACTAAATAACTTCCCCggttacctcttatctag
gtagagtgtagcacctatccGCAGCCTGACTTTAGAACCCTCTTCAGCATGGTCCTGCGCggttacctcttatcta
gtagagtgtagcacctatccGCTCTTTAGCCACATTACTAGATTGGACTTTATGACATTTggttacctcttatctagtg

gtagagtgtagcacctatccAAAGTCCTTTGTATTGTAAGTCTACCCTGAGTCGATGAGAggttacctcttatctagt
gtagagtgtagcacctatccCTTCATGAGACCGCAACACAAAGGGCAGTAGAGGAGAAATggttacctcttatcta
gtagagtgtagcacctatccCTTTTATGGAGAATGTCTAAAGTGAGTACAGTATGTGCCAggttacctcttatctagt
gtagagtgtagcacctatccTGGGAAAAGCATTTTGGAAGCTCTTTATTAGGAAGAAACCggttacctcttatctagt
gtagagtgtagcacctatccGTAGTAAACAGTGTTCTTTTGAGACATGGCCAGATATTGCggttacctcttatctagt
gtagagtgtagcacctatccGGTAATAAATGGAGACAGAACACTGCAACCTACCAATTTAggttacctcttatctag
gtagagtgtagcacctatccGCAGGAAGTACAGCTTGTGGTTGTATTTGAGGTTGGAAGCggttacctcttatctag
gtagagtgtagcacctatccAGAAATCTGGAGACATGAGTAGGGAGGTGGAGGTGCAGGGggttacctcttatcta
gtagagtgtagcacctatccACTGCATTTTTTTCTCCCTATTATTTTAGTTCTTTGGGGCggttacctcttatctagtgg
gtagagtgtagcacctatccCTCATCCTGGAGACCAATATGTCATTTATAATTTCTGGCTggttacctcttatctagtg
gtagagtgtagcacctatccCCTTCTAATAAACTTGAACTGGCAGAGATAGAATGTTAGCggttacctcttatctagt
gtagagtgtagcacctatccAGGAGGACTGAAGAATGAAAGATAGGAATCAAAGAACAAAggttacctcttatctag
gtagagtgtagcacctatccCCTTTGCATCCCAAGAGACCAATGTTGCCTTTTAATTTTCggttacctcttatctagtg
gtagagtgtagcacctatccCGTTCTGATGGTGGAGGAGGGCATGGATTCAGGAAAATGAggttacctcttatctag
gtagagtgtagcacctatccGAAACTAGCACTTCCAAAATGACTTCCAGGCCTATTAAAAggttacctcttatctagt
gtagagtgtagcacctatccAAGCCAAAATACTGTTTCAGAAATAAACTCCAGATGACAGggttacctcttatctagt
gtagagtgtagcacctatccTTTCTTCATATTCACAAAACCTAAGCTCCCATCTTCATAGggttacctcttatctagtg
gtagagtgtagcacctatccCAAGCATGTGACATTCATATAGAGACAGACAATCTAAAACggttacctcttatctagt
gtagagtgtagcacctatccTCTTAATCATTCGGTATCACTCTAGTCTTTCCCAATATGGggttacctcttatctagtg
gtagagtgtagcacctatccAAATAGAGGAAGAACACTTCTACAAGACAATTCTTTTCGGggttacctcttatctagt
gtagagtgtagcacctatccCCACTGGAATATAAAAGGCACACAAACACATACAAGCATGggttacctcttatctag
gtagagtgtagcacctatccAGCTAAGAACATATAGTAGTGAAGAAATCTGCTTGCAGAAggttacctcttatctagt
gtagagtgtagcacctatccAGCAGTTTTAGGCTCATAGCCAAATTGAAAGTACAGAGATggttacctcttatctagt
gtagagtgtagcacctatccCTCAGCTGTTCCCACATATGCGTAGCTTCCCCCATTATCAggttacctcttatctag
gtagagtgtagcacctatccTTCATTCTTGGTGTTGTACAATTTATGGGTCTGGACAAATggttacctcttatctagtg
gtagagtgtagcacctatccAAAATCCCCTCTGCTCCCTATTCACCTATTTTTAAGTATAggttacctcttatctagtg
gtagagtgtagcacctatccATAGTAAGGTTGGTATGAAATTGGATATGCAGTTTTGAGCggttacctcttatctagtg
gtagagtgtagcacctatccCCTTGTGGGTGCTTTCCTGAAGTTTTTACATTAGAAACTCggttacctcttatctagtg
gtagagtgtagcacctatccGCTGATGCTTTCACAAGAATGGGGGACACATGCACCTTCTggttacctcttatctag
gtagagtgtagcacctatccATTCCCTTGAGAGCAAAACTGAAGTTGAATCCTCTCCAAGggttacctcttatctag
gtagagtgtagcacctatccAGCTGCCAATGGGAGGTGTCCACAGAGATCCCTTATTCTAggttacctcttatctag
gtagagtgtagcacctatccGTATGGAACTAGAAACTGCATGTGCCAAGTTGTCAGGGACggttacctcttatctag
gtagagtgtagcacctatccGGTGTTATTGTCCCAGATAGACAGTTCCCAGACCATGTGTggttacctcttatctag
gtagagtgtagcacctatccAGATTTCTGTTTTACTAGTTCTATTTCCTTTCCTCTCCCTggttacctcttatctagtgg
gtagagtgtagcacctatccAGCAGTTTATGGTTTCTTTTCCTTGACAAATTGCCAAGTCggttacctcttatctagtg
gtagagtgtagcacctatccATGATCTGTCTTCCACCTTCATGACGTAAGCCCCTAAAGCggttacctcttatctag
gtagagtgtagcacctatccGACGTCTACTTTTTTTTGTCTGTTTGGTGAGGTGGAGAGAggttacctcttatctagtg
gtagagtgtagcacctatccTGCAAAGAACTGATAGGTTGAGCATTTTCAGAAAACACATggttacctcttatctagt
gtagagtgtagcacctatccCAATCTTGAAGTAGAAAAGCCCAGGAAACTTTCAAATCTTggttacctcttatctagt
gtagagtgtagcacctatccACCGTTCCCCTGGGTCATTCCTTTTTCCTTTGTCTCAGCCggttacctcttatctagt
gtagagtgtagcacctatccAAATCTCTCCCTTCCTGTCTAGCTTATATGATTTGTCCTCggttacctcttatctagtg
gtagagtgtagcacctatccCTGACCAGGTTCTTTCCCCCCTTCCCTTGTTCAAATCCCCggttacctcttatctag
gtagagtgtagcacctatccATGGCTCTCCCCAGACTGCACTGTTACATCTGACACCTGCggttacctcttatcta
gtagagtgtagcacctatccCTGCGCCGTACCCTCAGCTGGAGGCCTCTGCTAAGGTCAAggttacctcttatcta
gtagagtgtagcacctatccTCTTGCCTATTCTGACTCAATAAAACTTCATTGATTTGCCggttacctcttatctagtg
gtagagtgtagcacctatccTTGAGGAAAGGCTTGGTCTTATAATAAGAGGAAATGGAGCggttacctcttatctagt
gtagagtgtagcacctatccAATTTGTCTCCTTGTCCCTAGCTTACCAATAATATTAACCggttacctcttatctagtg
gtagagtgtagcacctatccCAGGATGAAGGTATGGAACAGAAGTCAAAAGGAGCCTCTGggttacctcttatcta
gtagagtgtagcacctatccCTGTTTCAGAGGAAGTGGCAACTCATCAGCTCTTGTAGGGggttacctcttatctag
gtagagtgtagcacctatccGAATCACCCTCCTCACTTCCAGCCTACCTTCTTCTATCCTggttacctcttatctag
gtagagtgtagcacctatccACGAGGTACTCTGGTGACTGGACTGGCTTCCCAAGAATCAggttacctcttatcta
gtagagtgtagcacctatccTGCCTTCCTCCCAGAGCTTTAATTTCTGCTTCTAGTTTCCggttacctcttatctagtg

gtagagtgtagcacctatccTCTCTGAGATCGTGATGTGTCTAGAAGTACTGATAATGCTggttacctcttatctagtg
gtagagtgtagcacctatccCTTGGAGCTGTGGGAATGAAAACTGAATATCCTGTCTCACggttacctcttatctagt
gtagagtgtagcacctatccCTTAAAGACAAGAAGCCTATTGATGAAAAGCCAATGACACggttacctcttatctagt
gtagagtgtagcacctatccTAAACTTTAACTAATCAACCTCTTCCAATCCCTTCTTTCCggttacctcttatctagtg
gtagagtgtagcacctatccGCCAAGAACATGCACATCCCTACCCCTTTCTAAAACTGTTggttacctcttatctagt
gtagagtgtagcacctatccTCCTTATGACTGACAGTCTCCAGGAAGTAATGGGGAACAGggttacctcttatctag
gtagagtgtagcacctatccGTGGTACCATTAAGTGAAGTGAGATATAGACAGGGTGAGGggttacctcttatctag
gtagagtgtagcacctatccGCAAAGCATGGTGGGGATAATTGGTTCAGGGCCCTGTGGTggttacctcttatctag
gtagagtgtagcacctatccTTTCAGAAAAGATATATGCAGCAAATAGCCCAGGCAGAGGggttacctcttatctag
gtagagtgtagcacctatccGGTGACTGATACATGAGAACCATTAAAATAAAGCTGTGATggttacctcttatctagtg
gtagagtgtagcacctatccAATAAGAATGAGGACAGAGGCTTAACATACAAAAATAGGGggttacctcttatctagt
gtagagtgtagcacctatccTGTAAGCAGCATAACAAAATCCACTCAAGAAAACAGACAAggttacctcttatctag
gtagagtgtagcacctatccAACAGTTTGAAGAAATACCAAGACCGTAAGCAACAGAAGAggttacctcttatctag
gtagagtgtagcacctatccTTACTAGACAGGAGAATTTAGAGATAGAGGAAAACTTGAGggttacctcttatctagtg
gtagagtgtagcacctatccCTGCCTGGGTTTGCATCCTGCCACATCCTTGCTAGCTGTGggttacctcttatctag
gtagagtgtagcacctatccAGTTCTTTACTCTACCTCTGCCTTAGTGCCTCATCTGTAAggttacctcttatctagtg
gtagagtgtagcacctatccCCTATTTCAGAGAGTTGGCATGAGGATCCTGTGAGTTACCggttacctcttatctagt
gtagagtgtagcacctatccACCACCACCTGACACAAGCTCTATGTGACTATTTGCTGCTggttacctcttatctagt
gtagagtgtagcacctatccGAAGATAAGCACAGCCATTTTTCCTTTTGACTTTCTCCCAggttacctcttatctagtg
gtagagtgtagcacctatccACTAAGTGTGGGAGGAAGAAATATCAAATTCAGGGATAGTggttacctcttatctagt
gtagagtgtagcacctatccCAGCTCTTGTCAGATACTGGAAAACTTCAAAGTCCTTACAggttacctcttatctagtg
gtagagtgtagcacctatccACTGTATCTGAAAACATGGAGTAAACATCCATCTGTCTGAggttacctcttatctagtg
gtagagtgtagcacctatccCTTCATCATCCATAACATGGCTTTCAGAAGTAAATATTGCggttacctcttatctagtg
gtagagtgtagcacctatccCACCACATGTCATTACTTTAAGAACAAATTATGCTTCCCCggttacctcttatctagtg
gtagagtgtagcacctatccGCTTTAACTCTGGCCATCAATCACCTTATAATTAGTAAAGggttacctcttatctagtg
gtagagtgtagcacctatccTTTAGTTTTCTCAGCATTTAGGATCATTCTGTTGGTATGGggttacctcttatctagtgg
gtagagtgtagcacctatccGCCTCACAAAGCTTCCATTCACATCAAAAGTATGCTCAAGggttacctcttatctagt
gtagagtgtagcacctatccGCTTAGCTGGCCATCTGTTACCAATTATTCTATTTAGCATggttacctcttatctagtg
gtagagtgtagcacctatccACTATATTGGCCTCCTCCACAAAGCCTTTCAAGCAAGGTTggttacctcttatctagt
gtagagtgtagcacctatccGAAAGTAAAACAACCAGGACTGTCACAAACAAGTTACCAAggttacctcttatctag
gtagagtgtagcacctatccACCGCATTCCAGTTTGTGCCCAGTTAGGTGTGCTAGGTACggttacctcttatctag
gtagagtgtagcacctatccGTTTGCCAATTTTTCTATGTCATGCTCCACCTGCCATCCTggttacctcttatctagtg
gtagagtgtagcacctatccCAATCTTAGTGTCTCCATGGCTTTCTTTTTAAAGTACATGggttacctcttatctagtgg
gtagagtgtagcacctatccGTAGATGAATCTGCTGTTTGATAACCTCTAATGTAAGGAAggttacctcttatctagtg
gtagagtgtagcacctatccCCTGTTTGTAAGAATATAGGCTTAACTTGGGAAGGAAAACggttacctcttatctagtg
gtagagtgtagcacctatccTGGAATGATTAGATTATCAGCAAAGTATGAAGCAGTGGGAggttacctcttatctagtg
gtagagtgtagcacctatccTCCATAAATGCAATGAGAGTGGAAAAAAGGGAGGTCAACCggttacctcttatctag
gtagagtgtagcacctatccGAAGGGCTTTCTATGGTAGGAAAACAACTTGAGAGGAAAAggttacctcttatctagt
gtagagtgtagcacctatccTTCAAGCAACATCCTTTAAATGGGTCACTTTCAGTCCACAggttacctcttatctagtg
gtagagtgtagcacctatccAAACTTAAAAACGTTTGTCCTCCTTAAAAGGAAATCCAGGggttacctcttatctagtg
gtagagtgtagcacctatccTGCTTCAGGTTTTATTTTCCTGGCATTATGCTGCACGTAAggttacctcttatctagtg
gtagagtgtagcacctatccCAGGGGTAAGAGAAGATTCTGCCTGCATAGTCTGGTCTGGggttacctcttatctag
gtagagtgtagcacctatccAGAACTCAAAGGGCATATACTTTCTAAGAGACTGCTGATGggttacctcttatctagt
gtagagtgtagcacctatccAGCTGACTTTAAATGACCTAGGTGGTAAGAGAGGGGGCTCggttacctcttatctag
gtagagtgtagcacctatccCTTTGAGAGTGGCGAGAAAGGAAGAAAAGAGGGTATTGAGggttacctcttatctag
gtagagtgtagcacctatccATTGGCACATTTCAACAATTGTTTTACCCCGAGAAAGTCAggttacctcttatctagtg
gtagagtgtagcacctatccTATAAAAAGTATATCAAGAAATGCTTACTCCCACCGTCTCggttacctcttatctagtg
gtagagtgtagcacctatccATCTTATTCTCTAGTCCCCCACAACTTTACTCATTTCCTAggttacctcttatctagtg
gtagagtgtagcacctatccTGGTTTAGAATAGATTGAACATCAGTAGAGAGAAAACCTCggttacctcttatctagtg
gtagagtgtagcacctatccCAGTGGATGAGGTTTTCTTGTTGGGAGCTGAGTTGGGGAGggttacctcttatctagt
gtagagtgtagcacctatccACTGCTCATGCGTTGTCTGAATTCCGAGAAATCTGTGCCAggttacctcttatctagt
gtagagtgtagcacctatccTCCTCCATGGTCCTCAAGCCTGTGACTCCAAACTCTACACggttacctcttatctag

gtagagtgtagcacctatccGACATTCCACAGATTATATTCTACTCCTATCAAACACCAGggttacctcttatctagtg
gtagagtgtagcacctatccTTCTTTTTTGAGATAAAGTCTTGCTATGTCGCCCAGGCTGggttacctcttatctagtg
gtagagtgtagcacctatccACTGCACCTGGCCCCATCCTATTACTTTTTGATTACTGCTggttacctcttatctagtg
gtagagtgtagcacctatccCAACTTTCTTCAGTCTTTCCTTCTATGACCCAACGGATGAggttacctcttatctagtg
gtagagtgtagcacctatccATCACAAATAAGTATGTTAACGTCTGAATGCAGTTTGCCAggttacctcttatctagtg
gtagagtgtagcacctatccACAAGTCCTTAATATGTTTCCTATAGTAACCTGTGGTACAggttacctcttatctagtg
gtagagtgtagcacctatccGTTCCTTCATGCAGAACAGGCACTAATGACTTACCAGAAAggttacctcttatctagt
gtagagtgtagcacctatccTCAGTTCATGCCCAGGTAACACAAATGCCTTTAAGTAAATggttacctcttatctagtg
gtagagtgtagcacctatccCAATCACACCCAGGAAATAACTTATCTAATCAATGAACACggttacctcttatctagt
gtagagtgtagcacctatccGTTTTACATTACTGACATGTTGAGGAAACATTTGACATGCggttacctcttatctagtg
gtagagtgtagcacctatccGATGTTCATCAGCAAACGTGTGGTCAAAATGCCTCAGAGTggttacctcttatctagt
gtagagtgtagcacctatccATTCTACCCTATCTGCAAATCTCTAGTACTTACTTTAGTCggttacctcttatctagtg
gtagagtgtagcacctatccGGTTAGTACTATTCTTCTGTCAAACTCACTACTTATATGGggttacctcttatctagtg
gtagagtgtagcacctatccCTTGCAGAAAAGCTAAAGTGTGGACAGTATCTATACCAAGggttacctcttatctagt
gtagagtgtagcacctatccTGTGTAACTGATATCAATCTTCACTGTTGTAAAGAAGGGTggttacctcttatctagtg
gtagagtgtagcacctatccAGCTTTCTCACCAATGACATCAACATGTACTATCTACGGTggttacctcttatctagtg
gtagagtgtagcacctatccCTTGGGGAACTGACACTTACAAAATCACTATGGCATCACTggttacctcttatctagt
gtagagtgtagcacctatccTTGATACAATTCTAGTCCTGGCTCTTTCCTTGATATGATTggttacctcttatctagtg
gtagagtgtagcacctatccCATATTCTGAAAATGAGGAGATTAGCTTACATATGCTTGCggttacctcttatctagtg
gtagagtgtagcacctatccCACACACACACGTCAACTTATGCTAGGTGTTGAATAACTAggttacctcttatctagt
gtagagtgtagcacctatccGACTACAGAGATTCAGAGTAAAGAGAAGTCACCACAGCCTggttacctcttatctag
gtagagtgtagcacctatccCCAGGAAGAACACAGGCTGGGTTGAATGATTGCAGCTGGAggttacctcttatctag
gtagagtgtagcacctatccCTAGTGAGATGGGCTGGTGCAGAGTCTAGGTGCAGATTCTggttacctcttatctag
gtagagtgtagcacctatccAGAGTGACTTGATGCTGACCCAGAAGCCCTTAAACCCAACggttacctcttatctag
gtagagtgtagcacctatccGGTACCGATGACTCCAGCCAAAGGGAAAGAATTATAGAAGggttacctcttatctag
gtagagtgtagcacctatccGCAGTTCTTAGCAGGTTATTTCCCTCACATCATGTTTACTggttacctcttatctagtg
gtagagtgtagcacctatccAACAAGATGAAGGAAATAAGCACATACGTATTCTTTCTCCggttacctcttatctagtg
gtagagtgtagcacctatccCAAATCTTTCCCTATAAAGCTGAGAATATGAGTATCAGGTggttacctcttatctagtg
gtagagtgtagcacctatccGGATTTCTGGGCCTCTAACCACCTCAGGTCAGCTCCAACAggttacctcttatctag
gtagagtgtagcacctatccCTGCATAAGGGTGAATATTGAATATTTCCAAGCTGCTGGTggttacctcttatctagtg
gtagagtgtagcacctatccGCTGATGAATGAGCCCTGCCACGTATAGGAGCTTTAGCCCggttacctcttatctag
gtagagtgtagcacctatccGATGTTATCCAGCTAATCCCAAGTAAGCCCCTAAGGATCTggttacctcttatctagt
gtagagtgtagcacctatccAGCCTTACTGTGTGTGCCTAGTTCCTGTGATAGAGCTTCTggttacctcttatctagtg
gtagagtgtagcacctatccTATTCCTAACTGACACTCAAATTCCCCCACTGACTCCTGAggttacctcttatctagt
gtagagtgtagcacctatccTGTCAGGAGTTATATTTAATCTGTGGCAATCCCCCATTTCggttacctcttatctagtg
gtagagtgtagcacctatccAAATATTTCTTCAGCCAATATCCATTGCTTCAATGACAGCggttacctcttatctagtg
gtagagtgtagcacctatccCATATGTTTATTCTGGCACCACCGGGAAGCAGCCTTTAGCggttacctcttatctag
gtagagtgtagcacctatccTTCTTGAATTGTTTCTTAGCTCTCAGCACCTGTCAGAATCggttacctcttatctagtg
gtagagtgtagcacctatccAAATCTGTTTGTGTCAGCAGCTCTTATAGGGTTTTGAATCggttacctcttatctagtg
gtagagtgtagcacctatccGGCTACTCACTCTATACCTTTCCGTAGTCATTTCCTACTGggttacctcttatctagtg
gtagagtgtagcacctatccCAACATTTATTAATTATGTCAAAGCCAAAAGCCCACATGGggttacctcttatctagtg
gtagagtgtagcacctatccCTGCAGAGGTAGAAATCAAGTTGTTTGGGCTGGACTCTTAggttacctcttatctagt
gtagagtgtagcacctatccCAAAATCATTCATGTTGTTTGCAGACCTCATTTCCTTGTAggttacctcttatctagtg
gtagagtgtagcacctatccGGGGTCCCTTTCCTTGCTGGCTGCCAACCAAAGTCCATCCggttacctcttatctag
gtagagtgtagcacctatccGCCTCTCCATCTTCAAGCCAACAAACAGCACACACATCCAggttacctcttatctag
gtagagtgtagcacctatccACTCATATGTCAAATCGCTCTGGCTTCCTCTTTTGCTACCggttacctcttatctagtg
gtagagtgtagcacctatccTTCTGTCCTTAAAGGCTCATGTGATCATATTAGGTCCACCggttacctcttatctagtg
gtagagtgtagcacctatccAGTTCTAACTCCCAGTACTTCTCAATGTGACTTCATTTGGggttacctcttatctagtg
gtagagtgtagcacctatccTTAAGGAGGAACCTAATTGAAAATGATGTCATAAGGGTGGggttacctcttatctagtg
gtagagtgtagcacctatccGGAAGATGATGTGAAGATACCCAGCAAGAACCCCATGTAAggttacctcttatctag
gtagagtgtagcacctatccGCTGTGACTGCAAGCCCGGGAATGCCAAGTACTGCCTGCCggttacctcttatcta
gtagagtgtagcacctatccCACCTTGATTTCAGATATCTTGCCTCCAGAACTATAAAGCggttacctcttatctagtg

gtagagtgtagcacctatccACAGCAGCCCTAGAAAACTGGTACACCATATAACATAACAggttacctcttatctag
gtagagtgtagcacctatccGAAGTGTTATCTCATCACATCCACAGATTCCAGGCATTGTggttacctcttatctagt
gtagagtgtagcacctatccGACATTCCAAGAAGTTCCGCCTATGATGCTGATGCTATTGggttacctcttatctagt
gtagagtgtagcacctatccTCCCCTTGACATTCCCTCCTCAGACCACACTTCTTTTTGCggttacctcttatctagt
gtagagtgtagcacctatccGGTCCTTCCTTGCCTTAGGGTATTTTTTCTTAACTTCCTTggttacctcttatctagtg
gtagagtgtagcacctatccACTTCTGTTTGGTTGAAATGTTGCTATTACTCACTCCCAGggttacctcttatctagtg
gtagagtgtagcacctatccCCAAGTATAGACACGGAAAAGGCTCCCAAATTCTTCAGGGggttacctcttatctag
gtagagtgtagcacctatccAACTATGTGGTTCAAGTACCAGGAGGACTTTAGATAGCTCggttacctcttatctagt
gtagagtgtagcacctatccAAAATAGCTTTGTGAATGAAACCCTAATGATGACTCTGGAggttacctcttatctagt
gtagagtgtagcacctatccAACACGTCTATGGCTAATACATTAGGAAACTAGTTCTCCTggttacctcttatctagt
gtagagtgtagcacctatccGTTATGAGTCACCTTGTTTATTTTGAACCAAAATGCTGAGggttacctcttatctagtg
gtagagtgtagcacctatccGTCTCGCATATCCCCAGACCTCACTGTAACTCTATCCTACggttacctcttatctag
gtagagtgtagcacctatccGGTTTCCCTGCAGCCATGTAAGCTCCTATAGTAAACAGAGggttacctcttatctag
gtagagtgtagcacctatccAGGTCGCTTTTGATCTCCAATCTTTTCATTTTCTCTCTAAggttacctcttatctagtg
gtagagtgtagcacctatccCACTCAACAACAATGCCCTCTCCTTCTTCCTCATGAATCTggttacctcttatctagt
gtagagtgtagcacctatccAGAACCACAATACTCACTCATGATCATAAATTCTGCGGGCggttacctcttatctag
gtagagtgtagcacctatccGCCCAAGGATCTGCCTCTGATCAAAAAGCCCTCCTCAATGggttacctcttatctag
gtagagtgtagcacctatccCCTTTGTGGGTCATCAGGCAGGAGAAGCTTGTCAAACACTggttacctcttatctag
gtagagtgtagcacctatccGGTTCTCCCAGGGACTCAGAATATATATAGTGTCCTGTTAggttacctcttatctagt
gtagagtgtagcacctatccCTGATTTTCTAAGGTGTAGGCTTTGATTTTACAGGTAGATggttacctcttatctagtg
gtagagtgtagcacctatccAGTGATCTCTATTTTAAAAAAATGTGTATGACTCAGCCGGggttacctcttatctagtg
gtagagtgtagcacctatccCCAGCACTTTGGGAGTCCGATGAGGGTGGATCACAAGGTCggttacctcttatcta
gtagagtgtagcacctatccAGGCAGGAGATTGGTGTGAACCCAGGAGGTGGAGTTGCAGggttacctcttatcta
gtagagtgtagcacctatccCTGTAGAAACAAAGCAATTGTCAGAACCAGACAGAGATATggttacctcttatctag
gtagagtgtagcacctatccTGGACATGGCCAGTGAAAGAATCAGTTAGCTTTCAGATATggttacctcttatctagt
gtagagtgtagcacctatccATCCAAGGACTATGAAGCGATATAAAAAGGTGTAACTTACggttacctcttatctagt
gtagagtgtagcacctatccTGGTATAAATATTTTTGTGCCTATCTCTTCACATACAGGCggttacctcttatctagtg
gtagagtgtagcacctatccTTTGTCCCAAAGTTGTATCAATTTACAGTTCCACCAGCCAggttacctcttatctagt
gtagagtgtagcacctatccCAACACTTGGTATTCTCATTCTCCTTGATTTTACCCTTCCggttacctcttatctagtg
gtagagtgtagcacctatccTTGGTGAAGTAACTGTTCAAGTCTTTTGCTTTCTATTTCGggttacctcttatctagtg
gtagagtgtagcacctatccCTATTTCTTATTCACCCTTACTTTGATGACAGTTTTGTCCggttacctcttatctagtg
gtagagtgtagcacctatccCTGTTTCTCAGACTCTCTGCTGCCTCCCTTGGAATCAACAggttacctcttatctagt
gtagagtgtagcacctatccGGATGTCTTCTTATTCTAGATCTTGACCCCATAATTCTTTggttacctcttatctagtg
gtagagtgtagcacctatccCTATTCTCAATGGTAGGGTTGCACCGAATTATTTTGTCAGggttacctcttatctagtg
gtagagtgtagcacctatccAGTCACTCTGTTTAACCTCTTTTCCACTCACGTATGAAGAggttacctcttatctagt
gtagagtgtagcacctatccGCTCAGCATGCAAACAGATGTAAACTTGACCCTAGTATTTggttacctcttatctagt
gtagagtgtagcacctatccCATTTTAACTTTTAGTTTACCTTTTCTGAGACAGAGTCGCggttacctcttatctagtg
gtagagtgtagcacctatccTTTGTATTTTTAGTATAGACGGAATTTCACCATCTTGGCCggttacctcttatctagtg
gtagagtgtagcacctatccTGAACCACCGCGCCCAGTCAACATTTAGTTTACCTTCTAAggttacctcttatctagt
gtagagtgtagcacctatccAGAGAAATTTCGGTGCTTATTCATCCAACTCTTGCCTAACggttacctcttatctagt
gtagagtgtagcacctatccAAAATAAGAATACCAAGATCCCTTCCCAAGGGATAGAGACggttacctcttatctag
gtagagtgtagcacctatccTACTGTGTGAGGTTAAAGAGTAATCATTTTAGAATGGGAGggttacctcttatctagtg
gtagagtgtagcacctatccTGATATTAAAAATTGAGATCTAGGTGGGGCACGGTGGATCggttacctcttatctagt
gtagagtgtagcacctatccCTCTGGGATGCCAAGGCAGGGAGAACACCTGATGTCAGGAggttacctcttatcta
gtagagtgtagcacctatccAGACTCTCTCAAAAAACAGATCTGAATAAGAAGAGCCCCAggttacctcttatctag
gtagagtgtagcacctatccGGATCACAGAGAAGAGTACTCCAGGTAAAGGGAAGAATTTggttacctcttatctag
gtagagtgtagcacctatccCAAGCTTAGTTTGTTTCCAGAAAAAGCAAAGGCTAGCATAggttacctcttatctagt
gtagagtgtagcacctatccCCAATTGGCGAAAGGGAACATGGTGTAAATGAGGCAAGAGggttacctcttatctag
gtagagtgtagcacctatccAAAAATATTTAAGTTTGGGAAAGGAAAGGGGAAGTGGAGGggttacctcttatctag
gtagagtgtagcacctatccTATTTCTCTGGCTGCTATGTGGAATATATATTGTGTATGGggttacctcttatctagtg
gtagagtgtagcacctatccTCTGCTAGCTTTGGGGATGTATTGCACTTGCTTCTCTAGTggttacctcttatctagtg
gtagagtgtagcacctatccTAGTGCTAAAAATTTCCCCCCCTAACACTGCCTTAGCTGTggttacctcttatctagt

gtagagtgtagcacctatccGGTTGTTTAATTTTCATGTCCTTGGATGATGTTGAGTGATggttacctcttatctagtgg
gtagagtgtagcacctatccTTGTGCTGTGGTCTGAGAATACGGGGTTTCTTTTTCTTTAggttacctcttatctagtgg
gtagagtgtagcacctatccCTGGCCTCTTTTGTTTGTTTAATTTGCCGAGAATTGCTTTggttacctcttatctagtgg
gtagagtgtagcacctatccTGATGGTGTGCTTGATTTCTGGGATGAATGGCTAGCAGCCggttacctcttatctagt
gtagagtgtagcacctatccATAAGTGGGAGTTAAACAATCAGAACATATGGACACACAGggttacctcttatctagt
gtagagtgtagcacctatccCTGTTGCAGGGTGGGAGGCGGGGGAGGGAACCTAGAGGACggttacctcttatct
gtagagtgtagcacctatccTGTAAAACACCTAAAGCTATAGAAACTCTGGAAGATAACCggttacctcttatctagt
gtagagtgtagcacctatccTGATGAAGATGGCAAAAGCAGTTGCAACTACAGCAAAAATggttacctcttatctagt
gtagagtgtagcacctatccGGGCAAAAGACACGAACAGAAACTTATCTTAAAAAGACACggttacctcttatctag
gtagagtgtagcacctatccTCAAAAAATAACAGACAGTGAGGTCAGGGAAAGGAAACACggttacctcttatctag
gtagagtgtagcacctatccTCCTCAAAGAACTAAAATCATAACTACCATTTGGCCCAGCggttacctcttatctagt
gtagagtgtagcacctatccATCAACCTAAGTGCTCTTCAGTGGTAGCGTGGATGAAGAAggttacctcttatctagt
gtagagtgtagcacctatccCAGGTGTGGTGGGGGACGCAGGTAGTTCCAGGTATGCAGGggttacctcttatcta
gtagagtgtagcacctatccTGCAGTCATAAAAAGAATATGATCATGTCTTTTGCGGCAAggttacctcttatctagtg
gtagagtgtagcacctatccCTAGAGGCCATTGCTGTAAGCAAACTAATGCAGAATAGAAggttacctcttatctagt
gtagagtgtagcacctatccAGGGAGAAGATCAGAAAAAAGACCTATGGGGTACTATGCTggttacctcttatctag
gtagagtgtagcacctatccAGAAGATTTAAGAGTTCAAACAGGGTATAATGAAGCAACCggttacctcttatctagt
gtagagtgtagcacctatccTTCTAAGAGGAAGGAAGTAATATGTGAGCAGGGCCGGGAAggttacctcttatctag
gtagagtgtagcacctatccACTTAGGTCTACAGAAGGTGGGAGTGTCTTGCAGGTGCTAggttacctcttatctag
gtagagtgtagcacctatccCAGAAAAAAAATGCAGCTGGTATACAAGACATCACAGGTAggttacctcttatctag
gtagagtgtagcacctatccTGGCTGAATCTATCTGGAAGGGCAAGAGAGTGTGGGAAATggttacctcttatctag
gtagagtgtagcacctatccTGGCTCAGTTGAACAGAAGATCAGATAACAAAAAGACAGTggttacctcttatctagt
gtagagtgtagcacctatccAACTCAGAGGACAAAAATAAAAGCAAGGAGATAAGGAGACggttacctcttatctag
gtagagtgtagcacctatccCAGTAGGTCAAATGAAAATTATGGAAACTTGGACTAGGGGggttacctcttatctagt
gtagagtgtagcacctatccTTGTTTGCTTTTTGATTTGTTTGGAGACAGATTCTTGCTCggttacctcttatctagtgg
gtagagtgtagcacctatccAAACTCAAGCAATCCTCCTGCTGCAGCCTTCCAAGTAGCTggttacctcttatctag
gtagagtgtagcacctatccACTGCACCTGGCCGTCAGTGTATTTGAAGTATAATCAACAggttacctcttatctagt
gtagagtgtagcacctatccAGGAACGAGATGTAACAAGAAAGAAAAATCACAAGCTTAGggttacctcttatctag
gtagagtgtagcacctatccTTTTAACTTGAGCAACTAGAAGGAGAGTGATGTCATTTACggttacctcttatctagtg
gtagagtgtagcacctatccAGAAAGACTAATCACGGGAGTTCAGGTGTGCAGGAATAAAggttacctcttatctag
gtagagtgtagcacctatccAAGTTGAAATTGCCAGTTAGACATCTATGTAGACATTTCCggttacctcttatctagtg
gtagagtgtagcacctatccGTCTGGAACTCTGGGGAACTATCAGGACTCAAAATACAGAggttacctcttatctag
gtagagtgtagcacctatccATTACCAGGAGAGAATACATAGACAAAAAAAGAATAGGCCggttacctcttatctag
gtagagtgtagcacctatccGCACTCCAATGTTGAATGATGCTAAGCAGTCAAGAAAGAGggttacctcttatctag
gtagagtgtagcacctatccCCAAATAGAGTGAGTTGTGGAGAGAATATAAAATGATGTGggttacctcttatctagt
gtagagtgtagcacctatccCCAAGGAGAGGAGAGAAACGGGACTGTAGCTATAGAGGATggttacctcttatcta
gtagagtgtagcacctatccACCAAGGGTTCATGTTAGCAAATTGAAAATGGGTGCCCACggttacctcttatctag
gtagagtgtagcacctatccACTATTCTCATTTGTGACATTCTATAAAAAGGATCCTCCCggttacctcttatctagtg
gtagagtgtagcacctatccTTCTTATCTAAAAGGAAGACACTGGGAGGCTATAAACACTggttacctcttatctagt
gtagagtgtagcacctatccCAGGATGCGTGACAACTCAGTTACTCTGCCTACATAAAACggttacctcttatctag
gtagagtgtagcacctatccTCTAAAATGTAAGATTAAGATCTCTAGTGGATATGAGGGCggttacctcttatctagtg
gtagagtgtagcacctatccCAACAGAAGACTATTGAAAGACTTGTCAGAAACCAGAAAAggttacctcttatctagt
gtagagtgtagcacctatccTCCTCAGTTTCAGAAAGCACATTGCAAAACACTCTCGTCTggttacctcttatctagt
gtagagtgtagcacctatccCAGTGTTTTAATACAAGTGCAGGGATTTCTTCTTCCTTTAggttacctcttatctagtgg
gtagagtgtagcacctatccCATCTATGAAGCTGGCTGTTATACTTGTACTGACCTTGGAggttacctcttatctagtg
gtagagtgtagcacctatccATACCCAATCAGAACAAATTTGGCATAAAAACGAAACAGCggttacctcttatctagt
gtagagtgtagcacctatccTTCAGGAGTCCATTGAGACTGAAATTTCCAATGAAGAACTggttacctcttatctagtg
gtagagtgtagcacctatccATCTCAAGGATCATCAAGACCAATCTCCTCATCAAACAAAggttacctcttatctagt
gtagagtgtagcacctatccAATCCCAGAAAGAGGAAGTGATTTGACAAGGTTCCATGGCggttacctcttatctag
gtagagtgtagcacctatccCATAACAGTGAGATGGCAATTAATTTAAAAGAGACAGACCggttacctcttatctagt
gtagagtgtagcacctatccATAAATTATAAGGCATCTCGGTAAGCAAATAGTCAGAGTCggttacctcttatctagtg
gtagagtgtagcacctatccAAGAAGCCAAGCTAATGAAGATAGGCAGAAATTAGTAAACggttacctcttatctagt

gtagagtgtagcacctatccATTCTTTCCCCCCACTTACTTACATTGTATGAGAGCTACAggttacctcttatctagtg
gtagagtgtagcacctatccCTGGAAAGGAACCAATCATAAATGCCACAACACAAGAGGGggttacctcttatctag
gtagagtgtagcacctatccTAAATTGGCAACCACACTTATTTGAGAAAACTTAGGCAAGggttacctcttatctagtg
gtagagtgtagcacctatccTGCTTTTTTCAAATCAGAAAAAAATGTGTGCAAGGCAGCTggttacctcttatctagtg
gtagagtgtagcacctatccTCTTTCTGGATGGTGTTGAAGAATATAGCTACAAAGTCAAggttacctcttatctagtg
gtagagtgtagcacctatccTTGCCGCAGTAGAAATCCTATCTAATCCCTTTAGATACTGggttacctcttatctagtg
gtagagtgtagcacctatccCAGTGGGATAACGGCTGCAATAAAAGAAATTGAATGTATGggttacctcttatctagt
gtagagtgtagcacctatccCCACCAGCAGAACTAGCCCACCCACACGTCAGTCTCTTAGggttacctcttatcta
gtagagtgtagcacctatccCCACGACTGAACTGACCATGAGGTGAACCACATATATGCCggttacctcttatctag
gtagagtgtagcacctatccGGGCCAACAAAATGATTCCAACATTACCATGATCTATCTGggttacctcttatctagt
gtagagtgtagcacctatccTTTCAAGACTTACTATGTAAAGCTACAGTTGTCAAGACAGggttacctcttatctagtg
gtagagtgtagcacctatccCATATATGTGCTCAGTTTGGGTTTGTTTCATTGGTTGTTTggttacctcttatctagtgg
gtagagtgtagcacctatccTGTTCACAAAGGACCAAGGTAATGTAATCGGGGAAAAAATggttacctcttatctagt
gtagagtgtagcacctatccAGTACTCGTATCAGTGTGGGATAAAAATGAATTTTGACCTggttacctcttatctagtg
gtagagtgtagcacctatccAAAATCAACTCCAAATGGAACATCCACTGATATGGCTTGGggttacctcttatctagt
gtagagtgtagcacctatccGTTTAGTACCATCCCCCCCAAGTGCTGTCTCAAGATAGAAggttacctcttatctag
gtagagtgtagcacctatccATCTGGTTGTTTAGAAGTATGTGGGCCAGGCACGGTGGCTggttacctcttatctag
gtagagtgtagcacctatccGAGAGGCCAAGGCAGGAGGATCAGGAGGTCAGGAAATAGAggttacctcttatcta
gtagagtgtagcacctatccAATAAACCATTTGTGTTGTATACTTACGATGGATACCACggttacctcttatctagtg
gtagagtgtagcacctatccAACTTATAGTGACAGATAACAAATTAGTGGTTGCCTAAGGggttacctcttatctagtg
gtagagtgtagcacctatccAGCTGAAATGCAAAGAAACATGAAGAGCTTTTTGGGGTGAggttacctcttatctagt
gtagagtgtagcacctatccCCAAGAAAACAAAAGTAAGGAAATGGAGAGGGAAAGTGGAggttacctcttatctag
gtagagtgtagcacctatccCTGGAATGTAGTGGCAAGATTTCCGCTCACTGCAAAACCTggttacctcttatctag
gtagagtgtagcacctatccGGACTACAGGCGCACGCTGCAACACCAGGCCAATTTTTTGggttacctcttatctag
gtagagtgtagcacctatccCACCACACCCGGCCCTGAGTCATATTTTTAAAGGACATTGggttacctcttatctag
gtagagtgtagcacctatccTCTATTTTGAGAAGAGACTGAAAGGTAGAAGCAAGTTTTCggttacctcttatctagtg
gtagagtgtagcacctatccGAAGGAAGTTTCTACAGTTTGGTTTTTTGTGTGGGTTTTCggttacctcttatctagtgg
gtagagtgtagcacctatccCCTGGTAGAAATTGGAAATCTAAACACCATATTAAAAGGCggttacctcttatctagt
gtagagtgtagcacctatccATACTTAAAGCTAGATACACAGGTACAGCAACTAAAATGGggttacctcttatctagt
gtagagtgtagcacctatccCCTGAAGTTTAGACAAGTAGGCCAATACAACCAATTAACAggttacctcttatctagt
gtagagtgtagcacctatccCCAGATAGCTAATTGGGAGAAAAAAAAAAATGAAACTGGGggttacctcttatctagt
gtagagtgtagcacctatccTTGAAGATTTCAATGAGAAACACCAAAGTTCCTAGACAAGggttacctcttatctagt
gtagagtgtagcacctatccTGTAAAAAGTTCTGTTGATAATGAAAAAGAATGAGGCCGGggttacctcttatctagtg
gtagagtgtagcacctatccCCAGCACATTGGGAAGCCGAGGCAGGTAGATCACTTGAGGggttacctcttatcta
gtagagtgtagcacctatccGGCAGGAGAATCGCTTAAATCAGGAGGCGGAGGTTGCAGCggttacctcttatcta
gtagagtgtagcacctatccAGAGACTCATAAGCTAGATTATAGTGAAATCAAGACTCTCggttacctcttatctagtg
gtagagtgtagcacctatccTGACAAAAAAGAACAAGAAGCAAGTTACAGAATTGGAGACggttacctcttatctag
gtagagtgtagcacctatccCTTATAACCAGCAAAAGACTCATACACAGACTACAGAGCTggttacctcttatctag
gtagagtgtagcacctatccTGAAGAGACAATACAATAGACTTGAACAGTCATTTCACCAggttacctcttatctagt
gtagagtgtagcacctatccTTATGAAAACTGCCCAACCCCAATAGTAAGCAAGGAAAGGggttacctcttatctag
gtagagtgtagcacctatccATCACAAATCAGTATCACTATTAACCCACCAGAATGACAAggttacctcttatctagt
gtagagtgtagcacctatccATGCACAAATGTCCAAACAATGCTTATAACAAGTAGAAGCggttacctcttatctagt
gtagagtgtagcacctatccTGAGGCAAGATAATTGCTTAAACTCAGGAGGCAAAGGTTGggttacctcttatctagt
gtagagtgtagcacctatccCCATCTCGGGGGGAAAAAAAAGTAGAAGCTTTTGTCAAAAggttacctcttatctagt
gtagagtgtagcacctatccAAAAAGAATGAAATTATCATTTCCTGGATGTAGCTGGAGGggttacctcttatctagtg
gtagagtgtagcacctatccTGGGAACAACAGATATTGTGATCTACTAGAGGGGGGCTAGggttacctcttatctag
gtagagtgtagcacctatccGGCTCATTTGTACCCTCAGTGTCACACAATACATCCACGTggttacctcttatctagt
gtagagtgtagcacctatccCATTTCTTTGGGTCTTCATTGCTTTATGAAGGCTCTCATGggttacctcttatctagtg
gtagagtgtagcacctatccGAACAAAGAAAAGTGGTAAGGGAATTGTATACCTTATGAGggttacctcttatctagt
gtagagtgtagcacctatccAAGTCACTTTAATCAAAACACTCTGGCACCAACACAAAAAggttacctcttatctagt
gtagagtgtagcacctatccCAGTGGGAAATAACTGGTTTATTAATAAATGATCCTGGCAggttacctcttatctagtg
gtagagtgtagcacctatccTCTGTATAGAAGAAAACAAGCTAGATCATCTACCTCACACggttacctcttatctagt

gtagagtgtagcacctatccGTTTTCACAGTCTTGAGGCAGAGAAAAACTTCCTATGTGAggttacctcttatctagt
gtagagtgtagcacctatccTGACAAAAGATCACTACATCTAGTAAGCAAAAAGAATGCCggttacctcttatctag
gtagagtgtagcacctatccTTTATGACATTTGGAACTGATGAAGATACAGGAAAATGGGggttacctcttatctagt
gtagagtgtagcacctatccTTTCTGGAACTACAAGTTGCCTAAGCTTCTTATAATAAGCggttacctcttatctagtg
gtagagtgtagcacctatccTACATACCTTTTGACCAAGCAATTTCACTTCACAAAATCCggttacctcttatctagt
gtagagtgtagcacctatccAGAAACAAAAGCACCTGTGTGTAATGGGATGACTATTGCAggttacctcttatctag
gtagagtgtagcacctatccGAGTATCCATTCTACAGATGTAACCCTTGAACAAACGAAAggttacctcttatctagt
gtagagtgtagcacctatccAGGAAAGGAAAGGAAAAGGACTTAGATCAATATCTAGTCTggttacctcttatctagt
gtagagtgtagcacctatccAAAATGTCTAAGATGTATTGCTAATCTGCCCATGAATGGGggttacctcttatctagt
gtagagtgtagcacctatccGCATAAGCACTGAGAAAGTCATAGAAGTTTTAACATCCAAggttacctcttatctagt
gtagagtgtagcacctatccAAGTACATCTCTTTGCATTGTTTTACCAACTACAATGAGCggttacctcttatctagtg
gtagagtgtagcacctatccATGCTTGATTGAAAATATACTTCTCATCTATGGCTGTTCCggttacctcttatctagtg
gtagagtgtagcacctatccTAAACCCTTTCTCTAACTTCTGTATCAATCTTCTTTCCTGggttacctcttatctagtg
gtagagtgtagcacctatccCCTTTTGTCCTCTTTCTATTCCTTCTCTGTCTCTCTTGTTggttacctcttatctagtgg
gtagagtgtagcacctatccAAAATCTCTGCAAATTCCTTCTCATAGTTATTTCCTTCCCggttacctcttatctagtg
gtagagtgtagcacctatccAAGAATTCACTTATTCACCTGCTCATGGTACTGAATCCCCggttacctcttatctagt
gtagagtgtagcacctatccAGGAGACAGAGATGGAAATCAAAAAATTAAAGTAGGGTGAggttacctcttatctag
gtagagtgtagcacctatccGTAATATATACTGAATCCCTAGAGGTAGGTGTGCAAAAGTggttacctcttatctagt
gtagagtgtagcacctatccCACCTGGAGCTAAACCTTTGGAGCAGAGTGCTGAAGAAAAggttacctcttatcta
gtagagtgtagcacctatccCCATGTTTTTGTGTTGGCATTCATTCTTCTTTTTTCCAGCggttacctcttatctagtgg
gtagagtgtagcacctatccAACTTTTTTGACAAACCACATGAACTGTAGAGGAGTACGTggttacctcttatctagt
gtagagtgtagcacctatccTAGTCAGTAAAAAGAATGAACACCCATACTCATGTGGGCAggttacctcttatctag
gtagagtgtagcacctatccGCATGCTCTGGTCATGCATAAACTTTTATTCTTTTGTTAGggttacctcttatctagtg
gtagagtgtagcacctatccTGCTATATATGTTGGTTCTCATAATTTGTTAGGGAGTAGGggttacctcttatctagtg
gtagagtgtagcacctatccCTTTTAGGAGAAATGAAGTTGTTTGAATTGGAGAGTAGAGggttacctcttatctagtg
gtagagtgtagcacctatccGGAAGCATTTTAAACGACTGTTACTTGCCAACTGTGCTGTggttacctcttatctagt
gtagagtgtagcacctatccAAGAATTCTGTAATAGCTCTGCTAGTTTCTCCTCATTGCAggttacctcttatctagtg
gtagagtgtagcacctatccCTACAGAGACAGCAAACAGAGGAAGGGGAGAATGGGTTACggttacctcttatcta
gtagagtgtagcacctatccCTAGTGGATAGAAATATGCTTGCACAATAGTACAATGAGAggttacctcttatctagt
gtagagtgtagcacctatccAACTGGAAACCATGTTAGAAATGTGTATCTCAGCTCCCTCggttacctcttatctagt
gtagagtgtagcacctatccACTAAGCAGTCATTCCTCTCTATAATTCTACCTCTGTAATggttacctcttatctagtg
gtagagtgtagcacctatccGAAAAAGCCACAAGCAAATATTTGTAGCATTTCTGGACAAggttacctcttatctagt
gtagagtgtagcacctatccACAATAGAAATGCCACTTTAAATACACAAAGGAGATGTGCggttacctcttatctagt
gtagagtgtagcacctatccCAATGTTCTTATCACAACAGCTAAAGAAATAACTGCAGTGggttacctcttatctagt
gtagagtgtagcacctatccTATTACTATTCCAATATATAGGGTGCCCAACTGTTTTCACggttacctcttatctagtg
gtagagtgtagcacctatccATCTGGAGATTACCCAACTAGAATAAAATCAAGCACCATTggttacctcttatctagt
gtagagtgtagcacctatccCCTTTTGCTTTTCTGACACCTGTAAACCTCTTTCCATACCggttacctcttatctagtg
gtagagtgtagcacctatccACTACACATTTTAGTCAGCTGCAGGCACCATCTGTTTCTAggttacctcttatctagt
gtagagtgtagcacctatccCATCCATGCCTGATAAACTGCTGTATAAATACTTTGGGGAggttacctcttatctagt
gtagagtgtagcacctatccTTTGTACCCGTTGTCAACATTTCCCCACTACATTTCCATCggttacctcttatctagt
gtagagtgtagcacctatccTCCAATTACAGTCTGAGAAAAGTCTCCAGGTTTATAAAAGggttacctcttatctagt
gtagagtgtagcacctatccAGAAGTCACATCATATTCTCCTCTTATCTTTATCCAAGGTggttacctcttatctagtg
gtagagtgtagcacctatccTTCAGAGGTTATAGCTCAATTTCAACAACATGCTCACACCggttacctcttatctagt
gtagagtgtagcacctatccCTGAAACATCATGGTATTGGGGGGTGGGGGTAGAAGTAGGggttacctcttatcta
gtagagtgtagcacctatccCCAAGGGGTTAATTTGATCAGGTATGCATGCTCTACTCTGggttacctcttatctagt
gtagagtgtagcacctatccATGAACCTGAAGGCCATGGTAGCTACTTTTAACTTTTCCAggttacctcttatctagt
gtagagtgtagcacctatccGTATGTATACCTAATCTAATCTCCAACTTCTTTGCCTGTCggttacctcttatctagtg
gtagagtgtagcacctatccCCACCAGCCAGTTCCTCCCATTCTTACAGACTCTAGTTTCggttacctcttatctag
gtagagtgtagcacctatccATTCACTTTAGAATCTAGAACCAGGATCAATCACCACGTGggttacctcttatctagt
gtagagtgtagcacctatccGTTCACCCCCAAACAATAACAGACATTAACACAAACAACAggttacctcttatctag
gtagagtgtagcacctatccTAAAAGATAGTAAAAGAGTCTGAGCCTATTAAGACAGGGCggttacctcttatctag
gtagagtgtagcacctatccAGTCTGACTAGGCGGATTTTCACAGGGTTTTTCAAGGTCAggttacctcttatctagt

gtagagtgtagcacctatccCTCCTGATTTGGGGTTTTTGTTGGAGTTTTGTTTTTGTTTggttacctcttatctagtgg
gtagagtgtagcacctatccTTGTTTTGAATGACTCCAATAGCTCCTTCCCACATCTTGGggttacctcttatctagtg
gtagagtgtagcacctatccGAAATCTATAGTTTTCTCATCATCCAGAGGCTACATTTAGggttacctcttatctagtg
gtagagtgtagcacctatccTTCTGTCTGGGAGAACTGAGTGGGGAGAAAAAGGTAAGCGggttacctcttatctag
gtagagtgtagcacctatccGTTCCATCTGAGTTCACCAAGTCCCAATATTCTGTGTGGGggttacctcttatctagt
gtagagtgtagcacctatccGAACCTTCCCTGTGTATATTTTCTAGCTTCTCTGTGGGTCggttacctcttatctagtg
gtagagtgtagcacctatccGGACCTCCCATCTATAGACCTCCTATTTCAGCCTGACCTGggttacctcttatctag
gtagagtgtagcacctatccAGTCTTTCTTGTTTTAGGCCCTTGACTGGTTACCTCATGCggttacctcttatctagtg
gtagagtgtagcacctatccTTTGACCATGTATTTAAGATCAGGTATATCTAAGGGCAAGggttacctcttatctagtg
gtagagtgtagcacctatccCAGACCTGCTTCCCACACAAGATGCAATGTTTTCAGACATggttacctcttatctag
gtagagtgtagcacctatccAGTATGCTTCCCAACTGAAACCTTACTAATTGTTCAACACggttacctcttatctagtg
gtagagtgtagcacctatccATGCTTGAAGTTAAATATGAGTCCTATACAGATAGGACTCggttacctcttatctagtg
gtagagtgtagcacctatccCATAAAGAAAATGATATGTGGTGCTTAAATGACGGATACCggttacctcttatctagt
gtagagtgtagcacctatccGCACATTTGATTCCTAAGGCATTGGCATTTAATTTGACACggttacctcttatctagtg
gtagagtgtagcacctatccTTTCCATTGTAGTAAACCTAAACCTCCATGTATACTCGAGggttacctcttatctagtg
gtagagtgtagcacctatccTGCACAGAAAGCCAATCATTTAGACAATCAGTACTGCCGAggttacctcttatctag
gtagagtgtagcacctatccCAGATGCTACAGCTGAGATGGGTGCTCAGCCTGAAATCCAggttacctcttatctag
gtagagtgtagcacctatccACAACATGTAAGAAAGCAGGAACTAATGAGGAACAAGGTGggttacctcttatctag
gtagagtgtagcacctatccCTGTCTATGGGACTACTGGGCCATTTTCGGTGGGAGACGAggttacctcttatctag
gtagagtgtagcacctatccCTACTAAGTGCCAAAATGAAGTAGTATGTTAAGTCCTACTggttacctcttatctagtg
gtagagtgtagcacctatccCATTTGAAAATCTAGTCATGGAAAGTCTGATAGGGCCTATggttacctcttatctagtg
gtagagtgtagcacctatccAATCTTTCTAGTTATTGCAGAGCAAGAGTTTTCCTCTATGggttacctcttatctagtg
gtagagtgtagcacctatccAATGCTTGGTCATAGATGTAAAGATGAGTAACAGTGTTTCggttacctcttatctagtg
gtagagtgtagcacctatccCAGTGGAGCTTTAGAAGAATACACAAAGAAACCTAAACAGggttacctcttatctag
gtagagtgtagcacctatccCATAAGCAGCATGGGTCTAAGATTGACTAAGAGGTTTGGAggttacctcttatctagt
gtagagtgtagcacctatccGTTTCAGTAAACATTCATCCTAAGACTCAACCACATTATGggttacctcttatctagtg
gtagagtgtagcacctatccCAGTTGTCATGAAGCTGTATGGGAAAACATATGGAAACAAggttacctcttatctagt
gtagagtgtagcacctatccACTGTGTCTTGTGTGAAAAAAGCTTTAAAAAAATACGGGCggttacctcttatctagtg
gtagagtgtagcacctatccTCTCTTTCCTCCTTGATATCTCATGTCATCTGTCAAATTTggttacctcttatctagtgg
gtagagtgtagcacctatccCTGGGAACATAATTGCTACTCTATGGGGCAGCCTCCACAGggttacctcttatctag
gtagagtgtagcacctatccACTGGTATCTTACTTCTTCCACTGTAAACAGTTTTCCAGAggttacctcttatctagtg
gtagagtgtagcacctatccGATAGACCTTTCATTATCACTTAACACTGACCACAACTTGggttacctcttatctagtg
gtagagtgtagcacctatccCAAATACTTGTTGCTGCAAAGGGATTTATTTCTGAACTGAggttacctcttatctagtg
gtagagtgtagcacctatccCCATTACGGTTCCTTTATCTTCTTGTTTATTAGGTCTTCCggttacctcttatctagtgg
gtagagtgtagcacctatccGTGGTTAACAGGACTTGCAGCAAAAGTTGAAAAATCCATTggttacctcttatctagt
gtagagtgtagcacctatccTCAGTTGAGAAGGGAAAAATTGAGAGCCTATGAACATTTGggttacctcttatctagt
gtagagtgtagcacctatccGAATTTTCTTATTAAACCCCAGATCTAGCTAGCAACAGGTggttacctcttatctagtg
gtagagtgtagcacctatccTTCTAGCTCTAAATTTCTATGAGTTGAAGCGGACAAGGTTggttacctcttatctagtg
gtagagtgtagcacctatccAAGAGAGCAGGTATTTTAATGATATGAGAAGTGGATCCGAggttacctcttatctagt
gtagagtgtagcacctatccTAATTTCAAGGCGCTGGTTTATGTTTTCAGTATTCAAAGCggttacctcttatctagtg
gtagagtgtagcacctatccTACTCTGTGGAGGTGAGCTTCTTTAAAGGAATGTGCCCCTggttacctcttatctagt
gtagagtgtagcacctatccCAGCATGCATTTTTTAACTAGGTTCACAGTATTGGGGTAAggttacctcttatctagtg
gtagagtgtagcacctatccTGAAGCTAAACAAATGTTCCCTAACCTTTTCATCCCAGCCggttacctcttatctagt
gtagagtgtagcacctatccCTTATTTGTCTTCAGTCTCAATTCTTAATCTAGGCTCATCggttacctcttatctagtgg
gtagagtgtagcacctatccTCTTTCCAACACAGATCCTTAAGCTGCTTTGAATAACTCAggttacctcttatctagtg
gtagagtgtagcacctatccACTATTCAGGTTTAAGTAGTCTTCTTTAAGCAGCCCAGGTggttacctcttatctagtg
gtagagtgtagcacctatccCTAAGGGCCTTACATCAACTCATTTAATCTTGTAATTACCggttacctcttatctagtg
gtagagtgtagcacctatccCGTGTTATCCTCGGAGAAAATTAAGGCATAGTAAGTTTCCggttacctcttatctagt
gtagagtgtagcacctatccATTAAGTGACAGAGACAAGTTACAAACCCAGGTTCCTAATggttacctcttatctagt
gtagagtgtagcacctatccCTTACAAGAGCTCTATATTAGACAGATTTACCTTGAACTGggttacctcttatctagtg
gtagagtgtagcacctatccATGGGATGGGATGAGATGGGGCTGGAGGCAGTTGTCTACAggttacctcttatctag
gtagagtgtagcacctatccGCTCATCCAGGCTCACAAGAACCAATTGTAAGATTTTCAGggttacctcttatctagt

gtagagtgtagcacctatccTGTTTGTTGCTTGAAATTGTTCATAATGGGAGTAGTTACGggttacctcttatctagtgg
gtagagtgtagcacctatccCTACAAGCTAGACCCTTCCTCTTTCCCCTACCCTCACATGggttacctcttatctag
gtagagtgtagcacctatccACTACAGGTATGCGCCACCACACTCAGCTTATTTTTTTGTggttacctcttatctagtg
gtagagtgtagcacctatccGTCACTGCACCTGACCTAGCTAACATTTTTAAATTCTGCTggttacctcttatctagtg
gtagagtgtagcacctatccCAAGGCAGTGGGGTTGATAACTAACTATTCCTGGTACATAggttacctcttatctagt
gtagagtgtagcacctatccCTGGTTTCTTTAAGGGGTGATATTAGAGGATACAGAAGGTggttacctcttatctagtg
gtagagtgtagcacctatccACTTATGGATATAGAAAAAGCAGTTGATGTTGTCCTTGAGggttacctcttatctagtg
gtagagtgtagcacctatccAGGAAAGGTCCTTAACCACAGTCTGTAGGCAAAAGTCAAGggttacctcttatctag
gtagagtgtagcacctatccTCAAAGAGTGAGTGATATAAACAGCCAAGAGTCAATTCTTggttacctcttatctagtg
gtagagtgtagcacctatccAGAGAATATTTCCAGGAAGACTTTGGGTGGCTCTAGTACCggttacctcttatctagt
gtagagtgtagcacctatccAATGGAAGAAACTAAGCAGAAAAGTCAGATACCAAAGAACggttacctcttatctag
gtagagtgtagcacctatccCAGAAGTGGAGTCCAAACCTGAGAAGATTATCAAGAATCCggttacctcttatctag
gtagagtgtagcacctatccCCTAGCAGGATCCGCAGATGTAAAATAATATGGACAGGTAggttacctcttatctag
gtagagtgtagcacctatccGCTCACTTCTTGAAACTAAAAGATGAAAGCCCAATTAACTggttacctcttatctagtg
gtagagtgtagcacctatccGCCTGACCACGAGCCTCTACCTGAGCTTGCCAAATGCCAGggttacctcttatcta
gtagagtgtagcacctatccCAGGAACAAAAATGATCTGGCTGCTAGGACTGAAATTCTGggttacctcttatctagt
gtagagtgtagcacctatccCACTTGAGCAATTGTGACACACTCTGAATTATAATCTTTCggttacctcttatctagtg
gtagagtgtagcacctatccGAACTAGTTACATAGTCAGAAAAAAATCAGATGCCATTGCggttacctcttatctagtg
gtagagtgtagcacctatccGACACACCATCCCCATTACACTCTCATTTTTACGTGACACggttacctcttatctagt
gtagagtgtagcacctatccTGGTCCAGCTTTGTGATTTTTAGCACAAATCAACTAGTGTggttacctcttatctagtg
gtagagtgtagcacctatccGGGATGTTTATCAGATAAAAGCTAAAATAGGCCCAACATAggttacctcttatctagtg
gtagagtgtagcacctatccGCCAGGAGTTTGAAAACAGTCTTGACAACATAGTGAGACCggttacctcttatctag
gtagagtgtagcacctatccGGCATGGTGGCCTGAGCCCGTTCCCAGCTACTAAGGAGGCggttacctcttatcta
gtagagtgtagcacctatccAAGCTAAAATATAAAGGGGAAGGTATGTGGAACCATTATGggttacctcttatctagtg
gtagagtgtagcacctatccGGAAACTTTTGAGAATGAAAATTGCTCCTAATACTGACTGggttacctcttatctagtg
gtagagtgtagcacctatccGGATACACACCCAAAAAAAAATACCAAATTGTGCTGGTGAggttacctcttatctagt
gtagagtgtagcacctatccGCCTACTCAGGGGCATCCTTTGAAAACAGTATCATCTTTTggttacctcttatctagtg
gtagagtgtagcacctatccTGATAGATCAGATACTGTCCACAAATCCTCAAATTCATGGggttacctcttatctagtg
gtagagtgtagcacctatccTACTGACTTGAAAGGATGGTGTGTAATGATTTGTACATTGggttacctcttatctagtg
gtagagtgtagcacctatccGCAGACTTGAGATTGCTAGGGTATATATTACAAAAGGAATggttacctcttatctagtg
gtagagtgtagcacctatccGGTCCAACAGAGTGTATTTTTGATGAAGGTATTTTAGTCAggttacctcttatctagtg
gtagagtgtagcacctatccGAAATAAAATAGGGGTCTTTCATAGGTTGCACTTAGAATGggttacctcttatctagtg
gtagagtgtagcacctatccTCGAGAAACTAAATACTTCAAAAATGTCGAGTCAACTTGGggttacctcttatctagtg
gtagagtgtagcacctatccCATTCTATATATGCTGGCATGTTTTTTGCCTCTTTAACCTggttacctcttatctagtgg
gtagagtgtagcacctatccTTATTCTAAATGTTTGCTGAGTATTCAGTTCTGTGGCTTGggttacctcttatctagtgg
gtagagtgtagcacctatccGACATGACTCATTCTTTTTTGCCTAGTAATCTCTCATTAGggttacctcttatctagtgg
gtagagtgtagcacctatccAATAGATGTTCTGGCATGGAGTTAAAATTGTTAGTGAGGCggttacctcttatctagtg
gtagagtgtagcacctatccGTTCCACCACCAAGACAGCTTCCATTCCAGCAGGGGCCTTggttacctcttatctag
gtagagtgtagcacctatccGCCAGGGCCAACTCACATATCTGTTAAGGACTTCACTGTGggttacctcttatctag
gtagagtgtagcacctatccACCTCTGTCATAAATTAACACAAATTCGGTAGCCTAACAGggttacctcttatctagtg
gtagagtgtagcacctatccTTCCATCATGGCTCTGGACGTCACAAGTCGGAAATCTTTTggttacctcttatctagtg
gtagagtgtagcacctatccCAAGTATTGGCAGTACAGTGCGCTAGAGGAAGATTAATTTggttacctcttatctagtg
gtagagtgtagcacctatccCATTTCTTGCATCCTGGCCCCATCTTTTATCTTCAAAACCggttacctcttatctagtg
gtagagtgtagcacctatccGGTCACATTGCCTTCTTTGTACCAAATCTCCCTCTGCTTCggttacctcttatctagtg
gtagagtgtagcacctatccGGATAACTCCCCCTTCTCGAAGTCCTTAACCAAATCACTTggttacctcttatctagt
gtagagtgtagcacctatccCACAGTTTTAAGAGATTAGGATCTGAATATGTTGAGAGGCggttacctcttatctagtg
gtagagtgtagcacctatccAAATGTCTATGCCTTTGAAGCTCAATAATTGAGCTTTCTGggttacctcttatctagtg
gtagagtgtagcacctatccATTTACTCTTTTCACTAAGATTTGAGTGGCTACATTGTGCggttacctcttatctagtgg
gtagagtgtagcacctatccGTATGACTAGTTTATTAAGGGCTCCATAGAAAAAGTTGACggttacctcttatctagtg
gtagagtgtagcacctatccCACTAACAGAAATAAATAATGCCCCTGATAAGTTCATCAGggttacctcttatctagtg
gtagagtgtagcacctatccCACAGCTAAAGAAAGAATCAGTGGAATTGAAGATAGATCAggttacctcttatctagt
gtagagtgtagcacctatccTCCCTAACATAGAAACAAAACTGTCAAGCAAAATTGAAGCggttacctcttatctagt

gtagagtgtagcacctatccTCAGAAACGTCCAGCTGACTTTTAACTAGAGACTTTACCAggttacctcttatctagt
gtagagtgtagcacctatccACCACCAACTAACCTCTAAGGACTTCCCATTTTAAAACCAggttacctcttatctag
gtagagtgtagcacctatccAAACAGAAGTTTCCCCTTGAGCCCCCTCAGTGGAGCCCTGggttacctcttatcta
gtagagtgtagcacctatccTTTGTGCCTTGCACTGCCCAACTCATGAATTGCTGAATGCggttacctcttatctagt
gtagagtgtagcacctatccTCCAAAATTAGTGACAGACATCACACCACAGATTTAGGAAggttacctcttatctag
gtagagtgtagcacctatccAACAAACAAAACCTATCCCTGGGCATGTCATATTTAAACTggttacctcttatctagt
gtagagtgtagcacctatccGGGACACCTTACCTTTAGACAGACAAAAATAGGAATTACAggttacctcttatctag
gtagagtgtagcacctatccAAGAAAAAAAAGGTTAAATAAGCACAAATCCCCACCAACGggttacctcttatctag
gtagagtgtagcacctatccAAAAACATGGAAAGCATCACCAAGAGAAGTTCCCTTTAAGggttacctcttatctag
gtagagtgtagcacctatccTTTAGGCAGAAGGAAAATAACATTGGTCAGAAGCTTGGATggttacctcttatctagt
gtagagtgtagcacctatccATGGTCAGTTTTATGCATCCACTTAGCTAGCTACAGTTATggttacctcttatctagtg
gtagagtgtagcacctatccCAAATACTGATCTTAGGGTAAATGGCCATCAATCAACAAGggttacctcttatctagt
gtagagtgtagcacctatccGCCATAAAAAAAGAATGAACTAACAGCCTTTACAATGACCggttacctcttatctag
gtagagtgtagcacctatccCTAAGTGAAGTAACTCACGATGGAAAACCAAACATGGCATggttacctcttatctag
gtagagtgtagcacctatccCTTTGGGGATTTGGGGGGAAAAGTGAGAGGGGGCGAGGGAggttacctcttatcta
gtagagtgtagcacctatccTACTCATGTAACCAAATATCCCTGTACCCACAAGAACTTAggttacctcttatctagt
gtagagtgtagcacctatccGTAAGGGAGGTTATACTAGATAATTCAAATGGGCTTAATCggttacctcttatctagt
gtagagtgtagcacctatccAAAAATTCTTAAGAGCAGAGCCAAGCTTTTCCTAAAATGGggttacctcttatctagt
gtagagtgtagcacctatccGCAGTTGCAGCTTCAGGTTTAACAGCCAGAGTTCCAGCCTggttacctcttatctag
gtagagtgtagcacctatccAGCTTGTCCTACAGATTTGGGGATTGTCTAGCCAGCTCTTggttacctcttatctagt
gtagagtgtagcacctatccGCCAATTGCTTACAAAAAAGTTCTGTGTCTTCTACTGTTGggttacctcttatctagtg
gtagagtgtagcacctatccACAACAATGTCACAAGGGATCGTAGGGTAGAATTGGGAATggttacctcttatctag
gtagagtgtagcacctatccTCATAAATTACTTGTACTACCTGTGATGTGCTATAGGCTTggttacctcttatctagtg
gtagagtgtagcacctatccTGGGATCATATTAAATGCTCAATTAAAGAGAGGAGGAAAGggttacctcttatctagt
gtagagtgtagcacctatccAAAGACTCAAATTGATATAGATATACAGCTCTCAAGGAGGggttacctcttatctagt
gtagagtgtagcacctatccCTTCATTGCAAAGAGTATATTGTGGAAAGGGGGGAGGGAAggttacctcttatctag
gtagagtgtagcacctatccACATCCAACATAATGTCTTATTAGCCCTTATCAAGCCTAAggttacctcttatctagt
gtagagtgtagcacctatccGCAGGAATAAGCCATGTTACCATTGATGTGATGTGATGAGggttacctcttatctagt
gtagagtgtagcacctatccCATCAACCGAATCTCAATTGTGAGACTTTCTAAAATTACCggttacctcttatctagt
gtagagtgtagcacctatccAACAAGGAAAGTCTGAGAATGTGCCATAGTATGAGACGCAggttacctcttatctag
gtagagtgtagcacctatccTGGAACAGAAATAGGACATTAGGCAAAAGCAAAGGAGATCggttacctcttatctag
gtagagtgtagcacctatccGAAGGAATGAAGGAAGGACATGCTGAAACAGGGAGTGGATggttacctcttatcta
gtagagtgtagcacctatccCAAGTACTGACCTGTTTATGAAGTATGTTCTAACTGCAGTggttacctcttatctagtg
gtagagtgtagcacctatccTAGCTATTGCTGCATAAAATTCACCCCAAAATGTAGTGGCggttacctcttatctagt
gtagagtgtagcacctatccTCTCCATGATATCAGTGAGTCAGGAATTCATACTGGGACCggttacctcttatctag
gtagagtgtagcacctatccGCTTGTCCCTACTTCATTACGTCTGGACCTCAGCTGGAAGggttacctcttatctag
gtagagtgtagcacctatccCTTACTGAAGGCTTTTTGAAATCATTTGAGTCCAGCTGTCggttacctcttatctagtg
gtagagtgtagcacctatccCCTGGTGGCTATGTTCTAAAAGTAAGCGAGCTAAGAGAAAggttacctcttatctag
gtagagtgtagcacctatccGGAATCCATACTGCTTAATTACAACCTGGACTTGAAAGTCggttacctcttatctagt
gtagagtgtagcacctatccGCAGTTCCAAATGTGTATTCAGTTTTAACGAGGAGACAACggttacctcttatctagt
gtagagtgtagcacctatccCTTTAACAAGCATATGGGATGGGATACATATTGGTGAAGCggttacctcttatctagt
gtagagtgtagcacctatccATGTTTCTTACAAATAAACTTTGTGGCTTCCAAGTTTGGCggttacctcttatctagtg
gtagagtgtagcacctatccATAGACAAACAGATAGGTAAACAGACTTCACCTGGTTAACggttacctcttatctag
gtagagtgtagcacctatccTTGCTTCTCCCACATATTGTCAACTACTCCCATATTCTACggttacctcttatctagtg
gtagagtgtagcacctatccTTTAGAGTCCATGTTCCGGTCACTGCACTAGGGTCTGGAGggttacctcttatctag
gtagagtgtagcacctatccCCTTAATGAGCTACAGGTTTATGGGCAACACAGAAAGCAAggttacctcttatctag
gtagagtgtagcacctatccTGTGTTTCATATGTTAACAATGGTAAGTATTGGGTGCTGTggttacctcttatctagtg
gtagagtgtagcacctatccCACTTGGAATAGACACATTACCTAGGCCTGGGAATTGGGAggttacctcttatctag
gtagagtgtagcacctatccCCCAGAAGTACCATATGACTTAGAATGAATAAGTAGGAGCggttacctcttatctag
gtagagtgtagcacctatccGAATGGGTAAAAGTATTCAAAACAGAGGGAAAAACAGGTTggttacctcttatctag
gtagagtgtagcacctatccTTCTAGGAGCAAGAGAAAATGTGGCATAGTCATCGATCAGggttacctcttatctag
gtagagtgtagcacctatccTCAAATGTACTATGCAAAGCAGCTGAGATTATAAGTCCAGggttacctcttatctagt

gtagagtgtagcacctatccACTTTTGTCAGGTTTTGTCCACTTATATATCCCCAGTGTCggttacctcttatctagtg
gtagagtgtagcacctatccGTATTTGTGGCTTCATACTTGGATTCCTGTCTCTGAATATggttacctcttatctagtgg
gtagagtgtagcacctatccCTGGACAGTCTTGCTATTCATTATTTCCATCCTAGATGCggttacctcttatctagtg
gtagagtgtagcacctatccCATAAATTGTGGCCTGAAGAAAAGACCTGCCGTCATAGGCggttacctcttatctag
gtagagtgtagcacctatccAGGCCAGCAACCAAAGCTACTGCTCCTCTGAAAGTTCTGTggttacctcttatctag
gtagagtgtagcacctatccGGCATTTGAAAATTCATGGTCCCCAGTGTCTTCAACTAAGggttacctcttatctagt
gtagagtgtagcacctatccCCATGGCAGTAGACCTGGAGTTGGTTATATCTAGAGAATGggttacctcttatctagt
gtagagtgtagcacctatccTGATGCTCACTGTCCTGATATACATGCCTGCAAGAAGATAggttacctcttatctagt
gtagagtgtagcacctatccGAGCTTATTGCACCAGCCTCAATCTAAAATTACATATATGggttacctcttatctagtg
gtagagtgtagcacctatccGTAAAACAAGCCAGACACTGCAAGACAAATACTGCGTGAAggttacctcttatctag
gtagagtgtagcacctatccTAGAAACAGAATAGAAGGGTCCTAGGGGTGAGGGGAGTAAggttacctcttatctag
gtagagtgtagcacctatccCATAAAATGAGTAAGTTCGGACACCTAATGTACAGCTTTAggttacctcttatctagtg
gtagagtgtagcacctatccGCCTCCCAAGTAGCTGTGTTTACAGGCATGCGCCACCATGggttacctcttatctag
gtagagtgtagcacctatccTGAGCCACTGTGCCAAGCAGATCTTATGTATTCTTATCAGggttacctcttatctagtg
gtagagtgtagcacctatccTTCGATTGAATGCAGCTATCCTTCATTTCACAATGTGTATggttacctcttatctagtgg
gtagagtgtagcacctatccAAAATACTCATCTATATTGTGGTTCATCTATCCCATCTGCggttacctcttatctagtg
gtagagtgtagcacctatccAATATTTTGCAAACCCCAATTATTACCACCCAGCTCACTTggttacctcttatctagtg
gtagagtgtagcacctatccGGACTTAGGATTCCATGACAGTAGAGAATTTCCAACCTCAggttacctcttatctagt
gtagagtgtagcacctatccTGATTCAATAAGATCATAGAAACAAGGCTCTCCAACACAGggttacctcttatctagt
gtagagtgtagcacctatccATCCCCGTGGTGTTTAATATTATGTGTTTGCAACATATCTggttacctcttatctagtgg
gtagagtgtagcacctatccAGTCAAATGGAGGAAATTTATGAAAGAAGGTAAATGTGGGggttacctcttatctagtg
gtagagtgtagcacctatccGATCTTTCATCCTCCTAAGACAATACTACCTTCCATTCTTggttacctcttatctagtg
gtagagtgtagcacctatccCAAATTTGACTTGGAGATCCGGCTAGATTTTGATGACTTAggttacctcttatctagtg
gtagagtgtagcacctatccTTATGTTAAGAATGTCGTAAATCACACTGCAGAAGCTAAGggttacctcttatctagtg
gtagagtgtagcacctatccGGTAATTACTTTGTGGCACCAATGGAAACCAAGACCTTTTggttacctcttatctagtg
gtagagtgtagcacctatccAAAGTATGTCAAGTCTTTGCCTCTGTGTACCTGGGTGTGGggttacctcttatctagtg
gtagagtgtagcacctatccCAAGTGGCTTTGATGGGTTCCTGTGTTAGCATTGTTCACGggttacctcttatctagtg
gtagagtgtagcacctatccGGCAGGCACACACTTATCAAATGACTTACCTTTATTAATTggttacctcttatctagtg
gtagagtgtagcacctatccATGCTACTGTCTAAGGCAGTTCTTATTCCCTCCAAATGTGggttacctcttatctagtg
gtagagtgtagcacctatccCAGCCACAGAATTTTCCAAAACCTGCTAATTAGCTTCAGCggttacctcttatctagt
gtagagtgtagcacctatccCTTCAGAAGGCCAAACAGTCAAATGCAAACTTCAGCATCAggttacctcttatctag
gtagagtgtagcacctatccTGCAGCTTATCAATTTTCTTGTGAATGTTTGGTTGTCATCggttacctcttatctagtgg
gtagagtgtagcacctatccACTTCTTGTTTGCATTCGTTATAGCCTTCATTATACCTGGggttacctcttatctagtgg
gtagagtgtagcacctatccTTGACACATAAGGCACATGAAAATATTATGTTGCTCTAGGggttacctcttatctagtg
gtagagtgtagcacctatccAAATGAGGTCCTATACTAAAAATTACTCCAAAAGCATGCCggttacctcttatctagtg
gtagagtgtagcacctatccGTCACTGACTTTTTGTGTTGACAAGATATGGTAGCAATCTggttacctcttatctagtg
gtagagtgtagcacctatccTGGGATGCCGGAAATGAAGTAAACCTTAAGTTATTTGGTTggttacctcttatctagtg
gtagagtgtagcacctatccCTCAGAGGATATGTGAACCGAACACTGTGGCCCACGTTGCggttacctcttatctag
gtagagtgtagcacctatccTGTAACAAGACACACAGGATTGAAAACCGCAGCTTTAGGAggttacctcttatctag
gtagagtgtagcacctatccACCTTTCCCAAAGATTCTCTTCATATTCTAGCTCTCAGGCggttacctcttatctagtg
gtagagtgtagcacctatccGGTAATTCCTTTGTAGTAGCTATTTTACTGCACCTTCCACggttacctcttatctagtg
gtagagtgtagcacctatccGGTAGGCTCACTCCCCTCTCATGGTTGTTTACATAAAATGggttacctcttatctagtg
gtagagtgtagcacctatccACGCTCAGGTTTCCAAGTGTCAGTTTTTTACATGGTCACGggttacctcttatctagtg
gtagagtgtagcacctatccATTTGTCTCTACTGTTAAATAATTCACGAACTCATCTGGGggttacctcttatctagtg
gtagagtgtagcacctatccGAATTCTTCCTTATCATTCACCCACACCCACACCCACTGCggttacctcttatctag
gtagagtgtagcacctatccGCTCAGCACTTTCACCAAATCCCTTCCAGTCTCTTCCTCAggttacctcttatctagt
gtagagtgtagcacctatccTTCAGTGTATAATGAGTATCTATAGGCTAACTATAGGCACggttacctcttatctagtg
gtagagtgtagcacctatccCTTGCATAACTGACACTTTAAGCCCCTTGAACAGCAACTTggttacctcttatctagt
gtagagtgtagcacctatccTTCCCCTTCCCCCAACTCCTGCAACCACCTACTCTCTGCTggttacctcttatctag
gtagagtgtagcacctatccCTGTGTTTGGCTATTTTGGATAACTCTTGAATCCTGCAGTggttacctcttatctagtg
gtagagtgtagcacctatccCCCTTCAGTGATTAGTCTAACATAATGTTGCTCACAGCTCggttacctcttatctagtg
gtagagtgtagcacctatccCCTTTTTTGGGGCTGGATAATGTTCCATTTATATACAACCggttacctcttatctagtg

gtagagtgtagcacctatccTCTCATGTAAGGGGAAAGAGAGAGCAGGGATATATGTTCTggttacctcttatctagt
gtagagtgtagcacctatccCACTCTCTGGACATGTTTTTATTTTATGGAGAAAGTCTGAggttacctcttatctagtg
gtagagtgtagcacctatccAGTCTATCCTAGCTGTATATTACTGAAAGGGTTGATAATGggttacctcttatctagtg
gtagagtgtagcacctatccTTTGAACAAGAGACATTTCCTACATCTTTCAGATCATCCCggttacctcttatctagtg
gtagagtgtagcacctatccTCTGAATGTTAGCTTTTCAATTGTAGAGGGTCAGGTATGGggttacctcttatctagtg
gtagagtgtagcacctatccGCCCCTTCTCCATGTTCTCATAGCATCCTGGCCATGTCCCggttacctcttatctag
gtagagtgtagcacctatccCATATTTCTATTTGTATGTTGATTTTCCCCAGGTCCAAGTggttacctcttatctagtg
gtagagtgtagcacctatccGGCCAAGAATCATAACCAATAGATGATAGAGATGGGATTTggttacctcttatctagt
gtagagtgtagcacctatccGCCTTTCTGATATCAGAGCCTATGATCTCTTGATTTTTACggttacctcttatctagtg
gtagagtgtagcacctatccGTGTGTCAAATTATCTGGAATCACTACTTCTGGGATAACAggttacctcttatctagtg
gtagagtgtagcacctatccGGTGAATTAGTATTATCATTTGTTTGGTTTGCTTAGGTCCggttacctcttatctagtg
gtagagtgtagcacctatccGTATACCATATAAAAATGTCATCAACACAGTCTTGGCATGggttacctcttatctagtg
gtagagtgtagcacctatccAGTGCTCAATAAATGTGGGAAAACTCCTGTACGAGAAGCAggttacctcttatctag
gtagagtgtagcacctatccGTTAACTTAGCTATCTCACTTTTCTTCTAGTTGTATCTGCggttacctcttatctagtg
gtagagtgtagcacctatccGACATTGGTCTTATTATTTGAAACTGTGGTGTTGAGTAGTggttacctcttatctagtg
gtagagtgtagcacctatccGCTGGCCCTGAACTACAAATCAAGCACACTTGCATAACCTggttacctcttatctag
gtagagtgtagcacctatccGGAATAGCCCAACAGTAGAACTACAGAAAGAAATATTTTGggttacctcttatctagt
gtagagtgtagcacctatccAATGCCTCAGTGAAAATAATACGTGAATGTAAAAACCCTGggttacctcttatctagt
gtagagtgtagcacctatccAAAGCATAAACTTCCTTTGATATCTGTCCGCCCTAGAAAAggttacctcttatctagt
gtagagtgtagcacctatccTGCTTGTTTGGAATTCCATTTATATTCTATTCCGTGCTACggttacctcttatctagtg
gtagagtgtagcacctatccATGAAAATGTTGTGGCTATTGGTAGGTTGGGGAAAGGGAGggttacctcttatctagt
gtagagtgtagcacctatccAAACAGATATCTTCCAAGGAAAAAGAGTATATCACGTAGGggttacctcttatctagt
gtagagtgtagcacctatccGTAAAGAGATAAACGTATAGATTTGCCACACTGCCAAAGCggttacctcttatctagt
gtagagtgtagcacctatccCTGAACAAATACCAGAGCCTCATTCTGCAGATTTAAATATggttacctcttatctagtg
gtagagtgtagcacctatccATTGTGTAAGTGTCTATTGTAAGCAGTTTTGCAAACTCTCggttacctcttatctagtg
gtagagtgtagcacctatccGCTACACTCAGCTGTGGCTAAGAAAACAATCAAAAGACATggttacctcttatctag
gtagagtgtagcacctatccGGTTTATTCTAATTTAGACCAGCATCCAATGAAACCTACAggttacctcttatctagtg
gtagagtgtagcacctatccAAAGTATTGAGCGTTCTTAAGTATTCCCAGCTGGATGCATggttacctcttatctagtg
gtagagtgtagcacctatccACCAGGTGGTCCTATAGGCTTAATGATGATTTGATTATAAggttacctcttatctagtg
gtagagtgtagcacctatccGTATTCTGGATACCTTTCTAACTTATCTCAATGTGCTAACggttacctcttatctagtg
gtagagtgtagcacctatccCTAAAGCTGCAACTATATTCCTAACAGAGACCAGGCTGAAggttacctcttatctag
gtagagtgtagcacctatccCAACTGTGTTGGAATTTCACTTTTGCCCCACTTTCCTAGGggttacctcttatctagtg
gtagagtgtagcacctatccAGAGATGAGTGCTAATAAACAGACAGGGCAGGCTACTTTCggttacctcttatctag
gtagagtgtagcacctatccCCAGACAGGTTGTTATGCCATCGAATAAAGTTTTTAAGATggttacctcttatctagtg
gtagagtgtagcacctatccGCTGATGTTTGGGAAAGGAGAGCTAATGGAATCAAATGTGggttacctcttatctagt
gtagagtgtagcacctatccATCCTCTTTGAATTAGACTAGAAAAACATTTGACCAGCTGggttacctcttatctagtg
gtagagtgtagcacctatccATTGTAGACCAGTGACCTTCCATTCTCCCCTTATCGTCTCggttacctcttatctagt
gtagagtgtagcacctatccGAGATGAAGCCTCCTTTGTTCTCTTGAGTACTTTTATGAGggttacctcttatctagtg
gtagagtgtagcacctatccATGCAAGTTTAGGGGAGGAGAGGCATTTGAGACCTACTCTggttacctcttatctag
gtagagtgtagcacctatccTCTGGTTGATGAAGGATAAAAGAGATGCAGAAAAATAGACggttacctcttatctagt
gtagagtgtagcacctatccAATCGTAAGGCATGGATTTAGCAAGGGTCTTATAAAACAAggttacctcttatctagt
gtagagtgtagcacctatccATCAGGAGCATGAAAGGATTTAATTTGCATCTAGGTTACAggttacctcttatctagtg
gtagagtgtagcacctatccAAATAAGCAATGGAGTATATGGGGTCTGAACACAATATTGggttacctcttatctagtg
gtagagtgtagcacctatccTTAATTGTGGTGAAATGTCATTGACTCTTTGGTCTAGAAGggttacctcttatctagtg
gtagagtgtagcacctatccCCCCAGATTATAACTTGGACATATTCAAACTATTACAAGCggttacctcttatctagtg
gtagagtgtagcacctatccTTCCTTCATGCACTGCCTTATATAAGAACCCATTGCCTCTggttacctcttatctagtg
gtagagtgtagcacctatccAGGGCAATATTTACTGAAGATCTACATAACTGAGTGATGCggttacctcttatctagtg
gtagagtgtagcacctatccGATGTACCCGCCATCTAAAAATCAACAGAACTTGTCAGATggttacctcttatctagt
gtagagtgtagcacctatccCAACAAGGAATCAGCAATTATATCAGGACTTTCAAATGCCggttacctcttatctagt
gtagagtgtagcacctatccATGGAAATATACTGGCCGAAAAGTAACTTGTCAGGGTCATggttacctcttatctagt
gtagagtgtagcacctatccCGCTGTCAATTAAAACGGGGCTGCCATGCTTTACTAAACCggttacctcttatctag
gtagagtgtagcacctatccCACATCATTAAAACTCAGTGACTATCAGGCTTGGAAGTAGggttacctcttatctagt

gtagagtgtagcacctatccTTCAAACAAGAGAATTCCATTTAGCCATCACAGTTAGTTCggttacctcttatctagt
gtagagtgtagcacctatccCCGGGAACACCTGGGCTCACGGAATCCTCTCACCTTAGCCggttacctcttatcta
gtagagtgtagcacctatccAATGCAGGCCAGTTTTTAAAAGGAGAGAATATAAAGGAACggttacctcttatctagt
gtagagtgtagcacctatccGTGCAACATTTACCATAATACTGAAATAGCCTGTTTGTTGggttacctcttatctagtg
gtagagtgtagcacctatccTGTTTTGTTGGGACACGGTGTCGCTCTGTGACCTAGGCTGggttacctcttatctag
gtagagtgtagcacctatccGCCTCCCAGGTTCAAGTGATTCCGGTGCCTCAGCTTCCTGggttacctcttatcta
gtagagtgtagcacctatccCGGGAGCCACGGTGTCCAGCCTGTTTGTTGGTTTTAAAAAggttacctcttatctag
gtagagtgtagcacctatccAAACAGTACACTTTTCATAGCTCTCAAATCTCTCTCATTCggttacctcttatctagtg
gtagagtgtagcacctatccTCACATATTACTGGGAATCGTCATATTCTGACTGAATGCTggttacctcttatctagtg
gtagagtgtagcacctatccTGGATTTGCACAGACCACACACCACAGAATAAAGACCTTTggttacctcttatctag
gtagagtgtagcacctatccGCTTTTGCAGTAACAGTCCATTCAGATGATCCTATTAATTggttacctcttatctagtg
gtagagtgtagcacctatccAAGTGATTTTGGAATGTTACGGAGTTCATCAGGTGAAAATggttacctcttatctagtg
gtagagtgtagcacctatccCTTTTTAAATGACTCCAAGCAGATAACCACTGGCACAATGggttacctcttatctagt
gtagagtgtagcacctatccTCTCCAACTGGGTTTAGATTCTAGCCTTGCACTGATCACCggttacctcttatctagt
gtagagtgtagcacctatccCCTCCCAACTAAGTCCAAGGATAGAAATGACCCATACAGGggttacctcttatcta
gtagagtgtagcacctatccCCAATGGAATCGACAGAGCACCAATCAAAAAGAGACGCTAggttacctcttatcta
gtagagtgtagcacctatccCTTACAGGAAGCATGACCTGGGGGTTCTCAGGAAACTTACggttacctcttatctag
gtagagtgtagcacctatccAAAACGCAAAGGGGAAGCCAGAAATGTCCTCTCATAGTGGggttacctcttatcta
gtagagtgtagcacctatccAAACCATCAGATCTCTTGAAGACTCACTATCCCAAGAGCAggttacctcttatctag
gtagagtgtagcacctatccAATTCGAGATGAGATTTGGCCCACCTGAGGACACAGAGCCggttacctcttatcta
gtagagtgtagcacctatccACCTCATATATCTTAGACTACTGAATCATGGTGGATTTTGggttacctcttatctagtg
gtagagtgtagcacctatccGCTGGTGCTGTTATCTTATCCTGAACTGTGTATATTAGTTggttacctcttatctagtg
gtagagtgtagcacctatccATTTCTAATTGATTGTGCTCCTCTCTCCTGGAAAAGCCCTggttacctcttatctagt
gtagagtgtagcacctatccTCTTTTTTGTTTGTCTTGGTTGGGTTTGATTTTGGCTTTGggttacctcttatctagtgg
gtagagtgtagcacctatccTTTTGGCTTACTTTATAATCCTCAGGTTTTCTGGTGTGCTggttacctcttatctagtg
gtagagtgtagcacctatccGAAAAATTCTATGAAGTAGCCATGAAAAGCCAAGAAGAGGggttacctcttatctag
gtagagtgtagcacctatccGGACCCAAACTGCCTTTCTTTTTAGCTTCAGAAGATAACGggttacctcttatctagt
gtagagtgtagcacctatccCCTTGTGTGTAAATAAATGCCCTCTACGACTGATGCAGGCggttacctcttatctag
gtagagtgtagcacctatccCCTAGAAGTGTATCATTGTCAAGGAAAGATTCAAACTCAAggttacctcttatctagt
gtagagtgtagcacctatccCCTAAAATACTAGACACCAGACTACATAGAAGATAAATCCggttacctcttatctag
gtagagtgtagcacctatccCTAGTTGAAGAGACAGAACATCAAGGGAAATAATAGGCACggttacctcttatctag
gtagagtgtagcacctatccCATGGTAAAATGAAAAGGCCTGGGGTTAGGGTTAGTCAGAggttacctcttatctag
gtagagtgtagcacctatccACTACAATCACTTTATCTTTCCATCTCTGGTTACATCTGTggttacctcttatctagtg
gtagagtgtagcacctatccCCTTCCTCCACGGGTACATTAAGGCTCAAACCACAAGAATggttacctcttatctag
gtagagtgtagcacctatccGGATCTTGAAAATCATCCTAATGGCTAGCAAGGAAAACCTggttacctcttatctag
gtagagtgtagcacctatccAGAGAACTCAATTACTTGTAACACCGGGCCCTGAAGATGTggttacctcttatctag
gtagagtgtagcacctatccCGCCATCAGAGTTGTTTGCATAATTTTGGAATATCCACTCggttacctcttatctagt
gtagagtgtagcacctatccTCCCCCCGCAAAAAAAAAACTCCGTTTTCAAATTCAGAAGggttacctcttatctag
gtagagtgtagcacctatccGTGATTGATTCGGTAATGGTCAAAGAAAAGTACAAAGATGggttacctcttatctagt
gtagagtgtagcacctatccCTCCCCGTGTTTAAGAGAAAGCTGCAAAATGGAATAATATggttacctcttatctagt
gtagagtgtagcacctatccCAGCATGGTCAGTTTCTGATGAGGTTTCTCTTCCGGGTTCggttacctcttatctagt
gtagagtgtagcacctatccCACATGGTGGAGAGCAGAGAAGGGAAATAAGCTCTCTTGTggttacctcttatctag
gtagagtgtagcacctatccTATGACCTTATCTAATCCTACTAACCTCTCAAAGGCCCAAggttacctcttatctagt
gtagagtgtagcacctatccACACTGGGGAATAGGGTTTCAACATGAATTCTGGGGAGACggttacctcttatctag
gtagagtgtagcacctatccCTTGTTCTTCTGGGTTTTGTCATATGTAGAAACTGCTATGggttacctcttatctagtg
gtagagtgtagcacctatccCATGAAGCTGACACTAAGAATTGGAGATCACAGATGGAAAggttacctcttatctag
gtagagtgtagcacctatccATGCCCTACCATTGATCTTCCTGTTACATGAGAAAAAAAAggttacctcttatctagt
gtagagtgtagcacctatccCATTTATTTAGCTCTTAATGTGAAGAAGTCCCCCATGCTAggttacctcttatctagtg
gtagagtgtagcacctatccTTTTATCCTCAAAACATTATCACTCAGAGATGACGAAACCggttacctcttatctagt
gtagagtgtagcacctatccAAAAGATTGAATGACTCGTCCAAAATCACACAAAGCCAGTggttacctcttatctag
gtagagtgtagcacctatccGCAGGCTGCCTCCTTCAACACCCATAAATTTTTTAACAAAggttacctcttatctagt
gtagagtgtagcacctatccGAAACTTCTCTTTTAAAAATGCACTTGATGAGGCTGGGTGggttacctcttatctagt

gtagagtgtagcacctatccACAAAATTAGCCGGGCATGGTAGTGCATTCCTGTAATCCCggttacctcttatctag
gtagagtgtagcacctatccGTTTTCCAATTTCCTGTTTATTTCACAAGTATGACTAGCCggttacctcttatctagtg
gtagagtgtagcacctatccAGATCCTACTTATCTCAGACTCTCACGAGCTTCCCCAAAGggttacctcttatctag
gtagagtgtagcacctatccTACTTCTATAATAGCACAGAACCTCTGAAGCCCCTAATCAggttacctcttatctagt
gtagagtgtagcacctatccCAGGAGTCTTATACCTGTTTTGTTGTAAAGGAGACAGAAAggttacctcttatctagt
gtagagtgtagcacctatccACAATATGTGAATTTTTCCTCTTGAAGGGGCAGCTGGTCTggttacctcttatctagt
gtagagtgtagcacctatccCAAGACTGGATACTTTCGGCTTGGATGAAAAACAGAAGAAggttacctcttatctag
gtagagtgtagcacctatccCTACTCAGAGCCAGGGGGACAGCCAGACCAAAAATTGTTGggttacctcttatcta
gtagagtgtagcacctatccGCCAAAAGATAGAAGCCAAGTGTCAAATCCAAGAACAAAAggttacctcttatctag
gtagagtgtagcacctatccACCTTGATGTCAAAATTCGAGCTGATCAGAGCCAACCCAGggttacctcttatctag
gtagagtgtagcacctatccAGAAGAACTGTCTCAGATATGTCTGGGCCTTCATGATCCCggttacctcttatctag
gtagagtgtagcacctatccGCAGACCACCAAAGATCCCAAAATAAAAGCAAAGGTTTAAggttacctcttatctag
gtagagtgtagcacctatccAGAATTTCTCTTGGTGCAGCAAAGGATCTCTAGTAAGAAAggttacctcttatctagt
gtagagtgtagcacctatccGATCGTTATAAAGGTTGATGATTCTTAAGTACATTCGAGGggttacctcttatctagtg
gtagagtgtagcacctatccGTTCTAACAATGTTCAGGTGCAGGGATCAAGTGTCTTTTAggttacctcttatctagt
gtagagtgtagcacctatccTCCACCAGAAGACATCAAGTTTAACTGAAATTCAAATCTGggttacctcttatctagt
gtagagtgtagcacctatccGGACAACCATCTGGAAAATAACGCTGCAAAACTACTACACggttacctcttatctag
gtagagtgtagcacctatccCAAAGCTGCAATGAAGTAAAACCGACTGAAGAGTATGTTAggttacctcttatctag
gtagagtgtagcacctatccGCTATACCAATCTGTGCATGTGAATAACTGTCAATATCCCggttacctcttatctagt
gtagagtgtagcacctatccATGGGAATAAACATTAAATATTTACCAGCTGGTGCACCACggttacctcttatctagt
gtagagtgtagcacctatccCCCTGAGTACACACAATAATAAATGAATAGACTTGGGCTTggttacctcttatctagt
gtagagtgtagcacctatccCACTATGGTTACATCAATAGGTGAAGCGATAAATGGAGGCggttacctcttatctag
gtagagtgtagcacctatccACAGAAGTCCTTGGCCATTTGTACTTTTTAAAAGAACATGggttacctcttatctagtg
gtagagtgtagcacctatccCCAAACTAACAATGTCTCCTAATATGAAATCCAGTTTCTGggttacctcttatctagt
gtagagtgtagcacctatccTGTCTGGTGCTAAGAAAATGTATGACAATTAAACCGATTGggttacctcttatctagt
gtagagtgtagcacctatccCAGGAGATGGGGTTACAGGACATAGTGTCATAGGAACAAGggttacctcttatctag
gtagagtgtagcacctatccAAATTTTGTGCCTTATAAGATATGCTGAACCAAGTGTTCCggttacctcttatctagtg
gtagagtgtagcacctatccAAATGGTTGATAGTAAGAGCTCCACTAAAATACCGAGTTGggttacctcttatctagt
gtagagtgtagcacctatccCCCTTCAACCTACTCAGTGAGGGACTGTCAGCCTTCCAGCggttacctcttatcta
gtagagtgtagcacctatccCCTCCATGGCTGTTATCCTCTTCATCCTAGTCAAGTGGCAggttacctcttatctag
gtagagtgtagcacctatccGAAATTTTTCCCCTGACTACTGCATTCTTACTATCTGAATggttacctcttatctagtg
gtagagtgtagcacctatccCTCTAAGGATGAGACAACAATAACAGATGGCATCAAGGGGggttacctcttatctag
gtagagtgtagcacctatccATGTCTCCCTTTTGAGAAAAGAATTGGCACTAATCAAAGGggttacctcttatctagt
gtagagtgtagcacctatccGGAGTCCTAGGAAGTAGAAGCCATTTCAGAGCCCTAAAAAggttacctcttatctag
gtagagtgtagcacctatccATTCAGTAGGATTCAGATAATTTGAAAGGAGCAAAAAGCCggttacctcttatctagt
gtagagtgtagcacctatccTCCTGGTAATGATTTTTGCACATTATCTATCATGGTCTAGggttacctcttatctagtg
gtagagtgtagcacctatccTGTAAAGCATTGTTTATATTATGGAGGCACACGAATGTCAggttacctcttatctagtg
gtagagtgtagcacctatccTTCTTGTCTTTCCCTTTCTTTGTTCTTCCAATCTATAGAGggttacctcttatctagtgg
gtagagtgtagcacctatccGAAAGGCCCAGTCAGTGTGTGTGTTTCCTGCAGCTTCTCAggttacctcttatctag
gtagagtgtagcacctatccTCATGTGTTGAACCTCTACTATGTGCTACTACTCCTAGTCggttacctcttatctagt
gtagagtgtagcacctatccCAGTAACTGGCGGAAAATTATGCAATCCCAAATGTTTTTAggttacctcttatctagt
gtagagtgtagcacctatccGGTCAAACGTGAAAACATGCTTGTAAAAATGTGGAGGTTGggttacctcttatctagt
gtagagtgtagcacctatccGGTATTTGGTTACATGGCTAGTTTCTTTAATGGTGATTTCggttacctcttatctagtg
gtagagtgtagcacctatccCCCATCACCCAAGCAGTATACATTATACCCAATATATAGTggttacctcttatctagt
gtagagtgtagcacctatccCCAGGTTGGAGCACAGTGGCATGGTCTCGGTTTACTGCAAggttacctcttatcta
gtagagtgtagcacctatccTTAGTAGAAACGGGGTTTCGCAGTGTTGGTCAGGCTGGTCggttacctcttatctag
gtagagtgtagcacctatccCCTCATCCCCCTCCCAGCCTTTCCTGCCACCCTGAGTCCCggttacctcttatcta
gtagagtgtagcacctatccTCCAGCTCCATTCAAGTTGCTACAAAAGACATTATTTCATggttacctcttatctagtg
gtagagtgtagcacctatccTCCTTGCAATTGTGAATTGAAGCATTGCACGTTTTTAGAAggttacctcttatctagtg
gtagagtgtagcacctatccGACGAGTCATGTTAGTGATATTTAAGGCACCTCAATGATTggttacctcttatctagt
gtagagtgtagcacctatccATTTTCTAAGGATCAGAGGCCAGTTAAATGTCCCCAGGTCggttacctcttatctag
gtagagtgtagcacctatccGACAAGCAAGCAAACCAGGGGTAGGAGTTATACACACTGCggttacctcttatcta

gtagagtgtagcacctatccGCAAAGGAGCTTGGAGACCTCAGGGAAAAGTCAGAAGGAGggttacctcttatcta
gtagagtgtagcacctatccCCAAAAGGAGAAGGGTGAGAATCTTGGTAGAAGTGAGGCTggttacctcttatcta
gtagagtgtagcacctatccTTTTCTATGTTTCATAATGTAGGTCTCCCATGTCTACATGggttacctcttatctagtg
gtagagtgtagcacctatccAGGAAAAAATGACCTTGCAAAATCCTAGCTGTTTATTGGAggttacctcttatctagt
gtagagtgtagcacctatccCAATGAATGTACAATGTTCCAAATCTGGGTTGATGCAGGCggttacctcttatctagt
gtagagtgtagcacctatccTGTGGGTTAGTTACATCTACAAAATCTACAACAGGACAGAggttacctcttatctagt
gtagagtgtagcacctatccGTTTGTAGGGCCTAAAGCTAACACAACTTGAGGTATTATCggttacctcttatctagt
gtagagtgtagcacctatccCCACAAAAATATAAAGTTAGCTACAGGGACTTGGAAGAGGggttacctcttatctag
gtagagtgtagcacctatccAGTGAGGGATACTGAAATCAAAGCTTCATTGGCGTCAAAGggttacctcttatctag
gtagagtgtagcacctatccGTGGAAACTTTTCCTAAAAGAAATACAAATGCGGCCGGGCggttacctcttatctag
gtagagtgtagcacctatccTTTTGGGAAGCCAAGGTGGGCAGAACATGAGGTCAAGAGAggttacctcttatcta
gtagagtgtagcacctatccAATACAAATGTTGGGCAAGATTTTCTTCACATAGGAGAACggttacctcttatctagt
gtagagtgtagcacctatccTCTGGATAGAGGGGAGAAGGGATAGGAATATTAAAAATAAggttacctcttatctagt
gtagagtgtagcacctatccTGTCATTACATTTAAATGCATGGTAATTCCCAGTGTCAAGggttacctcttatctagtg
gtagagtgtagcacctatccATAATTATGGGAGAAAATAGAATAGAGGAGAAATGGCAGCggttacctcttatctag
gtagagtgtagcacctatccCCTAGTGGATGTCTGGTGAGGAGCTTCATCAATAGCTGATggttacctcttatctag
gtagagtgtagcacctatccAATGCCTGTTCAGCTCTGTCTGAATGTAAATTGGACTTTTggttacctcttatctagtg
gtagagtgtagcacctatccGTCTGCTTTGACCATTGAACTTTCCTAACCTATCCCTTTTggttacctcttatctagtg
gtagagtgtagcacctatccCCATACAAAAACATCAGAAAAAGAACGCTAAAAACATGGCggttacctcttatctag
gtagagtgtagcacctatccGTTAATCTTAATACGGTCTTTCTACTGCATTCATGTTGACggttacctcttatctagtg
gtagagtgtagcacctatccTGGTTCTACTTTGCCACAGTACTCCCTTCCCCTCTTCTCTggttacctcttatctagt
gtagagtgtagcacctatccGGACCATACCTCTTCCCATCCCTCTACTCCTGTCTGTCTCggttacctcttatctag
gtagagtgtagcacctatccCCAATCGGACTTTCTTTCCTGGCAGACTGCATCTATCCATggttacctcttatctagt
gtagagtgtagcacctatccCCATTAGGGCTGAGTGTTTCCAGGGGCAGAGTCCAAAGAAggttacctcttatcta
gtagagtgtagcacctatccCTCTCCTTCCAGCAGCACAGTGATTCAGCAATTCTTTGGAggttacctcttatctag
gtagagtgtagcacctatccCTATCCAGAATGCCTCCAACAAAGTCCTCATATGCTTAAGggttacctcttatctag
gtagagtgtagcacctatccTGATAGAGGGTAGTTTATCACTAAGTTAATAGGGTCTCACggttacctcttatctagt
gtagagtgtagcacctatccTTTTTCCAGTCTTGTTCATTTCTCTGTATAGTCTTGAAGCggttacctcttatctagtg
gtagagtgtagcacctatccAAGCAGTCTTAAAAGTACAAATACCGCCCTACTCCCTCAAggttacctcttatctag
gtagagtgtagcacctatccTAGTTCCTTTTATACTTTACCAAATCTGCTCTACCATGTCggttacctcttatctagtg
gtagagtgtagcacctatccAGAATAGTAGTGTCTCAGGTGGTAACTCCCAAAAGGTCATggttacctcttatctag
gtagagtgtagcacctatccCTTCTGGTTTGAGACTTCACTGATATTTTTCTTTACACTCggttacctcttatctagtg
gtagagtgtagcacctatccTTGACAAATTGAAATTGGCAAGCCAGTTACCTCTAATAGCggttacctcttatctagt
gtagagtgtagcacctatccGAGAACCTCTGGCATTTTCCAAGCTCAGCATTTCCCTAATggttacctcttatctagt
gtagagtgtagcacctatccTCAGCAGCCTAAGAGGTTACCACAGGACACTCGAACATGCggttacctcttatcta
gtagagtgtagcacctatccAAAGATCTCCACTTGCCTTTCTATATTGTATCATACTGGAggttacctcttatctagtg
gtagagtgtagcacctatccCCAAGACAGGATTTACATTCAGCAACTGCCAAGCTATCTGggttacctcttatctag
gtagagtgtagcacctatccGAATGTATTTCTAGACTTGGAGTCGGAAGCACTTTTAAAAggttacctcttatctagtg
gtagagtgtagcacctatccAAAACCTTTGACTTTCTGCCTGTACTGAATCATTCCAATCggttacctcttatctagtg
gtagagtgtagcacctatccGGTCTTTTAGAGAGAACTTACCAATCCTACCTCAAGCCTGggttacctcttatctag
gtagagtgtagcacctatccCACCTATGAACTCTAGATAACATTTGCTGATACTGAGATCggttacctcttatctagt
gtagagtgtagcacctatccGGTAATGTGGACTACAGTGTCAAAGATAAATGCAAATGATggttacctcttatctagt
gtagagtgtagcacctatccGGTGACACTCATATTTGGGGAAAAGCAAAGTTAAAAAGAAggttacctcttatctagt
gtagagtgtagcacctatccACTAAAATGGCAATGTGAGTTATAGGGTGGGGAAATGACAggttacctcttatctag
gtagagtgtagcacctatccGAAGAATCTCAAGTTGTTTCCCTTTAGTCACCCAAGTAACggttacctcttatctagt
gtagagtgtagcacctatccTGTCCATTTTTGTTTGTATTCTGTAGATGAGGTTGAAGAGggttacctcttatctagtg
gtagagtgtagcacctatccCTGGTAAGTGTCTGGATCCTTAAAAGCGGGAGATTTATACggttacctcttatctagt
gtagagtgtagcacctatccACAGAGTTGAACAGGAATTCCTATAGCCCATTACATCAGCggttacctcttatctag
gtagagtgtagcacctatccGCTCCTCACCAGACATCCACTAGGAAAACAGCTGCCATTGggttacctcttatcta
gtagagtgtagcacctatccCGACTGGACTATCACACCCTCAATCGTATTCACATGGCAGggttacctcttatctag
gtagagtgtagcacctatccCACTGCTCTTCTCTATGTGGTCCCTCCAAGTCTGTTCTTCggttacctcttatctagt
gtagagtgtagcacctatccATGATCCATGCATACTCCTCCAGCACCACCCTGCAACGCTggttacctcttatcta

gtagagtgtagcacctatccGCTGAGGCTCTTCATCTGTTTTCTTTCCTGGTGTGTGTGTggttacctcttatctagtg
gtagagtgtagcacctatccTGTGTGTGTGTGTTGCCCTGCTTTACTTATCCCCTCATAAggttacctcttatctagt
gtagagtgtagcacctatccCTAACATCTTAGTGCACCCCATACAACTTCCCTTCCCAGAggttacctcttatctag
gtagagtgtagcacctatccAACATCTGGAATTTGTTTCTTCAGAATACTAGGCTCCCAAggttacctcttatctagt
gtagagtgtagcacctatccGAATCACTATTTGTTTTTTCAGGATGCTCTTCTTTTACCCggttacctcttatctagtg
gtagagtgtagcacctatccTACTGACCTCACTTACTACTACCCTACCTAGATTGTTCAGggttacctcttatctagt
gtagagtgtagcacctatccGTTGAAATGGCTTCAGGCAGTGGACAACTCTTTACCTAAAggttacctcttatctagt
gtagagtgtagcacctatccGTGAAACACCAACCCAAACTTCTGCCATTCTCTCTTCCTCggttacctcttatctag
gtagagtgtagcacctatccGAAAAAGTCGAAGGTTTCTTAGTTTCTGGTCCCTATGTGGggttacctcttatctagt
gtagagtgtagcacctatccACAGATCTCCCAAGCCCACTTTAATGTCCCGACTCCTAGCggttacctcttatcta
gtagagtgtagcacctatccGGTCCAATCCACCCCTTCAAAGCAAAAGCACTCCCATCATggttacctcttatcta
gtagagtgtagcacctatccCGCCTTGATATTTTGAAGTCGCAGTCTTTCCATGTCCATCggttacctcttatctagt
gtagagtgtagcacctatccGTCTCTGTCTGACCATGACACTTTCAACCATGCTCTTATGggttacctcttatctagt
gtagagtgtagcacctatccATTTTTGAAAATCTACTTCTTTGCTGCACACCGGTTTTCAggttacctcttatctagtg
gtagagtgtagcacctatccTGGTTTCAGACTCACTTGCTCCAATTCTTATCTAGGATTAggttacctcttatctagtg
gtagagtgtagcacctatccGTTTGTAGAGTAGATGTGCCTTACACAAGGATTCTTCAGAggttacctcttatctagt
gtagagtgtagcacctatccTGAATTTCAACAATCACAAAAGGTAAACATGCCTAGGTAGggttacctcttatctagt
gtagagtgtagcacctatccCATAACCTAATACTCTTACACCCCTTTTTTTCTATTCGTGggttacctcttatctagtg
gtagagtgtagcacctatccTCAGTCTTAATTGGTGCTGAAACCCCAAAATGATAAATGCggttacctcttatctagt
gtagagtgtagcacctatccCAGTGGAATTAATTAAAAAATCATGTGACCATTGGCCCCAggttacctcttatctagt
gtagagtgtagcacctatccAGGGTGGGTTCTTTCTCATTACCTCTAGTCAATTGAAGGGggttacctcttatctagt
gtagagtgtagcacctatccACCAGTCACAAAGTTTCTATGTACCTACAGGATTAAAAGGggttacctcttatctagt
gtagagtgtagcacctatccGAGATAAAACTTAAATGTCCCCACTACTTCAGATGACCTCggttacctcttatctagt
gtagagtgtagcacctatccCCCACAGGCTCCCGTAACACCATCCCTTGCTTTCTCCCTCggttacctcttatcta
gtagagtgtagcacctatccGATTACCTCTCCATCCTTTGTTTGGTTCTCAGCTCCTATGggttacctcttatctagtg
gtagagtgtagcacctatccACACTTGAAGTTGTCTCTGTTTTCGTAGTTCGGCCCTGACggttacctcttatctagt
gtagagtgtagcacctatccTACCTCCAACACACATGCACGTGGCCTCCCCCATCACTGGggttacctcttatcta
gtagagtgtagcacctatccCCCTCCATCCCGGGTCACTGGCGACACAAGCATCTTTCAGggttacctcttatcta
gtagagtgtagcacctatccGAACCTAAGATGAATCAACTTCCATGTCCTAGATCTGTTCggttacctcttatctagt
gtagagtgtagcacctatccTCACAGTTCTGCAGGCTCTGACATCAGCTTCTGGAGAGGCggttacctcttatcta
gtagagtgtagcacctatccGGAAGGTGAAGTAGGAGCAAGCACATCATATAGCCAGATCggttacctcttatcta
gtagagtgtagcacctatccAAGAGAGACAGAGAGGAAGATATAAAGAGAGAGGGAAGGGggttacctcttatcta
gtagagtgtagcacctatccGATCTCGTGAGAACCACTCATTATCCTGAAGACAACACCAggttacctcttatctag
gtagagtgtagcacctatccCCTAACTTCAGGCTGGAGATTACTATTCCACATGAGACTTggttacctcttatctagt
gtagagtgtagcacctatccTACATCATAGAAATTGCCAGTAGCAGCTCAATTCCTGTTTggttacctcttatctagt
gtagagtgtagcacctatccGGTTCCCCCAAAAGCAAACTCTAAGACAAAGCCTGTTGCAggttacctcttatcta
gtagagtgtagcacctatccTGCTCTATTCATTTCCTTTTACTGCTGCATCATATTACCAggttacctcttatctagtg
gtagagtgtagcacctatccGGTCAGAGAACTAAAATGTATCATATGGGCTAAAATCAAGggttacctcttatctagt
gtagagtgtagcacctatccGGGCTGGTTTTTTCTAGAAGCTCTGAAAGTAGAATCCATTggttacctcttatctagt
gtagagtgtagcacctatccCAGTGTTCCCTGGCCTGTGACATCTTTCTCACATCCCTCCggttacctcttatctag
gtagagtgtagcacctatccTGCTTCCATCATCACACCTTCTACTGTCTCTAACTCTACTggttacctcttatctagt
gtagagtgtagcacctatccCCAGCTGCATATCTCAGATACTCTACTTATCTCAAAATCCggttacctcttatctagt
gtagagtgtagcacctatccTGTAAGATAACACATGTACAGATTTCTGAGAGTAGGACATggttacctcttatctagt
gtagagtgtagcacctatccGTGTCATTATTCAGCTCACCACAAATGCATTGGCCACTGCggttacctcttatctag
gtagagtgtagcacctatccCAATTCATGCCCACAGAACATGCTAAGGAATCATAATGCAggttacctcttatctag
gtagagtgtagcacctatccCCTAGGGGAAAAGAGGGAGGCAACATTATCTGGCAGCACCggttacctcttatcta
gtagagtgtagcacctatccATTGATCAAGGTTGCCCCCAGGGTGTTAACTCCTCTGTGAggttacctcttatctag
gtagagtgtagcacctatccAGCCATGTGCATTAGTGCCAGCTGGGGTCCTATGGTAGACggttacctcttatcta
gtagagtgtagcacctatccACTGGAAGCACGAGGTTCAAAGTAAGGTACAATCAACATTggttacctcttatctag
gtagagtgtagcacctatccCCCAAATTTGGTTAAAGGTGAAACTGCATGGAACTGGATGggttacctcttatctag
gtagagtgtagcacctatccCGAAGCCTAAGGAATATGCAGAGATTCTATAGTGTTTAGTggttacctcttatctagt
gtagagtgtagcacctatccTGATGGTTTTCAGCTTCATAGATGTCCCTGCAAAGGACATggttacctcttatctagt

gtagagtgtagcacctatccTGTGAACAGTGCCGCAAAGTAGGAGTTGAACAATTAGAACggttacctcttatctag
gtagagtgtagcacctatccGGGCCTGTTGGGGGCTGGGGAGCTAAGGGAGGGATAGCATggttacctcttatcta
gtagagtgtagcacctatccAAAAAGAATATGCAGAGATTCGTAGTAGGGGTCCAATTCAggttacctcttatctagt
gtagagtgtagcacctatccCCACGGGCAACTTTAACTAGTTTGTATATCTCTCAGGACTggttacctcttatctagt
gtagagtgtagcacctatccATGCATTTCTCAAACATGTTTCTCAAATACATTGCTGCCCggttacctcttatctagtg
gtagagtgtagcacctatccTCCCCCTGCAAGTGAGTTATAATAGGAGTAAAGTAAATTTggttacctcttatctagt
gtagagtgtagcacctatccCAAAAATTAGACAAAAGCACCAAGAAAGGCAACTGGCTGGggttacctcttatcta
gtagagtgtagcacctatccCAGGAGGGGGAGCTTGCAATGAGCGGAAATCGCACCACTGggttacctcttatcta
gtagagtgtagcacctatccACTAAAGAGGTTATTCTGCCTATAGCTGCAGGCAATTATTggttacctcttatctagt
gtagagtgtagcacctatccCAAACTTCTTGGAGAGCAAAGGGAAAGCAGCACCAGGGAAggttacctcttatcta
gtagagtgtagcacctatccCTCAAATGCAACAAGAGTAGAACCAGGAGGCAGAAAAGATggttacctcttatcta
gtagagtgtagcacctatccTTCATGTTATCAGGACAACCACCCTTTAATGAACAAGCCTggttacctcttatctagt
gtagagtgtagcacctatccTCAGGACTGAGCTAGTGACTCTTCAACCATTTATCAGGAAggttacctcttatctagt
gtagagtgtagcacctatccGGAAAGTATCTGTGTGTGGAGACTAGAAGTACAGCAAAGAggttacctcttatctag
gtagagtgtagcacctatccGCTTATCAGAGAGGCAAAGACGAAAAGGACATACACACAGggttacctcttatcta
gtagagtgtagcacctatccACTTAATTGGTGGGAAAACATAAACTGACAGAAGCTTTTGggttacctcttatctagt
gtagagtgtagcacctatccAGGGTGCAGACTCTGATTCAGAACTCTACTTCTGGGGGAAggttacctcttatctag
gtagagtgtagcacctatccCTGACATGGAAGAGACTCACAATATGTAAAGAAAAATGACggttacctcttatctag
gtagagtgtagcacctatccATAGCTTTGTGTGTGAGGACTTACATATGATATGACTATCggttacctcttatctagtg
gtagagtgtagcacctatccAATCCTTGACATCACAGCCAAAGTTAAGTTCACACAGAGAggttacctcttatctag
gtagagtgtagcacctatccCAGAGCTGCTTCCTTTGTGTGACTTCCTGCTTTGTTTTTAggttacctcttatctagtg
gtagagtgtagcacctatccTGGGGAAAAATGGATGATGAGAATGATATGAATATGAGCTggttacctcttatctagt
gtagagtgtagcacctatccTACTAATATGCCTAGTGACATTTAGGTTTTGTTGCTGTTCggttacctcttatctagtg
gtagagtgtagcacctatccGTTCTAAACTCTCTGGCTCAAACCCCTTTTCCTATTAAAGggttacctcttatctagt
gtagagtgtagcacctatccATTCAGAAAAACCCTACAAAGCCTAAAAACTGAAAGCTTCggttacctcttatctag
gtagagtgtagcacctatccGATAAGAAGAACAAAATCAAGATATAAACCCACAAGGTCCggttacctcttatctag
gtagagtgtagcacctatccGTCAACACAGTTTCAATAGTGCAAAGAGACATTAATACACggttacctcttatctagt
gtagagtgtagcacctatccCCGAAAATTGGTAACGAACTGAAAAAGCTGAACACCAAGTggttacctcttatctag
gtagagtgtagcacctatccTTTCCATGAAACCTAGAAACACTTAGCCACTAGAGGTACCggttacctcttatctag
gtagagtgtagcacctatccCATCCTCTTCCCCATCTCATGCATCCAGACAACTACCCTTggttacctcttatctag
gtagagtgtagcacctatccCAAGAGATGAGACATTTATTCTGTGAGGGGGGTAGGGGGTggttacctcttatctag
gtagagtgtagcacctatccGGAGAAAATTTATCCCTAACACCCTAAAGCATTCATTGAAggttacctcttatctagt
gtagagtgtagcacctatccTACATCAATTTCTCTTGTAGGCTTCTAAAACAGTACCAACggttacctcttatctagtg
gtagagtgtagcacctatccATAATTTTCTGTAGGGCTTCAAACCCTCATAAAAGTTGGGggttacctcttatctagt
gtagagtgtagcacctatccGGCATAGAGGAGGGCCGGTTGGAAGGGGAGGGGATGAGAAggttacctcttatcta
gtagagtgtagcacctatccACAACACTGAAAGTCTCTATACTTAATGGACTCCTAGCCTggttacctcttatctagt
gtagagtgtagcacctatccTGCCCCCAGAATAGTGTTGACCAAGTATATTATCTTCCTTggttacctcttatctagt
gtagagtgtagcacctatccAGGAGATTAAAAGAGTCTTCCTTGAACAAACTGAAGTGCCggttacctcttatctag
gtagagtgtagcacctatccACCAATATGTAAGGTCTTCTGTGGGGGCCTGAGAAAATATggttacctcttatctagt
gtagagtgtagcacctatccTGGCCTATACACCCTCTCCTCCACCCATTCACACAGACCTggttacctcttatcta
gtagagtgtagcacctatccCCTTAAAACTTGAAAATCAGAGAACAAAGCACACAAACAGggttacctcttatctag
gtagagtgtagcacctatccTGGAACTCAGGAGTAGAGAGGAGTGGAGCAAAGAAACCCTggttacctcttatcta
gtagagtgtagcacctatccAATAGACATTCAACTATACAGAATTCTGGGCAGTGACTTTggttacctcttatctagt
gtagagtgtagcacctatccTAAAAGAAAAAATGAGTAATGCATTCAGGCATAGACAGGGggttacctcttatctag
gtagagtgtagcacctatccGGATAACCCTAGCTCCTTAGTTTCTTCCAGTTATTGAAAGggttacctcttatctagt
gtagagtgtagcacctatccTGAAGTAGCAGCATCAGTGAGAATCTAACAAGAACAGCCAggttacctcttatctag
gtagagtgtagcacctatccGTGTCACACTCTTGCAAGCCACTAAGGGGTACGGCGCTAAggttacctcttatcta
gtagagtgtagcacctatccAATAATAGCGATCTTTGCAATTCCTACTGATCTGAGCTCAggttacctcttatctagt
gtagagtgtagcacctatccAATAATAGCGATCTTTGCAACTCCTACTGATCTGAGCTCAggttacctcttatctagt
gtagagtgtagcacctatccAGGGAGTTATAATAGGAGTAAAATATTCCCTGAAAGGAGAggttacctcttatctagt
gtagagtgtagcacctatccAAAACTTGAGACTAGTGCCTGTATCTTTGTAATCCTGGGGggttacctcttatctagt
gtagagtgtagcacctatccATCCAAGAGATTAAAAAGGCAGAGGAAAAGCTTGAGACCTggttacctcttatctag

gtagagtgtagcacctatccGGAACCAGAATTTAGCTCCAAGACTGAGAAAACAAAGTTGggttacctcttatctag
gtagagtgtagcacctatccTGCCCCGCAAAGCAGCATCCCACCATTTTGAGAATACTTGggttacctcttatctag
gtagagtgtagcacctatccGGTTAAAGACATGAAATTTAAATTCCAGACTTGTACCCACggttacctcttatctagtg
gtagagtgtagcacctatccGAAATATCCACAGCAGCAAAGTCCTAGAGCAAATATGTGTggttacctcttatctagt
gtagagtgtagcacctatccTGTCCTTGAAATTAGAGAGAAGGAGACTTGAGACATAGTGggttacctcttatctagt
gtagagtgtagcacctatccACTCAATGTGCACAAGTCAACCAGCCATACCAAGGGTTCGggttacctcttatctag
gtagagtgtagcacctatccGAAATCACTTTCCATGCATGGGTTTTCTATTCAAATTCACggttacctcttatctagtg
gtagagtgtagcacctatccATAGTGGCTCCTATAAGAAATTTTCGCTAAAGCTGGTTACggttacctcttatctagtg
gtagagtgtagcacctatccCAGTCCCAAGAAGTCCCCCAATCTGAAAAATTTGACAAATggttacctcttatctagt
gtagagtgtagcacctatccTGATACCAGAATCCATCCTGATCCAGCCATATCAGTTACCggttacctcttatctagt
gtagagtgtagcacctatccCCACAATACTGCTACATAAATACAACCACAAAATCTCAGTggttacctcttatctagt
gtagagtgtagcacctatccATAAACATTTATTTTGCTCACAAGTTGACCGTACACCTGGggttacctcttatctagtg
gtagagtgtagcacctatccCGATGCTCTGGGACTTAACTGATCTTGGTTGGATTCATGCggttacctcttatctagtg
gtagagtgtagcacctatccGGTGGGTCTAGTACAGCCTCAGTATTCAGTTGGGATATTTggttacctcttatctagtg
gtagagtgtagcacctatccCTTTTTCTTGTGAAGGAGGCTGGGATCTAAGAGAATGAATggttacctcttatctagtg
gtagagtgtagcacctatccCCCAGGCTCAAAATGGGCATATCTTCTTATCTGCTGCCTTggttacctcttatctagt
gtagagtgtagcacctatccCCAAAGCAAATTACAGGCCAGCCCAGATTCCAGAGAGGGGggttacctcttatcta
gtagagtgtagcacctatccCACTAATACTTTGCATTGCAAGCAGCATGTTTACAAGGAGggttacctcttatctagtg
gtagagtgtagcacctatccTCAACCTTCCACACCAGCATCTTGATTTCAGAGAATTAGTggttacctcttatctagtg
gtagagtgtagcacctatccAGATTATTAGTGAGAAAACCACAGATATCTGAATCAAGCCggttacctcttatctagtg
gtagagtgtagcacctatccTTATATAAGATGTTAACACAAGGAGGAGCTGGCTGAAGGAggttacctcttatctagt
gtagagtgtagcacctatccGGGAAGTCTGTACTCTCTTTATAACTTTTCTGTGAAGCTAggttacctcttatctagtg
gtagagtgtagcacctatccAAGAACATGATAAAAAGGAATTCGCAGTATAGGCCATTTGggttacctcttatctagtg
gtagagtgtagcacctatccGATCCTCCATCCTCTATTCAGAAACCCCTGTGGCCTGGGAggttacctcttatctag
gtagagtgtagcacctatccGGGCCACCTCTCTGCAGACCTCCAAGAAAACAGCAACGCTggttacctcttatcta
gtagagtgtagcacctatccGGTGTTGACCCGGCTGGCTCTGGTGGTTAGGGGAGATGCCggttacctcttatcta
gtagagtgtagcacctatccCTTTATGAAGGGAGCGAAATCCAGACTTAGCCGTGTCCATggttacctcttatctag
gtagagtgtagcacctatccCTGAGACTGACAAATTGTGATCTTCACCATGTAGCAGCTCggttacctcttatctagt
gtagagtgtagcacctatccCAGTGTGATAGAGTCTGTATAGTGAACATATGTTTCTTGTggttacctcttatctagtgg
gtagagtgtagcacctatccTGTCTTCCTTCAGAAAAGCCCAACCTCCTTCTACCTTTGCggttacctcttatctagt
gtagagtgtagcacctatccGTTCCCAGAGGTAGATAGGCACAAGACATATGACCCAGACggttacctcttatctag
gtagagtgtagcacctatccCAGAAGAGCTCATCCCTCTGGCCACTGCAATTGATCAGGGggttacctcttatctag
gtagagtgtagcacctatccGAGCACCAATCAGACCTTCTTCATCATATTTGATATAAAGggttacctcttatctagtg
gtagagtgtagcacctatccTTTCTGAAGCTAAAGACCACCAGAGGGAAGAGCCAAGATAggttacctcttatctag
gtagagtgtagcacctatccGCCAGAGATACCCCTTAGACTTCCGTATTTCCAGCCATAAggttacctcttatctag
gtagagtgtagcacctatccGGGTTTCTCTCATTTGCACCCAAAACAATCCTGCCAAATGggttacctcttatctagt
gtagagtgtagcacctatccCCTGGAAGTGGGGTGTTGGAATTGACAGGCATTAAAATGAggttacctcttatctagt
gtagagtgtagcacctatccCCTCTTTACCTGGGATGTTAGTGGTGCCTCTTAAGTGGTAggttacctcttatctagtg
gtagagtgtagcacctatccTTTTCCTTGCTGTTCCTGGGACTCAACAAATTCACTTCTGggttacctcttatctagtg
gtagagtgtagcacctatccGCCGACCATATTTTGCAGCCTTTATTATCCTAGAAGCTTCggttacctcttatctagtg
gtagagtgtagcacctatccGCCTATGTAGCCTCACCAAAGAGATCATTTCTTCCTCCAGggttacctcttatctagt
gtagagtgtagcacctatccTCTGTTTTGAAAGGAGATAATACTGAATTGCAGGAAAAGCggttacctcttatctagtg
gtagagtgtagcacctatccCCTAATTTCCTAGGCCCTTACCAAAAACTTCTGGCTATGTggttacctcttatctagtg
gtagagtgtagcacctatccCTCATTTTAAAGAGACAGCCCCCCTCATTACGCCAGCCCAggttacctcttatctag
gtagagtgtagcacctatccAAGGTAAAGACATTATTCAGAGACAGAAACTCCAAAACCAggttacctcttatctagt
gtagagtgtagcacctatccAACATGCCATAGTAGATCCCTGTTCTAATTGACTAGCCTGggttacctcttatctagtg
gtagagtgtagcacctatccAATATCGAATAGTCTGGCCTCCCTCTGAGATTCTAAAGGTggttacctcttatctagtg
gtagagtgtagcacctatccCTTCTAAAGTAATCTCCAGACCCCAAAGCCTGCACCAAAAggttacctcttatctag
gtagagtgtagcacctatccGTAAATTCACGGGGAAGATTGCCATGGCCTCATTTTAGCCggttacctcttatctagt
gtagagtgtagcacctatccGCACTGCATTTTCATTCCCATCTGACTTATGTGTCCAATTggttacctcttatctagtg
gtagagtgtagcacctatccGATCCACAATGTGCAAAAGAAGAAAAGTGAGTAATTAAGCggttacctcttatctagt
gtagagtgtagcacctatccGGAGTTGGTATTTTATCCTGCAGGCCTTAAGGAACAGTGAggttacctcttatctagt

gtagagtgtagcacctatccCAGGTTTAATGCAGATAAGACTTGTGGTGTGTTTTGTAAAggttacctcttatctagtg
gtagagtgtagcacctatccAATGGTAGATTGGTAGGGAAAGGGTAGGACTGAAAATGAAggttacctcttatctag
gtagagtgtagcacctatccAATATAGGTTTGAACTAACAGAGTTGCAATGACTATGGCAggttacctcttatctagt
gtagagtgtagcacctatccCCAGAGAGTATGCAGACAATGGAAGAGGTTGCCAATATCAggttacctcttatctag
gtagagtgtagcacctatccGAACTCCTCCTATAAGATAGCACGAAGATTAAACTATGTAggttacctcttatctagt
gtagagtgtagcacctatccCTTGTTTTGTTTTTGTTGGAGGGACGGAAGAAGCAGACAGggttacctcttatctagt
gtagagtgtagcacctatccAAGGATTTGTACTGAAAATTGTCAAGATGTATAGGTGCTCggttacctcttatctagtg
gtagagtgtagcacctatccGCAACACATACACAAAAGAATGCCTTGGTTACCTACTGATggttacctcttatctag
gtagagtgtagcacctatccCTAGGTCCACTGCACAAGGTTTTAGGCTAGAAAATGAGGCggttacctcttatctag
gtagagtgtagcacctatccTCTTTCTGAGTTCCTTTTCACCTGGATGAGAGCATAGTGTggttacctcttatctagtg
gtagagtgtagcacctatccTAATTTATAAAAGCTAGAAGCGATGATTACTTGCCCAACGggttacctcttatctagt
gtagagtgtagcacctatccTTAGTCCTTCAACATCCCTTCCCTCACTAAGATTCTATGAggttacctcttatctagt
gtagagtgtagcacctatccTGTGTTCCTCTAAATGCTATACCTTTCCTTCTTTGCCACCggttacctcttatctagtg
gtagagtgtagcacctatccAATCCCTAAATAGTGTTCAATAAATGCCCATTGGTCTCTCggttacctcttatctagt
gtagagtgtagcacctatccAATGTTGATTCAATGTGGCAATTGTAGTCCATCTATCACAggttacctcttatctagtg
gtagagtgtagcacctatccAGGATTGAATGAAATTTTGCAGGTTAAGAGCTTAGCTTAGggttacctcttatctagtg
gtagagtgtagcacctatccGGCATATAATAAGCACTCGCAGATGATGGCTATCTTCAACggttacctcttatctag
gtagagtgtagcacctatccGTAACCCACCCATTTTCACTCCAGTTTCTACTTCTTTGTGggttacctcttatctagtg
gtagagtgtagcacctatccTACCATAATGATAATCAAAATCCCACCCAAGCATTCTTTGggttacctcttatctagt
gtagagtgtagcacctatccGAGCAAAACTTCTTTCTCATAATCAGGGCTAAGGCTTAAAggttacctcttatctagt
gtagagtgtagcacctatccAGTTCTTTCTTCAACTTGGCCAGCTGCACTTGAATGCGAGggttacctcttatctagt
gtagagtgtagcacctatccCCAAGTGGGATGAGTCATTGAATTCTAAGCACAAAATATTggttacctcttatctagt
gtagagtgtagcacctatccAGGAGCAGTGGAGAGTTCCTTTTTTATGGTGAAACAGTGCggttacctcttatctagt
gtagagtgtagcacctatccGGACGAAAGCCATCCCATCTCATCCCATTTCCTCTTTTCCggttacctcttatctag
gtagagtgtagcacctatccCAACTGTGAAAGGGAGGGAAGAATTGACAAGAAACTATTTggttacctcttatctag
gtagagtgtagcacctatccAAGAAAAAGTCCTTGAGTAATAGTATGGAACATTAAGCCCggttacctcttatctagt
gtagagtgtagcacctatccGGATGCTCATTAAATGCTTGCTGGCTGTGTGTAGAAGTCTggttacctcttatctagt
gtagagtgtagcacctatccGTCATGCTGCCCTATTGATAACCCTAGACTACATCACACCggttacctcttatctag
gtagagtgtagcacctatccTGCAACAGTTTATTAGCACTCCAGCATTAACTATATGAAGggttacctcttatctagt
gtagagtgtagcacctatccCTGAGTAGTTTGTTGAGAGAAGGACATTTTCAGATGGACAggttacctcttatctagt
gtagagtgtagcacctatccCGTGAATCACTCAGGTTTTCTTTTTGAAATAACATTGCTGggttacctcttatctagtg
gtagagtgtagcacctatccGCCCATTTCATCTTGTCAATGCTCTTCTTCATATTTAATCggttacctcttatctagtg
gtagagtgtagcacctatccATTCCAGTTGATACAATCATGTAACAGGCATTTAAGGAACggttacctcttatctagt
gtagagtgtagcacctatccATGCAAGCTACTCTACTAGGTACTCCAGGGAATACGAAGCggttacctcttatctag
gtagagtgtagcacctatccCTGTCCAGTGGCTTTTAATAGCAGAGGAAAGACAAAACAAggttacctcttatctag
gtagagtgtagcacctatccACAAACTGATGGGAAAGCATCAAAGGATTGTTGTATGCAGggttacctcttatctag
gtagagtgtagcacctatccGACATGATTATAACTGCATTTTAGAAAAATCACCCTGGCCggttacctcttatctagt
gtagagtgtagcacctatccGCTGAGGCAAGAGAGTCACTAGAACCTGGAAGACAGAGGTggttacctcttatcta
gtagagtgtagcacctatccCAGAAGGATGATGGGATAGATTGAAAGGGAGCAAGCAAGGggttacctcttatcta
gtagagtgtagcacctatccGAGGCCCAGGAAAGCCAGTAGGTAAGTTCTAGTCCAAGGAggttacctcttatcta
gtagagtgtagcacctatccGTATTCAACCTTCTCCCTTTTGTGTCCTATTTGGATCTCAggttacctcttatctagtg
gtagagtgtagcacctatccGGGACATCTGTTTTATTCAACCCACTAATTCAAATGCTAAggttacctcttatctagtg
gtagagtgtagcacctatccTATCTGGGCATCCCATGGCCTAATCAAGCTGACACATAAAggttacctcttatctag
gtagagtgtagcacctatccAGCTACTGTAATAATAGAGGTAAGATATGAAAGAGGCCTAggttacctcttatctagt
gtagagtgtagcacctatccTGTATTAAAAGATTTCAGTGAGGGAAAATGAATAGGCCTGggttacctcttatctagt
gtagagtgtagcacctatccAAATGATTGATTGATGGGTAGCAAGTGACTGATGAGGGATggttacctcttatctagt
gtagagtgtagcacctatccCAAGACCTCTACCTTTGGCAGCAGGGTGGGCTCACTGCAGggttacctcttatcta
gtagagtgtagcacctatccTCTGATGGAGAGTGGGGTATACATAGTAGGAGGATGGCAGggttacctcttatctag
gtagagtgtagcacctatccGGAAGCTGAGCACCTGCAATGGTCAAGTTTTGGAGTTAACggttacctcttatctag
gtagagtgtagcacctatccGGATACCTAGTTAGCTGGTAAAGCATTGCTTCTGAGGGTGggttacctcttatctag
gtagagtgtagcacctatccTGGCATGTGAGTCTGTGAACCGAGTGGGTAAGATCCCCCCggttacctcttatcta
gtagagtgtagcacctatccGGCACCATCTGATTAGTTAGGGGTCCAAGTGGAACAAAAAggttacctcttatctag

gtagagtgtagcacctatccTTTCTTTCTCCTGCAGATCCAGGATGTCCCTCTCCTGCCCggttacctcttatctag
gtagagtgtagcacctatccACTCCACATTCTCTGGCTTTTCGACTCCAGGATTCACACAggttacctcttatctag
gtagagtgtagcacctatccCTGAGAATTATACCATCAGCATCCCTGGTTATGAGGTTTTggttacctcttatctagt
gtagagtgtagcacctatccCCAGCTTGCAGAGGGCCTGTGTTGGTACTTCTCAGCCTTCggttacctcttatctag
gtagagtgtagcacctatccGAACCAATTCCCCCAGTAAATCCCTTCTCACATGTATATAggttacctcttatctagt
gtagagtgtagcacctatccCTGGAAAACTCTGACTATTACAGCACTTTGAATGCCAAGCggttacctcttatctag
gtagagtgtagcacctatccGGTTAGACTTTAATTCAGTGAGCAATGGAGAGGCTTTTAAggttacctcttatctagt
gtagagtgtagcacctatccGTTCTTTTGGAAAATTAATCTGATCAATGAAGAGGCCTAGggttacctcttatctagtg
gtagagtgtagcacctatccGGTTAATTGCTAAGACACCTAGTTAGTGCAGAGATGGGACggttacctcttatctag
gtagagtgtagcacctatccTCTTCAGACTCCTGGCCTATTTTGTGCAATGAGGAAGAACggttacctcttatctagt
gtagagtgtagcacctatccATTCTAAATATGTATTCCTCTCTCCAATAAGTCTTCCTCCggttacctcttatctagtg
gtagagtgtagcacctatccGCCTTCATGCTTATATTGGGCCAGACATTTAGCACACAAAggttacctcttatctagt
gtagagtgtagcacctatccGATGCTTCCAGTATTTTTTTATTAATGCCCAAAGAGCCCAggttacctcttatctagtg
gtagagtgtagcacctatccCTAAATGTAGATAAAGGCAATGGTTTTGTAAGACTCCTTCggttacctcttatctagtg
gtagagtgtagcacctatccGTTAAGTAACTTACTCAATACCACACAGTAAATGGTAGAGggttacctcttatctagt
gtagagtgtagcacctatccATTTAACCCCAAATCCAACTCCAGAATCCCAGCCCTTAAGggttacctcttatctag
gtagagtgtagcacctatccGAATGATTTTCTTGAGAAGGTTCAAAAGAAGCTAAGGCGAggttacctcttatctagt
gtagagtgtagcacctatccCAGTGATTTTCTTGCAGCTGTGTAGGCCATTATAGCAAACggttacctcttatctagt
gtagagtgtagcacctatccCTCAGCGCCCCAGAGAGAACCTACCAACTCAAATTGCAACggttacctcttatcta
gtagagtgtagcacctatccAACAACAAGTTCCTAACACACCGGAAATAAAGACTGGGGAggttacctcttatcta
gtagagtgtagcacctatccGGAACTGCCATTAACCTCACTTATTTTAAATAGCACTTAGggttacctcttatctagtg
gtagagtgtagcacctatccAAACCCAATTCTAATCTGTATGTGTGCTTTGTGCATTTGCggttacctcttatctagtg
gtagagtgtagcacctatccTCACAAAGAATGCGAGGGAAATACTGGAAAACCAAACAACggttacctcttatcta
gtagagtgtagcacctatccGCCTTACCTGTTAACTTCTGAGCTATTTTTGTGACATGAAggttacctcttatctagtg
gtagagtgtagcacctatccACTTTTTGTTTCGTGAAGGCCCTTGGGTAGATTTTTTATGggttacctcttatctagtg
gtagagtgtagcacctatccTCGCCAAAGTGAATTTAGTAGAGAAAACCCCAAGGGAGCAggttacctcttatcta
gtagagtgtagcacctatccTACATTAACATCTCTCTTTATCGTGTCTATTCATGACCAGggttacctcttatctagtg
gtagagtgtagcacctatccGAATTTTAAATCAGCTCCTTTTGTAATCCATCTCGAGAAGggttacctcttatctagtg
gtagagtgtagcacctatccCTGGGCTAAGCATTTATGTGAGTGGAGAAAGAACTATGTGggttacctcttatctagt
gtagagtgtagcacctatccTTGAAGCAATATGTGGAGAAGGGAATGAGAAAGAGGCAATggttacctcttatctag
gtagagtgtagcacctatccGTGTTGGACTATTGTGATATCAAATGCGGTAGCTATTCAGggttacctcttatctagt
gtagagtgtagcacctatccTCCAAAATAAATTAGTGTTCATTCAGAGTAAGAGTGGAGGggttacctcttatctagt
gtagagtgtagcacctatccGTAAAGGATAAGAATGAAAGAAAAAGGATAAGGAACAGGGggttacctcttatctag
gtagagtgtagcacctatccCCACTCATAAACACAAGAAATGGTTTCTTATTCAACAAGGggttacctcttatctagt
gtagagtgtagcacctatccTTAAGAAGGAAGAAGTAGTGAATCTCAAAGATGTTGTTGCggttacctcttatctagt
gtagagtgtagcacctatccATGCACTGACTCTTAAAATGTTGTAAGTCTACATGAAGACggttacctcttatctagt
gtagagtgtagcacctatccGGCAGCTGGTTCCAGAATATCCAAACCTGAAACAAAATTAggttacctcttatctag
gtagagtgtagcacctatccAAAAAGGCATAGGTCTAGGATAACCAACAATATTACCAGAggttacctcttatctag
gtagagtgtagcacctatccCAATTGTGTAAAAATCCGCATTTCTCCAGATAGTAGTGATggttacctcttatctagtg
gtagagtgtagcacctatccGCAAAACTTATTCCCGTTTAACTTTTTTTTTACCAGGCTCggttacctcttatctagtg
gtagagtgtagcacctatccATGCTATGATCCATCTTGAAAACGCCCATGAGTCAATTCTggttacctcttatctagt
gtagagtgtagcacctatccCAGACTCTAACAGGAGACCAACATAAATAAAATTATCCACggttacctcttatctag
gtagagtgtagcacctatccTTGCTGAAACCAGAAATTGTTTAGATCAAGTAAAGAGACCggttacctcttatctagt
gtagagtgtagcacctatccCAAAGAGGAGCCTGTGAGAGACCCAGACAAGCATCACCCTggttacctcttatcta
gtagagtgtagcacctatccCAAGGAGACATCAAATCAGATAGCAGCAACGAAGTGACTGggttacctcttatcta
gtagagtgtagcacctatccCATAGCTCATCTGTGTCTGGAAATGGCAATGCAAGAGTAAggttacctcttatctag
gtagagtgtagcacctatccCTCACTTGCATTATTGATAGCTGGATCTTCAGACCTAGACggttacctcttatctagt
gtagagtgtagcacctatccAAGGCTTTCCACTACTGTATTAATGGATTCTTGCACTGCTggttacctcttatctagtg
gtagagtgtagcacctatccTTTTAACTTGTGGTAAGAGCCATCTACTCAAATCTAAGAGggttacctcttatctagtg
gtagagtgtagcacctatccCATTTACGTTAGGTATGTCTCCTGATGCTATCCCTCTCCCggttacctcttatctagt
gtagagtgtagcacctatccCATGGCAGAAGGAGAAAGCAGAGCAGGGAAATCACACGACggttacctcttatcta
gtagagtgtagcacctatccGGAGCAAGAAAGAGAGAACATGGCAGGAAGGTGCTACACAggttacctcttatcta

gtagagtgtagcacctatccAACCATTCATGAGAAATGCATCACCATGATCCAAGCACCTggttacctcttatctag
gtagagtgtagcacctatccACATGAGATCTGGGCAGGGAAAAATACACAAACTGGACAAggttacctcttatctag
gtagagtgtagcacctatccGAAGTCTCCCTCCCTTCTTAATGCGCCTAAAGACTGCAGCggttacctcttatctag
gtagagtgtagcacctatccCAAGAGTTTGTACCCAGTAGAAGTCACAGCCAGTAACAGCggttacctcttatctag
gtagagtgtagcacctatccCACTCTGCCTCCTTTAAAGTTCTACCCCAGATCTCCCCACggttacctcttatctag
gtagagtgtagcacctatccCTGGGAATAGGCTGTCACTGAATATAGTATAAAAGCATTTggttacctcttatctagtg
gtagagtgtagcacctatccCTCACCAAAAAAATCTAAAACTGTACAGATAGAACGTCTCggttacctcttatctagt
gtagagtgtagcacctatccGTGAAATACAGAAACTGTCACAACCACAAGCCGCACACTAggttacctcttatctag
gtagagtgtagcacctatccGTACTGTGCGAGGATTGATTTAAGAAGCCTACCTAGGGGCggttacctcttatctag
gtagagtgtagcacctatccCCTCAGACCCTAGGCAGAATGAAGACATCTCCTACAATAAggttacctcttatctag
gtagagtgtagcacctatccAGTAATCTTTGAACTGAAACCTTGAATTCTTGTCGTGAGGggttacctcttatctagtg
gtagagtgtagcacctatccTTCTTGAACAGTCACAATGCAACATCACAAAACCAAGGAGggttacctcttatctag
gtagagtgtagcacctatccGAGCTATAGGAGAAAGCAGGACATCTTATCTTTTCTTGGGggttacctcttatctagt
gtagagtgtagcacctatccAGGGGAAGAACTGAGTGGAGGGAGGAAATGAAACAGCGGTggttacctcttatcta
gtagagtgtagcacctatccATGGCTTTGCTGTTAGCTATGCTATCTTCTGGTTCCTGAAggttacctcttatctagtg
gtagagtgtagcacctatccATTGTTCCCCCAGGCAAAAGGAGGATAGATTTGATGAAGGggttacctcttatctag
gtagagtgtagcacctatccCCAATGGATATATATGGCTCATTCACCACTACTACCACCTggttacctcttatctagt
gtagagtgtagcacctatccGCATCACCCCAGCACTACCTCTACACACACACGCATACACggttacctcttatcta
gtagagtgtagcacctatccACATATACAGACACACCAGTGACAGAAGTAAGGTCAGGCAggttacctcttatctag
gtagagtgtagcacctatccGCCTTGTTTTTGCTTCACCTTAATCTCTCTCTGTCCTGACggttacctcttatctagtg
gtagagtgtagcacctatccATAGGAAAGGGAAAAGATGAAGATAAGAGACAAATAGCTCggttacctcttatctag
gtagagtgtagcacctatccCAAATATTGGGCCTGGACCAACCTCTTACACCAAGGAGTTggttacctcttatctag
gtagagtgtagcacctatccCAAACTTTTCAGACTAAAACTTGCAGGGGGAGAAGGAAACggttacctcttatctag
gtagagtgtagcacctatccAAGTACTTCTCCAGGTATTGACTGCCATAGAGCAAATAAAggttacctcttatctagt
gtagagtgtagcacctatccGCCTTAGCTAATAGCAACTAAATTTACTGAGCAGCTACTAggttacctcttatctagt
gtagagtgtagcacctatccTTGAAAAGAGAGAATTGAGGCCCTAGGTAGTTATTTAATGggttacctcttatctagt
gtagagtgtagcacctatccCAATAGGGATAGCACTGTGTCTGAGAAGCCAAAGAAGAGAggttacctcttatctag
gtagagtgtagcacctatccCAGGGAATGATATACAGAGTTTATTGGGGGCACTTACACAggttacctcttatctagt
gtagagtgtagcacctatccAGTGGCAGCAGGCTGGATGAGAGAACCACTACCATTTGTAggttacctcttatctag
gtagagtgtagcacctatccCTCCATTTAACCCAAAACAAGGGCCTCCCAATCCCCTGTAggttacctcttatctag
gtagagtgtagcacctatccGGCAAGGGGTTCAATTGTCCTTCATAGATAAGATCAATGTggttacctcttatctagt
gtagagtgtagcacctatccAGCTCAGAATTCTGAACCAACGCTCTGCTTAGACCATAGGggttacctcttatctag
gtagagtgtagcacctatccTCAGGTGCCTCTGCCATACATTTTATTGGACAATTTCATAggttacctcttatctagtg
gtagagtgtagcacctatccAATAATTAAAAGATTAAGGATTCAGGGTAGGGTTTGGAGCggttacctcttatctagt
gtagagtgtagcacctatccCCTACAAGGTCCTAAAAAATGGCACAAACACCCAAATGAAggttacctcttatctag
gtagagtgtagcacctatccCAAGTGGAGCCAGAGGAGAATCTTTTGCAAACATGTGGGAggttacctcttatctag
gtagagtgtagcacctatccAGGTCACTGGGGTTTCCTCTCCTTCTCTGAATCAAGACTTggttacctcttatctagt
gtagagtgtagcacctatccTTAGATAAACAACTCAGAACCTTGACTAGCTGCTGGGCGCggttacctcttatctag
gtagagtgtagcacctatccTCAGACACTGATGCAATGGCAAGAGCAAGAGAAAAATGGCggttacctcttatctag
gtagagtgtagcacctatccTCTTCTGCAGACCCATTTCCAGGCCCATGCTTACCTCACAggttacctcttatctag
gtagagtgtagcacctatccGACTCTCTTACCAACATATGTCACAGATAGGAGAAAGGGGggttacctcttatctag
gtagagtgtagcacctatccGGCTTCCCAACAATCATATTAGCTACTTTCTTTCACTTGGggttacctcttatctagtg
gtagagtgtagcacctatccGTAGAGAGAAGCCAGAAGTTCCCTTCATAAGCCTTCCCCCggttacctcttatctag
gtagagtgtagcacctatccGCACACATTTGTTCCTTTTTCTACAAGCTAAAGCAGGAGCggttacctcttatctagt
gtagagtgtagcacctatccTGCACTGGGAAGAAAGCATGTGAGTTGGCAGGTTCAGAGCggttacctcttatctag
gtagagtgtagcacctatccGCCCGCCATATCCCCTCCACCCATGCAGAAAGTAACTATGggttacctcttatctag
gtagagtgtagcacctatccTCTTGTTTAAATGCCTTTCTATATCTCTTCACCTGTGTAGggttacctcttatctagtgg
gtagagtgtagcacctatccGGCTAGAAATTGCCCCAGAAGGAACCCTCAGTTAATGAAGggttacctcttatctag
gtagagtgtagcacctatccAGAGCAATAGGAGGGTTCTGAAATGTGTTCCATAGTCTTTggttacctcttatctagt
gtagagtgtagcacctatccACTTTGTAGTGTTTTTTTCCCTTCTTGTTTCTCTTCTGCggttacctcttatctagtgg
gtagagtgtagcacctatccGATCACATCCCAAAGAAACTGCTTGTACGCAAACCCTTGAggttacctcttatctag
gtagagtgtagcacctatccAGATATCACTTCCACACTGCTACCCCAAGCCTTCTTCATTggttacctcttatctagt

gtagagtgtagcacctatccCCTGAGTCAATAGAAGGTCAAGCAAGTCGTTACACTGTCTggttacctcttatctag
gtagagtgtagcacctatccAAGCTTTAGGATGAGGTGGCATTATGGCTTGAAGGTCTCCggttacctcttatctag
gtagagtgtagcacctatccCAGGTAGTGGCAATGGTAGGAAGTCTAGTGTTTCAATATAggttacctcttatctagt
gtagagtgtagcacctatccGTGACAAATTCTCAAAAGAAATCTTATGGACTTGAAAGGCggttacctcttatctagt
gtagagtgtagcacctatccACTTTATGTTTCATCCAAGCCCAGATAGTCAATGAATAAGggttacctcttatctagtg
gtagagtgtagcacctatccCTTAATTTCAGGGTTGTGACAAATAAATGAGATTGCAGTCggttacctcttatctagtg
gtagagtgtagcacctatccACTTCCATAAATGTGAGTATCCTTCCACCTCCCCAAGAAAggttacctcttatctag
gtagagtgtagcacctatccTTATTATTGTCACTATCAGTTTTAAGGACCACCTCTAGGggttacctcttatctagtg
gtagagtgtagcacctatccTTTAAAAATCAAGTCTACTGTGATTGATCCAGCTGTTTCCggttacctcttatctagtg
gtagagtgtagcacctatccCTACTTCTAGAGCTTGGAGAGTAGGGGTAAGCCAGATAATggttacctcttatctag
gtagagtgtagcacctatccGCTCCACTTCACAGAGAAGACTAGGCACAGCAAGAAGAGGggttacctcttatcta
gtagagtgtagcacctatccTTATTTCCATTGCTTCTCCTCTTATTTAGTGAAGACAGACggttacctcttatctagtg
gtagagtgtagcacctatccTCCTATAAGTCAATCTCATTTGCTAAGTCTAGTTCTTTGCggttacctcttatctagtg
gtagagtgtagcacctatccTTTTCCTTTCCTATTTACATGTTTCAAATTCCCATCCTGCggttacctcttatctagtgg
gtagagtgtagcacctatccCATTCATGTATTTCTCACACTGTTTCTCTCTCTCCCAAGAggttacctcttatctagtg
gtagagtgtagcacctatccCTGCCCTTTCTTTTCTTCCTATTAGCTTGCTACACCAAAGggttacctcttatctagtg
gtagagtgtagcacctatccGCTGTCTCTATAATGCCTTACTGAAAATGCAGTGAAAACCggttacctcttatctagt
gtagagtgtagcacctatccTGTCCCTGGATGAGACACAATGTCCCAACTGATGTTTTCAggttacctcttatctagt
gtagagtgtagcacctatccAAGAGGGTGTGCAAGCTTTCATAAGTGTGTTCTTTAATGGggttacctcttatctagt
gtagagtgtagcacctatccAGGAAAACTCTTGTGATGCTAGTAGAAGAATAATATAGGCggttacctcttatctagt
gtagagtgtagcacctatccTTCCTCATCATTTCACCAAATGAAATATTCAGCCCCGACTggttacctcttatctagt
gtagagtgtagcacctatccGAGGCTGAGGCTGGTGAATCACCTGAGGTTGGGAGTTCGAggttacctcttatcta
gtagagtgtagcacctatccTGCACTCGAGCCTGGGCAAGGGAGAAAGACTTCATCTCAAggttacctcttatcta
gtagagtgtagcacctatccGAAAGAAAGAAATATTCAGCCCCAAGTTGCTGAGCCCAGTggttacctcttatctag
gtagagtgtagcacctatccCCCCACTCCATAGGTGCAGGCTTACAGACTTCCCTAACAAggttacctcttatcta
gtagagtgtagcacctatccAGGCCACAACCTCAAACTCACAGCAAACAGGAAGCAGGAGggttacctcttatcta
gtagagtgtagcacctatccGTCACATGTAACACAGTCAGAGCTTCAAGCCCCCGGAAAAggttacctcttatcta
gtagagtgtagcacctatccAGATTTCATTCTCTGCCCCTGATGATGCCCCGCACTTTCTggttacctcttatctagt
gtagagtgtagcacctatccGGACACCATCTGCCCTACACTCACCCCAGGAAAGCATTGTggttacctcttatcta
gtagagtgtagcacctatccCATTCCTGAAATTTAGTCATCCAACCTCTGCTAGACTATTggttacctcttatctagtg
gtagagtgtagcacctatccAGTCTGCTCCCTCTTTCACATCTCATTTTGTCTCCTCTTCggttacctcttatctagtg
gtagagtgtagcacctatccAAACTACCCTGGCTCCTTCCCTCCTTCTCAGTTTAACACAggttacctcttatctag
gtagagtgtagcacctatccAGATTCTTCTCTTAGTTTCTCTACTGTTAGCACACTTGAGggttacctcttatctagtg
gtagagtgtagcacctatccGTTACAGATGTATTATTATTTGTCACTGGTCTCCATCATGggttacctcttatctagtg
gtagagtgtagcacctatccCCTCATTACATACTAGAATACCTATTGGCTACTATTTAGCggttacctcttatctagtg
gtagagtgtagcacctatccCACCATTACTGAAAAGAATTATCAGCAAGAACTAAAGAGCggttacctcttatctag
gtagagtgtagcacctatccTCAACAAGGACTGGAGGAAAATGCATATTAAAATACAAGGggttacctcttatctag
gtagagtgtagcacctatccGTGACACAACGGGCCAAACATTAAGGAAGATCACACTGAGggttacctcttatcta
gtagagtgtagcacctatccATGCACCCATGGAAGTAGATATTAAGAAAGCAGGAAACTCggttacctcttatctag
gtagagtgtagcacctatccCATATGAAGTGGGCAAGTACTAGACATTGGGAGGTCATTGggttacctcttatctag
gtagagtgtagcacctatccAGGAAGGAAAAGAGGAAGGATTAAATCAGCAACAGAACAAggttacctcttatcta
gtagagtgtagcacctatccCCTCATGCAACTACTGTTTCTGCCAAAAGAATAGAAAATGggttacctcttatctagt
gtagagtgtagcacctatccATCTCTGTTGTTTAGCCACTGGGATGATTTACACGGGAATggttacctcttatctagt
gtagagtgtagcacctatccCCCTCCCCATCCATGTTATCTGCTGTGGATTTGCAAAAGGggttacctcttatctag
gtagagtgtagcacctatccTATGCACACCCAAACCTACCATCGAACATAGAGAAATATAggttacctcttatctag
gtagagtgtagcacctatccGAATTATTCTCCCCCGATTTCAGAGTATGCCTAGTTTGGTggttacctcttatctagt
gtagagtgtagcacctatccGAGGGAGAGTTATCTATGGGTTTGTACTGTTCTGCTTTGTggttacctcttatctagtg
gtagagtgtagcacctatccTCTGTTTTTGTTTTGTGTTTCAGGAGATAGCATCTGTTGAggttacctcttatctagtgg
gtagagtgtagcacctatccAAATTGAAAGAATAGCTACCCCAGGTTTAGCAGAGACACAggttacctcttatctag
gtagagtgtagcacctatccCTTTCCTGAATCACAAACAGCAGTAGAAAAAAAAAACAGGggttacctcttatctag
gtagagtgtagcacctatccAGCATTCTGAGTTTGGGGTTTTCCTTTTTCCTTTGGCCACggttacctcttatctagtg
gtagagtgtagcacctatccGCAAATCAAACACAATTACTTGCCCGAGTTCCCCTAGCTTggttacctcttatctag

gtagagtgtagcacctatccTTGGGATGCTCCTTCAGGCTATCAACAAACTCTCTCCTATggttacctcttatctagt
gtagagtgtagcacctatccGAATTTATTTACTACTTTGGTGTTCCTACTCTCTTCACTCggttacctcttatctagtg
gtagagtgtagcacctatccTGGCATCTCTCCACCTTCCCTACCGTATGTAAAAGAGCACggttacctcttatctag
gtagagtgtagcacctatccCATTAGCACAAGAACAAAATGAGCAGAGGCATTTTAAAACggttacctcttatctag
gtagagtgtagcacctatccAACAGTCAACACATATTAAATTTACACCTACTACATGCCCggttacctcttatctagt
gtagagtgtagcacctatccGTGGCATTTTTAAGTACAGCACATGGTCCTTGTTCCTACAggttacctcttatctagt
gtagagtgtagcacctatccTTTAACTGAAACAATTTAGAGATCACTGAAGGGACCCACAggttacctcttatctagt
gtagagtgtagcacctatccCTTGAACAATTAGGATTCCTGGAGAGGCAAAAAGAGAAGAggttacctcttatctag
gtagagtgtagcacctatccGCAAGATTAAAAGCACGAGGGAGGGAACACTGGGAATTTCggttacctcttatcta
gtagagtgtagcacctatccCTACAGAGAAGACCTAAAATGAAGCAAGCAGAGAAAACCAggttacctcttatcta
gtagagtgtagcacctatccTCAATTCTATGAGAAGGTCATTTTTCCTCTCTTTAGGACAggttacctcttatctagtg
gtagagtgtagcacctatccCAGAGATTTCCCATATATATCCCCTGTCCCCCAATATACAggttacctcttatctagt
gtagagtgtagcacctatccATTAGGGTTCTCTCTTGGTGATATACATTCTGTGGGTTTGggttacctcttatctagtg
gtagagtgtagcacctatccTCACCATTATAATATCTATCATTAAGGGCATTTCACCACCggttacctcttatctagtg
gtagagtgtagcacctatccACTTGAAGTAGTGTTTCCTAATAACCAGTGCATGACCTCTggttacctcttatctagt
gtagagtgtagcacctatccAGAAACCGCATGACAGCAGCATATCTTTCTCACACAGGCCggttacctcttatcta
gtagagtgtagcacctatccACTGCTCAGCCTCATAATAGTGGCATGGGTGGTGACATGGggttacctcttatctag
gtagagtgtagcacctatccTTTGCTTCTTCCTCATTTTACTCTTGCCGCCTCCATGTAAggttacctcttatctagtg
gtagagtgtagcacctatccAGCTCTTTTTGTTCCCAGTTATCAGTATGTCTTCATCAGCggttacctcttatctagtg
gtagagtgtagcacctatccGAAGCAACTTTAGAACTAGAGAATAGACAGAGGTTGGAACggttacctcttatctag
gtagagtgtagcacctatccGGAGGGTTCAGAAGAAAACAAGAAAATGTGGGAAAGTTTGggttacctcttatctag
gtagagtgtagcacctatccTGGTCTCAGATGGAGATGAACTTCTTAGGAACTGAAGCAAggttacctcttatctag
gtagagtgtagcacctatccAAAGGGAAATAGAGGAGAAAAGTCTAGAAAATTTGAAGCCggttacctcttatctag
gtagagtgtagcacctatccTAATGTTAATTTCCAAGACCCTGAGGAAAATGTCTCCAGGggttacctcttatctagt
gtagagtgtagcacctatccGCCACTACAGCCATGACTGCAAGGGGCCAACATACAGCTCggttacctcttatcta
gtagagtgtagcacctatccCTTCAGAGCGGGGAAGCCCCAAGCCTTGACACCTTCCATGggttacctcttatcta
gtagagtgtagcacctatccGAGCCTGCAGGTGTACAGAAGTCAAGAGTTGAGGTTTGGGggttacctcttatcta
gtagagtgtagcacctatccCCAGAATGGTAGATCCACCTTCAGCTTGCACTTTGTGCCTggttacctcttatctag
gtagagtgtagcacctatccGCTTCCCAAGATCATGGGAGCCTACCTATTGCATCAGCGTggttacctcttatctag
gtagagtgtagcacctatccGTACCCCTATTGTATCTAAGGAGTTACTAGCTTGCTTTTGggttacctcttatctagtg
gtagagtgtagcacctatccCTCCCAGAATTCCCACGTTCCCCTCGGTCCCCTGGGAGGGggttacctcttatcta
gtagagtgtagcacctatccGTAATTGAATCATGTGGGCTACTCTTTCCTGTGCTACTCTggttacctcttatctagtg
gtagagtgtagcacctatccTTTCACCCTCCACTATGATTTTGAGGCCGCCCCAGCCATGggttacctcttatctag
gtagagtgtagcacctatccACTAATACAGGGGGTCAGGAGAAACTAGAAGAGCATGAAAggttacctcttatcta
gtagagtgtagcacctatccCAAAGATGGACTGGCCTTGAATTTTTCTGTTTTGTGAGTTggttacctcttatctagtg
gtagagtgtagcacctatccCATTGGAAAATAGAAATCATGCAAAAATCCATACACGGAGggttacctcttatctag
gtagagtgtagcacctatccAAAGTTTGGTAACATCCGTGAAAAAAATCTTACCCTTCCTggttacctcttatctagt
gtagagtgtagcacctatccCATCACTGCAGAGACGCTCTATTTTTGTGTTTTGAAATGGggttacctcttatctagtg
gtagagtgtagcacctatccACTGCTGGTTCTGTTGTTTACTATTGGAATTCCCCTGAAAggttacctcttatctagtg
gtagagtgtagcacctatccCTTCTTCAATTGAGCCAAAGTTCAGGAAAAGGTGTGACACggttacctcttatctag
gtagagtgtagcacctatccCATCTTGCATATGTCAGATCTCTACCAAATTATTTACCTGggttacctcttatctagtg
gtagagtgtagcacctatccTCTCATTTAATCGTTCCCACCATCTCACAAGATAGGCATTggttacctcttatctagt
gtagagtgtagcacctatccGGTAATTTTCCCAGAGTCACATTCTAGTGAGTGATGAAACggttacctcttatctagt
gtagagtgtagcacctatccTGAAGAACATGAAGGAGGATGAAGAACGTGAAAGAGAAGGggttacctcttatcta
gtagagtgtagcacctatccTGAAATTAAGTCATAAGGGAAGAACTGAAAAGGTGGCACTggttacctcttatctag
gtagagtgtagcacctatccCTGATCTAGCATTCTGAAGCAAGAGAGTCACTGAAAAAAAggttacctcttatctag
gtagagtgtagcacctatccGATGAGATACACCTAAAAAGAAAGTTAAAAGCTTGGTCTCggttacctcttatctagt
gtagagtgtagcacctatccATTCTCTACTTTAATTATAAATTAGCTGTGTGGGCTCTGCggttacctcttatctagtg
gtagagtgtagcacctatccAGTTAAGCTGGGAAAAGAGATCTTAGAGGAGCCAAAGACTggttacctcttatctag
gtagagtgtagcacctatccGAGAAAAAGGTCTCTATTTTTACACCTGGATTTGTCACCTggttacctcttatctagtg
gtagagtgtagcacctatccCTATGAAGAAGATCAACAGAAGAACCAAGGAGCAGAGCAGggttacctcttatcta
gtagagtgtagcacctatccAGAAATCATTTGGAAAGACAATACTTCATGCGTAATAGGGggttacctcttatctagt

gtagagtgtagcacctatccCTGGAAAGGAAATTGTATTGGATCAGAATGTGGATTTAGGggttacctcttatctagtg
gtagagtgtagcacctatccCACAAAAAGTGTCTATCCCTGTCACACGATGTTCATAGACggttacctcttatctagt
gtagagtgtagcacctatccCTAGCCACAGGTCAATAGGAAAGAACTTGAATCTTGATGGggttacctcttatctagt
gtagagtgtagcacctatccGCTAAGTCCCTAAGAGAAAAGCTTTGTGAATGCGGAGATAggttacctcttatctagt
gtagagtgtagcacctatccCCTGAGCATCCAAAGACAAAATGTTAGCAGGTACAGTGCAggttacctcttatctag
gtagagtgtagcacctatccGACACCTTACCCCTTATAACTCTGTCTCTTGGTAGTCCCCggttacctcttatctagt
gtagagtgtagcacctatccTTCCTTAAAGAACACTTGGGCATCTCTGTCTCATGGGATTggttacctcttatctagtg
gtagagtgtagcacctatccAATCCTAAATATTTGCTCTTGTTACCAATACTGTTCAGGCggttacctcttatctagtg
gtagagtgtagcacctatccAACTCTTCACATCTATTACCCTGTGATATGTTTTGGCTGTggttacctcttatctagtg
gtagagtgtagcacctatccACGTGTTGTGAGAGGGACATAACTGGACATAATTGAATCAggttacctcttatctagt
gtagagtgtagcacctatccAGATCTGATGCTTTTATAATGGGGTCCCTGTTTTGCTTGGggttacctcttatctagtg
gtagagtgtagcacctatccGGAACTCTGAGTCCGTTAAACCTCCTTGTCTTTATAAATTggttacctcttatctagtg
gtagagtgtagcacctatccTTTTGAAGGAAAATATCACTGTATCCAAATGCCATGAAGGggttacctcttatctagtg
gtagagtgtagcacctatccTGTGCCATTACCAGCTTCAGAAATCAATAATGCTTCAGAGggttacctcttatctagt
gtagagtgtagcacctatccCTCAAGCAAGGGAAAAACAAATAACATGTCATAATCCTCTggttacctcttatctagt
gtagagtgtagcacctatccCTACTGGGCATCTGTTGTCTTCTTCTCTTTATGTTCCATAggttacctcttatctagtg
gtagagtgtagcacctatccCTCTGGAAGCACTCACAAATTAGTACAATATTCACAACTGggttacctcttatctagt
gtagagtgtagcacctatccATACTCTTAAACTATACCTGAAGATACAGACATAGCATGCggttacctcttatctagt
gtagagtgtagcacctatccTTCTTCACACTAATAACCAGAGGGACACTGATGACTAGGCggttacctcttatctag
gtagagtgtagcacctatccTTGTTGTAGTAAAGGAATATTTGTCAAGTAGGAGGGTAAGggttacctcttatctagtg
gtagagtgtagcacctatccAATATTCTTATTCTGGGGTCCAAATATGTTGCAGGCAGGAggttacctcttatctagt
gtagagtgtagcacctatccATTCTTTAATATGCAAAGCTGTATCTCTAAGACCCTAAGCggttacctcttatctagtg
gtagagtgtagcacctatccCCACTTTGTCACTGGAAAGCTTGGTACTATAATTGAAACAggttacctcttatctagt
gtagagtgtagcacctatccTGTTATAAGACATCTCCCACCCAGCCACATGCCCAAGGAAggttacctcttatctag
gtagagtgtagcacctatccCTGGGCTCTAAAAGGGGAAGGTGTTTTCTTACTGACTGAAggttacctcttatctagt
gtagagtgtagcacctatccCACCCATGGCAACTCAGAGATAAACACTCCGGTCATTTGAggttacctcttatctag
gtagagtgtagcacctatccAGCACCAATAGCTGCTCTCCAGGCCTAAGACACCAAGAGTggttacctcttatcta
gtagagtgtagcacctatccCATCTATGAGGGCACCAGTGGCAACAAGAGCACAGGGACTggttacctcttatcta
gtagagtgtagcacctatccCTCCCAGACAGTGACTACTACTGCATACACGGGTCCCTCCggttacctcttatcta
gtagagtgtagcacctatccGTGACACAGCCACCATGACAAATCAGATCCTGCTCCTCACggttacctcttatcta
gtagagtgtagcacctatccGGGGCTGCTGATCCATTAGACACAAGAACTACAACTGTGCggttacctcttatctag
gtagagtgtagcacctatccAAGTGTATTAGTCTGTTCTCTATGCCGCTAATAAAGACATggttacctcttatctagtg
gtagagtgtagcacctatccCACAATCATGGTGGAAGGCAAAAGGCGTGTCTTACATGGCggttacctcttatctag
gtagagtgtagcacctatccGAGTGAGAATCAAGTGAAAGGGTTTTCCCCTTATGAAACCggttacctcttatctagt
gtagagtgtagcacctatccCCATATCACCAGGTAACTTCCACAAGTGAGTGGAATGTAGggttacctcttatctag
gtagagtgtagcacctatccCCCAAAGCTGCTCCTTAAGAAATAAATTATTATGCAAGACggttacctcttatctagt
gtagagtgtagcacctatccGAGAGAGAGCTAAAAATCTCAGGTTTTCAATTTTTTCTCCggttacctcttatctagtg
gtagagtgtagcacctatccAAGAGATAACTTGGTAAACTTCAGGTGGTTAGTTTCTTCAggttacctcttatctagtg
gtagagtgtagcacctatccTTTACTCTCCTAGGCTGTAAATATTATAGGCACCACTTAAggttacctcttatctagtg
gtagagtgtagcacctatccTGAGATAGATGTGGTCGGTTCTAAATCAAACATACTAAAGggttacctcttatctagt
gtagagtgtagcacctatccCCCAAAGAATCAAGAATCCCATTATAATTAGCTTTTCCTGggttacctcttatctagt
gtagagtgtagcacctatccGTCAACATTCCTTGAAAACAAATGGAAGTTTATCTGTCTCggttacctcttatctagtg
gtagagtgtagcacctatccCCACTTGGGAATTTATCTGTCTAGACATTGGTGAAAATATggttacctcttatctagtg
gtagagtgtagcacctatccTGCAGAACACATTCCATAAGTCTCCTGTCAAGAGGAACCTggttacctcttatctag
gtagagtgtagcacctatccGCTATGCTCAGGTCTTGCCATTTTCTCTGAAGTTCCACACggttacctcttatctagt
gtagagtgtagcacctatccACATCATGCGTTGGAATTGTCTGAATGGCTAATATTTTTCggttacctcttatctagtg
gtagagtgtagcacctatccCTTAGGGTGATTTTCCTCAGATCATTTAAATTGGCATAGTggttacctcttatctagtg
gtagagtgtagcacctatccATCCTGGAATTCAAACCAAGCATCTTAGTGTAACTGACGAggttacctcttatctagt
gtagagtgtagcacctatccAACACCAGTTTCCCACTTAGAACTTAGGCTGCGCAATTTTggttacctcttatctagt
gtagagtgtagcacctatccTCCTCTCCCCATTTTGTTGCCTCTATTATGTTAGTGGTGTggttacctcttatctagtg
gtagagtgtagcacctatccAAAGCCAGGAATCTTTACTTAAAGCTACTAAAAGCCAACAggttacctcttatctagt
gtagagtgtagcacctatccTGTCTGACATTGGGCTATAAGCTTCATGTGAATCATCTCAggttacctcttatctagt

gtagagtgtagcacctatccTCCTTTCACAACCCTGCAAGGCAGGAATTTTTTTTTTTTTggttacctcttatctagtg
gtagagtgtagcacctatccTTTTTGAGACAGTTTCATTCTCGTTGCCCAGGCTGGAAGGggttacctcttatctagt
gtagagtgtagcacctatccGCATGTGCCACCAGATCCAGCTAATTTCTATATTTTTAGTggttacctcttatctagtg
gtagagtgtagcacctatccCTGTGCCCAGCCTGAAGTAGGAACTATTTTTACCCTCATTggttacctcttatctagt
gtagagtgtagcacctatccAGAAAGATTAAATAAAGCCACCAAAAAGATCTGATCCCAGggttacctcttatctag
gtagagtgtagcacctatccTGGCAGTGGTGTTCTTTTTTAATGGTATGAAAACTAACACggttacctcttatctagtg
gtagagtgtagcacctatccGAATATTCATTTCTGTAACTTTATATAGCCTCCTGTCCTGggttacctcttatctagtg
gtagagtgtagcacctatccTCCATTCCCCATAGGATCTGACTCCATCCTGACACATAGTggttacctcttatctag
gtagagtgtagcacctatccCACATGATGGAAGGTGGTAGCTGTCTAAAATATGCATCTCggttacctcttatctagt
gtagagtgtagcacctatccTCTAGCTGGGTTCATGGCCTCTGGGAATAAAGACTATATTggttacctcttatctagt
gtagagtgtagcacctatccCTCAGTGCACTCAGGTAATTAAGTTATGACCAATCAAGGTggttacctcttatctagt
gtagagtgtagcacctatccCAGAGAGCATATCCTTCTTCACCCTTCCTTCATCCCACTGggttacctcttatctag
gtagagtgtagcacctatccCAGTGATGGCTAAAGCAGGAATGGCTCTTTTGGAACATGTggttacctcttatctag
gtagagtgtagcacctatccCCTTAGGAACTGTGGAGGTGAGAGCAGGAAGGGGCCCAAAggttacctcttatcta
gtagagtgtagcacctatccTACCAGCCAGGAACTGCCTCCCTCTCAACTTTTCACATGAggttacctcttatctag
gtagagtgtagcacctatccCATGTGTGTTTAAACGTATTTAAGTTCTCTGTCATGTGCAggttacctcttatctagtg
gtagagtgtagcacctatccAGGACTTAAAACCTGAAATTGGAGAGATTCGTGACAACATggttacctcttatctagt
gtagagtgtagcacctatccCAAATAGTTGGCATTGAGAATTTAAGCACTGAGGTCTGGAggttacctcttatctagt
gtagagtgtagcacctatccTTTTGCAGTGGTAAAACCCTTGGTCAAAATGAAACCCTCTggttacctcttatctagt
gtagagtgtagcacctatccAGATGACCTGCTGACTGAGGCTGCAGTGTAAGGAAAACGTggttacctcttatcta
gtagagtgtagcacctatccTGCTAGCTCCTTCTTGCCACCTTTTGCAAAATTACTCAAGggttacctcttatctagt
gtagagtgtagcacctatccCACTCTCTGCTTGTGGCCTCCCCTACAACTTGACAGAAAGggttacctcttatctag
gtagagtgtagcacctatccTGAAAAAGTCAACATATGTACCCCAAAGTGAAGAGGGATTggttacctcttatctag
gtagagtgtagcacctatccAAGAATGGTGTCCTGTCCTACAAATCTTATGAGCTTGCAAggttacctcttatctagt
gtagagtgtagcacctatccATAAATCCACTGTTAAAGCCATCATTAAGTTGGTAGAAGGggttacctcttatctagt
gtagagtgtagcacctatccGGAAAAGCAAAGCCAAGAGATCAGTAAATTAAAGGCCAGAggttacctcttatcta
gtagagtgtagcacctatccAATCTCTCAGGGCCAAGGCAAGGGTGAGAAAGAAGAAGGCggttacctcttatcta
gtagagtgtagcacctatccAAACAAAACAAAAAAGGAAGTGCCCAGCACACAAAACTAAggttacctcttatcta
gtagagtgtagcacctatccCACCTGACAGTGCATGCCTCCTTCCACTTTGTGCTTTGGGggttacctcttatctag
gtagagtgtagcacctatccCTAGTCCAGGCCTCTATCATCTCTACATAAGATATTTCACggttacctcttatctagt
gtagagtgtagcacctatccCTAAAGCAACCTCAAACCTAGCCTACAAAAGACAGTGACAggttacctcttatcta
gtagagtgtagcacctatccCCTCAAAGCAATGCTCAGGTCCTGGAACTACATCGTGGGGggttacctcttatcta
gtagagtgtagcacctatccCTCAGAATCAACATGGGGACAAATGAACAAGGAATTAAAAggttacctcttatctag
gtagagtgtagcacctatccGTGGCCACAAGAAACAACAAGCAAGAGAGAATATCCTCCAggttacctcttatcta
gtagagtgtagcacctatccTAGATTCAGGACTTCCCAAGGGAAAAACCATGGCGCTTCCggttacctcttatcta
gtagagtgtagcacctatccCCAAGGCAGAGAGTCCCACCCAGCAGGTAAGACAATTGCTggttacctcttatcta
gtagagtgtagcacctatccATCTCGGCTCAATGCAACCTCCAACTCCTGGGTTCAAGGGggttacctcttatcta
gtagagtgtagcacctatccCCTCCACTCTCTTTTTCTAAATGGGGCTTTTGTCATAATTggttacctcttatctagtg
gtagagtgtagcacctatccCTCTGCCTTGGTATATTGGATGAGTATGGGGGAAGGGACCggttacctcttatctag
gtagagtgtagcacctatccTGGCACACAGAGCTGCATCTAGAACTGATGACAAGGTTTTggttacctcttatctag
gtagagtgtagcacctatccCTCAGTGCAGTTATTGGATGGTCTTATGGGTTGAATTGTGggttacctcttatctagtg
gtagagtgtagcacctatccATTTGGAAATAGAGCCATTGCAGACGGAATTAGTTAAGACggttacctcttatctagt
gtagagtgtagcacctatccAAAAGGGGAAATTTGGACAAGACAGCATGCAAGGAGAACAggttacctcttatcta
gtagagtgtagcacctatccCCAAGGAAGGCTAAAGACTGCCACAAACCACCAGAAGCCAggttacctcttatcta
gtagagtgtagcacctatccACAGATTCTTCCTCACAGCCGGAAGAAGGCACCAACCCTGggttacctcttatcta
gtagagtgtagcacctatccTACTTGGTTATGGCAGCCCATGTGAATTAATACAGATAGGggttacctcttatctagt
gtagagtgtagcacctatccTCTGAGGAGAAGGTAAGGGTGTAAAGAAGAAATCATTGAGggttacctcttatctag
gtagagtgtagcacctatccTAGTGTTCCTAGTGTTCATCTCCCCTTTCTTAGCAGTGCTggttacctcttatctagtg
gtagagtgtagcacctatccCTAGATTCCTTTGCAACTAGGTTTGGCCATGTGACCACATggttacctcttatctagt
gtagagtgtagcacctatccGGTGTTTCTAAAGAGAATGGCAAGATACATCCTTCATTTTggttacctcttatctagtg
gtagagtgtagcacctatccCTGGAAGATACATGGAACAGGAGGAGCATAAACAGCCCCCggttacctcttatcta
gtagagtgtagcacctatccTAAATATGATAGAGCAACATAATAGAAGGACCCAGGATCCggttacctcttatctag

gtagagtgtagcacctatccAAGCCCACTACTATTCTCAAGTGCTTTTCTTTTAACTTCTggttacctcttatctagtg
gtagagtgtagcacctatccTCCTGTTTAAGCCATAATTACTTAGGATTTTCTGTCACTCggttacctcttatctagtg
gtagagtgtagcacctatccTAAATCGTGGCAGAGACAGTAATTATCATCCAGCACCCATggttacctcttatctag
gtagagtgtagcacctatccTGACCCCTTCAGCTAATTGTGTCCATATTAATATATCTGGggttacctcttatctagtg
gtagagtgtagcacctatccGCATGCCTTTAAAACATTGCCCATATACTCCCCACTCTTTggttacctcttatctagt
gtagagtgtagcacctatccGTGGATAGGGACATGGAGACGGTAGCACAAGAGAGATGAGggttacctcttatcta
gtagagtgtagcacctatccGTCTGTAGATAACCCTGACAATACATCTGCCTACCCATAGggttacctcttatctag
gtagagtgtagcacctatccTAAAGAAACTTAGCCACTGCCCTAATACATGATTTCAATCggttacctcttatctagt
gtagagtgtagcacctatccCTATAAACTTTGCTTCAAAATCATAGTGGGATGGAGGACTggttacctcttatctagt
gtagagtgtagcacctatccATTTGTTCCAAGTAACCTGAGAGAATGTAACTTTTAGAGGggttacctcttatctagtg
gtagagtgtagcacctatccCTGAGTCCTGAGGCCAATCTTTAACTTGTGACCTAATCTGggttacctcttatctagt
gtagagtgtagcacctatccAAGCTGGATTTATGAGCTACAATGGGGCTTGCTTCTGTCCggttacctcttatctagt
gtagagtgtagcacctatccCTGCCAGGATAGCCCCAGTAGCTTATTACACTAGTTTGTTggttacctcttatctagt
gtagagtgtagcacctatccTTGACAATATGGATGCCAGAAAGTTTAATATACACCATGCggttacctcttatctagt
gtagagtgtagcacctatccGGCAAAGGGGACAAAAGAACCAGGTGACATAACTGATGGAggttacctcttatcta
gtagagtgtagcacctatccGCATAACTGGTTTGCATGTCCCTTTTTGAAGCAATGGAGAggttacctcttatctagt
gtagagtgtagcacctatccTTAATTATACTTCTTTCCATCACCAGTTACTCGGCACTCCggttacctcttatctagtg
gtagagtgtagcacctatccGATTCTACATCTATCAGCAGTTGCAGATCCCTGGCACCATggttacctcttatctag
gtagagtgtagcacctatccCAAATGGGCTTGCACAGATGAATTCAGCTATTCCAGAATTggttacctcttatctagt
gtagagtgtagcacctatccCCCACAAAAAAAAAACTCCCCAGATGTCCCATCTCTTGGTggttacctcttatctag
gtagagtgtagcacctatccAGTATAACTATACTCAAATGAGAATGACCTGCAGTGCCTCggttacctcttatctagt
gtagagtgtagcacctatccCATGTCATAAAAATGTCTGAGGGACAAATAAGAGTCATTCggttacctcttatctagt
gtagagtgtagcacctatccGAGCAAGGTAGTAATGTATTCAATTCAGTTGAAGCCAAGAggttacctcttatctagt
gtagagtgtagcacctatccCTGTGTATATCTGACAGTGTATATCTGACATGTTCATGGGggttacctcttatctagtg
gtagagtgtagcacctatccCTTTAAAAGCCTACAATGCACACAATACCCCCGCACGTCCggttacctcttatctag
gtagagtgtagcacctatccTTCACATTTATGGAGATTATTGCCAGATGAAATTTCTGCCggttacctcttatctagtg
gtagagtgtagcacctatccAAATACAAAGAAGTCGACCAAAGCAAAACCTGTAAAAAGCggttacctcttatctag
gtagagtgtagcacctatccCTGAGGGGAAAAAGAAAAGGATTCAATCTCGTTTATAGAAggttacctcttatctagt
gtagagtgtagcacctatccAGACTTCTGCACCTGATATCACATTCCTGCCCCATTATTTggttacctcttatctagt
gtagagtgtagcacctatccACTTTCAGTCCTACGTCTTTTCTTACAGGCTCCTTTAATTggttacctcttatctagtg
gtagagtgtagcacctatccGTTTCCATCAACAACATCCCATCCTTTCCTGTCCCCGAAAggttacctcttatctag
gtagagtgtagcacctatccCATATGCTGTTGTTTCTGCCACATCTTCCCTGCTACCTACggttacctcttatctagt
gtagagtgtagcacctatccTCATTCCACCTTGGTTGAGACAGCCACTAATTTTTAAAATggttacctcttatctagtg
gtagagtgtagcacctatccGTGGAGATTCTGAGACATTGCCATATATTCAAAGTAGGTGggttacctcttatctagt
gtagagtgtagcacctatccACAAAAGGACTAAAACTGCCTAGTGTCTGACTGTAAATACggttacctcttatctagt
gtagagtgtagcacctatccGAACATCCCTATGCTAAAAATCAAACATTCAAAATGCTCCggttacctcttatctagt
gtagagtgtagcacctatccAATACAGGCACACGACACAGTTTATTCAGTCTCCCCCAAGggttacctcttatctag
gtagagtgtagcacctatccTATTTTCTGTGTGTGTGCAGATTCCCCCACTCAAGCATGCggttacctcttatctagt
gtagagtgtagcacctatccGTGCAGGCTGGACACATCAATGGCAGTTTCCCCACAATGCggttacctcttatctag
gtagagtgtagcacctatccGTGGTGTAAAGATATTATTGGAAATGTCAACAAGGCCTGCggttacctcttatctagt
gtagagtgtagcacctatccTCCCTATGGCTAACAGTAAAAAGAAGAAGAGGAAGCATTTggttacctcttatctag
gtagagtgtagcacctatccGCTTTTGGAGAAACTGGACAATGCTATTACATGTGAAACAggttacctcttatctagt
gtagagtgtagcacctatccCACCATATATGACCTGAAGGAACAGAAGGACAATCCACTGggttacctcttatctag
gtagagtgtagcacctatccGAAGATCTTGATTGTGTATAGAAAGAGTGGATCCATCAGTggttacctcttatctagt
gtagagtgtagcacctatccAACAAGCAAAGATCTATGACGGTGAACTAGAAATTGAAGGggttacctcttatctag
gtagagtgtagcacctatccTTTTTAGACACTTGTGATGATAAATCATCTGCTGGTCATGggttacctcttatctagtg
gtagagtgtagcacctatccTCTGACACAAGAACAAGTTTATAATGCCAACAAACATCAGggttacctcttatctagt
gtagagtgtagcacctatccTGGTGTTCTTGCCCTAGAAATATATTCACTACAACTAATGggttacctcttatctagtg
gtagagtgtagcacctatccCAGAATAACTGTGCTGGGACGTGCTAATGCAGCAGGCATGggttacctcttatctag
gtagagtgtagcacctatccTGCTGCTTTCAGGAAGTGAATTTCTTGTTATGACATAGCTggttacctcttatctagtg
gtagagtgtagcacctatccTCACCAGGGACATCTTTTCTTATTGGTTTCACAGATATTTggttacctcttatctagtg
gtagagtgtagcacctatccACCTTTTGTTATTCCTTGACGACTGTTCTACTCATCCTCCggttacctcttatctagtg

gtagagtgtagcacctatccAAATATAAAAACACTTTATAGAACAGCATGCTAGCCGCAGggttacctcttatctagt
gtagagtgtagcacctatccAAGGTTTTCAAAAGGAGCTGTCATAAAAGATGCCGTATATggttacctcttatctagt
gtagagtgtagcacctatccACAATGACTAAAGACATGGTTGTGCATGCCTAGCAAAGCCggttacctcttatctag
gtagagtgtagcacctatccAGGTGGTGACTTTGAAGGATGCTGAATGTCAAGTGAGAAAggttacctcttatctag
gtagagtgtagcacctatccACCTCCTTACATATGCAAAATCATACCTTCAGTCTGCCAGggttacctcttatctagt
gtagagtgtagcacctatccCATCAATGAGACTCCACTTGTTCTATCTTGACTTGAAATAggttacctcttatctagtg
gtagagtgtagcacctatccCAAGGTGATTGTGAAAATAGCGATAAGAAGATGACATGGTggttacctcttatctagt
gtagagtgtagcacctatccGTAGAAAAAGTGCCTATAGAAAACATGGTGAAAATGGGTGggttacctcttatctag
gtagagtgtagcacctatccATTGAAGGACATGACGAATCCATAACAGAACAAGAAATCAggttacctcttatctag
gtagagtgtagcacctatccAATGAAGTAGGCAATTCTGGAAGGAACATTTTTAAAAGCCggttacctcttatctagt
gtagagtgtagcacctatccCACTACTTGGTCCCTCAACTGCTTTTGCTATTTCTTCTCAggttacctcttatctagtg
gtagagtgtagcacctatccGTGGACAATAATCTTTTAATCAAAACATAGCACCACAGGTggttacctcttatctagt
gtagagtgtagcacctatccTGTTGTTGCTGTTGTTTATTCTGGTGATGGCACTGTGCTGggttacctcttatctagtg
gtagagtgtagcacctatccTGGTAGGTTATATTTTTCACTGGCAAATACTTATGTGTGCggttacctcttatctagtg
gtagagtgtagcacctatccTGATGGTGATGCCAGACAACAACAGACCGTCCACATGGGTggttacctcttatcta
gtagagtgtagcacctatccCTTTCTGATGGCTCAATGTAATCAAACTTTGTGTCATGCAggttacctcttatctagtg
gtagagtgtagcacctatccGGCAAATATTCCAAAATCTAGGATCCAAAACACTTCTGGTggttacctcttatctagt
gtagagtgtagcacctatccGATACTCAATCTGTACAAACCACTCACCTGTCACCTATAAggttacctcttatctagt
gtagagtgtagcacctatccGGACATGAGCTACCATAACCCACTGAAGCCTATACACAACggttacctcttatcta
gtagagtgtagcacctatccTGCCCTGGAAGTGGCAGCTGCACTACAAATCCCCATAGCTggttacctcttatcta
gtagagtgtagcacctatccTCCAGGTGAAGTTTCCTCCAGTAGCCCCAAGAACCCAAGCggttacctcttatcta
gtagagtgtagcacctatccAGACAAAACCAATTGGATGTATGACATAGCTAGAGATTTCggttacctcttatctagt
gtagagtgtagcacctatccGCAAGTCCAAATTTTGTAGAGCAAGCTGGCAGGCTGGAAAggttacctcttatctag
gtagagtgtagcacctatccTTGAGTCTGAAGATAGTCAAGAGGCAGAATTCCCTTCTCTggttacctcttatctagt
gtagagtgtagcacctatccAGTCTTTTCTCTGAAGGCTTTCGATGGGTTGGATGAGACCggttacctcttatctagt
gtagagtgtagcacctatccAACACCTTTATAGAATGAATTAATGGCATATGCAGCAACCggttacctcttatctagt
gtagagtgtagcacctatccAATGGAAAACCAAATATCATATGTTCTCACCCATAAGTGGggttacctcttatctagt
gtagagtgtagcacctatccTGCCACCTGTTTCCCCAAAAAAACTATGGAAACTTTAAAAggttacctcttatctagt
gtagagtgtagcacctatccTGGAAACATCTAGATTCATGTCTAATCAAGCAAACTAGGCggttacctcttatctagt
gtagagtgtagcacctatccGCCAAGCTGACTCATAAAATTCCCCATTGTAGGTCCTAAGggttacctcttatctag
gtagagtgtagcacctatccTAAATTACCCCCACGATGCTCCAAATACTATCCCCTGTTCggttacctcttatctag
gtagagtgtagcacctatccAGCTGATTCATCCTCCATAAGTACAAAAATCACCTCCCATggttacctcttatctagt
gtagagtgtagcacctatccCAGCCCAAACATCAGCGGAAAATCACAAAGCCCCAGATCTggttacctcttatcta
gtagagtgtagcacctatccGGTCTCTCCTCTCTAGAGCACAACACTCCATCTACACATGggttacctcttatctag
gtagagtgtagcacctatccTCTACACCCTTGCATAGATCATAACAGAAGCCACTCACCTggttacctcttatctag
gtagagtgtagcacctatccCTGCCCCTAGAGAGAACGTTGAACTAAACTAACCATATGGggttacctcttatctag
gtagagtgtagcacctatccGAACATTCAAACAAGACACACTCACACATTCATAGATCACggttacctcttatctag
gtagagtgtagcacctatccTCATTTTACTTAAAAGTTGGGCAATTTTGGCCGGGCATGGggttacctcttatctagt
gtagagtgtagcacctatccATTGCTTGAACCTGAGAGGTGGGGGTTGTGGTGAGCTGAGggttacctcttatctag
gtagagtgtagcacctatccCTTACTAGGAAATGTGTTTATTTAATCTCTAGGCTCACTCggttacctcttatctagtg
gtagagtgtagcacctatccGTAAGAAAATACAGATATCAGAAGATACATTCCAGGCTCAggttacctcttatctagt
gtagagtgtagcacctatccCCACAAGTTAAGTCAGGAGGCAGAAAATACCAAAAATACAggttacctcttatctag
gtagagtgtagcacctatccGCACTCACTTCCCTAAAATTTGCATCACTTATACACAGAGggttacctcttatctagt
gtagagtgtagcacctatccTGTCATTCAGCCGGTTTAAAAACAACATACAAACAACAGCggttacctcttatctag
gtagagtgtagcacctatccCATAAAATGAGTTATTGCAGATCCCAGTAGCACTATAGAAggttacctcttatctagt
gtagagtgtagcacctatccTGATTATTCAAATTGTGACCGAGAGCTGCAGACAGTGCCCggttacctcttatctag
gtagagtgtagcacctatccCCATTTCTGCTTCATGCTGGTCTGTTCCCTCCCCTAAGAGggttacctcttatctag
gtagagtgtagcacctatccTGCCTTGTTATCATCATTGGTAATTAGCATTTACTGAGCAggttacctcttatctagtg
gtagagtgtagcacctatccCAGAACTTAATGCTGCCCAGTGGGTTGTTGGTCTAATTTCggttacctcttatctagt
gtagagtgtagcacctatccCAGCACCTCAACAAAGACCAACCACCAAAAACCCAAAGCAggttacctcttatcta
gtagagtgtagcacctatccTCAGACTTGCCTTATCAGACCGAGAGTTGCTTGATTACACggttacctcttatctagt
gtagagtgtagcacctatccAAGTGGCCAGAAAAGAACTTGAAAGAGGAAAGCAGATCTTggttacctcttatctag

gtagagtgtagcacctatccGTGTCATTTCTTTCCTTCACCCATTGACCTCTTTTGTAGGggttacctcttatctagtg
gtagagtgtagcacctatccAAATGGTTCACAGCATGGTAGAGAAAAGACCCAAAAATTAggttacctcttatctagt
gtagagtgtagcacctatccGGTTTTGATTTCTGATGAAAGTAAAAGTTCTGGCCAGGCAggttacctcttatctagt
gtagagtgtagcacctatccCCAACATGGTGAAACCCCCTCTCTATACTTAAAAAAAAAAggttacctcttatctagt
gtagagtgtagcacctatccAAGTAAATAGTCTTCAACATTCAAGTTGTAGAGTGCAGAGggttacctcttatctagt
gtagagtgtagcacctatccAAAACCAAAAACAGCAATAGCCCCATATTTCCATCACATTggttacctcttatctagt
gtagagtgtagcacctatccGGAAAACCTAATTCCTGCATTACAGTTGGTCATTTTGGTCggttacctcttatctagt
gtagagtgtagcacctatccCTGTTGTACTAGTCAGGTGACCTTAAACTTAAACACTTGTggttacctcttatctagtg
gtagagtgtagcacctatccTCTCTAAAATAGGGATAGCAATGAAATCAACCTCAAAAGGggttacctcttatctagt
gtagagtgtagcacctatccAGTGAAATAACCCATATAAAACTGCTTGACATAGCTTAGCggttacctcttatctagt
gtagagtgtagcacctatccTCAGGTATGACACAAATGCAGGTTATTACATTGCCCTTGTggttacctcttatctagt
gtagagtgtagcacctatccCTTTTTTTTTTTTTGGATAGAGAGTCTCGCTCTGTCGCCCggttacctcttatctagtg
gtagagtgtagcacctatccATCTGCCTCCTGGTTCAAGCGATTCTCCAGTCTCAGCCTCggttacctcttatctag
gtagagtgtagcacctatccGCACCCGGCCCAAATAACATTTTTATGTAAGTAGACCCAggttacctcttatctag
gtagagtgtagcacctatccCAAAACAAGGCACGTCATGGTGAGTTTCCTCTTGACTAGAggttacctcttatctag
gtagagtgtagcacctatccAGCTGGGAGGAAGATGATGGAAGAGGTGCAGGTCTCAAAGggttacctcttatcta
gtagagtgtagcacctatccATCAATGGAGGATGCTTTTGTCTTAATAAGGGACTCTACAggttacctcttatctagt
gtagagtgtagcacctatccGAACTTACCTTCTATCCTCTTTTCCTCCTCTACTCACACAggttacctcttatctagt
gtagagtgtagcacctatccACTATGGTTAAGACTACGTGGATTCAAATGCTGCCTGACTggttacctcttatctagt
gtagagtgtagcacctatccATCTGTAAAATGCAGACAATGGAAATGGTATATAGACACCggttacctcttatctagt
gtagagtgtagcacctatccAACAATAGACACTGCAGACTGTTAGAGGGGAGAGGGAGGGggttacctcttatcta
gtagagtgtagcacctatccGGTGACAAGACCCTTACTCAAACCTCAGAATAAAGCAATAggttacctcttatctag
gtagagtgtagcacctatccACGTAACAAACCAGCACACTTATCCCTGTATCTAAAATGAggttacctcttatctagt
gtagagtgtagcacctatccACTAAACTGAAGAATCTTATTCCCTGGAATGTCATTTCTGggttacctcttatctagtg
gtagagtgtagcacctatccGGTACAGTCACAATTGTCATCTTGCAATGTATATTAACCCggttacctcttatctagt
gtagagtgtagcacctatccGAAATTTCAAAAACAATGTAATCCCCAACCTCCACCTTTCggttacctcttatctagt
gtagagtgtagcacctatccCTCAAATAGACTACATGGATTCAAATGCTGCGTGACTTTGggttacctcttatctagt
gtagagtgtagcacctatccGTTACGTCACCTTTCTGTGCCTAATCTGTGCATTATCTGTggttacctcttatctagtg
gtagagtgtagcacctatccGGAAAATGAATAAGAGATCAGAATCACCATTATCACCATCggttacctcttatctagt
gtagagtgtagcacctatccCATCTTCATTATCATTGTTACTACACATATTCCTCACTCCggttacctcttatctagtg
gtagagtgtagcacctatccTCAAGGAAACCAGTAACACTAAGTAACAGAATAAAAGCACggttacctcttatctag
gtagagtgtagcacctatccCCCTACACAAGAGGCCTGAATTAGAAGAGTTCGCAGCATGggttacctcttatcta
gtagagtgtagcacctatccATGAGTGTTGAGTGGGGAATGAAATATAATATCCTTATGCggttacctcttatctagtg
gtagagtgtagcacctatccTAGCACAAGGGAATATTTTTTAGAAATGACGCATCTGAGCggttacctcttatctagt
gtagagtgtagcacctatccAGGAAATCAGGACCCAGAGATAAACAACTTGCCTAATGACggttacctcttatctag
gtagagtgtagcacctatccGACAAGAACCCAAGTCTCCAAGTCCAGTGCTCTTCAGCCAggttacctcttatcta
gtagagtgtagcacctatccTTGTAAAACTCTGATGCAGATTTAGAGACCAAGGAAAGGTggttacctcttatctagt
gtagagtgtagcacctatccGCTATCCCAGTTGCATCCACCAAGAAGCTGATATTCTTCCggttacctcttatctag
gtagagtgtagcacctatccATTTCCTTCTGGTCAATTCTTCTTTCTCTCCTAGGTCACAggttacctcttatctagtg
gtagagtgtagcacctatccTTGTAAGAAGATGCTGCACTAACATGGCTTTAGCAATAAAggttacctcttatctagt
gtagagtgtagcacctatccGGTTAAAGCCCCTCTCCTTCCTCACCCATCTCCTCACCCAggttacctcttatcta
gtagagtgtagcacctatccCCCCTCTGTCCTGGTCCTCTCCATTTCAGGACTTTTTTTTggttacctcttatctagtg
gtagagtgtagcacctatccCTCACTGCAACCCCCATCTACTGGATTCAAGAGATTTTCCggttacctcttatctag
gtagagtgtagcacctatccCAACCCCCCACGTAGCTGGAAATATAGGCGCCTGACACCAggttacctcttatcta
gtagagtgtagcacctatccTGGGAGTACAGACAGGAGCCATCACATGGAAGCATCTCAAggttacctcttatcta
gtagagtgtagcacctatccGCCTTATTTCCTTGCTTCTCTTCCCTTCTACCCTCTCACCggttacctcttatctagt
gtagagtgtagcacctatccGAAATGGCTTTTTTCTTACTCACAACCCACATCACCAGGTggttacctcttatctagt
gtagagtgtagcacctatccCTCACTCCTTCTGTCAATCTCAATTCCAATATCCCTAGTTggttacctcttatctagtg
gtagagtgtagcacctatccCATCCTGCCACTTACCACACTTTACATTTCTACTTCCATAggttacctcttatctagt
gtagagtgtagcacctatccCCATCTGCTCCCCAGTCCCTGGTATATAGCATAGTTCCTGggttacctcttatctag
gtagagtgtagcacctatccTATCCAAAGTATTCCATTTAGCTGAAACTGCCATTGTTTGggttacctcttatctagtg
gtagagtgtagcacctatccCAACTATTCTACTTTTTGTTCTCCTGGTTGTTTCCATACCggttacctcttatctagtg

gtagagtgtagcacctatccACATTAGACATTCCTTCCCCTTTCATATTTTCCTTTCTTCggttacctcttatctagtgg
gtagagtgtagcacctatccAGAGAATTGAGGTCACAGGGTTGCATGAATTGCCAAGTCAggttacctcttatctag
gtagagtgtagcacctatccTACCAAACTAAGAGTAGAGACATCTCATTGCATAAAACACggttacctcttatctagt
gtagagtgtagcacctatccCATTCCTTCTTCTATGCTTTCACAGTTTCTTTTTCCACTTggttacctcttatctagtgg
gtagagtgtagcacctatccCTCCAGCCTCACCCACCTATTCTCATCTGTAGCAGCAATGggttacctcttatctag
gtagagtgtagcacctatccTGGAAAGATAGAAACCACCCAAATAGAACTCCCACAAGATggttacctcttatctag
gtagagtgtagcacctatccGGCCTCACCCATCTGCACCTTTAAACTCTGCTTCTCTCCTggttacctcttatctag
gtagagtgtagcacctatccCATTACCTGTGCACTAGATCCCAACCCTTCTTGCCTCCTCggttacctcttatctag
gtagagtgtagcacctatccCATTTCTTCAGTTATTCTCTCCTCTTTCCCATACCATAAAggttacctcttatctagtg
gtagagtgtagcacctatccCTGAATCATTTCCACCAGAACAAAAGACGTTCTTTCCTTCggttacctcttatctagt
gtagagtgtagcacctatccAACTTCTCCTGGTTTCTCTACATTTTCACCTTCCCTTTCTggttacctcttatctagtg
gtagagtgtagcacctatccCAATTTAACTCTTCCTATTGTCTTTTGAATCCACTCCAGTggttacctcttatctagtg
gtagagtgtagcacctatccAGTTCTCTCATACCCTCCAAACTGCTAAAGCTATGGTCAAggttacctcttatctagt
gtagagtgtagcacctatccAGATCATGACTATCTTCTTACAACCTCCCTCACCTACATTggttacctcttatctagtg
gtagagtgtagcacctatccGGCACTGTGTCCTGGTCTTTTCTGGTTCCTCTTTATCTCTggttacctcttatctagtg
gtagagtgtagcacctatccTCAGCCCCAAATTCACTTCTCTCTATATACTCACTCTCTTggttacctcttatctagtg
gtagagtgtagcacctatccGCTAAAAATGCCATTTTCATGCTGATGACTTCCAGAGATGggttacctcttatctagt
gtagagtgtagcacctatccTATCGTATCTAAAACTGAGTTCTCAATATATACCCCCAACggttacctcttatctagt
gtagagtgtagcacctatccGAGTCATCTTTAGCTCCATTTTCTCCTATACCCCACATCAggttacctcttatctagt
gtagagtgtagcacctatccGCTCCATTTATTTTTTGCCTTCATCAGCGTCTTCTATTATggttacctcttatctagtgg
gtagagtgtagcacctatccCTACAGGAGTTTAAAAGGAGCCACAAATACTATCAAGAATggttacctcttatctagt
gtagagtgtagcacctatccACACAAAGACTCCAAGAATGAGCCACAACTCTGCTGACCCggttacctcttatcta
gtagagtgtagcacctatccTCCCTCCTTCCCTGCTTCCCCTGACACCTCTGCCCTACCGggttacctcttatcta
gtagagtgtagcacctatccCAGAGAGGTCCTGCATACATTTCACCCAGATTCCTTCCATggttacctcttatctag
gtagagtgtagcacctatccCATCTCACATTTATAGCCCAATATCAAACCCAGGAAATTGggttacctcttatctagt
gtagagtgtagcacctatccGTATGTGTGTAGAGTTCTGTGTCACTTAATCCTATGTGTGggttacctcttatctagtg
gtagagtgtagcacctatccCTGCAATCAAGACACAGAAACTAACCCATCGCCACAAAGGggttacctcttatcta
gtagagtgtagcacctatccCTCATGCTACCTTTTTATGATCACACCTACCTCCCTCTGTggttacctcttatctagt
gtagagtgtagcacctatccTTTTTCACTCTGCATAATACCCTTGAGATCTACAGGTGTTggttacctcttatctagtg
gtagagtgtagcacctatccCTGTTTTCATTTCTTCAAGATAGGTTCCCAGGAGTTCAATggttacctcttatctagtg
gtagagtgtagcacctatccGTTGTATGGTCAATGTGTGGCTATTTCAGTTTAGTTTTCAggttacctcttatctagtgg
gtagagtgtagcacctatccGGATGTTTGCAAGCCCCCTGGTCCAAGTCACTGTCCTCCCggttacctcttatcta
gtagagtgtagcacctatccGGGACTCTTCTTTGATCTTTGTTCACCCCCTACCACTTCTggttacctcttatctagt
gtagagtgtagcacctatccTGATTCTGGCAAAATGAAAATCAGGTTGTGCTATTCCTCTggttacctcttatctagtg
gtagagtgtagcacctatccCTGCAAGGACCTCTATCTTTTGACTCTTCCCTGTTCTTTGggttacctcttatctagtg
gtagagtgtagcacctatccGTTTCCAATTATTCTCCCCAAAACTCACTCCTCTCCAGCTggttacctcttatctagt
gtagagtgtagcacctatccCATTCTCTTGCCTCTGAGTCTTTGTAATTGCTGTTCCCTCggttacctcttatctagtg
gtagagtgtagcacctatccGGTTGCTCTTCCCACATAGTTCATTTCCACACTCCTTCAGggttacctcttatctagt
gtagagtgtagcacctatccGGATTTCCTAGTCACCCTCTCACCCTACCAACCTTTTTGTggttacctcttatctagt
gtagagtgtagcacctatccTCCATAGTACATTGTCACCAACATGTCACATACTTTACTTggttacctcttatctagtg
gtagagtgtagcacctatccGTCTCACACGTTTTATCAGAATGCAGGTATCTTGAAACCAggttacctcttatctagt
gtagagtgtagcacctatccGTTCACTGCTGAATCCCCAATGATTGGAACAATTCCTGGCggttacctcttatctag
gtagagtgtagcacctatccTATAACTGCCAGCATTTCATATTTGTTCACAATGTCCATGggttacctcttatctagtg
gtagagtgtagcacctatccCATTGAAACAGAGGCCAAAGTACTAAAAGCATTTAAAAGGggttacctcttatctag
gtagagtgtagcacctatccGCAGTTTTTGCATTGTTGGGAATTTGTCATTTCATATTGGggttacctcttatctagtgg
gtagagtgtagcacctatccTGTAAAGTCGCAAACACAACTCACAATATCAACCATGCATggttacctcttatctag
gtagagtgtagcacctatccGCCAGCCATCGAAGTTCACAACCAATTGAGGACAACAATCggttacctcttatcta
gtagagtgtagcacctatccTCCTCTTACAACTACACAACTGCCAAAGAACTAAACGTCGggttacctcttatctag
gtagagtgtagcacctatccAGGTGAAAAAGCTCAGCTCAATAAGTCGGTGCCTCATGAGggttacctcttatctag
gtagagtgtagcacctatccGATTGTGACATGTGACAAAAGTGGATTTTATATGGCAATCggttacctcttatctagtg
gtagagtgtagcacctatccCCAAACTTCCACCAAAAAAAGCTTATGATCACTGTTTGGTggttacctcttatctagt
gtagagtgtagcacctatccTCTGAATCCCGGTGAAACGATTTACACCCGAGAAGTATGCggttacctcttatctag

gtagagtgtagcacctatccCTGCAGCCAGCATTGATCAATAGAAAGGGCCCAATTATTCggttacctcttatctag
gtagagtgtagcacctatccCACCTGACCTCTCGCCAACATATTACCACTTCAAGCATGTggttacctcttatctag
gtagagtgtagcacctatccTGCTTCCACAACCAGCAGATTGCAGAAAATCCTTTCCATGggttacctcttatctag
gtagagtgtagcacctatccCCAAAGCATGGATTTTTGTTCTACAGGAGTAAACAAACTTggttacctcttatctagt
gtagagtgtagcacctatccATTACTTTTGCACCGACCTATATTTGAAAGTGCACTGGGCggttacctcttatctagt
gtagagtgtagcacctatccCAAGAGTTCTTACATTTCGTCTTGCTCCAGAACAAAATGTggttacctcttatctagt
gtagagtgtagcacctatccCAGCTCCAATTCTGCAGTTTCATGTCAGAGCACAGAGCTCggttacctcttatctag
gtagagtgtagcacctatccGCCATGAGCTCATTCCCTGATAATCTGGAGAACAAGTCCAggttacctcttatctag
gtagagtgtagcacctatccTTACATTCCTGCTTTGATTGTCAATCCAAACCAAAAGATGggttacctcttatctagtg
gtagagtgtagcacctatccAAGAAGAAAACTTTACCTTATTACCCGCTGCATTACACTTggttacctcttatctagt
gtagagtgtagcacctatccGCCTCCCAAGTAGCATTACACTTTTGAACCAAAATTATGCggttacctcttatctagt
gtagagtgtagcacctatccTTATAGTAGGTATAAGAAAGTTCAACGCCAACAAGAAGAGggttacctcttatctagt
gtagagtgtagcacctatccGTATCAATACATGGATGTGCTAGAAAAACAAAGCCTAAGTggttacctcttatctagt
gtagagtgtagcacctatccTATCTCACTAATGCCAGTTGCTATTTTCTGAGTGCTTACTggttacctcttatctagtg
gtagagtgtagcacctatccAGTCTAACCACTTTACACACGTTAACTCACACACAGCTAAggttacctcttatctagt
gtagagtgtagcacctatccAACTTGGCTCTGGAGTTTGGGCTCTTTTGTCCCTTCATTGggttacctcttatctagt
gtagagtgtagcacctatccTAAATGTTTTAAACCACCCCAGGCGATAATAGAGGAGAAAggttacctcttatctagt
gtagagtgtagcacctatccACTAGCCTGAGAAGTGATAAGGGGGTTTCAAAGTGAGGTTggttacctcttatctag
gtagagtgtagcacctatccCAGGCCTCTTCTGTTTGCATCTGTCCCCTTCTCCCTCTGCggttacctcttatctag
gtagagtgtagcacctatccGCTGCTCCCATTGGCCATCCACCCTCACTGTTCCTCCCCAggttacctcttatcta
gtagagtgtagcacctatccCACATTGACTTACTGCTTCACTCCTCCTTCCAAATGCCTTggttacctcttatctagt
gtagagtgtagcacctatccTCTGTAGGGAAGAAAATGGGATTAGTCTATTACTGTAGCCggttacctcttatctagt
gtagagtgtagcacctatccTCTTTGGACCCTAAGATAAGAATCTCAGAGAGCATTAAGGggttacctcttatctagt
gtagagtgtagcacctatccCTATAAATAGTCTATAAATCACACCCCCTGAGTTCTTCGCggttacctcttatctagt
gtagagtgtagcacctatccACTACCGGTGTGATCTCTGACAAGTCCTGTGAGCTCTCTGggttacctcttatctag
gtagagtgtagcacctatccGTGGGAGGATTCATTGAGCTAAACTATGGAAAGCAGTGCTggttacctcttatctag
gtagagtgtagcacctatccATGTGAATGTTTGCTATTACTGTATATCCCCTGGTCATTGggttacctcttatctagtg
gtagagtgtagcacctatccAAATACGCTTTCTCCCTTCCTCCATCCCCATTCTTCTCTCggttacctcttatctagt
gtagagtgtagcacctatccTTCATAGAGTGGAGAATTGGGTTGCTTTAAAAAACTTGCTggttacctcttatctagtg
gtagagtgtagcacctatccGGACTTTTCTTCCAACCAGCATTTCTTTCCATGAGACTAAggttacctcttatctagt
gtagagtgtagcacctatccCTTCTAAGAGAAATTTTTGAAAGAATTGTCATGGGTCTCGggttacctcttatctagtg
gtagagtgtagcacctatccAGTCTGCTTACCATGAGAAATGTTCACAAACTACCACAAAggttacctcttatctagt
gtagagtgtagcacctatccGTGTGCACAGTACCTAAAAATGTACTTTGTTCAAATGCAAggttacctcttatctagt
gtagagtgtagcacctatccACATTTACCAGCATTGGAGGGTAACTTTGCATTATTCTGGggttacctcttatctagt
gtagagtgtagcacctatccAAGAAATTTAAGAGAAGGAGACAAAAAGAGAATGGCGCAAggttacctcttatctag
gtagagtgtagcacctatccCACCCTTTGCTTATGTAACTCCTGCCATGGCAATTTAAGGggttacctcttatctagt
gtagagtgtagcacctatccTGTACAATCCACTGACAATTTTCACCTCCCTTGATAGAAAggttacctcttatctagt
gtagagtgtagcacctatccGGGACTGTTAATCTTTCTCATTCACAATCTATTTCCCAAGggttacctcttatctagtg
gtagagtgtagcacctatccTTGTATACTGAGAAAAGCTTTGAAACATATAGGCAGAAGCggttacctcttatctagt
gtagagtgtagcacctatccTGCTACCTACGACTTCAGAAATACTTGAATGTATGTTTCCggttacctcttatctagtg
gtagagtgtagcacctatccCATGCTGCAACTTCAAATGAATGGATAATTGGTCAGGGAAggttacctcttatctagt
gtagagtgtagcacctatccGATAAATAGTGAAGTATCTGCATAGTCTGAGGTACACAGGggttacctcttatctagt
gtagagtgtagcacctatccTGGGATAGCATAAGACACGCCGCTTCCTGTGAGCATGAGGggttacctcttatcta
gtagagtgtagcacctatccCTGATGACCAACAGTGATTCTCCTCAACTTAACGGGGTTTggttacctcttatctagt
gtagagtgtagcacctatccGTGCAATTTTGGTATGTGCATATATTGCACTGTGGTGAAAggttacctcttatctagtg
gtagagtgtagcacctatccGTACCCATTAAGTATTTTTTATTCCTTACCCCCCCAATCCggttacctcttatctagtg
gtagagtgtagcacctatccAACACATGGTACCCGAAATTGTATTTCTGAGTTCTTTCAAggttacctcttatctagtg
gtagagtgtagcacctatccTGGGCTCCAATTCCATCTATGATGATGCAAAAGACATAATggttacctcttatctagt
gtagagtgtagcacctatccCTTTATCTAGTCATCTGTTGCTAAACTTAGGTTGATTCCAggttacctcttatctagtg
gtagagtgtagcacctatccTTTAGTTCTCTGAGAAATTGTTTTCCATAGAGGCTTGTACggttacctcttatctagtg
gtagagtgtagcacctatccGATGTTGAGTATTTTTCTCAGATGCTTTGTAGCCATTTGTggttacctcttatctagtg
gtagagtgtagcacctatccTATCCTATTCTACAGGTTCTGCATTCACTTTAACAGTTTGggttacctcttatctagtg

gtagagtgtagcacctatccCAGTTTTTGTTCCTACATGCCTTTCCTTTCCAGTGTCTGCggttacctcttatctagtg
gtagagtgtagcacctatccTCCTGTGATTCTGCCCTACAAGACATGCTTAGTTAATGGCggttacctcttatctagt
gtagagtgtagcacctatccGCTCTCACATGTGCCCTTGATCATTCCTTCTGTCTGCTTTggttacctcttatctagt
gtagagtgtagcacctatccCTGCCCTCCTTCAGCTCACAGTGCAGCAATTGTTTTGACAggttacctcttatctag
gtagagtgtagcacctatccTTATCACACATCCTATTCATAGGCACAGATAAGAAAAGCTggttacctcttatctagt
gtagagtgtagcacctatccAAGTTATCTAATTATGGGTATCCTTAAGTCCATGGTGGCCggttacctcttatctagt
gtagagtgtagcacctatccTTTTGTGTTTTATCCCTTTAGCTTTCCTTAGTTCCAAGAGggttacctcttatctagtgg
gtagagtgtagcacctatccTGGCTTTAATCTATCCATTTGTCAGTTAATGTCACGCAATggttacctcttatctagtg
gtagagtgtagcacctatccTCACAGACCAGACTAGTCCCAGGCAATTGTCTCAGAAATGggttacctcttatctag
gtagagtgtagcacctatccCTAAGTCAGCATAAACCTACCATCCACCCTACAAAATATGggttacctcttatctag
gtagagtgtagcacctatccCAGCAATGAATGTTGAAGAGTTCCATTAAGAGTGCCATTCggttacctcttatctagt
gtagagtgtagcacctatccTCATGAAAACAGCTCTAAACCAGTACATGTACTAATACCTggttacctcttatctagt
gtagagtgtagcacctatccTCTACAGTGTCAAAAGTTCAGATTCTAATGTTCATGCAAGggttacctcttatctagtg
gtagagtgtagcacctatccACACCACAACCACCAAAGCTTTCATGAATTTTTCTTTATGggttacctcttatctagtg
gtagagtgtagcacctatccTGCCCCAGAGTAATTGTGACAAAGTTTGGCATGCCATTAGggttacctcttatctag
gtagagtgtagcacctatccGGCATAAAGTCCAATCATCAATATACATTCTCTGAAGTCAggttacctcttatctagt
gtagagtgtagcacctatccCCCACATCAAAACAGCCAGAAGGGTGGATATCCTGGCCCAggttacctcttatcta
gtagagtgtagcacctatccAGTCCTACAACTAAGATGAATCAATCAATCTCCTTTTGCCggttacctcttatctagt
gtagagtgtagcacctatccGGATTATTCAAACCACACAATTAATTCCCCTGATTCAAAGggttacctcttatctagt
gtagagtgtagcacctatccAAATGCACTCATCCACTGATACTAGCACTCATTTATTCACggttacctcttatctagt
gtagagtgtagcacctatccTCAACAAACAGGCATTATCTGCTTCATTCAGGCTCAGGATggttacctcttatctagt
gtagagtgtagcacctatccCTCATGGAGCTTCAACTCTAGACAGATATTGATCAAATAAggttacctcttatctagt
gtagagtgtagcacctatccTCCCAGTCGTGATAAATGAAAAATAGAGATCTAAAAGTGCggttacctcttatctagt
gtagagtgtagcacctatccCAATCTGAACACTGCAGCATTGATTATCATTATACTCGAAggttacctcttatctagt
gtagagtgtagcacctatccTACATACCTAGATATAAGTAAGGATCACTTTACTCTGGTGggttacctcttatctagtg
gtagagtgtagcacctatccTCTCAGTAACTGAATATGCCAAAGGTTTATTTCTCACTTGggttacctcttatctagtg
gtagagtgtagcacctatccGGTAGCTCCCTTCAAGCAGTGAATCAGGAATCCAAGCCACggttacctcttatcta
gtagagtgtagcacctatccCTTCCAGGTCAGTGGCAACAAAAGTCCAAAGCAAGTGAATggttacctcttatctag
gtagagtgtagcacctatccCTCTTAAATGCTCCATCTCACCACAAGGGCACTACTAAATggttacctcttatctag
gtagagtgtagcacctatccGGGGAACATGGATATATTCAGGAAACATTTCTGCCAGGCAggttacctcttatctag
gtagagtgtagcacctatccTTCTCATAGTTCAAGTGGGAAGGGTAATTCAAGCTATTTCggttacctcttatctagtg
gtagagtgtagcacctatccAAAGGGGGTTAATGCGAGGATCTTTCAACACTTCAAGGAGggttacctcttatctag
gtagagtgtagcacctatccGGCTATTACTATTATTTCTACCACAACTACCAATAACACCggttacctcttatctagt
gtagagtgtagcacctatccCAGCTTCTGTCAATGGAAAAAGAGAGCCAGTCTAGGTGGGggttacctcttatcta
gtagagtgtagcacctatccGCATCCTCACCAAGTCACTTGCTGGTAGAATACAGAAGGCggttacctcttatcta
gtagagtgtagcacctatccCTGGCCCTCTCCTAATCTCCTGGTATGCTCAAGTTGACTTggttacctcttatctagt
gtagagtgtagcacctatccACTGGCCTAGTCTAAAATTGTCTAATTGGGTTAACAGACCggttacctcttatctagt
gtagagtgtagcacctatccGCTAAATATACCCAGTTACATAAGCACTTTCTCATCCTGTggttacctcttatctagt
gtagagtgtagcacctatccATCCTAGGTCAGAAATGAGGAATATTGCTTGTTTAACTGGggttacctcttatctagt
gtagagtgtagcacctatccTGATCTAAATTACAGACCCTCCTTTACCTAACTAAAAACCggttacctcttatctagt
gtagagtgtagcacctatccAGCAATTTCTGGTTATCATTATGGCTCCCAATCGTCTTTGggttacctcttatctagtg
gtagagtgtagcacctatccCAATCATGCTCAGCACCAATATATCTGCTTAATATGCTCCggttacctcttatctagt
gtagagtgtagcacctatccTGGGATCTTGTGCAAATTATACTGAACTCTTTATCATGTGggttacctcttatctagtg
gtagagtgtagcacctatccGCAAAGCAGCAATGTGCAGTAAAGTTAATGGCAATATCAAggttacctcttatctag
gtagagtgtagcacctatccAGAAAAAAAGTAACAGAGATTCCAGAGATCAAGGATAGACggttacctcttatctag
gtagagtgtagcacctatccAGAAAAACAACCTACAAGACCTACTGAAAATAAATGGCTGggttacctcttatctag
gtagagtgtagcacctatccAAGGATATAGTAAACTAAACTGCATTTAGTGGCTCAAACCggttacctcttatctagt
gtagagtgtagcacctatccAGCTGAAATGTATGTCATTATTAAAAGGTATACCGTGACCggttacctcttatctagt
gtagagtgtagcacctatccCGAATCAAAAAGAAAACAGAAACACTTTGCTGTTGTTGTCggttacctcttatctagt
gtagagtgtagcacctatccCTCCAAAACTAAAGTTTGAGAAATCACAAAAGAGTGACAGggttacctcttatctag
gtagagtgtagcacctatccAAATGTGAAATTTAGAAAAATAAGTAGGGTAGGGGCCAGAggttacctcttatctagt
gtagagtgtagcacctatccTCTGGTAATTGGTTTTGTTGTATAATAGTGACAAGAACGGggttacctcttatctagtg

gtagagtgtagcacctatccTGCCAGGAAAGTTTGGGAAGATGCTATCAATTTTACTTTAggttacctcttatctagtg
gtagagtgtagcacctatccACATTTTCTAAGTGTTTGGGGGGATATGGAGGAAGGTGGTggttacctcttatctagt
gtagagtgtagcacctatccATGTTGGGACATTTCAAAGAAGACTTGTCCACATGTTTTTggttacctcttatctagtg
gtagagtgtagcacctatccATATGGTGGGGGATATGAGTATGGGTGTGACTGCTTGTGTggttacctcttatctagt
gtagagtgtagcacctatccCATTTGTATGTCTTGTGAGTATATGTGTGCCTGTATGATTggttacctcttatctagtg
gtagagtgtagcacctatccTGTGAGTGTATGTGTAGGAGAGTGTAAGTGTGAGTGAATGggttacctcttatctagt
gtagagtgtagcacctatccTGTACGAAAATGTGTGGAAGTGAATATGAGTGTATATGTGggttacctcttatctagtg
gtagagtgtagcacctatccTGCTGACACAACACATCTAATAAACCAAAGGGGTAAGGCAggttacctcttatctag
gtagagtgtagcacctatccCTGGAAAGGGACAAAGAGTAGAGGTATTTATGGGTTAATTggttacctcttatctagt
gtagagtgtagcacctatccGCATTCCCTTAGCCAGACTGCATTCAACACCAAAGTCATTggttacctcttatctag
gtagagtgtagcacctatccGCTGGGGCCTGTCTTCTGTCTTCCTTTCACTGTTCTCTCTggttacctcttatctagt
gtagagtgtagcacctatccTTTGCCAAAGAAAACAGAAAGGACTCCAATAACATCCCCTggttacctcttatctag
gtagagtgtagcacctatccGACCATGTCTGTCTCAGTTCTAGGCTCTATCTCTGCAAGGggttacctcttatctag
gtagagtgtagcacctatccGCTGATGAATTCCTCCAAGCCTTGATTAAAAGCACTTTTTggttacctcttatctagtg
gtagagtgtagcacctatccTGTACCCAGCCTTCCCCCACATAGACCTCTCCCTCAACTCggttacctcttatcta
gtagagtgtagcacctatccCAGCAATTATCATTTGGAGTCACATATTCTAGCTATTTGCggttacctcttatctagtg
gtagagtgtagcacctatccGCCCCTTCCAGCTTTACAGTTCTATGATTCAGTGTTGTCAggttacctcttatctagt
gtagagtgtagcacctatccTGATTTAGAGTAAAAGTTTGTCAAAGGAAGGAGGCCAATCggttacctcttatctagt
gtagagtgtagcacctatccTGAATGGTTGGGGAAAATAATCTTTGATGAGAAAAATCGGggttacctcttatctagt
gtagagtgtagcacctatccACCTTGATTGTGGTGATAGGATCACAGGTGTTTACATATGggttacctcttatctagt
gtagagtgtagcacctatccATATAACCAATATTTTTCTCCCTCTGCTATGTACCAGATGggttacctcttatctagtg
gtagagtgtagcacctatccCCCTTTGATCATTTTATAGGTGGGAAACAGGCATCTCAAAggttacctcttatctagt
gtagagtgtagcacctatccGGATATGAATTTAACTCCTTATGTCGTTTCCACCAGACCAggttacctcttatctagt
gtagagtgtagcacctatccTTACTTCTCACTGGTGTACAGCTGGTGATTAATAAATGCTggttacctcttatctagtg
gtagagtgtagcacctatccGATGAAATCACATGGCTATCTACCCAGGGAAATTATAATAggttacctcttatctagt
gtagagtgtagcacctatccGAGATTAAATTTTATTGAGAACTATGTGCCTGGCATTGCAggttacctcttatctagtg
gtagagtgtagcacctatccGAAATAAAATCCAGACAATTTAAGTAACTTCTCCAAGGCCggttacctcttatctagt
gtagagtgtagcacctatccCAGCCTATTTTCTGTCTTATCCATAGTGAGTCTATTCTATggttacctcttatctagtg
gtagagtgtagcacctatccTGTCCTCAAATTCACTGGTTCAGTCCTCTATCATCTCCATggttacctcttatctagt
gtagagtgtagcacctatccAGCCCATCCAGCTAGGTTTTACTTTGGTTACTGAATTTTTggttacctcttatctagtg
gtagagtgtagcacctatccTGACTGCTATAAAATATTTGTCTGATCACTTCCACATCTGggttacctcttatctagtg
gtagagtgtagcacctatccATGTTGTGATCTTCCTGGTTTTTAGAATGACACATGACTTggttacctcttatctagtg
gtagagtgtagcacctatccATGAAACTCTACACCCTATTGAATCTTCTGTTTTTAGCAGggttacctcttatctagtg
gtagagtgtagcacctatccAAGTGCAGTGCAAGGGTTGGGTGGGTGTGTACTTCCAAGTggttacctcttatctag
gtagagtgtagcacctatccGGTTGTTCCTATTCTGCTTCCTCTTCGGCTCTGGTGACACggttacctcttatctagt
gtagagtgtagcacctatccACAGGACACCAACTCCCACCCCACTGGGCTCCACTGACACggttacctcttatct
gtagagtgtagcacctatccGCACCAACTAGCCTGCCTCCCATCACATAATTCTGTCTTGggttacctcttatctag
gtagagtgtagcacctatccGGATGCTCAGCTCCCCACTTGGCCCCACTAACACCAGAAAggttacctcttatcta
gtagagtgtagcacctatccGCATGTACTTAAGTCTCATTGCTGCTGGGTCTAGATGGAGggttacctcttatctagt
gtagagtgtagcacctatccAGGAGAATTGCACAGCCTGGTTTCATTTGGTCCTGCTAGCggttacctcttatctag
gtagagtgtagcacctatccCAGCAGGGGGTAAGGGGTAGAGGGGCATTTTTCCATTGGTggttacctcttatctag
gtagagtgtagcacctatccGTACTGCCAAAATGCTTCCTGGTCCTTTGCCTAGAAACAGggttacctcttatctag
gtagagtgtagcacctatccGAGCTTTTTATGTCTGTGCCTGTTTGTGGCTCTGGTTTAAggttacctcttatctagtg
gtagagtgtagcacctatccTGTGCAAGATACTCAGGAGGAAATAAGAGAGCCCAGGAGCggttacctcttatcta
gtagagtgtagcacctatccGCAGCTCGGCCTTCTTTCGCTTTTCAAAGACTTCCAATGCggttacctcttatctag
gtagagtgtagcacctatccGTTGTATTAAGCCCAAGATTTTTAGTTGTAAAGGACCTACggttacctcttatctagtg
gtagagtgtagcacctatccGGAGGAACCAGAACGATCTAAGGTCTTATAAACTCATAAAggttacctcttatctag
gtagagtgtagcacctatccTTTGAACCCAGGATGTCTAGTCACAGGAACTGTTTTTTGTggttacctcttatctagtg
gtagagtgtagcacctatccTGTTGGCAGGGTTGGTTTTTGCTGGGGCCTCTCTCCTTGGggttacctcttatctag
gtagagtgtagcacctatccCCTTCTTGCTATGTTCTTATAGGCTTTCCTCCATATGTGTggttacctcttatctagtg
gtagagtgtagcacctatccGCTAGTCACATTGGATTACAGCCCACACAAATAACCTCATggttacctcttatctag
gtagagtgtagcacctatccAAACCTCTTTCTAAATACAGTCTCCTTCTGAAGTACTGGAggttacctcttatctagt

gtagagtgtagcacctatccTGAATTTTTGGAGGACACCACTCAGTGTGTAGCAATATGTggttacctcttatctagtg
gtagagtgtagcacctatccATATATCATCTGAGAAAAGTAGCCTGAATGTACAATTGGCggttacctcttatctagtg
gtagagtgtagcacctatccATTAACCATGACGAACACCCATTTCCCTGGCCATTTTCCTggttacctcttatctagt
gtagagtgtagcacctatccAAAGAGTATAGCTGATTGCATGTATGGAAAAGGGAAAAGCggttacctcttatctagt
gtagagtgtagcacctatccCCCCCTCTAAGCTACTTGGGTTGACAGGAAATCCATTGGCggttacctcttatctag
gtagagtgtagcacctatccCATGGTGTGGCTTCTCCTCTCTTGCCCTGGGAAGGTGACAggttacctcttatctag
gtagagtgtagcacctatccCAGTGCCTCTTAGCTGCTTGTTGGGGTGATTGGTGATCTCggttacctcttatctagt
gtagagtgtagcacctatccTGCGTCATTCCCACAGTCACTCCAAAAGCTCAGTCACCCGggttacctcttatctag
gtagagtgtagcacctatccCAGTTTTCGCAATGTAATAGTAAGATTCAGAAGTAAGCACggttacctcttatctagtg
gtagagtgtagcacctatccGCATGCAGCATGCTACTCTTTTTATAAAGTCAAACATCATggttacctcttatctagtg
gtagagtgtagcacctatccAATATGGGATTCAGAGTGATGGCTGCCTTTGGTGGGTAAAggttacctcttatctagt
gtagagtgtagcacctatccAGACAATGAGGTGAGGACAAAGCCATGTAACAGGTGAGGCggttacctcttatctag
gtagagtgtagcacctatccGGTGGGTTCCAAGATAATCACTGCATTATTTGAAATAACAggttacctcttatctagtg
gtagagtgtagcacctatccTTAAATAAATAAAAACGGGGGAAACTGGGTTGCTAAGAGTggttacctcttatctagtg
gtagagtgtagcacctatccAATTGGGATTATGTATTACTTTCTTCTCTCCTTTTCCTCCggttacctcttatctagtgg
gtagagtgtagcacctatccGCTCAAGCCAACAAATAGCAGATCAGGAATCAGTTAGGCAggttacctcttatctag
gtagagtgtagcacctatccGAATTTTCAAGCCTAGAAACTCCCAGCCCTAATCTCAGAGggttacctcttatctag
gtagagtgtagcacctatccCAGGGATGAGGTCTGAGAATCTTTATTTTATGCTGAACTCggttacctcttatctagtg
gtagagtgtagcacctatccAGCCGTCCTGCCTTCAACGTCTTGATATAGTACATCTCATggttacctcttatctagtg
gtagagtgtagcacctatccAATCTTTACTGTTTCACTATTGCTTGCCAAACCTAAATCCggttacctcttatctagtg
gtagagtgtagcacctatccCAAAGTCCTCTGCAATCTGGCACCACCGTAATTACCCAAAggttacctcttatctag
gtagagtgtagcacctatccCTTACATGTCTTCCATAGTCATAGCTAGATACAATCAAGCggttacctcttatctagtg
gtagagtgtagcacctatccATGGGAGTTCATTAGCGCCTCTGGCCTATTTTCCTGTGGCggttacctcttatctagt
gtagagtgtagcacctatccAAATCCTTCTCTCTCCTACACCTCTTTAGATTTGGCAAATggttacctcttatctagtg
gtagagtgtagcacctatccGCCAGATTCAACTCTTTCCTTCCCATATGAGCCATTTCTGggttacctcttatctagtg
gtagagtgtagcacctatccCATGCTGCTAATAAAGACTACTCGGGACTGGGTAATTTATggttacctcttatctagtg
gtagagtgtagcacctatccGGCAGGGAAGAGAGCTTGTGCGGAGGAACTCCCATTTTTAggttacctcttatctag
gtagagtgtagcacctatccCACTACCAGGAGAACAATATAGGTGAAATTGCCCCCTTGAggttacctcttatctag
gtagagtgtagcacctatccCCAAACCATATCACCAGCCCAATAGCCCCCAAATGAACTGggttacctcttatctag
gtagagtgtagcacctatccACTTCTTTCTTTACAGATAGTAAGCCTTCATGGAATCACAggttacctcttatctagtg
gtagagtgtagcacctatccTTTTTCTTCCCCCTCATCTGATTGTGATTTGGATTATGTTggttacctcttatctagtgg
gtagagtgtagcacctatccCTGTTGGAAGTATGACACCCACCATTGATCCTATTCATTTggttacctcttatctagtg
gtagagtgtagcacctatccAAACTAGACCAGGAAGAGATTTGAGACAGTGAAGAAAAACggttacctcttatctag
gtagagtgtagcacctatccAGCAGAGAAAGCCAATGGTCAAATAAAGAAAAGACAACATggttacctcttatctag
gtagagtgtagcacctatccGACTGGAGAGAACATTCTGGTGCTGTATGAGTCCCTGCCTggttacctcttatctag
gtagagtgtagcacctatccCACCTCAGTCCCAGTGAATAAATTTCCCCCTTTCAGTTTAggttacctcttatctagtg
gtagagtgtagcacctatccGCTCCATTCTGCATAGGGCTGACTCACACACCTGGCCATTggttacctcttatctag
gtagagtgtagcacctatccCAGTTACTAGAAGGAATAATAGGAAGGTGATGACTCCCAAggttacctcttatctagt
gtagagtgtagcacctatccAGTGAGTGTCCTATGATGCTGGCTGCTAGAGATTTCAAACggttacctcttatctagt
gtagagtgtagcacctatccTGTTGCTCTCCACAGACATCCAAGGGAAAGGGGTTGATCTggttacctcttatctag
gtagagtgtagcacctatccGGCTCAGCTCACTCCCTCTTTGCATTCCCCAGATTCTCCCggttacctcttatctag
gtagagtgtagcacctatccGGACACTCCCAACATCCTCAGGACTACACTTTTTGATCTCggttacctcttatctagt
gtagagtgtagcacctatccAATGCGGGGTTATATTTTAACGTATTCTGTCAGGTTTGGCggttacctcttatctagtg
gtagagtgtagcacctatccAGAGAAAGAACTATCTCCTTAAAATCTGACTTCAAGGAACggttacctcttatctagtg
gtagagtgtagcacctatccATTCAAACACATATCAGGTCATGCTCTTCTTTCGCGATTTggttacctcttatctagtg
gtagagtgtagcacctatccATCAGCACATTTACATAGCATACAACCTCATTCTTAAAGGggttacctcttatctagtg
gtagagtgtagcacctatccTCATGGAGCAATGAGAATTCTGGGCAAATTTTAGATATCTggttacctcttatctagtg
gtagagtgtagcacctatccATTTGCTCTGTGACCAATTTAAGGATTTTACATTCGCACCggttacctcttatctagtg
gtagagtgtagcacctatccTGTCCCAAGCCACACTGAGATCGTAAAGAAATGTTCCTAGggttacctcttatctag
gtagagtgtagcacctatccCACTGAATCTACAAAGTGCACATATTTTTGCCTGCCAAGCggttacctcttatctagt
gtagagtgtagcacctatccGACTCCCCTTCTGTCAGTGCTATGAATTCTGTGGTGCCAGggttacctcttatctagt
gtagagtgtagcacctatccCCTGCAGGTCCCATGGAAAAACTAAAACAATAGGAAGTCAggttacctcttatctag

gtagagtgtagcacctatccACACATTCAGTTGTTGGCAGTCCTTATTTTCAGAACCACAggttacctcttatctagtg
gtagagtgtagcacctatccCAGCCAATGTTTTTACTGCACAGTACGTTTTCCTCTTTGTggttacctcttatctagtg
gtagagtgtagcacctatccCAGAGAGAAGAACCACAACTGAGGAGCAAAAAAGGGGGAAggttacctcttatcta
gtagagtgtagcacctatccATGCAAATTGAAGAAAAAAACAGCAGTCTTTGGACCTCAGggttacctcttatctagt
gtagagtgtagcacctatccTGGATTCAATGCCAAATGTGTATTTAGAGATACCCTGAAAggttacctcttatctagtg
gtagagtgtagcacctatccTTTTGCCTTTGTTAGAAGAGCTGTTTTGCAATATGGTTTGggttacctcttatctagtgg
gtagagtgtagcacctatccCTAATTCAGTTATTAAGGACAAAAGCTACCAGCCAATCCAggttacctcttatctagt
gtagagtgtagcacctatccCATTAAAAATATTTGCAGTGAAGGATGGGAAATGGAACTCggttacctcttatctagtg
gtagagtgtagcacctatccATGAATGTAGATGAAGCCTCAAGCCTTGTCTGAAGTCATTggttacctcttatctagtg
gtagagtgtagcacctatccGGGGGAGGTTTACTTGCTCTAATAGATTTGGCCTACTTGGggttacctcttatctagtg
gtagagtgtagcacctatccCATGTCACCAATATGATTACCTAGAGTACAAAGCCATCCTggttacctcttatctagt
gtagagtgtagcacctatccCACATGATGAAAGGAGAAATTAACTCAGACAGTGGTACAGggttacctcttatctag
gtagagtgtagcacctatccAAATCTTCGTACTCAATTGTTTTCATTGGAGAGGAGACGGggttacctcttatctagtg
gtagagtgtagcacctatccTCTGGAGATAGTTTTGTTGTCACCACTAGGACAAAGAAAGggttacctcttatctagtg
gtagagtgtagcacctatccTGGGTAGAAGCCAGAGATTTGGTTAAATATCTTACAATGCggttacctcttatctagtg
gtagagtgtagcacctatccGCCCCCCACAACAAAGAACTAACCAGGCAAAACTGCCAGCggttacctcttatcta
gtagagtgtagcacctatccATTGCCAGATCCTTTCAAAAATTGAGCTAACTAAAAGCAGggttacctcttatctagtg
gtagagtgtagcacctatccAATTGGTTTTATCTCATCTGAGGAAAGTGCTTAGGGTAAGggttacctcttatctagtg
gtagagtgtagcacctatccCTCAGAGGGCGAGAAGAAAATGAAAAGAGCAGTTACTGGAggttacctcttatctag
gtagagtgtagcacctatccCAAAAACCAATTACCCACAAGATCCTTATTAATGATGAGCggttacctcttatctagtg
gtagagtgtagcacctatccTCTTTGTTCTAAAAGATGACTTGGCTCCAGGATTAATATCggttacctcttatctagtg
gtagagtgtagcacctatccCAACAGGTATATAAAACAATACCCATCACTCATCAGAGAAggttacctcttatctagt
gtagagtgtagcacctatccGAAATATCATCTCACCCCAGCTAGAATGGCTATTATAAAAggttacctcttatctagtg
gtagagtgtagcacctatccAAACATACAATTACATATGATCCAGCAATTCCACTACTGGggttacctcttatctagtg
gtagagtgtagcacctatccGAATACTATTCAGCCATAATTGTCAGAACATGGATGAGCCggttacctcttatctagt
gtagagtgtagcacctatccAGTGGTTACTAAAGACTGTGAAAGATAGTGAGGAGGGGCTggttacctcttatctag
gtagagtgtagcacctatccAGAAAGATAGGAAAAATAAGTAGTTCTGCGTCCCTTAGCAggttacctcttatctagt
gtagagtgtagcacctatccAAGCTAGAATAGAGGATTTTAAATGTCCCCAACGCAAATAggttacctcttatctagtg
gtagagtgtagcacctatccAAGTGATTAAACATGAAAAAACAGATGGCTCCAATCCTCTggttacctcttatctagtg
gtagagtgtagcacctatccCTCTTGAGGCTTCTTTCTGGACATTTATATCCTAGAACTTggttacctcttatctagtgg
gtagagtgtagcacctatccAATTCTGTAATTTTTTTGCCCATGAGACAGCACTACGAGAggttacctcttatctagtg
gtagagtgtagcacctatccTCAGCCAATTTAGTAGAGAATTCACATGATATAAGCAGTCggttacctcttatctagtg
gtagagtgtagcacctatccGCCGGACTAGCATTTCTCATCATATTCCCTTAGTAGAGCAggttacctcttatctagt
gtagagtgtagcacctatccATGTGTTGCCCCAAAAAGTCATACAGTCAAATAAGATAACggttacctcttatctagtg
gtagagtgtagcacctatccATACTGACTATGATCCTGCACCTTGCCAAATGTTTTATTCggttacctcttatctagtg
gtagagtgtagcacctatccGCCAGTGAGATGGGTCTTATTAATCCCCTTGCACAGCTGGggttacctcttatctag
gtagagtgtagcacctatccCCTATTGTACAGGGAAGTAACTTGGCCAGGTTTCATTGCTggttacctcttatctagtg
gtagagtgtagcacctatccCCAAGTCTGTCCTGTATTCAAACCTGGTGCTCTTAAATACggttacctcttatctagtg
gtagagtgtagcacctatccACCCTCTGTCTTAGTCTATTCCTACTACTATAACAAAACAggttacctcttatctagtg
gtagagtgtagcacctatccCTGTGTCTGGTGGGGGCCTGATCTATAGATGGCATTGTTTggttacctcttatctagtg
gtagagtgtagcacctatccTCTTTACATAATGGAAAAGGCAGAAGGGACAAATGTTATGggttacctcttatctagtg
gtagagtgtagcacctatccACATGGAAGAAGAGGTGGAAGGACAGAAGGGGTGAACAGCggttacctcttatcta
gtagagtgtagcacctatccCTCCCAAAGGCCCCACACTTGAATACCATCAAGTTGGTGAggttacctcttatctag
gtagagtgtagcacctatccCATTCAGACCATAGCACCCTTTAACAGGCTGTTTCTAATTggttacctcttatctagtg
gtagagtgtagcacctatccACTTATTTCTCCCATGCACTCTTTTTATCATGAAACAGCCggttacctcttatctagtg
gtagagtgtagcacctatccGAAAAAAATCTGAAGAATATCTGACACCTCCTAGTTGTTCggttacctcttatctagtg
gtagagtgtagcacctatccATGTAAGTGTCATTGGGCTGATCAGAAAGGTACAAATCCTggttacctcttatctagtg
gtagagtgtagcacctatccAAACCAAGTACACAATTTTCTCCCTATAAGCTTCATGTTGggttacctcttatctagtg
gtagagtgtagcacctatccCTGGAGAGTATTAAAAGCTGTGGGACTATTTTCCTTCCTGggttacctcttatctagtg
gtagagtgtagcacctatccACAAAAGACATTAAAATAAACTGCTAAGGGAGGTTTGGTGggttacctcttatctagtg
gtagagtgtagcacctatccTTAAGGCCAAAAATGAAGTGTTACTGTTGAATGAATTCCCggttacctcttatctagtg
gtagagtgtagcacctatccATATTTAATTGGGAGCATAACTCTGAGAAGAATCAGACTCggttacctcttatctagtg

gtagagtgtagcacctatccGGAAAGTCAAGTAAAGAGCATTATATCCCATAGGGCACATggttacctcttatctag
gtagagtgtagcacctatccCAAGAGGAGAAACAGTCAGCAAAATAAAACCAGATGTGAAggttacctcttatctag
gtagagtgtagcacctatccTATGATTATACCATCCCTTCCAATATGAACAGCACTTTCAggttacctcttatctagtg
gtagagtgtagcacctatccTGAATCACTGAATCCTTTTCGAGTAAGTGAAACATTGAGCggttacctcttatctagt
gtagagtgtagcacctatccAGACCATGTATTCGAGAAGAAGAAATTATATAAACAGCCCggttacctcttatctagt
gtagagtgtagcacctatccCTTTCAGTCAAGTCTACTAGTTCTAATCCCAACATGCTAAggttacctcttatctagt
gtagagtgtagcacctatccTTTTCTGTAAAATAAGTTATACCCTTTTTGACCCTGCCACggttacctcttatctagtg
gtagagtgtagcacctatccGTGACCGACACATTTGGTTATTTCCATTGAATATATTCCTggttacctcttatctagtg
gtagagtgtagcacctatccTGGTCCCACAATAAGAGCGAACGTTTCTAACATATATTCAggttacctcttatctagt
gtagagtgtagcacctatccTTCAAATACAAATAGGGTAACAGTGCAAAGTTGTCTGATCggttacctcttatctagt
gtagagtgtagcacctatccTGCATAATGTCTGTACCTGAATATTGCAATGTGGTGAGAGggttacctcttatctagt
gtagagtgtagcacctatccCTATGCTGTCACAATTTGGGGAGAACATGAAGGGGAATTTggttacctcttatctagt
gtagagtgtagcacctatccCTCACTGTGCATGTGTGTTCTGTCACTATAGTGTGGTACTggttacctcttatctagt
gtagagtgtagcacctatccCAAACCACTTTCTCAAAATCACAAGTTCTCAGACTTTAAGggttacctcttatctagt
gtagagtgtagcacctatccAAGCCATCTAGTGCCTTCATCCAGCCCTCATCCTCTCCAAggttacctcttatctag
gtagagtgtagcacctatccACTTCTTAAATGTTGGGCAGAACAGCTCTCTCTAATGGGGggttacctcttatctagt
gtagagtgtagcacctatccAGAGGTAGTTGCATCTCACATTGTGTTTCTGCTCCTAGAAggttacctcttatctagt
gtagagtgtagcacctatccGTATCAAAAGGTACCTCTCCTCTGCTTTCTCTATCTCCAGggttacctcttatctagt
gtagagtgtagcacctatccGGAAGATGCTTGAATCATAATAAATCCTTTTTCCTCCCTGggttacctcttatctagtg
gtagagtgtagcacctatccGAATTCTGTTCCCTCTTCCCCATTTCCCTCCTACTGCCTCggttacctcttatctagt
gtagagtgtagcacctatccGATTGTGTAACTCTCCCCATTTTAGTCTCCCTGGCTCCAAggttacctcttatctagt
gtagagtgtagcacctatccTGCATGTCACATCCATTCTAATCACACTAAAAGACACCTAggttacctcttatctagt
gtagagtgtagcacctatccCCTATCATGTAAAAGGTACCCATACTTGGCCATATTTTGCggttacctcttatctagt
gtagagtgtagcacctatccTTTAATAAAAGAAGCAGTAGTCACAAAGAAGAACCGCAGCggttacctcttatctag
gtagagtgtagcacctatccGCTTATCAGTTCTAAGCTGTTAGTTGACAGTAAACATATGggttacctcttatctagtg
gtagagtgtagcacctatccGGCAAAGATTTAAAAGGCCCCATAAAAATACCCGTATATTggttacctcttatctagt
gtagagtgtagcacctatccCCTCATACCTGGGTAAAATCAAAGCACATGTTATTAGATGggttacctcttatctagt
gtagagtgtagcacctatccAGAAGTTAATCTGCCATTTCCCCCATCATTAATAAAGCATggttacctcttatctagt
gtagagtgtagcacctatccACTTTTATGGCTGTCTACCTACTCAGATGATTGGGGGTTTggttacctcttatctagtg
gtagagtgtagcacctatccAGAGTTTTCTAGTAAGGATAAATGCACTGGAGAGAAAAACggttacctcttatctagt
gtagagtgtagcacctatccCATATGAATTGTCTCTATCATGAACAGCATCATACTAGAGggttacctcttatctagt
gtagagtgtagcacctatccTGGATTCTGTTAGTCTTCAGTCACTTTTATGAAACAAAGCggttacctcttatctagtg
gtagagtgtagcacctatccCCAAATAAACTCTCCTTTTCTTGGCTCTTGCAATTTTCAAggttacctcttatctagtg
gtagagtgtagcacctatccAAGAGGTACAACTTTGGAGTTTCTTAACAAGTTCCCAAAAggttacctcttatctagt
gtagagtgtagcacctatccTGATATACCAGAGTACACATCAGCAGAGTATCTTAAACAAggttacctcttatctagt
gtagagtgtagcacctatccATCCATAGTACATTCTAATTTACAGTGCTGCAAAACAAGGggttacctcttatctagt
gtagagtgtagcacctatccTTATGAGTAACAAGTGATTCCAGTTAGGTCCCAGGCCTAAggttacctcttatctagt
gtagagtgtagcacctatccAGCTCAAAGAACTGACACATCCTCTAAAAACACAAACTATggttacctcttatctag
gtagagtgtagcacctatccTGCATGTTTATAAGTGTTCATGTAAGTGTGGAGCAATAAGggttacctcttatctagtg
gtagagtgtagcacctatccTTTGGTTCCAATTTGCATTTTTCATTGTCCTTCCACACTTggttacctcttatctagtgg
gtagagtgtagcacctatccGCCGTAGCTCAGTGTTGCCAGCAGCCCTTGAATAAGTTTTggttacctcttatctag
gtagagtgtagcacctatccACTTGGTGAAGGTTAAGAACATTCTTTTTAATGGGCCCATggttacctcttatctagtg
gtagagtgtagcacctatccAATCAAACAGAACACATATTTTGCTGGACATATCACAAGCggttacctcttatctagt
gtagagtgtagcacctatccCTTGTGAGGATAGATATGTGGACAAGTCACTGACCTGTTTggttacctcttatctagt
gtagagtgtagcacctatccAGTTTCATCCTCTGTTGTAAAATACAAAAAATGGACAGGGggttacctcttatctagt
gtagagtgtagcacctatccAGCTGCTCTGAAGGCTGAGGAAGGGAGGATTACTTAAGCCggttacctcttatcta
gtagagtgtagcacctatccTCTTCTACCTTCTGCCATAAAATTGTTTTTTCATCATCCCggttacctcttatctagtg
gtagagtgtagcacctatccTTCAGATACAGCAGAAAACTTGAACAGAATGAAATGGGGAggttacctcttatctag
gtagagtgtagcacctatccAGACATTCTTAAACATGGCATTTGAACATCTCTTTCCTACggttacctcttatctagtg
gtagagtgtagcacctatccTGCAACTGAATGAGGACAATCTTTCTATGGTAGATAAAGAggttacctcttatctagt
gtagagtgtagcacctatccCCTACAAAATGCTAGCAAGATAGGGGAGGAAATTATAAGAggttacctcttatctag
gtagagtgtagcacctatccGCATCTCTCCATACTGAACATAATAGTGCAACATTAGGCAggttacctcttatctagt

gtagagtgtagcacctatccGGTGTTGCCTTTATGAAAATCTACATGCATTTAGCCCTATggttacctcttatctagtg
gtagagtgtagcacctatccTACCAGGATCTGTAATTCATTGATTAAGGATTCATCTCTCggttacctcttatctagtg
gtagagtgtagcacctatccTGGTTTAAGGGTCCTAGTACAATATATGTCAATACAGGGCggttacctcttatctagtg
gtagagtgtagcacctatccTTAGGTCCTTCAATTATAAATGACATTTGCCACTCTAGTCggttacctcttatctagtg
gtagagtgtagcacctatccTGGGGTCCAAGGTCAAATACAAAAGATTCCTTATTCCAAAggttacctcttatctagtg
gtagagtgtagcacctatccGTGCAGTTTTGTTTCCACCTTTAGCAAATATTTCCATTGTggttacctcttatctagtgg
gtagagtgtagcacctatccTATTGAGTATGTTCTATGTACCAGATAGTGTGTTCAACTCggttacctcttatctagtgg
gtagagtgtagcacctatccTGACATACATAAAGGAGTATTTAAGCATGGACACCAAAGTggttacctcttatctagtg
gtagagtgtagcacctatccGCTCATGGGAAAATTACCTAACCTTGCTAAGCCTCAGATTggttacctcttatctagt
gtagagtgtagcacctatccAATATCTATAAGCTTGTTGTGGAAAAGATGAGAACCATGGggttacctcttatctagtg
gtagagtgtagcacctatccAGCAATTTACCAAGATCACACACAGCTAGACCACGGAAGAggttacctcttatctag
gtagagtgtagcacctatccACTCAAACCAAGTTCTTCTAATTCCAAGCTTTCTAATTCCggttacctcttatctagtg
gtagagtgtagcacctatccCATGATCTCTGAGGGTCACTAGGATGGCATGCAGATAGAAggttacctcttatctag
gtagagtgtagcacctatccGATGAAGACACCATCACAAACAGTGTTTCTATTCCTGGGAggttacctcttatctagt
gtagagtgtagcacctatccGGGTGACTTAAAGAAGACAAACCCTATTTAACTCTTAGAAggttacctcttatctagtg
gtagagtgtagcacctatccCTGATCAAGAGTTTCCAACCACCAGTGCATGACATACCACggttacctcttatctag
gtagagtgtagcacctatccATAACCCTACAGTTTATGACAGAAGATGAAGATACATTGCggttacctcttatctagtg
gtagagtgtagcacctatccAGTTGGAATGGGTTGTCCTGTCCCTTAGTTGTGAGCCTCAggttacctcttatctagt
gtagagtgtagcacctatccGTGACCACTCTATTTAAACCACTTGACTTATGCTGAGTTAggttacctcttatctagtg
gtagagtgtagcacctatccTGAATCAAATTGCCACAGTGACCTTCAACTATAACTCAGAggttacctcttatctagtg
gtagagtgtagcacctatccTTCACCCCTGCCTATTTTCTCCTAAACTCTCAACATCTAGggttacctcttatctagtg
gtagagtgtagcacctatccTTCAATTTGACATGCATTTGGATCCCACCCATATAAAGCAggttacctcttatctagtg
gtagagtgtagcacctatccCCCAAAGCCTCACAGATCAGACATGTCAACTCAAGGGAAAggttacctcttatctag
gtagagtgtagcacctatccCCACAGCAAAAGGAATGTTAGTTATTTTAAAAGGCTCTGTggttacctcttatctagtg
gtagagtgtagcacctatccTGACCCAGATTTAATGCAGACCTTCCACCTTCATCATAGAggttacctcttatctagt
gtagagtgtagcacctatccAAGTCTCATTTTTTGTAAGACGTTTGCTTGCATCAACAGCggttacctcttatctagtg
gtagagtgtagcacctatccACTGCAGTGTCTTCTCTAAAATTCTTGGTTGTTTGATCTCggttacctcttatctagtgg
gtagagtgtagcacctatccGGGGCTACTCTCCTAACTCTCTCCATGTTCTCACACCCTCggttacctcttatctag
gtagagtgtagcacctatccCACACATATACGCCACCCTGAAAGTCACTCTTCTCAATGGggttacctcttatctag
gtagagtgtagcacctatccCTAGAGAGTAGGGCTTGCACATGAAAAAGTAAAATGGCTAggttacctcttatctagt
gtagagtgtagcacctatccAGACCTCTCTTCCCACTCTACCTCTAATGTTGCAAATAATggttacctcttatctagtg
gtagagtgtagcacctatccCCTGAGGCTACTCCCACTTTAAAGCTGTCTCTGCCCTACAggttacctcttatctag
gtagagtgtagcacctatccATGCTTTCTTATTTTCTTCAGTGTGCATTTCCAAGGGACTggttacctcttatctagtgg
gtagagtgtagcacctatccTCCTCTCCATCCATCTCATTCACTTGCTGCCACTACAATTggttacctcttatctagtg
gtagagtgtagcacctatccCCTCACCTGGAACTCTGTGCCTTCACTCTTTACTTCCTCCggttacctcttatctagt
gtagagtgtagcacctatccGTCTATTGCTTTAACCACCTTTTGCCAACATTCTTGAATTggttacctcttatctagtgg
gtagagtgtagcacctatccCAGCCTCATATCAATCTGCCTTGTCTGTGCCTAGAACCAGggttacctcttatctag
gtagagtgtagcacctatccCAGATGCTTGTATTTGTACCTCTATGAGTCATTGCGTTATggttacctcttatctagtgg
gtagagtgtagcacctatccGTTCAGCCATCTCTTTTTTTCTAACTCAACAGCAGCTATTggttacctcttatctagtg
gtagagtgtagcacctatccTCTTCTCCTTTCTCCCTAAACCTCTGATCTGTTCCTTCCCggttacctcttatctagtg
gtagagtgtagcacctatccGTTCCTGCCACAAAACCTACTTACTTTTCCCCTTCTGCTAggttacctcttatctagtg
gtagagtgtagcacctatccATTTGGCAAATACTTAGATGGTGCTTACTACCTTCCAGGCggttacctcttatctagtg
gtagagtgtagcacctatccCTCCAAGACAACATTAGAAGCACTTTAAATACAGTAACTCggttacctcttatctagtg
gtagagtgtagcacctatccCAAGAAACTGAGGCACAAGAACATTTGCTAATGTATCCAAggttacctcttatctagt
gtagagtgtagcacctatccGACAAGCAACCTTCTCCAAAATATATGTTCTTAAGCATTCggttacctcttatctagtg
gtagagtgtagcacctatccGCATAGAGAAGCTCAGTAGTTTCCCCAGTTCATGCAGCTGggttacctcttatctag
gtagagtgtagcacctatccCCTATGCTTTTCCCACTAGGCTGCCACTGACTTCTCCACCggttacctcttatctag
gtagagtgtagcacctatccGTTTTTCAGGAACCTCTCTTCTATAGCATCAGCATATATCggttacctcttatctagtg
gtagagtgtagcacctatccCATCAGTCTCCCATCAAAAAATTGTTTTAGAAAGCTCTGCggttacctcttatctagtg
gtagagtgtagcacctatccTTCTGTGTGGATAGAACTCACGTTTGGTTTAGTCTTTGTTggttacctcttatctagtgg
gtagagtgtagcacctatccTGTAAAGTGAATGAAGGAGGGGACATGGTCTAAGGAGAAAggttacctcttatctag
gtagagtgtagcacctatccGGTGAACCTAAGGACTACAAATATTTTACACATGAGCACCggttacctcttatctagt

gtagagtgtagcacctatccTGATAAAGATGGTATAAAGTCAGAATTCCCCACCACCACCggttacctcttatctag
gtagagtgtagcacctatccCATTCCTGGAAATCAACTAGAAGTACCAAGGAGAATAAGAggttacctcttatctag
gtagagtgtagcacctatccCTTTCAAAAGCATAGAGAAGAGTAATAGGTCACTAAGACTggttacctcttatctagt
gtagagtgtagcacctatccAAAGTAGGCTTTTTCAAAGTGAAGGGTACATGAGGTATACggttacctcttatctagt
gtagagtgtagcacctatccGACAGCATTTTCTCTATACCTATTCTCATCTAAAAGAAGCggttacctcttatctagtg
gtagagtgtagcacctatccCCTATAATTGGAGACAGGGAGAAGAGAGGTGTTACATTATggttacctcttatctagt
gtagagtgtagcacctatccCCTATCTATCTGTTTTGGTCCAGAGTAAAGGCAAAATAAGggttacctcttatctagt
gtagagtgtagcacctatccTAATAAGAAAAAAATCCATCCAGACTAGAATGCAACACGCggttacctcttatctag
gtagagtgtagcacctatccTGTTGGCCCAGGAAAAATCCATCTCTTTCAACATGAGAAAggttacctcttatctagt
gtagagtgtagcacctatccATGGAATGCATACTAAGAGATAGGTTCACAGGTCATACATggttacctcttatctagt
gtagagtgtagcacctatccTTATGATTCACATACAAAAAGGATGTACAAAAGGTGCCTCggttacctcttatctagt
gtagagtgtagcacctatccGGAAAAATGATAGCAGAAAAAGAATGTCACCAGAAAAAGGggttacctcttatctag
gtagagtgtagcacctatccGGTAAAAATTTGGCCATAAAGTGTAATGCTCTATCAAGTGggttacctcttatctagt
gtagagtgtagcacctatccACAATAGTCGGAAGTGAAACAGGAGCTCTCAAATATCAGAggttacctcttatctag
gtagagtgtagcacctatccGAAAGTGACAACAAAACTGTAAAGAACACAAGGGTAAATCggttacctcttatctag
gtagagtgtagcacctatccAGGTAAAGGAGACTGCAATTATCTGAATGTTTGTGTACCCggttacctcttatctagt
gtagagtgtagcacctatccCAAGGTAATAGTATTAGGGTTTGGGACTTTTGGGTGTTCCggttacctcttatctagt
gtagagtgtagcacctatccCCACTATAAGAGGACTGTAAAAGATAGCCATCTATAAACCggttacctcttatctag
gtagagtgtagcacctatccCTGTGAGAAATAAACTTCTGGGGGTTTTGGGGGGGGGGGTggttacctcttatctag
gtagagtgtagcacctatccCCCAGGTTCACACCTTTCTCCTGCCTCATCCTCCCAAGTAggttacctcttatctag
gtagagtgtagcacctatccGCTACCCAATTCATGCTATTTTGTTATAAGACCCCAAATAggttacctcttatctagtg
gtagagtgtagcacctatccAAGATAACAGAAAAACTGGCAATGTTTATTATCCCCATGGggttacctcttatctagt
gtagagtgtagcacctatccATAATGTAGAGAATATAATGTCTAGGCTAGGAAGATGGCTggttacctcttatctagt
gtagagtgtagcacctatccAAAAACTCAGGACCGAAGAGACTAAAAGAATTGACCAAGTggttacctcttatctag
gtagagtgtagcacctatccTGAAACTCCAAGTGCTCATAATAATATAAGACTTGCCACGggttacctcttatctagt
gtagagtgtagcacctatccCACTAGAGATGTACTCTCTTTTGGATAGTGACATCTTTGCggttacctcttatctagtg
gtagagtgtagcacctatccGTTTCTGTGGGTCAAAAACATGAAAGTGGCTTTGCTGGGCggttacctcttatctag
gtagagtgtagcacctatccAGCTGTTAATAAGGGCCTTAGACATCTGAAAACCTGACGGggttacctcttatctag
gtagagtgtagcacctatccAGAATCCACTTTTAAGGCATTTTACTCACAGGTTGGTTCTggttacctcttatctagtg
gtagagtgtagcacctatccCAGTCTTGTTTCTTCCATATATGGGCATCTCCACAGGCTGggttacctcttatctagt
gtagagtgtagcacctatccCACCTGGCTTCCCACAGAATGTGTCCGAAAGAACAAGGTAggttacctcttatcta
gtagagtgtagcacctatccGCCCTTGCCTCAAAAGTCACATGCCATCGCTTCAGATCATggttacctcttatctag
gtagagtgtagcacctatccGCCACCTATGCCAGCTTTTTTCTATAAAGTGAAGTTTAGAggttacctcttatctagtg
gtagagtgtagcacctatccAGACTGTATAGTTTGAACTACCTTCTGTAAAGCAATTCAGggttacctcttatctagtg
gtagagtgtagcacctatccGTTAATTAAACCCATTGATGTTAAGATGCACCTTTTCCACggttacctcttatctagtg
gtagagtgtagcacctatccATGTCTTATAGTAGTTGGCATATTATTATCCCCATCAGCTggttacctcttatctagtg
gtagagtgtagcacctatccATCAGTTAAGTACAAAGGAGTTGTCATTACCATGCATGCAggttacctcttatctagt
gtagagtgtagcacctatccACATTGTTGGTACCACACATGTTGATTTAGCTGCTATTTAggttacctcttatctagtg
gtagagtgtagcacctatccCCACAAAGGTATAAAACTAGAACTCAGTAATAGGAGGAACggttacctcttatctag
gtagagtgtagcacctatccCAGAAACACAACATGCCAAATCTTATGAGCTGCAGCAGCAggttacctcttatcta
gtagagtgtagcacctatccAGATTTCAATTGCCCTAACTTCACACCTGCAAGGACTAGAggttacctcttatctag
gtagagtgtagcacctatccGAACAGAAATAAACGGCATAGAGACTAGAAAGATAATAGGggttacctcttatctag
gtagagtgtagcacctatccCTTTAGCCAACTTAACCAAGCAAAAAAGGAGAGAAGCCTCggttacctcttatctag
gtagagtgtagcacctatccAGAAACGTACAACCTACCAACATGCAATCATGAAGAAATAggttacctcttatctag
gtagagtgtagcacctatccAGCTTGGAACTATATGGCTTCAGGGGTGAATTCTACTAAGggttacctcttatctagt
gtagagtgtagcacctatccGAATACTTTCAAACTTACCATATGATACCAGCACTACCCTggttacctcttatctagt
gtagagtgtagcacctatccATCATGCACCCTGATCATGTGGGATTTATCCGAGATGCAAggttacctcttatctag
gtagagtgtagcacctatccCATATGCAGATCAATAAACACGATACGCCACATTAACAGAggttacctcttatctag
gtagagtgtagcacctatccAAACTAAAAGCTTTTTTTCTGAGACCCAAAACAAGACCAGggttacctcttatctagt
gtagagtgtagcacctatccCACTCTCAGCACTTCGGTTTGATATAGTACTAGAAATCCTggttacctcttatctagt
gtagagtgtagcacctatccTCAAATTGGAATGGAAGAAGCTAAGTGGTCTCTGTTTGCTggttacctcttatctagt
gtagagtgtagcacctatccCTAGTAAGCAAATTCAGTAAACTGACAGGATAGGAAATCAggttacctcttatctagt

gtagagtgtagcacctatccCCAAGGAGGTGAAAGATCTCTACACTAAAACTATAAAACAggttacctcttatctagt
gtagagtgtagcacctatccTTTGTATGGAACCACAAAATTCCCCCAAGTAGCTATCTTGggttacctcttatctagtg
gtagagtgtagcacctatccTTCTCTGACTTCAAAATCTCTTACAGAGCTACAGTAATCAggttacctcttatctagtg
gtagagtgtagcacctatccTGACCAAGATGCCAAGATCATACAATGAAGAAAAGAAAAGggttacctcttatctag
gtagagtgtagcacctatccCCACATGCAAAGTAATGAAATTAGACACTTGTCTTAAACCggttacctcttatctagtg
gtagagtgtagcacctatccGATTCCAATGAACTAAAAACTTCTGCAAAGGAAACAATCGggttacctcttatctagt
gtagagtgtagcacctatccCCAATTTACATATGGGTAAGGAGCCTGAATAGACATTTTTggttacctcttatctagtg
gtagagtgtagcacctatccCATTACTGGTCATCAGAGAAGGGCAAATTAAAACCACAATggttacctcttatctagt
gtagagtgtagcacctatccTGATTATTATGAAACAGACAACAGATAACAAGTGCTGGCAggttacctcttatctagtg
gtagagtgtagcacctatccGACCCAGCAGTCTCACTTCCGGCTATATATTCAAAGGAAAggttacctcttatctag
gtagagtgtagcacctatccAATAGCATTGATAGGAAATCGACCTAAGTGTTTATCAGTGggttacctcttatctagtg
gtagagtgtagcacctatccTTTGTAACAATATGAATGAACGTGGACGTTGCACCAAGTGggttacctcttatctagtg
gtagagtgtagcacctatccGCCAGACACAAAGACAAATAGTGAATGATCTCTTATCTGTggttacctcttatctagtg
gtagagtgtagcacctatccGTAGAGTGGTTACCAGGGACTAGTAGATGAGGGAAATGGAggttacctcttatctag
gtagagtgtagcacctatccACTTTCAGTTATGAGTTGAACAATCTCTGAGGATCTAATGggttacctcttatctagtg
gtagagtgtagcacctatccTAAAATGTCTTCAAATGTAACGCTGTGATTCAGCATTGAGggttacctcttatctagtg
gtagagtgtagcacctatccTGTACACACAAAAAAAGCATGTAAACAGAGACACAGTGGGggttacctcttatctag
gtagagtgtagcacctatccTGTTGATGAAGCAAATATTCACCATAGGACAAATGATCACggttacctcttatctagtg
gtagagtgtagcacctatccATGATGCTGTGCTACATTTTGTTTTCTGAGGATTATGCAAggttacctcttatctagtgg
gtagagtgtagcacctatccTATTGCCAAATCCCTGAACTAGATGAAAGAAATGTCAAAGggttacctcttatctagtg
gtagagtgtagcacctatccAGGCTGGTGTGACCAACAAACATATCATGCAAAGTTATCAggttacctcttatctagt
gtagagtgtagcacctatccTATGTAAAATAGTAAGTGCCCCAAAGTCAATGACATCTGAggttacctcttatctagtg
gtagagtgtagcacctatccTGTTAGAAGTCTTCACCACAATACAATACATGCATGTAAGggttacctcttatctagtg
gtagagtgtagcacctatccAAAACAAAAATGAAAATGTACCCTCCAGGCAACTTTAACCggttacctcttatctagt
gtagagtgtagcacctatccAGTCCGTTTGTATTTCAAATTCTTCCTTTTCTATGAGAGGggttacctcttatctagtgg
gtagagtgtagcacctatccCAAGCTTGTGTTCCTAGATTGGGCCAGTGCATGTCAACCCggttacctcttatctag
gtagagtgtagcacctatccTGATCTTCAATTATACTTTCTTGGCCTTTGCTATTCTAGCggttacctcttatctagtgg
gtagagtgtagcacctatccTCATCTTTCTTAATGGTTATACTTATTTCCAGTCACTGCCggttacctcttatctagtgg
gtagagtgtagcacctatccCCATTGTGGTATCTTTTGAGTTCTTTCCCATATGACAGAAggttacctcttatctagtg
gtagagtgtagcacctatccAAACATCTTTTCCTGGGTATATTAGTGATCAAAGGGCCTTggttacctcttatctagtg
gtagagtgtagcacctatccCACATTTTGAGCCATGGACTTTCAGATTATACCTTTCAGAggttacctcttatctagtg
gtagagtgtagcacctatccAGCAAAAGGTGATGATGGACAGAAATAGAAGACATGGGGTggttacctcttatctag
gtagagtgtagcacctatccCAGTCTTTTCTCTCTGCCCTGGATCCAAGTTTTACTGCATggttacctcttatctagtg
gtagagtgtagcacctatccTTTCTCCTCTTTTGTCTCTCTCACTCTTCTACTCTCAAAAggttacctcttatctagtgg
gtagagtgtagcacctatccGGGTTGCAGATGGGAAGCAGGGTGAACCCCAACATTGCCTggttacctcttatcta
gtagagtgtagcacctatccGCTGACAGCCCTCCCTCAAACCAGCATCTCTGTCTCTGCTggttacctcttatctag
gtagagtgtagcacctatccCACGTTTCTGCCCTTGTGCCTCATGCAGTCTTCTGTCACTggttacctcttatctagt
gtagagtgtagcacctatccTGGCCAGAGTATCAGAGTACCACCAATCGATTCCACAAATggttacctcttatctag
gtagagtgtagcacctatccGCGCTATCAATCAGCCAAGAGCCTCAATTAAAATTTGATTggttacctcttatctagtg
gtagagtgtagcacctatccCTGTTTTCACTTCAGTCCCTCCTACAAGTCAATTCATTTGggttacctcttatctagtg
gtagagtgtagcacctatccCATGTTCAACTCCCTATTTATGCCCTATCCCACCCCCAGCggttacctcttatctag
gtagagtgtagcacctatccAATAAATAAGACCTGGCATGAATGCTACTCATCTACAACTggttacctcttatctagtg
gtagagtgtagcacctatccGTTCAGGACTTATTTTTCTTGGAGCTAGAAGTTTTCCAGAggttacctcttatctagtg
gtagagtgtagcacctatccCATGGTGGAATAAGATAGGATGCAGAGGATGTCATAGAAAggttacctcttatctagt
gtagagtgtagcacctatccACTCGCCTCAAATCAGTGACACGACTCATCAGGTCTGCACggttacctcttatctag
gtagagtgtagcacctatccGCAGAGTATACAGAGTACAAATTTGTAGGAAGGGGGAGCAggttacctcttatctag
gtagagtgtagcacctatccAACACTATCTTGTCACTGTTACATTAAGACCTAGACCTTCggttacctcttatctagtg
gtagagtgtagcacctatccTAGTAAAATAAGGCTACATTTCATTGCTCCATCATGTGTGggttacctcttatctagtg
gtagagtgtagcacctatccAAAATTACTGGAGAAAGGAAGAAATCACATTCCAAGCTCAggttacctcttatctagt
gtagagtgtagcacctatccCACCAAGAGCTGCAAATTTGGCATTTCTGCACAGGAGGAAggttacctcttatctag
gtagagtgtagcacctatccGAACTTCCATCTTTATTGCTCTCTGTATCTTGTGAGTATTggttacctcttatctagtgg
gtagagtgtagcacctatccACTTACATAAATTCTCTTCTCAGCTAAAGTACTATGCTCCggttacctcttatctagtg

gtagagtgtagcacctatccGATTAATATACTCACAATCTGTAGAAATGGGAAAGAGGggttacctcttatctagt
gtagagtgtagcacctatccCTGGATTTCATCTGTGTCCCATTCTCTCAGCACTTTTCAGggttacctcttatctagtg
gtagagtgtagcacctatccCTGCCTCACAGCCACAGATGATACACAGGCCTTGGATCCTggttacctcttatctag
gtagagtgtagcacctatccTCATGTTTCCTACTTGGCAGGCTCCCAATTACCAATGGCAggttacctcttatctagt
gtagagtgtagcacctatccGAATACGTGAACTAAACAAAGCCAAGCACTTCAAATACTTggttacctcttatctagt
gtagagtgtagcacctatccAAGAAGGGACATACTAGACTGTTTTCAATTTTACCTCTTCggttacctcttatctagtg
gtagagtgtagcacctatccCCAACTGTGTGTCAAAGGAAGACAACGTGGGCTGGATAATggttacctcttatctag
gtagagtgtagcacctatccTTATGTAACTGAGTCACTTCCACTTGGGGGATGAGGGCTTggttacctcttatctagt
gtagagtgtagcacctatccCATGATAAATGCACGTTAATGCGAGAGGACAAAATGAAAAggttacctcttatctagt
gtagagtgtagcacctatccGCTTTGAATTGACTTTCTTTCACCTGCCTCTAAACTACGTggttacctcttatctagtg
gtagagtgtagcacctatccACTGCTGTTGTCGCTACTCATTCCAGGCTGATGTCTCTCAggttacctcttatctagt
gtagagtgtagcacctatccCAGCACCAAATCTTAGACCCTTTTCTTTTAATTCTCTATCggttacctcttatctagtg
gtagagtgtagcacctatccCATTCCACATTGTCCAGCTCCGACGGTTCCACTTCAAATGggttacctcttatctag
gtagagtgtagcacctatccACCATATTTTTGCAGAATACTCAATGATACAGGAAGTGTCggttacctcttatctagtg
gtagagtgtagcacctatccTTTTGAGACAGTCTGTCTCTGTCACGCAGGCTAGAGTGTAggttacctcttatctagt
gtagagtgtagcacctatccCAGGCACACACCACCATACCTAGCAAATTTTTGTATTTTTggttacctcttatctagtg
gtagagtgtagcacctatccCCACGCCTGGCCTCATCATATAATATTACATAAAAAACATggttacctcttatctagtg
gtagagtgtagcacctatccTTTTTGACACGGAGTCTCGCTCTCTTGCCCAGGCTGGAGTggttacctcttatctag
gtagagtgtagcacctatccAGCCTCCCAAGTATATGGGACTGCAAGTGCACACCACCACggttacctcttatcta
gtagagtgtagcacctatccCCACCATCCCTGGCCCAAAATATCGGTCTCAATCATTTATggttacctcttatctagt
gtagagtgtagcacctatccTCATCTTTTCCAACTACTATACAGATGTTGGATATTCCTCggttacctcttatctagtg
gtagagtgtagcacctatccCCTGCATTTTATCATCTATTTGTACCATATTCTACTCCACggttacctcttatctagtg
gtagagtgtagcacctatccCCAAATCCCACGTGGTTCAACATTGTTTCCGGCTTCCAGGggttacctcttatctag
gtagagtgtagcacctatccCACATCTTGTGGCAGGAAGCAGAATGAGCCAGGACTATCTggttacctcttatctag
gtagagtgtagcacctatccCCAGATATAATACAGGGACCTTGAAAATTCCTCTTAGCTCggttacctcttatctagtg
gtagagtgtagcacctatccGCCTCCTTCTCCCCATATAAAAACTGAAAGATTAGGATGAggttacctcttatctagt
gtagagtgtagcacctatccGCATAAAATTTCTACTTTTCATCAGCTGCTAGATCCGACTggttacctcttatctagtg
gtagagtgtagcacctatccGCAAGTTCTAATTGCCATGAGTGTGAGGTTACTGCAACGTggttacctcttatctagt
gtagagtgtagcacctatccGGAACTGGTGAAAGCTGAAAGTCATAAATATTCAGGAAATggttacctcttatctagtg
gtagagtgtagcacctatccCAGATGTATATGCAACCCAAATATCTGTAGTATGGAAAACggttacctcttatctagtg
gtagagtgtagcacctatccGAGGGGGAGAGTGCTGAAAAAAGAGAAAGGCAAAATGGGCggttacctcttatcta
gtagagtgtagcacctatccGTAATATACCAGACATAAATAAGGAGTTAGCTGTCTACCCggttacctcttatctagtg
gtagagtgtagcacctatccCATCTGAATCATCAATTGGCTCAAGTATAAATACCTGGAAggttacctcttatctagtg
gtagagtgtagcacctatccACAATTAATCTTAGAAAGTATAACTCAACCTGGGTGGAAGggttacctcttatctagtg
gtagagtgtagcacctatccATAAAAGTCAAACCATGCAGATGTACACCAAGTAAACATCggttacctcttatctagt
gtagagtgtagcacctatccGTTCCTCCTTATCTCATCCTCTCCCATCCCACTCTTCCAAggttacctcttatctagt
gtagagtgtagcacctatccTGTACATATCTTCACATTGTTTCTGTCTCTTTCACATGGCggttacctcttatctagtg
gtagagtgtagcacctatccTAGATGTCTTTGTCCTGTCAGATTTAACCAAAAACTTGACggttacctcttatctagtg
gtagagtgtagcacctatccTAAAGGAAAAAATGTTACTAGGACAGTAAGAACAGCACAGggttacctcttatctagt
gtagagtgtagcacctatccCCTTCTAAACATAGGCATCTGCAGACTTCGATTAGCCCAGggttacctcttatctag
gtagagtgtagcacctatccCCCCCAGGTTAATTATCATTTGAAGATATAATCACTCCAGggttacctcttatctagtg
gtagagtgtagcacctatccGCCTTGGGAATTAGCATCACAACTCTTCCTTCTCACTGCCggttacctcttatctagt
gtagagtgtagcacctatccCTGAGGGTTTGGAAAGCAAGAATCAGGATACTTTGGAGGTggttacctcttatctagt
gtagagtgtagcacctatccAGATGGCTCTTCAAAGATCAGACTCCTAAGGTTTAATGTTggttacctcttatctagtg
gtagagtgtagcacctatccAGTTTAAATGCTAACATGACTCTGTATTACTGGCAAAACCggttacctcttatctagtg
gtagagtgtagcacctatccGAAGATTTTGCAGAGATTGCTTGTATATATAGCCAATACGggttacctcttatctagtg
gtagagtgtagcacctatccTCAGATGGCTCAGTTTGAAAGAATTATTATATTGGCTCTGggttacctcttatctagtg
gtagagtgtagcacctatccGCTTGATATTCTAAGCACCTTACATGAGGTAACCAATTTAggttacctcttatctagtg
gtagagtgtagcacctatccGGAAACTAAAGAACAGAGAGCTTGTCAGCTTCATCTTCACggttacctcttatctagt
gtagagtgtagcacctatccAAATGCAGCAGTTTTAGCCTTCACACCTTTATGTCTCTTTggttacctcttatctagtg
gtagagtgtagcacctatccGAAAAAGAAAGTCTTCTCTGGATTCACATTCTCATTAGCAggttacctcttatctagtg
gtagagtgtagcacctatccCTGGGATATGACTAGAACAAGCCAATCAGAGCCTACACCTggttacctcttatctag

gtagagtgtagcacctatccGGCATCAACACCACCTAAGCTTCCTAGCTGAGAAGGGAGGggttacctcttatcta
gtagagtgtagcacctatccCCAAGGTGTTAGGAGAAGGGAAGTAATGGATACTGGGGAGggttacctcttatctag
gtagagtgtagcacctatccTCTGCTCTAGAACTCATTAGCCAGAAACAAAGAAACTGAAggttacctcttatctagt
gtagagtgtagcacctatccTTGGCATTATTTGTTCTGAAACAGTATGTCTTACATGCAGggttacctcttatctagtgg
gtagagtgtagcacctatccTTTAAAGAAAACCCAAATTTATAGTCTATCCTGTCTGCCCggttacctcttatctagtg
gtagagtgtagcacctatccAAACTACAACAGGTTATCATACATGCTAGGCACTGTGAGCggttacctcttatctagt
gtagagtgtagcacctatccAGATGGTCATTTTTGTCAGGAATGTTTATCATGCAGAACTggttacctcttatctagtg
gtagagtgtagcacctatccCTTTCTATGTAGTCGTCTTTTATGTATTTGTTACTTCGCCggttacctcttatctagtgg
gtagagtgtagcacctatccCTTACATGCAAGCAAAACACATGAGTGAAATGAAGAACTCggttacctcttatctagt
gtagagtgtagcacctatccGCTACCCAATTTAAAATATGTTCTCAAATGAAGATGCCACggttacctcttatctagtg
gtagagtgtagcacctatccGGGAACACACTTTAAAGTTTAATGACCTTCACTGGCAGCAggttacctcttatctagt
gtagagtgtagcacctatccACTCAATTTAATCCATGAAGATTTGTTAAGGTAGAGTGCCggttacctcttatctagtg
gtagagtgtagcacctatccGATAGAAGATATCCTCTACATTAGCCTTCACAAACAAGTAggttacctcttatctagtg
gtagagtgtagcacctatccTATGTATTTAAATAGCTGCACTAGGAAAAAATGGCATGCCggttacctcttatctagtg
gtagagtgtagcacctatccTGTAAACTCTATCCTCAACCTTACCTCTCCAGAATACATGggttacctcttatctagtg
gtagagtgtagcacctatccGCCTTCTGCTAGTTTTCCACAGCTATGATAAATCGATGTCggttacctcttatctagtg
gtagagtgtagcacctatccCCAGTGGACTCTAGAAAAATAGTGGTTTATCATGGGAAGCggttacctcttatctagt
gtagagtgtagcacctatccCCAGAAGATTTCACATTAAGCTGTGTTTGAATGGAAAAGAggttacctcttatctagtg
gtagagtgtagcacctatccCTGAAAGAGTCAACATAACAGATATCAACTCACAATCGGGggttacctcttatctag
gtagagtgtagcacctatccCTGCCCATAATCTAAATATACCAGCAACCATTCCCAGGCCggttacctcttatctag
gtagagtgtagcacctatccTGCTATTGAGAAGTATCATTCTATCAAAGCCTAAGGAGCCggttacctcttatctagtg
gtagagtgtagcacctatccAGGTCTGAGCCTGAACAAGCTGCAAAACATTTCAAACCTCggttacctcttatctag
gtagagtgtagcacctatccGGTGAATTAAAGCTCCAGAGAAGGCAAATTAGAGGGGTGAggttacctcttatctag
gtagagtgtagcacctatccATCCTCTCATTTAATGCAAGAATCCCTCTACAACAGTACAggttacctcttatctagtg
gtagagtgtagcacctatccGGCTGCTCCAACAATGCTTGTCAAGTTTTAAATACTGGGAggttacctcttatctagtg
gtagagtgtagcacctatccCTCCTCATACTAAGTCAAAACCTCCTCTCTGTCATTCCTAggttacctcttatctagtg
gtagagtgtagcacctatccAGGAAGACAGCTATCATGTTGCCTACTCAGTCATGTCATCggttacctcttatctagt
gtagagtgtagcacctatccGGACATCCTACAGCCTGAATATGTTTTGCATCTTTCTCCTggttacctcttatctagtg
gtagagtgtagcacctatccTCTAGAATTTCCAAAAAACATACCTGCTTCCTTTTCCTACggttacctcttatctagtg
gtagagtgtagcacctatccACAGGTGTTTATTCCACTTTGTAGTCTCTCAGAATTGAATggttacctcttatctagtgg
gtagagtgtagcacctatccCCATTTATACTGCTTTGTGTCTGTTTAAACTATGCTCAACggttacctcttatctagtgg
gtagagtgtagcacctatccCTCTACCCTTGAGTCTCCCTATATCCAATCTCCTCCATTCggttacctcttatctagtg
gtagagtgtagcacctatccCAAATGCCTCTCTATTCTACAACATCTAATTGATCCTGCCggttacctcttatctagtg
gtagagtgtagcacctatccCTGGAGAAGCAGCGTCATGTGAGATGGTAAAATACTATAGggttacctcttatctagt
gtagagtgtagcacctatccGTACATTCTGTGCTTCTCTTAACACCTGATCATTTTCTAAggttacctcttatctagtgg
gtagagtgtagcacctatccCATTTATTATGCCCTAGGTGACTAAGTGATGCTTTTACAAggttacctcttatctagtg
gtagagtgtagcacctatccGATGCTGTTTTGTTTTTGTGTAATTTTGTGGCAGTTTCACggttacctcttatctagtgg
gtagagtgtagcacctatccCGGAGAGATTGGAGTAATGATCAACTATGAAACACAAGACggttacctcttatctag
gtagagtgtagcacctatccCCAAACACCATGCTGCTCTGTCAGGACATTTCTATGTTGTggttacctcttatctagtg
gtagagtgtagcacctatccCAAAAAGTGGAATCCTAACCTCCTCTGAAAAATCTGAGTTggttacctcttatctagtg
gtagagtgtagcacctatccCCCAGGTCAGAACAAAGTGTCCAACATAGAGCTAGCTTCTggttacctcttatctag
gtagagtgtagcacctatccGCACCTTCGCATCTCTTAACTCAGTTATACTTCAGAACGTggttacctcttatctagtg
gtagagtgtagcacctatccACTGAGGCTTGGAGACTTGTGACGTGATGAAGCTCACCTAggttacctcttatctag
gtagagtgtagcacctatccCAACAATTGATTCAGTATACTACATTCTTCAGCGTTTGTCggttacctcttatctagtg
gtagagtgtagcacctatccAGTTAACTACTTTTTGTTTTGCTTCGGTTTTTGAGACAGGggttacctcttatctagtgg
gtagagtgtagcacctatccCTGTGGCACAAACACAGCTCCTTGAAGCCTGAACCTCCCAggttacctcttatctag
gtagagtgtagcacctatccTTTTTGTGGTGGGTGGAGGGGGTAGAGATGGGATCTCCCTggttacctcttatctag
gtagagtgtagcacctatccCCATGCCTGGCCATAAAGTTAACTTCTGCTTTTACTTGCCggttacctcttatctagtg
gtagagtgtagcacctatccTTTTGACAAGTGATATAAGCAGAAGAAACAGTGAGAGAGGggttacctcttatctagt
gtagagtgtagcacctatccTTCCTAAGATGGCTGCAAACATATGTCCTAAATGACACAGggttacctcttatctagt
gtagagtgtagcacctatccACTGTGGTAGTGAGAGGCCATTATATGCTGCTGATGACACggttacctcttatctagt
gtagagtgtagcacctatccGCCACACCTCCTTCTTCTTCTTGCTGCAGAACCCTGACTCggttacctcttatctag

gtagagtgtagcacctatccAGGACCCTCGAAATACTGGATCTTGAAATAATAACCAATTggttacctcttatctagtg
gtagagtgtagcacctatccTTTCTTGTTTTTTGAAAGCATGGGGAAGTTTCCCGCTGAAggttacctcttatctagtg
gtagagtgtagcacctatccAAAAAGAAAGGAAAAAAATGCATGGAGGTAGGGGGACAGggttacctcttatctag
gtagagtgtagcacctatccAGATATTTCTGAACCTTCATACCACTTACTCTGGGATTAAggttacctcttatctagtg
gtagagtgtagcacctatccTTCTAATCTCTGCTTTGCCATTTTCTTTATAATTGTGGGCggttacctcttatctagtgg
gtagagtgtagcacctatccAAAATGGTGATAATATTAGCCTACTCCAGGGAGATTCTGTggttacctcttatctagtg
gtagagtgtagcacctatccAGCCCAGCTTTAGCAACCACCCCCATAGACAGAGGCTTCTggttacctcttatctag
gtagagtgtagcacctatccGGAGTAAGAGGGAGCAGAGGGAGTGGAACGTGGCAGAAggttacctcttatcta
gtagagtgtagcacctatccAGAGAGAAGCCCTGCCCACTTTATCCAAGGTAAAACATCCggttacctcttatctag
gtagagtgtagcacctatccTATCTCTTTCTAACACAAGCTCTCTCCCATGATTAAGTCCggttacctcttatctagtg
gtagagtgtagcacctatccGTCCAGCAGTTTCTCCACCAGACCACATCCAGTTCCCAGAggttacctcttatctag
gtagagtgtagcacctatccTTTGTTTGAAAAGTCCACATTTCTATTTATGCACTGCTGCggttacctcttatctagtgg
gtagagtgtagcacctatccAAACTACACAGCAGGGCCAGGTGTGATGGCTCACACCTGTggttacctcttatctag
gtagagtgtagcacctatccATGGTGGCAGGTGCCTATAATTATCCCAGCTACTCGGGAGggttacctcttatctag
gtagagtgtagcacctatccCTGCACTCCAGCCTTGGCAAAAGAACAAAACTCCATCTCAggttacctcttatctag
gtagagtgtagcacctatccAAATGCCAGGTGCGGTGGCTCACACCGTTAATCCCAGCACggttacctcttatcta
gtagagtgtagcacctatccACAAACAAAAAGACTACACAACGTATGTTAATGCTATGGCggttacctcttatctagt
gtagagtgtagcacctatccTGGAGACCAAACACATTACTATTCACTTCCTGCTTTAGTTggttacctcttatctagtg
gtagagtgtagcacctatccCTCATAAAGCCCCGTATCACTAAGACAACAGCTGAGAGTAggttacctcttatctag
gtagagtgtagcacctatccGTGATAAAGTAATATTCACCATTGGACAACTGAGGAAAACggttacctcttatctagtg
gtagagtgtagcacctatccCTAACTTCAGCTGAACACTATCGTGAGAGAATTGTAAAAAggttacctcttatctagtg
gtagagtgtagcacctatccCTACTAGTAGCTTTGAAATGTCCACTCTTATACTAAAAGCggttacctcttatctagtg
gtagagtgtagcacctatccCAGAGAATGTATATCAGTAATAACAGATCAAGCCACTAACggttacctcttatctagt
gtagagtgtagcacctatccGGCAGGCCTCTTAAGTGTACTTGTATATCGACTTAAAATGggttacctcttatctagtg
gtagagtgtagcacctatccGACAGAGTTCCACTCTTTTGCCCCAGCTGGCGTGCAGTGGggttacctcttatctag
gtagagtgtagcacctatccCCCACGCCCAGCCTATTTCTATTGAATTTGATCAAAATGTggttacctcttatctagtg
gtagagtgtagcacctatccCCCCTTTTCCTTTCCAACTTCCTGTTTTGTATCATAAGCCggttacctcttatctagtg
gtagagtgtagcacctatccCCTCTACACTTGAGATCCCTTCCCTGAATATGTCCAAAGAggttacctcttatctagt
gtagagtgtagcacctatccACTGTGGAGTGGTTAAATCCAGCTGATTAGCATGTACATCggttacctcttatctagtg
gtagagtgtagcacctatccAGTTATGATTTTTGTGGTGAGAGCTGTAATTTCAACCCCCggttacctcttatctagtg
gtagagtgtagcacctatccCTTTCTCTCTCTATGCATTTTCATCCACCTTTTCAATCATggttacctcttatctagtgg
gtagagtgtagcacctatccAATTAATGATCTGTTTATGTCCAAGTCACTAGTGAAGCACggttacctcttatctagtg
gtagagtgtagcacctatccCAGATGTGTTTTTGATTCAGAGCATTACCCTCGTTCCTTCggttacctcttatctagtg
gtagagtgtagcacctatccCTCCACAAAATAATCTTCAAATAAGTGTTTCATGCCCTTCggttacctcttatctagtg
gtagagtgtagcacctatccCAAATCTAAGACTTTGATCATGAGGTATTCAGCACTATCAggttacctcttatctagtg
gtagagtgtagcacctatccAGATAGTATTTTAAAAATGGCTTAGGCTAGGCACGGTGGCggttacctcttatctagt
gtagagtgtagcacctatccTTTGGGAGGCAAAGACAGGTGGATCACGAGGTCAGGAGATggttacctcttatctag
gtagagtgtagcacctatccATGCCACTGAACTCCATCCTGCGCAACAGAGCCAGACTCTggttacctcttatctag
gtagagtgtagcacctatccGTTGGTGCAAAAGTAACTGGAGTTTGCCATTACTTTTAATggttacctcttatctagtg
gtagagtgtagcacctatccATTTGTGTTTAATATTTTGACAACTCAGCACCCACTTCTCggttacctcttatctagtgg
gtagagtgtagcacctatccATTAAATAGTCTTGCTGCAACAATAAATCCAAGTGCCTACggttacctcttatctagtg
gtagagtgtagcacctatccGGAGATTCCCTTCTTCTTAAATTACCTGTATTAGTCCATTggttacctcttatctagtgg
gtagagtgtagcacctatccTTCGTATGGTGACAGGAAGGTGAATGAGTGCTCAGTGAAGggttacctcttatctag
gtagagtgtagcacctatccCTGGTTCTTCCCACGACACCTGGAGATTACTGGAACTACAggttacctcttatctag
gtagagtgtagcacctatccCAGCAAAACCATATCATTACCCAAAGTCAGTTTCTGTTGCggttacctcttatctagtg
gtagagtgtagcacctatccCCCATGGACCCTGACTGTTACTGTTACTTATGAAATAAACggttacctcttatctagtg
gtagagtgtagcacctatccAGATGCATTCTCTTAATCAGTGAGTAAATAGGATCCAGGGggttacctcttatctagtg
gtagagtgtagcacctatccCAGGGAAATGGCAAGAGCTGAGAAGACATCATTTTAAGACggttacctcttatctag
gtagagtgtagcacctatccTGAATGACAAAGGGGACACAGAGAAGGAAGTTTCACAAGTggttacctcttatctag
gtagagtgtagcacctatccGCACAGCAGAGGTAGTAGAGATAGAGGGGCAATTATAATTggttacctcttatctag
gtagagtgtagcacctatccATTCAGGATTTTATTATGGGGAGAATGGCAGCAAACATATggttacctcttatctagtg
gtagagtgtagcacctatccACATAATCTCCAGACTTGGTGGTGTGTTCTGAAATTTGTTggttacctcttatctagtg

gtagagtgtagcacctatccCACAATCCTGGCTCACTACACCGTTGACCTCCCCAGGCTCggttacctcttatcta
gtagagtgtagcacctatccCCACACCCAGCCTATTGCTATTGAATTTAATCAAAATGTTggttacctcttatctagtg
gtagagtgtagcacctatccTTTAGTTTAGATTTGCAAGTCTTAAAATGACCTAGGCTGGggttacctcttatctagtg
gtagagtgtagcacctatccCCCCTTTTCCTTTCCAACTTGTTTTGTGTCATAAGCCAGAggttacctcttatctagtg
gtagagtgtagcacctatccGATCCCTACACTTGATATATATTTATAACATCTCCAGACCggttacctcttatctagtg
gtagagtgtagcacctatccAATATCGATTGAAATTGGCAACAGGGTTGTTCAGCAAGAAggttacctcttatctagt
gtagagtgtagcacctatccTCATCTCTCCACGTTGAAGAATTTTATATTCTATGCAGCAggttacctcttatctagtg
gtagagtgtagcacctatccGTGATCTTTTTTAATAGCTTCTAACTATCCTGATTTCCGCggttacctcttatctagtg
gtagagtgtagcacctatccCGAAGGAATTTCTTTGCTTAGTTGATACAGGAGCATTAACggttacctcttatctagtg
gtagagtgtagcacctatccAAACTTATAATATCATTTCTCTCCCTGTGCAGTAAGTCTGggttacctcttatctagtg
gtagagtgtagcacctatccTCCTCCCAGAAATCAGTAAAGGTACTATTTTTGTCTTTGCggttacctcttatctagtg
gtagagtgtagcacctatccGTTTTCTCAACCTCCTCAATTCTAAATATCAAAGGCAGCTggttacctcttatctagtg
gtagagtgtagcacctatccACTGCTGGAATGGTGTTAACAGGCTCCTTGGATTCAACTGggttacctcttatctagt
gtagagtgtagcacctatccAACCCACAAAATGCTAGTTCCTCTTCCCCTTCATGCTGCCggttacctcttatctag
gtagagtgtagcacctatccCTGCCACACCCTCCACATCGCTTTCAGAGTTCTTAACAGCggttacctcttatctag
gtagagtgtagcacctatccAAGAGGAGATATTTATAGGAATGTGGAGGGTGTATGAAACggttacctcttatctagt
gtagagtgtagcacctatccCCCCTTTCTAACAGGCTAAAATCTTTTTCAATCTTGCAGGggttacctcttatctagtg
gtagagtgtagcacctatccGCTGTTTGATTGAAGCCCTGCAGTATTGACAAGAAAGGCCggttacctcttatctag
gtagagtgtagcacctatccACGTCAGCATTTGAAAATGAAAAAGAAAAACATAGGCCCCggttacctcttatctag
gtagagtgtagcacctatccGGTTTTTTTAGACGGAGTTTTGCTCTTGGTGTCCAGGCTGggttacctcttatctagtg
gtagagtgtagcacctatccTCACTGCAACCTCCGCCCCTCCTGGTTTCAAGAGATTCTCggttacctcttatctag
gtagagtgtagcacctatccCCACCGTGCCCAGCATCAATACACATTTTTAAAGAAAATAggttacctcttatctagt
gtagagtgtagcacctatccTTTTGAGATGCCAGGAGAAAGAGGGAAGGATGAGAGAGACggttacctcttatcta
gtagagtgtagcacctatccGGAGCAATATTATACCAAAGTAACTTCCCCATCAGTAGACggttacctcttatctagt
gtagagtgtagcacctatccTTTCCATGATACCCACACAAGGCAAAACTGAAACATAGGCggttacctcttatctag
gtagagtgtagcacctatccGCCCAATTGCAACATCACCATCCTTCTAGCACTGTCACAAggttacctcttatctag
gtagagtgtagcacctatccAAATTTTTTCTCTCACCGCTCCAGAGCCTACACATCCAAAggttacctcttatctagt
gtagagtgtagcacctatccCTCTGAGGGAGCATTCATTCCAGGCCTCCCTTCTAGCTTCggttacctcttatctag
gtagagtgtagcacctatccCATCACCCCAATCTCTGTTTCCATCGTCTGTCTCTGTATTggttacctcttatctagtg
gtagagtgtagcacctatccACCTCTTTGTGTAAGGACAGCAGTCATTAAATTAGGGCCCggttacctcttatctagt
gtagagtgtagcacctatccGGGAGACACAATTCAAGCCACAGGACCCTCTATAATAACAggttacctcttatcta
gtagagtgtagcacctatccGATAGTCAGTGTTTATCTTTGCATATTCAGGCCTTTGTAGggttacctcttatctagtg
gtagagtgtagcacctatccCCTCATACTCTAGTACTTTTATAAACACTTCCTCAAGTCCggttacctcttatctagtg
gtagagtgtagcacctatccCCCTAAGAGTTATTCATATAAACTAGCAGGTGTAGAGAGGggttacctcttatctagt
gtagagtgtagcacctatccAGCCATGGGACTTGCTGGGTGTGCTGCCATATTTTGATTTggttacctcttatctagt
gtagagtgtagcacctatccCAGAGTAGAGTACAATTTGCTATGAATAAGAGGTGAGGAAggttacctcttatctagt
gtagagtgtagcacctatccTTTATCTGAATCCAAAGAAAGACTGTCTGTTCACCAGAACggttacctcttatctagt
gtagagtgtagcacctatccAGTTCAAAGAAGTCAATAGGTCTACTCAAAATGGACACCAggttacctcttatctagt
gtagagtgtagcacctatccTGGAAACAGAGAAGACTTTAATGGACACTGTCAAGCACTGggttacctcttatctag
gtagagtgtagcacctatccGTAATTGTGGCTTACTTGTTATGCTGTTCTGAGACTTTCAggttacctcttatctagtg
gtagagtgtagcacctatccAGTGTTGCTGTCAAATCTGTTGATACCTAGAACATGAAAAggttacctcttatctagtg
gtagagtgtagcacctatccTTTGTTTTGAGACAGGGTATGTCTCTGTTGCCTAGGCTGGggttacctcttatctagt
gtagagtgtagcacctatccTAGGACTAGAGGCACTCACCACCATCCCAGCTAATTTTTGggttacctcttatctag
gtagagtgtagcacctatccCCACCGTGCCTGGCTACACTAAGTATAGACCAAATGCCCCggttacctcttatcta
gtagagtgtagcacctatccGCACCACTCCCGCTAAATAATACCTCCAACATTTACAATTggttacctcttatctagt
gtagagtgtagcacctatccATTATTCAGAGTAGCAGAGAACTGTTTTTAATCTACCCAGggttacctcttatctagtg
gtagagtgtagcacctatccTTCACCAAGAAGACTGATCTCAAGAATTATGAAGCTGTCTggttacctcttatctagt
gtagagtgtagcacctatccAAATATTTTTGCCCAGGATATAACCCCCAACAGTTGCCACggttacctcttatctagt
gtagagtgtagcacctatccCTGTTCTGAAGATGTGTGAAGAGCTGTCTGATATAATGCCggttacctcttatctagt
gtagagtgtagcacctatccCCACTGCTACTCCCACACAACAGCTAATATTTTATAAAAAggttacctcttatctagt
gtagagtgtagcacctatccGTATTTTTCCAACCAAGAACACTAGAAATAAGCTCAGGAGggttacctcttatctagt
gtagagtgtagcacctatccTGCCCAAATCATGTCACCTCCTTTGCAGTTGTGCCTGGGGggttacctcttatctag

gtagagtgtagcacctatccCCCTGTGACTCTTCTCCTCGAACTTCCATACAACATGTCTggttacctcttatctagt
gtagagtgtagcacctatccTTGGAAGATGATGGGGGAAGCAACAAGGTAAGGGATCTGCggttacctcttatcta
gtagagtgtagcacctatccCCCAGAGAAAAAAGTCATGTTCAGTGAAGGGAAATGAGGGggttacctcttatcta
gtagagtgtagcacctatccAGCCATGTGTTCTTTTGCTCCAATGATCAACTTCATCCCTggttacctcttatctagt
gtagagtgtagcacctatccGGCCAAGTGAACATCAGGTCAAAAAATAGAATCCCAGGACggttacctcttatcta
gtagagtgtagcacctatccCCCACAGCAAAGCTCTTTAATAAGATAATTCCCCATGCCCggttacctcttatctag
gtagagtgtagcacctatccCTGGGTCAGATGCAGAAATATCATGAACTAAATGTCAATTggttacctcttatctagt
gtagagtgtagcacctatccTTTATAGTTGAGAGGAGGATTCTAGAAATGTAAATGGGGCggttacctcttatctagt
gtagagtgtagcacctatccTCTGCCCTAAACTTCACATCTCTCCTTCAGAAACTTGCTTggttacctcttatctagt
gtagagtgtagcacctatccTCTCCAAAAAGTATAACAGTCAGAACTTCCAGGCCTCTCCggttacctcttatctag
gtagagtgtagcacctatccGCTGCCAATTTAATTATGCTTTTCAGTATTTTTCAGCCTCggttacctcttatctagtg
gtagagtgtagcacctatccAGACTGGTAGCCTTCTCTCTTCTTTCCTGGACACACAGCTggttacctcttatctag
gtagagtgtagcacctatccGTGGTGAGGTACGGCCATGAGACTGAATTCCAACAGATGGggttacctcttatcta
gtagagtgtagcacctatccGTGTGCACCACTCTCAGGTCTGGCCCATGAAACTTCCATGggttacctcttatcta
gtagagtgtagcacctatccTCTTGCTTTTTTCCCATCTGCCAGCCTGGTGCAACAAACAggttacctcttatctagt
gtagagtgtagcacctatccCCTAAATCGTCACCAGGAAGCTCTAATCTAAAAAACAAACggttacctcttatctag
gtagagtgtagcacctatccCTCAAATTTTGTGTTACAGCAGTGATATGGTTTGGATGTGggttacctcttatctagtg
gtagagtgtagcacctatccTGTTTAAAAGTGTGACACCTTCCCCCTACTTCTCTCTTGCggttacctcttatctagt
gtagagtgtagcacctatccCTTTCACCATGTGAGATGCTGCTTCCATTTCGCTTTCTGCggttacctcttatctagt
gtagagtgtagcacctatccAGCCAGTTAAACCTCCTTTCAATATAAATTACCTAGTCCCggttacctcttatctagt
gtagagtgtagcacctatccTGTATTCATCTACATCAATCCACGTAAGTATCCAGGACCCggttacctcttatctagt
gtagagtgtagcacctatccAGTCTTTCTGGCTATGAGTACAACATGGAAATACATACTGggttacctcttatctagt
gtagagtgtagcacctatccAACATTGTTAGTTGCAGGATTAGGATTACATGACTTTCAGggttacctcttatctagtg
gtagagtgtagcacctatccCACCAGGAACATAGCTGTCTCTAGTGAATCTACCTTGACTggttacctcttatctag
gtagagtgtagcacctatccGTGGAGACCGTGGGCTTACTACACTGTTTTTCACTACAGCggttacctcttatctag
gtagagtgtagcacctatccAGTTATGCTTCTGAAAACACTGCAGATGATGTAAAAGAGCggttacctcttatctagt
gtagagtgtagcacctatccAAGGCACCTTTTTCATTGGCAGTCGTGTTTTTGATCCTTTggttacctcttatctagtg
gtagagtgtagcacctatccGGAAATCCTGTGTCACATTCTTCCCTGCTTAAGGCATGACggttacctcttatctag
gtagagtgtagcacctatccTGATTGTTTCTTGGAAACACACCCAGAAATGATTCATTTCggttacctcttatctagtg
gtagagtgtagcacctatccACGTCAAGTATGGCCAGTGGGTGATAAGTCACATTTCACTggttacctcttatctag
gtagagtgtagcacctatccATGAAAAACAGGAATTGGGAGGAGAAGGCAGAGCAAGAGCggttacctcttatcta
gtagagtgtagcacctatccGCATTGACATAGCTGAAGGAAATGTAGTTCAATACTCCGTggttacctcttatctagt
gtagagtgtagcacctatccCCTGATTAGATCTAAAGGAGTTTAAAAGAAGGTGTGGTTAggttacctcttatctagt
gtagagtgtagcacctatccGGCTAAGTCCTTCCTTTTGTAGGCTCTGATATTTGATGTGggttacctcttatctagtg
gtagagtgtagcacctatccCATGGGTGGATTCATCTCTTGGGAATTTAAAGTCAATAGAggttacctcttatctagt
gtagagtgtagcacctatccGGAGTAAACAAGAAAGCCAACATATTCATCTACATCAATCggttacctcttatctagt
gtagagtgtagcacctatccAAGTATCCAAGATCCTCATGTGTTTCTATAGCCAGCCCAGggttacctcttatctag
gtagagtgtagcacctatccTGGAAATACATATGACAGCAGAAGTCCACTTTGTGAGCCTggttacctcttatctag
gtagagtgtagcacctatccATTGCAAGACTGGGATTACCTGACTTTCAGTTCATTCATTggttacctcttatctagtg
gtagagtgtagcacctatccGTCTCTAGGAGTCAATGCATCAAGCTATTGAAGATGTGGGggttacctcttatctag
gtagagtgtagcacctatccTCTTTTGTTTTCCTTCATCCCACACTTATCTGTCCTTTACggttacctcttatctagtg
gtagagtgtagcacctatccCACCCATGCCTGAATGTACCATTTTTTCTCACTCTTCATAggttacctcttatctagtg
gtagagtgtagcacctatccCTTGAGTTGCCTGTTAATTGCATGTGAAATGCCTTATTTCggttacctcttatctagtg
gtagagtgtagcacctatccAAATCTCATGCTGAATTTTATTCCCAATGTTAGAGGAGGAggttacctcttatctagtg
gtagagtgtagcacctatccGGCCTCCCCAGACATGACTACTGTACAGCCTATGGAACTGggttacctcttatcta
gtagagtgtagcacctatccGAGAATGGACTGATACACTTCCTGATATCTGAGCTCAAGGggttacctcttatctag
gtagagtgtagcacctatccCTTTGCCTATAAAAGTTAGTGTTATTTTTCCCCCAGAGAAggttacctcttatctagtg
gtagagtgtagcacctatccTTTTGAAGCAAATGTCATTCATTTGATGGTCTCATCCCCAggttacctcttatctagtg
gtagagtgtagcacctatccAGTTGACTTTCTAAGCACAAATTTCTTGCCAAAGGAAAAGggttacctcttatctagt
gtagagtgtagcacctatccCCAATTAGTACTTGAGACACAATGAGATTGTTGAAAGAACggttacctcttatctagt
gtagagtgtagcacctatccAACACTAAGGTTCTTGACTCTGTTCATCCACTAAAGCACTggttacctcttatctagt
gtagagtgtagcacctatccGAGTTCCTTTTAGCTCTATGTTTTCATGATTCTAAGACACggttacctcttatctagtg

gtagagtgtagcacctatccCAGACATGCCATAGTACCAAAAACTCTTGCTAAGAAGAAAggttacctcttatctag
gtagagtgtagcacctatccAAGAGAAGACCTACCAGTGACTGAGCCTTTACCAAATTCTggttacctcttatctag
gtagagtgtagcacctatccTCACTCAATCCTTAAAACCACACTGTAAGTAGATCCTAGCggttacctcttatctagt
gtagagtgtagcacctatccATTTTACAGATGACAAAGGGAGGTACAGAGAAATTAAGTGggttacctcttatctagt
gtagagtgtagcacctatccTGTTATGGGCTGAATGATTTTGTCCCCCCCACCCACCCCAggttacctcttatctag
gtagagtgtagcacctatccTTGGAAGATAATTAAATTTAGAAGAGGTAATGAGAGGGGGggttacctcttatctagt
gtagagtgtagcacctatccCACTTTCATGCTCTCTCACTCTCTCCTCTCAATCTGTCTCggttacctcttatctagt
gtagagtgtagcacctatccCCATCTGCAGTCCAGGAAAAGATCCCTAACCAGAAACCAAggttacctcttatcta
gtagagtgtagcacctatccCTGTGAGAAATACATGTCTGCTGTCTAAACCACCCAGACTggttacctcttatctag
gtagagtgtagcacctatccTGAACTAAGACACCCAATACCCTACAGTTAGTACAAGGTCggttacctcttatctag
gtagagtgtagcacctatccTTGTACCAAAATATTTGAAGAAATTGTCCCCAGTGCAGCCggttacctcttatctagt
gtagagtgtagcacctatccAGTGCTACTACTCTGAGGGCAGGATTAGAAATTAATTTGTggttacctcttatctagt
gtagagtgtagcacctatccCAGCAAATGACACAGATGGTCTGGTTTGAAGAATAGGCACggttacctcttatctag
gtagagtgtagcacctatccCTTGATCAAAAGGATTTCACTATCCCAGGAGCAGGCAATGggttacctcttatctag
gtagagtgtagcacctatccCTTCCTGTGGCTTTCTTCAAACTGTAAATTCTACTGCATTggttacctcttatctagtg
gtagagtgtagcacctatccTGCTTCAACCTGGTCACAGATGCAACTGGTGAGGCAAGAGggttacctcttatcta
gtagagtgtagcacctatccCCTGGGGAGGGGATAATGATTGAGAAGAGAGAATTGACACggttacctcttatcta
gtagagtgtagcacctatccAGAAGGGAGAAGAAGATAAAGGCAGATGAAGGCAACAGCCggttacctcttatcta
gtagagtgtagcacctatccCCATTACTTACAGCTTACAACCTCCCAAGTCAGGCAGCAGggttacctcttatcta
gtagagtgtagcacctatccGGGAAGGTTATCAGAAGAAGCAGGGAGTGTGTCATGGTGGggttacctcttatcta
gtagagtgtagcacctatccTGGATTCTGTTCTGCTTTTATGGCATCTGTTTCCAATCTAggttacctcttatctagtg
gtagagtgtagcacctatccCTGCTCCTTCCAAATCCAAGTCAGTGCCCAGCTCAGCATCggttacctcttatcta
gtagagtgtagcacctatccCTTGGGACAGGCTGCATTGAGCTCGGAAAGAAAATTGGAAggttacctcttatctag
gtagagtgtagcacctatccAACCATCTAAGTTCACTTCTCTAACAGGAGCAAATTTGGCggttacctcttatctagt
gtagagtgtagcacctatccATGAGCAAAGGTATCATATTCCATAATCCTCCCTGTGTCTggttacctcttatctagt
gtagagtgtagcacctatccAAATGCACTGGAGTAAGAAAGAAATATCTGCACATGTCCTggttacctcttatctagt
gtagagtgtagcacctatccTGACTCCATCTGCTGCTTCACATGCCAAGTGCTAAGGATGggttacctcttatctag
gtagagtgtagcacctatccTATTCTGCACTTTTACCTCCATTGAACATATATCACTAGGggttacctcttatctagtg
gtagagtgtagcacctatccTGGACCACGAAATCTCATTGCCTTAGATGTATCTGTATGCggttacctcttatctagt
gtagagtgtagcacctatccTGGGAAATACAGATAACAAAGGTGCTACATTGATGCAGTTggttacctcttatctagt
gtagagtgtagcacctatccCTAATCCATTACTTCATTTTCCATGTGTTGACCCAGCTCTggttacctcttatctagtg
gtagagtgtagcacctatccTCTTGGGTATGAGATTTATTGTTCCAATGTGGACTGTGAAggttacctcttatctagtg
gtagagtgtagcacctatccTGAGATCATAGGTTCCACTAAGCCCCACTGTCCCAAAACAggttacctcttatctag
gtagagtgtagcacctatccCTAAGGAACCATGCCAAGCTTTGAGCCAACCATGCTGTGTggttacctcttatctag
gtagagtgtagcacctatccTTTCATGCAAGAGTCCTGAAGCAGAGTAGTGGAATTTTCTggttacctcttatctagt
gtagagtgtagcacctatccTAAAGCACCCCAATTCCTGTCCCAGAAACAATGTTTATTGggttacctcttatctagt
gtagagtgtagcacctatccAATGTAAAACTTTCTGCTGCCTCACCATCCCCACCCTCCCggttacctcttatctag
gtagagtgtagcacctatccGGGTTTCCAGTTCCTGCCGCAGACTCTGACTCCTCTCACCggttacctcttatcta
gtagagtgtagcacctatccCCTCAGGGGCTGTCAGGAGAGCAGAGGAACAGAAAAGCACggttacctcttatct
gtagagtgtagcacctatccCTTTGTGTGCAGAGCCACATGGGGCCAGAGTTCTACCTGCggttacctcttatcta
gtagagtgtagcacctatccAGGAACTTCTCTCTGCCTTCTTCGTTTGCTGGTGAGATCAggttacctcttatctagt
gtagagtgtagcacctatccGGGACCCAAAGATGACACAGGACAGAGCCAGGATTAGAATggttacctcttatcta
gtagagtgtagcacctatccATGACCCCATGCCTACTCCTCCTCACTGTCTCCTTGTGCGggttacctcttatctag
gtagagtgtagcacctatccGAGAAAGGAGGAACAGAGGATCTTGGGGTGGAAAAGTTTGggttacctcttatcta
gtagagtgtagcacctatccCAGCTTTCATCTGTTCTGCACACATTCCTCACAAGGCACCggttacctcttatctag
gtagagtgtagcacctatccGCCTCAGCTCACTCCTCTGGTGCAAAGTCCTCTGGATCCCggttacctcttatcta
gtagagtgtagcacctatccAGCTACTGTGGAACCAGGTGGAAACTTATAAAAGGGAAGCggttacctcttatctag
gtagagtgtagcacctatccTGAGCACTGGCAGACAATATGCAGAGATATAGTGTGGAGCggttacctcttatctag
gtagagtgtagcacctatccGGTTACCTACATCTGCAAAGATCCCTGTCCTGCTGCCTCCggttacctcttatctag
gtagagtgtagcacctatccGGGACATAGAGTTCTGTGCCTATGGCAGAAACAATCCAGAggttacctcttatctag
gtagagtgtagcacctatccGATTGTGGATCTACTTGCAGGACAAAGCCTAGATTTTAACggttacctcttatctagt
gtagagtgtagcacctatccTTCATGAAGAATCCAAGCCACCAGTCATTTTTTCCTCTTTggttacctcttatctagtg

gtagagtgtagcacctatccGATTTCTATCATCCCATCATCCAAATAGCGAATATAGTACggttacctcttatctagt
gtagagtgtagcacctatccGTTTTTAAACCTTGTCCCTTTGTCCCCTCCCCTATTTGGggttacctcttatctagtg
gtagagtgtagcacctatccCCGCGCCCGGCCAATCGTTGCCATTTTTATGTCTGTGAATggttacctcttatctag
gtagagtgtagcacctatccTAGTATTCTGTGGTGTATCTGTACCAATTTTATCCAATCCggttacctcttatctagtg
gtagagtgtagcacctatccATAGTGCTGCGATGAACAAAGGAGTGTGCGTGTCTTCTTGggttacctcttatctag
gtagagtgtagcacctatccTATAAGCATTCCCTTCTCTTCGAAGCCTGAATTACATCTGggttacctcttatctagt
gtagagtgtagcacctatccGATAATATCTCATTGCCAAGCCACCGAAGTTTTTGATGTGggttacctcttatctagt
gtagagtgtagcacctatccTTCAGATGTTGTTATTTGTCTTCTCCCTTCTTATGTACCAggttacctcttatctagtgg
gtagagtgtagcacctatccCCTTGTGGTACATGCTAGGGAACTTCAACACATCAGGCAAggttacctcttatctag
gtagagtgtagcacctatccGAAGTGAAAAAAATTCTGATTGCATGATTTGCCTCAGATCggttacctcttatctagt
gtagagtgtagcacctatccTGTTATTCAGACCCTGGCCTGACTCTTTAACCTGGATTCTggttacctcttatctagt
gtagagtgtagcacctatccTAGTACTCCTTCTACCCATCTCAAGTATAGCTGACCAAAAggttacctcttatctagt
gtagagtgtagcacctatccGCTCATTCATCCTCCCTCTAAAGCTGGACACATAGAAACTggttacctcttatctag
gtagagtgtagcacctatccTCTACACCCCGCAATACTCTTCTTAATCATCAAGACTCAGggttacctcttatctagt
gtagagtgtagcacctatccAGATCCTACTACTGTCCTGTATCCCTGAAATACTAACTAGggttacctcttatctagt
gtagagtgtagcacctatccCTTGGGATTTCTAAAGGTTTTCTTCCTGATACTCCAATCAggttacctcttatctagtg
gtagagtgtagcacctatccGCTACTCACACTAGCTGATATCAGGTTGGTTTTCTGTTTCggttacctcttatctagtg
gtagagtgtagcacctatccACCTATAAATGTATGTCCATATAAGTGTGTCTGTCTTCAGggttacctcttatctagtg
gtagagtgtagcacctatccATGAATGTTTGATTTCTACAGTGAAGATGATGCTGATGACggttacctcttatctagtg
gtagagtgtagcacctatccATGGTGACGGTAATGGCGATAATAATGGTTGGATAATGGCggttacctcttatctagt
gtagagtgtagcacctatccTGACCAACAAGCTGCAGTAGAGAACTTAGTAGACATGAGTggttacctcttatctag
gtagagtgtagcacctatccGGTACTGGTTTTCTCTGGTAGTGTATAGACGTGTTTACTGggttacctcttatctagtg
gtagagtgtagcacctatccTGAATGACATGGCAGAAGAGGCTCAGAAGCTGCAGGAGGCggttacctcttatcta
gtagagtgtagcacctatccCTTCAGGAAGAAGCACCTGTTAGCAAACCAAAGCATCGCCggttacctcttatcta
gtagagtgtagcacctatccTGAGTCATAATTCACCAAGCTCAGAGCCAAACTGACAGGCggttacctcttatcta
gtagagtgtagcacctatccGACTACTGGTTTCCCTTATGATGTCCTGAGCATGACTCCAggttacctcttatctagt
gtagagtgtagcacctatccGCTTCCCTGCCTTTGGTGTGTACTCTGTGTCCCATTGTCAggttacctcttatctagt
gtagagtgtagcacctatccTGAGCAGGGTCAGGGTGATAGTTTGGGGCTTCAGTCACTGggttacctcttatctag
gtagagtgtagcacctatccCATAGCCGATGGTCTGAGAATGAAACATGATGGAAATAATggttacctcttatctagt
gtagagtgtagcacctatccATGCACTCAGTCCTATAAATGTATCTTTTCAAATGCTCAGggttacctcttatctagtg
gtagagtgtagcacctatccATAGCTTATGTTATGCGTTTAGTCAATATCTTGCACATCCggttacctcttatctagtg
gtagagtgtagcacctatccCTGATTCCAGCTTGCATTTAGGACATTCATTTTTTATTGCggttacctcttatctagtg
gtagagtgtagcacctatccTGAGTGCTTTCAGGGAGAATAAATAAGGCTTTAAAGGAATggttacctcttatctagt
gtagagtgtagcacctatccACTTTAAGCTAACTTTACATGGTGTGCAATTGTATGTCAGggttacctcttatctagtg
gtagagtgtagcacctatccTCTGGCTTGCTTTAGAAAGGGGACACAGAAATCAATCCATggttacctcttatctag
gtagagtgtagcacctatccTAGCGATGAGAACTGGAAAAGAATAAAAATGTGCTTTTGCggttacctcttatctagt
gtagagtgtagcacctatccGTGAAGACATTTTACTACAACTCAGAAAGGAAACGGTTATggttacctcttatctagt
gtagagtgtagcacctatccGGAATGGGGACATCAAAAGATGTATGACACTTAAATTCAGggttacctcttatctagt
gtagagtgtagcacctatccACAAAAGATAAATAACACTCACATCATTCACTGCATCTGCggttacctcttatctagt
gtagagtgtagcacctatccCTGCTATCTTTTACAATAAAGTACGCATTAGTAACGATGCggttacctcttatctagtg
gtagagtgtagcacctatccATTTGAGGTCAAGTGAAAAGGTTAGTGTGCCAACTTCTAAggttacctcttatctagt
gtagagtgtagcacctatccTGAACTCTGACCATTCCTTTTCTCTGACATCCACATTCAAggttacctcttatctagt
gtagagtgtagcacctatccCCAGTCCTGCCCCATGGTCTTAGAGCTAAAGGAAGAGGAGggttacctcttatcta
gtagagtgtagcacctatccAATTCCAACTCCACCACTTATCAGCTCTATAAATTTGACAggttacctcttatctagt
gtagagtgtagcacctatccGTTCTGTTGGAAGGACACACTGAGTTATATGGAAAGCACTggttacctcttatctagt
gtagagtgtagcacctatccCTACATAAGCCTTAGCTCTTTGACAACAATGGCCTTTTTTggttacctcttatctagtg
gtagagtgtagcacctatccCTTCCTCAGTACCTCCACTTCCACTCCTCTCCAGACCCTAggttacctcttatctag
gtagagtgtagcacctatccCCTCAAGTCGACTGCAACAAATAGGGTTCCAGCTGGGCTCggttacctcttatcta
gtagagtgtagcacctatccATTTCAAATCAAATGTCCTCTCCTTCATGCTACCCACTAGggttacctcttatctagt
gtagagtgtagcacctatccACCTTGGTCTTTATCTATGTTTATGTGTAGTGCAGCTGAAggttacctcttatctagtg
gtagagtgtagcacctatccGCATTCAGGAGACCACTTAGGAAACTGTCATTGGTTAGGCggttacctcttatctag
gtagagtgtagcacctatccGAAAGTTCCAAAGACTGACTGAAAAGAGAAGCTAAGAGAAggttacctcttatctag

gtagagtgtagcacctatccCACGTGAATTTAGAGGGTGGGTTAAGCAGTGAAAAGTAAGggttacctcttatctagt
gtagagtgtagcacctatccATGGGGAAAAGTAGTCAAGCAACATAAAAAAAACTTCAGCggttacctcttatctagt
gtagagtgtagcacctatccCCTAGTGGCAGTGGATGAAGGAAGGACACCGAAAACAAGAggttacctcttatcta
gtagagtgtagcacctatccTTTCTAATATGTGAAAAGGACCCAAAACTGAAGTAATGCCggttacctcttatctagtg
gtagagtgtagcacctatccCTTGTGGTAAGTCGTTTGGGAAGAAATCAAGGAAAGAGGGggttacctcttatctag
gtagagtgtagcacctatccATGTAGAAGTCTAGATGAATGACAGGGAAGAGAGAGGCTGggttacctcttatctag
gtagagtgtagcacctatccAGCCAGAATGAAGACATCAGCGCTCTAGAACTTGCAGAAAggttacctcttatctag
gtagagtgtagcacctatccGGTTTCCCAATATGTCTGTAATAAAAGGCTCAAGGTAGTCggttacctcttatctagtg
gtagagtgtagcacctatccAAATCCCAAAGCTCAGGTGTGTTAGTTCAGGCTCTTCAAGggttacctcttatctagt
gtagagtgtagcacctatccTTGAAGGGTTATGGGATCAAGGATTAAGCTGGAGAAGGTGggttacctcttatctagt
gtagagtgtagcacctatccCTTTGAAAGGAGAGTGGGAAGAAAGAAAAATTGAGTAGACggttacctcttatctagt
gtagagtgtagcacctatccGAGCAGTTCTAAGAAAGATCCAGCCAGGCCAATGAAAAGCggttacctcttatctag
gtagagtgtagcacctatccCTGCATCTCACAGGAATGGGCCAGCAACATGATCAGTTATggttacctcttatctag
gtagagtgtagcacctatccCACTCGGCACAAACATGATGGCAAGCCCAGAAAGCCAGCTggttacctcttatcta
gtagagtgtagcacctatccCAGGACATTGGAGCTGCAGATTCACCACCACCCAGCGTCAggttacctcttatcta
gtagagtgtagcacctatccACAAATTGATCCACCCTACACACCAAATAATTATCCTAAGggttacctcttatctagtg
gtagagtgtagcacctatccAGAATTACAATAGGATCCAGCACCCAAAAGAAGTGAAAGCggttacctcttatctag
gtagagtgtagcacctatccCTCAACACATAAACATACACCCATGCCCATAGAAGGACTAggttacctcttatctag
gtagagtgtagcacctatccCACATGCTACAGCATTGTGCTAAGAGAATTAAACCAGTCAggttacctcttatctagt
gtagagtgtagcacctatccGGACAAATGCTGTATAAGTCTGCTTATATGAGTGTGTTAGggttacctcttatctagtg
gtagagtgtagcacctatccGTACAGGAAGCATAGCCCCTATATCCATTCTGGTGAGGGCggttacctcttatctag
gtagagtgtagcacctatccGCATGTCACACAGTGAGAGAGAAAGCAAGAACCTGGGGAGggttacctcttatcta
gtagagtgtagcacctatccATTCATGAAGGATCGGCCCCTATGACCCAAATGCCTCCCCggttacctcttatctag
gtagagtgtagcacctatccAGGGTACAAACGTCCAAACAATATCACTGAGGTTCCTAGAggttacctcttatctag
gtagagtgtagcacctatccCAGGGGATGGAAGGAGAGGGGAATAGGGAGTCAAGGTTTAggttacctcttatctag
gtagagtgtagcacctatccTAAAGAGTTTCAGTTTGCGAAGGAAAGAAAGGTTGGGAAGggttacctcttatctagt
gtagagtgtagcacctatccGAATATATTTGATAGAGGGGGAAAAAAATAGGAATGCTCCggttacctcttatctagtg
gtagagtgtagcacctatccCTGCTTCTAGATCCCTGATGAAGTCAGCACCGCTACTGACggttacctcttatctag
gtagagtgtagcacctatccCTGGTGGGTTGGTTCCAGGGCTTTCTCTGATTGTGCTCCTggttacctcttatctagt
gtagagtgtagcacctatccAGCAAGTTCTCAGACTCCACCCAGAAGCTTTGTCCTAGAGggttacctcttatctag
gtagagtgtagcacctatccGCCAGCAATTCCCACAAAAGGCCACAAAGTCCAGGTCCCTggttacctcttatcta
gtagagtgtagcacctatccTCCTAGACCTTGGAAACTGGCCCTGATCTTTTGCTCCTTTggttacctcttatctagtg
gtagagtgtagcacctatccATTTTGGTAAATGCATCCAGATGGCTTTCTCCCCTTTAGAggttacctcttatctagtg
gtagagtgtagcacctatccGGCAGAACAAAAAAGTGAGAAAGAAAGCCTGCGAAGCTGTggttacctcttatctag
gtagagtgtagcacctatccCTGCTAATAAAGACATACCCGGGTAATTTACAAAGGAAAGggttacctcttatctagt
gtagagtgtagcacctatccCACAACACATGGGATTTGTGGGAACTACAAGAAGAGATGTggttacctcttatctag
gtagagtgtagcacctatccCCAATTGAATCTTTGGCAAAATGTTGCTGTGGTGGACTGGggttacctcttatctagtg
gtagagtgtagcacctatccGACTCACAGTTCCATGTGGTGCAGGAGGCCTCACAATCATggttacctcttatctag
gtagagtgtagcacctatccTTATTTGTATCACAATAACAGCATAAGAAAGACCAGCCCCggttacctcttatctagt
gtagagtgtagcacctatccCTAGGGTAAGCAATGTGTGTTATCCACAAAAAGGACAGAGggttacctcttatctag
gtagagtgtagcacctatccTGATTCCCACTTCCCTCTCTGCTCCTGCTCTTGCTTCCCAggttacctcttatctagt
gtagagtgtagcacctatccGGACCTTTCTCGCTTTCCACTCTGCTTCAGGCACTAAACCggttacctcttatctag
gtagagtgtagcacctatccGCATTTCCATCTTCCTTTCTGCCTATCTGTCCACTCGCTTggttacctcttatctagtg
gtagagtgtagcacctatccTGAGTTTAAATTTCAGGCTCCACATACTGTGTTCAAGGTAggttacctcttatctagtg
gtagagtgtagcacctatccGGAGCCAGGGAAAAACAAGTCAGCAGTTAATTCTTCAAGCggttacctcttatctag
gtagagtgtagcacctatccTTCTCTCACTAATCTGGAAGCACTGGTTTTTATCTGAATAggttacctcttatctagtg
gtagagtgtagcacctatccGGTACCTTCTGAGACCTATAAATCCTTATGACTACTCCATggttacctcttatctagtg
gtagagtgtagcacctatccATTCTCATTTTGCAGATAGGAAGCCTGTGTAACTCAACTGggttacctcttatctagtg
gtagagtgtagcacctatccTGAACCCAGGCAGACTAGCTTCAGAGACAGTGCTCTTAACggttacctcttatctag
gtagagtgtagcacctatccACGGAGAATGGTTCAATGGAAGAAAGCTTATTAATACTTCggttacctcttatctagtg
gtagagtgtagcacctatccCATGGATGAAGCTAGGGCCCTTTAAGTGTTTTTCCTTTGTggttacctcttatctagtg
gtagagtgtagcacctatccCTGTCAGTAGAGGGCTCTGGAGAAGCATCACAGAAGGAAAggttacctcttatctag

gtagagtgtagcacctatccCTCCTTTAGACTTCATAGCTTCTCCAGGCTCCTCCTACAAggttacctcttatctagt
gtagagtgtagcacctatccGCTTCCCAGAGCCCAGCACCTGCAGTGGAGCAGCTTCTCCggttacctcttatcta
gtagagtgtagcacctatccGGCCAACATCACCCAGCAGCCAGCAGTACCCCCAACAACCggttacctcttatct
gtagagtgtagcacctatccGGTGAGATACCTCCTTGTGAACAGCTGTCTCCAACACCCTggttacctcttatctag
gtagagtgtagcacctatccCTAATTTCTAGATAATTTTGTCAGCACAGTATCACAGGGAggttacctcttatctagtg
gtagagtgtagcacctatccTGCCATCCAGTCAGCCACAGCTATATCCTCTCCAACAAGGggttacctcttatctag
gtagagtgtagcacctatccATCTCAGCCCTTTGATACACTTAGCGTATGTGCATTTTCTggttacctcttatctagtg
gtagagtgtagcacctatccGACCTGGACGTCTACTCAAATATGCTTTACCTCTTGGCTCggttacctcttatctagt
gtagagtgtagcacctatccAATGTAAAGTCTAGTCATGGAAATATGTATGCAGATGAGGggttacctcttatctagtg
gtagagtgtagcacctatccACCTATTTAACTTGCTATACTCCATTCCCAGAAAAGAACCggttacctcttatctagtg
gtagagtgtagcacctatccCAGTAAACCAATTCTTTCTGATGACAATAATGAGGCTGATggttacctcttatctagtg
gtagagtgtagcacctatccAGGTTAGACTTTCCTATAACTGAAATTAGACCCTTAGAAGggttacctcttatctagtg
gtagagtgtagcacctatccATGACAAAGCCATTAGAGACACCCAAATCAAATTCTGTTTggttacctcttatctagtg
gtagagtgtagcacctatccCAGAGAGATGGAGGGAGGTAGATGTGGAAAACTGGAATGAggttacctcttatctag
gtagagtgtagcacctatccTGACAATTTGAGATTAACACAGAAGAGGTTGGTGAGCATTggttacctcttatctagtg
gtagagtgtagcacctatccATTAAGAAATAAGGTCTCAGATGAAATTCCCAGGGTAGCCggttacctcttatctagt
gtagagtgtagcacctatccAACTTCCCAGGTCAACCATCAATTCCATTACCACCACCCAggttacctcttatctag
gtagagtgtagcacctatccTGGATTGCTGCAGTGTGGGGAATGTGCTGAGAGGAGCTCAggttacctcttatctag
gtagagtgtagcacctatccGGCTTTGATTTAAAACCTCAGGATCTGCTGTGTACTCATTggttacctcttatctagtg
gtagagtgtagcacctatccCCTCCCTGGGCTGTCATGATGATAAAAGGAAGCACTGAATggttacctcttatctag
gtagagtgtagcacctatccCTGGCACCTAGGAAGTCTTAGCAAATGTAATTACCTTCTTggttacctcttatctagtg
gtagagtgtagcacctatccCTAGCTGCACTAGGCTAGGTGGCACTGCTGCAACAAAGTGggttacctcttatctag
gtagagtgtagcacctatccCTGGGTGACTTAGAACAAGAGGAATCTATTGTCTCACAGTggttacctcttatctagt
gtagagtgtagcacctatccAAGGGGTCAGCTTGGCCGGGCTTGCTCTAAAACCTATAGAggttacctcttatctag
gtagagtgtagcacctatccGACCAATTCCAAGGAAGATAATTTTTCTATGGGCCAGAAAggttacctcttatctagtg
gtagagtgtagcacctatccGCAACTAGATGGTTCCATCTCGGGGTGATGAGAGACAGTGggttacctcttatctag
gtagagtgtagcacctatccTTAGATTCTCATAGGCAGGACTCAACCTAGATCCCTCGCAggttacctcttatctag
gtagagtgtagcacctatccAGGCGGCAATGTGAGTGATGGGAGGTGGCTGTACTTAACAggttacctcttatctag
gtagagtgtagcacctatccCACGGCCAGTACTAGCCAGTGGCCCGGGAGTTGGGGACCCggttacctcttatct
gtagagtgtagcacctatccGAAATTCCTTGACCCGGGGCCGCATCGCTCTAATCTCTACggttacctcttatctag
gtagagtgtagcacctatccCACCTTCCCTCTGTGCTCATCTGTCTCTGTGTCCATAGTTggttacctcttatctagtg
gtagagtgtagcacctatccGTTTAGCCACCCTCTTAATCTCATTTTAACCTGGTTCACTggttacctcttatctagtg
gtagagtgtagcacctatccCTCAATATATCTTTTTTGGGGGCGTGGCAGGGGGAACACAggttacctcttatctag
gtagagtgtagcacctatccCATAATATGTTTCCTTAGTTTTCTGCCCAGAGTAGCAACCggttacctcttatctagtg
gtagagtgtagcacctatccAGTGAACTATCAGTTAGCTATTTAAAGGACACTGTCTGCCggttacctcttatctagtg
gtagagtgtagcacctatccGGCTTAAAATATGCCTATATAGGTCACGCAACTTAGAGACggttacctcttatctagt
gtagagtgtagcacctatccGTGACTGCCTCTGCCCCCAGCCCCCACCGACCCCAGCCAAggttacctcttatct
gtagagtgtagcacctatccCCTTCTCTTCCCCTCCCTTCTTCATCTGATTTCTGAGCCTggttacctcttatctagtg
gtagagtgtagcacctatccCAGCTGCATTAAAATCTCTAGACTCTGCCAAACATGAACAggttacctcttatctagt
gtagagtgtagcacctatccAAAAATGGCAGAGACAGTGAGAAAAACATTCCAGTTCATCggttacctcttatctagt
gtagagtgtagcacctatccAGTCCCCACCCCCCACATTCCCATGCTGAAGCCCAAGTGCggttacctcttatcta
gtagagtgtagcacctatccGGAATACATTTTGCCACTAACACTTTCATATGAACTGCAGggttacctcttatctagtg
gtagagtgtagcacctatccTGGAACAAGAAGAGCACAGGCAAGGAGCATAGACAGGGGAggttacctcttatcta
gtagagtgtagcacctatccGGAAACTGAAAGGAGTGCGTACGTTTTCTCATAGGCCAAAggttacctcttatctag
gtagagtgtagcacctatccTGGCAAAAATAGATTCATGAGCATTTTCCTCCTCTCTCTCggttacctcttatctagtg
gtagagtgtagcacctatccCTCTCTCCCGTCCCTACTGTAAGCATCTTTGTTTCTCTTAggttacctcttatctagtg
gtagagtgtagcacctatccCATGTGAGAGGCAGGCCCACGGTAGACAGTTACCTGAATGggttacctcttatctag
gtagagtgtagcacctatccGGATAAAAGATTCCCTAGTCCAAAGTCACCTCCCCATCATggttacctcttatctag
gtagagtgtagcacctatccTCTTCCCAAGACAATTGATGAAAACAAAGCTCTTCGGGGGggttacctcttatctag
gtagagtgtagcacctatccTGGGATAGTATATTCACTGGTGGTCACCTTAGACAAATGCggttacctcttatctagtg
gtagagtgtagcacctatccGCTATAGTCATATGTTTGCTGTGTTCTCTGCTGTATCCTCggttacctcttatctagtg
gtagagtgtagcacctatccTGTGCAAATCCCTGGGCATAAAATGTGATATGAATCTGCTggttacctcttatctagtg

gtagagtgtagcacctatccCTAAAGGAGTTAACACATCTATAATGCTTAGAACAGTTCCggttacctcttatctagtg
gtagagtgtagcacctatccTGCTGCTATTGTTGTTGTTGACTGAGAAAACTCACAAGTAggttacctcttatctagtg
gtagagtgtagcacctatccAGAAAGGTCTGGAGAACTGCAACTGACAGCTGGGTGGGGTggttacctcttatctag
gtagagtgtagcacctatccGGAACTGAGAGCAAAACTAACCTTGCCCACTACACGAGCGggttacctcttatcta
gtagagtgtagcacctatccGCACATAGCGCTTACCAAGTCCCAAGTAGGCCCCTGGCACggttacctcttatcta
gtagagtgtagcacctatccTGTGGCCCTCGTTTCTCTCCCTGGTGGAGCTGGAAGCATTggttacctcttatctag
gtagagtgtagcacctatccAAGGCAGGAGAGTACAGTGGGACAAAGCTTCCATAAACTAggttacctcttatctag
gtagagtgtagcacctatccCCACTGCGTCCCGGAATCTCTGAAACTGAACCAGATACATggttacctcttatctag
gtagagtgtagcacctatccGATTCCTGACCTGGACCCACGGGACACCCTATAGCCTAAAggttacctcttatctag
gtagagtgtagcacctatccTGCTAGCACACAGTTTCCCAAAGTATGATACAGGTAGTTCggttacctcttatctagt
gtagagtgtagcacctatccGAATTTCTAACATACATAAATGCATTGTGAATCTCCAGGGggttacctcttatctagtg
gtagagtgtagcacctatccACTGTGCACCTATCCCAAATGATCAGAGAAGACTGAGTTTggttacctcttatctagt
gtagagtgtagcacctatccCCCTGGGGTTGTAGACGAATGCTAGATGGTGTCTCCTCAAggttacctcttatctag
gtagagtgtagcacctatccTCCTCATCTGAAAAATGAAAGAAGTCAGTCAGTGGTTATTggttacctcttatctagtg
gtagagtgtagcacctatccTTGTCAAATGCACTGTGTGAAGCTGGTGAAGGCAAAGGTGggttacctcttatctag
gtagagtgtagcacctatccGCGCATATTAAGCATCTGCCTGACCATTTTTTCATACAACggttacctcttatctagtg
gtagagtgtagcacctatccGAGTCACAAGTCCCTCAACTATAAGATGTTCTAAGATCCAggttacctcttatctagt
gtagagtgtagcacctatccGGTGAAGGTACCTGAATCAGAAGCATATAGAGGCAGATATggttacctcttatctag
gtagagtgtagcacctatccTTTCCTTTTCTGAAACTACCTCTTTCCCACTTCTCAGTCCggttacctcttatctagtg
gtagagtgtagcacctatccGTTAACCAACTCCTTCCAACTCCAGAAGTGGGCATGTCACggttacctcttatctag
gtagagtgtagcacctatccGACCAATCAGAACACCTCACTCGCCAGGCCATAGTGAAGGggttacctcttatcta
gtagagtgtagcacctatccGAAAGTGTTCTCCAGCATAGGACATACCGTAGAGCTATCAggttacctcttatctag
gtagagtgtagcacctatccAGGAAGGAGGGAAGAGGGAAGGGGGAGGGGGGAAGGAAGGggttacctcttatc
gtagagtgtagcacctatccGGAAGAAAGGAAGGAAGAAGGGGAGGGGAGGGGAGGGGAAggttacctcttatct
gtagagtgtagcacctatccAGGGAGGGAAGGGGAGGGGGAAAGGGGGAAGGAAGGAAGAggttacctcttatct
gtagagtgtagcacctatccGAGGGGGAGGGGAGGGAGGACCAAAGAAAGAAAAAAACCAggttacctcttatcta
gtagagtgtagcacctatccGCTGAATAAAAGCCTGCAGTCTTGGTAGCCATCTTTGTGTggttacctcttatctagtg
gtagagtgtagcacctatccTGATTCTTGATGACACTGGCTGACTCACCCCTGAACTCCAggttacctcttatctag
gtagagtgtagcacctatccGAGAGCCAGTAAATATGCCCTTCTATTTAAATTAACTTGCggttacctcttatctagtg
gtagagtgtagcacctatccTCAGATATCAATCTTGGGATTCCCAGAGCTACGGTGAGGCggttacctcttatctag
gtagagtgtagcacctatccCAGTGTTCATGCTGAGTCTACATTCAACAGAAAACCACCGggttacctcttatctag
gtagagtgtagcacctatccTGACTGTACATTGACAAGCACTAGCAGGACCTGACCGTTGggttacctcttatctag
gtagagtgtagcacctatccAAGGCCCCAGAGACTTCAAGTAATTTCTTCCACCTGCTGCggttacctcttatctag
gtagagtgtagcacctatccATCTGTGATGTTGGTAGGGGGTAGAGGGCAGGTGTGGAGGggttacctcttatctag
gtagagtgtagcacctatccAAAAAGATTTGCATGCAATTTCAGTGCTTTATACACTCCCggttacctcttatctagtg
gtagagtgtagcacctatccGGCTCCATCCCCTTGCTTTAGGTAGAGAAGAATTGCTGAAggttacctcttatctagt
gtagagtgtagcacctatccTGAAAGATCCCAATCTCTCAGAGGACTTCAGGTGGAAAGGggttacctcttatctag
gtagagtgtagcacctatccTGGCACCATTCCCCACAGAACTCTGGCACTCACACAAACAggttacctcttatctag
gtagagtgtagcacctatccTGCTGCCCCACCCCTGCATGTCCTTTTTGGAGTATTGGAGggttacctcttatctag
gtagagtgtagcacctatccGCCTAGCTAGAACCCCTCCTTTGTCAGATTTGGAATTAACggttacctcttatctagt
gtagagtgtagcacctatccGGCATGAGATTTTGATTCCTGTTAAAAGAGCTCCCTGTAGggttacctcttatctagtg
gtagagtgtagcacctatccACACTTCAAACTCCTATCAGTCATTCCTACCCGATTCCAGggttacctcttatctagt
gtagagtgtagcacctatccAGGAAAGTTAGGGACGTTTTAATAAAAGTAGAAGGAAGAGggttacctcttatctagt
gtagagtgtagcacctatccAGATGTCCACTGCAGTCTGGCTTTGTCTGGGCCCGCATGGggttacctcttatctag
gtagagtgtagcacctatccCAAATTATTGTTGTAAGTGGTTTTGCTCTTCCCAGTTGCAggttacctcttatctagtg
gtagagtgtagcacctatccGGCTTCCCAACCAGGAACTATGAAGTTGATAGTTTGCTTTggttacctcttatctagtg
gtagagtgtagcacctatccCAGCAAAGTTAAAGGTTAATGGTTTGACAAGTCCTTACCCggttacctcttatctagtg
gtagagtgtagcacctatccGTTGCATGAAAACTGGCCTCTCCCGCTGAAACCTAAATGTggttacctcttatctag
gtagagtgtagcacctatccGCCTGTGTTTGTGACCCGGTCCAAGCCTCATGTGTCCTCCggttacctcttatctag
gtagagtgtagcacctatccGCTCCCAGCCACGGCCGTTCCACCACACCAGTCTCGCAGAggttacctcttatcta
gtagagtgtagcacctatccTGCCCCAGAAGCTGTCAGATCTAAGGTCCCATGTTAGGAGggttacctcttatctag
gtagagtgtagcacctatccAAACCTATTTTACGAGTGAGAGCCTTTTGCCAAGAATAAGggttacctcttatctagtg

gtagagtgtagcacctatccTCCCCTGTTTGCCTGTTTCAATCTCAAAAGCAGATAACAAggttacctcttatctagt
gtagagtgtagcacctatccTTTCTGCCTGTTTGTTTGTTATACTTGGAATCCCAGAGAGggttacctcttatctagtg
gtagagtgtagcacctatccAACATGTTAAACCTTCTCCATAGCAACTGTATTCCTCTTAggttacctcttatctagtg
gtagagtgtagcacctatccGGGCCCAAGTAATCATTACCACCCTTACAGATGCAGCCTGggttacctcttatcta
gtagagtgtagcacctatccACTTCAAGGGTGTTGGTGCCAGGAGATGTGATATTTTCAAggttacctcttatctagt
gtagagtgtagcacctatccTGCCACCGTAAGTCCTTGGAAAAAGGTAAGAATTTATTTCggttacctcttatctagt
gtagagtgtagcacctatccTTAAATTAAGCCACCTCGTTTCATTTTGTTTCCAAAGACCggttacctcttatctagtg
gtagagtgtagcacctatccAGTCACTGGGCTTGGCGTGTTCCATTAGGAAGCTCATTAAggttacctcttatctag
gtagagtgtagcacctatccGGCAAGACACCATTGGCTCGTATGGCAAGGATCTGGATGGggttacctcttatcta
gtagagtgtagcacctatccTGAATGACTACGGACTGGCACAGCCTCAGATGCAGAGTGTggttacctcttatcta
gtagagtgtagcacctatccAAGTTGTATTTGATGTCTTCTGTTTTAGCATACTCTCCCAggttacctcttatctagtg
gtagagtgtagcacctatccTGGAATCTGTTGGAAGCTTCTCTATGTTGTCTGCTCACAGggttacctcttatctagt
gtagagtgtagcacctatccGTTTCGTGTTGCCTTCTGGTCACAGTCCTTGCTTCGGTAGggttacctcttatctagt
gtagagtgtagcacctatccGCCTGTCCATTTCAAAGGGCTCTTGGGCTCTGAACTGCCTggttacctcttatctag
gtagagtgtagcacctatccGTGCTGCAAGAACAAGCTCAGTCTGTTAAAAAAGAGTTCCggttacctcttatctag
gtagagtgtagcacctatccCAACTGCTCCCACATCCATGTCTCTGGTTACCTTGCCTGCggttacctcttatctag
gtagagtgtagcacctatccTTACCAGAACTTCCAGTCTCTAGCACACCTCCGATACCACggttacctcttatctag
gtagagtgtagcacctatccTCGGTTTGCAGCTTCCTCCCAGAACCTCTTGAAAGCGGGTggttacctcttatctag
gtagagtgtagcacctatccCTCCAGGATCACAGACTGACTCTCAGCACCAACCAAACCCggttacctcttatcta
gtagagtgtagcacctatccTGTCAAGCCCACAGTATCTATGAGTCAAAACCGCCAGCACggttacctcttatcta
gtagagtgtagcacctatccGTGTTAAACAATCTAATCGATCAACCAGTGAATCTCAATCggttacctcttatctagt
gtagagtgtagcacctatccAATCACCTGCAAGCCTTTAAAAAGACTAACCCCCGCGCCGggttacctcttatcta
gtagagtgtagcacctatccGGTCTGAGGTAGAGCAGTATTTTTTTTTTTACACTCCAAAggttacctcttatctagtg
gtagagtgtagcacctatccGGCGCAGTCAGGGTTATGAATCATTGATCTAATCCAAGAAggttacctcttatctag
gtagagtgtagcacctatccAAAAATTCTCACCCAATAATGACTGGACAGACACAAATACggttacctcttatctag
gtagagtgtagcacctatccAAGTGACCCCCATCTAATTTAAACAAGTAATTTCTCCAGAggttacctcttatctagt
gtagagtgtagcacctatccTTTTATGAAAGCCAACCCAATTTTTTTTCTTCCAGGACAGggttacctcttatctagtg
gtagagtgtagcacctatccGGACCTGCATCACTACCTACTGAACATAGACTTTGTCCATggttacctcttatctag
gtagagtgtagcacctatccATGCACCTTTTAAAACTATAACTCTTCAATGCCTCCAAAGggttacctcttatctagt
gtagagtgtagcacctatccCAAGAATGTCACTAACCATGTTTTCCAATCAGATCTTCACggttacctcttatctagt
gtagagtgtagcacctatccCGCTAGCAACATCTTAGACATCCAGAATATATGGCCCCAGggttacctcttatctag
gtagagtgtagcacctatccAATCTGCATTTTTAACCAGGTCCAAAGAGAACCCACGGCTggttacctcttatctag
gtagagtgtagcacctatccAGTACCCAGTAAACTGCTTTTACGAATCTTACTGTTAGAAggttacctcttatctagtg
gtagagtgtagcacctatccAGGCTCTCTTTTTGTGGCCTTAGCCCTTATTTTCCACTGAggttacctcttatctagtg
gtagagtgtagcacctatccGGATGCCCTGCTCCCACAAAACAGGTCCACCTCCAGCTTCggttacctcttatcta
gtagagtgtagcacctatccTGCCCACCTTCTGCAAAGCTCTCTCCCGTCCTCTCTGCCTggttacctcttatcta
gtagagtgtagcacctatccGAGCCTCTCCTCTCCAAAACCTCCCTAGATCTCTCCAGCCggttacctcttatcta
gtagagtgtagcacctatccGGTCTTGCCTGCTTATAATTTTATAGCGCTTATCCGTACAggttacctcttatctagtg
gtagagtgtagcacctatccGCTGTTTGTGTTTTATTGTTTAGCCCTTATTTTGCCCTTAggttacctcttatctagtgg
gtagagtgtagcacctatccTCAAATACTTACGCCTTCTCTCTTCCACTTATCCATCGTTggttacctcttatctagtg
gtagagtgtagcacctatccCCTGCCACGGAGCTTTGCACATAATAGACTCAACAAACATggttacctcttatctag
gtagagtgtagcacctatccCAGATCTCTGATGGCAAACTATACAACACATAAAACCTATggttacctcttatctagt
gtagagtgtagcacctatccGCTGGTATTAAAGCTTCCAATATGCTAGACACATGAGTTGggttacctcttatctagt
gtagagtgtagcacctatccGAAAGCAAGACAACATAATATCAGGTATCAACTAACAGTGggttacctcttatctag
gtagagtgtagcacctatccCTAATCAAATAGAAACCCCTTAACTACTAACTGCCACTGGggttacctcttatctag
gtagagtgtagcacctatccGCATTAGCAGCCAACCCCAGGCCTGTGACGTCCAAAGAAAggttacctcttatcta
gtagagtgtagcacctatccCCTCCTGACTAGGATTGTCTCTATTAGGGTATGGGCAGTCggttacctcttatctag
gtagagtgtagcacctatccCTGGATAGAATGACCCCATGTATTGCAACAGGAATATTCAggttacctcttatctagt
gtagagtgtagcacctatccAATATGAAAATGAGTAAGACACTGTCTTTGCCTCCGTAGGggttacctcttatctagt
gtagagtgtagcacctatccCCTAAGCAGGTTATTTGAATGCAATGCAATGAATATACAGggttacctcttatctagt
gtagagtgtagcacctatccTTAATTGATACAGTACTAGAGTACCATGTAATACACACGGggttacctcttatctagt
gtagagtgtagcacctatccCCTAACACAACCTGAGGCAATCAAGGAGACCCCTGTGGAGggttacctcttatcta

gtagagtgtagcacctatccGCAAAGAAGTTAGCCAAGTGAAAGGTGAAGAATCAATACTggttacctcttatctag
gtagagtgtagcacctatccTTGAGGGAAGCATGCAAAGAAAGGGGCGAAGTAACTGAAAggttacctcttatcta
gtagagtgtagcacctatccCACATTGTAGGAGGTAGGACGTGATGAGGTTTCTTCCACAggttacctcttatctag
gtagagtgtagcacctatccCTTGCTAGTTGATAAGCTTAGATGCAAAAGACCTTGGACTggttacctcttatctagt
gtagagtgtagcacctatccAGGCAAATGGAGAATGTCTAGGCTTGAAAGATTTAATGCAggttacctcttatctagt
gtagagtgtagcacctatccGAACAACTTAGATTCTACACAAACCAGAACATTAAATGCCggttacctcttatctagt
gtagagtgtagcacctatccCATAGCTATGTATACACTGATTCAAACATTCCATGGGTCAggttacctcttatctagt
gtagagtgtagcacctatccTCTCTGAGATAAATCCAGCCAAGATTCTATTTTCTGATTCggttacctcttatctagtg
gtagagtgtagcacctatccAATGAAGCATACCACAGGACAAATAGCACAGTGTTACCAAggttacctcttatctag
gtagagtgtagcacctatccTGGGCCTCATTCTCACTCGTCCAACTTCTCTTTCCAGCAAggttacctcttatctag
gtagagtgtagcacctatccACCTTTTCTGCAATTGTAGCTCATCATGTCAGGCAAGTTTggttacctcttatctagtg
gtagagtgtagcacctatccACAGCCAGTTCAGTTATTTGCTTTATACTTGTATACTTGCggttacctcttatctagtg
gtagagtgtagcacctatccTGTTCAAAACCATTAACACAGAAGAAAAACACTAGCAGGAggttacctcttatctag
gtagagtgtagcacctatccCCTAAAGTTATATATAGCTGGAATAATTCAACTGTCTGCCggttacctcttatctagtg
gtagagtgtagcacctatccCCTAATGGCTTCTAATCTGTACTGGACTTGGACTTTTCAAggttacctcttatctagtg
gtagagtgtagcacctatccGCAACTCTCTCCTGCCAACTTCACCATATTTGGATGCAGCggttacctcttatctag
gtagagtgtagcacctatccGGTATGATTTGCCTAGCAGTGATATTATTTGTTAGAGACAggttacctcttatctagtg
gtagagtgtagcacctatccGTCATTTGCCACAACACAGAAGGACCTGGAGGACATTATAggttacctcttatctag
gtagagtgtagcacctatccGGAATGTAAAAACAGAAGGTCAAATATACAAAGATAGGCCggttacctcttatctag
gtagagtgtagcacctatccGTGGCTCACAACTGTAAATCCAGCAGATTGGGAGGCTGAGggttacctcttatcta
gtagagtgtagcacctatccACATAGCAAGACCCTGACCCTACCAAAAACAGAAAAAAATggttacctcttatctag
gtagagtgtagcacctatccGAAAAAGCCAGGCCCTGTCTCTCTCTCCACATAGATAGATggttacctcttatctag
gtagagtgtagcacctatccGTACATACACACACACAAAGCTAGAGAATAAAACAATGGTggttacctcttatctag
gtagagtgtagcacctatccTGTAGTCAAGATCTTTGCTAAATGAGTAAACTATAGCTGCggttacctcttatctagtg
gtagagtgtagcacctatccACTATGTGAGTTGGGGGAGATGTTAGTTAATTTATTCCACggttacctcttatctagtg
gtagagtgtagcacctatccATTGTGATTTGGTTGCCCAGAAAAGAAAGAAAGAAAGAAGggttacctcttatctagt
gtagagtgtagcacctatccGAAAAAGAAACAGAAGAAGTTGCAGAGCAAAATAATGAGGggttacctcttatctag
gtagagtgtagcacctatccACTTGCTAGTGAGATGACTTTGGATAGGTGATTTAATCATggttacctcttatctagtg
gtagagtgtagcacctatccGACCAGCCTAGTCAAAATGCCAACATGCCCGTCTCTATTAggttacctcttatctag
gtagagtgtagcacctatccGAAAGAAAAAGAAAAGTCTCAGCTGCCCAATGTGTAAAATggttacctcttatctag
gtagagtgtagcacctatccTTATATGCAGCATGTAAATCAGAGTTGGGCACATAGGAAAggttacctcttatctagt
gtagagtgtagcacctatccCTCAAGGAATAATTTCATGGAATCACAGAGTAGTTCATACggttacctcttatctagt
gtagagtgtagcacctatccAGCACTTCTGGGAATGTTTGATGTTTGAAATATAGAGGGAggttacctcttatctagt
gtagagtgtagcacctatccTATATTACAAGGGTTAGTTATCAGAAAAGAATGCTGTGGCggttacctcttatctagt
gtagagtgtagcacctatccCTTGAATGATCCGACCCCACTTCCAGCCTCTTCCCTAGTTggttacctcttatctag
gtagagtgtagcacctatccGCAAGTGACACAGTTCAGCCAGAAAAGACATCTCTCTTCCggttacctcttatctag
gtagagtgtagcacctatccGAAACCTCCAAAGAATCAGAGACCTTGGTGCTAGCATTGCggttacctcttatctag
gtagagtgtagcacctatccCTGAATCAGTGCCAACAACAACCCACCTACATATTTTGCAggttacctcttatctag
gtagagtgtagcacctatccACCCCCTTTGAATTCAGGTTTTCTAACACATACAGCCAAAggttacctcttatctagt
gtagagtgtagcacctatccACCTGGCTGATACATTTATAGACTACATTGCTGTTGAGACggttacctcttatctagt
gtagagtgtagcacctatccGCTCATGTTTACCCATTATCACTTGTGCCTGCTTTCTTTTggttacctcttatctagtg
gtagagtgtagcacctatccGGTATAACGAGTTCCTTGGTGTGCTCCTGTTGCTATTTTCggttacctcttatctagtg
gtagagtgtagcacctatccCTTTCCTTCCAGCTCTTCCCCATACCTTCTGCTTGCTCCCggttacctcttatctag
gtagagtgtagcacctatccCACTTCCAGCTATTTATGCTGCAGGCTCTTCATTTCTTGCggttacctcttatctagt
gtagagtgtagcacctatccGTTACTGGTGCTACTAAATACCCCAACACAAACATCACAAggttacctcttatctag
gtagagtgtagcacctatccGGACATTCATCTTCCCATTAAAGACAAATTAACCACTTGCggttacctcttatctagt
gtagagtgtagcacctatccACCTTAGTCTCCACCGGTTCTCAAAAAGCCCTCCTGCCCAggttacctcttatcta
gtagagtgtagcacctatccTTCATCCCTGCCTGCTCTCTGTTGAAGTCTGTTCCGCTCCggttacctcttatctag
gtagagtgtagcacctatccGCTGCGCCATCTTTGAATAGTGTTTCCATTTATTATTTTGggttacctcttatctagtg
gtagagtgtagcacctatccAGTCATATGGGTTAAACCATAATGTCAGTAAAACAGCATCggttacctcttatctagt
gtagagtgtagcacctatccCAGCAGAAAGATGAATTTTGTTGGCCTTTGAAAGTCAGAGggttacctcttatctagt
gtagagtgtagcacctatccAGAACCTGTCTTATTTTCTGCTTCATGAACTAGTAACACAggttacctcttatctagtg

gtagagtgtagcacctatccGCTTTTGAGGTTGTGAGCCAAAGGAACAGCAAACACCATTggttacctcttatctag
gtagagtgtagcacctatccCTCCAGTATTGAGATTACCCGTGACAGAGTTAAAACTTGGggttacctcttatctagt
gtagagtgtagcacctatccGCACAACATAAGATCCTGCCATTGTAAAGGAATCACATGGggttacctcttatctag
gtagagtgtagcacctatccAAAAACTTCACTGGAATTTCCAAGTAGGAGGGGCTCGAGCggttacctcttatctag
gtagagtgtagcacctatccTACTATTTGAAAATTATTTAGGCCACAGGGAGCAAAATGCggttacctcttatctagt
gtagagtgtagcacctatccCATATATGAATCCAGGTAAATCAGAAACTCTTCCCAGATAggttacctcttatctagt
gtagagtgtagcacctatccCAAGTCATTTGCAGACACAGCTCAATAATGTACCAAGAATggttacctcttatctagt
gtagagtgtagcacctatccAGCTTGGGTGTGGATTCTGGCTCTTACTGTGTTAATTTGGggttacctcttatctagtg
gtagagtgtagcacctatccATAAGAGTGAAATGAGTTGGAATTGTACAGTGCTTAGAACggttacctcttatctagt
gtagagtgtagcacctatccCTGAGAAGCTTACATGAGGGGATAAATTTGGACTAATCAGggttacctcttatctagt
gtagagtgtagcacctatccAGGAAGCCCCTAAGAAACCTCTTGACACACAGTAAGAAGGggttacctcttatcta
gtagagtgtagcacctatccCTGCCTCCCTGTAATTCCTACCTTGAACCCTAAGTCTGAAggttacctcttatctagt
gtagagtgtagcacctatccACAAATAAATCCAAGTCTGCTTCTATGTGGCAAATCACCAggttacctcttatctagt
gtagagtgtagcacctatccGAACCTTCATTTCCTTCTCTTCTCTAAGCCAAGCATCCTCggttacctcttatctagt
gtagagtgtagcacctatccCTCTATAGGCACAGAGTTTGTCAGGTCTGCCATTAAGGATggttacctcttatctagt
gtagagtgtagcacctatccGAGGCTCTCATTGTGGACCCTTGACTTCTATTAACTGAGGggttacctcttatctagt
gtagagtgtagcacctatccGGTCACCCAATCACCTTCTACCTAGCCTACAGTGAACAAAggttacctcttatctag
gtagagtgtagcacctatccCCTCTTGATCTACTACAACCACAGCACTCAGCTTCATACTggttacctcttatctagt
gtagagtgtagcacctatccAGTACACCAAATCAGATTTTAAATGTCCTAGAGGAAATCGggttacctcttatctagt
gtagagtgtagcacctatccCAAAAATATTTGCATACATAAGAGAGTCAAGAGGCCATGAggttacctcttatctagt
gtagagtgtagcacctatccAAACTGCCTAGGTCTAAAATTTTCATAAAGTGCAATCCTCggttacctcttatctagt
gtagagtgtagcacctatccCAAAAGTTAATTCTTTGGTTTAAAAGTCCCCTCTTATCCCggttacctcttatctagtg
gtagagtgtagcacctatccGAGTGTTTCATGTTTATTCCGTTTCCATTGAATTTGACTCggttacctcttatctagtg
gtagagtgtagcacctatccGGATCACACTCTTCAAGAGCTCTCTCTATCAGGTAAGGGCggttacctcttatctag
gtagagtgtagcacctatccTGACTCCAAATTACCTCTCTTGCCTCACCTCTGATAACACggttacctcttatctagt
gtagagtgtagcacctatccGTTTCACCAACCCCACCCACCTACACTCACAGCACTACCTggttacctcttatcta
gtagagtgtagcacctatccAAACTGATACTGATGGAGGAGGAGGTTGATAAGGTCTCAGggttacctcttatctag
gtagagtgtagcacctatccGATAGGAAAACCAATATCTAGCATGCCTGGGTGAACCTTGggttacctcttatctag
gtagagtgtagcacctatccCACACTATGCTTGGCCTTCTAACTCTGTATCTGACTGAACggttacctcttatctagt
gtagagtgtagcacctatccTGGTGCCCCAAATAACTTTTCCAGCCCATTACCCTGTTATggttacctcttatctagt
gtagagtgtagcacctatccTTCCTGGGATCCTGCCATAAACCAAAAAGACCTCTAAGCAggttacctcttatctag
gtagagtgtagcacctatccATGCCAGCTCCCACCAGACTTTCAGTATCAGCACTCCTGCggttacctcttatcta
gtagagtgtagcacctatccTGGAGACTGGGCCAAAATTTACCATACATGTATTAACAACggttacctcttatctagt
gtagagtgtagcacctatccCTTGAAAGCAATGGCATGAGCTCAATGTAACCTCAATCTCggttacctcttatctagt
gtagagtgtagcacctatccCACACCTGGCCTAACATTACAATTTGAACTGCTAATACAAggttacctcttatctagt
gtagagtgtagcacctatccTGAAGAAGGAATGTGCAAACTCGTGCATAAAGAAAGCCTCggttacctcttatctag
gtagagtgtagcacctatccATGTGATCATTTCAATCCTTTCCAAGTTACTGGTGGTTTGggttacctcttatctagtg
gtagagtgtagcacctatccATCTCAAAATTTATGTAGAAGCAAAATGAGTGTGGATGGCggttacctcttatctagt
gtagagtgtagcacctatccCTTGTGAATGCCAAGCATATCAACAGTGTGGGAAAAGACCggttacctcttatctag
gtagagtgtagcacctatccCTAGTATATGTAAGGATTTACCATATGGTGTATGAGGTGTggttacctcttatctagtg
gtagagtgtagcacctatccAGCTTTGCTACATACCCTTATGAGATAAATCCTGTATACAggttacctcttatctagtg
gtagagtgtagcacctatccACAAAGTAGAAGGGAAGGTACAGAGATTTGTCTTATGGAGggttacctcttatctagt
gtagagtgtagcacctatccCAACTATGAGACAAATGGTTACTGTCCTTCATATGTAAACggttacctcttatctagt
gtagagtgtagcacctatccAATACAGTATATTTACAGGCAATTCTTCTTAGAGGAGGAGggttacctcttatctagt
gtagagtgtagcacctatccACACAATGGACTTTGGGAACACAGGGGGAAAGTGTTGGAGggttacctcttatcta
gtagagtgtagcacctatccATACTGCTCGGGGGATGGGTGCACCAAACTCTCACAAATCggttacctcttatcta
gtagagtgtagcacctatccTGAATACTAACTCTTACTATTTCTAGCTATGTGAGCTTGGggttacctcttatctagtg
gtagagtgtagcacctatccAGCTGCTTTGAATAGTAAATGAGATAAACAAGGTAAGGTGggttacctcttatctagt
gtagagtgtagcacctatccCTTAACAAACATTAGCCATATAGTCTTCCTTCAATACACGggttacctcttatctagt
gtagagtgtagcacctatccCATGCAAGAGATATTTATAGACATATTCGTGAGTGACAAGggttacctcttatctagt
gtagagtgtagcacctatccCTAGATGCTGAGGATACCATGGGAAAGAAAGATCAACACCggttacctcttatcta
gtagagtgtagcacctatccTCTACTAGTGCTGGCAGACAATTAAATGAACAGTTACAATggttacctcttatctagt

gtagagtgtagcacctatccAAACAATATTTCAATGCTGTTGGTAGGGAAGCTCACAGGGggttacctcttatctagt
gtagagtgtagcacctatccGCAAGGAGTCAAGGCCACAAAGCACTGAGAAGCCAGTTGAggttacctcttatcta
gtagagtgtagcacctatccCCAGTCTATTCGGTGGGGAAATTTTCTAATCAGCAAAACTggttacctcttatctagtg
gtagagtgtagcacctatccCAGGCCTGGGATTTTGTCAAGCAGTAGTGAAAGAAGTTACggttacctcttatctagt
gtagagtgtagcacctatccAAAAGAGTTGATAGTGACAAAAGAGAGACTGAAGTGATAGggttacctcttatctagt
gtagagtgtagcacctatccATTGAGAAATGGATTGAAGATGCAAGGTGGGTGACAGAGAggttacctcttatctag
gtagagtgtagcacctatccTCTAGAGAGCCTAAAGAGACAAAGAGTAAGATGGATATGAggttacctcttatctagt
gtagagtgtagcacctatccCAGAGTATGGCCATGGAAAAAGAAGAGGAACAGGGTTAGAggttacctcttatctag
gtagagtgtagcacctatccGTTACGAGTAAGTCAAAACCCTGAGAAGCCTGTGTCATTTggttacctcttatctagt
gtagagtgtagcacctatccCCACAGAAGGACAGAAGCCACATGCCAGAGTAGAACGTCTggttacctcttatcta
gtagagtgtagcacctatccTCTCTACTAAACATACAAAAAATGAGCCGGGCATGGTGGCggttacctcttatctag
gtagagtgtagcacctatccAAAAAGAATGTGGTAGAAGGATTGGCTCATTTAGGATGATggttacctcttatctagtg
gtagagtgtagcacctatccTTATAAAACATAATTCCCTGCTAGTAACAGGACCAGTCAGggttacctcttatctagt
gtagagtgtagcacctatccGGGTACCACAGCTTGAGTTTTCAGAGATGTATATTTCAACggttacctcttatctagtg
gtagagtgtagcacctatccGCCTTTGGCATGGTGAGAAAATGAACATCTATTAAAACTTggttacctcttatctagtg
gtagagtgtagcacctatccCCTTTGACATATTTATTTCATTAGTTTGGGGAGCTAGTTCggttacctcttatctagtgg
gtagagtgtagcacctatccATGGTGAAAAAAGGGAAATGAGATTGATTCTAGGATAGTGggttacctcttatctagtg
gtagagtgtagcacctatccGAAAATTCCAATTACTTGTAGCCTCCTTAGTACCACCCATggttacctcttatctagtg
gtagagtgtagcacctatccGGCAATTCCATTTCAGTATATCCATTCAAAGAAGACAAACggttacctcttatctagtg
gtagagtgtagcacctatccTGTAAAAGCAAAAGGCAAAGTTCTGTTGGTCTTGATTCCTggttacctcttatctagtg
gtagagtgtagcacctatccAGTTCCAACTTGTCCAGATGAAAGGCCATTACCTAAAAATggttacctcttatctagt
gtagagtgtagcacctatccGCTTTGGGGTCCTCAGCACTTGATCCAAGGGGAAAAAACCggttacctcttatctag
gtagagtgtagcacctatccTTTTCTTCTGCTCATTTTGTTCATTCATATCTCAGGTCCCggttacctcttatctagtgg
gtagagtgtagcacctatccGTCTTATACCATTCATGACATCCAAGTGAGAATACAGAAGggttacctcttatctagt
gtagagtgtagcacctatccATTAACATTTTACTACCTCTTCTACTCAGTTCAAGTCACCggttacctcttatctagtg
gtagagtgtagcacctatccGTTCTTCACCAACAGATTTAGTAGGACCAGTATTCTTGGCggttacctcttatctagt
gtagagtgtagcacctatccTCTGCCTCCTGGGTTCAAATGTGCCTCAGCCTTCCAAGTAggttacctcttatctag
gtagagtgtagcacctatccCCACCATGATCGGCCAGGAAATTAATTTTTAACTAAACATggttacctcttatctagtg
gtagagtgtagcacctatccAGATTCGTCTTTTAAACTTGCAGTTGGTACACAATGGAATggttacctcttatctagtg
gtagagtgtagcacctatccCAACATGGATGGAATTGGACAACATTATGATACTAAGTGAggttacctcttatctagt
gtagagtgtagcacctatccACGTGGAATCTACAAATAGAGAATTCACAAAAGCAAGAAGggttacctcttatctagt
gtagagtgtagcacctatccTGATGGTCAAAAGGTACAAAGGCACAGTTAGACAAGAGGAggttacctcttatctag
gtagagtgtagcacctatccTTCAAAATCGCTGAAAGTACATTTCGAATGCTCTCACCACggttacctcttatctagt
gtagagtgtagcacctatccCCACATTTTACTCATGAATCATAATTTCACTTTGTACCCCggttacctcttatctagtg
gtagagtgtagcacctatccTTGAGAACCAATTTTTGTTCACCCTGCTTATATACATTGGggttacctcttatctagtgg
gtagagtgtagcacctatccGTTGATCGCTCAACAATTAACAGGTAAGTCCAATTTCAATggttacctcttatctagtg
gtagagtgtagcacctatccTTTTATTCTTTCAAATCACAAAGTGACACAGATGCGATCGggttacctcttatctagtg
gtagagtgtagcacctatccTGAGTAGCATAAACATCTGCCAATTATTTGTCTCATATGGggttacctcttatctagtg
gtagagtgtagcacctatccTTTCAAAAACAGAGGTAAAACTGGGGCAAAAAACAGACCAggttacctcttatctag
gtagagtgtagcacctatccGATTATCCTCTCATTTTAAAACCAGAAACCTGAGCATAAGggttacctcttatctagtg
gtagagtgtagcacctatccTCTCTGAAATAAATAACATCCAGCAACGTTTCAACTTGACggttacctcttatctagtg
gtagagtgtagcacctatccTTCTGTCTTAATTTGACAACCTAATCTCCCAAGACTCGGTggttacctcttatctagtg
gtagagtgtagcacctatccAACATATCACCCTGAGTAATTTCTCTTTTTAGAACACCAGggttacctcttatctagtg
gtagagtgtagcacctatccTTGCAAAGATTCAATTAAACAGCAGTGTCCCTCTGGAGATggttacctcttatctagtg
gtagagtgtagcacctatccCAAAGCAATAAACATTCATTTGTCCTTTTCCTCCCTAAGGggttacctcttatctagtg
gtagagtgtagcacctatccCAGCATAAATGTAAGGTAAGCTGCTCCTTCTGTAGACAACggttacctcttatctagt
gtagagtgtagcacctatccGCCCTCATGCCGCTGTTCCTGCAAGCTTTTGCTCTAAGAAggttacctcttatctag
gtagagtgtagcacctatccGTGTGACTATGACTTGGCATTTTTTCCTTTAAAGTTACTCggttacctcttatctagtgg
gtagagtgtagcacctatccAATTTCCATCAAGAGAAGGACTATAACGTATTAGCCAAAGggttacctcttatctagt
gtagagtgtagcacctatccCCAAAAGAAGATGACTAGATAACAATCAGCAAAACTACTCggttacctcttatctagt
gtagagtgtagcacctatccCATTTTGTTGATTCGAATGACAGTTTCTCTGACAGAGTGAggttacctcttatctagtg
gtagagtgtagcacctatccCATTTTATGTGTTGTCCAGAAGGTAACGATATATCCCTTTggttacctcttatctagtgg

gtagagtgtagcacctatccTTAAACCAAAAATATGACTGGGGGAATCATAAAAAGCCAGggttacctcttatctagt
gtagagtgtagcacctatccTTTCATATGCACTTCGAAAAATACTGCTCTTACCTGGAATggttacctcttatctagtg
gtagagtgtagcacctatccAGAATTCCTATCTTACCTTTGTCCTTGCATCAATGAAGCAggttacctcttatctagtg
gtagagtgtagcacctatccTGTCTGAATTCCTTCAACAAGACCAATGCAAGTGATATTGggttacctcttatctagtg
gtagagtgtagcacctatccGTTCTTTAGGAATGTGAAACTTCGATGTCAAGTCTTAGCCggttacctcttatctagtg
gtagagtgtagcacctatccCCATGCACAACTAGACATCTATTGCCAAGGGGAGAGAGTCggttacctcttatctag
gtagagtgtagcacctatccCTACAATTGTTGCCCTTTTTCCAAGTTCAGAGCAAAGACCggttacctcttatctagt
gtagagtgtagcacctatccGAGAAAAGTAGATGAAAAATCAGGTCCAGGATAAATTGCAggttacctcttatctagt
gtagagtgtagcacctatccCTGTTTTTGTAGAGTGAGCATGCCTTTCTTTAAGATAGAAggttacctcttatctagtg
gtagagtgtagcacctatccCAGGATATATTGCTGGAATACGACCAAAGAGTTTAACATAggttacctcttatctagtg
gtagagtgtagcacctatccGTACAACTAGTTTATATTTCCCATGTATTAATGCCTCTCGggttacctcttatctagtg
gtagagtgtagcacctatccCTGCTTCCTGTCTCCTTTGCCTGATTTTCACTTCCTTTTCggttacctcttatctagtg
gtagagtgtagcacctatccCCATGATTCAATCTTCTTCCAATTTCTTTGTTCTCTTTGGggttacctcttatctagtgg
gtagagtgtagcacctatccATAGCCATCTTGTGACATAGTCCCAAAAGATATTATCTTCggttacctcttatctagtg
gtagagtgtagcacctatccTGATACACCTATTAAATGGCAATGTAAAATCTCACAAGGCggttacctcttatctagtg
gtagagtgtagcacctatccCAATTTTCTTCACCCTCTCCAAATAGACCTCTCTTTTTCAggttacctcttatctagtg
gtagagtgtagcacctatccGAGGCCCCAGCAGTTAGTATAGTCCCTAGCATATAATAGGggttacctcttatctag
gtagagtgtagcacctatccATGAACTCCTCTGAAAAACACGTTTCATGAATTAGATCAGggttacctcttatctagtg
gtagagtgtagcacctatccGTTCTCCATTTTACTGCTAAAGAAAGTGTGGCTCAGAAAAggttacctcttatctagtg
gtagagtgtagcacctatccGCATTCATAAATAGCCAAAAGTCTTTTTTTTCAGGCCAGCggttacctcttatctagtg
gtagagtgtagcacctatccCCATATGAACCTGTGGGCTATTCTTCACTACAAAGTTAAGggttacctcttatctagtg
gtagagtgtagcacctatccACTATTCTGCATAGTCTCAATAGTGTGTGTGTGCATATGTggttacctcttatctagtg
gtagagtgtagcacctatccCTTAGATCTGTTTGTTACATGTGCTAAGCTATTGGGATATggttacctcttatctagtg
gtagagtgtagcacctatccGATTGCCTCTAAGCAGTACATAAATTAAACCCAATTACTCggttacctcttatctagtg
gtagagtgtagcacctatccAATACAGAATTCACAGGTCAACGCTTTCTAATAACTAAGCggttacctcttatctagtg
gtagagtgtagcacctatccACATTGTAGAGCATGTCACTGTTCCATATTAACACTCCTTggttacctcttatctagtg
gtagagtgtagcacctatccGCAACATTGCTAACACACACAATTCACCCTTGCCTTTTGGggttacctcttatctagt
gtagagtgtagcacctatccTTCAAATCGAACGCCAAGATTCCTCTTTCCTAATTGCAGGggttacctcttatctagtg
gtagagtgtagcacctatccGCAGGAGCCGTTTGCAGCAGCTGTTGGCCCACAGTCAGTTggttacctcttatctag
gtagagtgtagcacctatccCTTTGTCTGCCTACATAGCTCATAGTCATAATCAGTTCTTggttacctcttatctagtg
gtagagtgtagcacctatccGAAAGAACAAGAATTATTATACCAGGTTAAACCGGTGATCggttacctcttatctagtg
gtagagtgtagcacctatccCAGCAGCTAGACTTATGGGAGAATTTGATTATCAAGATAGggttacctcttatctagtg
gtagagtgtagcacctatccAATAAAGAGTGTAGAATATACAAGACCTCAACCAAGTAGCggttacctcttatctagt
gtagagtgtagcacctatccGCTGGTGGAGTTGGGGGATACTAAAAAAAAAAAGTAAAAAggttacctcttatctagt
gtagagtgtagcacctatccGCTACTCGTGAGGCTGGGGCGGAAGAATCCCTTTAACCCAggttacctcttatctag
gtagagtgtagcacctatccAAAGAAAAAAGAAATGCAAGTCACTGTCTCCACGGAACATggttacctcttatctag
gtagagtgtagcacctatccCTTCCATCTAGCTAAAAAAGACATTCTCTGTCCTTTCAGGggttacctcttatctagtg
gtagagtgtagcacctatccGCCCTGCCTGCCTGTGTTGCCTTTCACTTCGCATCATTCCggttacctcttatctag
gtagagtgtagcacctatccCACTCGCCATCCTCTCCTCCTGCCTAGTGAATGCTAATTCggttacctcttatctag
gtagagtgtagcacctatccGCAACCTTTCCTAAACCCACCTGCCAACCACAGTAGCTGCggttacctcttatctag
gtagagtgtagcacctatccAGATGTTTTGATACAGGCAAGCAGTGTGAAATAAGCACATggttacctcttatctagtg
gtagagtgtagcacctatccCAAGCATTTAGCCTTTGAGTTACAAATAGTCCAGTTATACggttacctcttatctagtg
gtagagtgtagcacctatccACTATTGTCACCCTATGTACTATCAAATAGTAGGTTCTCCggttacctcttatctagtg
gtagagtgtagcacctatccTATTGTTGCATGTTTTAGTTATTTACTCTCCATCTCCCAGggttacctcttatctagtgg
gtagagtgtagcacctatccGGAGGTCCCCAATAAATGTTCCTGAGTAAAGAAATCAATCggttacctcttatctagt
gtagagtgtagcacctatccTTAAGTGCTAAGTGGTTAAAAACGCAAGTTCTGAGTCAGAggttacctcttatctagtg
gtagagtgtagcacctatccTGAGTAAACTTACGTACAGAGAATAGAGTCGCCTCAAGCAggttacctcttatctag
gtagagtgtagcacctatccGCTATGATGCCCAGTTGTTCAGCCAAACACCAGTCTAGATggttacctcttatctag
gtagagtgtagcacctatccTGATTAAAACAGATTACCCTGCATAATGTTATGGGCCTTAggttacctcttatctagtg
gtagagtgtagcacctatccAGGTATTAAGAAGAAAAACGGAGGTTTCCCAAAGAACGGAggttacctcttatctag
gtagagtgtagcacctatccAACATAGAAACCCTTCCTGGGACTGAGTTTCCAGCCTGCTggttacctcttatctag
gtagagtgtagcacctatccCTCTTACCTGAATTTCAAGAATCCAAGCCCTACAAATTTAggttacctcttatctagtg

gtagagtgtagcacctatccTTCTCTAGATAACTCTGGCTAACATAGATATATACTAGGGggttacctcttatctagtg
gtagagtgtagcacctatccGGAGAAGTCAACAGTCCTGCCTTACATAGAAAACAAATGCggttacctcttatctag
gtagagtgtagcacctatccGGAGACAGGACCAATCAGATACCCATTCACCTAGAACCTCggttacctcttatctag
gtagagtgtagcacctatccAATTCCTTTGAGTCTCTTTTATTTTCTTGGAGCTACAGACggttacctcttatctagtgg
gtagagtgtagcacctatccATGCCAATCTGTTTGTCCATTCTCTGCCTTGCATCTTCTGggttacctcttatctagtg
gtagagtgtagcacctatccGAAAGTTTCATTTGGTCTGTTGTGACCCTCATGGCCTAAAggttacctcttatctagtg
gtagagtgtagcacctatccGCTCTCGTGGCTGGCTGCTTCCCTATCATGATAATTCACAggttacctcttatctagt
gtagagtgtagcacctatccTACCCAGAGAAGTCAGTTGGCATAGTTCATTTTGGGGGCCggttacctcttatctag
gtagagtgtagcacctatccCATCTCTGGGACAATTAAAGACGAGAGTCATAGATGCCAAggttacctcttatctag
gtagagtgtagcacctatccACTGTTTCTTCCTGGAAAACTGTAAAAAGCTGCCTTGGAAggttacctcttatctagtg
gtagagtgtagcacctatccTGATGACATGTCTACAAAAGAGGAAAATGATACGGGCAGTggttacctcttatctagt
gtagagtgtagcacctatccGCAGCCACGAGATGATGACCAGCATAGGAGCTCTCAGCCCggttacctcttatcta
gtagagtgtagcacctatccCTGTAACAGCGACCACTTCTTACCTATACTCGGTTCTTCTggttacctcttatctagtg
gtagagtgtagcacctatccCAATAATGAGAGAGATTGAGTCAGATGATGGCAGTAGTCAggttacctcttatctagt
gtagagtgtagcacctatccACTAGTCAAGCATTCGTAGTATTCTGGTGAGCCCTATTATggttacctcttatctagtg
gtagagtgtagcacctatccTTATTTGTACCTTATTAGATTGGGGAGCCTGAAGAAATGCggttacctcttatctagtg
gtagagtgtagcacctatccGTGCAATGTTGTTTTACACCATAGACTAACTTCCCTTTCTggttacctcttatctagtg
gtagagtgtagcacctatccAGTAAGACTTCTAAATGATCTCAGTGGTATCAGTGTGGGCggttacctcttatctagtg
gtagagtgtagcacctatccTCACATCTGTTTGGGAAATATTGGGAAATACTGCTGGACTggttacctcttatctagtg
gtagagtgtagcacctatccTACCGAGATGTACCTTCCAGTACTGGAGCCACTCATTGTAggttacctcttatctagt
gtagagtgtagcacctatccCACATTGGGCAAACAGGCTATTATATTGAACTGTGCAGATggttacctcttatctagtg
gtagagtgtagcacctatccTCTATTGGACGGTACTAGTATGGCACGCTCACTGTGATATggttacctcttatctagtg
gtagagtgtagcacctatccCAGTCTTACCAGAGCTTCCCTTACAGAATGATGAACTTAGggttacctcttatctagt
gtagagtgtagcacctatccTAGGAACATACTTGTGGAGAGAGCTGCCAGGAAAGGGATTggttacctcttatctag
gtagagtgtagcacctatccCCCTCACCTCAGATACTGAATTTTTAAAGAAGCTGATCCAggttacctcttatctagtg
gtagagtgtagcacctatccTAGGAAGTGAAATGAATGTGCTGGAAGATATGGCCAAAGCggttacctcttatctag
gtagagtgtagcacctatccATGAAATCACCCTGCAGTCCCAAGCCCCATCGTGTCATGAggttacctcttatctag
gtagagtgtagcacctatccCAGATACTGGGGTACAAGTGGAGTGCATAATTCAACAGTCggttacctcttatctag
gtagagtgtagcacctatccTTCTAAAGAAAACCTGTACCAGTTCCTGAGAAACAGATTTggttacctcttatctagtg
gtagagtgtagcacctatccATACTTTAGATGAAGGATGGTGAGTTTAAGGGAAGAAAACggttacctcttatctagtg
gtagagtgtagcacctatccCATAGTTTGGGCGTCCTGTGGAATGCAGCCAGGTTCTTGGggttacctcttatctag
gtagagtgtagcacctatccGGATTCATACTCAGGTGTTGCATGCTGTGGGAAACTTAACggttacctcttatctagt
gtagagtgtagcacctatccAGGCAGCACAAAGAAAAAAGACTAGACACATTAGCATAAAggttacctcttatctag
gtagagtgtagcacctatccTTGACAGTTACAACAAGAACAATTACCCCAGGACTCCACCggttacctcttatctag
gtagagtgtagcacctatccGGTTCCCACATTTGAGCCTCAGACTTCACATTCTTTTCTTggttacctcttatctagtg
gtagagtgtagcacctatccTCAGTCAAGTTCTTTTAACCTATTGGATTAGAAACCAGTGggttacctcttatctagtg
gtagagtgtagcacctatccGTAAAGCCCATAATTACATACATTCCAGTTTGCTGCCAATggttacctcttatctagtg
gtagagtgtagcacctatccATGGTCAATAGTGCCCCATTTCACTCATAAAAGCCCAAGTggttacctcttatctagt
gtagagtgtagcacctatccGTCACCTACTCATAATGCTATTCCCTAAGTATGTTCCAAAggttacctcttatctagtg
gtagagtgtagcacctatccAAGGGCAAGATGTGTTGTTTTTCCAAGGCTGGCGGTTTTAggttacctcttatctagtg
gtagagtgtagcacctatccGTCAGCACTGCTCACCATTTTCAAAGCTACTCCATTTTTAggttacctcttatctagtg
gtagagtgtagcacctatccCAGTCTTCTTCCTGTCCATTCTTCAAGATTTTTATTTTCCggttacctcttatctagtgg
gtagagtgtagcacctatccCATTTACTGTAATGTGTGCTTTCTATTTGTGGCTTCTTTCggttacctcttatctagtgg
gtagagtgtagcacctatccGTATCCACTGGTAGAATGAGCAACATCTGATAACTAACCTggttacctcttatctagt
gtagagtgtagcacctatccCCTGATAGGTTAAACTTCCTAGTGCTTTCAAAAATAACTGggttacctcttatctagtg
gtagagtgtagcacctatccCTGATAAATAGCTATCCATCAGCCCCAGGGGAGTTAGGGCggttacctcttatctag
gtagagtgtagcacctatccGTATAAATACCCTCATCGTGGACAATTTTAGCCTACAACCggttacctcttatctagtg
gtagagtgtagcacctatccTGTGTAAAAATGGAACAGTTGTCTGTGATGTGCCCGTGTGggttacctcttatctagtg
gtagagtgtagcacctatccCAAAACCCTTGGCAGAGATATGGAACATAAAGTCCTAAAAggttacctcttatctagt
gtagagtgtagcacctatccCTATGTGAAAACTAGAAAAGTCAATAGGAGCTATTTCTGGggttacctcttatctagtg
gtagagtgtagcacctatccGGTTCTAATTCCCCTTGCTACCCACCAAAGATCATTCCCAggttacctcttatctagt
gtagagtgtagcacctatccCATTATTATGTGAGGATAGTACTCCAGGCTCCATGACTGCggttacctcttatctagt

gtagagtgtagcacctatccAAAAATCAGACACTTTCAGAGCTGTACATCAATGCACTAGggttacctcttatctagt
gtagagtgtagcacctatccCTGTTCTAAGAGATTGCAGAAGGCCTAAAGTTTACATTTGggttacctcttatctagtg
gtagagtgtagcacctatccCAGATCTAAGCACTATTTCTATTCAAAGCCAATTTACAGCggttacctcttatctagtg
gtagagtgtagcacctatccTATAATAGAGAATAAAGCAGGTTAGAAGTAGGAAAAGCCCggttacctcttatctagt
gtagagtgtagcacctatccTCTACACAGAATTCACTAACTAGACACACCAGTAAAAGGCggttacctcttatctag
gtagagtgtagcacctatccCTGTCGCAACACAGCATTACCTCCAAACTAAAAATATTTGggttacctcttatctagt
gtagagtgtagcacctatccACTCCAAATTTATTAAGGTAATCATAGGTGTCTCAGTCACggttacctcttatctagtg
gtagagtgtagcacctatccTTAAAAAAAAAATCCTTATGCTAAAATGTGGGCAGGGCACggttacctcttatctagt
gtagagtgtagcacctatccGGTGGCACGTGCCTGTAATCTAGCTACTCCGGAGGCTGAAggttacctcttatctag
gtagagtgtagcacctatccAAAACGTGGTATATACACACTATAGAATACCATTCAGCCTggttacctcttatctagt
gtagagtgtagcacctatccAAAGGAGGAAATCTGCCATTTGTGACATGGATAAATTTGAggttacctcttatctagtg
gtagagtgtagcacctatccAAAAAGACAAATACTGTTGGCCAGTGCAGTAGCCCACGGCggttacctcttatctag
gtagagtgtagcacctatccCACTCCAGTCTGGGTAACAGAGCAAGTCTCCATCTCAACAggttacctcttatctag
gtagagtgtagcacctatccAAAGTACATTCCACAGGGTGTGAGCTGGCTCAATCAAGCAggttacctcttatctag
gtagagtgtagcacctatccTATGTAAATAAAGTAGTTGCCCAAGATCAGTCTGATTGTGggttacctcttatctagtg
gtagagtgtagcacctatccAGTGAGAGGCTGACATGAAGTTACAAGTTATACCCTATGCggttacctcttatctagt
gtagagtgtagcacctatccGGAAGTGACCAATCAGAGACTGTAGTTACAAATGAAGACTggttacctcttatctag
gtagagtgtagcacctatccCAGCAGCCTGATTCATACTCCAATGCAAATGAAGACTTGGggttacctcttatctag
gtagagtgtagcacctatccAGGTATTTTCAGTTTTTCATTTGCCACACAGAAAAGGCAGggttacctcttatctagtg
gtagagtgtagcacctatccAAGTTCCTTGCCTCCAGACTTTATTCTCCTGCCCCAATACggttacctcttatctagt
gtagagtgtagcacctatccGAGAGTAGAATGGCAGTTATTGGGGTGTAAAGGGGACAGGggttacctcttatctag
gtagagtgtagcacctatccCAAAATTTCCATTAGATGGAAGGAGAAAGTTCAAGAGATCggttacctcttatctagt
gtagagtgtagcacctatccGAGGTAATGCATATAATAAGTAGCTCGATTTAGCCATTTCggttacctcttatctagtg
gtagagtgtagcacctatccTTTGTTTTTTGAGACAAGCTCCTGCTCTGTCGCCCAGACTggttacctcttatctagt
gtagagtgtagcacctatccGGCCAGGCTGGTCACGAACTCCTGACCTGAAGTGATCTGCggttacctcttatcta
gtagagtgtagcacctatccGCGCAGTGGCTCACGCCTTTTCCTTCCACATCTTTCCATTggttacctcttatctagt
gtagagtgtagcacctatccGTTCTCATGAAGGAAATGTTGGAACTAAAAAGGGTCAAAGggttacctcttatctagt
gtagagtgtagcacctatccTAACGGGAAAGGTCAGAAAAATGGGTAATAAGTTAGTACTggttacctcttatctagt
gtagagtgtagcacctatccATTTCTGAAACACACAGACAGGGAATGCCACAACTGATGCggttacctcttatctag
gtagagtgtagcacctatccGCCCAGGCCAGCTCCATCATCTCTGATCTCCCTCTCAGCCggttacctcttatcta
gtagagtgtagcacctatccAGAGTCATCTTCCCTTCCTACCTCCATTCCTGCCTCCCCAggttacctcttatctag
gtagagtgtagcacctatccTCTTTAACTCACCATAAACTTGCCTGTTCCCCTTAATTTCggttacctcttatctagtg
gtagagtgtagcacctatccCAAAAAAATGCCTTTGCCGCTGCCATCAATGTTGTCTCTGggttacctcttatctagt
gtagagtgtagcacctatccTCATACTCAACCCTTGTCTTATATGAATGACCCCCCAGCCggttacctcttatctagt
gtagagtgtagcacctatccCTCCTTGCACATCTCTTTCCCTGAATCCTGTGACACGCCCggttacctcttatctag
gtagagtgtagcacctatccATAAACTATCAATTGTCTACCCAGCAGGGGTTACCCCATGggttacctcttatctagt
gtagagtgtagcacctatccAAGTAGGCTTCTCCCCTAGCTCCAAGAGATGAACCATGTTggttacctcttatctag
gtagagtgtagcacctatccTCCTCATGTAATCTCATTCCTCATGGCTGTAATGGGTGCAggttacctcttatctagtg
gtagagtgtagcacctatccGATAAATTCCTCCCAATGCTCAATTCATGCTCTTGAACTCggttacctcttatctagtg
gtagagtgtagcacctatccGTTTGGCTCTGTGACAGCTTTTTCTGGGGGGGAGGGGAGGggttacctcttatctag
gtagagtgtagcacctatccACTGATGTGAAAGAAATGCTGAATTAACCCTGGAATCACTggttacctcttatctagtg
gtagagtgtagcacctatccTGTATGTGAGACAGTAAGTCCTTATTGACTAAGCCACTCTggttacctcttatctagtg
gtagagtgtagcacctatccCTGACATGTTCTCCCTTCTCAGCTTCCATTCTCACCCACCggttacctcttatctagt
gtagagtgtagcacctatccGGCCCCTACCTGATCTCTATACCCTTTGCATATTAGTAGTggttacctcttatctagtg
gtagagtgtagcacctatccTTATTCTATTCACTTTCTCTTTGTGCTTTCTCTCTTCCACggttacctcttatctagtgg
gtagagtgtagcacctatccGGACCTACTATGAGCCAGATCCTGTTCCATGTTCTATATTggttacctcttatctagtg
gtagagtgtagcacctatccACTGATAAATAAGACCACTGTTTCCCTGGACAAAAGAACAggttacctcttatctagt
gtagagtgtagcacctatccAGGGGAGGAGGAAAGGGAAGGGGTTGAGGTTGAAGAATTTggttacctcttatctag
gtagagtgtagcacctatccCCTCTTTTTCTATTCTCTAAGTGTGTACAGTATTGGACATggttacctcttatctagtgg
gtagagtgtagcacctatccATTTCACCTTGGGTCAGTTTGCGGGAGCTATACTTTTCTAggttacctcttatctagtg
gtagagtgtagcacctatccTTTCCCTCTGCATACAGCTTTGGTTACACTTTCTATATGTggttacctcttatctagtgg
gtagagtgtagcacctatccGTGTTTTGACTTCCATTATGGCCCAAGTTATATCAATTTGggttacctcttatctagtgg

gtagagtgtagcacctatccTCTTAGATTTTGCTGTTAGTTACTAAGAGAGACGTGTTCAggttacctcttatctagtg
gtagagtgtagcacctatccCTCATGATGATTGTTGACTTGTTTATTTCTCTCCCTAGTTggttacctcttatctagtgg
gtagagtgtagcacctatccCATGGCTCTAGAGAATTGAACATGTTATCCTTATCCCTAAggttacctcttatctagt
gtagagtgtagcacctatccTAATATAGGTATACTAGCTTTCTTTTTGGGGGAATTTGCCggttacctcttatctagtg
gtagagtgtagcacctatccTATCCTTTTAGTTCCAATCTTTCTTTATCATCTCCAGTGCggttacctcttatctagtg
gtagagtgtagcacctatccCTGTTACCACTGTAATGCTCTAATTATAGTTTTAGCTTCCggttacctcttatctagtg
gtagagtgtagcacctatccAGAACTGGCATTTCTGAAATTATGGTGTGTGGTTTTGAATggttacctcttatctagtg
gtagagtgtagcacctatccAACCAGTTCTTGTTTTTCCTGAGCTCATGGGTCCTGCTGGggttacctcttatctagt
gtagagtgtagcacctatccCGTGTGGGTGGATGGCAGGGCAGAAAGAGGGAAGATGTGGggttacctcttatcta
gtagagtgtagcacctatccGAGGTGGTGGTGGGTAAGCAATGCTAAGTGTAGGTCGTGTggttacctcttatctag
gtagagtgtagcacctatccAGCAATTATGTGAGACGAGAGAGAACTGGTGATTACTTGTggttacctcttatctagt
gtagagtgtagcacctatccCATCAGTTGTTACCCTGTTGAGACGTTGCAGATCTTTTTCggttacctcttatctagtg
gtagagtgtagcacctatccACCTGAATCTAGACCTGGATGGGAAATCTACTGAATTTGAggttacctcttatctagt
gtagagtgtagcacctatccAAATATACGATATCTGTTCCCTACATTTGCAAATTCTGGCggttacctcttatctagtg
gtagagtgtagcacctatccACAAATTCTGGCAAGGATCACACTTAAAGAAGGAGAAAAGggttacctcttatctag
gtagagtgtagcacctatccCCTATAATGGGCTTACTCACTCCTCTTTGTACAGGGAAATggttacctcttatctagt
gtagagtgtagcacctatccTTTCTTAGCTCTTGGGAAATGGGTTTTTGGGATAACTTGGggttacctcttatctagtg
gtagagtgtagcacctatccCTTCTCTGGAGGCAAAAATGGAAAATGGTAGGAAGTAAGAggttacctcttatctag
gtagagtgtagcacctatccGTAGCAATGAAAACAAAATACAAAAGTTTGTCTGGGGTGCggttacctcttatctagt
gtagagtgtagcacctatccATGCCAACATCCAAATTTGTACTAATCTCTCTAACCTCAGggttacctcttatctagt
gtagagtgtagcacctatccCCAGTCTGGAAACTAACTCATCCACCTTTCTCTATGCAGTggttacctcttatctagt
gtagagtgtagcacctatccAAGAAAATATGCCCCAAAATATATGATATGTACCTGCAGGggttacctcttatctagt
gtagagtgtagcacctatccGCACATACCAAGTCCTTAATTCTTCTGCTCTTTCGTTTTCggttacctcttatctagtg
gtagagtgtagcacctatccCTCACTGCAGGCTTGAACTCTTGGACTCAGGCAATCCTCCggttacctcttatcta
gtagagtgtagcacctatccAGCTGGGACTATAGGCAGGTGTTACCATGCCTGGCTAATTggttacctcttatctag
gtagagtgtagcacctatccTGTAATATCTCCAAGGAATCTTTAGAATATACACTGCCACggttacctcttatctagt
gtagagtgtagcacctatccTTCAATGACTGGCTGAACAAAAAGGTGTTCTTTAGAAAGAggttacctcttatctagt
gtagagtgtagcacctatccACTATGAGGTCCTTCCATTTTATAGTAAGAAAACTGAAGCggttacctcttatctagt
gtagagtgtagcacctatccCTCACCAAGGTCATAGATTTAATAAGGGTTAGTGCCAGGAggttacctcttatctag
gtagagtgtagcacctatccAATCCTGCATTTTCCACTCTATCACACCTTCTCTGGCTGAggttacctcttatctagt
gtagagtgtagcacctatccGGTTCTTAGATAGCCAGGACTTCTGACAAAGTGTGAGAGCggttacctcttatctag
gtagagtgtagcacctatccAGTGCCAGTGTCAAGAACTCAAAACAGTAGGAAGTCAGAAggttacctcttatctag
gtagagtgtagcacctatccCTCTGGAGGGTTGCAAGGAAATACTTACATGTGATTTCTAggttacctcttatctagt
gtagagtgtagcacctatccAATTGTTATTTCTCACCTTCCCTACAATCGATTCCGCCTGggttacctcttatctagt
gtagagtgtagcacctatccGGTATCCACATTACTTCTCTGAACACCAATTTCATTTTACggttacctcttatctagtg
gtagagtgtagcacctatccAATTTCTATTTCCCATCTTTCATTTCCTTTTCCACAACCCggttacctcttatctagtg
gtagagtgtagcacctatccTATGTATTGCCTCAGTTAAAGAAAATATACTCAGCTGGAGggttacctcttatctagt
gtagagtgtagcacctatccGCATATATGTCTACCATGCTATTCTGGCACATTTTTTGACggttacctcttatctagtg
gtagagtgtagcacctatccAAAATTTCTCAGGTAACTGTGATAGAAGAGATTTCACTGGggttacctcttatctagt
gtagagtgtagcacctatccAATCCATAATTCATTAGTCATCTCTCTGGTTAGTTCTCAGggttacctcttatctagtg
gtagagtgtagcacctatccCCCTTCAGAGGATCCTATAATTATTTTCATCATTTCCACTggttacctcttatctagtg
gtagagtgtagcacctatccTCAACATAATGAGAGGTTTGGGGGATCCGTTCTGAAGTGAggttacctcttatctag
gtagagtgtagcacctatccCAACTGCTTTGTATCTCTGAAACTCCATAAGGGCTTTAATggttacctcttatctagtg
gtagagtgtagcacctatccCATTGTCATATTTTCTTCAATCAGTACCTACTGCAGTGCCggttacctcttatctagtg
gtagagtgtagcacctatccGGTACTTCCCACAAACAAAATTAAGCATTCAGTGTACATTggttacctcttatctagt
gtagagtgtagcacctatccGGTTTGGACCAGTGTCTCCAGCAAATCTCACGTCAAATTGggttacctcttatctag
gtagagtgtagcacctatccGTTTAGCGCCATTCCCATTGGTACTGTCTTCACAATCATGggttacctcttatctagt
gtagagtgtagcacctatccCTGGAGGACAGTCATGCAATATCTTGGTGCACTGCAACCTggttacctcttatctag
gtagagtgtagcacctatccCTGGTTCTTTAAAATTGTGTGGCACCACCTCCCTCCTCTCggttacctcttatctagt
gtagagtgtagcacctatccTGTGCCTGCTCCCCCTTCGCTTTCTGACATGATTGGAAGCggttacctcttatctag
gtagagtgtagcacctatccAAGAACGGACTAATATACCTATCTTTGAAGAGAGTTCTCCggttacctcttatctagt
gtagagtgtagcacctatccTGCTTATTGATATCTTGTAGCAAACACATAGCCACATACAggttacctcttatctagt

gtagagtgtagcacctatccACTTGATTATAAAGTTCTGACTGTACCCTATAAGAAGCAGggttacctcttatctagtg
gtagagtgtagcacctatccCAAATTTCTAGCAGTTTGTTTCATCCATATTTCCAGGACCggttacctcttatctagtg
gtagagtgtagcacctatccTAAAATCCATTAGCTTTTCTGAAACATTTCACTGGGTACCggttacctcttatctagtg
gtagagtgtagcacctatccGCTCCCCTCAAAGCCCACATTATCCTCAGTGACATTTATAggttacctcttatctagt
gtagagtgtagcacctatccCCAGTGAATCATGGCCAAATAACAGCAGAGGAAGTAAAGGggttacctcttatctag
gtagagtgtagcacctatccGGATAAGGACACTCTGACTGTATTTTCCCCACTTTAGCCCggttacctcttatctagt
gtagagtgtagcacctatccTACAATATTCTGTAACTCATACCAGTAGCCACTCTAAAGAggttacctcttatctagtg
gtagagtgtagcacctatccGGTACCTTACTGAACACTATTTATGCTGATAAGATACAGggttacctcttatctagtg
gtagagtgtagcacctatccCAGAAATTGGAAAGGGGAGTAAAACAGTGAAATGAGAAATggttacctcttatctagt
gtagagtgtagcacctatccGAGAATATTCTCTCTCCTTCTTGGAATATTGAGTCTTCTAggttacctcttatctagtgg
gtagagtgtagcacctatccTGTAAACCACTTAAGCAGATCCAATGAACCTACACATTTTggttacctcttatctagtg
gtagagtgtagcacctatccCTCTACTAGTCATCAGTTGCCCAAACAGAAAATACTCAGTggttacctcttatctagt
gtagagtgtagcacctatccGTCCAAGAATGACAGGGTAAACTCTCCAAGACTATATTCAggttacctcttatctagt
gtagagtgtagcacctatccAATATATTCTCTCACAGTCCAGAGGGTGACCCAGATGTCCggttacctcttatctag
gtagagtgtagcacctatccTCTTACAGTTTCTGGTGGCTGCAGCGTTCTTTGGCTTACAggttacctcttatctagtg
gtagagtgtagcacctatccCTCCCTCTGCCTCTCCTTTGTAAGGACACTTCCAATGGCAggttacctcttatctag
gtagagtgtagcacctatccACCCAGAATAATCTCCCTATATCAAAATCCTTAACATCTGggttacctcttatctagtg
gtagagtgtagcacctatccCCACTTTATTCATGATTACCGGTTCCAGGAATTAGGATGTggttacctcttatctagtg
gtagagtgtagcacctatccCTACACAGAGCAAGACCACAGTTAAAGCATTTAAACGGAAggttacctcttatctag
gtagagtgtagcacctatccACCTGAGTTATGCACAAGCCTCACACAATGCCCTTCTCGTggttacctcttatctag
gtagagtgtagcacctatccCTGTGTAATGTCAGCTGTTTGGAAACCCATGCTCAGGGTCggttacctcttatctagt
gtagagtgtagcacctatccCCTTCAAGGCGATTACTGACCGATTAAGAAATTGATTCCCggttacctcttatctagt
gtagagtgtagcacctatccAGATATTTCTAATGTGAGGGGGCAATATTTTCACTTGACGggttacctcttatctagtg
gtagagtgtagcacctatccTTTTATAAAAGCATTTCAGCTCAAAAGTTGCCTAGACACCggttacctcttatctagtg
gtagagtgtagcacctatccAGCGTCGCTGACACAGATAGTTACAAATAGGAAGGCAGACggttacctcttatctag
gtagagtgtagcacctatccTACTCATTAACCAATCTCTCCTTACCCACCCCTCCACCCTggttacctcttatctag
gtagagtgtagcacctatccTAACCACCATTCTCTTCTCTACTTCTACGAGTTCAGCTTCggttacctcttatctagtg
gtagagtgtagcacctatccCTGTAACTGGCTTATTTACTAAACACAATGTTCTCCAAGggttacctcttatctagtg
gtagagtgtagcacctatccCTGATGGGCACTTAGAATGATTACATATCTTGTCTACAGTggttacctcttatctagtg
gtagagtgtagcacctatccCATGGGGGTGCAGATATTTCCTGGAGATACTGATTTTATTggttacctcttatctagtg
gtagagtgtagcacctatccGGTTTCCTCTTCCCCACACCTTTGCCAATACTTACCTCTTggttacctcttatctagtg
gtagagtgtagcacctatccTGCTACTGAGTTGAGTTCCTCCCACATTTTTTATATTAACggttacctcttatctagtgg
gtagagtgtagcacctatccAATACTTTCCCCCATTTTGTAGGCTGTCTCCTCACTCCGTggttacctcttatctagtg
gtagagtgtagcacctatccTTCCTTTGCCGTGACATCAATTTAATTTGATGTCATCTCAggttacctcttatctagtgg
gtagagtgtagcacctatccAGTCTGGCTTCATTTTTCTGGATGTAGATATCTTGTTTTCggttacctcttatctagtgg
gtagagtgtagcacctatccCCTTTCTCCATGGTGTGTTCTTGACACTTTTGCCAAAAATggttacctcttatctagtg
gtagagtgtagcacctatccGTGTCTGTTTTCATACCAGTCTCATGCTGTGTTGGTCACTggttacctcttatctagtg
gtagagtgtagcacctatccGGACATTTTAATATTAATTCGGAGGTTCATCAATGGGTACggttacctcttatctagtg
gtagagtgtagcacctatccAGAAATGATAAATGTGATGTGACAGACATGCTAATACTCCggttacctcttatctagtg
gtagagtgtagcacctatccTTAAAATGTTAGGAAGATATTTCAAAGAGGACAGCCATCCggttacctcttatctagtg
gtagagtgtagcacctatccTTTGTGCCACAGTCAAAAAGACAACACCAAGTTGGATAGAggttacctcttatctagt
gtagagtgtagcacctatccCCATATGAGGAAGTCACATATTCTCTATGCAAGTGTAAAAggttacctcttatctagtg
gtagagtgtagcacctatccAAAACTTGGATACAGAACAATGTACACATTATGCTTCCACggttacctcttatctagtg
gtagagtgtagcacctatccGCAACCTACATAAGAAACTAAATACGATGGTTAGTCATTGggttacctcttatctagtg
gtagagtgtagcacctatccAGGAGTGGGAAATGGATGAACAGGAGACCAGGCTGGAGGGggttacctcttatcta
gtagagtgtagcacctatccAAAAAGTCAGGTGAGGCTCTAGGAAAAATATCTGGCCAGAggttacctcttatctag
gtagagtgtagcacctatccCAGATTTCAGCTGCTAACATATGTAAATGATCCCATCCAGggttacctcttatctagt
gtagagtgtagcacctatccACTGAAATGTGCTGATGAAACTAAATTGGGATTCTAGAGAggttacctcttatctagtg
gtagagtgtagcacctatccATTACTGAAATATACCAGCCCTCCTTTGAAGCATTATAGGggttacctcttatctagtg
gtagagtgtagcacctatccCTGACAGGAATCATAAGCTAGGAAGGAAGTTGCATGTAAAggttacctcttatctagt
gtagagtgtagcacctatccGCCCTGCCAGAGACAAATGAAATCAGAACAGATTCTCACCggttacctcttatctag
gtagagtgtagcacctatccCTTACTCAGTCCCTTTAAATATGTCCCAGACAAAGAATGCggttacctcttatctagt

gtagagtgtagcacctatccGAAAGGAAGGTTTTCTTAGATCTGGAAAAGGCAAAAAAGAggttacctcttatctagt
gtagagtgtagcacctatccCCATTGTCTTTCCTTGGGCAACAGGCTGCCTTTACCAGCTggttacctcttatctag
gtagagtgtagcacctatccTAGAGAAAAGTGTGCACTTTCCAGCCAGGTGGCTCACACCggttacctcttatcta
gtagagtgtagcacctatccGCCAAGGCAGGCGAGTCACGAGGTCAAGAGATCAAGACCAggttacctcttatct
gtagagtgtagcacctatccGCACGTGCAAAATAGGACATTGTTTTTAAGTGTCTATGTAggttacctcttatctagtg
gtagagtgtagcacctatccTTTTCATTATAAGCTAGCACCTCAGCACTTTAATTTGGGGggttacctcttatctagtg
gtagagtgtagcacctatccGCATCTAAACCTCAGTCATTCATCATTAATAAGAAGTGAGggttacctcttatctagt
gtagagtgtagcacctatccCTGCCTAGTTCTGTTTATCATCAGGATGTACGCCTTGAAGggttacctcttatctagt
gtagagtgtagcacctatccTGTGCACCTCCCAGCCTCTGCCCACACACATCCCTGTAAAggttacctcttatcta
gtagagtgtagcacctatccACAAGCAGTCCCTAATCTTAGATGGTAGCATATTTCAGAGggttacctcttatctagt
gtagagtgtagcacctatccAGTCACCCTGTGTCCTTCTCCTAGACTTTATTGTCCTAAGggttacctcttatctagt
gtagagtgtagcacctatccTGTCTATAAACATATTTGAAGTCTAGAAACTGCCCCCAAAggttacctcttatctagt
gtagagtgtagcacctatccCCCAGGCATCAGTATTTGTTTTTAATTCTCAAAGTTACTCggttacctcttatctagtg
gtagagtgtagcacctatccACTTTTGTTGCATTTTAACTCCCATGACTGTTCAGACTTTggttacctcttatctagtg
gtagagtgtagcacctatccGGCCTACCCTATTTCATCTATCTAACGTCTCTTCTGCTTTggttacctcttatctagtg
gtagagtgtagcacctatccAGCTATGGTATTGTCTTGCTCGTCTCCAGACTTAACACAAggttacctcttatctagt
gtagagtgtagcacctatccGAATTTCTGTTCAACTCTATTCCACGTCCTGGCTTCCCTGggttacctcttatctagt
gtagagtgtagcacctatccATTACAAAAATACCCCTGAAGAGTCTGCCCACCCACCTGCggttacctcttatcta
gtagagtgtagcacctatccACGGCAATTTATGTAGTCTTTTTTTGTGCAATGCAGTAACggttacctcttatctagtg
gtagagtgtagcacctatccCCAAATCCCTTACAATGGCTATTTCTTTGCTCACCTATACggttacctcttatctagt
gtagagtgtagcacctatccCCTTTCTATCATAGGGCTTCATCACTTCAGAACTGTCACCggttacctcttatctagt
gtagagtgtagcacctatccGGGCCAAATAATTGTGGAGGCTGTGCTATTCATTAATATGggttacctcttatctagt
gtagagtgtagcacctatccTCACTAGATGCCAGAAGCACGCTTCTCCCAAGTTATGACAggttacctcttatctag
gtagagtgtagcacctatccGAAGAAGTTTCCAACCCTAACCATTTGAGAAGCACTGCTCggttacctcttatctag
gtagagtgtagcacctatccTATGGCTAGTCTCCCAAACTACTTCAAAGGGGTCGTCTCCggttacctcttatctag
gtagagtgtagcacctatccTTTTACCCAAGTTTACGATTGATTACTCTAGACCATCTTCggttacctcttatctagtg
gtagagtgtagcacctatccTTCCCAAAAGAAGCCAACACCAAAACAGCCGAGACAGACCggttacctcttatct
gtagagtgtagcacctatccCCTTAGGGGGATTCTACAGAACTTATCACATGTATGGCTTggttacctcttatctagt
gtagagtgtagcacctatccTCATGACTGTGATTTTTCCATTGTATACCTTTGCAGCCCAggttacctcttatctagtg
gtagagtgtagcacctatccCCATGCCTGGCTAATACTAGAACCTTTCTTAACATTAGAAggttacctcttatctagt
gtagagtgtagcacctatccGGTTCCTGACTCTAAATATTAATTCTTTCATTCCTTCCTCggttacctcttatctagtg
gtagagtgtagcacctatccTGTGGGATGGGGATTATAATGTTCTGGCAAAGCCCTTTGAggttacctcttatctagt
gtagagtgtagcacctatccCCAACTCTTAATTATTTGAGCTAATGGAGAAGGACAAAGCggttacctcttatctagt
gtagagtgtagcacctatccGGTTACCGAAAGTTAGTAGACTGATTTTATTGTGTGATTCggttacctcttatctagtg
gtagagtgtagcacctatccTTGAGCATTATGTTACTGGCATAGCATGACCAAAGGAATGggttacctcttatctagt
gtagagtgtagcacctatccATCTGGAGGTGATTTCAATGTTAACATTTTCTTGGCAAGTggttacctcttatctagtg
gtagagtgtagcacctatccACCATAAAATTTGCCATTCCTAGCAGTATTAAGTGTACAGggttacctcttatctagt
gtagagtgtagcacctatccACCATCACCACCACACATCTTCAGAGCTTTTATATCTTCCggttacctcttatctagt
gtagagtgtagcacctatccAGATAGCCTCTCCCTTCTATTGACTAATAATCTGCCAAGCggttacctcttatctagt
gtagagtgtagcacctatccTCCTGTAGCTCTTATTGAAATAAATGTAGTCTCCCAAAAGggttacctcttatctagtg
gtagagtgtagcacctatccTGACAAAGGTACCAAGTATCTTTGAAGAAAGAATACTGTCggttacctcttatctagt
gtagagtgtagcacctatccTGAGGAAACCAGATACTCACATGCAGAAAAATGGGATTAGggttacctcttatctag
gtagagtgtagcacctatccCTTCTAGAAGCATAAGGAAAACACTTTGTGACATGGGTATggttacctcttatctagt
gtagagtgtagcacctatccTTTTGGATAAGACCTCAACACACAGGAAACTTAAGCAAAAggttacctcttatctagt
gtagagtgtagcacctatccTAAAGGAAACAACCAACAGATTCAAGAGACAACCTACAGAggttacctcttatctag
gtagagtgtagcacctatccGAGATATCACCTCACACATTAGGATGCCTATTAGGAAAAAggttacctcttatctagt
gtagagtgtagcacctatccAGTATTGGCAAAGGTGTGGGGAAGAGGAAACCCTTGTACAggttacctcttatctag
gtagagtgtagcacctatccACCATATGACCCAGGAATCTGGCTCCGGGGTATATATCCAggttacctcttatctag
gtagagtgtagcacctatccATTCACAACAGTCAAGACAGGGAATCACCCTAAGTGTTCAggttacctcttatctag
gtagagtgtagcacctatccGAGGACATTATGTTTAGTGAGATAATCCAGGCACAGAAAGggttacctcttatctagt
gtagagtgtagcacctatccTTATTTGTGGAATCTGAAAAGGCTGAACTAGTGGAAGTAGggttacctcttatctagt
gtagagtgtagcacctatccTGGGGGATGGGGAGGGGTCGGTGGGTAGAAGACGTTTGTCggttacctcttatcta

gtagagtgtagcacctatccGAATACTCTAAGGGAATATTCATGCTGTGGACAAGAAATTggttacctcttatctagt
gtagagtgtagcacctatccGTATTCTCAGCATAAAAATGGTAACTACAAGGTAATGCACggttacctcttatctagt
gtagagtgtagcacctatccCGTTTTTGTCATTTCATAGTTATGCTTCATACATTCCCACggttacctcttatctagtg
gtagagtgtagcacctatccAATGCCTCAGTAACTCCATTCTTCCTCTATTTCTTTTTCAggttacctcttatctagtg
gtagagtgtagcacctatccCTCAAAGAGTACATGGAGAGAATATAAGGTTGTTGACTAGggttacctcttatctagt
gtagagtgtagcacctatccCTTAGCTTTATTAATAAACTCGATTTTAGCTGTCCACAGCggttacctcttatctagtg
gtagagtgtagcacctatccAGGACTACGGAAGATTCTTCAGAACATATCTTTAGCATTAggttacctcttatctagt
gtagagtgtagcacctatccCAAATAGCCTCTTTGTCCAACTCTCATAAAACTGCAAACTggttacctcttatctagt
gtagagtgtagcacctatccTATTTTTGAAAGACAAGGACTGAAGTGGATGGTAATGACAggttacctcttatctagt
gtagagtgtagcacctatccTAAATGTCTCCAATATCAATTTCCTTCACAGAACCGCATAggttacctcttatctagt
gtagagtgtagcacctatccAATCCCCATACAACTCCGAGCCCTAGTTTAAAAATGTCCCggttacctcttatctag
gtagagtgtagcacctatccCTCCACACAGTTTCTCTCCAAAGCCTGTAGCAATTTTAAAggttacctcttatctagt
gtagagtgtagcacctatccTTTTTCAAGATAAGGTGACTCCCTGTGTCTACTCTTTCTTggttacctcttatctagtg
gtagagtgtagcacctatccCCTATAACTAAATGAAAGCTAATACATTTGTCTGCTCCAGggttacctcttatctagt
gtagagtgtagcacctatccAGTTCTGGAAACTGGAGAGACAAGATCAAGGTGCCGGCAGggttacctcttatcta
gtagagtgtagcacctatccAGACCTCTTCTCTTGACTTGTCGGTGGCCACCAGCTCTCTggttacctcttatctag
gtagagtgtagcacctatccCACATGAACTCTTTTTTTGTAAGCAGGAGAGAGTGAACTCggttacctcttatctagt
gtagagtgtagcacctatccGAATCCACCCTTATGATGCCATTTCACCTCAATTACTTCCggttacctcttatctagt
gtagagtgtagcacctatccACACACATTCAGTTCATAGCCACTAGTATCAGCCTTTTAAggttacctcttatctagt
gtagagtgtagcacctatccCTGTGTCCAGATCCACAAAGTAAAATGAAGTTAAAGATGAggttacctcttatctagt
gtagagtgtagcacctatccAGAATTGAAGCCCCCCTAAAGCTCTTTTCAACTGTTAGAGggttacctcttatctagt
gtagagtgtagcacctatccTTTGCCTACTTATCCCCAAATTGACTCAAACCTAGAATTAggttacctcttatctagtg
gtagagtgtagcacctatccAACATGAAACTTCTTTGAGAGTTCTCCTTAGATACTGGGAggttacctcttatctagt
gtagagtgtagcacctatccCCAAAACTAGTGGGCAAAATTTTAACTAACAGCTCCGTTTggttacctcttatctagt
gtagagtgtagcacctatccGAACTTATACTTTGAACACATGGAGTTCTGGGCAGGAAAAggttacctcttatctagt
gtagagtgtagcacctatccAGTACTTGCAACTGTCATTTTTAAAACCCACAATTCTCACggttacctcttatctagtg
gtagagtgtagcacctatccAGGTCAAATTTCTTTCCTACAACTTTTAGAGGCACCTTTTggttacctcttatctagtg
gtagagtgtagcacctatccTTTTCAAAAATCTGTACCAAATAATAGCCCCAGAGGAAAGggttacctcttatctagt
gtagagtgtagcacctatccGAAACATTGCCGAGCATTTTCTCCTTTAGCCAGAAATTCAggttacctcttatctagt
gtagagtgtagcacctatccCTTTGGAGCAGCTGCCCTATAATAACTATTCAGAAAAAAAggttacctcttatctagt
gtagagtgtagcacctatccCAATTAGTTCTCATCCTAATCTTTCTGACTTTCAAGTCAGggttacctcttatctagtg
gtagagtgtagcacctatccCTTTCGTAGCATTATTTTAAAAAGCAAAGATTAAGGCCGGggttacctcttatctagt
gtagagtgtagcacctatccGCCTGGGCAATAAGCGTGAAACTCCATCTCAAAAGAAAAAggttacctcttatctag
gtagagtgtagcacctatccTTGATCCTCTATTACATGTTATGTACTATGTTGAGCCTTGggttacctcttatctagtg
gtagagtgtagcacctatccCTATAAATGGGCAAATAGATGTCGAGGGTGAGCTAAGGGCggttacctcttatctag
gtagagtgtagcacctatccCTATATAATAGAGGCAAGCAACCCAAGTCATCAATGAGCCggttacctcttatctag
gtagagtgtagcacctatccTCTTCGTTTCTCTTGCATTCCTCTTGCTTAAAAACTGTGAggttacctcttatctagtg
gtagagtgtagcacctatccAAAGAAGATATTTCCACTGCCCTTTTCTTTCTGTATACAGggttacctcttatctagtg
gtagagtgtagcacctatccAGCTTGCTCTAACTTTGGATTCTACTCTTCCCTCACCCACggttacctcttatctagt
gtagagtgtagcacctatccGGGACTACAGGCGCCCGCTACCACACCTAGCTAATTTTTTggttacctcttatctag
gtagagtgtagcacctatccCGCCCGGCCCCAGGATAATGAATTATTAAGAAATAAATGAggttacctcttatctag
gtagagtgtagcacctatccTGGGGAAATGTTTTCAGTTACTCTAACAAAAGAGAACTCCggttacctcttatctagt
gtagagtgtagcacctatccCTCTGAAAAAAAGGAGGGAGGGTGAAGCGGAGATCATTCTggttacctcttatcta
gtagagtgtagcacctatccTGAAATGTGTGTAAAGTCCAACGATGCTAAGTTTTTTGTCggttacctcttatctagtg
gtagagtgtagcacctatccCTTCTATAAGTACTTGGCCAGTTGACCATTTTGCTCTCTGggttacctcttatctagtg
gtagagtgtagcacctatccCGTCTCAAATATATTAGGGGCTCAATTAATGGTCGTTGAAggttacctcttatctagt
gtagagtgtagcacctatccGGGGACTTCGCATGCATAATAATAGGTGCTCAATACAGTAggttacctcttatctag
gtagagtgtagcacctatccCCTAGGGATTAACAGAACATCAGAGTCTCTCATTCAAACAggttacctcttatctag
gtagagtgtagcacctatccGTTTGAAGGGAGAATAAAGATGAACAAAACCTGACCCACTggttacctcttatctag
gtagagtgtagcacctatccTCCTCTCATCTGGACTCGCATAAACAGCCTCCTTCCTGGTggttacctcttatctag
gtagagtgtagcacctatccGCCATGCCTATCAAACCCATTCCTCGCAGAGCCCCCAAACggttacctcttatcta
gtagagtgtagcacctatccCTGCAGAATGGAATTCTCTGATTACTTTACACCTTGCTTAggttacctcttatctagtg

gtagagtgtagcacctatccAACCAAAGTTCAAATCTTGTAATGGGTTACGTGAGGCTCAggttacctcttatctagtg
gtagagtgtagcacctatccATCTGGCTCTTACTGCTTCTCTATGCTTTTGTTTGGTTGGggttacctcttatctagtgg
gtagagtgtagcacctatccTTTGGTTTGGTTGTTTTTTTGGTGGGCATAGAGGGCTTTTggttacctcttatctagtgg
gtagagtgtagcacctatccGCTGTTTCTAGTTCTTGCTGTCTCCCGATCATTCCTCACTggttacctcttatctagtg
gtagagtgtagcacctatccCTCCTCTTTCTCTTCTTCACTTTTAAGCCTTTGCTCTGGTggttacctcttatctagtgg
gtagagtgtagcacctatccGGTATCTCCCTTTCTGACTCCCTCAGTTGCTCAGACTATCggttacctcttatctagt
gtagagtgtagcacctatccCAAGTCATTCTATCATTATTTGGCTCTCTTACTGATCCTCggttacctcttatctagtgg
gtagagtgtagcacctatccTTCTGTGTCTTAGTCATCCCCGTTTCCCTATCAGATTGGCggttacctcttatctagtg
gtagagtgtagcacctatccAGAGATTTTGAATGAATGAATCCTCAATGGCTGCTCTACAggttacctcttatctagtg
gtagagtgtagcacctatccGCCTAAAACAGCCTCGATGACCTCTTTTCTACTTCGCCCCggttacctcttatctag
gtagagtgtagcacctatccCCAGAGGGACATTGAAACTTCTTACCTGAGCATAGTAAAGggttacctcttatctagt
gtagagtgtagcacctatccTTTCCCTAAACTATCTCTCAAAAATCAAGCCCAGTATATGggttacctcttatctagtg
gtagagtgtagcacctatccGTTCTCAACTCTACCACTACCACCATCGCCAAGTTCCTAAggttacctcttatctag
gtagagtgtagcacctatccTACAAGCTCAGAGTATCTTTAACATGTAGGGTTTTCTCTCggttacctcttatctagtg
gtagagtgtagcacctatccACTTACCAGTGAAAGGACCTAATTCCCCTATCCATAGGTAggttacctcttatctagt
gtagagtgtagcacctatccGGCACCCCATATTTCTTCTATAACTTCCCATCCTTTTCCAggttacctcttatctagtg
gtagagtgtagcacctatccGCCTCCTTAAGAAGTATGGGATATAACATGGCTCCTCTAGggttacctcttatctagt
gtagagtgtagcacctatccAGGGGCCTTGTTTCCAATATAAAAGCTACATAAAACTTCAggttacctcttatctagtg
gtagagtgtagcacctatccTGATTTCACACCACCCCCACCAATTCCTAAAAAGCTTCCAggttacctcttatctag
gtagagtgtagcacctatccCCTCCTGGTCATTTCTCTCTTGGCATCACTTCATACAAAAggttacctcttatctagtg
gtagagtgtagcacctatccTGCATATATGTAAAAAGACATTCAGTACAGCACTCTTTGGggttacctcttatctagtg
gtagagtgtagcacctatccCAATAAAGGACTTACAAAATAAACGACAGCACATCTAGACggttacctcttatctagt
gtagagtgtagcacctatccTGAGGTAGATCATTGCCACATGTTCATGCATTGCAGTAGCggttacctcttatctagt
gtagagtgtagcacctatccGATATGTAGTTAACAAACAGAAAGCAAGACACAGAACAGGggttacctcttatctag
gtagagtgtagcacctatccCTAGCGGTTACTGGAGGAGGGTTAATCTGTATTACATTTTggttacctcttatctagtg
gtagagtgtagcacctatccTTTAAAACAAAAGCTATCCAGAGGAAGTACAGGCAAAAGGggttacctcttatctag
gtagagtgtagcacctatccGCCTGGCTCACTCTACCCGCCTGACTCATGGCTTCCGGTGggttacctcttatcta
gtagagtgtagcacctatccAGACGACTTGTTTCTGTGATTTCTCAATCTTAGCAGCACTggttacctcttatctagtg
gtagagtgtagcacctatccTGCCCCCACATCAAAAAATGCAGCTCCTCTTTCCCAACATggttacctcttatctag
gtagagtgtagcacctatccAGCACATCTCGAAGCTTCTATTTCCCTGTCCCTATACACAggttacctcttatctagt
gtagagtgtagcacctatccTTTGTGAAAAACATTTCTTCCATATTAAGTACCAGACCGCggttacctcttatctagtg
gtagagtgtagcacctatccTTCTTGACAATAAACTCTGGGGAAAGAAGCTATTTGTGCTggttacctcttatctagtg
gtagagtgtagcacctatccCTGCTGGCCGGTTCCTTGGTGAAGAGTTCTGATGGCATAAggttacctcttatctag
gtagagtgtagcacctatccGGAAGTGTTGCCAGTATTTAGAGAACCCCGTAAATGAAATggttacctcttatctagt
gtagagtgtagcacctatccCCATGCCAAGGAGCCAAGAGGAATAATTAAAGGAGTGAATggttacctcttatctag
gtagagtgtagcacctatccACAAGTCACATAATCAATGATTCATTACTTCACACACAGGggttacctcttatctagtg
gtagagtgtagcacctatccCAACTGGTTGTGAACAGGAAGAACAAAAATCTTTATAGCAggttacctcttatctagtg
gtagagtgtagcacctatccTGGTTGAAGAAATTCTCAAAGTTTGGGCATGTCTGAAAAGggttacctcttatctagtg
gtagagtgtagcacctatccAAATCAGAATCCCAGATAACTCTGTGTGAATCTTACATTGggttacctcttatctagtg
gtagagtgtagcacctatccGATGTTTTAATCCTTTGTCACTAAATGGTTACTAACGCTGggttacctcttatctagtgg
gtagagtgtagcacctatccGCAAAATTTACCTGCCTATAAAGTCGCAAAAACAGTTGTAggttacctcttatctagtg
gtagagtgtagcacctatccATCTTGATTAACTAGTCCATCTTCCTACCACACATGATTAggttacctcttatctagtg
gtagagtgtagcacctatccCAAGTTTGAAGGGCCCCAGTAGAAAATCAAGTGTTAATACggttacctcttatctagt
gtagagtgtagcacctatccTGACAAAGAAAAGCTAACTTCAACATAACTTGTTTCTGGCggttacctcttatctagtg
gtagagtgtagcacctatccCCTATTTCAGGTGCTACAGATACTGAAAATAGGAGTATTTggttacctcttatctagtg
gtagagtgtagcacctatccAATTCCAAACGAAGGATTGACTCTGATAGGCTTACAACAAggttacctcttatctagt
gtagagtgtagcacctatccTTCTCAAGGAATTCAACCCCTCCAGTCAATACTTTTTCTTggttacctcttatctagtg
gtagagtgtagcacctatccTTTGCATGTGGGAGTACACAAATGAATTGAATATTGGATCggttacctcttatctagtg
gtagagtgtagcacctatccGTTTCACCTTACATGGCTTTATTGAAATAACATTCCATCGggttacctcttatctagtg
gtagagtgtagcacctatccTGATATTAACAGATTATGCATTTCCTCAGAGAAGCAGTAGggttacctcttatctagtg
gtagagtgtagcacctatccACAGACATATATATGGCTCTGGTTTGAAAGAGAAGACAATggttacctcttatctagtg
gtagagtgtagcacctatccTGGTCCACATGTATTAAAAAACATGTCAATTACTTGGTGCggttacctcttatctagtg

gtagagtgtagcacctatccTCATCAAATGGTAATTAGGGCAACATATGTAAATGCATGCggttacctcttatctagtg
gtagagtgtagcacctatccCTCCCAGGTTATTTGAATGGTATCTTTGGAGGGCTTACTCggttacctcttatctagtg
gtagagtgtagcacctatccACCCACAATACCTCCACTATTACAGCTTATAGGAAATTACggttacctcttatctagt
gtagagtgtagcacctatccCCATGGAGGGGAAGGGATCTAAAGGTGCTCATGTCAAGTTggttacctcttatctag
gtagagtgtagcacctatccTTACTGTCTACCCAATGTAAAATGAATTTTCTTCACAGCCggttacctcttatctagtg
gtagagtgtagcacctatccACAGTTTTCTTGTATTGACTTTAACTTTAGAGGGGACAGTggttacctcttatctagtg
gtagagtgtagcacctatccACTGTCCCTAACCATTAAATTCCCAGTCTTTCAAATAATGggttacctcttatctagtg
gtagagtgtagcacctatccGCGGGGGGTGCTGGGTGTGACAGGGTGCAGTGAATAAGATggttacctcttatcta
gtagagtgtagcacctatccAACATTTGTAAGACTTGAGTGACAGAGGCAAAATTTTAGCggttacctcttatctagtg
gtagagtgtagcacctatccAAGGGAAATCAGAATCGCCGAGCCTTGCCATCTGTCAGTGggttacctcttatctag
gtagagtgtagcacctatccGGTCTGTCCCAAGATGTTCTCAATCATGTGCTTCCTGTTGggttacctcttatctagtg
gtagagtgtagcacctatccACTTTTGGTCAGTGAATCAGAGTCCTAAATTATTTCAGGCggttacctcttatctagtg
gtagagtgtagcacctatccAGATGAATTCTGATGCATTACTTAGTCTGAGACCAAACTCggttacctcttatctagtg
gtagagtgtagcacctatccGCACAACTTCCTAGATTTCTGCTTCCATTTCGCCTAAAATggttacctcttatctagtg
gtagagtgtagcacctatccGTGGATTAAAAATACAGGTACCTTGACTTTAAGTAGACAGggttacctcttatctagtg
gtagagtgtagcacctatccTGTATTTTTTCCTAACCCAATTCCTCGATAAGGATGTAAGggttacctcttatctagtg
gtagagtgtagcacctatccCATCAAAACATGTTGCTTTAGTATATAAACCAGCTGATGCggttacctcttatctagtg
gtagagtgtagcacctatccGGATCACATGAGTCTAAGAATAAAACTGTCGATTTGTAACggttacctcttatctagtg
gtagagtgtagcacctatccCAGCTCTTACAAAATCCTTAAAAACGTACATCAGAGAAAGggttacctcttatctagt
gtagagtgtagcacctatccTGTCATCACAGAATATCCAAAGGACACCTCACAGCATGGAggttacctcttatctag
gtagagtgtagcacctatccTTTCCCTTCACTTATGGTTTACATAGGATATGCATTGGGAggttacctcttatctagtg
gtagagtgtagcacctatccAGGTCTCCTTCTGTTTCTGTTGCATTTACCTATTATTGGAggttacctcttatctagtgg
gtagagtgtagcacctatccAATATTCCATTTTACAGCTATCTACAGATAGGCAGCCCTTggttacctcttatctagtg
gtagagtgtagcacctatccGAGTTCTCTGCAGCTCGTCATTTGAAAGGATTCCACTTAAggttacctcttatctagtg
gtagagtgtagcacctatccGCATTTTCAGGTTTTCAGGTTTGCCCAGAAGTCAACGTTTggttacctcttatctagtg
gtagagtgtagcacctatccGGTCTAAGTAACCAAAATGAAACTTCTTTTGCGCAATAGAggttacctcttatctagtg
gtagagtgtagcacctatccCCCTACTGGGTAACTCTTTGGTTAGATTATATATAAGCATggttacctcttatctagtg
gtagagtgtagcacctatccTTAAGTAAACTGCCAGGTAAGAGTAGGTCTGTTATTAATCggttacctcttatctagtg
gtagagtgtagcacctatccAAATGTGCATATACTCAAACAGATGGTACAAACTTCCAACggttacctcttatctagtg
gtagagtgtagcacctatccCCTCCCCACCCCACCCCGCCCACCAAGTCTTACAGCTAAAggttacctcttatcta
gtagagtgtagcacctatccGGCTTTCCTAATACCCACGACGTGCACTATGACCCAGCTCggttacctcttatctag
gtagagtgtagcacctatccGCCTTGCACTCTTAGGCGGCAGCGTCCTTTCTCCCTCTTGggttacctcttatctag
gtagagtgtagcacctatccAACTGCTGCCATGTCAACCAGTGGAATCAAGGGCTTCTGGggttacctcttatctag
gtagagtgtagcacctatccTGCAGGCAGGCGGATCACTTTCTATAAGTTTAAGCTACAGggttacctcttatctagt
gtagagtgtagcacctatccGGATCCTTTACTGTCCCACGCACTAGATTCTTCTACTTTAggttacctcttatctagtg
gtagagtgtagcacctatccCCATGGGGTCCTAATTCCAGGCTCAGAATCTTCCATCCAGggttacctcttatctag
gtagagtgtagcacctatccTCAAAAAGACTGTGTAATGACCTTGTAAAACACTAACCCCggttacctcttatctagt
gtagagtgtagcacctatccGAACCAATCCTGATCTCTTAAACCCCCAGTGAGTCATAAGggttacctcttatctag
gtagagtgtagcacctatccGACACGGTCCAATCAGGTTCTGAACAATTCAATCATCCCCggttacctcttatctag
gtagagtgtagcacctatccTGGCCACACTCCCCCAATAGATCCAAACTGAAGTACCATCggttacctcttatctag
gtagagtgtagcacctatccGGTTTGAGGCAGGGAGTAGAGAAAGCATGGAGTGGTAGGTggttacctcttatctag
gtagagtgtagcacctatccTGGGGTGGGGGTTCCTCTGGGTTCTCTCGGTTCAGGAGGTggttacctcttatctag
gtagagtgtagcacctatccTGCAGGTTCTGTAGTGGAGGGAAGGTGGGCGGACAGGTCTggttacctcttatctag
gtagagtgtagcacctatccCCAGGGGACAAAGGTGGGACAAGAGGCTCTGTAATCATCTggttacctcttatctag
gtagagtgtagcacctatccGAGCTCACTCCCTGCAGAAGACGTACTCGGTGTAGCTGGTggttacctcttatctag
gtagagtgtagcacctatccTCTTGTCCTCGCTCTGGTCGGTGCTGCTGGCAAAGGCGCAggttacctcttatctag
gtagagtgtagcacctatccCTGCTCCAGGAAGTTGAACTTGAGGTAATAGCGCGAGGTGggttacctcttatctag
gtagagtgtagcacctatccGTGTCCCCAAACACCTCCTTGGCCAGCGACGTCTTTCCGCggttacctcttatcta
gtagagtgtagcacctatccGCCCGATGGTGTAGGAGCCGCGGTAGCCGATGGTGATGTAggttacctcttatcta
gtagagtgtagcacctatccCGCGCCCCCGGACGGACTGCGGCTAGCCAGCTCCAGCGTGggttacctcttatc
gtagagtgtagcacctatccTCGTCACCCAGCGAGCCCTCCTTGTGCACCCCGCGCCGCGggttacctcttatct
gtagagtgtagcacctatccGGGCTGCTGGGGCGCCCCGAGGCGGCGCACGAGCTCTGGCggttacctcttatc

gtagagtgtagcacctatccGCACGAGCTGCAAGTCCCGCAGGTAATCCAGGATGTAGCGggttacctcttatcta
gtagagtgtagcacctatccAGCCGTCCCGGTCCAGAAAGAAGCGGCCTTTGCTGTCCCGggttacctcttatcta
gtagagtgtagcacctatccGCACCGACACCACCGTGCAGCGCCGGGTCACGTACACCTGggttacctcttatct
gtagagtgtagcacctatccGAGGACGACGAGGAGCCACTGCCGCCCCCGCCGCCGCCCggttacctcttat
gtagagtgtagcacctatccCCAGAGCCATGACAGAGAGGTGGCCGGGCCGGGACAGTGGggttacctcttatct
gtagagtgtagcacctatccGCTGCACTCAGGAGCTGCAACCGCCTTCCCCGGAGCCCCGggttacctcttatct
gtagagtgtagcacctatccCCACCGCCACCGCCGCCACCTCCTAGAGCCGCGCGGAGCCggttacctcttat
gtagagtgtagcacctatccCGCCCCCTTGAGTTCCAGTGCGCTCCGCCCGCCTTGCCCGggttacctcttatct
gtagagtgtagcacctatccCGCGCTGCTTAAGTAGAGCCGCCTGCTCTGGTGCCGTGGCggttacctcttatcta
gtagagtgtagcacctatccGGTGGGGAGCGGGGACTCGATTTGCATCTTAAACGCGGACggttacctcttatcta
gtagagtgtagcacctatccGGCGCCCGGACTCCCACCCGCTCCCCACCCACCAGCCCCGggttacctcttat
gtagagtgtagcacctatccTTGGCGGCTTCGGAAGACAGCTCCTTGGAGGGACAGTCACggttacctcttatcta
gtagagtgtagcacctatccCCCCTCCCCTCTCGCCCCAGCCCGCACCCAAGGACTCTGGggttacctcttatct
gtagagtgtagcacctatccTTCGGAAGCCTGTCGCCCCGGGAGTGGAAATGCGTGGCCGggttacctcttatct
gtagagtgtagcacctatccCAGTGCAGCTGCGGCCTGCCATCGTTGGTTGTCGTGTCACggttacctcttatcta
gtagagtgtagcacctatccGTAGAGAGACGCGAAAAACCTGGAGAGACGAAAGAGCCGCggttacctcttatct
gtagagtgtagcacctatccATAAATAAACCTAGAAACAAAAAGCAAACCTCAGTGTCCCggttacctcttatctag
gtagagtgtagcacctatccCAGAGACAAAATGAGATGAAGCTTCAAATCTGCTCTTAATggttacctcttatctagt
gtagagtgtagcacctatccTTCGAAGTGAAGGGGTAAGACTCCGGAACGTTCGGTCTCCggttacctcttatcta
gtagagtgtagcacctatccTTGAATCGCGCCCTTAGACTCACAACAGTTGCAGATCTAAggttacctcttatctag
gtagagtgtagcacctatccCCTGCATGTTTATCCTTCCAAAAGTCCAGGCGCCCCTTCTggttacctcttatctag
gtagagtgtagcacctatccCGGTCACTCTTCGGCCCTGTCCAAGTTCCCGTCAGTCTCAggttacctcttatcta
gtagagtgtagcacctatccATATTTATATATCTGAAGGCCCTTGTCTGGATCCCATCTCggttacctcttatctagtg
gtagagtgtagcacctatccTCCTGCTGCCTACACGTTTGCTTTTCCCCCTCGGACTGCAggttacctcttatctag
gtagagtgtagcacctatccCCTCTGGATGGCTCCCTACACACTGTCTGCGCTGCTCCCAggttacctcttatcta
gtagagtgtagcacctatccCTGATTTGGGTATGAAAATAAGTAAGAGCTCCTGAATGCTggttacctcttatctagt
gtagagtgtagcacctatccAGAGAAGAGTGCAGAGGAGGGTGTCAAGAAATGATTGGGTggttacctcttatcta
gtagagtgtagcacctatccCGAATAATCTCCAAAGCGCTTTCAGTGTAGCCCAATAAGCggttacctcttatctag
gtagagtgtagcacctatccAGGGGTCTGCGCAGTTATCACATCAGTTTGAAGTATAGACggttacctcttatctag
gtagagtgtagcacctatccATAAAATTTATCCTTCTGTGTAACTACCCAGTGAGCTTCAggttacctcttatctagtg
gtagagtgtagcacctatccTGTTTATATCTGCTCAACAATTACCGCGCTCATGTCAGTGggttacctcttatctagt
gtagagtgtagcacctatccAAATGAGATCAACCCCCAAACCAGTTATCTTAATTGCAAGggttacctcttatctagt
gtagagtgtagcacctatccATAATAGATAACACAAAACTTCCAAGTCCCAGTGCAGGATggttacctcttatctag
gtagagtgtagcacctatccAGACCTGACAAAACCAGCCTCTTTTCTACAAATCCAAGAAggttacctcttatctag
gtagagtgtagcacctatccCAAATCCTGCCAGCTCTGTGTTAAAAATCTTTCTAGACTAggttacctcttatctagt
gtagagtgtagcacctatccTTTCACCATCTCCCTATCATCACCCTGGACCAAGCCACCAggttacctcttatcta
gtagagtgtagcacctatccCTTGCCTGGATTATTGCACTGGTCTCCTAACTGCTTCTACggttacctcttatctagt
gtagagtgtagcacctatccCTCCTACAGTCTTGTATCACCAGAACAAAAACAGGGACCCggttacctcttatcta
gtagagtgtagcacctatccCCAGTGGCTTCCTATCTCAGAAAGAGCAAAATCCAAAGTCggttacctcttatctag
gtagagtgtagcacctatccGAGTTATCCCACCAGCACCTCTCTGACACATCTCCTGCCCggttacctcttatcta
gtagagtgtagcacctatccGCTTTAGCCACATAGCCCCTTTCTTGTTTTCTCAGGGCCTggttacctcttatctagt
gtagagtgtagcacctatccTTGTAAAAGTCTCATTTCTCCTCCCTCCCGTGTTCCTCACggttacctcttatctagt
gtagagtgtagcacctatccAAGACCTTTCCTTGATTCCACTTCTTAATGCAGCACAGCCggttacctcttatctagt
gtagagtgtagcacctatccGCAGCTTCCTCTTCCTTGCCTTATTTGTTGACTCTTTTCCggttacctcttatctagtg
gtagagtgtagcacctatccCAGTTTTTGTCTGCTTGGTTCACTGCCACACCCCCACTGCggttacctcttatctag
gtagagtgtagcacctatccCTCACGCAAGCATTCAACCACTATTTATTCAGGGCCTACTggttacctcttatctag
gtagagtgtagcacctatccTTCCAGGCTGCATGTTAAAAAATAATCACTGCCCACTAGGggttacctcttatctag
gtagagtgtagcacctatccATAAAGATAAGTAAGCCTATTTCAGGTGGTGATAAATGCCggttacctcttatctagt
gtagagtgtagcacctatccAAAATTATAAGATGTGATAGAGGGACTGAGTGTGGAATACggttacctcttatctagt
gtagagtgtagcacctatccCTGGGGAAGGCATCTCTGTGAATGTGACATTTGAGCTGAAggttacctcttatctag
gtagagtgtagcacctatccAAGAGCATTCCAGGCACAGGGTCAAACAAGTTAAAGGGGCggttacctcttatcta
gtagagtgtagcacctatccGTTTATGTATTGTGTCATCTACTTGTATCCTGTGAATCTCggttacctcttatctagtg

gtagagtgtagcacctatccAGTACAGAGTCAAATGTGAGTGATACCGTGTGCTTATTCAggttacctcttatctagtg
gtagagtgtagcacctatccTGCAGCCTACTATCATCATTCTCTACATCTTGACATAAGGggttacctcttatctagtg
gtagagtgtagcacctatccCTTATGGAAGCCAGGTAATGCATGAGAGATATAATAATGAggttacctcttatctagtg
gtagagtgtagcacctatccCTAGTTAATTTTGTGTGTTCTGTTTCTTGCCTATTGGTTGggttacctcttatctagtgg
gtagagtgtagcacctatccTTCAAAAGGAAATACTACTGGGGAACTCTGATGCTGGGACggttacctcttatctag
gtagagtgtagcacctatccGAGTCATAACTCTTTAGCATCTCCAGTATGTTAAATATGCggttacctcttatctagtg
gtagagtgtagcacctatccGGAGGATTTGGAAACCCACTAGCCCTGTGTTTAGTCTCAAggttacctcttatctagt
gtagagtgtagcacctatccATGGTCATTGCTAATTGATGGTCATTGCTCCTTTTCAAGTggttacctcttatctagtgg
gtagagtgtagcacctatccAGTGTGCCTGCAAGTAACCTGTCCTTTGATCCTAAAGTGGggttacctcttatctagt
gtagagtgtagcacctatccGGAAACATCTCTGGACCTACATTCTCATCTCATTTGCATCggttacctcttatctagtg
gtagagtgtagcacctatccCCCCACCTACTGCCTTCTCTTCAAACCAGCTTCCCCTCCCggttacctcttatctag
gtagagtgtagcacctatccCTTATCCAAAGTTTCCATTACCATGGTTTCAGTTACCCATggttacctcttatctagtg
gtagagtgtagcacctatccTGAAATCTCCCACTGTCCTGCTTCGTCTTGCCCTGGACATggttacctcttatctagt
gtagagtgtagcacctatccTCCACACCTCTACACTCCCAACCCATGAGTCACTTAGTAGggttacctcttatctag
gtagagtgtagcacctatccCAGCTGTCAGATCAACTGTCTTAGTATCACCCTATCACAGggttacctcttatctagt
gtagagtgtagcacctatccCTTAATAAATGGCCACAAATTACAAGAGTAGTGATGCTGGggttacctcttatctagtg
gtagagtgtagcacctatccAGAGAAGCCATAAAATGCTGCCTTTAAGTGAAAAGGTGAAggttacctcttatctagt
gtagagtgtagcacctatccTTTGCTGTCACACCTCCAAACACAAAACGTATGGCACAGAggttacctcttatctag
gtagagtgtagcacctatccGAAAAGGCATTAAACTTGTGGATTGAAGACATGAAGAGAAggttacctcttatctagt
gtagagtgtagcacctatccAGCAATCAGGTTCAGTACTACCTACGGTTTCAGTCACCCAggttacctcttatctag
gtagagtgtagcacctatccGGAGACTACTGTATTCTGCATACCAAAGATTTAAGGAAAGggttacctcttatctagtg
gtagagtgtagcacctatccGTTAAAGGGGAGGAAGAACTGGTGAAATGAAGTGAGTGCCggttacctcttatctag
gtagagtgtagcacctatccTGATCTGTACATTGTAAAACACCATTCAAGTGTCAGAATCggttacctcttatctagtg
gtagagtgtagcacctatccCTTATCATGGTGCTTGACAAGTCTTCCCAATAAATACTGAggttacctcttatctagtg
gtagagtgtagcacctatccATTAAAATGAACACTATGGGGAAAGGGGAAGAGAGGCAGAggttacctcttatctag
gtagagtgtagcacctatccTTCCATTTGTCATCCTCCTAATATGTGTGAATCTTTTCTCggttacctcttatctagtgg
gtagagtgtagcacctatccGAAGGAGAGATTAATTGAATAACATTAAGTGAATCCGGAGggttacctcttatctagtg
gtagagtgtagcacctatccCCTGAAGAAAAGAGAACAAAATGTTATATAGGGTACACTGggttacctcttatctagt
gtagagtgtagcacctatccAAACAGGTCTAGACAGTAAAGAATATCCCAACATTGTATGggttacctcttatctagtg
gtagagtgtagcacctatccCAATGTTGATATTTCTATACCCTTAAGATGGTGTTTGGAGggttacctcttatctagtgg
gtagagtgtagcacctatccCAAGCATGTATAGAGTACTTACTATGAGCCAGCCCTATTTggttacctcttatctagtg
gtagagtgtagcacctatccTAATCCTAACAATAACTTTATGAGGTTGGTGCTATTACCCggttacctcttatctagtg
gtagagtgtagcacctatccGCACAGTGAACTAATTTTCCCAAGATCATACAGCAAATAGggttacctcttatctagt
gtagagtgtagcacctatccGCCTTAGAATCTGTGCTTTTAAACACTGCATTTTAAATCCggttacctcttatctagtg
gtagagtgtagcacctatccACAGATGAGAAACCCAAGGCAAAGAGAAGTAAGGTAACTTggttacctcttatctag
gtagagtgtagcacctatccACTCTAAAGTCTACGTGCTTGTGTTCCGCAGTACTGGTAAggttacctcttatctagtg
gtagagtgtagcacctatccCCTGAAAGAACTAGGGTTAAGAAAGAGTCAAGTCCTTGGCggttacctcttatctag
gtagagtgtagcacctatccTGCCTTCTTTTTTCCTTCCAACCTATTTTAGGTTTATGGGggttacctcttatctagtgg
gtagagtgtagcacctatccTGTTACACAGGTAAATTGCACGTCGCAGGGGTTTGGTGTAggttacctcttatctagt
gtagagtgtagcacctatccAGTTTTTGGATTCTCACCTTCCATCCAACCTCTACCCTCAggttacctcttatctagtg
gtagagtgtagcacctatccTCCTTGTATCCATGTGTACTTTTACCCAAATAAGAATCTCggttacctcttatctagtg
gtagagtgtagcacctatccACACTATTGTCCTAATCTTTCCCTAATTCAGTCCTGTACAggttacctcttatctagtg
gtagagtgtagcacctatccGATTTTATTTCTGGCTCTCTCAAATATCATTACGGTTTGCggttacctcttatctagtgg
gtagagtgtagcacctatccGCATGTTCATTTGGCTTTATTCTGATCTCATTAACTCTTGggttacctcttatctagtgg
gtagagtgtagcacctatccGCTGAGATCTAGGTTCCAGTTTCCCAAAGCAAGCAGTATCggttacctcttatctag
gtagagtgtagcacctatccGTCTTCTGGGGCTCATAAAATTGAACAAGTGCATTCTACTggttacctcttatctagtg
gtagagtgtagcacctatccTTGCATATCAAACTGAACACAAAAATAGCTGGCCCTCATCggttacctcttatctagt
gtagagtgtagcacctatccCACATTCTAATCACATTCTAGTGTTTTTCTGTTTCGAGTGggttacctcttatctagtgg
gtagagtgtagcacctatccGTGTTAACCAACCTGTGTTCCAAATGAAGAAAGAAACTCAggttacctcttatctagt
gtagagtgtagcacctatccGATGCTTCCTGATTTCATATCCCATGCTCTTTGAACTCTTggttacctcttatctagtg
gtagagtgtagcacctatccAACACCCAAATGAAGGTTGTCATTCAAAATAATCACCTTGggttacctcttatctagtg
gtagagtgtagcacctatccGATGCTGGAATCACTCACCATTTTTGGTACATTTGTTTGGggttacctcttatctagtg

gtagagtgtagcacctatccTAAAGTTGTAGTTGATATCTTTCTAGGGCTCTAAAACAGGggttacctcttatctagtg
gtagagtgtagcacctatccCAGACCCACATGCCTGAAAATAAATTAGGACAAAGCCAGTggttacctcttatctag
gtagagtgtagcacctatccTCATCGGGTATTCACCTTTTGACATTGGGTTTTAAAGATCggttacctcttatctagtg
gtagagtgtagcacctatccCTGAATAGCATTGAGTTTAGGTTACTTAGAAAGAAGGGCTggttacctcttatctagt
gtagagtgtagcacctatccATATGTGGTTCCAGAAAGTCTGCATATTTTATTGTGAGGTggttacctcttatctagtg
gtagagtgtagcacctatccCCACTGGCTAGACTGTAATGGTTGATTTCTTATGTAATTGggttacctcttatctagtg
gtagagtgtagcacctatccGGGGAAAAATATATTTCCCCAATTTGCTTTCCAGTTACTAggttacctcttatctagtg
gtagagtgtagcacctatccTAGATGGGAAAATCATACACTTTAGAGTTTGATGAACCTGggttacctcttatctagt
gtagagtgtagcacctatccTTTATCTTAAAATGGAGCAAAAATACCAACCTCATGGGACggttacctcttatctagt
gtagagtgtagcacctatccAATGAGATGATACAAAGCATTTTCTTCCACAATGTCTGGCggttacctcttatctagt
gtagagtgtagcacctatccATGTAGGTTTCTGCTCCCACCCTTTCATTCCCCCATTGCCggttacctcttatctag
gtagagtgtagcacctatccGAGTTTGCTTACCTCATTCTTCAATCTGTTTATCACTCTCggttacctcttatctagtg
gtagagtgtagcacctatccTCTCTTTAACAGTAACTAAAATGGACAGGGTAAATGGCACggttacctcttatctagt
gtagagtgtagcacctatccCAGCAGTTCCCCATTTCTTCCTAGGAGTCAGTCACATCTAggttacctcttatctag
gtagagtgtagcacctatccTCTCTGGTTGGAAAATTTAGTCAAAATCTAGACAGCTGGGggttacctcttatctagt
gtagagtgtagcacctatccTCAATTCCAGGGCTTATTTCTCATCCTAGACTTTGCAACTggttacctcttatctagtg
gtagagtgtagcacctatccACAGTTGGCATAGTTTTATCTCCGTAACTAAAAGTTCAGCggttacctcttatctagt
gtagagtgtagcacctatccCTCTAGGAGGTAAAGGCTGGGTAGGGGTGAGCAGAAGTGTggttacctcttatcta
gtagagtgtagcacctatccAGGGCCTTAAAAACTCCAGGTATCCTTAAAATCCAAAGACggttacctcttatctag
gtagagtgtagcacctatccCTGTTCTAAACCCAGGGACTTAAGCCATGAAATGTTGAGGggttacctcttatctag
gtagagtgtagcacctatccCCTAAAGAGGGCAAGAGCACCATCCCAAACCACAGCTATGggttacctcttatcta
gtagagtgtagcacctatccTCCCCCTAGCTTCCAGGCGACAGTGGCTCCACAACAGGAAggttacctcttatcta
gtagagtgtagcacctatccCTGGAAGAAGTATCAACAGTGCAAGTTTAGCTCCCCTTTAggttacctcttatctagt
gtagagtgtagcacctatccCACCTGCTATGTGCCTGATCTTGCCAGAAGTTATGGATGGggttacctcttatctag
gtagagtgtagcacctatccGTCCTTGGCCCATGAGAACCTGTACAATATCAAAGTAAGTggttacctcttatctag
gtagagtgtagcacctatccGGACATGCCACAGGTGAAGATCATAGGGAGGGAAGTTACCggttacctcttatcta
gtagagtgtagcacctatccTTCTAGAGAAACAGGAAAGATACTATACAAAGCTGACCTCggttacctcttatctag
gtagagtgtagcacctatccAGTCAGGTCAAACAAAACACTTTTCCTCAGCATTGCTAATggttacctcttatctagt
gtagagtgtagcacctatccATTATATTTTGGCAAGGTATCTGACAGCTCATTCATGATCggttacctcttatctagtg
gtagagtgtagcacctatccAGAGAAATGTGTGCTGGACAATAATATCAATTAGATGGAGggttacctcttatctagt
gtagagtgtagcacctatccGATTATTGATGAATGAAACAATGCGTAACTGAAGAGAACGggttacctcttatctagt
gtagagtgtagcacctatccTATGCTAATAAAATGTATAGTTACACAGAGCTTAGTGGGCggttacctcttatctagt
gtagagtgtagcacctatccGCAAGAAAGAGAAGCAAATAAGAGAGGATGTAGTAAGTATggttacctcttatctag
gtagagtgtagcacctatccGCTTATGACACAGTCGAGCAGTTTACATGAAAAGTGACCTggttacctcttatctag
gtagagtgtagcacctatccCTGTAACATGCACAGGTACAAACACCAAATACAATCGCTTggttacctcttatctag
gtagagtgtagcacctatccATTTATGTCAAATCTTTCCTCTTACAGATCCAAGCTGAGGggttacctcttatctagtg
gtagagtgtagcacctatccCTGACAGTAAAACTACACCTAGCATTCACATATCTTTAAGggttacctcttatctagt
gtagagtgtagcacctatccCTTAATAGGCAAGAGTATATTTGAAGGAGTATAAGCAAGCggttacctcttatctagt
gtagagtgtagcacctatccTTTAGCATAGAATAGCTTAGAAGAGAGTATTACTGGAGGCggttacctcttatctagt
gtagagtgtagcacctatccAGCTTGTCTTAGTCTGTTCACGTTGCTCTAACAAAATACCggttacctcttatctagt
gtagagtgtagcacctatccCTGAGAAGTCCAAAATCAAATCATCAGCAGATTAGGTGTCggttacctcttatctag
gtagagtgtagcacctatccAGGGTCTATTTCCTGGTGTGCAGATGGCTCCTTCTCTCTAggttacctcttatctagt
gtagagtgtagcacctatccCTAATCCCATTCATGAGAACCTGATCACTCCCAAGACCGCggttacctcttatcta
gtagagtgtagcacctatccCATTATATGGGGGATTTGGTTTCAGCATGGGAATTTGAAGggttacctcttatctagt
gtagagtgtagcacctatccGACCATAGCACATGTTTAATGAGTCTGAAAAGTCCTTAGGggttacctcttatctagt
gtagagtgtagcacctatccAGGCTGAATTAGGGACATTGTCTTTGGGGGTGAGTAGACTggttacctcttatctag
gtagagtgtagcacctatccAACTACAGGGTTCTAGGTGGCCGGAGCATTGTCCAGTATTggttacctcttatctag
gtagagtgtagcacctatccAAAGTGTTCTCATTGTCCAGCTCTTCTTCTTGTACAGCCAggttacctcttatctagt
gtagagtgtagcacctatccCTTGGGAGTTAGCAGCTGTATAAAAAGGGCAGTCCTAATCggttacctcttatctag
gtagagtgtagcacctatccAAATCCTAGTGCTTGCTTATCTTCCTTACTAAGAAATGTCggttacctcttatctagtg
gtagagtgtagcacctatccTGCTTCTTGGTTGGCAGATACAGCTTCCCTTGTTATCTTCggttacctcttatctagtg
gtagagtgtagcacctatccTCAAACCATCCTTTGCTGATATTAAGTTCTTCAGCTTTAGggttacctcttatctagtg

gtagagtgtagcacctatccGCCTTTCTTATAACAATTCTGTATCTACATGAAAAGCTGCggttacctcttatctagtg
gtagagtgtagcacctatccGCTGCAGACCTCAATCTATAATACACACCAAGCAATTTAAggttacctcttatctagt
gtagagtgtagcacctatccTAGTGGCAGTTTGTATGGGTGTCATGTGTTATTCAAGGTTggttacctcttatctagtg
gtagagtgtagcacctatccTGAGAGATCACTTTTCATTGTGACACATAATTTACCAGAGggttacctcttatctagtg
gtagagtgtagcacctatccGGCATTTTAAGTGGATACTTTTGACACTTGAGCTCACTACggttacctcttatctagtg
gtagagtgtagcacctatccGGAGGTGGCTATGAAATTATAGTAGTATAGTATGTGCTACggttacctcttatctagtg
gtagagtgtagcacctatccTGCATGTTTGTTTATAATTCTCTCAACTGCAGTGGTGCCAggttacctcttatctagtg
gtagagtgtagcacctatccTATTTCTAGGCCACATGAATGAAATGAGAGAACTCATTTCggttacctcttatctagtg
gtagagtgtagcacctatccACAAGGCAAGTTCCTTCCCCCCCAGGAGCCTGTAAAATCAggttacctcttatctag
gtagagtgtagcacctatccCTTGTTAGCTACAATAGGGATGCAGGCATTGAGTAAATACggttacctcttatctagt
gtagagtgtagcacctatccGGCTACAGGGCCCATTCAAGTCTGAAATCTAGCAGGGCAGggttacctcttatctag
gtagagtgtagcacctatccGGTTCCCATAGTCTTGGTCAGCTTCACCCCTGTGGCTTTGggttacctcttatctag
gtagagtgtagcacctatccCCCTAGCAGAGATTTTACATGAGGACCCCCTCCATAGCAAggttacctcttatctag
gtagagtgtagcacctatccTGGCTTCTGTGCACTCGCAGAATCAACACCACATGGAAGCggttacctcttatctag
gtagagtgtagcacctatccCTCTGTGTTGGCCCTTTTCACCCATAGCTGCAGTGGCTGGggttacctcttatctag
gtagagtgtagcacctatccCTGGCCCAAAAAAACACTTTTTCCCTCTAGGCTTCCAGGCggttacctcttatctag
gtagagtgtagcacctatccTGAATGTCTCCTCAGAAAATAGGATTGTCTTTTCTATCACggttacctcttatctagtg
gtagagtgtagcacctatccCTCTGCTTCCCTTATAAAAATAAATGCTTTTAACAGCACCggttacctcttatctagtg
gtagagtgtagcacctatccCAGTTCCCAAAAAGTTCCCCATTTCATCTGAGACCACCTCggttacctcttatctag
gtagagtgtagcacctatccCAAACTGTTCCAACTTTTGCGTATTACCCAGTTCTAAAGTggttacctcttatctagtg
gtagagtgtagcacctatccCACATTCTCAAGTATCTTTTCGGTAACACCCTACTCCTGGggttacctcttatctagt
gtagagtgtagcacctatccAAAAGAAAGAGTTTTACTGGATTTACGGTTCCACGTGGCTggttacctcttatctagt
gtagagtgtagcacctatccAGATCTCATGAGACCCATTTACCATCACAAGAACAGCATGggttacctcttatctag
gtagagtgtagcacctatccGAGCCAAAACATATCAGACAAGAACTGGGTTGACTTCACTggttacctcttatctag
gtagagtgtagcacctatccAGGGACTTATGATTTGGGCTGAGAAATTGCATTAGATGATggttacctcttatctagtg
gtagagtgtagcacctatccTCTTTCTAACGTAAGAGAACAATTTTCCTCACCAACTGCTggttacctcttatctagtg
gtagagtgtagcacctatccAAATAGGCATTTTTTTCTAATCTTCCCTCCCAATTTCAGGggttacctcttatctagtg
gtagagtgtagcacctatccTATAATCATAACCAATAGCTAGTATGTCCTGTATCTTCCCggttacctcttatctagtg
gtagagtgtagcacctatccGTGCCAAACAGTAGATCAATCATGTCTGAGATCCTTGAATggttacctcttatctagt
gtagagtgtagcacctatccGCACCATTCCACATAGTTTAAAGCATTTTATTGCACATACggttacctcttatctagtg
gtagagtgtagcacctatccGTTTCTCGCCTATTCATTCTACCAAAACCTTGTATCATTTggttacctcttatctagtg
gtagagtgtagcacctatccAACCCATAGAACTGTACAATGCAGAGTGATTTCTAATACAggttacctcttatctagt
gtagagtgtagcacctatccTACTAATAATAAAGGAAACTGCGGGGAAGAGGAGAAGAGTggttacctcttatctag
gtagagtgtagcacctatccGCAAACAGATTTTCATAAGCCCTCTAATTATTTACCCTTGggttacctcttatctagtg
gtagagtgtagcacctatccTTCTCCAAAGGAGTCTATCATCCTGCTTTCAAAATTGTTTggttacctcttatctagtg
gtagagtgtagcacctatccCTACAGATAGAAAGAAAAAGCTTTTGTTCATCTCAGGAACggttacctcttatctagt
gtagagtgtagcacctatccCTTTACCCTACCAAATTTAATATAAGACAAGATGCCACAGggttacctcttatctagt
gtagagtgtagcacctatccCGTGTTTAAATTATCTGATCCCTTTTACATCATCCTTCTGggttacctcttatctagtg
gtagagtgtagcacctatccAGTGATACTGTCCTATCCTCAACTGCTCTACATATTCAAGggttacctcttatctagtg
gtagagtgtagcacctatccAAGTAGCCATCTCCATTTCTTGAGCGCCTTCCACATGTAAggttacctcttatctagt
gtagagtgtagcacctatccTCTAAGGCAGACAGAGTTCCTATCTTCAGCAGAGGATGCAggttacctcttatctag
gtagagtgtagcacctatccAAACAAGCCTGGGTTTATCACATTCATAATCCACTTAACAggttacctcttatctagtg
gtagagtgtagcacctatccCACATAGGAAGACTGCCAAGGAGAAACTAGTACCATAGACggttacctcttatctag
gtagagtgtagcacctatccATAATATTTTCCCTATAGTTAGACAGCATTTCAGCTGACCggttacctcttatctagtg
gtagagtgtagcacctatccGCTGCTTCATCAGAGCACTGAAGGCCAAATATATATTAACggttacctcttatctagt
gtagagtgtagcacctatccAGATACTTAAATGGTTCTCATCAAACCTCCAGATGTTCAAggttacctcttatctagtg
gtagagtgtagcacctatccGAAAAGTGCATGAACTCTCTGATTGTCCTATCAGCAGGGTggttacctcttatctagt
gtagagtgtagcacctatccGGAAGACCAAAAGGCACATGTTTTTTTCTCCAGCTGTTTTggttacctcttatctagtg
gtagagtgtagcacctatccCCAGGTAGGGAGTTGAAGGGGTAGACAAGGGAATTATTAAggttacctcttatctag
gtagagtgtagcacctatccGCTGTGGAAGGAGGAAAATGAGGAGATACGTGAGTGGGGCggttacctcttatcta
gtagagtgtagcacctatccCTCAAAGGAGGGGTCCAGGAAGCCAATAGAGAATATAATGggttacctcttatctag
gtagagtgtagcacctatccCCTTAGCTGTCTCTGTCTCACTACCACTTTAACCTCCTTTggttacctcttatctagtg

gtagagtgtagcacctatccCACGTCTGAATCCTGTAGCTTAGAGTCCACTGTGAGAACAggttacctcttatctag
gtagagtgtagcacctatccTTTTCAGCTTCTCCTGGATTGATCAATACACCTTTCACTTggttacctcttatctagtg
gtagagtgtagcacctatccCCTTTATTGGTGACTTGTACTCTTGCTTGTGTTAAGGATGggttacctcttatctagtg
gtagagtgtagcacctatccTCATTATGTCATTTCTATCTATTTCTGTCTGGCTGAAGACggttacctcttatctagtg
gtagagtgtagcacctatccAAGCAGGTTCTTGTTTCTATCATCAGAATGGGTCCTTAAGggttacctcttatctagtg
gtagagtgtagcacctatccTGTTGGTTTTATTCCCACATTTCCCCTGACCCAATTAGAAggttacctcttatctagtg
gtagagtgtagcacctatccCAGTTATGGCATGTGCTGATGGTCAAGAGGAAAGCGGGCAggttacctcttatcta
gtagagtgtagcacctatccCCTGTCCCGGAAGTCCTGCCCAACACTGTCACATCAAAAGggttacctcttatcta
gtagagtgtagcacctatccCAATAGCATAACGAAGGTCAGCAGAGGGGAGCATTTTACCggttacctcttatcta
gtagagtgtagcacctatccCTAACAGGCAATGGATAGCAAAGTCATCAGTAGGTTTAGCggttacctcttatctag
gtagagtgtagcacctatccAGACGGTGAAACATTTTTGTCTACTTAGCTTCAAGGTGCAggttacctcttatctagtg
gtagagtgtagcacctatccCATCACCCCTAAATGAAAGACAAACCCATGAGCTACCACTggttacctcttatcta
gtagagtgtagcacctatccAAGTTTCTGTATATACGTACGACCAGCTCTCTTCCTCACCggttacctcttatctagt
gtagagtgtagcacctatccCAAGCTTCCACAAAATAGATACCATTTCATTTCCTTTCTGggttacctcttatctagtg
gtagagtgtagcacctatccAGTGACTGAATAATCAAACTGGATGGCTGAATTGAAGGGAggttacctcttatctag
gtagagtgtagcacctatccGAATAATGTGTTCTAAGCATCAATTTGGTCTGCTGGCTGCggttacctcttatctagtg
gtagagtgtagcacctatccGTTAAAAGTCTCAATGATTTAGAATTGGTGCGTGAAAGTCggttacctcttatctagtg
gtagagtgtagcacctatccGGAAAATGTATAAATCTCCTCTCCAAAGGGCCCAGTGGATggttacctcttatctag
gtagagtgtagcacctatccTTTTTACTATGTGTTCACAAGCAGAGGATAGAGAAATGCTggttacctcttatctagtg
gtagagtgtagcacctatccGGGATCTTTATCGCTCAGCTGCCTACCATCTATCTTAAACggttacctcttatctagt
gtagagtgtagcacctatccCCATACCTTGCCTCTTTCATAGATGTACACTGTATCAATAggttacctcttatctagtg
gtagagtgtagcacctatccCATGTACAATATGCTACCTTTTCAGGGAAAATCATGGGCAggttacctcttatctagt
gtagagtgtagcacctatccCTTCCTTAGGTAATAGAAACTGGGTTAATAGCAAATGGGGggttacctcttatctagt
gtagagtgtagcacctatccCAACTGGGGTATTGATAATCTTTAGTTTTCTGACATGCGGggttacctcttatctagtg
gtagagtgtagcacctatccTAAGTGTTAAGTGCTTAGCCAACCACTCTAAAAATACTCTggttacctcttatctagtg
gtagagtgtagcacctatccCATATTTTATGTTTAAATGGGAGCCAGGAGATAAGGAACTggttacctcttatctagtg
gtagagtgtagcacctatccCACTTTGCATTTGAGACTACAGATTCAACATGTGATGGAGggttacctcttatctagtg
gtagagtgtagcacctatccCTTCATCCTCATTTGTTGAAACGTCATTTGTGGCATTTTAggttacctcttatctagtg
gtagagtgtagcacctatccCTGAGATAGGTGTTTTAGTGTGTCTATTAAAGATTTGGAGggttacctcttatctagtg
gtagagtgtagcacctatccAAGATCCTAAACTTTTCAAACATGTCACAGGTAGTACTTGggttacctcttatctagtg
gtagagtgtagcacctatccGTAAAATGTACCGGTTAAAGCAGTATGTTTCTCAGATAGCggttacctcttatctagtg
gtagagtgtagcacctatccCATTTGAGCAAAAGAGTGTTGGGTTCATAAATAAGAAGTCggttacctcttatctagtg
gtagagtgtagcacctatccCTTGTAAGTACCTGTAAATAGCTACTCTGAAATTCCCTTTggttacctcttatctagtg
gtagagtgtagcacctatccGCCGTAAGTATAACACAGTGTTTTATAAACTTGGACCTGCggttacctcttatctagtg
gtagagtgtagcacctatccGCTAATAGTTTAGTTGATGTTGATAGTGGTAAGAAGGAACggttacctcttatctagtg
gtagagtgtagcacctatccAAATGTCAATTCAACTAAAATGTTACTCTCTTAGTGGGGGggttacctcttatctagtg
gtagagtgtagcacctatccATGAGTGTATCGTCTGTCCTTGAGACTTTGATGTAAAAAGggttacctcttatctagtg
gtagagtgtagcacctatccGTTTTTATTTCCCATGCTCCTTTATCTATTAGCAGACATCggttacctcttatctagtg
gtagagtgtagcacctatccGGCCCAGGCCAATCACTGTATCCCACCCCTTGGCTCTTGGggttacctcttatcta
gtagagtgtagcacctatccGTTGCTCAACCAAAGTGCAGCTTCCAGCAATGCTATCTTTggttacctcttatctagt
gtagagtgtagcacctatccCCCTTTGGTGGGCGGCGGTTAGGGGTGATGCTCATTCTCTggttacctcttatctag
gtagagtgtagcacctatccTGGTTCCTGTTGCTCTTCCCCAGTGTTCTCCCCATAAGCCggttacctcttatctag
gtagagtgtagcacctatccGGACTGACTCATGTTGCATTTTTAAAATAAGGCATAGTTCggttacctcttatctagtg
gtagagtgtagcacctatccCTGGTCAGAAAATCCTCATTGCTCTCCATTGCTTAAAAAAggttacctcttatctagtg
gtagagtgtagcacctatccCTCTAAACTCCTTTTCCAGCCTAATTTCACTCAACCTAGTggttacctcttatctagtg
gtagagtgtagcacctatccATTTCTTCAAAACACCCAGTACTTTCCCTTATTCATTCTGggttacctcttatctagtg
gtagagtgtagcacctatccATTATTTTCTGCCTCCAGCCATGCCAGTTTCCACCTTCTGggttacctcttatctagtg
gtagagtgtagcacctatccTCATCTTCCATAAATCCTTCTTCATTATGCCATGAATCAGggttacctcttatctagtg
gtagagtgtagcacctatccGAGAAATTTAATTCTTGCCTATATAAAAGAAGCTTGGGGGggttacctcttatctagtg
gtagagtgtagcacctatccTTTTCTATGCTGCATAACAACTTAGTACAAACCTAACAGCggttacctcttatctagtg
gtagagtgtagcacctatccCATAGGTCAAGATTCTGCATATGGCTTAGTTAGGTGGTCTggttacctcttatctagtg
gtagagtgtagcacctatccATGGACTGCATTCTCATCTAGAAACTAGGGAAGAATCTGCggttacctcttatctag

gtagagtgtagcacctatccTATTTCCTTGCAGCTGTAAAATTCGTGTGGTCTTGTTTCTggttacctcttatctagtgg
gtagagtgtagcacctatccTGATTTCAGAGAAGGTGTCAGTGCTCTTTTAAATGGCTCAggttacctcttatctagtg
gtagagtgtagcacctatccAGGAACTCAAAGTCAACTGCTTAGGAACTTTAATTACAGCggttacctcttatctagtg
gtagagtgtagcacctatccAACCTAATCATGGGACTGGTTTCACATCTCCTTTGCCATAggttacctcttatctagtg
gtagagtgtagcacctatccTAGAAGTAAGTCATAGTTTTCATCCCAACTCAACAAAAGGggttacctcttatctagtg
gtagagtgtagcacctatccAGATTGTGATTCAGTAGGGGGTCACCTTCATGTGCGTCCAggttacctcttatctagt
gtagagtgtagcacctatccAGTCCTCTTTTCCTGCTTTACTCTATCACTAGCCTCCATCggttacctcttatctagtg
gtagagtgtagcacctatccCTTTTAAAGCCCTAACCATCATATTCACCTAACATGCTGCggttacctcttatctagtg
gtagagtgtagcacctatccAGAAATAGGGTGAGAAGGATTATTCTTCCTAACTGGGTCAggttacctcttatctagtg
gtagagtgtagcacctatccCTCTTTAGCCTCAGGTGCAGACAATAACGAGACATATTACggttacctcttatctagt
gtagagtgtagcacctatccAAAGTCAGTCTACCTTTCATTGTCACTACCTGCCAGCAATggttacctcttatctagtg
gtagagtgtagcacctatccGTTAATGACAGGGATACGTCCTCACCTTCTTTCAAATGTGggttacctcttatctagtg
gtagagtgtagcacctatccTCACCTCCTTTCAAATAAGCAGAAGAGTGAGATAGACGAAggttacctcttatctagt
gtagagtgtagcacctatccATCCTCTTAACAGATCACATACTCTCTCCCAAATCCAATAggttacctcttatctagtg
gtagagtgtagcacctatccTGTTTATTTAATGGATTATCAGCACTCCCTATCCCACAAAggttacctcttatctagtg
gtagagtgtagcacctatccCAGATCTTTTTGTCTGGGCACTATGCGGCTCCCTTCTACTggttacctcttatctagtg
gtagagtgtagcacctatccGGTCCACCCATTTCAAAGTCAATACCACCGAGAAACTTCCggttacctcttatctag
gtagagtgtagcacctatccCCAGAGTGATGGATATCGTGGTATCTGTGTGTAATTTTACggttacctcttatctagtg
gtagagtgtagcacctatccCAGAAACTACCTGTGCTCCCTGGACTGTTGTTGGGCCCCCggttacctcttatctag
gtagagtgtagcacctatccCACTGGCGGTGACAAGGATGGGATCAATACATGTGAACATggttacctcttatctag
gtagagtgtagcacctatccAATCATTAAGTCAAATACAAATAGAGAAAGGGTACGGAGGggttacctcttatctagt
gtagagtgtagcacctatccAAATAGTGAGGATATGTCATGGATATTAACTCCAAACCTCggttacctcttatctagtg
gtagagtgtagcacctatccCTGTGCCGTGATCCACCATAGCTTGCCTACTTTCGCCATTggttacctcttatctagt
gtagagtgtagcacctatccCGTGCTGCTTCTTTGCGTTGCTGACCCACCTCCAAGTTCTggttacctcttatctagt
gtagagtgtagcacctatccACAACAGAGAACAGCTAAGCCATGTGAAAGTAGAGGACAAggttacctcttatctag
gtagagtgtagcacctatccGGTCTCCCTGTTCCTTCCAGAATCTCTCACAGTGGGTCGCggttacctcttatctag
gtagagtgtagcacctatccGTGCTCGCCAGATTCACCCTTTCAGAGGCCCCTAAAGACTggttacctcttatctag
gtagagtgtagcacctatccGTTTCGGAAACTGACCTTTCAAAGAGCCTGAGTCTGAGAAggttacctcttatctagt
gtagagtgtagcacctatccCACAGAGTAAAATCATGCTTTAATGGTGGGTTTAGGTCATggttacctcttatctagtg
gtagagtgtagcacctatccACAGAAATAAGTGGGAAAGGCAAGGAGGAAAGCTTCAGAGggttacctcttatctag
gtagagtgtagcacctatccGACGTCCATGGATAAAACAGCCAGTGAACTGCAGCCCTAGggttacctcttatctag
gtagagtgtagcacctatccGCAACAGCAACATGGAAGTTAAAATGGTTTACTCCACAAGggttacctcttatctagt
gtagagtgtagcacctatccAAATTAGCCGGGCGTGGTGGCAGATGCCTGCAGTCCCAGCggttacctcttatcta
gtagagtgtagcacctatccAAAAAGAAACCCTGAGGAAGTAAAATCAGCTTGGGAAAATggttacctcttatctagt
gtagagtgtagcacctatccAACCAAATGAAAGAGGAGCTCAAAGAATAAATTATAGCCCggttacctcttatctagt
gtagagtgtagcacctatccATTTTTTAATGTACTCTTCAACACCCCTTCTCTCTCTCTCggttacctcttatctagtgg
gtagagtgtagcacctatccACACACACACATTTTTCTACATCAACAGTAACCATCAAGGggttacctcttatctagt
gtagagtgtagcacctatccCAAGTGGATGTTCTTTCTCCTTTAGGGGACAAATTGTAAAggttacctcttatctagtg
gtagagtgtagcacctatccTGCACTCAGTCCAGCTATGAACCTCCCTGATTTTCTCCCTggttacctcttatctagt
gtagagtgtagcacctatccGGTAACCCCTCCTCTTTGTGTCCCATTTGCTATTCCCTTTggttacctcttatctagtg
gtagagtgtagcacctatccCCCTCTCTACTGGCTTCTTCTGGGGCTATAAATGTATTCAggttacctcttatctagtg
gtagagtgtagcacctatccATTTAAAAATACCCTCAGGCCAGATGCAGGCTTACACCTAggttacctcttatctagt
gtagagtgtagcacctatccTTTTTAAGGAGACAGAGAGACCAAGAAATACCTTCAAGTGggttacctcttatctagt
gtagagtgtagcacctatccTCACAATGAACTCCCTGAATAATCACATGCAAACTGTCCCggttacctcttatctagt
gtagagtgtagcacctatccGCTCCTGCCCCCTTAACTGTGCTGAAATGCTTTGACAAAGggttacctcttatctag
gtagagtgtagcacctatccACAAATCTAATGTCTTCTTTTTAGTCCTCAGCTTCCCTCAggttacctcttatctagtg
gtagagtgtagcacctatccTTTGTGACATGCTACTCTGTTGATCATTCTGTGTTTTTGAggttacctcttatctagtgg
gtagagtgtagcacctatccCATAATTCTGCTGTCTCACTTATGTTCCTTCTCAAATTCTggttacctcttatctagtgg
gtagagtgtagcacctatccGGTTAAATGTTGGTATTCCCCAGGGCTCTGTATTCATGTGggttacctcttatctagtg
gtagagtgtagcacctatccATCCCTGGGGCTCAAACTGTGGTCTCCAAACCAGCAACATggttacctcttatctag
gtagagtgtagcacctatccACTTGTTAGAAATGCAATTTAAGAAATACAGGACAGCCAGggttacctcttatctagtg
gtagagtgtagcacctatccCAGTGAGCCAAGATCACGCCACTGCCCTACAGCCTGGGTGggttacctcttatcta

gtagagtgtagcacctatccCTAAATCTGGCAACCCTAACTACAAATTACTAGTCCCTTTggttacctcttatctagtg
gtagagtgtagcacctatccCTGGCCTATAATTTGTGTTGTAATAAGTCCTCCAGGTGATggttacctcttatctagtg
gtagagtgtagcacctatccTGCTCTACATCCCACCCTTGACTGCCTTCATCTGCTCCCAggttacctcttatctag
gtagagtgtagcacctatccGTTCCTGAACCGCTGATTTGCATGCTCCTGAATTTTATTTggttacctcttatctagtg
gtagagtgtagcacctatccGTGTCCCACTTTTATGAGGAAAACCAGGAAGAATGCAAGAggttacctcttatctag
gtagagtgtagcacctatccTTCTGGGCTTTAGAACAATGTTTACCTGTCTTTGTATAAGggttacctcttatctagtgg
gtagagtgtagcacctatccAGAATCCATTTTTTTCTGCTCCTGCCTACCCACTCATGTAggttacctcttatctagtg
gtagagtgtagcacctatccCAGAGAATTAGGGAAAAATAATTAGTCACTGGAATTAGGCggttacctcttatctagtg
gtagagtgtagcacctatccTAAGAGAAGATGATAGAGGTGCCCCATATAATTCTTAGGGggttacctcttatctagt
gtagagtgtagcacctatccCAGTGCCCCTCAATCTATCTGGAGTTCCTCTGAGCCATTAggttacctcttatctagt
gtagagtgtagcacctatccGAGATGGAATGGTGGGAAGCAAAAGCAGATGGAATTGGGCggttacctcttatctag
gtagagtgtagcacctatccCCAAAGGACAGGAGCATAGATAGGACTTAGAGAACTGGCGggttacctcttatctag
gtagagtgtagcacctatccAAAAGGGTCCCACAGTCATGACAAGGGAGGCAGCCACACTggttacctcttatcta
gtagagtgtagcacctatccGGCAGTACCCAAGATCAGACAAGTCTACACATCAGCCAATggttacctcttatctag
gtagagtgtagcacctatccATGCCCTCAGACAAGAGCCCATTCCCACCACATTGCGACTggttacctcttatctag
gtagagtgtagcacctatccTATGACAAATGATTAACATGATTTAGCTGACTTGGAAGGGggttacctcttatctagtg
gtagagtgtagcacctatccTGCCCATGAAGCAGAATGAGAGGTTCAGTCAGTTACAGGAggttacctcttatctag
gtagagtgtagcacctatccGGGTAAACAAAGTCAGCTGCTGCAACACACCCTGAAATCTggttacctcttatctag
gtagagtgtagcacctatccTCTCACTCACACAAAGTTCAGTCAAGAATCAGCCACCCTCggttacctcttatctag
gtagagtgtagcacctatccCCTCAGGCACAGAGGCAGGGGAAGAAAGGGAAGGAGGCTGggttacctcttatct
gtagagtgtagcacctatccCCTGTTAACCTGGAAGGGACGCTTCTGCTCCTAGGCCATCggttacctcttatctag
gtagagtgtagcacctatccCAGTCTCACTAGAAAGAAGGCAGGAGATACAGAACAACACggttacctcttatctag
gtagagtgtagcacctatccCAAAATAGCATGAACTGGGTGGATCAAACAACAGAAACTTggttacctcttatctagt
gtagagtgtagcacctatccGGACACACACACATTCATTCCCTAACATTTGGCAAACATTggttacctcttatctagt
gtagagtgtagcacctatccCATTCAAGGTGATCTCTATGCTGATAGTTCCCAAATCTCTggttacctcttatctagtg
gtagagtgtagcacctatccCCCCAATCCGGACTCCTGATCCATATACCCACCTGCCTCCggttacctcttatcta
gtagagtgtagcacctatccATTTGTTACACAGGTATATTCCAGGATGCTGAAGTTTGAGggttacctcttatctagtg
gtagagtgtagcacctatccTTTAATATGTCTAAAGCTAAACCTATCCTCTTCCCCCAACggttacctcttatctagtg
gtagagtgtagcacctatccGTATTTCTTATCTCTAGACACCCACAACAGAAACCTGGAGggttacctcttatctagt
gtagagtgtagcacctatccGGACTATGCTCTCCTCTACCCTTTTACATTCTATCGGTCAggttacctcttatctagtg
gtagagtgtagcacctatccAGTAGATTCTGCTTCCTAACTGCCCTTATACCTGTTTCTTggttacctcttatctagtg
gtagagtgtagcacctatccTCTTGGCACTATTGACATTCCAGGCCAGATAACTCTTTGCggttacctcttatctagtg
gtagagtgtagcacctatccACCTAGTAGATGCTAAGGTTGCCCTCGCGAAACAAAAAGTggttacctcttatctag
gtagagtgtagcacctatccGCTGCAGGCCTCCAATTAAAAAATTACTGATCTGTTACTGggttacctcttatctagtg
gtagagtgtagcacctatccCAGGCCTGAGTTGTTTTAATAGTTGTTGGGTTGTATTTTTggttacctcttatctagtgg
gtagagtgtagcacctatccATTTTTAGTAGAGAAGGCGTTTTACCACATTGTCCAGGCTggttacctcttatctagtg
gtagagtgtagcacctatccCCACTGTGCCCGGCCCATTGGGTTGTATTATAATTGGCATggttacctcttatctagt
gtagagtgtagcacctatccCCAAAACATTGCACAGCCCACCCACTCACCTCCCACTCTAggttacctcttatcta
gtagagtgtagcacctatccGAACCCATTGTCTGCACTGCTACCTGAATGATCTTTGTGGggttacctcttatctagt
gtagagtgtagcacctatccCTGACCATTTCACTCTCCCATTAAAAGCCCCCTTTGCTTAggttacctcttatctagtg
gtagagtgtagcacctatccTTCTCCATGAACCTGACTTAACCAATTTCACTGCACATCCggttacctcttatctagtg
gtagagtgtagcacctatccCCACTATCCTCTACTTGTAATGCAGGAACATATTTTGGCCggttacctcttatctagtg
gtagagtgtagcacctatccGTTCAAGCTCATAACCTCCTGTGTGAAGTCATTCCTAAACggttacctcttatctagtg
gtagagtgtagcacctatccTTCTGGAAGACTTGGGTTTTTGTATGGCTACCTGTACTAAggttacctcttatctagtg
gtagagtgtagcacctatccTCATTAAGACAACCCTGACACATAGTAACCACCCAATTCAggttacctcttatctagt
gtagagtgtagcacctatccTTGATTCTGCAAAAAGGCTTGGACAGTAAATTGGGCTGCAggttacctcttatctagt
gtagagtgtagcacctatccTAAGGACTTAACTCTGAAATTTTGGAAGACAAAGTAGCCTggttacctcttatctagtg
gtagagtgtagcacctatccGGAAGCCAAGGTGTGAGGATCACTTGAGATCAGGAGTTTGggttacctcttatctag
gtagagtgtagcacctatccCTCCATCTCTACTAAAAAGATACAAAAACTAGCCAGGCGTggttacctcttatctagt
gtagagtgtagcacctatccCTGGGTGACACAACAAGACTCTGCCTGAAAAAAAATAAATggttacctcttatctagt
gtagagtgtagcacctatccCTTTTTACTCATTACAGTTTATTGAGCACACACTAAGTGCggttacctcttatctagtg
gtagagtgtagcacctatccCTGTGAGAATTACCAATAAAACATGAAAGAGTGAAAGAGCggttacctcttatctagt

gtagagtgtagcacctatccGGTGCGTCATTGTTATATGGAAATAAAAATCTCAACAGTGggttacctcttatctagtg
gtagagtgtagcacctatccGCAGCAGCAATTAAGCCTCATATGACTATATAACTTTAGAggttacctcttatctagtg
gtagagtgtagcacctatccTGTCAGGGAGGAGAAATGGAAAAATCATGCAATCAAGAGAggttacctcttatctag
gtagagtgtagcacctatccAATAATAAAGGACAGGACCTCTTTTTAAATGCTTGATGGCggttacctcttatctagtg
gtagagtgtagcacctatccGTCACACATATGGATTTTCTAATTCTATGAGAGGGTTTTCggttacctcttatctagtgg
gtagagtgtagcacctatccTCCAGTGTGTTGACAAGTTAATAAGTGATCTACTGACTTCggttacctcttatctagtg
gtagagtgtagcacctatccAACCAATTTCGGCATAGTAGGTGGCATTGCTTTCCCTGGGggttacctcttatctagt
gtagagtgtagcacctatccCTGCATGCCACCTCTCAGCCCACCTGCCAGCTACCACCCCggttacctcttatcta
gtagagtgtagcacctatccTTTCCAGCCCTGCCAAACCGCCCACCTCACCCTGAAAATTggttacctcttatctag
gtagagtgtagcacctatccGCTGTACCTAGCCCATGTCTACTCATCCTTTTCTCTGAAGggttacctcttatctagtg
gtagagtgtagcacctatccTGAAATGTCATCTCTCCTGTGAAGCCTACTTGGGTCTCCCggttacctcttatctagt
gtagagtgtagcacctatccGAATCTGTAATCCCTGGTTCACTGGCTCCTTGGTACTTAAggttacctcttatctagtg
gtagagtgtagcacctatccTGTCATGCCACACTAACCGTTTCTATTCATTTAGCTATTTggttacctcttatctagtgg
gtagagtgtagcacctatccTAGAAATAGAAAGATGAATGGTGCATGGGGTCTGCTAGAGggttacctcttatctagt
gtagagtgtagcacctatccAAGAACACCCAGGAAATTTATCGCAAAAAGATCTTCCCCTggttacctcttatctagt
gtagagtgtagcacctatccGATTTGGGCCCTATCTTCAGAAATTCAAAGAATATGGCTCggttacctcttatctagtg
gtagagtgtagcacctatccGGGTCCCATATAGATTGACTGTGCAGGATAAGACAAAAGTggttacctcttatctagt
gtagagtgtagcacctatccTGGACTACATAGGAGAAGTGAATTTAGGAAGAAAGTTGATggttacctcttatctagtg
gtagagtgtagcacctatccCACTGGATGCGTATGGACATTTGGGTAAAGATATCTAAGAggttacctcttatctagtg
gtagagtgtagcacctatccCTGAACGTCTGAGCCTGAAAGTTGGGGATGCAGAATGAGCggttacctcttatctag
gtagagtgtagcacctatccTGACTGGTAATCAGGGACCATGAGACCTTCAGAGAACCCCggttacctcttatctag
gtagagtgtagcacctatccGCTCAACGTGGGAAGAAAGCCTAGGCAGAGCTCTGGAGAAggttacctcttatcta
gtagagtgtagcacctatccCTGGCAGAGGAAAAGGAGTCATGGAGGACACTGAGAAGCAggttacctcttatcta
gtagagtgtagcacctatccGATGTGATTCGGTACAAAGGAAAGATATTACAGGAAAGAGggttacctcttatctagt
gtagagtgtagcacctatccAGATTAAGGGCATAGCTTCTGAAGTCAGACCACCTCCATGggttacctcttatctag
gtagagtgtagcacctatccGAGCAAAGTATTCTTTTACGCTTTATACCTCAGTTTTCTCggttacctcttatctagtgg
gtagagtgtagcacctatccTTATAGATTTCTCTCTTCCATTTTTGTCAACTTCCCCAAGggttacctcttatctagtgg
gtagagtgtagcacctatccAGTGATCTTCCCATCTTGGCCTTTCTAGTAGCTGGGACTAggttacctcttatctagtg
gtagagtgtagcacctatccCACCACACCCAGCCACAGAGACCACTTTTTAAAAATCTTCggttacctcttatctag
gtagagtgtagcacctatccGGAAATAGTTGTTGAATAAAATCACTCTACTAGGACCCTCggttacctcttatctagtg
gtagagtgtagcacctatccGCCAGAGCATCTTTTGTCATAATAGTTAGTCAGAGATTTGggttacctcttatctagtg
gtagagtgtagcacctatccTCTCCAAAGTCAGAAAAGTCTATTTTGGTTGTTTCCAAAGggttacctcttatctagtg
gtagagtgtagcacctatccAAATCCTAGAGTTGAAGAGACTTTAAAAGTTATCCAGTCCggttacctcttatctagtg
gtagagtgtagcacctatccTGACATCCATAAAACTCAACTTCCAATGACAGCCAACCCAggttacctcttatctag
gtagagtgtagcacctatccAGGTTTTATTTTATTCTTAGATCTTGAACACAAGGCGGCCggttacctcttatctagtg
gtagagtgtagcacctatccTTTTACCAACTGATCCTAATTCAATCTTATAGAGCTACCCggttacctcttatctagtg
gtagagtgtagcacctatccTCTATACCCCTGAGAAGATAACAATCCTTCAAGAAACTGAggttacctcttatctagt
gtagagtgtagcacctatccCTTATTTTATTGGAAGAAGGTACCACCTAGGACTTTCACAggttacctcttatctagtg
gtagagtgtagcacctatccCCAAAGCTTCAAAGAACAGGCAGAGCCTCCTGTTAGGGGCggttacctcttatcta
gtagagtgtagcacctatccTGACTTGAAGTTGAAGTCAACGCACATTGACCATTCTGAAggttacctcttatctagtg
gtagagtgtagcacctatccCAGAGCCTGGATGACAGCACCTGTGTACAGCATGGTTTACggttacctcttatctag
gtagagtgtagcacctatccCTGTTGAGACCCAGTGTTCAGAAAAAGATTACTTTCAAAAggttacctcttatctagtg
gtagagtgtagcacctatccACTTAAGAGTGCCGATGCAGACATATAAGGAAATTAATGTggttacctcttatctagtg
gtagagtgtagcacctatccTGCAGCCCAAGGATCAAGGACAAATTTTTAACTTTCGAGTggttacctcttatctagtg
gtagagtgtagcacctatccCCATAGATATTGATTTCTCTGACGGATCTGGGCAAAGTAAggttacctcttatctagtg
gtagagtgtagcacctatccTGGATGACTTGGAGAGGTTCCAGACTTCAGTGTAAGAAGTggttacctcttatctagt
gtagagtgtagcacctatccGTGACAGAAATAGCAAGAGACAGCAATTCCAAGAGCAATTggttacctcttatctag
gtagagtgtagcacctatccCTGCTCCTGTAGGACCCAGGTGAACCCTCAAAACTGTGAGggttacctcttatctag
gtagagtgtagcacctatccGCCCTGGGAGCTTGCTGGGTGCCTTAGCAGGCTACTCCTGggttacctcttatcta
gtagagtgtagcacctatccGCAAATCCAGTATGCAGACTCTACAGGCCTGGGAAAAATCggttacctcttatctag
gtagagtgtagcacctatccCCTGGGGCAGGTTTTCAAGCTGGTATTGCTCTCTGCCTGGggttacctcttatctag
gtagagtgtagcacctatccTCTGGGGGCTGATAAGGGACACGCAGGGAGTGAGACCAGCggttacctcttatcta

gtagagtgtagcacctatccAGGCAGCCATAATCCTCCCACGTACACAACTCCAGTGACCggttacctcttatcta
gtagagtgtagcacctatccTCCCTACCCACTCTGGTAGCCGAAGACAAAAGGCATATAAggttacctcttatctag
gtagagtgtagcacctatccGGCCCCGTCCACTGCCGGTTCTTCCCCGTAATACCACAGCggttacctcttatcta
gtagagtgtagcacctatccCAGAGTCCATTGTACTCCCTACCACCTCCACAGGAACAGGggttacctcttatctag
gtagagtgtagcacctatccCTACATCAAGGGAACATCCTATGGGACAAAAGAATATGAAggttacctcttatctagt
gtagagtgtagcacctatccGCCTTCAGTGCTAGACCTTCCTTCTGACAGAGCCTACCCAggttacctcttatctag
gtagagtgtagcacctatccAGGTGAAGCCCAATGCAAGGCAATACAAAATATGGTACCAggttacctcttatctag
gtagagtgtagcacctatccATTCAGGAAAGTTTGGACACACTTTTAGAAATGCAAAGTGggttacctcttatctagtg
gtagagtgtagcacctatccCCTAAGAATAATTGGTGTATCTGAGGAAGAAGAGAATCCTggttacctcttatctagtg
gtagagtgtagcacctatccAAAAACAAGAAGCACAAAGAACACCAAGGAAATTCATCACggttacctcttatctag
gtagagtgtagcacctatccATCAGCCAAGAATTTTGTAAATCCAGCAAAACTAAGCATCggttacctcttatctagtg
gtagagtgtagcacctatccTTCACCATTACCAAGCCACCTCTATAAGAACTGCTAAAAGggttacctcttatctagt
gtagagtgtagcacctatccATGAATGCAACAGTACCTCAGGTTTCAATACTAACATTGAggttacctcttatctagtg
gtagagtgtagcacctatccCGCAAAATGGATAAGAACTCAGCACAGCAAAAGGAACAGTggttacctcttatctag
gtagagtgtagcacctatccAAAGTAGGCTAAGGACATGAAAAGAAGATACACAAATGGCggttacctcttatctag
gtagagtgtagcacctatccAAACCACAATGAGGTACTACCTCACTCCTATAAGAATGGCggttacctcttatctag
gtagagtgtagcacctatccAACAGTAGATGCTGGCATGGATGCGGTAAACAGGGAACACggttacctcttatctag
gtagagtgtagcacctatccTACTTGCACACACATTTACAACAGTGCAATTCACAATTGCggttacctcttatctagtg
gtagagtgtagcacctatccAAGAAACTGTGGTATATACCTATATGGTGGAATACTATGCggttacctcttatctagtg
gtagagtgtagcacctatccGGAAAGAGTGGGAGGAGGGTCAGGGCTAGAAGACTACAAAggttacctcttatcta
gtagagtgtagcacctatccACTGAATTGCTGCAGTCCCATGATAAAACTTTAGCAGATAggttacctcttatctagtg
gtagagtgtagcacctatccGAATAACGAAAGTAGTTTCTTAAGATACAACCTACTCCTGggttacctcttatctagtg
gtagagtgtagcacctatccGTTGATAAAGCAATGGTAGGCTTCGAGAGGATTAACTCCAggttacctcttatctagt
gtagagtgtagcacctatccGTCAGCAGCCATCGACACTGTGGCAAGTCCCTCCACCAGAggttacctcttatcta
gtagagtgtagcacctatccCACTTACTGAAGGTCCAGGTGAATGTTAGCATTTTTTAGGggttacctcttatctagtg
gtagagtgtagcacctatccTGCACACCACCACCACATCCGACAAATTTTTCTATTTTTTggttacctcttatctagtg
gtagagtgtagcacctatccCATTGTGATATTTGCTTTATGCCAGTGATATGGAACTGACggttacctcttatctagtg
gtagagtgtagcacctatccAAAATTAACAGGACAAAAATCAACACTGTATCAGCTGGTGggttacctcttatctagt
gtagagtgtagcacctatccCATAGACTGGGTGACTTATATAACCGAAATTTATTGCTCAggttacctcttatctagtg
gtagagtgtagcacctatccCAATTCTGATGCAAGTGGATGGAGCAAATACAGTCTTAAGggttacctcttatctagt
gtagagtgtagcacctatccGAGTATCAACCCTCTGAGCTCTGTACCGGTCTGCACTAGGggttacctcttatctag
gtagagtgtagcacctatccCATTCACAGCCTAGCTTGCCCAGAAACAGATAATCAGGACggttacctcttatctag
gtagagtgtagcacctatccTGTAAAGAAAGTTACAGCAACAAGTTAAGAAGCCCTGCCAggttacctcttatctag
gtagagtgtagcacctatccTTGTGAACAACCAGCATTCAAGAGTGCACACCTATGCCTTggttacctcttatctagt
gtagagtgtagcacctatccATTCAAATATATGTCTTGGTTTTAGTGTAGCTCTTCAGCCggttacctcttatctagtgg
gtagagtgtagcacctatccGATTGACCATCTCAGCTATCTGTTTTAGCTTAGTATCATCggttacctcttatctagtg
gtagagtgtagcacctatccCAAAGCTCTGGAACAACATAGCAGCATCACTGGTTATCTAggttacctcttatctagt
gtagagtgtagcacctatccATCACCTCACCTCTTTGATCCTCCATGTATTTCCCTATAAggttacctcttatctagtg
gtagagtgtagcacctatccTTTGCATAAAACTTTAAGAAGCCTCCCTATGAAGTACACAggttacctcttatctagtg
gtagagtgtagcacctatccATCTATAGTTTTCATGAGATTCTTAAAAGGGTCCAAGGGAggttacctcttatctagtg
gtagagtgtagcacctatccCTAGCTGCCCACCTCACTGCACTATTTTAACACATTCCTCggttacctcttatctagt
gtagagtgtagcacctatccCATAAAGTAACTGTGAATTCTCCTCAAACTTCCTAAAACCggttacctcttatctagtg
gtagagtgtagcacctatccTCCTCCCAACTCAGGAAGCAGCTATGGTTTTCTATGTCTGggttacctcttatctagt
gtagagtgtagcacctatccCTTCAGAAGGGTTGCCATAGCAGAATAGAAAAAAACTAAAggttacctcttatctagt
gtagagtgtagcacctatccGCCTGAAGTATATGCCTTAACCACTGGATATAGATGTCTAggttacctcttatctagtg
gtagagtgtagcacctatccTCAAAGACAAGTGCAGTTCCATCCCAAAAATAGACAAGGAggttacctcttatctag
gtagagtgtagcacctatccCCAGTAAGGGGGAGTCAGAGGAAGAGAAGAGCTGAGATATggttacctcttatctag
gtagagtgtagcacctatccCATCACTGGTGGGAGAGGGTGAATAATAGAAACTCCAAATggttacctcttatctag
gtagagtgtagcacctatccGGGGACATCTGTTAACACTGAAAACATTCCAGCAAGATCCggttacctcttatctag
gtagagtgtagcacctatccCCAGAGTGGAAACAGTAGAAGAATCATCAATGTGTGGGCAggttacctcttatctag
gtagagtgtagcacctatccCTCCAAAATCCAGTGAGGCAAAGGAAGAGTTCATGGTATTggttacctcttatctagt
gtagagtgtagcacctatccGTAAATAACAGAAAAAGAGCAAAACCCATGAGGCCTCTGAggttacctcttatctag

gtagagtgtagcacctatccCTGACACAGCACAAACACCAGAGACCAGGGGCCAGAAGGAggttacctcttatct
gtagagtgtagcacctatccGGTTAGATAGATGTTGCCAACCTAAGGCCTGACACTTGATggttacctcttatctag
gtagagtgtagcacctatccCATAAGTGTCCTCCCCCTCCCCACAATGTACAGGAAGGTGggttacctcttatcta
gtagagtgtagcacctatccGATCCTGGGACCAACTAAGATTTAGTATTTGCCATTCAGCggttacctcttatctagt
gtagagtgtagcacctatccGGAGATTCAGTGAGAAAATGTGTTCAGTTACAGAAATTCTggttacctcttatctagt
gtagagtgtagcacctatccGATACACTTCTTAAATACTCAAGTTTATGGGTAGAGACGTggttacctcttatctagtg
gtagagtgtagcacctatccCAAACGATGACAAATGTTCTACTAAAATCACATAGCACCGggttacctcttatctag
gtagagtgtagcacctatccGGCATTACCCAGGTCCATACAGTCAAAGCAGCTTTTCAAGggttacctcttatctag
gtagagtgtagcacctatccAGTGTGACAACTCTGACGGTGGTGACCCTTTTCTCCTGCTggttacctcttatctag
gtagagtgtagcacctatccGTGCTGATTTAGACCAAGTGTTCACTCTTCCACCTGATGAggttacctcttatctagt
gtagagtgtagcacctatccCTGAAGCAGTCCATTGTCAACAAAAGCCCTTAAAATTTCggttacctcttatctagt
gtagagtgtagcacctatccCATGTTTTATGGTGCTAAAGTCTGCTGTCTTAACTAGTCTggttacctcttatctagtg
gtagagtgtagcacctatccCTTCTTTTTGAGGCAAATATTGATCCTACCCACGGGATATggttacctcttatctagt
gtagagtgtagcacctatccGATAGACATCAGGCTCTTGAACATTTTGACAATTCTGCATggttacctcttatctagt
gtagagtgtagcacctatccCAGTGAAACAGCATAGAACATAGATACTAAGATCCAAGTTggttacctcttatctagt
gtagagtgtagcacctatccAAAATCCAATGACAAATCGGTTTAATTCCAGCCTTTGCCAggttacctcttatctagt
gtagagtgtagcacctatccACTTCCAATATTTAAGGGAGTCCTCAGGGTCATGATACTAggttacctcttatctagt
gtagagtgtagcacctatccTCCATTGCCCCTCTTCCAGATATTGCCCCCAGCCATGGGTggttacctcttatcta
gtagagtgtagcacctatccCTTATATCATCTAAGCTCCCAGCAGAAATGACTTCCCTTAggttacctcttatctagt
gtagagtgtagcacctatccCGAAAAATATCCCTTTAAACAATGTTCCTGTGACTTTTGCggttacctcttatctagtg
gtagagtgtagcacctatccGTAAAGAGCCAGTTGATAAATTACTTCGTGATTCAGAGCAggttacctcttatctagt
gtagagtgtagcacctatccAGCTTGAGTTCCTGAAAGTGTTTGTCTCTGTCCTGCTGTCggttacctcttatctagt
gtagagtgtagcacctatccAGGAAACTCAGAAATAATTATTGTGCTCTCCTGCAGTAATggttacctcttatctagt
gtagagtgtagcacctatccAAATTGTGTATTGTGTTCGATTTTGTAAACAGCCTCAAGCggttacctcttatctagtg
gtagagtgtagcacctatccCATTGAAAGAAATGCATTTTAACTTCCCCGAGATTTGGTTggttacctcttatctagtg
gtagagtgtagcacctatccATGTAAGTTCTATGAGGGACAAGAGTACATCAAATTTGCTggttacctcttatctagt
gtagagtgtagcacctatccGTAACTAACAACCTGAAAACTTCCCCATGGCTTAAAGAACggttacctcttatctag
gtagagtgtagcacctatccGTTGGCTGTGACTTTACCTCTCATAACCACTGGAATGTTGggttacctcttatctagt
gtagagtgtagcacctatccTCCTGGCAAAGAGAAAAGAGATATGACAAACCACAAATCAggttacctcttatctag
gtagagtgtagcacctatccTGCAGGGAGGGTTTGGAATATCTGGAGACAATAATGCATCggttacctcttatctag
gtagagtgtagcacctatccTGGACTTACAGATTACATTTTCCTAAATTAATGCGGTGCTggttacctcttatctagtg
gtagagtgtagcacctatccCTTCCCATATTTCACCTTTGGTCTTTGAAACTTGTTTTAGggttacctcttatctagtg
gtagagtgtagcacctatccTGTACTATCAACTTCGTTATACCTAGGTCAAATACACATGggttacctcttatctagtg
gtagagtgtagcacctatccCCTGAATAGCTTAGGAAGTCTAGAGAGCAGGGTATTAACTggttacctcttatctag
gtagagtgtagcacctatccACAGAGGCCTCAGTGAATAGTTACAGAGCCCATGTCAGGGggttacctcttatcta
gtagagtgtagcacctatccACCTGGATCTTAATATCTATGTTCTATGCTGTTTACACTGggttacctcttatctagtg
gtagagtgtagcacctatccTTCCAGAAGAATGCAGAATTGGGAATATGTTCAAGAGCCTggttacctcttatctagt
gtagagtgtagcacctatccTCACTTGAACACTTTGAAACACCTGAAAACCCCAAACAAAggttacctcttatctag
gtagagtgtagcacctatccTGAATGTATGCCTAAGCTTGGAAGAAAATAAGAGAACTGCggttacctcttatctagt
gtagagtgtagcacctatccCAGGCAGCTAGAGAACAAAAAGTGCAACAGTCACGCTGGTggttacctcttatcta
gtagagtgtagcacctatccCCTGAGGCTGAGGTTTTAATGGCTTCATAAAAGCAGAAGAggttacctcttatctag
gtagagtgtagcacctatccAGTTACACTGAGGCCCCAAATAAAGCTATAATCCACAAAGggttacctcttatctag
gtagagtgtagcacctatccAGCCTAAGTAAAAGGTATACCCAGACTTGCCAGATTTAGCggttacctcttatctag
gtagagtgtagcacctatccGTGTGTCCAATGCATTATCTGGAACATACTTCTCCTAGAAggttacctcttatctagt
gtagagtgtagcacctatccAGAGACACTAGAAAAATTACCCACCAACCAGCACAAGGAGggttacctcttatcta
gtagagtgtagcacctatccTGCTTGCCTATCTCCATGTGGACTCTCAGTAGTGGGGGACggttacctcttatctag
gtagagtgtagcacctatccTGCCTTACACAGATTTATGGTTCAACTTTATACTGCATGAggttacctcttatctagtg
gtagagtgtagcacctatccGGCCCACCAGACCATGATACCTCAGCATTCCAAAGATAAGggttacctcttatcta
gtagagtgtagcacctatccGCAAAGCAGGCTCCTTCACTTGTCAGTTTAAAAATTCAAAggttacctcttatctagt
gtagagtgtagcacctatccATCAAACAAGCTGATTCCTTCTCTGTATTCTCGTAGGTAGggttacctcttatctagt
gtagagtgtagcacctatccGGTTCGATCCTGTGGCTTCACGCTGTGATTCTCATTTGGCggttacctcttatctagt
gtagagtgtagcacctatccAAAGGAAAAAACAAGCCAGTTCTAGCATTGGAAATGTAGCggttacctcttatctag

gtagagtgtagcacctatccCACACTGGACCTGTTGTGGCCGGGGCAGGGAGGGGGCGCggttacctcttatct
gtagagtgtagcacctatccTAACAAACCTGCACATTCTCAACATGTAACCTGGAACTAAggttacctcttatctagt
gtagagtgtagcacctatccAAAAGAATAGACCTTAAGGCACTTTGAAGCTTTTAGTTCCggttacctcttatctagtg
gtagagtgtagcacctatccAGCTATCTGATTTTAAAGGCTTAGTCTCAATGTGTCCCTTggttacctcttatctagtg
gtagagtgtagcacctatccAAATCCCAGTAGAGTAGCCAATTGTCCTGAAACCCGCTTGggttacctcttatctag
gtagagtgtagcacctatccTCTTCATCCTAATAAAAGGCAAATTAAGTACAGTTGACCCggttacctcttatctagtg
gtagagtgtagcacctatccCCATGCAGTAAAAAATCTGGGTATAACTTTCGATTCCACAggttacctcttatctagtg
gtagagtgtagcacctatccAAGAAAATGTATTCCTTTAGCTGCCATCCTGCCTCCCTGTggttacctcttatctagt
gtagagtgtagcacctatccCTCCCCAAGTGCCAACCCTAGTCCCCCACGCAGCCCCTTTggttacctcttatcta
gtagagtgtagcacctatccCAATCATGAACCAAAAACTCACCAAACGGCAAGCAGAAGTggttacctcttatctag
gtagagtgtagcacctatccTGCTCACAGGAAAAAGGTGGTTCACAGCATCTGAGACGCTggttacctcttatctag
gtagagtgtagcacctatccCTCATGAGGTTTTGAATGTCCAGGGACATTGGCCAATATCggttacctcttatctagt
gtagagtgtagcacctatccTTCTTGACTTCAGGGACAAAACAGCAATGGAACCAATCCAggttacctcttatctag
gtagagtgtagcacctatccCCTTCAAGGCTCAGAGGTTAACGGTTTTATACATAAGGGTggttacctcttatctagt
gtagagtgtagcacctatccCGACAGCAGAGGATAAAAAATTTCAGTTCTCCTTAAAGAAggttacctcttatctagt
gtagagtgtagcacctatccTGTTTACACACCAAGGAACAGTGATTCACTTTAACAACCCggttacctcttatctagt
gtagagtgtagcacctatccTCAGGCATCGCTGGCAGCAAACACTTTCACCATGACAGGCggttacctcttatctag
gtagagtgtagcacctatccGCATCGATCATAAACCAGTGCTGCAGATGGTCTGACTAGTggttacctcttatctag
gtagagtgtagcacctatccGCTACTGGAGAGGATGATGAAGTTCTAGATCTTGTGGAGAggttacctcttatctagt
gtagagtgtagcacctatccCTTCCAACAATGAGGCAAACTGAATTAAGTCAACTTCTGAggttacctcttatctagt
gtagagtgtagcacctatccAACTTGAAGTAGTTACTGAGAGCTGCTGTTTTATGTTATGggttacctcttatctagtg
gtagagtgtagcacctatccAGGTTAATAAAGCTCTTTGCCTAGTAAAATATGTGTGTGCggttacctcttatctagtg
gtagagtgtagcacctatccCTTTACATTGAGTAGGCTGAAAAGTAGGAGGAAGAGGAGGggttacctcttatctag
gtagagtgtagcacctatccGAGGCAGGAGAGGCAGGGACGTTACATGTGTATTAGTCCGggttacctcttatctag
gtagagtgtagcacctatccGGAGGGCAAAGGAGGAGCAAGGGCGTGTCTTACATGGCAAggttacctcttatcta
gtagagtgtagcacctatccATTCACTACCACGAGATCAGTATGGAGAAAACCGCTCCCAggttacctcttatctag
gtagagtgtagcacctatccTCAAGGTGAGATTTGCATGGTGACATGGCCACACTATATCggttacctcttatctagt
gtagagtgtagcacctatccTATATGGTACTAATCCTTCTACAACTAGGCCCCTTTAGATggttacctcttatctagtg
gtagagtgtagcacctatccCTGAAATCATGAAAATTCTGGTCAGGTCCTGTCTTGTCTTggttacctcttatctagtg
gtagagtgtagcacctatccCACATGTTAACCACTCCCTCTTTGACCAATATCCCTTTTTggttacctcttatctagtg
gtagagtgtagcacctatccTGCTGATGTACTATTTCTTTGTCAAGCAGAGGGAAATAACggttacctcttatctagtg
gtagagtgtagcacctatccTGTAGAAGAAATGAATGCCTTTAAGAATGTTTCAACCAGCggttacctcttatctagtg
gtagagtgtagcacctatccAAACTCCCCCTCCCTTACTCTCCAGCTGCATAAAAACTCTggttacctcttatctag
gtagagtgtagcacctatccGTTAAGATAGATTTGAGAGAGCTTGTTCTCCTGGCTTCTCggttacctcttatctagtg
gtagagtgtagcacctatccCAAGCACCGATGTCCCATTGTTTGGCTCCAGCTGCACATTggttacctcttatctag
gtagagtgtagcacctatccTTTGGGATTCTACAACACTTCCAATGTTTGCTTTAGTTTCggttacctcttatctagtgg
gtagagtgtagcacctatccGTTTTCTGGCCCTGCTTCTTTTGACCTCTATTCTTTATTGggttacctcttatctagtgg
gtagagtgtagcacctatccCAAGTCATCTCCTATTAATTCCTTTCATGTGACGTCTGCTggttacctcttatctagtg
gtagagtgtagcacctatccCTATATCTTGAAACCCTTTATTCCTACCCTTTTTGCCATAggttacctcttatctagtgg
gtagagtgtagcacctatccTGGGCACAGTTTTTTCCAAATGGAATTTATTATTTCAGGCggttacctcttatctagtgg
gtagagtgtagcacctatccCTTTTTATCCCAATAATAGCATCTTTAATGGCAGGTAAGCggttacctcttatctagtg
gtagagtgtagcacctatccACTTTCATAATGTTCTCTCTCTCAGATACTCTTCCATAGAggttacctcttatctagtgg
gtagagtgtagcacctatccCCATAGAGTACAGTGTGTAACGAGCCTTAAAGGTCTTTATggttacctcttatctagtg
gtagagtgtagcacctatccTGTTTGGGGGCAAGTAGACACCTTTGACACCTTCCATGTTggttacctcttatctagt
gtagagtgtagcacctatccTGGGGTTCTTAGTGGCCAGGGACATTGTCAAATATCAAAAggttacctcttatctagt
gtagagtgtagcacctatccCTGACTTCAGGGACAAACCATCAAAGGGACCAATCCAGAAggttacctcttatctag
gtagagtgtagcacctatccCTTCAAGACTCAAGGGTTAACAGTATTATAGTTAAGGGCAggttacctcttatctagtg
gtagagtgtagcacctatccTTTGTATAAAATGGTAGAGTCAGCCTATTCCTTCCTGCCTggttacctcttatctagtg
gtagagtgtagcacctatccCTGCATTTTAAAAACCTCTTCTTGAAGATATCCATTCTCCggttacctcttatctagtg
gtagagtgtagcacctatccTAATGGCATCTGGAAACTTATCTGCTGCCTCTTCATTGGCggttacctcttatctagtg
gtagagtgtagcacctatccCTTTGCTGACACTAAATTCTCTAGCATTAGACCCTTCACCggttacctcttatctagtg
gtagagtgtagcacctatccTTTTATAGCAATCCCGCATCCACATAAAAAGCTGCATTTTggttacctcttatctagtg

gtagagtgtagcacctatccGGGCCTGTTGGGGGGCCGGGGGCAAGGGGCAGGGAGAGCCggttacctcttatc
gtagagtgtagcacctatccTCACCAAAAGTGCAAGATTTCTGCACCCACTGGCATAGCTggttacctcttatctag
gtagagtgtagcacctatccATCTTGAAATGGAGGGCAACCACAGCTGCAGATCATGATCggttacctcttatctag
gtagagtgtagcacctatccATCACTAATGGCATTTTGAATGGGTCTCATGATGTTATCCggttacctcttatctagtg
gtagagtgtagcacctatccGTATTGCACTAAACACGACAAGAAATACATGAGAACTGTGggttacctcttatctagt
gtagagtgtagcacctatccACTGGAGAGATGAATTGCTCTTGTGAAGATGATTTTATGCggttacctcttatctagtg
gtagagtgtagcacctatccCATTTCTTTCAACTTCAAATGGCATAATGTGCGGTCTGCGggttacctcttatctagtg
gtagagtgtagcacctatccTATCAACATACATTTTATGCATGCATGACATGCCTAGCTTggttacctcttatctagtg
gtagagtgtagcacctatccACAGGGTTTATTTGTGAGTCTTTTCAAGTTGCCATAAAACggttacctcttatctagtg
gtagagtgtagcacctatccCACATATAAGTGGCCTTGCTCAGTTCAAACTCATGTTGTTggttacctcttatctagtg
gtagagtgtagcacctatccCTCTTTATCCTTCTACAAGCAACATTTGGGAAAAATAAGCggttacctcttatctagtg
gtagagtgtagcacctatccAATAAGTACATATGGATATACACTCACTCATAAGCTTCCGggttacctcttatctagtg
gtagagtgtagcacctatccTCCTTATCTGTAAAATACAACCAACCATAGAACCAAAAGGggttacctcttatctagt
gtagagtgtagcacctatccATTCAATGAGATTGTGCACAGTGTAAATAAAGCACTTAGCggttacctcttatctagtg
gtagagtgtagcacctatccAATGAAGCCATTTTCACATTTTTGGTTTGAAGTAGCAGCTggttacctcttatctagtg
gtagagtgtagcacctatccTGCTTGAAGATTCTCACACACAATATCTCAGAAACTGTTGggttacctcttatctagtg
gtagagtgtagcacctatccCACTATATGTGGACATTGATCATTCATCCTTTGACTTTCAggttacctcttatctagtgg
gtagagtgtagcacctatccCACACATTCTTATCAAGAATCAGGAAAAGTGACAATGCTTggttacctcttatctagtg
gtagagtgtagcacctatccAAGCTTGATACATTTTATCTTGTCTGCTTGTCCTTTTTCCggttacctcttatctagtgg
gtagagtgtagcacctatccTAAAGGTAATAGTGAAAATAGCCTTCCCAGGCCGGGCGCGggttacctcttatctag
gtagagtgtagcacctatccGCTGAGGCTAGAGAATTGCTCAAACCAGGACCCAGGAGGCggttacctcttatcta
gtagagtgtagcacctatccGGCTTCCCTTTTTCCCACTTTTTACCGTCAGTGACCCAGAggttacctcttatctagt
gtagagtgtagcacctatccTTAAAGTTCTTTTTTTCTTCCGGAATTAGCTGCATTGAGCggttacctcttatctagtgg
gtagagtgtagcacctatccAATTGTTATTAATATGAAGACATTACCCTTCCTCTGTCGGggttacctcttatctagtg
gtagagtgtagcacctatccTTTGCCCCAAATCACCCATATCATCTTCTTACATCTTTTTggttacctcttatctagtgg
gtagagtgtagcacctatccGTTTTCCTTTGTTTGACATTGAAAGTCCTGAATGGGATCAggttacctcttatctagtg
gtagagtgtagcacctatccAACTTGTTAAACATGGATTCTGATTCTGTAGGGGCTGATTggttacctcttatctagtg
gtagagtgtagcacctatccGCTAGTCTGGACCACATTTCAGCAGCACAGATCTAAAGCGggttacctcttatctag
gtagagtgtagcacctatccCTCAGCGTGTCGTCTTGGGACCAGCAGCCCAGGCATCACCggttacctcttatcta
gtagagtgtagcacctatccGGCACACCCCTGACATTCTGAATGAGAAACTTTGGGTTAAggttacctcttatctagt
gtagagtgtagcacctatccAGAACCACTGCTCTGAAAAATTAACACCCTGCTCTAAAGTggttacctcttatctagt
gtagagtgtagcacctatccCCTGCCAACATGGACCGATTTTATTGTTTACTATTAAGACggttacctcttatctagtg
gtagagtgtagcacctatccGACAGTCTGGCCATGGCTAAGTTAACTGCAGGAAAGGAGCggttacctcttatctag
gtagagtgtagcacctatccGAAATTTCATCAGGCAGGAGCATTCCCTACATCTGAGCATggttacctcttatctagt
gtagagtgtagcacctatccTCAGCAGGAGGGAAGTCTGACAGGGGTTAGTAGAGCAGTGggttacctcttatctag
gtagagtgtagcacctatccCACAGAGATAGGCATGTGCTTAAGGTACCCACTAAACAGGggttacctcttatctag
gtagagtgtagcacctatccAATGTTATTGGTTAGTATGACTTGGTGTAATCTCTAGAGCggttacctcttatctagtgg
gtagagtgtagcacctatccCCATGTAACAGCTAGATAACTTTGGAATTTACTCATCTTGggttacctcttatctagtg
gtagagtgtagcacctatccATTACAAGTGTGGGGAAAGAAAATAAGGGTTCCATATTTCggttacctcttatctagtg
gtagagtgtagcacctatccCTTGTGAGAGATTAGACAAAAGTCAGAAATGCAAGCAGATggttacctcttatctagt
gtagagtgtagcacctatccCTAGAGAAAGACCAAGCAAGAGTAGGATAGAACTAGGGAAggttacctcttatctag
gtagagtgtagcacctatccCTGCCCAAAGTCAGGTACGTATCAGAAAAAAGGAACACACggttacctcttatctag
gtagagtgtagcacctatccATGTATCATCATCAGTAACTCTCAGAACTTAAGGCCGCACggttacctcttatctagt
gtagagtgtagcacctatccGCTCTCACCTTAGCTGCAAATGGAATCACCTGGGGAGCTTggttacctcttatctag
gtagagtgtagcacctatccGGCATTACCTAGGCATCAGAGTTCCAAAATCTCCTCAAAGggttacctcttatctag
gtagagtgtagcacctatccTTATTAGTGTCCAGGGTTGAGTGTCTTGCATCTTTTAAGTggttacctcttatctagtgg
gtagagtgtagcacctatccGCCCAACTGCAGGTAAATGAGAAACTCAAGGATAGGAAAAggttacctcttatctag
gtagagtgtagcacctatccCACCACACCTGGCTAATGGATTTTATTTTTTTGGTGGAAAggttacctcttatctagtg
gtagagtgtagcacctatccCTGGCCCAGGTTCTTTATTCTTCATATTTATGCTCTAGACggttacctcttatctagtg
gtagagtgtagcacctatccGTTGCTAGTACATAGGGTGCCTTTGTGCAATTCAGAAGCAggttacctcttatctagt
gtagagtgtagcacctatccCTTCTTCAGGATGCTTTGTCAAGAACTGAGAACGGTCTGAggttacctcttatctagt
gtagagtgtagcacctatccCCTATCGCAGTTTCATGGATGGTGGCAGAAGACTTGAGACggttacctcttatctag

gtagagtgtagcacctatccCATCTCATTCCTCAGATGATTTCCTATTCATACTTCAACTggttacctcttatctagtg
gtagagtgtagcacctatccTTACCCGAAGATTTTGATACTTTAAGACATGATCAACTCCggttacctcttatctagtg
gtagagtgtagcacctatccGCCTAATCCTCGTTATTGGCACATTGAAGCAACATAAGCCggttacctcttatctag
gtagagtgtagcacctatccCTGAGAAATACAGATTAAACTAGCCAAATCCATGACAATGggttacctcttatctagt
gtagagtgtagcacctatccTGTGTACAAATGACCCATACCAAGTATTAGATGAAACAAGggttacctcttatctagt
gtagagtgtagcacctatccACAATGTGCCACTATGTCTGAAGATGTGCCATAACTGTTAggttacctcttatctagt
gtagagtgtagcacctatccGTTTAATCTCTTGCAAGAACTATACTGAAAAGATGGCCTAggttacctcttatctagt
gtagagtgtagcacctatccCAACAGGTGAATGATTAGTATGGAATTTTAAAAGGGGAACggttacctcttatctagt
gtagagtgtagcacctatccCAAAGAAGGAACAGAAGCTAACTAGCATAAACTGGGGAGAggttacctcttatcta
gtagagtgtagcacctatccGCCTTGGTAAATGTCCCTCCAATCCTGTGACGGTTAATTTggttacctcttatctagt
gtagagtgtagcacctatccAGACTGAGCAAAGAAGATCCGCCCTCACCAATGTGGGAGGggttacctcttatcta
gtagagtgtagcacctatccCTCTCTTCCTTCTTGAGCTGGCACTTGCATCTTCTCCTGCggttacctcttatctagt
gtagagtgtagcacctatccTTGGACTGAATTACACCATCTTTTAGTTCTCCAACTTACAggttacctcttatctagtg
gtagagtgtagcacctatccGAGAACCATGACTAATACACCCTCCCACTGCTTCATTCCAggttacctcttatctag
gtagagtgtagcacctatccACATTTGGCAAGCAACATTCCATGACTCATCTGGCTTCCCggttacctcttatctag
gtagagtgtagcacctatccGGAAGGACCCCAGATAATGAAAGAAACTAGCCTTGCAATAggttacctcttatctag
gtagagtgtagcacctatccGAACTAAAATGATACGTTGGCATTCTGATCTATGCATCAGggttacctcttatctagt
gtagagtgtagcacctatccCACCAGTTTGTTAGTGAGACTGTCATTTCCCCGGTGGCAGggttacctcttatctag
gtagagtgtagcacctatccGCCATGGACATGCAGCTGACGACATGAAGGCATTTTCCCAggttacctcttatcta
gtagagtgtagcacctatccCCACCTGGCCTCTCTAAACTCCCCCTCATTACTCCCTGATggttacctcttatctag
gtagagtgtagcacctatccTTTCATCCTATCCTAAGGTGAGTGCCTGAGCTGAAACTGGggttacctcttatctag
gtagagtgtagcacctatccAAAATTCCAGAAGTCTTTGGAGACAGGGCAATTCTAAACCggttacctcttatctag
gtagagtgtagcacctatccACATGCAAGCAGAATGTCACTCTCAAGACTGAAATATTACggttacctcttatctagt
gtagagtgtagcacctatccTATGCACCCAGGATCCATATTTATTCATTCAGCACCCAGAggttacctcttatctagt
gtagagtgtagcacctatccAGATACAGAGGAAAATGAGATGGAACATTAACCATGGGCTggttacctcttatctag
gtagagtgtagcacctatccTCACAGGACTAGTTATCAGCTTTGCAGGGCTTGAGTCAATggttacctcttatctagt
gtagagtgtagcacctatccAAGTTGACCAGGATCTTGGGGACTCACTCTCACTAACCCAggttacctcttatctag
gtagagtgtagcacctatccTTATCATTTCTAGTGTCTATAAGGAGTCTGCTATTAGACCggttacctcttatctagtg
gtagagtgtagcacctatccCTGGAAAGGCAAGTTATCAATGTTCACTCGGTTAGTTAAAggttacctcttatctagt
gtagagtgtagcacctatccGGCCACCTCACACACAACACGGGAATGCCTAACTTTGATGggttacctcttatcta
gtagagtgtagcacctatccGCAATCCTGTCACCATGATATGAAATTTGTTTTTCTCTAGggttacctcttatctagtg
gtagagtgtagcacctatccGCTAATACAGATGGTCACCCTAATTATCAAGTAAAGTTAGggttacctcttatctagt
gtagagtgtagcacctatccAAATGGTGCTGTCATGTCTTCTTGAGAACTAGTTATTGTTggttacctcttatctagtg
gtagagtgtagcacctatccCAGGTAGTGGGTTGTATCCTGCTTGGTTATATAAAAGGATggttacctcttatctagt
gtagagtgtagcacctatccTTGCCTGTGATGTGCATTACATTCTGGTTTAACGCTTATTggttacctcttatctagtg
gtagagtgtagcacctatccTCAATAGAGAGGATTTCTGGGCTGGGAGGAGATTTTGTTTggttacctcttatctagt
gtagagtgtagcacctatccTGTATTTCCCTACCATCTTATTCTTCCACATGCACAGTCCggttacctcttatctagt
gtagagtgtagcacctatccCTGTTGATTTCTACCATTCCTTTTCCTATTTTCACACATCggttacctcttatctagtg
gtagagtgtagcacctatccAACTAGATTTTATTAATAATGTTAAACGGTGAGGGTGGGGggttacctcttatctagtg
gtagagtgtagcacctatccAGTGGATACCAAGGGGTAAAAAAGAAGGGAAGGAGGAAGGggttacctcttatcta
gtagagtgtagcacctatccTGGAAGGCAGAACCCAAGTAGTCATACGGAAGGTACCAAGggttacctcttatcta
gtagagtgtagcacctatccGGAGGAAGTATTAAGATGTTGAAATGCATAGCTAATTTGCggttacctcttatctagt
gtagagtgtagcacctatccGCTTTTTGTATTAGCTGTCATCCCTATAATATTCAGAGACggttacctcttatctagtg
gtagagtgtagcacctatccACAAGGGTAATGTGTTGGGGTATAAATTTAAAATCCAGGAggttacctcttatctagt
gtagagtgtagcacctatccCATTGTGCTGTAAGTTCATGCTATGTTTTCACAAAGGCTAggttacctcttatctagtg
gtagagtgtagcacctatccAGAAATTGGAAAAGTACATATAGCAGAGAATCTTGTGGACggttacctcttatctagt
gtagagtgtagcacctatccATAGACGATATCATGGCTTCTCAGTGATCTGTGTGAGGGAggttacctcttatctagt
gtagagtgtagcacctatccCAGCACTCATTCTTTCCCTACTAACCAAGTGAGCATTGTTggttacctcttatctagt
gtagagtgtagcacctatccAGAAGGGCCTCAAGAACACACTCCTGTACATATTAGAAATggttacctcttatctag
gtagagtgtagcacctatccGATTGATTCTTCATTCTCATGTGACTTGGATTACTAGAGTggttacctcttatctagtg
gtagagtgtagcacctatccTCCCTGCAAAGCTGATAACTAGTCCTGTGAATATGTGGTCggttacctcttatctagt
gtagagtgtagcacctatccTGGGTAATGTTTCCATGGAGTCCAGACTCAGCAATCAGCTggttacctcttatctag

gtagagtgtagcacctatccTCTCCCTTCCTTACTCCCTCTCCTCTAATTTCACAAAGCCggttacctcttatctagt
gtagagtgtagcacctatccGGTGATCCCTCTTTCCCATCATAAAGGTTGTAGCCAATACggttacctcttatctagt
gtagagtgtagcacctatccATAGGTTTATGTGACATGGAAGTTTTCAGGATGGAGGCCTggttacctcttatctagt
gtagagtgtagcacctatccACGCTTAGGTTCTATGACGTAGGGACAACCATTTAGAAAGggttacctcttatctagt
gtagagtgtagcacctatccAAAAAGGGTATGAGCTAATGGGAATAGACTCAGTGGGGGAggttacctcttatctag
gtagagtgtagcacctatccCTTGGCTTCTCTGTGCAGCACTCCTTTCTCCTGGGTAGAAggttacctcttatctagt
gtagagtgtagcacctatccCTACAGGTGTGTGTCACCACCCAATTAACTTTTGTATTTTggttacctcttatctagtg
gtagagtgtagcacctatccGGCCAGTTGTTACACAGAAATGTGGGGGAAGGTTAGAGGAggttacctcttatcta
gtagagtgtagcacctatccAGGGAAAAGAGGTTCTAATTTCTATGCTTTGCCTCAGGGAggttacctcttatctagt
gtagagtgtagcacctatccGGGCGAGAGACAGGAGGGCAAAAGGTTGGAGACAGACTTTggttacctcttatcta
gtagagtgtagcacctatccGCTTCTGAGGCTTTCATTTGAAGGTATTGTTTTCTGAGCCggttacctcttatctagtg
gtagagtgtagcacctatccGCTTAATGATCTATGATTTACTAAATGGGACAGTCAAAGCggttacctcttatctagt
gtagagtgtagcacctatccGCACTCTGGGAGGCCAAGCTCAGGAAATTCTAGCACTTTGggttacctcttatcta
gtagagtgtagcacctatccGAAACCCCGTCTCTAACTAACATACAAAAGAAATTAGCTGggttacctcttatctagt
gtagagtgtagcacctatccGAAAGAAAAGAAAAGCAGATGATGACAGTTCTTAAGGGAAggttacctcttatctag
gtagagtgtagcacctatccTGGCATGCCACCGTCCCCTCTGAAAACGAAGTGACCACGGggttacctcttatcta
gtagagtgtagcacctatccCAGCCCTGACCACAGCCGATTGGACCCACAGCAGACACGGggttacctcttatct
gtagagtgtagcacctatccGTAATCCAAAATGAGATGACAGAGGCAGGGTGAGAGGCTCggttacctcttatcta
gtagagtgtagcacctatccACTTAAGCCCAGGAGTTTGGAGTTACAGCGAGCTATGACTggttacctcttatctag
gtagagtgtagcacctatccAGAGATAATAGTGAGTAAACCTGAAGCTCTCTCTGTAGGTggttacctcttatctagt
gtagagtgtagcacctatccTTAAATGTTAACACTGTAATATGAAAGCGTGGCTGAACAGggttacctcttatctagt
gtagagtgtagcacctatccCTTAAGAAATGACCAGCAAACTCCATGATCTCCAAGCCTAggttacctcttatctag
gtagagtgtagcacctatccGGTCATTTTAGCTTGGGGAATTGAGGACACAGAGGCAGGTggttacctcttatctag
gtagagtgtagcacctatccTCATGTCCCAACACTACAATCTGACTTAATCTGTATTTGTggttacctcttatctagtg
gtagagtgtagcacctatccATGAATAAAACCAACAGCATGAAAGAGAGGTGCCAAGATGggttacctcttatctag
gtagagtgtagcacctatccGATTCATAAAACAAGATATTACCGGCCATGCCAATGGATAggttacctcttatctagt
gtagagtgtagcacctatccAGTTCACACTTACGGGCACTAACTCCAACAGAAGCTTCCAggttacctcttatctag
gtagagtgtagcacctatccTGTTTATACAGAGGACTTCACACACCACCAACCATGCAATggttacctcttatctagt
gtagagtgtagcacctatccACGCACAGGGAATAATCAGAGGCAGGTAACAAACCACAGCggttacctcttatcta
gtagagtgtagcacctatccGTGGGTCCTGAGTCCTTGGTAGGGTCACTGCTTCTCTCTGggttacctcttatctag
gtagagtgtagcacctatccCCACCAAGACAGGAGAAGGTCTCAGACTTACCCCGGGCAGggttacctcttatcta
gtagagtgtagcacctatccGCTTTTATAATCCTCAGCCACTTTTCACTTTCTATTGGGTggttacctcttatctagtg
gtagagtgtagcacctatccGGTGCTATTACCTCTGTCCATTACATGTATGTTTATCACAggttacctcttatctagtg
gtagagtgtagcacctatccCTTAATGCAGCTTGACATGAGTGACTCCCTTTTGAGACATggttacctcttatctagt
gtagagtgtagcacctatccCAGAGAACAAGAGTTAGTATAAAAGAAAAACATGGTTGCCggttacctcttatctagt
gtagagtgtagcacctatccCTGAAAAGCCAAATGAGCACAACTGAGGAACAAATTAGTGggttacctcttatctag
gtagagtgtagcacctatccGACCAAGTCATACAATTCTACCAGAACACTGAATGGAAATggttacctcttatctagt
gtagagtgtagcacctatccTCAAGTGGAGACAAGGAGAAAAAATCCAGAGTCATAATAAggttacctcttatctagt
gtagagtgtagcacctatccGACTTGATATTCCAATAGAGTCTCTGGTTTGAAATGGGAAggttacctcttatctagt
gtagagtgtagcacctatccGTTTAAAAGAGCACAAACTACCTCAATTCCATTCTGTCTGggttacctcttatctagt
gtagagtgtagcacctatccTCAAATAAGGCATCAATCCATCTTACTCTGTTTTTAGCAGggttacctcttatctagtg
gtagagtgtagcacctatccAGCCCATGGATAATCACTCAATGTTTGCTAGTTAACTAAGggttacctcttatctagt
gtagagtgtagcacctatccTTTCTTCAAGGTAGAATGATTTGGAGAATGCATCATAACCggttacctcttatctagtg
gtagagtgtagcacctatccTGTTGAGACAGTCATATTTAAGCAATGCTTTTACAGCCACggttacctcttatctagt
gtagagtgtagcacctatccCATTCAATTGTACTGATTCCTCACTTTATAGTCCCTGCAAggttacctcttatctagtg
gtagagtgtagcacctatccATACATACAAAATAAACACTGACTCTATGAGGCATTCTCCggttacctcttatctagt
gtagagtgtagcacctatccATTTTACACCTTCCTTGTTAAAATGTGATGCTGTCGACCAggttacctcttatctagtg
gtagagtgtagcacctatccGGCGTGGTGGTGCGGGCCTGTATTCCCAACTACTCCAGAGggttacctcttatcta
gtagagtgtagcacctatccAGAGAAGAAAAGAAAAATGTGATGCTGTCCTAAGGGCTGGggttacctcttatctag
gtagagtgtagcacctatccGGAACCAGAACAATGGCATATAGATAATGGCCTAAGGTGGggttacctcttatctag
gtagagtgtagcacctatccTTTGCCCCAAGTTTCATCTTTTATAAAACCATTCTCATGGggttacctcttatctagtg
gtagagtgtagcacctatccGGAAATGAACCCTGATTGCTACCGCTGTATTTTCTCCTAAggttacctcttatctagt

gtagagtgtagcacctatccCAATAGTGGCAAAGAAGGGCAAGTCAGAAAAAAAGGAAACggttacctcttatctag
gtagagtgtagcacctatccTTCAAATATTTCCAGTTTACTCTGACAGTTCACATTCTCCggttacctcttatctagtgg
gtagagtgtagcacctatccGATAAAGATCTATTCATTGTATTCTATTTGCTGGCAGACCggttacctcttatctagtg
gtagagtgtagcacctatccCCAGAACAGGATAAAAGCCACAGCCATTAAAAAATAGATTggttacctcttatctag
gtagagtgtagcacctatccGTAATCCCAGGTACTCAGGAGGCTGGGGCACAAGAATTGCggttacctcttatctag
gtagagtgtagcacctatccAAAATAAACAATAGATTCTCCTGACACATTTCTGTTGGGGggttacctcttatctagtg
gtagagtgtagcacctatccGGCATGAGTTAATATGACTGACTAATTGAAAGCATCAGGTggttacctcttatctagtg
gtagagtgtagcacctatccACCTATGTGTGAAGTTGTGTGTGAGACCCTGGGATATTGTggttacctcttatctagtg
gtagagtgtagcacctatccGGTAGATCAGTTCCCTGCCTAAAAAGAATGTTCCTATCTAggttacctcttatctagtg
gtagagtgtagcacctatccCTAAACTTAGGATGTTGTGGTGGGCTCTTCGATGTAGTTCggttacctcttatctagtg
gtagagtgtagcacctatccAGATGAACTGGGAAAGAAGCGAGGTTTTTATTTCTGTAACggttacctcttatctagtg
gtagagtgtagcacctatccGCTTATATACCTTTTAAGCGATATGTCTGTGTGTAAGTGTggttacctcttatctagtgg
gtagagtgtagcacctatccTGCTGGAGTGATTACCCTTAATTTGTCTGATGGAGGCAAGggttacctcttatctagt
gtagagtgtagcacctatccGGAAGTCTGGAGAGTTCCTCTAGTCCCCAGTAAAGCTTGTggttacctcttatctag
gtagagtgtagcacctatccCTAAATGGGTCCTGTTAAGACTTCCTTCATTATTTTGTCAggttacctcttatctagtgg
gtagagtgtagcacctatccCTGCCTCTATCAGCTTTTAACATTTAACCTAACCACTCAGggttacctcttatctagtg
gtagagtgtagcacctatccGGGACTACAAGCATGAGCCCAGCCCTAGATGTAACTTCCTggttacctcttatctag
gtagagtgtagcacctatccGTTTTCTCAACTTTGGTGCTGTTAACATTTTGGGCCTGGTggttacctcttatctagtg
gtagagtgtagcacctatccTTTAGCAACATACCTGGACTCTACCCATAAGATGCCACAAggttacctcttatctag
gtagagtgtagcacctatccGGAACAAACTCATCCCCAGTTAAGACCACTAGCTTAGATGggttacctcttatctag
gtagagtgtagcacctatccTAAGAAAAGGGAAGTTGTAGAGTGAAGGAGAAGAAAAAGGggttacctcttatctag
gtagagtgtagcacctatccGGGTAGGGAGAGAAGACATAAGAAGTAAGAGGGAATCGTTggttacctcttatctag
gtagagtgtagcacctatccCATAATGGGTGGTGGGTGGCAGACGCTAAGCTTGATTTGAggttacctcttatctag
gtagagtgtagcacctatccGCTTGGAGCTTTATTAACTATCTATTGCTATGTGTGATGGggttacctcttatctagtgg
gtagagtgtagcacctatccACTCTGGGTGTTTCTGTGATAATGTTTTGGATGAGTCTAAggttacctcttatctagtgg
gtagagtgtagcacctatccTAGATTGAGTAAAACAGATTTTCCTCCCTAAGGTGAATGGggttacctcttatctagtg
gtagagtgtagcacctatccACAAAAAGGCTGACCCATCCCAGGATAAGAGGGAATTCCCggttacctcttatctag
gtagagtgtagcacctatccGAGCATTTTAACTCCTTTGTCCCAAACTCTTCCCTAATAGggttacctcttatctagtg
gtagagtgtagcacctatccGAGATTTCCTCCATAATTTAAATGCTGTCTTCTCCAAAGAggttacctcttatctagtg
gtagagtgtagcacctatccACAGGTTCAGATTTGAAATCACTTGTTTTTGGTACCCACTggttacctcttatctagtg
gtagagtgtagcacctatccCCCAGAGGAGCAGAGTTAATTCCAGGCAAGCCATATCATAggttacctcttatctag
gtagagtgtagcacctatccGCAAAGAGGCCTGGACAAGCTGTGTTGAAGCAGACTCCAGggttacctcttatcta
gtagagtgtagcacctatccGCACTACATTCCAGAAGCCAGGCAGAGGATCAAGAATCAGggttacctcttatctag
gtagagtgtagcacctatccATTCAGGAATGAGCAAGGGTACAAGCTGGGGAAAAGTTCAggttacctcttatctag
gtagagtgtagcacctatccCCCCAATGCCCTTCCAGGTTCTTAATCTCAAGTTGCCAGAggttacctcttatctag
gtagagtgtagcacctatccCCAACAGGGTGAAGTGGAAGGGGAAAATGGGTGTGTACATggttacctcttatctag
gtagagtgtagcacctatccCCTAACCGATTTGAGGGCATTTGATTGTTTTCTTTTCTTTggttacctcttatctagtgg
gtagagtgtagcacctatccTCAGCCTCCCAAGTAGTTATGACTGCAGGCACACGCCACCggttacctcttatcta
gtagagtgtagcacctatccCCAATGCACCCAGTCATGTTTCCAATTTTTGATCTTATAAggttacctcttatctagtg
gtagagtgtagcacctatccGGTATGAATTTTGTTTGTGTGCAGGTGGAGTTTTTTCCCAggttacctcttatctagtgg
gtagagtgtagcacctatccATTCTTCTAGTATTGACATAAGCTATGTAGGCACAAAAGGggttacctcttatctagtg
gtagagtgtagcacctatccATCATATGATCACGATTACTGTACATCCTAGCCTCATTGAggttacctcttatctagtg
gtagagtgtagcacctatccCTTAAAGAGGAAGTTAAAGTGAGTCTAGTCAGGAGGTGTTggttacctcttatctagt
gtagagtgtagcacctatccAAAATGAAGGGAAATGGGTTTGAGAGGAGTGAAGGGTATAggttacctcttatctag
gtagagtgtagcacctatccTTCACTCCTCCTGAACTGAACCTCTTCTTTACAACGAAATggttacctcttatctagtg
gtagagtgtagcacctatccCTTTTCTAGCAGACTTCTTCCTTCCTCAATTTCTTCAGTTggttacctcttatctagtgg
gtagagtgtagcacctatccCTTCTGCTTTAACTTTTCTTTCAACCTGTGGGCTACCTAAggttacctcttatctagtg
gtagagtgtagcacctatccGAGATACATGATTCTATTGTACAAAGTTAGGAAGGGAATGggttacctcttatctagtg
gtagagtgtagcacctatccTTGAATTTTAACCCCCTAGTTTTATGTGTCCATTATGACGggttacctcttatctagtgg
gtagagtgtagcacctatccGATTGCATTTCTGTTAATCTATTTACTGTCACCACCTTCTggttacctcttatctagtgg
gtagagtgtagcacctatccAGCTTTTTGAGGCAGGAACCATCCAGTATTGACTTGGCAGggttacctcttatctag
gtagagtgtagcacctatccGAACAGTGCTGGACTCATAGTAAGAACTTGTGGGCCGGGCggttacctcttatctag

gtagagtgtagcacctatccGATCAGGGGAGATCGACACCATGCTCGCTAACTCGGTAAAggttacctcttatcta
gtagagtgtagcacctatccTGTATTTAACTAAACAGAGATGCAAATGCCATGAGAAGGTggttacctcttatctagt
gtagagtgtagcacctatccCAGAAGATATTTTGGAAATAGATGAACGCAAGGACCACAGggttacctcttatctag
gtagagtgtagcacctatccTGAGCAGCTGTGTGTTTATTCCCCTCACGTTTCGTTCCTTggttacctcttatctagt
gtagagtgtagcacctatccGGCTCCACCTTCCTGTATCACCAGCAGCACGGTCTCATATggttacctcttatcta
gtagagtgtagcacctatccGCAAGTACCTCCAGGGCTCCATGTCACCACCCCACCTCCCggttacctcttatct
gtagagtgtagcacctatccCACATTCCTCCATAAGACACCCCTGGGTCAGATCTAGCCCggttacctcttatcta
gtagagtgtagcacctatccCAGCTGTGGGGTTTGTGGGGGTGAGAAGAGAGGTTTAAAAggttacctcttatctag
gtagagtgtagcacctatccAGTTTTGCTTAGAGTCAGGGATCCAATCAATATTTGAAACggttacctcttatctagtg
gtagagtgtagcacctatccCTTTCTTCCTCTCTCTTTCTGCCCCTAAAAATATTTTAACggttacctcttatctagtg
gtagagtgtagcacctatccCATTATTGTTATTCTTCCACCCTAATCTGAAACCTTTGTGggttacctcttatctagtg
gtagagtgtagcacctatccCCTTTCTCTCTCCCAGCCATCTTCTCTCCTGCTTTCGCCCggttacctcttatctag
gtagagtgtagcacctatccTTTCTATCAAAGCCCTCAGAGACATAAAAAATAGCCCATCggttacctcttatctagt
gtagagtgtagcacctatccAGTGTTCCTTGACTTGATAAACCTGACATGAGGCTCTGTAggttacctcttatctagt
gtagagtgtagcacctatccCTAGGCACGGGTTCCATCTCTGTAGCTCTGTCCTGGCACTggttacctcttatctag
gtagagtgtagcacctatccCAAAGAACCCCAGATACCCCAGGTCCCCTAATCCAATAGCggttacctcttatct
gtagagtgtagcacctatccTTGTGCAATGATGTAGTAACCAGAACAAGAAAGGGAAGCAggttacctcttatctag
gtagagtgtagcacctatccGGTGAGCAGATAGTGAAACCTGACTTAAGAGTGGATGTTTggttacctcttatctagt
gtagagtgtagcacctatccGAGTACTCCACTGCTTATGAAATTCAAGTAATCTTACATGggttacctcttatctagtg
gtagagtgtagcacctatccGAACACAGACACTTCATATTCAACAAGTTCACACTGAGCCggttacctcttatctag
gtagagtgtagcacctatccCATTGCTTGGAAGGCTGCTCCTCTTCCTGTGTTCTCTCTCggttacctcttatctagt
gtagagtgtagcacctatccAGTCTTTTCACTGCCCCTGTCTACCCCTCTCCAATCCATTggttacctcttatctagt
gtagagtgtagcacctatccGTTCTGACCATAACTCTACTTTTCTTAAGACAGCACTTTGggttacctcttatctagtg
gtagagtgtagcacctatccTGTTCATCATTCGGTCATAAGCTGCAAATCCATTCTCCTTggttacctcttatctagtg
gtagagtgtagcacctatccTCCCTCCACTCTCTCACACCTCCCAGCCTTGGCCCTTGCTggttacctcttatcta
gtagagtgtagcacctatccTTCTACCTTTACCCTCCATTCTCATCACCCTACACTATTAggttacctcttatctagtg
gtagagtgtagcacctatccGTTTTCATAACCAGATCAAGTATCGATGTCTCCACCAACAggttacctcttatctagt
gtagagtgtagcacctatccGAGCAGGCAGTGGTTCTCAAGTCACATATGTAATTTAAAAggttacctcttatctagt
gtagagtgtagcacctatccATTTTTTTTATTAAGATGAGAGGAGTCTCACTCTGTTGCCggttacctcttatctagtg
gtagagtgtagcacctatccCCACACCTGGTCAATGAATAGAACATTGATTCAGCATGCAggttacctcttatctag
gtagagtgtagcacctatccCTCAGTAGCCACATTAGGGAGTAGCTGCCATATTGTACGTggttacctcttatctag
gtagagtgtagcacctatccCACCAGAGCACTTGAGAGCCTCCTAGAGATCCTAATGAAAggttacctcttatcta
gtagagtgtagcacctatccCATTTCTGGGGAGAGTCTAATTTTGGTGGTCTGGAGTGGGggttacctcttatctagt
gtagagtgtagcacctatccCCCCAGGCAGTTCTGAGGTACATGTTCGTCTGACCATTTGggttacctcttatctag
gtagagtgtagcacctatccACCTATAGGGGTTTATGTTCAGATTCCACCACCTAAAAGCggttacctcttatctagt
gtagagtgtagcacctatccTCTCCTCACCTGCAAAAATAACTACCTAGAGACTTGTGGGggttacctcttatctag
gtagagtgtagcacctatccCCTGGCAGGTGTCTCATAATAGCAGCTGTGCACTGCACCCggttacctcttatcta
gtagagtgtagcacctatccACAACTCTCTGTGCTTCCCTCCACCACCACCCATCACGCAggttacctcttatcta
gtagagtgtagcacctatccGTGTCCTGTGCTTGCTATGCTGCCCCTCAACTCAACTGAGggttacctcttatctag
gtagagtgtagcacctatccAACTGGCTGCTGTGTGCTCCCACATTACTCTGCCTCACACggttacctcttatctag
gtagagtgtagcacctatccAAACTAAATCTGAGCAACTGGGTTTCTTGTAGTGGAGCAAggttacctcttatctagt
gtagagtgtagcacctatccAGGTGTCTTATAGAACAGAAGTGTGTAAGTAAAGTGGGGTggttacctcttatctagt
gtagagtgtagcacctatccCTGTCCTAACACTCTTCCCTTTAAAGTCTATCACAGAAAGggttacctcttatctagt
gtagagtgtagcacctatccCAGATCGTGTTATTCAGAGGAGCAAGAGGATGATTCTAGAggttacctcttatctag
gtagagtgtagcacctatccGTGTTTCACATAGCCAGCCAATATAGCAAGGTAAGAAGCTggttacctcttatctag
gtagagtgtagcacctatccGAGATAAAACCCAGTGCATACATATGTGTTACATTTACTCggttacctcttatctagt
gtagagtgtagcacctatccTCGTTTTATAGACAGCACTTGATGTACATAGCACTGTTCTggttacctcttatctagtg
gtagagtgtagcacctatccCTTTACTAAGTCACTGTCTTAAAACCATACACAGTGAGGCggttacctcttatctagt
gtagagtgtagcacctatccTGTAATCCCAGCACTACCAGAGGCCAAGGTGGGCAGACGGggttacctcttatct
gtagagtgtagcacctatccCCTTGGCAACATGGCAAAGACCCATTTCTACAAAAACAAAggttacctcttatctag
gtagagtgtagcacctatccCCTGAGAGGTTGGGGTTGTAGTAAGTCAAGATCATGCCACggttacctcttatctag
gtagagtgtagcacctatccATTATTATTATTCTCTGGAGAAAAGCCTGGTCAAGTACTCggttacctcttatctagtg

gtagagtgtagcacctatccCTCTCCCTAGGGTGTATTAGAGATAAACAGTTAGTCCATCggttacctcttatctagt
gtagagtgtagcacctatccCCCCAAGGAGTAAATAAGAGTAACTTCAGATGTGCTTCCTggttacctcttatctag
gtagagtgtagcacctatccGATCTAGGATCAAGCAACGACATCTCCTCCCCCTTTCACAggttacctcttatctag
gtagagtgtagcacctatccCAAGGATGAGAGGTATTCAATGACTAGGGCTTCTATCTGTggttacctcttatctagt
gtagagtgtagcacctatccGAATAACTCTTGAAATGGTGACTTGGTATACTGTATTCACggttacctcttatctagtg
gtagagtgtagcacctatccGATTTGCTTTTGTTCCAGATCTACAGTTCCAACATATCCAggttacctcttatctagtg
gtagagtgtagcacctatccCTGCTTTTGTGAACGGTGTGGCTGTAAGGAGGTTTTCACGggttacctcttatctagt
gtagagtgtagcacctatccAAGTCAGACAGATAAAGTGGCTTACCCTCTCTCCTATGTAggttacctcttatctagt
gtagagtgtagcacctatccCAGATTGTGTAGCAGAGGAAAGAAGATGGAAGGATTTAGAggttacctcttatctag
gtagagtgtagcacctatccTAAAACCCATATCCAGACCTAATATCTTCTGTTTTTCCTGggttacctcttatctagtg
gtagagtgtagcacctatccGGGCCAGTTTCTCTCAAGTAATGTGTTCTTTATTCATTCAggttacctcttatctagtg
gtagagtgtagcacctatccAGGAACAATGGCCTTATAGTGTCTGTATTTTCTGAACCAAggttacctcttatctagt
gtagagtgtagcacctatccACAGATGATTTTTATGCTTTTTCTGTGTGTCACGGGAAATggttacctcttatctagtg
gtagagtgtagcacctatccGACATGTTGGGATTTCCGAGATATTTTTATGATTTATGGGggttacctcttatctagtg
gtagagtgtagcacctatccCAGGAGATTGAAAGTTAAAGCCAACGGTGCATTCCAAAGCggttacctcttatctag
gtagagtgtagcacctatccAGTGGGAAATGCACCAGTGACAGCAAATAGATATAAAAATggttacctcttatctag
gtagagtgtagcacctatccAAGAAAATGGCTAGGAGAAAGAGGAATAAGATCTTAAAGCggttacctcttatctag
gtagagtgtagcacctatccTCTGAGTTTGACAGTTATTGTGCTATTCTTACAACTTCTCggttacctcttatctagtg
gtagagtgtagcacctatccAAAGGTACAAAAACGTATGGCCTCATATGGAATAAATGACggttacctcttatctagt
gtagagtgtagcacctatccGGAGGAATTGCCAGCCACCTGATTCTGCAGCTTTGTTTTTggttacctcttatctagt
gtagagtgtagcacctatccATATAGATCTTTCCAGGGGCCACAGCAGAGGGAGCTCAGGggttacctcttatcta
gtagagtgtagcacctatccGCTCACAAGCTGCTCTCCAAGGTACTCCCTTCCTCTGGCAggttacctcttatcta
gtagagtgtagcacctatccACAGCATTTCCCAGTGCCCGGCCATGTGATTAATAATCACggttacctcttatctag
gtagagtgtagcacctatccGGTGGGCACCTGTTATCCCAGCTGCTCCAGAGGCTGAGGCggttacctcttatcta
gtagagtgtagcacctatccACAAGGTTATTACTAAACTAAGTCCTTAGTGGTCCAAAAGggttacctcttatctagt
gtagagtgtagcacctatccTAAGGGAAAATTTAAGGCTTTTTGTTGCTGTTTGTGTCTGggttacctcttatctagtg
gtagagtgtagcacctatccAGATTCACTCCATGGATTTTGATGACACGTGGCACCCTGCggttacctcttatctag
gtagagtgtagcacctatccAGGAGTCCAAAGTTTTCTGGCCTTGACCTGCTGCTGGCTTggttacctcttatctag
gtagagtgtagcacctatccCAAAGAGGTATGGAGAGAATTTGCCCCATCAAAAGGTAGTggttacctcttatctag
gtagagtgtagcacctatccCCCCTTCATCTATCAATCATTATCTCCGTAGAGCTTTCTGggttacctcttatctagt
gtagagtgtagcacctatccGGTAGATAAGTCAATACACCAGTCACATACGAAACCCTCTggttacctcttatctag
gtagagtgtagcacctatccGTGACTTCAAGAGTGCAACCTGGACCGTGAGATTAATTGGggttacctcttatctag
gtagagtgtagcacctatccTGTCATTCTGGCATTCATGGGTGTACTGAGTATTCCTTATggttacctcttatctagtg
gtagagtgtagcacctatccAGAATATCCTTACTCTGACTGTTCATCCTGCAGATAAAAAggttacctcttatctagt
gtagagtgtagcacctatccGCTCAGTCTCAGATTTATGTTGCCACTTGCTCCAGCCTGCggttacctcttatctag
gtagagtgtagcacctatccAAGTTTCAGTTTCTTCAAGGCCCACAATTCTCACAGTGACggttacctcttatctagt
gtagagtgtagcacctatccTGGTACCATTTATAAAGAACTCACTATGTGCTAGCACTATggttacctcttatctagtg
gtagagtgtagcacctatccGCTCCTAAAACTCTAAGAGGTAGATGTTATTATTATCACCggttacctcttatctagtg
gtagagtgtagcacctatccGCGTGAGCCATCGTGCCCGGCCTTATTATTATTATTTTTAggttacctcttatctagtg
gtagagtgtagcacctatccGACTGCAGGCACCCACCACCACATCCGGCAAATTTTTGTAggttacctcttatcta
gtagagtgtagcacctatccGCATGAACCACCACACCTCGCCTATCACCATTTTGTGTAAggttacctcttatctag
gtagagtgtagcacctatccTCTACAGAGAAGTTGAATAATTTTCCCCAGATCATATAGCggttacctcttatctagt
gtagagtgtagcacctatccAATGTTCTTGATGACTACGCTATCCTGCCTCCTGGTGGTTggttacctcttatctagt
gtagagtgtagcacctatccTGTACATCTCCAACTCTGCTTCTTTGCTTTTCAGATTCCAggttacctcttatctagtg
gtagagtgtagcacctatccGCTGCATCCAAGTTTTTAGGACTTAGCTCGACAAAGTGCCggttacctcttatctag
gtagagtgtagcacctatccGCACCCATGGCCAATGCTGCCACCCAGACCAAGCCCCTCCggttacctcttatct
gtagagtgtagcacctatccGCATGGGATAGAAGCTGCATTTTTGGCAATGTTGGGTCTCggttacctcttatctagt
gtagagtgtagcacctatccGCCTTTTATGCCAACTATTCCCCAAAAGTCCTTCCAAGCAggttacctcttatctag
gtagagtgtagcacctatccCGTGGCCTTTAAGCGTTTTCCTGCACATTTATCTACCCACggttacctcttatctagt
gtagagtgtagcacctatccTGCATCTGTGAGAAAGCACCTTGTAGCAGAGAGAGCCCTGggttacctcttatcta
gtagagtgtagcacctatccCTGACTACATTAAGAGAATTGTGCTCAGGATACCAAATGTggttacctcttatctagt
gtagagtgtagcacctatccACAGGCCCTTTCCAGTTTCGGAGCATGAAGCCCGTCATTCggttacctcttatcta

gtagagtgtagcacctatccGCCATGCTGCTTGATGGTGGCATCACTGTCCCCTCATTCCggttacctcttatctag
gtagagtgtagcacctatccCCAGATCAACAGGCCACAGGTGAGAGAGCTGCTCAGTAAGggttacctcttatcta
gtagagtgtagcacctatccCCCTCCGGACAACTTGCCAAGCTTCAACATACTGTACTGTggttacctcttatctag
gtagagtgtagcacctatccTCGCTCTGATACCTTCTATGGGCACTGGAGAAAACCACTGggttacctcttatctag
gtagagtgtagcacctatccGAGGAAAAGTTCAGAGCCAATGCCTCCAAGATGCTGTCCTggttacctcttatcta
gtagagtgtagcacctatccGTGGAAAGCCTTATAAAGATAGTCAAAAATCTAGAAGACCggttacctcttatctagt
gtagagtgtagcacctatccCAAACTCTCCAGCATGTAATAATTCTATCACAAATCTCTCggttacctcttatctagt
gtagagtgtagcacctatccAATGACTTTGCATTTGGGGAGATTCAATGATTTGGTTTGTggttacctcttatctagtg
gtagagtgtagcacctatccAGGGTCTGCTGCTTGGTTTTCCCAGGAAAAATGAACAAAGggttacctcttatctag
gtagagtgtagcacctatccGACAGTTCCACTTACCTAAATCAAGATGAAACACACACACggttacctcttatctag
gtagagtgtagcacctatccATTAATCTCTTCATAGGAATTATGTGTGGACTTCATGAGCggttacctcttatctagtg
gtagagtgtagcacctatccAGGGATGTGAACCTGCATGTATATTTTCTGACAGTGGAGAggttacctcttatctagt
gtagagtgtagcacctatccGTGCATTGTGTCACCAACAGATCTCCTAGACCATGGCTTAggttacctcttatctag
gtagagtgtagcacctatccCAAGGGGTGCTACTGGGGAATGGGTGGGTTTAAATCCTGCggttacctcttatcta
gtagagtgtagcacctatccCCTTGAGCATGTTACCATTAGCCCTCTGCCTCAGTTTCCCggttacctcttatctag
gtagagtgtagcacctatccGTCAAGCCGAAGTAAAAAGCAGTCTGGAAAAATCGCATTTggttacctcttatctag
gtagagtgtagcacctatccAGAACCCATGGTCTTAAGCACTGCAATATATCACCTTTCAggttacctcttatctagt
gtagagtgtagcacctatccGTGTCATGGGGGTTAAAAATGCAACACTAACAATCATAGTggttacctcttatctagt
gtagagtgtagcacctatccGGGCAACCTATTAACTCTAGATGTTGATTAATAAGTATGCggttacctcttatctagtg
gtagagtgtagcacctatccACACGTCCTTCTTCATATGAAGGAGAAGTGCTTCCTTCATggttacctcttatctagt
gtagagtgtagcacctatccCCTATAAACCGTCAGATTTCAGGAGAACTTACTATCATAAggttacctcttatctagt
gtagagtgtagcacctatccTTGGGTGGGGACACGGTCATACCAAATCATTCTGCCTCTGggttacctcttatctag
gtagagtgtagcacctatccAACTCATTCCAGCATCAAGGCAAAAGTCTAACTCCAAAGTggttacctcttatctag
gtagagtgtagcacctatccCTAGATACAATGGGGGTAAAAGCACTGGGTAAATACATCCggttacctcttatctag
gtagagtgtagcacctatccGTCATTCTGATACAAGAGGTGGGACTCCCATGGCCTTGGGggttacctcttatcta
gtagagtgtagcacctatccCACAGCTTTTCCAGGCACATATTGCAAGCCGTCGGTGGATggttacctcttatctag
gtagagtgtagcacctatccGGGACTTTCTGTGGGGGCTGCAGTCCCACATTTCCCTTCTggttacctcttatctag
gtagagtgtagcacctatccAAGACTTGGGGCTTACACCCCTTGAAGCCATGGCCTGAGCggttacctcttatcta
gtagagtgtagcacctatccGGAACCAAGTAAGCCCTGAGACTGTAGAAAGCAGCAAGGCggttacctcttatcta
gtagagtgtagcacctatccTTGGGAGGCACTGCTGTGAAGAGTTCTCACATGTCCTGCAggttacctcttatctag
gtagagtgtagcacctatccACATTTGGCTCCGTGTTACTTATGTGAATTTCTACTGATGggttacctcttatctagtg
gtagagtgtagcacctatccTGGGTTTTTCTCTTCCATCACATCATCAAGATGCAAATTTggttacctcttatctagtg
gtagagtgtagcacctatccCTCTGCTTCTCTTTTAAACACAAGTTTCAATTCCAAACCAggttacctcttatctagtg
gtagagtgtagcacctatccCTCCCACCAGATACCCTAAATCTTCCCTCAAGTTCAAAGTggttacctcttatctag
gtagagtgtagcacctatccGTCTCTTTGCTAAAACGTAGCAACAGTCACCTTTATTCTAggttacctcttatctagtg
gtagagtgtagcacctatccTCCATCTGAGACCTGGATTCCATTGTCCATATCACTATCAggttacctcttatctagt
gtagagtgtagcacctatccTTGGTCAATGCCATTCAACATGTCTCTGGGAAGTTCCAAAggttacctcttatctagt
gtagagtgtagcacctatccTTCTGTCTTCTGAGCCCTTCAGACTTTTCCAACCTCTGCCggttacctcttatctagt
gtagagtgtagcacctatccCCAAAGTCTCTTCCACATGTTGGGGTATCTTTATAGCAGTggttacctcttatctagt
gtagagtgtagcacctatccTTCACATGATAGCAGGAAGAAGAAAAGCCAAGCAGAAGGGggttacctcttatcta
gtagagtgtagcacctatccTCATCAGATCTCAGGAGAATGCACTCACTAACATAAGAACggttacctcttatctag
gtagagtgtagcacctatccCAGCCAAACCATATTATATGTCCACTGAAATTTCTAGGATggttacctcttatctagt
gtagagtgtagcacctatccTCGAGAACAGCAGTAGAATGTACCACTTCCAGGTTAATAGggttacctcttatctag
gtagagtgtagcacctatccTGAGCCCAGGAGTTTGATGAGACAGTCTGGACAACATAGTggttacctcttatctag
gtagagtgtagcacctatccCTGCTACTCAGGAGGTTGAAGTTGGAGGATTGCTTGAGCCggttacctcttatctag
gtagagtgtagcacctatccGGACTAAATGTTCCAGTTTAAAAAGACCAAGATTTCCTGAggttacctcttatctagt
gtagagtgtagcacctatccAGTTGTGCATGCATCACCACAATTCAGTTTTGGAATACTTggttacctcttatctagtg
gtagagtgtagcacctatccCCCACCTCCAGCCCATAGTAACCAGTGATGTATCTCTATAggttacctcttatctag
gtagagtgtagcacctatccATATAATAAGCAGTCATATGTGTTTGCCTTCTTTCACTGGggttacctcttatctagtg
gtagagtgtagcacctatccGGTTTTGAGATTCTTCCACATATGTGTCAGCACTTTTTTGggttacctcttatctagtg
gtagagtgtagcacctatccATTGTAGGGATATGCCAATGTTTGTTTACCCATTCACTAAggttacctcttatctagtg
gtagagtgtagcacctatccTAGTGTTTCTATGAGCATTTACGTACAGGTCTTTGTGTGAggttacctcttatctagtg

gtagagtgtagcacctatccTGTTTCCATTTCTTTTTGATAGATACCTAGGAGTGACTCTggttacctcttatctagtg
gtagagtgtagcacctatccTTCCAAAGTAGCTCTATCATGTTACGTTTTCTCCAACAATggttacctcttatctagtg
gtagagtgtagcacctatccGGTTCCAGTTTCTCCTCATACTTACCAAACCTTAGAATTGggttacctcttatctagt
gtagagtgtagcacctatccTGACTCATTCTCCTCCATTTTTCCAGTTCTTTTTATTGTGggttacctcttatctagtgg
gtagagtgtagcacctatccCTTATGGTATGGACGTATACCTGGAATATATAAGAGCTCTggttacctcttatctagt
gtagagtgtagcacctatccAATTTATCATTCTGGCTTTGGATGTCATCAGAAGACTTTCggttacctcttatctagtg
gtagagtgtagcacctatccCAAATAAAAAATGCAGAAGGGTTGACGGTATAAAAGATGGggttacctcttatctag
gtagagtgtagcacctatccTTGGACCAAGCATTACTATAAATAGTGTCACTATGTAACGggttacctcttatctagt
gtagagtgtagcacctatccAGCTAACTGAAATAATCCTAACCCTACATGCACCTAAATCggttacctcttatctag
gtagagtgtagcacctatccAATCCATCTACCACCATACTGAGAGATTTTTAACATTCCTggttacctcttatctagtg
gtagagtgtagcacctatccACACAATCAACATGTTTATACAACAGGAATTCTGTACCCAggttacctcttatctagt
gtagagtgtagcacctatccTCTCATAGAACATTTACAAAAGTGGACTAACACTGAGCCAggttacctcttatctag
gtagagtgtagcacctatccCAGTCTCAACAACTTTTCAAAGTACTGGATCATACAGTTAggttacctcttatctagt
gtagagtgtagcacctatccTTTAAAAAAGATAATTAGAAAAGCTGCCAGCCGCAGTGGCggttacctcttatctag
gtagagtgtagcacctatccCCCCAGGAGATTGTGGCTACAATGAGATATGATGGCACCAggttacctcttatcta
gtagagtgtagcacctatccCAACAAAGTAAGACCCTGTCTGGCCATCAGAGAAATGCAAggttacctcttatcta
gtagagtgtagcacctatccGGATCTAGAACTGGAAATAGCATTTGACTCAGCCATCCCAggttacctcttatctag
gtagagtgtagcacctatccTAGTGGGTGCAGCACACCAGCATGCACATGTATACATATGggttacctcttatctag
gtagagtgtagcacctatccGGACTAACCAATGACAAAAATACTACAAATCAGAACTGGTggttacctcttatctag
gtagagtgtagcacctatccAGCAGTACTTTGAAATTACATGTAGAATCGCTGGCTCTCTggttacctcttatctagt
gtagagtgtagcacctatccAAATCCCTGATGAATTTGTCTGAACACAGCTGATCAAAGAggttacctcttatctagt
gtagagtgtagcacctatccTTGTAGCCCTGAGGTAAATTTAATTGTACTTTCTTGATGCggttacctcttatctagtg
gtagagtgtagcacctatccGACGTTTCTAGGGGCAACATCTAAGGACTATTAATAATATggttacctcttatctagt
gtagagtgtagcacctatccCCTCAGACCACCAGCAGGGGTTGGGGAAGCCACTCTCTCCggttacctcttatct
gtagagtgtagcacctatccACCTTGATGTTGGTCATTATGCCCACTGAGTAAACTGAAGggttacctcttatctagt
gtagagtgtagcacctatccATACATCAGGGAAATGTTGGCCTATAGAGAGAGCTGAAGGggttacctcttatctag
gtagagtgtagcacctatccGGCAGGCCACACATATAAATGGGAGAGACAGAGAGGAAGAggttacctcttatct
gtagagtgtagcacctatccGATCCACTCTCCACATCCTTTCCAGGTGATGATTTCTTTCggttacctcttatctagt
gtagagtgtagcacctatccGGCTGGAGTGCAAAGGCACAATCTCATCTCACTGCAACGTggttacctcttatcta
gtagagtgtagcacctatccATTACAAGCAAGAGCCACCGTGCCCGGCTTTTTTTTTTCggttacctcttatctagt
gtagagtgtagcacctatccATATTTCCTGGGCTGGTCTTCAACTTACAAAGGGCGAGTGggttacctcttatctag
gtagagtgtagcacctatccTGGAATGCTGGAGTTAGACCTGAGCCCTCTTGTCTCTTCCggttacctcttatctag
gtagagtgtagcacctatccGCTGGAGAACAAGCACTGTTTCTGAGCTGCCCTGCCCTGAggttacctcttatcta
gtagagtgtagcacctatccTGATTTAAAGTCTAGGAATTGATCTGACGTCTAGGAATTGggttacctcttatctagtg
gtagagtgtagcacctatccAGGGAGAACAACCAAAGATGCTGCTCAGTCCATTTTTATTggttacctcttatctagt
gtagagtgtagcacctatccAGGTACAAGCAAATTAGCCCTCCACATGGATCAGGCCAGAggttacctcttatcta
gtagagtgtagcacctatccTGGGACAAAGGCTTCAGGCTCACAGAGAAAGTTTCCATTTggttacctcttatctag
gtagagtgtagcacctatccCCCTGTACTTCTGCATCCTTGGCACACATCCATGGCCCCGggttacctcttatcta
gtagagtgtagcacctatccCAGAAACATTTTTGCCATCTTTAAAGAGCCCTAGTGCTGGggttacctcttatctagt
gtagagtgtagcacctatccGTGATTCATGAGAAGGACCCAGCATTTAGTAGCCCCGTGGggttacctcttatcta
gtagagtgtagcacctatccACTACAGTGCAAGGATACATGACACGATAAGCAACAGAAAggttacctcttatcta
gtagagtgtagcacctatccTCTGGAAAAATCCAAGCACAGGCTTTTAATGCTCTCTCCCggttacctcttatctag
gtagagtgtagcacctatccGAGACTCATGAAGCAGGCTCCTTTTGTCAGCAGCGAAAAAggttacctcttatctag
gtagagtgtagcacctatccTCTGCCCTAGAAAGAGACTCAGCATTCAAAGATTTTACATggttacctcttatctagt
gtagagtgtagcacctatccAGGCACCCTCTCCCTAGAAATAATCAGAATTCCAAACTCAggttacctcttatctag
gtagagtgtagcacctatccGCATAAACCAAGTTGCTTGCGCAGTCTAGATACAATGAATggttacctcttatctagt
gtagagtgtagcacctatccAGCTAGAGAGTGGGTTAAAGTGCTATGTTCCTAGACACCAggttacctcttatctag
gtagagtgtagcacctatccCCTCAGGTCTGCTATGTTATTTCTTTTCTGCGTGGCTTTTggttacctcttatctagtg
gtagagtgtagcacctatccGCTTTGCAGGGTAGGAAAATGTTATATAAAAGTTGGACCTggttacctcttatctagt
gtagagtgtagcacctatccCTCTGTTGCACTTGAGCCCTGTAAGCTGTGTGTTCTATCAggttacctcttatctagt
gtagagtgtagcacctatccTAACTAAAGCTGATATCCCCAGTAGGCTTAAAACTGTAGGggttacctcttatctagt
gtagagtgtagcacctatccATGTGTCTTGGGGTTGATCTACTCGTGGAGTGTCTTAGTGggttacctcttatctagt

gtagagtgtagcacctatccTGGTTCCGTTCTCCCATCTTCTTCGGGTACTCCAGTCAAggttacctcttatctagt
gtagagtgtagcacctatccTTCAGCAAGATGGTCTTCCAACTCTGATACCCTTTCTTCTggttacctcttatctagtg
gtagagtgtagcacctatccATTGATACTTGTGTATGTGTTACAAAGTTCTCATGCTGTGggttacctcttatctagtgg
gtagagtgtagcacctatccCTCCTGCAACCTTTTATCAAAGTTCTTATCTTCTTTGCATggttacctcttatctagtgg
gtagagtgtagcacctatccATTCTGGCCTTTCAGGTTTTCAGTGTTTTTTCGTTGATTCggttacctcttatctagtgg
gtagagtgtagcacctatccTCAGGCCCTATTCATCTAATTTGCTCCCACTCCTGAAGATggttacctcttatctagt
gtagagtgtagcacctatccGAGAACAGCAAGATGGGTGACTCCTCCTTCCTCTGGGATCggttacctcttatctag
gtagagtgtagcacctatccGGTGGCACAGAGAACAGGACCCGTTTAATATAGCACTTTGggttacctcttatctag
gtagagtgtagcacctatccGCTGGGGGAAATGCACTCGTCTGGGCTGTCTGGGTTCCTggttacctcttatctag
gtagagtgtagcacctatccGGAGGAAAGACTAAGTCAGCTAGTCCGTGGAGACTATGGCggttacctcttatctag
gtagagtgtagcacctatccGCTCAGACCCAGGGAGATCAGGGTTCAGTCCCTGAGCCCCggttacctcttatcta
gtagagtgtagcacctatccGTCAGGCCTGAAGAGGCACTATGGCTGCATCTGCCACAGCggttacctcttatcta
gtagagtgtagcacctatccCCTCTTGGGACCAACCCGTCCAACCTCCCTGGCTTCAGCGggttacctcttatcta
gtagagtgtagcacctatccCTTAGACAGCAGGCAGTGACAGCTGTGGTGCTCGCCACTCggttacctcttatcta
gtagagtgtagcacctatccTGAGTGGCTGTTGAGAATCTCCATGGCTCCGTGGCTGAGAggttacctcttatctag
gtagagtgtagcacctatccGAAAAAGCACGATTTCCCAGGCTAGGTAGCACGCTCACTCggttacctcttatctag
gtagagtgtagcacctatccCACCACACTGCTCTTCCTTGCTCTCCATGGATCATGCCAGggttacctcttatctag
gtagagtgtagcacctatccATTCACACACTGTTATGGTTTTTGCTGGGAGCCTCCGATCggttacctcttatctagt
gtagagtgtagcacctatccTCATGGCAGAAGATGAAGACCAAAGAGATGTTTTACATGGggttacctcttatctagt
gtagagtgtagcacctatccTCCATGTCTTTGCATTGTGAGTAGTATTGCAATGAACGTAggttacctcttatctagtg
gtagagtgtagcacctatccGATTGCTGAGTCAAATGGTATATTTGTCTTTAGGTCTTTGggttacctcttatctagtgg
gtagagtgtagcacctatccTTTCTCTAATGAAGATCTGTGATTATTTTAAGCCAGCTGCggttacctcttatctagtgg
gtagagtgtagcacctatccTTCGCTTGGAAGGTTTTATCTAAAAAACATCACACTCAGAggttacctcttatctagtg
gtagagtgtagcacctatccACCGTTAAGTACTGTGATAGCTTGTGTAAAGGAAACTCTTggttacctcttatctagtg
gtagagtgtagcacctatccCTGGGTAAATAATGGGCTTTCATTCAATTTAACACGTGTGggttacctcttatctagtg
gtagagtgtagcacctatccGGCTTACTATCTGCCAGGGGATATTCTCAGTGCAGGTAATggttacctcttatctagt
gtagagtgtagcacctatccGGCAAAACCTCCTGTTCTCATGAGCTTACGATCTATTATGggttacctcttatctagt
gtagagtgtagcacctatccAAAATAATTGAATAAGATGAGGAAAGTGCTCTGTGCCTGTggttacctcttatctagt
gtagagtgtagcacctatccCTGGTAGGAATGTAAGCTTCTGGTGCTGAGTTAGGTTCAGggttacctcttatctagt
gtagagtgtagcacctatccAGAAGAGTCCAAAGAATGTGAGCCTCCCTTTAATGGAATTggttacctcttatctagt
gtagagtgtagcacctatccGTTCTTTCTGGGCCATGATATAACCTATGACTGCAAAAGAggttacctcttatctagt
gtagagtgtagcacctatccAGGTAGCTTCCTTGTCGATTCTAGCAAGACTTCCCTGCACggttacctcttatctag
gtagagtgtagcacctatccAGCTATAGGGAATCAAGGTAGGGGGCAAAGGGGCAAAGTCggttacctcttatcta
gtagagtgtagcacctatccCATGTGTGTGCAATCTCTTCTCCCAACAAAAGATCTACCTggttacctcttatctagt
gtagagtgtagcacctatccTACTCATCCTCTGTTCTCTGACCATAAAAAAAAATAGCACggttacctcttatctagt
gtagagtgtagcacctatccGTTCACAGAAAGTAATAATTCAAGACATTCAAAGCAGGTGggttacctcttatctagt
gtagagtgtagcacctatccGCCCAGTAGTTTGCAGCTGTGTGGCACATCAAAAGAATTTggttacctcttatctagt
gtagagtgtagcacctatccTGGTAACTTTCTTAAAGCTTCATGGCTCAGCTGGGAATGGggttacctcttatctagt
gtagagtgtagcacctatccCTTAGGCAATATAGGGAGAAGCTGTTTCTACAAAAAAGTAggttacctcttatctagt
gtagagtgtagcacctatccTGGTGGCGTACACCTGTAACCCTAGCTTACTTGGGAGGCTggttacctcttatctag
gtagagtgtagcacctatccCCTGTCTCAAAAAAAGGAGGTGGGGGGGGGTCATAGTTTTggttacctcttatctag
gtagagtgtagcacctatccACTTTGGGAGGCAGAGGCGGGTGGCTCATCTGAGGTCGGGggttacctcttatcta
gtagagtgtagcacctatccTCTACTAAAAATCCAAAATTAGCTGGGCTTGGTGGCACATggttacctcttatctagt
gtagagtgtagcacctatccCCTTCCAAGAAATAACTAAAATTAGTTTAGTCTTCCAGCGggttacctcttatctagt
gtagagtgtagcacctatccTATCCTTTTTGTGCAGTTATTACAAACAAGTCAACCAACCggttacctcttatctagtg
gtagagtgtagcacctatccCCTGGAGGTTCAATAAAAACTTTAAAACAGCCCAATGCCCggttacctcttatctag
gtagagtgtagcacctatccGCATGGCATGATAGTGTTGCTTCTGACTCTAAAAAATAAAggttacctcttatctagt
gtagagtgtagcacctatccCTTATGAGTCAAGATCACAATTAAAGTACCTACCATCAAGggttacctcttatctagt
gtagagtgtagcacctatccCTGGTAGGAGCTAAACTATTTTTACCTGAACAATTTCTTCggttacctcttatctagtg
gtagagtgtagcacctatccGCACTATATCTCAAGCCTGCTGTAAAACATGTCCTACCATggttacctcttatctagt
gtagagtgtagcacctatccTAATCCACCAGAAATATCATGCAAAGAGCCAGGTGCAGTGggttacctcttatctag
gtagagtgtagcacctatccCAGGAAGCTGGCTTAAACCCAAAAGTTCAAGACTGTAGTGggttacctcttatctag

gtagagtgtagcacctatccGCAACATAGTGAGACTTTATGCCTAAAAAAAAAACCATGCggttacctcttatctagt
gtagagtgtagcacctatccCACATGTCAGTGCCATTTTGTTGGTAACATCCCCATAGGAggttacctcttatctagt
gtagagtgtagcacctatccCATCCATGTTTACAGCTAGGGCCCTGTACAGCAGTGTCTCggttacctcttatctag
gtagagtgtagcacctatccCATGAAGAGAAGTTATAACTGAGTGGGGAAAAAAATGTTCggttacctcttatctagt
gtagagtgtagcacctatccGGCAACATGACCGTGAAAAAGGTTAAAATTACTGGCTTACggttacctcttatctagt
gtagagtgtagcacctatccTGCCAAGTTCAGAAAAAAAGCTCACAGATATTATGCACCCggttacctcttatctag
gtagagtgtagcacctatccTCGGTCCCAACATGAGGCTTCACTTAAATACCCAGTTAGTggttacctcttatctag
gtagagtgtagcacctatccTGTACTAGGATCAATATGAAATAATTAAAGCAAGCGCCACggttacctcttatctagt
gtagagtgtagcacctatccTCTTACAGCTGAATGATATGATAACAGAATTGCCACGTTGggttacctcttatctagt
gtagagtgtagcacctatccTTCCATATGATTCAAGCCATCTAGTCTTATCACCTTAAACggttacctcttatctagtg
gtagagtgtagcacctatccCCCCTTATCACCTCCACCTCCACCCTAATGCCAATCACCAggttacctcttatcta
gtagagtgtagcacctatccGCAGCCCCGTAATGAATGATAATTTGTCTAGCACCCTGGGggttacctcttatctag
gtagagtgtagcacctatccGTCATGATCACGCCACTGCAGTGCACTGCACTGCCAGCCTggttacctcttatcta
gtagagtgtagcacctatccGTCAAGCTATTAGAAAAGTGGTCACGTCTATATTTATCTGggttacctcttatctagtg
gtagagtgtagcacctatccAGGCAGCTTTCTATCCTTTTCTTTCCCTCTGTCCACATTAggttacctcttatctagtg
gtagagtgtagcacctatccCTGCCAAGCCAGGGGAAATGCAGCAGCAGTAGCGATGGGCggttacctcttatct
gtagagtgtagcacctatccCCACAGGGGAAGGCTTGAAGTGTCCTTCACAGAACTGCCGggttacctcttatcta
gtagagtgtagcacctatccCTCTGAGCTACTCCAAAGCCACCAGCAAAGACGACGACCCggttacctcttatct
gtagagtgtagcacctatccTATTTAAGGCAGCTGACAAGAAGAATGGATGAACAAATCTggttacctcttatctagt
gtagagtgtagcacctatccAAAGAATGAGGGTATAGAAGAGCAACACATAACACACGTGggttacctcttatctag
gtagagtgtagcacctatccCTTTTTTTAAAAAGTCAGTACCGCAGCTGGGCGTGGTAGCggttacctcttatctag
gtagagtgtagcacctatccCCAAGGCAGGCAGATCAAGAGGACAGAAGTTTGAGACCAGggttacctcttatcta
gtagagtgtagcacctatccAAAAAGACTCTGTCTCAGTATCACAACCTATAATCGTCAAggttacctcttatctagt
gtagagtgtagcacctatccCATAGCTTCAATGTAGCCTTATAGCAAGTGGACTCAATAGggttacctcttatctagt
gtagagtgtagcacctatccTGAAATTCTGAAGATTCAAGCTGGGACACCTATCATGATGggttacctcttatctagt
gtagagtgtagcacctatccTACTACCTTAGATGTGAAAACACTAATATTAGCACCAGACggttacctcttatctagt
gtagagtgtagcacctatccGGGTTAACCGTATTAGCAGTAGAAGAAAAGAGTGAATTCTggttacctcttatctagt
gtagagtgtagcacctatccCACCCACAACTCAAATAAAATGAATCTCCCTAGCCCTTTCggttacctcttatctag
gtagagtgtagcacctatccTCTCCAAAGACAGTTATCCAATATCCATGCCTCCTGGTATggttacctcttatctagt
gtagagtgtagcacctatccGGGTCCCCTCTCCTTGAATCGAATCTCACTTTTGATCCATggttacctcttatctagt
gtagagtgtagcacctatccAAGTGATGCAGAGAGACTTCTTGGCTACATTGTAAGAGGAggttacctcttatctag
gtagagtgtagcacctatccTGCTAAAATTCTTAATTGGGAGGACATTAGGTTGAGAGAGggttacctcttatctagt
gtagagtgtagcacctatccCGGTCACGGTCATCCCAGGAAAACAACACTTAAGCTTAACggttacctcttatcta
gtagagtgtagcacctatccAATCAGAAATTGCCAATGAACCTCAAACTAGTGACTTTTCggttacctcttatctagt
gtagagtgtagcacctatccGCCCAATTCATGAGCTGTCTATTCAAATAAACTGGTTAAAggttacctcttatctagt
gtagagtgtagcacctatccTTGTCTTTTCACACTTAGAATCCTTGTTCTGGGGAATGCCggttacctcttatctagtg
gtagagtgtagcacctatccCCTAACCACCCTGAGGCCTACGAGTTATGCGGAAGCCCAGggttacctcttatcta
gtagagtgtagcacctatccCCTTCAGCTCTTCAGCCATCCTAGCCCAGGCACCAGACATggttacctcttatcta
gtagagtgtagcacctatccACTGTTGTGGATATCCAGCCAAGCCAAGCCTTTAGATAGCggttacctcttatctag
gtagagtgtagcacctatccCCCCAGCCTCTACCTCAATGCAACATTATTAGACACCCCAggttacctcttatcta
gtagagtgtagcacctatccGATCCAGACCCACCAAAAGGTTTCCTCCCCCTCTCTCTAAggttacctcttatcta
gtagagtgtagcacctatccGTCACCATACCTGATCAGACTCTTTCACAAGATAACGTGCggttacctcttatctag
gtagagtgtagcacctatccGTTCCCTGATCCATTCATTCATTGTCCCTAGAAATCCCCTggttacctcttatctagt
gtagagtgtagcacctatccCTGTTTCGCCAGGATGATGTATAAGCTTTTGAACCCCCTTggttacctcttatctagt
gtagagtgtagcacctatccACTCTGTGATTCGCCTGGTGTACACCTTAATTAAATTTGTggttacctcttatctagtg
gtagagtgtagcacctatccAAGCTTTCCTTTGACCTCTACGTTACTCAACCCACAGAACggttacctcttatctag
gtagagtgtagcacctatccGATTAAGCCATGAAATGTAAGCAATAAATGTCTGGAACACggttacctcttatctagt
gtagagtgtagcacctatccCAACAAGCAACACACACTCCTGATCCAATACAATGATTCCggttacctcttatctag
gtagagtgtagcacctatccGGACATCTCCAAAGTGTCTCCAAAAGCTTTAGAAACATTGggttacctcttatctagt
gtagagtgtagcacctatccCCAGGCTGGAGTGCGGTTGTGCAATCTTGGCTCATTGCAAggttacctcttatctag
gtagagtgtagcacctatccGTGCCACCACACCCAGGTAATTGTTTGTATTTTTAGTAGAggttacctcttatctagt
gtagagtgtagcacctatccCACCATGCCCAGCCTAGTAAATCAATCTTTTATTTCAATTggttacctcttatctagtg

gtagagtgtagcacctatccTTTGAAAAGGGAGCGTTATCAATCTAAAACTGATTTCTGCggttacctcttatctagtg
gtagagtgtagcacctatccGCAAAACCTCTACTGTAATTTTCAAAGTCTGAAGGCCTCAggttacctcttatctagtg
gtagagtgtagcacctatccAACACCCCACCTGTTGCCTAGAAGCCGAATCAGAATCTGCggttacctcttatctag
gtagagtgtagcacctatccTCCAGGTGAGTCATGCGTCCATTGAAGTTGAAGAAGTACTggttacctcttatctagt
gtagagtgtagcacctatccGAATCTTTTTAAAATTCCGAAATCCAGGCCATACCCTAGAggttacctcttatctagtg
gtagagtgtagcacctatccAAATCATGGGACACAGGCGTTGATGGTTTTGAAGGCTACCggttacctcttatctag
gtagagtgtagcacctatccCTAGATAGGGCACTGGGAAATCACTTGGGGAGCATAAACAggttacctcttatctag
gtagagtgtagcacctatccATTGGTTTCCACCCCCGGAGAGTCTCATGTCGCTGGTCTCggttacctcttatctag
gtagagtgtagcacctatccCCAGGTAATTCTCATGTACAGCACAGTTTGTACAGCCACAggttacctcttatctagt
gtagagtgtagcacctatccCAGCTGAGCCTGCACCACTCCTTGGCGAAGAAGGGTTTAAggttacctcttatctag
gtagagtgtagcacctatccACCTTAATTAAAATGAACAGTTGGCCTCACAGTTTTCAGCggttacctcttatctagtg
gtagagtgtagcacctatccACCAGGTAGAATGACAGCATCAAGCTTGCTCGAGTTCAGGggttacctcttatctag
gtagagtgtagcacctatccGCACATTTCTCTTCTTTCTGTTGGGCTCCCTTTCAAGTGAggttacctcttatctagtg
gtagagtgtagcacctatccGGCGCAAATACCTTCACCTTCAAAAATAGAAAGAAAAACAggttacctcttatctagt
gtagagtgtagcacctatccTGTCATTACCTCTATGAAAACAATCAAGCAACACTATCTCggttacctcttatctagtg
gtagagtgtagcacctatccCTAAACCAGGAAGCCAAGACCAAAAGCCGCGAAGTGATTCggttacctcttatcta
gtagagtgtagcacctatccCTAGAAATATTATAGCTCAACCACAAAGAAGAAAGTGCAGggttacctcttatctagt
gtagagtgtagcacctatccTCCAAAAGGAAGGGAGATCAAGTAGATTACAGAGAGGTGGggttacctcttatctag
gtagagtgtagcacctatccAGAGAGGATGTAATTTTATAGATAGGAACATACAGGACTCggttacctcttatctagtg
gtagagtgtagcacctatccTTAAATCAGCCCTTGAGATCCTTCAAAGCAGTGCTTCTCAggttacctcttatctagtg
gtagagtgtagcacctatccCCAACGAGGCCAATTCAGTGGAAATCTCTGGGAGTGAAGAggttacctcttatctag
gtagagtgtagcacctatccGCTCCAGCTTAAAATCTCCCCAGGCGATTCTGTTTGTCAGggttacctcttatctagt
gtagagtgtagcacctatccACCACTAATCTAAAGCAAATGGGAGATAAAACTCAGGGCTggttacctcttatctag
gtagagtgtagcacctatccGCTCTCGCTTGATCCACCTCCATGCCTCCAGCCCCACCAAggttacctcttatcta
gtagagtgtagcacctatccGAGGACATCACTCTAGTTTCCCTGTCTCCAACCCTGGGATggttacctcttatctag
gtagagtgtagcacctatccCCTGATCTGGGGAAACATGAGGACGTGGAGGCAAGGGAGGggttacctcttatcta
gtagagtgtagcacctatccCCACACCTTCTTCACTGGCTTCTATTCCAACCTAAAACACggttacctcttatctagt
gtagagtgtagcacctatccCAGGAGGTACACGAGGCTGGTCTGAATCAAATGTCCCTTGggttacctcttatctag
gtagagtgtagcacctatccGAGTCTCAGCCTCTTGTTACTAAACTTTGTATTTCACTAAggttacctcttatctagtg
gtagagtgtagcacctatccTGAGAAAAGACTTTGAATATTCTGAAGGGCAAGAAAGATGggttacctcttatctagtg
gtagagtgtagcacctatccATGGTGGCTCATACCTGTAATAGCAGCAGTTTGGAAGGCCggttacctcttatctag
gtagagtgtagcacctatccGCCAGGAATGGAGGTGTGTGCTGCAGTCCTACCTACTCAGggttacctcttatctag
gtagagtgtagcacctatccACATGGATTTGATTTTAAAGAATTCCCCAAACAAGTTGGCggttacctcttatctagtg
gtagagtgtagcacctatccGTTGTTTCACGGAATAATCAGATGAACATGCTCAAGGAGTggttacctcttatctagtg
gtagagtgtagcacctatccTGATAAGAAAGCACTGAGGCATTTACTGGGTTGTAATCTAggttacctcttatctagtg
gtagagtgtagcacctatccCCATGTACCTCTCCCGGACTTCATCTCAAGCACAGAGACCggttacctcttatctag
gtagagtgtagcacctatccAAATACTCACAAACCCCAGTGTGTAGGACATCGCCCTTCAggttacctcttatctag
gtagagtgtagcacctatccTCAGTAACTACTTGTTGAAAGGATGAATGAATAACACCCAggttacctcttatctagtg
gtagagtgtagcacctatccTCTTAATACCCTTCAAGTGAAAGAGGTGCTAGATATTTCCggttacctcttatctagtg
gtagagtgtagcacctatccTTTAGAGAAAAGAGAATTGGCTTCTAGGACTGACTTTGTCggttacctcttatctagtg
gtagagtgtagcacctatccAAAGAAGTCATTTTTCCTCCCAGGGAAACTCAGCCAGGAGggttacctcttatctag
gtagagtgtagcacctatccTGTCCTCCAGCTTCAAACTTCAAATCCTGCAGACAAAGAAggttacctcttatctagt
gtagagtgtagcacctatccCTTATTAAAAAGAAAGGTGCAGCAGCTGTGGCGGTTCCTCggttacctcttatctag
gtagagtgtagcacctatccCAACAGATGAATGGTACACAAAATGTGGTGTTCTACATACggttacctcttatctagtg
gtagagtgtagcacctatccCCCTAAGGACATAGGCCAAGTGAAATAAGGCAGTAACACAggttacctcttatctag
gtagagtgtagcacctatccCTTGTAGTAAGTACCTAGAGTAGACAAAGTCTTAGAGATAggttacctcttatctagtg
gtagagtgtagcacctatccCTGGAGCTGGGAGAAAGGAGAATGAGGAATTAGTGTTTCAggttacctcttatctag
gtagagtgtagcacctatccCGGTTGCAGTTTGGGATGATGAATATGGTCTGGGGACGGAggttacctcttatctag
gtagagtgtagcacctatccCACTTAATGCCAGTCGATCACACACTACTTAAAAATGGTTggttacctcttatctagtg
gtagagtgtagcacctatccGGTAAAGTTCATGTTACATATGTTTTACTCCCTCCCCCTCggttacctcttatctagtg
gtagagtgtagcacctatccAAAACAAAACACTATTGGCAAAGGGGTAGCAGGTGAGGGGggttacctcttatctag
gtagagtgtagcacctatccTCCTTGTGACTGCGAACATCCCTGCTGCCCGTTTTTGTCggttacctcttatctagt

gtagagtgtagcacctatccGCGGAGTGTGCGGAGGGAGGCGTGTTGGATACCTTTTAAAggttacctcttatctag
gtagagtgtagcacctatccGCGGTGACTCACGACTGTAAGTCGCGGCCCAACCCTTTGGggttacctcttatcta
gtagagtgtagcacctatccAAGCCTGGCCAAAATGGTGAACTCCTGTTTAAAAAAAAAAggttacctcttatctagt
gtagagtgtagcacctatccAACAAAAAAAAATTAGAAGCCCCTGGGGACGGAGAGCATTggttacctcttatctag
gtagagtgtagcacctatccATAAAAATAAAAATTAGAAGCCCGAGAGAGGGAAAACAGCggttacctcttatctag
gtagagtgtagcacctatccGCCCCCGCCACTCCGCAGGCTGCAGAGGGAAGAGAAAGCGggttacctcttatc
gtagagtgtagcacctatccGCAGCCCAGACCTAGCCCCGCAGCCCCACATACCCCAGCAggttacctcttatc
gtagagtgtagcacctatccCCGCTCGCTCCCACCTCCGGCTGCACCCGCGCGCCCCCTTggttacctcttatc
gtagagtgtagcacctatccGAGAGATCTGTCCGCTGAACCTGCGCATCCCGGCCCCTGCggttacctcttatcta
gtagagtgtagcacctatccTGTTCGGAGGGAAGAGAGAGGGCGCCAGGGAGACATAAGCggttacctcttatcta
gtagagtgtagcacctatccGATTCTATCCTTTGCCGCACTACTAAAAAGTAACTTCCATggttacctcttatctagtg
gtagagtgtagcacctatccTGTGTGCAATAATTTTTTTAACTGCTCAGTTCCAAAGCTGggttacctcttatctagtg
gtagagtgtagcacctatccCCATTGCATCTTCATCGGAGCTGGAGGACAAAAAAAAAATggttacctcttatctagt
gtagagtgtagcacctatccGGACACTTAACATGGAGAGTTTGAGATCCGGATACAGCGTggttacctcttatctag
gtagagtgtagcacctatccGCCGCTGCTCCTCATGACATTTTGTCTCACAAACTTAGCAggttacctcttatctagt
gtagagtgtagcacctatccACGCCATATGTGTTGCCTAATTTTCTTATATGATCACAGGggttacctcttatctagtg
gtagagtgtagcacctatccAAGTTACTTGTTAATAAAATTTCAGGTAGGATGCTCCCTGggttacctcttatctagtg
gtagagtgtagcacctatccAGACACCTTAGTTCATTATTCTAATAAGCTGTTGATCTGGggttacctcttatctagtg
gtagagtgtagcacctatccTGGAGAAGATAATTTGAAGGACCACAGGAAGTTATTTGCTggttacctcttatctagtg
gtagagtgtagcacctatccATGATGCCATATTTACCAAGCGTTCGAGCAATCAAAGCGTggttacctcttatctagt
gtagagtgtagcacctatccCTTCAGATTTGGGTACCCCCTGCAATATATGGTTCTACAAggttacctcttatctagtg
gtagagtgtagcacctatccAAACCTTGTTAAGCTTCACAAAGTTTCCACTGAAGATTTGggttacctcttatctagtg
gtagagtgtagcacctatccGCAGGTACATAGAAGTTGCCGGCTTTGCTTGCCATCCTTGggttacctcttatctag
gtagagtgtagcacctatccATCTTCAGCTCTGTGAAATTTCTTCGCTTTTTCTTAAGGGggttacctcttatctagtg
gtagagtgtagcacctatccAAACTTTTTCTTCTTCTCTTTGACACTCTCCATGGTTCCAggttacctcttatctagtg
gtagagtgtagcacctatccGATGAAATCTGCAATATCCAGCAAACATTACTTTCCCTACggttacctcttatctagtg
gtagagtgtagcacctatccTTATTAGTCCCATGTTCCTCTGATAAAAATTCACTAAGCCggttacctcttatctagtg
gtagagtgtagcacctatccGCTGAATAAACCCTGCATTTACTTTCAAGAAGAAACCTACggttacctcttatctagtg
gtagagtgtagcacctatccCCTGGAGGTAGATTGCTGCTCATAAGTGACAGATGGAGTAggttacctcttatctag
gtagagtgtagcacctatccGTAGGATATTTACTGTCAACGATGCCTAGATTTAGAAAGAggttacctcttatctagtg
gtagagtgtagcacctatccTGGAAAACACACATGCATCCGTCATCACTGCAGATGAGGAggttacctcttatctag
gtagagtgtagcacctatccCAAGGAGTGAATTTGAAAAGCAGGATAACTTAGTGTAGGTggttacctcttatctagtg
gtagagtgtagcacctatccTGCCCAAGATCAAACCTGACTCCACCCTCACTGCCTTATGggttacctcttatctag
gtagagtgtagcacctatccGAAATTACTCAAAATATCTGTACCAACGTTTCCTCAGGCTggttacctcttatctagtg
gtagagtgtagcacctatccATGTCATCTCTGTCAAAACCTTGATACTAGTGTATTTCTGggttacctcttatctagtg
gtagagtgtagcacctatccGGTTTGTACCCCTTTAGTTTCCTTGATATACGTTGATTAAggttacctcttatctagtg
gtagagtgtagcacctatccACTAAAACTACAAAAATTATGTGGGCCTGGTGGCACATGCggttacctcttatctagt
gtagagtgtagcacctatccATTGGCTTTTATTAGTGACTCATAAAACAGGCAGTTGCTAggttacctcttatctagtg
gtagagtgtagcacctatccCCAGTTGGTTTTCCTAAGGTAGCTTGAACAGAAATAGGGAggttacctcttatctagt
gtagagtgtagcacctatccACTTCAGGTTACTTTACTTATAAAGGGTTAAAGCAGAGAGggttacctcttatctagtg
gtagagtgtagcacctatccCTGTGAATCTCTTGTTTTTACGAAAAGCCAGTCTATTTTGggttacctcttatctagtg
gtagagtgtagcacctatccGTTTACGTTTCAGTTTGATTACAGTTCGATTACTTAGCACggttacctcttatctagtg
gtagagtgtagcacctatccGGGGCTCAGTTCAAAACCATGGCCTCTCATACATTTTATTggttacctcttatctagtg
gtagagtgtagcacctatccCCAATTTTCTTGCGGCTATTGATAAATCGATCATAACAGAggttacctcttatctagtg
gtagagtgtagcacctatccGTGTTCCTTATGGCTCCACAGTTTGTTTTTAAGTATTACCggttacctcttatctagtg
gtagagtgtagcacctatccAGTAGATGTCTCACTCTAATCTCTGCCTTCAGTGAATATTggttacctcttatctagtg
gtagagtgtagcacctatccCTCTGACTCTACCAAAAACATTTTATTTAATTAGCCAGGCggttacctcttatctagtg
gtagagtgtagcacctatccATAAATGATGCTTTAGGCAAGTGAAATTGTCGAAACTCTGggttacctcttatctagtg
gtagagtgtagcacctatccAGGATGTACGTATAGTTGATGTATTTGTTTAGCCCAGTTTggttacctcttatctagtg
gtagagtgtagcacctatccGTTTCAGAAACACTTTGGGGGTTCATTAATTTTTGGTTACggttacctcttatctagtg
gtagagtgtagcacctatccTTCTTAACCTCTTACATGAATATTTGTGTCAGGGCCAGGCggttacctcttatctagtg
gtagagtgtagcacctatccATTCGAGACCAGCTTGGGAACGTAGTGAGACTCCTTATCAggttacctcttatctag

gtagagtgtagcacctatccAATTATTTGGGCATGTTGGTACATTCCTGTAGTCCAGCTAggttacctcttatctagtg
gtagagtgtagcacctatccCACTTGAGCTAGGAGGCTATGGCTGCAGTGACTGTGATTGggttacctcttatctag
gtagagtgtagcacctatccAAAGAAAAAAAGGAGTAGGAAATTAGTGTGTCTAGGGCCGggttacctcttatctag
gtagagtgtagcacctatccATGAGAAACTAAAAATTAGCTTGGACTTTAATCTGCCCTCggttacctcttatctagtg
gtagagtgtagcacctatccCTGATAAAACCAGCCTCAAAATTACCAGAAAACAGTACTTggttacctcttatctagt
gtagagtgtagcacctatccACCACAGTAAATCATAAAGTGTCATAACAACCTCCTTTCTggttacctcttatctagtg
gtagagtgtagcacctatccCTTTATGGAAGTTGTTGTGCGTGTCTCTGTTGTCTTTGTGggttacctcttatctagtgg
gtagagtgtagcacctatccGCATAGGCTGATCTGAAGAAGACACTTCAATCATGGGTGAggttacctcttatctag
gtagagtgtagcacctatccCCTATGAAATTAGAGCATTGGGGAACAAAAGTAGGCAGAAggttacctcttatctag
gtagagtgtagcacctatccTGTGTGAGGTTCAGGTGGAAGGGTATAATTACAGTGGCATggttacctcttatctagt
gtagagtgtagcacctatccACCAATAAAAAAGATGCACAGAGCAATGCTGCCAGAGACTggttacctcttatctag
gtagagtgtagcacctatccCATCTCTGCCCCCACTTACTGATATTCCTGACACTACAGCggttacctcttatctag
gtagagtgtagcacctatccTTACCAACAACCATGGGAGGTCCTCTTCCGCCACATCTGGggttacctcttatctag
gtagagtgtagcacctatccGCTATGGTGTTCCTGGGACCAAGGAGCCAACTTGAAGGATggttacctcttatctag
gtagagtgtagcacctatccACAAGAAGTGCAAGCGACTCTAGAGTCAGAAGAAGTGGATggttacctcttatctag
gtagagtgtagcacctatccCTGATCACAACAGGAGCTTTACTGCAGAAATAACCATTTAggttacctcttatctagtg
gtagagtgtagcacctatccCAGTCCTGTGCCCTGTCACTAGTCAGACAACTCTACCATCggttacctcttatctag
gtagagtgtagcacctatccAGAAGGAAGGAGAGAGAGTGGAACCTTACAAAGTAAACCTggttacctcttatctag
gtagagtgtagcacctatccCATCATTTGAGAGCTAAATTTTGAGATTATGCCCCCGCCTggttacctcttatctagtg
gtagagtgtagcacctatccAACATTGGCAAATTGGCTCACTTTGAACCATCTCAGTGACggttacctcttatctagt
gtagagtgtagcacctatccAGTTCCTCGGTCACCTCCACAATCCAGATGGAATCCTTGGggttacctcttatctag
gtagagtgtagcacctatccGCGACATTGCTGAGGGACCTCTGGCTTTTGCTTCTCCAGAggttacctcttatctag
gtagagtgtagcacctatccTTGGACCTCAAGAATGAATTGGTGTACCAGTTGGAACAGGggttacctcttatctagt
gtagagtgtagcacctatccGAGATTCTGGAAGCAATCAGTCAAAATTCAGTTGTCATTAggttacctcttatctagtg
gtagagtgtagcacctatccATGACTTTATCCAGAATGACTGACAGCAGAGTGTAACATTggttacctcttatctagtg
gtagagtgtagcacctatccCCCAGAAGAATCAGTGTAGTTTCCATGGCAGAGTGAGTTGggttacctcttatctag
gtagagtgtagcacctatccAGTCTGTACTTCCTTGTCCTCATGCCAGTATAATGCTTTGggttacctcttatctagtg
gtagagtgtagcacctatccGTAGATGAAATACATGGAAGAGACATTAATACTGACTTCCggttacctcttatctagtg
gtagagtgtagcacctatccAAGTTCTCATTGTTCTTATGTCCACTTCTATTGATACCAGggttacctcttatctagtgg
gtagagtgtagcacctatccTCAATTGCCCCATCATTTAAGTGTATGGGAGGACTTACCCggttacctcttatctagt
gtagagtgtagcacctatccTTCACCGAAAACCAAGAAGAAGGACAAGGATGATGATGGTggttacctcttatctag
gtagagtgtagcacctatccCAAGGATGAGCATGTCTCAATTGAATGAAAAAGAAACTCCggttacctcttatctagt
gtagagtgtagcacctatccAGTACATTGAAACCCTTAATATTCCCAGAGCTGTGTTGGTggttacctcttatctagtg
gtagagtgtagcacctatccCACATTTTGGAAGCCATCAGTATCAGATTCTACCTCTGCAggttacctcttatctagtg
gtagagtgtagcacctatccCAAGCATTATCATAAATGATGTTTATGTCGTTGACTCCTGggttacctcttatctagtg
gtagagtgtagcacctatccAATATGACCAACTATGCTACAGTATGGGGATCAAAAACAAggttacctcttatctagt
gtagagtgtagcacctatccCATGACACCAGAGATGTTCCAAACACCATTGCATGAAGTTggttacctcttatctagt
gtagagtgtagcacctatccTAGCATAAAACTTCCATGTCTAGTTGGAATTGGCCAGTTTggttacctcttatctagtg
gtagagtgtagcacctatccCAATGATGATTTGACTCCATTGGGACGAATCCTGGCTAACggttacctcttatctagt
gtagagtgtagcacctatccTGGCAAAATGATGATAATGGTGTGTATTTTCTATGTGGGAggttacctcttatctagtg
gtagagtgtagcacctatccCTTTTATAAATGAAGGAAAGTGGCTGGGCTACAACCATTGggttacctcttatctagtg
gtagagtgtagcacctatccGAAACGGATTTTCTCATCATGTAGCCCTTTTATCAGTATTggttacctcttatctagtgg
gtagagtgtagcacctatccATGGGTGGAGAAGAAGCAGAAATACGTTTTTGTGCGCACAggttacctcttatctag
gtagagtgtagcacctatccAAGTTCAGCTCAAAGAGATTCTGATTAATTCTGGGTTTCCggttacctcttatctagtg
gtagagtgtagcacctatccAATTTGGATGTTGTTATCTCTCTTCCTGGCCTTTGGTGTGggttacctcttatctagtgg
gtagagtgtagcacctatccATGCTATCATAAGGAAAAGACAAAGATTCTCACCACTGAAggttacctcttatctagt
gtagagtgtagcacctatccGTACCCATCTCCCTTCTTTGCATTTGGTGAAAAGATTCGAggttacctcttatctagtg
gtagagtgtagcacctatccTGACTTTAGTCACCCCACTACAGTTGCTTCTCTTTGCCTCggttacctcttatctagtg
gtagagtgtagcacctatccACCGCTCTCCAGGCAGCCACTGAGGCTCTGGTTGTTGAAGggttacctcttatctag
gtagagtgtagcacctatccAGTTGGACCCCGTAAATGAAAGTATACTGAACATGATCCGggttacctcttatctagt
gtagagtgtagcacctatccTCTAGACCCTCAGCTGCTGATATCAACCTTATGATTGGCAggttacctcttatctagt
gtagagtgtagcacctatccGGTCCTCCTAAGATGGCCCAATATGACAATGGAAGCGGATggttacctcttatctag

gtagagtgtagcacctatccGGGAGGTTCTAGTTATGGCAGTGAAGGCTATGGCAGTGGCggttacctcttatctag
gtagagtgtagcacctatccTGTTGGTGGAGGCTATAGAGAAGTTTCCTGAGGTGGCTATggttacctcttatctagt
gtagagtgtagcacctatccTGGAGGAGATTACAGAGGGCCTAGTAGAGGCTACAGATGAggttacctcttatctag
gtagagtgtagcacctatccCTTTGGACAAGGAAGAGAAGGTGGCATCTGTTAAGGCTTGggttacctcttatctag
gtagagtgtagcacctatccGCCATGCTATGTGTGACATTTTTTCCAATGTTTATTTCCggttacctcttatctagtgg
gtagagtgtagcacctatccTGCATTTTAACCCATTCTGTGGTTCACTGTAGCTTAAGGAggttacctcttatctagtg
gtagagtgtagcacctatccATTTTTCTTAGAGACAGTATCTCATTATGTTACCCAGGCggttacctcttatctagtg
gtagagtgtagcacctatccACATGCCACCTTGCCTGGCTTCCTGCTTTCTTATTCACTAggttacctcttatctagtg
gtagagtgtagcacctatccTACATCAGTAACAATAAAATATCCCACCTTTGCTTGAAGGggttacctcttatctagtg
gtagagtgtagcacctatccTTTCCAGACACAAAACTCCATCGCTATTTTAACTTCATAGggttacctcttatctagtg
gtagagtgtagcacctatccAGGAAATAGGCCCTTCATCCACTTTGCAGTCTAGACCTAGggttacctcttatctag
gtagagtgtagcacctatccCCGTCTGGAGCAGCCACTACCATCATTGACATACAGCCTTggttacctcttatctag
gtagagtgtagcacctatccATACTGCTTATTTAAATTTCATCTCAACTCCACACTTCCCggttacctcttatctagtg
gtagagtgtagcacctatccTCCTATCACTCTATCTGGGGAGATGCCATGTGGTCCAGGCggttacctcttatctag
gtagagtgtagcacctatccGATTGAAATTAGTTCTGTCTGTGGCCCTCAAGATTTAAAggttacctcttatctagtg
gtagagtgtagcacctatccCCTTCAGCTTGGGGTGTGGGGTCATAGACAAAAAGAATGGggttacctcttatctag
gtagagtgtagcacctatccTCAGTGTTCCCATCCAAGGCAAAACCAGACCAGAAGAATGggttacctcttatctag
gtagagtgtagcacctatccCCAGGAGGCTGAGTTAAGACCAATACTAGGGCCAGGCATGggttacctcttatcta
gtagagtgtagcacctatccAAAAATTACAACTGGCACACACTTGTACCCAGCCACTAGGggttacctcttatctag
gtagagtgtagcacctatccTAAAAACCAATACTAGGAAGTGCTTTCTTACCATGAAAGCggttacctcttatctagtg
gtagagtgtagcacctatccCAGAATGAAGAGACAGATGGCAGAAGCAAGACCAGAGGCCggttacctcttatcta
gtagagtgtagcacctatccGGTCAACCTGAATTTGTTTTTGTAGCTCTGGGAAGTCCTGggttacctcttatctagtg
gtagagtgtagcacctatccATGGAACCTGGAACCATCTACTGTTAATTGTCCTTTTAGTggttacctcttatctagtg
gtagagtgtagcacctatccTTTGTATTTGGTGAAAAGATTTGAACTCGAGCCATCTCTGggttacctcttatctagtg
gtagagtgtagcacctatccACCGCTCTCCAGGCAGCCACGGAGGCTCTGGTTGTTGAGGggttacctcttatcta
gtagagtgtagcacctatccATACTGTGTATGGTTCCAGGATTTGTAAATGCTCTAATCAggttacctcttatctagtg
gtagagtgtagcacctatccACGGAATCCTTCAGGTGCCCTTCAAATCAGACATTACATTggttacctcttatctagt
gtagagtgtagcacctatccAGGGCTAGAATGCTTCCCCCTCTTATCTTGGGCAGGTCTTggttacctcttatctagt
gtagagtgtagcacctatccAATTTTGAATTTGGCCATAGTATACGGAGGAGAGCTAGGTggttacctcttatctagtg
gtagagtgtagcacctatccCGCTTAGACAAACGTATGGTAAAATGCAGCCCAGCCTGAAggttacctcttatctag
gtagagtgtagcacctatccCCTGGAGCAGCAAGTGGGCAAAGGAGTGTGTGTTTGGGGCggttacctcttatcta
gtagagtgtagcacctatccTGAGTCCCATATGTAATCAAGAGACCTGGAGGGCCCAAGGggttacctcttatctag
gtagagtgtagcacctatccGCTACATATCCTCCAATGGATTGTGGTGGTCATCTCCGGCggttacctcttatctag
gtagagtgtagcacctatccTTATTCCCATAGGCTTACAAACATTATGATGCTTCACACGggttacctcttatctagtg
gtagagtgtagcacctatccGTCTAAGGCCCTGCAAAACTGCTGTCTGGTAAAAACAAGTggttacctcttatctag
gtagagtgtagcacctatccGCAAGGCCCTGAATGTACTACGTATAGACTTAAGTCAGTAggttacctcttatctagt
gtagagtgtagcacctatccGTAATTGATGCCATTGTTTTCTTGACCTAGTTGAAGAGTCggttacctcttatctagtg
gtagagtgtagcacctatccTTCTTTCCAGTGAAGTGACAGATCCAAGCAAGGAACACCTggttacctcttatctag
gtagagtgtagcacctatccTGAGGGGAGCGGTAGGGGAGGGGGTAGATTCGGCAGCACTggttacctcttatcta
gtagagtgtagcacctatccGGCAGGGACCACATATTGTCGGTGTTAACCACTGTTATCAggttacctcttatctagt
gtagagtgtagcacctatccCCAGGTCGAGTGAGGACGAGTCATTTCCCCACTAAGCACTggttacctcttatctag
gtagagtgtagcacctatccGTGACGACGAGCCATTTCCGCACTATTGTCCCTTCTTCCTggttacctcttatctagt
gtagagtgtagcacctatccCTGCACCCTCAGCTCCAAATACACATTCTCGTTCACTCAAggttacctcttatctag
gtagagtgtagcacctatccCCCTGTGGGATGGGGTAAAAGTTCGGTGTTGGGAGGAGCCggttacctcttatctag
gtagagtgtagcacctatccGATTTTTGGTTTTGGTGGTGGCTTTTGGGCAAGGACAGGTggttacctcttatctagtg
gtagagtgtagcacctatccTGGATCGAATTCTCCTGGGAGCCAGACCAAAGCACCTTCCggttacctcttatctag
gtagagtgtagcacctatccCGCGGGGAGGTGTCATGCGCCGGAACCTGCGCTTGGGGCCggttacctcttatct
gtagagtgtagcacctatccGCGTCCCGGACTGGCGGCTCCGCGCATGCTCCTCCCACCGggttacctcttatct
gtagagtgtagcacctatccCGGAAGTACTGGGTGCAGCCTGATGGCGCAGGAGGTAGACggttacctcttatcta
gtagagtgtagcacctatccCTCGGGGACGCGCTTCCTGGTGCTGGGCCCTGGCGCTGCTggttacctcttatcta
gtagagtgtagcacctatccCGCGCGCACTGTCGGGGTTGGGGTCGGCGTTGACGATGGGggttacctcttatcta
gtagagtgtagcacctatccGACCCCTGCTCACCGTGGTTTGTGTCGGGTCACGAGATGGggttacctcttatctag

gtagagtgtagcacctatccCTCGGAGCGCACTTGAGAGCAAAGTTGAGGACTGGGGAGTggttacctcttatcta
gtagagtgtagcacctatccGACCTTTCGTCCCCTCTGGTCACTTGCGGCAGACTCAGGAggttacctcttatctag
gtagagtgtagcacctatccTCGTGGAAGAAGAAAAGGAGAGTAATGTTATTTAATGGAGggttacctcttatctagtg
gtagagtgtagcacctatccAGCTGATTTCTTGAGTAGTTTAAAAACACTGTCTATCGCAggttacctcttatctagtg
gtagagtgtagcacctatccTTATATCATGCACTGTCGTTAGTTATTACCGCGGAACATTggttacctcttatctagtg
gtagagtgtagcacctatccGTTTGAACCTGATCAGAACCTGATCCGAAGTCCTTATTGGggttacctcttatctagt
gtagagtgtagcacctatccCTGGTTAGACCGGGAAGATTGTCACCGCTTTTGTCCAGTCggttacctcttatctag
gtagagtgtagcacctatccGGACAGAGCCCGCACACCTGGGAGGTCGATTCCAGCCTTTggttacctcttatcta
gtagagtgtagcacctatccCTCCGGCGGTCAGAGATAGCTAAACACAGAGAAAGAAAACggttacctcttatctag
gtagagtgtagcacctatccGTCAACATCAGGCAGGAGAAAGACCAGCAGTTTTTGAAAAggttacctcttatctag
gtagagtgtagcacctatccCTGCACACAACAAACCTGAAAAAAACAGCTATTTTGTATGggttacctcttatctagtg
gtagagtgtagcacctatccGTGGGAAGGAGTTCTGCCTGCTGCGGGTCAAATCTGGCTCggttacctcttatctag
gtagagtgtagcacctatccAGAATCTCTGTGCCCACATGCCCTGAGTCAAAAAATAGGAggttacctcttatctag
gtagagtgtagcacctatccTGGAAGGATTAAATCTAGTAAAATGCCAGCTGTTGTCAAAggttacctcttatctagtg
gtagagtgtagcacctatccTTGTTTTGTGTGATATACTACGTGGAAAGAGAAGTTTGTGggttacctcttatctagtgg
gtagagtgtagcacctatccAAACGATTTGATTTAACATTAGGGCAAGGTGTTGACTATGggttacctcttatctagtg
gtagagtgtagcacctatccTTTAAGTCTCAGATATTAAGTACGTGCTCCCATCTAATTCggttacctcttatctagtg
gtagagtgtagcacctatccTTTGCTGTGTCAGTGAAATCTCCATGTGATTATGTATTTGggttacctcttatctagtgg
gtagagtgtagcacctatccGTGCCTATAAAACACTGTACCCAAGTAGCAGAAACTTTTGggttacctcttatctagt
gtagagtgtagcacctatccGTTGATTTGTTAAGATTTGTTAGATCTGGCTACAACCTTGggttacctcttatctagtgg
gtagagtgtagcacctatccACTTCTTTCTTTCCCAGTGCCAGATTATAGGTGACCAGTGggttacctcttatctagtg
gtagagtgtagcacctatccCTGCAGTGCCCCTCGAGTCCTATTGCTGTGGCCTTCTCCGggttacctcttatctag
gtagagtgtagcacctatccAGGTCACCTTCTAACTGATCTCTCATGTTTTATTTGGCCTggttacctcttatctagtg
gtagagtgtagcacctatccAAGGTTTACTTTATAGCAAATTCAGTCCTGTCAGATCTGCggttacctcttatctagtg
gtagagtgtagcacctatccAACATCCTTTAAGTTTAGCCCTCCCCAATTATAAGGCAGGggttacctcttatctagt
gtagagtgtagcacctatccGCCCCTGCCAGTCTCTTCCGTCTCCATTCCTGCGTCGAGGggttacctcttatctag
gtagagtgtagcacctatccCGTACCTACCGGAGACCAAATGTGCCAGGCTTGTCTTACAggttacctcttatctag
gtagagtgtagcacctatccTTCCTCTGCCTTTCTGTCTTAACCTTCTAATCAACTACATggttacctcttatctagtgg
gtagagtgtagcacctatccCATCTCTTTCTTTGCCCTCCCCACCCCATTCATCCCATACggttacctcttatctagt
gtagagtgtagcacctatccACTCATTCCTTTAGTCTAGCACATACCACACTCATTACTTggttacctcttatctagtg
gtagagtgtagcacctatccGTGTCTTTCCAGCTCCCACCCACAGTTCCATGAGAAAGGGggttacctcttatctag
gtagagtgtagcacctatccGTTTCCCTCCTTCTTACTATAGTTCACTATATATGTAGTGggttacctcttatctagtgg
gtagagtgtagcacctatccATTCTGTAAACATTTGCAAATGTCTACTATGATCTGGAGCggttacctcttatctagtg
gtagagtgtagcacctatccTGTCTTCAAAATTTTCACAGTGGGATCAGGGAACAGGGAAggttacctcttatctagt
gtagagtgtagcacctatccGTTTATATTTATTAACTATGGTCCGACCTTTGAATGGTCCggttacctcttatctagtgg
gtagagtgtagcacctatccCTAAGTAGATGGTTTCTTTTTCTTTATTAGGCGCTTTGACggttacctcttatctagtgg
gtagagtgtagcacctatccACCTGATCCTTATTGTCAAGCTAAGTATACTTTCTGTCCAggttacctcttatctagtg
gtagagtgtagcacctatccCAGTTATGGAGGGTGATGATGACATTGAAGTTTTTCGATTggttacctcttatctagtg
gtagagtgtagcacctatccCTCCTGGGACACTTGGTAAGGATGCATCTTGGTCTTATAAggttacctcttatctagtg
gtagagtgtagcacctatccAGTAATCACTATTTAGATGGCTGACTTTAGATCATGTTGGggttacctcttatctagtg
gtagagtgtagcacctatccCCATTATCCATTGTAATATTATGAATCTGAAACCAGCTCCggttacctcttatctagtg
gtagagtgtagcacctatccCACCTCTTGATTTCCATACTCTGTTGTAGTCACTTTTCCAggttacctcttatctagtg
gtagagtgtagcacctatccGAGACTGGCATGGGGGCTCCTCACACTTATTCTAAGCTCTggttacctcttatctag
gtagagtgtagcacctatccCTTGTCCACATAACATGCAGGCCTCCTATTTGAGATGAGCggttacctcttatctagt
gtagagtgtagcacctatccCACAGAGTGGATATGACAGTAGCAGCATGGAAGAACAGCTggttacctcttatctag
gtagagtgtagcacctatccTCTACTTGATATGCATTTGTGATGTGCCACATTTTCTCCTggttacctcttatctagtgg
gtagagtgtagcacctatccCTGGACCCATGGTTGTGTCACAGTTGCTTTTATACATGTAggttacctcttatctagtg
gtagagtgtagcacctatccCCATTGGATTCAGAAGTACATTAACTGGCAAGAACTACACggttacctcttatctagt
gtagagtgtagcacctatccCCAACTTGGCAACTGTACATTTCCCCATCTCCGACCTGAAggttacctcttatctag
gtagagtgtagcacctatccCTGCCTGCTTTTTTGAGGGAATTGATGATGTTCACTGGAAggttacctcttatctagtg
gtagagtgtagcacctatccATGGGACATTAGTTCAAGTAGCAACTATATCAGGTAAGTTggttacctcttatctagtg
gtagagtgtagcacctatccATTTGAATAGGTTTGTGGGGAACTGGTTATGTTAAGTTACggttacctcttatctagtgg

gtagagtgtagcacctatccTTTATCCCTCACCCCTCTCCGACCCTTTTCCCCGACTTCCggttacctcttatctag
gtagagtgtagcacctatccTTTGGTTTTCTATTCCTGAAGAGTTACTTCACTTGGATGGggttacctcttatctagtg
gtagagtgtagcacctatccTATGGCTGAGTAGTATTATTCCTTGGTATATATACACCACggttacctcttatctagtg
gtagagtgtagcacctatccTGCATGGGAAAGTATTTTTTGCGTATAATGAGTTCTCTTCggttacctcttatctagtg
gtagagtgtagcacctatccATTTACATTCCTAGCAGCAGAGAAGAAGTGTTCCCTTTCCggttacctcttatctagt
gtagagtgtagcacctatccCATTCTTGCAGGAGTGAGGCGGTATTACATTGTGGTTTTGggttacctcttatctagt
gtagagtgtagcacctatccCATTTCCCTGATCATTGCATTTCCCTGAGCATTTTCATGTggttacctcttatctagtg
gtagagtgtagcacctatccGTCTATTTGTGTCCTTAGCCACCCCCACCCCGCCTTTTTGggttacctcttatctag
gtagagtgtagcacctatccGTTTGTTTGAGGTGGAGTTTTATTCTTGTCGCTCAGGCTGggttacctcttatctagtg
gtagagtgtagcacctatccGGTTCAAGCGATTCTCCAGCCTTAGCGTCCCAAGTAGCTGggttacctcttatcta
gtagagtgtagcacctatccGTGAGCCACCACTCCCGGCCGTTAACCCACTTTTTGATGGggttacctcttatcta
gtagagtgtagcacctatccTGTTGCATTTGCTTTTGGTGGAACCACTAAACTCTGACCCggttacctcttatctagt
gtagagtgtagcacctatccGTAGCTTATCTCTGATCTTTGAATGTTTCCCTAAAAATGCggttacctcttatctagtg
gtagagtgtagcacctatccGAAAAATAATAGCTACTAATATAGACTGGTTCTGGCACTGggttacctcttatctagt
gtagagtgtagcacctatccTACTGTACATACATCACATCATTCATCATCACAACTTCTGggttacctcttatctagtg
gtagagtgtagcacctatccAGGCACAGAGAGGTTAAAGTTACACAGCTTAGTAAGTGGGggttacctcttatctag
gtagagtgtagcacctatccAGTTACGCCCCATTGCCCATTCTCCTAACTACTTTGATACggttacctcttatctagt
gtagagtgtagcacctatccCTCCCAAAATGTATATGATCTTATATTTGAGACCCTCTTGggttacctcttatctagtg
gtagagtgtagcacctatccTTTCTTCTTTTGGATGTATTGGAAAGTTGCTGAAAGTTGGggttacctcttatctagtg
gtagagtgtagcacctatccCAAGTGCTGCATTGTTGGTTAGTAATGAAAAGAGTTCAGCggttacctcttatctagt
gtagagtgtagcacctatccACTTGGGTCACATTTTATATCTTATTGGCCCGTTGCCTCAggttacctcttatctagtg
gtagagtgtagcacctatccTGGAGACAATATACCATAGCATTTGAGAAGTAAGGAGCTTggttacctcttatctagt
gtagagtgtagcacctatccTTCCCCAGAACAGTAGAAACATCTGAAATGAAAGCACTGTggttacctcttatctag
gtagagtgtagcacctatccTGATTTGGAGGTATGTTAACTCACGGTTCAGTGGAGAGGAggttacctcttatctagt
gtagagtgtagcacctatccACTCATTCTCTGCAAATAAGGAATTGAGGCTCAGCCAGGTggttacctcttatctag
gtagagtgtagcacctatccCTCTCAAGGTCAAAGAGCTGGATGTAGCAGAATCAGGAATggttacctcttatctag
gtagagtgtagcacctatccCCGAATGCACCATGTTGTTTTCTAATATTTCCTGCTACTTggttacctcttatctagtg
gtagagtgtagcacctatccTCAATGTGCCTTAAACCATACCAACAATACAAAAGTCCATggttacctcttatctagt
gtagagtgtagcacctatccGGCCAGTTCACACCTTTCCCCATTTCCACCTACATCGTTAggttacctcttatctag
gtagagtgtagcacctatccAATAGTGTTTTGAAGCATTCTTGTCTCATAAGGATAGTGGggttacctcttatctagtg
gtagagtgtagcacctatccTTTGAAAGTTGAAGAATGCTAAGAACAGTATCCCCAGTTCggttacctcttatctagt
gtagagtgtagcacctatccGATAAAGGCAGTCACTCTATTCATTTTACAGAACATGAAGggttacctcttatctagt
gtagagtgtagcacctatccGTGACAGGATGCAGTGTTTGGGGCCCAGTTCTATTAGGCAggttacctcttatctag
gtagagtgtagcacctatccAATGCCCAAGATGAGAGTTGTAACCATGCAGAGTAGTGTTggttacctcttatctag
gtagagtgtagcacctatccATATTTGGAAAGATTCTTCTGGTCTAGTACAAGATGGTAGggttacctcttatctagtg
gtagagtgtagcacctatccGTTTAGTTGATTTGTTGGAAGGACTTAACGTGGCAGGTGGggttacctcttatctagt
gtagagtgtagcacctatccCCAGCAACTCCAAGTACAGGGAAGAAAGAAACGTACCAAAggttacctcttatcta
gtagagtgtagcacctatccTGTTCTCATAACAGTAATAGCCAGCCTTCTAGAGCAGTGGggttacctcttatctag
gtagagtgtagcacctatccTGAAGCTCCACATTTCCCACACCTCCATTTTCCTTACACTggttacctcttatctagt
gtagagtgtagcacctatccCAATTCTTTTTGTGCTTTCCTTTTCTCTTTTGCCTCTGAAggttacctcttatctagtgg
gtagagtgtagcacctatccGAGAGCTTCTGCATTTGTATAATTTCGCTTTGACTATAGCggttacctcttatctagtg
gtagagtgtagcacctatccCAGAAATATCTTGTGCCTTGAAGAATGGAATAAAATTGGCggttacctcttatctagt
gtagagtgtagcacctatccAACCATTGTTTACTCAGGCATCCAGTCTTAATCAAGTTAGggttacctcttatctagt
gtagagtgtagcacctatccAAAATTAATATGATTAGGCCAGCCATGGTAGCTCAAACCTggttacctcttatctagt
gtagagtgtagcacctatccGGCAATACAGCAAGACCCCGTGTTGCCCAGGCTGGTCTCAggttacctcttatcta
gtagagtgtagcacctatccGTGAGCTGTGATGTTACCACCGCACTCTAGCCTCGGCAACggttacctcttatcta
gtagagtgtagcacctatccTTTTAAACTAGGAAACATGTTCAACCAAATGGCAAAGTGGggttacctcttatctagt
gtagagtgtagcacctatccCAGGACAATGAAACAGGAATTTATTATGAGACATGGAATGggttacctcttatctagt
gtagagtgtagcacctatccCATGGTTTGATTCCTACGACTGTTCCAAATTTGTGTTAAGggttacctcttatctagtg
gtagagtgtagcacctatccCTGAATTTGGAGCAGAGTTCAAGAACATAGAAACCAACTAggttacctcttatctag
gtagagtgtagcacctatccTTCTTTACAGTGGAGAACCTACTTATCTGGGAAATGAAACggttacctcttatctagt
gtagagtgtagcacctatccGACTCTTGGTTTAGCCATAAAAAGATTTTATTACCCCTTCggttacctcttatctagtg

gtagagtgtagcacctatccAGAATTTCTGTTGAGTCTCTTGCAAATTTTTGATGCAGTGggttacctcttatctagtgg
gtagagtgtagcacctatccATGAAGAAATCCCTTTACCTATCAGAAACAAAACACTCTCggttacctcttatctagtg
gtagagtgtagcacctatccAAAACACCTTAATTCTACTGCTCTTTTTTCTCCAATCACCggttacctcttatctagtg
gtagagtgtagcacctatccACTTTTGTGAATTCCTTAGCCTTTCTTCCTTGGTGCATAAggttacctcttatctagtgg
gtagagtgtagcacctatccAGAATTGCTGCATATTAACATCTCAGGACTCTTCTCTTGTggttacctcttatctagtg
gtagagtgtagcacctatccCGTACTATATTGGCCAAAGTGAGCGAGTTAGGTGATCTTGggttacctcttatctagt
gtagagtgtagcacctatccTCCGAGCCTTTGTTAATATGGAGAATTATGGTTCATATCAggttacctcttatctagtg
gtagagtgtagcacctatccACCCAGGGTCCTACAGATAGATATGGTGTGCCCAGATTTTggttacctcttatctagt
gtagagtgtagcacctatccACATTCAGTGACATTTTCATGGTGGGAGCTCTTCTTTCTGggttacctcttatctagtg
gtagagtgtagcacctatccGGAACCAAGTTTAGTATAGTCTGATTACATTCCATTCACCggttacctcttatctagtg
gtagagtgtagcacctatccCATAACTGGAGTGATTGTGCTTCTAGATGTGGCAAATCCAggttacctcttatctagtg
gtagagtgtagcacctatccATATTTCTGGCTGAGATTTTGGAACTAGCTAGTAACTGGCggttacctcttatctagtg
gtagagtgtagcacctatccTGTAAAGATGTAAGTTGTGTATGTTTGGCAGGTCACATTGggttacctcttatctagtg
gtagagtgtagcacctatccCAAATGTCATCCTTGATCTTGCCTAATGTAGTTTATGTGCggttacctcttatctagtg
gtagagtgtagcacctatccTTTCCAGGGTTTTGTAGTCACCTAGATTTTAAGCTGATAGggttacctcttatctagtg
gtagagtgtagcacctatccGGCACCTTCTGACTTCTTCATTGTTTCCTGTAGTTCTCCCggttacctcttatctagtg
gtagagtgtagcacctatccTGGAAGAGGTTGCACTTCTCCGAGAGAGGACAGGTTTCTGggttacctcttatctag
gtagagtgtagcacctatccCAAGCTTTCAAGGTCCTTTAAGTATTTGATGATCAACTGAggttacctcttatctagtg
gtagagtgtagcacctatccTTTCTATTCAAGGAGAAAACACCATTCAGTAAGAAGATGGggttacctcttatctagtg
gtagagtgtagcacctatccCCAATTATGTGATTCAGTTTCAACTTTCAAGTACTTCCTGggttacctcttatctagtgg
gtagagtgtagcacctatccAATTAACCTGATCCAAATGAGTAAAGTGGACCTTAGAAAGggttacctcttatctagtg
gtagagtgtagcacctatccATCCAGCTAGTAGTAATGAAGTCAGGTCTTGAACCCCGGTggttacctcttatctag
gtagagtgtagcacctatccGACTGAATTGGATGCACTATAGTACAGGCTTTTAGCACCGggttacctcttatctagt
gtagagtgtagcacctatccAGATGTTACTGGTCTGTGAGGAAATAAGTACAGATACTGAggttacctcttatctagtg
gtagagtgtagcacctatccGTCTTTGGTCTTATTTTGTCTAGGAATTCATTTTTGCTGGggttacctcttatctagtgg
gtagagtgtagcacctatccTGTAGTCCCACAACAGTCTGAAAATAACCCAGTCACCAGAggttacctcttatctag
gtagagtgtagcacctatccGAACTTTCTTTCACGCCTCCTTTAGCCTGTCGACTCCTTAggttacctcttatctagtg
gtagagtgtagcacctatccTGCCTAAATATTCCAAATTAATACTGCCTCTTTAACCTGCggttacctcttatctagtg
gtagagtgtagcacctatccTCACAATGTTACAAGAAAACTTGGATCTGTGCATTATCACggttacctcttatctagtg
gtagagtgtagcacctatccGCTAAACTCAGGCAACACGAACTTCTACCAAACTCAGGAAggttacctcttatctag
gtagagtgtagcacctatccACTACAGGATTTTGGATTTCCAGAACTGCCTTGAAGAGACggttacctcttatctagt
gtagagtgtagcacctatccCAAAGAACTTGGTTATGCAGGGAAAACAAATGAGGATATCggttacctcttatctagt
gtagagtgtagcacctatccAAGCATTCAAGATTTTTGAGTAGGCACAGTGGCTTACGCCggttacctcttatctagt
gtagagtgtagcacctatccTCACTTTGCGAAACTGAGGTGGTGGAAGGATCACTGAAGGggttacctcttatctag
gtagagtgtagcacctatccTTAAAAAAAAAAAAAAAAAGCTAGGGGTGGTGGTCCATGCggttacctcttatctagt
gtagagtgtagcacctatccTGAAGACAGAAAGCATCATAAAGTCAGGTAACTACAGCACggttacctcttatctag
gtagagtgtagcacctatccCTTAAGAATTAAACCTGTCAGCTCTACTTACAAGATGTAGggttacctcttatctagtg
gtagagtgtagcacctatccCAGGTGAAGATGATTCAAGAACATGCTTTCAAAATAGTAGggttacctcttatctagtg
gtagagtgtagcacctatccGCTGCCCTTTGAGCAATATGACTTAATGTGATTTACTTTAggttacctcttatctagtgg
gtagagtgtagcacctatccTGCAGTTTCTATTTTGCTATTGTTATAGATGTAGGCTGGGggttacctcttatctagtgg
gtagagtgtagcacctatccTTATGCCTGTAATCCTAACACATACTTTGGGAGGCCAAGGggttacctcttatctagt
gtagagtgtagcacctatccCAAAAATTAGCTGGTTGTGGTGGAGCATGCCTGTAGTCCCggttacctcttatctag
gtagagtgtagcacctatccCATTTTTAGTCTGACTAGGGCTTTAGAGTGAATATATCTCggttacctcttatctagtgg
gtagagtgtagcacctatccTCTGGCACAAAATTAGTTAAGTAAGTATACCCACAAAGTGggttacctcttatctagtg
gtagagtgtagcacctatccATATATTGTTGCATGTAGAGGAAAACACAGTGAAGAAACCggttacctcttatctagtg
gtagagtgtagcacctatccATTTGGAACCCTCAATATTGGTGGCTCCTTAAAAATTGGAggttacctcttatctagtg
gtagagtgtagcacctatccAGTCTTAGGAAGTCTTATCCAGTTTTATGGTCTGCTGTGGggttacctcttatctagtg
gtagagtgtagcacctatccTTTTTAAATAGTTGGCAGGCTATACAAAGAGTGATGTTCCggttacctcttatctagtg
gtagagtgtagcacctatccTGGCCAAAGTGTTTCCAGCATCTTCGTCATATTTACCCTTggttacctcttatctagtg
gtagagtgtagcacctatccTGGACTGATTATTCTAAGGATAAGCTCTTACTTCACTTTCggttacctcttatctagtgg
gtagagtgtagcacctatccGCTTTGTGCCCTTTTTAAAGTTGGGTGTCAGCAAGCAGCTggttacctcttatctagtg
gtagagtgtagcacctatccGTCTGTGCAAAGCTCATTACTATCTATGTGGGGTAGAAggttacctcttatctagtg

gtagagtgtagcacctatccAAAAATTTTAAAGGACCTAGTGGAGGCTTGAACCTTCTTCggttacctcttatctagt
gtagagtgtagcacctatccAACGTCTTCCTTTCAGCCTAAAGTTCTTAAAATGCACTAAggttacctcttatctagtg
gtagagtgtagcacctatccACCTATTTTTTCTGATATGGTTACAACAGCCCAAATTACCggttacctcttatctagtg
gtagagtgtagcacctatccAAAGACATTTATCTCTAAAAAGACTGGTTGGGTTCAAGGAggttacctcttatctagt
gtagagtgtagcacctatccTCTTTAGGCATCAGAGCTTATTTCAAGAATAGATTCCATGggttacctcttatctagtg
gtagagtgtagcacctatccCCTTGTATCTTCAAAAAGGCGAGTCTATCATAAACATTGTggttacctcttatctagtg
gtagagtgtagcacctatccCCAAATACAAGCTTTGGTTAGTAGAACTGATTATGTACTGggttacctcttatctagtg
gtagagtgtagcacctatccCAAGGAAAAACAACAACTTGACTAGCACAGGTAATTCTTGggttacctcttatctagt
gtagagtgtagcacctatccTCGAATCTAGATAAGCCTTTTTTCTTCTCATATGTTGCTGggttacctcttatctagtg
gtagagtgtagcacctatccGCTTTATTCTTAAGAAGGCCATCCGTTTTAAAAGTAGATCggttacctcttatctagtg
gtagagtgtagcacctatccTCAACCACTGAGACACAATCTAAAAGTCACACCACTGAAAggttacctcttatctag
gtagagtgtagcacctatccCGCTTTAAGAATTAATGTATTCCACCGGGGTTAAATCATTggttacctcttatctagtg
gtagagtgtagcacctatccGTAATATCTACTAGCACTAGTGCCCAGCATCAAAGTTTATggttacctcttatctagt
gtagagtgtagcacctatccCTGCAGACATTTTTCTATTAAAGTTTTCCAGTTCACTGAGggttacctcttatctagtg
gtagagtgtagcacctatccCCTCTTTAAAATTTGCTGTTTATGCTAGACTGTACAATGGggttacctcttatctagtg
gtagagtgtagcacctatccAGAAAAGTGAGTGAGCCATCACTAAAATTTACTGGAAACTggttacctcttatctagt
gtagagtgtagcacctatccCAAAGATAAAACCAAGGAGAACTAGTAAAGCATTCCTTACggttacctcttatctagt
gtagagtgtagcacctatccTGCCTCAAGTGCCACATACATAATAACATATCACAAAAGAggttacctcttatctagt
gtagagtgtagcacctatccTTTATACAAAGCTAAACACTGCCTCCCTTCTGTTCCATTTggttacctcttatctagtg
gtagagtgtagcacctatccTTATCCTTCTATAATACCACTGTGTGACATCTATCAGTCTggttacctcttatctagtg
gtagagtgtagcacctatccCAGCTTTGTAACATTTGAGAACATTCATCCAATCTTTTCCggttacctcttatctagtg
gtagagtgtagcacctatccCCTTATAACAGGTGTATGTGTAACTGAAGACCCTAAAATGggttacctcttatctagt
gtagagtgtagcacctatccACTGTTTAATACGTATAACTGGTTATTTCACATCCGAACCggttacctcttatctagtg
gtagagtgtagcacctatccAAATTCGGAGATATGACTGCTATTACACTAAGGAAACAAGggttacctcttatctagt
gtagagtgtagcacctatccCTCATTCATGGGTCTTCCGATAAACAATCTTTACATATACggttacctcttatctagtg
gtagagtgtagcacctatccTCTGTGCCACAAAATAGTAGAATTATATCAGAACTACAGCggttacctcttatctagt
gtagagtgtagcacctatccGGTGCTATTCATCATGCAGTTTTTACCAAGTGGGCATCATggttacctcttatctagt
gtagagtgtagcacctatccAAGATGCTTGGACACTTCCCAGTGAATCTGCTCCAAACTAggttacctcttatctag
gtagagtgtagcacctatccAATGGATAATTGAGATAGGCGGCCACCACACATCTTCACAggttacctcttatctag
gtagagtgtagcacctatccGGCACTACATGAGACTTCACATTCCCCTAGTCCAATAGCTggttacctcttatctag
gtagagtgtagcacctatccGTGGCCGTAATCCATTTGCACACACTACTAGTTCAACCAGggttacctcttatctag
gtagagtgtagcacctatccCAAAGTACAGATGGGTGGCAGGTATTTCATAGCGAAAGAAggttacctcttatctag
gtagagtgtagcacctatccACTTTGGGTGAATGTCTGATGAAAGCATCCCAGCTACTCTggttacctcttatctagt
gtagagtgtagcacctatccTCTGTCCAGGATTCTCACTGACTACATCAATGCGCAAATGggttacctcttatctag
gtagagtgtagcacctatccGTTTTTCAGAAGACAATGCAAGTAACAACTCATCACTCAAggttacctcttatctagt
gtagagtgtagcacctatccGCTAGTTCTCTTAAGCCGTGACACTGATCAGCCACACAAAggttacctcttatctag
gtagagtgtagcacctatccAAATTCAATCAAGTACATGTCCCACAAAAGTTTGTCACAGggttacctcttatctagt
gtagagtgtagcacctatccATTTAAAAAATACACCATCACTAAGGTTAGGGCATTCCAGggttacctcttatctagt
gtagagtgtagcacctatccACTAACACTAAACTTATAGGTTAGGTAATTACAGGACCACggttacctcttatctagt
gtagagtgtagcacctatccCGGAGGCCAAAGGACATTTTTTGAAGTTCAAAACCTAATCggttacctcttatctagt
gtagagtgtagcacctatccATTCATTATGACTTTCGAAGCTCTAACATACCTAGAAAGCggttacctcttatctagt
gtagagtgtagcacctatccGGTGCCATGCTTTATAACTTACCAAAACTATCATCCTAGCggttacctcttatctagt
gtagagtgtagcacctatccAGTTCTGTTTTTGTTTTCAGTCAGTCTCACCCTGTAGCCCggttacctcttatctagtg
gtagagtgtagcacctatccCCATGTTGGCCAGGATAGTATCGAACTCCGGACCTCAAGTggttacctcttatcta
gtagagtgtagcacctatccACTGCACCCGGCCTACTATAAGAATTCTACCCCAAAGAGGggttacctcttatcta
gtagagtgtagcacctatccTTTAGAAACAGGGATAGGAACAAGAAAACATACACCGTATggttacctcttatctag
gtagagtgtagcacctatccCAAGTATGATGTGCCAGGTACTAGATGTTATCCCACAAATggttacctcttatctagt
gtagagtgtagcacctatccTCTGGAAAAGTATTATTTCTACTTTGAAGTGGAGACAGCTggttacctcttatctagtg
gtagagtgtagcacctatccTGCAATACAAACACAGGCTAAGTGACTCTTATGTCCATGCggttacctcttatctag
gtagagtgtagcacctatccCCATAGCTTTATAACAGTGGGAAAAGATAGGCATTAGAATggttacctcttatctagt
gtagagtgtagcacctatccTGAATTACCAGAATGAGTCACCAGAAAAGAGAGTAATGTCggttacctcttatctag
gtagagtgtagcacctatccAGCAACCACAAAGTTTCTGTTTTCCGCAAAGTAGGTATATggttacctcttatctagt

gtagagtgtagcacctatccAATATACTCTAATTACTCAAGGTAACCTAAGCACATCTCCggttacctcttatctagtg
gtagagtgtagcacctatccGAAATAGGAACGCTATTACAGTTAGCGGCAGTGTAAATTAggttacctcttatctagtg
gtagagtgtagcacctatccTGCCCATCAATGATAGACTAAAGAAAGAAAACATGGCACAggttacctcttatctag
gtagagtgtagcacctatccGGGGAACCTCACACGCTGGTGCCTGTCGGGAGTTGGGGGGggttacctcttatcta
gtagagtgtagcacctatccAAAAACACACAGACATGGGAAATGGTAGCTAACCATTCAAggttacctcttatctag
gtagagtgtagcacctatccGCGTAATTCTCAAAATGGCCTCCAAGATCCCACACCCTAAggttacctcttatctag
gtagagtgtagcacctatccATATGGTACAGCTGAGCATATAAAAGATTATCTGGATGGTggttacctcttatctagtg
gtagagtgtagcacctatccTCTAATCATATGAGCCCCTTTTAAAGCAGTTTCCTCTGGCggttacctcttatctagtg
gtagagtgtagcacctatccGGAAGTCAGAGAGATTCAAAATGCAAAGGGGACTCAATGTggttacctcttatctag
gtagagtgtagcacctatccCTAAGCTAGAGGGGGCCAAAAAAGAAAAATCACAACCAGCggttacctcttatctag
gtagagtgtagcacctatccCAATAAGGAAACTGGATGTCAGTCCAACAATCACAAGGAAggttacctcttatctag
gtagagtgtagcacctatccAATAACTTGAATGAGCGTGGAGGTGGATTCTTCTGCAGAGggttacctcttatctagt
gtagagtgtagcacctatccTCAGCCTTTTAAGATTTTAAGCAGAGAAGCCACTTGAGCCggttacctcttatctagt
gtagagtgtagcacctatccCTGCTTTGTGTATAGCTGTCAGTTATGTACCAGTAAGAACggttacctcttatctagtg
gtagagtgtagcacctatccGTGACAAAAGTAAAATAAAACTAGGGCACTATTACGACTCggttacctcttatctagt
gtagagtgtagcacctatccAGACAAATAACCCAAATTAGAAAATAGGCAAAGGGGCTGGggttacctcttatctag
gtagagtgtagcacctatccGGGAGGCCAAAGCGGACGGATTGCCTGAGGTCAGAAGTTCggttacctcttatcta
gtagagtgtagcacctatccAGTGAGCCGAGATCAATGCCACTGCACTCTCCAGCCTGGGggttacctcttatcta
gtagagtgtagcacctatccTACAAATGGCCAAGAAGCACCATGAAAAGACAGTCAAGGTggttacctcttatctag
gtagagtgtagcacctatccATCTCTACAAAAATTACAAAGTATTAGCCGGGCATAATGGggttacctcttatctagtg
gtagagtgtagcacctatccAATCAAAACCACAATGAGTTTACACCCCCTAGCATGGCTAggttacctcttatctag
gtagagtgtagcacctatccTTGGAAAACAGCCTGGCATTACCTCAAATGGGGAGTATGAggttacctcttatctag
gtagagtgtagcacctatccGTATGTACCTAAGAGAAATTAAACCATATGTCCGCACAAAggttacctcttatctagtg
gtagagtgtagcacctatccAGATTAGTGGTTGTTGAGGGATGGCTGGGATGGAGGTTTTggttacctcttatctagtg
gtagagtgtagcacctatccGTGATGCATGCACTACTCTACCAATACACTAAAAATCAATggttacctcttatctagtg
gtagagtgtagcacctatccAAAAATATGGCAAAAAGCACAAAGACCTTTGTCCTAGGACggttacctcttatctagt
gtagagtgtagcacctatccAGAATGTACCCTAAGGAAGAAATTAATACAAGCAAACTGCggttacctcttatctagt
gtagagtgtagcacctatccCACAGCAAAAAAGGTAAAAACAAGTGTCCAACTTTAGAACggttacctcttatctagt
gtagagtgtagcacctatccGCAATGTTGGCTTTATAGGGAACTAAGTAACTAAATGTTCggttacctcttatctagtg
gtagagtgtagcacctatccCTAGAGGTCCAGTTACAGACAATTATAACAACTATAGGAAggttacctcttatctagtg
gtagagtgtagcacctatccCAGTCCCTAATTTCAACTTAGAGAAACTGTCCAAATATGCggttacctcttatctagtg
gtagagtgtagcacctatccCCAGGCTGGAGCACAGTGGTGCGACCTCGGCTCACTGCAAggttacctcttatcta
gtagagtgtagcacctatccGCCAACGTGCCCAGCTGTATTAAAAAGTATTCTTTCTCTTggttacctcttatctagtg
gtagagtgtagcacctatccTTCCTGAAGAACTAGTCCAAAAGTCATTATGAAACCAACTggttacctcttatctagtg
gtagagtgtagcacctatccTTCCTGGGCATAGGCCGAATTAACTTTGGGAAGGAATTCGggttacctcttatctagt
gtagagtgtagcacctatccAACTGGTAACCGCCGTTTCACAAAAAGACCCCTCCTTCTTggttacctcttatctag
gtagagtgtagcacctatccACAAGATTAGAAATTACAATGTGGGGGTCATGAAGCCTCTggttacctcttatctagt
gtagagtgtagcacctatccGACTGGTAATCTGGGCCAACTGGTTCTGCCATGGCACCTAggttacctcttatctag
gtagagtgtagcacctatccACATTAAGAAAACCTAACTCTGACCCCTTATAATTCCATCggttacctcttatctagtg
gtagagtgtagcacctatccTTGATAAATCGGTTCTGTCTGGGCAACGGACAAGGTGAACggttacctcttatctag
gtagagtgtagcacctatccAAGTGGGATCAAGTAAGCTTATACTATACTATCAGAACTGggttacctcttatctagtg
gtagagtgtagcacctatccCACAGAATATACCAGAATAAAGAGAACTATGAACTCACAGggttacctcttatctagt
gtagagtgtagcacctatccTCAGAGACATAAAGACTCATACAATGTGTTAAATGCTAGCggttacctcttatctagtg
gtagagtgtagcacctatccCTGGAAATACACAGGTAGGAAATCTCAACCAGAAAGGAGAggttacctcttatctag
gtagagtgtagcacctatccAGGTCAACAAAGATCACATCTGAGCTGAATTCTGAAGCCTggttacctcttatctagt
gtagagtgtagcacctatccCTGGAGTTAGCAAGGGGAAGTTAAAACATTTGAGCAAGTAggttacctcttatctagt
gtagagtgtagcacctatccACAAACAAAAATCTGTAGTTCCATAGGACCCAAGTAAAAGggttacctcttatctagt
gtagagtgtagcacctatccCAAAAGGACCTTGTACACTATGAAGCCAGGAAGCACAGACggttacctcttatctag
gtagagtgtagcacctatccAAATGTAATCAAGGTCATATAAGAACTAACTGCCAAAGCCggttacctcttatctagt
gtagagtgtagcacctatccGCTAAGAAAATACAATAGACCTAGCCATTCTCATCACCAGggttacctcttatctagt
gtagagtgtagcacctatccGTGGGAGAAAAGAGAATGAGCAAGGAGACTTGGGAGAATGggttacctcttatctag
gtagagtgtagcacctatccGCAATGGCAAGAAGGTTGGAAGTGTTTTTGTAAAGATCAAggttacctcttatctagtg

gtagagtgtagcacctatccCCTTAAAGAACAGGTTTCCCAAATAACACAGCTAAAGGATggttacctcttatctagt
gtagagtgtagcacctatccAACATGTCAATAGTGGGAGGAGTCAGAAATTAAACAGCAGggttacctcttatctag
gtagagtgtagcacctatccAGTAGGGTTTATACTCTCTTTTCAATACAAACCACCTGGAggttacctcttatctagtg
gtagagtgtagcacctatccAATGCTCAGAAAATCCTATATAAGAGAAACTGGAAAGAGCggttacctcttatctagt
gtagagtgtagcacctatccTGGGTTCTTGGCTCCAAAATTCAACTTAGGAAAAACATCAggttacctcttatctagtg
gtagagtgtagcacctatccTACAACTTAGTACAACTCTTCTCCCTCTACATAGTGAGAAggttacctcttatctagtg
gtagagtgtagcacctatccCAATTATGTGACTGCTCTGACTCAAAATCTACCCTATTAGggttacctcttatctagtg
gtagagtgtagcacctatccCTACAATGAGAAAAAACTCTACTGTAATTAATGGAGCACCggttacctcttatctagtg
gtagagtgtagcacctatccGACCTCATACTGTCCTTTCCTCTTTTTTTTCCTGTTCATTggttacctcttatctagtgg
gtagagtgtagcacctatccTGAAAGTCAGTGTCTAGGCTAGTAACCCTACAGAACTTATggttacctcttatctagtg
gtagagtgtagcacctatccTGAAGCAGTCATATTAATCCACCTTCAAGGTTAATGATCAggttacctcttatctagtg
gtagagtgtagcacctatccGTCTAGCAAAAAACACAACATACCTGCTGGAGAGACATGAggttacctcttatctag
gtagagtgtagcacctatccAACAGCTTGAGTGTATCACTATTGTTGGCCACTAGTACTTggttacctcttatctagtg
gtagagtgtagcacctatccGATGGATCATCTACTGGAGTATCATCTATCTTAAGCGAAGggttacctcttatctagtg
gtagagtgtagcacctatccTTTACGAACACAACTGTCAGTGCAGAGATAAAGTGAGACTggttacctcttatctagt
gtagagtgtagcacctatccTTTTTAGAAATGAGAATCAAGGGGTTTCCTAGTACTACAGggttacctcttatctagtg
gtagagtgtagcacctatccCCTTCACTCTGTTGTTGGGAATATGGTAAAAATAGGTGACggttacctcttatctagtg
gtagagtgtagcacctatccCTTTGAGAAGAGTCTATTTATGTGTGGGGGAAAAAAGCATggttacctcttatctagtg
gtagagtgtagcacctatccCTAAATCAGAGAAACCACTGGTAAGTAAAAAGAGGAATTCggttacctcttatctagt
gtagagtgtagcacctatccTCTCTCTCAGACACTTCATTTTGCCATATCTGGAAACCCTggttacctcttatctagtg
gtagagtgtagcacctatccGATCCACAAAAGCACCAATGAGTATCATTCATCATATATGggttacctcttatctagtg
gtagagtgtagcacctatccCACACACACACGTTATATATCTGTTGGACAGGATTTGATCggttacctcttatctagtg
gtagagtgtagcacctatccGAATCAATGTAGGCAAACAGGAATTAAAGCATTCTCAAAGggttacctcttatctagt
gtagagtgtagcacctatccTCAATCGGAGCTAAGATATGAGTTGAAACCACTGGGCAAAggttacctcttatctag
gtagagtgtagcacctatccGTAAAGAAAGGCTAATACATGGTCTAACACATTCCTCTCTggttacctcttatctagtg
gtagagtgtagcacctatccACCAACTTCATGCTCCAGTTAAGCTTTATAAATTCTCTACggttacctcttatctagtg
gtagagtgtagcacctatccTGAAAAAGGATAAGGGCAGGCTAAGAGCCAAGAAAGTATAggttacctcttatctag
gtagagtgtagcacctatccCCTGAACTTAAGCAATCCTGTTTCATCCTCCCTAGTAGCTggttacctcttatctagtg
gtagagtgtagcacctatccTCTTGAACTCCTGGCCACAACAGAACCTCCTGCCTCGGCTggttacctcttatctag
gtagagtgtagcacctatccAGCTACCACACCCAACCAAAACACTACCTTCTTGATCACAggttacctcttatctag
gtagagtgtagcacctatccGTGCATCACTGACGTCTTTCCCCATTTCCCCAATTTAGCTggttacctcttatctagtg
gtagagtgtagcacctatccACATTTCTTCTATGTTCTCTTCAGTTCTCTTGGGATTTCCggttacctcttatctagtgg
gtagagtgtagcacctatccATTGCAACGCCCTACTTCAATCTTCTACCAAAGCCATAAAggttacctcttatctagt
gtagagtgtagcacctatccTTAAGTATCCTAGCCTCACTTAATAAGGGAAGAAGCATTCggttacctcttatctagtg
gtagagtgtagcacctatccCCATTCATGGTTCCCAAATTAGTACTAGATATATGCATAGggttacctcttatctagtg
gtagagtgtagcacctatccATTCAGATTGTTTCATTTCCTATTTGAAGCTTGAAGGTGGggttacctcttatctagtgg
gtagagtgtagcacctatccTACCATGTAACTAGTATTTCTTAGCGTGAGTTCCTTTAACggttacctcttatctagtgg
gtagagtgtagcacctatccCAGGAAGAGAAGACTTAGAATCACCCCGAATTTGAAAAAGggttacctcttatctag
gtagagtgtagcacctatccTTTAGGAGGCTATTCAGAAGATTTTTACACATTAAGTCGGggttacctcttatctagtg
gtagagtgtagcacctatccCAGATTCTCCTGCCCATTTCCCTTATTTTATATGTGAAGAggttacctcttatctagtg
gtagagtgtagcacctatccTAAGACTTTGACATCAACTGGTGATTTATTTGGTACAGACggttacctcttatctagtg
gtagagtgtagcacctatccGTTATATTCCCTCCTCAATCATCCTAGTACATTAAACTACggttacctcttatctagtg
gtagagtgtagcacctatccAAAAGGGTGTCTATATTTTAGTTTTCAGTGTCTTCTGGGAggttacctcttatctagtgg
gtagagtgtagcacctatccACCTATTAACCATTAGCTAACATATTCAGACACCTTTACCggttacctcttatctagtg
gtagagtgtagcacctatccTTGTATTTTAAGAGAATAGTTGTGAGAGAGAGACACGATCggttacctcttatctagtg
gtagagtgtagcacctatccGTCTGTAAATTAATCTAAACTGTGCTGAGACTAACTTTGGggttacctcttatctagtg
gtagagtgtagcacctatccCTTTCCAGAACATTTGCCAAAAGATTTTAGAAGCTATACCggttacctcttatctagtg
gtagagtgtagcacctatccCAATATTGGCACTCTACTTTTCTTTGATTCTAACTTCACCggttacctcttatctagtgg
gtagagtgtagcacctatccGAAGCCCCAAAATGAGTATCTATGTGTCCCTTTTTTTCTAggttacctcttatctagtg
gtagagtgtagcacctatccCTTACTGAATTTCTTTTAATACTTCCTATAGACGCCTCTGggttacctcttatctagtgg
gtagagtgtagcacctatccACAAAAACAGCAGATACCTTTGGTAAATCCATAAAGCTTGggttacctcttatctagtg
gtagagtgtagcacctatccTAAGTCCAAGTGTTTTTAAAGAGCAATTCACAAGTTGCGAggttacctcttatctagtg

gtagagtgtagcacctatccGCTGATTCCTTGCTGCTGTCCACCTTAGAAAAGAAACACCggttacctcttatctag
gtagagtgtagcacctatccTTTACTTTTTTCCCCAAATACCCAGACTTCAAGATTACTCggttacctcttatctagtg
gtagagtgtagcacctatccGAATATTCTTTCAATCATCCACCCTCCACCCTCACCTCCAggttacctcttatctagt
gtagagtgtagcacctatccTTCTAAGTCTCTCTGAAAAAGTATAAGGGCAATTGTCAAGggttacctcttatctagtg
gtagagtgtagcacctatccGGTAGTGGATCACATAATAGATGTAAATACAGTGTCTATGggttacctcttatctagtg
gtagagtgtagcacctatccTTTACTCAAAATGTTCACGTGTATCACAACCTGGGATTCAggttacctcttatctagtg
gtagagtgtagcacctatccATTAATGGAAAGATGTGAAGTAAAATGAGTGTACTGGACCggttacctcttatctagtg
gtagagtgtagcacctatccTGTAGCTATTCTCACACATACTTCTGCAAAGACATTTCCTggttacctcttatctagtg
gtagagtgtagcacctatccGAGTTCATCCATCAAAAAGTTAGTTGTTTCCTGGGAAGCAggttacctcttatctagtg
gtagagtgtagcacctatccATTAGTTTCAGAGCTACTTATAGATAATATGTGGTCCCTGggttacctcttatctagtg
gtagagtgtagcacctatccACAAATGTTCATACCCAACCCAGACCATCTGAATCAGAAAggttacctcttatctag
gtagagtgtagcacctatccCCTTCAGGTGATTTTGATGCACATAAAGTTAGAGAACCACggttacctcttatctagtg
gtagagtgtagcacctatccAATTCTAACTATAAGCCTCCTTGGTCTATAATCAGTGAAGggttacctcttatctagtg
gtagagtgtagcacctatccGAGAGCCCAGAAGAATCCTGTGATGTGACTCTTTTTTGTAggttacctcttatctagtg
gtagagtgtagcacctatccTGCACAAATGATTTTATGAGGGTTATACTAGATACACCTGggttacctcttatctagtg
gtagagtgtagcacctatccGGCTATAAAAATGACCCAGCAAATCTACTAGGTGGAACTGggttacctcttatctag
gtagagtgtagcacctatccCAAAGATAACTAAAACCAAACAGAAACCTCTTTAGCCTTCggttacctcttatctagt
gtagagtgtagcacctatccGATGAGTCCTGAATTTTTGTTTGATCCCCCTACCTTTGCCggttacctcttatctagtg
gtagagtgtagcacctatccGGATCGCAAGATAATATACTAGACACCAATTAAGTTTCAGggttacctcttatctagtg
gtagagtgtagcacctatccTTTATTTGTAAAACCTGGCAATCAGTGGCCTAAGCGCCTTggttacctcttatctagtg
gtagagtgtagcacctatccGGAAAAAAAAGGCAAAAACATATCCAACCATTCCCACCACggttacctcttatctag
gtagagtgtagcacctatccTTCGAGGGAAGACAAGATTTTAGTTATGGCCAGAAGCTCTggttacctcttatctagt
gtagagtgtagcacctatccACTGTTCTGCCTAGGTTTAGCCTGGACTATTCAGCAACTGggttacctcttatctagt
gtagagtgtagcacctatccAGATCCAGTAAACATGTCCATATTTCATTATTCATCACCCggttacctcttatctagtg
gtagagtgtagcacctatccAAACATGACAAGCCTCACACAAACCACTGTGATTTTTTATggttacctcttatctagtg
gtagagtgtagcacctatccTCTGTCACCAAGGCTGTAGTACAGCTGCACAATCATAGCTggttacctcttatctag
gtagagtgtagcacctatccGGACTACAGGTACCCACATCCAGCTATGATTTCTTAAATTggttacctcttatctagtg
gtagagtgtagcacctatccATCGCAAAAAGCCACCCACTTCATTTATCTCATCACTGCAggttacctcttatctagt
gtagagtgtagcacctatccTCCAATACCTTCCTTCCAGCAACAAAAAGAAATAAACAGTggttacctcttatctagt
gtagagtgtagcacctatccACATAAAGTTTCTATGCTTCCTTTTCATCTGAAGATCCCAggttacctcttatctagtg
gtagagtgtagcacctatccAAAGCCATCAGTAAGTAACACTAAGTTCCTGTTAGATAGAggttacctcttatctagtg
gtagagtgtagcacctatccGCTTTATCAATGACTTATGCTTATTTAGTGGCTGCTTTTGggttacctcttatctagtgg
gtagagtgtagcacctatccGCAAATATTTTAAAGGCATGAGAAGATGTACTGGTTTAGGggttacctcttatctagtg
gtagagtgtagcacctatccATTACCTTGAAGCTGACATATTGTAGATGGTTTGAATGTCggttacctcttatctagtg
gtagagtgtagcacctatccCTTTCAAATAAGATGTAGCTGGCTGATTCAGTTCAAATTCggttacctcttatctagtg
gtagagtgtagcacctatccGGCATGTGAAATACCTGGTTCCAGTTGCGGCAAACTTGTGggttacctcttatctag
gtagagtgtagcacctatccCCCGGTCAAGAAGAGGCAAATATTTAAATACTTGGAGAATggttacctcttatctagt
gtagagtgtagcacctatccAATGCTCATTTGTAGTCCTCAGTTTCTTGGATTTCTCTGCggttacctcttatctagtgg
gtagagtgtagcacctatccCTTCTTCTGATGAATTACGGTCACTATCTCTTCCTCCTCGggttacctcttatctagtg
gtagagtgtagcacctatccTCCTATTCCGAAAAATGCATCAGTTTTTTTCCTACTCAACggttacctcttatctagtg
gtagagtgtagcacctatccCATTCTTCTGTCACTGAGATGTGTGTTTATATACACAAACggttacctcttatctagtg
gtagagtgtagcacctatccCCAAAACAAACTATTATGAAAAATGCCAATGAAGGCCAACggttacctcttatctagt
gtagagtgtagcacctatccATGAATATAAAAGGGAACCTACAGCGATACTTTGTAAAGGggttacctcttatctagtg
gtagagtgtagcacctatccAGTCATCTAAGGATATTTGTTCTTGAATTCCAGGCCAACCggttacctcttatctagtg
gtagagtgtagcacctatccTCCAGAGAGGATGTTCCACATTTAATAGTCTATACAGGGGggttacctcttatctagt
gtagagtgtagcacctatccAACTGGGCCCCATAGCAGGAAGTGAGCAAGCTTTACAGTCggttacctcttatctag
gtagagtgtagcacctatccTCATAGGAGTGAGAATCTTATTGTGAAATGTCCACATGAGggttacctcttatctagtg
gtagagtgtagcacctatccGGAACAGTTTCTTCCTGAAATCACCCCCTGCCACGGACAGggttacctcttatctag
gtagagtgtagcacctatccGTTACTGCCAGACATTTCTTAACTCCGCCTAAAATAAGCAggttacctcttatctagtg
gtagagtgtagcacctatccACCCCTACTATGAATTGTTTGGTCTGGAATGTAGCTGCTTggttacctcttatctagtg
gtagagtgtagcacctatccCACTGTTTTCCCCAAGTTGTGTTTCTGAATTATACCTATTggttacctcttatctagtgg
gtagagtgtagcacctatccTTTTTCATGGCAAAACAGTATTATGGCTAAAGAGAATGGGggttacctcttatctagtg

gtagagtgtagcacctatccGTTTGAATTCTGGCTCTACCTCTCACTGGCTATGTGGCTTggttacctcttatctagt
gtagagtgtagcacctatccAAATGCAATACTATTAAGGCTTTTCAAAGGGGAGATACGAggttacctcttatctagt
gtagagtgtagcacctatccTTAACGACATTAAAGTACACAACATGCTAAAAACCAGGGTggttacctcttatctagt
gtagagtgtagcacctatccATTCCGTTCACTCATTCATTCAAGATGGGGTCTCGCTCTGggttacctcttatctagt
gtagagtgtagcacctatccATGCACCACCACACCCGACTCTATTTTTGCTTTAAAATGGggttacctcttatctagt
gtagagtgtagcacctatccTGTACGAGAAAACACACCCTGGAAGTAGACTCACTGAATTggttacctcttatctag
gtagagtgtagcacctatccACCAACAGTAGACGGCAGATTACAGGTAATTTTTTTTTCTggttacctcttatctagtg
gtagagtgtagcacctatccCACATTTTCTTTTACTATCAATGCATTACCGTGTAATCGGggttacctcttatctagtg
gtagagtgtagcacctatccCTGGTAGTTACTTAGTTCAGAGCCTGTAATACTTTTGTTTggttacctcttatctagtg
gtagagtgtagcacctatccACAGTTCTGGCAGGGTAGTGGTTTCGACCTAAACTTGGGGggttacctcttatctag
gtagagtgtagcacctatccGCTCCAACACTCCTCTGGCCTATCAGCATAAAAGAATTTCggttacctcttatctag
gtagagtgtagcacctatccACAAATGTTTCTGCAGACTTTCTCCACCCAGTACCTCGCAggttacctcttatctag
gtagagtgtagcacctatccTGTGAAAGAGGATTGAGGGCTGGAGAGTAAAAAGAGAATGggttacctcttatctag
gtagagtgtagcacctatccGACCTTAATGCCATGAAAGTCAGTCAAAGACGTCTTAGCAggttacctcttatctag
gtagagtgtagcacctatccCAATAAACTGCCTATTCTTCACTGCTCCTATTCCCAACTAggttacctcttatctagt
gtagagtgtagcacctatccCTGATCAGGGAGAGAACAAGAGAGAAAATAGAATGACAGAggttacctcttatcta
gtagagtgtagcacctatccTCTGATGGAAGACTGGAAATTATGTGTGACTTGAGATTAGggttacctcttatctagt
gtagagtgtagcacctatccTCACTAGAGGCTGAGATAACACTGTTATTATGACCCATTTggttacctcttatctagt
gtagagtgtagcacctatccTAAAATTTGGGTCAAAGAACAAAGTGTTTGCTATCCAGAGggttacctcttatctagt
gtagagtgtagcacctatccATTTCTCTTCTGACCACCTAAAACTACTCACAATTTTGAGggttacctcttatctagtg
gtagagtgtagcacctatccTTCTACACCTTATTATAGAATACTGATTGCCTGTTTCTGGggttacctcttatctagtg
gtagagtgtagcacctatccATCTTCTAAAACCAGAACTAGACAACTTAAAGACGGAATGggttacctcttatctagt
gtagagtgtagcacctatccTTTTTAACAGTATGCAACACTAAGGGACAGTATCTCTTGAggttacctcttatctagtg
gtagagtgtagcacctatccCACTATATGATCACCAAAGCCTATTTTAAGAACTTACCAGggttacctcttatctagt
gtagagtgtagcacctatccTGTAAGGCCAAATTCCAATAAAAATATCCTCTGCACTATCggttacctcttatctagt
gtagagtgtagcacctatccCTGCTATACATAAAATTAAACATCTCAGAGCATCTTCCAGggttacctcttatctagt
gtagagtgtagcacctatccGTCAAAGTGAAAACACCTCCTATATTTGAAAAGAAGTGTGggttacctcttatctagt
gtagagtgtagcacctatccCCAAGCTGTACTATAAAACGTCAATGAGACATTCTAGACTggttacctcttatctagt
gtagagtgtagcacctatccCAGTTTAATTTCCTGCAAAGCTTCTAGGCCCCAAATAATGggttacctcttatctagt
gtagagtgtagcacctatccGCTTTTGGTGAGGGACTGATATGACTACAAACTGCTAAATggttacctcttatctagt
gtagagtgtagcacctatccTTAGTGAACTAACTCTATGTTCCATAAACTTTCCAGGTGGggttacctcttatctagtg
gtagagtgtagcacctatccCAGCCTGGGCAACACGATGAAACCTGTCTCTACAAAAATTggttacctcttatctag
gtagagtgtagcacctatccTTTGAAAGAAGCAACTTCATTAGGTGAAGACCGGAAGTGAggttacctcttatctag
gtagagtgtagcacctatccCATCGTTGTATTAGGCACTTTTACACTGGGCATGCAATAAggttacctcttatctagt
gtagagtgtagcacctatccGCTCATTTGTAGTTAACTCTAAAAAGCCTATTTTGGCATCggttacctcttatctagtg
gtagagtgtagcacctatccAGGTTAACTCACAACTAAATGCTTAGAGAGCGCACAGTGGggttacctcttatctag
gtagagtgtagcacctatccGAAAAGGAGGATTCTAATTAAAAAATCTGGGTCACTCAACggttacctcttatctagt
gtagagtgtagcacctatccCATTAGCAACGACAAAATAAAAATGATCTCTCTCACCTCTggttacctcttatctagt
gtagagtgtagcacctatccACCCCTGTCTGCAACCCCACCTAAAGGACTGCCCACGCCAggttacctcttatct
gtagagtgtagcacctatccAACACAAGTTGACCAAGTTTTTCGCCTGGCTAAGCGGCAGggttacctcttatctag
gtagagtgtagcacctatccCCGACTGGCTTCCCCTGACATTTCTCATGCGCCCCACCCTggttacctcttatcta
gtagagtgtagcacctatccGCTTTGACATTTCAGTGCCTCATTGCTCAAGCTACCAGCAggttacctcttatctag
gtagagtgtagcacctatccTGCCTGTCACCAGCCCCCGCCAAGGGTCATAAAGCCTGGCggttacctcttatct
gtagagtgtagcacctatccGCCGCCGCACACGCACCACGCCTTGTTGACATGTTTGGAAggttacctcttatcta
gtagagtgtagcacctatccGGAAACGCCCCCAAGCAAGCTCCAAGGTCTGGGTGCCGCGggttacctcttatct
gtagagtgtagcacctatccCGGCCCGCCCACAGACGGTCCGCGGTCCCTCCCCGGCTCCggttacctcttat
gtagagtgtagcacctatccGGGCTCGGCCCTGCCCCTCCGCCGTTACCTGCTGCCGGGGggttacctcttatc
gtagagtgtagcacctatccCTCACATGAGGCGCCCCTGGCCCCCCGCTTCGCGCTCCCGggttacctcttatc
gtagagtgtagcacctatccGCCCCCCGCGCTCCGCCCGCATCCGAGGCGCTGGTCCCGCggttacctcttat
gtagagtgtagcacctatccAAGCTTTCCTTCTCAGTCCCGCGCTGTCTGTGTTCAGGGAggttacctcttatctag
gtagagtgtagcacctatccGCTTCCTGCACCTGGGCCACAAACAGCGAGTCTAAAATGGggttacctcttatcta
gtagagtgtagcacctatccCGGCCGCGCGCTTCCCTGGCGCGTCCCCGCCCAAGGGGGCggttacctcttat

gtagagtgtagcacctatccCCAGTCTCCTTGCTGTGGACGAGCCTCTGCCTTCCTGCCggttacctcttatcta
gtagagtgtagcacctatccTTGTTGTTTTGAGATGGAATGTGGGAGGGCTGAAGCTAGCggttacctcttatctagt
gtagagtgtagcacctatccCTGGAAGGTTTAGGGAGGTGGAGCAGAAAAAAGAAAGGAAggttacctcttatcta
gtagagtgtagcacctatccGGATGCCTGAAACCGAGGAGTTAAATACAGTCTCCGAAGAggttacctcttatcta
gtagagtgtagcacctatccCTTAAAAGATGCCAAGGAAATGTACCAGAATAGTGAGAGTggttacctcttatctag
gtagagtgtagcacctatccAGAGTCCTGAAATTTGTAAATAGCTTTTGAAACGCAATGGggttacctcttatctagt
gtagagtgtagcacctatccCTGAAAAGAAAACCTCTGGTGAATCCTTTTGTAAATCAAGggttacctcttatctagt
gtagagtgtagcacctatccCAATATCATTAAGTTGAATCATGCGCCATTCCTCTCTACAggttacctcttatctagt
gtagagtgtagcacctatccGTAGTTTTCCATTCAAAGATGTCTTAGAACACTCAGACCCggttacctcttatctagt
gtagagtgtagcacctatccCAGTGGTGCCCAGGGAGCTCTCCGTTCTTCCCGTTTTCTCggttacctcttatctag
gtagagtgtagcacctatccCTAGGAAGCCATGTATAAAGTTACGTCGAAGTTCAGTGTGggttacctcttatctagt
gtagagtgtagcacctatccCAAGGCAGAAGGGTTGCTTGTGGTCAAGAATTGGAGACCAggttacctcttatcta
gtagagtgtagcacctatccATAATTTTAAAAAATCAGACGAGCTTGGTGTAATCCCGGCggttacctcttatctagt
gtagagtgtagcacctatccATAAAATGAATTCAATGTGGGGAAATGCCAGTGTTGTTTGggttacctcttatctagtg
gtagagtgtagcacctatccGATCACTGAATCAAACTGTACAAAGCCTCTAGTAGCATAAggttacctcttatctagt
gtagagtgtagcacctatccAGCTGCAACATTTGTTTAGGTCCCTCAGTGACTGGATGATggttacctcttatctagt
gtagagtgtagcacctatccTAAGTCAAATACAGGCCCATTTATTTAAAGTTGGGTGAACggttacctcttatctagt
gtagagtgtagcacctatccACATTAAACCAATTCAGTAAACATGATGAGCATAGGTTGCggttacctcttatctagt
gtagagtgtagcacctatccTCTTTTTTCGCCATTATGATCTATTCTCCTCCTAAACAAGggttacctcttatctagtg
gtagagtgtagcacctatccGACCTTGGCACACTTGACCTATTGAATACCCCTTTTCTTCggttacctcttatctagt
gtagagtgtagcacctatccACATTTATCAATTGTATCATCAAGATCAGTGGCATGGTAGggttacctcttatctagtg
gtagagtgtagcacctatccCTGGGAAAAATAATTTATTACTGACCCTGTTTGGAATTGCggttacctcttatctagtg
gtagagtgtagcacctatccATTGTTTTAAATTATCCACAGACAATGTACCCCCTCCACCggttacctcttatctagt
gtagagtgtagcacctatccCCAATGATCAGGATAGATTAGATTATGTTGCAGTAGTTCCggttacctcttatctagt
gtagagtgtagcacctatccGTTTTTAGGAGTACTGCTTTAGTATTTTTGTAGAACGTCCggttacctcttatctagtg
gtagagtgtagcacctatccTGGGGTTTGTCTTAGGTTTTTTTCTTATAATTACATGGGGggttacctcttatctagtgg
gtagagtgtagcacctatccAGAAGGAAAACTACAGAGGTATAAAGTACCATTCTCATCAggttacctcttatctagt
gtagagtgtagcacctatccAGGGTACATCTATTAATACGACTTATCCTTGTTGATGTCAggttacctcttatctagtg
gtagagtgtagcacctatccCAGAGGTTGAGATCAAGTTATTCTACTTAAAGTTACCCTTggttacctcttatctagtg
gtagagtgtagcacctatccAGCCAACACTGAAGAATGTGGATTTACCTGAACTGCAACAggttacctcttatctag
gtagagtgtagcacctatccTTAAAACTTTTCTGTCTTGGGAGGTGTATCTACATATCTCggttacctcttatctagtg
gtagagtgtagcacctatccACTTTTCTGTCTGTAATTTCTGTAGATCTGTCTTCAAGTCggttacctcttatctagtg
gtagagtgtagcacctatccAAGACTACCTTTGCCTGCGTTTTTCGTTTCTAAAAGTTTTggttacctcttatctagtg
gtagagtgtagcacctatccATCCTATGAAATAATTGGGGGTCTAATCATGATATGTGTCggttacctcttatctagtg
gtagagtgtagcacctatccGGATTGTGAGTTCATCTTCCGTACAACTTTAGCTGCAGGAggttacctcttatctagt
gtagagtgtagcacctatccCAGAAGCTGGAAGGGGAGGAGATTGTTAGTGAGTGCTGGAggttacctcttatcta
gtagagtgtagcacctatccAAGGAATTGTTAATGGAGTCCAGAGAAATCAGGATATCTTggttacctcttatctagt
gtagagtgtagcacctatccTCTTTCATATAATAATGGCAAGCCCAGGGAAACATTTGTCggttacctcttatctagt
gtagagtgtagcacctatccATTGTTGGACTTGGCTAAAGAAATTTCCAAGCAGGCTGGCggttacctcttatctag
gtagagtgtagcacctatccGGGAGGCCAAAGTAGGCGGATTGCTTGAGCTCAGGAGTTCggttacctcttatcta
gtagagtgtagcacctatccGACTGTGCCACTGCGCTGTATCTCTAGCCTGGGTGACTGAggttacctcttatcta
gtagagtgtagcacctatccGAAAGAAAAGGAAAGAAATCGCCAAGCAGAAGGTTGAGAGggttacctcttatcta
gtagagtgtagcacctatccTCTTTTGGTTGCATATGAAAAAGTTCAGATACCCAGCACTggttacctcttatctagt
gtagagtgtagcacctatccCTTAAACTCAGGAGTTTGAGGGCAACATGGGAAAACCCTGggttacctcttatctag
gtagagtgtagcacctatccGAAAAAGTTCAGATAGAGAGAACGTAAGTACAGAAGTAACggttacctcttatctag
gtagagtgtagcacctatccTTTAGTAGAATTTAGAGGAAAGATAAAGGGCCTGGAACAGggttacctcttatctagt
gtagagtgtagcacctatccAAAAAGACCTTTAGGGAAATGATAAATACCTGGGTACTACggttacctcttatctagt
gtagagtgtagcacctatccATTGGACTTTAGAACAATTTCAGGTGGTACCGCTTAGACCggttacctcttatctagt
gtagagtgtagcacctatccAAGAGGTCTTTTAAGGATTTTATGTGTATGCTTCCTAGACggttacctcttatctagtg
gtagagtgtagcacctatccCCATAGCATCTTGAATCTCAGCCCTACCATAAATCTCAAAggttacctcttatctagt
gtagagtgtagcacctatccGGAGTGAACCAAAGTAAGAGTCATAGGAAACATACAATATggttacctcttatctag
gtagagtgtagcacctatccTTGAATAGAAGATACAGAGACAGTTGAAAATGAAGAGTCCggttacctcttatctagt

gtagagtgtagcacctatccGCTCCCATCTTCTGTCATTTAAATATAGGTCCATAAAACGggttacctcttatctagtg
gtagagtgtagcacctatccCTTATAAAGAAGGAATGATGACTCAGAGTGCAGAGAAAGGggttacctcttatctag
gtagagtgtagcacctatccAAGCAGGACTGGGCCTTAATTAAGAAACTGGTTACATGTTggttacctcttatctagtg
gtagagtgtagcacctatccAGTGACCGCTATTGTTTCCTCTTCTCCCCTTTATAAATGGggttacctcttatctagtg
gtagagtgtagcacctatccTCCTACGCCTATTTCACCATTGTATATTGGTTCTACATGAggttacctcttatctagtg
gtagagtgtagcacctatccAAGTCTTTAAGAGGAAATGCACTTGAATATGGGCTGGACTggttacctcttatctagtg
gtagagtgtagcacctatccCTCCATCATGAAAATGTAGAGGAAGTAATATTCTAGGACTggttacctcttatctagtg
gtagagtgtagcacctatccGCTAACTTATGAAAAGCCTTGCAGTTTTTGTCTGGATGTCggttacctcttatctagtg
gtagagtgtagcacctatccGGAACCCAGTTGCCATGCTAATAAGAAGTCCAAGTCACATggttacctcttatctag
gtagagtgtagcacctatccAGCCATGTGGGTGAACCTTCTTGAGCATCTGTCCAACTTAggttacctcttatctagt
gtagagtgtagcacctatccAGATGACTCCGGCTCAGCTGCTATCTATTTGCAAACACAAggttacctcttatctagt
gtagagtgtagcacctatccAGCAAAGCTCGGTGAACCTACAGAACCCTGAGAGACAATAggttacctcttatctag
gtagagtgtagcacctatccCATTAAGTGTGGAGTGCTTTGTCATACAGTAGTAGATAACggttacctcttatctagtg
gtagagtgtagcacctatccAAACAGAAGCTGGTTTATACAAAAGGCAAATGAGGTAGACggttacctcttatctagt
gtagagtgtagcacctatccGGGGAACTAATATAGCCGCTACAGAGATTTAAAATACTAAggttacctcttatctagtg
gtagagtgtagcacctatccAGTAATTATCAAAGCCTTTTCCTCCCAAATAACCCTAAGCggttacctcttatctagtg
gtagagtgtagcacctatccGATGAGCCTTATCAGATTTCTAAGAGTTTAATCCTTTCCTggttacctcttatctagtgg
gtagagtgtagcacctatccATTTTTTTCTTGAGACAAGGCTATGTTGCTATGTTGCTCAggttacctcttatctagtgg
gtagagtgtagcacctatccAGCATGACCTGATCCCTTCTTATACAATTGAGAAAAAAGAggttacctcttatctagtg
gtagagtgtagcacctatccTCCAAGTCATCTTAATGAAGCTACTATAATCATGATTCCCggttacctcttatctagtg
gtagagtgtagcacctatccCCCCATGGCCCACAATGAAAATAAATGCAAAAATGCTAAAggttacctcttatctag
gtagagtgtagcacctatccGAAAATTCCACACCTGACCACAGTAAAAATGCAGTCAAGAggttacctcttatctag
gtagagtgtagcacctatccAAATTATCTTCTGGCTAAGTGTATAAAGTGTCTGTGCAACggttacctcttatctagtg
gtagagtgtagcacctatccTTTCATGTTTAGGTTTGGGGCCCATTCCTAAAATATCTCAggttacctcttatctagtg
gtagagtgtagcacctatccCTTCTAGTTCCAAGAAGTTCAGATAAGAGATATTTGACCTggttacctcttatctagtg
gtagagtgtagcacctatccAAACCAAAACAAAACAGTGATAATAACGTGCCTGATCCTCggttacctcttatctagt
gtagagtgtagcacctatccCCTCTTTTTCTTTTCCCACGGCATTTATCACTTCCTAGCTggttacctcttatctagtg
gtagagtgtagcacctatccGGTATTGTATGTCAGTTGTTTAGTGTCCATCTCCCACCTGggttacctcttatctagtg
gtagagtgtagcacctatccTAAGAATGGCAAAAACATGAATCTGGCCGGGTGTGGTGGCggttacctcttatctag
gtagagtgtagcacctatccTCCGGGAGGCAGAAGTTGCACCGAGCTGAGATTGAGCCACggttacctcttatcta
gtagagtgtagcacctatccCCAAAAAACAAAAACATGAATCTTAGTCTGCTTGGTTCACggttacctcttatctagtg
gtagagtgtagcacctatccAGGCTCCTAAACCAGTCTGTAAATGTTTCTTCCAAGGATGggttacctcttatctagtg
gtagagtgtagcacctatccTGTTTTATTACATGGCGTAGTTACCTGGACTTAACCCAAAggttacctcttatctagtg
gtagagtgtagcacctatccCCTGAGGAATAGATAGGGACTTAGATGTCACAGAAGAGGGggttacctcttatctag
gtagagtgtagcacctatccTCTTAAGAGGCAAATGGGAAGATGATGAGAGAACACAGTCggttacctcttatctag
gtagagtgtagcacctatccAACCCTGTTCTTTTGTGATTCACCAATTTCTTATGTTCCAggttacctcttatctagtgg
gtagagtgtagcacctatccGGTAAACCTCCATATAGAAACAGGGGCCATCCAAGCTGGGggttacctcttatctag
gtagagtgtagcacctatccCCCCTGGGAATTTGGAATTAATACCGGCAGAAAGAGAGTAggttacctcttatctag
gtagagtgtagcacctatccTCCTTAGACTTGTAACACATCAACCCAGGAGCTATAGGTGggttacctcttatctagt
gtagagtgtagcacctatccCTGAGAAAGAAGAAGAGAAAATAGACGCATAGAGAGGAACggttacctcttatctag
gtagagtgtagcacctatccCAGCCCTGGTTCTGCTCTATTCCCACAGCCTGGAAGAGGTggttacctcttatctag
gtagagtgtagcacctatccCAATCAAAACTCCTTACCATTGCAAACTTAATCTGGAAAGggttacctcttatctagtg
gtagagtgtagcacctatccCTGAAAGGTGATCATGGAATTTGGTCTTGCTGTGTCTTGTggttacctcttatctagtg
gtagagtgtagcacctatccGCCTTGCATCCTCTCCACAGCTGCCCTTTAACCAAAGATGggttacctcttatctag
gtagagtgtagcacctatccGCTGTGGATGGCCCTGTCTTGGATGGAGCCCCAGAAAAGTggttacctcttatctag
gtagagtgtagcacctatccGCATGACTACTGCCCCTTTTGAAAGATTTCCTGTTTTCCAggttacctcttatctagtg
gtagagtgtagcacctatccGGAACCTAAGACCGATTTATGCCGACTTCCTAGAACGAAAggttacctcttatctag
gtagagtgtagcacctatccGATGCATGCAGTTCTTTCCTGAGGGGCTGATACATCTCTAggttacctcttatctagt
gtagagtgtagcacctatccGGAACAGAACTTTTCACCCGAATTGAGTCATTCCTTTGAGggttacctcttatctagtg
gtagagtgtagcacctatccATGCTCCATGTGTACTTTCCCACTCTCTGACCTCTTTTTCggttacctcttatctagtg
gtagagtgtagcacctatccGCTCTCTTCTTTCAGGTCCTTTAGTCTGATGTCGCTTCAGggttacctcttatctagtg
gtagagtgtagcacctatccGCCTCAAGCCTCCACTGCTCAGTCAAAGCTGCTCTAGAGTggttacctcttatctag

gtagagtgtagcacctatccGAAGATGTGGATTGGGTTGGAGCTCAGGTCTCATTCTGCCggttacctcttatctag
gtagagtgtagcacctatccCAGGATACGCGAGCTTGCACAGATTACTTCTTCATGCTTCggttacctcttatctag
gtagagtgtagcacctatccATCTATAAAATGGGAATAAGGCTGTTCACAGTGGCTAACAggttacctcttatctagt
gtagagtgtagcacctatccATTGCTTGAGCACAGGAGTGCAAGGTGGCAGTGAGCTAAGggttacctcttatcta
gtagagtgtagcacctatccACATAGGGTTATCGTGAAAATTAAAGGTGTCAACATCCATggttacctcttatctagt
gtagagtgtagcacctatccAAAAAGGTCAGTAGCAGGCACACAGTAAATCCCCAGTAAAggttacctcttatcta
gtagagtgtagcacctatccCAGAGTGGGAGTGGGCCTGAACAAATGGCTGAAGAGCCCGggttacctcttatct
gtagagtgtagcacctatccGGTGTACACCCTATGTCAATGAAGTAGTGGTCTGTGATCAggttacctcttatctagt
gtagagtgtagcacctatccAGTTACAAAGTTATATACTCCCATACAAATGAGGACTTGGggttacctcttatctagt
gtagagtgtagcacctatccTTTCATCTGCCACGCAGAAAGGGGGGTTGCAAAAGGAGTAggttacctcttatcta
gtagagtgtagcacctatccTCCTTTTGATTTAGTTCTAGTAAGTCAGCCTGAATTGGCCggttacctcttatctagtg
gtagagtgtagcacctatccCTCCTGTCTCACTATTATTTTTGGCCCTAACTTGATAAGTggttacctcttatctagtg
gtagagtgtagcacctatccCAGTGGCTGTTTCAATTCTAGGCCTTCATAGCTGCAAGAGggttacctcttatctag
gtagagtgtagcacctatccCTTAAGCAGCAGTAACACCACCAAAGGGGCACTAACAGCCggttacctcttatct
gtagagtgtagcacctatccCTTTGTCCTTGAAAAGCAGTAACCCAGGCAAGCACTAAAAggttacctcttatctag
gtagagtgtagcacctatccGCATGCATGTCAAGAAACAGCACACCAGCTGCAAATAGTTggttacctcttatctag
gtagagtgtagcacctatccAAAAGATTCTATAAAATTTCAGATTGCCACCTCTACTCCCggttacctcttatctagt
gtagagtgtagcacctatccTGGTTTCCCATTCATTCTGCCATTGTAACACAGCATCCCTggttacctcttatctagt
gtagagtgtagcacctatccATGTCCATCAGCCTAGCCTCTGTTTGAATAGCCCCTGTGAggttacctcttatctag
gtagagtgtagcacctatccTAATATTTGTTTCATTGCCAAGGATATTTGTTCCACAGGGggttacctcttatctagtg
gtagagtgtagcacctatccGGGATTTCTTTGTTATTTTTACCCCTGGCCTTTACTTTTGggttacctcttatctagtgg
gtagagtgtagcacctatccCATGCTCCCTTTTTAGGCACGTGGCATTTTTCCTATATGAggttacctcttatctagt
gtagagtgtagcacctatccCAGCTCCTGTCATGTCCACTTTGCCAATCAGTTCCATTCTggttacctcttatctagt
gtagagtgtagcacctatccAAAAATCTCTTTAGCTGCCTGTTTTTCAAATGTACCCCATggttacctcttatctagtg
gtagagtgtagcacctatccACTTTTGATCAGGCCCCAAGTTGTGTGTAGGTTTTGAATGggttacctcttatctagt
gtagagtgtagcacctatccTTGCACACAGTTTACTTACTAAAAGGCATGGACATTGGCAggttacctcttatctagt
gtagagtgtagcacctatccGTTGGAGGAGAGGTCTGGTGAAAGAATACTGTATGTATCAggttacctcttatctagt
gtagagtgtagcacctatccTGGTTTAGCACCATTTCCCCCAGGTTGTTTAAAAGTGTGTggttacctcttatctagt
gtagagtgtagcacctatccCTCCCACCCCACTCAGTCTGTTTCTCCTGCTGGCCATTGAggttacctcttatctag
gtagagtgtagcacctatccGAACAGACTAATACAGACATCTTCAGCTTCAGCTGGAGTTggttacctcttatctag
gtagagtgtagcacctatccGGGCACTTTCACACAGCTCATTAATTGACCCCAGTTGACCggttacctcttatctag
gtagagtgtagcacctatccCAATTCCATCAGATTTTCTTATTTCTTGTTCACATCCTCCggttacctcttatctagtg
gtagagtgtagcacctatccGACATTTAGGAATTAAACACCATACACACTTGCAGCATTGggttacctcttatctagt
gtagagtgtagcacctatccTGCTGCCCCTTCTACTTGTCTGATAGATATGACTCTTTGAggttacctcttatctagt
gtagagtgtagcacctatccCACTTTCTAATTGGAGTCTTTTGGTGCTTGATTCTTTCATggttacctcttatctagtgg
gtagagtgtagcacctatccTTTTTGTGTTAAAATGTCCAGGTCACATTCTTATGCATGCggttacctcttatctagtg
gtagagtgtagcacctatccCCATTTTTCCATTAGTGCACTTGGGAATTACTGGACCCCTggttacctcttatctagt
gtagagtgtagcacctatccACTCTGTGTGTGAAGCCAGTGAATCTCCTACTAAATCCTCggttacctcttatctagt
gtagagtgtagcacctatccCATTCTGGTATTATATGCTGTGCTCCATTAAATCAAAGGTggttacctcttatctagtg
gtagagtgtagcacctatccCAATAATTGAGGGAAGTGACTGATGTATTTGCTGGGCCTTggttacctcttatctagt
gtagagtgtagcacctatccATAAAGCGCTGATTGCAATTATTCCTGTTGTTCGAAAGCAggttacctcttatctagt
gtagagtgtagcacctatccTGCTTCCGTCTTTGCAAGGTTGAAGGGAAATTCTTAGGCCggttacctcttatctagt
gtagagtgtagcacctatccCTAATGTATAATCTGGTCTGTATCCAAGTCATCTTCTATGggttacctcttatctagtg
gtagagtgtagcacctatccTTTTTGAGATGAAGTATCGCTCTTGGTGCCCAGGCTGGAGggttacctcttatctag
gtagagtgtagcacctatccGGCTTACCGCAACCTCTGCCTGCCAGGTTCCAGCAATTCTggttacctcttatcta
gtagagtgtagcacctatccCACCATGCCCTGCCGAGGTATAATATATAAACGGTGAAGTggttacctcttatctag
gtagagtgtagcacctatccTGATTAAGACAGAAAATACTCCGGTAACCCAGTGCTCCCTggttacctcttatctag
gtagagtgtagcacctatccATGGAATCATACTGTATGTGCTCTGCAAGTCAGCTTAACTggttacctcttatctagt
gtagagtgtagcacctatccCCTCTTTGTGTGTTAACTCTATCTAGGATTCGATTGTAAAggttacctcttatctagtg
gtagagtgtagcacctatccCAAAGGAAGGAAGTGTGTTTTGTGACTCTTGATTATAGCCggttacctcttatctagt
gtagagtgtagcacctatccTCCAAGATTACAGCCATAACTTCCCAACTGTCTTATCCAGggttacctcttatctagt
gtagagtgtagcacctatccGGCCTGCTTCCTGCCTCATCTTCCTAGAGCACACATTCGTggttacctcttatctag

gtagagtgtagcacctatccGATAAAATCAAACTGTTGAGAATGAGATTCTAGACCAGCAggttacctcttatctagt
gtagagtgtagcacctatccTTTTCTAGAAGCCACTAAAAAAAGGAACAGCTGGAATGAAggttacctcttatctagt
gtagagtgtagcacctatccTTGAAATACTTTCCATTCTTTTCTTGGTGAGACTCAGTGggttacctcttatctagtgg
gtagagtgtagcacctatccCTTGTGAAGCCCATTGTATTCCTTTCCTGTTGCTGCTGTAggttacctcttatctagtg
gtagagtgtagcacctatccTGGAGATCAGAATTTTGAAATGTACTAACAGCAGAGTGTAggttacctcttatctagtg
gtagagtgtagcacctatccGCCTCCAGGGGAGAACCTGTGTTCACCTTTGCCAGCTCCCggttacctcttatcta
gtagagtgtagcacctatccTTCCTCCATCTTCAAAGCCGACAGTGTAGCATCTTCAAGTggttacctcttatctagt
gtagagtgtagcacctatccCACATCTCCTTCTCTGACCGTGACCCTCTTGCCTCTCTTTggttacctcttatctagt
gtagagtgtagcacctatccGTCAATCCAGTGTACTCTTCCCATCACAAGATCTGTACCGggttacctcttatctag
gtagagtgtagcacctatccCAAGGGCTAAGACGTGGACAGATTTGGGGGGATGGTTATTggttacctcttatctag
gtagagtgtagcacctatccGCCCTCAATAAATGCTTTGTCTAATGGTTATAAAACCTGGggttacctcttatctagtg
gtagagtgtagcacctatccGGATTCTGTATTTGTGTTCCATGGCTGTCTCATCCTACCAggttacctcttatctagtg
gtagagtgtagcacctatccCAGAGCGCGCCACACATCTTGACAGACTCTCATGCAGTTCggttacctcttatctag
gtagagtgtagcacctatccTCTCTGTAGAAGTCGTGGTTACTAAACACCCTCATGAGAGggttacctcttatctag
gtagagtgtagcacctatccGCAAAACAGCTTACCTGGTGCTTCTGAGGAACACGTTAATggttacctcttatctag
gtagagtgtagcacctatccTTGCTTGTCAAAACTGATCTATCCTGAAAGACTATCTGCGggttacctcttatctagtg
gtagagtgtagcacctatccTGTTTTAAATGAAGATATCTCTGACCTTGCTTGTTACTGGggttacctcttatctagtgg
gtagagtgtagcacctatccTTGATGGCCTTTTCTTGAGCAATTTGATAACATTGTATGGggttacctcttatctagtgg
gtagagtgtagcacctatccGCGAGACACTAATCTTTAACCTTGGCTTTTTCTTTCATAAggttacctcttatctagtgg
gtagagtgtagcacctatccCTGTAACTCCCTCCAGCGTATATCTTTTTGTAGTCTTGTTggttacctcttatctagtgg
gtagagtgtagcacctatccCTAAGCCACCCTAAGACATTTGAGAAATGAAGCCATAAGTggttacctcttatctag
gtagagtgtagcacctatccCTCTAGGAAGCAGATGCCAAGATTAAGTTAGAAGTGCAGAggttacctcttatctag
gtagagtgtagcacctatccCCTATGAAAGATAAAGTGATGAGGGAGCAGGAGTGGGCAGggttacctcttatctag
gtagagtgtagcacctatccAAAGAAGAGAGAAGGAAAGATTGGGTAGGAAAAGCCTCCGggttacctcttatctag
gtagagtgtagcacctatccTCAGTCGGCCTGAGTTCTTACGCAAGACCGCGCACGCAGGggttacctcttatcta
gtagagtgtagcacctatccGCCAGGCCCGAGAACCCCATCTGAACTCTATCACAGGCTGggttacctcttatcta
gtagagtgtagcacctatccGTGACATTGGGTCATATGTGGGGGTGATTAAAGCTGGAGGggttacctcttatctag
gtagagtgtagcacctatccCTCCTGACGACACTGAACACTCTTGGAGGCATATGCTGTCggttacctcttatctag
gtagagtgtagcacctatccCCCTGTGCAACCTTGGCCATACGATAAGTGTCCTGAAATTggttacctcttatctag
gtagagtgtagcacctatccCTTTGAGTACTGACATGCCTGATACCCAAATAAAATGCTCggttacctcttatctagt
gtagagtgtagcacctatccTATTCCCAAACCTGAAAAAACCCGAAATCAGAAACACTTCggttacctcttatctag
gtagagtgtagcacctatccAGTAATGTAACAATAATAGCATGATGTGATAGAGCTTGGGggttacctcttatctagtg
gtagagtgtagcacctatccCTACAGCTGGAATCTAGTGCAAATAATTTCTTTAAACAGCggttacctcttatctagtg
gtagagtgtagcacctatccCTCTCAGAGTTGCCAGGAGGATTAAATAAAATGCCTAATTggttacctcttatctagtg
gtagagtgtagcacctatccAGTAAATGTCCTCTCCTTTGCTCTCTCTTGAATAGTTTATggttacctcttatctagtgg
gtagagtgtagcacctatccATAAATTGAGATTTTAAAAGGTATGTGGCTGACTCAAGGCggttacctcttatctagtg
gtagagtgtagcacctatccTGAATTCAAAGCTTCTGGTTTCCATCCCAGCACTCTATGCggttacctcttatctagt
gtagagtgtagcacctatccTGAGCCGCAGTGTTCCATCATACAGCCAATAGCAGAAACAggttacctcttatctag
gtagagtgtagcacctatccAGTGACACAGTGGGCAGGGAGATATATTGGGCTGAAAATCggttacctcttatctag
gtagagtgtagcacctatccCTAAACATTCTGAATGAAAAGTGAATAAAGGGAGCATTGCggttacctcttatctagt
gtagagtgtagcacctatccTTTTATCAAATAAGCTTATTGAGGTTGGGTGCGGTGGCTCggttacctcttatctagtg
gtagagtgtagcacctatccTGTAATCCTAGCTGAGGCAGGTAGATCGCTTCAGCTCAGGggttacctcttatctag
gtagagtgtagcacctatccATGGTGAAACCCTGTTTCAATTACAAAAATTAGCTGGGCAggttacctcttatctagtg
gtagagtgtagcacctatccGCTTATTGATAAAATAGGGAGATTTTAGGCCTAAACAGTCggttacctcttatctagtg
gtagagtgtagcacctatccAAAAACAATTTCTTTCCAGGAAGATTCACTGACATCTACCggttacctcttatctagtg
gtagagtgtagcacctatccGATTCTGCTTTCTGTCTCCTGCCTTGGATGTTACATGCAGggttacctcttatctagtg
gtagagtgtagcacctatccCTTGAGCTAAAAATGCCTGCCAAAGCCACTCCTTCCTGTGggttacctcttatctag
gtagagtgtagcacctatccTTTATTGCTAAGATGTTTATGTTCTCTGGTTTTGAGGTGCggttacctcttatctagtgg
gtagagtgtagcacctatccCACTACACAGCTTTGTTGATTTTATCCCTTTCATAACTTGggttacctcttatctagtgg
gtagagtgtagcacctatccTTGTTTCTTTTCCTCTACGATTTACAAACTGAGCTGACCAggttacctcttatctagtg
gtagagtgtagcacctatccTGGTATTCTTCGCAGCTATTAAGTGTCCATAAAGTCTCTTggttacctcttatctagtg
gtagagtgtagcacctatccAGGTTAGTTTCATTCCTGTGGCCTGTGAATTTAAGTCAAAggttacctcttatctagtg

gtagagtgtagcacctatccCTGCATTCTCCCCTCCAGTTTTCTTGTAACAGCTTTAGTCggttacctcttatctagtg
gtagagtgtagcacctatccGAAACATTTTCAGTCAATCCTAAGCTTGAACACTCATACTggttacctcttatctagtg
gtagagtgtagcacctatccGTATTTCTCATGCCTTTCTCCACATCTCTGCTCCTTGAAAggttacctcttatctagtg
gtagagtgtagcacctatccGCTTAAATGGCCTCTTAGCTACACTCCCAACGACAAAACAggttacctcttatctag
gtagagtgtagcacctatccGAACATTCTGTAAAATAGTTACCTGTGCTCAGATCTGATCggttacctcttatctagtg
gtagagtgtagcacctatccAAAATGTGATTAAAACTTTTGCTAGTCTTGGCCAGGCGCGggttacctcttatctagt
gtagagtgtagcacctatccCACGCCTGTAATACCGGAATTTTGGGAGGCCGGAGGCGAGggttacctcttatcta
gtagagtgtagcacctatccGTTCAAGACCAGCCTGGTGAAACCCCATCCCTACTAAAAAggttacctcttatctag
gtagagtgtagcacctatccGTCAACAATCAATAATTTGGCAATTTCATGACTCGAGTGTggttacctcttatctagtg
gtagagtgtagcacctatccTTTTATTACTTTATATGAACTTCAAAAAGGGGCTGGGTGCggttacctcttatctagtg
gtagagtgtagcacctatccTTTGGGAGGCCAAAGCAGTGGGGATTGCTTGAGTCTAGGAggttacctcttatctag
gtagagtgtagcacctatccAAATTAGCTGGGCATGGTGCTGGTGCATGCCTGTAGTCCCggttacctcttatctag
gtagagtgtagcacctatccGGCTGGAGGATCGCTTGAGCCCAGTTGGAGGCTGCAGTGAggttacctcttatcta
gtagagtgtagcacctatccTTTATGATCTTCTGCTTAGGGTTTAAGATTCCAACATTGCggttacctcttatctagtg
gtagagtgtagcacctatccTGTGTTGTGTTAGTAAATTTCTTTCTGAGGAAAACTGGACggttacctcttatctagtg
gtagagtgtagcacctatccGTTTTTTAAACAATACCTCCCACAGTGGTAGACTTTATCAggttacctcttatctagtg
gtagagtgtagcacctatccGGTTTTATTGACATGTACATTCCAATATGTTTACAGCTGCggttacctcttatctagtg
gtagagtgtagcacctatccCACACTCAGTATTGCACTTCATTAAAATTTCAGGCTCAAAggttacctcttatctagtg
gtagagtgtagcacctatccATAAACCAGAATATTCCTCTTTTTATCCCCACCCGTGATGggttacctcttatctagtg
gtagagtgtagcacctatccTTTCTTTTTTGCTTGAAAGCCAGCATCGTTCTTAGTCCATggttacctcttatctagtg
gtagagtgtagcacctatccCAATTATCTATAAATGTCCAATGCTTCACAGGCCAATGACggttacctcttatctagtg
gtagagtgtagcacctatccCAAACACCTCACAAGTTTGATGTCATTTGTTCAAAAGAAGggttacctcttatctagtg
gtagagtgtagcacctatccAAGAATTATGTACATGGTATTTGGTATCCTTCTTGCACTGggttacctcttatctagtg
gtagagtgtagcacctatccGGATGTAAAAATGGTCCCGTCCTTATCCTGAGCAGAGTTTggttacctcttatctagt
gtagagtgtagcacctatccCTCTTGGGGGATGAAGGCAATTTTTCTCTCTGTAGACAAAggttacctcttatctagtg
gtagagtgtagcacctatccCATTTCTGCACACTCCACATCCCAGAGCAAACTCTTCCCCggttacctcttatctag
gtagagtgtagcacctatccGTTCCTTCTAACTGCTTGCCTTTGGGACCTGTATAAATTTggttacctcttatctagtg
gtagagtgtagcacctatccAGATATGTACAATGGAAGAAGTGATGGGGTCCCTTACTCAggttacctcttatctagt
gtagagtgtagcacctatccCTAGAGAGGATCACTTCGAGCTACAGATCTTTTTCTATTAggttacctcttatctagtg
gtagagtgtagcacctatccGTCACAAAACCCCTCAATTTAGTCTGGCAGAACATTATTTggttacctcttatctagtg
gtagagtgtagcacctatccAGCCATGTTTTCCCTTTTATTTTTACTGATGAAAAAGCCCggttacctcttatctagtg
gtagagtgtagcacctatccAAGGCCAATGATTCCTGTAATTTTTAAAAGTACCTGAAGGggttacctcttatctagtg
gtagagtgtagcacctatccAATGTGCTTCACGTCATGGGTTTCATATCATTTAGAAAAGggttacctcttatctagtg
gtagagtgtagcacctatccAGGCTTACTTTGTCAGAAAAACACGCTAGGCTTATACTTGggttacctcttatctagtg
gtagagtgtagcacctatccTAGTAGATGAAGGAATAGGAACTCTGCATTAGTTTTCTACggttacctcttatctagtg
gtagagtgtagcacctatccAATTCTGATTGTGCTTGAGAGAAAAATATCACGAGATTGGggttacctcttatctagtg
gtagagtgtagcacctatccTGAGCTTACACTGATCAATGAGGAATGACGTCTCATTTACggttacctcttatctagtg
gtagagtgtagcacctatccTTTCTGACATAAGAGAAGGAAACCATGATGGATAAAAGGCggttacctcttatctagt
gtagagtgtagcacctatccTGGCTATTACTACTATTACTGAACAACCACAAACTGAGGCggttacctcttatctagt
gtagagtgtagcacctatccCTAGAGGAATAAAAAGTAAGTTCTGTACAAGAGTAAGACCggttacctcttatctagt
gtagagtgtagcacctatccGAGTGTCAAACTGAAGACAGCTAATCTCAGAGAAGTCAACggttacctcttatctag
gtagagtgtagcacctatccGCCACCTCAGACAAATGCAGGCACAGACCTCAGGATGGGTggttacctcttatcta
gtagagtgtagcacctatccAGGATGCCAGAGGTTTGCTAGAAAGTTAATTATTCCTGTTggttacctcttatctagtg
gtagagtgtagcacctatccGCTCCAAATCACACTGCAGATTGGTGCCCACCCCACCAGTggttacctcttatcta
gtagagtgtagcacctatccGTGCGCAGGATGGGGACACTCAATGGCAGATCTAGTTCAGggttacctcttatctag
gtagagtgtagcacctatccTACTCAACAAATATTTACCGCATGCCTAATATATGAAGGGggttacctcttatctagtg
gtagagtgtagcacctatccTTAGGTACTTGGGATTAAAGATGAAAAGTGCATGGCTCCCggttacctcttatctagt
gtagagtgtagcacctatccGGTGAGAAATAAAAATAGTGAAAACACACCTGCAAACATCggttacctcttatctagt
gtagagtgtagcacctatccAGTATATCAGCTACTTAACCATTTAACAGTCATAGGAGGCggttacctcttatctagtg
gtagagtgtagcacctatccGATACCTCATCTTTAAAATAGACACAGCAATACCTAACTCggttacctcttatctagtg
gtagagtgtagcacctatccTGTGAAGATTAAGTATCCTGGCACGAGAAGGTAGTCAATAggttacctcttatctagt
gtagagtgtagcacctatccAGAAAGAAATGGATAGAGGGATTATGCTATGGTCTGGATGggttacctcttatctagt

gtagagtgtagcacctatccGTGCTCCTCCAAATTTATATACTGACGTGTTAATGAAAGGggttacctcttatctagt
gtagagtgtagcacctatccGGTCCCCAGAGAGCCCCAGGTCCACAGGTCAGTGCCTCATggttacctcttatct
gtagagtgtagcacctatccCAGTTGGGGAAAAAAAGTCAAGCTGCAGGCACCGATGAGGggttacctcttatcta
gtagagtgtagcacctatccCCCACGGCTGCACAGACAGAAAAACCTTCAATCAGATAAAggttacctcttatcta
gtagagtgtagcacctatccTGTCCTTTGTATAGTCAGGGGGCTCTCTCCTTTTGTGGGCggttacctcttatctagt
gtagagtgtagcacctatccGCCAGGGTCCCGGGCCAAGCTGAGTCACGCTGCGGGCCAAggttacctcttatc
gtagagtgtagcacctatccGTCCATAAAAACCCTGGGCCCCAGCCTCACGGGAGCATCCggttacctcttatct
gtagagtgtagcacctatccCGCTCTAACCTCACCTTTGTATCTGTGCTCCTTAATCTTCggttacctcttatctagt
gtagagtgtagcacctatccAGACAATGAGAGACTGTTATATCTTGGTGCACTGCTGAGAggttacctcttatctagt
gtagagtgtagcacctatccCAATGTTATGGTGTTAGGAGGTGGAGCTGTGGGAGGTGATggttacctcttatctag
gtagagtgtagcacctatccGAATGGGATTAGTGCTCTTCCTATCAATGAGACCCAGGAGggttacctcttatctag
gtagagtgtagcacctatccCCCTGCCCTTCCACGACAGCAAGAAAGTGCCACCTATGAGggttacctcttatcta
gtagagtgtagcacctatccTCCTCACCAGACACCTCGACTGCTAATTCCTAGATCTTAGggttacctcttatctag
gtagagtgtagcacctatccACTGTGAGAATAAATTTCCACTGTTTATAAGCTACCCAGTggttacctcttatctagtg
gtagagtgtagcacctatccATATTAGAATTAACAGGGTTGAGAGTGGTTAGAGAATTGCggttacctcttatctagt
gtagagtgtagcacctatccCTGTCCTTCTAAGTAAACTGTATATGGCCATCTTCCCTACggttacctcttatctagt
gtagagtgtagcacctatccCCTGTGGGATCTCCTTTATTCTATTCCAGTTCTCTAAATGggttacctcttatctagtg
gtagagtgtagcacctatccATACCTCAATCTCAGTGTATGTTCTTAGGCCAGTTCATTAggttacctcttatctagtg
gtagagtgtagcacctatccTTTTCCATTTGATAACATAGGGATAATACCATCTACTCGCggttacctcttatctagtg
gtagagtgtagcacctatccAAATGGAGCAAATATGTGTGAAAATGCCCAGGACCGAGGTggttacctcttatctag
gtagagtgtagcacctatccGCTCCTCCCCCGCCTCCCTCCTGTGATTCCTTTGTGTTGGggttacctcttatctag
gtagagtgtagcacctatccAGGAAACCTGACCCTCCTCTCTCAGCTACTACCCACCACTggttacctcttatcta
gtagagtgtagcacctatccCTTTAAGTAAAATCTAGATTTCCCCTGAAGATACTGCTTGggttacctcttatctagtg
gtagagtgtagcacctatccTGGTTATCTCCCAAGCTCCTCCAGGTCATATGGCTACAACggttacctcttatctag
gtagagtgtagcacctatccACTGATTTTTCTCTTGGCTCCTATTTCTTTCCAGACCACAggttacctcttatctagtg
gtagagtgtagcacctatccCAACTTTCATTCAATAATTGCCCTTCCTCCAAGCTCAAGTggttacctcttatctagt
gtagagtgtagcacctatccAGTCATCTAGAATACTGCCCTTTCCACACTCCCTTCACTTggttacctcttatctagt
gtagagtgtagcacctatccATGGCTCACATCTCTGTTCTGTCCATTATCCTCCATAACCggttacctcttatctagt
gtagagtgtagcacctatccGGCCTTGCTTCTTTCAAAGCCTGCTACTTCTATTAAAAAAggttacctcttatctagtg
gtagagtgtagcacctatccAAAAATGATAACACAATTGGAAGCAACTTTCCCTAGTCTCggttacctcttatctagt
gtagagtgtagcacctatccTCAATAGTATTAGTTTCTTGAGTATCTTTCCAGGGGTTTCggttacctcttatctagtg
gtagagtgtagcacctatccACTAGATAGTCTCATTTTCCCTCTTACAAAAGGCAGCATAggttacctcttatctagt
gtagagtgtagcacctatccGCAATTTGCTTTTTTTTTTTGAAACAGCTCTGTCGCCCAGggttacctcttatctagtg
gtagagtgtagcacctatccTGATTCTCTGGCCTCAGCCTCGTGAGTAGCTCGGATTACAggttacctcttatctag
gtagagtgtagcacctatccTGTCAGTCACACCACTTGGCCTAATTTACTTTTTTCATCTggttacctcttatctagtg
gtagagtgtagcacctatccTTCCAAATCAGCACACAGAAACACCATCCTCACTCTTTTTggttacctcttatctagt
gtagagtgtagcacctatccCTGATGCGTAATATTAGCCACTACCCTACTGGACACTAGCggttacctcttatctag
gtagagtgtagcacctatccAACCTTGTTTATTTGCCCTTTTTTACTGTCCTCACATTTGggttacctcttatctagtgg
gtagagtgtagcacctatccGGGTTGATACGCTGGTAATTTTGAGAGGCACTATTAAGCTggttacctcttatctagt
gtagagtgtagcacctatccCAGATGGCAGTGTGTGTTTTCTAGCTTTGACAACACGGTAggttacctcttatctagt
gtagagtgtagcacctatccAGATATATCTTATGTAAAATTCTCTCCTATGGTTGCACGGggttacctcttatctagtg
gtagagtgtagcacctatccGTATTCCACCTCACCGTTGCCACTAGCCTCCTGTCCTCTTggttacctcttatctag
gtagagtgtagcacctatccATTTTCAGCTTCTTTTTGGTTTCATACTTCTTTCCCCTCAggttacctcttatctagtgg
gtagagtgtagcacctatccCTTTCCATAGACAATAATTTCTCCTGTATTCTGAGTGGTTggttacctcttatctagtg
gtagagtgtagcacctatccATTCCCTTGACTTTCTGATGCTCACTACACTACTGATTCCggttacctcttatctagt
gtagagtgtagcacctatccGATTAAATCACTTGGTTCATTTTATGGCAACGGTTCCAGGggttacctcttatctagt
gtagagtgtagcacctatccAACAGCCATCTTTTCACCAATCCTTAGTAGACTTCATTCTggttacctcttatctagtg
gtagagtgtagcacctatccGAGATCTGCTCCGTCACAGCTAACTTCTCACTCATGATGAggttacctcttatctag
gtagagtgtagcacctatccCTTCCTAATTCACCCAGAAAGTCATAAGCTTACAAGGATCggttacctcttatctagt
gtagagtgtagcacctatccTTCCTCCTGCTTCTTTAGGCTAGTTTGAATTGAGTTTCTGggttacctcttatctagtg
gtagagtgtagcacctatccTTAAAATATTAAAAGCATACCTGCAGGACAGTGTGGTAGTggttacctcttatctagt
gtagagtgtagcacctatccCAAAAATGTGTTGTTCCAAAACAGAAAAAAACAGCCACTGggttacctcttatctag

gtagagtgtagcacctatccGTTTGGGACACATCTATAGCAAAAGAGAAGAGACAGACACggttacctcttatctag
gtagagtgtagcacctatccTTTCAATCACACTTAAGATAGGCATATACATAGACAGCTCggttacctcttatctagt
gtagagtgtagcacctatccTGAAAACTAACAAATAAATATGTGGCCTGTCCAAAAGGTCggttacctcttatctagt
gtagagtgtagcacctatccTGAACATTTCACTGAAGAGGGGAAAACCTAAAAGTTTCACggttacctcttatctagt
gtagagtgtagcacctatccCAACACTGAACAAATATCAGAAAATACTGAGGAGAAAAGCggttacctcttatctag
gtagagtgtagcacctatccGTTGTTTCTAATAAAATGTTCTGCCTACCTGAGCCCTGGAggttacctcttatctagt
gtagagtgtagcacctatccGATCATAATCATCTCCCCGTCCAGCCTCAGCATCGCAGCGggttacctcttatcta
gtagagtgtagcacctatccGGAGAAAACACATCTAGATTACAGCAGGAAAAAAGAAAACggttacctcttatctag
gtagagtgtagcacctatccAGTATGGGCATAGTGAAAGTAAGATCTAGGGGTTTTAGAAggttacctcttatctagt
gtagagtgtagcacctatccCATGCCCATATTGCTTGCTGACTGTATGTAGGGTGGATCAggttacctcttatctagt
gtagagtgtagcacctatccTCTTCTCTTCTCCTCTTCCACCTCTCTCTCTGTTTCTTTGggttacctcttatctagtg
gtagagtgtagcacctatccGTGGCGCAATCATGGCTCCCGACAGCCTCAACCTCTTGGGggttacctcttatcta
gtagagtgtagcacctatccCCACATGCCTGGCCTAAAGCTATTTCGACTCTCAAATGGAggttacctcttatctag
gtagagtgtagcacctatccGCTTAAAGTAGGCAGTTACATTGTCCTGTCAAAACAAAAAggttacctcttatctagt
gtagagtgtagcacctatccATCCCCTTCTACTGTTTTAATAAATATGCTTAGCAGTTCCggttacctcttatctagtg
gtagagtgtagcacctatccTAAGCATCCTCTGATCACACATTTTTTAAGAAGAAAAGGCggttacctcttatctagt
gtagagtgtagcacctatccTTGGGAGGGTGAGGCAGACAGATTCCTTGAGCTCAGGAGTggttacctcttatcta
gtagagtgtagcacctatccCTCTATCAAAAATACAAACAAACACAGAAATTAGCCAGGCggttacctcttatctag
gtagagtgtagcacctatccGTTCAGAATTCTGCTCATAGTTATAAAGTTATGCAGTAGGggttacctcttatctagtg
gtagagtgtagcacctatccAAAAATATCATGATGGGTGATTTATCTGTATTTGCCCTGGggttacctcttatctagtg
gtagagtgtagcacctatccCAATTTGCATGCACTGATTGTCTCATAGTCAAAAAGCTATggttacctcttatctagtg
gtagagtgtagcacctatccCAAAGACAGGATCCATTGATGCAGAGGTTATAAACGGTCGggttacctcttatctag
gtagagtgtagcacctatccCTTACTCCAGCTGTTTCACTTTTCTATATCCTCTGCTTGGggttacctcttatctagtg
gtagagtgtagcacctatccAGCATTTGACTGCTACCCTGTAATAGCACAATATTCATTTggttacctcttatctagtg
gtagagtgtagcacctatccGTAACAATACAAGCTTCATAAATGTGCACTATTGGGTGTGggttacctcttatctagt
gtagagtgtagcacctatccCCTTTCTGGCTATCATCTCTCTACTTAGACATCCATTCATggttacctcttatctagtg
gtagagtgtagcacctatccGTATTTTCTTTTTGCTACATTGGGAGACTATGCACCTTTCggttacctcttatctagtg
gtagagtgtagcacctatccCATTCTCCACTTAACTTTGTATCTAGTAAGTGCAAAGTACggttacctcttatctagtg
gtagagtgtagcacctatccTATCAAGTAGGGCAGAGAGATAAGTGGTCCAGTAGTCATAggttacctcttatctag
gtagagtgtagcacctatccATCATGTGGTGGTGGTGTTGATGGAAGTGAGATGGGGGTAggttacctcttatctag
gtagagtgtagcacctatccCCTGAGCTGAAGCCCGAGGTATGACAAGACAGATTAGGCTggttacctcttatcta
gtagagtgtagcacctatccCAGGTCAGAGTAAAATATGCAGGGGGAGGTAGAGAAGAGCggttacctcttatcta
gtagagtgtagcacctatccTGGCATGGGGGTGTCAGTGGGTGAGCAGACAGGGCTGAGTggttacctcttatcta
gtagagtgtagcacctatccTTTGGTATGTGGCAGCAAATGGAGAGCAAATGGTAGCATCggttacctcttatctag
gtagagtgtagcacctatccGATGAGACTGGAGAGGGAGACTGGGCTACCGTATTATAAGggttacctcttatcta
gtagagtgtagcacctatccTTAGATCTAGAAGTATTAATTAGCAGCAGTGTAAGGGATGggttacctcttatctagt
gtagagtgtagcacctatccAGGAAGAAGGCTACTGTAATTACCAAGGAAAATGATTAGGggttacctcttatctag
gtagagtgtagcacctatccTGGAGATGTAATGGAAAGAAAAAAGGACTCAAGAGAGATCggttacctcttatctag
gtagagtgtagcacctatccGACTACAGGCAGATGCCAACCTACCTGTCTAATTTTTGTAggttacctcttatctagt
gtagagtgtagcacctatccACTGTGCCCGGCCAAGATTGATGAATATAATACCTGACATggttacctcttatctag
gtagagtgtagcacctatccCTCTGAAGTATGTGTGAGAGATCCAAATGTCTGGTACGCAggttacctcttatctag
gtagagtgtagcacctatccAAAAATTTTGGGGGGTCATAAACATATACATGGTAGGTGAggttacctcttatctagt
gtagagtgtagcacctatccCCCAAGGTAAGGAAGTATTTATTAAGTAGGCAAGAATGGAggttacctcttatctagt
gtagagtgtagcacctatccGAAGAAGTAGTTAAAGATAAATGACTAGAAAGGAAGGAGGggttacctcttatctag
gtagagtgtagcacctatccTTAAGGGTTTTAAGGAAGGAATGGTTAGCAGAGTCAAGTGggttacctcttatctagt
gtagagtgtagcacctatccCAGGTTATTACTGATGGGAAGCAGCTTGTATACAGAAATAggttacctcttatctagt
gtagagtgtagcacctatccGGGAAACATTGTACCAAGCTATCTTATTTTTAACTTCAGGggttacctcttatctagtg
gtagagtgtagcacctatccTATAGCATACCTTTCTGCATTTGTAGCACACATAATATGCggttacctcttatctagtg
gtagagtgtagcacctatccTCATTGCATAGCCAGCTGGGTCATTATAAAACCTCACACCggttacctcttatctag
gtagagtgtagcacctatccGCTTCACTCTTATGCAGACCTTCATATTCCAATCTCATTAggttacctcttatctagtg
gtagagtgtagcacctatccTCTTCTGACATCCTCATAGAGTTCTTTTATTGGATCAAGTggttacctcttatctagtg
gtagagtgtagcacctatccATCATTTATTAACCAACTCAACAAACACTTAGCATGCACCggttacctcttatctagt

gtagagtgtagcacctatccCTGTAAATTCTTTAGAAGCCCATATTTTTCCAAGAGCAGGggttacctcttatctagtg
gtagagtgtagcacctatccTTTCCTGCGTCTGAAATATATCTCCTGTTGCCCTAGCTATggttacctcttatctagtg
gtagagtgtagcacctatccAACAAAGTTACCTTGTTGCCACAAATAAGACACGCTCTGTggttacctcttatctagt
gtagagtgtagcacctatccGCAGAACGCCAGGTGACAGAAGTGAACAGGTAGTATTTGCggttacctcttatctag
gtagagtgtagcacctatccGTATCTATTAATATTTGTAGCACCATATGATAGGGGAGAGggttacctcttatctagtg
gtagagtgtagcacctatccGTGCTAAGTCTTTCCACATTCTACCCTTTATATTAGTTTGggttacctcttatctagtgg
gtagagtgtagcacctatccAGATAAGAAAATAGGGAGAGCTTAATAGAACAAAAAGGGGggttacctcttatctag
gtagagtgtagcacctatccATTAATAGGGAAAGTACTGCCACATTCTTCAAAAACAGCTggttacctcttatctagtg
gtagagtgtagcacctatccAGCTTGCCAAAACAACTTCCTACTTGCTGACTACCTTTGTggttacctcttatctagtg
gtagagtgtagcacctatccTGTCAAAATGCCAAGAACTTTCTCCTTGTTTTAACTAAGCggttacctcttatctagtg
gtagagtgtagcacctatccGTACTGAGGGTTCCAATTTCACTCTTTAATGGCAGGTATCggttacctcttatctagtg
gtagagtgtagcacctatccGAGGATCATTCTTCTTTTTGGTAGCCGTTGTCATGTTATGggttacctcttatctagtgg
gtagagtgtagcacctatccTCTCTAGTGGTCTGAGAGTAGAGATGCATGCCCTTTAATGggttacctcttatctagtg
gtagagtgtagcacctatccTGAGATGCATTATAACAGAGAGATTATATTCTCAGGTTCCggttacctcttatctagtg
gtagagtgtagcacctatccATGAGAAGGGAAGAAACTGCTAGGGGCTGGTGGGTAAAACggttacctcttatctag
gtagagtgtagcacctatccAAGTACATGATTTCAGATTCTTTCCATACCTTGCAAATGGggttacctcttatctagtg
gtagagtgtagcacctatccGCCAAGCCATCGATTTTCTAATACTCGCCGACAGCACTGTggttacctcttatctag
gtagagtgtagcacctatccTGACTGGGAGATTATTTAATGCAAGGCACTACTTAAAGCAggttacctcttatctagtg
gtagagtgtagcacctatccTCATTACAGTAAATTGGGATACTATTAACTAGGCCTTTGGggttacctcttatctagtg
gtagagtgtagcacctatccTATCTTGGAATCATGAGAACTGAAAATATAAGCAATGGGCggttacctcttatctagtg
gtagagtgtagcacctatccGAAATTAAGAAGAGCCTTAGCTACTGGGGGATAATCTAATggttacctcttatctagtg
gtagagtgtagcacctatccCCCCCTGGCTTCTGAATATTTTATACATTCTGTTGATTACggttacctcttatctagtgg
gtagagtgtagcacctatccTCACACAAATCACAAATTAAAGCTGAGCATGCAACATAGAggttacctcttatctagt
gtagagtgtagcacctatccCTCCTCATTAGCCTGGTACATGTATAGATTAAAAGAACACggttacctcttatctagtg
gtagagtgtagcacctatccCCATCTCAGATCACTGCAAGCTCCGTCTCCTGGCTTCATGggttacctcttatctag
gtagagtgtagcacctatccCCCGGCCTCAATCAATGTTTATTTGACAAATATATTCCCCggttacctcttatctagtg
gtagagtgtagcacctatccGTCTGGATACATTCAATACTTTAAAAGCTGAGGCAACCTGggttacctcttatctagtg
gtagagtgtagcacctatccCGAGAGCGCTTCGGTGAAACATATCATGCACATGTCATCGggttacctcttatctag
gtagagtgtagcacctatccCAGACAGTGCTCTTCGTTTTTAACACCCCCGCATGGATGGggttacctcttatctag
gtagagtgtagcacctatccCAGGCTATCTTAGCGTATTCCTGTTAAAGATGAATTACAAggttacctcttatctagtg
gtagagtgtagcacctatccCCCACAGGAAGTATCTATGCAGGCCAAGTAACAACAAAGTggttacctcttatctag
gtagagtgtagcacctatccTAAAAAATGCCACTCTCTATAAAGGGAATGCTGATAAGTGggttacctcttatctagtg
gtagagtgtagcacctatccTTTGTCATCATGCACAGCTATAAAGCAAAAAGTTCCTGGCggttacctcttatctagtg
gtagagtgtagcacctatccATAAAAAAAGTTCCCCAATGTATAATTTCCCAGTCACCACggttacctcttatctagtg
gtagagtgtagcacctatccAAGGTACCTTTGACTCTCTGAGCATATCCACATTCTTATTggttacctcttatctagtg
gtagagtgtagcacctatccTATGATCATGGTAACTGAGAAACATACAAAAAGAGTGTGGggttacctcttatctagtg
gtagagtgtagcacctatccCTTGAGAGCCTTATTTGACCCATTCTTCAACTTTGGTAAGggttacctcttatctagtg
gtagagtgtagcacctatccGAGTGACGCTGAGCTAAAACTCTTTTCTGTATGAAGTAAAggttacctcttatctagtg
gtagagtgtagcacctatccGTGTCACTATATACTCACCTGGCAATCTGCATCAGAACAAggttacctcttatctagt
gtagagtgtagcacctatccGCTACTCGTGGTGGGTTTGCCTAGGTTCAAGCAGAAAAATggttacctcttatctagt
gtagagtgtagcacctatccACTAGGTTTGTATAAGTTGCTATCCTTGGGAAATCAGCCTggttacctcttatctagtg
gtagagtgtagcacctatccAAGAAATTAAAGACCTATCTGAAGCCAAATACACTCTCGAggttacctcttatctagt
gtagagtgtagcacctatccCTGTTACAAGCCCCTGACTCCAAATGCACAATACCATAAAggttacctcttatctag
gtagagtgtagcacctatccGCATGGTATAAAGCCTATCATCAGAGTATGAAACTTGAAAggttacctcttatctagtg
gtagagtgtagcacctatccCCTACCTCAGGAACTCTGCAATTTGAAAAGGAATTTAAATggttacctcttatctagtg
gtagagtgtagcacctatccGAAAAAGAACGTGAAGATTTGAAAATTACAGACCAAGCTGggttacctcttatctagt
gtagagtgtagcacctatccTTTATTAATGCAGTATACAGCACTGACAGATTATAGGAGGggttacctcttatctagtg
gtagagtgtagcacctatccCAGCATGCAAAGTCACCCTAGAATTAAATTGATTCGGATTggttacctcttatctagtg
gtagagtgtagcacctatccCTAACTCCAACACTGGAAGTTCGTGTCAGTCAAATGGCAGggttacctcttatctag
gtagagtgtagcacctatccTTTTATTGCTATCAATGGAAACAGTACCATTCACAACACCggttacctcttatctagtg
gtagagtgtagcacctatccATCTTAGTGATCATCTAGTGCAGCAATTTTTATTGTGGGGggttacctcttatctagtg
gtagagtgtagcacctatccGGGAATTATCAATAGAAATGTATTTTATCTATGGAGCCCCggttacctcttatctagtg

gtagagtgtagcacctatccCTCACTTTATAAATGAAGTCAATGAGTTCCAGAGGTTAAGggttacctcttatctagtg
gtagagtgtagcacctatccCACAGCTAGTTAGTGGCAAATCAGGTTTCTTGACTTTTGGggttacctcttatctagtg
gtagagtgtagcacctatccTATCACTCAAAATTATTCACAGTTTAATCAGTTGGCACGGggttacctcttatctagtg
gtagagtgtagcacctatccAGGCATGGCTCAATCTTTAAACCTTTTACCTGTGTGTTCTggttacctcttatctagtg
gtagagtgtagcacctatccTCCACCATTGCCTTCATTGTTTTAGAATCTGCCAGTGCCAggttacctcttatctagtg
gtagagtgtagcacctatccGGTCAACCTTTATAGCTTCTTCTTCTTCTTTGAAGACCTAggttacctcttatctagtgg
gtagagtgtagcacctatccAGAACAGGGAAAAATATGCTTATTTATCACTCTGAAAGGGggttacctcttatctagtg
gtagagtgtagcacctatccTGAATATTTCCAATCTTCTCTTTAATCTTCACAGCAGTCCggttacctcttatctagtgg
gtagagtgtagcacctatccTATTTATAGATGGTAAGCGAGACAGCTGTGAACTCAACCAggttacctcttatctagt
gtagagtgtagcacctatccAGTCCACTTTTTCCCCCTACTACACAAAGCCTCTTTCACAggttacctcttatctagt
gtagagtgtagcacctatccAATCTTCACCTGTCTTTGATGAGTTTTGGTTCTTCGATGAggttacctcttatctagtgg
gtagagtgtagcacctatccCATTGCATAAAATGATTGCTGCAGTTTCACCATCATCATGggttacctcttatctagtg
gtagagtgtagcacctatccTAAGACATTTCCACGTTACTTAGATTAACAAGCAAGTCTCggttacctcttatctagtg
gtagagtgtagcacctatccATCTCTCAAAACTCCAAATCCAATCATTAATGTGCTTTGCggttacctcttatctagtg
gtagagtgtagcacctatccATGTCTGCCCCATTATATATCTAAACATTCTGTATGCCAAggttacctcttatctagtg
gtagagtgtagcacctatccAGTTTTCCTTTCTAGTGAACTTAAAGATACCCATCAAACCggttacctcttatctagtg
gtagagtgtagcacctatccGGTGGTGTGTCATGTTGCCTGTGTTATTGCTTTATTTCCTggttacctcttatctagtgg
gtagagtgtagcacctatccTTTTGATATGATGCAAACATGACTCAGAGAAGAAGAAAGCggttacctcttatctagtg
gtagagtgtagcacctatccAATAAAAGCCCCTAAAGTTTCCATATACTATGAAGTAGGGggttacctcttatctagtg
gtagagtgtagcacctatccCGAAGGGACACAATCTTTCAAGAGATCCACTAACTGTTTAggttacctcttatctagt
gtagagtgtagcacctatccATTCTGATCTTACCATTTGTTTTTGTTGTCCATCCTTTCCggttacctcttatctagtgg
gtagagtgtagcacctatccGCTGTTGAAAGGGAAGGCTGTTACACCACTGAATTTTTAGggttacctcttatctagtg
gtagagtgtagcacctatccAAATGCTGGCTGATTTTGAAAATACATAGATACTCAGTGGggttacctcttatctagtg
gtagagtgtagcacctatccTGATGTGGTAGCAGGCATACACGATCACTGTCCTATGTGCggttacctcttatctag
gtagagtgtagcacctatccAGTGACTGTTTCTCTGATGAACCACCCTGTCTTCTATCTGggttacctcttatctagtg
gtagagtgtagcacctatccATTTAGTATCATTAAATGCCAAAGAAGGTGGTACAGCAACggttacctcttatctagtg
gtagagtgtagcacctatccTGTACAGTCCCATGAGTAGTCAGGTACACATTAAATCCTTggttacctcttatctagtg
gtagagtgtagcacctatccAAGTTCCTCAGAAGTCTTTGTTGCCCCTTGTTGCCCCAGAggttacctcttatctagt
gtagagtgtagcacctatccACTATACTTGGTGCACTTTGTCAGAAATACATACATGTCTggttacctcttatctagtg
gtagagtgtagcacctatccTCACCTGAGGTCAGGGGGTTATGACCAGCCTGACTAACATggttacctcttatctag
gtagagtgtagcacctatccAGGTGTGGTGGTATATGCCTGTAATCCGAGCTACTTGGGAggttacctcttatctagt
gtagagtgtagcacctatccAGGGTAAGCTTCAGTTTTATATAGAATGATATATCCCCTCggttacctcttatctagtg
gtagagtgtagcacctatccGAATAGAAAATGACTTAACTAGCTCAACTCTAAGAAGTGCggttacctcttatctagtg
gtagagtgtagcacctatccAAATAGAAAAAGAAATAATGCATGCAAGGAGGAAGGGGAAggttacctcttatctag
gtagagtgtagcacctatccACATGAGTGGTAGGAAAGGATTACTGTCAAAAATACCTATggttacctcttatctagtg
gtagagtgtagcacctatccCAATAATAGCAAGATAGCATAAATTGACTTTTTCCGCCCTggttacctcttatctagtg
gtagagtgtagcacctatccGCAATTTTTACCAGCATTCACTTCTTCCTAACGTTGTCAGggttacctcttatctagtg
gtagagtgtagcacctatccCTTCAAGGAAATCTCAAATCTCTTCTGGAAGTATATGGGAggttacctcttatctagtg
gtagagtgtagcacctatccATCTCATTTCCTCAAAGATTTCCACCCTGACCCTGCCCTGggttacctcttatctagt
gtagagtgtagcacctatccGTAAGTTACAGGGCTCTACATTAGGAACAACCAGGCATAGggttacctcttatctag
gtagagtgtagcacctatccGTGGTAAACTACAGCAAACAGATCATTAGAAAGACAACACggttacctcttatctag
gtagagtgtagcacctatccCCAGTGTACCCTTGGCTTCCTAAACAATAACAAAAACAAGggttacctcttatctagt
gtagagtgtagcacctatccAACAGTGAGAAGTAACGACTAGTTCATAATATAAAAGCCCggttacctcttatctagt
gtagagtgtagcacctatccCAAAACAAAGCAAAACCTACTCTAGTTGTTGCAATACATCggttacctcttatctagtg
gtagagtgtagcacctatccGGCAATGATGGTTTCTGCAATAGCATTTTTTGTCACTCGGggttacctcttatctagtg
gtagagtgtagcacctatccTGACAGATGACCCTATTGAGCAGAACAGAGTATTTTAATAggttacctcttatctagtg
gtagagtgtagcacctatccCTCCTATCAAAGGAAATATTTGTTTTTAGTGAGACCATGGggttacctcttatctagtg
gtagagtgtagcacctatccATGGGCACAAACTTTTGAACTAAAATTTCTTCGTGATCCTggttacctcttatctagtg
gtagagtgtagcacctatccAAATTAAGTTCTCAATACTCTTTTCCTGGAAATCAAGGGCggttacctcttatctagtg
gtagagtgtagcacctatccGAGAAGTTTGTCACTTAGCATAAAATTTACACAGTGAAGCggttacctcttatctagtg
gtagagtgtagcacctatccTATAAACAGGTTCAAACTGCAACTACCTTTATTTGCTACCggttacctcttatctagtg
gtagagtgtagcacctatccTATTTTAATTCAATGAACACCCTGAGAAAGTCTGTTTGCCggttacctcttatctagtg

gtagagtgtagcacctatccGGTTGAGAAAATATCCACGCTGTTGGGAAAAGTGATAGAAggttacctcttatctagt
gtagagtgtagcacctatccTTCCCCCAGAAAATCCTCCCACTTCTCATATCCACATGATggttacctcttatctagt
gtagagtgtagcacctatccCAGTTCAATTCCTCCTCCTTCTCCAAGCTGCCTTCTATTCggttacctcttatctagt
gtagagtgtagcacctatccACTCTACATTCCATACTATCTTGTCCTTCTCAGGCTGCCCggttacctcttatctagt
gtagagtgtagcacctatccCAATGTGTGGAAAATGACAAGGAGCAACTAATCAGAGCCAggttacctcttatctag
gtagagtgtagcacctatccTGTAAAGTAACTCCCTGTCCTCCACACTCCCAGATCTCCTggttacctcttatctag
gtagagtgtagcacctatccCATGAGGACTGAGGAGAATGAGAAACTTCCACTTTCGAGAggttacctcttatctag
gtagagtgtagcacctatccCCTTTAGGTTTTTTGAATCAAGTTCTTTCTTAGGGTCTGAggttacctcttatctagtg
gtagagtgtagcacctatccGTCACGTAACAATTCCCACACAAATTAACTGGAAAATTTGggttacctcttatctagt
gtagagtgtagcacctatccGACAAATTATCACTCACTGAAAAGACAAACAGCATGATAGggttacctcttatctagt
gtagagtgtagcacctatccAGTTACACTAATCTACATCTCCTATCAATTCAGCAGAGTAggttacctcttatctagt
gtagagtgtagcacctatccCTTACTGCTGCCATATCCTTGATAAGTTTAATGATGATCTggttacctcttatctagtg
gtagagtgtagcacctatccCGTCGGAGATAACTACAAGAAAAAATTAGCACTTAAAAAGCggttacctcttatctagt
gtagagtgtagcacctatccAAAAGAAAATTCATCAAACCATTACCCCAAATCCCAAGACggttacctcttatctagt
gtagagtgtagcacctatccGAGTTCCCTTTCAAGTATTTCTCTGTTTGCACATTATTTCggttacctcttatctagtg
gtagagtgtagcacctatccTCCAGGGACCACACACGACTTATTTCTAGGGCACATGCGGggttacctcttatcta
gtagagtgtagcacctatccAATTCACTGTATCACTAAGCACACTACTATTTAGTTCAGCggttacctcttatctagtg
gtagagtgtagcacctatccCCTAGCAAGACTACAGGAAAGAAGTGATACAATGATTAACggttacctcttatctagt
gtagagtgtagcacctatccTGTTATCTGCATATTGTCATGGATTGGCTCTACAAGTAGCggttacctcttatctagtg
gtagagtgtagcacctatccGTTACACACAATTACAGAACATCCTTGCATATATACCTTGggttacctcttatctagt
gtagagtgtagcacctatccAACAGATATCCAAAAAGGCTGCAGCAATTTATATGAGATGggttacctcttatctagt
gtagagtgtagcacctatccGCCTATTTCCTCATAATGTTGCCAAAGCATCAAAGATATTggttacctcttatctagtg
gtagagtgtagcacctatccTTCTCTGATTAGTAGTGAGGTGTTTTCATTTGTTTACTGGggttacctcttatctagtgg
gtagagtgtagcacctatccCTGTGAACTGACTACTGGCATTTATAAGTTTTTTTCTGTGggttacctcttatctagtg
gtagagtgtagcacctatccCCTATATGATTTTGGGTTCTGTGTTTTGTTTAAGAAGGTCggttacctcttatctagtg
gtagagtgtagcacctatccTAAAGCTTTAGTTGACCTGAACTTTACTTTATTCCCCCATggttacctcttatctagtg
gtagagtgtagcacctatccGGATATCCAAATATTCTGACATCACATATTCAGCCTACTTggttacctcttatctagtg
gtagagtgtagcacctatccAGCCAGTTGTTTATGGATTTTATTTGGTGAGTAACCTAACggttacctcttatctagtg
gtagagtgtagcacctatccCTTAAATTTCTGAACAGTAGCTCACTCTATTCTGTTTTGGggttacctcttatctagtg
gtagagtgtagcacctatccATAACTAACCTGGAATTGAAAATCATCGGAAGAGTAACTGggttacctcttatctagt
gtagagtgtagcacctatccGGTGGTTCCTTTGGCTTTTAGCTGTACAGTGAGTGCTTTGggttacctcttatctagt
gtagagtgtagcacctatccGCATCCTAGAAACATGTCCAAGATACCCAGCTTATTCCAGggttacctcttatctag
gtagagtgtagcacctatccCTCCTATGATGCTGGCCAGACGACTAGGGGTTACTTCAGGggttacctcttatcta
gtagagtgtagcacctatccAGGCATAAGTCACTGTACCCCATCCCAGGCTTCATTTTTTggttacctcttatctagt
gtagagtgtagcacctatccATTCAAATGGTAATAGTAGTGGTAATTACTAACCCAAGGCggttacctcttatctagt
gtagagtgtagcacctatccATGGTAATATGGTAATCAGTAGCCAGAATACTCTCTTATCggttacctcttatctagtg
gtagagtgtagcacctatccCTCACTATAAACAGTAAAAAAGGGAGTAAAATACAGGCCCggttacctcttatctag
gtagagtgtagcacctatccATCTATACTTAGGGTCCTTAGATAATTCTCTTTGCATTGCggttacctcttatctagtg
gtagagtgtagcacctatccGTTTCAAGTAACAAAAGCAATGGAAAGGCTGTAGTGTGATggttacctcttatctagt
gtagagtgtagcacctatccTGTGTGAAGCAGCGAGAAGAGATCCTGAAGCAGGGTTAGCggttacctcttatcta
gtagagtgtagcacctatccGCCAACGTTAGAGCCACTCAGTGCTAAAACCATAGAGAGCggttacctcttatcta
gtagagtgtagcacctatccGGCATCTTCATCATTTGGCTGAGAATTGGCATTTTCTTTGggttacctcttatctagtg
gtagagtgtagcacctatccATGTTCCCATTCTTCACGGACTCTGGTAGCTTCCATGCGAggttacctcttatctagt
gtagagtgtagcacctatccTTGGACAATATGAGCACACACCTATTTAAAGTTAACACAGggttacctcttatctagt
gtagagtgtagcacctatccTATAGGGTTAGTTTGATTTACTACTCCTAGCAAAACAAGCggttacctcttatctagtg
gtagagtgtagcacctatccTACTATTCTCCCAGTTAATTCTCCATCATATGTCATTCTCggttacctcttatctagtg
gtagagtgtagcacctatccAAGAAACAAGTAAAGCTAAGGGAGCCACAATAAAAATGGAggttacctcttatctag
gtagagtgtagcacctatccGTTTAGCACGTCAAAAACCAATCACCATATTTTATCACTGggttacctcttatctagt
gtagagtgtagcacctatccTCTCATGGTTTACCATTATGTTCATAAGGTGCTTTATAGCggttacctcttatctagtg
gtagagtgtagcacctatccGCATCTAAGATTCAATGAAACACTGCTAGAAAGATAAAGCggttacctcttatctagt
gtagagtgtagcacctatccTGAAAAGAATAATAGTGTAGAAAGTTAGAATAGGGCTGGGggttacctcttatctagt
gtagagtgtagcacctatccAAATACAAAAAAATTACACCTGTAATCCCAGCAGTTTGGGggttacctcttatctagt

gtagagtgtagcacctatccAAATACAAAAATATTATGCCTGTAATCCCAGCACTTTGGGggttacctcttatctagtg
gtagagtgtagcacctatccAAAATTAGCCAGCCGTGGTGGCAGGCATCTGTAGTCCCAGggttacctcttatctag
gtagagtgtagcacctatccTTATGTTATGTAGCACCCCATTATATGAACAATATAGCGCggttacctcttatctagtg
gtagagtgtagcacctatccACTGGTATAGTTAACACTGTTAATTGCTCAGCCAACACACggttacctcttatctagtg
gtagagtgtagcacctatccCACCAAATAAAAAATCTACCCATAACAAATAGCAGCAAGGggttacctcttatctag
gtagagtgtagcacctatccACTCATTCCTCTCTAAAAAATGGCAGGCACATTCTGCAGGggttacctcttatctagt
gtagagtgtagcacctatccTTTTTCAAAATACTTTATACCAAGCTCATCCAACCTGCAGggttacctcttatctagtg
gtagagtgtagcacctatccGATTGGACATCCCTGCTTTATACCCTAATAACAAACAAACggttacctcttatctagtg
gtagagtgtagcacctatccGAAATAGCTGTTAACCAACCTGTTTTTGTAAGTTAGTCAGggttacctcttatctagtg
gtagagtgtagcacctatccTGAATATGATTCCAACCATATGTTCTTTCAGGTCAGCATCggttacctcttatctagtg
gtagagtgtagcacctatccCTCAGAACTTAGGAAATTCAGAGAAGTATATTCTCCTTTCggttacctcttatctagtg
gtagagtgtagcacctatccAGTTCACATCCCACATTTAATTGTTTGAAGAACAATCCAGggttacctcttatctagtg
gtagagtgtagcacctatccGAAAGTGTCCTAACAAACCTCTGTGATGATAGTGGTCTTAggttacctcttatctagtg
gtagagtgtagcacctatccAGCTATCGGAACACCAAGTCATCAAAGAAATGTATCATAAggttacctcttatctagt
gtagagtgtagcacctatccTTTATAGAGCTTATTATGTGCACGCTGTTCTTCTAAGTGCggttacctcttatctagtgg
gtagagtgtagcacctatccCACACACACACAAATCTTATCTAATCTTCGCAACAACCCCggttacctcttatctag
gtagagtgtagcacctatccAATGATGCTTGGTTTTACATTTCTTAAGATAGAGAGCCAGggttacctcttatctagtg
gtagagtgtagcacctatccTGTCCCTTTCGCCCTAACAATGAGGAAAAACCTGATCATCggttacctcttatctagt
gtagagtgtagcacctatccATTAGAATGTTGAGATCAGCTACCAGGTAGAATGAATTCCggttacctcttatctagtg
gtagagtgtagcacctatccCCAGTTAGTTGGATACTTCAAAACTCTTCTTTCAGTACTTggttacctcttatctagtgg
gtagagtgtagcacctatccACAAGCAAAAAACCAAAAACCAGTAAGGCTCTTAAAGACTggttacctcttatctag
gtagagtgtagcacctatccAACTGATATTTATAGAACCCCTCAAAGAAGAACAGCAGAGggttacctcttatctagt
gtagagtgtagcacctatccTGGACATTCACTAAAACAGACTATATTATAGGCCAGAATCggttacctcttatctagtg
gtagagtgtagcacctatccGGACTGAAATTATACGAAGTATACAATACAGTTTGACCACggttacctcttatctagtg
gtagagtgtagcacctatccTGGGAATTAAACAAGGTGCTTCTAAATTACCCAAGGGTCAggttacctcttatctagt
gtagagtgtagcacctatccGAAATCAAAATGGGGCCGGATACAGTGGCTCACGCCTGTAggttacctcttatctag
gtagagtgtagcacctatccAAACAGAATACCCCAAATTTGTGAAATGCAGCTAATGTAGggttacctcttatctagtg
gtagagtgtagcacctatccAAATACTTAGATGAGAAAAGAAAGGTCTTGGAGTCTACGAggttacctcttatctagtg
gtagagtgtagcacctatccCAGAAAAAGATGAACAAATGAATACAAGGCAAACATAGGGggttacctcttatctag
gtagagtgtagcacctatccTGAAACAAAAAGCTAGTTCTCTGAAGAGATCAAACTGATGggttacctcttatctagtg
gtagagtgtagcacctatccATGAAGACACAAATCACCAAGACACATCACTACAGGTTCTggttacctcttatctag
gtagagtgtagcacctatccAGTGTCAGAACTACTAAAGCTTACCCAAAAAGAAACAGATggttacctcttatctagt
gtagagtgtagcacctatccAATGGTTTTTAGTACATTCAGATATGTGCAACTATCGTCGggttacctcttatctagtg
gtagagtgtagcacctatccCTATATCATGTAGCTACCATCCCCTAGTTTTTTTTTTCACggttacctcttatctagtgg
gtagagtgtagcacctatccGAACTGTATACAAGCTAGTATCACATGTTGACTAAAATGCggttacctcttatctagtg
gtagagtgtagcacctatccGAGAAACATATAAAGGTACCACTGTGCTCTTGTCTGGGTGggttacctcttatctagt
gtagagtgtagcacctatccGTTAAGACCAGGTCTCTTAAAAAAGAATCCATTCTGATAGggttacctcttatctagtg
gtagagtgtagcacctatccGATTCACCCCCAAAAAATCTACAGTTCACATCACACTTAAggttacctcttatctagt
gtagagtgtagcacctatccGGTTTATACTTTCTTTTCAAGATCCAGAACAAGTCAGAGCggttacctcttatctagtg
gtagagtgtagcacctatccCATGGCGCTTAAGGGCTTAATCAATGCAATAACACAATAAggttacctcttatctagt
gtagagtgtagcacctatccTTGAAGACTACATGATTATCTAAGAAAACCCCAAAGAGTCggttacctcttatctagtg
gtagagtgtagcacctatccCTGCTGAGAGAAATTTTAAACAGATGTAAGAAAACCCCTGggttacctcttatctagt
gtagagtgtagcacctatccTTGACATGGCGATTCTCCCTAAACTATCAATTTGGTGTTAggttacctcttatctagtg
gtagagtgtagcacctatccCCCCCTTTTTTTTTTAAAGAAACGGTCAAGCTGATTCTAAggttacctcttatctagtg
gtagagtgtagcacctatccATAAAGCTACAGTAAGTAAGACACTTTGATGGATCAATGGggttacctcttatctagtg
gtagagtgtagcacctatccGTCCAGAAACAGACCTACCATTTATGGTAAATTGACTATTggttacctcttatctagtg
gtagagtgtagcacctatccCATCTATATGCAAAAGATAAGCCTCAATTCTTACACTGCAggttacctcttatctagtg
gtagagtgtagcacctatccAATATCTGTGTGACCTTGGGTTCTAGACCATACATAAAAAggttacctcttatctagtg
gtagagtgtagcacctatccAACTTCTGTTCTCTGAAAGATAAGACACAGACTAGGAGAAggttacctcttatctagt
gtagagtgtagcacctatccGTGTTCACAGCAGCTTTACTAATAGTAGCCAAACAGCAGAggttacctcttatctagt
gtagagtgtagcacctatccCATATAATGGAACACTTCTCAGCAACCAAAAGAATGAACCggttacctcttatctagt
gtagagtgtagcacctatccGTGAAAGAAACAAGACAAAAAGGCTACATTCTGAGTGGTTggttacctcttatctagt

gtagagtgtagcacctatccCAGGAAAAGGCAACACAACACTGGCAGAATATAGTTCAGTggttacctcttatctag
gtagagtgtagcacctatccAGAGAAATTTGGGGAGTGATGTGTTTTAAAACTTGGTTGTggttacctcttatctagtg
gtagagtgtagcacctatccGCAAAACCACATCTCTGCAGATATATTAAAATCAGCTGGGggttacctcttatctag
gtagagtgtagcacctatccTGGGAGGCTGAGACAGGAGGATCCATTGAGCCCGGGAGGTggttacctcttatct
gtagagtgtagcacctatccTGGATGACAGAGAGAGACCCGGTCTCAAGCAAAACAAAACggttacctcttatcta
gtagagtgtagcacctatccTCTCTTAAAAATCATGTTATTTCATGGGCACACACATGACggttacctcttatctagt
gtagagtgtagcacctatccAACCTTTCCTACTCAACCTTAACTCCTCAGCATACACTCTggttacctcttatctagt
gtagagtgtagcacctatccAGTCTCTCTGATCAGTCCTTGCTCCCATTCTCTCTACTAAggttacctcttatctagt
gtagagtgtagcacctatccGAATCCCTTCATTCTCTTTCAGATCTAACTCCAGTCCAATggttacctcttatctagt
gtagagtgtagcacctatccGGCACAATGATCTCAAGGTACTCTGAAAGCCAATGTCATCggttacctcttatctag
gtagagtgtagcacctatccCTATATAGTATCTTTGTTGATTTCTTTTTGGGGGGGGTGAggttacctcttatctagtg
gtagagtgtagcacctatccAGCTTTTTGAGGGCTTGCCATTCTCACAATTGTTTAGTACggttacctcttatctagtg
gtagagtgtagcacctatccCTGCTAAAATTCAATTGACTTAACAGGTAGTTACACAGTCggttacctcttatctagt
gtagagtgtagcacctatccCAAATGAAATCTCCAAAAGATGTTGAAAGAAGTTACCAGCggttacctcttatctagt
gtagagtgtagcacctatccTCAGACCTCAAACAAAAACAGGACAATGATTAGATACTACggttacctcttatctag
gtagagtgtagcacctatccCTGCCAAGGTGATTATTTGTAGCATTTAAAGACACTAGAAggttacctcttatctagt
gtagagtgtagcacctatccGAAGTACAAAAGTTTTAGGTATAGTGTTTTAGTCAGGACCggttacctcttatctagtg
gtagagtgtagcacctatccCATGTGTGTGATGCTACCATGATGGATACAAAGACAAATTggttacctcttatctagt
gtagagtgtagcacctatccCTTGGTCTTTAAGAAGCTATGGTTCAGTAGAAGAGAGAAAggttacctcttatctagt
gtagagtgtagcacctatccCAGAGAGAAAACTGCTTTTGTTTGGGGAAATCATAAACAAggttacctcttatctagt
gtagagtgtagcacctatccTGTGAAGGACAGGGACAATGGAGGAGGGAGAAGATAACTGggttacctcttatcta
gtagagtgtagcacctatccACAAATGAATGCAAAAACCCAAGGTATAGAGCAAATTTGCggttacctcttatctag
gtagagtgtagcacctatccCATTGTATACTTTTTCTTCTCCTTCAGGTGATGACAATAGggttacctcttatctagtg
gtagagtgtagcacctatccAGAGATGAGCTTTCCAAATACCTGATGAAGATATGCATGTggttacctcttatctagt
gtagagtgtagcacctatccGCATTAACACTATATTGGCCAGAGGACTCATTAGCTGATTggttacctcttatctagt
gtagagtgtagcacctatccCATGGAACTCAGGTGTTTAGGGCTATGAAAAGAAAAAATGggttacctcttatctagt
gtagagtgtagcacctatccTACATCAATTATCCTGTGCTAGAATATGTTGTAAGGACACggttacctcttatctagtg
gtagagtgtagcacctatccCAGAGTCTCAGCTTCACAGTTCCTCTGCTGGGCCACACTGggttacctcttatcta
gtagagtgtagcacctatccTGCTTTCACCATCTTTCCAGCCCAGGACTTTGACTTGTCGggttacctcttatctagt
gtagagtgtagcacctatccAATTTTTATGATGTGATTATCCCCTCCTGGAAGTTCACTTggttacctcttatctagtg
gtagagtgtagcacctatccCTGGTTGAGTAGAAAAGAGGCAGGAACCTGATTCAGCTGGggttacctcttatcta
gtagagtgtagcacctatccTGGACTTAGCATTGATACCAGCAAGCTCAAATAACAAAGAggttacctcttatctag
gtagagtgtagcacctatccTGAGGTTATTTCACACTGATTTTCTTATTTCAACCAGGTGggttacctcttatctagtg
gtagagtgtagcacctatccCTAAGGTTAATACAGGTTAAGGGATTTTCAGTAAATGGTGggttacctcttatctagtg
gtagagtgtagcacctatccGAATTAAAATCCAGGCCTGACTACGATATGGCTTCCGAGCggttacctcttatctag
gtagagtgtagcacctatccTGACTGTACTTTATCTATATCATCTGTAAGTGGAAACCTCggttacctcttatctagtg
gtagagtgtagcacctatccGTGCATCCTCACATTAAAGTACCATTTAAAGATAAGGGACggttacctcttatctagt
gtagagtgtagcacctatccAACTGTTCTTATGGCATTTAATAACCAAGAAATGACCTGGggttacctcttatctagt
gtagagtgtagcacctatccTTGATTCCTTCAATGTGATTTGAAAACAAGCAAGCAAGCAggttacctcttatctagt
gtagagtgtagcacctatccCAGTGAGGAACAGATTTGACAAAGTTTCATGAATACAAGTggttacctcttatctagt
gtagagtgtagcacctatccGCACAGTCTCAGACACATGCCTAGATAAAACTTAACTTGTggttacctcttatctagt
gtagagtgtagcacctatccTATAAGTGTTATAACAGACAGATTTTTCAGGCCAGAGGACggttacctcttatctagt
gtagagtgtagcacctatccACTCAAAGTATAGGTGAAGTAAGGTTTTAAGTGTGAATGGggttacctcttatctagt
gtagagtgtagcacctatccGTCTTTGTTGGGGAGTAGGGTAGGGAGACTGTAGGTACAAggttacctcttatctag
gtagagtgtagcacctatccTCATGACTGGGGAACATTTAGGAAAACAGGTGAGATGTCCggttacctcttatctag
gtagagtgtagcacctatccTGGTTACTATTCCTATGTTTTGGACCACTTGTCAATTTAGggttacctcttatctagtg
gtagagtgtagcacctatccTCCCTATAATACTTTGGTTGTGTGCCAAGGGTGGAATTCAggttacctcttatctagt
gtagagtgtagcacctatccTTTCAATTACCTGCTTTATTCTGCACAAATCTTCTACTGCggttacctcttatctagtg
gtagagtgtagcacctatccATATGCTATATTCCTGCTTCAGAACTTACAATGGTTTCTCggttacctcttatctagtg
gtagagtgtagcacctatccGTCCTTCCTGATTGACCCCACCCCATCCCTCCAACTAACCggttacctcttatcta
gtagagtgtagcacctatccACTTTAGTCTCATTCTCACAAACTCCATCTTAAATTGCACggttacctcttatctagtg
gtagagtgtagcacctatccGCTCTCTCACAGCTTGACATTCCCTCTCCTTATTCAAAGTggttacctcttatctagt

gtagagtgtagcacctatccAGCCTTTCAACAAATCTGCTTGTCACTTATGTCAGACTCTggttacctcttatctagtg
gtagagtgtagcacctatccTGGAAGTATAAAGAGGGAAGGTAACTGAACTAACAAATACggttacctcttatctagt
gtagagtgtagcacctatccCTTCCTTACAGATTTGAAGTAGGATATATGAGAGGAGGTTggttacctcttatctagtg
gtagagtgtagcacctatccGCCAAACTAGGGAAGGCAACATTAATACACAAAATAAATGggttacctcttatctagt
gtagagtgtagcacctatccGTGTCAAATATCTTAAGGCTGAATAAGTTTTACGCAAAGCggttacctcttatctagtg
gtagagtgtagcacctatccATTAACATTAATTCCTTGGAGAGCTGCATAATATGAGTGCggttacctcttatctagtg
gtagagtgtagcacctatccGCTGATACAACTCAAATCAACAAATGCTGTGTACAAGGCTggttacctcttatctagt
gtagagtgtagcacctatccTGAGGAACATGGAGGTACATAACTCAAGGAAAACATACATggttacctcttatctagt
gtagagtgtagcacctatccATTTGCATTATCCTTACTGGCTGGGCACCCCAAATCTGAAggttacctcttatctagt
gtagagtgtagcacctatccAAACCAGAAATGCTCCAATATGCATTTCCTTTAAGTGTCAggttacctcttatctagtg
gtagagtgtagcacctatccTTGGAATGCTCAACCTATAACATAGAGGAAGCCTGTTAAAggttacctcttatctagtg
gtagagtgtagcacctatccAGAAGTGTTAAATTGATTGCAATCATTGTTCTGAGGGAGAggttacctcttatctagtg
gtagagtgtagcacctatccCCTTGCTTAAAATGTACCATGAGCCAAAGATAGCTTTGCAggttacctcttatctagtg
gtagagtgtagcacctatccGTAAGATAAAGTAGTTACACCAGTGGAAATTCTTAGGTCTggttacctcttatctagtg
gtagagtgtagcacctatccTTATATTAGCTAAGCCTCTTTTACTTACTCAATACCAGGGggttacctcttatctagtg
gtagagtgtagcacctatccATGTACAGGAGTTGGAGGTTGCTCTGATTGCAGACTGGCCggttacctcttatctag
gtagagtgtagcacctatccGGTTATCTGAGACCCAATCCTACCACTGTGGCTCTTCCTGggttacctcttatctag
gtagagtgtagcacctatccGGAGAGCCATTAATACTGTATCTGCTAGCAAGCCACATCCggttacctcttatctag
gtagagtgtagcacctatccGAAAAAAATTCTAAATGAGGTAGAAGAAAGGCAGTCTGGCggttacctcttatctagt
gtagagtgtagcacctatccATACTCAAGCAAATAGGACTCTTTTTTCTTGTTTAGTGCCggttacctcttatctagtg
gtagagtgtagcacctatccTCAGTTGAAACACATGGATGAACTAGGCTGAAGAGAATATggttacctcttatctagt
gtagagtgtagcacctatccCTTATTTTTTATTTCATGAGGACTGTGGCAAAGTAGACACggttacctcttatctagtg
gtagagtgtagcacctatccCACTTGAGGCCAGGGGTTTGAGACCGGCCTGGACAACATGggttacctcttatcta
gtagagtgtagcacctatccCACTATAGGTCAAAGATTATTATTAAGTGCCACCAGAGTAggttacctcttatctagtg
gtagagtgtagcacctatccAAAGAATCAGCAGAGGGAAAGAGTAATTTTTTAGAGGAGTggttacctcttatctagtg
gtagagtgtagcacctatccCTGAGTTTGAGAGACCCTGGAAGAATTGATCAGCAGAGTTggttacctcttatctagt
gtagagtgtagcacctatccTCCAGGTCACGGGAAAATCTTAACCAAGGGAATGGCAGCAggttacctcttatctag
gtagagtgtagcacctatccCTGTAGCCATGGTATATTTGTAATCAGACCTCTTTTCAGAggttacctcttatctagtg
gtagagtgtagcacctatccAGCAATGATGTTTGGTTGGTGAAATTCCAGGTGATAAAGCggttacctcttatctagtg
gtagagtgtagcacctatccCAGAAATGAAAGTGCATGAATACTAAGATCTGAAAAGAGCggttacctcttatctagt
gtagagtgtagcacctatccTGAACAGACACATAGCGGGCATTGATTCCTTTTACACAGTggttacctcttatctagt
gtagagtgtagcacctatccTTGTTTTTATCTTCATCTCCTGATTCCCAAAAGGTTTCTGggttacctcttatctagtgg
gtagagtgtagcacctatccTCATGGCAGGTCGGCTTGAAGTTTTTATGTCAACAATGCTggttacctcttatctagtg
gtagagtgtagcacctatccTCTCCTTTGATGTATTATTAACTAACAGGACAACCAGGATggttacctcttatctagtg
gtagagtgtagcacctatccTAAAACTCTCTTCACTATCGCCATCATCTGACTTTGACAAggttacctcttatctagtg
gtagagtgtagcacctatccTGTGATGAAAGACGTTGCTCTGATGAAGGAACTGGTGATCggttacctcttatctagt
gtagagtgtagcacctatccGCTTGAATTTGAGACTCCAGCACCTAAAAAGGCGATATATggttacctcttatctagt
gtagagtgtagcacctatccTAAGGAACTCTTAGATGAGTACTCTGTAGCTAATATCTCAggttacctcttatctagtg
gtagagtgtagcacctatccCTTTGGAACAAGGTAAATTCTAAGGAAAGTTGACCCATTTggttacctcttatctagtg
gtagagtgtagcacctatccATAAAAGAATTAGACAGATGAAGATCAAGGCTACAGAAGCggttacctcttatctagt
gtagagtgtagcacctatccGTACACCAACTGCAACAACCCCTCAAATTTATTACTCTTAggttacctcttatctagtg
gtagagtgtagcacctatccCAGGGTATAGTTTATCTTGTAGTTCAAAAATCTTGTGAGGggttacctcttatctagtg
gtagagtgtagcacctatccAGAAATCTCTCTTCCACTGGTCTTATCATTAGTAACATGGggttacctcttatctagtg
gtagagtgtagcacctatccTTTATAAGTGTGAGTCCAGTATGTGAATCTAGTAACAAGGggttacctcttatctagtg
gtagagtgtagcacctatccCACATTTCAAATTGAAATTGGAATCTGTTGAGCATCTGTGggttacctcttatctagtg
gtagagtgtagcacctatccGTGCATACAACCAGGCTCTGTTTTCCCCAAGATTCTTTTCggttacctcttatctagtg
gtagagtgtagcacctatccCTCTCTCTTTTTGGGACATTTAAGTGATTTGAGGTAGGAGggttacctcttatctagtg
gtagagtgtagcacctatccGGAAAGTGAGGGAGATTGCTACAGTGATAATTTAGGGCACggttacctcttatctag
gtagagtgtagcacctatccCATTTTACAGGAGGGCAAACAAATGTTAACTAAGAGGAAGggttacctcttatctagt
gtagagtgtagcacctatccGAGGCACCATCTATATAATGTAGGGATTCTATCTTACTATggttacctcttatctagtg
gtagagtgtagcacctatccCTTTGAATATTCCATAAACTCTCTTGTCCAATGTACCTAGggttacctcttatctagtg
gtagagtgtagcacctatccGTGGATACCAAAATTACCTCAGGGCCATTTGCTGTAATGAggttacctcttatctagt

gtagagtgtagcacctatccGCAGCAAGCGGTGAAAGGTGTGTGGTAAAGGATGAGCAGCggttacctcttatcta
gtagagtgtagcacctatccGTGGATGTTCACATACTCTTGTATCTTTCTCCTAAGGCAGggttacctcttatctagtg
gtagagtgtagcacctatccCAAGTTTATCAGTTTGTGTGCTGGAAGGAATCCTCTTCACggttacctcttatctagtg
gtagagtgtagcacctatccTAGGGAAAAAAGCTGTAAGTTTAGTCACAAGCAAAATCAGggttacctcttatctagt
gtagagtgtagcacctatccAATGTATCAATTCTAGTGTACTCACAAAGATACCTGTAGGggttacctcttatctagtg
gtagagtgtagcacctatccACATATGTTTAGAAGTTGGGGTGGTCACTAACTCATACAGggttacctcttatctagtg
gtagagtgtagcacctatccTTCTACGCAAATGGACCACAGCTAGGAAAGAGAAGTAGGGggttacctcttatctag
gtagagtgtagcacctatccATTTCATTTCTCATTACAACCCTCAGATGGCTCCATTAATggttacctcttatctagtg
gtagagtgtagcacctatccATAAAAAAATACCATAAAAAGTCTCCCTGTCTCTACAGCCggttacctcttatctagtg
gtagagtgtagcacctatccCTGGCAATTTCTATTACCAGTTTCTTGGACTGTCTATAAAggttacctcttatctagtgg
gtagagtgtagcacctatccCACAAATGGCAGCACTGGAAACAAACTGATCTGTGTTCTTggttacctcttatctagt
gtagagtgtagcacctatccAATACCATAGTTTATTTCATCAGTCCACTTCAGAACAGTCggttacctcttatctagtg
gtagagtgtagcacctatccCTGCAATGAATCAACTTGTATATCTGTCTTTTCACACATGggttacctcttatctagtg
gtagagtgtagcacctatccTTTTGATAGATCTTACCAAACCATTCTCTAAGAAGCTGCTggttacctcttatctagtg
gtagagtgtagcacctatccCTACCAATGTAAGCCCATCAAACTTTTTAAACTATGTCACggttacctcttatctagtg
gtagagtgtagcacctatccGCTTTTTGCCTCCTTCTTTGTGCATATATTGATTACTGATggttacctcttatctagtgg
gtagagtgtagcacctatccAGAATGTAGTTTTGTTCTTACAGGGAGATAGCAGGCCATCggttacctcttatctagtg
gtagagtgtagcacctatccAAGGCTTGGCATTCGACTTTGTTAGACTGGGGTTATTTCAggttacctcttatctagtg
gtagagtgtagcacctatccATGGTCTGTATTCCTTGAGTATGGTCTTTCTGGGGTTTCAggttacctcttatctagtg
gtagagtgtagcacctatccTGCCAGGAGTATTCATCACGCTCACCCCACCTCTGGTTGGggttacctcttatctag
gtagagtgtagcacctatccAATCTGTTCAGTTCTCAGCTTACAATTCATTTCCACTGCTggttacctcttatctagtg
gtagagtgtagcacctatccCAGTCTAAGAGTGGCCAGGACCCTAGGGATATTTTTATGCggttacctcttatctagt
gtagagtgtagcacctatccTTAGACTCCCTTTCTGCAGCTCTGTACCACTAGTGCCCTGggttacctcttatctagt
gtagagtgtagcacctatccGTCCTAAACTTTGATCTCTGCCAACACAACTGACGTACTCggttacctcttatctagt
gtagagtgtagcacctatccGGCTTACCTCATGCCCTTTTCTTCTCTCAAGGATTACAGCggttacctcttatctagtg
gtagagtgtagcacctatccTCCTCAGAGCCTATGAACAATTGTTTTAAATATTTTGCCCggttacctcttatctagtg
gtagagtgtagcacctatccTTACTATAAGAACTAATGCTGACACTGGCTGTTTCATCGTggttacctcttatctagtg
gtagagtgtagcacctatccTAACTAGAACTCTGCCAAACAACAAGTTTTGAGATGTACTggttacctcttatctagtg
gtagagtgtagcacctatccTCTGTTATAATTCTTCTTTGATGCAACTCATTTGGAAGCGggttacctcttatctagtgg
gtagagtgtagcacctatccTCTTATATACTAGCAATAAAGACTGAGGACATGGTGGGAAggttacctcttatctagtg
gtagagtgtagcacctatccAACTTGCTTTATGTGTCCAGTAAACAGTCAATTTCCATAGggttacctcttatctagtg
gtagagtgtagcacctatccCTATCAAGACTGTCAATCATTCAAATCTGTGTACTTTCTGggttacctcttatctagtg
gtagagtgtagcacctatccTCTCTGCCTGACCTATCCAGTACTAAAAAGAGTGTTATGGggttacctcttatctagtg
gtagagtgtagcacctatccACACAAAACAGCCACAAATTCTCTGTCGCTCTTTCCATCAggttacctcttatctagt
gtagagtgtagcacctatccGTCTTTGAATTTGACCGGCCTTGTGAATTATACTAGCTAAggttacctcttatctagtg
gtagagtgtagcacctatccGCTTTGCAGCTTCTGTGCTCTCTTTCTTGAATGCTTCTGGggttacctcttatctagtg
gtagagtgtagcacctatccGGTACCACTTGAAGCACACTATGTAACCAAAACAGGAAAGggttacctcttatctag
gtagagtgtagcacctatccGCTGCAATTGCCAACCCACAGACTGTCAGCAAATAAACAAggttacctcttatctag
gtagagtgtagcacctatccAATCTCACATTGTAACTGTAGACTGTCCTATTTTTCCTTGggttacctcttatctagtgg
gtagagtgtagcacctatccAAGGTTACAGTATTAGGTAGATACATATCCAGGATTGATCggttacctcttatctagtg
gtagagtgtagcacctatccGCCCTCATATTTACACTTTCTTTTCTCTTCATTGATACTCggttacctcttatctagtgg
gtagagtgtagcacctatccCCTTCAACTGGGTTAACAAGTATTTTCTTTCAGCACGTTGggttacctcttatctagtg
gtagagtgtagcacctatccGCTACCAAAAATTAAGCAATTATTCTCACTGTCCCTCTTTggttacctcttatctagtg
gtagagtgtagcacctatccAGATCTCTTTCTTTGTTTTTGGTGTTCAGGTGATTCACTAggttacctcttatctagtgg
gtagagtgtagcacctatccTTATTCTGCTTGGGGTTTACTGGGCTTCTTGACTCAATGGggttacctcttatctagtg
gtagagtgtagcacctatccGTATTAACTCTTAAACACTGTCTCTCTTCCATTCTGTCTTggttacctcttatctagtgg
gtagagtgtagcacctatccACTCCAAGTAGGTAAGTTGTACATCATTTTGCCCTTCATTggttacctcttatctagtg
gtagagtgtagcacctatccCTTTTTCTGTGTTGAAGTCTAGATAATTTCTACAGGTCTCggttacctcttatctagtgg
gtagagtgtagcacctatccACCAATTTGTTTGCTGACATGCTATAAAATGTCCAACTGAggttacctcttatctagtg
gtagagtgtagcacctatccAGATAATTACAACCATTGTTGAGGGTCTATTTCTACTGTCggttacctcttatctagtg
gtagagtgtagcacctatccTTTTTGTTAGTTCTTGCTCATGGTGTCTTGTTTGTGAGTTggttacctcttatctagtgg
gtagagtgtagcacctatccGTTGCTTGATATCCTTGGAAATTAACCTAAAATGCCATTCggttacctcttatctagtg

gtagagtgtagcacctatccACATGGAAACCGAGGCCTACAAGTCATACAACTACAATAGggttacctcttatctag
gtagagtgtagcacctatccGCTCCTAGTGCCCTAATTCAATGCTCTCTCCATTATACAGggttacctcttatctagt
gtagagtgtagcacctatccAAAATCCAGGTCTCATTATTAATACTTTGCCCATAGCACTggttacctcttatctagtg
gtagagtgtagcacctatccCTTTTTGCTTTCAAGTTAGATCTGCGAATGATTTCATGCAggttacctcttatctagtg
gtagagtgtagcacctatccTTTGCTTGTTTTATTTTCTTCATCCTCTTCTTCCTCTGGTggttacctcttatctagtgg
gtagagtgtagcacctatccCACAGAATATCATGGAGAGAAGTGGTTTGGATGACATTACggttacctcttatctagt
gtagagtgtagcacctatccCAGTTGAAAGCCTGAAGAAATGACGGTTAAATGAGTTCAAggttacctcttatctagt
gtagagtgtagcacctatccGTACTCAGAAATAAAGTACCTCCATGGATTAAGACTTCAAggttacctcttatctagt
gtagagtgtagcacctatccCTCACTGCAGCCTCAAGCACCCAGGCACAAATGATCCTCCggttacctcttatcta
gtagagtgtagcacctatccCATGGGCTACTATGCTTGGCACCTTGAAAATCTGACCTTTggttacctcttatctagt
gtagagtgtagcacctatccCTGAAAGCACACGGAACTTGAAACAGTTAATAATTTGGGGggttacctcttatctagt
gtagagtgtagcacctatccTTATTCAAGCACATTTTTTCCAACTTGCATCTGGATGAACggttacctcttatctagtg
gtagagtgtagcacctatccGTGTATCTCAAGCTCTAGGCAAGCAACCATTTACTTAAATggttacctcttatctagt
gtagagtgtagcacctatccAATTCCCATCCATGAAGTGGTAGGTTATTTATCACATTTGggttacctcttatctagtg
gtagagtgtagcacctatccGTAATATACAGTAATCCTATTATTTCCAGAGACCAAAGGCggttacctcttatctagt
gtagagtgtagcacctatccGTCAACTTACCACTCAAAAGCCCTACTCCTTATCTTCTTTggttacctcttatctagt
gtagagtgtagcacctatccAATTTCACACAATTTCTGTTGGTAAAAAAGTGGCATCGTGggttacctcttatctagtg
gtagagtgtagcacctatccTGTCTTCATACCTCCATAGCAAAAACTCGACAAGCAGATTggttacctcttatctagt
gtagagtgtagcacctatccAGGGCCTGTTTCATTGCTATTTCAAGACCTTCTAGATCATggttacctcttatctagtg
gtagagtgtagcacctatccTGTATAACAAAAGCTAAAACTGGCCACTGGAAATTTCGTTggttacctcttatctagt
gtagagtgtagcacctatccACGGATGTCTCAAAAGGGAAGAACCAACTTCTATTAAAGAggttacctcttatctagt
gtagagtgtagcacctatccTCACTTAAAGCCTTTGTCATTGGTTTTTTGAACATGACACggttacctcttatctagtg
gtagagtgtagcacctatccAGTCATATAGCTATTAAGTTGTAGAGGTGGGTTTTGAATCggttacctcttatctagtg
gtagagtgtagcacctatccGAAATCTACATCTCATCTGCATCATTTTCTAAAAGATGGCggttacctcttatctagtg
gtagagtgtagcacctatccGATCTCTCTGATAATTGATCACTATATTACCCTGTTCTTCggttacctcttatctagtg
gtagagtgtagcacctatccTTAAAATTATTCTTGAGGTAGAATCCCCAGCTCCAATTTGggttacctcttatctagtg
gtagagtgtagcacctatccGTTGATACCAACTGGTCCAGATCATGTAACTGTTTTTTTTggttacctcttatctagtg
gtagagtgtagcacctatccTATCCCCAAAGTTTTATTATCTACTTAGAATGATGGCAGCggttacctcttatctagtg
gtagagtgtagcacctatccGTTGGGGTGGAGAGATAACTGTGAATGTGTACGTCATTGGggttacctcttatctagt
gtagagtgtagcacctatccGGTTGTAACATTTGGCTTAGATACCTAACAAGTAAGATTGggttacctcttatctagtg
gtagagtgtagcacctatccATCTCCAGATTTCCCTTCAACTAAGATGCAAAATTTAGAGggttacctcttatctagt
gtagagtgtagcacctatccATGTCAATCACACATTAGAAATTATCCCATGCCTTTTCCTggttacctcttatctagtg
gtagagtgtagcacctatccCTCAATATTCTTTTTTATCCTCATACTTACCAGTGTCTCCggttacctcttatctagtg
gtagagtgtagcacctatccCTCGTGCAGGTAAAATGGAGTGTTGTTTATCATATTCGGAggttacctcttatctagt
gtagagtgtagcacctatccAAGACAGTTCTCTTTGCAAGATCAGTATCAGTTATGGGACggttacctcttatctagt
gtagagtgtagcacctatccTGTACATGGAACAATGCAACAGATACAAACTCAGAAGAGCggttacctcttatctag
gtagagtgtagcacctatccTGTGACTCAATATTTCATGTGGAGAAACAAAATCAAAGCCggttacctcttatctagt
gtagagtgtagcacctatccTCAAATGCAGTAACATTTTGACTGATTTCAGCATTGTACCggttacctcttatctagtg
gtagagtgtagcacctatccAAAGGTTTCCAAGCTCCTTTTAGAATCAAAAGCACTGGGGggttacctcttatctagt
gtagagtgtagcacctatccATATTCAGGAAAAACAGTCAGATATAATCGTAAGTGCACCggttacctcttatctagt
gtagagtgtagcacctatccAGTTTTCTTTTGGTGTCTTCAGTGTTCTGTGAGGTTTCTCggttacctcttatctagtg
gtagagtgtagcacctatccATGGCTGCATCAAGGTGTCATGAGAGGTAACAAACTGATTggttacctcttatctagt
gtagagtgtagcacctatccGAAACTATTCTTTTCGTGAGCTCCTTTTTCTGTCACCTGAggttacctcttatctagtg
gtagagtgtagcacctatccTTGGAATATTTTTGCTCACCGTTGGTTCTGGAGTTTTTTCggttacctcttatctagtg
gtagagtgtagcacctatccCAGGTTTGAAATCCTGTGTCCCTTGGTAAAATTATCTATTggttacctcttatctagtg
gtagagtgtagcacctatccGTGTTTTCTTTACTCACCACTTCTCTAAGCTTATGTACATggttacctcttatctagtg
gtagagtgtagcacctatccTGATTGGCAGTAAATGTCCCTGGGTGTTAGTATAGGTTGCggttacctcttatctagt
gtagagtgtagcacctatccTGGTGTTACTATAAAGCACAGACATTTGTTCTGTTTTCAGggttacctcttatctagtg
gtagagtgtagcacctatccTCAAGCAACACAAGAAAATCCAAAGAAGAAAGCAAAGATGggttacctcttatctag
gtagagtgtagcacctatccGGTAATTCCTTGTTAGGCTGTCTTGCCTCCTGTAGGATGGggttacctcttatctagt
gtagagtgtagcacctatccCTAGCCCAGTTGTGACAAAGTTATTTCCAAATATTTCCAGggttacctcttatctagt
gtagagtgtagcacctatccAGGCAAATATCACCCTCAATTGAGAACCACTGATTTAATGggttacctcttatctagt

gtagagtgtagcacctatccGGTAATAGTAACCCAATGAGAATCACACTTGTCAGACTAGggttacctcttatctagt
gtagagtgtagcacctatccTTAAAGATATAAAAGGTTGCACTGCACTCTGGAGAAAATGggttacctcttatctagt
gtagagtgtagcacctatccCACACAGTTGGGGAGGGGAGAAGTTGCATGTATTTTTAATggttacctcttatctagt
gtagagtgtagcacctatccAAAAACCTCATTTGGATATATATGTCCCATTGCGCTCATTggttacctcttatctagtg
gtagagtgtagcacctatccGTTGTATAGTAGTAACACAGTGGCATGCTCTTTTAATAAGggttacctcttatctagtg
gtagagtgtagcacctatccATTCAGCAAATGTCTTCCTTTCCAATGATACCATGTTCACggttacctcttatctagtg
gtagagtgtagcacctatccTATCTCTTTAATCCATATGTCAGCTTCTCAATTGTCACTGggttacctcttatctagtg
gtagagtgtagcacctatccTTATCACATCATTCGTCCTCTAAAAAAGTTTCAGTGGTACggttacctcttatctagtg
gtagagtgtagcacctatccGGGATTACAGGTGTGTGTCACCATGCTCGGCTAATTTTTTggttacctcttatctagt
gtagagtgtagcacctatccCCACAGCGCCTGGCCTCTATTTTCTTGTGTATTACATTATggttacctcttatctagtg
gtagagtgtagcacctatccCTTTCCGAGAAGCTCTTTACTCTGATACTACCATACCTAAggttacctcttatctagt
gtagagtgtagcacctatccATCCAAGAAGTTTAAGATTTATTTTCAATGCCACTGCTCCggttacctcttatctagtg
gtagagtgtagcacctatccAGTAACCCTTCCCTACTCTGATTTACTACAGCATTAAGTAggttacctcttatctagt
gtagagtgtagcacctatccTGTAATCTTTAATTCCTTCACTAGTTGATCTCTTCTCTCCggttacctcttatctagtg
gtagagtgtagcacctatccGGACCTTACCTTAGTCATACATGTATGCTTTTTCTTACTGggttacctcttatctagtg
gtagagtgtagcacctatccTGGCTCACAGATGTATTAAAGAACTTCTACAGGACTCTTTggttacctcttatctagt
gtagagtgtagcacctatccAGTCTTTCTTCCACCAAACACTAGCTATTCAACAGACAGGggttacctcttatctagt
gtagagtgtagcacctatccGCATAGACTTCATATTGCTCTCTGGCTCTTCAAATACATTggttacctcttatctagtg
gtagagtgtagcacctatccACGGGGCAGGTACAGATGTTGAACGTAGATATCTCATTTCggttacctcttatctagt
gtagagtgtagcacctatccATCTTGATAGCCCACAAAATTTTCATGATGATGCCTGCATggttacctcttatctagt
gtagagtgtagcacctatccGGCAATACAGATCCTCTCCATTTTTAATCTCTTTCATAACggttacctcttatctagtg
gtagagtgtagcacctatccAAGGCAAAGCATTAAAATAATCGTAGATGTAGGTAGGTGAggttacctcttatctagt
gtagagtgtagcacctatccAAGTCTGATAATTTTATTCATCAACATTCCAGGTACCACGggttacctcttatctagtg
gtagagtgtagcacctatccAGTTTCTTTCCTTTTGAGAACTTTTCCATCTTCCAGTGTTggttacctcttatctagtgg
gtagagtgtagcacctatccAACCATCCAGTTGCTTAAGCTAAAATATTTCCAGTGTTGTggttacctcttatctagtg
gtagagtgtagcacctatccTCATCTTTCAAGCCCATGATATTAGGTTTCCTCTCAGAGCggttacctcttatctagt
gtagagtgtagcacctatccTCTCTTCTCATTCCCTATCATCACTGCCTTGGTTCATGCTggttacctcttatctagtg
gtagagtgtagcacctatccCAGACCACTGAAATACAAGCAGTTACTTTTTTGGGTAGGAggttacctcttatctagt
gtagagtgtagcacctatccTTAGATATATGAAATTTTCTGTGGTTCAGGGTGGAAGGGGggttacctcttatctagt
gtagagtgtagcacctatccCCTTTTCAAAATGAACAAATAGTTGCTAACCTAAGTCCACggttacctcttatctagt
gtagagtgtagcacctatccAAGATCTATTGTATTTTGGCATACTGATGATTTCCACCTCggttacctcttatctagtg
gtagagtgtagcacctatccTAAAAACCATACAACTCTGCTACACAAAGAGGACAATTTCggttacctcttatctagt
gtagagtgtagcacctatccCATGTCAGTGGGCAATATCTAGATTACCTTCATTTGTCAGggttacctcttatctagt
gtagagtgtagcacctatccTCTTTTGGAGTGTCAGTGCATGATGTGTGAAAGTACAGGCggttacctcttatctagt
gtagagtgtagcacctatccCTGAGGTCCAATATATGAAAACAGAATACTAACAGGTTTCggttacctcttatctagt
gtagagtgtagcacctatccAATGCACTTTTATCTAGTCTCCACATTAGCTTCCACATTCggttacctcttatctagtg
gtagagtgtagcacctatccTGTATTTCTCTGTTTAAAATTCTTCCACTGTTTCCTCTGCggttacctcttatctagtgg
gtagagtgtagcacctatccAATCACTTTGAACGGGCTTTCACATATTTCCTCCAGTAGCggttacctcttatctagt
gtagagtgtagcacctatccCTTTAACAACCCGTGATTCCTGAAATGTGTAATGAGATTTggttacctcttatctagtg
gtagagtgtagcacctatccCTCACCTGGGTTATTCTTCCAAGAGTCGCTTTCTTCCTGTggttacctcttatctagt
gtagagtgtagcacctatccGCCTGACAGATTTCTATTTATCCTTTGAGACTCTACTAAAggttacctcttatctagtg
gtagagtgtagcacctatccCTCTAACAGCATCTTACATACATACTTCTATTGTAGCACTggttacctcttatctagtg
gtagagtgtagcacctatccACATAGAAGCCAGGATTTTGTTATATTCATACATTCAGGGggttacctcttatctagtg
gtagagtgtagcacctatccCACAAGGCTGGTAGGTGCCCAGTTGGTATTCATTTAAATGggttacctcttatctagt
gtagagtgtagcacctatccAAACACACTTTCTTTACACTTCAGCATTTCGCACTAACAGggttacctcttatctagt
gtagagtgtagcacctatccAGTCAGCCTACCAAAGAGTAAAAGATAGCTATAGTCTTATggttacctcttatctagt
gtagagtgtagcacctatccATACCTATTACCATAAAGTCATTATGTCAGTCAAGGCAACggttacctcttatctagt
gtagagtgtagcacctatccGTCAAATACCGAACTACCCTAGATATCAATAAACAATCGCggttacctcttatctagt
gtagagtgtagcacctatccAACAACAGCAAAATTTTATTACCTAGCTAATGTCTGCAGGggttacctcttatctagt
gtagagtgtagcacctatccCCTTCTTTTACACTGGGCAACTGAGATTGGAATGGCTTTGggttacctcttatctagt
gtagagtgtagcacctatccGCTCAGGGACAGCAGGAAGAGTGCATTGGCTTTAATCACCggttacctcttatcta
gtagagtgtagcacctatccTCTTGACTTGCATTGGTTTTCTCCTAGATTGTTTGACCTGggttacctcttatctagtg

gtagagtgtagcacctatccTAAAAAACAACATTCATTTCATATTCATCCTGGGCTGACCggttacctcttatctagt
gtagagtgtagcacctatccAGTGCGTGTATTCAGTGAAATTTTTTAGTTAGGTGCCACTggttacctcttatctagtg
gtagagtgtagcacctatccAGAAAAGAGAGAGTAGATATGGAATAGGAAAACAAAGGAGggttacctcttatctag
gtagagtgtagcacctatccAAGAGGAAAAGAGAAAGAAGGTGAAAAGGAAAGCTCTCCAggttacctcttatcta
gtagagtgtagcacctatccATTTGTCAATAAGGCATATGTAGCAGAGGTTATCTAGTGTggttacctcttatctagtg
gtagagtgtagcacctatccAAAGTATAGCTCCAAATGGAATAAAATACCTTCTCCCTTGggttacctcttatctagt
gtagagtgtagcacctatccGTATTTTTCTCTACAGCGATCACATACCAGATACCAAGTGggttacctcttatctagt
gtagagtgtagcacctatccATGCCACCATCACCACAGTTACCAGCCCATCCTCCACAAAggttacctcttatcta
gtagagtgtagcacctatccGCTATTGTAACCTTGTCCACCAGCATATCGGCCACAACCTggttacctcttatctag
gtagagtgtagcacctatccCTCCAAGTGTCCTTAATAGTTTGATCCTTATTACTCATACggttacctcttatctagtg
gtagagtgtagcacctatccGCTTACTTGATCAAAAAGCAATCAATTAAATGATGCCCCAggttacctcttatctagt
gtagagtgtagcacctatccCTGAAAATGCAGTCTTCTGATAAAACAATATCACTTCCGTggttacctcttatctagt
gtagagtgtagcacctatccGCAATTCTGAACAGGAAAATAGTAAAATCAAAGGAGATGCggttacctcttatctag
gtagagtgtagcacctatccGCATGCCACTGTTTCTGCTGTGTGAGACAGACACATTAATggttacctcttatctagt
gtagagtgtagcacctatccGTGAAATAGCACTAAATGACCTGGCCATCATAATATTAGAggttacctcttatctagt
gtagagtgtagcacctatccAAAGCACTTATACGTACATTCAACAGTATCATTTACCTGGggttacctcttatctagt
gtagagtgtagcacctatccTGACTCCCCACACAGTTCACAGATGGTATCTTTCTCTGGTggttacctcttatctagt
gtagagtgtagcacctatccGCCATCTCTCCAAACAAATTACCCCTGGGCCTAACTTCTGggttacctcttatctag
gtagagtgtagcacctatccCCTTTTTGGACTTCTTATTCTCTTTCTCACACTCTTTCTTggttacctcttatctagtgg
gtagagtgtagcacctatccGACTTGGAAAAATCATATTCTGGGAAGCAGCTTTGATAGTggttacctcttatctagt
gtagagtgtagcacctatccAGAGATGTCTTCCCAAAAAGCCACTAAATGTTGTAAAGTTggttacctcttatctagt
gtagagtgtagcacctatccGGAGACTTAGACTTCATTGTGTTATGCATGGACATTTCAAggttacctcttatctagtg
gtagagtgtagcacctatccTGGTTCCTTAAGCATAGACAATACCTTGTTTTTGTTGATGggttacctcttatctagtg
gtagagtgtagcacctatccTATCTGGTCCATGGGGTGCAATATTAAACATATTCAGTGCggttacctcttatctagt
gtagagtgtagcacctatccTTAACTTGGTTTCTTTTTCCTCTGTAGGTTGTTGGCTATTggttacctcttatctagtgg
gtagagtgtagcacctatccCTTATAGGAGCATGTTCTTTGGAAAGTTCAGGATGAAACTggttacctcttatctagt
gtagagtgtagcacctatccAAGAAGAACAAGCCATTGCATCATGTACTATGCCTTCATGggttacctcttatctagt
gtagagtgtagcacctatccAGACATGGCTTGCTTGGCTGGCTCTAAAGGCTTGGATCCAggttacctcttatctag
gtagagtgtagcacctatccAAACTCGATTTTTCTTTTTTGGGCCTGGTGTGTCTATTAGggttacctcttatctagtg
gtagagtgtagcacctatccATGCTCTCATTTCCTGTTCATCTTTCAGTGGTGCTTTTCCggttacctcttatctagtg
gtagagtgtagcacctatccACTTTATTTGAACCTTCATCCACACTCTTAAATTCACTGCggttacctcttatctagtg
gtagagtgtagcacctatccCTGTTAGACTATTTGTTTTTAAAGTACTTGAGGTGCAGACggttacctcttatctagtg
gtagagtgtagcacctatccCCACATGGGGAGAGGGAGCTCTATTTTCAGATCCAGGGGAggttacctcttatcta
gtagagtgtagcacctatccAATCCTTTGGTACTGATTTTGGTTTTGGTGACGTTGACCGggttacctcttatctagtg
gtagagtgtagcacctatccAAAGGTTCTGTGAGGGGCATTTTCTTCTTGCGGAGGGTATggttacctcttatctagt
gtagagtgtagcacctatccCTTTGTCTTAACAGGAGCAGTACTTTTTTGAGGTAGATTTggttacctcttatctagtg
gtagagtgtagcacctatccATGTGGTGAGGAGGAGCGTGGAGAACTAGCACCCGATGGGggttacctcttatcta
gtagagtgtagcacctatccATTGGCTGGGAGAGTCTTTGGCTTAGGAGATGACAACCGAggttacctcttatctag
gtagagtgtagcacctatccAGATTTCAATGTCTGCAAGGTGTTATGGTTGGGGGATAATggttacctcttatctagt
gtagagtgtagcacctatccCAGATTTTAAGATGAGGGATTCACTAGCAGATAAACTTTCggttacctcttatctagt
gtagagtgtagcacctatccGCAGTAGTCCGGCTCATACGTCCATCAGGTTTAAGCGATCggttacctcttatctag
gtagagtgtagcacctatccTCTGATTTTCCTGATGTAGAGGCTGGTTTAGAAGATGTTGggttacctcttatctagtg
gtagagtgtagcacctatccTTTCCTCGATCACCTGTATGGTCCATTTTACAGTGAAACTggttacctcttatctagtg
gtagagtgtagcacctatccCATTACACAACATTACCACCAAAAACTATTTTCATGGACGggttacctcttatctagt
gtagagtgtagcacctatccGTGTGAACTATCAGTGATATCTCAACCTCTGCCCTAGTGCggttacctcttatctag
gtagagtgtagcacctatccGGGACTAGAAAGAGACAGAAATTAATCTTTTGGGGAATAAggttacctcttatctagt
gtagagtgtagcacctatccAAATGTAGAGTAGAGCTAAATTTCCAGCAGAGTGATACCTggttacctcttatctagt
gtagagtgtagcacctatccACGGTATGGGTTTTGTTGAATTAGAGAACTGAATTTATGCggttacctcttatctagtg
gtagagtgtagcacctatccTATGCATACAATTAGGCTAACAAAGAGTTGCTTAAGAGGCggttacctcttatctagt
gtagagtgtagcacctatccATGATCAGATTAGACTAGTTTACAATGAGCACATGAATGCggttacctcttatctagt
gtagagtgtagcacctatccTTTATAAACAGGTTACACTCCAAAAGTTCAGGAAGCTTAGggttacctcttatctagt
gtagagtgtagcacctatccTGAGTGGTAAAAGTCTGAACTGTGGACCCTCAAAACAAGGggttacctcttatctag

gtagagtgtagcacctatccAAAAGAAGAATCACCTCCCTTTTCTAAGGCAAAGCATTAGggttacctcttatctagt
gtagagtgtagcacctatccCACCCTGCCTTCATTCAGTGTCTCTCCTTCATTCTATTTggttacctcttatctagtg
gtagagtgtagcacctatccGTAAATGGTACAAACTTCAACATCACCTTTCCTCCTTAAAggttacctcttatctagt
gtagagtgtagcacctatccGTTAGTTGTTAGGAGAAAAGCAATCTGACTGAAGGGTTTCggttacctcttatctagt
gtagagtgtagcacctatccCAGTTCCTAGGAAATTAAGGTAGATTTGTTAACTGGCTATggttacctcttatctagtg
gtagagtgtagcacctatccATTCAGATGATGTAGGAAAATGTTTTTATGCTTCCGTGCggttacctcttatctagtg
gtagagtgtagcacctatccGTGTATAGTCACGCTCTATTTCTCTATGGCTCAGGACAGTggttacctcttatctagt
gtagagtgtagcacctatccTGTCTGTTACTCTTCACTGATGTAATACCAAGAATGAATCggttacctcttatctagtg
gtagagtgtagcacctatccCTTGATTTTTTTCTTCTGCTACTCCCAAGTATTCAGCTTCggttacctcttatctagtg
gtagagtgtagcacctatccGCTCCCAAGGCCACAGGACAGAGGTGACATTAGTTATTGAggttacctcttatcta
gtagagtgtagcacctatccGCAAACTGAGTAATTACATGGTGAAATAATAGAGTATGGCggttacctcttatctagt
gtagagtgtagcacctatccAGGGCACACTGATGAACTAAATGTTTAATTGCAACATGCCggttacctcttatctagt
ggttacctcttatctagtggTTCCTGGAGACTGCTTTCAGCAAGTAGGGGCCACCCACAGgaatcttaaacccaa
ggttacctcttatctagtggCATCTAGGAACATTTCCATGTGGTCTTTCCCAACATAACAgaatcttaaacccaactt
ggttacctcttatctagtggTCTCACCCACACTCAAGGGGGAAGATCATACAAAGGCATGgaatcttaaacccaa
ggttacctcttatctagtggGGCTTGGACAGCAGAGTACTTACCTCCAGAAAAAGGTAGCgaatcttaaacccaa
ggttacctcttatctagtggTTACGCTTCTCTTCTAAAAGTTGGCTATAGGTTTTGGTAGgaatcttaaacccaacttg
ggttacctcttatctagtggAGTTCACGACTTTAATGAGACTGAATTTAATGTTTCCTCCgaatcttaaacccaacttg
ggttacctcttatctagtggGGTTAAGCCATCTTTAATTACTGGAATAAATCACCCTTGGgaatcttaaacccaactt
ggttacctcttatctagtggTTCTCCTGCGAAACTATCTGGAGCTGGTGCTTTTCATGAGgaatcttaaacccaactt
ggttacctcttatctagtggTGCCTTTACTGAGGTCAATTTCAGTTAATCGTATTTTCTGgaatcttaaacccaacttgt
ggttacctcttatctagtggTCTCTGATAACTGCTTTGTCTCATAGAGAGAACGTCATGCgaatcttaaacccaactt
ggttacctcttatctagtggGCTCCATGTGTACTTTTTAAAAAAGCATCTGCCTTTTTGGgaatcttaaacccaacttg
ggttacctcttatctagtggCACACGCGCGCGCACACACACACAGAGATTACTTTATTGGgaatcttaaacccaa
ggttacctcttatctagtggAGATCCTCCTAGAGCCAACTCCAAGATAAGGTTTTGTGTGgaatcttaaacccaact
ggttacctcttatctagtggCTGGAAACAGGAGTGAAGGACTAGAAGGAGTGGATGAAGGgaatcttaaacccaa
ggttacctcttatctagtggGGTGCTACTTGAGGGAAATGGGGTTCAATGCTCACCAGCCgaatcttaaacccaa
ggttacctcttatctagtggCCCTTAGAGGCAGTGCTACCATGGAGTTAAGGAGGCAGTGgaatcttaaacccaa
ggttacctcttatctagtggCTGGGCTCTTACTAGCGTCTGGATGTAGAATCCCTGGGGCgaatcttaaacccaa
ggttacctcttatctagtggGTGTCAGCAGGAGTTGAGTCAAAGCCCATCTGGAACTGTGgaatcttaaacccaa
ggttacctcttatctagtggAGAGGGGAGATGGAGAGCTTTGAGTTGGGACACAAGTAGCgaatcttaaacccaa
ggttacctcttatctagtggTGTGTGTTAAAGTCTTCGGCTATTAGTCTATGTCTGTGTGgaatcttaaacccaacttg
ggttacctcttatctagtggTGGCCCTAATTGCTTTATAGTAAATAGTTGCCTTTAGCACgaatcttaaacccaacttg
ggttacctcttatctagtggTACCCCCTGACACTCCCCACCCCACCCCACCCAGACAGCCgaatcttaaaccca
ggttacctcttatctagtggTCACCCAGAGACCACCCAGGGATCACTCGTAGTGTTGTACgaatcttaaacccaa
ggttacctcttatctagtggGTTGCCCAGGCTTGCATGCAATGGCGTGATCTCGGCTCACgaatcttaaacccaa
ggttacctcttatctagtggGCGTGAGCCACTGCTCCTGGCCACAGAGTCGTTTTATTGCgaatcttaaacccaac
ggttacctcttatctagtggCTGCTATTTTAGCCTTCCCTACCCACAACCCCTTGACAACCgaatcttaaacccaact
ggttacctcttatctagtggGTCTCCGTAGTCTTGCCTTTTCCGGAACGTCATCTAGTCGgaatcttaaacccaact
ggttacctcttatctagtggTCTTTCCCTCATATTTTTACATCATACCTTAGTGTCTTCCgaatcttaaacccaacttgt
ggttacctcttatctagtggTTCCTCTATGACACCATTTAAGTTAGCTAGTACCCTCTTCgaatcttaaacccaacttg
ggttacctcttatctagtggTGCTTTCCCCATTTCCCCCCATTTTCAGATAATTGCACTGgaatcttaaacccaactt
ggttacctcttatctagtggGTCAGAACACATAGCCATTACCTGCTATGCTGTCTCAGTTgaatcttaaacccaact
ggttacctcttatctagtggCATGCACACACACACACACACCCCTATATGTACACACATAgaatcttaaacccaa
ggttacctcttatctagtggACGAGCATGTGGGGATGGATCTCTCTGCTCTTGGGTCTCCgaatcttaaacccaac
ggttacctcttatctagtggCCCGATTTCAGAAATTTCTTCTCTGTATCAGGTTGCCCGGgaatcttaaacccaact
ggttacctcttatctagtggGTTTAAGAGAATTCCTTCTAACCAACTAGGGTAAGAGCCCgaatcttaaacccaact
ggttacctcttatctagtggCCTCCCTCCTTAACTTAGGAGCTAAAATTCTCTATCACCTgaatcttaaacccaactt
ggttacctcttatctagtggCAGAGAAAGTCCTTTGATTCCTCACACTTGTTGTTCAAAGgaatcttaaacccaactt
ggttacctcttatctagtggTTGGCTTACACTGGAACATCAGCTTCTTGCACGATTACTGgaatcttaaacccaactt
ggttacctcttatctagtggGTCATGGCCGTCATAATTGCTAACTGCTAAACAAATCATGgaatcttaaacccaactt
ggttacctcttatctagtggAGATTATGAAAGAAAGTAATGTAAAAGAGAGCTGGCAGACgaatcttaaacccaact

ggttacctcttatctagtggTATTTGCTGTTTCGTTAGAGAGAAACTGATGCTGGGCGGCgaatcttaaacccaactt
ggttacctcttatctagtggCTTTAACGTGCCCTCATCAGACAGATGTGAATGCCAAGTCgaatcttaaacccaact
ggttacctcttatctagtggTCATGACACAATCTCACTGTCCTATCGTGGGAACATCCAAgaatcttaaacccaact
ggttacctcttatctagtggAGAGACCATTACTACGAAAAGCAAAATTTTCTTGGTGCTGgaatcttaaacccaactt
ggttacctcttatctagtggGCTACATTCAAGCCATCATCTGTCCAATATAGCACTTGTCgaatcttaaacccaactt
ggttacctcttatctagtggTCCCTTGTCCCCCTCTCAGGGCGCAAGATGTGGGTGTGGCgaatcttaaacccaa
ggttacctcttatctagtggCCCCACTACTCATACCTCTAGGGGAGCATGCAGACGGGCAgaatcttaaacccaa
ggttacctcttatctagtggCCCACAGCAGTGTCTAGGGGTGAATGTGTATAGCTGAAGCgaatcttaaacccaac
ggttacctcttatctagtggCCATAGGCGGCTTGTGTTAGTCAGCTCCATTAGACCCCTGgaatcttaaacccaac
ggttacctcttatctagtggGAAGGATTTCTGTAACCCGAGGTTCTTGCCTTGGTGTACCgaatcttaaacccaact
ggttacctcttatctagtggGAGTGGAAGTAGCTCTCAGCAGATTGGAGAGGAAGAAGGGgaatcttaaacccaa
ggttacctcttatctagtggGCTGGTGCCTGCTGGTACGCTCCTCTCGACGTCCAGCTGCgaatcttaaacccaa
ggttacctcttatctagtggGCCTGCTAGGGTCTGGGTGTTTCTATAGGCACAGGATGGGgaatcttaaacccaac
ggttacctcttatctagtggCAAAAACTCCTGTCCTTGCATAGGTCAGAGGGCACAGGCCgaatcttaaacccaa
ggttacctcttatctagtggGGGATCCCCGTCCTCCTCTACTCATCACTTCCCTTCCCCCgaatcttaaacccaa
ggttacctcttatctagtggAAGGTACCACTCTCTTCCCTTCCCAGCACTTCCCTTCCCCgaatcttaaacccaac
ggttacctcttatctagtggCACTATGAATTGTTTTCTTTACCACGGTCAAGGAATGCATgaatcttaaacccaacttg
ggttacctcttatctagtggGGGGGGTTTTCCTTTCGCTTCTAGGACAAAAGCCTCTGTCgaatcttaaacccaact
ggttacctcttatctagtggGCTGGCTTCATTTCTCACCGCATAACACCTGGTAGGGTCCgaatcttaaacccaac
ggttacctcttatctagtggCATCGTTCTATTCACTAGGCAACGGACACACCTTCCAAGGgaatcttaaacccaac
ggttacctcttatctagtggGCTGACTTTCCTAACTTACGTAGGACTTGCTGTTCTCTCTgaatcttaaacccaacttg
ggttacctcttatctagtggCATGAGTAATGCCTTCATTTGCTAGAAAGCTCACTGGGGGgaatcttaaacccaact
ggttacctcttatctagtggGACTTCGGGTTTACGCTTTCAGTCATAATTTGGTCCCAGCgaatcttaaacccaactt
ggttacctcttatctagtggTTCCATGCTACCTTTAGATACCAATGCTGCCATTAGCACCgaatcttaaacccaactt
ggttacctcttatctagtggACTCATGGATGTTCCCATTACATATGTGAAATTATCCACCgaatcttaaacccaacttg
ggttacctcttatctagtggCCTCCACCTCCCGGGTGCAAGTGATTCCCCTGCTTCAGCCgaatcttaaacccaa
ggttacctcttatctagtggCGTGAGCCACCACGCCTGGCCCTGCCAACCCATCTTATCAgaatcttaaacccaa
ggttacctcttatctagtggGGCATTGAACCCGGAGAGTCTACCCTTTTTGCTCTTGGCCgaatcttaaacccaac
ggttacctcttatctagtggCCACTGTGGGAAAGCACCATGCAGAGTGACTGGCACACAGgaatcttaaacccaa
ggttacctcttatctagtggGACTGAATTGATAGACAAAAAACCTGCAGGCTGAAGCATGgaatcttaaacccaac
ggttacctcttatctagtggCTGAATCAAAAACTGCGGGGACTGAGGCACGGGAAGGCTCgaatcttaaacccaa
ggttacctcttatctagtggTCTCTTCTGGCCAATGTTTTAAACACTAAAGCTTAAAGCCgaatcttaaacccaacttg
ggttacctcttatctagtggTGAGCTGGACATCAAAACATCTAGGACATGAATAGCTCTGgaatcttaaacccaact
ggttacctcttatctagtggGTAAAATGGTTTGTGGTGGGAAAGAGGAAGTAAGCAATACgaatcttaaacccaact
ggttacctcttatctagtggGTCAGTAACCAGTCTCTCTTACTCCCTGTGTCTTCTAGCCgaatcttaaacccaactt
ggttacctcttatctagtggTCCAAGCAATATCCAACCTAGCCATTGTACTTTTCCTGCCgaatcttaaacccaactt
ggttacctcttatctagtggCAGGGCCAAACATTTTTATAATGGCTTCCACAGGCAGAAGgaatcttaaacccaact
ggttacctcttatctagtggACACTAAAAGACCTTTCAGAATGTCTGAATTCATCCAGACgaatcttaaacccaactt
ggttacctcttatctagtggAGCTTATTAGCTCTGAGATAGAAAAGTGGCAGATCAGGTGgaatcttaaacccaact
ggttacctcttatctagtggTGGTGCTGAATTAGGCAGAGAAAGAGAAAGAGGAGGGACCgaatcttaaacccaa
ggttacctcttatctagtggGGAAGAAGGTAGTGTCCTAGTAAATTTGTAAGAAGGGTTGgaatcttaaacccaactt
ggttacctcttatctagtggGCTCCCATAATTCTTACAATGTTGTGGGAGGGACCCGGTGgaatcttaaacccaac
ggttacctcttatctagtggTTCTCGCCTGCTGCCGCCATGATTGTGAGGCTTCCCCACCgaatcttaaacccaa
ggttacctcttatctagtggTCAGCAGCGTGAAAGTGGACTAATACACACCACATCTATAgaatcttaaacccaact
ggttacctcttatctagtggGCTGTAAATATGAGCTTAGACCGGATGTATGCAGGAGCAGgaatcttaaacccaac
ggttacctcttatctagtggAACCCATGGAACAGTGGAAAAGAAGGGGAGCTCACCAGCCgaatcttaaacccaa
ggttacctcttatctagtggCCAAGCAACCCTGGTGATCCACAGGATGATGATAATACCCgaatcttaaacccaac
ggttacctcttatctagtggAGAGGGAGTTCCTCACATGGGAGGGTGTCCTTTGGATCACgaatcttaaacccaac
ggttacctcttatctagtggGTTGTTACTGAGTGAGGCTCTCTCAATCTCAGCGTCCTGGgaatcttaaacccaact
ggttacctcttatctagtggCTGGTCTCAGCACCCATTACCAGCAGAGGGCGCCAGAGCCgaatcttaaacccaa
ggttacctcttatctagtggCAAATTATTGCTTGGGGGTGGGTGGAGGGAAGTGTCACGGgaatcttaaacccaac
ggttacctcttatctagtggTCCCTCTTTCACTTTCTTAAGGAGGGAGAAAAACCTCTTTgaatcttaaacccaacttg

ggttacctcttatctagtggTGCCGTCACAGCGTTACTTGTTAATCAAAACAGGTTTCCCgaatcttaaacccaactt
ggttacctcttatctagtggCCGTTATCTCCTTCCTCCTGGGTGTGTATCCCCCTGTGGGgaatcttaaacccaact
ggttacctcttatctagtggGCAGGCCCGTGAATATGGCACCAGAGAAGACGGAGGAGGGgaatcttaaacccaa
ggttacctcttatctagtggCTGCTGCAGGGACAGAAACCTCTGCCAGGGAGACTCGCCCgaatcttaaacccaa
ggttacctcttatctagtggGGTCTCGGTCGAAAGAATCGCTAATCAAGAGCCACCAGCCgaatcttaaacccaac
ggttacctcttatctagtggGGAGGGAGCTATTACAAAGCAGTGCTGACATAGTGTGAAGgaatcttaaacccaact
ggttacctcttatctagtggCATCACCAGCTGCATTCATTAATACTTAGGCCTAACTGGGgaatcttaaacccaactt
ggttacctcttatctagtggGACTCAAATGAGTATTTCTGGGAAGAATTGTTGTTTGCGGgaatcttaaacccaacttg
ggttacctcttatctagtggGCTCTGCTCAGGACCAACGTAGAGATAAGTTGCCCTTCGCgaatcttaaacccaac
ggttacctcttatctagtggCCCCAGTTTCGGGAACACGTAGGGGGCCCAGGGCAGTGGGgaatcttaaacccaa
ggttacctcttatctagtggGCCAGGCATAGAGGGGTTTGGGACACATGTTCTGGACTAGgaatcttaaacccaac
ggttacctcttatctagtggGGCCCACTTTTCTGGGAAGGTGATGCCACGGAGAATGAGCgaatcttaaacccaac
ggttacctcttatctagtggTTCTGCTGAGGTGAAATCTAGTGGGGGACAGGAGACTCAGgaatcttaaacccaac
ggttacctcttatctagtggGGGGGCAATTTTCCCGCTTACCCGTAATGGCAGCCTATTTgaatcttaaacccaact
ggttacctcttatctagtggGACAGTAGGATGCCCTTTGCCAAGCCTGGGGACTTAGGCCgaatcttaaacccaac
ggttacctcttatctagtggGCCTTAAACACTTGAACAGCTCACTAGAAAAGAAAAAGTCgaatcttaaacccaactt
ggttacctcttatctagtggCCTGACTCCAGGGCATTGTGAATGCAGATGAAAGGGCACCgaatcttaaacccaac
ggttacctcttatctagtggGAGGAACTGGCGTGGGATGGAGGTAGGGAGGGTAAGGGCCgaatcttaaacccaa
ggttacctcttatctagtggCTGGAGGGAAGGGGTTCTGGTAATGTAAATTCCCTGGCTGgaatcttaaacccaact
ggttacctcttatctagtggCATGGAATTATGGGTGTCCACAACTCAAATCCTACCTGTGgaatcttaaacccaactt
ggttacctcttatctagtggAGCCCAGAGAGACTGAGAGGACAAAAGATTTTCCAGAACCgaatcttaaacccaac
ggttacctcttatctagtggTCATTCCTTGAGTTCTCCCTACACTCACAGCATGAAGGTCgaatcttaaacccaactt
ggttacctcttatctagtggGCTGTTTAATACACAGTCTCTTTAAGGGGGCACAGCACAGgaatcttaaacccaact
ggttacctcttatctagtggCAGTAATGAACTAGGTACTGTGGAAGGACACAAGGAGGCGgaatcttaaacccaac
ggttacctcttatctagtggGGCAACAGAAATGGCTTCACAGAGGCTGTTCTAGCAGAGCgaatcttaaacccaac
ggttacctcttatctagtggAAGAACATGAGAGAAGGAATGTAGGGAAGGACATTCCGGGgaatcttaaacccaac
ggttacctcttatctagtggGTTTGGAGGACTGGCAAGCAGCTTCACAGATGGGGGAAAGgaatcttaaacccaac
ggttacctcttatctagtggGAACCACGAACACAGACTGGGTCAGGAAGGAATTGGAGGGgaatcttaaacccaa
ggttacctcttatctagtggACCCCAACCGTGCACCGATTAAGCGTATGGAGTTGGCCCCgaatcttaaacccaa
ggttacctcttatctagtggTACCGTGTGAGTTCCACAACACCCTGGGTAGACTCTGTTTgaatcttaaacccaact
ggttacctcttatctagtggCACAGGCTGGTGCCCATCCGCAGACCATGCTTGGAGCAGCgaatcttaaacccaa
ggttacctcttatctagtggTGCCCTAGGAAGGTTACACTGGCAGCAAGATGGAGAAAGGgaatcttaaacccaac
ggttacctcttatctagtggTGCGAGGATCTGACCCGGGGCCCTGAATATCCTTAGGGGGgaatcttaaacccaa
ggttacctcttatctagtggGCGCTTGTTGAGTGTTGAGTATATGAGACGCGGGAAAGGGgaatcttaaacccaact
ggttacctcttatctagtggAAGAGGATCCTGAAGGCCAACCTAGGGTAGGGGAAAGCCCgaatcttaaacccaa
ggttacctcttatctagtggGAGTCAGAAGCAGCTCCAAACAGCCAAATTGGTGCATGTGgaatcttaaacccaac
ggttacctcttatctagtggGTTTTGCCGCAGAACATTCCTCTGTCCTAAAGTCATGGGCgaatcttaaacccaact
ggttacctcttatctagtggTCTGTCAGTGTAACCTTATCCCAGCTTCCTGTTTGAGCACgaatcttaaacccaactt
ggttacctcttatctagtggGGACCCGCTCTGCCACACAGAATGCAGTTCTGCTGTTGGGgaatcttaaacccaac
ggttacctcttatctagtggGGAAGGGAGGATTTTGGGGTGTAGGGGAGCTTGTAGCCTGgaatcttaaacccaac
ggttacctcttatctagtggCTTGAGTCGGCTGACACGTGCCCACTCCCCTGCCACCCCCgaatcttaaacccaa
ggttacctcttatctagtggCAGACACCTCTGCACATCACACAGCGCAGGTGTTCCAACCgaatcttaaacccaa
ggttacctcttatctagtggGGGCCCCCAGCCTCATGGAGACAAACTTCCCCAACTCCCAgaatcttaaacccaa
ggttacctcttatctagtggGGACTCCAAAGTGGACACTGAGCTAGGGGACTGCATCTGGgaatcttaaacccaa
ggttacctcttatctagtggGCTTGGCACACGGCAGAAGCTAAATGAATGGCTGAGGAGCgaatcttaaacccaa
ggttacctcttatctagtggCCCTTCTCCTTCCTTCCCATCTTTCCTACTTCTCTCTCCCgaatcttaaacccaactt
ggttacctcttatctagtggCCTTGATAAATAAACCAGACAGGATTCTGCTTTCTCCCTGgaatcttaaacccaactt
ggttacctcttatctagtggCCTTCAGTCCAGACCCACTTTCTGCCCACGTTATGGTTGCgaatcttaaacccaact
ggttacctcttatctagtggACCAGCCCCTATCCACTAGCCACCTGGTCTTTGCTTACATgaatcttaaacccaact
ggttacctcttatctagtggCGGAGAACTCACTACCTTTGAAAGCAATTACCCTTCACTGgaatcttaaacccaact
ggttacctcttatctagtggGTTGCCTTTGTGTGGTGGAGTGATGAAAGATTTGGGTTCCgaatcttaaacccaactt
ggttacctcttatctagtggCCTTGGGAATTTCTCAGACCGTGAGCTGAGAACTCTCTCCgaatcttaaacccaact

ggttacctcttatctagtggCAGATTGACCCCAGTTTTGTCCAGAGGAAGCAAGAATAAGgaatcttaaacccaac
ggttacctcttatctagtggTTTTGCTGCCAGTACTTCCAAAGTGTTTGGCTTACTCAGCgaatcttaaacccaactt
ggttacctcttatctagtggGATGTGGATCATTCAGTCTCTCCTTCACTGAAGATATTTGgaatcttaaacccaacttg
ggttacctcttatctagtggGGATGTTCTGAGAATTTTGGGAGGCAGGGTACCATATTGCgaatcttaaacccaact
ggttacctcttatctagtggTGGGTGCAAATCCAGGCTCTATTAGCTTTGTTGTTTTCAAgaatcttaaacccaacttg
ggttacctcttatctagtggGATGTTCGACTGACATAATTGGTGTGAGGATTCAACTTGCgaatcttaaacccaactt
ggttacctcttatctagtggCAGGGCCTACTTGAGGATGGAGGATCAGAGGAGGGTGAGGgaatcttaaacccaa
ggttacctcttatctagtggCCAAACTCCTGTAACACACAGTTTATCTATAGAACCAGCCgaatcttaaacccaact
ggttacctcttatctagtggCATGTGCCACAATTGGGTTTTACTTATTCGACAGAATTCCgaatcttaaacccaactt
ggttacctcttatctagtggCCAAGATCTCTGCTTGCTCCCGTCCTCCCAGGCATTAAAGgaatcttaaacccaac
ggttacctcttatctagtggCCTAACTGGTTGGTTAGGTTCTACCTGAGAGCAGACATCCgaatcttaaacccaact
ggttacctcttatctagtggCAGGAAGACAAGAATCATCCGAGAATCCAGGAATTCAGGCgaatcttaaacccaa
ggttacctcttatctagtggCACTAGTTAAGAAAAGGCCCCTGTAATTTTCAGAAGGGAGgaatcttaaacccaact
ggttacctcttatctagtggCTTCTGTCAGATTTGCACCATAGTATTTCACTTTCTTTGGgaatcttaaacccaacttg
ggttacctcttatctagtggTGGTATTGTGCTTTTAACTTGTTTCCAAATGTTTGCTGTCgaatcttaaacccaacttgt
ggttacctcttatctagtggTGATTTTCTATCTTGATCTTGTGCTCTGTGTGGTTCTAGGgaatcttaaacccaacttgt
ggttacctcttatctagtggGAGAGGGTGGATGTCTTGGCCTTGTTCTTGATATTAGGGGgaatcttaaacccaact
ggttacctcttatctagtggGCGTAGGCGTGTTGCAGATGTTCTTTCTCAAGGTGAGGTCgaatcttaaacccaact
ggttacctcttatctagtggGTCATTCTGTCATTTGCCATGAGTTGGATGTCAGAGTCCCgaatcttaaacccaactt
ggttacctcttatctagtggGAAGCTCTGTCGTTTGGCGCATATGTGTATGTATGTATACgaatcttaaacccaactt
ggttacctcttatctagtggAGGATTGCTACTTTTTCCCTGGCAAATTGAGCCTTTTTTCgaatcttaaacccaacttg
ggttacctcttatctagtggGGCTTACAAGAGCCACAAACTTAATGGAAATCAACTTAACgaatcttaaacccaact
ggttacctcttatctagtggCTGGGAAGTTTAAGAAGGTAGCTCTAGTTTTGAACATAGCgaatcttaaacccaactt
ggttacctcttatctagtggCTTTCTCTCTCCCTCTTTCCTCCTGCTTTTCTGTGTTAGCgaatcttaaacccaacttg
ggttacctcttatctagtggTCTCCATACACTGTCCAAGATTATAACACATGCTGTTAGCgaatcttaaacccaactt
ggttacctcttatctagtggACCAATCCCTATGGCCAGGGAAATGGGACACTGTGACCAGgaatcttaaacccaa
ggttacctcttatctagtggCCAACTCAAAAAGAACATGGTGGTGGTTGGTGGGGTGGGGgaatcttaaacccaa
ggttacctcttatctagtggCATCCTGAAGTCAAAAATTATACCACCCAATATGCAGCTCgaatcttaaacccaact
ggttacctcttatctagtggGCCAGTTTGTTTCAAAGCCCTGGTTATGTCACTTTCCTGGgaatcttaaacccaactt
ggttacctcttatctagtggGAAATAGGAACCGTCTTCTATATTAACTTCCTCAGCTACCgaatcttaaacccaactt
ggttacctcttatctagtggAGCAAAAACCTTTGCCACATGAGTTCAGGAGGGAGGTAGGgaatcttaaacccaac
ggttacctcttatctagtggGAGGTTTAATGGGCTCACCGTTCCTCATGGCTGGGGAGGCgaatcttaaacccaa
ggttacctcttatctagtggCATGGTGGCAGCCAAGAAGAGTGTGTGTAGGGGAACTCCCgaatcttaaacccaa
ggttacctcttatctagtggGATTCAGTTACCTCCTACCGGGTCCCTTCCACGACACGTGgaatcttaaacccaac
ggttacctcttatctagtggCCTGGGAGGTTGTTCCACACATCAGGAGTAACACAGACACgaatcttaaacccaa
ggttacctcttatctagtggGGCAGGAAGCATATGGCAAGAATGAGTAAAATCGGAATGGgaatcttaaacccaac
ggttacctcttatctagtggGTACCAATTAGCCTATGTTTAAGGAGCCGTGAAGGACCGCgaatcttaaacccaac
ggttacctcttatctagtggCTGCTGCTCTCTGGCCTCCAGTTCTCAGCCTCCCAGTCCCgaatcttaaacccaa
ggttacctcttatctagtggCCGTCCCATCTTCAGTGGCACCCACCTCCATTGCAGCACgaatcttaaacccaa
ggttacctcttatctagtggTTACCTGTTTGTGCGGCTTTCTCTACCATATTACAGGTGCgaatcttaaacccaactt
ggttacctcttatctagtggGTCAGTAACTAGACATTATTTATCTTTGTGACCGCGATATgaatcttaaacccaacttg
ggttacctcttatctagtggGTAATAGCAGAGGTGGTCATTTCCCCTGGTGCACTGTTCCgaatcttaaacccaact
ggttacctcttatctagtggGCAAATGCTCCAGTTTCAGGGAAAACTAAACTTTCCTTTTgaatcttaaacccaactt
ggttacctcttatctagtggAGTGCAGCAGGAAATTCATATCTGGGACCTCTCGATCCTCgaatcttaaacccaac
ggttacctcttatctagtggGTAGCTTGCCGTGCCACCAGTCAGGGTGAGAACTGCTGGCgaatcttaaacccaa
ggttacctcttatctagtggCGGCATTCTCTCCCCCACCCTACCTCTGAATCCCTAATTGgaatcttaaacccaac
ggttacctcttatctagtggCAGCTGTACCACACTGCTGCCCAGATCCTTCAAGACTTTAgaatcttaaacccaac
ggttacctcttatctagtggGCAGGACTCTCTGGCCTGCTAATTGTGCCTCTCCTGTTACgaatcttaaacccaact
ggttacctcttatctagtggCCCAGACCGCCCTAGGCTGATTTGCTGGGAGTTGGAAACCgaatcttaaacccaa
ggttacctcttatctagtggTGATATTCTCCCAGGAGCACTAGTCCCGGCCACCCTGCTGgaatcttaaacccaa
ggttacctcttatctagtggGGCTTCAGGCCCCTCCTGCGTCCTTTCCTAGCTTTTAGTAgaatcttaaacccaact
ggttacctcttatctagtggCTGCCTAAAGCCGTACAATTTCTCCGTGTTGTTCTTGTCCgaatcttaaacccaactt

ggttacctcttatctagtggGGAGCCTACCTTCCTTAAACTGCTTTCCCAATGTGAGGCCgaatcttaaacccaact
ggttacctcttatctagtggCCAGCTGTTACTGCTGGCTTCCCATGATCGTCAGTGACTTgaatcttaaacccaact
ggttacctcttatctagtggCATATGCACAAGTGGAGGGGAAGAAAAAAGCTTGCTCCTGgaatcttaaacccaac
ggttacctcttatctagtggCCCTGATGTGTCACTAGGTGCATACACATGATGAAACAGGgaatcttaaacccaact
ggttacctcttatctagtggCATTGCAAATGACTCCCTGGCCCCTCTGCTTCCTTTCCCCgaatcttaaacccaact
ggttacctcttatctagtggGAGCTCTGGAGGAAGCAGGAAGCGTGAGCAAGGCTGGGCCgaatcttaaacccaa
ggttacctcttatctagtggGGCCCGTGTCGCTGTTTTTATCAGTGCAATTGCTGCATTTgaatcttaaacccaacttg
ggttacctcttatctagtggGGGAGATGCCCTGCAGATGGAGTGGCTTCTTTCAGGGGGCgaatcttaaacccaac
ggttacctcttatctagtggGTGCGGGAATGAGGATTTGCAAAGGTAAATAGAGCTGTGAgaatcttaaacccaact
ggttacctcttatctagtggGGAGTGAAAAAGAGCCAGAAACACCTCTTTGAAGAGCAGCgaatcttaaacccaac
ggttacctcttatctagtggACAAAACCAGTTCCGGGCCGGGTACGGTGGCTCACACCTGgaatcttaaacccaa
ggttacctcttatctagtggCCCAGGAGGCAGAGATCGTGGTGAGCCAAGATCACGCCATgaatcttaaacccaa
ggttacctcttatctagtggTAGAGAAGACATGGCATCTGCCTCCTCAGCCACTCCGTGGgaatcttaaacccaac
ggttacctcttatctagtggGATTGTTGTCAATCCTGTGATAGAGATAAACCAGGGGCACgaatcttaaacccaact
ggttacctcttatctagtggGAGAGATCCGAGCAGTTTGAAGGAAGTACTCCCATAGCTCgaatcttaaacccaac
ggttacctcttatctagtggAGCACCTGACAGGTATCGAGGCCCCGCGTGTGTCAGGCAGgaatcttaaacccaa
ggttacctcttatctagtggGCTTTCCATGCACTGTCTTGTTTAACCATGGCCACCCCTCgaatcttaaacccaact
ggttacctcttatctagtggAGTACTGGTTGACTATCCCACATTCCAAAATCTGAACTCCgaatcttaaacccaactt
ggttacctcttatctagtggGACCGTAATCTCTCCAACTAGCATTATTGGGTTAATTGGCgaatcttaaacccaacttg
ggttacctcttatctagtggCCTCTGCCCTAGAAAGAGAGGGATGGTTAGAACACTGGCCgaatcttaaacccaac
ggttacctcttatctagtggGAAAGTTCAGGTCCCATCACTTATTAACTGTTTGACCACAgaatcttaaacccaacttg
ggttacctcttatctagtggCATTCCTGCCCCTGTAACTGAGGATAGTACTATCTACTGCgaatcttaaacccaactt
ggttacctcttatctagtggACCTGTGACCCTGCTTTGCAAACAGCTAATTGAAAACTGCgaatcttaaacccaact
ggttacctcttatctagtggCTTCCAGAGTTGAAAAGCAGAAGTGTTAAGGATTGGGTGGgaatcttaaacccaact
ggttacctcttatctagtggGGAGGTATCAATTTCTAATAGGGTCAGGTCTGGGAAAGAGgaatcttaaacccaact
ggttacctcttatctagtggGCTTCATAAGCCTGATCAACTTGTCCTTATGAAACTTTTAgaatcttaaacccaacttg
ggttacctcttatctagtggTCTTTCTGCTCTTCACCTAGTTAAGCCCCATTCAGTCTTCgaatcttaaacccaacttg
ggttacctcttatctagtggCCCTCCTCCCACAACCAAGTAATGCCCCGCTAACCCATGGgaatcttaaacccaa
ggttacctcttatctagtggGCTATCCCTCCCTAGCCCCACACTCCCTGACGGGCCCCAGgaatcttaaacccaa
ggttacctcttatctagtggCTTTGCAATTGTAAATAGTGCTGCAAGAAACATGCGTGTGgaatcttaaacccaacttg
ggttacctcttatctagtggGATTGCCATTTTAACTGGTATGAAATGGTATTTCATTGCGgaatcttaaacccaacttgt
ggttacctcttatctagtggTTAGATCCCATTTGTCAACTTGGGCTTTTGTTGCCATTGCgaatcttaaacccaacttg
ggttacctcttatctagtggCTTCACAGGGGCAGGAGCGGTATCTGTTTAGTTCGCCTCCgaatcttaaacccaac
ggttacctcttatctagtggTACCTATTGCAGTACCTGAGATGAGTTAGGTGCACAACAAgaatcttaaacccaactt
ggttacctcttatctagtggTCCTTAACCTTGATATGATCTTTCATACAGTAGCTCGTGCgaatcttaaacccaacttg
ggttacctcttatctagtggGGATGGTAACTTTGGAGAGGAAGTCCTAATTTAATGCCTGgaatcttaaacccaacttg
ggttacctcttatctagtggGTGCTAAAAGCTCAGAACTCAGTTTGCTGCCTCCTAGCTCgaatcttaaacccaact
ggttacctcttatctagtggGTTGGAGCATGTTGTGTGAATGGCATTTCCAGAGTTTCTGgaatcttaaacccaacttg
ggttacctcttatctagtggAAGTCAGGGTTAGCTTCAACTATATGCCAGGCAAAATCCCgaatcttaaacccaact
ggttacctcttatctagtggGATGGAGGTTTACTCCTCTCTTAGGTAAGGTCGGAAGGCGgaatcttaaacccaact
ggttacctcttatctagtggCTTCCTAGCTTTTACTCTTTCACGCGTGGCTTCCATTCCCgaatcttaaacccaacttg
ggttacctcttatctagtggTGGCACTTCCAGCCAGCAACCAAGGAGAAAGGGGAAGGCAgaatcttaaacccaa
ggttacctcttatctagtggCGTTGCTCACACCCCACTGGGCAGAATGTGTCATGTGGCCgaatcttaaacccaac
ggttacctcttatctagtggGGCAGACATGTGCCCAACTGAGAAAATAGACTTCTGTTACgaatcttaaacccaact
ggttacctcttatctagtggTGATCTCGGTTCACTGCAACATCCACTTCTTGGGTTCAAGgaatcttaaacccaactt
ggttacctcttatctagtggCCTCAGCCTTCCAAGTAGCGGGGATTATAGGTGGCGCCCAgaatcttaaacccaac
ggttacctcttatctagtggGGCCACGCTGGTCACCAACTCCAGACCTCAAGTAATCTGCgaatcttaaacccaac
ggttacctcttatctagtggGCGTGAGCCACCATGCCCAGCTGGAAGTTCTTCTTTTTAAgaatcttaaacccaact
ggttacctcttatctagtggAAGAAAATCGAAATACTTGGAAGTACTCCCATGACATGACgaatcttaaacccaactt
ggttacctcttatctagtggCCGTGAATTTTGCCTCTGGATTGACTCCCTTGTGGATTGGgaatcttaaacccaacttg
ggttacctcttatctagtggGACTGAACATGTATTGAGGGTCTGCTAGGTGCCAGGCGCGgaatcttaaacccaac
ggttacctcttatctagtggCCATGCTGTAACCAGGTGCTGACTGAGCTGTGCAGCAACCgaatcttaaacccaac

ggttacctcttatctagtggCTCAGCAATCTTAAACTTTAAAAGCTAAGTCTGGATTTCCgaatcttaaacccaacttg
ggttacctcttatctagtggCTTGCTTCCTTAGGGTGGGCCATATTGCAGGGCAAAGGACgaatcttaaacccaac
ggttacctcttatctagtggGTGTGCTTAACCTGGGATCTGGCACAAACAGGCTCACATGgaatcttaaacccaac
ggttacctcttatctagtggTCTTTGAGCACGTGCCAGGCACTTTGGTTTATGGAATCACgaatcttaaacccaactt
ggttacctcttatctagtggGTTGAGGAGCCTTGGGTCCCACGACCTCACACAGCCTACCgaatcttaaacccaa
ggttacctcttatctagtggCTTGCTACCTTCACAATTACAGACTCAAGGAAGATCCGCCgaatcttaaacccaact
ggttacctcttatctagtggCTAGTCCTGCCACTGCTGTGAACAGGTGTACCACTATGCCgaatcttaaacccaac
ggttacctcttatctagtggGTGACAAAGCCAGGATGCAAGCCTCCTGCCATAGGTCCAGgaatcttaaacccaac
ggttacctcttatctagtggGAATGCTCAGGCCTCGTGGGAGCCCCCTTGAGCTCTGTCCgaatcttaaacccaac
ggttacctcttatctagtggGTTTGGGCAGGATGCAGATTGCTAACATCTGTGCACAGCCgaatcttaaacccaact
ggttacctcttatctagtggCTGGGGCAATTAACCACTTGGAACTTTAGAGCAGAACCTCgaatcttaaacccaact
ggttacctcttatctagtggGACAAGAGCAAGACGATAGCCCAGATCTCCTACCCACTGCgaatcttaaacccaac
ggttacctcttatctagtggCCTCACTACATGGAGAATGAATAAAGAAGGGAGTGGATTCgaatcttaaacccaact
ggttacctcttatctagtggCATATGAGGATGTGGATCACATAAGCATGTGTAATCATGGgaatcttaaacccaacttg
ggttacctcttatctagtggTTAAGGCTTACTGTTAAGTGATGGGAGGAAGGACAGACACgaatcttaaacccaact
ggttacctcttatctagtggACCAGAAGTGGTTCACACCCTAGTTGTTATCAGTTGCACCgaatcttaaacccaactt
ggttacctcttatctagtggGAATGGAGGCAGGTGATGTCAGGAGGAGGATCATGGGGCCgaatcttaaacccaa
ggttacctcttatctagtggTTGGAATCCTGGTTCCATCATGTGCTACACTCACCCCAGGgaatcttaaacccaact
ggttacctcttatctagtggACCAGTGCTGTCTAGAAGATGCGTGACAGTAGGGACCCTGgaatcttaaacccaac
ggttacctcttatctagtggCAGTGTCCTCAACATTGAGTGCATACTGGACATAATAAGCgaatcttaaacccaactt
ggttacctcttatctagtggACACTCAGAATAAAAGGTACTGGCAGTGAGAGTGAGAGTCgaatcttaaacccaact
ggttacctcttatctagtggGAGAGTAAGGGAGGAAGAGACTCATTGCTCAGGGCAGAAGgaatcttaaacccaac
ggttacctcttatctagtggGACAGATGACAGTGAGAAAACAGAGCTGTATTCATCTGGTgaatcttaaacccaactt
ggttacctcttatctagtggGCACAGAGTTCAAGATGCGGAAGCTCCATGTGAGGTCAAGgaatcttaaacccaac
ggttacctcttatctagtggAGACACACAAGCACTGCATCAGGTATTTTAGGGATGCTACgaatcttaaacccaact
ggttacctcttatctagtggCATCCATTTTACAGCTGAGGAGACTGTGGTACAAAGAAAGgaatcttaaacccaactt
ggttacctcttatctagtggGCCCATGGTCACCCCCTTGAATACATTATTCAATAAGAGAgaatcttaaacccaactt
ggttacctcttatctagtggGTGAGTAATCTCAGAGGAACTGCAGAGATGAACTTCAGGGgaatcttaaacccaact
ggttacctcttatctagtggTGAATGCTACGGTGGACATCTAAGTCTGATATTTGTCCCCgaatcttaaacccaacttg
ggttacctcttatctagtggCTTAAAGAAGGAAGCAGTGATCAACAAGAAGTAGTATGGGgaatcttaaacccaact
ggttacctcttatctagtggCTGATGGAGTGTCAGGCTAGCAAAGCATCAGGTGCAACTGgaatcttaaacccaac
ggttacctcttatctagtggCAGTCAAGGATGTGGAAGTTTCCTTCAGAAGATTTTGGAGgaatcttaaacccaacttg
ggttacctcttatctagtggTGGAATGGGGCAGTCTTTAGCTACCCACCATATCTGTCTGgaatcttaaacccaactt
ggttacctcttatctagtggGGGTGCGCCCTTACAGGTGGAGCAATGGTGAGTGCACACTgaatcttaaacccaac
ggttacctcttatctagtggCGTGGCCCCTGCTGCTGTGTCATTTTCCTATTGGCTAGGGgaatcttaaacccaact
ggttacctcttatctagtggAAATAGAATGACTGGGTGAGTGCTTTGGCGGGAGTCAGGGgaatcttaaacccaac
ggttacctcttatctagtggGGGTGGAGCAGGTGATTGGAATGTAGTAGGGTGGAGCAGGgaatcttaaacccaac
ggttacctcttatctagtggGTGTCATGGGGTCAGGAGGATCAGCAACTAGGAGCCTGGGgaatcttaaacccaac
ggttacctcttatctagtggTAAGACAGGGACACACTCTAGAAGTGGTAACGAGAGTCGGgaatcttaaacccaac
ggttacctcttatctagtggGTGATGGATCAGGGCTGAAGCAGTCCGCAAGGTTAACTCCgaatcttaaacccaac
ggttacctcttatctagtggCCATCAGAGTTTTAAAGGATTGCTGAGCCTGCCAGTTGGCgaatcttaaacccaact
ggttacctcttatctagtggGTGTGTCTGGGGAGAGAGTCATGTGACTTGGGATATGGGCgaatcttaaacccaact
ggttacctcttatctagtggGGCAATAAGTTATTAGATCATCAGTGGGCTCTTATGTGGCgaatcttaaacccaacttg
ggttacctcttatctagtggATTAGAAATCCATCCTGCCAGACACTGTGGCTCATGCCTGgaatcttaaacccaact
ggttacctcttatctagtggTGGGAGGCTGAAGCAGGCCAGGATTTCTAGACTAGCCTGGgaatcttaaacccaac
ggttacctcttatctagtggGCTGGGCAAGCAGTGACCTCCCAACCCTGGAAGTATGCAGgaatcttaaacccaa
ggttacctcttatctagtggCACACACACACGCACACATACACACGCTCGCACTCTCCAGgaatcttaaacccaa
ggttacctcttatctagtggGAGTGGATTCATTTGGCCTACTAAAATGCCTTTGGTGCAGgaatcttaaacccaacttg
ggttacctcttatctagtggGGTTCCAGTCATTAATGTTCTGTGCTGCTAAACCCAAGCCgaatcttaaacccaactt
ggttacctcttatctagtggGGGACTATAGCTTAGTCGGACCTGTTTCTGAAACATGTAGgaatcttaaacccaactt
ggttacctcttatctagtggTACACAATGAGGCTTTCTCTCCAATGCCCCAGCCCCTTGCgaatcttaaacccaac
ggttacctcttatctagtggTCCCTGGTCCCTGCTTCCATCTGTGCAAACTGGACTGCCCgaatcttaaacccaac

ggttacctcttatctagtggGGGGACACTATAAAAGGCTACTGATGCGACTGGTGGAGGCgaatcttaaacccaac
ggttacctcttatctagtggGGCTCCCTGCTCCCTGTCCAACCAAGGACAAATCAAACAGgaatcttaaacccaac
ggttacctcttatctagtggAGTCATGGCTAAAACTTGGACTCTGCTATTTCGCCCAGAGgaatcttaaacccaact
ggttacctcttatctagtggCCCTTGGGGCAGACACCCTGGAGCAAGGCTGCCTAGGTCCgaatcttaaacccaa
ggttacctcttatctagtggGGGCCATTCACTCCTTGGCACAGCGGGAATGGTAACAGTGgaatcttaaacccaac
ggttacctcttatctagtggGGGGTTGTCCTGAGAACCTTCACGAGATCATGAAGCAGGCgaatcttaaacccaac
ggttacctcttatctagtggGACGGAGGCTAATGACATGGGCTCTGGAATCTGACCTGCCgaatcttaaacccaac
ggttacctcttatctagtggCAGTGCTCCGTAGTTAACCTTCTTTAGTCTCAGTCTTCCCgaatcttaaacccaacttg
ggttacctcttatctagtggCTCGGGAAAATTTGCATGCTGGGCTCTGAGGTTTTCCAGGgaatcttaaacccaact
ggttacctcttatctagtggACTTTCATTACCTAGCCTGGGGCTGGCACATAGTAGGACCgaatcttaaacccaact
ggttacctcttatctagtggACCTCGACTCTCTTGGCTGACAATCCTCCACGACCCTCCCgaatcttaaacccaac
ggttacctcttatctagtggGCACTGTGTGCTTGTCTGTACACCACACAATCACCAGGCAgaatcttaaacccaac
ggttacctcttatctagtggAACGCCATGCACACATACATCCACAATATACGCATGCCACgaatcttaaacccaac
ggttacctcttatctagtggGCATACACAAACACCATATTGGATACACCAAACACCAGACgaatcttaaacccaac
ggttacctcttatctagtggACACACCACATGAACATACCACACACACACACCACAAACCgaatcttaaacccaa
ggttacctcttatctagtggGCACACACGCGTATGCACCATTTACCTTTCAGCTGCTCTGgaatcttaaacccaact
ggttacctcttatctagtggCAAGCGTGAGGTCTTTCATCAGCTGTGGCCACCTCCCTGGgaatcttaaacccaac
ggttacctcttatctagtggTTCCCCTTCTGATTGTCAAGTAACACCATTAGCCCAGGCCgaatcttaaacccaact
ggttacctcttatctagtggGATGAGGCTGGTTATTCTTTCCAAAATTCCGCAAGGCTCCgaatcttaaacccaactt
ggttacctcttatctagtggGGCCAGAACGTTGGGCTGGAAGGTGTGCTAGAAGAGTGTCgaatcttaaacccaac
ggttacctcttatctagtggTCCCCAGAGCTCATGCTGTTCATACATATTACCAGGCCCCgaatcttaaacccaact
ggttacctcttatctagtggCAATAGAGGTCTAGGGTGGTGCCCAGAAATCTGCCTTTTGgaatcttaaacccaact
ggttacctcttatctagtggCACTTGGAGAAACACCAGAATCCCTTATTTCTTAGCTGGGgaatcttaaacccaactt
ggttacctcttatctagtggGTAAGTAGCAAGGCTGGGACTGGACCTCAGGTCTCCTTGCgaatcttaaacccaac
ggttacctcttatctagtggTGCCTTTAGTTCTCCAAGCATTATAGAGTTGGTTGGTGGCgaatcttaaacccaacttg
ggttacctcttatctagtggCTGAGAGCAGACAAGGCTTTTCTGTGGTCAGTCTTGTTCCgaatcttaaacccaactt
ggttacctcttatctagtggCTCCTCTTGACTGTCACCAGTTGTCACATAAGGCCACCGGgaatcttaaacccaac
ggttacctcttatctagtggCCCTGCTCAGAAGCCAGACCTATTGCTTAAAGCTACACGCgaatcttaaacccaac
ggttacctcttatctagtggCTGCTTAGACCAGCTGTTAGAAAACTCCAGCCTGTAGGCCgaatcttaaacccaac
ggttacctcttatctagtggGGCTGCCTTTGTACAAAGTTGCAATAGATCATATGGCCTGgaatcttaaacccaactt
ggttacctcttatctagtggCCCTCCCAGGGGCCATGGAGCATACATACGCAACAAAGAAgaatcttaaacccaa
ggttacctcttatctagtggGATATGTTCTACAAGGTCCCTTTCTACCCATAGCCTCGCCgaatcttaaacccaact
ggttacctcttatctagtggATCTCAGCAACCTAATGACTACCTTTCTGAACCGTCTTGGgaatcttaaacccaactt
ggttacctcttatctagtggAAGCACTGGGTATCTCTTCTCCCCTTGCCCTTCTGCATTCgaatcttaaacccaactt
ggttacctcttatctagtggGGCCAAGGGCTGAGGGCCGCTTCCAGAAGCAGATTCTTCTgaatcttaaacccaac
ggttacctcttatctagtggGGCCCCAGAGAGATTCCAAAGGCCTGAGGACTTGGCACAAgaatcttaaacccaa
ggttacctcttatctagtggTTCAATTTCACATATGCTTGTATACTCACATACTCCTGGGgaatcttaaacccaacttg
ggttacctcttatctagtggGACAAAAGGCTGTCATCATAATATGTGTTTCTGCGGGGGGgaatcttaaacccaact
ggttacctcttatctagtggCTGCTTTGATAGGAAGACTGATCTGATGCTTAAGCCCACGgaatcttaaacccaact
ggttacctcttatctagtggGCACCTGGCCTTGAAATAATACTACTGTCTATTTTATAGGgaatcttaaacccaacttg
ggttacctcttatctagtggGCTTAAGTCAGTACTCTAAGAAGGCTGTCATTGTTCATGGgaatcttaaacccaacttg
ggttacctcttatctagtggATTATGTCCAGAAGGATTAAATTAGGGCATTACACGTGTGgaatcttaaacccaacttg
ggttacctcttatctagtggAGTCTCTGCCCTCTGCCTTTGGACAGAGTTGTGCTTTTCCgaatcttaaacccaactt
ggttacctcttatctagtggGATCTTGAGTGTCTCTCCATCTCCTCTTGAGTGAGCTTGCgaatcttaaacccaacttg
ggttacctcttatctagtggGCCAACTACATGGAACCAAGCCTGGACTAGTGTGCTGGAGgaatcttaaacccaac
ggttacctcttatctagtggGGCACAGCCAAGGTCATCCAGCACCAGGCCACCCTCAAACgaatcttaaacccaa
ggttacctcttatctagtggGGTCAGCCACAGTGAGCATAACTGCCTACCTAACACTCAGgaatcttaaacccaac
ggttacctcttatctagtggTGCACGAGTGAGTCCAGTCAAGATCAACCAAACCTAGTCCgaatcttaaacccaac
ggttacctcttatctagtggTTCCCAGATGATCTATAAGTTCATGAGATGTAACACATGGgaatcttaaacccaacttg
ggttacctcttatctagtggTGCTTTTCTTCCCTGTACAAATGGTCATGGTCCAGAAGCCgaatcttaaacccaactt
ggttacctcttatctagtggGCTTCCTTTGTTATAAATACCAACTCAGTCCATAGAGATGgaatcttaaacccaacttg
ggttacctcttatctagtggCTGAGTATTTCCCACTGATTACAGAAATGAAGTCTCCTGGgaatcttaaacccaacttg

ggttacctcttatctagtggCTGGAAAACAGGACATGTGTTTGAAGGAAAATATTCAACCgaatcttaaacccaactt
ggttacctcttatctagtggTTCCTGAATGCTTGTGTCAGCTTTTTGGTGACAAGAAAGGgaatcttaaacccaactg
ggttacctcttatctagtggGGTGAGTGGGATGTTGTCATCAGCTTTGATGCTATGATGCgaatcttaaacccaactg
ggttacctcttatctagtggGCTCTGGATGATAGGGTTGAACAGTTCTGTACCTCCTTGGgaatcttaaacccaactt
ggttacctcttatctagtggGTGTGCTCTCACAGTAGACCTGCAGCAAAATTGTGAGGTGgaatcttaaacccaact
ggttacctcttatctagtggGGTAGACATCTTGTGAAAGGAATCAGAATATGTTGTAGGCgaatcttaaacccaactg
ggttacctcttatctagtggGCAGTGCTATTATAAGATATAGTATGAGCCATAAGGGTGGgaatcttaaacccaactg
ggttacctcttatctagtggACCAGCTAGACAGAAGAGACAGGGAAAGAAACTGGCGTTGgaatcttaaacccaac
ggttacctcttatctagtggCCTCTCTGTGAAGTCCTGAGTCCATGGAAAGGCCGAGGGGgaatcttaaacccaac
ggttacctcttatctagtggGCAACATGGCAGCTCTATTAGTGATGGAAACTGGATTGGGgaatcttaaacccaact
ggttacctcttatctagtggACCCAGCCTCTGGCAGAGTGGGACAGCTGAAGCTTCCTGGgaatcttaaacccaa
ggttacctcttatctagtggCCCGGTGGAAAGCAGATCGCAATGGATACTTCCTGCAGTCgaatcttaaacccaac
ggttacctcttatctagtggAGCAGCGGTCAAGATAAACTAAGCCCCTTATCCCCAGCCCgaatcttaaacccaac
ggttacctcttatctagtggTGCTTCAGTCTTTCTTCCATGGGAATGAAGTCAGGGATACgaatcttaaacccaactg
ggttacctcttatctagtggGGGTTCTCCCCTAATGTCTGAGATAACACAGCTGGAGGCCgaatcttaaacccaac
ggttacctcttatctagtggCTTCTGGCTCAGGAGGAGTTGTAGAGGTGAGAAAACAGTGgaatcttaaacccaact
ggttacctcttatctagtggCAGACAAGACAGTGATCAAGAATGGGTTCACAAAAGAGGCgaatcttaaacccaac
ggttacctcttatctagtggTGGCTTGGCAAGCTCATTCAGTGCGGAAAACAGTTACCTGgaatcttaaacccaact
ggttacctcttatctagtggGGAAAGCCCATGTCCTAGGAAAATGAGTGCCAAATTCAGCgaatcttaaacccaac
ggttacctcttatctagtggCTCTGTTTGGGGCCAGGTACTAGAGCATCTAGGATGCCAGgaatcttaaacccaac
ggttacctcttatctagtggCCCCACTGCAGGATTCCAGTGACAGGTGGTGAGGTCCTGGgaatcttaaacccaac
ggttacctcttatctagtggCTGGGGAGATGGACAGAGGAATACAAGTGCATAAAGCAGGgaatcttaaacccaac
ggttacctcttatctagtggGAAGACCAACAGTGAGACAGTAAGATGTTTGAGGAAAGGGgaatcttaaacccaac
ggttacctcttatctagtggTCAGTGCCATGGAGACTAGTCTTCAGTTCTCTGAATGCCCgaatcttaaacccaactt
ggttacctcttatctagtggTCAGAAGACAGAAAAGTGGGCAAGGCATAAAGATGTGTTGgaatcttaaacccaact
ggttacctcttatctagtggGTGGATGAAATCGCTCTCTCTCTCACACACACACACACGCgaatcttaaacccaac
ggttacctcttatctagtggCCCCTTCTTAAAGGGCAGGGGCAAGCAGAGCATCCCTCAAgaatcttaaacccaac
ggttacctcttatctagtggGAGAGATTTCCAGAGTGTAGTGTCACTGAAGTGATGGACGgaatcttaaacccaact
ggttacctcttatctagtggTTTACCTGCAGTGTTGGTATATTGGCATGGATACACCCTCgaatcttaaacccaactg
ggttacctcttatctagtggCTCCAATTTGGCTTTTCTTCTTATCTGCAAAATGGGAAGCgaatcttaaacccaacttg
ggttacctcttatctagtggTTCCAGTATTTGGTGACTGAGAACCTGAATATTTAAGGATgaatcttaaacccaacttg
ggttacctcttatctagtggCACAGGGAAGGGGCAGTGGAGGGATGGGTGCTCTCTGGGGgaatcttaaacccaa
ggttacctcttatctagtggGGGAAAGACTCCCTCTCTCTGAGCCCAGAGGACTGAGAAGgaatcttaaacccaac
ggttacctcttatctagtggGGCAGCCAGGGCTGTAACTACATCTCCTGCTCTCGAGGGGgaatcttaaacccaac
ggttacctcttatctagtggGGGGGACTATATGGGGCAAAGAGCTCTAACCCACCTCTCCgaatcttaaacccaac
ggttacctcttatctagtggCCAGAAGGCAGAGAAGCCAACATATGGAATCCCAGCATGGgaatcttaaacccaac
ggttacctcttatctagtggTGGACTGGCATAGAGAGAGGGTTTCATTAGCAGGCCTGAGgaatcttaaacccaac
ggttacctcttatctagtggAGGAAACTAGGTTTCCAGCCCTTGGGACTGCAATGGGCCCgaatcttaaacccaac
ggttacctcttatctagtggGACCTCTGGGGTCAGTGTTGCTGTGGGGATGTGGCATGACgaatcttaaacccaac
ggttacctcttatctagtggTTCATGGTGTCATACCACCTAGAAATAATTACTCCCTCTGgaatcttaaacccaactg
ggttacctcttatctagtggGATCCCTTGCTGTAAGACATAGGTTCAGCCTGATCCTGGCgaatcttaaacccaact
ggttacctcttatctagtggGAGCAGGAAGCACTGCGCAGAGCTAACAGCAGAGACCCAGgaatcttaaacccaa
ggttacctcttatctagtggCCTGGACTGGACGCCTTGTTTCATGTTTCCTTACTTTGTGgaatcttaaacccaactg
ggttacctcttatctagtggAATGGGGTAATATGAGTAGATAGCTCCACTCAAAGAGTTGgaatcttaaacccaactt
ggttacctcttatctagtggGATAACTAGCCATAAAGTTCTTAGCATTAGAGTGACTTGCgaatcttaaacccaactg
ggttacctcttatctagtggCAGAAGCTTCTACTACCCTGTTGGCATACTCAAGCCTAGGgaatcttaaacccaact
ggttacctcttatctagtggGGGCTAAAACGGGTGGTGGAGCCATTGGTCTCAAGTGGGCgaatcttaaacccaac
ggttacctcttatctagtggCAGGGATGTCAGCATCAACAGCACTGAACAGCCAGGGTTGgaatcttaaacccaac
ggttacctcttatctagtggGAGAAGCTGCCCTCCGCATGGGACCATGGGGTCATGCCAGgaatcttaaacccaa
ggttacctcttatctagtggGCACAGCCAGGGAAGTGCTGGTGAGAAAGAGACCAAGAAAgaatcttaaacccaa
ggttacctcttatctagtggGGAGTGAGGGAGAGTAAAACAGTAGGGAAATAAAAGAGCCgaatcttaaacccaac
ggttacctcttatctagtggCCCTAGCCCATTGCAATCTCATGATTAGGATGGTAAGCCCgaatcttaaacccaact

ggttacctcttatctagtggGGGTGCATTTTGTATTGTTTAGTGCTTAATTTGATGTCTGgaatcttaaacccaacttgt
ggttacctcttatctagtggCCTTCCACTGTTTAGCATATCCACTTTTAACCAAGGGCCCgaatcttaaacccaactt
ggttacctcttatctagtggGGGGTCATCTTCTTACTCCCTCTCCAATTCTATCCCTCAGgaatcttaaacccaactt
ggttacctcttatctagtggCTGCAATCAACTTTAAATTGGAAGAAAGGATCAGAGTCTCgaatcttaaacccaactt
ggttacctcttatctagtggCCTTGGCCTTGGGGTGGTGTTTCTGCCGTGGAGTGGGATCgaatcttaaacccaac
ggttacctcttatctagtggGCCTCAATGGCACTGAGTGATTCTGCTTAGGTGAATGTGCgaatcttaaacccaact
ggttacctcttatctagtggGGCAGACACAGACAACACTCTCTCTTAATCCCCAAGAAGGgaatcttaaacccaac
ggttacctcttatctagtggTTGCTTCCTTCTTGTGAATGAAAGATCACACAGGCATGGGgaatcttaaacccaactt
ggttacctcttatctagtggTGGAGCAATGAAATAGTGATACCTGCAAAGCCTGAATAACgaatcttaaacccaact
ggttacctcttatctagtggGAGCCAGTGAAGGTCACACGTTTTGCATAAAGTGCCTTGCgaatcttaaacccaact
ggttacctcttatctagtggGTGAACTTCGCCTGAGGAGAGGAAAGTGACTTATTAGAGCgaatcttaaacccaact
ggttacctcttatctagtggCATCCCACAGGCGGCTGCAAGAACTCGGTGAGGAGCTGGGgaatcttaaacccaa
ggttacctcttatctagtggGGCTGAACTGAATGAGAGTTGCTTAGGGAGCAGGGCCAGGgaatcttaaacccaa
ggttacctcttatctagtggAGGGCTGCCATGTTTATTACCTCCTTTTATTCTCAAAGTAgaatcttaaacccaacttg
ggttacctcttatctagtggGGTGTTGTAGCTGACATTCAAGCTCTGTTAGATCCTGAGGgaatcttaaacccaactt
ggttacctcttatctagtggGCTTTATGAAAGGATCGAGTCTTTGCACTTAGAGCAAGGGgaatcttaaacccaactt
ggttacctcttatctagtggGACTTCATAGCATTGCAAGTTCCCTGACCTCCAAACTCTGgaatcttaaacccaact
ggttacctcttatctagtggCCCTTGCTGTAAGACATAGGTCCAGTCTGATCCTGGCCTCgaatcttaaacccaact
ggttacctcttatctagtggGCCGTTCAGGAATCCAAATGCAATCTAGAGCAGGGAGCACgaatcttaaacccaac
ggttacctcttatctagtggGAGTAGGGACCAAAGACTCAGACTTCCTGGACTGGACCCCgaatcttaaacccaa
ggttacctcttatctagtggGGAGGTAATACAGATAGGTAGCTACCCTCAGAGTGGTGTGgaatcttaaacccaact
ggttacctcttatctagtggACTCCTACTACCCTGTTGGCACACTCAAGCCTAAGACTAGgaatcttaaacccaact
ggttacctcttatctagtggCCTGAACTGCTAGGTCAACAGGCAGAGGCTGAGTCCTGCCgaatcttaaacccaa
ggttacctcttatctagtggCTCCACCCAGGTGCCACCCCAGGCTAGCTAAAGCTGCCGGgaatcttaaacccaa
ggttacctcttatctagtggGGCAAGGACGTCAAGTTCAAGTCTTCACTGAGGAGCTTTCgaatcttaaacccaact
ggttacctcttatctagtggTGCAGATACATGACTGGGGCAACTGGAAGACCTTCCTGCCgaatcttaaacccaac
ggttacctcttatctagtggATACTTCACAGAATAATACAGGAGAGGATAGAGGTATTGGgaatcttaaacccaactt
ggttacctcttatctagtggGAAAGAGAGTAGGTTTTGTACTCCAGATGGTCTTCAGTTCgaatcttaaacccaactt
ggttacctcttatctagtggACTATGGAGGATGTTAACTGTTTGGCAGGATTGTTGTGAGgaatcttaaacccaactt
ggttacctcttatctagtggGTGGGCTGAAGTCTTCCAATATGGTTTCCTGGATGAGCTCgaatcttaaacccaactt
ggttacctcttatctagtggAACAGAGGAGAGAGAAGCAGAGGCTGTAACCAAGGCACTGgaatcttaaacccaa
ggttacctcttatctagtggCTGATGGGATAGAGGTGTTAGTTACCTGCTTCACACCCACgaatcttaaacccaact
ggttacctcttatctagtggAGATACAACAACTTTTGCAATGTGCTTGATCCCAGGTGACgaatcttaaacccaactt
ggttacctcttatctagtggTGGTTAGTTGAATTAAGTCATGGATACCTTTTCCACCTTTgaatcttaaacccaacttgt
ggttacctcttatctagtggCCCAGTTCAGACTAATGAGAGTTAAGGGGAAGTTGGCTGGgaatcttaaacccaact
ggttacctcttatctagtggGGGAACCACTCTAGGAACCATTCTATAACCATAAGGGGATgaatcttaaacccaact
ggttacctcttatctagtggTGGAAGGAGGAGGTGTTCTGATAAGTATCCTTGAGCTGCCgaatcttaaacccaact
ggttacctcttatctagtggGGATGATTTGGCTGACTACAGACAGCATAGATGACAGAGGgaatcttaaacccaact
ggttacctcttatctagtggGCTGCACGGGTCTAGGAGAAATAGCATATGCCATAGAGCCgaatcttaaacccaac
ggttacctcttatctagtggGAGGCTCTACCATCTGGATTTGTGTGAGCACACTCTATGAgaatcttaaacccaactt
ggttacctcttatctagtggAATCACCTGATGACACATTTTGCAGAATGTATTCCCATCCgaatcttaaacccaactt
ggttacctcttatctagtggCGCTTCCCTTCTCCTCTCTAAAGATCAGCGTAGTTCATCCgaatcttaaacccaactt
ggttacctcttatctagtggAGGCCTAGTAGGAAAAACCCTCTTTTGATGGAGAGTTGGGgaatcttaaacccaact
ggttacctcttatctagtggTCCCTAGGAAGCACCTCTCAAATGTGTCTATCTGTAATACgaatcttaaacccaactt
ggttacctcttatctagtggAGGCCACCGAAAATCTAGCCAAGTTAGAATGAACACTACCgaatcttaaacccaac
ggttacctcttatctagtggCTGGAAATCACAACCATGTCACCCAAAGGAAGAGAGCCAGgaatcttaaacccaa
ggttacctcttatctagtggAGCATTCCCAACAGTTTGCTCCGTGCCCTTCTCCAAACCCgaatcttaaacccaac
ggttacctcttatctagtggGGAAACCCAAGTGAGCTTGTGCACTCTGAGGGGATTGGGGgaatcttaaacccaa
ggttacctcttatctagtggCAGGTGTAAGCTATTGTACGTGAGCACCACTGTGTATGGCgaatcttaaacccaact
ggttacctcttatctagtggCTGCTTCTGTCAAGTTTCAGGGAATATAAGGGTATGAATCgaatcttaaacccaactt
ggttacctcttatctagtggACCACCATGAAACCACCATGTCTGGTGTAACACACTGCCCgaatcttaaacccaac
ggttacctcttatctagtggTCTCCTCCTGATCTAGGATTAGCCAGAACCATGTGTGCTGgaatcttaaacccaactt

ggttacctcttatctagtggGGCCTACAAAGACAGAAGTGCTCCAATTTTTCTCCCATCCgaatcttaaacccaact
ggttacctcttatctagtggGTGCTTGAAATGTCAGCATTCCTTTGCTTTTAGTTCTTCTgaatcttaaacccaacttgt
ggttacctcttatctagtggCCCAGATGAGTAGAGGCAGGAAGTTTTAGTCATACTACACgaatcttaaacccaact
ggttacctcttatctagtggTTGATCACTATGTTCTTGCATGGACTGTGCTTCCTCTCCCgaatcttaaacccaacttg
ggttacctcttatctagtggCCATCCCAGCCCTTGTGGAGTCTAGAAAGAGACATAGAGGgaatcttaaacccaac
ggttacctcttatctagtggATGAAGAGGCATTTGGTGCTATGAGAGGGTATCGTAGGGGgaatcttaaacccaact
ggttacctcttatctagtggGCAAGGGAAACAAGGGGTTAACTAGGCAGCCATAGTTGGCgaatcttaaacccaac
ggttacctcttatctagtggGACTGTGAATGCCTTTAGTAGGGTTCCCAGCACATAGTGCgaatcttaaacccaact
ggttacctcttatctagtggTGATTACTGGTATCAGTTTTATTGATTGTGAAATGCAGCCgaatcttaaacccaacttgt
ggttacctcttatctagtggCCTTTGTCAAGGTACGCTAAAGTCTTCTAGTTATAAAGTCgaatcttaaacccaacttg
ggttacctcttatctagtggTGGGGGGAACTCAATTTTTATGTCACGATATGGGAGGCTGgaatcttaaacccaactt
ggttacctcttatctagtggTCCCTACCTCCTAAGACATGGAGTTTTGGCTTCTGACCTCgaatcttaaacccaactt
ggttacctcttatctagtggGATGTGCATTATCATTTGACCCAGCAAGTCTCCTTGTAAGgaatcttaaacccaacttg
ggttacctcttatctagtggAGACAAGGGCAAGAGTGTTCATTTCAATATTTATAACAGCgaatcttaaacccaacttg
ggttacctcttatctagtggGACTGTGCAGGAGTTAAAAACCATGAGGTGGATCTTATCCgaatcttaaacccaactt
ggttacctcttatctagtggCTCAGTTCCCCCAGTCCTAATTGGTTAAGTTTGGGCAACCgaatcttaaacccaactt
ggttacctcttatctagtggAGTGCTTAGAACACTACTCTGCTCTAAGGGCTCACATGTGgaatcttaaacccaactt
ggttacctcttatctagtggGGCTCATTAACATATGCATCACCTCAAACACCATTTTTTGgaatcttaaacccaacttg
ggttacctcttatctagtggGTCTCCTTGACTTATGATGGGGTTATGCCCTGGTAAGCCCgaatcttaaacccaactt
ggttacctcttatctagtggGCCTATAATCTCAGCACTTTGGAAGGCAGTGGAGGGCAGAgaatcttaaacccaac
ggttacctcttatctagtggAGGAGTTTGAGACCGGGCTGGCCAACACAGTGAAACCCCAgaatcttaaacccaa
ggttacctcttatctagtggCTCAAGAGGCTGAGGCATTGAACCTGGGAGCTGGAGGTTGgaatcttaaacccaac
ggttacctcttatctagtggCATTGCACTCCAGCCTGGGCAAGAGATAGAGGCCATATCTgaatcttaaacccaac
ggttacctcttatctagtggCACAGCTTAGCTTGGACTGCCTTAAATGTGCTCAGAACACgaatcttaaacccaact
ggttacctcttatctagtggCAGACACGGTAAAGATTGGTGGTTTACCCCCGTGCATGGCgaatcttaaacccaac
ggttacctcttatctagtggCTCAGCACTGAGGGAGTATTGAACTGCAGGTCTCTAGCCCgaatcttaaacccaac
ggttacctcttatctagtggTCTACTGAATGTATGTGGTTTTTGGACTGTTGTCAAGTTGgaatcttaaacccaacttgt
ggttacctcttatctagtggCCCTTTGACCATGGTCTCCACATGTCTCCCATGCACCAGCgaatcttaaacccaac
ggttacctcttatctagtggCCATGTGTGAGTGTGAATATGCAGTATTTGTTTCTCTGTGgaatcttaaacccaacttg
ggttacctcttatctagtggTGTACCATATTTCCATTGTCCACTCTTTGGTGATGGACACgaatcttaaacccaacttg
ggttacctcttatctagtggATAACTTGGCTCCTGTGAATAGTGCTTCAGTGAACATGGGgaatcttaaacccaactt
ggttacctcttatctagtggACTGATTTAAAAGCTTTTGGGGCCAGGCATGGTGGCTCACgaatcttaaacccaact
ggttacctcttatctagtggAAGCTTTTGGGTAAATACTCAGAAGTGGGGTTGCTGGATCgaatcttaaacccaactt
ggttacctcttatctagtggCATCCCCACAAACAGTGTACAGGTGTTCCTTTATTGTTACgaatcttaaacccaacttg
ggttacctcttatctagtggGTGCATAGAAAAAGGGCTATAGAAAGACTGCAGATTTTCCgaatcttaaacccaactt
ggttacctcttatctagtggAGGGAGGCAAGCATGACCTCATGTTGCTCATCTTGTCACGgaatcttaaacccaact
ggttacctcttatctagtggCTCTCCACCCAGGTTGGGCTCTGCCATGTTGAGATGACAGgaatcttaaacccaac
ggttacctcttatctagtggGTGCCTTGTGGGAGAGCTTTATGATTCCACAACTATGGGGgaatcttaaacccaactt
ggttacctcttatctagtggTGGGTTTCAAGGGAGAGAGTTTGCAAACTTCAGGCTCTCCgaatcttaaacccaact
ggttacctcttatctagtggAGGCCACATTCTCCTGCCAGAGGAAGCAGAGTGGATACCCgaatcttaaacccaac
ggttacctcttatctagtggCACCTCATTCTGCCCCTATCCCTACCCCACACCACAGCTCgaatcttaaacccaac
ggttacctcttatctagtggTTTTCCTGGGTGATTTTGGCATTGAGTTTGACCCAAGGACgaatcttaaacccaacttg
ggttacctcttatctagtggGGCGGGGAGAGAAGTGGGGGCAGGAGTTGAAGAATTGGGGgaatcttaaacccaa
ggttacctcttatctagtggCCACAGAGGGAAACTTTCAATGGTTACTCCTTATACTTCTgaatcttaaacccaacttg
ggttacctcttatctagtggGCAATGAGAAAATATTCATGTAGTAGTCACCTGTGTGTGCgaatcttaaacccaacttg
ggttacctcttatctagtggTCATCTGTCTATGGAGGCTGACCTTTGGGTGGTGAACAGGgaatcttaaacccaact
ggttacctcttatctagtggGTCAAGTTCCACAGCTCCTTTTCTCATGATAACAGAATAAgaatcttaaacccaacttg
ggttacctcttatctagtggCAGAAATGTAGAGAGACTATTATAATGGACACATGCCTTGgaatcttaaacccaacttg
ggttacctcttatctagtggGCTGCATTTGCTTTATTTTTTGTTTGTCTGTTTTGCTCAGgaatcttaaacccaacttgt
ggttacctcttatctagtggTGCCCAGGCTGGGTTCAGTGGTGTGATCTCCGCTCACTGCgaatcttaaacccaac
ggttacctcttatctagtggTCAATAGTTGAAGTGGTTCTCCTGCCTCAGCCACCCCAGTgaatcttaaacccaact
ggttacctcttatctagtggGGCCAGACTGGTCTCGAACTGCTGACTCCAAGTGATCTGCgaatcttaaacccaac

ggttacctcttatctagtggACATGAGCCACTGCACCCGGCCTGCTTAGCATTTTAGAGTgaatcttaaacccaact
ggttacctcttatctagtggTCCTAGTGGCATGGAAGGCCCAATCCAAATTCAAATGTCCgaatcttaaacccaact
ggttacctcttatctagtggGGGGAGTCTTTCGTGGTTTTCTGTAAGGCCTCTGGAGGGCgaatcttaaacccaact
ggttacctcttatctagtggTGAGGGATGCCTTGGTCCTCAGCCGGAAAGAGGCAGTTGGgaatcttaaacccaac
ggttacctcttatctagtggCCTGGGTGATGGTTTCCCATGGCAGAAGCTAAGGCACCTGgaatcttaaacccaac
ggttacctcttatctagtggCAGGAACAAGCCTGTGATTACTCATTGACCTGAGCTGGGGgaatcttaaacccaac
ggttacctcttatctagtggGCAAATCACATGCAAATCTGAGAAAACTCCACCGACTCCCgaatcttaaacccaac
ggttacctcttatctagtggGTGAAGAGCATCTATTAGTTTATTGCATCTGTTAGGAAAGgaatcttaaacccaacttg
ggttacctcttatctagtggCCAGACAAGTGAATTTGGCAGGTTATCAACGACCCTCTGCgaatcttaaacccaact
ggttacctcttatctagtggGCACTGCCCTGATTGCCTTAGGACTCCCAAATTCAAGCCCgaatcttaaacccaac
ggttacctcttatctagtggTCACTCACATCCTCTGGAAAATGAAGGGGTTGACCTAATGgaatcttaaacccaact
ggttacctcttatctagtggCTGGTTCTAAAGCTCTAGGTATCCCTGAAAAGATCTGGCCgaatcttaaacccaact
ggttacctcttatctagtggTGGAAATTGAGGTCATAGATATTCCTTCAGTGTGCACTGCgaatcttaaacccaactt
ggttacctcttatctagtggAGGAAAACAGCAATCCTCATTCATTTTACTACCTATCTCCgaatcttaaacccaactt
ggttacctcttatctagtggTGAGAAAGATTATTAATAGGACACTTGGATGCATTCATCCgaatcttaaacccaactt
ggttacctcttatctagtggGAAAGTCTTGTGAGTGTTCCATGGCCGGACCTGATAACTCgaatcttaaacccaact
ggttacctcttatctagtggACCTGGGTGTTCAGAGGACCTGGGTGTCTCAGCTTCCTGCgaatcttaaacccaac
ggttacctcttatctagtggGGGAGAGTACATTCCTTGAGTCCTTGCCAAGAGCCAGGCCgaatcttaaacccaac
ggttacctcttatctagtggCCTGGCCCAGATGTGGAGCCAAAGCCCTCCCTGCTTCCCCgaatcttaaacccaa
ggttacctcttatctagtggGCTGGTCAAGACTAAGAGAATGTGGATACAACAGAAGTGGgaatcttaaacccaact
ggttacctcttatctagtggTCCAGGTGTATTTGTTGGATGCTTGATTTTTGAGCTGCAGgaatcttaaacccaacttg
ggttacctcttatctagtggTTCCAGGTCTTCACGCCCATACTTGTTCTACAGGGAGAGCgaatcttaaacccaact
ggttacctcttatctagtggGCCCCCTTCGAGAGCTGCCTCTGACTCCTCAGGCTAGGCCgaatcttaaacccaa
ggttacctcttatctagtggGGTGTGCACCCTCCTAGTTCTTGGGCCTTCTCCTTTTGAAgaatcttaaacccaactt
ggttacctcttatctagtggCTGAGTTCAGATCCCAGGCTCCCTTACTTACTGGAGAGCCgaatcttaaacccaac
ggttacctcttatctagtggACAGCTGTGGACCAGGTTACTTAACCCTCATGTGCCTCAGgaatcttaaacccaact
ggttacctcttatctagtggCATGTGAAGTCCTTAGAACAGTGCCTGGCGTGTAGCAAGCgaatcttaaacccaac
ggttacctcttatctagtggTTGTCTTGCTTTCCAGTAAACAATAGTTCTACTTGGGTGGgaatcttaaacccaacttg
ggttacctcttatctagtggATCTGTCTCATTTACTGTTGTATACCTGGTGTCTGCAGGCgaatcttaaacccaactt
ggttacctcttatctagtggGCCGCGTGCTAGGCCAGAATATCTGCCACATGCTAGGAGCgaatcttaaacccaa
ggttacctcttatctagtggGGTTCAGAGAAGCACACTGATTTGCTTTGGGTCACACAGCgaatcttaaacccaact
ggttacctcttatctagtggGATTCTGGGCCCAGTGCTCTTTTCCTTCGTCTAAGCTGCCgaatcttaaacccaact
ggttacctcttatctagtggTAGGAGGTTCTGATTTCATGAAACCAAAGCCAAAGCAAAGgaatcttaaacccaact
ggttacctcttatctagtggGGCCTCTTCCATGGAGTTCACAGATTTCTGGGTAGAGGACgaatcttaaacccaact
ggttacctcttatctagtggGGTGGACCTCTTGAGCACTCTGAACTGTGAGAATTCGCAGgaatcttaaacccaact
ggttacctcttatctagtggTTAAGCTAGAAGACCTCTGAAGAACTGTTGCTATTGTACCgaatcttaaacccaactt
ggttacctcttatctagtggAATGTGCACACGAATCACCTGGGAGCCGTATCACAAAGGCgaatcttaaacccaac
ggttacctcttatctagtggTGAGAAAGTTTGGGGAGGGACGTGAGATTTGACATTTTATgaatcttaaacccaactt
ggttacctcttatctagtggCGGGGTTTCACCGTGTTGGCCAGTCTGACCTGGAACTCCTgaatcttaaacccaac
ggttacctcttatctagtggGGGATTACAGGTGTGAGCCCTTGTGCTCAGTCAGATTTAGgaatcttaaacccaact
ggttacctcttatctagtggGCTGATGCTGATGCTGCTGGATCAAGGACCACGCTCTGAGgaatcttaaacccaac
ggttacctcttatctagtggTTATGGGGCTTTCTTGGACTGCACTCTGGCCAAACGCAGTgaatcttaaacccaact
ggttacctcttatctagtggCTGCTGTGGAGCCGCCTGCATCCAAGTCCAGCTCTGCTGCgaatcttaaacccaa
ggttacctcttatctagtggTGGGCCCTTGGGACAGATTCTTCTCTTGGAATCTGAGTGTgaatcttaaacccaactt
ggttacctcttatctagtggCGGGTGTTTTGGGTGTTGGGAAGAAGCAACGATTGGATGCgaatcttaaacccaact
ggttacctcttatctagtggGGGCCCTTTCGATTCTAAGATTGCATGACTGCCTGACTTGgaatcttaaacccaactt
ggttacctcttatctagtggGTGCATGTGTGTGTGTGTTTAGGAGGGAGTGGAGAGGAACgaatcttaaacccaact
ggttacctcttatctagtggGAAAGAGACTTCTAGGCAGTTGCTTTGGAAACCATGTTTGgaatcttaaacccaactt
ggttacctcttatctagtggCCTCCTTTGTTGTCTAGTCCCATTCCACCAGGTGGCCAAGgaatcttaaacccaact
ggttacctcttatctagtggAAATACTCCATCTCTCATGCTCCCCATCTCCCTGGATTCCgaatcttaaacccaact
ggttacctcttatctagtggGACAAAGGAAAGCAAGAGGAAAGGAAGGGAGGAAAGAAGGgaatcttaaacccaa
ggttacctcttatctagtggTCACACAGTGTTATTGGCATTGGACTTATTAGGTTGGTGCgaatcttaaacccaacttg

ggttacctcttatctagtggTTTGCACCAACCTAATATTTCCATCAGGTGGGATTTTAGGgaatcttaaacccaacttg
ggttacctcttatctagtggGATGTACCTTCACAAGTAGATAGAGCCTACCTCTCACAGGgaatcttaaacccaact
ggttacctcttatctagtggGTACGTGGAACCACAGCTCTCAGCATGGTCGGGTCCCCTCgaatcttaaacccaac
ggttacctcttatctagtggCCTTGCCCTCTCATGAGATAGTCAAGCATTTCTCCCTCCCgaatcttaaacccaact
ggttacctcttatctagtggCCTCATGCCCCTTGCTCCCTACATGGTATCCAACCTTGCCgaatcttaaacccaac
ggttacctcttatctagtggGGCCCTGGCCTGGTTCTCATCACTTAGTCCTGATATTCTGgaatcttaaacccaactt
ggttacctcttatctagtggCTCCTCCTTGACTTCAGTCAGATCTCGCTCTCTTTTGGGCgaatcttaaacccaactt
ggttacctcttatctagtggCCCTGGGGGCTCCTGCATCCTATAACCTCATGGATCAGTGgaatcttaaacccaac
ggttacctcttatctagtggCGGTTTGGAGATCAAAAGTCCAAAATGGTTCTCAGTGGGCgaatcttaaacccaact
ggttacctcttatctagtggGAGCCACCACGCCCGGCCAGAACTGGGTTCTTTTCGGAGGgaatcttaaacccaa
ggttacctcttatctagtggGGCCCTTTGCAGTTTCTGAAGGCTGCCTGCATTCCTTGGCgaatcttaaacccaact
ggttacctcttatctagtggAAGGGCCCAGTGAGTCTTTCTCATGGTGTCATCTCCCTGGgaatcttaaacccaact
ggttacctcttatctagtggAGGATCCTAGTGGTTACACTGGGCCCACTGCCTATTCTAGgaatcttaaacccaact
ggttacctcttatctagtggTAGCTCTGGGAGAAGATTCCACTCAGAGTTGGGCAGGAGGgaatcttaaacccaac
ggttacctcttatctagtggGCAACTGGCCAGGGAATGTAAAAGAGGGCTCAGAAGCATTgaatcttaaacccaac
ggttacctcttatctagtggACTATACATTTGCTCTAACCTACTTCACAACTTTGGCAAGgaatcttaaacccaacttg
ggttacctcttatctagtggCTCTGAGACTACAACAGAAAATTCAATATGTAGCAATTGCgaatcttaaacccaacttg
ggttacctcttatctagtggGTTCTGTAGGAATGAGCTTTATGTCAAACAATGAGCTGAGgaatcttaaacccaacttg
ggttacctcttatctagtggGGCCTCGATCTCCTGACTTCATGAGCTGCCTGCCTCGGCCgaatcttaaacccaac
ggttacctcttatctagtggGGCATGAGCCACCACTACTGGCCTGATATTTACATTTTTAgaatcttaaacccaacttg
ggttacctcttatctagtggTGATTCCATCACCCGAACCCAACCACTAGGCTCCACTGCCgaatcttaaacccaac
ggttacctcttatctagtggTTCAGCGGAGAGAACAATCAGTGAGACCTGAGTGGCTAAGgaatcttaaacccaac
ggttacctcttatctagtggGGTGGTGATATGTGAACACCCGTTTAAAAGGTTGCTCTGCgaatcttaaacccaactt
ggttacctcttatctagtggCCATGTCACCCCCAGGGCCTGCCTGTGCATTAGGGCAGTGgaatcttaaacccaa
ggttacctcttatctagtggGCTGCTTTTATTTCAACTTCCCACTTCCACCTTGGTGAGGgaatcttaaacccaacttg
ggttacctcttatctagtggAGAAATCACCTTTGTTTTCTTTAGGCCTGTAGATTGGCTTgaatcttaaacccaacttg
ggttacctcttatctagtggGGGTGTGCTTGGGTGTAGGGGAAGGACAGGTGGGACGTGCgaatcttaaacccaa
ggttacctcttatctagtggATATGCTCTCAGCATGCTTTAGGGGCTGTTTTAAGGAACAgaatcttaaacccaacttg
ggttacctcttatctagtggGAAATTACTTGTATCCCATTAGAGACTCTGCAGATCACGCgaatcttaaacccaactt
ggttacctcttatctagtggAGTAGGCACTATGAATATGGAAATCAGCAAGACATGGCCCgaatcttaaacccaact
ggttacctcttatctagtggGCACAGACAAAGCATCAGGTGGATTAAACATGTGGTGCTGgaatcttaaacccaact
ggttacctcttatctagtggGGGGTTGCCCAAAGCCCCACAGCAAGCAAGCAGATTAGTTgaatcttaaacccaac
ggttacctcttatctagtggACCATGTGATACCCCCTCATAGCCTTCTGGAGCAGGGATCgaatcttaaacccaac
ggttacctcttatctagtggGCTTAAGGAGCCAAACAGCAGATACTTTAGGCTTTGCCAGgaatcttaaacccaact
ggttacctcttatctagtggGCAAGGCAGCAAGAGTTTTCCTATAGACCACAGTTCCAGGgaatcttaaacccaac
ggttacctcttatctagtggCAGTATGTGTATATCTCCACATATACCTTATCTGACATTCgaatcttaaacccaacttg
ggttacctcttatctagtggAGATCATCTAAGCCACCTTTGCTTCTCTAGAGAGTGCTACgaatcttaaacccaactt
ggttacctcttatctagtggATCCATTATTATTTACCGAAAAGACTCCATGGAGAAGGCCgaatcttaaacccaactt
ggttacctcttatctagtggGATGTGGTTTTGCTGCTTGAAGTGGTTTTGGGAGAGGCTCgaatcttaaacccaacttg
ggttacctcttatctagtggGATTTCCAGGCATCGTGACTCCTGGGCTGAGCAGCACAACgaatcttaaacccaac
ggttacctcttatctagtggCTTCTGCTACTTTTCTTTCTTGAGAGGGCATGGTTCATAGgaatcttaaacccaacttg
ggttacctcttatctagtggCCTGTGGTCAGGAACTGGTACGGTAGCTTACATGTGTGAAgaatcttaaacccaact
ggttacctcttatctagtggCATTGCCCCCATTTGTTAGAAAAACAGAGGCACAGAGAGGgaatcttaaacccaac
ggttacctcttatctagtggACACAGCTAGGAAGTGTCAGAACGAGATTCAAACCAGATGgaatcttaaacccaac
ggttacctcttatctagtggCAAACCTTGGGTTTGGAACCTATGGTGGCTCTATCTTTGAgaatcttaaacccaacttg
ggttacctcttatctagtggTCTAAAATGATGAGCACGATAAAAGAGCCACATCATCTGCgaatcttaaacccaact
ggttacctcttatctagtggGAGCAAGTGGTAGATGCTGTTGGAGCGTGGAACTCCACACgaatcttaaacccaac
ggttacctcttatctagtggGTGAGAAACCATGGGGTGTTCAAAGGAGAGCCATTGTGGCgaatcttaaacccaac
ggttacctcttatctagtggGACTCAGGGCCCATGGCAGTGACCCCCAACCTCCTGGCAGgaatcttaaacccaa
ggttacctcttatctagtggCAGACCCAACTCCACACTCTGAGCTGTGTGCGTCAGGGGGgaatcttaaacccaa
ggttacctcttatctagtggAGCCTCACTTTGTTTATTAGCAAAATGGAGACACTAATACgaatcttaaacccaacttg
ggttacctcttatctagtggACTCATCTTTGACCTAAACTGGCATGAGACGGGAGACTGGgaatcttaaacccaact

ggttacctcttatctagtggGTTTGATTACAGCATGCTATCCCAGACCCTGCTGGTGGGgaatcttaaacccaac
ggttacctcttatctagtggAGTGGCCCTGTGCTAACTACCTCATTAGGTCTTGGGGTGGgaatcttaaacccaac
ggttacctcttatctagtggATGGAAAGGGCTTAGCAGAGTTAGTAGCACATCATAAATCgaatcttaaacccaact
ggttacctcttatctagtggATACAACTTTGTTTAGAGTCCTCCACTATTGTCTCCTGAGgaatcttaaacccaacttg
ggttacctcttatctagtggTCTTCCAAGATGCCTTCTGAATGTTTTGAATCTGGTGCTTgaatcttaaacccaacttg
ggttacctcttatctagtggGCATAAGCAACAACGTCGTGTCTGCTGCCCAGCCAAGCACgaatcttaaacccaa
ggttacctcttatctagtggGATGGAATTCAAAGAGACCACCTACAGCATGCTGTAAAACgaatcttaaacccaac
ggttacctcttatctagtggAACTTCCTCACAAGTAGATTTGTTCCAAAATGGATTGGGCgaatcttaaacccaactt
ggttacctcttatctagtggAGCATTGTGCTTGAGATGACCTTTGAGAACCTTCTAACTCgaatcttaaacccaactt
ggttacctcttatctagtggGAATGGGGAAACGCAGATGAAACATCCTTGACCTCCCAGGgaatcttaaacccaa
ggttacctcttatctagtggTCTCTTCGGAGGGGTTGGATTTGAAGGTCGAGTGGAGTTCgaatcttaaacccaact
ggttacctcttatctagtggGCCCTGCTTCTGTCCAAAACACAAAGTATAGGAAAAAAGCgaatcttaaacccaac
ggttacctcttatctagtggTAGATCAACTTGGCAACATATGCAGAAGCAGTGCGGAAGCgaatcttaaacccaac
ggttacctcttatctagtggCACGACACACCCCTTCACCTCTTTCCATGCCAGAGCTCTGgaatcttaaacccaac
ggttacctcttatctagtggGAGATGCAACGGGGCTGATAGTTGGAGGATAGAATGCCAGgaatcttaaacccaac
ggttacctcttatctagtggACTACAGTGGGCTTTCCAAGAGTCCCCAGAATTAGTCTGGgaatcttaaacccaac
ggttacctcttatctagtggGGACCTTTACCTACTGGTAGAGAACACAGGTGCTCTCTGCgaatcttaaacccaac
ggttacctcttatctagtggCTTTACAGTGAGATGACCTCGAAGGCCCTCTATGGAGAGGgaatcttaaacccaac
ggttacctcttatctagtggGGACAGCGGTATTTAGGGAAAATCTGTCTTTGTGGAGAAGgaatcttaaacccaact
ggttacctcttatctagtggGCCTTTTTTGGTGGGACCTGGGGGTGGAAGAGCCCATCTCgaatcttaaacccaac
ggttacctcttatctagtggTTGCACACTTTCACATACAAATTGTACAAAACCCTGACCCgaatcttaaacccaactt
ggttacctcttatctagtggGCACAATGATGTGACGGTGAAGCACCCCTAGAGAAGTGTCgaatcttaaacccaac
ggttacctcttatctagtggGCTGCGTGAACTGGGACAAGTCATGTCACCGCTCCTGGCCgaatcttaaacccaa
ggttacctcttatctagtggCAGTGGGGCACTGGAGGCTGCCTTGGTCCCTTCCAGCCCTgaatcttaaacccaa
ggttacctcttatctagtggCACACTCGGCTCTTCTGTTTCCAAGAACAATAATGACTGCgaatcttaaacccaact
ggttacctcttatctagtggTGCCTTAGATTGTGATCTTCGTGACTTCATCTGTGTGAACgaatcttaaacccaacttg
ggttacctcttatctagtggGGCCATGTCCAGGGGCCCTTCCTTAAAGCCTCCAGTTGAGgaatcttaaacccaa
ggttacctcttatctagtggGAGAGAGGTGGGAGAAGGTGGTACGGTATGAAGTGGAGTCgaatcttaaacccaa
ggttacctcttatctagtggTCTGAGCCGCTTCCTCCCTGAGCCTTTCATCCCTCCCTGCgaatcttaaacccaac
ggttacctcttatctagtggGCAGCCTCTGCACCCTTTGGCTAGATTCCTGCTCAGGAGAgaatcttaaacccaac
ggttacctcttatctagtggCCCCTGTGGAGGAAGGGATGGTTTGCTACATGGCCCTCACgaatcttaaacccaa
ggttacctcttatctagtggCCTTGCCTCACCCGGCTTCACTGCCCCTCTCTGGAGGCTGgaatcttaaacccaa
ggttacctcttatctagtggGTGTACATTTCCAGTCTACTTTCTTTGCCTGGAAGGGTGCgaatcttaaacccaactt
ggttacctcttatctagtggGGCCTGTATCTGATTTACTTTTGATTCCCTGGTCCGAGGCgaatcttaaacccaactt
ggttacctcttatctagtggGGGGTGGACGTCTGGCAGCCGTGTTTCAACTCTCTAAGACgaatcttaaacccaac
ggttacctcttatctagtggCGCTTAGCCTTTCCCACAGCAGGAACGGGGATGTGAGGTGgaatcttaaacccaa
ggttacctcttatctagtggAGCTCTGCAGAAGATATAGAACTGGGCCGGGGGCGGTGGCgaatcttaaacccaa
ggttacctcttatctagtggCGGGAGACTGAAGCAGGAGAGTGGCATGAACTCGGGAGGCgaatcttaaaccca
ggttacctcttatctagtggCAGACCACGCATACCCATATGATGGAGGAACCAATGTTCGgaatcttaaacccaac
ggttacctcttatctagtggGTAGGGGCTTATCCTACCAGATTAAAACAGTGTGGTAGTGgaatcttaaacccaact
ggttacctcttatctagtggCGGTATGTTGGCAGAACAGATAGGATCCCGCTACATATAGgaatcttaaacccaac
ggttacctcttatctagtggGATAATAAATGTGCCACTTCAGTGGGGAAATGACAGATTGgaatcttaaacccaact
ggttacctcttatctagtggATGGTACAGAGACAATTGATTCCACATTTGGGTGAAAAGCgaatcttaaacccaact
ggttacctcttatctagtggTGAGGTGGAGGCAGCTTTCTAAGCAAGATAGAAAACACAGgaatcttaaacccaac
ggttacctcttatctagtggCATATGATGCACAAGAGGATATTATCCCCAGGATGCAAGGgaatcttaaacccaac
ggttacctcttatctagtggCAGAATAATGAATGAGGGCATAGTTTTAAGTCTCCCTCCCgaatcttaaacccaact
ggttacctcttatctagtggTCTGTGTCCTCTATATGTAAAGTCCATAAAGTTCTGCGGGgaatcttaaacccaacttg
ggttacctcttatctagtggAATATGAGCTGCTCAATTTCGACACTGAATTCAGCTAACGgaatcttaaacccaactt
ggttacctcttatctagtggCCCACTCAAATTCATTTGACGGCTATTTGCCTTACCACGCgaatcttaaacccaact
ggttacctcttatctagtggCCCCAGGAACGGGGTGTGCAAACAGTGGTACCTAACTCAAgaatcttaaacccaa
ggttacctcttatctagtggGGGAGGAAAGTAGGAAGGAGGGGGTGCAGGCAGCCTGGCCgaatcttaaaccca
ggttacctcttatctagtggTTCTGTTCCTGGTGCGGTTCCTCCTCTAGCCACATGATGGgaatcttaaacccaact

ggttacctcttatctagtggGCCAGGGGCGCCGGGGTGTGTTTGTGTTTGTGCCTGCAAGgaatcttaaacccaa
ggttacctcttatctagtggGCCCAGACTTAGCACTCTCAGCTGCTCTGTAATGCAAGTCgaatcttaaacccaac
ggttacctcttatctagtggGGGCTTCACGGTCTTCTTTCTAGAAAAATCCCAACAAGGGgaatcttaaacccaact
ggttacctcttatctagtggTGCAGCATCTGCTCTGCCGGGTTTGCAATCTGTGCACAGGgaatcttaaacccaac
ggttacctcttatctagtggCGTGCCCCGGGTTGGAAAATGTGATTTGTGACAGAGAAGCgaatcttaaacccaac
ggttacctcttatctagtggTGCTCCAGGTTGGTTGCTCTCTTTCAATTGCAAAGCCGTCgaatcttaaacccaact
ggttacctcttatctagtggCAACTAGAGGCACACCCAGCAACTGTCCTTCTGCCCCCAGgaatcttaaacccaa
ggttacctcttatctagtggACTCCAGAGAAGAGCTGTCCCCCAAATTCAAGAAGAATCGgaatcttaaacccaac
ggttacctcttatctagtggGGATGCCTGGGGCAGCTTGTCTGTTCAAGCAGAATCTCCCgaatcttaaacccaac
ggttacctcttatctagtggTGAAACCTATTAGTCCCATCCTCTGATGCCTGGGACCAGGgaatcttaaacccaac
ggttacctcttatctagtggGAGGCAGAGGATGTTGACACAAACTTACAGACTGGGACTCgaatcttaaacccaac
ggttacctcttatctagtggTCCCAGCTCAGTGGTGGACAGCAGAGGACTCTAAGGTGGGgaatcttaaacccaa
ggttacctcttatctagtggTGGAGTCCCCCTTCAGCCACTTTAAGTTAACGGATGTCTCgaatcttaaacccaact
ggttacctcttatctagtggCTCATCTGTAAAATGAAGAGAAAGATGTCAATTGCTTGGGgaatcttaaacccaactt
ggttacctcttatctagtggGCGTGGGGCCCGGAACCTCTCCTGTGGCTGGGAGTGAGTGgaatcttaaacccaa
ggttacctcttatctagtggGAGAACCTCTTTACACAATTGCCCATCCTCCTAGCCCTGCgaatcttaaacccaac
ggttacctcttatctagtggTGCTGGCGGAGGCACCCTAACCAAAGGGGGACTGGATCTGgaatcttaaacccaa
ggttacctcttatctagtggGCATCCGCTCTTCATTTCCCCAAGGGTGATGGTTCCGAGAgaatcttaaacccaac
ggttacctcttatctagtggCTGAGCCACAGTTACTCCCAGGAATGGCCCAACACACAGCgaatcttaaacccaa
ggttacctcttatctagtggGACATGCCCTGACCCAGGCTCTTTATTTGGAAACCCAGGCgaatcttaaacccaac
ggttacctcttatctagtggGTGTCCATGGCAGGCAGGTGGAGGAAGGGGGAGTTTGGCCgaatcttaaacccaa
ggttacctcttatctagtggCCCATACATTGCTGGGGACCTCCAAGGGACAGACAGTGTCgaatcttaaacccaa
ggttacctcttatctagtggGATGGCATTAGTGCTGAAGCCCAATTGCCCCAGCAGCAGCgaatcttaaacccaa
ggttacctcttatctagtggGGCGGTGCCAGCAGATGTCTCCATGATGTGATTTAAGGGCgaatcttaaacccaac
ggttacctcttatctagtggGGCAGTCCTGGGTGAGTGCCATGTGCCCACCCCCACTTTAgaatcttaaacccaa
ggttacctcttatctagtggAGCACCACGCTAATACTTCTATGCCCATCAGTCCTTTCTCgaatcttaaacccaact
ggttacctcttatctagtggAAGTGCCAGTTGTAGTCCCCACTCCCCTCCAACCCCATCCgaatcttaaacccaa
ggttacctcttatctagtggGGTCTGAGGGTGTGGCTTGGACATCCATAAAGTTCCTCAGgaatcttaaacccaact
ggttacctcttatctagtggCCCAGTTGTCTTCTTTATCCTCACCCCGACCTTCCTAGGGgaatcttaaacccaact
ggttacctcttatctagtggACCCCTAATTGTTCAGGATGGAAGCAGAGAATGTCTTGGgaatcttaaacccaact
ggttacctcttatctagtggGGAGCAAGGTTTCAAATGCAGGTCTGACATGTCAGGCTCCgaatcttaaacccaac
ggttacctcttatctagtggGCCAAGGCAGAGTGTGTTCTTCCGAGAGAGGATATTAGAGgaatcttaaacccaac
ggttacctcttatctagtggCAGTCTGGAAATTTCTTGAAAGTCTCAAGCTTCATCTTAAgaatcttaaacccaacttg
ggttacctcttatctagtggGACAGGCTGGGGTCGTTCATACCACACTGTGCTTGCAGGGgaatcttaaacccaa
ggttacctcttatctagtggCACCGATGGACAGGAGAGGACTGGAGGACATGATGAGGGGgaatcttaaacccaa
ggttacctcttatctagtggCTTGGGGAGAAAGAGTGATGGGTATCTGGATGCCCTCAGGgaatcttaaacccaac
ggttacctcttatctagtggTGGAGGATGCAAGCATTCAAGATTGGTGTAAGAGCACAGCgaatcttaaacccaac
ggttacctcttatctagtggTTTTGAGATGGAATCCCACTCTTTCACCCAGGCTGGAGTGgaatcttaaacccaact
ggttacctcttatctagtggGGCCTCCCAAGACAAATATTTTTAATAAAGGGTTAGATAGgaatcttaaacccaactt
ggttacctcttatctagtggGGCTTGTGGCCCATGCGTTCTCCATCACAGCTATTCAACTgaatcttaaacccaact
ggttacctcttatctagtggGGCGTGTAGCCATGGACACTACAGAAACAGGCAGGTGGGAgaatcttaaacccaa
ggttacctcttatctagtggGGGCAGTAGTTTGCTGAACCCTGTGCTTAGTTTGCTTACTgaatcttaaacccaactt
ggttacctcttatctagtggGCTCTGGGAGTACACACGTGCACTTGTGTTGCTAGCCTCCgaatcttaaacccaac
ggttacctcttatctagtggCATGCCTGTAACTCCAGCGCTTTAGGAGGCCTAGGTGGGTgaatcttaaacccaa
ggttacctcttatctagtggCCAGCTTAGGCAACATAGCAAGACCTCCCCCTCTAGCAAAgaatcttaaacccaa
ggttacctcttatctagtggGGCTAAGGTGGGAGGAATGCTTGAGCCTGGAAAGTCTAGGgaatcttaaacccaa
ggttacctcttatctagtggTGTGCCACTGTACTCTAACCTGGGTGATGGAACAAAACCCgaatcttaaacccaac
ggttacctcttatctagtggCACATCTAACAAGCAGAAGGCCAGAATTTGCACCCAAGCCgaatcttaaacccaa
ggttacctcttatctagtggAGCCCAGATTCTTTTCCAGGCTATAACCATGTTTCTAAGCgaatcttaaacccaactt
ggttacctcttatctagtggACCTTCTCTAGCTGGCTCTTTGCCCAATATGTGACAGTGCgaatcttaaacccaact
ggttacctcttatctagtggAGGCAAATATCGGGAGACTGATCTCTTTTGGGGACCCACCgaatcttaaacccaac
ggttacctcttatctagtggCACTCCCAGAGTGCAATCCCACCAACAGCAGCCTGCCAAAgaatcttaaacccaa

ggttacctcttatctagtggGCCAGGAAGCCAACAATCACTGTGGCCCAGATGCCCACTCgaatcttaaacccaa
ggttacctcttatctagtggGCCCTCTGCCTCACCCTCTAGAAGCTGCCGTGCCAGGGCCgaatcttaaacccaa
ggttacctcttatctagtggGGTAGCGTCCCTTCTCCTCAGAGAGCTGGGTACGCTTGGCgaatcttaaacccaac
ggttacctcttatctagtggGAGGTTGTTGGTGGGTACACCCTCCGCACTTCCTCTGTGGgaatcttaaacccaac
ggttacctcttatctagtggCAGACAGTCAGATACCAATGGCTCTTAAGTCTCTGGGGTCgaatcttaaacccaact
ggttacctcttatctagtggCTCAGCACTTTGGGAGGCCTAGCCGTGCAGATCACCTGAGgaatcttaaacccaac
ggttacctcttatctagtggCCCTTGGAGCAGATGCTTCTGGAGCCGACCCCCACTACCCgaatcttaaacccaa
ggttacctcttatctagtggCCCCTTGCCCTTACCACAGTGCACACCAGCCCCAGTGTCCgaatcttaaacccaa
ggttacctcttatctagtggCCTGAAGTGATAAGGAATGAATGCCCCCCTGAGAAGAGCCgaatcttaaacccaac
ggttacctcttatctagtggACCCCAGCTCCCGCTCCCTTTAGTGAGGCCAATCCCAGGCgaatcttaaacccaa
ggttacctcttatctagtggGCTCCATTGCCCACACTGGTTGCTGGCTCAAGGGTTCGTCgaatcttaaacccaac
ggttacctcttatctagtggTACCTTCTTTTTCCCCACTTTCCTACCAGTGTACCCTTAAgaatcttaaacccaacttg
ggttacctcttatctagtggCAGAGACAGCTTCTGAAAAACCCAAACTAAAATAGCACTCgaatcttaaacccaact
ggttacctcttatctagtggACGCAGTTGCAAGCTCAGAGGTCTAGGGCAAGGTGGATGCgaatcttaaacccaac
ggttacctcttatctagtggGATGTCATTGGGTAAGACATTGGGCGTACATGGGACAGTGgaatcttaaacccaact
ggttacctcttatctagtggGGTCTCTATATGCAGCTCACACTCTGAAGAGAGGAGAGAGgaatcttaaacccaac
ggttacctcttatctagtggGGGGGGATCAGCAGTATAGCGAAGGGTAAGTGGTACAACCgaatcttaaacccaac
ggttacctcttatctagtggGTTGAGAGAAGTCCTCATGCTCCTGGGTTCTTCATCTCACgaatcttaaacccaactt
ggttacctcttatctagtggCTGTATTCTTAGGGAGTTTACAATTGCAGAAGAGCTAGGGgaatcttaaacccaactt
ggttacctcttatctagtggCCACTGAGGTTCCCTGAGAAGATTAAGTCCAGCTACTCTGgaatcttaaacccaact
ggttacctcttatctagtggCTATGCATCTAGAATGGTACTGGTTATGTACAAACCCTTGgaatcttaaacccaacttg
ggttacctcttatctagtggGTGTTGTGTTCTGTGATTCAGATGAGCAACACCAGTTGTGgaatcttaaacccaacttg
ggttacctcttatctagtggGCAGAGCTCTGAATGTCAAGAAGCCAGGCTGGGTGTGGTGgaatcttaaacccaac
ggttacctcttatctagtggGCCGGCCCTTGAGGGTCAGAGGATGGCAGGTATGGGGTAGgaatcttaaacccaa
ggttacctcttatctagtggGCCAGGTGAGGCTAGGCAGGTGGATGAGAGGAGTGGGAAAgaatcttaaacccaa
ggttacctcttatctagtggGCCAGAGAAGTAGGAGAAGGTAAATTATGGAGAACCTTATgaatcttaaacccaactt
ggttacctcttatctagtggTAGGAGGTGCGGTAGCTTTATCCAATGTGGTAGTGGAATCgaatcttaaacccaactt
ggttacctcttatctagtggTAACACCTGGCTTTATGAGTGGCAGATGAAGGGTAGAGCCgaatcttaaacccaact
ggttacctcttatctagtggGGTTATGTTACAATTGAGGTGACTACTTGAATCCAAGCAGgaatcttaaacccaacttg
ggttacctcttatctagtggGGAGAGGATGTGGTCACCTAAAATGACATTGACGGTCCTGgaatcttaaacccaact
ggttacctcttatctagtggTGCCTGGCTTAGCATTTTCACACTCTCAAGATGGTCCTGTgaatcttaaacccaacttg
ggttacctcttatctagtggAGGGAAGACCCCAGGACATCACAACACATAGAGGCTGAGGgaatcttaaacccaa
ggttacctcttatctagtggTGAGGTAGGAGGAAAACCAGAAAAACATGGTGTCACACAGgaatcttaaacccaac
ggttacctcttatctagtggGCTGGAAGTTCAGGCATGCAAAGGACATCCATGGGATCCGgaatcttaaacccaac
ggttacctcttatctagtggGGGACCATGACTCTCTATAGGACACAGAACAACCTGGGCCgaatcttaaacccaac
ggttacctcttatctagtggGGCTATAGCATCACAGATGAATTAAGGACGGTGTGTGGTGgaatcttaaacccaact
ggttacctcttatctagtggGGCCATGGAGTATTTCGTTTTAGCATGCACAGAACAACTGgaatcttaaacccaactt
ggttacctcttatctagtggAAGAACAATGCCTTGTCAACGACATTGGCTGATACTAAGCgaatcttaaacccaact
ggttacctcttatctagtggGCCTCCAAGGGCTCTGGCTTCCCAATAATGTTCCTACTACgaatcttaaacccaact
ggttacctcttatctagtggCTGCTTCTGCACATCATTTCTTCTAACTCCCAGGCCAGTGgaatcttaaacccaactt
ggttacctcttatctagtggGCTGTCTCAGCAAAGCTTCTTGGCTGGTTATATAAAAGGCgaatcttaaacccaactt
ggttacctcttatctagtggGCAACCCAAGAAGGCCTGCTGTGACTTGTTATCTGCAGAGgaatcttaaacccaac
ggttacctcttatctagtggTGCCAGTGGGGCCACTTAACACCTTCCCGTAAATGGTCTCgaatcttaaacccaac
ggttacctcttatctagtggCGCCTTCAGGAGTCAGAGCTGAGAGAAGGTCAAACCCGGGgaatcttaaacccaa
ggttacctcttatctagtggAACCTGACAGTTCCTTTCAAACTTCTCCCTGTGCAGCATGgaatcttaaacccaactt
ggttacctcttatctagtggCAGCAGAAAGGAAAGTGCATGGGCCTTGGAATCACAGACCgaatcttaaacccaac
ggttacctcttatctagtggGAACAACTCACTCTAATATGGCTTCTACTCTCTCCACTGGgaatcttaaacccaactt
ggttacctcttatctagtggCTACCACCCTGGTTACACTTGTTCAGTCTTCTGTGCTAGCgaatcttaaacccaactt
ggttacctcttatctagtggGCCTTCTTCCCTCTTCTTAAACACTCTCTCTAGATAATAGgaatcttaaacccaacttg
ggttacctcttatctagtggGTTTGTCCACCTGCCCACTTGCCATCTCCACTTGGATGCCgaatcttaaacccaact
ggttacctcttatctagtggCCATTCTACTCCAAGTCTACCACATCTCAGTAAATGACACgaatcttaaacccaactt
ggttacctcttatctagtggAAGCCTAAGGTTCATAGTAAATCCCTGTCTTCTGCCCTCCgaatcttaaacccaactt

ggttacctcttatctagtggGGATCCTATATCCAAGTGATATGATTTGGGTTTGTGTCCCgaatcttaaacccaacttg
ggttacctcttatctagtggGGAGGGGAGGGGTCAGGTGGGAGGTGATTAGATCATGGGGgaatcttaaacccaa
ggttacctcttatctagtggGTGTGTAGCACCTCCTCCTCTCTCTTTCTCCTTCTGTGACgaatcttaaacccaactt
ggttacctcttatctagtggACCCAGCCTCAGGTAGGTCTTTGTAGCAATGCGAGAACGGgaatcttaaacccaac
ggttacctcttatctagtggCAGCCTTCTCCAACCTGCATGATTCCTTGGCAGACTGAGCgaatcttaaacccaac
ggttacctcttatctagtggCTCCCCAGCCTTTGGAGCGGCTAATTGGATTTAGCTAGTGgaatcttaaacccaact
ggttacctcttatctagtggGCGGTTGGAGTGTGTGTTCCATGGGCTCCTCTCTTTCAGGgaatcttaaacccaact
ggttacctcttatctagtggCAGGGATCCTTCATAGAAGGTGTCATTCCCGTCAGGAGGCgaatcttaaacccaac
ggttacctcttatctagtggCTAGTTAGCCCTCCCTCTACTTGTGCCTCTGTGCCCTGTGgaatcttaaacccaact
ggttacctcttatctagtggAGGAACTGCACAGCTCTCTGTGTCTTTCTCACATCCTGCCgaatcttaaacccaact
ggttacctcttatctagtggCACCCCCGAGCTATCCTTCTTTGAAAATGCTCTCTATTTCgaatcttaaacccaactt
ggttacctcttatctagtggCACTGTCCTAATTAGCCTCCTACACTTACCAAAACCAGCCgaatcttaaacccaact
ggttacctcttatctagtggACTGCAGCAGAACTGAGTGGTCATTTCAAGACTTATATCAgaatcttaaacccaactt
ggttacctcttatctagtggTAAGTCCCTTCATGATCTGGCCCCTGCCTACTCCTCAAGCgaatcttaaacccaac
ggttacctcttatctagtggACACTCCAAGCTCTTTGCTGCCTTGTGGATTTCATGCAAGgaatcttaaacccaactt
ggttacctcttatctagtggGCCTAGACTCCTTTCCTCACAGAGAACATGCATATTTGCCgaatcttaaacccaact
ggttacctcttatctagtggGCTGAGGTTTCAGTTTAGTTGCCACCTTTTCAGAAACACCgaatcttaaacccaactt
ggttacctcttatctagtggCCCGTCCTAAAGTAGTTCTTTCTTCATCATTCTTTGCCATgaatcttaaacccaacttg
ggttacctcttatctagtggTCGGTACACTGTAAGCATAACTCCATTGTGGTGGTGGTGGgaatcttaaacccaact
ggttacctcttatctagtggGATTGAAGGCATTGGTCAGGTGCAAATACCTGGCTCATAGgaatcttaaacccaact
ggttacctcttatctagtggATTGAGCATGTACCATTTAGCAAGTGCTTTATGATCCCTGgaatcttaaacccaactt
ggttacctcttatctagtggGTCCACTCTTTCTTCAGGTACTTCTGGTTATCATGGATGGgaatcttaaacccaactt
ggttacctcttatctagtggGAAGCAAATTTACCCGCTTACAAGCTTGGGAAATAGGACCgaatcttaaacccaact
ggttacctcttatctagtggTTCTCCACATCCTCTCCAGCACCTGTACATATTGTTATATgaatcttaaacccaacttg
ggttacctcttatctagtggAAGGCTTTTGGATAGATTTACAATCTGAGACTTCTTCACCgaatcttaaacccaactt
ggttacctcttatctagtggGGAATAGTACTAGGAAAAAACCAACAAGCCTCTTGGACGCgaatcttaaacccaac
ggttacctcttatctagtggAAGTGACTCCAAAAGGACTTAGATTAAAGCAAGTGTTGGGgaatcttaaacccaact
ggttacctcttatctagtggGAGTGGTTGTCTTTAGTGGACTCATGGCATGTGGGGTGGGgaatcttaaacccaact
ggttacctcttatctagtggGTTGGAGATGAGGACGCGTACCTGAGTACCTGCCTCCTGGgaatcttaaacccaac
ggttacctcttatctagtggGTCAGATTCAGAAGTCCCCTCCAGGCCCACATTTCCATTTgaatcttaaacccaact
ggttacctcttatctagtggCCTAGGAGTCCTTTCATATCCCTATTCTTACCTTGGCCCCgaatcttaaacccaactt
ggttacctcttatctagtggGGGAATGTAAACTCGTACAACAGTATGGCAGACAGTACGGgaatcttaaacccaac
ggttacctcttatctagtggCCAAAAGTGGAACTACCATTCGATCCAGTAATTCCATTACgaatcttaaacccaactt
ggttacctcttatctagtggATGCACATGCATTTTTATGGCAGCACAATTTGCAATTTCAgaatcttaaacccaacttg
ggttacctcttatctagtggAAGAAAACATGGGATATATACACCATGGAATGCTACTCAGgaatcttaaacccaactt
ggttacctcttatctagtggATGGTCTTTGCAGAAACTTGGATGGCCATTATTGTAAGTGgaatcttaaacccaacttg
ggttacctcttatctagtggGACTTTGGGGACTCAGGGGAAGAGTGGAAGGTGGGTGAAGgaatcttaaacccaac
ggttacctcttatctagtggATCTATGTAAACAAAACCACCTGTACCCTGAAAACTACTGgaatcttaaacccaactt
ggttacctcttatctagtggCCTGTGGTCACACAACCAGAAAGTGACAAAATAAGGCCTCgaatcttaaacccaac
ggttacctcttatctagtggTGGTGTAAAGACAGTGAAAACAGCACAGGTCTCTGTTCTCgaatcttaaacccaact
ggttacctcttatctagtggGGCCGAAGGTTGGTGCCTTCCTCTGATGATTGCCACAGGCgaatcttaaacccaac
ggttacctcttatctagtggTATCACCTGCTTCAGTGTGGAGCCACTTACAGAATCATGAgaatcttaaacccaactt
ggttacctcttatctagtggGCCGAGAAGAGGGAAGTGAGAGGGAATGGAGCTTTTCCCCgaatcttaaacccaa
ggttacctcttatctagtggCTTGTTCTCAGGCCCAGAATTCAGGCTATGAGGTTTTGCGgaatcttaaacccaactt
ggttacctcttatctagtggGGAGGGCACAGACTTGGGTTCAAAGTACAGCTTCATCACCgaatcttaaacccaac
ggttacctcttatctagtggGTAGCCTTGGACACAAAGAACATCTTTGACTCTTGCTGCTgaatcttaaacccaactt
ggttacctcttatctagtggTTTCCTTCTCCTTCTCCATCCGATCGTGAGTAATACAGACgaatcttaaacccaactt
ggttacctcttatctagtggCCTCTTTTCATCTTGAGAGGCAGAAGGCATGCATGTAGGGgaatcttaaacccaact
ggttacctcttatctagtggAGGCCAGTGGGTTAATGCCCCCTGTCCTCAGTGAAGCTACgaatcttaaacccaac
ggttacctcttatctagtggGAGAACATTCTGTATGGTTTGAAGCTGAGAAGTTTCAAGCgaatcttaaacccaactt
ggttacctcttatctagtggGGATATGAGTCAGTTGAGATCCCAGACTTCTGGCCTAGACgaatcttaaacccaact
ggttacctcttatctagtggTTTTTAGGAATCCCTTTAGGCCACAACTTTTGCTCAGGTGgaatcttaaacccaactt

ggttacctcttatctagtggTGTCCATGGTCATTAGAGTAGATGGCAGGGCACTTCTAGGgaatcttaaacccaact
ggttacctcttatctagtggCTATCTTCATTCCAGTTGTGTATACATTAAATGTCTAGCCgaatcttaaacccaacttg
ggttacctcttatctagtggGAGTTTTCCAGAGATGAAATGGCTGCTGACTCATCTTCAGgaatcttaaacccaactt
ggttacctcttatctagtggCCATGGCATCGTGGCCTTCTCTACCAAACCTGTTTCTGCCgaatcttaaacccaact
ggttacctcttatctagtggGGGATTCCTTCTTACCACCTCGGTCTGTTTCACATGTCACgaatcttaaacccaactt
ggttacctcttatctagtggGCAGAGGTTGGAGCACTCCTGGATAGAGGTGTCAGAGAGGgaatcttaaacccaa
ggttacctcttatctagtggTTAGATGTGGAGCTTATTCAATGGGCTCTAAATTCCCCTCgaatcttaaacccaactt
ggttacctcttatctagtggTGTAACTCTCAAAAGGAGGTATTTAAGGGCTCAATGTGGCgaatcttaaacccaactt
ggttacctcttatctagtggGCACTCAGGCAGCTGTGGGCTGGACTATTTTGTGGAGCGGgaatcttaaacccaa
ggttacctcttatctagtggGCTACATAGGTATACTGTGCCATGGTAGTTTGCTGCACCCgaatcttaaacccaact
ggttacctcttatctagtggGCTATCCCTCCTCTAGCCCCCTACCCTAGGACGGGCCCCGgaatcttaaacccaa
ggttacctcttatctagtggTCTCATTGTTTAACTCCTGCTTATGAGTGAGAATATGCAGgaatcttaaacccaacttg
ggttacctcttatctagtggATTTTCTCCCATCCTGTAGGTTGCCTGATCACTCTGATGAgaatcttaaacccaactt
ggttacctcttatctagtggGCTGTGCGGAAGCTATTTAGTTAGTTAGATCCCATTTGTCgaatcttaaacccaactt
ggttacctcttatctagtggTGGTGTTTTAGTCATGAAGTCTATGTTCTGAATGGTATTGgaatcttaaacccaacttgt
ggttacctcttatctagtggTTCAGTTTTCTGCATATGGCTAGCGAATTTTCCCAACACCgaatcttaaacccaactt
ggttacctcttatctagtggGGAAATCCTTTCCCTGATGCTTGTTTTTGTCAGGTTTGTCgaatcttaaacccaacttg
ggttacctcttatctagtggGCTTATGATTGTCTTGGCTATACGGTCTCTTCTTTGGTTCgaatcttaaacccaacttg
ggttacctcttatctagtggCCTATCCATGAGCATTGGAATGTTTTTCCATTTGTTTGTGgaatcttaaacccaacttg
ggttacctcttatctagtggCACTTTGGCTAGCGTGCAGTCATTTGTAGATGACGTGCAGgaatcttaaacccaact
ggttacctcttatctagtggCTAACACAACGGCAAGCATCCGGAGTCTGATCTGTGTGGGgaatcttaaacccaa
ggttacctcttatctagtggCTGTCAGAAGTGTGCATGTGTCACCTTGGGCTCCTTAGCCgaatcttaaacccaac
ggttacctcttatctagtggCTGTCAAAAAGATGATTCCTACCTCACATAGTGAGTGGACgaatcttaaacccaact
ggttacctcttatctagtggGGATGGAGAGCACAAGAGAGACACGGTTCCTATCATTGGGgaatcttaaacccaa
ggttacctcttatctagtggCAGGTAAGACTCATCGGCACCTTTGCTATGCTAGGTAGTCgaatcttaaacccaact
ggttacctcttatctagtggGTCCTGTGGTAGACCTCCGAGGTAGACACTATCATCACCCgaatcttaaacccaa
ggttacctcttatctagtggCTAGTTGACCAAGTCCAGATTTGAACCCAGGCTGCTTGGCgaatcttaaacccaac
ggttacctcttatctagtggCCTATTATACTGACAGTGTCAAGAAGGCAAACAGTAGGGCgaatcttaaacccaac
ggttacctcttatctagtggAGGGATGCTGACATAGATTGATTAGGGTGTCTGGGAAGCCgaatcttaaacccaac
ggttacctcttatctagtggTGAGAGGGGGACCAGGGCAGGGGAAGCTGCAGAGACCAGCgaatcttaaaccca
ggttacctcttatctagtggAACAGGCAGCACTTTAGAGGCTGTGCCAGGGACTGTGGATgaatcttaaacccaa
ggttacctcttatctagtggGAGCAAAGAGATCATTGCAGATTTTAATCAGGTAATGACAgaatcttaaacccaactt
ggttacctcttatctagtggCTGAGATCTTGGTAGGTATTGGGTATTATGATATGCTGGGgaatcttaaacccaactt
ggttacctcttatctagtggCCTGTTTCCACAGCCTTAACCTTAACCGTGCATGGTCCTGgaatcttaaacccaact
ggttacctcttatctagtggTTCCATCCAGTCCTATTGCTTTCAGTCCCACTCTCATGCCgaatcttaaacccaactt
ggttacctcttatctagtggCCAGTTCCAGTCTTTCCTGGGAATTCCAACTCAGCAGTTGgaatcttaaacccaact
ggttacctcttatctagtggTCCATTTGTATGCTTATTGGCATCTCAAACTTAACATGTGgaatcttaaacccaacttg
ggttacctcttatctagtggTCCAGTTTTCAGTGGCGGCTTCATTATTCCATTGCCCGGGgaatcttaaacccaact
ggttacctcttatctagtggGACTGTCATCCTCCACTCCTCTCTTTAGTGCCCCAAAAGCgaatcttaaacccaac
ggttacctcttatctagtggGCCCCCTCCCCTTTCCGAATGCAAATCACCAGCATCTCTCgaatcttaaacccaa
ggttacctcttatctagtggCCTCCTCATAATTCACCTGGCTTCTACAGTTATAAAACTGgaatcttaaacccaactt
ggttacctcttatctagtggTTTCCTCTTAGAGTGAAAGAGAAAGTTTTCACCATGGTCTgaatcttaaacccaacttg
ggttacctcttatctagtggCATGCTATAACAATACAGTTGAGTGGTTGCAACAGAGACCgaatcttaaacccaact
ggttacctcttatctagtggCTCTGACCTCGTCTTCCAACACTCACACCCTTTCCTACTCgaatcttaaacccaact
ggttacctcttatctagtggGCAATGTCAGAAACTTTCTCAGAATGTCCTTACCTCAGGGgaatcttaaacccaact
ggttacctcttatctagtggCGGAGGACCCTTCACCCATGATCAAATGCATTCATCCATTgaatcttaaacccaact
ggttacctcttatctagtggGCTCCCGAGCTTCCTTTCAAGGTTTTTGCTCAAATGTCATgaatcttaaacccaactt
ggttacctcttatctagtggCAACACCCTCCCCTATTGGTCTCTTTCATTCTCATCTTGCgaatcttaaacccaactt
ggttacctcttatctagtggCCCCTTACTTCCACTGACAGTGTGTAACTTGTATATTCTCgaatcttaaacccaactt
ggttacctcttatctagtggCATTCCTTCATCATCTGTGACTTGAAGAAAGCCAGGTATGgaatcttaaacccaactt
ggttacctcttatctagtggAGTTCTAGAATATTCCTTGGGTGCCTGTCATGTGCTAGGCgaatcttaaacccaactt
ggttacctcttatctagtggACCTCTCACAAACCTTGGTAGGAAGGGGCCCTTGTTTCTCgaatcttaaacccaac

ggttacctcttatctagtggGCTGTTGTGTACTGAGGGGCAGGAAGAGGCCATGTCTCCGgaatcttaaacccaac
ggttacctcttatctagtggCAATTTCTGAGGTCTGAGCCACTTCTTCCCTCCAGGTGGCgaatcttaaacccaact
ggttacctcttatctagtggCCAGGATCCTTCCCCAGAAATCCAGTTCCCGCCACAATGGgaatcttaaacccaac
ggttacctcttatctagtggGCTCTCAATAATAGAGTCCTAACTGGAAGATGTCAGAGTGgaatcttaaacccaactt
ggttacctcttatctagtggACAGATATGCATCTCAAAACACTGCCTAGAGCCAACCTTCgaatcttaaacccaact
ggttacctcttatctagtggCATTCACATTGTTCTGAAACTGCCACTACTGTCCACTTCCgaatcttaaacccaactt
ggttacctcttatctagtggCCTGTTAAACCATAACTCACCATCCCTCCTTTCCCCAGTCgaatcttaaacccaact
ggttacctcttatctagtggCCATTTTACGTTCTGTCTCCATGAATTTGATTACTCTAGGgaatcttaaacccaacttg
ggttacctcttatctagtggAGCATTTAACCTCTTGTGACAGGCTTATTTCACTTAGCAGgaatcttaaacccaacttg
ggttacctcttatctagtggGCATTGCTTCCACCTTTGGACATATTGTGAATAATGCTGCgaatcttaaacccaacttg
ggttacctcttatctagtggCCAGAAGTGGAATTGCTGGGACATATGGTAATTCAGTGTTgaatcttaaacccaactt
ggttacctcttatctagtggCATGACTTAGTCACTGACTTCCCAGGTTGCCCTAAGGACCgaatcttaaacccaact
ggttacctcttatctagtggTCTTGTGTTTTGTTCCCCAAGTGAGTCAACTGAGGAAGCCgaatcttaaacccaactt
ggttacctcttatctagtggCACCAGCTCAATATTATCTGACCCAAGCCCGAGAGCTATGgaatcttaaacccaac
ggttacctcttatctagtggACCTCTGAAGGGGCATTAACAGACCCTGACAGACCAAGGCgaatcttaaacccaa
ggttacctcttatctagtggCTGCACTTTTTGGAGTCATGAGGAGAGCCCTGGAAATGGGgaatcttaaacccaact
ggttacctcttatctagtggTACGATCTTGGCCAAGATATTTAACCTTCTCTAAGAGTGGgaatcttaaacccaacttg
ggttacctcttatctagtggGTAGTGATATGGTTTGTCTCTGTATCCCCACTGAAATCTCgaatcttaaacccaacttg
ggttacctcttatctagtggCAAGGGAGGGACCTGGTGGGAGGTACTTGGATCACGGGGCgaatcttaaacccaa
ggttacctcttatctagtggCTCTCTCTCTCCTGCCACCATGTGAAGCAGGTCTTTGCTTgaatcttaaacccaactt
ggttacctcttatctagtggTAATACAGAAAATTGGTACCAGCAGAGCAGGGCACTTTTAgaatcttaaacccaactt
ggttacctcttatctagtggGGAATTGGGTAATGGGCAGAGATTGGAGCAGTTTGGAGGGgaatcttaaacccaac
ggttacctcttatctagtggATGGACAATGAAGTACAGGCTGATGTGGTCTCAGATGGAGgaatcttaaacccaact
ggttacctcttatctagtggGGAAAGGTCACTCTTGCTATGCCTTAGCAGATAGACTGGCgaatcttaaacccaact
ggttacctcttatctagtggGCAGCAAAACATTCAAAACGTGACCTGGCTTTTTCTGAAGgaatcttaaacccaactt
ggttacctcttatctagtggAAAGGGAAGCAGAACATAAAAAGTTTGGAAAATTTGCAGCgaatcttaaacccaactt
ggttacctcttatctagtggAAGAAGCTCCTCCCATCACAAGCCTGAAGGCCTAAGAGGGgaatcttaaacccaa
ggttacctcttatctagtggGGGACATGGCACACTGAGTCCCAACAGCTTCAGATCCAGCgaatcttaaacccaa
ggttacctcttatctagtggGCCCCAAACCTTGGTGGCTTTCATGTGGTATGGGCCTGTCgaatcttaaacccaact
ggttacctcttatctagtggCACCTGGATCTCTAGGCAGAAGTTTGTTGCAGGGGTGGAGgaatcttaaacccaac
ggttacctcttatctagtggAGGGCAATGCAGAGGGGAAATGTAGGATTTAAGCCCTCACgaatcttaaacccaac
ggttacctcttatctagtggGGGTCACCATCCTTCTGACCCCAGCATGGTAGAACCACAGgaatcttaaacccaa
ggttacctcttatctagtggCTGTGTACTTGGAAAACCCACAAGCACTCAATGCCAGCTGgaatcttaaacccaac
ggttacctcttatctagtggTTAATGAGGTCCCTGCCAAGTTTTGGACTTGCATGGGACCgaatcttaaacccaactt
ggttacctcttatctagtggGTGCCTGTACCCCCATTGTATCTTGGCAGTAACAAACCTGgaatcttaaacccaact
ggttacctcttatctagtggAATGAAAAGGGACAACAGATTTGGGAGGGGCTAGGGGCAGgaatcttaaacccaa
ggttacctcttatctagtggTTGTAATCCCTACGTGTCAAGGGAGGGTCCTGGTGGGAGGgaatcttaaacccaac
ggttacctcttatctagtggGTGTGGCACTCCTTCCTTTGATCTCTCTCTTTCTCCTGCCgaatcttaaacccaacttg
ggttacctcttatctagtggCACCTTTACTTTCCACTGTAACTGTAAGTTTCCTGAGGCCgaatcttaaacccaacttg
ggttacctcttatctagtggGGTGGATTACTTCAGCCCAGTAGTTTGAGACTAGCCTGGGgaatcttaaacccaact
ggttacctcttatctagtggCTGGGAATGTACTCAAAGGGTTATAAAGACTATTCTACTGgaatcttaaacccaacttg
ggttacctcttatctagtggAGGGGAACATCACACGCCAGGGCCTGTGGGGGGTTGGGGGgaatcttaaacccaa
ggttacctcttatctagtggTGTGTATACCTATGTAGCAAGCCTGCATGTTCTGCACATGgaatcttaaacccaacttg
ggttacctcttatctagtggGCAATGAGCCATGATTGTGCCACTGAATTACAACCTGTGTgaatcttaaacccaactt
ggttacctcttatctagtggAAAGAGGGGAAGAGATTGAGGGTCAGAACCTCATGGACTCgaatcttaaacccaac
ggttacctcttatctagtggCTTCTCTTCTTGGCCCAACTTTTAAAGACTTCTGTAGCTCgaatcttaaacccaacttg
ggttacctcttatctagtggCCAGCTGTAACATGCATTTCTTGAAGTTCTTAGAGACCCCgaatcttaaacccaactt
ggttacctcttatctagtggGATTCCTGCCATCTGTTGGTTGTGGTCAAAGCTGATGGGGgaatcttaaacccaactt
ggttacctcttatctagtggGGCCAACGCCAGCCTGTATGTGGAGAAGCCAGCATTTGATgaatcttaaacccaac
ggttacctcttatctagtggGTCCACCACCGTTGTCCCAATACCACCGTGAGAGGTGCCCgaatcttaaacccaa
ggttacctcttatctagtggCCCTGCAAAGAACCAAGTCTCCTGGGCCAGCCCTGTGCAGgaatcttaaacccaa
ggttacctcttatctagtggAGTTTCAGAGATTAAAGTGATTCAGTTAAAGGTTCAGAGCgaatcttaaacccaacttg

ggttacctcttatctagtggTGCCTCATAAGGGGCTTGTCCTTTTTAATTACACAGGTGCgaatcttaaacccaactt
ggttacctcttatctagtggTCCTTCCTCAGAGCTCCCTGCAGAGATTTTGCCACAGTTGgaatcttaaacccaact
ggttacctcttatctagtggGTTCGAGGATACAGATGTTGCTGAAATCAGAGCTCTCAGCgaatcttaaacccaact
ggttacctcttatctagtggAGAGAGATAAAATGGCCCAATGTCCCCTGCTCAAGAAAGTgaatcttaaacccaac
ggttacctcttatctagtggTACAAACCTTATGACCTTGGGCGAGCTTCTTACCATCCCCgaatcttaaacccaact
ggttacctcttatctagtggCAGGATTAAATGATGTATGTAAAGTGCTTTGCACAGCTGCgaatcttaaacccaactt
ggttacctcttatctagtggGTGCTTTTGTTTAACCATCACCAGTATCCTAAGTGATAGGgaatcttaaacccaactt
ggttacctcttatctagtggGCAAAGATATTACACAACTCTTGCTTGTTGCTAGTGCAACgaatcttaaacccaactt
ggttacctcttatctagtggGCCAGCTGAGGGGCAGAACTTGTGCATTTTGCAACCATCCgaatcttaaacccaac
ggttacctcttatctagtggGTTTGGCATTCAATCCTCATTACTGACTATGTATGGCTTCgaatcttaaacccaacttg
ggttacctcttatctagtggCAAATGGTGAATAATTGATTTAGCTGACCTGACAACACCCgaatcttaaacccaactt
ggttacctcttatctagtggTCTGGGCATCCTGGCTTCTAGCAATTCTCTATAGCGGCCCgaatcttaaacccaact
ggttacctcttatctagtggACTTAGATCATGAGCTCTGTCCTCCCAGTATCACCCTGGCgaatcttaaacccaact
ggttacctcttatctagtggATGGTTCACATTAGACATTCCCTGAAGAATGTACCGATCCgaatcttaaacccaactt
ggttacctcttatctagtggTGGCCTCTACATAGCAAAACTTGAGACATATTTAGTTAGCgaatcttaaacccaactt
ggttacctcttatctagtggACCATATTGTGTCATTGTTTAAAATGCCTTTGCTGGCCTCgaatcttaaacccaacttg
ggttacctcttatctagtggTTCATCTTAGAGCACATTAGTGTAGGAAGCATTTATTTCCgaatcttaaacccaacttg
ggttacctcttatctagtggCCTTGGGCATATCCACACCAAAGGCTATATGAATTCAGAGgaatcttaaacccaact
ggttacctcttatctagtggCAGAAGCAAATGAGGACATTTTGGCAGACATCTTTGTAGCgaatcttaaacccaactt
ggttacctcttatctagtggCTGGAATGTGACAGGGATTAGGCTGGGATTAGGTGATGGGgaatcttaaacccaac
ggttacctcttatctagtggTGGGTACACATGGACTGAGGCAAGTACCATGAGGTTAGTGgaatcttaaacccaac
ggttacctcttatctagtggGAAGGGGAGACAACTTATTCATATTTTTAGCTTGGCCAGCgaatcttaaacccaactt
ggttacctcttatctagtggCAGGCTGGAGAGCAGTGGAACAATCACAACTCTCTGCAGCgaatcttaaacccaa
ggttacctcttatctagtggTGGGTTCAAGTGATCGTCCTACCTCAACCTCTCAAGTAGCgaatcttaaacccaact
ggttacctcttatctagtggCCCCAGGTTGGTCTCTAGCTTCCAGGCTCAAGTGATCCACgaatcttaaacccaac
ggttacctcttatctagtggCCAAAGTGCTGGGGTTACAAGGATGAACCATCTCATTCAGgaatcttaaacccaac
ggttacctcttatctagtggGCCAAAATACTGTGCCTGAGTACTTGGGGATGCTGAATGCgaatcttaaacccaact
ggttacctcttatctagtggTGTGTTCCCTGCTCAGATAGACCTGTATGTAACTGGCCTGgaatcttaaacccaactt
ggttacctcttatctagtggCTGTTTTTGGTCCCTCCTACTTAAGTCGGAGAGGTGAGGCgaatcttaaacccaact
ggttacctcttatctagtggTTTAAAAGAATAGACTTCAGGCCAGGTGCAGTGGCTCATGgaatcttaaacccaact
ggttacctcttatctagtggGACAGGTGCCTGTCATGTAATCTCAGCTACTCGGGAGGCAgaatcttaaacccaac
ggttacctcttatctagtggGACTTCAAATAAATATGAGTTGTGCGGAATTTTGCTTGGCgaatcttaaacccaacttg
ggttacctcttatctagtggCGTGCAGTGGAATACTTTCCGGCCTGCAAATCAGGGGTTCgaatcttaaacccaac
ggttacctcttatctagtggGCTCTGCCATGGAAGTAGGTGTGTATCCATACTGGTAGGGgaatcttaaacccaact
ggttacctcttatctagtggCCTTGGTTTTCTTGGATGTGAAGTGAGGAAGTTACTCCTGgaatcttaaacccaactt
ggttacctcttatctagtggGACAAACTCATGACACGGGAAGGCTCTTTGGCCCTGTCACgaatcttaaacccaac
ggttacctcttatctagtggTGCTTCAGAAAGAGAGCCAGTTGTATTGCCTACAGGTGTCgaatcttaaacccaact
ggttacctcttatctagtggGTGTTTGAACAACCAACATTTGAAAGGCAAATCTTCCAACgaatcttaaacccaactt
ggttacctcttatctagtggGCAGTCTGCATGGGCTTTGTGCATGGGTTTGTCATTTGGGgaatcttaaacccaactt
ggttacctcttatctagtggCAGGACTGATTATAGTGTCTGCCTCATTGGGCTATTTAGCgaatcttaaacccaactt
ggttacctcttatctagtggGATGTGGGCTTTCCAGCTGACAGTGTTGTACCCTCTCTTGgaatcttaaacccaactt
ggttacctcttatctagtggCTCACATAGGTGGAGAATGGTAAGCCTGGGACTTGAACCCgaatcttaaacccaac
ggttacctcttatctagtggGGAGACTGTACTCCACTCACTTCACTGTCTCTCTGATTACgaatcttaaacccaactt
ggttacctcttatctagtggTGAGTCTTTGAAACTCTTTTCCTGGTGGACTTTATATACCgaatcttaaacccaacttg
ggttacctcttatctagtggACAATTCTTTTGCTTTGTCTTAGGACCAGAGTTATAGTCCgaatcttaaacccaacttg
ggttacctcttatctagtggTGGAAAGCTTTTCTATACCTGAAAGTACTGAGTCCTCATCgaatcttaaacccaactt
ggttacctcttatctagtggCAGAGGTCTCTGATTTCTTGTTTGCATAAAAACTAACTAGgaatcttaaacccaacttg
ggttacctcttatctagtggTAATAGAAGCTAGTGTTTGTTGAACACATGTACCAGGCACgaatcttaaacccaactt
ggttacctcttatctagtggCTCCAGAGCTCAGCAACTTTCACTTCCACTTTGCCAGCCCgaatcttaaacccaac
ggttacctcttatctagtggAGCAGCCTTGCTCCAACTGGCTAAAAAAGTGTGTGTGTGTgaatcttaaacccaact
ggttacctcttatctagtggACCGTGATGGTGTCTCTGATTTTGCCTAGCAGGGGTGTGAgaatcttaaacccaact
ggttacctcttatctagtggTGAGCCACTGCACCCAGCCTCATGTATAACTTTAGTAATCgaatcttaaacccaact

ggttacctcttatctagtggTCACAAGGGCACTGTGAAGAAGTCATCTTTTCATTGTCCCgaatcttaaacccaactt
ggttacctcttatctagtggGAGAACTTGAGAGCTTGCCAGACTTCAAACAACAAGACACgaatcttaaacccaac
ggttacctcttatctagtggGGTGGAGTAGGTTGAGGAGAGGGTCTCCAAGAGTGTTTGGgaatcttaaacccaac
ggttacctcttatctagtggCAGCACAGGCAATTATTTGCATAGCAATAATAAGCACAGAgaatcttaaacccaactt
ggttacctcttatctagtggCCAGATGTATCCCTTGCAGTTCACGATCTGATGTTTTTGCgaatcttaaacccaactt
ggttacctcttatctagtggTGCAAAAGTCATTGAGCATATATTATTCTCAATGGAGCTCgaatcttaaacccaacttg
ggttacctcttatctagtggGTTGTGGGAGAATGGATGACTTCCTTCCTCTTGAGGCTGGgaatcttaaacccaact
ggttacctcttatctagtggGTTGGAAAAATCTGTAGGCATTCACCAGAAATGCTTAGGGgaatcttaaacccaactt
ggttacctcttatctagtggGCTGAAGAGTGTAGAGAATATCCTATGAGCACTGGGAGGCgaatcttaaacccaac
ggttacctcttatctagtggGTTCAAAAAAGAAGTGGGATGACCAGATGGGAGCAGGAGAgaatcttaaacccaac
ggttacctcttatctagtggGACCTGGGGCAGTGGCAGGGGAGACAGAGAATACGCCCCCgaatcttaaaccca
ggttacctcttatctagtggGGATTTGTAGCTTCCATCTCACACTCCATATGTCTTCCTCgaatcttaaacccaactt
ggttacctcttatctagtggTTTTTTCTAGACAAATTTTTGCTCTGTCACCCAGGCTGGAgaatcttaaacccaacttg
ggttacctcttatctagtggCCTGAGTAGGTGGGATTACAGGTGTGCTCCCCAATGCTTGgaatcttaaacccaac
ggttacctcttatctagtggAGGCTGGTCTCGAATGCCTGACATCGTGATCCGTCCGCCTgaatcttaaacccaac
ggttacctcttatctagtggGAAACCAGCCCTCACTATAACCAGTGAAGCAGTTTGAAGCgaatcttaaacccaac
ggttacctcttatctagtggTAGCTAATAATGTTCCCCCACACTATCCTTTTCCCCCAGGgaatcttaaacccaactt
ggttacctcttatctagtggGCTTGATGGGTATCTCTGCATCTGTTTTCAATGTTCTTGCgaatcttaaacccaacttg
ggttacctcttatctagtggGGACATCCCTTGAGGGTAACAGATAGAAATAAAACTCACCgaatcttaaacccaac
ggttacctcttatctagtggTTCCTTGCATGGATATGCCACAATTTGTGCAGTCTTTTTCgaatcttaaacccaacttg
ggttacctcttatctagtggTTACACCTTGAAAGGACCTTACACATGTCAACCAACCTCCgaatcttaaacccaac
ggttacctcttatctagtggTGGTAGTTCCTCTGGAGCCAGAATCCAGTTTCCTGAGAGCgaatcttaaacccaac
ggttacctcttatctagtggAGGGAACAGAGATTTCGATTCCTAAATGGCTGGGGATCTGgaatcttaaacccaac
ggttacctcttatctagtggAGCCTCCTTTGCCAAAGCCAGGGCTAAACTCACTCCAGCCgaatcttaaacccaa
ggttacctcttatctagtggGTGTGTGGGGTTTCCTGGTTGGGGGACAGAAGGCACCTCCgaatcttaaacccaa
ggttacctcttatctagtggTGCATAGATCCCTTTGACCTTTTAAACATCCTTGGAGGCCgaatcttaaacccaactt
ggttacctcttatctagtggGCCTCCAGGCTTCTAGGAATAGGTAGAACAACAACAACAAgaatcttaaacccaac
ggttacctcttatctagtggGCTAAGTAGGAAAAGAGGCAGAAGACTGGTGACCTTTTCCgaatcttaaacccaac
ggttacctcttatctagtggCAGATTACTGATTTCAAGTATGACTTTCCAGGGGAGACACgaatcttaaacccaactt
ggttacctcttatctagtggGGAAGGATCACTTGAGGCCAGGAGTCCACGATCAGCCTGGgaatcttaaacccaa
ggttacctcttatctagtggCACACAACAATGGTCTTGTGGCAGAGCTGGGACTACTCCCgaatcttaaacccaac
ggttacctcttatctagtggGTCCCAGGGCAAAGTTCTTCCTTTCAATTGACAAACACAGgaatcttaaacccaact
ggttacctcttatctagtggCAGAGAAGATGATTTATCTTTGTCAAAATGCCATTGTGAGgaatcttaaacccaacttg
ggttacctcttatctagtggCGTAGCCAGAAGCCCAGGGACGTACGGGGGCCACAGGGTCgaatcttaaaccca
ggttacctcttatctagtggCAAAGGCTGGTGAACGGAGAAAGCTGTTGAGGTAGAGGTCgaatcttaaacccaac
ggttacctcttatctagtggCCAAGGGGGGAGCTGTGGGTCACTAAGAGGTTGCTGCTCAgaatcttaaacccaa
ggttacctcttatctagtggGCTCCCTCTGAAGTGGCAATCAGTCAGAGTGACCACTGCCgaatcttaaacccaa
ggttacctcttatctagtggTCAATGGTCCAGCAATTTCCCTCAATTTTAATCAATTTGGgaatcttaaacccaacttg
ggttacctcttatctagtggGGCAAGTGTTCTTGATCAGGATGCCCCCATGCTCCCTGGCgaatcttaaacccaac
ggttacctcttatctagtggGGAAGCCGTTTAGAGACCCTGTCCACTATTAGGCTGTGTGgaatcttaaacccaac
ggttacctcttatctagtggCCCAGCCCACTGTGATCAGCATGGTGACTGCTGGGGGTGCgaatcttaaacccaa
ggttacctcttatctagtggGGAAAGTAAGAGAGAGGTGGGTATAGGGAGAGCTATGGCAgaatcttaaacccaac
ggttacctcttatctagtggATGATTGGTACTATTGGTGTCTATTAGGGTTGCCAGTTACgaatcttaaacccaacttg
ggttacctcttatctagtggTCGGTGTCCTATATTTGTATTGGTTAGATGTGGCCACCTCgaatcttaaacccaactt
ggttacctcttatctagtggGCTGATGGGAAGTACAATAGTATCAGTGGTGACCTAGCACgaatcttaaacccaac
ggttacctcttatctagtggGCCAGTATTTTGATGGGGGTATTAAGATGAACAAAACAGGgaatcttaaacccaactt
ggttacctcttatctagtggGAAGATGTGTGAACCAATGTTATGATGCAGTATGATAAGCgaatcttaaacccaactt
ggttacctcttatctagtggGGCGCTTGTGCTTTTTTCCATTCATCACACTAGAGGGATGgaatcttaaacccaactt
ggttacctcttatctagtggGAATTGTAGCTCCCCATAATTTCCACATGTCCTGTGAGGGgaatcttaaacccaactt
ggttacctcttatctagtggTCCCCTACTCACGCTCTCTTATGTGCCACCATGTAAGACGgaatcttaaacccaac
ggttacctcttatctagtggACACAAGGGGTATGACGAACAATGGCTGAGAGCTCAAGCCgaatcttaaacccaa
ggttacctcttatctagtggCCACATTCAAGATAACTCTTCAGGAAGTAGGGTGAGAGGGgaatcttaaacccaac

ggttacctcttatctagtggACAAATTGGGTCATGACCTTCAGTATATTCCACAGGTGTGgaatcttaaacccaactt
ggttacctcttatctagtggAACTGGACCCAGACTGCTTGCTTGCCATTTTATTTTCTGCgaatcttaaacccaactt
ggttacctcttatctagtggGTGTGCTTCATTCTCTCTGTACTCCCCAATCCCACATCACgaatcttaaacccaact
ggttacctcttatctagtggTCCTTTTCAAACCAGAGTCAGAGTTACAAGAAATAAGCACgaatcttaaacccaact
ggttacctcttatctagtggGATTACAAGCATGAGCCACCAAGCCCAACCTCAATCTGATgaatcttaaacccaac
ggttacctcttatctagtggGTTGCTGCAGGCAATTCAGAGATGAACAAGGAACAGTTTGgaatcttaaacccaact
ggttacctcttatctagtggGGGGAGTCCTTGCTTCAGATGCTTGCCATGCTCCAGCTGCgaatcttaaacccaac
ggttacctcttatctagtggCTCACTAGTGCCAAGTGTGAACCCAACATCTCGGCCATGGgaatcttaaacccaac
ggttacctcttatctagtggGCCAGCAGACTGCAGGTGGCATAGAAAGACTGGCCAGTGGgaatcttaaacccaa
ggttacctcttatctagtggGTCCAGGCTCAGAGGCCCCACAGGAGAGGCACAGGCACGGgaatcttaaaccca
ggttacctcttatctagtggGACTGACCAGGTGTGCACACTGTCTATGCGAACAATGGCCgaatcttaaacccaac
ggttacctcttatctagtggCCTTCCTGGAGGCCTGACTGTGGAGATGCTCTGGGTTCCCgaatcttaaacccaac
ggttacctcttatctagtggCCAAGAGTGGCCTGTGCTTCTTACAAGCAAAAGGTCAGACgaatcttaaacccaac
ggttacctcttatctagtggCAGTGGAGTGGAAGGCCAGTTTCTGCTGGGAGACTTAGACgaatcttaaacccaac
ggttacctcttatctagtggAGATGCAGCATCTGAAATGGGCTCTAAAAGAGGGGTCAAGgaatcttaaacccaac
ggttacctcttatctagtggAAGCATTCCTCTGCCCGCAGAGTTGAAATCAGATGTTCAGgaatcttaaacccaact
ggttacctcttatctagtggTGTGCATGCACGCATACACACACACACCTGGCACTTACCCgaatcttaaacccaac
ggttacctcttatctagtggTTGCTCCCTGCAGAGAAAAGATATGAAGGAGAAAGGAAGCgaatcttaaacccaac
ggttacctcttatctagtggAGATGGTGATTGAGCTAATATCTCACCTTGTCCCAGGTGCgaatcttaaacccaact
ggttacctcttatctagtggAGGTGAGTTAACTGAATCAGAATAGATGCGTGATTTGCCCgaatcttaaacccaact
ggttacctcttatctagtggCCACCACTCCCGCCACCTGCCTTTATGCCCTGGACCCAGGgaatcttaaacccaa
ggttacctcttatctagtggGGCTTAGGACCCACCAGAGCCACACTTTACTCAGACTAAGgaatcttaaacccaac
ggttacctcttatctagtggCCCCATCTTAAAACTTACCTGCCCTATGCTTCATTCCCTGgaatcttaaacccaact
ggttacctcttatctagtggGAAGAATCCTTCAGAGCAGGCAAAGCCAATTAGACCTCACgaatcttaaacccaac
ggttacctcttatctagtggCATGCACCACCATGCCCGACAAATTTTATATTTTTAGTAGgaatcttaaacccaactt
ggttacctcttatctagtggCTGAAACTGAAGGCTAACCAGTCAGAAGCCACCAACTCACgaatcttaaacccaac
ggttacctcttatctagtggGCCTTTCCCTTGGATTCCCATGATGGACCCCACAAGTAACgaatcttaaacccaac
ggttacctcttatctagtggGCCTCAATTTACCTTTTAACACTTCTTGAGACTGTATTGCgaatcttaaacccaacttg
ggttacctcttatctagtggCTCAGCATATCCCTAAGAGTGAAATTCACCTTATCAGTCCgaatcttaaacccaact
ggttacctcttatctagtggAGGTCTGGCAGATACTCATTGGAGTCCAGCACTCAAAGTGgaatcttaaacccaac
ggttacctcttatctagtggCATTTCTCTTCAGCTTTTTCTTATAAATACTGGGACCCCCgaatcttaaacccaacttg
ggttacctcttatctagtggTCTGAGCCTGTCACATAGTCCAGCTTTATATACAGCACCCgaatcttaaacccaact
ggttacctcttatctagtggGCCTTTCTTCAGTGGTATAAAGGTACTGCCCCATACCAGCgaatcttaaacccaact
ggttacctcttatctagtggGTGACTGGTTGACTCCTTCCTCTCTTGCTTCCTTCCTTCCgaatcttaaacccaactt
ggttacctcttatctagtggCTCCCTTCCATCTTCCCTTCCTCTTTCCCTCCCTCTTACCgaatcttaaacccaact
ggttacctcttatctagtggCCCTTACTCCCTCTATTCCTCATTCCCTTCCTCTTTCCCCgaatcttaaacccaactt
ggttacctcttatctagtggCTGGGTATTTTGGTAGGCCCTTTACATAAAACCTCATTTCgaatcttaaacccaacttg
ggttacctcttatctagtggCAGAGCCTCAGAGCATGTAGTTAATGCCCCAAGAAAGCACgaatcttaaacccaac
ggttacctcttatctagtggGGAAGCTCATATTCATTCACAAATCTGTACTGAAACACACgaatcttaaacccaactt
ggttacctcttatctagtggAGCTAGGTTGGTGATTCCCAAACAGACTAGCTCTGCTTCCgaatcttaaacccaact
ggttacctcttatctagtggCTGGATGACAACATGTCTTCACATTCCCATCAGACTTCTGgaatcttaaacccaact
ggttacctcttatctagtggGGGGCAACCTGCTCTGGCATTGTCTGAGGATAAAAGACTGgaatcttaaacccaac
ggttacctcttatctagtggGCAGGGTGCTCCACAAGAGGTTGCTCTGTCTTGGTAGGAGgaatcttaaacccaac
ggttacctcttatctagtggCCGCCACATCCCTCACCAACTGTTGTCAAAATATCAACTGgaatcttaaacccaac
ggttacctcttatctagtggCCTTTTGTTTCTCTGAGGTTGCTTATTTCCCTCTCTGGACgaatcttaaacccaacttg
ggttacctcttatctagtggGCTCTATAACTCAGCTCTTAGAACACCGAACTCTAGTGAGgaatcttaaacccaact
ggttacctcttatctagtggGGGAAAATGTTTTTGGCAAATTTGGGGAGATTGTCAAGCCgaatcttaaacccaactt
ggttacctcttatctagtggGTGGCTGGCACCACCCTACATGCTGGGAAACAGTGCACAAgaatcttaaacccaa
ggttacctcttatctagtggCTAGACATGTCAGCAGCTTGAGAGAGAAGATAAATTCATAgaatcttaaacccaact
ggttacctcttatctagtggGGGTTGGGTATTTAGGGAGGGCCTCTCTCAGGAGGGAAGCgaatcttaaacccaac
ggttacctcttatctagtggCCTTCCACACTCTGCTTCATGTCTTTGAGTCAGCCTACCCgaatcttaaacccaact
ggttacctcttatctagtggCTCTCTCTGGCATGCTAAATCTCCACTCAGACTTGACTTCgaatcttaaacccaact

ggttacctcttatctagtggGGCATGACCTGGTTAAGATCTTGACTTTCCCTCTTAATAGgaatcttaaacccaacttg
ggttacctcttatctagtggCCTCCTTCCTACTTCTCTTTCTATCCCCAAAACCTGTAACgaatcttaaacccaacttg
ggttacctcttatctagtggTGGCCTGGTTGTAAAAGTGCCAAATTGTTTTCCAATGTGGgaatcttaaacccaacttg
ggttacctcttatctagtggTTCCACCAGGGGTGTATTAGAATTCTAGTTGCTCCTCATCgaatcttaaacccaacttg
ggttacctcttatctagtggTTAGGTATTCTAGTAAAGTTGTAGTGGCATCTTACCATGCgaatcttaaacccaacttg
ggttacctcttatctagtggTTACACCATTTTTCCTTAGGATTTTTGTTTCCCCATTGAGgaatcttaaacccaacttgt
ggttacctcttatctagtggTCTACCAGTTGTGGCTTGTCTTTTCACATTCTTATCAGGGgaatcttaaacccaacttg
ggttacctcttatctagtggTATGCATATGCCTATCCAATTATTCCATCACCATGTGGTGgaatcttaaacccaacttg
ggttacctcttatctagtggCTGTTGCACCTGACTACTTATTTTGAAATCAGTACCACACgaatcttaaacccaacttg
ggttacctcttatctagtggTAACCAGCTTGCCAATTTCTACAATTCTGGAACTTTCTGGgaatcttaaacccaacttg
ggttacctcttatctagtggCAGAAAATTAATATCTGCACAATGGAGCATCTTCCAATCCgaatcttaaacccaacttg
ggttacctcttatctagtggCAGCAATGTTGTCTAGTTTTTAATGTGCAGGAGGTCTTAGgaatcttaaacccaacttg
ggttacctcttatctagtggGCTGCGACCTTGCTAAATTAACCTCTTGGTGGAGGGAGGGgaatcttaaacccaac
ggttacctcttatctagtggGTTTTTCTGGCTCCTTTTCCTGTCTTACATTACCTGCTGGgaatcttaaacccaacttg
ggttacctcttatctagtggGTGAAAGGAGGGCATCACTGCTTTGTTTCTGATTTGAAGGgaatcttaaacccaacttg
ggttacctcttatctagtggATGAGTTGTGGTAGATGCTCATCAAATTGAGAAACTTTCCgaatcttaaacccaacttg
ggttacctcttatctagtggGCTGTTATTTTTAACCAACCAGGCAGACAGCACTGATGACgaatcttaaacccaactt
ggttacctcttatctagtggGGATATCAGAGTGATGATGGCCTCAGCATAAGTTGAGAACgaatcttaaacccaact
ggttacctcttatctagtggCTCCAGTAAAATCATCTGAGAGTAGCATTTACTTTGTGGGgaatcttaaacccaacttg
ggttacctcttatctagtggATAGGGCTATTCAGTTTCTTTCTTTTCAAGTGAGCTTTGGgaatcttaaacccaacttgt
ggttacctcttatctagtggTTGCCCTCATTCTTTCTTTTCTGATTAGCCTACCTAGTGGgaatcttaaacccaacttg
ggttacctcttatctagtggGCATGTGTTTTTGCTCTTTCTTTTCTAGTTTCTTTTGTGGgaatcttaaacccaacttgt
ggttacctcttatctagtggTATAGGCCTCTTGTGCTATGAATACCCGTCTAAGTGCTGCgaatcttaaacccaactt
ggttacctcttatctagtggCTCACAGATTTGATAGGGTGCTAACAATGTTTTCTTTAAGgaatcttaaacccaacttg
ggttacctcttatctagtggCTGTCGCCCAGAGCAAGAAAGTGCAGTGGCGCGATCTCGGgaatcttaaacccaa
ggttacctcttatctagtggGCGTGAGCCACTGCACCCAGCCCTCTAGAAATATTTTGATgaatcttaaacccaact
ggttacctcttatctagtggGAGCAGGTTTAGGTTTACAGCAAAACTGAGTGAAAGGTCCgaatcttaaacccaact
ggttacctcttatctagtggGTGTCCACCAGTACAGTATCATACAGAAGCGTTTCACTGCgaatcttaaacccaact
ggttacctcttatctagtggTCGCACCTCCCTTCCTCCTAACCCCTGGAAATCACTGATGgaatcttaaacccaac
ggttacctcttatctagtggGCATGTAAGGTTCCTCTATGTCTTTTTATGACTTGATAGCgaatcttaaacccaacttgt
ggttacctcttatctagtggGGCTGCTTCCAAGTTTGGGCAATTTTGAATCAAGCTGCTGgaatcttaaacccaactt
ggttacctcttatctagtggTCAACTCATTTGGCTAAGTACCAAGGAGTTTGACTGCTGGgaatcttaaacccaactt
ggttacctcttatctagtggATGAGTGAGAGTTTCTATTGCTCCACACCCTTGTCAGCATgaatcttaaacccaacttg
ggttacctcttatctagtggGCCTTTGGCCATTGAGATAGGTATGTAGTGATATTTCTTTgaatcttaaacccaacttgt
ggttacctcttatctagtggTGGTGATGTGTCTTTTCAAGTCTCCTTTTAGGACTTCTGCgaatcttaaacccaacttg
ggttacctcttatctagtggGAATTTTGAGGTTTTGGGGGGTTTATTTTAGATGATGGCCgaatcttaaacccaacttg
ggttacctcttatctagtggCTCCAAGTCCATGGCTTGACTTTTTATTCTCTTGACATTGgaatcttaaacccaacttg
ggttacctcttatctagtggATTGAAGGTCATGTAGATTTTCTCTGTTGTCTCCTTGGAGgaatcttaaacccaacttg
ggttacctcttatctagtggGCAGCATGTAAGGTCTGTGCCTAGATTCTTTTCATTTTGGgaatcttaaacccaacttg
ggttacctcttatctagtggGCATTGATCTATCGTTTATCCTTTTGCCCATACCACACTGgaatcttaaacccaacttg
ggttacctcttatctagtggGAATTCGCTATTCTGTGCCTCTTGCCTCTCCATATAAACCgaatcttaaacccaacttg
ggttacctcttatctagtggGACGACTGACTCTTTTATATTAAGAATGTACTCTGCAACCgaatcttaaacccaacttg
ggttacctcttatctagtggCTGCAGTCTCCGCCTCTGGGATTCATGCCATTCTCCTGCCgaatcttaaacccaac
ggttacctcttatctagtggCCACCACCACGCCCAGCTAATTTTCTTATCTTTTCTTTTTgaatcttaaacccaacttg
ggttacctcttatctagtggAGCCACCGCGCCCAGCCTGAATTTTCTATGTAGACAATTAgaatcttaaacccaact
ggttacctcttatctagtggGTGAGAGGGAACATTCTTGCTTAGTTCCTGATCTTTGTGGgaatcttaaacccaacttg
ggttacctcttatctagtggGATCAGTATCTAATCCCACTCCTCACCAACTCCCCTCTCCgaatcttaaacccaact
ggttacctcttatctagtggAATCAGAGTGAGAATTTGCTTCAGATTTAATACACTCAGGgaatcttaaacccaacttg
ggttacctcttatctagtggTCCTTTTAGGAAAAGACATTTGCCAGAGGAAAGGCTCTTCgaatcttaaacccaactt
ggttacctcttatctagtggCTAGGCATTCTACTACAAGATAACACTCCAAAAAGCAGTGgaatcttaaacccaact
ggttacctcttatctagtggACACCAGAGTCTATGAAATACAGAAGAAAAAATACCTGCCgaatcttaaacccaact
ggttacctcttatctagtggGCTAGGAATTATTCATGTGTTATGTCTTTAATCTTCCCAGgaatcttaaacccaacttgt

ggttacctcttatctagtggCCTCGTGTTCAGGTAGGATAGCTGAAACAGAATGGGTAAGgaatcttaaacccaact
ggttacctcttatctagtggAGGTCTCATAGCTAGAAAGTGGCAAAATCAGAAAACACAGgaatcttaaacccaact
ggttacctcttatctagtggGAAAGGTATGATTTTGTTTCGTCCTCGCATCAGCCCTGCGgaatcttaaacccaactt
ggttacctcttatctagtggCAGTGTCACGTGGCCAGCAAAGTCAGCATCAGGATTCATCgaatcttaaacccaac
ggttacctcttatctagtggGCCAGAGAGGCTTTGTGGGGGAGCTCACGTTTAGGCTGGAgaatcttaaacccaac
ggttacctcttatctagtggGCTGAGAAAGCCCAGGGATGACTGAGGAAGCTGCCGCATGgaatcttaaacccaa
ggttacctcttatctagtggGATGGCCAAATGGTTTGTCCAATATGCCTCCAGGTGAAGGgaatcttaaacccaact
ggttacctcttatctagtggCGGCTAGAGTAAATGGAGATCTTACTTGTCAGGGGTCTGGgaatcttaaacccaact
ggttacctcttatctagtggGCCTACCTGAGTAACTTAAGGCAAGTTATTAAAGCCCTCGgaatcttaaacccaactt
ggttacctcttatctagtggCATATTGGTTGAGAGGAGATTGGGTCACTCAGGGACTGGGgaatcttaaacccaact
ggttacctcttatctagtggAATCTTTGGATTGGATGCTATTTAGTGACCTTTCCAATTCgaatcttaaacccaacttgt
ggttacctcttatctagtggGCCTAGCATCTCTCATAAAACTTAACTTCCTAGGTAGCTGgaatcttaaacccaacttg
ggttacctcttatctagtggCTGCCAGGCAGTATTCCCTTCACTTTGTATTCATTATGGTgaatcttaaacccaacttg
ggttacctcttatctagtggCATCTGTCTGACTCCAAGGTGTTTGCTCACTATCCTGGGCgaatcttaaacccaactt
ggttacctcttatctagtggGATCAGTTTCTCTCCTCCAGGCACTCCTTGAACAGACCCCgaatcttaaacccaac
ggttacctcttatctagtggCTTCCCTTCTAAAGACGTCTGGCCTTCCAGCGGGAGGTGGgaatcttaaacccaac
ggttacctcttatctagtggCCTGCCCATATCCTGTCACTCCAGCTTCCTTCCCTTGCTGgaatcttaaacccaact
ggttacctcttatctagtggAGCAGCAGCCCTGATACGCACTTCACTTGCCTCTTTGCCCgaatcttaaacccaac
ggttacctcttatctagtggTGGGATTCCCATACTTCCAAAGAGGCACAATCCAGATGTGgaatcttaaacccaact
ggttacctcttatctagtggGAGGGAAGCCCAGATGTGAAGTAGAGTTTACTGACTCTCCgaatcttaaacccaact
ggttacctcttatctagtggGAGGGCAGCTGAAGTGCTTGGTCCACACCCTTTAGAGCAGgaatcttaaacccaac
ggttacctcttatctagtggCGCTTCCTTTTAAAGTGTGCCTGGAAGCATTGCAAAGGGGgaatcttaaacccaact
ggttacctcttatctagtggGTCAGGGGTGGTGCTATATTTAAAGGAAATTTAGGTCTTGgaatcttaaacccaacttg
ggttacctcttatctagtggCTCAAGAATTCCTGCACATTTTGTACCCTTCCCTTGGTGCgaatcttaaacccaacttg
ggttacctcttatctagtggACCAGTTTAGCCCCGATAATCTCAGAGCTGACTTTCTGACgaatcttaaacccaactt
ggttacctcttatctagtggGCTCCTAATCTCAGCCTTCTGGACTAACCGCTTATCCCTCgaatcttaaacccaact
ggttacctcttatctagtggGCTTCCAGGAATTTCCTCTTTAACATTCCCCAAAAAGGTGgaatcttaaacccaacttg
ggttacctcttatctagtggTGAAAGGAAAGGTGTTAGGAGTAAACTGGAGACCCGTGAGgaatcttaaacccaac
ggttacctcttatctagtggCCTGACAATAATTAGATGGAGTAGCCACTGCCCCCGTAGGgaatcttaaacccaac
ggttacctcttatctagtggGTCCTCTGTGTTTGTGTCACCAACATTGAGGCTCAAGGCCgaatcttaaacccaact
ggttacctcttatctagtggCAGCAAATGACAACCTCTTGAAGTGGTGTCATTGTCTGGGgaatcttaaacccaactt
ggttacctcttatctagtggCTCATGGCCACAGAAAACAAAACAGTGAGGTTCAGAGTGGgaatcttaaacccaac
ggttacctcttatctagtggGGTCCCAGAGACATGGGTTGCTGGTTCCCCAGTGGAATGCgaatcttaaacccaac
ggttacctcttatctagtggGGTGTGCCATTTATAGGGTATGAAAAGGCTGGCTACCCCCgaatcttaaacccaact
ggttacctcttatctagtggTGCAGATGAGTTCTCTATGTGGTCAGCACCGTGTTGCCTGgaatcttaaacccaactt
ggttacctcttatctagtggTGGAGCCTCCATGTTGAACATACTGGCCCCCAGGTAACCCgaatcttaaacccaac
ggttacctcttatctagtggCCAGCTTGCTTATCTATGTTTGCAGGTCAATTTTTCAGGCgaatcttaaacccaacttg
ggttacctcttatctagtggCACCACGCCCGGCTAATTTTTTCTATTGTTAGTATAGACGgaatcttaaacccaacttg
ggttacctcttatctagtggGCGTGAGCCACTGTGCCCGGCCGCCATTTTCCTTTTGTTAgaatcttaaacccaact
ggttacctcttatctagtggTTTGCTGAAGACTCTGTTGCCCTTACTATCCCCCCAAATAgaatcttaaacccaacttg
ggttacctcttatctagtggAGCTCCTGTATCATACTCTGAGACTCAAATTTCTCCTAAAgaatcttaaacccaacttg
ggttacctcttatctagtggTAGTAGTAAATATGTCCTTTGGGTGTTTTGAGGCTAAATGgaatcttaaacccaacttgt
ggttacctcttatctagtggGGGTTTCTAATGTAATCTCTGTCTGCCCGACTATATAGTGgaatcttaaacccaacttg
ggttacctcttatctagtggATGAGTGAGTATCTCATTACTTCATGATATTTTCTAGGGCgaatcttaaacccaacttgt
ggttacctcttatctagtggTATGTTGAGTACCTTGGTCTAGTGTTTTCCACACATGGCCgaatcttaaacccaacttg
ggttacctcttatctagtggAAGGGACTATTTTGGTCAATTCTCTGTCTGACTTTGCAACgaatcttaaacccaacttg
ggttacctcttatctagtggGGAGTGACCAAGTTACAGCTTTCTAGAAGAGTAATGCTTGgaatcttaaacccaactt
ggttacctcttatctagtggACCTTCAAAAGAGGAGAGAAGCATGTCTCAAAGGAAAGGTgaatcttaaacccaact
ggttacctcttatctagtggTATCCTTCTACTGCCTATTAACTGGGTTTGTGGCTCAAGGgaatcttaaacccaacttg
ggttacctcttatctagtggGGTAGCTCTTGACACCAGAGTTCATCTCATTCCAAGTGTGgaatcttaaacccaactt
ggttacctcttatctagtggGCTGGAAGGCACGTAATCCTCAAGGAGGAAGGCAGACCAAgaatcttaaacccaa
ggttacctcttatctagtggTCTCAGATGAGGAGTGAAACTACAGTGGTAGCAGGGAAGGgaatcttaaacccaac

ggttacctcttatctagtggGGATGTAACAGGCATCTAGCTGGCTCTTTTGGAATCAAGCgaatcttaaacccaact
ggttacctcttatctagtggACTTAGCAGCCCATTGTTATCTGAGTGGAAATGCAGCTGGgaatcttaaacccaact
ggttacctcttatctagtggCTGGAAGAGGTCAGGTCTAGAAATGTGATTTGAGAGTCAGgaatcttaaacccaact
ggttacctcttatctagtggTCATGATCTTGGATGAGATCTCTAAGAAGTGAGCACAGGGgaatcttaaacccaact
ggttacctcttatctagtggGGAACATCAGGAGGATCCAGCAAGGGAGAGAGAGGAAGAGgaatcttaaacccaa
ggttacctcttatctagtggTGGACGTATGGATGTCTTGTAGTGACAAACATGTTTTAAGgaatcttaaacccaacttg
ggttacctcttatctagtggGTTGCGTAAAAGACAACTGAAATGTCAAGGAAAATGATCGgaatcttaaacccaact
ggttacctcttatctagtggGTGCTGAGCACTTAGTTAAGGTTTTTAATCTGGAGGCCTGgaatcttaaacccaactt
ggttacctcttatctagtggTTTTTTCTTCCCCTCCACTTTCTCGGGCCAGTATTTCTGGgaatcttaaacccaacttg
ggttacctcttatctagtggTAAGTGAAAGAGTCGGGTGTGGTGGCATATGCCTGTAGTCgaatcttaaacccaact
ggttacctcttatctagtggACAGTGAGCTGTGATCATGCCAGTGTACCCCAGCCTGGGCgaatcttaaacccaa
ggttacctcttatctagtggCAGAGTCTGATAACTGATAAAGAGAAGAACCTGAGGGATCgaatcttaaacccaact
ggttacctcttatctagtggCCTCAGATCTGGGCTAATACGTAGGAGGGTGTGCACATCCgaatcttaaacccaac
ggttacctcttatctagtggTTTTGAAAGCTTAGCAGGAAGAGAAACATACCTGGAGGGGgaatcttaaacccaact
ggttacctcttatctagtggCCCAGAAGTTCTTCTGATCTCGGAGGAATTTGAAAGAAGGgaatcttaaacccaact
ggttacctcttatctagtggCCATGCCTGTTCCCTGCACATTCAGCTCTTGACTCTCACCgaatcttaaacccaact
ggttacctcttatctagtggTGAGTAGCTGGGACCACAGATACGTGCAACCAAGTCCAGCgaatcttaaacccaa
ggttacctcttatctagtggGGTCTCCATCTCCTGAGCTCAAGCAACCCACCCTCCTCAGgaatcttaaacccaa
ggttacctcttatctagtggTGCTGGGATCATAGGCGTGAGATACTTGATTCTCTTAATTgaatcttaaacccaacttg
ggttacctcttatctagtggTTATGTAACTCCTTCCACCATGTGCTAAGTACCTTGGTACgaatcttaaacccaactt
ggttacctcttatctagtggGACAGCCATGGGGTTGGTTGCAAATGGGGTATGGGACTGGgaatcttaaacccaac
ggttacctcttatctagtggAACGTTATGTGTAAAGGTGTAGGGAAGGGGAGAACTCTCCgaatcttaaacccaact
ggttacctcttatctagtggGAGCCACCACGCCCAGCCAAAAGGAACATTTCTGCCTCACgaatcttaaacccaa
ggttacctcttatctagtggTGCCTGCACAAGATCACAAACCCTGTCTATACCTGGTTCCgaatcttaaacccaact
ggttacctcttatctagtggAAGAAGAAAGTACTAGAGGCCAGGAGCGGTGGCTGACGCCgaatcttaaacccaa
ggttacctcttatctagtggCCTGGGAGACGGAGCTTGCAGTCGGCCGAAATCGTGCCACgaatcttaaacccaa
ggttacctcttatctagtggCAGATTTTGTCTCGTAGTTGCAGAAGAGCAGTTGTACCAGgaatcttaaacccaactt
ggttacctcttatctagtggCACATCCCCTCCCATCTGAACTTTACTACCCATGGCATCCgaatcttaaacccaact
ggttacctcttatctagtggCAGGGCCAGCTAATATTAAATGGGGTGGGCAGGGATCCAGgaatcttaaacccaac
ggttacctcttatctagtggAGATCAGATAGGCAAAGAGAAGACATGCTGTCATTGGGCCgaatcttaaacccaac
ggttacctcttatctagtggACCTCGTGATCTGCCTTCCTCGGCCTCGCAAAGGGCTGGGgaatcttaaacccaa
ggttacctcttatctagtggCCCATTGGGCCTTTTTCTTATGATTTTCATGACTTTTTGCgaatcttaaacccaacttgt
ggttacctcttatctagtggCTCAATATTCCCATCTTCTCCTTTACGAAAGTCTTCCTCCgaatcttaaacccaacttg
ggttacctcttatctagtggTGACTTCTGTTCTCCCAGGAGCAAGATGAGAGGTGGTCTCgaatcttaaacccaact
ggttacctcttatctagtggCAGATCCAGATGTGCCAAGCTTGGGTGGACTTTCTCTCCCgaatcttaaacccaact
ggttacctcttatctagtggGATGATGTGCCAGGTAAAGCTGTCTATAGCCATTAGCTGCgaatcttaaacccaact
ggttacctcttatctagtggACCTGGCAACCTCAACTCTGCATTCCTGGAAAATCTCAGGgaatcttaaacccaact
ggttacctcttatctagtggTAAGAATTGTGGGTCAAACATCTATCTTTCAGATGAGGCCgaatcttaaacccaacttg
ggttacctcttatctagtggTCAAAGCCACAGATACTCCAAGCTAAGGACCAATGCCAGCgaatcttaaacccaac
ggttacctcttatctagtggGGCTGGGACCGATGGATCTCAGGTTTTGACCCATACTCTGgaatcttaaacccaact
ggttacctcttatctagtggTGTCACTTTGGACAAATGTCTTCTCATCTCTCAGTCTTCTgaatcttaaacccaacttg
ggttacctcttatctagtggTGACTGGGGTTGAAAGACTTCTCTAGAAATGAGGGTGCCCgaatcttaaacccaact
ggttacctcttatctagtggGGTGACAGCATTCCAGTCTCCTCTAGAGAGCCCCATTCCCgaatcttaaacccaac
ggttacctcttatctagtggGTGTCCTCACAGTTTCTCAGACCTTGTGACTGGTTGAGGGgaatcttaaacccaact
ggttacctcttatctagtggGAGGTGTGTGGTTTCTACTGGAGTGTGAGAGTAAAAGTGGgaatcttaaacccaactt
ggttacctcttatctagtggATGGAGGCAGATAACAAGTGACAGAGTGTGTTGTGAGAGGgaatcttaaacccaact
ggttacctcttatctagtggGGAGCCAGTCTGCCCTGGTTGGAATCACAACTCTGCTACCgaatcttaaacccaac
ggttacctcttatctagtggGTGGCCTCCTTGAGCAAGCCACTAGACTTCTGTATGTCTCgaatcttaaacccaact
ggttacctcttatctagtggCCATACCCAAGTGCACCATATGCGTGTGTATGCACAGCGCgaatcttaaacccaac
ggttacctcttatctagtggGACACAGACACACACACACAGTTACCCTTATTATGACTGGgaatcttaaacccaact
ggttacctcttatctagtggGAGCCAGTACGTTCTATATTTTGCTGAACACAGTTTGATCgaatcttaaacccaacttg
ggttacctcttatctagtggGTTAGCAGCAGATCTTAAAACCAGGTATTCTGACCCACACgaatcttaaacccaact

ggttacctcttatctagtggGCATCAAATTAAGGCATTGGACTGTACAGAGAAGCAGTCCgaatcttaaacccaact
ggttacctcttatctagtggCTGGGAACTGCTGTGGATTTTGTTAGGTGTGATGCTACTGgaatcttaaacccaactt
ggttacctcttatctagtggAGTCGGGGTTCTCTAGAGTTGACACTCAGTATTAACCATCgaatcttaaacccaactt
ggttacctcttatctagtggATGACAAAATGCTAGTAACTGCCATTCCACTGAATGATGGgaatcttaaacccaactt
ggttacctcttatctagtggCTTCCCCATTTCAATGTAGCCACCATAAGGGCAAAGTTGCgaatcttaaacccaact
ggttacctcttatctagtggGAACAGGTTAGACCTTGTTAGCTGAGAGGAACTTCAGGCCgaatcttaaacccaact
ggttacctcttatctagtggTGTGGGTGCTCTGCCACATCCTCTCATCACCAGTCATTCCgaatcttaaacccaact
ggttacctcttatctagtggGCAGGCATCTGGGACTTCTGTCCTGAAGGCTGACCTCAGGgaatcttaaacccaac
ggttacctcttatctagtggGCAAGGGGCTGGCCAGATGTGACTTTCCCCAAGAGAAGCCgaatcttaaacccaa
ggttacctcttatctagtggAGCAGATGTTGGAGGACAGTCATTTTAGGTTTCTTGTCCCgaatcttaaacccaactt
ggttacctcttatctagtggTCCCAGGGCAATTGAGCTCAAATGGCCCACTGCATCTCCCgaatcttaaacccaac
ggttacctcttatctagtggCCCACTTTCCTCGGTTTACTTCTCTCCTCTTTCTTCCATCgaatcttaaacccaacttg
ggttacctcttatctagtggCCTGAGACAGAACGGTAAAGAAGATCAAGTCTGAGCAAGGgaatcttaaacccaac
ggttacctcttatctagtggTAGAGAGGAAGGATTGGCTGGAAAAACTTTCTGGATAGACgaatcttaaacccaact
ggttacctcttatctagtggGGACATTGGGAAAAACCCTGTGCTCATCACCAGCTCCACCgaatcttaaacccaac
ggttacctcttatctagtggCCCCCCTTTCTGCCTTCTCTGTGGAGACATTTCTTGGTGCgaatcttaaacccaact
ggttacctcttatctagtggATGCTCCTCCCTGAGAGCCAAACCTAAATTCCATCCTGGCgaatcttaaacccaac
ggttacctcttatctagtggGCCCTTGGATGAAAATAGACATTCCTTTCTCCTACAATCTgaatcttaaacccaactt
ggttacctcttatctagtggGCCTATGAAGCTCCTAAATTGTGTCCAACTCAAAGCCCCCgaatcttaaacccaact
ggttacctcttatctagtggGGGGTACGATATAATCTGCTTAGTCTGACAGTCAAGGACCgaatcttaaacccaact
ggttacctcttatctagtggCCCTCCTGGGTGGTTTTGAATCCATGCCGTTTCATTAGCTgaatcttaaacccaactt
ggttacctcttatctagtggGCTTGTTGCTCATAGGAGAATGATGTTAGAGGGATTAATGgaatcttaaacccaactt
ggttacctcttatctagtggCTTCTAGCTACAAGGTATGGGAATTCAGCGTACCTTTGACgaatcttaaacccaactt
ggttacctcttatctagtggCTGGTCTCCCTGGTTCCACACACACCCTCCTAGTAGCCCCgaatcttaaacccaac
ggttacctcttatctagtggCTGTTTTATCCTGCTGCCAGTGGATGGTTCAAAATGTGAGgaatcttaaacccaactt
ggttacctcttatctagtggTTGTCTCTGGTTTCCAAAGATTCAGTAGTTTCTCCACTGAgaatcttaaacccaacttg
ggttacctcttatctagtggGTTCCTTGGTAACTAGATTCAGCTTCATTTCTCTCTTCAGgaatcttaaacccaacttg
ggttacctcttatctagtggCACTGTGCCTTTGCTTAAGCTGGTTTCTTTTCTTGGTGCCgaatcttaaacccaacttg
ggttacctcttatctagtggAGGTGTTGCTCAAATTTTATTCCATTCCCCCAAAATCATGgaatcttaaacccaacttg
ggttacctcttatctagtggGTTTTCTTCATTTGCTGGTGAGTGTTTAAAAGGCAGTTGCgaatcttaaacccaacttg
ggttacctcttatctagtggGAAATCCCTGAGCATAACCGCGGGGTAGGGCGAGGTTTGGgaatcttaaacccaa
ggttacctcttatctagtggCACTTCTGACCCTTTCCAATCTCTCCTCTCAGAGACCAGGgaatcttaaacccaact
ggttacctcttatctagtggCTCTCAGTGAGCTCTTCCTCCTGGTGTTCATGTGCTTGGGgaatcttaaacccaact
ggttacctcttatctagtggCACAGGTCGTGATGTCCATCTTGCTAGCACTCTCTCTTTGgaatcttaaacccaactt
ggttacctcttatctagtggCCAACAGCCTGTGAGAAGCTGAATCCTGCCAACCACCGCGgaatcttaaacccaa
ggttacctcttatctagtggAGCCTTGAGGTGATTGCAGTTTTGTGAGAGACCCAGAGACgaatcttaaacccaact
ggttacctcttatctagtggAATGCTGTTGCTTAGAGCCACTATGGTTTGGGGTAATTTGgaatcttaaacccaactt
ggttacctcttatctagtggCAGGACATGCTTTCTTATTGATGAATGGCCAGTGGCATGGgaatcttaaacccaact
ggttacctcttatctagtggCCTTCTCTCTCTCTGTTTCCATTCCAGGGCTTTGTGTAAAgaatcttaaacccaacttg
ggttacctcttatctagtggAGCTTCACAAAGCTAGTACATAATTATGCAAAGCTAGGACgaatcttaaacccaactt
ggttacctcttatctagtggAACTGAAGACACTGGCTCAAAAGCACACATTCAGCAAGTGgaatcttaaacccaac
ggttacctcttatctagtggAGCAAGCTCTTTCCCCTAAACCAGAGTCCTCAACTGTGGCgaatcttaaacccaac
ggttacctcttatctagtggGGGCACAGCTTGGGCATTGGGATTTTAGAAACCACCCAGGgaatcttaaacccaac
ggttacctcttatctagtggAACCAATTTAAGAAGCACTAGGTACATCTCTGAGCCTCCCgaatcttaaacccaact
ggttacctcttatctagtggCATGTCCAATGGTACTTAATTACACTCTGTGTTTGTCACCgaatcttaaacccaacttg
ggttacctcttatctagtggAGGTGAGGAAGAAACAGAAGGGGAGGAATCAACTCTGACCgaatcttaaacccaac
ggttacctcttatctagtggCATTGAATGAGGAAGATTTTATTACAGACCAGGACACTGGgaatcttaaacccaactt
ggttacctcttatctagtggGACTCATATTTCACGTGTCCTTGAGTAATAGTCATTTCCCgaatcttaaacccaacttg
ggttacctcttatctagtggTGATAAAGGCCACACAGCAAATGAGTGACAGAACTGGATCgaatcttaaacccaac
ggttacctcttatctagtggACTTGGACTATGGATCTTTCCACCACACCAGAGCAAGCTCgaatcttaaacccaact
ggttacctcttatctagtggCCCATGCTCTGTTTCCTCCTGCAGAGTGCCAGATTTCAGGgaatcttaaacccaact
ggttacctcttatctagtggGCCTGGAAACTCTTTCCTTAGGCATACTTTATTGCCATCCgaatcttaaacccaactt

ggttacctcttatctagtggGGAACAGCAACCATCCCCCAGCACTCCCTTCTCTTTGCTCgaatcttaaacccaac
ggttacctcttatctagtggTCTTTTTCTACTGTACCCCTGAGGCCACCCTGAAGGAGGCgaatcttaaacccaac
ggttacctcttatctagtggCAGGTTCAGTTACCACCTAACAGAGATCGAGAAAGTGGTCgaatcttaaacccaac
ggttacctcttatctagtggAGTCTGAAGACTTCTAAAATCCCTTCCAAAACGGAGAGTCgaatcttaaacccaact
ggttacctcttatctagtggAGGATACTCATGGTTTGCTGGAATTTTCATGGATTCTTTAgaatcttaaacccaacttg
ggttacctcttatctagtggGTTCATTCAAGGGATATTCAGTGAGAAGCTGCTTTATATGgaatcttaaacccaacttg
ggttacctcttatctagtggAGTGGTGGCTCTCATGAGCTCAAACCTTCTTCTCCATTCCgaatcttaaacccaact
ggttacctcttatctagtggGAAAGTCAAATTCAGCCTAACTGGTGACCACTGGGAAAAGgaatcttaaacccaac
ggttacctcttatctagtggGAAAGGCAACCACGTACCATCACTCCCCAAGGTCTCGTTCgaatcttaaacccaac
ggttacctcttatctagtggCAGACCCCCTATTGTTTTATTTTTTGCCCTTGGGTGACTGgaatcttaaacccaacttg
ggttacctcttatctagtggCTCGGAGTTACTTCCATTGGAACCATTCTCCATTGCTCACgaatcttaaacccaact
ggttacctcttatctagtggACCACCAAACCTGACCCTTCTTAGGCATCTTCTCAGGGACgaatcttaaacccaac
ggttacctcttatctagtggTATCAATGTAACAAAGACTGTGTTGGACCTTGTGATTATGgaatcttaaacccaacttg
ggttacctcttatctagtggGCTTTACCAAGATACAACTCACATACCATAAATCTTACCCgaatcttaaacccaactt
ggttacctcttatctagtggACAGTTCCGAGTACGTTAACAGAGTTGTGCAACCATTACCgaatcttaaacccaact
ggttacctcttatctagtggCTTGTCACTCTTCCTATTATGCCTTCTCAGCAGTTGTTGGgaatcttaaacccaacttg
ggttacctcttatctagtggTGGAATTATAGATATGTGGGCTTTATGTCTTGTGTCTTTCgaatcttaaacccaacttgt
ggttacctcttatctagtggACCTAGGAGTGCAAATGCTGGGTCATATAGTAACTCTGTGgaatcttaaacccaact
ggttacctcttatctagtggAAGACTGAGCATTTGATGTTCTCACCAGCAATGTATGAGGgaatcttaaacccaact
ggttacctcttatctagtggGGGCGTGAAGTTGTATCTCATTGTTTTCATTTGCATTCCCgaatcttaaacccaacttg
ggttacctcttatctagtggTGTCCATGTTCACATCTCCTATGGAGAAGTGTCTGTTCAGgaatcttaaacccaactt
ggttacctcttatctagtggGAGTCTTGCTGTCACCCAGGGAGTGCAGGAGTGCAGTGGCgaatcttaaacccaa
ggttacctcttatctagtggGGCATGAGCCACCGCACCCGGCACTCCTTTGCCTATTTTTgaatcttaaacccaac
ggttacctcttatctagtggGAATTGCGACTGTTCTTTACATATTCTAGATACTAATCCGgaatcttaaacccaacttg
ggttacctcttatctagtggCATTCTGTGGGTAGTCTTTTCACTTTCCTGATGGTGATTTgaatcttaaacccaacttgt
ggttacctcttatctagtggGGTGTTATATCTAAGAGTTCTTGCCTAATTCACAATAACCgaatcttaaacccaacttg
ggttacctcttatctagtggGGTGTGAGGTAGGGGTCCAAGTTCACTTTTTCGTATGTGCgaatcttaaacccaact
ggttacctcttatctagtggGGGGTTGTTTCTGGACTGTTAGTTCTATAACATCAGTTTGgaatcttaaacccaacttg
ggttacctcttatctagtggCATTGTTTTGGCTATTCAGGGCTCCTTAAAATTCCATGTGgaatcttaaacccaacttg
ggttacctcttatctagtggTTGAATCCATAGATCACCTTGGGTTGTACCACTATCATAGgaatcttaaacccaactt
ggttacctcttatctagtggTCTTCCAACTGATAAAACTGGGATGCTCTTTCACTTTATAgaatcttaaacccaacttg
ggttacctcttatctagtggTTTGAATGTTAATCTTGTGTCCTGTGCACTAACTGAACTCgaatcttaaacccaacttg
ggttacctcttatctagtggACCCTAATGGTCCTGGATTGAACCTCCAGTAAAATGTTCCgaatcttaaacccaact
ggttacctcttatctagtggTTCAGTCTTTCACTATTAAGTATGATGCCAACTGTGGGTTgaatcttaaacccaacttg
ggttacctcttatctagtggGTTTCTGTCAAATTATTTTTGTGCCTCTTATGATCATGGGgaatcttaaacccaacttgt
ggttacctcttatctagtggGTTGCTCACTAGTAATTTTCTAGGGATTTTCGCATCTGTGgaatcttaaacccaacttg
ggttacctcttatctagtggTCCTAGCCTCAGAGAACGAGTCAACTAGTGTTCCTTCTCCgaatcttaaacccaact
ggttacctcttatctagtggTTAGTGCTAGGTCAATTCATATTGTCCAGCCAAGTGACAGgaatcttaaacccaactt
ggttacctcttatctagtggAATAGAGTCTCAGACTTGTGAGACACCATCAACTGTAACGgaatcttaaacccaact
ggttacctcttatctagtggCCCTACTAGCTTTGCATTTTCACCTAGCAGTAATTAACAGgaatcttaaacccaactt
ggttacctcttatctagtggGTTACTGCCTAGCTTCTGTGGGGACTTAGTTTGCAACCTCgaatcttaaacccaact
ggttacctcttatctagtggGGCTTCTTGAGTAAGAGTAAAGATGGATTTGGCTGAGGAGgaatcttaaacccaact
ggttacctcttatctagtggCATTTCTGGGAAGAATAGCTGGAGCTAGGCTGTGGAGAGCgaatcttaaacccaac
ggttacctcttatctagtggTTGATCTCTACTGGCACTCTGCAAATGAGATTTATGGTTGgaatcttaaacccaactt
ggttacctcttatctagtggGCCCAGGCTGGAATTCAGTGGCACAAACATGGCACACTGCgaatcttaaacccaa
ggttacctcttatctagtggCCTGAGCTCAAGGGATGTGTCTGCCTCAGCGTCCCAAAGCgaatcttaaacccaa
ggttacctcttatctagtggGTGGAGATTAAAGCTGAGAGTCGCATAGTCTGTCAATGGCgaatcttaaacccaact
ggttacctcttatctagtggCTCTCAGTCTAGCCCACTGCTGGTAGACAGACAGACTAGCgaatcttaaacccaac
ggttacctcttatctagtggGCTGTACATGGGCCATAACCTTGATTGTTTTGTTCTCTGCgaatcttaaacccaactt
ggttacctcttatctagtggGCACCTAGAATAACATGTAGCTCTTGGATGAAAGAAATAAgaatcttaaacccaactt
ggttacctcttatctagtggGGCTCTCTCAGGGGTTTACAGCACATGTAACTCCTGGAAGgaatcttaaacccaact
ggttacctcttatctagtggGGCTCACTTGTTGGGAATGTTCTGCTTCAGAAGATACCAGgaatcttaaacccaact

ggttacctcttatctagtggCCTTCAAGATGGTAGTAGACAGGCTGGCCATTGGGTGGGgaatcttaaacccaac
ggttacctcttatctagtggATGGTGAAAGGCAGTTGTCACTTGGAAAACATACTCTGTTgaatcttaaacccaactt
ggttacctcttatctagtggGCTTAGGAAGGAGAATTTGTCCTGAAATCTATTCCTTCTGgaatcttaaacccaactt
ggttacctcttatctagtggAGTAGCTGCGACCACAGGTGCACGCCAACATGCCCGGCTAgaatcttaaacccaa
ggttacctcttatctagtggACATCTGGGGTTGCAGGCATAAACCACTGCACCCGGCCTGgaatcttaaacccaa
ggttacctcttatctagtggTGAGGCTTTGTGTCAATGTATCAGATCCCCATGTTTATATgaatcttaaacccaacttg
ggttacctcttatctagtggGAGTGTGGCATTATAAAAGGCTATAGAATTTTCAAGGCCCgaatcttaaacccaactt
ggttacctcttatctagtggGGTTTCTTACACAAAACACCAGACAGAGAATTTAGAGGCCgaatcttaaacccaact
ggttacctcttatctagtggGACCCTTAGCTGGTCTTATGTCTGAGTTTCAGGGTTCTTCgaatcttaaacccaactt
ggttacctcttatctagtggCAGGGGTGGAGAATTCCTAAGGATCTTCCAGGTTTACAGTgaatcttaaacccaact
ggttacctcttatctagtggTGCAATGTTTCCTGAGAAGCTTCAAGTTCTATGATTGATCgaatcttaaacccaacttg
ggttacctcttatctagtggCAGGTTCCAACCAAATTTGATATTTGCTCTAGTGCTCTGGgaatcttaaacccaactt
ggttacctcttatctagtggCTACAGTAACTCTTAAGGAGGCCAGATAGGAGGTAACAAGgaatcttaaacccaac
ggttacctcttatctagtggATTTAACTAATTAATGGAAGAAGTGGAACCAGAACCTGGGgaatcttaaacccaactt
ggttacctcttatctagtggATGGGACCATAAGACATGTCCTAATTCTCCGGAAGCTTCCgaatcttaaacccaact
ggttacctcttatctagtggAGGGAACACAGAACAGAGAAGAGCACAGTCCCACCTGGGGgaatcttaaacccaa
ggttacctcttatctagtggGGTCTTAAAGAATGGATGAGTCGGCCAGGCACAGTGACTCgaatcttaaacccaac
ggttacctcttatctagtggCTACTAAGAAATACCAAAAAACTAGCTGGGCGCGGTGGCGgaatcttaaacccaac
ggttacctcttatctagtggGCATGAGTCAGCCAGGCATAGGATGGAAGAAGGAAATTTCgaatcttaaacccaac
ggttacctcttatctagtggTCTGTGCAGGGAAAGATAGGTGCTTCTTTCTGGAGCATAGgaatcttaaacccaact
ggttacctcttatctagtggGGGATTGGAGTGAAGGGAAACTAGAAAGGTAGCTGGAATCgaatcttaaacccaac
ggttacctcttatctagtggGCTGGCAGCTTTTCCTGAAGGATGTGGAGGGAAGCTATTGgaatcttaaacccaac
ggttacctcttatctagtggGTGGGTGCATAGAGGTTGTGAGAGGAGGCAAGAGAGGAGGgaatcttaaacccaa
ggttacctcttatctagtggAGTTTGGTGGCTTTAGCACAGGATGGAAAATGCTCAGATGgaatcttaaacccaact
ggttacctcttatctagtggGGAAAACCAGAGGTGATCATCTTTCCAGGGGAAAGTGAGCgaatcttaaacccaac
ggttacctcttatctagtggCAGTACTTGGTAGCTTAAGGTTGTGCACACTACTTCTCTTgaatcttaaacccaactt
ggttacctcttatctagtggACAATCAGATTTCGCATTGGAGTAAGCTAAAAATATGCTGgaatcttaaacccaactt
ggttacctcttatctagtggAATTGGTTGACTACTGTAGTTTAGGCAGTGAGCAAAGCCCgaatcttaaacccaact
ggttacctcttatctagtggCTCAGCAGTATTGCAAAGTAGGTATACGTATCACAGATGGgaatcttaaacccaact
ggttacctcttatctagtggGCACAAAGCCAATAAGTTGCCTGCTAAAGATCTCGGAGCTgaatcttaaacccaac
ggttacctcttatctagtggTTCAATTGTGGGTTGGTGGCTGGTTGCAGTGGCTCATGCCgaatcttaaacccaact
ggttacctcttatctagtggGGTGGATTGCTTGAGCTCAGGACTTGAAGACCAGCATGGGgaatcttaaacccaac
ggttacctcttatctagtggCAGAGTGAGACTCTGTTTCCCCAGTCCCCCTAAAAAATACgaatcttaaacccaac
ggttacctcttatctagtggGTTTCCTCAAATGTAGCTCTTCATTGCAACGGGTCTTGAAgaatcttaaacccaactt
ggttacctcttatctagtggGGGTGTGGTGCAACGTGTGTTTCCTCAGTGCAGCTCAGTGgaatcttaaacccaac
ggttacctcttatctagtggCCTACGGCTCACTGTAGTCAAAGATTCCAGTGTTCCCACCgaatcttaaacccaac
ggttacctcttatctagtggAGACAGACAAAGCTACAGTTCCTGTCTGAGTCCTACAGTGgaatcttaaacccaac
ggttacctcttatctagtggAGGCCAGTGAATGGGGCTGAGAAACTAACTCTGCCTCTTTgaatcttaaacccaac
ggttacctcttatctagtggCCTTTGTTATTTAGAACCTAGATTTGGCCAGGCGCGGTGGgaatcttaaacccaact
ggttacctcttatctagtggAACCCTAGATTTTGGGGAAACTTTCACAAATGCTTGTGTCgaatcttaaacccaactt
ggttacctcttatctagtggCTCTCCTCAGCCTGTTGCCTGGAAATGAGGGAAGGCAGAGgaatcttaaacccaa
ggttacctcttatctagtggGCTCCCTGAGGGCAAGTGATAGGTCTGTCTTGTTCACTGCgaatcttaaacccaac
ggttacctcttatctagtggGTGCATAGAATTGTGTCTGCTATATGATAGACATTTCAGAgaatcttaaacccaacttg
ggttacctcttatctagtggTCTCAAACAACCTTCTTCCTACATTCAGATGATAGTTGGCgaatcttaaacccaactt
ggttacctcttatctagtggAGTCAGAGAAGGGACCAGTTGATGCAGACTCCAAGTGGGCgaatcttaaacccaa
ggttacctcttatctagtggGTTTGACAGCTACAGTACAGCAATCACAGAATCATCTGCCgaatcttaaacccaact
ggttacctcttatctagtggGTCAAGGTGGTTCTAATGGGGCAAACGTCTAGTGAGGGCGgaatcttaaacccaac
ggttacctcttatctagtggTTGGGTAACATTTAGGATTCTGCTTTAAAGGGGAACAGAGgaatcttaaacccaactt
ggttacctcttatctagtggACTGGAAAAGAAGGGTGGGTTTAAGAGAGGAGTTTTGTTGgaatcttaaacccaact
ggttacctcttatctagtggGTTATGATTGTAGGCAGGTGGGAATAATTCATCAGAGAGGgaatcttaaacccaact
ggttacctcttatctagtggTTCCACTGCACAAGCTGAAGGGGTGGTCTTATCCTGGGGCgaatcttaaacccaac
ggttacctcttatctagtggGTCTAGATGCTGATGAGTGGGTGGATGTAGTGGCTGGCGGgaatcttaaacccaac

ggttacctcttatctagtggCCCATTGGGCTACAAACTCCTTTGGGACAGTGGCTTAGACgaatcttaaacccaact
ggttacctcttatctagtggCTGGAAGTCTCCGGGATATAGAGGTTGTCGGGGGTATGAGgaatcttaaacccaac
ggttacctcttatctagtggAGAATATGCATTGCAGCTGTGAAAAGGCTGTGACAGGAGCgaatcttaaacccaact
ggttacctcttatctagtggGAGCCTCCCAGGACCCAACCCTACACTCTGTACTGTTGGGgaatcttaaacccaac
ggttacctcttatctagtggACCTTACGTAACCAGGCAGAAGATCATATGGCCCTGACTCgaatcttaaacccaac
ggttacctcttatctagtggATGTCTGAGATTGAATCCAGGCTTTGCCAGGTTGTCATTGgaatcttaaacccaacttg
ggttacctcttatctagtggGAACTGGACACATAGAAGTAACCCCCGCATGTTAGTCACCgaatcttaaacccaac
ggttacctcttatctagtggCCACCCAGTGGCTATGTCCTTCTTACTTCTTGACCCCTGGgaatcttaaacccaact
ggttacctcttatctagtggAGGGACTTGTTGACTGAAGCCAGAACAATCAAACTCTCTCgaatcttaaacccaact
ggttacctcttatctagtggTGAGAGACACAGGAATTGATGTTAGTAGGAGCTGAGTCACgaatcttaaacccaact
ggttacctcttatctagtggGGCCACAGACTCCTGGTACTGAAGTCCTTGTGATTTCCTCgaatcttaaacccaact
ggttacctcttatctagtggAGTCTGTGACTACTCCATTTCCAATAAATCCTTTGTCACCgaatcttaaacccaacttg
ggttacctcttatctagtggTTGCATATGTAAATAGATTCCAGGCTCAGTGGAGCTCCCGgaatcttaaacccaactt
ggttacctcttatctagtggTCAGCCAGCATTTAGCGAGATCGTTCTGTTTTAGCATTGCgaatcttaaacccaacttg
ggttacctcttatctagtggGACTACATGGATGATTCAGATGGAATTTCTGCCTTCAAGGgaatcttaaacccaacttg
ggttacctcttatctagtggTCATGGGTTTAGGAGGAGTGAGTGGGTTTCTGGTGGCACCgaatcttaaacccaact
ggttacctcttatctagtggCACGATTGAGTACCCTTCCAGTGAGCTGTCAAACTCCTGCgaatcttaaacccaact
ggttacctcttatctagtggTCCCTTCAAAACGAAATTTGGAAGTCGAAATCTTTGGGGCgaatcttaaacccaactt
ggttacctcttatctagtggTCCATTCATTTGTCAATGGAGGTTAACATACATCTCTCCCgaatcttaaacccaacttg
ggttacctcttatctagtggTTTCTCCACTGTCTGCTCTTCTTTTGTGGTGGTTGGGGGGgaatcttaaacccaacttg
ggttacctcttatctagtggGTTTCATTTCTCACTGTTCCTCTATAAACACTGGGACTCCgaatcttaaacccaacttg
ggttacctcttatctagtggCCTCCATGCCTTTGTTAATACAGTCTCTCCCCACCCGCTGgaatcttaaacccaact
ggttacctcttatctagtggTACACTTGTCTTTTTCAGCAGAAATGTTGCATACTCCATGgaatcttaaacccaacttg
ggttacctcttatctagtggAATTCACTTTTTCGAATTCCAAACGTGCCGCCTGTACTTCgaatcttaaacccaacttg
ggttacctcttatctagtggTGCCTCATGTCATAGGCGTTATCTCCAACAGTTATCTGCCgaatcttaaacccaactt
ggttacctcttatctagtggTGTTGGAGGACATTGTTATCTGCCTCTCTAAAACTGTTGGgaatcttaaacccaacttg
ggttacctcttatctagtggCTCAAAACAGTAGCCCCTGCTCAAGGCGTCTTCACAGAGCgaatcttaaacccaac
ggttacctcttatctagtggCATGGGGGAGAGGGACAAGGAGGAAGAAGAGCAGGCTGGGgaatcttaaacccaa
ggttacctcttatctagtggTGTGTGATGGGTTGGGTTTCCCTGGAAACAGCTGATATGGgaatcttaaacccaactt
ggttacctcttatctagtggATACTTGTGAAAAGGAAGGAAAGGAAGCGGGACTGGGAAGgaatcttaaacccaac
ggttacctcttatctagtggGCTGTTATGCAGTCTCAATGGAATCCTCAGCTGATCTTGCgaatcttaaacccaactt
ggttacctcttatctagtggGAGGTAGGCTACCCCAGGAAGGTGCATGTCCTCTGGCAAGgaatcttaaacccaac
ggttacctcttatctagtggTTGGAAGCAGGCCGCCCCAAGAAGGTGCATGTCCTCTGGCgaatcttaaacccaa
ggttacctcttatctagtggGACTATGGGAGGAGGACCTGGGCAGTGCATCATACTGTCCgaatcttaaacccaac
ggttacctcttatctagtggTCAGCTTCTGAACAGTGGTTCTAAGTAGCCTCAGTAGGCCgaatcttaaacccaact
ggttacctcttatctagtggCCTATGAGTTGCAACCTCTATTCGGGTATGAAAATGCAACgaatcttaaacccaactt
ggttacctcttatctagtggTTGGAACAATACCAGACAATCTAGTGATATGTGCCATCACgaatcttaaacccaactt
ggttacctcttatctagtggCTGGAAAGCGGCATACTCTGTTCTATTTTATGAACCTCTCgaatcttaaacccaacttg
ggttacctcttatctagtggCTGGTTTCTACCAGGGCTGTCTAGTCTTCATGGTCTTATGgaatcttaaacccaacttg
ggttacctcttatctagtggTTGCTCCCAAGGGGTGTTTGTATGTGTGTTTGATCTCCAGgaatcttaaacccaacttg
ggttacctcttatctagtggCATCCCATCTTCACCACCCAGGCACAGACAGATCCTTGTGgaatcttaaacccaac
ggttacctcttatctagtggGCCCTTCCTGCTTGAATCAAAGCCTTGGGTGGTCTAGGGCgaatcttaaacccaac
ggttacctcttatctagtggCTTGAGGATCAAAACAGGTAGTTCTAGGCAGTCATGATCCgaatcttaaacccaactt
ggttacctcttatctagtggACCTTCTCTAATCTGCAGTTTCCTCATCTATCAAGTAGAGgaatcttaaacccaacttg
ggttacctcttatctagtggGACACTTAACATAGTATCTGTTGAGAAGTCAGTCTTTAACgaatcttaaacccaacttg
ggttacctcttatctagtggCAGATTGTTATTATTAGAATTCAAGGATGGGGAATACAGCgaatcttaaacccaacttg
ggttacctcttatctagtggGTAGCAATGACAAACTGGGCTTCAAGTCAGCAGAGGACAGgaatcttaaacccaac
ggttacctcttatctagtggGCTGGACTGGGACTCAGGATAACTGGCATCTAGGCAAGCCgaatcttaaacccaac
ggttacctcttatctagtggTGATTTTGGGCAAACACTTCAACTTTCCAAGGATTCATAGgaatcttaaacccaacttg
ggttacctcttatctagtggCCTCACAGTTCAGTCACAAATAGCACTGACCAATGAGCTGgaatcttaaacccaact
ggttacctcttatctagtggAGATGTGTGGCAAGGACAGAAAAGCAAACACAGCATGTTCgaatcttaaacccaac
ggttacctcttatctagtggCATCACACACACGGGCCTCTCGGGGGATGGGGGTTGGGGGgaatcttaaacccaa

ggttacctcttatctagtggGATGTGGTCTTCAGGTTCTTTCTTTCCCTGCTATGGTGgaatcttaaacccaacttg
ggttacctcttatctagtggAATACCTGAGTTACCACATGGAAGAGAGGTGCCCTAGGGCgaatcttaaacccaac
ggttacctcttatctagtggATGGCATAGGATTAGATTGAGCTTGTCCAACCCATGGCCCgaatcttaaacccaact
ggttacctcttatctagtggCTTCCAATGTGACCTGAGGAAGCCAAAAGATTGGACAGCCgaatcttaaacccaac
ggttacctcttatctagtggGCTGTAATAGCTCTCATTCTGACCATTAGGATGAATGGAGgaatcttaaacccaactt
ggttacctcttatctagtggAGCTCTGTCATATTTGTGCTCAATACAGAACTTAGGAGCCgaatcttaaacccaactt
ggttacctcttatctagtggCATAGATGTGATGATTGGGGCCATGCCAAGAGTTTTGTCCgaatcttaaacccaactt
ggttacctcttatctagtggTACGTGTGTGTACATGTATGTGTGTTATATTGATGTGAACgaatcttaaacccaacttg
ggttacctcttatctagtggTATACTCAAAGTGAGATTAACACACAATGGCTCATGGCACgaatcttaaacccaactt
ggttacctcttatctagtggAAAGTAATACCAACTTGGTGATAAGGAAAGGAAAGTGGGAgaatcttaaacccaact
ggttacctcttatctagtggTGCACAAGTTTCAACTGATGGCAAAGTTAATCTGAACTGCgaatcttaaacccaactt
ggttacctcttatctagtggCAAGAGCAATCACAATACATAGACATACACACATATCCTCgaatcttaaacccaact
ggttacctcttatctagtggAAGAGATTGAGTTCTGAAATAAAACATTTTACCACCTGGGgaatcttaaacccaactt
ggttacctcttatctagtggCTAGGACCTATCTCATGGGGTAATTCAGAGAGGATAATTGgaatcttaaacccaactt
ggttacctcttatctagtggTGAAAAGTGCCTAACATAATGCCTGGCATAGAGAAAGCACgaatcttaaacccaact
ggttacctcttatctagtggTGTTTATCTTTCTGGCTAGCAAGACCATGTGATCTGAAGGgaatcttaaacccaactt
ggttacctcttatctagtggGATGTGGTATTTTGTACCAGATTACTTTCTAACTGGAGAGgaatcttaaacccaacttg
ggttacctcttatctagtggAGTTCCTACCTTTCTCCACCCAGCCCTTCAGAGACATCACgaatcttaaacccaac
ggttacctcttatctagtggGGTCCAAAGGGGTGAAATGACTGCCTTATCTTCACTTGGGgaatcttaaacccaact
ggttacctcttatctagtggGTGTAAGAAATGATCTTTTACCTATTGCACACAGTAAGTGgaatcttaaacccaacttg
ggttacctcttatctagtggAAATTAACACTAATCAGCCAAGCACGGTGGTTCATGCCTGgaatcttaaacccaact
ggttacctcttatctagtggGAAAGATTGAGTGGCTATATGAATATCAGACAATGCAGACgaatcttaaacccaactt
ggttacctcttatctagtggTGCCACATGTGAAGAGAGGTATTTAGAAATAATAAATGGGgaatcttaaacccaactt
ggttacctcttatctagtggACAATTCTAAGTATATGTGCTCCTAAAACCAGAGCATTGGgaatcttaaacccaactt
ggttacctcttatctagtggCCTTGGGTTAAGCATGTTTGGTAGACAAGAGGCAAAAAGCgaatcttaaacccaact
ggttacctcttatctagtggATCCAGTCGTCAAAAGATCCCATTAAGTAAATGAATGTGCgaatcttaaacccaactt
ggttacctcttatctagtggGACAGTAGACTTGTATAAAAGCTAGATAAAGAGCTCCTACgaatcttaaacccaactt
ggttacctcttatctagtggAACAACCCAATAAAAATGGGCAAAAGTTGTGCACAGACACgaatcttaaacccaac
ggttacctcttatctagtggGGCTAGGACGTGGAGCAACTGGAATGCTTAGACATTGCTGgaatcttaaacccaac
ggttacctcttatctagtggGTCAGGCAGCCATTCTACTCCCTGATACTTACCTAGGAGGgaatcttaaacccaac
ggttacctcttatctagtggTAACCAAAACCCAGAAACAGATCAAGTATTCACATACAGGgaatcttaaacccaact
ggttacctcttatctagtggCCAAAGCACTAAATTGAGTGAAAGGAGGCAGATATAAGAGgaatcttaaacccaact
ggttacctcttatctagtggCTCCTAACAGTCAAATCTAATCCATGTTCCCAGAAATCAGgaatcttaaacccaactt
ggttacctcttatctagtggTACTGGTGATCTTACCACATACATTTTTCGTACAGCGGACgaatcttaaacccaactt
ggttacctcttatctagtggTTGTAGTTCCCATTTCAACTCCCTTCCTTTTTTAAGCCTGgaatcttaaacccaacttg
ggttacctcttatctagtggCTTATTGAAAGCTGCTATCTAATAGATCCTCAATCTATGCgaatcttaaacccaacttg
ggttacctcttatctagtggGCCTGAGTCAATCTTGCCTGTAGTGGGGCTGAGTCTGGGCgaatcttaaacccaac
ggttacctcttatctagtggTCTCAACTGGCTGGGGATCTGAGCACCAGGGAGGCTCTGGgaatcttaaacccaa
ggttacctcttatctagtggGGGAGTTCCCCCCTTTTCTCAAGGATGTCCCTGGCTTTCTgaatcttaaacccaact
ggttacctcttatctagtggGCCGGTTCGTGATTTGACGGCCTGGAGAGTCCCCAAAGGAgaatcttaaacccaa
ggttacctcttatctagtggCCCCAACTCCGCTTTCGGCATCTTCTGCCCTGCACTCTGCgaatcttaaacccaac
ggttacctcttatctagtggGGCCTGAAGTACTCAAAGAGGTTTCTCAGTTCTCACCCTGgaatcttaaacccaact
ggttacctcttatctagtggTGTCCAGACTCATTGTGTGGCCAGAGTTCCTTGAGATTGCgaatcttaaacccaactt
ggttacctcttatctagtggCTCCTTCTCCATCTATGGTGAATTCTGCCTTTTGCCAGACgaatcttaaacccaactt
ggttacctcttatctagtggAGACATTCTCATCCTCAGATTTCTAATGGGTTCTTTGATCgaatcttaaacccaacttg
ggttacctcttatctagtggGCAGTTTATTCTGCTTCTAGTGAGATTGACAGATTCTTACgaatcttaaacccaacttg
ggttacctcttatctagtggCTACCCAGGTTAGGTGTTGTCAAAGAATGGCCTGCATGACgaatcttaaacccaact
ggttacctcttatctagtggACCCCTATGTTTCTATATCCTTCAAGCTAAGTCTGGCCCTgaatcttaaacccaactt
ggttacctcttatctagtggTGTTCACTAACCCCCAACCTAGACTATGACATGTTCTGTGgaatcttaaacccaactt
ggttacctcttatctagtggTGTGTGAGCGGGACAAAGAGATTGTGCAGGCCACACATGCgaatcttaaacccaac
ggttacctcttatctagtggCAGCTTTCCTTTGGCTTCAAATTAAAGAGGCATGTAAGAGgaatcttaaacccaactt
ggttacctcttatctagtggCAGTTGTTTTGGCATTTGGACTGAGGATATAGTTTTGGCCgaatcttaaacccaacttg

ggttacctcttatctagtggCTTAGTCCTGAAAGCCCTTTCTCAGTAACAATGGTGCTACgaatcttaaacccaactt
ggttacctcttatctagtggCTGTGCTGCAGAGATGGGGTCCTGACAGACACAATGCCCCgaatcttaaacccaac
ggttacctcttatctagtggCCTGTCCCCTTCCTTAGTTATCTCTGCTCCAGATGACTAGgaatcttaaacccaactt
ggttacctcttatctagtggAGAGTAGAGAAGAGGTCACTGTCACTGTGGTCAGGAGCACgaatcttaaacccaac
ggttacctcttatctagtggGTGAAAGGGAAGCTTGAAATCTGTCAATTCCATATTCCTCgaatcttaaacccaacttg
ggttacctcttatctagtggGAGATGCCAGTGCACTATGATTGTGCCACTACACTCCCGCgaatcttaaacccaac
ggttacctcttatctagtggGAGCAAAACCCTGACTCTAAAACCAAAAAGAAAAGCGAATgaatcttaaacccaact
ggttacctcttatctagtggGTGCTAGGACTTCTGGGGGATGTTGCCTTCTAAACCTTCCgaatcttaaacccaact
ggttacctcttatctagtggGGCTGTTCATACACCTCTTTTTATACATCCTCATCGCATCgaatcttaaacccaacttg
ggttacctcttatctagtggGGATCCTGAATTCTGGATTCCAGGTTCTAGCTTCTGCACCgaatcttaaacccaactt
ggttacctcttatctagtggTTTTTTGGCCGGGCGGGGGGTGGGGTGGGGGGCAGGGAATgaatcttaaacccaa
ggttacctcttatctagtggATTGCAACCTTCAGCTCCCGGGTTCAATTCTTGAGCATACgaatcttaaacccaactt
ggttacctcttatctagtggCATGGCAGGGTTTTCACTTTTTATTGGCTGTGAGTATACCgaatcttaaacccaacttg
ggttacctcttatctagtggGCTTGTAGAAGCCCCTTATCTAACATCTGTCCTTTGGAGCgaatcttaaacccaactt
ggttacctcttatctagtggCTTGGAATACTGCAGGAAGGAAATAGGCCATATGATTTCCgaatcttaaacccaactt
ggttacctcttatctagtggTCAGCACAACATAGCACATAGATGAAAGAGTTGATGACTAgaatcttaaacccaactt
ggttacctcttatctagtggCTGGTAATATTAGGGACTCATTGAGTACTATGGATAAAAGgaatcttaaacccaacttg
ggttacctcttatctagtggGGTTTTTGGATAATGGATCTGAAATTCAGAGCAGTTAGGCgaatcttaaacccaacttg
ggttacctcttatctagtggGAGGAGACTTCATCCCTATTTTCTATCTCAAAGTCCATACgaatcttaaacccaacttg
ggttacctcttatctagtggCCACCACCCTGACTTAACTCCACGAGCATGTCCTTGCGTGgaatcttaaacccaac
ggttacctcttatctagtggGCTTTCTCCTTGGCTAATCTTCTATCTTCTATCTCTTGGCgaatcttaaacccaacttgt
ggttacctcttatctagtggAATCTAAGCCCAGTACTTCATTGCTGCTCAAAGAAACAGCgaatcttaaacccaact
ggttacctcttatctagtggGGCTTAGACATCTCAGAATTGAGAATTTTAGGTGCTCTGGgaatcttaaacccaacttg
ggttacctcttatctagtggCTTTGTATTGATTGCAGAATGAAATCTTTGCCTCTCCCTCgaatcttaaacccaacttg
ggttacctcttatctagtggCAACAGCCCCTTTCCTCCAGCAAAGTATGACTCAATCTTCgaatcttaaacccaact
ggttacctcttatctagtggGCTCCTTTCCTACACATAATGCTTTCTTCTCTCTCTGTGGgaatcttaaacccaacttg
ggttacctcttatctagtggATCCTTTTTTGCTTGTGGCTTATCTCCTCCACATCTAGACgaatcttaaacccaacttg
ggttacctcttatctagtggATGTCTCATTTCTGTAGTGTCTTGAGTACTCAAGGCAGCCgaatcttaaacccaacttg
ggttacctcttatctagtggTCCGTTAGAACACATATCTGGAAAGACACATCCCAATAGAgaatcttaaacccaact
ggttacctcttatctagtggGTGTGGAGCGGGCGTCTGATTGCACCTGCCTCGTGCCAGCgaatcttaaacccaa
ggttacctcttatctagtggGCCTCATCCGCTGATGGATCTGTGCCACTGCTGCTTGCTGgaatcttaaacccaac
ggttacctcttatctagtggCAGGCAAGCTTTTGCACAAGCCGTTCCCTCTGCCTGGGACgaatcttaaacccaac
ggttacctcttatctagtggCCATCTTCAAGCAAAATTTGCCTTTCTCTCCCCTGGGCTCgaatcttaaacccaactt
ggttacctcttatctagtggATTTCTTTCACTCCATCATAGCCTTCATCAGCCTTGGTGGgaatcttaaacccaacttg
ggttacctcttatctagtggGAGACTGTGTTCATTCAATCTCCCTACCACCAACTCTAGCgaatcttaaacccaact
ggttacctcttatctagtggCAGTTCAAACCTTCCACTGGTCCACAGGATGCCTCACTGCgaatcttaaacccaac
ggttacctcttatctagtggGACCTGGACATTTGGAGATTGAGAGGCCATATCTCTACTGgaatcttaaacccaact
ggttacctcttatctagtggCCCTGATGAGCTAGCTAGAGAGGATTCGTCTCTGAGGCACgaatcttaaacccaac
ggttacctcttatctagtggCCCTGATGAGCTAGCTAGAGAGGATTCATCTCTGAGGCACgaatcttaaacccaac
ggttacctcttatctagtggCCTAGGTCTACATCCCCAGGAGAGTTCCCTTGTACAGCCCgaatcttaaacccaac
ggttacctcttatctagtggGTGCACAGCTGGCTTTGGTGGCAGCAAGATGCCTTAGGCCgaatcttaaacccaac
ggttacctcttatctagtggAGGAACTGGCAGAGAAAGTGAATTCTCTATCGTAAAGCCCgaatcttaaacccaact
ggttacctcttatctagtggGGTGAATGATTCTGGCCTGCGGTCTATGGCAATGCCTAGGgaatcttaaacccaact
ggttacctcttatctagtggCCAGGGACCTATAGCTTTGAAACACTAGGGCTACAGGGAGgaatcttaaacccaac
ggttacctcttatctagtggCCAGTTTCCTCTGTCTCCTCTTTTCTTGTCTTCATTGCCCgaatcttaaacccaacttg
ggttacctcttatctagtggCTGAGCTCCCTGTCTCTGGTCTCTCCCTATATCCCCTTCCgaatcttaaacccaact
ggttacctcttatctagtggCCACACGCCTGCCATTTAGCTTCCCAAACTACAGCAGAATgaatcttaaacccaac
ggttacctcttatctagtggGAATCAGGAAATTTGCTTTACTGGGTACAAGTCCCTTGACgaatcttaaacccaacttg
ggttacctcttatctagtggCTTTAGGAGGCCAAGGTGGTGGGCGGATCATGAGGTCAAGgaatcttaaacccaac
ggttacctcttatctagtggACTAGCAGGATGTATGAAATAATGTGGATGCTCTGCTCCCgaatcttaaacccaactt
ggttacctcttatctagtggTGCTAACCATTCCCCTCCTTAAAAAACCACAATGACTCCCgaatcttaaacccaact
ggttacctcttatctagtggTACACACTTACACAGCTTTCTATATCCTGGCTACAACGACgaatcttaaacccaactt

ggttacctcttatctagtggGGATGCTTCACCCAAATCATACAACCCACTGAGCCTCAGGgaatcttaaacccaac
ggttacctcttatctagtggCAGCCCCATACTTCAAGGCCTCTTGCAGATTCCTCCTCCCgaatcttaaacccaac
ggttacctcttatctagtggTGCATACACATACCTGTTCGGTCCTTCAGAGCAGCTGCTGgaatcttaaacccaact
ggttacctcttatctagtggCTTCCTACCCTCAGAAGCAAGTATAGTGAATGGCACCAGCgaatcttaaacccaac
ggttacctcttatctagtggCACCAGTGGACCCTGCTGAGTGACCCTCCTCCCAATGGAGgaatcttaaacccaa
ggttacctcttatctagtggGCAGAACCACAGACTTATCTGCCATTTCACTCTGAGTGCAgaatcttaaacccaact
ggttacctcttatctagtggGACTGGTGCAGTAATAACAAGAATTTCCTAGAGGACTACCgaatcttaaacccaact
ggttacctcttatctagtggTGTCTGGTTTCTTACCTAAAACTTCACAATCCACTTAAGAgaatcttaaacccaacttg
ggttacctcttatctagtggCAGTCTGAGTTTGCTGGGAGGCTGGAATGATTGCCAAGCCgaatcttaaacccaac
ggttacctcttatctagtggGAAAGAAGTGGGGATGACAAATTCTGTTTTTTCTGGGGGCgaatcttaaacccaactt
ggttacctcttatctagtggGCAGGAGAGAATTTGGTTAAAAGTAAGGAAGATCTGTCAGgaatcttaaacccaactt
ggttacctcttatctagtggTGGCTGCATTATGAAAATGTGGAAATAGACCAGGAACCTCgaatcttaaacccaact
ggttacctcttatctagtggGTACAGTATAGTAGTCATAGTTTCCTCATCTGCAAAATGGgaatcttaaacccaacttg
ggttacctcttatctagtggGCACTCTCAATCTCAATATACTTAGTGTCTTAACTAAGGGgaatcttaaacccaacttg
ggttacctcttatctagtggATTGCGGATGAAGATGGAGGAGGAGCAGGAGGAGGAGAAGgaatcttaaacccaa
ggttacctcttatctagtggGGGACGTAGCTGGGGTCTCCTGTATGCAACATGGAATGAGgaatcttaaacccaac
ggttacctcttatctagtggGTCTGTAATTTTGGGGTGACATTATGTACCTCACAAGAGGgaatcttaaacccaacttg
ggttacctcttatctagtggCATAGCACAAGTTTTGGTACACACTAGGCACCTCTACATGgaatcttaaacccaact
ggttacctcttatctagtggGCCCAGACACAGTCCCAGTCCTACATTTGGAGCAAGTCACgaatcttaaacccaac
ggttacctcttatctagtggAGTTCCTTCCATGTCATCATGAAATCTCACTACTAGCAGCgaatcttaaacccaacttg
ggttacctcttatctagtggATTTTTGGTGCCTTAGGAAAGGAAAATTCTTTCAGTGAGCgaatcttaaacccaacttg
ggttacctcttatctagtggGTTAGAGGGGCTAAGCATTGAAACAGTAACGGGTGTTCCCgaatcttaaacccaact
ggttacctcttatctagtggCCGTGGCTTTGGGTGAGGCAAAAACATTTGGCACAGTGAGgaatcttaaacccaac
ggttacctcttatctagtggATCTATGGGCATACAACCCACTTAAGGTTTACTATTGCTCgaatcttaaacccaacttg
ggttacctcttatctagtggCGTGCATGCATGGAAGCACTCCTAACTATGCATGGCAGGCgaatcttaaacccaac
ggttacctcttatctagtggGCACTTGCCCGCCCACAGAAAGTTGCAGAGCTTCCGATCCgaatcttaaacccaac
ggttacctcttatctagtggCCCCCACCTCCAAGTGTTAACCTGCCCTACCAGAGCCAGGgaatcttaaacccaa
ggttacctcttatctagtggTGGGAGGGTGATGGCTGTGCCCAGATTTCAGCTTTTGTTCgaatcttaaacccaactt
ggttacctcttatctagtggCCTTAACTGCTTGCTCCATTAACTTTGAGGAATTGGAGACgaatcttaaacccaacttg
ggttacctcttatctagtggCGAGATCTTACCCACCAAAGACACCATTTAGGACCTGGCCgaatcttaaacccaac
ggttacctcttatctagtggCCAAAAGGGTTGGCCACAGCTCGACAATTAATTAGCTGGGgaatcttaaacccaac
ggttacctcttatctagtggCTGCCCAGTGAGAGGCCAGAGGGATCCAGGTATGTGCTGGgaatcttaaacccaa
ggttacctcttatctagtggTAGTGATGGTCAGGGAGGTTGAAATGAGCCACCAAGGGAGgaatcttaaacccaac
ggttacctcttatctagtggCCCTCCCTCCCTTCTTACTTTCTTCCCCTCTCTCCCTCCCgaatcttaaacccaact
ggttacctcttatctagtggTGCCTAAACCCCTGTTATGCTGTGGGCTGGACTAGGTGCCgaatcttaaacccaac
ggttacctcttatctagtggTTCTCCACCCTCACCCTATGTTAAGCTTCTTGATGGCAGGgaatcttaaacccaactt
ggttacctcttatctagtggGGATGGAAAGAAGCAAGGGATAGGATTTTGTGGAAGTTCCgaatcttaaacccaact
ggttacctcttatctagtggGTTCTGTATGTGACCCAGATGGCCCACATGGTCCCCTGACgaatcttaaacccaac
ggttacctcttatctagtggTCTCCCCTCAAAGGGCTTCCTTTGCTTCAAAGGGCTTTAGgaatcttaaacccaactt
ggttacctcttatctagtggGTTACAGGTAGGACTGCATTCCTCCTGGAGGCTCCATGGGgaatcttaaacccaac
ggttacctcttatctagtggTGCCTTTCCCAGCTTCTAGAGAATAGCTACATTCCTTGGCgaatcttaaacccaactt
ggttacctcttatctagtggGCCAGCAGTCTAAACTCTTTTTTATTGACCTCCTGCCTCCgaatcttaaacccaacttg
ggttacctcttatctagtggGTTGGACCCACACAGAGAATCAAGGATAATCCCAATATCCgaatcttaaacccaact
ggttacctcttatctagtggCCAGGGATTAGGATGTAGACATCATGGGGGCAGGGAGGGGgaatcttaaacccaa
ggttacctcttatctagtggCAGCAAACGAACCAAACTGAAAGGCCACAAAATGAGCAAGgaatcttaaacccaac
ggttacctcttatctagtggCTGTAAAGTATTTTGCTCAGAGAGCCCCATGATGATGAAGgaatcttaaacccaactt
ggttacctcttatctagtggTTTTGGCCCTGTCTCCTTCCTGTGGAAATCAGATAGATTCgaatcttaaacccaacttg
ggttacctcttatctagtggCTCTTGATTCCCTAGACATCGTAAGAGCAAACCACATACTgaatcttaaacccaactt
ggttacctcttatctagtggAGTGGACACTCTAGAAGGCCCTCACTTGGCTAACGACAGCgaatcttaaacccaac
ggttacctcttatctagtggTGTCTTCCTGGGCTCTCTACAGCACACATTCAGGGCACGGgaatcttaaacccaac
ggttacctcttatctagtggGTAATTCATGCAGAGCCGGCTGTGCTGGAGACCAGACTCTgaatcttaaacccaac
ggttacctcttatctagtggGGAAAGTGTGGTGGGTGTGGGCTCGGAAAGTGGGGAGTGCgaatcttaaacccaac

ggttacctcttatctagtggTCCTGGGTGGGATTGCAGAACCGATTGAGGCAGTTCCAGGgaatcttaaacccaac
ggttacctcttatctagtggTGCTGCATCGGAATGAAGGGTCTGCAAAATATCTCAAGCAgaatcttaaacccaact
ggttacctcttatctagtggGGGGAGGTTCAGACTCTTCAGCCAGAGGTTGCATGGCCCCgaatcttaaacccaa
ggttacctcttatctagtggGCAAAGGCAGATTGGTCCCCAGGCAAGGAAGGGGTTTTTCgaatcttaaacccaac
ggttacctcttatctagtggGGCTAGTTCAGCCTACAGTCAGGAATGAACAAAGTTTAGAgaatcttaaacccaact
ggttacctcttatctagtggAGGCAGTTTCAGTCTTAGATGCCCTTAGGAGCCAGACAGGgaatcttaaacccaac
ggttacctcttatctagtggACTGGCCAGGTGCAAGACACAGGGAGTGATAGGGACTGTGgaatcttaaacccaa
ggttacctcttatctagtggGGTCCAGCCATTTGATTCCAGTGGGAGGGTCTGGGTGAGGgaatcttaaacccaac
ggttacctcttatctagtggATCATCTCTGGCAGCTCCTTTTCTGCTGATTCCATTCTCCgaatcttaaacccaactt
ggttacctcttatctagtggTGCACTCATTTGAGGCAGAAACCTCTATGTCTTCATAATTgaatcttaaacccaacttg
ggttacctcttatctagtggCAGTGTCACAGTCATCGGCTAATACCCAAAGGGCTGTGGCgaatcttaaacccaac
ggttacctcttatctagtggGGCTGAAGTGTTGGTAATCTTAACCCTTACTTCTTTTCCGgaatcttaaacccaacttg
ggttacctcttatctagtggCCTAACAAGGAATTTTATCCAAAAGGAACAATGGCCAGATgaatcttaaacccaact
ggttacctcttatctagtggGATTCCCATGAGTTGAGGTTGATTGGCTGCAAGTGGGCCCgaatcttaaacccaac
ggttacctcttatctagtggGGCAGGGGCCAGACACTGTAGAATTACCTTCCACATAAACgaatcttaaacccaac
ggttacctcttatctagtggGCAGGGGACTTAAACTTTAAGACAAAGCAACACTGTTCACgaatcttaaacccaact
ggttacctcttatctagtggGGGCTCTGACAAATTACATTTACAGCCCAGTTTAAGGAGGgaatcttaaacccaact
ggttacctcttatctagtggGAGAAGGATGTGGTGAGTGGCAATGATATATCCCCATTCCgaatcttaaacccaact
ggttacctcttatctagtggTTCTAGTCCTTTCTGTATGGCAGGAACTGTTCTATGACCCgaatcttaaacccaactt
ggttacctcttatctagtggCCACCACTGCACTCATACAAGGCTAGATTCTTAACAGGGCgaatcttaaacccaac
ggttacctcttatctagtggGCCCTTAATCATTGTACCAGGTTCCAAGGGCTGCTGTAACgaatcttaaacccaact
ggttacctcttatctagtggTTCAAGAAATTCTCTTGCTTCAGCCTCGTGAGTAGCTGGGgaatcttaaacccaactt
ggttacctcttatctagtggCGAACTCCTGACCTGAGGTAATCCGCTGGCCTCGGCCTCCgaatcttaaacccaa
ggttacctcttatctagtggGGCATGAGCCACCGTGCCCGACCCAGAAATGTATTCTTTTgaatcttaaacccaac
ggttacctcttatctagtggGGTTGGTTCCTTCTGCGGCTCTGAGGAGAGAGTGCACCCCgaatcttaaacccaa
ggttacctcttatctagtggCCTAGTTTCTGGTGATTGTCAGTGAGCCTTGACATTCTTTgaatcttaaacccaacttg
ggttacctcttatctagtggCTCGGCCTTCATTTTCACATGATCTTCCCCTCTATGTCTCgaatcttaaacccaactt
ggttacctcttatctagtggTTCTTCAGTCTGAATTTAAGGCTCTTCCTAAAAAGCCAGGgaatcttaaacccaactt
ggttacctcttatctagtggAGGCACTGAGGGCTAACACTTGGACATATCTTTCTTGGAGgaatcttaaacccaact
ggttacctcttatctagtggGAAACAGACAACAGTCTTCAGGGGCAGTTGAATAACAGCCgaatcttaaacccaac
ggttacctcttatctagtggGCTGGGCTCAGCACATGATCATGGGGGCATGCATTTTTAGgaatcttaaacccaac
ggttacctcttatctagtggACCACTAACTGGAGACTGTCTTATTTCCATGGAAGACTTCgaatcttaaacccaactt
ggttacctcttatctagtggGATAAATTCCCATGATTGGCTAAAGCCTGCTGTTGTGAGCgaatcttaaacccaactt
ggttacctcttatctagtggCTGGGCTAAGGGATGCCCCAGTGGCTGGTAAAACGCTATTgaatcttaaacccaac
ggttacctcttatctagtggTGGAGGAGTATTATCCAATCCCTGAGGACCCCAATAGAACgaatcttaaacccaac
ggttacctcttatctagtggGCTGGGACATGCATCTTCTTTTGCCCTTGGTCATCGCAGCgaatcttaaacccaact
ggttacctcttatctagtggCCAGCGCCCCCACCACTTGTTCTCAGTTCTTTGGACTCAGgaatcttaaacccaac
ggttacctcttatctagtggGACTTCTCAGCCTCCATAATCACATGCGCCAATCCCCATAgaatcttaaacccaac
ggttacctcttatctagtggCTGTTTCTCTGGAGAACTCTGACTAAACAACAACATCAACgaatcttaaacccaactt
ggttacctcttatctagtggCCACACTCTCATTAGTATAATTGGGAGAAAAATATGCCCCgaatcttaaacccaactt
ggttacctcttatctagtggCTAATACAAGGAATGGCCATTTCCTCCAAAGGTCCTGTTGgaatcttaaacccaact
ggttacctcttatctagtggAGCTGGGCCTGGGCTGGGACTCAGACACACAGATTTCCCCgaatcttaaacccaa
ggttacctcttatctagtggCCCAAGGGTCTCTTTTATCCTGGATGCCTGACACCTTGCCgaatcttaaacccaac
ggttacctcttatctagtggTCATCCTGTGGCCTTGAATGTGCTTCCTCCATTTGACCCCgaatcttaaacccaact
ggttacctcttatctagtggCCTGCCTCCCACCTGAACAATGACATCAAGGCAATGACGGgaatcttaaacccaa
ggttacctcttatctagtggGCCTCCAGCAGCTTTGTCTTCCTGTGTGGCTTTATCTCTCgaatcttaaacccaactt
ggttacctcttatctagtggGACAGCAATGGCATAAGCTCTGGGGCCAGATGGATATGGGgaatcttaaacccaa
ggttacctcttatctagtggGTGGCCTTGGATGAGTTAATGTAATGTCTCTGAGCCACGGgaatcttaaacccaact
ggttacctcttatctagtggATAGTTTATTGTGTAAGTGTCTACTGAGTTGTAAGCCCTCgaatcttaaacccaacttg
ggttacctcttatctagtggTGAACAAGGCAATTCATATACCTGACGTGGTGCATGGCACgaatcttaaacccaac
ggttacctcttatctagtggTAGCGCACATCTTAGTCTGCACAAAATACTTAAGACTGGGgaatcttaaacccaact
ggttacctcttatctagtggGCTTCTAAGATGGTGCCTTGTTTCTGTGTCCCCCAGAGGGgaatcttaaacccaact

ggttacctcttatctagtggGATGGATGGGCAAAAGGGCCTAGCTAGTTCTCTGCAACCCgaatcttaaacccaac
ggttacctcttatctagtggGCCTAGTCATGTCTTAAAGGACCCACCTCAACGCTGTTGCgaatcttaaacccaac
ggttacctcttatctagtggAGAACACCACTTCATCCCAAGAATTGTCCAGGTCTCTTCCgaatcttaaacccaact
ggttacctcttatctagtggTCCTGGCTAAACCAAATTGGGTTGGATATCTAACCTTCCCgaatcttaaacccaactt
ggttacctcttatctagtggCCTCACAGGCATAATGAAGGTAATAATATGTGTTTCATACgaatcttaaacccaacttg
ggttacctcttatctagtggTTCAGTGAGTTTATTCATTCATTAGGCTTACAACAGTGCCgaatcttaaacccaacttg
ggttacctcttatctagtggGGAAGATTCCTCCAAAGGCATTCTCCTTGGATCTTCATTGgaatcttaaacccaactt
ggttacctcttatctagtggTCCTCCCTCTTTCCCTACTGCTCCTAGCATACTGCTGGGGgaatcttaaacccaact
ggttacctcttatctagtggTTGCGTCTACTTTTGAAAAGTGTGTGTTGCAGAATTGGCAgaatcttaaacccaacttg
ggttacctcttatctagtggACGACAAAGCTGTAAATCTTGCAAGACTTTAGCTATACTCgaatcttaaacccaactt
ggttacctcttatctagtggGAGGTTGGAACTTGACCACAGACACCTGGTTAGTTAGCAGgaatcttaaacccaact
ggttacctcttatctagtggAAAAAAGAGATTCCCAGCCTATGGATCCTAAAAACTTGCTgaatcttaaacccaactt
ggttacctcttatctagtggGGAGGATAAGAGCCATATTTTGCTGGATATTCACCTGAGCgaatcttaaacccaactt
ggttacctcttatctagtggGACCTGAGAGGACCTTTTGCTTTTCTTCTCCACCTGGTCCgaatcttaaacccaactt
ggttacctcttatctagtggAGGATGCATTTGCACGGGTCAGGAACTTGGCTGCTGCTACgaatcttaaacccaac
ggttacctcttatctagtggGTCCCTTGTCTACTTTCCATCTCAGGGACTCAGTTACTTCgaatcttaaacccaactt
ggttacctcttatctagtggCTGGCTGTGTTCCAAATCCTCTTCCCATTCTAGCATTCTGgaatcttaaacccaactt
ggttacctcttatctagtggACTATTACCTCATGCTCCAGCCGGGAGGTTCTGTATTGCCgaatcttaaacccaact
ggttacctcttatctagtggTCATACAATGCCTCTCTTTGAGTGACACATGGGTAACTGTgaatcttaaacccaactt
ggttacctcttatctagtggCGTAGGAACCGTCTCCCTGCAACACATTTCTTTCAGCTTGgaatcttaaacccaact
ggttacctcttatctagtggCTTCTTGGGGATTCACATGCTCTCTGAGGGTTCACCAAAGgaatcttaaacccaact
ggttacctcttatctagtggATATCTCAGCATGGTGGCAATGCCTCCCTTGTGTATTGCCgaatcttaaacccaactt
ggttacctcttatctagtggATGTGACTTTGAACACTTGAAAACACAGACAGCCTGGTGGgaatcttaaacccaact
ggttacctcttatctagtggCTCTGAACTGTTCAAGTTTTAAGCAATCAGTTCCAGTTAGgaatcttaaacccaacttg
ggttacctcttatctagtggGCGAGAGCTGCTGGTAGGCAATCAACATAAAATATCCATAgaatcttaaacccaact
ggttacctcttatctagtggGATCAGGAGGTTTTCTTTTGCCTACTGTCTATCAGCTGTGgaatcttaaacccaactt
ggttacctcttatctagtggGCTGTTTCTTTTATTAGACAAGATGGGAAAAGAGGCAAACgaatcttaaacccaactt
ggttacctcttatctagtggCTCCTTTGTATGGTACAAGAAGGTTATAAAGCCATAATAGgaatcttaaacccaacttg
ggttacctcttatctagtggCATGTGGCTGGGCTGAGCTATGTAACCCCAGGGGATCATGgaatcttaaacccaac
ggttacctcttatctagtggGTGTGGGATGGTGCCCTCAGAAATGTGCAATGGAAAAGCCgaatcttaaacccaac
ggttacctcttatctagtggGGAATGCAAACACGGGATGAAAGGGAATGTGAATCAACACgaatcttaaacccaac
ggttacctcttatctagtggGTATACAGACATGTTATATATTTTCTGATTAGGAGAGCCCgaatcttaaacccaacttg
ggttacctcttatctagtggGCTGAACTAAGCAGGGTGGCCTAACTTCTGTGACAAACCGgaatcttaaacccaac
ggttacctcttatctagtggGTCTGTTGGTCTGTTCCATGGCCTCAGAGGTCTGCAGAGGgaatcttaaacccaact
ggttacctcttatctagtggGCCCAAAGGGTCTTCTGAAAAAATCTGGGTAGAGCCAGGCgaatcttaaacccaac
ggttacctcttatctagtggGAATCTGCAGGACAGTCTCTGTGTGCCCAGGAGGAATGGGgaatcttaaacccaac
ggttacctcttatctagtggGGGAACATAGTGCCAGTGCATTGTCTTTGCTACTTGCTGCgaatcttaaacccaactt
ggttacctcttatctagtggTCCTTTTTCCACTCCACAAAGCTCTAGAATTCATCCTCTGgaatcttaaacccaactt
ggttacctcttatctagtggCACGGATTACAATAACAGCAGTGCTTGAGGTCAACAAGGGgaatcttaaacccaac
ggttacctcttatctagtggACTCAGACATGACCCAGCAGTTGCTTGGAGATATTTGGCCgaatcttaaacccaact
ggttacctcttatctagtggGTTCTTGTCTCTGCCCCTGTTGCTTTCTCTCTCTAAGGCCgaatcttaaacccaactt
ggttacctcttatctagtggGCCCGTTGTTTCAACTGTAAATGCCATGTGGTCTACTCTCgaatcttaaacccaactt
ggttacctcttatctagtggCTCAGCTAGAGGGCTCATATCTAACCACCTGCTGAGCACCgaatcttaaacccaac
ggttacctcttatctagtggGGCTTGAGCCACCGCGCCCGGCCCCCACATGCTCTTATTAgaatcttaaacccaa
ggttacctcttatctagtggATGGTGTACAACTTGATATATGTCACCATTTTGGAATGGCgaatcttaaacccaacttg
ggttacctcttatctagtggGTCACCATGTCATACAATAGATCCTCTAAACTTATTCCTCgaatcttaaacccaactt
ggttacctcttatctagtggCTCCTCAATCCCAATCCCCACTCCCCAGCTCCTGGTAGCCgaatcttaaacccaa
ggttacctcttatctagtggTTCTGTGCCTGGTTTCTTTCACTAAACAATGTCCTTCAGGgaatcttaaacccaacttg
ggttacctcttatctagtggGGCTAGCCAGTTTTCCCAGCATTTATTAAATAGGGAATCCgaatcttaaacccaactt
ggttacctcttatctagtggGGGATGGCATTGAATCTGCAAATTACCCTGGGCAGTATGGgaatcttaaacccaact
ggttacctcttatctagtggGCTCTCTGTTTGTGTGTTGTTGGTGTACAAGAATGCTTGTgaatcttaaacccaacttg
ggttacctcttatctagtggTTTGAAGGGTTTTTTGTGTTTGCTCTTGCTTTTCTAGTTCgaatcttaaacccaacttgt

ggttacctcttatctagtggCTTCTTGTTGAATTGATCGATTTACCATTATGTAATGGCCgaatcttaaacccaacttgt
ggttacctcttatctagtggGGCATGATTTTGCAGCAGCAGGTACTGGTTGTTCCTTTCCgaatcttaaacccaactt
ggttacctcttatctagtggTCTGAATGTTGGCCTGCCTTGCTTGTTTGGGGAAGTTCTCgaatcttaaacccaacttg
ggttacctcttatctagtggGGCAATGGCGGGCGCCCCTCCCCCAGATTTTTAAGGCTGgaatcttaaacccaac
ggttacctcttatctagtggTCCATTCGCTAGGTCAACATCAGGAAGGTTTTCCCTAATGgaatcttaaacccaactt
ggttacctcttatctagtggGGGGTGAGAGAAGGGTCTGGTTTCATGCTTCTGCATGTGGgaatcttaaacccaact
ggttacctcttatctagtggGAAGAGACTGTCCTGTCCTGATTGTATATGTATGTTTTGGgaatcttaaacccaacttg
ggttacctcttatctagtggCATTGACCTATATGTCTGCTTTGGTGCCAGTACCATGCTGgaatcttaaacccaacttg
ggttacctcttatctagtggCTCAGGATTGCTTTGGCTATTCAAGGTTTTGTGTGGTTTCgaatcttaaacccaacttg
ggttacctcttatctagtggCTGTAGATTGCTATTGGCAGTATGGGCATTTTAAAAATACgaatcttaaacccaacttg
ggttacctcttatctagtggAAACTTTTTATGGTGTTCCTTATAGAGATCTTTCACCTCCgaatcttaaacccaacttgt
ggttacctcttatctagtggATGGTACAAATTTTTGTCTGGTGATTTTGTATCCTGCAGCgaatcttaaacccaacttgt
ggttacctcttatctagtggCCAGTGCTATGTTGATTTGAAGTGGTAAGAGTGGGCATCCgaatcttaaacccaactt
ggttacctcttatctagtggGTGTTCTACTTTTCGCCATTGAGTGTGATATTAGCTGTGGgaatcttaaacccaacttg
ggttacctcttatctagtggAGAGAACATTCCTTCTATAGTTAATTTGTTGAGAGCTCTGgaatcttaaacccaacttg
ggttacctcttatctagtggCATCTATTGAGATGATCATGCAGTTTTGTCCTTCATTCTGgaatcttaaacccaacttgt
ggttacctcttatctagtggTGAACCATCCAAAACCCAGGACCTGATGACTTCATGACTGgaatcttaaacccaact
ggttacctcttatctagtggAACTAGATAGAAGTAATGCCAGCTTTTGTCAAACTTTCCCgaatcttaaacccaacttg
ggttacctcttatctagtggTGCAAGCCTCACCTTACCCTGATATCAAAGCCAATCAAGGgaatcttaaacccaact
ggttacctcttatctagtggTTCCTGATGAACATAAATGCCAAAATCCTCAACCAAACACgaatcttaaacccaactt
ggttacctcttatctagtggCACCATGATAAAGGGATATTTATGCCCGGCAAAACATGACgaatcttaaacccaact
ggttacctcttatctagtggGGGTTATCAACTTACGATTCAGGTTTCTTGGGTCCTAAGGgaatcttaaacccaacttg
ggttacctcttatctagtggTCCAGGTTTTGTGGGTCCTAAGGCTTCAACAATTAAAAACgaatcttaaacccaacttg
ggttacctcttatctagtggCAAACATCACATGTTTGTTAACTGTCTGACACAACTCAGCgaatcttaaacccaacttg
ggttacctcttatctagtggGTCTGTCTATCTATCTATCTTTCCTAGTCTTATTTCTAGCgaatcttaaacccaacttgt
ggttacctcttatctagtggTTGTCAAACCTGAAAAACTTCTGTCAAAGTTCTTTCACTGgaatcttaaacccaacttg
ggttacctcttatctagtggGCCCTTTCAATTGCAACACGTGTTCTTTTGGGGAGATGGGgaatcttaaacccaactt
ggttacctcttatctagtggCAATAGTTTTTGGGGGTAGGGATAGTTTTTGGTTACATGGgaatcttaaacccaacttg
ggttacctcttatctagtggCCCCTACCTCTCTCCCAACCTTCCTCTGCCCCAAGTACCCgaatcttaaacccaac
ggttacctcttatctagtggGCATCCTCATAGCTTAGTTCTCACTTAGAAGTTAGAATATgaatcttaaacccaacttg
ggttacctcttatctagtggAACGGCCTCCAGTTCCATCCAAGTTGCCTCAAAAGACATCgaatcttaaacccaac
ggttacctcttatctagtggTTTTATCCCTTTGTTGGTTGATGGGCACTTAGGATGGTGCgaatcttaaacccaacttg
ggttacctcttatctagtggGTTTTCCATAGCGGTTCCACTAATTTACATTCCCACCAGCgaatcttaaacccaacttg
ggttacctcttatctagtggTTGTGGTTTTCATTTGCATTTTCCCTGATGATTAGTGACGgaatcttaaacccaacttgt
ggttacctcttatctagtggTTCTTTTGAGAAATGTCCATTCATGTCCTCAGCCCACTTTgaatcttaaacccaacttg
ggttacctcttatctagtggTTGAAACACTTGTTCATACCAGTAGGAGCCCTGCCAGGGGgaatcttaaacccaac
ggttacctcttatctagtggTGCAAATGAGAAGCCCTGAAGCTTAGACATCATTAGTTTCgaatcttaaacccaacttg
ggttacctcttatctagtggCCCTTGTTTATCTGTGTTCACCCTCATGCTGTTTACCCACgaatcttaaacccaacttg
ggttacctcttatctagtggCATGTTCTTCATGGTGGATCTCTGTCTCCATTATGCTACCgaatcttaaacccaacttg
ggttacctcttatctagtggATTTCCCACTTGAAGTCTTTGTCCCCTTCTTGTCATTCTGgaatcttaaacccaacttg
ggttacctcttatctagtggTGGGTCATATGTGATCTGGACCAAACTATTCTTTCTAGTGgaatcttaaacccaacttg
ggttacctcttatctagtggTTCACTCTAGTCTAGAGATTTGGACTTCACCTAAGCAATGgaatcttaaacccaacttg
ggttacctcttatctagtggAGAGGGATACAATTCTCAGCATCTCCTTAGGGCCTGGACCgaatcttaaacccaac
ggttacctcttatctagtggGGAAGTGTCTTACACAGCTGTTGGAGTGTTCACATGTAGCgaatcttaaacccaactt
ggttacctcttatctagtggTCTGATCCCTTTTCCCTCCTTTGGTTTTCTCTGTGGTCCTgaatcttaaacccaacttg
ggttacctcttatctagtggTTGACATATCAGCATAACGGGAATAATGAGACTATCGTGCgaatcttaaacccaactt
ggttacctcttatctagtggGGATACCTCAGAGGCAGCTGGCTGAGTACGTCTGCTGTGCgaatcttaaacccaac
ggttacctcttatctagtggCAGTGTGGAGAGGCCACTGAGAATCTCTGATGACATGTGCgaatcttaaacccaac
ggttacctcttatctagtggCTCTAAAATGTTCCTACAGAACAACAACAGTGATTAGCAGgaatcttaaacccaactt
ggttacctcttatctagtggAATCTGAAGGTCATCATTTGAGCTAATTTCACCCGACAGGgaatcttaaacccaactt
ggttacctcttatctagtggCCTCGCCAACTTTACCCCTGGTATAGTCTAGCAAGACTTCgaatcttaaacccaact
ggttacctcttatctagtggGGGGTGGGTACGGAGTCTGTTTTTCACTCTGGGAAAGATGgaatcttaaacccaact

ggttacctcttatctagtggCCAGAAGGTAAATGAGGCCAGCTTCACTAGCGGTTTTAGCgaatcttaaacccaac
ggttacctcttatctagtggCATAGCCAAGGCTGGATCAGAACCAGAATAAGAATTTCTCgaatcttaaacccaact
ggttacctcttatctagtggGGGAAGCAGATGCAATTAGCACAGGAGCCATTCTGGAGGGgaatcttaaacccaa
ggttacctcttatctagtggCCAGATTCCAAAGAGGGGTCCTTGTCTCCAGTCCTGGCCCgaatcttaaacccaa
ggttacctcttatctagtggCGGTAACATCCCTACCCTTAGTACAAAAGGTGAGTTGACCgaatcttaaacccaac
ggttacctcttatctagtggGGGCATGGGTCTGGAGGTGGTTTGTCTGAAGCGAATCCTGgaatcttaaacccaac
ggttacctcttatctagtggGGTTGGTGCAAAATAATTAGCCGCTAATGGCAAAAACCACgaatcttaaacccaact
ggttacctcttatctagtggGAGCATAATTATTGCAGTACCTCACAGAGCTATTGTGAAGgaatcttaaacccaactt
ggttacctcttatctagtggCTTCAATCCTCACAGTTCTGAGGCAGGCTTCACTCTCCCCgaatcttaaacccaac
ggttacctcttatctagtggAGACTGAGAGGCAGCTCTAGCACAGAGTGGAGATGAGGAGgaatcttaaacccaa
ggttacctcttatctagtggCGTTTCCCCAAAACCAACCCCTACTGAGCCACTGACTCCGgaatcttaaacccaa
ggttacctcttatctagtggCTCCCCACCCCTCCCCCACCCTCTGGCATCCCTCTATCACgaatcttaaacccaa
ggttacctcttatctagtggAGAGCTCCTTCTTCCCTCGGGAAAGCTCTCTCTGTAAATGgaatcttaaacccaac
ggttacctcttatctagtggGAGGCTGGTTAGTGAACAATCGTCATAAGAGGGTCACTTGgaatcttaaacccaact
ggttacctcttatctagtggTGCAGGCTTTCTCGTTTGTTTCTAGCTATATTTCAAGAAGgaatcttaaacccaacttg
ggttacctcttatctagtggCAAGCTTCCATAGTTGACAGATGAAAAAGCTGAGGCCCAGgaatcttaaacccaac
ggttacctcttatctagtggCAGCTTTCACAGTTCCTTCAACAGCAGAGCTCAAGACTAGgaatcttaaacccaac
ggttacctcttatctagtggCACCACTTCCTTCCCTTTCATCAGACTGAACTCTTGGCAGgaatcttaaacccaac
ggttacctcttatctagtggCCCACTTTTCATGTTTTGGTTTTGCTTTGCCTAACACCTCgaatcttaaacccaacttg
ggttacctcttatctagtggCCAGGGAAACTCATCTCTTATTTGCTCTCACAATAGCCCCgaatcttaaacccaac
ggttacctcttatctagtggTGTCTTTGTGTATGTCTGCCTTTCCTGGAGGAAAAAGAACgaatcttaaacccaactt
ggttacctcttatctagtggGTTCGTCGTTCTATTCTTTGTCTCTTGGCATCATCATTGGgaatcttaaacccaacttg
ggttacctcttatctagtggGATGGATGGATGGATGGATGCTTGGGTGGTTGGGTAGATAgaatcttaaacccaac
ggttacctcttatctagtggTGGATGGGTGGGTGGGTGAGTGAATGGGTGGGTGGGTGGGgaatcttaaacccaa
ggttacctcttatctagtggGGACATGTAAATTTTCCTTTGAATGTGTAGGAAATTACCCgaatcttaaacccaacttg
ggttacctcttatctagtggGCCTAGCTTATGATCCCCTGATGTTTCTAAAAGCAGGAAGgaatcttaaacccaact
ggttacctcttatctagtggGCTCTAGTAGCAGGTGGAAATTAAATGCCCCCTCACTGTGgaatcttaaacccaac
ggttacctcttatctagtggCTCAACACCTTTACAGCCCATGGGTATATGTGTGTGCCTGgaatcttaaacccaac
ggttacctcttatctagtggCAAATGGCCCCTTGGACAGTCAAAAGTTTAGTGCCTTCGCgaatcttaaacccaac
ggttacctcttatctagtggTTATGGGGAGCCAATGAGGTCTGGCAAGCTCTAATGTCTGgaatcttaaacccaact
ggttacctcttatctagtggCAGGAATTCCATGTGATGTGTGCAGCAGCTCAGCCCTGCCgaatcttaaacccaa
ggttacctcttatctagtggGCAAGAAAGAAATGATGCTTCTGGGTTCTCTGAGTGACAGgaatcttaaacccaac
ggttacctcttatctagtggCTCTGACTACTTTCTTTTGTAGTCTCATCAGCTCAGAAGGgaatcttaaacccaactt
ggttacctcttatctagtggGTCAAAGTGTAGAACAATGAGCCTATGACAGAAGTGAGGCgaatcttaaacccaac
ggttacctcttatctagtggCATCCATCCCTCTGACAGTGGTAATAGGAATCCTGAGTGCgaatcttaaacccaac
ggttacctcttatctagtggACATGTGCTTCTACCGCATCTTATCAGCACTCAGGCAAGGgaatcttaaacccaac
ggttacctcttatctagtggGACACTTAGTGAGCATCTAATAAGTGTCAGGCATTAGGCCgaatcttaaacccaac
ggttacctcttatctagtggGTGGACTTTGCCAGTAAAAATAAGAAGTAGAGTTGTGTGCgaatcttaaacccaactt
ggttacctcttatctagtggGGCTAAATATGGACTTTTCCTAGAAGAGGGATTCAGATGGgaatcttaaacccaactt
ggttacctcttatctagtggGTACTATTTGTGTGACTTTGGGCAAGCCTGTTCTCCCCTCgaatcttaaacccaactt
ggttacctcttatctagtggCCTCATTCCTAAGCATAGCACATTTGAGCACAAATATTTCgaatcttaaacccaactt
ggttacctcttatctagtggCTGTTGGAAAAGTATCTGCTAAGGGTGAAACTGAGACCTGgaatcttaaacccaac
ggttacctcttatctagtggGCAGCACCCTTTATGTTGTCACCATTTCATGAGGGGCTCCgaatcttaaacccaact
ggttacctcttatctagtggGACCTTTACCCTATTCTTTGAAAACGAATTCAGCGGGGAGgaatcttaaacccaact
ggttacctcttatctagtggACCATCCCTCATCTCTTCATGGGTGTTTTCCCCTAGTGCCgaatcttaaacccaact
ggttacctcttatctagtggCCTTATCATGCACCAAGTGAATCCTTATAATCAATGGGTCgaatcttaaacccaactt
ggttacctcttatctagtggCTGTAGATAAGAGATTTCAAGTGCAGCAATCTTAAGAACCgaatcttaaacccaactt
ggttacctcttatctagtggAAGCATGCTCATGAGTCTCATCCCAGGGTTTTGTGAGGAGgaatcttaaacccaact
ggttacctcttatctagtggGGCTGGGGCATGAACTAAGAGTCAGACAGGCTTGGGTATGgaatcttaaacccaa
ggttacctcttatctagtggCCTGTATGGTAAGGACAACACTAGTACCTAGTTCGATGGGgaatcttaaacccaact
ggttacctcttatctagtggTCAACAGACAAAAAGCTTACACATGAGCTGACAATACAAGgaatcttaaacccaact
ggttacctcttatctagtggCCCGTTGCAGTCATTAACTGTTAAACTTTCCTTGCCCGGGgaatcttaaacccaact

ggttacctcttatctagtggTGGCAGTGATTTCCAATCCTGTTTTCTTCATCTGGAGAGCgaatcttaaacccaacttg
ggttacctcttatctagtggCTAAAGACAGAATCCTTTGACCTTCTGCTGTGTACCGTGGgaatcttaaacccaactt
ggttacctcttatctagtggGGCTGAAGATAAACATCTGGTCTTTTCCCGGCATGATCCCgaatcttaaacccaact
ggttacctcttatctagtggGTGCTGCTCCCAATGTGGTCCAGGAATGGTGGTAGTGAGCgaatcttaaacccaac
ggttacctcttatctagtggCCGGTGCCTCCTTCCCTCTCTGCCCCTTTTCAAAATCCCCgaatcttaaacccaac
ggttacctcttatctagtggGTATCCAATGTTCGGCTCCTACTTGTAAGTGAACACATACgaatcttaaacccaacttg
ggttacctcttatctagtggACTTAGGATAATGCCCTCCAGCTACATACAAGTTGCTGCAgaatcttaaacccaact
ggttacctcttatctagtggCCAATCCAGCATTGATGAGGACCTGGGTTGATTCCATGTCgaatcttaaacccaact
ggttacctcttatctagtggGTGAGCTGCCATTTGTTGTCAGTCCATGAGATAAGCCCAGgaatcttaaacccaact
ggttacctcttatctagtggACCTGATTATTTAACTCAGATAACAAGAAAAGCCCTGCTGgaatcttaaacccaacttg
ggttacctcttatctagtggGGCATGGTGGCTTATATCCCACGGTGGCTGTAATCCCAGCgaatcttaaacccaac
ggttacctcttatctagtggGGATTGCTTCAAGCCAAGAGTTCAAGATTAGCCTGGACAAgaatcttaaacccaact
ggttacctcttatctagtggACCAGGCCTGGTGATGTATGCCTATAGTCCTAACTACTTGgaatcttaaacccaactt
ggttacctcttatctagtggCAGTAAGCCATGATTATGCCATCGCACTCTAGCCTGGGTGgaatcttaaacccaact
ggttacctcttatctagtggCACTGTCTCAGTCGATCACTCAATTAAATAAAATCTTGTGgaatcttaaacccaacttg
ggttacctcttatctagtggCCCTGGTGCTGATATTTGGCTGAGTATCTTTGGGACTACCgaatcttaaacccaactt
ggttacctcttatctagtggGATATGCCCACCATCAATCATCATAGGAAGCCGAGGGTGCgaatcttaaacccaac
ggttacctcttatctagtggGCCCATACTTATACAAAAGGCCAGGTATGAGAGCTCCTACgaatcttaaacccaact
ggttacctcttatctagtggTCACCAGAGGCATAAGTCAGGACAAGGGTTCCAGTACCCCgaatcttaaacccaac
ggttacctcttatctagtggAGCCCAGCCCAGTGTGTTGATTTGCCGAAAGTCCAGCGAGgaatcttaaacccaac
ggttacctcttatctagtggCTAGTCCATGCCCACTGGACTGGAGCAGTGTTATCTGAAGgaatcttaaacccaact
ggttacctcttatctagtggGCAGGCTATTGGAGAGGCAGCATGATATATTAGAACATGGgaatcttaaacccaact
ggttacctcttatctagtggTATCCCTCACCTACTTCCCACCGTTTCCCCCCGAGTTTCCgaatcttaaacccaact
ggttacctcttatctagtggCTTTGCATCCTCATAGCTTAGCTTCCAGTTATGAGTGAGAgaatcttaaacccaacttg
ggttacctcttatctagtggGAATAATAATCTCCAGTTCCTTTGAGGCTTCTGTGAATGCgaatcttaaacccaacttg
ggttacctcttatctagtggGGCTGAGTAGTATTTCATGGTATATGTGTGTGTGTGTAGAgaatcttaaacccaacttg
ggttacctcttatctagtggTATACAGACTGTGCTATATATATATACACACACACTGCGGgaatcttaaacccaacttg
ggttacctcttatctagtggACACACACACACACACACAGTCGCTTGTTGATTGATGGACgaatcttaaacccaact
ggttacctcttatctagtggACACAGTAGTGGGATTGCAGGATCAAAAGGAAGAACAGGGgaatcttaaacccaac
ggttacctcttatctagtggATGATCCACCATGTTTTGACCTTGGACACATTAACCAACCgaatcttaaacccaactt
ggttacctcttatctagtggCTCATGTCTGGAATAGGTATTATAATTAGCGTTGGCAAAGgaatcttaaacccaacttg
ggttacctcttatctagtggAAGGACATGACCCAGAGTGAAGCTGACCCTGAATAAGCACgaatcttaaacccaac
ggttacctcttatctagtggAGCTCATCCACGGGGTTAGCAGAAGTCAGGCACAAAAAGCgaatcttaaacccaac
ggttacctcttatctagtggCCACATTCCCAATCTGGCCTTTCTGCAGTAGCATGCGACTgaatcttaaacccaact
ggttacctcttatctagtggAGCCAGAGCAGGGTAGGAATTTGGGGACACGGGGACACACgaatcttaaacccaa
ggttacctcttatctagtggGCCTCTTGTCACTAGAAAATGAAGGTAGGGCTTGTCTAGCgaatcttaaacccaact
ggttacctcttatctagtggTGTTTCACCTGGTTGATCTGCTACCAATAGGTCAAATCATgaatcttaaacccaacttg
ggttacctcttatctagtggGGCTTTCAGCAGCCCAGAAAGCATTGCATTAGCTTCATCCgaatcttaaacccaact
ggttacctcttatctagtggGCCACTTGTAGTACAATATGTAGACGTTGTCCATAGGTGCgaatcttaaacccaactt
ggttacctcttatctagtggTGGGGCAGGGGAGGGGTAGAGTAGGAAGAAGGTGGACTCGgaatcttaaacccaa
ggttacctcttatctagtggCCTCGCACATTCCAGGTCTCCTGCCTCTCACTGAGCCACCgaatcttaaacccaac
ggttacctcttatctagtggGAGAGCCACAGCTGGGAACATTGTTGGGTAGTTCACGATGgaatcttaaacccaac
ggttacctcttatctagtggGTGGCTGCTTCCTCTCATTCTCTCCACTATTGTTGGCAACgaatcttaaacccaacttg
ggttacctcttatctagtggCCTACTTATGTAGGATATCCTGTTATGTGCCCAGATTCTGgaatcttaaacccaacttg
ggttacctcttatctagtggTAGAGCTTACTTAGCACAGACCCTCAACACACATTATCTCgaatcttaaacccaactt
ggttacctcttatctagtggTCCCAAGGTGACATAGCTAATAAATGACTGAACCAGGAACgaatcttaaacccaact
ggttacctcttatctagtggGTATCCTAAGAGAGAAATTCCTTCTAGGTAGCAACAAGCGgaatcttaaacccaactt
ggttacctcttatctagtggAGGGGAAGTTTGAGGTGATAGAAGAACACACAGAAGGGGCgaatcttaaacccaac
ggttacctcttatctagtggTGAACAGAGGTCCTTAGGAAGCATAGGCATTACCCAGAAGgaatcttaaacccaac
ggttacctcttatctagtggACCTGGAGATAGTAGAGAACTGTGTATACTGGGGGAAGTGgaatcttaaacccaact
ggttacctcttatctagtggACCAGGTTCTGCAGAAATGAGTAAGCCACAGTGCCTCTCCgaatcttaaacccaac
ggttacctcttatctagtggGCCACAGTTAAAACTATCCCAGAGAGGCCAGGCGCGGTGGgaatcttaaacccaa

ggttacctcttatctagtggTGCCACTGCACTCCATCCAACCTGGGAGATAGAGCGAGACgaatcttaaacccaa
ggttacctcttatctagtggACAACTATCCCAGAGATAGGGAATACTAGAGTGAGAAGAGgaatcttaaacccaac
ggttacctcttatctagtggCATGTCAGGGAAGGGATACAGAATCAAAAAAGCTTCAGAGgaatcttaaacccaac
ggttacctcttatctagtggTGAAGGGTTGAAGAGGGACCCACCAAGGAGGGAAGCCAGGgaatcttaaacccaa
ggttacctcttatctagtggCCAGAAAGACATGGCATTTTCAAAGGGAGTCCATGTAGGAgaatcttaaacccaact
ggttacctcttatctagtggCTTCACCAGGCAGCAAAGAGGATACAAGATGACATCAGTGgaatcttaaacccaac
ggttacctcttatctagtggATTCTAGTCCTTTGAATGTTCCCATTTGACTGGAAGTGAGgaatcttaaacccaacttg
ggttacctcttatctagtggGGAAATAGGCCCTCTGACTCCACCAGCGTTGGACTTGGAGgaatcttaaacccaa
ggttacctcttatctagtggCACTTCTAGGTTCCCCACACCTAGCTTAGTGTCTGGCTCCgaatcttaaacccaac
ggttacctcttatctagtggCTGGACTCAGCCTGAATTTGAATACTGCTCCTTACCACTCgaatcttaaacccaact
ggttacctcttatctagtggGACCCTCATCTGTAAAATAGAGTTGACATTAGCTCCTCAGgaatcttaaacccaact
ggttacctcttatctagtggAATGACTTAGAACAGGGCTGGGTGCATATAAAGTTCTTAGgaatcttaaacccaactt
ggttacctcttatctagtggGAGACTAATGAGGTGTTACAATCAGTTCAAAGGAGATTCCgaatcttaaacccaact
ggttacctcttatctagtggGCAAAGGGAGGAAAGACTAACCTCTTGCACATTGGGGAGGgaatcttaaacccaa
ggttacctcttatctagtggCTGTCTATGGACAAGAATCATTTTGCATTGCCATCAGTTGgaatcttaaacccaactt
ggttacctcttatctagtggCAGCTCCCATAGAAACAAAATTAGATAACTGGGAAGAGTGgaatcttaaacccaact
ggttacctcttatctagtggGGAAAAGGTAAAATCAGTAATCCACCATCAAAAGGTAACCgaatcttaaacccaact
ggttacctcttatctagtggCTCTCATATATATATATATTTGGGAGCAGACTACACGTTCgaatcttaaacccaacttg
ggttacctcttatctagtggACCTCCGCCTCCAGGGTTCAAGCAAGTCTTCTGCCTCAGCgaatcttaaacccaa
ggttacctcttatctagtggGAGCCACTGCACCTGGCCATATTTTTACATTAGTGAAATGgaatcttaaacccaactt
ggttacctcttatctagtggTCAGTAGTAGCACAAGCTGATTTCTTTGTCCTGTACTGGCgaatcttaaacccaactt
ggttacctcttatctagtggTTGCATTTTGCCAGAGACTTGATCTATGTGCCTAGGCAAGgaatcttaaacccaactt
ggttacctcttatctagtggGGACTCGAGCAGCTTACAGATCAAAGATCCCACTATTGGCgaatcttaaacccaac
ggttacctcttatctagtggCTCTGGGTACCTGATGCTCCTTCAGCCATCAACAGATATGgaatcttaaacccaact
ggttacctcttatctagtggCCCAAGTTGATTTGTCTCCTCCACATGATGTGTCTGCCTCgaatcttaaacccaact
ggttacctcttatctagtggGTCAGCTAGCTCAAGAGGTCCTAAAGCAGAAACATCTGATgaatcttaaacccaac
ggttacctcttatctagtggTTCACTCATGATACAGCTCTCAAAGAACCTGACGGTCAGGgaatcttaaacccaac
ggttacctcttatctagtggGGAGCTTCCCTCCTCCACCTGCGGTCCCCAAACATATACCgaatcttaaacccaa
ggttacctcttatctagtggTGTGACCTGAACTCTTCCACTTTCTGCCACCTCAATTATCgaatcttaaacccaactt
ggttacctcttatctagtggCTGTTGGCTCTTTCTTTCCTTCTTCCTCTTTCCTCCACTCgaatcttaaacccaacttg
ggttacctcttatctagtggCCTCCAGTGCCCCTGCTTGCAATCTATTCTCTACAAGGGGgaatcttaaacccaac
ggttacctcttatctagtggCTGAAAAGCTCAGATGTGACTATGCTGTTTCCTGATTAACgaatcttaaacccaactt
ggttacctcttatctagtggTGCGTATGGCTATAGTTCTCAAACTTGTATCAGAATCACCgaatcttaaacccaactt
ggttacctcttatctagtggGCTCTCATAGTTTCTGACTCAGTAGGTCTGGGGTCAGGCCgaatcttaaacccaac
ggttacctcttatctagtggCTGACAAGTTCCTAGGTCCTGTGGATCTTTTGAGAACCACgaatcttaaacccaact
ggttacctcttatctagtggCTCTCACTGAGGCAAGCATGGTTTTCCTTCTAGACACACCgaatcttaaacccaact
ggttacctcttatctagtggATCGGTTCTGCTTGATATGTAATGTTCTTCCTCATCGACGgaatcttaaacccaacttg
ggttacctcttatctagtggAGCTACCTCTAGCTCCACACACCAAGATAGAATAACTGACgaatcttaaacccaac
ggttacctcttatctagtggGCCAATGTACTTAGCTCAGTTATAAAGTCCTTGAAGGCAGgaatcttaaacccaact
ggttacctcttatctagtggGAAACACTTTAGTTTAGGGATGAGCTATCAGATCAGCTCCgaatcttaaacccaact
ggttacctcttatctagtggCATGGGCATTCAGGGAAGAATTAGGTAAATAGTTGGGCAGgaatcttaaacccaact
ggttacctcttatctagtggGCAACACATGTGAAGACAGAAGTCATGACTGCCAACTGGCgaatcttaaacccaa
ggttacctcttatctagtggCTACGTTAGGATAGAGGGCAAAGTTAAATGCAACAAGTACgaatcttaaacccaac
ggttacctcttatctagtggAGCCCTGTGCTACATACAAAGTATAGGCCATTATATGTGGgaatcttaaacccaact
ggttacctcttatctagtggATGTTTCAAAAACATACCAGGAGTCACAAATTTTCCAGGGgaatcttaaacccaactt
ggttacctcttatctagtggCTTGAGAGTCAAATGAGTCACTTTTAAACTGTCTATGGTGgaatcttaaacccaacttg
ggttacctcttatctagtggTGCCTTTGGATAGTATTCAAGGAAGTCGCTTTCCTGCATCgaatcttaaacccaactt
ggttacctcttatctagtggTTGAATAAAGAGCAGAGTCCTTGCCTTTGATAATGAGCCCgaatcttaaacccaact
ggttacctcttatctagtggATGTGTCATCTTTCTTTTATAAGCCACACCCATAATACAGgaatcttaaacccaacttg
ggttacctcttatctagtggACAGCAATCAAAAGACAAAACCCCTCAAGAGTCTATGTTCgaatcttaaacccaact
ggttacctcttatctagtggGTTGCAATAGCACTTTTGGGAGCGTTATTTTTATCAACCCgaatcttaaacccaacttg
ggttacctcttatctagtggAATGAGCTTGGGAGTCAATATGACTGATTCTTGGCACACGgaatcttaaacccaact

ggttacctcttatctagtggTGTAGAACTCAGAGTGGCTTGGAGAGCAGTGCTGGAGGTCgaatcttaaacccaac
ggttacctcttatctagtggAGGCAGGAAATCCCAGGCTTATGAGGATTAAACTTTAGGGgaatcttaaacccaact
ggttacctcttatctagtggGTAGGCTACTGTGAGTGCTCTGGGTACATTTAAGGCAAGCgaatcttaaacccaact
ggttacctcttatctagtggCTTCACATATTAGCCTCAGGACATTCCTATGAGGTTGGTGgaatcttaaacccaactt
ggttacctcttatctagtggTGATTCCATTCTCTCCCTTTCAGTGTTTCTCAGTAGGCCCgaatcttaaacccaactt
ggttacctcttatctagtggTCTCTCCCCTATGTACTCACTCTACTCCAAGGGGCACTGGgaatcttaaacccaac
ggttacctcttatctagtggGCCAAGGAACCCTGAGCACATGCAGCCTAACTCTCTTCTCgaatcttaaacccaac
ggttacctcttatctagtggCCAGCCACCTCTTGGGATGCACACGTTTCTCTTCAACTGCgaatcttaaacccaac
ggttacctcttatctagtggCTGCGTCACATGGTAAGAGAGCTCAGCTATTGATCAATGGgaatcttaaacccaact
ggttacctcttatctagtggAGGAGCTGACAGCCTATTTTGTCTTGGGGCCTAAGCAGGGgaatcttaaacccaac
ggttacctcttatctagtggCAGATTTACCAGCAAAGGTCCATTTCTTTTCTTTCTTTCTgaatcttaaacccaacttgt
ggttacctcttatctagtggCTCCCAGGTTCAAGCAATTCTCCTCAACTTCCTGAGTAGCgaatcttaaacccaact
ggttacctcttatctagtggGTAGAGATAGGGTTTAACAATCTTGGCCAGGCTGGTCTTGgaatcttaaacccaact
ggttacctcttatctagtggCAGGCTTGAGCCACCATGCCCGGCCGGTCTGCTGTCTATTgaatcttaaacccaac
ggttacctcttatctagtggATTCTACCCACCATTTTCTTCATAGCTTAGGTGTATTATCgaatcttaaacccaacttg
ggttacctcttatctagtggTCCCAAAGAGGTGGACACTACTATTGTTTTAATTTTACAGgaatcttaaacccaacttg
ggttacctcttatctagtggATCCATAAGTCCACTTTGGTACTATACTTATGCTTGCTGGgaatcttaaacccaacttg
ggttacctcttatctagtggGCTGCAATAATGTCTTCAAAGACCATTCTCTTAATCACTGgaatcttaaacccaactt
ggttacctcttatctagtggATCTTGCCTTGACTTTGAGATTTTCTAGTGCTGGAATTAAgaatcttaaacccaacttg
ggttacctcttatctagtggAGCAGACCTGAAAAACACTTGCTTGGGAAGATCATGATGGgaatcttaaacccaac
ggttacctcttatctagtggCCTTGCTATTACTGGTCCCCATCCTACTATAGTATCATCCgaatcttaaacccaactt
ggttacctcttatctagtggCAGAATTGTTCCTAACATCTAGATCATGAAGATGAATTCCgaatcttaaacccaactt
ggttacctcttatctagtggTCCCAAAGGCTCGGGGAGGTTAAATAATCCATTGAAGATCgaatcttaaacccaact
ggttacctcttatctagtggCTGGGACAACAGGCTCCCAGTCCAAATAACTACAGTATCTgaatcttaaacccaac
ggttacctcttatctagtggTTTAGCTAAAGTCAGATCCACTCCTAAGAAGAAGTTGTGGgaatcttaaacccaactt
ggttacctcttatctagtggGAGGTAGATATTTAGTTTTACCCTTGGAGTAATTGCCCAGgaatcttaaacccaactt
ggttacctcttatctagtggATGTGATTTTTTCCCTCCTTTGTTGAAGTATAGAGAGTAGgaatcttaaacccaacttg
ggttacctcttatctagtggACTTCCTCATTTGCTCAGCTTATTCATTTGTGAAAAGAGGgaatcttaaacccaacttg
ggttacctcttatctagtggCTTGTGTAGGTGCGTTGTTGTTGTAGTGCTGTCTTCTCACgaatcttaaacccaacttg
ggttacctcttatctagtggACGAGGATGGGGGACAAGGAAGGAGGAAATGGTGCATACGgaatcttaaacccaa
ggttacctcttatctagtggGCTCCTGATGGGAATTCAATTGCACTTCAAAACCAACTTCgaatcttaaacccaactt
ggttacctcttatctagtggGAGTTGGTACACACAACAGTTAGAATTATCATAATTTGGCgaatcttaaacccaactt
ggttacctcttatctagtggGCCCTAGTCAGCTTCTCCTTTCTAGATGGTAGAACCAAGCgaatcttaaacccaact
ggttacctcttatctagtggCCCCTAGTCCATTCAAATTCTCTGGCCACCCTCGCCAACCgaatcttaaacccaac
ggttacctcttatctagtggGACTTCAGGCCCTGCTTGATGAAGGCAAAATGTTTTCCCCgaatcttaaacccaact
ggttacctcttatctagtggATCACTGTTCCTGAAGGTGGCCAAAGCACACTCCAAAGGGgaatcttaaacccaac
ggttacctcttatctagtggTACAGCCCAAGAATGGAGACCACAAGGGAGGCTAGACTCCgaatcttaaacccaa
ggttacctcttatctagtggGCATCAGCGAGTGTGGGTATATAGGAATGCTTAACAAGAGgaatcttaaacccaact
ggttacctcttatctagtggCCCTGCAGGAAATGTTGAGTCAGTTTCCACAGGCCCCCACgaatcttaaacccaac
ggttacctcttatctagtggGATCCCGAAACTCTGTTTTGTAGCATTCCTTCAGCAAAGCgaatcttaaacccaactt
ggttacctcttatctagtggTTTCAGACTCTCCATGCCTGAAACATACAGAGTCATCAACgaatcttaaacccaactt
ggttacctcttatctagtggGTTTTCTCTTTGAGGGTAGTGGGTTTGAAGAAATAAACAGgaatcttaaacccaacttg
ggttacctcttatctagtggGGAATGACTTGAAGGCAAAAAGCATCGTGGCTGTTGGGACgaatcttaaacccaac
ggttacctcttatctagtggGTTGCAGAAATAGGGGTCCATTCAAAGCACCTTGAATGCCgaatcttaaacccaac
ggttacctcttatctagtggCTAATCAACCAAAAAGACTGGCTTGATGTACATTAAGCTTgaatcttaaacccaactt
ggttacctcttatctagtggACTGGAACTTCTATATCCAGCTTCATTCTTCTAATACTCCgaatcttaaacccaacttg
ggttacctcttatctagtggCATCATGATATCACTGAGGACAAGGCACCATTGGTTTTTCgaatcttaaacccaactt
ggttacctcttatctagtggGACGTAGGATGTGGCATAGCATGTCTCCTTGAGATGTTCCgaatcttaaacccaact
ggttacctcttatctagtggTTATTTGGAGGTCTTTGAGTGTTTCTTTCGGGGACTCCCCgaatcttaaacccaactt
ggttacctcttatctagtggCCAGGCTCAGCTTGTGGTCACATACTCACTTTACCTCTCCgaatcttaaacccaact
ggttacctcttatctagtggACATAATCTACAAACCAACCTGATCTTCTGCATTGAGATCgaatcttaaacccaactt
ggttacctcttatctagtggGGTCCTTTTCATTTGATAAATTTCAGCCTGATGTGTCTGGgaatcttaaacccaacttg

ggttacctcttatctagtggTCCTATACCTATGGGACACCCTTGTAACCAACCATAAGGCgaatcttaaacccaact
ggttacctcttatctagtggGGGGCATTTCTCAGACCTTGGAGAGATGGAGAACACCGCCgaatcttaaacccaac
ggttacctcttatctagtggCCTCTGAGAAAGCCTGCCTGCCTTCACTCTGAAGAGAGCCgaatcttaaacccaac
ggttacctcttatctagtggGATGGAGAAAAGCAGAATCCTTTGTATTCATACTTAGAGCgaatcttaaacccaacttg
ggttacctcttatctagtggCCTCTTTGTGGTCTCCAGGGAACAGGAATTTGAGAGTTCCgaatcttaaacccaactt
ggttacctcttatctagtggACAAGGAAGAAGACAGACCTTTTGCTTGCCCTCAAGGATGgaatcttaaacccaact
ggttacctcttatctagtggCAGGAATAGAAGTTCTAAGAGGCATTGTTTGGTCAATGTCgaatcttaaacccaacttg
ggttacctcttatctagtggTGGGACTTTCCTTTGTTTCTATTTACCTCCCTTTAAGTTGgaatcttaaacccaacttgt
ggttacctcttatctagtggGGATTTTCCAAGTTGTCTTTCTATGAGAAAGTATGCTGTGgaatcttaaacccaacttg
ggttacctcttatctagtggCTTGGCTTTGTTTCTAGTTATGCAAAAGTAAATGATGAGCgaatcttaaacccaacttg
ggttacctcttatctagtggTCCTTCTAACCTCATCTCCCCCTAAATAAGGCAAATTTGCgaatcttaaacccaacttg
ggttacctcttatctagtggCATTGGCCTTCTTCATGGTTTTTGGATCCTGACAACTATCgaatcttaaacccaacttg
ggttacctcttatctagtggGCCTGGCAGGAAGTTATATTTAGTAAGTACTGTGTGAAGCgaatcttaaacccaacttg
ggttacctcttatctagtggTACTCAACCCTTGTTGCTCAGTTCCAATACTCATGCTTCCgaatcttaaacccaacttg
ggttacctcttatctagtggTCTCACCAAGTTCAGAATAAGCTGACACCAACTGGTTCCCgaatcttaaacccaact
ggttacctcttatctagtggGGGGGGTGGGGGTGGAGGAGAGAGAGAAAGAGAGAGAGAGgaatcttaaacccaa
ggttacctcttatctagtggCTGATGCGAGACTCAGCCAGTGTTTACTCAATTGTCTCACgaatcttaaacccaactt
ggttacctcttatctagtggTCTGATGTTGTCCCAGCATTCAATCATACTGTCAAATTTGgaatcttaaacccaacttg
ggttacctcttatctagtggTGTCTCCATACTTTCTTTCCTTTCAGATTTGCTTTGTTCCgaatcttaaacccaacttgt
ggttacctcttatctagtggTTCCAGCTCCTTCCCCTCCTCTTCTGATGTCTCTACTTTCgaatcttaaacccaacttg
ggttacctcttatctagtggGCATGAGCCACTGTGCCTGGCCTCTCTACTTTCTCAATAGgaatcttaaacccaact
ggttacctcttatctagtggCGATATTTCTGAACTTCAGATCCCTCATGTGTAATATGAGgaatcttaaacccaacttg
ggttacctcttatctagtggTATCTGCTATATCCATTTCAAAGGGTTGTTGTGAGGAATGgaatcttaaacccaacttg
ggttacctcttatctagtggTCCAATAGATGTACTGAATTCATACCTCGAGTAGAGCAAGgaatcttaaacccaactt
ggttacctcttatctagtggACACTGTAAACTAGGACACACTGGTAGTTCTTACATGAAGgaatcttaaacccaactt
ggttacctcttatctagtggCCATGAGGGAGATTCTTGCAGAAACATTTCCTGGGACTGGgaatcttaaacccaact
ggttacctcttatctagtggGTTGGGCTTTGCTTGTTGCTCCAAAATTGTGAGTAGACCCgaatcttaaacccaacttg
ggttacctcttatctagtggGCTGTTTCTGACAATCAGCCCGGGCACCTAGTCACTTCCCgaatcttaaacccaac
ggttacctcttatctagtggGGCAGCTTGCTCCACGCCGGTGCCTACCTGTGCAGATGGGgaatcttaaacccaa
ggttacctcttatctagtggAAGTGGCAAGGAGGCAGAGAAAGCATCTGTACCCTTACAAgaatcttaaacccaac
ggttacctcttatctagtggTGAGTACAAAGAGCTACCTATGGTGGTCTGGAGCCGGAGGgaatcttaaacccaac
ggttacctcttatctagtggCAAAGGATATAAGATGCATAGCCAACTGAGGAACCTGAGCgaatcttaaacccaac
ggttacctcttatctagtggAGCCCCGCAAGGCCTGAAGGAGAGCTGACATAGGCACCCCgaatcttaaacccaa
ggttacctcttatctagtggGGCCATGAACACCAGTGGAGAAGAGGCAGAAACACCAATGgaatcttaaacccaa
ggttacctcttatctagtggCTTCCCATGTGCCTCTAAGTGCCAGTGCAGGCCCCACAGGgaatcttaaacccaac
ggttacctcttatctagtggGGGAGAAAGGAGATGACGCAAAGGAACCTAACTGGACTTTgaatcttaaacccaac
ggttacctcttatctagtggTGGAACCTCCCAAGATAATGCCAAGGGTCAAAGGGTGCGCgaatcttaaacccaac
ggttacctcttatctagtggATAGTTGATGCAGCATGGAGTAGTGGGGATGTGGCCCAAGgaatcttaaacccaac
ggttacctcttatctagtggCAGCCACTCTGCCCAGACTTCAAGGTGGAAAGAACTCAGCgaatcttaaacccaac
ggttacctcttatctagtggCCCACTACCTGTTTCCGCATAATATGCCCTCTAATTGTTCgaatcttaaacccaacttg
ggttacctcttatctagtggAGAGTAAGAAGATGCCTGCACAAGACTGTCCATGTGTGGGgaatcttaaacccaac
ggttacctcttatctagtggTCATCGTCATCCTGCAGTGAAAGCTGTCCTTGGAGCTCCCgaatcttaaacccaact
ggttacctcttatctagtggTGCTGGTGCCCTGCCTTAAATGGAGAAATATGGAGTTATCgaatcttaaacccaactt
ggttacctcttatctagtggATTATAGGGAAGAAAAGCCCAGCCATTTCAAAATCACCAGgaatcttaaacccaact
ggttacctcttatctagtggGACAGATACTTTCTACTAGGATGTTGAAATGTCTTCTGAGgaatcttaaacccaacttg
ggttacctcttatctagtggGTGGTCCATACATCCTCTATCACAATGACATGACTATGCCgaatcttaaacccaactt
ggttacctcttatctagtggGCAGGTACCAGGTTGGATTTGACTCCTGCGCTATAGTTTCgaatcttaaacccaactt
ggttacctcttatctagtggTATTCCAAAGTAGAAGTTACATAAGCAAACATCCCTCCTCgaatcttaaacccaacttg
ggttacctcttatctagtggGGAAAAAATACTACAGGGAAGCACAGGCATACATTTTCCCgaatcttaaacccaact
ggttacctcttatctagtggCCATAGGTTGGTCTTTAAGAAAAGGAATTATTGAGTAGAGgaatcttaaacccaacttg
ggttacctcttatctagtggACTTTCTAAAGGGGCTCATTCTCATCTACATTCCCAACAGgaatcttaaacccaacttg
ggttacctcttatctagtggCCAAGCTTACTTTAGTGAATCCTCTGCATCATCAGGATTTgaatcttaaacccaacttg

ggttacctcttatctagtggTCCCTTTGGTGACTTTCATATTCATGACAACAGCATTTTCgaatcttaaacccaacttg
ggttacctcttatctagtggCCCTTTTTGTGCAGTCTGATATCCCATGTATGTTGAGTGGgaatcttaaacccaacttg
ggttacctcttatctagtggGCCCCAAATGTCTCTGTGTTTCCTCCCAATAACACTAGCCgaatcttaaacccaact
ggttacctcttatctagtggAGGCACTTAGGTCTAGGGAATGCCCCAAATATTGTATACAgaatcttaaacccaactt
ggttacctcttatctagtggGCTGGTGCACCTGCCTCCAAATCCCAACAGTAAGCACGGCgaatcttaaacccaa
ggttacctcttatctagtggTCCCTGTTCCCTGGGTTTTGGATTCTTGGTCCTGTACCCCgaatcttaaacccaactt
ggttacctcttatctagtggCCTGTTTGGGGATGCCTCATTGGGCTTCTTGGTCTTCAGCgaatcttaaacccaactt
ggttacctcttatctagtggCTTCCTCCTCTAGTTTTGCCATACATGTGCATGCACAGACgaatcttaaacccaactt
ggttacctcttatctagtggCCAGAATGCTTCCTACGGGAGAGGAGACACAGCCTTAAGCgaatcttaaacccaac
ggttacctcttatctagtggAAGATGGGTCCAGATTTAAACATTAACAGGGTGAGATCAGgaatcttaaacccaactt
ggttacctcttatctagtggAGATTGGTGCTCTAGTTTACATGTGTGAGGGCCTTTGTAGgaatcttaaacccaacttg
ggttacctcttatctagtggGCACTGCTGATCCCCAACCTTCGGATTCTCTTTTATGCCCgaatcttaaacccaactt
ggttacctcttatctagtggTGGCCCACGTCAAGAACCACATATTTGAATATCATGTAGGgaatcttaaacccaactt
ggttacctcttatctagtggTTGGTTGTGGCTTGTACTTAGAAAAACTGCTTGATACTGGgaatcttaaacccaacttg
ggttacctcttatctagtggGGACTCATGCTTACCAAATAGGATGTTGCTGAAGCATCCCgaatcttaaacccaact
ggttacctcttatctagtggGATCCCAGTGTTTTAAACGCCTGCCTAGACTAGAAGGTGGgaatcttaaacccaact
ggttacctcttatctagtggCACTTACCCAGACCCATTGCCTGTTTCCTCATAATATGCCgaatcttaaacccaactt
ggttacctcttatctagtggGGGCAAGACCATCCCATATCTGGATAACGCAACCACTAGGgaatcttaaacccaac
ggttacctcttatctagtggTCTCTCAGTGAAAGCTGTCCTTGGAATTTCCTTCTTGCACgaatcttaaacccaacttg
ggttacctcttatctagtggGATAACAGCAAATATTGCCAATGCCCTGGCTTAAATGGAGgaatcttaaacccaactt
ggttacctcttatctagtggGACCACCTTGAAATCTCCAGCATTTTTTCACTGGACAGGCgaatcttaaacccaactt
ggttacctcttatctagtggTTGGTCAAGCTGGTGGCCCCACGTTCACCTCCCTGGACCCgaatcttaaacccaac
ggttacctcttatctagtggTGAGGGATCTCTTCTTGAATCAGAACTACCATTCCTGGGGgaatcttaaacccaactt
ggttacctcttatctagtggTGCTTCTACCTAGAATAGAGTCTTACTCCCAGATCATGAGgaatcttaaacccaacttg
ggttacctcttatctagtggTGCTATGAAAGAGATCCTAGATTTACCCCCAAGTTCAGAGgaatcttaaacccaactt
ggttacctcttatctagtggTTGAGGCAACCACTGTTTTGCCCATGTGATTCTGCTTTCCgaatcttaaacccaacttg
ggttacctcttatctagtggGATCCCTTTATAGTGAGTGACACAATTTTTCTGATGGTAGgaatcttaaacccaacttg
ggttacctcttatctagtggGGTTCGAGTCCTGCTTATGCATTTTATAAGTCTTGCCTACgaatcttaaacccaacttg
ggttacctcttatctagtggAGCTCCAGTTTCCTGATGGAGATAATAACACATGTGTTGCgaatcttaaacccaacttg
ggttacctcttatctagtggGGAACACAATGGGGATTCATAATGTTATTTTCTCCCTGTCgaatcttaaacccaacttg
ggttacctcttatctagtggCGGAATCCAGGGAGGCAGTGCATTTATGGGCTAGGGTAGGgaatcttaaacccaac
ggttacctcttatctagtggGCAGTGCTCTCCAAAGACCTAAGAGAAGAGGCTGACGCTGgaatcttaaacccaac
ggttacctcttatctagtggTGCACTTCTCAGCAGAGTACTCCCACCCTTGGCTTCTCCCgaatcttaaacccaac
ggttacctcttatctagtggGTTCCCAGGACAAGCGACATCAGCATCACCTGAGAACTCGgaatcttaaacccaac
ggttacctcttatctagtggGCTTGGACCGCATTCTAGACCTACTAAATCAGAAACTCTGgaatcttaaacccaactt
ggttacctcttatctagtggTACTTCAGGTGTCCCCTCCTACACCAAGGAAAAGGCAGCCgaatcttaaacccaac
ggttacctcttatctagtggCAGGTGTCCTCCCATGTGAGCCTTAGGTTAGCAAATGCTCgaatcttaaacccaact
ggttacctcttatctagtggCAGTACCCTAGGCGAGGATCCTGATATAGGTCAGGGATGTgaatcttaaacccaac
ggttacctcttatctagtggGGCAGGCTTATAACTCCTGCTTCCCAGGTCATAGGATTCAgaatcttaaacccaact
ggttacctcttatctagtggCATACTGCTGCTGCATGCTTTCATCACTCTGCCCCATCCCgaatcttaaacccaact
ggttacctcttatctagtggGAATATGAGAAAATTTTAATGTCCACAACTGTCCTGCAAGgaatcttaaacccaacttg
ggttacctcttatctagtggTCACACTCCATGTCCCGAAGCTCGCTGCTGAGTTCACCTGgaatcttaaacccaac
ggttacctcttatctagtggCCAGAAGCTGGAAACTGCCAAGGGTAGCTCTCACACAGTGgaatcttaaacccaac
ggttacctcttatctagtggGACAGGGGATAAATATCCTAGCTGGCCTCTCCCTTGGCAGgaatcttaaacccaac
ggttacctcttatctagtggGGTCCCCAGAAGGATTGAACCCCTACAGTGGCAATGGCTCgaatcttaaacccaac
ggttacctcttatctagtggGCTTTCTCTCCCTTCTGTCTTACTTTCCTCAGTTCCTTCCgaatcttaaacccaacttg
ggttacctcttatctagtggCAGGATCTACTTCAGGAACCCAAAGCAAAGTAGTTGGATGgaatcttaaacccaact
ggttacctcttatctagtggCATGAGTGCAAAAGTAATTGCAGTGTTTGCATGGTGGGACgaatcttaaacccaactt
ggttacctcttatctagtggAGACCATGGAAATAATATCAGACAAAAAGCAGATTAGAGCgaatcttaaacccaactt
ggttacctcttatctagtggCGGAGACAAACCGCAACATCACCAACGCCTTTGGCCCAGGgaatcttaaacccaa
ggttacctcttatctagtggGAGGAGCATAGTGACCAGCCATTGGAAGTCGACAAAGACCgaatcttaaacccaac
ggttacctcttatctagtggACAACTACACGAGAAGTTGTCAAAGAACGCAATGTTGACCgaatcttaaacccaact

ggttacctcttatctagtggAGGTGAAAAACTTGATAAGTGGGTGCCTTGTGAGCTCAGCgaatcttaaacccaact
ggttacctcttatctagtggGTGCAATGAAAAGTGGATTTGGGGCCGGGCGCGGTGGCTCgaatcttaaacccaa
ggttacctcttatctagtggGGATTTTATATATGGCAACCAGCAATGACCAGCTCAGTGGgaatcttaaacccaac
ggttacctcttatctagtggTGGTGGTCTGCTGCTGGTCTGATCCACCGCTGCTCTCTGAgaatcttaaacccaac
ggttacctcttatctagtggTGCAGCATCTGCAGCTGGCATTGGTCAACATAACGGGTCCgaatcttaaacccaac
ggttacctcttatctagtggTCCTCATCCGCCATATTCACCTGACGTCTTGCCAACTAACgaatcttaaacccaact
ggttacctcttatctagtggGCAGGATGCAGAACACGCTTTCCAAGAGTTTGTCGAATCCgaatcttaaacccaac
ggttacctcttatctagtggTCAAGGTCTGAAACCGCAATTACCTTTGCAACAACCTAGGgaatcttaaacccaact
ggttacctcttatctagtggTTCTTGTCGATATTCCCCAAGGGCCTGGCATTGTGCAAGCgaatcttaaacccaact
ggttacctcttatctagtggATGGGGCCTTCATCCCATATTCTTTAATGCAAAGAAAGCAgaatcttaaacccaactt
ggttacctcttatctagtggTTCTTGTCGATACTCCCCAAGGGCCTGGCATTGTGCAAGCgaatcttaaacccaac
ggttacctcttatctagtggTTAGTAGTTGTTGAATTCTCACCTTCCCAAACCCCTCAGCgaatcttaaacccaactt
ggttacctcttatctagtggCTGTTGACTTTCCCAACATCTATATGACATGCACAGGACCgaatcttaaacccaact
ggttacctcttatctagtggACAACCCAGCAGAACTAAAAGCCCTGCATCTTCAGGACTGgaatcttaaacccaac
ggttacctcttatctagtggTTCCCAATGTTTTCCCATCACTGGAAGAAAACTTACATGGgaatcttaaacccaactt
ggttacctcttatctagtggTCTATTATTATGAGTACCACTCAAAGTAACTGTCCAGAGCgaatcttaaacccaactt
ggttacctcttatctagtggCTGGTACAGCCCACACTCAGAGCAGATTGTGACCAGAGGAgaatcttaaacccaa
ggttacctcttatctagtggGGGGGCTGTTAGATTCTTCAGGCCACACCACTCTGAAATGgaatcttaaacccaac
ggttacctcttatctagtggAAGCATGTTTTAGACATCTATTATGTGCAAAGCCCAAGGCgaatcttaaacccaactt
ggttacctcttatctagtggCTGCTACTATCCCAAATACAGGCCTCCATCCTAATGTGAGgaatcttaaacccaact
ggttacctcttatctagtggGCTCCTAATCTGAAGTTTGCATTAGGTTTTTGGTGAATACgaatcttaaacccaacttg
ggttacctcttatctagtggAGTTCTCTTTTGCAGTTTGTTGAGAGTTTGTCATGAATGGgaatcttaaacccaacttg
ggttacctcttatctagtggTGGTAAACCAGTCTTAATAAACTTGGGGGAATAAAAACCCgaatcttaaacccaact
ggttacctcttatctagtggCTACTGGAGTAGGTAGGCTGATATTTTGCTTAGGTTTGTGgaatcttaaacccaacttg
ggttacctcttatctagtggTCCGTAGTTCTTTTCGTTTTGTCAGGTTTTCTTATGAAGGgaatcttaaacccaacttgt
ggttacctcttatctagtggGGTAAAACGTATCCAGATAACCATCTGACCTAGAGTTTTCgaatcttaaacccaactt
ggttacctcttatctagtggCTTTCATGTTCCTTGATGTAAAGTTGTTCTTAATATCCCCgaatcttaaacccaacttgt
ggttacctcttatctagtggTTCCATAACTGCACTTACGTTCCTCTTTTTATTTTTGACCgaatcttaaacccaacttgt
ggttacctcttatctagtggTGTGTGTGTGAACCATCTTTTAGCTTTGTTGAATCTCACCgaatcttaaacccaacttg
ggttacctcttatctagtggCTGTGGAAGGGGCAGGAAGGGAGCAGAACTGTGCAGAGGGgaatcttaaacccaa
ggttacctcttatctagtggAACAGTGCTGGCCTCAGGAGAGACCCCACCAACCCATGGGgaatcttaaacccaa
ggttacctcttatctagtggGGCTGGGCCTGTGCACCCTGTATGGGCTGCCTCAGGAAGGgaatcttaaacccaa
ggttacctcttatctagtggGCCTGTGTCTTAGGCAATCTCGGAAGGTGATGCCAGCTATgaatcttaaacccaac
ggttacctcttatctagtggCCTAGCAGTCAGAGAAAATATCTTTCCTTGAAGCATCACCgaatcttaaacccaact
ggttacctcttatctagtggGGGGCTAAAATTTTGTCTAAATGCTACTTTAGCTGCATTGgaatcttaaacccaacttg
ggttacctcttatctagtggTCAAAGATTCAAACCCATTCAAGGGCTCCCCATTGCCTCCgaatcttaaacccaac
ggttacctcttatctagtggTATGGCTTGAACTTACTTCTCCCTCATCACTCCTGCACCCgaatcttaaacccaact
ggttacctcttatctagtggATGTCCTCCATCTAGCCAAGTGTGGACTTGCCTTCAGGCCgaatcttaaacccaac
ggttacctcttatctagtggGGTGCAATCCCCGCCTCTCCCGTTCAAGCAATTCTCCTGCgaatcttaaacccaac
ggttacctcttatctagtggTGACCTCGTGATTCACCTCCTTGGCCTCCCAGAGTGCTGGgaatcttaaacccaac
ggttacctcttatctagtggGTAGGCTTGCATAAAGCAGGTACCTTCCCTCTGTGTTTCCgaatcttaaacccaact
ggttacctcttatctagtggGCTCCCACACACCTTACCTCATCATATTGAAATTGCATATgaatcttaaacccaactt
ggttacctcttatctagtggAGGTTAGTGTGGTAGGCAGAATAATGGCTTCCCACAGAGGgaatcttaaacccaac
ggttacctcttatctagtggGGTTGCAGATGTAACTAAAGTTGCCAACTCATCAGCTGCCgaatcttaaacccaact
ggttacctcttatctagtggGGGAGGGGACAGAAGAGTCAGTGCTGGAGTGAGACAGCAGgaatcttaaacccaa
ggttacctcttatctagtggTGGAGAAATAGGAACACTTTTGCACTGATGGTGGGACTGTgaatcttaaacccaact
ggttacctcttatctagtggAAGACAAATGCACACATATGTTTATTACAGCAGTGTTCACgaatcttaaacccaactt
ggttacctcttatctagtggGGCTACCATAATCCAAGGAGGGTGAGCAGCCTCCAGAAGCgaatcttaaacccaa
ggttacctcttatctagtggAAAGGAATGGATTGACTCCTAGAGCCTCCAGAAAAACTGCgaatcttaaacccaac
ggttacctcttatctagtggTGGTGAGCCCACTGAGATCTGTTTTAGATTTCTAACCTCCgaatcttaaacccaactt
ggttacctcttatctagtggCCTAGTACAGATTCTGATAGCTAGAAGTGGGGTGCTGCAGgaatcttaaacccaac
ggttacctcttatctagtggGCCAAGGGGCAGGGGCTGGAAGAATTTTGAGTAGCATGTTgaatcttaaacccaac

ggttacctcttatctagtggCGATTCTGCTAATGAGGACTCAGAAGAAAGTGAAGGACACgaatcttaaacccaac
ggttacctcttatctagtggTGGAAAATGGAGAAAAGGGATCTTTGTTACGCACTAGCAGgaatcttaaacccaact
ggttacctcttatctagtggTATCTGAGGAAACTTCCAAACAAAATGTTGAAGCATTAGCgaatcttaaacccaactt
ggttacctcttatctagtggAAAGGTCTCCAGAACTTGATGATTTGAGAAATTCTCAGCCgaatcttaaacccaactt
ggttacctcttatctagtggGTTAAGAGACCTCAGGAAATAGTCTAATAGGTAAAAGAGTgaatcttaaacccaactt
ggttacctcttatctagtggTTACAGGGACAGTCTAAGTAGGTGGTTTTAAGGGGTCCAGgaatcttaaacccaact
ggttacctcttatctagtggGCTGGAAGCTTTGAACTGAGGAGAAATGGTAGTCAGCAGCgaatcttaaacccaac
ggttacctcttatctagtggGAGCTGGAATGACCCAGGGATACAGGAATTGAAGTATCTAgaatcttaaacccaac
ggttacctcttatctagtggCCTGCCCCAAAACTTCAGTCAAATATGCTGGAAGGAAGGGgaatcttaaacccaac
ggttacctcttatctagtggTCTGCAGGACCAAAAAGGGACACTATTTGAATTAGGTGAGgaatcttaaacccaact
ggttacctcttatctagtggTCCTGAAAATGAATTCCAAGATCTCCCTCTGAAGATCTTGgaatcttaaacccaactt
ggttacctcttatctagtggGAGTTTCCTTCTTTATGGGATATCAAGTCTTGAGGTGAAGgaatcttaaacccaacttg
ggttacctcttatctagtggGGGTGACATCAGTAACCTGTTATACTGAGGACACTCAGGCgaatcttaaacccaac
ggttacctcttatctagtggTGGATCCACTAGGAAAGTCACATCAAAAATGAGCATTTGTgaatcttaaacccaactt
ggttacctcttatctagtggCAATGGTGTCTGTTTCATGCACACAGATGTTCACTGCAGCgaatcttaaacccaact
ggttacctcttatctagtggAGCAATGACATAGAATCAACCTAAGTGTTCTAAATGACAGgaatcttaaacccaactt
ggttacctcttatctagtggATGCAGCCACAAAAAATAGTGAGATCGTGTCTTTTGAGGGgaatcttaaacccaact
ggttacctcttatctagtggACCAAATACTGCATAGTCTCACTTATAGTAGGAGCTAAATgaatcttaaacccaactt
ggttacctcttatctagtggGGTCTACTTGAGGATGGAGGGTGGGAGCAGGGAAAGGAGCgaatcttaaacccaa
ggttacctcttatctagtggGTACAACAAACCCCTCCCCATCACATTTGTTTGCCTATGCgaatcttaaacccaact
ggttacctcttatctagtggGTGTCTGTTTGGGATACTCCAGTTCCATTTCTCACTACTCgaatcttaaacccaacttg
ggttacctcttatctagtggCTTAGAACCCTCCCTATGTGGCAGTAAGCAAACTCTAATGgaatcttaaacccaact
ggttacctcttatctagtggCACTTTCCTCCAAGGTTACTAGAGGTTTGGACTGAGATGGgaatcttaaacccaact
ggttacctcttatctagtggGAGTGTGTTTTCCATCCCAATACAGAAGTAGTGGTTCCAGgaatcttaaacccaact
ggttacctcttatctagtggTCACAGGAAGAAGTCACTCATCCTCTCATCATCTTAACGCgaatcttaaacccaact
ggttacctcttatctagtggGCATTCAATCCCATTTTACATACTTGTGACCCTGGTACGCgaatcttaaacccaactt
ggttacctcttatctagtggCATAGTATACCATAATTTGCAGGCCACATAGGACCATCAGgaatcttaaacccaact
ggttacctcttatctagtggTTATGCCCTTACTGTTTCCTTGCTGTGATTTCCTTCGCACgaatcttaaacccaacttg
ggttacctcttatctagtggCAAGTCATCTTAACTACTGTTTCTTGATGGACAAGACAGGgaatcttaaacccaactt
ggttacctcttatctagtggCTTGGGTTTGCTAGTTATATTGCTTAATGCAAATGACTGCgaatcttaaacccaacttg
ggttacctcttatctagtggTCCAACCTTTGCTTTTAAGAAGTAGTCTGGAGTGGGTGACgaatcttaaacccaactt
ggttacctcttatctagtggGGTGACATTCCTTTAGAACTCAGATCATCCTCAAAATCGGgaatcttaaacccaact
ggttacctcttatctagtggAAATGGCTGGTATTGGAGACAGCTGTTAACTCTACTCTAGgaatcttaaacccaactt
ggttacctcttatctagtggGCTGAAGGAGTGTCATCTTTTCAGTTTTGTTGGTTGCCTGgaatcttaaacccaacttg
ggttacctcttatctagtggCATCGGTTTAGATGCTAGTTATTTACCACCCTTTGATGGCgaatcttaaacccaactt
ggttacctcttatctagtggGGTATTGGATGATCATCCATGGAGACCAGAATGAGATCCCgaatcttaaacccaac
ggttacctcttatctagtggCTGTAGTCTTTTCTCAGCAGCAGTTTCCCCAGATAGTCTCgaatcttaaacccaactt
ggttacctcttatctagtggCCCTTTTACTCATTGTATGCTTTCTGTGTTTACAGAGTTCgaatcttaaacccaacttgt
ggttacctcttatctagtggGTGCCTTAGCACGTAGCAAACCGTAGAAAAATTTTTCGAAgaatcttaaacccaact
ggttacctcttatctagtggAGAACTTTGCCTACATCAGTAGTAGTAACTAGCACTTCAAgaatcttaaacccaactt
ggttacctcttatctagtggCACTTTATTTTTTCAAAATGAGTTTGGTAGCCATCTGGGCgaatcttaaacccaacttg
ggttacctcttatctagtggATGCTTCCAGTCAGTTCTTACTGGACACGAATACCTGCTGgaatcttaaacccaact
ggttacctcttatctagtggACCTGAAATCAACAGGGAGTGCACAAAAACAGGCAGATCCgaatcttaaacccaa
ggttacctcttatctagtggCCTGGAGAAAGGGATGACATGGGGTGAGTTAGCCACTATGgaatcttaaacccaa
ggttacctcttatctagtggAGCATGGCAGAAGAGTGAATCAGTGCCACATAGTGTAGCTgaatcttaaacccaac
ggttacctcttatctagtggATACAGTCTTTCTGACAATATGACACAGTTCAGGGCAACCgaatcttaaacccaact
ggttacctcttatctagtggGACAGGCAGAGAGAGTGAGTCAAAAATGTTAGCTGGTGGCgaatcttaaacccaa
ggttacctcttatctagtggGCACTAAACAACATTGCATTCAAGAAACAGGCTTTGTAGCgaatcttaaacccaact
ggttacctcttatctagtggTGAAGGAATACCACAGAATTTAGAGTTCAGAACATGGCTCgaatcttaaacccaact
ggttacctcttatctagtggGAGAAAAGGCAATCAATGAAGGTAAAATCCAAGATAACTCgaatcttaaacccaact
ggttacctcttatctagtggAATGGACTTTAAGGTGGCTATTATTACTGTTCTCAGTGACgaatcttaaacccaacttg
ggttacctcttatctagtggAAATCAGTGAACTTAAAGATAGGTCTGCAGACATCCAAACgaatcttaaacccaact

ggttacctcttatctagtggGAACAGATTCTTAGGGACTTGTGGGACATACCACATGGTCgaatcttaaacccaact
ggttacctcttatctagtggAATTGGAGTCAAGGTGAAGAACGGGACAGAAATTTTATTTgaatcttaaacccaactt
ggttacctcttatctagtggGAGGGCCAACAATTTATAATGACAGCTGGCTTCTCCTCAGgaatcttaaacccaact
ggttacctcttatctagtggAGGAAGGTATTTCACACTAAAGAAAATGATACTAGGAGGGgaatcttaaacccaact
ggttacctcttatctagtggGACAATTCTGATAGAGAAAGAAATGGATCAATATGGTTGCgaatcttaaacccaactt
ggttacctcttatctagtggAAAGAGAAACCATTAGAAGGCAAGATCCATTAGAGAAGGGgaatcttaaacccaac
ggttacctcttatctagtggTGCTTTATTCCTACAATAGTGCCTGGTATAAAGCTGATGCgaatcttaaacccaactt
ggttacctcttatctagtggATGCCTCTGAAAATGCTACTTTTTACAACTTTATTGAGGCgaatcttaaacccaacttg
ggttacctcttatctagtggATCATCACAATCAAGCTGGCTAACATATCCAACACATCACgaatcttaaacccaact
ggttacctcttatctagtggCTGGAGTGTGGCGGGGTGATAAGAATATGGGTGATAAGAGgaatcttaaacccaac
ggttacctcttatctagtggCCTGAAATCAGACTGCTTGTGCTAGATTCTTGTCACTACCgaatcttaaacccaactt
ggttacctcttatctagtggGTGTATCTTGAGGCATGTTTGTTGTTCTCTCTGAACATCCgaatcttaaacccaacttg
ggttacctcttatctagtggATGAGAACAAAAGCACCTAAATCAAAAAGTTGTCAGCAGGgaatcttaaacccaact
ggttacctcttatctagtggAAGTGCTTGATATAGTTCCTGGCACATAGTAAGTACTCAGgaatcttaaacccaactt
ggttacctcttatctagtggGTTTTCCCCCAGGTTCTAAGTTTTCTGTAATGCATCATTTgaatcttaaacccaacttg
ggttacctcttatctagtggAGCTGCTCTATTTCTGAAAAACCCTCTTACGATCATACACgaatcttaaacccaactt
ggttacctcttatctagtggAGTTGTAGCATACAAGGTTAATATACAAAGGTCCAGTGGCgaatcttaaacccaact
ggttacctcttatctagtggGTATAAGGTCTATATGAAGAGCAAAAACTCTGATGAAAGGgaatcttaaacccaactt
ggttacctcttatctagtggCAAGATGTCAGTGCGTCCCAGCTTGTTCTAATGCAATTCCgaatcttaaacccaact
ggttacctcttatctagtggGCAAACCCAATATCGAAGGGAAAGAATAGAGCAAGAGGACgaatcttaaacccaac
ggttacctcttatctagtggACTATGAAGCCACAGTGATCAAGACAATGTGGTATTGGCCgaatcttaaacccaact
ggttacctcttatctagtggTGGACCTTCGCGTGCAGAAAAATGAGTCTAGACATAGCTCgaatcttaaacccaact
ggttacctcttatctagtggCGAGAAGGTAACAGAAGAAATTTAGATGATCTTGGGTGTGgaatcttaaacccaact
ggttacctcttatctagtggGGTAATGCCAAAAGAATGAGAATGCAAGCCACAGAATGGGgaatcttaaacccaac
ggttacctcttatctagtggGGGTAACCATATGAACAGATGATCCCTAATGTATATCAGAgaatcttaaacccaactt
ggttacctcttatctagtggGCACCAATGCTGGCAAGGATGTGGAGTTACAGGAACTCTCgaatcttaaacccaac
ggttacctcttatctagtggCAGCCACTTTAGAAGTTAAGGGGAGGGAAAGCATTAGGAGgaatcttaaacccaac
ggttacctcttatctagtggACATACTCTTACCCTGTGATCCAGCAAACATAATCTTTGGgaatcttaaacccaactt
ggttacctcttatctagtggTTGTCTGCACAAAACCCTTCACACCAATTTATATGAGAGGgaatcttaaacccaactt
ggttacctcttatctagtggTTCCTAAACTTAGAAACAAGCAAGCTATCCTTCAATAGACgaatcttaaacccaactt
ggttacctcttatctagtggAAGATAGATAGCTCTACCAAGCCATAAAAATACATGGAAGgaatcttaaacccaact
ggttacctcttatctagtggAGCCGATCTGAAAAGGCTACATATTGTATTAGTCCAACTAgaatcttaaacccaactt
ggttacctcttatctagtggCTCCTCTAGGAGTTAGTGGGGAGGAGGGATAAATAGGTAGgaatcttaaacccaac
ggttacctcttatctagtggTCTTTAGGGCATGAAACTATTCTGTACTACAGTAATACTGgaatcttaaacccaacttg
ggttacctcttatctagtggGGAACAACAGCGAGAATGAACCCTAAACTCTCAACTTTGGgaatcttaaacccaac
ggttacctcttatctagtggCAGTGGAGTAGGTCAGACGTGTGGGAACAAGGGGTACGGGgaatcttaaacccaa
ggttacctcttatctagtggTACAAGGGATACAGGGAAACTCTTCTGCTTAATTATTCTGgaatcttaaacccaactt
ggttacctcttatctagtggGTGGCTTGGAAAAAATGTGGGTCTTGTTTTCTACAAAAAGgaatcttaaacccaactt
ggttacctcttatctagtggTAGAGGAGCAGATGTCATGGAAGGCATATCCACACCTGTCgaatcttaaacccaac
ggttacctcttatctagtggAGGCAGTAGTTCTCAAAGTGTAGACCTGCACTAGCAGTGCgaatcttaaacccaac
ggttacctcttatctagtggCAGGAACTCCTTACAAATGCAAATTCTGAGGACCTGCTTCgaatcttaaacccaact
ggttacctcttatctagtggTAGTCCCAGCTATTAGTGTTTAAAACTCTCCCAGTGACTCgaatcttaaacccaactt
ggttacctcttatctagtggAGGGTCTGTGGTTGCATGCAACAGAAACTGTTCCAGACTGgaatcttaaacccaact
ggttacctcttatctagtggAAAGAGGGAAACAACTAGACCTCAGGAAGAGCAATCAAAGgaatcttaaacccaac
ggttacctcttatctagtggGGCAGAGAGAGGAGGCGCTGGCACTGTGACTGATAGTCCCgaatcttaaacccaa
ggttacctcttatctagtggGTGGCAGTTTTAGGGACATGGGCAGCAGGGCAGGCAAAACgaatcttaaacccaa
ggttacctcttatctagtggGGAGGCAGTGTGACAGAGTAGAAGGTATGTGGAGTTTGGCgaatcttaaacccaac
ggttacctcttatctagtggCAGTCTCTCAACTCTGCTGCTTATTACCATTGGACTTTGGgaatcttaaacccaactt
ggttacctcttatctagtggCAGCCTGCACATCAGTTGCAAAATTAGCTGATTTGACATGgaatcttaaacccaact
ggttacctcttatctagtggAGGAGCTGGCACATAGTAGGTAGTGACTTTCGTTGGGCACgaatcttaaacccaac
ggttacctcttatctagtggCACTTCCCACAGGCTTAGGGAGGGCTCCTTGGATACCATCgaatcttaaacccaac
ggttacctcttatctagtggACCCTAGGGGTCAGAATAGGCTAATACTCTTCCTAACATGgaatcttaaacccaact

ggttacctcttatctagtggCAGTTGTAGAAGATTTGTCTGAAGGTGGGGAAAGTAACCgaatcttaaacccaact
ggttacctcttatctagtggGGGGAATGAGTCACCATTTCCCTACCTACATGTATGTGTAgaatcttaaacccaactt
ggttacctcttatctagtggTCTCTGTGGCCCTAACCTGCTTCCATGGCAGCAGCTTGTGgaatcttaaacccaact
ggttacctcttatctagtggGGGCCTCTCTGTGTCATACATACACTCAGAGCCCTAGGGCgaatcttaaacccaac
ggttacctcttatctagtggGCCCTGTGGATGGGTGTGCCTGGGTGTGCGAGCCAGGGAGgaatcttaaacccaa
ggttacctcttatctagtggAGAGCCCTGCCCTACTCTTACTAAGCAGAAAAGGAAAGCCgaatcttaaacccaac
ggttacctcttatctagtggGGGTAGCTAGAATGTTCTCTGCTCCAGAGCGTCCTGTGAGgaatcttaaacccaact
ggttacctcttatctagtggACATCTACTGTATGTTAGAAAACAGGCTTGGTGATGTTGCgaatcttaaacccaacttg
ggttacctcttatctagtggGAGTGGTGGTGGCCAGCTTGTAACTGAGATCTGTGTGGCCgaatcttaaacccaac
ggttacctcttatctagtggGCACCTTGCTGTCAGTGAGAGAGAGACATGGTCAGGCTTCgaatcttaaacccaac
ggttacctcttatctagtggCTATCCCAGTGAGGTAGTGTGAGATGAGAGTCAGATTGCCgaatcttaaacccaact
ggttacctcttatctagtggCTTTCCACTGCATCTTTCTGGAGTAGAAAAAGATGTGCACgaatcttaaacccaacttg
ggttacctcttatctagtggACATGGAAGCTTCTCATGCTCATCTTTATCTTTGTGCAGGgaatcttaaacccaacttg
ggttacctcttatctagtggGAAGAAAATAGACTCAAAAGGCAGAGGACTTTGAAGGAAGgaatcttaaacccaact
ggttacctcttatctagtggGTCCTCTGAAGTCCAACATGTCAGTGTCCTATCAAGCCAGgaatcttaaacccaact
ggttacctcttatctagtggATTACATGCCAGACATATGCTCATTGCAGACATTCCAGAGgaatcttaaacccaactt
ggttacctcttatctagtggTGCAGGGCACTAGAGGACCGCAGAGAGTGGCTCCTCCAGGgaatcttaaacccaa
ggttacctcttatctagtggGGCAGAGAGGGACAGCAGCGGGGGACAGAGAAATGTATGGgaatcttaaacccaa
ggttacctcttatctagtggGCAGAGGCACCCTGTGCAAAAGCAAAGAGGCAGGAGAGAGgaatcttaaacccaa
ggttacctcttatctagtggGGTAAGTGGTAGGCATGGAATTCAACTCTTTTGGCTCACGgaatcttaaacccaactt
ggttacctcttatctagtggGACCTTCAAGGATGATTGAAAGCTGCTGAAAGAATAAGACgaatcttaaacccaact
ggttacctcttatctagtggCTGAGTGCTATACAATCCATATTGTTGACCAATAGGAAGCgaatcttaaacccaacttg
ggttacctcttatctagtggCCAAAGGCCATGACTTTCCCTTGCCCATCTACTATTTGCCgaatcttaaacccaact
ggttacctcttatctagtggGCCACATGCGCCCTACTATTTAGCTTCTCCTTCTCTCTGCgaatcttaaacccaactt
ggttacctcttatctagtggGAGGCTCAACTCACCATGTAAGAAGCCCAGTGACTTTGAGgaatcttaaacccaac
ggttacctcttatctagtggAGAAGCCCAGGGAGCCTTAGCTGTTGCAACCCTCCCAGCCgaatcttaaacccaa
ggttacctcttatctagtggGACGCCTGACTAAAGGGTCCCTTGAGATGACCATAGCCCCgaatcttaaacccaac
ggttacctcttatctagtggTGAGAACCCTCAGTCAGGAAAGCCCAGCCAAGCTGCTCCCgaatcttaaacccaa
ggttacctcttatctagtggAGAGTGACTTGATATGGAGAAATAAGTGATACAGACTGTGgaatcttaaacccaacttg
ggttacctcttatctagtggGCCTGCTCCTCTCTATCTTTGACCTGGCTGCTTGTCCTTCgaatcttaaacccaactt
ggttacctcttatctagtggTTGCCCATGCGTGAAGATTCCACTCAGATCCCACCTTCCCgaatcttaaacccaac
ggttacctcttatctagtggGTCTTCAACAAGGTCAAGATCAGGAGTCCACAGATCTGCCgaatcttaaacccaac
ggttacctcttatctagtggGGAGTTTGTTAAAGGGACGCATGCCCGGGAGCTAGCCCAGgaatcttaaacccaac
ggttacctcttatctagtggGGGGACCAAGCATCTTTACTGGCCTGGTTTCAGTTGGTGGgaatcttaaacccaact
ggttacctcttatctagtggCAGTTTTTGAAAACAAGTCATCTGGACCACAAATTTGAGCgaatcttaaacccaacttg
ggttacctcttatctagtggCACTGGGCATTATACTAGGTGCTGGCAACACAGAGATGAGgaatcttaaacccaac
ggttacctcttatctagtggGACTGGGCAGTGACTCAGAAAGGCAGCTGGAGGTGGGATGgaatcttaaacccaa
ggttacctcttatctagtggCAGCTAGGCAAACAGGTAAAACCAGGGTGAGGAGGCTGCCgaatcttaaacccaa
ggttacctcttatctagtggTCCTTCTACCCATTCCTATTCATGCACAAATTCTTTCCCCgaatcttaaacccaacttg
ggttacctcttatctagtggGTACCCTCCGAAAGATCACTTTGCCCACAGGAGAGTCGACgaatcttaaacccaac
ggttacctcttatctagtggCCCTCTTGCCCACCCACCTTTTCCATTACGGCTATACCTGgaatcttaaacccaact
ggttacctcttatctagtggACTCATCATCTATGCCTAGCCTGGACCCAGCTAACCCAGCgaatcttaaacccaac
ggttacctcttatctagtggTGTTGCAGTTCTGAAGGTAGGAAGTCAAAAATGGGTCTTGgaatcttaaacccaacttg
ggttacctcttatctagtggGGAGGCTCCAGAGAAGAATCCATACCCCTGCCTTTAGCAGgaatcttaaacccaac
ggttacctcttatctagtggTCCTTGACTCATTGTCTTTTCTTCCAACATCAAAGCCAGCgaatcttaaacccaacttg
ggttacctcttatctagtggCATTGGGCCCATATGGATAATCCAGGATAATCTCTCCATCgaatcttaaacccaactt
ggttacctcttatctagtggTGCAGGTTCTGGGGGAATTAGGATGTGGACATCACTAGAGgaatcttaaacccaact
ggttacctcttatctagtggCCAGAGTTCACACCAGAACTGCCATGTCTCCTTACCAGAGgaatcttaaacccaac
ggttacctcttatctagtggATGTCGTTCTCCTACTAATTCTAACTGTGCCCCCTGTTGCgaatcttaaacccaacttg
ggttacctcttatctagtggACCACTATGTCAGTCGTTCAGATAGAATTTCTCTCCTCACgaatcttaaacccaacttg
ggttacctcttatctagtggGAGTTTGCTCTCTCATCATCTACTCCTTTTTCTCACTGTGgaatcttaaacccaacttg
ggttacctcttatctagtggCATGCTAGCTTTGATCCCTCTCTTTGCCTACACTGACATCgaatcttaaacccaacttg

ggttacctcttatctagtggAATCCTCTAAATGCAGGATGCTGGTTGATCCTTTGCCCCCgaatcttaaacccaact
ggttacctcttatctagtggGACCTTCAATGGCATCCACCTTCTCCATGGTGGTTGGGTGgaatcttaaacccaact
ggttacctcttatctagtggGACTTTTCTGTGCCTATCCCCTATATCCTGAAAATACCAAgaatcttaaacccaacttg
ggttacctcttatctagtggCAAGATTACCAACAATCATTGGTTCCTCAACTTTGTTCCCgaatcttaaacccaacttg
ggttacctcttatctagtggAGGCCCAGCTCACTTTCTGTCTCCTTACCAAAGCAATGCCgaatcttaaacccaact
ggttacctcttatctagtggGCCCCATCACCATTGATTATCATAATCAGTGCCTCCCTCCgaatcttaaacccaact
ggttacctcttatctagtggTGTCTATACTCTTGGAGCATGTAAGCATTCTCCCTTGACCgaatcttaaacccaacttg
ggttacctcttatctagtggCTTTATCCCTTAGACACTAGTCTTACTACACAGTAGGTCCgaatcttaaacccaacttg
ggttacctcttatctagtggGTGATACCACCATGACAGGTTGGAAACTCATTCATCTGGGgaatcttaaacccaact
ggttacctcttatctagtggTTTTCTTCAACCAGATACTTTTCCCCAACAGGAGACTCAGgaatcttaaacccaactt
ggttacctcttatctagtggGTCACTTGTAGTGACAAGCTTTCTTTTTCAAGTTTGCAAAgaatcttaaacccaacttg
ggttacctcttatctagtggAGAAGGGTGGAAATAAAGTCAGTGGGCTGGGGATGTGGGGgaatcttaaacccaac
ggttacctcttatctagtggGGCTCCATGCAGCTTTTGACAGTGGTTCAAAGCTGTCACGgaatcttaaacccaact
ggttacctcttatctagtggTGCGTCAGCAGAGAAAGGTTTCAGTATTCCCACCGAAACCgaatcttaaacccaac
ggttacctcttatctagtggACATGTTCTGCTGGAGGCACTGAAAAGCTTAGCATCTCCCgaatcttaaacccaact
ggttacctcttatctagtggGGTTATTATGGGAGCTCACGTGGCTCAGGCAAGGCTGGGCgaatcttaaacccaac
ggttacctcttatctagtggTGGCTTTGGGCGCTTCAGTGGCTGGGCAGATCATGTGGTGgaatcttaaacccaac
ggttacctcttatctagtggACACTATGCCCTTGCCTAAGAGCCTCCACTCAGAGACAGGgaatcttaaacccaac
ggttacctcttatctagtggAGCTTTATCAAGTGCTCTGGGAAGCCAGTGATGATGGGTAgaatcttaaacccaact
ggttacctcttatctagtggGGCTAAAGGCCAAGTGGAAGCTAAGGTCATCAGAATAGAAgaatcttaaacccaac
ggttacctcttatctagtggTTGCTAAGATTCAGCTTCTGTCTATTGTCAAAGCTCATCAgaatcttaaacccaacttg
ggttacctcttatctagtggGGATACTCCAGTAAATGAGAAATCAAATGAATGTTCCTACgaatcttaaacccaacttg
ggttacctcttatctagtggCCCTCTGGGGGGCCTTAACAAATCTGCTGTGCAGCTTTGGgaatcttaaacccaac
ggttacctcttatctagtggCTCATATCTGCTCCTAAAGAAGACTTGCTTCCCCGCCCCCgaatcttaaacccaac
ggttacctcttatctagtggCAAGTTTTAGAGGGAAGAAACAATTTGGCCAACCACTGACgaatcttaaacccaact
ggttacctcttatctagtggGTGTAAACCAGGTTCTCATCCGTAGCTGCCCACAGGGTTCgaatcttaaacccaac
ggttacctcttatctagtggAATCCCAGAATCGTCGTTCCTTACCTGAGGATTATACTTCgaatcttaaacccaacttg
ggttacctcttatctagtggTGTAGTATCTTCTGCCTCTGGCACAAACACTGGCATGCAGgaatcttaaacccaact
ggttacctcttatctagtggCTGAGGTTGCTTTTTCTTCTTTGGAGTCAGCTTTCTCTGGgaatcttaaacccaacttg
ggttacctcttatctagtggTCAGTTTGTTGAATTTGCCATGTTCCCTCCCACCACAAGGgaatcttaaacccaactt
ggttacctcttatctagtggGGTGCCTTCATTGACATCTGCTGGGGAACCTATACCCCTCgaatcttaaacccaact
ggttacctcttatctagtggCCTCATCATACCCTTTCTTGAGACCATTATTCTTTCACTCgaatcttaaacccaacttg
ggttacctcttatctagtggGGTACAGGGTGTATGTGTGCACATTCCCACTTGCACCACCgaatcttaaacccaac
ggttacctcttatctagtggATGCTTACACACAGGAAGTGCTCAAGAATTATTTGCATCCgaatcttaaacccaacttg
ggttacctcttatctagtggGCATAAAGGCTGACAAGTACAGTTATCTAGCAGGTGCAGGgaatcttaaacccaact
ggttacctcttatctagtggCTATACTTCTTGTCAGAAAGACTTAGAACCAAAAGAATGGgaatcttaaacccaacttg
ggttacctcttatctagtggGGTATCATTTTTGTGGCATTAATAGCACCCATTCACACTGgaatcttaaacccaacttg
ggttacctcttatctagtggGTTTGGGTTTGTAATTCATGTTCTCTTTTCATCCCTTCACgaatcttaaacccaacttgt
ggttacctcttatctagtggGCAGGTTCTGGAGAAAAGCGGCTTGGGTTGGACTCCCAGCgaatcttaaacccaac
ggttacctcttatctagtggCTCTCTGTGCCTCAGATTCCTCATTTACCAACTGGGGATGgaatcttaaacccaactt
ggttacctcttatctagtggTGAAGTGCCTAGCCCACCGTTTGGCATGCTACCTTTGCACgaatcttaaacccaact
ggttacctcttatctagtggCACTCAACTAAATGTCAACAGAAGAAAGTGGGAAAGATCTgaatcttaaacccaact
ggttacctcttatctagtggCGAAAACCCCCTCAGAAAATCAACGAATCCAGGAGCTGGCgaatcttaaacccaa
ggttacctcttatctagtggATAGATAGACCATTAGCCAGACTCATAAAGAAGAAAAGAGgaatcttaaacccaact
ggttacctcttatctagtggGATCCCACAGAAATACAAACTACCGTCTGAGAATACTATAgaatcttaaacccaactt
ggttacctcttatctagtggGAGGAGCTGGTACCATTCCTTATGAAAACTTTCCAAACAAgaatcttaaacccaactt
ggttacctcttatctagtggTAGCTCAATATCCCTGACGAACATTGACGCGAAAATCCTCgaatcttaaacccaact
ggttacctcttatctagtggGCTTATCCACCACAGTCAAGTTGCCTTCATCCCTGGGATGgaatcttaaacccaact
ggttacctcttatctagtggGACAAACCCGCAGCCAGTATCATACAGAATGAGCAAAAGCgaatcttaaacccaac
ggttacctcttatctagtggGGGAAAGAAATAAGGCGTATTCAAATCAGAAGAGAGGAAGgaatcttaaacccaact
ggttacctcttatctagtggCTTAAGCTAATATGCAACTTCAGCAAATTCTCAGGGTACAgaatcttaaacccaacttg
ggttacctcttatctagtggATAATAGACAAACAAAGAGCCAAATCTTGAGTGAACTCCCgaatcttaaacccaact

ggttacctcttatctagtggAGGAATACAACTTACAAGGAAAGTGAAAGACCTCTTCAAGgaatcttaaacccaact
ggttacctcttatctagtggCAATGCTATCCCCGTCAAGTTACCACTGACGTTCTTCACGgaatcttaaacccaact
ggttacctcttatctagtggACAGTCACCAAAACAACATGGTACTGGAACCACAACAGATgaatcttaaacccaac
ggttacctcttatctagtggAAGACCTAAAACCATACAATCCCTAAAATAAAACCTAGGCgaatcttaaacccaact
ggttacctcttatctagtggAGTGGGCCACTGTGGGAATTTACGGACCCCGAACAGAGGGgaatcttaaacccaa
ggttacctcttatctagtggGTAAAAAGTGTGTGTTTGAATGACATGAAATCAGTGCACCgaatcttaaacccaacttg
ggttacctcttatctagtggCCATAAGGTCTGATTGTCTGCAGGGTTGGGCCAAAAAAGCgaatcttaaacccaact
ggttacctcttatctagtggGGAGGTCTATAAAAGGCCGCTCTGGGAATGTCTGTCTTGTgaatcttaaacccaact
ggttacctcttatctagtggAGGACTGAGATATGCCCTGGTATCCTGCAGTACCCTCAGGgaatcttaaacccaac
ggttacctcttatctagtggACATAGACCCTTATCAATGGTTCTGTTTTTGCCCTTTGCCgaatcttaaacccaacttg
ggttacctcttatctagtggGTGATCTTCGTACTTACTCCCTGTTCTTACAACCCCTCCCgaatcttaaacccaactt
ggttacctcttatctagtggGGTCCTAACAATATGTGATGTCACCCCCCAGTGGCCCAGCgaatcttaaacccaac
ggttacctcttatctagtggGGGGCGGATTCCCCCAATAGTAGGCGAAGGATATGAACAGgaatcttaaacccaac
ggttacctcttatctagtggCTACTATAAAGACACATGCTACATATGTTTATTGCAGCACgaatcttaaacccaacttg
ggttacctcttatctagtggGGATGAGTTCATGTCTTTTTCAAGGACATGGATGAAGCTGgaatcttaaacccaacttg
ggttacctcttatctagtggCCACCATGGCATGTGTATACATGTGGAACAAACCTGCATGgaatcttaaacccaact
ggttacctcttatctagtggCACCTACTACATGCCAAGAACTACCAAACATAATTCAGGCgaatcttaaacccaact
ggttacctcttatctagtggCTCATTTAATCTTCACAACGTCCTCAAACATGGTTAAAACgaatcttaaacccaacttg
ggttacctcttatctagtggAATTCTCCTACTTCGTGAACATTGGTTCTTTAACGGGCAGgaatcttaaacccaacttg
ggttacctcttatctagtggGGAGGGGGATGGCCATCATTCAGATTTCCCCATGTAGGACgaatcttaaacccaac
ggttacctcttatctagtggGAAAAGAAAGAGCCCAAAACAGCACTGGGACCTTAAACCGgaatcttaaacccaac
ggttacctcttatctagtggTGACTAGCATCTCTCCATTCTACTACCCACCCCAGCTCCCgaatcttaaacccaac
ggttacctcttatctagtggGCATCATCCATGCTATTGCAAAATGACAGGATTTCCTTCTgaatcttaaacccaacttg
ggttacctcttatctagtggCACATTTTCTTTGTCCATTATCTCTTGATGGACACTTAGGgaatcttaaacccaacttgt
ggttacctcttatctagtggCTTGGCTATTGTGAGTAATGCTGCAGTTAACACAGGACATgaatcttaaacccaacttg
ggttacctcttatctagtggGATCACCATCTACCCCTATAACCACCAGCCCCAACCCAGGgaatcttaaacccaa
ggttacctcttatctagtggCTGAAGGTGAGGCTAGAAATCTGTTTTTAAAATATCCTCCgaatcttaaacccaacttg
ggttacctcttatctagtggTTGCATGCTGAAGTATGAGAAAAACTGAATTGAATGATGCgaatcttaaacccaacttg
ggttacctcttatctagtggTGTCTCCCACACTACTCTTTCTCTTTCTCCTTCTACCTCCgaatcttaaacccaacttg
ggttacctcttatctagtggCTCCCACTGTCTCCTATCCATTTTGCCTCCTCTCCCCACCgaatcttaaacccaact
ggttacctcttatctagtggGGGCTTGTACCCTTTCCAATCTTCAGGGTTCTGTAAACAAgaatcttaaacccaacttg
ggttacctcttatctagtggATGAATGCATGGGCAAATGAACAAATTACAGAAGGTGTCCgaatcttaaacccaact
ggttacctcttatctagtggCTCAAATGTCTTCACGTGTGCTAGTTCATGCACCTTTGCCgaatcttaaacccaacttg
ggttacctcttatctagtggAGAGGCAGAAATCCCACCCCTATTCCCTAAATCCCAAACCgaatcttaaacccaac
ggttacctcttatctagtggTGGTGCTTGGATCAAAACATTCAAGGAGGACAAACCGGGGgaatcttaaacccaac
ggttacctcttatctagtggGGCAAGATCATCTCTGGCTGCCCTCTCCCCAGCAGTGGAGgaatcttaaacccaac
ggttacctcttatctagtggGTTACATCTCTTTCCTCATGTTCTTTTGGGCTGGTTCTTGgaatcttaaacccaacttgt
ggttacctcttatctagtggGCTCTTTGATCTGCGTTCCTTCAACAACTGGGTGAAGGGCgaatcttaaacccaact
ggttacctcttatctagtggCCAGGCCAATGTTCTCAACATGTGTTCCTTGCACTACCTGgaatcttaaacccaact
ggttacctcttatctagtggAGATGCAGACTCCTGAGCTCTACCCGCAAACTCCCCAGAGgaatcttaaacccaac
ggttacctcttatctagtggTCTAGAAATCTGTATTTTTAGCAAGCACTTCAGGTGAGCCgaatcttaaacccaacttg
ggttacctcttatctagtggGGGAAACTCCTAAAAAATACTGAAGCCCAGATTTCTCCCCgaatcttaaacccaact
ggttacctcttatctagtggGCTGGAGGAGTGGGTGAGCATTGGATAAAAGTTAGGGTGGgaatcttaaacccaac
ggttacctcttatctagtggTACATCCATTCACATTTAAACTTGTCTACTGTGGGAGGAGgaatcttaaacccaacttg
ggttacctcttatctagtggTCAAGAGCATGACATGAGCCATCAGTAGCATTTTAGAAAGgaatcttaaacccaactt
ggttacctcttatctagtggTAGGATTCACTGAAGCTGTAAAACCAAGATGAAGGCAAGGgaatcttaaacccaact
ggttacctcttatctagtggCTGCTGCACTGAGGAGCCCCTACCTAGCATTTCAACAGCCgaatcttaaacccaac
ggttacctcttatctagtggGTCAACTGTCAAATGCTTGGAAATCATTTAATATCTTCCCgaatcttaaacccaacttg
ggttacctcttatctagtggATAGGGTCTGCTGAGGGAAATACTTTTCACAGAGGGAGGGgaatcttaaacccaact
ggttacctcttatctagtggGGATTAGATGAAACAAGACTACTGTGCCCCGTTAATATTCgaatcttaaacccaacttg
ggttacctcttatctagtggAGCGGATATCATCTCCATAATTTTGCAGGCCGAAGTCTAGgaatcttaaacccaactt
ggttacctcttatctagtggGTGTCCTATGTGGGACAGTGACCTTTTGGAGGCAACCTCCgaatcttaaacccaact

ggttacctcttatctagtggGCTTCAAAGTGTGAAGAAAGGAGGAAGAACTGCTGCTCCCgaatcttaaacccaac
ggttacctcttatctagtggCCTGCACTTTACCACACTCATCTCCCCCAACAACCCCCGCgaatcttaaacccaa
ggttacctcttatctagtggAACTTCTGGTTCAGAGAGTTGAGTTAATCACACTCCTCAGgaatcttaaacccaactt
ggttacctcttatctagtggCCCTCCACACCCCATTCCCATTCCCTCCTCTATCCCATGGgaatcttaaacccaac
ggttacctcttatctagtggGATCCGTTGATATTCCTCTGTGTACCTCTAATGCACTGTGgaatcttaaacccaactt
ggttacctcttatctagtggGAGGGAATTGAGGCCTAGAAAGTAAATTGCCTGGCACAGCgaatcttaaacccaac
ggttacctcttatctagtggCTGGTCTCCAGCTCACTTTTACTCCAAAGTTAGTACTCTTgaatcttaaacccaactt
ggttacctcttatctagtggGCTAGTTAGTGGCAGAGCTTGGATGACAACTCAGGCCTGCgaatcttaaacccaac
ggttacctcttatctagtggATCCTGTGGGCTTCTCATTTTATGATCCAATGACCTGCCAgaatcttaaacccaactt
ggttacctcttatctagtggTGTGGGGCATACGGAGTCTTGTGGAGAGGCCTGATGTGTGgaatcttaaacccaac
ggttacctcttatctagtggGGGCCAAGGAAGGCTATGTGGGCAATCTGTCCTCTCAAGGgaatcttaaacccaa
ggttacctcttatctagtggCTCTAAGGCTGACATGATGCAATAGGTCTGCTGACAGGAGgaatcttaaacccaac
ggttacctcttatctagtggCCTTGTGTGCCTTGGCAGGTGGCCCTGGGAACTCAACTGCgaatcttaaacccaa
ggttacctcttatctagtggGTAAGCACTCAAGCAGGGTGAAATGAAGAGTCAATTCTGCgaatcttaaacccaac
ggttacctcttatctagtggCTACTTGGAGTCTAGTCTTGTGAACTAAGCTTATTGTGCCgaatcttaaacccaactt
ggttacctcttatctagtggCCAGTTTGGATGAGCACATTTCCTGATATGCAAAGGAGAGgaatcttaaacccaac
ggttacctcttatctagtggACAAATACCAAGCTGACTTCTTTCCAACATGACCTTGGGCgaatcttaaacccaac
ggttacctcttatctagtggCTCAAACATGAACTGAAGACAATGACATCCATTTCACAACgaatcttaaacccaact
ggttacctcttatctagtggGCCTGGCACATTATGAGACATGATTGAATGTTAGGTCACCgaatcttaaacccaact
ggttacctcttatctagtggGGTAGAGGACAGCTTAGCTGGGCCTTGAAGCACTTGGAAGgaatcttaaacccaa
ggttacctcttatctagtggGGCACGGGAGCCAAGTGAAACAAAACACTCCCTCCGAGGGgaatcttaaaccca
ggttacctcttatctagtggGCAGCTCTATGAGAACCACCTCCTTTCTGTCATATAAGGGgaatcttaaacccaac
ggttacctcttatctagtggGATGTTTTCCTGGCCCTGTCAATTTTATTCCCGGCTTGCCgaatcttaaacccaactt
ggttacctcttatctagtggCCACAAGACACTATCAGCTCAGACCCAGAAACAGGATGCCgaatcttaaacccaa
ggttacctcttatctagtggTAGACATGGTAACGTCTAGTCTGAGGCTTACAGGGGACTGgaatcttaaacccaac
ggttacctcttatctagtggTGGAGGGAAAGGTTTCTCTGCCTGTCTGACACTTGGTTTCgaatcttaaacccaact
ggttacctcttatctagtggCACCTGAGCTCACCTGTTCCCAGTAGCTGTACTTTTGATCgaatcttaaacccaact
ggttacctcttatctagtggTGCATTAAGCAGAGCCCCTTTCAAGTAGCCCAGGGATGAAgaatcttaaacccaac
ggttacctcttatctagtggGGGAGATGCAGTGACTCTGCTTTAGAAGAGAATTGAGATGgaatcttaaacccaac
ggttacctcttatctagtggCAGCAAGGCGACTTAGAATGAGGTTCGGAGGAGGTGAACCgaatcttaaacccaa
ggttacctcttatctagtggGCAACCCATGGTTACCTGGGGAGATTGTTCCAAATGCAGGgaatcttaaacccaac
ggttacctcttatctagtggCCAGCCTAGAGGTTCTGCACACCTTCCTCAAGGACCATACgaatcttaaacccaac
ggttacctcttatctagtggGGCAGCTGTGTTCTCAATTTTAACTTTTTCCCTGAGGGTGgaatcttaaacccaactt
ggttacctcttatctagtggTGGCCTCTTGAAGTTAAGTGTGGCCAGGTGGGGCTTTAGCgaatcttaaacccaac
ggttacctcttatctagtggCGGTGGTGTCTGATGGGGCTTTAGCAATAACATGTGAACGgaatcttaaacccaact
ggttacctcttatctagtggGAGGCTGATCTGTCAGTTGACATCCCAGAGTGACTGATGGgaatcttaaacccaac
ggttacctcttatctagtggCCCATATGGGACATGTAGAGTGAGTGAGAAATAAATGTTCgaatcttaaacccaact
ggttacctcttatctagtggGACTGCAGGCAAGGAGGTTAAGATCTCTCTCATTATCGCCgaatcttaaacccaac
ggttacctcttatctagtggTTCTTGTTCCCCACAGCCTTTCTCCCCAATGTGCGCATGCgaatcttaaacccaact
ggttacctcttatctagtggTTATTCTGTGACATCAAGAAGCTCAGTGAATCAGTCAGAGgaatcttaaacccaactt
ggttacctcttatctagtggGCCAAACACACAGATTGGCACCTGCAAAGATGAATATGAGgaatcttaaacccaac
ggttacctcttatctagtggCTCTGTCTTCAAGAAGCTAATAGTCTAGTTATGAACTAGCgaatcttaaacccaactt
ggttacctcttatctagtggGGAGAGAGTAGGATGAACTGTACGTAGGTAGCCAGTGATGgaatcttaaacccaac
ggttacctcttatctagtggGCTGAGCCTTAAGTGACAAGTAAATGCTGGGCAGATGCAGgaatcttaaacccaac
ggttacctcttatctagtggGGGTGTGTACAGGTAAGTTAGGTCAGATTGTGATGTTTGGgaatcttaaacccaactt
ggttacctcttatctagtggAGGACAGGATTACAGTTGTGTTTGGGAAGAAGACCTTGAGgaatcttaaacccaac
ggttacctcttatctagtggGGGGCACATTTGGTATGTTTTCAAAATTTCCAGTGGGAGCgaatcttaaacccaactt
ggttacctcttatctagtggTCAAGAGCTGGGCCTAGGTAGAATCAAATTGATTACATACgaatcttaaacccaact
ggttacctcttatctagtggACAGATGATTGACGTGGACATTGGCCAAAACAGGGAACACgaatcttaaacccaac
ggttacctcttatctagtggCCAAGAACGTAGATGTTAGCTGAATGCTTCTTTCCAATCCgaatcttaaacccaactt
ggttacctcttatctagtggGACTGACTATTCTACAGCTCATCCCCACTCCTGTTGATGCgaatcttaaacccaac
ggttacctcttatctagtggTGTTTCTCCTGGCTTTGATCATTTTGGTTTTGGGGAGCACgaatcttaaacccaacttg

ggttacctcttatctagtggGGTGACTCAGCATAAGTAGATGGAAGAGAGCCAAGCTCGGgaatcttaaacccaac
ggttacctcttatctagtggAGAGAAGAATCAGGAGGGTGTAATAGCAAGGGAGTCACTGgaatcttaaacccaac
ggttacctcttatctagtggGAATAGTGTAGTCTACACCTTCAAAGTCAGAAAAATTGGGgaatcttaaacccaactt
ggttacctcttatctagtggACCTCATCAGCAACTTAGGAAAACACAACATATCCAAGACgaatcttaaacccaact
ggttacctcttatctagtggGGCCAGAGTTGGACTGGGAGGTTGAGGAGATGCAGGTTCAgaatcttaaacccaac
ggttacctcttatctagtggGGAGAGACAAACGTGTTTATAAATGCAGGGGAGTGATTTGgaatcttaaacccaactt
ggttacctcttatctagtggCACAGAACATTTATCACTTGCTCTCCACAGACACTCTCCCgaatcttaaacccaact
ggttacctcttatctagtggCTCTTTCTGTCACAGCTCAACCTGCCATAGAGAACCCAGGgaatcttaaacccaac
ggttacctcttatctagtggGAAGGTTAGTTAGAAGGCTTTGCACAAATGCTATTGCACGgaatcttaaacccaactt
ggttacctcttatctagtggAGACAGGAAAAGTAGGATTGTTGCTGTAGCAGAAGTGGGCgaatcttaaacccaact
ggttacctcttatctagtggGGGAAAATAGTTTGGTGTGGAGGAAGGTAAGAAGTAAGGCgaatcttaaacccaact
ggttacctcttatctagtggGAGTAGGTAGCTGGATATGCAAATCTAGAGTTTGGATGAGgaatcttaaacccaactt
ggttacctcttatctagtggCATGAGACCAGAGGAGATCACCAAAGGTATTAGCCTAGAGgaatcttaaacccaac
ggttacctcttatctagtggGAAATTGAGCCCTGGTGTAGAAAAGATGAATAGAAACCAGgaatcttaaacccaact
ggttacctcttatctagtggAAGGAGAAGGCATCCAGGTGGAAAGTGAACCAGGGGAGTGgaatcttaaacccaa
ggttacctcttatctagtggAGTGAGGAAAGTATTGTACGGAAGAGGAAAAAATTAACTGgaatcttaaacccaactt
ggttacctcttatctagtggGGATTTAGTAGTGGCTAATTGGTATGTTGAACCAGGGAACgaatcttaaacccaactt
ggttacctcttatctagtggAGTCAACTCAGTGGAAGCCCTTTTTTCTCACATGAGTCAGgaatcttaaacccaactt
ggttacctcttatctagtggCCAAAAGGGTTGTTGTTTGCCAGTCTTTGGATGTACATCCgaatcttaaacccaactt
ggttacctcttatctagtggAGCCATCTGAGGTCAGAACATGTCTAGATTTTTGCAACCGgaatcttaaacccaactt
ggttacctcttatctagtggGCTATAGCTGCTTTTTGGCCCAGAGGGAATATGCCATGGCgaatcttaaacccaact
ggttacctcttatctagtggCATTTTCATAGTGGGGAGGAGGATATTATTAATTTCCCCCgaatcttaaacccaacttg
ggttacctcttatctagtggCTAAATTTGTATTCACTTGTAATTTGTCTGCTGACATCCCgaatcttaaacccaacttgt
ggttacctcttatctagtggCTTGTCACTGCCCTCCCATTACCTCATGAAAGCCAAACCCgaatcttaaacccaac
ggttacctcttatctagtggCCCTACAGGGACCTCCCATCTGCCAGTCACCCACCCTCTGgaatcttaaacccaa
ggttacctcttatctagtggGCTTCTGGAATGCAAAGTGCAGGTCTCTTTTTTCATAGTGgaatcttaaacccaacttg
ggttacctcttatctagtggTGTCCACAATGCATCGTAGTATATGAAATGTTCCCTGTCCgaatcttaaacccaactt
ggttacctcttatctagtggCCTAGTTGAAGGGGTCAACGAGGGCCTCTATGAGAAGGGAgaatcttaaacccaac
ggttacctcttatctagtggCAAGCAGACCTGCAGTAAAGAGGGGAACTTGTCAGGCCCCgaatcttaaacccaa
ggttacctcttatctagtggGGTGTTGCAGGGGAGTTGGGGATGGCAGGTCCAGGTAAGCgaatcttaaacccaa
ggttacctcttatctagtggGTGAAATGGGAGCCCTTGTGGAGTTTTAAGGAAAGGAGTGgaatcttaaacccaact
ggttacctcttatctagtggGATTCATCCTGGCTCTGATTGAAGAATAATAGAAGCCGGGgaatcttaaacccaact
ggttacctcttatctagtggGTGTTGAGCAAAAGTAGCGAGGCACAAAAGAAGCACAAGCgaatcttaaacccaac
ggttacctcttatctagtggGGGAGGAGATGTGGACTAGTGCCCGTAACAGTGGTGAGGGgaatcttaaacccaa
ggttacctcttatctagtggGTGTGTCTGGTCCTTGCTATAAACTGTTAAAGAGGAAAGGgaatcttaaacccaactt
ggttacctcttatctagtggGGGACGGCCAATGTCTTGTCCAAAATCACAAAGCTAGTGCgaatcttaaacccaac
ggttacctcttatctagtggCAGACATTACACTGAGTTAAAGAAGACAGACACAATGAGGgaatcttaaacccaact
ggttacctcttatctagtggCAACAAGCAAAGTGAATCTGTGGTCTATGGTGATAGAAGGgaatcttaaacccaact
ggttacctcttatctagtggCTGGATAGGGCCATGAGAGAACTATCTGGAGTGATGGACGgaatcttaaacccaac
ggttacctcttatctagtggCTTCGGCCTAAAGCTACTAGAGAATCCGGGTTGCTGACCCgaatcttaaacccaac
ggttacctcttatctagtggGCCATGACTCTAGTAGCCAGTGTAACCCTTTTCCCTTTGAgaatcttaaacccaactt
ggttacctcttatctagtggAAGTCTAGAGTTCAAAGTCATGTCCTGCATGGGCAAAGTGgaatcttaaacccaact
ggttacctcttatctagtggAGCTCTCAAGATCTAGAATGAGGCAGATAACCAAGACAGCgaatcttaaacccaac
ggttacctcttatctagtggCCCTGTAAATGGACTGGGGTAGCTTATCAGGATTTAGAGTgaatcttaaacccaactt
ggttacctcttatctagtggTAAAACTGATAACGGAAGGCGATGCCCTTACAGGAATGGCgaatcttaaacccaac
ggttacctcttatctagtggATTCATGGACTGCATACTTATACAATCTCATTAGTCTGCCgaatcttaaacccaacttg
ggttacctcttatctagtggTCTATGTCTGTATTCTGTAGTTTTCAAGACTTGAGTTTGCgaatcttaaacccaacttgt
ggttacctcttatctagtggGGTCGTTGGGTGAGCTAAAGCCATGATGGAGAGTCAGGTCgaatcttaaacccaac
ggttacctcttatctagtggGCTGGCTTTGAGGGTGATGAATTTGTGAGCACTCAGGCTCgaatcttaaacccaact
ggttacctcttatctagtggCTCCTCGTGAATCAGCTTATGCATAATAATGTCACACCTCgaatcttaaacccaactt
ggttacctcttatctagtggCCTCCACACAGGTATAACGTATCCTGGAAGACATAAAGTGgaatcttaaacccaact
ggttacctcttatctagtggTAAAAATGGATGGATATGGGCCGGGCGTGGTGGCTAACGCgaatcttaaacccaac

ggttacctcttatctagtggGAGGCTGAGGCAGGAGAACAGGAGAATGGTGTGAACTCGGgaatcttaaacccaa
ggttacctcttatctagtggAATGGATGGATATGAAGGTGGAGGCAGAGTTGGGTTAAGGgaatcttaaacccaact
ggttacctcttatctagtggGTATCCTAGTTACCAGAAAAAGGTCACTGGACTGAAGCAGgaatcttaaacccaact
ggttacctcttatctagtggGTTTTGGTTCTTTAAGGTATTCTACTCAGCATAACAGCTCgaatcttaaacccaacttg
ggttacctcttatctagtggTGCTAAGGGTGTCATAACAACATTCCACAAACCTAGTGGCgaatcttaaacccaact
ggttacctcttatctagtggCGATGTCGCTGGTGTTGGCTCCTCCTGAGGGCTGTGAGGGgaatcttaaacccaa
ggttacctcttatctagtggCTGGTTTGTGGATGCCCTTTTTCCTGTGTCTCCTCACACCgaatcttaaacccaactt
ggttacctcttatctagtggCCCCTTTGTATAAGAACACTAATCATGTTGGATTAAGGTCgaatcttaaacccaactt
ggttacctcttatctagtggACGGTCACACTCTGGGGCAGTGAAGGTTAGGACTACAACGgaatcttaaacccaa
ggttacctcttatctagtggCAACCGGTAACATATCCTGATCCTTGGCTCCCTCATCTGCgaatcttaaacccaact
ggttacctcttatctagtggACATAAGAACAAGGCAGTAAAAACAGTGAGCGATAACAGCgaatcttaaacccaac
ggttacctcttatctagtggAAGAGCTCTCTCTTACTGAGAACAGCCATGTGCCTGGCAGgaatcttaaacccaac
ggttacctcttatctagtggGAAGCCAGAGCCTGTAAGCTTAATCACGTCCCCTCCCCACgaatcttaaacccaa
ggttacctcttatctagtggTTCCTGTAGTGGAATCCCAATTAAAGCAATAATGTAGAGCgaatcttaaacccaactt
ggttacctcttatctagtggCTGAACTTTGTAACTTTGGTCGTGTTGCTTTCTGTGAATCgaatcttaaacccaacttg
ggttacctcttatctagtggGGCACAAAGTAAGCTATAGACTCAAAAATGGTGATTCCAGgaatcttaaacccaact
ggttacctcttatctagtggATCCCTGAGGTCCACTCCATAGCTTGATCAAAGAAGTGGGgaatcttaaacccaac
ggttacctcttatctagtggGACTAGTGTGGCAGCACCTTAAACCCACAAATAAGTTTGGgaatcttaaacccaact
ggttacctcttatctagtggGAGAAGGGAAACCAAAATTATTTCAGGACCTACTATGTGCgaatcttaaacccaactt
ggttacctcttatctagtggATACATAGGCATATGCTATTCCATCAGTGGTTCAGAGGGCgaatcttaaacccaactt
ggttacctcttatctagtggGCTATCTCAGGGATGATTAAATTGAGAGTTGCTCAGGGTCgaatcttaaacccaactt
ggttacctcttatctagtggCCCAGGGCTATTTCATTCCAAGGCCCATGTTACTTTCTGGgaatcttaaacccaactt
ggttacctcttatctagtggTTTTCTTTGAATCCCTAATAGTGGGTGCTATTATACCCCCgaatcttaaacccaacttg
ggttacctcttatctagtggCCTCCAAAGAGATGGCATCTTACAGTTTAATTACTCTGACgaatcttaaacccaactt
ggttacctcttatctagtggGGGTGCTGCTTGACAGTAACAGAATGGTCCATGGGGACAGgaatcttaaacccaac
ggttacctcttatctagtggCAATGATAGGAACACAGAGCTAAGCAGAGTGTCAAGAGACgaatcttaaacccaac
ggttacctcttatctagtggGTGGGGAGGAGACCCTGATGGAAGAAGAGAAGACTTCAGCgaatcttaaacccaa
ggttacctcttatctagtggGGGGTTAATGTTCCAGACTGATGACTTGAAGGAGGGTGTGgaatcttaaacccaact
ggttacctcttatctagtggGAGCAAAGAGAATCCCAATCACTCTCCCTCTCACCCTCCCgaatcttaaacccaa
ggttacctcttatctagtggGGCCAGCCCTTCTCTTCAGCCACATGGTATTTGGACAGCCgaatcttaaacccaac
ggttacctcttatctagtggCCCTGCAGGGAAAGCTCAATCAGCCAGCTGAGCCAGTCACgaatcttaaacccaa
ggttacctcttatctagtggGCCTATCGTGGCTGGAGGGCTAAAGCCTTGTTCTCAAATCgaatcttaaacccaact
ggttacctcttatctagtggAGGGTATATTTATCCTACCCTTCCCCCAGCCTCAACAGGTgaatcttaaacccaact
ggttacctcttatctagtggGGCTGGCCCATGTGGCTCCTGGCATTACAGGCACCAGAGGgaatcttaaacccaa
ggttacctcttatctagtggCTGGGGGAGTATGGAGGATGTGCTACTGTGTGGCAGGTGCgaatcttaaacccaa
ggttacctcttatctagtggACGGGATCCCATATACTATGATTCTGAAATTCACTTGCGCgaatcttaaacccaactt
ggttacctcttatctagtggTCCTAGCTTATCTATGGTTGTCATTATTTCAAGAGCCTAGgaatcttaaacccaacttg
ggttacctcttatctagtggAAAAGGAAGAGATGTGAGTAGGGTCTCACTGTGTTGCACGgaatcttaaacccaact
ggttacctcttatctagtggTTGATTGTCTAAGAGGTCCTGTGCTACAAGCTGGCAGGGCgaatcttaaacccaact
ggttacctcttatctagtggGCCAGGGGCAGAGACAGGTTTTCTGGGGGCATGAGACTTCgaatcttaaacccaa
ggttacctcttatctagtggGCTTCACAGGGAGCTCACTTCTGCCCAGAGCCAAGGACTGgaatcttaaacccaa
ggttacctcttatctagtggGCCTCCCCTGGGGCAGAAATGGTGGCAGCAAAGCAACTTCgaatcttaaacccaa
ggttacctcttatctagtggCCTTTGCAAAGGTTGCATTCCTCCCTTCCCTTGTGGCAGTgaatcttaaacccaactt
ggttacctcttatctagtggTTCAGTGTCTTGGAACACATGATTTAGAATCAGTGTTCAGgaatcttaaacccaacttg
ggttacctcttatctagtggTCCAGCATATGACTAACGGAAAGAAGTCAATCTGAAATGGgaatcttaaacccaact
ggttacctcttatctagtggGGTTGCCAGGGGTTAGAAGGAAGGGAGAGATGAATCGGGGgaatcttaaacccaa
ggttacctcttatctagtggTTTTAGGGCAGCAAAACTGTTCTGTATGATACTATAATGGgaatcttaaacccaacttg
ggttacctcttatctagtggGGTCCATTTAGGCTGCTCTAAGAAAATATCTTAGACTGGGgaatcttaaacccaactt
ggttacctcttatctagtggGGCACTGGCAGATTCAGTGTCTGGTGCAGGCGATCCTCTTgaatcttaaacccaac
ggttacctcttatctagtggCACTGGTGGAAGGGGCAAAGCAGCTGTCTAGGGCTTCTTAgaatcttaaacccaac
ggttacctcttatctagtggAGGTCCCACCTCTTAAAACCATCACACTGGGCTTTAGGTTgaatcttaaacccaact
ggttacctcttatctagtggTTGGGGAGACATAAACATTCAGACCACAGAAGCACAGCACgaatcttaaacccaa

ggttacctcttatctagtggGTGATGTCACGGAGAAAAACTTCTCCTCGACTCTCTTAGGgaatcttaaacccaact
ggttacctcttatctagtggGCATAGGAGTTCATGGAAAAGAAATGAAATTCAAAGATGCgaatcttaaacccaact
ggttacctcttatctagtggGGGGCTTCAAATGATGCTAAATTGTGGGGAAGTGACTAGGgaatcttaaacccaact
ggttacctcttatctagtggGAGTGGCTCTCCTCTTCCCAGTTCAGGAGAGGGAGACACCgaatcttaaacccaa
ggttacctcttatctagtggTGCCCTGATTTTAATCTGATAAGAAGAGGGCAGAGAACCCgaatcttaaacccaact
ggttacctcttatctagtggTCAATTGCCTTCAGCTCAAAATGGTCCTTACGTGGAAGTGgaatcttaaacccaact
ggttacctcttatctagtggGTAAGAGAGGTTGTACTTGCGTGGAGGCAGGATGAGTAGGgaatcttaaacccaac
ggttacctcttatctagtggCAGAGGGGGAAGGGACTCCAACTATAAAACAGCAAGCTTCgaatcttaaacccaa
ggttacctcttatctagtggGATCAGAGAACGTTTTACCCTCAGGCCAGCCGCTTCCTGGgaatcttaaacccaa
ggttacctcttatctagtggCTGGCTCGACTTAGAATGCATCAGAGTTCAAGAGTCTGGGgaatcttaaacccaact
ggttacctcttatctagtggGGTCAGGTTAATAGGCATTCTGTTCTAATAGATCACTGGGgaatcttaaacccaactt
ggttacctcttatctagtggATGAAGCCAAGAACAGGAGTTTGGAGCGTCTGTATCTGGCgaatcttaaacccaac
ggttacctcttatctagtggGCCGGCAATCCTTGAGCCTCATCTAACTCATATCGGGAAGgaatcttaaacccaac
ggttacctcttatctagtggTTGTGCCTTGGAAGGAAGGGGTTGGTGATGGTAAGTCTGGgaatcttaaacccaact
ggttacctcttatctagtggTCTGGTCCCTGAAGAGACATCCCATATGCTCTCGGGTGAGgaatcttaaacccaac
ggttacctcttatctagtggATGCTGGAAGAATGAAGTGGATGACTACCAAGCCCATCTTgaatcttaaacccaact
ggttacctcttatctagtggCAGACAGGGCACTGGGATAAGAGGCGCGAATTAGGATATGgaatcttaaacccaa
ggttacctcttatctagtggCTGTACTAATTTGCATGAATAGAAATGCAGAGAGAGCTCTgaatcttaaacccaactt
ggttacctcttatctagtggTCGTCATGTTAGGCAACGTATTTTCAATGTCATCTTATGGgaatcttaaacccaacttg
ggttacctcttatctagtggTTGGTCACCATCATTATTCACACGATTGTTATTCACCTGGgaatcttaaacccaacttg
ggttacctcttatctagtggTCTGGCTCTTTCTGGGGAAAAGGGCAGGCTTAATACAGGCgaatcttaaacccaac
ggttacctcttatctagtggTTAGTTAGCTTCACTAGCTTCACTTTATTGTGCCTCACAGgaatcttaaacccaacttg
ggttacctcttatctagtggGGTTTGTGGCAACCCTGCCTCAAGCAAACCTATCAGCACCgaatcttaaacccaa
ggttacctcttatctagtggCAACATGGGCTCATTTCCTGTTTCCACGTCACATTTGGGTgaatcttaaacccaactt
ggttacctcttatctagtggGGGGAACCATGAACCACAACCATATTAGGCAGCAAGCTTAgaatcttaaacccaac
ggttacctcttatctagtggTGGCCATTCCCCCATCTCTTTCTCTCTGCTCAGGCCTCCCgaatcttaaacccaac
ggttacctcttatctagtggGCCTCTAAGTGTTCAAATAACAAGAAGAGTCCCACGTCTCgaatcttaaacccaact
ggttacctcttatctagtggAAGGTTACTGAGGAGGGCATATAGAAAGCTGAGACAGACCgaatcttaaacccaac
ggttacctcttatctagtggATGCAGAACAGTTAACCATGTTGTGAATGCAAAAGAAACGgaatcttaaacccaact
ggttacctcttatctagtggGAGAGGTGAGGAAGCTGCAAAGGAGAAGTTGGAAGCTAGCgaatcttaaacccaa
ggttacctcttatctagtggGCCACCTCTATAATATAAAAGTACAAGGAGAAGCAGCAAGgaatcttaaacccaact
ggttacctcttatctagtggCTAGGACTTTCATACCTATAGAGGAGAAGTCAATGCCTGGgaatcttaaacccaact
ggttacctcttatctagtggTAAGCTGACTCTCTTGTTAGGGGCTATTGTATCTGGTGGCgaatcttaaacccaactt
ggttacctcttatctagtggGCCTGTGCTCTGTAAGCAGAACAAACTGGATAGCAGCACGgaatcttaaacccaa
ggttacctcttatctagtggGCATGGCTTACTCAATCTTTTAAGCCCACTGTTGAAAATTgaatcttaaacccaacttg
ggttacctcttatctagtggTGACAATGCACCTAGTTGCCCAAGAGCTCTGATGGAGTTGgaatcttaaacccaac
ggttacctcttatctagtggCATGCCTGCTAATACAACATTCATTCCAAAGCCTATGGACgaatcttaaacccaact
ggttacctcttatctagtggTTCCTCTGACGGATCTCAGCAAAGTACAGAACAACATGGCgaatcttaaacccaac
ggttacctcttatctagtggCTGCCACCCCTGAGACAGCAAGGCCCACACCTCCTCCTCCgaatcttaaacccaa
ggttacctcttatctagtggGGATGAAGACCTTTATGATGATCCACCCCACTTAATAAATgaatcttaaacccaactt
ggttacctcttatctagtggCGGTTTATGTTATCCATAAGGCTTCTGTTCTACATTAGGCgaatcttaaacccaacttg
ggttacctcttatctagtggGCAGGTGGCCCAAATCCCTAACTCCCACATTACTCAAGGGgaatcttaaacccaa
ggttacctcttatctagtggCATCACTGTAGATATTAGGGCTTTCATGATTCATGGGGGGgaatcttaaacccaactt
ggttacctcttatctagtggGTGGTAGAAATAGCAAGAGAGGCCTGGCGTGGTAGCTCACgaatcttaaacccaa
ggttacctcttatctagtggGGGAGGCCAACGTGGTGGATCACAAGGTCAGGAAATCGAGgaatcttaaacccaa
ggttacctcttatctagtggGATCACGCCACTGTTCTCCAGCCTGGCCACAGAGCAAGACgaatcttaaacccaa
ggttacctcttatctagtggAGAGCAAGAGAACTAGAAGTGGGTCCTGAAGATGTGACTGgaatcttaaacccaac
ggttacctcttatctagtggAGTGGTTGCCTGAGATGGAATCTACTTCTGGTGCACAGGCgaatcttaaacccaac
ggttacctcttatctagtggAATTGATAAAGTGGTAGCAGAGTTCAAGAGGATTGACTCCgaatcttaaacccaact
ggttacctcttatctagtggACGTTAAATAGCAGCACATTCTACAGAGAAACCTTTCAGGgaatcttaaacccaact
ggttacctcttatctagtggACTACCACCCTGTTCAGTCAGCAGCCATCAACACTGAGGCgaatcttaaacccaa
ggttacctcttatctagtggCAGCAAAATGAATATGAGTTGCTGAAGACTCAGACATCTTgaatcttaaacccaactt

ggttacctcttatctagtggCCGTGGTGATCTGGACACAGAATCCACAGTATCTCTGAGGgaatcttaaacccaac
ggttacctcttatctagtggGCTCTTGTATCTGTTTTGCCTTTCCCAATAGTGATTGCACgaatcttaaacccaacttg
ggttacctcttatctagtggAGTGGCTCTCTGAAAAGTATTTTTCTTCATTTCACTTCTGgaatcttaaacccaacttgt
ggttacctcttatctagtggCCATCAGCTGAGACATTTAGAAACATCTGTACTTTCACACgaatcttaaacccaactt
ggttacctcttatctagtggTTTCACCCTCAGTGTAAAACTTACAGATTTTGTTGCCGAGgaatcttaaacccaactt
ggttacctcttatctagtggGAAATGTTGGGCCAAATATAGAATGATTGAAAAATCACCCgaatcttaaacccaactt
ggttacctcttatctagtggATGTCTTGCTCATGGTAAGGACTTCGTGCTATCTTTGTACgaatcttaaacccaacttg
ggttacctcttatctagtggGGGCAAATTTCAATACCAATGACAGTAGATGTAATTTCACgaatcttaaacccaactt
ggttacctcttatctagtggGCTCAAAATATGGCTGATGGAGTGCTTGTGCCCTTCCATGgaatcttaaacccaact
ggttacctcttatctagtggTTTTCCAGGACATTTGCAAGACATTCGTCAAGTTTTGAGGgaatcttaaacccaactt
ggttacctcttatctagtggGTCACAGGATGCCAATAGTGACAGACGGAGTGCTGATGCCgaatcttaaacccaa
ggttacctcttatctagtggTCTTCACATTGTGGAGTTTCTCCTCTTCTCTTGGGATATCgaatcttaaacccaacttg
ggttacctcttatctagtggCTGTGACAGATGCCTATTTGCTAGATCAACTGAGAGCATCgaatcttaaacccaact
ggttacctcttatctagtggGAGTCGAGGCTTTTGGGAAGTGACAGGGTCATGAGAGTGGgaatcttaaacccaac
ggttacctcttatctagtggACGGATAGAAGCCACTATCCATGGGGAACAAGCCTCCACCgaatcttaaacccaa
ggttacctcttatctagtggGGACTTCCCAGCCTCCAGAACGCTGAACAACACATTTCTGgaatcttaaacccaac
ggttacctcttatctagtggCCTAATCTAAGACAGCAGCCTGAGTGGACTAGGACAATTGgaatcttaaacccaac
ggttacctcttatctagtggCAGAATTTCTACAGAGGCAACACACACAAAAATAGGTGTGgaatcttaaacccaact
ggttacctcttatctagtggAAATAACCTATGGGGTGGGAGAAAATATTTGTAAACCATGgaatcttaaacccaactt
ggttacctcttatctagtggTTCCTCCAAGTCAATAGCAAACAAACCAATAGAAAACCAGgaatcttaaacccaact
ggttacctcttatctagtggATAGACAAAGAAACTGAATAGACATCTCTCGAAAGAAGACgaatcttaaacccaact
ggttacctcttatctagtggGCAAGTTAATATCACAAGATACCACCTCACAGCTGTTAGGgaatcttaaacccaact
ggttacctcttatctagtggAGCGGCCCCTTGCTTTGTGTCTGCCAGAATCCATTATGGCgaatcttaaacccaact
ggttacctcttatctagtggCCGGCTGGGAATGTCTGGAGTCTGCAGGCCCCAGTCACTAgaatcttaaacccaa
ggttacctcttatctagtggTGGTTGCAGAGTGTGGTTGGGGGTTAGAGCGGCGCCCTGGgaatcttaaacccaa
ggttacctcttatctagtggGATGAGTGTTGGAAATGATGTGCAGAAAAGGGAACCCCAGgaatcttaaacccaac
ggttacctcttatctagtggGTGGAAATGTAAATTGGTGCAGCCTCTATGAAATAATGTAgaatcttaaacccaactt
ggttacctcttatctagtggGTGAAATGATGCATGTGTTAGCCTGTTTTAATCATTCCACgaatcttaaacccaacttg
ggttacctcttatctagtggGCTCCAGATGGTAGAGTTAATCCCTGAAGCTAGAAGATGCgaatcttaaacccaact
ggttacctcttatctagtggGTGGGATGCTGTTTGGGGATAATCAGGGACTCAGGATAGGgaatcttaaacccaact
ggttacctcttatctagtggGTTTAGGTTGTCTGCTGTATAGAGGAAACTAGAGGAGATAgaatcttaaacccaactt
ggttacctcttatctagtggAGTCACTGGGGCCCTGATGTCAAGGCACATGCTCTAGTGGgaatcttaaacccaac
ggttacctcttatctagtggGAGGCTGAGGAAGAGGATTCCCATCTCACCTGCTGCGGGCgaatcttaaacccaa
ggttacctcttatctagtggGGATGGTATTCATTAGGGATTATCACAGGAGGGATCAACTgaatcttaaacccaactt
ggttacctcttatctagtggGGAGCCAGCCCACAAAATGCTAATAGAGGCAGTGGGGAGGgaatcttaaacccaa
ggttacctcttatctagtggAGGTATGAGTCAAAGGAGAGAATTCCAGACAGATGAAAGCgaatcttaaacccaact
ggttacctcttatctagtggTAGTTTTACTGTAAAAAGGTCAAAGCCAGAAGTAAGGGGCgaatcttaaacccaactt
ggttacctcttatctagtggGCTCCTGTTCCAAGCCCATGCTCTGTGCGACTGCAGATGGgaatcttaaacccaac
ggttacctcttatctagtggTAGGCTATGGAAAAGGGAAGAGCAGTCCCCATGGTGGGCCgaatcttaaacccaa
ggttacctcttatctagtggGAAGCTAGCACCACACCAGGATCTCTGCTGGGCACACACCgaatcttaaacccaa
ggttacctcttatctagtggGGTCAGTATCATCCCCATTTTACAGATTTGAAAGCTGAGGgaatcttaaacccaactt
ggttacctcttatctagtggTGCAGAAGTCTTGCTTCAGTGTCTAACTTCCTCATCTCACgaatcttaaacccaactt
ggttacctcttatctagtggGCCATCTAAAGTGGCACCTTAGACACAGTAAGCAAACTGGgaatcttaaacccaac
ggttacctcttatctagtggAGAAGTCAGTCTCAAGATACTTGACAATGTCTCCTATTTGgaatcttaaacccaactt
ggttacctcttatctagtggTGTTGCTCACAACCAATCTGTCTGGTAGGAAAGGCAGATGgaatcttaaacccaact
ggttacctcttatctagtggAGGCCAAGGGGCTTGCTCAAGATTCCACAGTGAAGAATCTgaatcttaaacccaac
ggttacctcttatctagtggTTCAGAAGTCCACAGGATGACTGCCCTTGAAATCCAGTGGgaatcttaaacccaact
ggttacctcttatctagtggCTCCTCTTGCTCTCTGTGCGCCAACCTGAGCAGCCTCCTGgaatcttaaacccaac
ggttacctcttatctagtggCTTGCTTTAGGGCCACAGTTCATGTCACTCCTTCTTCCTGgaatcttaaacccaactt
ggttacctcttatctagtggGCTGACCACCCTGATGGAAGTGCCCTCTCCAGCTCACCTCgaatcttaaacccaa
ggttacctcttatctagtggCCCACTGCGATGGTTTGTGGAGGTGGGACTTTTAGGGGGTgaatcttaaacccaact
ggttacctcttatctagtggGCCAGGAAAAGGGACGTCACCAGGAACCGAGCCTGCTGACgaatcttaaacccaa

ggttacctcttatctagtggGCCACCCAGACTGTGATATTTTGCCATGATGCCTGAACTGgaatcttaaacccaact
ggttacctcttatctagtggGTTTGTTCCTCTAGATTATAAGCTCTCCAAGGCAGGGACCgaatcttaaacccaactt
ggttacctcttatctagtggGTCATGGCTATAAGTTTACTGCTATTTCCCCATCACATGGgaatcttaaacccaacttg
ggttacctcttatctagtggGTCAGCCATTAGAGTTTATTTACCTTAAGGGTGTCTGACGgaatcttaaacccaacttg
ggttacctcttatctagtggGGTCTCTCTTGAACTTTGGACATCTCTTCCACAAACTTCCgaatcttaaacccaacttg
ggttacctcttatctagtggCCCCACTTTAGCCACCTTCACCATGGCCTAACTTTTGCCCgaatcttaaacccaact
ggttacctcttatctagtggGTTTACTTCAACCTCCAACACTTACAGGTCCCTCAAATCCgaatcttaaacccaactt
ggttacctcttatctagtggCAGATCCCTTCATATCATGTTTTCAATCTGAGTTTGGCTCgaatcttaaacccaacttg
ggttacctcttatctagtggTACAAACTGTGGCCTGTGGAATAAATCTGGTCTGCACCTGgaatcttaaacccaact
ggttacctcttatctagtggGGCACAGGTGACTAGTGGCAACAGAATCTGTGTGGCTCACgaatcttaaacccaac
ggttacctcttatctagtggCTATCTGTCCTTTATAGAAAGAAGTTACCAACCCCTAATCgaatcttaaacccaacttg
ggttacctcttatctagtggCTATGAAGTAGAAAGTAAAAGATGATCTTTTGGTGCTGACgaatcttaaacccaacttg
ggttacctcttatctagtggGAAGGGTTTGGTGATCAGTATGCAATCATGAAACCATCACgaatcttaaacccaactt
ggttacctcttatctagtggCCATCACCCCCAAAGGTTTCCTCTGCCTTATTTATTTATCgaatcttaaacccaacttg
ggttacctcttatctagtggTGCTACCCTCTTAGCAAATTTAGCAACTCTTTGAGTATGCgaatcttaaacccaacttg
ggttacctcttatctagtggTGCCCAATACCTCCCTGTTTCCCCCTCCTTCATCCCCTGGgaatcttaaacccaac
ggttacctcttatctagtggATGAGTGGGATCATTTAGTATTTGTCCTTCGATGTCTGGCgaatcttaaacccaacttg
ggttacctcttatctagtggTCACCAGGTTTATCAATGTTGTTGCAAATGACAAGATTCCgaatcttaaacccaacttg
ggttacctcttatctagtggCAGGTTGCTTTCATGTCTTGGGTATTGCGAGTAATGTTGCgaatcttaaacccaacttg
ggttacctcttatctagtggGAACAGGTATCTCTTTGACATACTAATTTCATGCCTTTGGgaatcttaaacccaacttg
ggttacctcttatctagtggTCACATGGTAGTTCTATTTTTTGGTTTTTGAGGAGCCTCCgaatcttaaacccaacttg
ggttacctcttatctagtggCATTCTCACCAACAGTGTAAAAAGGTTCCTTTTTCTCCACgaatcttaaacccaacttg
ggttacctcttatctagtggGCATTTCCCCAATGATTAGTGATAGGGATATTGGGCTTTGgaatcttaaacccaacttg
ggttacctcttatctagtggTTCTCTCTGTATTTTGGATATTAGAGCCCCTTAACAGATGgaatcttaaacccaacttg
ggttacctcttatctagtggGGGTTGTCTCTTTATTTTGTAAATTGTTTCCTTTGCTGTGgaatcttaaacccaacttgt
ggttacctcttatctagtggAATCTTTGTCCAGATCAGTGTGTGGAGCTTTCCTCTTACGgaatcttaaacccaacttg
ggttacctcttatctagtggGTCTTCAACCCACTTTGAGTTGATTCTTGTATATGGGATGgaatcttaaacccaacttg
ggttacctcttatctagtggTTCATTCTTCTTCATGAGGACATCCAGTTTTTTCAACACCgaatcttaaacccaacttg
ggttacctcttatctagtggAATCAGTAGGTTTTTCATACACTGACAATGAACTATCTGGgaatcttaaacccaacttg
ggttacctcttatctagtggGTTGAAATGTCCATACTAAATGTGATCTAAAGAGTCCATGgaatcttaaacccaacttg
ggttacctcttatctagtggGCAATCTTGAGCCATAAGAACAAAGTTGGAGGCATCACACgaatcttaaacccaact
ggttacctcttatctagtggACCAATGGAATAGGATAGAGAGCCCAGGAATAAATCCACAgaatcttaaacccaac
ggttacctcttatctagtggTGACAAAGATGCCAAGAATGGGGAAGGGAGAGTTTCTTTCgaatcttaaacccaact
ggttacctcttatctagtggATATCCAGTTTCCTCAATTGTATATCCATTCACTCACCCGgaatcttaaacccaacttg
ggttacctcttatctagtggCTCAGCCTCCCACTCCCATATTTACAGTTTGATTAGGGAGgaatcttaaacccaactt
ggttacctcttatctagtggCAAGGTGATAGAATGCATGGTGACACCTAACCCAGACTGGgaatcttaaacccaac
ggttacctcttatctagtggGCCAAGGCCAGGGTGTGTAGTGGAGAGAATGGGCAGTGGCgaatcttaaacccaa
ggttacctcttatctagtggACAAGTTCCTTGTCCAAGAGCTTGGATTGTATCCTAACTGgaatcttaaacccaacttg
ggttacctcttatctagtggACACTAAGTGTTTGAAGAAGAGGGATGAAATGGTCAAGTTgaatcttaaacccaacttg
ggttacctcttatctagtggCTGACTGCTCCTTTCTGGAAATCTCAAATTTATATTTGCCgaatcttaaacccaacttg
ggttacctcttatctagtggGAATGGGTTAGGGAACTGAGTCAGAAAGTCCCTGTTGTACgaatcttaaacccaact
ggttacctcttatctagtggAGGAATGTCCAGCCCCACCTAATTTGGCAGCTGCCCCTCCgaatcttaaacccaac
ggttacctcttatctagtggAGGGATGTAGTGGATTCAAGAGGCTGATGATGCCATCTGGgaatcttaaacccaact
ggttacctcttatctagtggATCAGCTTTATTTCTGCAACATCCTGTGCCCACAGAGGGGgaatcttaaacccaact
ggttacctcttatctagtggCACCATGTGAAGATACAGATACACCCAGAACCTTAAAGGCgaatcttaaacccaac
ggttacctcttatctagtggATGTCCCCCAAAAGCCAATGTCTTGCCACCACCAGTGCCCgaatcttaaacccaac
ggttacctcttatctagtggGCTGGAGTAAGGTTTTCTCAATGCCCAAGAGATGAGCTGGgaatcttaaacccaact
ggttacctcttatctagtggTTCTTAGTGGGTCCAAGACCATGTGTTACATTTCCCTCCCgaatcttaaacccaacttg
ggttacctcttatctagtggCTGCTGTCCTCTTCGCCTCTCTGCCACCTCTTAACTGCACgaatcttaaacccaact
ggttacctcttatctagtggTACTTGTCTTTTCATCTACACTCTGGTCACTCTGACCCCCgaatcttaaacccaacttg
ggttacctcttatctagtggTCAGGATTTTCTCTTTTCCTGGCTTACTCTCCAGCACAAGgaatcttaaacccaacttg
ggttacctcttatctagtggTGTGCTTCATTGGAAAATGCATGTTGTTTGAATCACACTCgaatcttaaacccaacttg

ggttacctcttatctagtggGTCCAGGGGTGATTTAATTGACTTTTCTTCCCCATTGTGGgaatcttaaacccaactt
ggttacctcttatctagtggCCTGATCTGTAAATGCATTTCTGGGTTTTCCCTTGGACTCgaatcttaaacccaactt
ggttacctcttatctagtggGAGCTCCTACTACAATCCAAAGTGAACTAGTTGCAGTGCCgaatcttaaacccaac
ggttacctcttatctagtggGCAGATTCCTGGGCCTCTCCTTAGACCCACTGAATCAGACgaatcttaaacccaac
ggttacctcttatctagtggGAGGGTCAGAAATCTCCATCTTAGCAAGTTAATGTCCAGGgaatcttaaacccaac
ggttacctcttatctagtggGACGTATGATAAAAAGTAGAAGGAAGGACTTTCCAGCACCgaatcttaaacccaac
ggttacctcttatctagtggACATCAGGAGATGCATGTATAAACCAAATCCAAATAACCCgaatcttaaacccaac
ggttacctcttatctagtggATGAGTGAGGAGTTTGTCACTCACATCCCCCAGGTAAAACgaatcttaaacccaac
ggttacctcttatctagtggAGGAGGTTGTTAGGCCATGAGTGCCTCCCCCAAGAATGGGgaatcttaaacccaa
ggttacctcttatctagtggAGAGGCTTCTCACAGCATTGTGACCTCTAGCCCTCCCACCgaatcttaaacccaa
ggttacctcttatctagtggGCAGAGGAGCCATCACCAGACAACAGAACCAGCCAACAGCgaatcttaaaccca
ggttacctcttatctagtggCATCTCATTTTCTCCAGAACTGGGATAAAATAAATTCCTGgaatcttaaacccaactt
ggttacctcttatctagtggTCCCAATTTCAGGTATTTTGTGATAGCAGCAAAAACAGACgaatcttaaacccaactt
ggttacctcttatctagtggCCATCTCCACTGGGGATATGGCATTTAAGAGATCTCTTTGgaatcttaaacccaactt
ggttacctcttatctagtggTGTGGTGCTCACCAGCTTGCTTAAGTTACAGCACTTGCTGgaatcttaaacccaact
ggttacctcttatctagtggTGGCAAGTGTGAGTTAATGGAAGCCTCATAAAGCAAGAGGgaatcttaaacccaac
ggttacctcttatctagtggTGTATAACATCAAGTGCAAAAGTGTGTGCTAGGAAAGTTCgaatcttaaacccaactt
ggttacctcttatctagtggGACCAGAGATTGGGGAGTATATAGCTTTTGGGAATTCAGGgaatcttaaacccaact
ggttacctcttatctagtggGTGATCCTGAGCCACCTACTATGGAAAGGTGTCAGTGAGCgaatcttaaacccaac
ggttacctcttatctagtggGCTGAGCAGACTTCTACTGCGCATCGCCTATGTACAGGCCgaatcttaaacccaa
ggttacctcttatctagtggGGGTAGAGAGGGAAAAGTTTCACCCCAAATACCAGATGCCgaatcttaaacccaa
ggttacctcttatctagtggTGTTGGTTGTCAAGGCTGTCATTAACTTCTGGGAGTTCCCgaatcttaaacccaactt
ggttacctcttatctagtggGCAGTTCTGCCTGCAATCAGAAGCTGGCTTTGAGCCCCACgaatcttaaacccaa
ggttacctcttatctagtggCTTGGGATTTACTGTGGATTGAGTCTACGAAGCTGGAAGGgaatcttaaacccaact
ggttacctcttatctagtggATTCCACTGTCTCCTCATGTGGATTTTATCAAATTTCTCCgaatcttaaacccaacttg
ggttacctcttatctagtggGGGGAACCTAGAAGTATGATTGTGACGTGCATTGTTGAGAgaatcttaaacccaact
ggttacctcttatctagtggGCTGAAGACTGGATTTTCTACTCCTTCAGATGTTTTCCAGgaatcttaaacccaactt
ggttacctcttatctagtggCTCATAGCTAGGGTCTCCTGGTCCCCAAATGTAGACACACgaatcttaaacccaac
ggttacctcttatctagtggCCCAGAGCCATTTTGTATAAGAGCCACCATGGATTTAACCgaatcttaaacccaact
ggttacctcttatctagtggCAGCATCATCACCTGGCCACCAAATCAAAACTGAGCACCCgaatcttaaacccaa
ggttacctcttatctagtggGTTCTCTGCAGCCATGCAAGGTTATGAAATGGGCACAGACgaatcttaaacccaac
ggttacctcttatctagtggGTCTTTGTGTTTAAGAGGTTCATGGTCTACCTGAGAAATGgaatcttaaacccaactt
ggttacctcttatctagtggAGCATCAATGGTATATCAAGCACTGTACTGGCTTCTGGGCgaatcttaaacccaact
ggttacctcttatctagtggAAATGTCCTTGAGGAGCTACGGAAGATTCCTGCATGAAGCgaatcttaaacccaac
ggttacctcttatctagtggAGCTGAAGAAAAAACAACTGGCTTTGGGGGCTATATAAATgaatcttaaacccaact
ggttacctcttatctagtggACCCCAAAAGCTTGAGTGAAGAAAGACTTCTGTAAGTTACgaatcttaaacccaact
ggttacctcttatctagtggGCCAGGTGCTTTCTGTAGGACACACAGATACAGAATGAGGgaatcttaaacccaac
ggttacctcttatctagtggGAGGCATCACCATGTCTATCTCATCAGGGTTTAACCAGAGgaatcttaaacccaact
ggttacctcttatctagtggGCAGGCAGGAAGAAAACAGGCTGGGACTTGTAGGCACAGGgaatcttaaacccaa
ggttacctcttatctagtggGTGGATGCTGCTCTTCTTCAGGGAAGACTTGGCTCTGCTCgaatcttaaacccaact
ggttacctcttatctagtggTCAGGCCCACTCACATTATCTAAAATAATCTCCCTTACTCgaatcttaaacccaactt
ggttacctcttatctagtggATAACTGAGAGTTGTAACTGATATGGTTTGGCTATGTCCCgaatcttaaacccaactt
ggttacctcttatctagtggTTGAATCATAGGGGCAGTTTCTCCCATTCTGTTCTCATGAgaatcttaaacccaactt
ggttacctcttatctagtggCCCTCTTTGCTCCACTTTCATTCTCCTTCTTCCTGCTGCCgaatcttaaacccaactt
ggttacctcttatctagtggGTAAGTTTCCTGCGGCCTCCCCAGCCACGCTGAACTGTGAgaatcttaaacccaa
ggttacctcttatctagtggCATGAGAATGGACTAATACAGTAACTTTACCAAGTGGACAgaatcttaaacccaact
ggttacctcttatctagtggAACTGAATGAAAGAGGATGGTGATTCCTACTTCAGGGTGGgaatcttaaacccaact
ggttacctcttatctagtggGAACACTTTTCTATGATTGATCTAAGGTTGGCTTTTAGGGgaatcttaaacccaacttg
ggttacctcttatctagtggTGTTTTTGCTTGCATTTCTCTCTCTGCCCACTCTTGGGGCgaatcttaaacccaactt
ggttacctcttatctagtggCAGGGAACCCAGATGCCAGCCAAAGACATCCCTTTCCCTCgaatcttaaacccaa
ggttacctcttatctagtggCCCGGCCTTATATCTCAAAGCCCCTTTTCATCCAGGTTTCgaatcttaaacccaact
ggttacctcttatctagtggGCTACTAATGTCCTCTTCCATCCCAGACTCCCCTCACACCgaatcttaaacccaac

ggttacctcttatctagtggTGCAGGAGGACATACAACATTAACCTTTAAATAGAGATCCgaatcttaaacccaactt
ggttacctcttatctagtggAAAAGTCCTCATTGATCTCGGGGCCGTTTCACGCAGGTGCgaatcttaaacccaac
ggttacctcttatctagtggCAGCTTCACTTTCAGTCTCCTTCTTCGGCTCTGGCGCTGCgaatcttaaacccaact
ggttacctcttatctagtggTGATCTGAATTTCTTGGCAGTGATGGGCTCGGTGTTGGGCgaatcttaaacccaactt
ggttacctcttatctagtggGTGACTGAGACTGGATGGGGGATGGAGATGCTGGGGGCTTgaatcttaaacccaac
ggttacctcttatctagtggGGCTCAGGAGACCCAGAGACAGATCGGGAGGATGAGACCCgaatcttaaacccaa
ggttacctcttatctagtggGAGGTCTGAGGAGCTCGAGGCCGTGATGAGGGCGAATGCCgaatcttaaacccaa
ggttacctcttatctagtggGGGGAGCTTCTTTGGAAGGTGGAGAATGGGAGACCCGCCGgaatcttaaacccaa
ggttacctcttatctagtggGGTGGTGTGGGAGTCCTGAGCCGTCGAGGATGAGGGGCGGgaatcttaaacccaa
ggttacctcttatctagtggTCTCCTCCGTCTACCATGAGGGGAAGTCTCTTTTTGGTGCgaatcttaaacccaactt
ggttacctcttatctagtggCTCCGGGAAGGGGAATGTCTCCGCCGCTTGCCAACCTCACgaatcttaaacccaa
ggttacctcttatctagtggCTCTTGGGTCGTTCATCTTCTGAGGAGAAGGAGGAACCAGgaatcttaaacccaact
ggttacctcttatctagtggCGTCTGTATTGGCGTCTCTGCTGAGATCAATGAGGAAGGCgaatcttaaacccaact
ggttacctcttatctagtggTGACACTTTTATGGTTCCTTTAAATGCTCTGGCTATTCCCgaatcttaaacccaacttg
ggttacctcttatctagtggATGAATATTTTAATGCCACAGTCTTTCCTGTTGCCTGTGGgaatcttaaacccaacttg
ggttacctcttatctagtggATCAGGATTCGAGCTCTAGGATAAGACAGAAAATGAAAGCgaatcttaaacccaact
ggttacctcttatctagtggGATGTGTAACAACTTGATTTGGAACACTGGACTCTCATTCgaatcttaaacccaacttg
ggttacctcttatctagtggTGTAAAGGCCATGAACTAGTCCCAGAAAGGATTCCTTCAGgaatcttaaacccaact
ggttacctcttatctagtggTGTCTTGTTCTTCCAATTGGTGTATACAACTTTCAGAGCCgaatcttaaacccaacttg
ggttacctcttatctagtggAAGGGCCCAGACTTTGGAATAGTGTCTCGGTTTCACTGTTgaatcttaaacccaactt
ggttacctcttatctagtggAGTCCCAGACTCTTCTTCTCTTCCTGGACTGCCCATCTCCgaatcttaaacccaact
ggttacctcttatctagtggTTAATTTCTCTTCCTTCTCTCACCCTTCTTTTCAGGATTGgaatcttaaacccaacttgt
ggttacctcttatctagtggGTTTCTGCCTCCCTGGGTCTTTCTGTTTACTGACCGCAACgaatcttaaacccaactt
ggttacctcttatctagtggCCTCTTCTACTCCAAGTGCTTTCAAAGAAAGTCCTCTGCCgaatcttaaacccaactt
ggttacctcttatctagtggCATTACGTTTTTTCCCTCTACCTGTTTGCCTTTATTCTCTgaatcttaaacccaacttgt
ggttacctcttatctagtggTTCCTTTCCATTTCTACTCAGCATGAGGTGTCCATTGAACgaatcttaaacccaacttg
ggttacctcttatctagtggTCTTGCTAGAGGTGGGGAGAGAGGTTGGGGGCGGTTAAGGgaatcttaaacccaac
ggttacctcttatctagtggGCCTAGGAGATATTAGAGCTGCAGAGATTTGGCTATCTTGgaatcttaaacccaactt
ggttacctcttatctagtggCCCTAGGGATTAGTACATAGGCTTGCAAATAGCAGGTATGgaatcttaaacccaact
ggttacctcttatctagtggTACTTAGGCGGTATCTTCAGGCATATCTGTGTTTATGTGGgaatcttaaacccaacttg
ggttacctcttatctagtggCCCACAAATGTCTACATCCTAATTCCTAAGATCTGTAAACgaatcttaaacccaacttg
ggttacctcttatctagtggGGGTCCTTATAAGTAAAACAGAGAAGCAGGAAAATGAGGGgaatcttaaacccaact
ggttacctcttatctagtggGAGTGATGTTGGCTTTAGAGATGGAAGGAGTCACAAGCTGgaatcttaaacccaact
ggttacctcttatctagtggGTCTTAAAGGAATTGTTATAAAGGAATAGCTGAAGCTGGGgaatcttaaacccaacttg
ggttacctcttatctagtggATTGGGTAGCTGCATCTGGCAAGGGCCTCAGGCTGTTTCCgaatcttaaacccaac
ggttacctcttatctagtggAGGGGAGCTGGTGTGTTTAGAGATCACGTGGGGAGAGAGGgaatcttaaacccaac
ggttacctcttatctagtggGCAGAATGTGCAGGTTTGCTACATAGGAATACACATGCCAgaatcttaaacccaact
ggttacctcttatctagtggGCTATCCCTTCCTTAACCCCCAAACCCTGACAGGCCCTGGgaatcttaaacccaac
ggttacctcttatctagtggCACTTATGAATGATAACATGCAGTGTTGGGTTTTCTGTCCgaatcttaaacccaacttg
ggttacctcttatctagtggACGTGTGCATGTGTCTTTGTAGTACAGTGATTTATAATCCgaatcttaaacccaacttg
ggttacctcttatctagtggTGGTTCTAGATCCTTGAGGAATTGCAACATTGTCTTCCACgaatcttaaacccaacttg
ggttacctcttatctagtggAGTGAGAACTCACTTCCCTGGCCCCAGAGAGAGCAAAAGCgaatcttaaacccaac
ggttacctcttatctagtggCAACATGAGGTTTTTAGGGGACAAACATCCAAACTATGTCgaatcttaaacccaactt
ggttacctcttatctagtggGATTCTCCCCTAAAGCCCACAGAAGGAAAGCAGCCCTGCCgaatcttaaacccaa
ggttacctcttatctagtggAACCCAGTAAGACCCGTTTTGGACTTTGGACACCAAGAGCgaatcttaaacccaac
ggttacctcttatctagtggAGCCATTAAGCTTGAGGCAATTTGTTACAGCAGCAATTGGgaatcttaaacccaactt
ggttacctcttatctagtggGTATTGTTCCCAGTTCTCTGGACCTCAGATTTCTTTCCTGgaatcttaaacccaacttg
ggttacctcttatctagtggCACACGAGTGCAGAGTGCCTATTTACTAGAGAGATCAGCAgaatcttaaacccaact
ggttacctcttatctagtggTGCCAGAGGCAGGTCCTAAAAACCCACAAAGCAGGTGAACgaatcttaaacccaac
ggttacctcttatctagtggAGGTGTTTTTACCTTTGTCAGCATGCCACAAGTTACTTCCgaatcttaaacccaacttg
ggttacctcttatctagtggGTGAGGTCCCGTTTCAGCCAATGGAAAGTGGACACAGCAGgaatcttaaacccaac
ggttacctcttatctagtggGCTGGCAAGTGTACCCTTTCTGCAATAGGTAAAAACTGCCgaatcttaaacccaact

ggttacctcttatctagtggTTCAGGAACCTAAAAAAGTGGATGAAGAACTGAGGTTGCgaatcttaaacccaact
ggttacctcttatctagtggGGGATTATTTGGAATCCCTTGTCACCAGAAGCTCTTAAGGgaatcttaaacccaact
ggttacctcttatctagtggCTTCGTTGCAAATATGTTCTTGGCTTAGTTGAGGTAATGCgaatcttaaacccaacttg
ggttacctcttatctagtggCTTCCCTGAAGCTGTTCCTCCAGTCCCTGCCAGTGGGATCgaatcttaaacccaac
ggttacctcttatctagtggAGCCCCATCTTTTAAGCCTCCCTGGGACCTGCCCTCGCAGgaatcttaaacccaa
ggttacctcttatctagtggACCTACTCCACTTCCTCCTCATGGCCTCCTGCAGAATGCCgaatcttaaacccaac
ggttacctcttatctagtggAGGTTCTATTTTCCTCGCCCGCATTCTCTTTTGCAAAGCCgaatcttaaacccaactt
ggttacctcttatctagtggGCTTCTCTGCGCCTGCTTTATCTCTATTTTCTACACTCGCgaatcttaaacccaactt
ggttacctcttatctagtggAGGCGTCACAAAATTTGCATTTGTTGGAACCAAAATTTCCgaatcttaaacccaactt
ggttacctcttatctagtggGTCCCCAGGCCAGCCTCCCAGTCTGTGTGTTTCCCCCTGGgaatcttaaacccaa
ggttacctcttatctagtggCAGGACTCTGCTCTGGGGATTTACGGACAAGGGTATCAAGgaatcttaaacccaac
ggttacctcttatctagtggACTAACCCTCTCAAACTGAATGAGTGGCTTAAAATCTTCCgaatcttaaacccaactt
ggttacctcttatctagtggGAGCGAGCCAGCACCCCCACCCCCCAAATCCTGTGACAAGgaatcttaaaccca
ggttacctcttatctagtggTGGCAGCATTATGGGGGAAAAGCAATGATGATCTAATGAGgaatcttaaacccaac
ggttacctcttatctagtggCCCACCAACCTTAGCTGCGAACCTTACCTCAAGTGTTACCgaatcttaaacccaac
ggttacctcttatctagtggTGAGCAAGACATTCAATAAGCAAGTGTGATGGCTATGAGGgaatcttaaacccaac
ggttacctcttatctagtggCATTACTGGGGCTGTATCCTGCTACAATAATAAGGCTGACgaatcttaaacccaact
ggttacctcttatctagtggCAAGCCAAAATGTATTACTTCATCCACAAATAGTATCGTGgaatcttaaacccaactt
ggttacctcttatctagtggTACACAATAGAAAATTTGCTATATGGTGCATGGTGGCATGgaatcttaaacccaactt
ggttacctcttatctagtggGGAAGAAATGTGTAAGTATAGCCATTGTTGGAGCAGATGCgaatcttaaacccaact
ggttacctcttatctagtggATAGATCTCACCATGCTTATTTACTGCTTAAGTCATAGCCgaatcttaaacccaacttg
ggttacctcttatctagtggTAATGCAATTAATGAGGTTGATGTCCCTAAAACAAGAGAGgaatcttaaacccaactt
ggttacctcttatctagtggTGTGGAAATGATCTCATGGCAAAGTTTTTAAAGAGGCTGCgaatcttaaacccaactt
ggttacctcttatctagtggGCTGACCTTCTCTGTAAATAAGAAGTGACTGTCAGTAGCCgaatcttaaacccaact
ggttacctcttatctagtggCCCACTTGATCTGTCCCTGCAAGGTAAGAGTGGCATTTTCgaatcttaaacccaac
ggttacctcttatctagtggGCAACAGATCCTACAGTTAGACTCTTCTGATCACCCTGAGgaatcttaaacccaact
ggttacctcttatctagtggCTTTGGTTTCTTCTCTTTAACGTCCTGCATATATGCTGCCgaatcttaaacccaacttg
ggttacctcttatctagtggTGGGAGTTTCTTGATGTCAGAACACAGCAATTGTCCCACCgaatcttaaacccaac
ggttacctcttatctagtggAAAACAAATAATGGACCAATGGGCTCAAAAGCAAACTGGCgaatcttaaacccaac
ggttacctcttatctagtggGGGCTAACACTTAAACGGTTGTGTACTCACTATGTACTAGgaatcttaaacccaactt
ggttacctcttatctagtggGTCTGAGATGCAATCCATGTGATTTGTTCACAGAGCTCAGgaatcttaaacccaact
ggttacctcttatctagtggGCGGGTAAAAACAAAATTTGGCATCCAGTTTCAAAAGGAGgaatcttaaacccaac
ggttacctcttatctagtggCTTCCCACTCTGCATTTTCAAAGGCATATGGTATTGACTCgaatcttaaacccaactt
ggttacctcttatctagtggCTCAGTGCTGCACTGCTGGTACGTGGCGTGGTCCCCTCTCgaatcttaaacccaa
ggttacctcttatctagtggACACACTACCCATGCAGACTATCACTCTCAGTCTTGTTTAgaatcttaaacccaactt
ggttacctcttatctagtggGAGGCCAAAGTGTGGGGAAGCTCCTTAACCATGCTGCTGCgaatcttaaacccaa
ggttacctcttatctagtggGCTGGAGAGACTGAGAACATGAGGCCTAAAGAGGAGACTCgaatcttaaacccaa
ggttacctcttatctagtggGAGCAAATCCAAGAGTGTTGAGTGGAAGTGAAAGGGAGGCgaatcttaaacccaa
ggttacctcttatctagtggGGAAGAACTTTCTAGTATTTGGTACTACAATGGGAAAGACgaatcttaaacccaactt
ggttacctcttatctagtggATGTGTCTTTATAGCAGCATGACTTAAAATCCTTTGGGTAgaatcttaaacccaacttg
ggttacctcttatctagtggGGGTAGTCCCTACATGAGGGTTAGGTAGAATTGACCTGCTgaatcttaaacccaact
ggttacctcttatctagtggACGGGGTGGTGTTTGTTCCCTCAGTACTTTATCATCTATGgaatcttaaacccaactt
ggttacctcttatctagtggACAAATAGCAAATTATTGTGGGGGTGTGTGTGTGTGTGTGgaatcttaaacccaactt
ggttacctcttatctagtggCATCAGGTGATCCATCCGACTCATTTCCCAAAGTGCTGGGgaatcttaaacccaac
ggttacctcttatctagtggGGCCATCATGCCTGGCCCGCAAATTGTTGTTATTTATAACgaatcttaaacccaactt
ggttacctcttatctagtggATGGTACTGAGAATGATGCCTGAAACATAGTAACTACCCCgaatcttaaacccaact
ggttacctcttatctagtggACCATTATTGCTCTTGCATACCATATTGCTCTTGCATACCgaatcttaaacccaacttg
ggttacctcttatctagtggGCTACACACAGTATTTCATTTAGGCCTCACTATGTCCCTGgaatcttaaacccaactt
ggttacctcttatctagtggGCAAATGAGAAAACAGTCTGTACCTTGTATGCCATGCTGCgaatcttaaacccaact
ggttacctcttatctagtggTCTGCTTGATAATTGGTACCAAAATAAGCCTTTTTCTGGGgaatcttaaacccaacttg
ggttacctcttatctagtggGGCTGCTGCATTACTCCTGCTGCCCCATCCTACTGTGATCgaatcttaaacccaac
ggttacctcttatctagtggCATGAGATGGTGAGTTGTCCCCTTGCCAATGAGGGTTTGGgaatcttaaacccaact

ggttacctcttatctagtggGTGTGGGTTAGAGGGGGAGGCTGAATATGAAGGTCCTGGGgaatcttaaacccaac
ggttacctcttatctagtggCTTTGCAAAGGAAAAGGAGGTGAGTGACAGCCCAGGGTCCgaatcttaaacccaac
ggttacctcttatctagtggAGCCAGGCCATGGAGTTCCTCAGACCTGCTTTCTAAGGGCgaatcttaaacccaac
ggttacctcttatctagtggCCTCAGCAAGCCATTGAACTTCTCTGAGCTCAGTCCTTTCgaatcttaaacccaact
ggttacctcttatctagtggGGGGTGACAGTGCTCACATGCCAGGAATACAAAGGGATTGgaatcttaaacccaac
ggttacctcttatctagtggACACATCTGTCAGGGACCCCAATAAGGTCAGCTGTCTTCCgaatcttaaacccaac
ggttacctcttatctagtggCTCCCCCACCCCATGCCCAATACTGAACAGTGGAGGGACGgaatcttaaacccaa
ggttacctcttatctagtggACCCAAAGCAAGTACTGTTAGGATCTCTGGAAACTTTCCCgaatcttaaacccaact
ggttacctcttatctagtggTGAATGGGACAAGAAGAAGTTTTACAGATAGCCAATGGAGgaatcttaaacccaact
ggttacctcttatctagtggGGGGTTATGATAGAAACGAGAAAGTAAAACAAACCCATGCgaatcttaaacccaact
ggttacctcttatctagtggCCATTTCAGGTCCATATTTTCGCTTGAAAATTGAGATTCCgaatcttaaacccaacttg
ggttacctcttatctagtggAGTAGAGGAGTTGGTTGAAGGACAGGGTAATGCCAGGAGGgaatcttaaacccaac
ggttacctcttatctagtggGGGCACCTCAGGGCTCTACCAAGGACACGTAGAAGCTTTGgaatcttaaacccaac
ggttacctcttatctagtggACGAGCTACCTCAGCACGTAGTGGAGCCTTGGAAAACAGAgaatcttaaacccaac
ggttacctcttatctagtggGTCAAACATTTTCCAGCATTTCCTCTCCCATCATAGCTGGgaatcttaaacccaactt
ggttacctcttatctagtggGACACACACGTGCATACACATAAATACCTTGATAAGTTACgaatcttaaacccaactt
ggttacctcttatctagtggTAATTCATCAATGTTTCTAAGACTCATTCAGCGTGGCTGCgaatcttaaacccaacttg
ggttacctcttatctagtggCATCCACACGCCTCCTGACTGGCTACTCAGTGCCGATGACgaatcttaaacccaac
ggttacctcttatctagtggGGGCTGGTTTTAAGATACATTGAGGTGACATCAGGTGGCCgaatcttaaacccaact
ggttacctcttatctagtggTAGGATATCCTATTCAAAGACATCTGTGGAAGTGTGGACCgaatcttaaacccaactt
ggttacctcttatctagtggTGAATAACAGTGAAGGGGTTTCCACCAGCAAGTAACATAAgaatcttaaacccaact
ggttacctcttatctagtggGATGATGCTGAAGTAGAAAGAGTTTCTAGTCAGGGACTGGgaatcttaaacccaact
ggttacctcttatctagtggGGATTCTGAGGGGAAGAAAGTGTTCTGCAAAGCAATAAGCgaatcttaaacccaact
ggttacctcttatctagtggCATGTGCTAGGCAAATCTCTTTCACATACATGTGGGGAGGgaatcttaaacccaact
ggttacctcttatctagtggCCTGGGAGGCAGAGGTTGCATTGAGTTGAAATCACGCCACgaatcttaaacccaac
ggttacctcttatctagtggATTGTGTATTTTAGAGGAGACTGAGGATTGGAGGCTGAAGgaatcttaaacccaactt
ggttacctcttatctagtggAGCTAAGATAGCTCTTTGGGTTCTAAATTCTGTGATCTTCgaatcttaaacccaacttg
ggttacctcttatctagtggAACTGGGTAAGGGAGACCATTGAGAAGGACTAAGGACACCgaatcttaaacccaac
ggttacctcttatctagtggAGAGGAACTTGTTTGGCTTGAAACTTCTTAATGTTTAGGCgaatcttaaacccaacttg
ggttacctcttatctagtggATGACAGTGGTCATTGAGAAAACAAAATCCCCAAAGCAATgaatcttaaacccaact
ggttacctcttatctagtggGCTGGTCCGAGGCTACACAGCTAATCAGTGGGAGAGTTGGgaatcttaaacccaac
ggttacctcttatctagtggCAGAGGTCCAGTTTTGAAGTCCACACTTTTAGCCATTACAgaatcttaaacccaactt
ggttacctcttatctagtggTAGGCACATAGATTAGGGGTTAGTAGAATGCTCTGGGCAGgaatcttaaacccaact
ggttacctcttatctagtggTCTTAAAGGAGGTAAAGCTTGAATGAGACTGTTAGTAAGCgaatcttaaacccaactt
ggttacctcttatctagtggTGTGTAACATATCTCATATTCCAAGAGATCCTTTTGGTAGgaatcttaaacccaacttg
ggttacctcttatctagtggTATCTGCTCACAGTACTGGCTTCATTATTTGGAGTTAAAAgaatcttaaacccaacttg
ggttacctcttatctagtggGCGAATGGTCAGGTCCAAAGTAGATGCAGAATATGCCAGGgaatcttaaacccaac
ggttacctcttatctagtggTCCCTATTCAGCCCAGGACTATGTACCATAAGATTACTGCgaatcttaaacccaact
ggttacctcttatctagtggACGAAACAGTGACATACAGTAGGGAGAGCTGGATTGAGGCgaatcttaaacccaac
ggttacctcttatctagtggGGAAGTTTCCTGAAAGCATTTTGGGGAAACATCTTTTGGGgaatcttaaacccaactt
ggttacctcttatctagtggGAGGTTATGAGAGAATGACCCTCCCGTGATGGTGAGTGGGgaatcttaaacccaac
ggttacctcttatctagtggCCGTGCTGGACAGCATGCTTAGCCACCAGTCACATGTGGGgaatcttaaacccaac
ggttacctcttatctagtggTGACAATGTGGGTAGTCTGAACTGGGATATGCTGTAAGTGgaatcttaaacccaactt
ggttacctcttatctagtggTTTTTGAGGTGCCGTTTCACTCTTGTTGTCCAGGCTTGAGgaatcttaaacccaactt
ggttacctcttatctagtggGTGAGCCACCGTGCCTGGCCTATCTTTACCTTTTTAAGGTgaatcttaaacccaactt
ggttacctcttatctagtggGGCTCAGGAGGACTCACAGAATAACTGGCAACCCTGCCCCgaatcttaaacccaa
ggttacctcttatctagtggGACCCAGGGATCAACAACCAATGAGAGCAGCATGGAGGCCgaatcttaaacccaa
ggttacctcttatctagtggCAAAATGCTGATTCCAGGACTGGGATTCAGGATGATTTCCgaatcttaaacccaactt
ggttacctcttatctagtggCTGCTCCTTGAAAGGGTATCTAATTGGGCATTGCTTTTACgaatcttaaacccaactt
ggttacctcttatctagtggGGCAGGACTCCAAAAACAGACATGCTGATATGTACTGGGGgaatcttaaacccaac
ggttacctcttatctagtggCTCGTACCCAGCTCGGTACCTGGAACATGATAGGGATCCCgaatcttaaacccaac
ggttacctcttatctagtggGTAATCCTCAGGAGGACAAAGGCAGATAGCTTCCCTTCCCgaatcttaaacccaac

ggttacctcttatctagtggTCTTGGAGAATTTTGCCTTACAGAAGTCAAATCACATACCgaatcttaaacccaacttg
ggttacctcttatctagtggTATGATCAATGGAGGGGTTGTCACGAGCCAGGCTAAGAGCgaatcttaaacccaac
ggttacctcttatctagtggTCTCATTCTGTCTATGCCAGAGAATCAACTATGACATATGgaatcttaaacccaacttg
ggttacctcttatctagtggGGAACTGGGATTTGAATTCAGACAGTCTGACTCCAGACGCgaatcttaaacccaact
ggttacctcttatctagtggCCTTTTCATGAAAGCTTTGCTCGTTGTGTGGATGAGTGTGgaatcttaaacccaacttg
ggttacctcttatctagtggGTGTGCTATCACTGAGGTAGAAGACATCTCTCTCTCTCTCgaatcttaaacccaactt
ggttacctcttatctagtggCACACACACGCATGCAGACACATCCTTGAGCTCAAAGAGCgaatcttaaacccaac
ggttacctcttatctagtggCCTATTCAAATCTCAGCAAAACAGAGATACCAGGCTCTGCgaatcttaaacccaact
ggttacctcttatctagtggTTTTTTGCTGTAGACAAAGCTGTTTCTTGCACTGCTACATgaatcttaaacccaacttgt
ggttacctcttatctagtggCTTCTCACCCTGCATTTCCGAAGACATATGGTATTGGCTCgaatcttaaacccaactt
ggttacctcttatctagtggGAGTGCATATATCCATATACAAAATAGAGGTCTGTTCTTCgaatcttaaacccaacttg
ggttacctcttatctagtggGAAAGCCCAGGTATTGGTCAGAATTCATGCTCGGCTCAGGgaatcttaaacccaac
ggttacctcttatctagtggCCTCAAAAAAGAGGGAACCGAATCTTCAAATCTGAGCCACgaatcttaaacccaac
ggttacctcttatctagtggGGCACAAGATGGAGTCCTGGTGGCCAGAACTGAGTATGGGgaatcttaaacccaac
ggttacctcttatctagtggCCCAGAAGGCAGTTGGATAGGACAAGGGGATCGGAAGAGCgaatcttaaacccaa
ggttacctcttatctagtggATGGAGGGAGAAGGGATACCCAGAAGCAGGATTCAGGGGGgaatcttaaacccaa
ggttacctcttatctagtggGGCCATGCCCATGGACAGGAGAAGCATATGATGGGTGCACgaatcttaaacccaac
ggttacctcttatctagtggATCCACAGAGCAGAGGAGGTTGTGAAGAGCAGGCATAGGCgaatcttaaacccaac
ggttacctcttatctagtggTGCTCTCAAGACCTCTTAGTGTTGTAGGTAGATGACGCACgaatcttaaacccaactt
ggttacctcttatctagtggCCTTGATTTGAATTGGCATAAAAGGTCAAATCAGAAAGCCgaatcttaaacccaacttg
ggttacctcttatctagtggGTCCAAAGGCATCTCAGAGGTCATCTCAGGGAGAAAAGGGgaatcttaaacccaac
ggttacctcttatctagtggCCTCCACTCTACCTAAACAAGGCAGCCCTGCTGTGCTCCGgaatcttaaacccaac
ggttacctcttatctagtggGGCGACATCAGGGAAGATGTAAAATAGGATTGTGATAGGAgaatcttaaacccaact
ggttacctcttatctagtggAAGGGAAGCGTAGAACCCTTTTCCCCTTGTTAGCCAATTCgaatcttaaacccaact
ggttacctcttatctagtggGCAGCCTAACTATACCCGTGTCTTCAGAAAGAGCAGTGTCgaatcttaaacccaact
ggttacctcttatctagtggGGTCAGTGTGGGTCTCCTGAAGAGCCCGAACAACCACGGGgaatcttaaacccaa
ggttacctcttatctagtggGGGTATCATAGGAAGATCCTGCCTTTCTTCCAGTTCCAACgaatcttaaacccaactt
ggttacctcttatctagtggCTGTGAGAGAGCCTTAGGAAGTTCTCTGGGGAAGCCCTGCgaatcttaaacccaac
ggttacctcttatctagtggACTTTCAGGTGATTAAAAGTGGGGAAGAATGGTTTCAAGCgaatcttaaacccaacttg
ggttacctcttatctagtggGGAAAGAAAATGAAATAAGGCCGGCCTCGGTGGCTCACACgaatcttaaacccaac
ggttacctcttatctagtggCTGCAGTTAATATTTTGGCAAGCTTTCTTCACTTGTATGCgaatcttaaacccaacttgt
ggttacctcttatctagtggATTGTCAGATGGTACTTCATCATCCTTCTCTCCCATCTTCgaatcttaaacccaacttg
ggttacctcttatctagtggCATGCCAGATGTCTCTTATATGAAGCCGTCCCGCCCAAGGgaatcttaaacccaac
ggttacctcttatctagtggATACTGCAAAGCCCTTGGAAGGATGGGTCTGAACACTCACgaatcttaaacccaac
ggttacctcttatctagtggCTGCTGGCATGTTGCTTTGTCCCTGTGTCTTGTGCTACTAgaatcttaaacccaacttg
ggttacctcttatctagtggAGAGAACCAGGCACAAGTCAGCTGAGGCAGGAGACCCGCCgaatcttaaacccaa
ggttacctcttatctagtggAAGCCTGGGTGGGTGAGGGTCTCTGGGCTTCCGCCGTGCCgaatcttaaacccaa
ggttacctcttatctagtggCATTCTCCAGCACACCCTATAAACATGTCTAAGGCTGTCCgaatcttaaacccaact
ggttacctcttatctagtggGGAGATGACAGCAGGGAGAGAGAGAGGGGCAAAGAAGGGCgaatcttaaacccaa
ggttacctcttatctagtggGCAGGGTTTCTAAGCTCTCTAATCTCCCCTGTGCAGCTGGgaatcttaaacccaact
ggttacctcttatctagtggCAAATCCAGTGGTGATCTCTGTGCAACGTGGTATCACCTGgaatcttaaacccaact
ggttacctcttatctagtggGCCACCTCCTCCTTCCAGACGTTCCCAGCTAAAGTGGCACgaatcttaaacccaac
ggttacctcttatctagtggCATTACTCTCTATGTTTATTGCCTGCATAGCTCTTATTTGgaatcttaaacccaacttgt
ggttacctcttatctagtggCCACATGAGCATGGGGACACATCTCCTTTGTCACTGCCCCgaatcttaaacccaac
ggttacctcttatctagtggTCTATCAGACAGTGTAGATAGCAGCACATAAAGGAAAGGGgaatcttaaacccaact
ggttacctcttatctagtggATGACTACTAAATATAATAAAATCGGCTGGGCACGGTGGCgaatcttaaacccaact
ggttacctcttatctagtggAGTCAGAAATCTTGGGGATTATATTGCAAAGAATTTCCACgaatcttaaacccaacttg
ggttacctcttatctagtggTATTAAGAAATTGGCATCTACTCACCAGTTTGGACATCGCgaatcttaaacccaacttg
ggttacctcttatctagtggTCCTGCATAGGCTGAAAAAGGTCTCTTTCCATGCCTTTCCgaatcttaaacccaactt
ggttacctcttatctagtggCTCAGAGTCATTTCTTCCAATTCACCAATTAGGATGAGGGgaatcttaaacccaacttg
ggttacctcttatctagtggAAGAGGAAATGGTGCATGGGCAAGAAGAAATTTGGAAAGGgaatcttaaacccaact
ggttacctcttatctagtggGGAGGAATTATATTGAAAGGTGAAACAAGGGAGAAAAGATgaatcttaaacccaactt

ggttacctcttatctagtggCCAGCCCCTCTCTAATAACTACTACTCTTTGGAACAAGGGgaatcttaaacccaact
ggttacctcttatctagtggGTATACATGTGCCGTGCTGGTGTGCTACACCCATTAACTCgaatcttaaacccaact
ggttacctcttatctagtggTTGCTTATCAGCTTAAGGAGATACTGGCAAAAACCACATGgaatcttaaacccaactt
ggttacctcttatctagtggAGAGTCAGGGGCCACAGGTCTCAGGCTTCGACTGGATGTGgaatcttaaacccaac
ggttacctcttatctagtggGCTGAATCTCCTTCTAAGGGCGTTTGTGATATGAACTGAGgaatcttaaacccaactt
ggttacctcttatctagtggCCCCTCCCTCGAGCTTTACCCAGACTCAGAAGAAAGCCAGgaatcttaaacccaac
ggttacctcttatctagtggGTGGAAAACCTCACAAAAACAGTTAACTTCAACATTTGGGgaatcttaaacccaactt
ggttacctcttatctagtggAAGTGAATGATTCTACTTCCCAAATGCAGTAAACCCACTGgaatcttaaacccaactt
ggttacctcttatctagtggTTCCTCTAGATGATTGTGGCCCTTGAAAGTCATCAAGGTCgaatcttaaacccaactt
ggttacctcttatctagtggAGAGGGTGCTGTCAACAAAGAAATCAACTGAGTGGAAGTGgaatcttaaacccaact
ggttacctcttatctagtggTCCCATACGTAACTGAATAGCAAGAAAACAAGTAACCCAGgaatcttaaacccaact
ggttacctcttatctagtggGACAAGGGATAACAAGGTTGGCAATGATGTAGAGAAAAGGgaatcttaaacccaact
ggttacctcttatctagtggGTCACTATGGAAAACTGCATGGAGGAGCTTCAAAAAACTGgaatcttaaacccaact
ggttacctcttatctagtggCTGCATTCCCATGTTCGCTGCAGCCTTATTCACAATTGCCgaatcttaaacccaactt
ggttacctcttatctagtggGAGTGTCCATCAACAGATGAATGGCTATAGAAAACATATAgaatcttaaacccaactt
ggttacctcttatctagtggCCTGTAGGACATTGTGCTGAGTAAAATAAGCCTGTCACAGgaatcttaaacccaact
ggttacctcttatctagtggCCAGAGGGTGGGGTGGAGAGGGGTGGGGTACAGGGAATGGgaatcttaaacccaa
ggttacctcttatctagtggGACAAATAAGTTTTCAATGCTATTGCACAGTGTGGTGACCgaatcttaaacccaactt
ggttacctcttatctagtggATCATGTAGCCCCTGAGATAGATGGATATGTGAGCCTAGCgaatcttaaacccaact
ggttacctcttatctagtggCAGAACATATTGGGAGGAGATATTTGGCAGCTGAAGTGGCgaatcttaaacccaact
ggttacctcttatctagtggCTGCAGAGTCTCTAGCAGTGCTGGATCCAAGGGTATGCGGgaatcttaaacccaac
ggttacctcttatctagtggCACAGGCACATCCTGGCATGGGGAGCACCTCAGGAGCACAgaatcttaaacccaa
ggttacctcttatctagtggCCCACCCAACTGCAAGGGATACCCAGCCTACATGCAGAGGgaatcttaaacccaa
ggttacctcttatctagtggGCATGCAATAGGCCCCTCCCCACTAGGAGTCTATGGAGACgaatcttaaacccaac
ggttacctcttatctagtggCCAAGTGAGCTATGAAGCCAGATTTTCCCTCCTGGGCCACgaatcttaaacccaac
ggttacctcttatctagtggGAGACCAGCTACACTTGGTAGAGACAGCCATAGGAAAGTGgaatcttaaacccaac
ggttacctcttatctagtggAGGGTTTAGTAAAGCCAGCTGTTTCCACTTCTGAAAATAAgaatcttaaacccaactt
ggttacctcttatctagtggTGCTGGCTCTAGAAAGTGTGGGGTCTTTGTTGTGGAGCAGgaatcttaaacccaact
ggttacctcttatctagtggTGGAAGAAAGTGAACTGAACTTGGAAGTTTGAAGATCTCGgaatcttaaacccaactt
ggttacctcttatctagtggGCCTCAACTACTTGCATTAACTTGTACAAGTCATTCAACCgaatcttaaacccaactt
ggttacctcttatctagtggACTCTCCAAAGACAGGGAGACTGCTGATATAAGAGGGCACgaatcttaaacccaac
ggttacctcttatctagtggGTCAGCATAGGGCCCTGGGCTTACAACGGTGCCATGAGCCgaatcttaaacccaa
ggttacctcttatctagtggGGAAGGACAGCTATAGCAATGAAAGGACTAGTGCAGATTCgaatcttaaacccaact
ggttacctcttatctagtggACCAAGGTGTAGTAACATGTTTTAGTATGGAGGGGAAGGCgaatcttaaacccaact
ggttacctcttatctagtggGAAGGTAAAGATAGCAGCTAATGGAGACAGGACAGGACTGgaatcttaaacccaac
ggttacctcttatctagtggAGGGAGAGAAGAGATGTCAACTGCCTACTCTGGTAGCCTCgaatcttaaacccaac
ggttacctcttatctagtggCGCTCATAACTTTGTCTGGCTTAATCCTGCTCATCCCAGCgaatcttaaacccaactt
ggttacctcttatctagtggGTCTCCACGTTTTGGGAAGTCATCACTCACTTCTCTGGGCgaatcttaaacccaact
ggttacctcttatctagtggACATCTACCTTTCTCTATGGACTCTGAGCTTCTTGAGGGCgaatcttaaacccaactt
ggttacctcttatctagtggGAAGCACCCATACCTAGAACAAAGCTTAGCACATAGTAGGgaatcttaaacccaac
ggttacctcttatctagtggCCTCACAACAACCCTAAGCTGGAGACTCAAACAAGGCTGGgaatcttaaacccaac
ggttacctcttatctagtggGGAACACTGAACATTCACTGAGAACTGACAACTTGTGGGGgaatcttaaacccaact
ggttacctcttatctagtggGACTCTTGAATGCCAGTCTCTGTACCTGTACCATGTTGGCgaatcttaaacccaactt
ggttacctcttatctagtggGGGAGGTGGGACCTAGAGGGAAGTGTTTGGGTCCAGTGGGgaatcttaaacccaac
ggttacctcttatctagtggTGTCTTGTGGGAGTGGATGACTTCTGTCTTGCAGGACTGGgaatcttaaacccaactt
ggttacctcttatctagtggGTTGTGAAGCTTCTCCTCATGTTTTGCTGTGTTTGCACGCgaatcttaaacccaactt
ggttacctcttatctagtggGGCAGCACATAACCCTCTGCAAGCTGAGCAGATGCCAGTGgaatcttaaacccaac
ggttacctcttatctagtggGGATTTTCTAGCCACTAGAGTTGTGAGCCAAATAACATTTgaatcttaaacccaactt
ggttacctcttatctagtggACCCAGATCAGGTATTCTGTTAGAGCAACACTAAAGGGACgaatcttaaacccaact
ggttacctcttatctagtggCACACCTATTCATTTGCTCCAGAAAAGGTAGTTATTCTCCgaatcttaaacccaactt
ggttacctcttatctagtggCATCTTCCACTTCATGCTTATTCAATCCATTACCAATTCCgaatcttaaacccaacttg
ggttacctcttatctagtggAGCACATAGACCACTGCCAGGAACCCTTGACTTGCCGACCgaatcttaaacccaac

ggttacctcttatctagtggCTAGTTCTTCTTCCTACTGCAGCCGGAGCAATCTTTTTAAgaatcttaaacccaacttg
ggttacctcttatctagtggACTCTCTTGATGAAATCTCCTAGAAGACCTTTTGTAGCTCgaatcttaaacccaacttg
ggttacctcttatctagtggCCTCTCCAGCTTCCTCCCTCTGTGCTCCAACCACATGGGCgaatcttaaacccaac
ggttacctcttatctagtggCACCCAGCTTCAAATTCCCTTCTGGCTGCAGGGCCTTGGCgaatcttaaacccaac
ggttacctcttatctagtggTCTCTGCCTGATGTTTTCCACCCTTCCACCTACATTCATCgaatcttaaacccaacttg
ggttacctcttatctagtggTTGGAGCTCAGCTCAGACAAGGTTAGATCCTGCCATTGACgaatcttaaacccaact
ggttacctcttatctagtggGTGTGATTAACTGATAAATGACTGTCTGACCCCTCCTCCCgaatcttaaacccaactt
ggttacctcttatctagtggCTCAGTACTCTGTACCCAATGCCCACCCCAGCATCCTCACgaatcttaaacccaac
ggttacctcttatctagtggGTTCAGTACATGGGTGTTGAATAGATGCATGAATAATACAgaatcttaaacccaacttg
ggttacctcttatctagtggCTGTGATTCTGTGCTACTCCTCTGCGCTTCTCATGTACACgaatcttaaacccaacttg
ggttacctcttatctagtggCCACCTCCTGGCCTCAAGGGACTCTCCTGCCCCAGCCTCCgaatcttaaacccaa
ggttacctcttatctagtggAGATTACAGGCGCATGTCACTACACTCAGCTAATTTTTATgaatcttaaacccaacttg
ggttacctcttatctagtggTCTCAGGTGATCTACTGCCTGAGCCTCCCAAAGTGCTGGGgaatcttaaacccaac
ggttacctcttatctagtggGGATCCTTGGCAGGCCTTTAGCACACTGCACAGAAGGGACgaatcttaaacccaac
ggttacctcttatctagtggCTCTGGTTCCCTGCTTTGCCCAGACTGTTTCTATGTCCCCgaatcttaaacccaactt
ggttacctcttatctagtggCGGGGCATCTGCTTTGTGCTAAGTGTAAGCCCTGCCTCCAgaatcttaaacccaac
ggttacctcttatctagtggGCATAAGGAGAGACACACACATGAAAACCAACTAATTGTGgaatcttaaacccaact
ggttacctcttatctagtggGACAGCCCTGTAGAAGGAGTAGGCAAGTGTTCTAGTATGGgaatcttaaacccaact
ggttacctcttatctagtggTCGGAGGGATGAGTAAGAGTTTACCATACTTTTAAGGGGGgaatcttaaacccaactt
ggttacctcttatctagtggTGCAAATGCAGTATGAGGAGCCGTGGTACAGTTTAATGTGgaatcttaaacccaactt
ggttacctcttatctagtggGACGAAGGAGGAAGCTGGTGTGTTTGGAGAAGGCTGAGGGgaatcttaaacccaac
ggttacctcttatctagtggCCTTCCTTTAAAAGCCATGCCCTATTCTCCCTCTCCTGCCgaatcttaaacccaact
ggttacctcttatctagtggTGTCCCTGTGAGAACCCATCTTGCTGAGATTGTCGCTGGCgaatcttaaacccaact
ggttacctcttatctagtggGCAAGAATTTGTCTAACAGAATGAATGATCAACCTTGAGCgaatcttaaacccaacttg
ggttacctcttatctagtggCAGTTGGCTGCTCCCGCTGTTTGGCACCAACCAACCTGACgaatcttaaacccaac
ggttacctcttatctagtggGAAAAGCAGTTCTGCTTGGTGCATGCTGGCTGCTGCTGTGgaatcttaaacccaact
ggttacctcttatctagtggACCTGCTGTGTCCATTGGTCAAGCACATTCAGGACATCGCgaatcttaaacccaact
ggttacctcttatctagtggGACTCTGAGAGAGAGCAAGGGAGGTTTTATGACTGGGGACgaatcttaaacccaac
ggttacctcttatctagtggGCAGTACCCTCCAGACTGCTCCTCCTCTCACAAAAACACCgaatcttaaacccaac
ggttacctcttatctagtggCAGGGCTCATTCCTTCACTTCTTTCAGGTGCCTACTCAGGgaatcttaaacccaactt
ggttacctcttatctagtggGCATAAAGATTGAATCGGAGTAATTGGGATATCCATTGCCgaatcttaaacccaacttg
ggttacctcttatctagtggGAGAGCTGAGCATCTGTTTGGTTCACTGCCCAACACATGCgaatcttaaacccaact
ggttacctcttatctagtggAGATGAATGAATCATGGATGAATGAATCAAATCAGTCAGCgaatcttaaacccaacttg
ggttacctcttatctagtggTTTCTTCATTTCCTACCCCAATTTAGTTTCTACTCAGGACgaatcttaaacccaacttgt
ggttacctcttatctagtggGGCAAGGTCTTGCTCTGTTGCCAGGGCTAGAGTGCAGTGGgaatcttaaacccaac
ggttacctcttatctagtggCCCAAATAGCTGGGATGAAAGTGTACACCACAATGCCCGGgaatcttaaacccaac
ggttacctcttatctagtggCTCTGTTGACCAGGTTGCTTTCGAACTCCTGGCCTCAAGCgaatcttaaacccaact
ggttacctcttatctagtggGCGCGAGCCAACTTGCCCAGCCCTGGAATTTTTGAGCCTGgaatcttaaacccaac
ggttacctcttatctagtggGCATTATCTCCTTGCTTCATTGCACCATTAAAATCTTTCCgaatcttaaacccaacttg
ggttacctcttatctagtggATCAGATGCCTCTTCAAATACCTCCTATCAGAATGGTCTCgaatcttaaacccaacttg
ggttacctcttatctagtggGCTTCACCTCCTGCCTCCACCCCATCTGTACTGCTGGTCCgaatcttaaacccaac
ggttacctcttatctagtggCCTCTCCTGCTTCAGAGGCTTTGCCCTGCTCTTCTCTGCCgaatcttaaacccaact
ggttacctcttatctagtggCTGTCCTGCTCTTCTCTGCGTCTTGTGGTTGGAATGCCTGgaatcttaaacccaactt
ggttacctcttatctagtggACAGCAGAGAAATTCCTCCTCTGGCCTCTCCTCTTCTCTGgaatcttaaacccaact
ggttacctcttatctagtggTTTATTTATTTTCATTGAGATGGAGTCTCGTTCTGTCGCCgaatcttaaacccaacttgt
ggttacctcttatctagtggCAAGCGAACCTCTTGCTTCAGCCTCCCTCCTGTAGCTGGGgaatcttaaacccaac
ggttacctcttatctagtggCAGGCGTGAGTCACCGCACCCAGCCTCTCTCTTTATAATTgaatcttaaacccaact
ggttacctcttatctagtggTCACCTGCATCAAACTCCTGAATTCTGTCATGCAACTGGAgaatcttaaacccaactt
ggttacctcttatctagtggCATGGAGCTCAAAGAAAGATGTTGAGAAACGTAGAGTTGCgaatcttaaacccaact
ggttacctcttatctagtggGTCAGACAAGCGACAGAGACCTATTTACAAAGAGACCCAGgaatcttaaacccaac
ggttacctcttatctagtggCAAAAGTCACCTGCAATATTTAGGATAGTGTCTGAAACAGgaatcttaaacccaacttg
ggttacctcttatctagtggGTCAGTTGTCACTGAGGAAGCACTTTGGGGTAGAAGAAGCgaatcttaaacccaac

ggttacctcttatctagtggGTTTGAAGTCTGAGGCAGCAGGTGAGGTTTGATTTTATATgaatcttaaacccaacttg
ggttacctcttatctagtggGGATCTATCAGATGGTGGAGCTACCCTCATAAAAGATTTGgaatcttaaacccaactt
ggttacctcttatctagtggTCAAGTGTCATTTCTTCAGGGAAGATTGCCCTTCACCATGgaatcttaaacccaactt
ggttacctcttatctagtggCACAATGTGTGACAGTTCTTTGCTCACATATATATTTCTCgaatcttaaacccaacttg
ggttacctcttatctagtggCTGTTATCCAGGATGGGGTGCAATGGGGCAATCACGGCTCgaatcttaaacccaac
ggttacctcttatctagtggAGTAGCTGAGACTACAAGTGTGCACTATCAAGCCTGGCTAgaatcttaaacccaact
ggttacctcttatctagtggCCTCAAGTGATCCTCCCACCTTAGCCCAGAAGGAGTCTTAgaatcttaaacccaac
ggttacctcttatctagtggGTATCTACCTGTCATCAGGCCTATAGTAGATACTCAGTAGgaatcttaaacccaactt
ggttacctcttatctagtggGAGATAGCTCACCAAGTAGAACACTGGATGGTATTGGAACgaatcttaaacccaact
ggttacctcttatctagtggGACCCTGACCCCCAAGGTGCCGGTACTAAGAAGTGGGGCCgaatcttaaacccaa
ggttacctcttatctagtggGGGTGGAGCCCTCACAAATGGGATTTATGCCCTCATAAAAgaatcttaaacccaact
ggttacctcttatctagtggCTCTGAACCTGGATAGAGGACCTCACTAGACGCTGACCAGgaatcttaaacccaac
ggttacctcttatctagtggGTCTGAGACCATAATGAAGCCATTGGAATGGTGGGAAGGCgaatcttaaacccaac
ggttacctcttatctagtggGTCATGTTGCCCATAGAGCAGATCCTACTATTACAGTGCCgaatcttaaacccaact
ggttacctcttatctagtggAACTGATTAGTTTATGAATCAGTCTGCCAAAAACTAGGGCgaatcttaaacccaactt
ggttacctcttatctagtggGCCTTAAAGTTTGCAAGGAGACCCTATTAACTGCGCACTGgaatcttaaacccaact
ggttacctcttatctagtggGGGGGTTTCCCATATTAGCCAGCCTGGTCCTGAACTCCTGgaatcttaaacccaac
ggttacctcttatctagtggGCCACTGCACCTGGCTAGCCACTCACTGTTTTCATGTTAGgaatcttaaacccaact
ggttacctcttatctagtggCTCATGTCCACAGCTTGTTCTTTCATAGGCTGCCATGATGgaatcttaaacccaactt
ggttacctcttatctagtggTCACAGGTCTACATTTCGTCCTCTAAACTACTCTGTTGGGgaatcttaaacccaactt
ggttacctcttatctagtggATGTAGGATCACTTTTTTACTACAGGTTTTGTCATATGGGgaatcttaaacccaacttg
ggttacctcttatctagtggCTGCTGAAACAAATGGCTTTTAATCCTTAAAAGGGCAGGGgaatcttaaacccaactt
ggttacctcttatctagtggTCCTAGAGCTTAAGAAAGGGGAGGCCCAATAACCAGATGCgaatcttaaacccaac
ggttacctcttatctagtggCCAGTATCCAGACAATTGGCATGAAAATAAAGGAGCCTAGgaatcttaaacccaact
ggttacctcttatctagtggTTTGCAAAGCACTTCCACAAATGGCCACAAATGTCATCTCgaatcttaaacccaactt
ggttacctcttatctagtggGCTTAAGATATGGGTGTTTCTTGAACCTGGGAGGTGGAGGgaatcttaaacccaactt
ggttacctcttatctagtggCACTCCAGCCTAGATGACAGAGCGAGAGTTTGTCTCAAAAgaatcttaaacccaact
ggttacctcttatctagtggGTTGGAAGGAGTTTGAAATCGTTGAGGACTCAGCAGCATGgaatcttaaacccaact
ggttacctcttatctagtggATTCTAGAACTTTAGATCAAAAGTCAACTCAATATGGGGGgaatcttaaacccaactt
ggttacctcttatctagtggTGGGACAAGATTGTGGGATCCACAGAAAATTGGAACTTGGgaatcttaaacccaact
ggttacctcttatctagtggTCCACAGAGATAAGAAATACACTTGCTTTTATCTCACTTCgaatcttaaacccaacttg
ggttacctcttatctagtggGGGTTTTAGGCCCCAGAGAGAAATTGTAGCTGCAATCAATgaatcttaaacccaactt
ggttacctcttatctagtggGTCCTCAGAGGAAAGTGAGTAGGGAGCTGTCTGAAGGGACgaatcttaaacccaac
ggttacctcttatctagtggGAGAGGGATAATGGATTGCGTTTGCAAGTGCAGTTGGGGCgaatcttaaacccaact
ggttacctcttatctagtggGGATTCTTTTAGTGTCTTGGTTCTTACCTGTTGTGGTTGCgaatcttaaacccaacttg
ggttacctcttatctagtggGGGAGAGAAGTGCCTCCCACAGCACCACGACCAGAAAGGGgaatcttaaacccaa
ggttacctcttatctagtggGGCAATAGCCAAGGACTCCTGGCAGTGAACTCATGTCCACgaatcttaaacccaac
ggttacctcttatctagtggGGAAGCTGGAGCTCCTGCACTGACTATCAATGTGAGCCCCgaatcttaaacccaac
ggttacctcttatctagtggGATCTGGGTCAACAGACTCTGGTTCCTGTCTCAGCAGTGCgaatcttaaacccaact
ggttacctcttatctagtggCGGCAAACCAAAGCCTGCTTATCAAATCTGGCATACCACCgaatcttaaacccaac
ggttacctcttatctagtggGCTTTTGCGCTGCCTCAGGAGATCTGAGTAGTGGCCACAGgaatcttaaacccaac
ggttacctcttatctagtggTTACTTCTTGTTTTGGAGAGTTTGCTGACTGGCACCAGAGgaatcttaaacccaactt
ggttacctcttatctagtggGTGTGACCACACATTTCGAAATGCCGTCTTCCCAATCTAGgaatcttaaacccaactt
ggttacctcttatctagtggCTGGGTTTGGAATAAAGAAGTGTGAGGGGAGAAAAATTCCgaatcttaaacccaactt
ggttacctcttatctagtggAGACTACTGAGTCTTTTCTTGAGCAAAGGAGAGAGTGTGAgaatcttaaacccaactt
ggttacctcttatctagtggGAGGGTTTGATCCGATAGCCTTCTGCGTTAGAATGTAGTCgaatcttaaacccaactt
ggttacctcttatctagtggCTCTTCTGCATGTTCAGAGGGAGAAAATGAGTGATGAGAGgaatcttaaacccaactt
ggttacctcttatctagtggCCTTACATCGATAAGATAACAGAAGGCTTTTCACACTGGCgaatcttaaacccaactt
ggttacctcttatctagtggCAGAGGTGTTGATCAGCAAAGTCTTAGAGTGAGTCTAGGGgaatcttaaacccaact
ggttacctcttatctagtggTCTGGGTATTGGAACCATCAACAGAGCCTGTGCCCCAGACgaatcttaaacccaac
ggttacctcttatctagtggCTGTGCAGCCAGTTCGGGAACCACCAGTGAAATCCAACCCgaatcttaaacccaa
ggttacctcttatctagtggGATGTACCACTGTCCCAGCCACTTCACCTGTATTCCCAGCgaatcttaaacccaac

ggttacctcttatctagtggCAGAGAAAACTGCCAAGTCCCCACAGTGGCCTTGCATGGGgaatcttaaacccaa
ggttacctcttatctagtggGTCACCTCTCTGACCTCATCTCCTACTTTCCTTCACCCGGgaatcttaaacccaact
ggttacctcttatctagtggTCTCCAACACACCAAGCACACTCCTGCATCAGAGGCTGTGgaatcttaaacccaa
ggttacctcttatctagtggCCTGCTTTGTTCCTCCCCAATATTTGCATGAACCACTTTCgaatcttaaacccaactt
ggttacctcttatctagtggCTGTTCAGAAGTCACTTTCACAGAAAAGTCTTCCTCAGCCgaatcttaaacccaact
ggttacctcttatctagtggCCTACTCCATTCGGTAAGCTCTAATTGAGAACAGAAAAGCgaatcttaaacccaact
ggttacctcttatctagtggCACTGTTGTATCCCCAACATCTATATCTAGCACATCATTGgaatcttaaacccaactt
ggttacctcttatctagtggGAATAAATGGCTCTGCTAGATTTTTGTCACCTGGATTGCCgaatcttaaacccaactt
ggttacctcttatctagtggGAGCTTTGCAATCAGACAGCTCTGGCTTGCATTCTTCATGgaatcttaaacccaact
ggttacctcttatctagtggGTGTAAAGTGGAGGCATGGACACTGGAGGATCATGGTGAGgaatcttaaacccaac
ggttacctcttatctagtggGGCCAATAAGACCTCCTTGGGGAAAGAAGGCTAAAGTGGGgaatcttaaacccaa
ggttacctcttatctagtggCCCCACTGGGAGTTCCAAGGGGCTATGGAGTTGATAGGATgaatcttaaacccaac
ggttacctcttatctagtggCCTGAAGGAGTGGGGGATTGTGTAAGCCTCATTGGTGCCCgaatcttaaacccaac
ggttacctcttatctagtggGGGGAGCAGCAGGCAGCCTTCTCTGGCCAACATGTATCAGgaatcttaaacccaa
ggttacctcttatctagtggGGGCTGGCTGGAAACAGGCCGGTAAGCCCCATGGAACTACgaatcttaaacccaa
ggttacctcttatctagtggATGGCCATGTAGAACAGCCATGTAGAACAGGATGTCTCCCgaatcttaaacccaac
ggttacctcttatctagtggGCCCACGTTCTAGGTAACTGGCTACTCACTTTCACAGCCCgaatcttaaacccaac
ggttacctcttatctagtggTCCAGATAAGGAAACAAGACTTGGCAAATGTGTAGTGTAGgaatcttaaacccaact
ggttacctcttatctagtggGTTTAGCATGCACATGAACTTGCAAGACCTTGCTCTTCCCgaatcttaaacccaact
ggttacctcttatctagtggGTCCTTTACTAATTCTGTGATCTTACAAGCTGTGCCTTATgaatcttaaacccaacttg
ggttacctcttatctagtggAAGCTGTTGAGAGTCAGGCACTCAACCCAACAACATGCCCgaatcttaaacccaa
ggttacctcttatctagtggACCCAGCCAGAGGTAGAACCCAAGATTCCGACTCTAGTTGgaatcttaaacccaac
ggttacctcttatctagtggCGGCCCATTGATTTAGACAGCATGGAACTCATAGCCCAGGgaatcttaaacccaac
ggttacctcttatctagtggCCCTTCCAAACATACGACTCAGTAATTCCAGTAGGAGGGCgaatcttaaacccaac
ggttacctcttatctagtggTCGCACCTCCCTTCTCTTCAGGATGCTGAGGACCTCTGGCgaatcttaaacccaac
ggttacctcttatctagtggCTGGCACTGTCATCAAAGTGATCCCTCTCCAAGGACATGGgaatcttaaacccaac
ggttacctcttatctagtggCTCGAAGTACGATCAGGGCAGGCACAGCAAAGGAGGAAGGgaatcttaaacccaa
ggttacctcttatctagtggGGGGTCAACAATAAAAGCCAGCCTGCTGCCTGTCCCCACCgaatcttaaacccaa
ggttacctcttatctagtggCTGGGCTGTTCATGTCCCCAGTTATGCCCCCAAACATTCCgaatcttaaacccaac
ggttacctcttatctagtggCTCTCTCATTCTAATTCATAGCAGGGAAGTGGACTGGCTCgaatcttaaacccaact
ggttacctcttatctagtggGGGCAACTGTGTTTTCCCTCATCTCCAGACAGCATGGATGgaatcttaaacccaac
ggttacctcttatctagtggGGAGGTGAAGTCATAGCCTCGGCAAACCTTTGAAGCTGGGgaatcttaaacccaac
ggttacctcttatctagtggGGCCAAGGCCAGAGGAGGTAAACAGGGTTGCATGATGTAAgaatcttaaacccaa
ggttacctcttatctagtggGCTGTTTCTGCAATAGTGTTGTTTGGGTACACGACAGGACgaatcttaaacccaact
ggttacctcttatctagtggTCTTGTATAGGGATACATGCCAAGTGCCATTCCTGATTCCgaatcttaaacccaactt
ggttacctcttatctagtggGCAAAGATGAACATGGCCTTGTCCTTACCCTTAAAGAACCgaatcttaaacccaact
ggttacctcttatctagtggCAGGATAGTTAGTTTGTGATAGAGAAAACCAAATGGTCCCgaatcttaaacccaact
ggttacctcttatctagtggTCACCCCAAAATTCACATCTACCTTGAAGTGCAAAATGGCgaatcttaaacccaact
ggttacctcttatctagtggGGAGAGAGCATGCACTCAGACACACACAGGGAAAAAATCCgaatcttaaacccaa
ggttacctcttatctagtggGACTGAAGCCAAGAAAAGACAAGGATTGACAGGAGCCACCgaatcttaaacccaa
ggttacctcttatctagtggGAGCATGGCTCCACTGGCACTGTAATTTCAGGTTGCTAGCgaatcttaaacccaac
ggttacctcttatctagtggGTCATAGCAGCCCTAGGAAGCTAACATACAGGTGAAGGACgaatcttaaacccaac
ggttacctcttatctagtggGAGAAGAGATGTCCTAAGATGTTTCTTTGGTGACATCATGgaatcttaaacccaactt
ggttacctcttatctagtggGCTCCACTCATCAAGGGGTGGCCCATATCCAAGCTGGAGCgaatcttaaacccaa
ggttacctcttatctagtggCTCTTTCTGCAGTTTGGATATTTATGCTGATCATAGCTCCgaatcttaaacccaacttg
ggttacctcttatctagtggTTGGCCGTTTTGCTTCCTTGATTCCAGGAGCTGTCCCAGCgaatcttaaacccaact
ggttacctcttatctagtggCCTACAACTCAGTTCTTTGGCTTTTCCTTCTTTGTTTGCTgaatcttaaacccaacttgt
ggttacctcttatctagtggTTGCTTGCAGCAAAGAAGTCTAACTGATATAGGACCCATGgaatcttaaacccaact
ggttacctcttatctagtggGTTGCAAGAACAAATCTGCTATATTTATGGCAACTTGTGGgaatcttaaacccaactt
ggttacctcttatctagtggAACACCTTAATTCATCATGGTCTCTTCATCTGGAATCCCCgaatcttaaacccaactt
ggttacctcttatctagtggTCAGAGTTTAACTAAACTGGGCTTTAACTGAGGTGCCTGCgaatcttaaacccaactt
ggttacctcttatctagtggATGGGCTTCCCAGGTCATATCCACCAATGGGTCATTCTTGgaatcttaaacccaact

ggttacctcttatctagtggATGGCAGGTACTCAGCCCAGTACCCGACATATAGAGATGCgaatcttaaacccaac
ggttacctcttatctagtggTCCTGACCCTCCAAAGGTCATCCTGACTCTTCTCAAAGCCgaatcttaaacccaact
ggttacctcttatctagtggAGCCCCCTTCCTCTTTAGCTTGCTGGGGTCTCCTGACTGGgaatcttaaacccaact
ggttacctcttatctagtggGCTCAAGGAACCCCTCCCTCAACCTCAAAAAGAGAAGACAgaatcttaaacccaac
ggttacctcttatctagtggTGTCAATTCAATCCAATCTCAAAGTACGTTTTGCCTTCTGgaatcttaaacccaacttg
ggttacctcttatctagtggATGGGGAAAAAACAATACCTGGAGTCATTATATGTGCATCgaatcttaaacccaactt
ggttacctcttatctagtggTGCACAAGCATTTTACTTGAATATTTCCAGGTCCTGCAAGgaatcttaaacccaactt
ggttacctcttatctagtggGGCAGCCCAAACATTCTTCCAGCAGAATGGCCATTCAGACgaatcttaaacccaac
ggttacctcttatctagtggTACCCTTCCAAAAGCATACACTATTGATTCTGCCTCTCTCgaatcttaaacccaactt
ggttacctcttatctagtggGAAGAAAAACTGACTTCTCATGAACACAACTTGATATCTCgaatcttaaacccaactt
ggttacctcttatctagtggCAATGTTTGCTTGGACCAGACTATTAAAACAGCCACTTGGgaatcttaaacccaactt
ggttacctcttatctagtggTCCCCCAGGACTCCCATCCTATTCTATCTTACCTCTAGGGgaatcttaaacccaact
ggttacctcttatctagtggTTTTTCAAACTGGAACTTTCAAGAGTGAAAGGGAGGATGGgaatcttaaacccaactt
ggttacctcttatctagtggCAGTGCTGTGTCTCTTCCTCTAGACATTAAGCCCCTGGCCgaatcttaaacccaact
ggttacctcttatctagtggGGGAAGGATCTGCACTTCAGTCATGGTCCTGTGTCATGGCgaatcttaaacccaact
ggttacctcttatctagtggACAATCTCCCTCAAACTTACAACTCTTTTGTTGTAATCACgaatcttaaacccaacttg
ggttacctcttatctagtggGGCTAATTGCTTATCCTTCAAGACACCTGATGTCAACTGCgaatcttaaacccaactt
ggttacctcttatctagtggTGTGTAGCATTCTGTCTTGATATCTTTGTGCTTGATCATGgaatcttaaacccaacttgt
ggttacctcttatctagtggCAGCTGAGAACTGCTGTGAACTCCATTTCTTCTCTATACCgaatcttaaacccaactt
ggttacctcttatctagtggTCGGAGCCTGTCACAGAATAAGGAAATAGTACATAACAACgaatcttaaacccaact
ggttacctcttatctagtggGTCAACTGCTGTTTGGTTTGCCTGATACTGATTATTTGCCgaatcttaaacccaacttg
ggttacctcttatctagtggGGACCCACAGACTACTATAAACTAAGAGACAATTCCAGACgaatcttaaacccaact
ggttacctcttatctagtggTCTGATAGTAGCACTACCAATGGCATGATATATTTAAAGGgaatcttaaacccaactt
ggttacctcttatctagtggTTCAATTTCATGGGAGAACAAAAGAGTGATTAAGAGCTAGgaatcttaaacccaactt
ggttacctcttatctagtggAGCCCAGAGGAAAAAAGTATATGTAGCCATTCAGTGATTCgaatcttaaacccaactt
ggttacctcttatctagtggTTCAAACGATGGCATAAGTTAAACTGACAGCTATGTTTCCgaatcttaaacccaactt
ggttacctcttatctagtggGGCAGTAGAGGTAGTAGATCAGGTCAAAGTGGGTTGTGCCgaatcttaaacccaac
ggttacctcttatctagtggGAGGAACTATGCATGTCAGCGTCCATGCTAAGCACGAGGGgaatcttaaacccaac
ggttacctcttatctagtggTCAGTTAAGTTCCTCCCTTACCGAACTTAAGTGTGTTGGGgaatcttaaacccaactt
ggttacctcttatctagtggTTCTCCCATTATGTAGGTTGCCTGTATACTCTGATGATAGgaatcttaaacccaacttg
ggttacctcttatctagtggAAAACGTCAGGAAAAAACCGATGTTGGAAAGGATGTGGAGgaatcttaaacccaact
ggttacctcttatctagtggGGATATGTTCATGTCATTCACAGGGACATGGATGAAGCTGgaatcttaaacccaactt
ggttacctcttatctagtggCACTGGGGCCTGTCGGAGGGTGGGGGGTTAGGGAAGGGATgaatcttaaacccaa
ggttacctcttatctagtggGATGGGTGCAGCAAACGACCATGGCATATGTATAACTATGgaatcttaaacccaact
ggttacctcttatctagtggCACCCCCCAAAAAAGTCATACCCGTAGGTTAAGGCATCTCgaatcttaaacccaac
ggttacctcttatctagtggGTATTCTTATGTTGGGGGCAAGGAAGTAGCAAATTAAGCTgaatcttaaacccaactt
ggttacctcttatctagtggGTTTATGGAATTATTTTTCAGATGGGTTCCCATTGAAGAGgaatcttaaacccaacttg
ggttacctcttatctagtggGAGCTTTGAAGGATAAGAAAAAAGAAATGAGCTTTGAAGGgaatcttaaacccaactt
ggttacctcttatctagtggAAAGACAATCATGTTGGGAAAACTGGATATCCATGTACAGgaatcttaaacccaactt
ggttacctcttatctagtggCACCTGAATCTATAAAGCTACTAGAAGAAAATGTAGGGGGgaatcttaaacccaact
ggttacctcttatctagtggTGGTTTTAGGTATGACTTCAAAAGCACAGGCAGCAAAAGCgaatcttaaacccaact
ggttacctcttatctagtggGCATAGCACAAGAAACAATTACAGAGTGAAGAGATCACCCgaatcttaaacccaac
ggttacctcttatctagtggGTCATAACTGTTAAAGAGCTAATATCCCAATTAGACAGGGgaatcttaaacccaactt
ggttacctcttatctagtggAAACGGGCAAAGAACCGGAATAGACATCTCTCAAAAGATGgaatcttaaacccaac
ggttacctcttatctagtggGTGTTGGCGAGGATGTGGAGAAAAGTGAATCCATACACACgaatcttaaacccaact
ggttacctcttatctagtggAAATTAGTACAGGCATTTTGGAAAATAGTATGGAGGTTCCgaatcttaaacccaacttg
ggttacctcttatctagtggTAGAAGTCAGTATACCAAAGGAGATACCTGCACTCCCATGgaatcttaaacccaact
ggttacctcttatctagtggAAATGCAGTACATCTACACAATGGAATACTATTCTGCCTGgaatcttaaacccaactt
ggttacctcttatctagtggCGTTAAGTGAAATAAGCCAGGCACACTTACATGTGGAAGCgaatcttaaacccaact
ggttacctcttatctagtggTCAGAGGCAGGTGGGGGCTAGGAGGGTAGTGTATGGGGAGgaatcttaaacccaa
ggttacctcttatctagtggGCAGGAGATGTTGGTCAACAGGTCCAAAGTTACAATGTTAgaatcttaaacccaactt
ggttacctcttatctagtggCGCATTTATCCACTTGCTTCAAATCATAATTGCTAGGAGGgaatcttaaacccaactt

ggttacctcttatctagtggTCAAAATAGCTAAGAGATATGACTTTCGATGTTCTCACCCgaatcttaaacccaacttg
ggttacctcttatctagtggCCTTGATTTATCTTGCTATCTTGCAGTGCTTACATGCGTCgaatcttaaacccaacttg
ggttacctcttatctagtggCTGTGACTATACTTCCGGTCTCAGATATCTTACTATCTAGgaatcttaaacccaacttg
ggttacctcttatctagtggCCACCTACATCAAATCCACAGCTAAGCAGTCCATAGCTGCgaatcttaaacccaac
ggttacctcttatctagtggCCACAAAGTCATCAGAGACCTAGACTCCTGTCATTATGCTgaatcttaaacccaact
ggttacctcttatctagtggCGTGAGCCACCGCGCCTGGCCTATGTAGCTATACTGTTTAgaatcttaaacccaact
ggttacctcttatctagtggGGGTCTCCAGAATTTCTGCACATCTTGTAGACAGAGGCACgaatcttaaacccaact
ggttacctcttatctagtggGCCTTGGAAGCTAGAGATCATGTGTCCCTTCAAAGCAGAGgaatcttaaacccaact
ggttacctcttatctagtggGTCTGTCTCTCTGGGGCGAAGATTAGGCATGAGCACTGCCgaatcttaaacccaac
ggttacctcttatctagtggGGCCCTATTCATGTCACCCTATGGGAATTGAGGCCCAGGGgaatcttaaacccaac
ggttacctcttatctagtggCATGCCATGTGCCGATACCTATGACACTGCAGCAGGCTACgaatcttaaacccaac
ggttacctcttatctagtggTCAGACCCTTCACAGTTCTTGACAGTCATCCATAACTCGCgaatcttaaacccaact
ggttacctcttatctagtggAGGTGTCTAACTCCAAGGTGTCTAGAACAGAGTAGGTGACgaatcttaaacccaact
ggttacctcttatctagtggCCAGTTGACATAAAGTCCCAGATGATACCACTGAGGAGCCgaatcttaaacccaac
ggttacctcttatctagtggGGGCCTCATCCCTTCAGAGAAACTGTTCTTCCAGGGTCTCgaatcttaaacccaact
ggttacctcttatctagtggGGCACAGCCTGCTGACCCTTTCCACTCTGTATCTCACTCCgaatcttaaacccaac
ggttacctcttatctagtggCTCTTTCTTTTCTTGCCTTCCATGAAACTTTTCTCTCCTGgaatcttaaacccaacttgt
ggttacctcttatctagtggGCCATAAATGCCCACCCTCCCTCCAGCCCTCTTCTCACTCgaatcttaaacccaac
ggttacctcttatctagtggTAACCCCCAATTCGTTACCTCTACCTGGATGTGTTATTGGgaatcttaaacccaactt
ggttacctcttatctagtggCGCACCTCCCAAACTGGTAGAGCTGGCATCCCTTTCTCAGgaatcttaaacccaac
ggttacctcttatctagtggCGACTCTAAGGCCAATAACATTCCCATCCAGTACTATGGCgaatcttaaacccaact
ggttacctcttatctagtggGGCCCTCACTATCTATTACGCTGGCCCCATGTTCAAACCGgaatcttaaacccaac
ggttacctcttatctagtggGCTGCTGCCCTGATTAAAATCTTTCTTTGGCTTTTCTCGGgaatcttaaacccaacttg
ggttacctcttatctagtggCACTTCCTATTTCTCTCCATCTCACAGCCTGTGCCCCAGCgaatcttaaacccaact
ggttacctcttatctagtggTTTTCCAGCACGTGCCCTGTTAATATAGACCTTGCACCGCgaatcttaaacccaact
ggttacctcttatctagtggCTACCTGCTATTTGACATCACCCCCAAATCCAGTCTAGGGgaatcttaaacccaact
ggttacctcttatctagtggACCCCTCACTCTTAAGACTGGTGGATTACTGCCTACCTGGgaatcttaaacccaact
ggttacctcttatctagtggCATTGCTGTACCATACTAACTCGTTAACATACCTGAAACCgaatcttaaacccaactt
ggttacctcttatctagtggCCAACACAGAGCACAAATCCAGACGCAGAGTGGAGGCTCGgaatcttaaacccaa
ggttacctcttatctagtggAAGGCAGTTACTGAAGTGGAATGAAAGGAAGAAAGCACAGgaatcttaaacccaac
ggttacctcttatctagtggCTCTGGAATGTTCTTAGGCAATGATCACCCTCCTGGGCCCgaatcttaaacccaact
ggttacctcttatctagtggGTCCTTCACCTCCCAGTCGGTCCAAAGATAAGCCAGTGGGgaatcttaaacccaac
ggttacctcttatctagtggGCTACAGGGCAATGGTTTGTCTTGAACTACAAACCTAAAGgaatcttaaacccaactt
ggttacctcttatctagtggGGGTGCAGGTGCCTCTGCTAACCTCACCACTCCTGGGTCCgaatcttaaacccaa
ggttacctcttatctagtggTTCACTTCTCTAAACCTCTGTGTCCCCATATGCAAAATGGgaatcttaaacccaactt
ggttacctcttatctagtggGGAATGTTTTCACAGTGCCCAGAACAGGGTAAGTGCTCAGgaatcttaaacccaact
ggttacctcttatctagtggGGACTCAGCTCTGTTCCAGCTATTCTTCAAGAGGTGTGACgaatcttaaacccaact
ggttacctcttatctagtggGGCTCAAACACCTTTCCTCTTCAATGTGTAGTGGAGTTCCgaatcttaaacccaactt
ggttacctcttatctagtggATACACCCTGTGGAATGAAAATCAAAAATAGTTACTCTGGgaatcttaaacccaactt
ggttacctcttatctagtggCCAGGCCCTCCTTCATGAAATGTCTCTTTTCTGGCCTCCCgaatcttaaacccaact
ggttacctcttatctagtggCATGCACATCCTAAAAGTCCACTCCTGTCTTCTATACCCCgaatcttaaacccaact
ggttacctcttatctagtggTCACTTGCCTAATTGCTATTCCCACTGGGGTATTCCACAGgaatcttaaacccaactt
ggttacctcttatctagtggACACAAACCTCATACCCAGGATAGTACACACCAAAGCCAGgaatcttaaacccaa
ggttacctcttatctagtggTTAACTCCCAGATCTAATAGTCAGTTGCTGTGTGACCTGGgaatcttaaacccaactt
ggttacctcttatctagtggCCCTCTGTGCCTCCTTCTTCTCATCTGATATTGATATTACgaatcttaaacccaacttg
ggttacctcttatctagtggTATCACGTGTAAACACTCAGAATAGCTCCAAAAACATGGCgaatcttaaacccaact
ggttacctcttatctagtggTCTGAAAGCCATCTCTGAACTCCTACTTCCATTTAGTCACgaatcttaaacccaactt
ggttacctcttatctagtggGGAGAGAGTCTTGCTCTGTCGCCCGCACTGGAGTGCAGTGgaatcttaaacccaa
ggttacctcttatctagtggGCATGAGCCACCATGCCTAGCCACTTCTTTTCTAAATATTgaatcttaaacccaactt
ggttacctcttatctagtggGTGTAACAGGCTCTGAAATGGTTCACTCATTTGTTCATCCgaatcttaaacccaactt
ggttacctcttatctagtggTGGGATAGTCATGCTCTCCTCCTTCTGGTCTTGCTTCCCCgaatcttaaacccaact
ggttacctcttatctagtggCAGACAGTGCCATTGCCCCATGATCCCCAATGCCTTCATGgaatcttaaacccaac

ggttacctcttatctagtggGCCATGATCTGACCCTTATTTACCACTCCATCCTCTCTGCgaatcttaaacccaactt
ggttacctcttatctagtggGCTGTAAGCACTCGTCTCCTTGCTTTGTAGAGAAGGAGCCgaatcttaaacccaact
ggttacctcttatctagtggTTGGCGGTTTCAGCAAGACGTCTTCCCTGGTTCCCAGACCgaatcttaaacccaac
ggttacctcttatctagtggGTTTTCTGGTTTATTCATCTGCCCGGCATACTGTAAGGGGgaatcttaaacccaactt
ggttacctcttatctagtggTCTAACCAAAGGAACAAATAAACTCTGAAACTTCAGTGGCgaatcttaaacccaactt
ggttacctcttatctagtggGGAGTTGTCCAAGCAGAGATGACTTTTCCTAGCCTCCCTGgaatcttaaacccaact
ggttacctcttatctagtggGTCAGGGACTGTGTCTTGCTTATTGTTTGTTTTCTCTGCCgaatcttaaacccaacttg
ggttacctcttatctagtggACCTCTCTGGTTGCAGCTTGTCTTTTTGCTAATTACACAGgaatcttaaacccaacttg
ggttacctcttatctagtggCAAGTCTAGGGGCATGTCACTTTTCCATCAGCTTCCCTGCgaatcttaaacccaact
ggttacctcttatctagtggGTTCTCTTCTGCTTGGTGATACAAGAGGGCTTTTCCACTGgaatcttaaacccaactt
ggttacctcttatctagtggCCTGTGTGTTACCAGCAAAAACTTTTTTCACAAGCTGCTGgaatcttaaacccaactt
ggttacctcttatctagtggGCTCGTTCATTTCCAATTCAAGGACATCAACAATCCTCTCgaatcttaaacccaactt
ggttacctcttatctagtggCTTGTCCTATGTGATGTCAACACTTTAATCTGTCTATGGCgaatcttaaacccaacttg
ggttacctcttatctagtggTGACCCACAGCGTGTTACATTTGGCATTGCGGTGACTCAGgaatcttaaacccaact
ggttacctcttatctagtggGCAAAGCAAAACGCAAAATTTTGGGATTCTGATCCTCAGCgaatcttaaacccaactt
ggttacctcttatctagtggCCAGAAGCCTTGGTGTGTGGGGGGTGAGTGGATGTGGCTGgaatcttaaacccaac
ggttacctcttatctagtggAGTCTCACAACTTGGCTGCGGGCAAGCTCTGGTGTTTAGCgaatcttaaacccaact
ggttacctcttatctagtggGCTTGTCCTTTCACCTGGAATGTTGTCAGGAACCTGGGGCgaatcttaaacccaact
ggttacctcttatctagtggCCTTGGGCAAGTTGGCATGCTGCTCCGGGCCTCAGTTGCCgaatcttaaacccaac
ggttacctcttatctagtggTGGTGTCTTTAGGTCCAACCTCAGGTCTCCCTACATGTTGgaatcttaaacccaactt
ggttacctcttatctagtggGTGCACTACCAAGGTTTCTAACAGCCCAGTTCTGTCCTGGgaatcttaaacccaact
ggttacctcttatctagtggGTTTTCAGCTTGGGTGAGTCACTTTTCTCTCTGGACCTCCgaatcttaaacccaactt
ggttacctcttatctagtggGAATATCTATGATTCTGTGCCTCTCTGTATCTGAGTCCCCgaatcttaaacccaactt
ggttacctcttatctagtggGGGCTGCTTCCCAGGTTTATTTTGGGTGCAGTGGCCAGTCgaatcttaaacccaact
ggttacctcttatctagtggGACTTCACCCTACTGGCCCAGCAGAGCTATTGGGTGCTGCgaatcttaaacccaac
ggttacctcttatctagtggTGGGGTTGAGAGACATGTCCCTCAGGCATTCTCTTGTGCCgaatcttaaacccaact
ggttacctcttatctagtggGGCAAAACACAGCCTCCAACTCCAGTCAAACCACCACCACgaatcttaaacccaa
ggttacctcttatctagtggGCAGAAAAGAACTGTTGGGTGTAGTAGGGTGGAGGTGAAGgaatcttaaacccaac
ggttacctcttatctagtggGCCTTTATGAAGCATACATCACTTCGCACCAAGCATCACGgaatcttaaacccaact
ggttacctcttatctagtggTGTGGTCCCTGTTTTTAAGGAGATCACAGTTTTCTCAGGCgaatcttaaacccaactt
ggttacctcttatctagtggTCTTGGACATTCACCTAGAGATTAAAATGATGATTCTAGCgaatcttaaacccaacttg
ggttacctcttatctagtggGGTGGGAATAATGAATCTTTCAGAATTACTAAGTGGGGCCgaatcttaaacccaactt
ggttacctcttatctagtggTTCACCAGTGGTGACCTCCCCTCAAAGTCAGATACAGGGGgaatcttaaacccaac
ggttacctcttatctagtggGAGAGTCAAAACCTACCAGAAGGAGCATCACAAGGGAGACgaatcttaaacccaa
ggttacctcttatctagtggCCTTGCTCAGGTGCTCTGGGACTAGTTTATGTGCCAGCTGgaatcttaaacccaact
ggttacctcttatctagtggGTCTGCTGGACCTTCCATCTCCCAAACTGTAGCTCCCTGGgaatcttaaacccaac
ggttacctcttatctagtggGGCAAGGGAAGCTTCTTAACTCCTCTTAAATCAGCTAAGCgaatcttaaacccaact
ggttacctcttatctagtggCCAGTGAGGGCTTCTCTCCATCCTAAAACTTTCCCCCTGGgaatcttaaacccaact
ggttacctcttatctagtggTCTCTATGTTGCCTTGAAAGGGAAAAGAGAGCCGCAGGGTgaatcttaaacccaact
ggttacctcttatctagtggGGACCATGCCTAGAATGGCTTACATAATTTCCACCCACATgaatcttaaacccaact
ggttacctcttatctagtggACCAAAATGCTAGGGATTTGGTCTAGGCCCTGCTGCTCACgaatcttaaacccaact
ggttacctcttatctagtggTAATTGGATGCTGCAGCTGAGGAGTTAGGAGATCAGTCTCgaatcttaaacccaact
ggttacctcttatctagtggGGTGGAAAGTTAGGCAATCCTCATGGGTGAGGGGTCTGGCgaatcttaaacccaac
ggttacctcttatctagtggCCTCCTTGATACTGTCTGGGAGGTGTGATGGTTGGTTTCCgaatcttaaacccaactt
ggttacctcttatctagtggGACAAATGTAACTTTCTTGAGTTTCAAGACTGGGGGGATCgaatcttaaacccaactt
ggttacctcttatctagtggGGACCAGTTTCAGTCTCCCCTTTCTATTTTCAGTTCCTCGgaatcttaaacccaactt
ggttacctcttatctagtggGCTGCTTCATGCTGAGGAGGGGCATTACAAGCAGCTCCATgaatcttaaacccaac
ggttacctcttatctagtggGTCTCCTTCTATGACACAATCTTTCTCTCTCTAGTCTCCCgaatcttaaacccaactt
ggttacctcttatctagtggGCTTCAGTCCCATATACTTGGCCCGATTACCCACAGAAAGgaatcttaaacccaact
ggttacctcttatctagtggGGGTCTCCAAAATTGGCTTTCCTGGAACCATTCACAAGGCgaatcttaaacccaact
ggttacctcttatctagtggACCAGTTCCTCCAACTGTGTCTCATTGTAAAAGAAAACAGgaatcttaaacccaactt
ggttacctcttatctagtggCAAATTCTACAGAAATCAGGCAGAGAGAGAAATGTGCTTCgaatcttaaacccaact

ggttacctcttatctagtggGAGTACACTCTACTCAATTGCTAAAGGTTGTAAACAGCTCgaatcttaaacccaactt
ggttacctcttatctagtggCCTCCATTAGCTCAGTCCATGCAATCAACTCCCACTCTGCgaatcttaaacccaac
ggttacctcttatctagtggATCCTGGAAGTTTTCTCTCTAGCCCAATGGCAAAATCTCCgaatcttaaacccaact
ggttacctcttatctagtggGAATTCCCCAAAAGAAGCAAGTTCCAGACTGTAGCTGATGgaatcttaaacccaac
ggttacctcttatctagtggATTATGGATGAGAAAAGTCTCAACAGACACAATTGACAAGgaatcttaaacccaact
ggttacctcttatctagtggCCATCACAATGGCCCAGATACTGGTATTGCATCAGTGTGCgaatcttaaacccaac
ggttacctcttatctagtggTGCGCCCACATCTTCCACAATAGTTCCTGGGCCTACAGGGgaatcttaaacccaa
ggttacctcttatctagtggGCCCTGTGTCTCGGGCAAGTTGTTCATTTTGATTGTCACCgaatcttaaacccaactt
ggttacctcttatctagtggGACTACTGTCAATGTTCAAGATTTGGCATGAGTCTGTGCCgaatcttaaacccaactt
ggttacctcttatctagtggGCACTGTGAATTGGTGTCTAGTAGTTAATGCCTGCAGCACgaatcttaaacccaact
ggttacctcttatctagtggCTGTGACCCAAAGCATCAAAAATGTGATAGGTCCAATGCCgaatcttaaacccaac
ggttacctcttatctagtggACTATACAGTGAAACAGGGACAAAGTAAAAACAAGCACAGgaatcttaaacccaac
ggttacctcttatctagtggGAGCATCATCATACATTTCCAAGACTGGTTTCTAGATACGgaatcttaaacccaactt
ggttacctcttatctagtggGTTTTCAAGTACACACATTAAGGCCTAATAGTGATAACAGgaatcttaaacccaactt
ggttacctcttatctagtggCACTAGAAAGCCTGACATTTTTACTACTACTTAATCCAAGgaatcttaaacccaactt
ggttacctcttatctagtggAGTGGTAAATACAACTAACATAGTCTGAGAGAAATCCCAGgaatcttaaacccaact
ggttacctcttatctagtggCCTGAACATTAAAACTTTGTACCCCTATCACAATTTTTCCgaatcttaaacccaacttg
ggttacctcttatctagtggAAAACTGTCTAAACATTTCTTCTCTTACCTGCTTTTCCAGgaatcttaaacccaacttg
ggttacctcttatctagtggAGGGGTTCTTTTTTGACCCATAGCTTAAAGCTCTTGTAGCgaatcttaaacccaactt
ggttacctcttatctagtggTGGCTGGTGTCCCCTATCCATTTTTGCAGGCCTGACAAAGgaatcttaaacccaact
ggttacctcttatctagtggATGTGTATATACCAGTAAAACCCAAAGGAGAACAAATAGCgaatcttaaacccaact
ggttacctcttatctagtggAACAAACTGGAAACCAACCCCCAATTTTTCACCTACTCAGgaatcttaaacccaact
ggttacctcttatctagtggGGTTACAGTATAATCCAGAGCCTAAAACAAACATATGTTGgaatcttaaacccaactt
ggttacctcttatctagtggAATGTCTCTAAGTCCAGCAACATCACTATACCTTCTGTGCgaatcttaaacccaactt
ggttacctcttatctagtggTTTACTTCAAGCACGTGACCAGTAAGTACTTTAGTGCTAGgaatcttaaacccaactt
ggttacctcttatctagtggCAAAGCAATGCAAGCCTGTATGTAAAATTTGGCTCCATGCgaatcttaaacccaact
ggttacctcttatctagtggGATCTTTAGCTATGAGCAATGTTAATTAGCCAAATGTCTCgaatcttaaacccaacttg
ggttacctcttatctagtggTACACAAACCATTCAACATGCTTGGACTTCGTTTTTGTCCgaatcttaaacccaactt
ggttacctcttatctagtggTAGGACAAAAGTTTACCATGGTCCATAATTTGTATCAACCgaatcttaaacccaactt
ggttacctcttatctagtggAGTGAAAACCTAGGAAGTAAGACATTCTGAACTAGCTGTCgaatcttaaacccaact
ggttacctcttatctagtggCATGCTTCCCAACGTCATAACCCTTTCTTTCTCTCCTTTCgaatcttaaacccaactt
ggttacctcttatctagtggGGCCTGAGCCACCACATCCGGCCCCATAATCCTTTCTTAAgaatcttaaacccaac
ggttacctcttatctagtggCCCAGATGGCCAAAAAGCATCTATTATTCAATCCTAAATAgaatcttaaacccaactt
ggttacctcttatctagtggATAGCCTGCGAAGATCAGATGTTCACCTAAGTAAGAACACgaatcttaaacccaact
ggttacctcttatctagtggCCTTTTATGCTAAACCCCGACACCCCAAAATATCTAGCAGgaatcttaaacccaact
ggttacctcttatctagtggCCCAGACAAAAAGGTATGCTGGCAATTCCAAAGACATTTCgaatcttaaacccaact
ggttacctcttatctagtggTGTATACACACACAGACACAAAGATCCAATAGCATTTACCgaatcttaaacccaact
ggttacctcttatctagtggGCAATAGAAACTCACCAGTTTACAAACAGGTTCACATGGCgaatcttaaacccaact
ggttacctcttatctagtggGGCTGTGAACCAAAATTTGGGGTAGAGCAGTTCTCATGGCgaatcttaaacccaac
ggttacctcttatctagtggAGGCCATACCTTCCCAGATGCCAAAGAGCACTAGGTCCAGgaatcttaaacccaa
ggttacctcttatctagtggGCCCAACCCTGCTTAGAACAGCAGCGTAGGAGTCTGACTAgaatcttaaacccaa
ggttacctcttatctagtggGAATACTGGGGCCAGGCCAAGTGCAGTGGCTAACACCTGCgaatcttaaacccaa
ggttacctcttatctagtggAGAGACCAGCCTGGGCAACATGATGGGAACTTATCTCTCCgaatcttaaacccaac
ggttacctcttatctagtggGGCATGATGGCATGTACCTTTACTCCTAGCTACTTGGGAGgaatcttaaacccaact
ggttacctcttatctagtggACAGTGAGCCATGATGACACTATTGCACTCCAGCCTGGGCgaatcttaaacccaac
ggttacctcttatctagtggGCTCTAAGAGTATCCAAACTGAGGCAGTCAGGGTGCTTCCgaatcttaaacccaac
ggttacctcttatctagtggTCTTTAAAAAATTTCCCAGCCAGGTGTGGTAGCTCATGCCgaatcttaaacccaactt
ggttacctcttatctagtggCCCAAATTGAGAGAAGCAGATACCGTCTAGGCCCACAAAGgaatcttaaacccaa
ggttacctcttatctagtggAGGTAATCAGGAACATGGTTGGGGCCAGCAGTGGTGGGGCgaatcttaaacccaa
ggttacctcttatctagtggCCTCTGGCCAAAAAAGGGTCTGGCACCTGCTTAGGAGGGCgaatcttaaacccaa
ggttacctcttatctagtggTCAGCCTGTGGAAACAAACCCACAAGCAATGTGTTACTGGgaatcttaaacccaac
ggttacctcttatctagtggCTGCTTGCAGCGCAGAAAGCCGGTCACTGAGACAGTGAAGgaatcttaaacccaa

ggttacctcttatctagtggAAGGCTTTAATCAGGTGCTACAGCCAAGGAGATGACAGATgaatcttaaacccaact
ggttacctcttatctagtggGCAGGTGGTTGGTTACGCAATCATGATGGGTGAGGGGGTCgaatcttaaacccaac
ggttacctcttatctagtggTCTGGAAGGCCTGATGGTTGTTTTCCTGAGAAAGGAACTCgaatcttaaacccaactt
ggttacctcttatctagtggGACAACTGGGACAATTTCATAACCTGCATTAGAGGTTGGGgaatcttaaacccaact
ggttacctcttatctagtggGGTGCGGGTGCCAAAAGGCATTTGTCCCAGCCATACTCCCgaatcttaaacccaac
ggttacctcttatctagtggTCACTGCTACAATGCTCTGAACCCTATTTTCTGCAGATGCgaatcttaaacccaactt
ggttacctcttatctagtggTTGCCAAGGCTTTGATCGCCACTGCAATGTGCCAGTAAGGgaatcttaaacccaact
ggttacctcttatctagtggAGCTGGAGCTCTTCCTTGGAACCCCAAGCCATCATGTTTGgaatcttaaacccaact
ggttacctcttatctagtggGGAGCTGTGAACATCCTTGCAACTGGTGTTTGGCACAACCgaatcttaaacccaact
ggttacctcttatctagtggGACTAAGCCAGATGGACCACCTACAAATCACCTGGAGACCgaatcttaaacccaac
ggttacctcttatctagtggCTCCAGCTACTGTAGTCAAATGAAGAGGACACAGAAGCCCgaatcttaaacccaac
ggttacctcttatctagtggGGGACACATTGGTTGTATGACTAACTTTATAGGGAAAGACgaatcttaaacccaactt
ggttacctcttatctagtggGCAAAGAGAAGTTTCCATGGGGAAAATAACATGTAAAGTCgaatcttaaacccaactt
ggttacctcttatctagtggCCAAGTTGCATTTCAGAAGAGGGAGGAGAAAAAGCAAAGGgaatcttaaacccaac
ggttacctcttatctagtggCATATTCCTCACCCTTCAACAATAATCTGGAGGTCAAAAGgaatcttaaacccaactt
ggttacctcttatctagtggGTGGAGCATCTGGATCACTGAACAACGGGCTGGTTTGACCgaatcttaaacccaac
ggttacctcttatctagtggGACAAGCCCACACTTAGTTGCAAAGAGCCAGAAGGAAAGCgaatcttaaacccaac
ggttacctcttatctagtggCTTTGACCATTGGTGCAGAGCAAAGCTAAAGAGGAAGACCgaatcttaaacccaac
ggttacctcttatctagtggCTCTTCTTTCAAGTAACGTGAAATAAGGGCACCACTTTTCgaatcttaaacccaactt
ggttacctcttatctagtggCTTACCTATAAGGAAACCAGGGCTCAGAACAAGTGAAGGGgaatcttaaacccaac
ggttacctcttatctagtggCTCCCAGTTAGTCTGAAGCAGAGCAAAAATCACACACAAGgaatcttaaacccaac
ggttacctcttatctagtggATGAGCTCCTCTGTGCTCTGAGAGCCTGGCAAAGACTGGGgaatcttaaacccaac
ggttacctcttatctagtggTTCTCTTCCCTCAGGAAGGAATCGTGGATCATTCTCCAGCgaatcttaaacccaact
ggttacctcttatctagtggGACCATGTTCTAATAACCTTTCTCTCCACTTGGCTCCTGAgaatcttaaacccaactt
ggttacctcttatctagtggTGCCCATCACCCATCTGTCCCAGGAGTGTATCTTGAGGGCgaatcttaaacccaac
ggttacctcttatctagtggGTGGTGGAGGAAAGTTTAGCACTAGTTTCCTTTGGTAAGGgaatcttaaacccaactt
ggttacctcttatctagtggGCAGGTTACTAGTTCCTCTGCAGAATGGAACACTGCATAGgaatcttaaacccaact
ggttacctcttatctagtggCATTATTGTGAGGGGTCCACTCCGCAACAGAGCTGTGCTCgaatcttaaacccaac
ggttacctcttatctagtggGAGTAAAGTCAGTAAGTGAGTAGCATTCCATGGATGGGGGgaatcttaaacccaact
ggttacctcttatctagtggTCGTTCACCTTAGATGTCATTGAGGAATATCAGCAAGTCCgaatcttaaacccaactt
ggttacctcttatctagtggGAGTCTTTTACACATGTAGCTATTTTTCATGCATTGCATGgaatcttaaacccaacttgt
ggttacctcttatctagtggGTGTTAGCTTTGATACTGGTCACAGTTCTTTCCTGAGGACgaatcttaaacccaactt
ggttacctcttatctagtggGAGAATCTGTGGCCAGTGAATGTCATGTACTACTGTGGAGgaatcttaaacccaact
ggttacctcttatctagtggCATGCCCAAAGTCACACAAGAAGAACTGTGGATCCGTTTGgaatcttaaacccaact
ggttacctcttatctagtggCCAGAGATGTTAGTTTTGCATCCCTTTAAGTGTCATCTTGgaatcttaaacccaacttg
ggttacctcttatctagtggGGTGGCATAAACTCAACCAATTCCCCAGGTCTGTGTGAGCgaatcttaaacccaac
ggttacctcttatctagtggGCCCAGCCTCTTGGAATTTGGAGCAGTTTATCTTCTCCGGgaatcttaaacccaact
ggttacctcttatctagtggGAGCTATCGCTGCTTCTGCCCCAATGGTGTCTTCATCCTCgaatcttaaacccaact
ggttacctcttatctagtggGCCAAGCTGTTTTGATCTCTGTTGTTTGCCCCACAGTTCAgaatcttaaacccaactt
ggttacctcttatctagtggGTGGGGCTTGTCTCATTTGCTTCCCTTCTTTGGGGGTCTCgaatcttaaacccaactt
ggttacctcttatctagtggGCAGCAGACATTCTCCCAAGATGTAAAAATGTTCGTGGTGgaatcttaaacccaact
ggttacctcttatctagtggGAGATGCTTGATATCCCAATTATCATGATTTGATCACTACgaatcttaaacccaacttg
ggttacctcttatctagtggAAATGTCAATAGTAGTTTCTAGGTGGGTAAGATTGTTGCCgaatcttaaacccaacttg
ggttacctcttatctagtggTCTGAGAGGAAACCTACATTGCTTAGATAACATGAAAAGGgaatcttaaacccaactt
ggttacctcttatctagtggCATTAGAGCTGAGTGATGTCAATCAGAATCAGAGCTATCCgaatcttaaacccaactt
ggttacctcttatctagtggCACTGCTATTTTACTTAACCCCAATTTCTCTCTATCTCAGgaatcttaaacccaacttg
ggttacctcttatctagtggCCCCAAATGGTAGCAATGCCTCCACTTTACAAAGGGTTTCgaatcttaaacccaact
ggttacctcttatctagtggTCTTAACACCTTTGTCCCTCTGAGCTGTGTTGGTAATTTGgaatcttaaacccaacttg
ggttacctcttatctagtggGAGGAAGAAGGAAGTTGACGTTTACCACTATGGATCCAGCgaatcttaaacccaact
ggttacctcttatctagtggGAAAGTGGGGAGACAGTCTCAAAAGACAAAGCACTTGCCCgaatcttaaacccaac
ggttacctcttatctagtggCTAGTAAGCAACAAATCTAGAATTGAGCTCTGGTATCTAGgaatcttaaacccaactt
ggttacctcttatctagtggAGGAATTAGGTAGAATTTCCCCTCCCTTTATGTCCTCTCCgaatcttaaacccaactt

ggttacctcttatctagtggACATGTGGCTCTCTCCTCTTCCAACCATGTGCCTGGGCACgaatcttaaacccaac
ggttacctcttatctagtggCCAAAATAGTCTCTGAGCATATATATGCAGATTTGTGAGGgaatcttaaacccaactt
ggttacctcttatctagtggTTCAGTTATCATCTTCTCTCTACTTCCATAGCACTCAGTTgaatcttaaacccaacttg
ggttacctcttatctagtggCAGGGTGCTCAGAGTCTAATGTGAACACACATCCACTCAGgaatcttaaacccaac
ggttacctcttatctagtggGAAGTGGCCATGGAGTCCTACAGATGCATAAAGAAAGGGCgaatcttaaacccaac
ggttacctcttatctagtggATTTGCAGAGGAGAGGAGATCAGAAATGGCTTCTTGGAGAgaatcttaaacccaact
ggttacctcttatctagtggGGCAGGTGGGTAACATAGGAAAGAGGATTCAGGTGCTGGGgaatcttaaacccaac
ggttacctcttatctagtggGATGGCGGAATGGTGAGAAGGAAGATGGAAACTTAGACACgaatcttaaacccaac
ggttacctcttatctagtggTGTAATCAGATCTCCTCTTAGTGAAATCATGTAGGATACCgaatcttaaacccaactt
ggttacctcttatctagtggAGAGGGAATACTCCTTAAATTATTCTATAAGGCCAGTATCgaatcttaaacccaactt
ggttacctcttatctagtggCCTCAACAAAATGCTAGCTAACTGAGTCCAACAGCATATCgaatcttaaacccaact
ggttacctcttatctagtggTCATACCAGGAATGCAGGGATGGTTGAACATTTGCAAGTCgaatcttaaacccaact
ggttacctcttatctagtggAGCCTCAGCAAAATCAGCTTAGAAGGGACATGCCTTAAGGgaatcttaaacccaac
ggttacctcttatctagtggCCGAACAGGAAAAAGGTGAAAACATTCCCCTGAGAACTGGgaatcttaaacccaac
ggttacctcttatctagtggCAGTAAAGAGGAAGTCAAACTGCTGCTATTCACTGATGATgaatcttaaacccaactt
ggttacctcttatctagtggCTCATCCACAAAGCTCCTAGATTTGTTAAATGAATTCAGCgaatcttaaacccaactt
ggttacctcttatctagtggGCCAACAACCAAGCTGAGAATAAAATCGAGTACTCAACGCgaatcttaaacccaac
ggttacctcttatctagtggGGAATATACTTTACCAAGGAGGCAAAAGAGTTCTGCAAGGgaatcttaaacccaact
ggttacctcttatctagtggAACGTTGCTGAAAGAAATCATCCATGACACAAACAGATGGgaatcttaaacccaact
ggttacctcttatctagtggCCATGCTGCCAAAAACAGTCTATAAATTCAGTACAATTCCgaatcttaaacccaactt
ggttacctcttatctagtggGGAATCAACACAGAGCCCATGTAGCCAAAGCCTGACTAAGgaatcttaaacccaac
ggttacctcttatctagtggATACTACAAGGCTATCATTACCAGAACAGCATGGTACTGGgaatcttaaacccaact
ggttacctcttatctagtggGGGGACCTGATTAAACTGAAAACCTTCTGCACAGCAGAATgaatcttaaacccaact
ggttacctcttatctagtggAAAAGACAACCCGCAGAGTAAAAAGACAACCCACGGAGTGgaatcttaaacccaa
ggttacctcttatctagtggCACAAACTATGCATCCGACAAGGGACTAACATCCAGAATCgaatcttaaacccaac
ggttacctcttatctagtggCAAATAATCCCATCAAGAAGTGGGCAAAGGACGTGAATAGgaatcttaaacccaac
ggttacctcttatctagtggGGCATGGATGTGAGGCAAAAGGAACACTTTTACATGGCTGgaatcttaaacccaact
ggttacctcttatctagtggCACTATGGGAAACAGTATGGAGATTCCAAATAGAACTACTgaatcttaaacccaactt
ggttacctcttatctagtggGCAATCCCATATATATGGCATTCCTATATATACCATGGAAgaatcttaaacccaactt
ggttacctcttatctagtggATGGAAAACCAAGCATCGTATATTCTCACTTATAAGTGGGgaatcttaaacccaactt
ggttacctcttatctagtggCTTTGAAGACTTAGGGGGAAGGGTGGAAGTGAGGTGAGGGgaatcttaaacccaac
ggttacctcttatctagtggCTCTGTCGCCCAGGCTGGAGTGTCGTTGTGCAATCTCAGCgaatcttaaacccaac
ggttacctcttatctagtggCGTGAGCCACCGCGCCCGGCCCTTCACGTATTTTTGAGGGgaatcttaaacccaa
ggttacctcttatctagtggCCACAGTAGAAGGCATCAAGAAGGAGGAACTTGTAAACAGgaatcttaaacccaac
ggttacctcttatctagtggTGTTGTGTTTGGCAACAGTGGGGCCATTAGGACTTTAAGAgaatcttaaacccaactt
ggttacctcttatctagtggGGAGTAGGTTACTCCTGGGACACTGAGAACACTATGACACgaatcttaaacccaac
ggttacctcttatctagtggTGCCTTCAAGGGTCACAGAAAGCTTCCTGGAAGAGACGACgaatcttaaacccaac
ggttacctcttatctagtggGGGAGGGGGAAGCATATCAACCTAGAGGAACAGGGTGGTCgaatcttaaacccaa
ggttacctcttatctagtggGGCAGGCAGATTGCTTGAATTCAAGAGTTTGAGACCAGCCgaatcttaaacccaact
ggttacctcttatctagtggAAACTAGCCAGGGGTGATGGCAGACACCTGTGAACCCAGGgaatcttaaacccaa
ggttacctcttatctagtggGCTGAGGTAAGAGAATGACTTGAGCCTGGCAGGAATAGGTgaatcttaaacccaac
ggttacctcttatctagtggGGGAGGGTGGCAGGAGGTGGGACAGATAAATCACAGTTTGgaatcttaaacccaac
ggttacctcttatctagtggCAGCATGGCTAAAGAGTAGAGAGAGCAGTTAGTAGAGAGCgaatcttaaacccaac
ggttacctcttatctagtggTCCAGAGAAGGCATGAGCAACCAGGTTTCAAATGCAGGACgaatcttaaacccaac
ggttacctcttatctagtggGGGGGTCCCATTAGGTTCAGCTTTCAGTTGCTGGAAGAGCgaatcttaaacccaact
ggttacctcttatctagtggCTGTCAACATTCATCCCTACTCCACGCTCCCCACCTCTACgaatcttaaacccaac
ggttacctcttatctagtggTGAGAACGCCTGCCCAGATGTCCTGTTGGATGTCACTGGGgaatcttaaacccaac
ggttacctcttatctagtggAGCGGTGCTGAAGAGTGAGCCCCAGCCAACAGGATTTCCCgaatcttaaacccaa
ggttacctcttatctagtggATTCGATTCATTAACTAGAAAGAGGGAACAAGCAGCCAGCgaatcttaaacccaact
ggttacctcttatctagtggTGCCCCCATTCTTCAGTGAATCATTTGTTAGTTTGACCAGgaatcttaaacccaactt
ggttacctcttatctagtggCCTCATTCAAATACTTCTTGCGACATTAATTCCGAGGCAGgaatcttaaacccaactt
ggttacctcttatctagtggCAGCCTGAGGAGAAGAAATGCTCTCAATTTGGGTAGGGTGgaatcttaaacccaact

ggttacctcttatctagtggCTTGGCTCACAAGTGGATTTAAAAACAGGAAGAAAGAGAGgaatcttaaacccaact
ggttacctcttatctagtggCAAACCTAGTACAAGCTTCCAGTGAAAATTTGCGTGTTTGgaatcttaaacccaactt
ggttacctcttatctagtggTGACAGCCCAGTTGAGGTCCAGAGAGGGACACACAGTTTCgaatcttaaacccaac
ggttacctcttatctagtggAGCTACATCTTCTGACTTAAACTCCAACGTTCCTTTACCCgaatcttaaacccaactt
ggttacctcttatctagtggGGCTGATACCTGCACAGGTAGAAAGCAAACAGACCTCAACgaatcttaaacccaac
ggttacctcttatctagtggGGCTCCATCTCCTGCTTCCCTGGCTGGAACCACCTGATGGgaatcttaaacccaac
ggttacctcttatctagtggAGCACCTGGAATAGAGAAGACAGTCTACAACATACCCTCCgaatcttaaacccaac
ggttacctcttatctagtggTCATGCTTTCACTGACATCATCTCAATGGAGTCTCACAGCgaatcttaaacccaact
ggttacctcttatctagtggGTGAGTGGCAAAACCTAGCATGAAGCCCAGAGTATGTGGCgaatcttaaacccaac
ggttacctcttatctagtggACAGCCCACTGTAATGACTGTCAGGTTGCTTTTAAGATTAgaatcttaaacccaactt
ggttacctcttatctagtggAGAGTGTCCCTCCATGAAGCAGATTTGGGGTCTCATTCCGgaatcttaaacccaact
ggttacctcttatctagtggTGGACCATGTTAGGCAAGTCATTTAACACATTTGGACCCCgaatcttaaacccaact
ggttacctcttatctagtggGCCTTTAGTGCCCACTTTACTTGAGAGATTTTCAGAGCAGgaatcttaaacccaact
ggttacctcttatctagtggCCCCAGCATTGCCTGAAGCCCTGAGATTGCTTGATGGCCCgaatcttaaacccaac
ggttacctcttatctagtggCAGGCACTGCCCCAACTGGCCCTTACTACAACTCTAGAAAgaatcttaaacccaac
ggttacctcttatctagtggGGCCCAGAGTCAATATGGTATATGTTGACCTTGGAATTCGgaatcttaaacccaact
ggttacctcttatctagtggGTCACAAGAGCAGAAGCCACAAACTCAAATACTTTTAGGGgaatcttaaacccaact
ggttacctcttatctagtggGACCTGGTATTGGAGAGGCAATAGGGAGTGGTGGGGCCTGgaatcttaaacccaac
ggttacctcttatctagtggGGCTCAGTTAAAAGAGAGTGGCAGCAACTCAACTCCAGCCgaatcttaaacccaac
ggttacctcttatctagtggGCCCAGGGATTTCAGAACTTCTAGCTAGAAAAATAGAGGCgaatcttaaacccaact
ggttacctcttatctagtggCTGGTCCACCTAATAAGGGGACAGTTTGCTTCCTGCTCTCgaatcttaaacccaact
ggttacctcttatctagtggGACAAAACAGTGTATCAGTAATGCAGTGATTGTGAGCATGgaatcttaaacccaact
ggttacctcttatctagtggCCTGGTGTTGACATTAAGTAGTTTAATAACCTCGGGCAAGgaatcttaaacccaactt
ggttacctcttatctagtggGGAAGTAATAAGAGTACTTACTCCTTTCAGTGGTTGTGACgaatcttaaacccaactt
ggttacctcttatctagtggTGACATAGGTAAAACAATTAGAACAGTTCCTGACACACGGgaatcttaaacccaact
ggttacctcttatctagtggCAATGCTTATGAAAGCTCTCATCCCAGGATAAGCATCTCCgaatcttaaacccaact
ggttacctcttatctagtggCCTGACCTTGGAAAAGACAAAGCAGCTCATGTCCCCAGGGgaatcttaaacccaac
ggttacctcttatctagtggCTGGGAATGATCACCCAGTGTCTCCAAAACCTCAGCCAGCgaatcttaaacccaac
ggttacctcttatctagtggCACAGCTGCACTGAAGAGCTTCCTTCCTGGCTTGTGTCTCgaatcttaaacccaact
ggttacctcttatctagtggGGGGACCTTTTTAATTAGCTTTCTCTTTTGAATGACGCCAgaatcttaaacccaacttg
ggttacctcttatctagtggGTGAACTTGGCCCTCCACCATGACCCACGTTTAGGCCAGGgaatcttaaacccaac
ggttacctcttatctagtggGGTGGCAAAAGGTTTGCTTCCATGATTGGCAAGGAGAGGCgaatcttaaacccaac
ggttacctcttatctagtggGGGCTATCTGGAAACTTTTATGAAAGGGTCTCCTGTTTGCgaatcttaaacccaactt
ggttacctcttatctagtggCTGCTTTCAGGTCACCTTATGGTGTGTGTGTGTGTGTGTGgaatcttaaacccaactt
ggttacctcttatctagtggGTTCACAAGCCTGAGGCAGCCTACTAATTGCATGGCATGGgaatcttaaacccaac
ggttacctcttatctagtggAGCCAATGTGCCTCTTTGGAGACTGAAGAACAGGACTCAGgaatcttaaacccaac
ggttacctcttatctagtggGGAGGTAGCACCATTTCTCCGCTAGTTTTCATGTTGTAGGgaatcttaaacccaactt
ggttacctcttatctagtggGGAGAAACAGGCCCATAGAAGTCTACTAACTTGCCATAGCgaatcttaaacccaac
ggttacctcttatctagtggCTTCTTCAAAAGCCTATGCTCTTTTTACTGCCCCCATGCGgaatcttaaacccaactt
ggttacctcttatctagtggGGCTATTAACCATTCTCCAAACCATCACTTCCCTCCATGCgaatcttaaacccaact
ggttacctcttatctagtggGTGAAGGTTTCACTGACCTACCATTCCCTCAGCCAGATACgaatcttaaacccaact
ggttacctcttatctagtggGGGCTCAGCTGTGTCAGTGATTATACCAGATAATGGGCAGgaatcttaaacccaact
ggttacctcttatctagtggAGGCCAAGTCATGCTCATCATCGACGTCCAGTCCAGTGCCgaatcttaaacccaac
ggttacctcttatctagtggACCAAAGGAAGGGAGGGAGGGAGAGAGGCAGCAGGCATACgaatcttaaacccaa
ggttacctcttatctagtggGTATGTGTGGACCTCAGTTTCCCTCAACTGACCAGAGTACgaatcttaaacccaact
ggttacctcttatctagtggCTCTGGCAGCAAGGAAACCAACTCAGATGGGAGCAGATGGgaatcttaaacccaa
ggttacctcttatctagtggTGTATTTGTTTAGTTCTAAAGCCTTCCCCAACCCCACCTGgaatcttaaacccaactt
ggttacctcttatctagtggACCCTGGCTGCTTACAAGGCCCACCCTCTGCTCAGTCCTGgaatcttaaacccaa
ggttacctcttatctagtggGTCACTGTGTCAGTTGCCTGAACAGACCTCCAAGGAGCACgaatcttaaacccaac
ggttacctcttatctagtggCTGAATGCAACTGTAGACCTTACTACCCAGAAAAGGAGGCgaatcttaaacccaac
ggttacctcttatctagtggCCATTTGCAAGAGGTAGCATTAGTGGAGGAGTCTTGTTGCgaatcttaaacccaact
ggttacctcttatctagtggGGCTCAAGCCAGAGGAGCCCCCACAGCAAGTTGGCACAGGgaatcttaaacccaa

ggttacctcttatctagtggGATCATAGATCTTCCAGGACACCCAGAGTTATTCAGGAAGgaatcttaaacccaact
ggttacctcttatctagtggTCTCAGGCTGATGTGGCCGGGGAGATGTGGAATTGTGTAGgaatcttaaacccaac
ggttacctcttatctagtggGGACCAGAACAGACCAGGACATAAAACAGATTATTGTGTCgaatcttaaacccaact
ggttacctcttatctagtggGTTCCAGGCCAGATGATTTAGCTACATAGGGACCTAAAAGgaatcttaaacccaact
ggttacctcttatctagtggAGCACAAGTGGTCACCTGTCATGGCCCACAACTGGCACTCgaatcttaaacccaa
ggttacctcttatctagtggACTAACATTTATTGCCTCTTTGCTGGACCCTCTGTATGTAgaatcttaaacccaacttg
ggttacctcttatctagtggACCAGAAGGTTAAGACAATTGCCCAAGACCACACGGCTGGgaatcttaaacccaa
ggttacctcttatctagtggCAGGCAGTTGGGCTTGAGCTCCTACATGCTAAACCACCACgaatcttaaacccaa
ggttacctcttatctagtggTCTTATCATGCCAGGGGCTCATATCCTATGACACTTAAATgaatcttaaacccaactt
ggttacctcttatctagtggCTGCCAAAAGGCTGGAAAACTTGGGAGAGTCACTGCTCTGgaatcttaaacccaac
ggttacctcttatctagtggCACAGCAGCACTTAACTCTTCTGCCATACTCAACACAACCgaatcttaaacccaac
ggttacctcttatctagtggAGGTTTCATTGGAGGCTTCCCTGATCCTATTAAACATTCCgaatcttaaacccaactt
ggttacctcttatctagtggACTCATTTGCTCAGCAAACATTTCCTAAGGGTCTGCAATGgaatcttaaacccaact
ggttacctcttatctagtggCTTTCTCCTTAGGTTTCCTTTTTCTCTCTCTCTCTCTCTCgaatcttaaacccaacttgt
ggttacctcttatctagtggGCACGATCTCAGCTCATTGCACCCTCCTCCTCCTGGGTTCgaatcttaaacccaac
ggttacctcttatctagtggGAGCCACTGTGCCTGGCCCTCCCTTAGGTTTCTAAGAAAGgaatcttaaacccaac
ggttacctcttatctagtggGATCCTAGAATGCCCATTTTTAAAACGCCCCACTGAGGCgaatcttaaacccaact
ggttacctcttatctagtggGAGGCTGTACAGTCTGCTCACTGAGAGAGACTGCCTGAGCgaatcttaaacccaa
ggttacctcttatctagtggGCCTTGTCTCCTTGCGTGAGCTACTAAAGCACTCTGTACCgaatcttaaacccaact
ggttacctcttatctagtggATGTCAGCTATTATTATTTCCAGGCCTTGCATCATGTACgaatcttaaacccaacttg
ggttacctcttatctagtggCCAAAATACCCCCTGCCTCCATTCACGTATAGCCTCCCCCgaatcttaaacccaa
ggttacctcttatctagtggCCCTACCAGAGGGGTGCGTTGGTTACAGTCAATGAATCTAgaatcttaaacccaact
ggttacctcttatctagtggACGTTAGGTTTCACTCTTGTTGTACATTCTGTGGGTCTGGgaatcttaaacccaactt
ggttacctcttatctagtggCCTCTGTGCTCCTCATATTTATCCCTCTCTCTCCCCAACCgaatcttaaacccaact
ggttacctcttatctagtggGGAATCATACGGTATGTAGCCTTTTCCTGTTCAGATTGGCgaatcttaaacccaactt
ggttacctcttatctagtggAAGGTTCCTCCATGCTTTTTCATGAGGCTTTATAGCTCACgaatcttaaacccaactt
ggttacctcttatctagtggTGCTGTATAGTATTCCATTGTCCACCATACCACAGTTTATgaatcttaaacccaacttg
ggttacctcttatctagtggGGTTGCTTTCAAGTTTTGGCAATTATGGTTAAAACTGCTAgaatcttaaacccaacttg
ggttacctcttatctagtggCTCCTTTGGGCAAATACCAAGGAGCACGATTGGTGGATTGgaatcttaaacccaact
ggttacctcttatctagtggGTTTCATAAGACACGGCCAAACTGCCCTCTAAACTGGCTGgaatcttaaacccaac
ggttacctcttatctagtggGCTCCACATCCTTGCCGGCATTGGTGTTGTCAGTATTCTGgaatcttaaacccaact
ggttacctcttatctagtggTTATCTGAAGACATATGATATGGTGCATCATTTCACATGCgaatcttaaacccaacttg
ggttacctcttatctagtggTATCTTCTTTGGTGCAGTGTCTATTAAGATTGCAGGTCCGgaatcttaaacccaacttg
ggttacctcttatctagtggGGATTGCTGGTTGTATTATTGTTGAGTTCTTTGCATATTGgaatcttaaacccaacttgt
ggttacctcttatctagtggGGCTTTTCTGCAAATTCCTTGACATTGCCTTTCATGGGGCgaatcttaaacccaactt
ggttacctcttatctagtggACCCAAGGTCATCTAAGTTTTCTTTATGTTATCTTCTAGGgaatcttaaacccaacttg
ggttacctcttatctagtggAGGGTATAAGGTCTATGTCTAGATTGAGTTATTGCATGTGgaatcttaaacccaacttg
ggttacctcttatctagtggTGCTCCATTGTAGTGCCTTTGCTCATTTGTTGATTATCAGgaatcttaaacccaacttg
ggttacctcttatctagtggGTAGGTCTGTTTCTAGGCCATCTATTCTGTTCCATCTATCgaatcttaaacccaacttg
ggttacctcttatctagtggGTCCTGCAACTCCATTCTTCTTCATTATGGCTATTCTGTGgaatcttaaacccaacttg
ggttacctcttatctagtggGGGATTGCATTGCATCTATAGATCAAGTTGAGAAGAACTGgaatcttaaacccaactt
ggttacctcttatctagtggATATTGAGTCTTCTTATCCATGAGCATTGAATATCTCTCCgaatcttaaacccaacttg
ggttacctcttatctagtggCCCTCTTGTTCATTGCTTGTATATATGAAAGTAATTGACCgaatcttaaacccaacttg
ggttacctcttatctagtggGCATACATCTTTGACTTGTTCTTACTCTTACTGGGAAGGCgaatcttaaacccaactt
ggttacctcttatctagtggTTCTAATTTACTGGGCATTTTCATTGTGAATGGGTGCTGGgaatcttaaacccaacttg
ggttacctcttatctagtggGATGGATCACATGAATTGATTTTCAGATGTTCAGCGTGGCgaatcttaaacccaactt
ggttacctcttatctagtggCTGTACCTCTGGACTGTGAACTTCATGCCTGCTTCTCAGGgaatcttaaacccaact
ggttacctcttatctagtggGATAGCTGAAGTGGGCTGGAGTTGGGTTTTCTCCTCCTCCgaatcttaaacccaact
ggttacctcttatctagtggCTGGAGCTTGCTGGAGTTGAGTATTTCTTTTCCTCAGGTCgaatcttaaacccaactt
ggttacctcttatctagtggGTGGGTTAGGCTCTGGTTAGACAGCTTCCCCTGAGGGCAGgaatcttaaacccaa
ggttacctcttatctagtggTTTTTCTCCTCTTGCCAAAGCGTGAGGAAATTTTTCTCAGgaatcttaaacccaacttg
ggttacctcttatctagtggCCCATAAAAGTATGGTACTCCCTACTTCTGGATCCTCTGGgaatcttaaacccaact

ggttacctcttatctagtggTTTCCTACCCTGGCACTGGTTCCAATGGATGTTTCAGCCCgaatcttaaacccaact
ggttacctcttatctagtggCTGCCTATCTGTCCCTCCAATTCTGGGGGCAGTAGTTTCCgaatcttaaacccaact
ggttacctcttatctagtggACGATGGAGTGGGGACCTTCAAGCTGCTTCTACGTCAAACgaatcttaaacccaac
ggttacctcttatctagtggCCCTGGCATTTTTAATCATGATAACCATCCTATGAAGTAGgaatcttaaacccaactt
ggttacctcttatctagtggAATCAGGTACCAAGAAATTAAGTAATCAGCCCAAAGCCTCgaatcttaaacccaact
ggttacctcttatctagtggCTACTACAAATCCAAGCCAGTCCAACTCAGAAGCCAGGGCgaatcttaaacccaa
ggttacctcttatctagtggCCAGAAGTTGGGCAGAAATGAAGGGTTTCAAGTGGTCCTGgaatcttaaacccaac
ggttacctcttatctagtggCAGGAGAGGAGTACATGGGTAGAAAGTGGTGAACTTCAGGgaatcttaaacccaac
ggttacctcttatctagtggCTTCCTGGAAATAGCTCATAGCTTCCTGAGGGCCCCACCCgaatcttaaacccaac
ggttacctcttatctagtggCTATACCTGCTGTTGCCTCCTGTCTTTCTCCTACCACAGCgaatcttaaacccaact
ggttacctcttatctagtggCCAACTACCGCCATGCCTATATGACAACTAGGTACTAAGGgaatcttaaacccaac
ggttacctcttatctagtggTGGCCTATCTAATGTGATGACACCCTGGTCCAAGCCACTGgaatcttaaacccaac
ggttacctcttatctagtggCATTTCCTAGCTAGTTCCCCTCATTTACACTACTCCTCCCgaatcttaaacccaactt
ggttacctcttatctagtggTCCAACCTATAGACAAAGTGAGCTTCCAAATCATAAATCTgaatcttaaacccaactt
ggttacctcttatctagtggGGCATCCTGTGCAAGGTCTTTGTGGGGAGAGGGGACGTGGgaatcttaaacccaa
ggttacctcttatctagtggGGGACAGTGGAGGATATGAAATTGGAAAGTTACAGGGTCCgaatcttaaacccaac
ggttacctcttatctagtggAGCCTCCATTGAGAAATTCTTTGATTCATGTCAGTAAGGGgaatcttaaacccaactt
ggttacctcttatctagtggAGCTTTGGAGTCATCTTTGAGTTCTCTCTTCCTCTTACACgaatcttaaacccaacttg
ggttacctcttatctagtggCTTAACCAACATTGCTGTCAGCAGCCTGGTCCAAACCACCgaatcttaaacccaac
ggttacctcttatctagtggGCTGCTTCCCGCTGGCCTCATACAGTCTGGTCTCCATATGgaatcttaaacccaac
ggttacctcttatctagtggTAACACCTAAGTCAGACCACGTGACTCCTCTGTCCAGAACgaatcttaaacccaac
ggttacctcttatctagtggCACCAGGGCACACAGGGCTCTACTCAAAGCTGCCCACCGCgaatcttaaacccaa
ggttacctcttatctagtggGTCACTTCCCTTTTTCTCTCCACTCCAGTGCACTGCTCTCgaatcttaaacccaact
ggttacctcttatctagtggCCTGTTCCCAGATATCCACACAACTTTCTTTCCAGTTTCCgaatcttaaacccaactt
ggttacctcttatctagtggCCCTGATCATCCTGCCTAAAATCATCCCCCTCATTGCTCCgaatcttaaacccaac
ggttacctcttatctagtggGTGTCTGTTGCATACCTCCCCAACCTACCCCAGTACGATGgaatcttaaacccaac
ggttacctcttatctagtggAAGAATCATCCCTGTTATGGCCAGGTGTGGTGGCTCTGGCgaatcttaaacccaac
ggttacctcttatctagtggGTTGTTCTCCATTAGTCCCCAATGCATGGCAGAGTCTGGCgaatcttaaacccaact
ggttacctcttatctagtggCATGTAGCGAAAGGGTCTGGAAATGTGAGGCAGGGTGATCgaatcttaaacccaac
ggttacctcttatctagtggCTGGGCTGGAGGATAGACACAGGGTTGAGGGAACATCCTGgaatcttaaacccaa
ggttacctcttatctagtggGTGTGGCAAAGTTGGGCCTGTAGGAGTGAGCACTTGATGCgaatcttaaacccaac
ggttacctcttatctagtggAGCTGGAGAAGCTGAGCAGTGCAGAGGCTGGGAATGGCAGgaatcttaaacccaa
ggttacctcttatctagtggGGGTGGTCAGGGGGACTGTTAGTCCATGGAGAGTTTCCTGgaatcttaaacccaac
ggttacctcttatctagtggGCCTGCTTTGCTTCAGAGGTTGCCTCCCCTCCTACTCTGCgaatcttaaacccaac
ggttacctcttatctagtggTACTGAAGACACTAGGTTCCAAGGGACTGAATCTGAAGAGgaatcttaaacccaac
ggttacctcttatctagtggAATGAGGAAAAGTAAATGGAGGGTGGAGGAGATGCCATTCgaatcttaaacccaac
ggttacctcttatctagtggGAGAGGCAGCGTGATGGGAAGGAAAGAAAGCGAGCTGAGCgaatcttaaacccaa
ggttacctcttatctagtggGCCTTACTCCAGAACCTACAGAATGTGGGGCTTTGAAATGgaatcttaaacccaact
ggttacctcttatctagtggGCAGGAGCCTACCCCAGCCCTGGTGGTATTGTGGCAACCCgaatcttaaacccaa
ggttacctcttatctagtggGTCCTCCTCTCAGAGTTTTCTGCTGTGGAAACCATGCTCCgaatcttaaacccaact
ggttacctcttatctagtggGCAGGAATGTCTGAATCCTACCCCGTTTCTCAACAACCCCgaatcttaaacccaac
ggttacctcttatctagtggAGGTCTGGAGTTTTCTGAGCTAGATTGATGCCCCAAAGTCgaatcttaaacccaact
ggttacctcttatctagtggTGGCCCAACTAATAGACTAGAGGGGCTCTCCAATTCTCAGgaatcttaaacccaac
ggttacctcttatctagtggGCAAAAGTGTAGAAGGACACACACACATGCACGCACACACgaatcttaaacccaa
ggttacctcttatctagtggCACAGAGAGAGAGAGAGAGAGAGACTTCTAGAGGGTCTTTgaatcttaaacccaac
ggttacctcttatctagtggGCACTCCACAGGGCAAGGTGAGGGTGAGGCGAGCAAGGCCgaatcttaaaccca
ggttacctcttatctagtggCAGAGGCATGGATTCTCAGATTTGTGACACCTGCAGGGTCgaatcttaaacccaac
ggttacctcttatctagtggTGCACCCTGTGAGCCTCACTTACATTACCCTAGTCCCAGGgaatcttaaacccaac
ggttacctcttatctagtggGGATCTAGGTGAGAAGAGGTGTGGGAACCTGTATCTCCCCgaatcttaaacccaac
ggttacctcttatctagtggGCCTTGCATCTGCCATTCCTCTAGACTCTCAGTCCTTCTCgaatcttaaacccaact
ggttacctcttatctagtggGGATAGGGCCTAAGATTCTAAATTCTAACAAGCTTCCAGGgaatcttaaacccaact
ggttacctcttatctagtggCAGCAATGCCTAGAAATGATGGTTCTTCAAATCCAAGTGAgaatcttaaacccaact

ggttacctcttatctagtggCAGAACCTCTGCAGTGAATGGAGGAATTTCTGAGGGTCACgaatcttaaacccaac
ggttacctcttatctagtggCTCAAGAGCCTGAAAGAATAATAAATACATAAGCTTCTGGgaatcttaaacccaactt
ggttacctcttatctagtggTCCAAACGGCTTCTTTCCAGATAGATAGACCTGTCCTCCCgaatcttaaacccaact
ggttacctcttatctagtggGAATCAAATGGCAAATTGAAGCTTTTAGGTACTGTCCTGGgaatcttaaacccaactt
ggttacctcttatctagtggGACACCCTCAGGCTGCTTCCACTCACAGTACAAGGCAAGGgaatcttaaacccaa
ggttacctcttatctagtggGGTGAGAGAGGAAGCAAAAGGGAGGGGGGCAGGTGCCAGGgaatcttaaaccca
ggttacctcttatctagtggGTGAGAACTCACTCACTTCTCCTCCCACTCCAGGGAAGGCgaatcttaaacccaa
ggttacctcttatctagtggCCTGAACACCTCCCATTAGGCCCCCATCTTCAAAATTAGGgaatcttaaacccaac
ggttacctcttatctagtggAGGGGGACAAACATCCAAACTATAACAGACACCTTTGCACgaatcttaaacccaac
ggttacctcttatctagtggCCTGATATCTTACGGGTGCAAATAAAGTGACTCAACATTGgaatcttaaacccaactt
ggttacctcttatctagtggGGCAACCTAGAAGGACATGCAAAACACACACACACACCACgaatcttaaacccaa
ggttacctcttatctagtggGCACATACACACAGCAATACTCACATACACACACACACCCgaatcttaaacccaa
ggttacctcttatctagtggCACACACACACCCCACACACAAAAGCATATCTACAGCCACgaatcttaaacccaa
ggttacctcttatctagtggATTGACAAATAGGACAAAAGAGGATTGAGTAGCTGCTACGgaatcttaaacccaact
ggttacctcttatctagtggTCCCCAATTCCCAGGAAAATGTTTCATATACAGCATGTAGgaatcttaaacccaactt
ggttacctcttatctagtggAGGGGAGGAAAGACCAGGAACAATCTCTACCAGTGGGACCgaatcttaaacccaa
ggttacctcttatctagtggTGGCCAGGCCAGTTTGAAGGGCATCAAGGAAAACCTTCAGgaatcttaaacccaa
ggttacctcttatctagtggGTGTTGACACCAGATACAGCAGGAACTAGCCATGCGCAGGgaatcttaaacccaa
ggttacctcttatctagtggTGGCCATTGCTGGCTAGGGTTCCATGCAGACAGCTTAACCgaatcttaaacccaac
ggttacctcttatctagtggGACCTGGAGCTGCTAATATTCTCCCTTGTGCTATCTAGGTgaatcttaaacccaactt
ggttacctcttatctagtggGCATCAAGAAAAGTGCCTGGCAATGGAATATTATGCAGCCgaatcttaaacccaact
ggttacctcttatctagtggGAACTCCTGTAATTTTCAGTAACATGGGTAAGCTTGGGGGgaatcttaaacccaactt
ggttacctcttatctagtggGCCAGGCATAGAAAGATGAATATTGCACATTCTCACTCATgaatcttaaacccaactt
ggttacctcttatctagtggGGTGCTTACCAAAGGCTGATAGGGGAGGCGATAAAGAGGCgaatcttaaacccaa
ggttacctcttatctagtggCAAAATACCTGGAAGAGAAGACTTGGAATGTTTTCCATACgaatcttaaacccaactt
ggttacctcttatctagtggGCATGCATCAAAATATCACACGAACTTCATAAATATGCAGgaatcttaaacccaactt
ggttacctcttatctagtggCGCATGAATGAACTGAATGGACAGAAGCTTTGTGGGAGGCgaatcttaaacccaac
ggttacctcttatctagtggCAGGAATGATGATGCAGAGTATCAATATATGAAGTCCGGCgaatcttaaacccaact
ggttacctcttatctagtggTGTAGAAAAAGAAAGGACAGAGACATGTGGAGTGGCCACCgaatcttaaacccaa
ggttacctcttatctagtggCATGGATCATCAAGATCTTTTCCAAAGAGTCAAAATCATGgaatcttaaacccaactt
ggttacctcttatctagtggCCACAAGTGTTTATTGTCTGATAATGCTATCAACCTTGATgaatcttaaacccaacttg
ggttacctcttatctagtggGACATATGTCTGGGCCAGCCTAGCTGAAGATATTCCTGAGgaatcttaaacccaact
ggttacctcttatctagtggGCAGCACTCTCTCTCTGAGCCCAGGTCTCCTTACTTCTGGgaatcttaaacccaac
ggttacctcttatctagtggGCTTTGCTCTGTGTAAACACCTTGGGCTAAGTGACCCTCCgaatcttaaacccaact
ggttacctcttatctagtggGTTTTTGTAGGCAGCAATTGGCAGAAAGTGGGTCAGTTGGgaatcttaaacccaactt
ggttacctcttatctagtggCAGTCACCTCAGAGGCAGTGAAAATGTCTCATGCTCAACCgaatcttaaacccaac
ggttacctcttatctagtggGTCTCCAGGCATGTGTGATATCAACTTGGAAGAGGGGGTGgaatcttaaacccaac
ggttacctcttatctagtggACTTTTCCTTTAAAAAGGCACAGGGAAGCACAAAGACATTgaatcttaaacccaactt
ggttacctcttatctagtggGCTAATGACTACATCATCCTGATGTTTCCCTCTTCTATGGgaatcttaaacccaactt
ggttacctcttatctagtggGTTTCTTTCTTCTGATTAGGAGAAATTGGTCATGAGGACTgaatcttaaacccaacttg
ggttacctcttatctagtggACAAAGCTATGTTATTGTCCTCTCGGTTCTTCAGTGTGAGgaatcttaaacccaactt
ggttacctcttatctagtggGAATTACGCATTGCTAGAAAGTTTGAGAATCCTGTGATTGgaatcttaaacccaactt
ggttacctcttatctagtggCCATCAGGTTGGTCTAAAATTACACAACAGTACTCAACTCgaatcttaaacccaactt
ggttacctcttatctagtggAACAGCAAAACTTGGCTACAGAAGAATAGAAATAACCATGgaatcttaaacccaact
ggttacctcttatctagtggTGTAGAGTTTCACATACATATGCAGCTATTCCTGCACACCgaatcttaaacccaactt
ggttacctcttatctagtggGATAGCAAGTAAAAAGGCATATCAACCTTTTCAAGCTGCCgaatcttaaacccaactt
ggttacctcttatctagtggGATAGCGACCACAAGTGAGGCCCTCATTGCCACCTGGTGGgaatcttaaacccaa
ggttacctcttatctagtggACATGACACACATGAGATAGTAGAGGGTCACAACCCCTGGgaatcttaaacccaa
ggttacctcttatctagtggCTCCCTGGTCACCACGGAGCATACTTTGGTTCCAGGCTGGgaatcttaaacccaa
ggttacctcttatctagtggGAGCCACAAAGCTATTTTACCCATAAGGTCACCTGAAGCCgaatcttaaacccaact
ggttacctcttatctagtggTAACGTTGGTTATCCATCTAGATCAGCAGCGTGTTCTAGGgaatcttaaacccaactt
ggttacctcttatctagtggATCCCAACACCTAGGCCACAATTCAGACCAATTATACTAGgaatcttaaacccaact

ggttacctcttatctagtggTCCAGCCAGTGTTGAAATCATTGAACTGGAGTAAAGGCTGgaatcttaaacccaact
ggttacctcttatctagtggTTGAAGAGCAGTGGAAAGTAAACTAGCTTTAGCTGACAGCgaatcttaaacccaact
ggttacctcttatctagtggTCACCCATACGGTAAGCAGACTGTATTATCCCCGTGCCCCgaatcttaaacccaac
ggttacctcttatctagtggCTTGACTGTGTGACTCCATATCTTCAGTGACGGCTTGCTGgaatcttaaacccaactt
ggttacctcttatctagtggGAGAAACAGAATAGCAGCCATCAGACATCCAAGGATCTCCgaatcttaaacccaac
ggttacctcttatctagtggTACTTCTTTACAAGCCTTACAAACCACTAAGGCCAAGGGCgaatcttaaacccaact
ggttacctcttatctagtggACAGTAGCACTGTACCCTGATTCTAAGCTTGCAGTTGTAAgaatcttaaacccaactt
ggttacctcttatctagtggGACTTGCCCTGTGTGTTTTCTCCTTGTTTACCCGCTGGCCgaatcttaaacccaactt
ggttacctcttatctagtggCTACTTCAAATGCAAAAAAGGTAGAAGGTCAGCTGATTAGgaatcttaaacccaactt
ggttacctcttatctagtggTCCCCTAGACACCTGAGCTGTTTCAGTTTTGCCTCCTGGGgaatcttaaacccaact
ggttacctcttatctagtggTAGTAAAATAGGATTTACTCAGCTTCTACTCAGCCTCTACgaatcttaaacccaactt
ggttacctcttatctagtggCCCGAGTAGCTGGGAGTATAGGCGCCCCCCACCATACCTGgaatcttaaacccaa
ggttacctcttatctagtggGCATGAGCCACCGTGCCTGGCCAGCTTCTACTTATGTGTGgaatcttaaacccaac
ggttacctcttatctagtggTGCACATGTGCAAGTCTATTCATGACACTTCTGAGCTCAAgaatcttaaacccaactt
ggttacctcttatctagtggTGAACTCACAGCCGAGCATGCTAGCTCATGCCTATAATCCgaatcttaaacccaac
ggttacctcttatctagtggGTCCCAGCTAGTCAGGAGGCTGATTGCTTGAGCTCAGGAGgaatcttaaacccaac
ggttacctcttatctagtggGCATGACTGCAGCCTAGGCAATAAAGAAAGACTTTGTTCCgaatcttaaacccaact
ggttacctcttatctagtggCTAAGACACTGCTTTTAGACTTGGGCCTTGCCTAGCCCTCgaatcttaaacccaact
ggttacctcttatctagtggCTTCCCTCCTCAGGACCTTTGCCTATATCTTTCCCTCTTCgaatcttaaacccaactt
ggttacctcttatctagtggCAGGGTATCAATAACTGACTGCCATGTAACATGCCAACCCgaatcttaaacccaac
ggttacctcttatctagtggGCTCATGGTTCTATGGCCAGCAATTTGGGCTAGGATCAGCgaatcttaaacccaac
ggttacctcttatctagtggCGCTGCAGGCTGTCAACTGGTTGGACTCAGGGGACTCAGCgaatcttaaacccaa
ggttacctcttatctagtggCAGCTGGCAGTCTCAGAACAATATTCCAAGGGAGCAAGAGgaatcttaaacccaac
ggttacctcttatctagtggCACTGCAAGTCAAAGGCCAGTCCAGATTTAATGCATGGGGgaatcttaaacccaac
ggttacctcttatctagtggACCACACCTGGCTAACTAACACTGACACTTCAGATGTCAGgaatcttaaacccaac
ggttacctcttatctagtggTGCTCTCAAATCACCCTGTTCATTTTCCTTATAGCATTGGgaatcttaaacccaacttg
ggttacctcttatctagtggCTGGTTTCCCTCCTGAACTGTGAACTCCATGAGGGCAGGGgaatcttaaacccaac
ggttacctcttatctagtggTCAGTGTCTACCATAAAGTAGGGCCCAGCATCAGGATTCCgaatcttaaacccaac
ggttacctcttatctagtggGGTTTAGAAAGTTACTCTACCACTTCCTAGTTGTGGGACCgaatcttaaacccaactt
ggttacctcttatctagtggCACATAAGCAAGCTGAGTGTATTCACACATAGTAGGTGCCgaatcttaaacccaact
ggttacctcttatctagtggTGGTGCATGAGAAATCACAGAAAAACAGGGAATCGTTAACgaatcttaaacccaact
ggttacctcttatctagtggGACTTGCCCCATGATCAGTTGCCTCCCCCCAGGTTCCTCCgaatcttaaacccaac
ggttacctcttatctagtggCACAGCCAAACCATATCAGTCATGTATGTATAAACAGGGGgaatcttaaacccaact
ggttacctcttatctagtggAGATTCAATGCTACTCCCATCAAGCTAACATTGACTTTTTgaatcttaaacccaacttg
ggttacctcttatctagtggCAGATATATAGACCAATGAAACAGAGACCTCAGAAATAACgaatcttaaacccaact
ggttacctcttatctagtggCTAGCCATATGCAGTAAACAGAAACTGAACCCCTTCCTTAgaatcttaaacccaact
ggttacctcttatctagtggCTCAAGATGGATTAAAGCCTTGAATGTAAAACCCCAAACCgaatcttaaacccaact
ggttacctcttatctagtggGAGCTTCTGTTCAGCAAAAGAAATTATCACCATAGTGAACgaatcttaaacccaactt
ggttacctcttatctagtggAAAATCAAGCAACCCCATCAATAAATGGGCAAAGGATATGgaatcttaaacccaact
ggttacctcttatctagtggTGCACACGTATGCTTATTGCAGCACTAATTACAATAGAAAgaatcttaaacccaactt
ggttacctcttatctagtggAACCCAAATGCCCATCAATGATAAAGCAAACATGGCACATgaatcttaaacccaac
ggttacctcttatctagtggGGGACATGGATGAAGCTGCAAGCCATAATTCTCAGCCAACgaatcttaaacccaac
ggttacctcttatctagtggACACAAGGACACACAGAGAGGAACATCACACCCTGGGGCCgaatcttaaacccaa
ggttacctcttatctagtggGACAGGAGCCTGAGGTCAGCCACCAACAGTCTTTGTAACCgaatcttaaacccaac
ggttacctcttatctagtggCCCCTGATATTCTTTGTGCTGGAAAATCTGTCAGAATGACgaatcttaaacccaactt
ggttacctcttatctagtggACCACTGGGGTAGACTGAAATGAAATACAGTCATTAGTAGgaatcttaaacccaact
ggttacctcttatctagtggCTTCTGTTTTTCCAGTAGTGTCACTCTTCAGGGGCCCCTCgaatcttaaacccaactt
ggttacctcttatctagtggCTTTGACAGTCAGAAATATACCTACCCATTTCCAATTTCCgaatcttaaacccaactt
ggttacctcttatctagtggTTGGGGTAATCACTTCATCAGTAAAGTTGGCCTTCGTAAGgaatcttaaacccaactt
ggttacctcttatctagtggATGTTCTGTGGGTTACAAATGACTGCCCACCATGCTGTGGgaatcttaaacccaact
ggttacctcttatctagtggGTCTGTCCAAAAGGACTTCCCAAAAACCACTGTATGGCAGgaatcttaaacccaac
ggttacctcttatctagtggCTCCGGACTATGGCTAGGAACTATTACTATGGCTAACAAGgaatcttaaacccaact

ggttacctcttatctagtggGGGAGGCCAAGGCAGGCAGATCCCTTAAGTCTAGGAGTTTgaatcttaaacccaac
ggttacctcttatctagtggTGGGCAATGTGGTGAAACCCCCCTCTGCAAAAAATACAAAgaatcttaaacccaac
ggttacctcttatctagtggGCACACCTGTGGTCCCAGCTACTCATGGAGGCTAAGGAGGgaatcttaaacccaa
ggttacctcttatctagtggTCTCTGGACTATAAACTCTCATGAATCTAGCATTATTTGGgaatcttaaacccaacttg
ggttacctcttatctagtggGGATAGACTGTGATGACAAATAAATCAAATTTGGGATGCCgaatcttaaacccaactt
ggttacctcttatctagtggAACAGTCCAGGGAGGCTGAAAATAGAGTGTATGTTGCAGGgaatcttaaacccaac
ggttacctcttatctagtggGTGGCATCGCTGTCTGTCTGGAGGGTGCCATCACAGTCAGgaatcttaaacccaac
ggttacctcttatctagtggATGGAGTGGCCACATCACTTCCATTCACATTCCACTGCCCgaatcttaaacccaac
ggttacctcttatctagtggTACACGAAGGAAGAAGTGGAAGGTGGACTTTGGTAAACCGgaatcttaaacccaac
ggttacctcttatctagtggCCCAGCACTTTGGGAGGCTGAAGACGGTGGATCAACTGAGgaatcttaaacccaa
ggttacctcttatctagtggTCACTTCCCAAAGTGATGCTTAACTTCCTGCTAGACCAGCgaatcttaaacccaact
ggttacctcttatctagtggCAGTGTTCTTCCAGTCTGTCTCATGCTCTCCCATATCTGCgaatcttaaacccaactt
ggttacctcttatctagtggCACTCCCAGTTATTCCCTGGATAGAGTAATATGAGGCATGgaatcttaaacccaactt
ggttacctcttatctagtggATACACATCCCAACTCAGTGATACACAGTGTCAATGACAGgaatcttaaacccaact
ggttacctcttatctagtggCAATCACTTATGCAGTGCTTCCTATGTGCCAGAAACTGTGgaatcttaaacccaactt
ggttacctcttatctagtggGGTAAGAGGTTATGTCTACTCTGTTTAACTGTTCTGGGCCgaatcttaaacccaactt
ggttacctcttatctagtggCAGTCTCTAATGTTTCATCTGTAAAATGACGACAATTCCCgaatcttaaacccaactt
ggttacctcttatctagtggCAAGGGATATGAAAGTGTTTCACTCAATGCCCAGCACATGgaatcttaaacccaact
ggttacctcttatctagtggCTGCTGCAACAAATCACACAGCTCTCTCCTCACTACCAGCgaatcttaaacccaac
ggttacctcttatctagtggTCCTACACATAATTATAAGATCCTACATAGGTTGCTATCCgaatcttaaacccaacttg
ggttacctcttatctagtggCAGCATGCCTGAGCAGCTTACAAAGGAAGCTCTTTCCACGgaatcttaaacccaac
ggttacctcttatctagtggGTCCCATTTCTGAGAAACCATTCCAAACTGCGGTAAATGCgaatcttaaacccaact
ggttacctcttatctagtggTATAAGGACTTGGTACATTGGTAACAAAGCACATGCAAGAgaatcttaaacccaactt
ggttacctcttatctagtggCACTTGGTCATGAAATACCCCAGGGAGTGCACCAGGGCAGgaatcttaaacccaa
ggttacctcttatctagtggAACCCCAAGCGCTCTGTGGGTTTCTATAAGACCCCACTTTgaatcttaaacccaact
ggttacctcttatctagtggCCCTCTTGGGGTTTTGCTGATTGGTCCTTACTTAGAAGGTgaatcttaaacccaactt
ggttacctcttatctagtggGAGGCAGTGCATCTCTCTGCTTGAGGAAAACTCACTTTGGgaatcttaaacccaact
ggttacctcttatctagtggGGGAAGCTAGAGTGTTCTCCTGTTCTTTGCTGTCCTTGCCgaatcttaaacccaactt
ggttacctcttatctagtggCCTGAACTCACATCAGAAGCATGCCTTCAAGGGGAAAATGgaatcttaaacccaac
ggttacctcttatctagtggCACTTTTGTGCCAGTACTGTTCTCAGACAGGCAAACAGACgaatcttaaacccaact
ggttacctcttatctagtggGTCTTACAGTAAGCAGGTGGTACCAGCAGGGTTCTAACCCgaatcttaaacccaac
ggttacctcttatctagtggGTCACTTCACGGCTAGAGAAACTAAGACCAGAAGAGCTTCgaatcttaaacccaac
ggttacctcttatctagtggCTCAATGAGTGGGATGACCAGAACCCTGTACTGATCATGCgaatcttaaacccaac
ggttacctcttatctagtggGCTTGTGATATTGAGTGAGTTGTTTCACCTATACGACCCCgaatcttaaacccaactt
ggttacctcttatctagtggAAACATTTAGCACAGTTCCTGGTGCATAACAGGTCTGTAGgaatcttaaacccaactt
ggttacctcttatctagtggGCAGCAAAATCATCTCCCTCATTACTGCAGTTGATTTTCCgaatcttaaacccaactt
ggttacctcttatctagtggCCTACTAAATCACGCCTGTAATTCCAGCACTCTGGCAGGCgaatcttaaacccaac
ggttacctcttatctagtggGACTTTGAGCAAAGAAAAAGTCTGCTCTAACAGGAAGCTGgaatcttaaacccaact
ggttacctcttatctagtggTCCCTTCGAGGTGGATACTGAATGATTTGTGTGTGCGCACgaatcttaaacccaactt
ggttacctcttatctagtggTCCTTAGAGGAAACCACTGAAAGCCAATCATTTACAAAACgaatcttaaacccaactt
ggttacctcttatctagtggCATCTTGAAGAGTTCTTTGGTGCTCATTCATCAAACTTAGgaatcttaaacccaacttg
ggttacctcttatctagtggCTCCCACAAAATAAAAATGCCATAAAGCTTTACAGTACTGgaatcttaaacccaactt
ggttacctcttatctagtggGAGCAAGTGAGAGGAAAACCCGAAAATGAGCCTGTCCTGGgaatcttaaacccaa
ggttacctcttatctagtggGAAACAAACAAACAAGCAGCCCCTCCTCTCAGCTCTTGGGgaatcttaaacccaa
ggttacctcttatctagtggTGACAATATCTGCACTTCATAGTTGATTCATAGCATGACCgaatcttaaacccaacttg
ggttacctcttatctagtggCCAGCCCTTTTACAAATCAATCCTGTGAGGTGAACATCACgaatcttaaacccaact
ggttacctcttatctagtggGAGCCACAGAGAGGTTATATACTCATGATCAATAAGCTGGgaatcttaaacccaact
ggttacctcttatctagtggCCAGGAATGATACTGTCTTCCTTCTCCAAATATCCTGTTCgaatcttaaacccaactt
ggttacctcttatctagtggCAGCTCAACAATAGTAAGTAAATGTCCTCATGTGAAATCCgaatcttaaacccaactt
ggttacctcttatctagtggTCATTGAGCTCATTCTGTAAACAACCATTAGGCCAGGGGCgaatcttaaacccaact
ggttacctcttatctagtggTATGAGGCTCAATCTGTGATCACAGAGATGAAATATACAAgaatcttaaacccaactt
ggttacctcttatctagtggGCTGTAAGCTGTCTGCCTACTTCTGCCAGGTGTTAGTAATgaatcttaaacccaactt

ggttacctcttatctagtggAGTGGGGGAAGAAGAGTACTTCATGGTGGGAAGATATTCAgaatcttaaacccaact
ggttacctcttatctagtggACAATGCACAGTGGCCTATTGCAAGTTTTGAAAGCCCTGCgaatcttaaacccaact
ggttacctcttatctagtggAGGTTACTCACAAATCTTGTCGAGGAACTAATGTCCTAGGgaatcttaaacccaactt
ggttacctcttatctagtggGTGTGGGCTCCACTCCAGAAATTCTGATTGAAGTGGTCTCgaatcttaaacccaact
ggttacctcttatctagtggGTCAGATTCCTAGCTGATCTGATGTTTCAGGACTGAGAACgaatcttaaacccaactt
ggttacctcttatctagtggACACTCAGGTGATGGACACCCTTAGTATCCTGATTGGATCgaatcttaaacccaact
ggttacctcttatctagtggACATGTAACACATTTTCTCAAGTACCCCATAAATTTGCACgaatcttaaacccaactt
ggttacctcttatctagtggGTTACCTTCCTTTACATGATGCACCAGAAAGAATACCCACgaatcttaaacccaactt
ggttacctcttatctagtggGAGGTGGAGATATGCACAAAGCCACGTGGGAAATGTCATGgaatcttaaacccaac
ggttacctcttatctagtggACAAGCTATTTTGCTTGGTTGGAGATGAGGAGACAGAGGCgaatcttaaacccaact
ggttacctcttatctagtggTCGACCAAACTACTTTGAGCAATCAAGGGGATATATTGGCgaatcttaaacccaactt
ggttacctcttatctagtggGCTAGCTCTGTCCCCTCCAGAATACAGGCTCCAGGCAGGCgaatcttaaacccaa
ggttacctcttatctagtggCTCTTGGTGTCTAGCACGCTGCCTGTATTAAGGTAGGTCCgaatcttaaacccaact
ggttacctcttatctagtggTGCGGTTACTTTCAGTATCACTGCTTTCCTCATGGTTTCCgaatcttaaacccaacttg
ggttacctcttatctagtggGTCTAACTTGCCTTAGGGACCCAAGATGGCTGCTATTCACgaatcttaaacccaact
ggttacctcttatctagtggCTTCTCTTTTCATGGCTGTGAGAGACTAGCTACCTCTTCCgaatcttaaacccaactt
ggttacctcttatctagtggCTCTTCTTTCTGGGCACAATAGTCTCTATTTCCCAGCCTCgaatcttaaacccaactt
ggttacctcttatctagtggAGGTGTATCCATGCAACTCAGTTCTAGCCAATGGCGTGTGgaatcttaaacccaact
ggttacctcttatctagtggTCCACCACTTGCAGACCTACACATAAAGGCTCCCAGACACgaatcttaaacccaa
ggttacctcttatctagtggTACAGCAACCTTAGCAGCTACATATTGCAGTTGGCAGAGCgaatcttaaacccaact
ggttacctcttatctagtggGCAACTGGCTTAGTTCATATGATCATCCCCTGAGGTGAGGgaatcttaaacccaact
ggttacctcttatctagtggTCCCTAAATCACCCAGATATCCAAGTGGAAATCAGGCTCCgaatcttaaacccaac
ggttacctcttatctagtggGAATGTGGATGCTACCTGGTGGGCAACTGGAGGCCACTGGgaatcttaaacccaa
ggttacctcttatctagtggTGGGGTTTTCTAGATATACATGTCATCTGCAAACAGGGACgaatcttaaacccaactt
ggttacctcttatctagtggGTGCTGCTGGATTTGGTTTGACCACTGGCATATTTTAAGCgaatcttaaacccaactt
ggttacctcttatctagtggTGCACTCACTATCCTGCTTAATCAAATGTCACACTTTGCCgaatcttaaacccaactt
ggttacctcttatctagtggGCCCCACCGCTGATACATACCTCTCTTCCCAAGGGTAACCgaatcttaaacccaa
ggttacctcttatctagtggGATACTTATCATTCCCAGGCATGATTAAATGCTATTGCTGgaatcttaaacccaacttg
ggttacctcttatctagtggTTCATTGTATGACTATGCCATGGCTCACTTATCCATTCTGgaatcttaaacccaacttg
ggttacctcttatctagtggGGCATTGTGAACATTCTTGTGTGTATATCACTGTGTACACgaatcttaaacccaacttg
ggttacctcttatctagtggTTGCTGGGCCAAAGGCTATGTAAGTCTTCAACTTCCCTAGgaatcttaaacccaactt
ggttacctcttatctagtggGCTCTACATTGTCCAGATTCTACATGAAGCACTGGTGGCCgaatcttaaacccaact
ggttacctcttatctagtggGGTTATTGCCAAAGTCTATGTGAGAGAGACTGAGGGTCTGgaatcttaaacccaact
ggttacctcttatctagtggCAAGTCTCCAGATTGAACACAAGAGGAAGCTTATGTGCCGgaatcttaaacccaac
ggttacctcttatctagtggGAGGGCTTGGTAGTAAAAAAGAAAGCACTATCCATCAGGCgaatcttaaacccaac
ggttacctcttatctagtggGAATCATCGAGATGGGTAAAATCAGAAGAGTTGGATCAAGgaatcttaaacccaact
ggttacctcttatctagtggAGAATCAGTAGAGAAGTGAGTTAAGAAAATATCTCTGGGAgaatcttaaacccaactt
ggttacctcttatctagtggAGGAAGAGAAGACTGTGGAGAAGACAGAGACTGAAGAAGGgaatcttaaacccaa
ggttacctcttatctagtggGCCAGAGGAATAGAGCTTCAAAGAATGAGTGGTCAACCACgaatcttaaacccaac
ggttacctcttatctagtggGAGTCCATGTTAAAATGGTTTCAATGGAATAGCATCAATGgaatcttaaacccaactt
ggttacctcttatctagtggTAAAACAGTGAGATGGAATTCGAGTGCAAGATGTTTGTTGgaatcttaaacccaactt
ggttacctcttatctagtggACCCATTCTGGACCACTCTGATCACTAGTTTCACCCTCACgaatcttaaacccaact
ggttacctcttatctagtggCTCCCCTCACCTTTGCCCTCTTTGATTTTCTCCATTGTTGgaatcttaaacccaacttg
ggttacctcttatctagtggGAAAACTATGAAAGAAACTTCTGCCTGTAAATGCAGCTCCgaatcttaaacccaactt
ggttacctcttatctagtggGTAGAGACTGGGAAGGTAGGTGACTTTAGCTCACCAAGCCgaatcttaaacccaac
ggttacctcttatctagtggGTCCTGGGTGCAGCTCCTGGCACATGACATGGCCCACAAGgaatcttaaacccaa
ggttacctcttatctagtggGGTGCTGGGGCGAGAAGGGAGGGCAGCCTGCTCACTAGGGgaatcttaaacccaa
ggttacctcttatctagtggGAGGCTAGGAAGTACAGGGACCTGGCAAAGATGTCTCCTGgaatcttaaacccaa
ggttacctcttatctagtggCAAGGAACAGAGGCAGGTTGTTAGCTGGGCCCCTGGGAGGgaatcttaaacccaa
ggttacctcttatctagtggGCAGCCCTTCCCCCAGGTCAGGCAGCTGCACACCTCATGCgaatcttaaacccaa
ggttacctcttatctagtggGAGTAGGTGAGTCTACTTCTTTCCCAGGGGTGAGTAGCGGgaatcttaaacccaac
ggttacctcttatctagtggCAGCATGTAGTGGACATTTACATGTGCCGGATACCTCATAgaatcttaaacccaactt

ggttacctcttatctagtggCCTCACTGTGCAGATGAAGACACTGAGGCTCAGGGACTTGgaatcttaaacccaac
ggttacctcttatctagtggCATCTGGTCAATAGGGAGCCAGGATTCAAAATCACGTCAGgaatcttaaacccaact
ggttacctcttatctagtggGGTGTAACAGCATGAGTCTCCCATGAAAGGAAGTGGGGGCgaatcttaaacccaac
ggttacctcttatctagtggTGTGCTGCCTTCCTCCTCTGTGCAGCCTCAGCACTCACTGgaatcttaaacccaac
ggttacctcttatctagtggCTCTAGGAGTTTGCAATCTACGGGGAGTGATGGGCACATAgaatcttaaacccaact
ggttacctcttatctagtggATGCGCGAATGACTATGAAGAACAGAGGATTGGGGTCACCgaatcttaaacccaac
ggttacctcttatctagtggGAGACTCATGGAAGTCTCCTGAGGAGAAACCTCTGAGTCCgaatcttaaacccaac
ggttacctcttatctagtggTGGCCAGAAAAATTAAGAAGGGGCAGGGAATTCCAGAGAGgaatcttaaacccaac
ggttacctcttatctagtggAACAAAAGCAAAGAGGCATAAGATAGTCTGGAGTCTGGTCgaatcttaaacccaact
ggttacctcttatctagtggCAGGGTCAGCTTACAGAGGGTCCTATGCAGGCTAAATTCCgaatcttaaacccaac
ggttacctcttatctagtggCTCATGAAGCACTCCTGTGTTTGGAGGCAGCACTGGCAGCgaatcttaaacccaac
ggttacctcttatctagtggGAGCACCTACTATGTGGCAGGAACTCTCTTCATCAGCAAGgaatcttaaacccaact
ggttacctcttatctagtggCTAAATAAAACACAGAACACACCGCCCTCTTTGAAATGACgaatcttaaacccaact
ggttacctcttatctagtggTGGAAAGCATATCAGGGAAGGGGGTGTAAAGTGTCATGAGgaatcttaaacccaact
ggttacctcttatctagtggCTGAAGCCATGTTTAAACAGACACTTGGAGGAGAGGAGGGgaatcttaaacccaac
ggttacctcttatctagtggGGAAGGGGGATCAGCCAGTTCAAAGCCCTGGGGGGTGGGGgaatcttaaacccaa
ggttacctcttatctagtggTCCTGAGAGGGCAGAGGAGCCACGAGAAAGCTCTGAGATGgaatcttaaacccaa
ggttacctcttatctagtggGAGCAAGAGTGGGGGCAGGGAACCAGTGAGGAGACTGTGGgaatcttaaacccaa
ggttacctcttatctagtggGTGGTGCAGCGGCTTGGACCAGGGAGTTAGCAGTGGAAGCgaatcttaaacccaa
ggttacctcttatctagtggCTCTGCATATGGGAAAGGGTGGGCAGGAGAGAACAGTGCCgaatcttaaacccaa
ggttacctcttatctagtggCAGTGGAAGGGGTTTCAGGGAGTCCAGGTGTTCTGGTTGCgaatcttaaacccaac
ggttacctcttatctagtggTTGCCACAGTGCTTTCCTTAGCAAACTAAGGCAGGAACAGgaatcttaaacccaact
ggttacctcttatctagtggGAGGGGAACAAGAGATACTGAGGCCTACCTGAGGGTGGAGgaatcttaaacccaa
ggttacctcttatctagtggGACGTGAGTTTCCCTAGGTAACAAACTTGCACATGTGCCCgaatcttaaacccaact
ggttacctcttatctagtggGGCAGGCCATTGAAAAGCTCCTAGTTCTAATTACTGTTCCgaatcttaaacccaactt
ggttacctcttatctagtggCAGTCCGGCCTTTCCTCTTAGAGTTTGCAGTCTAAGGTAGgaatcttaaacccaactt
ggttacctcttatctagtggCTCACAACAACCCTCTGATGCAGGCACTATTATTTCCCCCgaatcttaaacccaact
ggttacctcttatctagtggTGAGGAAACTGAAGCACAGAGAGCTTGGTTGACTTGCCCAgaatcttaaacccaac
ggttacctcttatctagtggGTGGCTAAGCTGGGATTTGAACCAAGAAAACTCTCTTCCCgaatcttaaacccaact
ggttacctcttatctagtggGCCACCTCTCTGCTTGTGTTGCCTATTTCCACCATGTGAGgaatcttaaacccaactt
ggttacctcttatctagtggGGTGACAGAGAAGGGTAGATGTTTGGCATCTGTGAATTCTgaatcttaaacccaactt
ggttacctcttatctagtggTCTGTCCTCTGTATCCTGAATTGCCACTACCCTGAGCAGGgaatcttaaacccaact
ggttacctcttatctagtggACATTGGTCTTGAGGGGATGGGCAAACACTTTGGACTCTCgaatcttaaacccaact
ggttacctcttatctagtggTGCTCAGCGTAAGTTAGACAATGAATGAGACTTGTTGATGgaatcttaaacccaactg
ggttacctcttatctagtggTTCTGAGTAAACATGGGGCACCAACTCAAAGAGCAAGAGGgaatcttaaacccaac
ggttacctcttatctagtggGGGAATCTCATGGGCTCTGAAGGATTCTGGTCACATAGGGgaatcttaaacccaact
ggttacctcttatctagtggTGTTTTGGGACTTCATCCTGTCTTTTCCCTGCTCTGGGAGgaatcttaaacccaactg
ggttacctcttatctagtggGTTCTAATCCTAATTCTGTTGTTATCTACCTGAGTGACACgaatcttaaacccaactg
ggttacctcttatctagtggTCTCTGAGGCTCGACATTTACATCTACAAATAAAGAAGGCgaatcttaaacccaactt
ggttacctcttatctagtggAGCCTCTTCTATTTCTTCCAATTATTGATTGTATCGGCTCgaatcttaaacccaacttgt
ggttacctcttatctagtggGTTCTTATGGTGTTGCTGGACCTGTTGCTTTTCATTGCTGgaatcttaaacccaacttg
ggttacctcttatctagtggGTGATGGGTGTATAAGGCTCTGGCATTTTGCCCAGGTCTGgaatcttaaacccaact
ggttacctcttatctagtggTCCACTAATGGGTCTGCTGGGCATCTGAGACCACAGAGCCgaatcttaaacccaac
ggttacctcttatctagtggGGACCAGAGTTTGTCTAGAGAGAGAGCAGAGTACAAAGCCgaatcttaaacccaac
ggttacctcttatctagtggCCCAACCCAGCCACTTACTAGATATGTAGCCACAGCCACCgaatcttaaacccaac
ggttacctcttatctagtggCACTTTCTCAAACAGGCCTCCTAGGGTCTCCCACGTTGGCgaatcttaaacccaac
ggttacctcttatctagtggCCATCATTATGCACCAGTATTTAATAAATGCTGAGGAATGgaatcttaaacccaactg
ggttacctcttatctagtggCTCTGTCTCAGTCCTCACTTCAGTGAAGTGGGCAAAAGGGgaatcttaaacccaact
ggttacctcttatctagtggGGTCAAAGTGCCCTAGAAATAGCAAACACTGTCACAACCCgaatcttaaacccaac
ggttacctcttatctagtggGGGGGCTAAAGTGTGCTGCGTGTCTAGATGTGCCAGGCACgaatcttaaacccaac
ggttacctcttatctagtggCCAAGGTTTGCTCTAACTTTTCACACACTTTGGCAGACGGgaatcttaaacccaactt
ggttacctcttatctagtggCTCCTAAGTTGGAACTTTTGAGGCCATTACTGCCAGGAGGgaatcttaaacccaact

ggttacctcttatctagtggGAACTAGAAAATGGCCCCTTCTGGTCCCCAGTTTGGCCCCgaatcttaaacccaac
ggttacctcttatctagtggCACATAGATACTCATGTAGGTGAGATGGCGAACACAGCAGgaatcttaaacccaac
ggttacctcttatctagtggCCCATTTGGGATGACACACCTTCCTTAGAAGTGACAGTGGgaatcttaaacccaact
ggttacctcttatctagtggTGTGGGTTCTGCAGATTGCATCTGGAGTAAGAGTCCAGTGgaatcttaaacccaact
ggttacctcttatctagtggGGCCAAGGCCTCTTGCTTGCTAACATCCTCTGTCTGCCTCgaatcttaaacccaact
ggttacctcttatctagtggCATTCACTGAAGAAGCAAGCTGTCAAAAAGCAACTGCTACgaatcttaaacccaact
ggttacctcttatctagtggCAAATTGCTACCATCGAGGAGTCATAGTCTAATGAAAGAGgaatcttaaacccaactt
ggttacctcttatctagtggACAACATCAGACTCATCAGTGAATCTAGCAAGATCACTGGgaatcttaaacccaact
ggttacctcttatctagtggGTAGTTTCCAAGAGCTCCCGCTGGAAATTGGAAGCCATTGgaatcttaaacccaact
ggttacctcttatctagtggCCATGGTTATGGTTCGAAAGACTCTACATTGCTAAGATGCgaatcttaaacccaactt
ggttacctcttatctagtggTGTTGTTGTTGTTTTTGGAATAGACAAGCTGATTCTGAAGgaatcttaaacccaacttgt
ggttacctcttatctagtggATGCAAAGGACGTAAAACAGCTAAAGAAAATTTGCAGAAGgaatcttaaacccaact
ggttacctcttatctagtggGAATTACACTGTCAGATTTCGAGACCTACTACAAAGCTACgaatcttaaacccaactt
ggttacctcttatctagtggAACAGAAGACAGAGTCCAGAAACTGAACTACACATACACGgaatcttaaacccaac
ggttacctcttatctagtggGGTGCCAAGGCAATTGAGTAGGAAGAGAACGATGTTCTACgaatcttaaacccaact
ggttacctcttatctagtggCTTAAACAGGACACAAGAAGTGAAAAGGCAAGCCACAGCCgaatcttaaacccaac
ggttacctcttatctagtggCTATCTGAGAAAGTACTCATACCCAGAATACATTAAGAACgaatcttaaacccaactt
ggttacctcttatctagtggATCATGTGTATATGGACCATGCACATCATATGTATGTATCgaatcttaaacccaacttg
ggttacctcttatctagtggGCACACACACACACACACAAACTTTGATGAGGCACTTCACgaatcttaaacccaac
ggttacctcttatctagtggGAACTACTCTGAGTTACCAATACATACTCATCAGCATGGCgaatcttaaacccaactt
ggttacctcttatctagtggCATCTGCCTGCCTATACTATATGACCAGTAATCCCGTGACgaatcttaaacccaact
ggttacctcttatctagtggACACAACATGGACTAAACCAACAAGTACGTTGGGTGAAAGgaatcttaaacccaact
ggttacctcttatctagtggCTCTGCTGATAGCTGTAAGGAGAGTGGGCTCATCAGCAGGgaatcttaaacccaac
ggttacctcttatctagtggCTATCTTGACTTAGGTAGTGGTTACACAGGTGTACACAGGgaatcttaaacccaactt
ggttacctcttatctagtggTCTCTCTCCTCTCTCTTTCCCCTCTCCTGTTTCTCCCCTCgaatcttaaacccaactt
ggttacctcttatctagtggGGCAGATAAGTCAAACCTCAGAGAAACAAAGAAATCCACGgaatcttaaacccaac
ggttacctcttatctagtggCCATTAACCACCACAATGGAGACAGTAAAGTCAAGCTGGGgaatcttaaacccaac
ggttacctcttatctagtggCCCACTTTACTTTGTGGCACATACCTTCCTGGTTTCTGGCgaatcttaaacccaactt
ggttacctcttatctagtggTGCAGCAGCAGAGGAGAACAGATTTCCAATTGCAATAGTCgaatcttaaacccaact
ggttacctcttatctagtggTAGCAGAGTGGGCTCCAGGATGATAGCATAGCCCTCTGGGgaatcttaaacccaac
ggttacctcttatctagtggTCCTGAGTGCCTGCCACAAACCAGGTGTGTTACATAAGGGgaatcttaaacccaac
ggttacctcttatctagtggGGGATGACCTGTAAACCCTAGCAGTGGCGGGCGGGGGCCGgaatcttaaacccaa
ggttacctcttatctagtggGCAGGGGGATGGTAAGCAGGGGCACAATGGCAATGGCGACgaatcttaaacccaa
ggttacctcttatctagtggAAACCCAGACCTATTTCCATGACGGGGTTCTTACGTGTCCgaatcttaaacccaact
ggttacctcttatctagtggTGGGTCTTTTCATTTCCCTCCTCCCACAAAGAGAAAAAGCgaatcttaaacccaactt
ggttacctcttatctagtggCGCACACGCGCATGTACATTTGTATGTGTGTAAGGGTGTTgaatcttaaacccaactt
ggttacctcttatctagtggCTTAGTGTTTGCGCATAAGGATGTGGACATGTGTGGTCTGgaatcttaaacccaactt
ggttacctcttatctagtggATGTGTGTGGGAGGGAGGGTGTATATAAAGTCTGTTCTAGgaatcttaaacccaactt
ggttacctcttatctagtggAACATGAAAAATGTGAGTCAGAAAGCTGAAGGGTTAACTGgaatcttaaacccaactt
ggttacctcttatctagtggGAAAACCGTGGTTTGTAACTGGTGAGAGGAAGTATTGAGCgaatcttaaacccaact
ggttacctcttatctagtggGCCAGATCTGCAAAGGAAACCCCTCCCAGCCTGGAATTCCgaatcttaaacccaac
ggttacctcttatctagtggGAGCAGGCATTTAAAAGGGAAGGGAATGGGGGGAAACCTGgaatcttaaacccaac
ggttacctcttatctagtggGAATCCATCTGTCTTTCCCATCCACTACCCCATCCGCCTCgaatcttaaacccaact
ggttacctcttatctagtggCCTCATGCCTTCTCTTCTAGTGTGGCTCAGTGCTTAGTGCgaatcttaaacccaactt
ggttacctcttatctagtggAGTAACCCTGAATTGCAGCTCTTCCTTTCTCACTACATGAgaatcttaaacccaactt
ggttacctcttatctagtggCCTCACAGCTTATGTCTATTAGAACAGCACTTGGCACATGgaatcttaaacccaactt
ggttacctcttatctagtggGCAGGAAGGACTGAGGCCGTTGGTGCTTGGGGAGTGAGGGgaatcttaaacccaa
ggttacctcttatctagtggCCTCAAACACAAATCCCTCAATTGACCACAGGGGGCGCCCgaatcttaaacccaa
ggttacctcttatctagtggTTTGGCGCTGATAGCTGTGATCTGCCCAGCACAGTGGGGAgaatcttaaacccaac
ggttacctcttatctagtggCCGGGCTCAGATTCCTACTCCAGCACCTGGTGGCCAAGGGgaatcttaaacccaa
ggttacctcttatctagtggAGCCTGTTGGGAAAGAGAGAGGGGAGCGGGGAGGAATGGGgaatcttaaacccaa
ggttacctcttatctagtggGCGAGCCTTCCTACAGAAAGAATTGTGGGTGGCTGGGGAGgaatcttaaacccaac

ggttacctcttatctagtggGTGAGTGCAGAGAATACTCAGCCTTCTTCCAGATGGTGAGgaatcttaaacccaact
ggttacctcttatctagtggCACGTACACACCTTCTTTCTCATCCCAGGTCCTAGTCTGCgaatcttaaacccaact
ggttacctcttatctagtggTACGGCTTTCTGCCTGCTTGGGGTTTGGTCCACAATTCCCgaatcttaaacccaact
ggttacctcttatctagtggAGGCTTGGTTAGCAACTCAAGGCTCAAGCATAAATGGGACgaatcttaaacccaac
ggttacctcttatctagtggCCTTTTGAAACTCACCAATATAGTGATTGTAGCAACTAGCgaatcttaaacccaactt
ggttacctcttatctagtggCTAAGCACTATGCAAAGGCTTTAAAGCAGTGGTCCCAAGCgaatcttaaacccaac
ggttacctcttatctagtggGGAAGACCGTGGACTGGGATGGTTTGGGGATGATTCAAGCgaatcttaaacccaac
ggttacctcttatctagtggCCATCTAGGGGTGATGGAGACGGTGACAGGTCATTAGGCgaatcttaaacccaac
ggttacctcttatctagtggGATCCCTCGCATGTGCAGTTCATGACAGGGTTTGTGCTGCgaatcttaaacccaact
ggttacctcttatctagtggGGAGGTGGAGCTCAGGCAGTAAGGTGAGCAATGGGGAGCAgaatcttaaacccaa
ggttacctcttatctagtggCAGGCCACAAAATGGTACCTGTCTGTGTCCCCAGGGTTGGgaatcttaaacccaac
ggttacctcttatctagtggGGCCTTCATCTCATTCAGTTTTCATCAAAATTCTGTGTGGgaatcttaaacccaacttg
ggttacctcttatctagtggGATGAGAGGGCCAGGATTCAAGTTCAAGCAGTCTGACTCCgaatcttaaacccaac
ggttacctcttatctagtggAATGACTGCAGTAGGAAGGGAGCATTTCCCGTGTGCAGTGgaatcttaaacccaac
ggttacctcttatctagtggGAGAGGAGAAGGCATTCTAGGCAGAAAGGCCAGTGCTTACgaatcttaaacccaac
ggttacctcttatctagtggGTCAAGCACCCTCCTGCCTCCCAGTGGGGTTGAGAAGGGGgaatcttaaacccaa
ggttacctcttatctagtggGGCAGAAGTGAAGCTGGGAAGTTTGCACTTGCTAGTTGGGgaatcttaaacccaac
ggttacctcttatctagtggTCTTCTCAATGCCAAAGAGGACCTCAGTCTCTGTGTGCGCgaatcttaaacccaact
ggttacctcttatctagtggGTTGTTGTTGATTTGTGGCCCAGGCTAGAGTGCGGTGGTGgaatcttaaacccaact
ggttacctcttatctagtggAGGTTCAAGGGATTCTCCTGCCTCCAGACTCCCAAGTAGCgaatcttaaacccaac
ggttacctcttatctagtggTTTCACCGTGTTGGCCAGGCTGGTCTGTGTGTGCTTTGGAgaatcttaaacccaact
ggttacctcttatctagtggCCGTGCATTCAAAATCATAAGGCAAGTTTTCCTGTGTGGCgaatcttaaacccaact
ggttacctcttatctagtggAGAGCGTCAGGGTTGACAGTAGCTGTGATGCTCCAGATGGgaatcttaaacccaac
ggttacctcttatctagtggTCAGGGCAGTGGTTGAGGGGCTGTGGGAGGGTGGGGAGGGgaatcttaaacccaa
ggttacctcttatctagtggTCTCAACCATCTGTATTTGATTGGAATATTGTGTGACTTGgaatcttaaacccaacttg
ggttacctcttatctagtggGGGATTTTAGGAGAAAACGCTTAGCCATACAGAGCTGTGGgaatcttaaacccaac
ggttacctcttatctagtggACATCCTGTACTTAGAAAGGAATAACCTGCACATGACACGgaatcttaaacccaact
ggttacctcttatctagtggCACCTTTGCATTCTGAACCTAGCTGGGAAATTATGACAACgaatcttaaacccaact
ggttacctcttatctagtggGTGGGCGTCCTCACATTTTGTGTCTTAAAATGTCCAGCTCgaatcttaaacccaactt
ggttacctcttatctagtggTTTCCCCCATGAAATCCCCATTGAAAACAGAAACCAGAACgaatcttaaacccaact
ggttacctcttatctagtggGGTGACTGAAGCATAGCAAATAGAGTTTTGTTTCCTTCCCgaatcttaaacccaactt
ggttacctcttatctagtggCAAAGGCACTGGAGGAAATAAGGGTTGGGGGAGAGGGAGGgaatcttaaacccaa
ggttacctcttatctagtggGGGTCATGTGATCTGCGTGAGTTTCCTAAGGGAGAGAGGGgaatcttaaacccaac
ggttacctcttatctagtggCTGAATGGCAAGTGAGCAAGACCCTCGCTGCTGAAATCTGgaatcttaaacccaac
ggttacctcttatctagtggCCAGAGTTAGGAGCTGGAAAGAAGGTCAAGAGGACATCTGgaatcttaaacccaac
ggttacctcttatctagtggGGTCCAAAGGTCAGCTCCAGGGACCAGGCTGCAGGTCTCTgaatcttaaacccaa
ggttacctcttatctagtggTTCTCTTTCTGTATCACACATGCCGCCTCAACTAACCAGCgaatcttaaacccaact
ggttacctcttatctagtggAATGTGTGGCAGATCATCACCCCTGGGGCTTAGCTTTGGGgaatcttaaacccaac
ggttacctcttatctagtggCTCAGAGCAACGTAAGCACAACGGCCCCACTTTGCATACCgaatcttaaacccaac
ggttacctcttatctagtggGAACAAAATCATACTGGGCGTTTTGTTTTCGTTTGCAGCCgaatcttaaacccaactt
ggttacctcttatctagtggGAGCTAGATGGCAGATATTTTGGACTTTGAGGACCTAGGGgaatcttaaacccaact
ggttacctcttatctagtggGAGTGTGGCCATGTTCCAATTATTTACAAAAACAGGTGGCgaatcttaaacccaact
ggttacctcttatctagtggTCCTATGTTTATCACATCAGATATCCAACTGTGTATACTGgaatcttaaacccaacttg
ggttacctcttatctagtggGCAATATTTGCATGACCATTGACATTTAGATGCTTTCCAGgaatcttaaacccaactt
ggttacctcttatctagtggCCTTAGATTTTTGTCTTCATTTCGGGTAATTTCCATAGGGgaatcttaaacccaacttg
ggttacctcttatctagtggCTCAAAAGGCTTCACCCATTTACTCCCAGAGCTGTGGATGgaatcttaaacccaact
ggttacctcttatctagtggCCCTCCTCACTCAACTCCCCCAAACCCCACCCATGGCCCCgaatcttaaacccaa
ggttacctcttatctagtggCTGAACGTTCCTTCCCTCTTCCTGTCCACCTCACATCTGGgaatcttaaacccaact
ggttacctcttatctagtggAATCCCCTGACTTCCTGTTTCTCTCCCTACAATAGGTCCCgaatcttaaacccaact
ggttacctcttatctagtggTTTTTGCTTTTTGCTTCTCCCACTTCAACCCCAAACTCTCgaatcttaaacccaacttg
ggttacctcttatctagtggGCAGTGGTGTCATTACAGCTCACTGCAACCTTGATCTCTCgaatcttaaacccaact
ggttacctcttatctagtggCCTCAGCCTCCCAAGTATCCGGGACTAAAGGCTTGTACCAgaatcttaaacccaac

ggttacctcttatctagtggGTAGAGATGGGGTCTCACCAAGTTACCCAGGCTTGTCTGGgaatcttaaacccaac
ggttacctcttatctagtggGTGTGAGCCACCACACCCAGCACAAACTCTTTATTGTTGTgaatcttaaacccaact
ggttacctcttatctagtggGTTTCATAGAGCTTTGCCTATTTAAGAGTCTCCATTTGGCgaatcttaaacccaacttg
ggttacctcttatctagtggCACTAAGGCAGGCAGTTCACAGGAGTATGTGAAAAGACCCgaatcttaaacccaac
ggttacctcttatctagtggAGGGAATAGTGAGGTGTGTGGTACAGAGAGTCAAATGCAAgaatcttaaacccaact
ggttacctcttatctagtggGCTTCCTTCCTCCACACCCCACAGCCCACCCCCCGACTGCgaatcttaaacccaa
ggttacctcttatctagtggACCAGCTAAACTGAACTATTCCAACACACAGTTCACCCTGgaatcttaaacccaact
ggttacctcttatctagtggTGTTCATGATGATCCCTTCTTCCTCAAATACTCATTCCTTgaatcttaaacccaacttg
ggttacctcttatctagtggGCTATGTTGCTCAGGTTGATCTCAAATTTCTGGGCTCAAGgaatcttaaacccaactt
ggttacctcttatctagtggGTGTGAGCCACTGTGCCTGGCCTCACCCTCCCTTTCTAATgaatcttaaacccaact
ggttacctcttatctagtggACTGCTGTCTTGCTGCCTACCATCACCCTGTGCCTGGGGGgaatcttaaacccaac
ggttacctcttatctagtggTCTCCCTGCAGGCTTCTCAAAGGCACAGGAGTAGGCTACCgaatcttaaacccaac
ggttacctcttatctagtggCAGAGCAGATCACTCTCCAACATATAGGAATTTAGTACTCgaatcttaaacccaactt
ggttacctcttatctagtggGATCTGCCCTCAGATTACCTGTCCTGCCTCATCTGCTGCCgaatcttaaacccaact
ggttacctcttatctagtggCCTCTGCCTGAGAGATGTTCATTCAGCAACCCCCATCCCGgaatcttaaacccaac
ggttacctcttatctagtggTCATGGAAGCCCTACCTGATCTCTACTTTCCTGGTTCCCCgaatcttaaacccaact
ggttacctcttatctagtggGCCAGCAATCGTTAACCACTGTGGTACCCTTCTCATTCACgaatcttaaacccaact
ggttacctcttatctagtggGGGTACTGAGATTTAGAGTCAGCAGGTAACGTTCTCAAGGgaatcttaaacccaact
ggttacctcttatctagtggCACCCTAGGTCTTTCTGATTCCTTGTCACCAGCCTTCTCCgaatcttaaacccaactt
ggttacctcttatctagtggCACCATATTGAAATTGCCTATTTCCTCCTCCTGCTTCCCCgaatcttaaacccaactt
ggttacctcttatctagtggCGCCCATGCTTAGCACAGACTGGGTACCTAGAACAGACCCgaatcttaaacccaa
ggttacctcttatctagtggAATGCAGTCGGTGTGCTCAGGTGCTGAGATCCATGGCCAGgaatcttaaacccaac
ggttacctcttatctagtggTATGCCCAGAGGAGTTTGTACCTGCCTGGTTGCTTGAGGGgaatcttaaacccaact
ggttacctcttatctagtggCGCTTTCCAGATGAAGCCCCTGGAGAAGACGCAATCTTCCgaatcttaaacccaac
ggttacctcttatctagtggGGCTGAATTAGCTTATGACCTCAGGGCCCTGGTTGCATCCgaatcttaaacccaact
ggttacctcttatctagtggGTCCATGCCAGGTCTTATGCCCACACCCAGACTGGCCCAGgaatcttaaacccaa
ggttacctcttatctagtggCCTCCGATAAACAAAGGAGGCACCCGTGTGGAAGAAACAGgaatcttaaacccaa
ggttacctcttatctagtggGGGGTGAGGAAGTACAGAGTATGTGGGGGCATCTGACAGCgaatcttaaacccaa
ggttacctcttatctagtggCCTGAGACCCTGCTATGGGAGAGGGGGAGACATGCTCAGGgaatcttaaacccaa
ggttacctcttatctagtggGGAAGGAGCAGGAAGGACAAGAGAATTTCCACCCCTGGGGgaatcttaaacccaa
ggttacctcttatctagtggGCCCAAGTCAGATCATCCAGGGCCCTGGTGAAACAGTTTGgaatcttaaacccaac
ggttacctcttatctagtggCCCACCAGGGCTATCTGGGTCCTGTGGTCAAGGTGGCACCgaatcttaaacccaa
ggttacctcttatctagtggCCAGCACTTCGGGAGGCCAAGGCAGATCACTTGATATCAGgaatcttaaacccaac
ggttacctcttatctagtggGAGCCACTGTACTCCTTGCTGGGTGACAGAGCGAGAGTCCgaatcttaaacccaa
ggttacctcttatctagtggGGGAGTGGGGTGGGGGCAAAGTCATGGTGCTGAATGTTAGgaatcttaaacccaac
ggttacctcttatctagtggGAGACAGAGTTGGAGTCTCTCTCTGTCGCCCATGCTAGAGgaatcttaaacccaac
ggttacctcttatctagtggCTCCTGCCTCGGCCCCCCAAGTAACTGGGATTACAGACACgaatcttaaacccaa
ggttacctcttatctagtggGATGGGGTTTCGTCATGTTGGTTGGGCTGGTCTCAAACTCgaatcttaaacccaactt
ggttacctcttatctagtggCACTGCCATTATCTCCCTCCAGGCCTCAGTATCCTGCCTGgaatcttaaacccaac
ggttacctcttatctagtggAGACCAGGTAATGTTAAGGTCTTTCCAGCTGGGACAAAACgaatcttaaacccaact
ggttacctcttatctagtggAAATACAAAAATTAGACTGGCACACACCTGTAATCCCAGCgaatcttaaacccaact
ggttacctcttatctagtggTCAATGAAGAATAGTAAGGCCAAGTGCAGTGGCTCACACCgaatcttaaacccaac
ggttacctcttatctagtggGTTTGAAGCCAACCTGGGCAACGTAGCAAGACCCTATCCCgaatcttaaacccaac
ggttacctcttatctagtggCAAAAATTAGCAATGCATGGTGGTGTGCCTGTAGTTCCAGgaatcttaaacccaactt
ggttacctcttatctagtggACTTGAGCCCAGGCAGTCAAGGCCGCAATGAGCTATGATCgaatcttaaacccaac
ggttacctcttatctagtggCTCCAGCCTGGGTGATACAGTAAGACCGTGACTCAAAATAgaatcttaaacccaact
ggttacctcttatctagtggCGCTAGCATTTGTAGTGTGTTTTCTGTGTACCTAGTACTGgaatcttaaacccaacttg
ggttacctcttatctagtggTTCTCATTGAATCTTCATGACAACCACATGAAGAGGTAACgaatcttaaacccaactt
ggttacctcttatctagtggGAGCTGGGATTCAAGCCCGAGTCTGCTCTGACATCCAGCCgaatcttaaacccaac
ggttacctcttatctagtggGGAAGTTTCAAAACCTGAAAAATCCATGATGGCTTGTGCCgaatcttaaacccaactt
ggttacctcttatctagtggGAGGTGCCCTGTGTGACTCTGGACGGCTCCCAGATGAAGCgaatcttaaacccaa
ggttacctcttatctagtggGCGCCCCAGCCTCATGAACCCTCACCAGGTCTACTCAGGCgaatcttaaacccaa

ggttacctcttatctagtggCCAGGAGCCTGGCAGGGCCCTCTCTCTCTCCCTAACAAACgaatcttaaacccaa
ggttacctcttatctagtggTCCCCTCACCTTGTAAATAATCTCCCTCCCAGGCACCGAGgaatcttaaacccaac
ggttacctcttatctagtggGTGAACACTGTTGTGGAATGCCTGCCACACTCCCGTGGGCgaatcttaaacccaac
ggttacctcttatctagtggCTCAGTCTCCATTTGTGACCTTGGTGGCGAGAGGTTTGGGgaatcttaaacccaact
ggttacctcttatctagtggCAGAGCCGTGAGAGTTACCAGAGTCCAGAGAACAATCTGGgaatcttaaacccaac
ggttacctcttatctagtggGGGGAAAAAGCTGAGGTCCAGACAGGAGACAGACGGTACGgaatcttaaacccaa
ggttacctcttatctagtggGCAGGAATAGGTTGAGACAGCAGGTCAAGTTCATTTCCAGgaatcttaaacccaact
ggttacctcttatctagtggTGGCATTGGGACTGCTCAATGTTCTTTCCTCCTCTTTGGGgaatcttaaacccaacttg
ggttacctcttatctagtggGGGTGGATCATTAGAGGCCAGGAGTTCGAGATCAGCCGGGgaatcttaaacccaac
ggttacctcttatctagtggGGTGACACAGCGAGACCTTGTCTTAAATATATATACGTGTgaatcttaaacccaacttg
ggttacctcttatctagtggTCAGTAGAGTCAGGGTCTCACCATGTGGCCCAGGTTGGTCgaatcttaaacccaac
ggttacctcttatctagtggAGTGCTGGGATTACAAGCAAATGTCACCAGGCCCAACCTGgaatcttaaacccaac
ggttacctcttatctagtggCCCATTGTGGGTGGTCTTTGTAGGCCCAAGGAAGTGCCTTgaatcttaaacccaact
ggttacctcttatctagtggACAGAGATGTCCAGAAGAGATGGCACAGACTCTTTCTCTCgaatcttaaacccaact
ggttacctcttatctagtggGGTCTCGATCTCTTGACTGTGTGATCTGACCGCCTTGGCCgaatcttaaacccaact
ggttacctcttatctagtggAGGCGTGAACCACTGCACCCGGCCAGACTTTGAATCTTACgaatcttaaacccaac
ggttacctcttatctagtggCTGGTGCCTGTCCCTGGAGTGACTGAGTCATCCAGCTCTCgaatcttaaacccaac
ggttacctcttatctagtggCCTGGGTCAAACCTGTCTCTACCCTGGCTTCCCTATCTCCgaatcttaaacccaact
ggttacctcttatctagtggGCCAGTAGGAGCTTCTGCTTCTTGCAACTAAAAAGCCATGgaatcttaaacccaact
ggttacctcttatctagtggTCCCCACAATTTCATGCTGTCTTGCTCTGAGGCAACAACCgaatcttaaacccaact
ggttacctcttatctagtggCCAAACCCTGGGTAGTTGCATCTTTTCAGTCAATGGCTGCgaatcttaaacccaactt
ggttacctcttatctagtggCAGCCCCCACTCGCAACACAGCTGATAGGGTGGTCTACACgaatcttaaacccaa
ggttacctcttatctagtggCTCCACTCTTACTGTCTCGAAGTATCAAACCCAAATAAACgaatcttaaacccaactt
ggttacctcttatctagtggCGCTCTATCACCAAGTTGGAATGCAGTGGCGCGATCTCGGgaatcttaaacccaac
ggttacctcttatctagtggGCGTGAGCCACCAGGCCCGACCAAAACTGTCATTTAAGAAgaatcttaaacccaac
ggttacctcttatctagtggCCATCTAATCAGCTGAACAATTCTACAGATGGTGCATTACgaatcttaaacccaacttg
ggttacctcttatctagtggATGAGAAAACTGGCTGTCCAAATTGGGAAAGGGGCAGTTGgaatcttaaacccaact
ggttacctcttatctagtggGCTCACAGTGGTGGTCTTTAGGTATAATCCCAGCTCGATCgaatcttaaacccaactt
ggttacctcttatctagtggTGCCTGGAAATGACTCAGTGGTGCCCCAGTCAGGTGGTTCgaatcttaaacccaac
ggttacctcttatctagtggTGCCCCACCATAAGATAGTTGAGTATCAGAATCTCCAGAGgaatcttaaacccaact
ggttacctcttatctagtggCTTAATATTTGGAAAGTGAAAAGTAGGCTGGGAGCAGTGGgaatcttaaacccaactt
ggttacctcttatctagtggGCCAACATGGCAAAACCCTATCTCTATCAAAAGTACAAAAgaatcttaaacccaactt
ggttacctcttatctagtggCAGGAGGTGGAGGTTCCAGTGAGCCGATCATGCCACTGCAgaatcttaaacccaac
ggttacctcttatctagtggGCGTGAGCCACTGCGCCAGGCCCCCCAGTGTTTTTAATACgaatcttaaacccaac
ggttacctcttatctagtggAGTGGCTCCTGCGGTTGATTAGGTCTAGGTGTTCCCAGGCgaatcttaaacccaact
ggttacctcttatctagtggACAGGTGCAAGTAGGTGAAGATGGCATAGCCAGCAAGTGGgaatcttaaacccaac
ggttacctcttatctagtggAACACCCAACATTCCCCTGTGACTTTTCTGACAGCCCGAGgaatcttaaacccaac
ggttacctcttatctagtggAGCTTCTGCCAGGAAGCCCTTCCCAACTAACTCCAGCTCCgaatcttaaacccaac
ggttacctcttatctagtggTTGTCTTCTGACAGCCACCTTGCACTGTACTTGATGAATTgaatcttaaacccaacttg
ggttacctcttatctagtggTAGACTGTGGTTTTTCATGTGTGTGTGCACATGTGTAAACgaatcttaaacccaacttg
ggttacctcttatctagtggCTCATGTTAACTGAGACCTCACCTTGGTTACCATGGAACCgaatcttaaacccaactt
ggttacctcttatctagtggCACGGTTCTGACACAAATGAGTTTCCATGAATACAGTGCCgaatcttaaacccaactt
ggttacctcttatctagtggCTCCCACCTCAGCCTCCAGAGTAGCTATCACTACAGGCGAgaatcttaaacccaac
ggttacctcttatctagtggTTGGCCCCTCAGACTGCTGGGGTTACAGGTGTGAGCCGCCgaatcttaaacccaac
ggttacctcttatctagtggTCCTAGGATTGCCAAAACAGATTATGATGAATTGTGTGGCgaatcttaaacccaacttg
ggttacctcttatctagtggGTCAGCAGGGCTGTGCTCCCTCTGAAGATTCTAGAGGAAAgaatcttaaacccaac
ggttacctcttatctagtggACAATCTCTGCCTCTGACTTCACATGGCCTCCTCCCCTGCgaatcttaaacccaac
ggttacctcttatctagtggCCTCCCGGGTTCAAGCGAAATTCTCACGTGTCAGGCTCCCgaatcttaaacccaac
ggttacctcttatctagtggGTACAGACAGGGATTCACTACGTTGGCCAGGCTGGTCTCGgaatcttaaacccaac
ggttacctcttatctagtggGGCATGAGCCACCTTGCCCAGCGTGGATTTAAGGATTTAAgaatcttaaacccaact
ggttacctcttatctagtggACTCAGTTCACTACAGGTGGATAAGAACCTCCAGGGCTGGgaatcttaaacccaac
ggttacctcttatctagtggCTCCACTCCCAGTGCTAAACTAGCAAGTACACACAGCAGGgaatcttaaacccaac

ggttacctcttatctagtggGGGTTGGATCTGAATGAGGGCAATGAGCTCCGTTCACTGGgaatcttaaacccaac
ggttacctcttatctagtggCTCCTTCCAGCCTTCTGTACAGCCAACCACTCCCAAGCCCgaatcttaaacccaa
ggttacctcttatctagtggCATGGCTTTCTCTTCCTGCTCCCCACCCACCCTTGCTGCCgaatcttaaacccaac
ggttacctcttatctagtggCCTCCCACTTTCCCGCCTTGAATGACAGTTTAAGGGAAGCgaatcttaaacccaac
ggttacctcttatctagtggGGAAGCAAGGAAGCCCCTCCTCACTTCGAGCACCTTCACCgaatcttaaacccaa
ggttacctcttatctagtggTTCTCTGCTTTTAGGGTGCAAGGCCATTAGGGTTCCTCTGgaatcttaaacccaactt
ggttacctcttatctagtggCCCTTAGTCAAATGTCCTCTCCTAGCCACATACCCTTCCGgaatcttaaacccaac
ggttacctcttatctagtggGAAGTCCCAGGCGAAAAGGTTCCTCATTGTTAGCACTGCCgaatcttaaacccaac
ggttacctcttatctagtggACACTGTGTGCCAAGCAGTGTGCCAAGCCCTTCACACACAgaatcttaaacccaa
ggttacctcttatctagtggCATTGTACCGAGGAGAATACTGAGGGCTTAAGCAACAAGGgaatcttaaacccaac
ggttacctcttatctagtggCCTACGTTCATCAGATTCCAGAGTGCATTCTCTCCACCACgaatcttaaacccaact
ggttacctcttatctagtggGACTGCTTCCTAGCCAAGCTTCCTGAAAACACATAAGACCgaatcttaaacccaac
ggttacctcttatctagtggCCCTTCTCCTCTGAAATCACTGGAGTGGGAGTAGGGTTGCgaatcttaaacccaac
ggttacctcttatctagtggTCCCCTTGGAACTGGTTTTTCTCATAGCAGTTGCTTTGAGgaatcttaaacccaactt
ggttacctcttatctagtggCAGTCAGGGAACAAGTCACAAGATATCCCCAAATCAGAGGgaatcttaaacccaa
ggttacctcttatctagtggGCCTGACACCTCTCTCTGAACCCAACTATTTGCCAGTGGCgaatcttaaacccaac
ggttacctcttatctagtggGCCATCCTGGCTCTGCCCTGGAGGAGTCCTTATCTGAACCgaatcttaaacccaa
ggttacctcttatctagtggCCAGGCGCTTGGGAAGATGTCCTGGACCAAGGTTCCAAGCgaatcttaaacccaa
ggttacctcttatctagtggCCCTAGTGAGGAAGGCACCGTCATCATCCCTGTGCTGCAGgaatcttaaacccaa
ggttacctcttatctagtggGCGGCAGAACTGGCTCCTGCCTGCTGCACTCTCCTACTTCgaatcttaaacccaa
ggttacctcttatctagtggTCACTGCTGGATACCAATGGGCTAGAACAATGCCAGGTGGgaatcttaaacccaac
ggttacctcttatctagtggAATGCATAGAGAAAATCTCAACCCGGAATTAAGGTGTTGGgaatcttaaacccaact
ggttacctcttatctagtggCCATAAAAAGTCTATGTAGGCAGGGCGCAGTGGCTCACGCgaatcttaaacccaa
ggttacctcttatctagtggTCCACTAAAAATTCAAAAATTAGCCGGCCATGGTGGCGCGgaatcttaaacccaac
ggttacctcttatctagtggGAGGTTGAGGCATGAGAATCGCTTGATCTCAGAGGTGGAGgaatcttaaacccaac
ggttacctcttatctagtggCCATACTGGTATCTATACCCTAGTTTGGGAAGCCCTAAGCgaatcttaaacccaact
ggttacctcttatctagtggCTTTGTGTCCCTGTGGTTGAGGGTAAAGGCCAACAGTGCCgaatcttaaacccaac
ggttacctcttatctagtggCTTGCTAGAAAGAACTGGTTATAGATTTGATGGTTCCTTCgaatcttaaacccaacttg
ggttacctcttatctagtggGGCAAGAACAACAAGAGGAAGGAGGCCTGGACTGGAGCCCgaatcttaaacccaa
ggttacctcttatctagtggGGACTTCCAGGTTTGAGCTGATGAGCATCAGGACACCTGCgaatcttaaacccaac
ggttacctcttatctagtggTCATGTCCTGGAGACAGTCTACTACTTTGAACTGGTGGCCgaatcttaaacccaact
ggttacctcttatctagtggCTGCAGAACCTGCTTAGTAACCAGCACACAGAAGATGCTCgaatcttaaacccaac
ggttacctcttatctagtggCTCCCTCATTTTCTTCTGTTTCCACAGTCTCCTAAGGCTCgaatcttaaacccaactt
ggttacctcttatctagtggCGGGAGGCCAGGCATGGTCACTCACTCCTGTAATCCCAGCgaatcttaaacccaa
ggttacctcttatctagtggCAGTGAGGCAAGGTCGAAGCACTGCACTCCAGCCTGGCGAgaatcttaaacccaa
ggttacctcttatctagtggTGTCCAGTCCCACCCCAGATATCAGCCTCCAGCCCCCTGGgaatcttaaacccaa
ggttacctcttatctagtggTGGTGCAGTGACCTCACCAGAAGGCATGAGATCCAGCGAAgaatcttaaacccaa
ggttacctcttatctagtggTGCCAGTGAGAATATCACCAGCTAACTGCAGGCAGGGAGGgaatcttaaacccaa
ggttacctcttatctagtggATTAAACTTTGTTTCCTTGTACACAGCAGCCAACCGCAAGgaatcttaaacccaactt
ggttacctcttatctagtggTGCCTGAGATGGCACACACGCTTATGATGTATCAGGTCACgaatcttaaacccaac
ggttacctcttatctagtggCAAACTGAAAATGTCACCCCCATCTGGAGAGTGGGACACGgaatcttaaacccaa
ggttacctcttatctagtggCTGCACTAGTGGCTGGTTCAGTAATAAATCTGTAAGGGCCgaatcttaaacccaact
ggttacctcttatctagtggCTGCCTCCCATCAGTCATCCCAGGGGACGCCAGCTGCCACgaatcttaaacccaa
ggttacctcttatctagtggGGAGAGGCCCACAGAGGAGGAACCAATGTCTCCAGCCAACgaatcttaaacccaa
ggttacctcttatctagtggCCTGAGGCCTTTCCAAAGCCGTGTGAGTGACGCTGGGAGGgaatcttaaacccaa
ggttacctcttatctagtggCCAGAGAGACCAGGAGCAGAACCACCAACTAGGCTGAGCCgaatcttaaacccaa
ggttacctcttatctagtggGTTTTCGCTTGCTAACTTTTGGGGTAATTTGTCACACAGCgaatcttaaacccaacttg
ggttacctcttatctagtggGAGGATTGCTTGCGTTCAAGAGTTAGAGACCAGCCTGGGCgaatcttaaacccaac
ggttacctcttatctagtggAAATTATCTGGGCATGGTGGCGCGCTTCTATAATCCCAGCgaatcttaaacccaact
ggttacctcttatctagtggTGCAGTGAGCCGAGATCACACCTCTGCGCTCCAGCCTGGGgaatcttaaacccaa
ggttacctcttatctagtggGCTGGGCCTGGTGGTGCATGCTTGTGGTCCCAGTTTCTCGgaatcttaaacccaac
ggttacctcttatctagtggGTGGGAGAATCATCTGAGCTCGGAATGTCAAGGCTGTAGTgaatcttaaacccaac

ggttacctcttatctagtggCTGTACTCCAGCCTGGGCAAACAGAGTCTCTGGGACCCTGgaatcttaaacccaac
ggttacctcttatctagtggGCCCCCTCTGTTAGGTGTGGCCTTCTTATGTGCCTTGGCCgaatcttaaacccaact
ggttacctcttatctagtggCTGGGCTCCTGGATGAGCATGACACAGGACAGAACCACCCgaatcttaaacccaa
ggttacctcttatctagtggATAAGCCTTTGTTGAGGCCGGGCACTGTGGCTCAAGCCTGgaatcttaaacccaac
ggttacctcttatctagtggGCAGTGAACTGAGACTGCGCTACTTTACTCCAGCCTGGGCgaatcttaaacccaac
ggttacctcttatctagtggGTCACTGAGACTTTGGAGTTGTTTGTTACTACTAGCTTACgaatcttaaacccaacttg
ggttacctcttatctagtggCCTGCAGCTAATCAACAGACCACATTACCATTTTGTTCACgaatcttaaacccaactt
ggttacctcttatctagtggTTGCATCTGAACCAAGATTTACTATGGTCACTTAATAGGGgaatcttaaacccaacttg
ggttacctcttatctagtggCACTGGAGAAACCTCTATCTCAGTGCCAAGATAACTCTTCgaatcttaaacccaact
ggttacctcttatctagtggTCATTTTGAAACAGCGTCTCACTCACTCTGTTGCCTAGGCgaatcttaaacccaactt
ggttacctcttatctagtggCCCAAGTAGCTGGGTTTAAAGGGGCACCAACATGCTGGGCgaatcttaaacccaac
ggttacctcttatctagtggGGGATTTTACCATGTTGGGTAGGCTGATCTCCAACTCCTGgaatcttaaacccaactt
ggttacctcttatctagtggCAGGTGCGAGCCACCACGCCTGGCCTATAACCTTACTTTTgaatcttaaacccaac
ggttacctcttatctagtggTTGCCCTGGCTAGTAGGAAGCTCCAAGTCAAACTGCAGTCgaatcttaaacccaac
ggttacctcttatctagtggGTATGGCGCTTATTTTTAGTACTAACCTTGTCCTGGCTTCgaatcttaaacccaacttg
ggttacctcttatctagtggACTCATGAGGAACCTCACATCCATTTTCAAATGAGGAAAGgaatcttaaacccaactt
ggttacctcttatctagtggCACAGGCAGCATAACTTGTAAACATCAGTGACAGAACAAGgaatcttaaacccaact
ggttacctcttatctagtggTATCTGGGCTTCATCTGAGTTTGACTACGTAGTAGCTGTGgaatcttaaacccaacttg
ggttacctcttatctagtggGTGTCTCAGTTTCTTCCTTTATAGAAGTCAGGGTGAGAGGgaatcttaaacccaacttg
ggttacctcttatctagtggTACTGGGGAAAGTACTTGGCCCTTCATAAGAGCTCTTACCgaatcttaaacccaact
ggttacctcttatctagtggGCAATGCACCCTGGTGAGCTCAGAAACATCCCCTGCTGAGgaatcttaaacccaac
ggttacctcttatctagtggCAGCTGCATCAGGAGAAACTTAGGTCAGAAAAAAGATGGGgaatcttaaacccaac
ggttacctcttatctagtggGCTCATGCAATCACAGTGGGAACAAGCAGGGAGAAACTCCgaatcttaaacccaac
ggttacctcttatctagtggCTGGAAAGCAGGAGATAGGTGATTCCAGTTCACATTTTAAgaatcttaaacccaacttg
ggttacctcttatctagtggTGGGCTGGATAGCTCTCTGTTGTAGGGACCATTCTGTGTCgaatcttaaacccaactt
ggttacctcttatctagtggTTCCGAGATGGAGTCTCCCTCTGTCCATGGGCTGGAGTGCgaatcttaaacccaac
ggttacctcttatctagtggTCCTGGGTAGTTAGGATTACAAGTGTGTGCCACCTCGCCCgaatcttaaacccaact
ggttacctcttatctagtggGGTGTAAGCCACCGCACCTGGCCACCTGTTCTATGTCTTAgaatcttaaacccaact
ggttacctcttatctagtggCTGGCCTTTGCCCACCAGATCCCAGTAGCTCTCCCATCCCgaatcttaaacccaac
ggttacctcttatctagtggGTCCCTTGGGAGGCCAAATTGTCCCCTCTCCTCCTGTGTGgaatcttaaacccaac
ggttacctcttatctagtggGACCTGGGCTTGCTCCTTAAGCAACTCATAACTCACGCAGgaatcttaaacccaac
ggttacctcttatctagtggGTGAGGAAGGAAGTACAACCTGAGCTCAGCAGGGGAGAGGgaatcttaaacccaa
ggttacctcttatctagtggAACCACTGACTGCCCTCCCCAAAGTGGGCCCACTCCCTCCgaatcttaaacccaa
ggttacctcttatctagtggCTTCACAGGAAGTTCTATTATTTATTTGAGACAGAGTCTCgaatcttaaacccaacttg
ggttacctcttatctagtggTGGTGCTATCTCGGTTCGCTGCAACCTTTGACTCCGCGGTgaatcttaaacccaact
ggttacctcttatctagtggCTGGGATTACAGGCACGCCCCCACGCCAGGCTAATATTTGgaatcttaaacccaac
ggttacctcttatctagtggCTGGCCTCAATTGATCTGCCCACTCGACCTCCCAAAGTGCgaatcttaaacccaac
ggttacctcttatctagtggGCACCTGGCCAGCAAGGTCTGTAATCTAAAACAAACTTTTgaatcttaaacccaactt
ggttacctcttatctagtggATACATATTATACAGCATATATATGTGGCCTAGGGGTTACgaatcttaaacccaacttg
ggttacctcttatctagtggGAGTATCAGTTTCCTATATAATAGAAACTGGGCATGCCAGgaatcttaaacccaacttg
ggttacctcttatctagtggTGCCCAATTGGTAGATAAGCAAACTGTATCTAGTGGCAAGgaatcttaaacccaactt
ggttacctcttatctagtggCAAGTCCGGTACTCTTTCCATTTACCGCCTCGCTCCTGAGgaatcttaaacccaact
ggttacctcttatctagtggTACAAAGGACATGAATGCATCATTTTTACGGCTGCATAGTgaatcttaaacccaacttg
ggttacctcttatctagtggTGCCATTCTAACTGGTGTGAGATAGTGGTTTTGATTTGCAgaatcttaaacccaacttg
ggttacctcttatctagtggCCTTGGGCAGTAAGTCCATTTTCACAATATTGATTCTTCCgaatcttaaacccaacttg
ggttacctcttatctagtggTTGAAGAGGTCCTTCCCATCCCTTGTATTCCTAGGTATTTgaatcttaaacccaacttg
ggttacctcttatctagtggGGCTCCCTGTTTGTCTGCTATGGGTGTATAAGAATGCTTGgaatcttaaacccaacttg
ggttacctcttatctagtggTGTTTATTGATTTGCCTATGTTGAAGCAGCCTTGCATCCCgaatcttaaacccaacttg
ggttacctcttatctagtggAGTGCTGGAACTACACGCGCAAGCCACTGCACCCAGCCTGgaatcttaaacccaa
ggttacctcttatctagtggCCTTCACCAGCCCCTAGTGACAAAGACTACTTCTGTCTTGgaatcttaaacccaact
ggttacctcttatctagtggTCTCTGGCCTCCAGCATAAGGTAGAACCTACATCAGGGACgaatcttaaacccaac
ggttacctcttatctagtggGGGAGGGAGGGAAGGAAAAAGAAAGGGAGGGAGGGAGGGGgaatcttaaaccca

ggttacctcttatctagtggTCCCCAGGCAAGTGAGGTGGGAAGAGCATTCCAGGCAGGCgaatcttaaacccaa
ggttacctcttatctagtggGGTCAACAGGACTGACTACATAATTTGTAGGGCCCAGTGCgaatcttaaacccaac
ggttacctcttatctagtggGAGAGCGGGGGCCCTTTGTGATTGCACAGGTTACCTGCCCgaatcttaaacccaa
ggttacctcttatctagtggGCAGTCGGGTAGAGCTTGCCTCTCTCAATCCACACCTTGGgaatcttaaacccaac
ggttacctcttatctagtggCCTAACAGCTGAAGCCAGTTTCCCAGAAGAGAACAGAGCCgaatcttaaacccaa
ggttacctcttatctagtggCAGCAAGCCCCAAACAGAGCTGCAGTGTGTGGAGTTCTGGgaatcttaaacccaa
ggttacctcttatctagtggGCTGAAAGTAGAAAGTACTGAGTTTCTACTAATGAGGCCCgaatcttaaacccaact
ggttacctcttatctagtggCTGTGCCTGGCGAACACGGCAATTTGTTATCAACCCACAGgaatcttaaacccaac
ggttacctcttatctagtggGCTCTCTGATCCTATCTTTGGTAAAACCTGTAATTTACAGgaatcttaaacccaacttg
ggttacctcttatctagtggGGGCACCTGTACTTGGGTCTAGTTTTGACCTTGTACTTGGgaatcttaaacccaactt
ggttacctcttatctagtggTGTGAGTGTGTGTTCATACATACACACATACCCTTTTGTGgaatcttaaacccaacttg
ggttacctcttatctagtggTGCTTTGTTTCTGTTAAGCCTAGAGAAAGCTGGTAATCCCgaatcttaaacccaactt
ggttacctcttatctagtggTCCATATAGAGCCAGTAATCTCATTATTAGGTTTATACCCgaatcttaaacccaacttg
ggttacctcttatctagtggTCTACTAAAGACACATGCACGTATATGCTTATTACAGCACgaatcttaaacccaactt
ggttacctcttatctagtggCAGGGGGAGGGAAAAATCACATACTGGGGCCTGTCGGGGGgaatcttaaacccaa
ggttacctcttatctagtggAATGGTCTCCTTCAGGAATGGAAAAGAAAAGTAAAGAAAGgaatcttaaacccaact
ggttacctcttatctagtggCGTGATCTCATGAATTCTTCACAAGAACCTGAGAGATGGCgaatcttaaacccaact
ggttacctcttatctagtggGTGTCAACTTGACAGGGTCAGGGGATCTCCAGAGAGCTGGgaatcttaaacccaa
ggttacctcttatctagtggTTCTGGGTGTGTGGGTGAAAGTGTTCCTGGAGGTGATTTGgaatcttaaacccaactt
ggttacctcttatctagtggTCACCCTGTCCAGTGCAGGTGAGCATCACTTAGCCTACTGgaatcttaaacccaac
ggttacctcttatctagtggCCTGGCTGCTGGAGCTCAATTGAACTTCCCCTGCCCTCAGgaatcttaaacccaac
ggttacctcttatctagtggCCAAGACTCTCAGACTTTCCAGCGACACCACCAGCTTTCCgaatcttaaacccaac
ggttacctcttatctagtggGCTTGCAGACAGCAGATCGTGGGACTTTCCAGCTGCCATAgaatcttaaacccaac
ggttacctcttatctagtggACTCTAAAACCCATACAGGCCGGGCTTGATGGCTCACGCCgaatcttaaacccaa
ggttacctcttatctagtggGGGAGGCTAAGGTAGGTAGATCGCTTGAGCCTAGGAGTTCgaatcttaaacccaac
ggttacctcttatctagtggGAAATTAATGGGGCTTGGTGACACATGCTTATAGTCCCAGgaatcttaaacccaact
ggttacctcttatctagtggCTGTACTTACAGTCTGGGAGATAGAGAGAGACCCTGTCTCgaatcttaaacccaac
ggttacctcttatctagtggCCCATATGATTTTCATGAAACCACATTTCTTCTTGAAGAGgaatcttaaacccaacttg
ggttacctcttatctagtggACTATCCCATGGCTTAGGAATTAGCCTAGACTCAACTGCCgaatcttaaacccaact
ggttacctcttatctagtggCTTGTTCAACCTCAGAGAATCAATCAACAATCAGGCAGCCgaatcttaaacccaac
ggttacctcttatctagtggCAAAAGAATAATAGTTTGGACTGTGTATGGTTTCAGAAGGgaatcttaaacccaactt
ggttacctcttatctagtggTTCCCAGCACCTCCAGCAAACTATTGACTCCACACTAGCCgaatcttaaacccaac
ggttacctcttatctagtggAGGTCACTTGGGGCTCCACTTTCTTTTTCTTTTCTTTTTTgaatcttaaacccaacttgt
ggttacctcttatctagtggTGTTGGCCAGGATTGTCTCGATTTCTTGACCTTGTGATCCgaatcttaaacccaactt
ggttacctcttatctagtggCAGCTGGGAGCTGTGCTGGGACTGTGTGCTGGGAGCCTGGgaatcttaaacccaa
ggttacctcttatctagtggAGCTTTATTGGGTTTCCTTGCGGCATGGTAGCTCGGGTTTgaatcttaaacccaactt
ggttacctcttatctagtggTCATGGGGTCTCTCTCTGTGGCCCAGGCTGCAGTACAGTGgaatcttaaacccaac
ggttacctcttatctagtggGCCCAGGCCAGGTTAGTTCAGTTCTAAGAATGAGTGGCCCgaatcttaaacccaac
ggttacctcttatctagtggGAGAGGCACTTTGTTACGCTTTATGACCCAGCCCCAGAAGgaatcttaaacccaac
ggttacctcttatctagtggGAATCTACCTGTCAGTGGAAGGAAAACCAACCCACACTGCgaatcttaaacccaac
ggttacctcttatctagtggAGGCCATTTTTGGGAAGTACAGTTTGCCACAGAGTGGTAGgaatcttaaacccaact
ggttacctcttatctagtggGTACCTGGCATGTAGCAGAGACCCAATTTGTGGTGGCTGGgaatcttaaacccaac
ggttacctcttatctagtggCAGGGGAGGCCTTCAACAGGTCAACAGGTATTCAGGGTCCgaatcttaaacccaa
ggttacctcttatctagtggGTGTGACCGTCCTGACATTTTGCTTTCTGTATTGCTCTCCgaatcttaaacccaactt
ggttacctcttatctagtggGGTCGATCATGAGAATGACCAGAGACAGGCAACACTGTAGgaatcttaaacccaac
ggttacctcttatctagtggTGGCTGGGTCTATCCAAACTGTCTTGTATAGGTTGGCTCCgaatcttaaacccaact
ggttacctcttatctagtggGGAGGTTTTGGCATGGAGCGCTTCTGAGCCACTCCGTCTCgaatcttaaacccaac
ggttacctcttatctagtggGGGCCAGAGACAGAACGGCACTTGCCCAAGGTCATAGAGGgaatcttaaacccaa
ggttacctcttatctagtggCAGAGTCATGAATTAGAGCAGGGTTTCCCAGCCTCACCACgaatcttaaacccaac
ggttacctcttatctagtggAGATGGAGTCTTACTCTGTTGCTTATGCTGGAGTGCAGTGgaatcttaaacccaactt
ggttacctcttatctagtggGCGTGAGCCACCGCTCCCAGCCACTACTGACATTTAGACCgaatcttaaacccaa
ggttacctcttatctagtggGAGGGCTGTCCTGTGCATTGCAGAATAGTTAGAAGCATCCgaatcttaaacccaac

ggttacctcttatctagtggTTGGATACTGCCAAATGTCTCCTGGGGCCAGGGTTGGGGGgaatcttaaacccaac
ggttacctcttatctagtggGAGAACCACTGGACTAAAATTCAGGTCCTCTAGTTGCTTGgaatcttaaacccaact
ggttacctcttatctagtggGACCAAATTCAGATGCTGAATTGGTAGGGCTTTACTGTAGgaatcttaaacccaactt
ggttacctcttatctagtggGAATCGTCTACGAGATAACAGGCCTGATTTCTCCAAAAAGgaatcttaaacccaact
ggttacctcttatctagtggGGCAGGAATGACTCTTGGAGATTAAAAGAGACTCATGACCgaatcttaaacccaact
ggttacctcttatctagtggGGATATCTTGTTGTGGTTGAGAGAATGTCCTTGTTTTGGGgaatcttaaacccaacttg
ggttacctcttatctagtggGCGATAATCTGTAACTTACTCTGAAATAAATATCTCAGCCgaatcttaaacccaacttg
ggttacctcttatctagtggATTGAAGTTTTCTGAAGTTCACAAAGATGTTCATTGCAGCgaatcttaaacccaacttg
ggttacctcttatctagtggGCAACCCTAACTATTTATCCCTAAGCAACTCCTAAGCACGgaatcttaaacccaact
ggttacctcttatctagtggTCTGGTATTCTAGATGAGGTAGAATACATGCAACATTTACgaatcttaaacccaacttg
ggttacctcttatctagtggGCATGCACCACCACACTTGGCTAATTGAAAGATACAAACCgaatcttaaacccaact
ggttacctcttatctagtggTCCCTCATCCTGGGACATCAGTTTTCATGAGGGCAGCCGGgaatcttaaacccaac
ggttacctcttatctagtggAGCAGCTCTGTGGATGCTGAGCCATCCAGGATGAGACCATgaatcttaaacccaac
ggttacctcttatctagtggATCTCAGCCCAGTGATGATCTATACTGCAAGAAATACCACgaatcttaaacccaact
ggttacctcttatctagtggCCTTTTTCTCAGGATCTGTCTCTCTGGAGGAGATTTTGGGgaatcttaaacccaactt
ggttacctcttatctagtggAGAGTCTTGCTTTCTCACCCAGGCTTAAGTGCAGTGGTGCgaatcttaaacccaact
ggttacctcttatctagtggTCACCTCCTGGGCTCAAATAATCCTTGTGCCTCAGCCTCCgaatcttaaacccaact
ggttacctcttatctagtggGGCTGGCCTCCAACTCCTGTGCTCAAGTGATTCACCTTCTgaatcttaaacccaact
ggttacctcttatctagtggGGCATGAGCCACCACGCATGAAAAAATGGCCAGTATGTTTgaatcttaaacccaact
ggttacctcttatctagtggGTTCATTTGAATGGGGATGGGGAAGGGACTGAGGAGAGGGgaatcttaaacccaac
ggttacctcttatctagtggGTGTACATACATATGCAAACAGATAGGCTGCCGACCACTGgaatcttaaacccaact
ggttacctcttatctagtggGCTTATAATAATATAAGAAGGCTGGGCACAGTGGCTCACAgaatcttaaacccaact
ggttacctcttatctagtggGCCCAGGAGCTCAAGACCAGGTTGGGCAACGTGGCAAGACgaatcttaaacccaa
ggttacctcttatctagtggAATTAAAGAAAAGAAATTAGCCAAGCATGGTGGTGTTCACgaatcttaaacccaactt
ggttacctcttatctagtggGCTATGCTCTTCAGCTTGAATGATGGAGTTAGGCCCTGCCgaatcttaaacccaact
ggttacctcttatctagtggAGAAAGGCACACATGGAACAAGGCTATGTATTTTCTTGAGgaatcttaaacccaactt
ggttacctcttatctagtggAATCCAAATGGACCACAATGACTGCCTTGGGGAATCAGAGgaatcttaaacccaac
ggttacctcttatctagtggGCAGTGCTGCTCGCTGCCTTCCAGTCTGAGCTGACTTCTCgaatcttaaacccaac
ggttacctcttatctagtggCCCTTCTCCAGGGGCAGCCTACCTCCAGGAGTGCCAAGGCgaatcttaaacccaa
ggttacctcttatctagtggGGCAGAGCTCCCTGTGGGGTTGGCTAAGGCCTCTGAGACCgaatcttaaacccaa
ggttacctcttatctagtggGAGACGAGTCTCCCTCTGTCACCCAGGGGCTGGAGTGCAGgaatcttaaacccaa
ggttacctcttatctagtggTCCCAAGTAGCTAGGATTATAGGAGTGCGCCACCATGCCTgaatcttaaacccaac
ggttacctcttatctagtggTGTACTGCTGACTGGGCGTGGTGGCTCTCATCTGTAATCCgaatcttaaacccaact
ggttacctcttatctagtggGCTTGAACCGGGGAGGCAGAGGTTTTGGTGAGCTGTGATCgaatcttaaacccaac
ggttacctcttatctagtggATGCAGTCTTGTGCCTCACGCAACATTGTCCTCCAGCCCCgaatcttaaacccaac
ggttacctcttatctagtggCTCCTACTGGCATCATTGCTGCCCCTCCAGCCTGCTAGGCgaatcttaaacccaac
ggttacctcttatctagtggGGGCTTTTGCACTTACTCTTCCTTCCACCTGAAAGATTCTgaatcttaaacccaactt
ggttacctcttatctagtggATTCACATGACTCACTCCCTCACTTCGCCTAGACTTTAAAgaatcttaaacccaactt
ggttacctcttatctagtggCCCAGCACTTTGGGAGGCTAAGGCTAAAGGATCTCTTGAGgaatcttaaacccaac
ggttacctcttatctagtggTGAGACCAGCCTGGGCAGCATAGCTAGATCCTGTCTCTACgaatcttaaacccaac
ggttacctcttatctagtggAGCTTCTCAGGATGCTTAGGTGGGAGGTTCCCTGGAGCCCgaatcttaaacccaac
ggttacctcttatctagtggCCCATTTCACTGATGAGAAACTGCGACCTTGAGAGGCAGGgaatcttaaacccaac
ggttacctcttatctagtggCGCAGGAGGAGCAGAAAACAGATGTCCCTTTTCTTGGCACgaatcttaaacccaac
ggttacctcttatctagtggCAGCTTCTTTGTGTTTGACCAATGACGGATGAGCACATGGgaatcttaaacccaact
ggttacctcttatctagtggGACATCAAAGACTTCCTGCTGGATGGGCCAGTCCAACCGGgaatcttaaacccaac
ggttacctcttatctagtggAGAAACTGCTCCTGCTTGGGAGAACCTGTGCTTCTGGTTCgaatcttaaacccaact
ggttacctcttatctagtggGCTCAACCTGGACCTCATTAATAGGGCAAGAGCTCAGAAGgaatcttaaacccaac
ggttacctcttatctagtggTGGCTTGGATTACCTTCAAGGATGGAGAAAATGGAGCATTgaatcttaaacccaact
ggttacctcttatctagtggCTGAAAACACCCAGTATAATGCATGGTAACTAGTAAGCTCgaatcttaaacccaact
ggttacctcttatctagtggCTAAAATGTTTTGCTTAGGCTAGAGGCCGGGGCTCATGCCgaatcttaaacccaact
ggttacctcttatctagtggCAAAAGTCAGCTGTTCATGGTGGTTCACACCTGTAGTCCCgaatcttaaacccaact
ggttacctcttatctagtggCGCACCACTGCACTCCTGCCTGGGCTATGGAGCGAGACTCgaatcttaaacccaa

ggttacctcttatctagtggGAACTCTGGAATAAGCTGAGTGATCTTAAATAAACCACTGgaatcttaaacccaactt
ggttacctcttatctagtggGTGGTAACCCACAAAAAGCACTTAGACCAATGTCTGGCACgaatcttaaacccaac
ggttacctcttatctagtggACCCTACTTTGGGTGCTAAGCTCTGTGCTGGCTCTCAGGAgaatcttaaacccaact
ggttacctcttatctagtggGACATGGTTTCTGTGTTCCTGTAGCCCACAGCCATGGCAGgaatcttaaacccaact
ggttacctcttatctagtggCTGGGAATCAACAGACCATTACATTTGTGAGCACATGTTGgaatcttaaacccaactt
ggttacctcttatctagtggCTGTCTTCATTATTTGCTCAAAGGGATCCATGACCACTCCgaatcttaaacccaactt
ggttacctcttatctagtggTTGTTATAAATAAAGCAGAGGCCGGGCACGGTGGCTCACGgaatcttaaacccaac
ggttacctcttatctagtggAGCAGACACCTGTGATCCCGGCTACTAGGGAGGCTGAGGCgaatcttaaacccaa
ggttacctcttatctagtggGCAGTGAGCCAGGATCATGCCACTGGTCTCCAACCTGGAGgaatcttaaacccaa
ggttacctcttatctagtggGTGTGTCAGAGACTACCAGCTGTGCACCCATCTCTTTCCCgaatcttaaacccaact
ggttacctcttatctagtggCTTCCAGGTCTGGCCTATAAACCCCCACCCCACCCCATGCgaatcttaaacccaa
ggttacctcttatctagtggAGGCTGTTTGGAATGGTGACCCCAGGGGAACTTGGAAGCCgaatcttaaacccaa
ggttacctcttatctagtggCCTAGAGCCCTGAATGACTGCCTGGAAGAGCTGCCTCACCgaatcttaaacccaa
ggttacctcttatctagtggGGGAGAAGGAGGGATGGGGGATGGCAGGGCTTGAAATGAGgaatcttaaacccaa
ggttacctcttatctagtggGGCTTGGCCCATGTGACTGAGACTAGCACAGGACCCTATCgaatcttaaacccaa
ggttacctcttatctagtggCAGGGTGGGCTCAAATCGTGCTGGGGAGGCATCCTCGTGCgaatcttaaacccaa
ggttacctcttatctagtggCTCCCACCTCAACCTCCTGAGTACCTGACACCACAAGTGCgaatcttaaacccaa
ggttacctcttatctagtggGGGGTCTCCCTCTGTTTCTTGGGCTGGTCTTGAACTGCTGgaatcttaaacccaact
ggttacctcttatctagtggGGCGTGCACCACCATGCCCAGCCTGTCCAAGTCTTTTTAGgaatcttaaacccaac
ggttacctcttatctagtggGGTAAGAGATTGGTCAAGAAGAGAATCTTTGCTATCACCCgaatcttaaacccaact
ggttacctcttatctagtggTGAAGCCCAGGAAATACTAATGTAGATGGGTCCCAGCAAGgaatcttaaacccaac
ggttacctcttatctagtggCCAAACTCCCTACCACACCTACATCCACAATCTTCACACCgaatcttaaacccaac
ggttacctcttatctagtggATCTTCACACCTACATCCACAATACCACATCTACATCCACgaatcttaaacccaact
ggttacctcttatctagtggAAATTTTAAAAATCATTAGCCAGGTATGGTGGTGTGTTCCgaatcttaaacccaacttg
ggttacctcttatctagtggGCACTATGCACTCCAGTCCGGGTGACAGACTGAGACCCTGgaatcttaaacccaa
ggttacctcttatctagtggCTGGCCGACGCAGATACTGAACGTTGGTTAACACAATCCCgaatcttaaacccaac
ggttacctcttatctagtggGCAGCTAGCAGAGGCCATATGAGAGCTCTAGCCAATGAGGgaatcttaaacccaa
ggttacctcttatctagtggCCACGGGTTGGTTGGAAGGTTTTTGCTTTTTCTGGCAAACgaatcttaaacccaactt
ggttacctcttatctagtggCCATGGATTTGATGCCCAAGACCAAGAATATGCTGCTCTGgaatcttaaacccaact
ggttacctcttatctagtggCCTTTCACATGCCATTGGCCAGAATTGGATCAGCTGAGCCgaatcttaaacccaact
ggttacctcttatctagtggGGAAGTAGAATCACATGCTAGGCTTAGACTGATCACTGGGgaatcttaaacccaact
ggttacctcttatctagtggCTGGAGCATCAATCAAAGCATGCCCTATAGCAGGTAAATGgaatcttaaacccaact
ggttacctcttatctagtggCCACTCATAAACAAAAGGAAAAGGCAAATAAGACATCAGGgaatcttaaacccaact
ggttacctcttatctagtggGCAAGCAAGATGAATTCCTAGCAATAAGGTACTGGTTAGGgaatcttaaacccaact
ggttacctcttatctagtggCAATGGAATGCAACAGGACTATTGAAAAACAACGTTGCAGgaatcttaaacccaact
ggttacctcttatctagtggACCGTTATGAAAAGATGATTGTGATGTGTTATTAAGTGGCgaatcttaaacccaacttg
ggttacctcttatctagtggGGATGTGTATTCCAGAGTTAACAATGGCCATTTCTGATGGgaatcttaaacccaactt
ggttacctcttatctagtggAGGTTCAAGCGATTCTCCTGCCCTCAGTCTCTCAAGTAGCgaatcttaaacccaact
ggttacctcttatctagtggCCGCTCCCGGATATATTTGTTGATTTTTTAAACTAGTGAGgaatcttaaacccaacttg
ggttacctcttatctagtggAATGAGCATCTCAACAGCAAAGTGGTCAATGGGCCTAAGGgaatcttaaacccaac
ggttacctcttatctagtggCAATGAGCCAATGATCAGGTGTGTAAGAATTTATTCTTGGgaatcttaaacccaactt
ggttacctcttatctagtggCCCCGCATGAAAAGAATAGAGCCTGGTTGGGAGTACAGGGgaatcttaaacccaa
ggttacctcttatctagtggTCACAGGTAAAACTGTGCTGAAGAACAATTTGTCAATAGGgaatcttaaacccaactt
ggttacctcttatctagtggCACTCTGCTGCCCAGGCTAGAATGCAGGGGTGCCGTCTCGgaatcttaaacccaa
ggttacctcttatctagtggGGTGTGAGCCACTGCACCTGGCCCTGGTCTAGGCAATTATgaatcttaaacccaac
ggttacctcttatctagtggCTCCTTGGGCTCAGGTGATCCTCCTACTCAGACTCCCCAGgaatcttaaacccaac
ggttacctcttatctagtggGGTTTCAAAATCCTGGCTCAGGTGATCTTCCCGCCTCACCgaatcttaaacccaact
ggttacctcttatctagtggGCCCAGGCTAAAGTGCACTGGTGCGATATCAGCTCACTGCgaatcttaaacccaa
ggttacctcttatctagtggCCAGGCTGGTCTTACACTCCTAACCTCAGGAGATCCACCTgaatcttaaacccaac
ggttacctcttatctagtggCAATGTGGTTCCAGGAAGCACTGATTCATTAGGAGCTGGCgaatcttaaacccaact
ggttacctcttatctagtggCTAGTCTTGGCTTCCTGTCACCTCAGGGGCAGCTGTGCTGgaatcttaaacccaac
ggttacctcttatctagtggGCTTAGACAAGACTTCTTTGGGCCTCGGCTTGCCTGTGTGgaatcttaaacccaact

ggttacctcttatctagtggCCCTGCCCGGGAGCTGTTCAGAGGCTGAAGAGGAAGACCAgaatcttaaacccaa
ggttacctcttatctagtggCCTGGAGGCTGCCAGGTAGGGCTGGGGCCAGCATGACCTGgaatcttaaacccaa
ggttacctcttatctagtggCTGTGAAGTGCACATAGAAGAGAGACTGGGGCCAAGCCACgaatcttaaacccaac
ggttacctcttatctagtggAGCCAATGAGACAGGACAGCTAGATACTGTGTGGTTAGAGgaatcttaaacccaact
ggttacctcttatctagtggGGCAGAGAATTGAATTCAGAGGGGCCAGTTGTGCTGCTGCgaatcttaaacccaac
ggttacctcttatctagtggCAGAGTGAGACTCTGTCCCGAAGAGAGAGAGAGAGAGATTgaatcttaaacccaac
ggttacctcttatctagtggTCCAGAGCAATGACCGTGAGCATGAAGAAAGGTGGGTGGGgaatcttaaacccaac
ggttacctcttatctagtggTGGGAAGAGGAGAACTCAGGTTCATGGCCTCAGCAAGTGGgaatcttaaacccaac
ggttacctcttatctagtggGAACATTCGTCATGCACTCCAGCCATCCTGAATTGCTAGCgaatcttaaacccaact
ggttacctcttatctagtggCACACTGTTCCCTTTGCTAAAATGCTCCCCATTCTCTTGGgaatcttaaacccaactt
ggttacctcttatctagtggCCTATTTTTTGAGTTTCAGCTCTGGCCCCACCTCCCTGGGgaatcttaaacccaact
ggttacctcttatctagtggAGATACCTCTCTGTGCTTCCACCACACCTGGGGTTACCCCgaatcttaaacccaac
ggttacctcttatctagtggGTTCTCACAAGATCTAGGATGTCTATCCCTTCGTTCACTCgaatcttaaacccaactt
ggttacctcttatctagtggCCTGACCCAATAGAAAATGATGAGGGTGGCTGTTTGTCTGgaatcttaaacccaact
ggttacctcttatctagtggCTGATGGTTCTCTAGGCCTTTTTCTGCTGTTCTCTCTGCCgaatcttaaacccaactt
ggttacctcttatctagtggTAACACTCTATCTGGTTTTTATGGCTGGCTCCTTCTTTTTgaatcttaaacccaacttgt
ggttacctcttatctagtggGAGATCGAGCCTCTACCTGTCACCCGGTGGAGTGGAGTGGgaatcttaaacccaac
ggttacctcttatctagtggAGTAGCTGGGACTACAAGGTGCGTTGCCACCACACTCGGCgaatcttaaacccaac
ggttacctcttatctagtggAGCCACCACACCTGGACGGCTGGCTTCTTCTTGTTGGTCAgaatcttaaacccaac
ggttacctcttatctagtggGCCCCATTTTCTCTTTAGCTCTCTGCCGAGATTTACTGTCgaatcttaaacccaactt
ggttacctcttatctagtggTTGCACTCATAGTCCTGAGGACTAGCACAGTGCCTGGCACgaatcttaaacccaac
ggttacctcttatctagtggGTAGATAAGGATGCAGATATAGGGCCAGGCATGGTGGTTCgaatcttaaacccaact
ggttacctcttatctagtggCTGGTCAACATAGTGAAACCCCTGTCTACTGAAAATACAAgaatcttaaacccaact
ggttacctcttatctagtggGAGATTGCAGTGAGCCCAGATCGTGCCTGGGTGACAGAGCgaatcttaaacccaac
ggttacctcttatctagtggGAGGATGCAGATATAGAGAAGGTAAAGACTGGTACTGAGCgaatcttaaacccaact
ggttacctcttatctagtggGACAATAGCACAACTAGGCTAGCGCAGTGGCTCATGCCTGgaatcttaaacccaac
ggttacctcttatctagtggGCCGAGGCGGGTGAATCACTTGAGCCCGGGAGTAAGCCAGgaatcttaaacccaa
ggttacctcttatctagtggGGTGGAGGCTGCAGTGAGTCGAGTTTGCACCACTGCTCTTgaatcttaaacccaac
ggttacctcttatctagtggAACAACGAACAAATACAGGTGAGGTAAAGGGAGATGTGAGgaatcttaaacccaac
ggttacctcttatctagtggGGCCTACAAGTAAAACATCATCATGCCACTCTTTCCTTGCgaatcttaaacccaact
ggttacctcttatctagtggCCTTTGTAGCCTTATTTAGAAAGCAATGCTTCTCATGTGGgaatcttaaacccaactt
ggttacctcttatctagtggGCCTATATGTATCCATCTGTGCCCAAAAAGGCCTCATCTGgaatcttaaacccaact
ggttacctcttatctagtggCTGCAGAGTTTTAATGTTTTACAACGTGTATGTATGTGTGgaatcttaaacccaacttgt
ggttacctcttatctagtggTTAAAAAATGAAAGCAGGCCTGGGCACGGTGGCTCACGCCgaatcttaaacccaac
ggttacctcttatctagtggGCAGTGAGCCGAGATTGAGCTGCTGCATTCAAGCCCGGTAgaatcttaaacccaac
ggttacctcttatctagtggGGGCAAACAGCATCAAGGTCAAAGTTGAGTGAGTCCTGGGgaatcttaaacccaac
ggttacctcttatctagtggCAGCCAGGGGAGATTTTCCATGCAGAAGGGACTGGGCATCgaatcttaaacccaac
ggttacctcttatctagtggCCTCTCATAGCCCACCAGCATCCCCCAGGAGGACACTGGCgaatcttaaacccaa
ggttacctcttatctagtggGGTCTCACAGCTCCTATCGCATGATGTATAGATGCAGAGGgaatcttaaacccaact
ggttacctcttatctagtggCTGCAGCTCAGGCACCTGCCCAGTCCCTCTCAGAGCCGGGgaatcttaaacccaa
ggttacctcttatctagtggCACCTGTCCAGCCCCTCACAGAGAGCCAGGCACAAAGCGGgaatcttaaacccaa
ggttacctcttatctagtggGCAAGGAGCCAGGCACAAAAGAGATGCCCATGAGTGTAAAgaatcttaaacccaac
ggttacctcttatctagtggGTGAGTAGGATGTAGGGTAGCCGGCAGGGGAGAAGAGGGGgaatcttaaacccaa
ggttacctcttatctagtggGTAACCCCCACCCCATTGCCTAGCTGGGTGCCTGACTCCCgaatcttaaacccaa
ggttacctcttatctagtggCAGTAGACGTTGAACTGGACGGGCGCTGGCGTGGTGGCTCgaatcttaaacccaa
ggttacctcttatctagtggGGGAGGCCGAAGCGAGTGGATCATCTCAGGTTCGGAGTTCgaatcttaaacccaac
ggttacctcttatctagtggGAACTGAACAGGGGCAGCTCTTACATCTGTCCAGTCTCGGgaatcttaaacccaac
ggttacctcttatctagtggCCCACATGGGAGGCCAGTGTCCACTTGCTCCTTTGGAGGAgaatcttaaacccaac
ggttacctcttatctagtggGGTCAACTCTGCCCGGAGTGGAGGGAGCTGCGCAGACCGCgaatcttaaacccaa
ggttacctcttatctagtggACAGCGGGAGACGTCGAAGGAGGCGAGAAGGTGCGCGCGCgaatcttaaaccca
ggttacctcttatctagtggAGAGGCGCTCAGATCACTGGAGCTGGAGCTCAGTCGGCGGgaatcttaaacccaa
ggttacctcttatctagtggCCCGGAGCATCTCTGGCCCTCCTATTTCACTGAACTCGCCgaatcttaaacccaac

ggttacctcttatctagtggCCGCTTGCTTCCAGGACGGTGTGGGGAAGGGTAAGAGGGCgaatcttaaacccaa
ggttacctcttatctagtggGCTTGTAGCGAGGACCCCCAGCCCAGGGCAGAGCCTACCCgaatcttaaaccca
ggttacctcttatctagtggGGAGGGGCGACGACGAGCTGCGAGCAAAGATGTGCCCCGGgaatcttaaaccca
ggttacctcttatctagtggGGCGGTCCCTGTGACCTGTGGAGACACGGCCAGATCTGCCgaatcttaaacccaa
ggttacctcttatctagtggGTAGGGATGCGCTGTGAGTAGTCGGACGGAGGAAAGGGGCgaatcttaaacccaa
ggttacctcttatctagtggGGTGGCGAGTTTGGGCGTTTGGGGAGGGTGAGGGGAGGGGgaatcttaaacccaa
ggttacctcttatctagtggATGGCCGGTGCATAGAGGCAATTCTGGATTTATGAACTTGgaatcttaaacccaactt
ggttacctcttatctagtggCAGATTTGGAGCCATGATGTGGTTTGGAGGCCCCTCTGGCgaatcttaaacccaac
ggttacctcttatctagtggTTGACCATCACTCCCCGCTTCCTCTCACACCCCCTCAGGGgaatcttaaacccaa
ggttacctcttatctagtggTTGCATCGCATGTGAGGGTTTGTTGGGTGCTCCTAAACCCgaatcttaaacccaac
ggttacctcttatctagtggTGGCTGGGCTGTTGATTTAATTAGGTTTGCCTTCTCTTGCgaatcttaaacccaacttg
ggttacctcttatctagtggCAGGAAGCACAGCCATGAAGCTGCATGCGTCTGTGTGTGGgaatcttaaacccaa
ggttacctcttatctagtggCCTTTTCCTGGTGGTTTGCAGGCATGGAAGATTCTTCTGTgaatcttaaacccaactt
ggttacctcttatctagtggGGCCCTTCTAGAACATTAGAGAGGAGAAGAAATCACGTGCgaatcttaaacccaac
ggttacctcttatctagtggCTCTCGCCCACTCCACCACTGTGAATCCTGGCTACTGGGGgaatcttaaacccaa
ggttacctcttatctagtggCTAGAGAAATGTGTTTGTGTGTGGTGGGTGGGCGGGGGCCgaatcttaaacccaac
ggttacctcttatctagtggTGAGACTTGTTCAACTTGACACAGTCCTGTGGTCATAGCGgaatcttaaacccaact
ggttacctcttatctagtggCAGTCAAGAAGGGGGTTGTATCAATCCTCAGAACCCTGAGgaatcttaaacccaac
ggttacctcttatctagtggCAGAATCCGTGTCTGGGAGAAACCTCGTGGCCTGGTCTCCgaatcttaaacccaa
ggttacctcttatctagtggGAGGACGTGCCTCGTGGTCTAAAGCTTCGGCACAAGGGCCgaatcttaaacccaa
ggttacctcttatctagtggTCCACTTACGGGTATGACTGTGGGGTATATGCTGTACCCAgaatcttaaacccaact
ggttacctcttatctagtggAGAGGGATTTGTTAAGTGGTTGTGTTGCTGTTTTGCCTGGgaatcttaaacccaacttg
ggttacctcttatctagtggGCCTCATCTATCTTGAGTGCAGAATTTTTGCATAAACGGCgaatcttaaacccaactt
ggttacctcttatctagtggTCCCCCGCCCCTCGCCACCTCCATGCTGTGTTTCTGTGGCgaatcttaaacccaa
ggttacctcttatctagtggCTGCACTGGAAAGGAGGAGTTCTCCCCTCTGTTCTGACATgaatcttaaacccaac
ggttacctcttatctagtggGATTGCTGGTCTGAAAACCTGGTGAATTCTCACTGCCCACgaatcttaaacccaact
ggttacctcttatctagtggCTGGAGCCTGTGTTTGCCATGCCTGGGCCTTCTGCCTGCCgaatcttaaacccaa
ggttacctcttatctagtggGACACCCTGTTCTTTCCTCCTGTTTGTGTGAGTGGCCCTGgaatcttaaacccaact
ggttacctcttatctagtggCTGGGGGAAGTGGAAGACTTTGGATAGTTGCCGGTCCAGGgaatcttaaacccaa
ggttacctcttatctagtggAGGCAGTGGCAGGCCTGCTGATGGCTGCCCCTTTGCTATTgaatcttaaacccaac
ggttacctcttatctagtggCAGGTGGCCCTTCTGGATGTGTGGCCTTTTTATTTTGCATgaatcttaaacccaactt
ggttacctcttatctagtggTGGCTGCAGACATAAGATTCTTCTTTTTCTCTTGTTTTGCgaatcttaaacccaacttgt
ggttacctcttatctagtggCTCCAGATTGGTCAAGCTGGTCTCGAACATCTGACCTCAGgaatcttaaacccaac
ggttacctcttatctagtggCCCAAAGTACTGGGATTACAGGTGCCACAGTACCCGGCCTgaatcttaaacccaa
ggttacctcttatctagtggTGTGATCTCTTTGAGGGATCAGGACCACACCCCCCACCCCgaatcttaaacccaa
ggttacctcttatctagtggCCCAGTGTCCATGGTGGGACCCTGACTGAGGATGGGAACAgaatcttaaacccaa
ggttacctcttatctagtggCTGGAATCATTTGGACCATTGAGTCCTGTGGACAGTCACTgaatcttaaacccaact
ggttacctcttatctagtggGGTCCTTTGGCAGGAGCCCCCTTTGTCTCTTCTGTTTTCTgaatcttaaacccaactt
ggttacctcttatctagtggGCATGTTCTTCACCTTCTACTCCCTACAACTCCATGAGGCgaatcttaaacccaact
ggttacctcttatctagtggGTCTAGCAATGGTAGACTTCTTGTTGTGGGGCAGGGTTGGgaatcttaaacccaact
ggttacctcttatctagtggGTTGTGGTTTTGCAGATGAGGAACCCACAGCCCAGAGAGGgaatcttaaacccaa
ggttacctcttatctagtggCAGCTTGGGAGAGGCAGAGCTGTAGAGATTTGGACCACAGgaatcttaaacccaa
ggttacctcttatctagtggGAGGTTAAGAACAAGCAAAGGGCCGGGCGCGGTGGCTCACgaatcttaaaccca
ggttacctcttatctagtggGCGGAGGTTGCCGGGAGCCGAGATCGCACCATTGTATTCCgaatcttaaacccaa
ggttacctcttatctagtggGCAGGAGAACACAGTGTGTCCCCCACCCTCACCGTGCACCgaatcttaaaccca
ggttacctcttatctagtggCCTCATGCTCTGCCTGCCGACCTTTTCTGCAGCCACGGAGgaatcttaaacccaa
ggttacctcttatctagtggGAGGGCTGCCATGCTGGAAAGTTTGGGCTGTGGGTGTGGGgaatcttaaacccaa
ggttacctcttatctagtggGGGTAGAGGTCAGAACCCTTAGTCTTGGCTGGGTGCAGTAgaatcttaaacccaac
ggttacctcttatctagtggTGCAGTGAGCTGAGATTGCCAGTGCACTCCTGCCTGGGAGgaatcttaaacccaa
ggttacctcttatctagtggGCAAGAGGCTGCAATCAGGTATAGCACTGGATTTTGTGACgaatcttaaacccaact
ggttacctcttatctagtggCAATGTCAGGTCATAAAACACAAGACATATACATGATGTGgaatcttaaacccaactt
ggttacctcttatctagtggATCTTTTCCTTTTTGTGTGTGTGTGGAGTTAGGTGGGGGGgaatcttaaacccaacttg

ggttacctcttatctagtggGTGCCCAACACAGCCCTGGTACTTTGTTGAATCAGGTGGCgaatcttaaacccaac
ggttacctcttatctagtggGCCCGGCATGGTGGCTTATGCCTATAATTTTTGCACTTTGgaatcttaaacccaacttg
ggttacctcttatctagtggGGGCAGCAGAGTGAGACTCTGTCTCTGTAAAAAATAATAAgaatcttaaacccaactt
ggttacctcttatctagtggGAGACAGCTTGACATGGTGTGGAAGTGGGGGGCTTGCCTGgaatcttaaacccaac
ggttacctcttatctagtggCAGTAGCAGGAGGCAGTGGGGATGTGTCAGGGAGCTGAGGgaatcttaaacccaa
ggttacctcttatctagtggCAGAATGTGTCTGGCAGAGGACCTGGGCTCATAAGTGAGCgaatcttaaacccaac
ggttacctcttatctagtggCCTGTGGCTGCATGAAGTAGTGGTTTCTGAATAGAAAGGCgaatcttaaacccaact
ggttacctcttatctagtggTCACTGTAGTTCTCTGCAAGCATCAGAAACAAGCCCTGGCgaatcttaaacccaact
ggttacctcttatctagtggGCTGCAGAAGTGGGTCTCAGGAAGGACAGGAATCAAAGGGgaatcttaaacccaac
ggttacctcttatctagtggCAGGTCACTGTGCCTCCATTTCCTCAGCTGCTGAGCAGGGgaatcttaaacccaac
ggttacctcttatctagtggCCTCCCTGGGATGGTGCAGAGCACTTAGCAGAGCACTTAGgaatcttaaacccaac
ggttacctcttatctagtggGAGCAGTTGCTGTATGCCTGGCACCGTTTTAGCTGCTGGCgaatcttaaacccaact
ggttacctcttatctagtggCCCTGCCCTCCTGAAGCTTCTGTCCTAGTGGATGTCACACgaatcttaaacccaac
ggttacctcttatctagtggCCCAGAGACAGCTGGGGGACAGATGGAGGCAGTGCAGGCAgaatcttaaacccaa
ggttacctcttatctagtggACTGTGTTAACAGTAAAGTGCTTAGCAGAGGCTCCCACACgaatcttaaacccaact
ggttacctcttatctagtggCCTCCCTCATAAAATGGCAGCCTCCCTTTCTTCCATTCCCgaatcttaaacccaact
ggttacctcttatctagtggGGGCTTATCTTCCTAGTATACCATATGTCTCTCTTCTCCCgaatcttaaacccaacttg
ggttacctcttatctagtggGGGTGTCTACTCTGTGCCTGGCATTGCTGTAGATGTTGGGgaatcttaaacccaactt
ggttacctcttatctagtggAAGGACAGATTTCCATGCTAGAGCTTAGGTTACAGCTGGGgaatcttaaacccaact
ggttacctcttatctagtggCAGGCCTAGCAGGGTCAGTGTCTGGAGGGCTGGGGCGTGGgaatcttaaacccaa
ggttacctcttatctagtggAAGTTGTCCGGTCAGAGTCAGGGCAGGCTCCATGGAGGAGgaatcttaaacccaac
ggttacctcttatctagtggCCTCCACCTGAAGTAGGGGAAGGACAGGCGTTTCAGGCTCgaatcttaaacccaac
ggttacctcttatctagtggAGGCAGAAGTGAGCCTGTGGGGGTGAGGAGCAGCCAGGAGgaatcttaaacccaa
ggttacctcttatctagtggTGGCGTGAGCAAGGGAGGTAGGAGAACCCGAGCAGCATGCgaatcttaaacccaa
ggttacctcttatctagtggGATTCTGTCTTTTACTCTGTAAAATGGGGAGCCCTTGCGGgaatcttaaacccaacttg
ggttacctcttatctagtggGTTTTTTGTTTGTTATGAGGCAGAGTCTTGCTCTGTTGCCgaatcttaaacccaacttg
ggttacctcttatctagtggCTGCCTCCTGGGTTTAAGCAATCCTTACACCTCAGCCTCCgaatcttaaacccaact
ggttacctcttatctagtggGGTGTGAACCACTGTGACTGGCTTCCTTGCAGTGTTCTAAgaatcttaaacccaactt
ggttacctcttatctagtggTGATCTGACTTGGCTTTTGAAACGGTCAGTCTGGCTGCTGgaatcttaaacccaactt
ggttacctcttatctagtggCCAGGTCCAGGCTGGCAGTGGTGGATGGGGCCACAGGAATgaatcttaaacccaa
ggttacctcttatctagtggAACTGTGCTCTCCATATGGTAGCTGTGAGCCCTATGTGGGgaatcttaaacccaact
ggttacctcttatctagtggGTGTTAAACACACATTGGATTTCAGAGACTTAGTGAGAAGgaatcttaaacccaacttg
ggttacctcttatctagtggCCTCAGGTGATCCATCCCACTCAGCTTTCCAAAGTGCTGGgaatcttaaacccaac
ggttacctcttatctagtggGTGAGCCACTGCACCAGGCCTATTTGTTTATTTATATGAGgaatcttaaacccaacttg
ggttacctcttatctagtggGCAGTGATGTGTTCATAGATCATTATAGCCTCGACCTCCCgaatcttaaacccaactt
ggttacctcttatctagtggGCCTCCTGGGCAGCTGGGACCACAGGTCTGTGTCACCACGgaatcttaaacccaa
ggttacctcttatctagtggTGAGCTCCTGGGCTCCAGCAATCTGCCTTCCTCAGCCTCCgaatcttaaacccaac
ggttacctcttatctagtggGTGAGCCACTGCACCCTGCCAAGTTATCTCATGAACCAATgaatcttaaacccaact
ggttacctcttatctagtggCGCTGGATGGAACAATGCCTTGGACGAGGTCAGCATTTAGgaatcttaaacccaac
ggttacctcttatctagtggCCGTTGCCATCTTCGGGAAGTTTCCAGTGTGGGAGGGAGGgaatcttaaacccaac
ggttacctcttatctagtggCCAGGCCTGCTGTAGCCCAGAAGCTGGAAGAGACTCGCCCgaatcttaaacccaa
ggttacctcttatctagtggGCAGTGGGCTGGCAGTGCAGAGGTTCAAGGCCTTGGACTGgaatcttaaacccaac
ggttacctcttatctagtggGGGTGGAGGTGGAAAAGGCAGGTGGGGCGAGGGTGAGGGGgaatcttaaacccaa
ggttacctcttatctagtggTGGAGAGCCTGAATAACCAGGTATAAGTGATTTCTTTTTCgaatcttaaacccaacttg
ggttacctcttatctagtggCTGGGACTACAGGCACCCGCCCACCACGCCCGGCTAAGTTgaatcttaaacccaa
ggttacctcttatctagtggCGCCCAGCCCCAGGCGGAAATTATTTCAACTGGGTCCTGAgaatcttaaacccaac
ggttacctcttatctagtggCAAGGATTGTCAGCAAGGAGCAACTTCCCAAATATACATCgaatcttaaacccaact
ggttacctcttatctagtggGGTGGAAGAAGGAAATATGGGCCAGGCACTGTGGCTTATGgaatcttaaacccaac
ggttacctcttatctagtggGGTGGGAGGATTAGTTGAGCCCATCTGAGACCAGCCTGGGgaatcttaaacccaac
ggttacctcttatctagtggCAGCCTTCCGAGTAGCTGGAGTTACAGCCTTGTGCCACCAgaatcttaaacccaac
ggttacctcttatctagtggGGGCTTGGTGGCATGTACCTGTAGTTCCATCTTTGGTGGCgaatcttaaacccaactt
ggttacctcttatctagtggCTTGGGAGGCTGAGGCAGGAAGCTTGCTCGAACCCAGAAGgaatcttaaacccaa

ggttacctcttatctagtggAATAAATTTTTAAAAGCTGGGAGGTGGGGTTGGTAAATTCgaatcttaaacccaacttg
ggttacctcttatctagtggGACATTGGGCCTCACTCCTAGGCTGGTCTTTAACTCCCGGgaatcttaaacccaac
ggttacctcttatctagtggAGCCACTGTGCCCAGGCTGTGGTTTTTTATTTTTTAATTTgaatcttaaacccaacttgt
ggttacctcttatctagtggTTTTGAGACAGCATCTCATTCTGTTATCCAGGCTGGAGTGgaatcttaaacccaacttg
ggttacctcttatctagtggCTGCCTTGGCCTCCGAGTAGCTGGGATGCCACCGTGCCCGgaatcttaaacccaa
ggttacctcttatctagtggCCTGACCTCAAGTGATCCCACCTCGGCTTCCCAAAGTGCTgaatcttaaacccaac
ggttacctcttatctagtggTAGGACACAGGCATTTTTGAATATTATATGCACTTTGTCCgaatcttaaacccaacttg
ggttacctcttatctagtggCAGGAGTGAGCCACTGTGCCCAGCCGTCCATGTCATTTTTgaatcttaaacccaact
ggttacctcttatctagtggTCATTCCTTTGATTTGGATGCTTAGCTTCATAACAGCCTGgaatcttaaacccaacttg
ggttacctcttatctagtggAACAAATTCTCAGTCCTAGGATTATTGTGCATAATGTCTGgaatcttaaacccaacttg
ggttacctcttatctagtggGCACAGAAGCTGAATTTGCTGCTTTCTGGTGAGGCACGTGgaatcttaaacccaact
ggttacctcttatctagtggGGAATGCCTGTGGTGTGCCAAGTTCTGCTTGAGGCCTTGGgaatcttaaacccaact
ggttacctcttatctagtggACTAGTACCAGTTCTTAAGAGGCTTATTGTTAAGTGGGGGgaatcttaaacccaacttg
ggttacctcttatctagtggCTGTCTGAGGAATCAGGATTTGAGTGAGACACGAAGACTCgaatcttaaacccaact
ggttacctcttatctagtggCCAGAAGAACAAAGTGTTAGCAGAAAAGACAAGATTGATCgaatcttaaacccaact
ggttacctcttatctagtggGGCACATCTAAGCAACAAAAACTGCAATGGTATTAAAAGCgaatcttaaacccaact
ggttacctcttatctagtggGTGCTCATAGAAAGCAAAGTAGATGGCTGGGCACCGTGGCgaatcttaaacccaac
ggttacctcttatctagtggGCCACTGCATGCCAGCCTGGACACAGAGAACGACTCCGTCgaatcttaaacccaa
ggttacctcttatctagtggGCTAACAAAGTCAGGTTTTGGAGATCTAATTTCTAGTCAGgaatcttaaacccaacttg
ggttacctcttatctagtggCCCTCCTGTCCCTTCACAGCCAGTCTCCTTCTCCACCCCCgaatcttaaacccaac
ggttacctcttatctagtggCCCCTGAGTTGTTTGCTGACCTTACAGTGTTGCCTTTTTCgaatcttaaacccaacttg
ggttacctcttatctagtggTGCATTCGGGACCCTTCCATGTTGTTCTATGCATCAGGGGgaatcttaaacccaact
ggttacctcttatctagtggTGCTGACTGGGACTCTGTTGTATGGATATACCACTGTTTGgaatcttaaacccaacttg
ggttacctcttatctagtggGGCACAGGGTGTTCAGCAGCTCCGCTAGTGAGTGTGTGAGgaatcttaaacccaac
ggttacctcttatctagtggCCTTGCCAGCAGGTAGTGAAATTGGAAACATGTGCAGCCCgaatcttaaacccaac
ggttacctcttatctagtggAGATTGAGAATTCACAGTTCGTGCTGTATGTTACTTAAGGgaatcttaaacccaacttg
ggttacctcttatctagtggACTCACTGTCATCATCTGGGGAGAGAAGGGGAGTGGGCTGgaatcttaaacccaac
ggttacctcttatctagtggAGGTTGCTGGAGTGTGAGTGGTGTGATCGTGGCTCACTGCgaatcttaaacccaact
ggttacctcttatctagtggACTACAGGTGTGAGCCACCATGCCTGTATTTTTTGTAGAGgaatcttaaacccaacttg
ggttacctcttatctagtggGGCCTCAAGCCATCCTCCTGCCCAGTTCTCCCAAAGTGCTgaatcttaaacccaac
ggttacctcttatctagtggAGGTGGATACCTACACATATCACTTACACAGTTTGAAAATgaatcttaaacccaacttg
ggttacctcttatctagtggTACATATGACAAGATGTTCATGGTTGTCAATTCTAGGTTCgaatcttaaacccaacttg
ggttacctcttatctagtggGGACTACAGATGCCTGCTACCACGCCCGGTTAATTGTTTTgaatcttaaacccaact
ggttacctcttatctagtggAGCCACCGGGCCCGGCTGAGACAGTCTTGCTCTGTCACCCgaatcttaaacccaa
ggttacctcttatctagtggCCTCAGCCTCCCGAGTATTTGGGACTTCAGGCACAACCCCgaatcttaaacccaac
ggttacctcttatctagtggGCGTGAGCCACGGCGTCCAGCCTGTTCTTTAAATTTCTTTgaatcttaaacccaactt
ggttacctcttatctagtggGCATGTGTGAAGGCCCTGAGGCTGCAAAACTTTTGGTGTGgaatcttaaacccaact
ggttacctcttatctagtggGATTCGTGTGGCTGGAAAGGAGTGAACGTCCAGGGTCCAGgaatcttaaacccaac
ggttacctcttatctagtggGGGTAGATGGTGGATCAACGGAAGCAGGTGGCCCATTTGGgaatcttaaacccaac
ggttacctcttatctagtggGAGCTGGACTCGGCTCTAGCAGTGGAGAGGAAGAGAAGAGgaatcttaaacccaa
ggttacctcttatctagtggTATTTTGGAGACAGAAAGGACTTCTTGGGACCAGGCGTGGgaatcttaaacccaact
ggttacctcttatctagtggGTGAGCTGAGATTGAGCCATTGTATTCCAGCCTGGGCAACgaatcttaaacccaact
ggttacctcttatctagtggGGAGGGCAAAGTTTGAGGGAAGAGTTAGCTAGCTAGAGACgaatcttaaacccaac
ggttacctcttatctagtggGTGCTCAGGCATTTGGATTTTACCCTGTGGGTGATAGGGAgaatcttaaacccaactt
ggttacctcttatctagtggGCAGCCTGGACCCCCTGTGCTCAAGCAATCCTCCCACCTCgaatcttaaacccaa
ggttacctcttatctagtggGTGCATGGCACCACCCCCAGCTAATGTTTTGTTTTTAATTgaatcttaaacccaacttg
ggttacctcttatctagtggAACTCCTGGCTTTAAGCGATCCTCCCATCTTGACCTCCCAgaatcttaaacccaact
ggttacctcttatctagtggGCCATCATGCTCGGCTCTATGGAAGGTTTTGAGGTGGGGCgaatcttaaacccaac
ggttacctcttatctagtggTGCCAGAGAGATTTGTCCAGAGGAAGGGTGACCAGCCTGGgaatcttaaacccaac
ggttacctcttatctagtggGCAGAGATTGTGTCAGTTACCCTCTGGTGGGATGCTTGGCgaatcttaaacccaact
ggttacctcttatctagtggTATAGGAGAGTGGCTAACGAAGACAGAGTTCCCTTCCGGCgaatcttaaacccaac
ggttacctcttatctagtggATCCCACTGGGCCAGCAGCAGCATTGGGATCACGTCAGCCgaatcttaaacccaa

ggttacctcttatctagtggGTCGGTGAAAAGCCCTGGCCAGACCCAGCCGGTGGAGACCgaatcttaaacccaa
ggttacctcttatctagtggCTGGCAGGCTGGGCTTCCTATGGTCATTGCAGTCCCTCCCgaatcttaaacccaac
ggttacctcttatctagtggGCCCTCTGGAAAGGGAGAGGAGCAGGAGGGGGCTTCAGGCgaatcttaaacccaa
ggttacctcttatctagtggCCTGGCTGGTCTGAGCCCTCTTCTTTCCTCTCCCCAGGCCgaatcttaaacccaac
ggttacctcttatctagtggCTTTCTGCATTAGGAAGACATTCTCTGCATGCTGATCTCGgaatcttaaacccaactt
ggttacctcttatctagtggGGAGCATCCTGCACCTCCCTCCTGTAAGCGAGCTTTGCAAgaatcttaaacccaac
ggttacctcttatctagtggGCTTGAGTGCCTGGTGCGTCAGAGGCTCTCAGTGAGCGTGgaatcttaaacccaac
ggttacctcttatctagtggGGCGAGGAATCTGGCCCTCACTGTCCCAAGGCTGAGTGCCgaatcttaaacccaa
ggttacctcttatctagtggGGAGGAAAGCACTGCCTTGGGGTCAGAAATAAGGATTGGCgaatcttaaacccaac
ggttacctcttatctagtggAGTGATCATGGGATCAGAGGCAGCAGGTGAATGTGGGGTGgaatcttaaacccaac
ggttacctcttatctagtggGCCAGCCAGGGAAATGGGACAGGATTTTGTCTCTTTCCCCgaatcttaaacccaac
ggttacctcttatctagtggCGCTGGTTCTGAGTGTGTCGTTCCTGCTTGATCCCCTCCCgaatcttaaacccaact
ggttacctcttatctagtggGGAGGATGAGTAGATTGTACAACCACACCAGGCGGGCCTGgaatcttaaacccaa
ggttacctcttatctagtggGAAGTGTCCGGATTGGCAAACCAGCGGGTCCCTCTTCTAGgaatcttaaacccaac
ggttacctcttatctagtggGAGGCTGCTGTTAGATTCCAGCTCCCTCCTCCCCTGGTCCgaatcttaaacccaac
ggttacctcttatctagtggCTCATGCCAAAAGATAGACCCAGTGTTGCCAGAGCTTCCAgaatcttaaacccaac
ggttacctcttatctagtggCCCCCGGGGCCAGCAGTTTGCAGCTCCTTGGTTATAGGTGgaatcttaaacccaac
ggttacctcttatctagtggCCTGGGCCTAGCAGGGAGCTGGCATGTCACAGTGGGTGCTgaatcttaaacccaa
ggttacctcttatctagtggCTTGAATGCTAGACTGAGCAAGCACCTCCCTGCTCACAGGgaatcttaaacccaac
ggttacctcttatctagtggCAGTTCCTTACATACTTTAGTGATAGTCTGTATGTGCCTGgaatcttaaacccaacttg
ggttacctcttatctagtggTCTTCTGTTCTTTTCCTTTTTCCTTTCTCTCTCCGTCCCCgaatcttaaacccaacttgt
ggttacctcttatctagtggTCTGGCCGGTTAACACAGGGCTTCATAACATGGGGTCGGCgaatcttaaacccaac
ggttacctcttatctagtggGTGGTTTCCGGTAGCTCACTGACTTTGCCCTCTCTGCTGGgaatcttaaacccaact
ggttacctcttatctagtggGATTTCCATCAGGGTAACAAGCATGATTTTCAGATTTACCgaatcttaaacccaactt
ggttacctcttatctagtggGCAAAATTGCTTAAAGAAATGAACCTTCAGGTGTGGCGGCgaatcttaaacccaact
ggttacctcttatctagtggGGATCACTTGAGCCCAGGAGGTCTCTAGGCTGCAGTAAGCgaatcttaaacccaac
ggttacctcttatctagtggGGAGAGGGGAGGGGAGGGGAAGAGGGAGAGAGGGAGGGAGgaatcttaaaccca
ggttacctcttatctagtggGGCTGGGCGTGGTAGCTCTTGCCTGTTGTAGTCCTGGCACgaatcttaaacccaac
ggttacctcttatctagtggGCCAGTGGCATGCACCTGTGGTCCCAGGAGGCCAAAGCTGgaatcttaaacccaa
ggttacctcttatctagtggGAGGCTGCAGTGACCTATGTTCCCACCACTGTACTCCAGCgaatcttaaacccaac
ggttacctcttatctagtggAGAAAAGAAAGAAAAGGAGAGAGGGAGGGAAGGAAGGAAGgaatcttaaacccaa
ggttacctcttatctagtggAGGGAGGGAAGGGAGGAAGGTAGGAAAATGGAAGGAAGGGgaatcttaaacccaa
ggttacctcttatctagtggCCGGTCGGCTATCTTGACGAAAAAAGTCTCAGTCCACTCCgaatcttaaacccaac
ggttacctcttatctagtggGCTTAATATGCACTTGTAACTGTGTCAGGCTCTCACAGGGgaatcttaaacccaactt
ggttacctcttatctagtggGTCTGCAAGAGCCCAGTCCATGCATTTAGGAGGCACAGAGgaatcttaaacccaac
ggttacctcttatctagtggAGATATTAAGCCCTGACTTCGTGCCGGTTTGAACTTTTCCgaatcttaaacccaactt
ggttacctcttatctagtggAGATGTCATGGTGATGGGAAGACTGGGGGAGGAGAGGAGCgaatcttaaacccaa
ggttacctcttatctagtggGTAGGTCTCTTACCCAGTGTCTTCTCCTGCTCTCTGGCTCgaatcttaaacccaactt
ggttacctcttatctagtggGCCTCCCCTAGACCCAGCTCTACTTTAGGCCACGTGAGTGgaatcttaaacccaac
ggttacctcttatctagtggCTCCTACTGTCCCTGACCCATGTCCGCTTTTGTCTCCTCCgaatcttaaacccaact
ggttacctcttatctagtggCAGACCGTGTGGCTTGAACCTCAGCCCTGCTGCCAAGTAGgaatcttaaacccaa
ggttacctcttatctagtggCCCTCATCTGTAAAATGGGAATACAAATGGTCCCTGTGTGgaatcttaaacccaactt
ggttacctcttatctagtggTAAGTGAGTTGCTGCACACCTCACCTCTGGCTCCTTCAACgaatcttaaacccaact
ggttacctcttatctagtggGGGTGGTGACGCAAACCACATGGAAACGAGAGGCAGACCCgaatcttaaacccaa
ggttacctcttatctagtggTACAGGCATGAACCACCGCCCCCGGCCGCAGATGGGAAATgaatcttaaacccaa
ggttacctcttatctagtggTTCCTGGTGGGGAAGACCTCCATCCTCCTTCTCCAGCCCCgaatcttaaacccaac
ggttacctcttatctagtggGAGCTTCTGGGTTTCTGGGAGAGGCTGAGCAGGTGGAGACgaatcttaaacccaac
ggttacctcttatctagtggCCAGAATCCAGGAGCCTGCCTGACGCGTCAGCCGGGGAGGgaatcttaaacccaa
ggttacctcttatctagtggGAAGTCGGCTTGAGTGAGCCAGCCTTTCATGCATCCTTGTgaatcttaaacccaact
ggttacctcttatctagtggGGAGGGGCTGAGTTGTGTCTTAATGAGCAGTTCCAGCTGCgaatcttaaacccaact
ggttacctcttatctagtggAGCTCCTGCTGCATGCTGGGCAATGCTCTAAGTGCTCTGCgaatcttaaacccaac
ggttacctcttatctagtggTCTCCCTAGCAAACCCTTTGAGGGGACTGACAGGATGACCgaatcttaaacccaac

ggttacctcttatctagtggTACAGATGTTGGGATTGAGGCACAGACAGTTTATTTTTAAgaatcttaaacccaacttg
ggttacctcttatctagtggGGCATTAAGGCCATTGGTATTGTTGTGTAACCATCACCGCgaatcttaaacccaactt
ggttacctcttatctagtggACTCTCCTCTCCCCCTCCCCTCATCCCCTGGTATCCACAGgaatcttaaacccaac
ggttacctcttatctagtggACCCTAAGCATCCGTATAGGTGGAATCATAGTATTCATCCgaatcttaaacccaactt
ggttacctcttatctagtggGCTCACCACAACCTCTGCCTCCCAATTCAAGAGATTCTCCgaatcttaaacccaac
ggttacctcttatctagtggGGCATGAGCCACCGTGCCCCACCTTAGTGTAATCTTTCAAgaatcttaaacccaact
ggttacctcttatctagtggCCGTTGTAGCCTGTGTCACAATTTTATGCCCTTTTAAGACgaatcttaaacccaacttg
ggttacctcttatctagtggTGGCTTTGCTGCTCCCAGAGTCTTGACCCTGTGCTGGGGCgaatcttaaacccaac
ggttacctcttatctagtggCTTGTCAGTGCTTTCGGCAGCTTGATCAGGTAGATGCTGCgaatcttaaacccaact
ggttacctcttatctagtggGGGCATGGCAGAAATTCATCAGGGGCGGAGGAGGCACAGGgaatcttaaacccaa
ggttacctcttatctagtggGCTGCCCACAACATAAATGGCTAGCAAATAATTTGGATGCgaatcttaaacccaact
ggttacctcttatctagtggACCTACAAAATTATAGATGTAAGGCTGGGTGTGGTGGCTCgaatcttaaacccaactt
ggttacctcttatctagtggGGGAGGCCAAGGTGGGCAGATAATGAGTTCAGGAGTTCGAgaatcttaaacccaac
ggttacctcttatctagtggTACCCATGACCTCTCCACCCTGCCTAGGATCCAAACTATGgaatcttaaacccaac
ggttacctcttatctagtggGCCTACCCCCTGCCCCCGAGCATTCTCCCTCATTGGTGGCgaatcttaaacccaa
ggttacctcttatctagtggCCAGGCTGGTCCCAAACTCCTGGGCTCGAGAGATCCTCTTgaatcttaaacccaac
ggttacctcttatctagtggGCGTGAGCCGCTATGCCAGGCCAACTCTTATCTTGAATGTgaatcttaaacccaact
ggttacctcttatctagtggGGAGTGCAATAGCATGATCTCGGCTAACTGTAACCTCCACgaatcttaaacccaact
ggttacctcttatctagtggAGGTGATCCACCCGCCTCGGCCTCCCGTTTGTATTCTTAAgaatcttaaacccaact
ggttacctcttatctagtggTCCATGCTGCCTTCAGCTTCCTTTGCCGCTCTGCATCATGgaatcttaaacccaact
ggttacctcttatctagtggTCTGCTGATGCCTGCTGCTGTAGCTTATTTTTCACTGTGAgaatcttaaacccaacttg
ggttacctcttatctagtggGGACATGTGAGTTGTTCGCAGGTGTCCCCCCACCCCCTCCgaatcttaaacccaa
ggttacctcttatctagtggGCCTATGTCAGAATTTCCAGGGTATACAGCTAGGACGCAGgaatcttaaacccaact
ggttacctcttatctagtggGCGCTGTTCTGGAATAGGAGAGACACAGACCTGCTGGGCCgaatcttaaacccaa
ggttacctcttatctagtggCATCTGTCTACCTCAGTAGTTTCCCTGTCTGTGAATTGAGgaatcttaaacccaacttg
ggttacctcttatctagtggGGGTTAAATAAGATAGAGACCTGGTATGGTGGCTCATGCCgaatcttaaacccaact
ggttacctcttatctagtggGAGAGAAAGATCCGGCATGTGTAGAGATTAACTCGGAGCCgaatcttaaacccaac
ggttacctcttatctagtggCAGAGTAAGTGCTCCCTGCTGTTCGCCTTGTTCCTGTTGGgaatcttaaacccaact
ggttacctcttatctagtggCGTGGGCCAAGGACGCCAGGCAGGATCATAGTCCGGCTGGgaatcttaaacccaa
ggttacctcttatctagtggCCCAGGTGGGATTTCACCCCCACTTCCTTCCAGGCTTCCCgaatcttaaacccaac
ggttacctcttatctagtggGCCTGTTGGGGCCTCCTCCAGTAGATGTGTGCCCTTGAGCgaatcttaaacccaac
ggttacctcttatctagtggAGGGGTCCCACCCTGCTCCGGAAGTCTCAGTGAGGAATCAgaatcttaaacccaa
ggttacctcttatctagtggGTGCTGCCCCCTGCCTGACATTCCATCCACACTTGAGGCCgaatcttaaacccaac
ggttacctcttatctagtggGTCTCTCTTCCAGGAGAGGGACCCGGTTCATCAGCAGGATgaatcttaaacccaac
ggttacctcttatctagtggGGCTCAGCGCCTGGGCTTTTTATAAATGAGGGCTCTGCTGgaatcttaaacccaact
ggttacctcttatctagtggTTCGGTGAGAGACCACCAGAAGCCACAAAAAATTGCTGACgaatcttaaacccaac
ggttacctcttatctagtggGCAGTTCCAGCAGCATTGATTTGAATTCCCCCTCTGCCGCgaatcttaaacccaact
ggttacctcttatctagtggAGCGAGTTGCTTACCCTCTAGAGCCTCCACGTCCTCATCTgaatcttaaacccaact
ggttacctcttatctagtggGACGTGCATGTAGAGCTTTCCCAGAGCTGGCATGTGGTAGgaatcttaaacccaac
ggttacctcttatctagtggGCCTCTGTCTGCCAGGCTCAAGGGATACTCCTGCCTCAGCgaatcttaaacccaa
ggttacctcttatctagtggGCTGGGACTTGCAGGCAAGTGCTACCGCTCCTAGCTAATTgaatcttaaacccaac
ggttacctcttatctagtggACTGGCTGCGAGCCCCGGGGTAGGCACACTGGTACTGGGGgaatcttaaacccaa
ggttacctcttatctagtggATCCGTGCTCGGTAAATCATCATTGAAGGAGCAAGGGTGCgaatcttaaacccaact
ggttacctcttatctagtggCCAGCTCTGGAGTCCTGGGATCCAAGTGTGAGCCCCATGTgaatcttaaacccaac
ggttacctcttatctagtggTAGCTCTGACTGTCACCAGGCTTTGCTCTGAGCACTCTGCgaatcttaaacccaact
ggttacctcttatctagtggTCCCAAGGTTAACAGCTAACCAGTAGCAGGGCACGCTGGGgaatcttaaacccaa
ggttacctcttatctagtggGGTATGCTCTCCTGTCTAGAGGTGATCTCTTCCTCCAGGGgaatcttaaacccaact
ggttacctcttatctagtggGACACCCTCAGGCTACCCAGCCCTCGGGAGCTCACAGCAGgaatcttaaacccaa
ggttacctcttatctagtggGATGCCTTGCAAGGTTACCTGTGCCAGAGTTCCTGGGGTAgaatcttaaacccaact
ggttacctcttatctagtggAGGTAGGACCCAATACTACTTTGCATTTTCCTGGAATCCTgaatcttaaacccaacttg
ggttacctcttatctagtggCCTCTAAGAAATGCAGAGGTGGAATTAGAAGGTGCGTGTCgaatcttaaacccaact
ggttacctcttatctagtggGCACCACAGATTCTATCGAGTCTCATCTGGCAGCTTCAAGgaatcttaaacccaact

ggttacctcttatctagtggCCCGTGGGACTTGCCCTGCACCCAGTTCAGCTTGCTTGGGgaatcttaaacccaac
ggttacctcttatctagtggGTATGGGTTCAAGGCCAGCGGTCCAGCACTCCCACAGACCgaatcttaaacccaa
ggttacctcttatctagtggGGGAAGGGTTGCCTCTGTGTGGGTTGGGAGTTTAATCAGCgaatcttaaacccaact
ggttacctcttatctagtggGGTCACGGGGCCAAGGGCAAACATACACCCCTCCCAGCCCgaatcttaaaccca
ggttacctcttatctagtggAGCTGCAAAGTTGCACACCCTTCTCCCCAAATTGACAAATgaatcttaaacccaact
ggttacctcttatctagtggGTTGCCCAGCTGTTGAGTGGTTTAAACTTCCAGATGTTCCgaatcttaaacccaactt
ggttacctcttatctagtggCCCTCGAGGGAAGGAGGTACACATTTCCTAGGGTGCTTGGgaatcttaaacccaac
ggttacctcttatctagtggCCAGTTACTGGTCAGGCAGGGAAGCCCAAGTTCTGGCCCCgaatcttaaacccaa
ggttacctcttatctagtggGTGACTTTGGGCTGAGCAGTTGGCCTCTCTGAGTCTTAGGgaatcttaaacccaact
ggttacctcttatctagtggGGATTACAGACATGAGCCACTGCGCCGAGCCTGAGCCTCAgaatcttaaacccaa
ggttacctcttatctagtggCTTGCAACTGGGAGCATGTTTTGAGAGCTAGAGAAAGTATgaatcttaaacccaactt
ggttacctcttatctagtggGCCCGTCCGCATTTTTAACGTGGGGACACTGAGGCCCAGGgaatcttaaacccaa
ggttacctcttatctagtggGGCCTGTGCTGTCCCATATGGTAGCCGCTAGCCACTAGCCgaatcttaaacccaa
ggttacctcttatctagtggCTAGAAAATTAAAAATTACTGCCGGGTGCGGTGGCTCACGgaatcttaaacccaact
ggttacctcttatctagtggCCACTGGGCAGTGCTATGCTAAACTGTGACACAAACGGTGgaatcttaaacccaac
ggttacctcttatctagtggGTGGCAGGCCTACCAGGTGTGGATGAGGCTGTCTGGGTTCgaatcttaaacccaac
ggttacctcttatctagtggGAGTATTGCAATAGTGCATGCTTCATAGAGCTGTTGTAAGgaatcttaaacccaactt
ggttacctcttatctagtggGTGCCTAGAATAGTGTCTGGCCCGTTTTAAGAGCTCAGTGgaatcttaaacccaact
ggttacctcttatctagtggCCTGAGGACACAAGGATGGAAAGACTCAGTCTGTCTGTGGgaatcttaaacccaac
ggttacctcttatctagtggGGAAAGATTAAACAATTCAATGCAGGCTGGGTGCAGTGGGgaatcttaaacccaact
ggttacctcttatctagtggCACTTTGGGAGGCCAAGACGAGCGGATAGTTGAGATCAGAgaatcttaaacccaac
ggttacctcttatctagtggAAATTAGCCAGACGTAGTGGCGAGTGCCTGTAGTCCCAGCgaatcttaaacccaac
ggttacctcttatctagtggAGAGGGGTAAGTCCATCGGGACAAATCAGGGGGCATTCCCgaatcttaaacccaa
ggttacctcttatctagtggGGACATTGTGGGCGGGAGAAACAGCACCTACAGGGCTGAGgaatcttaaacccaa
ggttacctcttatctagtggGCCTGAGCAAACTGCAATGTGGCAAGTGGGAAATGAACCCgaatcttaaacccaac
ggttacctcttatctagtggCCCATGGAGAGCCAAGAAGGATTTTGAGCCAGGGAGTGACgaatcttaaacccaac
ggttacctcttatctagtggACTGCTTTGAGGAGGTTGTCTTGGAAGGAAAGCAAGAGTCgaatcttaaacccaact
ggttacctcttatctagtggCAAGAGAAGAAAGATTGCTATTTCCAAAGCTCAAGTAGTGgaatcttaaacccaactt
ggttacctcttatctagtggGTGGCACGTATGCTATGATCTCACACCTGTAATCCCTTCCgaatcttaaacccaactt
ggttacctcttatctagtggGTTGGGGAGGCTTTAGGCTGTGGCTGCCAGCCCATCCCACgaatcttaaacccaa
ggttacctcttatctagtggGGATGCAGAGCTTTTTAACCATATCAAAGAGAAGGATAGGgaatcttaaacccaactt
ggttacctcttatctagtggATGAAACTCCTGCTTTTGATATGCTTAAATTCTGGGTCTGgaatcttaaacccaacttg
ggttacctcttatctagtggCCAGATTCCCCTGCCCTTTTTCTGCCTGTCCTTAGCCCTGgaatcttaaacccaact
ggttacctcttatctagtggCCCGGGACTCACACTGGGGGAGGCAGGGAGACCAGCATGCgaatcttaaaccca
ggttacctcttatctagtggTCCCCCTGGCAAGGTGTGGAACATTTATACTCCCACGCCCgaatcttaaacccaac
ggttacctcttatctagtggTGGATGAACAGTGTTTACATTAAGAGGCTTGTCAATGGCCgaatcttaaacccaactt
ggttacctcttatctagtggCCCTTTAAGGGGCTATCTGAGGCTGGGTGAGGTGGCTCACgaatcttaaacccaac
ggttacctcttatctagtggCCAGGAGTTCAACACTAACCTGGGCAACAAAGCGAGAGCCgaatcttaaacccaa
ggttacctcttatctagtggCATGGTGGAGTGCGCCTGTGGTCCTAGCTGAGGTGAGAGGgaatcttaaacccaa
ggttacctcttatctagtggCTGGCTTCATCCTAAGAGTAGTTGCATTTTAAACTCTCAGgaatcttaaacccaactt
ggttacctcttatctagtggATTCCTAAAGTGATTATGAAGGCTGAACAGGTGATGCCTGgaatcttaaacccaactt
ggttacctcttatctagtggGGGCCCCTTCAGCCCCATGTGTCCCAGGCTAATCTCAAACgaatcttaaacccaa
ggttacctcttatctagtggTACAGACACGAGCCACCACTCCCAGCCAGGTTGCTATTAAgaatcttaaacccaac
ggttacctcttatctagtggTCCAGGAGCGTGGGTCATCATTTCTTTCAAATAATGCTTCgaatcttaaacccaactt
ggttacctcttatctagtggGTATAAGGAAGAAAATAAGTCGTGTTAACGCAATTGCTGCgaatcttaaacccaactt
ggttacctcttatctagtggAGTATTTCCATGTGTTGTAAAGAGCAAGCTTTTAGGTATCgaatcttaaacccaacttg
ggttacctcttatctagtggGTTGTAACATCCTATTGCAATGACAGCTTGTCTTTCTCCCgaatcttaaacccaactt
ggttacctcttatctagtggTTAAGAATATTTAATGTGGCCGGGCATGAAGGCTCCTGCCgaatcttaaacccaactt
ggttacctcttatctagtggACCAGCTTGGGCAACATGGTGAAACCGCATCTCCACTCAAgaatcttaaacccaac
ggttacctcttatctagtggCTCGGAAGGCTAAGGCTGGAAAATCATTTGAACCCAGGAGgaatcttaaacccaac
ggttacctcttatctagtggCAGTGAGTCGAGATCATGCACTGCACTTCAGCCTGGGTAAgaatcttaaacccaac
ggttacctcttatctagtggTGAGATGAGTGTTTTGGTGTATATGTCTGTGCATGTGTGCgaatcttaaacccaacttg

ggttacctcttatctagtggAGGCTCTCAATGCATGTTGTTCAGTTGTTTCCCCATTTGGgaatcttaaacccaacttg
ggttacctcttatctagtggCTGAGGCCAGGAAGACTTGGGTCTTGTTGAGGTGAAGGGGgaatcttaaacccaac
ggttacctcttatctagtggTGCTGCGTGGAAAATGAGAGTAGCTTTGAAGGGGCACAGCgaatcttaaacccaac
ggttacctcttatctagtggCATCTACTTCAGAACTCTGGCTGCTACGTGGGGTACGACCgaatcttaaacccaac
ggttacctcttatctagtggGTGGCTAGCTGTATGTTCCTAGCCGCGTCACTCAACCTTGgaatcttaaacccaact
ggttacctcttatctagtggTATGGCGTGCATCTCTTCCTCAGCTATATGGGTGTCCTCGgaatcttaaacccaact
ggttacctcttatctagtggGTCTTAGCCTCCTCAGTTGCTGGGACTACAGGAGTGTGCCgaatcttaaacccaac
ggttacctcttatctagtggCGAACTCCTTGGCTCAAACAATTCCCCCTCCTCTGCCTCCgaatcttaaacccaac
ggttacctcttatctagtggAGGCATGAGCTACCGCTACACCCAGCCTATCCTGTTTACCgaatcttaaacccaac
ggttacctcttatctagtggGGGCAAATTGCTTGGCCTCAGGATCTTGGTTTTGTTTTCTgaatcttaaacccaacttg
ggttacctcttatctagtggGTGCTTCCGCTAGATTGCTTGACCTGGTGCCTAATCCTGGgaatcttaaacccaact
ggttacctcttatctagtggCTCACTAGTGGAGTCCACCAGACCTGATGATTCCCCAGCGgaatcttaaacccaac
ggttacctcttatctagtggCCTGCACTTTAGTTCTTCAGGCTGGTTGTATATGGCTGGCgaatcttaaacccaactt
ggttacctcttatctagtggCAGAGTCAGAAGGGACCCATCCAAGTTTGTCACTCAATGGgaatcttaaacccaac
ggttacctcttatctagtggAGGCTCCAGATGGTGAGTGTCTATTTTGGAATTTCATAACgaatcttaaacccaacttg
ggttacctcttatctagtggCACACAAGCATATTTGCATAGGCTGGATTTGGAAGGGTGCgaatcttaaacccaact
ggttacctcttatctagtggGTGGCTGTTGCTAAGAGGCCACCCGGGAGCTTTCTGGGGCgaatcttaaacccaa
ggttacctcttatctagtggACTTAGCCTTGACTGTATGACTTTTTTACAGTCCTGTTGGgaatcttaaacccaacttg
ggttacctcttatctagtggAGCCAATCCTGCTTCTTAATGCAAGGAAAGAATCCCTGGGgaatcttaaacccaact
ggttacctcttatctagtggGTCTATCTGAACTCACGTGCCCTGGATCCTGGCCACAGGGgaatcttaaacccaac
ggttacctcttatctagtggCAGGTGATCTGCCTGCCTCAAGCCTCCAAAAGTGCTGGGAgaatcttaaacccaac
ggttacctcttatctagtggGGACTGTCTGCTGTCTCTTGCTGGCCTCCCAGTCTGTCTCgaatcttaaacccaact
ggttacctcttatctagtggAGCCAACTGCTCCTCTGGTTCACCCCCTGGGGCTCCCTGTgaatcttaaacccaac
ggttacctcttatctagtggCCTCCTAACCAGCCCCCTTCACGGTCCAGGAGCCTCCCAGgaatcttaaacccaa
ggttacctcttatctagtggGCACGTCAGGGTTGAGCTGCTCCCCCAGGCCCCATGCACGgaatcttaaacccaa
ggttacctcttatctagtggTTCTGGCAAATACGATGGATTCCCTGCATGGCCCCAGACCgaatcttaaacccaac
ggttacctcttatctagtggTCTTGCTGGGTGGCTTCAGCAGGGGCAGGGGTGGAAAGCCgaatcttaaacccaa
ggttacctcttatctagtggTCTGGCCCGAAGCCCCTGTGCCGTATTTCCCTCAGCACCGgaatcttaaacccaa
ggttacctcttatctagtggGGCTGTTAGGGCCAAGTGCCTGCTCTTTGGAGGGCATCTAgaatcttaaacccaac
ggttacctcttatctagtggCCCAGCGACCAATTCTGGAAACATCAAAGTGTTTATGAGCgaatcttaaacccaact
ggttacctcttatctagtggGGTGAGGGGTGGGCAGGGTGTGTATCTTCTTGCTGCTGGGgaatcttaaacccaac
ggttacctcttatctagtggCTGGCAGGTTACACCTGGGTACATTTCAGTCAAGCCGAGCgaatcttaaacccaac
ggttacctcttatctagtggGTACAGGTGAGGGAAACTGAGGCCCAGAGTGGATGTGACGgaatcttaaacccaa
ggttacctcttatctagtggGGCTCTGTAAAGTTCTGAGTAGGTGAGACTTATTATTAGCgaatcttaaacccaacttg
ggttacctcttatctagtggAGTGGCCTCTCCTTTTCTCTTCTTGGAAGCACAAGCCCTGgaatcttaaacccaact
ggttacctcttatctagtggAACCTCTTCACGGTGGTTTTGGGAGCCAAGCGAGGCCAGGgaatcttaaacccaa
ggttacctcttatctagtggACGTGGGGAGAGGGATGTTGAGGACAGTCCATCCTCAGAGgaatcttaaacccaa
ggttacctcttatctagtggCTCAGTGAATCCGTGAAAGGATGGTGCAGGTCCGCCCCCCgaatcttaaacccaa
ggttacctcttatctagtggCTTCCATGTGTGTCCTCGTGGCCAATTTGTCTTTGCACGGgaatcttaaacccaactt
ggttacctcttatctagtggACATGCCAGATGCTCGTTCCAAGTTCTGTGCCTCCACCAGgaatcttaaacccaac
ggttacctcttatctagtggGCTGAGAATGCTTGTTCAGAGGTAAACTTGTTAGCATCTGgaatcttaaacccaactt
ggttacctcttatctagtggGTGATGAGCAAAAAATAATGATTGAGCCTGGCGCGGTGGCgaatcttaaacccaac
ggttacctcttatctagtggGGGGTGGTCCTGTACATGACAGCTATGGTTTTAAGGCCCCgaatcttaaacccaac
ggttacctcttatctagtggGGCTTCCCAGAAATCCTTTCCAACCCAAGAGCGTCAGAGGgaatcttaaacccaac
ggttacctcttatctagtggGCAAGACTTTCAGGTGTTGCTTCTCTGGCCTCCCTCTTTCgaatcttaaacccaactt
ggttacctcttatctagtggGGCTGCTCAGCCCAATGTAAAGAAAGCGCTTTCTGAAGTTgaatcttaaacccaact
ggttacctcttatctagtggGCCAGTGTGGCAGTATGAACTTGAGAAAGAGACATCCCCTgaatcttaaacccaac
ggttacctcttatctagtggGATCCACAATTTTACCTTTGCGGGTTGGCACATGGACTGGgaatcttaaacccaact
ggttacctcttatctagtggCATTTCCTAGGGCTCACATTACAAATTAGCAATGGGCTCCgaatcttaaacccaact
ggttacctcttatctagtggTCAGGGTGTCAGGAGGGCCATGCTCCACATGAAGGCTCCGgaatcttaaacccaa
ggttacctcttatctagtggGCTGGCAATCTCTGGCTTGTGGCTGCAGCAATCTGCCTCCgaatcttaaacccaac
ggttacctcttatctagtggGCCTTCTTCCCTCTGTGTGTCTCTGAATTCTCTCTTCTTGgaatcttaaacccaacttg

ggttacctcttatctagtggGGGCCCACCCTGATCCAGCAGAACCTCTTCTTTACTTAACgaatcttaaacccaact
ggttacctcttatctagtggTGAATTCTTGGGAGCAATCAACCCAGTACAGATGTCTGGCgaatcttaaacccaact
ggttacctcttatctagtggGACAACATCTGGGCAGATGACAGCATCTGGTATTTTTATGgaatcttaaacccaactt
ggttacctcttatctagtggGGAGGGAGAGGTGTGGTTGGCTTGGCAGGTGGAGAGGGGCgaatcttaaacccaa
ggttacctcttatctagtggGCAAGGAGCTTGCTATGTGGGGGTGTGGGAGTGGGGTTGGgaatcttaaacccaac
ggttacctcttatctagtggTTTCAGGAGAGAGTGATGTGGCCTGTGTCTTGGTGGTAGCgaatcttaaacccaact
ggttacctcttatctagtggTGTGCCAATAGGACTTGCACACGGATTGGATATGGAGGTGgaatcttaaacccaact
ggttacctcttatctagtggGGGTGGTTCATGCCTACTATAGTCCTAGCACGTTGCGGGGgaatcttaaacccaac
ggttacctcttatctagtggGAAGGATGCAAAGTGTTGTTTCCAGGTGTGTCTGTGAGGGgaatcttaaacccaact
ggttacctcttatctagtggCATGAAAATGCAAGACTGGCCTAGCCTCTTAGCCTACATCgaatcttaaacccaact
ggttacctcttatctagtggGGGAACCCTGACTAATACAGAGGCTGAGGCAGGAGGGCCGgaatcttaaacccaa
ggttacctcttatctagtggGGAGTTCAAGACCAGTCTGGACAACTTAGAGAGACCTTGTgaatcttaaacccaact
ggttacctcttatctagtggTGGACACGGGGAAGATGGGCAGTAGATTGTGGGCGCAGAGgaatcttaaacccaa
ggttacctcttatctagtggTGCAGATGCCCATTAATGGCCACAGGAGATGCCAACTAGGgaatcttaaacccaac
ggttacctcttatctagtggTGGAAGATTTTGGCAGAGGTGGGATCCAAATCCATGGGAGgaatcttaaacccaact
ggttacctcttatctagtggGACATGGCCACTAGTGGTAGGGGTAGAAGTTGGGGAGCTGgaatcttaaacccaac
ggttacctcttatctagtggCAAGGAGCCACTGGTGAGGCAGAAAAACATGCCGGGAGCCgaatcttaaacccaa
ggttacctcttatctagtggGGAGGAGAGTCATCACTTGCAATGTTGCTAAGAGGTCGGGgaatcttaaacccaac
ggttacctcttatctagtggCTTGGCAAGAGCGCCTCATGGGAGCAATCAAGAGGAAGCCgaatcttaaacccaa
ggttacctcttatctagtggGAAGGTGACTGCTCTTGCGTTCTGCTCTCTCTTCATTGCCgaatcttaaacccaactt
ggttacctcttatctagtggGCTTAGGCAAATCAGTGTCTTCAGTAGGTTCCTTGAAGGCgaatcttaaacccaactt
ggttacctcttatctagtggAACAACATCTTTGCAGAAAAGGCTGGGCAGGGCAGGCAGGgaatcttaaacccaa
ggttacctcttatctagtggGTCAGGCACCCCTCCTCCACCCCCATCTCGGCATACATCCgaatcttaaacccaa
ggttacctcttatctagtggCCCAGTGAAGGGAGGATCTGAATGACATCTCTATATTCCCgaatcttaaacccaact
ggttacctcttatctagtggCTTGGGTACCCATGGCAGGCATCACTAGCTGATCATGACCgaatcttaaacccaac
ggttacctcttatctagtggTCTACCCAGCATTCCAGATAGCCTGTCAACCTGCTGACAGgaatcttaaacccaac
ggttacctcttatctagtggCTTAATGTCATGCCTGGTTCAGCAAGTGGTTTTGTGACTGgaatcttaaacccaactt
ggttacctcttatctagtggAGAAGAGATCAAATTCATGCACATGGCCGGGTGTGGTGGCgaatcttaaacccaac
ggttacctcttatctagtggTAGAATTCAGGTATTGGGTAAAAGGTTATCCAACTCTGAGgaatcttaaacccaactt
ggttacctcttatctagtggAGAGGAATCTCCCCCCACCTCCCCAGATATATAGGAGGGGgaatcttaaacccaa
ggttacctcttatctagtggTGGCCACTTGGGGCTCTGGTCTCAGCCTTTCTGGATTGCCgaatcttaaacccaact
ggttacctcttatctagtggGGCATTTCATAATCCTCTCTACCTCCTGCCCTGCCCTGGGgaatcttaaacccaact
ggttacctcttatctagtggGCTGCTCCTCTCTCCTGCCCTCTATCTTGGCCTCGCCCGGgaatcttaaacccaac
ggttacctcttatctagtggCAGTGGCACAATCTCACTGCAACCTCAGCTTCCTGGGCTTgaatcttaaacccaac
ggttacctcttatctagtggAGTAGTTGGGACTACAGGTGCATGCTATGATGCCCAGCTAgaatcttaaacccaact
ggttacctcttatctagtggAGGGCATTGGCTCTGCCTGGCTCCAGAGGGATATGAAGCCgaatcttaaacccaac
ggttacctcttatctagtggCCAGACAGCTGGACCCCTCCCTAGAGACTTAGGATGAGGCgaatcttaaacccaa
ggttacctcttatctagtggGCTCACAGGTGACACTGAAGTTGACGACCTTGTATCTCACgaatcttaaacccaact
ggttacctcttatctagtggGGGCCCTCTGCTTCCACTGTGGCCTGCTTACAACCTCACCgaatcttaaacccaac
ggttacctcttatctagtggGCAGAGTTCACTGAGTCTGCTTAAAACCTTAAACTTCAAAgaatcttaaacccaactt
ggttacctcttatctagtggATTCCCTATTTAATAATGGTGTCAGGAAAACCGGCTAGCCgaatcttaaacccaactt
ggttacctcttatctagtggAAAAACCCTAGATGAAAACCTAGGCAACACCATTCAGGACgaatcttaaacccaac
ggttacctcttatctagtggGAAACTACTATCAGAGTCAACAAGGAACCTACAGAATGGGgaatcttaaacccaact
ggttacctcttatctagtggTGCAATCTATCCATCTGACCAATGGCTAATATTCAGAATCgaatcttaaacccaactt
ggttacctcttatctagtggGTTTATAGCAGCACTTTTTACAATAGCAAAGACCTGGAACgaatcttaaacccaactt
ggttacctcttatctagtggCCAAACACAGCATGTTCTCATTCAGAAGTGGGAGTTGAACgaatcttaaacccaact
ggttacctcttatctagtggGAGGGATGGCATTAGGAGAAATACCCAGTGCATGCGGGGCgaatcttaaacccaa
ggttacctcttatctagtggAACCACCATGGCACACGCATATCTATGTAACAAACCTGCGgaatcttaaacccaac
ggttacctcttatctagtggACCTCCACAAAAACCCTATCTCCAGGCTTCCCATTGTATCgaatcttaaacccaact
ggttacctcttatctagtggAATCCTGAGCAGCCTGTTCCTGTTGATCGAGTGAATGAGCgaatcttaaacccaact
ggttacctcttatctagtggGCAGTGAGTGAGTTCTCCATCACTAGAGGTAGTCAAGAAGgaatcttaaacccaact
ggttacctcttatctagtggGTCTTCCAGCCTCATCAAGTGGTCCCTCCCTCTGTAGCCCgaatcttaaacccaac

ggttacctcttatctagtggGGGCACCACAGGACCTTTGCACTTGCTGGTCCCAGATCCTgaatcttaaacccaac
ggttacctcttatctagtggATCACCTCCTCAGACCACCCCTTCTAATAGACTCATTCCCgaatcttaaacccaact
ggttacctcttatctagtggTCTCATCACTGGATGCTGCTGAATTTCACTGTGGCACCAGgaatcttaaacccaact
ggttacctcttatctagtggTTGTCCTTGAACTCTGTGTTGTCTGACACTCAGTGGGTGGgaatcttaaacccaactt
ggttacctcttatctagtggATTTTTATTTTACATTCCGAGGTACATGTGCAGGATGTGCgaatcttaaacccaacttg
ggttacctcttatctagtggCAGGCTGCAGAGGGTTAGGTCAAGGCTGAACCATGTAGTCgaatcttaaacccaac
ggttacctcttatctagtggAGTGACTGCAGTGAAGCCCCTCCATGGTGGAAGAAAATGGgaatcttaaacccaac
ggttacctcttatctagtggCTGGGCAGATCCAGAGGCTGGAGGTAATGGCAATAACTCTgaatcttaaacccaac
ggttacctcttatctagtggAAGAACGTAAAAGAGTGCTTACAGTGTGTTACAGGCACTGgaatcttaaacccaact
ggttacctcttatctagtggGTTGTCTCCATGTTACAGAGGAGAGGACTGAGGCACAGAGgaatcttaaacccaac
ggttacctcttatctagtggCAGAGTCCACACGCTTGGCGCCTGCACTGCACCGAGAGGGgaatcttaaacccaa
ggttacctcttatctagtggCGGGAGTCCCCCACCTGCTCTCTGTTTTCAAACTCCCCATgaatcttaaacccaac
ggttacctcttatctagtggGCCAGGTGTGATGAAATCCTCTAGGTGATGGGCAGTGGACgaatcttaaacccaac
ggttacctcttatctagtggGGATTCTAGCCCATGGACCATAAGCCAGGCCCCCTGAGGgaatcttaaacccaa
ggttacctcttatctagtggTCACCTAGTCCCCAGAGAAGGGCCATGGGTGGGCTGACTGgaatcttaaacccaa
ggttacctcttatctagtggGGGCTCGCCTTCCATCCAGCTGAAGTCTTTCTGGCCGGGCgaatcttaaacccaac
ggttacctcttatctagtggGCCGGGGGTGGGCCTCAGCTGCTGCCACTGGCATTGTATCgaatcttaaacccaa
ggttacctcttatctagtggTCGGAAACCTCATAATGGCTTCATGAACCATTGCACTTGGgaatcttaaacccaactt
ggttacctcttatctagtggTGAAAAATTCCCTTGATTTATTGTTTTCGGAGGCTGTAGgaatcttaaacccaacttgt
ggttacctcttatctagtggGATTGAGGCCCTCCTGTGTGCCAGGCTAAATGGTCATCCCgaatcttaaacccaac
ggttacctcttatctagtggGCGATGGGGAAACGGAGCCAGGGGAGGAACTTGCCCCAGCgaatcttaaacccaa
ggttacctcttatctagtggTGTGTTGTCAGGTATGATCAGCCTATGTACTGCTTCCTTGgaatcttaaacccaacttg
ggttacctcttatctagtggGTGGGGTAGAGAATATCAGTTGTAGCTCTAAAGGGGTGTGgaatcttaaacccaact
ggttacctcttatctagtggACAGGCATGAACCACTGCACCTGGTCTCATTTATGAATTTgaatcttaaacccaacttg
ggttacctcttatctagtggGCAAGGCTATGGACAGATACTGGTAATGGAACCTGTGCTCgaatcttaaacccaact
ggttacctcttatctagtggCCCTATGGGGTCAAAAACAAGGTCTAGCCAGGTGCGGTGGgaatcttaaacccaac
ggttacctcttatctagtggGGTGCCACTGGACTCCAACCTGGTCAACAGAGTGAGCCTGgaatcttaaacccaac
ggttacctcttatctagtggGTGCCAGACACTATTCTAGGTGCTTTTCATCAATTAGCTCgaatcttaaacccaacttg
ggttacctcttatctagtggCCACATAATGATGTGTGGATCAGCAATGGACCGTATATACgaatcttaaacccaactt
ggttacctcttatctagtggCCGATGTTCAGTACAGTAACATGCCATTCAGGACTGCAGCgaatcttaaacccaact
ggttacctcttatctagtggAGGCTGTACCATATAGTCCAGATGTGTAGTGAGTAGGCTGgaatcttaaacccaact
ggttacctcttatctagtggCCCTCTGTAATGTTCGCACAACGAAATTGCCTAACGCCGCgaatcttaaacccaac
ggttacctcttatctagtggACCCTTAAGCAACGTGTGACTGTCCTATTATCATCACCCCgaatcttaaacccaact
ggttacctcttatctagtggGGCCCAAGGTCACCCATATAATAAAGGCGGGCTGGGTGCgaatcttaaacccaac
ggttacctcttatctagtggCCAGCACTTTGGGAGTCCAAGGGGAATAGGATCACTTCAGgaatcttaaacccaac
ggttacctcttatctagtggAGCCGGGCATGGTGGCATAGACTTGTGGTCCCAGCTGCCCgaatcttaaacccaa
ggttacctcttatctagtggGAGCTGGGACTTGAAGCCAGATAATAAGGCAGGCAGAGGGgaatcttaaacccaac
ggttacctcttatctagtggGAGCCCGTGCATCAGTCAGGGTCTAATGGCTCAGCTGAGGgaatcttaaacccaac
ggttacctcttatctagtggGGATCATGGGGGTGGTACAGTGTTGCCAGGGCTGGGGAGGgaatcttaaacccaa
ggttacctcttatctagtggCCCAGCCGCTGCCATGTGGAGAGGGACCTGTCACTCCTGCgaatcttaaacccaa
ggttacctcttatctagtggCCCTTCTTCTGCCCTCTGACCTCCTGCTGGTGCCTCCCATgaatcttaaacccaac
ggttacctcttatctagtggTGGCCAGTGCACTGGGGAGCTGTTCTTGGAGCCCACAGGCgaatcttaaacccaa
ggttacctcttatctagtggTGGCACAGAGCAGACAATGGATTGAGTCGGGAGGGAGGGGgaatcttaaacccaa
ggttacctcttatctagtggGGTGAAGTCAGAGAGGCCGGGGTCCACACTACACAGGGCCgaatcttaaacccaa
ggttacctcttatctagtggGCCAGTGGAGGGTTTGAGCAGATGTGTGCATCACGGCCACgaatcttaaacccaac
ggttacctcttatctagtggGTCACCCCTTCCTCTCCCTTTCTCAGGATGAGTGCTGAGGgaatcttaaacccaact
ggttacctcttatctagtggCGTCTGCCCAGTCCAGACTACGTGCCTCATGCCAGCCAGCgaatcttaaacccaa
ggttacctcttatctagtggGAGAAAGGAGCAGTGTCTCAAAAGAGGATATTCTGGAACGgaatcttaaacccaact
ggttacctcttatctagtggGTTCCAGTGGAAAACTTTTACAGGCCGTCCTGGGCAAGGCgaatcttaaacccaac
ggttacctcttatctagtggGACCTGTGCTGGCAGCCCTGGCCACACCTTGGTGGGACCGgaatcttaaacccaa
ggttacctcttatctagtggTTGGGGACCCTGCCACTCTCCTAGCTGTGCTTCCGCAGAGgaatcttaaacccaac
ggttacctcttatctagtggCTGCCCATGCAGGAAGCTGGTCCTGTCCAACCAGGGAATGgaatcttaaacccaac

ggttacctcttatctagtggGGATGCCCAGGGGAGGGAGAACACACTTGAGTGGGATGGCgaatcttaaacccaa
ggttacctcttatctagtggGTGACCAAGGCACCAAGTCATCTGGAAACCCACTGCAGGCgaatcttaaacccaa
ggttacctcttatctagtggCACCTTTGTGTGCCCCAAAGCTTGCTCCGGGAATCGATGGgaatcttaaacccaac
ggttacctcttatctagtggGGGCATTTGAGAGTAAGGCAAGAAGCGCTTGGCATGTGCCgaatcttaaacccaac
ggttacctcttatctagtggGATCTTCATCCACCTCTTGTTGGCCTGAAGGAAGGGGAGGgaatcttaaacccaac
ggttacctcttatctagtggTTTTTTATTTTTCTTGAGACAGTCTTGCTCTATCACCCAGgaatcttaaacccaacttgt
ggttacctcttatctagtggGACCTCAAGTGATCTGCCCACTTCAGCGAGATTATAGGCGgaatcttaaacccaac
ggttacctcttatctagtggGGCTGAGGTAAGATCTGTGGTAGCATCCAAGTCTTGACCCgaatcttaaacccaact
ggttacctcttatctagtggCACAGATGCAATTTTGTGCCTCATCATCTGACCCAGGAGCgaatcttaaacccaact
ggttacctcttatctagtggGCTGGGACCACAGGTGTGCGCCACTACCCCCAGCTAATTTgaatcttaaacccaac
ggttacctcttatctagtggAGCCACTGCGCCTGGCTAAAGTTGTTATTTTAAAATTATTgaatcttaaacccaacttg
ggttacctcttatctagtggAACTGGAAAACGGCTTTGCTCTTTTGTGGAACTGCTCTGGgaatcttaaacccaactt
ggttacctcttatctagtggCTGCAGGTTTCAACTCCTGGGATCCAGTGATCCTCTTGCCgaatcttaaacccaact
ggttacctcttatctagtggTTTTTGTAGAGATAGGGTCTCTCTTTGTTGCTCAGGCTGGgaatcttaaacccaacttg
ggttacctcttatctagtggCCTACCACCTCAGCCTCCCAGCGAGTTGAGATTAAAGGTGgaatcttaaacccaac
ggttacctcttatctagtggGGACTCCACAGAAAGGCCGAGTCTCTCCAAGACCCAGCCGgaatcttaaacccaa
ggttacctcttatctagtggTCATCTGTCCACTCCTCCAAAGAGGGTGAGACAGGGCCAAgaatcttaaacccaac
ggttacctcttatctagtggTCACTGTGCACCAACCCTCCCTCTTGCTTCAGAAGAGACCgaatcttaaacccaac
ggttacctcttatctagtggGGGTCACCAAGTCACAGAGGCCCTCAGCTCCTGTGCCGGCgaatcttaaacccaa
ggttacctcttatctagtggAGCATAGGGCTAGAAGGCCAGGTCACAATACCCTCTCTGGgaatcttaaacccaac
ggttacctcttatctagtggGCCCCAGTTTCCTGGTCTACAGAAGTGACAGATGGGACCCgaatcttaaacccaac
ggttacctcttatctagtggGAGGATAAAAATAGTGATATGGGCTGGGCACGGTGGCTCGgaatcttaaacccaac
ggttacctcttatctagtggTTGTAAATCAAACAGCTCTTGAGAGATGCCTGTTTGGCTGgaatcttaaacccaacttg
ggttacctcttatctagtggCCTGACCTTGTGATCCACCCTCTCCGGCCTCCCAAAGTGCgaatcttaaacccaac
ggttacctcttatctagtggAGGAGGTGGGAATCCCTTGTCTTCCTGGGCAGACTAGATGgaatcttaaacccaac
ggttacctcttatctagtggGGACGATGCCTTCCAAAGCTGCTAAGTCCTGTGGCAGGGGgaatcttaaacccaac
ggttacctcttatctagtggTTCTTCATGGTTTCTCTGGGGAGCCTCCTAGCCTCTGAATgaatcttaaacccaacttg
ggttacctcttatctagtggGTAGTGCCCTCTCGTGAATGAGAAAAGCCGCCCAGTCCTGgaatcttaaacccaac
ggttacctcttatctagtggGCAGTGGGCAGAGCCAAGACCACCTTTATTTGTGCATCCGgaatcttaaacccaac
ggttacctcttatctagtggCACACAGTTTGTTGTCCTTCTGTCCCTTGCCCATGAGGCCgaatcttaaacccaact
ggttacctcttatctagtggAGCTTGTCTCCACCCATTCCTCTACAACCCACACAACAACgaatcttaaacccaac
ggttacctcttatctagtggAGATGCAACTCATGCCACCTCCCCCAGGAAGTTCTCCCTGgaatcttaaacccaac
ggttacctcttatctagtggAGGCTGCCAGAGATCCCTGTGTTAACTGTGAACAGAGCCCgaatcttaaacccaac
ggttacctcttatctagtggCCTGTCTCATCCACTCGACTGAACTTCTTGGGTAGGAATGgaatcttaaacccaact
ggttacctcttatctagtggCGTGAGCTACCACGCCCGGCCGGAGGTGTGTTTTACATGCgaatcttaaacccaac
ggttacctcttatctagtggGGGACAGCGCCTGGAACCTAGCAGGTGTTTTATGTGTGGGgaatcttaaacccaac
ggttacctcttatctagtggAGGTGTGTGTTGTGACGAGGGATGGCACTGGGGGCCTGCCgaatcttaaacccaac
ggttacctcttatctagtggGCTTCATTGTGGGGGCTGAGGTGGCTTGTTTCCTTGGAGCgaatcttaaacccaact
ggttacctcttatctagtggGAGGAGCTGGGGACTGAGCTTGGATCTAGATTATTGTTGCgaatcttaaacccaact
ggttacctcttatctagtggTGGGACAGAGGGGAGAGGGTGAAAGTGGGGAGGAGGAGGAgaatcttaaacccaa
ggttacctcttatctagtggAAGCCTAGACCCATCGTACGCTGAAACAGAGAAGAATGGCgaatcttaaacccaac
ggttacctcttatctagtggGTGTGAGTCTAGGAACTTAAGAATGCTAGTCCTTGACCCCgaatcttaaacccaact
ggttacctcttatctagtggCTTCCACCTACCGGATGGGATTATGCAAGTGATATGACCCgaatcttaaacccaact
ggttacctcttatctagtggGATATGCAGATGAACTGCCCAGGGTGTCTCTGGTCTTGGGgaatcttaaacccaact
ggttacctcttatctagtggGAGTGTCCTGTTTTCCCATCCACAGCCTCCAGGCCTGGCGgaatcttaaacccaac
ggttacctcttatctagtggCCAAGGTGGACATCTCTCTGACACTGGCGATTTTTCTCTCgaatcttaaacccaactt
ggttacctcttatctagtggCTTAGCGGGGTCCCAGGAAGGCTGTGCTGGCCCTGTCAGCgaatcttaaacccaa
ggttacctcttatctagtggCTCACCCTGGAGCCTTTCCCAAGCACACATCGTGGACCCCgaatcttaaacccaa
ggttacctcttatctagtggCACATGTGCTTTGTAAGGGCTCTTCAGGTGACTGGAGCTTgaatcttaaacccaactt
ggttacctcttatctagtggCCCGGTTACTTGGCTTTTGTTGAGTCCCTGGTGGGTGTCCgaatcttaaacccaact
ggttacctcttatctagtggGGGGCACCTTTCCTGGTCTCCTCTCTCTCCAAGGAGCAGGgaatcttaaacccaac
ggttacctcttatctagtggACCCTTGCATTCCTACTGCATGTTCCCTCAGGCTGAAGCCgaatcttaaacccaact

ggttacctcttatctagtggGCACCCACCACCCACGGAGACTAGCTCATGGCCGAGTGGCgaatcttaaacccaa
ggttacctcttatctagtggGGGTCTGCACAATATAAAGACAGCTCACAGGAAAGGGGCCgaatcttaaacccaac
ggttacctcttatctagtggGGGGTTTCGCCATTTGGGACAGGCTGGTCTTGAACTCCTGgaatcttaaacccaact
ggttacctcttatctagtggGTTGTTGTTGTTGTTGAGATGGAGTATTGCTTTGTTGCCCgaatcttaaacccaacttg
ggttacctcttatctagtggAGCCTCTGGAGTGGCTAGGATTATAGGCGCCCACCACCACgaatcttaaacccaac
ggttacctcttatctagtggTTTCACCATGTTGGCCAGGATAGTCGAACTCTTGGCCCCAgaatcttaaacccaact
ggttacctcttatctagtggGTGTTGCTGGACGGAATGCAAGTGGATGTGGCCCTTTGGGgaatcttaaacccaac
ggttacctcttatctagtggTGTGCCCTTTGACCTGGCCATCTCATGTTGGGACATTGGGgaatcttaaacccaact
ggttacctcttatctagtggTTTGTAGGAAAAGAACAGTGCCCAAGGTTATTCATGGAACgaatcttaaacccaactt
ggttacctcttatctagtggGACTGGCAAATGATGACACATTGTGCTAGGCAAGATGCGGgaatcttaaacccaac
ggttacctcttatctagtggGACACAAAAAGCACAATGTGGAGCGGGGAGTATGGCATGCgaatcttaaacccaac
ggttacctcttatctagtggGGTGGCCTGTGCGTGGTGGGAGGAGGGAGATGCAGGGAGGgaatcttaaacccaa
ggttacctcttatctagtggTCTTTTCAGAGATAAGGTTTCACTCTGTGGCCCAGGCTGGgaatcttaaacccaactt
ggttacctcttatctagtggGTGATCATAGCTTGCTGCAACCTCAGACTATGAAGGAGACgaatcttaaacccaact
ggttacctcttatctagtggGCATGAATATATTGAAATCGGGAGCAGCTCAAACATTGAGgaatcttaaacccaactt
ggttacctcttatctagtggTGATTAGGCTGGGGTTATGCATTTTTAGGAGGGAGACCCCgaatcttaaacccaact
ggttacctcttatctagtggCCCGGTTGCATATCCCTATAATACACCTGACCAAAGCCAGgaatcttaaacccaac
ggttacctcttatctagtggAGCTACTGGGGAGGCTGAGGCAGGGGGTTGCTTGAGCCCAgaatcttaaacccaa
ggttacctcttatctagtggGACCTCATCTCTCCAAAACCACAAATTTGGCTGGGCGCAGgaatcttaaacccaac
ggttacctcttatctagtggGGGCAACAAGACCGAAACTCCGTCTCAGATAAAAAACAAAgaatcttaaacccaac
ggttacctcttatctagtggGGTGACCACCACCCCTAGTGCTCCAATAAAGGGCTGTGGGgaatcttaaacccaac
ggttacctcttatctagtggGCCCTCTGAGATATTGCTGGGCCTTAGGGGAGGAGGTCCAgaatcttaaacccaac
ggttacctcttatctagtggCCCCAGGGCATGGTGTGGACCACGCTGAAGGGAGGTGGGGgaatcttaaacccaa
ggttacctcttatctagtggCCCAGGGGCTCTCCTGTGTGATGCTGAATTGCTGGGAAGGgaatcttaaacccaac
ggttacctcttatctagtggCCCTGATTGGTTGGGTGGCTGCTGAGAGCCTTATGGTGCCgaatcttaaacccaac
ggttacctcttatctagtggCCGGCCTGGGGATCCTGGAGGTTTTAGGCTGGCAGACCCTgaatcttaaacccaac
ggttacctcttatctagtggCCATGGGTGAAAAACAGCTCTGGGGACCCTCATAGGCTTCgaatcttaaacccaac
ggttacctcttatctagtggGCAGCTTGGCCCAGCTTTCGAAACAAGGTGTTCCCAGGAGgaatcttaaacccaac
ggttacctcttatctagtggGGTAGGATCCTATGAAAAACTCCTAGGGCCTTGCCGTTCCgaatcttaaacccaact
ggttacctcttatctagtggTTCACACTCCTAGTTATGCCATCGAGGTAAAAAGAGGTGGgaatcttaaacccaact
ggttacctcttatctagtggAATAAGAAAAGAGAAAGACTGCGTGCATGTGCATGGGTGCgaatcttaaacccaact
ggttacctcttatctagtggGGGTGCACAGCCTGTACATATGTGTATTTGTGCTCCTGTGgaatcttaaacccaactt
ggttacctcttatctagtggGGGTGCACAGCCTGTACATATGTGTATGTGTGCTCCTGCTgaatcttaaacccaact
ggttacctcttatctagtggGAGCCTGTGGGTATGTGAGGGGAGTGAGGGACCCTGAGGGgaatcttaaacccaa
ggttacctcttatctagtggGGCAAAGGTGTGCCTCCCTTTAAGAGGACCTAGCGGTCGGgaatcttaaacccaac
ggttacctcttatctagtggAGGTTATCAGGGAGTAGTACTTTGGCCTGGAAAAGGAACCgaatcttaaacccaact
ggttacctcttatctagtggAACCTGACTGGAAACTCAGGCCACCAGCATCTGTTGGGGGgaatcttaaacccaac
ggttacctcttatctagtggCACTTTTGGTTATGATGCCACGGCCCTGGTTCAGGTATCCgaatcttaaacccaact
ggttacctcttatctagtggGACACAGTGATGGTGTGGACCCATGCACAGCCAGCTTTGCgaatcttaaacccaac
ggttacctcttatctagtggATAGCGACTTTGCCTGACATAATTGGTGTCTGCACATGGGgaatcttaaacccaactt
ggttacctcttatctagtggTTTAAGTTAAGAACATGTAGTTGTGATCAGGAGCGGTGGCgaatcttaaacccaactt
ggttacctcttatctagtggCTAAAAATACAAAAAGTAGGTGGGCGCCTGTAATTCCAGCgaatcttaaacccaact
ggttacctcttatctagtggCGTGCCATAGCACTCCACTTTGGGTGATAGAGGGAGACTCgaatcttaaacccaac
ggttacctcttatctagtggGGCAGGTGGGTGGTGAATCCGGGTGTTGTGAGTCATTTGCgaatcttaaacccaact
ggttacctcttatctagtggCGTTTGGAGTGGGTGCTCTTCTGAGGTTTTTGTTTCCCCCgaatcttaaacccaactt
ggttacctcttatctagtggCACTGTGTGTGCATGGTAAGAACAAGGGCCATGGAATTAGgaatcttaaacccaact
ggttacctcttatctagtggTGAGCCACCCCCGCCCATGGCCCTCATCTGTACCCTGAGGgaatcttaaacccaa
ggttacctcttatctagtggGAGATGATGTAGCAAAACACCCAGCCCAGAGCCTAGCACCgaatcttaaacccaa
ggttacctcttatctagtggCAGCCATTCTCTGGATGTGGCCTCTTCTGCCTCCACTGTGgaatcttaaacccaact
ggttacctcttatctagtggCCCAGCCTTTCAGGAGGGTGAATCTGCACCTGGGGTCCAGgaatcttaaacccaac
ggttacctcttatctagtggGAGACGTGAGCCAGGCGCTGATTCATCATGATGCAGGCTGgaatcttaaacccaac
ggttacctcttatctagtggGGGCTCTGTGCTCTGTGGCCCCTGCTGCTCCGGACTGGTTgaatcttaaacccaac

ggttacctcttatctagtggGACCGGTTGCCACGCCATGCTCACAGTCTCTGTTCTTGCCgaatcttaaacccaac
ggttacctcttatctagtggCTGCAGCTCAGCCAGCACTTTACAAGGGTGATCTGGAGCCgaatcttaaacccaac
ggttacctcttatctagtggGCAGTTGGGGAGCTGCCTCGATTTCTGGTTACAGAATTCCgaatcttaaacccaact
ggttacctcttatctagtggGTGTGGGATTCCAAACTGTAGCAGCAGTGACTACAGCTGGgaatcttaaacccaac
ggttacctcttatctagtggAGGCAGAGCCTGGCGTCAGAAAGCTGCATTGCGCTAATGCgaatcttaaacccaac
ggttacctcttatctagtggGGGAGCCTGTTGGAGAGACACTTGGATGTTTAGCGAGCTGgaatcttaaacccaac
ggttacctcttatctagtggGAAGTGGTCAGTGCGTCGACTCTGCCTCTAGCCTGCTGGGgaatcttaaacccaac
ggttacctcttatctagtggTGACCTTTCTGGCTCTCAGCATCCTCTTGTGTGAAATAGGgaatcttaaacccaactt
ggttacctcttatctagtggGCCTGGCTTCCTGGCAAGATTGCTATCAGGATTAAGGCAGgaatcttaaacccaact
ggttacctcttatctagtggGCCTGCCCAGCCCTCCTCCCTCTCCATCCACACAGCAGCCgaatcttaaacccaa
ggttacctcttatctagtggGGTCCTCACTGCCTCTGTGGCGCCTTGCACTGGGTCCAGCgaatcttaaacccaac
ggttacctcttatctagtggCAGTGGCTGAGTGTGACTTTGGCTGGCATAGTGGTTGCTGgaatcttaaacccaact
ggttacctcttatctagtggCGAGCATCTCCCATGGACAAGATCAGCTGTGGGTGGAGCCgaatcttaaacccaac
ggttacctcttatctagtggGGACAGCGAGTCTTCCCAGGCCAGGGCTTCAAGTGAGGAGgaatcttaaacccaa
ggttacctcttatctagtggAGAACCATACCCAGCAGGAGGCCGGGGAAAGTGACTCCTGgaatcttaaacccaa
ggttacctcttatctagtggAAGGCTTCCTGGAAGAGGTGGAACGTGAGCAGCATAGGATgaatcttaaacccaac
ggttacctcttatctagtggCAGGTTGACCTAAGGGCTGGCAGCAGTGGAGGCCCCCCACgaatcttaaacccaa
ggttacctcttatctagtggGAGGGGCCTCTTTAGCTCAGTCCAGTTGAGGCAGCAGAGCgaatcttaaacccaac
ggttacctcttatctagtggGGGTGTGGTGTGACCTGAATGTTGGGCACGTGGTCGTAACgaatcttaaacccaac
ggttacctcttatctagtggGAGAATCGCTCCAACCTGGGAGGCAGAGACTGTAATGAGCgaatcttaaacccaac
ggttacctcttatctagtggCTGCTCTACTCTAGCCTGTCTCAAAACAAAAAACAAGAAAgaatcttaaacccaactt
ggttacctcttatctagtggGAGAGAAGAAAGGCGGGGCACGATGCTTTTTAAGTCTACGgaatcttaaacccaac
ggttacctcttatctagtggCACTGACCCTTCCTTTACCCCCGCCCCCATCCCGTGGCCCgaatcttaaacccaa
ggttacctcttatctagtggCGGCGTTTTCTCTTCCCGGCTCCTCTGCCCCCAGCCCCACgaatcttaaacccaa
ggttacctcttatctagtggTCCCAGCCATTTCCTGCCCCAAGGCCGCCCACCCTCCTCCgaatcttaaacccaa
ggttacctcttatctagtggGCCGGGAAGCAGGGCTCCTGTTTTGCCTGTTTACCCTGGCgaatcttaaacccaac
ggttacctcttatctagtggCCAGCTGCGTAAGTCTCCGGCAGTGGGCACAGGCGGAGGGgaatcttaaacccaa
ggttacctcttatctagtggGCCTGCCTGGCACAAGACCCCCTGCTCCTGGAGGCAGCTCgaatcttaaacccaa
ggttacctcttatctagtggCGCTCAGCAGTGAGGCCTCTGAGTCACAGTGCAGCCCTGGgaatcttaaacccaa
ggttacctcttatctagtggGCCTTGTGGGACCCTCCTGGGCTTTCCTGTCCCTCAGAGCgaatcttaaacccaac
ggttacctcttatctagtggCTCTGATGCCAGAGAGACACTCTAGGTCTTTCGCGCCCTGgaatcttaaacccaac
ggttacctcttatctagtggGCCAGGCATTCAGAGATACATTCCTCAATCTCGGTCCGCCgaatcttaaacccaac
ggttacctcttatctagtggGTTCAAGTCAGCAGCCGAGTTACAGGAAAGCAGCCAGGGCgaatcttaaacccaa
ggttacctcttatctagtggTTTCGGGAGGAGAGGGCTGGGTCGCGTGTCCTGGCAGGGCgaatcttaaacccaa
ggttacctcttatctagtggGAAGCAGGAGCAGGAATACCATTTATCCATTGGCACAGGGgaatcttaaacccaac
ggttacctcttatctagtggCTTGTCCTGACCCAAAGATGCTGATGCTGTCCTTATACCCgaatcttaaacccaactt
ggttacctcttatctagtggCTGACTCAGAGGCACAGAGTTAGAAAGCAGCATGTCTGGGgaatcttaaacccaac
ggttacctcttatctagtggCTCCTCTCTCTCTGTCTCTCTTTCTCCCACGTGGCTGCTGgaatcttaaacccaactt
ggttacctcttatctagtggCATCGCTGAGGTGGATGGAGAAGCAGCTGGGGGCCTTGGCgaatcttaaacccaa
ggttacctcttatctagtggGGGTTAGTGTCTGGAGCTTGGGGGATGGGAATGGGTCCCGgaatcttaaacccaac
ggttacctcttatctagtggGGGGGGACATTTCTGCTGGCCCTGGCAGACACTGGACTAGgaatcttaaacccaac
ggttacctcttatctagtggCAGGCCTCAGCAGACTGAGCGGGAACCCCCACCAGCGAGGgaatcttaaaccca
ggttacctcttatctagtggCAGCGTTTCTCTTGGGCCTGTGTTTGGAACAAGTATCCTTgaatcttaaacccaactt
ggttacctcttatctagtggGGAGCGGAGGCATGAAAGGTGTGGCCGGTAGGGGTGGGGGgaatcttaaacccaa
ggttacctcttatctagtggCCGCCCTGGCCAGAGCCCACAAGCAGCGTCCCCTGGTCATgaatcttaaacccaa
ggttacctcttatctagtggTCTCCATAATCTCGCCCCTCCCTGACCAGGCCACCTGACGgaatcttaaacccaac
ggttacctcttatctagtggGAGGACTCTGTTTCTAGAAAAATGAGAGTGGGCATCCCTCgaatcttaaacccaact
ggttacctcttatctagtggGGGGGTATGCTGGGGTGGCTGGGAAAGGTGGTTGGGTGTCgaatcttaaacccaac
ggttacctcttatctagtggCTGCGTGGCAGGCCTAGGACTCTAAGTGCTTCCAGCCAGCgaatcttaaacccaac
ggttacctcttatctagtggACGGAGTCTTGCTCTGGAGTGCAATGGCATGAACTTGGCTgaatcttaaacccaact
ggttacctcttatctagtggCGTGAGCTACCACGCCCAGCCCCAGCCAGCTATATTAACTgaatcttaaacccaac
ggttacctcttatctagtggGGTTTAGAGAGTCACAGTAACTGGTCCTGGCTACACAGCCgaatcttaaacccaac

ggttacctcttatctagtggGGCCTAGAGTGCTCTCCTAACCGCTATGCCTGGGGTGAGGgaatcttaaacccaa
ggttacctcttatctagtggGCTGAAGCTAAAATAATCCGAGGGCTGGGCGAGGTGGCTCgaatcttaaacccaa
ggttacctcttatctagtggGCCACTGTACTCCACCCTGGGATACGGAGCAGGACTCTGCgaatcttaaacccaa
ggttacctcttatctagtggCCGAGAGTGCTTACCATGTGCCTACATTGTTCTGGGATACgaatcttaaacccaac
ggttacctcttatctagtggGAGGCAGCTCGGTTCCCCCAGGGACTCTCAACAGTTAGGGgaatcttaaacccaa
ggttacctcttatctagtggAGAGGCTGAGGCAGGAGGATGGAGTATGTCCAGGAGTTTGgaatcttaaacccaa
ggttacctcttatctagtggGCCTATAGTCCCAGCTACTCAGTAGGTGAAAGAGGAGGAGgaatcttaaacccaa
ggttacctcttatctagtggCACACCACTGCACTCAGTCTAGGCCACAGAGTGAGACTGCgaatcttaaacccaa
ggttacctcttatctagtggCCCTCATGTTCTCCCCTCTCCTGGCATTGCCTGGCACTTGgaatcttaaacccaac
ggttacctcttatctagtggATGAGTCTTCAAAGTGCTCTGATTCGTATGGTGGAGTGTGgaatcttaaacccaactt
ggttacctcttatctagtggATGGAGTTGAATGTTCAGAATCTTTGGAGTGCTGTCAGGGgaatcttaaacccaactt
ggttacctcttatctagtggCTTTGGTGTGAGAAATCCCCAAAGATTCAAACTTGTCACCgaatcttaaacccaact
ggttacctcttatctagtggTGAGGTCAGGAGTTTAAGACCAGCCGGTCCAACATGGTGAgaatcttaaacccaac
ggttacctcttatctagtggTGGTCTTGAGGTTTCGGTTTGGGCATAGGTTACCTTCTGCgaatcttaaacccaactt
ggttacctcttatctagtggTGAGTGATATGGAGACAAGTTTTAACATGGGCACTTTTTGgaatcttaaacccaacttg
ggttacctcttatctagtggAAGGCCAGATGTCAAAGTGGACTTTCTATTTTTGATGGAAgaatcttaaacccaactt
ggttacctcttatctagtggCAAGGATTTCTGGAACTGCAACTGGCCCATAGATGGTGGCgaatcttaaacccaac
ggttacctcttatctagtggTGTACAGCAAGCAGTTCTGTGACTTTCAGTCACCACTTGGgaatcttaaacccaact
ggttacctcttatctagtggTTTGAGACAGGGTCTTGCTTAGGCCGGAGAGCAGTGGTGCgaatcttaaacccaac
ggttacctcttatctagtggGCCTCCTGAATAGCTGGGACTTTGGGTGTGTGCCACCACGgaatcttaaacccaa
ggttacctcttatctagtggCCAGGCTGGTCTTCAACTCCAGAGCTCGAGTGATTCTCCCgaatcttaaacccaac
ggttacctcttatctagtggTGGGATTACCAGCATAAGCCACCACACCTGGCCTGAAGGCgaatcttaaacccaa
ggttacctcttatctagtggCCTCACAACCTTCTTATGCTTGTCTGTAGTTGGCTGTCTCgaatcttaaacccaactt
ggttacctcttatctagtggCAGAACAACCAGGTCTGGGCTTCTTATCTTGCGAGGCAGGgaatcttaaacccaac
ggttacctcttatctagtggTGGGTGCCCACCTCATACCGGCCATGACCTTCTATTGAGGgaatcttaaacccaac
ggttacctcttatctagtggTGGAGCTTGCTTCCTTCCTACGACCCTAGACTGCTCCTCCgaatcttaaacccaac
ggttacctcttatctagtggGAGTCCTGGCTGCCCTGGTCCTGTGGAATCCCGTCATGGCgaatcttaaacccaa
ggttacctcttatctagtggTTGGGCTTTAGAGTCACCTGGGGTTGAATCTCGGGCCTGCgaatcttaaacccaac
ggttacctcttatctagtggACCTGTCCTAGCCTCAATCGCCTTATCTGTACCATGGGGAgaatcttaaacccaac
ggttacctcttatctagtggCCCTCCTTTCGTGTGTCATCTCCACCTGGGTGTGGTGAGCgaatcttaaacccaac
ggttacctcttatctagtggGGCTGCGGCTGCCAGCGGTGATGAGAACTGATGACACTCCgaatcttaaacccaa
ggttacctcttatctagtggGCCTCTGGGCAGGGCCACTTCCTAGAACCATGGGGCTGCTgaatcttaaacccaa
ggttacctcttatctagtggTCGGCTTCCAGCTCTGGGCACTGGGGTGCGGTGTGGGACCgaatcttaaacccaa
ggttacctcttatctagtggTGCTCCTCTGCCAGGAACCGCCAGAACCAAAGTCTTCTCGgaatcttaaacccaa
ggttacctcttatctagtggAGCACAGGGCTGAGCACTCCGGAGAGGAGCCTTTGGGGGAgaatcttaaacccaa
ggttacctcttatctagtggGCTTCGGAGCTCAGGGAGTACTATGGGGGAGATTCACTAGgaatcttaaacccaac
ggttacctcttatctagtggGGCAGCCCTGTAGAGAAGACAAGGCAGCTGGGCATGGTGGgaatcttaaacccaa
ggttacctcttatctagtggAGCACTTGGTCAGGTGGCCATTTCTCTCCCTCCTACTCGGgaatcttaaacccaac
ggttacctcttatctagtggGGCGTGAACTTGGGATGCCACTGTGGACGTGTCCCCAGCCgaatcttaaacccaa
ggttacctcttatctagtggTCGTGTTGCCTTGGACCTATTACCTCACCTCTTCCTGGTGgaatcttaaacccaact
ggttacctcttatctagtggCCCCTTTCTCCTATGCTTCTGTTTTTCCTTCTACCCTTCCgaatcttaaacccaacttg
ggttacctcttatctagtggCCACCCAGCCACCCTTTCATCTAACCAGCCTTCCACCTAGgaatcttaaacccaa
ggttacctcttatctagtggTGGCACAGTTGGGCCACTTGTTTCTGATGGGAGCCACTTCgaatcttaaacccaac
ggttacctcttatctagtggAGATCACCCGTAATTATCACCCATCATTCTGACACTTAGCgaatcttaaacccaact
ggttacctcttatctagtggACATCTTGTTGTCTTGCTTTCAGATGGTTTTGTTTTTGACgaatcttaaacccaacttgt
ggttacctcttatctagtggTCATTATGCACCTGTACAGGAAAGGTACTCCATAAGTAGGgaatcttaaacccaact
ggttacctcttatctagtggGTACCAGGCACAGAGCAGGTACTGGGCGCATGTTGCCTGCgaatcttaaacccaa
ggttacctcttatctagtggCAGCCAAGTCAGTCCTGCGGGGCTTTCAGATTTCCTCTGCgaatcttaaacccaac
ggttacctcttatctagtggGTGTCTCTGATTGTTTTCTTGGCAGAGCCACCCACACCCCgaatcttaaacccaac
ggttacctcttatctagtggCACCACTCCACAGGCAATATTAGGGGTTGCTGAGGCAGCCgaatcttaaacccaa
ggttacctcttatctagtggAAAAATTTTAACTTTTCTGGCTGGGCGCGTTGGCTCACGCgaatcttaaacccaact
ggttacctcttatctagtggCCCCATCTTTATTAAATATACAAAGATTAGCCAGGCGTGGgaatcttaaacccaactt

ggttacctcttatctagtggGGGTCTCGCTTTTTCACCCAGGCTGGATTGCAATGATACGgaatcttaaacccaact
ggttacctcttatctagtggCCACCTCAGCCTCCCACAGGTGCACACCACTATGCCCAGCgaatcttaaacccaa
ggttacctcttatctagtggGTCTCAAACTCCTGGCCTCGGGCAATCCTGGACTCCCCAGgaatcttaaacccaa
ggttacctcttatctagtggGGGTTGAGCCATAGCCTGCAGCAAACTCTTAAAACATGAGgaatcttaaacccaac
ggttacctcttatctagtggTGACTACTGGTGTGGTTGGCCAGCTTTTCATGTTTATGGCgaatcttaaacccaactt
ggttacctcttatctagtggTTGTTAACTGGTCATTTCCTTTGGGGTGTTAGCAATTTTTgaatcttaaacccaacttgt
ggttacctcttatctagtggTTGCAGATGGTGGAACCGGGGTGCCCCACAGTAAGTCCCCgaatcttaaacccaa
ggttacctcttatctagtggACCAAACCATGCTTTGCAAGTGAATACAAGGATCTACCCCgaatcttaaacccaac
ggttacctcttatctagtggTATTTATTTTTCCGAGACCGTGTCTTGCTCTGTCACCCAGgaatcttaaacccaactt
ggttacctcttatctagtggGGGATTACAGGTGTCCGCCACTGCGCCAGCTAATTTTTTGgaatcttaaacccaact
ggttacctcttatctagtggCTCGAACTCCTGATCTCATGATCTTTCTGCCTCAGCCTCCgaatcttaaacccaac
ggttacctcttatctagtggGGCCAGTTGCCAGTTTTTAATGTTCTCAGGTGAGTGGCATgaatcttaaacccaactt
ggttacctcttatctagtggGCCCCACCTTAAACCCCTAAATCAGCTGTTGGCGTCGGGGgaatcttaaacccaa
ggttacctcttatctagtggGCACCAATTGCAATTTCTTACTTAGGTGTGTGCTTGTGGGgaatcttaaacccaactt
ggttacctcttatctagtggCAGCTTCTCTGCTAAGGACAGCGATCATGTCTCTTTTATCgaatcttaaacccaactt
ggttacctcttatctagtggTGAGGCCCCTCAGCTTGGACCTGGCAGATCTGGGACCCGGgaatcttaaacccaa
ggttacctcttatctagtggAGCTGGATCCCTCAGGACCCTGTGTACCTGTGGCCAGGGCgaatcttaaacccaa
ggttacctcttatctagtggGCCTGGATGAGAGTGGTAAGAGTTGCTGAGGTGCCAGGCCgaatcttaaacccaa
ggttacctcttatctagtggGCGCGGATGGAGCGAGATGTAGGGGAAAGGTTGGGACCGGgaatcttaaacccaa
ggttacctcttatctagtggGGCACATCCCTGCCCTAAACCCTGCATGGCTTTCCTGGGCgaatcttaaacccaa
ggttacctcttatctagtggGCTGCAGGTGGGGTGTGGTTTGCAGTGGGGGTGGGGCAGGgaatcttaaacccaa
ggttacctcttatctagtggGGGATGAAGGACACCTGGGGAGGGGAGAGAGCCTGCCGGCgaatcttaaaccca
ggttacctcttatctagtggGCCCTGCCAGTCCCCCGCTGCATGAGTCTGGGCATGCTTCgaatcttaaacccaa
ggttacctcttatctagtggGGCCTGCCGTCAGGGTTTGTGGGCAAGATTCAGTAACTGGgaatcttaaacccaac
ggttacctcttatctagtggCTCCCACCCCTCGCTCCTTCTCTTATTCTTTTCCTGTTGCgaatcttaaacccaactt
ggttacctcttatctagtggCAGTGGAGGCCGCGGGAACTAGAGATAGAGGTCCTCTTGCgaatcttaaacccaa
ggttacctcttatctagtggTGGAGGTGACAGAGGCCATCTTGTCCTAGAGGGACCTGGCgaatcttaaacccaa
ggttacctcttatctagtggCTGAGTGCTTGGGATTCATATGGGGCTGTAGACGGCAGGCgaatcttaaacccaac
ggttacctcttatctagtggTCCCACATCTCATTCTTAGAGCTGTGTGGCCCTGGGCGAGgaatcttaaacccaac
ggttacctcttatctagtggCAGGATGGGGAGGTGGTTGAAATCATCCAGGGGTCACTGGgaatcttaaacccaa
ggttacctcttatctagtggCGGCAGCCTGAGGATGGGGATAAACCTGGGTGTGGCAGACgaatcttaaacccaa
ggttacctcttatctagtggCCCCCAAAGAGGTTCACATCCTGATCTCTGGAACATGGGGgaatcttaaacccaac
ggttacctcttatctagtggCCAGAGGGCCCAGTCTCATCACAGAGTCCTTACAAGTGGGgaatcttaaacccaa
ggttacctcttatctagtggTGCTGGCTTTGAGGATGGAGGAAGGAGCCATGAGACAAGGgaatcttaaacccaa
ggttacctcttatctagtggGGAACTCCCCCAGAGCCTCCAGGAAGGAACACAGTCCTGCgaatcttaaacccaa
ggttacctcttatctagtggTGTTCCGGCAGCCCTAGGAAGTGAATGCAATGGGCGCAGCgaatcttaaacccaa
ggttacctcttatctagtggGGCATCCAGCCTGTGGGGAGTGGGAGCTGAGGGCAGTCCAgaatcttaaacccaa
ggttacctcttatctagtggCATCAGTGAGTTGGGCCACTGAAGCTGCCTGGTGACCTGGgaatcttaaacccaa
ggttacctcttatctagtggGCCCATGGCTTCTCCCTCTGACAGCGCAGCCCTGCCTCCGgaatcttaaacccaa
ggttacctcttatctagtggTGGCGTGTTGGTGCATTAGGGACCCCTTCCTCTCCTGCCCgaatcttaaacccaac
ggttacctcttatctagtggTGGCGTTCTCAGGAGCCCTTTACCAGTACCCTTAGGCCCCgaatcttaaacccaac
ggttacctcttatctagtggCTGCTGGCTGTGTCCCTGCTCTGGGCCCAGCACCCTCATGgaatcttaaacccaa
ggttacctcttatctagtggGGCTGCCCCTGGGTCTGCTTTCCTGCCCCTGACCCACAGGgaatcttaaacccaa
ggttacctcttatctagtggGCTGGTGCTCTCTGAGCTGAGGCCCACTGCTGACCTGCTGgaatcttaaacccaa
ggttacctcttatctagtggAGGCACTGGGACATCCTGCCTTCCTTCAGTGTCCCCTCCCgaatcttaaacccaa
ggttacctcttatctagtggGACAGGAGTGGGCTGGGTGAGCATGGGGTGGTCGCAGGGGgaatcttaaacccaa
ggttacctcttatctagtggCCTGCTATCTGGTGGGTTCCAGGATGGTGACAGACAGTGGgaatcttaaacccaac
ggttacctcttatctagtggTCACTGGTGAAGGCAGACATCACTGCAGGAGACCAGACTGgaatcttaaacccaa
ggttacctcttatctagtggCAGGGAAGATCTTGGGCTCCGAGACCTAGGCAGATCTTGCgaatcttaaacccaa
ggttacctcttatctagtggCCGACCCTCCGCAGATGCTACACATTATGCCAGGTAAACAgaatcttaaacccaac
ggttacctcttatctagtggAGGCTCCTTGTGCCTGTGAGGAGTTTGCACTAATGCTATGgaatcttaaacccaact
ggttacctcttatctagtggCCCAAGTCATCAGCTGGGCAGTGCATGGGCTGCTCGCCTGgaatcttaaacccaa

ggttacctcttatctagtggGCTCAAGACCTGCCTGCGGCCTTCCTCGGGCTTGCTTGCCgaatcttaaacccaa
ggttacctcttatctagtggCACTTGCCTGGGAGGGTGGAGGAGGAGGGGTCCAAGCCGCgaatcttaaaccca
ggttacctcttatctagtggTCAGCTCCTCTGCCTGGAACACCCCCATTTACTCCTCCGCgaatcttaaacccaa
ggttacctcttatctagtggCGGCAGTATCCCGGGCTTGGCCAGCTTGCTGACACCTGGCgaatcttaaacccaa
ggttacctcttatctagtggGCGATGCTTCTGGAGGATGCTCCGCTGGGGAGGGCTCTGCgaatcttaaacccaa
ggttacctcttatctagtggTCGATCCTCACTCTGCCTTCCTGTCTGACAAAGGCCGGGCgaatcttaaacccaa
ggttacctcttatctagtggGAGACATGGTGGCGGGTGGGGTGGGGAGAAGGGGACTGGGgaatcttaaaccca
ggttacctcttatctagtggAAATGTCCCAGAGGCCCTAGGTCACTCACCCTCTGCCCCCgaatcttaaacccaa
ggttacctcttatctagtggGCTGCCTCCTGCTGAATCTGAAGTATGCCTTCACTGAGGGgaatcttaaacccaac
ggttacctcttatctagtggCCCCCGGGGCATGAGGCTCCTTCCTCTGTGATCCCTCTGTgaatcttaaacccaa
ggttacctcttatctagtggTCTGTAGTCCTTCTAGAAGCTAGGTAACTCTTCTAAGTCCgaatcttaaacccaactt
ggttacctcttatctagtggCCTAGAGCAGTTCTGCGAGGTGGTGGGAAGACCTAGAGAGgaatcttaaacccaa
ggttacctcttatctagtggGCTCTCTGAGCTATAAGGCCTCAGGGTCTGTGGCATTCCCgaatcttaaacccaac
ggttacctcttatctagtggGAGTTCCACCACCCAGAAATGAAACAGTGTTCATTTCCAGgaatcttaaacccaact
ggttacctcttatctagtggTATGGTTGATTTTACCAATTTCCTGTCACTTGAATCACAGgaatcttaaacccaacttg
ggttacctcttatctagtggGCCTCGGCTTCCAGAGTAGCTGGGGTTACGGGTGCATGCCgaatcttaaacccaa
ggttacctcttatctagtggCCACCCTCAGATTATCCACCCTCCTACACCTCCCACAGTGgaatcttaaacccaa
ggttacctcttatctagtggGTCCTGTGTGTATAATTTTGCATCCTATTATTTCTCCTTCgaatcttaaacccaacttgt
ggttacctcttatctagtggCAGGCACACTTTGGTATTCAGGAGTTGCTTCTGCTGGACCgaatcttaaacccaact
ggttacctcttatctagtggTGACTGGAGGCTTAGGGGTCTGGTTCAGGGTGGACTCAACgaatcttaaacccaac
ggttacctcttatctagtggGCCCAGGCTGAAGCGCAATGGCATGAGTTCGGCTCACTGCgaatcttaaacccaa
ggttacctcttatctagtggGTGAGCCACCACGCTTGGCCTATTTTTTATTATTAAAAAAgaatcttaaacccaacttg
ggttacctcttatctagtggTGAGCTTTGCAGTGTTGGACTAATATCACAAGTCGAAGGGgaatcttaaacccaact
ggttacctcttatctagtggCTGAGACCTTAAGGGCCTGGTTGTTTCTACTGTTCCCTGCgaatcttaaacccaact
ggttacctcttatctagtggCCTGCAAGCTTGAGAATGTTTGTCATTTACAAATTTTACCgaatcttaaacccaacttg
ggttacctcttatctagtggTGTACCAGATTGAGTAGTGTCCCCCCAAAATTCATGCCTGgaatcttaaacccaact
ggttacctcttatctagtggTGGAGTCTCGCTCTGTCACTCAGGAAGGAGTGCAGTGGCGgaatcttaaacccaa
ggttacctcttatctagtggGGCACCCGCCACCAGGCTTGGCTATTTTTTGTACTTTTAGgaatcttaaacccaact
ggttacctcttatctagtggGCACGAGCCACTGTGCCTGGTGGACATTGGGTCTTTTTAGgaatcttaaacccaact
ggttacctcttatctagtggGGGTGGGGGCTAAAGCCAATGACCCATGTCCTCAGAGGAGgaatcttaaacccaa
ggttacctcttatctagtggTTGTAGAGATACAGGGAGAAGGCCATGTGACCACAGAGGCgaatcttaaacccaa
ggttacctcttatctagtggGCCAAGGAGCTCCAAGGATTGCAGGAAGCCACCAGATGCCgaatcttaaacccaa
ggttacctcttatctagtggAGGATCTTTCATGGAGCCTTCAGAGGGAGCACAGTCCCGCgaatcttaaacccaa
ggttacctcttatctagtggTGGACATCTAGTCCCCAGAACTGTGAGAGAGTGCATCTGCgaatcttaaacccaac
ggttacctcttatctagtggTGAGGGAAGTGGAGCGGGAGAATGAGCGGCACTCCCTCCCgaatcttaaaccca
ggttacctcttatctagtggAGAACTGGCCATTTCATTCTGCATTTAGAGCTCTGGGAGCgaatcttaaacccaact
ggttacctcttatctagtggCCTCTTCAGCTAGAGGAAGCTCCTTCTCCCACTTACTAGCgaatcttaaacccaact
ggttacctcttatctagtggGGGCAGATGGCTTAACATTTCTGGGCCTCAGTGACCTTACgaatcttaaacccaact
ggttacctcttatctagtggGCATGTAAAGCCCCAGCATGCAGCCTGAAAATGTGAGCCCgaatcttaaacccaa
ggttacctcttatctagtggGTGACGGCCACAGTCACGTCTCCTTTCTGCCTCCTCCTCCgaatcttaaacccaa
ggttacctcttatctagtggTGTCCAGTGACTAGGGAGGGCAGGAAGTTCAGAGGGTGAAgaatcttaaacccaa
ggttacctcttatctagtggACCCTGTGGGTGGGTTGAAAACCATGCCCTGTCTACCTCCgaatcttaaacccaac
ggttacctcttatctagtggTACTGCCTGCTGTGTGCCTCTCCAGGGAGCACCAGGAGGGgaatcttaaacccaa
ggttacctcttatctagtggGCCTTTTTGCTGACGCGATGCCCAGGAGCCCTCTCAGGGGgaatcttaaacccaa
ggttacctcttatctagtggGGATGCTTCACATCCTAGGTTTGTGTGATGTCGGCACAGCgaatcttaaacccaact
ggttacctcttatctagtggGCTGAGGAAGGGATGCGCTGGACTCCCAGTGTCCAGGGGCgaatcttaaaccca
ggttacctcttatctagtggAGGCAGAGTCCACCAGAATCGGGTAAAGTATGTGACTGCCgaatcttaaacccaa
ggttacctcttatctagtggGGGCATGACGTCAACTGTGAGGAGAAGAGATCAAGATACAgaatcttaaacccaac
ggttacctcttatctagtggCCCACACCCAGAGAGGAATATCTGCAAGTTCTGGTGCCAGgaatcttaaacccaa
ggttacctcttatctagtggGTACATTTAAAAATAACTGGCTGAGCGCAGTGGCTCACGCgaatcttaaacccaact
ggttacctcttatctagtggGAACCCTCAAAAAACTAAAAGAGGCCGGGTGTGGTGGCTCgaatcttaaacccaa
ggttacctcttatctagtggAGCCGGGCTTGGTGGTGGGTGCCTGTGACCCAAGCTACTCgaatcttaaacccaa

ggttacctcttatctagtggGCCTGGGTGGCAGAGAGAGTATCCGTCTCAAATAAATAAAgaatcttaaacccaact
ggttacctcttatctagtggCCCTGAAGGGATTATTATGCATTGTGTGCCTCTATCAAAAgaatcttaaacccaacttg
ggttacctcttatctagtggCAATGGAAGCATCCAGAAAACCCTGGCTCGGAATCTGAGCgaatcttaaacccaac
ggttacctcttatctagtggCAGGTCTCACATGGGGGACAGTCTTGCCACTGTCAAAGGCgaatcttaaacccaac
ggttacctcttatctagtggCTTTGTCCCCACGCCTGCCCCTTGCTTCAGATCTTCTAGGgaatcttaaacccaact
ggttacctcttatctagtggCCACTGAGCTGCTCTCTCCTACTGAGCCCCGCATGGTTCCgaatcttaaacccaac
ggttacctcttatctagtggCTGCCCTCTCACTTGCTGGTGTTGGCCTGCTCTGGCTGTTgaatcttaaacccaact
ggttacctcttatctagtggGACAAGGGCCTGCCTCTGGCCTCTGGGTGTGTCCTCCTGCgaatcttaaacccaac
ggttacctcttatctagtggCAGGCTCAGACTGAATGCCTGCTTCTCCTTGCAGCCTTCTgaatcttaaacccaact
ggttacctcttatctagtggCTTGCTGGGTAAGCACAGCCCTGGATTACTTGGCATCTGGgaatcttaaacccaact
ggttacctcttatctagtggAAGGAGACCCCTTCTAACTTCTGGAAGCACAAAAGTGTGTgaatcttaaacccaact
ggttacctcttatctagtggGGAGCATTCTTGGCTGCTGAAAAATCATCATGGCTAGCTTgaatcttaaacccaactt
ggttacctcttatctagtggTGTGTAGGAGGGACCCGGCGTGCAATTTGCATGGATTGTCgaatcttaaacccaact
ggttacctcttatctagtggCAGCAGGTAAAAGAGCTGGGAGGGAGTAAAGGGAGTTGTTgaatcttaaacccaac
ggttacctcttatctagtggCCCCTCTTGGTGTACACAGTATGCAATGGAGGCAGGGTGCgaatcttaaacccaac
ggttacctcttatctagtggGCTTCCCAGAGGAGGCCTTTGCCCCACAGAATCTTTGCTCgaatcttaaacccaac
ggttacctcttatctagtggCTTTGCTGCCTCTATGAATCAATGATTCTGTGTTGTCCATgaatcttaaacccaacttg
ggttacctcttatctagtggTGGTCTCCATCATTCTGGCCAAGCACAGTGACTAACATACgaatcttaaacccaact
ggttacctcttatctagtggCTGCCAAGCAGCCAAGCACCCTATATCTGGGAGATGTAGAgaatcttaaacccaac
ggttacctcttatctagtggGGGAACATGCAAAGCTCAGGGCAGCTCAATTTGGGGCGAGgaatcttaaacccaac
ggttacctcttatctagtggGCAGTTCTGCATCTTGGGGTGCAGTGCTAGAGTGAATGACgaatcttaaacccaact
ggttacctcttatctagtggGCCATGGGAGCACTGGAGAAGGGGCAGCAGTGGATTCAGCgaatcttaaacccaa
ggttacctcttatctagtggTGAGAAGATTGGGAAAGGCTGCACAGAGGAGGAGAAGGGCgaatcttaaacccaa
ggttacctcttatctagtggGGGTGGGTAACAGTTAGAGGAACAGCAGGTACAGAAGCCCgaatcttaaacccaac
ggttacctcttatctagtggGACACATCCACCATGTCCAGAGATCAGGGATGGTTGCAGCgaatcttaaacccaac
ggttacctcttatctagtggCAATCCCAGTTCATCTTATTCCCACCCATATGGCTGGTGCgaatcttaaacccaactt
ggttacctcttatctagtggGACTGGTACCTTCCTAGGAGGGGTGTTGAGAGTCTGAGCCgaatcttaaacccaac
ggttacctcttatctagtggTCTTGGTGTGCAGTGTCTCTGATCTGAAATGTGCACCGTGgaatcttaaacccaactt
ggttacctcttatctagtggGGCCTGATTGTGAAGCGGGTTTGAGACAGAAGTGAAGCAGgaatcttaaacccaac
ggttacctcttatctagtggGGCTTTTGCCCACCCTTCCCTTCCTTCACATCCCTGGTGGgaatcttaaacccaact
ggttacctcttatctagtggTGGCCTGCCACCTGGCCCTTCCCTCGCTCTGCAGTCTTGGgaatcttaaacccaac
ggttacctcttatctagtggGTGGGTGTGTGGCCCTGGGCTGCACTGTGCCTGCTGGAGCgaatcttaaacccaa
ggttacctcttatctagtggGCCCTTTCACCATGCGTGCTTCTTTGGATGTGACCCCTTGgaatcttaaacccaactt
ggttacctcttatctagtggCAGTTTTCTATGGTGTTTGGGGGAGAAATTGGGGTGCTGGgaatcttaaacccaactt
ggttacctcttatctagtggCGTCCCCTTCCTCACAGCTGGGGTCTGTTCTTCCTCCGGGgaatcttaaacccaac
ggttacctcttatctagtggGGGTACACAGAGGTGGCCTGATTTCTGGAGCCATAGCAAGgaatcttaaacccaac
ggttacctcttatctagtggCTGTCTCTGTCTCTGTCTCTGTCTCTGACTCTGTCTGTCTgaatcttaaacccaacttg
ggttacctcttatctagtggCTCTCTCTCTCTCTCTTCTTTCTCCCTCTCCCTTTTTCACgaatcttaaacccaacttg
ggttacctcttatctagtggGGAATACTTCATCATTAGACATATTTCGAGCCTCCGTTCCgaatcttaaacccaacttg
ggttacctcttatctagtggGGACTGAAGGATTTAAAGGGATGTGGCCAAGTGTGCAAAGgaatcttaaacccaact
ggttacctcttatctagtggGGCGCTCCACATCTCCGTGATATAGCGTGTACTCCTTTCCgaatcttaaacccaact
ggttacctcttatctagtggTCCTAGTTCAGTGACCTTGGACAAGGCACCCTGGGGAGCCgaatcttaaacccaac
ggttacctcttatctagtggGGGAGACCCTGAGTGTTTGGACCCCGCCTGTGCTCAGCACgaatcttaaacccaa
ggttacctcttatctagtggAGGGCAGGTCGTGGGCTCCTTCTTCTCCCTTTCAGCAACCgaatcttaaacccaac
ggttacctcttatctagtggTCTCTGGGGAGTTCTGTGGCACCCCTGCCTTACCTGGGGGgaatcttaaacccaac
ggttacctcttatctagtggACGGGGTTGGGCCTCCGGATCTGCATTGGAGCAAGTCCACgaatcttaaacccaac
ggttacctcttatctagtggGGCACACTCAGGTTTGAGAGCCACTGACCTAGAGGGGAAGgaatcttaaacccaac
ggttacctcttatctagtggAAGAGACCAGTACTGTGACTTGCACCATGGAGACACCTGGgaatcttaaacccaac
ggttacctcttatctagtggGAATGCTGTAATGCCACCCAGTTCCCACAGAGCTCTGGGGgaatcttaaacccaac
ggttacctcttatctagtggCCCAAGTCAGGGTTCTTTTCTGGAGTGCTATTCTGACAGGgaatcttaaacccaactt
ggttacctcttatctagtggTCCCCTTCTCTCCCTTTCCTCTCATCTCCTCCCGCTTTCGgaatcttaaacccaactt
ggttacctcttatctagtggCTTCCCATCTCCTGGCCCCTCCAGTTCTGAAGGGATGGGGgaatcttaaacccaac

ggttacctcttatctagtggGCAGGCGGAGGAGGGAACCATGTCAGCTCCTAGGTGCCCCgaatcttaaacccaa
ggttacctcttatctagtggCCTGTCCTCCGTGGCGCTTGGGTGACAGCAGAGCAGAGCCgaatcttaaacccaa
ggttacctcttatctagtggGGGCTGTTCGTGCTGTCATCGCTTTCATTTGCCCCATCCCgaatcttaaacccaact
ggttacctcttatctagtggCTTAGCGAGATGTGCCTGCGTGACTTGAGTAGAGACCTGCgaatcttaaacccaac
ggttacctcttatctagtggGTCAGGAGCGTGACCTCCCCGGAAGCCCAGCTAATCCAGGgaatcttaaacccaa
ggttacctcttatctagtggGCTGTGGGATGTGGGAGAGGACAGGTTTAGCAGAGTGGCGgaatcttaaacccaac
ggttacctcttatctagtggCTCGCCGAGGAGGACAACTCCCCAGCGCCTGAGACAGCGGgaatcttaaaccca
ggttacctcttatctagtggAAGTGAAGCAGGCTGTGGTGTGATGATGGCCCCCGAGGGGgaatcttaaacccaa
ggttacctcttatctagtggCAAGGCCACCCTCTTCAGCCACACAAACTGGGCAAGGGGCgaatcttaaacccaa
ggttacctcttatctagtggGATATGAGACCTGTCTGTGGGGTTGTGAGGATACATGGGGgaatcttaaacccaact
ggttacctcttatctagtggCCCCGAGGGTCTCTCTCTCTCTCTTCCCAACTTAGATTCCgaatcttaaacccaact
ggttacctcttatctagtggCAACCCAGAAAAGCAGCTCTGATCTCTCGTGGAGCTTCCCgaatcttaaacccaac
ggttacctcttatctagtggCCGCACATCTACTACTGTTTAGGATAACATTTTGACTTAGgaatcttaaacccaacttg
ggttacctcttatctagtggGTCTCTGTCAAACTTAGTGAACTTTCTTTCCAGCCTGGGCgaatcttaaacccaactt
ggttacctcttatctagtggACCCCCCTCCCCCACAAAAAACCTTACTGAACTTTCTTGGgaatcttaaacccaact
ggttacctcttatctagtggCTGTCACCTTGAGTCATTGTTTGACGGCTCAGGTCCTTGGgaatcttaaacccaactt
ggttacctcttatctagtggTTGGACCTTTGCTGACCTTTCCCTGAGGGATTTTGTGGGCgaatcttaaacccaactt
ggttacctcttatctagtggCCCCACAAATTTTGAAGGTCTGTCACTTGGAGAAGAACAGgaatcttaaacccaact
ggttacctcttatctagtggTCATCTTGAGCCTAAAGATGAAGGAGAGTGGAGGTGAGCCgaatcttaaacccaac
ggttacctcttatctagtggGGTGCAAGGACCTCACTTGGGTTATCTCTAGCCTTCCCCCgaatcttaaacccaac
ggttacctcttatctagtggGCTTCGCAGATTTGGAAGCCTCAGAGCTAGAATTCTCCCCgaatcttaaacccaact
ggttacctcttatctagtggCCCTTCCCTGCCACTGTCTGCCTGTCTTCCTGTCTGCCTCgaatcttaaacccaact
ggttacctcttatctagtggGAAGGCTATCGTGAGGTAACCCTGGGATCCTTATTGGCACgaatcttaaacccaact
ggttacctcttatctagtggGCGAGAAATACCCACAGCCTCCCTGGATGCTGCCCACAGGgaatcttaaacccaa
ggttacctcttatctagtggCTCACCTGCAAGGGGGTGGGGAGAAGTGGCCAGGGGTGGGgaatcttaaacccaa
ggttacctcttatctagtggCTCCCTGAGCCTAGGGGGAATGATATTATGGCTTCAGGCCgaatcttaaacccaac
ggttacctcttatctagtggCTACTCTGACAGGGAGCCAGTCTGGAGAAGGATGTCATGGgaatcttaaacccaac
ggttacctcttatctagtggCCCTATTGGGGCTGTTTTGGTCAACTCCTGGCAGTAGATGgaatcttaaacccaact
ggttacctcttatctagtggTCCTTGCATTTCCCAGCAAAGTGCCATTCCCATCTCTTTCgaatcttaaacccaactt
ggttacctcttatctagtggAGGCAAGGCAGACTAGTGGTTAAGAACAAAGACCCAGGGCgaatcttaaacccaac
ggttacctcttatctagtggCAGCACTTCGGGAGGCCAAGGCCGGGGTATCAGTTGAGGTgaatcttaaacccaa
ggttacctcttatctagtggGGCAGTGCCTGGGCTCGTATCCTAGCTCTGTCTCTTGCTGgaatcttaaacccaact
ggttacctcttatctagtggGGCTTGGGCATACTTGGGCTTTGCTGTGCACCAGTGTTCTgaatcttaaacccaact
ggttacctcttatctagtggGGTCATGCCACCTTCGACGGACAATGCGCCAGCACCCACCgaatcttaaacccaa
ggttacctcttatctagtggCCAGTTCCTCCTCTAGGCATTGTGCCCAGCGCACCCTTCTgaatcttaaacccaac
ggttacctcttatctagtggCTCACTCTGCATCTGCAGGTCTTCGCTGGGTGATCTTGGGgaatcttaaacccaact
ggttacctcttatctagtggTACTTCTCCAGGCCTTACTTCTCAGCTCTTCAGTGAGGACgaatcttaaacccaactt
ggttacctcttatctagtggCCTTTCCTCCTGTTCCTCAGCTTCTGCCAGCCCGCCAGGGgaatcttaaacccaac
ggttacctcttatctagtggATGTCCTCGCTTATGGACATGTCCTGCTTCTCTGGAAATCgaatcttaaacccaactt
ggttacctcttatctagtggCATGTTTGATATGATTTCTGGCTTCTAGCACTGTGCTAGGgaatcttaaacccaacttg
ggttacctcttatctagtggCACTGTCTGTGTAGCTGGAATTAATCACCCACTATTCCAGgaatcttaaacccaactt
ggttacctcttatctagtggGTGGCATGATCACTGCTCATTGCAACTTCCGCCTGCTGGGgaatcttaaacccaac
ggttacctcttatctagtggGCACTGTGCCGGACACGCATGGGTTGCTGGAGGCTGGCAGgaatcttaaacccaa
ggttacctcttatctagtggGCAGGGCTCTGCTGCATCATGCCCATGAAGGTGCCTGCTGgaatcttaaacccaac
ggttacctcttatctagtggGGTTCTATACTGGCTGCTTGCCTCCTCCTTGTGGAGACCCgaatcttaaacccaact
ggttacctcttatctagtggGGACTGGGTTCGGTTTGCATCTCCCGAGTCTGACTCGTGGgaatcttaaacccaac
ggttacctcttatctagtggCTGCTTGTGTTGTGAGCCCACTGTGTGCCGGAGCACTCTCgaatcttaaacccaac
ggttacctcttatctagtggGTCTTGTCCGCAGATGAACTCCAGTGGACACATGTTTCTAgaatcttaaacccaactt
ggttacctcttatctagtggGGTAGCGATGGAGTTTCAACATGTTAGTTAGGCCGGTCTCgaatcttaaacccaact
ggttacctcttatctagtggCACCTAGGCTGTTGGCTCACTGCAACCTCCACTGACTTCCgaatcttaaacccaac
ggttacctcttatctagtggTAGCTGGGACTACAGGTGCGCCACCACTGCCAGCTAATTTgaatcttaaacccaac
ggttacctcttatctagtggTCACACCTTGGTTCCTGTGGCTCCCATCCCCCCTCCCCTGgaatcttaaacccaac

ggttacctcttatctagtggCGGGTGAGTGTGACGGGAGCAGGCACTCGAAGGCTTGGGAgaatcttaaacccaa
ggttacctcttatctagtggCGAAGCAAAAATAGAGCCTGTCAGCGGGGGGACAGCTGCCgaatcttaaacccaa
ggttacctcttatctagtggGATCTCCCCGATTTATCTTCCAGGTTAGGAAAGCCCCAGCgaatcttaaacccaac
ggttacctcttatctagtggTGTGAGCACAGCCAGAGAAATTGCCACCTCTCCCTTCCTCgaatcttaaacccaac
ggttacctcttatctagtggAAGGAGACATCGTCTTTTTGGCATCAGTTTTGAAGAAATCgaatcttaaacccaactt
ggttacctcttatctagtggCGTGAGCCACCACACCCAGCCAGTTTTGAAGAAATCTTACgaatcttaaacccaac
ggttacctcttatctagtggCTGCTGCTACAACTGCTGCCCCATTGACCGGGTCTCAGAGgaatcttaaacccaa
ggttacctcttatctagtggGCACATACACGCACACACATGCCTGCACACACACACTTGCgaatcttaaacccaa
ggttacctcttatctagtggTGCGGAGTCCAGAAGGCTCAGAGACAGAGAGAGAGAGAGGgaatcttaaacccaa
ggttacctcttatctagtggCGGTGCTCCCCTCAAAGCTTTAGGACCAGGGACGCCTTCCgaatcttaaacccaa
ggttacctcttatctagtggCTGTACCTCATACCCCATTTACCAGCCTGTGCCAACCCTGgaatcttaaacccaac
ggttacctcttatctagtggGCCCGTCCTATATTGAGGGGGAGTGGATGCGCAGAGGTGGgaatcttaaacccaa
ggttacctcttatctagtggCGGCAGGGCCTCAGCTAGTGGAAATACCTGGTGGGGAGGGgaatcttaaacccaa
ggttacctcttatctagtggAATGACGAGTTAATGGGTGTAGCACACAAGCATGGCACATgaatcttaaacccaac
ggttacctcttatctagtggGGGGCTGGGGCCACGTTTTCTTGTGGGGCAATTCCGTGTCgaatcttaaacccaac
ggttacctcttatctagtggGTGTCCTCTGGGGAAGGGTCAGCTGTGAGGCAGTGGGGCGgaatcttaaacccaa
ggttacctcttatctagtggGGGAAACCTGGAAGCTGCCGATTCCCAGGCCTTGAAGGGGgaatcttaaacccaa
ggttacctcttatctagtggTCTTTGACATTCCAGCAAACATATTTTCACAGGAAAACTCgaatcttaaacccaactt
ggttacctcttatctagtggCGAATGTTCTCAAACTGAACCTGAGTGTGGAGCCAGCACCgaatcttaaacccaac
ggttacctcttatctagtggTCCAGCATTTTGGAAGCCCCCTCCCAGTCAGTCACTCAGGgaatcttaaacccaac
ggttacctcttatctagtggACATCATCACACACTCCCCTGTGTCTTGCTTCTTTCACTCgaatcttaaacccaactt
ggttacctcttatctagtggCATGTGGTACGTGCATTTTCAGATCATTCTCACTGGTCGCgaatcttaaacccaactt
ggttacctcttatctagtggCTACTGTTGATGGGCATTTGAGTAGTTTCCAGTGTTTGGCgaatcttaaacccaactt
ggttacctcttatctagtggAGAGCTGCTGTGATATCCTTAGACTTGTCTTCTGGTGAACgaatcttaaacccaactt
ggttacctcttatctagtggCACGGAGTGGAGCTGCAGGGTCATAGGGTTTGGGTATATTgaatcttaaacccaac
ggttacctcttatctagtggTCAGAGTGTCTGTTCACCAGCACGTATGAATTCCAGTTCCgaatcttaaacccaact
ggttacctcttatctagtggACAGCAAGTAATCAGGAGAGAAAAGAAAATGCTCCCTGAGgaatcttaaacccaac
ggttacctcttatctagtggGCCACACATTGTGTACTTTTGCAGGCGGAGGGCGAGGTGCgaatcttaaacccaa
ggttacctcttatctagtggGCAGTCCTTGCCATTGGTCAGGGACCACCACTTGACTGCAgaatcttaaacccaac
ggttacctcttatctagtggCCCCAACAGAACAAACTTGGCTTCAGACTTGCCTGTGGGCgaatcttaaacccaac
ggttacctcttatctagtggGACTGTGTCCCCTGTTCACTCCCCTGTCATTAGCTGAGGCgaatcttaaacccaac
ggttacctcttatctagtggGCTCCCCTTCCTGTTGTAAGTGGCCCCTCTCAGCTGACGGgaatcttaaacccaac
ggttacctcttatctagtggCTGGGCTGCTTCCCAGGGCTGGCAGGTCTTGGTGGTCACTgaatcttaaacccaa
ggttacctcttatctagtggGAGGAGTGAGTGTGTGCAGTCCTGGAGTCACTGCTGTGGGgaatcttaaacccaac
ggttacctcttatctagtggGGAGCCTCTTCCATTTGATGTGAGAATGATAGATCCGTCAgaatcttaaacccaactt
ggttacctcttatctagtggTGCAACCTGTCACAGTCCATTCAGGCTGGTATAACAAACCgaatcttaaacccaac
ggttacctcttatctagtggGGTTCTGGAGGTGGGGAAGTCCAAGATTAAGGCTCAGACAgaatcttaaacccaac
ggttacctcttatctagtggCTGGTGAGGGCCTGTTTCCTGGTTTAGCAATGATGCCTTCgaatcttaaacccaact
ggttacctcttatctagtggGCCCTCCAGCCTCATGACCGAATCACTTTTCAAAGGCCTCgaatcttaaacccaac
ggttacctcttatctagtggGGCAGTTAAGAGTTCAACGTATGAATTTAGGGGTCGGGGGgaatcttaaacccaac
ggttacctcttatctagtggCCTCACAGTAGTTCTATGAAATGAGTAGATGTTGTTTACCgaatcttaaacccaacttg
ggttacctcttatctagtggACATTTCCCCAGAGTAAGAAACGTAGTAAATTTGGGAGTCgaatcttaaacccaactt
ggttacctcttatctagtggGGGACAAGAAGGCCAGATGCAGTGGCTAACACCTCTAATCgaatcttaaacccaa
ggttacctcttatctagtggGGGACAAGAGGAAGTGACCTGCAGAGCACATATTGGGTTCgaatcttaaacccaac
ggttacctcttatctagtggGAAATTAGAACCACTGTTTTTCTCTGGCTCTTTTCATGACgaatcttaaacccaacttg
ggttacctcttatctagtggGTTGGTTTTCAGCAGTTCAACTATTATATACTTAGGTGTGgaatcttaaacccaacttg
ggttacctcttatctagtggCACTTTCATCCCTTCCCACCCCCGATATTGATCTTGCTCCgaatcttaaacccaact
ggttacctcttatctagtggCTGTTAGGCTTTTTGATATTGTCCTACAGGTCTCTGAGGCgaatcttaaacccaactt
ggttacctcttatctagtggTGGCCTCTCCTCAAATTGTACTGACTCTTGGTTCCTCTTCgaatcttaaacccaactt
ggttacctcttatctagtggTTGTAGAATTTCCATTTGGCTCTTTCTTTGCTAAGCTTTCgaatcttaaacccaacttgt
ggttacctcttatctagtggTCCAGAAACTGGGTCATCTCAGGATCATGGGGACGGTCCCgaatcttaaacccaa
ggttacctcttatctagtggCACAGACATTGTGAGTGATACTTGGTAAGGACTGATTCTGgaatcttaaacccaactt

ggttacctcttatctagtggATCTAAGACTTCAAATTCTGACTCCCTAGTATTGGGCAGCgaatcttaaacccaactt
ggttacctcttatctagtggGGCTTTAACTGGGCTACTCCATACAAACTTGGCTCAGGAGgaatcttaaacccaact
ggttacctcttatctagtggAGGGAACATTTGTACACAGAATTTGAACTCCCCTTTTTGGgaatcttaaacccaactt
ggttacctcttatctagtggGTGTTAGCCACTCACATGTTGCAGAACCGGGCCTACTCTCgaatcttaaacccaac
ggttacctcttatctagtggGACCAAAAGCAGGAACTCACTGGCTTGCTGAATGTCCAGGgaatcttaaacccaac
ggttacctcttatctagtggACTCCAGAATGTCCTACTTTTGGTCATCAGGTGCCTAGATgaatcttaaacccaactt
ggttacctcttatctagtggACCAGGGAACTGCTTCAATAGGAACTACTCAACCATTGCCgaatcttaaacccaact
ggttacctcttatctagtggTTTTTTAGACAAAATCTTACTCTGTGCTCTGCCACCCAGGgaatcttaaacccaactt
ggttacctcttatctagtggGGCATGAGCCACCACGCTTGGCCTAGAAACACTGTCTTATgaatcttaaacccaac
ggttacctcttatctagtggCAGCTTTGGAGTCAAACCCAACTTGGATGCAGAATCTCACgaatcttaaacccaact
ggttacctcttatctagtggGCCGCTGCAAGTGTGTGTGGCTCTGTGCATTGTGGGCAGGgaatcttaaacccaac
ggttacctcttatctagtggCGTCCCCCTAAATGTCCACATCCGAATCCCTAGAACCTGCgaatcttaaacccaac
ggttacctcttatctagtggTTGCAGATGCTAGTAAGGCTAAGGACCCTGAGGTGGACGGgaatcttaaacccaac
ggttacctcttatctagtggATCACATGCGTCCTTAGGAGTGGAAGAGGGAGGCAGAGAGgaatcttaaacccaac
ggttacctcttatctagtggCAGAGATATGGCCACAGAAGCAGAGTCAGAGATGCCTTGGgaatcttaaacccaac
ggttacctcttatctagtggGATGGAGGAAGAAGCCACAGCCAAGGCCTATGAAGCAGCCgaatcttaaacccaa
ggttacctcttatctagtggCCCTCTCAGCCACCAGAAAGAAATGCAGCCCTCCCAACAGgaatcttaaacccaa
ggttacctcttatctagtggGCTCAGTGAGACCCATGTCAGACTTCCAGAAATGTAAGATgaatcttaaacccaact
ggttacctcttatctagtggGCTTTGGAGCCAGTCTGCCAGTCCGCCTTGTTTCAAACCCgaatcttaaacccaac
ggttacctcttatctagtggCCTCCGTCTCCTTGGTTTAAGCGATTCTCATTCCTCGGCCgaatcttaaacccaact
ggttacctcttatctagtggCCTGGCCTCAAGGGATCCACCAGCCTCGGCATCACAAAGTgaatcttaaacccaa
ggttacctcttatctagtggGTGAGCCACCGCGCCCAGCCTCAAACCCTTTTCCTGTGATgaatcttaaacccaac
ggttacctcttatctagtggGCTGTGTGACCTTGTGGAAGTTGCTTCACTTCTCTGGGCCgaatcttaaacccaact
ggttacctcttatctagtggGAGCGGCGGATCAGGGCACAGACTTGGGAGACAGACTGTCgaatcttaaacccaa
ggttacctcttatctagtggCCACTGCTTCATCTGGAAAACAGGGGCAGGCATCCCAGGGgaatcttaaacccaa
ggttacctcttatctagtggGAGGCCAGAGTTTTTAGCCGATACATCATAGACATTCCCCgaatcttaaacccaact
ggttacctcttatctagtggCATTGGGTGGATGGGGAATATTGAGCCTCCAGCCCTAGGCgaatcttaaacccaac
ggttacctcttatctagtggCACTGGATTGGGAAGACAAAGCTATCACCCTCCTGATGGGgaatcttaaacccaac
ggttacctcttatctagtggCAGGTCCTAGGCCCTGCCCTGAAGAGGCTCACAGTGCAGGgaatcttaaacccaa
ggttacctcttatctagtggCAAAGAACTACTTCCTACCAGCTGACCCTAAACCCCTCAGgaatcttaaacccaac
ggttacctcttatctagtggGGCCTGGCCTTGCTGTCAGCTTTCTAAGCCAGTCACAAACgaatcttaaacccaac
ggttacctcttatctagtggTGGATGCCCCCCAGCCTCCTCTCCCCTCTCAGGGTGCTCCgaatcttaaacccaa
ggttacctcttatctagtggAGGCAGCTCTCACCTGAAGGCCTCCCTGGACACCGGTCTCgaatcttaaacccaa
ggttacctcttatctagtggCACTCTGTTAGGGCCAAACACAAGGAGGAAAGAGTGACTGgaatcttaaacccaac
ggttacctcttatctagtggGATGCCCCTCAAGATGGCTGTGCCTGTCAGCTGCATGGAGgaatcttaaacccaac
ggttacctcttatctagtggGGTGGAACCATGCACTGGAATGACACACGCCCGGCACATTgaatcttaaacccaa
ggttacctcttatctagtggCAGCTTGGATGTCCATGGGGGACGAAGGACTGCAGCTGGCgaatcttaaacccaa
ggttacctcttatctagtggCCGGCGCTCAGCTGGGCTTGAGGTGAGGCTGTCCTGGCTGgaatcttaaacccaa
ggttacctcttatctagtggGCACCCTCTGTGTGGTTGGCGTGATGCTGGGTGCCTGGGGgaatcttaaacccaac
ggttacctcttatctagtggGAGGTCCTTCCTGGAAGCCGTTACTGTTCCCTGCCCTGACgaatcttaaacccaac
ggttacctcttatctagtggCGTATCTGTCACCATCTACCTCCCTATAAAGTGCCTCTGCgaatcttaaacccaact
ggttacctcttatctagtggTGTGTGGCATTTGGCACAGTCTCTCATGCTGCAGAAAGGGgaatcttaaacccaact
ggttacctcttatctagtggGAAGACATGGGAGATTCAATGGAAAGATATGCATGGGGGGgaatcttaaacccaac
ggttacctcttatctagtggGGGTTGTATGTGGGCAGATGCATGTGGAGGCAGACAGGCCgaatcttaaacccaac
ggttacctcttatctagtggGCCTCCAGGCAGACTGATGGATTCCTTTCCCTGCAGTGAAgaatcttaaacccaac
ggttacctcttatctagtggCACACGTCACAGGACTGGAATTGAAGATCGCCAAGTAGACgaatcttaaacccaac
ggttacctcttatctagtggGACAGACACTCAGCATGTGGTGGGTGATGTTGGCTGGGGGgaatcttaaacccaac
ggttacctcttatctagtggCCAGGGAGGCAGTGCTGGACTCTGACCTGGGCACAGCTGGgaatcttaaacccaa
ggttacctcttatctagtggCCCTGCAGCCCATGGGAAGGCAGAGTGACAAGTCAGTGGCgaatcttaaacccaa
ggttacctcttatctagtggGCCTGGTGGCCTGGGAGGCTGCTAGCTCTGAGCGATGAGTgaatcttaaacccaa
ggttacctcttatctagtggGCCACCTGGCATGTGGCGGCCAAGACCACAGTTGGGGCGGgaatcttaaacccaa
ggttacctcttatctagtggTCGGCTCATGCCTGACAGCACTTTCTCAGACAGGCTTCCCgaatcttaaacccaac

ggttacctcttatctagtggGGGTGTTACATGCTGTTCTCACCCATAGCTCTTAGATGTGgaatcttaaacccaactt
ggttacctcttatctagtggCTTTCTGAGTTGAGGAAATGCATTTGCCTTGCACACAGGCgaatcttaaacccaact
ggttacctcttatctagtggGCATCACAGTTGCCTTGATCAAATATGGTGACTCTCCGTGgaatcttaaacccaact
ggttacctcttatctagtggGCGCCTGGAGGCTCTGAGGGGCCATGTACCTTCCCCAAGGgaatcttaaacccaa
ggttacctcttatctagtggGGGAAGGGTGTGTGTGACGGTGAGCTGCAGCAGGTGCTGGgaatcttaaacccaa
ggttacctcttatctagtggGTGGTCACGTAGGGGCAGGCAGAGGAGCCATTTCGGGGGGgaatcttaaacccaa
ggttacctcttatctagtggCTCACAGTGGTGAGCAAAGAGGTTCTGGGATGGGAGGGGAgaatcttaaacccaa
ggttacctcttatctagtggGCCTTCCCAGTCCTCTCTCACCCCTTCCTGCCCTCTCCCCgaatcttaaacccaa
ggttacctcttatctagtggGGCAGCTGTGCCTCACTTATCACATGGTCCATAGCTCTGTgaatcttaaacccaact
ggttacctcttatctagtggGTGGTGGTCTGGGTTATCTCCCACTCTTCACAGTAGGGCCgaatcttaaacccaac
ggttacctcttatctagtggGGCAATGAATAAGATTCCTTCTCTGTGTCAGCAAGGATGCgaatcttaaacccaact
ggttacctcttatctagtggAAGCCATGCTGTCTACAGGGCACATAGCACATGTTTTTCCgaatcttaaacccaact
ggttacctcttatctagtggAACAGAGCTATGAGATGCGCACCATTATGATGATTATCTGgaatcttaaacccaactt
ggttacctcttatctagtggGCCTGCCTCGGCCTCCTAAAGAGCTGCGATTACAAGCGTGgaatcttaaacccaa
ggttacctcttatctagtggGCAGCATTTGGACAGCCCTGTCTGACTCCAAAGCCACTGCgaatcttaaacccaa
ggttacctcttatctagtggCATTCTCCCTTCTTGGCCACTGGTGACTGTGACCTTGGGCgaatcttaaacccaac
ggttacctcttatctagtggCTCTTCTCAGCCTGGAGACTTGAGAGTGAAAGGGGCACTGgaatcttaaacccaac
ggttacctcttatctagtggTGACAGGTGTATACGAGGATGTAGCTGGGCAAGCCAGGCCgaatcttaaacccaa
ggttacctcttatctagtggGGTGGGTGCTGGTGCTGCTGGTGCATCATGCATACAAGGCgaatcttaaacccaac
ggttacctcttatctagtggTTGCCTGAACAGTTTGCATTTCTACTCGTGGTGCAGGAGCgaatcttaaacccaact
ggttacctcttatctagtggTTCAGCCACTTAGAAGAGCCCTGTTATCCCTGACTGCCGCgaatcttaaacccaac
ggttacctcttatctagtggGTCGTACGGGCTGCCCTGCCACCTACACTGACATCCTGGCgaatcttaaacccaa
ggttacctcttatctagtggCCCCACGGGGTGCCTGGAGCCTCTGATGGTCTACCCAGGGgaatcttaaacccaa
ggttacctcttatctagtggGTGACTTGATCATCTCCAGGGGTTCTTCGGTGAACAGCCCgaatcttaaacccaac
ggttacctcttatctagtggCAGCTGGTGTGTATGAATACCTGGAATGTGCCAGTTCACCgaatcttaaacccaact
ggttacctcttatctagtggGCAGTCCCATCTGTCAGCCTTGACACCCCTACTCCTGTCCgaatcttaaacccaac
ggttacctcttatctagtggTTCCAAAGCGTGCTGATGAGCAATGCAAGATGATTTCATGgaatcttaaacccaact
ggttacctcttatctagtggAGGCAACAATTGGTATGTGCGACCTGTGGTTTTGCAGATGgaatcttaaacccaact
ggttacctcttatctagtggGGGGCACCGGTAATTGTTTCTATTTGCGGTACTCTGGCTGgaatcttaaacccaact
ggttacctcttatctagtggAGGCAGTGTCTGAGCCTGTGACTTGAGTAGTAGCTCTGTGgaatcttaaacccaact
ggttacctcttatctagtggCCCCAAATCCTGAAGATTCATGATGAAGTGACTGGCCGGCgaatcttaaacccaac
ggttacctcttatctagtggGCTAGATTGAAAACAATAAGGATCCCAGAACGATAGCACTgaatcttaaacccaact
ggttacctcttatctagtggTGGCTCAAATTGCCTGCAGTTACTATCTCAGCCCTGCCTGgaatcttaaacccaact
ggttacctcttatctagtggTGAGCACCCAAAGTTTTTCAGTCAATTCCTGAGTTAATTAgaatcttaaacccaacttg
ggttacctcttatctagtggGTCTCAATAAATTGAAGATAAGGGCTGAGCACAGTGGCTCgaatcttaaacccaact
ggttacctcttatctagtggCCACTGCACTCCAGCCTGCGTGACAAACTGAGACTCCATCgaatcttaaacccaa
ggttacctcttatctagtggAGGTTTCATTCAGTGCCAGGCATATAATATGTACTCAGGGgaatcttaaacccaactt
ggttacctcttatctagtggTAAAATGGGATTAGAAGGTCAAAGCATATAGGCATAAAGGgaatcttaaacccaact
ggttacctcttatctagtggCATTACGTTTAGCTCTGAGCTTCCTAATTAGCACAACATGgaatcttaaacccaactt
ggttacctcttatctagtggTTGCAGGCCACACAGCTCTGTACTGCTTGTCAGTTCTGCCgaatcttaaacccaac
ggttacctcttatctagtggGAACTGGCAATAGGCCTTTAGGCTGTAGTTTGTCCTTGGCgaatcttaaacccaact
ggttacctcttatctagtggGAAACATGTTTGTCTTTCAAAGGCAGAAACTCCTCCTGGAgaatcttaaacccaact
ggttacctcttatctagtggTTCTGTGAGACTTGTTAAGCAGTATATGCGCAGTGCTTTCgaatcttaaacccaactt
ggttacctcttatctagtggGGCCTTCACATGGCTAGTTTCCTTACTGTTCATTCAGCAGgaatcttaaacccaactt
ggttacctcttatctagtggGGCAGGGGGTGCTGCATTGACAAGGAGTGGTCTCGGTGACgaatcttaaacccaa
ggttacctcttatctagtggGCTAACTTGAGTGGGCTTTTGGTCCAGCAGGCCTGGGCTCgaatcttaaacccaac
ggttacctcttatctagtggGCGTCCTGCCTCAAGAGTGGTTGTGTCATGAAATCTTGGCgaatcttaaacccaact
ggttacctcttatctagtggTGAAGCTCTGCTGTTTTATACCTGCTGTGTGATCTTGCTCgaatcttaaacccaacttg
ggttacctcttatctagtggGGTAGACGTGTGCCATGTGATTTGCTGCATCTATCACCCCgaatcttaaacccaact
ggttacctcttatctagtggCCCCCGCCCCCGACCCACAGGCCCCAATGTGCTCAAGTGTgaatcttaaacccaa
ggttacctcttatctagtggCTGAGCCTCACTTTCTTCATCTGCAAACCATAACACCTCTgaatcttaaacccaact
ggttacctcttatctagtggTGTTAGTTCCTCCACCTGGGGGCTGGGCGGGCGGTGGCTGgaatcttaaacccaa

ggttacctcttatctagtggGAGCCTGTGGGTATGAAGATTGCAGCCCCCTCCTCCTGGGgaatcttaaacccaa
ggttacctcttatctagtggTCCCCAACCCCGTTACCCCTGAGCGAGTTTGGGGGCAAGCgaatcttaaacccaa
ggttacctcttatctagtggGAAGAACTCTCACGCTGTGGTGCAGGACCGTGGCCTTGGCgaatcttaaacccaa
ggttacctcttatctagtggGGCCAGGTGGGTGCATTTGGGCCTCTTCCCTACAGCGCCTgaatcttaaacccaa
ggttacctcttatctagtggGCCAGGGCAGGCTATCTTAAGAGGAGGGAGCATATTTGATgaatcttaaacccaac
ggttacctcttatctagtggCTCAGCTGTCCATAGGAAGAGCAGAGTAAGGGGCTATGTGgaatcttaaacccaac
ggttacctcttatctagtggGATGGCCCAGTTCCAAAAATGAAGAAAGAAAGTAAGAAAGgaatcttaaacccaac
ggttacctcttatctagtggGGAAGGAAAGAAAGAAAGAAGAGAGGGAGAGGAAGGAAGGgaatcttaaacccaa
ggttacctcttatctagtggAAAGAAAGACAGACAAAGAAAGAAAGACAGGAAGGAAGGAgaatcttaaacccaa
ggttacctcttatctagtggAGGAAGGAAGGAAAGAAGGAAGGAGGGGAAGAAGGAAGAAgaatcttaaacccaa
ggttacctcttatctagtggCCTGAGTGAAGAAAAATACCCTGTCCTTTGTACCTGCGTGgaatcttaaacccaac
ggttacctcttatctagtggAGAAAGAAAGAAAAGAAAGACAGGAAGGAAGAAAGGAAAGgaatcttaaacccaac
ggttacctcttatctagtggGTCCTTTGTACCTGCGTGAAGAAAGAAAAGAAAAGAAAGAgaatcttaaacccaac
ggttacctcttatctagtggGTCCTTTGTACCTGCGTGAAGAAAGAAGAGAAAAGAAAGAgaatcttaaacccaact
ggttacctcttatctagtggAGAAAGAAAGAAACGAAGGAAGGAAGAAAGAAAGAGTGTAgaatcttaaacccaac
ggttacctcttatctagtggTGAGTGGAGTAGAACCCAGAGATCAGGCTTTTGAGGACGGgaatcttaaacccaac
ggttacctcttatctagtggGGGAAAAGGCTTTGGAGACCAAGCTAGGCAGGGAAGCGCCgaatcttaaacccaa
ggttacctcttatctagtggGACCAGACTGAGGCAGTGAAGTCACCATAATTGATGGGCCgaatcttaaacccaac
ggttacctcttatctagtggTTTCCCTGTGGTGGGGGTGAGGCGGTGGCGTGGGGGCTTCgaatcttaaacccaa
ggttacctcttatctagtggCCAGTGCAGTTGCTGGGGATGGGGGTCTTTCCCTCTCATCgaatcttaaacccaac
ggttacctcttatctagtggGGACACCCCCACCCCCCGTTGACTTATCTGGCTCCAACCGgaatcttaaacccaa
ggttacctcttatctagtggGGTGGTCAGGTCTCTGAGTCACTCAGTAGCTTGGGACTGGgaatcttaaacccaac
ggttacctcttatctagtggAGATGATGGCATAAACAAATAAACACTGGCTGGCCCCCAGgaatcttaaacccaac
ggttacctcttatctagtggTGGGGGGCCCCAGAAGCTTCCTGTTGCGTCTGCAGGGAAAgaatcttaaacccaa
ggttacctcttatctagtggGGCTCTGTGAGCTAAGGGAGGACACTTGGCCTCTCTGGGCgaatcttaaacccaa
ggttacctcttatctagtggGATCTGGGAGGTAAAACTGGCCAGTTGTCCATCTCCCCTCgaatcttaaacccaac
ggttacctcttatctagtggGTCTCTGGCTCCCCGTCTCTCAATGGGGTGACAGTCCTCGgaatcttaaacccaac
ggttacctcttatctagtggGAGTCACCTCTGGGAAACTGCCTGAGAGCTGGATGACACCgaatcttaaacccaa
ggttacctcttatctagtggCATCCACAGAGACATCTGGAAAAGGAGGCAGAGCAGAGCCgaatcttaaacccaa
ggttacctcttatctagtggAGCCTGGAGCTGGTGGAAGTGAGGGCATAGAGCACATTCCgaatcttaaacccaa
ggttacctcttatctagtggGGCACCGGTGGAGACCTCATGATCTGGTGGTCCGCCTCCCgaatcttaaacccaa
ggttacctcttatctagtggTGTGTGCCTGGAGCTCTCTGGCTTCCTTGGCTTTCCACTGgaatcttaaacccaact
ggttacctcttatctagtggGCCTCCATGTATTCTAGAATCTGCTATGGAGTTTCAGGGGgaatcttaaacccaact
ggttacctcttatctagtggTCCCACCCAAGGATTAGCACATGCTCATGAGGTGGCATGCgaatcttaaacccaac
ggttacctcttatctagtggGCATGAGCCACCACACCCAGCCTGGATCATTGTCTTTAAAgaatcttaaacccaac
ggttacctcttatctagtggTGGGAAGTCTATGAGCCTTGTCTCCCATTCCCCATCCTTCgaatcttaaacccaact
ggttacctcttatctagtggAGTACCTCTCCTGAACTTGGGGTATAACAGGTCATGCTCTgaatcttaaacccaact
ggttacctcttatctagtggCAAATCTATTTGAAGATCACTGGTAGCACCATTTTGTGTGgaatcttaaacccaacttg
ggttacctcttatctagtggGATGTCTGCACTTTCCTCTTTTCTCCCGGCGCTATGCTGGgaatcttaaacccaact
ggttacctcttatctagtggGCTTGTTCATTCATTCCCTGGGCCACAGAGTCTTGGGTTGgaatcttaaacccaact
ggttacctcttatctagtggTCCTAGCACAGGGCTGCAGGTGCATGTCTGGGTGTGTCCCgaatcttaaacccaa
ggttacctcttatctagtggGGCACAGAGCATGCACATTTGCAGCCTGACTGGAGGTTGCgaatcttaaacccaa
ggttacctcttatctagtggGGTTCCTTCCCTGAGAGCCTGGGAACTTCTGGAAGGCAGGgaatcttaaacccaa
ggttacctcttatctagtggACCTCTGCCAGTCAGCATGCCCCACACCATGTACCAGCTCgaatcttaaacccaa
ggttacctcttatctagtggAGGGGACAAGGCAAGTGCCAGGGCAGGTCCAGAGTCTGGGgaatcttaaaccca
ggttacctcttatctagtggGGGGCCTGGAGTGGGGGTGTGAATGAAGACATTGCTGTTGgaatcttaaacccaac
ggttacctcttatctagtggGAGGCTGGACATGGAGGTCCCAGGCACCATTGCTGTGGGCgaatcttaaacccaa
ggttacctcttatctagtggGTTGGGCTTGCTGCAGAGGGCTGCTGGAGGGGCTGGCTGGgaatcttaaacccaa
ggttacctcttatctagtggGGGTTATCCAAGTGACCCATTGCAGGAAGAGGGGTGGGGGgaatcttaaacccaa
ggttacctcttatctagtggGTAACCACTTTATGAGTCTGTCTCAGCCAGGATCTGGGGCgaatcttaaacccaac
ggttacctcttatctagtggAGGAAAGGCAAGATTGAGATGCCCTCGGTGGCCCAGAAGGgaatcttaaacccaa
ggttacctcttatctagtggCTTCTGGAAGGAAGTTGAATGTTGTGGGAAAAAGCTGACAgaatcttaaacccaact

ggttacctcttatctagtggGGCTGGTCTCAAACTTTTGAACTTGAGTGATCCTCCCACCgaatcttaaacccaact
ggttacctcttatctagtggGGAATTATAGGCATGAACCACCATTGTACAGCCACAAGGGgaatcttaaacccaac
ggttacctcttatctagtggGTTGGTGGTTGAAAGAGAGCCCCATAACCCTCCACTGAGGgaatcttaaacccaac
ggttacctcttatctagtggTGGCACGGAAAGTCCAACACTCTGGGAACCTCCTCAGTCCgaatcttaaacccaa
ggttacctcttatctagtggGGGATGGTCGTTCACCCTACCTGTGAATTGAATGATACCAgaatcttaaacccaact
ggttacctcttatctagtggACTTGGTTCCGGCTGTGTGCAAGGCATTGTGCGAGGCTGCgaatcttaaacccaac
ggttacctcttatctagtggGCGGTTGAAGTTTGGAGTATGATTGAACTCATCACGCAGGgaatcttaaacccaact
ggttacctcttatctagtggTCCTTGTCCCCCTCTTTTCCTCCCCTTTCTAGTAGCCCTCgaatcttaaacccaactt
ggttacctcttatctagtggAACATGCGGTATTTTGTTTTCTGTTTCAGCATTAGTTTGCgaatcttaaacccaacttgt
ggttacctcttatctagtggCCAGCTGCATCCATGTTGCTGCAGAGAGCGTGATTCCATTgaatcttaaacccaact
ggttacctcttatctagtggATCCAGTCCACCACTGACGGGCATTTATGTTGATTCCATGgaatcttaaacccaactt
ggttacctcttatctagtggATTTTCCTTTGAGTATACACCCAGGTATGAGATTGGTGGGgaatcttaaacccaactt
ggttacctcttatctagtggTTCTGAGAAATCTCTAAACTGCTCCTCACAGTGGCTGGACgaatcttaaacccaact
ggttacctcttatctagtggCCCACCAACAGTGTGTCAGTGTCTCCTTTTCTACACAGGCgaatcttaaacccaact
ggttacctcttatctagtggGAAAACCATTCTGATTGGCATGAAATGTATCTCATTGTGGgaatcttaaacccaactt
ggttacctcttatctagtggGTCTTCTTTTGAGCAGTGTCCGTCCTTGTGCTTTGCCCACgaatcttaaacccaactt
ggttacctcttatctagtggGTTCCTTATCGATTCTGGATATTGGAACTTTGTCAAATGCgaatcttaaacccaacttg
ggttacctcttatctagtggTTCTCCCATTCTGTAGCTTGTCTGTGTACTCCCTTGATAGgaatcttaaacccaacttg
ggttacctcttatctagtggATTCCTGTCAAGGCCTATATTGAGAGGGGTATTTCCTCGGgaatcttaaacccaactt
ggttacctcttatctagtggGTGTGGTGATAAATAGGGATTCTGTTTCATTTTCTTCTGCgaatcttaaacccaacttg
ggttacctcttatctagtggTGAATAGGAAGTCTTTTCCCCATTGCCTCTTTTTGTCAAGgaatcttaaacccaacttg
ggttacctcttatctagtggCTGTTCCGTTGGTCTGTGTATTTGTTTTTGCACCAGTACCgaatcttaaacccaacttg
ggttacctcttatctagtggTGAAGTCAGGTAATGTGAGGTCTCTAGCTTTGTTCTTATTgaatcttaaacccaacttg
ggttacctcttatctagtggTGTGTCATGTCTGATTTCTTTCAACAATGTTTTGTAGTCCgaatcttaaacccaacttgt
ggttacctcttatctagtggGTGAGAGTGGGGGTCTTACTGTAGCCTGGGCAATACCCTCgaatcttaaacccaac
ggttacctcttatctagtggTGACTGCCCCTGGCCCTGCTGTGCAGCACTTTCAGACCCCgaatcttaaacccaa
ggttacctcttatctagtggGACCCATTCTGGGGCCAGCTCAGTGTTTTACAGTGATGAAgaatcttaaacccaact
ggttacctcttatctagtggCTGAGGTGTGTGTCTATCTCTGGCCTCGTTATACCTGCAGgaatcttaaacccaact
ggttacctcttatctagtggCCCACTTGCCAGGCAGCAAACCGTGGCCCTCCGTTAGGGGgaatcttaaacccaa
ggttacctcttatctagtggCATGAAGTGTTGAGTGTATGGGTCCCTGGCCAGCTTTGGCgaatcttaaacccaact
ggttacctcttatctagtggCCACTGACAGACTGAGGCCGGTGATGGGGAGCACCTGGGCgaatcttaaacccaa
ggttacctcttatctagtggGGTAGGCCAGGCTGGCTACATAATATGTGGGGCTCAGTGCgaatcttaaacccaac
ggttacctcttatctagtggACCAAGCTCGGGCCCCTCTGAGCCAGGAGCCCTGTGTGCCgaatcttaaacccaa
ggttacctcttatctagtggAGGAGCTGGTCCTAGGGCAGGACTAAGGTTCAGGCTTCGTgaatcttaaacccaac
ggttacctcttatctagtggGCATCCTGGGTCCAGGCTGTGTTTTGTGTGTCTGCAGAGCgaatcttaaacccaact
ggttacctcttatctagtggCCTTGGAGAAAAGTTTCTCTCTCCCCTGCCCTTCCCCAGCgaatcttaaacccaac
ggttacctcttatctagtggCTCTCCCACTCAGGAACATGGGCAAATGCCATGGGCTGGGgaatcttaaacccaa
ggttacctcttatctagtggCCTGGGGGTTATGTCCACGGGGAGCCTCACATGAAGAAGAgaatcttaaacccaa
ggttacctcttatctagtggGGGATGTGTATTTCTTGTGGTCTTAGCATGCTGGCAATTCgaatcttaaacccaactt
ggttacctcttatctagtggGGTTTTCACATCCATTGGTCATTTCTGAGTTGAGCACCTCgaatcttaaacccaactt
ggttacctcttatctagtggGAATGTTTGTTGAGCACCTACTATGTGCAGGGCAGTGTAGgaatcttaaacccaactt
ggttacctcttatctagtggCTGTGAGGTTGTTGCTGTCACCCCCATTTCTCAGGCGAGGgaatcttaaacccaact
ggttacctcttatctagtggGCAAGAGTGCACTGGCTTTCGGGGCCTGAGCTGGCTGGCCgaatcttaaacccaa
ggttacctcttatctagtggCCTGGCTTTGCCATTTGCAGCTGTGGTCTGAGGCAAGTTAgaatcttaaacccaact
ggttacctcttatctagtggTGGGGCTATACTAGTTGCTCTGTGTAGGAAGGGTCTGAGGgaatcttaaacccaact
ggttacctcttatctagtggTGCACAAACCCCACACACACCAAGTATCCATGAAGCCTGGgaatcttaaacccaa
ggttacctcttatctagtggCGGTGTATGGCAGATCTTGGGCAGAGCTGGACTGGTCTTCgaatcttaaacccaac
ggttacctcttatctagtggCCCCACTCTGCTTTCATGGACAGACCCTAATGACCACCTGgaatcttaaacccaac
ggttacctcttatctagtggGCCTGGCTGCTGCTTGCGGGGTTAATTAAACCATTCCTGGgaatcttaaacccaact
ggttacctcttatctagtggGCACCAAAATAAATCCTCCAAATGGCTTTGAGCCTTCCAGgaatcttaaacccaact
ggttacctcttatctagtggTTGAATTCTGACAGCACCAGCATTTTACATCTTCCTGAGGgaatcttaaacccaactt
ggttacctcttatctagtggAAAGGAGAAACTGGGGGTAGTTCAGGCTGTGGCTGAGCAGgaatcttaaacccaa

ggttacctcttatctagtggCCCTGCTCCTAGGGGTTGAGAGGGGGATGGGGGGTGGACTgaatcttaaacccaa
ggttacctcttatctagtggTCCGGAGAATCCATTGCCTTCCCGATGCTTCTGGGGTCTCgaatcttaaacccaac
ggttacctcttatctagtggAGGTCTTGTGCATAATGGTTGAGGTGTCAATGTGCAGGCGgaatcttaaacccaac
ggttacctcttatctagtggTGTAATTGTTACCCTTTATTGAGTGCCTACTATGTGCCAGgaatcttaaacccaacttg
ggttacctcttatctagtggCACCTTCATTTTACAGATGTGAAAACAGGCTCGGAGAGGCgaatcttaaacccaac
ggttacctcttatctagtggGTGGCACAATATCAGCTCACTTCAACCTCCCTCTCCCGGGgaatcttaaacccaac
ggttacctcttatctagtggCCCGAGTAGCTGGATTTGCAGGCATGTGCCACCGCACCCGgaatcttaaacccaa
ggttacctcttatctagtggGCCCGGCCTTCCCTGGCACTCTTGAGCATGTTTCCAGACCgaatcttaaacccaac
ggttacctcttatctagtggGGCTCTGGTCCCAGATGGACAGCCCGGGCCCTTTGAAGACgaatcttaaacccaa
ggttacctcttatctagtggGTGCCTGTTCTCCTGCAGGGTCTCCGGAAGTCATCAGCCGgaatcttaaacccaac
ggttacctcttatctagtggGCTCTGTGGGCTGGCTGGGGTGCATGCTGGACAGGGAGCGgaatcttaaacccaa
ggttacctcttatctagtggGCTGGTAGGGAGGAAAACACCACTGAAATAGTTCAGACCGgaatcttaaacccaac
ggttacctcttatctagtggGCAGTGCTGCCAATGATCAGATTTGGGTCAGGAAGCCCAGgaatcttaaacccaac
ggttacctcttatctagtggTCCACGAAATCCTCACCGTTTCAGACCCTGTTTTCTAAGCgaatcttaaacccaact
ggttacctcttatctagtggGGAGGGGATGGTTGGGTTGGGGATTGGGATGGGAGGGGGGgaatcttaaacccaa
ggttacctcttatctagtggGTGCTGCAGAAAGGAGCACATTCCACACCCAGCTTCTTCCgaatcttaaacccaac
ggttacctcttatctagtggGGCTCAGTGGCCTCCCTGCCTGGAGTTGGGTTCCTATGCCgaatcttaaacccaac
ggttacctcttatctagtggTTCCATTTCAGTTAATCTGTGGCCGCGCATCACCCATGGCgaatcttaaacccaact
ggttacctcttatctagtggGTGTTGGGTGATAAGTCAAGGAGACATATGGCACAGCTCCgaatcttaaacccaac
ggttacctcttatctagtggCTGGGCATCCCAGGCTTTGTATTTTCACGTTGGTCGGCCGgaatcttaaacccaact
ggttacctcttatctagtggATGCATGTGCATCCCTTCATCCCTTCCTCTTCAGTTTCTTgaatcttaaacccaacttg
ggttacctcttatctagtggCAGTGGCGTGATATTGACTCGCTGCAACCTCCGCCTCCTGgaatcttaaacccaac
ggttacctcttatctagtggCCTTGATGGGACTCCCGGCTAAACACCCCCATTGTCTGGGgaatcttaaacccaac
ggttacctcttatctagtggCTCCTTGGAGTGCGGGAGGAAGGGCTGCGGAAGGTGTCTGgaatcttaaacccaa
ggttacctcttatctagtggGGTACGGTCACCCTCCGCCTCTGTCTCCCTCTCCAGTCCCgaatcttaaacccaa
ggttacctcttatctagtggACCCAGCTGCCGGGTCTTCAAAGCCTGTTCTGACTTACCCgaatcttaaacccaac
ggttacctcttatctagtggGGCCCCTGCCATTTAGGTCCATGTCTCTCGTGATCCTGTGgaatcttaaacccaact
ggttacctcttatctagtggGTGGAAGGACAAGGTTTGGCCCACTGTATATTTGAAGGCCgaatcttaaacccaact
ggttacctcttatctagtggCTGATGGATGAAGAACTCTGAGGAGATCTGGTTGAAAGAGgaatcttaaacccaact
ggttacctcttatctagtggCGCAGGCCGCGTTTTTGTCCTTGCCTATTGGCTACTTTTGgaatcttaaacccaactt
ggttacctcttatctagtggTCGCAGTGTGGTTACTGGGAAAACAGAGCTCTCTCTCCCCgaatcttaaacccaac
ggttacctcttatctagtggCGAGAGAGAGAGGGCCTGGGTCTGAATCATGGCTTTGTGCgaatcttaaacccaac
ggttacctcttatctagtggCTAGTCACTTTACCTTCTGAGCCTTGTTTTTCCCAGCTGCgaatcttaaacccaactt
ggttacctcttatctagtggTTGGCATATGTGCGGAGCTCTTTGGGGGTGACTCATGGGCgaatcttaaacccaac
ggttacctcttatctagtggGGGGATGTGGTGGTACAGGGTAGGGTGTAGGGGTCCCCAGgaatcttaaacccaa
ggttacctcttatctagtggGGGCAGAGTCCATGCAGGCGCTCTGGGGCTGACAGGGACCgaatcttaaaccca
ggttacctcttatctagtggCAGCCAACTGGAGTGGAGGGTTGTGGGGAAAATGTTTGGCgaatcttaaacccaac
ggttacctcttatctagtggTCTTGGTGGGGTGTTCATCAGGCCTGTAGCTGCCACAGTCgaatcttaaacccaac
ggttacctcttatctagtggGGTTGCAAGGTGCTGGGAGTGGGGTGCAGGGAAGGGCCTCgaatcttaaacccaa
ggttacctcttatctagtggCGTGGGTCTCTGCTGGCCACGCAAGGGGAAGATGGGACTGgaatcttaaacccaa
ggttacctcttatctagtggGGAAGCGGGGGCAGTTGTGTGCGAAAGGTACAGGGTCGTGgaatcttaaacccaa
ggttacctcttatctagtggGGCTTCTCCAAACTTTGGTGTCCTCATCTGCAAACTGGGGgaatcttaaacccaact
ggttacctcttatctagtggGCAGGGAATGTGAAATCCTGACCTGAAATGATCACATGTGgaatcttaaacccaact
ggttacctcttatctagtggCCTTCTTTTCTTTATGGCCTTTCCTTGGTAAATCTCTGTGgaatcttaaacccaacttgt
ggttacctcttatctagtggCTAGTGCTTATTAACTTCTGGGCAGTGCGAATTTGGTCTCgaatcttaaacccaactt
ggttacctcttatctagtggCCTGTAATAATCCTAGGAGGATGGTGCTATAATTAGCCCCgaatcttaaacccaact
ggttacctcttatctagtggGTCTAATGTCACACTGCTGAAAGTGGCTTGCTGGTGTTCCgaatcttaaacccaact
ggttacctcttatctagtggGGCTCCAGAGTTTGTGCTATTAACCATTATATCAGTCTGCgaatcttaaacccaactt
ggttacctcttatctagtggGTGAGAGAGAGAGAGAGAGAGATTGTGCATTGTACGGTGAgaatcttaaacccaac
ggttacctcttatctagtggAAGGCATTCCTGTGCTACAGTCTTGGTGGGTCAGGCTTACgaatcttaaacccaact
ggttacctcttatctagtggCCGTCCAGGGACTCAGAGGAGGTCAGACAGGTGGGACTCCgaatcttaaaccca
ggttacctcttatctagtggCCTGGGCCTGTGTCAGTTGAGTGCGTCCAGATGCAGCTGGgaatcttaaacccaa

ggttacctcttatctagtggCCGCAAGGGGAGTGCTCCTGTGTGCAGTTGCTCTGACCTGgaatcttaaacccaac
ggttacctcttatctagtggGGCCAGGCCCCAGCCGGTGAATCACTGAAACTGATCCCCGgaatcttaaacccaa
ggttacctcttatctagtggGCCTCTGTTTCTCATCCTCGCTACCACCCCTGCGTAGTCCgaatcttaaacccaac
ggttacctcttatctagtggACCGTGGCCCAGGGAAGGTCGGGAACTTGTCCAAAGGCACgaatcttaaacccaa
ggttacctcttatctagtggCCAGCTGTGTCTAATAGCATGCGTGCCCCCTCTGCTGTGCgaatcttaaacccaac
ggttacctcttatctagtggGGGGTCCGCACTATGGGTGACCTCCCTGCTCCCTTTGGCCgaatcttaaacccaac
ggttacctcttatctagtggGGCTCAGTGTGGGGAAGTTGGTGGATGGGGAGAGGGCAGCgaatcttaaacccaa
ggttacctcttatctagtggAGTGGAGACCTGGATTTGAGTGCCCTGAGTTAGGTGAGCCgaatcttaaacccaac
ggttacctcttatctagtggCCTGCAGTTTGAACATCCTGGCATTCAGCTCAGCCCACTGgaatcttaaacccaac
ggttacctcttatctagtggCATGTAGACAGTTTAGGAAGAGGGATTGTTTATTGTGTTGgaatcttaaacccaacttg
ggttacctcttatctagtggGATTTCTACGTGTTGACTCATGTAACACAGGCAACCACCCgaatcttaaacccaact
ggttacctcttatctagtggCCTTGGCAAGTAAGTGGCTAAGGGGGGATTGGAACCCAGGgaatcttaaacccaac
ggttacctcttatctagtggCCTAGGAGGAGATTCTGCCAAGGTACAGCCCTCCACGATGgaatcttaaacccaac
ggttacctcttatctagtggAGCCTCCTGTGAGCTGACTCACTTGAGTTATTCCACACCAgaatcttaaacccaact
ggttacctcttatctagtggTCCCTATTGTGATAGAGGAAGAAAGGTGAAGGAATTTGCCgaatcttaaacccaactt
ggttacctcttatctagtggATGGTGACATTCGCATTGAACACATGTCCACTGACCCCAAgaatcttaaacccaact
ggttacctcttatctagtggGGGATTACAGACACCTGCCACCAGGCCAGACGAATTTTTTgaatcttaaacccaact
ggttacctcttatctagtggCGTGAGCCACGCCCACCCACCCCCTTCACACCCCACCTTGgaatcttaaacccaa
ggttacctcttatctagtggGGTGTGAGCCACCACGCTTGGCCTCTTCCCTCCCTTCTTGgaatcttaaacccaac
ggttacctcttatctagtggGCATCCTCCCATCAGCCATGCATTCCTCAGGGTCAGGCGCgaatcttaaacccaac
ggttacctcttatctagtggGCACCTAGCCTGATCCATTGACCTGGGCCCAGTATAGGCCgaatcttaaacccaac
ggttacctcttatctagtggGGGAGGCCATGGGGTGTTAAGGTTAATGCTGGGCCACACAgaatcttaaacccaac
ggttacctcttatctagtggGGGCGCTGCTGCTTGGGTGGGTCCAGTCTGCCCCATGTGGgaatcttaaacccaa
ggttacctcttatctagtggGCACTTCCCTGTGCTCAGCCTCTCCCCGACTGTGCACATTgaatcttaaacccaac
ggttacctcttatctagtggCTCAGACACTCCAGGGTCTGTGTGGGATTAAGTCGTCCGCgaatcttaaacccaac
ggttacctcttatctagtggGGGGCTCCAGTGCCTCTCTAAGTGAAGGGTTTGGGGCCACgaatcttaaacccaac
ggttacctcttatctagtggCTCTCCAGGCCCACATGCAGAATCCAGCAGCCCCCTGGCCgaatcttaaacccaa
ggttacctcttatctagtggCTGCCAGATGAATAGGCTTGAGGCAGCATACGAGTTTGCTgaatcttaaacccaact
ggttacctcttatctagtggGACAACCCAGTTTCAGAATGGGCAGAAGACTTGAACACACgaatcttaaacccaac
ggttacctcttatctagtggACACATGAAAAGATGCTCAGTAGCCATGGTTGTTAGGGAGgaatcttaaacccaact
ggttacctcttatctagtggCCTACTAGGATGGCTGTAATTAAACACCACCACCACCAACgaatcttaaacccaac
ggttacctcttatctagtggGATGATACAGTCTCTGTGGAAGACAGTTTGGTGGTTCCTCgaatcttaaacccaactt
ggttacctcttatctagtggCCACTCCTAGGTGTATATCCAAGAGAGTTGAAGGCGTATGgaatcttaaacccaact
ggttacctcttatctagtggCAGGCATGTGCACTACTCACAACAGCCAGAAGGTGGGAACgaatcttaaacccaac
ggttacctcttatctagtggCCAATGTCTGTCCGCGGAAGAATAGCAAACAGTATTCAACgaatcttaaacccaact
ggttacctcttatctagtggTGAACCTCAAAAATGCTGTTCTAGGTGACAGAAGCTAGGCgaatcttaaacccaact
ggttacctcttatctagtggGGTTGCTGGGGGCCATGGGGAGTAGCTGCTTAATGGGTACgaatcttaaacccaac
ggttacctcttatctagtggGCCTCAGCCTCCTGAGCGGTTAGGGTTACAGTCATGTGCCgaatcttaaacccaac
ggttacctcttatctagtggGTGTGAGCCACCACACCTGGCCAATGAGTACAAAGTTTTAgaatcttaaacccaact
ggttacctcttatctagtggGGAACCAGATAGACATGGTGATTATACCACATTGTGAATGgaatcttaaacccaactt
ggttacctcttatctagtggCCCATCAGGTTAAAAGAGGACATGTGGGCCAGGCTCAGTGgaatcttaaacccaac
ggttacctcttatctagtggATGGAGAAATCCTTTCCCTACTAAAAATACGAAATTAGCCgaatcttaaacccaacttg
ggttacctcttatctagtggCTGGTGGATTCTATAGAGATGGGATGGGTTCATTAGCCCCgaatcttaaacccaact
ggttacctcttatctagtggGCTCCATTAGAGCACACAGTGTAGAAGTGGAACCAGGATTgaatcttaaacccaact
ggttacctcttatctagtggGATCACTTTTCTATGTTCCTCTTCCTCTACAGTGGACAGGgaatcttaaacccaacttg
ggttacctcttatctagtggGCTGTGTGCCATGGGTACATCACCTGACCCTGAAGAGCCCgaatcttaaacccaac
ggttacctcttatctagtggTTCACATCCTTCACAGTCAATCCCACCCTGGCCACTAGGCgaatcttaaacccaac
ggttacctcttatctagtggTGTGGTCATGGATGAGTGCCTTCAACCCTCAGTACCTCAGgaatcttaaacccaact
ggttacctcttatctagtggGGGAGAGCATGAGAACACTTATCTGATGTCTTCCCTGGGGgaatcttaaacccaact
ggttacctcttatctagtggACACACCATGTAGGGGCAAGTGCTTTGAGAATGGAAAAGCgaatcttaaacccaac
ggttacctcttatctagtggGCAGAGAACAGACCCAAACTCCCGAATCTAGTGCTGTGGCgaatcttaaacccaac
ggttacctcttatctagtggGTGTGTGCTCAGCCCAGGACTGAATGCACAGCAGGCTGTCgaatcttaaacccaac

ggttacctcttatctagtggCATGGATTGATGTGTCCGTATTCGCCATAACCAAGCAGGAgaatcttaaacccaact
ggttacctcttatctagtggAGCAGGGAGAACTGGCCTTCCTGGTGGGAAGTTGTGGGCCgaatcttaaacccaa
ggttacctcttatctagtggGTAGTGGTGGGAGACCTGTGTTCCTTCCTTCCTGAGCCCCgaatcttaaacccaac
ggttacctcttatctagtggGGCAGCACTTCTTGCACAAGCTAGGGGAGCGTGAGTATCGgaatcttaaacccaa
ggttacctcttatctagtggGCAGGATGGATTGTTATGTATCCTTTCTGATGGAGGCTACgaatcttaaacccaactt
ggttacctcttatctagtggGGATGGATTATCTCACCTTGCACAAGACAGTGAGTGCTAGgaatcttaaacccaact
ggttacctcttatctagtggGGCAGTGAGACTTAGAGCATCTCTCAAGCCCACTTTGGCCgaatcttaaacccaac
ggttacctcttatctagtggTCCACGCAACCACCCATCCACCCACCCACATTTCCTAGCCgaatcttaaacccaa
ggttacctcttatctagtggCAGCCCCTGGTCTTAAGGACTCACAGATCATGAGGAGGCGgaatcttaaacccaa
ggttacctcttatctagtggGAGTGCTGGGATTCAGAAAGGTGCAGGCTGCTCTGGTAGCgaatcttaaacccaac
ggttacctcttatctagtggTCCTGTGATGACCCAGGACAGACTGGTCTTGATCCAGGGGgaatcttaaacccaac
ggttacctcttatctagtggGTGACCTCAAGAGGCCCCTTTTAGCCACAGTAATCTATGAgaatcttaaacccaact
ggttacctcttatctagtggGGAGAAAGAGAGAAGGAAGAAATAGAGGGGCCGGTCAGCCgaatcttaaacccaa
ggttacctcttatctagtggCCTGGAGCTGCAGGCTGTATGTGTGGGAGGTGCACAGCAGgaatcttaaacccaa
ggttacctcttatctagtggCTGCCTCTGCAGTGGTGGCAGCTGGCCCAGGCAGACCTCCgaatcttaaacccaa
ggttacctcttatctagtggGCGTCTGGTGGCCAAGGGCAGAGTGGAGCCTGTGTTTCCGgaatcttaaacccaa
ggttacctcttatctagtggGGCAGCGCCGTGATTCTGAGGGTGATGCTGGCGTTCGCCCgaatcttaaacccaa
ggttacctcttatctagtggGCAGCCTCTGCTTGCCCCGTAGCAGATGAAAGACTGCGTGgaatcttaaacccaa
ggttacctcttatctagtggAGCCCTGCTCATCTCTGCCATGACTCGGGGGCGGCCTGCGgaatcttaaacccaa
ggttacctcttatctagtggCCCCTGAAGGCAGTGGATGGGGGAGTGGAGACAAGTCCTGgaatcttaaacccaa
ggttacctcttatctagtggGGAGGGGATGTTCCTGGGGGTGTGGAACTGTGTATCGGCCgaatcttaaacccaac
ggttacctcttatctagtggAAGGCAGCTCCTCCACGGACTTGCTCGGTGAATGGAGGGCgaatcttaaacccaa
ggttacctcttatctagtggCGACGATGGCTTTCACAACTGGAACCACTAGGCCCTCAGGgaatcttaaacccaa
ggttacctcttatctagtggCTGAGTTCCCCAGACAATGCTGGCACATAGTAGGCCCTCCgaatcttaaacccaac
ggttacctcttatctagtggAGGGACACTTTGCTCCTGACATCCATACAGCTCCTCTAGCgaatcttaaacccaac
ggttacctcttatctagtggAGCCCTAGAGAGAAAGAATGTGAAGGAACAAAGGGTCAGGgaatcttaaacccaac
ggttacctcttatctagtggTGTAATGCAGTGAGCAAATGCACAAGCCTTGAGTCGCTGCgaatcttaaacccaac
ggttacctcttatctagtggGTCAGCCCTGGAATACCTCCACTGACGGGGCACTCACTACgaatcttaaacccaa
ggttacctcttatctagtggCCTCTCGCGGGGTCGTGGGGGCTTTCTTCCAGTTCTCTGGgaatcttaaacccaac
ggttacctcttatctagtggTGACCGCTGTCAGGTTGGTGAAGGCAGTGCGCGTTGGGCCgaatcttaaacccaa
ggttacctcttatctagtggGATTGCACGTCGGGTACCAGGAGCCTGGCCAGCCCTCAGCgaatcttaaacccaa
ggttacctcttatctagtggCAGGGAGCAGTGAAGCCCGGTTTACTTTTGTTGGCTTGTTgaatcttaaacccaactt
ggttacctcttatctagtggTGGAGTGTGCACATTCTAACACAGCACCTTGTCAGCTGGCgaatcttaaacccaac
ggttacctcttatctagtggAAATTCTCGGCATTCCTCCTTGATGTCAATTACTTCAAGGgaatcttaaacccaactt
ggttacctcttatctagtggGGGCTTCAGGAGCAGGGCTTTGGAGACTGAAGGCTTCTCCgaatcttaaacccaac
ggttacctcttatctagtggCCATCCTGCTACCCACTCCGGCTCTATTGGCGACTGTGGCgaatcttaaacccaac
ggttacctcttatctagtggCCCTCTGCCTGGAAGAGCTGCCTGGCCTCTGATCAGGGTCgaatcttaaacccaa
ggttacctcttatctagtggGGCCCACCATCCAGCAGTGCAGGTGTCCTGGGGACTCGGGgaatcttaaacccaa
ggttacctcttatctagtggGTGGAAGGAAAGCCTCCAAGCCCTCGCATGCAGGCCCACCgaatcttaaacccaa
ggttacctcttatctagtggGGGGGTCTCCTGAATCTTATCCCTTCGCAGCGGTAGGCAGgaatcttaaacccaac
ggttacctcttatctagtggGGCCTGACACACCTGGCCTTGAAGTCTCCCCCAGCCTCTCgaatcttaaacccaa
ggttacctcttatctagtggTGAGGGGGTCCTGTGTTATCTGTGTTCCTGCAGACAGAGCgaatcttaaacccaact
ggttacctcttatctagtggATGCATGAATAAACTCACTGCCAACATTCAGCCCACCCACgaatcttaaacccaac
ggttacctcttatctagtggGCAAGAACAGCAGAAACTCGTAGCTGACCTCCATTTGGCCgaatcttaaacccaac
ggttacctcttatctagtggGAGTGAGGCCCCAGGCTCATTGAAGACTCTGTGGGTTACCgaatcttaaacccaac
ggttacctcttatctagtggGGCAAGTTAATTAACCTCTCTGTGCCTGCATGTTGCATCTgaatcttaaacccaactt
ggttacctcttatctagtggCTAATACTAGTACCCACCTTCTAAAGTGATTATGAGCATTgaatcttaaacccaactt
ggttacctcttatctagtggTGTTAGCTATTGATATTGGTGTGCCCAGAAGGCTTGTTACgaatcttaaacccaactt
ggttacctcttatctagtggGTGCTTGCCAAAAGTTGTTGTGTTGGTAATTAAGTACGACgaatcttaaacccaactt
ggttacctcttatctagtggGTATTCCATATGCTTTACAGGTGATTAACTTTGGCACTGCgaatcttaaacccaactt
ggttacctcttatctagtggCTGAAACCAGGAATTGTAGATGTTGACTTGTGAGTGTGTAgaatcttaaacccaactt
ggttacctcttatctagtggCAGTGGCAGAACTGAGGAACAGGGGTTTTTCTCTCCAAGGgaatcttaaacccaac

ggttacctcttatctagtggCCAGCTGCAGGGTTTCCTGGAATCAGCAAAGGCCTGTGAGgaatcttaaacccaac
ggttacctcttatctagtggAGCGCTGAGTAGTGCTGAGGATATAGCAATTGGGGAGCACgaatcttaaacccaac
ggttacctcttatctagtggGGGTTGTTAACCGTGGAGAAGAGAAAGGAAAGATGATGTCgaatcttaaacccaact
ggttacctcttatctagtggGAGAGGTCTTCGAAAGGTGTTTTGCAAAATGATTCCTTGGgaatcttaaacccaactt
ggttacctcttatctagtggTTTGTGAGAAAACGTGTGCTAAGGAAGGCCTCGACATTAGgaatcttaaacccaact
ggttacctcttatctagtggGTGGTGCATGCCTATCTTCCTGTCTACTTGGGAGGCTGTGgaatcttaaacccaact
ggttacctcttatctagtggCTCCAGCCTGGGTGAGACTGCGAGACTCTGTCTGGAAAATgaatcttaaacccaac
ggttacctcttatctagtggCCTGGGTTCAAGCAATTTACCTGCCTCTGCCTCCCGAGTAgaatcttaaacccaac
ggttacctcttatctagtggCCTGGCCAGTTCAGGTTTCAAAGTATAAGATCAATCTTATgaatcttaaacccaactt
ggttacctcttatctagtggCCCCCCACCAAGGAGGGCTAGGACAAGTGACAAAAGTCACgaatcttaaacccaa
ggttacctcttatctagtggAGCCAAATGGGTAGAGCTAGGAATCCGAAGTAGGTGATGAgaatcttaaacccaac
ggttacctcttatctagtggTTCAAGATGCTGCAACCCCTCTCCCCCATATGCTACATTAgaatcttaaacccaact
ggttacctcttatctagtggCTGCCTCAGTTTTCCATGGCTGCAGAGCTCCTGCTGTTCCgaatcttaaacccaact
ggttacctcttatctagtggTTGCCTTGACTGCACCTCTCTGAGCTACCTTCCTACCTCCgaatcttaaacccaact
ggttacctcttatctagtggGTGTCCAGCTCACTGTGGGACCCAGGACTCCTTGAAGAGGgaatcttaaacccaac
ggttacctcttatctagtggGGGAAGGCGGCAGGCTTAGGCGAGGTCCTGCAGGCTGGACgaatcttaaacccaa
ggttacctcttatctagtggAGCCCAGGCCCATCTCTTTACTTCCAGACCAACTGGTGTCgaatcttaaacccaac
ggttacctcttatctagtggCTGGCAACCTCTTACCGGATGCAGCTGGCAGGATAATCCCgaatcttaaacccaac
ggttacctcttatctagtggTCAGAGGTATCTGGGCCCCCAAAAGAGTCACGGGAGTCCCgaatcttaaacccaa
ggttacctcttatctagtggGCTGTATCAGCAAAACTGCTAAGCCCTGACCCATATTTTGgaatcttaaacccaact
ggttacctcttatctagtggCTGGGTAGACCAGTGAGGAGGGCTGGGACAAGTGACTGGAgaatcttaaacccaa
ggttacctcttatctagtggGTGGCGGAGTCACCGTTGAGTTCATCATAGCGATGCTGGCgaatcttaaacccaac
ggttacctcttatctagtggGCGAAGCAGACGGGCCATTCATTTGTGCTTCACAGTGTGGgaatcttaaacccaac
ggttacctcttatctagtggGGGGCCTGGTGTGGGACATGCTCAGAGGAGGACGAGGAGGgaatcttaaacccaa
ggttacctcttatctagtggCTGAGTGTGCTGCTGTCACTGCCTTGGTTTGGGTCCCCGCgaatcttaaacccaac
ggttacctcttatctagtggGGCAGAGGGTTCAATGCAGTGGGTGCCATGAAGGAAGATGgaatcttaaacccaac
ggttacctcttatctagtggCCCTAACCCAGGGGCACATGGAGAAGGAGGAAGGCAGCCCgaatcttaaacccaa
ggttacctcttatctagtggGGCCAGAGCCCTGGGAAGTGTCAGATCCTGGTGTGCCGCGgaatcttaaacccaa
ggttacctcttatctagtggATCAGCTGTCACTTCAGCATTTCCCTGCATCAGCCTTCCCgaatcttaaacccaact
ggttacctcttatctagtggGCCCCCTCTGGCATGGTGTTGGAGGCCGGGGGTCAGAGATgaatcttaaacccaa
ggttacctcttatctagtggGTGGGCAGCAGCTATTCTCTTCTTCCTTTCCAGGGGGATGgaatcttaaacccaact
ggttacctcttatctagtggTGGAGCTGTTAGGGCCTCTTCTAGAAGCCACCAGTTTGCGgaatcttaaacccaac
ggttacctcttatctagtggGTGGCTGGGATCCATCATTGCTGGTGTTATTCCTTCACATgaatcttaaacccaactt
ggttacctcttatctagtggCACATATAATTCACCTGAAACTCACCCATTTAAGGTATACgaatcttaaacccaactt
ggttacctcttatctagtggGGTGTGAACCACCGCACCAGACCCCAACTTTTGTATTTTTgaatcttaaacccaact
ggttacctcttatctagtggGGCAGGAGCCACGGTGCTCGGCCAACACAATTGTTCTTAAgaatcttaaacccaac
ggttacctcttatctagtggACCATCACCACAATCAATTTGAGAGCATTTTGATCACCCCgaatcttaaacccaact
ggttacctcttatctagtggGCAGTCACCTCTTTATTCCCACAATTCTCTGCGTGCTAAGgaatcttaaacccaact
ggttacctcttatctagtggATACCTATCCTGGAATGGTCTTTTGTGCCTGGCTTCTTTCgaatcttaaacccaactt
ggttacctcttatctagtggGCATGTCATTTCTCCTTAGGGCTGAATACTGTTCCCTAGCgaatcttaaacccaactt
ggttacctcttatctagtggCATCTGTTGATGAACATTTGGGGACAGCAAAGCTCTAGTTgaatcttaaacccaactt
ggttacctcttatctagtggCATGTGATTAAGGCCAAACTCAACTTCACAAAAGAGAGCCgaatcttaaacccaact
ggttacctcttatctagtggGGTGGATGTTACAATGTGTTCAGTGGACACGCCATTCAGCgaatcttaaacccaact
ggttacctcttatctagtggCACTTACCATATGGTGGGCACTATGGTAGGATTCATTAATgaatcttaaacccaactt
ggttacctcttatctagtggATATAGTCTTGTGAAAAGATTTATCAGAGGCTGTTCTGTGgaatcttaaacccaacttg
ggttacctcttatctagtggTGAGCCACCGTGCCCAGGTGAGTTTTAGACTTTTTTGGGGgaatcttaaacccaact
ggttacctcttatctagtggCTGCCCACTTCCCTCCAGTCATGGTCTCTGGCCGGCTCTCgaatcttaaacccaac
ggttacctcttatctagtggCCTGCCCCTGGTGTCCCTCTAAGACCCAGATGTTTCCAGCgaatcttaaacccaac
ggttacctcttatctagtggAGGCAGTGTCTGGCCTAGGGAAGGTGACTGCAGGAAATGTgaatcttaaacccaac
ggttacctcttatctagtggCAGGTCCCCCTCCCTGGACCGGTACACTCACCCCAGCTGCgaatcttaaacccaa
ggttacctcttatctagtggGCTTCATCCGGGAAGTTGCTCACCCTGCACCAGCCCTGGCgaatcttaaacccaa
ggttacctcttatctagtggCCCCAGCCGAGACCACCTCCTTCCCTTGCCTGGCTCTGCCgaatcttaaacccaa

ggttacctcttatctagtggTCCCTCCTCAGGATCTGTCCCCGGGTGCCCACCCTCAGACgaatcttaaacccaa
ggttacctcttatctagtggGTCCCCTCTGATGATGTATCTGACTTGGAGATCCTTGGCCgaatcttaaacccaac
ggttacctcttatctagtggTCAGTTCTGTTCTGGAGAATAGAAGCTTCCTGGGCTGGCCgaatcttaaacccaact
ggttacctcttatctagtggCACAGCCTTGGCACTGTTCCCCCCTTCCTTCCCTCCTCCGgaatcttaaacccaa
ggttacctcttatctagtggGCCGACAATGACCTCAGCAGAAAGGGGAAAAGGACGTGGGgaatcttaaacccaa
ggttacctcttatctagtggGCAAACAGCTCTGGGTTCAAGTCCGGGCCCTGTTGCTCAGgaatcttaaacccaa
ggttacctcttatctagtggCTCAGTTGTGGTGGCTTCTCTAACCGGGAACTTCTTCTATgaatcttaaacccaactt
ggttacctcttatctagtggTCTGGGTCAATGATGTGGTGCAGTGACTAGCATCGAGGGGgaatcttaaacccaac
ggttacctcttatctagtggGGGCTCCCTCACCACGCAGGGGCTCAGCCTCGCCAGATAGgaatcttaaaccca
ggttacctcttatctagtggCCATCCAGGCCCGTTTCTGGTCTCATTGCTTTTGTTCATGgaatcttaaacccaactt
ggttacctcttatctagtggGGGTTGCTCTTCCTTTTCTCTCTGACCTGTCAAAAAGCCCgaatcttaaacccaactt
ggttacctcttatctagtggGCTACCTCTTCCATGAAGCCCTTTTGGATGCCTCTTAGGGgaatcttaaacccaact
ggttacctcttatctagtggCCTAGCACTGCTTGCTGCATGGTCCAGGATCTGTGGGGCCgaatcttaaacccaa
ggttacctcttatctagtggCCACTTTGCTCACTTCATGAATGTGGGGGTGTATGAAGAGgaatcttaaacccaact
ggttacctcttatctagtggTCAGTACTGGGATTCTGGCCATACCCCCTCCTTAGCTGGGgaatcttaaacccaac
ggttacctcttatctagtggAGCCGTGCTGACCCCCCAACCACAGGACACTTGTGCCCGCgaatcttaaaccca
ggttacctcttatctagtggCTGTCAGCAGTGACTTCTCACTACCTGGCATCCTCTGAATgaatcttaaacccaact
ggttacctcttatctagtggCTGAGCAAAGCTCATAGGATTAAATACACACAGACCTGTGgaatcttaaacccaact
ggttacctcttatctagtggGTGCGGTTGGTGGGTGAGAAGGGCTGTCGTCCCCTCTGGCgaatcttaaacccaa
ggttacctcttatctagtggGTAGAGGGGCCATCTCCCACTACCTACTGACAAAGATTCTgaatcttaaacccaac
ggttacctcttatctagtggCAGACACTTAAGTCTTTTCCTAGGCCCATCTGTGTACTTCgaatcttaaacccaactt
ggttacctcttatctagtggCTCTGCTGAGTCAGTTTGGTAAGGACCACCGGCCGTCAATgaatcttaaacccaac
ggttacctcttatctagtggCTACCACCCTCCAGATGATGTCTGGTCACCCTGGCCTGCCgaatcttaaacccaa
ggttacctcttatctagtggCCTCTGACCCCCACCCTGCTTCTTGGCTGTGAATTTCCACgaatcttaaacccaac
ggttacctcttatctagtggCTCCTACACTGCAAATTCCCATTGCCGTGGTCCCTATACCgaatcttaaacccaac
ggttacctcttatctagtggCTGCCAGAGCTTAAGCCCAGGCAGCTGAGTTAAAAGTGTGgaatcttaaacccaac
ggttacctcttatctagtggCCATCTTGTCCCTGAAAATTGGGACCTAAGCTCGATTCCGgaatcttaaacccaact
ggttacctcttatctagtggCCCCAGAGGGGTGCGGTGAGTAAGGGGGTGGCAACTGGTTgaatcttaaacccaa
ggttacctcttatctagtggGGGACACAGTGCTGCTCACAGCCTGTCCTGCTGCGGTGGCgaatcttaaacccaa
ggttacctcttatctagtggATAAGCCTGTGTCCGAGGCAGCCCTCCAAATACAGCAGGGgaatcttaaacccaa
ggttacctcttatctagtggTAGATGGGCTCTGGGCCACTCAGTTCCTGGAAGTTTGAAGgaatcttaaacccaac
ggttacctcttatctagtggTCAAGTGATGCGACTTCCAACCTCAGGAAAGATCGCTGTGgaatcttaaacccaac
ggttacctcttatctagtggGCACAGCCAGTGGGAGCCAGCATGCCAGAATTAGGAGGGGgaatcttaaaccca
ggttacctcttatctagtggGCAAGGGGGGTTTAGCGGGTGGGAAAGGGAGAGTCCTGGCgaatcttaaacccaa
ggttacctcttatctagtggAGGTCTTTTTAAGCTGAACTCTGAGGGTTTTCAGTTTTTCgaatcttaaacccaacttg
ggttacctcttatctagtggGTAGTCCCTCCTCAGACAGGCAGACTGCTAGAACCCTCCCgaatcttaaacccaa
ggttacctcttatctagtggCACCGTTTTGCCTCGCCCAATTAAAAATGATTGGGTGTGGgaatcttaaacccaact
ggttacctcttatctagtggAGGAAATCTTCCTCCCAACTCTTGCTTTGCAATTAGGCAAgaatcttaaacccaactt
ggttacctcttatctagtggGTGTCAAGGTCATGAGTGTCAGGGAAAGACTGAGAAATGGgaatcttaaacccaac
ggttacctcttatctagtggTGATTCTGGACAATTGTCAAACCTTGGATGGCTGGGAGCGgaatcttaaacccaact
ggttacctcttatctagtggGTCAGCTTCCTGATCTTCCTATAGAACATGGTCCTAAGGGgaatcttaaacccaact
ggttacctcttatctagtggGGGCTTCAGGTCCACTACTCCCTCTCACATGCTGTAGGGGgaatcttaaacccaa
ggttacctcttatctagtggGCTGTGTGTTCCCAGCACGTTTGTGATTATTTCACAAATAgaatcttaaacccaacttg
ggttacctcttatctagtggACCAACCTCTGGTGGGGCGGTGCGGAATTATTATTATTTTgaatcttaaacccaactt
ggttacctcttatctagtggGTGGCATGATCTTAGCTCACTACAACCCTCAACTCCTGAGgaatcttaaacccaac
ggttacctcttatctagtggGCGTGAGTCATCATGCCTGGCAGTAATCTTTTCAGTGGAGgaatcttaaacccaact
ggttacctcttatctagtggCAGGGTTTGACTTCAGCCAGGGAACATAACCTTTCTGTGCgaatcttaaacccaact
ggttacctcttatctagtggCCCAGCAGGGTGTGTAAAGGTACCAGTTCCTGGGGCGTCAgaatcttaaacccaa
ggttacctcttatctagtggTCCCTTCGTACCCCCTCACTGACATGCCCAGGTCTGTCTGgaatcttaaacccaac
ggttacctcttatctagtggTGGGTGGATTTACAGACTCCGCTGAGAACCCCCTTATAGCgaatcttaaacccaac
ggttacctcttatctagtggGTTCGTGATGATATCATGGGCTGTGACCCTGGGAGTTTAAgaatcttaaacccaact
ggttacctcttatctagtggGGTTTTGTTAAACTTGGCTAGAATCTTCCTATATCTCCAGgaatcttaaacccaacttg

ggttacctcttatctagtggGCTGGAGTTCAGTGGTGCCGTCACTGCAGCCTCAACCTCCgaatcttaaacccaac
ggttacctcttatctagtggCCTGGGCTCAAGCCATCTGCCTACCTTAGCCTCATAAAGTgaatcttaaacccaact
ggttacctcttatctagtggGCCCAGCCACCAAACCCAGTGCAGACTCTGCTGAATGCAGgaatcttaaacccaa
ggttacctcttatctagtggAGGTGGGGGCACTATGTTTATCTCTAGGACCCTGACTGGCgaatcttaaacccaact
ggttacctcttatctagtggGTATGATTCTACTTCATCCTCTGTAACCCTAGCAGTCCGGgaatcttaaacccaactt
ggttacctcttatctagtggGGCAGCCTGGCCTCAGAGTCGGTGCTAACCACTGAGCGGCgaatcttaaacccaa
ggttacctcttatctagtggCAGCGCGAGCCAAGCAGAACTCCCGTCCAGACCTCTGTCCgaatcttaaacccaa
ggttacctcttatctagtggAGGCCCTGCGGCTGGTCCCCAGACAATGCCACTTCCTCGCgaatcttaaacccaa
ggttacctcttatctagtggGGCCTCTTTGCCCCATTTCCTCCTCCCACCCCATCCTCAGgaatcttaaacccaac
ggttacctcttatctagtggGGAGGGCCTGGCCTTTGGCGGTGGACAGTCTGTGGCTGGGgaatcttaaacccaa
ggttacctcttatctagtggGCCGTGCACTCCCAGCCCACTTGCCCCATGCCAGCACTCGgaatcttaaacccaa
ggttacctcttatctagtggCCTCAAGGAGGCAGGGGGTCCATGTTTCTGCTCCTCTCCCgaatcttaaacccaac
ggttacctcttatctagtggGCTTTAGGTTTGATCTCCCTCCCTTCTGCCCCTCTCCTGGgaatcttaaacccaact
ggttacctcttatctagtggTGGATTACCCCAAGGAGGCTCGAACATGGGAGGACATGCGgaatcttaaacccaac
ggttacctcttatctagtggCCTCCCAGCACCTGCCTTATCCAACTCCCCACGCTGTGGCgaatcttaaacccaa
ggttacctcttatctagtggGCATCACTGGACTCCCATTGAACTCTGTGCAGATTCGCTGgaatcttaaacccaact
ggttacctcttatctagtggGGTACCTGATGGACACCAAGCTACGTACAGCTTCAAGGCCgaatcttaaacccaac
ggttacctcttatctagtggCATCTCTGACTGTACATACATACACAAACACACACACACGgaatcttaaacccaact
ggttacctcttatctagtggTCTCCAATGCATGTGGAGTGTTGTGCCAGGTCATCCTCACgaatcttaaacccaact
ggttacctcttatctagtggAGGATTGTCAGGCATTAGGGATTTATCTATAGCAATTTGGgaatcttaaacccaacttg
ggttacctcttatctagtggTCCAACAACCTTGGAGGCTATAAAAGGGCATATTATTGCCgaatcttaaacccaactt
ggttacctcttatctagtggCATGCGCCACCATATCTGGCTAATTTTATATGTTTAGTAGgaatcttaaacccaacttg
ggttacctcttatctagtggGCGCGAGCCATTGTGCCTGGCCAGTATCATGGATATTTTTgaatcttaaacccaactt
ggttacctcttatctagtggTTCCCATGGAGAGAGGCACAAACCTGCGGGTATAGGTGGGgaatcttaaacccaac
ggttacctcttatctagtggCAGGGTGGGCAGGAGAGAGACTGGTGAGGGGATGATGCCCgaatcttaaacccaa
ggttacctcttatctagtggGGGGCAAGATGAAGCCTCTCTGGGTTTGGTGGGACCTTGGgaatcttaaacccaac
ggttacctcttatctagtggCTAGTGTGTGGTCACCCAGAGGCCAAGGATGTGCTGTTTCgaatcttaaacccaact
ggttacctcttatctagtggCCTGGGCATAGGAGCCAGGCAGAACTGTGTGGGGATGTTGgaatcttaaacccaac
ggttacctcttatctagtggCAGTTTCTCTACATCCTGGGTTCTTCATCCATGAAATGGGgaatcttaaacccaacttg
ggttacctcttatctagtggAGCCTTCAGCAGGGTTCCAGCCTCACAGCTAACTCTCAACgaatcttaaacccaac
ggttacctcttatctagtggGCTGGAGCCATCAACTCAGCTATAGGATACAACACTGAGCgaatcttaaacccaac
ggttacctcttatctagtggCCTTTGATGGGTCTCCTTTTCCCTAGTATGGGCCATCGGCgaatcttaaacccaact
ggttacctcttatctagtggGAGGTGACTCCATGCAGTTCTTTTTGGGGGAAGGATGCTGgaatcttaaacccaact
ggttacctcttatctagtggCCAGTCTGGTGGTGGTGACTGGGGTGGGGACTGCTGAGGCgaatcttaaacccaa
ggttacctcttatctagtggAGCCTTCCCTCTGGGTGTCTTCCTCCCCAGGATGGTCTGGgaatcttaaacccaac
ggttacctcttatctagtggGATGGGAGGAAGTTGGTCTGAAAGTGCCTCAGTGATCGCTgaatcttaaacccaact
ggttacctcttatctagtggAGAGGGGAAAGAGTTGGCCAGCCTTTGTTTAGGGACTCCCgaatcttaaacccaac
ggttacctcttatctagtggTCTGGGCTGGAGATTCCAAACTGACTCCTTCATGGCAGGCgaatcttaaacccaac
ggttacctcttatctagtggGGGTCTACCTCCAGGTGTGTGGGGAAAAGTTGAGAAGCTGgaatcttaaacccaac
ggttacctcttatctagtggTCAATGGTTTTTCGGAGGGCCTGGAGCTGGGCTGGGACTGgaatcttaaacccaac
ggttacctcttatctagtggAGGCAGCCTCCACTTTTGTTGCTCCTGGGGCCAGATGGCGgaatcttaaacccaac
ggttacctcttatctagtggCTGGGTGCTCTGTTATCCAACCAGGCCCTCTCCTAAGTTCgaatcttaaacccaact
ggttacctcttatctagtggCCTGCCTCAGCCGCCGTAGTAGCTGGGATTACAGGCATGCgaatcttaaacccaac
ggttacctcttatctagtggGCGTGAGCCCTGTAATCCCGGTGCCCAGCCAGTTCTCTTCgaatcttaaacccaac
ggttacctcttatctagtggTGCTGGCTGGCTCTACACAGCAGCAGGCATCACAAGGGGTgaatcttaaacccaac
ggttacctcttatctagtggGGTGAACCCACTATCTATTTAGAGAGGCCAGGTCAGGGTGgaatcttaaacccaac
ggttacctcttatctagtggGTGTCTTTTTCCCTAGGGTGGGTATTTTCATTCAGGAAGGgaatcttaaacccaacttg
ggttacctcttatctagtggTCCCCTGTTGTAGAAGAAGCAGTGAAGTTTTAGGGAGATGgaatcttaaacccaactt
ggttacctcttatctagtggCAGGGGTCTGGTCCAAAGCCTAGAAGAAGGTGAGCCATGCgaatcttaaacccaac
ggttacctcttatctagtggAGCATGAGCTCTGGTTGCACAATCGTGGGAGGTTACTTAAgaatcttaaacccaact
ggttacctcttatctagtggCGTATGGAGCGGAGAACAGTGTCCAGCATGCAGTACGTGCgaatcttaaacccaac
ggttacctcttatctagtggAGCTTCCTGATCTGAGAGGTGTCAGTGGGTGGTGTGGCCCgaatcttaaacccaac

ggttacctcttatctagtggCAGGGGTGGCCACGTGAGGGTGGAAGTGGTTTGAATGAGGgaatcttaaacccaac
ggttacctcttatctagtggAGGTAAGAGTTGGACAAACAGTGTCTTCTCTCTCTCTCTCgaatcttaaacccaactt
ggttacctcttatctagtggCTGGGATTACAGACATGCACCACCCACGCCCGGCTAATTTgaatcttaaacccaac
ggttacctcttatctagtggGCGTGAGCCACCCCACCTGGTCGACAGTGTCTTCTCTAAGgaatcttaaacccaac
ggttacctcttatctagtggGCCCTGAAGCGCCCAGGCTAGAACTCTCTATTGAATGAGGgaatcttaaacccaac
ggttacctcttatctagtggGGTGGTGCTGGGTGTGGCTTTGGCTTCCTATGGCCGCCCCgaatcttaaacccaac
ggttacctcttatctagtggTGGATCTGTGCATCCTACCGCTCCACTCCATGTCCTGGCCgaatcttaaacccaac
ggttacctcttatctagtggTCAGGGCTGAGCTGCTTCCTCTCCTGCTTCCAGTGAGGGGgaatcttaaacccaac
ggttacctcttatctagtggCCCTCCCGTGGGACTTGGCATGGCACCCACCTCTGCAGAGgaatcttaaacccaa
ggttacctcttatctagtggGTGTGTCCTTGGTTGGATTCTCTAGATGGGAGGGACCGGCgaatcttaaacccaac
ggttacctcttatctagtggGTTCACATATGTGCCTTGGAAGAGTTTTATACAGGAAAGGgaatcttaaacccaactt
ggttacctcttatctagtggAAGTCCAGACCAAGTCACCTAACCCCTCGGGCTTGCAGCCgaatcttaaacccaa
ggttacctcttatctagtggCCTTGAGTTATCTGGTTCAGTCCCTTTGCTGTGGGCACCCgaatcttaaacccaac
ggttacctcttatctagtggTCTGACAGCAGGGGCAAGGGAACTGACGAGTCTTGGGTGCgaatcttaaacccaa
ggttacctcttatctagtggCCTGGGTTCAAGTGATTTTTGTGCCTCTAGCCTCCTGAGTgaatcttaaacccaactt
ggttacctcttatctagtggGGGTTTTGCCATGTAGGTCAGGCTAGTCTAGAAGTCTGGCgaatcttaaacccaact
ggttacctcttatctagtggCAGGGTGCCACCTGCCTGAAAGAAAATGCCCACCCTAGGGgaatcttaaacccaa
ggttacctcttatctagtggGAGACCAACGTGGCCTCTCTAAATAGGTAGTGGGTTAACCgaatcttaaacccaac
ggttacctcttatctagtggAACCACTTGTGATACCTGCTGCTGCGTAGAGCCAGCCAGCgaatcttaaacccaac
ggttacctcttatctagtggCTGAAGGAAGACTCTAGGATTACAGGAGTGAGCCTGGCCCgaatcttaaacccaac
ggttacctcttatctagtggCCCATGGTTTACAGAAGTCAAACCCAAGTCTCAGAGAGGGgaatcttaaacccaac
ggttacctcttatctagtggGCTCCGTCTAGTTCAGAGCCTGTGTGGAGGTCGACACTGGgaatcttaaacccaac
ggttacctcttatctagtggTCCAAGATCAGTCTCTTTCCCTCAGCGGAAGCCAGCTGGGgaatcttaaacccaac
ggttacctcttatctagtggCTTGTAACCTCTTCAGGCGACCTCCTTGGGATGTTCACAAgaatcttaaacccaac
ggttacctcttatctagtggCCTAGGGCTGGGATGGGGGTGGTATCCCTCACATCATTCCgaatcttaaacccaac
ggttacctcttatctagtggCCTGCTTTTATCAGCGGGGGCCACAAAGAGGCGACAAGCCgaatcttaaacccaa
ggttacctcttatctagtggCCAGGGAAAGGATGCCGTCCCACATGGCTTCAGGCCTCCGgaatcttaaacccaa
ggttacctcttatctagtggCAGCTGTGTCCACGGCATGTTCCTCAAGGACAGCTGCTCCgaatcttaaacccaac
ggttacctcttatctagtggGCCTTGAGGGGCTGCTGATAGGAACCCAAGGCCTCACCCTgaatcttaaacccaa
ggttacctcttatctagtggAACAGATTACACAAGAACGCAAATGTAAGATCCCTTTGGGgaatcttaaacccaac
ggttacctcttatctagtggGCTGCAGCACCAGAGGACATGGAAGAGAAAGGCTGAAGTGgaatcttaaacccaa
ggttacctcttatctagtggGCGTTCCTCTGCCCCAGGGTCTCTGTGTGGCAGGCGAGTGgaatcttaaacccaa
ggttacctcttatctagtggGGTGCAGTCATCGTGTGGCGGAATGCATGCTCACACAGGGgaatcttaaacccaa
ggttacctcttatctagtggCCTTTGAGCAGTCCTTCCTCTGGTGAGGTGCACAGGCGGGgaatcttaaacccaac
ggttacctcttatctagtggACTGAGGCTGAGAGGGATGCTGCAATTTAGGACTTCTGGGgaatcttaaacccaac
ggttacctcttatctagtggGGACTATGGGGATTGGACCAAGGGGTCTTCTTGCTTCCGCgaatcttaaacccaac
ggttacctcttatctagtggGAGCTGGGGTCTGTCTTGGCAACCCTGTGACTCTTCTCCCgaatcttaaacccaac
ggttacctcttatctagtggGGCTGTGGCCCTTTCCCAACCAGCACCTTGGCCCTGTACCgaatcttaaacccaa
ggttacctcttatctagtggTGGGGGCCTACGACCGCAGATGGAGCAGAGGTGGACAGGGgaatcttaaacccaa
ggttacctcttatctagtggAAGCCGCGTTAGGCAGCACGGGAGCCAGGTGCGGGGTCTCgaatcttaaacccaa
ggttacctcttatctagtggGGCAGTGCTGTGCTCTGTGAGGACTGCTTTTGACTGCAGGgaatcttaaacccaac
ggttacctcttatctagtggCTCCCTGGAGCCTTCTCCCGAGGTGCGCGGGTGACCTTGGgaatcttaaacccaa
ggttacctcttatctagtggCCCCTTTGAGCTGCTTGGCACGTGGCTCCCTTGGTGTTCCgaatcttaaacccaac
ggttacctcttatctagtggGGTAAGTGGCTTACCTGAGGCCATGTGCTAACAGAGAAGAgaatcttaaacccaac
ggttacctcttatctagtggCCACATACTTTCCATATCTTTAGAGGCCTGACCAAAGCAGgaatcttaaacccaac
ggttacctcttatctagtggGGCACCTGCCTGCTGGGCCCACTGTTTTCTAACTTTCCTAgaatcttaaacccaact
ggttacctcttatctagtggGCGTCAGGCCTGAACCCTGCATTTTGTGGAGGTGGGTGGGgaatcttaaacccaac
ggttacctcttatctagtggCCTGGGGAACATGCCTAGACACGGGGGACAACAGTTGCCCgaatcttaaacccaa
ggttacctcttatctagtggGGGACCGCTTTCACAAAACCACTGCAGGTGAGTGAGTTCCgaatcttaaacccaac
ggttacctcttatctagtggGGCCTGGTGTCTCTAGACTCATTATTTCCCCCACCCAACCgaatcttaaacccaac
ggttacctcttatctagtggCGAGCCACATTTTTATTGCCATAATCCAGGCCTGGACAGGgaatcttaaacccaac
ggttacctcttatctagtggAGAGTCAGCCCAAAATACAGTCTTTGTGTTGCCATCTGGGgaatcttaaacccaac

ggttacctcttatctagtggGAACGGCAGGGCCAGAGGTCCCACGAGCTGGTCAGACCCGgaatcttaaaccca
ggttacctcttatctagtggTGCTGGCTGAGTGACCTTGGGCATTGTGATTCATATTCTCgaatcttaaacccaactt
ggttacctcttatctagtggAGTGTGGTTACGAATGCTTTTTATCCTGCTTATCAATTTCgaatcttaaacccaacttgt
ggttacctcttatctagtggGGGGGGCGGTGGAGGGGCAGTGTTGAGACAGAGTCTTGCTgaatcttaaacccaa
ggttacctcttatctagtggCCTGGCCCACTGAACCTTGACCTCCAGGGCTCAAGGGGTCgaatcttaaacccaa
ggttacctcttatctagtggCCTCCAGAGTAGCTGGGACTTCAGGCATGAGCTACCACACgaatcttaaacccaa
ggttacctcttatctagtggGCCACTATGTCCGGCTCACGTCCAGTTATCTAGGGCAGAGgaatcttaaacccaa
ggttacctcttatctagtggGGAACCTGTAGGCTGCCTTCCATTCTGGAGACTGCAGGGGgaatcttaaacccaa
ggttacctcttatctagtggCCTTGAGGTTGTAGGGCTGAGATCCCGTTTTCTTGCTGGCgaatcttaaacccaac
ggttacctcttatctagtggGGCTTGTGGCCTCCTCTCTCCATTTTCAAGTCCAGCAATGgaatcttaaacccaac
ggttacctcttatctagtggATCTTTTCTCCCTCTTTCTTTTCACCTCTGTGATCTCAGCgaatcttaaacccaacttg
ggttacctcttatctagtggGGATTCATGTGATTGGACTGGGGCCACCCTGATAATCCAGgaatcttaaacccaac
ggttacctcttatctagtggCCGACATGGGCATCTCTTTGATTGGAATGAGGGGTGGCAGgaatcttaaacccaac
ggttacctcttatctagtggCAGTACCTCAGAAGGGTTGCTGTGATGATGAAGTGAATGGgaatcttaaacccaact
ggttacctcttatctagtggTGTAGAGCCATGTGAATTTGGCAGCATTGAATTGTTGACCgaatcttaaacccaactt
ggttacctcttatctagtggAATGTTTGCTGTCATTACTCTAGTGGTAGAAATGTCATGCgaatcttaaacccaacttg
ggttacctcttatctagtggGTCTCCTGCTCTGGTGCCAGTGTCTCTGTGGCCTGAGGGGgaatcttaaacccaa
ggttacctcttatctagtggAGCACCAGCTCAGCTCTGGGCCATGTGTGTGGGGTGGGACgaatcttaaacccaa
ggttacctcttatctagtggCTCCTTCCTGAGACCATCCCCTTTGGCTTCCTCGGGGCCCgaatcttaaacccaa
ggttacctcttatctagtggCCTTCCCAAGGGCACCCTGAGTCCCAGGTCCCCAGGCCCAgaatcttaaaccca
ggttacctcttatctagtggGCCCTCAGAGTGTGATCCCATGTCCCCAGCTCTGTAACTCgaatcttaaacccaac
ggttacctcttatctagtggGCCTTGGCCAGACACTTTCGCCTGTCAGTTCTGCCAGTTGgaatcttaaacccaac
ggttacctcttatctagtggTTCCACGGCTTGCCTAGGGTCGTTGAGCAATGTGGAAGGGgaatcttaaacccaac
ggttacctcttatctagtggCCATGGCTTGCCTAAGGTCATCAAGCTGGCATCTCAACCCgaatcttaaacccaac
ggttacctcttatctagtggGACCCATAGCCACCTCTCTCATTAAGGCTTCTCCTGCAGGgaatcttaaacccaac
ggttacctcttatctagtggCACTCACCTCTCCCCCTTGACCTGGAAGAGCCCCTGAAGGgaatcttaaacccaa
ggttacctcttatctagtggCTGAGGCCCAGGGAGTGTCCAGAGCTTTCTGAGGTCCACCgaatcttaaacccaa
ggttacctcttatctagtggCCAGGATGAGTGATAGCCACCTGGTGACAATGTTGAAGGTgaatcttaaacccaac
ggttacctcttatctagtggGCTGGGTTCAATTCTGACTTGCCTGACCAGCTGGTGCCTCgaatcttaaacccaac
ggttacctcttatctagtggGTCCCTCCTTGAAGCTGCGCATCGACTCTCCTGTCCTGGGgaatcttaaacccaa
ggttacctcttatctagtggGTAGTGGCTGTTCTGAAGTGGCTTTGGAAACCCTGGAATGgaatcttaaacccaac
ggttacctcttatctagtggGACCTGCCTCAGGGCCTCCTCTCCAAGTTGAGTGACCCATgaatcttaaacccaa
ggttacctcttatctagtggGGGGGCCTGGAAATGTCCCATGGACCATCTCGTGACTCAGgaatcttaaacccaa
ggttacctcttatctagtggGGTCCTGCACAGAGTGTTCCTGCCCGTTGATGCCCTCCCCgaatcttaaacccaa
ggttacctcttatctagtggGCGAAGTCTGACACATAGTAGCCTCCCGGTAATCTTAATAgaatcttaaacccaac
ggttacctcttatctagtggTCCAGCCCAGCCCTTTATTCTGGCCAGCCCTGGTGGGTCCgaatcttaaacccaa
ggttacctcttatctagtggCATAGGTAGGGTCTACAGCAGAGTTTCTCAGCCTTGGCGCgaatcttaaacccaac
ggttacctcttatctagtggGGATGTTGCTCGGCATCTCTGGCCTCTACCCACTGGATGCgaatcttaaacccaac
ggttacctcttatctagtggTTGCCACATGTTCTTTTGTGGTGGTGTTGGTGGTGGTGCGgaatcttaaacccaactt
ggttacctcttatctagtggGACCACCTAATTAGGGCTGACGTGGGCTCTTCCACAGAGCgaatcttaaacccaa
ggttacctcttatctagtggCCCCAATGTGGAAGATGAGCGTTCCGTTCCTGGCTGTAGGgaatcttaaacccaac
ggttacctcttatctagtggGAGGCAAGATAGTCTAGTGCTTTTCATTCCATGCGCACTCgaatcttaaacccaact
ggttacctcttatctagtggGGAAAATTATTTTGGGCCCTTGTTCAAGGTTCAGTTGACCgaatcttaaacccaactt
ggttacctcttatctagtggCCAGGCTGGAGTAGAGTGGTGCCATCCTAGTTCACTGCATgaatcttaaacccaac
ggttacctcttatctagtggCTGGGCTCAATCGATCCTCCTGCTTCAACATCCCAAAGTGgaatcttaaacccaac
ggttacctcttatctagtggGGCATGAGCCATTACACCTGGCCCATTTGACTATTTTTTCgaatcttaaacccaactt
ggttacctcttatctagtggGGTTCTGCCACTTGCTAGCCATGGTGAGACCTTGCCCAGCgaatcttaaacccaa
ggttacctcttatctagtggCCGGGCTGTTGTGCTACTGGCTGTGCTGCCTGAATGGGCCgaatcttaaacccaa
ggttacctcttatctagtggGCAGCCAGCCAGCTCATCCCCTCTGCAGCTGATACCGAGGgaatcttaaacccaa
ggttacctcttatctagtggCTGGTAACAGTCAGCCCGGCCCCTTTGAAGTGGTTTTAAGgaatcttaaacccaac
ggttacctcttatctagtggGACTCCGAGGGCGCAGGCTGCTCATTAGCTGCTGTCTGGGgaatcttaaacccaa
ggttacctcttatctagtggGGTTCCTTCCCTCCCCGTGCTCATGGCTGAGGTCCAGGCCgaatcttaaacccaa

ggttacctcttatctagtggCCTCTCCTGGGCAAAGGACATAGAAAAGCTTTCCGGGGAGgaatcttaaacccaac
ggttacctcttatctagtggGTGAATTCTACTTTCCCCTGGGGGTCAAGGCAATAGGCTGgaatcttaaacccaact
ggttacctcttatctagtggACCTGAAGTGGGTGGGGTGAGTGTGTGTGTGGGAAGTTGGgaatcttaaacccaac
ggttacctcttatctagtggGGCCGAGTGTTTTTCAAGGGCCAGTTAACCATTCCCCTGTgaatcttaaacccaact
ggttacctcttatctagtggCACTGCTGGGAGGAACATCCGCACATGCTTTCTTGGTCACgaatcttaaacccaac
ggttacctcttatctagtggCCAGGATTTATACCCTGAGGCGACTTCACTGCCGTGGAGGgaatcttaaacccaac
ggttacctcttatctagtggACACTCCCACTGACTGTACACAAGAGCCCTGTTTTCCCCCgaatcttaaacccaac
ggttacctcttatctagtggTGCCAGGGGGATGAGCAGCCTGCCCTCAACTGCCCCATGCgaatcttaaacccaa
ggttacctcttatctagtggCCCAGCCTGTGCCCTGACAACCTAGGAGGACACGCTGGTCgaatcttaaacccaa
ggttacctcttatctagtggTGTTGCCTTGAGTGTCAGTTTTGGACCTCCCACTGAGCAGgaatcttaaacccaact
ggttacctcttatctagtggGACAGGACTGAGTTCTTTGGAAAATAATCACATCCCACTGgaatcttaaacccaact
ggttacctcttatctagtggGTGGAACCGAAGGCCTGGCTTTTGGGACACTCACCGTCCCgaatcttaaacccaa
ggttacctcttatctagtggTGTGTCCTAGGGAGAGACCCATGAGCTCTCTGGGCCTCCTgaatcttaaacccaac
ggttacctcttatctagtggGTAATAAAGGACTAAAGCTACCAGCCTCGCTGCGGGTTGGgaatcttaaacccaac
ggttacctcttatctagtggCATTGCACCAAGGCAGGGCTCGCTGGAGTGCAGACCCAGGgaatcttaaacccaa
ggttacctcttatctagtggTCTGGTCTGTCAGGGTACCAGCCACACATCCAAGGCAGGGgaatcttaaacccaa
ggttacctcttatctagtggTTGGCCTTTCTGCTCATGCTTACTCTCCCAAGCTTGTTGGgaatcttaaacccaactt
ggttacctcttatctagtggCCCTTGTGGGCTGTATAGTCTAGTGGAGGAAGGGGGAAGAgaatcttaaacccaac
ggttacctcttatctagtggGTAGGAAGGTGAACCCCATGTTAAGAAAGCATCTTCCATCgaatcttaaacccaact
ggttacctcttatctagtggCATGATAGCGAGGAAATGGTGGTCATCTCCATTGCTCTGGgaatcttaaacccaact
ggttacctcttatctagtggGAGAAAGTGCTGATTTCCAAGAAAAGGAAAGAGACCACGGgaatcttaaacccaac
ggttacctcttatctagtggAGCACCTACTATATGTGCTGGACCCTCTTGTAAGAGCTTCgaatcttaaacccaact
ggttacctcttatctagtggGGCAGACAGGCAATATTATTATCCCCATTTGATCTACAAGgaatcttaaacccaactt
ggttacctcttatctagtggGCTTTTCTGCAGAGCTTGTGCACAGGGGCCAAAGTTTGGGgaatcttaaacccaac
ggttacctcttatctagtggCTGTTGTTGGTGTCACTTGTTGGAATTCCTAGAATCCCAGgaatcttaaacccaactt
ggttacctcttatctagtggGACCCAAGGTCACGACTCCCTCCCTCCTCCCTTCCCTTCCgaatcttaaacccaa
ggttacctcttatctagtggTTTGAGTTCCTTATTCCCTAGCTTCCTCCAAGCAGCAGGCgaatcttaaacccaact
ggttacctcttatctagtggCAAGAGGTGGTCCCTTGTTGAAAGCCTGTCACATGGCAGGgaatcttaaacccaac
ggttacctcttatctagtggTTCCTTGGTTTGCACACGAGGAAATGGAGGCTCAGAGAGCgaatcttaaacccaac
ggttacctcttatctagtggTACTCAAAGCCACACAGCTGGGTTAAATGGGTTCAGCACCgaatcttaaacccaac
ggttacctcttatctagtggGCCGTTGAAAGTCCCCTCATCCTGGACCTGTGGTCTGTCCgaatcttaaacccaac
ggttacctcttatctagtggGGCATTCCCTCTGTCTCCTCTGCCCATCCCAATGCCAGGCgaatcttaaacccaac
ggttacctcttatctagtggAGGTGGGCCTCAGGTTTCTCCCCTCCCAGGTGGGCATAGGgaatcttaaacccaa
ggttacctcttatctagtggCCCGGCCCTGTTTCTTAGCTGCTTTGACACCTGGGGCAGGgaatcttaaacccaac
ggttacctcttatctagtggGGCCAGCTGTGTTGTGACTGCCCACCAGAGGCTGGGTCTGgaatcttaaacccaa
ggttacctcttatctagtggGGCGATGATCACAGTCCAGGGACAAGGTTTCTGAATCTTGgaatcttaaacccaact
ggttacctcttatctagtggACCCTCTTGTAAGAGCCTTGTACAAATTAGGCAGACAGACgaatcttaaacccaact
ggttacctcttatctagtggGTGCAAAGTGACAGCTTTTCTGCAGAGCCTGTGCACAGGGgaatcttaaacccaac
ggttacctcttatctagtggCTTCTGCCAGCCCCGGTCAGCCTGCCTTTGACCCAAGGCCgaatcttaaacccaa
ggttacctcttatctagtggGACAAAGGGATACAGTATTGTGCGGTGGCTGTGGATGGCGgaatcttaaacccaac
ggttacctcttatctagtggGCCCATGGTAGGTCCAGGGCGTTTGGGGGAAGGAGACCGAgaatcttaaacccaa
ggttacctcttatctagtggCTCCTAGCTGTCATCCAATTCTGTGTAGAAGGGGAAAGCCgaatcttaaacccaact
ggttacctcttatctagtggGGGAAGCTTGGTGACTATGACTGGATGAATTCTGTCCCCCgaatcttaaacccaact
ggttacctcttatctagtggGAGCTCACGTGGCGGGGCTCCAATATGGTCAGACTGGTGGgaatcttaaacccaa
ggttacctcttatctagtggCAGGGAGGAGGTGGCCATCTACAAGCCCAGGAGAGAGGCCgaatcttaaacccaa
ggttacctcttatctagtggCAACTCTGACCACACCTTGATCCTGGACTTCTGGCTTTCAgaatcttaaacccaact
ggttacctcttatctagtggGCAACCCAGCCTGTGAAAATTTGTTGTAGCAGCCCTGGCGgaatcttaaacccaac
ggttacctcttatctagtggGACCATGAGCAAGTCACTCACCCTCTCTGAACCCCCTTGGgaatcttaaacccaac
ggttacctcttatctagtggGCCTCACAGGCTGTGGTAAGAGGCATCAGAAATAGCCGACgaatcttaaacccaac
ggttacctcttatctagtggGCTCACTGCGAGCCAGGGCCTGTTTAAGCCTTTTCCATGTgaatcttaaacccaact
ggttacctcttatctagtggGTGTAGCCTTTGGCTGGGGAGGGGAGATACTATTGTTCCCgaatcttaaacccaact
ggttacctcttatctagtggACCTGTCCCATGTTAGTCAGCAAGGAAGGGGCTGGTCCATgaatcttaaacccaac

ggttacctcttatctagtggGGTGCCCACACACCAGATCACTATGCCGTCCCAGCTGTTGgaatcttaaacccaa
ggttacctcttatctagtggGGTGAAGGGCCCACCCAGTAGTGCCTGGCCTATGGGACACgaatcttaaacccaa
ggttacctcttatctagtggAGCCCCCTGTGACCAGAACCTCCCATTTGCTGCATTGAGTgaatcttaaacccaac
ggttacctcttatctagtggTCTGTCTGTCTCCCTCTCTCTCCTCTCTCCCCTACCTTCCgaatcttaaacccaact
ggttacctcttatctagtggGCCCATCAGCATCCTCTCCCTATTCAGCCACCCCCACCCCgaatcttaaacccaa
ggttacctcttatctagtggGCTCCTGTCCTGGAGCTCAGCCTGGCACCCAGACCTGCCGgaatcttaaacccaa
ggttacctcttatctagtggGCCCTTTCTCACTGGGCCCTCACTGCTCTAGAGTCCCCGCgaatcttaaacccaa
ggttacctcttatctagtggCCACCCAGGGACAGGGAGGCCCAGTGCCGGCTGTGGAGTTgaatcttaaacccaa
ggttacctcttatctagtggAAACGCATCACCCTGAATGGGGGCCAGGCCAGGCATCGGCgaatcttaaacccaa
ggttacctcttatctagtggGCTGTCCCCTGCCTTCCCTGGCGGCCCGGGTTCCTGGAAAgaatcttaaacccaa
ggttacctcttatctagtggGAGGACGGGAGTTTCCCTCCAACTTAAGCCATTAGAAAGCgaatcttaaacccaac
ggttacctcttatctagtggAGCACTTACTAAGTGTCAGAGACTGTGCTAGGCACTTTACgaatcttaaacccaact
ggttacctcttatctagtggGCCAAGTCCTGGGGTTAGCACCCAATAGCCCCGTGGCGGCgaatcttaaacccaa
ggttacctcttatctagtggGGGTGTCCCAGCTCCTGTATGCCATCTGCCTGTCCAGCCCgaatcttaaacccaa
ggttacctcttatctagtggGGGGAGATGCGGCCAGTTTCCCAGGTGAGGCTGGTCCTGCgaatcttaaacccaa
ggttacctcttatctagtggGGAGCTCCAACCACTGCTCCACGCTGCCTCCACTCCTGAAgaatcttaaacccaa
ggttacctcttatctagtggGCACCGGGGCCCAGGGGTCACCAGGCTCAGATGGAATTTGgaatcttaaacccaa
ggttacctcttatctagtggGGGAGAAAAGCTGATTGGGGGATGGGAAGAGACAGCTGGCgaatcttaaacccaa
ggttacctcttatctagtggCCTGGGAGGACTGACTTTCACCCAAAAGGGCCTGGGCTGGgaatcttaaacccaa
ggttacctcttatctagtggGGAGCACCTTCGAGTGAGGAGGTGTAGGAGGGAGGATGGCgaatcttaaacccaa
ggttacctcttatctagtggCTCACTCCATTGGGCAATCATATGGGAATGCAAGCCACTGgaatcttaaacccaac
ggttacctcttatctagtggCTTTTTTCCAAATGGAGTCCTGCTGTGTGGCCCAGGCTGGgaatcttaaacccaact
ggttacctcttatctagtggCCCTGTGCCCCTGTCTTACTGAAGAGGGAACCGAGGCTTCgaatcttaaacccaa
ggttacctcttatctagtggCCCTCCTGGGAAACTTTCATCCCACCTGCTCCTTCTACCCgaatcttaaacccaac
ggttacctcttatctagtggTCCTGAGGCAAAGCTCCCTGCACAGCAGTGTCCACAGCCGgaatcttaaacccaa
ggttacctcttatctagtggGGCCAGTGGGTAGAGTTGATGCAGGCCTCAGGCTGATCAAgaatcttaaacccaa
ggttacctcttatctagtggCTGACATCTGAGGGACGTCAGATCACAGAAAGGTCTTGAGgaatcttaaacccaac
ggttacctcttatctagtggCAGCAAGCCTCCCCCGGTCATTACATGCCTAGACTGTGCCgaatcttaaacccaa
ggttacctcttatctagtggGGTCCTTGGCACTGTCCTTCAAGGGAAGAACTGTTTGCCCgaatcttaaacccaac
ggttacctcttatctagtggCACATCCAGGAAGCAGCTGAGCTGGGAATTAAGCCCAGGGgaatcttaaacccaa
ggttacctcttatctagtggGCCTCCCTTCTTTCCATGCTGCTCTGCTCCTTACAAAAACgaatcttaaacccaact
ggttacctcttatctagtggGGGATTACAGGTATATGCCACCACGCTCGACTAATGTTTGgaatcttaaacccaac
ggttacctcttatctagtggCGTGAGCCACCGCGCCTGGCCAAAAAACAGAATTCTTTCAgaatcttaaacccaa
ggttacctcttatctagtggGCAGGTTCAGAAAGACTGCCCTGGCCTCCATGTGCCCAGGgaatcttaaacccaa
ggttacctcttatctagtggTCACTGTCTCAGCAGAGCTGGAGGAACCTGTTAAACTGTCgaatcttaaacccaac
ggttacctcttatctagtggGCTCCCTGAGCTCTGAGGCTGTCTTAACACTCACTGTCTCgaatcttaaacccaac
ggttacctcttatctagtggCTGCAAAGCCCCTTCCCTGTTTCCAGACCCCTGGTTTCCTgaatcttaaacccaac
ggttacctcttatctagtggAACACCCTATTCTAGAAAGAACTGGCATGTTTTGGGTACCgaatcttaaacccaact
ggttacctcttatctagtggTTCCGATGAGGCAACAGAGGCACAGTCTGGGGAGCTCAGCgaatcttaaacccaa
ggttacctcttatctagtggTTAGAATCACAAGCTGTGTCTGCACCTAGGTTGTGCCTGCgaatcttaaacccaac
ggttacctcttatctagtggGCGTGAGCCACCGCGTCCGGCCCGTGCCTGCATTTTTCAGgaatcttaaacccaa
ggttacctcttatctagtggCCAGTAGGCAGAAGCTTCTCTGTCCTTCCTCAATGGGAGAgaatcttaaacccaac
ggttacctcttatctagtggGGCCCCGATCTGCTGGGAGATCACCCTACTGCAACCTCCCgaatcttaaacccaa
ggttacctcttatctagtggTCAGCCTGTAGATTGCCTGAGAACCTGACTCAAACCCCACgaatcttaaacccaac
ggttacctcttatctagtggAAAATCAGGCTCTTTGGAATACCTTTATCAGGGGATGGCCgaatcttaaacccaact
ggttacctcttatctagtggAGTTTGCTCTGGTAGCAACAGCACTATCCACACCCCTCCCgaatcttaaacccaac
ggttacctcttatctagtggTGCAGAATTGGGCTGAAACCCAGATCTCCTGAGATTAGGGgaatcttaaacccaac
ggttacctcttatctagtggGTTGTGAGCTCATTTCTATGTGGACTTCCTAGATGACTGGgaatcttaaacccaactt
ggttacctcttatctagtggACAGCTCACGTAGGTGCCTCGTTCATAATTTCTCCTTCCCgaatcttaaacccaact
ggttacctcttatctagtggTGCAAAGCATGGTCTAGGCAGGTTGGCTCCAGGACTTTGGgaatcttaaacccaac
ggttacctcttatctagtggACCAAAACCATGGACTTTTGCCACTGCTTGACATTGCTCCgaatcttaaacccaac
ggttacctcttatctagtggTTGCCTTTGAGGGTGAAGTTGACCTCCGTTACTCTTAAGTgaatcttaaacccaactt

ggttacctcttatctagtggTTGAGAATCATTCTGAGGGATGACAGAGGCCTGCAATACCgaatcttaaacccaact
ggttacctcttatctagtggGTGGTTGTGTGACTGCTGTGACTGAGACTCAACAAGTACCgaatcttaaacccaact
ggttacctcttatctagtggCCTGTGGGTGTGGTTCAGCGCCCCCAGTGCCTCTCCCAGGgaatcttaaacccaa
ggttacctcttatctagtggGAGCAGGTGAACAAACTGAGGCCCAGGGAGTGGCTTCCCCgaatcttaaacccaa
ggttacctcttatctagtggGCTGCTGCTCTGCAACTGGTGCTCGGCTTGGTCCTGGGGGgaatcttaaacccaac
ggttacctcttatctagtggTCCTTTGCACCACCCTACAAGCCTGTCCTCCAGGTCCTCCgaatcttaaacccaac
ggttacctcttatctagtggGCCAAGGACCTCAGAATATGACTGGATTTGAAGGTAGGGCgaatcttaaacccaac
ggttacctcttatctagtggGGACAGAGACACCAGGATGCGCACACACAGAGGGAAGGCCgaatcttaaacccaa
ggttacctcttatctagtggGCCAAAGAGAGGCCTCAGGAGACACCCAACTGCCAACACCgaatcttaaacccaa
ggttacctcttatctagtggGGCAGCCCTAGCAGACCAATATACCGTGTGTCTGGGGCAGgaatcttaaacccaac
ggttacctcttatctagtggCCTCCCCTGCCCTCCCCTCCCCTTCCCGAGACAGTCTCTGgaatcttaaacccaa
ggttacctcttatctagtggGTGGTGCAATCTCGGCCCACTACAACCGCCACCTCCCAAGgaatcttaaacccaa
ggttacctcttatctagtggTCAGTAGGAATGGGATTTTTTCATGTTGCCCAGGCTGGTCgaatcttaaacccaactt
ggttacctcttatctagtggCAGGCGTGAACCACCACACCCGGCTGTAATATCATTTCTAgaatcttaaacccaact
ggttacctcttatctagtggGCCAAGCTCCAAGAGGCTTAGTAATTTGCCCAAGATGCCCgaatcttaaacccaac
ggttacctcttatctagtggGCAAGGGCTAAATTTGTAGCTGGGTAGTTGGCCTCTAGGCgaatcttaaacccaact
ggttacctcttatctagtggGAGATAGCCTGGGTTAGTGCCCTCTCAGAGCTTGGGGAGGgaatcttaaacccaac
ggttacctcttatctagtggAGGGCATCTGGGACCCAAGCTCAAGGGGAAGAGCCTTGGCgaatcttaaacccaa
ggttacctcttatctagtggGTGCTTGTTCCACAGGAAGGATGGTGCAGAGCCAGGCTGGgaatcttaaacccaac
ggttacctcttatctagtggGGGCCTGCCCCTCATCCATCCAGCACCATCAAAGGCCTTTgaatcttaaacccaac
ggttacctcttatctagtggGGGACAGATAGGGAGCTGTTGCAGGAGAAAGAACTGCCACgaatcttaaacccaac
ggttacctcttatctagtggGGTTCCGTGGTGAAGGGAAGACACAGCTCTGCAGTTTAATgaatcttaaacccaact
ggttacctcttatctagtggTGAAAACAGGGTCTGCTTCACACTTGAGCTAGAGCCCTCCgaatcttaaacccaac
ggttacctcttatctagtggGCCTGTCATAGGCCTCATTATTATGGTAGGCACTATTATTgaatcttaaacccaacttg
ggttacctcttatctagtggCCAGTCATCACTGGCCTAGCTTGCATGGTGGTTTAATGCTgaatcttaaacccaactt
ggttacctcttatctagtggTGCACAGCTTGGGGATGGCAGGATCCACACTCTGCTCCCGgaatcttaaacccaac
ggttacctcttatctagtggTTCCAGCACTATGCGAGACCGAGGTGGGCAGATCACGAGGgaatcttaaacccaa
ggttacctcttatctagtggGTGGTGGTGCAAGCCTGTACCGTGCAGCTACTCAGGAGGCgaatcttaaacccaac
ggttacctcttatctagtggACAGCTCGTTCCTCTTCAGGCCTTCCCGGTCCTGGGAGCGgaatcttaaacccaac
ggttacctcttatctagtggGGTGCACACAGGCATGCGCATGGCGGGGAGGTGTGGGTGCgaatcttaaacccaa
ggttacctcttatctagtggCCGTGCCTCGTTCTGCAGGGCAAGGCCGAGAAACAGCGGGgaatcttaaacccaa
ggttacctcttatctagtggGGCCATGGTTTCTAATTCCCTGGCACCAGAAGCCCTGCTGgaatcttaaacccaac
ggttacctcttatctagtggGGGAATGACAGGTGGTGAGGTGAAGGGAACACGGGCGTGGgaatcttaaacccaa
ggttacctcttatctagtggGGACAGAGTGGTTGGTTGGCTGATCTAAGTGTGGAGGAGGgaatcttaaacccaac
ggttacctcttatctagtggCAGCACCTACTATACCAACCTTAGAAAACTTCATACTTTCgaatcttaaacccaactt
ggttacctcttatctagtggGTCCTGCCTCAGCCCCCCGAGTAGGTGAGACTACAGGTGCgaatcttaaacccaa
ggttacctcttatctagtggTTGTATTTTATTGGAGCTGGGGTTTCACCGTGTTGCCCAGgaatcttaaacccaactt
ggttacctcttatctagtggGGGGAAGACAGGGCAGGTTGAAAGGTCTGCTTTCCAGGGGgaatcttaaacccaac
ggttacctcttatctagtggGGCTCAGAGAGGAGAAAGTCAGGCGCCAGAGGTAGATGAGgaatcttaaacccaa
ggttacctcttatctagtggCTGGGATCTGGATGTGTGCCTTCTGTGGCTCGGTGACTCCgaatcttaaacccaact
ggttacctcttatctagtggCCCATAGCTGACAGCCACGGGAGCCATGTGCCAGGCCTGGgaatcttaaacccaa
ggttacctcttatctagtggCGGTCACAGCACCAAGATGGGAACCCAGCAGGTGGCCTGGgaatcttaaacccaa
ggttacctcttatctagtggTGGGGTGGGGTGGGGTAGGTGGGCTCAGGCCTGCAGAGCTgaatcttaaacccaa
ggttacctcttatctagtggGAGCAGAAGGGAGATTTGGGATGTCTTCTGAGTGCTAGCCgaatcttaaacccaact
ggttacctcttatctagtggACTGATGACTATCAGTCCCCTGATTGCTTCCTGTCTTATGgaatcttaaacccaactt
ggttacctcttatctagtggGGGGATATAAAAGTTAAAAGACCCAGGCTGGGCGCGGTGGgaatcttaaacccaac
ggttacctcttatctagtggGTCCCAGCTACTCGGTAGGCTAAGGCAAGAGAATCGCTTGgaatcttaaacccaac
ggttacctcttatctagtggAGGAGGCAGGAAACAGACATCTTTAATGGGCCAGAGGCTGgaatcttaaacccaac
ggttacctcttatctagtggAGGGGGTTGGGACCTAGGTGAGCAGCCCGTCAGGAGCCCCgaatcttaaacccaa
ggttacctcttatctagtggGCACTCCTAGCCCCAAGTGCAATATACAGGTGAGGCGCCTgaatcttaaacccaac
ggttacctcttatctagtggGCAGGGAATTGGCATCCACTTAACCATCCCTAATGCCTGCgaatcttaaacccaac
ggttacctcttatctagtggCTGGCCTTCCCCGGATGCAGTATGTGATTCCTGGAAGGGGgaatcttaaacccaac

ggttacctcttatctagtggGTCGTAGTTTTGTCATCTTCAGGCAACCCAGGATGACATGgaatcttaaacccaactt
ggttacctcttatctagtggCACTCTCCCTCTTTTGCTAGAAGGGATCGGCTTAGGCAGCgaatcttaaacccaact
ggttacctcttatctagtggCAGCCTGTTACTGCAGTTTCCAAGTTGGAGGGTCCGTGAGgaatcttaaacccaact
ggttacctcttatctagtggCAGAGGGGTACATGGGGAAGCTGAGCCTCTGATGAATGGGgaatcttaaacccaac
ggttacctcttatctagtggGTGGCTCTCCTCCATTTGCCTCCCTGGTCTTGCCTCCTGGgaatcttaaacccaact
ggttacctcttatctagtggCAGCTTGGGGACATTACAGATAAGGCTGGGCGGGTGTGCCgaatcttaaacccaac
ggttacctcttatctagtggTCACCGAGAGAAGTTAGGAATCGACCTTCCTTGAGGTGGGgaatcttaaacccaac
ggttacctcttatctagtggGGATCTTTGTGGCCTTCTGGATGGATGCACCACGATAGGGgaatcttaaacccaact
ggttacctcttatctagtggGGCCTCTGCAGGGTTTGCTTTGAGGTACTTCCTTCCTGTCgaatcttaaacccaactt
ggttacctcttatctagtggGGAGTCTGTCTCCATTTTTCAGACACAAAAACTGAGGCATgaatcttaaacccaactt
ggttacctcttatctagtggTTGTGGGGAGAGAGTATTATTGTTTCAATTTCCTGGACACgaatcttaaacccaacttg
ggttacctcttatctagtggGAGCACATACTGCAGGCCAGGCATGGTGCTGCATGCATTCgaatcttaaacccaac
ggttacctcttatctagtggGGGGGGCGGGCGGGCACGAATAGGGGAGGAATTTAGATGAgaatcttaaacccaa
ggttacctcttatctagtggCAGCCAAGGTGCCTGCCTGGGGACACAGGGTGCCATAGGCgaatcttaaacccaa
ggttacctcttatctagtggGCCCAGGGTGTGGACTTAGGGTGGTCAGGGGAAGGAAGGAgaatcttaaacccaa
ggttacctcttatctagtggGGCAGGAGGGAGACAGGAAAGGTAAGCAGAGGCCAGATCCgaatcttaaacccaa
ggttacctcttatctagtggGTGCTGAGGGCGATGGTCTCATGATGCAGATTTTAAGCAGgaatcttaaacccaact
ggttacctcttatctagtggCAGAGGGAGGTCCAAGCTGGTGGCCAGTGGTGACGGGGCCgaatcttaaacccaa
ggttacctcttatctagtggGAGAGAAGTCCCTGAACGCGGTGCACGAAGAGCCTCTGGGgaatcttaaacccaa
ggttacctcttatctagtggGCGGAGGTGATGGGGATTGTTTGGGCTGGAGTCTGGAGGGgaatcttaaacccaac
ggttacctcttatctagtggCCCCAAGCTGGGTGAGCATCTAAGGTAAGTGAGGGCTGTGgaatcttaaacccaac
ggttacctcttatctagtggCCAGGTTCTCCTACCAGCAAGCTGCCTTGAAGACTCTGAGgaatcttaaacccaac
ggttacctcttatctagtggTAACCTTGTCCGTACCACTGAGCACCTGTTACCCATACACgaatcttaaacccaact
ggttacctcttatctagtggTCCAAACCACGGAACAACTTGAAGGATTGTTTTGCCAGACgaatcttaaacccaact
ggttacctcttatctagtggTAGGGAGTGGGGATGGCCTTCAGGAGCTCCAGAGGCCCCCgaatcttaaacccaa
ggttacctcttatctagtggTCTGACTCCTGGTTTCTGGCTTCAACAATGACTAACCTGTgaatcttaaacccaacttg
ggttacctcttatctagtggAGCACCTCTGCATTCTCCAGGTTTGTTGTGAGGACGAAGGgaatcttaaacccaact
ggttacctcttatctagtggCTGGGCCTGCAGGGGCACTCCCAGCTAATTTTTATATTTTgaatcttaaacccaacttg
ggttacctcttatctagtggGAGCCACCGCTCCCGGCCATGAATGTAGAATTCTGAGCCCgaatcttaaacccaac
ggttacctcttatctagtggCACTGGGAGTGGGCCACACATTTAGGGGGTTGTTGTTGTAgaatcttaaacccaact
ggttacctcttatctagtggCGGTGGCACTGCTTTGCAGCCTAATGGTTTTTGACACATGgaatcttaaacccaactt
ggttacctcttatctagtggAATCCTTAAGCCAGCACCTTGAGTTGGGTGTGATCAACTCgaatcttaaacccaactt
ggttacctcttatctagtggGACAGGGCTAGGGGTTACCATCCCTCTTAAGGCCTAGGGCgaatcttaaacccaac
ggttacctcttatctagtggTCACACCCAGGTTGCGGTTTTTGTCCTTGTCTTCTGGAGGgaatcttaaacccaactt
ggttacctcttatctagtggGACGAGGCTGGAGTGCGGGGGTGCTATCTCGGCTCACTGCgaatcttaaacccaa
ggttacctcttatctagtggTGAGCCTCCATGCCTGGCCTCATCTGCATTTTAGGGTCCCgaatcttaaacccaact
ggttacctcttatctagtggGGGCCCCAATGGACCTCTGGCCTGCGGGATTGAGATCGGGgaatcttaaacccaa
ggttacctcttatctagtggCTTTCCTCAACAGAGTCTGGGTCTCACCCTGCCCAGCAGCgaatcttaaacccaac
ggttacctcttatctagtggGCCTTCCTGCCCCTGCCCAGGCTGACTTCACACCATGCTCgaatcttaaacccaac
ggttacctcttatctagtggGACTGCCTGCCCTCTACCTTGGATGGTGCGGTCCACGGGGgaatcttaaacccaa
ggttacctcttatctagtggGTGTGTCCCTGGTGCTTCTCTCAGTGTCTGACACCTGGTGgaatcttaaacccaact
ggttacctcttatctagtggCCTGTTTGGTGCATGGGTGAGTGTGGTTGTTTACACTAGCgaatcttaaacccaacttg
ggttacctcttatctagtggGTTCTCACCCACTCCTGCCTCCTCTGAGAAAAAGTGTGGCgaatcttaaacccaac
ggttacctcttatctagtggCTTTTCCATAGAAAGGCAGCACCCCCCACCCCCCTCCACCgaatcttaaacccaa
ggttacctcttatctagtggTGAAACCAACTTTGTACATCTCCACTCTGACTTCTGGGCCgaatcttaaacccaactt
ggttacctcttatctagtggTGTGTCAGGCATGATTTTTGGCACTTGGGAAACATCAGGGgaatcttaaacccaactt
ggttacctcttatctagtggGGCCCAGGTGGGGTTGGAATGGAGGAATGGAGTCCTGCCCgaatcttaaacccaa
ggttacctcttatctagtggGTCCTGCGTGCCTTGGGACCTGGTCCTGGTGGTTCCCTCCgaatcttaaacccaac
ggttacctcttatctagtggCGTCCAGTAACGTTTGTCAGGGGTTGAATGTCGAGTGGAGgaatcttaaacccaact
ggttacctcttatctagtggGGAGTGCCATGACGACGATGACCCCCAGGAGCGTCCTTCCgaatcttaaacccaa
ggttacctcttatctagtggGGGAAGGCTGGGCAAAGAGGGGCAAGTCAGCCATGCCTGGgaatcttaaacccaa
ggttacctcttatctagtggGCAGCAGGCCTGCATGCACCTGGATGTCAGAGTGGGAGGCgaatcttaaacccaa

ggttacctcttatctagtggGGCGGATACAGCAGCGACCAACCGAAGCCACCAGTGGGAGgaatcttaaaccca
ggttacctcttatctagtggGCCACACTGGGCTTCAGACCCTCCAGAGTCAAAGCCTGGCgaatcttaaacccaa
ggttacctcttatctagtggTTTTGTGGCTGCAGTAACAAATTTCGACAGATTTGGTAGCgaatcttaaacccaactt
ggttacctcttatctagtggGTCAGCGAGGTTGACACCCTCTGAGGTGCTAAGGGACACCgaatcttaaacccaa
ggttacctcttatctagtggTCCAGCTGCTTGTGGCACTGGATGTTGCTCGGCTTGTGGCgaatcttaaacccaac
ggttacctcttatctagtggCTCTGCTCTTGTCTCCACGTACCTTCTCTATGTCCGTCTCgaatcttaaacccaactt
ggttacctcttatctagtggCTTGAGCTCAAGCCATTGTACTGCCTCAGCCTCCCACAGCgaatcttaaacccaac
ggttacctcttatctagtggGCGTGAGCCCCCACTCCTATCCAAGATTCTTAACTTCATTgaatcttaaacccaact
ggttacctcttatctagtggGGGTGAAGACAGCATTCAACCCACACTAGTAGGCCCCAGCgaatcttaaacccaa
ggttacctcttatctagtggAGACCTCATATCCAGATGCTCTTTTCTGTGGTTTTGTCACgaatcttaaacccaacttg
ggttacctcttatctagtggCACATTGAGTAAAAGCTGCAAGAAGAGTACAAAGAACTCCgaatcttaaacccaac
ggttacctcttatctagtggCCCAAGTTCACCAATGAAAAAATGTTGCCCCCTAAGTGGGgaatcttaaacccaac
ggttacctcttatctagtggGCCTCAGGCTCCCCCTACCCCTCACAGCGTCAGACAGAGAgaatcttaaaccca
ggttacctcttatctagtggCAGGCAGCTGCTCACTCTAAGGCTTGCTGTAGGCTCTGGCgaatcttaaacccaac
ggttacctcttatctagtggTGTGGCATCGTCCTGGCACAGTCACAGAGAGGCCAAAGACgaatcttaaacccaa
ggttacctcttatctagtggCCCCCCATTTAAAGCCAAATGGCCATCCTGGGTCCTGGGGgaatcttaaacccaa
ggttacctcttatctagtggTGTTCTCTCCACTCTGCCCTTGGATGTGTCAGTTGCAGGCgaatcttaaacccaac
ggttacctcttatctagtggCGGCTCCCTGCACCGCCTGCTGCCTTTTGGTCCATTTTGAgaatcttaaacccaac
ggttacctcttatctagtggTGGCAGGAGATGAGCAGGCTGTTCAGAGTCCGCCAGGAGCgaatcttaaacccaa
ggttacctcttatctagtggGCAGACTTTAGGTGAGGGGGAGAGGCAAGGAGGTTACAGGgaatcttaaacccaa
ggttacctcttatctagtggGCAGAATCAGGGATGGAAAACCAGTGAAAGAGAATGTCACgaatcttaaacccaac
ggttacctcttatctagtggGAGGTTTTGCCAGAGGCACATGGAGGGGGTAGCTTGCTTGgaatcttaaacccaac
ggttacctcttatctagtggAGAGTCCCTATAAATGCTGGTGGATCAATGTAGCTGATGGgaatcttaaacccaact
ggttacctcttatctagtggCCTGCCACTGTATGGTATTGAATTTACCTGGGTTATGGAGgaatcttaaacccaactt
ggttacctcttatctagtggGAAGGACTTTAACCAGGTACTCTCTGGGTCCTGTTTCTGCgaatcttaaacccaact
ggttacctcttatctagtggCTCAGGACACAGTGATAACAAGATGGGCTTCATTCCTGCCgaatcttaaacccaac
ggttacctcttatctagtggGCCTAGTGGAGAAAACAGATAATACCATTGGAAAACCCACgaatcttaaacccaac
ggttacctcttatctagtggACGCAGAGGGGGTGTTCAACTTAGACTGAGGGCCGGGGTGgaatcttaaacccaa
ggttacctcttatctagtggGTCATGGCCTAGATCATGGCTCATGGCCTGTAATCCCAGCgaatcttaaacccaac
ggttacctcttatctagtggATTCTAGACAAACCTGGGCAAGATGGTGACACCCTGTCTCgaatcttaaacccaac
ggttacctcttatctagtggTGGTGGTGCATGCCTGTGGTCCCACCAACGTGAGAGGCTGgaatcttaaacccaa
ggttacctcttatctagtggGGAGACTGAGGTTGAGAAAAGTTTGGGCTGTCTGAGAAACgaatcttaaacccaac
ggttacctcttatctagtggAGTGTCAGTAGAGTTCAGTGAGTGAGGGTGCAGATGGGGGgaatcttaaacccaac
ggttacctcttatctagtggGAGATGTGAAAGAGGGGCAGGTGCAGAAAGATCAAAATTGgaatcttaaacccaac
ggttacctcttatctagtggGGGCCAGCACAGAGGCAGGGAGAAAATTTAGGAGGCTTCCgaatcttaaacccaa
ggttacctcttatctagtggCAGTCAGAGATAATGGTGATCAGATGGAGATGGAAAGAGGgaatcttaaacccaac
ggttacctcttatctagtggTGAAGAGCAAGTCACTGTGTTAGGGGCTGGGTGAGATGAGgaatcttaaacccaac
ggttacctcttatctagtggAGAGAACCTGACCCTCTGGCCAAGACCTTCAGAAGCTCAGgaatcttaaacccaa
ggttacctcttatctagtggGGCCTGTGAGCTGGGCCATGAAGGGTGGGCTCTGATCTATgaatcttaaacccaac
ggttacctcttatctagtggGGGGGAAAGCCCTCAAAGCAGAAAGAGCAAAGCTGCATGGgaatcttaaacccaa
ggttacctcttatctagtggGCATGTGATTCAGGAACCGTGAGCAATGGTAGACTCGTGCgaatcttaaacccaac
ggttacctcttatctagtggGCTGAGGTCTGACTGTGAGAGTTTGAAGCCCCACTGAGAGgaatcttaaacccaac
ggttacctcttatctagtggCTGGTGCTAAGAACTAGTAAGGTATAAGATAAGCCTAATCgaatcttaaacccaactt
ggttacctcttatctagtggGAGGTGTGAGGCAGAGCCAGTACATAATGTCTACGTTCTGgaatcttaaacccaac
ggttacctcttatctagtggTGTAGAGCATTTACCTTGTGCCAGGCAACATTCTGGATGCgaatcttaaacccaact
ggttacctcttatctagtggACGCAGGGGTTTGAATCTTAGCTGTTTGAGGTCATTGTCCgaatcttaaacccaactt
ggttacctcttatctagtggCCAAACTCCAGGAGTAGACAGATACCCACCAGGACTAGGCgaatcttaaacccaa
ggttacctcttatctagtggGGCCAGGAACAAGGGTGAAGGCATGCACTCTGTGAGCAGCgaatcttaaacccaa
ggttacctcttatctagtggCAGTGGGTCATGGGACTGAAGGAACCATGTTACACATGCCgaatcttaaacccaac
ggttacctcttatctagtggCCCTTGGGGAGAAGTGAGTGAGCAGGTGGAGAGACGATAGgaatcttaaacccaa
ggttacctcttatctagtggGATGGAAACACTGAGGGCTAAGAAGAAGGGCCTCTCCTCCgaatcttaaacccaa
ggttacctcttatctagtggGCAAGGCTGGTTCTATTTAAGAATGATATATAGTCTAGTGgaatcttaaacccaacttg

ggttacctcttatctagtggCTGTGCCTGCCACATGCTCAGCATTATAATTAGAGCCACCgaatcttaaacccaact
ggttacctcttatctagtggATCCGTGATCCTTTATGCTGTCAAAGCCTAAATTCCCCACgaatcttaaacccaactt
ggttacctcttatctagtggTGCTAGCATCAGCACATTCCAAAACTGCCTGCCCCATCCCgaatcttaaacccaac
ggttacctcttatctagtggGTGTGATTGCTGTCACAAAACTAGCAGTTAGAATCTTTTGgaatcttaaacccaacttg
ggttacctcttatctagtggCTAAAGGGTGGAAACTTAGAAACTGAAGTGGATTTACCACgaatcttaaacccaactt
ggttacctcttatctagtggAACTAAATGTACCAACACAGGCTCAAAACACTCTAAGATAgaatcttaaacccaactt
ggttacctcttatctagtggCTGGGTAACATTGAATCACAAACAGTTGAAAACACTGGGGgaatcttaaacccaact
ggttacctcttatctagtggGGCAAATCACTGGAGATAAATCATTGTGTACAGATTTCCCgaatcttaaacccaactt
ggttacctcttatctagtggGATGTATCTCTAGATCGTGTGCTTGCCAGAATCTTTGCTGgaatcttaaacccaactt
ggttacctcttatctagtggACGGTGGCCAGGGAAGATCCACTGAATTAGGAGTCAGCAGgaatcttaaacccaac
ggttacctcttatctagtggGCTTCGTCAAGTCATTTAAGCTTGGTACCCGTCAGTTTCCgaatcttaaacccaactt
ggttacctcttatctagtggGCAGCTTCTTAAAGTCCAGCTCACACCTCTGTCAACTCCCgaatcttaaacccaact
ggttacctcttatctagtggTGGACATATATAGGTCCACTCTTCTCATATCCACAGCTGCgaatcttaaacccaactt
ggttacctcttatctagtggGGTCATTATCTCAATAATCCCTTAGTGTAGGGAAAAGGGCgaatcttaaacccaactt
ggttacctcttatctagtggTTCATTTTCTGTCTTGAGCATTAGCAAGCCACAGTCCATAgaatcttaaacccaacttg
ggttacctcttatctagtggTTGACCTGTGTCTGTTTCTCCAAGCCAAAGGGTAAGTCAGgaatcttaaacccaactt
ggttacctcttatctagtggGTTAATATCCAGTGCAATAATCTAAAACCTACTGGTCATCgaatcttaaacccaacttg
ggttacctcttatctagtggGCGATTTCCTTTTATGAAGATCTGGAAGTTACAAAGACAGgaatcttaaacccaacttg
ggttacctcttatctagtggGTCACTTGGACTTGAAAACTGGTGGTCCTGGGATAGAGCCgaatcttaaacccaact
ggttacctcttatctagtggTCCACCTCCCGGTTCAAGCAATTCTCCCAGCTTAGCCTCCgaatcttaaacccaac
ggttacctcttatctagtggGGCATGAGCCACCACCCCCGGCCAATCCTAATTAATATTGgaatcttaaacccaac
ggttacctcttatctagtggCATTCTTATACCAGCCGTTCTCAACAACATAGTGTCTGCCgaatcttaaacccaactt
ggttacctcttatctagtggATACTCCCTGTGTTCTTGTTGATGAAGGAAGTAGGATGAGgaatcttaaacccaacttg
ggttacctcttatctagtggGTCTACAGGAGGTATAGATTCATTTTTCTATGATATAGGGgaatcttaaacccaacttg
ggttacctcttatctagtggTGTAAAAAAGGCTGGGAATAATTGCTTACTGCTGGAAAGCgaatcttaaacccaactt
ggttacctcttatctagtggGACTGTCCAGGCCTTCTCCCACATCTCCAGAAACCACCCCgaatcttaaacccaac
ggttacctcttatctagtggTTGGTAACCCAAGCATAGAGAGCTAGACACACCTAGCCAGgaatcttaaacccaac
ggttacctcttatctagtggGCAAGTCAAGAGAAGTGCTGGACCTGCTGAAAATAACAGGgaatcttaaacccaac
ggttacctcttatctagtggAAGAACATCAGCAAGTGTTTAGTAAGGGAGGGCCAGTGCCgaatcttaaacccaac
ggttacctcttatctagtggTGTGTTCTCTATAGTGTCCTCTGTGCCTGTATATGAAGCCgaatcttaaacccaacttg
ggttacctcttatctagtggTGCCTGATTCCCAAGGCACGAGTCCATCCTCTGCCCAGGCgaatcttaaacccaac
ggttacctcttatctagtggTACCAGAGCACATTTTCCTGCTGACATCTGTTCCAGTGGGgaatcttaaacccaact
ggttacctcttatctagtggCTGCTCAAGAAATGGCTCTAGACCAGCGGTTCTCAAACTTgaatcttaaacccaact
ggttacctcttatctagtggGCCCACAGACCACACTGTGAGTGGCAAGGCTGTGAGCTCCgaatcttaaacccaa
ggttacctcttatctagtggCAGGTTGGGAGGGTGATTACTGGAGATGAAAGATGAAACCgaatcttaaacccaact
ggttacctcttatctagtggCCTAAAGCAAAACTACCTACTCATAGGGCTGCTCTGAGGGgaatcttaaacccaac
ggttacctcttatctagtggGTCACACAACGCTCCTAGCACTGTGCCTAACATCTAAACCgaatcttaaacccaac
ggttacctcttatctagtggCCATGCCAAATGATGGGTTCCATCACTCTTATTAGTAGTCgaatcttaaacccaacttg
ggttacctcttatctagtggCAGAAAGCCTGGTTTGTAGAAGTCAATGAGAAGAGTGGGGgaatcttaaacccaact
ggttacctcttatctagtggGCACAGCAGAATTCAGAGGAGGTCTACTCTAGTGCAGGGCgaatcttaaacccaac
ggttacctcttatctagtggCCCCCGATCAACTGAATAGGAATCTCCGGTAGTCAGGTCCgaatcttaaacccaac
ggttacctcttatctagtggCTAGAGCTCCCCGAGTTCTGAGGTGGACTCTCATCTCTGCgaatcttaaacccaac
ggttacctcttatctagtggCTGTGCCCCATTTTCCTAATTTGTAAAATTAGAGTAATGGgaatcttaaacccaacttg
ggttacctcttatctagtggACAGCTTAATTTATGACTATGCTACATTTGGCCCTACCAGgaatcttaaacccaacttg
ggttacctcttatctagtggGGTGCATGACTATTACTATCAATATCTAAGACCTAGTACCgaatcttaaacccaacttg
ggttacctcttatctagtggCTACGGAATTCTGTATTTAGACCTGTTTGTATTCAGGATGgaatcttaaacccaacttg
ggttacctcttatctagtggGTGGTTCTACCATTTACTACATATGTGACCTTGAGCAAGGgaatcttaaacccaacttg
ggttacctcttatctagtggATCAGCAGCCTATTTCCCTATCTATCAACAAGTATGATGGgaatcttaaacccaacttg
ggttacctcttatctagtggTGGCACTGAATAGGGGCTTACTAAGTGCATTTCTCCACCCgaatcttaaacccaact
ggttacctcttatctagtggCCCCCAGTTTCTATAGTCTACTGGAGAATACAGGAAACAGgaatcttaaacccaact
ggttacctcttatctagtggTCCTTGTATTAGAGACATACAGTGATAAAGAGCTAGAGTCgaatcttaaacccaacttg
ggttacctcttatctagtggTATGTATATGTGCTGGGTTCCTTGATATGGTACATCTGGGgaatcttaaacccaacttg

ggttacctcttatctagtggTTGTATGTGCATGGTTAGCCATCGTTTTGATTTTTGGTGGgaatcttaaacccaacttgt
ggttacctcttatctagtggTGATGTCTTTGAGAACACAGAGTCCTCGAGGGCTTGGATGgaatcttaaacccaact
ggttacctcttatctagtggCTCCAAGTTGCATAAGAGGCCATTCTACAGGAAGCAGAGGgaatcttaaacccaac
ggttacctcttatctagtggGTTCCATCCCACTCGACAAGGGGCAAAAGTGACTGAGAAGgaatcttaaacccaac
ggttacctcttatctagtggCGTAGAACTTGAAAGATGTATTCAACCTACCAATTCCCTGgaatcttaaacccaactt
ggttacctcttatctagtggCGAAACACTAATGCCTACTCAAGACTCAGAAGATGGGTTGgaatcttaaacccaact
ggttacctcttatctagtggCAGCAGCAGTACCTATGAGCCCCAGCCATGTCTCGTGTCCgaatcttaaacccaac
ggttacctcttatctagtggGGCCATGTTGGGCAATGTGTCTTGTTACTTGCCTGTGTGTgaatcttaaacccaactt
ggttacctcttatctagtggACCTACCACCCTCATTGAAGCCTACAAACCAGGCACTCTGgaatcttaaacccaac
ggttacctcttatctagtggACACTGCATTTCTAAAAGCCTCCTGGAACATTTTACTTCCgaatcttaaacccaactt
ggttacctcttatctagtggCTTGGGTTGAGTCCAAGCTTGAAAAATTTCATCTGAAAATgaatcttaaacccaacttg
ggttacctcttatctagtggGCCATGGGGCACTTTTCTCCCTAGCCAATCAGCCAGAGCTgaatcttaaacccaac
ggttacctcttatctagtggGGGTAGACCCAACCCCCTCTGTGAGACTGCTCCTGCTAACgaatcttaaacccaac
ggttacctcttatctagtggGGAAACATGGTCTCCTTAGCAAGGCTGTACTAAACCAGCCgaatcttaaacccaac
ggttacctcttatctagtggGGAACAAGTGAGCTCCATGCCAACTGGTTTTCAGCCCCAGgaatcttaaacccaac
ggttacctcttatctagtggCAGGCTGTGAACTCTTTCCACCACCTGCGTTCTTTAATACgaatcttaaacccaactt
ggttacctcttatctagtggGCCACAGTGATGGCTTAGGAGCTGCCAATACCAAACGTATgaatcttaaacccaac
ggttacctcttatctagtggGCTGCAACTGTACTGAACCACCTACAGATGCCCACACATGgaatcttaaacccaac
ggttacctcttatctagtggCCTAAGAGGCCCCTCCCTGAATTTTCTCCCTCATGGCTCCgaatcttaaacccaac
ggttacctcttatctagtggCACCTCTCTTTCTGGTTCAGGACATAGGTCCGATCCTTACgaatcttaaacccaactt
ggttacctcttatctagtggGTCCTTGGACTATTCTGCAAGATCTCAGGTCTGCACTCACgaatcttaaacccaact
ggttacctcttatctagtggGTTTGCTTCCATCTCAGCACTTCTCTTCCTATTACTTGCCgaatcttaaacccaacttg
ggttacctcttatctagtggTGACATACATACAAGCCATTCACTGTTGTTTAGGTGACAGgaatcttaaacccaactt
ggttacctcttatctagtggGCCAGGCACTGTGCTTGGTACTTCAAGTGAATTTTCTCATgaatcttaaacccaactt
ggttacctcttatctagtggCAAATATGCCGTGAAGGTGGTATTATTACTAACATCATCAgaatcttaaacccaactt
ggttacctcttatctagtggACCTTGTGCTCAGAGAAGCTAAGTAATTTAGAATTGTTCCgaatcttaaacccaactt
ggttacctcttatctagtggTCCAGTTCTGGAGTATTCTGAATGGAATCCCATGTCTGTCgaatcttaaacccaactt
ggttacctcttatctagtggCCATGCCATCATATTCCTTTCCTTACTTCTAGGGAATAGCgaatcttaaacccaacttg
ggttacctcttatctagtggCAGGGACCTGAAAATGCTCTGGTGTGTCATTTAGAACCCGgaatcttaaacccaact
ggttacctcttatctagtggGGAGACCACGAGTTTGCCCACAGTTACACTCCCTTCCCAGgaatcttaaacccaac
ggttacctcttatctagtggGCGGGGCCACCCTAGTGCTAGAGCTCCCTGTGGAATGGGCgaatcttaaacccaa
ggttacctcttatctagtggCTTCTCCCTCTGCCCAATCCTGTTTCCCTCACTCTCTCTCgaatcttaaacccaact
ggttacctcttatctagtggAGCTTCATCCATGTTCCTAGAAAGGACGTGAACTCATCCTgaatcttaaacccaact
ggttacctcttatctagtggGATCACCATTCTAACCGGTGTGAGATGCTATGTCACTGTGgaatcttaaacccaact
ggttacctcttatctagtggCTCTGAGGCCCTATCTCTCTTATAAGTGTTGTTCCTGAAGgaatcttaaacccaactt
ggttacctcttatctagtggGTCTCCATCTGAGACTCTGTTTCTCAAAAAACTCAACCATgaatcttaaacccaactt
ggttacctcttatctagtggAGGATAGAATGCATGTCTCTCCAAATGCACTCTCTGGCCCgaatcttaaacccaact
ggttacctcttatctagtggACTGGACCTGCTTTGCAATGAGAATGGTCAGCCCTTGAGCgaatcttaaacccaac
ggttacctcttatctagtggGCCTCTGTTTTGCATGGAACTCCACCAAGGCATCAGGCACgaatcttaaacccaac
ggttacctcttatctagtggGGCGCTGGCCACTCCAAAAAGTCACTCATCTCTTAATGGGgaatcttaaacccaac
ggttacctcttatctagtggGTGGCAGCCTCCACTTTAAGAAAAAGGAAAACCACTTTTCgaatcttaaacccaact
ggttacctcttatctagtggAGCCACTACTATGCATGCCAGTCAAGATTTTAAACATATGgaatcttaaacccaactt
ggttacctcttatctagtggTGACTATACTCAGATAAGAGTGGCCAATGATATAAGTAGGgaatcttaaacccaactt
ggttacctcttatctagtggACCATTCAGCCATCACACATTCCTTATTAGTCCTGATACCgaatcttaaacccaactt
ggttacctcttatctagtggTATCGTACCTCATTGTCACAGTTATGGTGAGATTATACTGgaatcttaaacccaacttg
ggttacctcttatctagtggATCTCAGGGAAGACATGCAATTGTAGCTCCCTTTATGGGGgaatcttaaacccaact
ggttacctcttatctagtggCTCTATGCCAGACCTGCATGGGGAGTTGTTGATCAATCGGgaatcttaaacccaact
ggttacctcttatctagtggTAGGTAAGATGGGTCAGTTCTAGTGATCTACTGTACAATGgaatcttaaacccaactt
ggttacctcttatctagtggAGCAATAGGTATTGTGTACTTTAAAACACGTTTAGGAGGAgaatcttaaacccaactt
ggttacctcttatctagtggAGTGTTCTAACTACGAAAGAAACACAAAGGAACAAAAGGGgaatcttaaacccaact
ggttacctcttatctagtggGGGGTCTCGTTCTGTCCCCCAGCCTGGAGTGTAGTGGTGCgaatcttaaacccaac
ggttacctcttatctagtggGAGGTGATGGGTACGTTTAGTACCTTGGTCTTGGACTGTGgaatcttaaacccaactt

ggttacctcttatctagtggCATGAGTGTATGCACATATCTGAACTTATCACATGTGTACgaatcttaaacccaactt
ggttacctcttatctagtggATCCAGAACTATGCAGAACAGGCACTCTTCCTCTCCATTCgaatcttaaacccaact
ggttacctcttatctagtggCCTTACTCTAGTCCACAGAGCTAGTAAATGCCTGAGCCACgaatcttaaacccaac
ggttacctcttatctagtggCATGAGATCAGCCTTCAGCAATTTTGAAGTGTATAGTCACgaatcttaaacccaactt
ggttacctcttatctagtggGGTACAGTGTCATAGAGCTGATATCTAGAGTTCATTCATCgaatcttaaacccaactt
ggttacctcttatctagtggGAGTAACTTCTCATTTCCTCCTTCCCTTTAGCTCCTGACCgaatcttaaacccaactt
ggttacctcttatctagtggAGCATAGTGTCCTCAAGGTTCATCTATGTTATCTATATTGgaatcttaaacccaacttg
ggttacctcttatctagtggTGTTTTAAGGCTGCATAGTATTCTATTGTGTGTAGAGACCgaatcttaaacccaacttg
ggttacctcttatctagtggCTTTCTTGCTTGCTTGCTTTCCTGCTTTCTCGCTTTCTTGgaatcttaaacccaacttgt
ggttacctcttatctagtggCAGGGTCTCACTCTGTCACCCAGACTTTACTGAAGTGCCAgaatcttaaacccaac
ggttacctcttatctagtggCCTCCTGGGTAGCTGGAACTACAGGCAGAGGCATGCGCCCgaatcttaaacccaa
ggttacctcttatctagtggGCATGCTTTGCTCACGAGGCTGGTCTCAAACTCCTGGGCTgaatcttaaacccaac
ggttacctcttatctagtggGGCATGAGCCACCACACCCAGCTACTGTATTTTCTTAATCgaatcttaaacccaact
ggttacctcttatctagtggTAGTGTGTTTCCACATCTTGGCTTTTGTGAACAGTGCTGCgaatcttaaacccaactt
ggttacctcttatctagtggCATGACAATGATGGCCACAAGAACATTTGGGCTCACAGCCgaatcttaaacccaac
ggttacctcttatctagtggGCAGTCCCAACAGAGAGAATAGTTACCCTTTCCTAATTGCgaatcttaaacccaact
ggttacctcttatctagtggGTTGGCTTTGATCACATACCCACTTCTGAACCAAGCACTGgaatcttaaacccaact
ggttacctcttatctagtggTGTCCAGAACCGAGGAATGTACTAACTTTCAAGCTAGAAGgaatcttaaacccaact
ggttacctcttatctagtggGGAGGGGTGAACCAAAGGAGAAGGAAAGAGGTTGGGATGGgaatcttaaacccaa
ggttacctcttatctagtggCTCCTCCACTCTTACAGTGTCTGCACCGAAGCCTGACTGCgaatcttaaacccaac
ggttacctcttatctagtggGGCAGAACAGCAATTCTGGCCTGCCACTTGCATGGAAGTTgaatcttaaacccaac
ggttacctcttatctagtggTGTGCAATCTGGTACATAAATACCCCGGCTCCCTTACCCCgaatcttaaacccaac
ggttacctcttatctagtggCTGAGATATATATGCTAGGAATATCGTCCTTAAAGTTCTCgaatcttaaacccaacttg
ggttacctcttatctagtggGCCGGGAGTGGTGGCTCATGCCTGTACCCCTAGCACTTTGgaatcttaaacccaac
ggttacctcttatctagtggTGGTGCATGTCTGCAGTCTCAGCTACTTGGGTGGCTAAGGgaatcttaaacccaac
ggttacctcttatctagtggCAGAGAAAGACCCTGTCTCAAAACATATATGTGCATATCCgaatcttaaacccaact
ggttacctcttatctagtggTTCTTCCATTCTCTGTGTCACTTCTCCATATCCCTATCTTgaatcttaaacccaacttg
ggttacctcttatctagtggTGAGATAGGCTGTTCCATGAAGTAGCAAGTTTCTCAGCAGgaatcttaaacccaact
ggttacctcttatctagtggTGCAGGACCTACATTTAGACCATGCAGTCCCTTCTAAGGCgaatcttaaacccaac
ggttacctcttatctagtggCAGAGTCATTCTTGCAGAAAAAGAGATGTATGTTCATGCCgaatcttaaacccaactt
ggttacctcttatctagtggGGTTATTTGAAGGTACGAGGGTTAGCTTCCTCTTTTGTTGgaatcttaaacccaacttg
ggttacctcttatctagtggGTGCTTCCTTTGCCTGGGAAGACCTCAACATTACAGGAGGgaatcttaaacccaac
ggttacctcttatctagtggAAATTTTGACACCCACCATCATCCAGGCAGAGAAAGAGCCgaatcttaaacccaac
ggttacctcttatctagtggACATGGCTGCCACATATGTTCAATAGGAGGTGAGAGTGTGgaatcttaaacccaact
ggttacctcttatctagtggCTTTGTGTTGGTTTGGACATGTGCCCAGGAGACAGAAGTGgaatcttaaacccaact
ggttacctcttatctagtggCCACCATGATGATTTTACTTCCCCAGTGGTAATTTTCTCCgaatcttaaacccaactt
ggttacctcttatctagtggGGACAAAATTTGTTGAGTGTTAACTATGTATTGGGCACTGgaatcttaaacccaacttg
ggttacctcttatctagtggCCCAAGAAATGGGTCCCATTAAGTTGTCCACATTTTACAGgaatcttaaacccaactt
ggttacctcttatctagtggGGCTTGCTGAAGTTGACACAATAGTGAGGTGCAAGTCTCCgaatcttaaacccaac
ggttacctcttatctagtggGCATCTGCAATCAGCCCTCAGTTGGGTAGGCCGCTGCTGCgaatcttaaacccaa
ggttacctcttatctagtggCACTCACATGTTCGGAGGTTAGCTGACAACTGGCAAATCTgaatcttaaacccaact
ggttacctcttatctagtggCTCCAGCTGGTCTCTCACCCTCCAAGTGGCTAGCCTAGGCgaatcttaaacccaa
ggttacctcttatctagtggGGCAGTGGCAGTGGTATGATGTCTAGGCCCCAAACGGACAgaatcttaaacccaa
ggttacctcttatctagtggGGCTGAAGTAAGTCACGTGGCCAGCCCAAATTTGAAAGGCgaatcttaaacccaac
ggttacctcttatctagtggGTCATGCTGCAAACACACGGGGAGGCATGCGGGATTGCAGgaatcttaaacccaa
ggttacctcttatctagtggCTACCACTGTGTCCTTCCTTGCATCCTGGCACCTCTCTGGgaatcttaaacccaac
ggttacctcttatctagtggGGTGTTCTTCACCTTGGTTTTGCCTGAATGAACATCTAAGgaatcttaaacccaactt
ggttacctcttatctagtggCCTCCCACAAACAGGCCTTCGGCACCTGCCAAGACACTTCgaatcttaaacccaa
ggttacctcttatctagtggTCCTTTTACAGTAAGTGCTTTGGAATAGGGCTCTCACCTCgaatcttaaacccaactt
ggttacctcttatctagtggGCAATTCCCCATCTCTGCTCTTTTTCTTGCTCCCTGGAGCgaatcttaaacccaactt
ggttacctcttatctagtggCTGTGGCTTCTCCTATATGACAACTATTTTATATCCAAAGgaatcttaaacccaacttg
ggttacctcttatctagtggTCTCCAGGTCTTTCCAGATTTAATTATTCAACTGCTCCTCgaatcttaaacccaacttg

ggttacctcttatctagtggTGTGGTACCCAGAGTCATCTACATCCTCTGAGTAAAACAGgaatcttaaacccaact
ggttacctcttatctagtggTTTCCTCCTCTACCTCTTCTTCCTTTTGTAGTTGCAGGTCgaatcttaaacccaacttg
ggttacctcttatctagtggCTCACTCACAGTAAGTAATTTGGAACAGGACTGAGAGCCCgaatcttaaacccaact
ggttacctcttatctagtggTGTTTTCGAACCAAAGGAGAGGCATCATCTGACTTTCTCCgaatcttaaacccaactt
ggttacctcttatctagtggTCCTCTGTGAGTTCAGTCCTTTCCTTCTAGTCAGTTGCTGgaatcttaaacccaacttg
ggttacctcttatctagtggGGGTGAGGCCGTGGTTTAAAGCAGTCTCTTTCGGGCAACCgaatcttaaacccaac
ggttacctcttatctagtggCCCTGCGGTGTCAGGGGAGCATTGTCTGGTTCCCATGCTGgaatcttaaacccaac
ggttacctcttatctagtggTGTGCCCTCATGTTGACTCCCGGCTTCCTCCAAGCTCAGGgaatcttaaacccaac
ggttacctcttatctagtggAGGGCCAGTGGCAATAATGCACTGTGGCTCTCTCTCTAGTgaatcttaaacccaact
ggttacctcttatctagtggGCCATGTCTTGGCTTCCCTATGACTTGCTCACTACAATGGgaatcttaaacccaactt
ggttacctcttatctagtggGACTATGGAACTTCTTTGAGGGGCTTGCAGACTCCCTAGGgaatcttaaacccaact
ggttacctcttatctagtggGGTTTCTGTACCCTCCAGCTTGACATTTTACACCAAGGGGgaatcttaaacccaact
ggttacctcttatctagtggCACTCAGACTCCTCCCAAGCACTCTGGTGACTCTCTGAGCgaatcttaaacccaac
ggttacctcttatctagtggTCTGCTCTGTAAGTGCCTTCAAGTTCTGCCTCAATGGAAGgaatcttaaacccaactt
ggttacctcttatctagtggGTCAACAAAACCAGCCATCTGTACATCAAAAGCATTTGCCgaatcttaaacccaact
ggttacctcttatctagtggTCCAGAACTCAAAGCAACAGAGTCTCTGTGTTCAGACTGGgaatcttaaacccaact
ggttacctcttatctagtggGGGGCAGAAAGAAAAGGGACAAGGACTTCGGGAACTAAAGgaatcttaaacccaac
ggttacctcttatctagtggCATGGGGATTTATGATACTATTCTCTCAATGATAGATCTGgaatcttaaacccaacttg
ggttacctcttatctagtggCAGGAACCAGAGTGGTGAGAATGATGTGATTTGAGTGAAGgaatcttaaacccaact
ggttacctcttatctagtggGGCTTAATAGGATTTCTTCCCTTTCTCTTATTTTCTGTACgaatcttaaacccaacttgt
ggttacctcttatctagtggACAGAGGACAGTTAACTTTCTGTTATTAGAGCTCTAATCCgaatcttaaacccaacttg
ggttacctcttatctagtggGGGAAGGGAGGGGAGAGGAGAGGAGGGAAGCGAGGGAGAGgaatcttaaaccca
ggttacctcttatctagtggGGGAGGGGTGAGATGGTGAGGTGTGGAGGAAAAGTTAAATgaatcttaaacccaac
ggttacctcttatctagtggCACCAAGTCCCAGAACAGGTTGTGTGTGCCCCCTTGAGGCGgaatcttaaacccaac
ggttacctcttatctagtggCCTTTTTGTGTTCCTTGAGCCAAAGGGCCCTCATGACTGGgaatcttaaacccaact
ggttacctcttatctagtggGGAGACTGCTCCTCGTCTGTGCACAGATGAGTGGCCAACTgaatcttaaacccaac
ggttacctcttatctagtggGCTTCCTAGTCTGGTGGTGAATCCTTCATAGTCTGGTGAGgaatcttaaacccaactt
ggttacctcttatctagtggTTTCTTCTCCCCTTGCACGTTTCCTGCACAGAACAGGAGGgaatcttaaacccaact
ggttacctcttatctagtggGAGGGGAGGGGAGGAAAAGGGAGGGGAGAGGAGGGGAGGGgaatcttaaaccca
ggttacctcttatctagtggAGTGACACACACCCATCCCCACCCCAAAAGCAAAAAGGACgaatcttaaacccaa
ggttacctcttatctagtggACAGACAGAAAAGAATGTGGAGACTGTGACATGGAATCCCgaatcttaaacccaac
ggttacctcttatctagtggACAGAGGGTTAGGTAAGTGTCTAAGACAGAGCCAGTGCACgaatcttaaacccaac
ggttacctcttatctagtggCTTCTGTCTCTTCTCATACATATCCCAATCCACAACAGCGgaatcttaaacccaactt
ggttacctcttatctagtggTTCATTCATTCAGTGACTCAGCAAATAGTAAGTGCCATGCgaatcttaaacccaacttg
ggttacctcttatctagtggATTGGGAGCAAAGCAGACAAATGTCCCTTGCCCTGATGGGgaatcttaaacccaac
ggttacctcttatctagtggAAAGCAGGTAAAAAGGAATGAAGGCCCTCTGTGAAGGAGGgaatcttaaacccaac
ggttacctcttatctagtggGGCAGGCATCTTGAGACAGTTGCATTTCAACAATGGCCTGgaatcttaaacccaact
ggttacctcttatctagtggAGGCCAATATGGATGGACCACAACAGGGAGGAGAGCCTGCgaatcttaaacccaa
ggttacctcttatctagtggTCATGAGTGAAAGGAAAAGAACTGAAGGAGTTGGGTGTGGgaatcttaaacccaact
ggttacctcttatctagtggAGTGAGCCAAAGGCAGAAGCAGAGAGACCAGGTATCAGGCgaatcttaaacccaa
ggttacctcttatctagtggGGACCAGGGTGGCATTCCAGGTGGTGAGTGGTGGATGGACgaatcttaaacccaac
ggttacctcttatctagtggTGAGGTCAAGACAGCAGAAGGAGCAAGTTGGGGAGGAAGGgaatcttaaacccaa
ggttacctcttatctagtggGCAGACGTCAAATGAGTAGTTGTATATGCAGCTCTGTTGCgaatcttaaacccaactt
ggttacctcttatctagtggGGGAGTCATCAGTGTATAGATGGTATCAAGGCTGTAAGACgaatcttaaacccaact
ggttacctcttatctagtggGCAAACAGTAGAAGTGACTGAGATGGAGCAGCCTGTAGGGgaatcttaaacccaac
ggttacctcttatctagtggTGCAGAGGGAGCTGTCACTAGGTCCAATGTTGCCACTAGGgaatcttaaacccaac
ggttacctcttatctagtggCTGCAACATGCAGGCTTTTGGAGACATTAGCAAGACACATgaatcttaaacccaact
ggttacctcttatctagtggAGGGAAGAATATAAGCCTAAGCAGAGTGAGTTCCAGAGAGgaatcttaaacccaac
ggttacctcttatctagtggGCTAAGAAGGGGAGCAGGGAAATGTGGTATCAAGGAAAGGgaatcttaaacccaac
ggttacctcttatctagtggGAGTGCTTTCTGAGGCACCATCCTTGAGTTGGTAAGAGGGgaatcttaaacccaact
ggttacctcttatctagtggGGTACAAAAATGCAGAGACCGGCCTTGCACACAAACATGGgaatcttaaacccaac
ggttacctcttatctagtggGGGCAGATACAGGCAAGGTAGACGTGGTGCAGGGATTGGGgaatcttaaacccaa

ggttacctcttatctagtggGGAAACAAGATCCAGACACTATAGGTCACTGTGCAGTCACgaatcttaaacccaac
ggttacctcttatctagtggGGTCCTGGAATGAGTGAGTTGGAAAGATGGAGTGTGGTGGgaatcttaaacccaact
ggttacctcttatctagtggCCGAGATTACAGGGAGTGGCAGTGATTTTAATGCCAAGACgaatcttaaacccaact
ggttacctcttatctagtggGAGAGAAGAGGTAAGAGAGTTAAGGTCAAGTTACTGGAAAgaatcttaaacccaact
ggttacctcttatctagtggACTGCTAGTGAAGTCACCAGTTTATGATATGAACAGTATGgaatcttaaacccaacttg
ggttacctcttatctagtggGGGTTTGTAGATGCCTGCAATAAACAGGGGCAACTGAAGGgaatcttaaacccaac
ggttacctcttatctagtggGAGAGCAGGCAGGGAGAAGCATCTGGAGGCTGGAATGAGGgaatcttaaacccaa
ggttacctcttatctagtggACACCTAGTCAACCTCCAGGCTAGTGATAGAAAGAAGCAGgaatcttaaacccaac
ggttacctcttatctagtggGAAGGACTTTATGAGGAGGCAGGGATCCCAGGAGTATCTGgaatcttaaacccaac
ggttacctcttatctagtggGCTGATGACTGATCTGGAATTGTACAAGGGAGGGTGAGGGgaatcttaaacccaac
ggttacctcttatctagtggGGGAGCTAAAGGATGAGCTGGGTGTCTGGGGCTTTTGTAGgaatcttaaacccaact
ggttacctcttatctagtggCGTGGAGATCAAGGGAGGATCTTCTGCAAGACACCAGTGGgaatcttaaacccaac
ggttacctcttatctagtggGGGTGTGGGGTCTTTCTTGGAGCACCCAGGGCTCTGAGGCgaatcttaaacccaac
ggttacctcttatctagtggTAAGCATGAGCCCCCTCCTCATTCTAGCCAATGTTTTTGCgaatcttaaacccaactt
ggttacctcttatctagtggGTGAGGAAAATGTCAGTTTCTTCTCTCATGTAGCCTTCAGgaatcttaaacccaacttg
ggttacctcttatctagtggAATCTGAGTATCTATGACCTCAGAGAAGGGATCTGGGCTGgaatcttaaacccaact
ggttacctcttatctagtggTATGGCCTAAGGAAAGAGTCTACACTCAGAAGAGAAGGACgaatcttaaacccaac
ggttacctcttatctagtggTGGGGGTAATTGGAGTGGTTCCAGAAAATTAAATATTGGGgaatcttaaacccaacttg
ggttacctcttatctagtggGGAGTGTGTTAGTTTTGCAGGGGTGGCATAACAAAGGACCgaatcttaaacccaact
ggttacctcttatctagtggGCCTGTTCCCTGTTTCTCTCCTAGCTTCTGGAGGTGCCCGgaatcttaaacccaact
ggttacctcttatctagtggAGATGCATCACTCCAATCTCTGCCGCTATCTTCACATAACgaatcttaaacccaactt
ggttacctcttatctagtggCCTCCCCTTTTAAGGACACTAGGCATTGGATTTACGGCCCgaatcttaaacccaact
ggttacctcttatctagtggGGCATAAGCCATCGCGCCTGACCTGCAAAGACCATATTTCgaatcttaaacccaac
ggttacctcttatctagtggTATGAATTTTGAGGTGGCACTATAGGGCCCACTACAGGGGgaatcttaaacccaact
ggttacctcttatctagtggCCCATGTGTCATTCCATGTTGATCATGTGCCAGTAACTTCgaatcttaaacccaacttg
ggttacctcttatctagtggGGTCATGTCTCTTGCCAGGTGATCATTTTCCCTAGGTCAGgaatcttaaacccaactt
ggttacctcttatctagtggCCCCTGGGGTTTCTGATTTAGTGGTTCAAGGATGCGGCCCgaatcttaaacccaact
ggttacctcttatctagtggAGTTCCCAGATGACGTTGGTGTTGATGTTGCTCATCCCAGgaatcttaaacccaactt
ggttacctcttatctagtggGACTCCCAGAAATCAAATTTCTCTGACTCTGTACTTAGAGgaatcttaaacccaacttg
ggttacctcttatctagtggTCCTGTGTGTGGTTGATGACTAGGTATGTGGAAAGAACCCgaatcttaaacccaactt
ggttacctcttatctagtggCTTTGTATCCCATTACAAGTTGCAAGGTATCTTCCCATACgaatcttaaacccaacttg
ggttacctcttatctagtggGAGACACATCATTGTGAGCTCATGGTAGAAGCTGGGTGGCgaatcttaaacccaac
ggttacctcttatctagtggCTTTTTATTCTAGAGCCTGTGTAATCTGCCACCCAGCAGCgaatcttaaacccaactt
ggttacctcttatctagtggTTGATGAGAACTATCTCGGTGGTGCTCTGAAAATCACTGCgaatcttaaacccaactt
ggttacctcttatctagtggGCTGCATCATCAAGGTGGTGACGATGGTGTTTCCCCTCGCgaatcttaaacccaac
ggttacctcttatctagtggGGCTTGCCAGCCCAGTTCATCCGAGTTCTTTGCTCCCTGTgaatcttaaacccaact
ggttacctcttatctagtggCTGAGGTTCATTGGTTAATGACCCTTTCTTCTCAGGAACCgaatcttaaacccaacttg
ggttacctcttatctagtggGGGTTGTGTTCTGTGCAGGTTGCTTCAACTTTTCTCCTCCgaatcttaaacccaacttg
ggttacctcttatctagtggGCCTGCCTCTAACCATTTCACATGGCCCTGCTCACTGCCCgaatcttaaacccaac
ggttacctcttatctagtggCCACCCAGGCTGGCTGTTCCTGGTGGAGGAGTTTCCAGCTgaatcttaaacccaac
ggttacctcttatctagtggTCCTACCCTCATGCAGGTGACAGGTGTCTGAGTAGCTGAAgaatcttaaacccaact
ggttacctcttatctagtggGGCAACTCTAAATATGACGCACCACTTTTTCTTCTTACTGgaatcttaaacccaacttg
ggttacctcttatctagtggAATGGGTACAAGAATATCAAGTGTGCGTAGTTGCTGTGAGgaatcttaaacccaactt
ggttacctcttatctagtggGTCATCCTATTGCCTCTGCCTAGAGCCCCTAATCTACTCCgaatcttaaacccaact
ggttacctcttatctagtggGGCTGGTCTTGAACTCTTGAACTCTTGCCCTCAAGCAATCgaatcttaaacccaactt
ggttacctcttatctagtggGGGGAGTCATACCTAGAGTTGCCTGGATCCAAAGCTCGTGgaatcttaaacccaac
ggttacctcttatctagtggCACACACACACACACACTCATCATCATCATCATCATCATCgaatcttaaacccaact
ggttacctcttatctagtggGCCACTTTTAGTCCTCATCAGTCCCAAGCCCTTCAACCAGgaatcttaaacccaact
ggttacctcttatctagtggCCCCTAGGTATGCATAGTCATCCCTCGGTATCTGTGGGGGgaatcttaaacccaac
ggttacctcttatctagtggGAACTCCCCCATGAATACCAAAATTCACAGATGCTTAAGTgaatcttaaacccaactt
ggttacctcttatctagtggGAAAAGGATGTCTGTGCGTATTCAGTACAAATGCAACCACgaatcttaaacccaact
ggttacctcttatctagtggGAATTCGTGAATGTGGAACCCACAGATAAGGAGGGCCCACgaatcttaaacccaac

ggttacctcttatctagtggGCTTTCCTTCTCCTCCCTCCTTGCTGCTTAACCCTGAGGCgaatcttaaacccaact
ggttacctcttatctagtggTCCTCCAGGGCTAGGTGAGGGGCTCACAAGCTGAGTCACCgaatcttaaacccaa
ggttacctcttatctagtggCTCTCCCAAATTAGGAGCCTCTTGAGAACAGGACCTGGGGgaatcttaaacccaac
ggttacctcttatctagtggTCTGCATGGATCCAGTATAGTCCCTGCATACAGCTGGCTCgaatcttaaacccaact
ggttacctcttatctagtggCCAGCCAGGCATTTTTCAGCTGGGTGTGGTGGCTCCTGCCgaatcttaaacccaac
ggttacctcttatctagtggGGCCAACATGGTGAAACCCACCCCCGCTCCCCGTCTCTGCgaatcttaaacccaa
ggttacctcttatctagtggGCCAGGTGTGGTGGCGTGCGTCTATAGTCTCAGGTACTCGgaatcttaaacccaac
ggttacctcttatctagtggGCCAAGCACCATGGTGAGCACATGGGTGACCCTGCATCACgaatcttaaacccaa
ggttacctcttatctagtggCTGGGGCTCCTGGTGAATGTCCTGACAGAGGTGGCTTGGCgaatcttaaacccaac
ggttacctcttatctagtggGTGCTACATGAAGGCACTGTGAGACAACTTATGCCAACCCgaatcttaaacccaac
ggttacctcttatctagtggGCATCTCACTGATGCCACCTTTCTTGCTTTCATTCACAGTgaatcttaaacccaactt
ggttacctcttatctagtggCTTGAGCTTCTCTCTGGCCAAGGGGAAAATTGCCTTCGGGgaatcttaaacccaact
ggttacctcttatctagtggGGGTGCGGGCAAACCTCACCTCTGGATTGTTATGGTTGTGgaatcttaaacccaact
ggttacctcttatctagtggCATCTGAGGCTTATCGCTTCCAGACATCACCCTTCTCTTCgaatcttaaacccaactt
ggttacctcttatctagtggCTTCCTTTCCATCCCTGCTGCTCTCCTCCACCCCCGCTGGgaatcttaaacccaac
ggttacctcttatctagtggGCAACACATCAGACCAGATCGACCCTGCTTCCAGCCAGCCgaatcttaaacccaa
ggttacctcttatctagtggCTCCTTCTTACCCACGTTCATGTTGGTAAGCCTGATGCAGgaatcttaaacccaactt
ggttacctcttatctagtggCCATTCCAGTCACTGAGGCAGCCATGCTTGGTGACAAGGCgaatcttaaacccaac
ggttacctcttatctagtggGCTGCAGGGGAAAAAGCAAAAAGAAAACAGAAGCATCAGCgaatcttaaacccaac
ggttacctcttatctagtggGCCCGCCCTTGCTGTGCTTATGGGCTGGTTGGGCACTGGAgaatcttaaacccaac
ggttacctcttatctagtggGTGAGCCTTCAGAAGGAACTCCCTCACCAGGCTTGCTATTgaatcttaaacccaact
ggttacctcttatctagtggGTCTGATGGAAGTACTGGTATCCAGCCTTTTTAGTACAAGgaatcttaaacccaactt
ggttacctcttatctagtggGTTGCTTTTGTCCAAGGATCCGATGTTTGGAAAGAATCTGgaatcttaaacccaactt
ggttacctcttatctagtggATGATCACCGTTTTCGTGAGTCCTTAATTCCTTGTATCAGgaatcttaaacccaacttg
ggttacctcttatctagtggGCTGATCAACAGGAATCAGAAAGTAAATCCAACTTGAAGCgaatcttaaacccaact
ggttacctcttatctagtggAGTATTAACAAATTAAGCTGGGTGTGGTGGCTCACCCCTGgaatcttaaacccaact
ggttacctcttatctagtggGGTGGGTGGATCACGTGGAGTCAGGAGTTCGAGACCAGCCgaatcttaaacccaa
ggttacctcttatctagtggGGCACAGGCCTGTAATGCCAGCTTCTTGGGAGGGTGAGGCgaatcttaaacccaa
ggttacctcttatctagtggCCTGGAAGGCAGAGGCTGCAGGGAGCGGGGATCATACCACgaatcttaaacccaa
ggttacctcttatctagtggGCTCATGACCTCACTGCTCTCAACTCCACAGTACCTTTGCgaatcttaaacccaact
ggttacctcttatctagtggGTGTGATATGAAGGAGGAGTGATGGTCTGTGCATTTGTTCgaatcttaaacccaactt
ggttacctcttatctagtggGCTGTCCCTTTGCTTCTTATGTGCAAGTGATTTTCAGTTCgaatcttaaacccaacttg
ggttacctcttatctagtggGCACTTCTTGGTGTTAACCTATTATGAGAATTCTTCAAGGgaatcttaaacccaacttg
ggttacctcttatctagtggCCTGTCTGTTACCAGCAATTACTGAGGTCTCTAATTCAGGgaatcttaaacccaactt
ggttacctcttatctagtggTTGCAAAGCTCCTACTGCAAATGGCTCATGTAACCAACACgaatcttaaacccaact
ggttacctcttatctagtggTTCTAGCATAGATTATACCCTTAAACAGAGGATATGAACCgaatcttaaacccaactt
ggttacctcttatctagtggGAGAGAGAGAGAGACAGAGAGGGGCAGGGGTGGGGGGAGGgaatcttaaaccca
ggttacctcttatctagtggTCCCCTTTTCCCCAAAACCTGTTTGAAATAGGCTTCTGTCgaatcttaaacccaactt
ggttacctcttatctagtggCTCTTCTTGCAGAGTATCAGTGCAGTACAAATTACACCTGgaatcttaaacccaactt
ggttacctcttatctagtggAGTTGGTAGACACATTCTTCCGGAAAGACAGTGAACTCAGgaatcttaaacccaact
ggttacctcttatctagtggGTGAGATAGGGTCTTGCTTGCTCTGCTGCCCAGACTGGAGgaatcttaaacccaac
ggttacctcttatctagtggCTCCCTCCGCATTCCTCCTGAGTTCTGGGACTACTGGCACgaatcttaaacccaac
ggttacctcttatctagtggTTGTAGAGATGGGAGTCTCGCTATATTGCCAAGGCTGGCCgaatcttaaacccaact
ggttacctcttatctagtggACAGATGTGAGCCACCATGCCCCACCTACTTTATATTTTTgaatcttaaacccaactt
ggttacctcttatctagtggCATAGACTGTTGACCATGTAGCCAAGCAGAGAGGATGGAGgaatcttaaacccaac
ggttacctcttatctagtggAGATGTATTCGTAAGGAGAGGGATTTCATTCCAAAGGCAAgaatcttaaacccaactt
ggttacctcttatctagtggGGCTTCACAGACAGGAAAATGCACAGAGCAAGCAAGACTGgaatcttaaacccaac
ggttacctcttatctagtggGGCAATTGCCTGTCATCTCCTTTGTCTATCTCAGATGAAAgaatcttaaacccaactt
ggttacctcttatctagtggTGGGACTTACCTTCCAGGGTTGCTGGGAAGATCCAGGAAGgaatcttaaacccaac
ggttacctcttatctagtggGCAATGTACCTGGTACTCCCAAGTTCTCAGTGAGGGAATGgaatcttaaacccaact
ggttacctcttatctagtggGACTGAAGGAGTCCACTGGCTTTTCTCAGAAAAGTCTATTgaatcttaaacccaactt
ggttacctcttatctagtggACTGCTTCCATCTGCCTATGTCCTTTCTCCCTGTGACCACgaatcttaaacccaactt

ggttacctcttatctagtggATGAAAAGAGTCCATGTTTACACACTACATTTGCTACAGCgaatcttaaacccaactt
ggttacctcttatctagtggCTTACATTCATCCAGTTTTCAGCAGTATGAATAGCGTGGCgaatcttaaacccaactt
ggttacctcttatctagtggTCACTGTAAATCCTTAGAACAGGCCCTTGGGATGGTAGACgaatcttaaacccaact
ggttacctcttatctagtggGGGGCTTCTCGTGGCATGTGAAGTGGTTATGTGGAGAGAGgaatcttaaacccaac
ggttacctcttatctagtggGGGAGGGAGCATTGGAATCACTGTGTGAAGTGGTTGTAGGgaatcttaaacccaac
ggttacctcttatctagtggAGGGATTACGAAGGAGTAAGAAGTGGCTGTTGGTCGGAGGgaatcttaaacccaa
ggttacctcttatctagtggGAGGATCGGTGTTACTCTGGCATGTGATTGGGCTGTGGGCgaatcttaaacccaac
ggttacctcttatctagtggGGGAGCATGGATAGGGGTTTATTGTGGAGTGGCTTTAGGGgaatcttaaacccaact
ggttacctcttatctagtggACTGCAGAAGGGCTGGGAGAGGGGAGAGCGTTCACTGTGGgaatcttaaacccaa
ggttacctcttatctagtggGTGGGCTTGAGGATGGGGGTCCTGTGGAGTGAGAAGCGGGgaatcttaaacccaa
ggttacctcttatctagtggGAAGCCGTGTGGGAGAGGAGGTTTGAGGATTCCTGTGGCGgaatcttaaacccaa
ggttacctcttatctagtggAGAGCTGCAGCTGTCCTGTGGAGTGTAAGGCATCTGAGGCgaatcttaaacccaa
ggttacctcttatctagtggGGGGTGTATTGTGGCGTGTGAAGTGGTTGATGAGAGGGATgaatcttaaacccaact
ggttacctcttatctagtggCTGAAGCTTTCCATGGGCGTGAAGCAGGTGGTTGTGGGGGgaatcttaaacccaa
ggttacctcttatctagtggACCTTCCATGGAGTGTGAAGTGCAGTGGAGGTGGGAGGCCgaatcttaaacccaa
ggttacctcttatctagtggGCTGTAGAGCAGTAAGGGGCTTTGTGGTGGGAGGGTAGGGgaatcttaaacccaa
ggttacctcttatctagtggGGGAGGACCGGGTTCTTCTTGGAACATGAGGTGGTGTGGGgaatcttaaacccaa
ggttacctcttatctagtggCTGTGGGGGAGGAGGTTAGGGCTTCCCACGGGAGGTGAGGgaatcttaaacccaa
ggttacctcttatctagtggTGCTGTGGACTGTAAAATGCCTGTTGGGTCAGACGATTGGgaatcttaaacccaact
ggttacctcttatctagtggAGTGTGAAATGGTGGGGAGAGGGTCTGAGGCTGACTGTGGgaatcttaaacccaa
ggttacctcttatctagtggGGCAGTGGGAGGCTTATGGCTGATTTTGGAGTGAGACATGgaatcttaaacccaac
ggttacctcttatctagtggCCTCCCAGGGAAGCCACTTAGAATCATGGGAACCCTTCCAgaatcttaaacccaa
ggttacctcttatctagtggTTGAGCACTTTAATGTACCTGGCCCAGTCCTAGGAGCTGGgaatcttaaacccaac
ggttacctcttatctagtggGCTCTATGCTTCAGAGAGAGTGGGCAAGAGCATCTTCAGGgaatcttaaacccaac
ggttacctcttatctagtggCCCAGCATTTGGGTGTGACATGCCCTTCTAGATAGTGAGCgaatcttaaacccaac
ggttacctcttatctagtggGAGGTACCATCTATCTGGAGATCCTAACCCTGGTGAGCCCgaatcttaaacccaac
ggttacctcttatctagtggGGATAGAGCAGGGACCCTGGAGTTGGCTAGATCTGGGCTCgaatcttaaacccaa
ggttacctcttatctagtggGATGGGCATCTGCCATTTAGCCTATGTGAACACCCTCTCCgaatcttaaacccaac
ggttacctcttatctagtggGGCCGACAGGTGACACAGGCCCTGCTCATCAGACAAGTTCgaatcttaaacccaa
ggttacctcttatctagtggTGAGGGGTCAATGATATTAATGGAAAGGGAGGCAGGATGCgaatcttaaacccaac
ggttacctcttatctagtggGCCTGGTTCTATAGTCTCCAGTTGGGGTGGCAGCTCCTGGgaatcttaaacccaac
ggttacctcttatctagtggGGCAGCTGCCCTCTGGGTGTATTAGTTCCAAGGTGTGGCTgaatcttaaacccaac
ggttacctcttatctagtggTGGTAGCTATCTACCACCTTTAATAGAACTCTTTCTGCTGgaatcttaaacccaacttg
ggttacctcttatctagtggCCCATAGGGGCAAGCCAGCTTAATAGTTACCGGTATAGACgaatcttaaacccaac
ggttacctcttatctagtggAGTTATCTCCTGGGGTGCCTGTTTTCTCCTCCATGAAGTGgaatcttaaacccaactt
ggttacctcttatctagtggGGGGTTGTTGTGAGAATTAAATGGACTAATTCATATCAGGgaatcttaaacccaactt
ggttacctcttatctagtggCTCTTTCTGAGCCTCAGTTTCCATTTATAGACTCAAAGCGgaatcttaaacccaactt
ggttacctcttatctagtggTCCTTACTTCCTTCCCTCCTTCCCTTCTCCCAAAATGAGGgaatcttaaacccaactt
ggttacctcttatctagtggGACAGGAATCTGAGGACCACCTAAGGCAAACTTGACCTACgaatcttaaacccaa
ggttacctcttatctagtggGAACTACACCCCAAGAAAAATGCATTGTCCAGGGAGTGACgaatcttaaacccaac
ggttacctcttatctagtggGAAGTAAAAGGTGATGTTGGCCAGGCATGGTGGCTTATGCgaatcttaaacccaac
ggttacctcttatctagtggACTGAAAATACTGGGCCTGGTGGTGCACACCTGTGATCCCgaatcttaaacccaac
ggttacctcttatctagtggCACTCCAGCCTGGATGACAGAGAGAAAGAGACCCTGTCTCgaatcttaaacccaa
ggttacctcttatctagtggCATCTCCCCATTCCCCCCACCCTAGCCTCTGTAATCGCCGgaatcttaaacccaa
ggttacctcttatctagtggCCACATTATAAGTGAGAACGTGCAGTATTTCTTCCCATGCgaatcttaaacccaactt
ggttacctcttatctagtggCTCCAAGTCCATCTATGTTATTGCAAGTAACAGAATTTCCgaatcttaaacccaactt
ggttacctcttatctagtggGGTGGATTCCATACCTTGCCTATCGTGAATAATGCTGCAGgaatcttaaacccaactt
ggttacctcttatctagtggGTGCAGACATCTCTATAACAAACATTTCAAATCTTTTGGGgaatcttaaacccaacttg
ggttacctcttatctagtggGGCTGTACTAATTGGCATTCCCATCAAAAGCGTGCAAAGGgaatcttaaacccaac
ggttacctcttatctagtggTCCACGTCTCTGTCAACACTTGTTATCTTCTGCCTTTTTGgaatcttaaacccaacttg
ggttacctcttatctagtggCAGAGGTATGGCTTGCAAATTTTTTCTCCCAGTCCATAGGgaatcttaaacccaactt
ggttacctcttatctagtggGTCCATTTTTGTTGTTGTTGCCTGAACTTTTGAAGTCTAGgaatcttaaacccaacttgt

ggttacctcttatctagtggCATTGCCCAGATTAATGTCATATAGTTTTTCTTCTAGAGGgaatcttaaacccaacttg
ggttacctcttatctagtggTCTGTGATGCTTATGAGTTGATTTTTGTACATGTTGTGAGgaatcttaaacccaacttgt
ggttacctcttatctagtggGGAAATCCAGTTTTCCCAATACCATTTCCCAAGAACATTTgaatcttaaacccaactt
ggttacctcttatctagtggGGTTAACCGTATCTGCATGGGTCCATTTATGGGATCTCTGgaatcttaaacccaactt
ggttacctcttatctagtggTGACATGTCCATTTTTTCTGCCAATACCATGCTGTTTTAAgaatcttaaacccaacttg
ggttacctcttatctagtggGTCAGGTAGTGTGATATCTCCAGCTTTTTACTTTTTGCTCgaatcttaaacccaacttg
ggttacctcttatctagtggGATTACATTGATGCTATAGACTAATTTGGGTAATATGGGCgaatcttaaacccaacttg
ggttacctcttatctagtggCAATATATAGCTTTGGGGGTACAGGCCTTTCACCTCCTTGgaatcttaaacccaactt
ggttacctcttatctagtggATGCTGTCAATTTTTGGTGGGGCACGGTGGCTCATGCCTGgaatcttaaacccaact
ggttacctcttatctagtggTGAGACCAGCCTGGCCAGCATAGCGAATCCCCATCTCTACgaatcttaaacccaa
ggttacctcttatctagtggGCTATTGCCAGCAATTTTTGTGTGAGATTTTTGTATCCTGgaatcttaaacccaacttg
ggttacctcttatctagtggCAGCAAACATCGACAATTTCACTTTTACTATTTGGATGGCgaatcttaaacccaactt
ggttacctcttatctagtggGGCAAGAGTGGGCATCCTTGTCTGGTTCCAGATCTTAGGGgaatcttaaacccaac
ggttacctcttatctagtggAGCTGTTGGCTTCTAATATATGACCTTTATTCTGTTATGGgaatcttaaacccaacttgt
ggttacctcttatctagtggCTCGAATGCTTTTTCTGCATCTCATGAGATGATCAAATGCgaatcttaaacccaactt
ggttacctcttatctagtggAAACATGAGCTTTTTCCTCAGGCAATTATTAGAACCACTGgaatcttaaacccaactt
ggttacctcttatctagtggGAAGAAGCTTTAGATCAGCAAAATAGTCGACACATGGTCCgaatcttaaacccaact
ggttacctcttatctagtggGGGCCAAGAGAAAGTTGGTGAGAGTGGCAATGTCAGCTGCgaatcttaaacccaa
ggttacctcttatctagtggAACTCATAGCTACAGCCCTAAGATCTGGAACCAGACAAGGgaatcttaaacccaac
ggttacctcttatctagtggGAAGAGAAAAGAGCTGCTTCGAACCACAGATGGATTTGCCgaatcttaaacccaac
ggttacctcttatctagtggCGACCTCCAGGGTCACCAAGGCAGAGACTAAGAGGGACAGgaatcttaaaccca
ggttacctcttatctagtggCAGCTCGCATGCATTACTTCCACTCAAGTCCACTGGCCTGgaatcttaaacccaac
ggttacctcttatctagtggGGGAGGCTGAGAAGTATAGGTAAGCAATGGCTGTTGGGAGgaatcttaaacccaac
ggttacctcttatctagtggGGCTATACCACAACAGTTATCTCTATCTTTTCTGGTTTTCgaatcttaaacccaacttg
ggttacctcttatctagtggGTCCTACATGGTGGGAGCAGGAGGAAGAGGGCAAAGGGGGgaatcttaaaccca
ggttacctcttatctagtggGGAAAATCCACCCTGATGATCCAATCACCTCCTACCAGGCgaatcttaaacccaa
ggttacctcttatctagtggGCACTGGGGATTACAGTATGACATGAGATTTGGGTAGGGAgaatcttaaacccaact
ggttacctcttatctagtggTCAGGAGGAATTTACTCTAAACCCTAAGGCCTCTCTCCACgaatcttaaacccaact
ggttacctcttatctagtggTGAGAGAAAGAGTGACTTAGCTACTGGGTGCCTTTGGAAGgaatcttaaacccaact
ggttacctcttatctagtggGCTTGTTGGTACAGCCTCTCCCTTTCTTTCATTCTTTGAAgaatcttaaacccaacttg
ggttacctcttatctagtggGATTGCGCTCTTCTCCTTCTTGCTTCAAGCTTTGCTGGAGgaatcttaaacccaactt
ggttacctcttatctagtggGGGAAACCCTGTAATGACCCAACAATGCTTGAGATTGGACgaatcttaaacccaac
ggttacctcttatctagtggAGAAAGCCATTACTGCTGTTTCCTTTAGCCGTAAACCCACgaatcttaaacccaact
ggttacctcttatctagtggAGTGGTGGTGATATTCTGATCCCCATCTGTCTCACTTCAGgaatcttaaacccaactt
ggttacctcttatctagtggCAGGCAGGGAACAAGGGCCATGTTGATAGACCCTAGAGCCgaatcttaaacccaa
ggttacctcttatctagtggAGCACTCAGTGAACGAATGCACCTGTAAGAAACCTGAAAGgaatcttaaacccaac
ggttacctcttatctagtggCACCGATATTGACAACTGACTCCCAAATCCCTGCTTCATCgaatcttaaacccaact
ggttacctcttatctagtggTGCTAATACCATGCCCAGATAGAACACAAAGCAATATTTAgaatcttaaacccaactt
ggttacctcttatctagtggCTCTCCATAATTTTAGAGAGTTTTCCCTAAGGAAAGAAAGgaatcttaaacccaactt
ggttacctcttatctagtggGGGGGTAAGTTCTTTGCCACACCCCACACCTACCACCCAAgaatcttaaacccaa
ggttacctcttatctagtggTGGGTGAATCATTCCCAATCTGAACTGATAATCCTTGCCGgaatcttaaacccaact
ggttacctcttatctagtggACCCAGTCCCTCCCTAGTCCTACAAATGAGCTCCAAGCCCgaatcttaaacccaa
ggttacctcttatctagtggGCCCTGCAGTTGCACAAGTCCCAACTTTGCTCCCGAGGGCgaatcttaaacccaa
ggttacctcttatctagtggCATCCAGAGGGTTCCCACACTCCACCCAGGCACGCTCCCGgaatcttaaaccca
ggttacctcttatctagtggCCACCACGAATTACTGATCGTTACAGAGGTGGCCGGAGTGgaatcttaaacccaac
ggttacctcttatctagtggAGAACATTGTGGCAGGTGAGAATCCAGGCTGAGGAGCAGCgaatcttaaacccaa
ggttacctcttatctagtggTGGGGATTGTGTACACCTAGATTCCAGTCTCAAACCTGCCgaatcttaaacccaact
ggttacctcttatctagtggGTCCATAAGACCTCAGCTGGCACGACCATCACCTGTGCTGgaatcttaaacccaa
ggttacctcttatctagtggGGGACAATGCCTGGTGCACAGTCAACATGCAGTCAACGGCgaatcttaaacccaa
ggttacctcttatctagtggGGGTCAGACACCCAGCAGCAACTCTGGAGAGCATCCCAGCgaatcttaaaccca
ggttacctcttatctagtggGCTTCAGCTCTCTTACCATCCCAGAGACTAGAGGAGCCCCgaatcttaaacccaa
ggttacctcttatctagtggCAGATTGGACCTCTCAGCTGGCATCATGGAAGTAGGAGTCgaatcttaaacccaac

ggttacctcttatctagtggGCTGCAAGGTCACTGGAGGTTTAGAAAAGGACAGATGTTCgaatcttaaacccaact
ggttacctcttatctagtggCCATCCCCACCGCTCCCAGCAAAGCCTGAATCCCTTGGTCgaatcttaaacccaa
ggttacctcttatctagtggGCTTCTCCCTAAACTAGACACTCCTCCAGGGCAGAGAGTGgaatcttaaacccaac
ggttacctcttatctagtggCTGACAAGGAAAAAAGCCAATAATTATTAAGTGCCTACTGgaatcttaaacccaactt
ggttacctcttatctagtggGTGGCACGATCTCAACTCACTGTAACTTCTGACTCCCTGGgaatcttaaacccaact
ggttacctcttatctagtggGGCCAGGATGGTCTTTATTTCCTGACCTCATGATCTGCCCgaatcttaaacccaact
ggttacctcttatctagtggGTGAGCCACTGCGCCTGGCCAACAAACACTTTAAAAATACgaatcttaaacccaac
ggttacctcttatctagtggCTTTCTTTTTCACTGAAGTTTGGAGTGATAGCCTTATTTCgaatcttaaacccaacttgt
ggttacctcttatctagtggGACTCCTAGAAGTCAGTATCACACTGTCTTGTGTATAGTGgaatcttaaacccaactt
ggttacctcttatctagtggGCATAGAGGTAATAAGCAATAACTCGGGAACTAAACCACAgaatcttaaacccaact
ggttacctcttatctagtggGTCTAACCTTAAGTAGGACATTTAACCTTTCTGGTTAGAGgaatcttaaacccaacttg
ggttacctcttatctagtggACCATAAGTTATCATAGTAACCAACTCATAGGGCTGTGGCgaatcttaaacccaact
ggttacctcttatctagtggTGTAAAGCACATAGAATAGTCCCTGCCACATACTAAGTGCgaatcttaaacccaact
ggttacctcttatctagtggGCCCCATAAGACTGGGAGACCCTTGAGGACAGTGGCCACGgaatcttaaacccaa
ggttacctcttatctagtggTCCACCCCCCTCAGCCTGGCACTGAGAAGGAGCTCTGTGCgaatcttaaacccaa
ggttacctcttatctagtggGATATGCGAATGGGGAAATCAAACATTGACCTGGTTGCGGgaatcttaaacccaact
ggttacctcttatctagtggTGAAACCTTACCAGTCTTTCACACCTTAGTTTGGAATGAGgaatcttaaacccaactt
ggttacctcttatctagtggGGTGCCCCAGGGAAAGAGCAGAATAGACACCACCAGGAGCgaatcttaaacccaa
ggttacctcttatctagtggACGTCAATGCCTCCTCAATTTCATCTAGCATTGGCTCTGCgaatcttaaacccaactt
ggttacctcttatctagtggGAGTCAGGCTCCTATTCACTTAACTGTGATTGATCATTTCgaatcttaaacccaacttg
ggttacctcttatctagtggCCCAGAATAGGGAAGCAAACAAGGACAGAAGCCAATGATTgaatcttaaacccaac
ggttacctcttatctagtggCACACAGACACCCAGCAGTTCTATATGATTATCCCCTTCCgaatcttaaacccaact
ggttacctcttatctagtggAAAATACTGAGATACAGCCATGCTTGGTCATGCAGCTAGAgaatcttaaacccaact
ggttacctcttatctagtggACCCTGCCCTCTTTACATCTTGGTTCTAACCTTTTCTTCCgaatcttaaacccaacttg
ggttacctcttatctagtggAGCAAGGCTTTGCGAGCTTAGCATCCTGGCATTCTCAACCgaatcttaaacccaact
ggttacctcttatctagtggGAAACAGGTCCAAATAAGGCCTCTCAGTCAACAAGTCGGGgaatcttaaacccaac
ggttacctcttatctagtggAGTTGGAGGGGATCCACCACAGGAGACTATGAGAGGCAGCgaatcttaaacccaa
ggttacctcttatctagtggTCTTCAAATCAGCAGCTTCTGCCTTGAATGTAAAAAGTGGgaatcttaaacccaactt
ggttacctcttatctagtggGCCTGAAATGATTTAATGTGATCCCTAGGAGGGTTGGGTGgaatcttaaacccaactt
ggttacctcttatctagtggTATGTGCTGCTTAGGCATATGAATTATGCATATGCTAAGGgaatcttaaacccaacttg
ggttacctcttatctagtggGTGTTATCATCTCTTAATGGGGAAATAAAATGGAGACCAGgaatcttaaacccaactt
ggttacctcttatctagtggACAGGTGAGAAAACTGAAGAACTAGGATTAAGTAACTCCCgaatcttaaacccaact
ggttacctcttatctagtggCATTTAGGGCCCTGGAATAACAGTAACATCTAACGTTGCCgaatcttaaacccaact
ggttacctcttatctagtggCCTTAGGGGACAGCCATCCAAATCCTGCTTCTGCACATCCgaatcttaaacccaac
ggttacctcttatctagtggAGGCACTGAACAGGCCTGGGAATGCTGGAATCTATCTGCAgaatcttaaacccaac
ggttacctcttatctagtggCCCAGGTGGTTAGTGATCAGTGCCATGGCTCAACTGTGGCgaatcttaaacccaac
ggttacctcttatctagtggTGTGTCTGGTTCAAATGCTGAGTCAAGACCTCACCCTTGCgaatcttaaacccaact
ggttacctcttatctagtggATAGGGATAGAATTTGCATGTGTGAGATAGTTAGAATGCCgaatcttaaacccaactt
ggttacctcttatctagtggAGCATTTCCAGTGCTTTTACTCATGTATTCTTACAGTCACgaatcttaaacccaacttg
ggttacctcttatctagtggGTCCCAGTGGGAAAGCGGAAGACCAAAGACACTATCAGTGgaatcttaaacccaac
ggttacctcttatctagtggGGATGGTGGTCTATCACAGGGGATGACTTCTGAACTTGGCgaatcttaaacccaact
ggttacctcttatctagtggAGGAACAACGTCAAGAAAAGACATGGGGCCTTGAAAGAAGgaatcttaaacccaac
ggttacctcttatctagtggGCTCACCTGACTCTCCAACCAGACTCTCTTCATTATGCCCgaatcttaaacccaact
ggttacctcttatctagtggGCCTGCTGGGAGATTAGCTGATGCCATTCCCATCACCATGgaatcttaaacccaac
ggttacctcttatctagtggGGGTTTCAGCAGCTAATCCTATATTTGGGACTTGCAGACCgaatcttaaacccaactt
ggttacctcttatctagtggGAATTGGAAAAGCTTAGGGATGGTCTTCTCTAGAGCTTCCgaatcttaaacccaactt
ggttacctcttatctagtggTGTACCATAGGTAGGCATGTATTCCTTGGTATTTTGTCTCgaatcttaaacccaacttg
ggttacctcttatctagtggTGTAGGATTGACCCCAACCACCAGCTCTAATTTGGGTCAGgaatcttaaacccaact
ggttacctcttatctagtggCTCTGGAACCATGATTGCTTTAGGGTAGGCAAGTGATCTCgaatcttaaacccaactt
ggttacctcttatctagtggTGAAGATGTGAAGGCTGGAACTGCCAGGAGCCATGGCTGCgaatcttaaacccaac
ggttacctcttatctagtggGTTCTAGCCTGAGGCCAAAGCTAGCACAGAAGGCAGATCCgaatcttaaacccaac
ggttacctcttatctagtggCCTGCTGGATCAAGCTGTTTCCTAAGCTAGTTGCCTCTGGgaatcttaaacccaact

ggttacctcttatctagtggTATGCTTCACCCAGTTTGAGTGGGATTTTCTGTTAGTTGCgaatcttaaacccaacttg
ggttacctcttatctagtggAACCCCATGATAAGACATAGATAAAACCAGGTTCTCCCAGgaatcttaaacccaact
ggttacctcttatctagtggGTCTCACCCACAATGTACTTTCAATTTCTAAAATGTGGCGgaatcttaaacccaacttg
ggttacctcttatctagtggATTGGCAATACACGTGAAGGAATACAAACATGGGTACAAGgaatcttaaacccaact
ggttacctcttatctagtggAAGCAGATTTCAGAAAATTGGACAGGATGAAGGGTTGCAGgaatcttaaacccaact
ggttacctcttatctagtggGGAAGCTTTTATCTAGAGAAATCCCAATTATGCAGGATTCgaatcttaaacccaacttg
ggttacctcttatctagtggCAATATAGTCACTTATGTAGTGTAAGCATGTTCCACAATGgaatcttaaacccaacttg
ggttacctcttatctagtggTGCTACTTTGACCATGCTTATAAACTTCTAAACTTGCCACgaatcttaaacccaacttg
ggttacctcttatctagtggATACACTTTTCCTAGGGAGGGGCTGGTTTTATATTATGCCgaatcttaaacccaacttg
ggttacctcttatctagtggAACCTCAGCTTTCTCTCTGGCAGACATCAGTGTCGCAGGGgaatcttaaacccaac
ggttacctcttatctagtggTGGGACACAATATAAAAGGCACATGGTAGACCGTTTTGCGgaatcttaaacccaact
ggttacctcttatctagtggTGTAAAACATCATGATCCACAGAGGTCAGAAGACTAACCAgaatcttaaacccaact
ggttacctcttatctagtggACTGGTCAAAGAGAAGGGAAAGTAGAAAATGTGTATGAATgaatcttaaacccaactt
ggttacctcttatctagtggTCTTCAGAGAGGTGCTAAATCCACAAACCACATTGCCTTCgaatcttaaacccaact
ggttacctcttatctagtggGTCTCTTATCTAAAGATCAGTCATTACATGACCCCTTTCCgaatcttaaacccaacttg
ggttacctcttatctagtggTGAGGCAGAATAAGTTCTGAAGGCAGGCAACCTCAGGCTGgaatcttaaacccaac
ggttacctcttatctagtggCCTAGAACTAAAGCAAAAGGAAAACCTCAACTTTCCATGCgaatcttaaacccaact
ggttacctcttatctagtggCAGAGGCTACTCCCTTTGCAACCCCCAGTTTTCTGTGCGAgaatcttaaacccaact
ggttacctcttatctagtggGGCTGCGGGCCAAGTCTTCCTTTGTATAGGAGTATAACTTgaatcttaaacccaactt
ggttacctcttatctagtggGTAACTTTGCTTCGGGCTCTGATTGGTTCCCTACCTCAACgaatcttaaacccaactt
ggttacctcttatctagtggGGGGGTACACAAAGTAACCAATAGGACATCAATGGGACACgaatcttaaacccaac
ggttacctcttatctagtggAGCCTCTTGCTTGGGCCACTCCTACCCTGTGGAGTGTGTTgaatcttaaacccaact
ggttacctcttatctagtggTCAAGGTGCTAGGAACCTGGACACCCTCCACCTGGACACCgaatcttaaacccaa
ggttacctcttatctagtggTATGGTTTGTGGCATAGTCACTGAAGTATTTGAGAAACACgaatcttaaacccaacttg
ggttacctcttatctagtggTGATTCGTTCACAAGAATCTGAGGAAGTTACCTTTCCCCCgaatcttaaacccaactt
ggttacctcttatctagtggACCAGAGTCATGGTTTATTAGAAATGCCAGCTTGGGTGGGgaatcttaaacccaact
ggttacctcttatctagtggCCAGAGAGGGAAACTGAGGGATCTACTATCACATGACAGCgaatcttaaacccaac
ggttacctcttatctagtggCTGTTTGAATTCCAACCTGCTCCTGAATATGTGTGACTTCgaatcttaaacccaacttg
ggttacctcttatctagtggAACTCTTAAGATGATTTCTGGCATATGGTAAACACTCAACgaatcttaaacccaacttg
ggttacctcttatctagtggGCCATGAATGGCTTCACCTCGAGGAAAGCTAATACAGAGAgaatcttaaacccaac
ggttacctcttatctagtggCAGGTCAGCCTGGATATTCACTGGTTGTGGTGGGCAGAATgaatcttaaacccaact
ggttacctcttatctagtggCCCTGTCCCTCTTGTTCAGAGCCTTTATAACCTCTTCCCCgaatcttaaacccaactt
ggttacctcttatctagtggAGAATCTTTCATGTGATGGGTTATTACTCTCATAGTTAGGgaatcttaaacccaacttgt
ggttacctcttatctagtggGGGCTTGACATAATCAGTGAGCTTTATAAGGGACTGGGCCgaatcttaaacccaact
ggttacctcttatctagtggATGTGAGAAGCTGCACTTAAGAGGAAGCTGCATGGCAAGGgaatcttaaacccaac
ggttacctcttatctagtggGGCTTGCAGGGAGCCACAAAATGGGAACCTCATTCCTACAgaatcttaaacccaac
ggttacctcttatctagtggCAAGGGACTCTTAGCTCGAGAGGAGACCACATCCCAGGTGgaatcttaaacccaa
ggttacctcttatctagtggGCAGAGAAACTGGGTAGATCTGCCTGGACTCCTGACCTACgaatcttaaacccaac
ggttacctcttatctagtggGATAATAAATGGCTGTTGCTTTAAACCACTATGCTTGTGGgaatcttaaacccaacttg
ggttacctcttatctagtggCCCTGGCTCATCAGAGGCTAGGAGAGTAGATAGTAACAGCgaatcttaaacccaac
ggttacctcttatctagtggGCACAACTCAGGGAATTTTGTGGTCCCAGAGGATGGCTGGgaatcttaaacccaac
ggttacctcttatctagtggTGACCCTCTGCAGGAAGATTACTGTCATCCCCAAATACCCgaatcttaaacccaact
ggttacctcttatctagtggGTCCCTAGAAAAGACTTCTTAAATTTTGTCATCCTTTAGGgaatcttaaacccaacttg
ggttacctcttatctagtggTGTTGTCCACCTCTCAGAGGCGCTTCATAGTGCTGGTCCGgaatcttaaacccaact
ggttacctcttatctagtggTTTTGCTCACTATTTTATCTCCAGAGCCAGTACAAGTGGCgaatcttaaacccaacttg
ggttacctcttatctagtggGCACTGTTATACGCTCATATACCCGGCATTCACACACAGCgaatcttaaacccaact
ggttacctcttatctagtggACAGGTATGAGCCACTTCACCTGGCCATAGCACTTTGTTCgaatcttaaacccaact
ggttacctcttatctagtggGATGCTAGTTTATGAGATTCTTGAGGACAGAGGTAATGTCgaatcttaaacccaacttg
ggttacctcttatctagtggAGTACCTTTGTGCTAAACACAGTGATCGATGTGTAGTCAAgaatcttaaacccaacttg
ggttacctcttatctagtggCCCAGTAAAATCCATCTTGGGAGTCTGGCCTTCTGAACTGgaatcttaaacccaact
ggttacctcttatctagtggAGAGGGAGGGACAGAGCTGTCTTAGGCAGATGTCTCCTCCgaatcttaaacccaac
ggttacctcttatctagtggCAGCTTGCATTGCAGGACATATTATCAACAGTTTTGGGTCgaatcttaaacccaacttg

ggttacctcttatctagtggCTCATCATCTCTCCGGCTTTCTCCCCAACCTTGTGCCTGCgaatcttaaacccaact
ggttacctcttatctagtggTCAGTAACATGACCTGGGACAGCTAGCTCCACCTCTCTGGgaatcttaaacccaac
ggttacctcttatctagtggTCACCTTACGTTCAGTATTGTTATAAGGGTTAGAAAGCAAgaatcttaaacccaacttg
ggttacctcttatctagtggCACAGGATTGCAAGGTGGCAGGGCCTGTGGAAAGCTGGGTgaatcttaaacccaac
ggttacctcttatctagtggCCAGGCTCAGTCCTGGGGTGAGGTAAAGGGCAGCAGAGACgaatcttaaacccaa
ggttacctcttatctagtggGGCAAAGTGCAACAAATTCTGTAATATGAACTTCCCTGGGgaatcttaaacccaact
ggttacctcttatctagtggAGATTCAGGCAGGAGGTAATCATGAGACGGTATAGCCCCCgaatcttaaacccaac
ggttacctcttatctagtggGAGCCGAGATTACCAAACTCCCTAAAAGATAAGGCCAGGGgaatcttaaacccaac
ggttacctcttatctagtggCAAATGGTGAGGACTTCAATTTGGGTGGCGTTTTGTATGCgaatcttaaacccaactt
ggttacctcttatctagtggGAAGTTGATCTAACAGGAAAACAGTGGGGATGGCAGGTGGgaatcttaaacccaac
ggttacctcttatctagtggGTCACCCCTCTGCCCTCTAGTCCAATGCTGTCTTCTGGGGgaatcttaaacccaac
ggttacctcttatctagtggACCCAAAGCCAAGCCCAAGGAACTCCCAGCTGCTCCACACgaatcttaaacccaa
ggttacctcttatctagtggAACATATATGCATATGTGTGTGTGTTGAGATGGTGGGGGGgaatcttaaacccaactt
ggttacctcttatctagtggCTGTTTCCTGCCTATGTTATAAGCCAGGATAAGAGCTTCCgaatcttaaacccaactt
ggttacctcttatctagtggGGAAGGAGACAGCCAAAGGCCAACAATATAGGGCGAGGCCgaatcttaaacccaa
ggttacctcttatctagtggCTTCTCAGCTCTGGGGAAGGTTAATGAATTGTACTTGGGGgaatcttaaacccaactt
ggttacctcttatctagtggCATAAGGCAGGAAATGTGGCCACTATCAACCCAGAGCTGGgaatcttaaacccaac
ggttacctcttatctagtggGGCAAGACACAGACAGAAATTCACATGCTTGACCTGCATCgaatcttaaacccaac
ggttacctcttatctagtggGAACAAGTCACCTTCACTGTGTTGGGTGTCACAAAGAGAGgaatcttaaacccaact
ggttacctcttatctagtggCCTTTGTCTCATGGAGCTCCTCTTCCAGTCATTGTTGTGGgaatcttaaacccaactt
ggttacctcttatctagtggCATCATCCTCACTGCAAACACCACTACCATCTCCCTTTTTgaatcttaaacccaactt
ggttacctcttatctagtggCAATTTAAAAAATTATGTGGCTGGGTGTGGTGGCTCATGCgaatcttaaacccaactt
ggttacctcttatctagtggTGGGAGGCTGAGATAGGAGAATCCCTAGAACCCTGGGGGCgaatcttaaacccaa
ggttacctcttatctagtggGTGAGCCGTGATTGCGCCATTGCACACTAGCCTGGGTGACgaatcttaaacccaac
ggttacctcttatctagtggGCCACGCTATGAATTAGAAAACCTTGAGGAGAGTCCCGGGgaatcttaaacccaac
ggttacctcttatctagtggTCTGAATATCACAACAGTCCAATAAGGTAGATAGAATGGGgaatcttaaacccaactt
ggttacctcttatctagtggTGTTCAAGCCTCTGTTTTAGAAGTTGCAGAGAAGAATGAGgaatcttaaacccaactt
ggttacctcttatctagtggGGAGGGAAAAGCAATGTATACAATGTGTCAGATGCTTTACgaatcttaaacccaactt
ggttacctcttatctagtggCACTTGCCTAATATTTATTGAGCACCAATTATGTGACTGGgaatcttaaacccaacttg
ggttacctcttatctagtggGCCTTCCTGGAGCTCATATTTTGGGGCAGAAACAGAGATCgaatcttaaacccaact
ggttacctcttatctagtggGGGGTGCATCCAGGAGTCCACACTCTAGATATAACAGTGCgaatcttaaacccaac
ggttacctcttatctagtggAGCACTATCCATGTCTACATCAGAGGGTTTGCTGATTAGCgaatcttaaacccaactt
ggttacctcttatctagtggGCAGATGACATTGACAAGCCCAGTAGCCTAACATCTTTGGgaatcttaaacccaact
ggttacctcttatctagtggGATCATCTTTCTCCTCTAGTTAATGGAGGTCTCTGTCATCgaatcttaaacccaacttg
ggttacctcttatctagtggGGCGTGAGCTACCGTGCCCGGCCCATCTTATCTTACTGGAgaatcttaaacccaac
ggttacctcttatctagtggACCTGTGTGCACCTCAGAGTAAGGATTGCATGCAAGGTGCgaatcttaaacccaac
ggttacctcttatctagtggGGATAGAGAGGGAGAAATCACCAAGTTTCTGGGAATAGGCgaatcttaaacccaac
ggttacctcttatctagtggGTTTGTACAAGTGGCAGAATTCTTTCTGAGACTTGCTCTCgaatcttaaacccaactt
ggttacctcttatctagtggGGGGAGCAAATTCACAACCCCACCTGCCTGTTATTGTCACgaatcttaaacccaac
ggttacctcttatctagtggGGGGTAGACAATCTTATTATCTCCATTTTACAAATGAGGCgaatcttaaacccaactt
ggttacctcttatctagtggACCTGATGCTGTTATTAGATATTGAGTAGCAGAGCAATGGgaatcttaaacccaactt
ggttacctcttatctagtggTTTCCACTAACCCATTCTGGTCTGAGAAGATCCTGGTGTCgaatcttaaacccaactt
ggttacctcttatctagtggGGCCTTGTGTTTCAGCTGGCCATTCTCTTTCACACCAGCCgaatcttaaacccaact
ggttacctcttatctagtggCCTTTGAGTTTCCTCTCCAACCTTCTGTCTATGAAGAGCCgaatcttaaacccaactt
ggttacctcttatctagtggTCCCTTCTGGAAGAAGTTCCCAGTAGTTGACGATCACTTTgaatcttaaacccaactt
ggttacctcttatctagtggCGATCTTTGTTGCCTCCATAGGTAAAATGGAACTGCACAGgaatcttaaacccaact
ggttacctcttatctagtggTCAGACTTTTCACAGGATTCTGGATGATATCGTGGATGTCgaatcttaaacccaactt
ggttacctcttatctagtggTAGATGATCAGAACTAGAAAACCAGTGAGAATTCCTTTTGgaatcttaaacccaactt
ggttacctcttatctagtggGTGGAGGGAGGTCCTGGATAGTGTGAGCAGAACTGAGATGgaatcttaaacccaac
ggttacctcttatctagtggGAAGGTAGTGCATGCACCATATATTAAGTTCAAAACAGCCgaatcttaaacccaact
ggttacctcttatctagtggGCACATTTCACATCTTGTCACTTCTTAATGATTAGTGTGCgaatcttaaacccaacttg
ggttacctcttatctagtggACAGATGGACTTTGCAATTATTGCTTAGTTACATGCAGACgaatcttaaacccaactt

ggttacctcttatctagtggTATGCATGCTCATGTATATAGCTGTGTGCAACTTGTTTCCgaatcttaaacccaacttg
ggttacctcttatctagtggTTGAGACTGTTGCCAAAATAAGAAGTACAGTCATTCTGTGgaatcttaaacccaactt
ggttacctcttatctagtggAGGAAAAATAACTCTATGAGCTTTATGCAAACTCAAAGGGgaatcttaaacccaactt
ggttacctcttatctagtggAGAAGTGCTTTTGTGATACACACATATATGGTAAATACCCgaatcttaaacccaactt
ggttacctcttatctagtggGTGATGTAGTCTTAGATATGGGTCCCTGTTCTACCAGTTCgaatcttaaacccaactt
ggttacctcttatctagtggTTCTGAGCTTTAGTTTTCTCCCCTTTAAACATGAGGAGGGgaatcttaaacccaactt
ggttacctcttatctagtggCCCATCACACAAGTGGCATGCAGTAAAGTTTACTTACCTCgaatcttaaacccaact
ggttacctcttatctagtggGGAAGGGCCATAAGATACATACAAGAGGATATAGGCAGGCgaatcttaaacccaac
ggttacctcttatctagtggGCAGAAAGTGGAGAGTGGATATTTTGAAAAGTGCTGGTTGgaatcttaaacccaactt
ggttacctcttatctagtggCTAGGCTATGGCCAGCAATTTAATTAGAGACAAGATCTCCgaatcttaaacccaactt
ggttacctcttatctagtggCCAGGCACATGCTGAAGGCCAAGGCAGTGCAGGGAGCCAGgaatcttaaacccaa
ggttacctcttatctagtggCCGAGTGAAAGGAACTAGCAAAGCACACCAGCAGGAGAGGgaatcttaaacccaa
ggttacctcttatctagtggGCACCTTCTCAGCATACCCAGGTGCCAAGCACAGGGCAGGgaatcttaaacccaa
ggttacctcttatctagtggGCACTGAACCAGGAGCCAAGAGACCCAGATTCAATCCCTCgaatcttaaacccaa
ggttacctcttatctagtggGCTCTGTAATATGGAGCAAGTAAATTCCTTCAGTGCCCTCgaatcttaaacccaactt
ggttacctcttatctagtggCAGCAACACCCAGTCTTGCCAAGTGTAGCTGTACCTCGGTgaatcttaaacccaac
ggttacctcttatctagtggCCAGTAGTACATGTGAAGTGTTTACAACAGAGCCAGACAGgaatcttaaacccaact
ggttacctcttatctagtggTGTGAACAGGTGCTGTAGGAAGACAATAGAACTTGGCTCCgaatcttaaacccaact
ggttacctcttatctagtggAGACTTAAGCTCTGATTCAAATTCCATAAATATGAGCTGGgaatcttaaacccaactt
ggttacctcttatctagtggGTACAGTTGTAGGGTGATAGATACGTTTAGAAGGCATTGGgaatcttaaacccaactt
ggttacctcttatctagtggGTGAACAAGGTGAGCCAGTGCAGCTGGGAAAGGCGAGAACgaatcttaaacccaa
ggttacctcttatctagtggTGGCCCAGCCCTCTCTCCTCTCTGGTCCCCTGTGTTGGTGgaatcttaaacccaac
ggttacctcttatctagtggTGGGACAGAAATGCCACAGGTAGCCTGAGGACATGAGCTCgaatcttaaacccaa
ggttacctcttatctagtggGCTGCCACTGTTCTCAGAAAGACTGGAATGTGGATGGAGGgaatcttaaacccaac
ggttacctcttatctagtggGCAGACCAAAGAGTTGCAAATGGGCCAGGAGGAAGGCTAGgaatcttaaacccaa
ggttacctcttatctagtggGGGCTCTTCCCAGTTCAACGCGTGCCAAATACCCAAAGGAgaatcttaaacccaa
ggttacctcttatctagtggAGCCATGGCTTAGAAGACCCGTGAAGGGGAAAATTGCCTCgaatcttaaacccaa
ggttacctcttatctagtggGGCAGGTGCTTTATACGTTGATCCCATTTGGTCCCCACAGgaatcttaaacccaact
ggttacctcttatctagtggATGCAGGTGCATTGTAACCCTTCAACCAATTTTCGTTATGgaatcttaaacccaactt
ggttacctcttatctagtggCAGCTGGTTAGCAACAAAGCTAGGACTCAAACTAGTGGCCgaatcttaaacccaac
ggttacctcttatctagtggGCCTACAGCCTAAATGACTCATTAAACACCTTCCCTGAACgaatcttaaacccaact
ggttacctcttatctagtggGCAAACTGTGGGACATAGCCTTTGTAGTTCATTAAGCTACgaatcttaaacccaactt
ggttacctcttatctagtggCAGGGTCTGCTCTGTCACCCAGGCTCTGTTGCCCAGGGAGgaatcttaaacccaa
ggttacctcttatctagtggTGGCTGCACTGTAGCCTCAATCTAAAAAATAGCTCATAAAgaatcttaaacccaactt
ggttacctcttatctagtggGCCAGCGAAAGACACAGAGATTTATGTGGGGCATTTCCTCgaatcttaaacccaac
ggttacctcttatctagtggACGGTATCTCAAGTGGAAGAGAAGGATTAGAATGCCCACGgaatcttaaacccaac
ggttacctcttatctagtggGGACTTCATAAAGCAGACTTCTGTAATCCACAGTGAACATgaatcttaaacccaactt
ggttacctcttatctagtggCAACAAACATTCAGCTAACATGTGTTTATCCTCTTCTGGGgaatcttaaacccaactt
ggttacctcttatctagtggGACAGACATATTATTGCCCTGTCCTCATAGAGCTCACAAGgaatcttaaacccaact
ggttacctcttatctagtggGAACACCCTGTATGCACTAGAATCTGTACAAGGATCCAGGgaatcttaaacccaac
ggttacctcttatctagtggCAATGTTGGAATATGAGGATCAAGAAAAAACATGTGCCCCgaatcttaaacccaact
ggttacctcttatctagtggTTGAGATGGAGTTTCCCTCTTGTCGCTCAGGCTGGAGTGCgaatcttaaacccaact
ggttacctcttatctagtggAGCTGGGATTACACGCACCAACCACCACGACTAATTTTTTgaatcttaaacccaact
ggttacctcttatctagtggGGCACTAGCCACCGTGCCTGGCCTATATATGTTTTAATATgaatcttaaacccaactt
ggttacctcttatctagtggCTGCGGTCATCATGCAGTGCAATAGAGTATTAAAATCTTTgaatcttaaacccaactt
ggttacctcttatctagtggCCCCATCCTTTCCCCTTTGACCCAGCCTCTGGTAACCACCgaatcttaaacccaac
ggttacctcttatctagtggATAAGTGAGAGCATATAATATTTGTGTTCCTGTGCCTGGCgaatcttaaacccaactt
ggttacctcttatctagtggTGTCCTCCAGTTCCATCCATGCTGTCACAAATATTAGAACgaatcttaaacccaactt
ggttacctcttatctagtggGGCTGGTGTGAAAAATGTTGAAATTAACATGGGAGTGCAGgaatcttaaacccaact
ggttacctcttatctagtggCTCCCCAGCAGTGTACAAGCGTTTCTTTCTCCACCTCCTCgaatcttaaacccaact
ggttacctcttatctagtggGCAGCTGTTCTAACAGGTTTGAAGTAATATCTCATTGTGGgaatcttaaacccaactt
ggttacctcttatctagtggAGCTTCTTGCTTATCTCATTTTCATCTGGGGAAATTGCTAgaatcttaaacccaacttg

ggttacctcttatctagtggGCCACCACGCCCGGCCACTAATTTTGATTTTTAAAAGCACgaatcttaaacccaact
ggttacctcttatctagtggAGGTGCTTCGCTCACCTCACCCTAGTCCCAGCACTGAAGAgaatcttaaacccaa
ggttacctcttatctagtggCAGAGCCCTTGTTCTCCAGCTTTTACAATCCAATGGGAGGgaatcttaaacccaact
ggttacctcttatctagtggAGACACATACTAGCTAACATAAATGCTCTTCTAAGTACACgaatcttaaacccaactt
ggttacctcttatctagtggGAGCCTGTAATTGAACCCAGACAGCCTGGCTCCAATCCTGgaatcttaaacccaac
ggttacctcttatctagtggCACTGAGTAGTCTCTAGATTAAGACAGGTGGTTGTAGTTTgaatcttaaacccaactt
ggttacctcttatctagtggAGATCATCAGCTCAGCTGGAGAAACAAAAAGAGAACATGAgaatcttaaacccaac
ggttacctcttatctagtggGCCCCCAGATAGTTCTCCTAAAAATGTGATCTTCCTTCCCgaatcttaaacccaact
ggttacctcttatctagtggCATTTCCAAGGAGGATAATAGTAAATCTCTCTACCAATGCgaatcttaaacccaactt
ggttacctcttatctagtggGCACAGTAAGTAGCCAAGTAGCCACACTGATGCCTACTACgaatcttaaacccaac
ggttacctcttatctagtggACTCTGAATTCTTTTGGAGACATCCCTGAGGAGGAAGCAGgaatcttaaacccaact
ggttacctcttatctagtggCTGTATATAACACACATCTGTATGATGGTTTATGGGTTACgaatcttaaacccaacttg
ggttacctcttatctagtggCATTTGCTATTTATCCTCTATAATGTCTCAGGAGATAAGCgaatcttaaacccaacttg
ggttacctcttatctagtggGTTGCCCCACTGAGGAAGAGAGTTTGAAAAAGTAGGTCACgaatcttaaacccaac
ggttacctcttatctagtggAAAATTCTGAAGCCCTGCCACTAACCGCTAACCACTTGGGgaatcttaaacccaac
ggttacctcttatctagtggAGTGAGTTTTCAGTCACATGCCAGTTCATCTCACAGAGTGgaatcttaaacccaact
ggttacctcttatctagtggCATCTCTGCAGTTCCCATGCCCAGTACAGGTCCTGGTATGgaatcttaaacccaac
ggttacctcttatctagtggCCAACATGTATGTTGAATAGATGGATGGATGGATGGATGGgaatcttaaacccaact
ggttacctcttatctagtggGCCAGTGGTGGGTGCTAGCTCAACTATGGCATTTGCCCAGgaatcttaaacccaac
ggttacctcttatctagtggTCAGAGGTTTTTCCCTGATCACCTCTTACTGTTACCTGCgaatcttaaacccaactt
ggttacctcttatctagtggCAGTCACCCTGCTTTACTATTTTCAGAGCTCTTAGCACTCgaatcttaaacccaactt
ggttacctcttatctagtggGGCCAGGGAATTAGTTTCTCTTATCCACTACTGAACCCTCgaatcttaaacccaact
ggttacctcttatctagtggCATGAATGAACAAATTAGCCTCGATCACTCTTTCAGGTACgaatcttaaacccaactt
ggttacctcttatctagtggGGCCTTATAATCTTGGTTGCAACATTTTCTCCAGGCCTACgaatcttaaacccaactt
ggttacctcttatctagtggCACCCATCTTCTCCAGTGCAAGTCCATAACCACTAACTCCgaatcttaaacccaac
ggttacctcttatctagtggTCCTCCTTCCTCTTTCCCATGAAACTTTCACTGATCAGGCgaatcttaaacccaactt
ggttacctcttatctagtggGAGAGGATCTAGAAATGGATGGAGGGATGGATGAATGGACgaatcttaaacccaac
ggttacctcttatctagtggTACCTTCAAGGTTTATGAGTTTTCACCAGTGGAGTACTTGgaatcttaaacccaacttg
ggttacctcttatctagtggCTTTCTCCAGTGAAAGAGAATTGCCATTATTCTAGGACCCgaatcttaaacccaactt
ggttacctcttatctagtggAGCTTTGTTCCAATTATCTACTGCTACATAACAGCCACCCgaatcttaaacccaactt
ggttacctcttatctagtggGTGGCATCAGCTGGGATGGCTCCACCAGGCAGGAGGATCGgaatcttaaacccaa
ggttacctcttatctagtggGGTAGGCTAGTTCTGGCTGCTACTGAGCCTGGCTGAGGGCgaatcttaaacccaa
ggttacctcttatctagtggAGCCTTGCATCCTCTCCATAGGCTACTTGAGCTTTCTTGCgaatcttaaacccaact
ggttacctcttatctagtggGGAGGAAGTGCATACCATTTTTATGCCCTAGTCTTGGAAGgaatcttaaacccaact
ggttacctcttatctagtggGTCACATACTGTACTGCCTAGATTCAAGGTTAGGGAAAACgaatcttaaacccaact
ggttacctcttatctagtggTGCCCTCAGCCAGTGGAAGTTGAGAAAGAGACTCCATTACgaatcttaaacccaac
ggttacctcttatctagtggGGTTCCAAAGAGTCTGTGAGATGGGTGGTATTGTTTGAACgaatcttaaacccaactt
ggttacctcttatctagtggTGCCACAGGCCTCCTAATACTGAGTCCCAGAATTAGAGGGgaatcttaaacccaac
ggttacctcttatctagtggGTCATAGGCAGCTCTGGTAAACCCACACTGATGCCTGCCCgaatcttaaacccaa
ggttacctcttatctagtggTAGGGCATAGAGCAACTGATGGGAGAGAAAAGTTACACTGgaatcttaaacccaac
ggttacctcttatctagtggCCCAATCTTAGCCAATTAAACACAAAGAATCATCTGCTAGgaatcttaaacccaact
ggttacctcttatctagtggTACTCTTTTACACAGGATTTGATGGTGTGAAGATGTGAGGgaatcttaaacccaactt
ggttacctcttatctagtggCCACACAGAGAAGTCAGAGACAAGAGATAATCCTAGTGGCgaatcttaaacccaac
ggttacctcttatctagtggCCACCTATCCTCCTACACTCTGCATGCCACTAGGAGACTGgaatcttaaacccaac
ggttacctcttatctagtggCCTCAGCAACATATGGTTTGGACCTGGGCAAACTCTCCCCgaatcttaaacccaac
ggttacctcttatctagtggCCTCCAACCCATTCCCCATCCCCACCATTCAGCAACCCCCgaatcttaaacccaa
ggttacctcttatctagtggCAGTTCTCCCATTTCCCACTTCTTTAAGTACAGAGTTCTCgaatcttaaacccaactt
ggttacctcttatctagtggTCCATCCCAACTTTATGACTCAGTTCTCTGACAAGAATGCgaatcttaaacccaactt
ggttacctcttatctagtggTGGCCACAGATCCCTTGCTATTTCTTTCATCAAGAGGGGCgaatcttaaacccaact
ggttacctcttatctagtggAGCACAAAGAAAGCACACATATTTAAGGTGATGGACATCCgaatcttaaacccaact
ggttacctcttatctagtggCCCGGGTTGGTCTTGGGGACATAATTGATCAACAGAATGCgaatcttaaacccaac
ggttacctcttatctagtggTCTGGGATTTCCAAGACTAGGTTACCAGAAGCCTTAAAGCgaatcttaaacccaact

ggttacctcttatctagtggTCCCAGCCACTGGAATTCATGCTGTCAAAAAGTCCACGCCgaatcttaaacccaac
ggttacctcttatctagtggGAAGCCACCTTGGAACCCTCAGATCAGCCCACCTGCCAGCgaatcttaaacccaa
ggttacctcttatctagtggTGCAAGCTGAGTACCACCGAGTTACCTTGCTTGATGTTCCgaatcttaaacccaact
ggttacctcttatctagtggGAGGCACACAGTAGAGCATCTGTGGCCACTGTAGTTAAGGgaatcttaaacccaac
ggttacctcttatctagtggACTAGAACACTCTTTATCAAAGTAGGGCTAGGTTCACCAGgaatcttaaacccaactt
ggttacctcttatctagtggGGACAAGAATCTATCATTGCAGTTTGATGGTGCTAGACAGgaatcttaaacccaactt
ggttacctcttatctagtggAGTTGAATGTGTACAGTTGGCAAGGTGTGGGCAAAAGAACgaatcttaaacccaact
ggttacctcttatctagtggATCACGTTAGAAGTCCAGGAGCAAGTATGAATGTTCCCCGgaatcttaaacccaact
ggttacctcttatctagtggCTGTCTGCCTTCAAATGACCCTGCTTCTAAAAGCCCATTTgaatcttaaacccaactt
ggttacctcttatctagtggGCGTACTCAGTTTTTCTGTTTAGACAAATTCTGTCTGGTGgaatcttaaacccaacttg
ggttacctcttatctagtggGACACAGTCAGCCCAATGAAGGAGGAGGAGGAAAAGGAGGgaatcttaaacccaa
ggttacctcttatctagtggGGAAGATGATCAGAGTGGAAATGTGGCCAGGCGAGGTGGCgaatcttaaacccaa
ggttacctcttatctagtggGGGAGGCTGAGGTGGGCGGATCACTTATTTATATACCTCCgaatcttaaacccaact
ggttacctcttatctagtggGGTCTTGGTCCTGTTGCTTGCTGCTTGGAGAGCTAGGCCCgaatcttaaacccaact
ggttacctcttatctagtggGAGGTGCCAGGTAAGAAGGCTTTATCTGGGTGCTGCAGCGgaatcttaaacccaac
ggttacctcttatctagtggGCAGAGAAGGAACGTAGCTACAAGCAGGAAGGAGTTCGGGgaatcttaaacccaa
ggttacctcttatctagtggGAGGGGTGTGAGCTCTCATTATCTGGATGTGGTGATCTGGgaatcttaaacccaact
ggttacctcttatctagtggGCCTGAGGGTCAGTTTCCTGAGAAGAGATTTCAGATGGGGgaatcttaaacccaact
ggttacctcttatctagtggTTTCTACGGAGCAATTGGGCTGGTTTCACTAGCACAGTGCgaatcttaaacccaact
ggttacctcttatctagtggGATGGCAGTGGCTGTGGAAGTGGTGCAGGTTAGGAGCCACgaatcttaaacccaa
ggttacctcttatctagtggCTGAGGGTGGTGGTGATGATACTGATGATAGAGGTAGCCCgaatcttaaacccaact
ggttacctcttatctagtggGGTGAGGTTGGTATTGTTGAGTCCTAAATCAGAGAAAAGAgaatcttaaacccaactt
ggttacctcttatctagtggGGTGGTAAGCCTGATTGCCAGTCTATGCAACAGTTAGCAGgaatcttaaacccaact
ggttacctcttatctagtggCAGCCTCTGGAGAGTCCTTCTAGCTGACTCCCTAAATGCCgaatcttaaacccaac
ggttacctcttatctagtggGTCACACCCAGAGTGACTTCCCATTCCTCATTGTCTCTTGgaatcttaaacccaactt
ggttacctcttatctagtggCCTGAGTATCCTGCTTGTGGATCCCCAATAGCCAGCACGCgaatcttaaacccaac
ggttacctcttatctagtggGGCATCCCATGGAGCAGTGCAATATGCCAGCCCTACCCCCgaatcttaaacccaa
ggttacctcttatctagtggCCAAATAAGATGAGGAATTATGGGTGTCTGATGTGCCAGGgaatcttaaacccaact
ggttacctcttatctagtggTCATCCCGTATTTCTCACAGTAGAACCATGAAGTAGAGACgaatcttaaacccaactt
ggttacctcttatctagtggCCTATGAGAAAATGGGACTGAGATGGTTAAGGTGGGATAGgaatcttaaacccaact
ggttacctcttatctagtggGCTTTGTCCGCTCCTTTGTTATTCCTTGAAGATGTTAGGGgaatcttaaacccaacttg
ggttacctcttatctagtggCGCCTATGATTAGAACACATAGGTTCAGACATCACAAGGCgaatcttaaacccaact
ggttacctcttatctagtggCGGGGCAATGCTAGTTATCTCTTGGTGATTTGTGCCTCTGgaatcttaaacccaactt
ggttacctcttatctagtggGTCTCTGCCTTAGATCCCTTGGAAAGAGAGAAACTAATTCgaatcttaaacccaactt
ggttacctcttatctagtggTGTTTATTTGCCATCCTATCAAAATATGACTACCTAGGGGgaatcttaaacccaacttg
ggttacctcttatctagtggGTATACCACCCTCAGTTGTCCCGCAGTAAGCTCACCCAGCgaatcttaaacccaac
ggttacctcttatctagtggCACACACATGATAGTTATGGCCAAACCTGTATCTCCTTCCgaatcttaaacccaactt
ggttacctcttatctagtggCCAGCTGACTCTAAGACTCCCACTCCTAAATGTCCTGCAGgaatcttaaacccaac
ggttacctcttatctagtggTCTCAGTGAATCAGACCACCACCCAAATAACCCAAGAGTCgaatcttaaacccaac
ggttacctcttatctagtggCATCTCCAAACCCCTCTACCCCACCTAGCCACTTCCAGTCgaatcttaaacccaa
ggttacctcttatctagtggTCAGGGGTCCAAGCTTCTTCATTCTGCCTCACCATCTCGCgaatcttaaacccaact
ggttacctcttatctagtggGGCTAGAAGGAAAAGGGCAATGGCCTGAAGAGGAAGGGAGgaatcttaaacccaa
ggttacctcttatctagtggACTCAGCTTCCTTTAAACAGCCTCCCCAAAAGTTTCACGCgaatcttaaacccaact
ggttacctcttatctagtggCTGATTCTCATTGGTCACAACTTAGTGCCATGGAAGACTAgaatcttaaacccaactt
ggttacctcttatctagtggAGACAGAAAGGGAGAGCAGATCTTGGGAGGCAGCTAGCTGgaatcttaaacccaa
ggttacctcttatctagtggTTAACCTATTTGCCAGTTTCTCTGCATCCAACCATCATGGgaatcttaaacccaactt
ggttacctcttatctagtggGGTCCAACCTCAGAGAAATAGCTACTTAAGTGATACTGGGgaatcttaaacccaact
ggttacctcttatctagtggTGCCACTTGGAATGGATTTTTCCAATTGTCCTGCCCCCACgaatcttaaacccaactt
ggttacctcttatctagtggGACCAATGACGTAAGAAGGACTAATGTTTCCACCTGCCTCgaatcttaaacccaact
ggttacctcttatctagtggAGCAAGAGATTCCACATGTCAGACAGGAAAGAAGGCTAGGgaatcttaaacccaac
ggttacctcttatctagtggAAGACAAACAATGGTGATAAATGAAATGGCAGAACCAAGGgaatcttaaacccaact
ggttacctcttatctagtggCAGAGCCCCTGGGCCACCAGTCTTCAATGTTTGCTACCAGgaatcttaaacccaac

ggttacctcttatctagtggCCCTTCTGTACCTCAGCATTTCCATCCTGCCCCCTTCACCgaatcttaaacccaact
ggttacctcttatctagtggTCATTATCTATCCCAGTGCCTGGCCTGTGATATCCCTAGAgaatcttaaacccaactt
ggttacctcttatctagtggCGAGTAGACGTTCAGGTCACAAAATCTAGCCTGTCTCCCCgaatcttaaacccaac
ggttacctcttatctagtggTGCCATTGTCTGAACAGATTAAATAAAGCAGAGCATCACCgaatcttaaacccaactt
ggttacctcttatctagtggTGGTTCAAATCTAGGTAGCTATACTTACCAGCTGTGAGGCgaatcttaaacccaactt
ggttacctcttatctagtggCCTTCTATTTCACTTTTCTGCTAAATGAGACAATATGGACgaatcttaaacccaacttg
ggttacctcttatctagtggACACACACACACACACACACAACACCCGATATAATACAATgaatcttaaacccaac
ggttacctcttatctagtggGACACTAGAAATATTTTAGTCAGGTCTAAAGGAACAGAAGgaatcttaaacccaactt
ggttacctcttatctagtggCCCTGAGAAAGGGGACTTAAGGTTGGCAGGATAACAAGTGgaatcttaaacccaac
ggttacctcttatctagtggGAGCAAGACTGAAGGAGAACGTTTAAGTTCCCTTTCCAACgaatcttaaacccaact
ggttacctcttatctagtggAAGATGGCCCCATGGAGAAACTGGCAGCATGGCATGATGGgaatcttaaacccaa
ggttacctcttatctagtggCTCAGACACATATTTGAGAGGTAGAGAGGGGAGAAGAGGGgaatcttaaacccaac
ggttacctcttatctagtggGCCCACTGCTTAAACACTTACCTTGTGCCAGACATTGCACgaatcttaaacccaact
ggttacctcttatctagtggACCTGCAACCCATCCTGCAAAACTGTGAAGATTTGATATGgaatcttaaacccaactt
ggttacctcttatctagtggGCCTGTCTTATTCTAAAGCCCCTTCTCTTTTCACAATCCCgaatcttaaacccaactt
ggttacctcttatctagtggGGACATCCAATATTTGAGTGGCAGAGGCAAAAACATGGCCgaatcttaaacccaac
ggttacctcttatctagtggGATAGTCCTCCTGCCAAATGGTGAAGGCTAATTACCCTCCgaatcttaaacccaact
ggttacctcttatctagtggGAACGCAGTGGCTTCTTCCCTGCTCTCTCTTTCTTAGGTCgaatcttaaacccaactt
ggttacctcttatctagtggAGCCCACATGGTGAGCAACTGAGGCCTCCTACCAACAGCCgaatcttaaacccaa
ggttacctcttatctagtggGCCCCCGTTAGGCTCCCAGATGACAACATCCCCTGCTCTCgaatcttaaacccaa
ggttacctcttatctagtggCCCTTATCCAGGATACCCAGTTAAGCCACACCACAATTGCgaatcttaaacccaac
ggttacctcttatctagtggTACACCCATCCAGTGCTGAAGAAAGGGGTTGTCAGGAGGGgaatcttaaacccaa
ggttacctcttatctagtggGAGTGCTATTTCCCGTCACTACAGCACACTTGTCACCAACgaatcttaaacccaact
ggttacctcttatctagtggGGTCAGGGAATTTCTTTGGTTGCAGCCCATTTTGCCTTTGgaatcttaaacccaactt
ggttacctcttatctagtggCACACCATTCCATAGCACAGAACACAAGTAGATGGCTCCCgaatcttaaacccaac
ggttacctcttatctagtggGGTATGGCAGAAGCAGAATGGACTCAGGAATCATGGTGCCgaatcttaaacccaac
ggttacctcttatctagtggTTTCAGATCGGCCACTGACACAATGGACACATTATTCAACgaatcttaaacccaactt
ggttacctcttatctagtggGAGCGTATGACACACACAGCCATACGGTCACAGAGCTTTCgaatcttaaacccaac
ggttacctcttatctagtggGCCGGGCTGGGTTGATCCTCTGACCTGGCTAAGCTTCCAAgaatcttaaacccaac
ggttacctcttatctagtggGGGATTCCCCCTTGCCCCCCGCCGTATGAGTTATGTGGGGgaatcttaaacccaa
ggttacctcttatctagtggGCCGTGAAGCTGGAGAAGGAGAAGCTAGAGCAAAACCCGGgaatcttaaacccaa
ggttacctcttatctagtggGGCCGATGTGGGGCTCATCCTGGGGGTTCTATTTGGGAAGgaatcttaaacccaac
ggttacctcttatctagtggGCAAAGCAGAAACCCTCATGCAGGCCCGAAAGAGAAAGCGgaatcttaaacccaa
ggttacctcttatctagtggCAGTGCCCGAAACCCACACTGCAGATCAGCCACATCGCCCgaatcttaaacccaa
ggttacctcttatctagtggGGGAAGTCTTTCCCCCAGTCTCCGTCATCACTCTGGGCTCgaatcttaaacccaac
ggttacctcttatctagtggCTGCCCTTCTAGGAATGGGGAACAGGGGAGGGGAGGAGCTgaatcttaaacccaa
ggttacctcttatctagtggGGAATTGGGAACACTAAGGGTGGGGGCAGGGGAGTTTGGGgaatcttaaacccaa
ggttacctcttatctagtggTTTTAAGTGTGGTAAAATGTGTGGCACGTAGTAGGTATTCgaatcttaaacccaacttg
ggttacctcttatctagtggCCTCCTTTTTACCTTTTCCTAGAGTGCTACTTGATGGTAGgaatcttaaacccaacttg
ggttacctcttatctagtggCCGCTAACGGCATAGTTCGGCCCTCACCATTCCCTCACATgaatcttaaacccaac
ggttacctcttatctagtggAATGTGAGCATTAACCATTCCCCATGATACAGATAATTCAgaatcttaaacccaactt
ggttacctcttatctagtggGGTAGTGCCTTCCTTTATAGCTCAGGCTGGAGTGCAGTGGgaatcttaaacccaact
ggttacctcttatctagtggCGTGAACCACCACGCCCGGCCCTGAAAATTCATTTTTAAAgaatcttaaacccaact
ggttacctcttatctagtggAGATGTGCACCTATTTACCCTGGACCAGCTCCTTGGAAGGgaatcttaaacccaact
ggttacctcttatctagtggGCTATGAAACTGCATCGCCAAAAACTGAGACTCATTTTCCgaatcttaaacccaactt
ggttacctcttatctagtggTGACAACAAAATCTCCAGGGCATTCTGAAACCCAACAGGGgaatcttaaacccaac
ggttacctcttatctagtggAGGCATACCAGTGGTAGTTAGGAGACTTGCCCAAATTTCCgaatcttaaacccaact
ggttacctcttatctagtggTTCATGGCTTGGCTTTCTGCTATTTGCTCATTGGTGACACgaatcttaaacccaacttg
ggttacctcttatctagtggCTATGGCTAGGAAATCCCAGTGGAGATCAAGAAACCTGGGgaatcttaaacccaac
ggttacctcttatctagtggTGGGGTGACCTTTGATTAGTCCTAACTCCCTTCCTAAGCCgaatcttaaacccaactt
ggttacctcttatctagtggCAATGGATTACATTGCCTTAGACTTTTTCCTAGATCGAAAgaatcttaaacccaacttg
ggttacctcttatctagtggGTGCCTTTCCAACAATAATTGATACCACTAGTGAACATTCgaatcttaaacccaactt

ggttacctcttatctagtggGTTCTGCTCTAACCCAAGTGATTCTGGATCCCAGACTCCCgaatcttaaacccaact
ggttacctcttatctagtggCCAGAAATACTCAGAAAAAGGAGGGTGCCGTATGTCGCACgaatcttaaacccaac
ggttacctcttatctagtggCAGAATCCAGCCAAAGGATTCCCAGAAATTGAGGCTCTTCgaatcttaaacccaact
ggttacctcttatctagtggTTTGCATCTTTGGATGTGTGTTTTTTACATTGCTCTATGCgaatcttaaacccaacttgt
ggttacctcttatctagtggCTCCTCCCATCCTACCACTGAGGAATCACAGGGAATAGACgaatcttaaacccaac
ggttacctcttatctagtggGTTGCCTAGGAGGACTGAACTTTTTTGGACCTGGGGCACCgaatcttaaacccaact
ggttacctcttatctagtggAGCTCTAGTATAATCAAAAGGTCAGGATCTCCACTGGACCgaatcttaaacccaact
ggttacctcttatctagtggCTGAATGCCTGCCATGTGTGAAGGGAAATGCCTGGACTTCgaatcttaaacccaact
ggttacctcttatctagtggGTCTCCCCACATCTACCCAGGAAGACAGGCACAGTGAGCCgaatcttaaacccaa
ggttacctcttatctagtggCCCTGGATCCTCTTTATTACACCAAGCCTGCCTCACAAGGgaatcttaaacccaact
ggttacctcttatctagtggGGGAAAGACCCACTCCTCCCATTCATGATAAGGTAATGCTgaatcttaaacccaact
ggttacctcttatctagtggGCTGTCTGCAGCAGGGAGAAGTTCCTGGCAACATGACCCTgaatcttaaacccaac
ggttacctcttatctagtggGAGTGACTGAAGCTAAGAAGGAAGAATCTTCCAAGGACTCgaatcttaaacccaact
ggttacctcttatctagtggTCATGGAGAAATAAAGATGCCAGTCTCCAGGAAGCACCAGgaatcttaaacccaac
ggttacctcttatctagtggTTCTGGAATTTAACTCCTGGCATTAACAGTTAGCTGGGAGgaatcttaaacccaacttg
ggttacctcttatctagtggAAATCAATGGACAGTGGTTTCATAGCAGATGGCAAAGAGGgaatcttaaacccaact
ggttacctcttatctagtggCGAGACCCATAATTCCAAGCAGCCTCCATAAGGTAGAGGCgaatcttaaacccaac
ggttacctcttatctagtggGTCATTTCTCTACCCTTCTGTACTTGGCACACACCCTAGGgaatcttaaacccaactt
ggttacctcttatctagtggCTACGTAAATGAGCTCCTTGCCTTCTGGCTCCTAGTTGGGgaatcttaaacccaact
ggttacctcttatctagtggGAGACCTATTAGCAGGAAGATCGGAAGGTAGGAGAATCTGgaatcttaaacccaac
ggttacctcttatctagtggGCATCTCCCCAGACTTCAGTAAATGCTTCCTCCCTGTGCCgaatcttaaacccaact
ggttacctcttatctagtggGGGTCTCTGCTGCCACTAACCCTGGGATGCTTCACCCTCCgaatcttaaacccaac
ggttacctcttatctagtggATGTTTGAGCTGGTTATGTTGGGAGGGGCCCCACTATGGGgaatcttaaacccaact
ggttacctcttatctagtggACTAAGACACGGAAAGGTTAAGTAGCTTGTGCAAGGTCATgaatcttaaacccaact
ggttacctcttatctagtggAGCAATGCTTGTTTCTATAACAAATAAACCCCCATATCTCgaatcttaaacccaacttg
ggttacctcttatctagtggCTTACGTCAGTCCAAAATAGGGTTCCTGAGGAGCAAGTGGgaatcttaaacccaact
ggttacctcttatctagtggGTGGTGATCCAGAGAAGAAGCTCCTTGCATTTTCTGGCTCgaatcttaaacccaact
ggttacctcttatctagtggGGAGGCTTGAACATGGCAGGTTGTGTGTCACGCCCTCCACgaatcttaaacccaac
ggttacctcttatctagtggGGCTGTCCAATTGCAAGACAGTCCCAGAGATGGTGAGCTCgaatcttaaacccaac
ggttacctcttatctagtggGAGTTTGGGAAAGCTCTTGACAATTTCTACCTCAGAAGTGgaatcttaaacccaacttg
ggttacctcttatctagtggCCAAAGCTTTTCACTCAAAACACAAGGCCTCCAGTTCATGgaatcttaaacccaact
ggttacctcttatctagtggCATCCATGTGTGTTCCTGAGGGGAAGAGACAGAACCAGAGgaatcttaaacccaac
ggttacctcttatctagtggCAGGGACTGCTGCTGCCTCCACCTATCACAGTAATCACTTgaatcttaaacccaact
ggttacctcttatctagtggCCGATAAAGTAATGTATAAGTAACTGCACATAAGTCTAGCgaatcttaaacccaacttg
ggttacctcttatctagtggCCATGATGATTTACAGTCCCAGTTAGAAATTCCAAGGGTCgaatcttaaacccaactt
ggttacctcttatctagtggCAAACTGTAGCTCTCTCTCTTTCCACTCACATCTCTGTGGgaatcttaaacccaacttg
ggttacctcttatctagtggAGTGTTAGTACATACTCTGAGAACACAGACACAGTGGGACgaatcttaaacccaact
ggttacctcttatctagtggCCTCTTGACTTTGCCTTTTTTCCACCTCATAAGCATTTCCgaatcttaaacccaacttg
ggttacctcttatctagtggCAAGCCCCAAGAAGGTAAATTTTCCCTGAGTAACCTGGGGgaatcttaaacccaac
ggttacctcttatctagtggAGCTCTCAGCCTGCTGATCGACTCTACAGCACTGTTTCTCgaatcttaaacccaact
ggttacctcttatctagtggCCAAATATAAGAATGTGAAGGAAAGAGAATTTCAAGCACCgaatcttaaacccaactt
ggttacctcttatctagtggCCTGCCATCAAGGATCTTACCGGGGGTAGACAGATAGTGCgaatcttaaacccaac
ggttacctcttatctagtggGCTGGCAATAAGTGGGGTAAAGGAAATCAAATTCAGGTGCgaatcttaaacccaact
ggttacctcttatctagtggTCCCCTAAGCCTCAGAGCCCTGACTGATGAGTGATCCAGGgaatcttaaacccaac
ggttacctcttatctagtggGGAAATTCGAGAACACCATGAGTCCATGTGGCTGCAATGGgaatcttaaacccaac
ggttacctcttatctagtggGCTATGGAGGTAGGTGGGTGAACAGAAGAGAACCATCATCgaatcttaaacccaac
ggttacctcttatctagtggTAGGCTCTTTGTGTGTGTGTGTAGTGGTGGGGAGGAGGAGgaatcttaaacccaact
ggttacctcttatctagtggACCACAAGGGTAGACTTGGAGGTGGTGAAAGTGGTCAGATgaatcttaaacccaac
ggttacctcttatctagtggGGAGTTGTCAGGACCTGCTGATGGCTATGGCATGGCTGTGgaatcttaaacccaac
ggttacctcttatctagtggCACGCCACTGGCTCAGCTGGATACTTCCAAAGGGCTGCTCgaatcttaaacccaac
ggttacctcttatctagtggGATGAGTACTTCTGCCCTATCCTCAAGGAGTCCCAGGCTGgaatcttaaacccaac
ggttacctcttatctagtggATGATCATGCTGAGAGAGAAAGCCCACAAGTTACTCTGGGgaatcttaaacccaact

ggttacctcttatctagtggGTGGACTCAGGAATGCTTTTTGGAGGAAATGACACCTAGCgaatcttaaacccaact
ggttacctcttatctagtggGCCAAAGATGTTGGGAGAAGCAGTCAGGCAGTGGAAACAGgaatcttaaacccaac
ggttacctcttatctagtggCGTCTCACCAGGGAAGTGGAGGGTGGAAGGGAGCACAGGGgaatcttaaacccaa
ggttacctcttatctagtggGCAGGTTCCAATCATAGATGGTTTTTATGCTGTTCCTCAGgaatcttaaacccaacttg
ggttacctcttatctagtggAGATAGGAGTGCCAGTGTCAGGAATTAGGATGACAAGCTCgaatcttaaacccaact
ggttacctcttatctagtggGATCCCGGTTTTCTTGGGTGTGTTCTTCTATTGTACGTCCgaatcttaaacccaacttg
ggttacctcttatctagtggAACGTCTAGCATGGAAGAAGAAATCTACAGAGTTCCCTGGgaatcttaaacccaact
ggttacctcttatctagtggCACTCCCAAACCCACCACCACTATCTGCATACCTGCTGCCgaatcttaaacccaac
ggttacctcttatctagtggAAGATAGACATCAAGTCAAGCACAAACTTCAAAACTCTCCgaatcttaaacccaact
ggttacctcttatctagtggGCTAACATTGGCTGAGTGTTTACTTATATACCAGGCATTCgaatcttaaacccaacttg
ggttacctcttatctagtggCAGGGATCTTGACCCTAATGCCTTTTCTGGGAAAACAGTCgaatcttaaacccaact
ggttacctcttatctagtggCTCCAAGTCCCACTAACCAAGTTCAATTCCTACCTCTCCCgaatcttaaacccaact
ggttacctcttatctagtggCTGCTTAATACGAGTCATTCCAATCTGATCTCCGATGGCAgaatcttaaacccaactt
ggttacctcttatctagtggTCCACTGCACTTATTCCCATAAAAACAAGAACTTGTCAACgaatcttaaacccaactt
ggttacctcttatctagtggAGTACTCCTCCACCAAATGCAACTACCAAGTACCCAGCACgaatcttaaacccaac
ggttacctcttatctagtggCACCAAATACACACATATAAGCACACATGTGAGTTTGGCCgaatcttaaacccaact
ggttacctcttatctagtggGTCTGTAGGCTGCAGACTACGTGGGCACAAACAAGATTCAgaatcttaaacccaac
ggttacctcttatctagtggTGTGTGACACTGAACAGCTTCATCATGCCTCAGTTTCCTCgaatcttaaacccaactt
ggttacctcttatctagtggGAGCAACAGTCACACTCTCAAGAAGATTCATTATGCTCTCgaatcttaaacccaactt
ggttacctcttatctagtggAGAGAATAAAGTATGTGGACCTGTCCAGAGCAGATGACCGgaatcttaaacccaac
ggttacctcttatctagtggGCTTGAGGCTCTTAATTCCAGCTCCATTGCCCATTGGTTGgaatcttaaacccaactt
ggttacctcttatctagtggCTCTCAATCCACCCTGGAGAAGGAAATTAAAGCCTAGAAGgaatcttaaacccaact
ggttacctcttatctagtggAAAGCCACAACCAGAAGTCAAAGCTTCCTGTACTCAGCCCgaatcttaaacccaac
ggttacctcttatctagtggCAGGGCCTGTGGATACAGAAATGAATAAAGCACAGTCCCCgaatcttaaacccaac
ggttacctcttatctagtggTCCTAACATGGCCCCTTGGAGAGTTGATTCCCAGCCACTCgaatcttaaacccaac
ggttacctcttatctagtggACTGGAAATTCACTAACCTGTAACCTGTACCAGTTCCCAGgaatcttaaacccaact
ggttacctcttatctagtggCCTCACATGGAAGATGAGGATGCTTCTGAGGAGGAAAATGgaatcttaaacccaact
ggttacctcttatctagtggGGAAGTGGGGGCTCTAAAGCTGATAGAATAGCTTGTGACGgaatcttaaacccaact
ggttacctcttatctagtggGTCTCTCACCTTCTCTCAGTCTTTCCCTCATGGGCTCAGGgaatcttaaacccaact
ggttacctcttatctagtggCACAAGGCAATCAAGATGGAGAGAGTGGTGTAGCAGGAGGgaatcttaaacccaac
ggttacctcttatctagtggGAGAGAGAGAGAGAGAGAGAGCACTTGCCATGTATCTCTCgaatcttaaacccaac
ggttacctcttatctagtggCCTTAAATCTCACTGGGCAGAGCTGGCTCACATAGCTGCCgaatcttaaacccaac
ggttacctcttatctagtggGGAGGCAGGCAAGCATGCATCTGGCTCATTCAACCTACATgaatcttaaacccaac
ggttacctcttatctagtggCTGTGTAGGTTTGTTGAAGGAATGCATGATCTGCCCTGGGgaatcttaaacccaactt
ggttacctcttatctagtggGCCAGCATCCAGGAGTTTGCAAATCTCTAGTCCTCGGGCCgaatcttaaacccaac
ggttacctcttatctagtggTCTGAACCAAGCTGGAAGGGGGAAGAAACAGGCCAGGCAGgaatcttaaacccaa
ggttacctcttatctagtggCACCTGCCATTCTGGAAGGGGGAAGAAAGAGTCTAGGCAGgaatcttaaacccaac
ggttacctcttatctagtggGCCAGATTTTGCCCTCCATGCCTGGGTCCTCAATGGCCTGgaatcttaaacccaac
ggttacctcttatctagtggTTCACTAGGGCATGGAATTCTGCCACTTACTACAGCTCAGgaatcttaaacccaact
ggttacctcttatctagtggTCTACCCCAAGCAGATCTTCCATTTAGCCATCATCCAGGGgaatcttaaacccaact
ggttacctcttatctagtggACCATTCCCAGAACCTCAATCCCCACCCCATTTATTTCTGgaatcttaaacccaact
ggttacctcttatctagtggGATTCCTAGTGAATAGTCAGTTGCCTTGTAACCTCCCAAGgaatcttaaacccaactt
ggttacctcttatctagtggGGGTCCACCCATTGCTAAGTCTGTCCCCTCCTCCCCCAGCgaatcttaaacccaa
ggttacctcttatctagtggCCCAAGACAATGTGCCATGAGATTCATCTTCCCTTCTACGgaatcttaaacccaact
ggttacctcttatctagtggGTCAGAAGTGCCCTGACAGGAAAGCCTGGTGACCTGCCAGgaatcttaaacccaa
ggttacctcttatctagtggTCCGCATGCTCTGCTCTGTTTCCCTTTTCCTCCTAAAGGGgaatcttaaacccaactt
ggttacctcttatctagtggGGCGTTCAGTTTAGTCACCCAGGGTAAGCCGTTTTGACTTgaatcttaaacccaactt
ggttacctcttatctagtggCTGCTCTGTCCTAACTCCCCGTGACTGACACACAAGCCTGgaatcttaaacccaac
ggttacctcttatctagtggCACTCTCTGAAACCTCCCTCCAGAAACCTCAAGGCAGTTGgaatcttaaacccaac
ggttacctcttatctagtggCCTGGACAAGGCATTCTGGTGTAGTGAAAGTCAACGTCCCgaatcttaaacccaac
ggttacctcttatctagtggCTTTCCAGCTCAGGGATATTGGCCGCTTATCCAGTGAGCCgaatcttaaacccaact
ggttacctcttatctagtggGCTGGTAAAAGGTTTACACCTGCAATGGTGCCTGGAACAGgaatcttaaacccaact

ggttacctcttatctagtggACACATAGAGGTGTGTGTATGTGTCTGTGGTGTATGTGTGgaatcttaaacccaacttg
ggttacctcttatctagtggCCTCCTGCCTACTAACTGCCTGGTCTTAATCTAGGTGTCGgaatcttaaacccaactt
ggttacctcttatctagtggTTGTCAAAGTCCCATCTATGTTTTAGAGTGGAAGACGGACgaatcttaaacccaacttg
ggttacctcttatctagtggGGTCATCGGGGCAAGGCTCTGAGAGGGTGGCATGGAAGGGgaatcttaaacccaa
ggttacctcttatctagtggGGGAGAAACCACATGTTTTCGTTCCTCTCCACACACTGCCgaatcttaaacccaact
ggttacctcttatctagtggCCTCACTGTCAGCCTTGACTCCTTCCTTCACACCCATGCCgaatcttaaacccaac
ggttacctcttatctagtggGCAGCCAGCTCTGTGGATATCTGGGAAGGAGGCTTCCAGGgaatcttaaacccaac
ggttacctcttatctagtggGCCTGCTGTGGACAGTGAGGAGGCCGAGATGCTAGAGAAGgaatcttaaacccaa
ggttacctcttatctagtggGGGAAGTCATTGGAGATGTGTGGACAGAGGAGTGTCATGCgaatcttaaacccaac
ggttacctcttatctagtggGAAGGCCAAGTGCAGAAGCAAGGGATGCATCAGGAGTCTCgaatcttaaacccaac
ggttacctcttatctagtggGCTGCGAAGCCATGGTTGCTTGGTCTAGAGTGGCCATAAGgaatcttaaacccaac
ggttacctcttatctagtggGAGGCCAGAGTTGGTTGCCATTGCTGTTGGAATAGGCTTGgaatcttaaacccaact
ggttacctcttatctagtggTTGTGGTGGTGAGATGGGAATGGGGCAGTGGAACCAAGAGgaatcttaaacccaac
ggttacctcttatctagtggCTGAGACTCTCTTGTGCCAGACGTCTTGCTTAGCACTGGGgaatcttaaacccaact
ggttacctcttatctagtggGACAGAAACAGGCTTTGCTCTCATGAAACACAAGTAGCACgaatcttaaacccaact
ggttacctcttatctagtggGTACTTTTTATTTAATGAAGCTGACTGTCTGGTAGGAGAGgaatcttaaacccaacttg
ggttacctcttatctagtggCTGCTCCCCAGGCTCCTTAGGGAAGGTGATACTGGGCTAGgaatcttaaacccaac
ggttacctcttatctagtggCAGCATTTTTGAGAATCTGGCCAATATGGAAAGATTTGCTgaatcttaaacccaacttg
ggttacctcttatctagtggCCCATGTAGCTGGGAATACAGGTGCCTACCCACAGGTCCGgaatcttaaacccaac
ggttacctcttatctagtggGCGTGAGCCACGGCACCCGGCCTGGAGACCACTTATTTTGgaatcttaaacccaac
ggttacctcttatctagtggCCCCTCTTATCTCTGGGAGTGGGGAGGACATGACCTTCTGgaatcttaaacccaac
ggttacctcttatctagtggGGCTATCCTCTCTGCTCCCTGCCTACCTATCCCACTTGGGgaatcttaaacccaac
ggttacctcttatctagtggCGACTTCTCCCTCTGACCACCACACTTCTACGTGAACCCCgaatcttaaacccaac
ggttacctcttatctagtggCCTCTCCAACCCCATTTAAATGTGTTCCATTGAGCCTCTGgaatcttaaacccaactt
ggttacctcttatctagtggCCCAGCAATACTGCAAACCCCTTGAAGACAGGGATGGGTGgaatcttaaacccaac
ggttacctcttatctagtggTGTATCCCCAACAGGCATCACAATGCAAACTGCTCGGGGCgaatcttaaacccaac
ggttacctcttatctagtggGAGACTGTGAGCCAAAGACCTTGACATCATTTAAGAGAGGgaatcttaaacccaact
ggttacctcttatctagtggCCTGACTCAGGCTAACCAACATGCCTGTGCAGCTAGGAGAgaatcttaaacccaac
ggttacctcttatctagtggGAGTTGGTGAGTTTCCTTCCCTGAAGCCAAGTCACAGGGGgaatcttaaacccaac
ggttacctcttatctagtggGAGCTCCTTATGAGGTAGTCTGTTAGAGGACTCATGGTAGgaatcttaaacccaactt
ggttacctcttatctagtggGGCATCCAAATCTCTCCTTTGCTGGAAACGTAACACCACCgaatcttaaacccaact
ggttacctcttatctagtggACACACAGACATGGGGTTTTTGCTTCACTAGACATAGCCCgaatcttaaacccaact
ggttacctcttatctagtggCCCTTTTCTTTTTGGCATATTTTGGGAGGCAACCCACCACgaatcttaaacccaacttg
ggttacctcttatctagtggCAGACTTAAGAAGGAGAAATTGAGGGCCTGGGTGTTCTACgaatcttaaacccaact
ggttacctcttatctagtggTCAGACAACTAGCTTCAGCGACTACTAGAAATATCTTCGGgaatcttaaacccaactt
ggttacctcttatctagtggACTTGAGCATTTAGAAACCATGATGATTTCAGGCATGCCCgaatcttaaacccaactt
ggttacctcttatctagtggCAGCACAAAAGGCAGGTGTTGAGTGTGTAGAACTGTTCAGgaatcttaaacccaact
ggttacctcttatctagtggCGTAGGGAAAGAAAACTTGCATTTTGTGCATCAGCTTTTGgaatcttaaacccaacttg
ggttacctcttatctagtggCAGAGTTGGGGGCTCTTGTCCAAGCTTCTCCCACAACCGCgaatcttaaacccaac
ggttacctcttatctagtggGCCACTCAGCAATCTGCATTCAGCTCTGCTGTTGCTGTGGgaatcttaaacccaact
ggttacctcttatctagtggCACCTCAGAGGTTCTGATTCTCTGGGTATGGCATGGGCCCgaatcttaaacccaac
ggttacctcttatctagtggTATTCCAGGTCCCTGCCTTCATACAAAGCACCATGAGGTGgaatcttaaacccaact
ggttacctcttatctagtggCATTTTCTTGTTCCTGGTCCTAGGGTTCATATTTAGCTTAgaatcttaaacccaacttgt
ggttacctcttatctagtggCGGACATTTAACATACCCTGCCTCATTTGATAGGGTTGTCgaatcttaaacccaacttg
ggttacctcttatctagtggGCCTCTCCGAGTTTCAGTGTCTCCATTATTGAAAGCAGGGgaatcttaaacccaactt
ggttacctcttatctagtggGAGCAGGAGATGAAGGGCACAGGTTCTATAGCCTGACTGCgaatcttaaacccaac
ggttacctcttatctagtggACTGTGTGATCTAAGGGAAGTTACTTAACTGCTCTGTACCgaatcttaaacccaacttg
ggttacctcttatctagtggAAAGGCTTTGGCTGCATCACAGAGAGTTTGGTAAGTTGTGgaatcttaaacccaactt
ggttacctcttatctagtggGCCATCCCATCCCCTGTGACTTGAATACCCAGATGGCCTGgaatcttaaacccaac
ggttacctcttatctagtggAATGGCCAGTCCTTGCCTTAAGTGGTGACATTACCTTGTGgaatcttaaacccaactt
ggttacctcttatctagtggGGTGCTGTGGCTCGGCTCGGGGGACCTCCTTTGGGAGATCgaatcttaaacccaac
ggttacctcttatctagtggCTTGGTGCCACACTTCAATGTCTCCCTTAATTTCAATTCCgaatcttaaacccaacttg

ggttacctcttatctagtggTCACGTTTCAAGGGTGTCAGACCACACGCAGGGACGCCTGgaatcttaaacccaa
ggttacctcttatctagtggGCAAGAACCCCCCAGTTGCTTATTTCCGCACCCCGACCTCgaatcttaaacccaa
ggttacctcttatctagtggACCCCAACCTCTTATCTCTGCACCCCACCCCTTTTCTGGAgaatcttaaacccaac
ggttacctcttatctagtggCTCTAGGCTTGCTTCCTTCAGTATAGGCAACCTTCCACCCgaatcttaaacccaact
ggttacctcttatctagtggTGTAAAATAGGCAAACAGTCCGAGGCGCCTGACGTCCAGGgaatcttaaacccaa
ggttacctcttatctagtggAGTCCCTTCCTAGTCTCTGTGCCCAGTACAACTCGTCCCAgaatcttaaacccaac
ggttacctcttatctagtggCAGACCCATCTGACCTCTCACCTCCTCGCCAGGCCGAGCTgaatcttaaacccaa
ggttacctcttatctagtggTAATCTTTTTATCGCCTCCGCTCCTCACACCTGGTCCGGCgaatcttaaacccaact
ggttacctcttatctagtggATCCAGCCCAGTTCATGACTTGTTTAGCAGCAATACTGAGgaatcttaaacccaact
ggttacctcttatctagtggGGCCGTCTTATTCTCAAAATGCACTTTATTACCCATTCTGgaatcttaaacccaacttg
ggttacctcttatctagtggGGCAGCCACTCCCAGAGCCCCTAGAACGCTGGTCCAAGGCgaatcttaaaccca
ggttacctcttatctagtggGACCATCACGGACGCCAAGCTTTGGGTTACTCTCACAGTGgaatcttaaacccaac
ggttacctcttatctagtggCCCAATCCAAAGCCTCCTTGCGTCCTCCTCTTGTATCCCCgaatcttaaacccaac
ggttacctcttatctagtggATACCTCTACTCCCTCCTTGGTGACCCCTTACCATCTCATgaatcttaaacccaact
ggttacctcttatctagtggACGTGGTCAGAATTCTTACACAACAACCAGGACCACACCGgaatcttaaacccaa
ggttacctcttatctagtggGTTTTAGCCTAGCCCTCATATCTTCATGCAGCGGCTGCCGgaatcttaaacccaac
ggttacctcttatctagtggACTTTCACTGGATAGGTAAAGTCCTTTCCTACAGGGTCTGgaatcttaaacccaact
ggttacctcttatctagtggCCGTTCTGTCAGACATAATTCCTCGGTTAAGCCTTCCCACgaatcttaaacccaac
ggttacctcttatctagtggCCCTTCTCAGAATTCTGGCCTGTCCTCGGAATGCTACAGGgaatcttaaacccaac
ggttacctcttatctagtggGGCACTCTCTAATCGGATATCCTGAGTTGTCCCAATTCTTgaatcttaaacccaact
ggttacctcttatctagtggGTTTCTCAATTCATCCAAAACCGTATTCAGGCCATCACCAgaatcttaaacccaactt
ggttacctcttatctagtggCACCCCCCAAAAATTTTAGCCACCCCAACACTTCAACACCgaatcttaaacccaa
ggttacctcttatctagtggTTGCCTTAATTTTGATGTTTGCCCAGGAGTTATTCAGGAGgaatcttaaacccaacttg
ggttacctcttatctagtggTTACTATTATTATTGCACTGTGATCCCAGAGTGTGCTTGGgaatcttaaacccaacttg
ggttacctcttatctagtggGAGACTTGCTTTATGATCAAGCATGTAATGGACCTTAGAAgaatcttaaacccaact
ggttacctcttatctagtggTGGGTGAAGTGCTGTGTAGATGTCTATGAGGTCCAATTGGgaatcttaaacccaactt
ggttacctcttatctagtggCTGCCTTGATAATCTGTCTAATACCGTCAGTGGGATGTTGgaatcttaaacccaact
ggttacctcttatctagtggCCAAAGCAGTGATAAGAGAAAGTTTATAACCCTGAATGCCgaatcttaaacccaac
ggttacctcttatctagtggTACAGCTGATCCCACAGATATGCAAAAGATCCTCAGAGACgaatcttaaacccaac
ggttacctcttatctagtggCAATCTCCCAAGATTGAATCCGGAAGAGATTAAAACCCTGgaatcttaaacccaact
ggttacctcttatctagtggAGATCCTAACAACAACAACAACAAAAAAGCTCTGGACCAGgaatcttaaacccaac
ggttacctcttatctagtggGCTGAATTCTACCAGGCATACAAAGAACTGATACTGATTGgaatcttaaacccaactt
ggttacctcttatctagtggTCTGTGAAGCCAGCATCAGCCTGATACAAAAATCTGGCAAgaatcttaaacccaac
ggttacctcttatctagtggAAGAAAACTCCAGGCCAATATTCCTCATGAACATAGATGCgaatcttaaacccaac
ggttacctcttatctagtggCCATAATCAAGCAGTCTTTATTCCTGGGATGCAAGGTTCGgaatcttaaacccaactt
ggttacctcttatctagtggAACATATTTTATGATGAAAACCCTCAACAGACTAGGCATCgaatcttaaacccaact
ggttacctcttatctagtggATGGGCAAAAGGAGGAAGGAATCCCCTTGAGAACAGAAACgaatcttaaacccaa
ggttacctcttatctagtggAGCAATCAGGCGGGAGAAAGAAATAAAAGACATTCAAATAgaatcttaaacccaact
ggttacctcttatctagtggCTATACCTAGAAAATTCTAAACACTCTACCAAAAGGCTCCgaatcttaaacccaactt
ggttacctcttatctagtggCCTGAAAATTCAGCTAACAAAGGGAGTAAAAGATCTCCACgaatcttaaacccaact
ggttacctcttatctagtggCAGTGCTATTCCCATCAAATACCAACATCATTCTTCACAGgaatcttaaacccaact
ggttacctcttatctagtggACCAAAAAAGACCCTAGCCAAGGCGATCCTAAGCAAAAAGgaatcttaaacccaa
ggttacctcttatctagtggCTACAGGGCTATAGTAACCCAAAGAGCATGCTACTGGTACgaatcttaaacccaac
ggttacctcttatctagtggGCAATTCAGAAAGGACTCACTCTTTAGTAAATGGTGCTGGgaatcttaaacccaactt
ggttacctcttatctagtggGCAGAAGATTGAAGCTGGAACCTCACCTTTTACCATATACgaatcttaaacccaact
ggttacctcttatctagtggAGACAACTTGGAAAATACTCCTCTTGACATCAGCCTTGGCgaatcttaaacccaac
ggttacctcttatctagtggCATAACCTACAGGGTAGGAGAAGATATTCAGAAACTATGCgaatcttaaacccaac
ggttacctcttatctagtggCAAAACCACAATAAAATACCATCTCTCACCAGTCAGAATGgaatcttaaacccaac
ggttacctcttatctagtggTGGTGGGAATGTAGATTAGTCCAGCCACTGTGGAAGGCAGgaatcttaaacccaac
ggttacctcttatctagtggTCCCAAAGAACTTGAAACAGAGCTGCCATTTGATCCAGCAgaatcttaaacccaac
ggttacctcttatctagtggGACGTGGAATCCACCCAGGTGGCCATCAATGGTAGAGTGGgaatcttaaacccaa
ggttacctcttatctagtggTGTGGTCCATACACACACAGTGGAATACTATGCTGTCATAgaatcttaaacccaactt

ggttacctcttatctagtggCAACATGGATGCAGTTGGAGGCCATAAACCTAAGCAAATTgaatcttaaacccaact
ggttacctcttatctagtggGTGTAGACACTAGAAGGGGTAGGGAAGGAGGGAGCATGGGgaatcttaaacccaa
ggttacctcttatctagtggGGTTACCATGCTCACTACCTGGGTGATGGAATCTGTACCCgaatcttaaacccaac
ggttacctcttatctagtggTGAGAGTATACTAGTATTTATTAAGGGCCCTGTGTCAGGCgaatcttaaacccaactt
ggttacctcttatctagtggTCCCCCCAACGCCTCTGAGTCAGGTAATATTAGTTTCCTCgaatcttaaacccaact
ggttacctcttatctagtggCCTTGCCCAAGGTCTGACAGCTAGCAAACGGGAGAGCTGGgaatcttaaacccaa
ggttacctcttatctagtggCAGGCAGGCTCCAGAACTTAAAACTACTATGCAATATTGCgaatcttaaacccaact
ggttacctcttatctagtggCTCATCTCTTCTTCTGGCTGTAAGCTTCTGAATAGGTGGGgaatcttaaacccaactt
ggttacctcttatctagtggGACAGGGAGGAATGGGATGACACCTGCATTTTGGCAGATGgaatcttaaacccaac
ggttacctcttatctagtggGGAGCAGTTGGCCAGTCACAGGCTGTGTGTGCACATGTGCgaatcttaaacccaa
ggttacctcttatctagtggCATGGAGCTCTGCTGTGGAAAGCCTAGAAATACATTCTACgaatcttaaacccaact
ggttacctcttatctagtggCGGGAGGCTGAGCCAGGAGAATTACTGGAACCTGGGAGGCgaatcttaaacccaa
ggttacctcttatctagtggCCAGTGTAAGAATGGTGACACTAAGATATCCAAGAGAGGGgaatcttaaacccaac
ggttacctcttatctagtggCAGTGGACGACCAAATGCTCACCACTGTTTTCTGGGGCTCgaatcttaaacccaac
ggttacctcttatctagtggAGCACCTTTGCTGCCTCCATCTAAAACTGGCCCTTCCAACgaatcttaaacccaac
ggttacctcttatctagtggGTCCCCATTTCAGAGAGATTTCTTTGAAACAGAGCCAGCCgaatcttaaacccaact
ggttacctcttatctagtggTGGTGATTTCAGAGAGTGGAGCCTGGTGGAAACAAAACCCgaatcttaaacccaac
ggttacctcttatctagtggGCCCACATTCCCTGCGTTTTCCCACTCTCTGCCTCACCCCgaatcttaaacccaa
ggttacctcttatctagtggCCAGCACTTACCACCAGAGTCCAAATTCCCTGGCAGCAGCgaatcttaaacccaa
ggttacctcttatctagtggAGACCTATGGGGCTTCCTAACACATTAAAAAGGCCCTCAAgaatcttaaacccaac
ggttacctcttatctagtggTCCCTGACTTGGGTAAGCACTGGGATGGAAAAAAGGCCTGgaatcttaaacccaac
ggttacctcttatctagtggAGGCAAGACCTCAATCTGGGGACTCAGATGTAATCAACCAgaatcttaaacccaac
ggttacctcttatctagtggGCATAGTGCTAGGTATTGGTGGGACGGAGATGACAGACTGgaatcttaaacccaac
ggttacctcttatctagtggTACAGATGTGAGCCACCGTGTCTGGCCATTAAGTTAAATTgaatcttaaacccaactt
ggttacctcttatctagtggGCAGATCTCTGATCTGGAGTGCCTCTCTTCCTTTTTGCCCgaatcttaaacccaact
ggttacctcttatctagtggGCAATGTTTGTCTCTTCAGGTAGAAGCTGAGAGCCAGTGCgaatcttaaacccaact
ggttacctcttatctagtggCTTTCCAAATGGTAGCGGCCCTATCAGCCTGGGTCCTGGCgaatcttaaacccaac
ggttacctcttatctagtggAACACTCTCCCAAATTACCCATAACAGACATGGAATACGGgaatcttaaacccaac
ggttacctcttatctagtggTGATCTCAAGTTCTGAGAGCTGTTTGTTACACAGCGTAGCgaatcttaaacccaactt
ggttacctcttatctagtggAGCTCTCTGCTTCATGGTTACTTCACAAGAGCTTCATAAGgaatcttaaacccaactt
ggttacctcttatctagtggATAGCCCATGATTGGTTATGAGCTCCTTGAGAGAAGAGTCgaatcttaaacccaact
ggttacctcttatctagtggGTGCCTGATATAAAGTGCTATGGTTTGAATGTATGTGCCCgaatcttaaacccaactt
ggttacctcttatctagtggCATTTTGAAACTTACACAACACTAGTAAAAGGTGAGGCCTgaatcttaaacccaactt
ggttacctcttatctagtggAAGACCAGAGGGAACTAGCTAGGCCCTTCTGTCCTACCTCgaatcttaaacccaac
ggttacctcttatctagtggTGTCCCCTTCCACTGCAATGAGGGCACAGTAAGAAAGCTCgaatcttaaacccaac
ggttacctcttatctagtggGGTGCTCTAATCTTGGACTTCCCAGACCGTATAACTGTGGgaatcttaaacccaact
ggttacctcttatctagtggTGTGGAAGTGGCTTGAAATTGGGTAATGTGTAGAGACTGGgaatcttaaacccaactt
ggttacctcttatctagtggTCTGGTGAGGGCTCAAATGAAAACAGCTGCAGAGAGAACCgaatcttaaacccaac
ggttacctcttatctagtggTCATGGAGATACAGACAGTGAAGGCCATCTTGATGAGGTCgaatcttaaacccaac
ggttacctcttatctagtggCGGAGGAAACGTCATCTTTGTTATAAGTTGGCAGAGAACTgaatcttaaacccaactt
ggttacctcttatctagtggGCCCATGTTCTAATGTTTTGTGAAAGGCAGAAGTTATAAGgaatcttaaacccaactt
ggttacctcttatctagtggGAGAGTGCTGCATGACTGTTTATAGGAAAAGACGAGAAGGgaatcttaaacccaac
ggttacctcttatctagtggGGGAAGCAGAACTTAAAGCGTTGGACAATTCTCAGCCCAGgaatcttaaacccaac
ggttacctcttatctagtggGGAGAGAATAATCCTAAGGGTGAGGCCAAGTGACTCTTTGgaatcttaaacccaac
ggttacctcttatctagtggGCCGCCACTCTCATAACAGGCTCAGAGTGCCAGGGCCCTGgaatcttaaacccaa
ggttacctcttatctagtggCCCAGGACTGCCTCAGGCTTCTGCTCCACTCATTCTGGCCgaatcttaaacccaa
ggttacctcttatctagtggCCTCAAGTGCACCCAGGTGTGGTTCAAAGTGCTAGTCCGGgaatcttaaacccaac
ggttacctcttatctagtggGCTCAGAGTGCGTGAGCTAAGCAGGCATGGCTACCTCCTCgaatcttaaacccaa
ggttacctcttatctagtggGCCTTGGGGCGTAGGCAGAGAACTGTCACAGTGGCAGGGCgaatcttaaacccaa
ggttacctcttatctagtggCAGGGCCACTCCTGAGACCCCCCAGCCTGTGCAGCCACCAgaatcttaaaccca
ggttacctcttatctagtggGCAGCATAAGCCGCGCCTAGCAAAGCCACGGGGATGAGGCgaatcttaaaccca
ggttacctcttatctagtggGGGGACCAACCCATATACCAGTGTGTCAAGTAGTCAGGAAgaatcttaaacccaac

ggttacctcttatctagtggGCCTTGTTGGGTTTCGAATGGGAGTGAGACCTATTACTCCgaatcttaaacccaactt
ggttacctcttatctagtggTCCTATGCCTGTCCCACCACTGCATTTGAAGCACATAATAgaatcttaaacccaactt
ggttacctcttatctagtggTGGGGGAATTCTTCTCCAGCAGTCTGCAAAATCTGACAAGgaatcttaaacccaact
ggttacctcttatctagtggCTGTTGAATTCATTCTCTAACCACTTCTATCAAGTCACTGgaatcttaaacccaacttg
ggttacctcttatctagtggAGACACACACACTTGCTGCTAGATTTTACTCTCTTGGTGGgaatcttaaacccaactt
ggttacctcttatctagtggCTTGCTGCTAGATAAGGGGTACCTAATGCAAGATGTACCTgaatcttaaacccaactt
ggttacctcttatctagtggACCTGAAACAATTAGGAAGTCACCGTTCAGCTTTGCAGGCgaatcttaaacccaact
ggttacctcttatctagtggGGCTGATAAGGGGTTGAATCTGTGAGGTTTTCCCATAGAGgaatcttaaacccaactt
ggttacctcttatctagtggCTTTAGCCAACAGCAGTTGTCTCTAACTCACCCAAACTGCgaatcttaaacccaact
ggttacctcttatctagtggATTTATTTATTTAGAGACAGAGTCACATTCTGTCACCCAGgaatcttaaacccaacttg
ggttacctcttatctagtggCCAGGCTGCTCTCAAACTCCTGGTCTCGTGATCCACCTTCgaatcttaaacccaac
ggttacctcttatctagtggTGAGCCACCGCACCCAGCCTTTAAGAGCTATTTTATTTCCgaatcttaaacccaactt
ggttacctcttatctagtggGGAGGCCCAAGATAGGAAAGAGACTTGCCCAAGGCCACACgaatcttaaacccaa
ggttacctcttatctagtggCCCAACCACCATGCCAGTGGGAGTTCCAGGTCCACCCTGCgaatcttaaacccaa
ggttacctcttatctagtggTGACCACCTCCAAGATGGCCCAACGCTGCCCTTCAAAAGGgaatcttaaacccaa
ggttacctcttatctagtggGGGCAATCTGATGGTGAATTCATTAGAGGTGAAATAAATGgaatcttaaacccaactt
ggttacctcttatctagtggCCTCTTGATTCTAAGAAATCCTTTCATTGGGTCTAGAGTGgaatcttaaacccaacttg
ggttacctcttatctagtggCAGCACAGTTTTCTATGAACAAGTGCAAAAACGGGGTCAGgaatcttaaacccaact
ggttacctcttatctagtggGATGGGATGTGACATTGCTGCTTCCCCAATCTTTACCTACgaatcttaaacccaactt
ggttacctcttatctagtggCACACTGGGGATATCAGAGCCAGAGAGAAGACGTGGAGTCgaatcttaaacccaa
ggttacctcttatctagtggCCATAGTGCAGAGGTTGAGATATCCTGCTCTGGGTTACAGgaatcttaaacccaact
ggttacctcttatctagtggAGAGCAATCTGCCATCTCCACAACTTTGTACCTTTGCCTTgaatcttaaacccaactt
ggttacctcttatctagtggGCCTCAGCCTCCCATCAACTGGCACCATTGGCATCCTGGCgaatcttaaacccaa
ggttacctcttatctagtggCTGCACAGTGCCTGCTACACACTTTCATTAAGTAACTCTCgaatcttaaacccaactt
ggttacctcttatctagtggCCAGGTTTTCACAGGGAGGAAGAGGAAGGGTCAGAATTCCgaatcttaaacccaac
ggttacctcttatctagtggACTAAACTATCACACTATCACACTTCAGACAGGAGTGCAGgaatcttaaacccaact
ggttacctcttatctagtggGTGGAGAGGAGGTATTCGGAGGCCACAACGTAGGGCAAGGgaatcttaaacccaa
ggttacctcttatctagtggTAGGCTGGGTATGATAGGTAGAATAATGGCTCCCCCAAAGgaatcttaaacccaact
ggttacctcttatctagtggCCCCTGTAACCTGTGACTATTACCTTACACCATGAAAGTGgaatcttaaacccaactt
ggttacctcttatctagtggAGAGGGAGGCAGGAGAGTCAGAGTCAGAGGAGGTGTGAGGgaatcttaaacccaa
ggttacctcttatctagtggGGATTCTCTCCCAGAGTCTCACAAGGAGTGCAGTCCTGTGgaatcttaaacccaac
ggttacctcttatctagtggGCTGATTTCCAGCTTCAGACCTCCAGGACTTTAATGATAAgaatcttaaacccaactt
ggttacctcttatctagtggCAGCCACCAAGTGTGCAATAATTTGTTATGGAAATCACAGgaatcttaaacccaactt
ggttacctcttatctagtggATGAGGATTGCTACATAAAGTATAGAACACACAGTCCAACgaatcttaaacccaactt
ggttacctcttatctagtggGCCCTAACCGGCTTCCCCCTCCCACTCAGCCCCTGCATCCgaatcttaaacccaa
ggttacctcttatctagtggTCCCAGGTCTGCTCTGTTACAGTGGTATCATTATACCTGCgaatcttaaacccaactt
ggttacctcttatctagtggTAGTCTATGACTCAACCGAGTGAAACTGTTAAGGCCTCAGgaatcttaaacccaact
ggttacctcttatctagtggCATCCAACTCATCAGAACAACCATTGGGGAGATGGATTGGgaatcttaaacccaact
ggttacctcttatctagtggGCAGTGCCTAGTAACAAGCTGTGTGCATTTTTAAACCTGGgaatcttaaacccaactt
ggttacctcttatctagtggGTTGATTTCCTGAGAAGCAGGGCAGGAACTAGGGCAAGGCgaatcttaaacccaa
ggttacctcttatctagtggAGGAGGCTCTCCATTTCAGGTGCCGACCCTGCACTTGCAGgaatcttaaacccaa
ggttacctcttatctagtggCTTGCACCCTGGGAGACTCACTTGCTTCACTCTAGTATCGgaatcttaaacccaact
ggttacctcttatctagtggAACCTAGCCTCTTTACATGGCTTATGGGAAACTGAGGCCCgaatcttaaacccaact
ggttacctcttatctagtggGCTTCTGCCAACACCTACCTACTGAGTTTCAGCCAGAGGCgaatcttaaacccaac
ggttacctcttatctagtggGCCCTGGGACAAGGCCCTGTCTCAACTGGATTAAGCAAATgaatcttaaacccaac
ggttacctcttatctagtggCCCTGGCCACAGTTGTTAGCTCCAGGATATACATGTGGGGgaatcttaaacccaac
ggttacctcttatctagtggCACGGAGAGAGAAAATGATTCCTCTTTCCTTTGGAGGGTGgaatcttaaacccaact
ggttacctcttatctagtggGTCTGGAACTGCTGCAACCATTTATTACCATGAGAAAAGCgaatcttaaacccaactt
ggttacctcttatctagtggCTTCACCGACTCCCATCTAAAGGACCCACTACCTCTGGACgaatcttaaacccaac
ggttacctcttatctagtggGCCAAGTTCTACAGCTACATTGCCCCATAGAACTTTCTGCgaatcttaaacccaact
ggttacctcttatctagtggTCTGTGCTGTTCGATACAGTAAGCCACTACCTGCGTGTGGgaatcttaaacccaact
ggttacctcttatctagtggGCCACACAGACCAGTGGTTACCATATTGGTCCATGCAGCCgaatcttaaacccaac

ggttacctcttatctagtggGCCTCCTCCAGAAACCAGCTCCAGATCCTTTCTTTCCTCCgaatcttaaacccaac
ggttacctcttatctagtggGGCTGATCTTTTCTGTGATCCTATAGCAACTGAGCATTCCgaatcttaaacccaactt
ggttacctcttatctagtggAGGCAAGGCCCATTCCTGTGTCCAAGCACTGAGCAGAAGCgaatcttaaacccaa
ggttacctcttatctagtggGTGGCCCAAATGTGTACCAGATGTTGAAAGGCTTTGTAGCgaatcttaaacccaac
ggttacctcttatctagtggGCCCAGCCTTGTGACATCTGTGACCATCAGTGTCAGGCTCgaatcttaaacccaac
ggttacctcttatctagtggTACCCCTCTGCTGCCTTGACTTTCTTTTTCTTTTCTTTCTgaatcttaaacccaacttgt
ggttacctcttatctagtggCTTGGCTCACTGCAACTTCCGACTCCCCTGTTCCCTGCTTgaatcttaaacccaac
ggttacctcttatctagtggGTATTTTTAATAGGCACAGAGTTTCACCATGATGGCCAGGgaatcttaaacccaactt
ggttacctcttatctagtggCGCCCACCGCGGCCTCCCAAAACTGCCTTGACTTTCTAGCgaatcttaaacccaa
ggttacctcttatctagtggGTGGGTGAGAGAGGAGGAGCTGGAAGACAATGCAGAGCTGgaatcttaaacccaa
ggttacctcttatctagtggCATTGCAATTGCCAGGGTGTGCTATGTGGCTTCGACATGCgaatcttaaacccaact
ggttacctcttatctagtggGAGGGACCCAGTAAGAGATAATTGAATTTTGAGGCCAGTTgaatcttaaacccaact
ggttacctcttatctagtggCACCTTCCACCATGACTGTGATGCCTCCCCAGCTCCCATGgaatcttaaacccaa
ggttacctcttatctagtggCAGAATCCTCCTCCTTGAGATGTGCTTCCTGATCCCAGGCgaatcttaaacccaac
ggttacctcttatctagtggATTGTCAAGCATGTGATGAGGCACTGGGGTTCTGTCAGTGgaatcttaaacccaac
ggttacctcttatctagtggCTGTCATACTGAAGGTCACGTTCTAATAGGGGAGACAGACgaatcttaaacccaact
ggttacctcttatctagtggAAGAACAATCGAGTGTGGGAATAGAGGGGAAGCAGGCAGGgaatcttaaacccaa
ggttacctcttatctagtggGGTGACAAAGTCCAGAGTGGTGGCAGGTGAGACAATGGAGgaatcttaaacccaa
ggttacctcttatctagtggGCTCTGCCAGATCAGATCTTGTCTTTACATGCCCCCACCCgaatcttaaacccaac
ggttacctcttatctagtggCCAGGGAATAACTTGTACTCAGCTTCTGGGGTTCCCTAGGgaatcttaaacccaact
ggttacctcttatctagtggTTTCTCCTAGCTGAGCCCACTGTCCAATTAAGGATCAGACgaatcttaaacccaac
ggttacctcttatctagtggGCAAGTCACTTCTCCTTTCTGGATCTGTTTCTTCCTGTGCgaatcttaaacccaactt
ggttacctcttatctagtggGTTGCACAAGATAATCTCTGGTGCCCCTTCCAATTTAATAgaatcttaaacccaactt
ggttacctcttatctagtggGGAGTGGGGCGGGGGAAGAAAGGGGAGGAAGAGGAGGAGGgaatcttaaaccca
ggttacctcttatctagtggAAGGGGAAGGGAGAGGGGGAAGGAAGAAAGGGAGGGAGGGgaatcttaaaccca
ggttacctcttatctagtggGCCACAATAAATACTTCGGTGTGAGGCTGTAGAAATTGGCgaatcttaaacccaac
ggttacctcttatctagtggGAAGCATCTTCTACCTCCCTCACTTCCTGATCCCCATCCCgaatcttaaacccaac
ggttacctcttatctagtggGCCTGGAAAGATGCTCCCTAGCAGGGTAGAGTCAGGAACCgaatcttaaacccaa
ggttacctcttatctagtggTGGAAAGATGCTCCCTAGCAGGGTGGAGTCAGGAGCCTGCgaatcttaaacccaa
ggttacctcttatctagtggCTGGTGCTTTGGGCTGTGGAGCGGATTTCTCTCTCCAGTCgaatcttaaacccaact
ggttacctcttatctagtggGGCAGGGATCTCAAGTGTGTCCTTGGCCTGCAGTGGGGACgaatcttaaacccaa
ggttacctcttatctagtggGGGCAGCCTGCTCCATAACGATGCCCTCATTCCCAGGAGCgaatcttaaacccaa
ggttacctcttatctagtggGCCAGTTACTCAAATCCCATTCCTGGCTTTCTGCTTCAGTgaatcttaaacccaactt
ggttacctcttatctagtggCAGGAGAAGCAACTCTGCCTTATACTCGAAAATTAACCAGgaatcttaaacccaact
ggttacctcttatctagtggTGCAGTTTGAATCAGGGGTATTTTCTGGCTTACAGGTTAGgaatcttaaacccaactt
ggttacctcttatctagtggAAGATTCCCTAAGGGAGAAGTGATCTATTCCAATATGGGCgaatcttaaacccaact
ggttacctcttatctagtggTGTGAGAAATGATACCGCCCTGGACTTTCATGCTGGGCCCgaatcttaaacccaac
ggttacctcttatctagtggGTCGCCAGCTTGCCTGCTCCTTAAGATGCTTGAAGTCCCCgaatcttaaacccaac
ggttacctcttatctagtggAAGACCTGCTGCAGGCCTCCCTCACGCGATCCCAAGTGAGgaatcttaaacccaa
ggttacctcttatctagtggGCCTCATTCTAAGGCTGACGAAGCTGAGGCTCAGTAACCCgaatcttaaacccaa
ggttacctcttatctagtggCTACCCAGGCCTTCTACAACAGGTCTCTACAACTAGTGTCgaatcttaaacccaac
ggttacctcttatctagtggGCTGCTTTCTGATTGGGAACATACCAAGTTCAAGGTCCCCgaatcttaaacccaact
ggttacctcttatctagtggTGCAACTGTTCTATATCTGTGCTGTTCAATACAGTACTATgaatcttaaacccaacttg
ggttacctcttatctagtggCACTTAGCATCCCACTTGACCCTTACAACAACTTCATGGGgaatcttaaacccaact
ggttacctcttatctagtggGCCTCCAAGAACACACAAAATCTGGTGGCTGTTCTGGGAGgaatcttaaacccaac
ggttacctcttatctagtggAGTCCTAACTCTGCCATTCTCCAACTTTGTGACCTCAGGCgaatcttaaacccaact
ggttacctcttatctagtggGCGATAACTAGAGGGAAAGAGTACTTGCCTCATACGGCAGgaatcttaaacccaac
ggttacctcttatctagtggGCCTCAGCATTTGTGGAGTTATGTTCTACTTATGGGAGACgaatcttaaacccaactt
ggttacctcttatctagtggCAGGGAAAGAGAATAGGGAGTATTGGAAGGAGGAGTTCAGgaatcttaaacccaac
ggttacctcttatctagtggGCAGTGAAGACTTCAAAAAGGAGTGAGGGGCCAGGCGCACgaatcttaaacccaa
ggttacctcttatctagtggGGCAGGCCAAACACTTGAGCTAAGGTGTTGGAGGCCAGCCgaatcttaaacccaa
ggttacctcttatctagtggCTCAGAAGGTGGGGATGGGAGAAACACTTGAGTCCAGGAGgaatcttaaacccaa

ggttacctcttatctagtggGTGAGACCCTGTCTCAAAAAATAAAGGGAGGAGTGGGCGGgaatcttaaacccaac
ggttacctcttatctagtggACAGGGAGGCCATGTGGCTGCACTGGAGTGAGCGAGGGAGgaatcttaaacccaa
ggttacctcttatctagtggGAGGAAGATCAGAGATGTAACTGATAGTGCGAGGATGGGAgaatcttaaacccaac
ggttacctcttatctagtggGCCATTGGAGGCTTTTGACGTAGGAGTGCCATGATCTGACgaatcttaaacccaact
ggttacctcttatctagtggACCGGAGGGATCAGGTGTGGGAGCCAGTTAGGGAGATGTTgaatcttaaacccaac
ggttacctcttatctagtggAATGGTAGCCATTGAGATGGAGAAATGGTAGGAATTGGGTgaatcttaaacccaactt
ggttacctcttatctagtggCGAGCCAACAGGGTTTGCTGATGAGTTATATGTGAGTGGGgaatcttaaacccaact
ggttacctcttatctagtggTGGGACTACGAGTTTCTTGGTCTGAGTAACTGCTTTTCACgaatcttaaacccaacttg
ggttacctcttatctagtggGTATAGAAGAAGCAGCTCTGCAAGAGTGAGAAAGAACTGAgaatcttaaacccaact
ggttacctcttatctagtggTTCTCCAGCTATACAGTGTCCAGCACAGAGTATGTGTTCCgaatcttaaacccaactt
ggttacctcttatctagtggGCCCTACATCCAAGCACTCTGAGAAAGCACATGGTCAGGCgaatcttaaacccaac
ggttacctcttatctagtggGCACAGGCTGATCTCCCTCTCCACCCACCCTTCCTTCCTCgaatcttaaacccaac
ggttacctcttatctagtggACAGGAAATGGACAAAGGCACAAACTGGAAGTCAGTGCAGgaatcttaaacccaac
ggttacctcttatctagtggCAGAGAGTATTCCTTAATGCTCACAGAAAATCTAGCCCAGgaatcttaaacccaact
ggttacctcttatctagtggAGCTGAGGACTAGAAAGATGAAGTGGCTTGCAGCTTCCCCgaatcttaaacccaac
ggttacctcttatctagtggCCCAAGGATTCTTCTAGCTCTCACATGCTTTCAGGTTTTCgaatcttaaacccaacttg
ggttacctcttatctagtggGCTTTCCTTCATTACCCTCATCTAAGCCACCAGCAAACTCgaatcttaaacccaact
ggttacctcttatctagtggGGGTTTCACCGTATTAGCCACGATGGTTTTGATCTCCTGAgaatcttaaacccaacttg
ggttacctcttatctagtggAGGCTTGAGCCACTGCGCCCGGCCACTGTCACTCTCTTTTgaatcttaaacccaac
ggttacctcttatctagtggGCCTCTAACTGGCTTCTTTCCCCTCCTTACCCTGGCCCTCgaatcttaaacccaact
ggttacctcttatctagtggTGCATGTGACAGTCTGAACGATCTGTTGAGAATTTGAACCgaatcttaaacccaacttg
ggttacctcttatctagtggGCCAATCAGGCCTTACATGATCTGGCCCCCATTTTTCTCTgaatcttaaacccaactt
ggttacctcttatctagtggGGCTCCTAAGAGGGCCACCCCACCCACAGTGATTCCCACCgaatcttaaacccaa
ggttacctcttatctagtggCACATCCTTTGTGTAGGCTGGACCTAGTGCTTTGCCTTTAgaatcttaaacccaacttg
ggttacctcttatctagtggGGTGATGGGATGTCACATCCACGATTAGGTTGCAAAAGACgaatcttaaacccaact
ggttacctcttatctagtggCTCACTCAGATGAAGCAAGTTTCCATGTTGTGAGCTGCCCgaatcttaaacccaact
ggttacctcttatctagtggGCCAACAGCCAGCAAGGAAGTGAAGCCCTCAGTCTAACAGgaatcttaaacccaa
ggttacctcttatctagtggGTGAGAGACCCTGAGACAGAAGACTCAGCTAAGCCAGGCCgaatcttaaacccaa
ggttacctcttatctagtggACACCCTCTCGGAACATTTTCTCTTCTTTTTCTCTCAGCCgaatcttaaacccaacttg
ggttacctcttatctagtggGGTTGGCTCTATCATGCAGCCTCTCAGCCCAGATGTCCCCgaatcttaaacccaac
ggttacctcttatctagtggTTCCCTAGCTAACTCTACACTCAGCCCATTACTCTGTTTCgaatcttaaacccaacttg
ggttacctcttatctagtggGGAACCTTCTCAGATATGTTCATGGCGGAATCTCAGCTCCgaatcttaaacccaact
ggttacctcttatctagtggAGGAGTTGGGGGTGGGGGAGGAATGGAGATGCTCTTCCACgaatcttaaacccaac
ggttacctcttatctagtggCTAGGACATTGGACCAGCATTTGCTCTCCTGATTTATCCCgaatcttaaacccaactt
ggttacctcttatctagtggGAAGGACCCCAGGTTGTTTGTTTCTTTCAAAGATTACCCCgaatcttaaacccaacttg
ggttacctcttatctagtggCAGGGTGGCAAAAACCCACATCCTCTTCCGCTGGCTCCACgaatcttaaacccaac
ggttacctcttatctagtggCCTCTTCTGCAACCCACCGACTAGTTTCAACACATCTTTCgaatcttaaacccaactt
ggttacctcttatctagtggCGGAGGAAGTTCTAGGCTCAGTGTGGATTGCCAAGTGGTGgaatcttaaacccaac
ggttacctcttatctagtggGTATCTTTTGCAAGGCTCCGTTTTCTTCCTTTCTACTGTCgaatcttaaacccaacttgt
ggttacctcttatctagtggTCCCATTTTTAAGGGCCCCTCCACCGAAAAATACCAGAGGgaatcttaaacccaac
ggttacctcttatctagtggTAAATGCGGCCCATAAATGAGAACATTCATCCAAATGTCCgaatcttaaacccaactt
ggttacctcttatctagtggTGAAAAGGGGTTTCATTGAGAATTTCATTAGTAATTGGGGgaatcttaaacccaacttg
ggttacctcttatctagtggGGGCCAGAATGTGGAACAATGAAAACTGTCATACTTCTGGgaatcttaaacccaact
ggttacctcttatctagtggCCTCAGAAATGTGTGCACATGTGGACCAAAAGACATGTACgaatcttaaacccaact
ggttacctcttatctagtggGATAGCCCCAATCCAGTAACAACAATAGTTCTAGTTCAACgaatcttaaacccaactt
ggttacctcttatctagtggCTTGGCTACCTTTGGGGAGAGGGGAGGGGCTGGATATGGGgaatcttaaacccaac
ggttacctcttatctagtggCGCTCCCAGAGTGGTAGCAATGTTCTGTTTCTTTGACCTGgaatcttaaacccaactt
ggttacctcttatctagtggTTAGTGACTTGTTCACTTAAGATTTGTTCACTTTTCTGCGgaatcttaaacccaacttgt
ggttacctcttatctagtggATACCAAAACATCAGTAACATGTCTGGCCAAAGGCAAGGCgaatcttaaacccaac
ggttacctcttatctagtggGGGTAACAAAGCAAAATCTTCCCTCTGCCGTTTTCAAGATgaatcttaaacccaactt
ggttacctcttatctagtggCTCGTGTTCTCAAAGACTTAAGTAAAATGCTCCTGGTATCgaatcttaaacccaacttg
ggttacctcttatctagtggGAAACATATAACCTCAGCCCTGTTTGTATGTGCTTACACCgaatcttaaacccaacttg

ggttacctcttatctagtggCTGTTTTCCTTTCTCCCTTCTGTGTAGCTTCTGTCCTCCCgaatcttaaacccaacttg
ggttacctcttatctagtggCAGCAGCCATGATTTCACGAGACAAGATTAGGAGAATGGGgaatcttaaacccaac
ggttacctcttatctagtggGGGTGTGGTTTCAGGTTCTCCTCTATTATGGAAATTCTAGgaatcttaaacccaacttg
ggttacctcttatctagtggCTGGGGACTACTAGATGGGGGATGGAGGGAGCTAGAGGGGgaatcttaaacccaa
ggttacctcttatctagtggACTATGCTCAGTACCTGGATGGTGGAATTAGCTGCACCCCgaatcttaaacccaac
ggttacctcttatctagtggGCAATATACCCAAGTTACAAACCTTCACATGTACCTCCTGgaatcttaaacccaact
ggttacctcttatctagtggGAGGTGAGGATAATTTGGGGTAGGTTGGAGAATCCAGAAGgaatcttaaacccaac
ggttacctcttatctagtggTTTTGGGTATGGAAAGGTAGGAGAGTAATAGAAACAAAGGgaatcttaaacccaact
ggttacctcttatctagtggACATTCCACTTAACATTATTTGAAGACCAAAGGTAAAGCCgaatcttaaacccaactt
ggttacctcttatctagtggGAGTAGCGATTTTCTGAGTATCCATGAACACATGAAGTCCgaatcttaaacccaact
ggttacctcttatctagtggTGAGGTCATCAGAAAAGCTTGTCATGAAAAAATCCAGCACgaatcttaaacccaac
ggttacctcttatctagtggGGGGTGGGGATGGGAGAGGAACAGCCGTGGAACCCCTCCCgaatcttaaaccca
ggttacctcttatctagtggTGGCCTTGTTCACAGGAGAGCAGTTTGCCCTAAGTAGTTTgaatcttaaacccaact
ggttacctcttatctagtggTAAAACAGCAACTTGCCATAGAAATAGTTATGACAGTCTGgaatcttaaacccaactt
ggttacctcttatctagtggCTCCTGCAGATGCTGCCAGGGGAGGAGCAGTGTGGTCTGGgaatcttaaacccaa
ggttacctcttatctagtggGACTTGGGTTCTCTGAAGTTAGACAAGCCTAGATTGGAATgaatcttaaacccaactt
ggttacctcttatctagtggTAATTGTTTTTGTGCCTCCGCTTCTCCATCTGTAACATGGgaatcttaaacccaacttg
ggttacctcttatctagtggGTCCATATCACCTGGGTGCTATGAACAATAATATTGAGAAgaatcttaaacccaactt
ggttacctcttatctagtggATAGCTCCCACCCCCCACCACATCTCTTCTGCCCTGCCCCgaatcttaaacccaa
ggttacctcttatctagtggGCAAGGGGGGCTGGGTGGGGATAGGAGGGGTTGGAGGCAGgaatcttaaaccca
ggttacctcttatctagtggTCCTGCCAAGTTCCCCACTTACTCCTGGATTTATTTTTCCgaatcttaaacccaactt
ggttacctcttatctagtggAAGGGTATGTGTTAGCGTATTCCTGTCTGAATTTAGAGGCgaatcttaaacccaactt
ggttacctcttatctagtggAGGAAGCATTCCAGAGTACTCATATGTCTTGGGATGTACCgaatcttaaacccaact
ggttacctcttatctagtggAGCAACTCTGATATAATCTTTATGAAGTGCCAGGAACTTGgaatcttaaacccaactt
ggttacctcttatctagtggTGCATGAGGAAATTGAGGCACAGAGAGGCTGAATAACCTGgaatcttaaacccaac
ggttacctcttatctagtggAAGCTCAGAGCAGAAGTCAAGAAAGTGCAGCTGGTGGGCCgaatcttaaacccaa
ggttacctcttatctagtggGCCAGCCTTCTCCTCCTCAAGGCCACCTCATCATCACTCCgaatcttaaacccaa
ggttacctcttatctagtggTAGCTCTCAGAACCCGATCCTATTTATGCACTGACTTGTGgaatcttaaacccaactt
ggttacctcttatctagtggGTCCCTGACTCACAATGGTTTGACTTACAATTCTTTGACTgaatcttaaacccaacttg
ggttacctcttatctagtggTGGAAGAAACAGTACTTCAAGTACTCATACAGCTATTCTGgaatcttaaacccaactt
ggttacctcttatctagtggGTATTCTGAATACTCTTAAGGTAGGCTAGGCTAAGCTGTGgaatcttaaacccaactt
ggttacctcttatctagtggGTTTCCTTTTGGCCGTCCAAACCTAAAATATGTACAACCCgaatcttaaacccaactt
ggttacctcttatctagtggCCCTCTCATTACATGCAGTTTCAAGCACAATCTGCAACCCgaatcttaaacccaact
ggttacctcttatctagtggCCCTTCCACTGTACAGTCGGTTTAGGGTAACACAAGCAGCgaatcttaaacccaac
ggttacctcttatctagtggGGCCTGCCAATCAGTGACACAAATGCTCTGGGCTCTCAGGgaatcttaaacccaa
ggttacctcttatctagtggCTAGGATAGGTGCTGAGGCAAATACAGTGGTAGATGACACgaatcttaaacccaac
ggttacctcttatctagtggAAGAGCTATCCAAGTCTTTAGGTGAACTAAACCTTCAGCCgaatcttaaacccaact
ggttacctcttatctagtggCACTCACTCTGTGCCCAATTTCCTCTTCCACACAATGGGGgaatcttaaacccaac
ggttacctcttatctagtggCATCTACCTCATGGAGTCATTGTAGGATTTGACTGGGACAgaatcttaaacccaact
ggttacctcttatctagtggAGCCCTCTGTAGAAAGGCTTCAGCAGGGTTATAGTCACCCgaatcttaaacccaac
ggttacctcttatctagtggTCTGGGGAGAGTTTCTGTAACGTACTTCAGATTCTCTGAGgaatcttaaacccaactt
ggttacctcttatctagtggGATGTGCGCATCCCCAGACATGTCCCCTCTGATGCACCACgaatcttaaacccaa
ggttacctcttatctagtggCCCTCCTCCTCCTCCTTCTCCTACACACACACACACACACgaatcttaaacccaa
ggttacctcttatctagtggCACACACACACACACTTCATCCTACTCTCCAGCATTCAGGgaatcttaaacccaac
ggttacctcttatctagtggGGGACTTCTGTAGCCAGTCTTCCTCCCTCAGTGCCCCCTCgaatcttaaacccaa
ggttacctcttatctagtggCCATTCTCTGTCAACTGATCCTAGTGTTCTTTTTTATAGCgaatcttaaacccaacttg
ggttacctcttatctagtggAGGCTCCATGAAGATAGGAACTTGGTCCGTCTTGCTCATGgaatcttaaacccaact
ggttacctcttatctagtggGAGACCCCAAAAATCCCCAGAAAATTTCAGTTCAGCTTGCgaatcttaaacccaac
ggttacctcttatctagtggGCCTGCTTAACCAAGGTTGTCCAAATTCAGCTAAGGGATGgaatcttaaacccaact
ggttacctcttatctagtggCACCATGAGTGTTTTGTGTGGAAACAGAGTGAGGGTTTGCgaatcttaaacccaact
ggttacctcttatctagtggGGGCTTCTTACATCTGACTGCTGCCACCTCGGGCCAGTACgaatcttaaacccaa
ggttacctcttatctagtggGGTAGGGTAGCTACGCTTATAGACCTGAAATTCTAGTATAgaatcttaaacccaactt

ggttacctcttatctagtggGGAGCCTTGTTATGCACTAGGCACTGGGAACTGAGCTGTGgaatcttaaacccaac
ggttacctcttatctagtggCCAAGTGGAGGACTAATGAGAAATCCTTCTCCTCCCCCAGgaatcttaaacccaac
ggttacctcttatctagtggTGCACAAGGCAGCACACCCACATGCCCCTCGTATATAGACgaatcttaaacccaa
ggttacctcttatctagtggTGGTTGACAGTAGTTGTTCTCAACCAAGGGCCATGTTGCCgaatcttaaacccaact
ggttacctcttatctagtggGAAGGGGGATGTGCTAATGGCACCAGTGGGGAATGCTGCCgaatcttaaacccaa
ggttacctcttatctagtggGGTCCCAGATTACCTAAGTGCTGAGGTTGAGATGCCCTGGgaatcttaaacccaac
ggttacctcttatctagtggTAGCCAGATCCAAGCAGCAGTTATACTTTCCCTGATTTCCgaatcttaaacccaactt
ggttacctcttatctagtggCCAGCATTAAAGAGCCATTGTAGACATTGGGAGTTTTCCCgaatcttaaacccaact
ggttacctcttatctagtggCACCCTTCCCTCATCTAAGAAACCTTTCACATCCCCAAGGgaatcttaaacccaac
ggttacctcttatctagtggCTCACTTTCCCATTCTCCTTTGTAATGCTCCACCTCAGCCgaatcttaaacccaactt
ggttacctcttatctagtggACTGAAAAGTGAGCAAGATGAGAGAGCCGATGCTGGGTAGgaatcttaaacccaac
ggttacctcttatctagtggGCCCAGAATGTGAGCTGGGGTCTCTGTCCTCATAGCATCGgaatcttaaacccaac
ggttacctcttatctagtggGAGCCTCCCAGTGTGACTCTCCAGCCATCTTGGAGACGTGgaatcttaaacccaa
ggttacctcttatctagtggGAAGGATGGACTCACAATTCACCTAGAACTCACTCACTGGgaatcttaaacccaac
ggttacctcttatctagtggCCACCTTGGCTACTAACACAGTAAAACACCCTAGTGTGCGgaatcttaaacccaac
ggttacctcttatctagtggCAAATAGCCAATGCCTTCATCCAGCCAGTTTTTCCCCAGCgaatcttaaacccaac
ggttacctcttatctagtggCACAGGAAATAGGAATCATAAGCCACCTGTGCTCCTTTCAgaatcttaaacccaact
ggttacctcttatctagtggCAGACTGCCCTTGAGCCAGCTTGAACGTTGGTTTAAGTGGgaatcttaaacccaac
ggttacctcttatctagtggTGCCACTGTGGGAGAAGACTGACCTGTTAGCCTTTTCTGCgaatcttaaacccaact
ggttacctcttatctagtggCTGAGTGCAGAATTCTTCACATTATCCATGCAGAACTAGGgaatcttaaacccaactt
ggttacctcttatctagtggGGGTCTGTACAGAGTTAGTGTCACAGTAAGAATCTCATTGgaatcttaaacccaactt
ggttacctcttatctagtggAGTAACAGCGACCACTTACCTCATGCCTCATATGTGCCAGgaatcttaaacccaac
ggttacctcttatctagtggGGAAACTGGGGCTCTAGGAGTATGCAAGATTTGTCCAAGGgaatcttaaacccaac
ggttacctcttatctagtggCTTCACTCTAACTCCATCCTAAAAACCACCTTCCCCATCCgaatcttaaacccaact
ggttacctcttatctagtggCACCATCTCTCTACCATCCTAGCAGCAACTGCAATGCTTGgaatcttaaacccaac
ggttacctcttatctagtggGAGCTTGTCAAATGAGACACAGTTTGTTGTTCTGGCCCTGgaatcttaaacccaactt
ggttacctcttatctagtggGACAAGGGTATTGGGCCAAAAGTCTAGTCTTAAAGGTTCCgaatcttaaacccaac
ggttacctcttatctagtggCAGGTAAATACAGACAAGCTCTATCTAATGGTGTTAACTGgaatcttaaacccaactt
ggttacctcttatctagtggGGTCACATGTTCTTTCTTTAGACAACTGCTCATCATGTCCgaatcttaaacccaactt
ggttacctcttatctagtggCATAAGCCAACAGTAAGAAGTTAAGAGTGCCAAGAAAAGGgaatcttaaacccaac
ggttacctcttatctagtggCCCCACCCACTGGCATGTGCGTACAGTGCCTCCATGAGGGgaatcttaaacccaa
ggttacctcttatctagtggGCTTTGGCCAATAGAATGTGAGTGGGCATACGATGTGTGCgaatcttaaacccaact
ggttacctcttatctagtggGAAGTCATGAAACAATTGCCTGGTTCTGATTTATCTCCTGgaatcttaaacccaactt
ggttacctcttatctagtggTGACCTGGTATTGAGAAGGCACCTGAGGCAAAACCAGAGCgaatcttaaacccaa
ggttacctcttatctagtggGGAAGATGTTTGTTACTGCAGCATAACCTATCAAAGGCTGgaatcttaaacccaactt
ggttacctcttatctagtggCCTAGGTAAAATAGAAGATTAGGTTCCCGTGAGCTCTGGGgaatcttaaacccaact
ggttacctcttatctagtggACTCATGGCCAGCAGCATTATAGCTCATGCCTAAATGAGGgaatcttaaacccaac
ggttacctcttatctagtggCACAGTCTTCTTGACTCAAGAACACTACACAGCACTTCAGgaatcttaaacccaact
ggttacctcttatctagtggCAGAAAAATCACCACCAACAAAAATTAGCTGGGAGTGGTGgaatcttaaacccaac
ggttacctcttatctagtggGGGCAACAGAGTGAGTCTCCATCTCAAGAAAGAAAACAAAgaatcttaaacccaac
ggttacctcttatctagtggGCAGAGGCTCACCTTGTTTACCCTCAGCTAGGATCATGTGgaatcttaaacccaac
ggttacctcttatctagtggGCCACCCACTCTAAGCATGTCCAAGAATCACTGTAAAAGCgaatcttaaacccaac
ggttacctcttatctagtggTTGGGGATTTAAGTATGTTTTAGCAAGTAGGTGAGTTCACgaatcttaaacccaacttg
ggttacctcttatctagtggGTGCCACATCATTTAACCCTCATAACAATCCCAAAAGGTGgaatcttaaacccaac
ggttacctcttatctagtggGTAGAGAAAAGTAAGATTCTGACAGTGAAATAATGTGTCCgaatcttaaacccaactt
ggttacctcttatctagtggGCATGGCTCCAAAATCACAGTATGGCTCTTTTCTCTGCCCgaatcttaaacccaac
ggttacctcttatctagtggGGCACCAAGACTCAACTTATCTGCCTGGATTTCAGGGCCCgaatcttaaacccaac
ggttacctcttatctagtggTTCCATCTCCATTCAATCCATGCTCACCAACACCCTCCTCgaatcttaaacccaact
ggttacctcttatctagtggGCTCTTCTGTCACGCATCTGGTCCACATTCTCAAGGCTGGgaatcttaaacccaac
ggttacctcttatctagtggTTGGTCCAATCTTGATGCTTCCAGCTCAGACCCAGAGAGGgaatcttaaacccaac
ggttacctcttatctagtggCTGAGTAAGGACTTGAACACAGATTTGCTGACCTTGGGGCgaatcttaaacccaac
ggttacctcttatctagtggCCTTAGAGCAAACAAATGGGCAGAGGAGGGGCGGGAGTGGgaatcttaaacccaa

ggttacctcttatctagtggGGAAGGAGACAGCTGATGACTTTGTCAGTCAAACAGCCTCgaatcttaaacccaac
ggttacctcttatctagtggGCCAACAAACCATTGAAGAATAATAGCAGAGACCTTCTGGgaatcttaaacccaact
ggttacctcttatctagtggGCTCAAGGCTCCTTCCGGATGTTATCTCATTCTTCTTCCCgaatcttaaacccaactt
ggttacctcttatctagtggTGTACCCATTTTGCTGATAGAAACATTAAGCCTCAAACGCgaatcttaaacccaactt
ggttacctcttatctagtggATCAACAACCTTATTCTGCAGAACCAGGGTTAAAGCTGAGgaatcttaaacccaact
ggttacctcttatctagtggCTAGAAATGAGACTGAAACAAGGGGGAAAAGAGGACTGGCgaatcttaaacccaac
ggttacctcttatctagtggGGTCCTTGGTATCCATCCTGTGAAGTAGAAATTGTTATCCgaatcttaaacccaactt
ggttacctcttatctagtggGCCTGGTATTTGCAGAAGACTATGCTAGGCCCAATGAGCGgaatcttaaacccaac
ggttacctcttatctagtggGCACCAGGCATCAGAGATCGGAAGATAACTAAGACATGCCgaatcttaaacccaac
ggttacctcttatctagtggCCCACTCTCTTGTTGAGTACTTTCTACGTGCCAGGCACTGgaatcttaaacccaact
ggttacctcttatctagtggCAGGGAACAAAAACACAGTTCTTGCCTTCGTGTAGCTCACgaatcttaaacccaact
ggttacctcttatctagtggGGGATGCTGCATACCCAAAGATTCGGGGCAGGCACAGGAGgaatcttaaacccaa
ggttacctcttatctagtggCAGTTGAGACTTCAGGGTACCATTCACATTGACTCTGGAGgaatcttaaacccaact
ggttacctcttatctagtggGTGCTGGCAAATAGGCACAGTCCTATTCTGTCGACACCACgaatcttaaacccaac
ggttacctcttatctagtggGTGTCTACTATGTGCTATTATGCAGTGTTCAGTGCTGGGGgaatcttaaacccaactt
ggttacctcttatctagtggACAAAAACAACCAAAACCAAAAACTGGGATAATGGAGAAGgaatcttaaacccaact
ggttacctcttatctagtggGCCAAGAGGGGCCACCCAAACCAGATTAATAAGTTCCAGGgaatcttaaacccaac
ggttacctcttatctagtggAGCACACAATAAATGTTTTTGGCTGATAGAGTCACATCAGgaatcttaaacccaactt
ggttacctcttatctagtggCACCCAGCCTTGCAGCTCCACGTGGGGGGCACAAAGAGGGgaatcttaaacccaa
ggttacctcttatctagtggGGGCCTCCAGCCTCCTACCCCACCCCCACCTTGGATGTGCgaatcttaaacccaa
ggttacctcttatctagtggCATTTACAGGAGTGGTCTCCCTCTCATTCCCCAAGCAAAGgaatcttaaacccaact
ggttacctcttatctagtggCATGGTGATCTATGAATGTGTCTGCTCATGACTTCTCTGGgaatcttaaacccaactt
ggttacctcttatctagtggTCCTCACGCCTCTGGGACATGGACCTCCTGGCAGTGACTCgaatcttaaacccaac
ggttacctcttatctagtggTAAAAATGAATCATTGGCCAGACGTAGTGGCTCATGCCTGgaatcttaaacccaact
ggttacctcttatctagtggCCCTCCTTGTTATGAACTAGCCCAGAAGCCTCAGACTTACgaatcttaaacccaact
ggttacctcttatctagtggAAACTAACATTTACTGAGAACCCATTGTGTGTCTCTCATGgaatcttaaacccaactt
ggttacctcttatctagtggGCCCTGGGAATTAAATGAGTGTTACCATTTTCAAGATGGGgaatcttaaacccaactt
ggttacctcttatctagtggGCCAAGTCTCCAAATTCCAGTTCTACTACTTTTCCCTCTCgaatcttaaacccaactt
ggttacctcttatctagtggGGATAGATGGATGGATGGATAGAGAAAGAGACAAAGCAGGgaatcttaaacccaac
ggttacctcttatctagtggCTGGGGGCCCACCCTGAAACTACTGAATCACAGTCTCTAGgaatcttaaacccaac
ggttacctcttatctagtggGCCCAGCTGTTTTTGCCAACTGGAATTTATGAGCCACCGCgaatcttaaacccaact
ggttacctcttatctagtggAGGAGTGCCTACGCCTACTCCTGGTCTCCATGAGTCTCCTgaatcttaaacccaac
ggttacctcttatctagtggTGTTGAGTTGCACGCCAGACACTATACTAGATGATGGGACgaatcttaaacccaact
ggttacctcttatctagtggGGGACTTCAATTGAGGAAGTGCCATTGGGGAGGTATGTAAgaatcttaaacccaact
ggttacctcttatctagtggGGTGGGGGTAGGGATTTCTGCCTGTGTTCTTCAGAATGGGgaatcttaaacccaact
ggttacctcttatctagtggGTTGTGTTGACTTTTCCTCATGCTGCTTCCCCTGGCATGGgaatcttaaacccaactt
ggttacctcttatctagtggGAGAGACTTCACAAGAACTGCAATGAATCAAGATGTGCTGgaatcttaaacccaact
ggttacctcttatctagtggGTTTTCAACCAGTCCTGATTGGACCGATTGACTCATCCTCgaatcttaaacccaactt
ggttacctcttatctagtggCCTTATTCCCAGAGGGCTAGGGTGAGTCCTTGCAGAGGGGgaatcttaaacccaac
ggttacctcttatctagtggGATGGCACCTGGCAGAAATAGACCATCTGTCTTTCCTCCAgaatcttaaacccaact
ggttacctcttatctagtggCCTGCACTTAGGGTCTCATTGCCCCTCCCCACCCTCCGACgaatcttaaacccaa
ggttacctcttatctagtggCTCCTTGACCTCTGACTCACTGTGGGGAGAACTCTGCCTCgaatcttaaacccaac
ggttacctcttatctagtggCTCCCTCTCTAGCTCCAACTGTCCCTTTGCCTCCATGGGGgaatcttaaacccaac
ggttacctcttatctagtggGGGCTCACTCAGACCACTCCACTCCACCTGCCACCAGTGGgaatcttaaacccaa
ggttacctcttatctagtggGGCCTCAACCCCCAGGTGACTCTTGGACCACGTACCATCCgaatcttaaacccaa
ggttacctcttatctagtggAGCCCATGCTTGCTTCTAAGTGCTCCATGGTTTGGTTTGGgaatcttaaacccaactt
ggttacctcttatctagtggCTCCCTATCAACACTGAGAGATCTCCTGCTGACCCTGTGCgaatcttaaacccaac
ggttacctcttatctagtggTGTCTGCCTTGTCCCTAGCATCCAGATCAACACCTAGCAAgaatcttaaacccaact
ggttacctcttatctagtggTGAATGAATCCTTGGAGAGCGCAGGCCTTCACTGTGAGGCgaatcttaaacccaac
ggttacctcttatctagtggCTCCTTCCTCTGGGCTAACAGCATGTGAAATATCCCTCTGgaatcttaaacccaact
ggttacctcttatctagtggACGATGTGGAATCTCAGGCTCCCTTCTTGGGCACATAGTCgaatcttaaacccaact
ggttacctcttatctagtggCCATCCACATTCCCCACATCTCCTCACCTCTTTCTACAGCgaatcttaaacccaact

ggttacctcttatctagtggTACTGCCTGAAATGCCCTCTCCTGCATCCTTTCCTGTGTGgaatcttaaacccaactt
ggttacctcttatctagtggGCCTCCCAGGCTTTCAAGAACATTCCTCCTGTGGCTCCCAgaatcttaaacccaac
ggttacctcttatctagtggTCTCTTTTTATTGTACTTCTCTTTACTGTGCTTTGCAGAGgaatcttaaacccaacttgt
ggttacctcttatctagtggGATTTGTGGCAACCCTGCATCTAGGAAATCTATCAGTGCCgaatcttaaacccaact
ggttacctcttatctagtggCAGTGTAGGCTTATTTCATTTCACTGTGTCCCATTTTGTTgaatcttaaacccaacttgt
ggttacctcttatctagtggGACTGCTCCACTCCAACCAGCCATGTCCCCCATCTCTCCCgaatcttaaacccaa
ggttacctcttatctagtggGAACTCTTAAGTATTCACATGAAAGGAAGGGTCACAAGTCgaatcttaaacccaactt
ggttacctcttatctagtggATACTAGAAATGATTCAGTGAGCCGAGATCGCACCACTGCgaatcttaaacccaact
ggttacctcttatctagtggGCTTAGTAAGTATGGCATGTCAAAAGCTCAGCCTCTTGTGgaatcttaaacccaactt
ggttacctcttatctagtggGTGCTATTCCCGTGAACGTATTAATGATAAGAAGCAAAACgaatcttaaacccaacttg
ggttacctcttatctagtggCCCTTAAGTCAAAGGCTAATTCAGAGCAAGCCCCTAACTCgaatcttaaacccaact
ggttacctcttatctagtggGGAAGCTGCAGAAAATGAGTTTGAAGCTAGCAGAGGCTGGgaatcttaaacccaac
ggttacctcttatctagtggGCAAGTGCTGATGGAGAGGCTGCAGTAAGTTAACCAGAAGgaatcttaaacccaac
ggttacctcttatctagtggTCATTTTAGATCAAGCAGCCTTATACTGCAAGAAGATGCCgaatcttaaacccaacttg
ggttacctcttatctagtggGCTTCACCAGGCAGGGTAACTCTCTTGTTAGGGGCCAATGgaatcttaaacccaac
ggttacctcttatctagtggGTGTTCTGTAAATAGAACAACAAAGCCTAGGTGACAGCACgaatcttaaacccaact
ggttacctcttatctagtggAGCCCGTGCTTGAGACCTGCTGCTGAAAATCAAAGATTCCgaatcttaaacccaact
ggttacctcttatctagtggAGCTACCCAAGAGTCTTGATAATGCTTTTAGTTATCCAAGgaatcttaaacccaacttg
ggttacctcttatctagtggTTTGTGCCTGTGAACACAGCATTCCTGCCATAGCCCATGGgaatcttaaacccaact
ggttacctcttatctagtggAGGCTACAGTTGCCATAGATAGTGAGTCTTCTGATGAATCgaatcttaaacccaacttg
ggttacctcttatctagtggAGAGGTCATTAAGACTATTTGTGATGCATGGGAAGAAGTCgaatcttaaacccaacttg
ggttacctcttatctagtggGCTTTAGTGGAGGCAAGAACTGTGGATGTGGTAGAAACAGgaatcttaaacccaact
ggttacctcttatctagtggGGAGCCTGAAGATGTGACAGAATTGTTACAATCTCGTGATgaatcttaaacccaactt
ggttacctcttatctagtggCTTAGTTGATAAAGCAGCAGTAGGGCTTGAGAAGATTGACgaatcttaaacccaactt
ggttacctcttatctagtggGGAAATTGTCATAGCTACCTAACCTTCAGCAACCACCACCgaatcttaaacccaact
ggttacctcttatctagtggAGACAAGACCCTCCACCAGCAAAAACAGATTATGATTTGCgaatcttaaacccaact
ggttacctcttatctagtggGGCCAGGCTAGAACTCCTGACTTCTGGTGATCCTCCTGCCgaatcttaaacccaac
ggttacctcttatctagtggTACAGGGATGAGCCACCACGCCCAGCTAGCAATAAAATATgaatcttaaacccaac
ggttacctcttatctagtggGGATGTGTACATTTAAGAATGGGCATGGTGGCTCACACCTgaatcttaaacccaact
ggttacctcttatctagtggAGCTGAGTGTGGTGGCATGCACCTCTGTAATCTCAGCTACgaatcttaaacccaact
ggttacctcttatctagtggAGACATAATGCTATTTCACACACAGTAGACTACAGCAGTGgaatcttaaacccaactt
ggttacctcttatctagtggGCACTGGAAAATCAAAAACTTCACGTGACTTGCTTTATGGgaatcttaaacccaacttg
ggttacctcttatctagtggGGTCTGTAACCAAACCCACAATATCTCAAGTTATGCTTGTgaatcttaaacccaacttg
ggttacctcttatctagtggTCTGTCTTAAAGCTGGGCAACTGGTAAATGATAGAGCCAGgaatcttaaacccaact
ggttacctcttatctagtggGCCCATGAGGCATTATTTCCAGACTTCAACACTTTGTGGTgaatcttaaacccaacttg
ggttacctcttatctagtggGAAGGCAGACAAAGGAGGAAAACCAGAAGGGATTACAGGCgaatcttaaacccaa
ggttacctcttatctagtggTTCCTAGGGAATAGCTTTCCTCCTCCAGGGCTCCAGGTTGgaatcttaaacccaact
ggttacctcttatctagtggCTGAACTCAGGAACAAGCGTCAAGACTTGTTTTATTGGTAgaatcttaaacccaacttg
ggttacctcttatctagtggCATTCCTCACGTCTAAGAGAGGGGAGAAGGAAAGACACGGgaatcttaaacccaa
ggttacctcttatctagtggTGGACTGTGGTGTATCTCTCTTTCCCATCTCCCAGGCAGGgaatcttaaacccaact
ggttacctcttatctagtggTGGGCTGTGTTCAATCAGGGTTAGTGCGTAATAAAGATCCgaatcttaaacccaactt
ggttacctcttatctagtggCCTGCTTCAGTGCCTAGAGAGGAACAAGGGAGAGAAGGGGgaatcttaaacccaa
ggttacctcttatctagtggCCAGTCCTGACGGTGCCTATACATCCTTGGGTCATCACTCgaatcttaaacccaact
ggttacctcttatctagtggCTCTGAGAAATGGAAATAATAATGCCTACTTCACCAGATCgaatcttaaacccaacttg
ggttacctcttatctagtggGGTAATGTATTGGGTAACATTTTGTAATTGTGTGCTACTGgaatcttaaacccaacttgt
ggttacctcttatctagtggGGGGTCATGTCCCTTTCCAAGATGCCTCAGTTGACCAGTAgaatcttaaacccaact
ggttacctcttatctagtggATACTCCAACCTACCATTGTCAACCGCAACACTGGTTGTGgaatcttaaacccaact
ggttacctcttatctagtggGTCACTTCCACCAACACTTTTTGAGCATTCGCTGCATGCGgaatcttaaacccaact
ggttacctcttatctagtggGACCTCACCCTCCGAGATGTTTACAGTTTACTTTGCTTCCgaatcttaaacccaacttg
ggttacctcttatctagtggCTCGGAAGAGATGTCTCCAAGACACTTTCTACCATGACCTgaatcttaaacccaact
ggttacctcttatctagtggCCTCCCACTGAATGATTCCTCTGAGAAAAAAGCTCTCTGCgaatcttaaacccaact
ggttacctcttatctagtggCTTCTGTTTGTGTCCCAGGAATCCCAAATTGATGTCACACgaatcttaaacccaacttg

ggttacctcttatctagtggCATTTGCTTAGTTGATTTTCATTAAAATCCTGCAGGGAGCgaatcttaaacccaacttg
ggttacctcttatctagtggTTGCCCAGGTCACACGGAAAGAAATTAACGTGGCCACGGCgaatcttaaacccaa
ggttacctcttatctagtggTCGCTGACCTCCACCCTGTAAGAGCTTTTATTACCAAGCGgaatcttaaacccaact
ggttacctcttatctagtggCAGGCTCAGGGACACTGAATTTGACCAAAGAAGCCAATAGgaatcttaaacccaac
ggttacctcttatctagtggGACTCAGAACGGGTTAAGTGCTCCCTGGCTCATTCCCAACgaatcttaaacccaac
ggttacctcttatctagtggGCTTGCTCTGAAATAAATCAGTGTCCCTTTCTGCTGCTGCgaatcttaaacccaact
ggttacctcttatctagtggGCAAATGCAATGGGCCTTTACTGACATTGTGCTTCCCTGGgaatcttaaacccaac
ggttacctcttatctagtggCCCAGCAAGGAGGAGGACATGTGTCTTGTCACTGGCCCGGgaatcttaaacccaa
ggttacctcttatctagtggGGCCTCTCATTGCTCCTTGGCTCACTGCATTGCAAGATTCgaatcttaaacccaac
ggttacctcttatctagtggCAAGTCATAGGACTTGCCGGTAACCTAGATTGGTTTTCTCgaatcttaaacccaact
ggttacctcttatctagtggCGGATGCTCAGAAGCTGAAGATAACTAGTGCTCCTGGTCCgaatcttaaacccaac
ggttacctcttatctagtggCCGCTGAGGACAGAAGGCATTGGTCTCCCCTGCAGGAAGCgaatcttaaacccaa
ggttacctcttatctagtggGTCAGTAGTCAGAGCATGCATCAGCCCCCGGCGCCACCAGgaatcttaaacccaa
ggttacctcttatctagtggAAGCCAGCAGACCTCCGAGCAGATGAAATCATTAGGAGGCgaatcttaaacccaa
ggttacctcttatctagtggGAGAGAGGAGGCGGGGATTTCTCTGCATGCTCCCTTTGCCgaatcttaaacccaac
ggttacctcttatctagtggCCAGCTGTCTGTGACCTAGTTATCCAAGAAAGGAAACACGgaatcttaaacccaac
ggttacctcttatctagtggGTTTGCTACATGAGAACCCCATTCTCCAAAGACATGCTGGgaatcttaaacccaact
ggttacctcttatctagtggTTGGTAGCAATAAAGACAAGCCCTATTACAGTAGCCTAAGgaatcttaaacccaact
ggttacctcttatctagtggGGGATAGCCCATGGATAGGAAGTAAAAATCTTGGTTCATGgaatcttaaacccaact
ggttacctcttatctagtggCACCTCTTTCTCTTGCTCTCTCTCTCTCATTACATGGCCCgaatcttaaacccaact
ggttacctcttatctagtggCTCTATGAAGGTCTCTCTTCTTGCTTTTAAGCATGAAGCCgaatcttaaacccaacttg
ggttacctcttatctagtggGTATTTAATGCTAACAGGTGCTGGCACTGCCCAAGATGCCgaatcttaaacccaac
ggttacctcttatctagtggTCCCTGCCCTCGTGAAATGTACACTGTGGTGAGGCAGGTGgaatcttaaacccaac
ggttacctcttatctagtggGAGGGCAGTGAGTGCTTGGAAGAAAAGATCAAGGGAGGAGgaatcttaaacccaa
ggttacctcttatctagtggGCTGAGGAGAGGGAATAGCAGGTGCAAAGGCCTGGGGTGgaatcttaaacccaa
ggttacctcttatctagtggGCAGAGAGGCCATTGAGGAAGGAGGTTAAGCAAGTGAGGGgaatcttaaacccaa
ggttacctcttatctagtggGAGGTGAAGTCATAGACCTGGGGGCACAGATCACAAAGCGgaatcttaaacccaa
ggttacctcttatctagtggCTCTGAGTGAGATCAGAAGATCCCAAAAGGCCACCACAAGgaatcttaaacccaa
ggttacctcttatctagtggCTTGCACACAGTCAAGCTCCCTTAGTCTCCAAAAGAGGAGgaatcttaaacccaac
ggttacctcttatctagtggAGCAGTCGGCTTCGAGAGTCATCCTGAGTTTTCAAGGCTGgaatcttaaacccaac
ggttacctcttatctagtggAGGAAGAGACCTAGTATGAGATAAAGTGTCCAGGCCCTGCgaatcttaaacccaac
ggttacctcttatctagtggCTCACAGAATTAACTGGCAAATGCTAGTAAGAGTATCAGGgaatcttaaacccaact
ggttacctcttatctagtggGGACCCACAAGAGTTGTCAGAGGCAGTTGTGAGGGGTTGCgaatcttaaacccaa
ggttacctcttatctagtggGGGTGGCACTGTGGCTGGAATACAGAGCCCACAGGACTTCgaatcttaaacccaa
ggttacctcttatctagtggCCTCCTTTTTGTGACCCTTGGTGGAAAAAGTATGAGAGCCgaatcttaaacccaact
ggttacctcttatctagtggAACTGCCTTTCCTCCCTAGATGCCTGCCACAAAAACAGACgaatcttaaacccaac
ggttacctcttatctagtggTCTCCCTAGTTCCAAGAACACTGAACTCATGGTAATGGCCgaatcttaaacccaact
ggttacctcttatctagtggACCATCTGTAGTTCAGCCATAGCACTTCACATGAAAGAGGgaatcttaaacccaact
ggttacctcttatctagtggGTACAATTACTTGATAGACTGTTTTGGTTCCTATCTTTCGgaatcttaaacccaacttgt
ggttacctcttatctagtggTGCACACTAGGATTTGGCCAGGAGAATCCACACTAACTGGgaatcttaaacccaac
ggttacctcttatctagtggGGCCCCCCTGATGCCTCTTTGCACCTCTGTCTGCACTGCCgaatcttaaacccaa
ggttacctcttatctagtggCCACTTTCAGTCCTCTGCACCAGCTGCTCCATGTGTCCAGgaatcttaaacccaac
ggttacctcttatctagtggCCCATCATCCCCCAGAACTCCTGATATCCCCTACACTCCCgaatcttaaacccaa
ggttacctcttatctagtggGTCTATAGCAACCACCCCTCACCACTTTATAACATTTATGgaatcttaaacccaact
ggttacctcttatctagtggCTGTGTCCACTCACTAGAATTCAAATATCACAAAAGCAGGgaatcttaaacccaact
ggttacctcttatctagtggCTCCAAATCCTAGAAGGAAGCTGGCATTTAATATGTGCTCgaatcttaaacccaact
ggttacctcttatctagtggGAGACCTGGGCTCAAATCCAGCATGGGAATAAGTAGGGAGgaatcttaaacccaa
ggttacctcttatctagtggTGTCCTTGAGCTCAGGTGACGCCCATCTGGCCAACTCTCCgaatcttaaacccaac
ggttacctcttatctagtggGGACAGTTCTTGGCAGTAGTAGTAGTGCTGGTGGGGAGGGgaatcttaaacccaac
ggttacctcttatctagtggGCTCCCTGTCACAAGGCCATGAAAGATTAGGCTCGCAGACgaatcttaaacccaa
ggttacctcttatctagtggATGGGGGCAGAACAGGGACGCAGCAGAGCCAGAGTCTTCCgaatcttaaacccaa
ggttacctcttatctagtggGAATCCCAGCTACCACCCAGGAATAGGAAGGGCCAGGCTCgaatcttaaacccaa

ggttacctcttatctagtggCCTCCCAGTATGCAGGCTGGTTGGAGTTTCTCTGTGGACCgaatcttaaacccaact
ggttacctcttatctagtggTGGCTGTCTCACATGCAAAATGGGAGTATTTGAATCTCCTgaatcttaaacccaactt
ggttacctcttatctagtggGGTCTAGGGAGGAAGCAAAAAGGAAGTAGAAGGATTCTCCgaatcttaaacccaac
ggttacctcttatctagtggGCCAGGAGCGACTATAAAACACAGCTCATCATTGCAGACCgaatcttaaacccaac
ggttacctcttatctagtggGAAAACTACATCTCCTTGGAGTAAAGTGCCACAGGTATCCgaatcttaaacccaact
ggttacctcttatctagtggTGGGCAGTTTGTGACAGATCAAGTCACATTTCTGACAGAGgaatcttaaacccaactt
ggttacctcttatctagtggAGGACAAGGAGATAATGTGAAAATAACAGAGAGGCAAAGGgaatcttaaacccaac
ggttacctcttatctagtggAGAAGGGGACTCACTCCAGAACCGAAACAAGATACCCCACgaatcttaaacccaa
ggttacctcttatctagtggGTTTGTAGCTTGTCATACGTGTGTGTGAATGTTCAAGGGGgaatcttaaacccaactt
ggttacctcttatctagtggAACGGTAAATAGTGTGGGTTCAGTTCTGCTCTGCTAATCCgaatcttaaacccaactt
ggttacctcttatctagtggGACAGTAGCAATCAAAATTCTATCGCCCACAAGGGTGTGGgaatcttaaacccaact
ggttacctcttatctagtggCTCCATCCACCTCTGTAATACAACTTCTGGGAGTGCAGTCgaatcttaaacccaact
ggttacctcttatctagtggCAAAGATCCTGGGAAAACAATGTTATTTTGCTCATAGGTGgaatcttaaacccaactt
ggttacctcttatctagtggCTGGGAAGAGTGAGGCAGGAGGAGCACGTGAGCCCGGGAGgaatcttaaaccca
ggttacctcttatctagtggCTCCAGCCTGGACAACATAATGAGACTTTATTTCTAAATTgaatcttaaacccaacttg
ggttacctcttatctagtggAATAGTCAATACTCACTTCCTGCCCCAGGGAATCCGCTGGgaatcttaaacccaac
ggttacctcttatctagtggCACAAACTTTTGTGTATGCCTTTTTTCACCTAGCATGATGgaatcttaaacccaacttg
ggttacctcttatctagtggATCCATGTTGTTATTGGGTAATATTCTACTGTATGGATGCgaatcttaaacccaacttgt
ggttacctcttatctagtggGTACTGAGAGGTCTTGTATACCCTTCACCAAGGTTCCTTCgaatcttaaacccaactt
ggttacctcttatctagtggCAGAGGTACAGTGTGTGGGGGTGGGGGGACAGGCATGGGCgaatcttaaacccaa
ggttacctcttatctagtggTGCATGGGTGTGCGTGTGTAGTTCTATGTCATTTTATCACgaatcttaaacccaacttg
ggttacctcttatctagtggTCTGTAAGTGCTTCTGCAAGATACAGAAGTATTACCTCCCgaatcttaaacccaactt
ggttacctcttatctagtggAGTTACACACCCCCCATGCTTATTTCCGTTTGCTTTCTCCCgaatcttaaacccaactt
ggttacctcttatctagtggTTCCTGCTTTGATATACTTTTTTAACCCCATTAGACAGTGgaatcttaaacccaacttgt
ggttacctcttatctagtggACTCTTTCTGTTGTTCTTCCTTCCTTCCTGATGACCCAAGgaatcttaaacccaacttg
ggttacctcttatctagtggGAGGAATTCTTTCTGAGCACCTACCTCCCTTGCTAGCAACgaatcttaaacccaact
ggttacctcttatctagtggTCATCGGAGAATATCTTTATTACACCTTCATCCTAGAAGGgaatcttaaacccaactt
ggttacctcttatctagtggTGGCCTCCATGGTTTCTGATGAGAAGTGTGTTGTCATTTGgaatcttaaacccaactt
ggttacctcttatctagtggTGAGCCACCGCGCCCGGCCCTAATTCACTGATTTCTAGAAgaatcttaaacccaac
ggttacctcttatctagtggTGTTTTGCTATGGTGTGTCTTGGCTATCTCTTATATCTTGgaatcttaaacccaacttgt
ggttacctcttatctagtggCTTTGGATTTCTTTGGGCTTATCGTGTTCAGATTTCTTGGgaatcttaaacccaacttgt
ggttacctcttatctagtggTTGAGGTTCACTCAGTATCTTGAATCTGTAGGTTTATGTGgaatcttaaacccaacttgt
ggttacctcttatctagtggTAAAGACATTTCAGCTATGCCTTCTTTCTCTTCTTCTGGGgaatcttaaacccaacttg
ggttacctcttatctagtggTCAGTCTTTTGTATTAGTCCCAAAGCTCTCTGAGGCTCTGgaatcttaaacccaactt
ggttacctcttatctagtggGTCTTCTGTTTCACTGATATTTTTTACTGTGTCCTTCCACgaatcttaaacccaacttgt
ggttacctcttatctagtggGCCAGACAATTCTAGAATCTCTGTCATCAAGGTAATGGTGgaatcttaaacccaactt
ggttacctcttatctagtggGAGGGCTTCCTCTGATACCAAGCCAGAGGGAAAAGAATGGgaatcttaaacccaac
ggttacctcttatctagtggCAGGCGAAGGTGGATGTCCAGATACTCCACTCAGCCTCCGgaatcttaaacccaa
ggttacctcttatctagtggGGCCCCTGCCAGGCCTCTGCTGGTACCACTCTGGCTGGTAgaatcttaaacccaa
ggttacctcttatctagtggTGGTGGTAGGGAGGTGAGGGGTGTGGGGGAGTTGGACTCCgaatcttaaacccaa
ggttacctcttatctagtggGGGGGTTGGGCACCTCCTTACATCTTGGTGAGGGTGAAAGgaatcttaaacccaac
ggttacctcttatctagtggAGCCCTGGCTGATAAGATGGAATGGGCTACTGTTTTTACAgaatcttaaacccaactt
ggttacctcttatctagtggTCTTGCTAGGCTGCCCTTTCCTGGTCCTCTGGCTGTAAGCgaatcttaaacccaact
ggttacctcttatctagtggCACTTCCAGGTTGCCAACTCCTCAAGAGGCCAGTCCTGGGgaatcttaaacccaa
ggttacctcttatctagtggCCCACTGTCATGTCATACTCTGAGTCCTGAGGCCTCCAACgaatcttaaacccaac
ggttacctcttatctagtggCTTTTCTCCACCTTTCAATGTCTTCTCCTGCTTATATTATgaatcttaaacccaacttgt
ggttacctcttatctagtggACATTTCAAAAGATGTTTATAAGACGATGTTGGAGCACAGgaatcttaaacccaactt
ggttacctcttatctagtggTAAAATGTTCTGCATACTACATGCAAAATCGCATCTCTACgaatcttaaacccaactt
ggttacctcttatctagtggCTCGGGAGGCTAAGAGATGAGAATTGCTTGAACCTGGGAGgaatcttaaacccaac
ggttacctcttatctagtggCTGCACTCCAGCCTGGGCGACAGACACGTGGTCTAAAAAAgaatcttaaacccaa
ggttacctcttatctagtggGCACTGAGTTAGTGAAGCAGAAAGAGGGATAAAGCAATTCgaatcttaaacccaact
ggttacctcttatctagtggGAGACCTGCAAAGAGATCCATTTCAGGATGGCTATGCTGCgaatcttaaacccaact

ggttacctcttatctagtggGGACAGGCATTGGGGTTGCTTTGTTGAACAAATCTAGCAGgaatcttaaacccaact
ggttacctcttatctagtggGGGATATTTTCTGCTGAAGCATGTATTCTCTGGCACAAGCgaatcttaaacccaactt
ggttacctcttatctagtggGCATTCTCTTGGACAGAACCACAAAGCATGGTCTGCATTGgaatcttaaacccaact
ggttacctcttatctagtggAGTCACCAGATGCAGTTTAAGCTACAATAACATAGTGAGGgaatcttaaacccaact
ggttacctcttatctagtggCATGCTCACTCCATGGGAAATGGTGGCAATTATTAAGAGCgaatcttaaacccaact
ggttacctcttatctagtggTGCTTACTGATAACTGTGCAGTTAACCAATGGCAGTGTGCgaatcttaaacccaact
ggttacctcttatctagtggCTGTGTTTCCCTGAGTTTTGCATGCTACATGCGATGCATGgaatcttaaacccaactt
ggttacctcttatctagtggTCAGCGTGTGAGTGTTGTTTGTGGTCCAATCTCCGTCCCCgaatcttaaacccaact
ggttacctcttatctagtggTGCATTCACAGAAGGTAACAAAAGATTTTTGTGACACGAGgaatcttaaacccaactt
ggttacctcttatctagtggGTGTTGGTGTTTCAGAAAAGAGCTTTGGCTTTGAATTCAGgaatcttaaacccaacttg
ggttacctcttatctagtggATGGGGCTACTATTTAAACAGTAGCAAATTCAGGCCTATCgaatcttaaacccaactt
ggttacctcttatctagtggAGGACCTCTCACTGATAGTAAGAAAGAGGTGGTAAGAGAGgaatcttaaacccaac
ggttacctcttatctagtggGGGCAAGGGAATGGCCTAAGAGGAATCCATAATGAGTGATgaatcttaaacccaac
ggttacctcttatctagtggGGACACAAAGGCTCAAATCTAGAACCAAATCTATGGCAGGgaatcttaaacccaac
ggttacctcttatctagtggAACGCTGTGGGATTTTTGTGAAGTTACGTACCCCTATGACgaatcttaaacccaactt
ggttacctcttatctagtggGACTCATATGGCATGAATAACGCACTCTCCAGTAATAAGGgaatcttaaacccaact
ggttacctcttatctagtggTGCCTCCAAGCCAAAACTGTGCATCTCTCCAAAGCTCTGCgaatcttaaacccaac
ggttacctcttatctagtggCTCTGGTTCTTAGGTTAATGGCGTTCAGGAAAAACACATCgaatcttaaacccaactt
ggttacctcttatctagtggGGCTCAGATCTTCTCTTTGATGATAGGTTTGGTCATTGACgaatcttaaacccaacttg
ggttacctcttatctagtggCTGTAAGCCATAACTCCTGGTTTTTCACTGGAGGGGAGTGgaatcttaaacccaact
ggttacctcttatctagtggAGGTTTTCCTCGAACTTTTACAGTTGTCTTCCTCACACCTgaatcttaaacccaacttg
ggttacctcttatctagtggGCGGAGCATTTATCTTATACTTCATCGTCTCTCCTTTTGCgaatcttaaacccaacttg
ggttacctcttatctagtggGTCCTGGGTCCCTACTTTTCCTTTTCTGACAAACACAGACgaatcttaaacccaactt
ggttacctcttatctagtggGCGAATTTTTAAAGTTCTTTCCATTTCCACTACCCTCTGCgaatcttaaacccaacttg
ggttacctcttatctagtggTTGTCACCATCAAAGATGAAGTCACTCTCTCTCTCCCCTGgaatcttaaacccaactt
ggttacctcttatctagtggACCCCCCATCAGGTTTCCCTTATAGTTAACAGTTGCTGGCgaatcttaaacccaact
ggttacctcttatctagtggAGGCATGTACACTTTTATTGTGTTGACCACCCAGGAACCCgaatcttaaacccaact
ggttacctcttatctagtggGGGGTGATTCCAGGGGTAGACATGGGGCTCAGGCCTGGCCgaatcttaaacccaa
ggttacctcttatctagtggCGAGTGGCTGGGGGAGATTGTTGCCACCTTGTGGGAGCAGgaatcttaaacccaa
ggttacctcttatctagtggAGCCAAAAATGATACAGTCCCTCCAGTGAAAGTCCTGAGCgaatcttaaacccaac
ggttacctcttatctagtggGTGTCACTTGCCCTGTTCCTCATCACAAGTTCATGGTTGCgaatcttaaacccaactt
ggttacctcttatctagtggTTGGCCACTTAAAGTTCATACACGCATAATCTGCACTGGCgaatcttaaacccaactt
ggttacctcttatctagtggGAAGAAGCACATGCAAAGACCTGGAAGCAAGAGAGGGGAGgaatcttaaacccaa
ggttacctcttatctagtggGGCAGGGAGGGTAAGCAGGGGGCCATGTTGCGCAGGCTGCgaatcttaaaccca
ggttacctcttatctagtggGGAACCAGGAAAGGATTCAAGACAGAAGGAGGGTCAAGACgaatcttaaacccaa
ggttacctcttatctagtggCAGGAAGCTGCCTCAGGAAGCAATGATCTTGAGTTGGGGCgaatcttaaacccaac
ggttacctcttatctagtggCCCTGGTCTCCTTTCCCTTTGCAGACCAAACCATTAGGGCgaatcttaaacccaac
ggttacctcttatctagtggGCCCTGCACAAATAAATACTTGTTCAATTAATGAGTCTCAgaatcttaaacccaactt
ggttacctcttatctagtggTTTGTCTTGGTTCCTAGGTTTTACGCATTCATGGGGGAAGgaatcttaaacccaactt
ggttacctcttatctagtggTGGCAGCCTCAATTGCTTGAGCAAAAGCTGAAGTCAAACCgaatcttaaacccaac
ggttacctcttatctagtggTGGTTCCGTTGATATTAGATAGAACTATTGACTTTCCTCCgaatcttaaacccaacttg
ggttacctcttatctagtggGGCAAGGCTTCCTGCTGACTCCACAGAACCCAGGACAAGGgaatcttaaacccaa
ggttacctcttatctagtggGTGTGCACGCACACATGTTCTGGAGAAAAGAGAGAAAGTGgaatcttaaacccaac
ggttacctcttatctagtggCTGCTAATTTCTGCTCAGAGCAATCACTGCCCATGAAACCgaatcttaaacccaact
ggttacctcttatctagtggATGGAGAGAGAAACCCTTGCCTATTTTCCCCTGTGCCTGCgaatcttaaacccaac
ggttacctcttatctagtggAGCTCAAGGCTTTGACTTTTACCCCTTAGACTAAAAGGCCgaatcttaaacccaactt
ggttacctcttatctagtggGTTTCAGTGTCATGAGATCATGAAAGAGGAAGGGACCTCTgaatcttaaacccaact
ggttacctcttatctagtggAGGCTTGAGTTCTGTCGGGGGCACCTCCTGGCTGTGAGGCgaatcttaaacccaa
ggttacctcttatctagtggTGGGGGCAGCAAATACTGAGGGTGACATCATCTTCACTGCgaatcttaaacccaac
ggttacctcttatctagtggGGGTCACATGAGATAGTGGCAAGAAAATGCACCTGTGAACgaatcttaaacccaac
ggttacctcttatctagtggCAGAGGTGCAGGACTTACCCAGAGCAGCAGCCTGGTGACCgaatcttaaacccaa
ggttacctcttatctagtggCCAGCTCCCAATTCAGAGCACCTTCACCTCTGCACATGACgaatcttaaacccaac

ggttacctcttatctagtggACTCAATCCTGAGGACAGCCACTCTGACCAGTAGTACAGCgaatcttaaacccaac
ggttacctcttatctagtggGATGGGAAATCTTAGTGTGTTTGGACTTATCATATCTTAGgaatcttaaacccaacttg
ggttacctcttatctagtggCAACCAAAGCATAAGAGTATTCCTTTTTCTGAGCCATCTCgaatcttaaacccaactt
ggttacctcttatctagtggGGCCTTCCCTTTTTGAACACGGAAAGTTCCTCATGCCAGGgaatcttaaacccaac
ggttacctcttatctagtggGTCCCAAGAAAACTGAGGCAGTCATCCATCCCAAATGTCCgaatcttaaacccaac
ggttacctcttatctagtggGAATGTCTACAATATGCCAGGCACTGTTCTGAGCTCTTTAgaatcttaaacccaactt
ggttacctcttatctagtggGGAGCCAGGACCCATGGCATTGTGAATCCAGAACTGTGTCgaatcttaaacccaac
ggttacctcttatctagtggCTCCCCGAATGCCCTGAACATTCTCAGGAATTGTGGGTGGgaatcttaaacccaac
ggttacctcttatctagtggACCTGCCAGTTGTAATGGTTCTGCATTGATTGGAAAGTGGgaatcttaaacccaactt
ggttacctcttatctagtggACTTGGGAGTGTGATAGTGAAAAGACCACTAAGAAAATGCgaatcttaaacccaac
ggttacctcttatctagtggGTTCCTACCACATTGTTTCTTGACTTTACATCTTGAAGTCgaatcttaaacccaacttg
ggttacctcttatctagtggTTTGTAAGCTGAGAAACCCAATATATTAGCAAAGTGGGGCgaatcttaaacccaactt
ggttacctcttatctagtggTGAGGATTTATATGGTGTCACTGTACAAGCTTAATGGCTCgaatcttaaacccaactt
ggttacctcttatctagtggCCAGACACCGTATAGCAGACATCTGTGGTTGCCTGCACAGgaatcttaaacccaac
ggttacctcttatctagtggTGAGATCCTGATTTCCACGTGGGAACATCATCCTGTCCCCgaatcttaaacccaac
ggttacctcttatctagtggGAACTTCTGCGAAACACAAGTTCTCGTCATCTTCCATGGGgaatcttaaacccaac
ggttacctcttatctagtggCCCCACTGGATGTGAACAATAATAGTTAAAGCCACTAAAGgaatcttaaacccaac
ggttacctcttatctagtggGTGAGGGAGAGAAGACGAATCCTCACTTCATGAAAGCAGAgaatcttaaacccaac
ggttacctcttatctagtggCCAGCTCTGCACTTTCAAGACTGAGTGAGCTTGGGTGTGGgaatcttaaacccaac
ggttacctcttatctagtggACCACAAGGCCTGGCAAATAAGTGCTCAGTGAGTGAGGGCgaatcttaaacccaa
ggttacctcttatctagtggCCCTAGTGGGGAAGACTGGGCAGTGACTATATAAAAAATGgaatcttaaacccaac
ggttacctcttatctagtggGCTCATGTTACTGTGTGTGATACAATAAAGGGAAAGGAGGgaatcttaaacccaact
ggttacctcttatctagtggTCACTTTCCAAGATGAACCCTGGACCAAATCCTGATGCAGgaatcttaaacccaac
ggttacctcttatctagtggCCTCCAGTTCCAGCCATCCTTAACCTCCTCATTCCCAGTCgaatcttaaacccaac
ggttacctcttatctagtggTATACTAAAGCTGTGCTCACTCAGTCTAGGATTGTCCCATgaatcttaaacccaactt
ggttacctcttatctagtggCCCAAAGATGCAGCTGGCCACTCTTCAACAACAGCAGCAGgaatcttaaacccaa
ggttacctcttatctagtggCCTCTCCAAGCCTACAGGAAACTCCAAGACTACACCTGGCgaatcttaaacccaa
ggttacctcttatctagtggAGGGAATAAGAAGCGGCTAGGTTTAAAGGCAGGAGCATAGgaatcttaaacccaac
ggttacctcttatctagtggTCAGCCACCTCTGCTTCAACTATGTATTTCCCACTCTGGGgaatcttaaacccaact
ggttacctcttatctagtggCAGGTCAGCTACATCCAGCACAATCCTGGAGTGTGAGCACgaatcttaaacccaa
ggttacctcttatctagtggGGATGGAGTGAACCCTGCAGGAGTACAATAAGTACGCCAGgaatcttaaacccaa
ggttacctcttatctagtggTCTTCCCAGCAAAATCCTTATCATCTCTGAGGCCCAGCTCgaatcttaaacccaact
ggttacctcttatctagtggGCTCTTAACCCCACAATGCCCTGTCCACAGACTCTGAAAGgaatcttaaacccaac
ggttacctcttatctagtggGTTGTGTCCCATGTCTGTTTCCCCAGCAGGTTGTGAGTTCgaatcttaaacccaactt
ggttacctcttatctagtggTGGAGTATCTACACCGGAGCAGCCCACAGTTCAGGGATGGgaatcttaaacccaa
ggttacctcttatctagtggGCTTCCCAAAGTGATGGCACCTCAAGGGTCAGGACCTCACgaatcttaaacccaa
ggttacctcttatctagtggCCATGACCCAGCATCTACTACAGCATCTGTCACAACTGGGgaatcttaaacccaac
ggttacctcttatctagtggCTGGACTAGGGGATATCTCAGAACAGTACTTAGCACCTAGgaatcttaaacccaac
ggttacctcttatctagtggTGCTGGACAGCTGAACTGAAAATTACTCACAGCCCCATTCgaatcttaaacccaac
ggttacctcttatctagtggTAGCACTTAAGATGTAAATACAGAGTGTCAGCCATGTGGCgaatcttaaacccaact
ggttacctcttatctagtggCAAGTATCATGTATCCAAAGACCAGGGTTCAGTTTGGCCCgaatcttaaacccaact
ggttacctcttatctagtggCCATCTACTAAGTGCTGAGTGCTAGAATCCATTGACTACCgaatcttaaacccaact
ggttacctcttatctagtggCACAGGGACATCTGTGGTAAAACAGTAAATTCTCTAACCTgaatcttaaacccaactt
ggttacctcttatctagtggGGTTCTGCCTTTGCCTACCTTCTATAGTCAGAGACTACTGgaatcttaaacccaactt
ggttacctcttatctagtggTAACTGAAAGTCACTCTAGTTCTCTGCACATGTGAAGCAGgaatcttaaacccaactt
ggttacctcttatctagtggGCTCTATCTAAATGTCATTCCCTCTTCTCCATCCCATCACgaatcttaaacccaactt
ggttacctcttatctagtggGGCAGGACACTGTCTGACTCATTGGCTCCTGAGTCCCTTGgaatcttaaacccaac
ggttacctcttatctagtggTGAGCTTGGCATATAGCAAGGGCTCAATAAATGTTTATTGgaatcttaaacccaactt
ggttacctcttatctagtggCTCTGCTGAGGGCTCCCCAAGTCTCTCCTCTGGGTCCAGCgaatcttaaacccaa
ggttacctcttatctagtggGTCTCCACTGCTCAATGGAGAGTTCCTCTAAGGCACAGGCgaatcttaaacccaac
ggttacctcttatctagtggTCCTGACTCTAGCAGAGGGATGTTCACAAAGGATGTGCTCgaatcttaaacccaac
ggttacctcttatctagtggTGGCCCTTGATCGGTTTCTAGGGATCCACTCTATCCCCTGgaatcttaaacccaact

ggttacctcttatctagtggGTTGATCATCAGAGGCCAAAAGTCCAGTCACAGCTGTGGGgaatcttaaacccaac
ggttacctcttatctagtggAGAATCAAGTGCATTCTCCAGCAGGCAATTGACACTGGCCgaatcttaaacccaac
ggttacctcttatctagtggCTGCTTATCCCAGAGAGGTGATTCATGGGCTCTTCCTGCCgaatcttaaacccaact
ggttacctcttatctagtggGGCAGGAATGGTAGAGGCAAATCTGAGGTGGGACCACTTGgaatcttaaacccaac
ggttacctcttatctagtggACCTGCCATCATCACCAATCAGACAACTTGGGGCTCTCCCgaatcttaaacccaac
ggttacctcttatctagtggCTGCCCTCAGGCTGCTCTGTCCTGACGGTAATTTTGGTCAgaatcttaaacccaact
ggttacctcttatctagtggACCCTTACTTTCTCCCCCATAATGGTACCTGATGGCCCTCgaatcttaaacccaact
ggttacctcttatctagtggAGCACACAATCATAGCAGCTATTAACTTGCTGACACATCCgaatcttaaacccaact
ggttacctcttatctagtggATCCCTAAGTGGATTGCAGAAGTCTCTGGGTGACCGTAGGgaatcttaaacccaact
ggttacctcttatctagtggGGTCTGCAGCAACCTTCCCGGCACTGGTTCCTCAGCCCACgaatcttaaacccaa
ggttacctcttatctagtggCTGTTCTCCACATTACCCAGACATGCATGCATTAGGGGAGgaatcttaaacccaact
ggttacctcttatctagtggCGGGGATAGTTGGGCCATCTCTCTGTATTCCACTTCCCTCgaatcttaaacccaact
ggttacctcttatctagtggAGTGGAGACTGTTAAACTCTCCAGTGGTGGAATAATGCGGgaatcttaaacccaact
ggttacctcttatctagtggGAGGTGAGGACACAGGCTCAGAGATAAAGTGATATGGCCCgaatcttaaacccaac
ggttacctcttatctagtggGTGGTGCCTGGGTTTGAATCCAAGTGGCCTAAGTTCAAAGgaatcttaaacccaact
ggttacctcttatctagtggTGAGGAGCTCCTCCCCAATGTTTTGATGTTGCAGGTGGGGgaatcttaaacccaact
ggttacctcttatctagtggCGAGTTGCCCAAGTTCTTTGCAATTCTGATGGCTAGGCCCgaatcttaaacccaact
ggttacctcttatctagtggCTCATACTTCACCATCAAAACTGATTCCTTTTCTGTGGCCgaatcttaaacccaacttg
ggttacctcttatctagtggAGGTTACATAAACCCACTGAGGCATATTCCTACAATGCACgaatcttaaacccaactt
ggttacctcttatctagtggCCTTAGTGGCATTCTGATGGCTGATAAAAGTAAGACTTACgaatcttaaacccaacttg
ggttacctcttatctagtggGGATGGTGTTTGGATTGGTCGCCACTGTATCTGTAATGCCgaatcttaaacccaactt
ggttacctcttatctagtggCACTGTTTGTATACATTTGGAGAATAAAGGAATGCATCACgaatcttaaacccaacttg
ggttacctcttatctagtggGGTAGGTGGGAAAATGTAGGATACAAATTTGCACATATGGgaatcttaaacccaactt
ggttacctcttatctagtggGCCCAGAAGGAAATACACAAAACACAGAGTGGTTTGCCTCgaatcttaaacccaac
ggttacctcttatctagtggGTAATGAGGAAATCATTTTTCTGCCCCATTCAGTTTCTTCgaatcttaaacccaacttg
ggttacctcttatctagtggGAGCAGCTTGGGTGTGGGGATCTCGGCTGTGAGCTACCCCgaatcttaaacccaac
ggttacctcttatctagtggACTTGCCTTCCCCAACCCTACAGCTGCACAGCAGACATGCgaatcttaaacccaac
ggttacctcttatctagtggGGCAAGACTTGCAGGTGCACAGCAGCAGAAACCCACAGCCgaatcttaaacccaa
ggttacctcttatctagtggGCTCCTTGTGCGGTCACTTTGTCTGTGTCTCCATAGCCAGgaatcttaaacccaactt
ggttacctcttatctagtggGTGTGGATGAGTGGAGAATACAGATATGCTCAGGCCGGGCgaatcttaaacccaac
ggttacctcttatctagtggGAGTGATGGAAGGTTGCAGGCCCCATCAGTCCATCAGGGGgaatcttaaacccaac
ggttacctcttatctagtggGTGAAGATCTGGACCCTTCTGCTCCACGTGGCCCGCTCTGgaatcttaaacccaac
ggttacctcttatctagtggCTGTCTCTGGTTGTCTATTAACGTGAGTTAGTGCCAGTACgaatcttaaacccaacttg
ggttacctcttatctagtggGCGAGCCCAGCAAGAGCAACAATGTATGGAACGAGGAGCGgaatcttaaacccaa
ggttacctcttatctagtggGGACACTTACTTTCTGCCCCTTGCCCTAATGTACAACTACgaatcttaaacccaactt
ggttacctcttatctagtggCAAGGGCCTAGTCTCCCTCACCTCTCTCCTGCACTGCTGGgaatcttaaacccaac
ggttacctcttatctagtggGCCCCCTCTCTTCCATTCTCCACTGGTAGCCAGAGTGAGCgaatcttaaacccaac
ggttacctcttatctagtggGCAAAGGGCTGTTGGACTTGAGCTTCCTTCCTGACACCCCgaatcttaaacccaac
ggttacctcttatctagtggCCTACCTAGCTTGTTCCTACCTTGGGACCCTGGCACTGACgaatcttaaacccaac
ggttacctcttatctagtggACCAAAAAGGCTCCCCACCTGAGTGTAAATTAGTTCAACCgaatcttaaacccaact
ggttacctcttatctagtggACTTGGAACCAGCCCAAGTGCCCATCCATGATAGACTGGAgaatcttaaacccaac
ggttacctcttatctagtggCACCAGGGCCTGTTGGGGGTGGGGGGTTAGGGAAGGGATAgaatcttaaacccaa
ggttacctcttatctagtggCAAACCTACACACTCTGCACATGTATACCAGAATTTAAAGgaatcttaaacccaactt
ggttacctcttatctagtggAGCTTTCAGCTCTTGACCACAAATAGGCTCTCCAGGGTCCgaatcttaaacccaact
ggttacctcttatctagtggGATTTATTCCTAAGAAGGAGGAAGGGAGTGGGGAAAGGGCgaatcttaaacccaac
ggttacctcttatctagtggGTAACTTGGTGACAGGGTCAATCATACCCAAAACCTCAGCgaatcttaaacccaact
ggttacctcttatctagtggATACCAATATAACAAACTTGCACATATACGCCCAAACTCAgaatcttaaacccaactt
ggttacctcttatctagtggGACAATTTAAAGCTGTATCCCCAGTACCTAGGACGTTGGCgaatcttaaacccaact
ggttacctcttatctagtggGTACATTACTAAGTGCAGTGCCTCCTTGACCCGTCTACCCgaatcttaaacccaact
ggttacctcttatctagtggAGAAAACCTTCAAGAAGTGATTCCGAATCCTAGTTTCACCgaatcttaaacccaactt
ggttacctcttatctagtggGATTCTAGACAAGTCTTAAATTCTCTGCACCTCAGTTTTCgaatcttaaacccaacttg
ggttacctcttatctagtggGGGCACTTTGAACTGTATCTGGCACATGCATTACCATGAGgaatcttaaacccaactt

ggttacctcttatctagtggCCCCCCACAGCTGACTGGTAATGAGGCTGGGCCAGGGGAGgaatcttaaacccaa
ggttacctcttatctagtggAGCAGGAAGCTGAGGATGCAGACCTGGCCACCCGCGATAGgaatcttaaacccaa
ggttacctcttatctagtggCTCACATCCACTCATAAGCCAACCCAGGAGATGCTCAGGGgaatcttaaacccaa
ggttacctcttatctagtggCCTGCCTTGAGGACGTGGGTGCAAAGAGGGGAACCAGCAGgaatcttaaacccaa
ggttacctcttatctagtggCCCTGGGGGTACGAACAGAGGACAATGTGGTCACAGAATGgaatcttaaacccaa
ggttacctcttatctagtggGGAGATTTGGATAGGCATCACAGAAAAAGTGACAGCCCGCgaatcttaaacccaac
ggttacctcttatctagtggGCACTCCTTACAGCATACCCGAAAGCATTGGTGAGGACACgaatcttaaacccaac
ggttacctcttatctagtggCCATTCCTGTCCTCTCCCCTGCAACTTCCCAATCCCTCACgaatcttaaacccaac
ggttacctcttatctagtggCCCGCCCACCCCTTCACTAGCTTCTGATTTCAGGCAACGTgaatcttaaacccaac
ggttacctcttatctagtggTCTCCTACACACCCCTGCCCCTGGACCCAGGAATTCTTCCgaatcttaaacccaa
ggttacctcttatctagtggGGCTATATTAAGTAATAAGCCCACATGCTTTCTGTTGAGAgaatcttaaacccaactt
ggttacctcttatctagtggCCCTCTAGTTATCTTCCCACAGGCCCATCAAGAATCAGGCgaatcttaaacccaac
ggttacctcttatctagtggTAGGAAGACCACCATGGGCCCTTAGGGAGTCAGCGAAGGCgaatcttaaacccaa
ggttacctcttatctagtggCCCTCTCTAGGATCCCCAAGAGAGCTTGACAGCGTCCCTCgaatcttaaacccaa
ggttacctcttatctagtggCTCTGCGGTTTCCTGCACATAATTGATCACAGTTGAGCTGgaatcttaaacccaact
ggttacctcttatctagtggGATCTTGGCATACATGCGTGGGAAATGGCCTCTCAAGGGGgaatcttaaacccaac
ggttacctcttatctagtggGAGAAAGTTCCTGCCTTGAGTAAATTGCTGTCTGTTAGGGgaatcttaaacccaactt
ggttacctcttatctagtggGGAGGTGGGTGGCTTGTGAAAAACCAGAGAGTCCAGGGCCgaatcttaaacccaa
ggttacctcttatctagtggCTCACACAGGTCTCCTTAGAGCAGAGTTTCTCATCTCCAGgaatcttaaacccaact
ggttacctcttatctagtggGATCTCTGGACTCTGCACACTAGATACCAGTAGCACCCCCgaatcttaaacccaa
ggttacctcttatctagtggTCCTGGGGAGCAAAATCATCTCCTATTCTCACCTCCTGAGgaatcttaaacccaact
ggttacctcttatctagtggCCAACACAGTCAGCTGCTATCTCCACAACAGGCGGGTGAGgaatcttaaacccaa
ggttacctcttatctagtggGAACAAATGCAAGCAAAGCCAAGTTGCCATTGAATGTGGCgaatcttaaacccaac
ggttacctcttatctagtggTCCTCTCACCTTAGACTCCCAAAGTGCACTCCGCCCTGCCgaatcttaaacccaa
ggttacctcttatctagtggGGCTGCGTTTTAACCAGCCTGACATCACGCCTCAAGCAGGgaatcttaaacccaa
ggttacctcttatctagtggGGACAGGATGATGGAGCTGAGGCTCAAGAGACAGTCAGCCgaatcttaaacccaa
ggttacctcttatctagtggTACCCTGTCCATGGTGTAGAAGATGGGCTGGCGAGAGGCCgaatcttaaacccaa
ggttacctcttatctagtggATGACAAGGGTCTGCACTTTAAGCAGTGGGAGTAGGGATCgaatcttaaacccaac
ggttacctcttatctagtggCTGGGGGCAATGAATCACGCCTGTAATCCCAGCCCTTTGAgaatcttaaacccaac
ggttacctcttatctagtggAAAGACTGAGGCGGGAGAATCTCTGGAACCTGGGAGGCGGgaatcttaaacccaa
ggttacctcttatctagtggGAGCCAAGACCAAGATCATGCCACCATGCTCCAGCTTAGGgaatcttaaacccaa
ggttacctcttatctagtggGACAAAATTGACAGGATTCATGATTGATTGAACTAGGGAGgaatcttaaacccaact
ggttacctcttatctagtggTATCAACTGGTTTCACAAACCAACTAACTATCACAGCAGCgaatcttaaacccaact
ggttacctcttatctagtggCTCACCCATGTTGGTCCAATTGCGTGTTTGGCCAGCAGTCgaatcttaaacccaac
ggttacctcttatctagtggTCCCTGCATCTTGTGGCATCGGCCTACTCTAGGTCAGGGGgaatcttaaacccaac
ggttacctcttatctagtggTCCAGCACTGCTTGTGGCCAAATCCAGCCACTGCCTGTCgaatcttaaacccaa
ggttacctcttatctagtggGGGACACAGTCATGCTCATTTGTTTACGTATTATCTATGGgaatcttaaacccaacttg
ggttacctcttatctagtggGGCCGTTTGCTGGAACAGTTTGCTGACCCTTGCCCTCAGCgaatcttaaacccaac
ggttacctcttatctagtggCACCAGCTGGACCTGACGCATGCCCTTCCTGTGGCCACACgaatcttaaacccaa
ggttacctcttatctagtggGACTGAGACTCACGGCCCCACTCACTGCAGGAAAGCTGGGgaatcttaaacccaa
ggttacctcttatctagtggCCCACAGTCTACGCCACACATCCATCCATGGCTCTGGAGGgaatcttaaacccaa
ggttacctcttatctagtggCTTGCAAAGTGCTATAAACAACCATATCTCACTGAGATGCgaatcttaaacccaact
ggttacctcttatctagtggCTCAGAGCTATATACTCTCTCTTAGTTCAAGTAAGCGTACgaatcttaaacccaactt
ggttacctcttatctagtggTACAGAGTGTCTACTTTGTGCTCAGCACTGTGCCAAGCAGgaatcttaaacccaact
ggttacctcttatctagtggAAGGACTGGGATAGTTGAGAGTCACTTCCTGGAGTGGGGGgaatcttaaacccaac
ggttacctcttatctagtggCACCTGAGACCTGGAACACACAGCCTGCGAGGAGAGCAGCgaatcttaaacccaa
ggttacctcttatctagtggCTCTCCTTACCCCCTCTTTCTGATTTTTACTCCAGGGCTCgaatcttaaacccaactt
ggttacctcttatctagtggCTCCAGATCTGTTCCAAAGCAAACAGCAGCCAGGAAATCCgaatcttaaacccaac
ggttacctcttatctagtggGGACACAGGTTGAATGCTGTCCCCGAAGAGTGCTCCTTCCgaatcttaaacccaac
ggttacctcttatctagtggCACAGCTATCCTGATAGGGGAAGGAAGTAGCACTGTGAGGgaatcttaaacccaac
ggttacctcttatctagtggCTGGCTTCTCTCTGGGTCATGCTCCCAGGTCCCCGTGGCCgaatcttaaacccaa
ggttacctcttatctagtggGTCATCTACACTCACAGGCCGCTGGGACGGTAAGCTCCTGgaatcttaaacccaa

ggttacctcttatctagtggGCAGGAGGTCTCCATGTTGTGGTAGAGTAGACACCTGCTGgaatcttaaacccaac
ggttacctcttatctagtggTGAGCCAAGCATGTCCATAGGGTGAACTGTAGCAGTCACCgaatcttaaacccaac
ggttacctcttatctagtggCCATGGAAATGGAGACAAAATGAAGTCATTTGCCTTGGCgaatcttaaacccaact
ggttacctcttatctagtggGTTCACATCCTTAGTGTGACATATCTGTGCCCTGTTCCCCgaatcttaaacccaactt
ggttacctcttatctagtggGTCAACTCCTGCTTTTCAGTGTCCCACCAATATGAGACGCgaatcttaaacccaact
ggttacctcttatctagtggTCCTGCTTCCTGGTGTGCCACTCCCACTTTCTCCACCTGGgaatcttaaacccaact
ggttacctcttatctagtggCCATGTGCCATACTTAGCTATTAATGTCAGTTCATATGTGgaatcttaaacccaacttg
ggttacctcttatctagtggGTGTCAGGAAGCCACTCTTTTTGCTTTGCAATTGTAACTCgaatcttaaacccaacttg
ggttacctcttatctagtggCTCTGTTTCTCCAACCAGTCTGGATAGTTGAATGGACAGGgaatcttaaacccaactt
ggttacctcttatctagtggCTGTTTTCTCCAGTCACCTATCAGAGTGTCTGACATGTAGgaatcttaaacccaacttg
ggttacctcttatctagtggGAAGCTTGAAGCATTCCGGCCCCACCACTCTGTGCCCTTGgaatcttaaacccaac
ggttacctcttatctagtggCACTTCAGCAAGTATCTATTTCTTCATTTAGAAAGTGGGAgaatcttaaacccaacttg
ggttacctcttatctagtggCAAAGTCAAGATTAGGCAATAAAAAGCTAGGAGGAAATAGgaatcttaaacccaact
ggttacctcttatctagtggATCAACTGGTAGGCTAAGAGGAGGGTCTAACCTTTTGGCCgaatcttaaacccaact
ggttacctcttatctagtggCATCCAAATGGTCATTTGGTCTCAGAATAATGAAGAACAGgaatcttaaacccaacttg
ggttacctcttatctagtggATCATGACTACTCTAACTCTCCTTATCCTTCCAGATGCTGgaatcttaaacccaacttg
ggttacctcttatctagtggGAGAGGCCCACACAGCTCACTCTCAGAGTCTGGCAGCTCCgaatcttaaacccaa
ggttacctcttatctagtggCCAGCATCATTGCCTTAGAAGGGGAGAGTGTTCCATCTGCgaatcttaaacccaact
ggttacctcttatctagtggGTAGCAATGGTGGTGACAGAAATAGGTAGCAATGGTTGTGgaatcttaaacccaact
ggttacctcttatctagtggCAGTGACCATGTACTGGCCAATCATTTAACTACATCTCAGgaatcttaaacccaactt
ggttacctcttatctagtggCACACCTTCTAGGCACTGGGGATATAGCACTGAACACACCgaatcttaaacccaac
ggttacctcttatctagtggTTCAGCCTCATGTCAATTATTGCTTGCTGGTGAGAGAAGAgaatcttaaacccaacttg
ggttacctcttatctagtggCTTTAAGTTCTAGGGAACATGTGCAAAATGTGCAGGTTTGgaatcttaaacccaacttg
ggttacctcttatctagtggGACAGAAAGTGACTGAAAGGTGAACTCTGTGTTAGGTAGGgaatcttaaacccaact
ggttacctcttatctagtggCTCCAAGATCACATAGCTACCCACGGACAGAGCAAGTCTGgaatcttaaacccaac
ggttacctcttatctagtggAGTCAGTGTTGGCTACATTGGCTAGAAATCTTCACGAAGGgaatcttaaacccaactt
ggttacctcttatctagtggTAAGCTTGTCTTGGAGCGTGGTTAGGATTTAGACAGGTAGgaatcttaaacccaacttg
ggttacctcttatctagtggAAGAGTTGTGAGCAGGAAGAAATGGGTATTTATCGAGGACgaatcttaaacccaact
ggttacctcttatctagtggAGATGCTGGGTGTTGGGCATGTAGGTAGCAACTGGCACGCgaatcttaaacccaac
ggttacctcttatctagtggAATGGCTCAGGTACCTGAGGAAGAAATTTACATTGGGGACgaatcttaaacccaact
ggttacctcttatctagtggTGAAGGAGAGGTGGTGGAAGCTGGAAGTGGCATAGGAGACgaatcttaaacccaac
ggttacctcttatctagtggGGAGTCATCACTCTCAAATATGGCTCCCCAGAGATGCTCCgaatcttaaacccaac
ggttacctcttatctagtggCAACCCCTCTATCCTCTTAGAATCTGGCATGGGATCAGGCgaatcttaaacccaact
ggttacctcttatctagtggTCAACTGGCAAGGCCATGGCAATCAGGCAGACAGGTTAGCgaatcttaaacccaac
ggttacctcttatctagtggGATTCACCTGAGGTGGCCCGAGGCAGCAGAGCCGGAGGGGgaatcttaaaccca
ggttacctcttatctagtggAGGGGAACTCAGAATCATCACTGTGAAGGTGAGAAGAGCCgaatcttaaacccaac
ggttacctcttatctagtggGGATGAAGGAGGATAAACTGATAGAAGAAAAATGAGCCAGgaatcttaaacccaact
ggttacctcttatctagtggCAAAGTGGAATACAGTGTGAAGATTGATTTGGGGCAAAAGgaatcttaaacccaactt
ggttacctcttatctagtggCCCTTTGTATGATGCCTAGACCCAGCAACAGGGCAGTTGCgaatcttaaacccaac
ggttacctcttatctagtggAGGTGAAAGAAAGGTAGGAAGGAAACGAGATAGATGTGAGgaatcttaaacccaac
ggttacctcttatctagtggAGGGAACTGAGGAAAGGGAAAAAGGAAATAATCCATGATAgaatcttaaacccaact
ggttacctcttatctagtggCAGGGATTGGTAAACTATGACCCTGAGACAAATTCAGTATgaatcttaaacccaactt
ggttacctcttatctagtggATCTATGCCTGCTTTCAAACCATAGTTGCAGGGTTGTTGCgaatcttaaacccaacttg
ggttacctcttatctagtggACTATCTAGCCCTTAACAGAAAAGGTTTGCCACCCTAGTCgaatcttaaacccaact
ggttacctcttatctagtggCAGGGCCAGAATTTTGAATGCCTTGCTAAGGTGTTAGAACgaatcttaaacccaactt
ggttacctcttatctagtggAACCAATCAAAGAACTCGACAGTGGCAATCAATGTGGCACgaatcttaaacccaac
ggttacctcttatctagtggGCCTAAGACTTAAGAACTGTATGCTGGTCACCAGTCTGCCgaatcttaaacccaact
ggttacctcttatctagtggGCCACTGTGGTCTTTAAATTTCTAATGTCCAAAATGGGGGgaatcttaaacccaacttg
ggttacctcttatctagtggTAGTGCCTGTTGTGCGCAGAGGAATCCAGAGATGAGGGAGgaatcttaaacccaac
ggttacctcttatctagtggAGACTAGAAGGAAAGATGGGATGCAAACGGCAGTGCCAAGgaatcttaaacccaac
ggttacctcttatctagtggGGGAGGAAACCACTCAGCTTGAGAAGAAGTAGGCTGTCAGgaatcttaaacccaac
ggttacctcttatctagtggAGGAAGGGTTTCCAGCAAGGAAGAAAAGATCATGATGCTGgaatcttaaacccaact

ggttacctcttatctagtggAGTTCAAAGTGAGTGGCACCTAGAAGCCAACTGTGGATGCgaatcttaaacccaac
ggttacctcttatctagtggTGGAGGGAGGATATGAAAGGTTTTCTATGCTATGCCAAGGgaatcttaaacccaact
ggttacctcttatctagtggGAAGGATTTGGCTTCCAGGAAAGACATGGTCAGAACTGGGgaatcttaaacccaac
ggttacctcttatctagtggCCCCTGCATGAATGAGGGGGGCAGAGCTTAAGACTGGTCAgaatcttaaacccaa
ggttacctcttatctagtggAGTCATCGTCCTAGCAGCCATTGGGGAAAAGCAGCTGGACgaatcttaaacccaa
ggttacctcttatctagtggTCCCCCTAAATTGACCCCCGGCCCTGGGACATCTGCCTACgaatcttaaacccaa
ggttacctcttatctagtggTCCCAGGCCCGTATCTTCTCCAAACGCAATCTGGACAGCCgaatcttaaacccaa
ggttacctcttatctagtggAGCAAGAGAGGAAACAATCAACATATTCAGGAGAGGATGGgaatcttaaacccaac
ggttacctcttatctagtggGGGTTTACACAGTGGATCAGGGTTTGTGTTGTTCTCTCACgaatcttaaacccaactt
ggttacctcttatctagtggCCCACGTGGGACAGTGCCAAAGCTCTCTCTCCTGATGGGAgaatcttaaacccaa
ggttacctcttatctagtggGAAAACAGGACCCGGTTGCAGTTTGACATGCACAGAAGCCgaatcttaaacccaa
ggttacctcttatctagtggCTGGGTCCCAAGCCATACCTCCTGTGTGACCTTAGGGCACgaatcttaaacccaa
ggttacctcttatctagtggAATTATGCATAGCTTAATGCTCAAGATGAGTTCCCAGGCCgaatcttaaacccaact
ggttacctcttatctagtggGGCAGATTTTAAGTCTATAAACCTGCATAAGACAAAGCAGgaatcttaaacccaact
ggttacctcttatctagtggACAATTGCCAAAGGATTGAAAGAACAACGCAGAGAAGAGGgaatcttaaacccaac
ggttacctcttatctagtggTACCAATAAGCCTTTGAGCAAACCAATGTTCAAACAATACgaatcttaaacccaactt
ggttacctcttatctagtggAACACAAATGCAGACTAAAGCATTATTCACTATCGAGTTGgaatcttaaacccaactt
ggttacctcttatctagtggAGCCTTTCTGGAAAGCCACTTTTCAGCACTACTACAAGCCgaatcttaaacccaact
ggttacctcttatctagtggTGCGTAGTTTGACTCAATAATTCTACCCCTAAAAGCCTCCgaatcttaaacccaactt
ggttacctcttatctagtggAGGAGAGCCATAGACAAGGATTTATCTATAGAAATGTTTGgaatcttaaacccaactt
ggttacctcttatctagtggAGGAGAAGGAGGAGGAAAGAGGAGAGGAAGAAGGAAAACGgaatcttaaacccaa
ggttacctcttatctagtggGCAAAAAATGGGAACCCAGGACATTCAGATGCTTCTTAGCgaatcttaaacccaact
ggttacctcttatctagtggACACCTCACTTCCTGAAGACCACAGCTTAGCCTAGCCTACgaatcttaaacccaac
ggttacctcttatctagtggCAGAGCGCTTATATTTGTCTACAGTTGAGGAAAATCTACCgaatcttaaacccaactt
ggttacctcttatctagtggCACCATCATGAAGTCCAAACATCTAAGTCAAACCATTGTGgaatcttaaacccaact
ggttacctcttatctagtggGCACAAAGAGGGGTAACTTGTGGGAACTAGAGGTTGACACgaatcttaaacccaac
ggttacctcttatctagtggGGGCAGCAATAGCAAGGGAGTTTGGATGAAAAATAATCAGgaatcttaaacccaact
ggttacctcttatctagtggTGATAGTCACACAGCATATAATTGCCAAATAAGTAGAGGGgaatcttaaacccaact
ggttacctcttatctagtggCCAAATATATCAGTGCCTAGTCTGCTCCAGCTGCCATAGCgaatcttaaacccaact
ggttacctcttatctagtggGGGTTCCAGTATGACCGAGCTTTGGTGAAGGGACTCTTCCgaatcttaaacccaac
ggttacctcttatctagtggCTGTGTGCTCACATGGCCTTCCCTCTGTGTGTGCATGGGGgaatcttaaacccaac
ggttacctcttatctagtggCTGAACAACAAAATAGGATGGCAAACAAATCAGTGCCTGGgaatcttaaacccaac
ggttacctcttatctagtggCACCCTTTTGCCAGTTCTACCATGGGTGTGATAACTGCCCgaatcttaaacccaact
ggttacctcttatctagtggTGTCCCTGCCAAGTCTCATGTGGAAATGTGATCTCCAGTGgaatcttaaacccaact
ggttacctcttatctagtggAGCCTGGTGGGAGGTGTTTGGGTCATTCAGATGAATCCCTgaatcttaaacccaact
ggttacctcttatctagtggGATGCCTTCATATTCTCACTCTGTGTTGCACGCAAGATCCgaatcttaaacccaact
ggttacctcttatctagtggTACCCTGCCTCCTCCTTTGCCTTCTGCCATGTGTCGAAGCgaatcttaaacccaact
ggttacctcttatctagtggCCTCACCAGAAGCATAGCTGGCCCTGTTTCATGTACAGCCgaatcttaaacccaac
ggttacctcttatctagtggGCCAAAATAAAACCTCTTTTCTTTATACATTACCCCGTCTgaatcttaaacccaacttg
ggttacctcttatctagtggGACTAACACAGTATAAGGATAGATATAATTGGAGCAAAGGgaatcttaaacccaactt
ggttacctcttatctagtggGGTCTAAAAAGGGTGAGAGACACATGGAACAGACCTGAGCgaatcttaaacccaa
ggttacctcttatctagtggACAAACAGCAGAATCCCACCTAGATCAGCCAAACCCAGCCgaatcttaaacccaa
ggttacctcttatctagtggCAGAGCTATATTTATAAACCTGTATCAGCCAGCCCCAGAGgaatcttaaacccaact
ggttacctcttatctagtggTATGACTGACAGTAAGTCACCAAGTGTATTGGGTAACAGCgaatcttaaacccaactt
ggttacctcttatctagtggAGATCCAAAAAGCTCAGCAAACCACAAGCTGGACAAACCCgaatcttaaacccaa
ggttacctcttatctagtggGGCAAGGTTCATTATAGACAAACTTCTGAAAATTAAGGACgaatcttaaacccaactt
ggttacctcttatctagtggCATCAGAAACCATGAAAGCCAGAAAGAAGTGACGTAACACgaatcttaaacccaac
ggttacctcttatctagtggTTTCCCTGCAATGAGGGGGATATCAAGACATTCTCAGATGgaatcttaaacccaactt
ggttacctcttatctagtggGACTAAAGGAATTTCTCCAAACAGCAAGGACATCTGGAAGgaatcttaaacccaact
ggttacctcttatctagtggGCTTCCCTTGAGTTTTCTAATTTACATTTGATAGTCGAAGgaatcttaaacccaacttg
ggttacctcttatctagtggTGTCTTACGTGGTTCTGAATGTATGTACAGAAAATACTTGgaatcttaaacccaacttg
ggttacctcttatctagtggCAGAGAACACAATGGGAAGCTTCATTCAATGTAAGGGTACgaatcttaaacccaact

ggttacctcttatctagtggGTTAGTTCAGTAAGGTCGTAAGCTATAAGACAAACAAAGGgaatcttaaacccaactt
ggttacctcttatctagtggACCCCCACAGTACTTGTAGGTGAAAACTATAAAATACTGAgaatcttaaacccaactt
ggttacctcttatctagtggGCATGTCCATGGATTGAAATACTCAACATAGTCAGTTCTCgaatcttaaacccaactt
ggttacctcttatctagtggGGCCAGAGTAGTCCAGACTGTGGTATTGGCAGAGGCATTGgaatcttaaacccaac
ggttacctcttatctagtggCACTGCTGTAAAGATACTACCTGAGACCGGATAATTTATAgaatcttaaacccaactt
ggttacctcttatctagtggGGCAGCAGGAGAGAGAGAGGGGCAGGGAAAGGAAACACTCgaatcttaaaccca
ggttacctcttatctagtggCCCTGTTAAGGGGTTAAAAAGATAAGCTACAGTTGTAGGAgaatcttaaacccaactt
ggttacctcttatctagtggGTTTCACCAAAGAGGGTGTGTGTTTGTGTGTGTGTTTGTGgaatcttaaacccaacttg
ggttacctcttatctagtggAAACCACATTGGCATATCACTACACATCCATCAGAATAGCgaatcttaaacccaact
ggttacctcttatctagtggACCAATAGGTAAATGGTTAAATCAACTGTGGTACATCCACgaatcttaaacccaactt
ggttacctcttatctagtggGCTGAGCAAAAACAGTGACTCCTAAAAGGCTACACACGGCgaatcttaaacccaa
ggttacctcttatctagtggGCTAGGGGTTATGGAAAGGGTGGGGGTCAGGAGAAGGGAGgaatcttaaacccaa
ggttacctcttatctagtggGCAACATAAGGGAAACTTCCGTGGTGAAAATGTTCTTATCgaatcttaaacccaactt
ggttacctcttatctagtggGGATTAGCAGAAAGTTAGTAGAGAATACATGGGATCTCTCgaatcttaaacccaactt
ggttacctcttatctagtggGGTATCCTAATATTTCACAAGATACCTCTTGCAGTTTTCCgaatcttaaacccaacttg
ggttacctcttatctagtggTGGCCAAGATGCTCCTGAGGATGTAAATCAAAGTTATGGGgaatcttaaacccaact
ggttacctcttatctagtggTAACTGTTACAGAGATTTAGGGATTAAGCCAGCGTGGTGCgaatcttaaacccaact
ggttacctcttatctagtggAGAGTGCCTATGAACAAACATATACCCAAAATGGAAACTGgaatcttaaacccaact
ggttacctcttatctagtggCATCGATATTTGATCCCAGCCAAAGATCTATTTCAAGTGGgaatcttaaacccaactt
ggttacctcttatctagtggGAACATAAGAGAATATATTTGTGGTTTTGGGGTGTAGAAGgaatcttaaacccaacttg
ggttacctcttatctagtggCTTCATCAAAAGACACAATAAAGAAGCAAAAAGCCTCAAGgaatcttaaacccaact
ggttacctcttatctagtggAAACAAGCATTGCACAGAGAAAAGGACACACAAATGGTCCgaatcttaaacccaac
ggttacctcttatctagtggCCCACAAACAGACAATTACCATTTCATGCATGACAGACTGgaatcttaaacccaact
ggttacctcttatctagtggCTATAACACATGGCTGGTGCGAAAGTACATTGGCACAGCCgaatcttaaacccaac
ggttacctcttatctagtggTGTTTTGGCACTGAGCAATAAACGTGAACATGTACAGTGGgaatcttaaacccaactt
ggttacctcttatctagtggAGGTCTGGAGCTGCCACAGACTTTTCTACCCCAACACCCCgaatcttaaacccaa
ggttacctcttatctagtggGTCCTCCATGTCCTGCAGGTAAACCCTACAGACAAGCACAgaatcttaaacccaa
ggttacctcttatctagtggCAACTTCTGGTGACCACTGCTTCACTCGTCAAAAGGAATGgaatcttaaacccaact
ggttacctcttatctagtggTGGCAGGAGAAAGAAGCAAGTCACAGAAAAACATATGTGGgaatcttaaacccaac
ggttacctcttatctagtggTTCACATAAAGCCCAAACAGACAGAACTAAGCCTTATATCgaatcttaaacccaact
ggttacctcttatctagtggTTTGGTCAAAGAAAAATGGGGGCCACGCTGGGCACGGTGGgaatcttaaacccaa
ggttacctcttatctagtggCCTGGGCGACAGAGACTCCCTCTCAAATAAATGAATATTAgaatcttaaacccaact
ggttacctcttatctagtggGCAGTGAGTCGAGATTGCGCCATTTCACTACAGCCTGGGCgaatcttaaacccaa
ggttacctcttatctagtggGGTGGGGGGCAAAGGGTAGAGGAAAGTCGGGGAGGGGTGGgaatcttaaaccca
ggttacctcttatctagtggAGGAGAGGGAGAGAGTAGAATAGATAATTATGCACACTGGgaatcttaaacccaac
ggttacctcttatctagtggGTTCTACATAAGCTCTAATCACCTTTGCCCTAGGCTCTGGgaatcttaaacccaactt
ggttacctcttatctagtggCTGCACCAAGCCATCCAAACATCCTGCCCCTTAAAAGGTGgaatcttaaacccaa
ggttacctcttatctagtggCAGATGAGGAAATGAACTTACAAGCAGATAGGGAAAGGGCgaatcttaaacccaac
ggttacctcttatctagtggCCGCCTCTTCCCATCGCTTTGCATTTCCATGATGTCTATCgaatcttaaacccaactt
ggttacctcttatctagtggCAGGGCCGTTAGATGCAAGATGTTAAATTGGAAGAAAGCTgaatcttaaacccaact
ggttacctcttatctagtggCACAACACGTCCTCAGAGGGAAGAATTGCCAGTTTACAGGgaatcttaaacccaac
ggttacctcttatctagtggGCACCCTGAAGTTAGGTCTATGCCAGAATTTTTAGTACTGgaatcttaaacccaactt
ggttacctcttatctagtggGGCCATAGCACAGTGCATGGGAAATAGAAGAAATGGGTGCgaatcttaaacccaa
ggttacctcttatctagtggAGCAACGGTGCTACTTTGGGCAAACCATTGACCTTAGCTCgaatcttaaacccaac
ggttacctcttatctagtggTGAACAGGTAGCATAAAAGCTCAGGCCAATTTAAATGGGCgaatcttaaacccaact
ggttacctcttatctagtggGGAGCCACCAAACAAAATTCCAGGTTGCGGTCCTTTTTCCgaatcttaaacccaac
ggttacctcttatctagtggCCCTTTTCCAAACCAAGGTCAGATTTGTACTTAGGCTGACgaatcttaaacccaactt
ggttacctcttatctagtggGGGAGAGGATAATGGAGGGAGGGTACAATGATGCAGATGCgaatcttaaacccaa
ggttacctcttatctagtggCTAGTGAGTCCAATTCTACAACTGTCTTAAGAATAAAACCgaatcttaaacccaactt
ggttacctcttatctagtggCTGAAGCAGGGCTTGTAATGCAAAAAGCGGGTGGGGTGGGgaatcttaaacccaa
ggttacctcttatctagtggCTGTGAACACCTATTATGTGCCAGCCACCACCTTGGACACgaatcttaaacccaac
ggttacctcttatctagtggTCCCCATTTTGCAGAGGAGGAAACAGGTTCTGAGATAGTAgaatcttaaacccaact

ggttacctcttatctagtggGAGTGTAGACATTGGGGAGGGCTTATAGGTGGGACCACAGgaatcttaaacccaac
ggttacctcttatctagtggGGGTACCATCTGTGTGCTCATCCACCTCCATTCTTGCCCCgaatcttaaacccaact
ggttacctcttatctagtggCAGTGGTCTCTCCACATCTCCCTCACTGCCAGGTGGCCACgaatcttaaacccaac
ggttacctcttatctagtggGTCCCTCAACCCAAGGTCTGAGCTTCCATCTCGAGGAGGCgaatcttaaacccaac
ggttacctcttatctagtggGGCAGTACTTCACTCATTCAGGCAAGGGGGTAATACGGCCgaatcttaaacccaac
ggttacctcttatctagtggCTGCATTATCCCCTGTGGCTTCTCTACATCCCAAACTCTCgaatcttaaacccaactt
ggttacctcttatctagtggCGCTGGGGGATGAATGTGGCCACTGATTATGGGGCTGAGGgaatcttaaacccaac
ggttacctcttatctagtggCCCAAGATCCCTACTCCAGAAGCTCAGTCAAAACACTGAGgaatcttaaacccaac
ggttacctcttatctagtggTAGTTCCCAGAAATAGAGGTGTCTTTCCTGCAGACCTCAGgaatcttaaacccaactt
ggttacctcttatctagtggCCCTGACTCACCAGCTGACCTTGGGCAAGCCATTCCTCAGgaatcttaaacccaac
ggttacctcttatctagtggTCAGGACCTTTGATGTCAGTTTCCACAGACAAAGGGCTGGgaatcttaaacccaact
ggttacctcttatctagtggTACAGTGGATCTGCCAGGGAGAAGAGTGGAGGGCTGGGGGgaatcttaaacccaa
ggttacctcttatctagtggGGTTTGAGGGGTGCCATGGTTGAATAAAGACAAACAGGACgaatcttaaacccaact
ggttacctcttatctagtggGCTTAAGCTTTCTGAGCCTGAGTTCTCTAATCTGCTGCTCgaatcttaaacccaactt
ggttacctcttatctagtggGGGGTCTCACTCTGTCACTCCCCAGGTTGGAGTGCAGGGGgaatcttaaacccaa
ggttacctcttatctagtggGTCTCCCAGGTAGTTGGAACCACAGGTGTGCGCTACCACGgaatcttaaacccaa
ggttacctcttatctagtggGCTCAAGTAATCCATCTGCCTGAAGTTCCCAAAGTGCTGGgaatcttaaacccaact
ggttacctcttatctagtggCTACCAGAATGCCTGGCACATTGTAGGTGCAAAAATGTCCgaatcttaaacccaact
ggttacctcttatctagtggGCTCCTGTATATCCCCTACCCCCACACATTCACAGCCTCCgaatcttaaacccaac
ggttacctcttatctagtggACATTTCCCACTAGAGTGGTGCATTTTGTACAATTGGGTCgaatcttaaacccaactt
ggttacctcttatctagtggTTCCATCAGGGTTCATTCTTGGCATTGCACATTATATGGGgaatcttaaacccaactt
ggttacctcttatctagtggAAACTCTCTGTGCCCTATCTATTTATCCATCCCTCTACCCgaatcttaaacccaactt
ggttacctcttatctagtggCTGATCTTTTTACTGTCTCTATATTTTGCCTATCCCAGAAgaatcttaaacccaacttgt
ggttacctcttatctagtggAGATTCTTCTGCGTGTTTGCATGGCATGATAGCTCATCGCgaatcttaaacccaactt
ggttacctcttatctagtggCCACTGTCTGGATATACCACAGTTGACTTATCTGTTCACCgaatcttaaacccaactt
ggttacctcttatctagtggAAGCCGCTATACACATACATGTGCAAGTTTTTGCGTAGACgaatcttaaacccaactt
ggttacctcttatctagtggACTCATTGGGTAAATATCAAGGAGGGCAATGGCTAGATTGgaatcttaaacccaactt
ggttacctcttatctagtggGTCTTTTAAAGTGGCTGTACCGTTTTGCATCCCCACCAGCgaatcttaaacccaactt
ggttacctcttatctagtggATTGCTCCACATCTCCATCAGCATTTGCTGTTGTTGGTGCgaatcttaaacccaactt
ggttacctcttatctagtggCAGCTATGACTTGGTGTTAAGCAGTCACGTAGAGAGCAGCgaatcttaaacccaact
ggttacctcttatctagtggTAAGACCCTAACCACTACATTTTGCACAGTATTTCTTTCCgaatcttaaacccaacttg
ggttacctcttatctagtggCGTGAGCCACCATGCCCGGCCCATTGTTTAATATATCATCgaatcttaaacccaact
ggttacctcttatctagtggTGTTTCAATGGCAATATAAACCAACACAGAATCTCTGCCCgaatcttaaacccaactt
ggttacctcttatctagtggCATTTTGGCCATATCTAGAGCCAAGAAAGTGAACATGAGCgaatcttaaacccaact
ggttacctcttatctagtggCCCTTAGAACAAGAGGAAGATTGGGATCAGGAGAGGCTGGgaatcttaaacccaac
ggttacctcttatctagtggCAAGCCAAGCTCCTGAGAAAAGCTTGTTTTGTGGGACAGGgaatcttaaacccaact
ggttacctcttatctagtggGAGGATAGCAAAGACTCCAGCTTATCTAGTTATGATCCAGgaatcttaaacccaactt
ggttacctcttatctagtggTGGTGATATTAGGGTCCATATTGCTGAGCAAAGATAGTTGgaatcttaaacccaactt
ggttacctcttatctagtggATTTGCATAAGGACACCTGCCGACATCTTAAGGAACAGCCgaatcttaaacccaact
ggttacctcttatctagtggTCGTTATAGTGTATTGCTTTTATTAGCCTGTGACAGATGCgaatcttaaacccaacttg
ggttacctcttatctagtggGGTCAATTTGGGTTGTCAAATAGCCAGTTATTTGGCACTCgaatcttaaacccaactt
ggttacctcttatctagtggATTGGAGATAACAAACTGTTAGGTCCTTGTTATTCTGTGCgaatcttaaacccaacttg
ggttacctcttatctagtggGGATTAGCATGTTACTCAAGCTCACAAAGACATAGCTGGGgaatcttaaacccaact
ggttacctcttatctagtggGTTCTTTGTTCAGAGTATTTGCCACATTGAGGACTCCTGGgaatcttaaacccaactt
ggttacctcttatctagtggATCGATGACTAAAACTCATTTTTCCTCAGACCATGAGAGCgaatcttaaacccaactt
ggttacctcttatctagtggGCTTTATTAATGGGCATCTTCATATCCTTAGCAAGCTTAAgaatcttaaacccaacttg
ggttacctcttatctagtggGAGCATTCTTTACCTATTGCCCTACCAATCTTTCAGCTCCgaatcttaaacccaactt
ggttacctcttatctagtggCTGGGTCAAACTATGTTGAGGGCCAGCCACAGGCAATCATgaatcttaaacccaac
ggttacctcttatctagtggATCATGCCAAGTGAAACAGCCTGGAAGAGTGACAAAAGCCgaatcttaaacccaac
ggttacctcttatctagtggTCAGAATACCTATGCTCTAAACATTTACTACTGTGGAAACgaatcttaaacccaactt
ggttacctcttatctagtggAAGAAGCCCAAAAGTAACAGAGGACTAAGCAAGACAGGAGgaatcttaaacccaac
ggttacctcttatctagtggGCAGCTCTGCTCCACGGAAATTTCAGAGCCTAGGTTCCTTgaatcttaaacccaact

ggttacctcttatctagtggCTCAAATCCTAGCCCTCATGCCCAAGTTCAAACCAGCAGGgaatcttaaacccaac
ggttacctcttatctagtggGAAACAGGGGCAAAGCATCTACACCAGATCTCTGTTAAGGgaatcttaaacccaac
ggttacctcttatctagtggTCCCACTGGAGAAGCTAGTCATATGGCCACATCTAGCTGCgaatcttaaacccaac
ggttacctcttatctagtggGAAGTGGTGAGTGGATGCCTGGCACGGTGGCTCATGCCTGgaatcttaaacccaa
ggttacctcttatctagtggAGAAGTGGGAAGTGGAAATCAGAAAACGCCTAGCTGTCTCgaatcttaaacccaac
ggttacctcttatctagtggCGGAGAGCATGGATATGAAATAGCTAGCAGAGTGTCTGGCgaatcttaaacccaac
ggttacctcttatctagtggCCTGAGAACCTTGACCCATTAGAGAAGAACGGGAGTAGGCgaatcttaaacccaa
ggttacctcttatctagtggTCCCCAGGGCAGAGATGAGACAGGGAATGGCTTTTCTCTGgaatcttaaacccaac
ggttacctcttatctagtggGTTCTAGTGTAAGGAGCCAAGACAAGCATCTCATTCCTCCgaatcttaaacccaact
ggttacctcttatctagtggCTCTCTCCAAATCTTCTTCTTGCCCATTTTTCACCTTGCCgaatcttaaacccaacttg
ggttacctcttatctagtggCCCCCCAGGCTGGGTTAGATGCCCTTTTGCTGAATTATCGgaatcttaaacccaac
ggttacctcttatctagtggGCTGTGTTGAAATTAGTCTGTTTGCACGTCTCTACCACTGgaatcttaaacccaactt
ggttacctcttatctagtggTGAGGAAAAATATAAAGCCTTAGATATCATCATCTTCCCCgaatcttaaacccaactt
ggttacctcttatctagtggTAGATCTCAATCCCTTATTTCTATGCAGGGAACTAGAATGgaatcttaaacccaactt
ggttacctcttatctagtggGGCACATATTAATTGGGTCCTTGGTCAATGCTTGTTCAACgaatcttaaacccaactt
ggttacctcttatctagtggGAAAGAGAGAAGGATGAGGCTAGGTGCAGTGGCTCATGCCgaatcttaaacccaa
ggttacctcttatctagtggGGCAGAAAGATTGCTTGAGTCCAGAAACTTGAGACCAGCCgaatcttaaacccaac
ggttacctcttatctagtggCTCTGTCTCCACGAATATATATTATATACATTAGCCAGGCgaatcttaaacccaactt
ggttacctcttatctagtggTGATCACACCACTGCACTCCAGACGGGATGACAGAGTGAGgaatcttaaacccaa
ggttacctcttatctagtggAACCTCCCGAAACATGAACTGAGAATAATTCCCATGATACgaatcttaaacccaac
ggttacctcttatctagtggCTGAAAGACTGAGAGCTCTGTTCCACCACTGACAAGCGTGgaatcttaaacccaac
ggttacctcttatctagtggCTCCTGGTCAATGTGTTGGGGTAATGGTGTGGGTTTTAGCgaatcttaaacccaactt
ggttacctcttatctagtggGCAGCATTCCCGGTCTCTGCCCACTAAATGCCAGTAGCACgaatcttaaacccaa
ggttacctcttatctagtggGTCCCCTAGGTGGGTAACAGTGCTCTGCCCCCTCCCAAACgaatcttaaacccaa
ggttacctcttatctagtggACCACAGTGTAGACTTAAATAAAATTACTAAAGACCGGTCgaatcttaaacccaactt
ggttacctcttatctagtggAGGCTGTTGCACTCTAGAAACTACTCTAAAACAACTACAGgaatcttaaacccaact
ggttacctcttatctagtggAAAGTCTGAAAGTTACAGTACATGAGGGGGGAAGGAACCCgaatcttaaacccaac
ggttacctcttatctagtggAGAATTATCTCAGTTAAGGTGACTGCATAATGAATCTGACgaatcttaaacccaactt
ggttacctcttatctagtggCCAGAAAGGTGGAAGAATCCTTATACCTTATGCTGCATGCgaatcttaaacccaact
ggttacctcttatctagtggGGAGAGCAGCCAGAATGAGGAGAGGCCAACAGACCAGGTCgaatcttaaacccaa
ggttacctcttatctagtggGAACATGTTGACTCCTATGTTCCAGGAGCTGTAGAGACGGgaatcttaaacccaac
ggttacctcttatctagtggCCCTCTAGAAACTTACATTCTAGTAGAGGGAGCCAGTGTGgaatcttaaacccaact
ggttacctcttatctagtggGTGGGACTGTTGCATAGCTAAGAGTTATGCAAGCACCAAGgaatcttaaacccaac
ggttacctcttatctagtggGCTTCTGAGTTGATAGTGCTGTTTTGTGCCTTTTCAGAGGgaatcttaaacccaacttg
ggttacctcttatctagtggAGTAAAAGAAAGCCACTGGGAACCCTTGCAGTAATTCCCGgaatcttaaacccaac
ggttacctcttatctagtggGAACTCGGTGACTGGATAAGTATGATGGGGAATGTAGAGGgaatcttaaacccaac
ggttacctcttatctagtggGACTCTAGCTTCAAATAAGAGAAAGGATTGAGGAACAAGGgaatcttaaacccaact
ggttacctcttatctagtggGACTACAGGGAGGCAAGGCTGTTGTTCCCATGTATCAAGGgaatcttaaacccaac
ggttacctcttatctagtggCTAGGTGTGTTCTATATGGTCCCAAAGCTTTAAACTGGAGgaatcttaaacccaactt
ggttacctcttatctagtggCGCAGGAGGTAGAAGCTCATCAAGACAGCGCCTAGGGAGAgaatcttaaacccaa
ggttacctcttatctagtggAGGACATCTAGAGATTTTCACTGCTCTCCTCTCAGCTTGCgaatcttaaacccaact
ggttacctcttatctagtggCCCCATCCTGATTGTTCTTCAGGGAACCAAAGCCTCCTTCgaatcttaaacccaac
ggttacctcttatctagtggCAGATGGAATCAAGGCTCTTCTCTCCTGCACCAAGGGTGGgaatcttaaacccaac
ggttacctcttatctagtggTCAGCCTGCACAGGAAAAAGTCTCAGAGAATGGCCCCAAGgaatcttaaacccaa
ggttacctcttatctagtggGGAAGGCTTGCTCTCTCTCTGTGTGGTGTAGCTAAGAGGGgaatcttaaacccaac
ggttacctcttatctagtggGAGAGAGAGAGAAAGACTCAAGGACATGATCAAGAGAGCCgaatcttaaacccaa
ggttacctcttatctagtggGCTGTCCCTCAGAGTAGTTACACTCCAAGATTTCATTATCgaatcttaaacccaactt
ggttacctcttatctagtggGTTATCTGCTCCCCAACCCCCAAAGAATTCCTCTTGTGGCgaatcttaaacccaac
ggttacctcttatctagtggCAGGCATAGAATGAGAAGCGACTCCCAGACAATAGGTCACgaatcttaaacccaa
ggttacctcttatctagtggGCCTTCAATGTATCAAAAGCTGGCACATTACTGGTTCTGCgaatcttaaacccaact
ggttacctcttatctagtggGACAATTGAAATTTCCAAATCTGGAACAACTGGATTGGAGgaatcttaaacccaactt
ggttacctcttatctagtggGACCTTACAAGCACCATAAACATTCTGTTGTGGCTCTTCCgaatcttaaacccaact

ggttacctcttatctagtggGCATGTTAATGCCGTTAGCACTTACCTCTAAGACCGGTAGgaatcttaaacccaact
ggttacctcttatctagtggAGGGGATACAATCTCATTACAAGCTCTAGTAGTCATCCAGgaatcttaaacccaact
ggttacctcttatctagtggAGCAAGGCTTCTCATCCTCAGACTACGCAGATTTTCCCCCgaatcttaaacccaac
ggttacctcttatctagtggTTTTAGGGAATACATGAAGTGGTTCACCCTCCTGCCTTCCgaatcttaaacccaact
ggttacctcttatctagtggCTGTGCTCATGTGCAAAGGAATCTTCACTTCCTACTACTGgaatcttaaacccaactt
ggttacctcttatctagtggGGTATCTGAGACTCAGTCTTTCTCATCCATGACATGGAACgaatcttaaacccaact
ggttacctcttatctagtggTCACTGAGTGCTTGCTGTGGGTGATGCACTGAGGCAATGCgaatcttaaacccaac
ggttacctcttatctagtggGGAGTGGATCTTAAGGACTCCCAAGGACCAGGCACATCCCgaatcttaaacccaa
ggttacctcttatctagtggCAAGCTGCCATGGTGGTATGGATTTGAACATGTAGCCTGGgaatcttaaacccaact
ggttacctcttatctagtggCCAAGAGTGAAGCTCAACCTTAGAACCTTGGACAGAATGGgaatcttaaacccaac
ggttacctcttatctagtggAGTGGAAACTACCCAGGAATCAATCATGGAAGTTAAATGAgaatcttaaacccaact
ggttacctcttatctagtggGTATTCAATATAGACTACAAAGATGAGGACCAGGAGCTGGgaatcttaaacccaact
ggttacctcttatctagtggAGTCACAGGTGCAAGTTACGTGAAATAGAACTCTGTAGTCgaatcttaaacccaact
ggttacctcttatctagtggCACACATACAGTTCTGGTCCATGTGATAACGTGCCTGCTGgaatcttaaacccaact
ggttacctcttatctagtggGCCATACTCTCCATTCGTCCTTGCCTCGCTTCTTTTACTCgaatcttaaacccaactt
ggttacctcttatctagtggAGAGCCCAACTTTTCCCATGACAGATCTGAAGATGCTAACgaatcttaaacccaact
ggttacctcttatctagtggGCAGTTGAGATGAGATTTGTGTTCTGAGCTCCCTGTGGGCgaatcttaaacccaact
ggttacctcttatctagtggCCATAGGGCAAAAGGAGATAGGTGTTTTCCTTGAACATACgaatcttaaacccaact
ggttacctcttatctagtggGATGGAAGAACAATGTTCTACAGCATCTCCAGAACAAAGGgaatcttaaacccaac
ggttacctcttatctagtggGGACTCTTTTCACCCAGTGTGTGGAGCCCCTCCCCACCCCgaatcttaaacccaa
ggttacctcttatctagtggCCAGGTTCAGCCTACACATCCATGCCTTCATCTCTCTTGCgaatcttaaacccaact
ggttacctcttatctagtggGAGAATACTGGAAATGCTCAAAGTGTGTCAACTCTCACTCgaatcttaaacccaact
ggttacctcttatctagtggTTCTGTTCTCAAGTCTTAGTCCACACTTGTGCTCTGAGACgaatcttaaacccaactt
ggttacctcttatctagtggCAGTCATGTGTGTTATCTTATTGAATTCTCTCCACAACTCgaatcttaaacccaacttg
ggttacctcttatctagtggACACAGATAGTGGTAGAATTTGGATAACAGAGCCCAAGTGgaatcttaaacccaact
ggttacctcttatctagtggACAACATGAGGAATGACACTAAGTGAAAGGCATGTTCATCgaatcttaaacccaact
ggttacctcttatctagtggGTGGAGTATTCATTGAGTGCCAGAAATTATGCTCTGTTCCgaatcttaaacccaactt
ggttacctcttatctagtggCAGTTGCCCTAAAAAACCCAGAGATCTGAACTGAGGCCCCgaatcttaaacccaa
ggttacctcttatctagtggGAGTCATTACCACTTTCTGTAGGTACCCATCCCAAGAGGGgaatcttaaacccaac
ggttacctcttatctagtggTTCAGTGATTGCGTGAACTTCAGTAACAATTACATATGAGgaatcttaaacccaacttg
ggttacctcttatctagtggCCCAATGCATAGATGGCAATGTGAGTAGTCAGGATTGACCgaatcttaaacccaac
ggttacctcttatctagtggTGGCATAGGAAACACAGAGCTGAGAAGACGGGCATGGGCCgaatcttaaacccaa
ggttacctcttatctagtggGGAGCTTATAGTGTGACCATGAAGACAAACACTAACAAGCgaatcttaaacccaac
ggttacctcttatctagtggGCTGTTGATAAAGCAGTAGGAGTAAAGGGGGAGTGGGTGCgaatcttaaacccaac
ggttacctcttatctagtggGGCCACAGTTGCAGCAGCAGGAGATGGCACCTGGTTGGAGgaatcttaaacccaa
ggttacctcttatctagtggTGAGAAACAGGGTAAATGCAATCAAGTTCCAACCTAGCCAgaatcttaaacccaac
ggttacctcttatctagtggCTTTATCCCCATTTAACAGATGAGAAAATGAAGACTCAGGgaatcttaaacccaactt
ggttacctcttatctagtggGCTAGAGCACAAACCAGGGAGTCTAGCTCCAGAAGCCATGgaatcttaaacccaa
ggttacctcttatctagtggTGTTGTGGCTATTTGGCCTGGACCCCACAGCCAGCCCCTCgaatcttaaacccaa
ggttacctcttatctagtggGGAGCTTAATTCTAAACCCACCAGTTTCTGAAACCAGGACgaatcttaaacccaact
ggttacctcttatctagtggTGGATACTCCAGAGGAACAGCTGGGGTGGGTGAGAGCAGCgaatcttaaacccaa
ggttacctcttatctagtggTGACAGCAGCTCCAGGCAGTCTTGACACAAAGGACATTTCgaatcttaaacccaac
ggttacctcttatctagtggTCTAGCATTGGTCAGCTGTCCCAGAAATCAAATTTGGGACgaatcttaaacccaact
ggttacctcttatctagtggCAGAGACCCCACAAACCTACCATTATTCCTCTCGGAGGCCgaatcttaaacccaa
ggttacctcttatctagtggTGTGTGCTCTCTGATTGAGTGCCCTGTTAGTTAAGTGGGGgaatcttaaacccaactt
ggttacctcttatctagtggTCCCCGTCAGAGAAGACCTGTCCCTTGAGTGTCAGGATGCgaatcttaaacccaa
ggttacctcttatctagtggGCAGGAGAGGGAGAAACCAAAGTTTGGCATTGAATTGTGCgaatcttaaacccaac
ggttacctcttatctagtggGGTGAACAAACCACAGCTCCTGCCCACAAGCAGCCGATGGgaatcttaaacccaa
ggttacctcttatctagtggGTGGATCTCCATCTGGCTGCACATCTCAGTAACCTAGAACgaatcttaaacccaac
ggttacctcttatctagtggAGACCCATTCCCAGTGCCTCCAACCCAGACTAATTAACAAgaatcttaaacccaac
ggttacctcttatctagtggGGTTCTGGTATCTAGAGAGGGTTGATAACCACTGGTCAGGgaatcttaaacccaact
ggttacctcttatctagtggCCCTCCTTCTCTCTAACCTTGGTACAGCCTTCATACTTTGgaatcttaaacccaactt

ggttacctcttatctagtggATCTTATCAGACCCTTAGTACCTGCAGGAAGACCCTTGTCgaatcttaaacccaactt
ggttacctcttatctagtggGAGAGTCAGATGTCATTTGCAGCTTCTGAGGCATGAATTGgaatcttaaacccaactt
ggttacctcttatctagtggGAAGCCTGGGTCACTCTTTGGCATTCATGACTCCATGTCCgaatcttaaacccaact
ggttacctcttatctagtggGGTTTAACATCAGTCGGTGCTTTCCCTGTGTATAGCCAAGgaatcttaaacccaactt
ggttacctcttatctagtggCCATGTTGTCTGGGTGGAGCTGCCTCAAACTCTTGGAAGCgaatcttaaacccaact
ggttacctcttatctagtggGTGAAACTGTATTCTCCTTGCCTAGGACACAGGCCAGGACgaatcttaaacccaac
ggttacctcttatctagtggGCCGTCATTCTTGCCCATGGCTATTAAGCTAGCATATGGCgaatcttaaacccaactt
ggttacctcttatctagtggGGTGGTCTTCTTAGCCACCAGCTGGGGAGAGACTGGCTAAgaatcttaaacccaac
ggttacctcttatctagtggATACCTGGATCCAGTTGAGACTGAAGATCTACCCCTGACCgaatcttaaacccaact
ggttacctcttatctagtggGTGGGCTAATCATCCTCCATTTTTGTTTTGTTTTGTTTTGgaatcttaaacccaacttgt
ggttacctcttatctagtggCGTGAGCCACCGTGCCCGGCCTCATCCTCCATTTTTGATTgaatcttaaacccaact
ggttacctcttatctagtggTTCTCTCTTGCAGTTAAAAGAATGTTCACTACCACACTGGgaatcttaaacccaacttg
ggttacctcttatctagtggACAAGCATAGCTTCTTCCCTTATCCCTCCAAGTTACCGAGgaatcttaaacccaactt
ggttacctcttatctagtggGCTGACATTTGTCTAATGCAATTTTCCCATGATGTGCTCCgaatcttaaacccaacttg
ggttacctcttatctagtggGTCCATGTCCAAGTAAGTTTAGGGTTAACTTGTTAAAAGCgaatcttaaacccaacttg
ggttacctcttatctagtggAGTGAGAACTGTTAATCTCTCATAAGAGGTGCAGTATACCgaatcttaaacccaacttg
ggttacctcttatctagtggTAACTCGTCATTTACATTACGTATATCCCCTAATGCTATCgaatcttaaacccaacttg
ggttacctcttatctagtggCATGTGGTGTTTGGTCTTCTGTCCTCGCGATAGTTTGCTGgaatcttaaacccaacttg
ggttacctcttatctagtggCCTTTGTCAGATGAGTCAATTGCAAAAGTTTTCTCCCATTgaatcttaaacccaacttg
ggttacctcttatctagtggAGCACAGGGCTCTTTTTTGTCCTGCATCTCCTGGGATTAGgaatcttaaacccaactt
ggttacctcttatctagtggTTCCTTTGCTTGGAATACTCTTCCCACACCATCCACTACCgaatcttaaacccaactt
ggttacctcttatctagtggCTACTCAACTATCATGTCTCAGCATGAATACTACTTTCTCgaatcttaaacccaacttg
ggttacctcttatctagtggGACACCACTACTTTAATGACATCACTACTGCCACCTTCCCgaatcttaaacccaact
ggttacctcttatctagtggACCCATGATTTGCTCTCTTACACTGTATACTTCCGCTTTCgaatcttaaacccaacttg
ggttacctcttatctagtggGTGGTGGCCAGTTCACAATATGTTCCCCAATGATTCCCACgaatcttaaacccaact
ggttacctcttatctagtggGGAGACACTTCCCATGTGAATGTATGGCCAGTGCTATGTGgaatcttaaacccaact
ggttacctcttatctagtggAATTGCCCACTCTAGGGAAACCAGACACCATGCCCTGAGGgaatcttaaacccaac
ggttacctcttatctagtggAGAGAGGAGCTAAGGACCCCAACAAACAGCTGGCACCAGCgaatcttaaacccaa
ggttacctcttatctagtggTGTGAATGAGCCAACTTGGAAGGTGTCCTCCAGCCCTGATgaatcttaaacccaact
ggttacctcttatctagtggCCACAATCCAGTGAGAGGCCACAAACAAGATCAGCCCAGCgaatcttaaacccaa
ggttacctcttatctagtggGCCAGTACTGTGAATTGCTTACTATTGTGGCATACTCCCCgaatcttaaacccaactt
ggttacctcttatctagtggGCATGTATATGCACACATACTACCAGGGTCATAATTATAGgaatcttaaacccaacttg
ggttacctcttatctagtggTTATGTATTTATATACCTCTACATACACACCCAGAGGTAGgaatcttaaacccaacttg
ggttacctcttatctagtggTAGACATATACAGGTATAGGGGGCTCGTTGGTAATTTGGGgaatcttaaacccaactt
ggttacctcttatctagtggGTGACTATAGCCCTGAACTGAAAAATCCAGGAGATCTGGCgaatcttaaacccaact
ggttacctcttatctagtggGAAGTGACATAACCCCCTGCCCCTGCCCCACCTCCAGTGCgaatcttaaacccaa
ggttacctcttatctagtggGGCAGGGAGGGAATGGAATTCAGCTGGTCTAACACAGGGCgaatcttaaacccaac
ggttacctcttatctagtggGAAGCCCACGTCACCTGCTGAAGTCCTGTCTTCTTTGCTGgaatcttaaacccaact
ggttacctcttatctagtggGTGGCAGAAGCACCCCATCCCAGGACTTTTTGGGTGGCGGgaatcttaaacccaac
ggttacctcttatctagtggTTCACAGCCCTTGGAGCACTAAGACCCAGGCTCAGTCCTCgaatcttaaacccaac
ggttacctcttatctagtggTCCTGCCAATGGTATCACATAGTCAACTCTTAACAGATACgaatcttaaacccaacttg
ggttacctcttatctagtggGCCACACACGGCACCTGAGAAAATTGTGTCTGCAGGACCCgaatcttaaacccaac
ggttacctcttatctagtggAGGGTGGAAAGAGCTGTTCTACAAGGAATTTGTGAGGGAGgaatcttaaacccaact
ggttacctcttatctagtggCGTTGGTGGGGAAGGGGGTGGGATGGGGATTAGAGCTTGGgaatcttaaacccaac
ggttacctcttatctagtggCTACTGAGTAGTTACCGTATGTCAGGTACATTACATTTGTgaatcttaaacccaacttg
ggttacctcttatctagtggCAGGTTGAGTACACAGAAAGGTCTTATCGCAGTCCAAGCCgaatcttaaacccaac
ggttacctcttatctagtggCAATGAGAATGTGAGTATGTGTGTGCATGTGGCATAGAGGgaatcttaaacccaactt
ggttacctcttatctagtggCATTCTAGTGCAAGGTTTGGCAAACTATAGCCTGAGTGCCgaatcttaaacccaactt
ggttacctcttatctagtggCACTGCCATGCTCATTTGTTTATATGTAGTCTATGGCTGCgaatcttaaacccaacttg
ggttacctcttatctagtggCCACATGGTCTGCAAGCCTAAAATATTTACCATCTTGCCCgaatcttaaacccaactt
ggttacctcttatctagtggTGCTGACTCCTGCTCTATATGACCACGGTAATATAAGGCGgaatcttaaacccaactt
ggttacctcttatctagtggCCTTCTATATATCTAGGATAACTACATATCCCAGACTAAGgaatcttaaacccaacttg

ggttacctcttatctagtggACATCCATTTGCAGGAAAAACTATAGCAATATTGTTCCTGgaatcttaaacccaacttg
ggttacctcttatctagtggGCTGTCCAAGGAGTTAGTCCCATACCATCCTCTGAAGGAGgaatcttaaacccaac
ggttacctcttatctagtggCTATGGGGACGTTGAACCAGTCACTCAGATAAGAACGATGgaatcttaaacccaact
ggttacctcttatctagtggACCTTTTCCCCTGTCCTTCTCAGTCCTCATTTCCACCCCCgaatcttaaacccaact
ggttacctcttatctagtggGGGATATCCCCAAGTAAGTTTTGGAGTAGTAAAAGACTCCgaatcttaaacccaactt
ggttacctcttatctagtggGCCCAAGGGATATGGCGGTGGTAGGGCAGGAGTTATCACCgaatcttaaacccaac
ggttacctcttatctagtggGTGCTGTCCATGGGCAAACAAGAAACTTCCCCGTTGGCACgaatcttaaacccaac
ggttacctcttatctagtggGGGGTTGCCCAAACCTCAGCCATTTCTGGAGCATCTTTACgaatcttaaacccaact
ggttacctcttatctagtggACTTTGCTGTACAAGAAAACTGTTGCATTGCAATAAATGGgaatcttaaacccaacttg
ggttacctcttatctagtggGACAGATAATGTTGCCCACCCAAAGCTTTGTTCCTGAGCCgaatcttaaacccaact
ggttacctcttatctagtggGGTGTTAAAGAGTGATACCAAACTGAGAATCTCTCCAGGCgaatcttaaacccaact
ggttacctcttatctagtggGAGAATTCCAAAGCCAGCCAAAGAGTTGGAGGAGGGGTGGgaatcttaaacccaac
ggttacctcttatctagtggGTTGGCATCTCACGCACAGCAGACACTTGAGTATCATTGCgaatcttaaacccaact
ggttacctcttatctagtggTTCCATTGTAGAGATAGAGACACCAAAGCTCAAGTCCAGGgaatcttaaacccaact
ggttacctcttatctagtggCAGGGTCACAGACTGTGGATAGAGTAGCTTCAGACAGAGGgaatcttaaacccaac
ggttacctcttatctagtggCAATGGTCCTGCCAGTGATCCAACCCCTGCTTACTGGATAgaatcttaaacccaact
ggttacctcttatctagtggGTGCCTGCCAGAGTCTGAGCACCTCACCTGGCTCATCCTCgaatcttaaacccaac
ggttacctcttatctagtggTCCCACCCTGGCAACCTTTTATCACTCTCTGCTCCTCTCCgaatcttaaacccaact
ggttacctcttatctagtggGTGGGATTTCCAGGACTGAAGACAGCCTGCCACAAGATCCgaatcttaaacccaac
ggttacctcttatctagtggGGACCAGTGCTAGTTAGGGGTTCAGGGTGATAGAAGGGGGgaatcttaaacccaac
ggttacctcttatctagtggAGCAGTGGCGAGAAAAGTAGTCAGGTAAGATGGCAAGTGCgaatcttaaacccaac
ggttacctcttatctagtggAGGCAGATTCCAGGTACAAAGATGCATAGAGTGGAGACACgaatcttaaacccaac
ggttacctcttatctagtggCCCTCAAGGGAGCTGCTGTCTAGCTCTTCCCTGCCCAACAgaatcttaaacccaac
ggttacctcttatctagtggAGCCACACATGGTTACTGAGCACGGAAATGTGTCCAGGCCgaatcttaaacccaac
ggttacctcttatctagtggGACTTGGTCTTCAAATGACAAAAAAGCATGTGAACATCACgaatcttaaacccaactt
ggttacctcttatctagtggTGGAGAACACTAGTCCATGCAGCAAGATGGGTATGGACCCgaatcttaaacccaac
ggttacctcttatctagtggGGGGGGATTCCAAGTGGGGGAAGCACAGCAATTACTTATTgaatcttaaacccaact
ggttacctcttatctagtggCATCGTTTTCACAATATCCATTGGCTACTGAGCTTTAAAAgaatcttaaacccaacttg
ggttacctcttatctagtggCCCTCAGTACCACCTCCCAGAAGTTACCTCAGTTCTAGGCgaatcttaaacccaac
ggttacctcttatctagtggATCTAGACTCTATTTCCGATGAGCCAATGTCCATATAGGCgaatcttaaacccaacttg
ggttacctcttatctagtggGCTATTGGCCAGAGCATGGAAAAGATAAATAGACACAGACgaatcttaaacccaact
ggttacctcttatctagtggCAGGAGCATTTATTGGCTGCTTACTATATACCAGGCTCTCgaatcttaaacccaacttg
ggttacctcttatctagtggGATGGGGAAACTGAGGCTCCTAAAAGGTTATATAACTGCCgaatcttaaacccaact
ggttacctcttatctagtggTGAAAGGAGAGATCATGTCCACATACTTGAAAGCTCAGGGgaatcttaaacccaact
ggttacctcttatctagtggATGGAGCACATCTGCATGAGGATTTCAGCTTCTGCTTTGAgaatcttaaacccaacttg
ggttacctcttatctagtggCAGGTGACATGACTTTGCTCTCTTTTGCATCCAGCTGCCGgaatcttaaacccaactt
ggttacctcttatctagtggCTCTTTTGGAACTCCAGATTGTGTCTTAGCCTGCAGGTTCgaatcttaaacccaacttg
ggttacctcttatctagtggCGTAAGACTTGGAAACCAGCCAGTGAAAGAGGGAGGGAAGgaatcttaaacccaac
ggttacctcttatctagtggGAGCTATTCTTGGAGCTAGGAGTGTTGGCACCAGGCTCTCgaatcttaaacccaact
ggttacctcttatctagtggGACAATATATACTAGATCCCACCACCTCCCCCCACCCCCGgaatcttaaacccaac
ggttacctcttatctagtggGTGACAGAGGCTCTGGTAGAGTAACTGAATACTGGCTGGCgaatcttaaacccaac
ggttacctcttatctagtggGGGGTAGTGATTGCCAGAATTCAGATCCCAGCTTTGCGGGgaatcttaaacccaac
ggttacctcttatctagtggGAGGCTCTCTGTACCTCAGCTTCCTCATCAACACAATAAGgaatcttaaacccaact
ggttacctcttatctagtggGGGCTATCACAACAGGTAGCTCAGTTCTAGTATCTTCAACgaatcttaaacccaactt
ggttacctcttatctagtggATGCTACTTAAAAAGACACTGTCTCCCCAACAATGCATGGgaatcttaaacccaact
ggttacctcttatctagtggTGTGTCTACATTTAGTGGGGAAAGAAGTAGAACAAGTTGGgaatcttaaacccaacttg
ggttacctcttatctagtggGATGAGCCCCCATATTTTCTAGGAAGGTTAAATCACAACCgaatcttaaacccaactt
ggttacctcttatctagtggCTTCCCTTTACACCATACTTTGAAATGATATGTCAGTAAGgaatcttaaacccaacttg
ggttacctcttatctagtggGTGTTATTTGGAGATGGAAGTAACTGCCCAAAAGAGGTCCgaatcttaaacccaact
ggttacctcttatctagtggTTGAGCTGATAAAATTCTACTCTGCAGGATGGTGTGGCAGgaatcttaaacccaactt
ggttacctcttatctagtggCAGACTGGAATGGAGTGTCAGGAACCAGTATATCCGGAGCgaatcttaaacccaac
ggttacctcttatctagtggGAGCTCAGACACAAAGCTCAGAGTAATTGCAAAGCTGGACgaatcttaaacccaac

ggttacctcttatctagtggCGATGACATTTTGACCCTAACTTGCAACTACTTAACATGCgaatcttaaacccaactt
ggttacctcttatctagtggGTTGTCATGATACTTACCTCCAAATATGTTTGTTCATGCCgaatcttaaacccaacttg
ggttacctcttatctagtggCTTAAATGAAACACCACCACCACCATCCATCACAGTCTAGgaatcttaaacccaac
ggttacctcttatctagtggGGCCTACACCTTCCATTTGAGTTCATTCTTCCCCTCACTGgaatcttaaacccaactt
ggttacctcttatctagtggGGCTTTGCAGCCAAACTTTGAGGTCTGGATCCCAGCTCCAgaatcttaaacccaac
ggttacctcttatctagtggTCTCTGTGACCCTGAGGAAATGACGTCTTTGAGCCTCAGTgaatcttaaacccaact
ggttacctcttatctagtggCCACAGATGATGTAGGCTGAGAGATTGGTACAGTGTCCAGgaatcttaaacccaac
ggttacctcttatctagtggGCATCTGAGGCATTGTATCCTTGGGCCCAGGGCAGATAGCgaatcttaaacccaa
ggttacctcttatctagtggGACAAAATTTTTCTCCCTTTGAAAGCCAAGGGTCACCAGGgaatcttaaacccaact
ggttacctcttatctagtggCCCCCAACTCCACCCAAGCCTCCTAATGAGCGCATAGAAGgaatcttaaacccaa
ggttacctcttatctagtggTTACCCTACTTGGTGAAACCTGAACTAGGATTCTGAGGCCgaatcttaaacccaact
ggttacctcttatctagtggGGATGTAAAGTGATGCAAAGTGAGCCCACGCTGTTGAGAGgaatcttaaacccaac
ggttacctcttatctagtggGGCTCTGGGTGCACATGGGTGGTGCAGAGTGCGTCCCAGGgaatcttaaacccaa
ggttacctcttatctagtggGATGGAGGCAGCTGGGAAATCTTCTCACGGCCCTCATGCCgaatcttaaacccaa
ggttacctcttatctagtggCCTGCAGAGAGTGGAGGGCACCCAGAAATTCCCCAACCCAgaatcttaaacccaa
ggttacctcttatctagtggAAGAGAACACAACTGTTGAGTTCAGGATGGATGGGCCCAGgaatcttaaacccaac
ggttacctcttatctagtggGCCTTCTTTCCTTCATCTTTTCAGTAAATATGTACTAAGGgaatcttaaacccaacttg
ggttacctcttatctagtggCAGCTCTAGATTAACAGGTACCAAAGGGCCTTGCACACAGgaatcttaaacccaac
ggttacctcttatctagtggTGTTCCCTGTTGAATCTCTGTGTCTATAAGAATCCCTGGCgaatcttaaacccaactt
ggttacctcttatctagtggTGAATGCTCTTTCCAGAATCCAACACTTCTCATCATCCCCgaatcttaaacccaactt
ggttacctcttatctagtggTGCCTTGATGGTTGTATTAGCCTCCCAACTGGAGTCTGGGgaatcttaaacccaact
ggttacctcttatctagtggCCCTCAGTACACCAGCCAGAGGAATCCATAAAAATATGAGgaatcttaaacccaac
ggttacctcttatctagtggGTGCCCTACATGGCCCCACGCTTCCTGCCCTCTTGTGTCCgaatcttaaacccaa
ggttacctcttatctagtggCTTGTCTCCCCTCACTCCCTCCTCCCCACACTGGCCTCTTgaatcttaaacccaac
ggttacctcttatctagtggGAGCCCACCAACTGCCCTTCCTCATGCTTTTAAATTTGTTgaatcttaaacccaactt
ggttacctcttatctagtggTCTTTTCCTAAATGTCATGTTCTCAGCGGGGTTTTCCCTGgaatcttaaacccaacttg
ggttacctcttatctagtggCTGCTTTATTTTTCCCTATAGCCCACTTATCTCCATCTAGgaatcttaaacccaacttg
ggttacctcttatctagtggGCCTGCCTCCCCCAAACTAATATATAAGCTCCCCATGGGCgaatcttaaacccaa
ggttacctcttatctagtggACTGCTAAATCTCCAGCACCTAGAACAGTGTCTGGAACAGgaatcttaaacccaac
ggttacctcttatctagtggCGTGCTCTATGAGAAGTGTACATTTTCTCAGCTAACACTGgaatcttaaacccaactt
ggttacctcttatctagtggGACCCATCAATTCTACTCATAGGTATTTGTACAAGAGAAGgaatcttaaacccaactt
ggttacctcttatctagtggATAAAGCCATAAAAGGGAAACAATCCAAATGTCCATCAACgaatcttaaacccaact
ggttacctcttatctagtggCATCCATGAAAAGGAATGAGGTATTACACATGCTGCATTGgaatcttaaacccaactt
ggttacctcttatctagtggGAGGCTGGTAACAATGAAGTATGAAGTGGAGAGTGACTGCgaatcttaaacccaac
ggttacctcttatctagtggGGAGGTGATTACATTTTTCTGGACTTATATAGTGGTGATGgaatcttaaacccaacttg
ggttacctcttatctagtggAACACCCTGACGTGTACATTGAAATGGGTTATTTCTATGGgaatcttaaacccaactt
ggttacctcttatctagtggTTACCCATGTCATCCCAGAGGTTAGATCTCAAGTCTGGGGgaatcttaaacccaact
ggttacctcttatctagtggGCCTGAGGCTCTGTTTTGTCTTAAGCACTGCTTGGTGGAGgaatcttaaacccaact
ggttacctcttatctagtggGTATGCATTTAATGTCCAGTTCATTTGGAGGACAGACAGCgaatcttaaacccaactt
ggttacctcttatctagtggAAAGATAGGTGGAAGCCAAGAGCACGAGAACACAGAAGGGgaatcttaaacccaa
ggttacctcttatctagtggAGGGATGGTTTCCTGGAGCCAGGCTGGAAAGAAGGTAGGGgaatcttaaacccaa
ggttacctcttatctagtggGACAGGAAAGGAAAGGAGGAGCATTCTGGCACACGGAGCTgaatcttaaacccaa
ggttacctcttatctagtggGAATGCAGGAGAGACAGGCTGAAGAGATGCACCCTGAGGGgaatcttaaacccaa
ggttacctcttatctagtggGCAGGCTGCATTTTCTCCTGTAGGCAACTGGCATTGGCAGgaatcttaaacccaac
ggttacctcttatctagtggCTGGCAAGGCTAAGCTTCCAGCCATGTGAACCAAGAGTTGgaatcttaaacccaac
ggttacctcttatctagtggGATATAGTTGGATCTGTGTCCCTGTCAACATCTTATGCTGgaatcttaaacccaactt
ggttacctcttatctagtggGTAATCTCCAATGTGGGAGGTAATTGGATCACGGGGCTGGgaatcttaaacccaac
ggttacctcttatctagtggGTGGCACCTCCCTGCCTCTCTCTCTTGCTACTGCTCTGGCgaatcttaaacccaac
ggttacctcttatctagtggGTGAATTTCCTGAGGTCTCCCCAGAAGCTAAGAAGATGCCgaatcttaaacccaac
ggttacctcttatctagtggTTCTTGCACAACCTGTAGAACCATGAGCTAATTAAACCTCgaatcttaaacccaactt
ggttacctcttatctagtggCGCAAGAGCTTGGGTTCAGCCTGTGGCAGTGGGAGTGGCCgaatcttaaacccaa
ggttacctcttatctagtggGTCCCAGAGGCATCCAGAAGGAGGATTTGGTGGCCAACTGgaatcttaaacccaa

ggttacctcttatctagtggCCAGAGGGAGCACAGGAGGTGTGGGAGCAGGTAAGATGGGgaatcttaaacccaa
ggttacctcttatctagtggGAGATTTTGGCTTTGAGTTGCCATCAGGGAATTGTCCAGCgaatcttaaacccaactt
ggttacctcttatctagtggTGAGCATACACTACATCCTACAGAATGTTCTCAGCATTTCgaatcttaaacccaactt
ggttacctcttatctagtggCCTGTCTGCCTTCTCCTCACCATAGCCTTCTCCCACCTCCgaatcttaaacccaac
ggttacctcttatctagtggCCCCATCCAAGCCTTCATCCCACCTCTACCTCCTCCCTGCgaatcttaaacccaa
ggttacctcttatctagtggCTCCTCTCCCATCCTTTCCCACTTCTGGCTTCTCCCTTCCgaatcttaaacccaact
ggttacctcttatctagtggCTGAGGCAATGGATACTTTTGAGAACTGAGAGGTAATCTGgaatcttaaacccaactt
ggttacctcttatctagtggGGTCCACCCTTACCCACAGGGGCTGTGTTCCAAGACCTTCgaatcttaaacccaa
ggttacctcttatctagtggCAGATAGCACCAAACCCTATATATACTGTGTTTATGTCTAgaatcttaaacccaacttg
ggttacctcttatctagtggGACAGCTGCTAAGTGACTGACAGGCAGGGAGCATCTACAGgaatcttaaacccaa
ggttacctcttatctagtggTCAGGACAAAGTGGGACAGTGCGAGATTTCATCACGCTGCgaatcttaaacccaac
ggttacctcttatctagtggTGTGAGAGGTGCTGTTCTAGGTGCTGAAGGCAGAGCTGTGgaatcttaaacccaac
ggttacctcttatctagtggCAAGGAGGAATGGGGTAGCTGCTGTGTCAAGCAGTGAGGCgaatcttaaacccaa
ggttacctcttatctagtggGAACAGAAAGCAAGTTTTCTCTTTCTGACCACTGCCCCCCgaatcttaaacccaac
ggttacctcttatctagtggTCCTTTGTCTCTTGGATGAACTTTCAGACAGTTTCCATGCgaatcttaaacccaacttg
ggttacctcttatctagtggGACCACCACTGCCATTTTCCCTTCAATTACTGTGCCTAAGgaatcttaaacccaact
ggttacctcttatctagtggAAACCAAACACCTAATGTTCTCACTCACAGGTGGGAATTCgaatcttaaacccaact
ggttacctcttatctagtggACACTGGGGCCTGTCATGGGGTGGGTGAGGGGGAGATAACgaatcttaaacccaa
ggttacctcttatctagtggGCTGTCCCCACTGTGTATGTGAGGGTCAGTCCACTGGGGCgaatcttaaacccaa
ggttacctcttatctagtggCCCGTGGCTCTGGAGTACCTGCCAGACTTGTGCCCTTAACgaatcttaaacccaac
ggttacctcttatctagtggGGTACACAACCTGTCTGAGCCTGTTTTCTCATCTGCAAAAgaatcttaaacccaactt
ggttacctcttatctagtggTGATGCCTTCCTTGCAGGTTTCCTGTGAGCATTCAATATGgaatcttaaacccaactt
ggttacctcttatctagtggCTGAGCACTGCTCCATCCACTGCCAGCGCTTAGTAAGGGGgaatcttaaacccaa
ggttacctcttatctagtggCCAAGAAATGGGAACTCCTGAATAGGATGATTGGCTTCTCgaatcttaaacccaact
ggttacctcttatctagtggGGCAGCCCCAGAACTTAGCATAGCATAGTATCGTACACTCgaatcttaaacccaac
ggttacctcttatctagtggAGGAATAAGGGAACCCCAGTTTGTATGCCTCCAGCCCCTGgaatcttaaacccaa
ggttacctcttatctagtggGGTGTTTGTCTGAGGGATCAAGAGGGCTGGGGGTGCTTCCgaatcttaaacccaac
ggttacctcttatctagtggACAGGCAGGTGCCAGCTGGTGATTTCTTGTCCTACACAGGgaatcttaaacccaac
ggttacctcttatctagtggATGTCTCAGCTGGACATTCACCTACTCATTCATTCACTCGgaatcttaaacccaactt
ggttacctcttatctagtggACCTGGGAGCAGAGGAGACAAAAGGCAATGAGACATGGGCgaatcttaaacccaa
ggttacctcttatctagtggGAAGCTCACAGTCTACCAACAGACACAGCCTGTAAATAAGgaatcttaaacccaac
ggttacctcttatctagtggGAGGTCAGAACTACTTTGGGGGCAAGAACAGGGGTCGGTGgaatcttaaacccaa
ggttacctcttatctagtggAGGCTGGAGCTGGCTGGGACCTCATTGAGTGAACAGGATGgaatcttaaacccaa
ggttacctcttatctagtggACCAGCTGAGAAAACAGACAAAGGCACAGAGGCATGGCTGgaatcttaaacccaa
ggttacctcttatctagtggAAGAAGACCACAAACAGCCAGGTGTGATGGTGCATGTCTGgaatcttaaacccaac
ggttacctcttatctagtggAGGCTACAGTGAGCAACAATTACACCACTGCACTCCAGCCgaatcttaaacccaa
ggttacctcttatctagtggCCTGGGAAGAGTTAGGAGCAATGCAGCTGGAGAAGGCAGCgaatcttaaacccaa
ggttacctcttatctagtggGCCTCATGGGCTTTATCCTGAAGGCAGCTGTGGACCACTGgaatcttaaacccaac
ggttacctcttatctagtggAGGTGAGCCTTTCTGATCCATGTATGAACTCATCAGGCCCgaatcttaaacccaact
ggttacctcttatctagtggTCTGACTCTCAGCAGCATTTGTCACTGGAGGCCATCTCCCgaatcttaaacccaac
ggttacctcttatctagtggGCGCGAGCTTGATTCTCCTCCCACCTCTCAGATCGTGTCTgaatcttaaacccaac
ggttacctcttatctagtggCCTGTTTCACCAGGAATTTCCAAACATGCATGTCTCCCAGgaatcttaaacccaact
ggttacctcttatctagtggCTCTGCTCTGTCTCTTGGGAATCTCAACCACACCTGCAGCgaatcttaaacccaac
ggttacctcttatctagtggGCGTGAGCCACCGCGCCCAGCCGCTTCAACATTTTAAAGTgaatcttaaacccaa
ggttacctcttatctagtggTCCAACAGGGTTGTATGTATGATAGGGAGTCCCACAATGCgaatcttaaacccaact
ggttacctcttatctagtggAGTCTTTCCAGCAGAGCATATTTGTAGCCATATGTGAGAGgaatcttaaacccaactt
ggttacctcttatctagtggGAGGTAAAAAGTAGCGTGGAGGCCGGGCGCGGTGGCTCACgaatcttaaacccaa
ggttacctcttatctagtggCAACCCTGTGCTGGGCACTATGGAATATACAAAGGCAACCgaatcttaaacccaac
ggttacctcttatctagtggCACCAGTGAGAACAGCAAGGGGCAATAGGACATTGTGACCgaatcttaaacccaa
ggttacctcttatctagtggCTTTGGGCTTGACAAACTTAGGTTTGCATTCTTGCTCCTCgaatcttaaacccaacttg
ggttacctcttatctagtggATGGGGAACATAGTACGTCCCTCATAGACTGGTTTCAAGGgaatcttaaacccaact
ggttacctcttatctagtggACATAATGCCTGGGATGTTGCAAGGACTCAATTCATGTGGgaatcttaaacccaactt

ggttacctcttatctagtggTCAGCCACTGTGAAAAAGTTTGGCAGTTCCTCATTAGAAGgaatcttaaacccaactt
ggttacctcttatctagtggGCAGCATTATTCACAGCAGTCAAAAGGTGACCCAACTGTCgaatcttaaacccaact
ggttacctcttatctagtggTGAAGTCCTGATATATGTCACAACATGGAGGGATCCCAGGgaatcttaaacccaact
ggttacctcttatctagtggGTCCAGATGGCAAATCCAGACTGGGGAGAACACCACAGGGgaatcttaaacccaa
ggttacctcttatctagtggGGAGACTAGAAGGCACTGCTTAATGGGGACAGGATCTCCTgaatcttaaacccaac
ggttacctcttatctagtggGAAGTGGTAGTTGCACAACATTGTGGGTGTACTAAATGCCgaatcttaaacccaactt
ggttacctcttatctagtggGCACATCATGCTTTTCCTGGTCCTAGGCTGGTCCTGGCTTgaatcttaaacccaact
ggttacctcttatctagtggGGGGGCAAGGCAGGACCCCTGAGCTCCTCTGTAGTCCTCAgaatcttaaacccaa
ggttacctcttatctagtggGCCTTTATGGAGCTTATAGATGGTAGGAGTGTTCCTAAGCgaatcttaaacccaactt
ggttacctcttatctagtggAGTTAACACCTTGTACCCACAATCTGCAACCTCTGTGCAAgaatcttaaacccaact
ggttacctcttatctagtggATGCCAGCCTCAGAGTTGAGGTGAGGATTAAAGAGGGATGgaatcttaaacccaac
ggttacctcttatctagtggCCACCACAAATTTCCAGCAAGTAGGTGGCTTACAGCAACAgaatcttaaacccaac
ggttacctcttatctagtggAATCCTTCCTGGCCTCTTCCAGCTTCTAGTGACTCCAGGCgaatcttaaacccaact
ggttacctcttatctagtggTGGCTGCAATCTTTGCTCTAACTGCACAAGGCCTCTCCTCgaatcttaaacccaact
ggttacctcttatctagtggCTCCTTCTACCCGTCTCTTTCACCCAGGCTGGTCTACCTGgaatcttaaacccaact
ggttacctcttatctagtggAGCACATTTATCATTGGATGTAAGGCCTAGCCAGATAATCgaatcttaaacccaactt
ggttacctcttatctagtggTGGGGGCGGGGAGCTACCATTCAATTCAACCCACTGTTGAgaatcttaaacccaac
ggttacctcttatctagtggGGACAGAGTCTAACTGAAATCTTCATCTCACTGGGGCTCCgaatcttaaacccaact
ggttacctcttatctagtggGCTGGAGCCTTTTCTCGTCTCTCCCATTCCCTCGACCCCGgaatcttaaacccaac
ggttacctcttatctagtggCACTACCAACCACCTGAAGCCCCCCATTTCTGCGCCCACCgaatcttaaacccaa
ggttacctcttatctagtggCAGGTGCCCCTACAGAGCTGACCCTCCGCAGCCCAGTGGCgaatcttaaaccca
ggttacctcttatctagtggGGGTCTTGCTCCCTCTTGGTTGAAGCCGAGGGAACAACACgaatcttaaacccaac
ggttacctcttatctagtggGGGAGCTGATAAAAAGCAAAGACAGGGCTGGGCGCAGTGGgaatcttaaacccaa
ggttacctcttatctagtggGCAGTGAGCCAGGATCATACCAACGCACTCCAGACTGGGCgaatcttaaacccaa
ggttacctcttatctagtggCCTAGCACTTTGAGAAGCCAAGGCTGGAGGATCACTGAAGgaatcttaaacccaac
ggttacctcttatctagtggGGGAGGCAGAGGCAGAAGTATAGCTTGAGCCCAGGAGTTCgaatcttaaacccaa
ggttacctcttatctagtggCCAAATCCAGTTAAATCAGAATTTCTGGGGATTGGGACCCgaatcttaaacccaact
ggttacctcttatctagtggAGATAGATATTTTATATCAAAATATGGGGTGGGTAGGCTCgaatcttaaacccaacttg
ggttacctcttatctagtggTCCAATGGGCAGCCAAGATAAAGAATGCCTACCCTGGGGCgaatcttaaacccaa
ggttacctcttatctagtggGGCATTGCTCCATTTCTAAGCACTGGAAATTTTCAAGATGgaatcttaaacccaactt
ggttacctcttatctagtggGGAGCAGAAAGAGTTTTATGGGGGAAAGCCTGGGTCTTGGgaatcttaaacccaac
ggttacctcttatctagtggGCTGTGTGGCCCTGACCACGTTCCTTAATCCTTCTGAAGCgaatcttaaacccaact
ggttacctcttatctagtggGGAAGACTCAAATTCCTTCTCCTTCCCCTCCTCCCAGAAGgaatcttaaacccaact
ggttacctcttatctagtggGTGTACCTTCTACCAAAGGTTTGTTGGTCCCAGGAAAGGGgaatcttaaacccaact
ggttacctcttatctagtggGTGTTCGCTAGATTAGAGGCAATTCCAGAAATCCACTCCCgaatcttaaacccaact
ggttacctcttatctagtggGCCGATTTGGCTGTGGCTTCGGGCTACTTGATGGTAATTTgaatcttaaacccaactt
ggttacctcttatctagtggAACTGCAGCACACAGAGATTTGGCAGGCTACAAAAGCTGGgaatcttaaacccaac
ggttacctcttatctagtggGTTCTGGTGAGCTTTTTGGCAGAATCCTCCCTGTTTTCCCgaatcttaaacccaactt
ggttacctcttatctagtggCCAGAAGGCCATGTGTCCAAGCTGCTTGATCTTCCTGAGCgaatcttaaacccaac
ggttacctcttatctagtggAGCAGACTCCCCTGTGACACCGGAAGCCAGAACCCAACAGgaatcttaaaccca
ggttacctcttatctagtggGCTCCCAGCATTGTCTCTGCATTTTGCCTTTTGCAGGAACgaatcttaaacccaactt
ggttacctcttatctagtggGCAGTTTGCAAACACTGCTATTGCAATTCTCCTCCCCGATgaatcttaaacccaactt
ggttacctcttatctagtggATTGACACGGTTTCACAGAACTGGGGAGGCCTCAGGAAACgaatcttaaacccaac
ggttacctcttatctagtggCTTTACGTGATGTCAGGAGAGAGAAATGCCAAGCAAAGGGgaatcttaaacccaac
ggttacctcttatctagtggCCTCCCACTGAATCCCTCCTATGACACTTGAGGATTATGGgaatcttaaacccaact
ggttacctcttatctagtggGGACACAGCCAAACCATATCACATGAGCTAATAAAGACAGgaatcttaaacccaac
ggttacctcttatctagtggTCCTCATCTCTTTGTCCCTGGTTCCCAGCAGAATGTCTGGgaatcttaaacccaactt
ggttacctcttatctagtggGGCACGGTACTTTACCCCTTGTGCATACTTAGTGCCTGCCgaatcttaaacccaact
ggttacctcttatctagtggATTAAGCTTATTTAGAGCCCACCCACCCTAATGAACTGGGgaatcttaaacccaact
ggttacctcttatctagtggGCAAAATTCCAGTGGGCCATGCATTCTCAGATATCCTGAGgaatcttaaacccaact
ggttacctcttatctagtggGGTGATTCTGATGCCAGAGTTGAAACCAGTCCAACCTCCCgaatcttaaacccaac
ggttacctcttatctagtggATACCCAGCCACCTTACCAGGACCAACAAGGACCCGTAGGgaatcttaaacccaa

ggttacctcttatctagtggGCCTAGTGCTCTGTCCCTCTGCTCACTACACCAGCCACACgaatcttaaacccaac
ggttacctcttatctagtggCTGCTCCTGAGTGCACCAAGCTACTTCCAACCTCCAGGCCgaatcttaaacccaac
ggttacctcttatctagtggCTGGTCCTTCTGATCACTCAGGTCACACCTCAGAGGTTATgaatcttaaacccaactt
ggttacctcttatctagtggCTAGCCACCCAGTAAAGCACAATCTATTACACACACACGCgaatcttaaacccaac
ggttacctcttatctagtggTACATTCCCTTTTATTGGGACAATGGCACCCATCACTAGCgaatcttaaacccaactt
ggttacctcttatctagtggTCCCGGCATCTTGTGAGTCCGAGGCAGGAGGATCACGAGGgaatcttaaacccaa
ggttacctcttatctagtggGAAAACAAGCAGAGATCTTTCATGTCTTTTTCACTACTGCgaatcttaaacccaacttg
ggttacctcttatctagtggGCCCCTAGTGATGGAGGGAGGCATCTCACCCCCGTTCAAGgaatcttaaacccaa
ggttacctcttatctagtggCCCAGTGCCAAGCTGTCACCTGCCTCCCTCTTGCCTCCTGgaatcttaaacccaac
ggttacctcttatctagtggAATTCCCTGTAGCTGCCCCCAACAAGCCTGGTCCTCCTGCgaatcttaaacccaac
ggttacctcttatctagtggTCCTCTCTGCCTTCTATTGTCCCAGTGAATGGCTTCCTGGgaatcttaaacccaactt
ggttacctcttatctagtggTCCAGGACAGCTGCCGACACACTGCCAGGGTGGGGCTTGCgaatcttaaacccaa
ggttacctcttatctagtggCAAGGGTTGTCTTTACAGGCCGTGAGAAATATGCATCTGGgaatcttaaacccaactt
ggttacctcttatctagtggGGTGTGGGAGAAGAAGGAAGTCCCTACAGCAGCTCCCTGCgaatcttaaacccaa
ggttacctcttatctagtggAATAGCACATACGTCCCGCCACCACCACATTGTCGGGCACgaatcttaaacccaac
ggttacctcttatctagtggGTTTCCAAGGCTTCATTTTCTTTACGGCTCATGCTATTTGgaatcttaaacccaacttgt
ggttacctcttatctagtggCAGGGAGCTGTCTGTCTTACATGGTATCTATCCTCAGTAGgaatcttaaacccaactt
ggttacctcttatctagtggACCTCAGTTTCCTCTTCTTTAGAATGGGATAGTGGGGAGGgaatcttaaacccaactt
ggttacctcttatctagtggCACTAGGTTGTTGTATGGATCAAATCTGAGGATACCTGGGgaatcttaaacccaactt
ggttacctcttatctagtggATTCCTGGTCATTGTGTCACACACTGACATAGGGCTGGACgaatcttaaacccaact
ggttacctcttatctagtggACTATCCCTTCAGTGTGACTTCAGGGGCCCAAATGACCCCgaatcttaaacccaac
ggttacctcttatctagtggGAACAGAAATGCTTTCCTCAGTTTGAAGATGGGAGGGCAGgaatcttaaacccaact
ggttacctcttatctagtggCGTCATCTCAGCTACTCTCCTTTCAGCCCCCGAGTCTTGCgaatcttaaacccaact
ggttacctcttatctagtggCCAGCCTGGCTTAGGACTTCGGAGACATTTTCCATCTGGCgaatcttaaacccaact
ggttacctcttatctagtggGCAGGGGAACTTCAATGTGCAAGTGTTTCCTGTCCTGGTCgaatcttaaacccaact
ggttacctcttatctagtggCTGGACTCCGAAGATGAGACTCAAGGCAAGGCTTTGGCCCgaatcttaaacccaac
ggttacctcttatctagtggGCAGAGGTGGGGTATCCTCCATGCCTTTAGGAGTGGCGGCgaatcttaaacccaac
ggttacctcttatctagtggCTCCTCACGGCAATGCCTACCCTCTGGCCAACGCTTCCACgaatcttaaacccaac
ggttacctcttatctagtggCCCTCCCTTCCTCTATCACATTCCAACCTTGATGCAGTTCgaatcttaaacccaactt
ggttacctcttatctagtggCAGGGGTTCAGCTCCTGCTCTGAACTGGAAGCATATGTAAgaatcttaaacccaact
ggttacctcttatctagtggAGGGTGCTGCACAGTGTTAGAGGCATCTGTAACCAGGGTGgaatcttaaacccaac
ggttacctcttatctagtggGGAATGTCTGGGAATGCTTTTGAGAGGAGGGGGTAAAGTGgaatcttaaacccaact
ggttacctcttatctagtggAACCCAGAACAAAAGGGAGGAAGGGCATTGAAACTTGGAGgaatcttaaacccaac
ggttacctcttatctagtggCAAGGAGGTTGCTTTGGCTGGAGGGGAGAATGAGTGAGCCgaatcttaaacccaac
ggttacctcttatctagtggAGCAAAAAGGGTCAGTTGATGACAGCCTAGTAGAGTAGGCgaatcttaaacccaac
ggttacctcttatctagtggTGTCATATCCCTTTTATCATAAAGAGAGAGCCACTGAAGGgaatcttaaacccaacttg
ggttacctcttatctagtggTAGAAGAGTGAAGAACAGGCTAGGCACAGTGGCTCACACCgaatcttaaacccaac
ggttacctcttatctagtggGCAGTGGCAGTTAAGGGGATGGGAGGGCGCTCAGTTTGGGgaatcttaaacccaac
ggttacctcttatctagtggGGCAGATGGACCTGGATCTTTACTTTGGCAGTGCCTTTGCgaatcttaaacccaactt
ggttacctcttatctagtggGGCCATGCAAGGCAACAATCTTTCAACCCAGTCACTTGTTgaatcttaaacccaact
ggttacctcttatctagtggACAGAGCCAGACTCACAGTAGGTCCTCAATGATGATAGCCgaatcttaaacccaac
ggttacctcttatctagtggTCCTGAGGGAGAGAGGACCATGATGCTGCATCTGCCAACCgaatcttaaacccaac
ggttacctcttatctagtggCCAATGCCGAGTACAGCGGTTATAGACTATCCAACGTTGTgaatcttaaacccaact
ggttacctcttatctagtggCAGGAGCCTGAATGACCGTCAGCCCACAGGTATTTGGGGGgaatcttaaacccaac
ggttacctcttatctagtggAGGTAACACTTGACCTCAAGAAGTTTCCCCTCTGGCTGGGgaatcttaaacccaact
ggttacctcttatctagtggCCCTAGTAACCTTACCCTTCTTCCTGAAACTCACCTCGCCgaatcttaaacccaact
ggttacctcttatctagtggCTGCACATAGCTCCCTGAGCCACACACCCAGATCCATGGCgaatcttaaacccaa
ggttacctcttatctagtggCCTTCCAGGTAACAGCTCCTGGGTACCCCCACACACAGGCgaatcttaaacccaa
ggttacctcttatctagtggCCCATTGTTTTCCCTTCAAACCTATTCCTCCTTCAGGCCGgaatcttaaacccaactt
ggttacctcttatctagtggGCTGCCCAAGTTCTGTTTGCTAAGTGATTGTAACACAGTGgaatcttaaacccaacttg
ggttacctcttatctagtggTCAGATTCCAGTTCTTCTCATTTACCCCGTGACCTGCTGGgaatcttaaacccaactt
ggttacctcttatctagtggCTCTGTGCTTCACTTTCCTCACCTATAAAACAAGTACTTCgaatcttaaacccaacttg

ggttacctcttatctagtggGCATCAAACAGTGTCTGGCACATCATAAGTAATATAAAGGgaatcttaaacccaactt
ggttacctcttatctagtggCTCCTACTCCCTCAAACCTCTGCAAATGGTTTGCCCTCACgaatcttaaacccaact
ggttacctcttatctagtggCAGGAATCTCACTTCTCCATCCCAATTGCTGCTGCCTGTGgaatcttaaacccaact
ggttacctcttatctagtggCCCAGTTCATGTCCCCAGAGTTGAGGCCCTGGTGACTTCCgaatcttaaacccaac
ggttacctcttatctagtggGCTCTCAGATTCTGTGGTTCCAGAACCCTGGTGTCCACTCgaatcttaaacccaact
ggttacctcttatctagtggGTCAGCGAAGTTTCCCCCTCCACACTCACACTCCCCACCCgaatcttaaacccaa
ggttacctcttatctagtggTTAGTCCACAGACAGGTAGGAAGCCACAAGCAGCCTGGCCgaatcttaaacccaa
ggttacctcttatctagtggGACCTCTGCACCCAGCTCCTGTCCCTCCACTCCCTACGGCgaatcttaaacccaa
ggttacctcttatctagtggAGTTCCAAAAACCCTGCCTCTGTGACCTGATTGTGTTATCgaatcttaaacccaactt
ggttacctcttatctagtggGCAGCCTTCTCCAAAGCCATGATTTCCTATAATGGGGGTGgaatcttaaacccaact
ggttacctcttatctagtggGAGCAGTGGTTCTCAACCAGAGGCGATCTTGCTCCCCAGGgaatcttaaacccaa
ggttacctcttatctagtggGGGGGATGAGTGCTGAAGGCATCCAGTGGATAGTGGCCAGgaatcttaaacccaa
ggttacctcttatctagtggGGCTCAACACATCAATAGTACTGAGGTTCAGAAACGCTGGgaatcttaaacccaac
ggttacctcttatctagtggATAAAGAGTGAGGCACCATGGCATTTTTGCTCTGCTGCTCgaatcttaaacccaactt
ggttacctcttatctagtggCGTATCCACACTTCAAAGAGATGTCAAAAATCCGTCATGGgaatcttaaacccaact
ggttacctcttatctagtggAGGTCCTGGCTGGGTAGCTAAGCAGCTGGCATGTTAGGGCgaatcttaaacccaac
ggttacctcttatctagtggGGGCTTATCTACAAAGTGGGCAGAGGGGCTTAGGGTTCCCgaatcttaaacccaac
ggttacctcttatctagtggAGCAAGAGCTGGAGGACCTCAGGGAGGAGGGCAGGTGGGCgaatcttaaaccca
ggttacctcttatctagtggAGAGATATCTGAGCCCAGGACAGCAAGGGGTGAACAGAATgaatcttaaacccaac
ggttacctcttatctagtggGGTGGGAAAGAGGGTTTTAAGAGAACCACACAAGCAAAGCgaatcttaaacccaac
ggttacctcttatctagtggGACCCATTGTAGTTGCTCATAAATGCCGGCTCCTTCACCCgaatcttaaacccaact
ggttacctcttatctagtggCTCTACTGGGACCCAACCCTAGTTAGACCACCTGACAGCGgaatcttaaacccaac
ggttacctcttatctagtggCGGAAACCTTAGGGCCTTCAGTGATACTTCAGAAGGGGAGgaatcttaaacccaac
ggttacctcttatctagtggGAGTGATTCAACACATTTTCTGGTAGGAAAACCCCTTTCCgaatcttaaacccaactt
ggttacctcttatctagtggCCCGTATACCCCCAGCCCCCACAAATTCCCAGCTTCCCCCgaatcttaaacccaa
ggttacctcttatctagtggCACATCATTATCACCCAGAGCCCATTATCACCGAGAACCCgaatcttaaacccaac
ggttacctcttatctagtggCTTCTGTCCTCTACCTCCCTCCCTCCTGACCCCTGCACACgaatcttaaacccaac
ggttacctcttatctagtggTTACTGTCTCCATAAGTTTTGCCTTTTCCAGAATGTCATAgaatcttaaacccaacttgt
ggttacctcttatctagtggTGGATTGTTTCACTTGGTAATATGCACTGAAGGTTCTTCCgaatcttaaacccaacttg
ggttacctcttatctagtggGCATTTTCTGGATGGACCACAGTTTATGTATCCATTCACCgaatcttaaacccaactt
ggttacctcttatctagtggGGCCATGGACATGGGAGGCAACGTGTGTTGTGGTCTCTGGgaatcttaaacccaac
ggttacctcttatctagtggCTGGAGAGATTCTGCAGGCTCCTTGGACTCTTGGCTCTTAgaatcttaaacccaact
ggttacctcttatctagtggGATTCAAGATGATAGCCCTGGGTCCCCACCTGCCTGCTGGgaatcttaaacccaac
ggttacctcttatctagtggAGGGGCACGTCACATCCACCCAGAAGTACCCATCAGTACGgaatcttaaacccaa
ggttacctcttatctagtggGCTAGACCTGAGGCAGTGAAGCATGATGACACAGACCCTGgaatcttaaacccaa
ggttacctcttatctagtggGTGAGTAGGGTGACTTATGACCCCTGATCTCCTGGCTGCCgaatcttaaacccaac
ggttacctcttatctagtggCAAGTGGCCTTGAAATATCATTCAACGTCTCTGAGTGTTAgaatcttaaacccaactt
ggttacctcttatctagtggTCTGCACATCTCTCAGTCGGCTGCCTGGGACATGGTGAGCgaatcttaaacccaac
ggttacctcttatctagtggTACCTACTGATGTCGCCAGGCTAAGGCAGAGCCCTGGGGCgaatcttaaacccaa
ggttacctcttatctagtggAAAACTCTGGGACAAGTGGATCTTTTGCCTGACCTCTACCgaatcttaaacccaact
ggttacctcttatctagtggGGTGCTTATCCACCAGTACATGCCCTTCCTTCTCCCAACCgaatcttaaacccaac
ggttacctcttatctagtggTCAAAATAAACCTTTTTCCTAGAAGCCAGCTGACCCTTCCgaatcttaaacccaactt
ggttacctcttatctagtggCAGTGGTCAGGACACCTGAGCCTCCTTAACTGCAGCTCAGgaatcttaaacccaa
ggttacctcttatctagtggGGCAGACCTACAAAGGGACTAAGATTTAGCCTGAAGCTGCgaatcttaaacccaac
ggttacctcttatctagtggACCTTTGCAATTGTTTGCGTTTCAAATTCAGTTTTCCAAGgaatcttaaacccaacttgt
ggttacctcttatctagtggGAGCAGTGGGTCTCCCAAGGCCTAGGCTAATGGAGCACTCgaatcttaaacccaa
ggttacctcttatctagtggGATGAAGTCTGCCCCCCTTTCTGCTTGAGATTCTTGTACCgaatcttaaacccaactt
ggttacctcttatctagtggGCCTCTGGGCTTCCTTTGTCATGGTACCAAGGTCCCCTGAgaatcttaaacccaac
ggttacctcttatctagtggCACCTGGTCAATGGTCCTTAGAAAGCCACTGAAGAGGCCCgaatcttaaacccaac
ggttacctcttatctagtggGAGGGCAAGCATCTTAATGGCAGCTGACAGTTTGGTTCTGgaatcttaaacccaact
ggttacctcttatctagtggCAGAGAAGGTTGAAGGACACTCTCAGCACATGTGAAGAGGgaatcttaaacccaac
ggttacctcttatctagtggCACGGCTTCCTAACCCCGGGAGAGGAAAGATAGAAGCCAGgaatcttaaacccaa

ggttacctcttatctagtggCCTTCACCTACAGGGTCAGGTCCAAATTCTTCAGCCTGGCgaatcttaaacccaac
ggttacctcttatctagtggTCAGAGTCTGACCCCAATGTAGGGAACAACTTCTGATGGCgaatcttaaacccaac
ggttacctcttatctagtggGCACCCAAAGTGGGGAAGCCAAGTTTGAAGTAAAACCTTCgaatcttaaacccaac
ggttacctcttatctagtggAGCATGGGCTCCTGGTATTATGCCACCCCTGCCTCTTCCAgaatcttaaacccaac
ggttacctcttatctagtggGAGACACACTCTGCAAGGCTCCTTTAATCACTTTATACACgaatcttaaacccaactt
ggttacctcttatctagtggGCTGAGATGCTGTGTCCCTCTTTGCATTCATCTGCTGTCCgaatcttaaacccaactt
ggttacctcttatctagtggCTTCCTCAGATCATCTTCCCTTTCTGTCCTCTCCTGAGAGgaatcttaaacccaactt
ggttacctcttatctagtggGCCTACTAAACAAGGCTCAGAGGGGATGTTACACACACACgaatcttaaacccaac
ggttacctcttatctagtggCATACAGAGCGAGAGAGGGCAAATCCCAAAGTCAGAGTGCgaatcttaaacccaa
ggttacctcttatctagtggAAGCCCGACCCCCTGACTTCCTGACTTGAAAGTCAACTCCgaatcttaaacccaac
ggttacctcttatctagtggGGAGCCTTCCATAGTCAACAGACTGGGAAAGTTGGGAAGCgaatcttaaacccaac
ggttacctcttatctagtggGCCTGGAGGTGGGGGGTAAATATTTGGGAAGGAATGAGGGgaatcttaaacccaac
ggttacctcttatctagtggCCTCAAAGACATCTCAGTTTGGGTGATCATACCTAGCTGCgaatcttaaacccaact
ggttacctcttatctagtggTAAAACAATGCAAGGGGCAGGGCGTGGTGGCTCAAGCCTGgaatcttaaacccaa
ggttacctcttatctagtggTGGGAGGCCAAGATAGGCAGATCACAATGTCAGGAGATCGgaatcttaaacccaac
ggttacctcttatctagtggCTGTACTGTATGTGATCTGTGTGTGATTCAGGGGCCCAGGgaatcttaaacccaact
ggttacctcttatctagtggTCCTTCACAAGGATCTTTGAACCCTCTGCACTCAGACTGCgaatcttaaacccaact
ggttacctcttatctagtggTGAAGAGGATTGCCTGGAGGGTTTTTCTGGGCTAGGCCTGgaatcttaaacccaact
ggttacctcttatctagtggCCTGGCCACACCTAATTGCCTGGAGACTCGGACAGGAAACgaatcttaaacccaa
ggttacctcttatctagtggGCCTGCCAAACACACTGAGCAGCCTTGAGGGGAGGCCAGGgaatcttaaacccaa
ggttacctcttatctagtggTGCCCGGAACAAGAGCAATTTAGGCCCCCAGGAGCCAGAGGgaatcttaaacccaa
ggttacctcttatctagtggAGACCAGCGAGGCCAGAGGCAGAAATCAAAGTTGAGTTGCgaatcttaaacccaa
ggttacctcttatctagtggGGGGCTATGGGAAAGGGTCTAAGAAAAATTGGCCCAGACCgaatcttaaacccaac
ggttacctcttatctagtggAGTATGTCTGTGGATTGGCTTGGCTTAAAGAGGCAGGAGCgaatcttaaacccaact
ggttacctcttatctagtggAGTCTTAACCCCCAGTACTCCAGAACGTGAACTTACCTGGgaatcttaaacccaact
ggttacctcttatctagtggCTGGAGTAAAATAGACCCCTAATTCAACATGACTGGTGTCgaatcttaaacccaactt
ggttacctcttatctagtggGGCAGAGGTTGGGGTGACGAGTCTACAAGCCACAGCATGCgaatcttaaacccaa
ggttacctcttatctagtggAGCAGAAGCTAGGCAGAGGGCATGGAATAGGTTCATCCTCgaatcttaaacccaac
ggttacctcttatctagtggGGAGCATAGCCATTCAACATCTTGCTCTTGGACTTCTGTCgaatcttaaacccaactt
ggttacctcttatctagtggGTCCAAGCCACCGAGTTTGCAGTACTTTATTCTACTAGCCgaatcttaaacccaactt
ggttacctcttatctagtggGAGGCAGCATCAAGGGGAATATTGCATCAAGGGACAATGGgaatcttaaacccaac
ggttacctcttatctagtggCTGTTCACACCCCATGAATCCAGCTTGTTCAGCACCCTGGgaatcttaaacccaac
ggttacctcttatctagtggTTATAATCTGGGCCTTTGAGTGAGTGAAAGGACACGTAACgaatcttaaacccaactt
ggttacctcttatctagtggAAGACACATGCACACGTGTTTTTATTGCAGCACTATTCACgaatcttaaacccaactt
ggttacctcttatctagtggGACTGGCTTAAGAAAATGTGGCACATGTACAGTGATAGACgaatcttaaacccaact
ggttacctcttatctagtggGTCCTTTGCAGGGGCATGGATGAAGCATCATTCTAAGCAAgaatcttaaacccaact
ggttacctcttatctagtggCGAGAACACATGGACACAGGGCGGTGGGGGGCTGGGGGAGgaatcttaaaccca
ggttacctcttatctagtggAGGAGAAATGCCAAATGTAAATGACCAGTTGATGGGTGCAgaatcttaaacccaact
ggttacctcttatctagtggGTTCCAAGGAGAGAAGGTTGGATGTGATCAGGGATGGAGGgaatcttaaacccaac
ggttacctcttatctagtggAGCCTGGACCTTGGAAGCTGAGTTCATCTTAGGAATGAAGgaatcttaaacccaact
ggttacctcttatctagtggCACCCCCATGGTGCTATGCAGTACCCCCTCACTCATGGAAgaatcttaaacccaac
ggttacctcttatctagtggAAGCCAGAGTGTTGAACAAGCTGGGCTCGTGGGGTGTGAGgaatcttaaacccaac
ggttacctcttatctagtggGGTCATCCTCTGGCCTCAAATGAAAACTTGTACAATCCAGgaatcttaaacccaact
ggttacctcttatctagtggAGTCAGTGTTCCTTTATATCATGAAGTATCCTGCATTCAGgaatcttaaacccaacttg
ggttacctcttatctagtggGATTCCAAGGTCAATCTTCTCCTGTTCCCTGGGTAAGGCCgaatcttaaacccaact
ggttacctcttatctagtggGCCTGGAGCTCAGACTTCTCCTTCATTGAGTTACTAAGTGgaatcttaaacccaactt
ggttacctcttatctagtggGGAGTTGGAGAAGGGTAGAAATTAAGGCAAGGGGCACATGgaatcttaaacccaac
ggttacctcttatctagtggTGCCTGGTATTCTTGTAAGCTGGCTCTATATTCTTTGAGCgaatcttaaacccaacttg
ggttacctcttatctagtggGGGGGTCCCTCCAAAATTGCTCTGACCTACAGCTCCCTTGgaatcttaaacccaac
ggttacctcttatctagtggCTCCCAGGAATGTGGCCATAGCTCCAGCTGAGATCTGGGCgaatcttaaacccaa
ggttacctcttatctagtggACGGAACGGACAGACTACAATCCCACTCTGACTCTCTTGCgaatcttaaacccaac
ggttacctcttatctagtggCCCTGAAAAGCTCTCAGAGGACAAGCTGCACAGTGTAGCCgaatcttaaacccaa

ggttacctcttatctagtggGTCTCGGGTGGGAGTCAGGCTTAAGTCCACTGCAACCTGCgaatcttaaacccaac
ggttacctcttatctagtggCACTCTGCTCTCTGAGTGGTCTTGCTGAAACCCTTCTAGGgaatcttaaacccaact
ggttacctcttatctagtggCACTGAGCTCAGTGGAGAAGCAACCTCTACCAGGAAGTCCgaatcttaaacccaac
ggttacctcttatctagtggCTTGCCTCCTGACACTTTCCTATCCTGGTGTAGGTGAGGCgaatcttaaacccaact
ggttacctcttatctagtggCTCTTGACCACTGACTTTTCACAAGTTCTTCTGGGTGGTCgaatcttaaacccaactt
ggttacctcttatctagtggGGTGTGTGATAATCTGTTGTCATAGGATCTCAGCCAAAACgaatcttaaacccaactt
ggttacctcttatctagtggTCTTTACTACTCTGCAACTTTCCCAAGAGTGAAATCCCTGgaatcttaaacccaactt
ggttacctcttatctagtggCACGTTAATAGCCCAGCCTCAGATACAGTATTCAAGAGTCgaatcttaaacccaact
ggttacctcttatctagtggTCAGCTTGACTGAAGAAGAGAGTTTGTTTAGGGAAGGATCgaatcttaaacccaact
ggttacctcttatctagtggTTCAGGGACAGATGGATCCAGGTGAACACTCAGCTCTGCCgaatcttaaacccaac
ggttacctcttatctagtggAGGTTCTTAACCTCTCTGGCACTCAGCATTCTCACTGTAAgaatcttaaacccaactt
ggttacctcttatctagtggCACCTGGAATAGGGCTTATCACACAGTGGGTGGCTGACAGgaatcttaaacccaac
ggttacctcttatctagtggATTTGAGTACATCTTGTGGGTCAGGGACTGTGTTAAATGCgaatcttaaacccaactt
ggttacctcttatctagtggTTTTACACCAAGCCAATGAGACAGGGAGCAAGTTAGCCTGgaatcttaaacccaact
ggttacctcttatctagtggGTCACAGAGCTGGTGACAGGCAAACCTGGGATCTGGACTCgaatcttaaacccaac
ggttacctcttatctagtggTCCTACAGAGCCTGCTGTATGTTACCCTGACATCTCATGCgaatcttaaacccaact
ggttacctcttatctagtggAGGATCATTGTGTAGCTGGCATTAAGAGACTCTAGGAAAGgaatcttaaacccaact
ggttacctcttatctagtggAGCAACATGCTTCAGTCTCACGCTAAAAGCAATTATAATTgaatcttaaacccaactt
ggttacctcttatctagtggACCCACAGTGAGGGTCAGCTGACAGAATGTGTGAAAACGCgaatcttaaacccaac
ggttacctcttatctagtggGGCTTGAATCCATCTCTCCCCACCAAGGTCTTCCTCCCACgaatcttaaacccaac
ggttacctcttatctagtggGTGGGAATGCACCTCACCAACTTTGAAAGGCCCAAATCCCgaatcttaaacccaac
ggttacctcttatctagtggGACAGAGAAGGCCTGGGGTAGACTTTAGGGTGATTCGGGGgaatcttaaacccaac
ggttacctcttatctagtggGACACTGGGACCAAAGCAAATGTGACCTGAATATTCAACCgaatcttaaacccaact
ggttacctcttatctagtggGGATAGGCAAATGCAGGGCTGGGACACCTTTGGGGCAGCCgaatcttaaacccaa
ggttacctcttatctagtggGGGGCAGGGGAACACCTCAAATACCTCCACTGGAGCCTCGgaatcttaaacccaa
ggttacctcttatctagtggGTGTGCCAGGCACTGTGCAGAGCATTTTCTACACATGACCgaatcttaaacccaact
ggttacctcttatctagtggCAGAAGGATGTGGTGACTTACTCAAGACCACACAGCCCCCgaatcttaaacccaac
ggttacctcttatctagtggACATATTAGGAGGTTGAGAGCTAAACTGGAGGCCAAGATGgaatcttaaacccaact
ggttacctcttatctagtggAGGAAACAGATACTGGGCTCTGAAAGTGGAGTGCCCCACCgaatcttaaacccaac
ggttacctcttatctagtggGTTAGCTCTATTCCTTCAGTAAGTTATGCATGTCCTCAGTgaatcttaaacccaacttg
ggttacctcttatctagtggACCCGCAAAGCCCTTAGAACAGTGTCTGGCATGTAAGAAGgaatcttaaacccaac
ggttacctcttatctagtggGCTACAATTACTCTGAAGTGTGGGTGGGCTAGGCTGTGTGgaatcttaaacccaact
ggttacctcttatctagtggTCTTCCTGACTGCCAAGACCAGATGGTTTCCCTAGGAGCTgaatcttaaacccaact
ggttacctcttatctagtggGGAAGAAACTAACATTGGTTTACCCACAGCAAGGCTTGGCgaatcttaaacccaact
ggttacctcttatctagtggGGGATTCAAACTGGGAATGTTCTAGGCATAACAGATGGTGgaatcttaaacccaact
ggttacctcttatctagtggGGCCTCCCTGGGCAAGCCACCCTCCGGGAACCTCCATGCAgaatcttaaacccaa
ggttacctcttatctagtggGAAGTTCTCTGAATCTTCTTCTCTTGGGTTTTTATAGAGGgaatcttaaacccaacttgt
ggttacctcttatctagtggCAACTTGACCTTCAGCCCCTCTTTCCTCCCCAGAGGTTGGgaatcttaaacccaac
ggttacctcttatctagtggGTCCCCACCCTCTAATCTTGCCTTGGTCTTTTGGGTGACCgaatcttaaacccaact
ggttacctcttatctagtggAGCAGTGTGGAGAACCCAGGGATTTTAAGAGATCTGAGCCgaatcttaaacccaac
ggttacctcttatctagtggCACCACAGTCCACTCCCTGGTTTTCGACATGGATCCCTTGgaatcttaaacccaact
ggttacctcttatctagtggTGCATGAATATCTTCCCAGGGTGAGGCCACTCAGGTTTGCgaatcttaaacccaact
ggttacctcttatctagtggCAAGCAGGAGAGGCCTGGGTAAATACACAGCTTCACCCTCgaatcttaaacccaac
ggttacctcttatctagtggGAGACAGTGTCATCTCTTGCTCTGAAATTCTTTTGAGTAGgaatcttaaacccaacttg
ggttacctcttatctagtggTACTCTCAGTTGCCATTTCTCCTTCTCTCCATGAATACCCgaatcttaaacccaactt
ggttacctcttatctagtggGGCAGCAATACAAGTCTACCAAGTACGTCCTTCTCAGTCCgaatcttaaacccaact
ggttacctcttatctagtggGACCAGTTTTGCTAGATGAAGGCACACGCCCATCAGGCCCgaatcttaaacccaac
ggttacctcttatctagtggATAAGGTTTAAAAACATAGGCCGGGCGTGGTGGCTTATGCgaatcttaaacccaact
ggttacctcttatctagtggGTTTGAGACCACCCTGGGAAACATGGTCAAGCCCCATCTCgaatcttaaacccaac
ggttacctcttatctagtggGCTGCTCGGAAGGCTGCTGCACGGGAATTGCTTGAGCCTGgaatcttaaacccaac
ggttacctcttatctagtggCACTCCAGCTTGGGCTACAAAGTGAGATTCCATCAAAAAAgaatcttaaacccaact
ggttacctcttatctagtggCTTGCAGCCCTGGTCCTAGCAGCACTGTGTCAGGTAGCAGgaatcttaaacccaac

ggttacctcttatctagtggTGATTTGGAGAAGAACAAAATTATAGCCAGGTGCAGTGGCgaatcttaaacccaact
ggttacctcttatctagtggCGGGAGGCTGGAGACACAAGAATCCCTTGAATTCGGGAGGgaatcttaaacccaa
ggttacctcttatctagtggGCTGAGATCACGTCATTGCATCATTGCACTCCAGCCTCGGgaatcttaaacccaac
ggttacctcttatctagtggGTATAGCCATACCATCATCCTCCCCAGCTCCTCTTCAACCgaatcttaaacccaac
ggttacctcttatctagtggCTAGTCCCACTCATGCTTTTATCTCCAGCATCCCCAGCCCgaatcttaaacccaac
ggttacctcttatctagtggTCTCCCCGTTTTAGACAACCAATGTTTCCATTGCCCAGGCgaatcttaaacccaact
ggttacctcttatctagtggACGATTACTCTCAGGAGGATATCTCTCTGCCCATTTTTGGgaatcttaaacccaactt
ggttacctcttatctagtggTGATGGTCAGCTGTCCAAGTTGTCCAAAGCCATGCTACACgaatcttaaacccaac
ggttacctcttatctagtggGCCTGACAAGGCCCATTTGTCACCCCCAGGGCTGCTAGCAgaatcttaaacccaa
ggttacctcttatctagtggCCTAGCTATGTTCAGGGCCTTCGCACCAGAGAATCTGCCCgaatcttaaacccaa
ggttacctcttatctagtggGCCCACATATCCACTCATCTTATGGTCCCAGGTGGCCACCgaatcttaaacccaa
ggttacctcttatctagtggAGTCACCTTCTGAACTTGGAAAGGGGTTCTTCTGACGGGCgaatcttaaacccaac
ggttacctcttatctagtggGCCCCTCTCAGATTTCCATAGGGCCATGCTCCATATGGGCgaatcttaaacccaa
ggttacctcttatctagtggGACCTTTTCCCTGACCATGGAGCCAGGTCATTGCTCACTGgaatcttaaacccaac
ggttacctcttatctagtggATAAAAACCCAAATACAGGCCAGGTGCTGTGGCTCACGCCgaatcttaaacccaa
ggttacctcttatctagtggAGGAGTTCGAGAGCAGCCTTGCCAACTCGGTGAAACCCCGgaatcttaaacccaa
ggttacctcttatctagtggAAAATTAGCTGGTCATGGTGGTGCCTGCCTATAATTCCAGgaatcttaaacccaact
ggttacctcttatctagtggGCTGAGGCAGGAGAACTACTTGAACTCGGGAGGCAGAAGTgaatcttaaacccaa
ggttacctcttatctagtggCAGCATGCAATTCAGCCCACTGAGCTGATCTGCCTTTACCgaatcttaaacccaac
ggttacctcttatctagtggGCTGTCCATTCACCTTGGAGCTGCCATCTGTAAACCACGCgaatcttaaacccaa
ggttacctcttatctagtggCTGGCAGCACCTCCCACAGTTCCAGAGCCAGTCCTGGGAGgaatcttaaacccaa
ggttacctcttatctagtggCGTGCTTGTGAGTATCTCCTCCTTGCATTCCCCTAGTTGCgaatcttaaacccaact
ggttacctcttatctagtggACTCTGGGCACTTGCATCCCCATTAGAGCATTTTTCTGACgaatcttaaacccaact
ggttacctcttatctagtggCAGCGATGCATAGGACAAGGTGTGGGGGAAGGAGCACGAAgaatcttaaacccaa
ggttacctcttatctagtggAAGAAAAGAAAAAGGCTAGAGGATGGGCTCCTTGGGGGTGgaatcttaaacccaa
ggttacctcttatctagtggATACAGCCATCAGAGAGAGTTGCACTAGGTGGCCTCACAGgaatcttaaacccaa
ggttacctcttatctagtggATGGTGACATGAGCTTTTCCCAAAAGGCAGCAACAATAAGgaatcttaaacccaac
ggttacctcttatctagtggCTAAAACAGCACTCTGCAAACTTGGCCTAATGCAAAGAGGgaatcttaaacccaac
ggttacctcttatctagtggCATGCCTATTATGTCCCAGTCACTGTTCCAGGTGCAGAGGgaatcttaaacccaac
ggttacctcttatctagtggTATCCTCTACCACTGCCCCTGGCTCCTCCTCCTTGGGGAGgaatcttaaacccaa
ggttacctcttatctagtggGGAGAGGCTGAGGATCCTACGGTGCTCTCACCTGAGGCAGgaatcttaaacccaa
ggttacctcttatctagtggGGTATATAATAAATACATATGTCAGGCTGTGTTAACCTCCgaatcttaaacccaacttg
ggttacctcttatctagtggTCAGTTGGGAGTTCACAGGAAGTCTTTCTCAGAGAGGGACgaatcttaaacccaac
ggttacctcttatctagtggGGCCCTGAGGAATGGCAGTGGAGCCCTGGAGCCCTGTGGCgaatcttaaacccaa
ggttacctcttatctagtggGAGCAGGGCAGACGCTGAGGGATCTGGTCAGAAGTGCAGCgaatcttaaacccaa
ggttacctcttatctagtggGGAGAATCAGAAGTCTATGTCAAATTTACAGACCACTATGgaatcttaaacccaactt
ggttacctcttatctagtggAGGCGTGCACCAACACCATGCCTGGGTAATACTTGTATTTgaatcttaaacccaac
ggttacctcttatctagtggTCTCAAACTCCTGACCTCTGCCCGCCTTGGCCTCCCAAAGgaatcttaaacccaa
ggttacctcttatctagtggTACTCTGAGTGAGATGAAAAGCCACTGAGAGGTTTGGAGCgaatcttaaacccaac
ggttacctcttatctagtggGGAAGCAGAAAGAAGAGTCCGCTGCAATAGATGAGGTGGGgaatcttaaacccaa
ggttacctcttatctagtggGAACAGCATGGTGGCAGTGGAAGTGGTGAGAAGTAGACAGgaatcttaaacccaa
ggttacctcttatctagtggAGATCCATCAGGACAGGAGAGGACAGGAGTCAGGGATGCCgaatcttaaacccaa
ggttacctcttatctagtggGATTGGGGTAGGAGTAATCTTGGGAGATACCAGCAGTTCCgaatcttaaacccaac
ggttacctcttatctagtggGAGATGCCTACTAAGCTGGTAGTTGAATGTAAGAGTCTGGgaatcttaaacccaact
ggttacctcttatctagtggGAGGTGGAGGAGGGCAACAGAGAGTGAGGTTCTGGCTGCCgaatcttaaacccaa
ggttacctcttatctagtggGTGAGGCCACTTGAAGATGGAGGATGGAGCTGAGGATGTCgaatcttaaacccaa
ggttacctcttatctagtggGGCCACTGGTGATCTGAATAGAAGGTGGTTTTAGGCTGGGgaatcttaaacccaac
ggttacctcttatctagtggAAATTAGCCGGACGTGGTGACGAATGCCTGTAATCCCAGCgaatcttaaacccaa
ggttacctcttatctagtggGTTGATGGAGAACGGGAAGAGAGTAAATGGTGACAAGTCCgaatcttaaacccaa
ggttacctcttatctagtggGAGAGATGGAGTGGTTGCTGGTGGGCATCTCAGATCAAGGgaatcttaaacccaa
ggttacctcttatctagtggGAGACACTTCCCTAGATGTGACACACTCGACTGCCCACAGgaatcttaaacccaa
ggttacctcttatctagtggGCACCCCATGCTGAGGTGGGCATAACACCAATTATCTACAgaatcttaaacccaac

ggttacctcttatctagtggAGAGGAAGCCAGGAGCCTTCCAAGTTTTAATGGAGCAAAgaatcttaaacccaact
ggttacctcttatctagtggCTACTTAGCAATCTCCATTCCTAACCTGGAAAATGGTTGGgaatcttaaacccaactt
ggttacctcttatctagtggCCCAGCAGTGATGTGGATTCCTGTTGCCCTTTGTCTACTGgaatcttaaacccaactt
ggttacctcttatctagtggCTATGGGTTAGGACACTCAGGGCACAGTGTGGAAAGGTGCgaatcttaaacccaac
ggttacctcttatctagtggGCCTCAGGAGAGAAGGCACAAATGACTGGCAGCAACTTGCgaatcttaaacccaa
ggttacctcttatctagtggGCTGGAACCATCTGCAGGCTCATTCACTCATATGCCTGGCgaatcttaaacccaac
ggttacctcttatctagtggGTTGATCAAAGGCCTCTCCCGGATCCTTAGGGTAGTCGCCgaatcttaaacccaac
ggttacctcttatctagtggGCACCTCAGGGTTCCGAGAACAAGTGTTCCAGCAACAGAAgaatcttaaacccaac
ggttacctcttatctagtggCATGGCTTTTTATGACCTCACTTCAGAAGTCCCAGTTTCAgaatcttaaacccaactt
ggttacctcttatctagtggTCCACTTAAATTCAAAGGAAGGGGACCCAGAGCCCACCTCgaatcttaaacccaac
ggttacctcttatctagtggCCACACCCCATCTGGCAGAGGGAAGAAAATTCACACAAGGgaatcttaaacccaa
ggttacctcttatctagtggTGCCCACGCCATACAGGTAGGTGGCAAAGGTGAAGTCTGGgaatcttaaacccaa
ggttacctcttatctagtggCCCTTCTCCCTCCTGACCACCCTTGATATAATGGCTCTCCgaatcttaaacccaact
ggttacctcttatctagtggCAATCTCCCCCTCTACAATGGAAGTTTTGTGAAGACCAGGgaatcttaaacccaact
ggttacctcttatctagtggGTCCCTATCCTCTGGGACAATGTGTGGCGTGTAATCTATAgaatcttaaacccaactt
ggttacctcttatctagtggGAGCCGTGATTAATTGTACCACTGTATTCCAGCCTTGGCGgaatcttaaacccaact
ggttacctcttatctagtggAAGCCTCAACCTCCAGGTCTCAAGCCATCCTCAGCTTCCCgaatcttaaacccaac
ggttacctcttatctagtggTTTTTTGTAGAGGTGTCTCCTTGTGTTACCCAGGCTGGTCgaatcttaaacccaactt
ggttacctcttatctagtggCAAGCATGAGCCACAATGCCTGGCCTGAGATCAACCTTCCgaatcttaaacccaac
ggttacctcttatctagtggAGCTTTTGCTCTCACAACACTGGGAGGCCACATATTTGGTgaatcttaaacccaact
ggttacctcttatctagtggCCAGGAGGCTGCTCAGGTTGCCCATCTTCACCTGCTCAGGgaatcttaaacccaac
ggttacctcttatctagtggCCCCAATCTCTCCTCCCTCCCCTGCATCCAGCCCCAGCCCgaatcttaaacccaa
ggttacctcttatctagtggGCCCAAGGCCCCAGTTGTCCCAAACTCCTTGCAGTTATTTgaatcttaaacccaact
ggttacctcttatctagtggGGTACCCCCCAGAGATATGATATCCTAAGCCCTGGACCTGgaatcttaaacccaac
ggttacctcttatctagtggGGGCAGAGGGACAATAGACACACAGAGGCAGAGACTGGAGgaatcttaaacccaa
ggttacctcttatctagtggGCCCTCAGAAGCTAGAAGAGGCAAAGGAAGAATCCGCCCCgaatcttaaacccaa
ggttacctcttatctagtggGTTAGCTCATTGAAACTGACTTTGAAATTCTGGCCTCCAGgaatcttaaacccaactt
ggttacctcttatctagtggGCCACCACAGGAAACTAGCATGCCTCCTCTAGAAAGCCTGgaatcttaaacccaac
ggttacctcttatctagtggGGCTAGAGAGTCCCAGCTGCAACTTGTGTTTCCTTCCAGCgaatcttaaacccaact
ggttacctcttatctagtggGTGGATGCAAACTCGTCCACTTCCAAGAACCAGCAAATAAgaatcttaaacccaact
ggttacctcttatctagtggGAGTAGAGACTGGTACAAACCAAAGGGCATATGTCGCTGCgaatcttaaacccaac
ggttacctcttatctagtggGAGATAGGAACCGCCACAGAAAGATGGGTGAAGGATATGGgaatcttaaacccaac
ggttacctcttatctagtggGGTAACCACCCAAAGCCCAATAGACATGTGAAGAGTGCTCgaatcttaaacccaac
ggttacctcttatctagtggCCAAGAGTTGGTGGGACTTGCAGATTTGGGAACATCTACAgaatcttaaacccaact
ggttacctcttatctagtggGTGTTCCATAAGGAATCCTCCCCAGTAGGTCCCTCAGGGGgaatcttaaacccaac
ggttacctcttatctagtggCAGACGGTGGCAGAGTAGGGAAGCATCCATCCACGTGACCgaatcttaaacccaa
ggttacctcttatctagtggGAGGGAATAAGACTACGCAATGGACATGAACGGAGTAGCCgaatcttaaacccaac
ggttacctcttatctagtggCACAACAACAGGACAGATCTTCAAGTACCAGGCTGAGTCCgaatcttaaacccaac
ggttacctcttatctagtggCTCTACACGTACAAAGCACTATGGATCTTACCAGAACACCgaatcttaaacccaact
ggttacctcttatctagtggCAATACATTAGAATCTTGCCTATAGGAAGAGGAAGATGGGgaatcttaaacccaactt
ggttacctcttatctagtggAGGGACTGATGGTGTTAGGATAACACTGTGTAATTGGCTGgaatcttaaacccaactt
ggttacctcttatctagtggCACCTATGCTGTTTGGAGATTTTGGAGAGAAAGAAGAGCCgaatcttaaacccaact
ggttacctcttatctagtggAGCAATAGAAGTAATGTTGCATGATATCAGAAGCTGGGCCgaatcttaaacccaact
ggttacctcttatctagtggGTTCCTCTCATACTCATCTGCCATGGCCCCTGGAGAGAGGgaatcttaaacccaac
ggttacctcttatctagtggGTCCCAACTGAGCCCTGCCTTCAATTCATCCCAGCTCAGGgaatcttaaacccaac
ggttacctcttatctagtggGAGTCTTCCTGGCTCAGGCCCCAGACATTGCAGAGCAGAGgaatcttaaacccaa
ggttacctcttatctagtggCCCTCTGTGTCCTATTCAAGTTCCTGCCCCATGAAATCTGgaatcttaaacccaactt
ggttacctcttatctagtggGGGGTAGTTTGCATCACTGCTGTAGGCACCAAATCATGCCgaatcttaaacccaact
ggttacctcttatctagtggGCTCGTGCCAACTGTTTCCCCATGGAAATAAGCACTCAGGgaatcttaaacccaac
ggttacctcttatctagtggTGGAAAAATCCAGATTGTCTTCTGATTTTTCAATGTCAGCgaatcttaaacccaacttg
ggttacctcttatctagtggACAAATCTCATTTGCACTTCCTAATTGGCCTTGGAGCCACgaatcttaaacccaactt
ggttacctcttatctagtggCTGTGTCCCTCCCACCTTCATAGCCTCAAGATGGGCCCCCgaatcttaaacccaac

ggttacctcttatctagtggCACTGGCTTGGGTCAGGAGTCCGCCACTAACTGGCTGCCCgaatcttaaacccaa
ggttacctcttatctagtggTGGGGGCACCAATATCTACCTCGGCTGCCTCAGTGGCTGGgaatcttaaacccaa
ggttacctcttatctagtggGGTGGTCCCCAAAAAGATATGTCCATGTCCAAATTCCTGGgaatcttaaacccaact
ggttacctcttatctagtggGTCCACATAAGAGACAAACAGAGGGGCCTGGCGCGGTGGCgaatcttaaacccaa
ggttacctcttatctagtggCCCAGCTACAGGGTGGCTGAGGCACAAGAGGCAGAGGTTGgaatcttaaacccaa
ggttacctcttatctagtggACTGCACTCCAGCCTGAGCAACAAAATAAGACTCCATCTCgaatcttaaacccaac
ggttacctcttatctagtggGCAGCCACAAACCAAGGAAGAGTGGAGCCACCAGAAGCTGgaatcttaaacccaa
ggttacctcttatctagtggGGAATGGAATCTCCCTTTGGAGCCTCTGGAGGGAGCACAGgaatcttaaacccaa
ggttacctcttatctagtggACCTTGATTTTGCACTTCTGGTCTTCAGAGTTGTGAAAGAgaatcttaaacccaacttg
ggttacctcttatctagtggCAGGCTGCTGGCTAATCTAGCTCGTAAAGACCCCTCCCTGgaatcttaaacccaa
ggttacctcttatctagtggACTGTGAAGTCACCACCATCCCGGAAGAGTTACTTAAATCgaatcttaaacccaact
ggttacctcttatctagtggTCCCAAAGGTGGTGAGCTCTTCACTCAGCAGCGGTCCTGGgaatcttaaacccaa
ggttacctcttatctagtggCTCAGGCCATGCCCTAGACTTAGTGAGTCAACAGCCAGGCgaatcttaaacccaa
ggttacctcttatctagtggAGACCGTCAACATGAGAAGGGGGTTAAGGGGCCTTGAGCCgaatcttaaacccaa
ggttacctcttatctagtggGCTGATTTTCAGCCCTGCTCTTCCTCATCAATTAGAATCTgaatcttaaacccaactt
ggttacctcttatctagtggGGTCGTGAGCATGGCTCATCAAGGAGTTGTGGTAAAAGAGgaatcttaaacccaac
ggttacctcttatctagtggGCATGCAAATCACCCAGCATAGTTGCATAATAAATTCTGGgaatcttaaacccaact
ggttacctcttatctagtggGCCCAAGGGCCCCCATATTTAGAACTATGTGAGCCTTCCGgaatcttaaacccaac
ggttacctcttatctagtggGAGGCACTCACTGCAAAATTCTAATCAGATCATGTTGAATgaatcttaaacccaactt
ggttacctcttatctagtggAGATAATGGTGACTATCTCTTTATTTGGGCCTTCCAAGACgaatcttaaacccaactt
ggttacctcttatctagtggGTGTCAGCATGGCACACAGTGGCCCACAAGACCCACGCCTgaatcttaaacccaa
ggttacctcttatctagtggGCATGCTGCTCCCTGTGTCTCCAATGTCCTTCCCCTCCCCgaatcttaaacccaac
ggttacctcttatctagtggCCTGCAAGACTCAGCCTAATCATCACCTCCTCTGTGAAGCgaatcttaaacccaac
ggttacctcttatctagtggCCCTTGCTTCACCACCATCGTTTCTATAGCAAAGTTCTTCgaatcttaaacccaactt
ggttacctcttatctagtggCCCTGCATAGCATAAATGTCAGGGACATCAGATGGCCTGGgaatcttaaacccaac
ggttacctcttatctagtggCTAGCTGTGTGACCTAATCTCTCTGGGTCTCAGTTCCCCCgaatcttaaacccaact
ggttacctcttatctagtggCATGGTTCCTGGAACTCAGTAAGCAACTAACAATATCCACgaatcttaaacccaact
ggttacctcttatctagtggCATGGGTGAACTATAGGGGCTCAGACTTGGGAGGAACCTCgaatcttaaacccaac
ggttacctcttatctagtggGCCAGAGGGCTCCGTTTATGTGACTGAGGGACCCAGAACAgaatcttaaacccaa
ggttacctcttatctagtggGTCCTTGAGAAACAGGTCGAAGCTAAGAACCCAGGGCCTCgaatcttaaacccaa
ggttacctcttatctagtggCCTGAGTCACAGGGCTACGTGACCAGCAAGGAGGGTCAGCgaatcttaaacccaa
ggttacctcttatctagtggCGCCAGACGAGGAAGCTGTTTGGGTCCCCAAAGTGCTGTCgaatcttaaacccaa
ggttacctcttatctagtggATGAATGTGAGGTGATAAGCGTTAGGTTATAGGAACGAGGgaatcttaaacccaact
ggttacctcttatctagtggGAGAGTAGGGCAAGTTTCAAATCGGTGGGGCCCAGGGCTCgaatcttaaacccaa
ggttacctcttatctagtggGGTAATGCCTCCCTTATGGGGCTGTCATCACCACCCTAGGgaatcttaaacccaac
ggttacctcttatctagtggCTCCTGGCCCATAATGAGCACTCTATAAATACTACTATGCgaatcttaaacccaact
ggttacctcttatctagtggACTCTCCAGGAGGAAGTGCAAAGTGTTTTCAGAGACCAGCgaatcttaaacccaac
ggttacctcttatctagtggTCCTTAGACACCTGCTTTGTACCAAGCCCTGCGCTAGCACgaatcttaaacccaac
ggttacctcttatctagtggAGGTACAGCACCTTGGAAGCAGTCAGTAAATACTTATTGGgaatcttaaacccaact
ggttacctcttatctagtggGGCCTCTCTTAACAATGTTTGTAAGGCCGGGCGCAGTGGCgaatcttaaacccaac
ggttacctcttatctagtggTGAATGGCATGAGGCAGGGTGGAGAGAAGGCCACAGTGGCgaatcttaaacccaa
ggttacctcttatctagtggAGCTCGGGGGCCAAGGTGAGCTCAGAGCACGCAGGGATTAgaatcttaaacccaa
ggttacctcttatctagtggTAAGGACTTTGGATTTTGTTCTAGGCACAATGGAAAGCTTgaatcttaaacccaacttg
ggttacctcttatctagtggTGGCCTGGAGGAGGGGGACAGGTAGGGAATCCAGAAGGTGgaatcttaaacccaa
ggttacctcttatctagtggAGCTGTGGCTTAGTCCCAGGCTGAGGCAGGTGAGATGTAGgaatcttaaacccaa
ggttacctcttatctagtggGTGCAGCCAGTAGACCACGCTGACAGATTGGTGTGGCAGGgaatcttaaacccaa
ggttacctcttatctagtggGGCTTGAGCCACTCCCTGAGACTTTAAAGCCGAGATCAGAgaatcttaaacccaac
ggttacctcttatctagtggAGGTGGAAAGGCTTGAGAGATATCGCAGTGGAGTGCCTGGgaatcttaaacccaa
ggttacctcttatctagtggGCTGTGGGAATGTATAAAATTACCATGGCTTCAAGTGGAGgaatcttaaacccaact
ggttacctcttatctagtggGTAGAGCAAGCCTGAGGAAAACCAATATTTAAAAGCTGGGgaatcttaaacccaac
ggttacctcttatctagtggTAAGAAGGAGTAATTAATGACCAAGGAAGAAAACCAGGAGgaatcttaaacccaac
ggttacctcttatctagtggCAAAGGACAGAGAAGCTTAGAATTAGGAGGTCAGGTCTGGgaatcttaaacccaac

ggttacctcttatctagtggGGGAAACTATAGCCCAGATAGGGGAAGTGACTCACCCAAGgaatcttaaacccaac
ggttacctcttatctagtggCTGCCCAGCAGTGAAATAGACTTTCTCCTCAGGCAGTGGAgaatcttaaacccaac
ggttacctcttatctagtggAGCTGAAAGGTACAGCTGGATGCTAGAGAATAATTGTGGGgaatcttaaacccaact
ggttacctcttatctagtggCTCACTATGTGGCAGAGGCTGGTCTTGAACTCTTGGGCTCgaatcttaaacccaact
ggttacctcttatctagtggTCTGTGTGTGTTTGGGGTGAGGGGCGGTAGCCGTGGAGGGgaatcttaaacccaac
ggttacctcttatctagtggCTAGACAGAAGGTTGTAGAAAAGGTGACCCTCCGGTATGGgaatcttaaacccaac
ggttacctcttatctagtggTCTCAGGATTCCCAAGGTCTATTTATTAGCACATGAATGGgaatcttaaacccaacttg
ggttacctcttatctagtggACATTGCAGAAGTTTCTTGCTGAGGCTCTCTAACTATGCCgaatcttaaacccaactt
ggttacctcttatctagtggTGTTATCAGCCGACTTCCATAGGGTTCCCTTCCTCTGTGGgaatcttaaacccaactt
ggttacctcttatctagtggGACCACAGTGTGAGTGCCAGATCCCCCCACCACCCCTGGGgaatcttaaacccaa
ggttacctcttatctagtggTCCCCACCAGGACCCATGGGCCAGATCTCACCACCACCCCgaatcttaaacccaa
ggttacctcttatctagtggCCACCACCCCTGGCAGCACAGGTCTACCTTCCCCACCAGGgaatcttaaacccaa
ggttacctcttatctagtggGGGTTTCTGTGTGGGCATTTGCACGACAGAGTGTGAAGCCgaatcttaaacccaact
ggttacctcttatctagtggGTAGAGCAATCTGCCCCCAGTGCTTGGAGGGAAGAGAGGAgaatcttaaacccaac
ggttacctcttatctagtggCACAGAGAGAGCACTCAGAGGCAAACCCACAAAAAACCCTgaatcttaaacccaa
ggttacctcttatctagtggGGGGCTATGTACTTCTCCCACTTTTATTCTCTCTGCCGCCgaatcttaaacccaactt
ggttacctcttatctagtggCCCTCTCCTTCCTCCCCTTACCCTCTCTCAGCCCTGCCTGgaatcttaaacccaac
ggttacctcttatctagtggGGCCACTCTGTTTAGTCCGAGATGCTGGGAGCCACTATGGgaatcttaaacccaac
ggttacctcttatctagtggCCCCTAACATGAAGGAAGGGGGTACAGGTGGGTACAGATGgaatcttaaacccaac
ggttacctcttatctagtggCACAAGGGTGCAAGAACCACAGGGGAATCACGCTAGGGGCgaatcttaaacccaa
ggttacctcttatctagtggACAGGATGTGGAGGAGGAGGAAACTGAGGAAGAATTGCCGgaatcttaaacccaac
ggttacctcttatctagtggGGAGGAGTCTCAGGTGGCACAGGGATTCGGGCTGAGGTGCgaatcttaaacccaa
ggttacctcttatctagtggAGTGCTCAGAAGCCGGAGTGCTGGAGGAGAGGGCTGTAGCgaatcttaaacccaa
ggttacctcttatctagtggGGGGTTTGGCTGAATGCTTCTGGCAGGAGGGTCAGAGCTCgaatcttaaacccaac
ggttacctcttatctagtggGGGGAGAAGATGGAGGGGAAAGCAGGAAGGAGGCAGACAGgaatcttaaacccaa
ggttacctcttatctagtggGAGTATGGCTACAGGGTAGGGTGGCTGAGGTGGAGCGGGGgaatcttaaacccaa
ggttacctcttatctagtggGATGCTACTGTGTACAGTCAATGTTCTCAGGGACACTGAGgaatcttaaacccaact
ggttacctcttatctagtggCTGAGCCAGGACCCAAGTCGCCAGGGAATGCTGGGGAAGGgaatcttaaacccaa
ggttacctcttatctagtggGGATGGCACAAGCCTCAGAGGAGAAGGCAGCACCATAGGGgaatcttaaacccaa
ggttacctcttatctagtggGGGAGCCAGTCAGCCCTGGAATCCACCCACAGCCTGGAGCgaatcttaaacccaa
ggttacctcttatctagtggGGGCCAGAAGGAGCAGGTCTCAGGAAAGAGTGCTGGCTGCgaatcttaaacccaa
ggttacctcttatctagtggGTGAGCACAGGTGAAAGGTTTGTGAGAAAGGGGGAGCAGCgaatcttaaacccaac
ggttacctcttatctagtggGGCAGAGGAAAATGAGGAGTGCACCAAACACACAGAGGTCgaatcttaaacccaa
ggttacctcttatctagtggGATAAAATCAAGATTGAAGTTCTGGTGTGGTGGCTCACACgaatcttaaacccaactt
ggttacctcttatctagtggCTGAACAACATGGTGAAACCCTCTCTCACTTAAAAATCAAgaatcttaaacccaactt
ggttacctcttatctagtggTCACAACCCAATTGTCCCATCCACCAATGAATGGATAAGCgaatcttaaacccaact
ggttacctcttatctagtggGTCACACAATGAAATTCTGATACCTGCCACGACATGGATGgaatcttaaacccaact
ggttacctcttatctagtggGACTGGTGGGTGCCGGGGCTGGGGGAGCAGAGAATGGGAGgaatcttaaacccaa
ggttacctcttatctagtggCTGGATGCAGGGTTTCTGTTTGGGATGATCAACAAGATCAgaatcttaaacccaactt
ggttacctcttatctagtggGGTGGTGATTGTTGTACATTGCAAATATACTTAATGCCAGgaatcttaaacccaacttg
ggttacctcttatctagtggATTGAAACCGAATCTGATCAAGCCTCTGAATTCTACTGCCgaatcttaaacccaactt
ggttacctcttatctagtggGGAGACAGAGTAACATGTTAGATGACACTACTGGGAATACgaatcttaaacccaact
ggttacctcttatctagtggTGACAATATTAAGAAAGTATTGTTGGCTGGGTGCAGTGGCgaatcttaaacccaactt
ggttacctcttatctagtggGGGGGGCCAAGGTGGGCAGATAGTTTGAGCTCAGGAGTTCgaatcttaaacccaac
ggttacctcttatctagtggCCCAGCTACTTGGGAGTCTGAGGCTGAACGATCACTTGAGgaatcttaaacccaac
ggttacctcttatctagtggCAGGGCAGGAGAGATGCAGCGTCATAAGTGAACAAGCGTCgaatcttaaacccaac
ggttacctcttatctagtggAGGTCGTCCATGAGCTGAAAATTGATGAATTTCATGAAGGgaatcttaaacccaactt
ggttacctcttatctagtggAAGTAGGGAGAAAATGGACTTTTGGTGGACTAAAACCAAGgaatcttaaacccaactt
ggttacctcttatctagtggATAAATCCTATGTTTGGGCTCAGAATGAAGAAGGGATACCgaatcttaaacccaactt
ggttacctcttatctagtggGATCTGGGATTTAGTTGGCCCTGAGATCTGCATGAACTAGgaatcttaaacccaactt
ggttacctcttatctagtggAATCCCACCTTAGGCCAAGAGCACAACAGGATCCTCTGTGgaatcttaaacccaac
ggttacctcttatctagtggCTAAGGAATGTCTGGAATAGTGTGCTCAATTCTCAACCCCgaatcttaaacccaactt

ggttacctcttatctagtggGACTTAGTTAAGGACACAGTGGTGAGCATTCCAGGGAGGCgaatcttaaacccaac
ggttacctcttatctagtggACAGGGATTTGTGCCATATCAGGAAGTAGCATGAGATATCgaatcttaaacccaact
ggttacctcttatctagtggCCCGAAGAGCAGAGAAGGAGGATAAATATGTCATCTGAGGgaatcttaaacccaac
ggttacctcttatctagtggTCTAGAAGTATGCAGTCGGTCAGGGGCAGTGGCTCATGCCgaatcttaaacccaac
ggttacctcttatctagtggTGCAGTGAGCCAAGTTTGTGCAACTGCACTTCAACCTGGGgaatcttaaacccaac
ggttacctcttatctagtggGCAAATAGAAGCATGCCGTCATCCTGTGGCTTCCTCTGCCgaatcttaaacccaac
ggttacctcttatctagtggCTCCAGGGAGAGACACATAAAGGGGTGAATAGAGCTCCAGgaatcttaaacccaa
ggttacctcttatctagtggTCTATGCCCCTCAGTGAGTGATTTTACCTCTTTGAGCCTCgaatcttaaacccaactt
ggttacctcttatctagtggCATGTACCTAATAGGTTTAATGTATGGATGTCAGATTCTCgaatcttaaacccaacttg
ggttacctcttatctagtggTGCTATGAATCCTATAAAGTCACTTCTGTTATGCTGCTTCgaatcttaaacccaacttg
ggttacctcttatctagtggGTTCCAGCACAATTGATACAATAGGAAACAATTTGAGCACgaatcttaaacccaact
ggttacctcttatctagtggCAAAACACACCTCAAACATCTCCCAGTCACCTCAGCTGGGgaatcttaaacccaa
ggttacctcttatctagtggGGTGTGTGTTATGAACCACACCCATCCACATCTGGTATAAgaatcttaaacccaact
ggttacctcttatctagtggCACAACTTCTCAATAACTCACAAGGTACAACCCTTCTGACgaatcttaaacccaac
ggttacctcttatctagtggTAAACTAATGTGGAAAACTGTGCTACCATTTTATTTGGGGgaatcttaaacccaacttg
ggttacctcttatctagtggGTATTTTCCCCTTACCCCATTCTCTCCATAAGCCCCGTGGgaatcttaaacccaac
ggttacctcttatctagtggAGGAACACATGTGAAGTACTATAGCAGAACTGGCTGTGCGgaatcttaaacccaac
ggttacctcttatctagtggATCCTCACATTGTAGGTACTCAGTAAATGACAGCTGCTGCgaatcttaaacccaact
ggttacctcttatctagtggGTCCAGCATTGGGGAAAGGCCTCTTGTCAGTTACTATGGCgaatcttaaacccaac
ggttacctcttatctagtggGGCCAGCCCCTCCCCGGCTGCCATTTACCCACCAGACAGGgaatcttaaaccca
ggttacctcttatctagtggCCATGCCCTGGCTGTTCCCTGTTCAGTCATGGGAGGTGTCgaatcttaaacccaac
ggttacctcttatctagtggTCCCTTTCCTCTGTGACTCAGTACCTTCACTGAAAGATCAgaatcttaaacccaactt
ggttacctcttatctagtggTTGCTCGAGAAGCTTATTACAGTTCGGCCAAATACTTTCTgaatcttaaacccaactt
ggttacctcttatctagtggGTCTGGAACATCTGTCACTTCCTGCTCTGTGTTCTCTGTGgaatcttaaacccaactt
ggttacctcttatctagtggCATGTCACGGATGAGGGTTTGCCTTAGATGTCCTCCCAGGgaatcttaaacccaac
ggttacctcttatctagtggGGTGTCTAATTCTACAGGTTTCCCTTCTGGATAGCTTGTCgaatcttaaacccaactt
ggttacctcttatctagtggGGGTGCAGCACAAGATACTGGGGATAGAGCTGGAGGCAGCgaatcttaaacccaa
ggttacctcttatctagtggCCACCATTCATCCACATGGAGCACACAGACAGTTGGATTCgaatcttaaacccaac
ggttacctcttatctagtggAGATGCCCACAATTCATCTCTTCACTCAGATTTCTAGGGCgaatcttaaacccaact
ggttacctcttatctagtggCCCTCTCTAAGCCTCAGTTTGGTCATCTGGACATGGGAACgaatcttaaacccaac
ggttacctcttatctagtggTGGAAGAGTTGCTACATGGGATTACAAAGCATATGTCATAgaatcttaaacccaactt
ggttacctcttatctagtggGCCCTTAGCCTTCAAGGTAGCTAGCGCTTAGCCCAGGGGGgaatcttaaacccaa
ggttacctcttatctagtggGGTCCTTGTGGTTTGGGTTTGAGGACACAGAGAAGTGGCCgaatcttaaacccaac
ggttacctcttatctagtggGAGGTAGCCTCACCACTCTGCAGCCCCTCAGATTGGCCCCgaatcttaaacccaa
ggttacctcttatctagtggGAGGCAACCCAAGCCACCCACCTTTCCCACCCACGTCTGCgaatcttaaaccca
ggttacctcttatctagtggAGTTGTTGGCTCATTTTTCGCCGGGGGCAAAGGGAGGGAGgaatcttaaacccaac
ggttacctcttatctagtggGGTGTGGCCCCACATAAGTGACTTTTCCCGAGGATTTGCCgaatcttaaacccaac
ggttacctcttatctagtggGACCTCTCAACATCCCCTGGAAATAAGAATTATTATCCCCgaatcttaaacccaact
ggttacctcttatctagtggGGCCCAGAGAGGTTAACAACAAAAAATCCTCACTGGCCCCgaatcttaaacccaa
ggttacctcttatctagtggTCTGACTTGGTTGAGTATCCATGGAATTTGGTTTTCATCGgaatcttaaacccaacttg
ggttacctcttatctagtggTGCCTATCTCTCCTCCTCCATCCACAACAGCCAAGAGCGCgaatcttaaacccaa
ggttacctcttatctagtggGTGAGTCCCCAGGGTGTTAAGCTCTAGGCCAGCTTCTAGGgaatcttaaacccaac
ggttacctcttatctagtggGCTTCAGGCTCTCAGGAGCTTACAGGCCAGCACTCAACCCgaatcttaaacccaa
ggttacctcttatctagtggGCTTCTAGCTGAGTGCCTGGCACACCACAAACATTCAATCgaatcttaaacccaac
ggttacctcttatctagtggGATCAGCCTACTCCTGAATCCTGATGCACCCAAGCCTCCCgaatcttaaacccaa
ggttacctcttatctagtggCACTGTCTCATCTGTCTCCTTTACAACCCAGACATGTGCCgaatcttaaacccaact
ggttacctcttatctagtggTGCCCTATCAGATCCTTTGAACCCACCCCACCTGGATGGCgaatcttaaacccaac
ggttacctcttatctagtggAGCCACCTCCTCGGAGAAGCTTTCCCCAACCCTCCATTGCgaatcttaaacccaa
ggttacctcttatctagtggTCATGCTGCCGTTGCAACTAGGCTGTGAGATCTTTGAGGGgaatcttaaacccaact
ggttacctcttatctagtggGGAAACTTCAACACCAAGTAATAGAAAACCCAACTAACAGgaatcttaaacccaac
ggttacctcttatctagtggCCTGCACTTTTCATTGTCCCACTCTGTCCTCCCCATTAGCgaatcttaaacccaact
ggttacctcttatctagtggGCAAAGGGAACATGAGGAGAATAGGGACACGTCCCCTCTCgaatcttaaacccaa

ggttacctcttatctagtggCTCGGTCATTCAAGCTCAATTCCCCTCATACCTATGAGCCgaatcttaaacccaact
ggttacctcttatctagtggCTTGAGCAAAGAGAATGAGACCACCTTGATGGCTGAGACCgaatcttaaacccaac
ggttacctcttatctagtggTTCCTCCAGAAACACAAGGCTTCGGTGGAGAGTGAACAAAgaatcttaaacccaac
ggttacctcttatctagtggGAAGTTGACAACCCAGTGACATCTGCAATTCCCTTGTGACgaatcttaaacccaact
ggttacctcttatctagtggGCAGACCCAATACCACCTAGATCCTTCTGCTCACCGTGGGgaatcttaaacccaac
ggttacctcttatctagtggCACCTACCAGCCCATCAAGTAGGTGCTAGGGATATAGCAAgaatcttaaacccaac
ggttacctcttatctagtggGATCCTATCCTCAAGTGCCTCTGTTATTGCTCAGTCTCATgaatcttaaacccaacttg
ggttacctcttatctagtggCCAAAATATGAGATAGGCAGAGAAGATGTTCCTTTTGTACgaatcttaaacccaacttg
ggttacctcttatctagtggCCTGCTAACACAGAAGCTCACCTCAGAAAGAAATGGTCAGgaatcttaaacccaac
ggttacctcttatctagtggCTCATTCCAGTCAGCTGCAGACCATGACCCCACTAGAGACgaatcttaaacccaac
ggttacctcttatctagtggGGCAAAACACTAGTGCCTGTGTTCTCCATTCCCCAAACCAgaatcttaaacccaact
ggttacctcttatctagtggGCCCTGCCCTCTCTCGGAGAGTCACAATTTGCTTCAATGCgaatcttaaacccaact
ggttacctcttatctagtggTTTAATCACTCACAAGGCTGGATGTGGTGGCTCACGCTTGgaatcttaaacccaact
ggttacctcttatctagtggCACTCATAAAACACTTTTTCATTGCAAAACCCCTCCCAACgaatcttaaacccaactt
ggttacctcttatctagtggGTTTTGAGCAAAACAAACACGGTCCCTACCCTCCATGGACgaatcttaaacccaac
ggttacctcttatctagtggGGAACTAAATTCCGTATCACCAAGAACAGTGCTCACTGGGgaatcttaaacccaact
ggttacctcttatctagtggGTTGATCTCCACAGGTATGGTCATCAACCCATCCATGGTGgaatcttaaacccaact
ggttacctcttatctagtggCAGGACTATCTTCCTGAGATTATTAGTATCTACATGAACCgaatcttaaacccaacttg
ggttacctcttatctagtggAGCCCAATTTAGATGCAATAACAAAGAGCGCCAAAATCCCgaatcttaaacccaact
ggttacctcttatctagtggGAGATCCAGCCTGGAGGAACACCCCTCTGTGGAATGTGGCgaatcttaaacccaa
ggttacctcttatctagtggGGGCACGGGGTATGTTTCATCTACTCACAGTCCACTGGCCgaatcttaaacccaac
ggttacctcttatctagtggCCCCAAGTCCATCCGTGAGTCAGGAGTTGTATCCTCCTCCgaatcttaaacccaac
ggttacctcttatctagtggTAAAAGGAAGGGACAGGCCGTCCGGGAGGTGAGGGGCGCCgaatcttaaacccaa
ggttacctcttatctagtggCTACTGGGAAGTGAGGACCCCTCTGCCAGGCCAGCCGCCCgaatcttaaacccaa
ggttacctcttatctagtggCACAATGACCTGGGGAAAGGAAGAACAGCCTTGTATGGGGgaatcttaaacccaac
ggttacctcttatctagtggGGACTCTGATGTCACACCCGTCTGAATTTGAATCCAGGCCgaatcttaaacccaact
ggttacctcttatctagtggGCTGAGTGGCCTTGGGGATATTAGAAAAACATTATAATCCgaatcttaaacccaacttg
ggttacctcttatctagtggATGGGAACATCAGTATCTACTTTACAAGGCCGCAGTGAAGgaatcttaaacccaact
ggttacctcttatctagtggATAATAATACCAGAGTGAGGCCGGGTGCAGTGGCTCATGCgaatcttaaacccaac
ggttacctcttatctagtggACCCCGTCTCTACTAAAGATGTGAAATTAGCTAGGCATGGgaatcttaaacccaact
ggttacctcttatctagtggCACCACTCCTGGTGCCATTCAGACCTTCCCTCGGTAAGACgaatcttaaacccaac
ggttacctcttatctagtggGGTAAGAGCTGACAGTTTCGACAAGCTGTTCCTTTGGAAGgaatcttaaacccaact
ggttacctcttatctagtggTGGACAAGTCAACTCTGGTGACACATGAAAATGATGTGCCgaatcttaaacccaact
ggttacctcttatctagtggGGTCTCCAGGCAGAGGATCAGTGTGGGAGGCTACAGGGTCgaatcttaaacccaa
ggttacctcttatctagtggGCCCTGTTCCCATGAGTTCTCACCCAAAGAGGCCAGGGCCgaatcttaaacccaa
ggttacctcttatctagtggAGGGGGACTAGTTTCAGCAACATGACCCCATGCCCTCTGCgaatcttaaacccaac
ggttacctcttatctagtggGAAGGGTTTCAGGGAGCCCTGGAACTCCTCCCCTGTTCTCgaatcttaaacccaac
ggttacctcttatctagtggAGAGCTTGGGTCTTAGAGTAGCTCTCCTGCTTCTCTCTAGgaatcttaaacccaactt
ggttacctcttatctagtggAGGCAGCCTTACTCCAGCCTTGCAGCTGTTACAGAGACTGgaatcttaaacccaac
ggttacctcttatctagtggGCCAACCTGGCTCAAATCCTATCTGTGTGACCTTGAGGAAgaatcttaaacccaact
ggttacctcttatctagtggTCGTCCTTCAGCCATTGTCACAAATCTCATGATCACCCTCgaatcttaaacccaactt
ggttacctcttatctagtggGACTGATACATCATTCACTACATTGAGTCTTCACCTGGGGgaatcttaaacccaactt
ggttacctcttatctagtggGGAACTGCAGATTATCAGAGAATTACTTCTTGATGTTTGCgaatcttaaacccaacttg
ggttacctcttatctagtggTAACGCATTTACCCAAATTGATTGTCACATCTGTGGATTGgaatcttaaacccaacttg
ggttacctcttatctagtggCACCACAACCTCTGCCTCCCGAATTCAGGCAATTCTCTTGgaatcttaaacccaact
ggttacctcttatctagtggGGCTGATCTCAAATTCCCAACTTCAGGTGACCCGCCCACCgaatcttaaacccaac
ggttacctcttatctagtggTTTTCTGAGACGGTGTCTTCCTCTATCACCCAGGCTGGAGgaatcttaaacccaact
ggttacctcttatctagtggGCATGAGCCACCGTGCCTGGACTGTTATATTATTGTTTAGgaatcttaaacccaacttg
ggttacctcttatctagtggGCCATGTCCTATTGTGCTTCTCCTGGTGCCCTCTCAATGGgaatcttaaacccaact
ggttacctcttatctagtggTCACCCCCTCCTCACCAGTGACCCTGAACTGAATCATGGGgaatcttaaacccaac
ggttacctcttatctagtggCCCGTGCCCAGATGCCACCTGCTCCTTCCAAAGAGACGAGgaatcttaaacccaa
ggttacctcttatctagtggAGAGTCATAGCTGGCCTCCAACCAACAAGTAACATGAATAgaatcttaaacccaact

ggttacctcttatctagtggACCGCTGAGATGTGGATGTGGTTTGTTAGGACAGCATCACgaatcttaaacccaact
ggttacctcttatctagtggCACTGCAAAGCTGGAGAATGAGAGCACAAGCCTCCTGCGCgaatcttaaacccaa
ggttacctcttatctagtggTCCAAACCTTGTACTCATCCTACTACGCTATGTGCTTCCCgaatcttaaacccaactt
ggttacctcttatctagtggAGGATCTCACTCTGTGGCCCAACCTGGAGTGCAGTGGTGCgaatcttaaacccaac
ggttacctcttatctagtggCTCGAAATCCTGGACTGAAGCAATCCTTTCACCTCAGCCTgaatcttaaacccaact
ggttacctcttatctagtggTGTCAGACTCACCCATCAGTCTAAGCTCCATGAAGGCAGGgaatcttaaacccaac
ggttacctcttatctagtggTTCTCAGTTCTGAGTACATATTAGACGCTATTCCTTGATGgaatcttaaacccaacttg
ggttacctcttatctagtggCTGCCCATTGAATAACTTGATTCTTTCATTTCTATGTTGGgaatcttaaacccaacttgt
ggttacctcttatctagtggGCATATGGGTGAAGTTTGGGCTTTTCGTGTAACCCTCATGgaatcttaaacccaacttg
ggttacctcttatctagtggCCCTCACTCTCCTCCAACCTTTCCACCTTTTGTTATCTCCgaatcttaaacccaacttg
ggttacctcttatctagtggTAGACCCAGTGGCCCAGCTGGAATCAGGAGGTGTGGATGGgaatcttaaacccaac
ggttacctcttatctagtggCTGTGCTGCACAGCTCCACCCATTGGACCCTCAAACTGGGgaatcttaaacccaac
ggttacctcttatctagtggCTTCCCCAAACCTGCCTCTCCACCTGCATTGCTGTCTTGGgaatcttaaacccaact
ggttacctcttatctagtggCCTTTCCATCCCTAACCTAGTTAGTCACCCAGTCCGACTGgaatcttaaacccaact
ggttacctcttatctagtggGGCAGCTGCACTTACTACTCTCTCCATCCCCATACCCTGGgaatcttaaacccaac
ggttacctcttatctagtggCACTTTCCATATAGCTGTAGGGAGACATTAAAAATAAACCgaatcttaaacccaacttg
ggttacctcttatctagtggCCTGCTTAGAAAAGGTAAGGGATGTTAGATATTATCTGCGgaatcttaaacccaacttg
ggttacctcttatctagtggGGGCCCCACCCTCAACCTACTTACTTGGAATCTTCCACGGgaatcttaaacccaac
ggttacctcttatctagtggTATTTCAGGCTGCGCTTATGATTGGGCCAGTTTAGAAAACgaatcttaaacccaacttg
ggttacctcttatctagtggGACGTAGCCTACTAGTTAGAATGAGATTCAGCTGTGAGTAgaatcttaaacccaactt
ggttacctcttatctagtggCCTGGTGTCAGGAACACAGGCTCCATGTCATCCACCGTCTgaatcttaaacccaac
ggttacctcttatctagtggGGCCCAAGATGGCATCATCCACATTCCTGGCTCAGGATGGgaatcttaaacccaac
ggttacctcttatctagtggCAGTCGCTAGTTACAAGAGAAACTGAGAAATGCAGTCTGCgaatcttaaacccaact
ggttacctcttatctagtggGGGCAAAAACAATATTGGGAAGTAACTGTGAAGACCAGTAgaatcttaaacccaact
ggttacctcttatctagtggGCCCAAAGTCACAGGACTAGGTAGTGTCCCCAAATACGGCgaatcttaaacccaac
ggttacctcttatctagtggGCTGATGGTTCACACAGGGTTGGCAATGCTGTCCAGATGGgaatcttaaacccaac
ggttacctcttatctagtggCTGAAGGTGGCCTCTGTCTTTCTAGGTATCTCTGTGAGCCgaatcttaaacccaactt
ggttacctcttatctagtggCAGGAGGTGACAATCGGACATGATACTTGCAGTAAGAGAGgaatcttaaacccaac
ggttacctcttatctagtggTCAGCTCTGAACCCTTCAGTCTATTAATGTCTACACCACCgaatcttaaacccaactt
ggttacctcttatctagtggCCACTGCACCACTGCCTCCTTTGCCTGATTCCCCTGATGCgaatcttaaacccaac
ggttacctcttatctagtggGCACCTCCTCTGTGCCAGACACGATGCTAAGCATTGGGAGgaatcttaaacccaac
ggttacctcttatctagtggAAACCCTCAGAGCATCCCCCAGCTAGTAGAAGAGACAGGCgaatcttaaacccaa
ggttacctcttatctagtggGCATTGGGAAGTCTGAGAGGCCTGAGGTCAAGTCTCAGCGgaatcttaaacccaac
ggttacctcttatctagtggCCTGTGTCATCCTGGGAAAATCTGCATTTGATTTCTCATCgaatcttaaacccaacttg
ggttacctcttatctagtggGGGTTTGGGTGAGGATTAGCAAGGTAATGTACAGGACTTGgaatcttaaacccaact
ggttacctcttatctagtggCGTATCTCAAACATCACATTATACCCCCTAAATGCAGATGgaatcttaaacccaactt
ggttacctcttatctagtggGAGCCAATGCAAGCTTTACCATATTCCTTTTCCTGCCTTGgaatcttaaacccaacttg
ggttacctcttatctagtggGAGTAGTCCTGCCAGCCTCCAAGGGACAGATAGCACAAGCgaatcttaaacccaa
ggttacctcttatctagtggCTGCCAAGCTGGAAAGAGAGTCCTATAGACCACACCAGCCgaatcttaaacccaac
ggttacctcttatctagtggCCTGGGAAACACCTGCTGGGAAGATGCACATAAACACCTAgaatcttaaacccaac
ggttacctcttatctagtggGGCATGACAGCTGGAGCAGGTCTCTCTCTTCACTACAGCCgaatcttaaacccaac
ggttacctcttatctagtggGCATTTCTCCCTCTGAAAGAAGCAAGTGGTACCCATGGGGgaatcttaaacccaac
ggttacctcttatctagtggTCTTTCCCCTGCTAATTCTCTCAAAATCCAGAAGTCGAGCgaatcttaaacccaactt
ggttacctcttatctagtggCCACGTACTAAGTGCTTATTATGTATGGAGGCCAGTGCTGgaatcttaaacccaactt
ggttacctcttatctagtggCCCAGCAACCCCAGGAGATTACTGCTGTGATGATCACTCCgaatcttaaacccaac
ggttacctcttatctagtggCACACAGATGGTAAGTGTGCAAGTTACGCTTCCAAGCAGGgaatcttaaacccaac
ggttacctcttatctagtggGCTACAAGGAACTGTGTCCAGGGGCCAAGGAGAGCTCCAGgaatcttaaacccaa
ggttacctcttatctagtggCCGCAGGTGCCTCTTTTTCCTAGGATTCTAGTAAGTCTACgaatcttaaacccaacttg
ggttacctcttatctagtggGATCACTAGGAAAAGGCACAAGTAGGCCGGGCGCGGTGGCgaatcttaaacccaa
ggttacctcttatctagtggGGGTTAGCACCCTCCCAAGCCTTTCTCATGCCCATACTGCgaatcttaaacccaac
ggttacctcttatctagtggGGGCAACAGCATCCAGTAAGATAATGTAGGCAGGCCCAACgaatcttaaacccaac
ggttacctcttatctagtggAGGTGCTTAGCCTCTGGGGATGTGCAGGAGGAGTCCAGACgaatcttaaacccaac

ggttacctcttatctagtggGGGTGAAAGATGACCCTGTGGTTGTCATTCACCATCCTGCgaatcttaaacccaact
ggttacctcttatctagtggTCTGTATGAAATGCTCTTCCCCACAACACCTGTCCTTGTCgaatcttaaacccaactt
ggttacctcttatctagtggCTGTCACCCCTCCTCAACTCAACAAATGTGAAGCACAGGGgaatcttaaacccaac
ggttacctcttatctagtggTCCCCTGAACCCAGCTCAGTGCCTAGCACACAGTGAGTACgaatcttaaacccaac
ggttacctcttatctagtggCCAAGTGGGGATCTTCATGTATCTACAGCATTTGACTCCCgaatcttaaacccaactt
ggttacctcttatctagtggTGCTTCTCAAAAAGTCTGTGAATCAAGCAAGGCAGATGTCgaatcttaaacccaactt
ggttacctcttatctagtggACTTGGTCAAGATCACCAGTGAGCAAGTGGTACAGCCAGAgaatcttaaacccaac
ggttacctcttatctagtggGGAGAGACAGGAAGAGAAAAGGAGAAATGGAGAGAGAGAGgaatcttaaacccaa
ggttacctcttatctagtggCTGCCAGTAATAATTTGGTGAGACTCACACTGAGAGGCTGgaatcttaaacccaact
ggttacctcttatctagtggTTGTATAGGCAGAGATAAGTGGACACATTTCGCGTTGAGGgaatcttaaacccaactt
ggttacctcttatctagtggCCTCTTTCTAATCATCATTCGTCCATCAGCCTTGGAGGTCgaatcttaaacccaacttg
ggttacctcttatctagtggTGAGAGAGGCATGTGTCTAGAGACTGGGAAGGCACGGTGGgaatcttaaacccaac
ggttacctcttatctagtggAAGGCCCATTGTAACACTGGTTTTCCTCAGATAGTCAGCCgaatcttaaacccaactt
ggttacctcttatctagtggAGGACTGTTTACATGGAGGTTTTCAGCAGCCTTAGCAGGCgaatcttaaacccaact
ggttacctcttatctagtggTATGTGTAAATAAACAGTGGGAGGAAAGGAGAAGGCATCCgaatcttaaacccaact
ggttacctcttatctagtggAGTGATTTCTCTGCTCAGCTCTCCTGGGAAAAGGAGTGGCgaatcttaaacccaact
ggttacctcttatctagtggGCACTTAACCTCTAGTATCAGATCAATGTTGGAAATGCACgaatcttaaacccaacttg
ggttacctcttatctagtggCCACAACCAACGTGCTTCAAAACTATTGGCTGAATTCAGGgaatcttaaacccaact
ggttacctcttatctagtggTGGACACTACCCAAGTGGATTCTCCTTGCACCAAAATGGCgaatcttaaacccaact
ggttacctcttatctagtggGCCTCCTCCAGTCACACAAGTAGTCTATCATGAATCTGCCgaatcttaaacccaact
ggttacctcttatctagtggTCCCAGTGAATGAAGTGTATTATGTTGGCAGAAGGATACTgaatcttaaacccaacttg
ggttacctcttatctagtggCCGGCTCCGGAAATGAAAGGGAAATGCACCAAGACAAATGgaatcttaaacccaac
ggttacctcttatctagtggCCAACTGGTGCCTGATTCAGCTCAGATGGCCCCTACTCTGgaatcttaaacccaac
ggttacctcttatctagtggCCCAGGTTGGTTAGTTGAGTTTTGGCAATCTCTACTAGAGgaatcttaaacccaacttg
ggttacctcttatctagtggTTATGCTAAATACACTTCCCCAGGTATTTATGCACGTTGGgaatcttaaacccaacttg
ggttacctcttatctagtggTTTGGGCAGCTTAAAAGCCAACATAGGTATAGCTCTTTTGgaatcttaaacccaacttg
ggttacctcttatctagtggCCCTCCTGGGGATGAATCCAAAAGCTTTACAAGGGGTATCgaatcttaaacccaact
ggttacctcttatctagtggGCAGTCCCAGAGGGAAGGGAGGAATTTGGCTGATGAGTGGgaatcttaaacccaac
ggttacctcttatctagtggTCTCCATGCTCCATCTGGGGAGACAGTTTGGTCTGTACTGgaatcttaaacccaactt
ggttacctcttatctagtggCCTGATACCTTATTCTGTATCTAACTGGCCTTAACATCTCgaatcttaaacccaacttg
ggttacctcttatctagtggGATCTTGTGAGAGAAAGAGAAAGGTTTCATGTCTTAGCCCgaatcttaaacccaactt
ggttacctcttatctagtggTCCACTCAGTGTGATGCATAACACTAGGCAGCCACTACAGgaatcttaaacccaac
ggttacctcttatctagtggATCAGAGCGTGCAGAGTGGTGCCATGGATCCTAGGAGGGGgaatcttaaacccaac
ggttacctcttatctagtggCCCTACTGGGACTCTGCCTACTCCCACAGTACCTTTACACgaatcttaaacccaac
ggttacctcttatctagtggAGCCGTAATGTAAAGTATGTGACAAGGTGTGAAACCAGACgaatcttaaacccaact
ggttacctcttatctagtggACCCAGGACACAAAAATCCACAGGCACAAAATGATGAGAGgaatcttaaacccaac
ggttacctcttatctagtggGGGTATTACTGCTAGTATCTGATTCTGTACAACTGCGGTCgaatcttaaacccaacttg
ggttacctcttatctagtggGAATTAGAGGGATGTTAAAATGCTGGGATTATGACTTCTGgaatcttaaacccaacttg
ggttacctcttatctagtggGTGCAAGTATTACACATTCTTAGGGAGTGAGTCCTTAGGCgaatcttaaacccaactt
ggttacctcttatctagtggTGTCTTTGGTTGAAACCTGAAATGAGGAAATTAAGGCCAGgaatcttaaacccaacttg
ggttacctcttatctagtggTGTCTACAGACACCACTAGACTGTAAGCTGCATTCTTCTGgaatcttaaacccaactt
ggttacctcttatctagtggATCACCACCCAGCCTAGTCCCTAATACTTAGTAGATACCCgaatcttaaacccaact
ggttacctcttatctagtggGAGTTCTACTCCATTGGGTTGTCCTCTGATTGCTTCAGAGgaatcttaaacccaacttg
ggttacctcttatctagtggTAGCATCTTCCCAACACACATAGCACAGGACTAGGTACACgaatcttaaacccaac
ggttacctcttatctagtggATCTTTCCTTCTCCTCTTCCTCCTCCCATCCTTCACCCTCgaatcttaaacccaacttg
ggttacctcttatctagtggTGGCTTCTTATGGGCCATTGTCCTTGAGCACTTTCAACACgaatcttaaacccaacttg
ggttacctcttatctagtggTTTACTGTGTGCCCAACATTCTGACATGCACTTTACATGCgaatcttaaacccaacttg
ggttacctcttatctagtggGAGACTGTGATTCAAATGCAGAGTGTCTGACCCCGGAGCCgaatcttaaacccaac
ggttacctcttatctagtggCCCCTATGTCATGCTGAATTTTTGCTAGTCTTCCCTCTGGgaatcttaaacccaacttg
ggttacctcttatctagtggATGACAGGAGTAAGAATTTTAACTACAGAGGGTCCTGGCAgaatcttaaacccaactt
ggttacctcttatctagtggCACAGTTTCCATCTACCATAACATCTGCCAATGTGCTGCCgaatcttaaacccaactt
ggttacctcttatctagtggTGACACAAAGAGAGAAGTCAGATCACAGAGGTAACCTGTCgaatcttaaacccaac

ggttacctcttatctagtggCCGATTCCAAAAATTCCTGGAACTGGCAAGAAGATGGCAGgaatcttaaacccaac
ggttacctcttatctagtggGAAAAGATTATAGCTCAGAAACTGATTTCTTGTGTTTCCCgaatcttaaacccaacttg
ggttacctcttatctagtggTGAGGAAGAGAGAAACCTGGAACGTTAATTATGGCTTAGCgaatcttaaacccaact
ggttacctcttatctagtggGCAGGTCTCCTCCATCCCCTCTGTGAGTGCTAACTCAGCCgaatcttaaacccaac
ggttacctcttatctagtggCCCCCAGGTGGTCTCTCTAACCTGACACCCTGCCCCTCCCgaatcttaaacccaa
ggttacctcttatctagtggCAGGGGTCTACCTGCTGCACTCCAAGCAGAGCCTATGCCCgaatcttaaacccaa
ggttacctcttatctagtggTCTTATCTCCTGCCAATCCTGGCCACCCAATCTCTGACTCgaatcttaaacccaact
ggttacctcttatctagtggCCAGGAGCAAAGCAGACATGATTCCTCCCTGCCTGAGGCCgaatcttaaacccaa
ggttacctcttatctagtggTTTTGGGCTCTGCCTAACACAGTGTCTGTTGCAGCTCCTCgaatcttaaacccaact
ggttacctcttatctagtggGTAGTGGGAAAGCAGCCACTGACATGACACAAATGGATGGgaatcttaaacccaac
ggttacctcttatctagtggGTCTGGACTTGGTCTATAGGCCACAGTTTGCTGACCCCTGgaatcttaaacccaact
ggttacctcttatctagtggAGAGGGCTAGGTTAGAAATTCGAAATCCAGCTACTCCACCgaatcttaaacccaact
ggttacctcttatctagtggTCGTCTTCAGCAAGTTTCTCTGCACCTCAGTGTCTTCATCgaatcttaaacccaactt
ggttacctcttatctagtggCCAGGCTCTGTTTTCCCAGTGAAATCCATCCCGTGGTTCCgaatcttaaacccaact
ggttacctcttatctagtggCACCATGACCTGGAGAGGCCTTCAATGATCTTCCTCCCACgaatcttaaacccaac
ggttacctcttatctagtggGGGACTACAGGCACACACCACCGTGTTGGGCTGATTTTTGgaatcttaaacccaac
ggttacctcttatctagtggGGCTGGTCTCGACCTCCTGACCTCAGACGATCCGCCCACCgaatcttaaacccaa
ggttacctcttatctagtggCCCCTGTAGGATTACAGTGGCCCCACTTACTCTTCTACCCgaatcttaaacccaact
ggttacctcttatctagtggCTTCCTTCCCACCCCAGGGCCATTGTGGATGCAGCACCAGgaatcttaaacccaa
ggttacctcttatctagtggGCCCTCCTGCTGTGGTCACATGTCAGCTCCTATCCTTCAGgaatcttaaacccaact
ggttacctcttatctagtggCACCCACACACACATGGAGGAACAGCCACCAGTCCTTCCCgaatcttaaacccaa
ggttacctcttatctagtggGAGTGCATGGACTTCACTGTACCTCTAGCATCCAGCCCAGgaatcttaaacccaac
ggttacctcttatctagtggGGTGGACCAATGCCTCATGTCTCTTCGTTTCCTCTGGCACgaatcttaaacccaact
ggttacctcttatctagtggGAGTATGTCCCAAATACCCTACGCTGAGATACCCCTAGGGgaatcttaaacccaac
ggttacctcttatctagtggCTCTGAAGAACCCGTTCAGACAAAAGACAGCAAGACCAAGgaatcttaaacccaac
ggttacctcttatctagtggCCAGGGGTTCTGGTCTCAGCTCCCTCAGGTTGGCTGTGTGgaatcttaaacccaac
ggttacctcttatctagtggCTCTGGAGCAGTCACCAGCTCTTGAGCCAATGGTGTTAGCgaatcttaaacccaac
ggttacctcttatctagtggCCAACGTTACCCCCTTTTGTGGAGCTCTCAGCCAGAGTGGgaatcttaaacccaac
ggttacctcttatctagtggGTAACAGCAGAGAGTTAAATGAAGAAGCGACAGATGGCCCgaatcttaaacccaac
ggttacctcttatctagtggAATTCCACATCTTGGAGGCTTGGGGCTCAGAACACCATGCgaatcttaaacccaac
ggttacctcttatctagtggAAATTGTGTGTGTTTGTGTGTCAGCCTCTCTGCAAACACCgaatcttaaacccaactt
ggttacctcttatctagtggATGTCGTGTCATGTTTTCTTTCCTCCATCTCCTTTTAATGgaatcttaaacccaacttgt
ggttacctcttatctagtggGGCTCCCCCCAGCTCAGCCCTCTCTGCTTAGATCAGGAGCgaatcttaaacccaa
ggttacctcttatctagtggGGGCACTGCGACGTCAGCACTGGCACGGGGAGGATTTGCCgaatcttaaacccaa
ggttacctcttatctagtggCGAGCTCAGGAGGCCTCTGCCACAGTTATCTGTCCTGCTGgaatcttaaacccaac
ggttacctcttatctagtggAAGCAGTAAGAGCTTTTCGAGGGATGGGGAGATAGGCGTGgaatcttaaacccaac
ggttacctcttatctagtggTGTACGATAGGGTTATGTCCCAGTAAACACGGAAAGTTGGgaatcttaaacccaact
ggttacctcttatctagtggACTTCACCTCACCTACCTGAAATGCACTCAGAACACTTCCgaatcttaaacccaact
ggttacctcttatctagtggATGGGTACTCAAAATATGGTTTCTACTGAGTGCGTATAGCgaatcttaaacccaactt
ggttacctcttatctagtggGTTTGTCTCACCTAAGTAGTCAAGAACTGTCTGTAGTTGGgaatcttaaacccaactt
ggttacctcttatctagtggGTTCTCATTACGAAATAATGTTAAGTTCTTGAGGTGATGGgaatcttaaacccaacttg
ggttacctcttatctagtggCCACACTGTATTCGTAAATTATAACATCACTTTGTGCTCCgaatcttaaacccaacttg
ggttacctcttatctagtggGGTTCATCAGCTGAGATGACAAGCAGAGGCCAGAACAGCCgaatcttaaacccaa
ggttacctcttatctagtggGTGGAAGCAAATCAGGGTAATGAGGTTGGTGTCAGAGGAGgaatcttaaacccaac
ggttacctcttatctagtggGCAAGTACTGTCTCCCCCCACCCTCTTATCAGTGAGGCTCgaatcttaaacccaac
ggttacctcttatctagtggGTGCTGTGTCGCAAATCCAGGTGCACCAGATTCCAGAGCCgaatcttaaacccaac
ggttacctcttatctagtggTATGTATATTTAAAAACTGTGCTAATGAAGCTGACAGGGGgaatcttaaacccaactt
ggttacctcttatctagtggGCTCTGTAGCTCGACAGTGGTGCTGGCTGATCAAAACTACgaatcttaaacccaact
ggttacctcttatctagtggCAGGCACGCGCACGCACACACACACACACAAACCCTGTGGgaatcttaaaccca
ggttacctcttatctagtggGGGGGGAGCTGGTGTTGTGTCCTTGGGACCACCTTGCACGgaatcttaaacccaa
ggttacctcttatctagtggATTTCAAAGTTAAAAGTTGAGGCCGGGCTCAGTGGCTCACgaatcttaaacccaact
ggttacctcttatctagtggACCCCAACAACCTCACAGCTAACCCTTCCAAGAGAACTTCgaatcttaaacccaac

ggttacctcttatctagtggGGTGTCAGCTTCTATTTATTCCTGCCCCCATTACTGACTCgaatcttaaacccaactt
ggttacctcttatctagtggAGAGGTGCATGGTCTTTGGTGGTAAGAATTACACAAAAAGgaatcttaaacccaactt
ggttacctcttatctagtggGTCAGACAATACCAAAAGTGAATGGATAGGGAAGAACTGGgaatcttaaacccaac
ggttacctcttatctagtggGGGGAGCCATTTGGCAGTACCTAGTCAACCTAAAGGTGCGgaatcttaaacccaa
ggttacctcttatctagtggACTCCCAGAGACACTCTCACAAGTGTGCACGAGGATGTGCgaatcttaaacccaa
ggttacctcttatctagtggGCCAGGTCTGAGTTCTTCATTTCAGAACCATTCTGCCAGGgaatcttaaacccaac
ggttacctcttatctagtggTGGCATCCCACAAAGCACTCAACTGGAACCAGGCTGCGGGgaatcttaaacccaa
ggttacctcttatctagtggGAGGCTCTAGTTCCTCTGATGCCAGCTCTAGGAAATTCAAgaatcttaaacccaac
ggttacctcttatctagtggGGCCACTGTCAAGACAGGGAACTGGGCTCAATCAGTGACCgaatcttaaacccaa
ggttacctcttatctagtggGGGCATGCTGAATCCCTGCGAAGCCAAGAATGCACCGGGGgaatcttaaacccaa
ggttacctcttatctagtggGCCAGAGACCAGCAAGATTCAAAGCAGATGAGCAGGCCACgaatcttaaacccaa
ggttacctcttatctagtggGGCAGCTTGTGGGGGGCATGATGGAAAGATGTGAAGTGGGgaatcttaaacccaa
ggttacctcttatctagtggTGAGCAGCCATTGAATGGGAAGCAGGGTGAAGGAGCCACGgaatcttaaacccaa
ggttacctcttatctagtggCCTTTGTGCTATGCTCTGTGCTAGGTGCTTTCTACAGGTTgaatcttaaacccaactt
ggttacctcttatctagtggCTGACGGATCCAGGTTTCAAACCCAGGCCAGTTGGATCCCgaatcttaaacccaa
ggttacctcttatctagtggTCTCCCCTGTGGCTTCACTGCCTTTACTTAAAGTGACCGCgaatcttaaacccaac
ggttacctcttatctagtggTCCCGATATTTTGAACCATGAACCCAGATTCAGAGCTGGCgaatcttaaacccaact
ggttacctcttatctagtggGCAGGTGGTTAGAGTGTGGCCTTTAGAGACAGGTGGACAGgaatcttaaacccaac
ggttacctcttatctagtggGGAAAGGCCCTTAAACTCCCTGAACCTCAATTTCTATGCTgaatcttaaacccaac
ggttacctcttatctagtggGACAGCGGGCTCCACCTGGTTCCTGGCACCAAACAAAAGCgaatcttaaacccaa
ggttacctcttatctagtggTTTCTCCAACTAGATTCTAAGTTCATGTTTGGAGACACACgaatcttaaacccaacttg
ggttacctcttatctagtggCTGTAAGGGTTAGTCCAGAGTGCACTGCACAAAAACAGTCgaatcttaaacccaac
ggttacctcttatctagtggGAGAAACTGGTGTTGTTTTAAAACCTCAGCCCCTGCTAGGgaatcttaaacccaac
ggttacctcttatctagtggGAGACCAGCCTGGGCATACTGGCAAGATCCCCATCTCTACgaatcttaaacccaa
ggttacctcttatctagtggTGGTTGACAGAACCAGACTCTGTCTCAGAAGAATGAAAGAgaatcttaaacccaact
ggttacctcttatctagtggTTAGAGCAGGCTAAACTGGCAGGGTTGTCACAAAAGCTGGgaatcttaaacccaac
ggttacctcttatctagtggCGTTTCAGAACTAGAAGGCACATCTCTTAGATGCTGCTTGgaatcttaaacccaactt
ggttacctcttatctagtggGACTTCTGCCTGTTCCTAAATTGTGAAACCCTAGTAACTCgaatcttaaacccaactt
ggttacctcttatctagtggGTCTCTCAAGAAACAAGCTGCAAAGGTGGCTGGTCTTCTGgaatcttaaacccaac
ggttacctcttatctagtggTGCCAGTGACCAAAGTACCCCATATTTGTCTAGCTCCAGCgaatcttaaacccaac
ggttacctcttatctagtggCCATAACACCGGAGAAATGTCCTTGTTAGACCCATCGTACgaatcttaaacccaac
ggttacctcttatctagtggCCACAAAAAGGAAGAGGCAGAATGTAAGCCCAAGACTACCgaatcttaaacccaa
ggttacctcttatctagtggTTCTGAAAGCACCTTTCCTATCTGACCGACGATTGATCTCgaatcttaaacccaactt
ggttacctcttatctagtggGCTTCCTTCTTAGAATACCCCAGGTGAGAGCAGGCACACGgaatcttaaacccaac
ggttacctcttatctagtggTCTGCTACACGGCTCACTTCTCCCTGCCTTTATCATCTCCgaatcttaaacccaac
ggttacctcttatctagtggCCCAACCCTTAACCATACCATCGGGTTGTTCTCAAACTGAgaatcttaaacccaact
ggttacctcttatctagtggACAAGCACTATACAAATGTAGGCATCATTATTACCAAGGCgaatcttaaacccaactt
ggttacctcttatctagtggGCCTCTGTAGTTTCCAAGAACAGGAGTAATTGCAGTTGGCgaatcttaaacccaac
ggttacctcttatctagtggACCAAACCCAGGAGTATTTGGGAAAAGGCACAAGTTGGCCgaatcttaaacccaac
ggttacctcttatctagtggCCAGTTTCGTCAATCCTTCCTGCTACTTGGAGCAATGTAGgaatcttaaacccaactt
ggttacctcttatctagtggACCCTTTGGATCCTTTTAATCCCAGCTGATCATTTTCAGCgaatcttaaacccaactt
ggttacctcttatctagtggCCCTGGTGGACTTTGTCCCAGATTGTACCAGTAGCCACAGgaatcttaaacccaac
ggttacctcttatctagtggAGCAGAGGCTCAGAGAAAGCCCTGGAAGCAGCAAGCCAAGgaatcttaaacccaa
ggttacctcttatctagtggCACACATCTGGATGGGGCTGGGAGGGGCAGCGAGGTGGCCgaatcttaaaccca
ggttacctcttatctagtggCCCCCCTTGGGTGTCTGTGATGCGAGGTGATTTGGTAAGCgaatcttaaacccaac
ggttacctcttatctagtggGGATGTATAGGCTCACATCCTGGTTCTGCAATTTACTGGCgaatcttaaacccaactt
ggttacctcttatctagtggGAGAATCTTACAGGTGTGAACAGGGAACATCGTTTCAGGGgaatcttaaacccaac
ggttacctcttatctagtggTCCTCATCTGGAAAATGGTGATCATAGTAGCATTGGCTTCgaatcttaaacccaactt
ggttacctcttatctagtggCCTGACTCCTTCCTCGCCCACAGCCATCTGCCCTCCACACgaatcttaaacccaa
ggttacctcttatctagtggACCCTTCCACCATTCCCCAACACCTACCGGATAGAGCCCGgaatcttaaacccaa
ggttacctcttatctagtggCTCCTCATCAATGTCCCCCGTCCCTGCCTCCCTTGTAGCCgaatcttaaacccaa
ggttacctcttatctagtggCCACTCATTGTTCACCAGGTCCCTAGTGCCTGCCACACAGgaatcttaaacccaa

ggttacctcttatctagtggCTGCTGCGTGGTAACAAGTTCCCTAACCCCTCTGGAATGGgaatcttaaacccaac
ggttacctcttatctagtggGAAAGATACAGAGTATTGATCCTGATTGTGTCTGTGAGGGgaatcttaaacccaactt
ggttacctcttatctagtggGGCAGACCCACCCTTAATCTGGGCACAATCTAATCAGCTGgaatcttaaacccaac
ggttacctcttatctagtggGGGACTCAGACTGGCTCTCCTTGCTCCATAGCCTGCAGACgaatcttaaacccaa
ggttacctcttatctagtggAGAGAACCCTGACTAATACACCCTCTGAGCTTCCATTTCCgaatcttaaacccaact
ggttacctcttatctagtggGCCTTGCACACAGTGGGAGCTCCAGAGCCACCAGAGGCCCgaatcttaaaccca
ggttacctcttatctagtggCCTCCCTGAAACCAGAAAGCAGAGCAATACAAGGCAAGACgaatcttaaacccaa
ggttacctcttatctagtggCTAGGGAGGCAAGTGAGGAACACAGACCAACTTCTCACTCgaatcttaaacccaa
ggttacctcttatctagtggGAAGGTGATCACTTAAGGAGGCAGGTTATAAGGGGAGGAGgaatcttaaacccaac
ggttacctcttatctagtggGCCAGGAGGCTCTGAAAGGTCATGAGTTTCAGAATTTAGCgaatcttaaacccaact
ggttacctcttatctagtggAGGCAGTGCAGTAGACGTGAAGCATGTAAGTGACCCCCTCgaatcttaaacccaa
ggttacctcttatctagtggGATAAGTGGAGAAAATAAAGCCACATTAACAGAGACCATCgaatcttaaacccaact
ggttacctcttatctagtggACAGAACAAAGTTGAAGGACTAATATTACCTGATCCTGAGgaatcttaaacccaactt
ggttacctcttatctagtggGACAGACAAATAGATCGATGGAAGAGAATACAGGGTCTAGgaatcttaaacccaac
ggttacctcttatctagtggGGTGCAGCGGAAAAAAGGATTGACTTTGCAACAAATGATGgaatcttaaacccaact
ggttacctcttatctagtggAAGAAAACAGGAGAAAATCTTCATGACCTTGGGTTTTGCAgaatcttaaacccaactt
ggttacctcttatctagtggTCCCAGCACTTTAGGAGGCCAGTCAGGTGGATCACTTCAGgaatcttaaacccaac
ggttacctcttatctagtggCCACTTATCAAAAGATACAAGCACAGACTGAAAGAAAGTCgaatcttaaacccaact
ggttacctcttatctagtggGCACATAAAATATGCTAGGCCAGGTGCAGTGACTCACACCgaatcttaaacccaac
ggttacctcttatctagtggTGGGAGGCCAAGGTGGGCAAATTGCTTAAGCTCAGGAGTTgaatcttaaacccaac
ggttacctcttatctagtggCTCCTGAGCCTGGGAGGTCAAGGTCACAGTGAGTCAAGACgaatcttaaacccaa
ggttacctcttatctagtggCACTCCAGCCTCATGACAGAGTAAGACTGTCTCAGAAAAAgaatcttaaacccaac
ggttacctcttatctagtggAGGAAATGTACGCTAACACAGTGAGATGCCGCTACACATGgaatcttaaacccaac
ggttacctcttatctagtggTGCTGCTGCTGGGAATGTTAAAACGGCACCATCACTTTGGgaatcttaaacccaact
ggttacctcttatctagtggAAGTTAAATACATACCTACCTCATCATCCAGCCATTCTTCgaatcttaaacccaactt
ggttacctcttatctagtggAAAAACTGAAAACAGGGCCAGACATGGTGGCTCATACCTGgaatcttaaacccaac
ggttacctcttatctagtggGGATCACCTGTCAGGAGTTTGAGACCACCGTGGCTAACATgaatcttaaacccaac
ggttacctcttatctagtggGAGGCTGAGACAGGAAAGTCACTGAAACCCCTGGAGGCGGgaatcttaaacccaa
ggttacctcttatctagtggAAGGAAACATATGTCCATGCAAAGACTTCTACGTGAATGCgaatcttaaacccaact
ggttacctcttatctagtggATTCAAAAGCTAAAAGCAACCACATGTCTATCCCTGGGCGgaatcttaaacccaac
ggttacctcttatctagtggGTAAAAGAGCCAGACAAAAAGAGTTCACTCTGTATGCTTCgaatcttaaacccaact
ggttacctcttatctagtggGGTGGGGAAAGGGTGCAACAACAGAAGTCACTACAAAGGGgaatcttaaacccaa
ggttacctcttatctagtggAGATATGTTCATTATCTTGATTGTGGGGATGGATTTGTGGgaatcttaaacccaacttg
ggttacctcttatctagtggGATGCGGCCGGGCATGGTGGCTCAACACTGTAATCCCAGCgaatcttaaacccaa
ggttacctcttatctagtggTACAAAATTAGCTGAGAGTGGTGGTGGGTGCCTGTAATCCgaatcttaaacccaact
ggttacctcttatctagtggGAAAGATTCCTGATATACTGAAAAAGTGTCAAGATGGGGGgaatcttaaacccaact
ggttacctcttatctagtggACTTGAATCCCAAGTGCTCTAAGAGCAGACAAATGCTGCTgaatcttaaacccaact
ggttacctcttatctagtggACTGGACTTCAGTCTACCAGAGTTCAGGCAGATGGAAAGTgaatcttaaacccaac
ggttacctcttatctagtggTTCAAAGTATTTGCACAAAGCTGTTCTACACAGAGAGCTGgaatcttaaacccaactt
ggttacctcttatctagtggCACACACACACACACACACGAGCTTAAAAACAGTGCCAGTgaatcttaaacccaa
ggttacctcttatctagtggCCAACCCAATGTCTGGCTGGAAGCAGTCATTCTGAGCTTCgaatcttaaacccaac
ggttacctcttatctagtggACATCAGACAGAGGCAGCAGGAAGAGTTCCACTGAGGAAGgaatcttaaacccaa
ggttacctcttatctagtggCAGCCCTTGAGAATGAGGGGAGATGAGACCCAGCAGAGGGgaatcttaaacccaa
ggttacctcttatctagtggCCTAGCAGCAAGGCTGAAACTCTTGACAGCTACAATTGCCgaatcttaaacccaac
ggttacctcttatctagtggACAGATCAACAGGTCTGAAAGGGTTCATCTGGTGTTCCCCgaatcttaaacccaac
ggttacctcttatctagtggGTGGTGATTTCTGAGATTTTGATATGCCCATCACCCGAGCgaatcttaaacccaactt
ggttacctcttatctagtggTCCTTCATCCCCCTTCCAACTTCCCTTCTACTGAGTCCCCgaatcttaaacccaact
ggttacctcttatctagtggCACTTAGAATAAGGGCCTCCAGCTTCATCCAAGTTGCTGCgaatcttaaacccaac
ggttacctcttatctagtggGGCTGAGTAGTATTACACAGTGTAGATATGCCACATTTTCgaatcttaaacccaactt
ggttacctcttatctagtggGCAAATGCAAACTATGCTGCTATAAACATGCATGTACATGgaatcttaaacccaactt
ggttacctcttatctagtggTCTTTTCCTTTGGATAGATACGCAGTAGTGGGATTGCTGGgaatcttaaacccaactt
ggttacctcttatctagtggCCACCCGCAGCATAAAAGTGTTCCCTTCTTACCACATCCGgaatcttaaacccaac

ggttacctcttatctagtggATTGTCTAGGTGTGACAAACGAGTTCACTAAAGTCTCAGGgaatcttaaacccaactt
ggttacctcttatctagtggGTAGCACTTCTCTACATGAACAATGATCAAGCTGAGAATCgaatcttaaacccaactt
ggttacctcttatctagtggCCAAGGAGGTGAATGACTGCTACAAGGAAAACTGCAAAACgaatcttaaacccaac
ggttacctcttatctagtggGGACGGGTAGAATCAATATTGTGAAGATGATCATACTGCCgaatcttaaacccaact
ggttacctcttatctagtggCCCCCTTGTTTTGAAAACAAAGAAACTTAAGGTTTGAGGGgaatcttaaacccaactt
ggttacctcttatctagtggCTCTGAGAGTCAGCTCAGGCGATGCTAGAGCCAGACTCCCgaatcttaaacccaa
ggttacctcttatctagtggTCCTGTGTCTTGTGTTTCCCCTACTCCTGAAAGTGCAACTgaatcttaaacccaactt
ggttacctcttatctagtggAGAACCCATAGGGCATGTGTGTGTGTGTGTGCGTGCACgaatcttaaacccaac
ggttacctcttatctagtggGGGTAGGTGGGCTGGCTGAAGACCCAGAGAAGAATCACAGgaatcttaaacccaa
ggttacctcttatctagtggGCTGTCTGATGGCAGAATTCCTTCTTCCTTCAGGGAAATCgaatcttaaacccaactt
ggttacctcttatctagtggCCCATGTTATTGAGGGTTATCCACTTTACACAACCTCCACgaatcttaaacccaactt
ggttacctcttatctagtggCAGTCACGGGGTGCTCTGAGACTAGCCCTCATCTTCTCCCgaatcttaaacccaac
ggttacctcttatctagtggCCCTGGCACCTCATCTTCCCCCTCCCCTTGTTTGCTCATCgaatcttaaacccaac
ggttacctcttatctagtggGGCTTCCAACTTTGGCTGCACATGAGAATCATCTGGGGGGgaatcttaaacccaac
ggttacctcttatctagtggCAGGCTGGAGTGAGGTGGTGCAATTATTCTAAGATGCTCAgaatcttaaacccaact
ggttacctcttatctagtggCATCTCTGACACTGGAATGCCACTTGCAATTCATAGTTTCgaatcttaaacccaactt
ggttacctcttatctagtggAGAATGTGGTCTATTCAAAAGGTCCAGGGCAACTCTAGGCgaatcttaaacccaac
ggttacctcttatctagtggGGCACGCGCCACCACATCTTGCTAGTTCTTTGTATCTTTAgaatcttaaacccaactt
ggttacctcttatctagtggGGAGTGAGCCACCACATCTGGCCCTGGGGAGCTTTATAACgaatcttaaacccaa
ggttacctcttatctagtggCTGCAGGTAAGTCCCATGCATTTAGTTTTAACTCCTTGGGgaatcttaaacccaactt
ggttacctcttatctagtggGTCCTGTGCCCTTTTCCTTCAAAGACTAATCCCACAGCAGgaatcttaaacccaact
ggttacctcttatctagtggATCTCATCTGCTTCTATCTAGTGATGGATTACATGTTTGTgaatcttaaacccaacttgt
ggttacctcttatctagtggCCCAAAAAATGAGAATCCCAGACAGACCAGTGGGTGAAGGgaatcttaaacccaa
ggttacctcttatctagtggCCCGCGAATTACATCTCACAGAAGCTCACCTGCTTGAAGGgaatcttaaacccaac
ggttacctcttatctagtggGAGTGCATATTCATGAGCCATGGATGTCAATGGCAGCAAGgaatcttaaacccaac
ggttacctcttatctagtggCACTCAACCAGTCGGATCTGCTGAGTCAATCATGGAATCCgaatcttaaacccaac
ggttacctcttatctagtggGGGTGTCTTGCCAGACCTAATGCTGTAGTGCTCAGTTTTCgaatcttaaacccaactt
ggttacctcttatctagtggGGAGGCCAGGCTGAGAAATGATAAGCAATGTGGCCAAGGCgaatcttaaacccaa
ggttacctcttatctagtggTTGACTCCAAAACGTCCTGACACACCTCTCTTTTCTCAGCgaatcttaaacccaactt
ggttacctcttatctagtggGGACTCATGCCTCAACTTGCCCATGCCAGCTCTCCTTCGCgaatcttaaacccaac
ggttacctcttatctagtggGGGCTGTCTTTCTATCCACTGACTTTTCAGAAAGTTGAACgaatcttaaacccaactt
ggttacctcttatctagtggACTCCCCTAACTTTCTTCACAATTCCTCCTCCCCCCATCCgaatcttaaacccaact
ggttacctcttatctagtggCTGGCTCCCAACGACTACATTCGTACAATATGATACCCCAgaatcttaaacccaac
ggttacctcttatctagtggCTCTGACTGAGAATGAACCAATCAGATTCTTTTCCTCTGCgaatcttaaacccaactt
ggttacctcttatctagtggTCTCCATGTGGTCCTAGCTGTAACACATGACTTGGATGCCgaatcttaaacccaact
ggttacctcttatctagtggGCCCAGCTCCCTGCCTTCTAGGAACCTGGTAGGATTGTCCgaatcttaaacccaa
ggttacctcttatctagtggGTGCCTAGTGCCTGCCAGTCACTAGGTTTGTGAGCAGATGgaatcttaaacccaac
ggttacctcttatctagtggGCCTGGGTCTCAGAGTCGTGAAGCAGGAACTGGAGTTCCCgaatcttaaacccaa
ggttacctcttatctagtggCCACTAAGGTTTGGGGGTTGTTCGTTAATATGGCATAGTCgaatcttaaacccaactt
ggttacctcttatctagtggAGGGTTTGGGAATTATAACTACAGCATAGTCTAGACCAAGgaatcttaaacccaactt
ggttacctcttatctagtggTTACACATTGACTATGGTTGCCTGCATGCTACAAGGGCAGgaatcttaaacccaact
ggttacctcttatctagtggAGTTTGCTGACCACTGACTTAGACCATTCTGACTGATTACgaatcttaaacccaactt
ggttacctcttatctagtggACCAGTTTCCCTGCCTCCGTGCTTGCATCCGCTGCCTTCCgaatcttaaacccaac
ggttacctcttatctagtggTCAATCATTCAGTGTCTCTTCATCACTTATCTCCTACCTGgaatcttaaacccaacttg
ggttacctcttatctagtggTCCTCATCTGTAAAGCAGGGGCGATATCAGTAACTACCTCgaatcttaaacccaact
ggttacctcttatctagtggGCTCATTCAGCCCTGACACTTGGCACAGTTCAATGCCTGGgaatcttaaacccaac
ggttacctcttatctagtggGATGTTTTGGTCAACAGACTGCATACACAACAGTGGTGCCgaatcttaaacccaact
ggttacctcttatctagtggGCTGCAAAGTTTCTGTTGCCTAGTGATTTGTTACAACTGCgaatcttaaacccaactt
ggttacctcttatctagtggGGGCTTTACCCTACAGCCTAGGTGCGTATTAAGCTATCCCgaatcttaaacccaac
ggttacctcttatctagtggTGTAAGTACCCTACTCTCTACGAGGTTCCCACAAAGACAGgaatcttaaacccaac
ggttacctcttatctagtggTTCTCAGAACAAGTCCCCATGATCAAGCAACACATGACTGgaatcttaaacccaac
ggttacctcttatctagtggAATAACAAAACCCACAACACTGAAAGCAGACTCAAGTATCgaatcttaaacccaac

ggttacctcttatctagtggACTGCCACATAATACCAATACATAGTAGGGACTCAATGAGgaatcttaaacccaact
ggttacctcttatctagtggCCTGACACTGACTTCTAGGATCCTTGCTAGGGGACATCCCgaatcttaaacccaac
ggttacctcttatctagtggCCTGCCAGATAAATGCAGTTGAGTGATTCCTTCTGAAGCCgaatcttaaacccaact
ggttacctcttatctagtggCCATGACGCTATAAAGAAGGTCATTCAGCAAACTAGATGGgaatcttaaacccaact
ggttacctcttatctagtggTTTTGAATTCCAGAGAAAATGGTGTTAGGTAAAGAGTGACgaatcttaaacccaacttg
ggttacctcttatctagtggGAACTTAGGTGTGTTTGAACTTGTCCCTGAACTCTTAGTCgaatcttaaacccaacttg
ggttacctcttatctagtggTGCCTCCTGTGAGACCCACAAGGCAGAGGGATTTGATGGGgaatcttaaacccaac
ggttacctcttatctagtggCTCCTTCTGTCACTGCTTAGCTCTGCTTAGCTGTACCTACgaatcttaaacccaacttg
ggttacctcttatctagtggCCTCTGTGAATCTCATCTTGCCCATTTGTTTAAGCTGCTGgaatcttaaacccaacttg
ggttacctcttatctagtggGCGTTTCAAGGGAAATACACCCTGAACAGCATCATCAAGCgaatcttaaacccaac
ggttacctcttatctagtggCAGCCTCTCCCCGATAACATCAACACACCAGCCTGATAGGgaatcttaaacccaac
ggttacctcttatctagtggCAGGAAGCGCCCTCACTCTCTTTGAACTCCAGGTTGGGGCgaatcttaaacccaac
ggttacctcttatctagtggGGCTGGCTGAAGAGTATCGCATCATTCCCAGAAGACCAGCgaatcttaaacccaac
ggttacctcttatctagtggCTGCCCACCTCGGCCTCTCAAAGTGTTATTGTTACTTCTAgaatcttaaacccaacttg
ggttacctcttatctagtggCCACTTAACTCTGTGCACCGGCCAAAATGATTTATCTGCGgaatcttaaacccaact
ggttacctcttatctagtggGTCCCCCAAAAAGACATGTGAAAGCTCTAACCCCGAGGACgaatcttaaacccaac
ggttacctcttatctagtggTGAGAAGCAGCGTAGAGACAGATACACAGGGAAGAGGGCCgaatcttaaacccaa
ggttacctcttatctagtggGGCAGAGACTGAAACGATGCATCCACAAGCCAAGGAAGGCgaatcttaaacccaa
ggttacctcttatctagtggTTCACCCCAAAGCCTTCAGAAGGAGCACAGCTGTAGGGCCgaatcttaaacccaac
ggttacctcttatctagtggGCCACCCAGTTTGTGGTACTTTGGTATGGCAGCCCCAGAGgaatcttaaacccaac
ggttacctcttatctagtggAGCCTCCAAGCTGCCATCTCCTGCCTCTTTTCCTTCACGTgaatcttaaacccaact
ggttacctcttatctagtggGAGCTCAGCATCTTCTCTCAAGCCTATCCCATATCCTCGGgaatcttaaacccaact
ggttacctcttatctagtggCCTGTAATCTCCGCTTCCCTCAACCTAACACTCTCCACGGgaatcttaaacccaac
ggttacctcttatctagtggGTCTCTCTCTACCCACAAGACTACATCTCAGTCATCTTTGgaatcttaaacccaacttg
ggttacctcttatctagtggGCCCCTTGCTCAGTAGACATACAACCAAAATGGAAAGATGgaatcttaaacccaact
ggttacctcttatctagtggGAAAGTGGTCAGATGAAGGAGATGGGTAATGTCTGAAGTCgaatcttaaacccaact
ggttacctcttatctagtggTCTAAGGGTACAAATGACTAAGGCTATAATCTAGTGAAGGgaatcttaaacccaacttg
ggttacctcttatctagtggGGTTTGGGATATAGAACACGAGAGGGGAAGCAAAGTCAGCgaatcttaaacccaac
ggttacctcttatctagtggCTGGCCTGGAGCACTTACCTTCCAGATTCAGCTATGAGCCgaatcttaaacccaac
ggttacctcttatctagtggCCCACACTCAAACTAGGGGACGCCACTAGGTGCCTCATCCgaatcttaaacccaa
ggttacctcttatctagtggGGGGAACCTAGGACAGACCTGCTGATACAAGTGATGGTGGgaatcttaaacccaac
ggttacctcttatctagtggCCCACTGGAATAGCAACGGCGTGAGGTGCACAGTGGTAGGgaatcttaaacccaa
ggttacctcttatctagtggTAGACCTTGCCCACCAGGTGACATAAGGGTCATTATCCCCgaatcttaaacccaac
ggttacctcttatctagtggGAGTGCTTACCTCATTACGTCTTGCTCTACATGTCCACCCgaatcttaaacccaactt
ggttacctcttatctagtggCATGCTCAGAACTTGCTAAGGCCAGAGGTGCTCAGTAGTGgaatcttaaacccaac
ggttacctcttatctagtggGCCTCTGTAAGTGCAGTGGATACCCCTCAAAAACCCACCCgaatcttaaacccaac
ggttacctcttatctagtggGCCTTCGCCTGGTCATACATGGCCCTTCATTGCTCTCCTCgaatcttaaacccaact
ggttacctcttatctagtggGCTTCCAGGAGCTCCCCCTACCCACTCCACCTCCCATGACgaatcttaaacccaa
ggttacctcttatctagtggGGACTCAGCTCAGAAGTCCCAGAGAACACTGTGGCTTCTGgaatcttaaacccaac
ggttacctcttatctagtggACCCCTAGTGCCTAGCACAGAGCCCAGCATGCAGGTTGCCgaatcttaaacccaa
ggttacctcttatctagtggACTGTGGCAGTCCATCCACTCTCTGATTTCTCTTCTCCCCgaatcttaaacccaactt
ggttacctcttatctagtggGGCAAGCTGCAGACCTTGCCTTATCTGCCATTCTTCTAGAgaatcttaaacccaactt
ggttacctcttatctagtggAGATCTGATGGAGACTGAGATGGGTAGGGTCCAGATCCTCgaatcttaaacccaac
ggttacctcttatctagtggGGGCCAGGCATTTCCCAGAGTTCTTCTGATCCCAGGGTCCgaatcttaaacccaac
ggttacctcttatctagtggAAGTAAAGCAAGTTAGAGAACCGTTCCAAGGGGCCTAGCCgaatcttaaacccaac
ggttacctcttatctagtggGCAACAGACAAGAAGAATCCAGACTTCTGGGCCAGAGAAGgaatcttaaacccaac
ggttacctcttatctagtggCACATCTGTGGAATAAGGGACAGTCACTGGGCTGACTTTTgaatcttaaacccaact
ggttacctcttatctagtggAGATGAATTATTAAAGCAGTGCTTGCCCTGAAGGAATCATgaatcttaaacccaacttg
ggttacctcttatctagtggCATAGGTCAACATCTTGACTCCATGCCCAATATGCTGTTCgaatcttaaacccaactt
ggttacctcttatctagtggCTTTGGGGCCTTTCAAATTTTTCACACTGTGGACCACTTCgaatcttaaacccaacttg
ggttacctcttatctagtggGCAGGGTGTAAAAGGGGAAATTACTAAGAAGTCAGGGATGgaatcttaaacccaact
ggttacctcttatctagtggACCAGTATGAAAAGAGGCCAACCCTGAGATCATCAACTCCgaatcttaaacccaac

ggttacctcttatctagtggCCTCCAGGAAGCCTCCTTGAGTCCCCGCCTTATCCAACAGgaatcttaaacccaac
ggttacctcttatctagtggACCTGGGCCTTCAGCATGCCACACAAACCTATCATTTCTGgaatcttaaacccaact
ggttacctcttatctagtggTGCACCATGAGCTAACTCAACAAGGACTGAGTTTTGCCCCgaatcttaaacccaact
ggttacctcttatctagtggGACTGGTACATATAGGTGCTTCATGAGTCATGAGGGTCTGgaatcttaaacccaactt
ggttacctcttatctagtggCACAAGTCTAGAAGGAGAAAGTCAGTCCGCAGGTTATCAGgaatcttaaacccaac
ggttacctcttatctagtggTGCCACTCTGCACTGCCAGACAAAATGACAGCCAGCCAGCgaatcttaaacccaa
ggttacctcttatctagtggCACTCTGAAGTCACTTACCCGTGCAAAATTAAATCCTCACgaatcttaaacccaactt
ggttacctcttatctagtggGATGACCCATCATATGAAGGTTCCTAATTCTCTCAGGGGGgaatcttaaacccaact
ggttacctcttatctagtggCTCAGCCAAGATCTCCCGCCTTCTCCTTGCCAGGGGCCTGgaatcttaaacccaac
ggttacctcttatctagtggAGGACACCTGCCAGCAGGGCTCTTGACCTGATTCCCAGGGgaatcttaaacccaa
ggttacctcttatctagtggGCCTAAGAGAACAAGACAGAGGCAGGGTGGAGGCACAGCCgaatcttaaacccaa
ggttacctcttatctagtggGCAGCTCTGGGCACACACATTGGCTGTTTACCTGGGATCCgaatcttaaacccaac
ggttacctcttatctagtggGCCTGAGGGGAAGCCACAGCCTTCACAAGGAAACCGGCCCgaatcttaaacccaa
ggttacctcttatctagtggCGCCCCGGGCCAGTTCCTATGGAGGCTGCCAAGAGAGCCCgaatcttaaacccaa
ggttacctcttatctagtggTGGCAGGGAGGTAGTGCTTGCCCGGGGTGCTGGGGAGCTGgaatcttaaacccaa
ggttacctcttatctagtggTGGGACTTGCTTCCTGTCCTTTCTCCATTGTCCACCTTTCgaatcttaaacccaacttg
ggttacctcttatctagtggTGCAACAGCCTTCTAGCCATCCTCTTCCTCAGCTGCTGTCgaatcttaaacccaact
ggttacctcttatctagtggGAGGACTCACCTGAGCTTCTCACTCACCTTCAGGAACCCCgaatcttaaacccaac
ggttacctcttatctagtggACAGGACATACACTGCCCCACCCAACCTTAGCCTCCACAGgaatcttaaacccaa
ggttacctcttatctagtggGCTCAAAGAGGGGGCCAAGGAGCAAGTTTAGGGAGTGGTGgaatcttaaacccaac
ggttacctcttatctagtggCCTCCATTAGCAGCACCCTCATTCTTTAATGCGCATTGTCgaatcttaaacccaactt
ggttacctcttatctagtggGGCACCCTGCCATGCCCTCATCACATTGCCAGCTTAGAGGgaatcttaaacccaac
ggttacctcttatctagtggCCCCAGAGGGAACGATGCCACTGAACCAGCAACTGTGCCTgaatcttaaacccaa
ggttacctcttatctagtggTTCCCTTCCTACTCAGCTTCCTACCTGTTGACCAAACTTCgaatcttaaacccaacttg
ggttacctcttatctagtggCAAGCCAATTGCCATTTGCTCCGTGAGATCATTCCTGGCCgaatcttaaacccaact
ggttacctcttatctagtggCCCTCTGGGCTCTCATCATCTCTCGGCGGACTTGCCTTGGgaatcttaaacccaac
ggttacctcttatctagtggGTGTCTCTCCCCATGCGAATGTAACCTCCTTGAGGACAGGgaatcttaaacccaac
ggttacctcttatctagtggGTGGACCCAGTCTGACCTGTAGCTCTGTCCAACCCTCCTGgaatcttaaacccaac
ggttacctcttatctagtggGCCACCTCCTGGCCTCCCTGCTTCCTGTTCAATTCCCTAGgaatcttaaacccaac
ggttacctcttatctagtggGTCCCTACCACAGCCCACCAGGCCCTACGTGACGTGGCCCgaatcttaaacccaa
ggttacctcttatctagtggGGTACCCCCTTTCCTCTTCCCCAACTCTCCACACCTCACGgaatcttaaacccaac
ggttacctcttatctagtggGCCCACCCAAAAGCATTCCCACCTCACCCCATTGCTCTGCgaatcttaaacccaac
ggttacctcttatctagtggCAACATGGTCACCCCTCATTTCTTCCCACTCTGCAGTCCCgaatcttaaacccaact
ggttacctcttatctagtggCAGCAAGCGTCTCCTGAGTAAGCTGCTCTGGGCGCACCCGgaatcttaaacccaa
ggttacctcttatctagtggCTTGGGGCAGATCCTGAAAGACCCAGGGTGAAGGGGATAGgaatcttaaacccaac
ggttacctcttatctagtggTTCTCCCAGTTCTGATCCATGTCTGGGCACAGAGGTGTGGgaatcttaaacccaact
ggttacctcttatctagtggGGCTGCAAGAGCCTGCACATGGCACGCCTAGTACCTGCTGgaatcttaaacccaa
ggttacctcttatctagtggTCCTGAGCCCTGTATCAGCACCCAGGACTGCTCGCAGAGGgaatcttaaacccaa
ggttacctcttatctagtggAGCCAGAAAGCATCCTGCCTTGGCTAAATCACACTCCTCAgaatcttaaacccaac
ggttacctcttatctagtggACCCTTGTCTGCACTCCAGCCATGACAATAGTCTGGTCACgaatcttaaacccaact
ggttacctcttatctagtggCCAGGGGCTACAGGATCTTTCCAGTCCTTTCCTGACTCTCgaatcttaaacccaact
ggttacctcttatctagtggCTCCCAGCATGCCATACCCTCTAAAGCCTCCACACCTCTGgaatcttaaacccaac
ggttacctcttatctagtggTCCTTCCTCCTGGACTGCCCTTTCCCCTCTCCTTTGATTAgaatcttaaacccaacttg
ggttacctcttatctagtggCCTCCTTCGAGTGGGTTCCAGTCCAGGCCAAGTGCTCTCCgaatcttaaacccaac
ggttacctcttatctagtggCCAGGGCTTTCTCCCCAACTAGGGCTATGAGCTCTTAGAGgaatcttaaacccaac
ggttacctcttatctagtggCAAAAATTAGCCATGCATGATGGTGCACCCTTGTAATCCCgaatcttaaacccaactt
ggttacctcttatctagtggGAGAGAAAGCTGAACCTTTATTCACCAGCACTAAAAGGCCgaatcttaaacccaact
ggttacctcttatctagtggCAGTTGGTGTCACTCCCTTGCTCAAAAAGCCCCCAGTGGCgaatcttaaacccaac
ggttacctcttatctagtggCCACCCATCAGCTCTCTGCCTCTACCTCCTGGGACCTGCCgaatcttaaacccaac
ggttacctcttatctagtggAGGGCCTATGCACTGCTGTCCTTGCTTCTGGAACACCCTGgaatcttaaacccaac
ggttacctcttatctagtggTGTGACCATCTCCCTCACCTCCTCCTATCCCTGCAGTGAGgaatcttaaacccaac
ggttacctcttatctagtggCCAGCCCACTTAAAACCAAAGCCCACCACACTCTAAGCACgaatcttaaacccaac

ggttacctcttatctagtggCCTATGTCCCTTCTCCACCTTGCTTTTCTCTATAGACTTCgaatcttaaacccaacttg
ggttacctcttatctagtggGGTTATTGTCCATATTGCCTCACTACAGCTTCAGAGGGGCgaatcttaaacccaact
ggttacctcttatctagtggACCTCGTTATTAGAGACTCAATGCTTTGAATAGTGCCTGGgaatcttaaacccaactt
ggttacctcttatctagtggCTCTAGCATCTAGCATATTTCCTGGTTCATAGTAGACATCgaatcttaaacccaactt
ggttacctcttatctagtggGACAGAAGGGAAGGAGAAAAGAAGAGAGGCAAAAGGGGAGgaatcttaaacccaa
ggttacctcttatctagtggGAGAAGAAGGAAGCAAGAAAGAAACAGATACAGTAGGGAGgaatcttaaacccaa
ggttacctcttatctagtggGCCTAAAGTCCTCAGACAATTGTGTGTGTTGTGGGCTTAGgaatcttaaacccaact
ggttacctcttatctagtggACCTATCTTAGATTTCCTGGGAAATTTCTCTGCCCTCAGCgaatcttaaacccaactt
ggttacctcttatctagtggTTTGCCAACACAGACCCACCCAGAGATGAAAACATGAATGgaatcttaaacccaac
ggttacctcttatctagtggTGCATGCATGCAGTTCCTCCAAACCTTTGGCCAAGAGCTCgaatcttaaacccaac
ggttacctcttatctagtggTATTTTTTGAAATGGATCTCGCTCTGTCGCCAGGCTGGAGgaatcttaaacccaactt
ggttacctcttatctagtggGCTGGGACTACAGACATGCACCACCCTGCCCAGCTAATTTgaatcttaaacccaac
ggttacctcttatctagtggTCTCCATGTTGGCAAGGTTGGTCTTGGTCTCTTGACCTCGgaatcttaaacccaact
ggttacctcttatctagtggGGCCTGAGCCACCGTGCCCAGCCTTAAAAAGAATGCATTTgaatcttaaacccaac
ggttacctcttatctagtggATATACATGTATATGTAGTTATCAAGCCAGGCGCGATGGCgaatcttaaacccaact
ggttacctcttatctagtggGCTGGATACGTGCTTAGGACATAGCCAGGGAAGCAGCAGGgaatcttaaacccaa
ggttacctcttatctagtggCAGACCAGTCCAAACTCTGATACTTGGTGGCTGAAGACCCgaatcttaaacccaac
ggttacctcttatctagtggGATGTTCCCCCATACTCTGCCAGGACTCAGGGACTGTCCGgaatcttaaacccaa
ggttacctcttatctagtggTGGTCCTTCTATCCTTCACCCTGTGAGCTTTGCTTTTCTCgaatcttaaacccaacttg
ggttacctcttatctagtggTCTTTGCTGTGTAACACAAACTACCCCCAAATTTAATGGCgaatcttaaacccaactt
ggttacctcttatctagtggGCATGGATGAGAGGGTGGTTCTGGCTTAGGGTTCCTCCTGgaatcttaaacccaac
ggttacctcttatctagtggCACCTAGGGCTGCAGTCATCTGAAGGCTGGATTGGGCTGGgaatcttaaacccaa
ggttacctcttatctagtggCCTGGCTGCTGGCTGGAGGCCTCAGTTTCTCACCATGTAGgaatcttaaacccaac
ggttacctcttatctagtggTTCTAATCTCAAAAGTGAGAACGCAGCCTGGTGCCGTGGCgaatcttaaacccaac
ggttacctcttatctagtggCCCAGCACTTTGGCAAGTCAAGACGGGAGGTTTGCTTGAGgaatcttaaacccaac
ggttacctcttatctagtggCTGGCAACATAGTGAGACCTCGTCTCTGCGAAAAAGAGAAgaatcttaaacccaac
ggttacctcttatctagtggCACCGCACTCCAGCCTGGATGAAAAAGCGAGACCCTGCCCgaatcttaaacccaa
ggttacctcttatctagtggAAAGTGTGAACACCAGGAGGGAGGCAGGATCACCGGGGCCgaatcttaaacccaa
ggttacctcttatctagtggACTGCTTGTAAAGTCCCTGGAACATAGCAGACATGTAGCCgaatcttaaacccaac
ggttacctcttatctagtggAAAATAATAGGCAGCCCTGGTAAGTTCTTCACAGCTAAGGgaatcttaaacccaact
ggttacctcttatctagtggTGTCCTTGAAGCCAACGGCAATTCACAACTAACTACTGAGgaatcttaaacccaact
ggttacctcttatctagtggTGTAATCTTAGCATGTCATTTGAGTGTCGCTGTCAGAAGCgaatcttaaacccaactt
ggttacctcttatctagtggGCCTGGTGTTTGAATATACATCATATTGAAGTCAAGGCTGgaatcttaaacccaactt
ggttacctcttatctagtggAGAATTCATCTAGCCCTATGGTGGACACAGTCGTTGATGCgaatcttaaacccaact
ggttacctcttatctagtggATTCACCCATGTGACCAAAAACCACTTGTACTCCAAGAGCgaatcttaaacccaac
ggttacctcttatctagtggCCTCCACTACCTGGGGCTGCAGTCATCTGATCCAAGTTACgaatcttaaacccaac
ggttacctcttatctagtggGCTATTTCCCTTTCCTTCTTCCTTTGCCCTCCACCCTACCgaatcttaaacccaactt
ggttacctcttatctagtggGGCTCTGCAACCCTCTGCCTCAGGGCCCTTTTCCTGGCTGgaatcttaaacccaa
ggttacctcttatctagtggTGGTCAGAGAGCCGGTGGATCAGTTTCCCAGGTCCTTCGGgaatcttaaacccaa
ggttacctcttatctagtggCACTGTCTCCCAGTGGTCCCTTGCAAGACAGAGTTGCAGTgaatcttaaacccaac
ggttacctcttatctagtggTCCTGTCTTACCTCTCCACACCTATTCTGGTGCTTCCTGGgaatcttaaacccaactt
ggttacctcttatctagtggATTATTGTCTCAGGATATTCCTCTGGGTGAACCTCCACCCgaatcttaaacccaactt
ggttacctcttatctagtggCTGTGCCGTGATCCATGAGGCCCTGCACACTCTGCCCTCCgaatcttaaacccaa
ggttacctcttatctagtggCCTCACGTTCACTACACTCTAGCCATCTTGGTTTCCTGGAgaatcttaaacccaact
ggttacctcttatctagtggGCATGCCTTCAACTGAGCAGCTTTCCTCCCTGATTTCCTGgaatcttaaacccaact
ggttacctcttatctagtggTGTTTATAGCCAACACTCTACCACCAGCCCCTCCAAGTAGgaatcttaaacccaac
ggttacctcttatctagtggCAGAAAGGAAAGTGACTTTGCTCTGGGTTTTTTCTTTTCTgaatcttaaacccaacttg
ggttacctcttatctagtggGGCTCACTGCAACTTCCGCCTCCAAGGCTCAAGCAATTCTgaatcttaaacccaac
ggttacctcttatctagtggGTAGAGATGGGGTTTTCATTATTTTGGCCAGGCTGGTCTCgaatcttaaacccaactt
ggttacctcttatctagtggAGGCACCGCACCTGGCCTGCTCTGTTTTAGTCACTGCATCgaatcttaaacccaac
ggttacctcttatctagtggAATAGTGCTGGATACATAGTAGATGTTCAACAGATACTTGgaatcttaaacccaactt
ggttacctcttatctagtggTACCTGCCAGGCCCCTAGCTGTTATTATTGGCGCCCCTGCgaatcttaaacccaac

ggttacctcttatctagtggGCATGTTGCCTAATATTACAGAGCTCCCCAGTGGCAGAGCgaatcttaaacccaac
ggttacctcttatctagtggCCATCTGGGTTGTGGAGGGATGAGAACCAGTCTGTCTCGGgaatcttaaacccaac
ggttacctcttatctagtggGAGAACCAGTTTTCTCATCGGTAGAATCAGATGAAGCACCgaatcttaaacccaact
ggttacctcttatctagtggAGAGCTTGGCACATAGTACGTATTTATTAGGTGCTCATTGgaatcttaaacccaacttg
ggttacctcttatctagtggAGTACTAACTTAAAAACATTCAGAAGAGCAGGCACATGCTgaatcttaaacccaactt
ggttacctcttatctagtggCATGCCTTATTTCATTCAATCCTTACAACACCCTCTGAGGgaatcttaaacccaacttg
ggttacctcttatctagtggCACTAAGACAAAGGCCATTGCTCTGACCCTGGACTCTGCGgaatcttaaacccaac
ggttacctcttatctagtggCTGTGCTATACTGCTTAGAGACGTGGCTCTTAGGTCCTTAgaatcttaaacccaacttg
ggttacctcttatctagtggCTGCTTCCCAGCAGGACCAGGGTAAAGGCATTTCCAAAGTgaatcttaaacccaac
ggttacctcttatctagtggCTGGGCTGCATTCATCGTAGGTTGTAGGATCTCACAGTGCgaatcttaaacccaact
ggttacctcttatctagtggTGAAGCCCACCTTCACCCTGGGTAGCAAGCTTCCCTTCGGgaatcttaaacccaac
ggttacctcttatctagtggTGCTTGCCTGGGTGTGTGTGTACCTCGGTATCTCTCACACgaatcttaaacccaactt
ggttacctcttatctagtggGCCCAGCGCTTGTCAGTGTGGTCTAGATCCTGACCTTCAGgaatcttaaacccaac
ggttacctcttatctagtggGGAAATTCTTTAGAATTGCAGACTATTTGCCATCTGTGGTgaatcttaaacccaacttg
ggttacctcttatctagtggTGAGACCAGCCTGTCCAATATGGTGTAACCGCGTCTCCACgaatcttaaacccaac
ggttacctcttatctagtggATTAGCCCTGGGTGTGCTGGTGCGTGCTGGTGAGTCCCAGgaatcttaaacccaac
ggttacctcttatctagtggGCAGTGAGCCGAGATCACACCCCTACATGCCAGTCTGGGCgaatcttaaacccaa
ggttacctcttatctagtggCAGCCCCAACCCATCTACCCCACTGTCCCCAACCAGACCGgaatcttaaacccaa
ggttacctcttatctagtggCTGGGGATGGGGCTCAGCACTTTATTTCAATGATCCCTCCgaatcttaaacccaact
ggttacctcttatctagtggGCTTTAGCCTGCAAAAAGCAATTTGTCCAATCTCCCCACCgaatcttaaacccaact
ggttacctcttatctagtggGCCCTGGGAGTGTTGGGGCCCCAGTCTCTGCCAGGCATTTgaatcttaaacccaac
ggttacctcttatctagtggCAAAAATCATAGCTTCTCGGAAAGCCACCTCCCTTTCACTgaatcttaaacccaactt
ggttacctcttatctagtggCCAAATAGAACGAGCTGGCTTTGGGGAGGGCCGGGGGTGCgaatcttaaacccaa
ggttacctcttatctagtggCCCTCCGCTCAGCGACCGCGATGGCCTCATTTGGTGCTGCgaatcttaaacccaa
ggttacctcttatctagtggCAGTCATATAAAGAGCGATAAAGAAAGACAAGGGCGGAATgaatcttaaacccaact
ggttacctcttatctagtggGTGTATTATGGCTTGTGGTTTTCTATTCAAACCCAGGGAGgaatcttaaacccaacttg
ggttacctcttatctagtggTGCGCTCCGCAGCTTGGATGGTTTGGGGAAGGCAGGAAAGgaatcttaaacccaac
ggttacctcttatctagtggTCCCTTTGGCCTGGGTTTTATTGCACCAGACGTTACCACCgaatcttaaacccaactt
ggttacctcttatctagtggGCATCCCGGGTCAGCCCTTCCTCTCAGGTTTATGTTTTCCgaatcttaaacccaactt
ggttacctcttatctagtggGGCTCACTGCTGGTGGCCCGGTGCAGCCTCAAAGCTCCTCgaatcttaaacccaa
ggttacctcttatctagtggGCGGGGTGATGCTTTGCAATTCAGACGGGGTAGCCTTTGCgaatcttaaacccaac
ggttacctcttatctagtggGGACTCAGGGTGGGGAAGGGAGAAAGGGGAAGGCTGGCCCgaatcttaaacccaa
ggttacctcttatctagtggGGCAGCTCAAGACAGTAAGGAGGTCTGGTTGTCCTGGAGGgaatcttaaacccaac
ggttacctcttatctagtggGGCCACCCTCATTCCCTGAGCAGTCACTAAATGTCCCCCCgaatcttaaacccaac
ggttacctcttatctagtggCCCCACTTCACAGACGTGGAAATCAAGGCATACAGAGCTGgaatcttaaacccaac
ggttacctcttatctagtggTCCAGGTCTCTTTGGCTTCAGAACTCACGCTATACCTGGCgaatcttaaacccaact
ggttacctcttatctagtggCTTGGAGCACTTTCCATGTGCCAAACACTGTTCTAAGTTCgaatcttaaacccaacttg
ggttacctcttatctagtggATCCTCACAGCAGCCGAGAATGTTTTAACCCAGTTGACAGgaatcttaaacccaact
ggttacctcttatctagtggGGATCATTTCTGCCACAAACCACACAGAGATGCCTGCAGGgaatcttaaacccaac
ggttacctcttatctagtggCACCTAACCAGGTGCTTGAGGGGGACAGAAGCCTGCAAAGgaatcttaaacccaa
ggttacctcttatctagtggGGTGCTTGCCTTCAGAGAGCCTAGAGTCTAGTGAGAAAGGgaatcttaaacccaac
ggttacctcttatctagtggAGGCCTCTGTAAGCCTGTCTCCTAGGAAGTTTTAGTGGGGgaatcttaaacccaact
ggttacctcttatctagtggCCCAGTGTCAACCAGCCTAATTCCCTGAGTCAGCAGCAGCgaatcttaaacccaac
ggttacctcttatctagtggGCCCAGTGTGTTTCAAATCCAGGAAAGGGCGTCGCAGGTCgaatcttaaacccaac
ggttacctcttatctagtggGGCCATCTGTGGGTACCCCCATCCCAGCCTGAATCACCCCgaatcttaaacccaa
ggttacctcttatctagtggGACCTGGGATGTCTCCTAGGCAGGACTTCCCCTCCCTAGCgaatcttaaacccaac
ggttacctcttatctagtggTGGGCAAGTAGTTTAACCTCTCTGTTACAGACGTTTTCTCgaatcttaaacccaacttg
ggttacctcttatctagtggCGTGGAAAGCTCTTAGAATAGCGCCTCATGTCTGATAAGCgaatcttaaacccaact
ggttacctcttatctagtggAGGGTCCATACAAAGCACTGTGCCTGATGCATTCCAGGGGgaatcttaaacccaac
ggttacctcttatctagtggGGAATGAGGAATCCCAGCATCTCAGCATGGGGAGGCTTCCgaatcttaaacccaac
ggttacctcttatctagtggGGGCTTAGTGGTGCACAGGCCCCAGATCCAGGTAGCCTAGgaatcttaaacccaa
ggttacctcttatctagtggCCACTCAAAGTTTCAGTGTCTTCGTCTGTCAGAGGGAAGGgaatcttaaacccaact

ggttacctcttatctagtggGTACCTACTTCCTAGGGTTTGTCGCAAGAGCTAATCCCTGgaatcttaaacccaact
ggttacctcttatctagtggAGTTCAGAGGACATAGCAAATGCTCCAAAAATGGAAGCTAgaatcttaaacccaact
ggttacctcttatctagtggATGGTGTCTCTTGCAGAGATTCTCCCAAGTTCTCTCCTGCgaatcttaaacccaactt
ggttacctcttatctagtggCCCATTGGACTATAGGTTTCTTCAGGTTAGAGATTTTGCCgaatcttaaacccaactt
ggttacctcttatctagtggAGTATGATCTGACCAACATTTGGCAAGTCTTAGATACCCAgaatcttaaacccaactt
ggttacctcttatctagtggGGAGAGGTAAGTGTAAGGCAGGTGGATACTGGGAAAAGGCgaatcttaaacccaa
ggttacctcttatctagtggGTGCAAGGGCATAGAACATGTTGGGGAATGATATGAGTGCgaatcttaaacccaac
ggttacctcttatctagtggTGCTTTTAGCTAACACTTTATGCCAGAAACTGCTCTGAAGgaatcttaaacccaactt
ggttacctcttatctagtggCAAGCGATTCTCATGCCGCAGCCTTCCAAGTAGCTGGGACgaatcttaaacccaa
ggttacctcttatctagtggGCCTAGGTTTCTCGAAGTGCTAGGATTATAGATGTGAGCCgaatcttaaacccaact
ggttacctcttatctagtggGTCATAAGAACCCACTGTGGTGGCTACTATTATCATCCTCgaatcttaaacccaactt
ggttacctcttatctagtggTGCTGTCAGCCACATGGCCAACAGGAGGTGGAGATGGGACgaatcttaaacccaa
ggttacctcttatctagtggAGCCCCAGCTTCTCTCCTCTGGCTCAGGGGTGGGCCCAGCgaatcttaaacccaa
ggttacctcttatctagtggTCCCCCCACCCCACCATGGCCAAAGACAAGAGTCCTCTTCgaatcttaaacccaa
ggttacctcttatctagtggGAAGCTCCATCCAGGTCGCTCCACAGGTACAGTCCTGCAGgaatcttaaacccaa
ggttacctcttatctagtggCCACTTCGTGCCAGCTTGGCTGAGCTCTACATACCTGGCCgaatcttaaacccaac
ggttacctcttatctagtggGGCACTTTCACTGGAACATCTGGAATCCTGGAAATGGGGCgaatcttaaacccaac
ggttacctcttatctagtggGCTCTGCTCAGCTGGGTGTCAGGCCAAGACCTACTGCTGGgaatcttaaacccaa
ggttacctcttatctagtggTCGAGACACAGGCAGCATTATTCTTTCTCCTTAGGCAGGCgaatcttaaacccaact
ggttacctcttatctagtggGTCACCCAGGCTGGAGTGCGTTAGTGTAATGTCAGTCCACgaatcttaaacccaac
ggttacctcttatctagtggAGCCTCCCAAGTAGCTGAAATTACTGGCATGCACCACCACgaatcttaaacccaa
ggttacctcttatctagtggCTGCCAGACTCAGACCTGTAAGCTCCCCTCTATCCCTGCCgaatcttaaacccaa
ggttacctcttatctagtggCTCCTAGGTTCTCATCAGAGGCAGCCCCTGACCTTGGGCCgaatcttaaacccaa
ggttacctcttatctagtggTCCACAAACCAGGGCCCCTGCTAGGTACTAGGGACAACCCgaatcttaaacccaa
ggttacctcttatctagtggCAAAAAGAAAGGCATGTGAGGAGTAGGGGGCCCCACAGGGgaatcttaaacccaa
ggttacctcttatctagtggCAGAAGGGACCGGTTTGCCTCGCTCTTCGAAAGGCCCTGCgaatcttaaacccaa
ggttacctcttatctagtggACCTCCCATAACCCCATGTGTTGCAGAGGACAGAGCAGACgaatcttaaacccaa
ggttacctcttatctagtggGGCTCTGTGCCTGCTCACTTATCCGGGAGAGTTACTCAGAgaatcttaaacccaac
ggttacctcttatctagtggGCATCCCTGGCAGCCCAGAACCTCGAATAGAGGGGACCGGgaatcttaaaccca
ggttacctcttatctagtggGTGAGAAGGATGACTCGTAGGTAGGGTGACCATATGACCCgaatcttaaacccaac
ggttacctcttatctagtggGATTCCCACACCCTTTTACTCTTGGAAGTGTCCCAAGTTGgaatcttaaacccaact
ggttacctcttatctagtggGGTGGGCTAGCAGCTCATGTGGCGGGTCCATTGTGTGCCCgaatcttaaacccaa
ggttacctcttatctagtggCTTCAGCTGTGAACACTCAGAAACACTAATGCAGATGGAGgaatcttaaacccaac
ggttacctcttatctagtggGCACGGATCCAAAACCAAGGTCCTCCAATGTCTTACTGACgaatcttaaacccaac
ggttacctcttatctagtggGTTCTACTTCCTTGCCAAGCTGCTGTCATAGCCAATGGGCgaatcttaaacccaact
ggttacctcttatctagtggACCTCCTAAACTGTTAAGTCCTAAACCAATACAAGGTGCTgaatcttaaacccaact
ggttacctcttatctagtggCCATCTGACTCCTCACCATGCCCCTCCTCCACTGCACTCCgaatcttaaacccaa
ggttacctcttatctagtggGCCTCCTCTGTTCCCCTGTGGCTTCCTTCAGGGCAGGAACgaatcttaaacccaac
ggttacctcttatctagtggCTGGTGCCACCTCCAGAAAGCCTGCTTCAACTTCCCTGGGgaatcttaaacccaa
ggttacctcttatctagtggAGCCAAGGCCTCTGAACAATTCCAGCTCCTTCTCTATGTTgaatcttaaacccaact
ggttacctcttatctagtggCTCCTACTCTTCTGGGGCTAATTTCTCATGTTGTGTCTCGgaatcttaaacccaactt
ggttacctcttatctagtggTGGGGTTTTCTGTACATGCTGTCCCCTCTGTTTGGTGTCCgaatcttaaacccaactt
ggttacctcttatctagtggGCACCCCTCCCCGCTAGGAGCCCTTGCTGACCTCCCTGACgaatcttaaaccca
ggttacctcttatctagtggGCATCGTGAACCTCTGTTTTCTAGGGCTCACCTTGGCAGGgaatcttaaacccaac
ggttacctcttatctagtggGGACTGTAAGCCAAGGTGGAACCAGAGAATAGAGCTCTGGgaatcttaaacccaa
ggttacctcttatctagtggCCCACAACCTAGCACAGTGCTGGAGGATAATACATACTGGgaatcttaaacccaac
ggttacctcttatctagtggGTCCTTTGGAAGATGAAGGTGGGCATTCTTCTGGCTCCCCgaatcttaaacccaac
ggttacctcttatctagtggTCATCACCCCCTCTATCAAATGATGATATGGGTCTGGCCCgaatcttaaacccaact
ggttacctcttatctagtggCCTCACAACAACTCTGAAAGAGGGGTCCCGTGTTACAGACgaatcttaaacccaac
ggttacctcttatctagtggAGCATGGATTTTTAGGCCTGAATATACCTGTGAATCATTCgaatcttaaacccaacttg
ggttacctcttatctagtggTGGGAAGGGAGGTATACATTTGAGAATGCATTGGGGGGACgaatcttaaacccaac
ggttacctcttatctagtggTGTATGAGGGTATAGGAGGGGGGATGGCAAAAAAGATCTGgaatcttaaacccaac

ggttacctcttatctagtggGGACACCCATCTAGACATTGCCCTGGGAAAATCACCGCTCgaatcttaaacccaac
ggttacctcttatctagtggTTAATCAGGATGGGGAAGTCGGTGCAGGCCCCTCCTAGGCgaatcttaaacccaac
ggttacctcttatctagtggCTCTCCACCAAGCTGCTGGTTATGTCATTTCCCTGCCTACgaatcttaaacccaact
ggttacctcttatctagtggCCTCCAGGACCGCTTCAAGCCCCATCTGCTCTGCAAAGTCgaatcttaaacccaac
ggttacctcttatctagtggTTCATTTACTTATTCATTTGACAAGGCTTTGGTGAGAGCCgaatcttaaacccaacttg
ggttacctcttatctagtggGACCTCGTGTGCCGGAAGGGGAGACTGGAGCTCAGACAGGgaatcttaaacccaa
ggttacctcttatctagtggGCCTATGGCATTTCCTCTCCTTTGTGCCGCTTCCCGCCACgaatcttaaacccaact
ggttacctcttatctagtggCAACAGAGAAAAAGACTCACCTGACTAATAACTCACAAGCgaatcttaaacccaact
ggttacctcttatctagtggAGGTCCCTCTCAGTGGCATCACCAAGGGCTTACTTGGGGGgaatcttaaacccaac
ggttacctcttatctagtggAGGCACTGCTTTACATGCCCCACAGCCTGACCCTCCTCCGgaatcttaaacccaac
ggttacctcttatctagtggGCTCTGTTCCTCTCCAATAAGAACTCAGAAACCCCAGTGCgaatcttaaacccaact
ggttacctcttatctagtggGAGATGCCTGCCCTGCCTCTTATTGGCTGCAAGACCCTGGgaatcttaaacccaac
ggttacctcttatctagtggGCTTCCCTGGGACTCTGTTTTCTCATCTGCAAATGAGAACgaatcttaaacccaactt
ggttacctcttatctagtggGCTTCTGTATGCACAGCCCCATGTTTGCACACAGGAAGGCgaatcttaaacccaac
ggttacctcttatctagtggCTAGGTGGCCAACCCTGGATGCTATCAGTAGGTATCGTGGgaatcttaaacccaac
ggttacctcttatctagtggGCACAAAGTGCTCACTGTCCACCAGATTAACTACCTTACAgaatcttaaacccaact
ggttacctcttatctagtggGGGCTACAGAGGGCTTTGCTGAGGAGCTATGGAAGTCTGCgaatcttaaacccaac
ggttacctcttatctagtggCCCTCCTGGGAGATCAGTGGAGCTCAAGTGTTACATGCCAgaatcttaaacccaac
ggttacctcttatctagtggGCACCAGTCACACAGGACTCAAGCTAACCAGAAGGCAGAGgaatcttaaacccaa
ggttacctcttatctagtggCGGTCAGCGCAGGAGAGAAGCTGAAGTCTCCACCAGCCAAgaatcttaaacccaa
ggttacctcttatctagtggGGCTGGAAAGAGAGAAATTTGACAACAAGTTTGGGCTGGCgaatcttaaacccaact
ggttacctcttatctagtggGGGCTCTCCGCTTCTCTCTGGCCTCACCCCTGACTCCCAGgaatcttaaacccaac
ggttacctcttatctagtggGCAACCCCAGTACTCTGCATGGAGCCCAGGACATAGGAGGgaatcttaaacccaa
ggttacctcttatctagtggAGGGAAGACTTTGCAGAGAAGATGGGGCCTGAAGTGGTCCgaatcttaaacccaac
ggttacctcttatctagtggCCTCCTGCCCGGAAAGCCTTAGCACCCCAACACTCTCTGCgaatcttaaacccaa
ggttacctcttatctagtggGCCTCTCCTGATTACTATCTCCATCCTCCCCAGACTGGGCgaatcttaaacccaac
ggttacctcttatctagtggGGGAGCAGCTCGCCTGGACAGTCCTGGCTTGAGGCCTCCCgaatcttaaacccaa
ggttacctcttatctagtggGTCAGCTTCAGGCATGGCTGAGACAAGAGGATCTACTTCCgaatcttaaacccaac
ggttacctcttatctagtggTGGTACTGGCCATGGATTGGAGGTATCAGTTCCTCACCACgaatcttaaacccaact
ggttacctcttatctagtggCTACAGGGTTGCTTGAGTGTCCTCATAACATGGTGCCTGAgaatcttaaacccaact
ggttacctcttatctagtggCCCACCTTGTAAGTCACACGCCTTCATTTCTGCAATACCCgaatcttaaacccaact
ggttacctcttatctagtggGACACACAAGGTGGCAAGGACCGTTGGGTGCAGTCTTGGGgaatcttaaacccaac
ggttacctcttatctagtggGGTGGCCTCTCTCCTCCGTCACAGCACTTTTCATGTGGACgaatcttaaacccaact
ggttacctcttatctagtggAGTAGCTGCTCAATAAATGTGTACAGACTGGGTGCAGTGGgaatcttaaacccaact
ggttacctcttatctagtggGGCAGGCAGCTCACTTGAGGTCCGGAGTTCGAGGCCAGCCgaatcttaaacccaa
ggttacctcttatctagtggTCCAGGTGTGGTGTCACATGCCTGTGATCTCAGCTACCCGgaatcttaaacccaac
ggttacctcttatctagtggAGGCCCCACACTGCCTGGCCGCATCACCTCTCTGCCTTGCgaatcttaaacccaa
ggttacctcttatctagtggCTCTGCCCCGTCACCTGGAGCTGAAGCCTGTGGTTCCTCGgaatcttaaacccaac
ggttacctcttatctagtggTTTGACGGACTGCTCCCTCTTCTTGACGGGTATCCCTAGGgaatcttaaacccaact
ggttacctcttatctagtggGAGGCATCCTCAGTAATAGCGCTTCTGATTCCCTTAACCCgaatcttaaacccaact
ggttacctcttatctagtggTCTGTCTCCTCACTCCAACACATAACCTCTCTGAGGGGAGgaatcttaaacccaact
ggttacctcttatctagtggGTTTGCTGATAAAGTTGTAGCACCTAGATCAAGGTAGGTGgaatcttaaacccaactt
ggttacctcttatctagtggGGGAGTGATAGTGTGCGCCTGTAATCCCAGCTGAAGCAGGgaatcttaaacccaac
ggttacctcttatctagtggTCCAGCCTGGGCAACAAGGTGAGACTCGGTCCAAAAAACAgaatcttaaacccaac
ggttacctcttatctagtggCCGCACCACTGCACTCCAGCTCCAGCCTCGGCGACAGAGCgaatcttaaacccaa
ggttacctcttatctagtggTGGTGCCCCACCCCTTGAGAATAGATGTCAGGAAACTGCCgaatcttaaacccaac
ggttacctcttatctagtggGATTCTCTTTTGGTTTAGACAGAAGGTTCTGGGGATGCCCgaatcttaaacccaactt
ggttacctcttatctagtggCATGAAAGAACTTCCCTGCCCCACAGCTGCAGGTATCCCCgaatcttaaacccaac
ggttacctcttatctagtggCTCACAGAAGAAACTTGACGTCAATCCCCAAGAAGCTTCCgaatcttaaacccaac
ggttacctcttatctagtggGCCTCAAGTGTCTCATCGGTAAAATGGGAATTGAAATAGGgaatcttaaacccaactt
ggttacctcttatctagtggGTGTGTTACTTACCACAGTGTCTGGCACAATCATGACAAGgaatcttaaacccaactt
ggttacctcttatctagtggAGCACTTATGCTGCAAGGCTGCAATGGAATATATTAAAAGgaatcttaaacccaactt

ggttacctcttatctagtggCTGTTCTAATTCCTGCTGTCTGGGTAATATGCCAGCCAGGgaatcttaaacccaact
ggttacctcttatctagtggAGAAATCAATGCATTTTTGTTCTCCTCTCTTATCCTTGGCgaatcttaaacccaacttg
ggttacctcttatctagtggCTGCCCTACATCTGACCGTCAACGTAAATACAGGCAGCCCgaatcttaaacccaac
ggttacctcttatctagtggGGCCCAATTGACCTGTTGACCCAGTTTGCAGAGACGGCTCgaatcttaaacccaac
ggttacctcttatctagtggTCACTGCAATTACTCCTTGAGCCCCCAGTGCTCCCGGCACgaatcttaaacccaac
ggttacctcttatctagtggCACCCACTTCAGTCTAGCACTGGTGGACTTGAGTCCTGCGgaatcttaaacccaac
ggttacctcttatctagtggTGGGAGGCCAAGGTGGGAGCCGAGGAGTTCAAGACCAGCCgaatcttaaacccaa
ggttacctcttatctagtggCCAACAATACAAAAATTAGACGGGCTTTGTGGCGCATGCCgaatcttaaacccaac
ggttacctcttatctagtggGAGACAGAAGAGAGGGAGAGAAGTGGCTGACCGTCATGCGgaatcttaaacccaa
ggttacctcttatctagtggCCTCTTCAGGATCCAGCTTTTCCTAGTGAAGTAGCAGATCgaatcttaaacccaact
ggttacctcttatctagtggCCTTTCTGTTGCCCTCATCTGCCGTATTATTCCAGCTGAGgaatcttaaacccaactt
ggttacctcttatctagtggAGAATGGGGGCTGTGGGCTGGGGGCTGAGTCCCGAGGGTCgaatcttaaacccaa
ggttacctcttatctagtggTCACGCACACAACGTTGTCTTGTGGAGGCTGGCAGTCAGGgaatcttaaacccaac
ggttacctcttatctagtggCTCTCTCTCCACTGCCATTGGAAATCAGAGTCCCATGGTCgaatcttaaacccaac
ggttacctcttatctagtggGTCAGTCTTCCTTCTAACTTTTTCCAGTTGACAAACAGCCgaatcttaaacccaactt
ggttacctcttatctagtggCATACGTCCCTCTGTCATGGGAACACTAAGAACTTGTGAGgaatcttaaacccaact
ggttacctcttatctagtggTGGCTTGCCAGGGGTCAAATCCAGGCTCTGCCCTCGGGCCgaatcttaaacccaa
ggttacctcttatctagtggGGCATGCCGCTAATAGTGCCCATCTCATAGGGTGACTGTGgaatcttaaacccaac
ggttacctcttatctagtggGCACACAGTGAGTGCCCATGAAGACTTCCAGGTTACGACCgaatcttaaacccaac
ggttacctcttatctagtggCCAGACTTTCCACTGAGGCAGAAGCTGCTACTTGTCCCCCgaatcttaaacccaac
ggttacctcttatctagtggTCCAGCCTTTCTGGCGGCAAGGTGTAAATACAAACTCAGGgaatcttaaacccaac
ggttacctcttatctagtggTGGATGAGACAGCCCATCTCTCCACCCTCCTGCTGGCCCCgaatcttaaacccaa
ggttacctcttatctagtggTCAGGATAGAGCCACCATGCCACACCACACCGTGGGCAGCgaatcttaaacccaa
ggttacctcttatctagtggTCCATCCATCTCGAAGCAGCCTCTGTCTCAGGACCAGAGCgaatcttaaacccaac
ggttacctcttatctagtggCAGCTGGTCTGCCTCTGCCACCAGATGGGCCCAGACAGGCgaatcttaaacccaa
ggttacctcttatctagtggGTCTGAGGCCCAAGGAGGAGCTGTGGCACTGTCCTGAGACgaatcttaaacccaa
ggttacctcttatctagtggCTCACGGGAGTCCCTGCCTCCAAACTCAGGAGCACCCCAGgaatcttaaacccaa
ggttacctcttatctagtggCCCAGCTCTTGAGCCCTGGGGTGTATAGATTTGAAGTTGCgaatcttaaacccaact
ggttacctcttatctagtggCCTAGGTGTGACTCCCAGCTTAGCAGCTGTGGGAACTTGGgaatcttaaacccaac
ggttacctcttatctagtggCTAACCTCTCTGAGTTTCAGCTTCCTTGTCTGCAGACCAGgaatcttaaacccaact
ggttacctcttatctagtggTCAGTGAACATGCACAATGCTGCAAATATCAAGTTATGAGgaatcttaaacccaactt
ggttacctcttatctagtggGGGGACTCTTGTGTAAGTGGTATATTGGGGCACTGTCTGGgaatcttaaacccaact
ggttacctcttatctagtggGAGAGGGGAGAAAGACCAAGGAGCTAAGCCACGTGGTGACgaatcttaaacccaa
ggttacctcttatctagtggGCCCGAGCTCTAGAATGTGAATGGCACCACAGAGCTGCCGgaatcttaaacccaa
ggttacctcttatctagtggGGCGAGAACACAGCCACTCATGGGGATGGGGAGTGGGGGCgaatcttaaacccaa
ggttacctcttatctagtggCCCTGGCAGCTGGACCCAGTGAAGGGGAACCGAGTGAGGCgaatcttaaacccaa
ggttacctcttatctagtggTGCCACCTCTTTCACAAGTTCTGTCCATCCTCTCAGCTAGgaatcttaaacccaact
ggttacctcttatctagtggCACATGCCCATAACATTTTCTCTCTACCATGCCTACTGAAgaatcttaaacccaactt
ggttacctcttatctagtggGCCCTTGAGAGCTGAGATGGTGCTGAAGATGGGGCTGTCCgaatcttaaacccaac
ggttacctcttatctagtggGGCTTTCAGCCAACCTGACCAGAGACTGAAGGATGCCCTTgaatcttaaacccaac
ggttacctcttatctagtggTGCCAAGACACGGGGTAATATAATGTCAGTTCTTGGAAATgaatcttaaacccaactt
ggttacctcttatctagtggCAGTAACTCAGAGTGTCCATCATGAGTGTCCTGACGCACCgaatcttaaacccaac
ggttacctcttatctagtggCTTTGAGAGGCCGAGGCAGGTGGATTTCACTTGAGGTCAGgaatcttaaacccaac
ggttacctcttatctagtggCGTGGGCGACAGAGTGAGACTCCGTCTTAGAAAAAATAAAgaatcttaaacccaac
ggttacctcttatctagtggGGTACCTCACTCATTTCAACCTAGTGTTAACCGTGACACCgaatcttaaacccaact
ggttacctcttatctagtggCCACATTTGGTCATTTCCTCAAGGAGCTGCCGGCATTTGGgaatcttaaacccaact
ggttacctcttatctagtggGCATCAGATGGAAGGCAGTGAGAAGGAAGGGTATGCCGGCgaatcttaaacccaa
ggttacctcttatctagtggGGTGGAGGTGATCACGGCAGGCTTCACAGAGAAGGTGACGgaatcttaaacccaa
ggttacctcttatctagtggTGGGTGGGAAAAGGGACAGGCTACATGGCTGGGGAAGGGGgaatcttaaacccaa
ggttacctcttatctagtggGAGCACAGCACACCTGGAGAGGACACAGAAGCTCCACACCgaatcttaaacccaa
ggttacctcttatctagtggAGCCGGGCACAGAGGCTCACAGCTGTAATCTCGGCACTTTgaatcttaaacccaac
ggttacctcttatctagtggCAGCTACTCGGGAGGTTGAGGTGGGATCACTTGAGTCCAGgaatcttaaacccaac

ggttacctcttatctagtggGCAACAGAGTGACACCTTGTCTCAAAACACACAGAGAGAAgaatcttaaacccaac
ggttacctcttatctagtggAGAAAGGAAAGGAAATAAGGAAGGAGGAAGAGGGAAAGGAgaatcttaaacccaa
ggttacctcttatctagtggGAAAAGGGGAAAATCGTCACTAAGGTAAATAGTAGCATTGgaatcttaaacccaact
ggttacctcttatctagtggGTTGAGAATCAACTACATTTCAGAACCTTTACAGTCATTCgaatcttaaacccaacttg
ggttacctcttatctagtggGAACTCATCCCCCCATCTGACAGGTAAGGGCGCCCAAAGGgaatcttaaacccaa
ggttacctcttatctagtggTGGCCTTTCTGACTCTCCAGAGAGGGAAAGCAGTTACCCCgaatcttaaacccaac
ggttacctcttatctagtggCCAGGCATCTAAAATCTGCATATCTGTGCCCAAGAGGTCCgaatcttaaacccaac
ggttacctcttatctagtggGAGTCCAATAAATTTCTGAATAGACTTCCTCCTATAGTTGgaatcttaaacccaacttg
ggttacctcttatctagtggCAGGTGTGAGCCACCATGTCCGGCCTTCTAGTTAACTGTTgaatcttaaacccaact
ggttacctcttatctagtggCCCCCACCTTAAAAGGATGACAGATTCATTAAGCTATGAGgaatcttaaacccaac
ggttacctcttatctagtggCTAATCTTACTCCCATGAATGCCAAAGTAAGACTTGGACCgaatcttaaacccaact
ggttacctcttatctagtggTCCGGCTCAATCAGGAAAATAAGCACAGCACCAGGTACCCgaatcttaaacccaa
ggttacctcttatctagtggCACCCGTCACTCATACACACACAAAGCCAAGCACCACAGCgaatcttaaacccaa
ggttacctcttatctagtggTGTAGAATCTGCCTGGCCCTGAACACCCCTGTAGATCGAGgaatcttaaacccaac
ggttacctcttatctagtggCTCAGGTCACTCCCCTCCCACTCCACCCCCAGACCTGCTGgaatcttaaacccaa
ggttacctcttatctagtggTCTGGGGGTGGATGATTCTTCCAGGTGATGCTAATACAGCgaatcttaaacccaac
ggttacctcttatctagtggGCCACTAGTGTATATGCATAAAATACCAAAGGAGCTAGGGgaatcttaaacccaact
ggttacctcttatctagtggATATGACCTAGGCTGATTATTTCACTTCTCTGAGCCTCACgaatcttaaacccaactt
ggttacctcttatctagtggGCTGGTGAGGTCACTGAGATGACCCAAGTGATGCCAGTGGgaatcttaaacccaa
ggttacctcttatctagtggGACCTCCATGGATTCGGCATCATCCCCTCTCCCTTCTAAGgaatcttaaacccaac
ggttacctcttatctagtggGAGGACTCGGCTAGACCCAGAAGGCACAGAGGAGGTGGGGgaatcttaaaccca
ggttacctcttatctagtggGCAGCCGGGCCTCTGTGGTTCACAGCAGCTTCCCACAGGGgaatcttaaacccaa
ggttacctcttatctagtggCCACCAGCTTTCCCACCATAGTCTAAGAACATTAGATTGGgaatcttaaacccaac
ggttacctcttatctagtggCTTCTGTTATGGCACTTCAGGATAAGATCAGCCATGGTGGgaatcttaaacccaact
ggttacctcttatctagtggAGCCCTTACTCCCTGGAATCTAGTGCACCTTCTGCTTTCCgaatcttaaacccaac
ggttacctcttatctagtggCCAGACTCAAGGGAAAGTCCTGGGATCAAAGAAATCCCCCgaatcttaaacccaa
ggttacctcttatctagtggGACCTTCCTTCAAAGAATCATACATGTTCACTGTCACATTgaatcttaaacccaactt
ggttacctcttatctagtggGATTTTGCTTCGTTGCAAGCCACCTGCCCCCATGGAAACCgaatcttaaacccaac
ggttacctcttatctagtggCAATTAACAGAAACTTGCTTAACAGTCATCATGCATTGGCgaatcttaaacccaactt
ggttacctcttatctagtggGTCACATAAGCCGTTTTTCTCCCCTCTTGCAGAAAATGACgaatcttaaacccaactt
ggttacctcttatctagtggCGAGATGCAGACAGGTCACACAGGTTGTTAGGAAGAGCAGgaatcttaaacccaa
ggttacctcttatctagtggAGTTCCCTGAAGGCACTAACAAATCCCAGGTCCTTGTTCCgaatcttaaacccaac
ggttacctcttatctagtggAGGTCACAGCTACAAAAGTGGTTCTCTCTCTCTCTCTCTCgaatcttaaacccaactt
ggttacctcttatctagtggAAGGTTTTTCTCAGAACATGGGCAGCTTTCCAGACTTCCGgaatcttaaacccaact
ggttacctcttatctagtggGACTTCTCAGTTTTCCAAGGCCATGGCTACCCAATTTGGCgaatcttaaacccaac
ggttacctcttatctagtggCTCTAAATCCGTTAAGCTCTGCAACGCAGAAGTTTTCCCGgaatcttaaacccaac
ggttacctcttatctagtggCACTGAGGATATGGAGGAGGAAAATATCCCAAGACAGTGGgaatcttaaacccaac
ggttacctcttatctagtggGGGTAAGAAAATATAACTGGTGATGAAAATGAAAGCTGAGgaatcttaaacccaactt
ggttacctcttatctagtggCTCTCCTGGGGTTAGATGGTCCCTCCAGGCAGCCTGCTTCgaatcttaaacccaa
ggttacctcttatctagtggGACCCTTCAAATGTCTCTCAGAGATGACCAGCCATGTCTGgaatcttaaacccaac
ggttacctcttatctagtggTGAAAAGAGCTGTTTCCTACCTAAAGGAAGGCTTGCTTTCgaatcttaaacccaactt
ggttacctcttatctagtggCCTTTGCTGCGTAACATATCATCCCAAAGCTCAGTGACTCgaatcttaaacccaac
ggttacctcttatctagtggACGCCTGCAGAATTCAGGAGCAACTTGGCTGGTGAGTTCCgaatcttaaacccaac
ggttacctcttatctagtggGTGGGCTGGAGGACCCACTCAAAGGTGGATTGTTCACACAgaatcttaaacccaac
ggttacctcttatctagtggGTCATCTCCATGGGGTTACTTGAGTGTCCTGGAGGCATGGgaatcttaaacccaac
ggttacctcttatctagtggGCTACAACACCTTTTAGGACCTACTCTCAGAGGTAATGCCgaatcttaaacccaact
ggttacctcttatctagtggAGAGGTCTGGCTAGCACAAATACGGACGCAGGCTGGGCACgaatcttaaacccaa
ggttacctcttatctagtggGGGCAGCATAGTGAGGCCCGCACTTCTAAAATAATAATAAgaatcttaaacccaact
ggttacctcttatctagtggCAACAGAGTGCTGGGCAAAGGATGTGGCATTTCATGGGCCgaatcttaaacccaac
ggttacctcttatctagtggCTCTCTGGGAGAGACCTGATGTGGAGTTACCTAATTGGAAgaatcttaaacccaac
ggttacctcttatctagtggGCCTGTACAGCCACACAGAGTCCTCCACTAAGAAGGACCCgaatcttaaacccaa
ggttacctcttatctagtggGGGCCCCACAAATTATGTAGCCATTCCTACCTTAGTAGCCgaatcttaaacccaac

ggttacctcttatctagtggCTGTCAAGAAACCATGGATGAGAAATCCCCAGGGATAGGGgaatcttaaacccaac
ggttacctcttatctagtggCATAAAAGCATCTAGAAGAAAGTAAGTTCTAAACTGCTGCgaatcttaaacccaactt
ggttacctcttatctagtggGCTAGATTTCTCCAATAGCCTGAGTTCCTCCAAAAAGCAGgaatcttaaacccaact
ggttacctcttatctagtggAGAGTTTGCATGCAGGTAGTTTGTTTTGGGAAATGACCGCgaatcttaaacccaactt
ggttacctcttatctagtggGCTCTGTGAGCAACTGGGGCTCAATCTCATCAGAGGGCTCgaatcttaaacccaac
ggttacctcttatctagtggGAAAGAGGAAGCATTTGTCCTCCAATTGCCACCCTGATGGgaatcttaaacccaact
ggttacctcttatctagtggAGGCTGAGTGGGTTCCTTTGGGATGCTGCAGGGTGGAAGCgaatcttaaacccaac
ggttacctcttatctagtggGTGGATGCGAGGGGCTCCCGTGGCACTACAATCTCCATGCgaatcttaaacccaa
ggttacctcttatctagtggCTCCTTCCCTGAGCTCCATCCCTGGGGAGATGGGGGAGGGgaatcttaaacccaa
ggttacctcttatctagtggCCATCTCTAGCCCTTCCCCAGTTGAGGCTTCCCTCTTGAGgaatcttaaacccaact
ggttacctcttatctagtggGCCTCAACTGTAATCAATTTGGAATTACATGAGAAAGGGGgaatcttaaacccaactt
ggttacctcttatctagtggGTCCTGGGGCCTGCCTGAGTTTTCTTCTAGGGCCATGGGGgaatcttaaacccaac
ggttacctcttatctagtggCCCCAGAGTGGACTTAAAAGGGGTTGGATGGTGTAAGCTGgaatcttaaacccaac
ggttacctcttatctagtggGTGATTACCTCCTCCAAGCTTGTTCCTCCTAATCGAAGACgaatcttaaacccaactt
ggttacctcttatctagtggGAACAAGAAGACAATGTTAAGGGGGTGAGGGGGAATGTGCgaatcttaaacccaac
ggttacctcttatctagtggTCCTTTCATGACACAAAAATGGAAGCCAGAACTCACGTCCgaatcttaaacccaact
ggttacctcttatctagtggTCACCTCCCCTATCATTTGTCCCCTCCTGAGACAGGGAGGgaatcttaaacccaac
ggttacctcttatctagtggGAGTACAGAGCGAGACCCTGTCTGTGGCTCTACACATACCgaatcttaaacccaac
ggttacctcttatctagtggGACAGGTGATAAGGGTACAGTTTTGGCTCTCAACTCTCAAgaatcttaaacccaactt
ggttacctcttatctagtggCCAGTCTCAGTTAGCATCCATAAGGCAAAACCCGCACCACgaatcttaaacccaac
ggttacctcttatctagtggGATTCTGAAGTCAGGTGGACAATTGCCTCTGGGGATCTCCgaatcttaaacccaact
ggttacctcttatctagtggGCCTCTCAAGTGGCTGGGATTACCATGTGCTACCATGCCCgaatcttaaacccaac
ggttacctcttatctagtggTTTTTAATAGAGACAGACTCCTGGCCTCAAGTGATCCACCgaatcttaaacccaactt
ggttacctcttatctagtggAGCCACCTCGCCTGGCTAACATCTTTTCTACAGGCAATTTgaatcttaaacccaactt
ggttacctcttatctagtggGAGCCACTGCACCCAGCTCAGGCACTTCTTATAATGTCCCgaatcttaaacccaac
ggttacctcttatctagtggCACCAAGACCCAGGCTGACTGCAGACACCTTCGGGCCACCgaatcttaaacccaa
ggttacctcttatctagtggGCTTTAATCCCCTTCATTGCACTCACCATTTCTCTTTGCAgaatcttaaacccaacttg
ggttacctcttatctagtggAGGGTCTCACTCTGTTGCCAATTCTGGAGTGCAGGGTTGCgaatcttaaacccaact
ggttacctcttatctagtggCCACCTCAGCCTCCTGAATACCTGGGAACATAGGTGTGTGgaatcttaaacccaac
ggttacctcttatctagtggGGCTAAAAGCCATGTCTTGTTTATGTCCGTGCCACCACTGgaatcttaaacccaact
ggttacctcttatctagtggGGGTTGGTATGTATACCACAACATATTTTCTGCATAAATGgaatcttaaacccaacttg
ggttacctcttatctagtggGCAGGAGACAAATGATCAGATGCCATTTCGTCTGGAGCTCgaatcttaaacccaact
ggttacctcttatctagtggGATCAGCCAGCCTCTAAGCTGCAAGAAGCAAGGTGGCACGgaatcttaaacccaa
ggttacctcttatctagtggTGTCATCAACGCCCATGTTGGTTACTTATTTCTCGTAGCCgaatcttaaacccaactt
ggttacctcttatctagtggGAGCCACCGCGCCCGGCCCCTTTTCTTCTAAGAATTCATCgaatcttaaacccaac
ggttacctcttatctagtggTTTCACTTAGATGGGCCTCTTTGGTTGGAGGAACAGCGTGgaatcttaaacccaactt
ggttacctcttatctagtggGCCTCCTGAGTAAAAAATTAGCATGTGCCACTGTGCCTGGgaatcttaaacccaact
ggttacctcttatctagtggACTCCGCCTCCAGAGGTCAAGCGATTCTCGTGCCTAAGCCgaatcttaaacccaa
ggttacctcttatctagtggCTGGGATTACCATGTGCCACCACACCTGACTAATTCTGGTgaatcttaaacccaact
ggttacctcttatctagtggGCCACCTCGCCTGGCTAACATCTTCTCTACGGGCAATTCCgaatcttaaacccaac
ggttacctcttatctagtggGTGCTAGGATTACAAGTGTGAGCAACCGTGCCTGGATAATgaatcttaaacccaact
ggttacctcttatctagtggGCCCTGGCTGGTCTCAAACTCCAGGTTACCCAAAGTACTGgaatcttaaacccaac
ggttacctcttatctagtggTCTGCAAAGGCTGGAGTTAGGAGAACCCGTGGTTGATGGCgaatcttaaacccaac
ggttacctcttatctagtggTTTTTTAAGATTCAGGTTCACACTTGTAATCCCAGCACATgaatcttaaacccaacttg
ggttacctcttatctagtggCCTGGGAGGGGAATGTTGAAGTGAACCAAGATTGTGCCACgaatcttaaacccaac
ggttacctcttatctagtggCACGATAATCAAGCAGCTCCTAACACAGTTCTCTAGTCACgaatcttaaacccaact
ggttacctcttatctagtggCCCTGTAGCTCTGTGAGATAAGCCTTCCATACCTCTCTTGgaatcttaaacccaactt
ggttacctcttatctagtggAGGATTCCTGAGATATGGAATGAAAGAACTAGCAAAAAGCgaatcttaaacccaactt
ggttacctcttatctagtggCTCCTCCCTCAGCTGGGATGGGATTCTTCTCCACGATGCCgaatcttaaacccaac
ggttacctcttatctagtggCCTTGCCTTGAGTGCAAAGGCGATGTGTATTTTTCAGCGGgaatcttaaacccaactt
ggttacctcttatctagtggCTCAGAACCAAAGCAACACACGCCTTTCAGAGCGCTGGAGgaatcttaaacccaa
ggttacctcttatctagtggTGAACAAGCTCTCATGGCTCAACAGTGGAGATCTGAACAAgaatcttaaacccaact

ggttacctcttatctagtggAAGTGCTCCTGTCAGCCTTCAATTTCTCTACCTGGGTCTCgaatcttaaacccaactt
ggttacctcttatctagtggCAGCATCCTTTTGGAAGACTTCAGCAATTTCTGTGGAGCCgaatcttaaacccaactt
ggttacctcttatctagtggGCAGTCAGGTAATTCCACGTGGGCTCACTGGTTACTTTCCgaatcttaaacccaact
ggttacctcttatctagtggCTCTTCCTTCTAGGGCGCCAATCTTTCCAAGGATAACCACgaatcttaaacccaact
ggttacctcttatctagtggCTCCTACCTGGCACACAGATCACTCCTGCCCACTCCTGCCgaatcttaaacccaa
ggttacctcttatctagtggATCTTGTACACGCCTTGCTGGTACATATGGAGGACCATGGgaatcttaaacccaact
ggttacctcttatctagtggTTTCCTCCTTCTCAAGGAATCCTGCTCCCTGAGTTTCTGCgaatcttaaacccaactt
ggttacctcttatctagtggCTCACCTTCAAACTCACTGAGGTTATTCTTGTTTGGCTTCgaatcttaaacccaacttg
ggttacctcttatctagtggTTGTTCTCAATTGGTCTGTTTATCAGGGTAACTGTTTCAGgaatcttaaacccaacttgt
ggttacctcttatctagtggCAGGTGTGGACTTAACTTTGCCCTCCACTCCCAGCAACCGgaatcttaaacccaac
ggttacctcttatctagtggCTTTACCAGTGACACCAACCTCAAAAGAGTCAAGGATTCCgaatcttaaacccaact
ggttacctcttatctagtggAGGTAGCAACAAAGGACACTAAGTGTGCACAATGGCTGGGgaatcttaaacccaac
ggttacctcttatctagtggTCTCATTTCATCCTTAAGACAACTTAGAGAGGTAAGAAACgaatcttaaacccaactt
ggttacctcttatctagtggGTGGAGAAAGATAAGAAGCTTGGGGAAAAGCAATGAATGGgaatcttaaacccaact
ggttacctcttatctagtggTCAGTAGAAGAGCTGTGATTCAAGCCAGGTGTGTTGCTCCgaatcttaaacccaact
ggttacctcttatctagtggAGCAGGAACTGGAAGGTGAATGAATATATACTGTGAGATGgaatcttaaacccaactt
ggttacctcttatctagtggTAAATCTCAGCCAGCAGCCATCCAAAGAATCTACCGGAGGgaatcttaaacccaac
ggttacctcttatctagtggGCTTTTATTGCTCAGGATAGGCTAAGCTATGCTGCAATAAgaatcttaaacccaactt
ggttacctcttatctagtggCACACAAGTCCACTGCTGGTCAAGCAACTCTGCAGTGCAGgaatcttaaacccaac
ggttacctcttatctagtggCCAGTATTCAGACTTCTTTCATCCTGTGGTTCCATCATCCgaatcttaaacccaactt
ggttacctcttatctagtggCATTGCCACTCACAGCCCATGGCCAGAACTAATCACTGGCgaatcttaaacccaac
ggttacctcttatctagtggCTGCAAGGGAAGCCAGGAGATATAGGCAGCCCCTTGGGTAgaatcttaaacccaa
ggttacctcttatctagtggGCTAAGTACAACTGTGCAACACTCCCCAACTTTCCTTGGCgaatcttaaacccaact
ggttacctcttatctagtggTTCTCGTGCCTCAGCCTCCTAAAATAGCTGGGACTACAGGgaatcttaaacccaact
ggttacctcttatctagtggAGGCTTTTTGGGAGTAGCAACAGGTTAAAGCAATTGCAGGgaatcttaaacccaactt
ggttacctcttatctagtggGGTATTCCTGAAAAATCTGGAGGTTACTGCTGCAGCCCTCgaatcttaaacccaact
ggttacctcttatctagtggCTGCAGGAGATCCTCCCACCTCCACATTTAAGAAAAACTGgaatcttaaacccaact
ggttacctcttatctagtggCCTTTTCGACTGGTTTATGATGGGTTATCTTCAGAACAAAgaatcttaaacccaacttg
ggttacctcttatctagtggAACACACAGGCCTTTTGATTCCCACAGGAGGAACCATCACgaatcttaaacccaac
ggttacctcttatctagtggGGACTTGAGCTTAGGCAGAGATGATCCGTGAGCTGGATCCgaatcttaaacccaac
ggttacctcttatctagtggCAGCCTCAGCTGCATTCATTTCATATACCTGCACCATGCCgaatcttaaacccaact
ggttacctcttatctagtggTTTGCAAGTGACAGAAACCCAACTCCCACTGGCTTCAAGCgaatcttaaacccaac
ggttacctcttatctagtggCCCAGAGGAAAGCTGGGCCAAGAGGAAAGGGCAGAGTTTTgaatcttaaacccaa
ggttacctcttatctagtggGCCCTTGATCCAACTGTACCTGAAGGAAACATGCAGTTGGgaatcttaaacccaact
ggttacctcttatctagtggGAGCCCCAGCCAATAAATATGCTAAGTAAAGGACACAGTGgaatcttaaacccaac
ggttacctcttatctagtggTGTCACTGACCTTGGTGTCTCTGTCTTTCCCAAGTCAGTGgaatcttaaacccaactt
ggttacctcttatctagtggCAAAGACTCTGCCATTGTGGAGTTTATATTCCAGTAGGGGgaatcttaaacccaactt
ggttacctcttatctagtggAGATAGGATGCCCAGTCTTAGGTCAAATGGGGAAGTTTGGgaatcttaaacccaact
ggttacctcttatctagtggGGCCTCATAATCTGACCCTAGAGAATGGTCAGTCTCTGGGgaatcttaaacccaact
ggttacctcttatctagtggGCCTTTCAAGTCTGACAGCCATTCCTTTTCATTAGCCAAGgaatcttaaacccaactt
ggttacctcttatctagtggGTCCCAGCTCTGTTACTGATAAGCAGCCCTTCCCCTGTCCgaatcttaaacccaac
ggttacctcttatctagtggGAACGGGTTGGATGATACAACATTTGCTTTCAGTGACTCCgaatcttaaacccaactt
ggttacctcttatctagtggTCTATTGTAGACATCGACCACAATTCCTGGTTCATGGTACgaatcttaaacccaactt
ggttacctcttatctagtggCTGAGATGCATGGGAGATTAGAACCTACCACCTCTACAACgaatcttaaacccaact
ggttacctcttatctagtggGACTAGGGATTTTGTCTGGTTCATCTTTGTGGCCCAGGGCgaatcttaaacccaactt
ggttacctcttatctagtggGTGAGCCAGGGAAAGCTTATGAAGATAGGGGCATTTGAGCgaatcttaaacccaac
ggttacctcttatctagtggCTTAGAATTTAAAAAGCGGGCCGGGTGCAGTGGCTCACGCgaatcttaaacccaac
ggttacctcttatctagtggTGGATTGTCAAAGCAAGAGATTAAAGTTGGGAAACAGTCCgaatcttaaacccaactt
ggttacctcttatctagtggGGCACGAACAGGAGGAATTGTCATAATAACCCACCCCAGCgaatcttaaacccaa
ggttacctcttatctagtggCCATCACCATTGATGTCAACTGGATATCATGTACCCCCTGgaatcttaaacccaactt
ggttacctcttatctagtggGCAAAGAAAGAAAACTGTCATGAAAGCAAGAAAGACTGAGgaatcttaaacccaact
ggttacctcttatctagtggTGACACTAAACGCATTGTGGTATCTTAGATTGGATCCTGGgaatcttaaacccaactt

ggttacctcttatctagtggTCTGCAGCTTAATTAGTAGTGACATGCAAATGCTGGTTTCgaatcttaaacccaacttg
ggttacctcttatctagtggTGTGATGATAAATTAGGAGTAATTGAAACCAGGTGACAGCgaatcttaaacccaacttg
ggttacctcttatctagtggCTTGGCTGGGCTACGGTCAGCTCAACAAGGCAGTTCTGCCgaatcttaaacccaac
ggttacctcttatctagtggCCCCAAAAATCTAGCCTGGTTTGTTTCCACAGTGCCTGGGgaatcttaaacccaact
ggttacctcttatctagtggGTGTCCAAAGCAAGTCACAAGGCCAACCCAGATCCAGGGGgaatcttaaacccaa
ggttacctcttatctagtggCCACAGAGGCATGTTGTAGGGTGCATGAATTTCTGCCATAgaatcttaaacccaact
ggttacctcttatctagtggGCAGCCACTGGCAGGGTAGAAGCTTGGGGGTGCAGAGGTCgaatcttaaacccaa
ggttacctcttatctagtggGAAACAACGACTTGCCATAGTTCTGAAACCCAGCTCCTCCgaatcttaaacccaac
ggttacctcttatctagtggATGGGGATAAGAGTAACAATGTTACGGTGTTATGAGGATGgaatcttaaacccaacttg
ggttacctcttatctagtggCCTGGCAGCCCCCTCCCCTTGCCCAGCCTAGATTTGCCTGgaatcttaaacccaa
ggttacctcttatctagtggGGTCAACTCAGAGCAGCTTCCAAGCCCACCTGGCCTCCTGgaatcttaaacccaa
ggttacctcttatctagtggGAAGAGCCTGAATTGGATAACAGTTAACTAGGCACCAGCCgaatcttaaacccaac
ggttacctcttatctagtggGGACCCCCAAGCCCAAGCTGGTAACTACTGTGTATTCCCCgaatcttaaacccaac
ggttacctcttatctagtggGATGGGGACTTTGGTGCTCCAGGGACAAACTCTGAGCAGCgaatcttaaacccaac
ggttacctcttatctagtggCCAAATACCCTTCACCTTCCCCAGCGGAACTATTTCCAGCgaatcttaaacccaac
ggttacctcttatctagtggGCTCCCCTCTCCTAAAACTACAGTTATCTAGAAGCAGAACgaatcttaaacccaact
ggttacctcttatctagtggCACAGCTTCCCTTCCTCTCCTGGTGACAACCAATGTCTCCgaatcttaaacccaact
ggttacctcttatctagtggCACTTTCAGAGCTGGAAGGGAGTTTAACGCAGCTAGTCCCgaatcttaaacccaac
ggttacctcttatctagtggTGAAGCCTGGAGAGAAAGGGATTTGTCCACAGAGCTAACCgaatcttaaacccaac
ggttacctcttatctagtggTGGATTTGATCCTGACAAGAACATTGTGGAGCAGCAGAGCgaatcttaaacccaact
ggttacctcttatctagtggCAGAGTGTCTGGAGCAACACTGAGACTAGAACCACCTTCCgaatcttaaacccaac
ggttacctcttatctagtggTTTTTTAATGAAGTCTGCTCTGTCACCCAGGCTGGAGTCCgaatcttaaacccaacttg
ggttacctcttatctagtggTCCGCCCGACTTAGCCTGCCAATGTGCTGAGATTACAGGCgaatcttaaacccaac
ggttacctcttatctagtggGCACCAGGACTGATTCTAATTCTTAACTCTGTGTCTTCAGgaatcttaaacccaacttg
ggttacctcttatctagtggGCATCAAGCTCTAGGTTTAGAATTAGCATGTGACACACACgaatcttaaacccaactt
ggttacctcttatctagtggGTCACCCAGGCTGGAGTGCAGGGGAATGATCTTGGCTCACgaatcttaaacccaac
ggttacctcttatctagtggGTAGAGACGGGGGTTTTCTATGTTGGCCAGGCTGGTCTTGgaatcttaaacccaact
ggttacctcttatctagtggCTCAAGCAATTAACCCACCTTAGCCTACCAAGGTGCTGGGgaatcttaaacccaac
ggttacctcttatctagtggTTACTTAATAGGATCAGGATCATGCAACACCACACCTGGCgaatcttaaacccaact
ggttacctcttatctagtggCCTCAAGCAATTAACCCACCTTAGCCTCCCAAGGTGCTGGgaatcttaaacccaac
ggttacctcttatctagtggAAAGCATTACTTAATAGGATCAGGCATGGTGGCTCACGCCgaatcttaaacccaact
ggttacctcttatctagtggGGGAGGCTTGAGATAGATGGGAGGATCACTTGAGTCTGGGgaatcttaaacccaac
ggttacctcttatctagtggGTGAGATGGAGTCTCATTCTGACCCCCCAGACTGGAGTGCgaatcttaaacccaac
ggttacctcttatctagtggCGACCTCCCCAGACTCAGGCGATCCTCCCAAACCTCAGCCgaatcttaaacccaa
ggttacctcttatctagtggGGTCTCGAACTCCTGGACTCGGAATATGCCTGCCTCGGCCgaatcttaaacccaac
ggttacctcttatctagtggATCACAGGAAAAGTGATTCAAAGTGTCACTTCAGAGACACgaatcttaaacccaactt
ggttacctcttatctagtggGAAGGTGATCTGCTAATTCAGGGACCTTGTTTTATGGCCCgaatcttaaacccaactt
ggttacctcttatctagtggTCTCAGAGACAGGGTAAATGTAGCTTTCTGACAGAGGTCCgaatcttaaacccaact
ggttacctcttatctagtggGGTTGTTTCCTTTTGTTCATCTTAGACAGCTAAATGGCACgaatcttaaacccaacttg
ggttacctcttatctagtggCCAGGAGTTCGAGATTAGCCGGGGCAATATAGTGAGACCCgaatcttaaacccaac
ggttacctcttatctagtggGCCTGGCAAGTTCAAGACTGCAGTGAGTCGTGATCACACCgaatcttaaacccaac
ggttacctcttatctagtggAACCTGGGTGAGAGTGAGAGATCCTGTCTCAAAAGAAATAgaatcttaaacccaact
ggttacctcttatctagtggAGACCTTGCTTACTCTGTTTTCTCAGTGGACTTTGTGGCCgaatcttaaacccaacttg
ggttacctcttatctagtggGAGCTATCTGTATCATAGGTAAGTTATACAGTCCTATCCCgaatcttaaacccaacttg
ggttacctcttatctagtggGTGGGTCATCCAAGACATTTCCACCAACCCAATAACGCACgaatcttaaacccaac
ggttacctcttatctagtggCATTGATTGAACACATAATATGCGCCAAGTACTATAAGCGgaatcttaaacccaacttg
ggttacctcttatctagtggTGGAACTTGATGTCCTCTCTTCTCTAACCACGACACTGCCgaatcttaaacccaactt
ggttacctcttatctagtggACCAATCTGAGGAAGAACAAGTGAATGGACAGTTAACAGGgaatcttaaacccaact
ggttacctcttatctagtggTGAGGGTCTCCTGACCTTCCCTGCCATCCATCTCTTCCGCgaatcttaaacccaac
ggttacctcttatctagtggTGCTTGATTTAATTGATATCTTTCAGGAAGCCAAGGTTGGgaatcttaaacccaacttg
ggttacctcttatctagtggCCCAGTGATAACCCACTTAAGCTTGGCTCATGCTCTCCAGgaatcttaaacccaact
ggttacctcttatctagtggCCATCCTGTCCTCTTCACACTGCCTGAATTTGAATCCCTGgaatcttaaacccaactt

ggttacctcttatctagtggGAGATAGTCTCACTCTGATGCCCAGGCTGGCGTACAGTGGgaatcttaaacccaac
ggttacctcttatctagtggGTAGAGACTGGCTTTTGACACATTACCCAGGCTGGTCTTGgaatcttaaacccaact
ggttacctcttatctagtggGCCCACCTTGGTACCATTTATTCTTGAATCCCAGGCAAGCgaatcttaaacccaact
ggttacctcttatctagtggATGGGCACTGATCTGATTGGGTTGCTGTGGGGACTAAATGgaatcttaaacccaact
ggttacctcttatctagtggATGTCTCTGGGACTGCATTAATGATCTCGCCTGGAGAGGCgaatcttaaacccaact
ggttacctcttatctagtggCCAAATCACCCTTCTGTTTTCCCCTGTCCATCCCCACCCCgaatcttaaacccaac
ggttacctcttatctagtggAAAGTAAACCAAATAAAGGCTGGGAGTGTTGGCTCATGCCgaatcttaaacccaact
ggttacctcttatctagtggAGCCCAGGAGGTTGAGGCTGCGTTGAGCCGCGATCATGCCgaatcttaaacccaa
ggttacctcttatctagtggAGCCAGGTGACAGAGGTGAGGGAGGTCTTGTCTCAAGAAAgaatcttaaacccaac
ggttacctcttatctagtggCTGCACCAGGAGCCATCTCCACAGCGCTTCCCATCCACCCgaatcttaaacccaa
ggttacctcttatctagtggCCTCTCTCTCCTGATCCTGAAAGCAGCCCAGGAAGTTGGGgaatcttaaacccaac
ggttacctcttatctagtggGAGCCCTTAGCAGACATAAACTTGGGCAGGAAGGGATTGGgaatcttaaacccaac
ggttacctcttatctagtggCGCTAGGTTTTCTCCATGTTCTGCTTCAGAAATAGACTGGgaatcttaaacccaactt
ggttacctcttatctagtggAAAGAGCAAGGCAGGGAACGGGCAGGGGCGGGCTGAACGCgaatcttaaaccca
ggttacctcttatctagtggCGAGGAAGGAGGCACAGATAGGGCGCACAGGAGAGCTGACgaatcttaaaccca
ggttacctcttatctagtggTCAGCATTTCTGGGGCAGCCAGGAGGTTATTCCTAAGGGGgaatcttaaacccaac
ggttacctcttatctagtggGCATTTCAGGCTGTGACTTTCCTGGGTCAGAGAAGCCTTGgaatcttaaacccaact
ggttacctcttatctagtggAAATCTGCCCCAGTGTACACGGCTGCTGCACTAAAAGGGCgaatcttaaacccaa
ggttacctcttatctagtggTATAGGGCCATTGTCTGAAACCATGTGACCAACCCCAGTCgaatcttaaacccaac
ggttacctcttatctagtggGGCATGCTTCCAGGGCCAGGATTAGGCTGTAAGCTGGTTCgaatcttaaacccaac
ggttacctcttatctagtggTCATGGGTTCTTTTTTGTTGTTTGCAGGGGAAGTGGGGACgaatcttaaacccaactt
ggttacctcttatctagtggTGCACACATCCATCCTCTGTGTCGCTAACTGGTGGAGACGgaatcttaaacccaac
ggttacctcttatctagtggTCTTGGTGTCTTTTGTTAACTGACCCATTCCCAACTTTATgaatcttaaacccaacttg
ggttacctcttatctagtggGTATGTTTAATATAAATGGAGTTTTGCCATGTTGGCCAGGgaatcttaaacccaacttg
ggttacctcttatctagtggTTACAGGCGCGAGCTACCGCACCCAGCTCCAACTTGATTAgaatcttaaacccaac
ggttacctcttatctagtggACTGTCTACTGGTGTGGACCAAGTCTGAGGAAACACTTGGgaatcttaaacccaac
ggttacctcttatctagtggGATGGCTGGCTTGCCTTGATGAAGCTGTCTGTTTAATGGGgaatcttaaacccaact
ggttacctcttatctagtggGTGAAGACACAGGCCTTGACCTTGGAGTATAGGCAGGGGCgaatcttaaacccaa
ggttacctcttatctagtggCCGGAAGGGCAGGTGTTCTTGGCAGTGGGGATGGCCAGTGgaatcttaaacccaa
ggttacctcttatctagtggGAAGGCTAGCTTCTGCCACCATGCCCTGCCTGCAAACTAGgaatcttaaacccaa
ggttacctcttatctagtggATGGTAAATGCATGGCAGGTGATTGGGACACAGAAGAGGGgaatcttaaacccaac
ggttacctcttatctagtggCCTGGAGGAGACACATTTGAGCCAGGTGAAAGGAGGCAGGgaatcttaaacccaa
ggttacctcttatctagtggGGCAAGGTCACAGAGGCAACAGGGATTTGGGGAGTGAGACgaatcttaaacccaa
ggttacctcttatctagtggCCGACTCTTATTGCAGGGGTAGGAAGCTAGGGAACTGTGGgaatcttaaacccaac
ggttacctcttatctagtggGGAGATGTGGAGAGAGTGGGAGCACTGGAGGGCCAATGGGgaatcttaaacccaa
ggttacctcttatctagtggACAAGGTCAAATTGAAATGTTTGGATGACAGCTTCGAGTGgaatcttaaacccaact
ggttacctcttatctagtggGGGGCAGCAGACTGGATTTGGCCCATGGGCTGTGGTTTTCgaatcttaaacccaac
ggttacctcttatctagtggTCCAGATGGGAGACTGGAGGTTCAGGGAAAAGCATATACTgaatcttaaacccaac
ggttacctcttatctagtggGTGAAACCATCTTGCTGGCATCTCTTTCTGTGCTGGTTGGgaatcttaaacccaact
ggttacctcttatctagtggCTAAGTTCCAGGTCTGTGCTAGGACCCAGAGATCGGTGAGgaatcttaaacccaac
ggttacctcttatctagtggCAGTGAGCCGGTGCATGTTGATTGTCCACTATGGGGCAGGgaatcttaaacccaac
ggttacctcttatctagtggGGCACGTCTATCCCTCCATCAGTCCATCCATCCACCCAGCgaatcttaaacccaa
ggttacctcttatctagtggGGCAGGACTCTGTGGGATTTATTGGTAACCCAGCCATGACgaatcttaaacccaac
ggttacctcttatctagtggCCAGCAAAGTACAGATGGGGGTGGGGTGGGTGCTCAGAGCgaatcttaaacccaa
ggttacctcttatctagtggGACTTTCAGCTGGAAGAGATCAGGAAGGCTGACAAGATGAgaatcttaaacccaac
ggttacctcttatctagtggCCTGTTAACCCTTTAAAGTGACATATGAGTCTCTGCAAGGgaatcttaaacccaact
ggttacctcttatctagtggGCTCTTCTGTACATGCTGTTAGCCCTTGGGTCCCATCAGGgaatcttaaacccaact
ggttacctcttatctagtggGCCTTGTGATTTCACCGTCCTGGAGATGGCTCCTTTCTCCgaatcttaaacccaact
ggttacctcttatctagtggGCTAGGAATTTTCATGTGTCAGCTCTTGAGTTGGGCACTGgaatcttaaacccaact
ggttacctcttatctagtggTGGGGAAGCTGAGGCTAAGAGGATTAAGTAGCTTGCCTGGgaatcttaaacccaac
ggttacctcttatctagtggGAGAAGGAAGCAAGCATGGAACCCATATCTCTTGACTCTCgaatcttaaacccaac
ggttacctcttatctagtggTTTGCTTTGACTTCTTCCTCCATTCAGGTCAGCTCCCTGCgaatcttaaacccaact

ggttacctcttatctagtggGCTCTTTCTGCTTTATTCCCACAACCACTGCACCAGCTCCgaatcttaaacccaact
ggttacctcttatctagtggCCTGGGTAATAAATTGGCCTTCTCACTTTTTACTTTCCCCgaatcttaaacccaacttg
ggttacctcttatctagtggATCTAACTTGTGGCAGGCATTGTCCTGGGTCCTGAGGAAGgaatcttaaacccaact
ggttacctcttatctagtggATAGCAAACTTCATCTGAGTTGGCCATAGGGAGTCTGGGCgaatcttaaacccaact
ggttacctcttatctagtggTGGCAGCCCCAGCCAACAGCACAAGAGGGAAGGGAGAAAGgaatcttaaacccaa
ggttacctcttatctagtggGGGAGGATGTCCTGTTACCTGAGCCCAAAGTTTAACACCCgaatcttaaacccaac
ggttacctcttatctagtggTCCGTCTAGAGAGATCGCTCAGCAAAGGTCATTGGTTATGgaatcttaaacccaact
ggttacctcttatctagtggTCAAGAAGGCAGGTTGACCTTTGGGAGAATTTCTAGCAGCgaatcttaaacccaact
ggttacctcttatctagtggCACGTGTCCTGAGAGGTGAATGAGAAATGTGTGCAGTTACgaatcttaaacccaact
ggttacctcttatctagtggTAGGGGAGAGTAGCATTGAGGAAACAAAAATGGCCCAGGGgaatcttaaacccaac
ggttacctcttatctagtggAGGGGATTGGGGAGGGATGTGGAAAGCATAGACAGAAGGCgaatcttaaacccaa
ggttacctcttatctagtggCTAAAAACTGCTTCTGTGGGCAGAGTTCATATGCTGGAGGgaatcttaaacccaact
ggttacctcttatctagtggTGACACCCTTTGGTATGACCCTGGGGAGAATAAATGTGACgaatcttaaacccaact
ggttacctcttatctagtggCACACTCTCAGGTTAGAAACTCACTTTAGGGGCCACAGCCgaatcttaaacccaac
ggttacctcttatctagtggCACTCATGCAAATTCTGTGCTTGCTTCCATGCTCCATTTCgaatcttaaacccaacttg
ggttacctcttatctagtggCTTGGGAGAAATAAAATTGCTGGATTAGATACAGGCCACCgaatcttaaacccaact
ggttacctcttatctagtggTGTGAGACTGAGATAGCTGATTAAGATGGTGATGCTTGAGgaatcttaaacccaactt
ggttacctcttatctagtggAATATGTAGGGAAGAGGGAGGCAAACTGCGGCCATGGGCCgaatcttaaacccaa
ggttacctcttatctagtggTGCTTATACAAGCACTGAGTGATAGCCTCAATTTTGCCTGgaatcttaaacccaacttg
ggttacctcttatctagtggGTCAGCCCCTGGGACAATAAATAGTAGATATAGCCTGGGCgaatcttaaacccaac
ggttacctcttatctagtggGGGCGTGGTGGCACGTGCCCGTAGCACCTGTAATCCCAGCgaatcttaaacccaa
ggttacctcttatctagtggTAGTCTCCTGGGTCAAAGAGATCTGAATTTGGATTTTGCCgaatcttaaacccaacttg
ggttacctcttatctagtggGGCTTCCAAGATGCCACCCTCTCCTAAATTTCCTCCTTCCgaatcttaaacccaact
ggttacctcttatctagtggGACCACAGCTCAGGTCTTGGCCCCACTCCTCTTTTCCGCCgaatcttaaacccaac
ggttacctcttatctagtggGGTGCTTGTGCCGAGAGTTCATTCTTCCCATTGTATTTACgaatcttaaacccaacttg
ggttacctcttatctagtggGCCCGGATCTCTCCCTTAAACTCTACAGTCGTAATGCAACgaatcttaaacccaact
ggttacctcttatctagtggACATGCACAAAATGGAGCTTCTGATCTTCATCCCCAAACCgaatcttaaacccaact
ggttacctcttatctagtggGTTTCTTCATTGGAAATGGGCATAGAACCTGTCTCAGGGCgaatcttaaacccaactt
ggttacctcttatctagtggGTGTTACCGCTACTCATAAGACCTACGACAGTCAGCATTCgaatcttaaacccaact
ggttacctcttatctagtggGGAGAGGACAGCATAACCTTTCCATTCTTTCATCTAATGGgaatcttaaacccaacttg
ggttacctcttatctagtggTCCTTCTCTTTCAGACCCTTACTATCTTATCCTAGAACCAgaatcttaaacccaacttg
ggttacctcttatctagtggAACAGCCAAGATGCCCTAGTCCCACCTCCTAGCAAGCACCgaatcttaaacccaa
ggttacctcttatctagtggGCCAGTGTTCTCCAGGCTCATCCTCTGCAACACAGCCTTCgaatcttaaacccaac
ggttacctcttatctagtggACAGGGTAACTGGACTATGGGAGACAAGAGTAGAGGTTCCgaatcttaaacccaac
ggttacctcttatctagtggCCTGAGCTTTGCACTAGAACTACTGAATCAGGTCGGGCCCgaatcttaaacccaac
ggttacctcttatctagtggGCTCCAAAGGCATTTTTGAGCAGTCAGATTTAGGCGCTCCgaatcttaaacccaact
ggttacctcttatctagtggTACAATCATCTACAACTCTCAAAGAGAGACGAAAAACCAGgaatcttaaacccaact
ggttacctcttatctagtggGGGCAAAGGGCATAAAAGGAGGATTTACAAAAGAGGAAGCgaatcttaaacccaac
ggttacctcttatctagtggTGCGAATACAGTATTCCTGTGAGTATGTGATGAGATGGGCgaatcttaaacccaactt
ggttacctcttatctagtggTGGAGGTACCACTTTGAAAAGCAATTTGGTAGCATGTACCgaatcttaaacccaactt
ggttacctcttatctagtggTATGCCGGGGGAAAAGTTTGAATGTCCTTACACCAAACTGgaatcttaaacccaact
ggttacctcttatctagtggGGTCTGTAAGTGGAGCATAATATTGTTCTACCCATATAAAgaatcttaaacccaacttg
ggttacctcttatctagtggTTCTTCTTTTTAGAAGGTTCTACCCTGCTTTCTATGCTGGgaatcttaaacccaacttgt
ggttacctcttatctagtggTCTAAGTGTTTTTTACCCAAGCAGTGATCCGTTTATCCCCgaatcttaaacccaacttg
ggttacctcttatctagtggGCCTGAAGGCAAGATCTGGGGGGTCTTACGTGGCCAGAAGgaatcttaaacccaa
ggttacctcttatctagtggCCATTTAGTGTCTGTGACTCCATCACAAAGAATGCTGTGCgaatcttaaacccaactt
ggttacctcttatctagtggTCAAAATGTATATGTGTATCAAGACATCAAGCTATGCACCgaatcttaaacccaacttg
ggttacctcttatctagtggGGCTTCTACTTAGGATATAGAAAGCAGGAAAGAGTAGCACgaatcttaaacccaact
ggttacctcttatctagtggACTAGAAAATAATGCTAAAGAATCTGCAGATTCACAGCTCgaatcttaaacccaacttg
ggttacctcttatctagtggTAAAAAGTAAGAAAAATGCCGGGGAGGAGCCAAGATGGCCgaatcttaaacccaac
ggttacctcttatctagtggCGTGAGCGACGCGGAAAATGGGTGATTTCTGCATTTCCATgaatcttaaacccaact
ggttacctcttatctagtggGGTCACTCCCACCTGAATATTGCGCTTCTCGGACCAGCTTgaatcttaaacccaac

ggttacctcttatctagtggGGGAAACTCTAAAACGCAGAGCACCCCTCCTCCTCCAAAGgaatcttaaacccaa
ggttacctcttatctagtggGCACCACACCACACCTGTTCCAAAATGGACCACATACTTGgaatcttaaacccaac
ggttacctcttatctagtggGAAGGCAAGAAATAACTAAAATCCGAGTAGAACTGAAGGAgaatcttaaacccaac
ggttacctcttatctagtggGGGTATGTGTCTGTGGCCCCAGCTATTTGGGAGGCCGGGGgaatcttaaacccaa
ggttacctcttatctagtggGCCCCGTCTGAGAAGTGACGAGCCTCTCCAACCGGCAGCCgaatcttaaaccca
ggttacctcttatctagtggCAAGAGTGCATACATTGGGGAAGAAATAGTTCACCAAAAAgaatcttaaacccaac
ggttacctcttatctagtggAATTGGTGAAAATAGGCTGAATATATGATGTGTGGCCTGAgaatcttaaacccaact
ggttacctcttatctagtggCAGCTCCCTCCCTTTGGATCAGGTTTATATGTTTGATGAGgaatcttaaacccaact
ggttacctcttatctagtggAACTTCATTCAAAACTAGTCCCATGAAAAGGCAAACTGAGgaatcttaaacccaact
ggttacctcttatctagtggATGGGCAAAAACGGGAAGCATTACCTTTGAAAACTGGCACgaatcttaaacccaac
ggttacctcttatctagtggATCCCCATAAAGCTACCAATGATTTTCCTCACAGAATTGGgaatcttaaacccaact
ggttacctcttatctagtggCAGAACAGAGCCATCAGAAATAACGCGGCATATCTACAACgaatcttaaacccaac
ggttacctcttatctagtggGAGCCAGGACATGAACACATGTTTTCTTAGGATAGACGCCgaatcttaaacccaac
ggttacctcttatctagtggATGGAATCCCTGCAAGCCACAGAGAAATGGGAGTTCACACgaatcttaaacccaac
ggttacctcttatctagtggCAGGTGTTCACAAGAATGATTGGAGGCAGGGTGAGAATCCgaatcttaaacccaac
ggttacctcttatctagtggGCCTGTGTAGCATTGGAATAAAGCCATAGATTCTTCACCCgaatcttaaacccaact
ggttacctcttatctagtggGACAATAAACAATGTTGGCCTTGGGACAGTGGTGAAGATCgaatcttaaacccaact
ggttacctcttatctagtggAGGAAAGGGCAGTATCATGGAGAAGGCACCACTATCAGGGgaatcttaaacccaa
ggttacctcttatctagtggAACACTCTGCTCCTCTCCTCAGTCAGCAGCAGGTTCTTGCgaatcttaaacccaac
ggttacctcttatctagtggCCTGATAGTGGAGCAAATCCTGAGGAACATCATCTTGCACgaatcttaaacccaac
ggttacctcttatctagtggGGACGGATCCCTCATACATGGCTTAGCACTGTCCCCTTGGgaatcttaaacccaac
ggttacctcttatctagtggGTGGCACCTCCTCCCTCTCTCTCTTGGTCCTGCTCTTGTCgaatcttaaacccaact
ggttacctcttatctagtggCTGGCTCCCTGTCACCTTCTGCCATAACTGTAAGCTCCCTgaatcttaaacccaac
ggttacctcttatctagtggAGCATCATGCTTTCTGTAAAGCCTGAAGAATCATGAGTCAgaatcttaaacccaact
ggttacctcttatctagtggGGAGTACAGTGGTACGATCTCGACTCACTGAAACCTCTGCgaatcttaaacccaac
ggttacctcttatctagtggACCTGAAAATGTAGGACCAACTTTGGAACTGGGTAATAGGgaatcttaaacccaac
ggttacctcttatctagtggGAAGAGTTTGAAGAATGCAGAAGAAAGTAGGAAGATGAGGgaatcttaaacccaac
ggttacctcttatctagtggCTTCTTAGAGGCTAGTTAAATGATTGAGACCAAAATGCTGgaatcttaaacccaact
ggttacctcttatctagtggTTGAGAACTGGGGGAAAGGTCACCCTTGTTATGCCTTAGCgaatcttaaacccaact
ggttacctcttatctagtggGCTGCATTGTGTTTATGCTGTAGAGATATTTGGAAGCCTGgaatcttaaacccaact
ggttacctcttatctagtggGTGACCCGACTGCTTCTAACAGCTTATACTCAGATGCAGGgaatcttaaacccaac
ggttacctcttatctagtggCAGGGGAAGAATCTAAGCAGGCGGCAGAGTGCCCACTTGCgaatcttaaacccaa
ggttacctcttatctagtggCAAGGCAACCCCTCTCATCACAGGCTCAGAGACCAAGGAGgaatcttaaacccaa
ggttacctcttatctagtggGGACACTGCTCTCTGCATCCAGGCCACTCCAGTTCCAGCCgaatcttaaacccaa
ggttacctcttatctagtggGGCCCCAGGTACAGTTCAGGTCACAGCTCTGGAAAGCACAgaatcttaaacccaa
ggttacctcttatctagtggGGCAGAAGCCTGTACAGGAGCAGAGCCCCCACAGAGAGCCgaatcttaaaccca
ggttacctcttatctagtggGCCAAGGAAAGATGTGGGATTGGAGCCTCCACACAAGTCCgaatcttaaacccaa
ggttacctcttatctagtggGCTTAGTGAAGCTGTGGGAATGGGGCCTCTACCTTACAGCgaatcttaaacccaac
ggttacctcttatctagtggTCAACTACAACCCATGAAAGCAGCAACACGGGCTTCGCCCgaatcttaaacccaa
ggttacctcttatctagtggCCCACCCCTCATTCAGGGCAAAGAATCAAAGGCGATTATCgaatcttaaacccaac
ggttacctcttatctagtggGCCCTGCTGGATTTCAGACTTGTGTGGAGCCTGTAGCCCCgaatcttaaacccaac
ggttacctcttatctagtggGGAAATGTTTAATCAGTGCCTGTATCTCCATTGTATCTTGgaatcttaaacccaacttg
ggttacctcttatctagtggCTTGATGTTGGAAAGAGCTAAGACTTTGGGGGACCTTGGGgaatcttaaacccaact
ggttacctcttatctagtggGCCATGTGAGAAGTACATGAGATTTGGGGGCCAGAGGGAGgaatcttaaacccaa
ggttacctcttatctagtggGGAGGTGGAATCTGGTGGGAGGTTCCTGAATCATCATGGGgaatcttaaacccaac
ggttacctcttatctagtggAGAGTGTGGTATCGCCCTCCACTCTTTCTCTTGCTCACTCgaatcttaaacccaact
ggttacctcttatctagtggGCTTCCTGTGACCTCACCAGAAACAGACACCAGCAAGTTGgaatcttaaacccaac
ggttacctcttatctagtggACAGCAATGCAAGAATGGCCAAATACCATAAGCATATCAGgaatcttaaacccaac
ggttacctcttatctagtggATGACTCAAACCAAAGACTATCTTCCCCCAAAATCCTGGCgaatcttaaacccaac
ggttacctcttatctagtggCATATTGTCAAGGGACAACGCGATCAATGGAACCAGACTCgaatcttaaacccaac
ggttacctcttatctagtggGATGTTGAAAACGTCAGGGAATTTAAAACAGCTGATTAAGgaatcttaaacccaact
ggttacctcttatctagtggAAGATACATAAATTGCATGATCAGGTGAATTAGCAGAGAGgaatcttaaacccaact

ggttacctcttatctagtggAACTGTAACAGAGATGAAGCATGCCTTTGATGGGCTCATCgaatcttaaacccaact
ggttacctcttatctagtggCAGCAAGAGTAGTGGAAAAAATAATAGGTCATCCAAGAGCgaatcttaaacccaact
ggttacctcttatctagtggACATTCAGTACATGTCTTTGGAATTCCAGAGAAGAGACAGgaatcttaaacccaactt
ggttacctcttatctagtggCAGGATAAACACACACACATACGCGCGCACACACACACACgaatcttaaacccaa
ggttacctcttatctagtggCTGCTGTAGAACAAAGGCAGAAATAAAATCTTTAAGGCAGgaatcttaaacccaactt
ggttacctcttatctagtggGACTTTTGTCAGAAGCTATACAAGCCAGAAGATAATGGAGgaatcttaaacccaactt
ggttacctcttatctagtggTGCCAGCATGCCTGCAGCACAAAGGAAGTTCTCTGGGCAGgaatcttaaacccaac
ggttacctcttatctagtggCCAGACAGAAATTTGGGTCTATGAAAAGAAATGGAAACTCgaatcttaaacccaactt
ggttacctcttatctagtggAGCACAAATGTTGGGAGGAAGGGAGCGGGAGTCACTCTCGgaatcttaaacccaac
ggttacctcttatctagtggTTATTTGAAACAGAATCTCATCTGTTGCTCACTCGGAGAGgaatcttaaacccaacttg
ggttacctcttatctagtggCTCATTGTGTTGCACAGGCTGGTCTGAAATTGCTGGCCTCgaatcttaaacccaactt
ggttacctcttatctagtggAAGAGCTGGGATTATAGATGTGAGCCAGCCCTGTTCGTATgaatcttaaacccaactt
ggttacctcttatctagtggTAGCCAAAGATGCATACTATAAACCCTACGGCTACCACTGgaatcttaaacccaact
ggttacctcttatctagtggAGAGATATAAGTACATAATCCAAAGTGGAGATACCATGGGgaatcttaaacccaactt
ggttacctcttatctagtggACACATAGGTGTATTAAAAGATGGAGATTATCAGACTGGAgaatcttaaacccaacttg
ggttacctcttatctagtggGATGCAATGTTATGTTTTCCATGTCTTCAGATAGAAAGACgaatcttaaacccaacttg
ggttacctcttatctagtggAGGCTTCGAAATAGAGAACATTGCCAGGGATAAAGAGGGGgaatcttaaacccaac
ggttacctcttatctagtggTCTACAACTAAAGTTGAAGTCCTCAACACTTGTCTCTCAGgaatcttaaacccaacttg
ggttacctcttatctagtggGACCCAATCGACCCTTACAGAGCAATCCACCCAGCAACAGgaatcttaaacccaa
ggttacctcttatctagtggGTGCATGGCAAATTTACCAAGACAGACCAAATTCTAGGCCgaatcttaaacccaact
ggttacctcttatctagtggTTGCTTTTGAAATACGTGCTTACAGGCAGATCAAGTTTGGgaatcttaaacccaacttg
ggttacctcttatctagtggCCCCACCAGCAGACACATTAGAGGAAGCTTTGATGACTACgaatcttaaacccaac
ggttacctcttatctagtggGTAGGAGAGTGACTTTTCTGACTTTTAGTACTATAAAACCgaatcttaaacccaacttg
ggttacctcttatctagtggGTGAGTCTAATTGATGTACCCTCAGACTCTACAGTGGGCCgaatcttaaacccaact
ggttacctcttatctagtggCTAAGACTCTACTTACCATGACTTTAAACTACCAGAAGAGgaatcttaaacccaacttg
ggttacctcttatctagtggTCATTCTCCATACCTCTAGCTGACTGTGTCCTCACTCTCCgaatcttaaacccaactt
ggttacctcttatctagtggGATAGGTCCTTGAATAAGTGTGCCTGTGTTCAATTTTCTCgaatcttaaacccaacttg
ggttacctcttatctagtggCTTTTCAGTCTTTGTTACTTGACATGAAAAGTGGGGATACgaatcttaaacccaacttg
ggttacctcttatctagtggTCAGTCATCAGCAGCTTTTTGGTGTGCAAATAGAAAGAGCgaatcttaaacccaactt
ggttacctcttatctagtggGGGTTCAAGTTCAGCACTGCCAATTAATAGCTCTGTGAGCgaatcttaaacccaact
ggttacctcttatctagtggGCTCCTGCCATGAGCTTGCTACTCTTCCTTGTTCCTTTTGgaatcttaaacccaacttg
ggttacctcttatctagtggGGATGCAAGTAAATTACTTGGGAGTGAATTGGAGGGTGCCgaatcttaaacccaact
ggttacctcttatctagtggGTATTGCCTAGGTCATCTCTAGGTCCTAGGTCTAAGATATgaatcttaaacccaacttg
ggttacctcttatctagtggATGTTCCTCTATCCCTTGAGCTTATAAGACAACCTGAGACgaatcttaaacccaactt
ggttacctcttatctagtggAGAGCATGTGGGAATATGATGTTCTTTGTGGGACCCAATGgaatcttaaacccaactt
ggttacctcttatctagtggGGACTCTAATACATATTGCAGTTCTTCACATCTCAAGACCgaatcttaaacccaacttg
ggttacctcttatctagtggTTGGATATGTACTTGCTAGAAAGAAAGTTGAAGAAGGTGCgaatcttaaacccaacttg
ggttacctcttatctagtggGGAACAGATCATGATGATGAAGTATAGCCAAAGACAACCCgaatcttaaacccaact
ggttacctcttatctagtggGCTACTGTGCAGAGATGAAGTTTGAGAGTCACCATTGGACgaatcttaaacccaact
ggttacctcttatctagtggGCCCCCACCTCCCCGCAAAAGATTATGTGTGCTGTCGCACgaatcttaaacccaac
ggttacctcttatctagtggCTTACCCTCGAGGCCTCTCCAGACTTCCTATTTTGGGCAGgaatcttaaacccaact
ggttacctcttatctagtggCTTGTCCCTCAACAGCAGACTTAAGATACAAACCTCCCTCgaatcttaaacccaact
ggttacctcttatctagtggTACCTTGAGTTAGGTGGATATCTCTTTGCATCTCTTATCCgaatcttaaacccaacttg
ggttacctcttatctagtggCCCCAAAAGTGCCATGTGCTCACTCTCATATCTGGGCAGCgaatcttaaacccaac
ggttacctcttatctagtggCTGCCCTGGCTAGTGAACGCTTCACATCCCTCACAACTTGgaatcttaaacccaac
ggttacctcttatctagtggCAGCGCTGGACTGCTTTGTCAACAGTCGCTGTGGTAGGCAgaatcttaaacccaac
ggttacctcttatctagtggCCTCCAAAGTCATCCAGGTCCAAATTCCAAGAACCTGTGAgaatcttaaacccaact
ggttacctcttatctagtggCAGAAGGGCCTTATAAAGCAAAGAAGTAGGCGAGAGAGAGgaatcttaaacccaac
ggttacctcttatctagtggGATGGAGGCAGGTGCCACACCATGAGCAGGAATGTTGCAGgaatcttaaacccaac
ggttacctcttatctagtggCCCTAGAGCCTCCAAAAGGAAAACAACTCTGCCAGCAGCCgaatcttaaacccaa
ggttacctcttatctagtggCCAGCAACATCAGTCATCTGACCTCCAGAACTAAGACCATgaatcttaaacccaac
ggttacctcttatctagtggCCCCCGATTGTTTTAGGGAATGAAGACCATTCTTGTGTGGgaatcttaaacccaactt

ggttacctcttatctagtggCCTTAACTATTAGTACAGGGTAAAATAAGGGCGGCTTAGCgaatcttaaacccaact
ggttacctcttatctagtggGAAGTTGGATGGAACAGTGGACGGTGCTACAGACTCAGGGgaatcttaaacccaa
ggttacctcttatctagtggAGCTCTGCCACCTGCATGGAGATGAATCATTTCACCTTTGgaatcttaaacccaact
ggttacctcttatctagtggGCCTGAACCGGGTCAGGGGTTGGACTGATGACCTCAGAGGgaatcttaaacccaa
ggttacctcttatctagtggTTTCTGAGTCTCCTGGGAGGAGGGGAAGTCCTCTGGGAGCgaatcttaaacccaa
ggttacctcttatctagtggACCAGTAGCTTTGGGTCAAGCCCTGCTTCAGATTCGGGCCgaatcttaaacccaac
ggttacctcttatctagtggGGGCAGAATCTGGCCCTGAAACAAGAAGAGGGATGGAGGGgaatcttaaacccaa
ggttacctcttatctagtggCAGATGGAGGCAGGGAATACTGACCTAGGCCTGATTCTGCgaatcttaaacccaa
ggttacctcttatctagtggTTGGGTGACCTTGGGATCATCATTTCACTTCACTGGGCTGgaatcttaaacccaact
ggttacctcttatctagtggGACCTCCCCTCTAGCCTCAAAATCTTACTACCTTCCAGCCgaatcttaaacccaac
ggttacctcttatctagtggGCAGGAAAGGAAGGAAGGGAGGGAGAAATGGAGGGGAGGGgaatcttaaaccca
ggttacctcttatctagtggTTAGCAATGACCACTATCCTTTCAAATGACCCTGCTCTAGgaatcttaaacccaactt
ggttacctcttatctagtggGGCGCCGGCCTCCCAAGCATGAAACCCCAATCCCAAGATAgaatcttaaacccaa
ggttacctcttatctagtggGGCTTTCCAGTGTGGGAATCCAGCAAGATGGAGCACCGAGgaatcttaaacccaa
ggttacctcttatctagtggCTACCGCATTTACTTCAGACCGTCTCCTATCTAAGCCAAGgaatcttaaacccaact
ggttacctcttatctagtggTAGGAGATGGTTTATATTTGCCAGATGTTAAAAGGTGTTCgaatcttaaacccaacttg
ggttacctcttatctagtggATTTTACCACGCCCAATCCTCCAGCGGCTCCCCAGATACCgaatcttaaacccaa
ggttacctcttatctagtggGCCAACTGTGCCCATGGGATCATTTTTAGTTCCTAGACACgaatcttaaacccaact
ggttacctcttatctagtggTACCAGGGAAGAATGGAGGAGAATTGTAAGGATAATAAAGgaatcttaaacccaac
ggttacctcttatctagtggTTCCATGTATTTGTAGAATGCTTTAAAGCAGAGTTTCCCCgaatcttaaacccaacttg
ggttacctcttatctagtggCCCTTGCTGATGGGCATTCCTGCTCTCAGAGTGTAATCCCgaatcttaaacccaac
ggttacctcttatctagtggTTGCTGGCTTACTCCGCTGCAGTAACACCATGACTCCATCgaatcttaaacccaac
ggttacctcttatctagtggCATCCCTAACTTTCCAAGCATCCCCTCATCCCCAAAGCAGgaatcttaaacccaac
ggttacctcttatctagtggCAGTCACGGCCTCTTTCTAGCTTCAGTGCCCCACCCTACCgaatcttaaacccaac
ggttacctcttatctagtggGCCGCCATCCCGTTCTACTCCAACCTTCCTTCTCAGATTGgaatcttaaacccaac
ggttacctcttatctagtggCTTTCAGTGAGTCAACCTGCTGCCTTGGATCTTCCTCTCCgaatcttaaacccaact
ggttacctcttatctagtggCTGGTCCCCAAGCCCAAGATCCATTCCTGTCTGCTCTTCCgaatcttaaacccaac
ggttacctcttatctagtggCTCTCTTAGGCCATGATCGTTTCTATAGTTTGCTCCTGTGgaatcttaaacccaacttg
ggttacctcttatctagtggACAGTGAAAGTGATACGTGGATGTGGTCAAAATTTGGATGgaatcttaaacccaactt
ggttacctcttatctagtggAATCTGTTTTTGGCTATCCTGTAACGGCAAGAAGGATGGGgaatcttaaacccaact
ggttacctcttatctagtggGAGAGCCTGCTGACTCCAAATCCAGTGACACTCTCCACGCgaatcttaaacccaa
ggttacctcttatctagtggGTTGAAGATATAGATGCCAGGCATAGACTCCAGGAAAGGGgaatcttaaacccaac
ggttacctcttatctagtggAGCAGTATGATCTATAGCCAAGGAAAAGATAGCAAATCACgaatcttaaacccaac
ggttacctcttatctagtggAGAATGTGGCAGAAATATGGAACACAAACGTAACTCCAGGgaatcttaaacccaac
ggttacctcttatctagtggGCCCCTTCGCCTACCTGCTGAGTGATGCTGGCAATAGGCCgaatcttaaacccaa
ggttacctcttatctagtggGCTCCTAGAGCTGCAGTGAGGATTGAGTGAGATAATTCTCgaatcttaaacccaac
ggttacctcttatctagtggTAGCACAGAAAATGCCCAGTTAGACATGGCGGCTTGAATCgaatcttaaacccaac
ggttacctcttatctagtggGAACAAAGAGGATTGGAAAGGTAATGACATCAGTTGTGAGgaatcttaaacccaac
ggttacctcttatctagtggAAGGATAGAGGTAGCCAACTGATGAGAGCAGAGCAAGCTGgaatcttaaacccaa
ggttacctcttatctagtggCATCAGAAGACCCAGAAAAGATAATGGCTCATTGGCATCCgaatcttaaacccaac
ggttacctcttatctagtggGGGGAGAAAAATGAGAGGGTTGATTGACAATATGTGTAAGgaatcttaaacccaact
ggttacctcttatctagtggCACTGTCTAGTCCTAATAGCCAAATAACTCCTTCCCCAACgaatcttaaacccaac
ggttacctcttatctagtggGACCAGAGAACATGGAGACACTAGGACTATCTGAGGGTGGgaatcttaaacccaa
ggttacctcttatctagtggTCCCTGACACTGTCATTCAGGTCTCTATTCTCCAACTGTGgaatcttaaacccaactt
ggttacctcttatctagtggCTTCTCCATGGAGACTATCCGTCTCAGGAGGAAAACAAACgaatcttaaacccaac
ggttacctcttatctagtggCACTCTATGGTGAGGTCAACCAGTTGACAGCCTCAGTCACgaatcttaaacccaac
ggttacctcttatctagtggTAACTTTCCAGTGCTCCATTCTAGTTATTAGTGGAGTGCCgaatcttaaacccaactt
ggttacctcttatctagtggAGGGAGGATAATTGGAATACTGCGTTGAAGAACTCTTCAGgaatcttaaacccaact
ggttacctcttatctagtggCTGGAGAAAGGTAAGAAAATTAGACAATCCTCTGGGGGTCgaatcttaaacccaac
ggttacctcttatctagtggTTCCTACGTCATGAAAGAAGAAAGATCCACATCATGGGGCgaatcttaaacccaac
ggttacctcttatctagtggAATTGCCTTTAAAGCATGAAAGGTTACAATGACATTGGGCgaatcttaaacccaactt
ggttacctcttatctagtggTCCAACCTAGAATCCTATACCCTTAGTCCAATGTGTGAGGgaatcttaaacccaact

ggttacctcttatctagtggGACACTCAAGTTTCCCAAAACTGTTGTCTATCAGACAGCCgaatcttaaacccaact
ggttacctcttatctagtggGCTGTTGGGAATCATATACCATCAAAGTAAGTCAGCTGGGgaatcttaaacccaact
ggttacctcttatctagtggAGGGGTACTGACAGTTCACTGCTATTGCTAACCTTTGGGGgaatcttaaacccaact
ggttacctcttatctagtggCTCTCCTCTGTTACCCAGTGTAAGTTTTGTATCTGTCTGCgaatcttaaacccaacttg
ggttacctcttatctagtggGCCAGTCCATAGGGAAGGCCTTCCAGGTGCCAGAAAGCCCgaatcttaaacccaa
ggttacctcttatctagtggCTCTCATGCTCCTAACTACCCAGAAACTCCATGTAGGCAAgaatcttaaacccaact
ggttacctcttatctagtggTGTGTTTTATTGCTCTAAGACAGACTCACACAAAGTTAGCgaatcttaaacccaacttg
ggttacctcttatctagtggACATGAGTCAGCAGTCTGAGCATGGCCTAGCTAGGCTCTGgaatcttaaacccaac
ggttacctcttatctagtggCAAGCTCATCTCATCTGGAACGAGGGATCCTCTCCCCAGCgaatcttaaacccaac
ggttacctcttatctagtggGAGGCTCCCATTTCCTTGCTGGCTGCCATCTGAAGCTGCCgaatcttaaacccaac
ggttacctcttatctagtggGCCACATGGCTCCCTCTGTCTTAGCCAGCAGTGGAGAGTCgaatcttaaacccaac
ggttacctcttatctagtggGGGTCAGGCCCACCTGCATAATCTCCCTTTCAGTGCACTCgaatcttaaacccaac
ggttacctcttatctagtggGGCCCCCAATTACCTCTATAAAATCCCTTTTGCCACATATgaatcttaaacccaactt
ggttacctcttatctagtggCAGGTCATTTCACACTTAACAGGAGGGGATAATAAAAGGGgaatcttaaacccaact
ggttacctcttatctagtggGGCTCCAACACCCAGGATCTAGGTGCCATACTGCCAGTGCgaatcttaaacccaa
ggttacctcttatctagtggGATGACTGGGCTGTGGGCATGGGCAATGGTTGAAGGCCTGgaatcttaaacccaac
ggttacctcttatctagtggTGCATAGCCCCCCTCCCTAAGACAAACTTACATAGTACCCgaatcttaaacccaac
ggttacctcttatctagtggCTGAGGCTCTACAGGATTGGGCCAATGTCCAAAGTTACCCgaatcttaaacccaac
ggttacctcttatctagtggCCAAAGCCCTGGCTCTCAGTGACTAAGGTCTAAGACCACCgaatcttaaacccaac
ggttacctcttatctagtggCCTACCTTCCCTTCAACAGTCCTTCATTGCCATTTGCCTGgaatcttaaacccaactt
ggttacctcttatctagtggACTTGACTGAACTTTTCCTTAGGATGAGTTCTTTCCCAGAgaatcttaaacccaacttg
ggttacctcttatctagtggTTGGAGAGTTGTGGATCTTTACACAGGGACACTGCTCAGCgaatcttaaacccaact
ggttacctcttatctagtggCACTTAAGACTGTCATTCATGCTCATCAGCGTGACCACACgaatcttaaacccaact
ggttacctcttatctagtggCCCAATAGAAAACTTTGCTTAGAATTGAACTCCACATCCCgaatcttaaacccaactt
ggttacctcttatctagtggAGAACCTAATGTATTCTGTAAAATTACCACAAGGCTCAAGgaatcttaaacccaactt
ggttacctcttatctagtggTTGAGCAAATTGTGAGTGTGTGCAGCATTGAATCAAGTAGgaatcttaaacccaactt
ggttacctcttatctagtggAGTGGGCCCCTCTTTAAAACTCTGAGAAAATACGAGATTGgaatcttaaacccaactt
ggttacctcttatctagtggTCATACCCTTAGTAAGGACAGAAGGATGCTTGTGGCTTGGgaatcttaaacccaact
ggttacctcttatctagtggCAAGCAAGACAGTCACCTGAGTGTTCCTCTTAGCTGTGCCgaatcttaaacccaact
ggttacctcttatctagtggGTCTGCCCTTGCGTGTGCCTCCCTGCTGTACTCCTCTCACgaatcttaaacccaac
ggttacctcttatctagtggCCATCGTTCTGGTGACTTCTCAGCCTCAATATTGCTGCTTgaatcttaaacccaactt
ggttacctcttatctagtggGGCATTGCCTCCTTTGTGCATGACTGTACATCAGCATTTAgaatcttaaacccaactt
ggttacctcttatctagtggCTCATCTCATGCCATGCTCTCCACTCAATCCATACCGGCCgaatcttaaacccaac
ggttacctcttatctagtggTGGCCTCAGATCGCCTACACATGCTGCTTCCCTGGACTGGgaatcttaaacccaac
ggttacctcttatctagtggCCCCGCATTGCACTTACTCTAACTCACCTTTCAGGTCTCAgaatcttaaacccaact
ggttacctcttatctagtggGTCCTTCAGTGACTACCTAGTCTTATGCAAGTCCCCACTGgaatcttaaacccaact
ggttacctcttatctagtggCATGACACCTTTTGCTTAATTCACAGTACTTACCACGATTgaatcttaaacccaacttg
ggttacctcttatctagtggTGTGCCATGCACAACTCTAAAGGTGGGATATTGTGGAGCAgaatcttaaacccaact
ggttacctcttatctagtggCAGCTATGAGGAGAGATAGATAATGAACATATACAGAGATgaatcttaaacccaactt
ggttacctcttatctagtggAGTTCAGAGAAGAAGTGAGCCAGGCAGAAATGAGGGGAGCgaatcttaaacccaa
ggttacctcttatctagtggGCAAAGCCCAGGGATAGGAATGAGTTTGAAATGTTCAAGGgaatcttaaacccaact
ggttacctcttatctagtggGAGCAGGGACAGAAGGGAGAGTGAGGACATCGAAGTCAGCgaatcttaaacccaa
ggttacctcttatctagtggCTGGGAACAGTGAGAGCTCTGGCAGATTTAAAGGAGGGGGgaatcttaaacccaac
ggttacctcttatctagtggTCAGGAGCTCAGGCAAGGAGAGACAGTGTCTTGGACTGTCgaatcttaaacccaac
ggttacctcttatctagtggGATGGTGAAAAGTGGTTGGATTTGGGATCTATTGGAGACAgaatcttaaacccaactt
ggttacctcttatctagtggGTGGAGAGGGAGATTCCTTTGGAAGACTTCAGGGGTTGGGgaatcttaaacccaac
ggttacctcttatctagtggTGTGGCCATGTGAAATGTGCTATGCCCTTTAGATTCCAAGgaatcttaaacccaactt
ggttacctcttatctagtggGGGGACAGAGAAAAGAAAGGGATTAGGACAAAGCCTTGGGgaatcttaaacccaac
ggttacctcttatctagtggTTGGGAAGAGCCAGTATGGGCATTGAGAAGGAGCAGCCAGgaatcttaaacccaac
ggttacctcttatctagtggGAAATTGAGCTATAGCTAGAAGATGACCTTACCAGGCAGGgaatcttaaacccaact
ggttacctcttatctagtggTGAAGATATGTGCAGGGCACGGGCAACCTTAAGAGATGGCgaatcttaaacccaac
ggttacctcttatctagtggTGCTATTCTCCTCTCTTCCTCCAATCTCCTGCAGAGGTGCgaatcttaaacccaactt

ggttacctcttatctagtggCCAGAGGAGCCAAGAGAAGACGTCCAGCATGGTGACCTGGgaatcttaaacccaa
ggttacctcttatctagtggGAGAGTACTGGGGAAAAGGGGGATAGTTGTAAGAATGAGGgaatcttaaacccaac
ggttacctcttatctagtggGAGATCCAAAGAGCAAGAACTGGCTTGTCTTAGAGAGGAGgaatcttaaacccaac
ggttacctcttatctagtggGGGAAGGAATGTACAGATGGAGCTAGGGGCATAGAATCAGgaatcttaaacccaac
ggttacctcttatctagtggCATCAGCTCACGGGGAGAGGTCCTGGAGACTTAAGGAGAGgaatcttaaacccaa
ggttacctcttatctagtggATCTGGTAACCAAGTGCACGTGCAAAGTGAGTCCTGCGAGgaatcttaaacccaac
ggttacctcttatctagtggCAACTTCTCAGGAGCAAGCTCAGACTAAGTGATTGTTTGGgaatcttaaacccaact
ggttacctcttatctagtggGGAGAGAAAGGAAAGAAAGATAAGTGGTTTTTCAAGGCACgaatcttaaacccaact
ggttacctcttatctagtggTAAGCTGGAAGGTGAGAGAAGTGAAGATGAGAAGGGAAACgaatcttaaacccaac
ggttacctcttatctagtggGGGTGAGTGGATTTGAGATAATGATGTCAGTCTCCAGAGGgaatcttaaacccaact
ggttacctcttatctagtggGAATAAGCTACTAGGACAGAAGAGTACTGTATGCAAGTGGgaatcttaaacccaact
ggttacctcttatctagtggGGGCATGACTATGTGAGTAGGTGGCAGAGGTGGGTGGAGGgaatcttaaacccaa
ggttacctcttatctagtggGGCCAAAGAGCTAAAGGGCCAGAGTCATAGAGATTCCACCgaatcttaaacccaa
ggttacctcttatctagtggCGGACAGAGAGGAGAAGAGGAAGAAGTAAGCCAGACATTCgaatcttaaacccaa
ggttacctcttatctagtggATACCAAACACCATTCAGTGTACTTCACGTGTATTACCTCgaatcttaaacccaactt
ggttacctcttatctagtggCTACCCTATCCTCATTCTAAAGATGAATAACCTGGGTCCCgaatcttaaacccaact
ggttacctcttatctagtggGGCAGAGACTGAATCCAACCTAGGCAATCTGATTCCAGGGgaatcttaaacccaac
ggttacctcttatctagtggTGACCATGAGTTTCCTCTACTGCTCTAAAAAACAAACAAAgaatcttaaacccaactt
ggttacctcttatctagtggGCCAAGTTCTTCTGGGTAATGAAGTCTAATGATGGGCTTCgaatcttaaacccaactt
ggttacctcttatctagtggGCTTCCTCACCCCCAGGCCTCAAGTTATCTGAGGTGTGAGgaatcttaaacccaac
ggttacctcttatctagtggACAGTCACCAATCACAGGGGAAGCTGCTTTATCAGGGAACgaatcttaaacccaac
ggttacctcttatctagtggAGGAGGTGATAGGACCAAGAAACTGTGGAACATATGGGGGgaatcttaaacccaac
ggttacctcttatctagtggCTGTCAATTCCAGGGGTGGGGAACAGGAGTTTAGAAGGGCgaatcttaaacccaac
ggttacctcttatctagtggGGGGATAGAGTCATATTTGGGAAGCCTAGAGGCATGTCCAgaatcttaaacccaac
ggttacctcttatctagtggGATGGGAGCCTTGACCTCTGATGGTGACTAAGGAGGGAGGgaatcttaaacccaa
ggttacctcttatctagtggGTCCCAGCAGTCCAGGAGCATCAGTGATAGAAGTCCTGGGgaatcttaaacccaa
ggttacctcttatctagtggCCTGAGGGGCTTGCCATTGCTGAGATACACTGACAACTGGgaatcttaaacccaac
ggttacctcttatctagtggGAACCCAGCAAGGTTCCCAGCACACAGTTGGTGCCTGTACgaatcttaaacccaa
ggttacctcttatctagtggTCCTCCATGCTCACCCAACTTGCCACACTGGGCTTGGTTGgaatcttaaacccaac
ggttacctcttatctagtggCCCATCTACCTGGACACCACCACTCCTACTCAGCCAGTTCgaatcttaaacccaa
ggttacctcttatctagtggGACTTGCTTGGCTTCTTATCTTCAGCATCACACCGCTGTGgaatcttaaacccaact
ggttacctcttatctagtggGTACATTTTCTGTCTGTCTGCACTAGAATGCCTGTGAAGGgaatcttaaacccaactt
ggttacctcttatctagtggATCTGTCTGCCTTATCTTCATGTCCTGACACTTAAAATGGgaatcttaaacccaacttg
ggttacctcttatctagtggGCCCCCATAGTTCATGAATAACCAGCCCTCTGTAGCCTGGgaatcttaaacccaac
ggttacctcttatctagtggCCCGAGACAGCTCATCACACTACATCTGCAACTAACCGTGgaatcttaaacccaac
ggttacctcttatctagtggGAATAGCTCACAAAGGGGAGTAAGCAAGTATTTGGCAGGGgaatcttaaacccaac
ggttacctcttatctagtggCATGTCCCTGGGCCACCTTTTCTGAATGACTCAGGCGGCCgaatcttaaacccaac
ggttacctcttatctagtggGACAACAAAGCTGATGAGATGGCAGGTCACAGGTAAGGAGgaatcttaaacccaa
ggttacctcttatctagtggTTAAAGAGCATGCCTTTCAAAGTAAAGGGAACAGAGGAAGgaatcttaaacccaact
ggttacctcttatctagtggCAGCACTGGGAAAACCAAGGAGAACCCGTGCAAAGTGTTGgaatcttaaacccaa
ggttacctcttatctagtggTGTATTAGTTTGAAAGCAAAGTGGGCTTTGTCTCTGCCACgaatcttaaacccaactt
ggttacctcttatctagtggCAAGTGATTTTTGACCTAGTTGTGGCATAGGGCCTGAAGCgaatcttaaacccaact
ggttacctcttatctagtggACATTGCTCAGCCACTATAAGATTCCTTCATGTTTTTCCCgaatcttaaacccaacttg
ggttacctcttatctagtggACGTTTTCTCACCTCAGCTAAACAAAACCGGTCTTTTCAGgaatcttaaacccaactt
ggttacctcttatctagtggCCTGGCTCAAGAGCACAGCTACCAGGTGGCAGGTTGCAGGgaatcttaaacccaa
ggttacctcttatctagtggCAGCTGTTGAGCCAAGTAGACCCACATTTGAATCTATCTCgaatcttaaacccaact
ggttacctcttatctagtggCCTGTATGAAGGGACCGTAGCTTCTGTATCTGCAACAAACgaatcttaaacccaact
ggttacctcttatctagtggAATGGAGAAGCACTATCCTGGAATCTAGGGATGTAGATACgaatcttaaacccaact
ggttacctcttatctagtggGAAAGCAGAACGCAGATATTCTGAGTAGGGGGTTTGGGGGgaatcttaaacccaac
ggttacctcttatctagtggAGAGGAAGTGTAGGCTTGGCCAGGTCCCTGTTCCAGGCCCgaatcttaaacccaa
ggttacctcttatctagtggGCTGTTAGCTCTCTCTCCTTCACACCTTCAGACTCTATACgaatcttaaacccaactt
ggttacctcttatctagtggCTCTGTGCCTTGGTGTGTAGTATTCTAGAATAATGGCTTCgaatcttaaacccaacttg

ggttacctcttatctagtggGGGTTTGACACAGACATTTTTCATGTTATAGCTCATACAGgaatcttaaacccaacttg
ggttacctcttatctagtggGCCCAACACACGGATAGGTAAATGTTTCTGTAAAGGCCAGgaatcttaaacccaact
ggttacctcttatctagtggACAGAAGCCATAGACAATGTGGCCAAGGACAAGGATGGTGgaatcttaaacccaac
ggttacctcttatctagtggTGGGATGGATTTGGCCCACTAACTGTTTGCCAACCCCTGGgaatcttaaacccaact
ggttacctcttatctagtggGAGCGCCTTCTATGTGCCAGGTACTATCCTTGGATAGAGGgaatcttaaacccaact
ggttacctcttatctagtggCAGTGACAGGACACAGTTCTTACCCTCATTGAGCTTACAGgaatcttaaacccaact
ggttacctcttatctagtggAATCAGAAACCCAGATCATCTGACATCAAAGCGTACATTCgaatcttaaacccaactt
ggttacctcttatctagtggACTGACTAGCACAGTTCTGGGCACACATTAATCACTTCACgaatcttaaacccaactt
ggttacctcttatctagtggATCATAGTAGGGGATTTGGGTTGTAACGGAGATGGAGAACgaatcttaaacccaactt
ggttacctcttatctagtggCTGATGTGAGTAATATGTCATTGAGCCCAAGGCCATGGGGgaatcttaaacccaact
ggttacctcttatctagtggGGAAATCATCCCAAGACCTTGTCTTAACCTAAATTCCCCCgaatcttaaacccaactt
ggttacctcttatctagtggTGGGGGGTGGGAGTTGTGGAGAATGAAACAGGGAATGGACgaatcttaaacccaac
ggttacctcttatctagtggGGGGCTCAGTCCCACTTACAGACCCTCTGAGAAGCCTTGTgaatcttaaacccaac
ggttacctcttatctagtggCCACCACTTCACCTTCACTGGTCAAGGGTTATCCCAGAAGgaatcttaaacccaac
ggttacctcttatctagtggCTCTGAGTTGCATATGCATCAGAATGACTGAGCAGGTTCCgaatcttaaacccaactt
ggttacctcttatctagtggGGCCCAGGGCAGAAAGCAAGACTTCAAGGGTGCGGCCAAGgaatcttaaacccaa
ggttacctcttatctagtggTGACCTGCGTGGTAACTGGTTATTTTAGCAATGGCTGGAGgaatcttaaacccaactt
ggttacctcttatctagtggACCAAGAAGGTGTAAGATGGAACGCAAGAGCATAACAGTAgaatcttaaacccaac
ggttacctcttatctagtggGGAGTGAACACCCAACCATGTGTCTACCACCAGTTCTCACgaatcttaaacccaac
ggttacctcttatctagtggATCCACCTTCATGTACCACGTGCATAGTACTTTTATGAACgaatcttaaacccaacttg
ggttacctcttatctagtggTCTTCCTTTTTTCTACATAAGGAGACCAAGCTCAGACAAGgaatcttaaacccaacttg
ggttacctcttatctagtggCCAACACAGTAGCCATTAGCCACATGCAGCCACTGAGCCCgaatcttaaacccaa
ggttacctcttatctagtggATGGCCACATGTAGTGAGTAGACAAAGCAGTGGCAGACACgaatcttaaacccaac
ggttacctcttatctagtggACTTTTCACTGTTGAAGTGCAATGAAAGAAATGTGTTGACgaatcttaaacccaacttg
ggttacctcttatctagtggACGATTTCCTTGTTTTCTTGCTATCACCTGGTTGCTTTCCgaatcttaaacccaacttgt
ggttacctcttatctagtggAAAAGGAGGGACTTACCCAGAAATAACACTGAATGCAACCgaatcttaaacccaac
ggttacctcttatctagtggACACAAAGGATGTCTACCCTTCCTCCCTCCACCCACTTGGgaatcttaaacccaac
ggttacctcttatctagtggCTAGTTTGCACCTGGACATAAGGCCATTTTGTACTTCCACgaatcttaaacccaacttg
ggttacctcttatctagtggCACAATTTGCATAAGGTGAGACTTGCATAACTTGGGGTTGgaatcttaaacccaacttg
ggttacctcttatctagtggATCTTCTGAGTTTTAAGTTTAGGACTCTTAAGCCTAGGACgaatcttaaacccaacttg
ggttacctcttatctagtggGAGACACCTCATATTCGTGTATTTTTCAGAAGACCGAAGCgaatcttaaacccaactt
ggttacctcttatctagtggTCCTATCCATTTTTGGGGGGAAATATTTCTGCAAGAAGTCgaatcttaaacccaacttg
ggttacctcttatctagtggGGGGTACAACATGATGTTATAATTTATGAGTGTGATGTGGgaatcttaaacccaacttg
ggttacctcttatctagtggGTACAATTCTCTATTATTAACTATGTTCAACACACGGTCCgaatcttaaacccaacttg
ggttacctcttatctagtggTCTCCATTCCCCCAGCCTCTGTAACCAGCATTCTACTCTGgaatcttaaacccaact
ggttacctcttatctagtggCTACCTAACATCAAGTTCTTCAGTACCATTCGTGTTGTCCgaatcttaaacccaacttg
ggttacctcttatctagtggCCATTCATCCATTGATGAACACTTAGCTTGATTCCATAACgaatcttaaacccaacttg
ggttacctcttatctagtggATGGGTTAATTCCCAGAAGTAGGATTGCTGGATCACATGGgaatcttaaacccaactt
ggttacctcttatctagtggGTCACCAAAAGTGTACAAAGTTTCCCTTTTCTCCCCATCCgaatcttaaacccaactt
ggttacctcttatctagtggTATGCCACTCTGACAAGTGTTGAGTGATATCTTCTTATGGgaatcttaaacccaacttg
ggttacctcttatctagtggTTTGTGGCCATTCGTCTTCTTTTAAGAAATGTCTATTCAGgaatcttaaacccaacttgt
ggttacctcttatctagtggGTGAGCTCCTTCCATATTTTGGATGTTAGCCCCTTTTTGGgaatcttaaacccaacttg
ggttacctcttatctagtggTTCTCCCATTCTGTGGGTTGTCTATGCATGCCAGTAATTGgaatcttaaacccaacttg
ggttacctcttatctagtggTGAACAAACTATCATTTTCTCATTGTGCACTCTTGGAACCgaatcttaaacccaacttg
ggttacctcttatctagtggCATTGGTCCATGTTTCTACTCTTTTGCCAATACCATGCTGgaatcttaaacccaacttg
ggttacctcttatctagtggCAGCTTTGTAGTATAGTTTGAAATCAGATGGTGTGATACCgaatcttaaacccaacttg
ggttacctcttatctagtggACTACCCATAGACCATGCACTGCAAAAGGGTCTGGGGCAGgaatcttaaacccaac
ggttacctcttatctagtggCTCAAAAGTCTTGAACAAAATACCAGTGTCCCATTCAGTCgaatcttaaacccaactt
ggttacctcttatctagtggAGAATCCAAATCTTCTGGAAGCCATGTCTTTAAAAGAGTAgaatcttaaacccaacttg
ggttacctcttatctagtggGTCATGTTTTGAGTGCTTAATAGCCACATGTGACTAGTGGgaatcttaaacccaacttg
ggttacctcttatctagtggGGACATTGCAATTTCAGAGCAAAACAAGTATTGGTGATTAgaatcttaaacccaacttg
ggttacctcttatctagtggCATATGGTTTCTCATTCTTCCGCCTCACTCTCAGGCAGCCgaatcttaaacccaactt

ggttacctcttatctagtggGTTGCAAATTTGCTTATGTCTAGCAAATGCCAGTGCCTCCgaatcttaaacccaactt
ggttacctcttatctagtggCAAAACAGAGTAAATGGGCTCTAAGTAGGGGCTAAAGAGGgaatcttaaacccaac
ggttacctcttatctagtggAGTGTTGTCCTCAGGCCCACAGCAGCAGCAGCACCCAGGGgaatcttaaacccaa
ggttacctcttatctagtggAGGCAAAGTCCTGGTCCCCACCCCAGACCCAGTGAATCAGgaatcttaaacccaa
ggttacctcttatctagtggGCCCTCTGGGTGATTCTGGTGCCCACTAAAGTTTGAGAACgaatcttaaacccaact
ggttacctcttatctagtggGTTCAGACTGCTATATACATGGACTTGAGATGAATAAGTGgaatcttaaacccaactt
ggttacctcttatctagtggAAGGTAACAAAGAGGTCTGTTTTTTCTCTTTGAAACATGGgaatcttaaacccaacttg
ggttacctcttatctagtggAATTTTCTTTTAGTTCTGGCAATTAGAAGCCCCAAAGGGGgaatcttaaacccaactt
ggttacctcttatctagtggGTCTTGGCAACTTCTAGAGGACACCCACATTCCTAGGCTCgaatcttaaacccaact
ggttacctcttatctagtggGTCAGGTCGCAGCTCTCTGGTTCTCCCTATTCTGCTTCCCgaatcttaaacccaact
ggttacctcttatctagtggCGGACCCACCTGGCTAATCCAAGAACACCACCTCATCTCGgaatcttaaacccaa
ggttacctcttatctagtggGCCTCTCTGGCATGGACCCTAACATATTTACAGGTTCCAGgaatcttaaacccaact
ggttacctcttatctagtggACCCTTGCACATTTTCAGGTTGACATCTAGCATTTTTCACgaatcttaaacccaacttg
ggttacctcttatctagtggTTTTTCCTAGCAAACATGTTCCCACTTTCTTTCTCCAGAGgaatcttaaacccaacttg
ggttacctcttatctagtggTTCCACACTTTAGGGTTTTTCTTTGATTACTCCAACACACgaatcttaaacccaacttg
ggttacctcttatctagtggAGACACCTCTTTGTCATATATGGGGGTGGAGAGAGAGGACgaatcttaaacccaact
ggttacctcttatctagtggCAAATAGTCCCTCTGAAGTTTTCACATCTCAGCACAACCCgaatcttaaacccaact
ggttacctcttatctagtggGCGGAGCGACTTTTCTCACAGGATGTGCAGACTGTGCAGCgaatcttaaacccaac
ggttacctcttatctagtggCTAGCAGTACTACAAATAGCATGACTGCCTGTGGGTGAGGgaatcttaaacccaact
ggttacctcttatctagtggCCAGTCAACCATGCTAGAGGAAAGACTGAGTTAACTTCTGgaatcttaaacccaact
ggttacctcttatctagtggCTGTCATCTGGCAGGGAGATCTTTGAAAAGACAGGCAGCCgaatcttaaacccaac
ggttacctcttatctagtggTCATTGTCAAGCCAAATTGAATGGAATTCACATTCACTTCgaatcttaaacccaacttg
ggttacctcttatctagtggCAGCTGGGCTCACTAAAACCCCCGCCTCCTGGTTCATGCGgaatcttaaacccaa
ggttacctcttatctagtggCAGGTGTGAGCTACCGTGCCTGGCCATGGAATTCATTTAAgaatcttaaacccaact
ggttacctcttatctagtggAACTGACTCCTGGTGTATGTCAGCCACTCTGTTTAGACTGgaatcttaaacccaactt
ggttacctcttatctagtggCTTCTCAAAAAACAAGTGACCAAGTACAAGCATGCAGAACgaatcttaaacccaact
ggttacctcttatctagtggCTGCCAAGGAAAAATTAAGTAGAATATTCAGTCTCACAGCgaatcttaaacccaactt
ggttacctcttatctagtggGTGGGAAAGAGAGCTCAGCATGGTTGCGGTGGCGATGGCGgaatcttaaacccaa
ggttacctcttatctagtggAGGGAGTTCAGGAAAGGCATCATCAATAGGGCAACGCTTGgaatcttaaacccaac
ggttacctcttatctagtggGACAAGAGGAATCAGTCATGTCAATATCTGAGGCGTGAGCgaatcttaaacccaact
ggttacctcttatctagtggGCTGAAGCTGGGGTTGGGTGTGGTTTGCAGCTCACAGAGGgaatcttaaacccaac
ggttacctcttatctagtggCTCTGAGCAAATTGCAAAGCTGCTGTGGGAGGATGCAGAGgaatcttaaacccaac
ggttacctcttatctagtggAGTTGGGGATGGTGGAGCAAGGGGAATCAGGTAGAAGGCCgaatcttaaacccaa
ggttacctcttatctagtggCCAAACAAGGTAGTGACAGCAGCAGAGGCGGTGAGAAATGgaatcttaaacccaa
ggttacctcttatctagtggCAGAGCCAGAGGATCTGCTGATGGTATCAGGGAGGTATGAgaatcttaaacccaac
ggttacctcttatctagtggATGGAGTTTCCATTTATTCCAGGGCAGTCATTGTGCTAGGgaatcttaaacccaactt
ggttacctcttatctagtggTGCCAACCCAACATTGCAGGCAAGATGGCAGAAGCAAGGGgaatcttaaacccaa
ggttacctcttatctagtggCTCTGCCTGTCTCAAAGTCCTCTGCATCACACTTAGCAGCgaatcttaaacccaact
ggttacctcttatctagtggGGGGCAATTATCACTTTTAAGAAGGATTTGGCATGTGCTAgaatcttaaacccaactt
ggttacctcttatctagtggGAGTTGCGTACAGTTGCACGGTGAGGAGGTAGCAGAGCTCgaatcttaaacccaa
ggttacctcttatctagtggGACTCCAGAGCCCAAACGCCTCGCCCTTACATTAGAACCCgaatcttaaacccaa
ggttacctcttatctagtggGCTGGAATTGCCTCATCTATACACTGGGAATTATATTTGCgaatcttaaacccaactt
ggttacctcttatctagtggCATAAATTAGCTGGCATACTGTAGACACCCAATATCAGTGgaatcttaaacccaactt
ggttacctcttatctagtggGCACATAATGCTGATCCGCTACTCCTAGTGACAAATCCAAgaatcttaaacccaact
ggttacctcttatctagtggCCAGTGGACGTGTTGCCTCTCCAAGACAAGGCCAAGCAGCgaatcttaaacccaa
ggttacctcttatctagtggGGGCCTAACCCTATCCACACAAGTAAAAGTCATAAGGGGTgaatcttaaacccaac
ggttacctcttatctagtggGACAATCAAGGACTTAGAATCCCAATCTGCCCATCTTCTCgaatcttaaacccaact
ggttacctcttatctagtggGAAGAAGGACAAGGAAAGGAAGACGCAGGAAGGAGCTTCCgaatcttaaacccaa
ggttacctcttatctagtggGAGAAGACCCAGAATTTCTCCTGGTCCACAGTTTCTCCACgaatcttaaacccaact
ggttacctcttatctagtggCAGAGGGGGAATGCCATGAGGATTTGAGATGAAAGATGCGgaatcttaaacccaac
ggttacctcttatctagtggAGGCATGATGTCTGCCTGAGAGGCCACACAAGCAGCAGTTgaatcttaaacccaac
ggttacctcttatctagtggCCATGGGCAGGAAGGCAGATGACACCTCAGGTCCAGGAGCgaatcttaaacccaa

ggttacctcttatctagtggTGACAGAAAGCATCTAGAATGTGTCAGCGAAAGGCACTGGgaatcttaaacccaac
ggttacctcttatctagtggGAGGAGGAGGAGCAGGAGGAAGAGGAGCAGGAGGAGCAGGgaatcttaaaccca
ggttacctcttatctagtggGGAAGTCTTCTTCCCTTGAGGAATTCTTAGTCTCCTGGGAgaatcttaaacccaactt
ggttacctcttatctagtggTGCCACATTTTCTTATGGGTCTTGATGGACATGCACAAAGgaatcttaaacccaactt
ggttacctcttatctagtggGATAGAGGGTGAAAGCAGACTTTACAGGGTACTGGATGCCgaatcttaaacccaac
ggttacctcttatctagtggGGAATGTGCCAAACACAAGAGAGAATCGTGGTGCCCTTCCgaatcttaaacccaa
ggttacctcttatctagtggACACGTGCAAGGGCAGAGGAAGGAAACCATCTGGCATAGGgaatcttaaacccaa
ggttacctcttatctagtggCACAAGTCCTCTCTGGATCCATCCACTGCGAAAAGGAGCCgaatcttaaacccaa
ggttacctcttatctagtggAGGGAAAATAGGGAACTACAAAAACGTTTTTGTCAGTAGCgaatcttaaacccaactt
ggttacctcttatctagtggGCCGAGTGTCAGCTGTGTTTCAGGCACTCAGGAAGACAGGgaatcttaaacccaa
ggttacctcttatctagtggGCGCACCAGAATGTTCTAGGAGGCTTCCATCTTGGTTGGCgaatcttaaacccaac
ggttacctcttatctagtggCTCCCTGAGTCCAGAGATGAATCAGATCCTGTCCTTGCTCgaatcttaaacccaac
ggttacctcttatctagtggGGATCTTTCTTGCAGCAGGAAGTTTGGAGACCAGCCCTGCgaatcttaaacccaac
ggttacctcttatctagtggCTGCGTAACTAGGAACAGCCACTGAGCCACTTACAAATGAgaatcttaaacccaac
ggttacctcttatctagtggGCCCTAGTCCTGAGCTGCTTGGCCCCCTGTTCTGTTCTGCgaatcttaaacccaac
ggttacctcttatctagtggCTTGTTTTTGGCTGGGCACTGTGGCTCATACCTGCAATCCgaatcttaaacccaact
ggttacctcttatctagtggGGGAGGCAGAGCTTGCAGTGAGCTGATGTCATGCGACTGCgaatcttaaacccaa
ggttacctcttatctagtggCAGACCAGCCTCATCATGCCCCTGGAGAGGTACCAGTGCCgaatcttaaacccaa
ggttacctcttatctagtggGCAGCACCTTCTCTTCACCAGTAGGGATCCCATCTTCATAgaatcttaaacccaact
ggttacctcttatctagtggGTCTTGAATCTGATAACCCCAATCTCTTATATTTGTAAGGgaatcttaaacccaacttg
ggttacctcttatctagtggTTTTTTACCTTTTCATTCCTCTTGTATTTAGCCAGTTCCCgaatcttaaacccaacttgt
ggttacctcttatctagtggTTTCTACACGAGAGCATGATGGATTTGGTGATGCTATGACgaatcttaaacccaactt
ggttacctcttatctagtggGTGTTCTGTCTTTGCTATGCTAAGATGCCATGCTAAGTGGgaatcttaaacccaactt
ggttacctcttatctagtggCTGACCTCAAGGATGCTCCTGAGCTAGAAAGACACATACCgaatcttaaacccaac
ggttacctcttatctagtggACTGCCAACAAAGTGCAGTGGCCACACACAGCGGGCAGTGgaatcttaaacccaa
ggttacctcttatctagtggCTGAAAGAGTCTTACTGGCTGACAACAGGGCTGCTCCATGgaatcttaaacccaac
ggttacctcttatctagtggGAAGAGGAGTGGGTGGACGCACAGGAAACAGCAAGAGCCCgaatcttaaaccca
ggttacctcttatctagtggTGGGAGCGTGTAAGAGCTGAGGCAGGAAGGGGGTGGTGGGgaatcttaaacccaa
ggttacctcttatctagtggGTGTCATGGTCAGATCTGCCTTTCCAAAGATCACTCTCGGgaatcttaaacccaac
ggttacctcttatctagtggGGACTAAAGTAGGGCATTGGAGGGGTGTGTGGAGTAAGGGgaatcttaaacccaa
ggttacctcttatctagtggGCCACTATGCTAACAGTAGCCACATCCCAAGTAGGCAGGCgaatcttaaacccaa
ggttacctcttatctagtggCAGGCTCTAGATCTTGCATACTTTTCTCTATTCCTCCTGGgaatcttaaacccaactt
ggttacctcttatctagtggTACAGTCTGGGCATTAACTCCTTCCAGAGCTCAAGCCCTGgaatcttaaacccaac
ggttacctcttatctagtggTGGAGTCCTACAGTGTGGATCAAGATTAAGCTTGGCTCTGgaatcttaaacccaact
ggttacctcttatctagtggCCCGTAGAATGGGAATCCCAGTACCTATCTCAAAGTAAGTgaatcttaaacccaact
ggttacctcttatctagtggGCTCAATCTACTTGAAAGCACGGTCAGAGCATTCTTTGGGgaatcttaaacccaac
ggttacctcttatctagtggTCTGTGTTCATCACTGTATATTTATTGACTGGCTAGCTCAgaatcttaaacccaacttg
ggttacctcttatctagtggGTTTAGTCTACGTGAACATAATGAGGGTCACTTCTGTTCCgaatcttaaacccaactt
ggttacctcttatctagtggCAGACTGGCCTTTTTCAGAAGCCAAGGGATTGTTATGTCCgaatcttaaacccaact
ggttacctcttatctagtggGACCCAAAGGCAGGGAATGATTTGAGCATGTTCACAAGGCgaatcttaaacccaac
ggttacctcttatctagtggGTGGTAGTAAGATGGCCAGGATTTGCTGTTGGGGATGGGGgaatcttaaacccaac
ggttacctcttatctagtggCAGGAAGGAGTTAGGGGTCCTGCACTCAAGCTAATGAAGGgaatcttaaacccaa
ggttacctcttatctagtggGCCTCCTTGCTTACAGCCTTCCTCTGTCAAAGGGTAGTGCgaatcttaaacccaac
ggttacctcttatctagtggGGAGATGCTTCCGCTTCATTTTCATTTTGTTTCCTTCCTTgaatcttaaacccaacttgt
ggttacctcttatctagtggTCCTGATGCACTTTATGGAATCCATTGTCCTTGTCATCTCgaatcttaaacccaacttg
ggttacctcttatctagtggTTGGTCCTATCCCTCCATGTGCCTGTGTTAGCTGCAGGGGgaatcttaaacccaac
ggttacctcttatctagtggAACCTGCCCACATTATGTACATAGGGAATTAAAATTCAACgaatcttaaacccaactt
ggttacctcttatctagtggACTGAAGCAAGGTGAGTGCTGAAGGCAAAACCATATCGACgaatcttaaacccaac
ggttacctcttatctagtggAGCAAGGGCATGGGGATGGAAGACAGCAAGACCCAAGTCCgaatcttaaacccaa
ggttacctcttatctagtggCACCACCTCCTTCCAAATCTGTGACTTGGTACAAATAACCgaatcttaaacccaact
ggttacctcttatctagtggGCTGATCCTGACATACTGACATATATGTTTTTCCCCTTGgaatcttaaacccaacttg
ggttacctcttatctagtggCCTTTTAGTTCTATCATGAAGTTCATTGGTTTGTCTGGGGgaatcttaaacccaacttg

ggttacctcttatctagtggCTCACTCGTTCCTGGGCATAGGACAAGCTAACCATGGGAGgaatcttaaacccaac
ggttacctcttatctagtggAGTTCTTCCCTAAAACTAACCCCCTCCTTGTTCAGGAACCgaatcttaaacccaact
ggttacctcttatctagtggAGCCGTGAAATTAGAATTATGAGAGGAACCTGAATTCTGCgaatcttaaacccaactt
ggttacctcttatctagtggGGCCAGAGGTCACAAGATTTCCAACTTCCCCAATTGCTCCgaatcttaaacccaac
ggttacctcttatctagtggGTGGCCCCCTCCTTGAGGCAGACTTAGTGCACAAGGACCGgaatcttaaacccaa
ggttacctcttatctagtggAACCAGTTAGCAGCACCCAATCCCTAGCCCCCTGCCCACCgaatcttaaacccaa
ggttacctcttatctagtggAAACCCCCAGCCTCTGAGTTCTCAGGGACATGGATTTGAGgaatcttaaacccaac
ggttacctcttatctagtggCCAAGTCTCAGTGAATTGGTTTGTCTGTGCAATGGGCAAGgaatcttaaacccaactt
ggttacctcttatctagtggCAGGTGGTTACATCTGCATCATTCCCAACTTATCCTCTGTgaatcttaaacccaactt
ggttacctcttatctagtggTCTCAGCATCCCTATTTTCCATGATTCTGGCTTTTTATGTgaatcttaaacccaacttgt
ggttacctcttatctagtggGTAACTACCACATAGCATTGTTATGAGGCCAAAATAAGGCgaatcttaaacccaactt
ggttacctcttatctagtggGTGTTCTCGTTTCCTCTCTGAAATGGAAGCATTGGCTGCCgaatcttaaacccaactt
ggttacctcttatctagtggCGGATATGTCAGATCTGAGGTGTGGGACTATGTAGTCAAGgaatcttaaacccaact
ggttacctcttatctagtggGCTGGGTGCCAAGTCAACAGTCACCTTACAGTCTTCAGGGgaatcttaaacccaac
ggttacctcttatctagtggATGCAGTGTTGGTGTTCAATAGTCCACGATCTTGGGCAAGgaatcttaaacccaactt
ggttacctcttatctagtggCTCTAAAGATTCGGTTCATTCATCTATAGAGCAGAAAGAAgaatcttaaacccaactt
ggttacctcttatctagtggTCCTTGTACTTATATTTGGCATCACGCTATCTATCTCACCgaatcttaaacccaacttg
ggttacctcttatctagtggGCAGATAAGGAAATTGAGGCTTAGAAATGTTTCATGACTCgaatcttaaacccaactt
ggttacctcttatctagtggGGGCAGTTACTTCAGGTTCAGTCCATGCCAAATTTGAGTCgaatcttaaacccaactt
ggttacctcttatctagtggTTGGAAGGCCATTAGGCTGAGATGGCTCCAGTGTTTTGGGgaatcttaaacccaact
ggttacctcttatctagtggGGCAAAGGGGAAGCAGGGACATCTTACATGGTTGGAGCAGgaatcttaaacccaa
ggttacctcttatctagtggGAGAGAAGGGGGAGGTGCCACAAATTTCTATGCAACCAGAgaatcttaaacccaac
ggttacctcttatctagtggGAGGGGATGATACTAAACCATTGATGAAGGATCCACCCCCgaatcttaaacccaac
ggttacctcttatctagtggCCCACCTCCAATACTGGGGATTACAACTGAAAAAACAGAAgaatcttaaacccaac
ggttacctcttatctagtggTCTGAAATCACCAACTAACCTCCAACTAGAGACTTCCCACgaatcttaaacccaac
ggttacctcttatctagtggCTTTACTTTGCTTTCATGTGCACTCTACAGAAGCCCGCTGgaatcttaaacccaactt
ggttacctcttatctagtggGCAGAGTTCTCTGAACCTTTTCCAAGTGGAGTTCTGCCTGgaatcttaaacccaactt
ggttacctcttatctagtggGGACTAGAATGAAATGTCCATCACCTAGCATAGCACCAAGgaatcttaaacccaac
ggttacctcttatctagtggTGGACATGCCCATTACCTTCCAGTTTTGTTATGTTGGCAGgaatcttaaacccaactt
ggttacctcttatctagtggCATGAGATCTTCCCTCTTAAATTTTCTGGTGCACAATATCgaatcttaaacccaacttg
ggttacctcttatctagtggCTCCATCTTCCCACTCCTTGGCAACCACTACTGTGATCCCgaatcttaaacccaac
ggttacctcttatctagtggCTGTGCCTGGCTTATGTCACTTAGCCTAATGTCTCCCAGGgaatcttaaacccaact
ggttacctcttatctagtggATGCTTCGATGAACATGGGAGTGCAGAGATAAATGAAGGCgaatcttaaacccaac
ggttacctcttatctagtggAACAATGGGTAAATGTAGAGCCGCACTCGGGAGACAAGCCgaatcttaaacccaa
ggttacctcttatctagtggGGCAGAGCATTGGATGGTGTTGAGATTGACTCAGCTTGGGgaatcttaaacccaac
ggttacctcttatctagtggCTAAAACAGATCTGGGCTCAAATCCTGGGTTTTGGTATTGgaatcttaaacccaactt
ggttacctcttatctagtggAGTAACCTCACTACTCCAGTCATGAACTTGAAGCCAGTTGgaatcttaaacccaact
ggttacctcttatctagtggCTTGTGAATGTAAGACAATTTTTCAAGGCAAGTACTGGAGgaatcttaaacccaactt
ggttacctcttatctagtggTGTGTTGTCTTACCAAGAAAGGACTAGTCCATTTTGCATCgaatcttaaacccaactt
ggttacctcttatctagtggTTATCCATTTCTATTCATCAAGGCACCACATTTTTCTCTCgaatcttaaacccaacttgt
ggttacctcttatctagtggGCAGAATGGATTCTAATACGAGGTACCCTGTAATGTCAGGgaatcttaaacccaact
ggttacctcttatctagtggAGGTTTGCATTGCCTTCATATACCCCCAAAACTCAGCATGgaatcttaaacccaactt
ggttacctcttatctagtggGGTACATCATGCACAATGATAAACTACCCCATGTGGTTTGgaatcttaaacccaactt
ggttacctcttatctagtggGGAAATGAAGAAGCTGTGATTTATGACTCCAGCTGATAGGgaatcttaaacccaactt
ggttacctcttatctagtggGAGTGGGGGAAATCTCCAAGACTGATGACTTTCTCCCTTGgaatcttaaacccaact
ggttacctcttatctagtggCTGAGTGAACCAGATCTAGGTCTGTGTCTATCTGCTACCAgaatcttaaacccaactt
ggttacctcttatctagtggGTTATTGTCCCTTTCTGTCCAGTCATTCATTTGCTCATCCgaatcttaaacccaacttg
ggttacctcttatctagtggCCACCTGCACTAGTTCAGTTGTAAAGATGATCTGATGTGGgaatcttaaacccaactt
ggttacctcttatctagtggCCAATGACATGACTGCTTTTGGCCTCTGATTTTCTCTCCCgaatcttaaacccaactt
ggttacctcttatctagtggGTACTTGGAGAGGAGTCCGCTGACCCACGGTCCAGTTTGGgaatcttaaacccaac
ggttacctcttatctagtggCTACCTCGAAGCCATTCACAAATTCCTTATTCTCTCTGGGgaatcttaaacccaactt
ggttacctcttatctagtggTGACTCCAAACTTCCTTCTGTCTCAACAACCCTATAGCCCgaatcttaaacccaact

ggttacctcttatctagtggATTCCCTCTGAGTTCCAGTTTTCCCAGGTATCATGTAGGGgaatcttaaacccaactt
ggttacctcttatctagtggGCGAGGAAAGGTGAATCTCACAAGCTTCAGAGAGAAGATGgaatcttaaacccaac
ggttacctcttatctagtggCTTGTTTAATAGCCATAGCAGTCAAGTCAGCAAAGCGGCGgaatcttaaacccaact
ggttacctcttatctagtggGACCCTGAACACTCACCGAAAGCTACCTAAGACCTCACACgaatcttaaacccaa
ggttacctcttatctagtggGCCACTTACTAGTGCTTGGTGTTGGGCAAGTCACATCACCgaatcttaaacccaact
ggttacctcttatctagtggCATCTACAAAATGCAGCAGTAATCCCTACTTTATATGGTGgaatcttaaacccaactt
ggttacctcttatctagtggTATTAAGCACTTTATTTTTTACCCAGTCGACAGCAAGTGGgaatcttaaacccaacttg
ggttacctcttatctagtggCTCTGTCTCTGCCAAAGCCACAGACACTCAGTTCCTGGTGgaatcttaaacccaac
ggttacctcttatctagtggGACAGGCTCCACTCCTTTGCACTTCTTCCACTTTCCCTGCgaatcttaaacccaact
ggttacctcttatctagtggCGCAAGTCCTTAGCTTGCTGAGAAGGGCTCTTGTCTCATTgaatcttaaacccaact
ggttacctcttatctagtggTGCCATATGGTTTCCCCGGAAATGCTATCACTCAACAAGGgaatcttaaacccaact
ggttacctcttatctagtggGGCAGACCTGCACTCTCCTGCTCCCCTTCACTGCTACAGCgaatcttaaacccaa
ggttacctcttatctagtggACAGTTCATCTCAGGCTTCTAGGTGTTTAGAGACCACCACgaatcttaaacccaact
ggttacctcttatctagtggACTCTTCTGTGTTAAAGACAGACAGCTCAGTTGGTGCTACgaatcttaaacccaactt
ggttacctcttatctagtggCATCTCTCTCGGCCAGTTTCTTAAAGGGCGTTGCAGCAGCgaatcttaaacccaac
ggttacctcttatctagtggGAGGTTCTGCACTTTGGATGGAGATGCATATGTAATGGGGgaatcttaaacccaact
ggttacctcttatctagtggTGTCTGACCTACTGACAGTTGTTGATGTGCTCTCGTGACCgaatcttaaacccaactt
ggttacctcttatctagtggGAGCAGGGAGGAAGAGATTGCTGGAGCACCTAGGAGCTGGgaatcttaaacccaa
ggttacctcttatctagtggACAGAAGGAGCACTGACCCATGTTATAGACAATCGCAGAAgaatcttaaacccaac
ggttacctcttatctagtggCCCTCAGACTTGGTCTGTGTGTTCTACGCAGTTGATCTGCgaatcttaaacccaact
ggttacctcttatctagtggGTCCTGACTTGGGTGGCTGGTGGAAATGGGCGTGTACTCCgaatcttaaacccaac
ggttacctcttatctagtggCACTCTCTTGGCCCTGTGGGATTTTTGAGGCTGAGATTCCgaatcttaaacccaact
ggttacctcttatctagtggCACACACACAGAAAAGAATGATTTCAGGCCCTTCCTTGAGgaatcttaaacccaac
ggttacctcttatctagtggCTGCACAGTGAAGTCAGCCCACATCTATCGATATGGCTGCgaatcttaaacccaac
ggttacctcttatctagtggACCCGTTACAATACCCTGACTTTGGTGGTAGCTGGAATTGgaatcttaaacccaact
ggttacctcttatctagtggGGGCAGGGTTTTCAGTAGTGTCACCTGATTGCTGGTGACCgaatcttaaacccaac
ggttacctcttatctagtggCCTTGGAAATCTTATTCCTCCAGAGCCAACAGAGTAGGAGgaatcttaaacccaac
ggttacctcttatctagtggACTATAATGCTCTATACCAGGGGCAGTGATCTATGGCCACgaatcttaaacccaact
ggttacctcttatctagtggTGCCTGGGCCTTGCTACTACTACAGATTCTAAGGCTCCACgaatcttaaacccaac
ggttacctcttatctagtggCTGAAACAGAATCTTACGGTGGGAGAAGAATGCCAGACTCgaatcttaaacccaac
ggttacctcttatctagtggTGAGAGCTTCTACCCTAGGGATAGCCCCCTCTTTCTGTACgaatcttaaacccaact
ggttacctcttatctagtggGGAGACAGGAAATTCGTGTCCAACACAATGCGTGAGAGGCgaatcttaaacccaa
ggttacctcttatctagtggGTCAGAGAAAACCCTTCAGGCAAGAGAGACCTAAGCTGGCgaatcttaaacccaa
ggttacctcttatctagtggACCCCAAGAGTTCCTCCTCTGGACAGCTGCCTCATGAGGGgaatcttaaacccaa
ggttacctcttatctagtggGTCCAGATGGACTCCCGCCTGCCATTCATTACATATGTGGgaatcttaaacccaac
ggttacctcttatctagtggTAGTTCCTCTCTTACGGGTCCTCTTTTAATGGATGAGGGCgaatcttaaacccaactt
ggttacctcttatctagtggTGAGAACTCTGAACACACAGAATTCATAGGGCTTAGTCATgaatcttaaacccaactt
ggttacctcttatctagtggTTCCCCTGCCACTGGGTGAAAATTCAGGTTGTTCCAAGGCgaatcttaaacccaac
ggttacctcttatctagtggAGACCTTCCTTTCCCCTGGTCCACCCAAACTGCTAGAGCCgaatcttaaacccaac
ggttacctcttatctagtggAGGTAGCCCGCCTTGGGGAAGAATCATGTCTCACGTTCTCgaatcttaaacccaac
ggttacctcttatctagtggTTCAGATCAGGGTGGACGAGGGGAGAAAAAGAGGTAAGGGgaatcttaaacccaa
ggttacctcttatctagtggACTGCTGTGAATCAGCTCTGCTCTCTCTGGGCCTAACGGGgaatcttaaacccaac
ggttacctcttatctagtggAGACTGAACACTTTTCAGCAGGCCAGCCTCAAAATGAGCTgaatcttaaacccaac
ggttacctcttatctagtggGCAACCTCAGTTATACAGGACGCGGATTCACCCAGGTCCGgaatcttaaacccaa
ggttacctcttatctagtggGGAGCTTGAGGCTAGCTCGCTGTCTCTCCTGTGTGCCCACgaatcttaaacccaac
ggttacctcttatctagtggCACTCAATGCATTTGCTGTGTGCCAGGAGAAGGGGTCACCgaatcttaaacccaac
ggttacctcttatctagtggGCAGCTCCCCAACCTTTCTCTCTTGCTCAGTAGAGTCCCCgaatcttaaacccaac
ggttacctcttatctagtggGCACCTCTCCCACTTGGCTTGAAGTGAGGGACAGACATCCgaatcttaaacccaa
ggttacctcttatctagtggCCTTAGAGAAGGACTCAAAGCCTGTCATCAGCTCCTCCAAgaatcttaaacccaac
ggttacctcttatctagtggTCTGTCTGTTGGCTTCTGACCTTTTTATATTCTGCATGGGgaatcttaaacccaacttg
ggttacctcttatctagtggGGCAATTAGCTCAGGCTCCCAGCTTAGGCTTAACCTCAGCgaatcttaaacccaac
ggttacctcttatctagtggCAGGAGGCTAGCTCTGTGGTGGATGAAATGCTCCTCCCACgaatcttaaacccaac

ggttacctcttatctagtggCTTAACATCTTGCGGTGTGACTGTTTGCTGCCATTTCCCCgaatcttaaacccaactt
ggttacctcttatctagtggCACACTTCTGTCTTATTCGTCTCTATTTCTCCAATGCTAGgaatcttaaacccaacttg
ggttacctcttatctagtggGCATGGTCAGCCATATTGAGGGAGAGATGACTGATGGCAGgaatcttaaacccaac
ggttacctcttatctagtggTCCAGGCCTATCTGTGAACAGATCGAAGCATTAAGGAAACgaatcttaaacccaact
ggttacctcttatctagtggGAGTGGGTTATGCCAACAAAGAAGGCTCTTCCACCTCCTGgaatcttaaacccaac
ggttacctcttatctagtggGGTCCAGGTATATCATGATTCCTTCCAGGTGCACCCCTGGgaatcttaaacccaact
ggttacctcttatctagtggGCATGACCTAGTCAGAGATCTGTTTGAGGCACAAATTTCCgaatcttaaacccaactt
ggttacctcttatctagtggAGAGCACCTCCTTCATGCTAGTGTTCTCCCTAACTTGCTCgaatcttaaacccaactt
ggttacctcttatctagtggGAGATGCACCAAAGCAAGTGGGATAAAGGAAGCCTAGGGGgaatcttaaacccaa
ggttacctcttatctagtggAGAAGCCCCAGAGAATAGAATCTATACCTGTGGACTGAGCgaatcttaaacccaac
ggttacctcttatctagtggCGAATATCCATCCTTTCAAGACTCAGAGCTCCTTGCAGGGgaatcttaaacccaact
ggttacctcttatctagtggGTTGGATGACAGATAGGTGGATGGATGGGTGGAAGAATGGgaatcttaaacccaact
ggttacctcttatctagtggCGGTGTCTCTGGATTCTCTCAAAGCCAAACAGTGATCAGGgaatcttaaacccaact
ggttacctcttatctagtggTTCTCTATCATTAACAGCAGCAGCAATATCTCATAACGGCgaatcttaaacccaactt
ggttacctcttatctagtggTGCTACCAGCATCCAGCAGTTCTTCTTCAGATAACCACGCgaatcttaaacccaact
ggttacctcttatctagtggCCAGAAAACCACCTTTCCCAACAGCCTCAGCCCATGTGCCgaatcttaaacccaa
ggttacctcttatctagtggCACCATTGGCCAAAGCTACTCACAGTATTTATTCTTCCTGgaatcttaaacccaactt
ggttacctcttatctagtggGACTTCTCGGAAACTCTTTCTGCTTACCTAAACCCGAGAGgaatcttaaacccaact
ggttacctcttatctagtggTCTGGTTCCTCAGCAGCTATCTTTCTACCATAAGGAGGGCgaatcttaaacccaact
ggttacctcttatctagtggGATTACCTTAGACTAGCAAACGAAAGACAGAGAGACTGGGgaatcttaaacccaac
ggttacctcttatctagtggATCTTAACACTACTTCTGTACTATCTTTTTCTGTGTGTGCgaatcttaaacccaacttgt
ggttacctcttatctagtggTTTGCATTTGCAACCCTAAGTCCTAACAGATTAATTTCTCgaatcttaaacccaacttg
ggttacctcttatctagtggATGCTGGTATAAGTTTTTGCAACTCCTACTATACACATGGgaatcttaaacccaacttg
ggttacctcttatctagtggTGAACTTACTCATCCAGGTCCTGTGCCAGATTCTTGAAAGgaatcttaaacccaactt
ggttacctcttatctagtggCATTCCGTATCTCAAGGTGCTCACACTTTTAATAGGAATTgaatcttaaacccaacttg
ggttacctcttatctagtggGTGTAGCGAACACTTAAATGACAAATCAATTACTCCTGATgaatcttaaacccaactt
ggttacctcttatctagtggGTTAGTATGGTAAGTGGGTGGATGAGTAAGAAGGAAGATCgaatcttaaacccaactt
ggttacctcttatctagtggCCAGAGCAATTTAAGGCATACTCTCTGAGGGACAGGACCCgaatcttaaacccaac
ggttacctcttatctagtggGAAGCTCCAGAGGTGATTCTAATTTTCAGCCAGGGTTGCCgaatcttaaacccaact
ggttacctcttatctagtggGTGATGCTCAACCTTCTTGTGCATCCAAATCACCCTGGGCgaatcttaaacccaact
ggttacctcttatctagtggCTGATTCAGCTGGTCTAGGGTGGGAAACAGGTGTCAGGGGgaatcttaaacccaac
ggttacctcttatctagtggCCCCAGGTGACTCTAATATTCAGCCACGATTAAAAACCACgaatcttaaacccaact
ggttacctcttatctagtggTTGGAGGTGCCAACAGAGTCCTATGTATCATGCTGAGTACgaatcttaaacccaact
ggttacctcttatctagtggCAAACAGCAGCATTATACAGATAAGAAAATAGACACTCTGgaatcttaaacccaactt
ggttacctcttatctagtggTGACAAGAGCTGGAATTTGAACCCAGTTTTGTCTGCTTTCgaatcttaaacccaactt
ggttacctcttatctagtggACCTCAGCATTCATCTATTCGTTTATTGTTAGTCAGTCTGgaatcttaaacccaacttg
ggttacctcttatctagtggTCTCCACAAAGCGGCCACCTACCTCCTTAGGAACACAGCCgaatcttaaacccaa
ggttacctcttatctagtggAATGAACCAGCTTTAAGCCACCCCTAGTACCTCAGGCTGCgaatcttaaacccaac
ggttacctcttatctagtggGCTCTGGACACAACACAGTCCCAGTGGCTTCAAAGAGGGGgaatcttaaacccaa
ggttacctcttatctagtggGTGGCAGGTAGGGTTTTCTGTTGGAGAAAGAGCACCCGGGgaatcttaaacccaac
ggttacctcttatctagtggGTCTTGGTGTGAGAAGCAAGATTTGCCTCCTGGCTACAAGgaatcttaaacccaact
ggttacctcttatctagtggGTCCCAAGAACAACCTGCCCTTTGTGATCAAACCTGCCAGgaatcttaaacccaac
ggttacctcttatctagtggGCCTGTCATCTCAGCAGACAAAGGAGGCCTCCAGCCCCAGgaatcttaaacccaa
ggttacctcttatctagtggCCACTCACCTTGATGGCCTCCTGTTACAGGTGCCTCCCTGgaatcttaaacccaac
ggttacctcttatctagtggGATGCATGCCATCCTGCCATCTGCTCTCAGAGTGTTAGAAgaatcttaaacccaact
ggttacctcttatctagtggCTTGTTGTCCACACACCGAATCCTTAGCCCAGCTTGGAGCgaatcttaaacccaac
ggttacctcttatctagtggTCAGCTTCATCACCTGCTGCTACCCACAGCTTAATGACCCgaatcttaaacccaact
ggttacctcttatctagtggTCCTGCTGCCGAAATTCTCTTCTCCCTTCCTTCCTCTTGGgaatcttaaacccaactt
ggttacctcttatctagtggCTACCGTACTAGGAAGACTTCCTGCTGTACCGGTGAACAGgaatcttaaacccaac
ggttacctcttatctagtggCATACCTGGCTCAGTTGTGACCTCCTCCAGGAAGTCATCCgaatcttaaacccaac
ggttacctcttatctagtggAGCCCTCCTTCATGCTCTCATAATGCTATACTAGTAGCTCgaatcttaaacccaactt
ggttacctcttatctagtggGCAGGTTTTGGGGCCTCATTCCCAGTTACTGACTTAGTAGgaatcttaaacccaactt

ggttacctcttatctagtggCAGTGATGTTGATGCTGCTGGCCCAGGGAACACCCTTGAGgaatcttaaacccaac
ggttacctcttatctagtggATGATTCCACTTGCCCCACTCACTACATCAGTCAATAAGGgaatcttaaacccaact
ggttacctcttatctagtggGTTGGAAAGGGCTGCAGCCTCATCTGAAAGCTCAACTGGGgaatcttaaacccaac
ggttacctcttatctagtggACCGGTGGTTGTTAGTAGGGCTCAATTTTCCACTGGTGGTgaatcttaaacccaactt
ggttacctcttatctagtggGGTAGCTGGCTTTCCCCACAGCAAGTAACCCAAAAGAGACgaatcttaaacccaac
ggttacctcttatctagtggCTGCCATGTGCAGGTGGTAAAACAAACCAACCCTAGTTCAgaatcttaaacccaact
ggttacctcttatctagtggCGAAGGTAAGACCATGAATGCTGGCTATCAATCACATGGGgaatcttaaacccaact
ggttacctcttatctagtggGTAGGGTGTCTAGGGAAAACCTTTTTGGGGAGATGCTATGgaatcttaaacccaactt
ggttacctcttatctagtggCTTAGAAGTAGAGAATTCCAGGCAGAGGAAAAGCAAAGGCgaatcttaaacccaac
ggttacctcttatctagtggCTGGTAGAAGAGCCAGCAAGGAGCAGGAGGGGCAGCTGTTgaatcttaaacccaa
ggttacctcttatctagtggAAGTTCTGGGATACATGTGCCAAATGTGCAGGTTTGTTACgaatcttaaacccaacttg
ggttacctcttatctagtggGCTATCCCTCCCCTTGCCCCCCACCCAACAACAGGCCCCAgaatcttaaacccaa
ggttacctcttatctagtggGAACAGTGCTGCAAGAAACATACGTGTCCATGTGTCTTTAgaatcttaaacccaactt
ggttacctcttatctagtggCATTCTAACTGGTTTGAGATGGTATTGCATTGTGGTTTTGgaatcttaaacccaacttgt
ggttacctcttatctagtggGCATCTACTATGCTCAGGGTCTTATCAGCCATATAAATAAgaatcttaaacccaacttg
ggttacctcttatctagtggCGATCTCGGCTCACTACAGGCTCTGCCCACTGGTTTCACGgaatcttaaacccaac
ggttacctcttatctagtggCCGAGTAGCTGGGACTACAGGCGTGGAAAAATACATCATTgaatcttaaacccaact
ggttacctcttatctagtggAGGAACTGATCTTCCAGTTGAAGAAACCTTTCATGCTTTTgaatcttaaacccaacttg
ggttacctcttatctagtggCTAGATAATTTAGCTTCAATACATTCCCATTACCACACCCgaatcttaaacccaacttg
ggttacctcttatctagtggTAATCAACTCAGGAGCAAGAGTTTGAGGGAAAAAAGTACCgaatcttaaacccaact
ggttacctcttatctagtggGAGCCAGAACCTTTCCAATCTTGGAGAATATGCAACCTGGgaatcttaaacccaact
ggttacctcttatctagtggAAGCTGGAAGAGACTTCAGAGATCATTCCATCCTTCTGTCgaatcttaaacccaactt
ggttacctcttatctagtggCCAGCCCAAGATCTCAAAGACTGTTGGACAGCCCTCCATCgaatcttaaacccaac
ggttacctcttatctagtggTGAGAACCCGTCCCATGCAAGGTGCTCAGGGTAGCAGAGGgaatcttaaacccaa
ggttacctcttatctagtggCTCCCATCTCAGGATTTCTAGGTCAGCCTGTGTTCCATGGgaatcttaaacccaact
ggttacctcttatctagtggCCAGAAGCTGCCTTAGGTGTGGGAAAGAGCAGAGAAAGGGgaatcttaaacccaa
ggttacctcttatctagtggGCCTCCAGGCTCTCCTCTGCTCAACCACAGCAGACACCTTgaatcttaaacccaac
ggttacctcttatctagtggGGAGTTTATAAATGTCTCTGCTAAAAGAAGATTGAAAGCCgaatcttaaacccaacttg
ggttacctcttatctagtggAGAGGTCAGCAAACTTCATCTCCTAAACCCAAACTAGCCCgaatcttaaacccaac
ggttacctcttatctagtggCTCAGTGTCCATAAGTAAAGTTTAGAACACAGCCACACCCgaatcttaaacccaact
ggttacctcttatctagtggGTTGCAGAGCTGGGTAGTTGTGACAGACCATCTGGCCTGCgaatcttaaacccaac
ggttacctcttatctagtggCCCCTGGTACAGCCAAGGGTCCAATAGATGCCTAAATCCCgaatcttaaacccaac
ggttacctcttatctagtggTCTTCCTTCTCTTTTTATTCCTTCTGCTTCCCAGCAAAACgaatcttaaacccaacttg
ggttacctcttatctagtggGGGATCCACAGGTCTGCCTTTGATGCTAGTGTATTTTGGGgaatcttaaacccaactt
ggttacctcttatctagtggAGGGTTTCAGATTCCTGATTCAAACCATCCAAACCCCATCgaatcttaaacccaact
ggttacctcttatctagtggTACCTCCCTACTCTTATCCCCCAGCCAATGAAGCATCTGCgaatcttaaacccaact
ggttacctcttatctagtggAGCTGTTCTTTCCTGTCCCTGTGGTCCATAAAAGCCTGGGgaatcttaaacccaactt
ggttacctcttatctagtggACGAAGTCGAAGTGCATAGACAACTCCACCACCCACAGCTgaatcttaaacccaac
ggttacctcttatctagtggTCTCTCTAGAAGGGCAGATATGTGAAGAGGAGTCTGGCAGgaatcttaaacccaact
ggttacctcttatctagtggTCATTGTCAGAGATAACCTAGGTCTGAGGAAGCAGAGGCCgaatcttaaacccaac
ggttacctcttatctagtggGTGTAAAAGTGTTTGTTCTCACTGCTAACCAAAGACAGGCgaatcttaaacccaactt
ggttacctcttatctagtggAGCTCAGCGTTAGTAAGACTGCAGGGAAAAAGAAACACGCgaatcttaaacccaac
ggttacctcttatctagtggGCCCTTTGACAAAGGAATTCTCCTGGATTCTTCTAAAGGCgaatcttaaacccaactt
ggttacctcttatctagtggAAGTAGAAAGGTAGAACAGAGAACATTGTATACATACTGCgaatcttaaacccaactt
ggttacctcttatctagtggAGGAAGTTTCTGACAGTTCTTTTCTGTTAGCCATGGGGAGgaatcttaaacccaacttg
ggttacctcttatctagtggCCCACAGTGGGTACCTCTCAGGACCGTAATGCAGGGATCCgaatcttaaacccaac
ggttacctcttatctagtggGGCAGAGATAGGGCCACTGAGAAAGTGACTCAGCAGGGCCgaatcttaaacccaa
ggttacctcttatctagtggTAGCAGGGATCCTGACAAGGACATGCGATACCTCATTTCCgaatcttaaacccaact
ggttacctcttatctagtggGGCTCAGAGGTGGCCAAGCCCTGAGCAGAGCCCAGCTTCCgaatcttaaacccaa
ggttacctcttatctagtggTGGGTTGAACTGTATCTCCCACCCCCGCCCTCCCCTCCCCgaatcttaaacccaa
ggttacctcttatctagtggGAGGTTGGACGGGTGTGGGCCCCTAACCCAATACAACTGGgaatcttaaacccaac
ggttacctcttatctagtggGGAAGAGGAGAGCACACAGAGACACACACAGGGAGAGCACgaatcttaaacccaa

ggttacctcttatctagtggGGGAGTGTCCTGTGAGAGAGAGGCACAAATGGAGGGATGCgaatcttaaacccaa
ggttacctcttatctagtggACGTCAAAGATTGCTGGCCATTACCAGAAGCTAAGAAGGGgaatcttaaacccaac
ggttacctcttatctagtggGACCCTGTTGAGTGCCTGATTCTAGACTTTCCGTCTCCAGgaatcttaaacccaact
ggttacctcttatctagtggACCCTGAAAGTCTCTGTGGTCAGCGGAGTGAGTCTGAGCCgaatcttaaacccaac
ggttacctcttatctagtggGCTGCAGCTGGTTGCAACTTGGGTCTTACTGATACAGACCgaatcttaaacccaact
ggttacctcttatctagtggAGTGTGGGTTCTGAGCCAGGCACCTTCTCCCCACAGCAGCgaatcttaaacccaa
ggttacctcttatctagtggTGGGAGGACAGGAGTAAAATGGGGGAAGCGTGGGAAGGCGgaatcttaaacccaa
ggttacctcttatctagtggGTGAGTGAGGAAGCACCTGGATTCAGAGTTCCAATCTCAGgaatcttaaacccaact
ggttacctcttatctagtggGCGGTGATCCTTTAAGTTACTTAACCTCTCTGTGGCTCAGgaatcttaaacccaactt
ggttacctcttatctagtggGGGACTAATAATAGTTTCTCCCTCACAGGGCTATTGTGAGgaatcttaaacccaactt
ggttacctcttatctagtggAACTATAACTTGCATAGAACGGAGCCCAGTGCATAGTGTGgaatcttaaacccaact
ggttacctcttatctagtggCATTACCTCCACCTTTCCATGTATGCCTTTACAGTTATTCgaatcttaaacccaacttg
ggttacctcttatctagtggTATGAGTAGAGAGAGAAACTCAGGCTTAAGAGGTTAAAGGgaatcttaaacccaact
ggttacctcttatctagtggTGGCAAGAAGGAGCAAGGGAGGACAAAGCAGGAATGTGAGgaatcttaaacccaa
ggttacctcttatctagtggCCAGATTACCTGGCGCCATAGTGCATCCTTTGTTCAGTTCgaatcttaaacccaactt
ggttacctcttatctagtggCCTAAGTTCTGGTTGATCTGCCATCAATTTGCTGTACAGCgaatcttaaacccaactt
ggttacctcttatctagtggCTCAACCTCAGCACTGTTGATATTTCAGGTCAGGTTATTCgaatcttaaacccaactt
ggttacctcttatctagtggCTGGCCCTTGTACACTAGATACCAATAGCATCTTTCCCACgaatcttaaacccaact
ggttacctcttatctagtggCTACCAAATATCCTCCAGAAGGGAAAATTGTCCCCAGCTGgaatcttaaacccaact
ggttacctcttatctagtggGTTACCTGTGAAACTGCCTAATTGAGCCATTCTGGGGAGCgaatcttaaacccaact
ggttacctcttatctagtggGAGCATTGTTGAGGCCAAAAGGCTTAGGAAGAAATTACCCgaatcttaaacccaact
ggttacctcttatctagtggGGACATGAGCCACTTATTGAAGGATGGGAACGATTTCTGCgaatcttaaacccaact
ggttacctcttatctagtggCACATCAGGAGCCAAGTGAGAAACCAAGAAGAGACTAAGCgaatcttaaacccaa
ggttacctcttatctagtggTCCTCGTCCTTCAAGGCTCAGTTCAGATGCTACCTCCTCCgaatcttaaacccaact
ggttacctcttatctagtggTGGCAAGATACCGCTTTGCATTGTTCCTCTTTGCTCCCAGgaatcttaaacccaactt
ggttacctcttatctagtggTGCCATTGGCAGTAGGGTGGGTTGGTTCTTCATTGAGTAAgaatcttaaacccaactt
ggttacctcttatctagtggCCTTATCATCGTGATGACCAAAAATCTCCATGACACTATCgaatcttaaacccaactt
ggttacctcttatctagtggGCGCCTGTCTCCTCCTGTCGACAGTAAGCCCCTGGAGGACgaatcttaaacccaa
ggttacctcttatctagtggTCTCAATCATATCTGCATCACTAGGACCTACTCCAGTTCCgaatcttaaacccaactt
ggttacctcttatctagtggGGGGAAAATTGTGAATTCAGAATTGGAACATAAATTGGGGgaatcttaaacccaactt
ggttacctcttatctagtggGTAAGTCAAAATGGTCGATTTTCAGCAGCTGCATATAACCgaatcttaaacccaactt
ggttacctcttatctagtggTGGACACAGGAAAGGGAATATCACAATCTGGGGACTGTGGgaatcttaaacccaac
ggttacctcttatctagtggCTGGCTTAAATACCTACCTTTGGTCTTATCCTAATTAGGCgaatcttaaacccaacttg
ggttacctcttatctagtggGTGCCGTAAGCAATGGGGATGGATCCACTGCAGTCTGAGGgaatcttaaacccaa
ggttacctcttatctagtggGGGAGGGGAGTGTTTTTAAAAGAAGGGGGAAGAAGAGAGGgaatcttaaacccaac
ggttacctcttatctagtggTACAGAGAAGAAGGAAAACGTGGAGAAAAAAGAGAAGGAAgaatcttaaacccaac
ggttacctcttatctagtggGTTCCACTCCAATGCTGCAGTACAATCAGCTCAGAGCCTGgaatcttaaacccaac
ggttacctcttatctagtggCTGACATAGCCTAGCGCCTGCTTATTTTTAGATGAAATTAgaatcttaaacccaacttg
ggttacctcttatctagtggGGTTGCATCTCTCCTTGGCAGGTTCTATATTCAGCTCTGCgaatcttaaacccaactt
ggttacctcttatctagtggCTGCTTCTCCTGCAGCCCGTGAGAGATTGCAGAGAGGGGCgaatcttaaacccaa
ggttacctcttatctagtggGAGTAGCTGGATGGAAGGGAGCTGGAACTGCTTTGATTCCgaatcttaaacccaact
ggttacctcttatctagtggTGGCATTTGCACTGGATCATTCTTTGTTGGGGCCGGGGGCgaatcttaaacccaact
ggttacctcttatctagtggCTGGGGAGATACTCCAAGACACCCCCAGTTGAGAGCCACTgaatcttaaacccaa
ggttacctcttatctagtggCCGTAATCGAGTAACCCAAATTACGGTGGAACAGACTCCCgaatcttaaacccaac
ggttacctcttatctagtggGTCTCCTCCTGCCTCCGGGTCACTACTGGGTATCCTTGAGgaatcttaaacccaac
ggttacctcttatctagtggCCACTGGTAGCCACAGCTGGTACTGCCCTGTCTGGCCTTGgaatcttaaacccaa
ggttacctcttatctagtggCAGCTCACTCCTTCCTTGCTTCATACCAGGCATGCACCATgaatcttaaacccaact
ggttacctcttatctagtggACTCCACACAGCAGAAGAGAGAACCAAATGTAATTGTTTCgaatcttaaacccaact
ggttacctcttatctagtggGCGCTGGGCTGGCTTGTCATCGCACTCCTAGCTTCTCTCCgaatcttaaacccaac
ggttacctcttatctagtggCGCAGCCAAACCATGACTGCTCCTTGCACCCAATACCCACgaatcttaaacccaa
ggttacctcttatctagtggCCCAAGTCCTCACTTGCCACTCCAGGACTCACCTCCTCCCgaatcttaaacccaa
ggttacctcttatctagtggCCCCCCGAAACTGGGCACATGGGACACAAATTACTCAGAAgaatcttaaacccaa

ggttacctcttatctagtggCTCCTCATCAAGCCCAAAGGGAAACATCTGAAGACAACTCgaatcttaaacccaac
ggttacctcttatctagtggAGCCAAGGCCAAGACAGGAAAGAAGAATTGGTGGCAGAGGgaatcttaaacccaa
ggttacctcttatctagtggGCCCAGGAGTCAGCCAGATAAAGTTCACAGGGAGCTGGGCgaatcttaaacccaa
ggttacctcttatctagtggGCCCCCCATGGTGGAAAACAGCTTTGGGAGGCCAAGGTGGgaatcttaaacccaa
ggttacctcttatctagtggATCTCTACTAAAAATACAAGAAATTAGAGGGCATGGTGGCgaatcttaaacccaactt
ggttacctcttatctagtggGGCCAGGTCAGAGTCTGATCTAAGAAGACTAGAGGAGACCgaatcttaaacccaac
ggttacctcttatctagtggCCTGGCTGGGCTGCCCTACGATGGTCCCTCTGACAGCTGGgaatcttaaacccaa
ggttacctcttatctagtggGTGCCAGCCATACTACCCTTTAAAAATAAGTACCAACTCCgaatcttaaacccaact
ggttacctcttatctagtggCACCCTGATCCCCAGGCAGCTTGTTCCCTTGGGACGCCTCgaatcttaaacccaa
ggttacctcttatctagtggCACCCCTGACTTACAGACACAAACGCACACACACTCAGGCgaatcttaaacccaa
ggttacctcttatctagtggGAGGCCTTTGGAAGACGATTGGGTCATGAGGCTGGAGCCCgaatcttaaacccaa
ggttacctcttatctagtggGAGGCCCCATGAGAGCTCTCTAGTTCTCCACCCTGCAGAGgaatcttaaacccaa
ggttacctcttatctagtggGGGTCCCTTACCAGAACTTGCTCATGCTGGTACCCTGATCgaatcttaaacccaac
ggttacctcttatctagtggCTGTGACCAATAATAGTTTATTGTTTGCTGGGCGCGGTGGgaatcttaaacccaactt
ggttacctcttatctagtggGACTACTAGTTTCCACCTTCTCTGTCCGGTACTGAACGTGgaatcttaaacccaact
ggttacctcttatctagtggCCAGTGCTGATGCTGCTGTCTCCTCCTAGCCACAGCCTTGgaatcttaaacccaac
ggttacctcttatctagtggTGGTTTTCAGGTAGAATTTCCTAATCCAGATCCAAATATCgaatcttaaacccaacttg
ggttacctcttatctagtggGAAAAGTCAGATTGGGCTATCCCAGTAAGCTTGCCAGTAGgaatcttaaacccaac
ggttacctcttatctagtggCCATTAGGTTGGGAAGGATGGCTGTTTCTACATGTGCCAGgaatcttaaacccaact
ggttacctcttatctagtggTTCATAGGATGGGCTGTTACTCAAGATTCTGTAAGTTTCAgaatcttaaacccaacttg
ggttacctcttatctagtggTGTGGAGAGATGGGTCTAAGCTGTATGATGGGTTTCTGCCgaatcttaaacccaact
ggttacctcttatctagtggGGCATCTTCCATGCAACACAGACCATGCCAGAGGCTGATCgaatcttaaacccaa
ggttacctcttatctagtggTTCTAAGGTAGAGATGAACACATCTCCTCCTTCAGCTCCTgaatcttaaacccaact
ggttacctcttatctagtggGCCTGGTTTAGGCCATGTGCTCATCCCTGGACAAATCAGAgaatcttaaacccaac
ggttacctcttatctagtggTTGGCCCAGGCTGAATGGAGCCGTTTAGATCTACAGTGAGgaatcttaaacccaac
ggttacctcttatctagtggCAGTTACCGCTCCTATCGAAGCCAGGTACTGTATCCTCTCgaatcttaaacccaac
ggttacctcttatctagtggCGAATAATCTCATAGGTCTCGTATCAATCGTTCCTCCCTCgaatcttaaacccaactt
ggttacctcttatctagtggCATATAGGTGCATACGTCTGAATGTTTGGAAAGGCAGGGGgaatcttaaacccaact
ggttacctcttatctagtggTCTCCTACTACAGATAAGATCAATAAAGCCCCTTTAAATCgaatcttaaacccaactt
ggttacctcttatctagtggCCTGTTAAATCAGCGATTTTTCCCTCTTGTTAGATCATTCgaatcttaaacccaacttg
ggttacctcttatctagtggACTTCCTTAACCTCATGCCCCCCTTCAACAACTGCCTTAGgaatcttaaacccaact
ggttacctcttatctagtggATTGAGGGTTGACTATGCTGTTTCCAACTCTTCACCTCTCgaatcttaaacccaactt
ggttacctcttatctagtggGAAGCTCATCAACATCCTCCATCTGGCCCAATCCAATGGGgaatcttaaacccaac
ggttacctcttatctagtggCTGACTTCTAAGCCTTATTTGACAATTGTGCACACCTCCCgaatcttaaacccaactt
ggttacctcttatctagtggGCCTGCCTGGCTACTCCTTCAGAATCTCCTTGGCTGGGTCgaatcttaaacccaac
ggttacctcttatctagtggTACGCACTCTGCACTCTCTCTGGGGAAACACATCCATGGCgaatcttaaacccaac
ggttacctcttatctagtggATCCGTGTGCTGACTACTCTAGTTGAGAACTCTCCCTTGAgaatcttaaacccaact
ggttacctcttatctagtggAGGCCAGAGCAGAACTTTCCCACCTCCACTTCACATTTGCgaatcttaaacccaac
ggttacctcttatctagtggCTCAGTTGCCCAGCCATCAGCAAGTACTATTGATTCTGCCgaatcttaaacccaact
ggttacctcttatctagtggCTCCTGGACCACTGCAAGAGCCTCCTAGCCATCTCCCTGCgaatcttaaacccaa
ggttacctcttatctagtggTCTCCTTGACAGTTCATTCTCCATCCAGCCACAAGAGTGAgaatcttaaacccaact
ggttacctcttatctagtggTCATGTTTCTCTCTTGCTTAATATTGAACAAGCTCTCCACgaatcttaaacccaacttg
ggttacctcttatctagtggTCCTCTCTGGCATTTTTTCTGTTCCTCCTCCATACCAAGCgaatcttaaacccaactt
ggttacctcttatctagtggGTCCTGTACCACAGTTCTTTACCCAGCTGTCTTTTTTGCCgaatcttaaacccaactt
ggttacctcttatctagtggACCGGAACACTTAGCTGGTACCTCAGGAAATCTTTATCCCgaatcttaaacccaact
ggttacctcttatctagtggTCCTCGGTAGCACTCTTTTAAATGATGCCATTTATTTATGgaatcttaaacccaacttg
ggttacctcttatctagtggCCCCCCACTGGAGTGTGAACAGGTACCATATCTGTCTTATgaatcttaaacccaact
ggttacctcttatctagtggCATTTGCCTGTGTGTCCACAACTCTTTGATCAATATCTGGgaatcttaaacccaactt
ggttacctcttatctagtggAATGCGAAAGAATAAACTTGTGTCTCCCATTGCCCTTTTCgaatcttaaacccaactt
ggttacctcttatctagtggGGATGGTGGGTGGAAGAGTGGCTGGCTGGCTGGCCGGTTGgaatcttaaacccaa
ggttacctcttatctagtggATATCTCCCCATTACTTGCAAATATAGGATAGTGAGCAGGgaatcttaaacccaactt
ggttacctcttatctagtggATCTATAGCTGCATTAACAACCCGCCCAAAATGTCATAGCgaatcttaaacccaact

ggttacctcttatctagtggCTTCGGGTCCACATGGTATTGGCTGGAATGCTGACATGGCgaatcttaaacccaact
ggttacctcttatctagtggGTGGCTGTTGTGGCAGAGTTCAGCTGTTCATCTGGGACTGgaatcttaaacccaact
ggttacctcttatctagtggGGGTTTCTCCTGGCATGGCAGGTGGGTTCCGAGGCTTCCCgaatcttaaacccaac
ggttacctcttatctagtggTGACCTAGCCTTGAAAGTCCCAGGACGTCATTTCTGCCACgaatcttaaacccaact
ggttacctcttatctagtggGACTCCACCTCTTGAAGTGAGCAGTGTTGGTGAAGCCATCgaatcttaaacccaact
ggttacctcttatctagtggACCATACCTGTAAAGAGACAGCCCACTGGGTTCTGCAGGGgaatcttaaacccaac
ggttacctcttatctagtggACCACTTTGGGAATATGCATCTATGTGTTTTACTTTCAGCgaatcttaaacccaacttg

**Primers for probe pro**

| FISH probe region |
| --- |
| Ch13 region 1 |
| Ch13 region 2 |
| Ch13 region 3 |
| Myc (AF647N) |
| Myc (AF488) |

**Fluorescently labeled**

| FISH probe region |
| --- |
| Ch13 region 1 |
| Ch13 region 2 |
| Ch13 region 3 |
| Myc (AF647N) |
| Myc (AF488) |

**›duction**

| Forward primer (5'->3') |
|---|
| TCATAGGTTCATTATAGGAC GCAGTCATATAGAGCAGACG |
| GAATCTTAAACCCAACTTGT GACTTATCAGTCTTGTGTAC |
| GCTGATCTACCACTTAGTTA GTAGAGTGTAGCACCTATCC |
| GAATCTTAAACCCAACTTGT GGTTACCTCTTATCTAGTGG |
| CAGAGTACGATATGTCACGG GGTTACCTCTTATCTAGTGG |

**l detection oligos**

| Readout sequence |
|---|
| GTCCTATAATGAACCTATGA |
| ACAAGTTGGGTTTAAGATTC |
| TAACTAAGTGGTAGATCAGC |
| ACAAGTTGGGTTTAAGATTC |
| CCGTGACATATCGTACTCTG |

| **Reverse primer (5'->3')** |
|---|
| CGATTGAGGCCGGTAATACGACTCACTATAGGG GTCACAGGGCAGTATAACTT |
| CGATTGAGGCCGGTAATACGACTCACTATAGGG GTTTAGGACAAGTAACTATA |
| CGATTGAGGCCGGTAATACGACTCACTATAGGG CCACTAGATAAGAGGTAACC |
| CGATTGAGGCCGGTAATACGACTCACTATAGGG ACAAGTTGGGTTTAAGATTC |
| CGATTGAGGCCGGTAATACGACTCACTATAGGG ACAAGTTGGGTTTAAGATTC |

| **Fluorophore** |
|---|
| Alexa Fluor 790 |
| Alexa Fluor 647N |
| Alexa Fluor 555 |
| Alexa Fluor 647N |
| Alexa Fluor 488 |

| **RT primer (5'->3')** |
|---|
| TCATAGGTTCATTATAGGAC |
| GAATCTTAAACCCAACTTGT |
| GCTGATCTACCACTTAGTTA |
| GAATCTTAAACCCAACTTGT |
| CAGAGTACGATATGTCACGG |

| **Fluorophore position** |
|---|
| 5' |
| 5', 3' |
| 5', 3' |
| 5', 3' |
| 5', 3' |
